## Supplementary Table 4 for "Distinct roles for MNK1 and MNK2 in social and cognitive behavior through kinase-specific regulation of the synaptic proteome and phosphoproteome"

**RNA Sequencing results**

all genes

MNK1KO over WT RNA-Seq

| ensembl | baseMean | log2FoldCh | pvalue | padj | symbol |
|---|---|---|---|---|---|
| ENSMUSG( | 38.39162 | 0.03113 | 0.919564 | 0.990247 | 0610009E02Rik |
| ENSMUSG( | 149.0358 | -0.00179 | 0.992594 | 0.999327 | 0610009L18Rik |
| ENSMUSG( | 47.15189 | -0.02013 | 0.942084 | 0.993415 | 0610010K14Rik |
| ENSMUSG( | 488.1113 | 0.13535 | 0.34004 | 0.777959 | 0610030E20Rik |
| ENSMUSG( | 11.25929 | 0.069201 | 0.903901 | NA | 0610038B21Rik |
| ENSMUSG( | 60.55498 | -0.40319 | 0.104439 | 0.490485 | 0610040B10Rik |
| ENSMUSG( | 31.329 | 0.387663 | 0.25834 | 0.709574 | 0610040F04Rik |
| ENSMUSG( | 454.5432 | 0.114848 | 0.368115 | 0.795774 | 0610040J01Rik |
| ENSMUSG( | 12.93891 | -0.27539 | 0.63303 | NA | 0610043K17Rik |
| ENSMUSG( | 83.84037 | -0.0189 | 0.933713 | 0.993206 | 1110002E22Rik |
| ENSMUSG( | 490.9043 | -0.04483 | 0.694324 | 0.9343 | 1110002L01Rik |
| ENSMUSG( | 1620.367 | 0.019025 | 0.867815 | 0.976631 | 1110004F10Rik |
| ENSMUSG( | 39.51538 | -0.11302 | 0.753949 | 0.950981 | 1110006O24Rik |
| ENSMUSG( | 21.62515 | 0.227461 | 0.6132 | 0.908902 | 1110015O18Rik |
| ENSMUSG( | 97.8957 | -0.06884 | 0.75242 | 0.950536 | 1110019D14Rik |
| ENSMUSG( | 26.83199 | 0.2094 | 0.580434 | 0.896974 | 1110020A21Rik |
| ENSMUSG( | 366.2282 | -0.09857 | 0.56216 | 0.889236 | 1110032F04Rik |
| ENSMUSG( | 368.4378 | -0.05936 | 0.634895 | 0.914078 | 1110038B12Rik |
| ENSMUSG( | 433.8837 | 0.111593 | 0.419305 | 0.821987 | 1110038F14Rik |
| ENSMUSG( | 128.2734 | 0.196891 | 0.260311 | 0.710245 | 1110046J04Rik |
| ENSMUSG( | 180.5493 | 0.000117 | 0.999402 | 0.99989 | 1110059E24Rik |
| ENSMUSG( | 211.9859 | 0.073487 | 0.665913 | 0.927054 | 1110059G10Rik |
| ENSMUSG( | 1735.153 | 0.092117 | 0.304765 | 0.752574 | 1110065P20Rik |
| ENSMUSG( | 107.4967 | 0.045985 | 0.817805 | 0.965785 | 1190005I06Rik |
| ENSMUSG( | 166.822 | 0.07032 | 0.701521 | 0.937441 | 1300002E11Rik |
| ENSMUSG( | 1371.961 | -0.09744 | 0.327935 | 0.772842 | 1500004A13Rik |
| ENSMUSG( | 353.9453 | 0.102733 | 0.446508 | 0.833726 | 1500009L16Rik |
| ENSMUSG( | 3432.101 | -0.10769 | 0.175179 | 0.610399 | 1500011B03Rik |
| ENSMUSG( | 8.071107 | 0.079886 | 0.920831 | NA | 1500012K07Rik |
| ENSMUSG( | 100.1464 | -0.10999 | 0.587938 | 0.900678 | 1500015A07Rik |
| ENSMUSG( | 50.2518 | -0.36705 | 0.175159 | 0.610399 | 1500015L24Rik |
| ENSMUSG( | 87.37377 | 0.116377 | 0.615473 | 0.910481 | 1500026H17Rik |
| ENSMUSG( | 67.73877 | -0.26932 | 0.266307 | 0.716441 | 1500035N22Rik |
| ENSMUSG( | 523.3249 | -0.45892 | 0.005532 | 0.099488 | 1600012H06Rik |
| ENSMUSG( | 664.3217 | -0.02915 | 0.802225 | 0.962119 | 1600014C10Rik |
| ENSMUSG( | 12.29313 | -0.09479 | 0.854077 | NA | 1600014C23Rik |
| ENSMUSG( | 322.4489 | 0.097286 | 0.459215 | 0.840244 | 1600020E01Rik |
| ENSMUSG( | 13.69109 | -0.08988 | 0.883477 | NA | 1600029I14Rik |
| ENSMUSG( | 10.04207 | 0.108021 | 0.865564 | NA | 1700001G11Rik |
| ENSMUSG( | 10.15388 | -0.49842 | 0.386025 | NA | 1700001G17Rik |
| ENSMUSG( | 246.4802 | 0.230029 | 0.138133 | 0.553165 | 1700001O22Rik |
| ENSMUSG( | 27.60484 | -0.83378 | 0.058968 | 0.375231 | 1700003D09Rik |
| ENSMUSG( | 1827.387 | 0.208802 | 0.032103 | 0.273791 | 1700003E16Rik |
| ENSMUSG( | 21.04411 | -0.39439 | 0.350443 | NA | 1700003F12Rik |
| ENSMUSG( | 1340.842 | -1.49752 | 4.90E-42 | 1.99E-38 | 1700003M07Rik |
| ENSMUSG( | 21.71344 | -0.14299 | 0.722569 | 0.943752 | 1700007L15Rik |
| ENSMUSG( | 15.19319 | -0.02893 | 0.950678 | NA | 1700007P06Rik |
| ENSMUSG( | 185.9767 | -0.02727 | 0.880466 | 0.981469 | 1700008J07Rik |
| ENSMUSG( | 46.80159 | -0.22244 | 0.458811 | 0.840244 | 1700008O03Rik |
| ENSMUSG( | 31.27039 | 0.270674 | 0.427375 | 0.82549 | 1700010I14Rik |
| ENSMUSG( | 159.1381 | -0.28103 | 0.095989 | 0.475984 | 1700012D01Rik |
| ENSMUSG( | 14.99817 | -0.31811 | 0.531718 | NA | 1700016A09Rik |
| ENSMUSG( | 428.156 | 0.075244 | 0.666976 | 0.92729 | 1700017B05Rik |
| ENSMUSG( | 8.592125 | -0.17551 | 0.797795 | NA | 1700018A04Rik |
| ENSMUSG( | 277.4782 | 0.27747 | 0.049599 | 0.340062 | 1700019D03Rik |
| ENSMUSG( | 19.04458 | -0.1778 | 0.700867 | NA | 1700022N22Rik |
| ENSMUSG( | 3970.909 | 0.06695 | 0.459161 | 0.840244 | 1700025G04Rik |
| ENSMUSG( | 68.21311 | 0.153302 | 0.535365 | 0.88087 | 1700028E10Rik |
| ENSMUSG( | 31.25923 | -0.27954 | 0.396689 | 0.811742 | 1700028J19Rik |
| ENSMUSG( | 11.29171 | -0.75919 | 0.164429 | NA | 1700028K03Rik |
| ENSMUSG( | 8.358 | 1.042186 | 0.124902 | NA | 1700030C12Rik |
| ENSMUSG( | 135.3942 | 0.110903 | 0.569973 | 0.893041 | 1700030J22Rik |
| ENSMUSG( | 246.6124 | 0.223925 | 0.131987 | 0.544244 | 1700030K09Rik |
| ENSMUSG( | 87.30485 | 0.170163 | 0.442158 | 0.832774 | 1700034H15Rik |
| ENSMUSG( | 155.4663 | -0.07216 | 0.676618 | 0.930542 | 1700037C18Rik |
| ENSMUSG( | 15.17532 | -0.43842 | 0.356194 | NA | 1700040D17Rik |
| ENSMUSG( | 608.8266 | -0.3529 | 0.010099 | 0.143118 | 1700047M11Rik |
| ENSMUSG( | 111.1947 | 0.809543 | 0.001844 | 0.048742 | 1700048O20Rik |
| ENSMUSG( | 17.06533 | -0.23858 | 0.64356 | NA | 1700055D18Rik |
| ENSMUSG( | 35.99608 | 0.095708 | 0.77343 | 0.955447 | 1700063D05Rik |
| ENSMUSG( | 615.4602 | -0.17141 | 0.218859 | 0.664602 | 1700066M21Rik |
| ENSMUSG( | 176.4968 | 0.033026 | 0.845037 | 0.972023 | 1700086O06Rik |
| ENSMUSG( | 215.3781 | -0.06467 | 0.669544 | 0.928147 | 1700088E04Rik |
| ENSMUSG( | 13.4377 | 0.366858 | 0.460147 | NA | 1700093K21Rik |
| ENSMUSG( | 103.9199 | -0.03329 | 0.869318 | 0.97729 | 1700096K18Rik |
| ENSMUSG( | 67.96453 | -0.24501 | 0.292466 | 0.742521 | 1700101I11Rik |
| ENSMUSG( | 13.08954 | -0.57176 | 0.279298 | NA | 1700102H20Rik |
| ENSMUSG( | 24.99193 | 0.36558 | 0.340942 | 0.778775 | 1700102P08Rik |
| ENSMUSG( | 43.94872 | -0.42243 | 0.182543 | 0.619943 | 1700109H08Rik |
| ENSMUSG( | 27.0994 | 0.093364 | 0.797718 | 0.960507 | 1700110C19Rik |
| ENSMUSG( | 19.8335 | 0.184923 | 0.666202 | NA | 1700112D23Rik |
| ENSMUSG( | 22.81732 | 0.165016 | 0.679653 | 0.930983 | 1700112J16Rik |
| ENSMUSG( | 82.55037 | -0.22427 | 0.302605 | 0.750911 | 1700113A16Rik |
| ENSMUSG( | 80.4181 | -0.00177 | 0.993555 | 0.999369 | 1700120C14Rik |
| ENSMUSG( | 507.0953 | 0.064419 | 0.606991 | 0.906685 | 1700123O20Rik |
| ENSMUSG( | 44.45752 | -0.19816 | 0.49527 | 0.860831 | 1700124L16Rik |
| ENSMUSG( | 125.7838 | -0.38383 | 0.04842 | 0.336526 | 1810010H24Rik |
| ENSMUSG( | 198.6125 | -0.23323 | 0.177839 | 0.615225 | 1810014B01Rik |
| ENSMUSG( | 73.42792 | 0.085875 | 0.717711 | 0.941697 | 1810021B22Rik |
| ENSMUSG( | 41.00202 | -0.24963 | 0.428211 | 0.825894 | 1810024B03Rik |
| ENSMUSG( | 819.3509 | 0.08811 | 0.406249 | 0.816525 | 1810030O07Rik |
| ENSMUSG( | 13.0314 | -0.41396 | 0.434718 | NA | 1810034E14Rik |
| ENSMUSG( | 2142.997 | -0.00896 | 0.930173 | 0.991848 | 1810037I17Rik |
| ENSMUSG( | 33.03679 | -0.06439 | 0.854082 | 0.973491 | 1810044D09Rik |
| ENSMUSG( | 817.0139 | -0.19819 | 0.1609 | 0.590706 | 1810055G02Rik |
| ENSMUSG( | 55.30763 | 0.000449 | 0.998746 | 0.99989 | 1810062O18Rik |
| ENSMUSG( | 277.5323 | 0.048001 | 0.740568 | 0.947823 | 2010001A14Rik |
| ENSMUSG( | 424.6456 | -0.07337 | 0.660074 | 0.924627 | 2010315B03Rik |
| ENSMUSG( | 10.74893 | -0.36561 | 0.532879 | NA | 2200002D01Rik |
| ENSMUSG( | 2355.575 | 0.042761 | 0.729124 | 0.944738 | 2210016L21Rik |
| ENSMUSG( | 76.8944 | -0.22404 | 0.346141 | 0.779153 | 2210406O10Rik |
| ENSMUSG( | 471.4707 | 0.047959 | 0.707187 | 0.93949 | 2210408F21Rik |
| ENSMUSG( | 201.633 | 0.202756 | 0.175172 | 0.610399 | 2210408I21Rik |
| ENSMUSG( | 1942.748 | -0.03333 | 0.696943 | 0.935486 | 2210416O15Rik |
| ENSMUSG( | 372.4004 | 0.01845 | 0.891549 | 0.983955 | 2300009A05Rik |
| ENSMUSG( | 27.21316 | 0.623507 | 0.136904 | 0.55149 | 2310002F09Rik |
| ENSMUSG( | 276.0805 | -0.00613 | 0.966124 | 0.995377 | 2310009B15Rik |
| ENSMUSG( | 253.5511 | -0.18415 | 0.215349 | 0.659922 | 2310010J17Rik |
| ENSMUSG( | 942.6112 | 0.061469 | 0.53893 | 0.882849 | 2310011J03Rik |
| ENSMUSG( | 10.77132 | -0.04791 | 0.93025 | NA | 2310016G11Rik |
| ENSMUSG( | 518.8809 | 0.265965 | 0.059609 | 0.377385 | 2310022A10Rik |
| ENSMUSG( | 8453.174 | 0.214528 | 0.042128 | 0.316282 | 2310022B05Rik |
| ENSMUSG( | 27.90817 | -0.99631 | 0.009418 | 0.137123 | 2310030G06Rik |
| ENSMUSG( | 403.2968 | 0.124867 | 0.342626 | 0.778971 | 2310033P09Rik |
| ENSMUSG( | 106.6143 | -0.11243 | 0.602855 | 0.904887 | 2310039H08Rik |
| ENSMUSG( | 38.86285 | -0.04429 | 0.893517 | 0.984512 | 2310040G24Rik |
| ENSMUSG( | 346.7212 | -0.04104 | 0.777744 | 0.956168 | 2310057M21Rik |
| ENSMUSG( | 1245.316 | 0.034074 | 0.82236 | 0.966352 | 2310061I04Rik |
| ENSMUSG( | 1279.588 | -0.27618 | 0.021933 | 0.225738 | 2410002F23Rik |
| ENSMUSG( | 785.1762 | 0.145343 | 0.222479 | 0.668434 | 2410004B18Rik |
| ENSMUSG( | 35.66245 | 0.383255 | 0.271925 | 0.722255 | 2410004I01Rik |
| ENSMUSG( | 70.79305 | -0.65223 | 0.009399 | 0.13706 | 2410004P03Rik |
| ENSMUSG( | 797.9374 | 0.028957 | 0.776886 | 0.955828 | 2410006H16Rik |
| ENSMUSG( | 38.16585 | 0.642475 | 0.041394 | 0.314189 | 2410018L13Rik |
| ENSMUSG( | 107.6258 | -0.02857 | 0.901115 | 0.985633 | 2410022M11Rik |
| ENSMUSG( | 38.4241 | -0.10129 | 0.740177 | 0.947823 | 2500002B13Rik |
| ENSMUSG( | 247.4807 | -0.06959 | 0.621626 | 0.911432 | 2500004C02Rik |
| ENSMUSG( | 340.0288 | -0.02921 | 0.829111 | 0.969286 | 2510002D24Rik |
| ENSMUSG( | 2851.599 | -0.01081 | 0.915651 | 0.988634 | 2510009E07Rik |
| ENSMUSG( | 931.9018 | -0.24762 | 0.079258 | 0.435105 | 2510039O18Rik |
| ENSMUSG( | 41.7959 | -0.39425 | 0.244302 | 0.691435 | 2610005L07Rik |
| ENSMUSG( | 945.8837 | -0.12825 | 0.254493 | 0.704536 | 2610008E11Rik |
| ENSMUSG( | 20.65981 | -0.04776 | 0.906663 | NA | 2610020C07Rik |
| ENSMUSG( | 393.9043 | 0.267993 | 0.066101 | 0.397686 | 2610035D17Rik |
| ENSMUSG( | 17.06571 | -0.06994 | 0.8767 | NA | 2610035F20Rik |
| ENSMUSG( | 203.4525 | 0.132193 | 0.375214 | 0.800786 | 2610306M01Rik |
| ENSMUSG( | 10.94114 | 0.60619 | 0.276323 | NA | 2610307P16Rik |
| ENSMUSG( | 91.80845 | 0.040125 | 0.84613 | 0.972226 | 2610316D01Rik |
| ENSMUSG( | 33.43534 | 0.079489 | 0.823817 | 0.966603 | 2610318N02Rik |
| ENSMUSG( | 821.6239 | -0.03245 | 0.739573 | 0.947768 | 2610507I01Rik |
| ENSMUSG( | 24.69747 | 0.099582 | 0.809439 | 0.963674 | 2700038G22Rik |
| ENSMUSG( | 51.04799 | -0.17149 | 0.523569 | 0.875628 | 2700046A07Rik |
| ENSMUSG( | 217.6566 | 0.092619 | 0.542346 | 0.883182 | 2700049A03Rik |
| ENSMUSG( | 98.85096 | 0.187977 | 0.350444 | 0.782193 | 2700054A10Rik |
| ENSMUSG( | 216.8738 | -0.03938 | 0.80307 | 0.962169 | 2700062C07Rik |
| ENSMUSG( | 198.8965 | -0.14827 | 0.407603 | 0.817728 | 2700097O09Rik |
| ENSMUSG( | 159.2743 | -0.22714 | 0.203226 | 0.643631 | 2810001A02Rik |
| ENSMUSG( | 298.3334 | 0.050565 | 0.723877 | 0.944096 | 2810001G20Rik |
| ENSMUSG( | 942.597 | 0.162033 | 0.133921 | 0.547197 | 2810004N23Rik |
| ENSMUSG( | 784.8646 | -0.05982 | 0.630902 | 0.914073 | 2810013P06Rik |
| ENSMUSG( | 210.3032 | 0.040221 | 0.819914 | 0.965888 | 2810021J22Rik |
| ENSMUSG( | 290.8432 | -0.12373 | 0.538688 | 0.882755 | 2810029C07Rik |
| ENSMUSG( | 273.8348 | -0.07533 | 0.629844 | 0.913965 | 2810032G03Rik |
| ENSMUSG( | 118.9463 | 0.209745 | 0.410289 | 0.818258 | 2810049E08Rik |

MNK2KO over WT RNA-Seq

| ensembl | baseMean | log2FoldCh | pvalue | padj | symbol |
|---|---|---|---|---|---|
| ENSMUSG( | 35.90888 | 0.130089 | 0.705771 | 0.96315 | 0610009E02Rik |
| ENSMUSG( | 125.9145 | -0.23425 | 0.253251 | 0.88018 | 0610009L18Rik |
| ENSMUSG( | 46.10229 | 0.196235 | 0.517417 | 0.930939 | 0610010K14Rik |
| ENSMUSG( | 438.3463 | 0.100556 | 0.516244 | 0.930939 | 0610030E20Rik |
| ENSMUSG( | 7.173468 | 0.314774 | 0.688986 | NA | 0610039K10Rik |
| ENSMUSG( | 55.0151 | -0.42266 | 0.133171 | 0.822912 | 0610040B10Rik |
| ENSMUSG( | 24.40681 | -0.03304 | 0.934413 | 0.99306 | 0610040F04Rik |
| ENSMUSG( | 383.0719 | -0.11593 | 0.429176 | 0.917257 | 0610040J01Rik |
| ENSMUSG( | 11.97634 | -0.23715 | 0.68306 | 0.958678 | 0610043K17Rik |
| ENSMUSG( | 72.78895 | -0.15916 | 0.527299 | 0.934466 | 1110002E22Rik |
| ENSMUSG( | 434.9818 | -0.13139 | 0.322756 | 0.904023 | 1110002L01Rik |
| ENSMUSG( | 1450.717 | -0.03622 | 0.815309 | 0.979198 | 1110004F10Rik |
| ENSMUSG( | 35.60279 | -0.1574 | 0.6844 | 0.958678 | 1110006O24Rik |
| ENSMUSG( | 17.81553 | -0.07112 | 0.881873 | 0.989644 | 1110015O18Rik |
| ENSMUSG( | 77.15497 | -0.54839 | 0.029892 | 0.61992 | 1110019D14Rik |
| ENSMUSG( | 24.51742 | 0.219821 | 0.59259 | 0.945133 | 1110020A21Rik |
| ENSMUSG( | 392.5864 | 0.358477 | 0.074547 | 0.767242 | 1110032F04Rik |
| ENSMUSG( | 347.6255 | 0.045781 | 0.745128 | 0.96951 | 1110038B12Rik |
| ENSMUSG( | 372.67 | -0.05542 | 0.7328 | 0.967228 | 1110038F14Rik |
| ENSMUSG( | 101.0602 | -0.2266 | 0.281462 | 0.892412 | 1110046J04Rik |
| ENSMUSG( | 157.6877 | -0.13123 | 0.472229 | 0.923329 | 1110059E24Rik |
| ENSMUSG( | 192.5406 | 0.056611 | 0.785782 | 0.976285 | 1110059G10Rik |
| ENSMUSG( | 1424.541 | -0.21891 | 0.072334 | 0.763629 | 1110065P20Rik |
| ENSMUSG( | 96.74233 | -0.01377 | 0.956475 | 0.994798 | 1190005I06Rik |
| ENSMUSG( | 134.7715 | -0.29867 | 0.174129 | 0.844317 | 1300002E11Rik |
| ENSMUSG( | 1265.5 | -0.06245 | 0.606768 | 0.949772 | 1500004A13Rik |
| ENSMUSG( | 311.3448 | -0.01075 | 0.953223 | 0.994798 | 1500009L16Rik |
| ENSMUSG( | 3187.423 | -0.05279 | 0.64388 | 0.953903 | 1500011B03Rik |
| ENSMUSG( | 91.83442 | -0.07539 | 0.736999 | 0.967405 | 1500015A07Rik |
| ENSMUSG( | 47.96237 | -0.20769 | 0.478661 | 0.923329 | 1500015L24Rik |
| ENSMUSG( | 70.56823 | -0.2644 | 0.331422 | 0.907861 | 1500026H17Rik |
| ENSMUSG( | 56.87262 | -0.57139 | 0.045481 | 0.693959 | 1500035N22Rik |
| ENSMUSG( | 590.2032 | 0.192508 | 0.296235 | 0.898538 | 1600012H06Rik |
| ENSMUSG( | 580.4727 | -0.15263 | 0.241937 | 0.876528 | 1600014C10Rik |
| ENSMUSG( | 9.537566 | -0.58498 | 0.36444 | NA | 1600014C23Rik |
| ENSMUSG( | 300.9245 | 0.170474 | 0.298158 | 0.898538 | 1600020E01Rik |
| ENSMUSG( | 17.57484 | 0.772409 | 0.138678 | 0.829085 | 1600029I14Rik |
| ENSMUSG( | 10.7404 | -0.0136 | 0.981405 | NA | 1700001G17Rik |
| ENSMUSG( | 216.3209 | 0.133769 | 0.429799 | 0.917257 | 1700001O22Rik |
| ENSMUSG( | 27.44115 | -0.5079 | 0.227355 | 0.868507 | 1700003D09Rik |
| ENSMUSG( | 1583.564 | 0.073915 | 0.545979 | 0.938063 | 1700003E16Rik |
| ENSMUSG( | 18.82029 | -0.45084 | 0.313063 | 0.90048 | 1700003F12Rik |
| ENSMUSG( | 1627.938 | -0.31693 | 0.023333 | 0.585577 | 1700003M07Rik |
| ENSMUSG( | 17.0035 | -0.67912 | 0.159572 | 0.837724 | 1700007L15Rik |
| ENSMUSG( | 11.68001 | -0.57616 | 0.305375 | 0.898538 | 1700007P06Rik |
| ENSMUSG( | 167.227 | -0.08593 | 0.682466 | 0.958678 | 1700008J07Rik |
| ENSMUSG( | 43.01096 | -0.20896 | 0.536121 | 0.936967 | 1700008O03Rik |
| ENSMUSG( | 25.52759 | -0.06202 | 0.88306 | 0.989644 | 1700010I14Rik |
| ENSMUSG( | 14.19932 | 0.681598 | 0.204791 | 0.864412 | 1700012B09Rik |
| ENSMUSG( | 166.1212 | 0.136813 | 0.48919 | 0.925673 | 1700012D01Rik |
| ENSMUSG( | 13.87786 | -0.25645 | 0.646831 | 0.953988 | 1700016A09Rik |
| ENSMUSG( | 399.4197 | 0.147293 | 0.395387 | 0.917257 | 1700017B05Rik |
| ENSMUSG( | 7.186055 | -0.38525 | 0.609881 | NA | 1700018A04Rik |
| ENSMUSG( | 247.3941 | 0.204577 | 0.294319 | 0.898538 | 1700019D03Rik |
| ENSMUSG( | 17.23679 | -0.1548 | 0.767124 | 0.974236 | 1700022N22Rik |
| ENSMUSG( | 3601.136 | 0.054054 | 0.605134 | 0.949082 | 1700025G04Rik |
| ENSMUSG( | 64.8251 | 0.258942 | 0.333886 | 0.907907 | 1700028E10Rik |
| ENSMUSG( | 29.6517 | -0.16776 | 0.642012 | 0.953903 | 1700028J19Rik |
| ENSMUSG( | 13.85917 | 0.228289 | 0.664798 | 0.956766 | 1700028K03Rik |
| ENSMUSG( | 114.9337 | -0.08562 | 0.703035 | 0.962428 | 1700030J22Rik |
| ENSMUSG( | 223.9139 | 0.211933 | 0.228296 | 0.868507 | 1700030K09Rik |
| ENSMUSG( | 78.22543 | 0.124898 | 0.609717 | 0.950397 | 1700034H15Rik |
| ENSMUSG( | 138.7174 | -0.13573 | 0.46957 | 0.923329 | 1700037C18Rik |
| ENSMUSG( | 13.44451 | -0.47379 | 0.425373 | 0.917257 | 1700040D17Rik |
| ENSMUSG( | 654.5671 | 0.148615 | 0.532819 | 0.934795 | 1700047M11Rik |
| ENSMUSG( | 91.12527 | 0.561021 | 0.035587 | 0.664751 | 1700048O20Rik |
| ENSMUSG( | 15.28063 | -0.3177 | 0.53592 | 0.936952 | 1700055D18Rik |
| ENSMUSG( | 25.52508 | -0.68046 | 0.119299 | 0.813384 | 1700063D05Rik |
| ENSMUSG( | 620.1114 | 0.118839 | 0.442009 | 0.921554 | 1700066M21Rik |
| ENSMUSG( | 157.9695 | -0.01137 | 0.951754 | 0.994798 | 1700086O06Rik |
| ENSMUSG( | 191.8293 | -0.13315 | 0.430963 | 0.917257 | 1700088E04Rik |
| ENSMUSG( | 17.6471 | 1.187465 | 0.012999 | 0.458156 | 1700093K21Rik |
| ENSMUSG( | 90.32504 | -0.19052 | 0.445321 | 0.921734 | 1700096K18Rik |
| ENSMUSG( | 52.17276 | -0.8399 | 0.003046 | 0.219421 | 1700101I11Rik |
| ENSMUSG( | 12.11559 | -0.47128 | 0.418245 | 0.917257 | 1700102H20Rik |
| ENSMUSG( | 26.42654 | 0.701991 | 0.083575 | 0.77681 | 1700102P08Rik |
| ENSMUSG( | 48.1451 | 0.175767 | 0.631732 | 0.953251 | 1700109H08Rik |
| ENSMUSG( | 25.45028 | 0.158112 | 0.693325 | 0.96076 | 1700110C19Rik |
| ENSMUSG( | 19.24091 | 0.285449 | 0.573995 | 0.942912 | 1700112D23Rik |
| ENSMUSG( | 19.38641 | -0.06727 | 0.88675 | 0.990118 | 1700112J16Rik |
| ENSMUSG( | 74.75288 | -0.24265 | 0.330858 | 0.907513 | 1700113A16Rik |
| ENSMUSG( | 74.81177 | 0.06017 | 0.813441 | 0.979198 | 1700120C14Rik |
| ENSMUSG( | 429.8281 | -0.14469 | 0.295766 | 0.898538 | 1700123O20Rik |
| ENSMUSG( | 41.84819 | -0.11052 | 0.725947 | 0.966887 | 1700124L16Rik |
| ENSMUSG( | 10.37446 | 0.326925 | 0.624013 | NA | 1810009A15Rik |
| ENSMUSG( | 130.6986 | 0.035713 | 0.867357 | 0.987764 | 1810010H24Rik |
| ENSMUSG( | 164.0668 | -0.56458 | 0.002192 | 0.183242 | 1810014B01Rik |
| ENSMUSG( | 61.27309 | -0.17132 | 0.513453 | 0.930482 | 1810021B22Rik |
| ENSMUSG( | 36.06555 | -0.36749 | 0.343589 | 0.910026 | 1810024B03Rik |
| ENSMUSG( | 757.844 | 0.125472 | 0.329233 | 0.907083 | 1810030O07Rik |
| ENSMUSG( | 12.35341 | -0.34934 | 0.544682 | 0.938063 | 1810034E14Rik |
| ENSMUSG( | 1773.886 | -0.30633 | 0.025052 | 0.594627 | 1810037I17Rik |
| ENSMUSG( | 27.59754 | -0.34476 | 0.38562 | 0.917257 | 1810044D09Rik |
| ENSMUSG( | 853.8696 | 0.203838 | 0.205703 | 0.864412 | 1810055G02Rik |
| ENSMUSG( | 50.90192 | 0.029086 | 0.920697 | 0.992644 | 1810062O18Rik |
| ENSMUSG( | 217.6476 | -0.43062 | 0.028438 | 0.613486 | 2010001A14Rik |
| ENSMUSG( | 395.6973 | -0.01155 | 0.94853 | 0.994798 | 2010315B03Rik |
| ENSMUSG( | 10.79615 | -0.04633 | 0.937755 | NA | 2200002D01Rik |
| ENSMUSG( | 2016.18 | -0.14348 | 0.315282 | 0.901306 | 2210016L21Rik |
| ENSMUSG( | 65.81789 | -0.4389 | 0.103948 | 0.804969 | 2210406O10Rik |
| ENSMUSG( | 398.5079 | -0.17679 | 0.198762 | 0.862972 | 2210408F21Rik |
| ENSMUSG( | 184.8759 | 0.215527 | 0.265263 | 0.886134 | 2210408I21Rik |
| ENSMUSG( | 1770.373 | -0.03519 | 0.755219 | 0.971098 | 2210416O15Rik |
| ENSMUSG( | 330.6781 | -0.06088 | 0.688552 | 0.959742 | 2300009A05Rik |
| ENSMUSG( | 10.22416 | 0.306203 | 0.612133 | NA | 2310001H17Rik |
| ENSMUSG( | 19.3654 | 0.000911 | 0.998541 | 0.999396 | 2310002F09Rik |
| ENSMUSG( | 239.7113 | -0.15841 | 0.359964 | 0.914223 | 2310009B15Rik |
| ENSMUSG( | 221.5409 | -0.3232 | 0.079421 | 0.77458 | 2310010J17Rik |
| ENSMUSG( | 781.1584 | -0.22507 | 0.076374 | 0.770504 | 2310011J03Rik |
| ENSMUSG( | 10.65363 | 0.168702 | 0.77243 | NA | 2310016G11Rik |
| ENSMUSG( | 438.1073 | 0.05542 | 0.735859 | 0.967344 | 2310022A10Rik |
| ENSMUSG( | 6812.553 | -0.1376 | 0.305974 | 0.898538 | 2310022B05Rik |
| ENSMUSG( | 29.28496 | -0.48692 | 0.208176 | 0.864412 | 2310030G06Rik |
| ENSMUSG( | 346.7143 | -0.04486 | 0.789904 | 0.976798 | 2310033P09Rik |
| ENSMUSG( | 95.31919 | -0.18623 | 0.462601 | 0.923329 | 2310039H08Rik |
| ENSMUSG( | 32.7073 | -0.3249 | 0.419958 | 0.917257 | 2310040G24Rik |
| ENSMUSG( | 316.1109 | -0.03357 | 0.8379 | 0.981376 | 2310057M21Rik |
| ENSMUSG( | 1130.758 | 0.019494 | 0.90422 | 0.991089 | 2310061I04Rik |
| ENSMUSG( | 1229.378 | -0.1144 | 0.38458 | 0.917257 | 2410002F23Rik |
| ENSMUSG( | 652.4927 | -0.12546 | 0.39055 | 0.917257 | 2410004B18Rik |
| ENSMUSG( | 24.6624 | -0.41527 | 0.301557 | 0.898538 | 2410004I01Rik |
| ENSMUSG( | 83.4108 | 0.177572 | 0.526545 | 0.934374 | 2410004P03Rik |
| ENSMUSG( | 700.2255 | -0.08112 | 0.510222 | 0.929089 | 2410006H16Rik |
| ENSMUSG( | 28.06423 | 0.075107 | 0.843211 | 0.983281 | 2410018L13Rik |
| ENSMUSG( | 95.84499 | -0.10163 | 0.690086 | 0.960071 | 2410022M11Rik |
| ENSMUSG( | 36.59478 | 0.004916 | 0.98842 | 0.997862 | 2500002B13Rik |
| ENSMUSG( | 238.1865 | 0.07621 | 0.698119 | 0.960867 | 2500004C02Rik |
| ENSMUSG( | 296.0093 | -0.17165 | 0.34522 | 0.91045 | 2510002D24Rik |
| ENSMUSG( | 2578.569 | -0.03597 | 0.808173 | 0.978622 | 2510009E07Rik |
| ENSMUSG( | 1012.153 | 0.26506 | 0.119156 | 0.813384 | 2510039O18Rik |
| ENSMUSG( | 42.44941 | -0.04709 | 0.899616 | 0.990558 | 2610005L07Rik |
| ENSMUSG( | 952.002 | 0.160599 | 0.201128 | 0.86439 | 2610008E11Rik |
| ENSMUSG( | 22.39752 | 0.40473 | 0.363255 | 0.914223 | 2610020C07Rik |
| ENSMUSG( | 339.4235 | 0.114245 | 0.482235 | 0.924578 | 2610035D17Rik |
| ENSMUSG( | 15.15181 | -0.13457 | 0.786451 | 0.976303 | 2610035F20Rik |
| ENSMUSG( | 170.0158 | -0.13291 | 0.481676 | 0.924578 | 2610306M01Rik |
| ENSMUSG( | 101.0581 | 0.548872 | 0.0189 | 0.548832 | 2610316D01Rik |
| ENSMUSG( | 27.65501 | -0.24022 | 0.559147 | 0.939587 | 2610318N02Rik |
| ENSMUSG( | 726.8102 | -0.11994 | 0.310845 | 0.899422 | 2610507I01Rik |
| ENSMUSG( | 22.54625 | 0.126733 | 0.78603 | 0.976285 | 2700038G22Rik |
| ENSMUSG( | 53.64821 | 0.224957 | 0.425318 | 0.917257 | 2700046A07Rik |
| ENSMUSG( | 198.7571 | 0.099973 | 0.590908 | 0.944975 | 2700049A03Rik |
| ENSMUSG( | 91.59647 | 0.210828 | 0.418857 | 0.917257 | 2700054A10Rik |
| ENSMUSG( | 198.9707 | -0.02287 | 0.898211 | 0.990558 | 2700062C07Rik |
| ENSMUSG( | 205.2055 | 0.218042 | 0.273557 | 0.889848 | 2700097O09Rik |
| ENSMUSG( | 147.3632 | -0.17377 | 0.377977 | 0.916784 | 2810001A02Rik |
| ENSMUSG( | 257.8732 | -0.11309 | 0.54018 | 0.937383 | 2810001G20Rik |
| ENSMUSG( | 811.3475 | -0.00267 | 0.984073 | 0.996963 | 2810004N23Rik |
| ENSMUSG( | 699.4747 | -0.12922 | 0.352188 | 0.912093 | 2810013P06Rik |
| ENSMUSG( | 192.0801 | 0.047587 | 0.806083 | 0.977674 | 2810021J22Rik |
| ENSMUSG( | 255.999 | -0.22976 | 0.248031 | 0.877904 | 2810029C07Rik |
| ENSMUSG( | 233.9105 | -0.27628 | 0.1205 | 0.813384 | 2810032G03Rik |
| ENSMUSG( | 107.054 | 0.174549 | 0.490723 | 0.92646 | 2810049E08Rik |
| ENSMUSG( | 29.69274 | -0.11202 | 0.777609 | 0.976024 | 2810403D21Rik |

genes present in both MNK1KO and MNK2KO

| MNK1KO over WT RNA-Seq | | | | | | MNK2KO over WT RNA-Seq | | | | | |
|---|---|---|---|---|---|---|---|---|---|---|---|
| ensembl | symbol | baseMean | log2FoldCh | pvalue | padj | ensembl | symbol | baseMean | log2FoldCh | pvalue | padj |
| ENSMUSG( | 0610009E0 | 38.39162 | 0.03113 | 0.919564 | 0.990247 | ENSMUSG( | 0610009E0 | 35.90888 | 0.130089 | 0.705771 | 0.96315 |
| ENSMUSG( | 0610009L1 | 149.0358 | -0.00179 | 0.992594 | 0.999327 | ENSMUSG( | 0610009L1 | 125.9145 | -0.23425 | 0.253251 | 0.88018 |
| ENSMUSG( | 0610010K1 | 47.15189 | -0.02013 | 0.942084 | 0.993415 | ENSMUSG( | 0610010K1 | 46.10229 | 0.196235 | 0.517417 | 0.930939 |
| ENSMUSG( | 0610030E2 | 488.1113 | 0.13535 | 0.34004 | 0.777959 | ENSMUSG( | 0610030E2 | 438.3463 | 0.100556 | 0.516244 | 0.930939 |
| ENSMUSG( | 0610040B1 | 60.55498 | -0.40319 | 0.104439 | 0.490485 | ENSMUSG( | 0610040B1 | 55.0151 | -0.42266 | 0.133171 | 0.822912 |
| ENSMUSG( | 0610040F0 | 31.329 | 0.387663 | 0.25834 | 0.709574 | ENSMUSG( | 0610040F0 | 24.40681 | -0.03304 | 0.934413 | 0.99306 |
| ENSMUSG( | 0610040J0 | 454.5432 | 0.114848 | 0.368115 | 0.795774 | ENSMUSG( | 0610040J0 | 383.0719 | -0.11593 | 0.429176 | 0.917257 |
| ENSMUSG( | 0610043K1 | 12.93891 | -0.27539 | 0.63303 | NA | ENSMUSG( | 0610043K1 | 11.97634 | -0.23715 | 0.68306 | 0.958678 |
| ENSMUSG( | 1110002E2 | 83.84037 | -0.0189 | 0.933713 | 0.993206 | ENSMUSG( | 1110002E2 | 72.78895 | -0.15916 | 0.527299 | 0.934466 |
| ENSMUSG( | 1110002L0 | 490.9043 | -0.04483 | 0.694324 | 0.9343 | ENSMUSG( | 1110002L0 | 434.9818 | -0.13139 | 0.322756 | 0.904023 |
| ENSMUSG( | 1110004F1 | 1620.367 | 0.019025 | 0.867815 | 0.976631 | ENSMUSG( | 1110004F1 | 1450.717 | -0.03622 | 0.815309 | 0.979198 |
| ENSMUSG( | 1110006O2 | 39.51538 | -0.11302 | 0.753949 | 0.950981 | ENSMUSG( | 1110006O2 | 35.60279 | -0.1574 | 0.6844 | 0.958678 |
| ENSMUSG( | 1110015O1 | 21.62515 | 0.227461 | 0.6132 | 0.908902 | ENSMUSG( | 1110015O1 | 17.81553 | -0.07112 | 0.881873 | 0.989644 |
| ENSMUSG( | 1110019D1 | 97.8957 | -0.06884 | 0.75242 | 0.950536 | ENSMUSG( | 1110019D1 | 77.15497 | -0.54839 | 0.029892 | 0.61992 |
| ENSMUSG( | 1110020A2 | 26.83199 | 0.2094 | 0.580434 | 0.896974 | ENSMUSG( | 1110020A2 | 24.51742 | 0.219821 | 0.59259 | 0.945133 |
| ENSMUSG( | 1110032F0 | 366.2282 | -0.09857 | 0.56216 | 0.889236 | ENSMUSG( | 1110032F0 | 392.5864 | 0.358477 | 0.074547 | 0.767242 |
| ENSMUSG( | 1110038B1 | 368.4378 | -0.05936 | 0.634895 | 0.914078 | ENSMUSG( | 1110038B1 | 347.6255 | 0.045781 | 0.745128 | 0.96951 |
| ENSMUSG( | 1110038F1 | 433.8837 | 0.111593 | 0.419305 | 0.821987 | ENSMUSG( | 1110038F1 | 372.67 | -0.05542 | 0.7328 | 0.967228 |
| ENSMUSG( | 1110046J0 | 128.2734 | 0.196891 | 0.260311 | 0.710245 | ENSMUSG( | 1110046J0 | 101.0602 | -0.2266 | 0.281462 | 0.892412 |
| ENSMUSG( | 1110059E2 | 180.5493 | 0.000117 | 0.999402 | 0.99989 | ENSMUSG( | 1110059E2 | 157.6877 | -0.13123 | 0.472229 | 0.923329 |
| ENSMUSG( | 1110059G1 | 211.9859 | 0.073487 | 0.665913 | 0.927054 | ENSMUSG( | 1110059G1 | 192.5406 | 0.056611 | 0.785782 | 0.976285 |
| ENSMUSG( | 1110065P2 | 1735.153 | 0.092117 | 0.304765 | 0.752574 | ENSMUSG( | 1110065P2 | 1424.541 | -0.21891 | 0.072334 | 0.763629 |
| ENSMUSG( | 1190005I0 | 107.4967 | 0.045985 | 0.817805 | 0.965785 | ENSMUSG( | 1190005I0 | 96.74233 | -0.01377 | 0.956475 | 0.994798 |
| ENSMUSG( | 1300002E1 | 166.822 | 0.07032 | 0.701521 | 0.937441 | ENSMUSG( | 1300002E1 | 134.7715 | -0.29867 | 0.174129 | 0.844317 |
| ENSMUSG( | 1500004A1 | 1371.961 | -0.09744 | 0.327935 | 0.772842 | ENSMUSG( | 1500004A1 | 1265.5 | -0.06245 | 0.606768 | 0.949772 |
| ENSMUSG( | 1500009L1 | 353.9453 | 0.102733 | 0.446508 | 0.833726 | ENSMUSG( | 1500009L1 | 311.3448 | -0.01075 | 0.953223 | 0.994798 |
| ENSMUSG( | 1500011B0 | 3432.101 | -0.10769 | 0.175179 | 0.610399 | ENSMUSG( | 1500011B0 | 3187.423 | -0.05279 | 0.64388 | 0.953903 |
| ENSMUSG( | 1500015A0 | 100.1464 | -0.10999 | 0.587938 | 0.900678 | ENSMUSG( | 1500015A0 | 91.83442 | -0.07539 | 0.736999 | 0.967405 |
| ENSMUSG( | 1500015L2 | 50.2518 | -0.36705 | 0.175159 | 0.610399 | ENSMUSG( | 1500015L2 | 47.96237 | -0.20769 | 0.478661 | 0.923329 |
| ENSMUSG( | 1500026H1 | 87.37377 | 0.116377 | 0.615473 | 0.910481 | ENSMUSG( | 1500026H1 | 70.56823 | -0.2644 | 0.331422 | 0.907861 |
| ENSMUSG( | 1500035N2 | 67.73877 | -0.26932 | 0.266307 | 0.716441 | ENSMUSG( | 1500035N2 | 56.87262 | -0.57139 | 0.045481 | 0.693959 |
| ENSMUSG( | 1600012H0 | 523.3249 | -0.45892 | 0.005532 | 0.099488 | ENSMUSG( | 1600012H0 | 590.2032 | 0.192508 | 0.296235 | 0.898538 |
| ENSMUSG( | 1600014C1 | 664.3217 | -0.02915 | 0.802225 | 0.962119 | ENSMUSG( | 1600014C1 | 580.4727 | -0.15263 | 0.241937 | 0.876528 |
| ENSMUSG( | 1600014C2 | 12.29313 | -0.09479 | 0.854077 | NA | ENSMUSG( | 1600014C2 | 9.537566 | -0.58498 | 0.36444 | NA |
| ENSMUSG( | 1600020E0 | 322.4489 | 0.097286 | 0.459215 | 0.840244 | ENSMUSG( | 1600020E0 | 300.9245 | 0.170474 | 0.298158 | 0.898538 |
| ENSMUSG( | 1600029I1 | 13.69109 | -0.08988 | 0.883477 | NA | ENSMUSG( | 1600029I1 | 17.57484 | 0.772409 | 0.138678 | 0.829085 |
| ENSMUSG( | 1700001G1 | 10.15388 | -0.49842 | 0.386025 | NA | ENSMUSG( | 1700001G1 | 10.7404 | -0.0136 | 0.981405 | NA |
| ENSMUSG( | 1700001O2 | 246.4802 | 0.230029 | 0.138133 | 0.553165 | ENSMUSG( | 1700001O2 | 216.3209 | 0.133769 | 0.429799 | 0.917257 |
| ENSMUSG( | 1700003D0 | 27.60484 | -0.83378 | 0.058968 | 0.375231 | ENSMUSG( | 1700003D0 | 27.44115 | -0.5079 | 0.227355 | 0.868507 |
| ENSMUSG( | 1700003E1 | 1827.387 | 0.208802 | 0.032103 | 0.273791 | ENSMUSG( | 1700003E1 | 1583.564 | 0.073915 | 0.545979 | 0.938063 |
| ENSMUSG( | 1700003F1 | 21.04411 | -0.39439 | 0.350443 | NA | ENSMUSG( | 1700003F1 | 18.82029 | -0.45084 | 0.313063 | 0.90048 |
| ENSMUSG( | 1700003M | 1340.842 | -1.49752 | 4.90E-42 | 1.99E-38 | ENSMUSG( | 1700003M | 1627.938 | -0.31693 | 0.023333 | 0.585577 |
| ENSMUSG( | 1700007L1 | 21.71344 | -0.14299 | 0.722569 | 0.943752 | ENSMUSG( | 1700007L1 | 17.0035 | -0.67912 | 0.159572 | 0.837724 |
| ENSMUSG( | 1700007P0 | 15.19319 | -0.02893 | 0.950678 | NA | ENSMUSG( | 1700007P0 | 11.68001 | -0.57616 | 0.305375 | 0.898538 |
| ENSMUSG( | 1700008J0 | 185.9767 | -0.02727 | 0.880466 | 0.981469 | ENSMUSG( | 1700008J0 | 167.227 | -0.08593 | 0.682466 | 0.958678 |
| ENSMUSG( | 1700008O0 | 46.80159 | -0.22244 | 0.458811 | 0.840244 | ENSMUSG( | 1700008O0 | 43.01096 | -0.20896 | 0.536121 | 0.936967 |
| ENSMUSG( | 1700010I1 | 31.27039 | 0.270674 | 0.427375 | 0.82549 | ENSMUSG( | 1700010I1 | 25.52759 | -0.06202 | 0.88306 | 0.989644 |
| ENSMUSG( | 1700012D0 | 159.1381 | -0.28103 | 0.095989 | 0.475984 | ENSMUSG( | 1700012D0 | 166.1212 | 0.136813 | 0.48919 | 0.925673 |
| ENSMUSG( | 1700016A0 | 14.99817 | -0.31811 | 0.531718 | NA | ENSMUSG( | 1700016A0 | 13.87786 | -0.25645 | 0.646831 | 0.953988 |
| ENSMUSG( | 1700017B0 | 428.156 | 0.075244 | 0.666976 | 0.92729 | ENSMUSG( | 1700017B0 | 399.4197 | 0.147293 | 0.395387 | 0.917257 |
| ENSMUSG( | 1700018A0 | 8.592125 | -0.17551 | 0.797795 | NA | ENSMUSG( | 1700018A0 | 7.186055 | -0.38525 | 0.609881 | NA |
| ENSMUSG( | 1700019D0 | 277.4782 | 0.27747 | 0.049599 | 0.340062 | ENSMUSG( | 1700019D0 | 247.3941 | 0.204577 | 0.294319 | 0.898538 |
| ENSMUSG( | 1700022N2 | 19.04458 | -0.1778 | 0.700867 | NA | ENSMUSG( | 1700022N2 | 17.23679 | -0.1548 | 0.767124 | 0.974236 |
| ENSMUSG( | 1700025G0 | 3970.909 | 0.06695 | 0.459161 | 0.840244 | ENSMUSG( | 1700025G0 | 3601.136 | 0.054054 | 0.605134 | 0.949082 |
| ENSMUSG( | 1700028E1 | 68.21311 | 0.153302 | 0.535365 | 0.88087 | ENSMUSG( | 1700028E1 | 64.8251 | 0.258942 | 0.333886 | 0.907907 |
| ENSMUSG( | 1700028J1 | 31.25923 | -0.27954 | 0.396689 | 0.811742 | ENSMUSG( | 1700028J1 | 29.6517 | -0.16776 | 0.642012 | 0.953903 |
| ENSMUSG( | 1700028K0 | 11.29171 | -0.75919 | 0.164429 | NA | ENSMUSG( | 1700028K0 | 13.85917 | 0.228289 | 0.664798 | 0.956766 |
| ENSMUSG( | 1700030J2 | 135.3942 | 0.110903 | 0.569973 | 0.893041 | ENSMUSG( | 1700030J2 | 114.9337 | -0.08562 | 0.703035 | 0.962428 |
| ENSMUSG( | 1700030K0 | 246.6124 | 0.223925 | 0.131987 | 0.544244 | ENSMUSG( | 1700030K0 | 223.9139 | 0.211933 | 0.228296 | 0.868507 |
| ENSMUSG( | 1700034H1 | 87.30485 | 0.170163 | 0.442158 | 0.832774 | ENSMUSG( | 1700034H1 | 78.22543 | 0.124898 | 0.609717 | 0.950397 |
| ENSMUSG( | 1700037C1 | 155.4663 | -0.07216 | 0.676618 | 0.930542 | ENSMUSG( | 1700037C1 | 138.7174 | -0.13573 | 0.46957 | 0.923329 |
| ENSMUSG( | 1700040D1 | 15.17532 | -0.43842 | 0.356194 | NA | ENSMUSG( | 1700040D1 | 13.44451 | -0.47379 | 0.425373 | 0.917257 |
| ENSMUSG( | 1700047M | 608.8266 | -0.3529 | 0.010099 | 0.143118 | ENSMUSG( | 1700047M | 654.5671 | 0.148615 | 0.532819 | 0.934795 |
| ENSMUSG( | 1700048O2 | 111.1947 | 0.809543 | 0.001844 | 0.048742 | ENSMUSG( | 1700048O2 | 91.12527 | 0.561021 | 0.035587 | 0.664751 |
| ENSMUSG( | 1700055D1 | 17.06533 | -0.23858 | 0.64356 | NA | ENSMUSG( | 1700055D1 | 15.28063 | -0.3177 | 0.53592 | 0.936952 |
| ENSMUSG( | 1700063D0 | 35.99608 | 0.095708 | 0.77343 | 0.955447 | ENSMUSG( | 1700063D0 | 25.52508 | -0.68046 | 0.119299 | 0.813384 |
| ENSMUSG( | 1700066M | 615.4602 | -0.17141 | 0.218859 | 0.664602 | ENSMUSG( | 1700066M | 620.1114 | 0.118839 | 0.442009 | 0.921554 |
| ENSMUSG( | 1700086O0 | 176.4968 | 0.033026 | 0.845037 | 0.972023 | ENSMUSG( | 1700086O0 | 157.9695 | -0.01137 | 0.951754 | 0.994798 |
| ENSMUSG( | 1700088E0 | 215.3781 | -0.06467 | 0.669544 | 0.928147 | ENSMUSG( | 1700088E0 | 191.8293 | -0.13315 | 0.430963 | 0.917257 |
| ENSMUSG( | 1700093K2 | 13.4377 | 0.366858 | 0.460147 | NA | ENSMUSG( | 1700093K2 | 17.6471 | 1.187465 | 0.012999 | 0.458156 |
| ENSMUSG( | 1700096K1 | 103.9199 | -0.03329 | 0.869318 | 0.97729 | ENSMUSG( | 1700096K1 | 90.32504 | -0.19052 | 0.445321 | 0.921734 |
| ENSMUSG( | 1700101I1 | 67.96453 | -0.24501 | 0.292466 | 0.742521 | ENSMUSG( | 1700101I1 | 52.17276 | -0.8399 | 0.003046 | 0.219421 |
| ENSMUSG( | 1700102H2 | 13.08954 | -0.57176 | 0.279298 | NA | ENSMUSG( | 1700102H2 | 12.11559 | -0.47128 | 0.418245 | 0.917257 |
| ENSMUSG( | 1700102P0 | 24.99193 | 0.36558 | 0.340942 | 0.778775 | ENSMUSG( | 1700102P0 | 26.42654 | 0.701991 | 0.083575 | 0.77681 |
| ENSMUSG( | 1700109H0 | 43.94872 | -0.42243 | 0.182543 | 0.619943 | ENSMUSG( | 1700109H0 | 48.1451 | 0.175767 | 0.631732 | 0.953251 |
| ENSMUSG( | 1700110C1 | 27.0994 | 0.093364 | 0.797718 | 0.960507 | ENSMUSG( | 1700110C1 | 25.45028 | 0.158112 | 0.693325 | 0.96076 |
| ENSMUSG( | 1700112D2 | 19.8335 | 0.184923 | 0.666202 | NA | ENSMUSG( | 1700112D2 | 19.24091 | 0.285449 | 0.573995 | 0.942912 |
| ENSMUSG( | 1700112J1 | 22.81732 | 0.165016 | 0.679653 | 0.930983 | ENSMUSG( | 1700112J1 | 19.38641 | -0.06727 | 0.88675 | 0.990118 |
| ENSMUSG( | 1700113A1 | 82.55037 | -0.22427 | 0.302605 | 0.750911 | ENSMUSG( | 1700113A1 | 74.75288 | -0.24265 | 0.330858 | 0.907513 |
| ENSMUSG( | 1700120C1 | 80.4181 | -0.00177 | 0.993555 | 0.999369 | ENSMUSG( | 1700120C1 | 74.81177 | 0.06017 | 0.813441 | 0.979198 |
| ENSMUSG( | 1700123O2 | 507.0953 | 0.064419 | 0.606991 | 0.906685 | ENSMUSG( | 1700123O2 | 429.8281 | -0.14469 | 0.295766 | 0.898538 |
| ENSMUSG( | 1700124L1 | 44.45752 | -0.19816 | 0.49527 | 0.860831 | ENSMUSG( | 1700124L1 | 41.84819 | -0.11052 | 0.725947 | 0.966887 |
| ENSMUSG( | 1810010H2 | 125.7838 | -0.38383 | 0.04842 | 0.336526 | ENSMUSG( | 1810010H2 | 130.6986 | 0.035713 | 0.867357 | 0.987764 |
| ENSMUSG( | 1810014B0 | 198.6125 | -0.23323 | 0.177839 | 0.615225 | ENSMUSG( | 1810014B0 | 164.0668 | -0.56458 | 0.002192 | 0.183242 |
| ENSMUSG( | 1810021B2 | 73.42792 | 0.085875 | 0.717711 | 0.941697 | ENSMUSG( | 1810021B2 | 61.27309 | -0.17132 | 0.513453 | 0.930482 |
| ENSMUSG( | 1810024B0 | 41.00202 | -0.24963 | 0.428211 | 0.825894 | ENSMUSG( | 1810024B0 | 36.06555 | -0.36749 | 0.343589 | 0.910026 |
| ENSMUSG( | 1810030O0 | 819.3509 | 0.08811 | 0.406249 | 0.816525 | ENSMUSG( | 1810030O0 | 757.844 | 0.125472 | 0.329233 | 0.907083 |
| ENSMUSG( | 1810034E1 | 13.0314 | -0.41396 | 0.434718 | NA | ENSMUSG( | 1810034E1 | 12.35341 | -0.34934 | 0.544682 | 0.938063 |
| ENSMUSG( | 1810037I1 | 2142.997 | -0.00896 | 0.930173 | 0.991848 | ENSMUSG( | 1810037I1 | 1773.886 | -0.30633 | 0.025052 | 0.594627 |
| ENSMUSG( | 1810044D0 | 33.03679 | -0.06439 | 0.854082 | 0.973491 | ENSMUSG( | 1810044D0 | 27.59754 | -0.34476 | 0.38562 | 0.917257 |
| ENSMUSG( | 1810055G0 | 817.0139 | -0.19819 | 0.1609 | 0.590706 | ENSMUSG( | 1810055G0 | 853.8696 | 0.203838 | 0.205703 | 0.864412 |
| ENSMUSG( | 1810062O1 | 55.30763 | 0.000449 | 0.998746 | 0.99989 | ENSMUSG( | 1810062O1 | 50.90192 | 0.029086 | 0.920697 | 0.992644 |
| ENSMUSG( | 2010001A1 | 277.5323 | 0.048001 | 0.740568 | 0.947823 | ENSMUSG( | 2010001A1 | 217.6476 | -0.43062 | 0.028438 | 0.613486 |
| ENSMUSG( | 2010315B0 | 424.6456 | -0.07337 | 0.660074 | 0.924627 | ENSMUSG( | 2010315B0 | 395.6973 | -0.01155 | 0.94853 | 0.994798 |
| ENSMUSG( | 2200002D0 | 10.74893 | -0.36561 | 0.532879 | NA | ENSMUSG( | 2200002D0 | 10.79615 | -0.04633 | 0.937755 | NA |
| ENSMUSG( | 2210016L2 | 2355.575 | 0.042761 | 0.729124 | 0.944738 | ENSMUSG( | 2210016L2 | 2016.18 | -0.14348 | 0.315282 | 0.901306 |
| ENSMUSG( | 2210406O1 | 76.8944 | -0.22404 | 0.346141 | 0.779153 | ENSMUSG( | 2210406O1 | 65.81789 | -0.4389 | 0.103948 | 0.804969 |
| ENSMUSG( | 2210408F2 | 471.4707 | 0.047959 | 0.707187 | 0.93949 | ENSMUSG( | 2210408F2 | 398.5079 | -0.17679 | 0.198762 | 0.862972 |
| ENSMUSG( | 2210408I2 | 201.633 | 0.202756 | 0.175172 | 0.610399 | ENSMUSG( | 2210408I2 | 184.8759 | 0.215527 | 0.265263 | 0.886134 |
| ENSMUSG( | 2210416O1 | 1942.748 | -0.03333 | 0.696943 | 0.935486 | ENSMUSG( | 2210416O1 | 1770.373 | -0.03519 | 0.755219 | 0.971098 |
| ENSMUSG( | 2300009A0 | 372.4004 | 0.01845 | 0.891549 | 0.983955 | ENSMUSG( | 2300009A0 | 330.6781 | -0.06088 | 0.688552 | 0.959742 |
| ENSMUSG( | 2310002F0 | 27.21316 | 0.623507 | 0.136904 | 0.55149 | ENSMUSG( | 2310002F0 | 19.3654 | 0.000911 | 0.998541 | 0.999396 |
| ENSMUSG( | 2310009B1 | 276.0805 | -0.00613 | 0.966124 | 0.995377 | ENSMUSG( | 2310009B1 | 239.7113 | -0.15841 | 0.359964 | 0.914223 |
| ENSMUSG( | 2310010J1 | 253.5511 | -0.18415 | 0.215349 | 0.659922 | ENSMUSG( | 2310010J1 | 221.5409 | -0.3232 | 0.079421 | 0.77458 |
| ENSMUSG( | 2310011J0 | 942.6112 | 0.061469 | 0.53893 | 0.882849 | ENSMUSG( | 2310011J0 | 781.1584 | -0.22507 | 0.076374 | 0.770504 |
| ENSMUSG( | 2310016G1 | 10.77132 | -0.04791 | 0.93025 | NA | ENSMUSG( | 2310016G1 | 10.65363 | 0.168702 | 0.77243 | NA |
| ENSMUSG( | 2310022A1 | 518.8809 | 0.265965 | 0.059609 | 0.377385 | ENSMUSG( | 2310022A1 | 438.1073 | 0.05542 | 0.735859 | 0.967344 |
| ENSMUSG( | 2310022B0 | 8453.174 | 0.214528 | 0.042128 | 0.316282 | ENSMUSG( | 2310022B0 | 6812.553 | -0.1376 | 0.305974 | 0.898538 |
| ENSMUSG( | 2310030G0 | 27.90817 | -0.99631 | 0.009418 | 0.137123 | ENSMUSG( | 2310030G0 | 29.28496 | -0.48692 | 0.208176 | 0.864412 |
| ENSMUSG( | 2310033P0 | 403.2968 | 0.124867 | 0.342626 | 0.778971 | ENSMUSG( | 2310033P0 | 346.7143 | -0.04486 | 0.789904 | 0.976798 |
| ENSMUSG( | 2310039H0 | 106.6143 | -0.11243 | 0.602855 | 0.904887 | ENSMUSG( | 2310039H0 | 95.31919 | -0.18623 | 0.462601 | 0.923329 |
| ENSMUSG( | 2310040G2 | 38.86285 | -0.04429 | 0.893517 | 0.984512 | ENSMUSG( | 2310040G2 | 32.7073 | -0.3249 | 0.419958 | 0.917257 |
| ENSMUSG( | 2310057M | 346.7212 | -0.04104 | 0.777744 | 0.956168 | ENSMUSG( | 2310057M | 316.1109 | -0.03357 | 0.8379 | 0.981376 |
| ENSMUSG( | 2310061I0 | 1245.316 | 0.034074 | 0.82236 | 0.966352 | ENSMUSG( | 2310061I0 | 1130.758 | 0.019494 | 0.90422 | 0.991089 |
| ENSMUSG( | 2410002F2 | 1279.588 | -0.27618 | 0.021933 | 0.225738 | ENSMUSG( | 2410002F2 | 1229.378 | -0.1144 | 0.38458 | 0.917257 |
| ENSMUSG( | 2410004B1 | 785.1762 | 0.145343 | 0.222479 | 0.668434 | ENSMUSG( | 2410004B1 | 652.4927 | -0.12546 | 0.39055 | 0.917257 |
| ENSMUSG( | 2410004I0 | 35.66245 | 0.383255 | 0.271925 | 0.722255 | ENSMUSG( | 2410004I0 | 24.6624 | -0.41527 | 0.301557 | 0.898538 |
| ENSMUSG( | 2410004P0 | 70.79305 | -0.65223 | 0.009399 | 0.13706 | ENSMUSG( | 2410004P0 | 83.4108 | 0.177572 | 0.526545 | 0.934374 |
| ENSMUSG( | 2410006H1 | 797.9374 | 0.028957 | 0.776886 | 0.955828 | ENSMUSG( | 2410006H1 | 700.2255 | -0.08112 | 0.510222 | 0.929089 |
| ENSMUSG( | 2410018L1 | 38.16585 | 0.642475 | 0.041394 | 0.314189 | ENSMUSG( | 2410018L1 | 28.06423 | 0.075107 | 0.843211 | 0.983281 |
| ENSMUSG( | 2410022M | 107.6258 | -0.02857 | 0.901115 | 0.985633 | ENSMUSG( | 2410022M | 95.84499 | -0.10163 | 0.690086 | 0.960071 |
| ENSMUSG( | 2500002B1 | 38.4241 | -0.10129 | 0.740177 | 0.947823 | ENSMUSG( | 2500002B1 | 36.59478 | 0.004916 | 0.98842 | 0.997862 |
| ENSMUSG( | 2500004C0 | 247.4807 | -0.06959 | 0.621626 | 0.911432 | ENSMUSG( | 2500004C0 | 238.1865 | 0.07621 | 0.698119 | 0.960867 |
| ENSMUSG( | 2510002D2 | 340.0288 | -0.02921 | 0.829111 | 0.969286 | ENSMUSG( | 2510002D2 | 296.0093 | -0.17165 | 0.34522 | 0.91045 |
| ENSMUSG( | 2510009E0 | 2851.599 | -0.01081 | 0.915651 | 0.988634 | ENSMUSG( | 2510009E0 | 2578.569 | -0.03597 | 0.808173 | 0.978622 |
| ENSMUSG( | 2510039O1 | 931.9018 | -0.24762 | 0.079258 | 0.435105 | ENSMUSG( | 2510039O1 | 1012.153 | 0.26506 | 0.119156 | 0.813384 |
| ENSMUSG( | 2610005L0 | 41.7959 | -0.39425 | 0.244302 | 0.691435 | ENSMUSG( | 2610005L0 | 42.44941 | -0.04709 | 0.899616 | 0.990558 |
| ENSMUSG( | 2610008E1 | 945.8837 | -0.12825 | 0.254493 | 0.704536 | ENSMUSG( | 2610008E1 | 952.002 | 0.160599 | 0.201128 | 0.86439 |
| ENSMUSG( | 2610020C0 | 20.65981 | -0.04776 | 0.906663 | NA | ENSMUSG( | 2610020C0 | 22.39752 | 0.40473 | 0.363255 | 0.914223 |
| ENSMUSG( | 2610035D1 | 393.9043 | 0.267993 | 0.066101 | 0.397686 | ENSMUSG( | 2610035D1 | 339.4235 | 0.114245 | 0.482235 | 0.924578 |
| ENSMUSG( | 2610035F2 | 17.06571 | -0.06994 | 0.8767 | NA | ENSMUSG( | 2610035F2 | 15.15181 | -0.13457 | 0.786451 | 0.976303 |
| ENSMUSG( | 2610306M | 203.4525 | 0.132193 | 0.375214 | 0.800786 | ENSMUSG( | 2610306M | 170.0158 | -0.13291 | 0.481676 | 0.924578 |
| ENSMUSG( | 2610316D0 | 91.80845 | 0.040125 | 0.84613 | 0.972226 | ENSMUSG( | 2610316D0 | 101.0581 | 0.548872 | 0.0189 | 0.548832 |
| ENSMUSG( | 2610318N0 | 33.43534 | 0.079489 | 0.823817 | 0.966603 | ENSMUSG( | 2610318N0 | 27.65501 | -0.24022 | 0.559147 | 0.939587 |
| ENSMUSG( | 2610507I0 | 821.6239 | -0.03245 | 0.739573 | 0.947768 | ENSMUSG( | 2610507I0 | 726.8102 | -0.11994 | 0.310845 | 0.899422 |
| ENSMUSG( | 2700038G2 | 24.69747 | 0.099582 | 0.809439 | 0.963674 | ENSMUSG( | 2700038G2 | 22.54625 | 0.126733 | 0.78603 | 0.976285 |
| ENSMUSG( | 2700046A0 | 51.04799 | -0.17149 | 0.523569 | 0.875628 | ENSMUSG( | 2700046A0 | 53.64821 | 0.224957 | 0.425318 | 0.917257 |
| ENSMUSG( | 2700049A0 | 217.6566 | 0.092619 | 0.542346 | 0.883182 | ENSMUSG( | 2700049A0 | 198.7571 | 0.099973 | 0.590908 | 0.944975 |
| ENSMUSG( | 2700054A1 | 98.85096 | 0.187977 | 0.350444 | 0.782193 | ENSMUSG( | 2700054A1 | 91.59647 | 0.210828 | 0.418857 | 0.917257 |
| ENSMUSG( | 2700062C0 | 216.8738 | -0.03938 | 0.80307 | 0.962169 | ENSMUSG( | 2700062C0 | 198.9707 | -0.02287 | 0.898211 | 0.990558 |
| ENSMUSG( | 2700097O0 | 198.8965 | -0.14827 | 0.407603 | 0.817728 | ENSMUSG( | 2700097O0 | 205.2055 | 0.218042 | 0.273557 | 0.889848 |
| ENSMUSG( | 2810001A0 | 159.2743 | -0.22714 | 0.203226 | 0.643631 | ENSMUSG( | 2810001A0 | 147.3632 | -0.17377 | 0.377977 | 0.916784 |
| ENSMUSG( | 2810001G2 | 298.3334 | 0.050565 | 0.723877 | 0.944096 | ENSMUSG( | 2810001G2 | 257.8732 | -0.11309 | 0.54018 | 0.937383 |
| ENSMUSG( | 2810004N2 | 942.597 | 0.162033 | 0.133921 | 0.547197 | ENSMUSG( | 2810004N2 | 811.3475 | -0.00267 | 0.984073 | 0.996963 |
| ENSMUSG( | 2810013P0 | 784.8646 | -0.05982 | 0.630902 | 0.914073 | ENSMUSG( | 2810013P0 | 699.4747 | -0.12922 | 0.352188 | 0.912093 |
| ENSMUSG( | 2810021J2 | 210.3032 | 0.040221 | 0.819914 | 0.965888 | ENSMUSG( | 2810021J2 | 192.0801 | 0.047587 | 0.806083 | 0.977674 |
| ENSMUSG( | 2810029C0 | 290.8432 | -0.12373 | 0.538688 | 0.882755 | ENSMUSG( | 2810029C0 | 255.999 | -0.22976 | 0.248031 | 0.877904 |
| ENSMUSG( | 2810032G0 | 273.8348 | -0.07533 | 0.629844 | 0.913965 | ENSMUSG( | 2810032G0 | 233.9105 | -0.27628 | 0.1205 | 0.813384 |
| ENSMUSG( | 2810049E0 | 118.9463 | 0.209745 | 0.410289 | 0.818258 | ENSMUSG( | 2810049E0 | 107.054 | 0.174549 | 0.490723 | 0.92646 |
| ENSMUSG( | 2810403D2 | 29.81542 | -0.35296 | 0.319852 | 0.765305 | ENSMUSG( | 2810403D2 | 29.69274 | -0.11202 | 0.777609 | 0.976024 |
| ENSMUSG( | 2810408A1 | 29.37231 | 0.25621 | 0.472261 | 0.848011 | ENSMUSG( | 2810408A1 | 28.77169 | 0.405258 | 0.338376 | 0.907972 |
| ENSMUSG( | 2810414N0 | 55.54007 | 0.35161 | 0.201784 | 0.6415 | ENSMUSG( | 2810414N0 | 47.20227 | 0.154026 | 0.618466 | 0.952254 |
| ENSMUSG( | 2810429I0 | 93.46402 | -0.38881 | 0.103109 | 0.48855 | ENSMUSG( | 2810429I0 | 96.40555 | 0.004667 | 0.984634 | 0.996963 |
| ENSMUSG( | 2810454H0 | 13.28403 | -0.30153 | 0.577727 | NA | ENSMUSG( | 2810454H0 | 11.29906 | -0.5138 | 0.398968 | NA |

| | | | | | |
|---|---|---|---|---|---|
| ENSMUSG( | 29.81542 | -0.35296 | 0.319852 | 0.765305 | 2810403D21Rik |
| ENSMUSG( | 29.37231 | 0.25621 | 0.472261 | 0.848011 | 2810408A11Rik |
| ENSMUSG( | 55.54007 | 0.35161 | 0.201784 | 0.6415 | 2810414N06Rik |
| ENSMUSG( | 93.46402 | -0.38881 | 0.103109 | 0.48855 | 2810429I04Rik |
| ENSMUSG( | 13.28403 | -0.30153 | 0.577727 | NA | 2810454H06Rik |
| ENSMUSG( | 963.3291 | -0.11698 | 0.364897 | 0.79407 | 2810459M11Rik |
| ENSMUSG( | 44.96157 | 0.201968 | 0.507477 | 0.86795 | 2900005J15Rik |
| ENSMUSG( | 35.03033 | -0.08415 | 0.805728 | 0.962616 | 2900009J06Rik |
| ENSMUSG( | 2483.819 | 0.152014 | 0.09201 | 0.465937 | 2900026A02Rik |
| ENSMUSG( | 20.42786 | -0.28302 | 0.646459 | NA | 2900040C04Rik |
| ENSMUSG( | 7890.468 | 0.053723 | 0.71493 | 0.941144 | 2900052N01Rik |
| ENSMUSG( | 112.6438 | 0.235065 | 0.255499 | 0.706195 | 2900060N12Rik |
| ENSMUSG( | 192.8946 | -0.05298 | 0.778889 | 0.956442 | 2900076A07Rik |
| ENSMUSG( | 255.8629 | -0.01097 | 0.940208 | 0.993415 | 2900079G21Rik |
| ENSMUSG( | 50.09456 | -0.14591 | 0.6058 | 0.905919 | 2900093K20Rik |
| ENSMUSG( | 33.29159 | 0.105904 | 0.748268 | 0.949604 | 3010001F23Rik |
| ENSMUSG( | 10.65345 | -0.19221 | 0.753626 | NA | 3110006O06Rik |
| ENSMUSG( | 30.22536 | -0.0264 | 0.937103 | 0.993415 | 3110009E18Rik |
| ENSMUSG( | 62.25087 | -0.00535 | 0.984153 | 0.99781 | 3110039M20Rik |
| ENSMUSG( | 161.7531 | 0.056235 | 0.749832 | 0.949664 | 3110040N11Rik |
| ENSMUSG( | 29.98093 | -0.40677 | 0.281921 | 0.73186 | 3110045C21Rik |
| ENSMUSG( | 50.78185 | -0.15847 | 0.640658 | 0.916434 | 3110070M22Rik |
| ENSMUSG( | 226.9625 | 0.053178 | 0.708503 | 0.939519 | 3110082I17Rik |
| ENSMUSG( | 197.3728 | 0.153249 | 0.416542 | 0.8209 | 3110082J24Rik |
| ENSMUSG( | 27.38438 | 0.172952 | 0.624605 | 0.911971 | 3110099E03Rik |
| ENSMUSG( | 36.50079 | -0.05513 | 0.865004 | 0.975565 | 3300002I08Rik |
| ENSMUSG( | 19.12394 | -0.45749 | 0.309266 | NA | 3300002P13Rik |
| ENSMUSG( | 54.58843 | -0.00014 | 0.999628 | 0.99989 | 3425401B19Rik |
| ENSMUSG( | 12.68203 | -0.1739 | 0.742468 | NA | 3632454L22Rik |
| ENSMUSG( | 192.2017 | -0.00597 | 0.970371 | 0.996369 | 3830408C21Rik |
| ENSMUSG( | 20.26804 | -0.09127 | 0.835042 | NA | 3930402G23Rik |
| ENSMUSG( | 574.0225 | -0.14742 | 0.195186 | 0.6346 | 4632404H12Rik |
| ENSMUSG( | 35.04697 | 0.014487 | 0.966167 | 0.995377 | 4632427E13Rik |
| ENSMUSG( | 32.76017 | -0.08988 | 0.804203 | 0.962169 | 4632428C04Rik |
| ENSMUSG( | 35.57095 | 0.064093 | 0.858845 | 0.974707 | 4732414G09Rik |
| ENSMUSG( | 324.2219 | -0.05841 | 0.640963 | 0.91663 | 4732471J01Rik |
| ENSMUSG( | 82.50977 | -0.37744 | 0.090227 | 0.461412 | 4732491K20Rik |
| ENSMUSG( | 21.48453 | -0.11912 | 0.809769 | 0.96372 | 4833411C07Rik |
| ENSMUSG( | 21.19607 | -0.14991 | 0.718741 | NA | 4833412C05Rik |
| ENSMUSG( | 36.72951 | -0.37524 | 0.226141 | 0.672905 | 4833418N02Rik |
| ENSMUSG( | 986.6284 | 0.070682 | 0.631407 | 0.914073 | 4833420G17Rik |
| ENSMUSG( | 119.6319 | 0.080094 | 0.669712 | 0.928147 | 4833422C13Rik |
| ENSMUSG( | 14.25634 | 0.034739 | 0.942909 | NA | 4833422M21Rik |
| ENSMUSG( | 57.9741 | -0.09843 | 0.720904 | 0.942684 | 4833438C02Rik |
| ENSMUSG( | 9383.077 | 0.099841 | 0.325157 | 0.770608 | 4833439L19Rik |
| ENSMUSG( | 36.11508 | 0.077762 | 0.810773 | 0.964272 | 4921504A21Rik |
| ENSMUSG( | 157.0545 | 0.023358 | 0.894784 | 0.984512 | 4921524J17Rik |
| ENSMUSG( | 126.1788 | -0.08918 | 0.631776 | 0.914073 | 4921531C22Rik |
| ENSMUSG( | 30.45987 | -0.26836 | 0.429679 | 0.826446 | 4921536K21Rik |
| ENSMUSG( | 10.77221 | -0.14886 | 0.797138 | NA | 4930405A21Rik |
| ENSMUSG( | 25.77967 | -0.15004 | 0.711472 | 0.940028 | 4930426D05Rik |
| ENSMUSG( | 28.2245 | 0.2558 | 0.479364 | 0.854046 | 4930426L09Rik |
| ENSMUSG( | 14.62602 | 0.091036 | 0.870854 | NA | 4930444P10Rik |
| ENSMUSG( | 126.5975 | -0.45381 | 0.166543 | 0.599201 | 4930447C04Rik |
| ENSMUSG( | 12.38235 | -0.01733 | 0.974802 | NA | 4930447K03Rik |
| ENSMUSG( | 53.1558 | 0.02216 | 0.93745 | 0.993415 | 4930452G13Rik |
| ENSMUSG( | 583.5645 | -0.05656 | 0.602924 | 0.904887 | 4930453N24Rik |
| ENSMUSG( | 41.13564 | -0.1774 | 0.621247 | 0.911432 | 4930473A02Rik |
| ENSMUSG( | 14.03703 | 0.223754 | 0.663524 | NA | 4930474G06Rik |
| ENSMUSG( | 24.10223 | -0.24074 | 0.560317 | 0.888934 | 4930479D17Rik |
| ENSMUSG( | 106.3184 | 0.196772 | 0.3882 | 0.805915 | 4930481A15Rik |
| ENSMUSG( | 279.2801 | -0.03596 | 0.810953 | 0.964318 | 4930503L19Rik |
| ENSMUSG( | 53.53292 | -0.25071 | 0.333444 | 0.775602 | 4930506C21Rik |
| ENSMUSG( | 21.03148 | -0.59537 | 0.149847 | NA | 4930507D05Rik |
| ENSMUSG( | 17.62854 | 0.139268 | 0.80423 | NA | 4930513N10Rik |
| ENSMUSG( | 20.67142 | 0.034248 | 0.94564 | NA | 4930517G24Rik |
| ENSMUSG( | 56.87815 | -0.00044 | 0.99862 | 0.99989 | 4930519K11Rik |
| ENSMUSG( | 15.17476 | 0.106942 | 0.836704 | NA | 4930520O04Rik |
| ENSMUSG( | 30.60369 | -8.27E-05 | 0.999829 | 0.99989 | 4930522L14Rik |
| ENSMUSG( | 113.7812 | 0.010513 | 0.955415 | 0.993874 | 4930523C07Rik |
| ENSMUSG( | 29.40442 | 0.106898 | 0.754187 | 0.950981 | 4930525G20Rik |
| ENSMUSG( | 42.83531 | -0.114 | 0.708792 | 0.939519 | 4930539J05Rik |
| ENSMUSG( | 25.13263 | 0.120135 | 0.750993 | 0.950053 | 4930550C14Rik |
| ENSMUSG( | 19.7764 | 0.589054 | 0.156278 | NA | 4930553P18Rik |
| ENSMUSG( | 19.24142 | -0.08008 | 0.855078 | NA | 4930556M19Rik |
| ENSMUSG( | 53.49038 | -0.00861 | 0.973389 | 0.99675 | 4930563E22Rik |
| ENSMUSG( | 246.4156 | 0.194132 | 0.207727 | 0.64962 | 4930570G19Rik |
| ENSMUSG( | 23.62456 | -0.39179 | 0.32515 | 0.770608 | 4930577N17Rik |
| ENSMUSG( | 83.84711 | 0.113872 | 0.618793 | 0.911432 | 4930578M01Rik |
| ENSMUSG( | 38.49625 | 0.107746 | 0.724233 | 0.944096 | 4930579G24Rik |
| ENSMUSG( | 184.0921 | 0.107551 | 0.566831 | 0.891644 | 4930581F22Rik |
| ENSMUSG( | 35.97769 | -0.12171 | 0.70496 | 0.938303 | 4930590J08Rik |
| ENSMUSG( | 35.96302 | -0.10742 | 0.751022 | 0.950053 | 4930594M22Rik |
| ENSMUSG( | 11.93012 | -0.38703 | 0.506629 | NA | 4930599N23Rik |
| ENSMUSG( | 17.44727 | -0.34067 | 0.483101 | NA | 4931403E22Rik |
| ENSMUSG( | 1930.201 | 0.161341 | 0.141532 | 0.558606 | 4931406C07Rik |
| ENSMUSG( | 9.997274 | -0.37006 | 0.526288 | NA | 4931408D14Rik |
| ENSMUSG( | 113.363 | 0.002829 | 0.988596 | 0.999109 | 4931414P19Rik |
| ENSMUSG( | 30.14398 | -0.25696 | 0.477636 | 0.852702 | 4931415C17Rik |
| ENSMUSG( | 582.8918 | 0.169097 | 0.122224 | 0.527707 | 4932441J04Rik |
| ENSMUSG( | 11.60774 | -0.42952 | 0.424225 | NA | 4932702P03Rik |
| ENSMUSG( | 558.219 | 0.201052 | 0.11499 | 0.51132 | 4933404O12Rik |
| ENSMUSG( | 7.958368 | 0.022925 | 0.972852 | NA | 4933406B17Rik |
| ENSMUSG( | 41.81532 | 0.043802 | 0.903003 | 0.986055 | 4933406C10Rik |
| ENSMUSG( | 18.9339 | -0.21363 | 0.616911 | NA | 4933406I18Rik |
| ENSMUSG( | 89.4612 | -0.09795 | 0.653541 | 0.922819 | 4933407K13Rik |
| ENSMUSG( | 148.1669 | 0.005034 | 0.978222 | 0.997428 | 4933407L21Rik |
| ENSMUSG( | 110.2248 | -0.10925 | 0.607299 | 0.906826 | 4933412E12Rik |
| ENSMUSG( | 25.26575 | -0.04419 | 0.907243 | 0.98755 | 4933412O06Rik |
| ENSMUSG( | 319.3208 | -0.023 | 0.855975 | 0.973813 | 4933421O10Rik |
| ENSMUSG( | 15.10154 | -0.41738 | 0.397775 | NA | 4933424M12Rik |
| ENSMUSG( | 114.7523 | 0.140627 | 0.504458 | 0.866883 | 4933427D14Rik |
| ENSMUSG( | 26.79078 | 0.081943 | 0.822391 | 0.966352 | 4933427I22Rik |
| ENSMUSG( | 8013.95 | 0.170085 | 0.043053 | 0.31819 | 4933431E20Rik |
| ENSMUSG( | 72.49951 | 0.513653 | 0.065316 | 0.395951 | 4933432I09Rik |
| ENSMUSG( | 484.8386 | -0.16842 | 0.185724 | 0.623185 | 4933434E20Rik |
| ENSMUSG( | 885.2266 | 0.370597 | 0.065423 | 0.396102 | 4933439C10Rik |
| ENSMUSG( | 379.3905 | -0.05272 | 0.666201 | 0.927057 | 5031425E22Rik |
| ENSMUSG( | 8698.88 | 0.106398 | 0.154575 | 0.581712 | 5031439G07Rik |
| ENSMUSG( | 22.27376 | -0.39247 | 0.331418 | 0.77519 | 5033404E19Rik |
| ENSMUSG( | 94.29948 | 0.241008 | 0.27733 | 0.727292 | 5033417F24Rik |
| ENSMUSG( | 25.05707 | -0.30376 | 0.429311 | 0.826446 | 5033421B08Rik |
| ENSMUSG( | 10.46605 | -0.21048 | 0.710879 | NA | 5033426O07Rik |
| ENSMUSG( | 128.5463 | -0.07206 | 0.720727 | 0.942638 | 5033430I15Rik |
| ENSMUSG( | 39.70299 | -0.07838 | 0.804467 | 0.962346 | 5330413P13Rik |
| ENSMUSG( | 24.95027 | 0.081982 | 0.835104 | 0.970426 | 5330416C01Rik |
| ENSMUSG( | 143.2226 | -0.20062 | 0.264429 | 0.715323 | 5330434G04Rik |
| ENSMUSG( | 48.39146 | 0.040347 | 0.893148 | 0.984512 | 5430402O13Rik |
| ENSMUSG( | 177.2658 | 0.010587 | 0.9498 | 0.993821 | 5430405H02Rik |
| ENSMUSG( | 325.891 | -0.08512 | 0.550931 | 0.886038 | 5430416N02Rik |
| ENSMUSG( | 354.5768 | -0.01296 | 0.917403 | 0.989182 | 5530601H04Rik |
| ENSMUSG( | 15.56929 | -0.34434 | 0.463204 | NA | 5730405O15Rik |
| ENSMUSG( | 2253.945 | 0.132551 | 0.22059 | 0.666079 | 5730409E04Rik |
| ENSMUSG( | 2138.661 | 0.073522 | 0.393717 | 0.809643 | 5730455P16Rik |
| ENSMUSG( | 16.51529 | 0.11175 | 0.820984 | NA | 5830418P13Rik |
| ENSMUSG( | 10.60412 | 0.295789 | 0.618324 | NA | 5830432E09Rik |
| ENSMUSG( | 26.73011 | 0.062729 | 0.896452 | 0.984868 | 5830444B04Rik |
| ENSMUSG( | 26.37695 | -0.12248 | 0.736552 | 0.946437 | 5830454E08Rik |
| ENSMUSG( | 30.96575 | -0.19182 | 0.575051 | 0.894572 | 5930430L01Rik |
| ENSMUSG( | 17.87902 | 0.511708 | 0.269215 | NA | 5930438M14Rik |
| ENSMUSG( | 33.0876 | 0.445461 | 0.194285 | 0.633391 | 6030407O03Rik |
| ENSMUSG( | 11.67214 | 0.528249 | 0.32475 | NA | 6030442K20Rik |
| ENSMUSG( | 28.64404 | -0.56001 | 0.133741 | 0.547197 | 6030443J06Rik |
| ENSMUSG( | 847.3164 | -0.02482 | 0.819654 | 0.965888 | 6030458C11Rik |
| ENSMUSG( | 41.50835 | 0.02176 | 0.940539 | 0.993415 | 6230400D17Rik |
| ENSMUSG( | 4045.549 | 0.012182 | 0.911248 | 0.988483 | 6330403K07Rik |
| ENSMUSG( | 30.10373 | 0.200808 | 0.552089 | 0.886141 | 6330415G19Rik |
| ENSMUSG( | 136.5107 | 0.20327 | 0.25374 | 0.704011 | 6330418K02Rik |
| ENSMUSG( | 19.58027 | -0.23332 | 0.583261 | NA | 6430503K07Rik |
| ENSMUSG( | 10798.6 | 0.15606 | 0.059417 | 0.376611 | 6430548M08Rik |
| ENSMUSG( | 41.30744 | -0.14932 | 0.608687 | 0.907185 | 6430550D23Rik |
| ENSMUSG( | 114.9906 | -0.62477 | 0.00078 | 0.027489 | 6430571L13Rik |
| ENSMUSG( | 25.22923 | 0.482515 | 0.224375 | 0.67048 | 6430573P05Rik |
| ENSMUSG( | 107.6971 | -0.05169 | 0.806289 | 0.962804 | 6430584L05Rik |
| ENSMUSG( | 663.1571 | 0.050526 | 0.671574 | 0.928781 | 6530402F18Rik |
| ENSMUSG( | 35.53178 | -0.06141 | 0.862415 | 0.975055 | 6530411M01Rik |
| ENSMUSG( | 10.30468 | 0.281158 | 0.627597 | NA | 6720483E21Rik |
| ENSMUSG( | 79.23309 | -0.15205 | 0.498462 | 0.862866 | 6820408C15Rik |
| ENSMUSG( | 107.5652 | -0.32902 | 0.096506 | 0.476986 | 6820431F20Rik |
| ENSMUSG( | 89.71032 | 0.4627 | 0.033915 | 0.282701 | 8030453O22Rik |
| ENSMUSG( | 195.4316 | 0.143868 | 0.399221 | 0.813048 | 8430429K09Rik |
| ENSMUSG( | 47.86621 | -0.02996 | 0.920573 | 0.990307 | 9030612E09Rik |
| ENSMUSG( | 12.16967 | 0.293454 | 0.585313 | NA | 9030624G23Rik |
| ENSMUSG( | 17.59965 | 0.276802 | 0.536079 | NA | 9130019P16Rik |
| ENSMUSG( | 96.0002 | 0.201224 | 0.325545 | 0.770741 | 9130023H24Rik |
| ENSMUSG( | 472.182 | -0.13892 | 0.301942 | 0.750166 | 9130024F11Rik |
| ENSMUSG( | 47.04095 | 0.439898 | 0.125246 | 0.532314 | 9130213A22Rik |
| ENSMUSG( | 226.9369 | 0.120453 | 0.467544 | 0.844819 | 9130221H12Rik |

| | | | | | |
|---|---|---|---|---|---|
| ENSMUSG( | 28.77169 | 0.405258 | 0.338376 | 0.907972 | 2810408A11Rik |
| ENSMUSG( | 47.20227 | 0.154026 | 0.618466 | 0.952254 | 2810414N06Rik |
| ENSMUSG( | 96.40555 | 0.004667 | 0.984634 | 0.996963 | 2810429I04Rik |
| ENSMUSG( | 11.29906 | -0.5138 | 0.398968 | NA | 2810454H06Rik |
| ENSMUSG( | 870.7487 | -0.1443 | 0.330542 | 0.907398 | 2810459M11Rik |
| ENSMUSG( | 43.03243 | 0.339146 | 0.303363 | 0.898538 | 2900005J15Rik |
| ENSMUSG( | 29.02641 | -0.41873 | 0.30193 | 0.898538 | 2900009J06Rik |
| ENSMUSG( | 2217.076 | 0.097325 | 0.380062 | 0.916995 | 2900026A02Rik |
| ENSMUSG( | 18.56999 | -0.2946 | 0.639384 | 0.953903 | 2900040C04Rik |
| ENSMUSG( | 6642.536 | -0.18306 | 0.307678 | 0.898538 | 2900052N01Rik |
| ENSMUSG( | 107.4605 | 0.338402 | 0.180908 | 0.850372 | 2900060N12Rik |
| ENSMUSG( | 204.018 | 0.366145 | 0.092953 | 0.79479 | 2900076A07Rik |
| ENSMUSG( | 228.3648 | -0.07728 | 0.644999 | 0.953988 | 2900079G21Rik |
| ENSMUSG( | 50.38885 | 0.130039 | 0.669139 | 0.956766 | 2900093K20Rik |
| ENSMUSG( | 30.02641 | 0.064107 | 0.859634 | 0.985947 | 3010001F23Rik |
| ENSMUSG( | 8.671255 | -0.47252 | 0.510844 | NA | 3110006O06Rik |
| ENSMUSG( | 26.12148 | -0.2112 | 0.591894 | 0.944975 | 3110009E18Rik |
| ENSMUSG( | 62.72844 | 0.243108 | 0.409096 | 0.917257 | 3110039M20Rik |
| ENSMUSG( | 134.8087 | -0.20318 | 0.327919 | 0.906255 | 3110040N11Rik |
| ENSMUSG( | 25.53585 | -0.65264 | 0.123731 | 0.814691 | 3110045C21Rik |
| ENSMUSG( | 38.56993 | -0.76944 | 0.043223 | 0.690304 | 3110070M22Rik |
| ENSMUSG( | 201.4689 | -0.03064 | 0.857471 | 0.985884 | 3110082I17Rik |
| ENSMUSG( | 163.4001 | -0.13235 | 0.545103 | 0.938063 | 3110082J24Rik |
| ENSMUSG( | 19.92358 | -0.54527 | 0.216641 | 0.865167 | 3110099E03Rik |
| ENSMUSG( | 30.08499 | -0.32597 | 0.398701 | 0.917257 | 3300002I08Rik |
| ENSMUSG( | 17.06843 | -0.49985 | 0.30867 | 0.898538 | 3300002P13Rik |
| ENSMUSG( | 39.91774 | -0.73016 | 0.023864 | 0.588226 | 3425401B19Rik |
| ENSMUSG( | 14.1899 | 0.419163 | 0.43429 | 0.91869 | 3632454L22Rik |
| ENSMUSG( | 171.136 | -0.07613 | 0.675613 | 0.95694 | 3830408C21Rik |
| ENSMUSG( | 14.24672 | -0.93915 | 0.100341 | 0.801311 | 3930402G23Rik |
| ENSMUSG( | 524.9156 | -0.13494 | 0.29078 | 0.897019 | 4632404H12Rik |
| ENSMUSG( | 38.55911 | 0.541968 | 0.140686 | 0.830678 | 4632427E13Rik |
| ENSMUSG( | 30.97181 | 0.011541 | 0.975938 | 0.996416 | 4632428C04Rik |
| ENSMUSG( | 27.79807 | -0.4372 | 0.304165 | 0.898538 | 4732414G09Rik |
| ENSMUSG( | 338.3409 | 0.315784 | 0.040139 | 0.681706 | 4732471J01Rik |
| ENSMUSG( | 76.2026 | -0.33337 | 0.17172 | 0.842999 | 4732491K20Rik |
| ENSMUSG( | 20.5873 | 0.062166 | 0.89339 | 0.990558 | 4833411C07Rik |
| ENSMUSG( | 16.80491 | -0.57965 | 0.232875 | 0.868507 | 4833412C05Rik |
| ENSMUSG( | 35.95653 | -0.1563 | 0.636876 | 0.953903 | 4833418N02Rik |
| ENSMUSG( | 939.6415 | 0.191554 | 0.274928 | 0.889874 | 4833420G17Rik |
| ENSMUSG( | 102.9612 | -0.0851 | 0.686362 | 0.95899 | 4833422C13Rik |
| ENSMUSG( | 13.74006 | 0.177845 | 0.733253 | 0.967228 | 4833422M21Rik |
| ENSMUSG( | 60.19828 | 0.265541 | 0.341963 | 0.909146 | 4833438C02Rik |
| ENSMUSG( | 7767.433 | -0.18268 | 0.158758 | 0.837724 | 4833439L19Rik |
| ENSMUSG( | 34.75391 | 0.250665 | 0.503118 | 0.927825 | 4921504A21Rik |
| ENSMUSG( | 140.2434 | -0.03785 | 0.848208 | 0.985321 | 4921524J17Rik |
| ENSMUSG( | 109.6436 | -0.25167 | 0.257477 | 0.883401 | 4921531C22Rik |
| ENSMUSG( | 25.95719 | -0.48326 | 0.213048 | 0.864412 | 4921536K21Rik |
| ENSMUSG( | 8.673234 | -0.51073 | 0.4603 | NA | 4930405A21Rik |
| ENSMUSG( | 30.03579 | 0.498911 | 0.266947 | 0.886134 | 4930426D05Rik |
| ENSMUSG( | 25.74877 | 0.242865 | 0.563668 | 0.941206 | 4930426L09Rik |
| ENSMUSG( | 12.45064 | -0.15169 | 0.808095 | 0.978622 | 4930444P10Rik |
| ENSMUSG( | 145.6472 | 0.238906 | 0.443699 | 0.921734 | 4930447C04Rik |
| ENSMUSG( | 9.55463 | -0.55566 | 0.384988 | NA | 4930447K03Rik |
| ENSMUSG( | 44.95755 | -0.18682 | 0.554955 | 0.939565 | 4930452G13Rik |
| ENSMUSG( | 544.5012 | 0.009021 | 0.945105 | 0.994798 | 4930453N24Rik |
| ENSMUSG( | 37.73596 | -0.13741 | 0.706394 | 0.963475 | 4930473A02Rik |
| ENSMUSG( | 11.42699 | -0.10416 | 0.863289 | NA | 4930474G06Rik |
| ENSMUSG( | 20.22823 | -0.5065 | 0.271608 | 0.888062 | 4930479D17Rik |
| ENSMUSG( | 88.26161 | -0.07338 | 0.747866 | 0.96984 | 4930481A15Rik |
| ENSMUSG( | 300.1189 | 0.420887 | 0.036157 | 0.668395 | 4930503L19Rik |
| ENSMUSG( | 44.35553 | -0.55604 | 0.076774 | 0.770558 | 4930506C21Rik |
| ENSMUSG( | 21.78222 | -0.18434 | 0.669914 | 0.95694 | 4930507D05Rik |
| ENSMUSG( | 16.70202 | 0.190176 | 0.755991 | 0.971371 | 4930513N10Rik |
| ENSMUSG( | 15.62041 | -0.51493 | 0.353252 | 0.912263 | 4930517G24Rik |
| ENSMUSG( | 47.40361 | -0.27193 | 0.362654 | 0.914223 | 4930519K11Rik |
| ENSMUSG( | 10.99134 | -0.58192 | 0.371722 | NA | 4930520O04Rik |
| ENSMUSG( | 32.27056 | 0.376796 | 0.342372 | 0.909646 | 4930522L14Rik |
| ENSMUSG( | 109.0103 | 0.165256 | 0.499823 | 0.927825 | 4930523C07Rik |
| ENSMUSG( | 26.25167 | 0.031451 | 0.935207 | 0.993078 | 4930525G20Rik |
| ENSMUSG( | 38.99867 | -0.1065 | 0.746982 | 0.969763 | 4930539J05Rik |
| ENSMUSG( | 19.14108 | -0.45617 | 0.309224 | 0.898538 | 4930550C14Rik |
| ENSMUSG( | 17.73384 | 0.544544 | 0.238053 | 0.87259 | 4930553P18Rik |
| ENSMUSG( | 20.17144 | 0.302758 | 0.499994 | 0.927825 | 4930556M19Rik |
| ENSMUSG( | 46.11528 | -0.17762 | 0.548824 | 0.938075 | 4930563E22Rik |
| ENSMUSG( | 230.973 | 0.270087 | 0.125679 | 0.81516 | 4930570G19Rik |
| ENSMUSG( | 23.43136 | -0.11596 | 0.787388 | 0.97637 | 4930577N17Rik |
| ENSMUSG( | 78.05976 | 0.186566 | 0.453178 | 0.922316 | 4930578M01Rik |
| ENSMUSG( | 31.18177 | -0.21805 | 0.558513 | 0.93957 | 4930579G24Rik |
| ENSMUSG( | 163.7748 | 0.049572 | 0.837303 | 0.981335 | 4930581F22Rik |
| ENSMUSG( | 33.53511 | -0.04324 | 0.900033 | 0.990558 | 4930590J08Rik |
| ENSMUSG( | 32.75553 | -0.13787 | 0.725195 | 0.966887 | 4930594M22Rik |
| ENSMUSG( | 11.73985 | -0.18319 | 0.758673 | 0.971858 | 4930599N23Rik |
| ENSMUSG( | 14.86934 | -0.57951 | 0.306537 | 0.898538 | 4931403E22Rik |
| ENSMUSG( | 1620.995 | -0.07283 | 0.554386 | 0.939565 | 4931406C07Rik |
| ENSMUSG( | 8.074322 | -0.87416 | 0.212163 | NA | 4931408D14Rik |
| ENSMUSG( | 94.41538 | -0.26799 | 0.228511 | 0.868507 | 4931414P19Rik |
| ENSMUSG( | 32.71908 | 0.234248 | 0.514825 | 0.930939 | 4931415C17Rik |
| ENSMUSG( | 483.6129 | -0.09089 | 0.567399 | 0.941777 | 4932441J04Rik |
| ENSMUSG( | 480.395 | 0.044171 | 0.747698 | 0.96984 | 4933404O12Rik |
| ENSMUSG( | 8.248966 | -0.69444 | 0.362689 | NA | 4933405L10Rik |
| ENSMUSG( | 43.30341 | 0.408702 | 0.274455 | 0.889874 | 4933406C10Rik |
| ENSMUSG( | 19.4448 | 0.105976 | 0.820445 | 0.979198 | 4933406I18Rik |
| ENSMUSG( | 87.51091 | 0.113777 | 0.633913 | 0.953891 | 4933407K13Rik |
| ENSMUSG( | 121.2219 | -0.32996 | 0.123484 | 0.814691 | 4933407L21Rik |
| ENSMUSG( | 97.04881 | -0.22242 | 0.363726 | 0.914223 | 4933412E12Rik |
| ENSMUSG( | 24.34071 | 0.103605 | 0.800913 | 0.97681 | 4933412O06Rik |
| ENSMUSG( | 278.8935 | -0.14397 | 0.343215 | 0.909646 | 4933421O10Rik |
| ENSMUSG( | 12.19519 | -0.85514 | 0.143134 | 0.831311 | 4933424M12Rik |
| ENSMUSG( | 96.46578 | -0.06169 | 0.823663 | 0.979198 | 4933427D14Rik |
| ENSMUSG( | 19.87191 | -0.54058 | 0.230331 | 0.868507 | 4933427I22Rik |
| ENSMUSG( | 6936.179 | 0.026554 | 0.807941 | 0.978622 | 4933431E20Rik |
| ENSMUSG( | 54.64301 | 0.017917 | 0.950255 | 0.994798 | 4933432I09Rik |
| ENSMUSG( | 492.805 | 0.142438 | 0.344873 | 0.91045 | 4933434E20Rik |
| ENSMUSG( | 722.6119 | 0.074785 | 0.751291 | 0.970317 | 4933439C10Rik |
| ENSMUSG( | 359.4357 | 0.065348 | 0.662472 | 0.956205 | 5031425E22Rik |
| ENSMUSG( | 7426.369 | -0.08144 | 0.403146 | 0.917257 | 5031439G07Rik |
| ENSMUSG( | 20.69002 | -0.27107 | 0.571551 | 0.942116 | 5033404E19Rik |
| ENSMUSG( | 76.08388 | -0.08025 | 0.74832 | 0.96984 | 5033417F24Rik |
| ENSMUSG( | 21.39604 | -0.55007 | 0.206131 | 0.864412 | 5033421B08Rik |
| ENSMUSG( | 9.887613 | -0.13121 | 0.832372 | NA | 5033426O07Rik |
| ENSMUSG( | 112.0993 | -0.20432 | 0.356062 | 0.913331 | 5033430I15Rik |
| ENSMUSG( | 35.14164 | -0.14424 | 0.671097 | 0.95694 | 5330413P13Rik |
| ENSMUSG( | 24.6419 | 0.289761 | 0.476511 | 0.923329 | 5330416C01Rik |
| ENSMUSG( | 137.5371 | -0.05398 | 0.786484 | 0.976303 | 5330434G04Rik |
| ENSMUSG( | 42.92926 | -0.0603 | 0.850944 | 0.985541 | 5430402O13Rik |
| ENSMUSG( | 174.4795 | 0.232122 | 0.233288 | 0.868507 | 5430405H02Rik |
| ENSMUSG( | 303.0333 | -0.02541 | 0.861315 | 0.986281 | 5430416N02Rik |
| ENSMUSG( | 328.1173 | 0.026956 | 0.863522 | 0.986547 | 5530601H04Rik |
| ENSMUSG( | 14.40311 | -0.23341 | 0.681111 | 0.958395 | 5730405O15Rik |
| ENSMUSG( | 1863.381 | -0.14999 | 0.243098 | 0.877399 | 5730409E04Rik |
| ENSMUSG( | 1876.909 | -0.03549 | 0.747946 | 0.96984 | 5730455P16Rik |
| ENSMUSG( | 11.99911 | -0.56965 | 0.388516 | 0.917257 | 5830418P13Rik |
| ENSMUSG( | 26.26241 | 0.235328 | 0.636583 | 0.953903 | 5830444B04Rik |
| ENSMUSG( | 24.04399 | -0.12054 | 0.762702 | 0.973566 | 5830454E08Rik |
| ENSMUSG( | 36.58598 | 0.553077 | 0.153036 | 0.837724 | 5930430L01Rik |
| ENSMUSG( | 25.0339 | -0.09696 | 0.818242 | 0.979198 | 6030407O03Rik |
| ENSMUSG( | 8.255128 | -0.10314 | 0.88392 | NA | 6030442K20Rik |
| ENSMUSG( | 27.24114 | -0.39883 | 0.31803 | 0.902849 | 6030443J06Rik |
| ENSMUSG( | 774.9558 | -0.01419 | 0.910867 | 0.992196 | 6030458C11Rik |
| ENSMUSG( | 32.23376 | -0.45525 | 0.204937 | 0.864412 | 6230400D17Rik |
| ENSMUSG( | 3523.992 | -0.12306 | 0.424928 | 0.917257 | 6330403K07Rik |
| ENSMUSG( | 25.10634 | -0.03721 | 0.923641 | 0.992815 | 6330415G19Rik |
| ENSMUSG( | 110.4373 | -0.13444 | 0.512601 | 0.930164 | 6330418K02Rik |
| ENSMUSG( | 16.95365 | -0.39695 | 0.406876 | 0.917257 | 6430503K07Rik |
| ENSMUSG( | 9663.649 | 0.107796 | 0.34957 | 0.911652 | 6430548M08Rik |
| ENSMUSG( | 40.50197 | 0.088411 | 0.795862 | 0.97681 | 6430550D23Rik |
| ENSMUSG( | 120.2076 | -0.18038 | 0.383759 | 0.917257 | 6430571L13Rik |
| ENSMUSG( | 19.37869 | 0.050535 | 0.908262 | 0.991958 | 6430573P05Rik |
| ENSMUSG( | 93.55701 | -0.20068 | 0.383551 | 0.917257 | 6430584L05Rik |
| ENSMUSG( | 592.2464 | 0.00053 | 0.997198 | 0.999268 | 6530402F18Rik |
| ENSMUSG( | 31.20432 | -0.20801 | 0.583461 | 0.944266 | 6530411M01Rik |
| ENSMUSG( | 11.8479 | 0.867097 | 0.123268 | 0.814691 | 6720483E21Rik |
| ENSMUSG( | 79.6593 | 0.122728 | 0.616383 | 0.951413 | 6820408C15Rik |
| ENSMUSG( | 106.5273 | -0.08014 | 0.716289 | 0.965692 | 6820431F20Rik |
| ENSMUSG( | 74.89298 | 0.210752 | 0.439336 | 0.920721 | 8030453O22Rik |
| ENSMUSG( | 167.9081 | -0.02928 | 0.877015 | 0.989644 | 8430429K09Rik |
| ENSMUSG( | 44.0405 | 0.02605 | 0.938506 | 0.993883 | 9030612E09Rik |
| ENSMUSG( | 12.26503 | 0.510058 | 0.368661 | 0.915491 | 9030624G23Rik |
| ENSMUSG( | 15.14167 | 0.164325 | 0.752549 | 0.970521 | 9130019P16Rik |
| ENSMUSG( | 86.14009 | 0.171912 | 0.461174 | 0.923329 | 9130023H24Rik |
| ENSMUSG( | 460.0231 | 0.056396 | 0.681962 | 0.958678 | 9130024F11Rik |
| ENSMUSG( | 38.17844 | 0.14899 | 0.645498 | 0.953988 | 9130213A22Rik |
| ENSMUSG( | 202.9951 | 0.076632 | 0.675784 | 0.95694 | 9130221H12Rik |
| ENSMUSG( | 282.6939 | 0.070009 | 0.643126 | 0.953903 | 9130401M01Rik |
| ENSMUSG( | 17.40126 | -0.33137 | 0.497854 | 0.927825 | 9230112J17Rik |
| ENSMUSG( | 60.69726 | -0.37588 | 0.167977 | 0.840222 | 9230114K14Rik |
| ENSMUSG( | 11.52922 | -0.14649 | 0.797191 | 0.97681 | 9230116N13Rik |

| | | | | | | | | | | | |
|---|---|---|---|---|---|---|---|---|---|---|---|
| ENSMUSG( | 2810459M | 963.3291 | -0.11698 | 0.364897 | 0.79407 | ENSMUSG( | 2810459M | 870.7487 | -0.1443 | 0.330542 | 0.907398 |
| ENSMUSG( | 2900005J1 | 44.96157 | 0.201968 | 0.507477 | 0.86795 | ENSMUSG( | 2900005J1 | 43.03243 | 0.339146 | 0.303363 | 0.898538 |
| ENSMUSG( | 2900009J0 | 35.03033 | -0.08415 | 0.805728 | 0.962616 | ENSMUSG( | 2900009J0 | 29.02641 | -0.41873 | 0.30193 | 0.898538 |
| ENSMUSG( | 2900026A | 2483.819 | 0.152014 | 0.09201 | 0.465937 | ENSMUSG( | 2900026A | 2217.076 | 0.097325 | 0.380062 | 0.916995 |
| ENSMUSG( | 2900040C | 20.42786 | -0.28302 | 0.646459 | NA | ENSMUSG( | 2900040C | 18.56999 | -0.2946 | 0.639384 | 0.953903 |
| ENSMUSG( | 2900052N | 7890.468 | 0.053723 | 0.71493 | 0.941144 | ENSMUSG( | 2900052N | 6642.536 | -0.18306 | 0.307678 | 0.898538 |
| ENSMUSG( | 2900060N1 | 112.6438 | 0.235065 | 0.255499 | 0.706195 | ENSMUSG( | 2900060N1 | 107.4605 | 0.338402 | 0.180908 | 0.850372 |
| ENSMUSG( | 2900076A | 192.8946 | -0.05298 | 0.778889 | 0.956442 | ENSMUSG( | 2900076A | 204.018 | 0.366145 | 0.092953 | 0.79479 |
| ENSMUSG( | 2900079G | 255.8629 | -0.01097 | 0.940208 | 0.993415 | ENSMUSG( | 2900079G | 228.3648 | -0.07728 | 0.644999 | 0.953988 |
| ENSMUSG( | 2900093K2 | 50.09456 | -0.14591 | 0.6058 | 0.905919 | ENSMUSG( | 2900093K2 | 50.38885 | 0.130039 | 0.669139 | 0.956766 |
| ENSMUSG( | 3010001F2 | 33.29159 | 0.105904 | 0.748268 | 0.949604 | ENSMUSG( | 3010001F2 | 30.02641 | 0.064107 | 0.859634 | 0.985947 |
| ENSMUSG( | 3110006O | 10.65345 | -0.19221 | 0.753626 | NA | ENSMUSG( | 3110006O | 8.671255 | -0.47252 | 0.510844 | NA |
| ENSMUSG( | 3110009E1 | 30.22536 | -0.0264 | 0.937103 | 0.993415 | ENSMUSG( | 3110009E1 | 26.12148 | -0.2112 | 0.591894 | 0.944975 |
| ENSMUSG( | 3110039M | 62.25087 | -0.00535 | 0.984153 | 0.99781 | ENSMUSG( | 3110039M | 62.72844 | 0.243108 | 0.409096 | 0.917257 |
| ENSMUSG( | 3110040N1 | 161.7531 | 0.056235 | 0.749832 | 0.949664 | ENSMUSG( | 3110040N1 | 134.8087 | -0.20318 | 0.327919 | 0.906255 |
| ENSMUSG( | 3110045C2 | 29.98093 | -0.40677 | 0.281921 | 0.73186 | ENSMUSG( | 3110045C2 | 25.53585 | -0.65264 | 0.123731 | 0.814691 |
| ENSMUSG( | 3110070M | 50.78185 | -0.15847 | 0.640658 | 0.916434 | ENSMUSG( | 3110070M | 38.56993 | -0.76944 | 0.043223 | 0.690304 |
| ENSMUSG( | 3110082I1 | 226.9625 | 0.053178 | 0.708503 | 0.939519 | ENSMUSG( | 3110082I1 | 201.4689 | -0.03064 | 0.857471 | 0.985884 |
| ENSMUSG( | 3110082J2 | 197.3728 | 0.153249 | 0.416542 | 0.8209 | ENSMUSG( | 3110082J2 | 163.4001 | -0.13235 | 0.545103 | 0.938063 |
| ENSMUSG( | 3110099E0 | 27.38438 | 0.172952 | 0.624605 | 0.911971 | ENSMUSG( | 3110099E0 | 19.92358 | -0.54527 | 0.216641 | 0.865167 |
| ENSMUSG( | 3300002I0 | 36.50079 | -0.05513 | 0.865004 | 0.975565 | ENSMUSG( | 3300002I0 | 30.08499 | -0.32597 | 0.398701 | 0.917257 |
| ENSMUSG( | 3300002P1 | 19.12394 | -0.45749 | 0.309266 | NA | ENSMUSG( | 3300002P1 | 17.06843 | -0.49985 | 0.30867 | 0.898538 |
| ENSMUSG( | 3425401B1 | 54.58843 | -0.00014 | 0.999628 | 0.99989 | ENSMUSG( | 3425401B1 | 39.91774 | -0.73016 | 0.023864 | 0.588226 |
| ENSMUSG( | 3632454L2 | 12.68203 | -0.1739 | 0.742468 | NA | ENSMUSG( | 3632454L2 | 14.1899 | 0.419163 | 0.43429 | 0.91869 |
| ENSMUSG( | 3830408C2 | 192.2017 | -0.00597 | 0.970371 | 0.996369 | ENSMUSG( | 3830408C2 | 171.136 | -0.07613 | 0.675613 | 0.95694 |
| ENSMUSG( | 3930402G | 20.26804 | -0.09127 | 0.835042 | NA | ENSMUSG( | 3930402G | 14.24672 | -0.93915 | 0.100341 | 0.801311 |
| ENSMUSG( | 4632404H1 | 574.0225 | -0.14742 | 0.195186 | 0.6346 | ENSMUSG( | 4632404H1 | 524.9156 | -0.13494 | 0.29078 | 0.897019 |
| ENSMUSG( | 4632427E1 | 35.04697 | 0.014487 | 0.966167 | 0.995377 | ENSMUSG( | 4632427E1 | 38.55911 | 0.541968 | 0.140686 | 0.830678 |
| ENSMUSG( | 4632428C | 32.76017 | -0.08988 | 0.804203 | 0.962169 | ENSMUSG( | 4632428C | 30.97181 | 0.011541 | 0.975938 | 0.996416 |
| ENSMUSG( | 4732414G | 35.57095 | 0.064093 | 0.858845 | 0.974707 | ENSMUSG( | 4732414G | 27.79807 | -0.4372 | 0.304165 | 0.898538 |
| ENSMUSG( | 4732471J0 | 324.2219 | -0.05841 | 0.640963 | 0.91663 | ENSMUSG( | 4732471J0 | 338.3409 | 0.315784 | 0.040139 | 0.681706 |
| ENSMUSG( | 4732491K2 | 82.50977 | -0.37744 | 0.090227 | 0.461412 | ENSMUSG( | 4732491K2 | 76.2026 | -0.33337 | 0.17172 | 0.842999 |
| ENSMUSG( | 4833411C | 21.48453 | -0.11912 | 0.809769 | 0.96372 | ENSMUSG( | 4833411C | 20.5873 | 0.062166 | 0.89339 | 0.990558 |
| ENSMUSG( | 4833412C | 21.19607 | -0.14991 | 0.718741 | NA | ENSMUSG( | 4833412C | 16.80491 | -0.57965 | 0.232875 | 0.868507 |
| ENSMUSG( | 4833418N | 36.72951 | -0.37524 | 0.226141 | 0.672905 | ENSMUSG( | 4833418N | 35.95653 | -0.1563 | 0.636876 | 0.953903 |
| ENSMUSG( | 4833420G1 | 986.6284 | 0.070682 | 0.631407 | 0.914073 | ENSMUSG( | 4833420G1 | 939.6415 | 0.191554 | 0.274928 | 0.889874 |
| ENSMUSG( | 4833422C1 | 119.6319 | 0.080094 | 0.669712 | 0.928147 | ENSMUSG( | 4833422C1 | 102.9612 | -0.0851 | 0.686362 | 0.95899 |
| ENSMUSG( | 4833422M | 14.25634 | 0.034739 | 0.942909 | NA | ENSMUSG( | 4833422M | 13.74006 | 0.177845 | 0.733253 | 0.967228 |
| ENSMUSG( | 4833438C | 57.9741 | -0.09843 | 0.720904 | 0.942684 | ENSMUSG( | 4833438C | 60.19828 | 0.265541 | 0.341963 | 0.909146 |
| ENSMUSG( | 4833439L1 | 9383.077 | 0.099841 | 0.325157 | 0.770608 | ENSMUSG( | 4833439L1 | 7767.433 | -0.18268 | 0.158758 | 0.837724 |
| ENSMUSG( | 4921504A | 36.11508 | 0.077762 | 0.810773 | 0.964272 | ENSMUSG( | 4921504A | 34.75391 | 0.250665 | 0.503118 | 0.927825 |
| ENSMUSG( | 4921524J1 | 157.0545 | 0.023358 | 0.894784 | 0.984512 | ENSMUSG( | 4921524J1 | 140.2434 | -0.03785 | 0.848208 | 0.985321 |
| ENSMUSG( | 4921531C2 | 126.1788 | -0.08918 | 0.631776 | 0.914073 | ENSMUSG( | 4921531C2 | 109.6436 | -0.25167 | 0.257477 | 0.883401 |
| ENSMUSG( | 4921536K2 | 30.45987 | -0.26836 | 0.429679 | 0.826446 | ENSMUSG( | 4921536K2 | 25.95719 | -0.48326 | 0.213048 | 0.864412 |
| ENSMUSG( | 4930405A | 10.77221 | -0.14886 | 0.797138 | NA | ENSMUSG( | 4930405A | 8.673234 | -0.51073 | 0.4603 | NA |
| ENSMUSG( | 4930426D | 25.77967 | -0.15004 | 0.711472 | 0.940028 | ENSMUSG( | 4930426D | 30.03579 | 0.498911 | 0.266947 | 0.886134 |
| ENSMUSG( | 4930426L0 | 28.2245 | 0.2558 | 0.479364 | 0.854046 | ENSMUSG( | 4930426L0 | 25.74877 | 0.242865 | 0.563668 | 0.941206 |
| ENSMUSG( | 4930444P1 | 14.62602 | 0.091036 | 0.870854 | NA | ENSMUSG( | 4930444P1 | 12.45064 | -0.15169 | 0.808095 | 0.978622 |
| ENSMUSG( | 4930447C | 126.5975 | -0.45381 | 0.166543 | 0.599201 | ENSMUSG( | 4930447C | 145.6472 | 0.238906 | 0.443699 | 0.921734 |
| ENSMUSG( | 4930447K | 12.38235 | -0.01733 | 0.974802 | NA | ENSMUSG( | 4930447K | 9.55463 | -0.55566 | 0.384988 | NA |
| ENSMUSG( | 4930452G1 | 53.1558 | 0.02216 | 0.93745 | 0.993415 | ENSMUSG( | 4930452G1 | 44.95755 | -0.18682 | 0.554955 | 0.939565 |
| ENSMUSG( | 4930453N | 583.5645 | -0.05656 | 0.602924 | 0.904887 | ENSMUSG( | 4930453N | 544.5012 | 0.009021 | 0.945105 | 0.994798 |
| ENSMUSG( | 4930473A | 41.13564 | -0.1774 | 0.621247 | 0.911432 | ENSMUSG( | 4930473A | 37.73596 | -0.13741 | 0.706394 | 0.963475 |
| ENSMUSG( | 4930474G | 14.03703 | 0.223754 | 0.663524 | NA | ENSMUSG( | 4930474G | 11.42699 | -0.10416 | 0.863289 | NA |
| ENSMUSG( | 4930479D1 | 24.10223 | -0.24074 | 0.560317 | 0.888934 | ENSMUSG( | 4930479D1 | 20.22823 | -0.5065 | 0.271608 | 0.888062 |
| ENSMUSG( | 4930481A1 | 106.3184 | 0.196772 | 0.3882 | 0.805915 | ENSMUSG( | 4930481A1 | 88.26161 | -0.07338 | 0.747866 | 0.96984 |
| ENSMUSG( | 4930503L1 | 279.2801 | -0.03596 | 0.810953 | 0.964318 | ENSMUSG( | 4930503L1 | 300.1189 | 0.420887 | 0.036157 | 0.668395 |
| ENSMUSG( | 4930506C2 | 53.53292 | -0.25071 | 0.333444 | 0.775602 | ENSMUSG( | 4930506C2 | 44.35553 | -0.55604 | 0.076774 | 0.770558 |
| ENSMUSG( | 4930507D | 21.03148 | -0.59537 | 0.149847 | NA | ENSMUSG( | 4930507D | 21.78222 | -0.18434 | 0.669914 | 0.95694 |
| ENSMUSG( | 4930513N1 | 17.62854 | 0.139268 | 0.80423 | NA | ENSMUSG( | 4930513N1 | 16.70202 | 0.190176 | 0.755991 | 0.971371 |
| ENSMUSG( | 4930517G | 20.67142 | 0.034248 | 0.94564 | NA | ENSMUSG( | 4930517G | 15.62041 | -0.51493 | 0.353252 | 0.912263 |
| ENSMUSG( | 4930519K1 | 56.87815 | -0.00044 | 0.99862 | 0.99989 | ENSMUSG( | 4930519K1 | 47.40361 | -0.27193 | 0.362654 | 0.914223 |
| ENSMUSG( | 4930520O | 15.17476 | 0.106942 | 0.836704 | NA | ENSMUSG( | 4930520O | 10.99134 | -0.58192 | 0.371722 | NA |
| ENSMUSG( | 4930522L1 | 30.60369 | -8.27E-05 | 0.999829 | 0.99989 | ENSMUSG( | 4930522L1 | 32.27056 | 0.376796 | 0.342372 | 0.909646 |
| ENSMUSG( | 4930523C | 113.7812 | 0.010513 | 0.955415 | 0.993874 | ENSMUSG( | 4930523C | 109.0103 | 0.165256 | 0.499823 | 0.927825 |
| ENSMUSG( | 4930525G | 29.40442 | 0.106898 | 0.754187 | 0.950981 | ENSMUSG( | 4930525G | 26.25167 | 0.031451 | 0.935207 | 0.993078 |
| ENSMUSG( | 4930539J0 | 42.83531 | -0.114 | 0.708792 | 0.939519 | ENSMUSG( | 4930539J0 | 38.99867 | -0.1065 | 0.746982 | 0.969763 |
| ENSMUSG( | 4930550C1 | 25.13263 | 0.120135 | 0.750993 | 0.950053 | ENSMUSG( | 4930550C1 | 19.14108 | -0.45617 | 0.309224 | 0.898538 |
| ENSMUSG( | 4930553P1 | 19.7764 | 0.589054 | 0.156278 | NA | ENSMUSG( | 4930553P1 | 17.73384 | 0.544544 | 0.238053 | 0.87259 |
| ENSMUSG( | 4930556M | 19.24142 | -0.08008 | 0.855078 | NA | ENSMUSG( | 4930556M | 20.17144 | 0.302758 | 0.499994 | 0.927825 |
| ENSMUSG( | 4930563E2 | 53.49038 | -0.00861 | 0.973389 | 0.99675 | ENSMUSG( | 4930563E2 | 46.11528 | -0.17762 | 0.548824 | 0.938075 |
| ENSMUSG( | 4930570G1 | 246.4156 | 0.194132 | 0.207727 | 0.64962 | ENSMUSG( | 4930570G1 | 230.973 | 0.270087 | 0.125679 | 0.81516 |
| ENSMUSG( | 4930577N1 | 23.62456 | -0.39179 | 0.32515 | 0.770608 | ENSMUSG( | 4930577N1 | 23.43136 | -0.11596 | 0.787388 | 0.97637 |
| ENSMUSG( | 4930578M | 83.84711 | 0.113872 | 0.618793 | 0.911432 | ENSMUSG( | 4930578M | 78.05976 | 0.186566 | 0.453178 | 0.922316 |
| ENSMUSG( | 4930579G | 38.49625 | 0.107746 | 0.724233 | 0.944096 | ENSMUSG( | 4930579G | 31.18177 | -0.21805 | 0.558513 | 0.93957 |
| ENSMUSG( | 4930581F2 | 184.0921 | 0.107551 | 0.566831 | 0.891644 | ENSMUSG( | 4930581F2 | 163.7748 | 0.049572 | 0.837303 | 0.981335 |
| ENSMUSG( | 4930590J0 | 35.97769 | -0.12171 | 0.70496 | 0.938303 | ENSMUSG( | 4930590J0 | 33.53511 | -0.04324 | 0.900033 | 0.990558 |
| ENSMUSG( | 4930594M | 35.96302 | -0.10742 | 0.751022 | 0.950053 | ENSMUSG( | 4930594M | 32.75553 | -0.13787 | 0.725195 | 0.966887 |
| ENSMUSG( | 4930599N | 11.93012 | -0.38703 | 0.506629 | NA | ENSMUSG( | 4930599N | 11.73985 | -0.18319 | 0.758673 | 0.971858 |
| ENSMUSG( | 4931403E2 | 17.44727 | -0.34067 | 0.483101 | NA | ENSMUSG( | 4931403E2 | 14.86934 | -0.57951 | 0.306537 | 0.898538 |
| ENSMUSG( | 4931406C | 1930.201 | 0.161341 | 0.141532 | 0.558606 | ENSMUSG( | 4931406C | 1620.995 | -0.07283 | 0.554386 | 0.939565 |
| ENSMUSG( | 4931408D1 | 9.997274 | -0.37006 | 0.526288 | NA | ENSMUSG( | 4931408D1 | 8.074322 | -0.87416 | 0.212163 | NA |
| ENSMUSG( | 4931414P1 | 113.363 | 0.002829 | 0.988596 | 0.999109 | ENSMUSG( | 4931414P1 | 94.41538 | -0.26799 | 0.228511 | 0.868507 |
| ENSMUSG( | 4931415C1 | 30.14398 | -0.25696 | 0.477636 | 0.852702 | ENSMUSG( | 4931415C1 | 32.71908 | 0.234248 | 0.514825 | 0.930939 |
| ENSMUSG( | 4932441J0 | 582.8918 | 0.169097 | 0.122224 | 0.527707 | ENSMUSG( | 4932441J0 | 483.6129 | -0.09089 | 0.567399 | 0.941777 |
| ENSMUSG( | 4933404O1 | 558.219 | 0.201052 | 0.11499 | 0.51132 | ENSMUSG( | 4933404O1 | 480.395 | 0.044171 | 0.747698 | 0.96984 |
| ENSMUSG( | 4933406C1 | 41.81532 | 0.043802 | 0.903003 | 0.986055 | ENSMUSG( | 4933406C1 | 43.30341 | 0.408702 | 0.274455 | 0.889874 |
| ENSMUSG( | 4933406I1 | 18.9339 | -0.21363 | 0.616911 | NA | ENSMUSG( | 4933406I1 | 19.4448 | 0.105976 | 0.820445 | 0.979198 |
| ENSMUSG( | 4933407K1 | 89.4612 | -0.09795 | 0.653541 | 0.922819 | ENSMUSG( | 4933407K1 | 87.51091 | 0.113777 | 0.633913 | 0.953891 |
| ENSMUSG( | 4933407L2 | 148.1669 | 0.005034 | 0.978222 | 0.997428 | ENSMUSG( | 4933407L2 | 121.2219 | -0.32996 | 0.123484 | 0.814691 |
| ENSMUSG( | 4933412E1 | 110.2248 | -0.10925 | 0.607299 | 0.906826 | ENSMUSG( | 4933412E1 | 97.04881 | -0.22242 | 0.363726 | 0.914223 |
| ENSMUSG( | 4933412O | 25.26575 | -0.04419 | 0.907243 | 0.98755 | ENSMUSG( | 4933412O | 24.34071 | 0.103605 | 0.800913 | 0.97681 |
| ENSMUSG( | 4933421O1 | 319.3208 | -0.023 | 0.855975 | 0.973813 | ENSMUSG( | 4933421O1 | 278.8935 | -0.14397 | 0.343215 | 0.909646 |
| ENSMUSG( | 4933424M | 15.10154 | -0.41738 | 0.397775 | NA | ENSMUSG( | 4933424M | 12.19519 | -0.85514 | 0.143134 | 0.831311 |
| ENSMUSG( | 4933427D1 | 114.7523 | 0.140627 | 0.504458 | 0.866883 | ENSMUSG( | 4933427D1 | 96.46578 | -0.06169 | 0.823663 | 0.979198 |
| ENSMUSG( | 4933427I2 | 26.79078 | 0.081943 | 0.822391 | 0.966352 | ENSMUSG( | 4933427I2 | 19.87191 | -0.54058 | 0.230331 | 0.868507 |
| ENSMUSG( | 4933431E2 | 8013.95 | 0.170085 | 0.043053 | 0.31819 | ENSMUSG( | 4933431E2 | 6936.179 | 0.026554 | 0.807941 | 0.978622 |
| ENSMUSG( | 4933432I0 | 72.49951 | 0.513653 | 0.065316 | 0.395951 | ENSMUSG( | 4933432I0 | 54.64301 | 0.017917 | 0.950255 | 0.994798 |
| ENSMUSG( | 4933434E2 | 484.8386 | -0.16842 | 0.185724 | 0.623185 | ENSMUSG( | 4933434E2 | 492.805 | 0.142438 | 0.344873 | 0.91045 |
| ENSMUSG( | 4933439C1 | 885.2266 | 0.370597 | 0.065423 | 0.396102 | ENSMUSG( | 4933439C1 | 722.6119 | 0.074785 | 0.751291 | 0.970317 |
| ENSMUSG( | 5031425E2 | 379.3905 | -0.05272 | 0.666201 | 0.927057 | ENSMUSG( | 5031425E2 | 359.4357 | 0.065348 | 0.662472 | 0.956205 |
| ENSMUSG( | 5031439G | 8698.88 | 0.106398 | 0.154575 | 0.581712 | ENSMUSG( | 5031439G | 7426.369 | -0.08144 | 0.403146 | 0.917257 |
| ENSMUSG( | 5033404E1 | 22.27376 | -0.39247 | 0.331418 | 0.77519 | ENSMUSG( | 5033404E1 | 20.69002 | -0.27107 | 0.571551 | 0.942116 |
| ENSMUSG( | 5033417F2 | 94.29948 | 0.241008 | 0.27733 | 0.727292 | ENSMUSG( | 5033417F2 | 76.08388 | -0.08025 | 0.74832 | 0.96984 |
| ENSMUSG( | 5033421B | 25.05707 | -0.30376 | 0.429311 | 0.826446 | ENSMUSG( | 5033421B | 21.39604 | -0.55007 | 0.206131 | 0.864412 |
| ENSMUSG( | 5033426O | 10.46605 | -0.21048 | 0.710879 | NA | ENSMUSG( | 5033426O | 9.887613 | -0.13121 | 0.832372 | NA |
| ENSMUSG( | 5033430I1 | 128.5463 | -0.07206 | 0.720727 | 0.942638 | ENSMUSG( | 5033430I1 | 112.0993 | -0.20432 | 0.356062 | 0.913331 |
| ENSMUSG( | 5330413P1 | 39.70299 | -0.07838 | 0.804467 | 0.962346 | ENSMUSG( | 5330413P1 | 35.14164 | -0.14424 | 0.671097 | 0.95694 |
| ENSMUSG( | 5330416C | 24.95027 | 0.081982 | 0.835104 | 0.970426 | ENSMUSG( | 5330416C | 24.6419 | 0.289761 | 0.476511 | 0.923329 |
| ENSMUSG( | 5330434G | 143.2226 | -0.20062 | 0.264429 | 0.715323 | ENSMUSG( | 5330434G | 137.5371 | -0.05398 | 0.786484 | 0.976303 |
| ENSMUSG( | 5430402O1 | 48.39146 | 0.040347 | 0.893148 | 0.984512 | ENSMUSG( | 5430402O1 | 42.92926 | -0.0603 | 0.850944 | 0.985541 |
| ENSMUSG( | 5430405H | 177.2658 | 0.010587 | 0.9498 | 0.993821 | ENSMUSG( | 5430405H | 174.4795 | 0.232122 | 0.233288 | 0.868507 |
| ENSMUSG( | 5430416N | 325.891 | -0.08512 | 0.550931 | 0.886038 | ENSMUSG( | 5430416N | 303.0333 | -0.02541 | 0.861315 | 0.986281 |
| ENSMUSG( | 5530601H | 354.5768 | -0.01296 | 0.917403 | 0.989182 | ENSMUSG( | 5530601H | 328.1173 | 0.026956 | 0.863522 | 0.986547 |
| ENSMUSG( | 5730405O1 | 15.56929 | -0.34434 | 0.463204 | NA | ENSMUSG( | 5730405O1 | 14.40311 | -0.23341 | 0.681111 | 0.958395 |
| ENSMUSG( | 5730409E0 | 2253.945 | 0.132551 | 0.22059 | 0.666079 | ENSMUSG( | 5730409E0 | 1863.381 | -0.14999 | 0.243098 | 0.877399 |
| ENSMUSG( | 5730455P1 | 2138.661 | 0.073522 | 0.393717 | 0.809643 | ENSMUSG( | 5730455P1 | 1876.909 | -0.03549 | 0.747946 | 0.96984 |
| ENSMUSG( | 5830418P1 | 16.51529 | 0.11175 | 0.820984 | NA | ENSMUSG( | 5830418P1 | 11.99911 | -0.56965 | 0.388516 | 0.917257 |
| ENSMUSG( | 5830444B | 26.73011 | 0.062729 | 0.896452 | 0.984868 | ENSMUSG( | 5830444B | 26.26241 | 0.235328 | 0.636583 | 0.953903 |
| ENSMUSG( | 5830454E0 | 26.37695 | -0.12248 | 0.736552 | 0.946437 | ENSMUSG( | 5830454E0 | 24.04399 | -0.12054 | 0.762702 | 0.973566 |
| ENSMUSG( | 5930430L0 | 30.96575 | -0.19182 | 0.575051 | 0.894572 | ENSMUSG( | 5930430L0 | 36.58598 | 0.553077 | 0.153036 | 0.837724 |
| ENSMUSG( | 6030407O | 33.0876 | 0.445461 | 0.194285 | 0.633391 | ENSMUSG( | 6030407O | 25.0339 | -0.09696 | 0.818242 | 0.979198 |
| ENSMUSG( | 6030442K2 | 11.67214 | 0.528249 | 0.32475 | NA | ENSMUSG( | 6030442K2 | 8.255128 | -0.10314 | 0.88392 | NA |
| ENSMUSG( | 6030443J0 | 28.64404 | -0.56001 | 0.133741 | 0.547197 | ENSMUSG( | 6030443J0 | 27.24114 | -0.39883 | 0.31803 | 0.902849 |
| ENSMUSG( | 6030458C1 | 847.3164 | -0.02482 | 0.819654 | 0.965888 | ENSMUSG( | 6030458C1 | 774.9558 | -0.01419 | 0.910867 | 0.992196 |
| ENSMUSG( | 6230400D1 | 41.50835 | 0.02176 | 0.940539 | 0.993415 | ENSMUSG( | 6230400D1 | 32.23376 | -0.45525 | 0.204937 | 0.864412 |
| ENSMUSG( | 6330403K | 4045.549 | 0.012182 | 0.911248 | 0.988483 | ENSMUSG( | 6330403K | 3523.992 | -0.12306 | 0.424928 | 0.917257 |
| ENSMUSG( | 6330415G1 | 30.10373 | 0.200808 | 0.552089 | 0.886141 | ENSMUSG( | 6330415G1 | 25.10634 | -0.03721 | 0.923641 | 0.992815 |
| ENSMUSG( | 6330418K | 136.5107 | 0.20327 | 0.25374 | 0.704011 | ENSMUSG( | 6330418K | 110.4373 | -0.13444 | 0.512601 | 0.930164 |
| ENSMUSG( | 6430503K | 19.58027 | -0.23332 | 0.583261 | NA | ENSMUSG( | 6430503K | 16.95365 | -0.39695 | 0.406876 | 0.917257 |
| ENSMUSG( | 6430548M | 10798.6 | 0.15606 | 0.059417 | 0.376611 | ENSMUSG( | 6430548M | 9663.649 | 0.107796 | 0.34957 | 0.911652 |
| ENSMUSG( | 6430550D | 41.30744 | -0.14932 | 0.608687 | 0.907185 | ENSMUSG( | 6430550D | 40.50197 | 0.088411 | 0.795862 | 0.97681 |
| ENSMUSG( | 6430571L1 | 114.9906 | -0.62477 | 0.00078 | 0.027489 | ENSMUSG( | 6430571L1 | 120.2076 | -0.18038 | 0.383759 | 0.917257 |
| ENSMUSG( | 6430573P | 25.22923 | 0.482515 | 0.224375 | 0.67048 | ENSMUSG( | 6430573P | 19.37869 | 0.050535 | 0.908262 | 0.991958 |
| ENSMUSG( | 6430584L0 | 107.6971 | -0.05169 | 0.806289 | 0.962804 | ENSMUSG( | 6430584L0 | 93.55701 | -0.20068 | 0.383551 | 0.917257 |
| ENSMUSG( | 6530402F1 | 663.1571 | 0.050526 | 0.671574 | 0.928781 | ENSMUSG( | 6530402F1 | 592.2464 | 0.00053 | 0.997198 | 0.999268 |
| ENSMUSG( | 6530411M | 35.53178 | -0.06141 | 0.862415 | 0.975055 | ENSMUSG( | 6530411M | 31.20432 | -0.20801 | 0.583461 | 0.944266 |
| ENSMUSG( | 6720483E2 | 10.30468 | 0.281158 | 0.627597 | NA | ENSMUSG( | 6720483E2 | 11.8479 | 0.867097 | 0.123268 | 0.814691 |
| ENSMUSG( | 6820408C1 | 79.23309 | -0.15205 | 0.498462 | 0.862866 | ENSMUSG( | 6820408C1 | 79.6593 | 0.122728 | 0.616383 | 0.951413 |
| ENSMUSG( | 6820431F2 | 107.5652 | -0.32902 | 0.096506 | 0.476986 | ENSMUSG( | 6820431F2 | 106.5273 | -0.08014 | 0.716289 | 0.965692 |
| ENSMUSG( | 8030453O | 89.71032 | 0.4627 | 0.033915 | 0.282701 | ENSMUSG( | 8030453O | 74.89298 | 0.210752 | 0.439336 | 0.920721 |
| ENSMUSG( | 8430429K | 195.4316 | 0.143868 | 0.399221 | 0.813048 | ENSMUSG( | 8430429K | 167.9081 | -0.02928 | 0.877015 | 0.989644 |
| ENSMUSG( | 9030612E0 | 47.86621 | -0.02996 | 0.920573 | 0.990307 | ENSMUSG( | 9030612E0 | 44.0405 | 0.02605 | 0.938506 | 0.993883 |
| ENSMUSG( | 9030624G | 12.16967 | 0.293454 | 0.585313 | NA | ENSMUSG( | 9030624G | 12.26503 | 0.510058 | 0.368661 | 0.915491 |
| ENSMUSG( | 9130019P1 | 17.59965 | 0.276802 | 0.536079 | NA | ENSMUSG( | 9130019P1 | 15.14167 | 0.164325 | 0.752549 | 0.970521 |
| ENSMUSG( | 9130023H | 96.0002 | 0.201224 | 0.325545 | 0.770741 | ENSMUSG( | 9130023H | 86.14009 | 0.171912 | 0.461174 | 0.923329 |
| ENSMUSG( | 9130024F1 | 472.182 | -0.13892 | 0.301942 | 0.750166 | ENSMUSG( | 9130024F1 | 460.0231 | 0.056396 | 0.681962 | 0.958678 |
| ENSMUSG( | 9130213A | 47.04095 | 0.439898 | 0.125246 | 0.532314 | ENSMUSG( | 9130213A | 38.17844 | 0.14899 | 0.645498 | 0.953988 |
| ENSMUSG( | 9130221H1 | 226.9369 | 0.120453 | 0.467544 | 0.844819 | ENSMUSG( | 9130221H1 | 202.9951 | 0.076632 | 0.675784 | 0.95694 |
| ENSMUSG( | 9130401M | 318.3216 | 0.139556 | 0.306484 | 0.754123 | ENSMUSG( | 9130401M | 282.6939 | 0.070009 | 0.643126 | 0.953903 |
| ENSMUSG( | 9230112J1 | 23.22172 | 0.204506 | 0.64641 | 0.918612 | ENSMUSG( | 9230112J1 | 17.40126 | -0.33137 | 0.497854 | 0.927825 |
| ENSMUSG( | 9230114K1 | 73.57 | -0.04399 | 0.846352 | 0.972226 | ENSMUSG( | 9230114K1 | 60.69726 | -0.37588 | 0.167977 | 0.840222 |
| ENSMUSG( | 9230116N1 | 11.66867 | -0.41374 | 0.467445 | NA | ENSMUSG( | 9230116N1 | 11.52922 | -0.14649 | 0.797191 | 0.97681 |
| ENSMUSG( | 9330102E0 | 1417.876 | 0.113533 | 0.224884 | 0.671259 | ENSMUSG( | 9330102E0 | 1209.333 | -0.07349 | 0.534779 | 0.936303 |
| ENSMUSG( | 9330104G | 128.2266 | -0.01101 | 0.957808 | 0.993874 | ENSMUSG( | 9330104G | 120.9448 | 0.094065 | 0.641119 | 0.953903 |
| ENSMUSG( | 9330111N | 9.621292 | 0.296261 | 0.628149 | NA | ENSMUSG( | 9330111N | 8.774966 | 0.315213 | 0.631444 | NA |
| ENSMUSG( | 9330117O1 | 33.6231 | -0.30951 | 0.366448 | 0.794761 | ENSMUSG( | 9330117O1 | 28.9509 | -0.47232 | 0.240242 | 0.875274 |
| ENSMUSG( | 9330133O1 | 215.7032 | -0.12357 | 0.441698 | 0.832577 | ENSMUSG( | 9330133O1 | 199.4707 | -0.06482 | 0.733732 | 0.967228 |

| | | | | | |
|---|---|---|---|---|---|
| ENSMUSG( | 318.3216 | 0.139556 | 0.306484 | 0.754123 | 9130401M01Rik |
| ENSMUSG( | 23.22172 | 0.204506 | 0.64641 | 0.918612 | 9230112J17Rik |
| ENSMUSG( | 73.57 | -0.04399 | 0.846352 | 0.972226 | 9230114K14Rik |
| ENSMUSG( | 11.66867 | -0.41374 | 0.467445 | NA | 9230116N13Rik |
| ENSMUSG( | 1417.876 | 0.113533 | 0.224884 | 0.671259 | 9330102E08Rik |
| ENSMUSG( | 128.2266 | -0.01101 | 0.957808 | 0.993874 | 9330104G04Rik |
| ENSMUSG( | 9.621292 | 0.296261 | 0.628149 | NA | 9330111N05Rik |
| ENSMUSG( | 33.6231 | -0.30951 | 0.366448 | 0.794761 | 9330117O12Rik |
| ENSMUSG( | 215.7032 | -0.12357 | 0.441698 | 0.832577 | 9330133O14Rik |
| ENSMUSG( | 16.61318 | -0.24131 | 0.624196 | NA | 9330136K24Rik |
| ENSMUSG( | 718.6265 | -0.04572 | 0.708983 | 0.939519 | 9330151L19Rik |
| ENSMUSG( | 17.72769 | -0.37601 | 0.398644 | NA | 9330158H04Rik |
| ENSMUSG( | 1962.26 | 0.225632 | 0.011732 | 0.156231 | 9330159F19Rik |
| ENSMUSG( | 93.12834 | -0.09336 | 0.648999 | 0.920025 | 9330159M07Rik |
| ENSMUSG( | 58.3475 | -0.35397 | 0.167332 | 0.599971 | 9330179D12Rik |
| ENSMUSG( | 95.86535 | -0.1087 | 0.617021 | 0.910866 | 9330188P03Rik |
| ENSMUSG( | 654.2415 | 0.034192 | 0.779224 | 0.956632 | 9430015G10Rik |
| ENSMUSG( | 281.3182 | 0.104012 | 0.555977 | 0.887457 | 9430021M05Rik |
| ENSMUSG( | 36.63167 | 0.309913 | 0.348243 | 0.78132 | 9430037G07Rik |
| ENSMUSG( | 62.60016 | 0.023832 | 0.921521 | 0.990436 | 9430038I01Rik |
| ENSMUSG( | 392.4759 | 0.096749 | 0.488941 | 0.859092 | 9430041J12Rik |
| ENSMUSG( | 34.95979 | -0.23348 | 0.461711 | 0.841564 | 9430065F17Rik |
| ENSMUSG( | 61.22051 | 0.441738 | 0.080159 | 0.437771 | 9530026P05Rik |
| ENSMUSG( | 40.82685 | 0.606243 | 0.042371 | 0.316636 | 9530036O11Rik |
| ENSMUSG( | 34.61174 | -0.17226 | 0.594614 | 0.902971 | 9530051G07Rik |
| ENSMUSG( | 34.20013 | 0.744402 | 0.031094 | 0.268324 | 9530052E02Rik |
| ENSMUSG( | 15.07777 | 0.586387 | 0.301502 | NA | 9530059O14Rik |
| ENSMUSG( | 27.83883 | -0.03284 | 0.927797 | 0.991229 | 9530062K07Rik |
| ENSMUSG( | 4282.209 | -0.28462 | 0.021644 | 0.225182 | 9530068E07Rik |
| ENSMUSG( | 25.35497 | -0.03697 | 0.924851 | 0.991152 | 9530080O11Rik |
| ENSMUSG( | 139.3877 | -0.06241 | 0.73467 | 0.945584 | 9530082P21Rik |
| ENSMUSG( | 53.29593 | -0.43954 | 0.152208 | 0.578042 | 9630001P10Rik |
| ENSMUSG( | 408.0715 | 0.022938 | 0.846934 | 0.972374 | 9630013A20Rik |
| ENSMUSG( | 14.66481 | -0.15231 | 0.792093 | NA | 9630028B13Rik |
| ENSMUSG( | 43.38124 | 0.082944 | 0.780635 | 0.957491 | 9830166K06Rik |
| ENSMUSG( | 39.10202 | -0.02474 | 0.937379 | 0.993415 | 9930012K11Rik |
| ENSMUSG( | 21.56466 | -0.66252 | 0.100129 | 0.483045 | 9930111J21Rik1 |
| ENSMUSG( | 58.97537 | -0.16523 | 0.528295 | 0.878236 | 9930111J21Rik2 |
| ENSMUSG( | 160.8609 | 0.015125 | 0.929657 | 0.991848 | A130010J15Rik |
| ENSMUSG( | 57.2012 | -0.95937 | 0.000386 | 0.017375 | A230001M10Rik |
| ENSMUSG( | 13.65426 | -0.43711 | 0.422264 | NA | A230009B12Rik |
| ENSMUSG( | 23.52068 | -0.06348 | 0.876471 | 0.980777 | A230028O05Rik |
| ENSMUSG( | 50.80421 | -0.04663 | 0.889563 | 0.98353 | A230051N06Rik |
| ENSMUSG( | 787.2146 | 0.075609 | 0.667042 | 0.92729 | A230057D06Rik |
| ENSMUSG( | 246.8252 | 0.034161 | 0.82018 | 0.965888 | A230072C01Rik |
| ENSMUSG( | 34.63884 | -0.23677 | 0.460338 | 0.840676 | A230072E10Rik |
| ENSMUSG( | 193.2963 | 0.341936 | 0.068364 | 0.405144 | A230077H06Rik |
| ENSMUSG( | 20.59494 | 0.633637 | 0.116676 | NA | A230083G16Rik |
| ENSMUSG( | 138.9117 | -0.01962 | 0.910192 | 0.9881 | A230103J11Rik |
| ENSMUSG( | 19.87375 | -0.07496 | 0.857246 | NA | A230107N01Rik |
| ENSMUSG( | 248.005 | 0.187422 | 0.288737 | 0.739578 | A2m |
| ENSMUSG( | 177.9602 | 0.011197 | 0.951361 | 0.993821 | A2ml1 |
| ENSMUSG( | 51.15378 | -0.04037 | 0.890536 | 0.983756 | A330008L17Rik |
| ENSMUSG( | 317.2648 | 0.104695 | 0.414983 | 0.820184 | A330009N23Rik |
| ENSMUSG( | 31.81295 | -0.58547 | 0.115392 | 0.512362 | A330015K06Rik |
| ENSMUSG( | 1343.695 | 0.3893 | 0.001321 | 0.039614 | A330023F24Rik |
| ENSMUSG( | 12.20139 | -0.30286 | 0.575108 | NA | A330032P22Rik |
| ENSMUSG( | 20.22469 | 0.29613 | 0.487196 | NA | A330033J07Rik |
| ENSMUSG( | 11.22131 | -0.05796 | 0.916617 | NA | A330040F15Rik |
| ENSMUSG( | 78.7202 | 0.213643 | 0.348664 | 0.781338 | A330041J22Rik |
| ENSMUSG( | 141.7293 | 0.277469 | 0.14614 | 0.567605 | A330048O09Rik |
| ENSMUSG( | 17.10601 | -0.78835 | 0.106361 | NA | A330049N07Rik |
| ENSMUSG( | 28.25776 | -0.28797 | 0.432333 | 0.82768 | A330069E16Rik |
| ENSMUSG( | 184.2759 | -0.12572 | 0.430137 | 0.826789 | A330074K22Rik |
| ENSMUSG( | 45.38979 | -0.19075 | 0.535579 | 0.881043 | A330076C08Rik |
| ENSMUSG( | 97.29644 | -0.21 | 0.376974 | 0.801213 | A330076H08Rik |
| ENSMUSG( | 248.5221 | 0.23499 | 0.15109 | 0.576088 | A330102I10Rik |
| ENSMUSG( | 16.20218 | 0.413087 | 0.396821 | NA | A3galt2 |
| ENSMUSG( | 295.806 | -0.00394 | 0.979594 | 0.997428 | A430005L14Rik |
| ENSMUSG( | 47.07079 | -0.29221 | 0.308099 | 0.75626 | A430018G15Rik |
| ENSMUSG( | 295.505 | 0.226626 | 0.125692 | 0.53257 | A430033K04Rik |
| ENSMUSG( | 138.716 | -0.26951 | 0.157465 | 0.585653 | A430035B10Rik |
| ENSMUSG( | 409.924 | 0.078952 | 0.633965 | 0.914073 | A430046D13Rik |
| ENSMUSG( | 24.82521 | -0.49408 | 0.194607 | 0.633457 | A430057M04Rik |
| ENSMUSG( | 33.96265 | -0.26746 | 0.447594 | 0.834051 | A430108G06Rik |
| ENSMUSG( | 69.35609 | -0.19043 | 0.488973 | 0.859092 | A4galt |
| ENSMUSG( | 9.6766 | -0.41566 | 0.494052 | NA | A530021J07Rik |
| ENSMUSG( | 57.33669 | -0.07333 | 0.782492 | 0.957558 | A530072M11Rik |
| ENSMUSG( | 47.18343 | -0.24157 | 0.388426 | 0.806278 | A630001G21Rik |
| ENSMUSG( | 37.58645 | 0.888641 | 0.006997 | 0.115211 | A630023P12Rik |
| ENSMUSG( | 32.24702 | -0.17761 | 0.621101 | 0.911432 | A630066F11Rik |
| ENSMUSG( | 284.9878 | 0.118773 | 0.393869 | 0.809643 | A630072M18Rik |
| ENSMUSG( | 71.343 | 0.143254 | 0.558181 | 0.888427 | A630089N07Rik |
| ENSMUSG( | 1707.048 | 0.05738 | 0.547607 | 0.885153 | A730008H23Rik |
| ENSMUSG( | 26.07894 | 0.744294 | 0.0532 | 0.354231 | A730020M07Rik |
| ENSMUSG( | 112.587 | 0.057929 | 0.766776 | 0.954331 | A730036I17Rik |
| ENSMUSG( | 255.5309 | 0.367157 | 0.016229 | 0.189102 | A730056A06Rik |
| ENSMUSG( | 50.78019 | -0.12999 | 0.634685 | 0.914073 | A730081D07Rik |
| ENSMUSG( | 13.4544 | 0.15243 | 0.7616 | NA | A730098A19Rik |
| ENSMUSG( | 11.45946 | 0.259709 | 0.642189 | NA | A830005F24Rik |
| ENSMUSG( | 57.04016 | 0.078819 | 0.783843 | 0.957645 | A830008E24Rik |
| ENSMUSG( | 201.2509 | 0.182797 | 0.286068 | 0.735242 | A830009L08Rik |
| ENSMUSG( | 67.98895 | 0.030924 | 0.894861 | 0.984512 | A830011K09Rik |
| ENSMUSG( | 1076.131 | 0.056317 | 0.554846 | 0.887009 | A830018L16Rik |
| ENSMUSG( | 91.78798 | 0.271879 | 0.2544 | 0.704536 | A830019P07Rik |
| ENSMUSG( | 51.42271 | 0.283621 | 0.288059 | 0.738882 | A830021F12Rik |
| ENSMUSG( | 32.85206 | -0.50535 | 0.208314 | 0.650319 | A830035O19Rik |
| ENSMUSG( | 1820.067 | -0.09624 | 0.594513 | 0.902971 | A830036E02Rik |
| ENSMUSG( | 737.5554 | 0.062647 | 0.617502 | 0.910872 | A830082K12Rik |
| ENSMUSG( | 704.1457 | -0.01098 | 0.93884 | 0.993415 | A830082N09Rik |
| ENSMUSG( | 26.65192 | 0.19091 | 0.607656 | 0.906878 | A930001A20Rik |
| ENSMUSG( | 13.01021 | 0.238338 | 0.693878 | NA | A930001C03Rik |
| ENSMUSG( | 8.887422 | -0.59337 | 0.377498 | NA | A930003A15Rik |
| ENSMUSG( | 27.47452 | -0.05083 | 0.887354 | 0.983389 | A930003O13Rik |
| ENSMUSG( | 118.1077 | -0.1925 | 0.331487 | 0.77519 | A930004D18Rik |
| ENSMUSG( | 389.1821 | 0.187403 | 0.130486 | 0.541757 | A930005H10Rik |
| ENSMUSG( | 28.26604 | -0.40955 | 0.274145 | 0.724254 | A930006K02Rik |
| ENSMUSG( | 40.29636 | -0.08373 | 0.784042 | 0.957689 | A930007A09Rik |
| ENSMUSG( | 54.43652 | -0.33743 | 0.191876 | 0.630811 | A930007I19Rik |
| ENSMUSG( | 27.55391 | -0.1992 | 0.595377 | 0.903058 | A930009A15Rik |
| ENSMUSG( | 1132.139 | -0.04035 | 0.737588 | 0.947329 | A930011G23Rik |
| ENSMUSG( | 94.09326 | -0.1281 | 0.572941 | 0.893584 | A930012L18Rik |
| ENSMUSG( | 22.54035 | 0.293899 | 0.483956 | 0.856501 | A930016O22Rik |
| ENSMUSG( | 363.9197 | -0.11479 | 0.373819 | 0.80044 | A930017M01Rik |
| ENSMUSG( | 75.32247 | 0.017624 | 0.939441 | 0.993415 | A930024E05Rik |
| ENSMUSG( | 80.0296 | 0.013716 | 0.95351 | 0.993874 | A930033H14Rik |
| ENSMUSG( | 73.82956 | -0.11495 | 0.625757 | 0.912274 | A930037H05Rik |
| ENSMUSG( | 258.276 | 0.138963 | 0.345477 | 0.779153 | AA387883 |
| ENSMUSG( | 11.44604 | -0.46924 | 0.384108 | NA | AA413626 |
| ENSMUSG( | 109.9538 | -0.20581 | 0.323935 | 0.769037 | AA414768 |
| ENSMUSG( | 39.77855 | 0.045235 | 0.880408 | 0.981469 | AA465934 |
| ENSMUSG( | 134.39 | 0.153598 | 0.448268 | 0.834051 | AA986860 |
| ENSMUSG( | 368.7304 | 0.038195 | 0.800877 | 0.96188 | Aaas |
| ENSMUSG( | 1606.891 | 0.320437 | 0.000489 | 0.02028 | Aacs |
| ENSMUSG( | 2071.231 | 0.055674 | 0.535734 | 0.881209 | Aagab |
| ENSMUSG( | 9681.118 | 0.176504 | 0.043896 | 0.321154 | Aak1 |
| ENSMUSG( | 1015.643 | 0.070316 | 0.511481 | 0.870901 | Aamdc |
| ENSMUSG( | 4012.551 | 0.093903 | 0.282961 | 0.73186 | Aamp |
| ENSMUSG( | 15.45261 | 0.887046 | 0.06647 | NA | Aanat |
| ENSMUSG( | 1045.459 | 0.124035 | 0.199704 | 0.640239 | Aar2 |
| ENSMUSG( | 69.66609 | 0.061134 | 0.79452 | 0.959799 | Aard |
| ENSMUSG( | 3796.49 | 0.041495 | 0.672928 | 0.929007 | Aars1 |
| ENSMUSG( | 1056.862 | 0.226117 | 0.064427 | 0.393453 | Aars2 |
| ENSMUSG( | 13.03604 | 0.074172 | 0.895325 | NA | Aarsd1 |
| ENSMUSG( | 287.785 | -0.04103 | 0.782283 | 0.957558 | Aasdh |
| ENSMUSG( | 450.5583 | 0.02327 | 0.846694 | 0.972371 | Aasdhppt |
| ENSMUSG( | 574.3672 | 0.240911 | 0.120367 | 0.524495 | Aass |
| ENSMUSG( | 249.2508 | 0.102116 | 0.558478 | 0.888521 | Aatf |
| ENSMUSG( | 10274.98 | 0.086738 | 0.416148 | 0.82084 | Aatk |
| ENSMUSG( | 34.44937 | -1.00417 | 0.014531 | 0.177481 | AB124611 |
| ENSMUSG( | 29858.55 | 0.082556 | 0.467068 | 0.844819 | Abat |
| ENSMUSG( | 1954.28 | -0.19552 | 0.170513 | 0.605271 | Abca1 |
| ENSMUSG( | 9.766431 | -0.90658 | 0.131659 | NA | Abca13 |
| ENSMUSG( | 41.54773 | 0.207511 | 0.536033 | 0.88142 | Abca17 |
| ENSMUSG( | 9875.743 | -0.13867 | 0.278259 | 0.728302 | Abca2 |
| ENSMUSG( | 3837.629 | -0.16286 | 0.236979 | 0.684317 | Abca3 |
| ENSMUSG( | 131.6827 | -0.14735 | 0.619483 | 0.911432 | Abca4 |
| ENSMUSG( | 1523.374 | -0.12119 | 0.411258 | 0.819183 | Abca5 |
| ENSMUSG( | 59.55413 | -0.32372 | 0.216881 | 0.661816 | Abca6 |
| ENSMUSG( | 839.1562 | 0.052992 | 0.66726 | 0.92729 | Abca7 |
| ENSMUSG( | 125.4349 | -0.29285 | 0.226788 | 0.67384 | Abca8a |
| ENSMUSG( | 637.5161 | -0.31746 | 0.018736 | 0.205889 | Abca8b |
| ENSMUSG( | 644.3575 | 0.157664 | 0.318559 | 0.764234 | Abca9 |
| ENSMUSG( | 1322.757 | -0.05909 | 0.59925 | 0.903801 | Abcb10 |
| ENSMUSG( | 1149.777 | 0.111023 | 0.590486 | 0.90113 | Abcb1a |
| ENSMUSG( | 196.6681 | -0.04322 | 0.780292 | 0.957491 | Abcb1b |
| ENSMUSG( | 302.8013 | 0.048011 | 0.76687 | 0.954331 | Abcb4 |
| ENSMUSG( | 68.98276 | -0.19961 | 0.386338 | 0.804623 | Abcb6 |

| | | | | | |
|---|---|---|---|---|---|
| ENSMUSG( | 1209.333 | -0.07349 | 0.534779 | 0.936303 | 9330102E08Rik |
| ENSMUSG( | 120.9448 | 0.094065 | 0.641119 | 0.953903 | 9330104G04Rik |
| ENSMUSG( | 8.774966 | 0.315213 | 0.631444 | NA | 9330111N05Rik |
| ENSMUSG( | 28.9509 | -0.47232 | 0.240242 | 0.875274 | 9330117O12Rik |
| ENSMUSG( | 199.4707 | -0.06482 | 0.733732 | 0.967228 | 9330133O14Rik |
| ENSMUSG( | 13.50192 | -0.61183 | 0.295116 | 0.898538 | 9330136K24Rik |
| ENSMUSG( | 699.7531 | 0.137871 | 0.362026 | 0.914223 | 9330151L19Rik |
| ENSMUSG( | 16.88484 | -0.31434 | 0.568063 | 0.941777 | 9330158H04Rik |
| ENSMUSG( | 1729.319 | 0.132402 | 0.302006 | 0.898538 | 9330159F19Rik |
| ENSMUSG( | 83.04267 | -0.16469 | 0.476532 | 0.923329 | 9330159M07Rik |
| ENSMUSG( | 55.77004 | -0.22034 | 0.461689 | 0.923329 | 9330179D12Rik |
| ENSMUSG( | 82.2987 | -0.29266 | 0.220091 | 0.866104 | 9330188P03Rik |
| ENSMUSG( | 638.6872 | 0.231331 | 0.123827 | 0.814691 | 9430015G10Rik |
| ENSMUSG( | 255.7979 | 0.095212 | 0.576135 | 0.942912 | 9430021M05Rik |
| ENSMUSG( | 30.14665 | 0.025643 | 0.947479 | 0.994798 | 9430037G07Rik |
| ENSMUSG( | 59.10064 | 0.10006 | 0.725517 | 0.966887 | 9430038I01Rik |
| ENSMUSG( | 321.0907 | -0.21663 | 0.151378 | 0.837724 | 9430041J12Rik |
| ENSMUSG( | 32.57035 | -0.15976 | 0.648341 | 0.95422 | 9430065F17Rik |
| ENSMUSG( | 50.42757 | 0.1946 | 0.50405 | 0.927825 | 9530026P05Rik |
| ENSMUSG( | 31.93674 | 0.219268 | 0.535176 | 0.936508 | 9530036O11Rik |
| ENSMUSG( | 31.41211 | -0.20692 | 0.569751 | 0.941777 | 9530051G07Rik |
| ENSMUSG( | 21.79909 | -0.24357 | 0.57622 | 0.942912 | 9530052E02Rik |
| ENSMUSG( | 13.06296 | 0.466938 | 0.437669 | 0.920339 | 9530059O14Rik |
| ENSMUSG( | 24.88371 | -0.0995 | 0.803646 | 0.976978 | 9530062K07Rik |
| ENSMUSG( | 4308.232 | 0.017658 | 0.899893 | 0.990558 | 9530068E07Rik |
| ENSMUSG( | 21.45403 | -0.26215 | 0.557352 | 0.939565 | 9530080O11Rik |
| ENSMUSG( | 132.1078 | 0.056851 | 0.796166 | 0.97681 | 9530082P21Rik |
| ENSMUSG( | 52.28861 | -0.20167 | 0.508974 | 0.929089 | 9630001P10Rik |
| ENSMUSG( | 369.1243 | -0.00453 | 0.977278 | 0.996416 | 9630013A20Rik |
| ENSMUSG( | 13.40335 | -0.11677 | 0.845598 | 0.984209 | 9630028B13Rik |
| ENSMUSG( | 43.24084 | 0.312825 | 0.31601 | 0.901868 | 9830166K06Rik |
| ENSMUSG( | 42.42726 | 0.472565 | 0.161881 | 0.837724 | 9930012K11Rik |
| ENSMUSG( | 23.71109 | -0.01264 | 0.975789 | 0.996416 | 9930111J21Rik1 |
| ENSMUSG( | 67.96375 | 0.500077 | 0.07073 | 0.758714 | 9930111J21Rik2 |
| ENSMUSG( | 153.3439 | 0.133766 | 0.471977 | 0.923329 | A130010J15Rik |
| ENSMUSG( | 52.57514 | -0.91488 | 0.001663 | 0.150176 | A230001M10Rik |
| ENSMUSG( | 11.3826 | -0.70202 | 0.291857 | NA | A230009B12Rik |
| ENSMUSG( | 21.7795 | -0.0052 | 0.990241 | 0.998104 | A230028O05Rik |
| ENSMUSG( | 46 | -0.07544 | 0.80581 | 0.977674 | A230051N06Rik |
| ENSMUSG( | 727.4838 | 0.108372 | 0.595214 | 0.945701 | A230057D06Rik |
| ENSMUSG( | 216.9001 | -0.07505 | 0.678329 | 0.957435 | A230072C01Rik |
| ENSMUSG( | 30.57092 | -0.38317 | 0.305758 | 0.898538 | A230072E10Rik |
| ENSMUSG( | 154.4903 | -0.01667 | 0.933806 | 0.99306 | A230077H06Rik |
| ENSMUSG( | 16.83994 | 0.37102 | 0.42566 | 0.917257 | A230083G16Rik |
| ENSMUSG( | 124.7209 | -0.06126 | 0.758651 | 0.971858 | A230103J11Rik |
| ENSMUSG( | 17.34406 | -0.17918 | 0.701236 | 0.961756 | A230107N01Rik |
| ENSMUSG( | 237.8227 | 0.328699 | 0.057162 | 0.723362 | A2m |
| ENSMUSG( | 155.2157 | -0.13542 | 0.526216 | 0.934374 | A2ml1 |
| ENSMUSG( | 53.58426 | 0.317613 | 0.318918 | 0.903344 | A330008L17Rik |
| ENSMUSG( | 259.1606 | -0.21116 | 0.170107 | 0.841265 | A330009N23Rik |
| ENSMUSG( | 33.37577 | -0.13375 | 0.72284 | 0.966887 | A330015K06Rik |
| ENSMUSG( | 1242.795 | 0.429595 | 0.011892 | 0.445105 | A330023F24Rik |
| ENSMUSG( | 9.694195 | -0.75843 | 0.231856 | NA | A330032P22Rik |
| ENSMUSG( | 21.45599 | 0.681949 | 0.120485 | 0.813384 | A330033J07Rik |
| ENSMUSG( | 64.53832 | -0.10862 | 0.69288 | 0.960524 | A330041J22Rik |
| ENSMUSG( | 116.8976 | -0.02331 | 0.922917 | 0.992644 | A330048O09Rik |
| ENSMUSG( | 20.71383 | 0.176178 | 0.736765 | 0.967344 | A330049N07Rik |
| ENSMUSG( | 24.68324 | -0.41681 | 0.303206 | 0.898538 | A330069E16Rik |
| ENSMUSG( | 160.4577 | -0.26197 | 0.145364 | 0.8336 | A330074K22Rik |
| ENSMUSG( | 36.6196 | -0.61322 | 0.082099 | 0.77681 | A330076C08Rik |
| ENSMUSG( | 106.202 | 0.310363 | 0.239972 | 0.8752 | A330076H08Rik |
| ENSMUSG( | 206.9341 | -0.0131 | 0.946801 | 0.994798 | A330102I10Rik |
| ENSMUSG( | 15.0092 | 0.50079 | 0.35226 | 0.912093 | A3galt2 |
| ENSMUSG( | 264.1974 | -0.06477 | 0.69882 | 0.960925 | A430005L14Rik |
| ENSMUSG( | 42.57266 | -0.3051 | 0.374839 | 0.915491 | A430018G15Rik |
| ENSMUSG( | 263.7303 | 0.163769 | 0.302036 | 0.898538 | A430033K04Rik |
| ENSMUSG( | 128.202 | -0.24409 | 0.258305 | 0.883971 | A430035B10Rik |
| ENSMUSG( | 373.3407 | 0.082015 | 0.643787 | 0.953903 | A430046D13Rik |
| ENSMUSG( | 24.80698 | -0.17697 | 0.662391 | 0.956205 | A430057M04Rik |
| ENSMUSG( | 32.86609 | -0.10704 | 0.776792 | 0.975717 | A430108G06Rik |
| ENSMUSG( | 73.97643 | 0.279659 | 0.350503 | 0.911652 | A4galt |
| ENSMUSG( | 9.320874 | -0.22386 | 0.721602 | NA | A530021J07Rik |
| ENSMUSG( | 47.31611 | -0.41041 | 0.172877 | 0.843702 | A530072M11Rik |
| ENSMUSG( | 48.03297 | 0.116716 | 0.732422 | 0.967228 | A630001G21Rik |
| ENSMUSG( | 26.39427 | 0.309884 | 0.464107 | 0.923329 | A630023P12Rik |
| ENSMUSG( | 25.5768 | -0.61634 | 0.147589 | 0.836241 | A630066F11Rik |
| ENSMUSG( | 248.2841 | -0.01433 | 0.932428 | 0.99306 | A630072M18Rik |
| ENSMUSG( | 59.24272 | -0.11822 | 0.67603 | 0.95694 | A630089N07Rik |
| ENSMUSG( | 1500.361 | -0.04312 | 0.759129 | 0.971909 | A730008H23Rik |
| ENSMUSG( | 21.44426 | 0.529675 | 0.219536 | 0.866104 | A730020M07Rik |
| ENSMUSG( | 94.37031 | -0.20812 | 0.376503 | 0.915491 | A730036I17Rik |
| ENSMUSG( | 213.6514 | 0.124727 | 0.516198 | 0.930939 | A730056A06Rik |
| ENSMUSG( | 48.95784 | 0.016434 | 0.956746 | 0.994798 | A730081D07Rik |
| ENSMUSG( | 10.53642 | -0.27631 | 0.637279 | NA | A730098A19Rik |
| ENSMUSG( | 10.76152 | 0.295058 | 0.636242 | NA | A830005F24Rik |
| ENSMUSG( | 51.68835 | 0.060086 | 0.83558 | 0.981237 | A830008E24Rik |
| ENSMUSG( | 170.9807 | -0.00718 | 0.969723 | 0.99594 | A830009L08Rik |
| ENSMUSG( | 58.57666 | -0.15416 | 0.600723 | 0.947749 | A830011K09Rik |
| ENSMUSG( | 985.9156 | 0.06841 | 0.601183 | 0.947749 | A830018L16Rik |
| ENSMUSG( | 66.90436 | -0.40236 | 0.155353 | 0.837724 | A830019P07Rik |
| ENSMUSG( | 36.60233 | -0.4862 | 0.200149 | 0.863016 | A830021F12Rik |
| ENSMUSG( | 27.55659 | -0.82404 | 0.061431 | 0.736111 | A830035O19Rik |
| ENSMUSG( | 1708.138 | -0.01349 | 0.948717 | 0.994798 | A830036E02Rik |
| ENSMUSG( | 625.4051 | -0.15368 | 0.297408 | 0.898538 | A830082K12Rik |
| ENSMUSG( | 630.7223 | -0.06608 | 0.671743 | 0.95694 | A830082N09Rik |
| ENSMUSG( | 19.46235 | -0.54302 | 0.260507 | 0.884533 | A930001A20Rik |
| ENSMUSG( | 11.10807 | 0.11391 | 0.851292 | NA | A930001C03Rik |
| ENSMUSG( | 20.68289 | -0.65147 | 0.173995 | 0.844317 | A930003O13Rik |
| ENSMUSG( | 109.6181 | -0.14002 | 0.529747 | 0.934795 | A930004D18Rik |
| ENSMUSG( | 329.8353 | -0.02036 | 0.890881 | 0.990558 | A930005H10Rik |
| ENSMUSG( | 25.17112 | -0.45563 | 0.277649 | 0.891468 | A930006K02Rik |
| ENSMUSG( | 40.8137 | 0.219364 | 0.50026 | 0.927825 | A930007A09Rik |
| ENSMUSG( | 53.89403 | -0.06787 | 0.80908 | 0.97873 | A930007I19Rik |
| ENSMUSG( | 29.18958 | 0.220601 | 0.583522 | 0.944266 | A930009A15Rik |
| ENSMUSG( | 1016.971 | -0.08603 | 0.542728 | 0.938034 | A930011G23Rik |
| ENSMUSG( | 89.54764 | 0.011824 | 0.95894 | 0.994798 | A930012L18Rik |
| ENSMUSG( | 18.77609 | 0.0252 | 0.958374 | 0.994798 | A930016O22Rik |
| ENSMUSG( | 324.1223 | -0.19204 | 0.202062 | 0.864412 | A930017M01Rik |
| ENSMUSG( | 75.73572 | 0.312287 | 0.234854 | 0.871042 | A930024E05Rik |
| ENSMUSG( | 73.08648 | 0.024032 | 0.929507 | 0.99306 | A930033H14Rik |
| ENSMUSG( | 66.74081 | -0.14085 | 0.605962 | 0.949497 | A930037H05Rik |
| ENSMUSG( | 212.0544 | -0.16751 | 0.325001 | 0.905273 | AA387883 |
| ENSMUSG( | 11.93873 | -0.12337 | 0.842206 | 0.983044 | AA413626 |
| ENSMUSG( | 93.01186 | -0.46524 | 0.058324 | 0.728256 | AA414768 |
| ENSMUSG( | 38.49648 | 0.222701 | 0.488526 | 0.92554 | AA465934 |
| ENSMUSG( | 135.2235 | 0.418108 | 0.04247 | 0.690304 | AA986860 |
| ENSMUSG( | 335.3592 | 0.030491 | 0.85304 | 0.985598 | Aaas |
| ENSMUSG( | 1318.177 | 0.035792 | 0.745519 | 0.96951 | Aacs |
| ENSMUSG( | 1804.519 | -0.07572 | 0.517864 | 0.930939 | Aagab |
| ENSMUSG( | 8622.578 | 0.114456 | 0.244811 | 0.877904 | Aak1 |
| ENSMUSG( | 877.9044 | -0.08788 | 0.53421 | 0.935936 | Aamdc |
| ENSMUSG( | 3397.412 | -0.12046 | 0.287092 | 0.894091 | Aamp |
| ENSMUSG( | 12.64003 | 0.631276 | 0.26883 | 0.886755 | Aanat |
| ENSMUSG( | 896.7165 | -0.0473 | 0.685433 | 0.958678 | Aar2 |
| ENSMUSG( | 64.70536 | 0.092495 | 0.751048 | 0.970317 | Aard |
| ENSMUSG( | 3232.421 | -0.15817 | 0.172904 | 0.843702 | Aars1 |
| ENSMUSG( | 898.567 | 0.037383 | 0.78653 | 0.976303 | Aars2 |
| ENSMUSG( | 14.84791 | 0.652756 | 0.251214 | 0.879321 | Aarsd1 |
| ENSMUSG( | 275.9538 | 0.116993 | 0.579614 | 0.944209 | Aasdh |
| ENSMUSG( | 417.0212 | 0.059264 | 0.700917 | 0.96162 | Aasdhppt |
| ENSMUSG( | 463.596 | -0.10992 | 0.550926 | 0.939232 | Aass |
| ENSMUSG( | 208.0876 | -0.16377 | 0.423049 | 0.917257 | Aatf |
| ENSMUSG( | 9939.049 | 0.252134 | 0.086684 | 0.782073 | Aatk |
| ENSMUSG( | 34.84916 | -0.59827 | 0.158874 | 0.837724 | AB124611 |
| ENSMUSG( | 24490.91 | -0.23032 | 0.086337 | 0.781091 | Abat |
| ENSMUSG( | 2129.842 | 0.315161 | 0.056994 | 0.723362 | Abca1 |
| ENSMUSG( | 11.4156 | -0.09201 | 0.876015 | NA | Abca13 |
| ENSMUSG( | 31.67504 | -0.31781 | 0.396941 | 0.917257 | Abca17 |
| ENSMUSG( | 10207.4 | 0.223935 | 0.168426 | 0.840222 | Abca2 |
| ENSMUSG( | 3910.104 | 0.164842 | 0.316999 | 0.902088 | Abca3 |
| ENSMUSG( | 139.9299 | 0.279673 | 0.299797 | 0.898538 | Abca4 |
| ENSMUSG( | 1560.547 | 0.215121 | 0.185584 | 0.855305 | Abca5 |
| ENSMUSG( | 71.88728 | 0.489627 | 0.074297 | 0.766789 | Abca6 |
| ENSMUSG( | 787.6195 | 0.14061 | 0.380302 | 0.917202 | Abca7 |
| ENSMUSG( | 141.8778 | 0.336352 | 0.13321 | 0.822912 | Abca8a |
| ENSMUSG( | 682.3428 | 0.167517 | 0.291655 | 0.897854 | Abca8b |
| ENSMUSG( | 662.3695 | 0.478739 | 0.004928 | 0.28966 | Abca9 |
| ENSMUSG( | 1238.732 | 0.023562 | 0.864219 | 0.986552 | Abcb10 |
| ENSMUSG( | 1366.393 | 0.784582 | 0.084118 | 0.777886 | Abcb1a |
| ENSMUSG( | 177.5818 | -0.07149 | 0.695049 | 0.960867 | Abcb1b |
| ENSMUSG( | 274.1786 | 0.038402 | 0.823511 | 0.979198 | Abcb4 |
| ENSMUSG( | 67.35853 | -0.01155 | 0.964319 | 0.995433 | Abcb6 |
| ENSMUSG( | 1258.256 | 0.058185 | 0.689486 | 0.960071 | Abcb7 |
| ENSMUSG( | 1805.382 | 0.073336 | 0.592057 | 0.944975 | Abcb8 |
| ENSMUSG( | 2232.036 | -0.06642 | 0.671898 | 0.95694 | Abcb9 |
| ENSMUSG( | 812.8169 | 0.082153 | 0.543258 | 0.938063 | Abcc1 |
| ENSMUSG( | 360.3059 | 0.222437 | 0.159119 | 0.837724 | Abcc10 |
| ENSMUSG( | 19.1853 | -0.63662 | 0.170813 | 0.842157 | Abcc12 |

| | | | | | | | | | | | |
|---|---|---|---|---|---|---|---|---|---|---|---|
| ENSMUSG( | 9330136K2 | 16.61318 | -0.24131 | 0.624196 | NA | ENSMUSG( | 9330136K2 | 13.50192 | -0.61183 | 0.295116 | 0.898538 |
| ENSMUSG( | 9330151L1 | 718.6265 | -0.04572 | 0.708983 | 0.939519 | ENSMUSG( | 9330151L1 | 699.7531 | 0.137871 | 0.362026 | 0.914223 |
| ENSMUSG( | 9330158H( | 17.72769 | -0.37601 | 0.398644 | NA | ENSMUSG( | 9330158H( | 16.88484 | -0.31434 | 0.568063 | 0.941777 |
| ENSMUSG( | 9330159F1 | 1962.26 | 0.225632 | 0.011732 | 0.156231 | ENSMUSG( | 9330159F1 | 1729.319 | 0.132402 | 0.302006 | 0.898538 |
| ENSMUSG( | 9330159M | 93.12834 | -0.09336 | 0.648999 | 0.920025 | ENSMUSG( | 9330159M | 83.04267 | -0.16469 | 0.476532 | 0.923329 |
| ENSMUSG( | 9330179D1 | 58.3475 | -0.35397 | 0.167332 | 0.599971 | ENSMUSG( | 9330179D1 | 55.77004 | -0.22034 | 0.461689 | 0.923329 |
| ENSMUSG( | 9330188P( | 95.86535 | -0.1087 | 0.617021 | 0.910866 | ENSMUSG( | 9330188P( | 82.2987 | -0.29266 | 0.220091 | 0.866104 |
| ENSMUSG( | 9430015G1 | 654.2415 | 0.034192 | 0.779224 | 0.956632 | ENSMUSG( | 9430015G1 | 638.6872 | 0.231331 | 0.123827 | 0.814691 |
| ENSMUSG( | 9430021M | 281.3182 | 0.104012 | 0.555977 | 0.887457 | ENSMUSG( | 9430021M | 255.7979 | 0.095212 | 0.576135 | 0.942912 |
| ENSMUSG( | 9430037G( | 36.63167 | 0.309913 | 0.348243 | 0.78132 | ENSMUSG( | 9430037G( | 30.14665 | 0.025643 | 0.947479 | 0.994798 |
| ENSMUSG( | 9430038I0: | 62.60016 | 0.023832 | 0.921521 | 0.990436 | ENSMUSG( | 9430038I0: | 59.10064 | 0.10006 | 0.725517 | 0.966887 |
| ENSMUSG( | 9430041J1 | 392.4759 | 0.096749 | 0.488941 | 0.859092 | ENSMUSG( | 9430041J1 | 321.0907 | -0.21663 | 0.151378 | 0.837724 |
| ENSMUSG( | 9430065F1 | 34.95979 | -0.23348 | 0.461711 | 0.841564 | ENSMUSG( | 9430065F1 | 32.57035 | -0.15976 | 0.648341 | 0.95422 |
| ENSMUSG( | 9530026P( | 61.22051 | 0.441738 | 0.080159 | 0.437771 | ENSMUSG( | 9530026P( | 50.42757 | 0.1946 | 0.50405 | 0.927825 |
| ENSMUSG( | 9530036O1 | 40.82685 | 0.606243 | 0.042371 | 0.316636 | ENSMUSG( | 9530036O1 | 31.93674 | 0.219268 | 0.535176 | 0.936508 |
| ENSMUSG( | 9530051G( | 34.61174 | -0.17226 | 0.594614 | 0.902971 | ENSMUSG( | 9530051G( | 31.41211 | -0.20692 | 0.569751 | 0.941777 |
| ENSMUSG( | 9530052E( | 34.20013 | 0.744402 | 0.031094 | 0.268324 | ENSMUSG( | 9530052E( | 21.79909 | -0.24357 | 0.57622 | 0.942912 |
| ENSMUSG( | 9530059O1 | 15.07777 | 0.586387 | 0.301502 | NA | ENSMUSG( | 9530059O1 | 13.06296 | 0.466938 | 0.437669 | 0.920339 |
| ENSMUSG( | 9530062K( | 27.83883 | -0.03284 | 0.927797 | 0.991229 | ENSMUSG( | 9530062K( | 24.88371 | -0.0995 | 0.803646 | 0.976978 |
| ENSMUSG( | 9530068E( | 4282.209 | -0.28462 | 0.021644 | 0.225182 | ENSMUSG( | 9530068E( | 4308.232 | 0.017658 | 0.899893 | 0.990558 |
| ENSMUSG( | 9530080O1 | 25.35497 | -0.03697 | 0.924851 | 0.991152 | ENSMUSG( | 9530080O1 | 21.45403 | -0.26215 | 0.557352 | 0.939565 |
| ENSMUSG( | 9530082P2 | 139.3877 | -0.06241 | 0.73467 | 0.945584 | ENSMUSG( | 9530082P2 | 132.1078 | 0.056851 | 0.796166 | 0.97681 |
| ENSMUSG( | 9630001P1 | 53.29593 | -0.43954 | 0.152208 | 0.578042 | ENSMUSG( | 9630001P1 | 52.28861 | -0.20167 | 0.508974 | 0.929089 |
| ENSMUSG( | 9630013A2 | 408.0715 | 0.022938 | 0.846934 | 0.972374 | ENSMUSG( | 9630013A2 | 369.1243 | -0.00453 | 0.977278 | 0.996416 |
| ENSMUSG( | 9630028B1 | 14.66481 | -0.15231 | 0.792093 | NA | ENSMUSG( | 9630028B1 | 13.40335 | -0.11677 | 0.845598 | 0.984209 |
| ENSMUSG( | 9830166K( | 43.38124 | 0.082944 | 0.780635 | 0.957491 | ENSMUSG( | 9830166K( | 43.24084 | 0.312825 | 0.31601 | 0.901868 |
| ENSMUSG( | 9930012K1 | 39.10202 | -0.02474 | 0.937379 | 0.993415 | ENSMUSG( | 9930012K1 | 42.42726 | 0.472565 | 0.161881 | 0.837724 |
| ENSMUSG( | 9930111J2 | 21.56466 | -0.66252 | 0.100129 | 0.483045 | ENSMUSG( | 9930111J2 | 23.71109 | -0.01264 | 0.975789 | 0.996416 |
| ENSMUSG( | 9930111J2 | 58.97537 | -0.16523 | 0.528295 | 0.878236 | ENSMUSG( | 9930111J2 | 67.96375 | 0.500077 | 0.07073 | 0.758714 |
| ENSMUSG( | A130010J1 | 160.8609 | 0.015125 | 0.929657 | 0.991848 | ENSMUSG( | A130010J1 | 153.3439 | 0.133766 | 0.471977 | 0.923329 |
| ENSMUSG( | A230001M | 57.2012 | -0.95937 | 0.000386 | 0.017375 | ENSMUSG( | A230001M | 52.57514 | -0.91488 | 0.001663 | 0.150176 |
| ENSMUSG( | A230009B1 | 13.65426 | -0.43711 | 0.422264 | NA | ENSMUSG( | A230009B1 | 11.3826 | -0.70202 | 0.291857 | NA |
| ENSMUSG( | A230028O( | 23.52068 | -0.06348 | 0.876471 | 0.980777 | ENSMUSG( | A230028O( | 21.7795 | -0.0052 | 0.990241 | 0.998104 |
| ENSMUSG( | A230051N( | 50.80421 | -0.04663 | 0.889563 | 0.98353 | ENSMUSG( | A230051N( | 46 | -0.07544 | 0.80581 | 0.977674 |
| ENSMUSG( | A230057D( | 787.2146 | 0.075609 | 0.667042 | 0.92729 | ENSMUSG( | A230057D( | 727.4838 | 0.108372 | 0.595214 | 0.945701 |
| ENSMUSG( | A230072C( | 246.8252 | 0.034161 | 0.82018 | 0.965888 | ENSMUSG( | A230072C( | 216.9001 | -0.07505 | 0.678329 | 0.957435 |
| ENSMUSG( | A230072E1 | 34.63884 | -0.23677 | 0.460338 | 0.840676 | ENSMUSG( | A230072E1 | 30.57092 | -0.38317 | 0.305758 | 0.898538 |
| ENSMUSG( | A230077H( | 193.2963 | 0.341936 | 0.068364 | 0.405144 | ENSMUSG( | A230077H( | 154.4903 | -0.01667 | 0.933806 | 0.99306 |
| ENSMUSG( | A230083G: | 20.59494 | 0.633637 | 0.116676 | NA | ENSMUSG( | A230083G: | 16.83994 | 0.37102 | 0.42566 | 0.917257 |
| ENSMUSG( | A230103J1 | 138.9117 | -0.01962 | 0.910192 | 0.9881 | ENSMUSG( | A230103J1 | 124.7209 | -0.06126 | 0.758651 | 0.971858 |
| ENSMUSG( | A230107N( | 19.87375 | -0.07496 | 0.857246 | NA | ENSMUSG( | A230107N( | 17.34406 | -0.17918 | 0.701236 | 0.961756 |
| ENSMUSG( | A2m | 248.005 | 0.187422 | 0.288737 | 0.739578 | ENSMUSG( | A2m | 237.8227 | 0.328699 | 0.057162 | 0.723362 |
| ENSMUSG( | A2ml1 | 177.9602 | 0.011197 | 0.951361 | 0.993821 | ENSMUSG( | A2ml1 | 155.2157 | -0.13542 | 0.526216 | 0.934374 |
| ENSMUSG( | A330008L1 | 51.15378 | -0.04037 | 0.890536 | 0.983756 | ENSMUSG( | A330008L1 | 53.58426 | 0.317613 | 0.318918 | 0.903344 |
| ENSMUSG( | A330009N: | 317.2648 | 0.104695 | 0.414983 | 0.820184 | ENSMUSG( | A330009N: | 259.1606 | -0.21116 | 0.170107 | 0.841265 |
| ENSMUSG( | A330015K( | 31.81295 | -0.58547 | 0.115392 | 0.512362 | ENSMUSG( | A330015K( | 33.37577 | -0.13375 | 0.72284 | 0.966887 |
| ENSMUSG( | A330023F2 | 1343.695 | 0.3893 | 0.001321 | 0.039614 | ENSMUSG( | A330023F2 | 1242.795 | 0.429595 | 0.011892 | 0.445105 |
| ENSMUSG( | A330032P2 | 12.20139 | -0.30286 | 0.575108 | NA | ENSMUSG( | A330032P2 | 9.694195 | -0.75843 | 0.231856 | NA |
| ENSMUSG( | A330033J0 | 20.22469 | 0.29613 | 0.487196 | NA | ENSMUSG( | A330033J0 | 21.45599 | 0.681949 | 0.120485 | 0.813384 |
| ENSMUSG( | A330041J2 | 78.7202 | 0.213643 | 0.348664 | 0.781338 | ENSMUSG( | A330041J2 | 64.53832 | -0.10862 | 0.69288 | 0.960524 |
| ENSMUSG( | A330048O( | 141.7293 | 0.277469 | 0.14614 | 0.567605 | ENSMUSG( | A330048O( | 116.8976 | -0.02331 | 0.922917 | 0.992644 |
| ENSMUSG( | A330049N( | 17.10601 | -0.78835 | 0.106361 | NA | ENSMUSG( | A330049N( | 20.71383 | 0.176178 | 0.736765 | 0.967344 |
| ENSMUSG( | A330069E1 | 28.25776 | -0.28797 | 0.432333 | 0.82768 | ENSMUSG( | A330069E1 | 24.68324 | -0.41681 | 0.303206 | 0.898538 |
| ENSMUSG( | A330074K2 | 184.2759 | -0.12572 | 0.430137 | 0.826789 | ENSMUSG( | A330074K2 | 160.4577 | -0.26197 | 0.145364 | 0.8336 |
| ENSMUSG( | A330076C( | 45.38979 | -0.19075 | 0.535579 | 0.881043 | ENSMUSG( | A330076C( | 36.6196 | -0.61322 | 0.082099 | 0.77681 |
| ENSMUSG( | A330076H( | 97.29644 | -0.21 | 0.376974 | 0.801213 | ENSMUSG( | A330076H( | 106.202 | 0.310363 | 0.239972 | 0.8752 |
| ENSMUSG( | A330102I1 | 248.5221 | 0.23499 | 0.15109 | 0.576088 | ENSMUSG( | A330102I1 | 206.9341 | -0.0131 | 0.946801 | 0.994798 |
| ENSMUSG( | A3galt2 | 16.20218 | 0.413087 | 0.396821 | NA | ENSMUSG( | A3galt2 | 15.0092 | 0.50079 | 0.35226 | 0.912093 |
| ENSMUSG( | A430005L1 | 295.806 | -0.00394 | 0.979594 | 0.997428 | ENSMUSG( | A430005L1 | 264.1974 | -0.06477 | 0.69882 | 0.960925 |
| ENSMUSG( | A430018G: | 47.07079 | -0.29221 | 0.308099 | 0.75626 | ENSMUSG( | A430018G: | 42.57266 | -0.3051 | 0.374839 | 0.915491 |
| ENSMUSG( | A430033K( | 295.505 | 0.226626 | 0.125692 | 0.53257 | ENSMUSG( | A430033K( | 263.7303 | 0.163769 | 0.302036 | 0.898538 |
| ENSMUSG( | A430035B1 | 138.716 | -0.26951 | 0.157465 | 0.585653 | ENSMUSG( | A430035B1 | 128.202 | -0.24409 | 0.258305 | 0.883971 |
| ENSMUSG( | A430046D: | 409.924 | 0.078952 | 0.633965 | 0.914073 | ENSMUSG( | A430046D: | 373.3407 | 0.082015 | 0.643787 | 0.953903 |
| ENSMUSG( | A430057M | 24.82521 | -0.49408 | 0.194607 | 0.633457 | ENSMUSG( | A430057M | 24.80698 | -0.17697 | 0.662391 | 0.956205 |
| ENSMUSG( | A430108G( | 33.96265 | -0.26746 | 0.447594 | 0.834051 | ENSMUSG( | A430108G( | 32.86609 | -0.10704 | 0.776792 | 0.975717 |
| ENSMUSG( | A4galt | 69.35609 | -0.19043 | 0.488973 | 0.859092 | ENSMUSG( | A4galt | 73.97643 | 0.279659 | 0.350503 | 0.911652 |
| ENSMUSG( | A530021J0 | 9.6766 | -0.41566 | 0.494052 | NA | ENSMUSG( | A530021J0 | 9.320874 | -0.22386 | 0.721602 | NA |
| ENSMUSG( | A530072M | 57.33669 | -0.07333 | 0.782492 | 0.957558 | ENSMUSG( | A530072M | 47.31611 | -0.41041 | 0.172877 | 0.843702 |
| ENSMUSG( | A630001G: | 47.18343 | -0.24157 | 0.388426 | 0.806278 | ENSMUSG( | A630001G: | 48.03297 | 0.116716 | 0.732422 | 0.967228 |
| ENSMUSG( | A630023P1 | 37.58645 | 0.888641 | 0.006997 | 0.115211 | ENSMUSG( | A630023P1 | 26.39427 | 0.309884 | 0.464107 | 0.923329 |
| ENSMUSG( | A630066F1 | 32.24702 | -0.17761 | 0.621101 | 0.911432 | ENSMUSG( | A630066F1 | 25.5768 | -0.61634 | 0.147589 | 0.836241 |
| ENSMUSG( | A630072M | 284.9878 | 0.118773 | 0.393869 | 0.809643 | ENSMUSG( | A630072M | 248.2841 | -0.01433 | 0.932428 | 0.99306 |
| ENSMUSG( | A630089N( | 71.343 | 0.143254 | 0.558181 | 0.888427 | ENSMUSG( | A630089N( | 59.24272 | -0.11822 | 0.67603 | 0.95694 |
| ENSMUSG( | A730008H: | 1707.048 | 0.05738 | 0.547607 | 0.885153 | ENSMUSG( | A730008H: | 1500.361 | -0.04312 | 0.759129 | 0.971909 |
| ENSMUSG( | A730020M | 26.07894 | 0.744294 | 0.0532 | 0.354231 | ENSMUSG( | A730020M | 21.44426 | 0.529675 | 0.219536 | 0.866104 |
| ENSMUSG( | A730036I1 | 112.587 | 0.057929 | 0.766776 | 0.954331 | ENSMUSG( | A730036I1 | 94.37031 | -0.20812 | 0.376503 | 0.915491 |
| ENSMUSG( | A730056A( | 255.5309 | 0.367157 | 0.016229 | 0.189102 | ENSMUSG( | A730056A( | 213.6514 | 0.124727 | 0.516198 | 0.930939 |
| ENSMUSG( | A730081D( | 50.78019 | -0.12999 | 0.634685 | 0.914073 | ENSMUSG( | A730081D( | 48.95784 | 0.016434 | 0.956746 | 0.994798 |
| ENSMUSG( | A730098A1 | 13.4544 | 0.15243 | 0.7616 | NA | ENSMUSG( | A730098A1 | 10.53642 | -0.27631 | 0.637279 | NA |
| ENSMUSG( | A830005F2 | 11.45946 | 0.259709 | 0.642189 | NA | ENSMUSG( | A830005F2 | 10.76152 | 0.295058 | 0.636242 | NA |
| ENSMUSG( | A830008E2 | 57.04016 | 0.078819 | 0.783843 | 0.957645 | ENSMUSG( | A830008E2 | 51.68835 | 0.060086 | 0.83558 | 0.981237 |
| ENSMUSG( | A830009L0 | 201.2509 | 0.182797 | 0.286068 | 0.735242 | ENSMUSG( | A830009L0 | 170.9807 | -0.00718 | 0.969723 | 0.99594 |
| ENSMUSG( | A830011K( | 67.98895 | 0.030924 | 0.894861 | 0.984512 | ENSMUSG( | A830011K( | 58.57666 | -0.15416 | 0.600723 | 0.947749 |
| ENSMUSG( | A830018L1 | 1076.131 | 0.056317 | 0.554846 | 0.887009 | ENSMUSG( | A830018L1 | 985.9156 | 0.06841 | 0.601183 | 0.947749 |
| ENSMUSG( | A830019P( | 91.78798 | 0.271879 | 0.2544 | 0.704536 | ENSMUSG( | A830019P( | 66.90436 | -0.40236 | 0.155353 | 0.837724 |
| ENSMUSG( | A830021F1 | 51.42271 | 0.283621 | 0.288059 | 0.738882 | ENSMUSG( | A830021F1 | 36.60233 | -0.4862 | 0.200149 | 0.863016 |
| ENSMUSG( | A830035O: | 32.85206 | -0.50535 | 0.208314 | 0.650319 | ENSMUSG( | A830035O: | 27.55659 | -0.82404 | 0.061431 | 0.736111 |
| ENSMUSG( | A830036E( | 1820.067 | -0.09624 | 0.594513 | 0.902971 | ENSMUSG( | A830036E( | 1708.138 | -0.01349 | 0.948717 | 0.994798 |
| ENSMUSG( | A830082K1 | 737.5554 | 0.062647 | 0.617502 | 0.910872 | ENSMUSG( | A830082K1 | 625.4051 | -0.15368 | 0.297408 | 0.898538 |
| ENSMUSG( | A830082N( | 704.1457 | -0.01098 | 0.93884 | 0.993415 | ENSMUSG( | A830082N( | 630.7223 | -0.06608 | 0.671743 | 0.95694 |
| ENSMUSG( | A930001A: | 26.65192 | 0.19091 | 0.607656 | 0.906878 | ENSMUSG( | A930001A: | 19.46235 | -0.54302 | 0.260507 | 0.884533 |
| ENSMUSG( | A930001C( | 13.01021 | 0.238338 | 0.693878 | NA | ENSMUSG( | A930001C( | 11.10807 | 0.11391 | 0.851292 | NA |
| ENSMUSG( | A930003O: | 27.47452 | -0.05083 | 0.887354 | 0.983389 | ENSMUSG( | A930003O: | 20.68289 | -0.65147 | 0.173995 | 0.844317 |
| ENSMUSG( | A930004D: | 118.1077 | -0.1925 | 0.331487 | 0.77519 | ENSMUSG( | A930004D: | 109.6181 | -0.14002 | 0.529747 | 0.934795 |
| ENSMUSG( | A930005H: | 389.1821 | 0.187403 | 0.130486 | 0.541757 | ENSMUSG( | A930005H: | 329.8353 | -0.02036 | 0.890881 | 0.990558 |
| ENSMUSG( | A930006K( | 28.26604 | -0.40955 | 0.274145 | 0.724254 | ENSMUSG( | A930006K( | 25.17112 | -0.45563 | 0.277649 | 0.891468 |
| ENSMUSG( | A930007A( | 40.29636 | -0.08373 | 0.784042 | 0.957689 | ENSMUSG( | A930007A( | 40.8137 | 0.219364 | 0.50026 | 0.927825 |
| ENSMUSG( | A930007I1 | 54.43652 | -0.33743 | 0.191876 | 0.630811 | ENSMUSG( | A930007I1 | 53.89403 | -0.06787 | 0.80908 | 0.97873 |
| ENSMUSG( | A930009A1 | 27.55391 | -0.1992 | 0.595377 | 0.903058 | ENSMUSG( | A930009A1 | 29.18958 | 0.220601 | 0.583522 | 0.944266 |
| ENSMUSG( | A930011G: | 1132.139 | -0.04035 | 0.737588 | 0.947329 | ENSMUSG( | A930011G: | 1016.971 | -0.08603 | 0.542728 | 0.938034 |
| ENSMUSG( | A930012L1 | 94.09326 | -0.1281 | 0.572941 | 0.893584 | ENSMUSG( | A930012L1 | 89.54764 | 0.011824 | 0.95894 | 0.994798 |
| ENSMUSG( | A930016O: | 22.54035 | 0.293899 | 0.483956 | 0.856501 | ENSMUSG( | A930016O: | 18.77609 | 0.0252 | 0.958374 | 0.994798 |
| ENSMUSG( | A930017M | 363.9197 | -0.11479 | 0.373819 | 0.80044 | ENSMUSG( | A930017M | 324.1223 | -0.19204 | 0.202062 | 0.864412 |
| ENSMUSG( | A930024E( | 75.32247 | 0.017624 | 0.939441 | 0.993415 | ENSMUSG( | A930024E( | 75.73572 | 0.312287 | 0.234854 | 0.871042 |
| ENSMUSG( | A930033H: | 80.0296 | 0.013716 | 0.95351 | 0.993874 | ENSMUSG( | A930033H: | 73.08648 | 0.024032 | 0.929507 | 0.99306 |
| ENSMUSG( | A930037H( | 73.82956 | -0.11495 | 0.625757 | 0.912274 | ENSMUSG( | A930037H( | 66.74081 | -0.14085 | 0.605962 | 0.949497 |
| ENSMUSG( | AA387883 | 258.276 | 0.138963 | 0.345477 | 0.779153 | ENSMUSG( | AA387883 | 212.0544 | -0.16751 | 0.325001 | 0.905273 |
| ENSMUSG( | AA413626 | 11.44604 | -0.46924 | 0.384108 | NA | ENSMUSG( | AA413626 | 11.93873 | -0.12337 | 0.842206 | 0.983044 |
| ENSMUSG( | AA414768 | 109.9538 | -0.20581 | 0.323935 | 0.769037 | ENSMUSG( | AA414768 | 93.01186 | -0.46524 | 0.058324 | 0.728256 |
| ENSMUSG( | AA465934 | 39.77855 | 0.045235 | 0.880408 | 0.981469 | ENSMUSG( | AA465934 | 38.49648 | 0.222701 | 0.488526 | 0.92554 |
| ENSMUSG( | AA986860 | 134.39 | 0.153598 | 0.448268 | 0.834051 | ENSMUSG( | AA986860 | 135.2235 | 0.418108 | 0.04247 | 0.690304 |
| ENSMUSG( | Aaas | 368.7304 | 0.038195 | 0.800877 | 0.96188 | ENSMUSG( | Aaas | 335.3592 | 0.030491 | 0.85304 | 0.985598 |
| ENSMUSG( | Aacs | 1606.891 | 0.320437 | 0.000489 | 0.02028 | ENSMUSG( | Aacs | 1318.177 | 0.035792 | 0.745519 | 0.96951 |
| ENSMUSG( | Aagab | 2071.231 | 0.055674 | 0.535734 | 0.881209 | ENSMUSG( | Aagab | 1804.519 | -0.07572 | 0.517864 | 0.930939 |
| ENSMUSG( | Aak1 | 9681.118 | 0.176504 | 0.043896 | 0.321154 | ENSMUSG( | Aak1 | 8622.578 | 0.114456 | 0.244811 | 0.877904 |
| ENSMUSG( | Aamdc | 1015.643 | 0.070316 | 0.511481 | 0.870901 | ENSMUSG( | Aamdc | 877.9044 | -0.08788 | 0.53421 | 0.935936 |
| ENSMUSG( | Aamp | 4012.551 | 0.093903 | 0.282961 | 0.73186 | ENSMUSG( | Aamp | 3397.412 | -0.12046 | 0.287092 | 0.894091 |
| ENSMUSG( | Aanat | 15.45261 | 0.887046 | 0.06647 | NA | ENSMUSG( | Aanat | 12.64003 | 0.631276 | 0.26883 | 0.886755 |
| ENSMUSG( | Aar2 | 1045.459 | 0.124035 | 0.199704 | 0.640239 | ENSMUSG( | Aar2 | 896.7165 | -0.0473 | 0.685433 | 0.958678 |
| ENSMUSG( | Aard | 69.66609 | 0.061134 | 0.79452 | 0.959799 | ENSMUSG( | Aard | 64.70536 | 0.092495 | 0.751048 | 0.970317 |
| ENSMUSG( | Aars1 | 3796.49 | 0.041495 | 0.672928 | 0.929007 | ENSMUSG( | Aars1 | 3232.421 | -0.15817 | 0.172904 | 0.843702 |
| ENSMUSG( | Aars2 | 1056.862 | 0.226117 | 0.064427 | 0.393453 | ENSMUSG( | Aars2 | 898.567 | 0.037383 | 0.78653 | 0.976303 |
| ENSMUSG( | Aarsd1 | 13.03604 | 0.074172 | 0.895325 | NA | ENSMUSG( | Aarsd1 | 14.84791 | 0.652756 | 0.251214 | 0.879321 |
| ENSMUSG( | Aasdh | 287.785 | -0.04103 | 0.782283 | 0.957558 | ENSMUSG( | Aasdh | 275.9538 | 0.116993 | 0.579614 | 0.944209 |
| ENSMUSG( | Aasdhppt | 450.5583 | 0.02327 | 0.846694 | 0.972371 | ENSMUSG( | Aasdhppt | 417.0212 | 0.059264 | 0.700917 | 0.96162 |
| ENSMUSG( | Aass | 574.3672 | 0.240911 | 0.120367 | 0.524495 | ENSMUSG( | Aass | 463.596 | -0.10992 | 0.550926 | 0.939232 |
| ENSMUSG( | Aatf | 249.2508 | 0.102116 | 0.558478 | 0.888521 | ENSMUSG( | Aatf | 208.0876 | -0.16377 | 0.423049 | 0.917257 |
| ENSMUSG( | Aatk | 10274.98 | 0.086738 | 0.416148 | 0.82084 | ENSMUSG( | Aatk | 9939.049 | 0.252134 | 0.086684 | 0.782073 |
| ENSMUSG( | AB124611 | 34.44937 | -1.00417 | 0.014531 | 0.177481 | ENSMUSG( | AB124611 | 34.84916 | -0.59827 | 0.158874 | 0.837724 |
| ENSMUSG( | Abat | 29858.55 | 0.082556 | 0.467068 | 0.844819 | ENSMUSG( | Abat | 24490.91 | -0.23032 | 0.086337 | 0.781091 |
| ENSMUSG( | Abca1 | 1954.28 | -0.19552 | 0.170513 | 0.605271 | ENSMUSG( | Abca1 | 2129.842 | 0.315161 | 0.056994 | 0.723362 |
| ENSMUSG( | Abca13 | 9.766431 | -0.90658 | 0.131659 | NA | ENSMUSG( | Abca13 | 11.4156 | -0.09201 | 0.876015 | NA |
| ENSMUSG( | Abca17 | 41.54773 | 0.207511 | 0.536033 | 0.88142 | ENSMUSG( | Abca17 | 31.67504 | -0.31781 | 0.396941 | 0.917257 |
| ENSMUSG( | Abca2 | 9875.743 | -0.13867 | 0.278259 | 0.728302 | ENSMUSG( | Abca2 | 10207.4 | 0.223935 | 0.168426 | 0.840222 |
| ENSMUSG( | Abca3 | 3837.629 | -0.16286 | 0.236979 | 0.684317 | ENSMUSG( | Abca3 | 3910.104 | 0.164842 | 0.316999 | 0.902088 |
| ENSMUSG( | Abca4 | 131.6827 | -0.14735 | 0.619483 | 0.911432 | ENSMUSG( | Abca4 | 139.9299 | 0.279673 | 0.299797 | 0.898538 |
| ENSMUSG( | Abca5 | 1523.374 | -0.12119 | 0.411258 | 0.819183 | ENSMUSG( | Abca5 | 1560.547 | 0.215121 | 0.185584 | 0.855305 |
| ENSMUSG( | Abca6 | 59.55413 | -0.32372 | 0.216881 | 0.661816 | ENSMUSG( | Abca6 | 71.88728 | 0.489627 | 0.074297 | 0.766789 |
| ENSMUSG( | Abca7 | 839.1562 | 0.052992 | 0.66726 | 0.92729 | ENSMUSG( | Abca7 | 787.6195 | 0.14061 | 0.380302 | 0.917202 |
| ENSMUSG( | Abca8a | 125.4349 | -0.29285 | 0.226788 | 0.67384 | ENSMUSG( | Abca8a | 141.8778 | 0.336352 | 0.13321 | 0.822912 |
| ENSMUSG( | Abca8b | 637.5161 | -0.31746 | 0.018736 | 0.205889 | ENSMUSG( | Abca8b | 682.3428 | 0.167517 | 0.291655 | 0.897854 |
| ENSMUSG( | Abca9 | 644.3575 | 0.157664 | 0.318559 | 0.764234 | ENSMUSG( | Abca9 | 662.3695 | 0.478739 | 0.004928 | 0.28966 |
| ENSMUSG( | Abcb10 | 1322.757 | -0.05909 | 0.59925 | 0.903801 | ENSMUSG( | Abcb10 | 1238.732 | 0.023562 | 0.864219 | 0.986552 |
| ENSMUSG( | Abcb1a | 1149.777 | 0.111023 | 0.590486 | 0.90113 | ENSMUSG( | Abcb1a | 1366.393 | 0.784582 | 0.084118 | 0.777886 |
| ENSMUSG( | Abcb1b | 196.6681 | -0.04322 | 0.780292 | 0.957491 | ENSMUSG( | Abcb1b | 177.5818 | -0.07149 | 0.695049 | 0.960867 |
| ENSMUSG( | Abcb4 | 302.8013 | 0.048011 | 0.76687 | 0.954331 | ENSMUSG( | Abcb4 | 274.1786 | 0.038402 | 0.823511 | 0.979198 |
| ENSMUSG( | Abcb6 | 68.98276 | -0.19961 | 0.386338 | 0.804623 | ENSMUSG( | Abcb6 | 67.35853 | -0.01155 | 0.964319 | 0.995433 |
| ENSMUSG( | Abcb7 | 1346.31 | -0.01482 | 0.91889 | 0.990194 | ENSMUSG( | Abcb7 | 1258.256 | 0.058185 | 0.689486 | 0.960071 |
| ENSMUSG( | Abcb8 | 1896.952 | -0.05634 | 0.603994 | 0.905187 | ENSMUSG( | Abcb8 | 1805.382 | 0.073336 | 0.592057 | 0.944975 |
| ENSMUSG( | Abcb9 | 2248.243 | -0.33682 | 0.013582 | 0.170246 | ENSMUSG( | Abcb9 | 2232.036 | -0.06642 | 0.671898 | 0.95694 |
| ENSMUSG( | Abcc1 | 789.019 | -0.29111 | 0.011889 | 0.157801 | ENSMUSG( | Abcc1 | 812.8169 | 0.082153 | 0.543258 | 0.938063 |
| ENSMUSG( | Abcc10 | 348.7343 | -0.14312 | 0.32311 | 0.768415 | ENSMUSG( | Abcc10 | 360.3059 | 0.222437 | 0.159119 | 0.837724 |
| ENSMUSG( | Abcc12 | 18.74705 | -1.04296 | 0.016026 | NA | ENSMUSG( | Abcc12 | 19.1853 | -0.63662 | 0.170813 | 0.842157 |
| ENSMUSG( | Abcc3 | 174.7684 | -0.26292 | 0.180019 | 0.617634 | ENSMUSG( | Abcc3 | 192.8253 | 0.301738 | 0.178998 | 0.849041 |
| ENSMUSG( | Abcc4 | 604.9932 | 0.235316 | 0.079442 | 0.43547 | ENSMUSG( | Abcc4 | 631.984 | 0.589376 | 0.002004 | 0.169624 |
| ENSMUSG( | Abcc5 | 5565.852 | 0.103559 | 0.354272 | 0.786661 | ENSMUSG( | Abcc5 | 5263.539 | 0.206501 | 0.140374 | 0.830678 |
| ENSMUSG( | Abcc6 | 85.8307 | 0.016186 | 0.950982 | 0.993821 | ENSMUSG( | Abcc6 | 89.77457 | 0.40281 | 0.140748 | 0.830678 |
| ENSMUSG( | Abcc8 | 980.9976 | 0.017665 | 0.90833 | 0.987809 | ENSMUSG( | Abcc8 | 991.9264 | 0.309486 | 0.082144 | 0.77681 |

| | | | | | |
|---|---|---|---|---|---|
| ENSMUSG( | 1346.31 | -0.01482 | 0.91889 | 0.990194 | Abcb7 |
| ENSMUSG( | 1896.952 | -0.05634 | 0.603994 | 0.905187 | Abcb8 |
| ENSMUSG( | 2248.243 | -0.33682 | 0.013582 | 0.170246 | Abcb9 |
| ENSMUSG( | 789.019 | -0.29111 | 0.011889 | 0.157801 | Abcc1 |
| ENSMUSG( | 348.7343 | -0.14312 | 0.32311 | 0.768415 | Abcc10 |
| ENSMUSG( | 18.74705 | -1.04296 | 0.016026 | NA | Abcc12 |
| ENSMUSG( | 174.7684 | -0.26292 | 0.180019 | 0.617634 | Abcc3 |
| ENSMUSG( | 604.9932 | 0.235316 | 0.079442 | 0.43547 | Abcc4 |
| ENSMUSG( | 5565.852 | 0.103559 | 0.354272 | 0.786661 | Abcc5 |
| ENSMUSG( | 85.8307 | 0.016186 | 0.950982 | 0.993821 | Abcc6 |
| ENSMUSG( | 980.9976 | 0.017665 | 0.90833 | 0.987809 | Abcc8 |
| ENSMUSG( | 354.4775 | -0.05712 | 0.887182 | 0.983326 | Abcc9 |
| ENSMUSG( | 664.6232 | -0.1057 | 0.345817 | 0.779153 | Abcd1 |
| ENSMUSG( | 2903.536 | -0.07937 | 0.517469 | 0.873306 | Abcd2 |
| ENSMUSG( | 4341.029 | 0.083013 | 0.41708 | 0.8209 | Abcd3 |
| ENSMUSG( | 748.2222 | 0.106357 | 0.382089 | 0.803095 | Abcd4 |
| ENSMUSG( | 2120.011 | 0.117191 | 0.301703 | 0.749979 | Abce1 |
| ENSMUSG( | 2107.72 | 0.219137 | 0.009706 | 0.140314 | Abcf1 |
| ENSMUSG( | 2639.687 | 0.0949 | 0.277528 | 0.727457 | Abcf2 |
| ENSMUSG( | 1403.11 | 0.067417 | 0.486316 | 0.858575 | Abcf3 |
| ENSMUSG( | 1864.487 | -0.45031 | 0.002135 | 0.053442 | Abcg1 |
| ENSMUSG( | 800.4094 | 0.05431 | 0.752982 | 0.950667 | Abcg2 |
| ENSMUSG( | 18.05099 | -1.01444 | 0.040236 | NA | Abcg3 |
| ENSMUSG( | 1947.76 | -0.16076 | 0.185232 | 0.623185 | Abcg4 |
| ENSMUSG( | 183.3324 | 0.000291 | 0.998678 | 0.99989 | Abhd1 |
| ENSMUSG( | 437.4568 | -0.16932 | 0.144505 | 0.564381 | Abhd10 |
| ENSMUSG( | 1029.503 | -0.03069 | 0.775475 | 0.955606 | Abhd11 |
| ENSMUSG( | 14690.36 | -0.20273 | 0.067503 | 0.402091 | Abhd12 |
| ENSMUSG( | 14.09475 | -1.40211 | 0.041183 | NA | Abhd12b |
| ENSMUSG( | 1096.767 | -0.34665 | 0.014918 | 0.179376 | Abhd13 |
| ENSMUSG( | 1022.837 | -0.14313 | 0.120521 | 0.524506 | Abhd14a |
| ENSMUSG( | 459.2044 | 0.180464 | 0.258351 | 0.709574 | Abhd14b |
| ENSMUSG( | 55.02149 | -0.09022 | 0.72297 | 0.943761 | Abhd15 |
| ENSMUSG( | 3031.261 | -0.29767 | 0.022757 | 0.230554 | Abhd16a |
| ENSMUSG( | 10712.54 | 0.138597 | 0.099604 | 0.48212 | Abhd17a |
| ENSMUSG( | 788.7259 | 0.09667 | 0.363089 | 0.793211 | Abhd17b |
| ENSMUSG( | 1110.18 | 0.069652 | 0.517994 | 0.873754 | Abhd17c |
| ENSMUSG( | 396.7756 | 0.005896 | 0.962262 | 0.994543 | Abhd18 |
| ENSMUSG( | 6160.265 | -0.17244 | 0.233566 | 0.681508 | Abhd2 |
| ENSMUSG( | 4801.503 | 0.095206 | 0.400927 | 0.813897 | Abhd3 |
| ENSMUSG( | 8340.646 | 0.084518 | 0.434677 | 0.828978 | Abhd4 |
| ENSMUSG( | 1156.975 | 0.122364 | 0.265119 | 0.715962 | Abhd5 |
| ENSMUSG( | 2134.364 | -0.21677 | 0.139349 | 0.555477 | Abhd6 |
| ENSMUSG( | 15216.72 | 0.229203 | 0.00341 | 0.073541 | Abhd8 |
| ENSMUSG( | 3406.944 | 0.123908 | 0.187951 | 0.626616 | Abi1 |
| ENSMUSG( | 5540.778 | -0.14774 | 0.090234 | 0.461412 | Abi2 |
| ENSMUSG( | 281.9643 | -0.06195 | 0.688513 | 0.933441 | Abi3 |
| ENSMUSG( | 251.7913 | -0.9771 | 2.97E-09 | 1.27E-06 | Abi3bp |
| ENSMUSG( | 738.9172 | 0.018525 | 0.857129 | 0.974393 | Abitram |
| ENSMUSG( | 2108.757 | -0.19209 | 0.106578 | 0.494174 | Abl1 |
| ENSMUSG( | 2054.786 | -0.019 | 0.843571 | 0.971847 | Abl2 |
| ENSMUSG( | 3711.223 | 0.019562 | 0.849811 | 0.972638 | Ablim1 |
| ENSMUSG( | 4083.382 | -0.05303 | 0.594003 | 0.902869 | Ablim2 |
| ENSMUSG( | 990.5706 | -0.02016 | 0.850802 | 0.972638 | Ablim3 |
| ENSMUSG( | 18284.94 | 0.164206 | 0.027745 | 0.255176 | Abr |
| ENSMUSG( | 122.798 | 0.140643 | 0.522118 | 0.875186 | Abracl |
| ENSMUSG( | 102.9059 | -0.10891 | 0.611116 | 0.908551 | Abraxas1 |
| ENSMUSG( | 741.464 | 0.06396 | 0.555822 | 0.887457 | Abraxas2 |
| ENSMUSG( | 237.9013 | 0.057993 | 0.700857 | 0.93743 | Abt1 |
| ENSMUSG( | 1089.062 | 0.037058 | 0.738333 | 0.947329 | Abtb1 |
| ENSMUSG( | 876.9864 | 0.021747 | 0.82644 | 0.968349 | Abtb2 |
| ENSMUSG( | 598.4285 | 0.112266 | 0.331435 | 0.77519 | Abtb3 |
| ENSMUSG( | 2432.815 | 0.020037 | 0.807453 | 0.963257 | Acaa1a |
| ENSMUSG( | 18.01139 | 0.09691 | 0.823802 | NA | Acaa1b |
| ENSMUSG( | 1909.16 | 0.019284 | 0.846306 | 0.972226 | Acaa2 |
| ENSMUSG( | 3876.386 | 0.170544 | 0.050488 | 0.343938 | Acaca |
| ENSMUSG( | 279.9885 | 0.0569 | 0.691193 | 0.933881 | Acacb |
| ENSMUSG( | 165.6642 | -0.01423 | 0.929764 | 0.991848 | Acad10 |
| ENSMUSG( | 1685.688 | 0.048889 | 0.664007 | 0.926511 | Acad11 |
| ENSMUSG( | 266.3842 | 0.018609 | 0.901393 | 0.985633 | Acad12 |
| ENSMUSG( | 938.419 | -0.00677 | 0.95139 | 0.993821 | Acad8 |
| ENSMUSG( | 1014.825 | 0.024734 | 0.814037 | 0.964574 | Acad9 |
| ENSMUSG( | 3295.353 | 0.121531 | 0.311861 | 0.759517 | Acadl |
| ENSMUSG( | 2532.542 | 0.106522 | 0.363233 | 0.793211 | Acadm |
| ENSMUSG( | 612.2225 | 0.088603 | 0.418158 | 0.821966 | Acads |
| ENSMUSG( | 2259.664 | 0.1083 | 0.29508 | 0.743798 | Acadsb |
| ENSMUSG( | 2538.039 | 0.182115 | 0.089283 | 0.458563 | Acadvl |
| ENSMUSG( | 132.3793 | -0.40783 | 0.038842 | 0.30362 | Acan |
| ENSMUSG( | 10.07824 | -0.32472 | 0.595034 | NA | Acap1 |
| ENSMUSG( | 3470.524 | 0.080767 | 0.337113 | 0.777008 | Acap2 |
| ENSMUSG( | 5973.572 | 0.207016 | 0.029569 | 0.260563 | Acap3 |
| ENSMUSG( | 3344.52 | 0.009607 | 0.921096 | 0.990436 | Acat1 |
| ENSMUSG( | 1076.572 | 0.042764 | 0.645286 | 0.917957 | Acat2 |
| ENSMUSG( | 270.0648 | 0.075263 | 0.629031 | 0.913828 | Acat3 |
| ENSMUSG( | 990.2469 | 0.108396 | 0.280386 | 0.729291 | Acbd3 |
| ENSMUSG( | 981.0817 | 0.029461 | 0.790636 | 0.959461 | Acbd4 |
| ENSMUSG( | 4118.972 | 0.08199 | 0.505661 | 0.866932 | Acbd5 |
| ENSMUSG( | 1319.87 | -0.01518 | 0.884297 | 0.982647 | Acbd6 |
| ENSMUSG( | 172.3435 | 0.078253 | 0.670261 | 0.928525 | Accs |
| ENSMUSG( | 1209.225 | 0.063252 | 0.564964 | 0.890735 | Acd |
| ENSMUSG( | 713.7026 | -0.2306 | 0.200526 | 0.640661 | Ace |
| ENSMUSG( | 84.15947 | -0.28043 | 0.479157 | 0.854046 | Ace2 |
| ENSMUSG( | 238.062 | 0.403821 | 0.044918 | 0.325112 | Acer2 |
| ENSMUSG( | 579.5805 | -0.26352 | 0.077444 | 0.429743 | Acer3 |
| ENSMUSG( | 2125.862 | -0.33224 | 0.000236 | 0.012098 | Ache |
| ENSMUSG( | 4331.975 | 0.030148 | 0.769289 | 0.954674 | Acin1 |
| ENSMUSG( | 1594.498 | -0.3241 | 0.005632 | 0.100551 | Ackr1 |
| ENSMUSG( | 64.86934 | 0.049456 | 0.854465 | 0.973547 | Ackr2 |
| ENSMUSG( | 332.4315 | -0.43871 | 0.028915 | 0.258972 | Ackr3 |
| ENSMUSG( | 13.52785 | -0.22582 | 0.68076 | NA | Ackr4 |
| ENSMUSG( | 4593.659 | 0.106222 | 0.222568 | 0.668434 | Acly |
| ENSMUSG( | 1677.121 | 0.142279 | 0.200151 | 0.640302 | Aco1 |
| ENSMUSG( | 30436.36 | 0.086852 | 0.289929 | 0.739839 | Aco2 |
| ENSMUSG( | 467.4169 | 0.159896 | 0.295065 | 0.743798 | Acot1 |
| ENSMUSG( | 6599.033 | 0.239798 | 0.009547 | 0.138264 | Acot11 |
| ENSMUSG( | 1598.466 | 0.00738 | 0.934074 | 0.99323 | Acot13 |
| ENSMUSG( | 232.8933 | 0.024012 | 0.879962 | 0.981469 | Acot2 |
| ENSMUSG( | 23.31191 | -0.07155 | 0.853797 | 0.973491 | Acot3 |
| ENSMUSG( | 81.76732 | 0.058482 | 0.78282 | 0.957558 | Acot4 |
| ENSMUSG( | 50.39883 | -0.2124 | 0.445453 | 0.833239 | Acot5 |
| ENSMUSG( | 191.0906 | 0.009922 | 0.951921 | 0.993821 | Acot6 |
| ENSMUSG( | 3832.37 | -0.00199 | 0.988688 | 0.999109 | Acot7 |
| ENSMUSG( | 316.124 | -0.15017 | 0.304834 | 0.752574 | Acot8 |
| ENSMUSG( | 303.4877 | 0.019157 | 0.889683 | 0.983551 | Acot9 |
| ENSMUSG( | 13037.52 | 0.129577 | 0.166437 | 0.599114 | Acox1 |
| ENSMUSG( | 1609.38 | -0.01001 | 0.927634 | 0.991229 | Acox3 |
| ENSMUSG( | 1401.669 | -0.03675 | 0.70124 | 0.93743 | Acp1 |
| ENSMUSG( | 3040.411 | -0.30668 | 0.040294 | 0.31004 | Acp2 |
| ENSMUSG( | 40.09068 | -1.16122 | 0.000377 | 0.017069 | Acp3 |
| ENSMUSG( | 18.71082 | -0.1217 | 0.794528 | NA | Acp5 |
| ENSMUSG( | 390.3801 | 0.06125 | 0.665699 | 0.927054 | Acp6 |
| ENSMUSG( | 29.47335 | -0.20425 | 0.558789 | 0.888638 | Acp7 |
| ENSMUSG( | 56.24335 | -0.19583 | 0.476611 | 0.851717 | Acr |
| ENSMUSG( | 147.8458 | 0.569566 | 0.008964 | 0.132538 | Acrbp |
| ENSMUSG( | 22556.33 | 0.076182 | 0.493146 | 0.860552 | Acsbg1 |
| ENSMUSG( | 2133.475 | 0.034887 | 0.756431 | 0.951479 | Acsf2 |
| ENSMUSG( | 579.4882 | 0.031012 | 0.794823 | 0.959799 | Acsf3 |
| ENSMUSG( | 1222.82 | -0.23161 | 0.021839 | 0.225699 | Acsl1 |
| ENSMUSG( | 10791.05 | 0.033793 | 0.79619 | 0.960088 | Acsl3 |
| ENSMUSG( | 1612.926 | -0.46267 | 0.005071 | 0.094652 | Acsl4 |
| ENSMUSG( | 2014.739 | -0.13081 | 0.335973 | 0.777 | Acsl5 |
| ENSMUSG( | 19165.59 | -0.012 | 0.913983 | 0.988483 | Acsl6 |
| ENSMUSG( | 18.85329 | -0.35512 | 0.447994 | NA | Acsm5 |
| ENSMUSG( | 2920.782 | 0.173678 | 0.144932 | 0.565187 | Acss1 |
| ENSMUSG( | 3505.757 | 0.08762 | 0.468802 | 0.845636 | Acss2 |
| ENSMUSG( | 53.37884 | -0.00483 | 0.986229 | 0.998326 | Acss3 |
| ENSMUSG( | 282.5231 | 0.12615 | 0.401815 | 0.814965 | Acta1 |
| ENSMUSG( | 52.71579 | -0.74139 | 0.021324 | 0.223 | Acta2 |
| ENSMUSG( | 24522.23 | -0.07918 | 0.386043 | 0.804623 | Actb |
| ENSMUSG( | 9938.316 | -0.00941 | 0.9152 | 0.988634 | Actg1 |
| ENSMUSG( | 432.8853 | 0.007278 | 0.949825 | 0.993821 | Actl6a |
| ENSMUSG( | 2320.087 | 0.100825 | 0.435785 | 0.829093 | Actl6b |
| ENSMUSG( | 594.3063 | 0.057013 | 0.594559 | 0.902971 | Actmap |
| ENSMUSG( | 2374.816 | -0.14416 | 0.19246 | 0.631514 | Actn1 |
| ENSMUSG( | 635.7893 | -0.72395 | 4.10E-06 | 0.000549 | Actn2 |
| ENSMUSG( | 46.73538 | -0.07389 | 0.803763 | 0.962169 | Actn3 |
| ENSMUSG( | 5709.115 | -0.01535 | 0.880396 | 0.981469 | Actn4 |
| ENSMUSG( | 2672.009 | 0.106962 | 0.318502 | 0.764234 | Actr10 |
| ENSMUSG( | 5618.237 | 0.026242 | 0.751415 | 0.950237 | Actr1a |
| ENSMUSG( | 9191.111 | 0.102904 | 0.259067 | 0.709814 | Actr1b |
| ENSMUSG( | 6445.827 | 0.016881 | 0.830829 | 0.969798 | Actr2 |
| ENSMUSG( | 4292.658 | 0.04128 | 0.64232 | 0.916865 | Actr3 |
| ENSMUSG( | 2922.622 | 0.090888 | 0.37473 | 0.800466 | Actr3b |
| ENSMUSG( | 14.42824 | -0.43657 | 0.369295 | NA | Actr3-ps |
| ENSMUSG( | 190.653 | -0.0626 | 0.709604 | 0.939669 | Actr5 |
| ENSMUSG( | 322.0231 | 0.127702 | 0.342214 | 0.778828 | Actr6 |
| ENSMUSG( | 1064.125 | 0.161175 | 0.18332 | 0.62047 | Actr8 |
| ENSMUSG( | 13.94302 | -0.95326 | 0.087748 | NA | Actrt3 |

| | | | | | |
|---|---|---|---|---|---|
| ENSMUSG( | 192.8253 | 0.301738 | 0.178998 | 0.849041 | Abcc3 |
| ENSMUSG( | 631.984 | 0.589376 | 0.002004 | 0.169624 | Abcc4 |
| ENSMUSG( | 5263.539 | 0.206501 | 0.140374 | 0.830678 | Abcc5 |
| ENSMUSG( | 89.77457 | 0.40281 | 0.140748 | 0.830678 | Abcc6 |
| ENSMUSG( | 991.9264 | 0.309486 | 0.082144 | 0.77681 | Abcc8 |
| ENSMUSG( | 474.4263 | 0.92131 | 0.10475 | 0.804969 | Abcc9 |
| ENSMUSG( | 631.9996 | 0.024185 | 0.851198 | 0.985541 | Abcd1 |
| ENSMUSG( | 2565.736 | -0.17104 | 0.205016 | 0.864412 | Abcd2 |
| ENSMUSG( | 3711.35 | -0.10257 | 0.393745 | 0.917257 | Abcd3 |
| ENSMUSG( | 709.7543 | 0.218549 | 0.103452 | 0.804969 | Abcd4 |
| ENSMUSG( | 1830.05 | -0.04144 | 0.768496 | 0.974236 | Abce1 |
| ENSMUSG( | 1812.048 | 0.058685 | 0.578226 | 0.943353 | Abcf1 |
| ENSMUSG( | 2341.36 | 0.017446 | 0.876145 | 0.989644 | Abcf2 |
| ENSMUSG( | 1241.323 | -0.01655 | 0.894337 | 0.990558 | Abcf3 |
| ENSMUSG( | 2114 | 0.213133 | 0.224288 | 0.867489 | Abcg1 |
| ENSMUSG( | 848.0058 | 0.46328 | 0.050343 | 0.708648 | Abcg2 |
| ENSMUSG( | 21.62434 | -0.05748 | 0.908335 | 0.991958 | Abcg3 |
| ENSMUSG( | 1942.233 | 0.105676 | 0.408021 | 0.917257 | Abcg4 |
| ENSMUSG( | 165.7379 | -0.01897 | 0.925363 | 0.99306 | Abhd1 |
| ENSMUSG( | 404.9657 | -0.12535 | 0.379376 | 0.916995 | Abhd10 |
| ENSMUSG( | 918.3412 | -0.09529 | 0.452811 | 0.922316 | Abhd11 |
| ENSMUSG( | 14495.37 | 0.036744 | 0.781335 | 0.976106 | Abhd12 |
| ENSMUSG( | 1236.879 | 0.279522 | 0.090597 | 0.791074 | Abhd13 |
| ENSMUSG( | 971.7539 | -0.0181 | 0.877103 | 0.989644 | Abhd14a |
| ENSMUSG( | 418.8303 | 0.185887 | 0.353925 | 0.912411 | Abhd14b |
| ENSMUSG( | 56.12335 | 0.263106 | 0.49245 | 0.927315 | Abhd15 |
| ENSMUSG( | 3148.688 | 0.098647 | 0.517588 | 0.930939 | Abhd16a |
| ENSMUSG( | 9058.792 | -0.07405 | 0.463671 | 0.923329 | Abhd17a |
| ENSMUSG( | 709.2867 | 0.058659 | 0.685919 | 0.958956 | Abhd17b |
| ENSMUSG( | 993.9011 | 0.016916 | 0.902013 | 0.991003 | Abhd17c |
| ENSMUSG( | 378.631 | 0.138502 | 0.327856 | 0.906255 | Abhd18 |
| ENSMUSG( | 6187.295 | 0.116296 | 0.473485 | 0.923329 | Abhd2 |
| ENSMUSG( | 4174.208 | -0.03771 | 0.757902 | 0.971858 | Abhd3 |
| ENSMUSG( | 6847.858 | -0.22558 | 0.081462 | 0.77681 | Abhd4 |
| ENSMUSG( | 975.0743 | -0.10517 | 0.399656 | 0.917257 | Abhd5 |
| ENSMUSG( | 2241.806 | 0.199015 | 0.216658 | 0.865167 | Abhd6 |
| ENSMUSG( | 12851.21 | 0.019511 | 0.835954 | 0.981237 | Abhd8 |
| ENSMUSG( | 3052.646 | 0.074236 | 0.598529 | 0.946859 | Abi1 |
| ENSMUSG( | 5402.407 | 0.050627 | 0.675386 | 0.95694 | Abi2 |
| ENSMUSG( | 251.7072 | -0.11343 | 0.519741 | 0.931729 | Abi3 |
| ENSMUSG( | 280.9922 | -0.2432 | 0.270114 | 0.88756 | Abi3bp |
| ENSMUSG( | 656.3404 | -0.05106 | 0.682412 | 0.958678 | Abitram |
| ENSMUSG( | 2120.831 | 0.102208 | 0.486947 | 0.92541 | Abl1 |
| ENSMUSG( | 1909.271 | 0.038624 | 0.73183 | 0.967228 | Abl2 |
| ENSMUSG( | 3537.285 | 0.149502 | 0.263232 | 0.885994 | Ablim1 |
| ENSMUSG( | 3733.579 | -0.04344 | 0.719097 | 0.966799 | Ablim2 |
| ENSMUSG( | 944.1912 | 0.105958 | 0.404789 | 0.917257 | Ablim3 |
| ENSMUSG( | 15433.5 | -0.05283 | 0.589097 | 0.944975 | Abr |
| ENSMUSG( | 118.8262 | 0.285227 | 0.253234 | 0.88018 | Abracl |
| ENSMUSG( | 102.6802 | 0.174488 | 0.476551 | 0.923329 | Abraxas1 |
| ENSMUSG( | 664.567 | 0.021612 | 0.863321 | 0.986547 | Abraxas2 |
| ENSMUSG( | 206.2258 | -0.09057 | 0.606487 | 0.949743 | Abt1 |
| ENSMUSG( | 1008.836 | 0.088327 | 0.525749 | 0.934374 | Abtb1 |
| ENSMUSG( | 755.7341 | -0.14521 | 0.267515 | 0.886134 | Abtb2 |
| ENSMUSG( | 528.6435 | 0.032716 | 0.8473 | 0.985161 | Abtb3 |
| ENSMUSG( | 2180.729 | -0.02596 | 0.8256 | 0.979537 | Acaa1a |
| ENSMUSG( | 16.3758 | 0.111583 | 0.816495 | 0.979198 | Acaa1b |
| ENSMUSG( | 1598.744 | -0.23738 | 0.092015 | 0.791889 | Acaa2 |
| ENSMUSG( | 3327.005 | 0.003711 | 0.970543 | 0.99594 | Acaca |
| ENSMUSG( | 254.3093 | 0.060043 | 0.730359 | 0.967228 | Acacb |
| ENSMUSG( | 153.6766 | 0.050736 | 0.817774 | 0.979198 | Acad10 |
| ENSMUSG( | 1363.292 | -0.31479 | 0.009567 | 0.406458 | Acad11 |
| ENSMUSG( | 227.1564 | -0.19267 | 0.293494 | 0.898538 | Acad12 |
| ENSMUSG( | 808.9117 | -0.17326 | 0.16669 | 0.840183 | Acad8 |
| ENSMUSG( | 879.8566 | -0.12391 | 0.316247 | 0.901868 | Acad9 |
| ENSMUSG( | 2775.286 | -0.10852 | 0.458953 | 0.923329 | Acadl |
| ENSMUSG( | 2143.631 | -0.10966 | 0.400672 | 0.917257 | Acadm |
| ENSMUSG( | 520.3235 | -0.11585 | 0.374104 | 0.915491 | Acads |
| ENSMUSG( | 1947.583 | -0.05236 | 0.65628 | 0.955533 | Acadsb |
| ENSMUSG( | 2076.214 | -0.12966 | 0.308219 | 0.898538 | Acadvl |
| ENSMUSG( | 154.24 | 0.326189 | 0.118867 | 0.813384 | Acan |
| ENSMUSG( | 3235.593 | 0.144682 | 0.170362 | 0.841579 | Acap2 |
| ENSMUSG( | 5213.952 | 0.089812 | 0.405106 | 0.917257 | Acap3 |
| ENSMUSG( | 2753.614 | -0.30058 | 0.008241 | 0.388416 | Acat1 |
| ENSMUSG( | 902.2057 | -0.20926 | 0.09438 | 0.795735 | Acat2 |
| ENSMUSG( | 217.9741 | -0.28978 | 0.137708 | 0.828492 | Acat3 |
| ENSMUSG( | 837.3432 | -0.11146 | 0.397408 | 0.917257 | Acbd3 |
| ENSMUSG( | 888.7482 | 0.011356 | 0.928336 | 0.99306 | Acbd4 |
| ENSMUSG( | 3509.042 | -0.11665 | 0.417592 | 0.917257 | Acbd5 |
| ENSMUSG( | 1153.91 | -0.14309 | 0.33521 | 0.907907 | Acbd6 |
| ENSMUSG( | 154.0747 | 0.032817 | 0.868688 | 0.988097 | Accs |
| ENSMUSG( | 9.838647 | 0.194334 | 0.765258 | NA | Accsl |
| ENSMUSG( | 1038.165 | -0.11019 | 0.394132 | 0.917257 | Acd |
| ENSMUSG( | 790.0869 | 0.319301 | 0.046338 | 0.697814 | Ace |
| ENSMUSG( | 98.04069 | 0.432058 | 0.520312 | 0.931964 | Ace2 |
| ENSMUSG( | 195.3052 | 0.143234 | 0.51967 | 0.931729 | Acer2 |
| ENSMUSG( | 601.2593 | 0.126105 | 0.428331 | 0.917257 | Acer3 |
| ENSMUSG( | 2179.818 | 0.02857 | 0.800609 | 0.97681 | Ache |
| ENSMUSG( | 3979.143 | 0.053557 | 0.63811 | 0.953903 | Acin1 |
| ENSMUSG( | 1638.001 | 0.040478 | 0.767359 | 0.974236 | Ackr1 |
| ENSMUSG( | 68.18265 | 0.45358 | 0.117861 | 0.813384 | Ackr2 |
| ENSMUSG( | 394.2298 | 0.347043 | 0.162679 | 0.837724 | Ackr3 |
| ENSMUSG( | 3921.06 | -0.08226 | 0.433342 | 0.918434 | Acly |
| ENSMUSG( | 1417.025 | -0.07411 | 0.517654 | 0.930939 | Aco1 |
| ENSMUSG( | 25598.22 | -0.14576 | 0.144205 | 0.83148 | Aco2 |
| ENSMUSG( | 358.8787 | -0.35901 | 0.019523 | 0.556192 | Acot1 |
| ENSMUSG( | 5181.213 | -0.18623 | 0.103747 | 0.804969 | Acot11 |
| ENSMUSG( | 1319.309 | -0.29731 | 0.024767 | 0.592568 | Acot13 |
| ENSMUSG( | 188.3519 | -0.34902 | 0.071358 | 0.761942 | Acot2 |
| ENSMUSG( | 20.37848 | -0.21984 | 0.620827 | 0.953114 | Acot3 |
| ENSMUSG( | 75.0064 | 0.076762 | 0.749643 | 0.96998 | Acot4 |
| ENSMUSG( | 45.35423 | -0.2273 | 0.450484 | 0.922202 | Acot5 |
| ENSMUSG( | 175.328 | 0.027795 | 0.876793 | 0.989644 | Acot6 |
| ENSMUSG( | 3184.814 | -0.28475 | 0.096959 | 0.79645 | Acot7 |
| ENSMUSG( | 291.2987 | -0.10801 | 0.501258 | 0.927825 | Acot8 |
| ENSMUSG( | 243.2484 | -0.36108 | 0.040233 | 0.681706 | Acot9 |
| ENSMUSG( | 11131.04 | -0.05652 | 0.624077 | 0.953234 | Acox1 |
| ENSMUSG( | 9.826724 | 0.34156 | 0.596208 | NA | Acox2 |
| ENSMUSG( | 1465.59 | -0.00686 | 0.957883 | 0.994798 | Acox3 |
| ENSMUSG( | 1255.835 | -0.08902 | 0.456837 | 0.923329 | Acp1 |
| ENSMUSG( | 3248.181 | 0.169107 | 0.323497 | 0.904501 | Acp2 |
| ENSMUSG( | 48.71314 | -0.09588 | 0.743579 | 0.96951 | Acp3 |
| ENSMUSG( | 19.38088 | 0.22804 | 0.626452 | 0.953234 | Acp5 |
| ENSMUSG( | 376.9074 | 0.230049 | 0.22807 | 0.868507 | Acp6 |
| ENSMUSG( | 27.43368 | -0.13453 | 0.717023 | 0.965864 | Acp7 |
| ENSMUSG( | 62.26818 | 0.344943 | 0.216674 | 0.865167 | Acr |
| ENSMUSG( | 142.3553 | 0.702966 | 0.004571 | 0.277991 | Acrbp |
| ENSMUSG( | 18184.1 | -0.29239 | 0.023966 | 0.588226 | Acsbg1 |
| ENSMUSG( | 1854.33 | -0.10482 | 0.428103 | 0.917257 | Acsf2 |
| ENSMUSG( | 482.5576 | -0.236 | 0.089307 | 0.788502 | Acsf3 |
| ENSMUSG( | 1240.245 | 0.08961 | 0.469404 | 0.923329 | Acsl1 |
| ENSMUSG( | 9846.19 | 0.041524 | 0.789765 | 0.976792 | Acsl3 |
| ENSMUSG( | 1899.944 | 0.302441 | 0.115616 | 0.813384 | Acsl4 |
| ENSMUSG( | 2039.649 | 0.176066 | 0.266072 | 0.886134 | Acsl5 |
| ENSMUSG( | 17506.55 | -0.00303 | 0.980939 | 0.9967 | Acsl6 |
| ENSMUSG( | 17.29171 | -0.37826 | 0.485347 | 0.92541 | Acsm5 |
| ENSMUSG( | 2372.42 | -0.15801 | 0.15102 | 0.837173 | Acss1 |
| ENSMUSG( | 2944.752 | -0.14907 | 0.208926 | 0.864412 | Acss2 |
| ENSMUSG( | 49.44789 | 0.040826 | 0.891945 | 0.990558 | Acss3 |
| ENSMUSG( | 227.7369 | -0.23835 | 0.16756 | 0.840222 | Acta1 |
| ENSMUSG( | 68.13562 | 0.367293 | 0.278868 | 0.891737 | Acta2 |
| ENSMUSG( | 21227.21 | -0.23791 | 0.080045 | 0.776379 | Actb |
| ENSMUSG( | 8515.782 | -0.19357 | 0.082701 | 0.77681 | Actg1 |
| ENSMUSG( | 403.6748 | 0.076954 | 0.573599 | 0.942912 | Actl6a |
| ENSMUSG( | 1989.215 | -0.07596 | 0.586445 | 0.944975 | Actl6b |
| ENSMUSG( | 528.1155 | -0.01744 | 0.8929 | 0.990558 | Actmap |
| ENSMUSG( | 2381.77 | 0.13596 | 0.265209 | 0.886134 | Actn1 |
| ENSMUSG( | 695.8915 | -0.10788 | 0.425487 | 0.917257 | Actn2 |
| ENSMUSG( | 41.12359 | -0.17017 | 0.602779 | 0.94789 | Actn3 |
| ENSMUSG( | 5413.845 | 0.10076 | 0.36941 | 0.915491 | Actn4 |
| ENSMUSG( | 2249.267 | -0.12537 | 0.368721 | 0.915491 | Actr10 |
| ENSMUSG( | 4842.191 | -0.13799 | 0.190961 | 0.858148 | Actr1a |
| ENSMUSG( | 7817.066 | -0.09646 | 0.379053 | 0.916995 | Actr1b |
| ENSMUSG( | 5728.772 | -0.05717 | 0.624789 | 0.953234 | Actr2 |
| ENSMUSG( | 3799.275 | -0.04486 | 0.70482 | 0.962677 | Actr3 |
| ENSMUSG( | 2555.515 | -0.02823 | 0.820867 | 0.979198 | Actr3b |
| ENSMUSG( | 15.35673 | -0.03098 | 0.957263 | 0.994798 | Actr3-ps |
| ENSMUSG( | 171.5471 | -0.09227 | 0.620104 | 0.952896 | Actr5 |
| ENSMUSG( | 296.2382 | 0.145364 | 0.366782 | 0.914839 | Actr6 |
| ENSMUSG( | 926.5554 | 0.033829 | 0.812776 | 0.979148 | Actr8 |
| ENSMUSG( | 15.10261 | -0.30865 | 0.633066 | 0.953844 | Actrt3 |
| ENSMUSG( | 1010.051 | 0.21172 | 0.191362 | 0.858412 | Acvr1 |
| ENSMUSG( | 4675.926 | 0.180862 | 0.255817 | 0.883082 | Acvr1b |
| ENSMUSG( | 361.7163 | -0.1959 | 0.314248 | 0.900949 | Acvr1c |
| ENSMUSG( | 1330.906 | 0.26966 | 0.117429 | 0.813384 | Acvr2a |
| ENSMUSG( | 121.6847 | 0.236061 | 0.247982 | 0.877904 | Acvr2b |
| ENSMUSG( | 779.5022 | 0.462774 | 0.036776 | 0.671756 | Acvrl1 |
| ENSMUSG( | 323.1619 | -0.22138 | 0.141478 | 0.830678 | Acy1 |

| | | | | | | | | | | | |
|---|---|---|---|---|---|---|---|---|---|---|---|
| ENSMUSG( | Abcc9 | 354.4775 | -0.05712 | 0.887182 | 0.983326 | ENSMUSG( | Abcc9 | 474.4263 | 0.92131 | 0.10475 | 0.804969 |
| ENSMUSG( | Abcd1 | 664.6232 | -0.1057 | 0.345817 | 0.779153 | ENSMUSG( | Abcd1 | 631.9996 | 0.024185 | 0.851198 | 0.985541 |
| ENSMUSG( | Abcd2 | 2903.536 | -0.07937 | 0.517469 | 0.873306 | ENSMUSG( | Abcd2 | 2565.736 | -0.17104 | 0.205016 | 0.864412 |
| ENSMUSG( | Abcd3 | 4341.029 | 0.083013 | 0.41708 | 0.8209 | ENSMUSG( | Abcd3 | 3711.35 | -0.10257 | 0.393745 | 0.917257 |
| ENSMUSG( | Abcd4 | 748.2222 | 0.106357 | 0.382089 | 0.803095 | ENSMUSG( | Abcd4 | 709.7543 | 0.218549 | 0.103452 | 0.804969 |
| ENSMUSG( | Abce1 | 2120.011 | 0.117191 | 0.301703 | 0.749979 | ENSMUSG( | Abce1 | 1830.05 | -0.04144 | 0.768496 | 0.974236 |
| ENSMUSG( | Abcf1 | 2107.72 | 0.219137 | 0.009706 | 0.140314 | ENSMUSG( | Abcf1 | 1812.048 | 0.058685 | 0.578226 | 0.943353 |
| ENSMUSG( | Abcf2 | 2639.687 | 0.0949 | 0.277528 | 0.727457 | ENSMUSG( | Abcf2 | 2341.36 | 0.017446 | 0.876145 | 0.989644 |
| ENSMUSG( | Abcf3 | 1403.11 | 0.067417 | 0.486316 | 0.858575 | ENSMUSG( | Abcf3 | 1241.323 | -0.01655 | 0.894337 | 0.990558 |
| ENSMUSG( | Abcg1 | 1864.487 | -0.45031 | 0.002135 | 0.053442 | ENSMUSG( | Abcg1 | 2114 | 0.213133 | 0.224288 | 0.867489 |
| ENSMUSG( | Abcg2 | 800.4094 | 0.05431 | 0.752982 | 0.950667 | ENSMUSG( | Abcg2 | 848.0058 | 0.46328 | 0.050343 | 0.708648 |
| ENSMUSG( | Abcg3 | 18.05099 | -1.01444 | 0.040236 | NA | ENSMUSG( | Abcg3 | 21.62434 | -0.05748 | 0.908335 | 0.991958 |
| ENSMUSG( | Abcg4 | 1947.76 | -0.16076 | 0.185232 | 0.623185 | ENSMUSG( | Abcg4 | 1942.233 | 0.105676 | 0.408021 | 0.917257 |
| ENSMUSG( | Abhd1 | 183.3324 | 0.000291 | 0.998678 | 0.99989 | ENSMUSG( | Abhd1 | 165.7379 | -0.01897 | 0.925363 | 0.99306 |
| ENSMUSG( | Abhd10 | 437.4568 | -0.16932 | 0.144505 | 0.564381 | ENSMUSG( | Abhd10 | 404.9657 | -0.12535 | 0.379376 | 0.916995 |
| ENSMUSG( | Abhd11 | 1029.503 | -0.03069 | 0.775475 | 0.955606 | ENSMUSG( | Abhd11 | 918.3412 | -0.09529 | 0.452811 | 0.922316 |
| ENSMUSG( | Abhd12 | 14690.36 | -0.20273 | 0.067503 | 0.402091 | ENSMUSG( | Abhd12 | 14495.37 | 0.036744 | 0.781335 | 0.976106 |
| ENSMUSG( | Abhd13 | 1096.767 | -0.34665 | 0.014918 | 0.179376 | ENSMUSG( | Abhd13 | 1236.879 | 0.279522 | 0.090597 | 0.791074 |
| ENSMUSG( | Abhd14a | 1022.837 | -0.14313 | 0.120521 | 0.524506 | ENSMUSG( | Abhd14a | 971.7539 | -0.0181 | 0.877103 | 0.989644 |
| ENSMUSG( | Abhd14b | 459.2044 | 0.180464 | 0.258351 | 0.709574 | ENSMUSG( | Abhd14b | 418.8303 | 0.185887 | 0.353925 | 0.912411 |
| ENSMUSG( | Abhd15 | 55.02149 | -0.09022 | 0.72297 | 0.943761 | ENSMUSG( | Abhd15 | 56.12335 | 0.263106 | 0.49245 | 0.927315 |
| ENSMUSG( | Abhd16a | 3031.261 | -0.29767 | 0.022757 | 0.230554 | ENSMUSG( | Abhd16a | 3148.688 | 0.098647 | 0.517588 | 0.930939 |
| ENSMUSG( | Abhd17a | 10712.54 | 0.138597 | 0.099604 | 0.48212 | ENSMUSG( | Abhd17a | 9058.792 | -0.07405 | 0.463671 | 0.923329 |
| ENSMUSG( | Abhd17b | 788.7259 | 0.09667 | 0.363089 | 0.793211 | ENSMUSG( | Abhd17b | 709.2867 | 0.058659 | 0.685919 | 0.958956 |
| ENSMUSG( | Abhd17c | 1110.18 | 0.069652 | 0.517994 | 0.873754 | ENSMUSG( | Abhd17c | 993.9011 | 0.016916 | 0.902013 | 0.991003 |
| ENSMUSG( | Abhd18 | 396.7756 | 0.005896 | 0.962262 | 0.994543 | ENSMUSG( | Abhd18 | 378.631 | 0.138502 | 0.327856 | 0.906255 |
| ENSMUSG( | Abhd2 | 6160.265 | -0.17244 | 0.233566 | 0.681508 | ENSMUSG( | Abhd2 | 6187.295 | 0.116296 | 0.473485 | 0.923329 |
| ENSMUSG( | Abhd3 | 4801.503 | 0.095206 | 0.400927 | 0.813897 | ENSMUSG( | Abhd3 | 4174.208 | -0.03771 | 0.757902 | 0.971858 |
| ENSMUSG( | Abhd4 | 8340.646 | 0.084518 | 0.434677 | 0.828978 | ENSMUSG( | Abhd4 | 6847.858 | -0.22558 | 0.081462 | 0.77681 |
| ENSMUSG( | Abhd5 | 1156.975 | 0.122364 | 0.265119 | 0.715962 | ENSMUSG( | Abhd5 | 975.0743 | -0.10517 | 0.399656 | 0.917257 |
| ENSMUSG( | Abhd6 | 2134.364 | -0.21677 | 0.139349 | 0.555477 | ENSMUSG( | Abhd6 | 2241.806 | 0.199015 | 0.216658 | 0.865167 |
| ENSMUSG( | Abhd8 | 15216.72 | 0.229203 | 0.00341 | 0.073541 | ENSMUSG( | Abhd8 | 12851.21 | 0.019511 | 0.835954 | 0.981237 |
| ENSMUSG( | Abi1 | 3406.944 | 0.123908 | 0.187951 | 0.626616 | ENSMUSG( | Abi1 | 3052.646 | 0.074236 | 0.598529 | 0.946859 |
| ENSMUSG( | Abi2 | 5540.778 | -0.14774 | 0.090234 | 0.461412 | ENSMUSG( | Abi2 | 5402.407 | 0.050627 | 0.675386 | 0.95694 |
| ENSMUSG( | Abi3 | 281.9643 | -0.06195 | 0.688513 | 0.933441 | ENSMUSG( | Abi3 | 251.7072 | -0.11343 | 0.519741 | 0.931729 |
| ENSMUSG( | Abi3bp | 251.7913 | -0.9771 | 2.97E-09 | 1.27E-06 | ENSMUSG( | Abi3bp | 280.9922 | -0.2432 | 0.270114 | 0.88756 |
| ENSMUSG( | Abitram | 738.9172 | 0.018525 | 0.857129 | 0.974393 | ENSMUSG( | Abitram | 656.3404 | -0.05106 | 0.682412 | 0.958678 |
| ENSMUSG( | Abl1 | 2108.757 | -0.19209 | 0.106578 | 0.494174 | ENSMUSG( | Abl1 | 2120.831 | 0.102208 | 0.486947 | 0.92541 |
| ENSMUSG( | Abl2 | 2054.786 | -0.019 | 0.843571 | 0.971847 | ENSMUSG( | Abl2 | 1909.271 | 0.038624 | 0.73183 | 0.967228 |
| ENSMUSG( | Ablim1 | 3711.223 | 0.019562 | 0.849811 | 0.972638 | ENSMUSG( | Ablim1 | 3537.285 | 0.149502 | 0.263232 | 0.885994 |
| ENSMUSG( | Ablim2 | 4083.382 | -0.05303 | 0.594003 | 0.902869 | ENSMUSG( | Ablim2 | 3733.579 | -0.04344 | 0.719097 | 0.966799 |
| ENSMUSG( | Ablim3 | 990.5706 | -0.02016 | 0.850802 | 0.972638 | ENSMUSG( | Ablim3 | 944.1912 | 0.105958 | 0.404789 | 0.917257 |
| ENSMUSG( | Abr | 18284.94 | 0.164206 | 0.027745 | 0.255176 | ENSMUSG( | Abr | 15433.5 | -0.05283 | 0.589097 | 0.944975 |
| ENSMUSG( | Abracl | 122.798 | 0.140643 | 0.522118 | 0.875186 | ENSMUSG( | Abracl | 118.8262 | 0.285227 | 0.253234 | 0.88018 |
| ENSMUSG( | Abraxas1 | 102.9059 | -0.10891 | 0.611116 | 0.908551 | ENSMUSG( | Abraxas1 | 102.6802 | 0.174488 | 0.476551 | 0.923329 |
| ENSMUSG( | Abraxas2 | 741.464 | 0.06396 | 0.555822 | 0.887457 | ENSMUSG( | Abraxas2 | 664.567 | 0.021612 | 0.863321 | 0.986547 |
| ENSMUSG( | Abt1 | 237.9013 | 0.057993 | 0.700857 | 0.93743 | ENSMUSG( | Abt1 | 206.2258 | -0.09057 | 0.606487 | 0.949743 |
| ENSMUSG( | Abtb1 | 1089.062 | 0.037058 | 0.738333 | 0.947329 | ENSMUSG( | Abtb1 | 1008.836 | 0.088327 | 0.525749 | 0.934374 |
| ENSMUSG( | Abtb2 | 876.9864 | 0.021747 | 0.82644 | 0.968349 | ENSMUSG( | Abtb2 | 755.7341 | -0.14521 | 0.267515 | 0.886134 |
| ENSMUSG( | Abtb3 | 598.4285 | 0.112266 | 0.331435 | 0.77519 | ENSMUSG( | Abtb3 | 528.6435 | 0.032716 | 0.8473 | 0.985161 |
| ENSMUSG( | Acaa1a | 2432.815 | 0.020037 | 0.807453 | 0.963257 | ENSMUSG( | Acaa1a | 2180.729 | -0.02596 | 0.8256 | 0.979537 |
| ENSMUSG( | Acaa1b | 18.01139 | 0.09691 | 0.823802 | NA | ENSMUSG( | Acaa1b | 16.3758 | 0.111583 | 0.816495 | 0.979198 |
| ENSMUSG( | Acaa2 | 1909.16 | 0.019284 | 0.846306 | 0.972226 | ENSMUSG( | Acaa2 | 1598.744 | -0.23738 | 0.092015 | 0.791889 |
| ENSMUSG( | Acaca | 3876.386 | 0.170544 | 0.050488 | 0.343938 | ENSMUSG( | Acaca | 3327.005 | 0.003711 | 0.970543 | 0.99594 |
| ENSMUSG( | Acacb | 279.9885 | 0.0569 | 0.691193 | 0.933881 | ENSMUSG( | Acacb | 254.3093 | 0.060043 | 0.730359 | 0.967228 |
| ENSMUSG( | Acad10 | 165.6642 | -0.01423 | 0.929764 | 0.991848 | ENSMUSG( | Acad10 | 153.6766 | 0.050736 | 0.817774 | 0.979198 |
| ENSMUSG( | Acad11 | 1685.688 | 0.048889 | 0.664007 | 0.926511 | ENSMUSG( | Acad11 | 1363.292 | -0.31479 | 0.009567 | 0.406458 |
| ENSMUSG( | Acad12 | 266.3842 | 0.018609 | 0.901393 | 0.985633 | ENSMUSG( | Acad12 | 227.1564 | -0.19267 | 0.293494 | 0.898538 |
| ENSMUSG( | Acad8 | 938.419 | -0.00677 | 0.95139 | 0.993821 | ENSMUSG( | Acad8 | 808.9117 | -0.17326 | 0.16669 | 0.840183 |
| ENSMUSG( | Acad9 | 1014.825 | 0.024734 | 0.814037 | 0.964574 | ENSMUSG( | Acad9 | 879.8566 | -0.12391 | 0.316247 | 0.901868 |
| ENSMUSG( | Acadl | 3295.353 | 0.121531 | 0.311861 | 0.759517 | ENSMUSG( | Acadl | 2775.286 | -0.10852 | 0.458953 | 0.923329 |
| ENSMUSG( | Acadm | 2532.542 | 0.106522 | 0.363233 | 0.793211 | ENSMUSG( | Acadm | 2143.631 | -0.10966 | 0.400672 | 0.917257 |
| ENSMUSG( | Acads | 612.2225 | 0.088603 | 0.418158 | 0.821966 | ENSMUSG( | Acads | 520.3235 | -0.11585 | 0.374104 | 0.915491 |
| ENSMUSG( | Acadsb | 2259.664 | 0.1083 | 0.29508 | 0.743798 | ENSMUSG( | Acadsb | 1947.583 | -0.05236 | 0.65628 | 0.955533 |
| ENSMUSG( | Acadvl | 2538.039 | 0.182115 | 0.089283 | 0.458563 | ENSMUSG( | Acadvl | 2076.214 | -0.12966 | 0.308219 | 0.898538 |
| ENSMUSG( | Acan | 132.3793 | -0.40783 | 0.038842 | 0.30362 | ENSMUSG( | Acan | 154.24 | 0.326189 | 0.118867 | 0.813384 |
| ENSMUSG( | Acap2 | 3470.524 | 0.080767 | 0.337113 | 0.777008 | ENSMUSG( | Acap2 | 3235.593 | 0.144682 | 0.170362 | 0.841579 |
| ENSMUSG( | Acap3 | 5973.572 | 0.207016 | 0.029569 | 0.260563 | ENSMUSG( | Acap3 | 5213.952 | 0.089812 | 0.405106 | 0.917257 |
| ENSMUSG( | Acat1 | 3344.52 | 0.009607 | 0.921096 | 0.990436 | ENSMUSG( | Acat1 | 2753.614 | -0.30058 | 0.008241 | 0.388416 |
| ENSMUSG( | Acat2 | 1076.572 | 0.042764 | 0.645286 | 0.917957 | ENSMUSG( | Acat2 | 902.2057 | -0.20926 | 0.09438 | 0.795735 |
| ENSMUSG( | Acat3 | 270.0648 | 0.075263 | 0.629031 | 0.913828 | ENSMUSG( | Acat3 | 217.9741 | -0.28978 | 0.137708 | 0.828492 |
| ENSMUSG( | Acbd3 | 990.2469 | 0.108396 | 0.280386 | 0.729291 | ENSMUSG( | Acbd3 | 837.3432 | -0.11146 | 0.397408 | 0.917257 |
| ENSMUSG( | Acbd4 | 981.0817 | 0.029461 | 0.790636 | 0.959461 | ENSMUSG( | Acbd4 | 888.7482 | 0.011356 | 0.928336 | 0.99306 |
| ENSMUSG( | Acbd5 | 4118.972 | 0.08199 | 0.505661 | 0.866932 | ENSMUSG( | Acbd5 | 3509.042 | -0.11665 | 0.417592 | 0.917257 |
| ENSMUSG( | Acbd6 | 1319.87 | -0.01518 | 0.884297 | 0.982647 | ENSMUSG( | Acbd6 | 1153.91 | -0.14309 | 0.33521 | 0.907907 |
| ENSMUSG( | Accs | 172.3435 | 0.078253 | 0.670261 | 0.928525 | ENSMUSG( | Accs | 154.0747 | 0.032817 | 0.868688 | 0.988097 |
| ENSMUSG( | Acd | 1209.225 | 0.063252 | 0.564964 | 0.890735 | ENSMUSG( | Acd | 1038.165 | -0.11019 | 0.394132 | 0.917257 |
| ENSMUSG( | Ace | 713.7026 | -0.2306 | 0.200526 | 0.640661 | ENSMUSG( | Ace | 790.0869 | 0.319301 | 0.046338 | 0.697814 |
| ENSMUSG( | Ace2 | 84.15947 | -0.28043 | 0.479157 | 0.854046 | ENSMUSG( | Ace2 | 98.04069 | 0.432058 | 0.520312 | 0.931964 |
| ENSMUSG( | Acer2 | 238.062 | 0.403821 | 0.044918 | 0.325112 | ENSMUSG( | Acer2 | 195.3052 | 0.143234 | 0.51967 | 0.931729 |
| ENSMUSG( | Acer3 | 579.5805 | -0.26352 | 0.077444 | 0.429743 | ENSMUSG( | Acer3 | 601.2593 | 0.126105 | 0.428331 | 0.917257 |
| ENSMUSG( | Ache | 2125.862 | -0.33224 | 0.000236 | 0.012098 | ENSMUSG( | Ache | 2179.818 | 0.02857 | 0.800609 | 0.97681 |
| ENSMUSG( | Acin1 | 4331.975 | 0.030148 | 0.769289 | 0.954674 | ENSMUSG( | Acin1 | 3979.143 | 0.053557 | 0.63811 | 0.953903 |
| ENSMUSG( | Ackr1 | 1594.498 | -0.3241 | 0.005632 | 0.100551 | ENSMUSG( | Ackr1 | 1638.001 | 0.040478 | 0.767359 | 0.974236 |
| ENSMUSG( | Ackr2 | 64.86934 | 0.049456 | 0.854465 | 0.973547 | ENSMUSG( | Ackr2 | 68.18265 | 0.45358 | 0.117861 | 0.813384 |
| ENSMUSG( | Ackr3 | 332.4315 | -0.43871 | 0.028915 | 0.258972 | ENSMUSG( | Ackr3 | 394.2298 | 0.347043 | 0.162679 | 0.837724 |
| ENSMUSG( | Acly | 4593.659 | 0.106222 | 0.222568 | 0.668434 | ENSMUSG( | Acly | 3921.06 | -0.08226 | 0.433342 | 0.918434 |
| ENSMUSG( | Aco1 | 1677.121 | 0.142279 | 0.200151 | 0.640302 | ENSMUSG( | Aco1 | 1417.025 | -0.07411 | 0.517654 | 0.930939 |
| ENSMUSG( | Aco2 | 30436.36 | 0.086852 | 0.289929 | 0.739839 | ENSMUSG( | Aco2 | 25598.22 | -0.14576 | 0.144205 | 0.83148 |
| ENSMUSG( | Acot1 | 467.4169 | 0.159896 | 0.295065 | 0.743798 | ENSMUSG( | Acot1 | 358.8787 | -0.35901 | 0.019523 | 0.556192 |
| ENSMUSG( | Acot11 | 6599.033 | 0.239798 | 0.009547 | 0.138264 | ENSMUSG( | Acot11 | 5181.213 | -0.18623 | 0.103747 | 0.804969 |
| ENSMUSG( | Acot13 | 1598.466 | 0.00738 | 0.934074 | 0.99323 | ENSMUSG( | Acot13 | 1319.309 | -0.29731 | 0.024767 | 0.592568 |
| ENSMUSG( | Acot2 | 232.8933 | 0.024012 | 0.879962 | 0.981469 | ENSMUSG( | Acot2 | 188.3519 | -0.34902 | 0.071358 | 0.761942 |
| ENSMUSG( | Acot3 | 23.31191 | -0.07155 | 0.853797 | 0.973491 | ENSMUSG( | Acot3 | 20.37848 | -0.21984 | 0.620827 | 0.953114 |
| ENSMUSG( | Acot4 | 81.76732 | 0.058482 | 0.78282 | 0.957558 | ENSMUSG( | Acot4 | 75.0064 | 0.076762 | 0.749643 | 0.96998 |
| ENSMUSG( | Acot5 | 50.39883 | -0.2124 | 0.445453 | 0.833239 | ENSMUSG( | Acot5 | 45.35423 | -0.2273 | 0.450484 | 0.922202 |
| ENSMUSG( | Acot6 | 191.0906 | 0.009922 | 0.951921 | 0.993821 | ENSMUSG( | Acot6 | 175.328 | 0.027795 | 0.876793 | 0.989644 |
| ENSMUSG( | Acot7 | 3832.37 | -0.00199 | 0.988688 | 0.999109 | ENSMUSG( | Acot7 | 3184.814 | -0.28475 | 0.096959 | 0.79645 |
| ENSMUSG( | Acot8 | 316.124 | -0.15017 | 0.304834 | 0.752574 | ENSMUSG( | Acot8 | 291.2987 | -0.10801 | 0.501258 | 0.927825 |
| ENSMUSG( | Acot9 | 303.4877 | 0.019157 | 0.889683 | 0.983551 | ENSMUSG( | Acot9 | 243.2484 | -0.36108 | 0.040233 | 0.681706 |
| ENSMUSG( | Acox1 | 13037.52 | 0.129577 | 0.166437 | 0.599114 | ENSMUSG( | Acox1 | 11131.04 | -0.05652 | 0.624077 | 0.953234 |
| ENSMUSG( | Acox3 | 1609.38 | -0.01001 | 0.927634 | 0.991229 | ENSMUSG( | Acox3 | 1465.59 | -0.00686 | 0.957883 | 0.994798 |
| ENSMUSG( | Acp1 | 1401.669 | -0.03675 | 0.70124 | 0.93743 | ENSMUSG( | Acp1 | 1255.835 | -0.08902 | 0.456837 | 0.923329 |
| ENSMUSG( | Acp2 | 3040.411 | -0.30668 | 0.040294 | 0.31004 | ENSMUSG( | Acp2 | 3248.181 | 0.169107 | 0.323497 | 0.904501 |
| ENSMUSG( | Acp3 | 40.09068 | -1.16122 | 0.000377 | 0.017069 | ENSMUSG( | Acp3 | 48.71314 | -0.09588 | 0.743579 | 0.96951 |
| ENSMUSG( | Acp5 | 18.71082 | -0.1217 | 0.794528 | NA | ENSMUSG( | Acp5 | 19.38088 | 0.22804 | 0.626452 | 0.953234 |
| ENSMUSG( | Acp6 | 390.3801 | 0.06125 | 0.665699 | 0.927054 | ENSMUSG( | Acp6 | 376.9074 | 0.230049 | 0.22807 | 0.868507 |
| ENSMUSG( | Acp7 | 29.47335 | -0.20425 | 0.558789 | 0.888638 | ENSMUSG( | Acp7 | 27.43368 | -0.13453 | 0.717023 | 0.965864 |
| ENSMUSG( | Acr | 56.24335 | -0.19583 | 0.476611 | 0.851717 | ENSMUSG( | Acr | 62.26818 | 0.344943 | 0.216674 | 0.865167 |
| ENSMUSG( | Acrbp | 147.8458 | 0.569566 | 0.008964 | 0.132538 | ENSMUSG( | Acrbp | 142.3553 | 0.702966 | 0.004571 | 0.277991 |
| ENSMUSG( | Acsbg1 | 22556.33 | 0.076182 | 0.493146 | 0.860552 | ENSMUSG( | Acsbg1 | 18184.1 | -0.29239 | 0.023966 | 0.588226 |
| ENSMUSG( | Acsf2 | 2133.475 | 0.034887 | 0.756431 | 0.951479 | ENSMUSG( | Acsf2 | 1854.33 | -0.10482 | 0.428103 | 0.917257 |
| ENSMUSG( | Acsf3 | 579.4882 | 0.031012 | 0.794823 | 0.959799 | ENSMUSG( | Acsf3 | 482.5576 | -0.236 | 0.089307 | 0.788502 |
| ENSMUSG( | Acsl1 | 1222.82 | -0.23161 | 0.021839 | 0.225699 | ENSMUSG( | Acsl1 | 1240.245 | 0.08961 | 0.469404 | 0.923329 |
| ENSMUSG( | Acsl3 | 10791.05 | 0.033793 | 0.79619 | 0.960088 | ENSMUSG( | Acsl3 | 9846.19 | 0.041524 | 0.789765 | 0.976792 |
| ENSMUSG( | Acsl4 | 1612.926 | -0.46267 | 0.005071 | 0.094652 | ENSMUSG( | Acsl4 | 1899.944 | 0.302441 | 0.115616 | 0.813384 |
| ENSMUSG( | Acsl5 | 2014.739 | -0.13081 | 0.335973 | 0.777 | ENSMUSG( | Acsl5 | 2039.649 | 0.176066 | 0.266072 | 0.886134 |
| ENSMUSG( | Acsl6 | 19165.59 | -0.012 | 0.913983 | 0.988483 | ENSMUSG( | Acsl6 | 17506.55 | -0.00303 | 0.980939 | 0.9967 |
| ENSMUSG( | Acsm5 | 18.85329 | -0.35512 | 0.447994 | NA | ENSMUSG( | Acsm5 | 17.29171 | -0.37826 | 0.485347 | 0.92541 |
| ENSMUSG( | Acss1 | 2920.782 | 0.173678 | 0.144932 | 0.565187 | ENSMUSG( | Acss1 | 2372.42 | -0.15801 | 0.15102 | 0.837173 |
| ENSMUSG( | Acss2 | 3505.757 | 0.08762 | 0.468802 | 0.845636 | ENSMUSG( | Acss2 | 2944.752 | -0.14907 | 0.208926 | 0.864412 |
| ENSMUSG( | Acss3 | 53.37884 | -0.00483 | 0.986229 | 0.998326 | ENSMUSG( | Acss3 | 49.44789 | 0.040826 | 0.891945 | 0.990558 |
| ENSMUSG( | Acta1 | 282.5231 | 0.12615 | 0.401815 | 0.814965 | ENSMUSG( | Acta1 | 227.7369 | -0.23835 | 0.16756 | 0.840222 |
| ENSMUSG( | Acta2 | 52.71579 | -0.74139 | 0.021324 | 0.223 | ENSMUSG( | Acta2 | 68.13562 | 0.367293 | 0.278868 | 0.891737 |
| ENSMUSG( | Actb | 24522.23 | -0.07918 | 0.386043 | 0.804623 | ENSMUSG( | Actb | 21227.21 | -0.23791 | 0.080045 | 0.776379 |
| ENSMUSG( | Actg1 | 9938.316 | -0.00941 | 0.9152 | 0.988634 | ENSMUSG( | Actg1 | 8515.782 | -0.19357 | 0.082701 | 0.77681 |
| ENSMUSG( | Actl6a | 432.8853 | 0.007278 | 0.949825 | 0.993821 | ENSMUSG( | Actl6a | 403.6748 | 0.076954 | 0.573599 | 0.942912 |
| ENSMUSG( | Actl6b | 2320.087 | 0.100825 | 0.435785 | 0.829093 | ENSMUSG( | Actl6b | 1989.215 | -0.07596 | 0.586445 | 0.944975 |
| ENSMUSG( | Actmap | 594.3063 | 0.057013 | 0.594559 | 0.902971 | ENSMUSG( | Actmap | 528.1155 | -0.01744 | 0.8929 | 0.990558 |
| ENSMUSG( | Actn1 | 2374.816 | -0.14416 | 0.19246 | 0.631514 | ENSMUSG( | Actn1 | 2381.77 | 0.13596 | 0.265209 | 0.886134 |
| ENSMUSG( | Actn2 | 635.7893 | -0.72395 | 4.10E-06 | 0.000549 | ENSMUSG( | Actn2 | 695.8915 | -0.10788 | 0.425487 | 0.917257 |
| ENSMUSG( | Actn3 | 46.73538 | -0.07389 | 0.803763 | 0.962169 | ENSMUSG( | Actn3 | 41.12359 | -0.17017 | 0.602779 | 0.94789 |
| ENSMUSG( | Actn4 | 5709.115 | -0.01535 | 0.880396 | 0.981469 | ENSMUSG( | Actn4 | 5413.845 | 0.10076 | 0.36941 | 0.915491 |
| ENSMUSG( | Actr10 | 2672.009 | 0.106962 | 0.318502 | 0.764234 | ENSMUSG( | Actr10 | 2249.267 | -0.12537 | 0.368721 | 0.915491 |
| ENSMUSG( | Actr1a | 5618.237 | 0.026242 | 0.751415 | 0.950237 | ENSMUSG( | Actr1a | 4842.191 | -0.13799 | 0.190961 | 0.858148 |
| ENSMUSG( | Actr1b | 9191.111 | 0.102904 | 0.259067 | 0.709814 | ENSMUSG( | Actr1b | 7817.066 | -0.09646 | 0.379053 | 0.916995 |
| ENSMUSG( | Actr2 | 6445.827 | 0.016881 | 0.830829 | 0.969798 | ENSMUSG( | Actr2 | 5728.772 | -0.05717 | 0.624789 | 0.953234 |
| ENSMUSG( | Actr3 | 4292.658 | 0.04128 | 0.64232 | 0.916865 | ENSMUSG( | Actr3 | 3799.275 | -0.04486 | 0.70482 | 0.962677 |
| ENSMUSG( | Actr3b | 2922.622 | 0.090888 | 0.37473 | 0.800466 | ENSMUSG( | Actr3b | 2555.515 | -0.02823 | 0.820867 | 0.979198 |
| ENSMUSG( | Actr3-ps | 14.42824 | -0.43657 | 0.369295 | NA | ENSMUSG( | Actr3-ps | 15.35673 | -0.03098 | 0.957263 | 0.994798 |
| ENSMUSG( | Actr5 | 190.653 | -0.0626 | 0.709604 | 0.939669 | ENSMUSG( | Actr5 | 171.5471 | -0.09227 | 0.620104 | 0.952896 |
| ENSMUSG( | Actr6 | 322.0231 | 0.127702 | 0.342214 | 0.778828 | ENSMUSG( | Actr6 | 296.2382 | 0.145364 | 0.366782 | 0.914839 |
| ENSMUSG( | Actr8 | 1064.125 | 0.161175 | 0.18332 | 0.62047 | ENSMUSG( | Actr8 | 926.5554 | 0.033829 | 0.812776 | 0.979148 |
| ENSMUSG( | Actrt3 | 13.94302 | -0.95326 | 0.087748 | NA | ENSMUSG( | Actrt3 | 15.10261 | -0.30865 | 0.633066 | 0.953844 |
| ENSMUSG( | Acvr1 | 924.2595 | -0.32792 | 0.020415 | 0.215588 | ENSMUSG( | Acvr1 | 1010.051 | 0.21172 | 0.191362 | 0.858412 |
| ENSMUSG( | Acvr1b | 4377.367 | -0.29177 | 0.03222 | 0.274192 | ENSMUSG( | Acvr1b | 4675.926 | 0.180862 | 0.255817 | 0.883082 |
| ENSMUSG( | Acvr1c | 330.1769 | -0.85181 | 2.92E-06 | 0.000423 | ENSMUSG( | Acvr1c | 361.7163 | -0.1959 | 0.314248 | 0.900949 |
| ENSMUSG( | Acvr2a | 1187.519 | -0.33789 | 0.025773 | 0.247138 | ENSMUSG( | Acvr2a | 1330.906 | 0.26966 | 0.117429 | 0.813384 |
| ENSMUSG( | Acvr2b | 115.9211 | -0.1783 | 0.396484 | 0.811742 | ENSMUSG( | Acvr2b | 121.6847 | 0.236061 | 0.247982 | 0.877904 |
| ENSMUSG( | Acvrl1 | 664.9407 | -0.24864 | 0.178895 | 0.616298 | ENSMUSG( | Acvrl1 | 779.5022 | 0.462774 | 0.036776 | 0.671756 |
| ENSMUSG( | Acy1 | 356.6677 | -0.20178 | 0.135221 | 0.548658 | ENSMUSG( | Acy1 | 323.1619 | -0.22138 | 0.141478 | 0.830678 |
| ENSMUSG( | Acy3 | 74.96037 | -0.0636 | 0.77782 | 0.956168 | ENSMUSG( | Acy3 | 67.01315 | -0.09818 | 0.710259 | 0.964363 |
| ENSMUSG( | Acyp1 | 294.3612 | 0.043209 | 0.820645 | 0.965888 | ENSMUSG( | Acyp1 | 259.471 | -0.06346 | 0.769828 | 0.974236 |
| ENSMUSG( | Acyp2 | 474.4918 | 0.076266 | 0.540302 | 0.88315 | ENSMUSG( | Acyp2 | 388.8494 | -0.24638 | 0.126858 | 0.816972 |
| ENSMUSG( | Ada | 25.6018 | -0.13847 | 0.73078 | 0.944956 | ENSMUSG( | Ada | 22.45376 | -0.23015 | 0.60566 | 0.949298 |
| ENSMUSG( | Adal | 423.8852 | 0.149029 | 0.238981 | 0.686574 | ENSMUSG( | Adal | 377.5423 | 0.088051 | 0.517636 | 0.930939 |
| ENSMUSG( | Adam10 | 2454.158 | -0.27451 | 0.103159 | 0.48855 | ENSMUSG( | Adam10 | 2724.276 | 0.297183 | 0.108553 | 0.804969 |
| ENSMUSG( | Adam11 | 6660.318 | -0.0149 | 0.900177 | 0.985633 | ENSMUSG( | Adam11 | 6315.061 | 0.100156 | 0.385562 | 0.917257 |

| | | | | | |
|---|---|---|---|---|---|
| ENSMUSG( | 924.2595 | -0.32792 | 0.020415 | 0.215588 | Acvr1 |
| ENSMUSG( | 4377.367 | -0.29177 | 0.03222 | 0.274192 | Acvr1b |
| ENSMUSG( | 330.1769 | -0.85181 | 2.92E-06 | 0.000423 | Acvr1c |
| ENSMUSG( | 1187.519 | -0.33789 | 0.025773 | 0.247138 | Acvr2a |
| ENSMUSG( | 115.9211 | -0.1783 | 0.396484 | 0.811742 | Acvr2b |
| ENSMUSG( | 664.9407 | -0.24864 | 0.178895 | 0.616298 | Acvrl1 |
| ENSMUSG( | 356.6677 | -0.20178 | 0.135221 | 0.548658 | Acy1 |
| ENSMUSG( | 74.96037 | -0.0636 | 0.77782 | 0.956168 | Acy3 |
| ENSMUSG( | 294.3612 | 0.043209 | 0.820645 | 0.965888 | Acyp1 |
| ENSMUSG( | 474.4918 | 0.076266 | 0.540302 | 0.88315 | Acyp2 |
| ENSMUSG( | 25.6018 | -0.13847 | 0.73078 | 0.944956 | Ada |
| ENSMUSG( | 423.8852 | 0.149029 | 0.238981 | 0.686574 | Adal |
| ENSMUSG( | 2454.158 | -0.27451 | 0.103159 | 0.48855 | Adam10 |
| ENSMUSG( | 6660.318 | -0.0149 | 0.900177 | 0.985633 | Adam11 |
| ENSMUSG( | 224.6166 | -0.25607 | 0.134615 | 0.547918 | Adam12 |
| ENSMUSG( | 3968.131 | 0.005964 | 0.956152 | 0.993874 | Adam15 |
| ENSMUSG( | 1061.033 | -0.029 | 0.855049 | 0.973665 | Adam17 |
| ENSMUSG( | 39.07732 | -0.33169 | 0.319518 | 0.765296 | Adam18 |
| ENSMUSG( | 820.625 | -0.1266 | 0.468321 | 0.845336 | Adam19 |
| ENSMUSG( | 168.0321 | -0.40663 | 0.014588 | 0.177737 | Adam1a |
| ENSMUSG( | 9.752875 | 0.371382 | 0.544373 | NA | Adam21 |
| ENSMUSG( | 6960.81 | -0.29842 | 0.01263 | 0.162076 | Adam22 |
| ENSMUSG( | 5101.284 | -0.50526 | 6.92E-05 | 0.005054 | Adam23 |
| ENSMUSG( | 12.2272 | -0.24655 | 0.689444 | NA | Adam3 |
| ENSMUSG( | 15.30772 | 0.24768 | 0.629941 | NA | Adam32 |
| ENSMUSG( | 222.9974 | 0.231194 | 0.289618 | 0.739839 | Adam33 |
| ENSMUSG( | 13.52862 | 0.925175 | 0.068845 | NA | Adam4 |
| ENSMUSG( | 12.17634 | -0.20867 | 0.697731 | NA | Adam5 |
| ENSMUSG( | 129.1882 | 0.987664 | 1.85E-05 | 0.001854 | Adam8 |
| ENSMUSG( | 3121.341 | -0.07159 | 0.572875 | 0.893584 | Adam9 |
| ENSMUSG( | 391.9244 | -0.80754 | 2.24E-05 | 0.002129 | Adamts1 |
| ENSMUSG( | 1007.411 | 0.280147 | 0.066327 | 0.398419 | Adamts10 |
| ENSMUSG( | 33.04246 | -0.56902 | 0.137898 | 0.552903 | Adamts12 |
| ENSMUSG( | 35.25834 | -0.27632 | 0.424579 | 0.824675 | Adamts13 |
| ENSMUSG( | 22.49746 | 0.340911 | 0.400474 | 0.813717 | Adamts14 |
| ENSMUSG( | 190.9671 | -0.25926 | 0.142609 | 0.561026 | Adamts15 |
| ENSMUSG( | 201.3151 | -0.12937 | 0.423358 | 0.823662 | Adamts16 |
| ENSMUSG( | 164.0531 | 0.162577 | 0.382651 | 0.803227 | Adamts17 |
| ENSMUSG( | 74.73357 | -0.33154 | 0.167342 | 0.599971 | Adamts18 |
| ENSMUSG( | 19.47796 | -0.64546 | 0.283552 | NA | Adamts19 |
| ENSMUSG( | 484.4419 | -0.09912 | 0.482778 | 0.856184 | Adamts2 |
| ENSMUSG( | 586.5466 | 0.088253 | 0.447986 | 0.834051 | Adamts20 |
| ENSMUSG( | 298.6378 | -0.49705 | 0.008449 | 0.128676 | Adamts3 |
| ENSMUSG( | 2396.513 | -0.2526 | 0.011726 | 0.156231 | Adamts4 |
| ENSMUSG( | 148.5951 | -0.321 | 0.08282 | 0.444372 | Adamts5 |
| ENSMUSG( | 204.0094 | -0.67529 | 0.000135 | 0.007928 | Adamts6 |
| ENSMUSG( | 24.37573 | -0.15022 | 0.707526 | 0.93949 | Adamts7 |
| ENSMUSG( | 128.7961 | -0.50551 | 0.05871 | 0.373994 | Adamts8 |
| ENSMUSG( | 258.3729 | -0.00557 | 0.973832 | 0.99675 | Adamts9 |
| ENSMUSG( | 181.3646 | -0.69883 | 0.000664 | 0.024862 | Adamtsl1 |
| ENSMUSG( | 308.0532 | -0.10955 | 0.43901 | 0.830714 | Adamtsl2 |
| ENSMUSG( | 178.8523 | -0.14033 | 0.386161 | 0.804623 | Adamtsl3 |
| ENSMUSG( | 265.9548 | -0.70565 | 0.000101 | 0.006438 | Adamtsl4 |
| ENSMUSG( | 336.1072 | -0.19591 | 0.137348 | 0.552389 | Adamtsl5 |
| ENSMUSG( | 10863.74 | 0.154362 | 0.056084 | 0.36519 | Adap1 |
| ENSMUSG( | 211.6809 | -0.13726 | 0.429255 | 0.826446 | Adap2 |
| ENSMUSG( | 4115.5 | 0.143971 | 0.156812 | 0.5851 | Adar |
| ENSMUSG( | 7798.919 | 0.009341 | 0.904378 | 0.986246 | Adarb1 |
| ENSMUSG( | 322.1745 | 0.061455 | 0.653415 | 0.922819 | Adarb2 |
| ENSMUSG( | 138.1582 | 0.171891 | 0.43158 | 0.827432 | Adat1 |
| ENSMUSG( | 100.9222 | 0.051283 | 0.807115 | 0.9631 | Adat2 |
| ENSMUSG( | 976.2986 | 0.08474 | 0.420496 | 0.822898 | Adck1 |
| ENSMUSG( | 410.3938 | -0.15487 | 0.42326 | 0.823662 | Adck2 |
| ENSMUSG( | 455.9965 | 0.057803 | 0.644711 | 0.917782 | Adck5 |
| ENSMUSG( | 28924.51 | 0.204825 | 0.144235 | 0.563869 | Adcy1 |
| ENSMUSG( | 8388.587 | -0.0932 | 0.488643 | 0.859092 | Adcy2 |
| ENSMUSG( | 1742.122 | -0.1176 | 0.448787 | 0.83469 | Adcy3 |
| ENSMUSG( | 135.0212 | 0.399319 | 0.103077 | 0.48855 | Adcy4 |
| ENSMUSG( | 10001.58 | -0.31818 | 0.008717 | 0.131277 | Adcy5 |
| ENSMUSG( | 1993.807 | 0.225564 | 0.038743 | 0.303142 | Adcy6 |
| ENSMUSG( | 282.6723 | -0.28727 | 0.069552 | 0.408118 | Adcy7 |
| ENSMUSG( | 975.7224 | -0.32707 | 0.022429 | 0.228899 | Adcy8 |
| ENSMUSG( | 777.7899 | -0.00497 | 0.979903 | 0.997428 | Adcy9 |
| ENSMUSG( | 516.5098 | -0.56194 | 0.001623 | 0.045245 | Adcyap1 |
| ENSMUSG( | 13527.1 | -0.01008 | 0.92804 | 0.991229 | Adcyap1r1 |
| ENSMUSG( | 13687.67 | 0.101606 | 0.186637 | 0.623917 | Add1 |
| ENSMUSG( | 7464.56 | 0.142605 | 0.160839 | 0.590628 | Add2 |
| ENSMUSG( | 9165.123 | 0.10424 | 0.338705 | 0.777518 | Add3 |
| ENSMUSG( | 34.86822 | -0.12084 | 0.742136 | 0.948478 | Adgb |
| ENSMUSG( | 2314.174 | -0.08323 | 0.590188 | 0.90113 | Adgra1 |
| ENSMUSG( | 616.3918 | 0.154809 | 0.250225 | 0.700009 | Adgra2 |
| ENSMUSG( | 1408.503 | 0.034688 | 0.784731 | 0.958171 | Adgra3 |
| ENSMUSG( | 18451.99 | -0.00775 | 0.94207 | 0.993415 | Adgrb1 |
| ENSMUSG( | 19382.09 | 0.166663 | 0.153196 | 0.57989 | Adgrb2 |
| ENSMUSG( | 2821.066 | -0.17514 | 0.246548 | 0.694958 | Adgrb3 |
| ENSMUSG( | 141.7045 | -0.28351 | 0.167378 | 0.599971 | Adgrd1 |
| ENSMUSG( | 461.505 | -0.29452 | 0.049828 | 0.340731 | Adgre1 |
| ENSMUSG( | 27.96203 | -1.9407 | 0.001881 | 0.049538 | Adgre4 |
| ENSMUSG( | 673.983 | 0.059346 | 0.63818 | 0.916124 | Adgre5 |
| ENSMUSG( | 28.32831 | -0.09056 | 0.805744 | 0.962616 | Adgrf2 |
| ENSMUSG( | 58.11818 | 0.10131 | 0.711116 | 0.940028 | Adgrf4 |
| ENSMUSG( | 1431.915 | 0.179322 | 0.403801 | 0.816132 | Adgrf5 |
| ENSMUSG( | 4485.042 | -0.38834 | 0.010467 | 0.146428 | Adgrg1 |
| ENSMUSG( | 33.88062 | 0.11147 | 0.750188 | 0.949664 | Adgrg2 |
| ENSMUSG( | 9.620186 | -0.37505 | 0.538641 | NA | Adgrg3 |
| ENSMUSG( | 93.5599 | -0.45904 | 0.042746 | 0.316989 | Adgrg6 |
| ENSMUSG( | 22001.62 | 0.004285 | 0.97119 | 0.996369 | Adgrl1 |
| ENSMUSG( | 2899.468 | -0.23746 | 0.061782 | 0.384389 | Adgrl2 |
| ENSMUSG( | 3427.532 | 0.043699 | 0.704092 | 0.938172 | Adgrl3 |
| ENSMUSG( | 752.9256 | 0.288989 | 0.4446 | 0.833239 | Adgrl4 |
| ENSMUSG( | 112.3443 | -0.26475 | 0.249989 | 0.69959 | Adgrv1 |
| ENSMUSG( | 1755.783 | 0.011509 | 0.905787 | 0.986694 | Adh5 |
| ENSMUSG( | 2639.707 | 0.147673 | 0.175579 | 0.611004 | Adhfe1 |
| ENSMUSG( | 1137.59 | 0.037293 | 0.713973 | 0.940903 | Adi1 |
| ENSMUSG( | 4866.745 | -0.12295 | 0.193141 | 0.632494 | Adipor1 |
| ENSMUSG( | 3519.088 | 0.053522 | 0.645533 | 0.918025 | Adipor2 |
| ENSMUSG( | 1015.167 | 0.092562 | 0.463919 | 0.842812 | Adissp |
| ENSMUSG( | 3561.42 | 0.184697 | 0.116474 | 0.514944 | Adk |
| ENSMUSG( | 89.51888 | -0.57498 | 0.017366 | 0.197592 | Adm |
| ENSMUSG( | 98.90447 | 0.111351 | 0.584663 | 0.898884 | Adnp |
| ENSMUSG( | 696.7506 | 0.045987 | 0.693187 | 0.934202 | Adnp2 |
| ENSMUSG( | 1680.949 | -0.00857 | 0.926666 | 0.991229 | Ado |
| ENSMUSG( | 12180.04 | -0.01237 | 0.907412 | 0.987566 | Adora1 |
| ENSMUSG( | 1220.907 | -1.29293 | 0.00076 | 0.027082 | Adora2a |
| ENSMUSG( | 204.6506 | -0.46073 | 0.039387 | 0.305596 | Adora2b |
| ENSMUSG( | 39.71769 | -0.04629 | 0.908845 | 0.98794 | Adora3 |
| ENSMUSG( | 658.1774 | -0.30488 | 0.026508 | 0.248968 | Adpgk |
| ENSMUSG( | 2715.144 | -0.30721 | 0.001441 | 0.041882 | Adprh |
| ENSMUSG( | 273.4923 | 0.183756 | 0.210173 | 0.652567 | Adprm |
| ENSMUSG( | 544.5488 | -0.06345 | 0.577562 | 0.895864 | Adprs |
| ENSMUSG( | 251.3194 | -0.20868 | 0.290244 | 0.740097 | Adra1a |
| ENSMUSG( | 808.1531 | -0.2331 | 0.109035 | 0.498536 | Adra1b |
| ENSMUSG( | 825.4306 | -0.18394 | 0.339133 | 0.777518 | Adra1d |
| ENSMUSG( | 748.4734 | -0.12433 | 0.369741 | 0.797289 | Adra2a |
| ENSMUSG( | 13.55628 | -1.32818 | 0.022593 | NA | Adra2b |
| ENSMUSG( | 835.8157 | -0.08896 | 0.390925 | 0.80826 | Adra2c |
| ENSMUSG( | 1009.305 | -0.66608 | 8.92E-05 | 0.005955 | Adrb1 |
| ENSMUSG( | 83.59689 | -0.59007 | 0.009458 | 0.137493 | Adrb2 |
| ENSMUSG( | 41.90078 | 0.259954 | 0.391808 | 0.809274 | Adrb3 |
| ENSMUSG( | 1239.711 | -0.06667 | 0.574955 | 0.894572 | Adrm1 |
| ENSMUSG( | 21.89963 | 0.072214 | 0.881634 | 0.981948 | Adrm1b |
| ENSMUSG( | 701.1157 | 0.007883 | 0.942043 | 0.993415 | Adsl |
| ENSMUSG( | 427.5436 | -0.15877 | 0.193589 | 0.633067 | Adss1 |
| ENSMUSG( | 1791.278 | -0.08368 | 0.357363 | 0.788775 | Adss2 |
| ENSMUSG( | 1001.887 | -0.00441 | 0.973141 | 0.99675 | Aebp1 |
| ENSMUSG( | 1530.431 | -0.0203 | 0.816842 | 0.96566 | Aebp2 |
| ENSMUSG( | 678.8831 | 0.065226 | 0.538466 | 0.88259 | Aen |
| ENSMUSG( | 901.2851 | -0.01722 | 0.875265 | 0.979998 | Afap1 |
| ENSMUSG( | 1104.781 | -0.06865 | 0.597948 | 0.90372 | Afap1l1 |
| ENSMUSG( | 574.871 | -0.00906 | 0.944193 | 0.993503 | Afap1l2 |
| ENSMUSG( | 2322.677 | 0.075928 | 0.380033 | 0.801757 | Afdn |
| ENSMUSG( | 761.62 | 0.164764 | 0.199785 | 0.640239 | Aff1 |
| ENSMUSG( | 88.76147 | -0.03595 | 0.873154 | 0.978753 | Aff2 |
| ENSMUSG( | 1272.382 | -0.06883 | 0.483118 | 0.856216 | Aff3 |
| ENSMUSG( | 4023.695 | 0.109278 | 0.204495 | 0.645438 | Aff4 |
| ENSMUSG( | 694.782 | 0.159631 | 0.13482 | 0.54834 | Afg1l |
| ENSMUSG( | 194.7582 | 0.217815 | 0.163471 | 0.594869 | Afg2a |
| ENSMUSG( | 238.8492 | 0.15335 | 0.272264 | 0.7223 | Afg2b |
| ENSMUSG( | 1726.67 | 0.136922 | 0.147036 | 0.569057 | Afg3l1 |
| ENSMUSG( | 2211.549 | -0.0448 | 0.66499 | 0.926824 | Afg3l2 |
| ENSMUSG( | 24.85805 | -0.05966 | 0.871558 | 0.978114 | Afmid |
| ENSMUSG( | 1693.516 | 0.061649 | 0.491636 | 0.860231 | Aftph |
| ENSMUSG( | 297.9345 | -0.14057 | 0.354769 | 0.78738 | Aga |
| ENSMUSG( | 8727.004 | 0.064397 | 0.408647 | 0.818173 | Agap1 |
| ENSMUSG( | 64538.43 | 0.250075 | 0.003874 | 0.079851 | Agap2 |
| ENSMUSG( | 12667.86 | 0.167992 | 0.044473 | 0.32391 | Agap3 |
| ENSMUSG( | 89.14478 | -0.1532 | 0.512597 | 0.871762 | Agbl2 |
| ENSMUSG( | 253.834 | 0.025958 | 0.878351 | 0.981431 | Agbl3 |

| | | | | | |
|---|---|---|---|---|---|
| ENSMUSG( | 67.01315 | -0.09818 | 0.710259 | 0.964363 | Acy3 |
| ENSMUSG( | 259.471 | -0.06346 | 0.769828 | 0.974236 | Acyp1 |
| ENSMUSG( | 388.8494 | -0.24638 | 0.126858 | 0.816972 | Acyp2 |
| ENSMUSG( | 22.45376 | -0.23015 | 0.60566 | 0.949298 | Ada |
| ENSMUSG( | 377.5423 | 0.088051 | 0.517636 | 0.930939 | Adal |
| ENSMUSG( | 2724.276 | 0.297183 | 0.108553 | 0.804969 | Adam10 |
| ENSMUSG( | 6315.061 | 0.100156 | 0.385562 | 0.917257 | Adam11 |
| ENSMUSG( | 230.1365 | 0.101576 | 0.570076 | 0.941777 | Adam12 |
| ENSMUSG( | 3866.901 | 0.196086 | 0.140003 | 0.830678 | Adam15 |
| ENSMUSG( | 1040.507 | 0.183899 | 0.293348 | 0.898538 | Adam17 |
| ENSMUSG( | 36.30321 | -0.25441 | 0.488955 | 0.925573 | Adam18 |
| ENSMUSG( | 834.5238 | 0.19481 | 0.31443 | 0.900949 | Adam19 |
| ENSMUSG( | 165.205 | -0.14799 | 0.537235 | 0.937216 | Adam1a |
| ENSMUSG( | 9.787104 | 0.573847 | 0.366302 | NA | Adam21 |
| ENSMUSG( | 7298.94 | 0.122089 | 0.362031 | 0.914223 | Adam22 |
| ENSMUSG( | 5610.917 | 0.083869 | 0.567676 | 0.941777 | Adam23 |
| ENSMUSG( | 12.49573 | 0.148539 | 0.822802 | 0.979198 | Adam3 |
| ENSMUSG( | 12.59357 | -0.01612 | 0.976178 | 0.996416 | Adam32 |
| ENSMUSG( | 172.1941 | -0.24904 | 0.380823 | 0.917202 | Adam33 |
| ENSMUSG( | 11.50135 | -0.06923 | 0.912204 | 0.992225 | Adam5 |
| ENSMUSG( | 106.5392 | 0.757284 | 0.008348 | 0.388416 | Adam8 |
| ENSMUSG( | 3057.5 | 0.136329 | 0.338883 | 0.907972 | Adam9 |
| ENSMUSG( | 437.0466 | -0.10556 | 0.611383 | 0.950771 | Adamts1 |
| ENSMUSG( | 942.4591 | 0.348664 | 0.1044 | 0.804969 | Adamts10 |
| ENSMUSG( | 30.97554 | -0.46918 | 0.231034 | 0.868507 | Adamts12 |
| ENSMUSG( | 34.92681 | -0.05147 | 0.891452 | 0.990558 | Adamts13 |
| ENSMUSG( | 20.72388 | 0.361429 | 0.409181 | 0.917257 | Adamts14 |
| ENSMUSG( | 194.8147 | 0.07714 | 0.684628 | 0.958678 | Adamts15 |
| ENSMUSG( | 180.5775 | -0.17548 | 0.325538 | 0.905339 | Adamts16 |
| ENSMUSG( | 165.2752 | 0.432719 | 0.029851 | 0.61992 | Adamts17 |
| ENSMUSG( | 80.70148 | 0.190451 | 0.452997 | 0.922316 | Adamts18 |
| ENSMUSG( | 22.67749 | 0.070476 | 0.911987 | 0.992196 | Adamts19 |
| ENSMUSG( | 513.9972 | 0.332195 | 0.043684 | 0.690304 | Adamts2 |
| ENSMUSG( | 554.6731 | 0.196112 | 0.156917 | 0.837724 | Adamts20 |
| ENSMUSG( | 342.5764 | 0.213772 | 0.295343 | 0.898538 | Adamts3 |
| ENSMUSG( | 2318.613 | -0.06671 | 0.625667 | 0.953234 | Adamts4 |
| ENSMUSG( | 138.639 | -0.25775 | 0.22989 | 0.868507 | Adamts5 |
| ENSMUSG( | 211.35 | -0.22055 | 0.263936 | 0.886134 | Adamts6 |
| ENSMUSG( | 26.56672 | 0.360039 | 0.366675 | 0.914703 | Adamts7 |
| ENSMUSG( | 138.4796 | 0.02813 | 0.920615 | 0.992644 | Adamts8 |
| ENSMUSG( | 227.0556 | -0.09546 | 0.666842 | 0.956766 | Adamts9 |
| ENSMUSG( | 195.0946 | -0.13366 | 0.511448 | 0.92971 | Adamtsl1 |
| ENSMUSG( | 300.3905 | 0.09449 | 0.557256 | 0.939565 | Adamtsl2 |
| ENSMUSG( | 175.4666 | 0.077826 | 0.652519 | 0.954799 | Adamtsl3 |
| ENSMUSG( | 260.4578 | -0.42937 | 0.043913 | 0.690976 | Adamtsl4 |
| ENSMUSG( | 331.0723 | 0.03642 | 0.80563 | 0.977674 | Adamtsl5 |
| ENSMUSG( | 9345.322 | -0.00662 | 0.947465 | 0.994798 | Adap1 |
| ENSMUSG( | 220.3696 | 0.258949 | 0.28029 | 0.892356 | Adap2 |
| ENSMUSG( | 3537.187 | -0.02133 | 0.855225 | 0.985649 | Adar |
| ENSMUSG( | 6907.909 | -0.07343 | 0.488634 | 0.92554 | Adarb1 |
| ENSMUSG( | 277.2426 | -0.10806 | 0.508808 | 0.929089 | Adarb2 |
| ENSMUSG( | 123.9346 | 0.135534 | 0.556006 | 0.939565 | Adat1 |
| ENSMUSG( | 82.43137 | -0.27783 | 0.241031 | 0.876146 | Adat2 |
| ENSMUSG( | 845.4748 | -0.05806 | 0.634916 | 0.953903 | Adck1 |
| ENSMUSG( | 435.8671 | 0.290117 | 0.210425 | 0.864412 | Adck2 |
| ENSMUSG( | 420.2767 | 0.096696 | 0.504498 | 0.927825 | Adck5 |
| ENSMUSG( | 26422.15 | 0.215111 | 0.220287 | 0.866104 | Adcy1 |
| ENSMUSG( | 7755.368 | -0.04681 | 0.751195 | 0.970317 | Adcy2 |
| ENSMUSG( | 1868.063 | 0.341018 | 0.048092 | 0.704961 | Adcy3 |
| ENSMUSG( | 155.4815 | 0.965388 | 0.000796 | 0.094221 | Adcy4 |
| ENSMUSG( | 10613.3 | 0.138006 | 0.262547 | 0.885357 | Adcy5 |
| ENSMUSG( | 1889.784 | 0.331773 | 0.006631 | 0.338858 | Adcy6 |
| ENSMUSG( | 289.1882 | 0.070758 | 0.73355 | 0.967228 | Adcy7 |
| ENSMUSG( | 927.2292 | -0.18695 | 0.235485 | 0.871807 | Adcy8 |
| ENSMUSG( | 872.787 | 0.553339 | 0.210814 | 0.864412 | Adcy9 |
| ENSMUSG( | 534.8912 | -0.13763 | 0.48766 | 0.925427 | Adcyap1 |
| ENSMUSG( | 12698.17 | 0.077104 | 0.518499 | 0.930939 | Adcyap1r1 |
| ENSMUSG( | 11498.53 | -0.13415 | 0.16466 | 0.838743 | Add1 |
| ENSMUSG( | 6647.805 | 0.08139 | 0.502677 | 0.927825 | Add2 |
| ENSMUSG( | 7574.441 | -0.18291 | 0.144608 | 0.833258 | Add3 |
| ENSMUSG( | 33.86159 | 0.102085 | 0.787964 | 0.97637 | Adgb |
| ENSMUSG( | 2344.084 | 0.220777 | 0.188302 | 0.85698 | Adgra1 |
| ENSMUSG( | 623.7421 | 0.436652 | 0.008336 | 0.388416 | Adgra2 |
| ENSMUSG( | 1315.405 | 0.107278 | 0.458534 | 0.923329 | Adgra3 |
| ENSMUSG( | 18160.38 | 0.209216 | 0.068784 | 0.753052 | Adgrb1 |
| ENSMUSG( | 18347.55 | 0.271995 | 0.059553 | 0.729747 | Adgrb2 |
| ENSMUSG( | 2954.3 | 0.228942 | 0.192605 | 0.859764 | Adgrb3 |
| ENSMUSG( | 142.0862 | 0.018141 | 0.938175 | 0.993731 | Adgrd1 |
| ENSMUSG( | 454.417 | -0.04661 | 0.798209 | 0.97681 | Adgre1 |
| ENSMUSG( | 27.74625 | -1.50057 | 0.016259 | 0.513155 | Adgre4 |
| ENSMUSG( | 693.9556 | 0.390477 | 0.034132 | 0.651297 | Adgre5 |
| ENSMUSG( | 24.9758 | -0.19386 | 0.621299 | 0.953199 | Adgrf2 |
| ENSMUSG( | 51.41056 | 0.024863 | 0.932673 | 0.99306 | Adgrf4 |
| ENSMUSG( | 1637.045 | 0.752671 | 0.103452 | 0.804969 | Adgrf5 |
| ENSMUSG( | 4591.84 | -0.02058 | 0.903424 | 0.991089 | Adgrg1 |
| ENSMUSG( | 35.41448 | 0.456388 | 0.18892 | 0.85698 | Adgrg2 |
| ENSMUSG( | 10.83168 | 0.28054 | 0.630751 | NA | Adgrg3 |
| ENSMUSG( | 105.3004 | 0.181033 | 0.44773 | 0.922202 | Adgrg6 |
| ENSMUSG( | 22136.35 | 0.279884 | 0.056412 | 0.723362 | Adgrl1 |
| ENSMUSG( | 3067.209 | 0.200178 | 0.141392 | 0.830678 | Adgrl2 |
| ENSMUSG( | 3189.55 | 0.104688 | 0.406912 | 0.917257 | Adgrl3 |
| ENSMUSG( | 867.2267 | 0.863089 | 0.05945 | 0.729747 | Adgrl4 |
| ENSMUSG( | 114.4848 | 0.085497 | 0.732799 | 0.967228 | Adgrv1 |
| ENSMUSG( | 1524.008 | -0.13406 | 0.268771 | 0.886755 | Adh5 |
| ENSMUSG( | 2135.7 | -0.20162 | 0.132271 | 0.822912 | Adhfe1 |
| ENSMUSG( | 964.6474 | -0.17992 | 0.169033 | 0.840223 | Adi1 |
| ENSMUSG( | 4637.813 | 0.009875 | 0.929272 | 0.99306 | Adipor1 |
| ENSMUSG( | 3465.145 | 0.26966 | 0.048418 | 0.706142 | Adipor2 |
| ENSMUSG( | 891.8869 | -0.01329 | 0.927094 | 0.99306 | Adissp |
| ENSMUSG( | 2834.136 | -0.21156 | 0.1341 | 0.822912 | Adk |
| ENSMUSG( | 99.35277 | 0.072638 | 0.77724 | 0.975838 | Adm |
| ENSMUSG( | 83.70176 | -0.09442 | 0.698338 | 0.960908 | Adnp |
| ENSMUSG( | 602.7527 | -0.10442 | 0.463884 | 0.923329 | Adnp2 |
| ENSMUSG( | 1526.635 | -0.0219 | 0.87108 | 0.988771 | Ado |
| ENSMUSG( | 11510.7 | 0.093931 | 0.445417 | 0.921734 | Adora1 |
| ENSMUSG( | 1514.321 | -0.12574 | 0.388143 | 0.917257 | Adora2a |
| ENSMUSG( | 232.1107 | 0.211799 | 0.379934 | 0.916995 | Adora2b |
| ENSMUSG( | 46.63 | 0.648754 | 0.121621 | 0.813384 | Adora3 |
| ENSMUSG( | 717.2868 | 0.224492 | 0.113476 | 0.809623 | Adpgk |
| ENSMUSG( | 2639.569 | -0.1037 | 0.368321 | 0.915491 | Adprh |
| ENSMUSG( | 249.0707 | 0.186011 | 0.272972 | 0.889112 | Adprm |
| ENSMUSG( | 502.5834 | -0.0197 | 0.896747 | 0.990558 | Adprs |
| ENSMUSG( | 229.5823 | -0.18793 | 0.413218 | 0.917257 | Adra1a |
| ENSMUSG( | 828.0639 | 0.119474 | 0.476012 | 0.923329 | Adra1b |
| ENSMUSG( | 884.6583 | 0.285654 | 0.17362 | 0.844317 | Adra1d |
| ENSMUSG( | 799.3684 | 0.324133 | 0.041873 | 0.690304 | Adra2a |
| ENSMUSG( | 18.22796 | 0.035932 | 0.942798 | 0.994774 | Adra2b |
| ENSMUSG( | 875.5437 | 0.302947 | 0.013877 | 0.46927 | Adra2c |
| ENSMUSG( | 1170.376 | 0.110681 | 0.547546 | 0.938063 | Adrb1 |
| ENSMUSG( | 106.6103 | 0.429264 | 0.09746 | 0.796721 | Adrb2 |
| ENSMUSG( | 42.26897 | 0.509164 | 0.113331 | 0.809623 | Adrb3 |
| ENSMUSG( | 1158.08 | 0.006294 | 0.961379 | 0.99511 | Adrm1 |
| ENSMUSG( | 16.04527 | -0.6318 | 0.284917 | 0.892412 | Adrm1b |
| ENSMUSG( | 607.281 | -0.14153 | 0.257781 | 0.883401 | Adsl |
| ENSMUSG( | 408.3852 | -0.01992 | 0.88807 | 0.990442 | Adss1 |
| ENSMUSG( | 1661.334 | -0.03244 | 0.773596 | 0.974672 | Adss2 |
| ENSMUSG( | 8.896529 | 0.140376 | 0.837651 | NA | Adtrp |
| ENSMUSG( | 993.613 | 0.236048 | 0.096706 | 0.796245 | Aebp1 |
| ENSMUSG( | 1386.088 | -0.04136 | 0.740005 | 0.968811 | Aebp2 |
| ENSMUSG( | 620.1345 | 0.069313 | 0.583415 | 0.944266 | Aen |
| ENSMUSG( | 790.2347 | -0.12844 | 0.302425 | 0.898538 | Afap1 |
| ENSMUSG( | 1101.881 | 0.193696 | 0.266344 | 0.886134 | Afap1l1 |
| ENSMUSG( | 523.9529 | -0.00232 | 0.986807 | 0.997455 | Afap1l2 |
| ENSMUSG( | 2158.231 | 0.130872 | 0.221428 | 0.866104 | Afdn |
| ENSMUSG( | 672.7913 | 0.08699 | 0.579902 | 0.944209 | Aff1 |
| ENSMUSG( | 79.89998 | -0.05469 | 0.834691 | 0.981237 | Aff2 |
| ENSMUSG( | 1187.54 | 0.003688 | 0.97406 | 0.996416 | Aff3 |
| ENSMUSG( | 3636.1 | 0.08655 | 0.393389 | 0.917257 | Aff4 |
| ENSMUSG( | 573.9834 | -0.12326 | 0.332356 | 0.907907 | Afg1l |
| ENSMUSG( | 164.0038 | -0.00983 | 0.960154 | 0.994798 | Afg2a |
| ENSMUSG( | 229.5074 | 0.289091 | 0.078999 | 0.77458 | Afg2b |
| ENSMUSG( | 1460.521 | -0.07617 | 0.497234 | 0.927825 | Afg3l1 |
| ENSMUSG( | 2152.966 | 0.146684 | 0.25422 | 0.881345 | Afg3l2 |
| ENSMUSG( | 23.88699 | 0.108978 | 0.801646 | 0.97681 | Afmid |
| ENSMUSG( | 1522.106 | 0.022538 | 0.834136 | 0.981237 | Aftph |
| ENSMUSG( | 290.2128 | 0.060965 | 0.736677 | 0.967344 | Aga |
| ENSMUSG( | 7944.291 | 0.061502 | 0.548355 | 0.938063 | Agap1 |
| ENSMUSG( | 57920.07 | 0.210848 | 0.053673 | 0.720445 | Agap2 |
| ENSMUSG( | 11070.93 | 0.05169 | 0.61757 | 0.951952 | Agap3 |
| ENSMUSG( | 81.03074 | -0.15718 | 0.527367 | 0.934466 | Agbl2 |
| ENSMUSG( | 236.0068 | 0.077013 | 0.648257 | 0.95422 | Agbl3 |
| ENSMUSG( | 167.1918 | -0.05326 | 0.809577 | 0.978779 | Agbl4 |
| ENSMUSG( | 387.0585 | -0.06043 | 0.705215 | 0.962916 | Agbl5 |
| ENSMUSG( | 32.5999 | 0.133407 | 0.734372 | 0.967228 | Ager |
| ENSMUSG( | 1632.975 | 0.133683 | 0.229141 | 0.868507 | Agfg1 |
| ENSMUSG( | 841.8501 | -0.03607 | 0.793151 | 0.97681 | Agfg2 |
| ENSMUSG( | 2047.245 | -0.02907 | 0.819759 | 0.979198 | Aggf1 |
| ENSMUSG( | 639.2629 | -0.22601 | 0.09173 | 0.791889 | Agk |

| | | | | | | | | | | |
|---|---|---|---|---|---|---|---|---|---|---|
| ENSMUSG( | Adam12 | 224.6166 | -0.25607 | 0.134615 | 0.547918 | ENSMUSG( | Adam12 | 230.1365 | 0.101576 | 0.570076 | 0.941777 |
| ENSMUSG( | Adam15 | 3968.131 | 0.005964 | 0.956152 | 0.993874 | ENSMUSG( | Adam15 | 3866.901 | 0.196086 | 0.140003 | 0.830678 |
| ENSMUSG( | Adam17 | 1061.033 | -0.029 | 0.855049 | 0.973665 | ENSMUSG( | Adam17 | 1040.507 | 0.183899 | 0.293348 | 0.898538 |
| ENSMUSG( | Adam18 | 39.07732 | -0.33169 | 0.319518 | 0.765296 | ENSMUSG( | Adam18 | 36.30321 | -0.25441 | 0.488955 | 0.925573 |
| ENSMUSG( | Adam19 | 820.625 | -0.1266 | 0.468321 | 0.845336 | ENSMUSG( | Adam19 | 834.5238 | 0.19481 | 0.31443 | 0.900949 |
| ENSMUSG( | Adam1a | 168.0321 | -0.40663 | 0.014588 | 0.177737 | ENSMUSG( | Adam1a | 165.205 | -0.14799 | 0.537235 | 0.937216 |
| ENSMUSG( | Adam21 | 9.752875 | 0.371382 | 0.544373 | NA | ENSMUSG( | Adam21 | 9.787104 | 0.573847 | 0.366302 | NA |
| ENSMUSG( | Adam22 | 6960.81 | -0.29842 | 0.01263 | 0.162076 | ENSMUSG( | Adam22 | 7298.94 | 0.122089 | 0.362031 | 0.914223 |
| ENSMUSG( | Adam23 | 5101.284 | -0.50526 | 6.92E-05 | 0.005054 | ENSMUSG( | Adam23 | 5610.917 | 0.083869 | 0.567676 | 0.941777 |
| ENSMUSG( | Adam3 | 12.2272 | -0.24655 | 0.689444 | NA | ENSMUSG( | Adam3 | 12.49573 | 0.148539 | 0.822802 | 0.979198 |
| ENSMUSG( | Adam32 | 15.30772 | 0.24768 | 0.629941 | NA | ENSMUSG( | Adam32 | 12.59357 | -0.01612 | 0.976178 | 0.996416 |
| ENSMUSG( | Adam33 | 222.9974 | 0.231194 | 0.289618 | 0.739839 | ENSMUSG( | Adam33 | 172.1941 | -0.24904 | 0.380823 | 0.917202 |
| ENSMUSG( | Adam5 | 12.17634 | -0.20867 | 0.697731 | NA | ENSMUSG( | Adam5 | 11.50135 | -0.06923 | 0.912204 | 0.992225 |
| ENSMUSG( | Adam8 | 129.1882 | 0.987664 | 1.85E-05 | 0.001854 | ENSMUSG( | Adam8 | 106.5392 | 0.757284 | 0.008348 | 0.388416 |
| ENSMUSG( | Adam9 | 3121.341 | -0.07159 | 0.572875 | 0.893584 | ENSMUSG( | Adam9 | 3057.5 | 0.136329 | 0.338883 | 0.907972 |
| ENSMUSG( | Adamts1 | 391.9244 | -0.80754 | 2.24E-05 | 0.002129 | ENSMUSG( | Adamts1 | 437.0466 | -0.10556 | 0.611383 | 0.950771 |
| ENSMUSG( | Adamts10 | 1007.411 | 0.280147 | 0.066327 | 0.398419 | ENSMUSG( | Adamts10 | 942.4591 | 0.348664 | 0.1044 | 0.804969 |
| ENSMUSG( | Adamts12 | 33.04246 | -0.56902 | 0.137898 | 0.552903 | ENSMUSG( | Adamts12 | 30.97554 | -0.46918 | 0.231034 | 0.868507 |
| ENSMUSG( | Adamts13 | 35.25834 | -0.27632 | 0.424579 | 0.824675 | ENSMUSG( | Adamts13 | 34.92681 | -0.05147 | 0.891452 | 0.990558 |
| ENSMUSG( | Adamts14 | 22.49746 | 0.340911 | 0.400474 | 0.813717 | ENSMUSG( | Adamts14 | 20.72388 | 0.361429 | 0.409181 | 0.917257 |
| ENSMUSG( | Adamts15 | 190.9671 | -0.25926 | 0.142609 | 0.561026 | ENSMUSG( | Adamts15 | 194.8147 | 0.07714 | 0.684628 | 0.958678 |
| ENSMUSG( | Adamts16 | 201.3151 | -0.12937 | 0.423358 | 0.823662 | ENSMUSG( | Adamts16 | 180.5775 | -0.17548 | 0.325538 | 0.905339 |
| ENSMUSG( | Adamts17 | 164.0531 | 0.162577 | 0.382651 | 0.803227 | ENSMUSG( | Adamts17 | 165.2752 | 0.432719 | 0.029851 | 0.61992 |
| ENSMUSG( | Adamts18 | 74.73357 | -0.33154 | 0.167342 | 0.599971 | ENSMUSG( | Adamts18 | 80.70148 | 0.190451 | 0.452997 | 0.922316 |
| ENSMUSG( | Adamts19 | 19.47796 | -0.64546 | 0.283552 | NA | ENSMUSG( | Adamts19 | 22.67749 | 0.070476 | 0.911987 | 0.992196 |
| ENSMUSG( | Adamts2 | 484.4419 | -0.09912 | 0.482778 | 0.856184 | ENSMUSG( | Adamts2 | 513.9972 | 0.332195 | 0.043684 | 0.690304 |
| ENSMUSG( | Adamts20 | 586.5466 | 0.088253 | 0.447986 | 0.834051 | ENSMUSG( | Adamts20 | 554.6731 | 0.196112 | 0.156917 | 0.837724 |
| ENSMUSG( | Adamts3 | 298.6378 | -0.49705 | 0.008449 | 0.128676 | ENSMUSG( | Adamts3 | 342.5764 | 0.213772 | 0.295343 | 0.898538 |
| ENSMUSG( | Adamts4 | 2396.513 | -0.2526 | 0.011726 | 0.156231 | ENSMUSG( | Adamts4 | 2318.613 | -0.06671 | 0.625667 | 0.953234 |
| ENSMUSG( | Adamts5 | 148.5951 | -0.321 | 0.08282 | 0.444372 | ENSMUSG( | Adamts5 | 138.639 | -0.25775 | 0.22989 | 0.868507 |
| ENSMUSG( | Adamts6 | 204.0094 | -0.67529 | 0.000135 | 0.007928 | ENSMUSG( | Adamts6 | 211.35 | -0.22055 | 0.263936 | 0.886134 |
| ENSMUSG( | Adamts7 | 24.37573 | -0.15022 | 0.707526 | 0.93949 | ENSMUSG( | Adamts7 | 26.56672 | 0.360039 | 0.366675 | 0.914703 |
| ENSMUSG( | Adamts8 | 128.7961 | -0.50551 | 0.05871 | 0.373994 | ENSMUSG( | Adamts8 | 138.4796 | 0.02813 | 0.920615 | 0.992644 |
| ENSMUSG( | Adamts9 | 258.3729 | -0.00557 | 0.973832 | 0.99675 | ENSMUSG( | Adamts9 | 227.0556 | -0.09546 | 0.666842 | 0.956766 |
| ENSMUSG( | Adamtsl1 | 181.3646 | -0.69883 | 0.000664 | 0.024862 | ENSMUSG( | Adamtsl1 | 195.0946 | -0.13366 | 0.511448 | 0.92971 |
| ENSMUSG( | Adamtsl2 | 308.0532 | -0.10955 | 0.43901 | 0.830714 | ENSMUSG( | Adamtsl2 | 300.3905 | 0.09449 | 0.557256 | 0.939565 |
| ENSMUSG( | Adamtsl3 | 178.8523 | -0.14033 | 0.386161 | 0.804623 | ENSMUSG( | Adamtsl3 | 175.4666 | 0.077826 | 0.652519 | 0.954799 |
| ENSMUSG( | Adamtsl4 | 265.9548 | -0.70565 | 0.000101 | 0.006438 | ENSMUSG( | Adamtsl4 | 260.4578 | -0.42937 | 0.043913 | 0.690976 |
| ENSMUSG( | Adamtsl5 | 336.1072 | -0.19591 | 0.137348 | 0.552389 | ENSMUSG( | Adamtsl5 | 331.0723 | 0.03642 | 0.80563 | 0.977674 |
| ENSMUSG( | Adap1 | 10863.74 | 0.154362 | 0.056084 | 0.36519 | ENSMUSG( | Adap1 | 9345.322 | -0.00662 | 0.947465 | 0.994798 |
| ENSMUSG( | Adap2 | 211.6809 | -0.13726 | 0.429255 | 0.826446 | ENSMUSG( | Adap2 | 220.3696 | 0.258949 | 0.28029 | 0.892356 |
| ENSMUSG( | Adar | 4115.5 | 0.143971 | 0.156812 | 0.5851 | ENSMUSG( | Adar | 3537.187 | -0.02133 | 0.855225 | 0.985649 |
| ENSMUSG( | Adarb1 | 7798.919 | 0.009341 | 0.904378 | 0.986246 | ENSMUSG( | Adarb1 | 6907.909 | -0.07343 | 0.488634 | 0.92554 |
| ENSMUSG( | Adarb2 | 322.1745 | 0.061455 | 0.653415 | 0.922819 | ENSMUSG( | Adarb2 | 277.2426 | -0.10806 | 0.508808 | 0.929089 |
| ENSMUSG( | Adat1 | 138.1582 | 0.171891 | 0.43158 | 0.827432 | ENSMUSG( | Adat1 | 123.9346 | 0.135534 | 0.556006 | 0.939565 |
| ENSMUSG( | Adat2 | 100.9222 | 0.051283 | 0.807115 | 0.9631 | ENSMUSG( | Adat2 | 82.43137 | -0.27783 | 0.241031 | 0.876146 |
| ENSMUSG( | Adck1 | 976.2986 | 0.08474 | 0.420496 | 0.822898 | ENSMUSG( | Adck1 | 845.4748 | -0.05806 | 0.634916 | 0.953903 |
| ENSMUSG( | Adck2 | 410.3938 | -0.15487 | 0.42326 | 0.823662 | ENSMUSG( | Adck2 | 435.8671 | 0.290117 | 0.210425 | 0.864412 |
| ENSMUSG( | Adck5 | 455.9965 | 0.057803 | 0.644711 | 0.917782 | ENSMUSG( | Adck5 | 420.2767 | 0.096696 | 0.504498 | 0.927825 |
| ENSMUSG( | Adcy1 | 28924.51 | 0.204825 | 0.144235 | 0.563869 | ENSMUSG( | Adcy1 | 26422.15 | 0.215111 | 0.220287 | 0.866104 |
| ENSMUSG( | Adcy2 | 8388.587 | -0.0932 | 0.488643 | 0.859092 | ENSMUSG( | Adcy2 | 7755.368 | -0.04681 | 0.751195 | 0.970317 |
| ENSMUSG( | Adcy3 | 1742.122 | -0.1176 | 0.448787 | 0.83469 | ENSMUSG( | Adcy3 | 1868.063 | 0.341018 | 0.048092 | 0.704961 |
| ENSMUSG( | Adcy4 | 135.0212 | 0.399319 | 0.103077 | 0.48855 | ENSMUSG( | Adcy4 | 155.4815 | 0.965388 | 0.000796 | 0.094221 |
| ENSMUSG( | Adcy5 | 10001.58 | -0.31818 | 0.008717 | 0.131277 | ENSMUSG( | Adcy5 | 10613.3 | 0.138006 | 0.262547 | 0.885357 |
| ENSMUSG( | Adcy6 | 1993.807 | 0.225564 | 0.038743 | 0.303142 | ENSMUSG( | Adcy6 | 1889.784 | 0.331773 | 0.006631 | 0.338858 |
| ENSMUSG( | Adcy7 | 282.6723 | -0.28727 | 0.069552 | 0.408118 | ENSMUSG( | Adcy7 | 289.1882 | 0.070758 | 0.73355 | 0.967228 |
| ENSMUSG( | Adcy8 | 975.7224 | -0.32707 | 0.022429 | 0.228899 | ENSMUSG( | Adcy8 | 927.2292 | -0.18695 | 0.235485 | 0.871807 |
| ENSMUSG( | Adcy9 | 777.7899 | -0.00497 | 0.979903 | 0.997428 | ENSMUSG( | Adcy9 | 872.787 | 0.553339 | 0.210814 | 0.864412 |
| ENSMUSG( | Adcyap1 | 516.5098 | -0.56194 | 0.001623 | 0.045245 | ENSMUSG( | Adcyap1 | 534.8912 | -0.13763 | 0.48766 | 0.925427 |
| ENSMUSG( | Adcyap1r1 | 13527.1 | -0.01008 | 0.92804 | 0.991229 | ENSMUSG( | Adcyap1r1 | 12698.17 | 0.077104 | 0.518499 | 0.930939 |
| ENSMUSG( | Add1 | 13687.67 | 0.101606 | 0.186637 | 0.623917 | ENSMUSG( | Add1 | 11498.53 | -0.13415 | 0.16466 | 0.838743 |
| ENSMUSG( | Add2 | 7464.56 | 0.142605 | 0.160839 | 0.590628 | ENSMUSG( | Add2 | 6647.805 | 0.08139 | 0.502677 | 0.927825 |
| ENSMUSG( | Add3 | 9165.123 | 0.10424 | 0.338705 | 0.777518 | ENSMUSG( | Add3 | 7574.441 | -0.18291 | 0.144608 | 0.833258 |
| ENSMUSG( | Adgb | 34.86822 | -0.12084 | 0.742136 | 0.948478 | ENSMUSG( | Adgb | 33.86159 | 0.102085 | 0.787964 | 0.97637 |
| ENSMUSG( | Adgra1 | 2314.174 | -0.08323 | 0.590188 | 0.90113 | ENSMUSG( | Adgra1 | 2344.084 | 0.220777 | 0.188302 | 0.85698 |
| ENSMUSG( | Adgra2 | 616.3918 | 0.154809 | 0.250225 | 0.700009 | ENSMUSG( | Adgra2 | 623.7421 | 0.436652 | 0.008336 | 0.388416 |
| ENSMUSG( | Adgra3 | 1408.503 | 0.034688 | 0.784731 | 0.958171 | ENSMUSG( | Adgra3 | 1315.405 | 0.107278 | 0.458534 | 0.923329 |
| ENSMUSG( | Adgrb1 | 18451.99 | -0.00775 | 0.94207 | 0.993415 | ENSMUSG( | Adgrb1 | 18160.38 | 0.209216 | 0.068784 | 0.753052 |
| ENSMUSG( | Adgrb2 | 19382.09 | 0.166663 | 0.153196 | 0.57989 | ENSMUSG( | Adgrb2 | 18347.55 | 0.271995 | 0.059553 | 0.729747 |
| ENSMUSG( | Adgrb3 | 2821.066 | -0.17514 | 0.246548 | 0.694958 | ENSMUSG( | Adgrb3 | 2954.3 | 0.228942 | 0.192605 | 0.859764 |
| ENSMUSG( | Adgrd1 | 141.7045 | -0.28351 | 0.167378 | 0.599971 | ENSMUSG( | Adgrd1 | 142.0862 | 0.018141 | 0.938175 | 0.993731 |
| ENSMUSG( | Adgre1 | 461.505 | -0.29452 | 0.049828 | 0.340731 | ENSMUSG( | Adgre1 | 454.417 | -0.04661 | 0.798209 | 0.97681 |
| ENSMUSG( | Adgre4 | 27.96203 | -1.9407 | 0.001881 | 0.049538 | ENSMUSG( | Adgre4 | 27.74625 | -1.50057 | 0.016259 | 0.513155 |
| ENSMUSG( | Adgre5 | 673.983 | 0.059346 | 0.63818 | 0.916124 | ENSMUSG( | Adgre5 | 693.9556 | 0.390477 | 0.034132 | 0.651297 |
| ENSMUSG( | Adgrf2 | 28.32831 | -0.09056 | 0.805744 | 0.962616 | ENSMUSG( | Adgrf2 | 24.9758 | -0.19386 | 0.621299 | 0.953199 |
| ENSMUSG( | Adgrf4 | 58.11818 | 0.10131 | 0.711116 | 0.940028 | ENSMUSG( | Adgrf4 | 51.41056 | 0.024863 | 0.932673 | 0.99306 |
| ENSMUSG( | Adgrf5 | 1431.915 | 0.179322 | 0.403801 | 0.816132 | ENSMUSG( | Adgrf5 | 1637.045 | 0.752671 | 0.103452 | 0.804969 |
| ENSMUSG( | Adgrg1 | 4485.042 | -0.38834 | 0.010467 | 0.146428 | ENSMUSG( | Adgrg1 | 4591.84 | -0.02058 | 0.903424 | 0.991089 |
| ENSMUSG( | Adgrg2 | 33.88062 | 0.11147 | 0.750188 | 0.949664 | ENSMUSG( | Adgrg2 | 35.41448 | 0.456388 | 0.18892 | 0.85698 |
| ENSMUSG( | Adgrg3 | 9.620186 | -0.37505 | 0.538641 | NA | ENSMUSG( | Adgrg3 | 10.83168 | 0.28054 | 0.630751 | NA |
| ENSMUSG( | Adgrg6 | 93.5599 | -0.45904 | 0.042746 | 0.316989 | ENSMUSG( | Adgrg6 | 105.3004 | 0.181033 | 0.44773 | 0.922202 |
| ENSMUSG( | Adgrl1 | 22001.62 | 0.004285 | 0.97119 | 0.996369 | ENSMUSG( | Adgrl1 | 22136.35 | 0.279884 | 0.056412 | 0.723362 |
| ENSMUSG( | Adgrl2 | 2899.468 | -0.23746 | 0.061782 | 0.384389 | ENSMUSG( | Adgrl2 | 3067.209 | 0.200178 | 0.141392 | 0.830678 |
| ENSMUSG( | Adgrl3 | 3427.532 | 0.043699 | 0.704092 | 0.938172 | ENSMUSG( | Adgrl3 | 3189.55 | 0.104688 | 0.406912 | 0.917257 |
| ENSMUSG( | Adgrl4 | 752.9256 | 0.288989 | 0.4446 | 0.833239 | ENSMUSG( | Adgrl4 | 867.2267 | 0.863089 | 0.05945 | 0.729747 |
| ENSMUSG( | Adgrv1 | 112.3443 | -0.26475 | 0.249989 | 0.69959 | ENSMUSG( | Adgrv1 | 114.4848 | 0.085497 | 0.732799 | 0.967228 |
| ENSMUSG( | Adh5 | 1755.783 | 0.011509 | 0.905787 | 0.986694 | ENSMUSG( | Adh5 | 1524.008 | -0.13406 | 0.268771 | 0.886755 |
| ENSMUSG( | Adhfe1 | 2639.707 | 0.147673 | 0.175579 | 0.611004 | ENSMUSG( | Adhfe1 | 2135.7 | -0.20162 | 0.132271 | 0.822912 |
| ENSMUSG( | Adi1 | 1137.59 | 0.037293 | 0.713973 | 0.940903 | ENSMUSG( | Adi1 | 964.6474 | -0.17992 | 0.169033 | 0.840223 |
| ENSMUSG( | Adipor1 | 4866.745 | -0.12295 | 0.193141 | 0.632494 | ENSMUSG( | Adipor1 | 4637.813 | 0.009875 | 0.929272 | 0.99306 |
| ENSMUSG( | Adipor2 | 3519.088 | 0.053522 | 0.645533 | 0.918025 | ENSMUSG( | Adipor2 | 3465.145 | 0.26966 | 0.048418 | 0.706142 |
| ENSMUSG( | Adissp | 1015.167 | 0.092562 | 0.463919 | 0.842812 | ENSMUSG( | Adissp | 891.8869 | -0.01329 | 0.927094 | 0.99306 |
| ENSMUSG( | Adk | 3561.42 | 0.184697 | 0.116474 | 0.514944 | ENSMUSG( | Adk | 2834.136 | -0.21156 | 0.1341 | 0.822912 |
| ENSMUSG( | Adm | 89.51888 | -0.57498 | 0.017366 | 0.197592 | ENSMUSG( | Adm | 99.35277 | 0.072638 | 0.77724 | 0.975838 |
| ENSMUSG( | Adnp | 98.90447 | 0.111351 | 0.584663 | 0.898884 | ENSMUSG( | Adnp | 83.70176 | -0.09442 | 0.698338 | 0.960908 |
| ENSMUSG( | Adnp2 | 696.7506 | 0.045987 | 0.693187 | 0.934202 | ENSMUSG( | Adnp2 | 602.7527 | -0.10442 | 0.463884 | 0.923329 |
| ENSMUSG( | Ado | 1680.949 | -0.00857 | 0.926666 | 0.991229 | ENSMUSG( | Ado | 1526.635 | -0.0219 | 0.87108 | 0.988771 |
| ENSMUSG( | Adora1 | 12180.04 | -0.01237 | 0.907412 | 0.987566 | ENSMUSG( | Adora1 | 11510.7 | 0.093931 | 0.445417 | 0.921734 |
| ENSMUSG( | Adora2a | 1220.907 | -1.29293 | 0.00076 | 0.027082 | ENSMUSG( | Adora2a | 1514.321 | -0.12574 | 0.388143 | 0.917257 |
| ENSMUSG( | Adora2b | 204.6506 | -0.46073 | 0.039387 | 0.305596 | ENSMUSG( | Adora2b | 232.1107 | 0.211799 | 0.379934 | 0.916995 |
| ENSMUSG( | Adora3 | 39.71769 | -0.04629 | 0.908845 | 0.98794 | ENSMUSG( | Adora3 | 46.63 | 0.648754 | 0.121621 | 0.813384 |
| ENSMUSG( | Adpgk | 658.1774 | -0.30488 | 0.026508 | 0.248968 | ENSMUSG( | Adpgk | 717.2868 | 0.224492 | 0.113476 | 0.809623 |
| ENSMUSG( | Adprh | 2715.144 | -0.30721 | 0.001441 | 0.041882 | ENSMUSG( | Adprh | 2639.569 | -0.1037 | 0.368321 | 0.915491 |
| ENSMUSG( | Adprm | 273.4923 | 0.183756 | 0.210173 | 0.652567 | ENSMUSG( | Adprm | 249.0707 | 0.186011 | 0.272972 | 0.889112 |
| ENSMUSG( | Adprs | 544.5488 | -0.06345 | 0.577562 | 0.895864 | ENSMUSG( | Adprs | 502.5834 | -0.0197 | 0.896747 | 0.990558 |
| ENSMUSG( | Adra1a | 251.3194 | -0.20868 | 0.290244 | 0.740097 | ENSMUSG( | Adra1a | 229.5823 | -0.18793 | 0.413218 | 0.917257 |
| ENSMUSG( | Adra1b | 808.1531 | -0.2331 | 0.109035 | 0.498536 | ENSMUSG( | Adra1b | 828.0639 | 0.119474 | 0.476012 | 0.923329 |
| ENSMUSG( | Adra1d | 825.4306 | -0.18394 | 0.339133 | 0.777518 | ENSMUSG( | Adra1d | 884.6583 | 0.285654 | 0.17362 | 0.844317 |
| ENSMUSG( | Adra2a | 748.4734 | -0.12433 | 0.369741 | 0.797289 | ENSMUSG( | Adra2a | 799.3684 | 0.324133 | 0.041873 | 0.690304 |
| ENSMUSG( | Adra2b | 13.55628 | -1.32818 | 0.022593 | NA | ENSMUSG( | Adra2b | 18.22796 | 0.035932 | 0.942798 | 0.994774 |
| ENSMUSG( | Adra2c | 835.8157 | -0.08896 | 0.390925 | 0.80826 | ENSMUSG( | Adra2c | 875.5437 | 0.302947 | 0.013877 | 0.46927 |
| ENSMUSG( | Adrb1 | 1009.305 | -0.66608 | 8.92E-05 | 0.005955 | ENSMUSG( | Adrb1 | 1170.376 | 0.110681 | 0.547546 | 0.938063 |
| ENSMUSG( | Adrb2 | 83.59689 | -0.59007 | 0.009458 | 0.137493 | ENSMUSG( | Adrb2 | 106.6103 | 0.429264 | 0.09746 | 0.796721 |
| ENSMUSG( | Adrb3 | 41.90078 | 0.259954 | 0.391808 | 0.809274 | ENSMUSG( | Adrb3 | 42.26897 | 0.509164 | 0.113331 | 0.809623 |
| ENSMUSG( | Adrm1 | 1239.711 | -0.06667 | 0.574955 | 0.894572 | ENSMUSG( | Adrm1 | 1158.08 | 0.006294 | 0.961379 | 0.99511 |
| ENSMUSG( | Adrm1b | 21.89963 | 0.072214 | 0.881634 | 0.981948 | ENSMUSG( | Adrm1b | 16.04527 | -0.6318 | 0.284917 | 0.892412 |
| ENSMUSG( | Adsl | 701.1157 | 0.007883 | 0.942043 | 0.993415 | ENSMUSG( | Adsl | 607.281 | -0.14153 | 0.257781 | 0.883401 |
| ENSMUSG( | Adss1 | 427.5436 | -0.15877 | 0.193589 | 0.633067 | ENSMUSG( | Adss1 | 408.3852 | -0.01992 | 0.88807 | 0.990442 |
| ENSMUSG( | Adss2 | 1791.278 | -0.08368 | 0.357363 | 0.788775 | ENSMUSG( | Adss2 | 1661.334 | -0.03244 | 0.773596 | 0.974672 |
| ENSMUSG( | Aebp1 | 1001.887 | -0.00441 | 0.973141 | 0.99675 | ENSMUSG( | Aebp1 | 993.613 | 0.236048 | 0.096706 | 0.796245 |
| ENSMUSG( | Aebp2 | 1530.431 | -0.0203 | 0.816842 | 0.96566 | ENSMUSG( | Aebp2 | 1386.088 | -0.04136 | 0.740005 | 0.968811 |
| ENSMUSG( | Aen | 678.8831 | 0.065226 | 0.538466 | 0.88259 | ENSMUSG( | Aen | 620.1345 | 0.069313 | 0.583415 | 0.944266 |
| ENSMUSG( | Afap1 | 901.2851 | -0.01722 | 0.875265 | 0.979998 | ENSMUSG( | Afap1 | 790.2347 | -0.12844 | 0.302425 | 0.898538 |
| ENSMUSG( | Afap1l1 | 1104.781 | -0.06865 | 0.597948 | 0.90372 | ENSMUSG( | Afap1l1 | 1101.881 | 0.193696 | 0.266344 | 0.886134 |
| ENSMUSG( | Afap1l2 | 574.871 | -0.00906 | 0.944193 | 0.993503 | ENSMUSG( | Afap1l2 | 523.9529 | -0.00232 | 0.986807 | 0.997455 |
| ENSMUSG( | Afdn | 2322.677 | 0.075928 | 0.380033 | 0.801757 | ENSMUSG( | Afdn | 2158.231 | 0.130872 | 0.221428 | 0.866104 |
| ENSMUSG( | Aff1 | 761.62 | 0.164764 | 0.199785 | 0.640239 | ENSMUSG( | Aff1 | 672.7913 | 0.08699 | 0.579902 | 0.944209 |
| ENSMUSG( | Aff2 | 88.76147 | -0.03595 | 0.873154 | 0.978753 | ENSMUSG( | Aff2 | 79.89998 | -0.05469 | 0.834691 | 0.981237 |
| ENSMUSG( | Aff3 | 1272.382 | -0.06883 | 0.483118 | 0.856216 | ENSMUSG( | Aff3 | 1187.54 | 0.003688 | 0.97406 | 0.996416 |
| ENSMUSG( | Aff4 | 4023.695 | 0.109278 | 0.204495 | 0.645438 | ENSMUSG( | Aff4 | 3636.1 | 0.08655 | 0.393389 | 0.917257 |
| ENSMUSG( | Afg1l | 694.782 | 0.159631 | 0.13482 | 0.54834 | ENSMUSG( | Afg1l | 573.9834 | -0.12326 | 0.332356 | 0.907907 |
| ENSMUSG( | Afg2a | 194.7582 | 0.217815 | 0.163471 | 0.594869 | ENSMUSG( | Afg2a | 164.0038 | -0.00983 | 0.960154 | 0.994798 |
| ENSMUSG( | Afg2b | 238.8492 | 0.15335 | 0.272264 | 0.7223 | ENSMUSG( | Afg2b | 229.5074 | 0.289091 | 0.078999 | 0.77458 |
| ENSMUSG( | Afg3l1 | 1726.67 | 0.136922 | 0.147036 | 0.569057 | ENSMUSG( | Afg3l1 | 1460.521 | -0.07617 | 0.497234 | 0.927825 |
| ENSMUSG( | Afg3l2 | 2211.549 | -0.0448 | 0.66499 | 0.926824 | ENSMUSG( | Afg3l2 | 2152.966 | 0.146684 | 0.25422 | 0.881345 |
| ENSMUSG( | Afmid | 24.85805 | -0.05966 | 0.871558 | 0.978114 | ENSMUSG( | Afmid | 23.88699 | 0.108978 | 0.801646 | 0.97681 |
| ENSMUSG( | Aftph | 1693.516 | 0.061649 | 0.491636 | 0.860231 | ENSMUSG( | Aftph | 1522.106 | 0.022538 | 0.834136 | 0.981237 |
| ENSMUSG( | Aga | 297.9345 | -0.14057 | 0.354769 | 0.78738 | ENSMUSG( | Aga | 290.2128 | 0.060965 | 0.736677 | 0.967344 |
| ENSMUSG( | Agap1 | 8727.004 | 0.064397 | 0.408647 | 0.818173 | ENSMUSG( | Agap1 | 7944.291 | 0.061502 | 0.548355 | 0.938063 |
| ENSMUSG( | Agap2 | 64538.43 | 0.250075 | 0.003874 | 0.079851 | ENSMUSG( | Agap2 | 57920.07 | 0.210848 | 0.053673 | 0.720445 |
| ENSMUSG( | Agap3 | 12667.86 | 0.167992 | 0.044473 | 0.32391 | ENSMUSG( | Agap3 | 11070.93 | 0.05169 | 0.61757 | 0.951952 |
| ENSMUSG( | Agbl2 | 89.14478 | -0.1532 | 0.512597 | 0.871762 | ENSMUSG( | Agbl2 | 81.03074 | -0.15718 | 0.527367 | 0.934466 |
| ENSMUSG( | Agbl3 | 253.834 | 0.025958 | 0.878351 | 0.981431 | ENSMUSG( | Agbl3 | 236.0068 | 0.077013 | 0.648257 | 0.95422 |
| ENSMUSG( | Agbl4 | 225.9885 | 0.505463 | 0.010952 | 0.150804 | ENSMUSG( | Agbl4 | 167.1918 | -0.05326 | 0.809577 | 0.978779 |
| ENSMUSG( | Agbl5 | 446.9081 | 0.083978 | 0.520906 | 0.875186 | ENSMUSG( | Agbl5 | 387.0585 | -0.06043 | 0.705215 | 0.962916 |
| ENSMUSG( | Ager | 32.02174 | -0.1896 | 0.616652 | 0.910866 | ENSMUSG( | Ager | 32.5999 | 0.133407 | 0.734372 | 0.967228 |
| ENSMUSG( | Agfg1 | 1761.861 | 0.088072 | 0.323972 | 0.769037 | ENSMUSG( | Agfg1 | 1632.975 | 0.133683 | 0.229141 | 0.868507 |
| ENSMUSG( | Agfg2 | 964.4598 | 0.087173 | 0.442026 | 0.832762 | ENSMUSG( | Agfg2 | 841.8501 | -0.03607 | 0.793151 | 0.97681 |
| ENSMUSG( | Aggf1 | 2380.536 | 0.137845 | 0.145343 | 0.566152 | ENSMUSG( | Aggf1 | 2047.245 | -0.02907 | 0.819759 | 0.979198 |
| ENSMUSG( | Agk | 780.928 | 0.093117 | 0.393958 | 0.809681 | ENSMUSG( | Agk | 639.2629 | -0.22601 | 0.09173 | 0.791889 |
| ENSMUSG( | Agl | 4950.267 | 0.058313 | 0.663006 | 0.9265 | ENSMUSG( | Agl | 4028.189 | -0.28274 | 0.035839 | 0.668395 |
| ENSMUSG( | Agmat | 14.47941 | 0.15299 | 0.76197 | NA | ENSMUSG( | Agmat | 10.43548 | -0.5969 | 0.360001 | NA |
| ENSMUSG( | Agmo | 400.3897 | -0.04954 | 0.752634 | 0.950536 | ENSMUSG( | Agmo | 397.6842 | 0.197163 | 0.29053 | 0.897019 |
| ENSMUSG( | Ago1 | 3162.291 | 0.081185 | 0.331814 | 0.775288 | ENSMUSG( | Ago1 | 2738.419 | -0.0646 | 0.518101 | 0.930939 |
| ENSMUSG( | Ago2 | 3345.524 | 0.152843 | 0.118673 | 0.521258 | ENSMUSG( | Ago2 | 3034.854 | 0.141202 | 0.205656 | 0.864412 |
| ENSMUSG( | Ago3 | 1639.789 | -0.1912 | 0.054685 | 0.360128 | ENSMUSG( | Ago3 | 1664.822 | 0.126107 | 0.279996 | 0.892356 |
| ENSMUSG( | Ago4 | 540.7666 | -0.1538 | 0.191158 | 0.630245 | ENSMUSG( | Ago4 | 506.74 | -0.0759 | 0.591507 | 0.944975 |
| ENSMUSG( | Agpat1 | 6297.062 | -0.24669 | 0.038405 | 0.302389 | ENSMUSG( | Agpat1 | 6196.567 | -0.01153 | 0.931524 | 0.99306 |

| | | | | | |
|---|---|---|---|---|---|
| ENSMUSG( | 225.9885 | 0.505463 | 0.010952 | 0.150804 | Agbl4 |
| ENSMUSG( | 446.9081 | 0.083978 | 0.520906 | 0.875186 | Agbl5 |
| ENSMUSG( | 32.02174 | -0.1896 | 0.616652 | 0.910866 | Ager |
| ENSMUSG( | 1761.861 | 0.088072 | 0.323972 | 0.769037 | Agfg1 |
| ENSMUSG( | 964.4598 | 0.087173 | 0.442026 | 0.832762 | Agfg2 |
| ENSMUSG( | 2380.536 | 0.137845 | 0.145343 | 0.566152 | Aggf1 |
| ENSMUSG( | 780.928 | 0.093117 | 0.393958 | 0.809681 | Agk |
| ENSMUSG( | 4950.267 | 0.058313 | 0.663006 | 0.9265 | Agl |
| ENSMUSG( | 14.47941 | 0.15299 | 0.76197 | NA | Agmat |
| ENSMUSG( | 400.3897 | -0.04954 | 0.752634 | 0.950536 | Agmo |
| ENSMUSG( | 3162.291 | 0.081185 | 0.331814 | 0.775288 | Ago1 |
| ENSMUSG( | 3345.524 | 0.152843 | 0.118673 | 0.521258 | Ago2 |
| ENSMUSG( | 1639.789 | -0.1912 | 0.054685 | 0.360128 | Ago3 |
| ENSMUSG( | 540.7666 | -0.1538 | 0.191158 | 0.630245 | Ago4 |
| ENSMUSG( | 6297.062 | -0.24669 | 0.038405 | 0.302389 | Agpat1 |
| ENSMUSG( | 66.39721 | -0.20645 | 0.376738 | 0.801213 | Agpat2 |
| ENSMUSG( | 24657.59 | 0.044085 | 0.668965 | 0.927963 | Agpat3 |
| ENSMUSG( | 2451.736 | -0.40328 | 0.001129 | 0.035751 | Agpat4 |
| ENSMUSG( | 3657.701 | -0.05856 | 0.655908 | 0.923929 | Agpat5 |
| ENSMUSG( | 1337.591 | 0.051021 | 0.588213 | 0.900712 | Agps |
| ENSMUSG( | 7001.688 | 0.087436 | 0.442267 | 0.832774 | Agrn |
| ENSMUSG( | 1813.162 | 0.084565 | 0.546135 | 0.884974 | Agt |
| ENSMUSG( | 5250.796 | 0.174625 | 0.022815 | 0.230858 | Agtpbp1 |
| ENSMUSG( | 18.23423 | -0.82541 | 0.078008 | NA | Agtr1a |
| ENSMUSG( | 13.15575 | 0.116219 | 0.823386 | NA | Agtr2 |
| ENSMUSG( | 1062.71 | -0.07143 | 0.566702 | 0.891644 | Agtrap |
| ENSMUSG( | 1240.988 | -0.0009 | 0.994136 | 0.999369 | Ahctf1 |
| ENSMUSG( | 408.5938 | 0.010342 | 0.930126 | 0.991848 | Ahcy |
| ENSMUSG( | 30728.28 | 0.07737 | 0.508224 | 0.867999 | Ahcyl1 |
| ENSMUSG( | 9828.154 | 0.118214 | 0.201642 | 0.641362 | Ahcyl2 |
| ENSMUSG( | 1560.571 | 0.105618 | 0.307371 | 0.755324 | Ahdc1 |
| ENSMUSG( | 3920.711 | 0.156404 | 0.234405 | 0.681922 | Ahi1 |
| ENSMUSG( | 1316.606 | 0.259872 | 0.035139 | 0.288296 | Ahnak |
| ENSMUSG( | 78.02105 | -0.24202 | 0.32763 | 0.772692 | Ahnak2 |
| ENSMUSG( | 156.874 | 0.241989 | 0.175054 | 0.610399 | Ahr |
| ENSMUSG( | 37.26796 | -0.33028 | 0.33382 | 0.775602 | Ahrr |
| ENSMUSG( | 2308.552 | 0.13632 | 0.222453 | 0.668434 | Ahsa1 |
| ENSMUSG( | 848.0786 | 0.105891 | 0.317481 | 0.763823 | Ahsa2 |
| ENSMUSG( | 51.53926 | -0.10518 | 0.729135 | 0.944738 | AI115009 |
| ENSMUSG( | 12.81575 | 0.642273 | 0.228046 | NA | AI182371 |
| ENSMUSG( | 20.14847 | 0.271427 | 0.564578 | NA | AI197445 |
| ENSMUSG( | 64.92554 | 0.09227 | 0.713477 | 0.940865 | AI429214 |
| ENSMUSG( | 76.74703 | -0.32585 | 0.220981 | 0.666476 | AI467606 |
| ENSMUSG( | 529.1535 | -0.61584 | 0.000587 | 0.022791 | AI504432 |
| ENSMUSG( | 4646.03 | -0.68355 | 5.38E-08 | 1.71E-05 | AI593442 |
| ENSMUSG( | 1351.163 | 0.049571 | 0.685785 | 0.932981 | AI597479 |
| ENSMUSG( | 18.52078 | 0.371946 | 0.381638 | NA | AI661453 |
| ENSMUSG( | 10.55806 | -0.40497 | 0.480224 | NA | AI662270 |
| ENSMUSG( | 1165.132 | 0.005311 | 0.968524 | 0.996369 | AI837181 |
| ENSMUSG( | 11.46745 | 0.088528 | 0.874396 | NA | AI839979 |
| ENSMUSG( | 816.7793 | 0.106287 | 0.294008 | 0.743459 | AI854703 |
| ENSMUSG( | 399.708 | 0.037251 | 0.826099 | 0.968159 | AI987944 |
| ENSMUSG( | 1801.03 | 0.047758 | 0.635372 | 0.914296 | Aida |
| ENSMUSG( | 312.8991 | 0.108904 | 0.462974 | 0.842422 | Aif1 |
| ENSMUSG( | 353.9725 | -0.28419 | 0.032801 | 0.277278 | Aif1l |
| ENSMUSG( | 867.6499 | 0.079037 | 0.480335 | 0.854748 | Aifm1 |
| ENSMUSG( | 485.453 | 0.027151 | 0.828364 | 0.968994 | Aifm2 |
| ENSMUSG( | 3370.851 | 0.299769 | 0.00126 | 0.038624 | Aifm3 |
| ENSMUSG( | 1615.508 | -0.35586 | 0.008904 | 0.132538 | Aig1 |
| ENSMUSG( | 147.8928 | -0.27977 | 0.146171 | 0.567605 | Aim2 |
| ENSMUSG( | 1077.516 | 0.019359 | 0.86031 | 0.974842 | Aimp1 |
| ENSMUSG( | 375.9235 | -0.02619 | 0.851736 | 0.973104 | Aimp2 |
| ENSMUSG( | 2579.787 | 0.037566 | 0.645441 | 0.918017 | Aip |
| ENSMUSG( | 502.6432 | 0.067919 | 0.563815 | 0.890032 | Airim |
| ENSMUSG( | 26.82693 | -0.34188 | 0.377319 | 0.801213 | Airn |
| ENSMUSG( | 1366.297 | -0.259 | 0.052686 | 0.352396 | Ajap1 |
| ENSMUSG( | 6068.282 | -0.28278 | 0.000598 | 0.022994 | Ajm1 |
| ENSMUSG( | 87.28489 | -0.27758 | 0.264346 | 0.715323 | Ajuba |
| ENSMUSG( | 3555.185 | 0.046712 | 0.653145 | 0.922819 | Ak1 |
| ENSMUSG( | 42.7078 | -0.10867 | 0.731653 | 0.944956 | AK157302 |
| ENSMUSG( | 749.0713 | 0.161549 | 0.103499 | 0.489288 | Ak2 |
| ENSMUSG( | 6921.987 | 0.15767 | 0.155808 | 0.58375 | Ak3 |
| ENSMUSG( | 2313.68 | 0.113114 | 0.320675 | 0.765984 | Ak4 |
| ENSMUSG( | 2716.069 | 0.177477 | 0.135923 | 0.549949 | Ak5 |
| ENSMUSG( | 86.17276 | -0.43213 | 0.088607 | 0.45666 | Ak7 |
| ENSMUSG( | 45.90541 | -0.08102 | 0.775858 | 0.955606 | Ak8 |
| ENSMUSG( | 18.40391 | -0.10573 | 0.835719 | NA | Ak9 |
| ENSMUSG( | 117.5406 | 0.068706 | 0.724617 | 0.944096 | Akain1 |
| ENSMUSG( | 1623.885 | -0.16072 | 0.183424 | 0.62047 | Akap1 |
| ENSMUSG( | 593.8777 | 0.063375 | 0.586849 | 0.900235 | Akap10 |
| ENSMUSG( | 5615.38 | -0.14257 | 0.277131 | 0.727175 | Akap11 |
| ENSMUSG( | 1546.42 | 0.055559 | 0.721685 | 0.94325 | Akap12 |
| ENSMUSG( | 2111.393 | -0.23822 | 0.021203 | 0.222025 | Akap13 |
| ENSMUSG( | 579.7761 | 0.133771 | 0.252257 | 0.702302 | Akap17b |
| ENSMUSG( | 1121.773 | -0.06917 | 0.550859 | 0.886038 | Akap5 |
| ENSMUSG( | 7660.221 | -0.12897 | 0.334818 | 0.776386 | Akap6 |
| ENSMUSG( | 1136.919 | 0.059679 | 0.541971 | 0.883182 | Akap7 |
| ENSMUSG( | 2641.725 | 0.13645 | 0.229155 | 0.676908 | Akap8 |
| ENSMUSG( | 4038.49 | 0.260046 | 0.066455 | 0.39868 | Akap8l |
| ENSMUSG( | 1282.833 | -0.19282 | 0.053002 | 0.353344 | Akap9 |
| ENSMUSG( | 118.8133 | -0.10662 | 0.59077 | 0.90113 | Akip1 |
| ENSMUSG( | 2072.94 | 0.028549 | 0.775494 | 0.955606 | Akirin1 |
| ENSMUSG( | 1289.809 | 0.090843 | 0.471926 | 0.847528 | Akirin2 |
| ENSMUSG( | 298.4003 | -0.16824 | 0.223038 | 0.668948 | Akna |
| ENSMUSG( | 8383.875 | -0.0085 | 0.921852 | 0.990436 | Akr1a1 |
| ENSMUSG( | 639.8076 | -0.00496 | 0.9647 | 0.995155 | Akr1b1 |
| ENSMUSG( | 1109.423 | 0.166926 | 0.169191 | 0.603271 | Akr1b10 |
| ENSMUSG( | 25.47016 | 0.134233 | 0.74395 | 0.948497 | Akr1b8 |
| ENSMUSG( | 61.88533 | -0.0591 | 0.843724 | 0.971847 | Akr1c14 |
| ENSMUSG( | 19.94027 | 0.105268 | 0.817256 | NA | Akr1c18 |
| ENSMUSG( | 698.587 | 0.08252 | 0.468623 | 0.845503 | Akr1e1 |
| ENSMUSG( | 748.5374 | 0.066437 | 0.508824 | 0.868744 | Akr7a5 |
| ENSMUSG( | 8373.518 | 0.094032 | 0.21878 | 0.664559 | Akt1 |
| ENSMUSG( | 2042.394 | 0.137946 | 0.131248 | 0.543347 | Akt1s1 |
| ENSMUSG( | 10365.42 | 0.133569 | 0.192498 | 0.631514 | Akt2 |
| ENSMUSG( | 5536.7 | 0.163209 | 0.096725 | 0.477375 | Akt3 |
| ENSMUSG( | 5151.634 | -0.07167 | 0.441324 | 0.832449 | Aktip |
| ENSMUSG( | 539.6214 | -0.02495 | 0.837073 | 0.971111 | Alad |
| ENSMUSG( | 1323.606 | -0.02115 | 0.830376 | 0.96974 | Alas1 |
| ENSMUSG( | 1058.663 | -0.71855 | 0.086514 | 0.452045 | Alas2 |
| ENSMUSG( | 3806.14 | -0.30732 | 0.028396 | 0.257525 | Alcam |
| ENSMUSG( | 249.4545 | -0.01681 | 0.904074 | 0.986246 | Aldh16a1 |
| ENSMUSG( | 736.634 | 0.065921 | 0.561409 | 0.889052 | Aldh18a1 |
| ENSMUSG( | 11167.9 | 0.338948 | 0.003588 | 0.075902 | Aldh1a1 |
| ENSMUSG( | 246.7612 | 0.216188 | 0.289828 | 0.739839 | Aldh1a2 |
| ENSMUSG( | 26.79296 | -0.31023 | 0.444091 | 0.833239 | Aldh1a3 |
| ENSMUSG( | 77.79549 | -0.08788 | 0.732905 | 0.944956 | Aldh1a7 |
| ENSMUSG( | 195.5808 | -0.06859 | 0.709704 | 0.939669 | Aldh1b1 |
| ENSMUSG( | 7954.063 | 0.093748 | 0.398549 | 0.812878 | Aldh1l1 |
| ENSMUSG( | 281.3571 | 0.018862 | 0.902329 | 0.985707 | Aldh1l2 |
| ENSMUSG( | 5982.284 | 0.081937 | 0.416972 | 0.8209 | Aldh2 |
| ENSMUSG( | 40.18778 | -0.09118 | 0.776978 | 0.955828 | Aldh3a1 |
| ENSMUSG( | 1638.527 | 0.073636 | 0.426707 | 0.825155 | Aldh3a2 |
| ENSMUSG( | 257.2718 | -0.04893 | 0.758873 | 0.952113 | Aldh3b1 |
| ENSMUSG( | 45.01767 | 0.144662 | 0.620873 | 0.911432 | Aldh3b2 |
| ENSMUSG( | 1055.409 | 0.020782 | 0.84067 | 0.971179 | Aldh4a1 |
| ENSMUSG( | 7929.781 | 0.09658 | 0.383505 | 0.803428 | Aldh5a1 |
| ENSMUSG( | 5130.421 | 0.132326 | 0.238732 | 0.686489 | Aldh6a1 |
| ENSMUSG( | 9301.366 | 0.126738 | 0.220775 | 0.666475 | Aldh7a1 |
| ENSMUSG( | 1982.618 | 0.09149 | 0.445327 | 0.833239 | Aldh9a1 |
| ENSMUSG( | 32654.36 | -0.01003 | 0.923324 | 0.990694 | Aldoa |
| ENSMUSG( | 10.04508 | 0.055888 | 0.93105 | NA | Aldoart1 |
| ENSMUSG( | 82.00166 | 0.062832 | 0.80924 | 0.963595 | Aldob |
| ENSMUSG( | 122575.2 | 0.210732 | 0.026484 | 0.24888 | Aldoc |
| ENSMUSG( | 1051.65 | -0.29419 | 0.035206 | 0.288548 | Alg1 |
| ENSMUSG( | 1018.604 | -0.39208 | 0.013734 | 0.171117 | Alg10b |
| ENSMUSG( | 698.9897 | -0.40077 | 0.021387 | 0.22352 | Alg11 |
| ENSMUSG( | 391.9259 | -0.16059 | 0.304993 | 0.75285 | Alg12 |
| ENSMUSG( | 274.8999 | 0.027382 | 0.840634 | 0.971179 | Alg13 |
| ENSMUSG( | 759.5222 | -0.04099 | 0.745656 | 0.949242 | Alg14 |
| ENSMUSG( | 4774.114 | -0.51759 | 5.36E-05 | 0.004202 | Alg2 |
| ENSMUSG( | 408.2849 | -0.35756 | 0.019637 | 0.21122 | Alg3 |
| ENSMUSG( | 643.7274 | -0.18541 | 0.112931 | 0.506708 | Alg5 |
| ENSMUSG( | 260.6337 | -0.35496 | 0.030447 | 0.265225 | Alg6 |
| ENSMUSG( | 245.5674 | -0.21815 | 0.172977 | 0.608361 | Alg8 |
| ENSMUSG( | 683.8922 | -0.1997 | 0.183386 | 0.62047 | Alg9 |
| ENSMUSG( | 211.9287 | -0.39341 | 0.035799 | 0.290916 | Alk |
| ENSMUSG( | 47.54825 | -0.39224 | 0.154046 | 0.580901 | Alkal2 |
| ENSMUSG( | 305.3942 | -0.0579 | 0.657109 | 0.924142 | Alkbh1 |
| ENSMUSG( | 102.1854 | -0.08362 | 0.674943 | 0.929651 | Alkbh2 |
| ENSMUSG( | 283.6402 | -0.27734 | 0.041945 | 0.315416 | Alkbh3 |
| ENSMUSG( | 152.3405 | 0.043014 | 0.806095 | 0.962802 | Alkbh4 |
| ENSMUSG( | 2224.493 | 0.117308 | 0.180306 | 0.617778 | Alkbh5 |
| ENSMUSG( | 1012.809 | -0.03045 | 0.745695 | 0.949242 | Alkbh6 |
| ENSMUSG( | 308.8664 | -0.01454 | 0.91502 | 0.988634 | Alkbh7 |
| ENSMUSG( | 846.2912 | 0.13668 | 0.171784 | 0.606517 | Alkbh8 |
| ENSMUSG( | 311.0237 | 0.028611 | 0.841351 | 0.971179 | Alms1 |
| ENSMUSG( | 18.81754 | -0.52481 | 0.277266 | NA | Alox12 |

| | | | | | |
|---|---|---|---|---|---|
| ENSMUSG( | 4028.189 | -0.28274 | 0.035839 | 0.668395 | Agl |
| ENSMUSG( | 10.43548 | -0.5969 | 0.360001 | NA | Agmat |
| ENSMUSG( | 397.6842 | 0.197163 | 0.29053 | 0.897019 | Agmo |
| ENSMUSG( | 2738.419 | -0.0646 | 0.518101 | 0.930939 | Ago1 |
| ENSMUSG( | 3034.854 | 0.141202 | 0.205656 | 0.864412 | Ago2 |
| ENSMUSG( | 1664.822 | 0.126107 | 0.279996 | 0.892356 | Ago3 |
| ENSMUSG( | 506.74 | -0.0759 | 0.591507 | 0.944975 | Ago4 |
| ENSMUSG( | 6196.567 | -0.01153 | 0.931524 | 0.99306 | Agpat1 |
| ENSMUSG( | 70.68048 | 0.246939 | 0.330634 | 0.907508 | Agpat2 |
| ENSMUSG( | 22019.24 | -0.0123 | 0.916579 | 0.992529 | Agpat3 |
| ENSMUSG( | 2673.105 | 0.141663 | 0.318735 | 0.903297 | Agpat4 |
| ENSMUSG( | 3480.264 | 0.06777 | 0.641578 | 0.953903 | Agpat5 |
| ENSMUSG( | 1190.509 | -0.01788 | 0.875727 | 0.989644 | Agps |
| ENSMUSG( | 6652.388 | 0.203258 | 0.19973 | 0.863016 | Agrn |
| ENSMUSG( | 1681.77 | 0.132009 | 0.366607 | 0.914703 | Agt |
| ENSMUSG( | 4435.107 | -0.03894 | 0.688795 | 0.959742 | Agtpbp1 |
| ENSMUSG( | 25.25431 | 0.472494 | 0.26463 | 0.886134 | Agtr1a |
| ENSMUSG( | 12.61589 | 0.24767 | 0.648445 | 0.95422 | Agtr2 |
| ENSMUSG( | 906.3044 | -0.27179 | 0.037906 | 0.6763 | Agtrap |
| ENSMUSG( | 1269.856 | 0.320924 | 0.01827 | 0.542424 | Ahctf1 |
| ENSMUSG( | 363.2935 | -0.06482 | 0.648608 | 0.95422 | Ahcy |
| ENSMUSG( | 24992.4 | -0.26303 | 0.049238 | 0.708275 | Ahcyl1 |
| ENSMUSG( | 8233.629 | -0.12566 | 0.295703 | 0.898538 | Ahcyl2 |
| ENSMUSG( | 1414.866 | 0.089339 | 0.449706 | 0.922202 | Ahdc1 |
| ENSMUSG( | 3409.95 | 0.022491 | 0.88674 | 0.990118 | Ahi1 |
| ENSMUSG( | 1338.287 | 0.541856 | 0.000413 | 0.058872 | Ahnak |
| ENSMUSG( | 73.2642 | -0.11445 | 0.753015 | 0.970622 | Ahnak2 |
| ENSMUSG( | 158.538 | 0.503242 | 0.005576 | 0.303106 | Ahr |
| ENSMUSG( | 36.70989 | -0.07916 | 0.823893 | 0.979198 | Ahrr |
| ENSMUSG( | 1922.062 | -0.12667 | 0.371191 | 0.915491 | Ahsa1 |
| ENSMUSG( | 764.5511 | 0.076351 | 0.550806 | 0.939206 | Ahsa2 |
| ENSMUSG( | 46.71218 | -0.1296 | 0.668765 | 0.956766 | AI115009 |
| ENSMUSG( | 12.49158 | 0.806662 | 0.159404 | 0.837724 | AI182371 |
| ENSMUSG( | 16.42945 | -0.02259 | 0.962046 | 0.995237 | AI197445 |
| ENSMUSG( | 61.33087 | 0.199093 | 0.464477 | 0.923329 | AI429214 |
| ENSMUSG( | 82.64421 | 0.197821 | 0.532048 | 0.934795 | AI467606 |
| ENSMUSG( | 644.889 | 0.274797 | 0.187959 | 0.85698 | AI504432 |
| ENSMUSG( | 5150.045 | -0.03561 | 0.824576 | 0.979198 | AI593442 |
| ENSMUSG( | 1171.926 | -0.09766 | 0.47814 | 0.923329 | AI597479 |
| ENSMUSG( | 18.92881 | 0.684633 | 0.160948 | 0.837724 | AI661453 |
| ENSMUSG( | 1028.371 | -0.09009 | 0.549348 | 0.938519 | AI837181 |
| ENSMUSG( | 722.9063 | 0.026695 | 0.822261 | 0.979198 | AI854703 |
| ENSMUSG( | 371.3193 | 0.088388 | 0.62879 | 0.953251 | AI987944 |
| ENSMUSG( | 1587.561 | -0.05112 | 0.697286 | 0.960867 | Aida |
| ENSMUSG( | 260.8746 | -0.14824 | 0.372296 | 0.915491 | Aif1 |
| ENSMUSG( | 343.5736 | -0.08201 | 0.59694 | 0.946555 | Aif1l |
| ENSMUSG( | 754.3894 | -0.05162 | 0.686814 | 0.959098 | Aifm1 |
| ENSMUSG( | 408.9956 | -0.21141 | 0.148326 | 0.836241 | Aifm2 |
| ENSMUSG( | 2760.942 | 0.007797 | 0.956213 | 0.994798 | Aifm3 |
| ENSMUSG( | 1726.743 | 0.127039 | 0.426879 | 0.917257 | Aig1 |
| ENSMUSG( | 151.1704 | 0.079249 | 0.723488 | 0.966887 | Aim2 |
| ENSMUSG( | 929.0094 | -0.15025 | 0.335985 | 0.907913 | Aimp1 |
| ENSMUSG( | 332.773 | -0.11342 | 0.460522 | 0.923329 | Aimp2 |
| ENSMUSG( | 2215.61 | -0.13771 | 0.208287 | 0.864412 | Aip |
| ENSMUSG( | 458.6686 | 0.0709 | 0.610616 | 0.950771 | Airim |
| ENSMUSG( | 31.00026 | 0.361322 | 0.346613 | 0.911108 | Airn |
| ENSMUSG( | 1516.942 | 0.311337 | 0.04332 | 0.690304 | Ajap1 |
| ENSMUSG( | 6222.237 | 0.071477 | 0.459326 | 0.923329 | Ajm1 |
| ENSMUSG( | 93.46298 | 0.213412 | 0.399525 | 0.917257 | Ajuba |
| ENSMUSG( | 2970.826 | -0.21392 | 0.116312 | 0.813384 | Ak1 |
| ENSMUSG( | 37.74508 | -0.2404 | 0.515739 | 0.930939 | AK157302 |
| ENSMUSG( | 646.5605 | 0.006493 | 0.95902 | 0.994798 | Ak2 |
| ENSMUSG( | 5890.426 | -0.03739 | 0.79106 | 0.97681 | Ak3 |
| ENSMUSG( | 2003.063 | -0.03585 | 0.795911 | 0.97681 | Ak4 |
| ENSMUSG( | 2383.402 | 0.074805 | 0.551929 | 0.939565 | Ak5 |
| ENSMUSG( | 112.7194 | 0.59993 | 0.047315 | 0.700936 | Ak7 |
| ENSMUSG( | 44.3269 | 0.086575 | 0.797671 | 0.97681 | Ak8 |
| ENSMUSG( | 18.73463 | 0.252927 | 0.641141 | 0.953903 | Ak9 |
| ENSMUSG( | 102.9345 | -0.06745 | 0.778858 | 0.976106 | Akain1 |
| ENSMUSG( | 1640.733 | 0.141915 | 0.278533 | 0.891628 | Akap1 |
| ENSMUSG( | 538.7819 | 0.048918 | 0.711048 | 0.964363 | Akap10 |
| ENSMUSG( | 5736.135 | 0.18813 | 0.208557 | 0.864412 | Akap11 |
| ENSMUSG( | 1401.462 | 0.04519 | 0.775662 | 0.975466 | Akap12 |
| ENSMUSG( | 2134.013 | 0.067445 | 0.603972 | 0.948139 | Akap13 |
| ENSMUSG( | 504.9756 | 0.000173 | 0.999064 | 0.999503 | Akap17b |
| ENSMUSG( | 989.6576 | -0.16944 | 0.252212 | 0.879902 | Akap5 |
| ENSMUSG( | 7479.649 | 0.076716 | 0.615261 | 0.951177 | Akap6 |
| ENSMUSG( | 986.146 | -0.08616 | 0.473301 | 0.923329 | Akap7 |
| ENSMUSG( | 2434.884 | 0.167067 | 0.20655 | 0.864412 | Akap8 |
| ENSMUSG( | 3586.486 | 0.189152 | 0.298669 | 0.898538 | Akap8l |
| ENSMUSG( | 1286.523 | 0.092346 | 0.441954 | 0.921554 | Akap9 |
| ENSMUSG( | 110.6464 | -0.02587 | 0.904724 | 0.991089 | Akip1 |
| ENSMUSG( | 1795.68 | -0.12367 | 0.382545 | 0.917257 | Akirin1 |
| ENSMUSG( | 1142.964 | 0.005726 | 0.969652 | 0.99594 | Akirin2 |
| ENSMUSG( | 308.1452 | 0.199719 | 0.241655 | 0.876528 | Akna |
| ENSMUSG( | 7270.225 | -0.15689 | 0.165765 | 0.839705 | Akr1a1 |
| ENSMUSG( | 543.6647 | -0.21571 | 0.109749 | 0.80637 | Akr1b1 |
| ENSMUSG( | 927.7362 | -0.08192 | 0.572378 | 0.942515 | Akr1b10 |
| ENSMUSG( | 18.86841 | -0.49653 | 0.317309 | 0.902322 | Akr1b8 |
| ENSMUSG( | 71.7232 | 0.603496 | 0.073467 | 0.763629 | Akr1c14 |
| ENSMUSG( | 18.60617 | 0.141681 | 0.764234 | 0.974 | Akr1c18 |
| ENSMUSG( | 578.9868 | -0.19916 | 0.147946 | 0.836241 | Akr1e1 |
| ENSMUSG( | 635.8024 | -0.14404 | 0.281183 | 0.892356 | Akr7a5 |
| ENSMUSG( | 7400.354 | 0.007323 | 0.940992 | 0.994558 | Akt1 |
| ENSMUSG( | 1748.596 | -0.04176 | 0.724367 | 0.966887 | Akt1s1 |
| ENSMUSG( | 8023.68 | -0.35643 | 0.00178 | 0.156702 | Akt2 |
| ENSMUSG( | 4747.779 | -0.00701 | 0.94825 | 0.994798 | Akt3 |
| ENSMUSG( | 4434.236 | -0.24741 | 0.050688 | 0.709454 | Aktip |
| ENSMUSG( | 479.7599 | -0.0976 | 0.490769 | 0.92646 | Alad |
| ENSMUSG( | 1227.652 | 0.032669 | 0.785921 | 0.976285 | Alas1 |
| ENSMUSG( | 1077.312 | -0.33603 | 0.47898 | 0.923329 | Alas2 |
| ENSMUSG( | 4146.633 | 0.220016 | 0.146548 | 0.835973 | Alcam |
| ENSMUSG( | 227.316 | -0.01763 | 0.915354 | 0.992525 | Aldh16a1 |
| ENSMUSG( | 620.2399 | -0.16981 | 0.230929 | 0.868507 | Aldh18a1 |
| ENSMUSG( | 8428.974 | -0.19353 | 0.167508 | 0.840222 | Aldh1a1 |
| ENSMUSG( | 174.7311 | -0.57253 | 0.018011 | 0.539314 | Aldh1a2 |
| ENSMUSG( | 25.43007 | -0.18962 | 0.634891 | 0.953903 | Aldh1a3 |
| ENSMUSG( | 65.54038 | -0.36764 | 0.209747 | 0.864412 | Aldh1a7 |
| ENSMUSG( | 172.7024 | -0.18011 | 0.440391 | 0.921166 | Aldh1b1 |
| ENSMUSG( | 6470.242 | -0.24386 | 0.048564 | 0.706142 | Aldh1l1 |
| ENSMUSG( | 254.8601 | -0.0051 | 0.975664 | 0.996416 | Aldh1l2 |
| ENSMUSG( | 5114.661 | -0.10304 | 0.33725 | 0.907972 | Aldh2 |
| ENSMUSG( | 33.28468 | -0.41598 | 0.284086 | 0.892412 | Aldh3a1 |
| ENSMUSG( | 1375.25 | -0.16937 | 0.144088 | 0.831355 | Aldh3a2 |
| ENSMUSG( | 212.3926 | -0.36597 | 0.043943 | 0.690976 | Aldh3b1 |
| ENSMUSG( | 40.12419 | 0.061373 | 0.852917 | 0.985598 | Aldh3b2 |
| ENSMUSG( | 915.2415 | -0.12587 | 0.29854 | 0.898538 | Aldh4a1 |
| ENSMUSG( | 6653.679 | -0.14466 | 0.283494 | 0.892412 | Aldh5a1 |
| ENSMUSG( | 4388.964 | -0.04896 | 0.733988 | 0.967228 | Aldh6a1 |
| ENSMUSG( | 7666.664 | -0.16651 | 0.206905 | 0.864412 | Aldh7a1 |
| ENSMUSG( | 1640.948 | -0.19327 | 0.120744 | 0.813384 | Aldh9a1 |
| ENSMUSG( | 27650.05 | -0.23255 | 0.081918 | 0.77681 | Aldoa |
| ENSMUSG( | 72.91958 | -0.00342 | 0.989681 | 0.997928 | Aldob |
| ENSMUSG( | 99702.55 | -0.11259 | 0.389449 | 0.917257 | Aldoc |
| ENSMUSG( | 1090.657 | 0.098581 | 0.539862 | 0.937378 | Alg1 |
| ENSMUSG( | 1212.995 | 0.387182 | 0.036478 | 0.66949 | Alg10b |
| ENSMUSG( | 805.3293 | 0.296032 | 0.129052 | 0.820358 | Alg11 |
| ENSMUSG( | 399.8865 | 0.171343 | 0.305732 | 0.898538 | Alg12 |
| ENSMUSG( | 234.3888 | -0.17084 | 0.282774 | 0.892412 | Alg13 |
| ENSMUSG( | 686.6701 | -0.06957 | 0.625893 | 0.953234 | Alg14 |
| ENSMUSG( | 5285.752 | 0.092527 | 0.538282 | 0.937216 | Alg2 |
| ENSMUSG( | 409.4585 | -0.05295 | 0.753806 | 0.970671 | Alg3 |
| ENSMUSG( | 655.0178 | 0.13856 | 0.33409 | 0.907907 | Alg5 |
| ENSMUSG( | 297.4072 | 0.306737 | 0.073175 | 0.763629 | Alg6 |
| ENSMUSG( | 243.0439 | 0.038777 | 0.832991 | 0.981118 | Alg8 |
| ENSMUSG( | 693.7669 | 0.120117 | 0.476595 | 0.923329 | Alg9 |
| ENSMUSG( | 214.6848 | -0.05558 | 0.775155 | 0.975109 | Alk |
| ENSMUSG( | 48.90026 | -0.01503 | 0.961189 | 0.99511 | Alkal2 |
| ENSMUSG( | 290.0944 | 0.062707 | 0.677331 | 0.957106 | Alkbh1 |
| ENSMUSG( | 94.51091 | -0.03576 | 0.878237 | 0.989644 | Alkbh2 |
| ENSMUSG( | 267.3402 | -0.16925 | 0.290511 | 0.897019 | Alkbh3 |
| ENSMUSG( | 140.662 | 0.091889 | 0.652177 | 0.954766 | Alkbh4 |
| ENSMUSG( | 2019.589 | 0.105589 | 0.354063 | 0.912411 | Alkbh5 |
| ENSMUSG( | 902.4547 | -0.09598 | 0.402412 | 0.917257 | Alkbh6 |
| ENSMUSG( | 267.0751 | -0.18131 | 0.28353 | 0.892412 | Alkbh7 |
| ENSMUSG( | 756.5563 | 0.083004 | 0.476751 | 0.923329 | Alkbh8 |
| ENSMUSG( | 283.0628 | 0.034212 | 0.834408 | 0.981237 | Alms1 |
| ENSMUSG( | 26.38375 | 0.668311 | 0.130933 | 0.822884 | Alox12 |
| ENSMUSG( | 54.48009 | -0.38231 | 0.22566 | 0.868468 | Alox12b |
| ENSMUSG( | 69.83196 | 0.265854 | 0.333817 | 0.907907 | Alox5 |
| ENSMUSG( | 230.6119 | 0.024156 | 0.905935 | 0.991089 | Alox5ap |
| ENSMUSG( | 66.70176 | 0.006775 | 0.979752 | 0.996479 | Alox8 |
| ENSMUSG( | 136.2273 | 0.105037 | 0.624968 | 0.953234 | Aloxe3 |
| ENSMUSG( | 306.5889 | 0.347557 | 0.05026 | 0.708648 | Alpk1 |
| ENSMUSG( | 23.06133 | 0.183183 | 0.650658 | 0.95422 | Alpk3 |
| ENSMUSG( | 680.4737 | 0.364163 | 0.007255 | 0.357969 | Alpl |
| ENSMUSG( | 1109.317 | 0.147661 | 0.248211 | 0.877904 | Als2 |
| ENSMUSG( | 692.931 | 0.320269 | 0.047936 | 0.704961 | Als2cl |

| | | | | | | | | | | | |
|---|---|---|---|---|---|---|---|---|---|---|---|
| ENSMUSG( | Agpat2 | 66.39721 | -0.20645 | 0.376738 | 0.801213 | ENSMUSG( | Agpat2 | 70.68048 | 0.246939 | 0.330634 | 0.907508 |
| ENSMUSG( | Agpat3 | 24657.59 | 0.044085 | 0.668965 | 0.927963 | ENSMUSG( | Agpat3 | 22019.24 | -0.0123 | 0.916579 | 0.992529 |
| ENSMUSG( | Agpat4 | 2451.736 | -0.40328 | 0.001129 | 0.035751 | ENSMUSG( | Agpat4 | 2673.105 | 0.141663 | 0.318735 | 0.903297 |
| ENSMUSG( | Agpat5 | 3657.701 | -0.05856 | 0.655908 | 0.923929 | ENSMUSG( | Agpat5 | 3480.264 | 0.06777 | 0.641578 | 0.953903 |
| ENSMUSG( | Agps | 1337.591 | 0.051021 | 0.588213 | 0.900712 | ENSMUSG( | Agps | 1190.509 | -0.01788 | 0.875727 | 0.989644 |
| ENSMUSG( | Agrn | 7001.688 | 0.087436 | 0.442267 | 0.832774 | ENSMUSG( | Agrn | 6652.388 | 0.203258 | 0.19973 | 0.863016 |
| ENSMUSG( | Agt | 1813.162 | 0.084565 | 0.546135 | 0.884974 | ENSMUSG( | Agt | 1681.77 | 0.132009 | 0.366607 | 0.914703 |
| ENSMUSG( | Agtpbp1 | 5250.796 | 0.174625 | 0.022815 | 0.230858 | ENSMUSG( | Agtpbp1 | 4435.107 | -0.03894 | 0.688795 | 0.959742 |
| ENSMUSG( | Agtr1a | 18.23423 | -0.82541 | 0.078008 | NA | ENSMUSG( | Agtr1a | 25.25431 | 0.472494 | 0.26463 | 0.886134 |
| ENSMUSG( | Agtr2 | 13.15575 | 0.116219 | 0.823386 | NA | ENSMUSG( | Agtr2 | 12.61589 | 0.24767 | 0.648445 | 0.95422 |
| ENSMUSG( | Agtrap | 1062.71 | -0.07143 | 0.566702 | 0.891644 | ENSMUSG( | Agtrap | 906.3044 | -0.27179 | 0.037906 | 0.6763 |
| ENSMUSG( | Ahctf1 | 1240.988 | -0.0009 | 0.994136 | 0.999369 | ENSMUSG( | Ahctf1 | 1269.856 | 0.320924 | 0.01827 | 0.542424 |
| ENSMUSG( | Ahcy | 408.5938 | 0.010342 | 0.930126 | 0.991848 | ENSMUSG( | Ahcy | 363.2935 | -0.06482 | 0.648608 | 0.95422 |
| ENSMUSG( | Ahcyl1 | 30728.28 | 0.07737 | 0.508224 | 0.867999 | ENSMUSG( | Ahcyl1 | 24992.4 | -0.26303 | 0.049238 | 0.708275 |
| ENSMUSG( | Ahcyl2 | 9828.154 | 0.118214 | 0.201642 | 0.641362 | ENSMUSG( | Ahcyl2 | 8233.629 | -0.12566 | 0.295703 | 0.898538 |
| ENSMUSG( | Ahdc1 | 1560.571 | 0.105618 | 0.307371 | 0.755324 | ENSMUSG( | Ahdc1 | 1414.866 | 0.089339 | 0.449706 | 0.922202 |
| ENSMUSG( | Ahi1 | 3920.711 | 0.156404 | 0.234405 | 0.681922 | ENSMUSG( | Ahi1 | 3409.95 | 0.022491 | 0.88674 | 0.990118 |
| ENSMUSG( | Ahnak | 1316.606 | 0.259872 | 0.035139 | 0.288296 | ENSMUSG( | Ahnak | 1338.287 | 0.541856 | 0.000413 | 0.058872 |
| ENSMUSG( | Ahnak2 | 78.02105 | -0.24202 | 0.32763 | 0.772692 | ENSMUSG( | Ahnak2 | 73.2642 | -0.11445 | 0.753015 | 0.970622 |
| ENSMUSG( | Ahr | 156.874 | 0.241989 | 0.175054 | 0.610399 | ENSMUSG( | Ahr | 158.538 | 0.503242 | 0.005576 | 0.303106 |
| ENSMUSG( | Ahrr | 37.26796 | -0.33028 | 0.33382 | 0.775602 | ENSMUSG( | Ahrr | 36.70989 | -0.07916 | 0.823893 | 0.979198 |
| ENSMUSG( | Ahsa1 | 2308.552 | 0.13632 | 0.222453 | 0.668434 | ENSMUSG( | Ahsa1 | 1922.062 | -0.12667 | 0.371191 | 0.915491 |
| ENSMUSG( | Ahsa2 | 848.0786 | 0.105891 | 0.317481 | 0.763823 | ENSMUSG( | Ahsa2 | 764.5511 | 0.076351 | 0.550806 | 0.939206 |
| ENSMUSG( | AI115009 | 51.53926 | -0.10518 | 0.729135 | 0.944738 | ENSMUSG( | AI115009 | 46.71218 | -0.1296 | 0.668765 | 0.956766 |
| ENSMUSG( | AI182371 | 12.81575 | 0.642273 | 0.228046 | NA | ENSMUSG( | AI182371 | 12.49158 | 0.806662 | 0.159404 | 0.837724 |
| ENSMUSG( | AI197445 | 20.14847 | 0.271427 | 0.564578 | NA | ENSMUSG( | AI197445 | 16.42945 | -0.02259 | 0.962046 | 0.995237 |
| ENSMUSG( | AI429214 | 64.92554 | 0.09227 | 0.713477 | 0.940865 | ENSMUSG( | AI429214 | 61.33087 | 0.199093 | 0.464477 | 0.923329 |
| ENSMUSG( | AI467606 | 76.74703 | -0.32585 | 0.220981 | 0.666476 | ENSMUSG( | AI467606 | 82.64421 | 0.197821 | 0.532048 | 0.934795 |
| ENSMUSG( | AI504432 | 529.1535 | -0.61584 | 0.000587 | 0.022791 | ENSMUSG( | AI504432 | 644.889 | 0.274797 | 0.187959 | 0.85698 |
| ENSMUSG( | AI593442 | 4646.03 | -0.68355 | 5.38E-08 | 1.71E-05 | ENSMUSG( | AI593442 | 5150.045 | -0.03561 | 0.824576 | 0.979198 |
| ENSMUSG( | AI597479 | 1351.163 | 0.049571 | 0.685785 | 0.932981 | ENSMUSG( | AI597479 | 1171.926 | -0.09766 | 0.47814 | 0.923329 |
| ENSMUSG( | AI661453 | 18.52078 | 0.371946 | 0.381638 | NA | ENSMUSG( | AI661453 | 18.92881 | 0.684633 | 0.160948 | 0.837724 |
| ENSMUSG( | AI837181 | 1165.132 | 0.005311 | 0.968524 | 0.996369 | ENSMUSG( | AI837181 | 1028.371 | -0.09009 | 0.549348 | 0.938519 |
| ENSMUSG( | AI854703 | 816.7793 | 0.106287 | 0.294008 | 0.743459 | ENSMUSG( | AI854703 | 722.9063 | 0.026695 | 0.822261 | 0.979198 |
| ENSMUSG( | AI987944 | 399.708 | 0.037251 | 0.826099 | 0.968159 | ENSMUSG( | AI987944 | 371.3193 | 0.088388 | 0.62879 | 0.953251 |
| ENSMUSG( | Aida | 1801.03 | 0.047758 | 0.635372 | 0.914296 | ENSMUSG( | Aida | 1587.561 | -0.05112 | 0.697286 | 0.960867 |
| ENSMUSG( | Aif1 | 312.8991 | 0.108904 | 0.462974 | 0.842422 | ENSMUSG( | Aif1 | 260.8746 | -0.14824 | 0.372296 | 0.915491 |
| ENSMUSG( | Aif1l | 353.9725 | -0.28419 | 0.032801 | 0.277278 | ENSMUSG( | Aif1l | 343.5736 | -0.08201 | 0.59694 | 0.946555 |
| ENSMUSG( | Aifm1 | 867.6499 | 0.079037 | 0.480335 | 0.854748 | ENSMUSG( | Aifm1 | 754.3894 | -0.05162 | 0.686814 | 0.959098 |
| ENSMUSG( | Aifm2 | 485.453 | 0.027151 | 0.828364 | 0.968994 | ENSMUSG( | Aifm2 | 408.9956 | -0.21141 | 0.148326 | 0.836241 |
| ENSMUSG( | Aifm3 | 3370.851 | 0.299769 | 0.00126 | 0.038624 | ENSMUSG( | Aifm3 | 2760.942 | 0.007797 | 0.956213 | 0.994798 |
| ENSMUSG( | Aig1 | 1615.508 | -0.35586 | 0.008904 | 0.132538 | ENSMUSG( | Aig1 | 1726.743 | 0.127039 | 0.426879 | 0.917257 |
| ENSMUSG( | Aim2 | 147.8928 | -0.27977 | 0.146171 | 0.567605 | ENSMUSG( | Aim2 | 151.1704 | 0.079249 | 0.723488 | 0.966887 |
| ENSMUSG( | Aimp1 | 1077.516 | 0.019359 | 0.86031 | 0.974842 | ENSMUSG( | Aimp1 | 929.0094 | -0.15025 | 0.335985 | 0.907913 |
| ENSMUSG( | Aimp2 | 375.9235 | -0.02619 | 0.851736 | 0.973104 | ENSMUSG( | Aimp2 | 332.773 | -0.11342 | 0.460522 | 0.923329 |
| ENSMUSG( | Aip | 2579.787 | 0.037566 | 0.645441 | 0.918017 | ENSMUSG( | Aip | 2215.61 | -0.13771 | 0.208287 | 0.864412 |
| ENSMUSG( | Airim | 502.6432 | 0.067919 | 0.563815 | 0.890032 | ENSMUSG( | Airim | 458.6686 | 0.0709 | 0.610616 | 0.950771 |
| ENSMUSG( | Airn | 26.82693 | -0.34188 | 0.377319 | 0.801213 | ENSMUSG( | Airn | 31.00026 | 0.361322 | 0.346613 | 0.911108 |
| ENSMUSG( | Ajap1 | 1366.297 | -0.259 | 0.052686 | 0.352396 | ENSMUSG( | Ajap1 | 1516.942 | 0.311337 | 0.04332 | 0.690304 |
| ENSMUSG( | Ajm1 | 6068.282 | -0.28278 | 0.000598 | 0.022994 | ENSMUSG( | Ajm1 | 6222.237 | 0.071477 | 0.459326 | 0.923329 |
| ENSMUSG( | Ajuba | 87.28489 | -0.27758 | 0.264346 | 0.715323 | ENSMUSG( | Ajuba | 93.46298 | 0.213412 | 0.399525 | 0.917257 |
| ENSMUSG( | Ak1 | 3555.185 | 0.046712 | 0.653145 | 0.922819 | ENSMUSG( | Ak1 | 2970.826 | -0.21392 | 0.116312 | 0.813384 |
| ENSMUSG( | AK157302 | 42.7078 | -0.10867 | 0.731653 | 0.944956 | ENSMUSG( | AK157302 | 37.74508 | -0.2404 | 0.515739 | 0.930939 |
| ENSMUSG( | Ak2 | 749.0713 | 0.161549 | 0.103499 | 0.489288 | ENSMUSG( | Ak2 | 646.5605 | 0.006493 | 0.95902 | 0.994798 |
| ENSMUSG( | Ak3 | 6921.987 | 0.15767 | 0.155808 | 0.58375 | ENSMUSG( | Ak3 | 5890.426 | -0.03739 | 0.79106 | 0.97681 |
| ENSMUSG( | Ak4 | 2313.68 | 0.113114 | 0.320675 | 0.765984 | ENSMUSG( | Ak4 | 2003.063 | -0.03585 | 0.795911 | 0.97681 |
| ENSMUSG( | Ak5 | 2716.069 | 0.177477 | 0.135923 | 0.549949 | ENSMUSG( | Ak5 | 2383.402 | 0.074805 | 0.551929 | 0.939565 |
| ENSMUSG( | Ak7 | 86.17276 | -0.43213 | 0.088607 | 0.45666 | ENSMUSG( | Ak7 | 112.7194 | 0.59993 | 0.047315 | 0.700936 |
| ENSMUSG( | Ak8 | 45.90541 | -0.08102 | 0.775858 | 0.955606 | ENSMUSG( | Ak8 | 44.3269 | 0.086575 | 0.797671 | 0.97681 |
| ENSMUSG( | Ak9 | 18.40391 | -0.10573 | 0.835719 | NA | ENSMUSG( | Ak9 | 18.73463 | 0.252927 | 0.641141 | 0.953903 |
| ENSMUSG( | Akain1 | 117.5406 | 0.068706 | 0.724617 | 0.944096 | ENSMUSG( | Akain1 | 102.9345 | -0.06745 | 0.778858 | 0.976106 |
| ENSMUSG( | Akap1 | 1623.885 | -0.16072 | 0.183424 | 0.62047 | ENSMUSG( | Akap1 | 1640.733 | 0.141915 | 0.278533 | 0.891628 |
| ENSMUSG( | Akap10 | 593.8777 | 0.063375 | 0.586849 | 0.900235 | ENSMUSG( | Akap10 | 538.7819 | 0.048918 | 0.711048 | 0.964363 |
| ENSMUSG( | Akap11 | 5615.38 | -0.14257 | 0.277131 | 0.727175 | ENSMUSG( | Akap11 | 5736.135 | 0.18813 | 0.208557 | 0.864412 |
| ENSMUSG( | Akap12 | 1546.42 | 0.055559 | 0.721685 | 0.94325 | ENSMUSG( | Akap12 | 1401.462 | 0.04519 | 0.775662 | 0.975466 |
| ENSMUSG( | Akap13 | 2111.393 | -0.23822 | 0.021203 | 0.222025 | ENSMUSG( | Akap13 | 2134.013 | 0.067445 | 0.603972 | 0.948139 |
| ENSMUSG( | Akap17b | 579.7761 | 0.133771 | 0.252257 | 0.702302 | ENSMUSG( | Akap17b | 504.9756 | 0.000173 | 0.999064 | 0.999503 |
| ENSMUSG( | Akap5 | 1121.773 | -0.06917 | 0.550859 | 0.886038 | ENSMUSG( | Akap5 | 989.6576 | -0.16944 | 0.252212 | 0.879902 |
| ENSMUSG( | Akap6 | 7660.221 | -0.12897 | 0.334818 | 0.776386 | ENSMUSG( | Akap6 | 7479.649 | 0.076716 | 0.615261 | 0.951177 |
| ENSMUSG( | Akap7 | 1136.919 | 0.059679 | 0.541971 | 0.883182 | ENSMUSG( | Akap7 | 986.146 | -0.08616 | 0.473301 | 0.923329 |
| ENSMUSG( | Akap8 | 2641.725 | 0.13645 | 0.229155 | 0.676908 | ENSMUSG( | Akap8 | 2434.884 | 0.167067 | 0.20655 | 0.864412 |
| ENSMUSG( | Akap8l | 4038.49 | 0.260046 | 0.066455 | 0.39868 | ENSMUSG( | Akap8l | 3586.486 | 0.189152 | 0.298669 | 0.898538 |
| ENSMUSG( | Akap9 | 1282.833 | -0.19282 | 0.053002 | 0.353344 | ENSMUSG( | Akap9 | 1286.523 | 0.092346 | 0.441954 | 0.921554 |
| ENSMUSG( | Akip1 | 118.8133 | -0.10662 | 0.59077 | 0.90113 | ENSMUSG( | Akip1 | 110.6464 | -0.02587 | 0.904724 | 0.991089 |
| ENSMUSG( | Akirin1 | 2072.94 | 0.028549 | 0.775494 | 0.955606 | ENSMUSG( | Akirin1 | 1795.68 | -0.12367 | 0.382545 | 0.917257 |
| ENSMUSG( | Akirin2 | 1289.809 | 0.090843 | 0.471926 | 0.847528 | ENSMUSG( | Akirin2 | 1142.964 | 0.005726 | 0.969652 | 0.99594 |
| ENSMUSG( | Akna | 298.4003 | -0.16824 | 0.223038 | 0.668948 | ENSMUSG( | Akna | 308.1452 | 0.199719 | 0.241655 | 0.876528 |
| ENSMUSG( | Akr1a1 | 8383.875 | -0.0085 | 0.921852 | 0.990436 | ENSMUSG( | Akr1a1 | 7270.225 | -0.15689 | 0.165765 | 0.839705 |
| ENSMUSG( | Akr1b1 | 639.8076 | -0.00496 | 0.9647 | 0.995155 | ENSMUSG( | Akr1b1 | 543.6647 | -0.21571 | 0.109749 | 0.80637 |
| ENSMUSG( | Akr1b10 | 1109.423 | 0.166926 | 0.169191 | 0.603271 | ENSMUSG( | Akr1b10 | 927.7362 | -0.08192 | 0.572378 | 0.942515 |
| ENSMUSG( | Akr1b8 | 25.47016 | 0.134233 | 0.74395 | 0.948497 | ENSMUSG( | Akr1b8 | 18.86841 | -0.49653 | 0.317309 | 0.902322 |
| ENSMUSG( | Akr1c14 | 61.88533 | -0.0591 | 0.843724 | 0.971847 | ENSMUSG( | Akr1c14 | 71.7232 | 0.603496 | 0.073467 | 0.763629 |
| ENSMUSG( | Akr1c18 | 19.94027 | 0.105268 | 0.817256 | NA | ENSMUSG( | Akr1c18 | 18.60617 | 0.141681 | 0.764234 | 0.974 |
| ENSMUSG( | Akr1e1 | 698.587 | 0.08252 | 0.468623 | 0.845503 | ENSMUSG( | Akr1e1 | 578.9868 | -0.19916 | 0.147946 | 0.836241 |
| ENSMUSG( | Akr7a5 | 748.5374 | 0.066437 | 0.508824 | 0.868744 | ENSMUSG( | Akr7a5 | 635.8024 | -0.14404 | 0.281183 | 0.892356 |
| ENSMUSG( | Akt1 | 8373.518 | 0.094032 | 0.21878 | 0.664559 | ENSMUSG( | Akt1 | 7400.354 | 0.007323 | 0.940992 | 0.994558 |
| ENSMUSG( | Akt1s1 | 2042.394 | 0.137946 | 0.131248 | 0.543347 | ENSMUSG( | Akt1s1 | 1748.596 | -0.04176 | 0.724367 | 0.966887 |
| ENSMUSG( | Akt2 | 10365.42 | 0.133569 | 0.192498 | 0.631514 | ENSMUSG( | Akt2 | 8023.68 | -0.35643 | 0.00178 | 0.156702 |
| ENSMUSG( | Akt3 | 5536.7 | 0.163209 | 0.096725 | 0.477375 | ENSMUSG( | Akt3 | 4747.779 | -0.00701 | 0.94825 | 0.994798 |
| ENSMUSG( | Aktip | 5151.634 | -0.07167 | 0.441324 | 0.832449 | ENSMUSG( | Aktip | 4434.236 | -0.24741 | 0.050688 | 0.709454 |
| ENSMUSG( | Alad | 539.6214 | -0.02495 | 0.837073 | 0.971111 | ENSMUSG( | Alad | 479.7599 | -0.0976 | 0.490769 | 0.92646 |
| ENSMUSG( | Alas1 | 1323.606 | -0.02115 | 0.830376 | 0.96974 | ENSMUSG( | Alas1 | 1227.652 | 0.032669 | 0.785921 | 0.976285 |
| ENSMUSG( | Alas2 | 1058.663 | -0.71855 | 0.086514 | 0.452045 | ENSMUSG( | Alas2 | 1077.312 | -0.33603 | 0.47898 | 0.923329 |
| ENSMUSG( | Alcam | 3806.14 | -0.30732 | 0.028396 | 0.257525 | ENSMUSG( | Alcam | 4146.633 | 0.220016 | 0.146548 | 0.835973 |
| ENSMUSG( | Aldh16a1 | 249.4545 | -0.01681 | 0.904074 | 0.986246 | ENSMUSG( | Aldh16a1 | 227.316 | -0.01763 | 0.915354 | 0.992525 |
| ENSMUSG( | Aldh18a1 | 736.634 | 0.065921 | 0.561409 | 0.889052 | ENSMUSG( | Aldh18a1 | 620.2399 | -0.16981 | 0.230929 | 0.868507 |
| ENSMUSG( | Aldh1a1 | 11167.9 | 0.338948 | 0.003588 | 0.075902 | ENSMUSG( | Aldh1a1 | 8428.974 | -0.19353 | 0.167508 | 0.840222 |
| ENSMUSG( | Aldh1a2 | 246.7612 | 0.216188 | 0.289828 | 0.739839 | ENSMUSG( | Aldh1a2 | 174.7311 | -0.57253 | 0.018011 | 0.539314 |
| ENSMUSG( | Aldh1a3 | 26.79296 | -0.31023 | 0.444091 | 0.833239 | ENSMUSG( | Aldh1a3 | 25.43007 | -0.18962 | 0.634891 | 0.953903 |
| ENSMUSG( | Aldh1a7 | 77.79549 | -0.08788 | 0.732905 | 0.944956 | ENSMUSG( | Aldh1a7 | 65.54038 | -0.36764 | 0.209747 | 0.864412 |
| ENSMUSG( | Aldh1b1 | 195.5808 | -0.06859 | 0.709704 | 0.939669 | ENSMUSG( | Aldh1b1 | 172.7024 | -0.18011 | 0.440391 | 0.921166 |
| ENSMUSG( | Aldh1l1 | 7954.063 | 0.093748 | 0.398549 | 0.812878 | ENSMUSG( | Aldh1l1 | 6470.242 | -0.24386 | 0.048564 | 0.706142 |
| ENSMUSG( | Aldh1l2 | 281.3571 | 0.018862 | 0.902329 | 0.985707 | ENSMUSG( | Aldh1l2 | 254.8601 | -0.0051 | 0.975664 | 0.996416 |
| ENSMUSG( | Aldh2 | 5982.284 | 0.081937 | 0.416972 | 0.8209 | ENSMUSG( | Aldh2 | 5114.661 | -0.10304 | 0.33725 | 0.907972 |
| ENSMUSG( | Aldh3a1 | 40.18778 | -0.09118 | 0.776978 | 0.955828 | ENSMUSG( | Aldh3a1 | 33.28468 | -0.41598 | 0.284086 | 0.892412 |
| ENSMUSG( | Aldh3a2 | 1638.527 | 0.073636 | 0.426707 | 0.825155 | ENSMUSG( | Aldh3a2 | 1375.25 | -0.16937 | 0.144088 | 0.831355 |
| ENSMUSG( | Aldh3b1 | 257.2718 | -0.04893 | 0.758873 | 0.952113 | ENSMUSG( | Aldh3b1 | 212.3926 | -0.36597 | 0.043943 | 0.690976 |
| ENSMUSG( | Aldh3b2 | 45.01767 | 0.144662 | 0.620873 | 0.911432 | ENSMUSG( | Aldh3b2 | 40.12419 | 0.061373 | 0.852917 | 0.985598 |
| ENSMUSG( | Aldh4a1 | 1055.409 | 0.020782 | 0.84067 | 0.971179 | ENSMUSG( | Aldh4a1 | 915.2415 | -0.12587 | 0.29854 | 0.898538 |
| ENSMUSG( | Aldh5a1 | 7929.781 | 0.09658 | 0.383505 | 0.803428 | ENSMUSG( | Aldh5a1 | 6653.679 | -0.14466 | 0.283494 | 0.892412 |
| ENSMUSG( | Aldh6a1 | 5130.421 | 0.132326 | 0.238732 | 0.686489 | ENSMUSG( | Aldh6a1 | 4388.964 | -0.04896 | 0.733988 | 0.967228 |
| ENSMUSG( | Aldh7a1 | 9301.366 | 0.126738 | 0.220775 | 0.666475 | ENSMUSG( | Aldh7a1 | 7666.664 | -0.16651 | 0.206905 | 0.864412 |
| ENSMUSG( | Aldh9a1 | 1982.618 | 0.09149 | 0.445327 | 0.833239 | ENSMUSG( | Aldh9a1 | 1640.948 | -0.19327 | 0.120744 | 0.813384 |
| ENSMUSG( | Aldoa | 32654.36 | -0.01003 | 0.923324 | 0.990694 | ENSMUSG( | Aldoa | 27650.05 | -0.23255 | 0.081918 | 0.77681 |
| ENSMUSG( | Aldob | 82.00166 | 0.062832 | 0.80924 | 0.963595 | ENSMUSG( | Aldob | 72.91958 | -0.00342 | 0.989681 | 0.997928 |
| ENSMUSG( | Aldoc | 122575.2 | 0.210732 | 0.026484 | 0.24888 | ENSMUSG( | Aldoc | 99702.55 | -0.11259 | 0.389449 | 0.917257 |
| ENSMUSG( | Alg1 | 1051.65 | -0.29419 | 0.035206 | 0.288548 | ENSMUSG( | Alg1 | 1090.657 | 0.098581 | 0.539862 | 0.937378 |
| ENSMUSG( | Alg10b | 1018.604 | -0.39208 | 0.013734 | 0.171117 | ENSMUSG( | Alg10b | 1212.995 | 0.387182 | 0.036478 | 0.66949 |
| ENSMUSG( | Alg11 | 698.9897 | -0.40077 | 0.021387 | 0.22352 | ENSMUSG( | Alg11 | 805.3293 | 0.296032 | 0.129052 | 0.820358 |
| ENSMUSG( | Alg12 | 391.9259 | -0.16059 | 0.304993 | 0.75285 | ENSMUSG( | Alg12 | 399.8865 | 0.171343 | 0.305732 | 0.898538 |
| ENSMUSG( | Alg13 | 274.8999 | 0.027382 | 0.840634 | 0.971179 | ENSMUSG( | Alg13 | 234.3888 | -0.17084 | 0.282774 | 0.892412 |
| ENSMUSG( | Alg14 | 759.5222 | -0.04099 | 0.745656 | 0.949242 | ENSMUSG( | Alg14 | 686.6701 | -0.06957 | 0.625893 | 0.953234 |
| ENSMUSG( | Alg2 | 4774.114 | -0.51759 | 5.36E-05 | 0.004202 | ENSMUSG( | Alg2 | 5285.752 | 0.092527 | 0.538282 | 0.937216 |
| ENSMUSG( | Alg3 | 408.2849 | -0.35756 | 0.019637 | 0.21122 | ENSMUSG( | Alg3 | 409.4585 | -0.05295 | 0.753806 | 0.970671 |
| ENSMUSG( | Alg5 | 643.7274 | -0.18541 | 0.112931 | 0.506708 | ENSMUSG( | Alg5 | 655.0178 | 0.13856 | 0.33409 | 0.907907 |
| ENSMUSG( | Alg6 | 260.6337 | -0.35496 | 0.030447 | 0.265225 | ENSMUSG( | Alg6 | 297.4072 | 0.306737 | 0.073175 | 0.763629 |
| ENSMUSG( | Alg8 | 245.5674 | -0.21815 | 0.172977 | 0.608361 | ENSMUSG( | Alg8 | 243.0439 | 0.038777 | 0.832991 | 0.981118 |
| ENSMUSG( | Alg9 | 683.8922 | -0.1997 | 0.183386 | 0.62047 | ENSMUSG( | Alg9 | 693.7669 | 0.120117 | 0.476595 | 0.923329 |
| ENSMUSG( | Alk | 211.9287 | -0.39341 | 0.035799 | 0.290916 | ENSMUSG( | Alk | 214.6848 | -0.05558 | 0.775155 | 0.975109 |
| ENSMUSG( | Alkal2 | 47.54825 | -0.39224 | 0.154046 | 0.580901 | ENSMUSG( | Alkal2 | 48.90026 | -0.01503 | 0.961189 | 0.99511 |
| ENSMUSG( | Alkbh1 | 305.3942 | -0.0579 | 0.657109 | 0.924142 | ENSMUSG( | Alkbh1 | 290.0944 | 0.062707 | 0.677331 | 0.957106 |
| ENSMUSG( | Alkbh2 | 102.1854 | -0.08362 | 0.674943 | 0.929651 | ENSMUSG( | Alkbh2 | 94.51091 | -0.03576 | 0.878237 | 0.989644 |
| ENSMUSG( | Alkbh3 | 283.6402 | -0.27734 | 0.041945 | 0.315416 | ENSMUSG( | Alkbh3 | 267.3402 | -0.16925 | 0.290511 | 0.897019 |
| ENSMUSG( | Alkbh4 | 152.3405 | 0.043014 | 0.806095 | 0.962802 | ENSMUSG( | Alkbh4 | 140.662 | 0.091889 | 0.652177 | 0.954766 |
| ENSMUSG( | Alkbh5 | 2224.493 | 0.117308 | 0.180306 | 0.617778 | ENSMUSG( | Alkbh5 | 2019.589 | 0.105589 | 0.354063 | 0.912411 |
| ENSMUSG( | Alkbh6 | 1012.809 | -0.03045 | 0.745695 | 0.949242 | ENSMUSG( | Alkbh6 | 902.4547 | -0.09598 | 0.402412 | 0.917257 |
| ENSMUSG( | Alkbh7 | 308.8664 | -0.01454 | 0.91502 | 0.988634 | ENSMUSG( | Alkbh7 | 267.0751 | -0.18131 | 0.28353 | 0.892412 |
| ENSMUSG( | Alkbh8 | 846.2912 | 0.13668 | 0.171784 | 0.606517 | ENSMUSG( | Alkbh8 | 756.5563 | 0.083004 | 0.476751 | 0.923329 |
| ENSMUSG( | Alms1 | 311.0237 | 0.028611 | 0.841351 | 0.971179 | ENSMUSG( | Alms1 | 283.0628 | 0.034212 | 0.834408 | 0.981237 |
| ENSMUSG( | Alox12 | 18.81754 | -0.52481 | 0.277266 | NA | ENSMUSG( | Alox12 | 26.38375 | 0.668311 | 0.130933 | 0.822884 |
| ENSMUSG( | Alox12b | 72.62439 | 0.183433 | 0.525574 | 0.876728 | ENSMUSG( | Alox12b | 54.48009 | -0.38231 | 0.22566 | 0.868468 |
| ENSMUSG( | Alox5 | 71.23522 | 0.039915 | 0.865048 | 0.975565 | ENSMUSG( | Alox5 | 69.83196 | 0.265854 | 0.333817 | 0.907907 |
| ENSMUSG( | Alox5ap | 255.332 | 0.050384 | 0.784655 | 0.958149 | ENSMUSG( | Alox5ap | 230.6119 | 0.024156 | 0.905935 | 0.991089 |
| ENSMUSG( | Alox8 | 77.12517 | 0.15137 | 0.532487 | 0.879905 | ENSMUSG( | Alox8 | 66.70176 | 0.006775 | 0.979752 | 0.996479 |
| ENSMUSG( | Aloxe3 | 158.6789 | 0.262657 | 0.166511 | 0.599201 | ENSMUSG( | Aloxe3 | 136.2273 | 0.105037 | 0.624968 | 0.953234 |
| ENSMUSG( | Alpk1 | 278.2003 | -0.1945 | 0.195857 | 0.635235 | ENSMUSG( | Alpk1 | 306.5889 | 0.347557 | 0.05026 | 0.708648 |
| ENSMUSG( | Alpk3 | 25.30893 | 0.193634 | 0.596741 | 0.903526 | ENSMUSG( | Alpk3 | 23.06133 | 0.183183 | 0.650658 | 0.95422 |
| ENSMUSG( | Alpl | 587.5964 | -0.32758 | 0.01003 | 0.142701 | ENSMUSG( | Alpl | 680.4737 | 0.364163 | 0.007255 | 0.357969 |
| ENSMUSG( | Als2 | 1280.359 | 0.280819 | 0.009878 | 0.141868 | ENSMUSG( | Als2 | 1109.317 | 0.147661 | 0.248211 | 0.877904 |
| ENSMUSG( | Als2cl | 743.2464 | 0.253724 | 0.031781 | 0.272023 | ENSMUSG( | Als2cl | 692.931 | 0.320269 | 0.047936 | 0.704961 |
| ENSMUSG( | Alx3 | 29.24748 | 0.026253 | 0.944604 | 0.993583 | ENSMUSG( | Alx3 | 34.53562 | 0.668366 | 0.108544 | 0.804969 |
| ENSMUSG( | Alx4 | 90.28787 | 0.285182 | 0.239714 | 0.687561 | ENSMUSG( | Alx4 | 85.58242 | 0.364647 | 0.219537 | 0.866104 |
| ENSMUSG( | Alyref | 537.8451 | 0.054384 | 0.625138 | 0.912249 | ENSMUSG( | Alyref | 481.1657 | -0.00192 | 0.988673 | 0.997862 |
| ENSMUSG( | Alyref2 | 74.72471 | -0.24114 | 0.29484 | 0.743798 | ENSMUSG( | Alyref2 | 64.37208 | -0.45484 | 0.105801 | 0.804969 |
| ENSMUSG( | Amacr | 240.4968 | 0.09347 | 0.528204 | 0.878236 | ENSMUSG( | Amacr | 209.9721 | -0.0339 | 0.844606 | 0.983642 |
| ENSMUSG( | Ambra1 | 1810.855 | -0.02918 | 0.79627 | 0.960088 | ENSMUSG( | Ambra1 | 1722.787 | 0.09643 | 0.399357 | 0.917257 |
| ENSMUSG( | Amd1 | 2210.788 | 0.232299 | 0.014962 | 0.179768 | ENSMUSG( | Amd1 | 1887.023 | 0.050569 | 0.660702 | 0.956205 |
| ENSMUSG( | Amdhd1 | 12.09965 | -0.13863 | 0.794206 | NA | ENSMUSG( | Amdhd1 | 11.36145 | -0.01638 | 0.977098 | NA |

| | | | | | |
|---|---|---|---|---|---|
| ENSMUSG( | 72.62439 | 0.183433 | 0.525574 | 0.876728 | Alox12b |
| ENSMUSG( | 71.23522 | 0.039915 | 0.865048 | 0.975565 | Alox5 |
| ENSMUSG( | 255.332 | 0.050384 | 0.784655 | 0.958149 | Alox5ap |
| ENSMUSG( | 77.12517 | 0.15137 | 0.532487 | 0.879905 | Alox8 |
| ENSMUSG( | 158.6789 | 0.262657 | 0.166511 | 0.599201 | Aloxe3 |
| ENSMUSG( | 278.2003 | -0.1945 | 0.195857 | 0.635235 | Alpk1 |
| ENSMUSG( | 25.30893 | 0.193634 | 0.596741 | 0.903526 | Alpk3 |
| ENSMUSG( | 587.5964 | -0.32758 | 0.01003 | 0.142701 | Alpl |
| ENSMUSG( | 1280.359 | 0.280819 | 0.009878 | 0.141868 | Als2 |
| ENSMUSG( | 743.2464 | 0.253724 | 0.031781 | 0.272023 | Als2cl |
| ENSMUSG( | 29.24748 | 0.026253 | 0.944604 | 0.993583 | Alx3 |
| ENSMUSG( | 90.28787 | 0.285182 | 0.239714 | 0.687561 | Alx4 |
| ENSMUSG( | 537.8451 | 0.054384 | 0.625138 | 0.912249 | Alyref |
| ENSMUSG( | 74.72471 | -0.24114 | 0.29484 | 0.743798 | Alyref2 |
| ENSMUSG( | 240.4968 | 0.09347 | 0.528204 | 0.878236 | Amacr |
| ENSMUSG( | 1810.855 | -0.02918 | 0.79627 | 0.960088 | Ambra1 |
| ENSMUSG( | 2210.788 | 0.232299 | 0.014962 | 0.179768 | Amd1 |
| ENSMUSG( | 12.09965 | -0.13863 | 0.794206 | NA | Amdhd1 |
| ENSMUSG( | 504.8616 | 0.001482 | 0.991514 | 0.999232 | Amdhd2 |
| ENSMUSG( | 221.8823 | -0.15015 | 0.323572 | 0.768536 | Amer1 |
| ENSMUSG( | 4084.967 | -0.24063 | 0.043707 | 0.320345 | Amer2 |
| ENSMUSG( | 514.2843 | 0.074238 | 0.638506 | 0.916319 | Amer3 |
| ENSMUSG( | 9894.297 | -0.12843 | 0.260709 | 0.710468 | Amfr |
| ENSMUSG( | 32.22257 | 0.488256 | 0.161508 | 0.59148 | Amh |
| ENSMUSG( | 44.12747 | 0.100777 | 0.730865 | 0.944956 | Amhr2 |
| ENSMUSG( | 3407.515 | -0.19469 | 0.174827 | 0.610304 | Amigo1 |
| ENSMUSG( | 452.6997 | 0.093772 | 0.470506 | 0.84674 | Amigo2 |
| ENSMUSG( | 55.19027 | -0.14045 | 0.592591 | 0.902094 | Amigo3 |
| ENSMUSG( | 119.7014 | -0.25886 | 0.160117 | 0.589846 | Ammecr1 |
| ENSMUSG( | 938.0911 | -0.05594 | 0.608055 | 0.907154 | Ammecr1l |
| ENSMUSG( | 301.2149 | 0.221856 | 0.10911 | 0.498553 | Amn |
| ENSMUSG( | 360.4731 | -0.33214 | 0.010925 | 0.150598 | Amn1 |
| ENSMUSG( | 4218.253 | 0.22349 | 0.141612 | 0.558606 | Amot |
| ENSMUSG( | 2272.901 | 0.082856 | 0.500042 | 0.863455 | Amotl1 |
| ENSMUSG( | 630.6307 | -0.1254 | 0.357473 | 0.788872 | Amotl2 |
| ENSMUSG( | 1887.683 | 0.101428 | 0.262372 | 0.713005 | Ampd2 |
| ENSMUSG( | 1399.894 | 0.00392 | 0.969864 | 0.996369 | Ampd3 |
| ENSMUSG( | 4289.791 | 0.219367 | 0.028982 | 0.259144 | Amph |
| ENSMUSG( | 1617.323 | 0.06818 | 0.482078 | 0.855778 | Amt |
| ENSMUSG( | 496.9374 | 0.211749 | 0.364577 | 0.793796 | Amy1 |
| ENSMUSG( | 402.2282 | -0.12079 | 0.333908 | 0.775602 | Amz1 |
| ENSMUSG( | 1395.235 | 0.118204 | 0.191948 | 0.630811 | Amz2 |
| ENSMUSG( | 2394.491 | -0.0003 | 0.997691 | 0.999848 | Anapc1 |
| ENSMUSG( | 244.9993 | -0.04979 | 0.733451 | 0.944956 | Anapc10 |
| ENSMUSG( | 2146.533 | -0.03227 | 0.728236 | 0.944355 | Anapc11 |
| ENSMUSG( | 988.8642 | -0.01563 | 0.885886 | 0.983008 | Anapc13 |
| ENSMUSG( | 881.0147 | 0.059076 | 0.586201 | 0.899709 | Anapc15 |
| ENSMUSG( | 2013.372 | 0.064987 | 0.50149 | 0.864477 | Anapc16 |
| ENSMUSG( | 2788.673 | 0.083356 | 0.316896 | 0.763754 | Anapc2 |
| ENSMUSG( | 1479.193 | 0.149545 | 0.13253 | 0.544627 | Anapc4 |
| ENSMUSG( | 7645.66 | 0.126337 | 0.139607 | 0.555959 | Anapc5 |
| ENSMUSG( | 859.5383 | -0.05069 | 0.61943 | 0.911432 | Anapc7 |
| ENSMUSG( | 102.6516 | 0.094874 | 0.657494 | 0.92425 | Ang |
| ENSMUSG( | 437.9975 | -0.44156 | 0.006729 | 0.11299 | Angel1 |
| ENSMUSG( | 2139.559 | 0.038822 | 0.656609 | 0.923929 | Angel2 |
| ENSMUSG( | 330.5529 | 0.106359 | 0.58418 | 0.898738 | Angpt1 |
| ENSMUSG( | 129.3183 | 0.040143 | 0.839831 | 0.971179 | Angpt2 |
| ENSMUSG( | 8.792941 | -0.59207 | 0.393504 | NA | Angpt4 |
| ENSMUSG( | 28.26469 | -0.09834 | 0.79028 | 0.959461 | Angptl1 |
| ENSMUSG( | 200.9817 | -0.14587 | 0.461044 | 0.84109 | Angptl2 |
| ENSMUSG( | 12.65588 | 0.271611 | 0.59341 | NA | Angptl3 |
| ENSMUSG( | 191.2143 | -0.18164 | 0.255558 | 0.706195 | Angptl4 |
| ENSMUSG( | 221.5052 | -0.09012 | 0.592625 | 0.902094 | Angptl6 |
| ENSMUSG( | 12.4651 | 0.572899 | 0.313855 | NA | Angptl7 |
| ENSMUSG( | 9505.403 | -0.18502 | 0.108167 | 0.497094 | Ank |
| ENSMUSG( | 2241.53 | 0.100039 | 0.237806 | 0.685809 | Ank1 |
| ENSMUSG( | 36616.85 | 0.037954 | 0.752573 | 0.950536 | Ank2 |
| ENSMUSG( | 16751.03 | -0.27586 | 0.052366 | 0.35132 | Ank3 |
| ENSMUSG( | 15.37761 | -0.1293 | 0.794964 | NA | Ankdd1a |
| ENSMUSG( | 14.15283 | 0.112663 | 0.819196 | NA | Ankef1 |
| ENSMUSG( | 45.1639 | -0.16703 | 0.595982 | 0.903152 | Ankfn1 |
| ENSMUSG( | 2033.62 | 0.070887 | 0.426269 | 0.824898 | Ankfy1 |
| ENSMUSG( | 806.6844 | -0.05371 | 0.654983 | 0.92381 | Ankhd1 |
| ENSMUSG( | 2068.232 | 0.032899 | 0.749045 | 0.949604 | Ankib1 |
| ENSMUSG( | 12.37411 | 0.416485 | 0.485882 | NA | Ankle1 |
| ENSMUSG( | 1186.092 | -0.20023 | 0.164514 | 0.595363 | Ankle2 |
| ENSMUSG( | 23.07367 | -0.06282 | 0.871875 | 0.978266 | Ankmy1 |
| ENSMUSG( | 985.9789 | -0.02746 | 0.783888 | 0.957645 | Ankmy2 |
| ENSMUSG( | 990.6892 | 0.093346 | 0.320043 | 0.765523 | Ankra2 |
| ENSMUSG( | 1342.662 | 0.061427 | 0.566982 | 0.891644 | Ankrd10 |
| ENSMUSG( | 9454.6 | -0.02686 | 0.775103 | 0.955606 | Ankrd11 |
| ENSMUSG( | 1451.847 | -0.07486 | 0.457461 | 0.839558 | Ankrd12 |
| ENSMUSG( | 2010.197 | 0.022831 | 0.787421 | 0.959147 | Ankrd13a |
| ENSMUSG( | 4843.403 | 0.245313 | 0.003817 | 0.079179 | Ankrd13b |
| ENSMUSG( | 3588.793 | 0.029699 | 0.786129 | 0.958774 | Ankrd13c |
| ENSMUSG( | 1483.675 | 0.190132 | 0.192025 | 0.630811 | Ankrd13d |
| ENSMUSG( | 825.5051 | -0.12761 | 0.281243 | 0.730583 | Ankrd16 |
| ENSMUSG( | 3069.475 | 0.126871 | 0.107274 | 0.495579 | Ankrd17 |
| ENSMUSG( | 24.54449 | -0.4916 | 0.228421 | 0.675849 | Ankrd23 |
| ENSMUSG( | 1054.309 | 0.311916 | 0.078661 | 0.433892 | Ankrd24 |
| ENSMUSG( | 614.4775 | 0.171238 | 0.132972 | 0.545614 | Ankrd26 |
| ENSMUSG( | 1643.831 | 0.196694 | 0.027469 | 0.254456 | Ankrd27 |
| ENSMUSG( | 2300.574 | -0.06617 | 0.492258 | 0.860297 | Ankrd28 |
| ENSMUSG( | 982.5277 | 0.076386 | 0.431559 | 0.827432 | Ankrd29 |
| ENSMUSG( | 13.28665 | 0.309508 | 0.53679 | NA | Ankrd31 |
| ENSMUSG( | 1684.731 | 0.082265 | 0.48749 | 0.858812 | Ankrd33b |
| ENSMUSG( | 1280.453 | 0.093373 | 0.539034 | 0.882877 | Ankrd34a |
| ENSMUSG( | 217.8747 | 0.126219 | 0.474503 | 0.849425 | Ankrd34b |
| ENSMUSG( | 118.876 | 0.300763 | 0.18885 | 0.627678 | Ankrd34c |
| ENSMUSG( | 425.4321 | -0.02385 | 0.861594 | 0.975026 | Ankrd35 |
| ENSMUSG( | 90.90947 | -0.23037 | 0.293066 | 0.742763 | Ankrd37 |
| ENSMUSG( | 278.8746 | 0.017516 | 0.899493 | 0.985633 | Ankrd39 |
| ENSMUSG( | 15275.43 | -0.00629 | 0.946277 | 0.99369 | Ankrd40 |
| ENSMUSG( | 258.9999 | 0.079497 | 0.579087 | 0.896692 | Ankrd42 |
| ENSMUSG( | 416.8205 | -0.11002 | 0.424674 | 0.824675 | Ankrd44 |
| ENSMUSG( | 1225.627 | -0.03147 | 0.749452 | 0.949664 | Ankrd45 |
| ENSMUSG( | 4361.687 | 0.127952 | 0.207392 | 0.649561 | Ankrd46 |
| ENSMUSG( | 265.2726 | 0.043132 | 0.764841 | 0.954127 | Ankrd49 |
| ENSMUSG( | 1509.997 | 0.156817 | 0.110207 | 0.500785 | Ankrd50 |
| ENSMUSG( | 1814.977 | 0.214539 | 0.044731 | 0.324745 | Ankrd52 |
| ENSMUSG( | 506.4571 | 0.044184 | 0.728143 | 0.944355 | Ankrd54 |
| ENSMUSG( | 54.05335 | -0.11063 | 0.689756 | 0.933609 | Ankrd55 |
| ENSMUSG( | 1448.788 | -0.11327 | 0.307541 | 0.755462 | Ankrd6 |
| ENSMUSG( | 12.79722 | -0.23369 | 0.658233 | NA | Ankrd61 |
| ENSMUSG( | 502.5407 | -0.22696 | 0.193525 | 0.633027 | Ankrd63 |
| ENSMUSG( | 23.10994 | -0.22264 | 0.573819 | 0.893584 | Ankrd66 |
| ENSMUSG( | 252.9269 | -0.10737 | 0.55752 | 0.888279 | Ankrd9 |
| ENSMUSG( | 1655.81 | 0.183271 | 0.051198 | 0.347172 | Anks1 |
| ENSMUSG( | 9123.799 | 0.040709 | 0.617899 | 0.910872 | Anks1b |
| ENSMUSG( | 1178.124 | 0.07344 | 0.422568 | 0.823508 | Anks3 |
| ENSMUSG( | 513.7227 | 0.015664 | 0.912025 | 0.988483 | Anks6 |
| ENSMUSG( | 42.25443 | -0.32453 | 0.298484 | 0.74713 | Ankub1 |
| ENSMUSG( | 832.2366 | 0.126097 | 0.306178 | 0.753712 | Ankzf1 |
| ENSMUSG( | 758.974 | -0.07721 | 0.43963 | 0.83129 | Anln |
| ENSMUSG( | 90.28403 | -0.23617 | 0.374407 | 0.800445 | Ano1 |
| ENSMUSG( | 848.83 | -0.15555 | 0.281818 | 0.731842 | Ano10 |
| ENSMUSG( | 80.12672 | -0.69976 | 0.010006 | 0.142653 | Ano2 |
| ENSMUSG( | 2306.737 | -0.88127 | 5.12E-10 | 2.52E-07 | Ano3 |
| ENSMUSG( | 518.543 | -0.4142 | 0.00156 | 0.044321 | Ano4 |
| ENSMUSG( | 31.70647 | -0.09396 | 0.80878 | 0.963455 | Ano5 |
| ENSMUSG( | 947.3334 | 0.053729 | 0.693395 | 0.934202 | Ano6 |
| ENSMUSG( | 8.714241 | 0.099472 | 0.885807 | NA | Ano7 |
| ENSMUSG( | 2656.921 | -0.10666 | 0.309345 | 0.75737 | Ano8 |
| ENSMUSG( | 4587.976 | 0.104119 | 0.271505 | 0.722082 | Anp32a |
| ENSMUSG( | 2371.29 | 0.045545 | 0.621868 | 0.911432 | Anp32b |
| ENSMUSG( | 4710.675 | 0.018621 | 0.861414 | 0.975026 | Anp32e |
| ENSMUSG( | 322.0514 | 0.107117 | 0.455911 | 0.838611 | Anpep |
| ENSMUSG( | 1573.623 | 0.005117 | 0.956228 | 0.993874 | Antkmt |
| ENSMUSG( | 1506.304 | -0.07811 | 0.500844 | 0.863671 | Antxr1 |
| ENSMUSG( | 102.0805 | -0.12146 | 0.578998 | 0.896692 | Antxr2 |
| ENSMUSG( | 105.6158 | -0.05003 | 0.825433 | 0.967868 | Anxa1 |
| ENSMUSG( | 770.0136 | 0.070266 | 0.651286 | 0.921451 | Anxa11 |
| ENSMUSG( | 518.9985 | -0.00946 | 0.94016 | 0.993415 | Anxa2 |
| ENSMUSG( | 585.4998 | 0.120598 | 0.315231 | 0.762288 | Anxa3 |
| ENSMUSG( | 160.4106 | 0.180287 | 0.27514 | 0.72509 | Anxa4 |
| ENSMUSG( | 2977.424 | 0.026253 | 0.823666 | 0.966603 | Anxa5 |
| ENSMUSG( | 4186.227 | -0.04389 | 0.642915 | 0.916865 | Anxa6 |
| ENSMUSG( | 1465.944 | 0.083779 | 0.426644 | 0.825155 | Anxa7 |
| ENSMUSG( | 18.11178 | -0.50249 | 0.252055 | NA | Anxa8 |
| ENSMUSG( | 40.93526 | 0.397295 | 0.233429 | 0.681508 | Anxa9 |
| ENSMUSG( | 17.84049 | -0.23394 | 0.592803 | NA | Aoah |
| ENSMUSG( | 115.9853 | 0.087518 | 0.681828 | 0.931619 | Aoc2 |
| ENSMUSG( | 62.70768 | -0.51927 | 0.062269 | 0.386085 | Aoc3 |
| ENSMUSG( | 924.7392 | 0.109256 | 0.394978 | 0.810135 | Aopep |
| ENSMUSG( | 413.233 | -0.16094 | 0.181138 | 0.618139 | Aox1 |
| ENSMUSG( | 87.4804 | -0.00439 | 0.983826 | 0.99779 | Aox3 |
| ENSMUSG( | 37.83271 | 0.336581 | 0.362347 | 0.792943 | Aox4 |
| ENSMUSG( | 3414.234 | 0.092078 | 0.276797 | 0.726951 | Ap1ar |

| | | | | | |
|---|---|---|---|---|---|
| ENSMUSG( | 34.53562 | 0.668366 | 0.108544 | 0.804969 | Alx3 |
| ENSMUSG( | 85.58242 | 0.364647 | 0.219537 | 0.866104 | Alx4 |
| ENSMUSG( | 481.1657 | -0.00192 | 0.988673 | 0.997862 | Alyref |
| ENSMUSG( | 64.37208 | -0.45484 | 0.105801 | 0.804969 | Alyref2 |
| ENSMUSG( | 209.9721 | -0.0339 | 0.844606 | 0.983642 | Amacr |
| ENSMUSG( | 1722.787 | 0.09643 | 0.399357 | 0.917257 | Ambra1 |
| ENSMUSG( | 1887.023 | 0.050569 | 0.660702 | 0.956205 | Amd1 |
| ENSMUSG( | 11.36145 | -0.01638 | 0.977098 | NA | Amdhd1 |
| ENSMUSG( | 446.1972 | -0.08733 | 0.565226 | 0.94131 | Amdhd2 |
| ENSMUSG( | 226.6195 | 0.175004 | 0.286159 | 0.893906 | Amer1 |
| ENSMUSG( | 3732.704 | -0.2341 | 0.114164 | 0.811037 | Amer2 |
| ENSMUSG( | 453.9898 | -0.00902 | 0.956174 | 0.994798 | Amer3 |
| ENSMUSG( | 9566.597 | 0.047761 | 0.715522 | 0.965104 | Amfr |
| ENSMUSG( | 28.85787 | 0.43564 | 0.261895 | 0.884926 | Amh |
| ENSMUSG( | 37.49265 | -0.10244 | 0.754151 | 0.970715 | Amhr2 |
| ENSMUSG( | 3493.17 | 0.152776 | 0.352641 | 0.912093 | Amigo1 |
| ENSMUSG( | 484.4988 | 0.51495 | 0.000504 | 0.067381 | Amigo2 |
| ENSMUSG( | 47.61043 | -0.29139 | 0.324783 | 0.905257 | Amigo3 |
| ENSMUSG( | 106.5471 | -0.34972 | 0.106944 | 0.804969 | Ammecr1 |
| ENSMUSG( | 863.6824 | -0.02284 | 0.850468 | 0.985541 | Ammecr1l |
| ENSMUSG( | 293.4278 | 0.399947 | 0.009711 | 0.406458 | Amn |
| ENSMUSG( | 388.2211 | 0.162909 | 0.279387 | 0.892356 | Amn1 |
| ENSMUSG( | 3347.094 | -0.17522 | 0.301009 | 0.898538 | Amot |
| ENSMUSG( | 1903.427 | -0.16548 | 0.161387 | 0.837724 | Amotl1 |
| ENSMUSG( | 673.9052 | 0.327926 | 0.087098 | 0.782809 | Amotl2 |
| ENSMUSG( | 1673.176 | 0.024405 | 0.851917 | 0.985598 | Ampd2 |
| ENSMUSG( | 1222.776 | -0.12288 | 0.284427 | 0.892412 | Ampd3 |
| ENSMUSG( | 3576.818 | -0.02936 | 0.801529 | 0.97681 | Amph |
| ENSMUSG( | 1336.92 | -0.22378 | 0.065622 | 0.74546 | Amt |
| ENSMUSG( | 454.371 | 0.217297 | 0.619893 | 0.952896 | Amy1 |
| ENSMUSG( | 392.2212 | 0.078144 | 0.579765 | 0.944209 | Amz1 |
| ENSMUSG( | 1186.051 | -0.08171 | 0.469688 | 0.923329 | Amz2 |
| ENSMUSG( | 2262.717 | 0.106514 | 0.429605 | 0.917257 | Anapc1 |
| ENSMUSG( | 217.4255 | -0.1429 | 0.428723 | 0.917257 | Anapc10 |
| ENSMUSG( | 1787.328 | -0.31185 | 0.009263 | 0.402642 | Anapc11 |
| ENSMUSG( | 814.781 | -0.32944 | 0.010757 | 0.421367 | Anapc13 |
| ENSMUSG( | 765.8095 | -0.07954 | 0.516982 | 0.930939 | Anapc15 |
| ENSMUSG( | 1694.265 | -0.17215 | 0.188285 | 0.85698 | Anapc16 |
| ENSMUSG( | 2402.722 | -0.07898 | 0.462692 | 0.923329 | Anapc2 |
| ENSMUSG( | 1292.228 | 0.028196 | 0.825458 | 0.979445 | Anapc4 |
| ENSMUSG( | 6654.523 | -0.00322 | 0.975985 | 0.996416 | Anapc5 |
| ENSMUSG( | 771.7387 | -0.09158 | 0.443033 | 0.921554 | Anapc7 |
| ENSMUSG( | 105.2108 | 0.414213 | 0.06169 | 0.736228 | Ang |
| ENSMUSG( | 441.0757 | -0.11135 | 0.549221 | 0.938514 | Angel1 |
| ENSMUSG( | 1978.841 | 0.080849 | 0.442174 | 0.921554 | Angel2 |
| ENSMUSG( | 280.0938 | -0.1081 | 0.61142 | 0.950771 | Angpt1 |
| ENSMUSG( | 103.9883 | -0.31691 | 0.209283 | 0.864412 | Angpt2 |
| ENSMUSG( | 31.16115 | 0.455498 | 0.257471 | 0.883401 | Angptl1 |
| ENSMUSG( | 268.8376 | 0.855235 | 0.000304 | 0.047107 | Angptl2 |
| ENSMUSG( | 9.124513 | -0.52326 | 0.444441 | NA | Angptl3 |
| ENSMUSG( | 153.5417 | -0.58663 | 0.00293 | 0.217424 | Angptl4 |
| ENSMUSG( | 186.2775 | -0.34565 | 0.073866 | 0.765627 | Angptl6 |
| ENSMUSG( | 14.74956 | 1.175964 | 0.035917 | 0.668395 | Angptl7 |
| ENSMUSG( | 9529.206 | 0.097715 | 0.476956 | 0.923329 | Ank |
| ENSMUSG( | 1877.49 | -0.14262 | 0.261987 | 0.884926 | Ank1 |
| ENSMUSG( | 34249.06 | 0.11429 | 0.391797 | 0.917257 | Ank2 |
| ENSMUSG( | 17238.92 | 0.091737 | 0.559532 | 0.939807 | Ank3 |
| ENSMUSG( | 15.40491 | 0.165657 | 0.753407 | 0.970671 | Ankdd1a |
| ENSMUSG( | 13.52938 | 0.203602 | 0.703619 | 0.962481 | Ankef1 |
| ENSMUSG( | 43.66401 | 0.037222 | 0.91612 | 0.992529 | Ankfn1 |
| ENSMUSG( | 1758.294 | -0.08014 | 0.445935 | 0.921734 | Ankfy1 |
| ENSMUSG( | 770.6895 | 0.088785 | 0.547882 | 0.938063 | Ankhd1 |
| ENSMUSG( | 1935.268 | 0.10922 | 0.360882 | 0.914223 | Ankib1 |
| ENSMUSG( | 10.96331 | 0.344409 | 0.588415 | NA | Ankle1 |
| ENSMUSG( | 1266.359 | 0.258179 | 0.118174 | 0.813384 | Ankle2 |
| ENSMUSG( | 21.07973 | -0.02105 | 0.962356 | 0.995237 | Ankmy1 |
| ENSMUSG( | 847.114 | -0.2053 | 0.087704 | 0.785431 | Ankmy2 |
| ENSMUSG( | 877.1084 | 0.008847 | 0.940948 | 0.994558 | Ankra2 |
| ENSMUSG( | 1303.041 | 0.237023 | 0.049289 | 0.708275 | Ankrd10 |
| ENSMUSG( | 8993.985 | 0.0982 | 0.392664 | 0.917257 | Ankrd11 |
| ENSMUSG( | 1435.351 | 0.157219 | 0.165135 | 0.838743 | Ankrd12 |
| ENSMUSG( | 1889.913 | 0.111741 | 0.28861 | 0.894794 | Ankrd13a |
| ENSMUSG( | 4305.566 | 0.179256 | 0.113542 | 0.809623 | Ankrd13b |
| ENSMUSG( | 3153.732 | -0.07824 | 0.577025 | 0.942912 | Ankrd13c |
| ENSMUSG( | 1231.669 | -0.07508 | 0.63226 | 0.953452 | Ankrd13d |
| ENSMUSG( | 798.0158 | 0.051247 | 0.760625 | 0.972688 | Ankrd16 |
| ENSMUSG( | 2748.392 | 0.078342 | 0.456015 | 0.923329 | Ankrd17 |
| ENSMUSG( | 26.9662 | 0.134363 | 0.767333 | 0.974236 | Ankrd23 |
| ENSMUSG( | 867.3367 | 0.029345 | 0.885588 | 0.98977 | Ankrd24 |
| ENSMUSG( | 562.4623 | 0.177898 | 0.199737 | 0.863016 | Ankrd26 |
| ENSMUSG( | 1412.709 | 0.033527 | 0.777769 | 0.976084 | Ankrd27 |
| ENSMUSG( | 2133.269 | -0.01222 | 0.922245 | 0.992644 | Ankrd28 |
| ENSMUSG( | 852.8746 | -0.06999 | 0.656358 | 0.955533 | Ankrd29 |
| ENSMUSG( | 9.175977 | -0.53411 | 0.401975 | NA | Ankrd31 |
| ENSMUSG( | 1564.533 | 0.137016 | 0.280859 | 0.892356 | Ankrd33b |
| ENSMUSG( | 1110.804 | -0.05 | 0.763392 | 0.973951 | Ankrd34a |
| ENSMUSG( | 198.917 | 0.130101 | 0.506138 | 0.927825 | Ankrd34b |
| ENSMUSG( | 96.31108 | -0.02654 | 0.920864 | 0.992644 | Ankrd34c |
| ENSMUSG( | 353.1926 | -0.31692 | 0.041423 | 0.688699 | Ankrd35 |
| ENSMUSG( | 91.59668 | 0.06092 | 0.786067 | 0.976285 | Ankrd37 |
| ENSMUSG( | 265.1936 | 0.135163 | 0.373366 | 0.915491 | Ankrd39 |
| ENSMUSG( | 13018.38 | -0.20818 | 0.099295 | 0.800296 | Ankrd40 |
| ENSMUSG( | 237.7077 | 0.097679 | 0.542364 | 0.938034 | Ankrd42 |
| ENSMUSG( | 399.6478 | 0.04352 | 0.780313 | 0.976106 | Ankrd44 |
| ENSMUSG( | 1093.084 | -0.09973 | 0.453378 | 0.922316 | Ankrd45 |
| ENSMUSG( | 3706.923 | -0.07293 | 0.556932 | 0.939565 | Ankrd46 |
| ENSMUSG( | 243.8034 | 0.058165 | 0.744011 | 0.96951 | Ankrd49 |
| ENSMUSG( | 1359.711 | 0.127531 | 0.295334 | 0.898538 | Ankrd50 |
| ENSMUSG( | 1552.8 | 0.044012 | 0.742664 | 0.969415 | Ankrd52 |
| ENSMUSG( | 482.0976 | 0.170737 | 0.222405 | 0.866791 | Ankrd54 |
| ENSMUSG( | 43.10531 | -0.54153 | 0.093975 | 0.795735 | Ankrd55 |
| ENSMUSG( | 1349.568 | -0.05158 | 0.705659 | 0.96315 | Ankrd6 |
| ENSMUSG( | 9.437839 | -0.96224 | 0.159134 | NA | Ankrd61 |
| ENSMUSG( | 487.516 | -0.0414 | 0.833789 | 0.981218 | Ankrd63 |
| ENSMUSG( | 20.4063 | -0.32187 | 0.471173 | 0.923329 | Ankrd66 |
| ENSMUSG( | 246.4628 | 0.085943 | 0.645759 | 0.953988 | Ankrd9 |
| ENSMUSG( | 1498.18 | 0.167243 | 0.175304 | 0.845305 | Anks1 |
| ENSMUSG( | 8162.05 | -0.01215 | 0.898746 | 0.990558 | Anks1b |
| ENSMUSG( | 1061.306 | 0.043532 | 0.750229 | 0.970113 | Anks3 |
| ENSMUSG( | 467.4513 | 0.016161 | 0.912855 | 0.992298 | Anks6 |
| ENSMUSG( | 48.88776 | 0.392328 | 0.263912 | 0.886134 | Ankub1 |
| ENSMUSG( | 740.0105 | 0.057294 | 0.736828 | 0.967344 | Ankzf1 |
| ENSMUSG( | 741.6499 | 0.125968 | 0.333121 | 0.907907 | Anln |
| ENSMUSG( | 108.62 | 0.544318 | 0.050672 | 0.709454 | Ano1 |
| ENSMUSG( | 862.1299 | 0.164356 | 0.322507 | 0.904023 | Ano10 |
| ENSMUSG( | 118.8308 | 0.6991 | 0.00966 | 0.406458 | Ano2 |
| ENSMUSG( | 2777.152 | 0.057946 | 0.727883 | 0.966887 | Ano3 |
| ENSMUSG( | 567.3914 | 0.147755 | 0.340635 | 0.907972 | Ano4 |
| ENSMUSG( | 28.38883 | -0.13674 | 0.735262 | 0.967344 | Ano5 |
| ENSMUSG( | 926.5655 | 0.253463 | 0.088492 | 0.786756 | Ano6 |
| ENSMUSG( | 10.16133 | 0.713971 | 0.263708 | NA | Ano7 |
| ENSMUSG( | 2647.961 | 0.150669 | 0.175701 | 0.845305 | Ano8 |
| ENSMUSG( | 3828.874 | -0.15423 | 0.28703 | 0.894091 | Anp32a |
| ENSMUSG( | 2147.562 | 0.025116 | 0.865641 | 0.987041 | Anp32b |
| ENSMUSG( | 4135.533 | -0.09276 | 0.543777 | 0.938063 | Anp32e |
| ENSMUSG( | 352.0266 | 0.58809 | 0.001466 | 0.140378 | Anpep |
| ENSMUSG( | 1406.303 | -0.05391 | 0.642136 | 0.953903 | Antkmt |
| ENSMUSG( | 1358.339 | -0.10855 | 0.356615 | 0.913358 | Antxr1 |
| ENSMUSG( | 103.2156 | 0.188791 | 0.398231 | 0.917257 | Antxr2 |
| ENSMUSG( | 97.21988 | -0.04256 | 0.878923 | 0.989644 | Anxa1 |
| ENSMUSG( | 695.919 | 0.048272 | 0.7708 | 0.974236 | Anxa11 |
| ENSMUSG( | 504.4138 | 0.17942 | 0.253452 | 0.88018 | Anxa2 |
| ENSMUSG( | 585.856 | 0.376636 | 0.013037 | 0.458202 | Anxa3 |
| ENSMUSG( | 155.4874 | 0.352436 | 0.055019 | 0.723362 | Anxa4 |
| ENSMUSG( | 2509.88 | -0.20624 | 0.08497 | 0.779799 | Anxa5 |
| ENSMUSG( | 3614.468 | -0.2076 | 0.047504 | 0.701031 | Anxa6 |
| ENSMUSG( | 1341.587 | 0.095961 | 0.43369 | 0.918564 | Anxa7 |
| ENSMUSG( | 24.31525 | 0.625006 | 0.141757 | 0.830709 | Anxa8 |
| ENSMUSG( | 38.18597 | 0.448267 | 0.191163 | 0.858412 | Anxa9 |
| ENSMUSG( | 16.34055 | -0.21747 | 0.645449 | 0.953988 | Aoah |
| ENSMUSG( | 99.12798 | -0.07922 | 0.725229 | 0.966887 | Aoc2 |
| ENSMUSG( | 68.9079 | 0.083172 | 0.759501 | 0.972031 | Aoc3 |
| ENSMUSG( | 877.6626 | 0.225092 | 0.147506 | 0.836241 | Aopep |
| ENSMUSG( | 402.2376 | 0.039762 | 0.789068 | 0.976593 | Aox1 |
| ENSMUSG( | 85.41095 | 0.179426 | 0.46529 | 0.923329 | Aox3 |
| ENSMUSG( | 36.35416 | 0.458389 | 0.182542 | 0.850579 | Aox4 |
| ENSMUSG( | 2995.89 | -0.01764 | 0.881086 | 0.989644 | Ap1ar |
| ENSMUSG( | 5903.804 | 0.05785 | 0.598908 | 0.947045 | Ap1b1 |
| ENSMUSG( | 2282.782 | 0.019737 | 0.860868 | 0.986226 | Ap1g1 |
| ENSMUSG( | 237.0424 | -0.05998 | 0.835353 | 0.981237 | Ap1g2 |
| ENSMUSG( | 1373.783 | -0.07266 | 0.626892 | 0.953234 | Ap1m1 |
| ENSMUSG( | 18.9363 | -0.38225 | 0.399948 | 0.917257 | Ap1m2 |
| ENSMUSG( | 6158.149 | -0.2968 | 0.012699 | 0.455854 | Ap1s1 |
| ENSMUSG( | 630.4046 | 0.05511 | 0.757363 | 0.971858 | Ap1s2 |
| ENSMUSG( | 34.16363 | 0.080059 | 0.830747 | 0.98051 | Ap1s3 |
| ENSMUSG( | 7811.275 | -0.07955 | 0.515467 | 0.930939 | Ap2a1 |
| ENSMUSG( | 5335.884 | -0.10262 | 0.346566 | 0.911108 | Ap2a2 |
| ENSMUSG( | 8215.767 | -0.14219 | 0.165182 | 0.838743 | Ap2b1 |

| | | | | | | | | | | | |
|---|---|---|---|---|---|---|---|---|---|---|---|
| ENSMUSG( | Amdhd2 | 504.8616 | 0.001482 | 0.991514 | 0.999232 | ENSMUSG( | Amdhd2 | 446.1972 | -0.08733 | 0.565226 | 0.94131 |
| ENSMUSG( | Amer1 | 221.8823 | -0.15015 | 0.323572 | 0.768536 | ENSMUSG( | Amer1 | 226.6195 | 0.175004 | 0.286159 | 0.893906 |
| ENSMUSG( | Amer2 | 4084.967 | -0.24063 | 0.043707 | 0.320345 | ENSMUSG( | Amer2 | 3732.704 | -0.2341 | 0.114164 | 0.811037 |
| ENSMUSG( | Amer3 | 514.2843 | 0.074238 | 0.638506 | 0.916319 | ENSMUSG( | Amer3 | 453.9898 | -0.00902 | 0.956174 | 0.994798 |
| ENSMUSG( | Amfr | 9894.297 | -0.12843 | 0.260709 | 0.710468 | ENSMUSG( | Amfr | 9566.597 | 0.047761 | 0.715522 | 0.965104 |
| ENSMUSG( | Amh | 32.22257 | 0.488256 | 0.161508 | 0.59148 | ENSMUSG( | Amh | 28.85787 | 0.43564 | 0.261895 | 0.884926 |
| ENSMUSG( | Amhr2 | 44.12747 | 0.100777 | 0.730865 | 0.944956 | ENSMUSG( | Amhr2 | 37.49265 | -0.10244 | 0.754151 | 0.970715 |
| ENSMUSG( | Amigo1 | 3407.515 | -0.19469 | 0.174827 | 0.610304 | ENSMUSG( | Amigo1 | 3493.17 | 0.152776 | 0.352641 | 0.912093 |
| ENSMUSG( | Amigo2 | 452.6997 | 0.093772 | 0.470506 | 0.84674 | ENSMUSG( | Amigo2 | 484.4988 | 0.51495 | 0.000504 | 0.067381 |
| ENSMUSG( | Amigo3 | 55.19027 | -0.14045 | 0.592591 | 0.902094 | ENSMUSG( | Amigo3 | 47.61043 | -0.29139 | 0.324783 | 0.905257 |
| ENSMUSG( | Ammecr1 | 119.7014 | -0.25886 | 0.160117 | 0.589846 | ENSMUSG( | Ammecr1 | 106.5471 | -0.34972 | 0.106944 | 0.804969 |
| ENSMUSG( | Ammecr1l | 938.0911 | -0.05594 | 0.608055 | 0.907154 | ENSMUSG( | Ammecr1l | 863.6824 | -0.02284 | 0.850468 | 0.985541 |
| ENSMUSG( | Amn | 301.2149 | 0.221856 | 0.10911 | 0.498553 | ENSMUSG( | Amn | 293.4278 | 0.399947 | 0.009711 | 0.406458 |
| ENSMUSG( | Amn1 | 360.4731 | -0.33214 | 0.010925 | 0.150598 | ENSMUSG( | Amn1 | 388.2211 | 0.162909 | 0.279387 | 0.892356 |
| ENSMUSG( | Amot | 4218.253 | 0.22349 | 0.141612 | 0.558606 | ENSMUSG( | Amot | 3347.094 | -0.17522 | 0.301009 | 0.898538 |
| ENSMUSG( | Amotl1 | 2272.901 | 0.082856 | 0.500042 | 0.863455 | ENSMUSG( | Amotl1 | 1903.427 | -0.16548 | 0.161387 | 0.837724 |
| ENSMUSG( | Amotl2 | 630.6307 | -0.1254 | 0.357473 | 0.788872 | ENSMUSG( | Amotl2 | 673.9052 | 0.327926 | 0.087098 | 0.782809 |
| ENSMUSG( | Ampd2 | 1887.683 | 0.101428 | 0.262372 | 0.713005 | ENSMUSG( | Ampd2 | 1673.176 | 0.024405 | 0.851917 | 0.985598 |
| ENSMUSG( | Ampd3 | 1399.894 | 0.00392 | 0.969864 | 0.996369 | ENSMUSG( | Ampd3 | 1222.776 | -0.12288 | 0.284427 | 0.892412 |
| ENSMUSG( | Amph | 4289.791 | 0.219367 | 0.028982 | 0.259144 | ENSMUSG( | Amph | 3576.818 | -0.02936 | 0.801529 | 0.97681 |
| ENSMUSG( | Amt | 1617.323 | 0.06818 | 0.482078 | 0.855778 | ENSMUSG( | Amt | 1336.92 | -0.22378 | 0.065622 | 0.74546 |
| ENSMUSG( | Amy1 | 496.9374 | 0.211749 | 0.364577 | 0.793796 | ENSMUSG( | Amy1 | 454.371 | 0.217297 | 0.619893 | 0.952896 |
| ENSMUSG( | Amz1 | 402.2282 | -0.12079 | 0.333908 | 0.775602 | ENSMUSG( | Amz1 | 392.2212 | 0.078144 | 0.579765 | 0.944209 |
| ENSMUSG( | Amz2 | 1395.235 | 0.118204 | 0.191948 | 0.630811 | ENSMUSG( | Amz2 | 1186.051 | -0.08171 | 0.469688 | 0.923329 |
| ENSMUSG( | Anapc1 | 2394.491 | -0.0003 | 0.997691 | 0.999848 | ENSMUSG( | Anapc1 | 2262.717 | 0.106514 | 0.429605 | 0.917257 |
| ENSMUSG( | Anapc10 | 244.9993 | -0.04979 | 0.733451 | 0.944956 | ENSMUSG( | Anapc10 | 217.4255 | -0.1429 | 0.428723 | 0.917257 |
| ENSMUSG( | Anapc11 | 2146.533 | -0.03227 | 0.728236 | 0.944355 | ENSMUSG( | Anapc11 | 1787.328 | -0.31185 | 0.009263 | 0.402642 |
| ENSMUSG( | Anapc13 | 988.8642 | -0.01563 | 0.885886 | 0.983008 | ENSMUSG( | Anapc13 | 814.781 | -0.32944 | 0.010757 | 0.421367 |
| ENSMUSG( | Anapc15 | 881.0147 | 0.059076 | 0.586201 | 0.899709 | ENSMUSG( | Anapc15 | 765.8095 | -0.07954 | 0.516982 | 0.930939 |
| ENSMUSG( | Anapc16 | 2013.372 | 0.064987 | 0.50149 | 0.864477 | ENSMUSG( | Anapc16 | 1694.265 | -0.17215 | 0.188285 | 0.85698 |
| ENSMUSG( | Anapc2 | 2788.673 | 0.083356 | 0.316896 | 0.763754 | ENSMUSG( | Anapc2 | 2402.722 | -0.07898 | 0.462692 | 0.923329 |
| ENSMUSG( | Anapc4 | 1479.193 | 0.149545 | 0.13253 | 0.544627 | ENSMUSG( | Anapc4 | 1292.228 | 0.028196 | 0.825458 | 0.979445 |
| ENSMUSG( | Anapc5 | 7645.66 | 0.126337 | 0.139607 | 0.555959 | ENSMUSG( | Anapc5 | 6654.523 | -0.00322 | 0.975985 | 0.996416 |
| ENSMUSG( | Anapc7 | 859.5383 | -0.05069 | 0.61943 | 0.911432 | ENSMUSG( | Anapc7 | 771.7387 | -0.09158 | 0.443033 | 0.921554 |
| ENSMUSG( | Ang | 102.6516 | 0.094874 | 0.657494 | 0.92425 | ENSMUSG( | Ang | 105.2108 | 0.414213 | 0.06169 | 0.736228 |
| ENSMUSG( | Angel1 | 437.9975 | -0.44156 | 0.006729 | 0.11299 | ENSMUSG( | Angel1 | 441.0757 | -0.11135 | 0.549221 | 0.938514 |
| ENSMUSG( | Angel2 | 2139.559 | 0.038822 | 0.656609 | 0.923929 | ENSMUSG( | Angel2 | 1978.841 | 0.080849 | 0.442174 | 0.921554 |
| ENSMUSG( | Angpt1 | 330.5529 | 0.106359 | 0.58418 | 0.898738 | ENSMUSG( | Angpt1 | 280.0938 | -0.1081 | 0.61142 | 0.950771 |
| ENSMUSG( | Angpt2 | 129.3183 | 0.040143 | 0.839831 | 0.971179 | ENSMUSG( | Angpt2 | 103.9883 | -0.31691 | 0.209283 | 0.864412 |
| ENSMUSG( | Angptl1 | 28.26469 | -0.09834 | 0.79028 | 0.959461 | ENSMUSG( | Angptl1 | 31.16115 | 0.455498 | 0.257471 | 0.883401 |
| ENSMUSG( | Angptl2 | 200.9817 | -0.14587 | 0.461044 | 0.84109 | ENSMUSG( | Angptl2 | 268.8376 | 0.855235 | 0.000304 | 0.047107 |
| ENSMUSG( | Angptl3 | 12.65588 | 0.271611 | 0.59341 | NA | ENSMUSG( | Angptl3 | 9.124513 | -0.52326 | 0.444441 | NA |
| ENSMUSG( | Angptl4 | 191.2143 | -0.18164 | 0.255558 | 0.706195 | ENSMUSG( | Angptl4 | 153.5417 | -0.58663 | 0.00293 | 0.217424 |
| ENSMUSG( | Angptl6 | 221.5052 | -0.09012 | 0.592625 | 0.902094 | ENSMUSG( | Angptl6 | 186.2775 | -0.34565 | 0.073866 | 0.765627 |
| ENSMUSG( | Angptl7 | 12.4651 | 0.572899 | 0.313855 | NA | ENSMUSG( | Angptl7 | 14.74956 | 1.175964 | 0.035917 | 0.668395 |
| ENSMUSG( | Ank | 9505.403 | -0.18502 | 0.108167 | 0.497094 | ENSMUSG( | Ank | 9529.206 | 0.097715 | 0.476956 | 0.923329 |
| ENSMUSG( | Ank1 | 2241.53 | 0.100039 | 0.237806 | 0.685809 | ENSMUSG( | Ank1 | 1877.49 | -0.14262 | 0.261987 | 0.884926 |
| ENSMUSG( | Ank2 | 36616.85 | 0.037954 | 0.752573 | 0.950536 | ENSMUSG( | Ank2 | 34249.06 | 0.11429 | 0.391797 | 0.917257 |
| ENSMUSG( | Ank3 | 16751.03 | -0.27586 | 0.052366 | 0.35132 | ENSMUSG( | Ank3 | 17238.92 | 0.091737 | 0.559532 | 0.939807 |
| ENSMUSG( | Ankdd1a | 15.37761 | -0.1293 | 0.794964 | NA | ENSMUSG( | Ankdd1a | 15.40491 | 0.165657 | 0.753407 | 0.970671 |
| ENSMUSG( | Ankef1 | 14.15283 | 0.112663 | 0.819196 | NA | ENSMUSG( | Ankef1 | 13.52938 | 0.203602 | 0.703619 | 0.962481 |
| ENSMUSG( | Ankfn1 | 45.1639 | -0.16703 | 0.595982 | 0.903152 | ENSMUSG( | Ankfn1 | 43.66401 | 0.037222 | 0.91612 | 0.992529 |
| ENSMUSG( | Ankfy1 | 2033.62 | 0.070887 | 0.426269 | 0.824898 | ENSMUSG( | Ankfy1 | 1758.294 | -0.08014 | 0.445935 | 0.921734 |
| ENSMUSG( | Ankhd1 | 806.6844 | -0.05371 | 0.654983 | 0.92381 | ENSMUSG( | Ankhd1 | 770.6895 | 0.088785 | 0.547882 | 0.938063 |
| ENSMUSG( | Ankib1 | 2068.232 | 0.032899 | 0.749045 | 0.949604 | ENSMUSG( | Ankib1 | 1935.268 | 0.10922 | 0.360882 | 0.914223 |
| ENSMUSG( | Ankle1 | 12.37411 | 0.416485 | 0.485882 | NA | ENSMUSG( | Ankle1 | 10.96331 | 0.344409 | 0.588415 | NA |
| ENSMUSG( | Ankle2 | 1186.092 | -0.20023 | 0.164514 | 0.595363 | ENSMUSG( | Ankle2 | 1266.359 | 0.258179 | 0.118174 | 0.813384 |
| ENSMUSG( | Ankmy1 | 23.07367 | -0.06282 | 0.871875 | 0.978266 | ENSMUSG( | Ankmy1 | 21.07973 | -0.02105 | 0.962356 | 0.995237 |
| ENSMUSG( | Ankmy2 | 985.9789 | -0.02746 | 0.783888 | 0.957645 | ENSMUSG( | Ankmy2 | 847.114 | -0.2053 | 0.087704 | 0.785431 |
| ENSMUSG( | Ankra2 | 990.6892 | 0.093346 | 0.320043 | 0.765523 | ENSMUSG( | Ankra2 | 877.1084 | 0.008847 | 0.940948 | 0.994558 |
| ENSMUSG( | Ankrd10 | 1342.662 | 0.061427 | 0.566982 | 0.891644 | ENSMUSG( | Ankrd10 | 1303.041 | 0.237023 | 0.049289 | 0.708275 |
| ENSMUSG( | Ankrd11 | 9454.6 | -0.02686 | 0.775103 | 0.955606 | ENSMUSG( | Ankrd11 | 8993.985 | 0.0982 | 0.392664 | 0.917257 |
| ENSMUSG( | Ankrd12 | 1451.847 | -0.07486 | 0.457461 | 0.839558 | ENSMUSG( | Ankrd12 | 1435.351 | 0.157219 | 0.165135 | 0.838743 |
| ENSMUSG( | Ankrd13a | 2010.197 | 0.022831 | 0.787421 | 0.959147 | ENSMUSG( | Ankrd13a | 1889.913 | 0.111741 | 0.28861 | 0.894794 |
| ENSMUSG( | Ankrd13b | 4843.403 | 0.245313 | 0.003817 | 0.079179 | ENSMUSG( | Ankrd13b | 4305.566 | 0.179256 | 0.113542 | 0.809623 |
| ENSMUSG( | Ankrd13c | 3588.793 | 0.029699 | 0.786129 | 0.958774 | ENSMUSG( | Ankrd13c | 3153.732 | -0.07824 | 0.577025 | 0.942912 |
| ENSMUSG( | Ankrd13d | 1483.675 | 0.190132 | 0.192025 | 0.630811 | ENSMUSG( | Ankrd13d | 1231.669 | -0.07508 | 0.63226 | 0.953452 |
| ENSMUSG( | Ankrd16 | 825.5051 | -0.12761 | 0.281243 | 0.730583 | ENSMUSG( | Ankrd16 | 798.0158 | 0.051247 | 0.760625 | 0.972688 |
| ENSMUSG( | Ankrd17 | 3069.475 | 0.126871 | 0.107274 | 0.495579 | ENSMUSG( | Ankrd17 | 2748.392 | 0.078342 | 0.456015 | 0.923329 |
| ENSMUSG( | Ankrd23 | 24.54449 | -0.4916 | 0.228421 | 0.675849 | ENSMUSG( | Ankrd23 | 26.9662 | 0.134363 | 0.767333 | 0.974236 |
| ENSMUSG( | Ankrd24 | 1054.309 | 0.311916 | 0.078661 | 0.433892 | ENSMUSG( | Ankrd24 | 867.3367 | 0.029345 | 0.885588 | 0.98977 |
| ENSMUSG( | Ankrd26 | 614.4775 | 0.171238 | 0.132972 | 0.545614 | ENSMUSG( | Ankrd26 | 562.4623 | 0.177898 | 0.199737 | 0.863016 |
| ENSMUSG( | Ankrd27 | 1643.831 | 0.196694 | 0.027469 | 0.254456 | ENSMUSG( | Ankrd27 | 1412.709 | 0.033527 | 0.777769 | 0.976084 |
| ENSMUSG( | Ankrd28 | 2300.574 | -0.06617 | 0.492258 | 0.860297 | ENSMUSG( | Ankrd28 | 2133.269 | -0.01222 | 0.922245 | 0.992644 |
| ENSMUSG( | Ankrd29 | 982.5277 | 0.076386 | 0.431559 | 0.827432 | ENSMUSG( | Ankrd29 | 852.8746 | -0.06999 | 0.656358 | 0.955533 |
| ENSMUSG( | Ankrd31 | 13.28665 | 0.309508 | 0.53679 | NA | ENSMUSG( | Ankrd31 | 9.175977 | -0.53411 | 0.401975 | NA |
| ENSMUSG( | Ankrd33b | 1684.731 | 0.082265 | 0.48749 | 0.858812 | ENSMUSG( | Ankrd33b | 1564.533 | 0.137016 | 0.280859 | 0.892356 |
| ENSMUSG( | Ankrd34a | 1280.453 | 0.093373 | 0.539034 | 0.882877 | ENSMUSG( | Ankrd34a | 1110.804 | -0.05 | 0.763392 | 0.973951 |
| ENSMUSG( | Ankrd34b | 217.8747 | 0.126219 | 0.474503 | 0.849425 | ENSMUSG( | Ankrd34b | 198.917 | 0.130101 | 0.506138 | 0.927825 |
| ENSMUSG( | Ankrd34c | 118.876 | 0.300763 | 0.18885 | 0.627678 | ENSMUSG( | Ankrd34c | 96.31108 | -0.02654 | 0.920864 | 0.992644 |
| ENSMUSG( | Ankrd35 | 425.4321 | -0.02385 | 0.861594 | 0.975026 | ENSMUSG( | Ankrd35 | 353.1926 | -0.31692 | 0.041423 | 0.688699 |
| ENSMUSG( | Ankrd37 | 90.90947 | -0.23037 | 0.293066 | 0.742763 | ENSMUSG( | Ankrd37 | 91.59668 | 0.06092 | 0.786067 | 0.976285 |
| ENSMUSG( | Ankrd39 | 278.8746 | 0.017516 | 0.899493 | 0.985633 | ENSMUSG( | Ankrd39 | 265.1936 | 0.135163 | 0.373366 | 0.915491 |
| ENSMUSG( | Ankrd40 | 15275.43 | -0.00629 | 0.946277 | 0.99369 | ENSMUSG( | Ankrd40 | 13018.38 | -0.20818 | 0.099295 | 0.800296 |
| ENSMUSG( | Ankrd42 | 258.9999 | 0.079497 | 0.579087 | 0.896692 | ENSMUSG( | Ankrd42 | 237.7077 | 0.097679 | 0.542364 | 0.938034 |
| ENSMUSG( | Ankrd44 | 416.8205 | -0.11002 | 0.424674 | 0.824675 | ENSMUSG( | Ankrd44 | 399.6478 | 0.04352 | 0.780313 | 0.976106 |
| ENSMUSG( | Ankrd45 | 1225.627 | -0.03147 | 0.749452 | 0.949664 | ENSMUSG( | Ankrd45 | 1093.084 | -0.09973 | 0.453378 | 0.922316 |
| ENSMUSG( | Ankrd46 | 4361.687 | 0.127952 | 0.207392 | 0.649561 | ENSMUSG( | Ankrd46 | 3706.923 | -0.07293 | 0.556932 | 0.939565 |
| ENSMUSG( | Ankrd49 | 265.2726 | 0.043132 | 0.764841 | 0.954127 | ENSMUSG( | Ankrd49 | 243.8034 | 0.058165 | 0.744011 | 0.96951 |
| ENSMUSG( | Ankrd50 | 1509.997 | 0.156817 | 0.110207 | 0.500785 | ENSMUSG( | Ankrd50 | 1359.711 | 0.127531 | 0.295334 | 0.898538 |
| ENSMUSG( | Ankrd52 | 1814.977 | 0.214539 | 0.044731 | 0.324745 | ENSMUSG( | Ankrd52 | 1552.8 | 0.044012 | 0.742664 | 0.969415 |
| ENSMUSG( | Ankrd54 | 506.4571 | 0.044184 | 0.728143 | 0.944355 | ENSMUSG( | Ankrd54 | 482.0976 | 0.170737 | 0.222405 | 0.866791 |
| ENSMUSG( | Ankrd55 | 54.05335 | -0.11063 | 0.689756 | 0.933609 | ENSMUSG( | Ankrd55 | 43.10531 | -0.54153 | 0.093975 | 0.795735 |
| ENSMUSG( | Ankrd6 | 1448.788 | -0.11327 | 0.307541 | 0.755462 | ENSMUSG( | Ankrd6 | 1349.568 | -0.05158 | 0.705659 | 0.96315 |
| ENSMUSG( | Ankrd61 | 12.79722 | -0.23369 | 0.658233 | NA | ENSMUSG( | Ankrd61 | 9.437839 | -0.96224 | 0.159134 | NA |
| ENSMUSG( | Ankrd63 | 502.5407 | -0.22696 | 0.193525 | 0.633027 | ENSMUSG( | Ankrd63 | 487.516 | -0.0414 | 0.833789 | 0.981218 |
| ENSMUSG( | Ankrd66 | 23.10994 | -0.22264 | 0.573819 | 0.893584 | ENSMUSG( | Ankrd66 | 20.4063 | -0.32187 | 0.471173 | 0.923329 |
| ENSMUSG( | Ankrd9 | 252.9269 | -0.10737 | 0.55752 | 0.888279 | ENSMUSG( | Ankrd9 | 246.4628 | 0.085943 | 0.645759 | 0.953988 |
| ENSMUSG( | Anks1 | 1655.81 | 0.183271 | 0.051198 | 0.347172 | ENSMUSG( | Anks1 | 1498.18 | 0.167243 | 0.175304 | 0.845305 |
| ENSMUSG( | Anks1b | 9123.799 | 0.040709 | 0.617899 | 0.910872 | ENSMUSG( | Anks1b | 8162.05 | -0.01215 | 0.898746 | 0.990558 |
| ENSMUSG( | Anks3 | 1178.124 | 0.07344 | 0.422568 | 0.823508 | ENSMUSG( | Anks3 | 1061.306 | 0.043532 | 0.750229 | 0.970113 |
| ENSMUSG( | Anks6 | 513.7227 | 0.015664 | 0.912025 | 0.988483 | ENSMUSG( | Anks6 | 467.4513 | 0.016161 | 0.912855 | 0.992298 |
| ENSMUSG( | Ankub1 | 42.25443 | -0.32453 | 0.298484 | 0.74713 | ENSMUSG( | Ankub1 | 48.88776 | 0.392328 | 0.263912 | 0.886134 |
| ENSMUSG( | Ankzf1 | 832.2366 | 0.126097 | 0.306178 | 0.753712 | ENSMUSG( | Ankzf1 | 740.0105 | 0.057294 | 0.736828 | 0.967344 |
| ENSMUSG( | Anln | 758.974 | -0.07721 | 0.43963 | 0.83129 | ENSMUSG( | Anln | 741.6499 | 0.125968 | 0.333121 | 0.907907 |
| ENSMUSG( | Ano1 | 90.28403 | -0.23617 | 0.374407 | 0.800445 | ENSMUSG( | Ano1 | 108.62 | 0.544318 | 0.050672 | 0.709454 |
| ENSMUSG( | Ano10 | 848.83 | -0.15555 | 0.281818 | 0.731842 | ENSMUSG( | Ano10 | 862.1299 | 0.164356 | 0.322507 | 0.904023 |
| ENSMUSG( | Ano2 | 80.12672 | -0.69976 | 0.010006 | 0.142653 | ENSMUSG( | Ano2 | 118.8308 | 0.6991 | 0.00966 | 0.406458 |
| ENSMUSG( | Ano3 | 2306.737 | -0.88127 | 5.12E-10 | 2.52E-07 | ENSMUSG( | Ano3 | 2777.152 | 0.057946 | 0.727883 | 0.966887 |
| ENSMUSG( | Ano4 | 518.543 | -0.4142 | 0.00156 | 0.044321 | ENSMUSG( | Ano4 | 567.3914 | 0.147755 | 0.340635 | 0.907972 |
| ENSMUSG( | Ano5 | 31.70647 | -0.09396 | 0.80878 | 0.963455 | ENSMUSG( | Ano5 | 28.38883 | -0.13674 | 0.735262 | 0.967344 |
| ENSMUSG( | Ano6 | 947.3334 | 0.053729 | 0.693395 | 0.934202 | ENSMUSG( | Ano6 | 926.5655 | 0.253463 | 0.088492 | 0.786756 |
| ENSMUSG( | Ano7 | 8.714241 | 0.099472 | 0.885807 | NA | ENSMUSG( | Ano7 | 10.16133 | 0.713971 | 0.263708 | NA |
| ENSMUSG( | Ano8 | 2656.921 | -0.10666 | 0.309345 | 0.75737 | ENSMUSG( | Ano8 | 2647.961 | 0.150669 | 0.175701 | 0.845305 |
| ENSMUSG( | Anp32a | 4587.976 | 0.104119 | 0.271505 | 0.722082 | ENSMUSG( | Anp32a | 3828.874 | -0.15423 | 0.28703 | 0.894091 |
| ENSMUSG( | Anp32b | 2371.29 | 0.045545 | 0.621868 | 0.911432 | ENSMUSG( | Anp32b | 2147.562 | 0.025116 | 0.865641 | 0.987041 |
| ENSMUSG( | Anp32e | 4710.675 | 0.018621 | 0.861414 | 0.975026 | ENSMUSG( | Anp32e | 4135.533 | -0.09276 | 0.543777 | 0.938063 |
| ENSMUSG( | Anpep | 322.0514 | 0.107117 | 0.455911 | 0.838611 | ENSMUSG( | Anpep | 352.0266 | 0.58809 | 0.001466 | 0.140378 |
| ENSMUSG( | Antkmt | 1573.623 | 0.005117 | 0.956228 | 0.993874 | ENSMUSG( | Antkmt | 1406.303 | -0.05391 | 0.642136 | 0.953903 |
| ENSMUSG( | Antxr1 | 1506.304 | -0.07811 | 0.500844 | 0.863671 | ENSMUSG( | Antxr1 | 1358.339 | -0.10855 | 0.356615 | 0.913358 |
| ENSMUSG( | Antxr2 | 102.0805 | -0.12146 | 0.578998 | 0.896692 | ENSMUSG( | Antxr2 | 103.2156 | 0.188791 | 0.398231 | 0.917257 |
| ENSMUSG( | Anxa1 | 105.6158 | -0.05003 | 0.825433 | 0.967868 | ENSMUSG( | Anxa1 | 97.21988 | -0.04256 | 0.878923 | 0.989644 |
| ENSMUSG( | Anxa11 | 770.0136 | 0.070266 | 0.651286 | 0.921451 | ENSMUSG( | Anxa11 | 695.919 | 0.048272 | 0.7708 | 0.974236 |
| ENSMUSG( | Anxa2 | 518.9985 | -0.00946 | 0.94016 | 0.993415 | ENSMUSG( | Anxa2 | 504.4138 | 0.17942 | 0.253452 | 0.88018 |
| ENSMUSG( | Anxa3 | 585.4998 | 0.120598 | 0.315231 | 0.762288 | ENSMUSG( | Anxa3 | 585.856 | 0.376636 | 0.013037 | 0.458202 |
| ENSMUSG( | Anxa4 | 160.4106 | 0.180287 | 0.27514 | 0.72509 | ENSMUSG( | Anxa4 | 155.4874 | 0.352436 | 0.055019 | 0.723362 |
| ENSMUSG( | Anxa5 | 2977.424 | 0.026253 | 0.823666 | 0.966603 | ENSMUSG( | Anxa5 | 2509.88 | -0.20624 | 0.08497 | 0.779799 |
| ENSMUSG( | Anxa6 | 4186.227 | -0.04389 | 0.642915 | 0.916865 | ENSMUSG( | Anxa6 | 3614.468 | -0.2076 | 0.047504 | 0.701031 |
| ENSMUSG( | Anxa7 | 1465.944 | 0.083779 | 0.426644 | 0.825155 | ENSMUSG( | Anxa7 | 1341.587 | 0.095961 | 0.43369 | 0.918564 |
| ENSMUSG( | Anxa8 | 18.11178 | -0.50249 | 0.252055 | NA | ENSMUSG( | Anxa8 | 24.31525 | 0.625006 | 0.141757 | 0.830709 |
| ENSMUSG( | Anxa9 | 40.93526 | 0.397295 | 0.233429 | 0.681508 | ENSMUSG( | Anxa9 | 38.18597 | 0.448267 | 0.191163 | 0.858412 |
| ENSMUSG( | Aoah | 17.84049 | -0.23394 | 0.592803 | NA | ENSMUSG( | Aoah | 16.34055 | -0.21747 | 0.645449 | 0.953988 |
| ENSMUSG( | Aoc2 | 115.9853 | 0.087518 | 0.681828 | 0.931619 | ENSMUSG( | Aoc2 | 99.12798 | -0.07922 | 0.725229 | 0.966887 |
| ENSMUSG( | Aoc3 | 62.70768 | -0.51927 | 0.062269 | 0.386085 | ENSMUSG( | Aoc3 | 68.9079 | 0.083172 | 0.759501 | 0.972031 |
| ENSMUSG( | Aopep | 924.7392 | 0.109256 | 0.394978 | 0.810135 | ENSMUSG( | Aopep | 877.6626 | 0.225092 | 0.147506 | 0.836241 |
| ENSMUSG( | Aox1 | 413.233 | -0.16094 | 0.181138 | 0.618139 | ENSMUSG( | Aox1 | 402.2376 | 0.039762 | 0.789068 | 0.976593 |
| ENSMUSG( | Aox3 | 87.4804 | -0.00439 | 0.983826 | 0.99779 | ENSMUSG( | Aox3 | 85.41095 | 0.179426 | 0.46529 | 0.923329 |
| ENSMUSG( | Aox4 | 37.83271 | 0.336581 | 0.362347 | 0.792943 | ENSMUSG( | Aox4 | 36.35416 | 0.458389 | 0.182542 | 0.850579 |
| ENSMUSG( | Ap1ar | 3414.234 | 0.092078 | 0.276797 | 0.726951 | ENSMUSG( | Ap1ar | 2995.89 | -0.01764 | 0.881086 | 0.989644 |
| ENSMUSG( | Ap1b1 | 6743.051 | 0.16832 | 0.091088 | 0.463586 | ENSMUSG( | Ap1b1 | 5903.804 | 0.05785 | 0.598908 | 0.947045 |
| ENSMUSG( | Ap1g1 | 2652.178 | 0.180707 | 0.05701 | 0.368322 | ENSMUSG( | Ap1g1 | 2282.782 | 0.019737 | 0.860868 | 0.986226 |
| ENSMUSG( | Ap1g2 | 289.3977 | 0.24428 | 0.337816 | 0.777422 | ENSMUSG( | Ap1g2 | 237.0424 | -0.05998 | 0.835353 | 0.981237 |
| ENSMUSG( | Ap1m1 | 1604.523 | 0.106655 | 0.39363 | 0.809643 | ENSMUSG( | Ap1m1 | 1373.783 | -0.07266 | 0.626892 | 0.953234 |
| ENSMUSG( | Ap1m2 | 22.97846 | -0.06138 | 0.872849 | 0.978753 | ENSMUSG( | Ap1m2 | 18.9363 | -0.38225 | 0.399948 | 0.917257 |
| ENSMUSG( | Ap1s1 | 7209.545 | -0.09523 | 0.23991 | 0.687848 | ENSMUSG( | Ap1s1 | 6158.149 | -0.2968 | 0.012699 | 0.455854 |
| ENSMUSG( | Ap1s2 | 641.2126 | -0.15915 | 0.252468 | 0.702649 | ENSMUSG( | Ap1s2 | 630.4046 | 0.05511 | 0.757363 | 0.971858 |
| ENSMUSG( | Ap1s3 | 36.45648 | 0.043095 | 0.889416 | 0.98353 | ENSMUSG( | Ap1s3 | 34.16363 | 0.080059 | 0.830747 | 0.98051 |
| ENSMUSG( | Ap2a1 | 9191.293 | 0.121373 | 0.236933 | 0.684303 | ENSMUSG( | Ap2a1 | 7811.275 | -0.07955 | 0.515467 | 0.930939 |
| ENSMUSG( | Ap2a2 | 6483.729 | 0.186948 | 0.049572 | 0.340062 | ENSMUSG( | Ap2a2 | 5335.884 | -0.10262 | 0.346566 | 0.911108 |
| ENSMUSG( | Ap2b1 | 10010.8 | 0.158703 | 0.059447 | 0.376652 | ENSMUSG( | Ap2b1 | 8215.767 | -0.14219 | 0.165182 | 0.838743 |
| ENSMUSG( | Ap2m1 | 10223.53 | 0.133386 | 0.242146 | 0.689699 | ENSMUSG( | Ap2m1 | 8492.475 | -0.13563 | 0.341831 | 0.909146 |
| ENSMUSG( | Ap2s1 | 4148.582 | 0.006408 | 0.958374 | 0.993874 | ENSMUSG( | Ap2s1 | 3476.174 | -0.24965 | 0.110781 | 0.807582 |
| ENSMUSG( | Ap3b1 | 1447.812 | 0.207297 | 0.052319 | 0.35132 | ENSMUSG( | Ap3b1 | 1227.909 | 0.004164 | 0.974889 | 0.996416 |
| ENSMUSG( | Ap3b2 | 3173.942 | 0.257558 | 0.029342 | 0.259647 | ENSMUSG( | Ap3b2 | 2576.505 | -0.06735 | 0.628034 | 0.953234 |
| ENSMUSG( | Ap3d1 | 5954.325 | 0.258795 | 0.005576 | 0.099928 | ENSMUSG( | Ap3d1 | 4827.339 | -0.06854 | 0.553514 | 0.939565 |
| ENSMUSG( | Ap3m1 | 1680.752 | 0.002692 | 0.977078 | 0.997428 | ENSMUSG( | Ap3m1 | 1519.861 | -0.02067 | 0.854786 | 0.985649 |
| ENSMUSG( | Ap3m2 | 3384.445 | 0.076667 | 0.433508 | 0.828346 | ENSMUSG( | Ap3m2 | 2918.151 | -0.0842 | 0.468567 | 0.923329 |
| ENSMUSG( | Ap3s1 | 1795.812 | 0.020332 | 0.86237 | 0.975055 | ENSMUSG( | Ap3s1 | 1608.365 | -0.03482 | 0.830097 | 0.980224 |

| | | | | | |
|---|---|---|---|---|---|
| ENSMUSG( | 6743.051 | 0.16832 | 0.091088 | 0.463586 | Ap1b1 |
| ENSMUSG( | 2652.178 | 0.180707 | 0.05701 | 0.368322 | Ap1g1 |
| ENSMUSG( | 289.3977 | 0.24428 | 0.337816 | 0.777422 | Ap1g2 |
| ENSMUSG( | 1604.523 | 0.106655 | 0.39363 | 0.809643 | Ap1m1 |
| ENSMUSG( | 22.97846 | -0.06138 | 0.872849 | 0.978753 | Ap1m2 |
| ENSMUSG( | 7209.545 | -0.09523 | 0.23991 | 0.687848 | Ap1s1 |
| ENSMUSG( | 641.2126 | -0.15915 | 0.252468 | 0.702649 | Ap1s2 |
| ENSMUSG( | 36.45648 | 0.043095 | 0.889416 | 0.98353 | Ap1s3 |
| ENSMUSG( | 9191.293 | 0.121373 | 0.236933 | 0.684303 | Ap2a1 |
| ENSMUSG( | 6483.729 | 0.186948 | 0.049572 | 0.340062 | Ap2a2 |
| ENSMUSG( | 10010.8 | 0.158703 | 0.059447 | 0.376652 | Ap2b1 |
| ENSMUSG( | 10223.53 | 0.133386 | 0.242146 | 0.689699 | Ap2m1 |
| ENSMUSG( | 4148.582 | 0.006408 | 0.958374 | 0.993874 | Ap2s1 |
| ENSMUSG( | 1447.812 | 0.207297 | 0.052319 | 0.35132 | Ap3b1 |
| ENSMUSG( | 3173.942 | 0.257558 | 0.029342 | 0.259647 | Ap3b2 |
| ENSMUSG( | 5954.325 | 0.258795 | 0.005576 | 0.099928 | Ap3d1 |
| ENSMUSG( | 1680.752 | 0.002692 | 0.977078 | 0.997428 | Ap3m1 |
| ENSMUSG( | 3384.445 | 0.076667 | 0.433508 | 0.828346 | Ap3m2 |
| ENSMUSG( | 1795.812 | 0.020332 | 0.86237 | 0.975055 | Ap3s1 |
| ENSMUSG( | 3178.293 | 0.082925 | 0.295181 | 0.743798 | Ap3s2 |
| ENSMUSG( | 69.31028 | 0.248975 | 0.313282 | 0.76033 | Ap4b1 |
| ENSMUSG( | 368.5337 | -0.00654 | 0.9611 | 0.994009 | Ap4e1 |
| ENSMUSG( | 764.5979 | 0.131305 | 0.233553 | 0.681508 | Ap4m1 |
| ENSMUSG( | 1095.358 | 0.055566 | 0.584799 | 0.898923 | Ap4s1 |
| ENSMUSG( | 70.52815 | 0.111948 | 0.640128 | 0.916434 | Ap5b1 |
| ENSMUSG( | 434.1468 | 0.046629 | 0.700963 | 0.93743 | Ap5m1 |
| ENSMUSG( | 393.0522 | -0.12956 | 0.283405 | 0.73186 | Ap5s1 |
| ENSMUSG( | 811.6986 | 0.106853 | 0.340016 | 0.777959 | Ap5z1 |
| ENSMUSG( | 347.5928 | 0.335233 | 0.018244 | 0.20352 | Apaf1 |
| ENSMUSG( | 10845.88 | 0.189484 | 0.029686 | 0.261211 | Apba1 |
| ENSMUSG( | 15435.05 | -0.22362 | 0.233648 | 0.681508 | Apba2 |
| ENSMUSG( | 872.2373 | 0.187659 | 0.058136 | 0.371733 | Apba3 |
| ENSMUSG( | 14225.44 | 0.191575 | 0.034746 | 0.286661 | Apbb1 |
| ENSMUSG( | 311.8282 | -0.07755 | 0.631247 | 0.914073 | Apbb1ip |
| ENSMUSG( | 869.0755 | -0.09645 | 0.330095 | 0.774055 | Apbb2 |
| ENSMUSG( | 888.4038 | -0.06191 | 0.608983 | 0.907377 | Apbb3 |
| ENSMUSG( | 74805.14 | 0.076581 | 0.48945 | 0.859092 | Apc |
| ENSMUSG( | 6510.915 | 0.016919 | 0.843149 | 0.971847 | Apc2 |
| ENSMUSG( | 2327.503 | 0.111127 | 0.483681 | 0.85636 | Apcdd1 |
| ENSMUSG( | 541.8325 | 0.045498 | 0.680152 | 0.930983 | Apeh |
| ENSMUSG( | 582.6408 | 0.034307 | 0.774634 | 0.955606 | Apex1 |
| ENSMUSG( | 137.3925 | 0.242315 | 0.201561 | 0.641362 | Apex2 |
| ENSMUSG( | 1112.489 | -0.23575 | 0.034056 | 0.283276 | Aph1a |
| ENSMUSG( | 1013.927 | -0.1941 | 0.182765 | 0.620307 | Aph1b |
| ENSMUSG( | 232.6637 | -0.36556 | 0.029476 | 0.260274 | Aph1c |
| ENSMUSG( | 1990.679 | 0.058061 | 0.515958 | 0.872701 | Api5 |
| ENSMUSG( | 323.9112 | 0.018136 | 0.889513 | 0.98353 | Apip |
| ENSMUSG( | 181.8846 | 0.097404 | 0.537898 | 0.882548 | Aplf |
| ENSMUSG( | 4059.6 | -0.05401 | 0.707348 | 0.93949 | Apln |
| ENSMUSG( | 56.79774 | -0.49724 | 0.140984 | 0.557902 | Aplnr |
| ENSMUSG( | 65065.95 | -0.3542 | 0.001716 | 0.046884 | Aplp1 |
| ENSMUSG( | 23983.04 | -0.21872 | 0.075738 | 0.426666 | Aplp2 |
| ENSMUSG( | 1753.506 | -0.30358 | 0.028809 | 0.258592 | Apmap |
| ENSMUSG( | 9.83239 | 0.422057 | 0.469743 | NA | Apoa1 |
| ENSMUSG( | 37.84375 | -0.01035 | 0.973757 | 0.99675 | Apoa2 |
| ENSMUSG( | 127.6655 | -0.22813 | 0.207735 | 0.64962 | Apobec1 |
| ENSMUSG( | 172.8904 | -0.16248 | 0.297092 | 0.745805 | Apobec3 |
| ENSMUSG( | 93.70331 | -0.15226 | 0.505714 | 0.866932 | Apobr |
| ENSMUSG( | 70.00903 | 0.36862 | 0.169419 | 0.603554 | Apoc1 |
| ENSMUSG( | 42.83764 | 0.151888 | 0.616746 | 0.910866 | Apoc3 |
| ENSMUSG( | 5580.729 | -0.34565 | 0.004234 | 0.084545 | Apod |
| ENSMUSG( | 173435.2 | 0.018098 | 0.85038 | 0.972638 | Apoe |
| ENSMUSG( | 11.0228 | 0.087835 | 0.882232 | NA | Apoh |
| ENSMUSG( | 14.38931 | -0.3089 | 0.571202 | NA | Apol11b |
| ENSMUSG( | 16.11078 | 0.23102 | 0.6667 | NA | Apol7a |
| ENSMUSG( | 72.31612 | -0.00621 | 0.979631 | 0.997428 | Apol8 |
| ENSMUSG( | 140.5494 | -0.56256 | 0.008457 | 0.128676 | Apold1 |
| ENSMUSG( | 774.8368 | -0.05195 | 0.680976 | 0.931244 | Apoo |
| ENSMUSG( | 96.23116 | -0.0613 | 0.78722 | 0.959147 | Apool |
| ENSMUSG( | 48628.28 | -0.47878 | 0.000184 | 0.009888 | App |
| ENSMUSG( | 2812.099 | 0.003832 | 0.967823 | 0.996237 | Appbp2 |
| ENSMUSG( | 10.63586 | 0.853328 | 0.181131 | NA | Appbp2os |
| ENSMUSG( | 2098.114 | 0.064779 | 0.531161 | 0.879305 | Appl1 |
| ENSMUSG( | 6330.283 | 0.084864 | 0.278372 | 0.728374 | Appl2 |
| ENSMUSG( | 616.5274 | -0.08215 | 0.491482 | 0.860053 | Aprt |
| ENSMUSG( | 956.0281 | 0.010903 | 0.907946 | 0.9878 | Aptx |
| ENSMUSG( | 113.8999 | -0.14636 | 0.840482 | 0.971179 | Aqp1 |
| ENSMUSG( | 1053.606 | 0.022873 | 0.85044 | 0.972638 | Aqp11 |
| ENSMUSG( | 9.474804 | 0.458045 | 0.443437 | NA | Aqp2 |
| ENSMUSG( | 10985.49 | -0.19353 | 0.257565 | 0.708686 | Aqp4 |
| ENSMUSG( | 15.50281 | -0.10356 | 0.83445 | NA | Aqp5 |
| ENSMUSG( | 17.02559 | -0.78051 | 0.137982 | NA | Aqp6 |
| ENSMUSG( | 233.2137 | 0.065566 | 0.714335 | 0.941045 | Aqp9 |
| ENSMUSG( | 1188.361 | 0.156488 | 0.102791 | 0.488221 | Aqr |
| ENSMUSG( | 365.3475 | 0.073215 | 0.640564 | 0.916434 | Ar |
| ENSMUSG( | 3514.759 | -0.08496 | 0.279554 | 0.729114 | Araf |
| ENSMUSG( | 1169.531 | -0.09304 | 0.346341 | 0.779153 | Arap1 |
| ENSMUSG( | 3326.911 | -0.00267 | 0.977959 | 0.997428 | Arap2 |
| ENSMUSG( | 540.7238 | 0.365719 | 0.007395 | 0.118765 | Arap3 |
| ENSMUSG( | 2272.723 | 0.061079 | 0.609091 | 0.907455 | Arb2a |
| ENSMUSG( | 7430.516 | -1.16974 | 3.02E-16 | 3.50E-13 | Arc |
| ENSMUSG( | 3121.741 | 0.13013 | 0.137935 | 0.552903 | Arcn1 |
| ENSMUSG( | 3898.286 | -0.11009 | 0.333355 | 0.775602 | Arel1 |
| ENSMUSG( | 9882.402 | -0.0116 | 0.905408 | 0.986465 | Arf1 |
| ENSMUSG( | 1129.313 | -0.02594 | 0.802369 | 0.962119 | Arf2 |
| ENSMUSG( | 51782.73 | 0.044914 | 0.562978 | 0.889738 | Arf3 |
| ENSMUSG( | 2494.905 | -0.0742 | 0.494157 | 0.860611 | Arf4 |
| ENSMUSG( | 3862.153 | -0.04786 | 0.658029 | 0.92425 | Arf5 |
| ENSMUSG( | 902.5488 | 0.046818 | 0.657191 | 0.924142 | Arf6 |
| ENSMUSG( | 2178.705 | -0.08934 | 0.396719 | 0.811742 | Arfgap1 |
| ENSMUSG( | 1756.777 | 0.052277 | 0.569316 | 0.893041 | Arfgap2 |
| ENSMUSG( | 460.2811 | -0.00596 | 0.960843 | 0.993991 | Arfgap3 |
| ENSMUSG( | 2372.469 | 0.157717 | 0.068277 | 0.405144 | Arfgef1 |
| ENSMUSG( | 2008.8 | 0.033206 | 0.705053 | 0.938303 | Arfgef2 |
| ENSMUSG( | 1830.697 | -0.07373 | 0.482297 | 0.855884 | Arfgef3 |
| ENSMUSG( | 151.305 | 0.126751 | 0.540707 | 0.88315 | Arfip1 |
| ENSMUSG( | 2785.973 | 0.153199 | 0.10834 | 0.497337 | Arfip2 |
| ENSMUSG( | 894.389 | 0.00797 | 0.937039 | 0.993415 | Arfrp1 |
| ENSMUSG( | 44.42652 | -0.46543 | 0.135601 | 0.549038 | Arg2 |
| ENSMUSG( | 2884.401 | -0.13471 | 0.289541 | 0.739839 | Arglu1 |
| ENSMUSG( | 2175.091 | 0.124889 | 0.18528 | 0.623185 | Arhgap1 |
| ENSMUSG( | 626.515 | 0.073351 | 0.581786 | 0.897862 | Arhgap10 |
| ENSMUSG( | 328.0915 | 0.084362 | 0.525451 | 0.876728 | Arhgap11a |
| ENSMUSG( | 2097.418 | 0.162942 | 0.171554 | 0.606099 | Arhgap12 |
| ENSMUSG( | 136.2941 | 0.132452 | 0.492116 | 0.860297 | Arhgap15 |
| ENSMUSG( | 625.7393 | -0.08337 | 0.493512 | 0.860611 | Arhgap17 |
| ENSMUSG( | 501.3161 | -0.09691 | 0.437888 | 0.830072 | Arhgap18 |
| ENSMUSG( | 135.0343 | -0.213 | 0.221342 | 0.667193 | Arhgap19 |
| ENSMUSG( | 1407.967 | -0.14808 | 0.103761 | 0.489428 | Arhgap20 |
| ENSMUSG( | 6904.072 | 0.087879 | 0.306077 | 0.753712 | Arhgap21 |
| ENSMUSG( | 226.3544 | -0.02232 | 0.899291 | 0.985633 | Arhgap22 |
| ENSMUSG( | 6716.117 | -0.04928 | 0.552827 | 0.886141 | Arhgap23 |
| ENSMUSG( | 363.183 | -0.07395 | 0.592097 | 0.901851 | Arhgap24 |
| ENSMUSG( | 232.3727 | 0.269219 | 0.085596 | 0.451059 | Arhgap25 |
| ENSMUSG( | 2528.321 | 0.221556 | 0.013363 | 0.16868 | Arhgap26 |
| ENSMUSG( | 679.2899 | -0.05697 | 0.631772 | 0.914073 | Arhgap27 |
| ENSMUSG( | 25.66316 | -0.25709 | 0.489928 | 0.859319 | Arhgap28 |
| ENSMUSG( | 711.7408 | 0.141176 | 0.372475 | 0.798768 | Arhgap29 |
| ENSMUSG( | 330.0185 | 0.238891 | 0.101014 | 0.484746 | Arhgap30 |
| ENSMUSG( | 1508.005 | 0.189269 | 0.083427 | 0.445566 | Arhgap31 |
| ENSMUSG( | 7073.198 | 0.051993 | 0.520271 | 0.875186 | Arhgap32 |
| ENSMUSG( | 13319.43 | 0.17296 | 0.080521 | 0.438901 | Arhgap33 |
| ENSMUSG( | 7262.825 | 0.203547 | 0.015861 | 0.186421 | Arhgap35 |
| ENSMUSG( | 51.40395 | 0.44818 | 0.30072 | 0.74995 | Arhgap36 |
| ENSMUSG( | 2351.567 | 0.394031 | 0.000868 | 0.029574 | Arhgap39 |
| ENSMUSG( | 248.5967 | -0.06555 | 0.746365 | 0.949289 | Arhgap4 |
| ENSMUSG( | 9.237093 | 0.440177 | 0.489287 | NA | Arhgap40 |
| ENSMUSG( | 584.7651 | 0.068439 | 0.522229 | 0.875186 | Arhgap42 |
| ENSMUSG( | 5150.749 | 0.164583 | 0.068382 | 0.405144 | Arhgap44 |
| ENSMUSG( | 539.226 | -0.11076 | 0.372081 | 0.798768 | Arhgap45 |
| ENSMUSG( | 6545.325 | 0.052974 | 0.736164 | 0.946267 | Arhgap5 |
| ENSMUSG( | 157.158 | -0.02718 | 0.899895 | 0.985633 | Arhgap6 |
| ENSMUSG( | 59.4204 | -0.02102 | 0.933615 | 0.993194 | Arhgap9 |
| ENSMUSG( | 14963.9 | 0.078456 | 0.282376 | 0.73186 | Arhgdia |
| ENSMUSG( | 489.9738 | 0.06773 | 0.57981 | 0.89687 | Arhgdib |
| ENSMUSG( | 2587.425 | -0.07514 | 0.415679 | 0.82084 | Arhgdig |
| ENSMUSG( | 2249.606 | 0.310919 | 0.067715 | 0.40256 | Arhgef1 |
| ENSMUSG( | 2504.654 | 0.03715 | 0.660527 | 0.924795 | Arhgef10 |
| ENSMUSG( | 2976.108 | 0.06975 | 0.442579 | 0.83278 | Arhgef10l |
| ENSMUSG( | 3247.462 | 0.142932 | 0.124678 | 0.531714 | Arhgef11 |
| ENSMUSG( | 13854.99 | 0.025511 | 0.811757 | 0.964426 | Arhgef12 |
| ENSMUSG( | 558.4659 | -0.01464 | 0.898181 | 0.985633 | Arhgef15 |
| ENSMUSG( | 21.24606 | 0.050632 | 0.918088 | 0.989645 | Arhgef16 |
| ENSMUSG( | 7117.948 | 0.172794 | 0.0806 | 0.438981 | Arhgef17 |
| ENSMUSG( | 1564.615 | 0.150291 | 0.096319 | 0.476986 | Arhgef18 |
| ENSMUSG( | 1388.625 | 0.057855 | 0.568466 | 0.892252 | Arhgef19 |
| ENSMUSG( | 4320.833 | -0.05798 | 0.515951 | 0.872701 | Arhgef2 |
| ENSMUSG( | 5382.054 | 0.300915 | 0.00795 | 0.124138 | Arhgef25 |

| | | | | | |
|---|---|---|---|---|---|
| ENSMUSG( | 8492.475 | -0.13563 | 0.341831 | 0.909146 | Ap2m1 |
| ENSMUSG( | 3476.174 | -0.24965 | 0.110781 | 0.807582 | Ap2s1 |
| ENSMUSG( | 1227.909 | 0.004164 | 0.974889 | 0.996416 | Ap3b1 |
| ENSMUSG( | 2576.505 | -0.06735 | 0.628034 | 0.953234 | Ap3b2 |
| ENSMUSG( | 4827.339 | -0.06854 | 0.553514 | 0.939565 | Ap3d1 |
| ENSMUSG( | 1519.861 | -0.02067 | 0.854786 | 0.985649 | Ap3m1 |
| ENSMUSG( | 2918.151 | -0.0842 | 0.468567 | 0.923329 | Ap3m2 |
| ENSMUSG( | 1608.365 | -0.03482 | 0.830097 | 0.980224 | Ap3s1 |
| ENSMUSG( | 2815.658 | 0.002399 | 0.981118 | 0.996756 | Ap3s2 |
| ENSMUSG( | 52.64144 | -0.29378 | 0.329793 | 0.907083 | Ap4b1 |
| ENSMUSG( | 343.9417 | 0.069896 | 0.685961 | 0.958956 | Ap4e1 |
| ENSMUSG( | 665.7702 | 0.00576 | 0.967301 | 0.995606 | Ap4m1 |
| ENSMUSG( | 941.64 | -0.11907 | 0.398498 | 0.917257 | Ap4s1 |
| ENSMUSG( | 69.49371 | 0.346744 | 0.246409 | 0.877904 | Ap5b1 |
| ENSMUSG( | 431.1737 | 0.284796 | 0.039622 | 0.68111 | Ap5m1 |
| ENSMUSG( | 341.9728 | -0.26566 | 0.067364 | 0.75212 | Ap5s1 |
| ENSMUSG( | 743.7728 | 0.125702 | 0.400811 | 0.917257 | Ap5z1 |
| ENSMUSG( | 294.2541 | 0.138775 | 0.352555 | 0.912093 | Apaf1 |
| ENSMUSG( | 9293.84 | 0.020033 | 0.850736 | 0.985541 | Apba1 |
| ENSMUSG( | 15449.13 | 0.060953 | 0.725875 | 0.966887 | Apba2 |
| ENSMUSG( | 786.1053 | 0.160494 | 0.206439 | 0.864412 | Apba3 |
| ENSMUSG( | 12086.99 | -0.00322 | 0.973991 | 0.996416 | Apbb1 |
| ENSMUSG( | 289.9382 | -0.02425 | 0.878968 | 0.989644 | Apbb1ip |
| ENSMUSG( | 807.3333 | -0.03483 | 0.7824 | 0.976106 | Apbb2 |
| ENSMUSG( | 859.0915 | 0.110669 | 0.474215 | 0.923329 | Apbb3 |
| ENSMUSG( | 64608.71 | -0.07829 | 0.539833 | 0.937378 | Apc |
| ENSMUSG( | 5855.112 | -0.01896 | 0.866961 | 0.987487 | Apc2 |
| ENSMUSG( | 2711.328 | 0.736391 | 0.000262 | 0.04337 | Apcdd1 |
| ENSMUSG( | 483.0726 | -0.01298 | 0.927332 | 0.99306 | Apeh |
| ENSMUSG( | 506.8326 | -0.10848 | 0.493088 | 0.927358 | Apex1 |
| ENSMUSG( | 118.0838 | 0.090702 | 0.658644 | 0.956205 | Apex2 |
| ENSMUSG( | 1173.532 | 0.193144 | 0.161753 | 0.837724 | Aph1a |
| ENSMUSG( | 1030.834 | 0.129352 | 0.421067 | 0.917257 | Aph1b |
| ENSMUSG( | 251.8418 | 0.158192 | 0.373354 | 0.915491 | Aph1c |
| ENSMUSG( | 1766.55 | -0.02118 | 0.868736 | 0.988097 | Api5 |
| ENSMUSG( | 280.212 | -0.13443 | 0.393686 | 0.917257 | Apip |
| ENSMUSG( | 172.3364 | 0.210102 | 0.238904 | 0.873304 | Aplf |
| ENSMUSG( | 3266.257 | -0.44712 | 0.004762 | 0.284671 | Apln |
| ENSMUSG( | 54.1964 | -0.3027 | 0.423 | 0.917257 | Aplnr |
| ENSMUSG( | 65827.33 | -0.02812 | 0.832436 | 0.98075 | Aplp1 |
| ENSMUSG( | 24133.06 | 0.07781 | 0.576476 | 0.942912 | Aplp2 |
| ENSMUSG( | 1884.658 | 0.188198 | 0.232091 | 0.868507 | Apmap |
| ENSMUSG( | 35.02765 | 0.022862 | 0.948001 | 0.994798 | Apoa2 |
| ENSMUSG( | 127.2332 | 0.027699 | 0.893194 | 0.990558 | Apobec1 |
| ENSMUSG( | 158.26 | -0.13017 | 0.522902 | 0.932848 | Apobec3 |
| ENSMUSG( | 84.40631 | -0.19822 | 0.431087 | 0.917257 | Apobr |
| ENSMUSG( | 65.61307 | 0.427159 | 0.128571 | 0.81925 | Apoc1 |
| ENSMUSG( | 29.02652 | -0.80775 | 0.030671 | 0.628144 | Apoc3 |
| ENSMUSG( | 6104.075 | 0.197996 | 0.13313 | 0.822912 | Apod |
| ENSMUSG( | 158798.3 | 0.032444 | 0.777742 | 0.976084 | Apoe |
| ENSMUSG( | 9.279901 | -0.15074 | 0.813775 | NA | Apoh |
| ENSMUSG( | 14.38149 | -0.00703 | 0.989815 | 0.997928 | Apol11b |
| ENSMUSG( | 12.49158 | -0.16151 | 0.788384 | 0.97637 | Apol7a |
| ENSMUSG( | 72.04269 | 0.262195 | 0.328594 | 0.906727 | Apol8 |
| ENSMUSG( | 143.9188 | -0.17049 | 0.451524 | 0.922202 | Apold1 |
| ENSMUSG( | 655.433 | -0.28758 | 0.061947 | 0.736482 | Apoo |
| ENSMUSG( | 85.21084 | -0.14558 | 0.525967 | 0.934374 | Apool |
| ENSMUSG( | 54846.98 | 0.174357 | 0.259564 | 0.883971 | App |
| ENSMUSG( | 2490.206 | -0.08278 | 0.528703 | 0.934795 | Appbp2 |
| ENSMUSG( | 1864.798 | -0.00809 | 0.945999 | 0.994798 | Appl1 |
| ENSMUSG( | 5634.131 | 0.018422 | 0.869187 | 0.988097 | Appl2 |
| ENSMUSG( | 558.9057 | -0.10282 | 0.472723 | 0.923329 | Aprt |
| ENSMUSG( | 817.2149 | -0.18247 | 0.149096 | 0.836241 | Aptx |
| ENSMUSG( | 119.7891 | 0.246691 | 0.690346 | 0.960071 | Aqp1 |
| ENSMUSG( | 957.3558 | 0.018229 | 0.897233 | 0.990558 | Aqp11 |
| ENSMUSG( | 10114.88 | -0.15997 | 0.337976 | 0.907972 | Aqp4 |
| ENSMUSG( | 14.40706 | -0.06669 | 0.895252 | 0.990558 | Aqp5 |
| ENSMUSG( | 16.71296 | -0.46456 | 0.354092 | 0.912411 | Aqp6 |
| ENSMUSG( | 182.2689 | -0.39291 | 0.034299 | 0.652355 | Aqp9 |
| ENSMUSG( | 1032.848 | 0.025449 | 0.819018 | 0.979198 | Aqr |
| ENSMUSG( | 319.2164 | -0.04145 | 0.801445 | 0.97681 | Ar |
| ENSMUSG( | 3255.701 | -0.0363 | 0.725454 | 0.966887 | Araf |
| ENSMUSG( | 1052.496 | -0.12682 | 0.274726 | 0.889874 | Arap1 |
| ENSMUSG( | 3019.677 | -0.01516 | 0.901132 | 0.990558 | Arap2 |
| ENSMUSG( | 548.9139 | 0.639291 | 0.000177 | 0.032278 | Arap3 |
| ENSMUSG( | 1973.613 | -0.08156 | 0.581204 | 0.944266 | Arb2a |
| ENSMUSG( | 7027.272 | -0.99766 | 0.096056 | 0.796133 | Arc |
| ENSMUSG( | 2683.012 | -0.03714 | 0.738627 | 0.968018 | Arcn1 |
| ENSMUSG( | 3846.4 | 0.12331 | 0.38601 | 0.917257 | Arel1 |
| ENSMUSG( | 8792.662 | -0.08328 | 0.515167 | 0.930939 | Arf1 |
| ENSMUSG( | 999.1264 | -0.1152 | 0.343397 | 0.909753 | Arf2 |
| ENSMUSG( | 45702.97 | -0.04622 | 0.6586 | 0.956205 | Arf3 |
| ENSMUSG( | 2281.106 | -0.06784 | 0.651585 | 0.954492 | Arf4 |
| ENSMUSG( | 3444.938 | -0.11365 | 0.423608 | 0.917257 | Arf5 |
| ENSMUSG( | 826.2419 | 0.056719 | 0.668797 | 0.956766 | Arf6 |
| ENSMUSG( | 1978.709 | -0.09646 | 0.399559 | 0.917257 | Arfgap1 |
| ENSMUSG( | 1571.816 | 0.003357 | 0.97634 | 0.996416 | Arfgap2 |
| ENSMUSG( | 393.4276 | -0.19199 | 0.186125 | 0.855305 | Arfgap3 |
| ENSMUSG( | 2174.719 | 0.1736 | 0.118486 | 0.813384 | Arfgef1 |
| ENSMUSG( | 1827.312 | 0.029815 | 0.786166 | 0.976285 | Arfgef2 |
| ENSMUSG( | 1770.963 | 0.100092 | 0.365284 | 0.914449 | Arfgef3 |
| ENSMUSG( | 132.9229 | 0.012442 | 0.959126 | 0.994798 | Arfip1 |
| ENSMUSG( | 2440.106 | 0.044919 | 0.688921 | 0.959742 | Arfip2 |
| ENSMUSG( | 789.1544 | -0.08546 | 0.492537 | 0.927315 | Arfrp1 |
| ENSMUSG( | 39.30619 | -0.5996 | 0.086393 | 0.781091 | Arg2 |
| ENSMUSG( | 2953.903 | 0.197271 | 0.213519 | 0.864421 | Arglu1 |
| ENSMUSG( | 1830.434 | -0.10352 | 0.38265 | 0.917257 | Arhgap1 |
| ENSMUSG( | 533.58 | -0.12431 | 0.418077 | 0.917257 | Arhgap10 |
| ENSMUSG( | 296.8981 | 0.057774 | 0.753846 | 0.970671 | Arhgap11a |
| ENSMUSG( | 1750.019 | -0.09034 | 0.510171 | 0.929089 | Arhgap12 |
| ENSMUSG( | 108.1796 | -0.28339 | 0.206212 | 0.864412 | Arhgap15 |
| ENSMUSG( | 584.4282 | -0.00637 | 0.961425 | 0.99511 | Arhgap17 |
| ENSMUSG( | 458.1103 | -0.08981 | 0.520725 | 0.932121 | Arhgap18 |
| ENSMUSG( | 123.8552 | -0.18207 | 0.374117 | 0.915491 | Arhgap19 |
| ENSMUSG( | 1349.148 | 0.001684 | 0.987554 | 0.997519 | Arhgap20 |
| ENSMUSG( | 6662.839 | 0.246468 | 0.02688 | 0.603041 | Arhgap21 |
| ENSMUSG( | 207.1931 | -0.00587 | 0.974842 | 0.996416 | Arhgap22 |
| ENSMUSG( | 6039.199 | -0.0871 | 0.450803 | 0.922202 | Arhgap23 |
| ENSMUSG( | 314.2603 | -0.23324 | 0.112633 | 0.809623 | Arhgap24 |
| ENSMUSG( | 177.0378 | -0.23023 | 0.214205 | 0.864421 | Arhgap25 |
| ENSMUSG( | 2192.245 | 0.084234 | 0.442311 | 0.921554 | Arhgap26 |
| ENSMUSG( | 671.5862 | 0.179345 | 0.209927 | 0.864412 | Arhgap27 |
| ENSMUSG( | 23.45353 | -0.26787 | 0.508164 | 0.929089 | Arhgap28 |
| ENSMUSG( | 696.8424 | 0.338799 | 0.057124 | 0.723362 | Arhgap29 |
| ENSMUSG( | 277.6932 | 0.020363 | 0.899688 | 0.990558 | Arhgap30 |
| ENSMUSG( | 1404.557 | 0.252317 | 0.056659 | 0.723362 | Arhgap31 |
| ENSMUSG( | 6692.774 | 0.157755 | 0.108134 | 0.804969 | Arhgap32 |
| ENSMUSG( | 11737.56 | 0.081258 | 0.431214 | 0.917257 | Arhgap33 |
| ENSMUSG( | 6434.593 | 0.128736 | 0.215895 | 0.865167 | Arhgap35 |
| ENSMUSG( | 36.56677 | -0.28959 | 0.495639 | 0.927774 | Arhgap36 |
| ENSMUSG( | 2020.135 | 0.243643 | 0.080798 | 0.77681 | Arhgap39 |
| ENSMUSG( | 248.181 | 0.194147 | 0.370019 | 0.915491 | Arhgap4 |
| ENSMUSG( | 542.3111 | 0.116512 | 0.364153 | 0.914223 | Arhgap42 |
| ENSMUSG( | 4490.251 | 0.041483 | 0.694221 | 0.960867 | Arhgap44 |
| ENSMUSG( | 546.723 | 0.194579 | 0.199632 | 0.863016 | Arhgap45 |
| ENSMUSG( | 5692.614 | -0.08383 | 0.652054 | 0.9547 | Arhgap5 |
| ENSMUSG( | 157.9353 | 0.226958 | 0.371125 | 0.915491 | Arhgap6 |
| ENSMUSG( | 58.49338 | 0.203849 | 0.474535 | 0.923329 | Arhgap9 |
| ENSMUSG( | 12582.98 | -0.15677 | 0.118272 | 0.813384 | Arhgdia |
| ENSMUSG( | 421.9361 | -0.09869 | 0.478171 | 0.923329 | Arhgdib |
| ENSMUSG( | 2285.049 | -0.16932 | 0.132515 | 0.822912 | Arhgdig |
| ENSMUSG( | 1904.375 | 0.109272 | 0.572146 | 0.942475 | Arhgef1 |
| ENSMUSG( | 2205.125 | -0.06331 | 0.557743 | 0.939565 | Arhgef10 |
| ENSMUSG( | 2700.343 | 0.059222 | 0.590398 | 0.944975 | Arhgef10l |
| ENSMUSG( | 2849.179 | 0.038005 | 0.725134 | 0.966887 | Arhgef11 |
| ENSMUSG( | 11449.04 | -0.26872 | 0.019874 | 0.558472 | Arhgef12 |
| ENSMUSG( | 559.1709 | 0.253693 | 0.087492 | 0.784337 | Arhgef15 |
| ENSMUSG( | 23.62908 | 0.523609 | 0.307277 | 0.898538 | Arhgef16 |
| ENSMUSG( | 6507.103 | 0.183402 | 0.130217 | 0.821488 | Arhgef17 |
| ENSMUSG( | 1427.841 | 0.154372 | 0.159279 | 0.837724 | Arhgef18 |
| ENSMUSG( | 1207.362 | -0.07831 | 0.526062 | 0.934374 | Arhgef19 |
| ENSMUSG( | 4044.839 | 0.020476 | 0.844824 | 0.983831 | Arhgef2 |
| ENSMUSG( | 4476.138 | 0.051418 | 0.676074 | 0.95694 | Arhgef25 |
| ENSMUSG( | 3079.158 | 0.046693 | 0.762569 | 0.973539 | Arhgef26 |
| ENSMUSG( | 1098.846 | 0.129508 | 0.279082 | 0.892257 | Arhgef28 |
| ENSMUSG( | 1065.622 | 0.078735 | 0.550014 | 0.938699 | Arhgef3 |
| ENSMUSG( | 59.66613 | 0.191905 | 0.474132 | 0.923329 | Arhgef33 |
| ENSMUSG( | 172.9717 | 0.158399 | 0.375736 | 0.915491 | Arhgef37 |
| ENSMUSG( | 20.86715 | 0.213971 | 0.637438 | 0.953903 | Arhgef39 |
| ENSMUSG( | 9158.824 | 0.075805 | 0.478329 | 0.923329 | Arhgef4 |
| ENSMUSG( | 1201.306 | 0.067015 | 0.575329 | 0.942912 | Arhgef40 |
| ENSMUSG( | 183.6884 | 0.300634 | 0.150956 | 0.837173 | Arhgef5 |
| ENSMUSG( | 377.5175 | -0.00705 | 0.958989 | 0.994798 | Arhgef6 |
| ENSMUSG( | 4272.509 | 0.008408 | 0.932386 | 0.99306 | Arhgef7 |
| ENSMUSG( | 9150.982 | -0.05702 | 0.563767 | 0.941206 | Arhgef9 |
| ENSMUSG( | 3554.431 | 0.086501 | 0.464096 | 0.923329 | Arid1a |
| ENSMUSG( | 2524.457 | 0.138731 | 0.244319 | 0.877904 | Arid1b |
| ENSMUSG( | 1121.975 | 0.249382 | 0.030811 | 0.629149 | Arid2 |

| | | | | | | | | | | | |
|---|---|---|---|---|---|---|---|---|---|---|---|
| ENSMUSG( | Ap3s2 | 3178.293 | 0.082925 | 0.295181 | 0.743798 | ENSMUSG( | Ap3s2 | 2815.658 | 0.002399 | 0.981118 | 0.996756 |
| ENSMUSG( | Ap4b1 | 69.31028 | 0.248975 | 0.313282 | 0.76033 | ENSMUSG( | Ap4b1 | 52.64144 | -0.29378 | 0.329793 | 0.907083 |
| ENSMUSG( | Ap4e1 | 368.5337 | -0.00654 | 0.9611 | 0.994009 | ENSMUSG( | Ap4e1 | 343.9417 | 0.069896 | 0.685961 | 0.958956 |
| ENSMUSG( | Ap4m1 | 764.5979 | 0.131305 | 0.233553 | 0.681508 | ENSMUSG( | Ap4m1 | 665.7702 | 0.00576 | 0.967301 | 0.995606 |
| ENSMUSG( | Ap4s1 | 1095.358 | 0.055566 | 0.584799 | 0.898923 | ENSMUSG( | Ap4s1 | 941.64 | -0.11907 | 0.398498 | 0.917257 |
| ENSMUSG( | Ap5b1 | 70.52815 | 0.111948 | 0.640128 | 0.916434 | ENSMUSG( | Ap5b1 | 69.49371 | 0.346744 | 0.246409 | 0.877904 |
| ENSMUSG( | Ap5m1 | 434.1468 | 0.046629 | 0.700963 | 0.93743 | ENSMUSG( | Ap5m1 | 431.1737 | 0.284796 | 0.039622 | 0.68111 |
| ENSMUSG( | Ap5s1 | 393.0522 | -0.12956 | 0.283405 | 0.73186 | ENSMUSG( | Ap5s1 | 341.9728 | -0.26566 | 0.067364 | 0.75212 |
| ENSMUSG( | Ap5z1 | 811.6986 | 0.106853 | 0.340016 | 0.777959 | ENSMUSG( | Ap5z1 | 743.7728 | 0.125702 | 0.400811 | 0.917257 |
| ENSMUSG( | Apaf1 | 347.5928 | 0.335233 | 0.018244 | 0.20352 | ENSMUSG( | Apaf1 | 294.2541 | 0.138775 | 0.352555 | 0.912093 |
| ENSMUSG( | Apba1 | 10845.88 | 0.189484 | 0.029686 | 0.261211 | ENSMUSG( | Apba1 | 9293.84 | 0.020033 | 0.850736 | 0.985541 |
| ENSMUSG( | Apba2 | 15435.05 | -0.22362 | 0.233648 | 0.681508 | ENSMUSG( | Apba2 | 15449.13 | 0.060953 | 0.725875 | 0.966887 |
| ENSMUSG( | Apba3 | 872.2373 | 0.187659 | 0.058136 | 0.371733 | ENSMUSG( | Apba3 | 786.1053 | 0.160494 | 0.206439 | 0.864412 |
| ENSMUSG( | Apbb1 | 14225.44 | 0.191575 | 0.034746 | 0.286661 | ENSMUSG( | Apbb1 | 12086.99 | -0.00322 | 0.973991 | 0.996416 |
| ENSMUSG( | Apbb1ip | 311.8282 | -0.07755 | 0.631247 | 0.914073 | ENSMUSG( | Apbb1ip | 289.9382 | -0.02425 | 0.878968 | 0.989644 |
| ENSMUSG( | Apbb2 | 869.0755 | -0.09645 | 0.330095 | 0.774055 | ENSMUSG( | Apbb2 | 807.3333 | -0.03483 | 0.7824 | 0.976106 |
| ENSMUSG( | Apbb3 | 888.4038 | -0.06191 | 0.608983 | 0.907377 | ENSMUSG( | Apbb3 | 859.0915 | 0.110669 | 0.474215 | 0.923329 |
| ENSMUSG( | Apc | 74805.14 | 0.076581 | 0.48945 | 0.859092 | ENSMUSG( | Apc | 64608.71 | -0.07829 | 0.539833 | 0.937378 |
| ENSMUSG( | Apc2 | 6510.915 | 0.016919 | 0.843149 | 0.971847 | ENSMUSG( | Apc2 | 5855.112 | -0.01896 | 0.866961 | 0.987487 |
| ENSMUSG( | Apcdd1 | 2327.503 | 0.111127 | 0.483681 | 0.85636 | ENSMUSG( | Apcdd1 | 2711.328 | 0.736391 | 0.000262 | 0.04337 |
| ENSMUSG( | Apeh | 541.8325 | 0.045498 | 0.680152 | 0.930983 | ENSMUSG( | Apeh | 483.0726 | -0.01298 | 0.927332 | 0.99306 |
| ENSMUSG( | Apex1 | 582.6408 | 0.034307 | 0.774634 | 0.955606 | ENSMUSG( | Apex1 | 506.8326 | -0.10848 | 0.493088 | 0.927358 |
| ENSMUSG( | Apex2 | 137.3925 | 0.242315 | 0.201561 | 0.641362 | ENSMUSG( | Apex2 | 118.0838 | 0.090702 | 0.658644 | 0.956205 |
| ENSMUSG( | Aph1a | 1112.489 | -0.23575 | 0.034056 | 0.283276 | ENSMUSG( | Aph1a | 1173.532 | 0.193144 | 0.161753 | 0.837724 |
| ENSMUSG( | Aph1b | 1013.927 | -0.1941 | 0.182765 | 0.620307 | ENSMUSG( | Aph1b | 1030.834 | 0.129352 | 0.421067 | 0.917257 |
| ENSMUSG( | Aph1c | 232.6637 | -0.36556 | 0.029476 | 0.260274 | ENSMUSG( | Aph1c | 251.8418 | 0.158192 | 0.373354 | 0.915491 |
| ENSMUSG( | Api5 | 1990.679 | 0.058061 | 0.515958 | 0.872701 | ENSMUSG( | Api5 | 1766.55 | -0.02118 | 0.868736 | 0.988097 |
| ENSMUSG( | Apip | 323.9112 | 0.018136 | 0.889513 | 0.98353 | ENSMUSG( | Apip | 280.212 | -0.13443 | 0.393686 | 0.917257 |
| ENSMUSG( | Aplf | 181.8846 | 0.097404 | 0.537898 | 0.882548 | ENSMUSG( | Aplf | 172.3364 | 0.210102 | 0.238904 | 0.873304 |
| ENSMUSG( | Apln | 4059.6 | -0.05401 | 0.707348 | 0.93949 | ENSMUSG( | Apln | 3266.257 | -0.44712 | 0.004762 | 0.284671 |
| ENSMUSG( | Aplnr | 56.79774 | -0.49724 | 0.140984 | 0.557902 | ENSMUSG( | Aplnr | 54.1964 | -0.3027 | 0.423 | 0.917257 |
| ENSMUSG( | Aplp1 | 65065.95 | -0.3542 | 0.001716 | 0.046884 | ENSMUSG( | Aplp1 | 65827.33 | -0.02812 | 0.832436 | 0.98075 |
| ENSMUSG( | Aplp2 | 23983.04 | -0.21872 | 0.075738 | 0.426666 | ENSMUSG( | Aplp2 | 24133.06 | 0.07781 | 0.576476 | 0.942912 |
| ENSMUSG( | Apmap | 1753.506 | -0.30358 | 0.028809 | 0.258592 | ENSMUSG( | Apmap | 1884.658 | 0.188198 | 0.232091 | 0.868507 |
| ENSMUSG( | Apoa2 | 37.84375 | -0.01035 | 0.973757 | 0.99675 | ENSMUSG( | Apoa2 | 35.02765 | 0.022862 | 0.948001 | 0.994798 |
| ENSMUSG( | Apobec1 | 127.6655 | -0.22813 | 0.207735 | 0.64962 | ENSMUSG( | Apobec1 | 127.2332 | 0.027699 | 0.893194 | 0.990558 |
| ENSMUSG( | Apobec3 | 172.8904 | -0.16248 | 0.297092 | 0.745805 | ENSMUSG( | Apobec3 | 158.26 | -0.13017 | 0.522902 | 0.932848 |
| ENSMUSG( | Apobr | 93.70331 | -0.15226 | 0.505714 | 0.866932 | ENSMUSG( | Apobr | 84.40631 | -0.19822 | 0.431087 | 0.917257 |
| ENSMUSG( | Apoc1 | 70.00903 | 0.36862 | 0.169419 | 0.603554 | ENSMUSG( | Apoc1 | 65.61307 | 0.427159 | 0.128571 | 0.81925 |
| ENSMUSG( | Apoc3 | 42.83764 | 0.151888 | 0.616746 | 0.910866 | ENSMUSG( | Apoc3 | 29.02652 | -0.80775 | 0.030671 | 0.628144 |
| ENSMUSG( | Apod | 5580.729 | -0.34565 | 0.004234 | 0.084545 | ENSMUSG( | Apod | 6104.075 | 0.197996 | 0.13313 | 0.822912 |
| ENSMUSG( | Apoe | 173435.2 | 0.018098 | 0.85038 | 0.972638 | ENSMUSG( | Apoe | 158798.3 | 0.032444 | 0.777742 | 0.976084 |
| ENSMUSG( | Apoh | 11.0228 | 0.087835 | 0.882232 | NA | ENSMUSG( | Apoh | 9.279901 | -0.15074 | 0.813775 | NA |
| ENSMUSG( | Apol11b | 14.38931 | -0.3089 | 0.571202 | NA | ENSMUSG( | Apol11b | 14.38149 | -0.00703 | 0.989815 | 0.997928 |
| ENSMUSG( | Apol7a | 16.11078 | 0.23102 | 0.6667 | NA | ENSMUSG( | Apol7a | 12.49158 | -0.16151 | 0.788384 | 0.97637 |
| ENSMUSG( | Apol8 | 72.31612 | -0.00621 | 0.979631 | 0.997428 | ENSMUSG( | Apol8 | 72.04269 | 0.262195 | 0.328594 | 0.906727 |
| ENSMUSG( | Apold1 | 140.5494 | -0.56256 | 0.008457 | 0.128676 | ENSMUSG( | Apold1 | 143.9188 | -0.17049 | 0.451524 | 0.922202 |
| ENSMUSG( | Apoo | 774.8368 | -0.05195 | 0.680976 | 0.931244 | ENSMUSG( | Apoo | 655.433 | -0.28758 | 0.061947 | 0.736482 |
| ENSMUSG( | Apool | 96.23116 | -0.0613 | 0.78722 | 0.959147 | ENSMUSG( | Apool | 85.21084 | -0.14558 | 0.525967 | 0.934374 |
| ENSMUSG( | App | 48628.28 | -0.47878 | 0.000184 | 0.009888 | ENSMUSG( | App | 54846.98 | 0.174357 | 0.259564 | 0.883971 |
| ENSMUSG( | Appbp2 | 2812.099 | 0.003832 | 0.967823 | 0.996237 | ENSMUSG( | Appbp2 | 2490.206 | -0.08278 | 0.528703 | 0.934795 |
| ENSMUSG( | Appl1 | 2098.114 | 0.064779 | 0.531161 | 0.879305 | ENSMUSG( | Appl1 | 1864.798 | -0.00809 | 0.945999 | 0.994798 |
| ENSMUSG( | Appl2 | 6330.283 | 0.084864 | 0.278372 | 0.728374 | ENSMUSG( | Appl2 | 5634.131 | 0.018422 | 0.869187 | 0.988097 |
| ENSMUSG( | Aprt | 616.5274 | -0.08215 | 0.491482 | 0.860053 | ENSMUSG( | Aprt | 558.9057 | -0.10282 | 0.472723 | 0.923329 |
| ENSMUSG( | Aptx | 956.0281 | 0.010903 | 0.907946 | 0.9878 | ENSMUSG( | Aptx | 817.2149 | -0.18247 | 0.149096 | 0.836241 |
| ENSMUSG( | Aqp1 | 113.8999 | -0.14636 | 0.840482 | 0.971179 | ENSMUSG( | Aqp1 | 119.7891 | 0.246691 | 0.690346 | 0.960071 |
| ENSMUSG( | Aqp11 | 1053.606 | 0.022873 | 0.85044 | 0.972638 | ENSMUSG( | Aqp11 | 957.3558 | 0.018229 | 0.897233 | 0.990558 |
| ENSMUSG( | Aqp4 | 10985.49 | -0.19353 | 0.257565 | 0.708686 | ENSMUSG( | Aqp4 | 10114.88 | -0.15997 | 0.337976 | 0.907972 |
| ENSMUSG( | Aqp5 | 15.50281 | -0.10356 | 0.83445 | NA | ENSMUSG( | Aqp5 | 14.40706 | -0.06669 | 0.895252 | 0.990558 |
| ENSMUSG( | Aqp6 | 17.02559 | -0.78051 | 0.137982 | NA | ENSMUSG( | Aqp6 | 16.71296 | -0.46456 | 0.354092 | 0.912411 |
| ENSMUSG( | Aqp9 | 233.2137 | 0.065566 | 0.714335 | 0.941045 | ENSMUSG( | Aqp9 | 182.2689 | -0.39291 | 0.034299 | 0.652355 |
| ENSMUSG( | Aqr | 1188.361 | 0.156488 | 0.102791 | 0.488221 | ENSMUSG( | Aqr | 1032.848 | 0.025449 | 0.819018 | 0.979198 |
| ENSMUSG( | Ar | 365.3475 | 0.073215 | 0.640564 | 0.916434 | ENSMUSG( | Ar | 319.2164 | -0.04145 | 0.801445 | 0.97681 |
| ENSMUSG( | Araf | 3514.759 | -0.08496 | 0.279554 | 0.729114 | ENSMUSG( | Araf | 3255.701 | -0.0363 | 0.725454 | 0.966887 |
| ENSMUSG( | Arap1 | 1169.531 | -0.09304 | 0.346341 | 0.779153 | ENSMUSG( | Arap1 | 1052.496 | -0.12682 | 0.274726 | 0.889874 |
| ENSMUSG( | Arap2 | 3326.911 | -0.00267 | 0.977959 | 0.997428 | ENSMUSG( | Arap2 | 3019.677 | -0.01516 | 0.901132 | 0.990558 |
| ENSMUSG( | Arap3 | 540.7238 | 0.365719 | 0.007395 | 0.118765 | ENSMUSG( | Arap3 | 548.9139 | 0.639291 | 0.000177 | 0.032278 |
| ENSMUSG( | Arb2a | 2272.723 | 0.061079 | 0.609091 | 0.907455 | ENSMUSG( | Arb2a | 1973.613 | -0.08156 | 0.581204 | 0.944266 |
| ENSMUSG( | Arc | 7430.516 | -1.16974 | 3.02E-16 | 3.50E-13 | ENSMUSG( | Arc | 7027.272 | -0.99766 | 0.096056 | 0.796133 |
| ENSMUSG( | Arcn1 | 3121.741 | 0.13013 | 0.137935 | 0.552903 | ENSMUSG( | Arcn1 | 2683.012 | -0.03714 | 0.738627 | 0.968018 |
| ENSMUSG( | Arel1 | 3898.286 | -0.11009 | 0.333355 | 0.775602 | ENSMUSG( | Arel1 | 3846.4 | 0.12331 | 0.38601 | 0.917257 |
| ENSMUSG( | Arf1 | 9882.402 | -0.0116 | 0.905408 | 0.986465 | ENSMUSG( | Arf1 | 8792.662 | -0.08328 | 0.515167 | 0.930939 |
| ENSMUSG( | Arf2 | 1129.313 | -0.02594 | 0.802369 | 0.962119 | ENSMUSG( | Arf2 | 999.1264 | -0.1152 | 0.343397 | 0.909753 |
| ENSMUSG( | Arf3 | 51782.73 | 0.044914 | 0.562978 | 0.889738 | ENSMUSG( | Arf3 | 45702.97 | -0.04622 | 0.6586 | 0.956205 |
| ENSMUSG( | Arf4 | 2494.905 | -0.0742 | 0.494157 | 0.860611 | ENSMUSG( | Arf4 | 2281.106 | -0.06784 | 0.651585 | 0.954492 |
| ENSMUSG( | Arf5 | 3862.153 | -0.04786 | 0.658029 | 0.92425 | ENSMUSG( | Arf5 | 3444.938 | -0.11365 | 0.423608 | 0.917257 |
| ENSMUSG( | Arf6 | 902.5488 | 0.046818 | 0.657191 | 0.924142 | ENSMUSG( | Arf6 | 826.2419 | 0.056719 | 0.668797 | 0.956766 |
| ENSMUSG( | Arfgap1 | 2178.705 | -0.08934 | 0.396719 | 0.811742 | ENSMUSG( | Arfgap1 | 1978.709 | -0.09646 | 0.399559 | 0.917257 |
| ENSMUSG( | Arfgap2 | 1756.777 | 0.052277 | 0.569316 | 0.893041 | ENSMUSG( | Arfgap2 | 1571.816 | 0.003357 | 0.97634 | 0.996416 |
| ENSMUSG( | Arfgap3 | 460.2811 | -0.00596 | 0.960843 | 0.993991 | ENSMUSG( | Arfgap3 | 393.4276 | -0.19199 | 0.186125 | 0.855305 |
| ENSMUSG( | Arfgef1 | 2372.469 | 0.157717 | 0.068277 | 0.405144 | ENSMUSG( | Arfgef1 | 2174.719 | 0.1736 | 0.118486 | 0.813384 |
| ENSMUSG( | Arfgef2 | 2008.8 | 0.033206 | 0.705053 | 0.938303 | ENSMUSG( | Arfgef2 | 1827.312 | 0.029815 | 0.786166 | 0.976285 |
| ENSMUSG( | Arfgef3 | 1830.697 | -0.07373 | 0.482297 | 0.855884 | ENSMUSG( | Arfgef3 | 1770.963 | 0.100092 | 0.365284 | 0.914449 |
| ENSMUSG( | Arfip1 | 151.305 | 0.126751 | 0.540707 | 0.88315 | ENSMUSG( | Arfip1 | 132.9229 | 0.012442 | 0.959126 | 0.994798 |
| ENSMUSG( | Arfip2 | 2785.973 | 0.153199 | 0.10834 | 0.497337 | ENSMUSG( | Arfip2 | 2440.106 | 0.044919 | 0.688921 | 0.959742 |
| ENSMUSG( | Arfrp1 | 894.389 | 0.00797 | 0.937039 | 0.993415 | ENSMUSG( | Arfrp1 | 789.1544 | -0.08546 | 0.492537 | 0.927315 |
| ENSMUSG( | Arg2 | 44.42652 | -0.46543 | 0.135601 | 0.549038 | ENSMUSG( | Arg2 | 39.30619 | -0.5996 | 0.086393 | 0.781091 |
| ENSMUSG( | Arglu1 | 2884.401 | -0.13471 | 0.289541 | 0.739839 | ENSMUSG( | Arglu1 | 2953.903 | 0.197271 | 0.213519 | 0.864421 |
| ENSMUSG( | Arhgap1 | 2175.091 | 0.124889 | 0.18528 | 0.623185 | ENSMUSG( | Arhgap1 | 1830.434 | -0.10352 | 0.38265 | 0.917257 |
| ENSMUSG( | Arhgap10 | 626.515 | 0.073351 | 0.581786 | 0.897862 | ENSMUSG( | Arhgap10 | 533.58 | -0.12431 | 0.418077 | 0.917257 |
| ENSMUSG( | Arhgap11a | 328.0915 | 0.084362 | 0.525451 | 0.876728 | ENSMUSG( | Arhgap11a | 296.8981 | 0.057774 | 0.753846 | 0.970671 |
| ENSMUSG( | Arhgap12 | 2097.418 | 0.162942 | 0.171554 | 0.606099 | ENSMUSG( | Arhgap12 | 1750.019 | -0.09034 | 0.510171 | 0.929089 |
| ENSMUSG( | Arhgap15 | 136.2941 | 0.132452 | 0.492116 | 0.860297 | ENSMUSG( | Arhgap15 | 108.1796 | -0.28339 | 0.206212 | 0.864412 |
| ENSMUSG( | Arhgap17 | 625.7393 | -0.08337 | 0.493512 | 0.860611 | ENSMUSG( | Arhgap17 | 584.4282 | -0.00637 | 0.961425 | 0.99511 |
| ENSMUSG( | Arhgap18 | 501.3161 | -0.09691 | 0.437888 | 0.830072 | ENSMUSG( | Arhgap18 | 458.1103 | -0.08981 | 0.520725 | 0.932121 |
| ENSMUSG( | Arhgap19 | 135.0343 | -0.213 | 0.221342 | 0.667193 | ENSMUSG( | Arhgap19 | 123.8552 | -0.18207 | 0.374117 | 0.915491 |
| ENSMUSG( | Arhgap20 | 1407.967 | -0.14808 | 0.103761 | 0.489428 | ENSMUSG( | Arhgap20 | 1349.148 | 0.001684 | 0.987554 | 0.997519 |
| ENSMUSG( | Arhgap21 | 6904.072 | 0.087879 | 0.306077 | 0.753712 | ENSMUSG( | Arhgap21 | 6662.839 | 0.246468 | 0.02688 | 0.603041 |
| ENSMUSG( | Arhgap22 | 226.3544 | -0.02232 | 0.899291 | 0.985633 | ENSMUSG( | Arhgap22 | 207.1931 | -0.00587 | 0.974842 | 0.996416 |
| ENSMUSG( | Arhgap23 | 6716.117 | -0.04928 | 0.552827 | 0.886141 | ENSMUSG( | Arhgap23 | 6039.199 | -0.0871 | 0.450803 | 0.922202 |
| ENSMUSG( | Arhgap24 | 363.183 | -0.07395 | 0.592097 | 0.901851 | ENSMUSG( | Arhgap24 | 314.2603 | -0.23324 | 0.112633 | 0.809623 |
| ENSMUSG( | Arhgap25 | 232.3727 | 0.269219 | 0.085596 | 0.451059 | ENSMUSG( | Arhgap25 | 177.0378 | -0.23023 | 0.214205 | 0.864421 |
| ENSMUSG( | Arhgap26 | 2528.321 | 0.221556 | 0.013363 | 0.16868 | ENSMUSG( | Arhgap26 | 2192.245 | 0.084234 | 0.442311 | 0.921554 |
| ENSMUSG( | Arhgap27 | 679.2899 | -0.05697 | 0.631772 | 0.914073 | ENSMUSG( | Arhgap27 | 671.5862 | 0.179345 | 0.209927 | 0.864412 |
| ENSMUSG( | Arhgap28 | 25.66316 | -0.25709 | 0.489928 | 0.859319 | ENSMUSG( | Arhgap28 | 23.45353 | -0.26787 | 0.508164 | 0.929089 |
| ENSMUSG( | Arhgap29 | 711.7408 | 0.141176 | 0.372475 | 0.798768 | ENSMUSG( | Arhgap29 | 696.8424 | 0.338799 | 0.057124 | 0.723362 |
| ENSMUSG( | Arhgap30 | 330.0185 | 0.238891 | 0.101014 | 0.484746 | ENSMUSG( | Arhgap30 | 277.6932 | 0.020363 | 0.899688 | 0.990558 |
| ENSMUSG( | Arhgap31 | 1508.005 | 0.189269 | 0.083427 | 0.445566 | ENSMUSG( | Arhgap31 | 1404.557 | 0.252317 | 0.056659 | 0.723362 |
| ENSMUSG( | Arhgap32 | 7073.198 | 0.051993 | 0.520271 | 0.875186 | ENSMUSG( | Arhgap32 | 6692.774 | 0.157755 | 0.108134 | 0.804969 |
| ENSMUSG( | Arhgap33 | 13319.43 | 0.17296 | 0.080521 | 0.438901 | ENSMUSG( | Arhgap33 | 11737.56 | 0.081258 | 0.431214 | 0.917257 |
| ENSMUSG( | Arhgap35 | 7262.825 | 0.203547 | 0.015861 | 0.186421 | ENSMUSG( | Arhgap35 | 6434.593 | 0.128736 | 0.215895 | 0.865167 |
| ENSMUSG( | Arhgap36 | 51.40395 | 0.44818 | 0.30072 | 0.74995 | ENSMUSG( | Arhgap36 | 36.56677 | -0.28959 | 0.495639 | 0.927774 |
| ENSMUSG( | Arhgap39 | 2351.567 | 0.394031 | 0.000868 | 0.029574 | ENSMUSG( | Arhgap39 | 2020.135 | 0.243643 | 0.080798 | 0.77681 |
| ENSMUSG( | Arhgap4 | 248.5967 | -0.06555 | 0.746365 | 0.949289 | ENSMUSG( | Arhgap4 | 248.181 | 0.194147 | 0.370019 | 0.915491 |
| ENSMUSG( | Arhgap42 | 584.7651 | 0.068439 | 0.522229 | 0.875186 | ENSMUSG( | Arhgap42 | 542.3111 | 0.116512 | 0.364153 | 0.914223 |
| ENSMUSG( | Arhgap44 | 5150.749 | 0.164583 | 0.068382 | 0.405144 | ENSMUSG( | Arhgap44 | 4490.251 | 0.041483 | 0.694221 | 0.960867 |
| ENSMUSG( | Arhgap45 | 539.226 | -0.11076 | 0.372081 | 0.798768 | ENSMUSG( | Arhgap45 | 546.723 | 0.194579 | 0.199632 | 0.863016 |
| ENSMUSG( | Arhgap5 | 6545.325 | 0.052974 | 0.736164 | 0.946267 | ENSMUSG( | Arhgap5 | 5692.614 | -0.08383 | 0.652054 | 0.9547 |
| ENSMUSG( | Arhgap6 | 157.158 | -0.02718 | 0.899895 | 0.985633 | ENSMUSG( | Arhgap6 | 157.9353 | 0.226958 | 0.371125 | 0.915491 |
| ENSMUSG( | Arhgap9 | 59.4204 | -0.02102 | 0.933615 | 0.993194 | ENSMUSG( | Arhgap9 | 58.49338 | 0.203849 | 0.474535 | 0.923329 |
| ENSMUSG( | Arhgdia | 14963.9 | 0.078456 | 0.282376 | 0.73186 | ENSMUSG( | Arhgdia | 12582.98 | -0.15677 | 0.118272 | 0.813384 |
| ENSMUSG( | Arhgdib | 489.9738 | 0.06773 | 0.57981 | 0.89687 | ENSMUSG( | Arhgdib | 421.9361 | -0.09869 | 0.478171 | 0.923329 |
| ENSMUSG( | Arhgdig | 2587.425 | -0.07514 | 0.415679 | 0.82084 | ENSMUSG( | Arhgdig | 2285.049 | -0.16932 | 0.132515 | 0.822912 |
| ENSMUSG( | Arhgef1 | 2249.606 | 0.310919 | 0.067715 | 0.40256 | ENSMUSG( | Arhgef1 | 1904.375 | 0.109272 | 0.572146 | 0.942475 |
| ENSMUSG( | Arhgef10 | 2504.654 | 0.03715 | 0.660527 | 0.924795 | ENSMUSG( | Arhgef10 | 2205.125 | -0.06331 | 0.557743 | 0.939565 |
| ENSMUSG( | Arhgef10l | 2976.108 | 0.06975 | 0.442579 | 0.83278 | ENSMUSG( | Arhgef10l | 2700.343 | 0.059222 | 0.590398 | 0.944975 |
| ENSMUSG( | Arhgef11 | 3247.462 | 0.142932 | 0.124678 | 0.531714 | ENSMUSG( | Arhgef11 | 2849.179 | 0.038005 | 0.725134 | 0.966887 |
| ENSMUSG( | Arhgef12 | 13854.99 | 0.025511 | 0.811757 | 0.964426 | ENSMUSG( | Arhgef12 | 11449.04 | -0.26872 | 0.019874 | 0.558472 |
| ENSMUSG( | Arhgef15 | 558.4659 | -0.01464 | 0.898181 | 0.985633 | ENSMUSG( | Arhgef15 | 559.1709 | 0.253693 | 0.087492 | 0.784337 |
| ENSMUSG( | Arhgef16 | 21.24606 | 0.050632 | 0.918088 | 0.989645 | ENSMUSG( | Arhgef16 | 23.62908 | 0.523609 | 0.307277 | 0.898538 |
| ENSMUSG( | Arhgef17 | 7117.948 | 0.172794 | 0.0806 | 0.438981 | ENSMUSG( | Arhgef17 | 6507.103 | 0.183402 | 0.130217 | 0.821488 |
| ENSMUSG( | Arhgef18 | 1564.615 | 0.150291 | 0.096319 | 0.476986 | ENSMUSG( | Arhgef18 | 1427.841 | 0.154372 | 0.159279 | 0.837724 |
| ENSMUSG( | Arhgef19 | 1388.625 | 0.057855 | 0.568466 | 0.892252 | ENSMUSG( | Arhgef19 | 1207.362 | -0.07831 | 0.526062 | 0.934374 |
| ENSMUSG( | Arhgef2 | 4320.833 | -0.05798 | 0.515951 | 0.872701 | ENSMUSG( | Arhgef2 | 4044.839 | 0.020476 | 0.844824 | 0.983831 |
| ENSMUSG( | Arhgef25 | 5382.054 | 0.300915 | 0.00795 | 0.124138 | ENSMUSG( | Arhgef25 | 4476.138 | 0.051418 | 0.676074 | 0.95694 |
| ENSMUSG( | Arhgef26 | 3436.848 | 0.096606 | 0.423082 | 0.823662 | ENSMUSG( | Arhgef26 | 3079.158 | 0.046693 | 0.762569 | 0.973539 |
| ENSMUSG( | Arhgef28 | 1187.045 | 0.08315 | 0.399494 | 0.813178 | ENSMUSG( | Arhgef28 | 1098.846 | 0.129508 | 0.279082 | 0.892257 |
| ENSMUSG( | Arhgef3 | 1177.483 | 0.094279 | 0.400024 | 0.813641 | ENSMUSG( | Arhgef3 | 1065.622 | 0.078735 | 0.550014 | 0.938699 |
| ENSMUSG( | Arhgef33 | 54.44623 | -0.35698 | 0.173707 | 0.609084 | ENSMUSG( | Arhgef33 | 59.66613 | 0.191905 | 0.474132 | 0.923329 |
| ENSMUSG( | Arhgef37 | 194.2365 | 0.215333 | 0.258671 | 0.709814 | ENSMUSG( | Arhgef37 | 172.9717 | 0.158399 | 0.375736 | 0.915491 |
| ENSMUSG( | Arhgef39 | 18.15531 | -0.54407 | 0.230211 | NA | ENSMUSG( | Arhgef39 | 20.86715 | 0.213971 | 0.637438 | 0.953903 |
| ENSMUSG( | Arhgef4 | 10353.37 | 0.156755 | 0.040555 | 0.311208 | ENSMUSG( | Arhgef4 | 9158.824 | 0.075805 | 0.478329 | 0.923329 |
| ENSMUSG( | Arhgef40 | 1307.677 | 0.041284 | 0.68015 | 0.930983 | ENSMUSG( | Arhgef40 | 1201.306 | 0.067015 | 0.575329 | 0.942912 |
| ENSMUSG( | Arhgef5 | 200.3944 | 0.274159 | 0.077842 | 0.431216 | ENSMUSG( | Arhgef5 | 183.6884 | 0.300634 | 0.150956 | 0.837173 |
| ENSMUSG( | Arhgef6 | 416.6119 | 0.007932 | 0.947465 | 0.993821 | ENSMUSG( | Arhgef6 | 377.5175 | -0.00705 | 0.958989 | 0.994798 |
| ENSMUSG( | Arhgef7 | 4939.218 | 0.153116 | 0.063722 | 0.390618 | ENSMUSG( | Arhgef7 | 4272.509 | 0.008408 | 0.932386 | 0.99306 |
| ENSMUSG( | Arhgef9 | 10192.32 | -0.01455 | 0.862451 | 0.975055 | ENSMUSG( | Arhgef9 | 9150.982 | -0.05702 | 0.563767 | 0.941206 |
| ENSMUSG( | Arid1a | 3892.201 | 0.0773 | 0.457781 | 0.839765 | ENSMUSG( | Arid1a | 3554.431 | 0.086501 | 0.464096 | 0.923329 |
| ENSMUSG( | Arid1b | 2677.405 | 0.040339 | 0.690573 | 0.933881 | ENSMUSG( | Arid1b | 2524.457 | 0.138731 | 0.244319 | 0.877904 |
| ENSMUSG( | Arid2 | 1145.203 | 0.05254 | 0.573653 | 0.893584 | ENSMUSG( | Arid2 | 1121.975 | 0.249382 | 0.030811 | 0.629149 |
| ENSMUSG( | Arid3a | 168.0756 | -0.09924 | 0.569825 | 0.893041 | ENSMUSG( | Arid3a | 170.0559 | 0.194924 | 0.263356 | 0.886133 |
| ENSMUSG( | Arid3b | 179.3913 | -0.10855 | 0.560756 | 0.888934 | ENSMUSG( | Arid3b | 175.7251 | 0.108452 | 0.614313 | 0.951177 |
| ENSMUSG( | Arid4a | 895.8549 | -0.07801 | 0.443411 | 0.833108 | ENSMUSG( | Arid4a | 927.0142 | 0.278749 | 0.019089 | 0.549166 |
| ENSMUSG( | Arid4b | 1127.126 | -0.16081 | 0.120436 | 0.524495 | ENSMUSG( | Arid4b | 1144.317 | 0.152911 | 0.225251 | 0.868468 |
| ENSMUSG( | Arid5a | 505.1359 | -0.02763 | 0.808487 | 0.963455 | ENSMUSG( | Arid5a | 435.0895 | -0.19233 | 0.162762 | 0.837724 |
| ENSMUSG( | Arid5b | 612.723 | 0.065279 | 0.601942 | 0.904887 | ENSMUSG( | Arid5b | 614.0939 | 0.323769 | 0.009244 | 0.402642 |
| ENSMUSG( | Arih1 | 2401.398 | 0.083704 | 0.348526 | 0.78132 | ENSMUSG( | Arih1 | 2173.088 | 0.061327 | 0.64441 | 0.953903 |
| ENSMUSG( | Arih2 | 1493.7 | -0.02986 | 0.730408 | 0.944956 | ENSMUSG( | Arih2 | 1396.651 | 0.044368 | 0.678283 | 0.957435 |

| | | | | | |
|---|---|---|---|---|---|
| ENSMUSG( | 3436.848 | 0.096606 | 0.423082 | 0.823662 | Arhgef26 |
| ENSMUSG( | 1187.045 | 0.08315 | 0.399494 | 0.813178 | Arhgef28 |
| ENSMUSG( | 1177.483 | 0.094279 | 0.400024 | 0.813641 | Arhgef3 |
| ENSMUSG( | 54.44623 | -0.35698 | 0.173707 | 0.609084 | Arhgef33 |
| ENSMUSG( | 194.2365 | 0.215333 | 0.258671 | 0.709814 | Arhgef37 |
| ENSMUSG( | 18.15531 | -0.54407 | 0.230211 | NA | Arhgef39 |
| ENSMUSG( | 10353.37 | 0.156755 | 0.040555 | 0.311208 | Arhgef4 |
| ENSMUSG( | 1307.677 | 0.041284 | 0.68015 | 0.930983 | Arhgef40 |
| ENSMUSG( | 200.3944 | 0.274159 | 0.077842 | 0.431216 | Arhgef5 |
| ENSMUSG( | 416.6119 | 0.007932 | 0.947465 | 0.993821 | Arhgef6 |
| ENSMUSG( | 4939.218 | 0.153116 | 0.063722 | 0.390618 | Arhgef7 |
| ENSMUSG( | 10192.32 | -0.01455 | 0.862451 | 0.975055 | Arhgef9 |
| ENSMUSG( | 3892.201 | 0.0773 | 0.457781 | 0.839765 | Arid1a |
| ENSMUSG( | 2677.405 | 0.040339 | 0.690573 | 0.933881 | Arid1b |
| ENSMUSG( | 1145.203 | 0.05254 | 0.573653 | 0.893584 | Arid2 |
| ENSMUSG( | 168.0756 | -0.09924 | 0.569825 | 0.893041 | Arid3a |
| ENSMUSG( | 179.3913 | -0.10855 | 0.560756 | 0.888934 | Arid3b |
| ENSMUSG( | 895.8549 | -0.07801 | 0.443411 | 0.833108 | Arid4a |
| ENSMUSG( | 1127.126 | -0.16081 | 0.120436 | 0.524495 | Arid4b |
| ENSMUSG( | 505.1359 | -0.02763 | 0.808487 | 0.963455 | Arid5a |
| ENSMUSG( | 612.723 | 0.065279 | 0.601942 | 0.904887 | Arid5b |
| ENSMUSG( | 2401.398 | 0.083704 | 0.348526 | 0.78132 | Arih1 |
| ENSMUSG( | 1493.7 | -0.02986 | 0.730408 | 0.944956 | Arih2 |
| ENSMUSG( | 2678.356 | 0.347334 | 0.000598 | 0.022994 | Ark2c |
| ENSMUSG( | 829.9453 | 0.090583 | 0.381939 | 0.802983 | Ark2n |
| ENSMUSG( | 2301.776 | -0.06077 | 0.514495 | 0.872055 | Arl1 |
| ENSMUSG( | 313.202 | -0.3422 | 0.104847 | 0.491651 | Arl10 |
| ENSMUSG( | 26.09119 | -0.24828 | 0.495778 | 0.861294 | Arl11 |
| ENSMUSG( | 342.9785 | -0.01231 | 0.927984 | 0.991229 | Arl13b |
| ENSMUSG( | 842.2687 | -0.01065 | 0.918719 | 0.990075 | Arl14ep |
| ENSMUSG( | 585.0882 | 0.018139 | 0.886581 | 0.983204 | Arl15 |
| ENSMUSG( | 646.812 | -0.00332 | 0.97807 | 0.997428 | Arl16 |
| ENSMUSG( | 1346.916 | -0.03531 | 0.740527 | 0.947823 | Arl2 |
| ENSMUSG( | 9193.912 | 0.003014 | 0.976287 | 0.997428 | Arl2bp |
| ENSMUSG( | 2915.706 | -0.01859 | 0.858978 | 0.974707 | Arl3 |
| ENSMUSG( | 838.1206 | -0.08155 | 0.470666 | 0.84674 | Arl4a |
| ENSMUSG( | 1402.957 | -0.02698 | 0.801652 | 0.962077 | Arl4c |
| ENSMUSG( | 422.6032 | -0.79544 | 3.94E-06 | 0.000532 | Arl4d |
| ENSMUSG( | 2600.7 | 0.0668 | 0.541461 | 0.883182 | Arl5a |
| ENSMUSG( | 445.9243 | -0.06104 | 0.629704 | 0.913965 | Arl5b |
| ENSMUSG( | 70.04178 | -0.6493 | 0.010372 | 0.145282 | Arl5c |
| ENSMUSG( | 416.0229 | -0.00351 | 0.980732 | 0.997512 | Arl6 |
| ENSMUSG( | 10862.41 | -0.33944 | 0.005646 | 0.100551 | Arl6ip1 |
| ENSMUSG( | 898.7957 | 0.115251 | 0.298947 | 0.747827 | Arl6ip4 |
| ENSMUSG( | 5567.012 | -0.4416 | 0.000164 | 0.009126 | Arl6ip5 |
| ENSMUSG( | 208.5542 | -0.36994 | 0.016527 | 0.191072 | Arl6ip6 |
| ENSMUSG( | 7725.216 | -0.04801 | 0.552034 | 0.886141 | Arl8a |
| ENSMUSG( | 4861.888 | -0.07841 | 0.416851 | 0.8209 | Arl8b |
| ENSMUSG( | 1744.138 | 0.088754 | 0.363236 | 0.793211 | Armc1 |
| ENSMUSG( | 473.2154 | -0.10882 | 0.367271 | 0.795221 | Armc10 |
| ENSMUSG( | 219.4921 | 0.011749 | 0.942601 | 0.993415 | Armc2 |
| ENSMUSG( | 33.11806 | -0.13642 | 0.719789 | 0.941986 | Armc3 |
| ENSMUSG( | 498.1949 | -0.02384 | 0.839751 | 0.971179 | Armc5 |
| ENSMUSG( | 607.858 | -0.07692 | 0.570624 | 0.893192 | Armc6 |
| ENSMUSG( | 215.3333 | -0.10662 | 0.489087 | 0.859092 | Armc7 |
| ENSMUSG( | 1621.625 | 0.087459 | 0.341729 | 0.778828 | Armc8 |
| ENSMUSG( | 683.0373 | 0.101145 | 0.471621 | 0.847331 | Armc9 |
| ENSMUSG( | 1242.309 | -0.00639 | 0.955156 | 0.993874 | Armcx1 |
| ENSMUSG( | 2759.29 | 0.110807 | 0.203346 | 0.643631 | Armcx2 |
| ENSMUSG( | 1117.233 | 0.077982 | 0.483628 | 0.85636 | Armcx3 |
| ENSMUSG( | 1621.166 | -0.01076 | 0.931228 | 0.992282 | Armcx4 |
| ENSMUSG( | 392.4482 | -0.14724 | 0.250078 | 0.699718 | Armcx5 |
| ENSMUSG( | 83.0496 | -0.06388 | 0.794192 | 0.959799 | Armcx6 |
| ENSMUSG( | 20.3687 | -0.2082 | 0.658169 | NA | Armh1 |
| ENSMUSG( | 648.6517 | -0.06248 | 0.634866 | 0.914078 | Armh3 |
| ENSMUSG( | 901.6714 | -0.44469 | 0.012592 | 0.161972 | Armh4 |
| ENSMUSG( | 361.8934 | -0.01248 | 0.921652 | 0.990436 | Armt1 |
| ENSMUSG( | 1230.911 | 0.175301 | 0.10094 | 0.484681 | Arnt |
| ENSMUSG( | 15978.06 | 0.089036 | 0.219768 | 0.665168 | Arnt2 |
| ENSMUSG( | 4929.023 | -0.00862 | 0.931109 | 0.992271 | Arpc1a |
| ENSMUSG( | 1851.501 | 0.058908 | 0.592847 | 0.902275 | Arpc1b |
| ENSMUSG( | 4909.658 | 0.073899 | 0.348523 | 0.78132 | Arpc2 |
| ENSMUSG( | 2536.39 | -0.04516 | 0.672516 | 0.929007 | Arpc3 |
| ENSMUSG( | 2943.194 | 0.036139 | 0.71524 | 0.941288 | Arpc4 |
| ENSMUSG( | 2236.532 | -0.03755 | 0.688823 | 0.933441 | Arpc5 |
| ENSMUSG( | 2172.609 | 0.000442 | 0.996269 | 0.99943 | Arpc5l |
| ENSMUSG( | 556.081 | 0.063056 | 0.591666 | 0.901449 | Arpin |
| ENSMUSG( | 5272.992 | -0.00681 | 0.948144 | 0.993821 | Arpp19 |
| ENSMUSG( | 11513.03 | -0.11911 | 0.243215 | 0.69052 | Arpp21 |
| ENSMUSG( | 27.07158 | 0.329869 | 0.376258 | 0.801127 | Arr3 |
| ENSMUSG( | 21237.13 | 0.145148 | 0.11002 | 0.500285 | Arrb1 |
| ENSMUSG( | 1365.943 | 0.130361 | 0.281927 | 0.73186 | Arrb2 |
| ENSMUSG( | 239.9236 | -0.14997 | 0.366172 | 0.794603 | Arrdc1 |
| ENSMUSG( | 413.7135 | -0.35048 | 0.03679 | 0.296095 | Arrdc2 |
| ENSMUSG( | 776.0378 | -0.1722 | 0.110471 | 0.50108 | Arrdc3 |
| ENSMUSG( | 335.0688 | 0.038006 | 0.786029 | 0.958774 | Arrdc4 |
| ENSMUSG( | 1268.657 | -0.14443 | 0.339102 | 0.777518 | Arsa |
| ENSMUSG( | 5712.083 | -0.28303 | 0.04086 | 0.312615 | Arsb |
| ENSMUSG( | 661.2293 | -0.64894 | 2.70E-06 | 0.000405 | Arsg |
| ENSMUSG( | 51.60967 | 0.095074 | 0.747484 | 0.949604 | Arsi |
| ENSMUSG( | 40.50148 | 0.201742 | 0.514325 | 0.872055 | Arsj |
| ENSMUSG( | 374.3907 | -0.16444 | 0.233454 | 0.681508 | Arsk |
| ENSMUSG( | 155.6424 | -0.17872 | 0.473131 | 0.848727 | Art3 |
| ENSMUSG( | 13.18839 | -0.02177 | 0.965475 | NA | Artn |
| ENSMUSG( | 147.1788 | -0.15729 | 0.454378 | 0.837311 | Arv1 |
| ENSMUSG( | 933.1993 | 0.141357 | 0.245172 | 0.692812 | Arvcf |
| ENSMUSG( | 544.1914 | -0.03187 | 0.806427 | 0.962804 | Arx |
| ENSMUSG( | 653.9716 | -0.56392 | 3.39E-05 | 0.002956 | Arxes1 |
| ENSMUSG( | 1037.125 | -0.61808 | 0.002869 | 0.065631 | Arxes2 |
| ENSMUSG( | 846.1425 | 0.045683 | 0.756487 | 0.951479 | As3mt |
| ENSMUSG( | 2069.226 | -0.31292 | 0.044358 | 0.323404 | Asah1 |
| ENSMUSG( | 502.8808 | -0.25236 | 0.064731 | 0.394417 | Asah2 |
| ENSMUSG( | 3616.828 | 0.224393 | 0.023678 | 0.23542 | Asap1 |
| ENSMUSG( | 1804.098 | 0.302737 | 0.001727 | 0.046912 | Asap2 |
| ENSMUSG( | 304.325 | 0.015515 | 0.909976 | 0.9881 | Asap3 |
| ENSMUSG( | 1526.018 | 0.08494 | 0.361283 | 0.792797 | Asb1 |
| ENSMUSG( | 13.58617 | -0.3939 | 0.502732 | NA | Asb10 |
| ENSMUSG( | 80.31326 | -0.44837 | 0.052734 | 0.352577 | Asb11 |
| ENSMUSG( | 1307.711 | 0.033383 | 0.726964 | 0.944223 | Asb13 |
| ENSMUSG( | 78.39892 | 0.112266 | 0.611784 | 0.908586 | Asb16 |
| ENSMUSG( | 50.42365 | -0.12766 | 0.630807 | 0.914073 | Asb18 |
| ENSMUSG( | 84.00435 | -0.21246 | 0.372484 | 0.798768 | Asb2 |
| ENSMUSG( | 237.7625 | 0.272506 | 0.072341 | 0.415061 | Asb3 |
| ENSMUSG( | 27.60489 | -0.27276 | 0.470312 | 0.84674 | Asb4 |
| ENSMUSG( | 30.2437 | -0.18683 | 0.582211 | 0.897918 | Asb5 |
| ENSMUSG( | 743.3453 | 0.052139 | 0.687841 | 0.93324 | Asb6 |
| ENSMUSG( | 536.5787 | 0.119782 | 0.369021 | 0.796581 | Asb7 |
| ENSMUSG( | 735.7309 | 0.080491 | 0.52177 | 0.875186 | Asb8 |
| ENSMUSG( | 442.8406 | -0.06455 | 0.665269 | 0.926953 | Ascc1 |
| ENSMUSG( | 1180.017 | 0.115488 | 0.24189 | 0.689179 | Ascc2 |
| ENSMUSG( | 429.5571 | 0.089607 | 0.50016 | 0.863455 | Ascc3 |
| ENSMUSG( | 176.9079 | 0.058921 | 0.718135 | 0.941721 | Ascl1 |
| ENSMUSG( | 10.45237 | -0.33207 | 0.617925 | NA | Ascl5 |
| ENSMUSG( | 307.7903 | 0.050845 | 0.701044 | 0.93743 | Asf1a |
| ENSMUSG( | 15.89693 | -0.58612 | 0.315145 | NA | Asf1b |
| ENSMUSG( | 42.04607 | -0.14339 | 0.635779 | 0.9147 | Asgr1 |
| ENSMUSG( | 4227.921 | 0.000327 | 0.997386 | 0.999605 | Ash1l |
| ENSMUSG( | 1159.352 | -0.03284 | 0.744728 | 0.94902 | Ash2l |
| ENSMUSG( | 2608.171 | -0.19339 | 0.139143 | 0.555202 | Asic1 |
| ENSMUSG( | 2431.738 | 0.066541 | 0.571921 | 0.893523 | Asic2 |
| ENSMUSG( | 10.11714 | -0.32478 | 0.580065 | NA | Asic3 |
| ENSMUSG( | 735.8545 | -0.936 | 1.48E-07 | 3.75E-05 | Asic4 |
| ENSMUSG( | 459.4449 | 0.189262 | 0.150311 | 0.574875 | Asl |
| ENSMUSG( | 1481.194 | 0.03254 | 0.820577 | 0.965888 | Asns |
| ENSMUSG( | 110.1426 | -0.08752 | 0.673293 | 0.929007 | Asnsd1 |
| ENSMUSG( | 314.8092 | -0.16802 | 0.220603 | 0.666079 | Aspa |
| ENSMUSG( | 95.57404 | 0.056433 | 0.821266 | 0.966248 | Aspdh |
| ENSMUSG( | 166.0118 | 0.166235 | 0.357668 | 0.789196 | Aspg |
| ENSMUSG( | 3943.292 | -0.39167 | 0.001342 | 0.039921 | Asph |
| ENSMUSG( | 867.0701 | -0.29198 | 0.010268 | 0.144326 | Asphd1 |
| ENSMUSG( | 4393.913 | -0.03254 | 0.757363 | 0.951513 | Asphd2 |
| ENSMUSG( | 48.06855 | -0.14178 | 0.680157 | 0.930983 | Aspm |
| ENSMUSG( | 31.16933 | -0.62812 | 0.12927 | 0.538528 | Aspn |
| ENSMUSG( | 56.48502 | 0.119909 | 0.711572 | 0.940028 | Asprv1 |
| ENSMUSG( | 1112.495 | 0.023923 | 0.80419 | 0.962169 | Aspscr1 |
| ENSMUSG( | 30895.14 | 0.094149 | 0.422073 | 0.823495 | Asrgl1 |
| ENSMUSG( | 824.6885 | 0.070528 | 0.480258 | 0.854705 | Ass1 |
| ENSMUSG( | 139.9762 | -0.07619 | 0.679017 | 0.930914 | Aste1 |
| ENSMUSG( | 9416.757 | -0.24495 | 0.098233 | 0.479534 | Astn1 |
| ENSMUSG( | 1981.039 | -0.2531 | 0.086992 | 0.453311 | Astn2 |
| ENSMUSG( | 1094.89 | 0.071309 | 0.4903 | 0.859319 | Asxl1 |
| ENSMUSG( | 946.6428 | -0.04644 | 0.697681 | 0.93578 | Asxl2 |
| ENSMUSG( | 234.8943 | -0.42881 | 0.00415 | 0.083415 | Asxl3 |
| ENSMUSG( | 3523.747 | -0.36952 | 0.008884 | 0.132538 | Atad1 |
| ENSMUSG( | 221.7608 | -0.03289 | 0.856679 | 0.974154 | Atad2 |
| ENSMUSG( | 703.8302 | 0.020586 | 0.860792 | 0.975003 | Atad2b |
| ENSMUSG( | 1115.117 | 0.178902 | 0.096355 | 0.476986 | Atad3a |
| ENSMUSG( | 22.99835 | 0.15797 | 0.686392 | 0.933041 | Atad3aos |

| | | | | | |
|---|---|---|---|---|---|
| ENSMUSG( | 170.0559 | 0.194924 | 0.263356 | 0.886133 | Arid3a |
| ENSMUSG( | 175.7251 | 0.108452 | 0.614313 | 0.951177 | Arid3b |
| ENSMUSG( | 927.0142 | 0.278749 | 0.019089 | 0.549166 | Arid4a |
| ENSMUSG( | 1144.317 | 0.152911 | 0.225251 | 0.868468 | Arid4b |
| ENSMUSG( | 435.0895 | -0.19233 | 0.162762 | 0.837724 | Arid5a |
| ENSMUSG( | 614.0939 | 0.323769 | 0.009244 | 0.402642 | Arid5b |
| ENSMUSG( | 2173.088 | 0.061327 | 0.64441 | 0.953903 | Arih1 |
| ENSMUSG( | 1396.651 | 0.044368 | 0.678283 | 0.957435 | Arih2 |
| ENSMUSG( | 2472.578 | 0.38319 | 0.000923 | 0.102962 | Ark2c |
| ENSMUSG( | 763.4488 | 0.112127 | 0.396769 | 0.917257 | Ark2n |
| ENSMUSG( | 2125.46 | -0.02303 | 0.845018 | 0.983863 | Arl1 |
| ENSMUSG( | 338.8805 | 0.18213 | 0.454598 | 0.92252 | Arl10 |
| ENSMUSG( | 26.27268 | 0.013037 | 0.974776 | 0.996416 | Arl11 |
| ENSMUSG( | 327.3698 | 0.121 | 0.430331 | 0.917257 | Arl13b |
| ENSMUSG( | 766.5836 | -0.01499 | 0.899101 | 0.990558 | Arl14ep |
| ENSMUSG( | 523.9147 | -0.03375 | 0.82178 | 0.979198 | Arl15 |
| ENSMUSG( | 568.3338 | -0.11245 | 0.420391 | 0.917257 | Arl16 |
| ENSMUSG( | 1171.66 | -0.17816 | 0.190043 | 0.857355 | Arl2 |
| ENSMUSG( | 7884.993 | -0.17908 | 0.173436 | 0.844317 | Arl2bp |
| ENSMUSG( | 2503.861 | -0.19957 | 0.126265 | 0.81516 | Arl3 |
| ENSMUSG( | 823.4527 | 0.13224 | 0.30736 | 0.898538 | Arl4a |
| ENSMUSG( | 1338.633 | 0.106794 | 0.365266 | 0.914449 | Arl4c |
| ENSMUSG( | 390.8717 | -0.73426 | 0.000225 | 0.039463 | Arl4d |
| ENSMUSG( | 2345.659 | 0.036002 | 0.791245 | 0.97681 | Arl5a |
| ENSMUSG( | 425.7908 | 0.07511 | 0.60103 | 0.947749 | Arl5b |
| ENSMUSG( | 65.02911 | -0.61043 | 0.032732 | 0.637512 | Arl5c |
| ENSMUSG( | 369.6793 | -0.08279 | 0.631626 | 0.953251 | Arl6 |
| ENSMUSG( | 11395.32 | 0.087097 | 0.557168 | 0.939565 | Arl6ip1 |
| ENSMUSG( | 810.6408 | 0.082543 | 0.575708 | 0.942912 | Arl6ip4 |
| ENSMUSG( | 5922.877 | 0.040392 | 0.768175 | 0.974236 | Arl6ip5 |
| ENSMUSG( | 229.8659 | 0.201558 | 0.212649 | 0.864412 | Arl6ip6 |
| ENSMUSG( | 7080.937 | -0.03197 | 0.795069 | 0.97681 | Arl8a |
| ENSMUSG( | 4561.91 | 0.007741 | 0.946451 | 0.994798 | Arl8b |
| ENSMUSG( | 1539.785 | -0.00443 | 0.97314 | 0.996416 | Armc1 |
| ENSMUSG( | 443.3077 | -0.02533 | 0.857485 | 0.985884 | Armc10 |
| ENSMUSG( | 204.9489 | 0.079368 | 0.704686 | 0.962677 | Armc2 |
| ENSMUSG( | 34.9094 | 0.302795 | 0.457858 | 0.923329 | Armc3 |
| ENSMUSG( | 470.4634 | 0.085777 | 0.565917 | 0.941499 | Armc5 |
| ENSMUSG( | 556.5466 | -0.06228 | 0.680258 | 0.958115 | Armc6 |
| ENSMUSG( | 193.7644 | -0.14438 | 0.418625 | 0.917257 | Armc7 |
| ENSMUSG( | 1387.146 | -0.09757 | 0.40693 | 0.917257 | Armc8 |
| ENSMUSG( | 601.4259 | 0.002115 | 0.988943 | 0.997887 | Armc9 |
| ENSMUSG( | 1121.017 | -0.03918 | 0.776646 | 0.975717 | Armcx1 |
| ENSMUSG( | 2461.347 | 0.052838 | 0.621944 | 0.953234 | Armcx2 |
| ENSMUSG( | 1027.413 | 0.101496 | 0.42913 | 0.917257 | Armcx3 |
| ENSMUSG( | 1552.71 | 0.133583 | 0.374955 | 0.915491 | Armcx4 |
| ENSMUSG( | 366.7925 | -0.06968 | 0.637947 | 0.953903 | Armcx5 |
| ENSMUSG( | 80.47958 | 0.133461 | 0.614179 | 0.951177 | Armcx6 |
| ENSMUSG( | 19.88016 | 0.026378 | 0.957066 | 0.994798 | Armh1 |
| ENSMUSG( | 592.2048 | -0.05665 | 0.696639 | 0.960867 | Armh3 |
| ENSMUSG( | 1021.169 | 0.215478 | 0.247337 | 0.877904 | Armh4 |
| ENSMUSG( | 323.7774 | -0.06296 | 0.663117 | 0.956349 | Armt1 |
| ENSMUSG( | 1107.395 | 0.142063 | 0.242629 | 0.877399 | Arnt |
| ENSMUSG( | 13560.27 | -0.11672 | 0.214141 | 0.864421 | Arnt2 |
| ENSMUSG( | 4282.773 | -0.15165 | 0.223677 | 0.867404 | Arpc1a |
| ENSMUSG( | 1592.903 | -0.11066 | 0.380043 | 0.916995 | Arpc1b |
| ENSMUSG( | 4257.12 | -0.07099 | 0.575092 | 0.942912 | Arpc2 |
| ENSMUSG( | 2201.881 | -0.19379 | 0.170571 | 0.841739 | Arpc3 |
| ENSMUSG( | 2569.23 | -0.08842 | 0.401956 | 0.917257 | Arpc4 |
| ENSMUSG( | 2005.911 | -0.08742 | 0.489079 | 0.925673 | Arpc5 |
| ENSMUSG( | 1897.513 | -0.12693 | 0.29448 | 0.898538 | Arpc5l |
| ENSMUSG( | 490.7609 | -0.02615 | 0.839253 | 0.981912 | Arpin |
| ENSMUSG( | 4622.892 | -0.12386 | 0.405563 | 0.917257 | Arpp19 |
| ENSMUSG( | 10761.2 | -0.04484 | 0.734267 | 0.967228 | Arpp21 |
| ENSMUSG( | 18.54645 | -0.5004 | 0.321512 | 0.904019 | Arr3 |
| ENSMUSG( | 17950.14 | -0.06786 | 0.502769 | 0.927825 | Arrb1 |
| ENSMUSG( | 1240.462 | 0.121117 | 0.403759 | 0.917257 | Arrb2 |
| ENSMUSG( | 228.6398 | -0.0088 | 0.969605 | 0.99594 | Arrdc1 |
| ENSMUSG( | 403.7429 | -0.12297 | 0.510962 | 0.929546 | Arrdc2 |
| ENSMUSG( | 817.1949 | 0.245897 | 0.108175 | 0.804969 | Arrdc3 |
| ENSMUSG( | 298.3974 | -0.02911 | 0.840685 | 0.982538 | Arrdc4 |
| ENSMUSG( | 1279.821 | 0.154934 | 0.370477 | 0.915491 | Arsa |
| ENSMUSG( | 6032.275 | 0.156575 | 0.333978 | 0.907907 | Arsb |
| ENSMUSG( | 713.0862 | -0.08323 | 0.625013 | 0.953234 | Arsg |
| ENSMUSG( | 44.83621 | -0.01651 | 0.957854 | 0.994798 | Arsi |
| ENSMUSG( | 41.01615 | 0.497813 | 0.144941 | 0.83339 | Arsj |
| ENSMUSG( | 374.8986 | 0.111981 | 0.464292 | 0.923329 | Arsk |
| ENSMUSG( | 192.8512 | 0.659882 | 0.035596 | 0.664751 | Art3 |
| ENSMUSG( | 12.2858 | 0.033403 | 0.95137 | 0.994798 | Artn |
| ENSMUSG( | 142.7341 | 0.038668 | 0.853978 | 0.98563 | Arv1 |
| ENSMUSG( | 854.3158 | 0.158703 | 0.237782 | 0.87259 | Arvcf |
| ENSMUSG( | 529.9678 | 0.152825 | 0.265372 | 0.886134 | Arx |
| ENSMUSG( | 712.4161 | 0.004908 | 0.977572 | 0.996416 | Arxes1 |
| ENSMUSG( | 1157.254 | 0.029486 | 0.942982 | 0.994774 | Arxes2 |
| ENSMUSG( | 720.8883 | -0.15624 | 0.304289 | 0.898538 | As3mt |
| ENSMUSG( | 2222.227 | 0.177885 | 0.315166 | 0.901306 | Asah1 |
| ENSMUSG( | 496.681 | -0.00827 | 0.95916 | 0.994798 | Asah2 |
| ENSMUSG( | 3068.076 | 0.027142 | 0.812737 | 0.979148 | Asap1 |
| ENSMUSG( | 1581.085 | 0.197037 | 0.106781 | 0.804969 | Asap2 |
| ENSMUSG( | 307.9181 | 0.305105 | 0.052824 | 0.71525 | Asap3 |
| ENSMUSG( | 1378.365 | 0.061544 | 0.580188 | 0.944209 | Asb1 |
| ENSMUSG( | 9.46021 | -1.5584 | 0.019267 | NA | Asb10 |
| ENSMUSG( | 87.21789 | 0.099771 | 0.67781 | 0.95713 | Asb11 |
| ENSMUSG( | 1159.85 | -0.04863 | 0.705719 | 0.96315 | Asb13 |
| ENSMUSG( | 68.87621 | -0.01417 | 0.958677 | 0.994798 | Asb16 |
| ENSMUSG( | 44.75931 | -0.18477 | 0.544923 | 0.938063 | Asb18 |
| ENSMUSG( | 78.43492 | -0.12274 | 0.622274 | 0.953234 | Asb2 |
| ENSMUSG( | 205.3749 | 0.129487 | 0.44689 | 0.921891 | Asb3 |
| ENSMUSG( | 20.45999 | -1.0358 | 0.019693 | 0.556192 | Asb4 |
| ENSMUSG( | 26.61879 | -0.33396 | 0.404535 | 0.917257 | Asb5 |
| ENSMUSG( | 658.4344 | -0.02644 | 0.859288 | 0.985884 | Asb6 |
| ENSMUSG( | 473.2101 | 0.024154 | 0.869979 | 0.988097 | Asb7 |
| ENSMUSG( | 657.3812 | 0.023616 | 0.86352 | 0.986547 | Asb8 |
| ENSMUSG( | 398.4433 | -0.10521 | 0.539811 | 0.937378 | Ascc1 |
| ENSMUSG( | 1006.73 | -0.0746 | 0.598881 | 0.947045 | Ascc2 |
| ENSMUSG( | 391.0394 | 0.094263 | 0.513277 | 0.930482 | Ascc3 |
| ENSMUSG( | 167.6183 | 0.161301 | 0.374305 | 0.915491 | Ascl1 |
| ENSMUSG( | 11.90118 | 0.316 | 0.611702 | 0.950771 | Ascl5 |
| ENSMUSG( | 292.2714 | 0.156461 | 0.326823 | 0.905419 | Asf1a |
| ENSMUSG( | 16.30037 | -0.19042 | 0.705595 | 0.96315 | Asf1b |
| ENSMUSG( | 29.19663 | -1.14991 | 0.002273 | 0.185073 | Asgr1 |
| ENSMUSG( | 3877.613 | 0.021203 | 0.849755 | 0.985522 | Ash1l |
| ENSMUSG( | 1037.355 | -0.08427 | 0.494708 | 0.927372 | Ash2l |
| ENSMUSG( | 2667.658 | 0.146865 | 0.334681 | 0.907907 | Asic1 |
| ENSMUSG( | 2467.766 | 0.359596 | 0.008264 | 0.388416 | Asic2 |
| ENSMUSG( | 11.03847 | 0.208464 | 0.729389 | NA | Asic3 |
| ENSMUSG( | 890.8242 | 0.036632 | 0.827344 | 0.980224 | Asic4 |
| ENSMUSG( | 372.5673 | -0.14469 | 0.345092 | 0.91045 | Asl |
| ENSMUSG( | 1227.987 | -0.25233 | 0.092022 | 0.791889 | Asns |
| ENSMUSG( | 106.0224 | 0.059373 | 0.78565 | 0.976285 | Asnsd1 |
| ENSMUSG( | 331.8692 | 0.253828 | 0.133864 | 0.822912 | Aspa |
| ENSMUSG( | 94.79988 | 0.295346 | 0.182392 | 0.850579 | Aspdh |
| ENSMUSG( | 136.5585 | -0.10933 | 0.665936 | 0.956766 | Aspg |
| ENSMUSG( | 3962.758 | -0.08278 | 0.566396 | 0.941745 | Asph |
| ENSMUSG( | 848.4037 | -0.0734 | 0.583058 | 0.944266 | Asphd1 |
| ENSMUSG( | 4165.436 | 0.083984 | 0.500274 | 0.927825 | Asphd2 |
| ENSMUSG( | 44.97684 | -0.04391 | 0.907684 | 0.991958 | Aspm |
| ENSMUSG( | 42.98121 | 0.59296 | 0.098224 | 0.798438 | Aspn |
| ENSMUSG( | 53.96523 | 0.230538 | 0.472341 | 0.923329 | Asprv1 |
| ENSMUSG( | 1006.766 | 0.005674 | 0.961056 | 0.99511 | Aspscr1 |
| ENSMUSG( | 25233.22 | -0.23079 | 0.119049 | 0.813384 | Asrgl1 |
| ENSMUSG( | 721.1693 | -0.05098 | 0.690217 | 0.960071 | Ass1 |
| ENSMUSG( | 141.2902 | 0.208931 | 0.283411 | 0.892412 | Aste1 |
| ENSMUSG( | 9742.469 | 0.133424 | 0.419593 | 0.917257 | Astn1 |
| ENSMUSG( | 2037.746 | 0.11044 | 0.504927 | 0.927825 | Astn2 |
| ENSMUSG( | 995.3706 | 0.067384 | 0.56688 | 0.941777 | Asxl1 |
| ENSMUSG( | 900.2086 | 0.080288 | 0.564188 | 0.941206 | Asxl2 |
| ENSMUSG( | 227.3386 | -0.23218 | 0.155458 | 0.837724 | Asxl3 |
| ENSMUSG( | 3783.142 | 0.127722 | 0.424393 | 0.917257 | Atad1 |
| ENSMUSG( | 223.6431 | 0.259797 | 0.247901 | 0.877904 | Atad2 |
| ENSMUSG( | 675.6879 | 0.168486 | 0.207364 | 0.864412 | Atad2b |
| ENSMUSG( | 974.1318 | 0.064773 | 0.615098 | 0.951177 | Atad3a |
| ENSMUSG( | 21.3255 | 0.184737 | 0.674242 | 0.95694 | Atad3aos |
| ENSMUSG( | 144.4172 | 0.285097 | 0.136125 | 0.827092 | Atad5 |
| ENSMUSG( | 1556.7 | -0.09635 | 0.430928 | 0.917257 | Atat1 |
| ENSMUSG( | 6432.244 | 0.108884 | 0.430353 | 0.917257 | Atcay |
| ENSMUSG( | 1182.572 | 0.017145 | 0.884998 | 0.989688 | Ate1 |
| ENSMUSG( | 342.5935 | -0.01592 | 0.928437 | 0.99306 | Atf1 |
| ENSMUSG( | 1743.832 | -0.0389 | 0.794701 | 0.97681 | Atf2 |
| ENSMUSG( | 21.88771 | -0.14445 | 0.770932 | 0.974236 | Atf3 |
| ENSMUSG( | 2783.17 | -0.10904 | 0.442689 | 0.921554 | Atf4 |
| ENSMUSG( | 2006.023 | -0.13831 | 0.339031 | 0.907972 | Atf5 |
| ENSMUSG( | 2388.901 | 0.17874 | 0.318181 | 0.902849 | Atf6 |
| ENSMUSG( | 2748.328 | 0.195071 | 0.186178 | 0.855305 | Atf6b |
| ENSMUSG( | 773.5149 | 0.045631 | 0.71934 | 0.966876 | Atf7 |
| ENSMUSG( | 1961.336 | 0.040973 | 0.712601 | 0.964363 | Atf7ip |
| ENSMUSG( | 181.9689 | -0.21602 | 0.257022 | 0.883082 | Atg10 |
| ENSMUSG( | 432.9959 | -0.09898 | 0.540778 | 0.937439 | Atg101 |

| | | | | | |
|---|---|---|---|---|---|
| ENSMUSG( | Ark2c | 2678.356 | 0.347334 | 0.000598 | 0.022994 |
| ENSMUSG( | Ark2n | 829.9453 | 0.090583 | 0.381939 | 0.802983 |
| ENSMUSG( | Arl1 | 2301.776 | -0.06077 | 0.514495 | 0.872055 |
| ENSMUSG( | Arl10 | 313.202 | -0.3422 | 0.104847 | 0.491651 |
| ENSMUSG( | Arl11 | 26.09119 | -0.24828 | 0.495778 | 0.861294 |
| ENSMUSG( | Arl13b | 342.9785 | -0.01231 | 0.927984 | 0.991229 |
| ENSMUSG( | Arl14ep | 842.2687 | -0.01065 | 0.918719 | 0.990075 |
| ENSMUSG( | Arl15 | 585.0882 | 0.018139 | 0.886581 | 0.983204 |
| ENSMUSG( | Arl16 | 646.812 | -0.00332 | 0.97807 | 0.997428 |
| ENSMUSG( | Arl2 | 1346.916 | -0.03531 | 0.740527 | 0.947823 |
| ENSMUSG( | Arl2bp | 9193.912 | 0.003014 | 0.976287 | 0.997428 |
| ENSMUSG( | Arl3 | 2915.706 | -0.01859 | 0.858978 | 0.974707 |
| ENSMUSG( | Arl4a | 838.1206 | -0.08155 | 0.470666 | 0.84674 |
| ENSMUSG( | Arl4c | 1402.957 | -0.02698 | 0.801652 | 0.962077 |
| ENSMUSG( | Arl4d | 422.6032 | -0.79544 | 3.94E-06 | 0.000532 |
| ENSMUSG( | Arl5a | 2600.7 | 0.0668 | 0.541461 | 0.883182 |
| ENSMUSG( | Arl5b | 445.9243 | -0.06104 | 0.629704 | 0.913965 |
| ENSMUSG( | Arl5c | 70.04178 | -0.6493 | 0.010372 | 0.145282 |
| ENSMUSG( | Arl6 | 416.0229 | -0.00351 | 0.980732 | 0.997512 |
| ENSMUSG( | Arl6ip1 | 10862.41 | -0.33944 | 0.005646 | 0.100551 |
| ENSMUSG( | Arl6ip4 | 898.7957 | 0.115251 | 0.298947 | 0.747827 |
| ENSMUSG( | Arl6ip5 | 5567.012 | -0.4416 | 0.000164 | 0.009126 |
| ENSMUSG( | Arl6ip6 | 208.5542 | -0.36994 | 0.016527 | 0.191072 |
| ENSMUSG( | Arl8a | 7725.216 | -0.04801 | 0.552034 | 0.886141 |
| ENSMUSG( | Arl8b | 4861.888 | -0.07841 | 0.416851 | 0.8209 |
| ENSMUSG( | Armc1 | 1744.138 | 0.088754 | 0.363236 | 0.793211 |
| ENSMUSG( | Armc10 | 473.2154 | -0.10882 | 0.367271 | 0.795221 |
| ENSMUSG( | Armc2 | 219.4921 | 0.011749 | 0.942601 | 0.993415 |
| ENSMUSG( | Armc3 | 33.11806 | -0.13642 | 0.719789 | 0.941986 |
| ENSMUSG( | Armc5 | 498.1949 | -0.02384 | 0.839751 | 0.971179 |
| ENSMUSG( | Armc6 | 607.858 | -0.07692 | 0.570624 | 0.893192 |
| ENSMUSG( | Armc7 | 215.3333 | -0.10662 | 0.489087 | 0.859092 |
| ENSMUSG( | Armc8 | 1621.625 | 0.087459 | 0.341729 | 0.778828 |
| ENSMUSG( | Armc9 | 683.0373 | 0.101145 | 0.471621 | 0.847331 |
| ENSMUSG( | Armcx1 | 1242.309 | -0.00639 | 0.955156 | 0.993874 |
| ENSMUSG( | Armcx2 | 2759.29 | 0.110807 | 0.203346 | 0.643631 |
| ENSMUSG( | Armcx3 | 1117.233 | 0.077982 | 0.483628 | 0.85636 |
| ENSMUSG( | Armcx4 | 1621.166 | -0.01076 | 0.931228 | 0.992282 |
| ENSMUSG( | Armcx5 | 392.4482 | -0.14724 | 0.250078 | 0.699718 |
| ENSMUSG( | Armcx6 | 83.0496 | -0.06388 | 0.794192 | 0.959799 |
| ENSMUSG( | Armh1 | 20.3687 | -0.2082 | 0.658169 | NA |
| ENSMUSG( | Armh3 | 648.6517 | -0.06248 | 0.634866 | 0.914078 |
| ENSMUSG( | Armh4 | 901.6714 | -0.44469 | 0.012592 | 0.161972 |
| ENSMUSG( | Armt1 | 361.8934 | -0.01248 | 0.921652 | 0.990436 |
| ENSMUSG( | Arnt | 1230.911 | 0.175301 | 0.10094 | 0.484681 |
| ENSMUSG( | Arnt2 | 15978.06 | 0.089036 | 0.219768 | 0.665168 |
| ENSMUSG( | Arpc1a | 4929.023 | -0.00862 | 0.931109 | 0.992271 |
| ENSMUSG( | Arpc1b | 1851.501 | 0.058908 | 0.592847 | 0.902275 |
| ENSMUSG( | Arpc2 | 4909.658 | 0.073899 | 0.348523 | 0.78132 |
| ENSMUSG( | Arpc3 | 2536.39 | -0.04516 | 0.672516 | 0.929007 |
| ENSMUSG( | Arpc4 | 2943.194 | 0.036139 | 0.71524 | 0.941288 |
| ENSMUSG( | Arpc5 | 2236.532 | -0.03755 | 0.688823 | 0.933441 |
| ENSMUSG( | Arpc5l | 2172.609 | 0.000442 | 0.996269 | 0.99943 |
| ENSMUSG( | Arpin | 556.081 | 0.063056 | 0.591666 | 0.901449 |
| ENSMUSG( | Arpp19 | 5272.992 | -0.00681 | 0.948144 | 0.993821 |
| ENSMUSG( | Arpp21 | 11513.03 | -0.11911 | 0.243215 | 0.69052 |
| ENSMUSG( | Arr3 | 27.07158 | 0.329869 | 0.376258 | 0.801127 |
| ENSMUSG( | Arrb1 | 21237.13 | 0.145148 | 0.11002 | 0.500285 |
| ENSMUSG( | Arrb2 | 1365.943 | 0.130361 | 0.281927 | 0.73186 |
| ENSMUSG( | Arrdc1 | 239.9236 | -0.14997 | 0.366172 | 0.794603 |
| ENSMUSG( | Arrdc2 | 413.7135 | -0.35048 | 0.03679 | 0.296095 |
| ENSMUSG( | Arrdc3 | 776.0378 | -0.1722 | 0.110471 | 0.50108 |
| ENSMUSG( | Arrdc4 | 335.0688 | 0.038006 | 0.786029 | 0.958774 |
| ENSMUSG( | Arsa | 1268.657 | -0.14443 | 0.339102 | 0.777518 |
| ENSMUSG( | Arsb | 5712.083 | -0.28303 | 0.04086 | 0.312615 |
| ENSMUSG( | Arsg | 661.2293 | -0.64894 | 2.70E-06 | 0.000405 |
| ENSMUSG( | Arsi | 51.60967 | 0.095074 | 0.747484 | 0.949604 |
| ENSMUSG( | Arsj | 40.50148 | 0.201742 | 0.514325 | 0.872055 |
| ENSMUSG( | Arsk | 374.3907 | -0.16444 | 0.233454 | 0.681508 |
| ENSMUSG( | Art3 | 155.6424 | -0.17872 | 0.473131 | 0.848727 |
| ENSMUSG( | Artn | 13.18839 | -0.02177 | 0.965475 | NA |
| ENSMUSG( | Arv1 | 147.1788 | -0.15729 | 0.454378 | 0.837311 |
| ENSMUSG( | Arvcf | 933.1993 | 0.141357 | 0.245172 | 0.692812 |
| ENSMUSG( | Arx | 544.1914 | -0.03187 | 0.806427 | 0.962804 |
| ENSMUSG( | Arxes1 | 653.9716 | -0.56392 | 3.39E-05 | 0.002956 |
| ENSMUSG( | Arxes2 | 1037.125 | -0.61808 | 0.002869 | 0.065631 |
| ENSMUSG( | As3mt | 846.1425 | 0.045683 | 0.756487 | 0.951479 |
| ENSMUSG( | Asah1 | 2069.226 | -0.31292 | 0.044358 | 0.323404 |
| ENSMUSG( | Asah2 | 502.8808 | -0.25236 | 0.064731 | 0.394417 |
| ENSMUSG( | Asap1 | 3616.828 | 0.224393 | 0.023678 | 0.23542 |
| ENSMUSG( | Asap2 | 1804.098 | 0.302737 | 0.001727 | 0.046912 |
| ENSMUSG( | Asap3 | 304.325 | 0.015515 | 0.909976 | 0.9881 |
| ENSMUSG( | Asb1 | 1526.018 | 0.08494 | 0.361283 | 0.792797 |
| ENSMUSG( | Asb10 | 13.58617 | -0.3939 | 0.502732 | NA |
| ENSMUSG( | Asb11 | 80.31326 | -0.44837 | 0.052734 | 0.352577 |
| ENSMUSG( | Asb13 | 1307.711 | 0.033383 | 0.726964 | 0.944223 |
| ENSMUSG( | Asb16 | 78.39892 | 0.112266 | 0.611784 | 0.908586 |
| ENSMUSG( | Asb18 | 50.42365 | -0.12766 | 0.630807 | 0.914073 |
| ENSMUSG( | Asb2 | 84.00435 | -0.21246 | 0.372484 | 0.798768 |
| ENSMUSG( | Asb3 | 237.7625 | 0.272506 | 0.072341 | 0.415061 |
| ENSMUSG( | Asb4 | 27.60489 | -0.27276 | 0.470312 | 0.84674 |
| ENSMUSG( | Asb5 | 30.2437 | -0.18683 | 0.582211 | 0.897918 |
| ENSMUSG( | Asb6 | 743.3453 | 0.052139 | 0.687841 | 0.93324 |
| ENSMUSG( | Asb7 | 536.5787 | 0.119782 | 0.369021 | 0.796581 |
| ENSMUSG( | Asb8 | 735.7309 | 0.080491 | 0.52177 | 0.875186 |
| ENSMUSG( | Ascc1 | 442.8406 | -0.06455 | 0.665269 | 0.926953 |
| ENSMUSG( | Ascc2 | 1180.017 | 0.115488 | 0.24189 | 0.689179 |
| ENSMUSG( | Ascc3 | 429.5571 | 0.089607 | 0.50016 | 0.863455 |
| ENSMUSG( | Ascl1 | 176.9079 | 0.058921 | 0.718135 | 0.941721 |
| ENSMUSG( | Ascl5 | 10.45237 | -0.33207 | 0.617925 | NA |
| ENSMUSG( | Asf1a | 307.7903 | 0.050845 | 0.701044 | 0.93743 |
| ENSMUSG( | Asf1b | 15.89693 | -0.58612 | 0.315145 | NA |
| ENSMUSG( | Asgr1 | 42.04607 | -0.14339 | 0.635779 | 0.9147 |
| ENSMUSG( | Ash1l | 4227.921 | 0.000327 | 0.997386 | 0.999605 |
| ENSMUSG( | Ash2l | 1159.352 | -0.03284 | 0.744728 | 0.94902 |
| ENSMUSG( | Asic1 | 2608.171 | -0.19339 | 0.139143 | 0.555202 |
| ENSMUSG( | Asic2 | 2431.738 | 0.066541 | 0.571921 | 0.893523 |
| ENSMUSG( | Asic3 | 10.11714 | -0.32478 | 0.580065 | NA |
| ENSMUSG( | Asic4 | 735.8545 | -0.936 | 1.48E-07 | 3.75E-05 |
| ENSMUSG( | Asl | 459.4449 | 0.189262 | 0.150311 | 0.574875 |
| ENSMUSG( | Asns | 1481.194 | 0.03254 | 0.820577 | 0.965888 |
| ENSMUSG( | Asnsd1 | 110.1426 | -0.08752 | 0.673293 | 0.929007 |
| ENSMUSG( | Aspa | 314.8092 | -0.16802 | 0.220603 | 0.666079 |
| ENSMUSG( | Aspdh | 95.57404 | 0.056433 | 0.821266 | 0.966248 |
| ENSMUSG( | Aspg | 166.0118 | 0.166235 | 0.357668 | 0.789196 |
| ENSMUSG( | Asph | 3943.292 | -0.39167 | 0.001342 | 0.039921 |
| ENSMUSG( | Asphd1 | 867.0701 | -0.29198 | 0.010268 | 0.144326 |
| ENSMUSG( | Asphd2 | 4393.913 | -0.03254 | 0.757363 | 0.951513 |
| ENSMUSG( | Aspm | 48.06855 | -0.14178 | 0.680157 | 0.930983 |
| ENSMUSG( | Aspn | 31.16933 | -0.62812 | 0.12927 | 0.538528 |
| ENSMUSG( | Asprv1 | 56.48502 | 0.119909 | 0.711572 | 0.940028 |
| ENSMUSG( | Aspscr1 | 1112.495 | 0.023923 | 0.80419 | 0.962169 |
| ENSMUSG( | Asrgl1 | 30895.14 | 0.094149 | 0.422073 | 0.823495 |
| ENSMUSG( | Ass1 | 824.6885 | 0.070528 | 0.480258 | 0.854705 |
| ENSMUSG( | Aste1 | 139.9762 | -0.07619 | 0.679017 | 0.930914 |
| ENSMUSG( | Astn1 | 9416.757 | -0.24495 | 0.098233 | 0.479534 |
| ENSMUSG( | Astn2 | 1981.039 | -0.2531 | 0.086992 | 0.453311 |
| ENSMUSG( | Asxl1 | 1094.89 | 0.071309 | 0.4903 | 0.859319 |
| ENSMUSG( | Asxl2 | 946.6428 | -0.04644 | 0.697681 | 0.93578 |
| ENSMUSG( | Asxl3 | 234.8943 | -0.42881 | 0.00415 | 0.083415 |
| ENSMUSG( | Atad1 | 3523.747 | -0.36952 | 0.008884 | 0.132538 |
| ENSMUSG( | Atad2 | 221.7608 | -0.03289 | 0.856679 | 0.974154 |
| ENSMUSG( | Atad2b | 703.8302 | 0.020586 | 0.860792 | 0.975003 |
| ENSMUSG( | Atad3a | 1115.117 | 0.178902 | 0.096355 | 0.476986 |
| ENSMUSG( | Atad3aos | 22.99835 | 0.15797 | 0.686392 | 0.933041 |
| ENSMUSG( | Atad5 | 144.7688 | 0.031019 | 0.858349 | 0.974683 |
| ENSMUSG( | Atat1 | 1750.587 | -0.02288 | 0.814444 | 0.964574 |
| ENSMUSG( | Atcay | 6474.726 | -0.14623 | 0.238989 | 0.686574 |
| ENSMUSG( | Ate1 | 1318.697 | 0.059728 | 0.559985 | 0.888917 |
| ENSMUSG( | Atf1 | 369.0616 | -0.0595 | 0.676967 | 0.930542 |
| ENSMUSG( | Atf2 | 1976.098 | 0.057424 | 0.571363 | 0.893319 |
| ENSMUSG( | Atf3 | 23.07155 | -0.3297 | 0.418706 | 0.821966 |
| ENSMUSG( | Atf4 | 3127.19 | -0.03836 | 0.745426 | 0.949158 |
| ENSMUSG( | Atf5 | 2366.198 | 0.076011 | 0.459709 | 0.840338 |
| ENSMUSG( | Atf6 | 2278.341 | -0.23674 | 0.110876 | 0.501952 |
| ENSMUSG( | Atf6b | 2723.178 | -0.09868 | 0.429125 | 0.826446 |
| ENSMUSG( | Atf7 | 861.0564 | 0.082797 | 0.446224 | 0.833554 |
| ENSMUSG( | Atf7ip | 2193.008 | 0.093622 | 0.266596 | 0.716517 |
| ENSMUSG( | Atg10 | 209.446 | -0.06491 | 0.704415 | 0.938303 |
| ENSMUSG( | Atg101 | 499.6858 | 0.053695 | 0.690111 | 0.933812 |
| ENSMUSG( | Atg12 | 2864.179 | 0.068517 | 0.532311 | 0.879905 |
| ENSMUSG( | Atg13 | 2465.532 | 0.142061 | 0.208441 | 0.650319 |
| ENSMUSG( | Atg14 | 509.9897 | 0.108457 | 0.365841 | 0.794583 |
| ENSMUSG( | Atg16l1 | 1298.472 | 0.033283 | 0.799949 | 0.961622 |
| ENSMUSG( | Atg16l2 | 660.0769 | 0.321249 | 0.012627 | 0.162076 |
| ENSMUSG( | Atg2a | 1891.74 | 0.250103 | 0.011837 | 0.157374 |
| ENSMUSG( | Atg2b | 956.8492 | 0.108533 | 0.336782 | 0.777 |
| ENSMUSG( | Atg3 | 1494.493 | 0.112373 | 0.294763 | 0.743798 |

| | | | | | |
|---|---|---|---|---|---|
| ENSMUSG( | Ark2c | 2472.578 | 0.38319 | 0.000923 | 0.102962 |
| ENSMUSG( | Ark2n | 763.4488 | 0.112127 | 0.396769 | 0.917257 |
| ENSMUSG( | Arl1 | 2125.46 | -0.02303 | 0.845018 | 0.983863 |
| ENSMUSG( | Arl10 | 338.8805 | 0.18213 | 0.454598 | 0.92252 |
| ENSMUSG( | Arl11 | 26.27268 | 0.013037 | 0.974776 | 0.996416 |
| ENSMUSG( | Arl13b | 327.3698 | 0.121 | 0.430331 | 0.917257 |
| ENSMUSG( | Arl14ep | 766.5836 | -0.01499 | 0.899101 | 0.990558 |
| ENSMUSG( | Arl15 | 523.9147 | -0.03375 | 0.82178 | 0.979198 |
| ENSMUSG( | Arl16 | 568.3338 | -0.11245 | 0.420391 | 0.917257 |
| ENSMUSG( | Arl2 | 1171.66 | -0.17816 | 0.190043 | 0.857355 |
| ENSMUSG( | Arl2bp | 7884.993 | -0.17908 | 0.173436 | 0.844317 |
| ENSMUSG( | Arl3 | 2503.861 | -0.19957 | 0.126265 | 0.81516 |
| ENSMUSG( | Arl4a | 823.4527 | 0.13224 | 0.30736 | 0.898538 |
| ENSMUSG( | Arl4c | 1338.633 | 0.106794 | 0.365266 | 0.914449 |
| ENSMUSG( | Arl4d | 390.8717 | -0.73426 | 0.000225 | 0.039463 |
| ENSMUSG( | Arl5a | 2345.659 | 0.036002 | 0.791245 | 0.97681 |
| ENSMUSG( | Arl5b | 425.7908 | 0.07511 | 0.60103 | 0.947749 |
| ENSMUSG( | Arl5c | 65.02911 | -0.61043 | 0.032732 | 0.637512 |
| ENSMUSG( | Arl6 | 369.6793 | -0.08279 | 0.631626 | 0.953251 |
| ENSMUSG( | Arl6ip1 | 11395.32 | 0.087097 | 0.557168 | 0.939565 |
| ENSMUSG( | Arl6ip4 | 810.6408 | 0.082543 | 0.575708 | 0.942912 |
| ENSMUSG( | Arl6ip5 | 5922.877 | 0.040392 | 0.768175 | 0.974236 |
| ENSMUSG( | Arl6ip6 | 229.8659 | 0.201558 | 0.212649 | 0.864412 |
| ENSMUSG( | Arl8a | 7080.937 | -0.03197 | 0.795069 | 0.97681 |
| ENSMUSG( | Arl8b | 4561.91 | 0.007741 | 0.946451 | 0.994798 |
| ENSMUSG( | Armc1 | 1539.785 | -0.00443 | 0.97314 | 0.996416 |
| ENSMUSG( | Armc10 | 443.3077 | -0.02533 | 0.857485 | 0.985884 |
| ENSMUSG( | Armc2 | 204.9489 | 0.079368 | 0.704686 | 0.962677 |
| ENSMUSG( | Armc3 | 34.9094 | 0.302795 | 0.457858 | 0.923329 |
| ENSMUSG( | Armc5 | 470.4634 | 0.085777 | 0.565917 | 0.941499 |
| ENSMUSG( | Armc6 | 556.5466 | -0.06228 | 0.680258 | 0.958115 |
| ENSMUSG( | Armc7 | 193.7644 | -0.14438 | 0.418625 | 0.917257 |
| ENSMUSG( | Armc8 | 1387.146 | -0.09757 | 0.40693 | 0.917257 |
| ENSMUSG( | Armc9 | 601.4259 | 0.002115 | 0.988943 | 0.997887 |
| ENSMUSG( | Armcx1 | 1121.017 | -0.03918 | 0.776646 | 0.975717 |
| ENSMUSG( | Armcx2 | 2461.347 | 0.052838 | 0.621944 | 0.953234 |
| ENSMUSG( | Armcx3 | 1027.413 | 0.101496 | 0.42913 | 0.917257 |
| ENSMUSG( | Armcx4 | 1552.71 | 0.133583 | 0.374955 | 0.915491 |
| ENSMUSG( | Armcx5 | 366.7925 | -0.06968 | 0.637947 | 0.953903 |
| ENSMUSG( | Armcx6 | 80.47958 | 0.133461 | 0.614179 | 0.951177 |
| ENSMUSG( | Armh1 | 19.88016 | 0.026378 | 0.957066 | 0.994798 |
| ENSMUSG( | Armh3 | 592.2048 | -0.05665 | 0.696639 | 0.960867 |
| ENSMUSG( | Armh4 | 1021.169 | 0.215478 | 0.247337 | 0.877904 |
| ENSMUSG( | Armt1 | 323.7774 | -0.06296 | 0.663117 | 0.956349 |
| ENSMUSG( | Arnt | 1107.395 | 0.142063 | 0.242629 | 0.877399 |
| ENSMUSG( | Arnt2 | 13560.27 | -0.11672 | 0.214141 | 0.864421 |
| ENSMUSG( | Arpc1a | 4282.773 | -0.15165 | 0.223677 | 0.867404 |
| ENSMUSG( | Arpc1b | 1592.903 | -0.11066 | 0.380043 | 0.916995 |
| ENSMUSG( | Arpc2 | 4257.12 | -0.07099 | 0.575092 | 0.942912 |
| ENSMUSG( | Arpc3 | 2201.881 | -0.19379 | 0.170571 | 0.841739 |
| ENSMUSG( | Arpc4 | 2569.23 | -0.08842 | 0.401956 | 0.917257 |
| ENSMUSG( | Arpc5 | 2005.911 | -0.08742 | 0.489079 | 0.925673 |
| ENSMUSG( | Arpc5l | 1897.513 | -0.12693 | 0.29448 | 0.898538 |
| ENSMUSG( | Arpin | 490.7609 | -0.02615 | 0.839253 | 0.981912 |
| ENSMUSG( | Arpp19 | 4622.892 | -0.12386 | 0.405563 | 0.917257 |
| ENSMUSG( | Arpp21 | 10761.2 | -0.04484 | 0.734267 | 0.967228 |
| ENSMUSG( | Arr3 | 18.54645 | -0.5004 | 0.321512 | 0.904019 |
| ENSMUSG( | Arrb1 | 17950.14 | -0.06786 | 0.502769 | 0.927825 |
| ENSMUSG( | Arrb2 | 1240.462 | 0.121117 | 0.403759 | 0.917257 |
| ENSMUSG( | Arrdc1 | 228.6398 | -0.0088 | 0.969605 | 0.99594 |
| ENSMUSG( | Arrdc2 | 403.7429 | -0.12297 | 0.510962 | 0.929546 |
| ENSMUSG( | Arrdc3 | 817.1949 | 0.245897 | 0.108175 | 0.804969 |
| ENSMUSG( | Arrdc4 | 298.3974 | -0.02911 | 0.840685 | 0.982538 |
| ENSMUSG( | Arsa | 1279.821 | 0.154934 | 0.370477 | 0.915491 |
| ENSMUSG( | Arsb | 6032.275 | 0.156575 | 0.333978 | 0.907907 |
| ENSMUSG( | Arsg | 713.0862 | -0.08323 | 0.625013 | 0.953234 |
| ENSMUSG( | Arsi | 44.83621 | -0.01651 | 0.957854 | 0.994798 |
| ENSMUSG( | Arsj | 41.01615 | 0.497813 | 0.144941 | 0.83339 |
| ENSMUSG( | Arsk | 374.8986 | 0.111981 | 0.464292 | 0.923329 |
| ENSMUSG( | Art3 | 192.8512 | 0.659882 | 0.035596 | 0.664751 |
| ENSMUSG( | Artn | 12.2858 | 0.033403 | 0.95137 | 0.994798 |
| ENSMUSG( | Arv1 | 142.7341 | 0.038668 | 0.853978 | 0.98563 |
| ENSMUSG( | Arvcf | 854.3158 | 0.158703 | 0.237782 | 0.87259 |
| ENSMUSG( | Arx | 529.9678 | 0.152825 | 0.265372 | 0.886134 |
| ENSMUSG( | Arxes1 | 712.4161 | 0.004908 | 0.977572 | 0.996416 |
| ENSMUSG( | Arxes2 | 1157.254 | 0.029486 | 0.942982 | 0.994774 |
| ENSMUSG( | As3mt | 720.8883 | -0.15624 | 0.304289 | 0.898538 |
| ENSMUSG( | Asah1 | 2222.227 | 0.177885 | 0.315166 | 0.901306 |
| ENSMUSG( | Asah2 | 496.681 | -0.00827 | 0.95916 | 0.994798 |
| ENSMUSG( | Asap1 | 3068.076 | 0.027142 | 0.812737 | 0.979148 |
| ENSMUSG( | Asap2 | 1581.085 | 0.197037 | 0.106781 | 0.804969 |
| ENSMUSG( | Asap3 | 307.9181 | 0.305105 | 0.052824 | 0.71525 |
| ENSMUSG( | Asb1 | 1378.365 | 0.061544 | 0.580188 | 0.944209 |
| ENSMUSG( | Asb10 | 9.46021 | -1.5584 | 0.019267 | NA |
| ENSMUSG( | Asb11 | 87.21789 | 0.099771 | 0.67781 | 0.95713 |
| ENSMUSG( | Asb13 | 1159.85 | -0.04863 | 0.705719 | 0.96315 |
| ENSMUSG( | Asb16 | 68.87621 | -0.01417 | 0.958677 | 0.994798 |
| ENSMUSG( | Asb18 | 44.75931 | -0.18477 | 0.544923 | 0.938063 |
| ENSMUSG( | Asb2 | 78.43492 | -0.12274 | 0.622274 | 0.953234 |
| ENSMUSG( | Asb3 | 205.3749 | 0.129487 | 0.44689 | 0.921891 |
| ENSMUSG( | Asb4 | 20.45999 | -1.0358 | 0.019693 | 0.556192 |
| ENSMUSG( | Asb5 | 26.61879 | -0.33396 | 0.404535 | 0.917257 |
| ENSMUSG( | Asb6 | 658.4344 | -0.02644 | 0.859288 | 0.985884 |
| ENSMUSG( | Asb7 | 473.2101 | 0.024154 | 0.869979 | 0.988097 |
| ENSMUSG( | Asb8 | 657.3812 | 0.023616 | 0.86352 | 0.986547 |
| ENSMUSG( | Ascc1 | 398.4433 | -0.10521 | 0.539811 | 0.937378 |
| ENSMUSG( | Ascc2 | 1006.73 | -0.0746 | 0.598881 | 0.947045 |
| ENSMUSG( | Ascc3 | 391.0394 | 0.094263 | 0.513277 | 0.930482 |
| ENSMUSG( | Ascl1 | 167.6183 | 0.161301 | 0.374305 | 0.915491 |
| ENSMUSG( | Ascl5 | 11.90118 | 0.316 | 0.611702 | 0.950771 |
| ENSMUSG( | Asf1a | 292.2714 | 0.156461 | 0.326823 | 0.905419 |
| ENSMUSG( | Asf1b | 16.30037 | -0.19042 | 0.705595 | 0.96315 |
| ENSMUSG( | Asgr1 | 29.19663 | -1.14991 | 0.002273 | 0.185073 |
| ENSMUSG( | Ash1l | 3877.613 | 0.021203 | 0.849755 | 0.985522 |
| ENSMUSG( | Ash2l | 1037.355 | -0.08427 | 0.494708 | 0.927372 |
| ENSMUSG( | Asic1 | 2667.658 | 0.146865 | 0.334681 | 0.907907 |
| ENSMUSG( | Asic2 | 2467.766 | 0.359596 | 0.008264 | 0.388416 |
| ENSMUSG( | Asic3 | 11.03847 | 0.208464 | 0.729389 | NA |
| ENSMUSG( | Asic4 | 890.8242 | 0.036632 | 0.827344 | 0.980224 |
| ENSMUSG( | Asl | 372.5673 | -0.14469 | 0.345092 | 0.91045 |
| ENSMUSG( | Asns | 1227.987 | -0.25233 | 0.092022 | 0.791889 |
| ENSMUSG( | Asnsd1 | 106.0224 | 0.059373 | 0.78565 | 0.976285 |
| ENSMUSG( | Aspa | 331.8692 | 0.253828 | 0.133864 | 0.822912 |
| ENSMUSG( | Aspdh | 94.79988 | 0.295346 | 0.182392 | 0.850579 |
| ENSMUSG( | Aspg | 136.5585 | -0.10933 | 0.665936 | 0.956766 |
| ENSMUSG( | Asph | 3962.758 | -0.08278 | 0.566396 | 0.941745 |
| ENSMUSG( | Asphd1 | 848.4037 | -0.0734 | 0.583058 | 0.944266 |
| ENSMUSG( | Asphd2 | 4165.436 | 0.083984 | 0.500274 | 0.927825 |
| ENSMUSG( | Aspm | 44.97684 | -0.04391 | 0.907684 | 0.991958 |
| ENSMUSG( | Aspn | 42.98121 | 0.59296 | 0.098224 | 0.798438 |
| ENSMUSG( | Asprv1 | 53.96523 | 0.230538 | 0.472341 | 0.923329 |
| ENSMUSG( | Aspscr1 | 1006.766 | 0.005674 | 0.961056 | 0.99511 |
| ENSMUSG( | Asrgl1 | 25233.22 | -0.23079 | 0.119049 | 0.813384 |
| ENSMUSG( | Ass1 | 721.1693 | -0.05098 | 0.690217 | 0.960071 |
| ENSMUSG( | Aste1 | 141.2902 | 0.208931 | 0.283411 | 0.892412 |
| ENSMUSG( | Astn1 | 9742.469 | 0.133424 | 0.419593 | 0.917257 |
| ENSMUSG( | Astn2 | 2037.746 | 0.11044 | 0.504927 | 0.927825 |
| ENSMUSG( | Asxl1 | 995.3706 | 0.067384 | 0.56688 | 0.941777 |
| ENSMUSG( | Asxl2 | 900.2086 | 0.080288 | 0.564188 | 0.941206 |
| ENSMUSG( | Asxl3 | 227.3386 | -0.23218 | 0.155458 | 0.837724 |
| ENSMUSG( | Atad1 | 3783.142 | 0.127722 | 0.424393 | 0.917257 |
| ENSMUSG( | Atad2 | 223.6431 | 0.259797 | 0.247901 | 0.877904 |
| ENSMUSG( | Atad2b | 675.6879 | 0.168486 | 0.207364 | 0.864412 |
| ENSMUSG( | Atad3a | 974.1318 | 0.064773 | 0.615098 | 0.951177 |
| ENSMUSG( | Atad3aos | 21.3255 | 0.184737 | 0.674242 | 0.95694 |
| ENSMUSG( | Atad5 | 144.4172 | 0.285097 | 0.136125 | 0.827092 |
| ENSMUSG( | Atat1 | 1556.7 | -0.09635 | 0.430928 | 0.917257 |
| ENSMUSG( | Atcay | 6432.244 | 0.108884 | 0.430353 | 0.917257 |
| ENSMUSG( | Ate1 | 1182.572 | 0.017145 | 0.884998 | 0.989688 |
| ENSMUSG( | Atf1 | 342.5935 | -0.01592 | 0.928437 | 0.99306 |
| ENSMUSG( | Atf2 | 1743.832 | -0.0389 | 0.794701 | 0.97681 |
| ENSMUSG( | Atf3 | 21.88771 | -0.14445 | 0.770932 | 0.974236 |
| ENSMUSG( | Atf4 | 2783.17 | -0.10904 | 0.442689 | 0.921554 |
| ENSMUSG( | Atf5 | 2006.023 | -0.13831 | 0.339031 | 0.907972 |
| ENSMUSG( | Atf6 | 2388.901 | 0.17874 | 0.318181 | 0.902849 |
| ENSMUSG( | Atf6b | 2748.328 | 0.195071 | 0.186178 | 0.855305 |
| ENSMUSG( | Atf7 | 773.5149 | 0.045631 | 0.71934 | 0.966876 |
| ENSMUSG( | Atf7ip | 1961.336 | 0.040973 | 0.712601 | 0.964363 |
| ENSMUSG( | Atg10 | 181.9689 | -0.21602 | 0.257022 | 0.883082 |
| ENSMUSG( | Atg101 | 432.9959 | -0.09898 | 0.540778 | 0.937439 |
| ENSMUSG( | Atg12 | 2391.189 | -0.19291 | 0.142864 | 0.831221 |
| ENSMUSG( | Atg13 | 2147.549 | 0.016864 | 0.893137 | 0.990558 |
| ENSMUSG( | Atg14 | 448.4844 | 0.010612 | 0.937313 | 0.993685 |
| ENSMUSG( | Atg16l1 | 1219.451 | 0.124787 | 0.429315 | 0.917257 |
| ENSMUSG( | Atg16l2 | 606.4172 | 0.343937 | 0.014923 | 0.489529 |
| ENSMUSG( | Atg2a | 1620.891 | 0.084704 | 0.524002 | 0.933009 |
| ENSMUSG( | Atg2b | 910.6371 | 0.233231 | 0.105967 | 0.804969 |
| ENSMUSG( | Atg3 | 1303.773 | -0.01538 | 0.904425 | 0.991089 |

| | | | | | |
|---|---|---|---|---|---|
| ENSMUSG( | 144.7688 | 0.031019 | 0.858349 | 0.974683 | Atad5 |
| ENSMUSG( | 1750.587 | -0.02288 | 0.814444 | 0.964574 | Atat1 |
| ENSMUSG( | 6474.726 | -0.14623 | 0.238989 | 0.686574 | Atcay |
| ENSMUSG( | 1318.697 | 0.059728 | 0.559985 | 0.888917 | Ate1 |
| ENSMUSG( | 369.0616 | -0.0595 | 0.676967 | 0.930542 | Atf1 |
| ENSMUSG( | 1976.098 | 0.057424 | 0.571363 | 0.893319 | Atf2 |
| ENSMUSG( | 23.07155 | -0.3297 | 0.418706 | 0.821966 | Atf3 |
| ENSMUSG( | 3127.19 | -0.03836 | 0.745426 | 0.949158 | Atf4 |
| ENSMUSG( | 2366.198 | 0.076011 | 0.459709 | 0.840338 | Atf5 |
| ENSMUSG( | 2278.341 | -0.23674 | 0.110876 | 0.501952 | Atf6 |
| ENSMUSG( | 2723.178 | -0.09868 | 0.429125 | 0.826446 | Atf6b |
| ENSMUSG( | 861.0564 | 0.082797 | 0.446224 | 0.833554 | Atf7 |
| ENSMUSG( | 2193.008 | 0.093622 | 0.266596 | 0.716517 | Atf7ip |
| ENSMUSG( | 209.446 | -0.06491 | 0.704415 | 0.938303 | Atg10 |
| ENSMUSG( | 499.6858 | 0.053695 | 0.690111 | 0.933812 | Atg101 |
| ENSMUSG( | 2864.179 | 0.068517 | 0.532311 | 0.879905 | Atg12 |
| ENSMUSG( | 2465.532 | 0.142061 | 0.208441 | 0.650319 | Atg13 |
| ENSMUSG( | 509.9897 | 0.108457 | 0.365841 | 0.794583 | Atg14 |
| ENSMUSG( | 1298.472 | 0.033283 | 0.799949 | 0.961622 | Atg16l1 |
| ENSMUSG( | 660.0769 | 0.321249 | 0.012627 | 0.162076 | Atg16l2 |
| ENSMUSG( | 1891.74 | 0.250103 | 0.011837 | 0.157374 | Atg2a |
| ENSMUSG( | 956.8492 | 0.108533 | 0.336782 | 0.777 | Atg2b |
| ENSMUSG( | 1494.493 | 0.112373 | 0.294763 | 0.743798 | Atg3 |
| ENSMUSG( | 265.2207 | -0.09058 | 0.527893 | 0.878017 | Atg4a |
| ENSMUSG( | 1478.398 | 0.142607 | 0.191337 | 0.630245 | Atg4b |
| ENSMUSG( | 781.5211 | 0.203676 | 0.071381 | 0.412911 | Atg4c |
| ENSMUSG( | 830.1384 | 0.104181 | 0.29039 | 0.740353 | Atg4d |
| ENSMUSG( | 574.0935 | 0.174798 | 0.168241 | 0.601572 | Atg5 |
| ENSMUSG( | 870.9553 | 0.009649 | 0.920079 | 0.990247 | Atg7 |
| ENSMUSG( | 1623.378 | 0.000193 | 0.999535 | 0.99989 | Atg9a |
| ENSMUSG( | 143.9387 | -0.17792 | 0.308452 | 0.756439 | Atg9b |
| ENSMUSG( | 933.981 | 0.08718 | 0.374768 | 0.800466 | Atic |
| ENSMUSG( | 2318.406 | -0.41673 | 0.000656 | 0.024688 | Atl1 |
| ENSMUSG( | 2099.579 | -0.33258 | 0.010289 | 0.144496 | Atl2 |
| ENSMUSG( | 1266.631 | -0.14793 | 0.21146 | 0.654808 | Atl3 |
| ENSMUSG( | 444.8898 | 0.07392 | 0.582746 | 0.89798 | Atm |
| ENSMUSG( | 1873.326 | 0.051485 | 0.596125 | 0.903152 | Atmin |
| ENSMUSG( | 698.424 | -0.02843 | 0.786613 | 0.958856 | Atn1 |
| ENSMUSG( | 36.65941 | 0.12607 | 0.719177 | 0.941986 | Atoh7 |
| ENSMUSG( | 68.03479 | -0.08154 | 0.756792 | 0.951513 | Atoh8 |
| ENSMUSG( | 1240.885 | 0.085869 | 0.449756 | 0.835283 | Atosa |
| ENSMUSG( | 1287.077 | 0.004132 | 0.970112 | 0.996369 | Atosb |
| ENSMUSG( | 1155.001 | 0.05826 | 0.530965 | 0.879289 | Atox1 |
| ENSMUSG( | 447.1729 | 0.817071 | 1.14E-05 | 0.00123 | Atp10a |
| ENSMUSG( | 105.267 | -0.82312 | 0.000712 | 0.025833 | Atp10b |
| ENSMUSG( | 148.6147 | 0.028466 | 0.870211 | 0.97782 | Atp10d |
| ENSMUSG( | 2451.902 | -0.16683 | 0.174851 | 0.610304 | Atp11a |
| ENSMUSG( | 2556.38 | -0.18916 | 0.082997 | 0.44459 | Atp11b |
| ENSMUSG( | 312.9997 | -0.36088 | 0.037056 | 0.297106 | Atp11c |
| ENSMUSG( | 1848.86 | -0.06098 | 0.637506 | 0.915642 | Atp13a1 |
| ENSMUSG( | 6712.63 | 0.061267 | 0.63699 | 0.915191 | Atp13a2 |
| ENSMUSG( | 1703.406 | -0.20418 | 0.157291 | 0.585553 | Atp13a3 |
| ENSMUSG( | 1102.495 | 0.055662 | 0.628892 | 0.913816 | Atp13a4 |
| ENSMUSG( | 517.8025 | -0.2747 | 0.149372 | 0.573313 | Atp13a5 |
| ENSMUSG( | 30988.14 | -0.24241 | 0.070388 | 0.40908 | Atp1a1 |
| ENSMUSG( | 138067.9 | 0.043285 | 0.707629 | 0.93949 | Atp1a2 |
| ENSMUSG( | 80666 | -0.42456 | 0.000278 | 0.013625 | Atp1a3 |
| ENSMUSG( | 51031.18 | -0.55058 | 1.38E-05 | 0.001449 | Atp1b1 |
| ENSMUSG( | 44568.21 | -0.16443 | 0.22096 | 0.666476 | Atp1b2 |
| ENSMUSG( | 5041.471 | -0.12127 | 0.179627 | 0.61741 | Atp1b3 |
| ENSMUSG( | 171.3736 | 0.02205 | 0.895318 | 0.984512 | Atp23 |
| ENSMUSG( | 19.66935 | -1.48952 | 0.001426 | NA | Atp2a1 |
| ENSMUSG( | 25008.39 | -0.32832 | 0.014721 | 0.178407 | Atp2a2 |
| ENSMUSG( | 393.8346 | -0.02093 | 0.915404 | 0.988634 | Atp2a3 |
| ENSMUSG( | 13825.74 | -0.52239 | 0.001106 | 0.035255 | Atp2b1 |
| ENSMUSG( | 25916.45 | -0.3755 | 0.002279 | 0.055785 | Atp2b2 |
| ENSMUSG( | 1473.666 | -0.16847 | 0.366353 | 0.794736 | Atp2b3 |
| ENSMUSG( | 6670.777 | -0.33681 | 0.011117 | 0.152038 | Atp2b4 |
| ENSMUSG( | 5024.887 | -0.12519 | 0.340275 | 0.777962 | Atp2c1 |
| ENSMUSG( | 39.75186 | -0.28743 | 0.363395 | 0.793339 | Atp2c2 |
| ENSMUSG( | 28.38907 | 0.035809 | 0.919315 | 0.990247 | Atp4a |
| ENSMUSG( | 24672.74 | 0.156167 | 0.065935 | 0.397572 | Atp5f1a |
| ENSMUSG( | 33341.08 | 0.090638 | 0.25561 | 0.706195 | Atp5f1b |
| ENSMUSG( | 6396.159 | 0.055126 | 0.53301 | 0.879905 | Atp5f1c |
| ENSMUSG( | 6934.009 | 0.009974 | 0.907837 | 0.987798 | Atp5f1d |
| ENSMUSG( | 3575.22 | 0.078443 | 0.452254 | 0.836145 | Atp5f1e |
| ENSMUSG( | 4087.651 | -0.06242 | 0.482345 | 0.855884 | Atp5if1 |
| ENSMUSG( | 3799.038 | -0.03042 | 0.714832 | 0.941144 | Atp5mc1 |
| ENSMUSG( | 1745.929 | -0.02187 | 0.811141 | 0.964318 | Atp5mc2 |
| ENSMUSG( | 6562.236 | 0.004232 | 0.965174 | 0.995303 | Atp5mc3 |
| ENSMUSG( | 2898.238 | 0.025534 | 0.812621 | 0.964574 | Atp5me |
| ENSMUSG( | 4943.986 | -0.00194 | 0.981838 | 0.997512 | Atp5mf |
| ENSMUSG( | 2552.317 | -0.03262 | 0.692484 | 0.934101 | Atp5mg |
| ENSMUSG( | 2908.78 | 0.064728 | 0.505326 | 0.866883 | Atp5mj |
| ENSMUSG( | 4903.9 | -0.026 | 0.768127 | 0.954641 | Atp5mk |
| ENSMUSG( | 6753.571 | 0.026481 | 0.774945 | 0.955606 | Atp5pb |
| ENSMUSG( | 6429.809 | 0.025266 | 0.773019 | 0.955272 | Atp5pd |
| ENSMUSG( | 5722.676 | 0.027815 | 0.749679 | 0.949664 | Atp5pf |
| ENSMUSG( | 2977.91 | 0.083017 | 0.426849 | 0.825231 | Atp5po |
| ENSMUSG( | 3285.924 | -0.08434 | 0.666106 | 0.927057 | ATP6 |
| ENSMUSG( | 15422.67 | -0.26105 | 0.023387 | 0.233882 | Atp6ap1 |
| ENSMUSG( | 267.8709 | -0.36818 | 0.009245 | 0.135575 | Atp6ap1l |
| ENSMUSG( | 5883.516 | -0.53764 | 7.78E-05 | 0.005438 | Atp6ap2 |
| ENSMUSG( | 11637.8 | -0.03791 | 0.851034 | 0.972638 | Atp6-ps |
| ENSMUSG( | 19111.13 | -0.26618 | 0.025021 | 0.243138 | Atp6v0a1 |
| ENSMUSG( | 5425.241 | 0.016832 | 0.881255 | 0.981798 | Atp6v0a2 |
| ENSMUSG( | 7733.874 | -0.23809 | 0.022692 | 0.23004 | Atp6v0b |
| ENSMUSG( | 12369.8 | -0.54801 | 8.50E-06 | 0.001033 | Atp6v0c |
| ENSMUSG( | 5838.962 | 0.019101 | 0.846043 | 0.972226 | Atp6v0d1 |
| ENSMUSG( | 1034.76 | -0.16925 | 0.238472 | 0.686308 | Atp6v0e |
| ENSMUSG( | 14633.74 | -0.36093 | 0.003525 | 0.075237 | Atp6v0e2 |
| ENSMUSG( | 7274.631 | 0.039209 | 0.663939 | 0.926511 | Atp6v1a |
| ENSMUSG( | 10281.01 | 0.144032 | 0.266416 | 0.716441 | Atp6v1b2 |
| ENSMUSG( | 4710.743 | 0.068681 | 0.589714 | 0.90113 | Atp6v1c1 |
| ENSMUSG( | 31.94076 | -0.11102 | 0.791344 | 0.959461 | Atp6v1c2 |
| ENSMUSG( | 3588.372 | -0.10796 | 0.282216 | 0.73186 | Atp6v1d |
| ENSMUSG( | 6073.559 | 0.029162 | 0.813967 | 0.964574 | Atp6v1e1 |
| ENSMUSG( | 2470.618 | -0.02742 | 0.774967 | 0.955606 | Atp6v1f |
| ENSMUSG( | 1798.429 | 0.018501 | 0.843501 | 0.971847 | Atp6v1g1 |
| ENSMUSG( | 11269.17 | 0.023745 | 0.848153 | 0.972374 | Atp6v1g2 |
| ENSMUSG( | 3309.376 | -0.11156 | 0.249342 | 0.698744 | Atp6v1h |
| ENSMUSG( | 133.24 | -0.0986 | 0.67779 | 0.930542 | Atp7a |
| ENSMUSG( | 119.7529 | -0.2148 | 0.231914 | 0.679732 | Atp7b |
| ENSMUSG( | 162.0326 | -0.78229 | 0.160673 | 0.590555 | ATP8 |
| ENSMUSG( | 6283.884 | -0.27896 | 0.049064 | 0.338072 | Atp8a1 |
| ENSMUSG( | 1710.432 | -0.33168 | 0.023375 | 0.233882 | Atp8a2 |
| ENSMUSG( | 131.2751 | -0.59503 | 0.020541 | 0.216203 | Atp8b1 |
| ENSMUSG( | 2405.974 | -0.14022 | 0.191893 | 0.630811 | Atp8b2 |
| ENSMUSG( | 18.53972 | -0.13822 | 0.766227 | NA | Atp8b5 |
| ENSMUSG( | 14468.33 | -0.11122 | 0.413892 | 0.819998 | Atp9a |
| ENSMUSG( | 2242.344 | -0.00752 | 0.949954 | 0.993821 | Atp9b |
| ENSMUSG( | 709.4036 | -0.12447 | 0.346116 | 0.779153 | Atpaf1 |
| ENSMUSG( | 597.4821 | 0.022145 | 0.843745 | 0.971847 | Atpaf2 |
| ENSMUSG( | 285.5588 | 0.076663 | 0.678195 | 0.930542 | Atpsckmt |
| ENSMUSG( | 458.3816 | -0.07617 | 0.57089 | 0.8933 | Atr |
| ENSMUSG( | 2315.878 | -0.19023 | 0.080209 | 0.437895 | Atraid |
| ENSMUSG( | 549.7555 | 0.166505 | 0.154685 | 0.581863 | Atrip |
| ENSMUSG( | 5818.899 | -0.2847 | 0.063184 | 0.388743 | Atrn |
| ENSMUSG( | 2869.692 | -0.40686 | 0.007417 | 0.118765 | Atrnl1 |
| ENSMUSG( | 4084.291 | -0.08127 | 0.434456 | 0.828978 | Atrx |
| ENSMUSG( | 2478.108 | 0.064089 | 0.456945 | 0.839183 | Atxn1 |
| ENSMUSG( | 5778.544 | 0.070457 | 0.495999 | 0.861543 | Atxn10 |
| ENSMUSG( | 1857.251 | 0.014101 | 0.890025 | 0.983687 | Atxn1l |
| ENSMUSG( | 2706.117 | 0.036134 | 0.68802 | 0.93324 | Atxn2 |
| ENSMUSG( | 4348.075 | 0.120573 | 0.331901 | 0.775288 | Atxn2l |
| ENSMUSG( | 598.3097 | 0.082849 | 0.446165 | 0.833546 | Atxn3 |
| ENSMUSG( | 844.7948 | -0.07767 | 0.44829 | 0.834051 | Atxn7 |
| ENSMUSG( | 597.559 | -0.02705 | 0.816265 | 0.965455 | Atxn7l1 |
| ENSMUSG( | 705.6652 | 0.140063 | 0.321451 | 0.766644 | Atxn7l2 |
| ENSMUSG( | 15310.19 | 0.2014 | 0.007215 | 0.117261 | Atxn7l3 |
| ENSMUSG( | 7641.906 | 0.0874 | 0.337718 | 0.777422 | Atxn7l3b |
| ENSMUSG( | 150.7472 | -0.63262 | 0.002359 | 0.057092 | AU020206 |
| ENSMUSG( | 168.7503 | -0.31262 | 0.191173 | 0.630245 | AU021092 |
| ENSMUSG( | 1814.471 | 0.114033 | 0.26036 | 0.710245 | AU022252 |
| ENSMUSG( | 55.54078 | 0.345286 | 0.18943 | 0.62795 | AU022754 |
| ENSMUSG( | 40.65683 | -0.28212 | 0.374873 | 0.800479 | AU023762 |
| ENSMUSG( | 2521.207 | -0.22435 | 0.10341 | 0.489156 | AU040320 |
| ENSMUSG( | 12.76267 | -0.12268 | 0.81275 | NA | AU040972 |
| ENSMUSG( | 87.53681 | -0.10976 | 0.638719 | 0.91636 | AU041133 |
| ENSMUSG( | 1186.65 | 0.07838 | 0.515497 | 0.872701 | Auh |
| ENSMUSG( | 2422.39 | -0.09959 | 0.312541 | 0.760241 | Aup1 |
| ENSMUSG( | 29.26297 | -0.1943 | 0.570009 | 0.893041 | Aurka |
| ENSMUSG( | 1562.867 | -0.01041 | 0.906138 | 0.986921 | Aurkaip1 |
| ENSMUSG( | 19.4542 | -0.74706 | 0.080518 | NA | Aurkb |
| ENSMUSG( | 1970.249 | 0.216656 | 0.01104 | 0.151501 | Auts2 |
| ENSMUSG( | 13.75814 | -0.39947 | 0.425356 | NA | AV039307 |
| ENSMUSG( | 251.1361 | 0.061272 | 0.678046 | 0.930542 | Aven |
| ENSMUSG( | 17.80261 | 0.055761 | 0.903637 | NA | Avil |

| | | | | | |
|---|---|---|---|---|---|
| ENSMUSG( | 2391.189 | -0.19291 | 0.142864 | 0.831221 | Atg12 |
| ENSMUSG( | 2147.549 | 0.016864 | 0.893137 | 0.990558 | Atg13 |
| ENSMUSG( | 448.4844 | 0.010612 | 0.937313 | 0.993685 | Atg14 |
| ENSMUSG( | 1219.451 | 0.124787 | 0.429315 | 0.917257 | Atg16l1 |
| ENSMUSG( | 606.4172 | 0.343937 | 0.014923 | 0.489529 | Atg16l2 |
| ENSMUSG( | 1620.891 | 0.084704 | 0.524002 | 0.933009 | Atg2a |
| ENSMUSG( | 910.6371 | 0.233231 | 0.105967 | 0.804969 | Atg2b |
| ENSMUSG( | 1303.773 | -0.01538 | 0.904425 | 0.991089 | Atg3 |
| ENSMUSG( | 262.1743 | 0.141168 | 0.382696 | 0.917257 | Atg4a |
| ENSMUSG( | 1292.908 | 0.029723 | 0.80978 | 0.978779 | Atg4b |
| ENSMUSG( | 705.9476 | 0.177285 | 0.16687 | 0.840183 | Atg4c |
| ENSMUSG( | 760.8943 | 0.125177 | 0.357821 | 0.913759 | Atg4d |
| ENSMUSG( | 489.0254 | -0.01705 | 0.903089 | 0.991089 | Atg5 |
| ENSMUSG( | 743.7287 | -0.18714 | 0.129271 | 0.820358 | Atg7 |
| ENSMUSG( | 1708.173 | 0.398549 | 0.326601 | 0.905371 | Atg9a |
| ENSMUSG( | 146.8082 | 0.138315 | 0.49113 | 0.926604 | Atg9b |
| ENSMUSG( | 796.1403 | -0.10827 | 0.391556 | 0.917257 | Atic |
| ENSMUSG( | 2509.898 | 0.113369 | 0.440646 | 0.921166 | Atl1 |
| ENSMUSG( | 2236.095 | 0.1359 | 0.330333 | 0.907251 | Atl2 |
| ENSMUSG( | 1279.56 | 0.152747 | 0.270816 | 0.888062 | Atl3 |
| ENSMUSG( | 403.6553 | 0.072904 | 0.663963 | 0.956766 | Atm |
| ENSMUSG( | 1620.803 | -0.09604 | 0.446632 | 0.921734 | Atmin |
| ENSMUSG( | 621.6319 | -0.1053 | 0.55031 | 0.938699 | Atn1 |
| ENSMUSG( | 31.84109 | -0.01185 | 0.9756 | 0.996416 | Atoh7 |
| ENSMUSG( | 69.62257 | 0.27041 | 0.356 | 0.913331 | Atoh8 |
| ENSMUSG( | 1187.615 | 0.225977 | 0.110069 | 0.806572 | Atosa |
| ENSMUSG( | 1177.726 | 0.016324 | 0.886375 | 0.990011 | Atosb |
| ENSMUSG( | 951.4445 | -0.24812 | 0.068249 | 0.752208 | Atox1 |
| ENSMUSG( | 488.9212 | 1.220674 | 4.79E-06 | 0.002153 | Atp10a |
| ENSMUSG( | 124.4412 | 0.061615 | 0.810074 | 0.978779 | Atp10b |
| ENSMUSG( | 149.8076 | 0.317902 | 0.154673 | 0.837724 | Atp10d |
| ENSMUSG( | 2560.641 | 0.227251 | 0.129569 | 0.820557 | Atp11a |
| ENSMUSG( | 2691.033 | 0.227283 | 0.132967 | 0.822912 | Atp11b |
| ENSMUSG( | 364.9415 | 0.355347 | 0.047063 | 0.700664 | Atp11c |
| ENSMUSG( | 1825.689 | 0.17213 | 0.265197 | 0.886134 | Atp13a1 |
| ENSMUSG( | 6533.429 | 0.246284 | 0.136937 | 0.828492 | Atp13a2 |
| ENSMUSG( | 1854.001 | 0.302584 | 0.072603 | 0.763629 | Atp13a3 |
| ENSMUSG( | 953.9593 | -0.0893 | 0.503261 | 0.927825 | Atp13a4 |
| ENSMUSG( | 626.9266 | 0.523329 | 0.040047 | 0.681706 | Atp13a5 |
| ENSMUSG( | 31528.49 | 0.089026 | 0.565248 | 0.94131 | Atp1a1 |
| ENSMUSG( | 124331.7 | 0.010942 | 0.930718 | 0.99306 | Atp1a2 |
| ENSMUSG( | 86408.74 | 0.075186 | 0.589341 | 0.944975 | Atp1a3 |
| ENSMUSG( | 57623.26 | 0.1197 | 0.450551 | 0.922202 | Atp1b1 |
| ENSMUSG( | 43427.12 | 0.036384 | 0.802657 | 0.97681 | Atp1b2 |
| ENSMUSG( | 4830.505 | 0.026252 | 0.820281 | 0.979198 | Atp1b3 |
| ENSMUSG( | 151.1122 | -0.05907 | 0.764617 | 0.974094 | Atp23 |
| ENSMUSG( | 19.1685 | -1.08482 | 0.028042 | 0.61023 | Atp2a1 |
| ENSMUSG( | 26542.83 | 0.130696 | 0.410458 | 0.917257 | Atp2a2 |
| ENSMUSG( | 361.2865 | 0.012117 | 0.963153 | 0.995433 | Atp2a3 |
| ENSMUSG( | 15938.63 | 0.201334 | 0.265934 | 0.886134 | Atp2b1 |
| ENSMUSG( | 27150.79 | 0.053946 | 0.715183 | 0.965063 | Atp2b2 |
| ENSMUSG( | 1618.404 | 0.361317 | 0.09741 | 0.796721 | Atp2b3 |
| ENSMUSG( | 7073.971 | 0.121318 | 0.402028 | 0.917257 | Atp2b4 |
| ENSMUSG( | 5063.261 | 0.165592 | 0.281557 | 0.892412 | Atp2c1 |
| ENSMUSG( | 49.78024 | 0.58575 | 0.055243 | 0.723362 | Atp2c2 |
| ENSMUSG( | 26.89793 | 0.123496 | 0.746377 | 0.969687 | Atp4a |
| ENSMUSG( | 20332.13 | -0.13374 | 0.249917 | 0.877904 | Atp5f1a |
| ENSMUSG( | 27677.24 | -0.18259 | 0.070541 | 0.758714 | Atp5f1b |
| ENSMUSG( | 5351.598 | -0.199 | 0.091724 | 0.791889 | Atp5f1c |
| ENSMUSG( | 5860.891 | -0.21651 | 0.052636 | 0.71525 | Atp5f1d |
| ENSMUSG( | 2909.199 | -0.26217 | 0.04024 | 0.681706 | Atp5f1e |
| ENSMUSG( | 3567.686 | -0.19475 | 0.112135 | 0.808673 | Atp5if1 |
| ENSMUSG( | 3253.622 | -0.21819 | 0.04188 | 0.690304 | Atp5mc1 |
| ENSMUSG( | 1524.249 | -0.15329 | 0.245524 | 0.877904 | Atp5mc2 |
| ENSMUSG( | 5524.738 | -0.23614 | 0.083714 | 0.77681 | Atp5mc3 |
| ENSMUSG( | 2411.308 | -0.25154 | 0.07408 | 0.766789 | Atp5me |
| ENSMUSG( | 4103.434 | -0.28697 | 0.009708 | 0.406458 | Atp5mf |
| ENSMUSG( | 2096.402 | -0.35673 | 0.003866 | 0.248067 | Atp5mg |
| ENSMUSG( | 2427.529 | -0.19818 | 0.123241 | 0.814691 | Atp5mj |
| ENSMUSG( | 4150.049 | -0.25256 | 0.054716 | 0.723362 | Atp5mk |
| ENSMUSG( | 5613.926 | -0.25175 | 0.056552 | 0.723362 | Atp5pb |
| ENSMUSG( | 5402.493 | -0.21903 | 0.091064 | 0.791889 | Atp5pd |
| ENSMUSG( | 4801.863 | -0.22043 | 0.06287 | 0.736482 | Atp5pf |
| ENSMUSG( | 2551.064 | -0.09871 | 0.440982 | 0.921166 | Atp5po |
| ENSMUSG( | 2846.044 | -0.24517 | 0.57245 | 0.942533 | ATP6 |
| ENSMUSG( | 15413.39 | 0.019832 | 0.88143 | 0.989644 | Atp6ap1 |
| ENSMUSG( | 278.6533 | 0.043072 | 0.782275 | 0.976106 | Atp6ap1l |
| ENSMUSG( | 6744.812 | 0.170726 | 0.297835 | 0.898538 | Atp6ap2 |
| ENSMUSG( | 10168.19 | -0.16922 | 0.720007 | 0.966887 | Atp6-ps |
| ENSMUSG( | 19695.3 | 0.10304 | 0.450124 | 0.922202 | Atp6v0a1 |
| ENSMUSG( | 4801.167 | -0.0664 | 0.590903 | 0.944975 | Atp6v0a2 |
| ENSMUSG( | 14.83654 | 0.60631 | 0.277839 | 0.891468 | Atp6v0a4 |
| ENSMUSG( | 7758.909 | 0.051034 | 0.649357 | 0.95422 | Atp6v0b |
| ENSMUSG( | 13340.39 | -0.00784 | 0.95611 | 0.994798 | Atp6v0c |
| ENSMUSG( | 4956.578 | -0.19233 | 0.101253 | 0.801456 | Atp6v0d1 |
| ENSMUSG( | 1042.867 | 0.129062 | 0.415496 | 0.917257 | Atp6v0e |
| ENSMUSG( | 14800.59 | -0.034 | 0.810754 | 0.978779 | Atp6v0e2 |
| ENSMUSG( | 6370.019 | -0.07775 | 0.514606 | 0.930939 | Atp6v1a |
| ENSMUSG( | 8617.492 | -0.09644 | 0.499455 | 0.927825 | Atp6v1b2 |
| ENSMUSG( | 3962.375 | -0.16859 | 0.252037 | 0.879902 | Atp6v1c1 |
| ENSMUSG( | 26.75412 | -0.42222 | 0.362611 | 0.914223 | Atp6v1c2 |
| ENSMUSG( | 3262.507 | -0.11706 | 0.363663 | 0.914223 | Atp6v1d |
| ENSMUSG( | 5166.468 | -0.17729 | 0.245448 | 0.877904 | Atp6v1e1 |
| ENSMUSG( | 2023.562 | -0.35938 | 0.003011 | 0.219289 | Atp6v1f |
| ENSMUSG( | 1595.94 | -0.06268 | 0.675692 | 0.95694 | Atp6v1g1 |
| ENSMUSG( | 9595.53 | -0.1798 | 0.268811 | 0.886755 | Atp6v1g2 |
| ENSMUSG( | 3034.904 | -0.09428 | 0.478268 | 0.923329 | Atp6v1h |
| ENSMUSG( | 136.0408 | 0.239602 | 0.379346 | 0.916995 | Atp7a |
| ENSMUSG( | 125.6087 | 0.203688 | 0.309104 | 0.898538 | Atp7b |
| ENSMUSG( | 202.1891 | 0.208166 | 0.815051 | 0.979198 | ATP8 |
| ENSMUSG( | 6728.672 | 0.197826 | 0.226426 | 0.868507 | Atp8a1 |
| ENSMUSG( | 1863.398 | 0.201586 | 0.225344 | 0.868468 | Atp8a2 |
| ENSMUSG( | 141.1861 | -0.04482 | 0.861991 | 0.98632 | Atp8b1 |
| ENSMUSG( | 2276.276 | -0.02403 | 0.875585 | 0.989644 | Atp8b2 |
| ENSMUSG( | 14.87222 | -0.5276 | 0.362524 | 0.914223 | Atp8b5 |
| ENSMUSG( | 14494.53 | 0.164656 | 0.316686 | 0.901868 | Atp9a |
| ENSMUSG( | 2146.759 | 0.136993 | 0.3516 | 0.912093 | Atp9b |
| ENSMUSG( | 653.1735 | -0.09724 | 0.516195 | 0.930939 | Atpaf1 |
| ENSMUSG( | 533.2328 | -0.04027 | 0.757185 | 0.971858 | Atpaf2 |
| ENSMUSG( | 258.5751 | 0.06658 | 0.733793 | 0.967228 | Atpsckmt |
| ENSMUSG( | 444.3769 | 0.110522 | 0.494717 | 0.927372 | Atr |
| ENSMUSG( | 2341.911 | 0.117453 | 0.358808 | 0.913759 | Atraid |
| ENSMUSG( | 494.027 | 0.135363 | 0.340824 | 0.907972 | Atrip |
| ENSMUSG( | 6155.334 | 0.160917 | 0.334089 | 0.907907 | Atrn |
| ENSMUSG( | 3164.819 | 0.171628 | 0.312531 | 0.900037 | Atrnl1 |
| ENSMUSG( | 3958.579 | 0.09585 | 0.430321 | 0.917257 | Atrx |
| ENSMUSG( | 2279.647 | 0.092108 | 0.383766 | 0.917257 | Atxn1 |
| ENSMUSG( | 4795.045 | -0.20814 | 0.131544 | 0.822912 | Atxn10 |
| ENSMUSG( | 1636.258 | -0.08556 | 0.452187 | 0.922316 | Atxn1l |
| ENSMUSG( | 2487.326 | 0.061389 | 0.548745 | 0.938075 | Atxn2 |
| ENSMUSG( | 3995.349 | 0.143065 | 0.342317 | 0.909642 | Atxn2l |
| ENSMUSG( | 548.0867 | 0.094888 | 0.466545 | 0.923329 | Atxn3 |
| ENSMUSG( | 838.468 | 0.166656 | 0.162206 | 0.837724 | Atxn7 |
| ENSMUSG( | 529.0285 | -0.1194 | 0.407315 | 0.917257 | Atxn7l1 |
| ENSMUSG( | 622.7066 | 0.049382 | 0.74965 | 0.96998 | Atxn7l2 |
| ENSMUSG( | 12988.26 | 0.002423 | 0.979988 | 0.996479 | Atxn7l3 |
| ENSMUSG( | 6526.953 | -0.10182 | 0.413774 | 0.917257 | Atxn7l3b |
| ENSMUSG( | 156.1359 | -0.18873 | 0.374591 | 0.915491 | AU020206 |
| ENSMUSG( | 195.3527 | 0.387341 | 0.153473 | 0.837724 | AU021092 |
| ENSMUSG( | 1549.527 | -0.0744 | 0.527369 | 0.934466 | AU022252 |
| ENSMUSG( | 50.54635 | 0.34086 | 0.236514 | 0.872173 | AU022754 |
| ENSMUSG( | 34.43982 | -0.52513 | 0.17017 | 0.841342 | AU023762 |
| ENSMUSG( | 2631.073 | 0.176239 | 0.282223 | 0.892412 | AU040320 |
| ENSMUSG( | 15.70758 | 0.658265 | 0.18681 | 0.856099 | AU040972 |
| ENSMUSG( | 82.45138 | -0.01848 | 0.936682 | 0.993573 | AU041133 |
| ENSMUSG( | 1003.289 | -0.14376 | 0.312134 | 0.900037 | Auh |
| ENSMUSG( | 2366.174 | 0.102772 | 0.374239 | 0.915491 | Aup1 |
| ENSMUSG( | 24.60318 | -0.43301 | 0.272309 | 0.888771 | Aurka |
| ENSMUSG( | 1316.335 | -0.25006 | 0.033206 | 0.641315 | Aurkaip1 |
| ENSMUSG( | 23.0779 | 0.145868 | 0.744667 | 0.96951 | Aurkb |
| ENSMUSG( | 1828.775 | 0.263545 | 0.028155 | 0.611882 | Auts2 |
| ENSMUSG( | 12.61649 | -0.37421 | 0.488598 | 0.92554 | AV039307 |
| ENSMUSG( | 219.8232 | -0.04996 | 0.773482 | 0.974672 | Aven |
| ENSMUSG( | 14.72991 | -0.20015 | 0.722227 | 0.966887 | Avil |
| ENSMUSG( | 1291.555 | 0.118521 | 0.352663 | 0.912093 | Avl9 |
| ENSMUSG( | 325.983 | -0.15968 | 0.395928 | 0.917257 | Avpi1 |
| ENSMUSG( | 10.16365 | 0.796525 | 0.212524 | NA | Avpr1a |
| ENSMUSG( | 402.8171 | 0.170598 | 0.228313 | 0.868507 | AW011738 |
| ENSMUSG( | 42.72594 | 0.362466 | 0.326142 | 0.905339 | AW112010 |
| ENSMUSG( | 138.0614 | -0.17983 | 0.34406 | 0.910256 | AW146154 |
| ENSMUSG( | 260.5827 | -0.09711 | 0.580367 | 0.944209 | AW209491 |
| ENSMUSG( | 109.2651 | -0.5132 | 0.028024 | 0.61023 | AW495222 |
| ENSMUSG( | 92.80892 | -0.05559 | 0.83101 | 0.98051 | AW551984 |
| ENSMUSG( | 422.0466 | -0.07096 | 0.641363 | 0.953903 | AW554918 |
| ENSMUSG( | 57.90532 | -0.07429 | 0.797588 | 0.97681 | AW822252 |
| ENSMUSG( | 23.53121 | 0.314196 | 0.439443 | 0.920721 | Axdnd1 |
| ENSMUSG( | 795.9907 | -0.05487 | 0.653049 | 0.955015 | Axin1 |
| ENSMUSG( | 1051.257 | 0.358846 | 0.005524 | 0.303106 | Axin2 |

| | | | | | | | | | | | |
|---|---|---|---|---|---|---|---|---|---|---|---|
| ENSMUSG( | Atg4a | 265.2207 | -0.09058 | 0.527893 | 0.878017 | ENSMUSG( | Atg4a | 262.1743 | 0.141168 | 0.382696 | 0.917257 |
| ENSMUSG( | Atg4b | 1478.398 | 0.142607 | 0.191337 | 0.630245 | ENSMUSG( | Atg4b | 1292.908 | 0.029723 | 0.80978 | 0.978779 |
| ENSMUSG( | Atg4c | 781.5211 | 0.203676 | 0.071381 | 0.412911 | ENSMUSG( | Atg4c | 705.9476 | 0.177285 | 0.16687 | 0.840183 |
| ENSMUSG( | Atg4d | 830.1384 | 0.104181 | 0.29039 | 0.740353 | ENSMUSG( | Atg4d | 760.8943 | 0.125177 | 0.357821 | 0.913759 |
| ENSMUSG( | Atg5 | 574.0935 | 0.174798 | 0.168241 | 0.601572 | ENSMUSG( | Atg5 | 489.0254 | -0.01705 | 0.903089 | 0.991089 |
| ENSMUSG( | Atg7 | 870.9553 | 0.009649 | 0.920079 | 0.990247 | ENSMUSG( | Atg7 | 743.7287 | -0.18714 | 0.129271 | 0.820358 |
| ENSMUSG( | Atg9a | 1623.378 | 0.000193 | 0.999535 | 0.99989 | ENSMUSG( | Atg9a | 1708.173 | 0.398549 | 0.326601 | 0.905371 |
| ENSMUSG( | Atg9b | 143.9387 | -0.17792 | 0.308452 | 0.756439 | ENSMUSG( | Atg9b | 146.8082 | 0.138315 | 0.49113 | 0.926604 |
| ENSMUSG( | Atic | 933.981 | 0.08718 | 0.374768 | 0.800466 | ENSMUSG( | Atic | 796.1403 | -0.10827 | 0.391556 | 0.917257 |
| ENSMUSG( | Atl1 | 2318.406 | -0.41673 | 0.000656 | 0.024688 | ENSMUSG( | Atl1 | 2509.898 | 0.113369 | 0.440646 | 0.921166 |
| ENSMUSG( | Atl2 | 2099.579 | -0.33258 | 0.010289 | 0.144496 | ENSMUSG( | Atl2 | 2236.095 | 0.1359 | 0.330333 | 0.907251 |
| ENSMUSG( | Atl3 | 1266.631 | -0.14793 | 0.21146 | 0.654808 | ENSMUSG( | Atl3 | 1279.56 | 0.152747 | 0.270816 | 0.888062 |
| ENSMUSG( | Atm | 444.8898 | 0.07392 | 0.582746 | 0.89798 | ENSMUSG( | Atm | 403.6553 | 0.072904 | 0.663963 | 0.956766 |
| ENSMUSG( | Atmin | 1873.326 | 0.051485 | 0.596125 | 0.903152 | ENSMUSG( | Atmin | 1620.803 | -0.09604 | 0.446632 | 0.921734 |
| ENSMUSG( | Atn1 | 698.424 | -0.02843 | 0.786613 | 0.958856 | ENSMUSG( | Atn1 | 621.6319 | -0.1053 | 0.55031 | 0.938699 |
| ENSMUSG( | Atoh7 | 36.65941 | 0.12607 | 0.719177 | 0.941986 | ENSMUSG( | Atoh7 | 31.84109 | -0.01185 | 0.9756 | 0.996416 |
| ENSMUSG( | Atoh8 | 68.03479 | -0.08154 | 0.756792 | 0.951513 | ENSMUSG( | Atoh8 | 69.62257 | 0.27041 | 0.356 | 0.913331 |
| ENSMUSG( | Atosa | 1240.885 | 0.085869 | 0.449756 | 0.835283 | ENSMUSG( | Atosa | 1187.615 | 0.225977 | 0.110069 | 0.806572 |
| ENSMUSG( | Atosb | 1287.077 | 0.004132 | 0.970112 | 0.996369 | ENSMUSG( | Atosb | 1177.726 | 0.016324 | 0.886375 | 0.990011 |
| ENSMUSG( | Atox1 | 1155.001 | 0.05826 | 0.530965 | 0.879289 | ENSMUSG( | Atox1 | 951.4445 | -0.24812 | 0.068249 | 0.752208 |
| ENSMUSG( | Atp10a | 447.1729 | 0.817071 | 1.14E-05 | 0.00123 | ENSMUSG( | Atp10a | 488.9212 | 1.220674 | 4.79E-06 | 0.002153 |
| ENSMUSG( | Atp10b | 105.267 | -0.82312 | 0.000712 | 0.025833 | ENSMUSG( | Atp10b | 124.4412 | 0.061615 | 0.810074 | 0.978779 |
| ENSMUSG( | Atp10d | 148.6147 | 0.028466 | 0.870211 | 0.97782 | ENSMUSG( | Atp10d | 149.8076 | 0.317902 | 0.154673 | 0.837724 |
| ENSMUSG( | Atp11a | 2451.902 | -0.16683 | 0.174851 | 0.610304 | ENSMUSG( | Atp11a | 2560.641 | 0.227251 | 0.129569 | 0.820557 |
| ENSMUSG( | Atp11b | 2556.38 | -0.18916 | 0.082997 | 0.44459 | ENSMUSG( | Atp11b | 2691.033 | 0.227283 | 0.132967 | 0.822912 |
| ENSMUSG( | Atp11c | 312.9997 | -0.36088 | 0.037056 | 0.297106 | ENSMUSG( | Atp11c | 364.9415 | 0.355347 | 0.047063 | 0.700664 |
| ENSMUSG( | Atp13a1 | 1848.86 | -0.06098 | 0.637506 | 0.915642 | ENSMUSG( | Atp13a1 | 1825.689 | 0.17213 | 0.265197 | 0.886134 |
| ENSMUSG( | Atp13a2 | 6712.63 | 0.061267 | 0.63699 | 0.915191 | ENSMUSG( | Atp13a2 | 6533.429 | 0.246284 | 0.136937 | 0.828492 |
| ENSMUSG( | Atp13a3 | 1703.406 | -0.20418 | 0.157291 | 0.585553 | ENSMUSG( | Atp13a3 | 1854.001 | 0.302584 | 0.072603 | 0.763629 |
| ENSMUSG( | Atp13a4 | 1102.495 | 0.055662 | 0.628892 | 0.913816 | ENSMUSG( | Atp13a4 | 953.9593 | -0.0893 | 0.503261 | 0.927825 |
| ENSMUSG( | Atp13a5 | 517.8025 | -0.2747 | 0.149372 | 0.573313 | ENSMUSG( | Atp13a5 | 626.9266 | 0.523329 | 0.040047 | 0.681706 |
| ENSMUSG( | Atp1a1 | 30988.14 | -0.24241 | 0.070388 | 0.40908 | ENSMUSG( | Atp1a1 | 31528.49 | 0.089026 | 0.565248 | 0.94131 |
| ENSMUSG( | Atp1a2 | 138067.9 | 0.043285 | 0.707629 | 0.93949 | ENSMUSG( | Atp1a2 | 124331.7 | 0.010942 | 0.930718 | 0.99306 |
| ENSMUSG( | Atp1a3 | 80666 | -0.42456 | 0.000278 | 0.013625 | ENSMUSG( | Atp1a3 | 86408.74 | 0.075186 | 0.589341 | 0.944975 |
| ENSMUSG( | Atp1b1 | 51031.18 | -0.55058 | 1.38E-05 | 0.001449 | ENSMUSG( | Atp1b1 | 57623.26 | 0.1197 | 0.450551 | 0.922202 |
| ENSMUSG( | Atp1b2 | 44568.21 | -0.16443 | 0.22096 | 0.666476 | ENSMUSG( | Atp1b2 | 43427.12 | 0.036384 | 0.802657 | 0.97681 |
| ENSMUSG( | Atp1b3 | 5041.471 | -0.12127 | 0.179627 | 0.61741 | ENSMUSG( | Atp1b3 | 4830.505 | 0.026252 | 0.820281 | 0.979198 |
| ENSMUSG( | Atp23 | 171.3736 | 0.02205 | 0.895318 | 0.984512 | ENSMUSG( | Atp23 | 151.1122 | -0.05907 | 0.764617 | 0.974094 |
| ENSMUSG( | Atp2a1 | 19.66935 | -1.48952 | 0.001426 | NA | ENSMUSG( | Atp2a1 | 19.1685 | -1.08482 | 0.028042 | 0.61023 |
| ENSMUSG( | Atp2a2 | 25008.39 | -0.32832 | 0.014721 | 0.178407 | ENSMUSG( | Atp2a2 | 26542.83 | 0.130696 | 0.410458 | 0.917257 |
| ENSMUSG( | Atp2a3 | 393.8346 | -0.02093 | 0.915404 | 0.988634 | ENSMUSG( | Atp2a3 | 361.2865 | 0.012117 | 0.963153 | 0.995433 |
| ENSMUSG( | Atp2b1 | 13825.74 | -0.52239 | 0.001106 | 0.035255 | ENSMUSG( | Atp2b1 | 15938.63 | 0.201334 | 0.265934 | 0.886134 |
| ENSMUSG( | Atp2b2 | 25916.45 | -0.3755 | 0.002279 | 0.055785 | ENSMUSG( | Atp2b2 | 27150.79 | 0.053946 | 0.715183 | 0.965063 |
| ENSMUSG( | Atp2b3 | 1473.666 | -0.16847 | 0.366353 | 0.794736 | ENSMUSG( | Atp2b3 | 1618.404 | 0.361317 | 0.09741 | 0.796721 |
| ENSMUSG( | Atp2b4 | 6670.777 | -0.33681 | 0.011117 | 0.152038 | ENSMUSG( | Atp2b4 | 7073.971 | 0.121318 | 0.402028 | 0.917257 |
| ENSMUSG( | Atp2c1 | 5024.887 | -0.12519 | 0.340275 | 0.777962 | ENSMUSG( | Atp2c1 | 5063.261 | 0.165592 | 0.281557 | 0.892412 |
| ENSMUSG( | Atp2c2 | 39.75186 | -0.28743 | 0.363395 | 0.793339 | ENSMUSG( | Atp2c2 | 49.78024 | 0.58575 | 0.055243 | 0.723362 |
| ENSMUSG( | Atp4a | 28.38907 | 0.035809 | 0.919315 | 0.990247 | ENSMUSG( | Atp4a | 26.89793 | 0.123496 | 0.746377 | 0.969687 |
| ENSMUSG( | Atp5f1a | 24672.74 | 0.156167 | 0.065935 | 0.397572 | ENSMUSG( | Atp5f1a | 20332.13 | -0.13374 | 0.249917 | 0.877904 |
| ENSMUSG( | Atp5f1b | 33341.08 | 0.090638 | 0.25561 | 0.706195 | ENSMUSG( | Atp5f1b | 27677.24 | -0.18259 | 0.070541 | 0.758714 |
| ENSMUSG( | Atp5f1c | 6396.159 | 0.055126 | 0.53301 | 0.879905 | ENSMUSG( | Atp5f1c | 5351.598 | -0.199 | 0.091724 | 0.791889 |
| ENSMUSG( | Atp5f1d | 6934.009 | 0.009974 | 0.907837 | 0.987798 | ENSMUSG( | Atp5f1d | 5860.891 | -0.21651 | 0.052636 | 0.71525 |
| ENSMUSG( | Atp5f1e | 3575.22 | 0.078443 | 0.452254 | 0.836145 | ENSMUSG( | Atp5f1e | 2909.199 | -0.26217 | 0.04024 | 0.681706 |
| ENSMUSG( | Atp5if1 | 4087.651 | -0.06242 | 0.482345 | 0.855884 | ENSMUSG( | Atp5if1 | 3567.686 | -0.19475 | 0.112135 | 0.808673 |
| ENSMUSG( | Atp5mc1 | 3799.038 | -0.03042 | 0.714832 | 0.941144 | ENSMUSG( | Atp5mc1 | 3253.622 | -0.21819 | 0.04188 | 0.690304 |
| ENSMUSG( | Atp5mc2 | 1745.929 | -0.02187 | 0.811141 | 0.964318 | ENSMUSG( | Atp5mc2 | 1524.249 | -0.15329 | 0.245524 | 0.877904 |
| ENSMUSG( | Atp5mc3 | 6562.236 | 0.004232 | 0.965174 | 0.995303 | ENSMUSG( | Atp5mc3 | 5524.738 | -0.23614 | 0.083714 | 0.77681 |
| ENSMUSG( | Atp5me | 2898.238 | 0.025534 | 0.812621 | 0.964574 | ENSMUSG( | Atp5me | 2411.308 | -0.25154 | 0.07408 | 0.766789 |
| ENSMUSG( | Atp5mf | 4943.986 | -0.00194 | 0.981838 | 0.997512 | ENSMUSG( | Atp5mf | 4103.434 | -0.28697 | 0.009708 | 0.406458 |
| ENSMUSG( | Atp5mg | 2552.317 | -0.03262 | 0.692484 | 0.934101 | ENSMUSG( | Atp5mg | 2096.402 | -0.35673 | 0.003866 | 0.248067 |
| ENSMUSG( | Atp5mj | 2908.78 | 0.064728 | 0.505326 | 0.866883 | ENSMUSG( | Atp5mj | 2427.529 | -0.19818 | 0.123241 | 0.814691 |
| ENSMUSG( | Atp5mk | 4903.9 | -0.026 | 0.768127 | 0.954641 | ENSMUSG( | Atp5mk | 4150.049 | -0.25256 | 0.054716 | 0.723362 |
| ENSMUSG( | Atp5pb | 6753.571 | 0.026481 | 0.774945 | 0.955606 | ENSMUSG( | Atp5pb | 5613.926 | -0.25175 | 0.056552 | 0.723362 |
| ENSMUSG( | Atp5pd | 6429.809 | 0.025266 | 0.773019 | 0.955272 | ENSMUSG( | Atp5pd | 5402.493 | -0.21903 | 0.091064 | 0.791889 |
| ENSMUSG( | Atp5pf | 5722.676 | 0.027815 | 0.749679 | 0.949664 | ENSMUSG( | Atp5pf | 4801.863 | -0.22043 | 0.06287 | 0.736482 |
| ENSMUSG( | Atp5po | 2977.91 | 0.083017 | 0.426849 | 0.825231 | ENSMUSG( | Atp5po | 2551.064 | -0.09871 | 0.440982 | 0.921166 |
| ENSMUSG( | ATP6 | 3285.924 | -0.08434 | 0.666106 | 0.927057 | ENSMUSG( | ATP6 | 2846.044 | -0.24517 | 0.57245 | 0.942533 |
| ENSMUSG( | Atp6ap1 | 15422.67 | -0.26105 | 0.023387 | 0.233882 | ENSMUSG( | Atp6ap1 | 15413.39 | 0.019832 | 0.88143 | 0.989644 |
| ENSMUSG( | Atp6ap1l | 267.8709 | -0.36818 | 0.009245 | 0.135575 | ENSMUSG( | Atp6ap1l | 278.6533 | 0.043072 | 0.782275 | 0.976106 |
| ENSMUSG( | Atp6ap2 | 5883.516 | -0.53764 | 7.78E-05 | 0.005438 | ENSMUSG( | Atp6ap2 | 6744.812 | 0.170726 | 0.297835 | 0.898538 |
| ENSMUSG( | Atp6-ps | 11637.8 | -0.03791 | 0.851034 | 0.972638 | ENSMUSG( | Atp6-ps | 10168.19 | -0.16922 | 0.720007 | 0.966887 |
| ENSMUSG( | Atp6v0a1 | 19111.13 | -0.26618 | 0.025021 | 0.243138 | ENSMUSG( | Atp6v0a1 | 19695.3 | 0.10304 | 0.450124 | 0.922202 |
| ENSMUSG( | Atp6v0a2 | 5425.241 | 0.016832 | 0.881255 | 0.981798 | ENSMUSG( | Atp6v0a2 | 4801.167 | -0.0664 | 0.590903 | 0.944975 |
| ENSMUSG( | Atp6v0b | 7733.874 | -0.23809 | 0.022692 | 0.23004 | ENSMUSG( | Atp6v0b | 7758.909 | 0.051034 | 0.649357 | 0.95422 |
| ENSMUSG( | Atp6v0c | 12369.8 | -0.54801 | 8.50E-06 | 0.001033 | ENSMUSG( | Atp6v0c | 13340.39 | -0.00784 | 0.95611 | 0.994798 |
| ENSMUSG( | Atp6v0d1 | 5838.962 | 0.019101 | 0.846043 | 0.972226 | ENSMUSG( | Atp6v0d1 | 4956.578 | -0.19233 | 0.101253 | 0.801456 |
| ENSMUSG( | Atp6v0e | 1034.76 | -0.16925 | 0.238472 | 0.686308 | ENSMUSG( | Atp6v0e | 1042.867 | 0.129062 | 0.415496 | 0.917257 |
| ENSMUSG( | Atp6v0e2 | 14633.74 | -0.36093 | 0.003525 | 0.075237 | ENSMUSG( | Atp6v0e2 | 14800.59 | -0.034 | 0.810754 | 0.978779 |
| ENSMUSG( | Atp6v1a | 7274.631 | 0.039209 | 0.663939 | 0.926511 | ENSMUSG( | Atp6v1a | 6370.019 | -0.07775 | 0.514606 | 0.930939 |
| ENSMUSG( | Atp6v1b2 | 10281.01 | 0.144032 | 0.266416 | 0.716441 | ENSMUSG( | Atp6v1b2 | 8617.492 | -0.09644 | 0.499455 | 0.927825 |
| ENSMUSG( | Atp6v1c1 | 4710.743 | 0.068681 | 0.589714 | 0.90113 | ENSMUSG( | Atp6v1c1 | 3962.375 | -0.16859 | 0.252037 | 0.879902 |
| ENSMUSG( | Atp6v1c2 | 31.94076 | -0.11102 | 0.791344 | 0.959461 | ENSMUSG( | Atp6v1c2 | 26.75412 | -0.42222 | 0.362611 | 0.914223 |
| ENSMUSG( | Atp6v1d | 3588.372 | -0.10796 | 0.282216 | 0.73186 | ENSMUSG( | Atp6v1d | 3262.507 | -0.11706 | 0.363663 | 0.914223 |
| ENSMUSG( | Atp6v1e1 | 6073.559 | 0.029162 | 0.813967 | 0.964574 | ENSMUSG( | Atp6v1e1 | 5166.468 | -0.17729 | 0.245448 | 0.877904 |
| ENSMUSG( | Atp6v1f | 2470.618 | -0.02742 | 0.774967 | 0.955606 | ENSMUSG( | Atp6v1f | 2023.562 | -0.35938 | 0.003011 | 0.219289 |
| ENSMUSG( | Atp6v1g1 | 1798.429 | 0.018501 | 0.843501 | 0.971847 | ENSMUSG( | Atp6v1g1 | 1595.94 | -0.06268 | 0.675692 | 0.95694 |
| ENSMUSG( | Atp6v1g2 | 11269.17 | 0.023745 | 0.848153 | 0.972374 | ENSMUSG( | Atp6v1g2 | 9595.53 | -0.1798 | 0.268811 | 0.886755 |
| ENSMUSG( | Atp6v1h | 3309.376 | -0.11156 | 0.249342 | 0.698744 | ENSMUSG( | Atp6v1h | 3034.904 | -0.09428 | 0.478268 | 0.923329 |
| ENSMUSG( | Atp7a | 133.24 | -0.0986 | 0.67779 | 0.930542 | ENSMUSG( | Atp7a | 136.0408 | 0.239602 | 0.379346 | 0.916995 |
| ENSMUSG( | Atp7b | 119.7529 | -0.2148 | 0.231914 | 0.679732 | ENSMUSG( | Atp7b | 125.6087 | 0.203688 | 0.309104 | 0.898538 |
| ENSMUSG( | ATP8 | 162.0326 | -0.78229 | 0.160673 | 0.590555 | ENSMUSG( | ATP8 | 202.1891 | 0.208166 | 0.815051 | 0.979198 |
| ENSMUSG( | Atp8a1 | 6283.884 | -0.27896 | 0.049064 | 0.338072 | ENSMUSG( | Atp8a1 | 6728.672 | 0.197826 | 0.226426 | 0.868507 |
| ENSMUSG( | Atp8a2 | 1710.432 | -0.33168 | 0.023375 | 0.233882 | ENSMUSG( | Atp8a2 | 1863.398 | 0.201586 | 0.225344 | 0.868468 |
| ENSMUSG( | Atp8b1 | 131.2751 | -0.59503 | 0.020541 | 0.216203 | ENSMUSG( | Atp8b1 | 141.1861 | -0.04482 | 0.861991 | 0.98632 |
| ENSMUSG( | Atp8b2 | 2405.974 | -0.14022 | 0.191893 | 0.630811 | ENSMUSG( | Atp8b2 | 2276.276 | -0.02403 | 0.875585 | 0.989644 |
| ENSMUSG( | Atp8b5 | 18.53972 | -0.13822 | 0.766227 | NA | ENSMUSG( | Atp8b5 | 14.87222 | -0.5276 | 0.362524 | 0.914223 |
| ENSMUSG( | Atp9a | 14468.33 | -0.11122 | 0.413892 | 0.819998 | ENSMUSG( | Atp9a | 14494.53 | 0.164656 | 0.316686 | 0.901868 |
| ENSMUSG( | Atp9b | 2242.344 | -0.00752 | 0.949954 | 0.993821 | ENSMUSG( | Atp9b | 2146.759 | 0.136993 | 0.3516 | 0.912093 |
| ENSMUSG( | Atpaf1 | 709.4036 | -0.12447 | 0.346116 | 0.779153 | ENSMUSG( | Atpaf1 | 653.1735 | -0.09724 | 0.516195 | 0.930939 |
| ENSMUSG( | Atpaf2 | 597.4821 | 0.022145 | 0.843745 | 0.971847 | ENSMUSG( | Atpaf2 | 533.2328 | -0.04027 | 0.757185 | 0.971858 |
| ENSMUSG( | Atpsckmt | 285.5588 | 0.076663 | 0.678195 | 0.930542 | ENSMUSG( | Atpsckmt | 258.5751 | 0.06658 | 0.733793 | 0.967228 |
| ENSMUSG( | Atr | 458.3816 | -0.07617 | 0.57089 | 0.8933 | ENSMUSG( | Atr | 444.3769 | 0.110522 | 0.494717 | 0.927372 |
| ENSMUSG( | Atraid | 2315.878 | -0.19023 | 0.080209 | 0.437895 | ENSMUSG( | Atraid | 2341.911 | 0.117453 | 0.358808 | 0.913759 |
| ENSMUSG( | Atrip | 549.7555 | 0.166505 | 0.154685 | 0.581863 | ENSMUSG( | Atrip | 494.027 | 0.135363 | 0.340824 | 0.907972 |
| ENSMUSG( | Atrn | 5818.899 | -0.2847 | 0.063184 | 0.388743 | ENSMUSG( | Atrn | 6155.334 | 0.160917 | 0.334089 | 0.907907 |
| ENSMUSG( | Atrnl1 | 2869.692 | -0.40686 | 0.007417 | 0.118765 | ENSMUSG( | Atrnl1 | 3164.819 | 0.171628 | 0.312531 | 0.900037 |
| ENSMUSG( | Atrx | 4084.291 | -0.08127 | 0.434456 | 0.828978 | ENSMUSG( | Atrx | 3958.579 | 0.09585 | 0.430321 | 0.917257 |
| ENSMUSG( | Atxn1 | 2478.108 | 0.064089 | 0.456945 | 0.839183 | ENSMUSG( | Atxn1 | 2279.647 | 0.092108 | 0.383766 | 0.917257 |
| ENSMUSG( | Atxn10 | 5778.544 | 0.070457 | 0.495999 | 0.861543 | ENSMUSG( | Atxn10 | 4795.045 | -0.20814 | 0.131544 | 0.822912 |
| ENSMUSG( | Atxn1l | 1857.251 | 0.014101 | 0.890025 | 0.983687 | ENSMUSG( | Atxn1l | 1636.258 | -0.08556 | 0.452187 | 0.922316 |
| ENSMUSG( | Atxn2 | 2706.117 | 0.036134 | 0.68802 | 0.93324 | ENSMUSG( | Atxn2 | 2487.326 | 0.061389 | 0.548745 | 0.938075 |
| ENSMUSG( | Atxn2l | 4348.075 | 0.120573 | 0.331901 | 0.775288 | ENSMUSG( | Atxn2l | 3995.349 | 0.143065 | 0.342317 | 0.909642 |
| ENSMUSG( | Atxn3 | 598.3097 | 0.082849 | 0.446165 | 0.833546 | ENSMUSG( | Atxn3 | 548.0867 | 0.094888 | 0.466545 | 0.923329 |
| ENSMUSG( | Atxn7 | 844.7948 | -0.07767 | 0.44829 | 0.834051 | ENSMUSG( | Atxn7 | 838.468 | 0.166656 | 0.162206 | 0.837724 |
| ENSMUSG( | Atxn7l1 | 597.559 | -0.02705 | 0.816265 | 0.965455 | ENSMUSG( | Atxn7l1 | 529.0285 | -0.1194 | 0.407315 | 0.917257 |
| ENSMUSG( | Atxn7l2 | 705.6652 | 0.140063 | 0.321451 | 0.766644 | ENSMUSG( | Atxn7l2 | 622.7066 | 0.049382 | 0.74965 | 0.96998 |
| ENSMUSG( | Atxn7l3 | 15310.19 | 0.2014 | 0.007215 | 0.117261 | ENSMUSG( | Atxn7l3 | 12988.26 | 0.002423 | 0.979988 | 0.996479 |
| ENSMUSG( | Atxn7l3b | 7641.906 | 0.0874 | 0.337718 | 0.777422 | ENSMUSG( | Atxn7l3b | 6526.953 | -0.10182 | 0.413774 | 0.917257 |
| ENSMUSG( | AU020206 | 150.7472 | -0.63262 | 0.002359 | 0.057092 | ENSMUSG( | AU020206 | 156.1359 | -0.18873 | 0.374591 | 0.915491 |
| ENSMUSG( | AU021092 | 168.7503 | -0.31262 | 0.191173 | 0.630245 | ENSMUSG( | AU021092 | 195.3527 | 0.387341 | 0.153473 | 0.837724 |
| ENSMUSG( | AU022252 | 1814.471 | 0.114033 | 0.26036 | 0.710245 | ENSMUSG( | AU022252 | 1549.527 | -0.0744 | 0.527369 | 0.934466 |
| ENSMUSG( | AU022754 | 55.54078 | 0.345286 | 0.18943 | 0.62795 | ENSMUSG( | AU022754 | 50.54635 | 0.34086 | 0.236514 | 0.872173 |
| ENSMUSG( | AU023762 | 40.65683 | -0.28212 | 0.374873 | 0.800479 | ENSMUSG( | AU023762 | 34.43982 | -0.52513 | 0.17017 | 0.841342 |
| ENSMUSG( | AU040320 | 2521.207 | -0.22435 | 0.10341 | 0.489156 | ENSMUSG( | AU040320 | 2631.073 | 0.176239 | 0.282223 | 0.892412 |
| ENSMUSG( | AU040972 | 12.76267 | -0.12268 | 0.81275 | NA | ENSMUSG( | AU040972 | 15.70758 | 0.658265 | 0.18681 | 0.856099 |
| ENSMUSG( | AU041133 | 87.53681 | -0.10976 | 0.638719 | 0.91636 | ENSMUSG( | AU041133 | 82.45138 | -0.01848 | 0.936682 | 0.993573 |
| ENSMUSG( | Auh | 1186.65 | 0.07838 | 0.515497 | 0.872701 | ENSMUSG( | Auh | 1003.289 | -0.14376 | 0.312134 | 0.900037 |
| ENSMUSG( | Aup1 | 2422.39 | -0.09959 | 0.312541 | 0.760241 | ENSMUSG( | Aup1 | 2366.174 | 0.102772 | 0.374239 | 0.915491 |
| ENSMUSG( | Aurka | 29.26297 | -0.1943 | 0.570009 | 0.893041 | ENSMUSG( | Aurka | 24.60318 | -0.43301 | 0.272309 | 0.888771 |
| ENSMUSG( | Aurkaip1 | 1562.867 | -0.01041 | 0.906138 | 0.986921 | ENSMUSG( | Aurkaip1 | 1316.335 | -0.25006 | 0.033206 | 0.641315 |
| ENSMUSG( | Aurkb | 19.4542 | -0.74706 | 0.080518 | NA | ENSMUSG( | Aurkb | 23.0779 | 0.145868 | 0.744667 | 0.96951 |
| ENSMUSG( | Auts2 | 1970.249 | 0.216656 | 0.01104 | 0.151501 | ENSMUSG( | Auts2 | 1828.775 | 0.263545 | 0.028155 | 0.611882 |
| ENSMUSG( | AV039307 | 13.75814 | -0.39947 | 0.425356 | NA | ENSMUSG( | AV039307 | 12.61649 | -0.37421 | 0.488598 | 0.92554 |
| ENSMUSG( | Aven | 251.1361 | 0.061272 | 0.678046 | 0.930542 | ENSMUSG( | Aven | 219.8232 | -0.04996 | 0.773482 | 0.974672 |
| ENSMUSG( | Avil | 17.80261 | 0.055761 | 0.903637 | NA | ENSMUSG( | Avil | 14.72991 | -0.20015 | 0.722227 | 0.966887 |
| ENSMUSG( | Avl9 | 1362.706 | 0.007748 | 0.944973 | 0.993612 | ENSMUSG( | Avl9 | 1291.555 | 0.118521 | 0.352663 | 0.912093 |
| ENSMUSG( | Avpi1 | 357.2255 | -0.15402 | 0.31526 | 0.762288 | ENSMUSG( | Avpi1 | 325.983 | -0.15968 | 0.395928 | 0.917257 |
| ENSMUSG( | AW011738 | 407.0272 | -0.05973 | 0.632668 | 0.914073 | ENSMUSG( | AW011738 | 402.8171 | 0.170598 | 0.228313 | 0.868507 |
| ENSMUSG( | AW112010 | 33.29527 | -0.72474 | 0.035621 | 0.290384 | ENSMUSG( | AW112010 | 42.72594 | 0.362466 | 0.326142 | 0.905339 |
| ENSMUSG( | AW146154 | 160.7045 | -0.0032 | 0.984427 | 0.99781 | ENSMUSG( | AW146154 | 138.0614 | -0.17983 | 0.34406 | 0.910256 |
| ENSMUSG( | AW209491 | 308.8942 | 0.127486 | 0.398631 | 0.812878 | ENSMUSG( | AW209491 | 260.5827 | -0.09711 | 0.580367 | 0.944209 |
| ENSMUSG( | AW495222 | 127.8109 | -0.27811 | 0.12971 | 0.539528 | ENSMUSG( | AW495222 | 109.2651 | -0.5132 | 0.028024 | 0.61023 |
| ENSMUSG( | AW551984 | 105.1304 | 0.053585 | 0.835618 | 0.970557 | ENSMUSG( | AW551984 | 92.80892 | -0.05559 | 0.83101 | 0.98051 |
| ENSMUSG( | AW554918 | 465.2372 | -0.04927 | 0.67977 | 0.930983 | ENSMUSG( | AW554918 | 422.0466 | -0.07096 | 0.641363 | 0.953903 |
| ENSMUSG( | AW822252 | 65.95333 | 0.046318 | 0.859787 | 0.974818 | ENSMUSG( | AW822252 | 57.90532 | -0.07429 | 0.797588 | 0.97681 |
| ENSMUSG( | Axdnd1 | 24.08541 | 0.125873 | 0.742156 | 0.948478 | ENSMUSG( | Axdnd1 | 23.53121 | 0.314196 | 0.439443 | 0.920721 |
| ENSMUSG( | Axin1 | 897.1385 | 0.021462 | 0.838171 | 0.971111 | ENSMUSG( | Axin1 | 795.9907 | -0.05487 | 0.653049 | 0.955015 |
| ENSMUSG( | Axin2 | 1060.642 | 0.133367 | 0.161322 | 0.591331 | ENSMUSG( | Axin2 | 1051.257 | 0.358846 | 0.005524 | 0.303106 |
| ENSMUSG( | Axl | 7177.045 | -0.01381 | 0.895486 | 0.984512 | ENSMUSG( | Axl | 6602.419 | 0.014286 | 0.904575 | 0.991089 |
| ENSMUSG( | Azi2 | 1673.762 | 0.033115 | 0.73083 | 0.944956 | ENSMUSG( | Azi2 | 1515.356 | 0.014829 | 0.892759 | 0.990558 |
| ENSMUSG( | Azin1 | 5125.471 | 0.259749 | 0.008513 | 0.129175 | ENSMUSG( | Azin1 | 4621.874 | 0.230244 | 0.077847 | 0.77321 |
| ENSMUSG( | Azin2 | 945.8367 | 0.124292 | 0.35178 | 0.783451 | ENSMUSG( | Azin2 | 804.632 | -0.07244 | 0.602868 | 0.94789 |
| ENSMUSG( | B130024G1 | 16.99551 | 0.325356 | 0.523101 | NA | ENSMUSG( | B130024G1 | 16.9537 | 0.574615 | 0.252936 | 0.88018 |
| ENSMUSG( | B130034C1 | 48.08935 | -0.41902 | 0.184402 | 0.621442 | ENSMUSG( | B130034C1 | 46.62823 | -0.25281 | 0.477625 | 0.923329 |
| ENSMUSG( | B130055M | 102.0477 | 0.216255 | 0.28257 | 0.73186 | ENSMUSG( | B130055M | 93.04471 | 0.204223 | 0.383735 | 0.917257 |
| ENSMUSG( | B230110C( | 9.947126 | -0.58143 | 0.316767 | NA | ENSMUSG( | B230110C( | 11.19607 | 0.085246 | 0.888807 | NA |
| ENSMUSG( | B230206H( | 76.32039 | -0.47079 | 0.049485 | 0.339723 | ENSMUSG( | B230206H( | 91.35542 | 0.359386 | 0.205026 | 0.864412 |
| ENSMUSG( | B230206L( | 73.2217 | 0.088905 | 0.715299 | 0.941288 | ENSMUSG( | B230206L( | 61.23852 | -0.18951 | 0.503426 | 0.927825 |

| | | | | | |
|---|---|---|---|---|---|
| ENSMUSG( | 1362.706 | 0.007748 | 0.944973 | 0.993612 | Avl9 |
| ENSMUSG( | 12.48738 | -1.58972 | 0.313163 | NA | Avp |
| ENSMUSG( | 357.2255 | -0.15402 | 0.31526 | 0.762288 | Avpi1 |
| ENSMUSG( | 407.0272 | -0.05973 | 0.632668 | 0.914073 | AW011738 |
| ENSMUSG( | 33.29527 | -0.72474 | 0.035621 | 0.290384 | AW112010 |
| ENSMUSG( | 160.7045 | -0.0032 | 0.984427 | 0.99781 | AW146154 |
| ENSMUSG( | 308.8942 | 0.127486 | 0.398631 | 0.812878 | AW209491 |
| ENSMUSG( | 127.8109 | -0.27811 | 0.12971 | 0.539528 | AW495222 |
| ENSMUSG( | 105.1304 | 0.053585 | 0.835618 | 0.970557 | AW551984 |
| ENSMUSG( | 465.2372 | -0.04927 | 0.67977 | 0.930983 | AW554918 |
| ENSMUSG( | 65.95333 | 0.046318 | 0.859787 | 0.974818 | AW822252 |
| ENSMUSG( | 24.08541 | 0.125873 | 0.742156 | 0.948478 | Axdnd1 |
| ENSMUSG( | 897.1385 | 0.021462 | 0.838171 | 0.971111 | Axin1 |
| ENSMUSG( | 1060.642 | 0.133367 | 0.161322 | 0.591331 | Axin2 |
| ENSMUSG( | 7177.045 | -0.01381 | 0.895486 | 0.984512 | Axl |
| ENSMUSG( | 1673.762 | 0.033115 | 0.73083 | 0.944956 | Azi2 |
| ENSMUSG( | 5125.471 | 0.259749 | 0.008513 | 0.129175 | Azin1 |
| ENSMUSG( | 945.8367 | 0.124292 | 0.35178 | 0.783451 | Azin2 |
| ENSMUSG( | 16.99551 | 0.325356 | 0.523101 | NA | B130024G19Rik |
| ENSMUSG( | 48.08935 | -0.41902 | 0.184402 | 0.621442 | B130034C11Rik |
| ENSMUSG( | 102.0477 | 0.216255 | 0.28257 | 0.73186 | B130055M24Rik |
| ENSMUSG( | 9.947126 | -0.58143 | 0.316767 | NA | B230110C06Rik |
| ENSMUSG( | 11.68766 | -0.60408 | 0.276232 | NA | B230112J18Rik |
| ENSMUSG( | 76.32039 | -0.47079 | 0.049485 | 0.339723 | B230206H07Rik |
| ENSMUSG( | 73.2217 | 0.088905 | 0.715299 | 0.941288 | B230206L02Rik |
| ENSMUSG( | 11.47238 | 0.157567 | 0.770179 | NA | B230208H11Rik |
| ENSMUSG( | 204.4928 | -0.20879 | 0.183888 | 0.62087 | B230216N24Rik |
| ENSMUSG( | 237.829 | 0.044195 | 0.801818 | 0.962116 | B230217O12Rik |
| ENSMUSG( | 2476.761 | -0.0592 | 0.518197 | 0.873754 | B230219D22Rik |
| ENSMUSG( | 92.39584 | 0.312473 | 0.120453 | 0.524495 | B230307C23Rik |
| ENSMUSG( | 100.445 | -0.02697 | 0.90807 | 0.9878 | B230311B06Rik |
| ENSMUSG( | 17.63207 | 0.166428 | 0.703576 | NA | B230319C09Rik |
| ENSMUSG( | 780.3731 | 0.319608 | 0.013507 | 0.169727 | B230354K17Rik |
| ENSMUSG( | 4545.912 | -0.32723 | 0.011147 | 0.152217 | B2m |
| ENSMUSG( | 1236.49 | -0.45884 | 0.005038 | 0.094254 | B3galnt1 |
| ENSMUSG( | 437.5117 | -0.07662 | 0.565332 | 0.890779 | B3galnt2 |
| ENSMUSG( | 1403.428 | -0.45874 | 0.005886 | 0.103768 | B3galt1 |
| ENSMUSG( | 1014.531 | -0.58118 | 0.000469 | 0.01982 | B3galt2 |
| ENSMUSG( | 140.1902 | -0.28423 | 0.126566 | 0.534155 | B3galt4 |
| ENSMUSG( | 1392.682 | -0.31116 | 0.010237 | 0.144259 | B3galt5 |
| ENSMUSG( | 403.1987 | -0.43126 | 0.011445 | 0.153933 | B3galt6 |
| ENSMUSG( | 12155.01 | -0.08058 | 0.441154 | 0.832445 | B3gat1 |
| ENSMUSG( | 1179.959 | -0.04479 | 0.710221 | 0.9397 | B3gat2 |
| ENSMUSG( | 7915.867 | 0.010794 | 0.894128 | 0.984512 | B3gat3 |
| ENSMUSG( | 949.1037 | -0.28244 | 0.061129 | 0.381794 | B3glct |
| ENSMUSG( | 36.0286 | -0.43911 | 0.279925 | 0.729155 | B3gnt2 |
| ENSMUSG( | 26.61696 | 0.144109 | 0.685951 | 0.932981 | B3gnt3 |
| ENSMUSG( | 22.49013 | -0.20427 | 0.608596 | 0.907154 | B3gnt4 |
| ENSMUSG( | 26.83439 | -0.2489 | 0.516552 | 0.872788 | B3gnt5 |
| ENSMUSG( | 49.8599 | -0.60737 | 0.04333 | 0.319117 | B3gnt7 |
| ENSMUSG( | 42.08969 | -0.3781 | 0.19507 | 0.634458 | B3gnt8 |
| ENSMUSG( | 144.1388 | -0.74447 | 0.003674 | 0.077033 | B3gnt9 |
| ENSMUSG( | 224.5581 | 0.049221 | 0.739355 | 0.947768 | B3gntl1 |
| ENSMUSG( | 13.45648 | -0.06128 | 0.902933 | NA | B430212C06Rik |
| ENSMUSG( | 40.2164 | -0.49082 | 0.153338 | 0.57989 | B430319G15Rik |
| ENSMUSG( | 4221.81 | -0.14009 | 0.260095 | 0.710107 | B4galnt1 |
| ENSMUSG( | 21.92969 | -0.37202 | 0.421099 | 0.823126 | B4galnt3 |
| ENSMUSG( | 2691.425 | -0.03171 | 0.76115 | 0.952907 | B4galnt4 |
| ENSMUSG( | 350.2137 | -0.18071 | 0.232967 | 0.681135 | B4galt1 |
| ENSMUSG( | 6186.448 | 0.196027 | 0.041414 | 0.314189 | B4galt2 |
| ENSMUSG( | 1333.304 | -0.1301 | 0.327557 | 0.772692 | B4galt3 |
| ENSMUSG( | 1007.922 | -0.04734 | 0.695811 | 0.934979 | B4galt4 |
| ENSMUSG( | 1814.584 | -0.07534 | 0.586983 | 0.900235 | B4galt5 |
| ENSMUSG( | 4033.578 | -0.19763 | 0.136325 | 0.550595 | B4galt6 |
| ENSMUSG( | 726.9811 | -0.25631 | 0.07198 | 0.414057 | B4galt7 |
| ENSMUSG( | 29.4922 | -0.18978 | 0.697536 | 0.935662 | B4gat1 |
| ENSMUSG( | 619.9937 | 0.004406 | 0.970276 | 0.996369 | B630019K06Rik |
| ENSMUSG( | 11.91605 | 0.618283 | 0.252055 | NA | B930025P03Rik |
| ENSMUSG( | 371.1205 | -0.06374 | 0.631672 | 0.914073 | B9d1 |
| ENSMUSG( | 265.4226 | -0.16398 | 0.304801 | 0.752574 | B9d2 |
| ENSMUSG( | 24487.1 | 0.04501 | 0.685658 | 0.932981 | Baalc |
| ENSMUSG( | 1831.012 | 0.05982 | 0.504782 | 0.866883 | Babam1 |
| ENSMUSG( | 726.5192 | 0.046908 | 0.694028 | 0.934202 | Babam2 |
| ENSMUSG( | 3892.629 | -0.31476 | 0.032934 | 0.277503 | Bace1 |
| ENSMUSG( | 213.1326 | -0.44441 | 0.013877 | 0.172084 | Bace2 |
| ENSMUSG( | 474.1376 | 0.121309 | 0.308087 | 0.75626 | Bach1 |
| ENSMUSG( | 360.6406 | 0.14756 | 0.255042 | 0.705501 | Bach2 |
| ENSMUSG( | 900.013 | 0.079304 | 0.445346 | 0.833239 | Bad |
| ENSMUSG( | 5997.505 | 0.052207 | 0.566723 | 0.891644 | Bag1 |
| ENSMUSG( | 192.1527 | -0.01952 | 0.901874 | 0.985633 | Bag2 |
| ENSMUSG( | 347.9899 | -0.08376 | 0.519523 | 0.874979 | Bag3 |
| ENSMUSG( | 1002.012 | 0.021216 | 0.832999 | 0.970333 | Bag4 |
| ENSMUSG( | 800.3036 | 0.012102 | 0.925681 | 0.991204 | Bag5 |
| ENSMUSG( | 7247.288 | 0.171248 | 0.03546 | 0.290048 | Bag6 |
| ENSMUSG( | 846.2354 | -0.06144 | 0.552002 | 0.886141 | Bahcc1 |
| ENSMUSG( | 1018.382 | -0.02006 | 0.853624 | 0.973491 | Bahd1 |
| ENSMUSG( | 4901.49 | 0.1177 | 0.318167 | 0.764234 | Baiap2 |
| ENSMUSG( | 23.68821 | 0.325871 | 0.480909 | 0.855113 | Baiap2l1 |
| ENSMUSG( | 79.69022 | -0.28565 | 0.232825 | 0.680928 | Baiap2l2 |
| ENSMUSG( | 385.0804 | 0.349322 | 0.519703 | 0.874979 | Baiap3 |
| ENSMUSG( | 743.6679 | 0.080372 | 0.460618 | 0.840787 | Bak1 |
| ENSMUSG( | 134.9903 | -0.34861 | 0.121324 | 0.525713 | Bambi |
| ENSMUSG( | 1163.402 | -0.07247 | 0.481192 | 0.855242 | Banf1 |
| ENSMUSG( | 56.98939 | -0.24923 | 0.339768 | 0.777939 | Bank1 |
| ENSMUSG( | 814.4719 | 0.466862 | 0.000527 | 0.021317 | Banp |
| ENSMUSG( | 6509.552 | 0.210561 | 0.006284 | 0.108547 | Bap1 |
| ENSMUSG( | 22.72024 | -0.1142 | 0.802313 | 0.962119 | Bard1 |
| ENSMUSG( | 102.8889 | 0.136101 | 0.567067 | 0.891644 | Barx2 |
| ENSMUSG( | 15721.16 | 0.016592 | 0.831325 | 0.969889 | Basp1 |
| ENSMUSG( | 16.98472 | -0.53354 | 0.246266 | NA | Batf2 |
| ENSMUSG( | 23.8375 | 0.016063 | 0.967329 | 0.996006 | Batf3 |
| ENSMUSG( | 1036.849 | -0.01366 | 0.884155 | 0.982647 | Bax |
| ENSMUSG( | 161.2635 | 0.002267 | 0.990698 | 0.999232 | Baz1a |
| ENSMUSG( | 4319.169 | 0.202799 | 0.025906 | 0.247245 | Baz1b |
| ENSMUSG( | 1998.51 | 0.128019 | 0.214703 | 0.659193 | Baz2a |
| ENSMUSG( | 1121.756 | 0.010678 | 0.910426 | 0.9881 | Baz2b |
| ENSMUSG( | 41.31229 | 0.051972 | 0.860946 | 0.975025 | BB218582 |
| ENSMUSG( | 191.8904 | -0.22186 | 0.220344 | 0.665918 | Bbc3 |
| ENSMUSG( | 1636.866 | 0.067874 | 0.5328 | 0.879905 | Bbip1 |
| ENSMUSG( | 4801.524 | 0.117566 | 0.252941 | 0.703255 | Bbln |
| ENSMUSG( | 169.1649 | 0.298647 | 0.085456 | 0.450845 | Bbof1 |
| ENSMUSG( | 93.15177 | -0.29719 | 0.236023 | 0.683136 | Bbox1 |
| ENSMUSG( | 1797.421 | 0.200919 | 0.019542 | 0.210473 | Bbs1 |
| ENSMUSG( | 99.26509 | -0.29985 | 0.158437 | 0.587906 | Bbs10 |
| ENSMUSG( | 82.29624 | -0.04518 | 0.833316 | 0.970333 | Bbs12 |
| ENSMUSG( | 705.6213 | -0.03471 | 0.753438 | 0.950753 | Bbs2 |
| ENSMUSG( | 1114.03 | 0.007265 | 0.954915 | 0.993874 | Bbs4 |
| ENSMUSG( | 303.1228 | 0.012814 | 0.941625 | 0.993415 | Bbs5 |
| ENSMUSG( | 332.851 | 0.146154 | 0.275353 | 0.72513 | Bbs7 |
| ENSMUSG( | 336.9276 | 0.033765 | 0.823031 | 0.966449 | Bbs9 |
| ENSMUSG( | 709.0589 | -0.01603 | 0.876373 | 0.980735 | Bbx |
| ENSMUSG( | 207.788 | 0.130842 | 0.401997 | 0.814965 | BC002059 |
| ENSMUSG( | 2216.554 | -0.29166 | 0.00724 | 0.117427 | BC004004 |
| ENSMUSG( | 8143.268 | 0.043957 | 0.612112 | 0.908586 | BC005537 |
| ENSMUSG( | 2170.791 | -0.0276 | 0.779048 | 0.956487 | BC005624 |
| ENSMUSG( | 142.3003 | 0.154688 | 0.356566 | 0.788187 | BC006965 |
| ENSMUSG( | 13.27344 | -0.0736 | 0.898433 | NA | BC016548 |
| ENSMUSG( | 13.65991 | 0.322933 | 0.588706 | NA | BC016579 |
| ENSMUSG( | 37.51892 | -0.51005 | 0.105504 | 0.492417 | BC024063 |
| ENSMUSG( | 96.14751 | 0.254622 | 0.25289 | 0.703255 | BC024139 |
| ENSMUSG( | 140.4536 | -0.0129 | 0.953196 | 0.993874 | BC025920 |
| ENSMUSG( | 129.3672 | -0.24424 | 0.293132 | 0.742763 | BC028528 |
| ENSMUSG( | 168.6447 | 0.206647 | 0.226987 | 0.674187 | BC030500 |
| ENSMUSG( | 3951.15 | -0.21196 | 0.024126 | 0.237752 | BC031181 |
| ENSMUSG( | 478.261 | -0.09297 | 0.498772 | 0.862973 | BC034090 |
| ENSMUSG( | 43.21814 | -0.28596 | 0.345207 | 0.779153 | BC035044 |
| ENSMUSG( | 24.97761 | 0.303375 | 0.470731 | 0.846741 | BC037704 |
| ENSMUSG( | 31.79678 | -0.13308 | 0.710561 | 0.939748 | BC046251 |
| ENSMUSG( | 23.7395 | -0.40041 | 0.305851 | 0.753712 | BC049715 |
| ENSMUSG( | 101.7384 | 0.047017 | 0.833254 | 0.970333 | BC051226 |
| ENSMUSG( | 40.70029 | 0.186129 | 0.555992 | 0.887457 | BC055402 |
| ENSMUSG( | 67.65118 | 0.00763 | 0.976325 | 0.997428 | BC065397 |
| ENSMUSG( | 54223.3 | 0.190462 | 0.715844 | 0.941484 | Bc1 |
| ENSMUSG( | 24.2265 | -0.27155 | 0.496925 | 0.861754 | BC106179 |
| ENSMUSG( | 585.8519 | -0.08443 | 0.596755 | 0.903526 | Bcam |
| ENSMUSG( | 24570.47 | -0.08353 | 0.429061 | 0.826446 | Bcan |
| ENSMUSG( | 1301.839 | -0.35379 | 0.007582 | 0.120453 | Bcap29 |
| ENSMUSG( | 2376.898 | -0.36585 | 0.00228 | 0.055785 | Bcap31 |
| ENSMUSG( | 1261.403 | 0.051235 | 0.638314 | 0.916154 | Bcar1 |
| ENSMUSG( | 1128.461 | 0.109453 | 0.309826 | 0.757746 | Bcar3 |
| ENSMUSG( | 31037.84 | -0.10598 | 0.240634 | 0.688247 | Bcas1 |
| ENSMUSG( | 74.80579 | 0.15124 | 0.506694 | 0.86758 | Bcas1os2 |
| ENSMUSG( | 1371.056 | -0.01857 | 0.841622 | 0.971179 | Bcas2 |
| ENSMUSG( | 1379.515 | 0.039813 | 0.674602 | 0.929638 | Bcas3 |
| ENSMUSG( | 1488.263 | 0.206569 | 0.090042 | 0.460864 | Bcat1 |
| ENSMUSG( | 419.2167 | 0.113961 | 0.389275 | 0.807113 | Bcat2 |
| ENSMUSG( | 953.0468 | 0.00017 | 0.998669 | 0.99989 | Bccip |
| ENSMUSG( | 142.0992 | -0.12143 | 0.546142 | 0.884974 | Bcdin3d |

| | | | | | |
|---|---|---|---|---|---|
| ENSMUSG( | 6602.419 | 0.014286 | 0.904575 | 0.991089 | Axl |
| ENSMUSG( | 1515.356 | 0.014829 | 0.892759 | 0.990558 | Azi2 |
| ENSMUSG( | 4621.874 | 0.230244 | 0.077847 | 0.77321 | Azin1 |
| ENSMUSG( | 804.632 | -0.07244 | 0.602868 | 0.94789 | Azin2 |
| ENSMUSG( | 16.9537 | 0.574615 | 0.252936 | 0.88018 | B130024G19Rik |
| ENSMUSG( | 46.62823 | -0.25281 | 0.477625 | 0.923329 | B130034C11Rik |
| ENSMUSG( | 93.04471 | 0.204223 | 0.383735 | 0.917257 | B130055M24Rik |
| ENSMUSG( | 11.19607 | 0.085246 | 0.888807 | NA | B230110C06Rik |
| ENSMUSG( | 91.35542 | 0.359386 | 0.205026 | 0.864412 | B230206H07Rik |
| ENSMUSG( | 61.23852 | -0.18951 | 0.503426 | 0.927825 | B230206L02Rik |
| ENSMUSG( | 10.28926 | 0.132382 | 0.823429 | NA | B230208H11Rik |
| ENSMUSG( | 175.1298 | -0.41986 | 0.029995 | 0.620326 | B230216N24Rik |
| ENSMUSG( | 214.8261 | 0.02009 | 0.91122 | 0.992196 | B230217O12Rik |
| ENSMUSG( | 2307.774 | 0.003404 | 0.978216 | 0.996416 | B230219D22Rik |
| ENSMUSG( | 82.44791 | 0.263611 | 0.263918 | 0.886134 | B230307C23Rik |
| ENSMUSG( | 80.48764 | -0.44073 | 0.096332 | 0.796133 | B230311B06Rik |
| ENSMUSG( | 12.30149 | -0.65663 | 0.2357 | 0.871888 | B230319C09Rik |
| ENSMUSG( | 591.272 | -0.19907 | 0.169314 | 0.840806 | B230354K17Rik |
| ENSMUSG( | 4463.729 | -0.09244 | 0.502274 | 0.927825 | B2m |
| ENSMUSG( | 1365.83 | 0.134986 | 0.468219 | 0.923329 | B3galnt1 |
| ENSMUSG( | 427.1132 | 0.131612 | 0.456783 | 0.923329 | B3galnt2 |
| ENSMUSG( | 1496.77 | 0.035373 | 0.852221 | 0.985598 | B3galt1 |
| ENSMUSG( | 1238.92 | 0.311354 | 0.08985 | 0.79071 | B3galt2 |
| ENSMUSG( | 145.7424 | 0.120056 | 0.585941 | 0.944975 | B3galt4 |
| ENSMUSG( | 1485.295 | 0.159377 | 0.309977 | 0.898538 | B3galt5 |
| ENSMUSG( | 430.5053 | 0.063038 | 0.733077 | 0.967228 | B3galt6 |
| ENSMUSG( | 11765.01 | 0.095287 | 0.398519 | 0.917257 | B3gat1 |
| ENSMUSG( | 1099.003 | 0.017757 | 0.896331 | 0.990558 | B3gat2 |
| ENSMUSG( | 7111.434 | -0.0295 | 0.773947 | 0.97478 | B3gat3 |
| ENSMUSG( | 951.4903 | 0.010969 | 0.946712 | 0.994798 | B3glct |
| ENSMUSG( | 42.27207 | 0.340173 | 0.427142 | 0.917257 | B3gnt2 |
| ENSMUSG( | 29.09023 | 0.616147 | 0.09337 | 0.795507 | B3gnt3 |
| ENSMUSG( | 21.63286 | -0.01931 | 0.964998 | 0.995433 | B3gnt4 |
| ENSMUSG( | 34.55616 | 0.669893 | 0.067269 | 0.75212 | B3gnt5 |
| ENSMUSG( | 57.29609 | 0.144493 | 0.636544 | 0.953903 | B3gnt7 |
| ENSMUSG( | 42.78966 | -0.03187 | 0.916745 | 0.992529 | B3gnt8 |
| ENSMUSG( | 160.7657 | -0.05197 | 0.839618 | 0.982077 | B3gnt9 |
| ENSMUSG( | 199.9851 | -0.02024 | 0.903941 | 0.991089 | B3gntl1 |
| ENSMUSG( | 13.15263 | 0.135209 | 0.797183 | 0.97681 | B430212C06Rik |
| ENSMUSG( | 35.84421 | -0.60386 | 0.087034 | 0.782639 | B430319G15Rik |
| ENSMUSG( | 4299.684 | 0.182832 | 0.209571 | 0.864412 | B4galnt1 |
| ENSMUSG( | 20.36775 | -0.35277 | 0.448389 | 0.922202 | B4galnt3 |
| ENSMUSG( | 2716.475 | 0.256077 | 0.07046 | 0.758605 | B4galnt4 |
| ENSMUSG( | 349.6524 | 0.099869 | 0.656 | 0.955533 | B4galt1 |
| ENSMUSG( | 5637.777 | 0.197368 | 0.076759 | 0.770558 | B4galt2 |
| ENSMUSG( | 1381.85 | 0.240372 | 0.109281 | 0.805353 | B4galt3 |
| ENSMUSG( | 992.1285 | 0.175873 | 0.250194 | 0.877904 | B4galt4 |
| ENSMUSG( | 1848.661 | 0.242894 | 0.125201 | 0.81516 | B4galt5 |
| ENSMUSG( | 4298.673 | 0.253405 | 0.083843 | 0.777053 | B4galt6 |
| ENSMUSG( | 750.0241 | 0.117373 | 0.455644 | 0.923329 | B4galt7 |
| ENSMUSG( | 28.29374 | -0.02978 | 0.948116 | 0.994798 | B4gat1 |
| ENSMUSG( | 550.2385 | -0.08137 | 0.631048 | 0.953251 | B630019K06Rik |
| ENSMUSG( | 9.483009 | 0.297889 | 0.634133 | NA | B930025P03Rik |
| ENSMUSG( | 330.658 | -0.13278 | 0.420727 | 0.917257 | B9d1 |
| ENSMUSG( | 247.1961 | -0.10843 | 0.567045 | 0.941777 | B9d2 |
| ENSMUSG( | 20131.87 | -0.26431 | 0.047119 | 0.700664 | Baalc |
| ENSMUSG( | 1557.049 | -0.14275 | 0.193723 | 0.860096 | Babam1 |
| ENSMUSG( | 626.1692 | -0.11939 | 0.391175 | 0.917257 | Babam2 |
| ENSMUSG( | 4246.301 | 0.218711 | 0.208274 | 0.864412 | Bace1 |
| ENSMUSG( | 231.3333 | 0.104127 | 0.611016 | 0.950771 | Bace2 |
| ENSMUSG( | 421.8258 | 0.056831 | 0.688829 | 0.959742 | Bach1 |
| ENSMUSG( | 324.0548 | 0.110448 | 0.46999 | 0.923329 | Bach2 |
| ENSMUSG( | 790.5515 | -0.02988 | 0.82496 | 0.979297 | Bad |
| ENSMUSG( | 5089.176 | -0.15907 | 0.187547 | 0.85698 | Bag1 |
| ENSMUSG( | 167.1963 | -0.15134 | 0.400483 | 0.917257 | Bag2 |
| ENSMUSG( | 311.6333 | -0.12797 | 0.415308 | 0.917257 | Bag3 |
| ENSMUSG( | 910.6789 | 0.012235 | 0.920417 | 0.992644 | Bag4 |
| ENSMUSG( | 728.6968 | 0.00996 | 0.945391 | 0.994798 | Bag5 |
| ENSMUSG( | 6079.606 | -0.06298 | 0.538446 | 0.937216 | Bag6 |
| ENSMUSG( | 846.6238 | 0.207158 | 0.131699 | 0.822912 | Bahcc1 |
| ENSMUSG( | 913.5098 | -0.06184 | 0.618436 | 0.952254 | Bahd1 |
| ENSMUSG( | 4505.187 | 0.141207 | 0.310307 | 0.89905 | Baiap2 |
| ENSMUSG( | 25.62202 | 0.754583 | 0.089873 | 0.79071 | Baiap2l1 |
| ENSMUSG( | 79.02316 | -0.03385 | 0.901045 | 0.990558 | Baiap2l2 |
| ENSMUSG( | 311.4893 | 0.015909 | 0.976386 | 0.996416 | Baiap3 |
| ENSMUSG( | 647.1573 | -0.05524 | 0.668636 | 0.956766 | Bak1 |
| ENSMUSG( | 153.4242 | 0.307135 | 0.178384 | 0.848538 | Bambi |
| ENSMUSG( | 1025.538 | -0.17654 | 0.188806 | 0.85698 | Banf1 |
| ENSMUSG( | 44.29037 | -0.7925 | 0.009492 | 0.406458 | Bank1 |
| ENSMUSG( | 689.8288 | 0.274211 | 0.107589 | 0.804969 | Banp |
| ENSMUSG( | 5557.423 | 0.031143 | 0.755938 | 0.971371 | Bap1 |
| ENSMUSG( | 24.69507 | 0.348256 | 0.435527 | 0.919062 | Bard1 |
| ENSMUSG( | 7.267365 | -0.26908 | 0.788118 | NA | Barhl2 |
| ENSMUSG( | 87.21012 | -0.0741 | 0.793951 | 0.97681 | Barx2 |
| ENSMUSG( | 14164.47 | -0.01715 | 0.92017 | 0.992644 | Basp1 |
| ENSMUSG( | 14.40958 | -0.73985 | 0.159704 | 0.837724 | Batf2 |
| ENSMUSG( | 23.94488 | 0.270746 | 0.510565 | 0.929395 | Batf3 |
| ENSMUSG( | 950.7933 | 0.004511 | 0.968908 | 0.99594 | Bax |
| ENSMUSG( | 155.7474 | 0.1521 | 0.490767 | 0.92646 | Baz1a |
| ENSMUSG( | 3889.05 | 0.169032 | 0.130813 | 0.822884 | Baz1b |
| ENSMUSG( | 1795.442 | 0.089603 | 0.459133 | 0.923329 | Baz2a |
| ENSMUSG( | 1077.115 | 0.161639 | 0.197796 | 0.862161 | Baz2b |
| ENSMUSG( | 37.96161 | 0.051565 | 0.878613 | 0.989644 | BB218582 |
| ENSMUSG( | 166.8544 | -0.37101 | 0.052998 | 0.71525 | Bbc3 |
| ENSMUSG( | 1375.353 | -0.17529 | 0.207355 | 0.864412 | Bbip1 |
| ENSMUSG( | 3845.194 | -0.26741 | 0.024233 | 0.589324 | Bbln |
| ENSMUSG( | 160.4766 | 0.399103 | 0.045634 | 0.693959 | Bbof1 |
| ENSMUSG( | 78.20162 | -0.58299 | 0.017222 | 0.52741 | Bbox1 |
| ENSMUSG( | 1520.697 | -0.00504 | 0.96286 | 0.995304 | Bbs1 |
| ENSMUSG( | 103.8106 | 0.136488 | 0.61058 | 0.950771 | Bbs10 |
| ENSMUSG( | 70.87989 | -0.20505 | 0.403598 | 0.917257 | Bbs12 |
| ENSMUSG( | 662.0815 | 0.049634 | 0.732236 | 0.967228 | Bbs2 |
| ENSMUSG( | 940.4544 | -0.22518 | 0.132813 | 0.822912 | Bbs4 |
| ENSMUSG( | 303.0858 | 0.267324 | 0.190106 | 0.857355 | Bbs5 |
| ENSMUSG( | 296.6979 | 0.091726 | 0.573654 | 0.942912 | Bbs7 |
| ENSMUSG( | 310.2445 | 0.069935 | 0.684245 | 0.958678 | Bbs9 |
| ENSMUSG( | 631.5232 | -0.08699 | 0.495265 | 0.927372 | Bbx |
| ENSMUSG( | 184.4745 | 0.064959 | 0.719433 | 0.966876 | BC002059 |
| ENSMUSG( | 2218.636 | -0.00302 | 0.9819 | 0.996756 | BC004004 |
| ENSMUSG( | 7182.469 | -0.05083 | 0.649716 | 0.95422 | BC005537 |
| ENSMUSG( | 1859.273 | -0.21749 | 0.088371 | 0.786756 | BC005624 |
| ENSMUSG( | 134.3734 | 0.234226 | 0.265149 | 0.886134 | BC006965 |
| ENSMUSG( | 10.79584 | -0.04659 | 0.939895 | NA | BC016579 |
| ENSMUSG( | 42.64679 | 0.15637 | 0.623334 | 0.953234 | BC024063 |
| ENSMUSG( | 77.48913 | -0.08794 | 0.728377 | 0.966913 | BC024139 |
| ENSMUSG( | 122.1935 | -0.16069 | 0.48826 | 0.925453 | BC025920 |
| ENSMUSG( | 160.8132 | 0.615594 | 0.012544 | 0.451781 | BC028528 |
| ENSMUSG( | 146.4543 | 0.067607 | 0.735264 | 0.967344 | BC030500 |
| ENSMUSG( | 3715.448 | -0.11686 | 0.35141 | 0.912093 | BC031181 |
| ENSMUSG( | 424.5244 | -0.16112 | 0.344861 | 0.91045 | BC034090 |
| ENSMUSG( | 41.43686 | -0.12768 | 0.692684 | 0.96043 | BC035044 |
| ENSMUSG( | 24.48908 | 0.512518 | 0.233191 | 0.868507 | BC037704 |
| ENSMUSG( | 35.44804 | 0.42515 | 0.224696 | 0.868299 | BC046251 |
| ENSMUSG( | 27.75291 | 0.336544 | 0.369145 | 0.915491 | BC049715 |
| ENSMUSG( | 90.10805 | -0.02764 | 0.913235 | 0.992298 | BC051226 |
| ENSMUSG( | 36.02995 | 0.118067 | 0.744124 | 0.96951 | BC055402 |
| ENSMUSG( | 62.75588 | 0.066208 | 0.808809 | 0.97873 | BC065397 |
| ENSMUSG( | 48081.68 | 0.121058 | 0.860583 | 0.986137 | Bc1 |
| ENSMUSG( | 23.71968 | -0.03522 | 0.932425 | 0.99306 | BC106179 |
| ENSMUSG( | 669.1914 | 0.530799 | 0.008371 | 0.388416 | Bcam |
| ENSMUSG( | 23877.84 | 0.103916 | 0.410223 | 0.917257 | Bcan |
| ENSMUSG( | 1362.733 | 0.069154 | 0.665737 | 0.956766 | Bcap29 |
| ENSMUSG( | 2385.837 | -0.06142 | 0.665851 | 0.956766 | Bcap31 |
| ENSMUSG( | 1126.728 | -0.00277 | 0.982083 | 0.996756 | Bcar1 |
| ENSMUSG( | 931.9538 | -0.18273 | 0.172902 | 0.843702 | Bcar3 |
| ENSMUSG( | 26971.85 | -0.25418 | 0.05949 | 0.729747 | Bcas1 |
| ENSMUSG( | 63.50779 | -0.05449 | 0.832875 | 0.981069 | Bcas1os2 |
| ENSMUSG( | 1251.334 | -0.01549 | 0.894314 | 0.990558 | Bcas2 |
| ENSMUSG( | 1208.481 | -0.07573 | 0.543423 | 0.938063 | Bcas3 |
| ENSMUSG( | 1267.74 | 0.020076 | 0.875844 | 0.989644 | Bcat1 |
| ENSMUSG( | 403.1694 | 0.264936 | 0.090805 | 0.791074 | Bcat2 |
| ENSMUSG( | 845.3149 | -0.08166 | 0.526689 | 0.934374 | Bccip |
| ENSMUSG( | 139.7584 | 0.099305 | 0.673532 | 0.95694 | Bcdin3d |
| ENSMUSG( | 321.9317 | 0.067926 | 0.71212 | 0.964363 | Bche |
| ENSMUSG( | 2035.031 | -0.18971 | 0.125134 | 0.81516 | Bckdha |
| ENSMUSG( | 506.9615 | -0.08722 | 0.559331 | 0.939748 | Bckdhb |
| ENSMUSG( | 980.1228 | -0.1065 | 0.366415 | 0.914703 | Bckdk |
| ENSMUSG( | 446.4948 | -0.0214 | 0.873803 | 0.989644 | Bcl10 |
| ENSMUSG( | 1569.056 | 0.022607 | 0.881836 | 0.989644 | Bcl11a |
| ENSMUSG( | 1637.311 | 0.057469 | 0.696064 | 0.960867 | Bcl11b |
| ENSMUSG( | 1097.371 | -0.01769 | 0.909409 | 0.991958 | Bcl2 |
| ENSMUSG( | 15.05115 | 0.262723 | 0.598914 | 0.947045 | Bcl2a1b |
| ENSMUSG( | 2710.98 | -0.00594 | 0.956895 | 0.994798 | Bcl2l1 |
| ENSMUSG( | 177.5228 | 0.312861 | 0.110343 | 0.807051 | Bcl2l11 |
| ENSMUSG( | 90.38825 | 0.260036 | 0.258984 | 0.883971 | Bcl2l12 |
| ENSMUSG( | 1648.183 | -0.06881 | 0.543722 | 0.938063 | Bcl2l13 |
| ENSMUSG( | 9.229688 | -0.25929 | 0.737274 | NA | Bcl2l15 |
| ENSMUSG( | 7178.261 | -0.08003 | 0.44692 | 0.921891 | Bcl2l2 |

| | | | | | | | | | | | |
|---|---|---|---|---|---|---|---|---|---|---|---|
| ENSMUSG( | B230208H1 | 11.47238 | 0.157567 | 0.770179 | NA | ENSMUSG( | B230208H1 | 10.28926 | 0.132382 | 0.823429 | NA |
| ENSMUSG( | B230216N2 | 204.4928 | -0.20879 | 0.183888 | 0.62087 | ENSMUSG( | B230216N2 | 175.1298 | -0.41986 | 0.029995 | 0.620326 |
| ENSMUSG( | B230217O1 | 237.829 | 0.044195 | 0.801818 | 0.962116 | ENSMUSG( | B230217O1 | 214.8261 | 0.02009 | 0.91122 | 0.992196 |
| ENSMUSG( | B230219D2 | 2476.761 | -0.0592 | 0.518197 | 0.873754 | ENSMUSG( | B230219D2 | 2307.774 | 0.003404 | 0.978216 | 0.996416 |
| ENSMUSG( | B230307C2 | 92.39584 | 0.312473 | 0.120453 | 0.524495 | ENSMUSG( | B230307C2 | 82.44791 | 0.263611 | 0.263918 | 0.886134 |
| ENSMUSG( | B230311B( | 100.445 | -0.02697 | 0.90807 | 0.9878 | ENSMUSG( | B230311B( | 80.48764 | -0.44073 | 0.096332 | 0.796133 |
| ENSMUSG( | B230319C( | 17.63207 | 0.166428 | 0.703576 | NA | ENSMUSG( | B230319C( | 12.30149 | -0.65663 | 0.2357 | 0.871888 |
| ENSMUSG( | B230354K1 | 780.3731 | 0.319608 | 0.013507 | 0.169727 | ENSMUSG( | B230354K1 | 591.272 | -0.19907 | 0.169314 | 0.840806 |
| ENSMUSG( | B2m | 4545.912 | -0.32723 | 0.011147 | 0.152217 | ENSMUSG( | B2m | 4463.729 | -0.09244 | 0.502274 | 0.927825 |
| ENSMUSG( | B3galnt1 | 1236.49 | -0.45884 | 0.005038 | 0.094254 | ENSMUSG( | B3galnt1 | 1365.83 | 0.134986 | 0.468219 | 0.923329 |
| ENSMUSG( | B3galnt2 | 437.5117 | -0.07662 | 0.565332 | 0.890779 | ENSMUSG( | B3galnt2 | 427.1132 | 0.131612 | 0.456783 | 0.923329 |
| ENSMUSG( | B3galt1 | 1403.428 | -0.45874 | 0.005886 | 0.103768 | ENSMUSG( | B3galt1 | 1496.77 | 0.035373 | 0.852221 | 0.985598 |
| ENSMUSG( | B3galt2 | 1014.531 | -0.58118 | 0.000469 | 0.01982 | ENSMUSG( | B3galt2 | 1238.92 | 0.311354 | 0.08985 | 0.79071 |
| ENSMUSG( | B3galt4 | 140.1902 | -0.28423 | 0.126566 | 0.534155 | ENSMUSG( | B3galt4 | 145.7424 | 0.120056 | 0.585941 | 0.944975 |
| ENSMUSG( | B3galt5 | 1392.682 | -0.31116 | 0.010237 | 0.144259 | ENSMUSG( | B3galt5 | 1485.295 | 0.159377 | 0.309977 | 0.898538 |
| ENSMUSG( | B3galt6 | 403.1987 | -0.43126 | 0.011445 | 0.153933 | ENSMUSG( | B3galt6 | 430.5053 | 0.063038 | 0.733077 | 0.967228 |
| ENSMUSG( | B3gat1 | 12155.01 | -0.08058 | 0.441154 | 0.832445 | ENSMUSG( | B3gat1 | 11765.01 | 0.095287 | 0.398519 | 0.917257 |
| ENSMUSG( | B3gat2 | 1179.959 | -0.04479 | 0.710221 | 0.9397 | ENSMUSG( | B3gat2 | 1099.003 | 0.017757 | 0.896331 | 0.990558 |
| ENSMUSG( | B3gat3 | 7915.867 | 0.010794 | 0.894128 | 0.984512 | ENSMUSG( | B3gat3 | 7111.434 | -0.0295 | 0.773947 | 0.97478 |
| ENSMUSG( | B3glct | 949.1037 | -0.28244 | 0.061129 | 0.381794 | ENSMUSG( | B3glct | 951.4903 | 0.010969 | 0.946712 | 0.994798 |
| ENSMUSG( | B3gnt2 | 36.0286 | -0.43911 | 0.279925 | 0.729155 | ENSMUSG( | B3gnt2 | 42.27207 | 0.340173 | 0.427142 | 0.917257 |
| ENSMUSG( | B3gnt3 | 26.61696 | 0.144109 | 0.685951 | 0.932981 | ENSMUSG( | B3gnt3 | 29.09023 | 0.616147 | 0.09337 | 0.795507 |
| ENSMUSG( | B3gnt4 | 22.49013 | -0.20427 | 0.608596 | 0.907154 | ENSMUSG( | B3gnt4 | 21.63286 | -0.01931 | 0.964998 | 0.995433 |
| ENSMUSG( | B3gnt5 | 26.83439 | -0.2489 | 0.516552 | 0.872788 | ENSMUSG( | B3gnt5 | 34.55616 | 0.669893 | 0.067269 | 0.75212 |
| ENSMUSG( | B3gnt7 | 49.8599 | -0.60737 | 0.04333 | 0.319117 | ENSMUSG( | B3gnt7 | 57.29609 | 0.144493 | 0.636544 | 0.953903 |
| ENSMUSG( | B3gnt8 | 42.08969 | -0.3781 | 0.19507 | 0.634458 | ENSMUSG( | B3gnt8 | 42.78966 | -0.03187 | 0.916745 | 0.992529 |
| ENSMUSG( | B3gnt9 | 144.1388 | -0.74447 | 0.003674 | 0.077033 | ENSMUSG( | B3gnt9 | 160.7657 | -0.05197 | 0.839618 | 0.982077 |
| ENSMUSG( | B3gntl1 | 224.5581 | 0.049221 | 0.739355 | 0.947768 | ENSMUSG( | B3gntl1 | 199.9851 | -0.02024 | 0.903941 | 0.991089 |
| ENSMUSG( | B430212C( | 13.45648 | -0.06128 | 0.902933 | NA | ENSMUSG( | B430212C( | 13.15263 | 0.135209 | 0.797183 | 0.97681 |
| ENSMUSG( | B430319G1 | 40.2164 | -0.49082 | 0.153338 | 0.57989 | ENSMUSG( | B430319G1 | 35.84421 | -0.60386 | 0.087034 | 0.782639 |
| ENSMUSG( | B4galnt1 | 4221.81 | -0.14009 | 0.260095 | 0.710107 | ENSMUSG( | B4galnt1 | 4299.684 | 0.182832 | 0.209571 | 0.864412 |
| ENSMUSG( | B4galnt3 | 21.92969 | -0.37202 | 0.421099 | 0.823126 | ENSMUSG( | B4galnt3 | 20.36775 | -0.35277 | 0.448389 | 0.922202 |
| ENSMUSG( | B4galnt4 | 2691.425 | -0.03171 | 0.76115 | 0.952907 | ENSMUSG( | B4galnt4 | 2716.475 | 0.256077 | 0.07046 | 0.758605 |
| ENSMUSG( | B4galt1 | 350.2137 | -0.18071 | 0.232967 | 0.681135 | ENSMUSG( | B4galt1 | 349.6524 | 0.099869 | 0.656 | 0.955533 |
| ENSMUSG( | B4galt2 | 6186.448 | 0.196027 | 0.041414 | 0.314189 | ENSMUSG( | B4galt2 | 5637.777 | 0.197368 | 0.076759 | 0.770558 |
| ENSMUSG( | B4galt3 | 1333.304 | -0.1301 | 0.327557 | 0.772692 | ENSMUSG( | B4galt3 | 1381.85 | 0.240372 | 0.109281 | 0.805353 |
| ENSMUSG( | B4galt4 | 1007.922 | -0.04734 | 0.695811 | 0.934979 | ENSMUSG( | B4galt4 | 992.1285 | 0.175873 | 0.250194 | 0.877904 |
| ENSMUSG( | B4galt5 | 1814.584 | -0.07534 | 0.586983 | 0.900235 | ENSMUSG( | B4galt5 | 1848.661 | 0.242894 | 0.125201 | 0.81516 |
| ENSMUSG( | B4galt6 | 4033.578 | -0.19763 | 0.136325 | 0.550595 | ENSMUSG( | B4galt6 | 4298.673 | 0.253405 | 0.083843 | 0.777053 |
| ENSMUSG( | B4galt7 | 726.9811 | -0.25631 | 0.07198 | 0.414057 | ENSMUSG( | B4galt7 | 750.0241 | 0.117373 | 0.455644 | 0.923329 |
| ENSMUSG( | B4gat1 | 29.4922 | -0.18978 | 0.697536 | 0.935662 | ENSMUSG( | B4gat1 | 28.29374 | -0.02978 | 0.948116 | 0.994798 |
| ENSMUSG( | B630019K( | 619.9937 | 0.004406 | 0.970276 | 0.996369 | ENSMUSG( | B630019K( | 550.2385 | -0.08137 | 0.631048 | 0.953251 |
| ENSMUSG( | B930025P( | 11.91605 | 0.618283 | 0.252055 | NA | ENSMUSG( | B930025P( | 9.483009 | 0.297889 | 0.634133 | NA |
| ENSMUSG( | B9d1 | 371.1205 | -0.06374 | 0.631672 | 0.914073 | ENSMUSG( | B9d1 | 330.658 | -0.13278 | 0.420727 | 0.917257 |
| ENSMUSG( | B9d2 | 265.4226 | -0.16398 | 0.304801 | 0.752574 | ENSMUSG( | B9d2 | 247.1961 | -0.10843 | 0.567045 | 0.941777 |
| ENSMUSG( | Baalc | 24487.1 | 0.04501 | 0.685658 | 0.932981 | ENSMUSG( | Baalc | 20131.87 | -0.26431 | 0.047119 | 0.700664 |
| ENSMUSG( | Babam1 | 1831.012 | 0.05982 | 0.504782 | 0.866883 | ENSMUSG( | Babam1 | 1557.049 | -0.14275 | 0.193723 | 0.860096 |
| ENSMUSG( | Babam2 | 726.5192 | 0.046908 | 0.694028 | 0.934202 | ENSMUSG( | Babam2 | 626.1692 | -0.11939 | 0.391175 | 0.917257 |
| ENSMUSG( | Bace1 | 3892.629 | -0.31476 | 0.032934 | 0.277503 | ENSMUSG( | Bace1 | 4246.301 | 0.218711 | 0.208274 | 0.864412 |
| ENSMUSG( | Bace2 | 213.1326 | -0.44441 | 0.013877 | 0.172084 | ENSMUSG( | Bace2 | 231.3333 | 0.104127 | 0.611016 | 0.950771 |
| ENSMUSG( | Bach1 | 474.1376 | 0.121309 | 0.308087 | 0.75626 | ENSMUSG( | Bach1 | 421.8258 | 0.056831 | 0.688829 | 0.959742 |
| ENSMUSG( | Bach2 | 360.6406 | 0.14756 | 0.255042 | 0.705501 | ENSMUSG( | Bach2 | 324.0548 | 0.110448 | 0.46999 | 0.923329 |
| ENSMUSG( | Bad | 900.013 | 0.079304 | 0.445346 | 0.833239 | ENSMUSG( | Bad | 790.5515 | -0.02988 | 0.82496 | 0.979297 |
| ENSMUSG( | Bag1 | 5997.505 | 0.052207 | 0.566723 | 0.891644 | ENSMUSG( | Bag1 | 5089.176 | -0.15907 | 0.187547 | 0.85698 |
| ENSMUSG( | Bag2 | 192.1527 | -0.01952 | 0.901874 | 0.985633 | ENSMUSG( | Bag2 | 167.1963 | -0.15134 | 0.400483 | 0.917257 |
| ENSMUSG( | Bag3 | 347.9899 | -0.08376 | 0.519523 | 0.874979 | ENSMUSG( | Bag3 | 311.6333 | -0.12797 | 0.415308 | 0.917257 |
| ENSMUSG( | Bag4 | 1002.012 | 0.021216 | 0.832999 | 0.970333 | ENSMUSG( | Bag4 | 910.6789 | 0.012235 | 0.920417 | 0.992644 |
| ENSMUSG( | Bag5 | 800.3036 | 0.012102 | 0.925681 | 0.991204 | ENSMUSG( | Bag5 | 728.6968 | 0.00996 | 0.945391 | 0.994798 |
| ENSMUSG( | Bag6 | 7247.288 | 0.171248 | 0.03546 | 0.290048 | ENSMUSG( | Bag6 | 6079.606 | -0.06298 | 0.538446 | 0.937216 |
| ENSMUSG( | Bahcc1 | 846.2354 | -0.06144 | 0.552002 | 0.886141 | ENSMUSG( | Bahcc1 | 846.6238 | 0.207158 | 0.131699 | 0.822912 |
| ENSMUSG( | Bahd1 | 1018.382 | -0.02006 | 0.853624 | 0.973491 | ENSMUSG( | Bahd1 | 913.5098 | -0.06184 | 0.618436 | 0.952254 |
| ENSMUSG( | Baiap2 | 4901.49 | 0.1177 | 0.318167 | 0.764234 | ENSMUSG( | Baiap2 | 4505.187 | 0.141207 | 0.310307 | 0.89905 |
| ENSMUSG( | Baiap2l1 | 23.68821 | 0.325871 | 0.480909 | 0.855113 | ENSMUSG( | Baiap2l1 | 25.62202 | 0.754583 | 0.089873 | 0.79071 |
| ENSMUSG( | Baiap2l2 | 79.69022 | -0.28565 | 0.232825 | 0.680928 | ENSMUSG( | Baiap2l2 | 79.02316 | -0.03385 | 0.901045 | 0.990558 |
| ENSMUSG( | Baiap3 | 385.0804 | 0.349322 | 0.519703 | 0.874979 | ENSMUSG( | Baiap3 | 311.4893 | 0.015909 | 0.976386 | 0.996416 |
| ENSMUSG( | Bak1 | 743.6679 | 0.080372 | 0.460618 | 0.840787 | ENSMUSG( | Bak1 | 647.1573 | -0.05524 | 0.668636 | 0.956766 |
| ENSMUSG( | Bambi | 134.9903 | -0.34861 | 0.121324 | 0.525713 | ENSMUSG( | Bambi | 153.4242 | 0.307135 | 0.178384 | 0.848538 |
| ENSMUSG( | Banf1 | 1163.402 | -0.07247 | 0.481192 | 0.855242 | ENSMUSG( | Banf1 | 1025.538 | -0.17654 | 0.188806 | 0.85698 |
| ENSMUSG( | Bank1 | 56.98939 | -0.24923 | 0.339768 | 0.777939 | ENSMUSG( | Bank1 | 44.29037 | -0.7925 | 0.009492 | 0.406458 |
| ENSMUSG( | Banp | 814.4719 | 0.466862 | 0.000527 | 0.021317 | ENSMUSG( | Banp | 689.8288 | 0.274211 | 0.107589 | 0.804969 |
| ENSMUSG( | Bap1 | 6509.552 | 0.210561 | 0.006284 | 0.108547 | ENSMUSG( | Bap1 | 5557.423 | 0.031143 | 0.755938 | 0.971371 |
| ENSMUSG( | Bard1 | 22.72024 | -0.1142 | 0.802313 | 0.962119 | ENSMUSG( | Bard1 | 24.69507 | 0.348256 | 0.435527 | 0.919062 |
| ENSMUSG( | Barx2 | 102.8889 | 0.136101 | 0.567067 | 0.891644 | ENSMUSG( | Barx2 | 87.21012 | -0.0741 | 0.793951 | 0.97681 |
| ENSMUSG( | Basp1 | 15721.16 | 0.016592 | 0.831325 | 0.969889 | ENSMUSG( | Basp1 | 14164.47 | -0.01715 | 0.92017 | 0.992644 |
| ENSMUSG( | Batf2 | 16.98472 | -0.53354 | 0.246266 | NA | ENSMUSG( | Batf2 | 14.40958 | -0.73985 | 0.159704 | 0.837724 |
| ENSMUSG( | Batf3 | 23.8375 | 0.016063 | 0.967329 | 0.996006 | ENSMUSG( | Batf3 | 23.94488 | 0.270746 | 0.510565 | 0.929395 |
| ENSMUSG( | Bax | 1036.849 | -0.01366 | 0.884155 | 0.982647 | ENSMUSG( | Bax | 950.7933 | 0.004511 | 0.968908 | 0.99594 |
| ENSMUSG( | Baz1a | 161.2635 | 0.002267 | 0.990698 | 0.999232 | ENSMUSG( | Baz1a | 155.7474 | 0.1521 | 0.490767 | 0.92646 |
| ENSMUSG( | Baz1b | 4319.169 | 0.202799 | 0.025906 | 0.247245 | ENSMUSG( | Baz1b | 3889.05 | 0.169032 | 0.130813 | 0.822884 |
| ENSMUSG( | Baz2a | 1998.51 | 0.128019 | 0.214703 | 0.659193 | ENSMUSG( | Baz2a | 1795.442 | 0.089603 | 0.459133 | 0.923329 |
| ENSMUSG( | Baz2b | 1121.756 | 0.010678 | 0.910426 | 0.9881 | ENSMUSG( | Baz2b | 1077.115 | 0.161639 | 0.197796 | 0.862161 |
| ENSMUSG( | BB218582 | 41.31229 | 0.051972 | 0.860946 | 0.975025 | ENSMUSG( | BB218582 | 37.96161 | 0.051565 | 0.878613 | 0.989644 |
| ENSMUSG( | Bbc3 | 191.8904 | -0.22186 | 0.220344 | 0.665918 | ENSMUSG( | Bbc3 | 166.8544 | -0.37101 | 0.052998 | 0.71525 |
| ENSMUSG( | Bbip1 | 1636.866 | 0.067874 | 0.5328 | 0.879905 | ENSMUSG( | Bbip1 | 1375.353 | -0.17529 | 0.207355 | 0.864412 |
| ENSMUSG( | Bbln | 4801.524 | 0.117566 | 0.252941 | 0.703255 | ENSMUSG( | Bbln | 3845.194 | -0.26741 | 0.024233 | 0.589324 |
| ENSMUSG( | Bbof1 | 169.1649 | 0.298647 | 0.085456 | 0.450845 | ENSMUSG( | Bbof1 | 160.4766 | 0.399103 | 0.045634 | 0.693959 |
| ENSMUSG( | Bbox1 | 93.15177 | -0.29719 | 0.236023 | 0.683136 | ENSMUSG( | Bbox1 | 78.20162 | -0.58299 | 0.017222 | 0.52741 |
| ENSMUSG( | Bbs1 | 1797.421 | 0.200919 | 0.019542 | 0.210473 | ENSMUSG( | Bbs1 | 1520.697 | -0.00504 | 0.96286 | 0.995304 |
| ENSMUSG( | Bbs10 | 99.26509 | -0.29985 | 0.158437 | 0.587906 | ENSMUSG( | Bbs10 | 103.8106 | 0.136488 | 0.61058 | 0.950771 |
| ENSMUSG( | Bbs12 | 82.29624 | -0.04518 | 0.833316 | 0.970333 | ENSMUSG( | Bbs12 | 70.87989 | -0.20505 | 0.403598 | 0.917257 |
| ENSMUSG( | Bbs2 | 705.6213 | -0.03471 | 0.753438 | 0.950753 | ENSMUSG( | Bbs2 | 662.0815 | 0.049634 | 0.732236 | 0.967228 |
| ENSMUSG( | Bbs4 | 1114.03 | 0.007265 | 0.954915 | 0.993874 | ENSMUSG( | Bbs4 | 940.4544 | -0.22518 | 0.132813 | 0.822912 |
| ENSMUSG( | Bbs5 | 303.1228 | 0.012814 | 0.941625 | 0.993415 | ENSMUSG( | Bbs5 | 303.0858 | 0.267324 | 0.190106 | 0.857355 |
| ENSMUSG( | Bbs7 | 332.851 | 0.146154 | 0.275353 | 0.72513 | ENSMUSG( | Bbs7 | 296.6979 | 0.091726 | 0.573654 | 0.942912 |
| ENSMUSG( | Bbs9 | 336.9276 | 0.033765 | 0.823031 | 0.966449 | ENSMUSG( | Bbs9 | 310.2445 | 0.069935 | 0.684245 | 0.958678 |
| ENSMUSG( | Bbx | 709.0589 | -0.01603 | 0.876373 | 0.980735 | ENSMUSG( | Bbx | 631.5232 | -0.08699 | 0.495265 | 0.927372 |
| ENSMUSG( | BC002059 | 207.788 | 0.130842 | 0.401997 | 0.814965 | ENSMUSG( | BC002059 | 184.4745 | 0.064959 | 0.719433 | 0.966876 |
| ENSMUSG( | BC004004 | 2216.554 | -0.29166 | 0.00724 | 0.117427 | ENSMUSG( | BC004004 | 2218.636 | -0.00302 | 0.9819 | 0.996756 |
| ENSMUSG( | BC005537 | 8143.268 | 0.043957 | 0.612112 | 0.908586 | ENSMUSG( | BC005537 | 7182.469 | -0.05083 | 0.649716 | 0.95422 |
| ENSMUSG( | BC005624 | 2170.791 | -0.0276 | 0.779048 | 0.956487 | ENSMUSG( | BC005624 | 1859.273 | -0.21749 | 0.088371 | 0.786756 |
| ENSMUSG( | BC006965 | 142.3003 | 0.154688 | 0.356566 | 0.788187 | ENSMUSG( | BC006965 | 134.3734 | 0.234226 | 0.265149 | 0.886134 |
| ENSMUSG( | BC016579 | 13.65991 | 0.322933 | 0.588706 | NA | ENSMUSG( | BC016579 | 10.79584 | -0.04659 | 0.939895 | NA |
| ENSMUSG( | BC024063 | 37.51892 | -0.51005 | 0.105504 | 0.492417 | ENSMUSG( | BC024063 | 42.64679 | 0.15637 | 0.623334 | 0.953234 |
| ENSMUSG( | BC024139 | 96.14751 | 0.254622 | 0.25289 | 0.703255 | ENSMUSG( | BC024139 | 77.48913 | -0.08794 | 0.728377 | 0.966913 |
| ENSMUSG( | BC025920 | 140.4536 | -0.0129 | 0.953196 | 0.993874 | ENSMUSG( | BC025920 | 122.1935 | -0.16069 | 0.48826 | 0.925453 |
| ENSMUSG( | BC028528 | 129.3672 | -0.24424 | 0.293132 | 0.742763 | ENSMUSG( | BC028528 | 160.8132 | 0.615594 | 0.012544 | 0.451781 |
| ENSMUSG( | BC030500 | 168.6447 | 0.206647 | 0.226987 | 0.674187 | ENSMUSG( | BC030500 | 146.4543 | 0.067607 | 0.735264 | 0.967344 |
| ENSMUSG( | BC031181 | 3951.15 | -0.21196 | 0.024126 | 0.237752 | ENSMUSG( | BC031181 | 3715.448 | -0.11686 | 0.35141 | 0.912093 |
| ENSMUSG( | BC034090 | 478.261 | -0.09297 | 0.498772 | 0.862973 | ENSMUSG( | BC034090 | 424.5244 | -0.16112 | 0.344861 | 0.91045 |
| ENSMUSG( | BC035044 | 43.21814 | -0.28596 | 0.345207 | 0.779153 | ENSMUSG( | BC035044 | 41.43686 | -0.12768 | 0.692684 | 0.96043 |
| ENSMUSG( | BC037704 | 24.97761 | 0.303375 | 0.470731 | 0.846741 | ENSMUSG( | BC037704 | 24.48908 | 0.512518 | 0.233191 | 0.868507 |
| ENSMUSG( | BC046251 | 31.79678 | -0.13308 | 0.710561 | 0.939748 | ENSMUSG( | BC046251 | 35.44804 | 0.42515 | 0.224696 | 0.868299 |
| ENSMUSG( | BC049715 | 23.7395 | -0.40041 | 0.305851 | 0.753712 | ENSMUSG( | BC049715 | 27.75291 | 0.336544 | 0.369145 | 0.915491 |
| ENSMUSG( | BC051226 | 101.7384 | 0.047017 | 0.833254 | 0.970333 | ENSMUSG( | BC051226 | 90.10805 | -0.02764 | 0.913235 | 0.992298 |
| ENSMUSG( | BC055402 | 40.70029 | 0.186129 | 0.555992 | 0.887457 | ENSMUSG( | BC055402 | 36.02995 | 0.118067 | 0.744124 | 0.96951 |
| ENSMUSG( | BC065397 | 67.65118 | 0.00763 | 0.976325 | 0.997428 | ENSMUSG( | BC065397 | 62.75588 | 0.066208 | 0.808809 | 0.97873 |
| ENSMUSG( | Bc1 | 54223.3 | 0.190462 | 0.715844 | 0.941484 | ENSMUSG( | Bc1 | 48081.68 | 0.121058 | 0.860583 | 0.986137 |
| ENSMUSG( | BC106179 | 24.2265 | -0.27155 | 0.496925 | 0.861754 | ENSMUSG( | BC106179 | 23.71968 | -0.03522 | 0.932425 | 0.99306 |
| ENSMUSG( | Bcam | 585.8519 | -0.08443 | 0.596755 | 0.903526 | ENSMUSG( | Bcam | 669.1914 | 0.530799 | 0.008371 | 0.388416 |
| ENSMUSG( | Bcan | 24570.47 | -0.08353 | 0.429061 | 0.826446 | ENSMUSG( | Bcan | 23877.84 | 0.103916 | 0.410223 | 0.917257 |
| ENSMUSG( | Bcap29 | 1301.839 | -0.35379 | 0.007582 | 0.120453 | ENSMUSG( | Bcap29 | 1362.733 | 0.069154 | 0.665737 | 0.956766 |
| ENSMUSG( | Bcap31 | 2376.898 | -0.36585 | 0.00228 | 0.055785 | ENSMUSG( | Bcap31 | 2385.837 | -0.06142 | 0.665851 | 0.956766 |
| ENSMUSG( | Bcar1 | 1261.403 | 0.051235 | 0.638314 | 0.916154 | ENSMUSG( | Bcar1 | 1126.728 | -0.00277 | 0.982083 | 0.996756 |
| ENSMUSG( | Bcar3 | 1128.461 | 0.109453 | 0.309826 | 0.757746 | ENSMUSG( | Bcar3 | 931.9538 | -0.18273 | 0.172902 | 0.843702 |
| ENSMUSG( | Bcas1 | 31037.84 | -0.10598 | 0.240634 | 0.688247 | ENSMUSG( | Bcas1 | 26971.85 | -0.25418 | 0.05949 | 0.729747 |
| ENSMUSG( | Bcas1os2 | 74.80579 | 0.15124 | 0.506694 | 0.86758 | ENSMUSG( | Bcas1os2 | 63.50779 | -0.05449 | 0.832875 | 0.981069 |
| ENSMUSG( | Bcas2 | 1371.056 | -0.01857 | 0.841622 | 0.971179 | ENSMUSG( | Bcas2 | 1251.334 | -0.01549 | 0.894314 | 0.990558 |
| ENSMUSG( | Bcas3 | 1379.515 | 0.039813 | 0.674602 | 0.929638 | ENSMUSG( | Bcas3 | 1208.481 | -0.07573 | 0.543423 | 0.938063 |
| ENSMUSG( | Bcat1 | 1488.263 | 0.206569 | 0.090042 | 0.460864 | ENSMUSG( | Bcat1 | 1267.74 | 0.020076 | 0.875844 | 0.989644 |
| ENSMUSG( | Bcat2 | 419.2167 | 0.113961 | 0.389275 | 0.807113 | ENSMUSG( | Bcat2 | 403.1694 | 0.264936 | 0.090805 | 0.791074 |
| ENSMUSG( | Bccip | 953.0468 | 0.00017 | 0.998669 | 0.99989 | ENSMUSG( | Bccip | 845.3149 | -0.08166 | 0.526689 | 0.934374 |
| ENSMUSG( | Bcdin3d | 142.0992 | -0.12143 | 0.546142 | 0.884974 | ENSMUSG( | Bcdin3d | 139.7584 | 0.099305 | 0.673532 | 0.95694 |
| ENSMUSG( | Bche | 300.2851 | -0.44599 | 0.006556 | 0.111119 | ENSMUSG( | Bche | 321.9317 | 0.067926 | 0.71212 | 0.964363 |
| ENSMUSG( | Bckdha | 2450.17 | 0.085534 | 0.355929 | 0.788009 | ENSMUSG( | Bckdha | 2035.031 | -0.18971 | 0.125134 | 0.81516 |
| ENSMUSG( | Bckdhb | 612.4107 | 0.187841 | 0.120732 | 0.524807 | ENSMUSG( | Bckdhb | 506.9615 | -0.08722 | 0.559331 | 0.939748 |
| ENSMUSG( | Bckdk | 1182.619 | 0.165589 | 0.106589 | 0.494174 | ENSMUSG( | Bckdk | 980.1228 | -0.1065 | 0.366415 | 0.914703 |
| ENSMUSG( | Bcl10 | 491.4792 | -0.01192 | 0.915919 | 0.988634 | ENSMUSG( | Bcl10 | 446.4948 | -0.0214 | 0.873803 | 0.989644 |
| ENSMUSG( | Bcl11a | 1714.595 | 0.013791 | 0.919577 | 0.990247 | ENSMUSG( | Bcl11a | 1569.056 | 0.022607 | 0.881836 | 0.989644 |
| ENSMUSG( | Bcl11b | 1670.948 | -0.1527 | 0.180598 | 0.617841 | ENSMUSG( | Bcl11b | 1637.311 | 0.057469 | 0.696064 | 0.960867 |
| ENSMUSG( | Bcl2 | 1271.661 | 0.141207 | 0.282729 | 0.73186 | ENSMUSG( | Bcl2 | 1097.371 | -0.01769 | 0.909409 | 0.991958 |
| ENSMUSG( | Bcl2a1b | 15.37751 | 0.077556 | 0.867358 | NA | ENSMUSG( | Bcl2a1b | 15.05115 | 0.262723 | 0.598914 | 0.947045 |
| ENSMUSG( | Bcl2l1 | 3147.91 | 0.151719 | 0.13219 | 0.544244 | ENSMUSG( | Bcl2l1 | 2710.98 | -0.00594 | 0.956895 | 0.994798 |
| ENSMUSG( | Bcl2l11 | 184.6315 | 0.170139 | 0.37364 | 0.800375 | ENSMUSG( | Bcl2l11 | 177.5228 | 0.312861 | 0.110343 | 0.807051 |
| ENSMUSG( | Bcl2l12 | 86.45892 | -0.13073 | 0.544913 | 0.884712 | ENSMUSG( | Bcl2l12 | 90.38825 | 0.260036 | 0.258984 | 0.883971 |
| ENSMUSG( | Bcl2l13 | 1895.194 | 0.064878 | 0.49032 | 0.859319 | ENSMUSG( | Bcl2l13 | 1648.183 | -0.06881 | 0.543722 | 0.938063 |
| ENSMUSG( | Bcl2l15 | 12.34819 | 0.292211 | 0.646173 | NA | ENSMUSG( | Bcl2l15 | 9.229688 | -0.25929 | 0.737274 | NA |
| ENSMUSG( | Bcl2l2 | 8378.924 | 0.098421 | 0.2108 | 0.653564 | ENSMUSG( | Bcl2l2 | 7178.261 | -0.08003 | 0.44692 | 0.921891 |
| ENSMUSG( | Bcl3 | 16.27096 | 0.50593 | 0.2941 | NA | ENSMUSG( | Bcl3 | 15.3884 | 0.580627 | 0.236987 | 0.872173 |
| ENSMUSG( | Bcl6 | 951.6595 | 0.163367 | 0.190054 | 0.629259 | ENSMUSG( | Bcl6 | 820.5567 | 0.01238 | 0.925904 | 0.99306 |
| ENSMUSG( | Bcl6b | 103.8737 | -0.18504 | 0.435209 | 0.829093 | ENSMUSG( | Bcl6b | 97.98464 | -0.07733 | 0.757525 | 0.971858 |
| ENSMUSG( | Bcl7a | 2386.307 | 0.089175 | 0.406964 | 0.817163 | ENSMUSG( | Bcl7a | 2105.483 | -0.00384 | 0.975286 | 0.996416 |
| ENSMUSG( | Bcl7b | 1289.233 | 0.01373 | 0.909732 | 0.988084 | ENSMUSG( | Bcl7b | 1128.259 | -0.10776 | 0.464026 | 0.923329 |
| ENSMUSG( | Bcl7c | 1164.228 | 0.069 | 0.464379 | 0.843364 | ENSMUSG( | Bcl7c | 991.7131 | -0.1317 | 0.302672 | 0.898538 |
| ENSMUSG( | Bcl9 | 1455.137 | 0.045324 | 0.65347 | 0.922819 | ENSMUSG( | Bcl9 | 1328.655 | 0.052508 | 0.626334 | 0.953234 |
| ENSMUSG( | Bcl9l | 2073.034 | 0.138305 | 0.157763 | 0.585927 | ENSMUSG( | Bcl9l | 1976.601 | 0.263498 | 0.022598 | 0.581156 |
| ENSMUSG( | Bclaf1 | 2367.014 | 0.07645 | 0.396827 | 0.811742 | ENSMUSG( | Bclaf1 | 2189.69 | 0.119821 | 0.26152 | 0.884926 |
| ENSMUSG( | Bclaf3 | 171.097 | -0.13739 | 0.414272 | 0.819998 | ENSMUSG( | Bclaf3 | 174.8486 | 0.186443 | 0.307459 | 0.898538 |
| ENSMUSG( | Bco2 | 378.7376 | 0.05594 | 0.674666 | 0.929638 | ENSMUSG( | Bco2 | 338.0306 | -0.00567 | 0.969239 | 0.99594 |

| | | | | | |
|---|---|---|---|---|---|
| ENSMUSG( | 300.2851 | -0.44599 | 0.006556 | 0.111119 | Bche |
| ENSMUSG( | 2450.17 | 0.085534 | 0.355929 | 0.788009 | Bckdha |
| ENSMUSG( | 612.4107 | 0.187841 | 0.120732 | 0.524807 | Bckdhb |
| ENSMUSG( | 1182.619 | 0.165589 | 0.106589 | 0.494174 | Bckdk |
| ENSMUSG( | 491.4792 | -0.01192 | 0.915919 | 0.988634 | Bcl10 |
| ENSMUSG( | 1714.595 | 0.013791 | 0.919577 | 0.990247 | Bcl11a |
| ENSMUSG( | 1670.948 | -0.1527 | 0.180598 | 0.617841 | Bcl11b |
| ENSMUSG( | 1271.661 | 0.141207 | 0.282729 | 0.73186 | Bcl2 |
| ENSMUSG( | 15.37751 | 0.077556 | 0.867358 | NA | Bcl2a1b |
| ENSMUSG( | 3147.91 | 0.151719 | 0.13219 | 0.544244 | Bcl2l1 |
| ENSMUSG( | 184.6315 | 0.170139 | 0.37364 | 0.800375 | Bcl2l11 |
| ENSMUSG( | 86.45892 | -0.13073 | 0.544913 | 0.884712 | Bcl2l12 |
| ENSMUSG( | 1895.194 | 0.064878 | 0.49032 | 0.859319 | Bcl2l13 |
| ENSMUSG( | 12.34819 | 0.292211 | 0.646173 | NA | Bcl2l15 |
| ENSMUSG( | 8378.924 | 0.098421 | 0.2108 | 0.653564 | Bcl2l2 |
| ENSMUSG( | 16.27096 | 0.50593 | 0.2941 | NA | Bcl3 |
| ENSMUSG( | 951.6595 | 0.163367 | 0.190054 | 0.629259 | Bcl6 |
| ENSMUSG( | 103.8737 | -0.18504 | 0.435209 | 0.829093 | Bcl6b |
| ENSMUSG( | 2386.307 | 0.089175 | 0.406964 | 0.817163 | Bcl7a |
| ENSMUSG( | 1289.233 | 0.01373 | 0.909732 | 0.988084 | Bcl7b |
| ENSMUSG( | 1164.228 | 0.069 | 0.464379 | 0.843364 | Bcl7c |
| ENSMUSG( | 1455.137 | 0.045324 | 0.65347 | 0.922819 | Bcl9 |
| ENSMUSG( | 2073.034 | 0.138305 | 0.157763 | 0.585927 | Bcl9l |
| ENSMUSG( | 2367.014 | 0.07645 | 0.396827 | 0.811742 | Bclaf1 |
| ENSMUSG( | 171.097 | -0.13739 | 0.414272 | 0.819998 | Bclaf3 |
| ENSMUSG( | 378.7376 | 0.05594 | 0.674666 | 0.929638 | Bco2 |
| ENSMUSG( | 962.7409 | 0.172055 | 0.085252 | 0.450128 | Bcor |
| ENSMUSG( | 596.9967 | 0.00502 | 0.963357 | 0.994911 | Bcorl1 |
| ENSMUSG( | 6483.715 | 0.033301 | 0.716837 | 0.941697 | Bcr |
| ENSMUSG( | 596.4804 | 0.058834 | 0.585612 | 0.899171 | Bcs1l |
| ENSMUSG( | 2243.334 | 0.071847 | 0.463661 | 0.842812 | Bdh1 |
| ENSMUSG( | 83.66288 | -0.02524 | 0.927403 | 0.991229 | Bdh2 |
| ENSMUSG( | 28.59354 | -0.31581 | 0.378574 | 0.801688 | Bdkrb2 |
| ENSMUSG( | 607.2912 | -0.5613 | 0.000762 | 0.027082 | Bdnf |
| ENSMUSG( | 845.8237 | 0.194794 | 0.076152 | 0.42702 | Bdp1 |
| ENSMUSG( | 647.9405 | 0.281045 | 0.027822 | 0.255388 | Bean1 |
| ENSMUSG( | 3803.14 | 0.06894 | 0.396647 | 0.811742 | Becn1 |
| ENSMUSG( | 10068.13 | 0.155396 | 0.069079 | 0.407145 | Begain |
| ENSMUSG( | 294.0992 | -0.03803 | 0.766177 | 0.954173 | Bend3 |
| ENSMUSG( | 199.0277 | -0.0552 | 0.732314 | 0.944956 | Bend4 |
| ENSMUSG( | 233.3879 | -0.17137 | 0.306436 | 0.754119 | Bend5 |
| ENSMUSG( | 5523.502 | 0.091146 | 0.268369 | 0.718663 | Bend6 |
| ENSMUSG( | 95.18193 | 0.10286 | 0.676396 | 0.930542 | Bend7 |
| ENSMUSG( | 31.52565 | -0.07399 | 0.828046 | 0.968966 | Best1 |
| ENSMUSG( | 629.8123 | 0.069668 | 0.627614 | 0.912961 | Bet1 |
| ENSMUSG( | 511.6652 | 0.081446 | 0.506551 | 0.867425 | Bet1l |
| ENSMUSG( | 662.1128 | -0.10851 | 0.505837 | 0.866936 | Bex1 |
| ENSMUSG( | 3498.304 | -0.13737 | 0.400388 | 0.813717 | Bex2 |
| ENSMUSG( | 2677.803 | -0.03512 | 0.80839 | 0.963455 | Bex3 |
| ENSMUSG( | 225.8902 | -0.01661 | 0.924431 | 0.99114 | Bex4 |
| ENSMUSG( | 1478.846 | -0.18222 | 0.155661 | 0.583641 | Bfar |
| ENSMUSG( | 51.21766 | 0.092002 | 0.735477 | 0.946109 | Bfsp1 |
| ENSMUSG( | 250.6486 | -0.01911 | 0.891182 | 0.983819 | Bfsp2 |
| ENSMUSG( | 26.62442 | -0.20637 | 0.577801 | 0.895986 | Bglap3 |
| ENSMUSG( | 924.0989 | -0.23147 | 0.092086 | 0.466032 | Bgn |
| ENSMUSG( | 43.0568 | 0.17073 | 0.554069 | 0.886397 | Bhlha9 |
| ENSMUSG( | 1320.164 | 0.184406 | 0.069268 | 0.407813 | Bhlhb9 |
| ENSMUSG( | 696.4398 | 0.206637 | 0.049238 | 0.33882 | Bhlhe22 |
| ENSMUSG( | 1905.101 | -0.03728 | 0.790847 | 0.959461 | Bhlhe40 |
| ENSMUSG( | 1318.929 | -0.0104 | 0.92084 | 0.990409 | Bhlhe41 |
| ENSMUSG( | 13.04313 | 0.998772 | 0.105696 | NA | Bhmt2 |
| ENSMUSG( | 301.9257 | -0.0063 | 0.96421 | 0.995155 | Bicc1 |
| ENSMUSG( | 545.6614 | -0.05522 | 0.612837 | 0.908902 | Bicd1 |
| ENSMUSG( | 2212.063 | 0.01579 | 0.864637 | 0.975475 | Bicd2 |
| ENSMUSG( | 2568.829 | 0.300621 | 0.001692 | 0.046443 | Bicdl1 |
| ENSMUSG( | 791.7156 | 0.154789 | 0.127251 | 0.534725 | Bicra |
| ENSMUSG( | 1944.173 | 0.094278 | 0.373798 | 0.80044 | Bicral |
| ENSMUSG( | 168.962 | 0.047674 | 0.794006 | 0.959799 | Bid |
| ENSMUSG( | 6150.622 | 0.037203 | 0.723097 | 0.943761 | Bin1 |
| ENSMUSG( | 425.921 | -0.1242 | 0.36741 | 0.795221 | Bin2 |
| ENSMUSG( | 274.784 | -0.1332 | 0.363824 | 0.793343 | Bin3 |
| ENSMUSG( | 1042.493 | -0.03257 | 0.722758 | 0.943761 | Birc2 |
| ENSMUSG( | 98.16606 | -0.04378 | 0.836528 | 0.971111 | Birc3 |
| ENSMUSG( | 37.92165 | -0.135 | 0.665113 | 0.926824 | Birc5 |
| ENSMUSG( | 2655.17 | -0.24133 | 0.031052 | 0.268255 | Birc6 |
| ENSMUSG( | 470.2538 | -0.14059 | 0.35715 | 0.788702 | Bivm |
| ENSMUSG( | 2197.511 | -0.26246 | 0.016257 | 0.189161 | Blcap |
| ENSMUSG( | 84.05588 | -0.28306 | 0.197481 | 0.637186 | Blm |
| ENSMUSG( | 1495.028 | 0.028735 | 0.776477 | 0.955658 | Blmh |
| ENSMUSG( | 210.3362 | 0.128266 | 0.433208 | 0.828207 | Blnk |
| ENSMUSG( | 260.4856 | 0.110694 | 0.42227 | 0.823508 | Bloc1s1 |
| ENSMUSG( | 626.9454 | -0.12478 | 0.224715 | 0.671058 | Bloc1s2 |
| ENSMUSG( | 256.1736 | 0.046006 | 0.76564 | 0.954173 | Bloc1s3 |
| ENSMUSG( | 259.625 | -0.14964 | 0.343694 | 0.779153 | Bloc1s4 |
| ENSMUSG( | 677.9456 | 0.071601 | 0.561753 | 0.889052 | Bloc1s5 |
| ENSMUSG( | 723.917 | 0.055804 | 0.605827 | 0.905919 | Bloc1s6 |
| ENSMUSG( | 2979.676 | -0.28848 | 0.024802 | 0.242192 | Bltp1 |
| ENSMUSG( | 16570.06 | 0.10551 | 0.283124 | 0.73186 | Bltp2 |
| ENSMUSG( | 964.6642 | 0.043795 | 0.678665 | 0.930611 | Bltp3a |
| ENSMUSG( | 2405.965 | 0.009772 | 0.91292 | 0.988483 | Bltp3b |
| ENSMUSG( | 334.4423 | -0.08349 | 0.580286 | 0.896952 | Blvra |
| ENSMUSG( | 373.601 | -0.06528 | 0.597925 | 0.90372 | Blvrb |
| ENSMUSG( | 645.2024 | 0.031851 | 0.790799 | 0.959461 | Blzf1 |
| ENSMUSG( | 872.133 | -0.01712 | 0.85972 | 0.97481 | Bmal1 |
| ENSMUSG( | 209.5844 | 0.136814 | 0.4044 | 0.816132 | Bmal2 |
| ENSMUSG( | 7099.364 | 0.10324 | 0.275003 | 0.72509 | Bmerb1 |
| ENSMUSG( | 206.776 | -0.34154 | 0.022365 | 0.228729 | Bmf |
| ENSMUSG( | 1055.634 | -0.05844 | 0.667081 | 0.92729 | Bmi1 |
| ENSMUSG( | 2858.696 | 0.197627 | 0.185918 | 0.623352 | Bmp1 |
| ENSMUSG( | 9.20583 | -0.67018 | 0.285982 | NA | Bmp15 |
| ENSMUSG( | 132.2666 | -0.11755 | 0.556295 | 0.887663 | Bmp2 |
| ENSMUSG( | 786.0931 | -0.1901 | 0.106103 | 0.493403 | Bmp2k |
| ENSMUSG( | 511.0903 | -0.66251 | 0.000337 | 0.015675 | Bmp3 |
| ENSMUSG( | 271.5096 | -0.31279 | 0.056169 | 0.36528 | Bmp4 |
| ENSMUSG( | 110.1544 | -0.56518 | 0.010217 | 0.144102 | Bmp5 |
| ENSMUSG( | 503.9148 | -0.1846 | 0.159487 | 0.588863 | Bmp6 |
| ENSMUSG( | 724.5855 | -0.04505 | 0.711639 | 0.940028 | Bmp7 |
| ENSMUSG( | 294.8971 | -0.33352 | 0.040925 | 0.312769 | Bmper |
| ENSMUSG( | 4431.556 | -0.02947 | 0.819275 | 0.965888 | Bmpr1a |
| ENSMUSG( | 2464.838 | -0.08363 | 0.539337 | 0.883094 | Bmpr1b |
| ENSMUSG( | 5193.963 | -0.15741 | 0.266753 | 0.716789 | Bmpr2 |
| ENSMUSG( | 1266.686 | 0.228463 | 0.041631 | 0.315097 | Bms1 |
| ENSMUSG( | 387.7062 | 0.023105 | 0.858886 | 0.974707 | Bmt2 |
| ENSMUSG( | 1610.489 | -0.00119 | 0.991393 | 0.999232 | Bmyc |
| ENSMUSG( | 28.31439 | -0.60697 | 0.10671 | 0.49424 | Bnc2 |
| ENSMUSG( | 436.9298 | 0.039592 | 0.735199 | 0.945972 | Bnip1 |
| ENSMUSG( | 1323.775 | 0.001551 | 0.988475 | 0.999109 | Bnip2 |
| ENSMUSG( | 1713.701 | 0.030823 | 0.736245 | 0.946267 | Bnip3 |
| ENSMUSG( | 3576.927 | 0.040916 | 0.678139 | 0.930542 | Bnip3l |
| ENSMUSG( | 282.3765 | 7.93E-05 | 0.999593 | 0.99989 | Bnip5 |
| ENSMUSG( | 794.8225 | -0.30352 | 0.097011 | 0.477375 | Boc |
| ENSMUSG( | 2656.831 | 0.013628 | 0.878293 | 0.981431 | Bod1 |
| ENSMUSG( | 1778.749 | 0.064701 | 0.439863 | 0.831633 | Bod1l |
| ENSMUSG( | 2035.444 | -0.00359 | 0.970046 | 0.996369 | Bok |
| ENSMUSG( | 230.8782 | -0.06129 | 0.682983 | 0.93205 | Bola1 |
| ENSMUSG( | 1249.904 | 0.09787 | 0.290069 | 0.739883 | Bola2 |
| ENSMUSG( | 1067.274 | 0.073086 | 0.451268 | 0.835567 | Bola3 |
| ENSMUSG( | 1191.709 | 0.105029 | 0.326155 | 0.771128 | Bop1 |
| ENSMUSG( | 113.9314 | -0.05659 | 0.782048 | 0.957558 | Bora |
| ENSMUSG( | 610.696 | 0.053309 | 0.661518 | 0.925271 | Borcs5 |
| ENSMUSG( | 1110.322 | 0.049891 | 0.644942 | 0.91787 | Borcs6 |
| ENSMUSG( | 721.587 | 0.095236 | 0.357201 | 0.788702 | Borcs7 |
| ENSMUSG( | 904.0409 | -0.00832 | 0.938825 | 0.993415 | Borcs8 |
| ENSMUSG( | 4149.796 | -0.05829 | 0.589099 | 0.90113 | Bpgm |
| ENSMUSG( | 710.9752 | 0.026658 | 0.830483 | 0.96974 | Bphl |
| ENSMUSG( | 1576.978 | 0.089287 | 0.362396 | 0.792943 | Bpnt1 |
| ENSMUSG( | 3505.112 | -0.38322 | 0.0078 | 0.122469 | Bpnt2 |
| ENSMUSG( | 2874.657 | 0.04375 | 0.657918 | 0.92425 | Bptf |
| ENSMUSG( | 2456.407 | -0.00738 | 0.927482 | 0.991229 | Braf |
| ENSMUSG( | 1989.117 | 0.043513 | 0.625338 | 0.912249 | Brap |
| ENSMUSG( | 686.0425 | -0.03654 | 0.76536 | 0.954127 | Brat1 |
| ENSMUSG( | 180.3369 | -0.01956 | 0.919722 | 0.990247 | Brca1 |
| ENSMUSG( | 139.2145 | -0.49693 | 0.008807 | 0.132321 | Brca2 |
| ENSMUSG( | 675.4068 | 0.066359 | 0.518222 | 0.873754 | Brcc3 |
| ENSMUSG( | 1533.45 | -0.01444 | 0.870826 | 0.977951 | Brd1 |
| ENSMUSG( | 1861.437 | 0.053072 | 0.566515 | 0.891644 | Brd10 |
| ENSMUSG( | 10493.99 | 0.139106 | 0.062263 | 0.386085 | Brd2 |
| ENSMUSG( | 1800.564 | 0.055707 | 0.52904 | 0.878551 | Brd3 |
| ENSMUSG( | 513.3902 | -0.08117 | 0.618107 | 0.910929 | Brd3os |
| ENSMUSG( | 3760.635 | -0.0951 | 0.554135 | 0.886397 | Brd4 |
| ENSMUSG( | 1841.311 | 0.127619 | 0.14343 | 0.562473 | Brd7 |
| ENSMUSG( | 1320.84 | -0.04423 | 0.672761 | 0.929007 | Brd8 |
| ENSMUSG( | 29.3119 | 0.011512 | 0.973579 | 0.99675 | Brd8dc |
| ENSMUSG( | 2717.879 | 0.310536 | 0.003705 | 0.07745 | Brd9 |
| ENSMUSG( | 436.9257 | 0.078881 | 0.545249 | 0.884748 | Brdt |
| ENSMUSG( | 1860.057 | 0.07904 | 0.384501 | 0.804007 | Brf1 |
| ENSMUSG( | 339.3397 | 0.037915 | 0.790432 | 0.959461 | Brf2 |
| ENSMUSG( | 1627.424 | 0.021183 | 0.84995 | 0.972638 | Bri3 |

| | | | | | |
|---|---|---|---|---|---|
| ENSMUSG( | 15.3884 | 0.580627 | 0.236987 | 0.872173 | Bcl3 |
| ENSMUSG( | 820.5567 | 0.01238 | 0.925904 | 0.99306 | Bcl6 |
| ENSMUSG( | 97.98464 | -0.07733 | 0.757525 | 0.971858 | Bcl6b |
| ENSMUSG( | 2105.483 | -0.00384 | 0.975286 | 0.996416 | Bcl7a |
| ENSMUSG( | 1128.259 | -0.10776 | 0.464026 | 0.923329 | Bcl7b |
| ENSMUSG( | 991.7131 | -0.1317 | 0.302672 | 0.898538 | Bcl7c |
| ENSMUSG( | 1328.655 | 0.052508 | 0.626334 | 0.953234 | Bcl9 |
| ENSMUSG( | 1976.601 | 0.263498 | 0.022598 | 0.581156 | Bcl9l |
| ENSMUSG( | 2189.69 | 0.119821 | 0.26152 | 0.884926 | Bclaf1 |
| ENSMUSG( | 174.8486 | 0.186443 | 0.307459 | 0.898538 | Bclaf3 |
| ENSMUSG( | 338.0306 | -0.00567 | 0.969239 | 0.99594 | Bco2 |
| ENSMUSG( | 898.2424 | 0.238475 | 0.04292 | 0.690304 | Bcor |
| ENSMUSG( | 553.0341 | 0.051066 | 0.69223 | 0.960404 | Bcorl1 |
| ENSMUSG( | 5715.583 | -0.0623 | 0.531936 | 0.934795 | Bcr |
| ENSMUSG( | 507.6107 | -0.14611 | 0.280271 | 0.892356 | Bcs1l |
| ENSMUSG( | 1985.353 | -0.01443 | 0.913356 | 0.992298 | Bdh1 |
| ENSMUSG( | 65.9111 | -0.49289 | 0.060664 | 0.734317 | Bdh2 |
| ENSMUSG( | 25.80624 | -0.34543 | 0.383928 | 0.917257 | Bdkrb2 |
| ENSMUSG( | 680.3447 | 0.093249 | 0.596821 | 0.946541 | Bdnf |
| ENSMUSG( | 765.1284 | 0.176995 | 0.155658 | 0.837724 | Bdp1 |
| ENSMUSG( | 581.8964 | 0.243098 | 0.072395 | 0.763629 | Bean1 |
| ENSMUSG( | 3308.695 | -0.06599 | 0.557735 | 0.939565 | Becn1 |
| ENSMUSG( | 8819.221 | 0.044829 | 0.646296 | 0.953988 | Begain |
| ENSMUSG( | 261.186 | -0.11602 | 0.451647 | 0.922202 | Bend3 |
| ENSMUSG( | 180.5308 | -0.06135 | 0.730974 | 0.967228 | Bend4 |
| ENSMUSG( | 220.0617 | -0.07504 | 0.694456 | 0.960867 | Bend5 |
| ENSMUSG( | 4760.255 | -0.07116 | 0.575578 | 0.942912 | Bend6 |
| ENSMUSG( | 89.4574 | 0.17576 | 0.47365 | 0.923329 | Bend7 |
| ENSMUSG( | 25.53465 | -0.43751 | 0.275769 | 0.890277 | Best1 |
| ENSMUSG( | 10.05298 | -0.22262 | 0.744328 | NA | Best3 |
| ENSMUSG( | 541.6324 | -0.10773 | 0.558543 | 0.93957 | Bet1 |
| ENSMUSG( | 463.904 | 0.063064 | 0.644832 | 0.953903 | Bet1l |
| ENSMUSG( | 584.544 | -0.21331 | 0.325922 | 0.905339 | Bex1 |
| ENSMUSG( | 3065.772 | -0.26297 | 0.211292 | 0.864412 | Bex2 |
| ENSMUSG( | 2285.103 | -0.23827 | 0.20249 | 0.864412 | Bex3 |
| ENSMUSG( | 188.1586 | -0.31104 | 0.174376 | 0.844317 | Bex4 |
| ENSMUSG( | 1526.847 | 0.179719 | 0.204526 | 0.864412 | Bfar |
| ENSMUSG( | 41.72794 | -0.25076 | 0.434992 | 0.918876 | Bfsp1 |
| ENSMUSG( | 224.471 | -0.07879 | 0.659202 | 0.956205 | Bfsp2 |
| ENSMUSG( | 11.13234 | 0.742072 | 0.313848 | NA | Bglap |
| ENSMUSG( | 10.19607 | 0.748445 | 0.319423 | NA | Bglap2 |
| ENSMUSG( | 24.37072 | -0.15088 | 0.742429 | 0.969415 | Bglap3 |
| ENSMUSG( | 1026.027 | 0.333744 | 0.019092 | 0.549166 | Bgn |
| ENSMUSG( | 34.38345 | -0.21834 | 0.523395 | 0.932848 | Bhlha9 |
| ENSMUSG( | 1154.025 | 0.067533 | 0.577753 | 0.943131 | Bhlhb9 |
| ENSMUSG( | 615.4284 | 0.119831 | 0.376902 | 0.915593 | Bhlhe22 |
| ENSMUSG( | 1835.809 | 0.119553 | 0.448732 | 0.922202 | Bhlhe40 |
| ENSMUSG( | 1410.259 | 0.419401 | 0.00057 | 0.072448 | Bhlhe41 |
| ENSMUSG( | 291.0231 | 0.147765 | 0.34508 | 0.91045 | Bicc1 |
| ENSMUSG( | 568.5338 | 0.311389 | 0.024775 | 0.592568 | Bicd1 |
| ENSMUSG( | 2104.371 | 0.139251 | 0.257908 | 0.883466 | Bicd2 |
| ENSMUSG( | 2273.365 | 0.225841 | 0.076453 | 0.770555 | Bicdl1 |
| ENSMUSG( | 743.0734 | 0.237412 | 0.059745 | 0.729747 | Bicra |
| ENSMUSG( | 1725.163 | 0.016379 | 0.90079 | 0.990558 | Bicral |
| ENSMUSG( | 154.3676 | 0.060025 | 0.769399 | 0.974236 | Bid |
| ENSMUSG( | 5505.478 | -0.013 | 0.904414 | 0.991089 | Bin1 |
| ENSMUSG( | 389.557 | -0.10781 | 0.465653 | 0.923329 | Bin2 |
| ENSMUSG( | 258.9628 | -0.0246 | 0.879329 | 0.989644 | Bin3 |
| ENSMUSG( | 959.6806 | -0.00231 | 0.983397 | 0.996963 | Birc2 |
| ENSMUSG( | 85.97012 | -0.14353 | 0.58835 | 0.944975 | Birc3 |
| ENSMUSG( | 37.17232 | 0.086001 | 0.795043 | 0.97681 | Birc5 |
| ENSMUSG( | 2728.988 | 0.118571 | 0.409657 | 0.917257 | Birc6 |
| ENSMUSG( | 483.2366 | 0.209493 | 0.186818 | 0.856099 | Bivm |
| ENSMUSG( | 2159.867 | -0.02922 | 0.814306 | 0.979198 | Blcap |
| ENSMUSG( | 78.63286 | -0.18122 | 0.458967 | 0.923329 | Blm |
| ENSMUSG( | 1282.192 | -0.15027 | 0.220223 | 0.866104 | Blmh |
| ENSMUSG( | 185.6918 | 0.0397 | 0.824537 | 0.979198 | Blnk |
| ENSMUSG( | 211.0536 | -0.24496 | 0.158131 | 0.837724 | Bloc1s1 |
| ENSMUSG( | 555.7495 | -0.21098 | 0.10188 | 0.801719 | Bloc1s2 |
| ENSMUSG( | 223.8508 | -0.07056 | 0.698708 | 0.960908 | Bloc1s3 |
| ENSMUSG( | 250.1935 | 0.006057 | 0.973561 | 0.996416 | Bloc1s4 |
| ENSMUSG( | 551.4805 | -0.27117 | 0.04466 | 0.693697 | Bloc1s5 |
| ENSMUSG( | 631.4965 | -0.07716 | 0.589082 | 0.944975 | Bloc1s6 |
| ENSMUSG( | 3251.548 | 0.241579 | 0.165819 | 0.839724 | Bltp1 |
| ENSMUSG( | 14648.86 | 0.022161 | 0.846239 | 0.984578 | Bltp2 |
| ENSMUSG( | 881.1193 | 0.051721 | 0.666663 | 0.956766 | Bltp3a |
| ENSMUSG( | 2267.953 | 0.10801 | 0.339308 | 0.907972 | Bltp3b |
| ENSMUSG( | 281.0541 | -0.34444 | 0.05126 | 0.711023 | Blvra |
| ENSMUSG( | 331.3094 | -0.1551 | 0.309628 | 0.898538 | Blvrb |
| ENSMUSG( | 560.2575 | -0.10993 | 0.403425 | 0.917257 | Blzf1 |
| ENSMUSG( | 620.9489 | -0.85308 | 1.72E-11 | 2.01E-08 | Bmal1 |
| ENSMUSG( | 199.2951 | 0.264778 | 0.160442 | 0.837724 | Bmal2 |
| ENSMUSG( | 5867.679 | -0.18297 | 0.131608 | 0.822912 | Bmerb1 |
| ENSMUSG( | 200.9872 | -0.12395 | 0.527487 | 0.934466 | Bmf |
| ENSMUSG( | 1019.06 | 0.104555 | 0.510065 | 0.929089 | Bmi1 |
| ENSMUSG( | 2624.363 | 0.218442 | 0.170587 | 0.841739 | Bmp1 |
| ENSMUSG( | 10.56377 | 0.107694 | 0.854711 | NA | Bmp15 |
| ENSMUSG( | 152.4564 | 0.530017 | 0.023775 | 0.588226 | Bmp2 |
| ENSMUSG( | 756.5152 | -0.02411 | 0.853039 | 0.985598 | Bmp2k |
| ENSMUSG( | 666.2651 | 0.422525 | 0.031294 | 0.631113 | Bmp3 |
| ENSMUSG( | 326.5928 | 0.477586 | 0.008771 | 0.393969 | Bmp4 |
| ENSMUSG( | 130.0557 | 0.239693 | 0.259366 | 0.883971 | Bmp5 |
| ENSMUSG( | 459.0648 | -0.17693 | 0.237352 | 0.872173 | Bmp6 |
| ENSMUSG( | 703.1051 | 0.135553 | 0.29484 | 0.898538 | Bmp7 |
| ENSMUSG( | 332.8819 | 0.298134 | 0.112704 | 0.809623 | Bmper |
| ENSMUSG( | 4234.362 | 0.105392 | 0.474593 | 0.923329 | Bmpr1a |
| ENSMUSG( | 2154.85 | -0.2091 | 0.159746 | 0.837724 | Bmpr1b |
| ENSMUSG( | 5374.915 | 0.208866 | 0.181932 | 0.850579 | Bmpr2 |
| ENSMUSG( | 1121.637 | 0.148573 | 0.261385 | 0.884926 | Bms1 |
| ENSMUSG( | 381.4906 | 0.228792 | 0.147515 | 0.836241 | Bmt2 |
| ENSMUSG( | 17.31576 | 1.010633 | 0.047355 | 0.700936 | Bmx |
| ENSMUSG( | 1394.153 | -0.158 | 0.2754 | 0.889984 | Bmyc |
| ENSMUSG( | 27.93344 | -0.38219 | 0.360251 | 0.914223 | Bnc2 |
| ENSMUSG( | 379.1056 | -0.10818 | 0.45029 | 0.922202 | Bnip1 |
| ENSMUSG( | 1243.142 | 0.088778 | 0.451205 | 0.922202 | Bnip2 |
| ENSMUSG( | 1501.629 | -0.08733 | 0.556866 | 0.939565 | Bnip3 |
| ENSMUSG( | 3133.094 | -0.07657 | 0.575798 | 0.942912 | Bnip3l |
| ENSMUSG( | 236.7794 | -0.23846 | 0.172269 | 0.843466 | Bnip5 |
| ENSMUSG( | 845.6296 | 0.163184 | 0.403074 | 0.917257 | Boc |
| ENSMUSG( | 2301.384 | -0.13814 | 0.265399 | 0.886134 | Bod1 |
| ENSMUSG( | 1613.538 | 0.051984 | 0.619745 | 0.952896 | Bod1l |
| ENSMUSG( | 1785.506 | -0.1169 | 0.275614 | 0.890209 | Bok |
| ENSMUSG( | 204.7216 | -0.15593 | 0.402839 | 0.917257 | Bola1 |
| ENSMUSG( | 1062.154 | -0.1093 | 0.446546 | 0.921734 | Bola2 |
| ENSMUSG( | 869.607 | -0.26326 | 0.028926 | 0.614171 | Bola3 |
| ENSMUSG( | 1062.283 | 0.045413 | 0.715912 | 0.965424 | Bop1 |
| ENSMUSG( | 104.2704 | -0.05149 | 0.816314 | 0.979198 | Bora |
| ENSMUSG( | 540.3588 | -0.03262 | 0.81581 | 0.979198 | Borcs5 |
| ENSMUSG( | 971.1365 | -0.07263 | 0.616878 | 0.951666 | Borcs6 |
| ENSMUSG( | 613.6407 | -0.11109 | 0.46551 | 0.923329 | Borcs7 |
| ENSMUSG( | 784.607 | -0.15573 | 0.22326 | 0.867404 | Borcs8 |
| ENSMUSG( | 3472.733 | -0.32427 | 0.020299 | 0.563507 | Bpgm |
| ENSMUSG( | 578.6247 | -0.32283 | 0.037356 | 0.673514 | Bphl |
| ENSMUSG( | 1352.305 | -0.08957 | 0.466816 | 0.923329 | Bpnt1 |
| ENSMUSG( | 3791.955 | 0.136939 | 0.406177 | 0.917257 | Bpnt2 |
| ENSMUSG( | 2727.86 | 0.159234 | 0.145128 | 0.83339 | Bptf |
| ENSMUSG( | 2298.008 | 0.068017 | 0.499604 | 0.927825 | Braf |
| ENSMUSG( | 1758.946 | -0.0428 | 0.686162 | 0.958956 | Brap |
| ENSMUSG( | 627.3684 | -0.02038 | 0.883748 | 0.989644 | Brat1 |
| ENSMUSG( | 151.7123 | -0.27723 | 0.155769 | 0.837724 | Brca1 |
| ENSMUSG( | 145.6906 | -0.04903 | 0.819843 | 0.979198 | Brca2 |
| ENSMUSG( | 583.5978 | -0.09173 | 0.512503 | 0.930164 | Brcc3 |
| ENSMUSG( | 1443.711 | 0.076722 | 0.503894 | 0.927825 | Brd1 |
| ENSMUSG( | 1706.864 | 0.068885 | 0.563366 | 0.941206 | Brd10 |
| ENSMUSG( | 9515.738 | 0.126213 | 0.204957 | 0.864412 | Brd2 |
| ENSMUSG( | 1619.884 | 0.016995 | 0.887419 | 0.990266 | Brd3 |
| ENSMUSG( | 456.2554 | -0.15528 | 0.370543 | 0.915491 | Brd3os |
| ENSMUSG( | 3630.234 | 0.074607 | 0.627927 | 0.953234 | Brd4 |
| ENSMUSG( | 1601.806 | -0.00487 | 0.965607 | 0.995433 | Brd7 |
| ENSMUSG( | 1204.59 | -0.0366 | 0.778752 | 0.976106 | Brd8 |
| ENSMUSG( | 29.24447 | 0.259123 | 0.481325 | 0.924564 | Brd8dc |
| ENSMUSG( | 2298.615 | 0.110001 | 0.433718 | 0.918564 | Brd9 |
| ENSMUSG( | 372.1521 | -0.12597 | 0.42569 | 0.917257 | Brdt |
| ENSMUSG( | 1606.914 | -0.0749 | 0.503408 | 0.927825 | Brf1 |
| ENSMUSG( | 303.955 | -0.0067 | 0.967882 | 0.995852 | Brf2 |
| ENSMUSG( | 1464.707 | -0.01956 | 0.894747 | 0.990558 | Bri3 |
| ENSMUSG( | 4142.145 | 0.180072 | 0.271065 | 0.888062 | Bri3bp |
| ENSMUSG( | 25.81943 | -0.01539 | 0.968156 | 0.995887 | Bricd5 |
| ENSMUSG( | 6528.292 | 0.217874 | 0.226614 | 0.868507 | Brinp1 |
| ENSMUSG( | 3643.037 | 0.389954 | 0.014766 | 0.487333 | Brinp2 |
| ENSMUSG( | 830.462 | 0.188815 | 0.387404 | 0.917257 | Brinp3 |
| ENSMUSG( | 11.83776 | 0.846507 | 0.164766 | 0.838743 | Brip1 |
| ENSMUSG( | 728.9738 | 0.214396 | 0.146653 | 0.836044 | Brip1os |
| ENSMUSG( | 388.4606 | -0.00117 | 0.993479 | 0.998813 | Brix1 |
| ENSMUSG( | 4123.856 | -0.16694 | 0.133234 | 0.822912 | Brk1 |
| ENSMUSG( | 34.75197 | -0.05905 | 0.863063 | 0.986547 | Brme1 |
| ENSMUSG( | 1091.948 | -0.05582 | 0.683794 | 0.958678 | Brms1l |
| ENSMUSG( | 1642.987 | -0.13755 | 0.215485 | 0.865167 | Brox |

| | | | | | | | | | | | |
|---|---|---|---|---|---|---|---|---|---|---|---|
| ENSMUSG( | Bcor | 962.7409 | 0.172055 | 0.085252 | 0.450128 | ENSMUSG( | Bcor | 898.2424 | 0.238475 | 0.04292 | 0.690304 |
| ENSMUSG( | Bcorl1 | 596.9967 | 0.00502 | 0.963357 | 0.994911 | ENSMUSG( | Bcorl1 | 553.0341 | 0.051066 | 0.69223 | 0.960404 |
| ENSMUSG( | Bcr | 6483.715 | 0.033301 | 0.716837 | 0.941697 | ENSMUSG( | Bcr | 5715.583 | -0.0623 | 0.531936 | 0.934795 |
| ENSMUSG( | Bcs1l | 596.4804 | 0.058834 | 0.585612 | 0.899171 | ENSMUSG( | Bcs1l | 507.6107 | -0.14611 | 0.280271 | 0.892356 |
| ENSMUSG( | Bdh1 | 2243.334 | 0.071847 | 0.463661 | 0.842812 | ENSMUSG( | Bdh1 | 1985.353 | -0.01443 | 0.913356 | 0.992298 |
| ENSMUSG( | Bdh2 | 83.66288 | -0.02524 | 0.927403 | 0.991229 | ENSMUSG( | Bdh2 | 65.9111 | -0.49289 | 0.060664 | 0.734317 |
| ENSMUSG( | Bdkrb2 | 28.59354 | -0.31581 | 0.378574 | 0.801688 | ENSMUSG( | Bdkrb2 | 25.80624 | -0.34543 | 0.383928 | 0.917257 |
| ENSMUSG( | Bdnf | 607.2912 | -0.5613 | 0.000762 | 0.027082 | ENSMUSG( | Bdnf | 680.3447 | 0.093249 | 0.596821 | 0.946541 |
| ENSMUSG( | Bdp1 | 845.8237 | 0.194794 | 0.076152 | 0.42702 | ENSMUSG( | Bdp1 | 765.1284 | 0.176995 | 0.155658 | 0.837724 |
| ENSMUSG( | Bean1 | 647.9405 | 0.281045 | 0.027822 | 0.255388 | ENSMUSG( | Bean1 | 581.8964 | 0.243098 | 0.072395 | 0.763629 |
| ENSMUSG( | Becn1 | 3803.14 | 0.06894 | 0.396647 | 0.811742 | ENSMUSG( | Becn1 | 3308.695 | -0.06599 | 0.557735 | 0.939565 |
| ENSMUSG( | Begain | 10068.13 | 0.155396 | 0.069079 | 0.407145 | ENSMUSG( | Begain | 8819.221 | 0.044829 | 0.646296 | 0.953988 |
| ENSMUSG( | Bend3 | 294.0992 | -0.03803 | 0.766177 | 0.954173 | ENSMUSG( | Bend3 | 261.186 | -0.11602 | 0.451647 | 0.922202 |
| ENSMUSG( | Bend4 | 199.0277 | -0.0552 | 0.732314 | 0.944956 | ENSMUSG( | Bend4 | 180.5308 | -0.06135 | 0.730974 | 0.967228 |
| ENSMUSG( | Bend5 | 233.3879 | -0.17137 | 0.306436 | 0.754119 | ENSMUSG( | Bend5 | 220.0617 | -0.07504 | 0.694456 | 0.960867 |
| ENSMUSG( | Bend6 | 5523.502 | 0.091146 | 0.268369 | 0.718663 | ENSMUSG( | Bend6 | 4760.255 | -0.07116 | 0.575578 | 0.942912 |
| ENSMUSG( | Bend7 | 95.18193 | 0.10286 | 0.676396 | 0.930542 | ENSMUSG( | Bend7 | 89.4574 | 0.17576 | 0.47365 | 0.923329 |
| ENSMUSG( | Best1 | 31.52565 | -0.07399 | 0.828046 | 0.968966 | ENSMUSG( | Best1 | 25.53465 | -0.43751 | 0.275769 | 0.890277 |
| ENSMUSG( | Bet1 | 629.8123 | 0.069668 | 0.627614 | 0.912961 | ENSMUSG( | Bet1 | 541.6324 | -0.10773 | 0.558543 | 0.93957 |
| ENSMUSG( | Bet1l | 511.6652 | 0.081446 | 0.506551 | 0.867425 | ENSMUSG( | Bet1l | 463.904 | 0.063064 | 0.644832 | 0.953903 |
| ENSMUSG( | Bex1 | 662.1128 | -0.10851 | 0.505837 | 0.866936 | ENSMUSG( | Bex1 | 584.544 | -0.21331 | 0.325922 | 0.905339 |
| ENSMUSG( | Bex2 | 3498.304 | -0.13737 | 0.400388 | 0.813717 | ENSMUSG( | Bex2 | 3065.772 | -0.26297 | 0.211292 | 0.864412 |
| ENSMUSG( | Bex3 | 2677.803 | -0.03512 | 0.80839 | 0.963455 | ENSMUSG( | Bex3 | 2285.103 | -0.23827 | 0.20249 | 0.864412 |
| ENSMUSG( | Bex4 | 225.8902 | -0.01661 | 0.924431 | 0.99114 | ENSMUSG( | Bex4 | 188.1586 | -0.31104 | 0.174376 | 0.844317 |
| ENSMUSG( | Bfar | 1478.846 | -0.18222 | 0.155661 | 0.583641 | ENSMUSG( | Bfar | 1526.847 | 0.179719 | 0.204526 | 0.864412 |
| ENSMUSG( | Bfsp1 | 51.21766 | 0.092002 | 0.735477 | 0.946109 | ENSMUSG( | Bfsp1 | 41.72794 | -0.25076 | 0.434992 | 0.918876 |
| ENSMUSG( | Bfsp2 | 250.6486 | -0.01911 | 0.891182 | 0.983819 | ENSMUSG( | Bfsp2 | 224.471 | -0.07879 | 0.659202 | 0.956205 |
| ENSMUSG( | Bglap3 | 26.62442 | -0.20637 | 0.577801 | 0.895986 | ENSMUSG( | Bglap3 | 24.37072 | -0.15088 | 0.742429 | 0.969415 |
| ENSMUSG( | Bgn | 924.0989 | -0.23147 | 0.092086 | 0.466032 | ENSMUSG( | Bgn | 1026.027 | 0.333744 | 0.019092 | 0.549166 |
| ENSMUSG( | Bhlha9 | 43.0568 | 0.17073 | 0.554069 | 0.886397 | ENSMUSG( | Bhlha9 | 34.38345 | -0.21834 | 0.523395 | 0.932848 |
| ENSMUSG( | Bhlhb9 | 1320.164 | 0.184406 | 0.069268 | 0.407813 | ENSMUSG( | Bhlhb9 | 1154.025 | 0.067533 | 0.577753 | 0.943131 |
| ENSMUSG( | Bhlhe22 | 696.4398 | 0.206637 | 0.049238 | 0.33882 | ENSMUSG( | Bhlhe22 | 615.4284 | 0.119831 | 0.376902 | 0.915593 |
| ENSMUSG( | Bhlhe40 | 1905.101 | -0.03728 | 0.790847 | 0.959461 | ENSMUSG( | Bhlhe40 | 1835.809 | 0.119553 | 0.448732 | 0.922202 |
| ENSMUSG( | Bhlhe41 | 1318.929 | -0.0104 | 0.92084 | 0.990409 | ENSMUSG( | Bhlhe41 | 1410.259 | 0.419401 | 0.00057 | 0.072448 |
| ENSMUSG( | Bicc1 | 301.9257 | -0.0063 | 0.96421 | 0.995155 | ENSMUSG( | Bicc1 | 291.0231 | 0.147765 | 0.34508 | 0.91045 |
| ENSMUSG( | Bicd1 | 545.6614 | -0.05522 | 0.612837 | 0.908902 | ENSMUSG( | Bicd1 | 568.5338 | 0.311389 | 0.024775 | 0.592568 |
| ENSMUSG( | Bicd2 | 2212.063 | 0.01579 | 0.864637 | 0.975475 | ENSMUSG( | Bicd2 | 2104.371 | 0.139251 | 0.257908 | 0.883466 |
| ENSMUSG( | Bicdl1 | 2568.829 | 0.300621 | 0.001692 | 0.046443 | ENSMUSG( | Bicdl1 | 2273.365 | 0.225841 | 0.076453 | 0.770555 |
| ENSMUSG( | Bicra | 791.7156 | 0.154789 | 0.127251 | 0.534725 | ENSMUSG( | Bicra | 743.0734 | 0.237412 | 0.059745 | 0.729747 |
| ENSMUSG( | Bicral | 1944.173 | 0.094278 | 0.373798 | 0.80044 | ENSMUSG( | Bicral | 1725.163 | 0.016379 | 0.90079 | 0.990558 |
| ENSMUSG( | Bid | 168.962 | 0.047674 | 0.794006 | 0.959799 | ENSMUSG( | Bid | 154.3676 | 0.060025 | 0.769399 | 0.974236 |
| ENSMUSG( | Bin1 | 6150.622 | 0.037203 | 0.723097 | 0.943761 | ENSMUSG( | Bin1 | 5505.478 | -0.013 | 0.904414 | 0.991089 |
| ENSMUSG( | Bin2 | 425.921 | -0.1242 | 0.36741 | 0.795221 | ENSMUSG( | Bin2 | 389.557 | -0.10781 | 0.465653 | 0.923329 |
| ENSMUSG( | Bin3 | 274.784 | -0.1332 | 0.363824 | 0.793343 | ENSMUSG( | Bin3 | 258.9628 | -0.0246 | 0.879329 | 0.989644 |
| ENSMUSG( | Birc2 | 1042.493 | -0.03257 | 0.722758 | 0.943761 | ENSMUSG( | Birc2 | 959.6806 | -0.00231 | 0.983397 | 0.996963 |
| ENSMUSG( | Birc3 | 98.16606 | -0.04378 | 0.836528 | 0.971111 | ENSMUSG( | Birc3 | 85.97012 | -0.14353 | 0.58835 | 0.944975 |
| ENSMUSG( | Birc5 | 37.92165 | -0.135 | 0.665113 | 0.926824 | ENSMUSG( | Birc5 | 37.17232 | 0.086001 | 0.795043 | 0.97681 |
| ENSMUSG( | Birc6 | 2655.17 | -0.24133 | 0.031052 | 0.268255 | ENSMUSG( | Birc6 | 2728.988 | 0.118571 | 0.409657 | 0.917257 |
| ENSMUSG( | Bivm | 470.2538 | -0.14059 | 0.35715 | 0.788702 | ENSMUSG( | Bivm | 483.2366 | 0.209493 | 0.186818 | 0.856099 |
| ENSMUSG( | Blcap | 2197.511 | -0.26246 | 0.016257 | 0.189161 | ENSMUSG( | Blcap | 2159.867 | -0.02922 | 0.814306 | 0.979198 |
| ENSMUSG( | Blm | 84.05588 | -0.28306 | 0.197481 | 0.637186 | ENSMUSG( | Blm | 78.63286 | -0.18122 | 0.458967 | 0.923329 |
| ENSMUSG( | Blmh | 1495.028 | 0.028735 | 0.776477 | 0.955658 | ENSMUSG( | Blmh | 1282.192 | -0.15027 | 0.220223 | 0.866104 |
| ENSMUSG( | Blnk | 210.3362 | 0.128266 | 0.433208 | 0.828207 | ENSMUSG( | Blnk | 185.6918 | 0.0397 | 0.824537 | 0.979198 |
| ENSMUSG( | Bloc1s1 | 260.4856 | 0.110694 | 0.42227 | 0.823508 | ENSMUSG( | Bloc1s1 | 211.0536 | -0.24496 | 0.158131 | 0.837724 |
| ENSMUSG( | Bloc1s2 | 626.9454 | -0.12478 | 0.224715 | 0.671058 | ENSMUSG( | Bloc1s2 | 555.7495 | -0.21098 | 0.10188 | 0.801719 |
| ENSMUSG( | Bloc1s3 | 256.1736 | 0.046006 | 0.76564 | 0.954173 | ENSMUSG( | Bloc1s3 | 223.8508 | -0.07056 | 0.698708 | 0.960908 |
| ENSMUSG( | Bloc1s4 | 259.625 | -0.14964 | 0.343694 | 0.779153 | ENSMUSG( | Bloc1s4 | 250.1935 | 0.006057 | 0.973561 | 0.996416 |
| ENSMUSG( | Bloc1s5 | 677.9456 | 0.071601 | 0.561753 | 0.889052 | ENSMUSG( | Bloc1s5 | 551.4805 | -0.27117 | 0.04466 | 0.693697 |
| ENSMUSG( | Bloc1s6 | 723.917 | 0.055804 | 0.605827 | 0.905919 | ENSMUSG( | Bloc1s6 | 631.4965 | -0.07716 | 0.589082 | 0.944975 |
| ENSMUSG( | Bltp1 | 2979.676 | -0.28848 | 0.024802 | 0.242192 | ENSMUSG( | Bltp1 | 3251.548 | 0.241579 | 0.165819 | 0.839724 |
| ENSMUSG( | Bltp2 | 16570.06 | 0.10551 | 0.283124 | 0.73186 | ENSMUSG( | Bltp2 | 14648.86 | 0.022161 | 0.846239 | 0.984578 |
| ENSMUSG( | Bltp3a | 964.6642 | 0.043795 | 0.678665 | 0.930611 | ENSMUSG( | Bltp3a | 881.1193 | 0.051721 | 0.666663 | 0.956766 |
| ENSMUSG( | Bltp3b | 2405.965 | 0.009772 | 0.91292 | 0.988483 | ENSMUSG( | Bltp3b | 2267.953 | 0.10801 | 0.339308 | 0.907972 |
| ENSMUSG( | Blvra | 334.4423 | -0.08349 | 0.580286 | 0.896952 | ENSMUSG( | Blvra | 281.0541 | -0.34444 | 0.05126 | 0.711023 |
| ENSMUSG( | Blvrb | 373.601 | -0.06528 | 0.597925 | 0.90372 | ENSMUSG( | Blvrb | 331.3094 | -0.1551 | 0.309628 | 0.898538 |
| ENSMUSG( | Blzf1 | 645.2024 | 0.031851 | 0.790799 | 0.959461 | ENSMUSG( | Blzf1 | 560.2575 | -0.10993 | 0.403425 | 0.917257 |
| ENSMUSG( | Bmal1 | 872.133 | -0.01712 | 0.85972 | 0.97481 | ENSMUSG( | Bmal1 | 620.9489 | -0.85308 | 1.72E-11 | 2.01E-08 |
| ENSMUSG( | Bmal2 | 209.5844 | 0.136814 | 0.4044 | 0.816132 | ENSMUSG( | Bmal2 | 199.2951 | 0.264778 | 0.160442 | 0.837724 |
| ENSMUSG( | Bmerb1 | 7099.364 | 0.10324 | 0.275003 | 0.72509 | ENSMUSG( | Bmerb1 | 5867.679 | -0.18297 | 0.131608 | 0.822912 |
| ENSMUSG( | Bmf | 206.776 | -0.34154 | 0.022365 | 0.228729 | ENSMUSG( | Bmf | 200.9872 | -0.12395 | 0.527487 | 0.934466 |
| ENSMUSG( | Bmi1 | 1055.634 | -0.05844 | 0.667081 | 0.92729 | ENSMUSG( | Bmi1 | 1019.06 | 0.104555 | 0.510065 | 0.929089 |
| ENSMUSG( | Bmp1 | 2858.696 | 0.197627 | 0.185918 | 0.623352 | ENSMUSG( | Bmp1 | 2624.363 | 0.218442 | 0.170587 | 0.841739 |
| ENSMUSG( | Bmp15 | 9.20583 | -0.67018 | 0.285982 | NA | ENSMUSG( | Bmp15 | 10.56377 | 0.107694 | 0.854711 | NA |
| ENSMUSG( | Bmp2 | 132.2666 | -0.11755 | 0.556295 | 0.887663 | ENSMUSG( | Bmp2 | 152.4564 | 0.530017 | 0.023775 | 0.588226 |
| ENSMUSG( | Bmp2k | 786.0931 | -0.1901 | 0.106103 | 0.493403 | ENSMUSG( | Bmp2k | 756.5152 | -0.02411 | 0.853039 | 0.985598 |
| ENSMUSG( | Bmp3 | 511.0903 | -0.66251 | 0.000337 | 0.015675 | ENSMUSG( | Bmp3 | 666.2651 | 0.422525 | 0.031294 | 0.631113 |
| ENSMUSG( | Bmp4 | 271.5096 | -0.31279 | 0.056169 | 0.36528 | ENSMUSG( | Bmp4 | 326.5928 | 0.477586 | 0.008771 | 0.393969 |
| ENSMUSG( | Bmp5 | 110.1544 | -0.56518 | 0.010217 | 0.144102 | ENSMUSG( | Bmp5 | 130.0557 | 0.239693 | 0.259366 | 0.883971 |
| ENSMUSG( | Bmp6 | 503.9148 | -0.1846 | 0.159487 | 0.588863 | ENSMUSG( | Bmp6 | 459.0648 | -0.17693 | 0.237352 | 0.872173 |
| ENSMUSG( | Bmp7 | 724.5855 | -0.04505 | 0.711639 | 0.940028 | ENSMUSG( | Bmp7 | 703.1051 | 0.135553 | 0.29484 | 0.898538 |
| ENSMUSG( | Bmper | 294.8971 | -0.33352 | 0.040925 | 0.312769 | ENSMUSG( | Bmper | 332.8819 | 0.298134 | 0.112704 | 0.809623 |
| ENSMUSG( | Bmpr1a | 4431.556 | -0.02947 | 0.819275 | 0.965888 | ENSMUSG( | Bmpr1a | 4234.362 | 0.105392 | 0.474593 | 0.923329 |
| ENSMUSG( | Bmpr1b | 2464.838 | -0.08363 | 0.539337 | 0.883094 | ENSMUSG( | Bmpr1b | 2154.85 | -0.2091 | 0.159746 | 0.837724 |
| ENSMUSG( | Bmpr2 | 5193.963 | -0.15741 | 0.266753 | 0.716789 | ENSMUSG( | Bmpr2 | 5374.915 | 0.208866 | 0.181932 | 0.850579 |
| ENSMUSG( | Bms1 | 1266.686 | 0.228463 | 0.041631 | 0.315097 | ENSMUSG( | Bms1 | 1121.637 | 0.148573 | 0.261385 | 0.884926 |
| ENSMUSG( | Bmt2 | 387.7062 | 0.023105 | 0.858886 | 0.974707 | ENSMUSG( | Bmt2 | 381.4906 | 0.228792 | 0.147515 | 0.836241 |
| ENSMUSG( | Bmyc | 1610.489 | -0.00119 | 0.991393 | 0.999232 | ENSMUSG( | Bmyc | 1394.153 | -0.158 | 0.2754 | 0.889984 |
| ENSMUSG( | Bnc2 | 28.31439 | -0.60697 | 0.10671 | 0.49424 | ENSMUSG( | Bnc2 | 27.93344 | -0.38219 | 0.360251 | 0.914223 |
| ENSMUSG( | Bnip1 | 436.9298 | 0.039592 | 0.735199 | 0.945972 | ENSMUSG( | Bnip1 | 379.1056 | -0.10818 | 0.45029 | 0.922202 |
| ENSMUSG( | Bnip2 | 1323.775 | 0.001551 | 0.988475 | 0.999109 | ENSMUSG( | Bnip2 | 1243.142 | 0.088778 | 0.451205 | 0.922202 |
| ENSMUSG( | Bnip3 | 1713.701 | 0.030823 | 0.736245 | 0.946267 | ENSMUSG( | Bnip3 | 1501.629 | -0.08733 | 0.556866 | 0.939565 |
| ENSMUSG( | Bnip3l | 3576.927 | 0.040916 | 0.678139 | 0.930542 | ENSMUSG( | Bnip3l | 3133.094 | -0.07657 | 0.575798 | 0.942912 |
| ENSMUSG( | Bnip5 | 282.3765 | 7.93E-05 | 0.999593 | 0.99989 | ENSMUSG( | Bnip5 | 236.7794 | -0.23846 | 0.172269 | 0.843466 |
| ENSMUSG( | Boc | 794.8225 | -0.30352 | 0.097011 | 0.477375 | ENSMUSG( | Boc | 845.6296 | 0.163184 | 0.403074 | 0.917257 |
| ENSMUSG( | Bod1 | 2656.831 | 0.013628 | 0.878293 | 0.981431 | ENSMUSG( | Bod1 | 2301.384 | -0.13814 | 0.265399 | 0.886134 |
| ENSMUSG( | Bod1l | 1778.749 | 0.064701 | 0.439863 | 0.831633 | ENSMUSG( | Bod1l | 1613.538 | 0.051984 | 0.619745 | 0.952896 |
| ENSMUSG( | Bok | 2035.444 | -0.00359 | 0.970046 | 0.996369 | ENSMUSG( | Bok | 1785.506 | -0.1169 | 0.275614 | 0.890209 |
| ENSMUSG( | Bola1 | 230.8782 | -0.06129 | 0.682983 | 0.93205 | ENSMUSG( | Bola1 | 204.7216 | -0.15593 | 0.402839 | 0.917257 |
| ENSMUSG( | Bola2 | 1249.904 | 0.09787 | 0.290069 | 0.739883 | ENSMUSG( | Bola2 | 1062.154 | -0.1093 | 0.446546 | 0.921734 |
| ENSMUSG( | Bola3 | 1067.274 | 0.073086 | 0.451268 | 0.835567 | ENSMUSG( | Bola3 | 869.607 | -0.26326 | 0.028926 | 0.614171 |
| ENSMUSG( | Bop1 | 1191.709 | 0.105029 | 0.326155 | 0.771128 | ENSMUSG( | Bop1 | 1062.283 | 0.045413 | 0.715912 | 0.965424 |
| ENSMUSG( | Bora | 113.9314 | -0.05659 | 0.782048 | 0.957558 | ENSMUSG( | Bora | 104.2704 | -0.05149 | 0.816314 | 0.979198 |
| ENSMUSG( | Borcs5 | 610.696 | 0.053309 | 0.661518 | 0.925271 | ENSMUSG( | Borcs5 | 540.3588 | -0.03262 | 0.81581 | 0.979198 |
| ENSMUSG( | Borcs6 | 1110.322 | 0.049891 | 0.644942 | 0.91787 | ENSMUSG( | Borcs6 | 971.1365 | -0.07263 | 0.616878 | 0.951666 |
| ENSMUSG( | Borcs7 | 721.587 | 0.095236 | 0.357201 | 0.788702 | ENSMUSG( | Borcs7 | 613.6407 | -0.11109 | 0.46551 | 0.923329 |
| ENSMUSG( | Borcs8 | 904.0409 | -0.00832 | 0.938825 | 0.993415 | ENSMUSG( | Borcs8 | 784.607 | -0.15573 | 0.22326 | 0.867404 |
| ENSMUSG( | Bpgm | 4149.796 | -0.05829 | 0.589099 | 0.90113 | ENSMUSG( | Bpgm | 3472.733 | -0.32427 | 0.020299 | 0.563507 |
| ENSMUSG( | Bphl | 710.9752 | 0.026658 | 0.830483 | 0.96974 | ENSMUSG( | Bphl | 578.6247 | -0.32283 | 0.037356 | 0.673514 |
| ENSMUSG( | Bpnt1 | 1576.978 | 0.089287 | 0.362396 | 0.792943 | ENSMUSG( | Bpnt1 | 1352.305 | -0.08957 | 0.466816 | 0.923329 |
| ENSMUSG( | Bpnt2 | 3505.112 | -0.38322 | 0.0078 | 0.122469 | ENSMUSG( | Bpnt2 | 3791.955 | 0.136939 | 0.406177 | 0.917257 |
| ENSMUSG( | Bptf | 2874.657 | 0.04375 | 0.657918 | 0.92425 | ENSMUSG( | Bptf | 2727.86 | 0.159234 | 0.145128 | 0.83339 |
| ENSMUSG( | Braf | 2456.407 | -0.00738 | 0.927482 | 0.991229 | ENSMUSG( | Braf | 2298.008 | 0.068017 | 0.499604 | 0.927825 |
| ENSMUSG( | Brap | 1989.117 | 0.043513 | 0.625338 | 0.912249 | ENSMUSG( | Brap | 1758.946 | -0.0428 | 0.686162 | 0.958956 |
| ENSMUSG( | Brat1 | 686.0425 | -0.03654 | 0.76536 | 0.954127 | ENSMUSG( | Brat1 | 627.3684 | -0.02038 | 0.883748 | 0.989644 |
| ENSMUSG( | Brca1 | 180.3369 | -0.01956 | 0.919722 | 0.990247 | ENSMUSG( | Brca1 | 151.7123 | -0.27723 | 0.155769 | 0.837724 |
| ENSMUSG( | Brca2 | 139.2145 | -0.49693 | 0.008807 | 0.132321 | ENSMUSG( | Brca2 | 145.6906 | -0.04903 | 0.819843 | 0.979198 |
| ENSMUSG( | Brcc3 | 675.4068 | 0.066359 | 0.518222 | 0.873754 | ENSMUSG( | Brcc3 | 583.5978 | -0.09173 | 0.512503 | 0.930164 |
| ENSMUSG( | Brd1 | 1533.45 | -0.01444 | 0.870826 | 0.977951 | ENSMUSG( | Brd1 | 1443.711 | 0.076722 | 0.503894 | 0.927825 |
| ENSMUSG( | Brd10 | 1861.437 | 0.053072 | 0.566515 | 0.891644 | ENSMUSG( | Brd10 | 1706.864 | 0.068885 | 0.563366 | 0.941206 |
| ENSMUSG( | Brd2 | 10493.99 | 0.139106 | 0.062263 | 0.386085 | ENSMUSG( | Brd2 | 9515.738 | 0.126213 | 0.204957 | 0.864412 |
| ENSMUSG( | Brd3 | 1800.564 | 0.055707 | 0.52904 | 0.878551 | ENSMUSG( | Brd3 | 1619.884 | 0.016995 | 0.887419 | 0.990266 |
| ENSMUSG( | Brd3os | 513.3902 | -0.08117 | 0.618107 | 0.910929 | ENSMUSG( | Brd3os | 456.2554 | -0.15528 | 0.370543 | 0.915491 |
| ENSMUSG( | Brd4 | 3760.635 | -0.0951 | 0.554135 | 0.886397 | ENSMUSG( | Brd4 | 3630.234 | 0.074607 | 0.627927 | 0.953234 |
| ENSMUSG( | Brd7 | 1841.311 | 0.127619 | 0.14343 | 0.562473 | ENSMUSG( | Brd7 | 1601.806 | -0.00487 | 0.965607 | 0.995433 |
| ENSMUSG( | Brd8 | 1320.84 | -0.04423 | 0.672761 | 0.929007 | ENSMUSG( | Brd8 | 1204.59 | -0.0366 | 0.778752 | 0.976106 |
| ENSMUSG( | Brd8dc | 29.3119 | 0.011512 | 0.973579 | 0.99675 | ENSMUSG( | Brd8dc | 29.24447 | 0.259123 | 0.481325 | 0.924564 |
| ENSMUSG( | Brd9 | 2717.879 | 0.310536 | 0.003705 | 0.07745 | ENSMUSG( | Brd9 | 2298.615 | 0.110001 | 0.433718 | 0.918564 |
| ENSMUSG( | Brdt | 436.9257 | 0.078881 | 0.545249 | 0.884748 | ENSMUSG( | Brdt | 372.1521 | -0.12597 | 0.42569 | 0.917257 |
| ENSMUSG( | Brf1 | 1860.057 | 0.07904 | 0.384501 | 0.804007 | ENSMUSG( | Brf1 | 1606.914 | -0.0749 | 0.503408 | 0.927825 |
| ENSMUSG( | Brf2 | 339.3397 | 0.037915 | 0.790432 | 0.959461 | ENSMUSG( | Brf2 | 303.955 | -0.0067 | 0.967882 | 0.995852 |
| ENSMUSG( | Bri3 | 1627.424 | 0.021183 | 0.84995 | 0.972638 | ENSMUSG( | Bri3 | 1464.707 | -0.01956 | 0.894747 | 0.990558 |
| ENSMUSG( | Bri3bp | 3954.915 | -0.2293 | 0.103762 | 0.489428 | ENSMUSG( | Bri3bp | 4142.145 | 0.180072 | 0.271065 | 0.888062 |
| ENSMUSG( | Bricd5 | 27.33258 | -0.129 | 0.72508 | 0.944096 | ENSMUSG( | Bricd5 | 25.81943 | -0.01539 | 0.968156 | 0.995887 |
| ENSMUSG( | Brinp1 | 5950.482 | -0.33408 | 0.020167 | 0.214666 | ENSMUSG( | Brinp1 | 6528.292 | 0.217874 | 0.226614 | 0.868507 |
| ENSMUSG( | Brinp2 | 3402.531 | -0.05475 | 0.688846 | 0.933441 | ENSMUSG( | Brinp2 | 3643.037 | 0.389954 | 0.014766 | 0.487333 |
| ENSMUSG( | Brinp3 | 790.8256 | -0.2331 | 0.21061 | 0.653372 | ENSMUSG( | Brinp3 | 830.462 | 0.188815 | 0.387404 | 0.917257 |
| ENSMUSG( | Brip1os | 727.4697 | -0.05686 | 0.699218 | 0.936886 | ENSMUSG( | Brip1os | 728.9738 | 0.214396 | 0.146653 | 0.836044 |
| ENSMUSG( | Brix1 | 448.9996 | 0.150921 | 0.210029 | 0.652413 | ENSMUSG( | Brix1 | 388.4606 | -0.00117 | 0.993479 | 0.998813 |
| ENSMUSG( | Brk1 | 4903.735 | 0.070281 | 0.408851 | 0.818173 | ENSMUSG( | Brk1 | 4123.856 | -0.16694 | 0.133234 | 0.822912 |
| ENSMUSG( | Brme1 | 42.06768 | 0.193285 | 0.506368 | 0.867425 | ENSMUSG( | Brme1 | 34.75197 | -0.05905 | 0.863063 | 0.986547 |
| ENSMUSG( | Brms1l | 1241.71 | 0.051059 | 0.623106 | 0.911589 | ENSMUSG( | Brms1l | 1091.948 | -0.05582 | 0.683794 | 0.958678 |
| ENSMUSG( | Brox | 1870.152 | -0.02753 | 0.764854 | 0.954127 | ENSMUSG( | Brox | 1642.987 | -0.13755 | 0.215485 | 0.865167 |
| ENSMUSG( | Brpf1 | 835.5544 | 0.080831 | 0.529459 | 0.878883 | ENSMUSG( | Brpf1 | 776.6218 | 0.139877 | 0.305533 | 0.898538 |
| ENSMUSG( | Brpf3 | 1428.889 | 0.028627 | 0.800677 | 0.961745 | ENSMUSG( | Brpf3 | 1330.045 | 0.089055 | 0.453813 | 0.922316 |
| ENSMUSG( | Brsk1 | 31199.81 | 0.230042 | 0.004035 | 0.081715 | ENSMUSG( | Brsk1 | 27019.28 | 0.09117 | 0.364285 | 0.914223 |
| ENSMUSG( | Brsk2 | 7140.637 | 0.097008 | 0.314507 | 0.761558 | ENSMUSG( | Brsk2 | 6333.891 | 0.022048 | 0.825648 | 0.979537 |
| ENSMUSG( | Brwd1 | 3130.278 | -0.14814 | 0.110809 | 0.501952 | ENSMUSG( | Brwd1 | 2948.082 | -0.0497 | 0.64301 | 0.953903 |
| ENSMUSG( | Brwd3 | 292.8804 | -0.08158 | 0.610396 | 0.908189 | ENSMUSG( | Brwd3 | 293.6991 | 0.180417 | 0.372599 | 0.915491 |
| ENSMUSG( | Bscl2 | 2784.859 | -0.21452 | 0.08776 | 0.454663 | ENSMUSG( | Bscl2 | 2884.625 | 0.163003 | 0.276625 | 0.890889 |
| ENSMUSG( | Bsdc1 | 2093.05 | -0.10047 | 0.231664 | 0.67949 | ENSMUSG( | Bsdc1 | 1934.189 | -0.05828 | 0.577148 | 0.942912 |
| ENSMUSG( | Bsg | 24114.48 | -0.24002 | 0.068353 | 0.405144 | ENSMUSG( | Bsg | 24205.82 | 0.052239 | 0.712391 | 0.964363 |
| ENSMUSG( | Bsn | 15019.81 | 0.130882 | 0.292966 | 0.742763 | ENSMUSG( | Bsn | 14344.2 | 0.26141 | 0.096264 | 0.796133 |
| ENSMUSG( | Bspry | 16.64628 | 0.117294 | 0.796579 | NA | ENSMUSG( | Bspry | 13.15984 | -0.27226 | 0.611512 | 0.950771 |
| ENSMUSG( | Bst2 | 226.6864 | -0.25943 | 0.121451 | 0.525713 | ENSMUSG( | Bst2 | 216.9095 | -0.09496 | 0.608517 | 0.950113 |
| ENSMUSG( | Bsx | 9.686582 | -0.60334 | 0.324049 | NA | ENSMUSG( | Bsx | 9.61276 | -0.29785 | 0.635859 | NA |
| ENSMUSG( | Btaf1 | 1144.29 | 0.050355 | 0.665012 | 0.926824 | ENSMUSG( | Btaf1 | 1043.227 | 0.051608 | 0.693209 | 0.96076 |
| ENSMUSG( | Btbd1 | 7024.352 | 0.070458 | 0.526789 | 0.877267 | ENSMUSG( | Btbd1 | 5839.311 | -0.20313 | 0.179825 | 0.850372 |
| ENSMUSG( | Btbd10 | 1188.719 | 0.187759 | 0.143199 | 0.56185 | ENSMUSG( | Btbd10 | 1035.234 | 0.060666 | 0.656509 | 0.955533 |

| | | | | | |
|---|---|---|---|---|---|
| ENSMUSG( | 3954.915 | -0.2293 | 0.103762 | 0.489428 | Bri3bp |
| ENSMUSG( | 27.33258 | -0.129 | 0.72508 | 0.944096 | Bricd5 |
| ENSMUSG( | 5950.482 | -0.33408 | 0.020167 | 0.214666 | Brinp1 |
| ENSMUSG( | 3402.531 | -0.05475 | 0.688846 | 0.933441 | Brinp2 |
| ENSMUSG( | 790.8256 | -0.2331 | 0.21061 | 0.653372 | Brinp3 |
| ENSMUSG( | 727.4697 | -0.05686 | 0.699218 | 0.936886 | Brip1os |
| ENSMUSG( | 448.9996 | 0.150921 | 0.210029 | 0.652413 | Brix1 |
| ENSMUSG( | 4903.735 | 0.070281 | 0.408851 | 0.818173 | Brk1 |
| ENSMUSG( | 42.06768 | 0.193285 | 0.506368 | 0.867425 | Brme1 |
| ENSMUSG( | 1241.71 | 0.051059 | 0.623106 | 0.911589 | Brms1l |
| ENSMUSG( | 1870.152 | -0.02753 | 0.764854 | 0.954127 | Brox |
| ENSMUSG( | 835.5544 | 0.080831 | 0.529459 | 0.878883 | Brpf1 |
| ENSMUSG( | 1428.889 | 0.028627 | 0.800677 | 0.961745 | Brpf3 |
| ENSMUSG( | 31199.81 | 0.230042 | 0.004035 | 0.081715 | Brsk1 |
| ENSMUSG( | 7140.637 | 0.097008 | 0.314507 | 0.761558 | Brsk2 |
| ENSMUSG( | 3130.278 | -0.14814 | 0.110809 | 0.501952 | Brwd1 |
| ENSMUSG( | 292.8804 | -0.08158 | 0.610396 | 0.908189 | Brwd3 |
| ENSMUSG( | 2784.859 | -0.21452 | 0.08776 | 0.454663 | Bscl2 |
| ENSMUSG( | 2093.05 | -0.10047 | 0.231664 | 0.67949 | Bsdc1 |
| ENSMUSG( | 24114.48 | -0.24002 | 0.068353 | 0.405144 | Bsg |
| ENSMUSG( | 15019.81 | 0.130882 | 0.292966 | 0.742763 | Bsn |
| ENSMUSG( | 16.64628 | 0.117294 | 0.796579 | NA | Bspry |
| ENSMUSG( | 10.33688 | 0.233162 | 0.680456 | NA | Bst1 |
| ENSMUSG( | 226.6864 | -0.25943 | 0.121451 | 0.525713 | Bst2 |
| ENSMUSG( | 9.686582 | -0.60334 | 0.324049 | NA | Bsx |
| ENSMUSG( | 1144.29 | 0.050355 | 0.665012 | 0.926824 | Btaf1 |
| ENSMUSG( | 7024.352 | 0.070458 | 0.526789 | 0.877267 | Btbd1 |
| ENSMUSG( | 1188.719 | 0.187759 | 0.143199 | 0.56185 | Btbd10 |
| ENSMUSG( | 84.33916 | -0.15266 | 0.488657 | 0.859092 | Btbd16 |
| ENSMUSG( | 2044.334 | -0.2647 | 0.040513 | 0.311135 | Btbd17 |
| ENSMUSG( | 112.9148 | -0.02482 | 0.923201 | 0.990693 | Btbd19 |
| ENSMUSG( | 3053.169 | 0.115542 | 0.167806 | 0.60061 | Btbd2 |
| ENSMUSG( | 5079.528 | 0.066198 | 0.523534 | 0.875628 | Btbd3 |
| ENSMUSG( | 1906.665 | 0.112108 | 0.263607 | 0.714284 | Btbd6 |
| ENSMUSG( | 1117.41 | -0.07617 | 0.505092 | 0.866883 | Btbd7 |
| ENSMUSG( | 1061.832 | -0.10826 | 0.391557 | 0.809154 | Btbd8 |
| ENSMUSG( | 1754.182 | 0.160522 | 0.120353 | 0.524495 | Btbd9 |
| ENSMUSG( | 580.9594 | -0.47235 | 0.001289 | 0.039144 | Btd |
| ENSMUSG( | 2223.178 | -0.01977 | 0.83146 | 0.969889 | Btf3 |
| ENSMUSG( | 778.2974 | -0.2159 | 0.038797 | 0.303414 | Btf3l4 |
| ENSMUSG( | 55.77767 | 4.479622 | 3.64E-23 | 5.91E-20 | Btf3l4b |
| ENSMUSG( | 1045.442 | -0.16552 | 0.124822 | 0.531813 | Btg1 |
| ENSMUSG( | 461.001 | -0.7984 | 0.00126 | 0.038624 | Btg2 |
| ENSMUSG( | 119.2653 | -0.18563 | 0.344962 | 0.779153 | Btg3 |
| ENSMUSG( | 69.97366 | -0.04813 | 0.867949 | 0.976631 | Btk |
| ENSMUSG( | 25.95297 | -0.02131 | 0.955238 | 0.993874 | Btla |
| ENSMUSG( | 24.88298 | -0.34829 | 0.400382 | 0.813717 | Btn2a2 |
| ENSMUSG( | 1445.504 | 0.137684 | 0.280126 | 0.729155 | Btrc |
| ENSMUSG( | 15.23505 | -0.57618 | 0.232751 | NA | Bub1 |
| ENSMUSG( | 120.0572 | -0.26005 | 0.207088 | 0.649503 | Bub1b |
| ENSMUSG( | 1254.465 | -0.0105 | 0.924026 | 0.991077 | Bub3 |
| ENSMUSG( | 224.4843 | 0.332338 | 0.031651 | 0.271198 | Bud13 |
| ENSMUSG( | 700.7006 | 0.098117 | 0.392466 | 0.809274 | Bud23 |
| ENSMUSG( | 902.5369 | 0.065495 | 0.512693 | 0.871762 | Bud31 |
| ENSMUSG( | 67.72327 | -0.21546 | 0.413252 | 0.819541 | Bves |
| ENSMUSG( | 616.8054 | 0.224815 | 0.04662 | 0.330354 | Bysl |
| ENSMUSG( | 5494.548 | 0.012709 | 0.905761 | 0.986694 | Bzw1 |
| ENSMUSG( | 762.1943 | 0.071029 | 0.52708 | 0.877386 | Bzw2 |
| ENSMUSG( | 765.7579 | -0.05307 | 0.598049 | 0.903724 | C030006K11Rik |
| ENSMUSG( | 31.04554 | -0.21025 | 0.590205 | 0.90113 | C030013G03Rik |
| ENSMUSG( | 279.3142 | -0.03032 | 0.845757 | 0.972191 | C030018K13Rik |
| ENSMUSG( | 105.7652 | -0.58699 | 0.004024 | 0.081586 | C030029H02Rik |
| ENSMUSG( | 108.5211 | 0.164859 | 0.406487 | 0.816818 | C130026L21Rik |
| ENSMUSG( | 108.7582 | 0.293151 | 0.133575 | 0.546843 | C130046K22Rik |
| ENSMUSG( | 10.51045 | 0.148445 | 0.806609 | NA | C130050O18Rik |
| ENSMUSG( | 1386.082 | -0.03389 | 0.775592 | 0.955606 | C130074G19Rik |
| ENSMUSG( | 577.5213 | 0.012085 | 0.920804 | 0.990409 | C1d |
| ENSMUSG( | 687.804 | -0.0275 | 0.839584 | 0.971179 | C1galt1 |
| ENSMUSG( | 526.5545 | -0.34468 | 0.007417 | 0.118765 | C1galt1c1 |
| ENSMUSG( | 3698.841 | -0.05792 | 0.673404 | 0.929031 | C1qa |
| ENSMUSG( | 4076.99 | -0.05668 | 0.648055 | 0.919798 | C1qb |
| ENSMUSG( | 1399.791 | -0.0603 | 0.555587 | 0.887457 | C1qbp |
| ENSMUSG( | 3874.267 | -0.16004 | 0.21764 | 0.662824 | C1qc |
| ENSMUSG( | 860.102 | -0.43259 | 0.002322 | 0.05646 | C1ql1 |
| ENSMUSG( | 492.601 | -0.35188 | 0.056359 | 0.366006 | C1ql2 |
| ENSMUSG( | 1307.459 | -0.39554 | 0.026528 | 0.249008 | C1ql3 |
| ENSMUSG( | 125.2449 | -0.39386 | 0.108583 | 0.497529 | C1qtnf1 |
| ENSMUSG( | 387.1884 | -0.20987 | 0.119859 | 0.523965 | C1qtnf12 |
| ENSMUSG( | 51.90828 | -0.18941 | 0.491957 | 0.860297 | C1qtnf2 |
| ENSMUSG( | 7891.826 | -0.3313 | 0.00027 | 0.013409 | C1qtnf4 |
| ENSMUSG( | 258.7766 | -0.29493 | 0.071723 | 0.413564 | C1qtnf6 |
| ENSMUSG( | 32.12204 | 0.043985 | 0.902177 | 0.985633 | C1qtnf7 |
| ENSMUSG( | 66.83072 | -0.29147 | 0.212096 | 0.655777 | C1ra |
| ENSMUSG( | 23.04258 | -0.24142 | 0.551761 | 0.886141 | C1rl |
| ENSMUSG( | 38.39586 | -0.14865 | 0.710533 | 0.939748 | C1s1 |
| ENSMUSG( | 12.76947 | 0.364955 | 0.568559 | NA | C1s2 |
| ENSMUSG( | 51.46536 | -0.18023 | 0.510862 | 0.870755 | C2 |
| ENSMUSG( | 12.73244 | -0.10548 | 0.856879 | NA | C230014O12Rik |
| ENSMUSG( | 43.33326 | 0.198427 | 0.511424 | 0.870898 | C230035I16Rik |
| ENSMUSG( | 15.466 | 0.350362 | 0.518109 | NA | C230072F16Rik |
| ENSMUSG( | 424.0269 | -0.04552 | 0.780732 | 0.957491 | C2cd2 |
| ENSMUSG( | 6400.42 | -0.29603 | 0.01793 | 0.200989 | C2cd2l |
| ENSMUSG( | 906.5645 | 0.198467 | 0.052094 | 0.350316 | C2cd3 |
| ENSMUSG( | 58.12102 | 0.290144 | 0.325151 | 0.770608 | C2cd4b |
| ENSMUSG( | 467.4771 | 0.253868 | 0.02927 | 0.259574 | C2cd4c |
| ENSMUSG( | 11.62342 | -0.18674 | 0.732335 | NA | C2cd4d |
| ENSMUSG( | 2089.696 | 0.218698 | 0.09595 | 0.475936 | C2cd5 |
| ENSMUSG( | 38.59524 | -0.59891 | 0.192995 | 0.632208 | C3 |
| ENSMUSG( | 34.45003 | -0.29445 | 0.373937 | 0.800445 | C330013E15Rik |
| ENSMUSG( | 441.3964 | 0.197062 | 0.161902 | 0.592398 | C330018D20Rik |
| ENSMUSG( | 104.9212 | -0.31787 | 0.172646 | 0.608104 | C3ar1 |
| ENSMUSG( | 53.87437 | 0.511094 | 0.10804 | 0.497059 | C4a |
| ENSMUSG( | 560.3643 | 1.532107 | 6.23E-27 | 1.26E-23 | C4b |
| ENSMUSG( | 150.1769 | -0.13323 | 0.470646 | 0.84674 | C530005A16Rik |
| ENSMUSG( | 10.5206 | -0.31484 | 0.585646 | NA | C530044C16Rik |
| ENSMUSG( | 37.83762 | 0.023032 | 0.94969 | 0.993821 | C5ar1 |
| ENSMUSG( | 40.48409 | -0.29718 | 0.430674 | 0.826896 | C5ar2 |
| ENSMUSG( | 361.477 | -0.00855 | 0.95907 | 0.993937 | C630043F03Rik |
| ENSMUSG( | 28.91667 | 0.592631 | 0.183599 | 0.62067 | C7 |
| ENSMUSG( | 66.5705 | -0.00038 | 0.99873 | 0.99989 | C730002L08Rik |
| ENSMUSG( | 428.8045 | -0.02715 | 0.847442 | 0.972374 | C87436 |
| ENSMUSG( | 36.58809 | -0.04423 | 0.895869 | 0.984551 | C8g |
| ENSMUSG( | 100.7196 | -0.20478 | 0.300934 | 0.749979 | C920006O11Rik |
| ENSMUSG( | 70.33252 | 0.179148 | 0.441582 | 0.832555 | C920021L13Rik |
| ENSMUSG( | 519.8258 | 0.070847 | 0.570642 | 0.893192 | C9orf72 |
| ENSMUSG( | 123.7638 | 0.110852 | 0.563793 | 0.890032 | Caap1 |
| ENSMUSG( | 5179.314 | 0.071359 | 0.404573 | 0.816132 | Cab39 |
| ENSMUSG( | 769.9301 | 0.063713 | 0.563982 | 0.890139 | Cab39l |
| ENSMUSG( | 33.49923 | 0.097174 | 0.771065 | 0.954726 | Cabcoco1 |
| ENSMUSG( | 3209.557 | 0.133893 | 0.229973 | 0.677964 | Cabin1 |
| ENSMUSG( | 1382.743 | 0.773546 | 4.31E-12 | 3.49E-09 | Cables1 |
| ENSMUSG( | 1018.633 | 0.033349 | 0.754274 | 0.950981 | Cables2 |
| ENSMUSG( | 5144.593 | 0.09903 | 0.366166 | 0.794603 | Cabp1 |
| ENSMUSG( | 3059.425 | 0.42238 | 6.53E-05 | 0.004839 | Cabp7 |
| ENSMUSG( | 25.08696 | 0.009018 | 0.983891 | 0.997794 | Cabyr |
| ENSMUSG( | 1710.801 | -0.1489 | 0.127174 | 0.534725 | Cacfd1 |
| ENSMUSG( | 845.9754 | 0.218721 | 0.053595 | 0.35565 | Cachd1 |
| ENSMUSG( | 5771.441 | -0.02332 | 0.858731 | 0.974707 | Cacna1a |
| ENSMUSG( | 3271.678 | -0.06707 | 0.577209 | 0.895659 | Cacna1b |
| ENSMUSG( | 2370.624 | -0.35408 | 0.001614 | 0.045046 | Cacna1c |
| ENSMUSG( | 1996.976 | 0.050839 | 0.593972 | 0.902869 | Cacna1d |
| ENSMUSG( | 5750.693 | -0.27428 | 0.04483 | 0.324745 | Cacna1e |
| ENSMUSG( | 2794.807 | 0.056094 | 0.623007 | 0.911589 | Cacna1g |
| ENSMUSG( | 2271.04 | 0.023705 | 0.81258 | 0.964574 | Cacna1h |
| ENSMUSG( | 4036.775 | -0.08589 | 0.50499 | 0.866883 | Cacna1i |
| ENSMUSG( | 57.27179 | -0.42924 | 0.144826 | 0.564979 | Cacna1s |
| ENSMUSG( | 4685.632 | -0.4992 | 0.000794 | 0.02774 | Cacna2d1 |
| ENSMUSG( | 1743.304 | -0.37895 | 0.000468 | 0.01982 | Cacna2d2 |
| ENSMUSG( | 2721.097 | -0.19306 | 0.173504 | 0.60901 | Cacna2d3 |
| ENSMUSG( | 267.0449 | -0.37056 | 0.015032 | 0.180217 | Cacna2d4 |
| ENSMUSG( | 3423.557 | 0.108818 | 0.376306 | 0.801127 | Cacnb1 |
| ENSMUSG( | 733.2355 | 0.091123 | 0.470558 | 0.84674 | Cacnb2 |
| ENSMUSG( | 15015.85 | 0.181828 | 0.021671 | 0.225182 | Cacnb3 |
| ENSMUSG( | 3381.728 | -0.05254 | 0.548064 | 0.885153 | Cacnb4 |
| ENSMUSG( | 3021.319 | -0.06287 | 0.590499 | 0.90113 | Cacng2 |
| ENSMUSG( | 3697.407 | -0.30972 | 0.006932 | 0.114527 | Cacng3 |
| ENSMUSG( | 3573.795 | -0.32864 | 0.005037 | 0.094254 | Cacng4 |
| ENSMUSG( | 414.8407 | -0.7748 | 8.98E-07 | 0.000178 | Cacng5 |
| ENSMUSG( | 39.29945 | -0.52342 | 0.091035 | 0.463586 | Cacng6 |
| ENSMUSG( | 14530.68 | 0.010774 | 0.893185 | 0.984512 | Cacng7 |
| ENSMUSG( | 2237.915 | -0.11533 | 0.382519 | 0.803227 | Cacng8 |
| ENSMUSG( | 930.8849 | 0.07269 | 0.445404 | 0.833239 | Cactin |
| ENSMUSG( | 1701.192 | 0.082814 | 0.353919 | 0.786371 | Cacul1 |
| ENSMUSG( | 1302.865 | 0.054008 | 0.638852 | 0.916418 | Cacybp |
| ENSMUSG( | 849.7194 | 0.220377 | 0.047937 | 0.334426 | Cad |
| ENSMUSG( | 7854.368 | -0.2047 | 0.134665 | 0.547984 | Cadm1 |
| ENSMUSG( | 5069.966 | -0.46477 | 0.003204 | 0.070583 | Cadm2 |
| ENSMUSG( | 14140.32 | -0.27636 | 0.057339 | 0.369652 | Cadm3 |

| | | | | | |
|---|---|---|---|---|---|
| ENSMUSG( | 776.6218 | 0.139877 | 0.305533 | 0.898538 | Brpf1 |
| ENSMUSG( | 1330.045 | 0.089055 | 0.453813 | 0.922316 | Brpf3 |
| ENSMUSG( | 27019.28 | 0.09117 | 0.364285 | 0.914223 | Brsk1 |
| ENSMUSG( | 6333.891 | 0.022048 | 0.825648 | 0.979537 | Brsk2 |
| ENSMUSG( | 2948.082 | -0.0497 | 0.64301 | 0.953903 | Brwd1 |
| ENSMUSG( | 293.6991 | 0.180417 | 0.372599 | 0.915491 | Brwd3 |
| ENSMUSG( | 2884.625 | 0.163003 | 0.276625 | 0.890889 | Bscl2 |
| ENSMUSG( | 1934.189 | -0.05828 | 0.577148 | 0.942912 | Bsdc1 |
| ENSMUSG( | 24205.82 | 0.052239 | 0.712391 | 0.964363 | Bsg |
| ENSMUSG( | 14344.2 | 0.26141 | 0.096264 | 0.796133 | Bsn |
| ENSMUSG( | 13.15984 | -0.27226 | 0.611512 | 0.950771 | Bspry |
| ENSMUSG( | 216.9095 | -0.09496 | 0.608517 | 0.950113 | Bst2 |
| ENSMUSG( | 9.61276 | -0.29785 | 0.635859 | NA | Bsx |
| ENSMUSG( | 1043.227 | 0.051608 | 0.693209 | 0.96076 | Btaf1 |
| ENSMUSG( | 5839.311 | -0.20313 | 0.179825 | 0.850372 | Btbd1 |
| ENSMUSG( | 1035.234 | 0.060666 | 0.656509 | 0.955533 | Btbd10 |
| ENSMUSG( | 75.97803 | -0.19782 | 0.426798 | 0.917257 | Btbd16 |
| ENSMUSG( | 1952.478 | -0.11675 | 0.408955 | 0.917257 | Btbd17 |
| ENSMUSG( | 109.5816 | 0.150041 | 0.579171 | 0.943928 | Btbd19 |
| ENSMUSG( | 2608.291 | -0.06797 | 0.498886 | 0.927825 | Btbd2 |
| ENSMUSG( | 4807.323 | 0.173175 | 0.159609 | 0.837724 | Btbd3 |
| ENSMUSG( | 1741.828 | 0.119793 | 0.285234 | 0.89286 | Btbd6 |
| ENSMUSG( | 1027.99 | -0.04902 | 0.705063 | 0.96281 | Btbd7 |
| ENSMUSG( | 992.3819 | -0.03439 | 0.798782 | 0.97681 | Btbd8 |
| ENSMUSG( | 1562.913 | 0.102535 | 0.434365 | 0.91869 | Btbd9 |
| ENSMUSG( | 613.3956 | -0.00202 | 0.989952 | 0.997928 | Btd |
| ENSMUSG( | 1908.602 | -0.20092 | 0.100213 | 0.801311 | Btf3 |
| ENSMUSG( | 750.4989 | -0.04799 | 0.725687 | 0.966887 | Btf3l4 |
| ENSMUSG( | 1097.842 | 0.239138 | 0.046299 | 0.697814 | Btg1 |
| ENSMUSG( | 488.4474 | -0.26062 | 0.329707 | 0.907083 | Btg2 |
| ENSMUSG( | 116.3078 | 0.010742 | 0.959039 | 0.994798 | Btg3 |
| ENSMUSG( | 69.37523 | 0.195641 | 0.478905 | 0.923329 | Btk |
| ENSMUSG( | 23.76072 | -0.00911 | 0.982368 | 0.996756 | Btla |
| ENSMUSG( | 27.93557 | 0.290102 | 0.480969 | 0.924548 | Btn2a2 |
| ENSMUSG( | 1256.906 | 0.007658 | 0.957728 | 0.994798 | Btrc |
| ENSMUSG( | 15.77968 | -0.14088 | 0.774933 | 0.974931 | Bub1 |
| ENSMUSG( | 119.746 | -0.00088 | 0.996965 | 0.999268 | Bub1b |
| ENSMUSG( | 1096.6 | -0.13444 | 0.294643 | 0.898538 | Bub3 |
| ENSMUSG( | 203.2142 | 0.308883 | 0.069615 | 0.75601 | Bud13 |
| ENSMUSG( | 620.0392 | 0.012738 | 0.928541 | 0.99306 | Bud23 |
| ENSMUSG( | 778.5232 | -0.09906 | 0.455696 | 0.923329 | Bud31 |
| ENSMUSG( | 70.93695 | 0.193294 | 0.472373 | 0.923329 | Bves |
| ENSMUSG( | 512.3579 | -0.03551 | 0.790734 | 0.97681 | Bysl |
| ENSMUSG( | 4830.74 | -0.09475 | 0.51688 | 0.930939 | Bzw1 |
| ENSMUSG( | 684.1658 | 0.025466 | 0.851381 | 0.985541 | Bzw2 |
| ENSMUSG( | 683.2867 | -0.11245 | 0.364656 | 0.914223 | C030006K11Rik |
| ENSMUSG( | 26.59682 | -0.45545 | 0.304741 | 0.898538 | C030013G03Rik |
| ENSMUSG( | 227.6962 | -0.38842 | 0.038797 | 0.679972 | C030018K13Rik |
| ENSMUSG( | 104.3541 | -0.31285 | 0.157726 | 0.837724 | C030029H02Rik |
| ENSMUSG( | 90.47491 | -0.07328 | 0.770444 | 0.974236 | C130026L21Rik |
| ENSMUSG( | 89.24779 | 0.00974 | 0.966227 | 0.995475 | C130046K22Rik |
| ENSMUSG( | 11.45781 | 0.678155 | 0.343708 | 0.910026 | C130050O18Rik |
| ENSMUSG( | 1271.4 | -0.00991 | 0.938431 | 0.993863 | C130074G19Rik |
| ENSMUSG( | 513.5101 | -0.0669 | 0.685234 | 0.958678 | C1d |
| ENSMUSG( | 722.5327 | 0.36126 | 0.017857 | 0.536449 | C1galt1 |
| ENSMUSG( | 570.7285 | 0.17703 | 0.261511 | 0.884926 | C1galt1c1 |
| ENSMUSG( | 3651.931 | 0.174004 | 0.299928 | 0.898538 | C1qa |
| ENSMUSG( | 3951.273 | 0.122762 | 0.396757 | 0.917257 | C1qb |
| ENSMUSG( | 1263.156 | -0.09331 | 0.488945 | 0.925573 | C1qbp |
| ENSMUSG( | 3896.786 | 0.130792 | 0.395089 | 0.917257 | C1qc |
| ENSMUSG( | 902.0381 | 0.006065 | 0.970696 | 0.99594 | C1ql1 |
| ENSMUSG( | 521.2064 | 0.106948 | 0.570064 | 0.941777 | C1ql2 |
| ENSMUSG( | 1584.383 | 0.429569 | 0.020687 | 0.569765 | C1ql3 |
| ENSMUSG( | 138.2704 | 0.200406 | 0.479949 | 0.924179 | C1qtnf1 |
| ENSMUSG( | 411.5661 | 0.237027 | 0.091262 | 0.791889 | C1qtnf12 |
| ENSMUSG( | 45.1347 | -0.30811 | 0.338477 | 0.907972 | C1qtnf2 |
| ENSMUSG( | 7920.143 | -0.03416 | 0.786774 | 0.976303 | C1qtnf4 |
| ENSMUSG( | 254.9988 | -0.04662 | 0.789418 | 0.976593 | C1qtnf6 |
| ENSMUSG( | 26.85346 | -0.19006 | 0.631314 | 0.953251 | C1qtnf7 |
| ENSMUSG( | 8.415866 | 0.424837 | 0.551543 | NA | C1qtnf9 |
| ENSMUSG( | 65.91216 | -0.02196 | 0.936377 | 0.993573 | C1ra |
| ENSMUSG( | 20.74246 | -0.25674 | 0.547597 | 0.938063 | C1rl |
| ENSMUSG( | 39.05865 | 0.192992 | 0.568994 | 0.941777 | C1s1 |
| ENSMUSG( | 51.95573 | 0.102796 | 0.725973 | 0.966887 | C2 |
| ENSMUSG( | 11.54094 | -0.15693 | 0.810543 | 0.978779 | C230014O12Rik |
| ENSMUSG( | 38.18246 | 0.116777 | 0.732849 | 0.967228 | C230035I16Rik |
| ENSMUSG( | 14.49344 | 0.451494 | 0.405693 | 0.917257 | C230072F16Rik |
| ENSMUSG( | 466.8261 | 0.473718 | 0.02276 | 0.582823 | C2cd2 |
| ENSMUSG( | 6959.859 | 0.224897 | 0.116058 | 0.813384 | C2cd2l |
| ENSMUSG( | 808.9763 | 0.147082 | 0.296432 | 0.898538 | C2cd3 |
| ENSMUSG( | 41.73846 | -0.42719 | 0.198922 | 0.863016 | C2cd4b |
| ENSMUSG( | 388.5549 | 0.00182 | 0.989723 | 0.997928 | C2cd4c |
| ENSMUSG( | 11.69856 | 0.114458 | 0.842276 | 0.983044 | C2cd4d |
| ENSMUSG( | 1721.875 | -0.06839 | 0.667175 | 0.956766 | C2cd5 |
| ENSMUSG( | 34.36815 | -0.70221 | 0.149098 | 0.836241 | C3 |
| ENSMUSG( | 30.29999 | -0.3753 | 0.32099 | 0.904019 | C330013E15Rik |
| ENSMUSG( | 350.1339 | -0.21491 | 0.211734 | 0.864412 | C330018D20Rik |
| ENSMUSG( | 109.5002 | 0.10875 | 0.701429 | 0.961945 | C3ar1 |
| ENSMUSG( | 47.83648 | 0.454684 | 0.213195 | 0.864412 | C4a |
| ENSMUSG( | 695.6347 | 2.10844 | 7.36E-46 | 6.45E-42 | C4b |
| ENSMUSG( | 131.2693 | -0.2685 | 0.173787 | 0.844317 | C530005A16Rik |
| ENSMUSG( | 10.74605 | 0.092806 | 0.878442 | NA | C530044C16Rik |
| ENSMUSG( | 43.57808 | 0.651755 | 0.084325 | 0.778664 | C5ar1 |
| ENSMUSG( | 47.97678 | 0.468221 | 0.229676 | 0.868507 | C5ar2 |
| ENSMUSG( | 304.7038 | -0.24558 | 0.152575 | 0.837724 | C630043F03Rik |
| ENSMUSG( | 19.94579 | -0.18628 | 0.702439 | 0.962215 | C7 |
| ENSMUSG( | 75.54433 | 0.560046 | 0.02875 | 0.614171 | C730002L08Rik |
| ENSMUSG( | 388.0243 | -0.047 | 0.767559 | 0.974236 | C87436 |
| ENSMUSG( | 33.92481 | 0.019502 | 0.956729 | 0.994798 | C8g |
| ENSMUSG( | 91.3734 | -0.2013 | 0.405385 | 0.917257 | C920006O11Rik |
| ENSMUSG( | 66.5766 | 0.269228 | 0.316355 | 0.901868 | C920021L13Rik |
| ENSMUSG( | 468.755 | 0.042074 | 0.759712 | 0.972189 | C9orf72 |
| ENSMUSG( | 113.1393 | 0.122836 | 0.564857 | 0.941206 | Caap1 |
| ENSMUSG( | 4565.946 | -0.02509 | 0.850103 | 0.985538 | Cab39 |
| ENSMUSG( | 677.4717 | -0.03784 | 0.769312 | 0.974236 | Cab39l |
| ENSMUSG( | 35.94151 | 0.516852 | 0.122193 | 0.813384 | Cabcoco1 |
| ENSMUSG( | 2770.24 | -0.01786 | 0.876854 | 0.989644 | Cabin1 |
| ENSMUSG( | 1133.2 | 0.523385 | 7.16E-05 | 0.01665 | Cables1 |
| ENSMUSG( | 918.2876 | 0.006184 | 0.962787 | 0.995287 | Cables2 |
| ENSMUSG( | 4461.937 | -0.04302 | 0.70974 | 0.96416 | Cabp1 |
| ENSMUSG( | 2593.658 | 0.23898 | 0.079579 | 0.774721 | Cabp7 |
| ENSMUSG( | 21.19851 | -0.21753 | 0.642452 | 0.953903 | Cabyr |
| ENSMUSG( | 1688.621 | 0.084828 | 0.428793 | 0.917257 | Cacfd1 |
| ENSMUSG( | 886.7048 | 0.578282 | 9.81E-05 | 0.02045 | Cachd1 |
| ENSMUSG( | 5780.608 | 0.243726 | 0.121677 | 0.813384 | Cacna1a |
| ENSMUSG( | 3376.961 | 0.283437 | 0.038399 | 0.679434 | Cacna1b |
| ENSMUSG( | 2554.193 | 0.151244 | 0.249792 | 0.877904 | Cacna1c |
| ENSMUSG( | 1901.367 | 0.173788 | 0.116894 | 0.813384 | Cacna1d |
| ENSMUSG( | 6176.594 | 0.209655 | 0.175377 | 0.845305 | Cacna1e |
| ENSMUSG( | 2613.086 | 0.130235 | 0.308175 | 0.898538 | Cacna1g |
| ENSMUSG( | 2240.249 | 0.244688 | 0.024431 | 0.590702 | Cacna1h |
| ENSMUSG( | 4037.103 | 0.181957 | 0.251501 | 0.879838 | Cacna1i |
| ENSMUSG( | 51.35895 | -0.4993 | 0.082098 | 0.77681 | Cacna1s |
| ENSMUSG( | 5377.648 | 0.203789 | 0.245514 | 0.877904 | Cacna2d1 |
| ENSMUSG( | 1794.138 | -0.00093 | 0.994055 | 0.998813 | Cacna2d2 |
| ENSMUSG( | 2796.196 | 0.161822 | 0.37238 | 0.915491 | Cacna2d3 |
| ENSMUSG( | 240.1972 | -0.40017 | 0.027179 | 0.604183 | Cacna2d4 |
| ENSMUSG( | 2886.354 | -0.11664 | 0.410852 | 0.917257 | Cacnb1 |
| ENSMUSG( | 714.339 | 0.271025 | 0.062374 | 0.736482 | Cacnb2 |
| ENSMUSG( | 13033.79 | 0.047877 | 0.613441 | 0.951177 | Cacnb3 |
| ENSMUSG( | 3145.239 | 0.005199 | 0.965187 | 0.995433 | Cacnb4 |
| ENSMUSG( | 3088.533 | 0.259981 | 0.070608 | 0.758714 | Cacng2 |
| ENSMUSG( | 3884.998 | 0.119004 | 0.344699 | 0.91045 | Cacng3 |
| ENSMUSG( | 3651.164 | 0.020032 | 0.884055 | 0.989644 | Cacng4 |
| ENSMUSG( | 501.1664 | 0.136672 | 0.361522 | 0.914223 | Cacng5 |
| ENSMUSG( | 40.6606 | -0.12753 | 0.689923 | 0.960071 | Cacng6 |
| ENSMUSG( | 13133.63 | -0.01327 | 0.905283 | 0.991089 | Cacng7 |
| ENSMUSG( | 2287.231 | 0.214793 | 0.136405 | 0.827092 | Cacng8 |
| ENSMUSG( | 842.3573 | 0.055259 | 0.640283 | 0.953903 | Cactin |
| ENSMUSG( | 1515.227 | 0.014207 | 0.923017 | 0.992644 | Cacul1 |
| ENSMUSG( | 1182.529 | 0.037484 | 0.813259 | 0.979198 | Cacybp |
| ENSMUSG( | 725.44 | 0.043683 | 0.733835 | 0.967228 | Cad |
| ENSMUSG( | 7921.725 | 0.096942 | 0.525861 | 0.934374 | Cadm1 |
| ENSMUSG( | 5912.282 | 0.273517 | 0.146981 | 0.836241 | Cadm2 |
| ENSMUSG( | 15139.25 | 0.199963 | 0.218975 | 0.866104 | Cadm3 |
| ENSMUSG( | 17659.77 | -0.05797 | 0.665996 | 0.956766 | Cadm4 |
| ENSMUSG( | 3769.28 | -0.04643 | 0.719216 | 0.966844 | Cadps |
| ENSMUSG( | 1375.244 | 0.099353 | 0.43313 | 0.918434 | Cadps2 |
| ENSMUSG( | 49.43768 | 0.157863 | 0.587501 | 0.944975 | Cage1 |
| ENSMUSG( | 1152.775 | 0.011357 | 0.942748 | 0.994774 | Calb1 |
| ENSMUSG( | 417.3599 | -0.05879 | 0.900405 | 0.990558 | Calb2 |
| ENSMUSG( | 35.97885 | -0.1742 | 0.628392 | 0.953251 | Calca |
| ENSMUSG( | 1754.096 | -0.03492 | 0.779846 | 0.976106 | Calcoco1 |
| ENSMUSG( | 175.4831 | 0.658246 | 0.037878 | 0.6763 | Calcrl |
| ENSMUSG( | 833.0781 | 0.209445 | 0.200492 | 0.864176 | Cald1 |
| ENSMUSG( | 182.5108 | -0.12355 | 0.56436 | 0.941206 | Calhm2 |
| ENSMUSG( | 238.0816 | 0.066421 | 0.711389 | 0.964363 | Calhm5 |
| ENSMUSG( | 9.097958 | -0.83949 | 0.251266 | NA | Calhm6 |

| | | | | | | | | | | | |
|---|---|---|---|---|---|---|---|---|---|---|---|
| ENSMUSG( | Btbd16 | 84.33916 | -0.15266 | 0.488657 | 0.859092 | ENSMUSG( | Btbd16 | 75.97803 | -0.19782 | 0.426798 | 0.917257 |
| ENSMUSG( | Btbd17 | 2044.334 | -0.2647 | 0.040513 | 0.311135 | ENSMUSG( | Btbd17 | 1952.478 | -0.11675 | 0.408955 | 0.917257 |
| ENSMUSG( | Btbd19 | 112.9148 | -0.02482 | 0.923201 | 0.990693 | ENSMUSG( | Btbd19 | 109.5816 | 0.150041 | 0.579171 | 0.943928 |
| ENSMUSG( | Btbd2 | 3053.169 | 0.115542 | 0.167806 | 0.60061 | ENSMUSG( | Btbd2 | 2608.291 | -0.06797 | 0.498886 | 0.927825 |
| ENSMUSG( | Btbd3 | 5079.528 | 0.066198 | 0.523534 | 0.875628 | ENSMUSG( | Btbd3 | 4807.323 | 0.173175 | 0.159609 | 0.837724 |
| ENSMUSG( | Btbd6 | 1906.665 | 0.112108 | 0.263607 | 0.714284 | ENSMUSG( | Btbd6 | 1741.828 | 0.119793 | 0.285234 | 0.89286 |
| ENSMUSG( | Btbd7 | 1117.41 | -0.07617 | 0.505092 | 0.866883 | ENSMUSG( | Btbd7 | 1027.99 | -0.04902 | 0.705063 | 0.96281 |
| ENSMUSG( | Btbd8 | 1061.832 | -0.10826 | 0.391557 | 0.809154 | ENSMUSG( | Btbd8 | 992.3819 | -0.03439 | 0.798782 | 0.97681 |
| ENSMUSG( | Btbd9 | 1754.182 | 0.160522 | 0.120353 | 0.524495 | ENSMUSG( | Btbd9 | 1562.913 | 0.102535 | 0.434365 | 0.91869 |
| ENSMUSG( | Btd | 580.9594 | -0.47235 | 0.001289 | 0.039144 | ENSMUSG( | Btd | 613.3956 | -0.00202 | 0.989952 | 0.997928 |
| ENSMUSG( | Btf3 | 2223.178 | -0.01977 | 0.83146 | 0.969889 | ENSMUSG( | Btf3 | 1908.602 | -0.20092 | 0.100213 | 0.801311 |
| ENSMUSG( | Btf3l4 | 778.2974 | -0.2159 | 0.038797 | 0.303414 | ENSMUSG( | Btf3l4 | 750.4989 | -0.04799 | 0.725687 | 0.966887 |
| ENSMUSG( | Btg1 | 1045.442 | -0.16552 | 0.124822 | 0.531813 | ENSMUSG( | Btg1 | 1097.842 | 0.239138 | 0.046299 | 0.697814 |
| ENSMUSG( | Btg2 | 461.001 | -0.7984 | 0.00126 | 0.038624 | ENSMUSG( | Btg2 | 488.4474 | -0.26062 | 0.329707 | 0.907083 |
| ENSMUSG( | Btg3 | 119.2653 | -0.18563 | 0.344962 | 0.779153 | ENSMUSG( | Btg3 | 116.3078 | 0.010742 | 0.959039 | 0.994798 |
| ENSMUSG( | Btk | 69.97366 | -0.04813 | 0.867949 | 0.976631 | ENSMUSG( | Btk | 69.37523 | 0.195641 | 0.478905 | 0.923329 |
| ENSMUSG( | Btla | 25.95297 | -0.02131 | 0.955238 | 0.993874 | ENSMUSG( | Btla | 23.76072 | -0.00911 | 0.982368 | 0.996756 |
| ENSMUSG( | Btn2a2 | 24.88298 | -0.34829 | 0.400382 | 0.813717 | ENSMUSG( | Btn2a2 | 27.93557 | 0.290102 | 0.480969 | 0.924548 |
| ENSMUSG( | Btrc | 1445.504 | 0.137684 | 0.280126 | 0.729155 | ENSMUSG( | Btrc | 1256.906 | 0.007658 | 0.957728 | 0.994798 |
| ENSMUSG( | Bub1 | 15.23505 | -0.57618 | 0.232751 | NA | ENSMUSG( | Bub1 | 15.77968 | -0.14088 | 0.774933 | 0.974931 |
| ENSMUSG( | Bub1b | 120.0572 | -0.26005 | 0.207088 | 0.649503 | ENSMUSG( | Bub1b | 119.746 | -0.00088 | 0.996965 | 0.999268 |
| ENSMUSG( | Bub3 | 1254.465 | -0.0105 | 0.924026 | 0.991077 | ENSMUSG( | Bub3 | 1096.6 | -0.13444 | 0.294643 | 0.898538 |
| ENSMUSG( | Bud13 | 224.4843 | 0.332338 | 0.031651 | 0.271198 | ENSMUSG( | Bud13 | 203.2142 | 0.308883 | 0.069615 | 0.75601 |
| ENSMUSG( | Bud23 | 700.7006 | 0.098117 | 0.392466 | 0.809274 | ENSMUSG( | Bud23 | 620.0392 | 0.012738 | 0.928541 | 0.99306 |
| ENSMUSG( | Bud31 | 902.5369 | 0.065495 | 0.512693 | 0.871762 | ENSMUSG( | Bud31 | 778.5232 | -0.09906 | 0.455696 | 0.923329 |
| ENSMUSG( | Bves | 67.72327 | -0.21546 | 0.413252 | 0.819541 | ENSMUSG( | Bves | 70.93695 | 0.193294 | 0.472373 | 0.923329 |
| ENSMUSG( | Bysl | 616.8054 | 0.224815 | 0.04662 | 0.330354 | ENSMUSG( | Bysl | 512.3579 | -0.03551 | 0.790734 | 0.97681 |
| ENSMUSG( | Bzw1 | 5494.548 | 0.012709 | 0.905761 | 0.986694 | ENSMUSG( | Bzw1 | 4830.74 | -0.09475 | 0.51688 | 0.930939 |
| ENSMUSG( | Bzw2 | 762.1943 | 0.071029 | 0.52708 | 0.877386 | ENSMUSG( | Bzw2 | 684.1658 | 0.025466 | 0.851381 | 0.985541 |
| ENSMUSG( | C030006K1 | 765.7579 | -0.05307 | 0.598049 | 0.903724 | ENSMUSG( | C030006K1 | 683.2867 | -0.11245 | 0.364656 | 0.914223 |
| ENSMUSG( | C030013G( | 31.04554 | -0.21025 | 0.590205 | 0.90113 | ENSMUSG( | C030013G( | 26.59682 | -0.45545 | 0.304741 | 0.898538 |
| ENSMUSG( | C030018K1 | 279.3142 | -0.03032 | 0.845757 | 0.972191 | ENSMUSG( | C030018K1 | 227.6962 | -0.38842 | 0.038797 | 0.679972 |
| ENSMUSG( | C030029H( | 105.7652 | -0.58699 | 0.004024 | 0.081586 | ENSMUSG( | C030029H( | 104.3541 | -0.31285 | 0.157726 | 0.837724 |
| ENSMUSG( | C130026L2 | 108.5211 | 0.164859 | 0.406487 | 0.816818 | ENSMUSG( | C130026L2 | 90.47491 | -0.07328 | 0.770444 | 0.974236 |
| ENSMUSG( | C130046K2 | 108.7582 | 0.293151 | 0.133575 | 0.546843 | ENSMUSG( | C130046K2 | 89.24779 | 0.00974 | 0.966227 | 0.995475 |
| ENSMUSG( | C130050O1 | 10.51045 | 0.148445 | 0.806609 | NA | ENSMUSG( | C130050O1 | 11.45781 | 0.678155 | 0.343708 | 0.910026 |
| ENSMUSG( | C130074G1 | 1386.082 | -0.03389 | 0.775592 | 0.955606 | ENSMUSG( | C130074G1 | 1271.4 | -0.00991 | 0.938431 | 0.993863 |
| ENSMUSG( | C1d | 577.5213 | 0.012085 | 0.920804 | 0.990409 | ENSMUSG( | C1d | 513.5101 | -0.0669 | 0.685234 | 0.958678 |
| ENSMUSG( | C1galt1 | 687.804 | -0.0275 | 0.839584 | 0.971179 | ENSMUSG( | C1galt1 | 722.5327 | 0.36126 | 0.017857 | 0.536449 |
| ENSMUSG( | C1galt1c1 | 526.5545 | -0.34468 | 0.007417 | 0.118765 | ENSMUSG( | C1galt1c1 | 570.7285 | 0.17703 | 0.261511 | 0.884926 |
| ENSMUSG( | C1qa | 3698.841 | -0.05792 | 0.673404 | 0.929031 | ENSMUSG( | C1qa | 3651.931 | 0.174004 | 0.299928 | 0.898538 |
| ENSMUSG( | C1qb | 4076.99 | -0.05668 | 0.648055 | 0.919798 | ENSMUSG( | C1qb | 3951.273 | 0.122762 | 0.396757 | 0.917257 |
| ENSMUSG( | C1qbp | 1399.791 | -0.0603 | 0.555587 | 0.887457 | ENSMUSG( | C1qbp | 1263.156 | -0.09331 | 0.488945 | 0.925573 |
| ENSMUSG( | C1qc | 3874.267 | -0.16004 | 0.21764 | 0.662824 | ENSMUSG( | C1qc | 3896.786 | 0.130792 | 0.395089 | 0.917257 |
| ENSMUSG( | C1ql1 | 860.102 | -0.43259 | 0.002322 | 0.05646 | ENSMUSG( | C1ql1 | 902.0381 | 0.006065 | 0.970696 | 0.99594 |
| ENSMUSG( | C1ql2 | 492.601 | -0.35188 | 0.056359 | 0.366006 | ENSMUSG( | C1ql2 | 521.2064 | 0.106948 | 0.570064 | 0.941777 |
| ENSMUSG( | C1ql3 | 1307.459 | -0.39554 | 0.026528 | 0.249008 | ENSMUSG( | C1ql3 | 1584.383 | 0.429569 | 0.020687 | 0.569765 |
| ENSMUSG( | C1qtnf1 | 125.2449 | -0.39386 | 0.108583 | 0.497529 | ENSMUSG( | C1qtnf1 | 138.2704 | 0.200406 | 0.479949 | 0.924179 |
| ENSMUSG( | C1qtnf12 | 387.1884 | -0.20987 | 0.119859 | 0.523965 | ENSMUSG( | C1qtnf12 | 411.5661 | 0.237027 | 0.091262 | 0.791889 |
| ENSMUSG( | C1qtnf2 | 51.90828 | -0.18941 | 0.491957 | 0.860297 | ENSMUSG( | C1qtnf2 | 45.1347 | -0.30811 | 0.338477 | 0.907972 |
| ENSMUSG( | C1qtnf4 | 7891.826 | -0.3313 | 0.00027 | 0.013409 | ENSMUSG( | C1qtnf4 | 7920.143 | -0.03416 | 0.786774 | 0.976303 |
| ENSMUSG( | C1qtnf6 | 258.7766 | -0.29493 | 0.071723 | 0.413564 | ENSMUSG( | C1qtnf6 | 254.9988 | -0.04662 | 0.789418 | 0.976593 |
| ENSMUSG( | C1qtnf7 | 32.12204 | 0.043985 | 0.902177 | 0.985633 | ENSMUSG( | C1qtnf7 | 26.85346 | -0.19006 | 0.631314 | 0.953251 |
| ENSMUSG( | C1ra | 66.83072 | -0.29147 | 0.212096 | 0.655777 | ENSMUSG( | C1ra | 65.91216 | -0.02196 | 0.936377 | 0.993573 |
| ENSMUSG( | C1rl | 23.04258 | -0.24142 | 0.551761 | 0.886141 | ENSMUSG( | C1rl | 20.74246 | -0.25674 | 0.547597 | 0.938063 |
| ENSMUSG( | C1s1 | 38.39586 | -0.14865 | 0.710533 | 0.939748 | ENSMUSG( | C1s1 | 39.05865 | 0.192992 | 0.568994 | 0.941777 |
| ENSMUSG( | C2 | 51.46536 | -0.18023 | 0.510862 | 0.870755 | ENSMUSG( | C2 | 51.95573 | 0.102796 | 0.725973 | 0.966887 |
| ENSMUSG( | C230014O1 | 12.73244 | -0.10548 | 0.856879 | NA | ENSMUSG( | C230014O1 | 11.54094 | -0.15693 | 0.810543 | 0.978779 |
| ENSMUSG( | C230035I1( | 43.33326 | 0.198427 | 0.511424 | 0.870898 | ENSMUSG( | C230035I1( | 38.18246 | 0.116777 | 0.732849 | 0.967228 |
| ENSMUSG( | C230072F1 | 15.466 | 0.350362 | 0.518109 | NA | ENSMUSG( | C230072F1 | 14.49344 | 0.451494 | 0.405693 | 0.917257 |
| ENSMUSG( | C2cd2 | 424.0269 | -0.04552 | 0.780732 | 0.957491 | ENSMUSG( | C2cd2 | 466.8261 | 0.473718 | 0.02276 | 0.582823 |
| ENSMUSG( | C2cd2l | 6400.42 | -0.29603 | 0.01793 | 0.200989 | ENSMUSG( | C2cd2l | 6959.859 | 0.224897 | 0.116058 | 0.813384 |
| ENSMUSG( | C2cd3 | 906.5645 | 0.198467 | 0.052094 | 0.350316 | ENSMUSG( | C2cd3 | 808.9763 | 0.147082 | 0.296432 | 0.898538 |
| ENSMUSG( | C2cd4b | 58.12102 | 0.290144 | 0.325151 | 0.770608 | ENSMUSG( | C2cd4b | 41.73846 | -0.42719 | 0.198922 | 0.863016 |
| ENSMUSG( | C2cd4c | 467.4771 | 0.253868 | 0.02927 | 0.259574 | ENSMUSG( | C2cd4c | 388.5549 | 0.00182 | 0.989723 | 0.997928 |
| ENSMUSG( | C2cd4d | 11.62342 | -0.18674 | 0.732335 | NA | ENSMUSG( | C2cd4d | 11.69856 | 0.114458 | 0.842276 | 0.983044 |
| ENSMUSG( | C2cd5 | 2089.696 | 0.218698 | 0.09595 | 0.475936 | ENSMUSG( | C2cd5 | 1721.875 | -0.06839 | 0.667175 | 0.956766 |
| ENSMUSG( | C3 | 38.59524 | -0.59891 | 0.192995 | 0.632208 | ENSMUSG( | C3 | 34.36815 | -0.70221 | 0.149098 | 0.836241 |
| ENSMUSG( | C330013E1 | 34.45003 | -0.29445 | 0.373937 | 0.800445 | ENSMUSG( | C330013E1 | 30.29999 | -0.3753 | 0.32099 | 0.904019 |
| ENSMUSG( | C330018D2 | 441.3964 | 0.197062 | 0.161902 | 0.592398 | ENSMUSG( | C330018D2 | 350.1339 | -0.21491 | 0.211734 | 0.864412 |
| ENSMUSG( | C3ar1 | 104.9212 | -0.31787 | 0.172646 | 0.608104 | ENSMUSG( | C3ar1 | 109.5002 | 0.10875 | 0.701429 | 0.961945 |
| ENSMUSG( | C4a | 53.87437 | 0.511094 | 0.10804 | 0.497059 | ENSMUSG( | C4a | 47.83648 | 0.454684 | 0.213195 | 0.864412 |
| ENSMUSG( | C4b | 560.3643 | 1.532107 | 6.23E-27 | 1.26E-23 | ENSMUSG( | C4b | 695.6347 | 2.10844 | 7.36E-46 | 6.45E-42 |
| ENSMUSG( | C530005A1 | 150.1769 | -0.13323 | 0.470646 | 0.84674 | ENSMUSG( | C530005A1 | 131.2693 | -0.2685 | 0.173787 | 0.844317 |
| ENSMUSG( | C530044C1 | 10.5206 | -0.31484 | 0.585646 | NA | ENSMUSG( | C530044C1 | 10.74605 | 0.092806 | 0.878442 | NA |
| ENSMUSG( | C5ar1 | 37.83762 | 0.023032 | 0.94969 | 0.993821 | ENSMUSG( | C5ar1 | 43.57808 | 0.651755 | 0.084325 | 0.778664 |
| ENSMUSG( | C5ar2 | 40.48409 | -0.29718 | 0.430674 | 0.826896 | ENSMUSG( | C5ar2 | 47.97678 | 0.468221 | 0.229676 | 0.868507 |
| ENSMUSG( | C630043F0 | 361.477 | -0.00855 | 0.95907 | 0.993937 | ENSMUSG( | C630043F0 | 304.7038 | -0.24558 | 0.152575 | 0.837724 |
| ENSMUSG( | C7 | 28.91667 | 0.592631 | 0.183599 | 0.62067 | ENSMUSG( | C7 | 19.94579 | -0.18628 | 0.702439 | 0.962215 |
| ENSMUSG( | C730002L0 | 66.5705 | -0.00038 | 0.99873 | 0.99989 | ENSMUSG( | C730002L0 | 75.54433 | 0.560046 | 0.02875 | 0.614171 |
| ENSMUSG( | C87436 | 428.8045 | -0.02715 | 0.847442 | 0.972374 | ENSMUSG( | C87436 | 388.0243 | -0.047 | 0.767559 | 0.974236 |
| ENSMUSG( | C8g | 36.58809 | -0.04423 | 0.895869 | 0.984551 | ENSMUSG( | C8g | 33.92481 | 0.019502 | 0.956729 | 0.994798 |
| ENSMUSG( | C920006O1 | 100.7196 | -0.20478 | 0.300934 | 0.749979 | ENSMUSG( | C920006O1 | 91.3734 | -0.2013 | 0.405385 | 0.917257 |
| ENSMUSG( | C920021L1 | 70.33252 | 0.179148 | 0.441582 | 0.832555 | ENSMUSG( | C920021L1 | 66.5766 | 0.269228 | 0.316355 | 0.901868 |
| ENSMUSG( | C9orf72 | 519.8258 | 0.070847 | 0.570642 | 0.893192 | ENSMUSG( | C9orf72 | 468.755 | 0.042074 | 0.759712 | 0.972189 |
| ENSMUSG( | Caap1 | 123.7638 | 0.110852 | 0.563793 | 0.890032 | ENSMUSG( | Caap1 | 113.1393 | 0.122836 | 0.564857 | 0.941206 |
| ENSMUSG( | Cab39 | 5179.314 | 0.071359 | 0.404573 | 0.816132 | ENSMUSG( | Cab39 | 4565.946 | -0.02509 | 0.850103 | 0.985538 |
| ENSMUSG( | Cab39l | 769.9301 | 0.063713 | 0.563982 | 0.890139 | ENSMUSG( | Cab39l | 677.4717 | -0.03784 | 0.769312 | 0.974236 |
| ENSMUSG( | Cabcoco1 | 33.49923 | 0.097174 | 0.771065 | 0.954726 | ENSMUSG( | Cabcoco1 | 35.94151 | 0.516852 | 0.122193 | 0.813384 |
| ENSMUSG( | Cabin1 | 3209.557 | 0.133893 | 0.229973 | 0.677964 | ENSMUSG( | Cabin1 | 2770.24 | -0.01786 | 0.876854 | 0.989644 |
| ENSMUSG( | Cables1 | 1382.743 | 0.773546 | 4.31E-12 | 3.49E-09 | ENSMUSG( | Cables1 | 1133.2 | 0.523385 | 7.16E-05 | 0.01665 |
| ENSMUSG( | Cables2 | 1018.633 | 0.033349 | 0.754274 | 0.950981 | ENSMUSG( | Cables2 | 918.2876 | 0.006184 | 0.962787 | 0.995287 |
| ENSMUSG( | Cabp1 | 5144.593 | 0.09903 | 0.366166 | 0.794603 | ENSMUSG( | Cabp1 | 4461.937 | -0.04302 | 0.70974 | 0.96416 |
| ENSMUSG( | Cabp7 | 3059.425 | 0.42238 | 6.53E-05 | 0.004839 | ENSMUSG( | Cabp7 | 2593.658 | 0.23898 | 0.079579 | 0.774721 |
| ENSMUSG( | Cabyr | 25.08696 | 0.009018 | 0.983891 | 0.997794 | ENSMUSG( | Cabyr | 21.19851 | -0.21753 | 0.642452 | 0.953903 |
| ENSMUSG( | Cacfd1 | 1710.801 | -0.1489 | 0.127174 | 0.534725 | ENSMUSG( | Cacfd1 | 1688.621 | 0.084828 | 0.428793 | 0.917257 |
| ENSMUSG( | Cachd1 | 845.9754 | 0.218721 | 0.053595 | 0.35565 | ENSMUSG( | Cachd1 | 886.7048 | 0.578282 | 9.81E-05 | 0.02045 |
| ENSMUSG( | Cacna1a | 5771.441 | -0.02332 | 0.858731 | 0.974707 | ENSMUSG( | Cacna1a | 5780.608 | 0.243726 | 0.121677 | 0.813384 |
| ENSMUSG( | Cacna1b | 3271.678 | -0.06707 | 0.577209 | 0.895659 | ENSMUSG( | Cacna1b | 3376.961 | 0.283437 | 0.038399 | 0.679434 |
| ENSMUSG( | Cacna1c | 2370.624 | -0.35408 | 0.001614 | 0.045046 | ENSMUSG( | Cacna1c | 2554.193 | 0.151244 | 0.249792 | 0.877904 |
| ENSMUSG( | Cacna1d | 1996.976 | 0.050839 | 0.593972 | 0.902869 | ENSMUSG( | Cacna1d | 1901.367 | 0.173788 | 0.116894 | 0.813384 |
| ENSMUSG( | Cacna1e | 5750.693 | -0.27428 | 0.04483 | 0.324745 | ENSMUSG( | Cacna1e | 6176.594 | 0.209655 | 0.175377 | 0.845305 |
| ENSMUSG( | Cacna1g | 2794.807 | 0.056094 | 0.623007 | 0.911589 | ENSMUSG( | Cacna1g | 2613.086 | 0.130235 | 0.308175 | 0.898538 |
| ENSMUSG( | Cacna1h | 2271.04 | 0.023705 | 0.81258 | 0.964574 | ENSMUSG( | Cacna1h | 2240.249 | 0.244688 | 0.024431 | 0.590702 |
| ENSMUSG( | Cacna1i | 4036.775 | -0.08589 | 0.50499 | 0.866883 | ENSMUSG( | Cacna1i | 4037.103 | 0.181957 | 0.251501 | 0.879838 |
| ENSMUSG( | Cacna1s | 57.27179 | -0.42924 | 0.144826 | 0.564979 | ENSMUSG( | Cacna1s | 51.35895 | -0.4993 | 0.082098 | 0.77681 |
| ENSMUSG( | Cacna2d1 | 4685.632 | -0.4992 | 0.000794 | 0.02774 | ENSMUSG( | Cacna2d1 | 5377.648 | 0.203789 | 0.245514 | 0.877904 |
| ENSMUSG( | Cacna2d2 | 1743.304 | -0.37895 | 0.000468 | 0.01982 | ENSMUSG( | Cacna2d2 | 1794.138 | -0.00093 | 0.994055 | 0.998813 |
| ENSMUSG( | Cacna2d3 | 2721.097 | -0.19306 | 0.173504 | 0.60901 | ENSMUSG( | Cacna2d3 | 2796.196 | 0.161822 | 0.37238 | 0.915491 |
| ENSMUSG( | Cacna2d4 | 267.0449 | -0.37056 | 0.015032 | 0.180217 | ENSMUSG( | Cacna2d4 | 240.1972 | -0.40017 | 0.027179 | 0.604183 |
| ENSMUSG( | Cacnb1 | 3423.557 | 0.108818 | 0.376306 | 0.801127 | ENSMUSG( | Cacnb1 | 2886.354 | -0.11664 | 0.410852 | 0.917257 |
| ENSMUSG( | Cacnb2 | 733.2355 | 0.091123 | 0.470558 | 0.84674 | ENSMUSG( | Cacnb2 | 714.339 | 0.271025 | 0.062374 | 0.736482 |
| ENSMUSG( | Cacnb3 | 15015.85 | 0.181828 | 0.021671 | 0.225182 | ENSMUSG( | Cacnb3 | 13033.79 | 0.047877 | 0.613441 | 0.951177 |
| ENSMUSG( | Cacnb4 | 3381.728 | -0.05254 | 0.548064 | 0.885153 | ENSMUSG( | Cacnb4 | 3145.239 | 0.005199 | 0.965187 | 0.995433 |
| ENSMUSG( | Cacng2 | 3021.319 | -0.06287 | 0.590499 | 0.90113 | ENSMUSG( | Cacng2 | 3088.533 | 0.259981 | 0.070608 | 0.758714 |
| ENSMUSG( | Cacng3 | 3697.407 | -0.30972 | 0.006932 | 0.114527 | ENSMUSG( | Cacng3 | 3884.998 | 0.119004 | 0.344699 | 0.91045 |
| ENSMUSG( | Cacng4 | 3573.795 | -0.32864 | 0.005037 | 0.094254 | ENSMUSG( | Cacng4 | 3651.164 | 0.020032 | 0.884055 | 0.989644 |
| ENSMUSG( | Cacng5 | 414.8407 | -0.7748 | 8.98E-07 | 0.000178 | ENSMUSG( | Cacng5 | 501.1664 | 0.136672 | 0.361522 | 0.914223 |
| ENSMUSG( | Cacng6 | 39.29945 | -0.52342 | 0.091035 | 0.463586 | ENSMUSG( | Cacng6 | 40.6606 | -0.12753 | 0.689923 | 0.960071 |
| ENSMUSG( | Cacng7 | 14530.68 | 0.010774 | 0.893185 | 0.984512 | ENSMUSG( | Cacng7 | 13133.63 | -0.01327 | 0.905283 | 0.991089 |
| ENSMUSG( | Cacng8 | 2237.915 | -0.11533 | 0.382519 | 0.803227 | ENSMUSG( | Cacng8 | 2287.231 | 0.214793 | 0.136405 | 0.827092 |
| ENSMUSG( | Cactin | 930.8849 | 0.07269 | 0.445404 | 0.833239 | ENSMUSG( | Cactin | 842.3573 | 0.055259 | 0.640283 | 0.953903 |
| ENSMUSG( | Cacul1 | 1701.192 | 0.082814 | 0.353919 | 0.786371 | ENSMUSG( | Cacul1 | 1515.227 | 0.014207 | 0.923017 | 0.992644 |
| ENSMUSG( | Cacybp | 1302.865 | 0.054008 | 0.638852 | 0.916418 | ENSMUSG( | Cacybp | 1182.529 | 0.037484 | 0.813259 | 0.979198 |
| ENSMUSG( | Cad | 849.7194 | 0.220377 | 0.047937 | 0.334426 | ENSMUSG( | Cad | 725.44 | 0.043683 | 0.733835 | 0.967228 |
| ENSMUSG( | Cadm1 | 7854.368 | -0.2047 | 0.134665 | 0.547984 | ENSMUSG( | Cadm1 | 7921.725 | 0.096942 | 0.525861 | 0.934374 |
| ENSMUSG( | Cadm2 | 5069.966 | -0.46477 | 0.003204 | 0.070583 | ENSMUSG( | Cadm2 | 5912.282 | 0.273517 | 0.146981 | 0.836241 |
| ENSMUSG( | Cadm3 | 14140.32 | -0.27636 | 0.057339 | 0.369652 | ENSMUSG( | Cadm3 | 15139.25 | 0.199963 | 0.218975 | 0.866104 |
| ENSMUSG( | Cadm4 | 17684.78 | -0.34309 | 0.001235 | 0.038239 | ENSMUSG( | Cadm4 | 17659.77 | -0.05797 | 0.665996 | 0.956766 |
| ENSMUSG( | Cadps | 4321.981 | 0.081543 | 0.443766 | 0.833239 | ENSMUSG( | Cadps | 3769.28 | -0.04643 | 0.719216 | 0.966844 |
| ENSMUSG( | Cadps2 | 1687.474 | 0.394236 | 0.000149 | 0.008522 | ENSMUSG( | Cadps2 | 1375.244 | 0.099353 | 0.43313 | 0.918434 |
| ENSMUSG( | Cage1 | 55.09677 | 0.206744 | 0.419993 | 0.822442 | ENSMUSG( | Cage1 | 49.43768 | 0.157863 | 0.587501 | 0.944975 |
| ENSMUSG( | Calb1 | 1255.38 | -0.00662 | 0.958069 | 0.993874 | ENSMUSG( | Calb1 | 1152.775 | 0.011357 | 0.942748 | 0.994774 |
| ENSMUSG( | Calb2 | 441.1399 | -0.16414 | 0.436065 | 0.829093 | ENSMUSG( | Calb2 | 417.3599 | -0.05879 | 0.900405 | 0.990558 |
| ENSMUSG( | Calca | 33.65182 | -0.74412 | 0.033386 | 0.279991 | ENSMUSG( | Calca | 35.97885 | -0.1742 | 0.628392 | 0.953251 |
| ENSMUSG( | Calcoco1 | 1974.959 | 0.040019 | 0.711711 | 0.940028 | ENSMUSG( | Calcoco1 | 1754.096 | -0.03492 | 0.779846 | 0.976106 |
| ENSMUSG( | Calcrl | 134.5151 | -0.34213 | 0.240073 | 0.687899 | ENSMUSG( | Calcrl | 175.4831 | 0.658246 | 0.037878 | 0.6763 |
| ENSMUSG( | Cald1 | 838.4443 | -0.0395 | 0.787892 | 0.959226 | ENSMUSG( | Cald1 | 833.0781 | 0.209445 | 0.200492 | 0.864176 |
| ENSMUSG( | Calhm2 | 166.7363 | -0.75974 | 0.000198 | 0.010508 | ENSMUSG( | Calhm2 | 182.5108 | -0.12355 | 0.56436 | 0.941206 |
| ENSMUSG( | Calhm5 | 219.2793 | -0.48856 | 0.004244 | 0.084545 | ENSMUSG( | Calhm5 | 238.0816 | 0.066421 | 0.711389 | 0.964363 |
| ENSMUSG( | Calhm6 | 9.941745 | -0.86212 | 0.189638 | NA | ENSMUSG( | Calhm6 | 9.097958 | -0.83949 | 0.251266 | NA |
| ENSMUSG( | Calm1 | 102299.6 | 0.038693 | 0.646299 | 0.918612 | ENSMUSG( | Calm1 | 90250.76 | -0.05439 | 0.609227 | 0.95038 |
| ENSMUSG( | Calm2 | 19205.85 | -0.08498 | 0.559934 | 0.888917 | ENSMUSG( | Calm2 | 16636.84 | -0.24375 | 0.215302 | 0.865167 |
| ENSMUSG( | Calm3 | 57830.19 | 0.130313 | 0.098754 | 0.480598 | ENSMUSG( | Calm3 | 47582.84 | -0.1669 | 0.12541 | 0.81516 |
| ENSMUSG( | Calml4 | 36.80415 | -0.53464 | 0.259862 | 0.709951 | ENSMUSG( | Calml4 | 39.14562 | -0.04862 | 0.903991 | 0.991089 |
| ENSMUSG( | Caln1 | 5136.638 | 0.08586 | 0.406495 | 0.816818 | ENSMUSG( | Caln1 | 4930.57 | 0.23054 | 0.037952 | 0.6763 |
| ENSMUSG( | Calr | 16931.29 | -0.15911 | 0.256196 | 0.706924 | ENSMUSG( | Calr | 16959.03 | 0.118689 | 0.436672 | 0.919879 |
| ENSMUSG( | Calr3 | 44.82494 | -0.05858 | 0.8506 | 0.972638 | ENSMUSG( | Calr3 | 43.12162 | 0.110464 | 0.737817 | 0.967709 |
| ENSMUSG( | Calr4 | 88.59303 | -0.00098 | 0.996559 | 0.99943 | ENSMUSG( | Calr4 | 73.06791 | -0.3263 | 0.227691 | 0.868507 |
| ENSMUSG( | Calu | 5641.362 | -0.16224 | 0.207351 | 0.649561 | ENSMUSG( | Calu | 5502.845 | 0.041064 | 0.772817 | 0.974463 |
| ENSMUSG( | Caly | 4977.821 | -0.37363 | 9.32E-06 | 0.001095 | ENSMUSG( | Caly | 5104.449 | -0.00839 | 0.941619 | 0.994578 |
| ENSMUSG( | Camk1 | 1727.028 | 0.023832 | 0.801458 | 0.961947 | ENSMUSG( | Camk1 | 1562.803 | 0.004894 | 0.96573 | 0.995433 |
| ENSMUSG( | Camk1d | 3348.267 | 0.020518 | 0.802667 | 0.962169 | ENSMUSG( | Camk1d | 2979.687 | -0.04683 | 0.648825 | 0.95422 |
| ENSMUSG( | Camk1g | 842.7267 | 0.291615 | 0.076875 | 0.429306 | ENSMUSG( | Camk1g | 711.334 | 0.080553 | 0.650235 | 0.95422 |
| ENSMUSG( | Camk2a | 213368.9 | 0.195676 | 0.053767 | 0.356103 | ENSMUSG( | Camk2a | 188471.5 | 0.112775 | 0.366007 | 0.914623 |
| ENSMUSG( | Camk2b | 18823.93 | 0.09917 | 0.43902 | 0.830714 | ENSMUSG( | Camk2b | 16360.87 | -0.03634 | 0.794352 | 0.97681 |
| ENSMUSG( | Camk2d | 1882.153 | 0.049598 | 0.651487 | 0.921525 | ENSMUSG( | Camk2d | 1648.856 | -0.06882 | 0.669276 | 0.956766 |
| ENSMUSG( | Camk2g | 8221.592 | 0.040853 | 0.62208 | 0.911432 | ENSMUSG( | Camk2g | 7230.097 | -0.06143 | 0.541436 | 0.937841 |

| | | | | | |
|---|---|---|---|---|---|
| ENSMUSG( | 17684.78 | -0.34309 | 0.001235 | 0.038239 | Cadm4 |
| ENSMUSG( | 4321.981 | 0.081543 | 0.443766 | 0.833239 | Cadps |
| ENSMUSG( | 1687.474 | 0.394236 | 0.000149 | 0.008522 | Cadps2 |
| ENSMUSG( | 55.09677 | 0.206744 | 0.419993 | 0.822442 | Cage1 |
| ENSMUSG( | 1255.38 | -0.00662 | 0.958069 | 0.993874 | Calb1 |
| ENSMUSG( | 441.1399 | -0.16414 | 0.436065 | 0.829093 | Calb2 |
| ENSMUSG( | 33.65182 | -0.74412 | 0.033386 | 0.279991 | Calca |
| ENSMUSG( | 1974.959 | 0.040019 | 0.711711 | 0.940028 | Calcoco1 |
| ENSMUSG( | 134.5151 | -0.34213 | 0.240073 | 0.687899 | Calcrl |
| ENSMUSG( | 838.4443 | -0.0395 | 0.787892 | 0.959226 | Cald1 |
| ENSMUSG( | 166.7363 | -0.75974 | 0.000198 | 0.010508 | Calhm2 |
| ENSMUSG( | 219.2793 | -0.48856 | 0.004244 | 0.084545 | Calhm5 |
| ENSMUSG( | 9.941745 | -0.86212 | 0.189638 | NA | Calhm6 |
| ENSMUSG( | 102299.6 | 0.038693 | 0.646299 | 0.918612 | Calm1 |
| ENSMUSG( | 19205.85 | -0.08498 | 0.559934 | 0.888917 | Calm2 |
| ENSMUSG( | 57830.19 | 0.130313 | 0.098754 | 0.480598 | Calm3 |
| ENSMUSG( | 36.80415 | -0.53464 | 0.259862 | 0.709951 | Calml4 |
| ENSMUSG( | 5136.638 | 0.08586 | 0.406495 | 0.816818 | Caln1 |
| ENSMUSG( | 16931.29 | -0.15911 | 0.256196 | 0.706924 | Calr |
| ENSMUSG( | 44.82494 | -0.05858 | 0.8506 | 0.972638 | Calr3 |
| ENSMUSG( | 88.59303 | -0.00098 | 0.996559 | 0.99943 | Calr4 |
| ENSMUSG( | 5641.362 | -0.16224 | 0.207351 | 0.649561 | Calu |
| ENSMUSG( | 4977.821 | -0.37363 | 9.32E-06 | 0.001095 | Caly |
| ENSMUSG( | 1727.028 | 0.023832 | 0.801458 | 0.961947 | Camk1 |
| ENSMUSG( | 3348.267 | 0.020518 | 0.802667 | 0.962169 | Camk1d |
| ENSMUSG( | 842.7267 | 0.291615 | 0.076875 | 0.429306 | Camk1g |
| ENSMUSG( | 213368.9 | 0.195676 | 0.053767 | 0.356103 | Camk2a |
| ENSMUSG( | 18823.93 | 0.09917 | 0.43902 | 0.830714 | Camk2b |
| ENSMUSG( | 1882.153 | 0.049598 | 0.651487 | 0.921525 | Camk2d |
| ENSMUSG( | 8221.592 | 0.040853 | 0.62208 | 0.911432 | Camk2g |
| ENSMUSG( | 164398.7 | 0.140012 | 0.146941 | 0.568875 | Camk2n1 |
| ENSMUSG( | 7665.766 | 0.124182 | 0.112751 | 0.506698 | Camk2n2 |
| ENSMUSG( | 687.3438 | -0.02859 | 0.818008 | 0.965795 | Camk4 |
| ENSMUSG( | 2596.777 | 0.039312 | 0.749973 | 0.949664 | Camkk1 |
| ENSMUSG( | 9832.342 | 0.123198 | 0.132337 | 0.544251 | Camkk2 |
| ENSMUSG( | 215.2044 | 0.062219 | 0.686875 | 0.933041 | Camkmt |
| ENSMUSG( | 14890.07 | 0.204661 | 0.032472 | 0.275758 | Camkv |
| ENSMUSG( | 744.0924 | -0.2341 | 0.146954 | 0.568875 | Caml |
| ENSMUSG( | 29.88576 | -1.36231 | 0.240305 | 0.687899 | Camp |
| ENSMUSG( | 2559.669 | 0.13183 | 0.170279 | 0.605271 | Camsap1 |
| ENSMUSG( | 4182.221 | 0.107828 | 0.166586 | 0.599201 | Camsap2 |
| ENSMUSG( | 930.5956 | 0.235034 | 0.044345 | 0.323404 | Camsap3 |
| ENSMUSG( | 5324.49 | -0.01407 | 0.874133 | 0.979336 | Camta1 |
| ENSMUSG( | 15464.73 | 0.266018 | 0.001574 | 0.04447 | Camta2 |
| ENSMUSG( | 2717.538 | 0.150229 | 0.07484 | 0.424133 | Cand1 |
| ENSMUSG( | 188.9063 | -0.02665 | 0.862926 | 0.975055 | Cand2 |
| ENSMUSG( | 1701.421 | -0.15695 | 0.270048 | 0.719904 | Cant1 |
| ENSMUSG( | 15047.7 | -0.1423 | 0.309306 | 0.75737 | Canx |
| ENSMUSG( | 6313.74 | 0.238776 | 0.007535 | 0.120197 | Cap1 |
| ENSMUSG( | 4737.47 | 0.022882 | 0.831319 | 0.969889 | Cap2 |
| ENSMUSG( | 86.07161 | -0.19446 | 0.386869 | 0.805056 | Capg |
| ENSMUSG( | 542.363 | -0.0566 | 0.644708 | 0.917782 | Capn1 |
| ENSMUSG( | 1016.393 | 0.087056 | 0.422568 | 0.823508 | Capn10 |
| ENSMUSG( | 54.00918 | 1.963999 | 5.03E-06 | 0.000663 | Capn11 |
| ENSMUSG( | 36.94822 | -0.09187 | 0.776605 | 0.955658 | Capn12 |
| ENSMUSG( | 2434.831 | 0.183811 | 0.055462 | 0.362449 | Capn15 |
| ENSMUSG( | 4354.123 | -0.05482 | 0.61019 | 0.908174 | Capn2 |
| ENSMUSG( | 42.98332 | -0.08165 | 0.792613 | 0.95947 | Capn3 |
| ENSMUSG( | 2152.943 | 0.227935 | 0.011282 | 0.153004 | Capn5 |
| ENSMUSG( | 118.666 | 0.064252 | 0.755709 | 0.95146 | Capn6 |
| ENSMUSG( | 1219.308 | 0.034893 | 0.711555 | 0.940028 | Capn7 |
| ENSMUSG( | 15.11449 | -0.66164 | 0.193346 | NA | Capn9 |
| ENSMUSG( | 4583.241 | 0.035009 | 0.694508 | 0.9343 | Capns1 |
| ENSMUSG( | 4900.466 | 0.118105 | 0.170608 | 0.605271 | Caprin1 |
| ENSMUSG( | 276.1663 | -0.0315 | 0.835052 | 0.970426 | Caprin2 |
| ENSMUSG( | 33.71288 | -0.43443 | 0.208952 | 0.650854 | Capsl |
| ENSMUSG( | 731.7152 | 0.172286 | 0.117719 | 0.518015 | Capza1 |
| ENSMUSG( | 6248.976 | 0.084051 | 0.392487 | 0.809274 | Capza2 |
| ENSMUSG( | 6673.776 | 0.074645 | 0.376186 | 0.801127 | Capzb |
| ENSMUSG( | 3420.745 | -0.24179 | 0.049895 | 0.341042 | Car10 |
| ENSMUSG( | 6307.74 | -0.11121 | 0.160752 | 0.590628 | Car11 |
| ENSMUSG( | 486.6326 | -0.66527 | 1.05E-05 | 0.001196 | Car12 |
| ENSMUSG( | 121.7363 | 0.278558 | 0.160204 | 0.589853 | Car13 |
| ENSMUSG( | 570.5012 | -0.48692 | 0.00037 | 0.016819 | Car14 |
| ENSMUSG( | 70.93035 | 0.01181 | 0.962038 | 0.994538 | Car15 |
| ENSMUSG( | 12289.05 | 0.072549 | 0.473213 | 0.84878 | Car2 |
| ENSMUSG( | 21.3548 | -0.77975 | 0.072823 | 0.416499 | Car3 |
| ENSMUSG( | 1662.935 | -0.06672 | 0.633109 | 0.914073 | Car4 |
| ENSMUSG( | 17.0948 | -0.2234 | 0.617548 | NA | Car5a |
| ENSMUSG( | 434.5505 | 0.185877 | 0.211321 | 0.654503 | Car5b |
| ENSMUSG( | 575.3656 | 0.098302 | 0.552665 | 0.886141 | Car7 |
| ENSMUSG( | 1950.493 | 0.010277 | 0.934821 | 0.993231 | Car8 |
| ENSMUSG( | 26.39125 | -0.34949 | 0.444772 | 0.833239 | Car9 |
| ENSMUSG( | 197.6762 | 0.390097 | 0.023523 | 0.234655 | Card10 |
| ENSMUSG( | 13.21745 | -0.40389 | 0.443777 | NA | Card11 |
| ENSMUSG( | 37.58294 | -0.5083 | 0.098581 | 0.480543 | Card14 |
| ENSMUSG( | 285.4312 | -0.33485 | 0.012917 | 0.164193 | Card19 |
| ENSMUSG( | 78.69359 | -0.08654 | 0.693885 | 0.934202 | Card6 |
| ENSMUSG( | 105.0905 | 0.02056 | 0.923167 | 0.990693 | Card9 |
| ENSMUSG( | 359.627 | -0.0171 | 0.898315 | 0.985633 | Carf |
| ENSMUSG( | 4471.925 | -0.00296 | 0.976699 | 0.997428 | Carhsp1 |
| ENSMUSG( | 25.17354 | -0.05372 | 0.887045 | 0.983326 | Carlr |
| ENSMUSG( | 2523.006 | 0.027851 | 0.797922 | 0.960538 | Carm1 |
| ENSMUSG( | 1796.595 | 0.145488 | 0.188542 | 0.627572 | Carmil1 |
| ENSMUSG( | 3657.139 | 0.088371 | 0.343585 | 0.779153 | Carmil2 |
| ENSMUSG( | 1608.104 | 0.49072 | 0.000534 | 0.021434 | Carmil3 |
| ENSMUSG( | 257.7587 | 0.122624 | 0.39847 | 0.812876 | Carnmt1 |
| ENSMUSG( | 463.6033 | -0.28991 | 0.012565 | 0.161972 | Carns1 |
| ENSMUSG( | 1109.364 | 0.121184 | 0.251926 | 0.701908 | Cars1 |
| ENSMUSG( | 383.4367 | 0.145082 | 0.261056 | 0.710889 | Cars2 |
| ENSMUSG( | 282.2909 | -0.5223 | 0.005537 | 0.099488 | Cartpt |
| ENSMUSG( | 1713.044 | 0.107653 | 0.253899 | 0.704011 | Casc3 |
| ENSMUSG( | 1831.686 | -0.47051 | 0.001691 | 0.046443 | Casd1 |
| ENSMUSG( | 3167.146 | -0.07984 | 0.416154 | 0.82084 | Cask |
| ENSMUSG( | 8181.896 | 0.125865 | 0.102621 | 0.488148 | Caskin1 |
| ENSMUSG( | 1952.8 | -0.0945 | 0.350219 | 0.781943 | Caskin2 |
| ENSMUSG( | 46.42063 | -0.24653 | 0.441689 | 0.832577 | Casp1 |
| ENSMUSG( | 22.02305 | -0.01871 | 0.963658 | 0.994922 | Casp12 |
| ENSMUSG( | 321.3919 | -0.16139 | 0.207104 | 0.649503 | Casp2 |
| ENSMUSG( | 561.9068 | 0.018689 | 0.878527 | 0.981431 | Casp3 |
| ENSMUSG( | 12.29211 | -0.7792 | 0.151716 | NA | Casp4 |
| ENSMUSG( | 183.3604 | 0.021609 | 0.89021 | 0.983687 | Casp6 |
| ENSMUSG( | 120.4299 | -0.04836 | 0.805816 | 0.962616 | Casp7 |
| ENSMUSG( | 137.4956 | -0.07334 | 0.670846 | 0.928741 | Casp8 |
| ENSMUSG( | 342.4549 | 0.043095 | 0.728177 | 0.944355 | Casp8ap2 |
| ENSMUSG( | 850.8353 | 0.028215 | 0.772179 | 0.954994 | Casp9 |
| ENSMUSG( | 26.50583 | -0.11685 | 0.754277 | 0.950981 | Casq1 |
| ENSMUSG( | 111.0136 | -0.35894 | 0.254296 | 0.704536 | Casq2 |
| ENSMUSG( | 22.24224 | -0.35894 | 0.408935 | 0.818173 | Casr |
| ENSMUSG( | 20.04611 | 0.095619 | 0.823719 | NA | Cass4 |
| ENSMUSG( | 224.5648 | 0.009404 | 0.950536 | 0.993821 | Cast |
| ENSMUSG( | 179.7947 | 0.393895 | 0.01105 | 0.151501 | Castor1 |
| ENSMUSG( | 7364.441 | 0.248096 | 0.010667 | 0.148645 | Castor2 |
| ENSMUSG( | 68.6252 | -0.29707 | 0.222634 | 0.668434 | Casz1 |
| ENSMUSG( | 3175.723 | 0.076351 | 0.484589 | 0.856665 | Cat |
| ENSMUSG( | 106.0384 | 0.14341 | 0.521448 | 0.875186 | Catip |
| ENSMUSG( | 147.5151 | -0.00335 | 0.986217 | 0.998326 | Catsper2 |
| ENSMUSG( | 28.88632 | 0.860084 | 0.016663 | 0.192198 | Catsperd |
| ENSMUSG( | 79.35792 | 0.339715 | 0.15921 | 0.588305 | Catspere2 |
| ENSMUSG( | 28.91055 | 0.626997 | 0.079843 | 0.43634 | Catsperg1 |
| ENSMUSG( | 143.5347 | 0.068208 | 0.722861 | 0.943761 | Catsperz |
| ENSMUSG( | 696.2232 | -0.23052 | 0.228239 | 0.675678 | Cav1 |
| ENSMUSG( | 1454.625 | -0.17434 | 0.279047 | 0.728965 | Cav2 |
| ENSMUSG( | 883.0136 | 0.194511 | 0.125782 | 0.53257 | Cavin1 |
| ENSMUSG( | 297.3525 | 0.187667 | 0.306546 | 0.754161 | Cavin2 |
| ENSMUSG( | 1131.427 | 0.185487 | 0.104017 | 0.489802 | Cavin3 |
| ENSMUSG( | 23.37268 | 0.148557 | 0.717642 | 0.941697 | Cavin4 |
| ENSMUSG( | 10015.73 | -0.13805 | 0.24724 | 0.696222 | Cbarp |
| ENSMUSG( | 1044.25 | 0.118923 | 0.289277 | 0.739725 | Cbfa2t2 |
| ENSMUSG( | 1300.786 | 0.365355 | 0.000647 | 0.024524 | Cbfa2t3 |
| ENSMUSG( | 1054.145 | 0.020302 | 0.875387 | 0.980037 | Cbfb |
| ENSMUSG( | 2457.373 | 0.169054 | 0.087649 | 0.454663 | Cbl |
| ENSMUSG( | 516.5752 | -0.0054 | 0.9637 | 0.994922 | Cblb |
| ENSMUSG( | 313.1533 | -0.15903 | 0.259995 | 0.709951 | Cbll1 |
| ENSMUSG( | 417.2756 | -0.17739 | 0.208809 | 0.650823 | Cbln1 |
| ENSMUSG( | 931.7955 | -0.24655 | 0.133168 | 0.545727 | Cbln2 |
| ENSMUSG( | 17.14338 | 0.225674 | 0.632237 | NA | Cbln3 |
| ENSMUSG( | 381.4232 | -0.19628 | 0.226768 | 0.67384 | Cbln4 |
| ENSMUSG( | 1074.166 | -0.0863 | 0.489063 | 0.859092 | Cbr1 |
| ENSMUSG( | 70.54433 | 0.260841 | 0.316682 | 0.763648 | Cbr2 |
| ENSMUSG( | 421.6322 | -0.04961 | 0.703129 | 0.937735 | Cbr3 |
| ENSMUSG( | 353.4891 | 0.082546 | 0.50778 | 0.86795 | Cbr4 |
| ENSMUSG( | 3517.124 | 0.246583 | 0.005286 | 0.096877 | Cbs |
| ENSMUSG( | 981.3271 | 0.07364 | 0.455596 | 0.838412 | Cbx1 |
| ENSMUSG( | 105.773 | 0.005473 | 0.980289 | 0.997512 | Cbx2 |
| ENSMUSG( | 1490.41 | 0.018461 | 0.863302 | 0.975055 | Cbx3 |
| ENSMUSG( | 1311.216 | -0.14422 | 0.219619 | 0.664984 | Cbx4 |
| ENSMUSG( | 6703.149 | 0.061067 | 0.551895 | 0.886141 | Cbx5 |

| | | | | | |
|---|---|---|---|---|---|
| ENSMUSG( | 90250.76 | -0.05439 | 0.609227 | 0.95038 | Calm1 |
| ENSMUSG( | 16636.84 | -0.24375 | 0.215302 | 0.865167 | Calm2 |
| ENSMUSG( | 47582.84 | -0.1669 | 0.12541 | 0.81516 | Calm3 |
| ENSMUSG( | 39.14562 | -0.04862 | 0.903991 | 0.991089 | Calml4 |
| ENSMUSG( | 4930.57 | 0.23054 | 0.037952 | 0.6763 | Caln1 |
| ENSMUSG( | 16959.03 | 0.118689 | 0.436672 | 0.919879 | Calr |
| ENSMUSG( | 43.12162 | 0.110464 | 0.737817 | 0.967709 | Calr3 |
| ENSMUSG( | 73.06791 | -0.3263 | 0.227691 | 0.868507 | Calr4 |
| ENSMUSG( | 5502.845 | 0.041064 | 0.772817 | 0.974463 | Calu |
| ENSMUSG( | 5104.449 | -0.00839 | 0.941619 | 0.994578 | Caly |
| ENSMUSG( | 1562.803 | 0.004894 | 0.96573 | 0.995433 | Camk1 |
| ENSMUSG( | 2979.687 | -0.04683 | 0.648825 | 0.95422 | Camk1d |
| ENSMUSG( | 711.334 | 0.080553 | 0.650235 | 0.95422 | Camk1g |
| ENSMUSG( | 188471.5 | 0.112775 | 0.366007 | 0.914623 | Camk2a |
| ENSMUSG( | 16360.87 | -0.03634 | 0.794352 | 0.97681 | Camk2b |
| ENSMUSG( | 1648.856 | -0.06882 | 0.669276 | 0.956766 | Camk2d |
| ENSMUSG( | 7230.097 | -0.06143 | 0.541436 | 0.937841 | Camk2g |
| ENSMUSG( | 139849.5 | -0.05605 | 0.673573 | 0.95694 | Camk2n1 |
| ENSMUSG( | 6453.059 | -0.10461 | 0.324249 | 0.905257 | Camk2n2 |
| ENSMUSG( | 589.891 | -0.21044 | 0.123503 | 0.814691 | Camk4 |
| ENSMUSG( | 2325.678 | -0.00996 | 0.93882 | 0.993898 | Camkk1 |
| ENSMUSG( | 8496.687 | -0.02682 | 0.778986 | 0.976106 | Camkk2 |
| ENSMUSG( | 179.5983 | -0.20799 | 0.256663 | 0.883082 | Camkmt |
| ENSMUSG( | 12603.81 | -0.00075 | 0.994703 | 0.998813 | Camkv |
| ENSMUSG( | 792.8792 | 0.224733 | 0.181667 | 0.850579 | Caml |
| ENSMUSG( | 41.29437 | 0.121216 | 0.907123 | 0.991642 | Camp |
| ENSMUSG( | 2242.246 | 0.02106 | 0.853272 | 0.985598 | Camsap1 |
| ENSMUSG( | 3815.552 | 0.110551 | 0.293707 | 0.898538 | Camsap2 |
| ENSMUSG( | 791.703 | 0.04447 | 0.709083 | 0.964042 | Camsap3 |
| ENSMUSG( | 4752.582 | -0.0765 | 0.505961 | 0.927825 | Camta1 |
| ENSMUSG( | 13235.94 | 0.097149 | 0.351005 | 0.911707 | Camta2 |
| ENSMUSG( | 2328.836 | -0.02518 | 0.831953 | 0.980738 | Cand1 |
| ENSMUSG( | 159.9694 | -0.24192 | 0.178942 | 0.849041 | Cand2 |
| ENSMUSG( | 1771.076 | 0.22831 | 0.163541 | 0.838291 | Cant1 |
| ENSMUSG( | 14945.28 | 0.109554 | 0.487143 | 0.92541 | Canx |
| ENSMUSG( | 4934.664 | -0.20177 | 0.069542 | 0.75601 | Cap1 |
| ENSMUSG( | 4012.135 | -0.19758 | 0.164767 | 0.838743 | Cap2 |
| ENSMUSG( | 76.8293 | -0.23503 | 0.36452 | 0.914223 | Capg |
| ENSMUSG( | 484.0792 | -0.11731 | 0.452215 | 0.922316 | Capn1 |
| ENSMUSG( | 930.9209 | 0.103104 | 0.386364 | 0.917257 | Capn10 |
| ENSMUSG( | 23.69333 | 0.415587 | 0.482813 | 0.924615 | Capn11 |
| ENSMUSG( | 33.51204 | -0.09731 | 0.782891 | 0.976106 | Capn12 |
| ENSMUSG( | 2188.716 | 0.148919 | 0.210147 | 0.864412 | Capn15 |
| ENSMUSG( | 3837.22 | -0.15508 | 0.151342 | 0.837724 | Capn2 |
| ENSMUSG( | 36.68983 | -0.24964 | 0.498155 | 0.927825 | Capn3 |
| ENSMUSG( | 1774.522 | -0.05178 | 0.617336 | 0.951952 | Capn5 |
| ENSMUSG( | 110.8408 | 0.113557 | 0.674436 | 0.95694 | Capn6 |
| ENSMUSG( | 1129.983 | 0.081115 | 0.507045 | 0.928624 | Capn7 |
| ENSMUSG( | 14.46626 | -0.46561 | 0.369079 | 0.915491 | Capn9 |
| ENSMUSG( | 3920.954 | -0.15252 | 0.199004 | 0.863016 | Capns1 |
| ENSMUSG( | 4311.492 | 0.018349 | 0.879014 | 0.989644 | Caprin1 |
| ENSMUSG( | 248.8639 | -0.05167 | 0.798596 | 0.97681 | Caprin2 |
| ENSMUSG( | 12.07759 | -0.33368 | 0.580335 | 0.944209 | Caps2 |
| ENSMUSG( | 37.17672 | 0.124587 | 0.718812 | 0.966668 | Capsl |
| ENSMUSG( | 642.3914 | 0.063798 | 0.650893 | 0.95422 | Capza1 |
| ENSMUSG( | 5379.415 | -0.08154 | 0.526321 | 0.934374 | Capza2 |
| ENSMUSG( | 5684.935 | -0.12317 | 0.312569 | 0.900037 | Capzb |
| ENSMUSG( | 3475.313 | 0.083251 | 0.548375 | 0.938063 | Car10 |
| ENSMUSG( | 5816.931 | -0.07518 | 0.434177 | 0.91869 | Car11 |
| ENSMUSG( | 518.3722 | -0.1473 | 0.306042 | 0.898538 | Car12 |
| ENSMUSG( | 97.38063 | -0.09346 | 0.676878 | 0.957106 | Car13 |
| ENSMUSG( | 681.1482 | 0.323543 | 0.037463 | 0.673565 | Car14 |
| ENSMUSG( | 72.98111 | 0.3633 | 0.22642 | 0.868507 | Car15 |
| ENSMUSG( | 10125.36 | -0.22805 | 0.088482 | 0.786756 | Car2 |
| ENSMUSG( | 21.97051 | -0.40298 | 0.390799 | 0.917257 | Car3 |
| ENSMUSG( | 1621.741 | 0.132685 | 0.40031 | 0.917257 | Car4 |
| ENSMUSG( | 18.02299 | 0.251283 | 0.615267 | 0.951177 | Car5a |
| ENSMUSG( | 363.6433 | -0.06359 | 0.714731 | 0.964986 | Car5b |
| ENSMUSG( | 475.3741 | -0.18627 | 0.280273 | 0.892356 | Car7 |
| ENSMUSG( | 1617.976 | -0.2758 | 0.047308 | 0.700936 | Car8 |
| ENSMUSG( | 27.14847 | 0.068507 | 0.881099 | 0.989644 | Car9 |
| ENSMUSG( | 187.2865 | 0.499429 | 0.012039 | 0.445877 | Card10 |
| ENSMUSG( | 11.25165 | -0.61582 | 0.292872 | NA | Card11 |
| ENSMUSG( | 45.74515 | 0.350388 | 0.237373 | 0.872173 | Card14 |
| ENSMUSG( | 280.9543 | -0.08614 | 0.582928 | 0.944266 | Card19 |
| ENSMUSG( | 79.3419 | 0.219278 | 0.393187 | 0.917257 | Card6 |
| ENSMUSG( | 104.1987 | 0.267406 | 0.314127 | 0.900949 | Card9 |
| ENSMUSG( | 327.0511 | -0.0143 | 0.930337 | 0.99306 | Carf |
| ENSMUSG( | 3836.817 | -0.18439 | 0.130232 | 0.821488 | Carhsp1 |
| ENSMUSG( | 20.12578 | -0.47764 | 0.271298 | 0.888062 | Carlr |
| ENSMUSG( | 2246.398 | -0.04007 | 0.744878 | 0.96951 | Carm1 |
| ENSMUSG( | 1504.659 | -0.09822 | 0.401042 | 0.917257 | Carmil1 |
| ENSMUSG( | 3126.169 | -0.09787 | 0.401335 | 0.917257 | Carmil2 |
| ENSMUSG( | 1284.704 | 0.145729 | 0.417002 | 0.917257 | Carmil3 |
| ENSMUSG( | 235.1941 | 0.11491 | 0.517159 | 0.930939 | Carnmt1 |
| ENSMUSG( | 477.5786 | 0.084994 | 0.617518 | 0.951952 | Carns1 |
| ENSMUSG( | 1019.684 | 0.147664 | 0.221501 | 0.866104 | Cars1 |
| ENSMUSG( | 324.7946 | -0.06245 | 0.679771 | 0.957966 | Cars2 |
| ENSMUSG( | 285.9277 | -0.17544 | 0.271016 | 0.888062 | Cartpt |
| ENSMUSG( | 1538.025 | 0.06673 | 0.55097 | 0.939232 | Casc3 |
| ENSMUSG( | 2063.644 | 0.178103 | 0.286232 | 0.893906 | Casd1 |
| ENSMUSG( | 2944.789 | -0.01831 | 0.881989 | 0.989644 | Cask |
| ENSMUSG( | 7305.049 | 0.068858 | 0.480949 | 0.924548 | Caskin1 |
| ENSMUSG( | 1898.813 | 0.092648 | 0.425886 | 0.917257 | Caskin2 |
| ENSMUSG( | 42.03857 | -0.29211 | 0.408493 | 0.917257 | Casp1 |
| ENSMUSG( | 25.50493 | 0.64236 | 0.14331 | 0.831311 | Casp12 |
| ENSMUSG( | 319.9768 | 0.098934 | 0.489647 | 0.925858 | Casp2 |
| ENSMUSG( | 511.4136 | 0.008851 | 0.952179 | 0.994798 | Casp3 |
| ENSMUSG( | 12.50378 | -0.32387 | 0.582266 | 0.944266 | Casp4 |
| ENSMUSG( | 164.2253 | -0.02623 | 0.887262 | 0.990263 | Casp6 |
| ENSMUSG( | 113.2552 | 0.045909 | 0.828687 | 0.980224 | Casp7 |
| ENSMUSG( | 136.9638 | 0.188184 | 0.33221 | 0.907907 | Casp8 |
| ENSMUSG( | 344.8298 | 0.319231 | 0.031894 | 0.632006 | Casp8ap2 |
| ENSMUSG( | 746.2574 | -0.08316 | 0.479712 | 0.924021 | Casp9 |
| ENSMUSG( | 25.88317 | 0.081237 | 0.833432 | 0.981196 | Casq1 |
| ENSMUSG( | 101.2392 | -0.33289 | 0.286972 | 0.894091 | Casq2 |
| ENSMUSG( | 25.86038 | 0.382723 | 0.384849 | 0.917257 | Casr |
| ENSMUSG( | 18.04512 | 0.08395 | 0.859611 | 0.985947 | Cass4 |
| ENSMUSG( | 223.0249 | 0.261724 | 0.165506 | 0.838878 | Cast |
| ENSMUSG( | 172.0771 | 0.528659 | 0.00559 | 0.303106 | Castor1 |
| ENSMUSG( | 6534.221 | 0.17773 | 0.083248 | 0.77681 | Castor2 |
| ENSMUSG( | 63.42714 | -0.25459 | 0.340093 | 0.907972 | Casz1 |
| ENSMUSG( | 2635.531 | -0.20189 | 0.106099 | 0.804969 | Cat |
| ENSMUSG( | 96.83255 | 0.151233 | 0.535344 | 0.936508 | Catip |
| ENSMUSG( | 141.1367 | 0.135086 | 0.498434 | 0.927825 | Catsper2 |
| ENSMUSG( | 22.4508 | 0.452511 | 0.306471 | 0.898538 | Catsperd |
| ENSMUSG( | 74.38867 | 0.412647 | 0.115854 | 0.813384 | Catspere2 |
| ENSMUSG( | 26.44234 | 0.614426 | 0.199841 | 0.863016 | Catsperg1 |
| ENSMUSG( | 129.7868 | 0.044663 | 0.820397 | 0.979198 | Catsperz |
| ENSMUSG( | 762.5431 | 0.300868 | 0.128002 | 0.81845 | Cav1 |
| ENSMUSG( | 1445.216 | 0.079 | 0.676583 | 0.95694 | Cav2 |
| ENSMUSG( | 838.2533 | 0.309833 | 0.039378 | 0.679972 | Cavin1 |
| ENSMUSG( | 294.2342 | 0.415956 | 0.072834 | 0.763629 | Cavin2 |
| ENSMUSG( | 896.5543 | -0.22596 | 0.096239 | 0.796133 | Cavin3 |
| ENSMUSG( | 20.19002 | 0.043775 | 0.926101 | 0.99306 | Cavin4 |
| ENSMUSG( | 10306.05 | 0.212076 | 0.137046 | 0.828492 | Cbarp |
| ENSMUSG( | 932.1227 | 0.06361 | 0.62658 | 0.953234 | Cbfa2t2 |
| ENSMUSG( | 1116.145 | 0.208549 | 0.063147 | 0.736482 | Cbfa2t3 |
| ENSMUSG( | 994.4978 | 0.113906 | 0.480748 | 0.924548 | Cbfb |
| ENSMUSG( | 2178.675 | 0.094858 | 0.377201 | 0.915912 | Cbl |
| ENSMUSG( | 476.7192 | 0.036612 | 0.786743 | 0.976303 | Cblb |
| ENSMUSG( | 301.8814 | -0.00123 | 0.994082 | 0.998813 | Cbll1 |
| ENSMUSG( | 449.641 | 0.300065 | 0.039057 | 0.679972 | Cbln1 |
| ENSMUSG( | 967.1222 | 0.142552 | 0.414908 | 0.917257 | Cbln2 |
| ENSMUSG( | 38.00398 | 2.105481 | 0.043107 | 0.690304 | Cbln3 |
| ENSMUSG( | 411.1003 | 0.285756 | 0.108073 | 0.804969 | Cbln4 |
| ENSMUSG( | 922.3966 | -0.2736 | 0.081721 | 0.77681 | Cbr1 |
| ENSMUSG( | 65.12924 | 0.287112 | 0.314381 | 0.900949 | Cbr2 |
| ENSMUSG( | 352.3979 | -0.32252 | 0.030225 | 0.623534 | Cbr3 |
| ENSMUSG( | 300.7039 | -0.11639 | 0.430863 | 0.917257 | Cbr4 |
| ENSMUSG( | 2771.31 | -0.16664 | 0.109869 | 0.806572 | Cbs |
| ENSMUSG( | 869.3698 | -0.01061 | 0.934451 | 0.99306 | Cbx1 |
| ENSMUSG( | 101.5328 | 0.168248 | 0.468443 | 0.923329 | Cbx2 |
| ENSMUSG( | 1347.506 | -0.00826 | 0.957046 | 0.994798 | Cbx3 |
| ENSMUSG( | 1295.388 | 0.089232 | 0.511217 | 0.929615 | Cbx4 |
| ENSMUSG( | 5828.763 | -0.07441 | 0.550425 | 0.93874 | Cbx5 |
| ENSMUSG( | 25531.23 | 0.097686 | 0.322801 | 0.904023 | Cbx6 |
| ENSMUSG( | 1497.317 | 0.058607 | 0.66918 | 0.956766 | Cbx7 |
| ENSMUSG( | 203.537 | -0.01236 | 0.955244 | 0.994798 | Cbx8 |
| ENSMUSG( | 968.4267 | -0.13579 | 0.311677 | 0.900031 | Cby1 |
| ENSMUSG( | 1036.576 | 0.056294 | 0.673882 | 0.95694 | Cc2d1a |
| ENSMUSG( | 970.9668 | -0.07526 | 0.571446 | 0.942116 | Cc2d1b |
| ENSMUSG( | 699.1787 | -0.13136 | 0.350461 | 0.911652 | Cc2d2a |
| ENSMUSG( | 9.891919 | -0.11582 | 0.855264 | NA | Cc2d2b |
| ENSMUSG( | 1500.89 | 0.227704 | 0.083409 | 0.77681 | Ccar1 |
| ENSMUSG( | 2080.397 | -0.02142 | 0.862708 | 0.986547 | Ccar2 |
| ENSMUSG( | 265.5626 | 0.377258 | 0.067632 | 0.75212 | Ccbe1 |
| ENSMUSG( | 102.4485 | 0.010469 | 0.963892 | 0.995433 | Ccdc102a |

| | | | | | | | | | |
|---|---|---|---|---|---|---|---|---|---|
| ENSMUSG( | Camk2n1 | 164398.7 | 0.140012 | 0.146941 | 0.568875 | ENSMUSG( | Camk2n1 | 139849.5 | -0.05605 | 0.673573 | 0.95694 |
| ENSMUSG( | Camk2n2 | 7665.766 | 0.124182 | 0.112751 | 0.506698 | ENSMUSG( | Camk2n2 | 6453.059 | -0.10461 | 0.324249 | 0.905257 |
| ENSMUSG( | Camk4 | 687.3438 | -0.02859 | 0.818008 | 0.965795 | ENSMUSG( | Camk4 | 589.891 | -0.21044 | 0.123503 | 0.814691 |
| ENSMUSG( | Camkk1 | 2596.777 | 0.039312 | 0.749973 | 0.949664 | ENSMUSG( | Camkk1 | 2325.678 | -0.00996 | 0.93882 | 0.993898 |
| ENSMUSG( | Camkk2 | 9832.342 | 0.123198 | 0.132337 | 0.544251 | ENSMUSG( | Camkk2 | 8496.687 | -0.02682 | 0.778986 | 0.976106 |
| ENSMUSG( | Camkmt | 215.2044 | 0.062219 | 0.686875 | 0.933041 | ENSMUSG( | Camkmt | 179.5983 | -0.20799 | 0.256663 | 0.883082 |
| ENSMUSG( | Camkv | 14890.07 | 0.204661 | 0.032472 | 0.275758 | ENSMUSG( | Camkv | 12603.81 | -0.00075 | 0.994703 | 0.998813 |
| ENSMUSG( | Caml | 744.0924 | -0.2341 | 0.146954 | 0.568875 | ENSMUSG( | Caml | 792.8792 | 0.224733 | 0.181667 | 0.850579 |
| ENSMUSG( | Camp | 29.88576 | -1.36231 | 0.240305 | 0.687899 | ENSMUSG( | Camp | 41.29437 | 0.121216 | 0.907123 | 0.991642 |
| ENSMUSG( | Camsap1 | 2559.669 | 0.13183 | 0.170279 | 0.605271 | ENSMUSG( | Camsap1 | 2242.246 | 0.02106 | 0.853272 | 0.985598 |
| ENSMUSG( | Camsap2 | 4182.221 | 0.107828 | 0.166586 | 0.599201 | ENSMUSG( | Camsap2 | 3815.552 | 0.110551 | 0.293707 | 0.898538 |
| ENSMUSG( | Camsap3 | 930.5956 | 0.235034 | 0.044345 | 0.323404 | ENSMUSG( | Camsap3 | 791.703 | 0.04447 | 0.709083 | 0.964042 |
| ENSMUSG( | Camta1 | 5324.49 | -0.01407 | 0.874133 | 0.979336 | ENSMUSG( | Camta1 | 4752.582 | -0.0765 | 0.505961 | 0.927825 |
| ENSMUSG( | Camta2 | 15464.73 | 0.266018 | 0.001574 | 0.04447 | ENSMUSG( | Camta2 | 13235.94 | 0.097149 | 0.351005 | 0.911707 |
| ENSMUSG( | Cand1 | 2717.538 | 0.150229 | 0.07484 | 0.424133 | ENSMUSG( | Cand1 | 2328.836 | -0.02518 | 0.831953 | 0.980738 |
| ENSMUSG( | Cand2 | 188.9063 | -0.02665 | 0.862926 | 0.975055 | ENSMUSG( | Cand2 | 159.9694 | -0.24192 | 0.178942 | 0.849041 |
| ENSMUSG( | Cant1 | 1701.421 | -0.15695 | 0.270048 | 0.719904 | ENSMUSG( | Cant1 | 1771.076 | 0.22831 | 0.163541 | 0.838291 |
| ENSMUSG( | Canx | 15047.7 | -0.1423 | 0.309306 | 0.75737 | ENSMUSG( | Canx | 14945.28 | 0.109554 | 0.487143 | 0.92541 |
| ENSMUSG( | Cap1 | 6313.74 | 0.238776 | 0.007535 | 0.120197 | ENSMUSG( | Cap1 | 4934.664 | -0.20177 | 0.069542 | 0.75601 |
| ENSMUSG( | Cap2 | 4737.47 | 0.022882 | 0.831319 | 0.969889 | ENSMUSG( | Cap2 | 4012.135 | -0.19758 | 0.164767 | 0.838743 |
| ENSMUSG( | Capg | 86.07161 | -0.19446 | 0.386869 | 0.805056 | ENSMUSG( | Capg | 76.8293 | -0.23503 | 0.36452 | 0.914223 |
| ENSMUSG( | Capn1 | 542.363 | -0.0566 | 0.644708 | 0.917782 | ENSMUSG( | Capn1 | 484.0792 | -0.11731 | 0.452215 | 0.922316 |
| ENSMUSG( | Capn10 | 1016.393 | 0.087056 | 0.422568 | 0.823508 | ENSMUSG( | Capn10 | 930.9209 | 0.103104 | 0.386364 | 0.917257 |
| ENSMUSG( | Capn11 | 54.00918 | 1.963999 | 5.03E-06 | 0.000663 | ENSMUSG( | Capn11 | 23.69333 | 0.415587 | 0.482813 | 0.924615 |
| ENSMUSG( | Capn12 | 36.94822 | -0.09187 | 0.776605 | 0.955658 | ENSMUSG( | Capn12 | 33.51204 | -0.09731 | 0.782891 | 0.976106 |
| ENSMUSG( | Capn15 | 2434.831 | 0.183811 | 0.055462 | 0.362449 | ENSMUSG( | Capn15 | 2188.716 | 0.148919 | 0.210147 | 0.864412 |
| ENSMUSG( | Capn2 | 4354.123 | -0.05482 | 0.61019 | 0.908174 | ENSMUSG( | Capn2 | 3837.22 | -0.15508 | 0.151342 | 0.837724 |
| ENSMUSG( | Capn3 | 42.98332 | -0.08165 | 0.792613 | 0.95947 | ENSMUSG( | Capn3 | 36.68983 | -0.24964 | 0.498155 | 0.927825 |
| ENSMUSG( | Capn5 | 2152.943 | 0.227935 | 0.011282 | 0.153004 | ENSMUSG( | Capn5 | 1774.522 | -0.05178 | 0.617336 | 0.951952 |
| ENSMUSG( | Capn6 | 118.666 | 0.064252 | 0.755709 | 0.95146 | ENSMUSG( | Capn6 | 110.8408 | 0.113557 | 0.674436 | 0.95694 |
| ENSMUSG( | Capn7 | 1219.308 | 0.034893 | 0.711555 | 0.940028 | ENSMUSG( | Capn7 | 1129.983 | 0.081115 | 0.507045 | 0.928624 |
| ENSMUSG( | Capn9 | 15.11449 | -0.66164 | 0.193346 | NA | ENSMUSG( | Capn9 | 14.46626 | -0.46561 | 0.369079 | 0.915491 |
| ENSMUSG( | Capns1 | 4583.241 | 0.035009 | 0.694508 | 0.9343 | ENSMUSG( | Capns1 | 3920.954 | -0.15252 | 0.199004 | 0.863016 |
| ENSMUSG( | Caprin1 | 4900.466 | 0.118105 | 0.170608 | 0.605271 | ENSMUSG( | Caprin1 | 4311.492 | 0.018349 | 0.879014 | 0.989644 |
| ENSMUSG( | Caprin2 | 276.1663 | -0.0315 | 0.835052 | 0.970426 | ENSMUSG( | Caprin2 | 248.8639 | -0.05167 | 0.798596 | 0.97681 |
| ENSMUSG( | Capsl | 33.71288 | -0.43443 | 0.208952 | 0.650854 | ENSMUSG( | Capsl | 37.17672 | 0.124587 | 0.718812 | 0.966668 |
| ENSMUSG( | Capza1 | 731.7152 | 0.172286 | 0.117719 | 0.518015 | ENSMUSG( | Capza1 | 642.3914 | 0.063798 | 0.650893 | 0.95422 |
| ENSMUSG( | Capza2 | 6248.976 | 0.084051 | 0.392487 | 0.809274 | ENSMUSG( | Capza2 | 5379.415 | -0.08154 | 0.526321 | 0.934374 |
| ENSMUSG( | Capzb | 6673.776 | 0.074645 | 0.376186 | 0.801127 | ENSMUSG( | Capzb | 5684.935 | -0.12317 | 0.312569 | 0.900037 |
| ENSMUSG( | Car10 | 3420.745 | -0.24179 | 0.049895 | 0.341042 | ENSMUSG( | Car10 | 3475.313 | 0.083251 | 0.548375 | 0.938063 |
| ENSMUSG( | Car11 | 6307.74 | -0.11121 | 0.160752 | 0.590628 | ENSMUSG( | Car11 | 5816.931 | -0.07518 | 0.434177 | 0.91869 |
| ENSMUSG( | Car12 | 486.6326 | -0.66527 | 1.05E-05 | 0.001196 | ENSMUSG( | Car12 | 518.3722 | -0.1473 | 0.306042 | 0.898538 |
| ENSMUSG( | Car13 | 121.7363 | 0.278558 | 0.160204 | 0.589853 | ENSMUSG( | Car13 | 97.38063 | -0.09346 | 0.676878 | 0.957106 |
| ENSMUSG( | Car14 | 570.5012 | -0.48692 | 0.00037 | 0.016819 | ENSMUSG( | Car14 | 681.1482 | 0.323543 | 0.037463 | 0.673565 |
| ENSMUSG( | Car15 | 70.93035 | 0.01181 | 0.962038 | 0.994538 | ENSMUSG( | Car15 | 72.98111 | 0.3633 | 0.22642 | 0.868507 |
| ENSMUSG( | Car2 | 12289.05 | 0.072549 | 0.473213 | 0.84878 | ENSMUSG( | Car2 | 10125.36 | -0.22805 | 0.088482 | 0.786756 |
| ENSMUSG( | Car3 | 21.3548 | -0.77975 | 0.072823 | 0.416499 | ENSMUSG( | Car3 | 21.97051 | -0.40298 | 0.390799 | 0.917257 |
| ENSMUSG( | Car4 | 1662.935 | -0.06672 | 0.633109 | 0.914073 | ENSMUSG( | Car4 | 1621.741 | 0.132685 | 0.40031 | 0.917257 |
| ENSMUSG( | Car5a | 17.0948 | -0.2234 | 0.617548 | NA | ENSMUSG( | Car5a | 18.02299 | 0.251283 | 0.615267 | 0.951177 |
| ENSMUSG( | Car5b | 434.5505 | 0.185877 | 0.211321 | 0.654503 | ENSMUSG( | Car5b | 363.6433 | -0.06359 | 0.714731 | 0.964986 |
| ENSMUSG( | Car7 | 575.3656 | 0.098302 | 0.552665 | 0.886141 | ENSMUSG( | Car7 | 475.3741 | -0.18627 | 0.280273 | 0.892356 |
| ENSMUSG( | Car8 | 1950.493 | 0.010277 | 0.934821 | 0.993231 | ENSMUSG( | Car8 | 1617.976 | -0.2758 | 0.047308 | 0.700936 |
| ENSMUSG( | Car9 | 26.39125 | -0.34949 | 0.444772 | 0.833239 | ENSMUSG( | Car9 | 27.14847 | 0.068507 | 0.881099 | 0.989644 |
| ENSMUSG( | Card10 | 197.6762 | 0.390097 | 0.023523 | 0.234655 | ENSMUSG( | Card10 | 187.2865 | 0.499429 | 0.012039 | 0.445877 |
| ENSMUSG( | Card11 | 13.21745 | -0.40389 | 0.443777 | NA | ENSMUSG( | Card11 | 11.25165 | -0.61582 | 0.292872 | NA |
| ENSMUSG( | Card14 | 37.58294 | -0.5083 | 0.098581 | 0.480543 | ENSMUSG( | Card14 | 45.74515 | 0.350388 | 0.237373 | 0.872173 |
| ENSMUSG( | Card19 | 285.4312 | -0.33485 | 0.012917 | 0.164193 | ENSMUSG( | Card19 | 280.9543 | -0.08614 | 0.582928 | 0.944266 |
| ENSMUSG( | Card6 | 78.69359 | -0.08654 | 0.693885 | 0.934202 | ENSMUSG( | Card6 | 79.3419 | 0.219278 | 0.393187 | 0.917257 |
| ENSMUSG( | Card9 | 105.0905 | 0.02056 | 0.923167 | 0.990693 | ENSMUSG( | Card9 | 104.1987 | 0.267406 | 0.314127 | 0.900949 |
| ENSMUSG( | Carf | 359.627 | -0.0171 | 0.898315 | 0.985633 | ENSMUSG( | Carf | 327.0511 | -0.0143 | 0.930337 | 0.99306 |
| ENSMUSG( | Carhsp1 | 4471.925 | -0.00296 | 0.976699 | 0.997428 | ENSMUSG( | Carhsp1 | 3836.817 | -0.18439 | 0.130232 | 0.821488 |
| ENSMUSG( | Carlr | 25.17354 | -0.05372 | 0.887045 | 0.983326 | ENSMUSG( | Carlr | 20.12578 | -0.47764 | 0.271298 | 0.888062 |
| ENSMUSG( | Carm1 | 2523.006 | 0.027851 | 0.797922 | 0.960538 | ENSMUSG( | Carm1 | 2246.398 | -0.04007 | 0.744878 | 0.96951 |
| ENSMUSG( | Carmil1 | 1796.595 | 0.145488 | 0.188542 | 0.627572 | ENSMUSG( | Carmil1 | 1504.659 | -0.09822 | 0.401042 | 0.917257 |
| ENSMUSG( | Carmil2 | 3657.139 | 0.088371 | 0.343585 | 0.779153 | ENSMUSG( | Carmil2 | 3126.169 | -0.09787 | 0.401335 | 0.917257 |
| ENSMUSG( | Carmil3 | 1608.104 | 0.49072 | 0.000534 | 0.021434 | ENSMUSG( | Carmil3 | 1284.704 | 0.145729 | 0.417002 | 0.917257 |
| ENSMUSG( | Carnmt1 | 257.7587 | 0.122624 | 0.39847 | 0.812876 | ENSMUSG( | Carnmt1 | 235.1941 | 0.11491 | 0.517159 | 0.930939 |
| ENSMUSG( | Carns1 | 463.6033 | -0.28991 | 0.012565 | 0.161972 | ENSMUSG( | Carns1 | 477.5786 | 0.084994 | 0.617518 | 0.951952 |
| ENSMUSG( | Cars1 | 1109.364 | 0.121184 | 0.251926 | 0.701908 | ENSMUSG( | Cars1 | 1019.684 | 0.147664 | 0.221501 | 0.866104 |
| ENSMUSG( | Cars2 | 383.4367 | 0.145082 | 0.261056 | 0.710889 | ENSMUSG( | Cars2 | 324.7946 | -0.06245 | 0.679771 | 0.957966 |
| ENSMUSG( | Cartpt | 282.2909 | -0.5223 | 0.005537 | 0.099488 | ENSMUSG( | Cartpt | 285.9277 | -0.17544 | 0.271016 | 0.888062 |
| ENSMUSG( | Casc3 | 1713.044 | 0.107653 | 0.253899 | 0.704011 | ENSMUSG( | Casc3 | 1538.025 | 0.06673 | 0.55097 | 0.939232 |
| ENSMUSG( | Casd1 | 1831.686 | -0.47051 | 0.001691 | 0.046443 | ENSMUSG( | Casd1 | 2063.644 | 0.178103 | 0.286232 | 0.893906 |
| ENSMUSG( | Cask | 3167.146 | -0.07984 | 0.416154 | 0.82084 | ENSMUSG( | Cask | 2944.789 | -0.01831 | 0.881989 | 0.989644 |
| ENSMUSG( | Caskin1 | 8181.896 | 0.125865 | 0.102621 | 0.488148 | ENSMUSG( | Caskin1 | 7305.049 | 0.068858 | 0.480949 | 0.924548 |
| ENSMUSG( | Caskin2 | 1952.8 | -0.0945 | 0.350219 | 0.781943 | ENSMUSG( | Caskin2 | 1898.813 | 0.092648 | 0.425886 | 0.917257 |
| ENSMUSG( | Casp1 | 46.42063 | -0.24653 | 0.441689 | 0.832577 | ENSMUSG( | Casp1 | 42.03857 | -0.29211 | 0.408493 | 0.917257 |
| ENSMUSG( | Casp12 | 22.02305 | -0.01871 | 0.963658 | 0.994922 | ENSMUSG( | Casp12 | 25.50493 | 0.64236 | 0.14331 | 0.831311 |
| ENSMUSG( | Casp2 | 321.3919 | -0.16139 | 0.207104 | 0.649503 | ENSMUSG( | Casp2 | 319.9768 | 0.098934 | 0.489647 | 0.925858 |
| ENSMUSG( | Casp3 | 561.9068 | 0.018689 | 0.878527 | 0.981431 | ENSMUSG( | Casp3 | 511.4136 | 0.008851 | 0.952179 | 0.994798 |
| ENSMUSG( | Casp4 | 12.29211 | -0.7792 | 0.151716 | NA | ENSMUSG( | Casp4 | 12.50378 | -0.32387 | 0.582266 | 0.944266 |
| ENSMUSG( | Casp6 | 183.3604 | 0.021609 | 0.89021 | 0.983687 | ENSMUSG( | Casp6 | 164.2253 | -0.02623 | 0.887262 | 0.990263 |
| ENSMUSG( | Casp7 | 120.4299 | -0.04836 | 0.805816 | 0.962616 | ENSMUSG( | Casp7 | 113.2552 | 0.045909 | 0.828687 | 0.980224 |
| ENSMUSG( | Casp8 | 137.4956 | -0.07334 | 0.670846 | 0.928741 | ENSMUSG( | Casp8 | 136.9638 | 0.188184 | 0.33221 | 0.907907 |
| ENSMUSG( | Casp8ap2 | 342.4549 | 0.043095 | 0.728177 | 0.944355 | ENSMUSG( | Casp8ap2 | 344.8298 | 0.319231 | 0.031894 | 0.632006 |
| ENSMUSG( | Casp9 | 850.8353 | 0.028215 | 0.772179 | 0.954994 | ENSMUSG( | Casp9 | 746.2574 | -0.08316 | 0.479712 | 0.924021 |
| ENSMUSG( | Casq1 | 26.50583 | -0.11685 | 0.754277 | 0.950981 | ENSMUSG( | Casq1 | 25.88317 | 0.081237 | 0.833432 | 0.981196 |
| ENSMUSG( | Casq2 | 111.0136 | -0.35894 | 0.254296 | 0.704536 | ENSMUSG( | Casq2 | 101.2392 | -0.33289 | 0.286972 | 0.894091 |
| ENSMUSG( | Casr | 22.24224 | -0.35894 | 0.408935 | 0.818173 | ENSMUSG( | Casr | 25.86038 | 0.382723 | 0.384849 | 0.917257 |
| ENSMUSG( | Cass4 | 20.04611 | 0.095619 | 0.823719 | NA | ENSMUSG( | Cass4 | 18.04512 | 0.08395 | 0.859611 | 0.985947 |
| ENSMUSG( | Cast | 224.5648 | 0.009404 | 0.950536 | 0.993821 | ENSMUSG( | Cast | 223.0249 | 0.261724 | 0.165506 | 0.838878 |
| ENSMUSG( | Castor1 | 179.7947 | 0.393895 | 0.01105 | 0.151501 | ENSMUSG( | Castor1 | 172.0771 | 0.528659 | 0.00559 | 0.303106 |
| ENSMUSG( | Castor2 | 7364.441 | 0.248096 | 0.010667 | 0.148645 | ENSMUSG( | Castor2 | 6534.221 | 0.17773 | 0.083248 | 0.77681 |
| ENSMUSG( | Casz1 | 68.6252 | -0.29707 | 0.222634 | 0.668434 | ENSMUSG( | Casz1 | 63.42714 | -0.25459 | 0.340093 | 0.907972 |
| ENSMUSG( | Cat | 3175.723 | 0.076351 | 0.484589 | 0.856665 | ENSMUSG( | Cat | 2635.531 | -0.20189 | 0.106099 | 0.804969 |
| ENSMUSG( | Catip | 106.0384 | 0.14341 | 0.521448 | 0.875186 | ENSMUSG( | Catip | 96.83255 | 0.151233 | 0.535344 | 0.936508 |
| ENSMUSG( | Catsper2 | 147.5151 | -0.00335 | 0.986217 | 0.998326 | ENSMUSG( | Catsper2 | 141.1367 | 0.135086 | 0.498434 | 0.927825 |
| ENSMUSG( | Catsperd | 28.88632 | 0.860084 | 0.016663 | 0.192198 | ENSMUSG( | Catsperd | 22.4508 | 0.452511 | 0.306471 | 0.898538 |
| ENSMUSG( | Catspere2 | 79.35792 | 0.339715 | 0.15921 | 0.588305 | ENSMUSG( | Catspere2 | 74.38867 | 0.412647 | 0.115854 | 0.813384 |
| ENSMUSG( | Catsperg1 | 28.91055 | 0.626997 | 0.079843 | 0.43634 | ENSMUSG( | Catsperg1 | 26.44234 | 0.614426 | 0.199841 | 0.863016 |
| ENSMUSG( | Catsperz | 143.5347 | 0.068208 | 0.722861 | 0.943761 | ENSMUSG( | Catsperz | 129.7868 | 0.044663 | 0.820397 | 0.979198 |
| ENSMUSG( | Cav1 | 696.2232 | -0.23052 | 0.228239 | 0.675678 | ENSMUSG( | Cav1 | 762.5431 | 0.300868 | 0.128002 | 0.81845 |
| ENSMUSG( | Cav2 | 1454.625 | -0.17434 | 0.279047 | 0.728965 | ENSMUSG( | Cav2 | 1445.216 | 0.079 | 0.676583 | 0.95694 |
| ENSMUSG( | Cavin1 | 883.0136 | 0.194511 | 0.125782 | 0.53257 | ENSMUSG( | Cavin1 | 838.2533 | 0.309833 | 0.039378 | 0.679972 |
| ENSMUSG( | Cavin2 | 297.3525 | 0.187667 | 0.306546 | 0.754161 | ENSMUSG( | Cavin2 | 294.2342 | 0.415956 | 0.072834 | 0.763629 |
| ENSMUSG( | Cavin3 | 1131.427 | 0.185487 | 0.104017 | 0.489802 | ENSMUSG( | Cavin3 | 896.5543 | -0.22596 | 0.096239 | 0.796133 |
| ENSMUSG( | Cavin4 | 23.37268 | 0.148557 | 0.717642 | 0.941697 | ENSMUSG( | Cavin4 | 20.19002 | 0.043775 | 0.926101 | 0.99306 |
| ENSMUSG( | Cbarp | 10015.73 | -0.13805 | 0.24724 | 0.696222 | ENSMUSG( | Cbarp | 10306.05 | 0.212076 | 0.137046 | 0.828492 |
| ENSMUSG( | Cbfa2t2 | 1044.25 | 0.118923 | 0.289277 | 0.739725 | ENSMUSG( | Cbfa2t2 | 932.1227 | 0.06361 | 0.62658 | 0.953234 |
| ENSMUSG( | Cbfa2t3 | 1300.786 | 0.365355 | 0.000647 | 0.024524 | ENSMUSG( | Cbfa2t3 | 1116.145 | 0.208549 | 0.063147 | 0.736482 |
| ENSMUSG( | Cbfb | 1054.145 | 0.020302 | 0.875387 | 0.980037 | ENSMUSG( | Cbfb | 994.4978 | 0.113906 | 0.480748 | 0.924548 |
| ENSMUSG( | Cbl | 2457.373 | 0.169054 | 0.087649 | 0.454663 | ENSMUSG( | Cbl | 2178.675 | 0.094858 | 0.377201 | 0.915912 |
| ENSMUSG( | Cblb | 516.5752 | -0.0054 | 0.9637 | 0.994922 | ENSMUSG( | Cblb | 476.7192 | 0.036612 | 0.786743 | 0.976303 |
| ENSMUSG( | Cbll1 | 313.1533 | -0.15903 | 0.259995 | 0.709951 | ENSMUSG( | Cbll1 | 301.8814 | -0.00123 | 0.994082 | 0.998813 |
| ENSMUSG( | Cbln1 | 417.2756 | -0.17739 | 0.208809 | 0.650823 | ENSMUSG( | Cbln1 | 449.641 | 0.300065 | 0.039057 | 0.679972 |
| ENSMUSG( | Cbln2 | 931.7955 | -0.24655 | 0.133168 | 0.545727 | ENSMUSG( | Cbln2 | 967.1222 | 0.142552 | 0.414908 | 0.917257 |
| ENSMUSG( | Cbln3 | 17.14338 | 0.225674 | 0.632237 | NA | ENSMUSG( | Cbln3 | 38.00398 | 2.105481 | 0.043107 | 0.690304 |
| ENSMUSG( | Cbln4 | 381.4232 | -0.19628 | 0.226768 | 0.67384 | ENSMUSG( | Cbln4 | 411.1003 | 0.285756 | 0.108073 | 0.804969 |
| ENSMUSG( | Cbr1 | 1074.166 | -0.0863 | 0.489063 | 0.859092 | ENSMUSG( | Cbr1 | 922.3966 | -0.2736 | 0.081721 | 0.77681 |
| ENSMUSG( | Cbr2 | 70.54433 | 0.260841 | 0.316682 | 0.763648 | ENSMUSG( | Cbr2 | 65.12924 | 0.287112 | 0.314381 | 0.900949 |
| ENSMUSG( | Cbr3 | 421.6322 | -0.04961 | 0.703129 | 0.937735 | ENSMUSG( | Cbr3 | 352.3979 | -0.32252 | 0.030225 | 0.623534 |
| ENSMUSG( | Cbr4 | 353.4891 | 0.082546 | 0.50778 | 0.86795 | ENSMUSG( | Cbr4 | 300.7039 | -0.11639 | 0.430863 | 0.917257 |
| ENSMUSG( | Cbs | 3517.124 | 0.246583 | 0.005286 | 0.096877 | ENSMUSG( | Cbs | 2771.31 | -0.16664 | 0.109869 | 0.806572 |
| ENSMUSG( | Cbx1 | 981.3271 | 0.07364 | 0.455596 | 0.838412 | ENSMUSG( | Cbx1 | 869.3698 | -0.01061 | 0.934451 | 0.99306 |
| ENSMUSG( | Cbx2 | 105.773 | 0.005473 | 0.980289 | 0.997512 | ENSMUSG( | Cbx2 | 101.5328 | 0.168248 | 0.468443 | 0.923329 |
| ENSMUSG( | Cbx3 | 1490.41 | 0.018461 | 0.863302 | 0.975055 | ENSMUSG( | Cbx3 | 1347.506 | -0.00826 | 0.957046 | 0.994798 |
| ENSMUSG( | Cbx4 | 1311.216 | -0.14422 | 0.219619 | 0.664984 | ENSMUSG( | Cbx4 | 1295.388 | 0.089232 | 0.511217 | 0.929615 |
| ENSMUSG( | Cbx5 | 6703.149 | 0.061067 | 0.551895 | 0.886141 | ENSMUSG( | Cbx5 | 5828.763 | -0.07441 | 0.550425 | 0.93874 |
| ENSMUSG( | Cbx6 | 29194.38 | 0.209194 | 0.011206 | 0.152581 | ENSMUSG( | Cbx6 | 25531.23 | 0.097686 | 0.322801 | 0.904023 |
| ENSMUSG( | Cbx7 | 1638.129 | 0.047084 | 0.662127 | 0.925836 | ENSMUSG( | Cbx7 | 1497.317 | 0.058607 | 0.66918 | 0.956766 |
| ENSMUSG( | Cbx8 | 223.4684 | -0.01652 | 0.935609 | 0.993363 | ENSMUSG( | Cbx8 | 203.537 | -0.01236 | 0.955244 | 0.994798 |
| ENSMUSG( | Cby1 | 1115.45 | 0.011318 | 0.910944 | 0.988395 | ENSMUSG( | Cby1 | 968.4267 | -0.13579 | 0.311677 | 0.900031 |
| ENSMUSG( | Cc2d1a | 1195.49 | 0.192663 | 0.082889 | 0.444486 | ENSMUSG( | Cc2d1a | 1036.576 | 0.056294 | 0.673882 | 0.95694 |
| ENSMUSG( | Cc2d1b | 1117.032 | 0.06105 | 0.580957 | 0.89745 | ENSMUSG( | Cc2d1b | 970.9668 | -0.07526 | 0.571446 | 0.942116 |
| ENSMUSG( | Cc2d2a | 770.1511 | -0.12505 | 0.221484 | 0.667258 | ENSMUSG( | Cc2d2a | 699.1787 | -0.13136 | 0.350461 | 0.911652 |
| ENSMUSG( | Cc2d2b | 11.70991 | 0.113228 | 0.832569 | NA | ENSMUSG( | Cc2d2b | 9.891919 | -0.11582 | 0.855264 | NA |
| ENSMUSG( | Ccar1 | 1644.006 | 0.22543 | 0.02022 | 0.214698 | ENSMUSG( | Ccar1 | 1500.89 | 0.227704 | 0.083409 | 0.77681 |
| ENSMUSG( | Ccar2 | 2378.342 | 0.094619 | 0.387204 | 0.805442 | ENSMUSG( | Ccar2 | 2080.397 | -0.02142 | 0.862708 | 0.986547 |
| ENSMUSG( | Ccbe1 | 233.301 | -0.2614 | 0.18334 | 0.62047 | ENSMUSG( | Ccbe1 | 265.5626 | 0.377258 | 0.067632 | 0.75212 |
| ENSMUSG( | Ccdc102a | 117.075 | 0.108045 | 0.569893 | 0.893041 | ENSMUSG( | Ccdc102a | 102.4485 | 0.010469 | 0.963892 | 0.995433 |
| ENSMUSG( | Ccdc103 | 69.26551 | -0.28685 | 0.235115 | 0.682364 | ENSMUSG( | Ccdc103 | 63.74941 | -0.25243 | 0.341889 | 0.909146 |
| ENSMUSG( | Ccdc106 | 1129.198 | 0.078601 | 0.50513 | 0.866883 | ENSMUSG( | Ccdc106 | 989.8915 | -0.03436 | 0.803877 | 0.977 |
| ENSMUSG( | Ccdc107 | 855.6583 | -0.31844 | 0.001968 | 0.051087 | ENSMUSG( | Ccdc107 | 855.2396 | -0.0378 | 0.772064 | 0.974236 |
| ENSMUSG( | Ccdc110 | 15.87328 | -0.06815 | 0.887698 | NA | ENSMUSG( | Ccdc110 | 13.75386 | -0.20532 | 0.697545 | 0.960867 |
| ENSMUSG( | Ccdc112 | 184.688 | -0.32606 | 0.082237 | 0.442642 | ENSMUSG( | Ccdc112 | 191.5299 | 0.075867 | 0.697651 | 0.960867 |
| ENSMUSG( | Ccdc113 | 40.86127 | -0.63451 | 0.03555 | 0.2902 | ENSMUSG( | Ccdc113 | 46.35319 | 0.071872 | 0.81826 | 0.979198 |
| ENSMUSG( | Ccdc115 | 401.1321 | -0.03816 | 0.798226 | 0.96069 | ENSMUSG( | Ccdc115 | 352.6683 | -0.14345 | 0.350619 | 0.911652 |
| ENSMUSG( | Ccdc116 | 96.11975 | -0.14512 | 0.481061 | 0.855135 | ENSMUSG( | Ccdc116 | 110.8419 | 0.485196 | 0.027963 | 0.61023 |
| ENSMUSG( | Ccdc117 | 540.3906 | 0.15183 | 0.2312 | 0.679229 | ENSMUSG( | Ccdc117 | 489.2401 | 0.132993 | 0.34549 | 0.910581 |
| ENSMUSG( | Ccdc12 | 780.2502 | -0.06705 | 0.551099 | 0.886038 | ENSMUSG( | Ccdc12 | 668.4572 | -0.2573 | 0.045621 | 0.693959 |
| ENSMUSG( | Ccdc120 | 357.1136 | 0.181477 | 0.2059 | 0.647605 | ENSMUSG( | Ccdc120 | 314.6981 | 0.087984 | 0.603722 | 0.948137 |
| ENSMUSG( | Ccdc121rt1 | 21.28079 | 0.160227 | 0.701818 | 0.937441 | ENSMUSG( | Ccdc121rt1 | 23.65411 | 0.674446 | 0.145059 | 0.83339 |
| ENSMUSG( | Ccdc122 | 108.9544 | -0.26409 | 0.198414 | 0.638387 | ENSMUSG( | Ccdc122 | 100.7667 | -0.24302 | 0.330005 | 0.907251 |
| ENSMUSG( | Ccdc124 | 2547.887 | -0.05658 | 0.556654 | 0.887713 | ENSMUSG( | Ccdc124 | 2129.171 | -0.32725 | 0.011492 | 0.436431 |
| ENSMUSG( | Ccdc125 | 24.01832 | -1.07418 | 0.01005 | 0.142735 | ENSMUSG( | Ccdc125 | 24.74039 | -0.53159 | 0.246978 | 0.877904 |
| ENSMUSG( | Ccdc126 | 224.3287 | -0.45502 | 0.018764 | 0.206056 | ENSMUSG( | Ccdc126 | 262.6873 | 0.295247 | 0.151118 | 0.837173 |
| ENSMUSG( | Ccdc127 | 2877.963 | -0.08889 | 0.423243 | 0.823662 | ENSMUSG( | Ccdc127 | 2722.455 | 0.022078 | 0.864642 | 0.986591 |
| ENSMUSG( | Ccdc13 | 468.3598 | 0.026797 | 0.822829 | 0.966352 | ENSMUSG( | Ccdc13 | 391.5235 | -0.23818 | 0.105698 | 0.804969 |

| | | | | | |
|---|---|---|---|---|---|
| ENSMUSG( | 29194.38 | 0.209194 | 0.011206 | 0.152581 | Cbx6 |
| ENSMUSG( | 1638.129 | 0.047084 | 0.662127 | 0.925836 | Cbx7 |
| ENSMUSG( | 223.4684 | -0.01652 | 0.935609 | 0.993363 | Cbx8 |
| ENSMUSG( | 1115.45 | 0.011318 | 0.910944 | 0.988395 | Cby1 |
| ENSMUSG( | 1195.49 | 0.192663 | 0.082889 | 0.444486 | Cc2d1a |
| ENSMUSG( | 1117.032 | 0.06105 | 0.580957 | 0.89745 | Cc2d1b |
| ENSMUSG( | 770.1511 | -0.12505 | 0.221484 | 0.667258 | Cc2d2a |
| ENSMUSG( | 11.70991 | 0.113228 | 0.832569 | NA | Cc2d2b |
| ENSMUSG( | 1644.006 | 0.22543 | 0.02022 | 0.214698 | Ccar1 |
| ENSMUSG( | 2378.342 | 0.094619 | 0.387204 | 0.805442 | Ccar2 |
| ENSMUSG( | 233.301 | -0.2614 | 0.18334 | 0.62047 | Ccbe1 |
| ENSMUSG( | 117.075 | 0.108045 | 0.569893 | 0.893041 | Ccdc102a |
| ENSMUSG( | 69.26551 | -0.28685 | 0.235115 | 0.682364 | Ccdc103 |
| ENSMUSG( | 1129.198 | 0.078601 | 0.50513 | 0.866883 | Ccdc106 |
| ENSMUSG( | 855.6583 | -0.31844 | 0.001968 | 0.051087 | Ccdc107 |
| ENSMUSG( | 15.87328 | -0.06815 | 0.887698 | NA | Ccdc110 |
| ENSMUSG( | 184.688 | -0.32606 | 0.082237 | 0.442642 | Ccdc112 |
| ENSMUSG( | 40.86127 | -0.63451 | 0.03555 | 0.2902 | Ccdc113 |
| ENSMUSG( | 401.1321 | -0.03816 | 0.798226 | 0.96069 | Ccdc115 |
| ENSMUSG( | 96.11975 | -0.14512 | 0.481061 | 0.855135 | Ccdc116 |
| ENSMUSG( | 540.3906 | 0.15183 | 0.2312 | 0.679229 | Ccdc117 |
| ENSMUSG( | 780.2502 | -0.06705 | 0.551099 | 0.886038 | Ccdc12 |
| ENSMUSG( | 357.1136 | 0.181477 | 0.2059 | 0.647605 | Ccdc120 |
| ENSMUSG( | 21.28079 | 0.160227 | 0.701818 | 0.937441 | Ccdc121rt1 |
| ENSMUSG( | 108.9544 | -0.26409 | 0.198414 | 0.638387 | Ccdc122 |
| ENSMUSG( | 2547.887 | -0.05658 | 0.556654 | 0.887713 | Ccdc124 |
| ENSMUSG( | 24.01832 | -1.07418 | 0.01005 | 0.142735 | Ccdc125 |
| ENSMUSG( | 224.3287 | -0.45502 | 0.018764 | 0.206056 | Ccdc126 |
| ENSMUSG( | 2877.963 | -0.08889 | 0.423243 | 0.823662 | Ccdc127 |
| ENSMUSG( | 468.3598 | 0.026797 | 0.822829 | 0.966352 | Ccdc13 |
| ENSMUSG( | 465.5222 | -0.17081 | 0.261448 | 0.711405 | Ccdc134 |
| ENSMUSG( | 1976.93 | 0.135362 | 0.181142 | 0.618139 | Ccdc136 |
| ENSMUSG( | 762.7908 | 0.122392 | 0.252072 | 0.701908 | Ccdc137 |
| ENSMUSG( | 144.3181 | 0.108297 | 0.534098 | 0.880394 | Ccdc138 |
| ENSMUSG( | 80.41699 | -0.38837 | 0.098786 | 0.480598 | Ccdc14 |
| ENSMUSG( | 1362.622 | 0.466723 | 2.29E-06 | 0.000361 | Ccdc141 |
| ENSMUSG( | 16.13853 | -0.19432 | 0.700734 | NA | Ccdc146 |
| ENSMUSG( | 313.2138 | 0.205941 | 0.130578 | 0.541841 | Ccdc148 |
| ENSMUSG( | 1550.925 | 0.082361 | 0.404689 | 0.816221 | Ccdc149 |
| ENSMUSG( | 45.24568 | -0.3542 | 0.208105 | 0.650319 | Ccdc15 |
| ENSMUSG( | 13.83369 | 0.403415 | 0.456558 | NA | Ccdc152 |
| ENSMUSG( | 75.70257 | -0.70062 | 0.012962 | 0.164641 | Ccdc153 |
| ENSMUSG( | 1590.818 | 0.054178 | 0.536014 | 0.88142 | Ccdc157 |
| ENSMUSG( | 8.404453 | 0.937954 | 0.215684 | NA | Ccdc158 |
| ENSMUSG( | 176.3784 | 0.198623 | 0.270366 | 0.720226 | Ccdc159 |
| ENSMUSG( | 27.02668 | -0.35696 | 0.338463 | 0.777489 | Ccdc160 |
| ENSMUSG( | 58.20129 | -0.25967 | 0.319581 | 0.765305 | Ccdc162 |
| ENSMUSG( | 88.66125 | -0.18944 | 0.38795 | 0.805835 | Ccdc163 |
| ENSMUSG( | 133.3302 | 0.073071 | 0.696645 | 0.935318 | Ccdc166 |
| ENSMUSG( | 866.9853 | 0.091366 | 0.366608 | 0.794842 | Ccdc167 |
| ENSMUSG( | 67.55411 | 0.070724 | 0.798753 | 0.960896 | Ccdc17 |
| ENSMUSG( | 26.57946 | 0.214113 | 0.596083 | 0.903152 | Ccdc170 |
| ENSMUSG( | 72.34036 | -0.03573 | 0.884136 | 0.982647 | Ccdc171 |
| ENSMUSG( | 391.8385 | 0.156224 | 0.207443 | 0.649561 | Ccdc174 |
| ENSMUSG( | 349.7355 | 0.328187 | 0.027912 | 0.255775 | Ccdc177 |
| ENSMUSG( | 97.17836 | -0.01921 | 0.93195 | 0.9924 | Ccdc18 |
| ENSMUSG( | 23.21963 | 0.172517 | 0.65927 | 0.924506 | Ccdc180 |
| ENSMUSG( | 504.6752 | 0.190823 | 0.18914 | 0.627756 | Ccdc181 |
| ENSMUSG( | 391.0051 | -0.1386 | 0.440176 | 0.831851 | Ccdc184 |
| ENSMUSG( | 914.6008 | 0.19778 | 0.04818 | 0.335655 | Ccdc186 |
| ENSMUSG( | 302.062 | -1.0235 | 3.55E-06 | 0.000492 | Ccdc187 |
| ENSMUSG( | 1609.411 | 0.181555 | 0.141254 | 0.558134 | Ccdc190 |
| ENSMUSG( | 213.2318 | 0.001126 | 0.994665 | 0.999369 | Ccdc191 |
| ENSMUSG( | 98.30035 | 0.38112 | 0.082377 | 0.442878 | Ccdc22 |
| ENSMUSG( | 28.92148 | -0.37389 | 0.29604 | 0.744386 | Ccdc24 |
| ENSMUSG( | 919.7235 | 0.123115 | 0.353923 | 0.786371 | Ccdc25 |
| ENSMUSG( | 341.9904 | -0.14332 | 0.36277 | 0.793121 | Ccdc28a |
| ENSMUSG( | 683.9272 | -0.39156 | 0.000498 | 0.020389 | Ccdc28b |
| ENSMUSG( | 612.168 | 0.029053 | 0.814474 | 0.964574 | Ccdc3 |
| ENSMUSG( | 311.8328 | 0.003831 | 0.981487 | 0.997512 | Ccdc30 |
| ENSMUSG( | 889.5311 | -0.00506 | 0.969845 | 0.996369 | Ccdc32 |
| ENSMUSG( | 35.15837 | -0.25354 | 0.418309 | 0.821966 | Ccdc33 |
| ENSMUSG( | 864.7589 | -0.00977 | 0.937071 | 0.993415 | Ccdc34 |
| ENSMUSG( | 506.2057 | 0.09664 | 0.476189 | 0.851294 | Ccdc39 |
| ENSMUSG( | 57.43391 | -0.35942 | 0.151083 | 0.576088 | Ccdc40 |
| ENSMUSG( | 579.3912 | 0.131441 | 0.220296 | 0.665898 | Ccdc43 |
| ENSMUSG( | 4579.202 | -0.16501 | 0.183283 | 0.62047 | Ccdc47 |
| ENSMUSG( | 2311.102 | 0.051761 | 0.633051 | 0.914073 | Ccdc50 |
| ENSMUSG( | 143.728 | 0.213668 | 0.227421 | 0.67424 | Ccdc51 |
| ENSMUSG( | 277.6952 | -0.00592 | 0.971413 | 0.996415 | Ccdc57 |
| ENSMUSG( | 534.3719 | -0.19333 | 0.125788 | 0.53257 | Ccdc59 |
| ENSMUSG( | 1707.558 | 0.310936 | 0.001093 | 0.034959 | Ccdc6 |
| ENSMUSG( | 36.80361 | -0.0304 | 0.930315 | 0.991848 | Ccdc60 |
| ENSMUSG( | 244.9751 | 0.070378 | 0.614156 | 0.909405 | Ccdc61 |
| ENSMUSG( | 146.8893 | -0.00961 | 0.954928 | 0.993874 | Ccdc62 |
| ENSMUSG( | 9.357191 | 0.265816 | 0.673908 | NA | Ccdc63 |
| ENSMUSG( | 129.1756 | 0.022818 | 0.920048 | 0.990247 | Ccdc65 |
| ENSMUSG( | 419.5341 | 0.075952 | 0.560312 | 0.888934 | Ccdc66 |
| ENSMUSG( | 18.26185 | 0.039362 | 0.9304 | NA | Ccdc69 |
| ENSMUSG( | 609.4953 | 0.040201 | 0.749659 | 0.949664 | Ccdc71 |
| ENSMUSG( | 357.0813 | 0.06687 | 0.621825 | 0.911432 | Ccdc71l |
| ENSMUSG( | 21.60178 | -0.11187 | 0.785898 | 0.958774 | Ccdc73 |
| ENSMUSG( | 1308.833 | 0.169232 | 0.061243 | 0.381822 | Ccdc74a |
| ENSMUSG( | 168.7631 | 0.103195 | 0.526251 | 0.877265 | Ccdc77 |
| ENSMUSG( | 11.2613 | -0.31764 | 0.635486 | NA | Ccdc78 |
| ENSMUSG( | 289.3015 | 0.106161 | 0.45712 | 0.839312 | Ccdc8 |
| ENSMUSG( | 596.8122 | -0.07189 | 0.564093 | 0.890139 | Ccdc80 |
| ENSMUSG( | 35.67722 | -0.24161 | 0.523576 | 0.875628 | Ccdc81 |
| ENSMUSG( | 652.6497 | -0.0249 | 0.838606 | 0.971111 | Ccdc82 |
| ENSMUSG( | 1698.891 | 0.293605 | 0.002906 | 0.066108 | Ccdc85a |
| ENSMUSG( | 3922.149 | 0.087963 | 0.333785 | 0.775602 | Ccdc85b |
| ENSMUSG( | 5702.611 | 0.183755 | 0.043795 | 0.320589 | Ccdc85c |
| ENSMUSG( | 351.1588 | -0.19379 | 0.296715 | 0.745349 | Ccdc86 |
| ENSMUSG( | 60.09915 | 0.086451 | 0.726471 | 0.944223 | Ccdc87 |
| ENSMUSG( | 2157.988 | 0.105409 | 0.398811 | 0.812878 | Ccdc88a |
| ENSMUSG( | 165.4069 | -0.07491 | 0.668869 | 0.927963 | Ccdc88b |
| ENSMUSG( | 1490.382 | -0.24772 | 0.012803 | 0.163512 | Ccdc88c |
| ENSMUSG( | 14.81235 | -0.24387 | 0.634973 | NA | Ccdc89 |
| ENSMUSG( | 1226.968 | 0.101872 | 0.316288 | 0.763453 | Ccdc9 |
| ENSMUSG( | 494.1461 | 0.088093 | 0.478628 | 0.853679 | Ccdc90b |
| ENSMUSG( | 1466.007 | 0.095391 | 0.325283 | 0.77061 | Ccdc91 |
| ENSMUSG( | 2060.597 | 0.094061 | 0.451205 | 0.835567 | Ccdc92 |
| ENSMUSG( | 940.7118 | 0.072252 | 0.52303 | 0.875628 | Ccdc92b |
| ENSMUSG( | 1148.848 | 0.199166 | 0.041133 | 0.313521 | Ccdc93 |
| ENSMUSG( | 62.93376 | -0.15493 | 0.572216 | 0.893584 | Ccdc96 |
| ENSMUSG( | 1707.615 | 0.05645 | 0.524042 | 0.87583 | Ccdc97 |
| ENSMUSG( | 395.738 | -0.01767 | 0.893644 | 0.984512 | Ccdc9b |
| ENSMUSG( | 35.7532 | -0.79071 | 0.02192 | 0.225738 | Ccer2 |
| ENSMUSG( | 212.6441 | -0.04864 | 0.742243 | 0.948478 | Cchcr1 |
| ENSMUSG( | 12177.75 | -0.4974 | 8.51E-05 | 0.005781 | Cck |
| ENSMUSG( | 2318.008 | -0.36756 | 0.012256 | 0.160458 | Cckbr |
| ENSMUSG( | 19.49735 | -0.40965 | 0.351946 | NA | Ccl12 |
| ENSMUSG( | 21.13305 | -0.31873 | 0.481515 | NA | Ccl17 |
| ENSMUSG( | 13.14004 | -0.37256 | 0.483993 | NA | Ccl22 |
| ENSMUSG( | 10.0078 | -0.0301 | 0.958842 | NA | Ccl24 |
| ENSMUSG( | 243.4767 | -0.05538 | 0.707287 | 0.93949 | Ccl25 |
| ENSMUSG( | 1934.219 | 0.122053 | 0.285884 | 0.734989 | Ccl27a |
| ENSMUSG( | 157.3381 | 0.517013 | 0.012294 | 0.160458 | Ccl27al |
| ENSMUSG( | 69.78579 | -0.39951 | 0.220296 | 0.665898 | Ccl28 |
| ENSMUSG( | 9.131405 | 0.193585 | 0.761235 | NA | Ccl3 |
| ENSMUSG( | 24.32439 | -0.98118 | 0.126207 | 0.53365 | Ccl5 |
| ENSMUSG( | 162.323 | -0.08154 | 0.69584 | 0.934979 | Ccl6 |
| ENSMUSG( | 149.4797 | 0.161308 | 0.43591 | 0.829093 | Ccl9 |
| ENSMUSG( | 2302.647 | 0.012662 | 0.877952 | 0.981431 | Ccm2 |
| ENSMUSG( | 130.4633 | 0.651478 | 0.006802 | 0.113745 | Ccm2l |
| ENSMUSG( | 379.1657 | -1.38242 | 5.09E-13 | 4.59E-10 | Ccn1 |
| ENSMUSG( | 1181.618 | -0.60864 | 0.000538 | 0.021456 | Ccn2 |
| ENSMUSG( | 4100.122 | -0.31128 | 0.017231 | 0.196681 | Ccn3 |
| ENSMUSG( | 78.98489 | -0.93311 | 0.000115 | 0.007087 | Ccn4 |
| ENSMUSG( | 12.42181 | -1.61397 | 0.008857 | NA | Ccn5 |
| ENSMUSG( | 30.98754 | 0.369369 | 0.267152 | 0.717525 | Ccna1 |
| ENSMUSG( | 113.3168 | -0.21561 | 0.276671 | 0.726854 | Ccna2 |
| ENSMUSG( | 42.28966 | -0.25787 | 0.373566 | 0.800375 | Ccnb1 |
| ENSMUSG( | 34.62937 | -0.19329 | 0.550701 | 0.886038 | Ccnb2 |
| ENSMUSG( | 655.9043 | -0.0467 | 0.679575 | 0.930983 | Ccnc |
| ENSMUSG( | 2610.096 | -0.10731 | 0.345592 | 0.779153 | Ccnd1 |
| ENSMUSG( | 4520.522 | 0.07385 | 0.464795 | 0.843568 | Ccnd2 |
| ENSMUSG( | 1306.774 | -0.01821 | 0.841695 | 0.971179 | Ccnd3 |
| ENSMUSG( | 1312.455 | 0.039142 | 0.706936 | 0.93949 | Ccndbp1 |
| ENSMUSG( | 305.8667 | -0.11479 | 0.486645 | 0.858713 | Ccne1 |
| ENSMUSG( | 55.24553 | -0.19862 | 0.448887 | 0.83478 | Ccne2 |
| ENSMUSG( | 139.3245 | -0.04129 | 0.817207 | 0.965693 | Ccnf |
| ENSMUSG( | 2818.867 | 0.073546 | 0.513033 | 0.871762 | Ccng1 |
| ENSMUSG( | 999.458 | 0.016703 | 0.859712 | 0.97481 | Ccng2 |
| ENSMUSG( | 895.1982 | -0.06976 | 0.510344 | 0.870063 | Ccnh |
| ENSMUSG( | 15197.45 | 0.086021 | 0.411419 | 0.819301 | Ccni |
| ENSMUSG( | 280.896 | -0.17907 | 0.198934 | 0.639205 | Ccnj |
| ENSMUSG( | 120.3736 | -0.07556 | 0.691302 | 0.933881 | Ccnjl |

| | | | | | |
|---|---|---|---|---|---|
| ENSMUSG( | 63.74941 | -0.25243 | 0.341889 | 0.909146 | Ccdc103 |
| ENSMUSG( | 989.8915 | -0.03436 | 0.803877 | 0.977 | Ccdc106 |
| ENSMUSG( | 855.2396 | -0.0378 | 0.772064 | 0.974236 | Ccdc107 |
| ENSMUSG( | 13.75386 | -0.20532 | 0.697545 | 0.960867 | Ccdc110 |
| ENSMUSG( | 191.5299 | 0.075867 | 0.697651 | 0.960867 | Ccdc112 |
| ENSMUSG( | 46.35319 | 0.071872 | 0.81826 | 0.979198 | Ccdc113 |
| ENSMUSG( | 352.6683 | -0.14345 | 0.350619 | 0.911652 | Ccdc115 |
| ENSMUSG( | 110.8419 | 0.485196 | 0.027963 | 0.61023 | Ccdc116 |
| ENSMUSG( | 489.2401 | 0.132993 | 0.34549 | 0.910581 | Ccdc117 |
| ENSMUSG( | 668.4572 | -0.2573 | 0.045621 | 0.693959 | Ccdc12 |
| ENSMUSG( | 314.6981 | 0.087984 | 0.603722 | 0.948137 | Ccdc120 |
| ENSMUSG( | 23.65411 | 0.674446 | 0.145059 | 0.83339 | Ccdc121rt1 |
| ENSMUSG( | 100.7667 | -0.24302 | 0.330005 | 0.907251 | Ccdc122 |
| ENSMUSG( | 2129.171 | -0.32725 | 0.011492 | 0.436431 | Ccdc124 |
| ENSMUSG( | 24.74039 | -0.53159 | 0.246978 | 0.877904 | Ccdc125 |
| ENSMUSG( | 262.6873 | 0.295247 | 0.151118 | 0.837173 | Ccdc126 |
| ENSMUSG( | 2722.455 | 0.022078 | 0.864642 | 0.986591 | Ccdc127 |
| ENSMUSG( | 391.5235 | -0.23818 | 0.105698 | 0.804969 | Ccdc13 |
| ENSMUSG( | 514.7622 | 0.378086 | 0.041688 | 0.690304 | Ccdc134 |
| ENSMUSG( | 1661.752 | -0.09689 | 0.412959 | 0.917257 | Ccdc136 |
| ENSMUSG( | 657.6874 | -0.03702 | 0.768245 | 0.974236 | Ccdc137 |
| ENSMUSG( | 123.1034 | -0.0802 | 0.693622 | 0.960767 | Ccdc138 |
| ENSMUSG( | 81.23935 | -0.04065 | 0.876784 | 0.989644 | Ccdc14 |
| ENSMUSG( | 1102.098 | 0.159891 | 0.246501 | 0.877904 | Ccdc141 |
| ENSMUSG( | 20.51165 | 0.724388 | 0.145877 | 0.835074 | Ccdc146 |
| ENSMUSG( | 278.1428 | 0.130567 | 0.394774 | 0.917257 | Ccdc148 |
| ENSMUSG( | 1308.234 | -0.14466 | 0.233031 | 0.868507 | Ccdc149 |
| ENSMUSG( | 42.83681 | -0.19735 | 0.553242 | 0.939565 | Ccdc15 |
| ENSMUSG( | 11.38098 | 0.120045 | 0.839788 | NA | Ccdc152 |
| ENSMUSG( | 102.3746 | 0.501414 | 0.115752 | 0.813384 | Ccdc153 |
| ENSMUSG( | 1472.943 | 0.100435 | 0.358353 | 0.913759 | Ccdc157 |
| ENSMUSG( | 170.255 | 0.358921 | 0.153968 | 0.837724 | Ccdc159 |
| ENSMUSG( | 26.02727 | -0.16375 | 0.680805 | 0.958395 | Ccdc160 |
| ENSMUSG( | 60.87495 | 0.169721 | 0.562413 | 0.940761 | Ccdc162 |
| ENSMUSG( | 87.46889 | 0.05901 | 0.839952 | 0.98221 | Ccdc163 |
| ENSMUSG( | 115.3565 | -0.08927 | 0.697975 | 0.960867 | Ccdc166 |
| ENSMUSG( | 755.4196 | -0.03537 | 0.771487 | 0.974236 | Ccdc167 |
| ENSMUSG( | 64.72633 | 0.226258 | 0.506063 | 0.927825 | Ccdc17 |
| ENSMUSG( | 25.20145 | 0.364853 | 0.440739 | 0.921166 | Ccdc170 |
| ENSMUSG( | 63.47102 | -0.12655 | 0.662139 | 0.956205 | Ccdc171 |
| ENSMUSG( | 349.5648 | 0.092471 | 0.528591 | 0.934795 | Ccdc174 |
| ENSMUSG( | 290.9296 | 0.085932 | 0.591538 | 0.944975 | Ccdc177 |
| ENSMUSG( | 80.84886 | -0.30918 | 0.213287 | 0.864412 | Ccdc18 |
| ENSMUSG( | 27.65864 | 0.870575 | 0.044905 | 0.693959 | Ccdc180 |
| ENSMUSG( | 422.9135 | -0.04999 | 0.766297 | 0.974236 | Ccdc181 |
| ENSMUSG( | 333.9933 | -0.35176 | 0.076698 | 0.770558 | Ccdc184 |
| ENSMUSG( | 797.1834 | 0.076625 | 0.539293 | 0.937216 | Ccdc186 |
| ENSMUSG( | 334.7798 | -0.30333 | 0.104449 | 0.804969 | Ccdc187 |
| ENSMUSG( | 1222.914 | -0.36237 | 0.010801 | 0.421367 | Ccdc190 |
| ENSMUSG( | 185.9047 | -0.13124 | 0.45842 | 0.923329 | Ccdc191 |
| ENSMUSG( | 77.05871 | -0.03335 | 0.898457 | 0.990558 | Ccdc22 |
| ENSMUSG( | 24.62852 | -0.60863 | 0.141001 | 0.830678 | Ccdc24 |
| ENSMUSG( | 791.7853 | -0.04515 | 0.788429 | 0.97637 | Ccdc25 |
| ENSMUSG( | 321.9934 | -0.04816 | 0.795104 | 0.97681 | Ccdc28a |
| ENSMUSG( | 711.5171 | 0.019698 | 0.882217 | 0.989644 | Ccdc28b |
| ENSMUSG( | 626.9297 | 0.351666 | 0.026798 | 0.603041 | Ccdc3 |
| ENSMUSG( | 282.2799 | -0.02567 | 0.894909 | 0.990558 | Ccdc30 |
| ENSMUSG( | 790.1481 | -0.08501 | 0.623965 | 0.953234 | Ccdc32 |
| ENSMUSG( | 36.08022 | 0.132486 | 0.717601 | 0.965964 | Ccdc33 |
| ENSMUSG( | 726.4604 | -0.26209 | 0.068174 | 0.75212 | Ccdc34 |
| ENSMUSG( | 500.4135 | 0.315395 | 0.05735 | 0.723362 | Ccdc39 |
| ENSMUSG( | 65.09 | 0.300574 | 0.30737 | 0.898538 | Ccdc40 |
| ENSMUSG( | 508.6064 | 0.022561 | 0.863961 | 0.986552 | Ccdc43 |
| ENSMUSG( | 4572.652 | 0.103685 | 0.450505 | 0.922202 | Ccdc47 |
| ENSMUSG( | 2185.927 | 0.155621 | 0.2245 | 0.867924 | Ccdc50 |
| ENSMUSG( | 126.5713 | 0.112011 | 0.586121 | 0.944975 | Ccdc51 |
| ENSMUSG( | 243.1086 | -0.12436 | 0.481094 | 0.924548 | Ccdc57 |
| ENSMUSG( | 537.6863 | 0.096982 | 0.472714 | 0.923329 | Ccdc59 |
| ENSMUSG( | 1445.269 | 0.109853 | 0.399188 | 0.917257 | Ccdc6 |
| ENSMUSG( | 32.27756 | -0.14847 | 0.689736 | 0.960071 | Ccdc60 |
| ENSMUSG( | 201.8892 | -0.21356 | 0.247991 | 0.877904 | Ccdc61 |
| ENSMUSG( | 132.6554 | -0.02537 | 0.909686 | 0.991958 | Ccdc62 |
| ENSMUSG( | 7.684662 | -0.06935 | 0.924852 | NA | Ccdc63 |
| ENSMUSG( | 107.3016 | -0.24971 | 0.309735 | 0.898538 | Ccdc65 |
| ENSMUSG( | 383.8916 | 0.080803 | 0.592133 | 0.944975 | Ccdc66 |
| ENSMUSG( | 13.66966 | -0.60721 | 0.266005 | 0.886134 | Ccdc69 |
| ENSMUSG( | 548.8161 | 0.007788 | 0.959816 | 0.994798 | Ccdc71 |
| ENSMUSG( | 341.1062 | 0.194832 | 0.221091 | 0.866104 | Ccdc71l |
| ENSMUSG( | 20.6691 | 0.014651 | 0.972805 | 0.996416 | Ccdc73 |
| ENSMUSG( | 1176.455 | 0.133531 | 0.235295 | 0.871741 | Ccdc74a |
| ENSMUSG( | 153.2292 | 0.093233 | 0.62489 | 0.953234 | Ccdc77 |
| ENSMUSG( | 13.31029 | 0.454021 | 0.421838 | 0.917257 | Ccdc78 |
| ENSMUSG( | 248.8488 | -0.0552 | 0.750246 | 0.970113 | Ccdc8 |
| ENSMUSG( | 541.5305 | -0.08363 | 0.543523 | 0.938063 | Ccdc80 |
| ENSMUSG( | 43.38287 | 0.569588 | 0.077977 | 0.77321 | Ccdc81 |
| ENSMUSG( | 616.6332 | 0.08062 | 0.556639 | 0.939565 | Ccdc82 |
| ENSMUSG( | 1516.833 | 0.235889 | 0.086772 | 0.782073 | Ccdc85a |
| ENSMUSG( | 3614.364 | 0.117431 | 0.307554 | 0.898538 | Ccdc85b |
| ENSMUSG( | 5005.768 | 0.080626 | 0.471933 | 0.923329 | Ccdc85c |
| ENSMUSG( | 302.8906 | -0.37496 | 0.017449 | 0.531273 | Ccdc86 |
| ENSMUSG( | 45.1391 | -0.48115 | 0.124181 | 0.814691 | Ccdc87 |
| ENSMUSG( | 1961.805 | 0.098376 | 0.474363 | 0.923329 | Ccdc88a |
| ENSMUSG( | 159.2566 | 0.094643 | 0.662397 | 0.956205 | Ccdc88b |
| ENSMUSG( | 1388.275 | -0.17969 | 0.127953 | 0.81845 | Ccdc88c |
| ENSMUSG( | 13.609 | -0.18805 | 0.720287 | 0.966887 | Ccdc89 |
| ENSMUSG( | 1055.172 | -0.06445 | 0.580543 | 0.944266 | Ccdc9 |
| ENSMUSG( | 420.5442 | -0.11517 | 0.416728 | 0.917257 | Ccdc90b |
| ENSMUSG( | 1228.753 | -0.15187 | 0.221412 | 0.866104 | Ccdc91 |
| ENSMUSG( | 1742.24 | -0.12487 | 0.365066 | 0.914449 | Ccdc92 |
| ENSMUSG( | 849.9016 | 0.051027 | 0.678884 | 0.957617 | Ccdc92b |
| ENSMUSG( | 997.9239 | 0.068706 | 0.548857 | 0.938075 | Ccdc93 |
| ENSMUSG( | 64.00389 | 0.159547 | 0.545104 | 0.938063 | Ccdc96 |
| ENSMUSG( | 1481.295 | -0.08776 | 0.456837 | 0.923329 | Ccdc97 |
| ENSMUSG( | 369.9106 | 0.058195 | 0.672614 | 0.95694 | Ccdc9b |
| ENSMUSG( | 38.45591 | -0.1737 | 0.60806 | 0.950113 | Ccer2 |
| ENSMUSG( | 182.1268 | -0.22736 | 0.233451 | 0.868507 | Cchcr1 |
| ENSMUSG( | 12668.37 | -0.07105 | 0.614054 | 0.951177 | Cck |
| ENSMUSG( | 2462.27 | 0.100292 | 0.520083 | 0.931904 | Cckbr |
| ENSMUSG( | 18.13028 | -0.31867 | 0.490812 | 0.92646 | Ccl12 |
| ENSMUSG( | 26.42926 | 0.563937 | 0.142133 | 0.830709 | Ccl17 |
| ENSMUSG( | 14.63473 | 0.231714 | 0.650291 | 0.95422 | Ccl22 |
| ENSMUSG( | 11.75981 | 0.662455 | 0.27291 | 0.889077 | Ccl24 |
| ENSMUSG( | 216.2022 | -0.12439 | 0.450034 | 0.922202 | Ccl25 |
| ENSMUSG( | 1660.395 | -0.04836 | 0.733006 | 0.967228 | Ccl27a |
| ENSMUSG( | 130.343 | 0.272525 | 0.171782 | 0.842999 | Ccl27al |
| ENSMUSG( | 49.5389 | -1.42313 | 7.66E-05 | 0.017018 | Ccl28 |
| ENSMUSG( | 12.09222 | 1.031511 | 0.097636 | 0.796743 | Ccl3 |
| ENSMUSG( | 19.0225 | -1.77959 | 0.012383 | 0.451463 | Ccl5 |
| ENSMUSG( | 144.632 | -0.15223 | 0.523675 | 0.932848 | Ccl6 |
| ENSMUSG( | 119.381 | -0.22311 | 0.388329 | 0.917257 | Ccl9 |
| ENSMUSG( | 2000.932 | -0.12691 | 0.22021 | 0.866104 | Ccm2 |
| ENSMUSG( | 119.9197 | 0.685597 | 0.009757 | 0.406458 | Ccm2l |
| ENSMUSG( | 420.0405 | -0.53918 | 0.231136 | 0.868507 | Ccn1 |
| ENSMUSG( | 1368.231 | 0.150408 | 0.433252 | 0.918434 | Ccn2 |
| ENSMUSG( | 4434.265 | 0.196514 | 0.155702 | 0.837724 | Ccn3 |
| ENSMUSG( | 80.63943 | -0.5064 | 0.048042 | 0.704961 | Ccn4 |
| ENSMUSG( | 21.47146 | 0.614804 | 0.169839 | 0.840806 | Ccn5 |
| ENSMUSG( | 27.21049 | 0.28109 | 0.456488 | 0.923329 | Ccna1 |
| ENSMUSG( | 107.7331 | -0.08259 | 0.71171 | 0.964363 | Ccna2 |
| ENSMUSG( | 40.82709 | -0.11042 | 0.735609 | 0.967344 | Ccnb1 |
| ENSMUSG( | 32.19173 | -0.10574 | 0.767864 | 0.974236 | Ccnb2 |
| ENSMUSG( | 599.536 | -0.04256 | 0.759906 | 0.972335 | Ccnc |
| ENSMUSG( | 2230.198 | -0.30927 | 0.023663 | 0.588226 | Ccnd1 |
| ENSMUSG( | 4106.792 | 0.063958 | 0.631024 | 0.953251 | Ccnd2 |
| ENSMUSG( | 1179.354 | -0.0475 | 0.667299 | 0.956766 | Ccnd3 |
| ENSMUSG( | 1127.185 | -0.13874 | 0.308683 | 0.898538 | Ccndbp1 |
| ENSMUSG( | 295.4995 | 0.070619 | 0.71537 | 0.965063 | Ccne1 |
| ENSMUSG( | 58.6334 | 0.239408 | 0.386886 | 0.917257 | Ccne2 |
| ENSMUSG( | 133.1814 | 0.085586 | 0.682026 | 0.958678 | Ccnf |
| ENSMUSG( | 2501.079 | -0.00532 | 0.969918 | 0.99594 | Ccng1 |
| ENSMUSG( | 975.0815 | 0.206469 | 0.063848 | 0.736482 | Ccng2 |
| ENSMUSG( | 814.0528 | -0.07712 | 0.526483 | 0.934374 | Ccnh |
| ENSMUSG( | 13391.77 | -0.01084 | 0.939729 | 0.994276 | Ccni |
| ENSMUSG( | 281.3726 | 0.089738 | 0.605461 | 0.94924 | Ccnj |
| ENSMUSG( | 103.726 | -0.24006 | 0.306397 | 0.898538 | Ccnjl |
| ENSMUSG( | 796.8282 | 0.090995 | 0.486696 | 0.92541 | Ccnk |
| ENSMUSG( | 1206.41 | 0.172768 | 0.264874 | 0.886134 | Ccnl1 |
| ENSMUSG( | 3127.031 | 0.290703 | 0.061103 | 0.736111 | Ccnl2 |
| ENSMUSG( | 32.08644 | -0.109 | 0.782967 | 0.976106 | Ccno |
| ENSMUSG( | 284.1003 | -0.15288 | 0.310976 | 0.899422 | Ccnq |
| ENSMUSG( | 858.3643 | 0.099075 | 0.401522 | 0.917257 | Ccnt1 |
| ENSMUSG( | 2457.383 | 0.16932 | 0.143561 | 0.831311 | Ccnt2 |
| ENSMUSG( | 4635.969 | 0.059774 | 0.575115 | 0.942912 | Ccny |
| ENSMUSG( | 558.7805 | 0.159142 | 0.204333 | 0.864412 | Ccnyl1 |
| ENSMUSG( | 2951.117 | -0.0273 | 0.838578 | 0.981622 | Ccp110 |
| ENSMUSG( | 455.6929 | 0.093903 | 0.591539 | 0.944975 | Ccpg1 |
| ENSMUSG( | 26.95611 | -0.07345 | 0.860386 | 0.986137 | Ccr1 |
| ENSMUSG( | 35.87246 | -0.19721 | 0.589329 | 0.944975 | Ccr2 |

| | | | | | | | | | | | |
|---|---|---|---|---|---|---|---|---|---|---|---|
| ENSMUSG( | Ccdc134 | 465.5222 | -0.17081 | 0.261448 | 0.711405 | ENSMUSG( | Ccdc134 | 514.7622 | 0.378086 | 0.041688 | 0.690304 |
| ENSMUSG( | Ccdc136 | 1976.93 | 0.135362 | 0.181142 | 0.618139 | ENSMUSG( | Ccdc136 | 1661.752 | -0.09689 | 0.412959 | 0.917257 |
| ENSMUSG( | Ccdc137 | 762.7908 | 0.122392 | 0.252072 | 0.701908 | ENSMUSG( | Ccdc137 | 657.6874 | -0.03702 | 0.768245 | 0.974236 |
| ENSMUSG( | Ccdc138 | 144.3181 | 0.108297 | 0.534098 | 0.880394 | ENSMUSG( | Ccdc138 | 123.1034 | -0.0802 | 0.693622 | 0.960767 |
| ENSMUSG( | Ccdc14 | 80.41699 | -0.38837 | 0.098786 | 0.480598 | ENSMUSG( | Ccdc14 | 81.23935 | -0.04065 | 0.876784 | 0.989644 |
| ENSMUSG( | Ccdc141 | 1362.622 | 0.466723 | 2.29E-06 | 0.000361 | ENSMUSG( | Ccdc141 | 1102.098 | 0.159891 | 0.246501 | 0.877904 |
| ENSMUSG( | Ccdc146 | 16.13853 | -0.19432 | 0.700734 | NA | ENSMUSG( | Ccdc146 | 20.51165 | 0.724388 | 0.145877 | 0.835074 |
| ENSMUSG( | Ccdc148 | 313.2138 | 0.205941 | 0.130578 | 0.541841 | ENSMUSG( | Ccdc148 | 278.1428 | 0.130567 | 0.394774 | 0.917257 |
| ENSMUSG( | Ccdc149 | 1550.925 | 0.082361 | 0.404689 | 0.816221 | ENSMUSG( | Ccdc149 | 1308.234 | -0.14466 | 0.233031 | 0.868507 |
| ENSMUSG( | Ccdc15 | 45.24568 | -0.3542 | 0.208105 | 0.650319 | ENSMUSG( | Ccdc15 | 42.83681 | -0.19735 | 0.553242 | 0.939565 |
| ENSMUSG( | Ccdc152 | 13.83369 | 0.403415 | 0.456558 | NA | ENSMUSG( | Ccdc152 | 11.38098 | 0.120045 | 0.839788 | NA |
| ENSMUSG( | Ccdc153 | 75.70257 | -0.70062 | 0.012962 | 0.164641 | ENSMUSG( | Ccdc153 | 102.3746 | 0.501414 | 0.115752 | 0.813384 |
| ENSMUSG( | Ccdc157 | 1590.818 | 0.054178 | 0.536014 | 0.88142 | ENSMUSG( | Ccdc157 | 1472.943 | 0.100435 | 0.358353 | 0.913759 |
| ENSMUSG( | Ccdc159 | 176.3784 | 0.198623 | 0.270366 | 0.720226 | ENSMUSG( | Ccdc159 | 170.255 | 0.358921 | 0.153968 | 0.837724 |
| ENSMUSG( | Ccdc160 | 27.02668 | -0.35696 | 0.338463 | 0.777489 | ENSMUSG( | Ccdc160 | 26.02727 | -0.16375 | 0.680805 | 0.958395 |
| ENSMUSG( | Ccdc162 | 58.20129 | -0.25967 | 0.319581 | 0.765305 | ENSMUSG( | Ccdc162 | 60.87495 | 0.169721 | 0.562413 | 0.940761 |
| ENSMUSG( | Ccdc163 | 88.66125 | -0.18944 | 0.38795 | 0.805835 | ENSMUSG( | Ccdc163 | 87.46889 | 0.05901 | 0.839952 | 0.98221 |
| ENSMUSG( | Ccdc166 | 133.3302 | 0.073071 | 0.696645 | 0.935318 | ENSMUSG( | Ccdc166 | 115.3565 | -0.08927 | 0.697975 | 0.960867 |
| ENSMUSG( | Ccdc167 | 866.9853 | 0.091366 | 0.366608 | 0.794842 | ENSMUSG( | Ccdc167 | 755.4196 | -0.03537 | 0.771487 | 0.974236 |
| ENSMUSG( | Ccdc17 | 67.55411 | 0.070724 | 0.798753 | 0.960896 | ENSMUSG( | Ccdc17 | 64.72633 | 0.226258 | 0.506063 | 0.927825 |
| ENSMUSG( | Ccdc170 | 26.57946 | 0.214113 | 0.596083 | 0.903152 | ENSMUSG( | Ccdc170 | 25.20145 | 0.364853 | 0.440739 | 0.921166 |
| ENSMUSG( | Ccdc171 | 72.34036 | -0.03573 | 0.884136 | 0.982647 | ENSMUSG( | Ccdc171 | 63.47102 | -0.12655 | 0.662139 | 0.956205 |
| ENSMUSG( | Ccdc174 | 391.8385 | 0.156224 | 0.207443 | 0.649561 | ENSMUSG( | Ccdc174 | 349.5648 | 0.092471 | 0.528591 | 0.934795 |
| ENSMUSG( | Ccdc177 | 349.7355 | 0.328187 | 0.027912 | 0.255775 | ENSMUSG( | Ccdc177 | 290.9296 | 0.085932 | 0.591538 | 0.944975 |
| ENSMUSG( | Ccdc18 | 97.17836 | -0.01921 | 0.93195 | 0.9924 | ENSMUSG( | Ccdc18 | 80.84886 | -0.30918 | 0.213287 | 0.864412 |
| ENSMUSG( | Ccdc180 | 23.21963 | 0.172517 | 0.65927 | 0.924506 | ENSMUSG( | Ccdc180 | 27.65864 | 0.870575 | 0.044905 | 0.693959 |
| ENSMUSG( | Ccdc181 | 504.6752 | 0.190823 | 0.18914 | 0.627756 | ENSMUSG( | Ccdc181 | 422.9135 | -0.04999 | 0.766297 | 0.974236 |
| ENSMUSG( | Ccdc184 | 391.0051 | -0.1386 | 0.440176 | 0.831851 | ENSMUSG( | Ccdc184 | 333.9933 | -0.35176 | 0.076698 | 0.770558 |
| ENSMUSG( | Ccdc186 | 914.6008 | 0.19778 | 0.04818 | 0.335655 | ENSMUSG( | Ccdc186 | 797.1834 | 0.076625 | 0.539293 | 0.937216 |
| ENSMUSG( | Ccdc187 | 302.062 | -1.0235 | 3.55E-06 | 0.000492 | ENSMUSG( | Ccdc187 | 334.7798 | -0.30333 | 0.104449 | 0.804969 |
| ENSMUSG( | Ccdc190 | 1609.411 | 0.181555 | 0.141254 | 0.558134 | ENSMUSG( | Ccdc190 | 1222.914 | -0.36237 | 0.010801 | 0.421367 |
| ENSMUSG( | Ccdc191 | 213.2318 | 0.001126 | 0.994665 | 0.999369 | ENSMUSG( | Ccdc191 | 185.9047 | -0.13124 | 0.45842 | 0.923329 |
| ENSMUSG( | Ccdc22 | 98.30035 | 0.38112 | 0.082377 | 0.442878 | ENSMUSG( | Ccdc22 | 77.05871 | -0.03335 | 0.898457 | 0.990558 |
| ENSMUSG( | Ccdc24 | 28.92148 | -0.37389 | 0.29604 | 0.744386 | ENSMUSG( | Ccdc24 | 24.62852 | -0.60863 | 0.141001 | 0.830678 |
| ENSMUSG( | Ccdc25 | 919.7235 | 0.123115 | 0.353923 | 0.786371 | ENSMUSG( | Ccdc25 | 791.7853 | -0.04515 | 0.788429 | 0.97637 |
| ENSMUSG( | Ccdc28a | 341.9904 | -0.14332 | 0.36277 | 0.793121 | ENSMUSG( | Ccdc28a | 321.9934 | -0.04816 | 0.795104 | 0.97681 |
| ENSMUSG( | Ccdc28b | 683.9272 | -0.39156 | 0.000498 | 0.020389 | ENSMUSG( | Ccdc28b | 711.5171 | 0.019698 | 0.882217 | 0.989644 |
| ENSMUSG( | Ccdc3 | 612.168 | 0.029053 | 0.814474 | 0.964574 | ENSMUSG( | Ccdc3 | 626.9297 | 0.351666 | 0.026798 | 0.603041 |
| ENSMUSG( | Ccdc30 | 311.8328 | 0.003831 | 0.981487 | 0.997512 | ENSMUSG( | Ccdc30 | 282.2799 | -0.02567 | 0.894909 | 0.990558 |
| ENSMUSG( | Ccdc32 | 889.5311 | -0.00506 | 0.969845 | 0.996369 | ENSMUSG( | Ccdc32 | 790.1481 | -0.08501 | 0.623965 | 0.953234 |
| ENSMUSG( | Ccdc33 | 35.15837 | -0.25354 | 0.418309 | 0.821966 | ENSMUSG( | Ccdc33 | 36.08022 | 0.132486 | 0.717601 | 0.965964 |
| ENSMUSG( | Ccdc34 | 864.7589 | -0.00977 | 0.937071 | 0.993415 | ENSMUSG( | Ccdc34 | 726.4604 | -0.26209 | 0.068174 | 0.75212 |
| ENSMUSG( | Ccdc39 | 506.2057 | 0.09664 | 0.476189 | 0.851294 | ENSMUSG( | Ccdc39 | 500.4135 | 0.315395 | 0.05735 | 0.723362 |
| ENSMUSG( | Ccdc40 | 57.43391 | -0.35942 | 0.151083 | 0.576088 | ENSMUSG( | Ccdc40 | 65.09 | 0.300574 | 0.30737 | 0.898538 |
| ENSMUSG( | Ccdc43 | 579.3912 | 0.131441 | 0.220296 | 0.665898 | ENSMUSG( | Ccdc43 | 508.6064 | 0.022561 | 0.863961 | 0.986552 |
| ENSMUSG( | Ccdc47 | 4579.202 | -0.16501 | 0.183283 | 0.62047 | ENSMUSG( | Ccdc47 | 4572.652 | 0.103685 | 0.450505 | 0.922202 |
| ENSMUSG( | Ccdc50 | 2311.102 | 0.051761 | 0.633051 | 0.914073 | ENSMUSG( | Ccdc50 | 2185.927 | 0.155621 | 0.2245 | 0.867924 |
| ENSMUSG( | Ccdc51 | 143.728 | 0.213668 | 0.227421 | 0.67424 | ENSMUSG( | Ccdc51 | 126.5713 | 0.112011 | 0.586121 | 0.944975 |
| ENSMUSG( | Ccdc57 | 277.6952 | -0.00592 | 0.971413 | 0.996415 | ENSMUSG( | Ccdc57 | 243.1086 | -0.12436 | 0.481094 | 0.924548 |
| ENSMUSG( | Ccdc59 | 534.3719 | -0.19333 | 0.125788 | 0.53257 | ENSMUSG( | Ccdc59 | 537.6863 | 0.096982 | 0.472714 | 0.923329 |
| ENSMUSG( | Ccdc6 | 1707.558 | 0.310936 | 0.001093 | 0.034959 | ENSMUSG( | Ccdc6 | 1445.269 | 0.109853 | 0.399188 | 0.917257 |
| ENSMUSG( | Ccdc60 | 36.80361 | -0.0304 | 0.930315 | 0.991848 | ENSMUSG( | Ccdc60 | 32.27756 | -0.14847 | 0.689736 | 0.960071 |
| ENSMUSG( | Ccdc61 | 244.9751 | 0.070378 | 0.614156 | 0.909405 | ENSMUSG( | Ccdc61 | 201.8892 | -0.21356 | 0.247991 | 0.877904 |
| ENSMUSG( | Ccdc62 | 146.8893 | -0.00961 | 0.954928 | 0.993874 | ENSMUSG( | Ccdc62 | 132.6554 | -0.02537 | 0.909686 | 0.991958 |
| ENSMUSG( | Ccdc63 | 9.357191 | 0.265816 | 0.673908 | NA | ENSMUSG( | Ccdc63 | 7.684662 | -0.06935 | 0.924852 | NA |
| ENSMUSG( | Ccdc65 | 129.1756 | 0.022818 | 0.920048 | 0.990247 | ENSMUSG( | Ccdc65 | 107.3016 | -0.24971 | 0.309735 | 0.898538 |
| ENSMUSG( | Ccdc66 | 419.5341 | 0.075952 | 0.560312 | 0.888934 | ENSMUSG( | Ccdc66 | 383.8916 | 0.080803 | 0.592133 | 0.944975 |
| ENSMUSG( | Ccdc69 | 18.26185 | 0.039362 | 0.9304 | NA | ENSMUSG( | Ccdc69 | 13.66966 | -0.60721 | 0.266005 | 0.886134 |
| ENSMUSG( | Ccdc71 | 609.4953 | 0.040201 | 0.749659 | 0.949664 | ENSMUSG( | Ccdc71 | 548.8161 | 0.007788 | 0.959816 | 0.994798 |
| ENSMUSG( | Ccdc71l | 357.0813 | 0.06687 | 0.621825 | 0.911432 | ENSMUSG( | Ccdc71l | 341.1062 | 0.194832 | 0.221091 | 0.866104 |
| ENSMUSG( | Ccdc73 | 21.60178 | -0.11187 | 0.785898 | 0.958774 | ENSMUSG( | Ccdc73 | 20.6691 | 0.014651 | 0.972805 | 0.996416 |
| ENSMUSG( | Ccdc74a | 1308.833 | 0.169232 | 0.061243 | 0.381822 | ENSMUSG( | Ccdc74a | 1176.455 | 0.133531 | 0.235295 | 0.871741 |
| ENSMUSG( | Ccdc77 | 168.7631 | 0.103195 | 0.526251 | 0.877265 | ENSMUSG( | Ccdc77 | 153.2292 | 0.093233 | 0.62489 | 0.953234 |
| ENSMUSG( | Ccdc78 | 11.2613 | -0.31764 | 0.635486 | NA | ENSMUSG( | Ccdc78 | 13.31029 | 0.454021 | 0.421838 | 0.917257 |
| ENSMUSG( | Ccdc8 | 289.3015 | 0.106161 | 0.45712 | 0.839312 | ENSMUSG( | Ccdc8 | 248.8488 | -0.0552 | 0.750246 | 0.970113 |
| ENSMUSG( | Ccdc80 | 596.8122 | -0.07189 | 0.564093 | 0.890139 | ENSMUSG( | Ccdc80 | 541.5305 | -0.08363 | 0.543523 | 0.938063 |
| ENSMUSG( | Ccdc81 | 35.67722 | -0.24161 | 0.523576 | 0.875628 | ENSMUSG( | Ccdc81 | 43.38287 | 0.569588 | 0.077977 | 0.77321 |
| ENSMUSG( | Ccdc82 | 652.6497 | -0.0249 | 0.838606 | 0.971111 | ENSMUSG( | Ccdc82 | 616.6332 | 0.08062 | 0.556639 | 0.939565 |
| ENSMUSG( | Ccdc85a | 1698.891 | 0.293605 | 0.002906 | 0.066108 | ENSMUSG( | Ccdc85a | 1516.833 | 0.235889 | 0.086772 | 0.782073 |
| ENSMUSG( | Ccdc85b | 3922.149 | 0.087963 | 0.333785 | 0.775602 | ENSMUSG( | Ccdc85b | 3614.364 | 0.117431 | 0.307554 | 0.898538 |
| ENSMUSG( | Ccdc85c | 5702.611 | 0.183755 | 0.043795 | 0.320589 | ENSMUSG( | Ccdc85c | 5005.768 | 0.080626 | 0.471933 | 0.923329 |
| ENSMUSG( | Ccdc86 | 351.1588 | -0.19379 | 0.296715 | 0.745349 | ENSMUSG( | Ccdc86 | 302.8906 | -0.37496 | 0.017449 | 0.531273 |
| ENSMUSG( | Ccdc87 | 60.09915 | 0.086451 | 0.726471 | 0.944223 | ENSMUSG( | Ccdc87 | 45.1391 | -0.48115 | 0.124181 | 0.814691 |
| ENSMUSG( | Ccdc88a | 2157.988 | 0.105409 | 0.398811 | 0.812878 | ENSMUSG( | Ccdc88a | 1961.805 | 0.098376 | 0.474363 | 0.923329 |
| ENSMUSG( | Ccdc88b | 165.4069 | -0.07491 | 0.668869 | 0.927963 | ENSMUSG( | Ccdc88b | 159.2566 | 0.094643 | 0.662397 | 0.956205 |
| ENSMUSG( | Ccdc88c | 1490.382 | -0.24772 | 0.012803 | 0.163512 | ENSMUSG( | Ccdc88c | 1388.275 | -0.17969 | 0.127953 | 0.81845 |
| ENSMUSG( | Ccdc89 | 14.81235 | -0.24387 | 0.634973 | NA | ENSMUSG( | Ccdc89 | 13.609 | -0.18805 | 0.720287 | 0.966887 |
| ENSMUSG( | Ccdc9 | 1226.968 | 0.101872 | 0.316288 | 0.763453 | ENSMUSG( | Ccdc9 | 1055.172 | -0.06445 | 0.580543 | 0.944266 |
| ENSMUSG( | Ccdc90b | 494.1461 | 0.088093 | 0.478628 | 0.853679 | ENSMUSG( | Ccdc90b | 420.5442 | -0.11517 | 0.416728 | 0.917257 |
| ENSMUSG( | Ccdc91 | 1466.007 | 0.095391 | 0.325283 | 0.77061 | ENSMUSG( | Ccdc91 | 1228.753 | -0.15187 | 0.221412 | 0.866104 |
| ENSMUSG( | Ccdc92 | 2060.597 | 0.094061 | 0.451205 | 0.835567 | ENSMUSG( | Ccdc92 | 1742.24 | -0.12487 | 0.365066 | 0.914449 |
| ENSMUSG( | Ccdc92b | 940.7118 | 0.072252 | 0.52303 | 0.875628 | ENSMUSG( | Ccdc92b | 849.9016 | 0.051027 | 0.678884 | 0.957617 |
| ENSMUSG( | Ccdc93 | 1148.848 | 0.199166 | 0.041133 | 0.313521 | ENSMUSG( | Ccdc93 | 997.9239 | 0.068706 | 0.548857 | 0.938075 |
| ENSMUSG( | Ccdc96 | 62.93376 | -0.15493 | 0.572216 | 0.893584 | ENSMUSG( | Ccdc96 | 64.00389 | 0.159547 | 0.545104 | 0.938063 |
| ENSMUSG( | Ccdc97 | 1707.615 | 0.05645 | 0.524042 | 0.87583 | ENSMUSG( | Ccdc97 | 1481.295 | -0.08776 | 0.456837 | 0.923329 |
| ENSMUSG( | Ccdc9b | 395.738 | -0.01767 | 0.893644 | 0.984512 | ENSMUSG( | Ccdc9b | 369.9106 | 0.058195 | 0.672614 | 0.95694 |
| ENSMUSG( | Ccer2 | 35.7532 | -0.79071 | 0.02192 | 0.225738 | ENSMUSG( | Ccer2 | 38.45591 | -0.1737 | 0.60806 | 0.950113 |
| ENSMUSG( | Cchcr1 | 212.6441 | -0.04864 | 0.742243 | 0.948478 | ENSMUSG( | Cchcr1 | 182.1268 | -0.22736 | 0.233451 | 0.868507 |
| ENSMUSG( | Cck | 12177.75 | -0.4974 | 8.51E-05 | 0.005781 | ENSMUSG( | Cck | 12668.37 | -0.07105 | 0.614054 | 0.951177 |
| ENSMUSG( | Cckbr | 2318.008 | -0.36756 | 0.012256 | 0.160458 | ENSMUSG( | Cckbr | 2462.27 | 0.100292 | 0.520083 | 0.931904 |
| ENSMUSG( | Ccl12 | 19.49735 | -0.40965 | 0.351946 | NA | ENSMUSG( | Ccl12 | 18.13028 | -0.31867 | 0.490812 | 0.92646 |
| ENSMUSG( | Ccl17 | 21.13305 | -0.31873 | 0.481515 | NA | ENSMUSG( | Ccl17 | 26.42926 | 0.563937 | 0.142133 | 0.830709 |
| ENSMUSG( | Ccl22 | 13.14004 | -0.37256 | 0.483993 | NA | ENSMUSG( | Ccl22 | 14.63473 | 0.231714 | 0.650291 | 0.95422 |
| ENSMUSG( | Ccl24 | 10.0078 | -0.0301 | 0.958842 | NA | ENSMUSG( | Ccl24 | 11.75981 | 0.662455 | 0.27291 | 0.889077 |
| ENSMUSG( | Ccl25 | 243.4767 | -0.05538 | 0.707287 | 0.93949 | ENSMUSG( | Ccl25 | 216.2022 | -0.12439 | 0.450034 | 0.922202 |
| ENSMUSG( | Ccl27a | 1934.219 | 0.122053 | 0.285884 | 0.734989 | ENSMUSG( | Ccl27a | 1660.395 | -0.04836 | 0.733006 | 0.967228 |
| ENSMUSG( | Ccl27al | 157.3381 | 0.517013 | 0.012294 | 0.160458 | ENSMUSG( | Ccl27al | 130.343 | 0.272525 | 0.171782 | 0.842999 |
| ENSMUSG( | Ccl28 | 69.78579 | -0.39951 | 0.220296 | 0.665898 | ENSMUSG( | Ccl28 | 49.5389 | -1.42313 | 7.66E-05 | 0.017018 |
| ENSMUSG( | Ccl3 | 9.131405 | 0.193585 | 0.761235 | NA | ENSMUSG( | Ccl3 | 12.09222 | 1.031511 | 0.097636 | 0.796743 |
| ENSMUSG( | Ccl5 | 24.32439 | -0.98118 | 0.126207 | 0.53365 | ENSMUSG( | Ccl5 | 19.0225 | -1.77959 | 0.012383 | 0.451463 |
| ENSMUSG( | Ccl6 | 162.323 | -0.08154 | 0.69584 | 0.934979 | ENSMUSG( | Ccl6 | 144.632 | -0.15223 | 0.523675 | 0.932848 |
| ENSMUSG( | Ccl9 | 149.4797 | 0.161308 | 0.43591 | 0.829093 | ENSMUSG( | Ccl9 | 119.381 | -0.22311 | 0.388329 | 0.917257 |
| ENSMUSG( | Ccm2 | 2302.647 | 0.012662 | 0.877952 | 0.981431 | ENSMUSG( | Ccm2 | 2000.932 | -0.12691 | 0.22021 | 0.866104 |
| ENSMUSG( | Ccm2l | 130.4633 | 0.651478 | 0.006802 | 0.113745 | ENSMUSG( | Ccm2l | 119.9197 | 0.685597 | 0.009757 | 0.406458 |
| ENSMUSG( | Ccn1 | 379.1657 | -1.38242 | 5.09E-13 | 4.59E-10 | ENSMUSG( | Ccn1 | 420.0405 | -0.53918 | 0.231136 | 0.868507 |
| ENSMUSG( | Ccn2 | 1181.618 | -0.60864 | 0.000538 | 0.021456 | ENSMUSG( | Ccn2 | 1368.231 | 0.150408 | 0.433252 | 0.918434 |
| ENSMUSG( | Ccn3 | 4100.122 | -0.31128 | 0.017231 | 0.196681 | ENSMUSG( | Ccn3 | 4434.265 | 0.196514 | 0.155702 | 0.837724 |
| ENSMUSG( | Ccn4 | 78.98489 | -0.93311 | 0.000115 | 0.007087 | ENSMUSG( | Ccn4 | 80.63943 | -0.5064 | 0.048042 | 0.704961 |
| ENSMUSG( | Ccn5 | 12.42181 | -1.61397 | 0.008857 | NA | ENSMUSG( | Ccn5 | 21.47146 | 0.614804 | 0.169839 | 0.840806 |
| ENSMUSG( | Ccna1 | 30.98754 | 0.369369 | 0.267152 | 0.717525 | ENSMUSG( | Ccna1 | 27.21049 | 0.28109 | 0.456488 | 0.923329 |
| ENSMUSG( | Ccna2 | 113.3168 | -0.21561 | 0.276671 | 0.726854 | ENSMUSG( | Ccna2 | 107.7331 | -0.08259 | 0.71171 | 0.964363 |
| ENSMUSG( | Ccnb1 | 42.28966 | -0.25787 | 0.373566 | 0.800375 | ENSMUSG( | Ccnb1 | 40.82709 | -0.11042 | 0.735609 | 0.967344 |
| ENSMUSG( | Ccnb2 | 34.62937 | -0.19329 | 0.550701 | 0.886038 | ENSMUSG( | Ccnb2 | 32.19173 | -0.10574 | 0.767864 | 0.974236 |
| ENSMUSG( | Ccnc | 655.9043 | -0.0467 | 0.679575 | 0.930983 | ENSMUSG( | Ccnc | 599.536 | -0.04256 | 0.759906 | 0.972335 |
| ENSMUSG( | Ccnd1 | 2610.096 | -0.10731 | 0.345592 | 0.779153 | ENSMUSG( | Ccnd1 | 2230.198 | -0.30927 | 0.023663 | 0.588226 |
| ENSMUSG( | Ccnd2 | 4520.522 | 0.07385 | 0.464795 | 0.843568 | ENSMUSG( | Ccnd2 | 4106.792 | 0.063958 | 0.631024 | 0.953251 |
| ENSMUSG( | Ccnd3 | 1306.774 | -0.01821 | 0.841695 | 0.971179 | ENSMUSG( | Ccnd3 | 1179.354 | -0.0475 | 0.667299 | 0.956766 |
| ENSMUSG( | Ccndbp1 | 1312.455 | 0.039142 | 0.706936 | 0.93949 | ENSMUSG( | Ccndbp1 | 1127.185 | -0.13874 | 0.308683 | 0.898538 |
| ENSMUSG( | Ccne1 | 305.8667 | -0.11479 | 0.486645 | 0.858713 | ENSMUSG( | Ccne1 | 295.4995 | 0.070619 | 0.71537 | 0.965063 |
| ENSMUSG( | Ccne2 | 55.24553 | -0.19862 | 0.448887 | 0.83478 | ENSMUSG( | Ccne2 | 58.6334 | 0.239408 | 0.386886 | 0.917257 |
| ENSMUSG( | Ccnf | 139.3245 | -0.04129 | 0.817207 | 0.965693 | ENSMUSG( | Ccnf | 133.1814 | 0.085586 | 0.682026 | 0.958678 |
| ENSMUSG( | Ccng1 | 2818.867 | 0.073546 | 0.513033 | 0.871762 | ENSMUSG( | Ccng1 | 2501.079 | -0.00532 | 0.969918 | 0.99594 |
| ENSMUSG( | Ccng2 | 999.458 | 0.016703 | 0.859712 | 0.97481 | ENSMUSG( | Ccng2 | 975.0815 | 0.206469 | 0.063848 | 0.736482 |
| ENSMUSG( | Ccnh | 895.1982 | -0.06976 | 0.510344 | 0.870063 | ENSMUSG( | Ccnh | 814.0528 | -0.07712 | 0.526483 | 0.934374 |
| ENSMUSG( | Ccni | 15197.45 | 0.086021 | 0.411419 | 0.819301 | ENSMUSG( | Ccni | 13391.77 | -0.01084 | 0.939729 | 0.994276 |
| ENSMUSG( | Ccnj | 280.896 | -0.17907 | 0.198934 | 0.639205 | ENSMUSG( | Ccnj | 281.3726 | 0.089738 | 0.605461 | 0.94924 |
| ENSMUSG( | Ccnjl | 120.3736 | -0.07556 | 0.691302 | 0.933881 | ENSMUSG( | Ccnjl | 103.726 | -0.24006 | 0.306397 | 0.898538 |
| ENSMUSG( | Ccnk | 885.2894 | 0.128137 | 0.193915 | 0.633085 | ENSMUSG( | Ccnk | 796.8282 | 0.090995 | 0.486696 | 0.92541 |
| ENSMUSG( | Ccnl1 | 1165.733 | -0.20247 | 0.093467 | 0.469798 | ENSMUSG( | Ccnl1 | 1206.41 | 0.172768 | 0.264874 | 0.886134 |
| ENSMUSG( | Ccnl2 | 3155.063 | 0.061186 | 0.528379 | 0.878285 | ENSMUSG( | Ccnl2 | 3127.031 | 0.290703 | 0.061103 | 0.736111 |
| ENSMUSG( | Ccno | 32.71538 | -0.3505 | 0.362593 | 0.792943 | ENSMUSG( | Ccno | 32.08644 | -0.109 | 0.782967 | 0.976106 |
| ENSMUSG( | Ccnq | 295.5937 | -0.32369 | 0.023797 | 0.235641 | ENSMUSG( | Ccnq | 284.1003 | -0.15288 | 0.310976 | 0.899422 |
| ENSMUSG( | Ccnt1 | 893.2474 | -0.0539 | 0.598389 | 0.903801 | ENSMUSG( | Ccnt1 | 858.3643 | 0.099075 | 0.401522 | 0.917257 |
| ENSMUSG( | Ccnt2 | 2489.367 | -0.05755 | 0.557034 | 0.888198 | ENSMUSG( | Ccnt2 | 2457.383 | 0.16932 | 0.143561 | 0.831311 |
| ENSMUSG( | Ccny | 5214.445 | 0.129403 | 0.125927 | 0.532882 | ENSMUSG( | Ccny | 4635.969 | 0.059774 | 0.575115 | 0.942912 |
| ENSMUSG( | Ccnyl1 | 577.2126 | -0.01195 | 0.910314 | 0.9881 | ENSMUSG( | Ccnyl1 | 558.7805 | 0.159142 | 0.204333 | 0.864412 |
| ENSMUSG( | Ccp110 | 3008.117 | -0.25469 | 0.009869 | 0.141868 | ENSMUSG( | Ccp110 | 2951.117 | -0.0273 | 0.838578 | 0.981622 |
| ENSMUSG( | Ccpg1 | 469.2784 | -0.09137 | 0.552573 | 0.886141 | ENSMUSG( | Ccpg1 | 455.6929 | 0.093903 | 0.591539 | 0.944975 |
| ENSMUSG( | Ccr1 | 28.3765 | -0.22025 | 0.571295 | 0.893319 | ENSMUSG( | Ccr1 | 26.95611 | -0.07345 | 0.860386 | 0.986137 |
| ENSMUSG( | Ccr2 | 37.67693 | -0.31267 | 0.354827 | 0.78738 | ENSMUSG( | Ccr2 | 35.87246 | -0.19721 | 0.589329 | 0.944975 |
| ENSMUSG( | Ccr5 | 235.0308 | -0.29544 | 0.225313 | 0.672044 | ENSMUSG( | Ccr5 | 297.5152 | 0.616914 | 0.021953 | 0.575341 |
| ENSMUSG( | Ccrl2 | 30.42285 | -0.33624 | 0.38471 | 0.804031 | ENSMUSG( | Ccrl2 | 36.36337 | 0.453616 | 0.22489 | 0.86838 |
| ENSMUSG( | Ccs | 619.2863 | -0.00326 | 0.981236 | 0.997512 | ENSMUSG( | Ccs | 515.433 | -0.28031 | 0.053962 | 0.722223 |
| ENSMUSG( | Ccsap | 711.03 | 0.074675 | 0.60686 | 0.906685 | ENSMUSG( | Ccsap | 667.1981 | 0.157619 | 0.308268 | 0.898538 |
| ENSMUSG( | Ccser1 | 151.0622 | 0.166829 | 0.345917 | 0.779153 | ENSMUSG( | Ccser1 | 137.6495 | 0.165788 | 0.404575 | 0.917257 |
| ENSMUSG( | Ccser2 | 4114.435 | -0.00238 | 0.982148 | 0.997512 | ENSMUSG( | Ccser2 | 3701.677 | -0.04029 | 0.745859 | 0.96951 |
| ENSMUSG( | Cct2 | 3982.35 | 0.015257 | 0.872937 | 0.978753 | ENSMUSG( | Cct2 | 3374.976 | -0.20317 | 0.101158 | 0.801456 |
| ENSMUSG( | Cct3 | 2556.892 | 0.083168 | 0.476656 | 0.851717 | ENSMUSG( | Cct3 | 2119.867 | -0.19664 | 0.145371 | 0.8336 |
| ENSMUSG( | Cct4 | 2038.288 | 0.14525 | 0.136484 | 0.550651 | ENSMUSG( | Cct4 | 1712.915 | -0.08881 | 0.480917 | 0.924548 |
| ENSMUSG( | Cct5 | 3225.502 | 0.118716 | 0.293246 | 0.742763 | ENSMUSG( | Cct5 | 2729.585 | -0.09608 | 0.482949 | 0.924671 |
| ENSMUSG( | Cct6a | 3032.207 | 0.128995 | 0.220549 | 0.666079 | ENSMUSG( | Cct6a | 2563.501 | -0.08747 | 0.520853 | 0.932234 |
| ENSMUSG( | Cct7 | 4215.626 | 0.188948 | 0.155951 | 0.58375 | ENSMUSG( | Cct7 | 3323.735 | -0.23517 | 0.150514 | 0.837173 |
| ENSMUSG( | Cct8 | 3621.252 | 0.117251 | 0.382399 | 0.803227 | ENSMUSG( | Cct8 | 2962.374 | -0.20321 | 0.229694 | 0.868507 |
| ENSMUSG( | Ccz1 | 968.1054 | 0.035593 | 0.718957 | 0.941986 | ENSMUSG( | Ccz1 | 849.682 | -0.07581 | 0.521999 | 0.932571 |
| ENSMUSG( | Cd101 | 16.30192 | 0.272256 | 0.594656 | NA | ENSMUSG( | Cd101 | 13.09038 | -0.06204 | 0.91487 | 0.992407 |
| ENSMUSG( | Cd109 | 84.89947 | 0.042215 | 0.854123 | 0.973491 | ENSMUSG( | Cd109 | 91.54279 | 0.507531 | 0.060624 | 0.734317 |
| ENSMUSG( | Cd14 | 132.2252 | 0.039341 | 0.859669 | 0.97481 | ENSMUSG( | Cd14 | 155.1546 | 0.702734 | 0.009799 | 0.406458 |
| ENSMUSG( | Cd151 | 1187.507 | -0.1637 | 0.261409 | 0.711405 | ENSMUSG( | Cd151 | 1198.355 | 0.137593 | 0.422183 | 0.917257 |

| | | | | | |
|---|---|---|---|---|---|
| ENSMUSG( | 885.2894 | 0.128137 | 0.193915 | 0.633085 | Ccnk |
| ENSMUSG( | 1165.733 | -0.20247 | 0.093467 | 0.469798 | Ccnl1 |
| ENSMUSG( | 3155.063 | 0.061186 | 0.528379 | 0.878285 | Ccnl2 |
| ENSMUSG( | 32.71538 | -0.3505 | 0.362593 | 0.792943 | Ccno |
| ENSMUSG( | 295.5937 | -0.32369 | 0.023797 | 0.235641 | Ccnq |
| ENSMUSG( | 893.2474 | -0.0539 | 0.598389 | 0.903801 | Ccnt1 |
| ENSMUSG( | 2489.367 | -0.05755 | 0.557034 | 0.888198 | Ccnt2 |
| ENSMUSG( | 5214.445 | 0.129403 | 0.125927 | 0.532882 | Ccny |
| ENSMUSG( | 577.2126 | -0.01195 | 0.910314 | 0.9881 | Ccnyl1 |
| ENSMUSG( | 3008.117 | -0.25469 | 0.009869 | 0.141868 | Ccp110 |
| ENSMUSG( | 469.2784 | -0.09137 | 0.552573 | 0.886141 | Ccpg1 |
| ENSMUSG( | 28.3765 | -0.22025 | 0.571295 | 0.893319 | Ccr1 |
| ENSMUSG( | 37.67693 | -0.31267 | 0.354827 | 0.78738 | Ccr2 |
| ENSMUSG( | 235.0308 | -0.29544 | 0.225313 | 0.672044 | Ccr5 |
| ENSMUSG( | 30.42285 | -0.33624 | 0.38471 | 0.804031 | Ccrl2 |
| ENSMUSG( | 619.2863 | -0.00326 | 0.981236 | 0.997512 | Ccs |
| ENSMUSG( | 711.03 | 0.074675 | 0.60686 | 0.906685 | Ccsap |
| ENSMUSG( | 151.0622 | 0.166829 | 0.345917 | 0.779153 | Ccser1 |
| ENSMUSG( | 4114.435 | -0.00238 | 0.982148 | 0.997512 | Ccser2 |
| ENSMUSG( | 3982.35 | 0.015257 | 0.872937 | 0.978753 | Cct2 |
| ENSMUSG( | 2556.892 | 0.083168 | 0.476656 | 0.851717 | Cct3 |
| ENSMUSG( | 2038.288 | 0.14525 | 0.136484 | 0.550651 | Cct4 |
| ENSMUSG( | 3225.502 | 0.118716 | 0.293246 | 0.742763 | Cct5 |
| ENSMUSG( | 3032.207 | 0.128995 | 0.220549 | 0.666079 | Cct6a |
| ENSMUSG( | 4215.626 | 0.188948 | 0.155951 | 0.58375 | Cct7 |
| ENSMUSG( | 3621.252 | 0.117251 | 0.382399 | 0.803227 | Cct8 |
| ENSMUSG( | 968.1054 | 0.035593 | 0.718957 | 0.941986 | Ccz1 |
| ENSMUSG( | 16.30192 | 0.272256 | 0.594656 | NA | Cd101 |
| ENSMUSG( | 84.89947 | 0.042215 | 0.854123 | 0.973491 | Cd109 |
| ENSMUSG( | 132.2252 | 0.039341 | 0.859669 | 0.97481 | Cd14 |
| ENSMUSG( | 1187.507 | -0.1637 | 0.261409 | 0.711405 | Cd151 |
| ENSMUSG( | 107.4806 | 0.497403 | 0.015857 | 0.186421 | Cd163 |
| ENSMUSG( | 2359.164 | -0.30733 | 0.06839 | 0.405144 | Cd164 |
| ENSMUSG( | 164.9926 | 0.029298 | 0.856019 | 0.973813 | Cd164l2 |
| ENSMUSG( | 11.36167 | -0.83623 | 0.269792 | NA | Cd177 |
| ENSMUSG( | 159.3131 | -0.88874 | 3.57E-05 | 0.003067 | Cd180 |
| ENSMUSG( | 43.95301 | 0.127263 | 0.676587 | 0.930542 | Cd1d1 |
| ENSMUSG( | 2348.987 | -0.39504 | 0.015143 | 0.181303 | Cd200 |
| ENSMUSG( | 14.73041 | -0.06247 | 0.908571 | NA | Cd200r1 |
| ENSMUSG( | 10.00699 | 0.17978 | 0.7778 | NA | Cd209a |
| ENSMUSG( | 18.48335 | -0.02912 | 0.948238 | NA | Cd209f |
| ENSMUSG( | 16.81965 | -0.19554 | 0.66171 | NA | Cd22 |
| ENSMUSG( | 229.9394 | -0.43195 | 0.043035 | 0.31819 | Cd248 |
| ENSMUSG( | 482.9938 | -0.36046 | 0.021902 | 0.225738 | Cd24a |
| ENSMUSG( | 67.46751 | -0.16953 | 0.486682 | 0.858713 | Cd274 |
| ENSMUSG( | 465.457 | -0.39252 | 0.010317 | 0.144633 | Cd276 |
| ENSMUSG( | 761.9627 | -0.20676 | 0.107342 | 0.495755 | Cd2ap |
| ENSMUSG( | 2648.642 | 0.041631 | 0.602113 | 0.904887 | Cd2bp2 |
| ENSMUSG( | 107.8052 | -0.3669 | 0.187695 | 0.626295 | Cd300a |
| ENSMUSG( | 220.0389 | -0.13478 | 0.418512 | 0.821966 | Cd300c2 |
| ENSMUSG( | 8.27189 | 0.406949 | 0.613995 | NA | Cd300lb |
| ENSMUSG( | 21.40428 | -1.42124 | 0.012156 | 0.159821 | Cd300ld |
| ENSMUSG( | 10.70734 | -0.34045 | 0.601973 | NA | Cd300lf |
| ENSMUSG( | 695.6807 | -0.05925 | 0.640409 | 0.916434 | Cd302 |
| ENSMUSG( | 497.5454 | -0.18793 | 0.240759 | 0.688247 | Cd320 |
| ENSMUSG( | 439.1057 | -0.38584 | 0.014778 | 0.178616 | Cd33 |
| ENSMUSG( | 2637.213 | -0.23993 | 0.131936 | 0.544244 | Cd34 |
| ENSMUSG( | 59.96084 | -0.52183 | 0.112457 | 0.505997 | Cd36 |
| ENSMUSG( | 296.5726 | -0.23581 | 0.12116 | 0.525683 | Cd37 |
| ENSMUSG( | 352.5242 | -0.31944 | 0.046133 | 0.329204 | Cd38 |
| ENSMUSG( | 162.5858 | -1.1698 | 0.018966 | 0.207297 | Cd4 |
| ENSMUSG( | 34.49739 | -0.19773 | 0.539309 | 0.883094 | Cd40 |
| ENSMUSG( | 201.6448 | -0.12956 | 0.546672 | 0.884974 | Cd44 |
| ENSMUSG( | 256.3437 | -0.28672 | 0.121037 | 0.525487 | Cd46 |
| ENSMUSG( | 5045.727 | -0.52786 | 0.00021 | 0.011071 | Cd47 |
| ENSMUSG( | 38.60133 | -0.76855 | 0.040916 | 0.312769 | Cd48 |
| ENSMUSG( | 133.0533 | -0.65901 | 0.000688 | 0.025203 | Cd52 |
| ENSMUSG( | 512.0747 | -0.35437 | 0.054877 | 0.360515 | Cd53 |
| ENSMUSG( | 174.9969 | -0.45199 | 0.015473 | 0.18373 | Cd55 |
| ENSMUSG( | 228.4049 | 0.423793 | 0.033804 | 0.282338 | Cd59a |
| ENSMUSG( | 19.85417 | -0.00272 | 0.994877 | NA | Cd59b |
| ENSMUSG( | 102.0231 | -0.59913 | 0.012814 | 0.163518 | Cd6 |
| ENSMUSG( | 4154.78 | -0.12403 | 0.275751 | 0.725669 | Cd63 |
| ENSMUSG( | 638.1372 | -0.2719 | 0.041889 | 0.315367 | Cd68 |
| ENSMUSG( | 42.88195 | -0.64984 | 0.030968 | 0.267891 | Cd7 |
| ENSMUSG( | 60.45545 | -0.50352 | 0.04278 | 0.316989 | Cd72 |
| ENSMUSG( | 386.0178 | -0.55306 | 2.31E-05 | 0.002143 | Cd74 |
| ENSMUSG( | 25.50045 | -0.18413 | 0.633488 | 0.914073 | Cd79a |
| ENSMUSG( | 29.93713 | -0.59824 | 0.082119 | 0.442642 | Cd79b |
| ENSMUSG( | 29.41299 | -0.5059 | 0.175404 | 0.610788 | Cd80 |
| ENSMUSG( | 17152.06 | -0.42079 | 0.001195 | 0.037142 | Cd81 |
| ENSMUSG( | 1185.836 | -0.6592 | 3.24E-07 | 7.29E-05 | Cd82 |
| ENSMUSG( | 344.4012 | -0.2158 | 0.19949 | 0.640054 | Cd83 |
| ENSMUSG( | 198.033 | -0.31414 | 0.093456 | 0.469798 | Cd84 |
| ENSMUSG( | 161.8977 | -0.25722 | 0.173264 | 0.608563 | Cd86 |
| ENSMUSG( | 3060.016 | -0.66211 | 1.16E-06 | 0.000211 | Cd9 |
| ENSMUSG( | 499.5409 | -0.23359 | 0.240715 | 0.688247 | Cd93 |
| ENSMUSG( | 2918.518 | -0.24327 | 0.090923 | 0.463326 | Cd99l2 |
| ENSMUSG( | 24.91423 | 0.023954 | 0.949396 | 0.993821 | Cda |
| ENSMUSG( | 1507.741 | -0.1027 | 0.275229 | 0.72509 | Cdadc1 |
| ENSMUSG( | 830.7874 | 0.05814 | 0.57253 | 0.893584 | Cdan1 |
| ENSMUSG( | 1595.286 | 0.017253 | 0.840421 | 0.971179 | Cdc123 |
| ENSMUSG( | 194.9762 | -0.14716 | 0.438867 | 0.830714 | Cdc14a |
| ENSMUSG( | 296.0637 | -0.05523 | 0.717799 | 0.941697 | Cdc14b |
| ENSMUSG( | 1213.431 | 0.006351 | 0.944951 | 0.993612 | Cdc16 |
| ENSMUSG( | 36.56862 | -0.0256 | 0.94284 | 0.993415 | Cdc20 |
| ENSMUSG( | 1586.553 | 0.041052 | 0.658631 | 0.924506 | Cdc23 |
| ENSMUSG( | 159.1718 | 0.111078 | 0.503115 | 0.865929 | Cdc25a |
| ENSMUSG( | 611.1459 | 0.310641 | 0.058514 | 0.373218 | Cdc25b |
| ENSMUSG( | 15.96368 | -0.17586 | 0.709672 | NA | Cdc25c |
| ENSMUSG( | 522.3082 | -0.0246 | 0.848699 | 0.972374 | Cdc26 |
| ENSMUSG( | 1423.751 | 0.027361 | 0.770243 | 0.954689 | Cdc27 |
| ENSMUSG( | 1849.075 | 0.119234 | 0.195231 | 0.6346 | Cdc34 |
| ENSMUSG( | 2769.161 | 0.040503 | 0.649012 | 0.920025 | Cdc37 |
| ENSMUSG( | 3653.743 | 0.046207 | 0.616863 | 0.910866 | Cdc37l1 |
| ENSMUSG( | 904.6025 | 0.055792 | 0.657287 | 0.924144 | Cdc40 |
| ENSMUSG( | 8175.899 | 0.017378 | 0.845584 | 0.97218 | Cdc42 |
| ENSMUSG( | 14746.18 | 0.170842 | 0.0235 | 0.234568 | Cdc42bpa |
| ENSMUSG( | 6708.185 | 0.16621 | 0.070211 | 0.40908 | Cdc42bpb |
| ENSMUSG( | 737.931 | 0.086188 | 0.413287 | 0.819541 | Cdc42bpg |
| ENSMUSG( | 684.9908 | -0.31211 | 0.003366 | 0.073204 | Cdc42ep1 |
| ENSMUSG( | 215.8356 | -0.15102 | 0.357028 | 0.788702 | Cdc42ep2 |
| ENSMUSG( | 216.7843 | -0.40267 | 0.008619 | 0.130286 | Cdc42ep3 |
| ENSMUSG( | 2311.189 | -0.01009 | 0.914637 | 0.988501 | Cdc42ep4 |
| ENSMUSG( | 39.83126 | -0.33072 | 0.289677 | 0.739839 | Cdc42ep5 |
| ENSMUSG( | 3022.368 | 0.123269 | 0.14167 | 0.558688 | Cdc42se1 |
| ENSMUSG( | 4859.635 | 0.005472 | 0.945499 | 0.99365 | Cdc42se2 |
| ENSMUSG( | 64.75764 | 0.457907 | 0.056994 | 0.368322 | Cdc45 |
| ENSMUSG( | 899.1096 | 0.190653 | 0.059914 | 0.378725 | Cdc5l |
| ENSMUSG( | 24.44506 | 0.383513 | 0.329987 | 0.774026 | Cdc6 |
| ENSMUSG( | 337.2502 | 0.186647 | 0.17434 | 0.610304 | Cdc7 |
| ENSMUSG( | 905.3489 | 0.126601 | 0.259319 | 0.709948 | Cdc73 |
| ENSMUSG( | 18.07812 | -0.11887 | 0.803646 | NA | Cdca2 |
| ENSMUSG( | 55.11637 | -0.44181 | 0.09733 | 0.477629 | Cdca3 |
| ENSMUSG( | 141.1076 | 0.034209 | 0.861114 | 0.975026 | Cdca4 |
| ENSMUSG( | 14.4693 | 0.304976 | 0.550301 | NA | Cdca5 |
| ENSMUSG( | 162.8998 | -0.14332 | 0.436153 | 0.829093 | Cdca7 |
| ENSMUSG( | 38.00336 | 0.107916 | 0.731383 | 0.944956 | Cdca7l |
| ENSMUSG( | 49.02776 | -0.11969 | 0.660818 | 0.924962 | Cdca8 |
| ENSMUSG( | 13.33476 | -0.67315 | 0.191965 | NA | Cdcp1 |
| ENSMUSG( | 212.5003 | -0.14863 | 0.358991 | 0.790503 | Cdh1 |
| ENSMUSG( | 1536.099 | -0.45536 | 0.008077 | 0.125133 | Cdh10 |
| ENSMUSG( | 2059.303 | -0.46702 | 0.001641 | 0.045581 | Cdh11 |
| ENSMUSG( | 159.1818 | -0.78042 | 0.000111 | 0.00695 | Cdh12 |
| ENSMUSG( | 2347.406 | -0.4328 | 0.001719 | 0.046884 | Cdh13 |
| ENSMUSG( | 40.96653 | -0.29004 | 0.361173 | 0.792797 | Cdh15 |
| ENSMUSG( | 402.568 | -0.47328 | 0.015314 | 0.18239 | Cdh18 |
| ENSMUSG( | 706.2037 | 0.058678 | 0.713118 | 0.940734 | Cdh19 |
| ENSMUSG( | 3651.572 | -0.15512 | 0.29536 | 0.743814 | Cdh2 |
| ENSMUSG( | 1268.691 | -0.09601 | 0.487817 | 0.858824 | Cdh20 |
| ENSMUSG( | 1467.41 | 0.023041 | 0.833063 | 0.970333 | Cdh22 |
| ENSMUSG( | 58.64411 | -0.01898 | 0.943386 | 0.993415 | Cdh23 |
| ENSMUSG( | 441.1926 | 0.1495 | 0.317908 | 0.764145 | Cdh24 |
| ENSMUSG( | 28.73335 | -0.03808 | 0.920055 | 0.990247 | Cdh26 |
| ENSMUSG( | 15.81102 | -0.53819 | 0.348991 | NA | Cdh3 |
| ENSMUSG( | 857.4603 | -0.42833 | 0.003211 | 0.070583 | Cdh4 |
| ENSMUSG( | 723.951 | -0.05229 | 0.807776 | 0.963257 | Cdh5 |
| ENSMUSG( | 706.1736 | -0.67505 | 8.55E-05 | 0.005781 | Cdh6 |
| ENSMUSG( | 280.9809 | -0.27801 | 0.109423 | 0.499391 | Cdh7 |
| ENSMUSG( | 853.2092 | -0.50703 | 0.001039 | 0.033844 | Cdh8 |
| ENSMUSG( | 495.5066 | -0.47007 | 0.008452 | 0.128676 | Cdh9 |
| ENSMUSG( | 168.1625 | -0.68835 | 0.011509 | 0.154494 | Cdhr1 |
| ENSMUSG( | 79.85213 | -0.12487 | 0.632873 | 0.914073 | Cdhr3 |
| ENSMUSG( | 38.95063 | 0.22296 | 0.51657 | 0.872788 | Cdhr4 |
| ENSMUSG( | 172.983 | -0.01719 | 0.914044 | 0.988483 | Cdin1 |
| ENSMUSG( | 6348.749 | 0.115865 | 0.162923 | 0.593778 | Cdip1 |
| ENSMUSG( | 5175.236 | -0.32842 | 0.012552 | 0.161972 | Cdipt |
| ENSMUSG( | 33.5482 | -0.28241 | 0.408448 | 0.818167 | Cdk1 |

| | | | | | |
|---|---|---|---|---|---|
| ENSMUSG( | 297.5152 | 0.616914 | 0.021953 | 0.575341 | Ccr5 |
| ENSMUSG( | 36.36337 | 0.453616 | 0.22489 | 0.86838 | Ccrl2 |
| ENSMUSG( | 515.433 | -0.28031 | 0.053962 | 0.722223 | Ccs |
| ENSMUSG( | 667.1981 | 0.157619 | 0.308268 | 0.898538 | Ccsap |
| ENSMUSG( | 137.6495 | 0.165788 | 0.404575 | 0.917257 | Ccser1 |
| ENSMUSG( | 3701.677 | -0.04029 | 0.745859 | 0.96951 | Ccser2 |
| ENSMUSG( | 3374.976 | -0.20317 | 0.101158 | 0.801456 | Cct2 |
| ENSMUSG( | 2119.867 | -0.19664 | 0.145371 | 0.8336 | Cct3 |
| ENSMUSG( | 1712.915 | -0.08881 | 0.480917 | 0.924548 | Cct4 |
| ENSMUSG( | 2729.585 | -0.09608 | 0.482949 | 0.924671 | Cct5 |
| ENSMUSG( | 2563.501 | -0.08747 | 0.520853 | 0.932234 | Cct6a |
| ENSMUSG( | 3323.735 | -0.23517 | 0.150514 | 0.837173 | Cct7 |
| ENSMUSG( | 2962.374 | -0.20321 | 0.229694 | 0.868507 | Cct8 |
| ENSMUSG( | 849.682 | -0.07581 | 0.521999 | 0.932571 | Ccz1 |
| ENSMUSG( | 13.09038 | -0.06204 | 0.91487 | 0.992407 | Cd101 |
| ENSMUSG( | 91.54279 | 0.507531 | 0.060624 | 0.734317 | Cd109 |
| ENSMUSG( | 155.1546 | 0.702734 | 0.009799 | 0.406458 | Cd14 |
| ENSMUSG( | 1198.355 | 0.137593 | 0.422183 | 0.917257 | Cd151 |
| ENSMUSG( | 111.6262 | 0.813907 | 0.000217 | 0.038388 | Cd163 |
| ENSMUSG( | 2776.594 | 0.41912 | 0.027374 | 0.606201 | Cd164 |
| ENSMUSG( | 141.219 | -0.16141 | 0.413742 | 0.917257 | Cd164l2 |
| ENSMUSG( | 12.32479 | -0.27123 | 0.690313 | 0.960071 | Cd177 |
| ENSMUSG( | 173.096 | -0.24042 | 0.389874 | 0.917257 | Cd180 |
| ENSMUSG( | 43.02298 | 0.350522 | 0.287559 | 0.894091 | Cd1d1 |
| ENSMUSG( | 2781.642 | 0.370065 | 0.033419 | 0.643993 | Cd200 |
| ENSMUSG( | 16.86139 | 0.569461 | 0.265327 | 0.886134 | Cd200r1 |
| ENSMUSG( | 8.715729 | 0.063815 | 0.93168 | NA | Cd209a |
| ENSMUSG( | 16.56553 | -0.12322 | 0.81409 | 0.979198 | Cd209f |
| ENSMUSG( | 15.57934 | -0.11423 | 0.824998 | 0.979297 | Cd22 |
| ENSMUSG( | 277.5264 | 0.398523 | 0.117634 | 0.813384 | Cd248 |
| ENSMUSG( | 538.3435 | 0.238267 | 0.16628 | 0.840082 | Cd24a |
| ENSMUSG( | 60.65086 | -0.18826 | 0.504565 | 0.927825 | Cd274 |
| ENSMUSG( | 457.7099 | -0.14586 | 0.36477 | 0.914379 | Cd276 |
| ENSMUSG( | 10.17836 | -0.16398 | 0.799866 | NA | Cd28 |
| ENSMUSG( | 803.1372 | 0.217914 | 0.11958 | 0.813384 | Cd2ap |
| ENSMUSG( | 2281.343 | -0.12449 | 0.243674 | 0.877904 | Cd2bp2 |
| ENSMUSG( | 104.7574 | -0.16598 | 0.536301 | 0.93705 | Cd300a |
| ENSMUSG( | 211.6975 | 0.039934 | 0.834605 | 0.981237 | Cd300c2 |
| ENSMUSG( | 19.40935 | -1.54766 | 0.014916 | 0.489529 | Cd300ld |
| ENSMUSG( | 9.309095 | -0.48003 | 0.481299 | NA | Cd300lf |
| ENSMUSG( | 666.0116 | 0.085537 | 0.528595 | 0.934795 | Cd302 |
| ENSMUSG( | 502.9003 | 0.121395 | 0.478117 | 0.923329 | Cd320 |
| ENSMUSG( | 470.1727 | 0.113937 | 0.556219 | 0.939565 | Cd33 |
| ENSMUSG( | 2778.929 | 0.189543 | 0.318116 | 0.902849 | Cd34 |
| ENSMUSG( | 55.58241 | -0.47378 | 0.15892 | 0.837724 | Cd36 |
| ENSMUSG( | 315.1376 | 0.219618 | 0.188895 | 0.85698 | Cd37 |
| ENSMUSG( | 360.1853 | 0.03415 | 0.842194 | 0.983044 | Cd38 |
| ENSMUSG( | 182.6515 | -0.37323 | 0.138211 | 0.828492 | Cd4 |
| ENSMUSG( | 38.5441 | 0.383548 | 0.231274 | 0.868507 | Cd40 |
| ENSMUSG( | 197.4177 | 0.08216 | 0.725358 | 0.966887 | Cd44 |
| ENSMUSG( | 258.4165 | 0.029691 | 0.87981 | 0.989644 | Cd46 |
| ENSMUSG( | 5692.078 | 0.135214 | 0.418681 | 0.917257 | Cd47 |
| ENSMUSG( | 40.19891 | -0.26186 | 0.525285 | 0.934374 | Cd48 |
| ENSMUSG( | 124.9349 | -0.56484 | 0.006407 | 0.335013 | Cd52 |
| ENSMUSG( | 527.6949 | 0.027528 | 0.897333 | 0.990558 | Cd53 |
| ENSMUSG( | 194.0822 | 0.143347 | 0.462459 | 0.923329 | Cd55 |
| ENSMUSG( | 253.3777 | 0.894032 | 2.19E-07 | 0.000135 | Cd59a |
| ENSMUSG( | 19.79568 | 0.244665 | 0.570084 | 0.941777 | Cd59b |
| ENSMUSG( | 89.74412 | -0.73094 | 0.004213 | 0.263542 | Cd6 |
| ENSMUSG( | 3977.402 | 0.022145 | 0.858946 | 0.985884 | Cd63 |
| ENSMUSG( | 681.2038 | 0.198631 | 0.241771 | 0.876528 | Cd68 |
| ENSMUSG( | 41.64138 | -0.41443 | 0.190509 | 0.857801 | Cd7 |
| ENSMUSG( | 57.17602 | -0.36325 | 0.185618 | 0.855305 | Cd72 |
| ENSMUSG( | 372.176 | -0.35812 | 0.012309 | 0.450129 | Cd74 |
| ENSMUSG( | 19.48936 | -0.80955 | 0.080672 | 0.77681 | Cd79a |
| ENSMUSG( | 29.11797 | -0.36084 | 0.321321 | 0.904019 | Cd79b |
| ENSMUSG( | 30.79274 | -0.01526 | 0.972667 | 0.996416 | Cd80 |
| ENSMUSG( | 18037.16 | 0.025899 | 0.862032 | 0.98632 | Cd81 |
| ENSMUSG( | 1279.668 | -0.09475 | 0.543143 | 0.938063 | Cd82 |
| ENSMUSG( | 378.6972 | 0.327444 | 0.077344 | 0.771983 | Cd83 |
| ENSMUSG( | 205.2927 | 0.084791 | 0.68233 | 0.958678 | Cd84 |
| ENSMUSG( | 167.3332 | 0.126796 | 0.5501 | 0.938699 | Cd86 |
| ENSMUSG( | 3384.698 | -0.02608 | 0.869805 | 0.988097 | Cd9 |
| ENSMUSG( | 479.9521 | -0.06072 | 0.796005 | 0.97681 | Cd93 |
| ENSMUSG( | 2901.725 | 0.023574 | 0.884362 | 0.989688 | Cd99l2 |
| ENSMUSG( | 20.94284 | -0.23717 | 0.581489 | 0.944266 | Cda |
| ENSMUSG( | 1416.251 | -0.01137 | 0.91888 | 0.992529 | Cdadc1 |
| ENSMUSG( | 768.1664 | 0.103939 | 0.487333 | 0.92541 | Cdan1 |
| ENSMUSG( | 1411.702 | -0.07066 | 0.572761 | 0.94262 | Cdc123 |
| ENSMUSG( | 177.5794 | -0.13282 | 0.5059 | 0.927825 | Cdc14a |
| ENSMUSG( | 280.1919 | 0.057341 | 0.704772 | 0.962677 | Cdc14b |
| ENSMUSG( | 1077.659 | -0.06789 | 0.558087 | 0.939565 | Cdc16 |
| ENSMUSG( | 36.82939 | 0.240543 | 0.516432 | 0.930939 | Cdc20 |
| ENSMUSG( | 1424.19 | -0.00097 | 0.993352 | 0.998813 | Cdc23 |
| ENSMUSG( | 148.6586 | 0.176825 | 0.350088 | 0.911652 | Cdc25a |
| ENSMUSG( | 517.2929 | 0.113889 | 0.524761 | 0.933907 | Cdc25b |
| ENSMUSG( | 16.85583 | 0.219896 | 0.65445 | 0.955015 | Cdc25c |
| ENSMUSG( | 470.6862 | -0.05846 | 0.66508 | 0.956766 | Cdc26 |
| ENSMUSG( | 1343.456 | 0.123115 | 0.316356 | 0.901868 | Cdc27 |
| ENSMUSG( | 1558.311 | -0.10719 | 0.331436 | 0.907861 | Cdc34 |
| ENSMUSG( | 2398.874 | -0.10761 | 0.319947 | 0.903771 | Cdc37 |
| ENSMUSG( | 3550.744 | 0.225083 | 0.045351 | 0.693959 | Cdc37l1 |
| ENSMUSG( | 818.9759 | 0.037822 | 0.782792 | 0.976106 | Cdc40 |
| ENSMUSG( | 7049.006 | -0.14812 | 0.254235 | 0.881345 | Cdc42 |
| ENSMUSG( | 12434.03 | -0.04809 | 0.611665 | 0.950771 | Cdc42bpa |
| ENSMUSG( | 6102.056 | 0.163417 | 0.15906 | 0.837724 | Cdc42bpb |
| ENSMUSG( | 644.0176 | -0.04326 | 0.751596 | 0.97035 | Cdc42bpg |
| ENSMUSG( | 695.3918 | 0.019583 | 0.895195 | 0.990558 | Cdc42ep1 |
| ENSMUSG( | 210.0483 | 0.035888 | 0.842311 | 0.983044 | Cdc42ep2 |
| ENSMUSG( | 220.9452 | -0.05471 | 0.745852 | 0.96951 | Cdc42ep3 |
| ENSMUSG( | 2004.912 | -0.1567 | 0.175638 | 0.845305 | Cdc42ep4 |
| ENSMUSG( | 35.2593 | -0.41145 | 0.231622 | 0.868507 | Cdc42ep5 |
| ENSMUSG( | 2614.668 | -0.02492 | 0.810551 | 0.978779 | Cdc42se1 |
| ENSMUSG( | 4291.877 | -0.08668 | 0.425591 | 0.917257 | Cdc42se2 |
| ENSMUSG( | 58.54605 | 0.435856 | 0.123742 | 0.814691 | Cdc45 |
| ENSMUSG( | 767.8944 | 0.007912 | 0.94947 | 0.994798 | Cdc5l |
| ENSMUSG( | 22.99383 | 0.498808 | 0.236658 | 0.872173 | Cdc6 |
| ENSMUSG( | 325.4717 | 0.336611 | 0.075633 | 0.768524 | Cdc7 |
| ENSMUSG( | 857.52 | 0.226765 | 0.140893 | 0.830678 | Cdc73 |
| ENSMUSG( | 21.24134 | 0.574119 | 0.207977 | 0.864412 | Cdca2 |
| ENSMUSG( | 56.28092 | -0.04627 | 0.881289 | 0.989644 | Cdca3 |
| ENSMUSG( | 133.1651 | 0.146987 | 0.515874 | 0.930939 | Cdca4 |
| ENSMUSG( | 10.53946 | -0.31385 | 0.623266 | NA | Cdca5 |
| ENSMUSG( | 170.4487 | 0.255856 | 0.192471 | 0.859726 | Cdca7 |
| ENSMUSG( | 31.86552 | -0.12913 | 0.737762 | 0.967709 | Cdca7l |
| ENSMUSG( | 42.80122 | -0.25571 | 0.403686 | 0.917257 | Cdca8 |
| ENSMUSG( | 12.65203 | -0.5153 | 0.35484 | 0.912411 | Cdcp1 |
| ENSMUSG( | 193.159 | -0.15317 | 0.365767 | 0.914623 | Cdh1 |
| ENSMUSG( | 1732.275 | 0.195238 | 0.284938 | 0.892412 | Cdh10 |
| ENSMUSG( | 2294.5 | 0.150475 | 0.377973 | 0.916784 | Cdh11 |
| ENSMUSG( | 194.322 | 0.170016 | 0.4172 | 0.917257 | Cdh12 |
| ENSMUSG( | 2726.998 | 0.290515 | 0.067359 | 0.75212 | Cdh13 |
| ENSMUSG( | 42.34129 | 0.086904 | 0.792432 | 0.97681 | Cdh15 |
| ENSMUSG( | 458.1384 | 0.21125 | 0.336179 | 0.907972 | Cdh18 |
| ENSMUSG( | 682.5463 | 0.222759 | 0.192713 | 0.859764 | Cdh19 |
| ENSMUSG( | 3844.739 | 0.258499 | 0.120093 | 0.813384 | Cdh2 |
| ENSMUSG( | 1228.607 | 0.084892 | 0.624574 | 0.953234 | Cdh20 |
| ENSMUSG( | 1453.536 | 0.258131 | 0.092009 | 0.791889 | Cdh22 |
| ENSMUSG( | 58.32903 | 0.243599 | 0.367454 | 0.915217 | Cdh23 |
| ENSMUSG( | 405.3516 | 0.177783 | 0.233567 | 0.868507 | Cdh24 |
| ENSMUSG( | 23.14981 | -0.39704 | 0.36461 | 0.914223 | Cdh26 |
| ENSMUSG( | 15.15046 | -0.44665 | 0.405909 | 0.917257 | Cdh3 |
| ENSMUSG( | 905.6389 | 0.033243 | 0.815969 | 0.979198 | Cdh4 |
| ENSMUSG( | 754.5078 | 0.327349 | 0.425186 | 0.917257 | Cdh5 |
| ENSMUSG( | 825.0668 | 0.123767 | 0.518732 | 0.930939 | Cdh6 |
| ENSMUSG( | 302.2578 | 0.208221 | 0.219162 | 0.866104 | Cdh7 |
| ENSMUSG( | 999.0669 | 0.255888 | 0.15911 | 0.837724 | Cdh8 |
| ENSMUSG( | 564.2662 | 0.212848 | 0.283192 | 0.892412 | Cdh9 |
| ENSMUSG( | 232.5245 | 0.53587 | 0.026517 | 0.603041 | Cdhr1 |
| ENSMUSG( | 87.14408 | 0.388035 | 0.18171 | 0.850579 | Cdhr3 |
| ENSMUSG( | 37.47484 | 0.372256 | 0.260905 | 0.884853 | Cdhr4 |
| ENSMUSG( | 152.3479 | -0.12339 | 0.502749 | 0.927825 | Cdin1 |
| ENSMUSG( | 5402.895 | -0.08061 | 0.453903 | 0.922392 | Cdip1 |
| ENSMUSG( | 5374.495 | 0.070327 | 0.63296 | 0.953844 | Cdipt |
| ENSMUSG( | 31.62808 | -0.14789 | 0.68894 | 0.959742 | Cdk1 |
| ENSMUSG( | 1851.802 | -0.10695 | 0.389158 | 0.917257 | Cdk10 |
| ENSMUSG( | 2493.293 | -0.06219 | 0.633337 | 0.953844 | Cdk11b |
| ENSMUSG( | 498.9059 | 0.076804 | 0.591587 | 0.944975 | Cdk12 |
| ENSMUSG( | 1229.695 | -0.0147 | 0.894518 | 0.990558 | Cdk13 |
| ENSMUSG( | 5084.217 | 0.012164 | 0.905802 | 0.991089 | Cdk14 |
| ENSMUSG( | 10.93428 | -0.38422 | 0.540846 | NA | Cdk15 |
| ENSMUSG( | 20726.35 | -0.01197 | 0.901211 | 0.990558 | Cdk16 |
| ENSMUSG( | 2134.583 | -0.01566 | 0.89916 | 0.990558 | Cdk17 |
| ENSMUSG( | 699.2625 | -0.05533 | 0.691457 | 0.960071 | Cdk18 |
| ENSMUSG( | 3431.333 | -0.01332 | 0.897775 | 0.990558 | Cdk19 |
| ENSMUSG( | 145.6008 | 0.323504 | 0.163135 | 0.838291 | Cdk2 |
| ENSMUSG( | 237.3741 | 0.077512 | 0.64891 | 0.95422 | Cdk20 |
| ENSMUSG( | 1511.21 | -0.0758 | 0.545605 | 0.938063 | Cdk2ap1 |

| | | | | | | | | | | | |
|---|---|---|---|---|---|---|---|---|---|---|---|
| ENSMUSG( | Cd163 | 107.4806 | 0.497403 | 0.015857 | 0.186421 | ENSMUSG( | Cd163 | 111.6262 | 0.813907 | 0.000217 | 0.038388 |
| ENSMUSG( | Cd164 | 2359.164 | -0.30733 | 0.06839 | 0.405144 | ENSMUSG( | Cd164 | 2776.594 | 0.41912 | 0.027374 | 0.606201 |
| ENSMUSG( | Cd164l2 | 164.9926 | 0.029298 | 0.856019 | 0.973813 | ENSMUSG( | Cd164l2 | 141.219 | -0.16141 | 0.413742 | 0.917257 |
| ENSMUSG( | Cd177 | 11.36167 | -0.83623 | 0.269792 | NA | ENSMUSG( | Cd177 | 12.32479 | -0.27123 | 0.690313 | 0.960071 |
| ENSMUSG( | Cd180 | 159.3131 | -0.88874 | 3.57E-05 | 0.003067 | ENSMUSG( | Cd180 | 173.096 | -0.24042 | 0.389874 | 0.917257 |
| ENSMUSG( | Cd1d1 | 43.95301 | 0.127263 | 0.676587 | 0.930542 | ENSMUSG( | Cd1d1 | 43.02298 | 0.350522 | 0.287559 | 0.894091 |
| ENSMUSG( | Cd200 | 2348.987 | -0.39504 | 0.015143 | 0.181303 | ENSMUSG( | Cd200 | 2781.642 | 0.370065 | 0.033419 | 0.643993 |
| ENSMUSG( | Cd200r1 | 14.73041 | -0.06247 | 0.908571 | NA | ENSMUSG( | Cd200r1 | 16.86139 | 0.569461 | 0.265327 | 0.886134 |
| ENSMUSG( | Cd209a | 10.00699 | 0.17978 | 0.7778 | NA | ENSMUSG( | Cd209a | 8.715729 | 0.063815 | 0.93168 | NA |
| ENSMUSG( | Cd209f | 18.48335 | -0.02912 | 0.948238 | NA | ENSMUSG( | Cd209f | 16.56553 | -0.12322 | 0.81409 | 0.979198 |
| ENSMUSG( | Cd22 | 16.81965 | -0.19554 | 0.66171 | NA | ENSMUSG( | Cd22 | 15.57934 | -0.11423 | 0.824998 | 0.979297 |
| ENSMUSG( | Cd248 | 229.9394 | -0.43195 | 0.043035 | 0.31819 | ENSMUSG( | Cd248 | 277.5264 | 0.398523 | 0.117634 | 0.813384 |
| ENSMUSG( | Cd24a | 482.9938 | -0.36046 | 0.021902 | 0.225738 | ENSMUSG( | Cd24a | 538.3435 | 0.238267 | 0.16628 | 0.840082 |
| ENSMUSG( | Cd274 | 67.46751 | -0.16953 | 0.486682 | 0.858713 | ENSMUSG( | Cd274 | 60.65086 | -0.18826 | 0.504565 | 0.927825 |
| ENSMUSG( | Cd276 | 465.457 | -0.39252 | 0.010317 | 0.144633 | ENSMUSG( | Cd276 | 457.7099 | -0.14586 | 0.36477 | 0.914379 |
| ENSMUSG( | Cd2ap | 761.9627 | -0.20676 | 0.107342 | 0.495755 | ENSMUSG( | Cd2ap | 803.1372 | 0.217914 | 0.11958 | 0.813384 |
| ENSMUSG( | Cd2bp2 | 2648.642 | 0.041631 | 0.602113 | 0.904887 | ENSMUSG( | Cd2bp2 | 2281.343 | -0.12449 | 0.243674 | 0.877904 |
| ENSMUSG( | Cd300a | 107.8052 | -0.3669 | 0.187695 | 0.626295 | ENSMUSG( | Cd300a | 104.7574 | -0.16598 | 0.536301 | 0.93705 |
| ENSMUSG( | Cd300c2 | 220.0389 | -0.13478 | 0.418512 | 0.821966 | ENSMUSG( | Cd300c2 | 211.6975 | 0.039934 | 0.834605 | 0.981237 |
| ENSMUSG( | Cd300ld | 21.40428 | -1.42124 | 0.012156 | 0.159821 | ENSMUSG( | Cd300ld | 19.40935 | -1.54766 | 0.014916 | 0.489529 |
| ENSMUSG( | Cd300lf | 10.70734 | -0.34045 | 0.601973 | NA | ENSMUSG( | Cd300lf | 9.309095 | -0.48003 | 0.481299 | NA |
| ENSMUSG( | Cd302 | 695.6807 | -0.05925 | 0.640409 | 0.916434 | ENSMUSG( | Cd302 | 666.0116 | 0.085537 | 0.528595 | 0.934795 |
| ENSMUSG( | Cd320 | 497.5454 | -0.18793 | 0.240759 | 0.688247 | ENSMUSG( | Cd320 | 502.9003 | 0.121395 | 0.478117 | 0.923329 |
| ENSMUSG( | Cd33 | 439.1057 | -0.38584 | 0.014778 | 0.178616 | ENSMUSG( | Cd33 | 470.1727 | 0.113937 | 0.556219 | 0.939565 |
| ENSMUSG( | Cd34 | 2637.213 | -0.23993 | 0.131936 | 0.544244 | ENSMUSG( | Cd34 | 2778.929 | 0.189543 | 0.318116 | 0.902849 |
| ENSMUSG( | Cd36 | 59.96084 | -0.52183 | 0.112457 | 0.505997 | ENSMUSG( | Cd36 | 55.58241 | -0.47378 | 0.15892 | 0.837724 |
| ENSMUSG( | Cd37 | 296.5726 | -0.23581 | 0.12116 | 0.525683 | ENSMUSG( | Cd37 | 315.1376 | 0.219618 | 0.188895 | 0.85698 |
| ENSMUSG( | Cd38 | 352.5242 | -0.31944 | 0.046133 | 0.329204 | ENSMUSG( | Cd38 | 360.1853 | 0.03415 | 0.842194 | 0.983044 |
| ENSMUSG( | Cd4 | 162.5858 | -1.1698 | 0.018966 | 0.207297 | ENSMUSG( | Cd4 | 182.6515 | -0.37323 | 0.138211 | 0.828492 |
| ENSMUSG( | Cd40 | 34.49739 | -0.19773 | 0.539309 | 0.883094 | ENSMUSG( | Cd40 | 38.5441 | 0.383548 | 0.231274 | 0.868507 |
| ENSMUSG( | Cd44 | 201.6448 | -0.12956 | 0.546672 | 0.884974 | ENSMUSG( | Cd44 | 197.4177 | 0.08216 | 0.725358 | 0.966887 |
| ENSMUSG( | Cd46 | 256.3437 | -0.28672 | 0.121037 | 0.525487 | ENSMUSG( | Cd46 | 258.4165 | 0.029691 | 0.87981 | 0.989644 |
| ENSMUSG( | Cd47 | 5045.727 | -0.52786 | 0.00021 | 0.011071 | ENSMUSG( | Cd47 | 5692.078 | 0.135214 | 0.418681 | 0.917257 |
| ENSMUSG( | Cd48 | 38.60133 | -0.76855 | 0.040916 | 0.312769 | ENSMUSG( | Cd48 | 40.19891 | -0.26186 | 0.525285 | 0.934374 |
| ENSMUSG( | Cd52 | 133.0533 | -0.65901 | 0.000688 | 0.025203 | ENSMUSG( | Cd52 | 124.9349 | -0.56484 | 0.006407 | 0.335013 |
| ENSMUSG( | Cd53 | 512.0747 | -0.35437 | 0.054877 | 0.360515 | ENSMUSG( | Cd53 | 527.6949 | 0.027528 | 0.897333 | 0.990558 |
| ENSMUSG( | Cd55 | 174.9969 | -0.45199 | 0.015473 | 0.18373 | ENSMUSG( | Cd55 | 194.0822 | 0.143347 | 0.462459 | 0.923329 |
| ENSMUSG( | Cd59a | 228.4049 | 0.423793 | 0.033804 | 0.282338 | ENSMUSG( | Cd59a | 253.3777 | 0.894032 | 2.19E-07 | 0.000135 |
| ENSMUSG( | Cd59b | 19.85417 | -0.00272 | 0.994877 | NA | ENSMUSG( | Cd59b | 19.79568 | 0.244665 | 0.570084 | 0.941777 |
| ENSMUSG( | Cd6 | 102.0231 | -0.59913 | 0.012814 | 0.163518 | ENSMUSG( | Cd6 | 89.74412 | -0.73094 | 0.004213 | 0.263542 |
| ENSMUSG( | Cd63 | 4154.78 | -0.12403 | 0.275751 | 0.725669 | ENSMUSG( | Cd63 | 3977.402 | 0.022145 | 0.858946 | 0.985884 |
| ENSMUSG( | Cd68 | 638.1372 | -0.2719 | 0.041889 | 0.315367 | ENSMUSG( | Cd68 | 681.2038 | 0.198631 | 0.241771 | 0.876528 |
| ENSMUSG( | Cd7 | 42.88195 | -0.64984 | 0.030968 | 0.267891 | ENSMUSG( | Cd7 | 41.64138 | -0.41443 | 0.190509 | 0.857801 |
| ENSMUSG( | Cd72 | 60.45545 | -0.50352 | 0.04278 | 0.316989 | ENSMUSG( | Cd72 | 57.17602 | -0.36325 | 0.185618 | 0.855305 |
| ENSMUSG( | Cd74 | 386.0178 | -0.55306 | 2.31E-05 | 0.002143 | ENSMUSG( | Cd74 | 372.176 | -0.35812 | 0.012309 | 0.450129 |
| ENSMUSG( | Cd79a | 25.50045 | -0.18413 | 0.633488 | 0.914073 | ENSMUSG( | Cd79a | 19.48936 | -0.80955 | 0.080672 | 0.77681 |
| ENSMUSG( | Cd79b | 29.93713 | -0.59824 | 0.082119 | 0.442642 | ENSMUSG( | Cd79b | 29.11797 | -0.36084 | 0.321321 | 0.904019 |
| ENSMUSG( | Cd80 | 29.41299 | -0.5059 | 0.175404 | 0.610788 | ENSMUSG( | Cd80 | 30.79274 | -0.01526 | 0.972667 | 0.996416 |
| ENSMUSG( | Cd81 | 17152.06 | -0.42079 | 0.001195 | 0.037142 | ENSMUSG( | Cd81 | 18037.16 | 0.025899 | 0.862032 | 0.98632 |
| ENSMUSG( | Cd82 | 1185.836 | -0.6592 | 3.24E-07 | 7.29E-05 | ENSMUSG( | Cd82 | 1279.668 | -0.09475 | 0.543143 | 0.938063 |
| ENSMUSG( | Cd83 | 344.4012 | -0.2158 | 0.19949 | 0.640054 | ENSMUSG( | Cd83 | 378.6972 | 0.327444 | 0.077344 | 0.771983 |
| ENSMUSG( | Cd84 | 198.033 | -0.31414 | 0.093456 | 0.469798 | ENSMUSG( | Cd84 | 205.2927 | 0.084791 | 0.68233 | 0.958678 |
| ENSMUSG( | Cd86 | 161.8977 | -0.25722 | 0.173264 | 0.608563 | ENSMUSG( | Cd86 | 167.3332 | 0.126796 | 0.5501 | 0.938699 |
| ENSMUSG( | Cd9 | 3060.016 | -0.66211 | 1.16E-06 | 0.000211 | ENSMUSG( | Cd9 | 3384.698 | -0.02608 | 0.869805 | 0.988097 |
| ENSMUSG( | Cd93 | 499.5409 | -0.23359 | 0.240715 | 0.688247 | ENSMUSG( | Cd93 | 479.9521 | -0.06072 | 0.796005 | 0.97681 |
| ENSMUSG( | Cd99l2 | 2918.518 | -0.24327 | 0.090923 | 0.463326 | ENSMUSG( | Cd99l2 | 2901.725 | 0.023574 | 0.884362 | 0.989688 |
| ENSMUSG( | Cda | 24.91423 | 0.023954 | 0.949396 | 0.993821 | ENSMUSG( | Cda | 20.94284 | -0.23717 | 0.581489 | 0.944266 |
| ENSMUSG( | Cdadc1 | 1507.741 | -0.1027 | 0.275229 | 0.72509 | ENSMUSG( | Cdadc1 | 1416.251 | -0.01137 | 0.91888 | 0.992529 |
| ENSMUSG( | Cdan1 | 830.7874 | 0.05814 | 0.57253 | 0.893584 | ENSMUSG( | Cdan1 | 768.1664 | 0.103939 | 0.487333 | 0.92541 |
| ENSMUSG( | Cdc123 | 1595.286 | 0.017253 | 0.840421 | 0.971179 | ENSMUSG( | Cdc123 | 1411.702 | -0.07066 | 0.572761 | 0.94262 |
| ENSMUSG( | Cdc14a | 194.9762 | -0.14716 | 0.438867 | 0.830714 | ENSMUSG( | Cdc14a | 177.5794 | -0.13282 | 0.5059 | 0.927825 |
| ENSMUSG( | Cdc14b | 296.0637 | -0.05523 | 0.717799 | 0.941697 | ENSMUSG( | Cdc14b | 280.1919 | 0.057341 | 0.704772 | 0.962677 |
| ENSMUSG( | Cdc16 | 1213.431 | 0.006351 | 0.944951 | 0.993612 | ENSMUSG( | Cdc16 | 1077.659 | -0.06789 | 0.558087 | 0.939565 |
| ENSMUSG( | Cdc20 | 36.56862 | -0.0256 | 0.94284 | 0.993415 | ENSMUSG( | Cdc20 | 36.82939 | 0.240543 | 0.516432 | 0.930939 |
| ENSMUSG( | Cdc23 | 1586.553 | 0.041052 | 0.658631 | 0.924506 | ENSMUSG( | Cdc23 | 1424.19 | -0.00097 | 0.993352 | 0.998813 |
| ENSMUSG( | Cdc25a | 159.1718 | 0.111078 | 0.503115 | 0.865929 | ENSMUSG( | Cdc25a | 148.6586 | 0.176825 | 0.350088 | 0.911652 |
| ENSMUSG( | Cdc25b | 611.1459 | 0.310641 | 0.058514 | 0.373218 | ENSMUSG( | Cdc25b | 517.2929 | 0.113889 | 0.524761 | 0.933907 |
| ENSMUSG( | Cdc25c | 15.96368 | -0.17586 | 0.709672 | NA | ENSMUSG( | Cdc25c | 16.85583 | 0.219896 | 0.65445 | 0.955015 |
| ENSMUSG( | Cdc26 | 522.3082 | -0.0246 | 0.848699 | 0.972374 | ENSMUSG( | Cdc26 | 470.6862 | -0.05846 | 0.66508 | 0.956766 |
| ENSMUSG( | Cdc27 | 1423.751 | 0.027361 | 0.770243 | 0.954689 | ENSMUSG( | Cdc27 | 1343.456 | 0.123115 | 0.316356 | 0.901868 |
| ENSMUSG( | Cdc34 | 1849.075 | 0.119234 | 0.195231 | 0.6346 | ENSMUSG( | Cdc34 | 1558.311 | -0.10719 | 0.331436 | 0.907861 |
| ENSMUSG( | Cdc37 | 2769.161 | 0.040503 | 0.649012 | 0.920025 | ENSMUSG( | Cdc37 | 2398.874 | -0.10761 | 0.319947 | 0.903771 |
| ENSMUSG( | Cdc37l1 | 3653.743 | 0.046207 | 0.616863 | 0.910866 | ENSMUSG( | Cdc37l1 | 3550.744 | 0.225083 | 0.045351 | 0.693959 |
| ENSMUSG( | Cdc40 | 904.6025 | 0.055792 | 0.657287 | 0.924144 | ENSMUSG( | Cdc40 | 818.9759 | 0.037822 | 0.782792 | 0.976106 |
| ENSMUSG( | Cdc42 | 8175.899 | 0.017378 | 0.845584 | 0.97218 | ENSMUSG( | Cdc42 | 7049.006 | -0.14812 | 0.254235 | 0.881345 |
| ENSMUSG( | Cdc42bpa | 14746.18 | 0.170842 | 0.0235 | 0.234568 | ENSMUSG( | Cdc42bpa | 12434.03 | -0.04809 | 0.611665 | 0.950771 |
| ENSMUSG( | Cdc42bpb | 6708.185 | 0.16621 | 0.070211 | 0.40908 | ENSMUSG( | Cdc42bpb | 6102.056 | 0.163417 | 0.15906 | 0.837724 |
| ENSMUSG( | Cdc42bpg | 737.931 | 0.086188 | 0.413287 | 0.819541 | ENSMUSG( | Cdc42bpg | 644.0176 | -0.04326 | 0.751596 | 0.97035 |
| ENSMUSG( | Cdc42ep1 | 684.9908 | -0.31211 | 0.003366 | 0.073204 | ENSMUSG( | Cdc42ep1 | 695.3918 | 0.019583 | 0.895195 | 0.990558 |
| ENSMUSG( | Cdc42ep2 | 215.8356 | -0.15102 | 0.357028 | 0.788702 | ENSMUSG( | Cdc42ep2 | 210.0483 | 0.035888 | 0.842311 | 0.983044 |
| ENSMUSG( | Cdc42ep3 | 216.7843 | -0.40267 | 0.008619 | 0.130286 | ENSMUSG( | Cdc42ep3 | 220.9452 | -0.05471 | 0.745852 | 0.96951 |
| ENSMUSG( | Cdc42ep4 | 2311.189 | -0.01009 | 0.914637 | 0.988501 | ENSMUSG( | Cdc42ep4 | 2004.912 | -0.1567 | 0.175638 | 0.845305 |
| ENSMUSG( | Cdc42ep5 | 39.83126 | -0.33072 | 0.289677 | 0.739839 | ENSMUSG( | Cdc42ep5 | 35.2593 | -0.41145 | 0.231622 | 0.868507 |
| ENSMUSG( | Cdc42se1 | 3022.368 | 0.123269 | 0.14167 | 0.558688 | ENSMUSG( | Cdc42se1 | 2614.668 | -0.02492 | 0.810551 | 0.978779 |
| ENSMUSG( | Cdc42se2 | 4859.635 | 0.005472 | 0.945499 | 0.99365 | ENSMUSG( | Cdc42se2 | 4291.877 | -0.08668 | 0.425591 | 0.917257 |
| ENSMUSG( | Cdc45 | 64.75764 | 0.457907 | 0.056994 | 0.368322 | ENSMUSG( | Cdc45 | 58.54605 | 0.435856 | 0.123742 | 0.814691 |
| ENSMUSG( | Cdc5l | 899.1096 | 0.190653 | 0.059914 | 0.378725 | ENSMUSG( | Cdc5l | 767.8944 | 0.007912 | 0.94947 | 0.994798 |
| ENSMUSG( | Cdc6 | 24.44506 | 0.383513 | 0.329987 | 0.774026 | ENSMUSG( | Cdc6 | 22.99383 | 0.498808 | 0.236658 | 0.872173 |
| ENSMUSG( | Cdc7 | 337.2502 | 0.186647 | 0.17434 | 0.610304 | ENSMUSG( | Cdc7 | 325.4717 | 0.336611 | 0.075633 | 0.768524 |
| ENSMUSG( | Cdc73 | 905.3489 | 0.126601 | 0.259319 | 0.709948 | ENSMUSG( | Cdc73 | 857.52 | 0.226765 | 0.140893 | 0.830678 |
| ENSMUSG( | Cdca2 | 18.07812 | -0.11887 | 0.803646 | NA | ENSMUSG( | Cdca2 | 21.24134 | 0.574119 | 0.207977 | 0.864412 |
| ENSMUSG( | Cdca3 | 55.11637 | -0.44181 | 0.09733 | 0.477629 | ENSMUSG( | Cdca3 | 56.28092 | -0.04627 | 0.881289 | 0.989644 |
| ENSMUSG( | Cdca4 | 141.1076 | 0.034209 | 0.861114 | 0.975026 | ENSMUSG( | Cdca4 | 133.1651 | 0.146987 | 0.515874 | 0.930939 |
| ENSMUSG( | Cdca5 | 14.4693 | 0.304976 | 0.550301 | NA | ENSMUSG( | Cdca5 | 10.53946 | -0.31385 | 0.623266 | NA |
| ENSMUSG( | Cdca7 | 162.8998 | -0.14332 | 0.436153 | 0.829093 | ENSMUSG( | Cdca7 | 170.4487 | 0.255856 | 0.192471 | 0.859726 |
| ENSMUSG( | Cdca7l | 38.00336 | 0.107916 | 0.731383 | 0.944956 | ENSMUSG( | Cdca7l | 31.86552 | -0.12913 | 0.737762 | 0.967709 |
| ENSMUSG( | Cdca8 | 49.02776 | -0.11969 | 0.660818 | 0.924962 | ENSMUSG( | Cdca8 | 42.80122 | -0.25571 | 0.403686 | 0.917257 |
| ENSMUSG( | Cdcp1 | 13.33476 | -0.67315 | 0.191965 | NA | ENSMUSG( | Cdcp1 | 12.65203 | -0.5153 | 0.35484 | 0.912411 |
| ENSMUSG( | Cdh1 | 212.5003 | -0.14863 | 0.358991 | 0.790503 | ENSMUSG( | Cdh1 | 193.159 | -0.15317 | 0.365767 | 0.914623 |
| ENSMUSG( | Cdh10 | 1536.099 | -0.45536 | 0.008077 | 0.125133 | ENSMUSG( | Cdh10 | 1732.275 | 0.195238 | 0.284938 | 0.892412 |
| ENSMUSG( | Cdh11 | 2059.303 | -0.46702 | 0.001641 | 0.045581 | ENSMUSG( | Cdh11 | 2294.5 | 0.150475 | 0.377973 | 0.916784 |
| ENSMUSG( | Cdh12 | 159.1818 | -0.78042 | 0.000111 | 0.00695 | ENSMUSG( | Cdh12 | 194.322 | 0.170016 | 0.4172 | 0.917257 |
| ENSMUSG( | Cdh13 | 2347.406 | -0.4328 | 0.001719 | 0.046884 | ENSMUSG( | Cdh13 | 2726.998 | 0.290515 | 0.067359 | 0.75212 |
| ENSMUSG( | Cdh15 | 40.96653 | -0.29004 | 0.361173 | 0.792797 | ENSMUSG( | Cdh15 | 42.34129 | 0.086904 | 0.792432 | 0.97681 |
| ENSMUSG( | Cdh18 | 402.568 | -0.47328 | 0.015314 | 0.18239 | ENSMUSG( | Cdh18 | 458.1384 | 0.21125 | 0.336179 | 0.907972 |
| ENSMUSG( | Cdh19 | 706.2037 | 0.058678 | 0.713118 | 0.940734 | ENSMUSG( | Cdh19 | 682.5463 | 0.222759 | 0.192713 | 0.859764 |
| ENSMUSG( | Cdh2 | 3651.572 | -0.15512 | 0.29536 | 0.743814 | ENSMUSG( | Cdh2 | 3844.739 | 0.258499 | 0.120093 | 0.813384 |
| ENSMUSG( | Cdh20 | 1268.691 | -0.09601 | 0.487817 | 0.858824 | ENSMUSG( | Cdh20 | 1228.607 | 0.084892 | 0.624574 | 0.953234 |
| ENSMUSG( | Cdh22 | 1467.41 | 0.023041 | 0.833063 | 0.970333 | ENSMUSG( | Cdh22 | 1453.536 | 0.258131 | 0.092009 | 0.791889 |
| ENSMUSG( | Cdh23 | 58.64411 | -0.01898 | 0.943386 | 0.993415 | ENSMUSG( | Cdh23 | 58.32903 | 0.243599 | 0.367454 | 0.915217 |
| ENSMUSG( | Cdh24 | 441.1926 | 0.1495 | 0.317908 | 0.764145 | ENSMUSG( | Cdh24 | 405.3516 | 0.177783 | 0.233567 | 0.868507 |
| ENSMUSG( | Cdh26 | 28.73335 | -0.03808 | 0.920055 | 0.990247 | ENSMUSG( | Cdh26 | 23.14981 | -0.39704 | 0.36461 | 0.914223 |
| ENSMUSG( | Cdh3 | 15.81102 | -0.53819 | 0.348991 | NA | ENSMUSG( | Cdh3 | 15.15046 | -0.44665 | 0.405909 | 0.917257 |
| ENSMUSG( | Cdh4 | 857.4603 | -0.42833 | 0.003211 | 0.070583 | ENSMUSG( | Cdh4 | 905.6389 | 0.033243 | 0.815969 | 0.979198 |
| ENSMUSG( | Cdh5 | 723.951 | -0.05229 | 0.807776 | 0.963257 | ENSMUSG( | Cdh5 | 754.5078 | 0.327349 | 0.425186 | 0.917257 |
| ENSMUSG( | Cdh6 | 706.1736 | -0.67505 | 8.55E-05 | 0.005781 | ENSMUSG( | Cdh6 | 825.0668 | 0.123767 | 0.518732 | 0.930939 |
| ENSMUSG( | Cdh7 | 280.9809 | -0.27801 | 0.109423 | 0.499391 | ENSMUSG( | Cdh7 | 302.2578 | 0.208221 | 0.219162 | 0.866104 |
| ENSMUSG( | Cdh8 | 853.2092 | -0.50703 | 0.001039 | 0.033844 | ENSMUSG( | Cdh8 | 999.0669 | 0.255888 | 0.15911 | 0.837724 |
| ENSMUSG( | Cdh9 | 495.5066 | -0.47007 | 0.008452 | 0.128676 | ENSMUSG( | Cdh9 | 564.2662 | 0.212848 | 0.283192 | 0.892412 |
| ENSMUSG( | Cdhr1 | 168.1625 | -0.68835 | 0.011509 | 0.154494 | ENSMUSG( | Cdhr1 | 232.5245 | 0.53587 | 0.026517 | 0.603041 |
| ENSMUSG( | Cdhr3 | 79.85213 | -0.12487 | 0.632873 | 0.914073 | ENSMUSG( | Cdhr3 | 87.14408 | 0.388035 | 0.18171 | 0.850579 |
| ENSMUSG( | Cdhr4 | 38.95063 | 0.22296 | 0.51657 | 0.872788 | ENSMUSG( | Cdhr4 | 37.47484 | 0.372256 | 0.260905 | 0.884853 |
| ENSMUSG( | Cdin1 | 172.983 | -0.01719 | 0.914044 | 0.988483 | ENSMUSG( | Cdin1 | 152.3479 | -0.12339 | 0.502749 | 0.927825 |
| ENSMUSG( | Cdip1 | 6348.749 | 0.115865 | 0.162923 | 0.593778 | ENSMUSG( | Cdip1 | 5402.895 | -0.08061 | 0.453903 | 0.922392 |
| ENSMUSG( | Cdipt | 5175.236 | -0.32842 | 0.012552 | 0.161972 | ENSMUSG( | Cdipt | 5374.495 | 0.070327 | 0.63296 | 0.953844 |
| ENSMUSG( | Cdk1 | 33.5482 | -0.28241 | 0.408448 | 0.818167 | ENSMUSG( | Cdk1 | 31.62808 | -0.14789 | 0.68894 | 0.959742 |
| ENSMUSG( | Cdk10 | 2171.692 | 0.085865 | 0.436571 | 0.829373 | ENSMUSG( | Cdk10 | 1851.802 | -0.10695 | 0.389158 | 0.917257 |
| ENSMUSG( | Cdk11b | 2880.573 | 0.088418 | 0.316549 | 0.763648 | ENSMUSG( | Cdk11b | 2493.293 | -0.06219 | 0.633337 | 0.953844 |
| ENSMUSG( | Cdk12 | 534.9482 | 0.005307 | 0.966661 | 0.995784 | ENSMUSG( | Cdk12 | 498.9059 | 0.076804 | 0.591587 | 0.944975 |
| ENSMUSG( | Cdk13 | 1393.418 | 0.077064 | 0.402574 | 0.815505 | ENSMUSG( | Cdk13 | 1229.695 | -0.0147 | 0.894518 | 0.990558 |
| ENSMUSG( | Cdk14 | 5780.323 | 0.111464 | 0.203084 | 0.643631 | ENSMUSG( | Cdk14 | 5084.217 | 0.012164 | 0.905802 | 0.991089 |
| ENSMUSG( | Cdk15 | 13.49778 | -0.00453 | 0.993196 | NA | ENSMUSG( | Cdk15 | 10.93428 | -0.38422 | 0.540846 | NA |
| ENSMUSG( | Cdk16 | 24199.06 | 0.162034 | 0.035502 | 0.290108 | ENSMUSG( | Cdk16 | 20726.35 | -0.01197 | 0.901211 | 0.990558 |
| ENSMUSG( | Cdk17 | 2414.991 | 0.073686 | 0.466843 | 0.844819 | ENSMUSG( | Cdk17 | 2134.583 | -0.01566 | 0.89916 | 0.990558 |
| ENSMUSG( | Cdk18 | 779.0262 | -0.01002 | 0.931392 | 0.992327 | ENSMUSG( | Cdk18 | 699.2625 | -0.05533 | 0.691457 | 0.960071 |
| ENSMUSG( | Cdk19 | 3865.2 | 0.060586 | 0.487683 | 0.858824 | ENSMUSG( | Cdk19 | 3431.333 | -0.01332 | 0.897775 | 0.990558 |
| ENSMUSG( | Cdk2 | 136.6539 | -0.13628 | 0.504606 | 0.866883 | ENSMUSG( | Cdk2 | 145.6008 | 0.323504 | 0.163135 | 0.838291 |
| ENSMUSG( | Cdk20 | 270.4517 | 0.188244 | 0.265941 | 0.716441 | ENSMUSG( | Cdk20 | 237.3741 | 0.077512 | 0.64891 | 0.95422 |
| ENSMUSG( | Cdk2ap1 | 1737.418 | 0.062452 | 0.49383 | 0.860611 | ENSMUSG( | Cdk2ap1 | 1511.21 | -0.0758 | 0.545605 | 0.938063 |
| ENSMUSG( | Cdk2ap1rt | 30.35773 | -0.33832 | 0.349791 | 0.781943 | ENSMUSG( | Cdk2ap1rt | 24.89727 | -0.71672 | 0.073476 | 0.763629 |
| ENSMUSG( | Cdk2ap2 | 605.3322 | 0.137914 | 0.216423 | 0.661421 | ENSMUSG( | Cdk2ap2 | 522.8146 | -0.01913 | 0.896613 | 0.990558 |
| ENSMUSG( | Cdk3 | 51.07271 | 0.190295 | 0.529201 | 0.878662 | ENSMUSG( | Cdk3 | 41.51064 | -0.12095 | 0.748649 | 0.96984 |
| ENSMUSG( | Cdk4 | 1563.696 | -0.08003 | 0.378403 | 0.801688 | ENSMUSG( | Cdk4 | 1422.09 | -0.08717 | 0.434886 | 0.918876 |
| ENSMUSG( | Cdk5 | 2994.266 | 0.055894 | 0.602698 | 0.904887 | ENSMUSG( | Cdk5 | 2658.455 | -0.01958 | 0.879592 | 0.989644 |
| ENSMUSG( | Cdk5r1 | 5659.866 | -0.03273 | 0.730713 | 0.944956 | ENSMUSG( | Cdk5r1 | 5312.271 | 0.051817 | 0.636572 | 0.953903 |
| ENSMUSG( | Cdk5r2 | 21496.94 | 0.119222 | 0.160825 | 0.590628 | ENSMUSG( | Cdk5r2 | 19029.54 | 0.039542 | 0.71366 | 0.96497 |
| ENSMUSG( | Cdk5rap1 | 191.5699 | 0.263828 | 0.124314 | 0.531714 | ENSMUSG( | Cdk5rap1 | 165.695 | 0.119676 | 0.630849 | 0.953251 |
| ENSMUSG( | Cdk5rap2 | 1180.194 | 0.059708 | 0.592538 | 0.902094 | ENSMUSG( | Cdk5rap2 | 1006.022 | -0.13087 | 0.289603 | 0.896726 |
| ENSMUSG( | Cdk5rap3 | 1228.072 | 0.073151 | 0.452864 | 0.836327 | ENSMUSG( | Cdk5rap3 | 1054.954 | -0.09842 | 0.388904 | 0.917257 |
| ENSMUSG( | Cdk6 | 143.8747 | -0.02674 | 0.881356 | 0.981798 | ENSMUSG( | Cdk6 | 141.1649 | 0.186135 | 0.335296 | 0.907907 |
| ENSMUSG( | Cdk7 | 312.5339 | -0.07083 | 0.614334 | 0.909419 | ENSMUSG( | Cdk7 | 291.1125 | -0.00319 | 0.984609 | 0.996963 |
| ENSMUSG( | Cdk8 | 773.8844 | 0.085753 | 0.408459 | 0.818167 | ENSMUSG( | Cdk8 | 703.6322 | 0.074035 | 0.59689 | 0.946555 |
| ENSMUSG( | Cdk9 | 2396.34 | 0.244503 | 0.130496 | 0.541757 | ENSMUSG( | Cdk9 | 1961.512 | -0.05855 | 0.725362 | 0.966887 |
| ENSMUSG( | Cdkal1 | 259.478 | -0.04773 | 0.729362 | 0.944738 | ENSMUSG( | Cdkal1 | 231.2559 | -0.12069 | 0.464446 | 0.923329 |
| ENSMUSG( | Cdkl1 | 175.989 | -0.12655 | 0.483103 | 0.856216 | ENSMUSG( | Cdkl1 | 163.7866 | -0.06303 | 0.748709 | 0.96984 |
| ENSMUSG( | Cdkl2 | 866.5811 | 0.051763 | 0.645835 | 0.918224 | ENSMUSG( | Cdkl2 | 804.4571 | 0.099268 | 0.475681 | 0.923329 |
| ENSMUSG( | Cdkl3 | 413.9491 | -0.16354 | 0.178827 | 0.616298 | ENSMUSG( | Cdkl3 | 384.1847 | -0.11727 | 0.463019 | 0.923329 |
| ENSMUSG( | Cdkl4 | 377.4057 | 0.140712 | 0.243628 | 0.690987 | ENSMUSG( | Cdkl4 | 338.9401 | 0.101032 | 0.475201 | 0.923329 |

| | | | | | |
|---|---|---|---|---|---|
| ENSMUSG( | 2171.692 | 0.085865 | 0.436571 | 0.829373 | Cdk10 |
| ENSMUSG( | 2880.573 | 0.088418 | 0.316549 | 0.763648 | Cdk11b |
| ENSMUSG( | 534.9482 | 0.005307 | 0.966661 | 0.995784 | Cdk12 |
| ENSMUSG( | 1393.418 | 0.077064 | 0.402574 | 0.815505 | Cdk13 |
| ENSMUSG( | 5780.323 | 0.111464 | 0.203084 | 0.643631 | Cdk14 |
| ENSMUSG( | 13.49778 | -0.00453 | 0.993196 | NA | Cdk15 |
| ENSMUSG( | 24199.06 | 0.162034 | 0.035502 | 0.290108 | Cdk16 |
| ENSMUSG( | 2414.991 | 0.073686 | 0.466843 | 0.844819 | Cdk17 |
| ENSMUSG( | 779.0262 | -0.01002 | 0.931392 | 0.992327 | Cdk18 |
| ENSMUSG( | 3865.2 | 0.060586 | 0.487683 | 0.858824 | Cdk19 |
| ENSMUSG( | 136.6539 | -0.13628 | 0.504606 | 0.866883 | Cdk2 |
| ENSMUSG( | 270.4517 | 0.188244 | 0.265941 | 0.716441 | Cdk20 |
| ENSMUSG( | 1737.418 | 0.062452 | 0.49383 | 0.860611 | Cdk2ap1 |
| ENSMUSG( | 30.35773 | -0.33832 | 0.349791 | 0.781943 | Cdk2ap1rt |
| ENSMUSG( | 605.3322 | 0.137914 | 0.216423 | 0.661421 | Cdk2ap2 |
| ENSMUSG( | 51.07271 | 0.190295 | 0.529201 | 0.878662 | Cdk3 |
| ENSMUSG( | 1563.696 | -0.08003 | 0.378403 | 0.801688 | Cdk4 |
| ENSMUSG( | 2994.266 | 0.055894 | 0.602698 | 0.904887 | Cdk5 |
| ENSMUSG( | 5659.866 | -0.03273 | 0.730713 | 0.944956 | Cdk5r1 |
| ENSMUSG( | 21496.94 | 0.119222 | 0.160825 | 0.590628 | Cdk5r2 |
| ENSMUSG( | 191.5699 | 0.263828 | 0.124314 | 0.531714 | Cdk5rap1 |
| ENSMUSG( | 1180.194 | 0.059708 | 0.592538 | 0.902094 | Cdk5rap2 |
| ENSMUSG( | 1228.072 | 0.073151 | 0.452864 | 0.836327 | Cdk5rap3 |
| ENSMUSG( | 143.8747 | -0.02674 | 0.881356 | 0.981798 | Cdk6 |
| ENSMUSG( | 312.5339 | -0.07083 | 0.614334 | 0.909419 | Cdk7 |
| ENSMUSG( | 773.8844 | 0.085753 | 0.408459 | 0.818167 | Cdk8 |
| ENSMUSG( | 2396.34 | 0.244503 | 0.130496 | 0.541757 | Cdk9 |
| ENSMUSG( | 259.478 | -0.04773 | 0.729362 | 0.944738 | Cdkal1 |
| ENSMUSG( | 175.989 | -0.12655 | 0.483103 | 0.856216 | Cdkl1 |
| ENSMUSG( | 866.5811 | 0.051763 | 0.645835 | 0.918224 | Cdkl2 |
| ENSMUSG( | 413.9491 | -0.16354 | 0.178827 | 0.616298 | Cdkl3 |
| ENSMUSG( | 377.4057 | 0.140712 | 0.243628 | 0.690987 | Cdkl4 |
| ENSMUSG( | 2442.347 | 0.004006 | 0.965071 | 0.995303 | Cdkl5 |
| ENSMUSG( | 596.6308 | -0.04716 | 0.74412 | 0.948497 | Cdkn1a |
| ENSMUSG( | 1176.845 | 0.106467 | 0.2781 | 0.728086 | Cdkn1b |
| ENSMUSG( | 302.5252 | -0.09115 | 0.534788 | 0.880725 | Cdkn1c |
| ENSMUSG( | 364.646 | -0.12114 | 0.379471 | 0.801757 | Cdkn2aip |
| ENSMUSG( | 784.2941 | 0.147032 | 0.149628 | 0.573617 | Cdkn2aipnl |
| ENSMUSG( | 40.5918 | 0.280756 | 0.394273 | 0.810014 | Cdkn2b |
| ENSMUSG( | 280.3766 | -0.04818 | 0.742289 | 0.948478 | Cdkn2c |
| ENSMUSG( | 655.0878 | -0.06869 | 0.65928 | 0.924506 | Cdkn2d |
| ENSMUSG( | 55.01681 | -0.29424 | 0.251091 | 0.700723 | Cdnf |
| ENSMUSG( | 963.8321 | 0.204267 | 0.056841 | 0.3679 | Cdo1 |
| ENSMUSG( | 536.1413 | -0.42526 | 0.000274 | 0.013445 | Cdon |
| ENSMUSG( | 302.5155 | 0.002116 | 0.98825 | 0.999109 | Cdpf1 |
| ENSMUSG( | 1029.042 | -0.10286 | 0.350054 | 0.781943 | Cdr1os |
| ENSMUSG( | 1237.296 | 0.098665 | 0.447596 | 0.834051 | Cdr2 |
| ENSMUSG( | 800.0107 | -0.04218 | 0.702945 | 0.937735 | Cdr2l |
| ENSMUSG( | 3480.769 | -0.20753 | 0.093396 | 0.469793 | Cds1 |
| ENSMUSG( | 17161.63 | -0.11045 | 0.405786 | 0.81652 | Cds2 |
| ENSMUSG( | 37.28933 | -0.27227 | 0.379252 | 0.801757 | Cdsn |
| ENSMUSG( | 55.08836 | -0.37565 | 0.1481 | 0.571232 | Cdt1 |
| ENSMUSG( | 2731.673 | -0.01493 | 0.88634 | 0.983139 | Cdv3 |
| ENSMUSG( | 279.8211 | 0.067816 | 0.613814 | 0.90923 | Cdyl |
| ENSMUSG( | 67.84651 | 0.586644 | 0.013854 | 0.172062 | Cdyl2 |
| ENSMUSG( | 160.9958 | -0.00214 | 0.992253 | 0.999327 | Ceacam1 |
| ENSMUSG( | 617.7419 | 0.461193 | 0.0015 | 0.042984 | Cebpa |
| ENSMUSG( | 203.1253 | -0.21119 | 0.337889 | 0.777422 | Cebpb |
| ENSMUSG( | 175.8623 | -0.3864 | 0.055043 | 0.36102 | Cebpd |
| ENSMUSG( | 1361.996 | 0.063914 | 0.518019 | 0.873754 | Cebpg |
| ENSMUSG( | 1135.082 | 0.234387 | 0.016474 | 0.190587 | Cebpz |
| ENSMUSG( | 467.201 | -0.08824 | 0.559468 | 0.888669 | Cebpzos |
| ENSMUSG( | 260.0443 | -0.22172 | 0.182363 | 0.619591 | Cecr2 |
| ENSMUSG( | 124.7841 | 0.13441 | 0.45954 | 0.840338 | Cela1 |
| ENSMUSG( | 4278.427 | -0.00673 | 0.936306 | 0.993415 | Celf1 |
| ENSMUSG( | 7745.759 | 0.048782 | 0.555874 | 0.887457 | Celf2 |
| ENSMUSG( | 4531.55 | 0.211541 | 0.096739 | 0.477375 | Celf3 |
| ENSMUSG( | 18445.01 | 0.162107 | 0.045728 | 0.328047 | Celf4 |
| ENSMUSG( | 8917.826 | 0.256635 | 0.025497 | 0.246501 | Celf5 |
| ENSMUSG( | 502.3164 | 0.225633 | 0.065174 | 0.395697 | Celf6 |
| ENSMUSG( | 340.5891 | 0.116016 | 0.348986 | 0.781661 | Celrr |
| ENSMUSG( | 609.8126 | -0.12085 | 0.438294 | 0.830406 | Celsr1 |
| ENSMUSG( | 8184.915 | -0.28547 | 0.025223 | 0.244689 | Celsr2 |
| ENSMUSG( | 3269.322 | -0.04522 | 0.709847 | 0.939669 | Celsr3 |
| ENSMUSG( | 774.7387 | 0.168326 | 0.294972 | 0.743798 | Cemip |
| ENSMUSG( | 508.1397 | -0.20516 | 0.200847 | 0.640988 | Cemip2 |
| ENSMUSG( | 340.5539 | -0.10406 | 0.49494 | 0.860798 | Cenatac |
| ENSMUSG( | 6125.12 | 0.041826 | 0.732946 | 0.944956 | Cend1 |
| ENSMUSG( | 80.08411 | 0.074434 | 0.727046 | 0.944223 | Cenpa |
| ENSMUSG( | 17478.21 | 0.10763 | 0.165847 | 0.597653 | Cenpb |
| ENSMUSG( | 447.9202 | 0.121847 | 0.289992 | 0.739839 | Cenpc1 |
| ENSMUSG( | 60.03787 | -0.3254 | 0.242862 | 0.690275 | Cenpe |
| ENSMUSG( | 109.0788 | -0.07388 | 0.743641 | 0.948478 | Cenpf |
| ENSMUSG( | 14.13697 | -0.25258 | 0.608737 | NA | Cenph |
| ENSMUSG( | 20.21363 | -0.1785 | 0.659708 | NA | Cenpi |
| ENSMUSG( | 109.8522 | -0.19103 | 0.327153 | 0.772056 | Cenpj |
| ENSMUSG( | 15.00024 | -0.54348 | 0.252641 | NA | Cenpk |
| ENSMUSG( | 39.65971 | 0.018678 | 0.950536 | 0.993821 | Cenpl |
| ENSMUSG( | 98.53577 | -0.09985 | 0.617749 | 0.910872 | Cenpm |
| ENSMUSG( | 30.95619 | -0.1867 | 0.596366 | 0.903264 | Cenpn |
| ENSMUSG( | 110.3093 | -0.09447 | 0.61999 | 0.911432 | Cenpo |
| ENSMUSG( | 17.4057 | -0.16138 | 0.716658 | NA | Cenpp |
| ENSMUSG( | 181.7023 | 0.043373 | 0.81653 | 0.965455 | Cenpq |
| ENSMUSG( | 45.40985 | 0.202975 | 0.55494 | 0.887073 | Cenps |
| ENSMUSG( | 405.0179 | 0.045298 | 0.732193 | 0.944956 | Cenpt |
| ENSMUSG( | 43.87946 | -0.25657 | 0.404479 | 0.816132 | Cenpu |
| ENSMUSG( | 586.9386 | 0.080588 | 0.487478 | 0.858812 | Cenpv |
| ENSMUSG( | 54.65292 | -0.02586 | 0.923783 | 0.991046 | Cenpw |
| ENSMUSG( | 486.4657 | 0.080146 | 0.48986 | 0.859319 | Cenpx |
| ENSMUSG( | 1626.418 | 0.124467 | 0.162941 | 0.593778 | Cep104 |
| ENSMUSG( | 101.6155 | 0.056757 | 0.788649 | 0.959226 | Cep112 |
| ENSMUSG( | 1624.19 | 0.094795 | 0.296275 | 0.744386 | Cep120 |
| ENSMUSG( | 261.8692 | 0.068368 | 0.631442 | 0.914073 | Cep126 |
| ENSMUSG( | 469.2317 | -0.0008 | 0.995963 | 0.99943 | Cep128 |
| ENSMUSG( | 929.0851 | 0.180315 | 0.27862 | 0.728705 | Cep131 |
| ENSMUSG( | 256.4295 | 0.078508 | 0.579387 | 0.896722 | Cep135 |
| ENSMUSG( | 652.4974 | -0.09071 | 0.52671 | 0.877267 | Cep15 |
| ENSMUSG( | 155.5141 | 0.297325 | 0.087825 | 0.454663 | Cep152 |
| ENSMUSG( | 406.139 | 0.16597 | 0.169461 | 0.603569 | Cep162 |
| ENSMUSG( | 701.1045 | -0.04664 | 0.702441 | 0.937668 | Cep164 |
| ENSMUSG( | 1463.811 | -0.04832 | 0.615441 | 0.910481 | Cep170 |
| ENSMUSG( | 11111.04 | 0.289093 | 0.001145 | 0.036093 | Cep170b |
| ENSMUSG( | 1144.455 | 0.085552 | 0.405167 | 0.81652 | Cep19 |
| ENSMUSG( | 367.2806 | -0.07663 | 0.585135 | 0.899171 | Cep192 |
| ENSMUSG( | 1258.824 | -0.0295 | 0.773964 | 0.955606 | Cep20 |
| ENSMUSG( | 1854.867 | 0.108527 | 0.23544 | 0.682423 | Cep250 |
| ENSMUSG( | 375.2022 | 0.109569 | 0.360979 | 0.792797 | Cep290 |
| ENSMUSG( | 730.0359 | -0.16822 | 0.103911 | 0.489523 | Cep295 |
| ENSMUSG( | 1567.931 | -0.05656 | 0.617396 | 0.910872 | Cep350 |
| ENSMUSG( | 188.9717 | -0.07135 | 0.660714 | 0.924951 | Cep41 |
| ENSMUSG( | 408.228 | 0.170266 | 0.156244 | 0.583988 | Cep43 |
| ENSMUSG( | 242.5508 | 0.008168 | 0.956547 | 0.993874 | Cep44 |
| ENSMUSG( | 11.58667 | -0.46485 | 0.4222 | NA | Cep55 |
| ENSMUSG( | 324.4399 | -0.03399 | 0.821369 | 0.966248 | Cep57 |
| ENSMUSG( | 100.6047 | 0.039497 | 0.857126 | 0.974393 | Cep57l1 |
| ENSMUSG( | 1010.708 | 0.029375 | 0.766618 | 0.954302 | Cep63 |
| ENSMUSG( | 944.1391 | 0.145444 | 0.225942 | 0.672685 | Cep68 |
| ENSMUSG( | 388.7787 | -0.04175 | 0.722421 | 0.943752 | Cep70 |
| ENSMUSG( | 64.2762 | 0.144251 | 0.582643 | 0.897918 | Cep72 |
| ENSMUSG( | 156.1403 | 0.017938 | 0.916479 | 0.988844 | Cep76 |
| ENSMUSG( | 323.1567 | 0.136338 | 0.315671 | 0.762671 | Cep78 |
| ENSMUSG( | 1254.791 | 0.067604 | 0.52658 | 0.877267 | Cep83 |
| ENSMUSG( | 347.9155 | 0.071557 | 0.587481 | 0.900403 | Cep85 |
| ENSMUSG( | 521.8139 | -0.04071 | 0.754511 | 0.951007 | Cep85l |
| ENSMUSG( | 593.7013 | 0.090473 | 0.433136 | 0.828207 | Cep89 |
| ENSMUSG( | 398.4515 | -0.01857 | 0.884685 | 0.982647 | Cep95 |
| ENSMUSG( | 468.7667 | -0.16676 | 0.188741 | 0.627678 | Cep97 |
| ENSMUSG( | 1007.215 | -0.33453 | 0.05454 | 0.35941 | Cept1 |
| ENSMUSG( | 367.722 | -0.14618 | 0.303843 | 0.752148 | Cercam |
| ENSMUSG( | 3126.714 | 0.070124 | 0.38451 | 0.804007 | Cerk |
| ENSMUSG( | 96.06295 | -0.04539 | 0.838912 | 0.971179 | Cerkl |
| ENSMUSG( | 1410.416 | -0.04718 | 0.636586 | 0.914969 | Cerox1 |
| ENSMUSG( | 2954.351 | -0.43375 | 0.002 | 0.051341 | Cers2 |
| ENSMUSG( | 14.45031 | 0.772173 | 0.123019 | NA | Cers3 |
| ENSMUSG( | 1928.563 | -0.12428 | 0.292958 | 0.742763 | Cers4 |
| ENSMUSG( | 1304.291 | -0.13406 | 0.325347 | 0.77061 | Cers5 |
| ENSMUSG( | 1322.119 | -0.08015 | 0.634189 | 0.914073 | Cers6 |
| ENSMUSG( | 1940.219 | 0.04871 | 0.580334 | 0.896952 | Cert1 |
| ENSMUSG( | 20.69065 | -0.44288 | 0.290962 | NA | Ces2g |
| ENSMUSG( | 38.96894 | 0.233634 | 0.456199 | 0.838832 | Ces5a |
| ENSMUSG( | 813.7126 | 0.059195 | 0.629468 | 0.913965 | Cetn2 |
| ENSMUSG( | 915.8001 | 0.014868 | 0.88245 | 0.982244 | Cetn3 |
| ENSMUSG( | 146.4 | -0.02702 | 0.87801 | 0.981431 | Cetn4 |
| ENSMUSG( | 176.8597 | 0.172787 | 0.326571 | 0.771368 | Cfap100 |
| ENSMUSG( | 235.1101 | -0.06065 | 0.694438 | 0.9343 | Cfap119 |
| ENSMUSG( | 63.56006 | -0.35869 | 0.137432 | 0.552453 | Cfap126 |
| ENSMUSG( | 26.56829 | -0.30008 | 0.452503 | 0.836145 | Cfap144 |
| ENSMUSG( | 10.00197 | 0.07164 | 0.913709 | NA | Cfap157 |

| | | | | | |
|---|---|---|---|---|---|
| ENSMUSG( | 24.89727 | -0.71672 | 0.073476 | 0.763629 | Cdk2ap1rt |
| ENSMUSG( | 522.8146 | -0.01913 | 0.896613 | 0.990558 | Cdk2ap2 |
| ENSMUSG( | 41.51064 | -0.12095 | 0.748649 | 0.96984 | Cdk3 |
| ENSMUSG( | 1422.09 | -0.08717 | 0.434886 | 0.918876 | Cdk4 |
| ENSMUSG( | 2658.455 | -0.01958 | 0.879592 | 0.989644 | Cdk5 |
| ENSMUSG( | 5312.271 | 0.051817 | 0.636572 | 0.953903 | Cdk5r1 |
| ENSMUSG( | 19029.54 | 0.039542 | 0.71366 | 0.96497 | Cdk5r2 |
| ENSMUSG( | 165.695 | 0.119676 | 0.630849 | 0.953251 | Cdk5rap1 |
| ENSMUSG( | 1006.022 | -0.13087 | 0.289603 | 0.896726 | Cdk5rap2 |
| ENSMUSG( | 1054.954 | -0.09842 | 0.388904 | 0.917257 | Cdk5rap3 |
| ENSMUSG( | 141.1649 | 0.186135 | 0.335296 | 0.907907 | Cdk6 |
| ENSMUSG( | 291.1125 | -0.00319 | 0.984609 | 0.996963 | Cdk7 |
| ENSMUSG( | 703.6322 | 0.074035 | 0.59689 | 0.946555 | Cdk8 |
| ENSMUSG( | 1961.512 | -0.05855 | 0.725362 | 0.966887 | Cdk9 |
| ENSMUSG( | 231.2559 | -0.12069 | 0.464446 | 0.923329 | Cdkal1 |
| ENSMUSG( | 163.7866 | -0.06303 | 0.748709 | 0.96984 | Cdkl1 |
| ENSMUSG( | 804.4571 | 0.099268 | 0.475681 | 0.923329 | Cdkl2 |
| ENSMUSG( | 384.1847 | -0.11727 | 0.463019 | 0.923329 | Cdkl3 |
| ENSMUSG( | 338.9401 | 0.101032 | 0.475201 | 0.923329 | Cdkl4 |
| ENSMUSG( | 2170.44 | -0.06921 | 0.528475 | 0.934795 | Cdkl5 |
| ENSMUSG( | 467.8581 | -0.52407 | 0.000571 | 0.072448 | Cdkn1a |
| ENSMUSG( | 1092.264 | 0.15556 | 0.204801 | 0.864412 | Cdkn1b |
| ENSMUSG( | 297.6334 | 0.124223 | 0.404901 | 0.917257 | Cdkn1c |
| ENSMUSG( | 356.7031 | 0.090702 | 0.54183 | 0.937841 | Cdkn2aip |
| ENSMUSG( | 675.3028 | -0.01622 | 0.900579 | 0.990558 | Cdkn2aipnl |
| ENSMUSG( | 39.0088 | 0.442017 | 0.225467 | 0.868468 | Cdkn2b |
| ENSMUSG( | 243.2693 | -0.20242 | 0.228981 | 0.868507 | Cdkn2c |
| ENSMUSG( | 594.5021 | -0.08468 | 0.614008 | 0.951177 | Cdkn2d |
| ENSMUSG( | 54.05067 | -0.04637 | 0.86739 | 0.987764 | Cdnf |
| ENSMUSG( | 790.7215 | -0.09913 | 0.512754 | 0.930164 | Cdo1 |
| ENSMUSG( | 533.2194 | -0.13772 | 0.322322 | 0.904023 | Cdon |
| ENSMUSG( | 257.7682 | -0.19742 | 0.221653 | 0.866104 | Cdpf1 |
| ENSMUSG( | 1069.149 | 0.267993 | 0.057254 | 0.723362 | Cdr1os |
| ENSMUSG( | 1073.364 | -0.04345 | 0.78192 | 0.976106 | Cdr2 |
| ENSMUSG( | 721.1619 | -0.07328 | 0.550214 | 0.938699 | Cdr2l |
| ENSMUSG( | 3479.522 | 0.06892 | 0.608965 | 0.950226 | Cds1 |
| ENSMUSG( | 17138.83 | 0.15629 | 0.323676 | 0.904856 | Cds2 |
| ENSMUSG( | 34.60524 | -0.23779 | 0.493852 | 0.927372 | Cdsn |
| ENSMUSG( | 49.55925 | -0.39797 | 0.170271 | 0.841364 | Cdt1 |
| ENSMUSG( | 2420.752 | -0.0999 | 0.478751 | 0.923329 | Cdv3 |
| ENSMUSG( | 252.5282 | 0.03479 | 0.840363 | 0.982367 | Cdyl |
| ENSMUSG( | 55.05745 | 0.302166 | 0.27152 | 0.888062 | Cdyl2 |
| ENSMUSG( | 151.0877 | 0.096843 | 0.713098 | 0.964595 | Ceacam1 |
| ENSMUSG( | 516.3705 | 0.240871 | 0.114715 | 0.811703 | Cebpa |
| ENSMUSG( | 193.0959 | -0.08951 | 0.682033 | 0.958678 | Cebpb |
| ENSMUSG( | 181.1301 | 0.011165 | 0.962153 | 0.995237 | Cebpd |
| ENSMUSG( | 1221.779 | 0.018189 | 0.875372 | 0.989644 | Cebpg |
| ENSMUSG( | 976.0141 | 0.073421 | 0.577495 | 0.943127 | Cebpz |
| ENSMUSG( | 406.0932 | -0.23981 | 0.147452 | 0.836241 | Cebpzos |
| ENSMUSG( | 251.8297 | -0.03514 | 0.83841 | 0.981622 | Cecr2 |
| ENSMUSG( | 100.4663 | -0.23343 | 0.27568 | 0.890209 | Cela1 |
| ENSMUSG( | 4046.68 | 0.100242 | 0.333661 | 0.907907 | Celf1 |
| ENSMUSG( | 7237.66 | 0.118866 | 0.267859 | 0.886134 | Celf2 |
| ENSMUSG( | 3849.817 | 0.015481 | 0.91333 | 0.992298 | Celf3 |
| ENSMUSG( | 16844.3 | 0.168018 | 0.162696 | 0.837724 | Celf4 |
| ENSMUSG( | 7523.59 | 0.043308 | 0.74521 | 0.96951 | Celf5 |
| ENSMUSG( | 421.6392 | -0.00412 | 0.977104 | 0.996416 | Celf6 |
| ENSMUSG( | 277.2631 | -0.21967 | 0.145826 | 0.835057 | Celrr |
| ENSMUSG( | 610.1364 | 0.155468 | 0.370517 | 0.915491 | Celsr1 |
| ENSMUSG( | 8859.651 | 0.22046 | 0.128841 | 0.820095 | Celsr2 |
| ENSMUSG( | 3190.366 | 0.148316 | 0.265891 | 0.886134 | Celsr3 |
| ENSMUSG( | 762.7192 | 0.380241 | 0.031309 | 0.631113 | Cemip |
| ENSMUSG( | 548.1853 | 0.28482 | 0.136234 | 0.827092 | Cemip2 |
| ENSMUSG( | 358.6714 | 0.310847 | 0.102659 | 0.803503 | Cenatac |
| ENSMUSG( | 5114.947 | -0.21943 | 0.104946 | 0.804969 | Cend1 |
| ENSMUSG( | 77.3739 | 0.243499 | 0.301936 | 0.898538 | Cenpa |
| ENSMUSG( | 15066.56 | -0.05161 | 0.608566 | 0.950113 | Cenpb |
| ENSMUSG( | 423.1054 | 0.221338 | 0.113763 | 0.809746 | Cenpc1 |
| ENSMUSG( | 58.45402 | -0.09431 | 0.760371 | 0.972575 | Cenpe |
| ENSMUSG( | 98.66257 | -0.10129 | 0.714884 | 0.964986 | Cenpf |
| ENSMUSG( | 14.89956 | 0.159061 | 0.752162 | 0.970461 | Cenph |
| ENSMUSG( | 18.34251 | -0.14789 | 0.753971 | 0.970671 | Cenpi |
| ENSMUSG( | 112.06 | 0.132878 | 0.532824 | 0.934795 | Cenpj |
| ENSMUSG( | 16.46591 | 0.070899 | 0.882102 | 0.989644 | Cenpk |
| ENSMUSG( | 40.07637 | 0.292638 | 0.358633 | 0.913759 | Cenpl |
| ENSMUSG( | 95.27053 | 0.065116 | 0.774958 | 0.974931 | Cenpm |
| ENSMUSG( | 32.93059 | 0.2775 | 0.456604 | 0.923329 | Cenpn |
| ENSMUSG( | 110.7087 | 0.186904 | 0.39747 | 0.917257 | Cenpo |
| ENSMUSG( | 14.592 | -0.46979 | 0.363858 | 0.914223 | Cenpp |
| ENSMUSG( | 150.7936 | -0.24155 | 0.262863 | 0.885357 | Cenpq |
| ENSMUSG( | 35.72621 | -0.23301 | 0.553528 | 0.939565 | Cenps |
| ENSMUSG( | 355.6451 | -0.06193 | 0.696738 | 0.960867 | Cenpt |
| ENSMUSG( | 44.91883 | 0.109641 | 0.717267 | 0.965964 | Cenpu |
| ENSMUSG( | 496.7393 | -0.14113 | 0.353886 | 0.912411 | Cenpv |
| ENSMUSG( | 48.31303 | -0.08594 | 0.794128 | 0.97681 | Cenpw |
| ENSMUSG( | 424.1916 | -0.05161 | 0.718562 | 0.966633 | Cenpx |
| ENSMUSG( | 1464.559 | 0.092708 | 0.414036 | 0.917257 | Cep104 |
| ENSMUSG( | 86.18103 | -0.17394 | 0.482325 | 0.924578 | Cep112 |
| ENSMUSG( | 1426.951 | -0.01102 | 0.927453 | 0.99306 | Cep120 |
| ENSMUSG( | 252.2158 | 0.228147 | 0.178408 | 0.848538 | Cep126 |
| ENSMUSG( | 411.1867 | -0.12596 | 0.463472 | 0.923329 | Cep128 |
| ENSMUSG( | 776.9529 | -0.06691 | 0.70668 | 0.963529 | Cep131 |
| ENSMUSG( | 243.1845 | 0.195861 | 0.223472 | 0.867404 | Cep135 |
| ENSMUSG( | 583.8289 | -0.15499 | 0.37319 | 0.915491 | Cep15 |
| ENSMUSG( | 145.5224 | 0.378578 | 0.050233 | 0.708648 | Cep152 |
| ENSMUSG( | 382.1575 | 0.252659 | 0.076234 | 0.769674 | Cep162 |
| ENSMUSG( | 630.1924 | -0.08581 | 0.540086 | 0.937383 | Cep164 |
| ENSMUSG( | 1387.115 | 0.063909 | 0.570488 | 0.941872 | Cep170 |
| ENSMUSG( | 9462.119 | 0.107999 | 0.335703 | 0.907907 | Cep170b |
| ENSMUSG( | 1010.101 | -0.00653 | 0.957807 | 0.994798 | Cep19 |
| ENSMUSG( | 342.1033 | -0.00548 | 0.971439 | 0.996236 | Cep192 |
| ENSMUSG( | 1081.611 | -0.20954 | 0.08596 | 0.780185 | Cep20 |
| ENSMUSG( | 1608.735 | -0.0336 | 0.774549 | 0.974931 | Cep250 |
| ENSMUSG( | 343.503 | 0.131688 | 0.421345 | 0.917257 | Cep290 |
| ENSMUSG( | 760.7509 | 0.218547 | 0.080496 | 0.77681 | Cep295 |
| ENSMUSG( | 1544.048 | 0.166416 | 0.194257 | 0.860096 | Cep350 |
| ENSMUSG( | 176.991 | 0.005967 | 0.973157 | 0.996416 | Cep41 |
| ENSMUSG( | 367.7441 | 0.135591 | 0.339433 | 0.907972 | Cep43 |
| ENSMUSG( | 223.279 | 0.048693 | 0.797897 | 0.97681 | Cep44 |
| ENSMUSG( | 11.76176 | -0.08447 | 0.885865 | 0.989776 | Cep55 |
| ENSMUSG( | 319.5501 | 0.185592 | 0.250379 | 0.87792 | Cep57 |
| ENSMUSG( | 96.89557 | 0.19318 | 0.407739 | 0.917257 | Cep57l1 |
| ENSMUSG( | 896.8641 | -0.04698 | 0.691748 | 0.960247 | Cep63 |
| ENSMUSG( | 843.1954 | 0.0921 | 0.493652 | 0.927372 | Cep68 |
| ENSMUSG( | 380.8041 | 0.167521 | 0.300763 | 0.898538 | Cep70 |
| ENSMUSG( | 63.41372 | 0.380949 | 0.204215 | 0.864412 | Cep72 |
| ENSMUSG( | 154.0422 | 0.238985 | 0.190295 | 0.857355 | Cep76 |
| ENSMUSG( | 289.4236 | 0.088686 | 0.55409 | 0.939565 | Cep78 |
| ENSMUSG( | 1129.613 | 0.030006 | 0.812101 | 0.979148 | Cep83 |
| ENSMUSG( | 322.0297 | 0.118147 | 0.414195 | 0.917257 | Cep85 |
| ENSMUSG( | 493.2019 | 0.062828 | 0.672487 | 0.95694 | Cep85l |
| ENSMUSG( | 540.2451 | 0.090226 | 0.500709 | 0.927825 | Cep89 |
| ENSMUSG( | 374.863 | 0.076144 | 0.642596 | 0.953903 | Cep95 |
| ENSMUSG( | 458.5539 | 0.044864 | 0.74929 | 0.96998 | Cep97 |
| ENSMUSG( | 1120.15 | 0.25257 | 0.204123 | 0.864412 | Cept1 |
| ENSMUSG( | 362.1184 | 0.090723 | 0.665583 | 0.956766 | Cercam |
| ENSMUSG( | 2707.422 | -0.07568 | 0.468934 | 0.923329 | Cerk |
| ENSMUSG( | 80.45424 | -0.29721 | 0.261829 | 0.884926 | Cerkl |
| ENSMUSG( | 1316.963 | 0.027266 | 0.845124 | 0.983878 | Cerox1 |
| ENSMUSG( | 3389.831 | 0.257801 | 0.151919 | 0.837724 | Cers2 |
| ENSMUSG( | 15.36566 | 1.118017 | 0.027093 | 0.603041 | Cers3 |
| ENSMUSG( | 1924.94 | 0.140928 | 0.269795 | 0.887508 | Cers4 |
| ENSMUSG( | 1383.591 | 0.297904 | 0.049667 | 0.708648 | Cers5 |
| ENSMUSG( | 1456.198 | 0.441906 | 0.021042 | 0.573245 | Cers6 |
| ENSMUSG( | 1830.902 | 0.144379 | 0.21098 | 0.864412 | Cert1 |
| ENSMUSG( | 21.65547 | 0.009842 | 0.981971 | 0.996756 | Ces2g |
| ENSMUSG( | 32.673 | 0.021194 | 0.952007 | 0.994798 | Ces5a |
| ENSMUSG( | 692.6412 | -0.14615 | 0.324684 | 0.905257 | Cetn2 |
| ENSMUSG( | 852.6865 | 0.074487 | 0.527284 | 0.934466 | Cetn3 |
| ENSMUSG( | 127.8813 | -0.16795 | 0.426852 | 0.917257 | Cetn4 |
| ENSMUSG( | 163.6404 | 0.224866 | 0.345083 | 0.91045 | Cfap100 |
| ENSMUSG( | 203.534 | -0.21327 | 0.236075 | 0.872173 | Cfap119 |
| ENSMUSG( | 68.88089 | 0.152498 | 0.586696 | 0.944975 | Cfap126 |
| ENSMUSG( | 33.11424 | 0.576743 | 0.100655 | 0.801444 | Cfap144 |
| ENSMUSG( | 10.53048 | 0.504833 | 0.428587 | NA | Cfap157 |
| ENSMUSG( | 26.92271 | 0.095915 | 0.831916 | 0.980738 | Cfap161 |
| ENSMUSG( | 768.5366 | 0.049697 | 0.70685 | 0.963529 | Cfap20 |
| ENSMUSG( | 60.87339 | 0.317108 | 0.328672 | 0.906727 | Cfap206 |
| ENSMUSG( | 252.6417 | -0.06376 | 0.711335 | 0.964363 | Cfap20dc |
| ENSMUSG( | 80.37971 | 0.115443 | 0.665638 | 0.956766 | Cfap210 |
| ENSMUSG( | 12.28328 | 0.551648 | 0.369229 | 0.915491 | Cfap221 |
| ENSMUSG( | 181.7887 | -0.0071 | 0.969264 | 0.99594 | Cfap251 |
| ENSMUSG( | 22.25034 | 0.19956 | 0.629098 | 0.953251 | Cfap276 |
| ENSMUSG( | 480.8082 | -0.10677 | 0.532345 | 0.934795 | Cfap298 |
| ENSMUSG( | 14.65856 | 0.822197 | 0.105808 | 0.804969 | Cfap299 |
| ENSMUSG( | 155.1462 | 0.064192 | 0.749821 | 0.969988 | Cfap300 |
| ENSMUSG( | 2254.213 | -0.1401 | 0.297797 | 0.898538 | Cfap36 |
| ENSMUSG( | 921.725 | -0.00114 | 0.992637 | 0.99874 | Cfap410 |

| | | | | | | | | | | | |
|---|---|---|---|---|---|---|---|---|---|---|---|
| ENSMUSG( | Cdkl5 | 2442.347 | 0.004006 | 0.965071 | 0.995303 | ENSMUSG( | Cdkl5 | 2170.44 | -0.06921 | 0.528475 | 0.934795 |
| ENSMUSG( | Cdkn1a | 596.6308 | -0.04716 | 0.74412 | 0.948497 | ENSMUSG( | Cdkn1a | 467.8581 | -0.52407 | 0.000571 | 0.072448 |
| ENSMUSG( | Cdkn1b | 1176.845 | 0.106467 | 0.2781 | 0.728086 | ENSMUSG( | Cdkn1b | 1092.264 | 0.15556 | 0.204801 | 0.864412 |
| ENSMUSG( | Cdkn1c | 302.5252 | -0.09115 | 0.534788 | 0.880725 | ENSMUSG( | Cdkn1c | 297.6334 | 0.124223 | 0.404901 | 0.917257 |
| ENSMUSG( | Cdkn2aip | 364.646 | -0.12114 | 0.379471 | 0.801757 | ENSMUSG( | Cdkn2aip | 356.7031 | 0.090702 | 0.54183 | 0.937841 |
| ENSMUSG( | Cdkn2aipnl | 784.2941 | 0.147032 | 0.149628 | 0.573617 | ENSMUSG( | Cdkn2aipnl | 675.3028 | -0.01622 | 0.900579 | 0.990558 |
| ENSMUSG( | Cdkn2b | 40.5918 | 0.280756 | 0.394273 | 0.810014 | ENSMUSG( | Cdkn2b | 39.0088 | 0.442017 | 0.225467 | 0.868468 |
| ENSMUSG( | Cdkn2c | 280.3766 | -0.04818 | 0.742289 | 0.948478 | ENSMUSG( | Cdkn2c | 243.2693 | -0.20242 | 0.228981 | 0.868507 |
| ENSMUSG( | Cdkn2d | 655.0878 | -0.06869 | 0.65928 | 0.924506 | ENSMUSG( | Cdkn2d | 594.5021 | -0.08468 | 0.614008 | 0.951177 |
| ENSMUSG( | Cdnf | 55.01681 | -0.29424 | 0.251091 | 0.700723 | ENSMUSG( | Cdnf | 54.05067 | -0.04637 | 0.86739 | 0.987764 |
| ENSMUSG( | Cdo1 | 963.8321 | 0.204267 | 0.056841 | 0.3679 | ENSMUSG( | Cdo1 | 790.7215 | -0.09913 | 0.512754 | 0.930164 |
| ENSMUSG( | Cdon | 536.1413 | -0.42526 | 0.000274 | 0.013445 | ENSMUSG( | Cdon | 533.2194 | -0.13772 | 0.322322 | 0.904023 |
| ENSMUSG( | Cdpf1 | 302.5155 | 0.002116 | 0.98825 | 0.999109 | ENSMUSG( | Cdpf1 | 257.7682 | -0.19742 | 0.221653 | 0.866104 |
| ENSMUSG( | Cdr1os | 1029.042 | -0.10286 | 0.350054 | 0.781943 | ENSMUSG( | Cdr1os | 1069.149 | 0.267993 | 0.057254 | 0.723362 |
| ENSMUSG( | Cdr2 | 1237.296 | 0.098665 | 0.447596 | 0.834051 | ENSMUSG( | Cdr2 | 1073.364 | -0.04345 | 0.78192 | 0.976106 |
| ENSMUSG( | Cdr2l | 800.0107 | -0.04218 | 0.702945 | 0.937735 | ENSMUSG( | Cdr2l | 721.1619 | -0.07328 | 0.550214 | 0.938699 |
| ENSMUSG( | Cds1 | 3480.769 | -0.20753 | 0.093396 | 0.469793 | ENSMUSG( | Cds1 | 3479.522 | 0.06892 | 0.608965 | 0.950226 |
| ENSMUSG( | Cds2 | 17161.63 | -0.11045 | 0.405786 | 0.81652 | ENSMUSG( | Cds2 | 17138.83 | 0.15629 | 0.323676 | 0.904856 |
| ENSMUSG( | Cdsn | 37.28933 | -0.27227 | 0.379252 | 0.801757 | ENSMUSG( | Cdsn | 34.60524 | -0.23779 | 0.493852 | 0.927372 |
| ENSMUSG( | Cdt1 | 55.08836 | -0.37565 | 0.1481 | 0.571232 | ENSMUSG( | Cdt1 | 49.55925 | -0.39797 | 0.170271 | 0.841364 |
| ENSMUSG( | Cdv3 | 2731.673 | -0.01493 | 0.88634 | 0.983139 | ENSMUSG( | Cdv3 | 2420.752 | -0.0999 | 0.478751 | 0.923329 |
| ENSMUSG( | Cdyl | 279.8211 | 0.067816 | 0.613814 | 0.90923 | ENSMUSG( | Cdyl | 252.5282 | 0.03479 | 0.840363 | 0.982367 |
| ENSMUSG( | Cdyl2 | 67.84651 | 0.586644 | 0.013854 | 0.172062 | ENSMUSG( | Cdyl2 | 55.05745 | 0.302166 | 0.27152 | 0.888062 |
| ENSMUSG( | Ceacam1 | 160.9958 | -0.00214 | 0.992253 | 0.999327 | ENSMUSG( | Ceacam1 | 151.0877 | 0.096843 | 0.713098 | 0.964595 |
| ENSMUSG( | Cebpa | 617.7419 | 0.461193 | 0.0015 | 0.042984 | ENSMUSG( | Cebpa | 516.3705 | 0.240871 | 0.114715 | 0.811703 |
| ENSMUSG( | Cebpb | 203.1253 | -0.21119 | 0.337889 | 0.777422 | ENSMUSG( | Cebpb | 193.0959 | -0.08951 | 0.682033 | 0.958678 |
| ENSMUSG( | Cebpd | 175.8623 | -0.3864 | 0.055043 | 0.36102 | ENSMUSG( | Cebpd | 181.1301 | 0.011165 | 0.962153 | 0.995237 |
| ENSMUSG( | Cebpg | 1361.996 | 0.063914 | 0.518019 | 0.873754 | ENSMUSG( | Cebpg | 1221.779 | 0.018189 | 0.875372 | 0.989644 |
| ENSMUSG( | Cebpz | 1135.082 | 0.234387 | 0.016474 | 0.190587 | ENSMUSG( | Cebpz | 976.0141 | 0.073421 | 0.577495 | 0.943127 |
| ENSMUSG( | Cebpzos | 467.201 | -0.08824 | 0.559468 | 0.888669 | ENSMUSG( | Cebpzos | 406.0932 | -0.23981 | 0.147452 | 0.836241 |
| ENSMUSG( | Cecr2 | 260.0443 | -0.22172 | 0.182363 | 0.619591 | ENSMUSG( | Cecr2 | 251.8297 | -0.03514 | 0.83841 | 0.981622 |
| ENSMUSG( | Cela1 | 124.7841 | 0.13441 | 0.45954 | 0.840338 | ENSMUSG( | Cela1 | 100.4663 | -0.23343 | 0.27568 | 0.890209 |
| ENSMUSG( | Celf1 | 4278.427 | -0.00673 | 0.936306 | 0.993415 | ENSMUSG( | Celf1 | 4046.68 | 0.100242 | 0.333661 | 0.907907 |
| ENSMUSG( | Celf2 | 7745.759 | 0.048782 | 0.555874 | 0.887457 | ENSMUSG( | Celf2 | 7237.66 | 0.118866 | 0.267859 | 0.886134 |
| ENSMUSG( | Celf3 | 4531.55 | 0.211541 | 0.096739 | 0.477375 | ENSMUSG( | Celf3 | 3849.817 | 0.015481 | 0.91333 | 0.992298 |
| ENSMUSG( | Celf4 | 18445.01 | 0.162107 | 0.045728 | 0.328047 | ENSMUSG( | Celf4 | 16844.3 | 0.168018 | 0.162696 | 0.837724 |
| ENSMUSG( | Celf5 | 8917.826 | 0.256635 | 0.025497 | 0.246501 | ENSMUSG( | Celf5 | 7523.59 | 0.043308 | 0.74521 | 0.96951 |
| ENSMUSG( | Celf6 | 502.3164 | 0.225633 | 0.065174 | 0.395697 | ENSMUSG( | Celf6 | 421.6392 | -0.00412 | 0.977104 | 0.996416 |
| ENSMUSG( | Celrr | 340.5891 | 0.116016 | 0.348986 | 0.781661 | ENSMUSG( | Celrr | 277.2631 | -0.21967 | 0.145826 | 0.835057 |
| ENSMUSG( | Celsr1 | 609.8126 | -0.12085 | 0.438294 | 0.830406 | ENSMUSG( | Celsr1 | 610.1364 | 0.155468 | 0.370517 | 0.915491 |
| ENSMUSG( | Celsr2 | 8184.915 | -0.28547 | 0.025223 | 0.244689 | ENSMUSG( | Celsr2 | 8859.651 | 0.22046 | 0.128841 | 0.820095 |
| ENSMUSG( | Celsr3 | 3269.322 | -0.04522 | 0.709847 | 0.939669 | ENSMUSG( | Celsr3 | 3190.366 | 0.148316 | 0.265891 | 0.886134 |
| ENSMUSG( | Cemip | 774.7387 | 0.168326 | 0.294972 | 0.743798 | ENSMUSG( | Cemip | 762.7192 | 0.380241 | 0.031309 | 0.631113 |
| ENSMUSG( | Cemip2 | 508.1397 | -0.20516 | 0.200847 | 0.640988 | ENSMUSG( | Cemip2 | 548.1853 | 0.28482 | 0.136234 | 0.827092 |
| ENSMUSG( | Cenatac | 340.5539 | -0.10406 | 0.49494 | 0.860798 | ENSMUSG( | Cenatac | 358.6714 | 0.310847 | 0.102659 | 0.803503 |
| ENSMUSG( | Cend1 | 6125.12 | 0.041826 | 0.732946 | 0.944956 | ENSMUSG( | Cend1 | 5114.947 | -0.21943 | 0.104946 | 0.804969 |
| ENSMUSG( | Cenpa | 80.08411 | 0.074434 | 0.727046 | 0.944223 | ENSMUSG( | Cenpa | 77.3739 | 0.243499 | 0.301936 | 0.898538 |
| ENSMUSG( | Cenpb | 17478.21 | 0.10763 | 0.165847 | 0.597653 | ENSMUSG( | Cenpb | 15066.56 | -0.05161 | 0.608566 | 0.950113 |
| ENSMUSG( | Cenpc1 | 447.9202 | 0.121847 | 0.289992 | 0.739839 | ENSMUSG( | Cenpc1 | 423.1054 | 0.221338 | 0.113763 | 0.809746 |
| ENSMUSG( | Cenpe | 60.03787 | -0.3254 | 0.242862 | 0.690275 | ENSMUSG( | Cenpe | 58.45402 | -0.09431 | 0.760371 | 0.972575 |
| ENSMUSG( | Cenpf | 109.0788 | -0.07388 | 0.743641 | 0.948478 | ENSMUSG( | Cenpf | 98.66257 | -0.10129 | 0.714884 | 0.964986 |
| ENSMUSG( | Cenph | 14.13697 | -0.25258 | 0.608737 | NA | ENSMUSG( | Cenph | 14.89956 | 0.159061 | 0.752162 | 0.970461 |
| ENSMUSG( | Cenpi | 20.21363 | -0.1785 | 0.659708 | NA | ENSMUSG( | Cenpi | 18.34251 | -0.14789 | 0.753971 | 0.970671 |
| ENSMUSG( | Cenpj | 109.8522 | -0.19103 | 0.327153 | 0.772056 | ENSMUSG( | Cenpj | 112.06 | 0.132878 | 0.532824 | 0.934795 |
| ENSMUSG( | Cenpk | 15.00024 | -0.54348 | 0.252641 | NA | ENSMUSG( | Cenpk | 16.46591 | 0.070899 | 0.882102 | 0.989644 |
| ENSMUSG( | Cenpl | 39.65971 | 0.018678 | 0.950536 | 0.993821 | ENSMUSG( | Cenpl | 40.07637 | 0.292638 | 0.358633 | 0.913759 |
| ENSMUSG( | Cenpm | 98.53577 | -0.09985 | 0.617749 | 0.910872 | ENSMUSG( | Cenpm | 95.27053 | 0.065116 | 0.774958 | 0.974931 |
| ENSMUSG( | Cenpn | 30.95619 | -0.1867 | 0.596366 | 0.903264 | ENSMUSG( | Cenpn | 32.93059 | 0.2775 | 0.456604 | 0.923329 |
| ENSMUSG( | Cenpo | 110.3093 | -0.09447 | 0.61999 | 0.911432 | ENSMUSG( | Cenpo | 110.7087 | 0.186904 | 0.39747 | 0.917257 |
| ENSMUSG( | Cenpp | 17.4057 | -0.16138 | 0.716658 | NA | ENSMUSG( | Cenpp | 14.592 | -0.46979 | 0.363858 | 0.914223 |
| ENSMUSG( | Cenpq | 181.7023 | 0.043373 | 0.81653 | 0.965455 | ENSMUSG( | Cenpq | 150.7936 | -0.24155 | 0.262863 | 0.885357 |
| ENSMUSG( | Cenps | 45.40985 | 0.202975 | 0.55494 | 0.887073 | ENSMUSG( | Cenps | 35.72621 | -0.23301 | 0.553528 | 0.939565 |
| ENSMUSG( | Cenpt | 405.0179 | 0.045298 | 0.732193 | 0.944956 | ENSMUSG( | Cenpt | 355.6451 | -0.06193 | 0.696738 | 0.960867 |
| ENSMUSG( | Cenpu | 43.87946 | -0.25657 | 0.404479 | 0.816132 | ENSMUSG( | Cenpu | 44.91883 | 0.109641 | 0.717267 | 0.965964 |
| ENSMUSG( | Cenpv | 586.9386 | 0.080588 | 0.487478 | 0.858812 | ENSMUSG( | Cenpv | 496.7393 | -0.14113 | 0.353886 | 0.912411 |
| ENSMUSG( | Cenpw | 54.65292 | -0.02586 | 0.923783 | 0.991046 | ENSMUSG( | Cenpw | 48.31303 | -0.08594 | 0.794128 | 0.97681 |
| ENSMUSG( | Cenpx | 486.4657 | 0.080146 | 0.48986 | 0.859319 | ENSMUSG( | Cenpx | 424.1916 | -0.05161 | 0.718562 | 0.966633 |
| ENSMUSG( | Cep104 | 1626.418 | 0.124467 | 0.162941 | 0.593778 | ENSMUSG( | Cep104 | 1464.559 | 0.092708 | 0.414036 | 0.917257 |
| ENSMUSG( | Cep112 | 101.6155 | 0.056757 | 0.788649 | 0.959226 | ENSMUSG( | Cep112 | 86.18103 | -0.17394 | 0.482325 | 0.924578 |
| ENSMUSG( | Cep120 | 1624.19 | 0.094795 | 0.296275 | 0.744386 | ENSMUSG( | Cep120 | 1426.951 | -0.01102 | 0.927453 | 0.99306 |
| ENSMUSG( | Cep126 | 261.8692 | 0.068368 | 0.631442 | 0.914073 | ENSMUSG( | Cep126 | 252.2158 | 0.228147 | 0.178408 | 0.848538 |
| ENSMUSG( | Cep128 | 469.2317 | -0.0008 | 0.995963 | 0.99943 | ENSMUSG( | Cep128 | 411.1867 | -0.12596 | 0.463472 | 0.923329 |
| ENSMUSG( | Cep131 | 929.0851 | 0.180315 | 0.27862 | 0.728705 | ENSMUSG( | Cep131 | 776.9529 | -0.06691 | 0.70668 | 0.963529 |
| ENSMUSG( | Cep135 | 256.4295 | 0.078508 | 0.579387 | 0.896722 | ENSMUSG( | Cep135 | 243.1845 | 0.195861 | 0.223472 | 0.867404 |
| ENSMUSG( | Cep15 | 652.4974 | -0.09071 | 0.52671 | 0.877267 | ENSMUSG( | Cep15 | 583.8289 | -0.15499 | 0.37319 | 0.915491 |
| ENSMUSG( | Cep152 | 155.5141 | 0.297325 | 0.087825 | 0.454663 | ENSMUSG( | Cep152 | 145.5224 | 0.378578 | 0.050233 | 0.708648 |
| ENSMUSG( | Cep162 | 406.139 | 0.16597 | 0.169461 | 0.603569 | ENSMUSG( | Cep162 | 382.1575 | 0.252659 | 0.076234 | 0.769674 |
| ENSMUSG( | Cep164 | 701.1045 | -0.04664 | 0.702441 | 0.937668 | ENSMUSG( | Cep164 | 630.1924 | -0.08581 | 0.540086 | 0.937383 |
| ENSMUSG( | Cep170 | 1463.811 | -0.04832 | 0.615441 | 0.910481 | ENSMUSG( | Cep170 | 1387.115 | 0.063909 | 0.570488 | 0.941872 |
| ENSMUSG( | Cep170b | 11111.04 | 0.289093 | 0.001145 | 0.036093 | ENSMUSG( | Cep170b | 9462.119 | 0.107999 | 0.335703 | 0.907907 |
| ENSMUSG( | Cep19 | 1144.455 | 0.085552 | 0.405167 | 0.81652 | ENSMUSG( | Cep19 | 1010.101 | -0.00653 | 0.957807 | 0.994798 |
| ENSMUSG( | Cep192 | 367.2806 | -0.07663 | 0.585135 | 0.899171 | ENSMUSG( | Cep192 | 342.1033 | -0.00548 | 0.971439 | 0.996236 |
| ENSMUSG( | Cep20 | 1258.824 | -0.0295 | 0.773964 | 0.955606 | ENSMUSG( | Cep20 | 1081.611 | -0.20954 | 0.08596 | 0.780185 |
| ENSMUSG( | Cep250 | 1854.867 | 0.108527 | 0.23544 | 0.682423 | ENSMUSG( | Cep250 | 1608.735 | -0.0336 | 0.774549 | 0.974931 |
| ENSMUSG( | Cep290 | 375.2022 | 0.109569 | 0.360979 | 0.792797 | ENSMUSG( | Cep290 | 343.503 | 0.131688 | 0.421345 | 0.917257 |
| ENSMUSG( | Cep295 | 730.0359 | -0.16822 | 0.103911 | 0.489523 | ENSMUSG( | Cep295 | 760.7509 | 0.218547 | 0.080496 | 0.77681 |
| ENSMUSG( | Cep350 | 1567.931 | -0.05656 | 0.617396 | 0.910872 | ENSMUSG( | Cep350 | 1544.048 | 0.166416 | 0.194257 | 0.860096 |
| ENSMUSG( | Cep41 | 188.9717 | -0.07135 | 0.660714 | 0.924951 | ENSMUSG( | Cep41 | 176.991 | 0.005967 | 0.973157 | 0.996416 |
| ENSMUSG( | Cep43 | 408.228 | 0.170266 | 0.156244 | 0.583988 | ENSMUSG( | Cep43 | 367.7441 | 0.135591 | 0.339433 | 0.907972 |
| ENSMUSG( | Cep44 | 242.5508 | 0.008168 | 0.956547 | 0.993874 | ENSMUSG( | Cep44 | 223.279 | 0.048693 | 0.797897 | 0.97681 |
| ENSMUSG( | Cep55 | 11.58667 | -0.46485 | 0.4222 | NA | ENSMUSG( | Cep55 | 11.76176 | -0.08447 | 0.885865 | 0.989776 |
| ENSMUSG( | Cep57 | 324.4399 | -0.03399 | 0.821369 | 0.966248 | ENSMUSG( | Cep57 | 319.5501 | 0.185592 | 0.250379 | 0.87792 |
| ENSMUSG( | Cep57l1 | 100.6047 | 0.039497 | 0.857126 | 0.974393 | ENSMUSG( | Cep57l1 | 96.89557 | 0.19318 | 0.407739 | 0.917257 |
| ENSMUSG( | Cep63 | 1010.708 | 0.029375 | 0.766618 | 0.954302 | ENSMUSG( | Cep63 | 896.8641 | -0.04698 | 0.691748 | 0.960247 |
| ENSMUSG( | Cep68 | 944.1391 | 0.145444 | 0.225942 | 0.672685 | ENSMUSG( | Cep68 | 843.1954 | 0.0921 | 0.493652 | 0.927372 |
| ENSMUSG( | Cep70 | 388.7787 | -0.04175 | 0.722421 | 0.943752 | ENSMUSG( | Cep70 | 380.8041 | 0.167521 | 0.300763 | 0.898538 |
| ENSMUSG( | Cep72 | 64.2762 | 0.144251 | 0.582643 | 0.897918 | ENSMUSG( | Cep72 | 63.41372 | 0.380949 | 0.204215 | 0.864412 |
| ENSMUSG( | Cep76 | 156.1403 | 0.017938 | 0.916479 | 0.988844 | ENSMUSG( | Cep76 | 154.0422 | 0.238985 | 0.190295 | 0.857355 |
| ENSMUSG( | Cep78 | 323.1567 | 0.136338 | 0.315671 | 0.762671 | ENSMUSG( | Cep78 | 289.4236 | 0.088686 | 0.55409 | 0.939565 |
| ENSMUSG( | Cep83 | 1254.791 | 0.067604 | 0.52658 | 0.877267 | ENSMUSG( | Cep83 | 1129.613 | 0.030006 | 0.812101 | 0.979148 |
| ENSMUSG( | Cep85 | 347.9155 | 0.071557 | 0.587481 | 0.900403 | ENSMUSG( | Cep85 | 322.0297 | 0.118147 | 0.414195 | 0.917257 |
| ENSMUSG( | Cep85l | 521.8139 | -0.04071 | 0.754511 | 0.951007 | ENSMUSG( | Cep85l | 493.2019 | 0.062828 | 0.672487 | 0.95694 |
| ENSMUSG( | Cep89 | 593.7013 | 0.090473 | 0.433136 | 0.828207 | ENSMUSG( | Cep89 | 540.2451 | 0.090226 | 0.500709 | 0.927825 |
| ENSMUSG( | Cep95 | 398.4515 | -0.01857 | 0.884685 | 0.982647 | ENSMUSG( | Cep95 | 374.863 | 0.076144 | 0.642596 | 0.953903 |
| ENSMUSG( | Cep97 | 468.7667 | -0.16676 | 0.188741 | 0.627678 | ENSMUSG( | Cep97 | 458.5539 | 0.044864 | 0.74929 | 0.96998 |
| ENSMUSG( | Cept1 | 1007.215 | -0.33453 | 0.05454 | 0.35941 | ENSMUSG( | Cept1 | 1120.15 | 0.25257 | 0.204123 | 0.864412 |
| ENSMUSG( | Cercam | 367.722 | -0.14618 | 0.303843 | 0.752148 | ENSMUSG( | Cercam | 362.1184 | 0.090723 | 0.665583 | 0.956766 |
| ENSMUSG( | Cerk | 3126.714 | 0.070124 | 0.38451 | 0.804007 | ENSMUSG( | Cerk | 2707.422 | -0.07568 | 0.468934 | 0.923329 |
| ENSMUSG( | Cerkl | 96.06295 | -0.04539 | 0.838912 | 0.971179 | ENSMUSG( | Cerkl | 80.45424 | -0.29721 | 0.261829 | 0.884926 |
| ENSMUSG( | Cerox1 | 1410.416 | -0.04718 | 0.636586 | 0.914969 | ENSMUSG( | Cerox1 | 1316.963 | 0.027266 | 0.845124 | 0.983878 |
| ENSMUSG( | Cers2 | 2954.351 | -0.43375 | 0.002 | 0.051341 | ENSMUSG( | Cers2 | 3389.831 | 0.257801 | 0.151919 | 0.837724 |
| ENSMUSG( | Cers3 | 14.45031 | 0.772173 | 0.123019 | NA | ENSMUSG( | Cers3 | 15.36566 | 1.118017 | 0.027093 | 0.603041 |
| ENSMUSG( | Cers4 | 1928.563 | -0.12428 | 0.292958 | 0.742763 | ENSMUSG( | Cers4 | 1924.94 | 0.140928 | 0.269795 | 0.887508 |
| ENSMUSG( | Cers5 | 1304.291 | -0.13406 | 0.325347 | 0.77061 | ENSMUSG( | Cers5 | 1383.591 | 0.297904 | 0.049667 | 0.708648 |
| ENSMUSG( | Cers6 | 1322.119 | -0.08015 | 0.634189 | 0.914073 | ENSMUSG( | Cers6 | 1456.198 | 0.441906 | 0.021042 | 0.573245 |
| ENSMUSG( | Cert1 | 1940.219 | 0.04871 | 0.580334 | 0.896952 | ENSMUSG( | Cert1 | 1830.902 | 0.144379 | 0.21098 | 0.864412 |
| ENSMUSG( | Ces2g | 20.69065 | -0.44288 | 0.290962 | NA | ENSMUSG( | Ces2g | 21.65547 | 0.009842 | 0.981971 | 0.996756 |
| ENSMUSG( | Ces5a | 38.96894 | 0.233634 | 0.456199 | 0.838832 | ENSMUSG( | Ces5a | 32.673 | 0.021194 | 0.952007 | 0.994798 |
| ENSMUSG( | Cetn2 | 813.7126 | 0.059195 | 0.629468 | 0.913965 | ENSMUSG( | Cetn2 | 692.6412 | -0.14615 | 0.324684 | 0.905257 |
| ENSMUSG( | Cetn3 | 915.8001 | 0.014868 | 0.88245 | 0.982244 | ENSMUSG( | Cetn3 | 852.6865 | 0.074487 | 0.527284 | 0.934466 |
| ENSMUSG( | Cetn4 | 146.4 | -0.02702 | 0.87801 | 0.981431 | ENSMUSG( | Cetn4 | 127.8813 | -0.16795 | 0.426852 | 0.917257 |
| ENSMUSG( | Cfap100 | 176.8597 | 0.172787 | 0.326571 | 0.771368 | ENSMUSG( | Cfap100 | 163.6404 | 0.224866 | 0.345083 | 0.91045 |
| ENSMUSG( | Cfap119 | 235.1101 | -0.06065 | 0.694438 | 0.9343 | ENSMUSG( | Cfap119 | 203.534 | -0.21327 | 0.236075 | 0.872173 |
| ENSMUSG( | Cfap126 | 63.56006 | -0.35869 | 0.137432 | 0.552453 | ENSMUSG( | Cfap126 | 68.88089 | 0.152498 | 0.586696 | 0.944975 |
| ENSMUSG( | Cfap144 | 26.56829 | -0.30008 | 0.452503 | 0.836145 | ENSMUSG( | Cfap144 | 33.11424 | 0.576743 | 0.100655 | 0.801444 |
| ENSMUSG( | Cfap157 | 10.00197 | 0.07164 | 0.913709 | NA | ENSMUSG( | Cfap157 | 10.53048 | 0.504833 | 0.428587 | NA |
| ENSMUSG( | Cfap161 | 22.54179 | -0.84585 | 0.041045 | 0.313149 | ENSMUSG( | Cfap161 | 26.92271 | 0.095915 | 0.831916 | 0.980738 |
| ENSMUSG( | Cfap20 | 786.8982 | -0.15874 | 0.148403 | 0.571935 | ENSMUSG( | Cfap20 | 768.5366 | 0.049697 | 0.70685 | 0.963529 |
| ENSMUSG( | Cfap206 | 57.64306 | -0.11542 | 0.708777 | 0.939519 | ENSMUSG( | Cfap206 | 60.87339 | 0.317108 | 0.328672 | 0.906727 |
| ENSMUSG( | Cfap20dc | 296.7192 | 0.129228 | 0.39957 | 0.813178 | ENSMUSG( | Cfap20dc | 252.6417 | -0.06376 | 0.711335 | 0.964363 |
| ENSMUSG( | Cfap210 | 85.56593 | 0.015683 | 0.942873 | 0.993415 | ENSMUSG( | Cfap210 | 80.37971 | 0.115443 | 0.665638 | 0.956766 |
| ENSMUSG( | Cfap221 | 9.726751 | -0.42044 | 0.528423 | NA | ENSMUSG( | Cfap221 | 12.28328 | 0.551648 | 0.369229 | 0.915491 |
| ENSMUSG( | Cfap251 | 213.6191 | 0.179843 | 0.288139 | 0.738889 | ENSMUSG( | Cfap251 | 181.7887 | -0.0071 | 0.969264 | 0.99594 |
| ENSMUSG( | Cfap276 | 23.52863 | 0.091047 | 0.819074 | 0.965888 | ENSMUSG( | Cfap276 | 22.25034 | 0.19956 | 0.629098 | 0.953251 |
| ENSMUSG( | Cfap298 | 539.3327 | -0.03488 | 0.791894 | 0.959461 | ENSMUSG( | Cfap298 | 480.8082 | -0.10677 | 0.532345 | 0.934795 |
| ENSMUSG( | Cfap299 | 11.46549 | -0.05663 | 0.914848 | NA | ENSMUSG( | Cfap299 | 14.65856 | 0.822197 | 0.105808 | 0.804969 |
| ENSMUSG( | Cfap300 | 177.9544 | 0.198999 | 0.219249 | 0.664715 | ENSMUSG( | Cfap300 | 155.1462 | 0.064192 | 0.749821 | 0.969988 |
| ENSMUSG( | Cfap36 | 2649.433 | 0.063545 | 0.541689 | 0.883182 | ENSMUSG( | Cfap36 | 2254.213 | -0.1401 | 0.297797 | 0.898538 |
| ENSMUSG( | Cfap410 | 1014.289 | 0.005384 | 0.960347 | 0.993991 | ENSMUSG( | Cfap410 | 921.725 | -0.00114 | 0.992637 | 0.99874 |
| ENSMUSG( | Cfap418 | 845.9292 | 0.105805 | 0.381986 | 0.802983 | ENSMUSG( | Cfap418 | 753.6 | 0.03606 | 0.816746 | 0.979198 |
| ENSMUSG( | Cfap43 | 65.3778 | 0.179294 | 0.523999 | 0.87583 | ENSMUSG( | Cfap43 | 65.25205 | 0.440887 | 0.147858 | 0.836241 |
| ENSMUSG( | Cfap44 | 147.976 | 0.072368 | 0.711837 | 0.940028 | ENSMUSG( | Cfap44 | 155.7281 | 0.457493 | 0.026102 | 0.603041 |
| ENSMUSG( | Cfap45 | 62.8959 | -0.11925 | 0.660752 | 0.924951 | ENSMUSG( | Cfap45 | 65.67098 | 0.285127 | 0.328095 | 0.906255 |
| ENSMUSG( | Cfap46 | 136.3282 | 0.028313 | 0.880981 | 0.981692 | ENSMUSG( | Cfap46 | 119.5665 | -0.08251 | 0.693768 | 0.960767 |
| ENSMUSG( | Cfap47 | 26.21201 | 0.270435 | 0.45799 | 0.839769 | ENSMUSG( | Cfap47 | 23.79877 | 0.265999 | 0.518241 | 0.930939 |
| ENSMUSG( | Cfap52 | 40.71363 | -0.43101 | 0.191322 | 0.630245 | ENSMUSG( | Cfap52 | 49.04983 | 0.401678 | 0.243948 | 0.877904 |
| ENSMUSG( | Cfap53 | 10.66431 | -0.19731 | 0.742391 | NA | ENSMUSG( | Cfap53 | 13.12132 | 0.632263 | 0.255523 | 0.883043 |
| ENSMUSG( | Cfap54 | 191.9297 | 0.088912 | 0.616719 | 0.910866 | ENSMUSG( | Cfap54 | 228.9947 | 0.760141 | 0.001331 | 0.132494 |
| ENSMUSG( | Cfap57 | 24.95002 | -0.68366 | 0.093737 | 0.470609 | ENSMUSG( | Cfap57 | 32.27856 | 0.405957 | 0.302226 | 0.898538 |
| ENSMUSG( | Cfap61 | 53.9437 | -0.17402 | 0.511157 | 0.870859 | ENSMUSG( | Cfap61 | 55.91998 | 0.2121 | 0.471889 | 0.923329 |
| ENSMUSG( | Cfap65 | 216.2927 | 0.444565 | 0.109663 | 0.500208 | ENSMUSG( | Cfap65 | 174.1707 | 0.11942 | 0.631271 | 0.953251 |
| ENSMUSG( | Cfap68 | 1369.923 | 0.02537 | 0.790958 | 0.959461 | ENSMUSG( | Cfap68 | 1158.092 | -0.20149 | 0.118817 | 0.813384 |
| ENSMUSG( | Cfap69 | 461.4481 | 0.18566 | 0.234258 | 0.68192 | ENSMUSG( | Cfap69 | 409.3169 | 0.109859 | 0.540571 | 0.937383 |
| ENSMUSG( | Cfap70 | 40.1651 | -0.09387 | 0.81375 | 0.964574 | ENSMUSG( | Cfap70 | 38.27095 | 0.05624 | 0.896741 | 0.990558 |
| ENSMUSG( | Cfap74 | 530.8344 | 0.150642 | 0.202903 | 0.64342 | ENSMUSG( | Cfap74 | 480.5824 | 0.136375 | 0.361733 | 0.914223 |
| ENSMUSG( | Cfap90 | 195.8097 | 0.106737 | 0.54056 | 0.88315 | ENSMUSG( | Cfap90 | 159.7333 | -0.2241 | 0.271263 | 0.888062 |
| ENSMUSG( | Cfap91 | 48.70352 | 0.018066 | 0.953272 | 0.993874 | ENSMUSG( | Cfap91 | 51.30069 | 0.427954 | 0.189558 | 0.857355 |
| ENSMUSG( | Cfap92 | 85.13198 | -0.2517 | 0.363009 | 0.793211 | ENSMUSG( | Cfap92 | 78.50307 | -0.23466 | 0.442378 | 0.921554 |

| | | | | | |
|---|---|---|---|---|---|
| ENSMUSG( | 22.54179 | -0.84585 | 0.041045 | 0.313149 | Cfap161 |
| ENSMUSG( | 786.8982 | -0.15874 | 0.148403 | 0.571935 | Cfap20 |
| ENSMUSG( | 57.64306 | -0.11542 | 0.708777 | 0.939519 | Cfap206 |
| ENSMUSG( | 296.7192 | 0.129228 | 0.39957 | 0.813178 | Cfap20dc |
| ENSMUSG( | 85.56593 | 0.015683 | 0.942873 | 0.993415 | Cfap210 |
| ENSMUSG( | 9.726751 | -0.42044 | 0.528423 | NA | Cfap221 |
| ENSMUSG( | 213.6191 | 0.179843 | 0.288139 | 0.738889 | Cfap251 |
| ENSMUSG( | 23.52863 | 0.091047 | 0.819074 | 0.965888 | Cfap276 |
| ENSMUSG( | 539.3327 | -0.03488 | 0.791894 | 0.959461 | Cfap298 |
| ENSMUSG( | 11.46549 | -0.05663 | 0.914848 | NA | Cfap299 |
| ENSMUSG( | 177.9544 | 0.198999 | 0.219249 | 0.664715 | Cfap300 |
| ENSMUSG( | 2649.433 | 0.063545 | 0.541689 | 0.883182 | Cfap36 |
| ENSMUSG( | 1014.289 | 0.005384 | 0.960347 | 0.993991 | Cfap410 |
| ENSMUSG( | 845.9292 | 0.105805 | 0.381986 | 0.802983 | Cfap418 |
| ENSMUSG( | 65.3778 | 0.179294 | 0.523999 | 0.87583 | Cfap43 |
| ENSMUSG( | 147.976 | 0.072368 | 0.711837 | 0.940028 | Cfap44 |
| ENSMUSG( | 62.8959 | -0.11925 | 0.660752 | 0.924951 | Cfap45 |
| ENSMUSG( | 136.3282 | 0.028313 | 0.880981 | 0.981692 | Cfap46 |
| ENSMUSG( | 26.21201 | 0.270435 | 0.45799 | 0.839769 | Cfap47 |
| ENSMUSG( | 40.71363 | -0.43101 | 0.191322 | 0.630245 | Cfap52 |
| ENSMUSG( | 10.66431 | -0.19731 | 0.742391 | NA | Cfap53 |
| ENSMUSG( | 191.9297 | 0.088912 | 0.616719 | 0.910866 | Cfap54 |
| ENSMUSG( | 24.95002 | -0.68366 | 0.093737 | 0.470609 | Cfap57 |
| ENSMUSG( | 53.9437 | -0.17402 | 0.511157 | 0.870859 | Cfap61 |
| ENSMUSG( | 216.2927 | 0.444565 | 0.109663 | 0.500208 | Cfap65 |
| ENSMUSG( | 1369.923 | 0.02537 | 0.790958 | 0.959461 | Cfap68 |
| ENSMUSG( | 461.4481 | 0.18566 | 0.234258 | 0.68192 | Cfap69 |
| ENSMUSG( | 40.1651 | -0.09387 | 0.81375 | 0.964574 | Cfap70 |
| ENSMUSG( | 530.8344 | 0.150642 | 0.202903 | 0.64342 | Cfap74 |
| ENSMUSG( | 195.8097 | 0.106737 | 0.54056 | 0.88315 | Cfap90 |
| ENSMUSG( | 48.70352 | 0.018066 | 0.953272 | 0.993874 | Cfap91 |
| ENSMUSG( | 85.13198 | -0.2517 | 0.363009 | 0.793211 | Cfap92 |
| ENSMUSG( | 24.86386 | 0.231223 | 0.570663 | 0.893192 | Cfap95 |
| ENSMUSG( | 161.472 | -0.04696 | 0.804257 | 0.962169 | Cfap96 |
| ENSMUSG( | 462.7853 | 0.038596 | 0.765295 | 0.954127 | Cfap97 |
| ENSMUSG( | 43.85818 | -0.18886 | 0.505728 | 0.866932 | Cfap97d2 |
| ENSMUSG( | 16.67842 | 0.357069 | 0.481878 | NA | Cfap99 |
| ENSMUSG( | 2000.111 | 0.03182 | 0.769558 | 0.954674 | Cfdp1 |
| ENSMUSG( | 1279.438 | 0.073486 | 0.612989 | 0.908902 | Cfh |
| ENSMUSG( | 28263.12 | 0.028545 | 0.74802 | 0.949604 | Cfl1 |
| ENSMUSG( | 2175.214 | -0.01712 | 0.863788 | 0.975244 | Cfl2 |
| ENSMUSG( | 1405.737 | 0.06704 | 0.596855 | 0.903526 | Cflar |
| ENSMUSG( | 419.6352 | 0.455737 | 0.00048 | 0.020057 | Cfp |
| ENSMUSG( | 31.35387 | -0.41828 | 0.225566 | 0.672427 | Cftr |
| ENSMUSG( | 43.65127 | -0.16449 | 0.611246 | 0.908586 | Cgas |
| ENSMUSG( | 1732.263 | 0.086703 | 0.447304 | 0.834051 | Cggbp1 |
| ENSMUSG( | 146.8513 | 0.31037 | 0.080901 | 0.43975 | Cgn |
| ENSMUSG( | 780.8791 | 0.021021 | 0.889214 | 0.983434 | Cgnl1 |
| ENSMUSG( | 691.4307 | -0.34549 | 0.00316 | 0.06974 | Cgref1 |
| ENSMUSG( | 779.587 | -0.07715 | 0.497894 | 0.862343 | Cgrrf1 |
| ENSMUSG( | 112.9006 | 0.096202 | 0.730578 | 0.944956 | Chac1 |
| ENSMUSG( | 137.4279 | -0.12154 | 0.49546 | 0.861026 | Chac2 |
| ENSMUSG( | 9.222676 | -0.678 | 0.277221 | NA | Chad |
| ENSMUSG( | 793.1797 | -0.32161 | 0.012621 | 0.162076 | Chadl |
| ENSMUSG( | 262.7217 | -0.12373 | 0.405954 | 0.81652 | Chaf1a |
| ENSMUSG( | 24.73444 | -0.55215 | 0.155927 | 0.58375 | Chaf1b |
| ENSMUSG( | 778.7327 | 0.117148 | 0.377179 | 0.801213 | Champ1 |
| ENSMUSG( | 328.0561 | -0.24184 | 0.085738 | 0.451367 | Chaserr |
| ENSMUSG( | 22.97106 | -0.51776 | 0.302633 | 0.750911 | Chat |
| ENSMUSG( | 638.3237 | 0.00396 | 0.973659 | 0.99675 | Chchd1 |
| ENSMUSG( | 5463.022 | -0.02657 | 0.757065 | 0.951513 | Chchd10 |
| ENSMUSG( | 4751.031 | 0.053424 | 0.559251 | 0.888638 | Chchd2 |
| ENSMUSG( | 2386.711 | 0.031686 | 0.761118 | 0.952907 | Chchd3 |
| ENSMUSG( | 644.3137 | 0.022399 | 0.871635 | 0.97812 | Chchd4 |
| ENSMUSG( | 290.9424 | -0.03545 | 0.80521 | 0.962616 | Chchd5 |
| ENSMUSG( | 1295.595 | -0.15641 | 0.090781 | 0.462752 | Chchd6 |
| ENSMUSG( | 415.0877 | -0.04489 | 0.739738 | 0.947823 | Chchd7 |
| ENSMUSG( | 723.6682 | -0.02016 | 0.865243 | 0.975565 | Chd1 |
| ENSMUSG( | 429.8812 | 0.053116 | 0.677808 | 0.930542 | Chd1l |
| ENSMUSG( | 1545.72 | 0.192545 | 0.049624 | 0.340062 | Chd2 |
| ENSMUSG( | 40500.87 | 0.224204 | 0.004875 | 0.092488 | Chd3 |
| ENSMUSG( | 7488.854 | -0.05899 | 0.493311 | 0.860611 | Chd4 |
| ENSMUSG( | 8707.346 | 0.293669 | 0.00172 | 0.046884 | Chd5 |
| ENSMUSG( | 2417.592 | -0.21318 | 0.064532 | 0.393651 | Chd6 |
| ENSMUSG( | 883.9128 | -0.28809 | 0.01205 | 0.158771 | Chd7 |
| ENSMUSG( | 2907.171 | 0.063762 | 0.538342 | 0.88259 | Chd8 |
| ENSMUSG( | 2059.982 | 0.083194 | 0.400525 | 0.813717 | Chd9 |
| ENSMUSG( | 128.1281 | -0.17773 | 0.317138 | 0.763823 | Chdh |
| ENSMUSG( | 48.09572 | -0.05511 | 0.839197 | 0.971179 | Chek1 |
| ENSMUSG( | 61.9095 | 0.012846 | 0.958587 | 0.993874 | Chek2 |
| ENSMUSG( | 1740.657 | 0.138595 | 0.226667 | 0.673729 | Cherp |
| ENSMUSG( | 1160.577 | 0.041278 | 0.691431 | 0.933881 | Chfr |
| ENSMUSG( | 6933.921 | -0.4715 | 1.06E-06 | 0.000197 | Chga |
| ENSMUSG( | 15754.11 | -0.29143 | 0.02946 | 0.260274 | Chgb |
| ENSMUSG( | 899.2009 | -0.34727 | 0.001974 | 0.051135 | Chi3l1 |
| ENSMUSG( | 71.09129 | -0.23106 | 0.361654 | 0.792797 | Chia1 |
| ENSMUSG( | 637.3378 | -0.03091 | 0.766472 | 0.954193 | Chic1 |
| ENSMUSG( | 434.4897 | -0.16103 | 0.175272 | 0.61046 | Chic2 |
| ENSMUSG( | 2014.874 | -0.21622 | 0.038722 | 0.303123 | Chid1 |
| ENSMUSG( | 28.34672 | -0.28987 | 0.613187 | 0.908902 | Chil3 |
| ENSMUSG( | 2376.375 | 0.179283 | 0.048637 | 0.337278 | Chka |
| ENSMUSG( | 98.85496 | 0.051276 | 0.808439 | 0.963455 | Chkb |
| ENSMUSG( | 2917.438 | -0.32858 | 0.05997 | 0.378891 | Chl1 |
| ENSMUSG( | 871.4919 | -0.07375 | 0.555366 | 0.887457 | Chm |
| ENSMUSG( | 268.4034 | -0.3714 | 0.008466 | 0.128691 | Chml |
| ENSMUSG( | 4311.355 | 0.043327 | 0.576866 | 0.89546 | Chmp1a |
| ENSMUSG( | 608.5694 | -0.02045 | 0.843692 | 0.971847 | Chmp1b |
| ENSMUSG( | 1059.942 | 0.068357 | 0.574282 | 0.894186 | Chmp1b2 |
| ENSMUSG( | 1975.763 | 0.063176 | 0.547971 | 0.885153 | Chmp2a |
| ENSMUSG( | 995.3262 | -0.08268 | 0.461428 | 0.841432 | Chmp2b |
| ENSMUSG( | 3142.358 | 0.051547 | 0.538478 | 0.88259 | Chmp3 |
| ENSMUSG( | 7286.668 | 0.012525 | 0.884494 | 0.982647 | Chmp4b |
| ENSMUSG( | 3537.498 | 0.057862 | 0.602707 | 0.904887 | Chmp5 |
| ENSMUSG( | 1057.373 | 0.073386 | 0.432115 | 0.82768 | Chmp6 |
| ENSMUSG( | 2305.18 | 0.109253 | 0.268504 | 0.718807 | Chmp7 |
| ENSMUSG( | 21928.38 | 0.03717 | 0.7402 | 0.947823 | Chn1 |
| ENSMUSG( | 48.17524 | -0.30257 | 0.338344 | 0.777489 | Chn1os3 |
| ENSMUSG( | 1216.473 | 0.0167 | 0.887128 | 0.983326 | Chn2 |
| ENSMUSG( | 20.83484 | -0.09127 | 0.834525 | NA | Chodl |
| ENSMUSG( | 836.0822 | 0.110906 | 0.27761 | 0.72746 | Chordc1 |
| ENSMUSG( | 4348.728 | 0.057593 | 0.490079 | 0.859319 | Chp1 |
| ENSMUSG( | 170.1941 | -0.38678 | 0.042594 | 0.316734 | Chp2 |
| ENSMUSG( | 2507.615 | -0.02905 | 0.853638 | 0.973491 | Chpf |
| ENSMUSG( | 1765.212 | -0.06887 | 0.612536 | 0.908747 | Chpf2 |
| ENSMUSG( | 4493.269 | -0.04052 | 0.778472 | 0.956215 | Chpt1 |
| ENSMUSG( | 265.7696 | -0.14161 | 0.297973 | 0.746543 | Chrac1 |
| ENSMUSG( | 1336.088 | 0.516387 | 0.000866 | 0.029559 | Chrd |
| ENSMUSG( | 1707.796 | -0.23002 | 0.152593 | 0.578962 | Chrdl1 |
| ENSMUSG( | 9.195718 | 0.638238 | 0.338097 | NA | Chrdl2 |
| ENSMUSG( | 6307.047 | -0.28518 | 0.046764 | 0.330815 | Chrm1 |
| ENSMUSG( | 282.9 | -0.54127 | 0.002541 | 0.059739 | Chrm2 |
| ENSMUSG( | 1285.87 | -0.27864 | 0.154075 | 0.580901 | Chrm3 |
| ENSMUSG( | 672.0598 | -0.76924 | 4.59E-05 | 0.003685 | Chrm4 |
| ENSMUSG( | 10.28635 | 0.519772 | 0.417029 | NA | Chrm5 |
| ENSMUSG( | 12.61649 | 0.263967 | 0.635885 | NA | Chrna1 |
| ENSMUSG( | 11.21461 | -0.37541 | 0.591504 | NA | Chrna10 |
| ENSMUSG( | 30.71477 | -0.0517 | 0.882045 | 0.982069 | Chrna1os |
| ENSMUSG( | 22.38008 | -0.14021 | 0.747683 | 0.949604 | Chrna2 |
| ENSMUSG( | 34.23598 | -0.3913 | 0.222364 | 0.668412 | Chrna3 |
| ENSMUSG( | 1276.168 | -0.37133 | 0.010908 | 0.150598 | Chrna4 |
| ENSMUSG( | 66.01763 | -0.64865 | 0.032681 | 0.27685 | Chrna5 |
| ENSMUSG( | 12.55841 | 0.027256 | 0.959315 | NA | Chrna6 |
| ENSMUSG( | 195.7092 | -0.53616 | 0.041926 | 0.315414 | Chrna7 |
| ENSMUSG( | 29.19181 | -0.05893 | 0.86274 | 0.975055 | Chrnb1 |
| ENSMUSG( | 1498.572 | -0.26661 | 0.077948 | 0.431652 | Chrnb2 |
| ENSMUSG( | 54.4742 | -0.7257 | 0.019257 | 0.20921 | Chrnb3 |
| ENSMUSG( | 9.83904 | -0.24967 | 0.676381 | NA | Chrnb4 |
| ENSMUSG( | 9506.718 | -0.13026 | 0.234236 | 0.68192 | Chst1 |
| ENSMUSG( | 3450.898 | -0.15707 | 0.183057 | 0.62047 | Chst10 |
| ENSMUSG( | 3402.617 | -0.22664 | 0.070166 | 0.40908 | Chst11 |
| ENSMUSG( | 724.4478 | -0.36482 | 0.02578 | 0.247138 | Chst12 |
| ENSMUSG( | 86.92535 | -0.61112 | 0.022257 | 0.227763 | Chst14 |
| ENSMUSG( | 1006.331 | -0.40306 | 0.003992 | 0.081133 | Chst15 |
| ENSMUSG( | 17235.33 | 0.008371 | 0.941591 | 0.993415 | Chst2 |
| ENSMUSG( | 240.0084 | -0.43578 | 0.028876 | 0.258911 | Chst3 |
| ENSMUSG( | 39.85374 | -0.78447 | 0.052972 | 0.353293 | Chst5 |
| ENSMUSG( | 106.88 | -0.11934 | 0.586253 | 0.899709 | Chst7 |
| ENSMUSG( | 453.4536 | -0.03816 | 0.777261 | 0.955886 | Chst8 |
| ENSMUSG( | 11.56353 | -0.62883 | 0.260431 | NA | Chst9 |
| ENSMUSG( | 735.0255 | -0.24324 | 0.094538 | 0.471713 | Chsy1 |
| ENSMUSG( | 308.8091 | -0.65573 | 0.000801 | 0.027945 | Chsy3 |
| ENSMUSG( | 71.30314 | 0.060369 | 0.804072 | 0.962169 | Chtf18 |
| ENSMUSG( | 30.68867 | -0.11758 | 0.74631 | 0.949289 | Chtf8 |
| ENSMUSG( | 4654.681 | 0.006172 | 0.947498 | 0.993821 | Chtop |
| ENSMUSG( | 1188.605 | -0.00046 | 0.996535 | 0.99943 | Chuk |
| ENSMUSG( | 203.6615 | 0.022015 | 0.881739 | 0.98196 | Churc1 |
| ENSMUSG( | 937.5554 | -0.0165 | 0.904613 | 0.986275 | Ciao1 |
| ENSMUSG( | 784.6979 | -0.00177 | 0.989324 | 0.999111 | Ciao2a |

| | | | | | |
|---|---|---|---|---|---|
| ENSMUSG( | 753.6 | 0.03606 | 0.816746 | 0.979198 | Cfap418 |
| ENSMUSG( | 65.25205 | 0.440887 | 0.147858 | 0.836241 | Cfap43 |
| ENSMUSG( | 155.7281 | 0.457493 | 0.026102 | 0.603041 | Cfap44 |
| ENSMUSG( | 65.67098 | 0.285127 | 0.328095 | 0.906255 | Cfap45 |
| ENSMUSG( | 119.5665 | -0.08251 | 0.693768 | 0.960767 | Cfap46 |
| ENSMUSG( | 23.79877 | 0.265999 | 0.518241 | 0.930939 | Cfap47 |
| ENSMUSG( | 49.04983 | 0.401678 | 0.243948 | 0.877904 | Cfap52 |
| ENSMUSG( | 13.12132 | 0.632263 | 0.255523 | 0.883043 | Cfap53 |
| ENSMUSG( | 228.9947 | 0.760141 | 0.001331 | 0.132494 | Cfap54 |
| ENSMUSG( | 32.27856 | 0.405957 | 0.302226 | 0.898538 | Cfap57 |
| ENSMUSG( | 8.366763 | -0.00797 | 0.990937 | NA | Cfap58 |
| ENSMUSG( | 55.91998 | 0.2121 | 0.471889 | 0.923329 | Cfap61 |
| ENSMUSG( | 174.1707 | 0.11942 | 0.631271 | 0.953251 | Cfap65 |
| ENSMUSG( | 1158.092 | -0.20149 | 0.118817 | 0.813384 | Cfap68 |
| ENSMUSG( | 409.3169 | 0.109859 | 0.540571 | 0.937383 | Cfap69 |
| ENSMUSG( | 38.27095 | 0.05624 | 0.896741 | 0.990558 | Cfap70 |
| ENSMUSG( | 8.169091 | 0.777893 | 0.285136 | NA | Cfap73 |
| ENSMUSG( | 480.5824 | 0.136375 | 0.361733 | 0.914223 | Cfap74 |
| ENSMUSG( | 14.39655 | 1.082716 | 0.052175 | 0.712974 | Cfap77 |
| ENSMUSG( | 159.7333 | -0.2241 | 0.271263 | 0.888062 | Cfap90 |
| ENSMUSG( | 51.30069 | 0.427954 | 0.189558 | 0.857355 | Cfap91 |
| ENSMUSG( | 78.50307 | -0.23466 | 0.442378 | 0.921554 | Cfap92 |
| ENSMUSG( | 21.56872 | 0.153031 | 0.751711 | 0.970386 | Cfap95 |
| ENSMUSG( | 150.8896 | 0.038989 | 0.846853 | 0.985027 | Cfap96 |
| ENSMUSG( | 413.6872 | -0.02153 | 0.888263 | 0.990496 | Cfap97 |
| ENSMUSG( | 35.9032 | -0.51497 | 0.12798 | 0.81845 | Cfap97d2 |
| ENSMUSG( | 15.43278 | 0.41384 | 0.438049 | 0.920503 | Cfap99 |
| ENSMUSG( | 1655.168 | -0.26205 | 0.061371 | 0.736111 | Cfdp1 |
| ENSMUSG( | 1252.755 | 0.274657 | 0.102814 | 0.803962 | Cfh |
| ENSMUSG( | 23992.35 | -0.18338 | 0.204673 | 0.864412 | Cfl1 |
| ENSMUSG( | 1965.407 | -0.04567 | 0.757953 | 0.971858 | Cfl2 |
| ENSMUSG( | 1229.657 | -0.05238 | 0.698074 | 0.960867 | Cflar |
| ENSMUSG( | 361.7625 | 0.316068 | 0.082436 | 0.77681 | Cfp |
| ENSMUSG( | 32.57561 | 0.00511 | 0.988691 | 0.997862 | Cftr |
| ENSMUSG( | 43.45829 | 0.117798 | 0.723504 | 0.966887 | Cgas |
| ENSMUSG( | 1577.078 | 0.081617 | 0.586232 | 0.944975 | Cggbp1 |
| ENSMUSG( | 125.2234 | 0.129371 | 0.514488 | 0.930939 | Cgn |
| ENSMUSG( | 759.1172 | 0.208131 | 0.21419 | 0.864421 | Cgnl1 |
| ENSMUSG( | 670.3642 | -0.1465 | 0.268168 | 0.886493 | Cgref1 |
| ENSMUSG( | 776.7274 | 0.179418 | 0.203062 | 0.864412 | Cgrrf1 |
| ENSMUSG( | 7.025431 | -0.00816 | 0.99215 | NA | Ch25h |
| ENSMUSG( | 106.4361 | 0.174101 | 0.560962 | 0.940254 | Chac1 |
| ENSMUSG( | 128.9589 | -0.0455 | 0.823035 | 0.979198 | Chac2 |
| ENSMUSG( | 14.52755 | 0.870194 | 0.088986 | 0.788051 | Chad |
| ENSMUSG( | 829.0752 | 0.096039 | 0.531994 | 0.934795 | Chadl |
| ENSMUSG( | 249.6011 | 0.000456 | 0.997799 | 0.999268 | Chaf1a |
| ENSMUSG( | 23.72715 | -0.33853 | 0.445182 | 0.921734 | Chaf1b |
| ENSMUSG( | 658.8043 | -0.09673 | 0.505655 | 0.927825 | Champ1 |
| ENSMUSG( | 340.2708 | 0.143225 | 0.363578 | 0.914223 | Chaserr |
| ENSMUSG( | 22.2368 | -0.34216 | 0.437458 | 0.920144 | Chat |
| ENSMUSG( | 559.12 | -0.11363 | 0.39573 | 0.917257 | Chchd1 |
| ENSMUSG( | 4585.701 | -0.2789 | 0.02736 | 0.606201 | Chchd10 |
| ENSMUSG( | 3893.296 | -0.26661 | 0.037491 | 0.673565 | Chchd2 |
| ENSMUSG( | 2029.961 | -0.17553 | 0.197498 | 0.862161 | Chchd3 |
| ENSMUSG( | 566.9919 | -0.07988 | 0.60486 | 0.948977 | Chchd4 |
| ENSMUSG( | 246.2167 | -0.27386 | 0.124444 | 0.814913 | Chchd5 |
| ENSMUSG( | 1148.015 | -0.245 | 0.04579 | 0.695584 | Chchd6 |
| ENSMUSG( | 407.2819 | 0.166044 | 0.267847 | 0.886134 | Chchd7 |
| ENSMUSG( | 683.6453 | 0.084486 | 0.521985 | 0.932571 | Chd1 |
| ENSMUSG( | 368.0756 | -0.12141 | 0.426685 | 0.917257 | Chd1l |
| ENSMUSG( | 1356.823 | 0.088738 | 0.428766 | 0.917257 | Chd2 |
| ENSMUSG( | 34404.33 | 0.031094 | 0.744661 | 0.96951 | Chd3 |
| ENSMUSG( | 6819.805 | -0.05948 | 0.592229 | 0.944975 | Chd4 |
| ENSMUSG( | 7217.302 | 0.036044 | 0.732555 | 0.967228 | Chd5 |
| ENSMUSG( | 2387.414 | 0.030158 | 0.811742 | 0.979148 | Chd6 |
| ENSMUSG( | 875.6477 | -0.03044 | 0.797904 | 0.97681 | Chd7 |
| ENSMUSG( | 2721.464 | 0.141562 | 0.210085 | 0.864412 | Chd8 |
| ENSMUSG( | 1940.351 | 0.175063 | 0.112084 | 0.808673 | Chd9 |
| ENSMUSG( | 139.4568 | 0.32753 | 0.081542 | 0.77681 | Chdh |
| ENSMUSG( | 44.36251 | 0.000278 | 0.999275 | 0.999503 | Chek1 |
| ENSMUSG( | 60.10573 | 0.202476 | 0.448982 | 0.922202 | Chek2 |
| ENSMUSG( | 1527.119 | 0.030854 | 0.823674 | 0.979198 | Cherp |
| ENSMUSG( | 1068.458 | 0.07144 | 0.610643 | 0.950771 | Chfr |
| ENSMUSG( | 7340.062 | -0.00038 | 0.997683 | 0.999268 | Chga |
| ENSMUSG( | 16599.39 | 0.143017 | 0.363172 | 0.914223 | Chgb |
| ENSMUSG( | 831.4301 | -0.30286 | 0.02315 | 0.585577 | Chi3l1 |
| ENSMUSG( | 64.74209 | -0.24173 | 0.412202 | 0.917257 | Chia1 |
| ENSMUSG( | 589.4602 | 0.005328 | 0.970338 | 0.99594 | Chic1 |
| ENSMUSG( | 435.5287 | 0.116697 | 0.389592 | 0.917257 | Chic2 |
| ENSMUSG( | 1963.32 | -0.01134 | 0.931947 | 0.99306 | Chid1 |
| ENSMUSG( | 31.04082 | 0.197039 | 0.716454 | 0.965773 | Chil3 |
| ENSMUSG( | 2077.718 | 0.064396 | 0.565007 | 0.941248 | Chka |
| ENSMUSG( | 93.60255 | 0.167129 | 0.476786 | 0.923329 | Chkb |
| ENSMUSG( | 3216.988 | 0.23547 | 0.231788 | 0.868507 | Chl1 |
| ENSMUSG( | 820.919 | 0.022652 | 0.857459 | 0.985884 | Chm |
| ENSMUSG( | 269.7516 | -0.07559 | 0.659709 | 0.956205 | Chml |
| ENSMUSG( | 3761.804 | -0.08257 | 0.413595 | 0.917257 | Chmp1a |
| ENSMUSG( | 557.4592 | -0.00725 | 0.956032 | 0.994798 | Chmp1b |
| ENSMUSG( | 889.3957 | -0.18046 | 0.265152 | 0.886134 | Chmp1b2 |
| ENSMUSG( | 1669.091 | -0.16264 | 0.231536 | 0.868507 | Chmp2a |
| ENSMUSG( | 926.6077 | -0.02615 | 0.871882 | 0.988779 | Chmp2b |
| ENSMUSG( | 2688.516 | -0.1348 | 0.244327 | 0.877904 | Chmp3 |
| ENSMUSG( | 6112.503 | -0.23752 | 0.051534 | 0.711358 | Chmp4b |
| ENSMUSG( | 2966.333 | -0.19108 | 0.188809 | 0.85698 | Chmp5 |
| ENSMUSG( | 929.5863 | -0.03019 | 0.793011 | 0.97681 | Chmp6 |
| ENSMUSG( | 1994.06 | -0.03728 | 0.738423 | 0.967967 | Chmp7 |
| ENSMUSG( | 18907.33 | -0.1267 | 0.35606 | 0.913331 | Chn1 |
| ENSMUSG( | 43.44466 | -0.33683 | 0.326036 | 0.905339 | Chn1os3 |
| ENSMUSG( | 1124.202 | 0.057677 | 0.670003 | 0.95694 | Chn2 |
| ENSMUSG( | 22.88803 | 0.399458 | 0.338673 | 0.907972 | Chodl |
| ENSMUSG( | 801.1033 | 0.245214 | 0.044434 | 0.693697 | Chordc1 |
| ENSMUSG( | 3731.984 | -0.11886 | 0.301846 | 0.898538 | Chp1 |
| ENSMUSG( | 152.0943 | -0.46249 | 0.017178 | 0.52741 | Chp2 |
| ENSMUSG( | 2521.776 | 0.251987 | 0.18457 | 0.853513 | Chpf |
| ENSMUSG( | 1769.744 | 0.205816 | 0.199367 | 0.863016 | Chpf2 |
| ENSMUSG( | 4150.986 | -0.00304 | 0.985939 | 0.997188 | Chpt1 |
| ENSMUSG( | 243.1753 | -0.13296 | 0.401058 | 0.917257 | Chrac1 |
| ENSMUSG( | 1096.11 | 0.249078 | 0.126163 | 0.81516 | Chrd |
| ENSMUSG( | 1585.254 | -0.16997 | 0.323943 | 0.905257 | Chrdl1 |
| ENSMUSG( | 8.908951 | 0.722858 | 0.331046 | NA | Chrdl2 |
| ENSMUSG( | 6931.903 | 0.262831 | 0.10459 | 0.804969 | Chrm1 |
| ENSMUSG( | 355.14 | 0.412117 | 0.035291 | 0.661159 | Chrm2 |
| ENSMUSG( | 1392.659 | 0.23343 | 0.307358 | 0.898538 | Chrm3 |
| ENSMUSG( | 764.2852 | -0.01997 | 0.917113 | 0.992529 | Chrm4 |
| ENSMUSG( | 10.14248 | 0.744651 | 0.255729 | NA | Chrm5 |
| ENSMUSG( | 10.69201 | 0.098725 | 0.874376 | NA | Chrna1 |
| ENSMUSG( | 13.36018 | 0.336186 | 0.569695 | 0.941777 | Chrna10 |
| ENSMUSG( | 25.02062 | -0.38222 | 0.334157 | 0.907907 | Chrna1os |
| ENSMUSG( | 22.29013 | 0.103889 | 0.826935 | 0.980159 | Chrna2 |
| ENSMUSG( | 33.97584 | -0.10989 | 0.746184 | 0.969649 | Chrna3 |
| ENSMUSG( | 1355.582 | 0.099237 | 0.57787 | 0.943131 | Chrna4 |
| ENSMUSG( | 77.03398 | 0.144274 | 0.612365 | 0.951071 | Chrna5 |
| ENSMUSG( | 15.38266 | 0.765016 | 0.165436 | 0.838866 | Chrna6 |
| ENSMUSG( | 211.7688 | 0.026527 | 0.923669 | 0.992815 | Chrna7 |
| ENSMUSG( | 32.83879 | 0.524026 | 0.194152 | 0.860096 | Chrnb1 |
| ENSMUSG( | 1614.407 | 0.226793 | 0.228532 | 0.868507 | Chrnb2 |
| ENSMUSG( | 62.92889 | 0.065705 | 0.837556 | 0.981335 | Chrnb3 |
| ENSMUSG( | 10.28441 | 0.074274 | 0.910598 | NA | Chrnb4 |
| ENSMUSG( | 9801.251 | 0.223237 | 0.069782 | 0.756483 | Chst1 |
| ENSMUSG( | 3287.6 | -0.02101 | 0.876766 | 0.989644 | Chst10 |
| ENSMUSG( | 3540.964 | 0.163792 | 0.262774 | 0.885357 | Chst11 |
| ENSMUSG( | 802.3127 | 0.220696 | 0.233313 | 0.868507 | Chst12 |
| ENSMUSG( | 98.76907 | 0.107167 | 0.69513 | 0.960867 | Chst14 |
| ENSMUSG( | 1092.573 | 0.133979 | 0.405448 | 0.917257 | Chst15 |
| ENSMUSG( | 15915.23 | 0.046876 | 0.729059 | 0.966913 | Chst2 |
| ENSMUSG( | 246.2604 | -0.04492 | 0.86671 | 0.987458 | Chst3 |
| ENSMUSG( | 52.32307 | 0.367537 | 0.353563 | 0.912263 | Chst5 |
| ENSMUSG( | 116.7479 | 0.397196 | 0.095566 | 0.796133 | Chst7 |
| ENSMUSG( | 444.3966 | 0.172285 | 0.30673 | 0.898538 | Chst8 |
| ENSMUSG( | 14.54754 | 0.365442 | 0.483479 | 0.92499 | Chst9 |
| ENSMUSG( | 746.2275 | 0.079583 | 0.609396 | 0.950397 | Chsy1 |
| ENSMUSG( | 356.3445 | 0.109597 | 0.618053 | 0.952232 | Chsy3 |
| ENSMUSG( | 63.16991 | -0.0195 | 0.94557 | 0.994798 | Chtf18 |
| ENSMUSG( | 28.65254 | -0.01754 | 0.9644 | 0.995433 | Chtf8 |
| ENSMUSG( | 4090.232 | -0.10261 | 0.467638 | 0.923329 | Chtop |
| ENSMUSG( | 1119.436 | 0.09641 | 0.475586 | 0.923329 | Chuk |
| ENSMUSG( | 174.3875 | -0.17551 | 0.346946 | 0.911434 | Churc1 |
| ENSMUSG( | 857.857 | -0.00725 | 0.96233 | 0.995237 | Ciao1 |
| ENSMUSG( | 675.3064 | -0.17608 | 0.21835 | 0.866104 | Ciao2a |
| ENSMUSG( | 448.2729 | -0.22358 | 0.089387 | 0.788575 | Ciao2b |
| ENSMUSG( | 633.5425 | 0.04586 | 0.72257 | 0.966887 | Ciao3 |
| ENSMUSG( | 883.0255 | -0.02341 | 0.846309 | 0.984578 | Ciapin1 |
| ENSMUSG( | 356.9848 | 0.781088 | 1.95E-05 | 0.00609 | Ciart |
| ENSMUSG( | 683.4871 | -0.19438 | 0.124988 | 0.81516 | Cib1 |
| ENSMUSG( | 716.6671 | -0.15759 | 0.258992 | 0.883971 | Cib2 |
| ENSMUSG( | 917.0574 | 0.100698 | 0.417266 | 0.917257 | Cibar1 |
| ENSMUSG( | 15.03866 | -0.24287 | 0.640157 | 0.953903 | Cibar2 |
| ENSMUSG( | 8300.277 | 0.131624 | 0.22856 | 0.868507 | Cic |

| | | | | | | | | | | | |
|---|---|---|---|---|---|---|---|---|---|---|---|
| ENSMUSG( | Cfap95 | 24.86386 | 0.231223 | 0.570663 | 0.893192 | ENSMUSG( | Cfap95 | 21.56872 | 0.153031 | 0.751711 | 0.970386 |
| ENSMUSG( | Cfap96 | 161.472 | -0.04696 | 0.804257 | 0.962169 | ENSMUSG( | Cfap96 | 150.8896 | 0.038989 | 0.846853 | 0.985027 |
| ENSMUSG( | Cfap97 | 462.7853 | 0.038596 | 0.765295 | 0.954127 | ENSMUSG( | Cfap97 | 413.6872 | -0.02153 | 0.888263 | 0.990496 |
| ENSMUSG( | Cfap97d2 | 43.85818 | -0.18886 | 0.505728 | 0.866932 | ENSMUSG( | Cfap97d2 | 35.9032 | -0.51497 | 0.12798 | 0.81845 |
| ENSMUSG( | Cfap99 | 16.67842 | 0.357069 | 0.481878 | NA | ENSMUSG( | Cfap99 | 15.43278 | 0.41384 | 0.438049 | 0.920503 |
| ENSMUSG( | Cfdp1 | 2000.111 | 0.03182 | 0.769558 | 0.954674 | ENSMUSG( | Cfdp1 | 1655.168 | -0.26205 | 0.061371 | 0.736111 |
| ENSMUSG( | Cfh | 1279.438 | 0.073486 | 0.612989 | 0.908902 | ENSMUSG( | Cfh | 1252.755 | 0.274657 | 0.102814 | 0.803962 |
| ENSMUSG( | Cfl1 | 28263.12 | 0.028545 | 0.74802 | 0.949604 | ENSMUSG( | Cfl1 | 23992.35 | -0.18338 | 0.204673 | 0.864412 |
| ENSMUSG( | Cfl2 | 2175.214 | -0.01712 | 0.863788 | 0.975244 | ENSMUSG( | Cfl2 | 1965.407 | -0.04567 | 0.757953 | 0.971858 |
| ENSMUSG( | Cflar | 1405.737 | 0.06704 | 0.596855 | 0.903526 | ENSMUSG( | Cflar | 1229.657 | -0.05238 | 0.698074 | 0.960867 |
| ENSMUSG( | Cfp | 419.6352 | 0.455737 | 0.00048 | 0.020057 | ENSMUSG( | Cfp | 361.7625 | 0.316068 | 0.082436 | 0.77681 |
| ENSMUSG( | Cftr | 31.35387 | -0.41828 | 0.225566 | 0.672427 | ENSMUSG( | Cftr | 32.57561 | 0.00511 | 0.988691 | 0.997862 |
| ENSMUSG( | Cgas | 43.65127 | -0.16449 | 0.611246 | 0.908586 | ENSMUSG( | Cgas | 43.45829 | 0.117798 | 0.723504 | 0.966887 |
| ENSMUSG( | Cggbp1 | 1732.263 | 0.086703 | 0.447304 | 0.834051 | ENSMUSG( | Cggbp1 | 1577.078 | 0.081617 | 0.586232 | 0.944975 |
| ENSMUSG( | Cgn | 146.8513 | 0.31037 | 0.080901 | 0.43975 | ENSMUSG( | Cgn | 125.2234 | 0.129371 | 0.514488 | 0.930939 |
| ENSMUSG( | Cgnl1 | 780.8791 | 0.021021 | 0.889214 | 0.983434 | ENSMUSG( | Cgnl1 | 759.1172 | 0.208131 | 0.21419 | 0.864421 |
| ENSMUSG( | Cgref1 | 691.4307 | -0.34549 | 0.00316 | 0.06974 | ENSMUSG( | Cgref1 | 670.3642 | -0.1465 | 0.268168 | 0.886493 |
| ENSMUSG( | Cgrrf1 | 779.587 | -0.07715 | 0.497894 | 0.862343 | ENSMUSG( | Cgrrf1 | 776.7274 | 0.179418 | 0.203062 | 0.864412 |
| ENSMUSG( | Chac1 | 112.9006 | 0.096202 | 0.730578 | 0.944956 | ENSMUSG( | Chac1 | 106.4361 | 0.174101 | 0.560962 | 0.940254 |
| ENSMUSG( | Chac2 | 137.4279 | -0.12154 | 0.49546 | 0.861026 | ENSMUSG( | Chac2 | 128.9589 | -0.0455 | 0.823035 | 0.979198 |
| ENSMUSG( | Chad | 9.222676 | -0.678 | 0.277221 | NA | ENSMUSG( | Chad | 14.52755 | 0.870194 | 0.088986 | 0.788051 |
| ENSMUSG( | Chadl | 793.1797 | -0.32161 | 0.012621 | 0.162076 | ENSMUSG( | Chadl | 829.0752 | 0.096039 | 0.531994 | 0.934795 |
| ENSMUSG( | Chaf1a | 262.7217 | -0.12373 | 0.405954 | 0.81652 | ENSMUSG( | Chaf1a | 249.6011 | 0.000456 | 0.997799 | 0.999268 |
| ENSMUSG( | Chaf1b | 24.73444 | -0.55215 | 0.155927 | 0.58375 | ENSMUSG( | Chaf1b | 23.72715 | -0.33853 | 0.445182 | 0.921734 |
| ENSMUSG( | Champ1 | 778.7327 | 0.117148 | 0.377179 | 0.801213 | ENSMUSG( | Champ1 | 658.8043 | -0.09673 | 0.505655 | 0.927825 |
| ENSMUSG( | Chaserr | 328.0561 | -0.24184 | 0.085738 | 0.451367 | ENSMUSG( | Chaserr | 340.2708 | 0.143225 | 0.363578 | 0.914223 |
| ENSMUSG( | Chat | 22.97106 | -0.51776 | 0.302633 | 0.750911 | ENSMUSG( | Chat | 22.2368 | -0.34216 | 0.437458 | 0.920144 |
| ENSMUSG( | Chchd1 | 638.3237 | 0.00396 | 0.973659 | 0.99675 | ENSMUSG( | Chchd1 | 559.12 | -0.11363 | 0.39573 | 0.917257 |
| ENSMUSG( | Chchd10 | 5463.022 | -0.02657 | 0.757065 | 0.951513 | ENSMUSG( | Chchd10 | 4585.701 | -0.2789 | 0.02736 | 0.606201 |
| ENSMUSG( | Chchd2 | 4751.031 | 0.053424 | 0.559251 | 0.888638 | ENSMUSG( | Chchd2 | 3893.296 | -0.26661 | 0.037491 | 0.673565 |
| ENSMUSG( | Chchd3 | 2386.711 | 0.031686 | 0.761118 | 0.952907 | ENSMUSG( | Chchd3 | 2029.961 | -0.17553 | 0.197498 | 0.862161 |
| ENSMUSG( | Chchd4 | 644.3137 | 0.022399 | 0.871635 | 0.97812 | ENSMUSG( | Chchd4 | 566.9919 | -0.07988 | 0.60486 | 0.948977 |
| ENSMUSG( | Chchd5 | 290.9424 | -0.03545 | 0.80521 | 0.962616 | ENSMUSG( | Chchd5 | 246.2167 | -0.27386 | 0.124444 | 0.814913 |
| ENSMUSG( | Chchd6 | 1295.595 | -0.15641 | 0.090781 | 0.462752 | ENSMUSG( | Chchd6 | 1148.015 | -0.245 | 0.04579 | 0.695584 |
| ENSMUSG( | Chchd7 | 415.0877 | -0.04489 | 0.739738 | 0.947823 | ENSMUSG( | Chchd7 | 407.2819 | 0.166044 | 0.267847 | 0.886134 |
| ENSMUSG( | Chd1 | 723.6682 | -0.02016 | 0.865243 | 0.975565 | ENSMUSG( | Chd1 | 683.6453 | 0.084486 | 0.521985 | 0.932571 |
| ENSMUSG( | Chd1l | 429.8812 | 0.053116 | 0.677808 | 0.930542 | ENSMUSG( | Chd1l | 368.0756 | -0.12141 | 0.426685 | 0.917257 |
| ENSMUSG( | Chd2 | 1545.72 | 0.192545 | 0.049624 | 0.340062 | ENSMUSG( | Chd2 | 1356.823 | 0.088738 | 0.428766 | 0.917257 |
| ENSMUSG( | Chd3 | 40500.87 | 0.224204 | 0.004875 | 0.092488 | ENSMUSG( | Chd3 | 34404.33 | 0.031094 | 0.744661 | 0.96951 |
| ENSMUSG( | Chd4 | 7488.854 | -0.05899 | 0.493311 | 0.860611 | ENSMUSG( | Chd4 | 6819.805 | -0.05948 | 0.592229 | 0.944975 |
| ENSMUSG( | Chd5 | 8707.346 | 0.293669 | 0.00172 | 0.046884 | ENSMUSG( | Chd5 | 7217.302 | 0.036044 | 0.732555 | 0.967228 |
| ENSMUSG( | Chd6 | 2417.592 | -0.21318 | 0.064532 | 0.393651 | ENSMUSG( | Chd6 | 2387.414 | 0.030158 | 0.811742 | 0.979148 |
| ENSMUSG( | Chd7 | 883.9128 | -0.28809 | 0.01205 | 0.158771 | ENSMUSG( | Chd7 | 875.6477 | -0.03044 | 0.797904 | 0.97681 |
| ENSMUSG( | Chd8 | 2907.171 | 0.063762 | 0.538342 | 0.88259 | ENSMUSG( | Chd8 | 2721.464 | 0.141562 | 0.210085 | 0.864412 |
| ENSMUSG( | Chd9 | 2059.982 | 0.083194 | 0.400525 | 0.813717 | ENSMUSG( | Chd9 | 1940.351 | 0.175063 | 0.112084 | 0.808673 |
| ENSMUSG( | Chdh | 128.1281 | -0.17773 | 0.317138 | 0.763823 | ENSMUSG( | Chdh | 139.4568 | 0.32753 | 0.081542 | 0.77681 |
| ENSMUSG( | Chek1 | 48.09572 | -0.05511 | 0.839197 | 0.971179 | ENSMUSG( | Chek1 | 44.36251 | 0.000278 | 0.999275 | 0.999503 |
| ENSMUSG( | Chek2 | 61.9095 | 0.012846 | 0.958587 | 0.993874 | ENSMUSG( | Chek2 | 60.10573 | 0.202476 | 0.448982 | 0.922202 |
| ENSMUSG( | Cherp | 1740.657 | 0.138595 | 0.226667 | 0.673729 | ENSMUSG( | Cherp | 1527.119 | 0.030854 | 0.823674 | 0.979198 |
| ENSMUSG( | Chfr | 1160.577 | 0.041278 | 0.691431 | 0.933881 | ENSMUSG( | Chfr | 1068.458 | 0.07144 | 0.610643 | 0.950771 |
| ENSMUSG( | Chga | 6933.921 | -0.4715 | 1.06E-06 | 0.000197 | ENSMUSG( | Chga | 7340.062 | -0.00038 | 0.997683 | 0.999268 |
| ENSMUSG( | Chgb | 15754.11 | -0.29143 | 0.02946 | 0.260274 | ENSMUSG( | Chgb | 16599.39 | 0.143017 | 0.363172 | 0.914223 |
| ENSMUSG( | Chi3l1 | 899.2009 | -0.34727 | 0.001974 | 0.051135 | ENSMUSG( | Chi3l1 | 831.4301 | -0.30286 | 0.02315 | 0.585577 |
| ENSMUSG( | Chia1 | 71.09129 | -0.23106 | 0.361654 | 0.792797 | ENSMUSG( | Chia1 | 64.74209 | -0.24173 | 0.412202 | 0.917257 |
| ENSMUSG( | Chic1 | 637.3378 | -0.03091 | 0.766472 | 0.954193 | ENSMUSG( | Chic1 | 589.4602 | 0.005328 | 0.970338 | 0.99594 |
| ENSMUSG( | Chic2 | 434.4897 | -0.16103 | 0.175272 | 0.61046 | ENSMUSG( | Chic2 | 435.5287 | 0.116697 | 0.389592 | 0.917257 |
| ENSMUSG( | Chid1 | 2014.874 | -0.21622 | 0.038722 | 0.303123 | ENSMUSG( | Chid1 | 1963.32 | -0.01134 | 0.931947 | 0.99306 |
| ENSMUSG( | Chil3 | 28.34672 | -0.28987 | 0.613187 | 0.908902 | ENSMUSG( | Chil3 | 31.04082 | 0.197039 | 0.716454 | 0.965773 |
| ENSMUSG( | Chka | 2376.375 | 0.179283 | 0.048637 | 0.337278 | ENSMUSG( | Chka | 2077.718 | 0.064396 | 0.565007 | 0.941248 |
| ENSMUSG( | Chkb | 98.85496 | 0.051276 | 0.808439 | 0.963455 | ENSMUSG( | Chkb | 93.60255 | 0.167129 | 0.476786 | 0.923329 |
| ENSMUSG( | Chl1 | 2917.438 | -0.32858 | 0.05997 | 0.378891 | ENSMUSG( | Chl1 | 3216.988 | 0.23547 | 0.231788 | 0.868507 |
| ENSMUSG( | Chm | 871.4919 | -0.07375 | 0.555366 | 0.887457 | ENSMUSG( | Chm | 820.919 | 0.022652 | 0.857459 | 0.985884 |
| ENSMUSG( | Chml | 268.4034 | -0.3714 | 0.008466 | 0.128691 | ENSMUSG( | Chml | 269.7516 | -0.07559 | 0.659709 | 0.956205 |
| ENSMUSG( | Chmp1a | 4311.355 | 0.043327 | 0.576866 | 0.89546 | ENSMUSG( | Chmp1a | 3761.804 | -0.08257 | 0.413595 | 0.917257 |
| ENSMUSG( | Chmp1b | 608.5694 | -0.02045 | 0.843692 | 0.971847 | ENSMUSG( | Chmp1b | 557.4592 | -0.00725 | 0.956032 | 0.994798 |
| ENSMUSG( | Chmp1b2 | 1059.942 | 0.068357 | 0.574282 | 0.894186 | ENSMUSG( | Chmp1b2 | 889.3957 | -0.18046 | 0.265152 | 0.886134 |
| ENSMUSG( | Chmp2a | 1975.763 | 0.063176 | 0.547971 | 0.885153 | ENSMUSG( | Chmp2a | 1669.091 | -0.16264 | 0.231536 | 0.868507 |
| ENSMUSG( | Chmp2b | 995.3262 | -0.08268 | 0.461428 | 0.841432 | ENSMUSG( | Chmp2b | 926.6077 | -0.02615 | 0.871882 | 0.988779 |
| ENSMUSG( | Chmp3 | 3142.358 | 0.051547 | 0.538478 | 0.88259 | ENSMUSG( | Chmp3 | 2688.516 | -0.1348 | 0.244327 | 0.877904 |
| ENSMUSG( | Chmp4b | 7286.668 | 0.012525 | 0.884494 | 0.982647 | ENSMUSG( | Chmp4b | 6112.503 | -0.23752 | 0.051534 | 0.711358 |
| ENSMUSG( | Chmp5 | 3537.498 | 0.057862 | 0.602707 | 0.904887 | ENSMUSG( | Chmp5 | 2966.333 | -0.19108 | 0.188809 | 0.85698 |
| ENSMUSG( | Chmp6 | 1057.373 | 0.073386 | 0.432115 | 0.82768 | ENSMUSG( | Chmp6 | 929.5863 | -0.03019 | 0.793011 | 0.97681 |
| ENSMUSG( | Chmp7 | 2305.18 | 0.109253 | 0.268504 | 0.718807 | ENSMUSG( | Chmp7 | 1994.06 | -0.03728 | 0.738423 | 0.967967 |
| ENSMUSG( | Chn1 | 21928.38 | 0.03717 | 0.7402 | 0.947823 | ENSMUSG( | Chn1 | 18907.33 | -0.1267 | 0.35606 | 0.913331 |
| ENSMUSG( | Chn1os3 | 48.17524 | -0.30257 | 0.338344 | 0.777489 | ENSMUSG( | Chn1os3 | 43.44466 | -0.33683 | 0.326036 | 0.905339 |
| ENSMUSG( | Chn2 | 1216.473 | 0.0167 | 0.887128 | 0.983326 | ENSMUSG( | Chn2 | 1124.202 | 0.057677 | 0.670003 | 0.95694 |
| ENSMUSG( | Chodl | 20.83484 | -0.09127 | 0.834525 | NA | ENSMUSG( | Chodl | 22.88803 | 0.399458 | 0.338673 | 0.907972 |
| ENSMUSG( | Chordc1 | 836.0822 | 0.110906 | 0.27761 | 0.72746 | ENSMUSG( | Chordc1 | 801.1033 | 0.245214 | 0.044434 | 0.693697 |
| ENSMUSG( | Chp1 | 4348.728 | 0.057593 | 0.490079 | 0.859319 | ENSMUSG( | Chp1 | 3731.984 | -0.11886 | 0.301846 | 0.898538 |
| ENSMUSG( | Chp2 | 170.1941 | -0.38678 | 0.042594 | 0.316734 | ENSMUSG( | Chp2 | 152.0943 | -0.46249 | 0.017178 | 0.52741 |
| ENSMUSG( | Chpf | 2507.615 | -0.02905 | 0.853638 | 0.973491 | ENSMUSG( | Chpf | 2521.776 | 0.251987 | 0.18457 | 0.853513 |
| ENSMUSG( | Chpf2 | 1765.212 | -0.06887 | 0.612536 | 0.908747 | ENSMUSG( | Chpf2 | 1769.744 | 0.205816 | 0.199367 | 0.863016 |
| ENSMUSG( | Chpt1 | 4493.269 | -0.04052 | 0.778472 | 0.956215 | ENSMUSG( | Chpt1 | 4150.986 | -0.00304 | 0.985939 | 0.997188 |
| ENSMUSG( | Chrac1 | 265.7696 | -0.14161 | 0.297973 | 0.746543 | ENSMUSG( | Chrac1 | 243.1753 | -0.13296 | 0.401058 | 0.917257 |
| ENSMUSG( | Chrd | 1336.088 | 0.516387 | 0.000866 | 0.029559 | ENSMUSG( | Chrd | 1096.11 | 0.249078 | 0.126163 | 0.81516 |
| ENSMUSG( | Chrdl1 | 1707.796 | -0.23002 | 0.152593 | 0.578962 | ENSMUSG( | Chrdl1 | 1585.254 | -0.16997 | 0.323943 | 0.905257 |
| ENSMUSG( | Chrdl2 | 9.195718 | 0.638238 | 0.338097 | NA | ENSMUSG( | Chrdl2 | 8.908951 | 0.722858 | 0.331046 | NA |
| ENSMUSG( | Chrm1 | 6307.047 | -0.28518 | 0.046764 | 0.330815 | ENSMUSG( | Chrm1 | 6931.903 | 0.262831 | 0.10459 | 0.804969 |
| ENSMUSG( | Chrm2 | 282.9 | -0.54127 | 0.002541 | 0.059739 | ENSMUSG( | Chrm2 | 355.14 | 0.412117 | 0.035291 | 0.661159 |
| ENSMUSG( | Chrm3 | 1285.87 | -0.27864 | 0.154075 | 0.580901 | ENSMUSG( | Chrm3 | 1392.659 | 0.23343 | 0.307358 | 0.898538 |
| ENSMUSG( | Chrm4 | 672.0598 | -0.76924 | 4.59E-05 | 0.003685 | ENSMUSG( | Chrm4 | 764.2852 | -0.01997 | 0.917113 | 0.992529 |
| ENSMUSG( | Chrm5 | 10.28635 | 0.519772 | 0.417029 | NA | ENSMUSG( | Chrm5 | 10.14248 | 0.744651 | 0.255729 | NA |
| ENSMUSG( | Chrna1 | 12.61649 | 0.263967 | 0.635885 | NA | ENSMUSG( | Chrna1 | 10.69201 | 0.098725 | 0.874376 | NA |
| ENSMUSG( | Chrna10 | 11.21461 | -0.37541 | 0.591504 | NA | ENSMUSG( | Chrna10 | 13.36018 | 0.336186 | 0.569695 | 0.941777 |
| ENSMUSG( | Chrna1os | 30.71477 | -0.0517 | 0.882045 | 0.982069 | ENSMUSG( | Chrna1os | 25.02062 | -0.38222 | 0.334157 | 0.907907 |
| ENSMUSG( | Chrna2 | 22.38008 | -0.14021 | 0.747683 | 0.949604 | ENSMUSG( | Chrna2 | 22.29013 | 0.103889 | 0.826935 | 0.980159 |
| ENSMUSG( | Chrna3 | 34.23598 | -0.3913 | 0.222364 | 0.668412 | ENSMUSG( | Chrna3 | 33.97584 | -0.10989 | 0.746184 | 0.969649 |
| ENSMUSG( | Chrna4 | 1276.168 | -0.37133 | 0.010908 | 0.150598 | ENSMUSG( | Chrna4 | 1355.582 | 0.099237 | 0.57787 | 0.943131 |
| ENSMUSG( | Chrna5 | 66.01763 | -0.64865 | 0.032681 | 0.27685 | ENSMUSG( | Chrna5 | 77.03398 | 0.144274 | 0.612365 | 0.951071 |
| ENSMUSG( | Chrna6 | 12.55841 | 0.027256 | 0.959315 | NA | ENSMUSG( | Chrna6 | 15.38266 | 0.765016 | 0.165436 | 0.838866 |
| ENSMUSG( | Chrna7 | 195.7092 | -0.53616 | 0.041926 | 0.315414 | ENSMUSG( | Chrna7 | 211.7688 | 0.026527 | 0.923669 | 0.992815 |
| ENSMUSG( | Chrnb1 | 29.19181 | -0.05893 | 0.86274 | 0.975055 | ENSMUSG( | Chrnb1 | 32.83879 | 0.524026 | 0.194152 | 0.860096 |
| ENSMUSG( | Chrnb2 | 1498.572 | -0.26661 | 0.077948 | 0.431652 | ENSMUSG( | Chrnb2 | 1614.407 | 0.226793 | 0.228532 | 0.868507 |
| ENSMUSG( | Chrnb3 | 54.4742 | -0.7257 | 0.019257 | 0.20921 | ENSMUSG( | Chrnb3 | 62.92889 | 0.065705 | 0.837556 | 0.981335 |
| ENSMUSG( | Chrnb4 | 9.83904 | -0.24967 | 0.676381 | NA | ENSMUSG( | Chrnb4 | 10.28441 | 0.074274 | 0.910598 | NA |
| ENSMUSG( | Chst1 | 9506.718 | -0.13026 | 0.234236 | 0.68192 | ENSMUSG( | Chst1 | 9801.251 | 0.223237 | 0.069782 | 0.756483 |
| ENSMUSG( | Chst10 | 3450.898 | -0.15707 | 0.183057 | 0.62047 | ENSMUSG( | Chst10 | 3287.6 | -0.02101 | 0.876766 | 0.989644 |
| ENSMUSG( | Chst11 | 3402.617 | -0.22664 | 0.070166 | 0.40908 | ENSMUSG( | Chst11 | 3540.964 | 0.163792 | 0.262774 | 0.885357 |
| ENSMUSG( | Chst12 | 724.4478 | -0.36482 | 0.02578 | 0.247138 | ENSMUSG( | Chst12 | 802.3127 | 0.220696 | 0.233313 | 0.868507 |
| ENSMUSG( | Chst14 | 86.92535 | -0.61112 | 0.022257 | 0.227763 | ENSMUSG( | Chst14 | 98.76907 | 0.107167 | 0.69513 | 0.960867 |
| ENSMUSG( | Chst15 | 1006.331 | -0.40306 | 0.003992 | 0.081133 | ENSMUSG( | Chst15 | 1092.573 | 0.133979 | 0.405448 | 0.917257 |
| ENSMUSG( | Chst2 | 17235.33 | 0.008371 | 0.941591 | 0.993415 | ENSMUSG( | Chst2 | 15915.23 | 0.046876 | 0.729059 | 0.966913 |
| ENSMUSG( | Chst3 | 240.0084 | -0.43578 | 0.028876 | 0.258911 | ENSMUSG( | Chst3 | 246.2604 | -0.04492 | 0.86671 | 0.987458 |
| ENSMUSG( | Chst5 | 39.85374 | -0.78447 | 0.052972 | 0.353293 | ENSMUSG( | Chst5 | 52.32307 | 0.367537 | 0.353563 | 0.912263 |
| ENSMUSG( | Chst7 | 106.88 | -0.11934 | 0.586253 | 0.899709 | ENSMUSG( | Chst7 | 116.7479 | 0.397196 | 0.095566 | 0.796133 |
| ENSMUSG( | Chst8 | 453.4536 | -0.03816 | 0.777261 | 0.955886 | ENSMUSG( | Chst8 | 444.3966 | 0.172285 | 0.30673 | 0.898538 |
| ENSMUSG( | Chst9 | 11.56353 | -0.62883 | 0.260431 | NA | ENSMUSG( | Chst9 | 14.54754 | 0.365442 | 0.483479 | 0.92499 |
| ENSMUSG( | Chsy1 | 735.0255 | -0.24324 | 0.094538 | 0.471713 | ENSMUSG( | Chsy1 | 746.2275 | 0.079583 | 0.609396 | 0.950397 |
| ENSMUSG( | Chsy3 | 308.8091 | -0.65573 | 0.000801 | 0.027945 | ENSMUSG( | Chsy3 | 356.3445 | 0.109597 | 0.618053 | 0.952232 |
| ENSMUSG( | Chtf18 | 71.30314 | 0.060369 | 0.804072 | 0.962169 | ENSMUSG( | Chtf18 | 63.16991 | -0.0195 | 0.94557 | 0.994798 |
| ENSMUSG( | Chtf8 | 30.68867 | -0.11758 | 0.74631 | 0.949289 | ENSMUSG( | Chtf8 | 28.65254 | -0.01754 | 0.9644 | 0.995433 |
| ENSMUSG( | Chtop | 4654.681 | 0.006172 | 0.947498 | 0.993821 | ENSMUSG( | Chtop | 4090.232 | -0.10261 | 0.467638 | 0.923329 |
| ENSMUSG( | Chuk | 1188.605 | -0.00046 | 0.996535 | 0.99943 | ENSMUSG( | Chuk | 1119.436 | 0.09641 | 0.475586 | 0.923329 |
| ENSMUSG( | Churc1 | 203.6615 | 0.022015 | 0.881739 | 0.98196 | ENSMUSG( | Churc1 | 174.3875 | -0.17551 | 0.346946 | 0.911434 |
| ENSMUSG( | Ciao1 | 937.5554 | -0.0165 | 0.904613 | 0.986275 | ENSMUSG( | Ciao1 | 857.857 | -0.00725 | 0.96233 | 0.995237 |
| ENSMUSG( | Ciao2a | 784.6979 | -0.00177 | 0.989324 | 0.999111 | ENSMUSG( | Ciao2a | 675.3064 | -0.17608 | 0.21835 | 0.866104 |
| ENSMUSG( | Ciao2b | 513.9519 | -0.08895 | 0.41874 | 0.821966 | ENSMUSG( | Ciao2b | 448.2729 | -0.22358 | 0.089387 | 0.788575 |
| ENSMUSG( | Ciao3 | 722.7597 | 0.15119 | 0.215403 | 0.659961 | ENSMUSG( | Ciao3 | 633.5425 | 0.04586 | 0.72257 | 0.966887 |
| ENSMUSG( | Ciapin1 | 970.3781 | -0.02076 | 0.849329 | 0.972638 | ENSMUSG( | Ciapin1 | 883.0255 | -0.02341 | 0.846309 | 0.984578 |
| ENSMUSG( | Ciart | 261.6563 | -0.30024 | 0.111532 | 0.503511 | ENSMUSG( | Ciart | 356.9848 | 0.781088 | 1.95E-05 | 0.00609 |
| ENSMUSG( | Cib1 | 841.9794 | 0.145211 | 0.185475 | 0.623185 | ENSMUSG( | Cib1 | 683.4871 | -0.19438 | 0.124988 | 0.81516 |
| ENSMUSG( | Cib2 | 842.6548 | 0.051072 | 0.604572 | 0.905556 | ENSMUSG( | Cib2 | 716.6671 | -0.15759 | 0.258992 | 0.883971 |
| ENSMUSG( | Cibar1 | 954.3187 | -0.04908 | 0.649674 | 0.920645 | ENSMUSG( | Cibar1 | 917.0574 | 0.100698 | 0.417266 | 0.917257 |
| ENSMUSG( | Cibar2 | 16.76652 | -0.18821 | 0.69489 | NA | ENSMUSG( | Cibar2 | 15.03866 | -0.24287 | 0.640157 | 0.953903 |
| ENSMUSG( | Cic | 9246.038 | 0.171032 | 0.041375 | 0.314189 | ENSMUSG( | Cic | 8300.277 | 0.131624 | 0.22856 | 0.868507 |
| ENSMUSG( | Cidea | 85.9235 | -0.23446 | 0.284287 | 0.73274 | ENSMUSG( | Cidea | 89.80358 | 0.168341 | 0.469415 | 0.923329 |
| ENSMUSG( | Cideb | 401.8572 | -0.08647 | 0.604759 | 0.905556 | ENSMUSG( | Cideb | 340.4701 | -0.30385 | 0.099377 | 0.800296 |
| ENSMUSG( | Ciita | 8.252032 | -0.73411 | 0.283799 | NA | ENSMUSG( | Ciita | 7.986195 | -0.46432 | 0.504736 | NA |
| ENSMUSG( | Cilk1 | 1414.319 | 0.001562 | 0.986238 | 0.998326 | ENSMUSG( | Cilk1 | 1259.357 | -0.06369 | 0.585272 | 0.944975 |
| ENSMUSG( | Cilp2 | 94.79361 | -0.25936 | 0.209009 | 0.650854 | ENSMUSG( | Cilp2 | 86.70189 | -0.24891 | 0.274216 | 0.889874 |
| ENSMUSG( | Cimap1b | 45.55698 | -0.02902 | 0.925181 | 0.991204 | ENSMUSG( | Cimap1b | 49.45644 | 0.457348 | 0.165288 | 0.838743 |
| ENSMUSG( | Cimap1d | 14.85456 | -0.12645 | 0.792022 | NA | ENSMUSG( | Cimap1d | 14.01804 | -0.03128 | 0.951633 | 0.994798 |
| ENSMUSG( | Cimap3 | 13.15676 | -0.1146 | 0.836052 | NA | ENSMUSG( | Cimap3 | 14.41689 | 0.38238 | 0.465328 | 0.923329 |
| ENSMUSG( | Cimip2b | 18.61803 | -0.61369 | 0.157052 | NA | ENSMUSG( | Cimip2b | 21.97191 | 0.238747 | 0.598488 | 0.946859 |
| ENSMUSG( | Cimip2c | 11.17068 | -0.15503 | 0.781983 | NA | ENSMUSG( | Cimip2c | 14.77069 | 0.829051 | 0.150035 | 0.837086 |
| ENSMUSG( | Cinp | 1459.592 | 0.04263 | 0.640634 | 0.916434 | ENSMUSG( | Cinp | 1256.609 | -0.12357 | 0.271686 | 0.888062 |
| ENSMUSG( | Cip2a | 91.597 | -0.1376 | 0.525723 | 0.876835 | ENSMUSG( | Cip2a | 87.79238 | 0.02105 | 0.928709 | 0.99306 |
| ENSMUSG( | Cipc | 2907.628 | 0.041834 | 0.626987 | 0.912825 | ENSMUSG( | Cipc | 2632.902 | 0.02267 | 0.841067 | 0.9826 |
| ENSMUSG( | Cir1 | 432.5043 | 0.027025 | 0.833317 | 0.970333 | ENSMUSG( | Cir1 | 399.0708 | 0.057063 | 0.719596 | 0.966876 |
| ENSMUSG( | Cirbp | 1509.628 | -0.27487 | 0.025183 | 0.244448 | ENSMUSG( | Cirbp | 1442.278 | -0.12925 | 0.335965 | 0.907913 |
| ENSMUSG( | Cisd1 | 1993.657 | -0.10826 | 0.201526 | 0.641362 | ENSMUSG( | Cisd1 | 1831 | -0.08712 | 0.466345 | 0.923329 |
| ENSMUSG( | Cisd2 | 1512.466 | -0.35672 | 0.001495 | 0.042956 | ENSMUSG( | Cisd2 | 1581.677 | 0.065305 | 0.628304 | 0.953251 |
| ENSMUSG( | Cisd3 | 644.1316 | -0.00144 | 0.98998 | 0.999144 | ENSMUSG( | Cisd3 | 565.7674 | -0.11745 | 0.448176 | 0.922202 |
| ENSMUSG( | Cish | 144.6814 | -0.42185 | 0.029922 | 0.262341 | ENSMUSG( | Cish | 141.1368 | -0.20299 | 0.334073 | 0.907907 |
| ENSMUSG( | Cit | 5765.086 | 0.149736 | 0.05251 | 0.35151 | ENSMUSG( | Cit | 5192.742 | 0.118206 | 0.240978 | 0.876146 |
| ENSMUSG( | Cited1 | 28.67159 | -0.26774 | 0.466706 | 0.844819 | ENSMUSG( | Cited1 | 29.69695 | 0.071132 | 0.859307 | 0.985884 |
| ENSMUSG( | Cited2 | 482.9094 | 0.050722 | 0.766236 | 0.954173 | ENSMUSG( | Cited2 | 480.6729 | 0.286764 | 0.133809 | 0.822912 |
| ENSMUSG( | Cited4 | 130.1074 | -0.33165 | 0.198201 | 0.638387 | ENSMUSG( | Cited4 | 132.2501 | -0.00056 | 0.998315 | 0.999313 |

| | | | | | |
|---|---|---|---|---|---|
| ENSMUSG( | 513.9519 | -0.08895 | 0.41874 | 0.821966 | Ciao2b |
| ENSMUSG( | 722.7597 | 0.15119 | 0.215403 | 0.659961 | Ciao3 |
| ENSMUSG( | 970.3781 | -0.02076 | 0.849329 | 0.972638 | Ciapin1 |
| ENSMUSG( | 261.6563 | -0.30024 | 0.111532 | 0.503511 | Ciart |
| ENSMUSG( | 841.9794 | 0.145211 | 0.185475 | 0.623185 | Cib1 |
| ENSMUSG( | 842.6548 | 0.051072 | 0.604572 | 0.905556 | Cib2 |
| ENSMUSG( | 954.3187 | -0.04908 | 0.649674 | 0.920645 | Cibar1 |
| ENSMUSG( | 16.76652 | -0.18821 | 0.69489 | NA | Cibar2 |
| ENSMUSG( | 9246.038 | 0.171032 | 0.041375 | 0.314189 | Cic |
| ENSMUSG( | 85.9235 | -0.23446 | 0.284287 | 0.73274 | Cidea |
| ENSMUSG( | 401.8572 | -0.08647 | 0.604759 | 0.905556 | Cideb |
| ENSMUSG( | 8.252032 | -0.73411 | 0.283799 | NA | Ciita |
| ENSMUSG( | 1414.319 | 0.001562 | 0.986238 | 0.998326 | Cilk1 |
| ENSMUSG( | 94.79361 | -0.25936 | 0.209009 | 0.650854 | Cilp2 |
| ENSMUSG( | 45.55698 | -0.02902 | 0.925181 | 0.991204 | Cimap1b |
| ENSMUSG( | 14.85456 | -0.12645 | 0.792022 | NA | Cimap1d |
| ENSMUSG( | 13.15676 | -0.1146 | 0.836052 | NA | Cimap3 |
| ENSMUSG( | 18.61803 | -0.61369 | 0.157052 | NA | Cimip2b |
| ENSMUSG( | 11.17068 | -0.15503 | 0.781983 | NA | Cimip2c |
| ENSMUSG( | 1459.592 | 0.04263 | 0.640634 | 0.916434 | Cinp |
| ENSMUSG( | 91.597 | -0.1376 | 0.525723 | 0.876835 | Cip2a |
| ENSMUSG( | 2907.628 | 0.041834 | 0.626987 | 0.912825 | Cipc |
| ENSMUSG( | 432.5043 | 0.027025 | 0.833317 | 0.970333 | Cir1 |
| ENSMUSG( | 1509.628 | -0.27487 | 0.025183 | 0.244448 | Cirbp |
| ENSMUSG( | 1993.657 | -0.10826 | 0.201526 | 0.641362 | Cisd1 |
| ENSMUSG( | 1512.466 | -0.35672 | 0.001495 | 0.042956 | Cisd2 |
| ENSMUSG( | 644.1316 | -0.00144 | 0.98998 | 0.999144 | Cisd3 |
| ENSMUSG( | 144.6814 | -0.42185 | 0.029922 | 0.262341 | Cish |
| ENSMUSG( | 5765.086 | 0.149736 | 0.05251 | 0.35151 | Cit |
| ENSMUSG( | 28.67159 | -0.26774 | 0.466706 | 0.844819 | Cited1 |
| ENSMUSG( | 482.9094 | 0.050722 | 0.766236 | 0.954173 | Cited2 |
| ENSMUSG( | 130.1074 | -0.33165 | 0.198201 | 0.638387 | Cited4 |
| ENSMUSG( | 1709.12 | 0.270081 | 0.012865 | 0.163922 | Ciz1 |
| ENSMUSG( | 59.89473 | -0.65159 | 0.00796 | 0.124138 | CK137956 |
| ENSMUSG( | 42.34878 | -0.48369 | 0.126723 | 0.534201 | Ckap2 |
| ENSMUSG( | 54.63984 | -0.3187 | 0.225285 | 0.672044 | Ckap2l |
| ENSMUSG( | 1120.252 | -0.19277 | 0.145429 | 0.566269 | Ckap4 |
| ENSMUSG( | 8679.452 | 0.160891 | 0.123577 | 0.530552 | Ckap5 |
| ENSMUSG( | 63859.18 | 0.03711 | 0.6929 | 0.934202 | Ckb |
| ENSMUSG( | 112.5718 | -0.26391 | 0.184183 | 0.621101 | Cklf |
| ENSMUSG( | 3378.526 | -0.06319 | 0.634024 | 0.914073 | Ckmt1 |
| ENSMUSG( | 57.40085 | 0.076234 | 0.762555 | 0.953519 | Cks1b |
| ENSMUSG( | 22.51579 | -0.19301 | 0.668481 | 0.927864 | Cks2 |
| ENSMUSG( | 3006.529 | 0.114918 | 0.201158 | 0.641362 | Clasp1 |
| ENSMUSG( | 9274.888 | 0.076988 | 0.364498 | 0.793796 | Clasp2 |
| ENSMUSG( | 2222.08 | 0.079002 | 0.599345 | 0.903801 | Clasrp |
| ENSMUSG( | 303.3331 | -0.05415 | 0.694354 | 0.9343 | Clba1 |
| ENSMUSG( | 18.93111 | -0.40051 | 0.368343 | NA | Clca3a1 |
| ENSMUSG( | 1141.686 | -0.1674 | 0.171472 | 0.605943 | Clcc1 |
| ENSMUSG( | 68.74442 | 0.00563 | 0.981356 | 0.997512 | Clcf1 |
| ENSMUSG( | 119.3782 | 0.108307 | 0.563236 | 0.889897 | Clcn1 |
| ENSMUSG( | 2620.6 | 0.176226 | 0.110019 | 0.500285 | Clcn2 |
| ENSMUSG( | 7771.304 | -0.22972 | 0.096899 | 0.477375 | Clcn3 |
| ENSMUSG( | 8813.379 | -0.22456 | 0.067209 | 0.400638 | Clcn4 |
| ENSMUSG( | 356.3565 | -0.46479 | 0.019307 | 0.209368 | Clcn5 |
| ENSMUSG( | 3408.13 | -0.28077 | 0.041748 | 0.315339 | Clcn6 |
| ENSMUSG( | 1937.952 | -0.14982 | 0.229705 | 0.677538 | Clcn7 |
| ENSMUSG( | 85.37629 | -0.38288 | 0.134489 | 0.54768 | Cldn1 |
| ENSMUSG( | 2779.512 | -0.15633 | 0.284827 | 0.733433 | Cldn10 |
| ENSMUSG( | 10667.46 | -0.78267 | 1.59E-07 | 3.86E-05 | Cldn11 |
| ENSMUSG( | 1140.824 | -0.43743 | 0.003369 | 0.073204 | Cldn12 |
| ENSMUSG( | 95.64746 | -0.35891 | 0.10131 | 0.484996 | Cldn14 |
| ENSMUSG( | 6.88576 | -0.63247 | 0.412427 | NA | Cldn15 |
| ENSMUSG( | 51.43726 | -0.85047 | 0.002315 | 0.056456 | Cldn19 |
| ENSMUSG( | 62.10573 | 0.033771 | 0.962822 | 0.994824 | Cldn2 |
| ENSMUSG( | 8.706004 | 0.049878 | 0.943414 | NA | Cldn20 |
| ENSMUSG( | 15.78845 | -0.54401 | 0.285869 | NA | Cldn22 |
| ENSMUSG( | 16.06602 | -0.90828 | 0.055464 | NA | Cldn23 |
| ENSMUSG( | 13.10486 | -0.66237 | 0.216084 | NA | Cldn3 |
| ENSMUSG( | 65.84915 | -0.31801 | 0.196546 | 0.635942 | Cldn34c1 |
| ENSMUSG( | 1950.177 | -0.14878 | 0.442315 | 0.832774 | Cldn5 |
| ENSMUSG( | 3419.752 | -0.20045 | 0.071253 | 0.412315 | Cldnd1 |
| ENSMUSG( | 49.07152 | 0.242964 | 0.380414 | 0.801757 | Clec10a |
| ENSMUSG( | 271.4226 | 0.175852 | 0.217119 | 0.661995 | Clec11a |
| ENSMUSG( | 23.34332 | -1.93732 | 0.000124 | 0.007462 | Clec12a |
| ENSMUSG( | 101.5122 | -0.40026 | 0.109777 | 0.500285 | Clec14a |
| ENSMUSG( | 2927.491 | 0.176006 | 0.043244 | 0.318825 | Clec16a |
| ENSMUSG( | 136.1264 | -0.14676 | 0.483426 | 0.856304 | Clec18a |
| ENSMUSG( | 26.41011 | -0.83103 | 0.039269 | 0.305097 | Clec1a |
| ENSMUSG( | 184.1652 | -0.01088 | 0.955031 | 0.993874 | Clec2d |
| ENSMUSG( | 3097.045 | 0.009079 | 0.919876 | 0.990247 | Clec2l |
| ENSMUSG( | 130.2098 | -0.17934 | 0.316601 | 0.763648 | Clec3b |
| ENSMUSG( | 46.38507 | -0.2533 | 0.416341 | 0.82084 | Clec4a1 |
| ENSMUSG( | 55.7492 | -0.16448 | 0.55823 | 0.888427 | Clec4a2 |
| ENSMUSG( | 72.06554 | -0.50239 | 0.118295 | 0.520125 | Clec4a3 |
| ENSMUSG( | 12.2945 | -0.49158 | 0.397227 | NA | Clec4n |
| ENSMUSG( | 132.6719 | 0.202758 | 0.304061 | 0.752269 | Clec5a |
| ENSMUSG( | 24.66307 | -0.60273 | 0.128239 | 0.536641 | Clec7a |
| ENSMUSG( | 511.1746 | -0.37284 | 0.004211 | 0.084333 | Clgn |
| ENSMUSG( | 648.2984 | -0.05019 | 0.718479 | 0.941911 | Clic1 |
| ENSMUSG( | 11.73031 | -0.31053 | 0.584137 | NA | Clic3 |
| ENSMUSG( | 7786.698 | -0.02379 | 0.828808 | 0.969097 | Clic4 |
| ENSMUSG( | 293.2385 | 0.295875 | 0.216141 | 0.661017 | Clic5 |
| ENSMUSG( | 214.7135 | -0.1665 | 0.849645 | 0.972638 | Clic6 |
| ENSMUSG( | 1425.135 | 0.026208 | 0.783067 | 0.957558 | Clint1 |
| ENSMUSG( | 2775.797 | 0.06699 | 0.522109 | 0.875186 | Clip1 |
| ENSMUSG( | 4402.548 | -0.06643 | 0.44351 | 0.833108 | Clip2 |
| ENSMUSG( | 13384.6 | 0.116666 | 0.258614 | 0.709814 | Clip3 |
| ENSMUSG( | 1126.926 | -0.02789 | 0.769567 | 0.954674 | Clip4 |
| ENSMUSG( | 3322.513 | -0.13204 | 0.215646 | 0.660333 | Clk1 |
| ENSMUSG( | 1000.602 | 0.148885 | 0.201747 | 0.6415 | Clk2 |
| ENSMUSG( | 2113.634 | -0.04691 | 0.622172 | 0.911432 | Clk3 |
| ENSMUSG( | 1831.656 | -0.10534 | 0.369196 | 0.796641 | Clk4 |
| ENSMUSG( | 5372.511 | 0.060601 | 0.487886 | 0.858824 | Clmn |
| ENSMUSG( | 726.1943 | -0.53434 | 3.41E-05 | 0.002956 | Clmp |
| ENSMUSG( | 560.2289 | -0.34657 | 0.002456 | 0.058231 | Cln3 |
| ENSMUSG( | 459.4054 | -0.26553 | 0.087733 | 0.454663 | Cln5 |
| ENSMUSG( | 573.6429 | -0.39114 | 0.00316 | 0.06974 | Cln6 |
| ENSMUSG( | 1136.804 | -0.31324 | 0.04943 | 0.339723 | Cln8 |
| ENSMUSG( | 1487.518 | 0.043842 | 0.640427 | 0.916434 | Clns1a |
| ENSMUSG( | 1878.215 | 0.07876 | 0.372492 | 0.798768 | Clock |
| ENSMUSG( | 113.5385 | 0.035519 | 0.878498 | 0.981431 | Clp1 |
| ENSMUSG( | 2166.507 | 0.097513 | 0.329718 | 0.773619 | Clpb |
| ENSMUSG( | 1477.551 | 0.061297 | 0.50743 | 0.86795 | Clpp |
| ENSMUSG( | 8390.923 | -0.23755 | 0.072111 | 0.414057 | Clptm1 |
| ENSMUSG( | 4181.742 | -0.23927 | 0.060385 | 0.3797 | Clptm1l |
| ENSMUSG( | 843.5138 | 0.117667 | 0.221438 | 0.667258 | Clpx |
| ENSMUSG( | 12.80944 | -0.34275 | 0.49962 | NA | Clrn1 |
| ENSMUSG( | 61.23392 | -0.20854 | 0.408872 | 0.818173 | Clspn |
| ENSMUSG( | 48799.67 | -0.18029 | 0.20609 | 0.647953 | Clstn1 |
| ENSMUSG( | 2546.227 | -0.3306 | 0.019214 | 0.208885 | Clstn2 |
| ENSMUSG( | 12964.2 | -0.29252 | 0.035623 | 0.290384 | Clstn3 |
| ENSMUSG( | 4326.456 | 0.020874 | 0.83452 | 0.970344 | Clta |
| ENSMUSG( | 5193.757 | -0.07568 | 0.424779 | 0.824675 | Cltb |
| ENSMUSG( | 12271.9 | 0.030306 | 0.733179 | 0.944956 | Cltc |
| ENSMUSG( | 94301.22 | 0.112925 | 0.342183 | 0.778828 | Clu |
| ENSMUSG( | 510.0621 | 0.030787 | 0.80487 | 0.962473 | Cluap1 |
| ENSMUSG( | 3118.734 | 0.113212 | 0.158832 | 0.588112 | Cluh |
| ENSMUSG( | 601.4293 | -0.12567 | 0.39432 | 0.810014 | Clvs1 |
| ENSMUSG( | 346.1323 | 0.199937 | 0.188849 | 0.627678 | Clvs2 |
| ENSMUSG( | 129.2021 | -0.21799 | 0.252802 | 0.703255 | Clxn |
| ENSMUSG( | 612.8338 | 0.012596 | 0.922108 | 0.99045 | Clybl |
| ENSMUSG( | 2251.04 | -0.10154 | 0.235778 | 0.682914 | Cmas |
| ENSMUSG( | 429.7682 | 0.014679 | 0.922487 | 0.99045 | Cmbl |
| ENSMUSG( | 259.2244 | -0.09035 | 0.557685 | 0.888397 | Cmc1 |
| ENSMUSG( | 283.6745 | 0.05177 | 0.724782 | 0.944096 | Cmc2 |
| ENSMUSG( | 83.2918 | 0.081284 | 0.743305 | 0.948478 | Cmc4 |
| ENSMUSG( | 4110.876 | 0.265413 | 0.009291 | 0.136006 | Cmip |
| ENSMUSG( | 195.5101 | -0.07532 | 0.686621 | 0.933041 | Cmklr1 |
| ENSMUSG( | 18.16081 | -0.00864 | 0.985164 | NA | Cmklr2 |
| ENSMUSG( | 2238.799 | 0.006048 | 0.958234 | 0.993874 | Cmpk1 |
| ENSMUSG( | 826.7804 | -0.13438 | 0.259413 | 0.709948 | Cmpk2 |
| ENSMUSG( | 170.8966 | 0.043528 | 0.78343 | 0.957558 | Cmss1 |
| ENSMUSG( | 785.2533 | 0.038455 | 0.776588 | 0.955658 | Cmtm3 |
| ENSMUSG( | 6236.231 | 0.026851 | 0.813465 | 0.964574 | Cmtm4 |
| ENSMUSG( | 3296.773 | -0.22244 | 0.050064 | 0.342055 | Cmtm5 |
| ENSMUSG( | 2193.253 | -0.12047 | 0.300157 | 0.749354 | Cmtm6 |
| ENSMUSG( | 165.6229 | 0.017704 | 0.919925 | 0.990247 | Cmtm7 |
| ENSMUSG( | 60.16929 | -0.34889 | 0.174992 | 0.610399 | Cmtm8 |
| ENSMUSG( | 1033.966 | 0.012115 | 0.927847 | 0.991229 | Cmtr1 |
| ENSMUSG( | 88.17447 | -0.02675 | 0.907402 | 0.987566 | Cmtr2 |
| ENSMUSG( | 342.8856 | 0.216093 | 0.134139 | 0.547472 | Cmya5 |
| ENSMUSG( | 259.3255 | 0.178157 | 0.217391 | 0.662547 | Cnbd2 |
| ENSMUSG( | 4576.044 | 0.004022 | 0.967151 | 0.995949 | Cnbp |
| ENSMUSG( | 1398.87 | -0.00359 | 0.973556 | 0.99675 | Cndp2 |
| ENSMUSG( | 764.8124 | -0.22375 | 0.06718 | 0.400611 | Cnep1r1 |
| ENSMUSG( | 14.69898 | 0.24951 | 0.61817 | NA | Cnga2 |
| ENSMUSG( | 31.02103 | 0.24465 | 0.483396 | 0.856304 | Cnga4 |

| | | | | | |
|---|---|---|---|---|---|
| ENSMUSG( | 89.80358 | 0.168341 | 0.469415 | 0.923329 | Cidea |
| ENSMUSG( | 340.4701 | -0.30385 | 0.099377 | 0.800296 | Cideb |
| ENSMUSG( | 7.986195 | -0.46432 | 0.504736 | NA | Ciita |
| ENSMUSG( | 1259.357 | -0.06369 | 0.585272 | 0.944975 | Cilk1 |
| ENSMUSG( | 86.70189 | -0.24891 | 0.274216 | 0.889874 | Cilp2 |
| ENSMUSG( | 49.45644 | 0.457348 | 0.165288 | 0.838743 | Cimap1b |
| ENSMUSG( | 14.01804 | -0.03128 | 0.951633 | 0.994798 | Cimap1d |
| ENSMUSG( | 14.41689 | 0.38238 | 0.465328 | 0.923329 | Cimap3 |
| ENSMUSG( | 21.97191 | 0.238747 | 0.598488 | 0.946859 | Cimip2b |
| ENSMUSG( | 14.77069 | 0.829051 | 0.150035 | 0.837086 | Cimip2c |
| ENSMUSG( | 1256.609 | -0.12357 | 0.271686 | 0.888062 | Cinp |
| ENSMUSG( | 87.79238 | 0.02105 | 0.928709 | 0.99306 | Cip2a |
| ENSMUSG( | 2632.902 | 0.02267 | 0.841067 | 0.9826 | Cipc |
| ENSMUSG( | 399.0708 | 0.057063 | 0.719596 | 0.966876 | Cir1 |
| ENSMUSG( | 1442.278 | -0.12925 | 0.335965 | 0.907913 | Cirbp |
| ENSMUSG( | 1831 | -0.08712 | 0.466345 | 0.923329 | Cisd1 |
| ENSMUSG( | 1581.677 | 0.065305 | 0.628304 | 0.953251 | Cisd2 |
| ENSMUSG( | 565.7674 | -0.11745 | 0.448176 | 0.922202 | Cisd3 |
| ENSMUSG( | 141.1368 | -0.20299 | 0.334073 | 0.907907 | Cish |
| ENSMUSG( | 5192.742 | 0.118206 | 0.240978 | 0.876146 | Cit |
| ENSMUSG( | 29.69695 | 0.071132 | 0.859307 | 0.985884 | Cited1 |
| ENSMUSG( | 480.6729 | 0.286764 | 0.133809 | 0.822912 | Cited2 |
| ENSMUSG( | 132.2501 | -0.00056 | 0.998315 | 0.999313 | Cited4 |
| ENSMUSG( | 1476.705 | 0.124782 | 0.355156 | 0.912411 | Ciz1 |
| ENSMUSG( | 67.3723 | 0.010779 | 0.969866 | 0.99594 | CK137956 |
| ENSMUSG( | 44.8509 | -0.01877 | 0.951591 | 0.994798 | Ckap2 |
| ENSMUSG( | 53.80888 | -0.07231 | 0.801535 | 0.97681 | Ckap2l |
| ENSMUSG( | 1150.869 | 0.160898 | 0.308769 | 0.898538 | Ckap4 |
| ENSMUSG( | 7149.62 | -0.12942 | 0.28568 | 0.893906 | Ckap5 |
| ENSMUSG( | 53059.11 | -0.24029 | 0.044258 | 0.693437 | Ckb |
| ENSMUSG( | 123.8536 | 0.299252 | 0.310859 | 0.899422 | Cklf |
| ENSMUSG( | 2877.041 | -0.27374 | 0.075312 | 0.767994 | Ckmt1 |
| ENSMUSG( | 47.23892 | -0.24055 | 0.414212 | 0.917257 | Cks1b |
| ENSMUSG( | 21.19199 | -0.13853 | 0.772604 | 0.974463 | Cks2 |
| ENSMUSG( | 2562.623 | -0.07716 | 0.50991 | 0.929089 | Clasp1 |
| ENSMUSG( | 7978.208 | -0.09012 | 0.383235 | 0.917257 | Clasp2 |
| ENSMUSG( | 2049.536 | 0.111096 | 0.537526 | 0.937216 | Clasrp |
| ENSMUSG( | 268.4834 | -0.14274 | 0.386031 | 0.917257 | Clba1 |
| ENSMUSG( | 21.12989 | 0.139761 | 0.778011 | 0.976106 | Clca3a1 |
| ENSMUSG( | 1123.55 | 0.061214 | 0.668517 | 0.956766 | Clcc1 |
| ENSMUSG( | 62.36628 | -0.00381 | 0.989435 | 0.997928 | Clcf1 |
| ENSMUSG( | 110.2557 | 0.157315 | 0.486832 | 0.92541 | Clcn1 |
| ENSMUSG( | 2457.161 | 0.255766 | 0.06199 | 0.736482 | Clcn2 |
| ENSMUSG( | 7769.795 | 0.04899 | 0.751193 | 0.970317 | Clcn3 |
| ENSMUSG( | 8844.505 | 0.064195 | 0.642465 | 0.953903 | Clcn4 |
| ENSMUSG( | 409.2985 | 0.241698 | 0.282617 | 0.892412 | Clcn5 |
| ENSMUSG( | 3465.133 | 0.053561 | 0.732749 | 0.967228 | Clcn6 |
| ENSMUSG( | 1971.967 | 0.172109 | 0.248125 | 0.877904 | Clcn7 |
| ENSMUSG( | 97.57006 | 0.283191 | 0.240611 | 0.875927 | Cldn1 |
| ENSMUSG( | 2524.563 | -0.16391 | 0.297097 | 0.898538 | Cldn10 |
| ENSMUSG( | 13071.56 | 0.175112 | 0.362299 | 0.914223 | Cldn11 |
| ENSMUSG( | 1306.01 | 0.247272 | 0.143069 | 0.831221 | Cldn12 |
| ENSMUSG( | 90.98513 | -0.20648 | 0.448421 | 0.922202 | Cldn14 |
| ENSMUSG( | 49.07594 | -0.71346 | 0.026761 | 0.603041 | Cldn19 |
| ENSMUSG( | 73.03097 | 0.679173 | 0.105218 | 0.804969 | Cldn2 |
| ENSMUSG( | 9.422801 | 0.477485 | 0.487435 | NA | Cldn20 |
| ENSMUSG( | 13.86273 | -0.67791 | 0.238378 | 0.872841 | Cldn22 |
| ENSMUSG( | 19.27774 | 0.04347 | 0.933283 | 0.99306 | Cldn23 |
| ENSMUSG( | 12.498 | -0.51978 | 0.356266 | 0.913358 | Cldn3 |
| ENSMUSG( | 61.88525 | -0.20913 | 0.451012 | 0.922202 | Cldn34c1 |
| ENSMUSG( | 1973.324 | 0.161367 | 0.449615 | 0.922202 | Cldn5 |
| ENSMUSG( | 3332.983 | 0.001009 | 0.9942 | 0.998813 | Cldnd1 |
| ENSMUSG( | 44.50817 | 0.215933 | 0.487045 | 0.92541 | Clec10a |
| ENSMUSG( | 219.4121 | -0.16581 | 0.310404 | 0.899182 | Clec11a |
| ENSMUSG( | 31.27681 | -0.26595 | 0.529774 | 0.934795 | Clec12a |
| ENSMUSG( | 125.4205 | 0.486339 | 0.062623 | 0.736482 | Clec14a |
| ENSMUSG( | 2544.09 | 0.045744 | 0.655295 | 0.955294 | Clec16a |
| ENSMUSG( | 112.6919 | -0.44004 | 0.05844 | 0.728256 | Clec18a |
| ENSMUSG( | 30.13148 | -0.04249 | 0.918479 | 0.992529 | Clec1a |
| ENSMUSG( | 231.6481 | 0.822438 | 0.000721 | 0.08668 | Clec2d |
| ENSMUSG( | 2767.693 | -0.04626 | 0.673165 | 0.95694 | Clec2l |
| ENSMUSG( | 128.7917 | 0.06614 | 0.742903 | 0.969415 | Clec3b |
| ENSMUSG( | 38.44845 | -0.59223 | 0.092553 | 0.793565 | Clec4a1 |
| ENSMUSG( | 60.91488 | 0.357603 | 0.196304 | 0.862161 | Clec4a2 |
| ENSMUSG( | 78.2013 | 0.044906 | 0.891569 | 0.990558 | Clec4a3 |
| ENSMUSG( | 11.21554 | -0.41723 | 0.50085 | NA | Clec4n |
| ENSMUSG( | 117.0433 | 0.115468 | 0.584944 | 0.944975 | Clec5a |
| ENSMUSG( | 24.6428 | -0.31225 | 0.442055 | 0.921554 | Clec7a |
| ENSMUSG( | 521.4776 | -0.0208 | 0.884369 | 0.989688 | Clgn |
| ENSMUSG( | 620.2892 | 0.097247 | 0.513734 | 0.930586 | Clic1 |
| ENSMUSG( | 11.5744 | -0.10887 | 0.859541 | 0.985947 | Clic3 |
| ENSMUSG( | 6739.997 | -0.1779 | 0.119246 | 0.813384 | Clic4 |
| ENSMUSG( | 345.3325 | 0.91966 | 0.060008 | 0.730482 | Clic5 |
| ENSMUSG( | 210.8219 | 0.042567 | 0.959498 | 0.994798 | Clic6 |
| ENSMUSG( | 1299.533 | 0.027816 | 0.798109 | 0.97681 | Clint1 |
| ENSMUSG( | 2679.5 | 0.22782 | 0.099121 | 0.799802 | Clip1 |
| ENSMUSG( | 3965.691 | -0.09895 | 0.347942 | 0.911652 | Clip2 |
| ENSMUSG( | 11473.63 | -0.05875 | 0.675063 | 0.95694 | Clip3 |
| ENSMUSG( | 1055.295 | 0.048022 | 0.70412 | 0.962677 | Clip4 |
| ENSMUSG( | 3633.813 | 0.374902 | 0.023254 | 0.585577 | Clk1 |
| ENSMUSG( | 905.3952 | 0.12953 | 0.388874 | 0.917257 | Clk2 |
| ENSMUSG( | 2034.226 | 0.111842 | 0.426203 | 0.917257 | Clk3 |
| ENSMUSG( | 1874.411 | 0.225855 | 0.089099 | 0.788502 | Clk4 |
| ENSMUSG( | 4783.572 | -0.00328 | 0.978072 | 0.996416 | Clmn |
| ENSMUSG( | 770.5585 | -0.04579 | 0.745506 | 0.96951 | Clmp |
| ENSMUSG( | 588.6217 | 0.087696 | 0.518425 | 0.930939 | Cln3 |
| ENSMUSG( | 480.3507 | 0.147527 | 0.380105 | 0.916995 | Cln5 |
| ENSMUSG( | 600.489 | 0.038979 | 0.794036 | 0.97681 | Cln6 |
| ENSMUSG( | 1196.254 | 0.122059 | 0.481484 | 0.924578 | Cln8 |
| ENSMUSG( | 1280.115 | -0.12743 | 0.326338 | 0.905371 | Clns1a |
| ENSMUSG( | 1640.652 | -0.0444 | 0.709924 | 0.96416 | Clock |
| ENSMUSG( | 98.86327 | -0.0806 | 0.76391 | 0.974 | Clp1 |
| ENSMUSG( | 1882.505 | -0.03649 | 0.751431 | 0.970317 | Clpb |
| ENSMUSG( | 1291.438 | -0.06264 | 0.627452 | 0.953234 | Clpp |
| ENSMUSG( | 8457.377 | 0.066691 | 0.662277 | 0.956205 | Clptm1 |
| ENSMUSG( | 4170.311 | 0.03438 | 0.812518 | 0.979148 | Clptm1l |
| ENSMUSG( | 730.2945 | -0.03178 | 0.80803 | 0.978622 | Clpx |
| ENSMUSG( | 13.89743 | 0.165134 | 0.74915 | 0.96998 | Clrn1 |
| ENSMUSG( | 57.88365 | -0.07573 | 0.794351 | 0.97681 | Clspn |
| ENSMUSG( | 50066.74 | 0.167694 | 0.314537 | 0.900949 | Clstn1 |
| ENSMUSG( | 2604.272 | 0.025503 | 0.878943 | 0.989644 | Clstn2 |
| ENSMUSG( | 13560.13 | 0.122489 | 0.438945 | 0.920721 | Clstn3 |
| ENSMUSG( | 3653.786 | -0.20908 | 0.127338 | 0.817359 | Clta |
| ENSMUSG( | 4426.408 | -0.284 | 0.020844 | 0.571913 | Cltb |
| ENSMUSG( | 10615.52 | -0.12262 | 0.247607 | 0.877904 | Cltc |
| ENSMUSG( | 85690.89 | 0.107748 | 0.416512 | 0.917257 | Clu |
| ENSMUSG( | 467.6884 | 0.042584 | 0.792714 | 0.97681 | Cluap1 |
| ENSMUSG( | 2785.022 | 0.057564 | 0.564798 | 0.941206 | Cluh |
| ENSMUSG( | 544.9856 | -0.14912 | 0.408664 | 0.917257 | Clvs1 |
| ENSMUSG( | 301.7574 | 0.074315 | 0.647795 | 0.95422 | Clvs2 |
| ENSMUSG( | 117.6788 | -0.21017 | 0.343611 | 0.910026 | Clxn |
| ENSMUSG( | 517.7802 | -0.21817 | 0.102008 | 0.801851 | Clybl |
| ENSMUSG( | 2064.067 | -0.08398 | 0.439678 | 0.920721 | Cmas |
| ENSMUSG( | 346.6934 | -0.36848 | 0.017125 | 0.52741 | Cmbl |
| ENSMUSG( | 223.3343 | -0.26984 | 0.117081 | 0.813384 | Cmc1 |
| ENSMUSG( | 241.514 | -0.1589 | 0.370312 | 0.915491 | Cmc2 |
| ENSMUSG( | 79.85659 | 0.201319 | 0.466308 | 0.923329 | Cmc4 |
| ENSMUSG( | 3514.716 | 0.093801 | 0.388893 | 0.917257 | Cmip |
| ENSMUSG( | 212.3923 | 0.418658 | 0.073129 | 0.763629 | Cmklr1 |
| ENSMUSG( | 20.6085 | 0.590888 | 0.201275 | 0.864412 | Cmklr2 |
| ENSMUSG( | 2021.081 | -0.02603 | 0.882421 | 0.989644 | Cmpk1 |
| ENSMUSG( | 691.8477 | -0.41181 | 0.002971 | 0.218646 | Cmpk2 |
| ENSMUSG( | 140.3149 | -0.28028 | 0.157248 | 0.837724 | Cmss1 |
| ENSMUSG( | 749.1783 | 0.173072 | 0.249541 | 0.877904 | Cmtm3 |
| ENSMUSG( | 5731.765 | 0.053383 | 0.677477 | 0.957106 | Cmtm4 |
| ENSMUSG( | 3061.748 | -0.16287 | 0.192779 | 0.859764 | Cmtm5 |
| ENSMUSG( | 2055.367 | -0.03498 | 0.790578 | 0.97681 | Cmtm6 |
| ENSMUSG( | 147.7835 | -0.02676 | 0.899453 | 0.990558 | Cmtm7 |
| ENSMUSG( | 73.17677 | 0.497332 | 0.103202 | 0.804311 | Cmtm8 |
| ENSMUSG( | 990.4986 | 0.158386 | 0.295732 | 0.898538 | Cmtr1 |
| ENSMUSG( | 87.55248 | 0.216601 | 0.360082 | 0.914223 | Cmtr2 |
| ENSMUSG( | 276.4102 | -0.12247 | 0.451529 | 0.922202 | Cmya5 |
| ENSMUSG( | 223.8446 | 0.034096 | 0.838589 | 0.981622 | Cnbd2 |
| ENSMUSG( | 4172.485 | 0.003795 | 0.976856 | 0.996416 | Cnbp |
| ENSMUSG( | 1228.697 | -0.11414 | 0.369009 | 0.915491 | Cndp2 |
| ENSMUSG( | 767.3733 | 0.065368 | 0.640075 | 0.953903 | Cnep1r1 |
| ENSMUSG( | 12.18578 | -0.06282 | 0.916625 | 0.992529 | Cnga2 |
| ENSMUSG( | 32.24419 | 0.573063 | 0.128911 | 0.820245 | Cnga4 |
| ENSMUSG( | 13.8211 | 0.258788 | 0.632467 | 0.953682 | Cngb1 |
| ENSMUSG( | 2308.287 | 0.008783 | 0.954065 | 0.994798 | Cnih1 |
| ENSMUSG( | 15195.01 | 0.014625 | 0.881732 | 0.989644 | Cnih2 |
| ENSMUSG( | 2214.298 | 0.096071 | 0.444537 | 0.921734 | Cnih3 |
| ENSMUSG( | 2320.141 | -0.01995 | 0.878872 | 0.989644 | Cnih4 |
| ENSMUSG( | 2773.719 | 0.286555 | 0.011865 | 0.445064 | Cnksr2 |
| ENSMUSG( | 371.4595 | 0.04147 | 0.790823 | 0.97681 | Cnksr3 |
| ENSMUSG( | 55.78725 | -0.04577 | 0.895409 | 0.990558 | Cnmd |
| ENSMUSG( | 275.2789 | 0.313273 | 0.045128 | 0.693959 | Cnn2 |
| ENSMUSG( | 4591.276 | -0.07042 | 0.587946 | 0.944975 | Cnn3 |

| | | | | | | | | | | | |
|---|---|---|---|---|---|---|---|---|---|---|---|
| ENSMUSG( | Ciz1 | 1709.12 | 0.270081 | 0.012865 | 0.163922 | ENSMUSG( | Ciz1 | 1476.705 | 0.124782 | 0.355156 | 0.912411 |
| ENSMUSG( | CK137956 | 59.89473 | -0.65159 | 0.00796 | 0.124138 | ENSMUSG( | CK137956 | 67.3723 | 0.010779 | 0.969866 | 0.99594 |
| ENSMUSG( | Ckap2 | 42.34878 | -0.48369 | 0.126723 | 0.534201 | ENSMUSG( | Ckap2 | 44.8509 | -0.01877 | 0.951591 | 0.994798 |
| ENSMUSG( | Ckap2l | 54.63984 | -0.3187 | 0.225285 | 0.672044 | ENSMUSG( | Ckap2l | 53.80888 | -0.07231 | 0.801535 | 0.97681 |
| ENSMUSG( | Ckap4 | 1120.252 | -0.19277 | 0.145429 | 0.566269 | ENSMUSG( | Ckap4 | 1150.869 | 0.160898 | 0.308769 | 0.898538 |
| ENSMUSG( | Ckap5 | 8679.452 | 0.160891 | 0.123577 | 0.530552 | ENSMUSG( | Ckap5 | 7149.62 | -0.12942 | 0.28568 | 0.893906 |
| ENSMUSG( | Ckb | 63859.18 | 0.03711 | 0.6929 | 0.934202 | ENSMUSG( | Ckb | 53059.11 | -0.24029 | 0.044258 | 0.693437 |
| ENSMUSG( | Cklf | 112.5718 | -0.26391 | 0.184183 | 0.621101 | ENSMUSG( | Cklf | 123.8536 | 0.299252 | 0.310859 | 0.899422 |
| ENSMUSG( | Ckmt1 | 3378.526 | -0.06319 | 0.634024 | 0.914073 | ENSMUSG( | Ckmt1 | 2877.041 | -0.27374 | 0.075312 | 0.767994 |
| ENSMUSG( | Cks1b | 57.40085 | 0.076234 | 0.762555 | 0.953519 | ENSMUSG( | Cks1b | 47.23892 | -0.24055 | 0.414212 | 0.917257 |
| ENSMUSG( | Cks2 | 22.51579 | -0.19301 | 0.668481 | 0.927864 | ENSMUSG( | Cks2 | 21.19199 | -0.13853 | 0.772604 | 0.974463 |
| ENSMUSG( | Clasp1 | 3006.529 | 0.114918 | 0.201158 | 0.641362 | ENSMUSG( | Clasp1 | 2562.623 | -0.07716 | 0.50991 | 0.929089 |
| ENSMUSG( | Clasp2 | 9274.888 | 0.076988 | 0.364498 | 0.793796 | ENSMUSG( | Clasp2 | 7978.208 | -0.09012 | 0.383235 | 0.917257 |
| ENSMUSG( | Clasrp | 2222.08 | 0.079002 | 0.599345 | 0.903801 | ENSMUSG( | Clasrp | 2049.536 | 0.111096 | 0.537526 | 0.937216 |
| ENSMUSG( | Clba1 | 303.3331 | -0.05415 | 0.694354 | 0.9343 | ENSMUSG( | Clba1 | 268.4834 | -0.14274 | 0.386031 | 0.917257 |
| ENSMUSG( | Clca3a1 | 18.93111 | -0.40051 | 0.368343 | NA | ENSMUSG( | Clca3a1 | 21.12989 | 0.139761 | 0.778011 | 0.976106 |
| ENSMUSG( | Clcc1 | 1141.686 | -0.1674 | 0.171472 | 0.605943 | ENSMUSG( | Clcc1 | 1123.55 | 0.061214 | 0.668517 | 0.956766 |
| ENSMUSG( | Clcf1 | 68.74442 | 0.00563 | 0.981356 | 0.997512 | ENSMUSG( | Clcf1 | 62.36628 | -0.00381 | 0.989435 | 0.997928 |
| ENSMUSG( | Clcn1 | 119.3782 | 0.108307 | 0.563236 | 0.889897 | ENSMUSG( | Clcn1 | 110.2557 | 0.157315 | 0.486832 | 0.92541 |
| ENSMUSG( | Clcn2 | 2620.6 | 0.176226 | 0.110019 | 0.500285 | ENSMUSG( | Clcn2 | 2457.161 | 0.255766 | 0.06199 | 0.736482 |
| ENSMUSG( | Clcn3 | 7771.304 | -0.22972 | 0.096899 | 0.477375 | ENSMUSG( | Clcn3 | 7769.795 | 0.04899 | 0.751193 | 0.970317 |
| ENSMUSG( | Clcn4 | 8813.379 | -0.22456 | 0.067209 | 0.400638 | ENSMUSG( | Clcn4 | 8844.505 | 0.064195 | 0.642465 | 0.953903 |
| ENSMUSG( | Clcn5 | 356.3565 | -0.46479 | 0.019307 | 0.209368 | ENSMUSG( | Clcn5 | 409.2985 | 0.241698 | 0.282617 | 0.892412 |
| ENSMUSG( | Clcn6 | 3408.13 | -0.28077 | 0.041748 | 0.315339 | ENSMUSG( | Clcn6 | 3465.133 | 0.053561 | 0.732749 | 0.967228 |
| ENSMUSG( | Clcn7 | 1937.952 | -0.14982 | 0.229705 | 0.677538 | ENSMUSG( | Clcn7 | 1971.967 | 0.172109 | 0.248125 | 0.877904 |
| ENSMUSG( | Cldn1 | 85.37629 | -0.38288 | 0.134489 | 0.54768 | ENSMUSG( | Cldn1 | 97.57006 | 0.283191 | 0.240611 | 0.875927 |
| ENSMUSG( | Cldn10 | 2779.512 | -0.15633 | 0.284827 | 0.733433 | ENSMUSG( | Cldn10 | 2524.563 | -0.16391 | 0.297097 | 0.898538 |
| ENSMUSG( | Cldn11 | 10667.46 | -0.78267 | 1.59E-07 | 3.86E-05 | ENSMUSG( | Cldn11 | 13071.56 | 0.175112 | 0.362299 | 0.914223 |
| ENSMUSG( | Cldn12 | 1140.824 | -0.43743 | 0.003369 | 0.073204 | ENSMUSG( | Cldn12 | 1306.01 | 0.247272 | 0.143069 | 0.831221 |
| ENSMUSG( | Cldn14 | 95.64746 | -0.35891 | 0.10131 | 0.484996 | ENSMUSG( | Cldn14 | 90.98513 | -0.20648 | 0.448421 | 0.922202 |
| ENSMUSG( | Cldn19 | 51.43726 | -0.85047 | 0.002315 | 0.056456 | ENSMUSG( | Cldn19 | 49.07594 | -0.71346 | 0.026761 | 0.603041 |
| ENSMUSG( | Cldn2 | 62.10573 | 0.033771 | 0.962822 | 0.994824 | ENSMUSG( | Cldn2 | 73.03097 | 0.679173 | 0.105218 | 0.804969 |
| ENSMUSG( | Cldn20 | 8.706004 | 0.049878 | 0.943414 | NA | ENSMUSG( | Cldn20 | 9.422801 | 0.477485 | 0.487435 | NA |
| ENSMUSG( | Cldn22 | 15.78845 | -0.54401 | 0.285869 | NA | ENSMUSG( | Cldn22 | 13.86273 | -0.67791 | 0.238378 | 0.872841 |
| ENSMUSG( | Cldn23 | 16.06602 | -0.90828 | 0.055464 | NA | ENSMUSG( | Cldn23 | 19.27774 | 0.04347 | 0.933283 | 0.99306 |
| ENSMUSG( | Cldn3 | 13.10486 | -0.66237 | 0.216084 | NA | ENSMUSG( | Cldn3 | 12.498 | -0.51978 | 0.356266 | 0.913358 |
| ENSMUSG( | Cldn34c1 | 65.84915 | -0.31801 | 0.196546 | 0.635942 | ENSMUSG( | Cldn34c1 | 61.88525 | -0.20913 | 0.451012 | 0.922202 |
| ENSMUSG( | Cldn5 | 1950.177 | -0.14878 | 0.442315 | 0.832774 | ENSMUSG( | Cldn5 | 1973.324 | 0.161367 | 0.449615 | 0.922202 |
| ENSMUSG( | Cldnd1 | 3419.752 | -0.20045 | 0.071253 | 0.412315 | ENSMUSG( | Cldnd1 | 3332.983 | 0.001009 | 0.9942 | 0.998813 |
| ENSMUSG( | Clec10a | 49.07152 | 0.242964 | 0.380414 | 0.801757 | ENSMUSG( | Clec10a | 44.50817 | 0.215933 | 0.487045 | 0.92541 |
| ENSMUSG( | Clec11a | 271.4226 | 0.175852 | 0.217119 | 0.661995 | ENSMUSG( | Clec11a | 219.4121 | -0.16581 | 0.310404 | 0.899182 |
| ENSMUSG( | Clec12a | 23.34332 | -1.93732 | 0.000124 | 0.007462 | ENSMUSG( | Clec12a | 31.27681 | -0.26595 | 0.529774 | 0.934795 |
| ENSMUSG( | Clec14a | 101.5122 | -0.40026 | 0.109777 | 0.500285 | ENSMUSG( | Clec14a | 125.4205 | 0.486339 | 0.062623 | 0.736482 |
| ENSMUSG( | Clec16a | 2927.491 | 0.176006 | 0.043244 | 0.318825 | ENSMUSG( | Clec16a | 2544.09 | 0.045744 | 0.655295 | 0.955294 |
| ENSMUSG( | Clec18a | 136.1264 | -0.14676 | 0.483426 | 0.856304 | ENSMUSG( | Clec18a | 112.6919 | -0.44004 | 0.05844 | 0.728256 |
| ENSMUSG( | Clec1a | 26.41011 | -0.83103 | 0.039269 | 0.305097 | ENSMUSG( | Clec1a | 30.13148 | -0.04249 | 0.918479 | 0.992529 |
| ENSMUSG( | Clec2d | 184.1652 | -0.01088 | 0.955031 | 0.993874 | ENSMUSG( | Clec2d | 231.6481 | 0.822438 | 0.000721 | 0.08668 |
| ENSMUSG( | Clec2l | 3097.045 | 0.009079 | 0.919876 | 0.990247 | ENSMUSG( | Clec2l | 2767.693 | -0.04626 | 0.673165 | 0.95694 |
| ENSMUSG( | Clec3b | 130.2098 | -0.17934 | 0.316601 | 0.763648 | ENSMUSG( | Clec3b | 128.7917 | 0.06614 | 0.742903 | 0.969415 |
| ENSMUSG( | Clec4a1 | 46.38507 | -0.2533 | 0.416341 | 0.82084 | ENSMUSG( | Clec4a1 | 38.44845 | -0.59223 | 0.092553 | 0.793565 |
| ENSMUSG( | Clec4a2 | 55.7492 | -0.16448 | 0.55823 | 0.888427 | ENSMUSG( | Clec4a2 | 60.91488 | 0.357603 | 0.196304 | 0.862161 |
| ENSMUSG( | Clec4a3 | 72.06554 | -0.50239 | 0.118295 | 0.520125 | ENSMUSG( | Clec4a3 | 78.2013 | 0.044906 | 0.891569 | 0.990558 |
| ENSMUSG( | Clec4n | 12.2945 | -0.49158 | 0.397227 | NA | ENSMUSG( | Clec4n | 11.21554 | -0.41723 | 0.50085 | NA |
| ENSMUSG( | Clec5a | 132.6719 | 0.202758 | 0.304061 | 0.752269 | ENSMUSG( | Clec5a | 117.0433 | 0.115468 | 0.584944 | 0.944975 |
| ENSMUSG( | Clec7a | 24.66307 | -0.60273 | 0.128239 | 0.536641 | ENSMUSG( | Clec7a | 24.6428 | -0.31225 | 0.442055 | 0.921554 |
| ENSMUSG( | Clgn | 511.1746 | -0.37284 | 0.004211 | 0.084333 | ENSMUSG( | Clgn | 521.4776 | -0.0208 | 0.884369 | 0.989688 |
| ENSMUSG( | Clic1 | 648.2984 | -0.05019 | 0.718479 | 0.941911 | ENSMUSG( | Clic1 | 620.2892 | 0.097247 | 0.513734 | 0.930586 |
| ENSMUSG( | Clic3 | 11.73031 | -0.31053 | 0.584137 | NA | ENSMUSG( | Clic3 | 11.5744 | -0.10887 | 0.859541 | 0.985947 |
| ENSMUSG( | Clic4 | 7786.698 | -0.02379 | 0.828808 | 0.969097 | ENSMUSG( | Clic4 | 6739.997 | -0.1779 | 0.119246 | 0.813384 |
| ENSMUSG( | Clic5 | 293.2385 | 0.295875 | 0.216141 | 0.661017 | ENSMUSG( | Clic5 | 345.3325 | 0.91966 | 0.060008 | 0.730482 |
| ENSMUSG( | Clic6 | 214.7135 | -0.1665 | 0.849645 | 0.972638 | ENSMUSG( | Clic6 | 210.8219 | 0.042567 | 0.959498 | 0.994798 |
| ENSMUSG( | Clint1 | 1425.135 | 0.026208 | 0.783067 | 0.957558 | ENSMUSG( | Clint1 | 1299.533 | 0.027816 | 0.798109 | 0.97681 |
| ENSMUSG( | Clip1 | 2775.797 | 0.06699 | 0.522109 | 0.875186 | ENSMUSG( | Clip1 | 2679.5 | 0.22782 | 0.099121 | 0.799802 |
| ENSMUSG( | Clip2 | 4402.548 | -0.06643 | 0.44351 | 0.833108 | ENSMUSG( | Clip2 | 3965.691 | -0.09895 | 0.347942 | 0.911652 |
| ENSMUSG( | Clip3 | 13384.6 | 0.116666 | 0.258614 | 0.709814 | ENSMUSG( | Clip3 | 11473.63 | -0.05875 | 0.675063 | 0.95694 |
| ENSMUSG( | Clip4 | 1126.926 | -0.02789 | 0.769567 | 0.954674 | ENSMUSG( | Clip4 | 1055.295 | 0.048022 | 0.70412 | 0.962677 |
| ENSMUSG( | Clk1 | 3322.513 | -0.13204 | 0.215646 | 0.660333 | ENSMUSG( | Clk1 | 3633.813 | 0.374902 | 0.023254 | 0.585577 |
| ENSMUSG( | Clk2 | 1000.602 | 0.148885 | 0.201747 | 0.6415 | ENSMUSG( | Clk2 | 905.3952 | 0.12953 | 0.388874 | 0.917257 |
| ENSMUSG( | Clk3 | 2113.634 | -0.04691 | 0.622172 | 0.911432 | ENSMUSG( | Clk3 | 2034.226 | 0.111842 | 0.426203 | 0.917257 |
| ENSMUSG( | Clk4 | 1831.656 | -0.10534 | 0.369196 | 0.796641 | ENSMUSG( | Clk4 | 1874.411 | 0.225855 | 0.089099 | 0.788502 |
| ENSMUSG( | Clmn | 5372.511 | 0.060601 | 0.487886 | 0.858824 | ENSMUSG( | Clmn | 4783.572 | -0.00328 | 0.978072 | 0.996416 |
| ENSMUSG( | Clmp | 726.1943 | -0.53434 | 3.41E-05 | 0.002956 | ENSMUSG( | Clmp | 770.5585 | -0.04579 | 0.745506 | 0.96951 |
| ENSMUSG( | Cln3 | 560.2289 | -0.34657 | 0.002456 | 0.058231 | ENSMUSG( | Cln3 | 588.6217 | 0.087696 | 0.518425 | 0.930939 |
| ENSMUSG( | Cln5 | 459.4054 | -0.26553 | 0.087733 | 0.454663 | ENSMUSG( | Cln5 | 480.3507 | 0.147527 | 0.380105 | 0.916995 |
| ENSMUSG( | Cln6 | 573.6429 | -0.39114 | 0.00316 | 0.06974 | ENSMUSG( | Cln6 | 600.489 | 0.038979 | 0.794036 | 0.97681 |
| ENSMUSG( | Cln8 | 1136.804 | -0.31324 | 0.04943 | 0.339723 | ENSMUSG( | Cln8 | 1196.254 | 0.122059 | 0.481484 | 0.924578 |
| ENSMUSG( | Clns1a | 1487.518 | 0.043842 | 0.640427 | 0.916434 | ENSMUSG( | Clns1a | 1280.115 | -0.12743 | 0.326338 | 0.905371 |
| ENSMUSG( | Clock | 1878.215 | 0.07876 | 0.372492 | 0.798768 | ENSMUSG( | Clock | 1640.652 | -0.0444 | 0.709924 | 0.96416 |
| ENSMUSG( | Clp1 | 113.5385 | 0.035519 | 0.878498 | 0.981431 | ENSMUSG( | Clp1 | 98.86327 | -0.0806 | 0.76391 | 0.974 |
| ENSMUSG( | Clpb | 2166.507 | 0.097513 | 0.329718 | 0.773619 | ENSMUSG( | Clpb | 1882.505 | -0.03649 | 0.751431 | 0.970317 |
| ENSMUSG( | Clpp | 1477.551 | 0.061297 | 0.50743 | 0.86795 | ENSMUSG( | Clpp | 1291.438 | -0.06264 | 0.627452 | 0.953234 |
| ENSMUSG( | Clptm1 | 8390.923 | -0.23755 | 0.072111 | 0.414057 | ENSMUSG( | Clptm1 | 8457.377 | 0.066691 | 0.662277 | 0.956205 |
| ENSMUSG( | Clptm1l | 4181.742 | -0.23927 | 0.060385 | 0.3797 | ENSMUSG( | Clptm1l | 4170.311 | 0.03438 | 0.812518 | 0.979148 |
| ENSMUSG( | Clpx | 843.5138 | 0.117667 | 0.221438 | 0.667258 | ENSMUSG( | Clpx | 730.2945 | -0.03178 | 0.80803 | 0.978622 |
| ENSMUSG( | Clrn1 | 12.80944 | -0.34275 | 0.49962 | NA | ENSMUSG( | Clrn1 | 13.89743 | 0.165134 | 0.74915 | 0.96998 |
| ENSMUSG( | Clspn | 61.23392 | -0.20854 | 0.408872 | 0.818173 | ENSMUSG( | Clspn | 57.88365 | -0.07573 | 0.794351 | 0.97681 |
| ENSMUSG( | Clstn1 | 48799.67 | -0.18029 | 0.20609 | 0.647953 | ENSMUSG( | Clstn1 | 50066.74 | 0.167694 | 0.314537 | 0.900949 |
| ENSMUSG( | Clstn2 | 2546.227 | -0.3306 | 0.019214 | 0.208885 | ENSMUSG( | Clstn2 | 2604.272 | 0.025503 | 0.878943 | 0.989644 |
| ENSMUSG( | Clstn3 | 12964.2 | -0.29252 | 0.035623 | 0.290384 | ENSMUSG( | Clstn3 | 13560.13 | 0.122489 | 0.438945 | 0.920721 |
| ENSMUSG( | Clta | 4326.456 | 0.020874 | 0.83452 | 0.970344 | ENSMUSG( | Clta | 3653.786 | -0.20908 | 0.127338 | 0.817359 |
| ENSMUSG( | Cltb | 5193.757 | -0.07568 | 0.424779 | 0.824675 | ENSMUSG( | Cltb | 4426.408 | -0.284 | 0.020844 | 0.571913 |
| ENSMUSG( | Cltc | 12271.9 | 0.030306 | 0.733179 | 0.944956 | ENSMUSG( | Cltc | 10615.52 | -0.12262 | 0.247607 | 0.877904 |
| ENSMUSG( | Clu | 94301.22 | 0.112925 | 0.342183 | 0.778828 | ENSMUSG( | Clu | 85690.89 | 0.107748 | 0.416512 | 0.917257 |
| ENSMUSG( | Cluap1 | 510.0621 | 0.030787 | 0.80487 | 0.962473 | ENSMUSG( | Cluap1 | 467.6884 | 0.042584 | 0.792714 | 0.97681 |
| ENSMUSG( | Cluh | 3118.734 | 0.113212 | 0.158832 | 0.588112 | ENSMUSG( | Cluh | 2785.022 | 0.057564 | 0.564798 | 0.941206 |
| ENSMUSG( | Clvs1 | 601.4293 | -0.12567 | 0.39432 | 0.810014 | ENSMUSG( | Clvs1 | 544.9856 | -0.14912 | 0.408664 | 0.917257 |
| ENSMUSG( | Clvs2 | 346.1323 | 0.199937 | 0.188849 | 0.627678 | ENSMUSG( | Clvs2 | 301.7574 | 0.074315 | 0.647795 | 0.95422 |
| ENSMUSG( | Clxn | 129.2021 | -0.21799 | 0.252802 | 0.703255 | ENSMUSG( | Clxn | 117.6788 | -0.21017 | 0.343611 | 0.910026 |
| ENSMUSG( | Clybl | 612.8338 | 0.012596 | 0.922108 | 0.99045 | ENSMUSG( | Clybl | 517.7802 | -0.21817 | 0.102008 | 0.801851 |
| ENSMUSG( | Cmas | 2251.04 | -0.10154 | 0.235778 | 0.682914 | ENSMUSG( | Cmas | 2064.067 | -0.08398 | 0.439678 | 0.920721 |
| ENSMUSG( | Cmbl | 429.7682 | 0.014679 | 0.922487 | 0.99045 | ENSMUSG( | Cmbl | 346.6934 | -0.36848 | 0.017125 | 0.52741 |
| ENSMUSG( | Cmc1 | 259.2244 | -0.09035 | 0.557685 | 0.888397 | ENSMUSG( | Cmc1 | 223.3343 | -0.26984 | 0.117081 | 0.813384 |
| ENSMUSG( | Cmc2 | 283.6745 | 0.05177 | 0.724782 | 0.944096 | ENSMUSG( | Cmc2 | 241.514 | -0.1589 | 0.370312 | 0.915491 |
| ENSMUSG( | Cmc4 | 83.2918 | 0.081284 | 0.743305 | 0.948478 | ENSMUSG( | Cmc4 | 79.85659 | 0.201319 | 0.466308 | 0.923329 |
| ENSMUSG( | Cmip | 4110.876 | 0.265413 | 0.009291 | 0.136006 | ENSMUSG( | Cmip | 3514.716 | 0.093801 | 0.388893 | 0.917257 |
| ENSMUSG( | Cmklr1 | 195.5101 | -0.07532 | 0.686621 | 0.933041 | ENSMUSG( | Cmklr1 | 212.3923 | 0.418658 | 0.073129 | 0.763629 |
| ENSMUSG( | Cmklr2 | 18.16081 | -0.00864 | 0.985164 | NA | ENSMUSG( | Cmklr2 | 20.6085 | 0.590888 | 0.201275 | 0.864412 |
| ENSMUSG( | Cmpk1 | 2238.799 | 0.006048 | 0.958234 | 0.993874 | ENSMUSG( | Cmpk1 | 2021.081 | -0.02603 | 0.882421 | 0.989644 |
| ENSMUSG( | Cmpk2 | 826.7804 | -0.13438 | 0.259413 | 0.709948 | ENSMUSG( | Cmpk2 | 691.8477 | -0.41181 | 0.002971 | 0.218646 |
| ENSMUSG( | Cmss1 | 170.8966 | 0.043528 | 0.78343 | 0.957558 | ENSMUSG( | Cmss1 | 140.3149 | -0.28028 | 0.157248 | 0.837724 |
| ENSMUSG( | Cmtm3 | 785.2533 | 0.038455 | 0.776588 | 0.955658 | ENSMUSG( | Cmtm3 | 749.1783 | 0.173072 | 0.249541 | 0.877904 |
| ENSMUSG( | Cmtm4 | 6236.231 | 0.026851 | 0.813465 | 0.964574 | ENSMUSG( | Cmtm4 | 5731.765 | 0.053383 | 0.677477 | 0.957106 |
| ENSMUSG( | Cmtm5 | 3296.773 | -0.22244 | 0.050064 | 0.342055 | ENSMUSG( | Cmtm5 | 3061.748 | -0.16287 | 0.192779 | 0.859764 |
| ENSMUSG( | Cmtm6 | 2193.253 | -0.12047 | 0.300157 | 0.749354 | ENSMUSG( | Cmtm6 | 2055.367 | -0.03498 | 0.790578 | 0.97681 |
| ENSMUSG( | Cmtm7 | 165.6229 | 0.017704 | 0.919925 | 0.990247 | ENSMUSG( | Cmtm7 | 147.7835 | -0.02676 | 0.899453 | 0.990558 |
| ENSMUSG( | Cmtm8 | 60.16929 | -0.34889 | 0.174992 | 0.610399 | ENSMUSG( | Cmtm8 | 73.17677 | 0.497332 | 0.103202 | 0.804311 |
| ENSMUSG( | Cmtr1 | 1033.966 | 0.012115 | 0.927847 | 0.991229 | ENSMUSG( | Cmtr1 | 990.4986 | 0.158386 | 0.295732 | 0.898538 |
| ENSMUSG( | Cmtr2 | 88.17447 | -0.02675 | 0.907402 | 0.987566 | ENSMUSG( | Cmtr2 | 87.55248 | 0.216601 | 0.360082 | 0.914223 |
| ENSMUSG( | Cmya5 | 342.8856 | 0.216093 | 0.134139 | 0.547472 | ENSMUSG( | Cmya5 | 276.4102 | -0.12247 | 0.451529 | 0.922202 |
| ENSMUSG( | Cnbd2 | 259.3255 | 0.178157 | 0.217391 | 0.662547 | ENSMUSG( | Cnbd2 | 223.8446 | 0.034096 | 0.838589 | 0.981622 |
| ENSMUSG( | Cnbp | 4576.044 | 0.004022 | 0.967151 | 0.995949 | ENSMUSG( | Cnbp | 4172.485 | 0.003795 | 0.976856 | 0.996416 |
| ENSMUSG( | Cndp2 | 1398.87 | -0.00359 | 0.973556 | 0.99675 | ENSMUSG( | Cndp2 | 1228.697 | -0.11414 | 0.369009 | 0.915491 |
| ENSMUSG( | Cnep1r1 | 764.8124 | -0.22375 | 0.06718 | 0.400611 | ENSMUSG( | Cnep1r1 | 767.3733 | 0.065368 | 0.640075 | 0.953903 |
| ENSMUSG( | Cnga2 | 14.69898 | 0.24951 | 0.61817 | NA | ENSMUSG( | Cnga2 | 12.18578 | -0.06282 | 0.916625 | 0.992529 |
| ENSMUSG( | Cnga4 | 31.02103 | 0.24465 | 0.483396 | 0.856304 | ENSMUSG( | Cnga4 | 32.24419 | 0.573063 | 0.128911 | 0.820245 |
| ENSMUSG( | Cngb1 | 13.05847 | -0.16478 | 0.76767 | NA | ENSMUSG( | Cngb1 | 13.8211 | 0.258788 | 0.632467 | 0.953682 |
| ENSMUSG( | Cnih1 | 2254.812 | -0.35088 | 0.011441 | 0.153933 | ENSMUSG( | Cnih1 | 2308.287 | 0.008783 | 0.954065 | 0.994798 |
| ENSMUSG( | Cnih2 | 17107.37 | 0.086256 | 0.279911 | 0.729155 | ENSMUSG( | Cnih2 | 15195.01 | 0.014625 | 0.881732 | 0.989644 |
| ENSMUSG( | Cnih3 | 2243.591 | -0.13874 | 0.197376 | 0.637004 | ENSMUSG( | Cnih3 | 2214.298 | 0.096071 | 0.444537 | 0.921734 |
| ENSMUSG( | Cnih4 | 2490.484 | -0.08381 | 0.429769 | 0.826519 | ENSMUSG( | Cnih4 | 2320.141 | -0.01995 | 0.878872 | 0.989644 |
| ENSMUSG( | Cnksr2 | 2906.925 | 0.162446 | 0.110726 | 0.501952 | ENSMUSG( | Cnksr2 | 2773.719 | 0.286555 | 0.011865 | 0.445064 |
| ENSMUSG( | Cnksr3 | 417.1675 | 0.109953 | 0.346321 | 0.779153 | ENSMUSG( | Cnksr3 | 371.4595 | 0.04147 | 0.790823 | 0.97681 |
| ENSMUSG( | Cnmd | 55.88709 | -0.34437 | 0.301998 | 0.750166 | ENSMUSG( | Cnmd | 55.78725 | -0.04577 | 0.895409 | 0.990558 |
| ENSMUSG( | Cnn2 | 279.1611 | 0.092848 | 0.513201 | 0.871762 | ENSMUSG( | Cnn2 | 275.2789 | 0.313273 | 0.045128 | 0.693959 |
| ENSMUSG( | Cnn3 | 5444.404 | 0.152046 | 0.16052 | 0.590124 | ENSMUSG( | Cnn3 | 4591.276 | -0.07042 | 0.587946 | 0.944975 |
| ENSMUSG( | Cnnm1 | 3740.487 | -0.29517 | 0.010111 | 0.143118 | ENSMUSG( | Cnnm1 | 3853.054 | 0.076213 | 0.555107 | 0.939565 |
| ENSMUSG( | Cnnm2 | 894.8336 | -0.21939 | 0.130657 | 0.54201 | ENSMUSG( | Cnnm2 | 952.7644 | 0.235765 | 0.172441 | 0.843608 |
| ENSMUSG( | Cnnm3 | 1493.825 | -0.23866 | 0.060797 | 0.38089 | ENSMUSG( | Cnnm3 | 1500.008 | 0.055303 | 0.712124 | 0.964363 |
| ENSMUSG( | Cnnm4 | 492.5008 | -0.23725 | 0.139136 | 0.555202 | ENSMUSG( | Cnnm4 | 530.9946 | 0.253998 | 0.142604 | 0.831221 |
| ENSMUSG( | Cnot1 | 2561.992 | 0.217356 | 0.007841 | 0.123 | ENSMUSG( | Cnot1 | 2195.477 | 0.049144 | 0.631864 | 0.953251 |
| ENSMUSG( | Cnot10 | 1407.048 | 0.032932 | 0.786353 | 0.958843 | ENSMUSG( | Cnot10 | 1337.956 | 0.154059 | 0.262547 | 0.885357 |
| ENSMUSG( | Cnot11 | 666.6512 | 0.00975 | 0.934935 | 0.993231 | ENSMUSG( | Cnot11 | 607.3188 | 0.009506 | 0.942729 | 0.994774 |
| ENSMUSG( | Cnot2 | 1177.852 | 0.137539 | 0.15548 | 0.583591 | ENSMUSG( | Cnot2 | 1079.679 | 0.151097 | 0.221604 | 0.866104 |
| ENSMUSG( | Cnot3 | 971.811 | 0.09467 | 0.393465 | 0.809501 | ENSMUSG( | Cnot3 | 914.1597 | 0.185312 | 0.227032 | 0.868507 |
| ENSMUSG( | Cnot4 | 672.9098 | 0.032057 | 0.787302 | 0.959147 | ENSMUSG( | Cnot4 | 599.5417 | -0.03744 | 0.788552 | 0.976395 |
| ENSMUSG( | Cnot6 | 1885.983 | 0.008707 | 0.939942 | 0.993415 | ENSMUSG( | Cnot6 | 1743.42 | 0.048075 | 0.712352 | 0.964363 |
| ENSMUSG( | Cnot6l | 825.9561 | 0.049633 | 0.620552 | 0.911432 | ENSMUSG( | Cnot6l | 790.9838 | 0.188006 | 0.141562 | 0.830678 |
| ENSMUSG( | Cnot7 | 1878.854 | 0.001949 | 0.985025 | 0.997831 | ENSMUSG( | Cnot7 | 1745.134 | 0.052743 | 0.700159 | 0.961335 |
| ENSMUSG( | Cnot8 | 1276.987 | 0.098129 | 0.360667 | 0.79247 | ENSMUSG( | Cnot8 | 1122.216 | -0.0067 | 0.95939 | 0.994798 |
| ENSMUSG( | Cnot9 | 658.6529 | -0.03078 | 0.791845 | 0.959461 | ENSMUSG( | Cnot9 | 606.8436 | 0.002003 | 0.987329 | 0.99751 |
| ENSMUSG( | Cnp | 19610.45 | 0.031551 | 0.72419 | 0.944096 | ENSMUSG( | Cnp | 16472.1 | -0.2127 | 0.083071 | 0.77681 |
| ENSMUSG( | Cnppd1 | 1082.258 | -0.10821 | 0.303873 | 0.752148 | ENSMUSG( | Cnppd1 | 1065.024 | 0.118411 | 0.351405 | 0.912093 |
| ENSMUSG( | Cnpy2 | 1342.095 | -0.25829 | 0.038045 | 0.300991 | ENSMUSG( | Cnpy2 | 1309.698 | -0.04676 | 0.733326 | 0.967228 |
| ENSMUSG( | Cnpy3 | 2909.019 | -0.33582 | 0.003428 | 0.073753 | ENSMUSG( | Cnpy3 | 2938.535 | -0.01653 | 0.89896 | 0.990558 |
| ENSMUSG( | Cnpy4 | 490.0882 | -0.16511 | 0.245517 | 0.693175 | ENSMUSG( | Cnpy4 | 498.5628 | 0.154731 | 0.320594 | 0.903887 |
| ENSMUSG( | Cnr1 | 3459.649 | -0.599 | 3.23E-05 | 0.002865 | ENSMUSG( | Cnr1 | 4191.5 | 0.275786 | 0.108512 | 0.804969 |
| ENSMUSG( | Cnr2 | 9.584655 | -0.16015 | 0.813929 | NA | ENSMUSG( | Cnr2 | 10.28948 | 0.346584 | 0.628484 | NA |
| ENSMUSG( | Cnrip1 | 1487.599 | 0.044192 | 0.740896 | 0.948028 | ENSMUSG( | Cnrip1 | 1293.548 | -0.09718 | 0.576733 | 0.942912 |

| | | | | | |
|---|---|---|---|---|---|
| ENSMUSG( | 13.05847 | -0.16478 | 0.76767 | NA | Cngb1 |
| ENSMUSG( | 2254.812 | -0.35088 | 0.011441 | 0.153933 | Cnih1 |
| ENSMUSG( | 17107.37 | 0.086256 | 0.279911 | 0.729155 | Cnih2 |
| ENSMUSG( | 2243.591 | -0.13874 | 0.197376 | 0.637004 | Cnih3 |
| ENSMUSG( | 2490.484 | -0.08381 | 0.429769 | 0.826519 | Cnih4 |
| ENSMUSG( | 2906.925 | 0.162446 | 0.110726 | 0.501952 | Cnksr2 |
| ENSMUSG( | 417.1675 | 0.109953 | 0.346321 | 0.779153 | Cnksr3 |
| ENSMUSG( | 55.88709 | -0.34437 | 0.301998 | 0.750166 | Cnmd |
| ENSMUSG( | 279.1611 | 0.092848 | 0.513201 | 0.871762 | Cnn2 |
| ENSMUSG( | 5444.404 | 0.152046 | 0.16052 | 0.590124 | Cnn3 |
| ENSMUSG( | 3740.487 | -0.29517 | 0.010111 | 0.143118 | Cnnm1 |
| ENSMUSG( | 894.8336 | -0.21939 | 0.130657 | 0.54201 | Cnnm2 |
| ENSMUSG( | 1493.825 | -0.23866 | 0.060797 | 0.38089 | Cnnm3 |
| ENSMUSG( | 492.5008 | -0.23725 | 0.139136 | 0.555202 | Cnnm4 |
| ENSMUSG( | 2561.992 | 0.217356 | 0.007841 | 0.123 | Cnot1 |
| ENSMUSG( | 1407.048 | 0.032932 | 0.786353 | 0.958843 | Cnot10 |
| ENSMUSG( | 666.6512 | 0.00975 | 0.934935 | 0.993231 | Cnot11 |
| ENSMUSG( | 1177.852 | 0.137539 | 0.15548 | 0.583591 | Cnot2 |
| ENSMUSG( | 971.811 | 0.09467 | 0.393465 | 0.809501 | Cnot3 |
| ENSMUSG( | 672.9098 | 0.032057 | 0.787302 | 0.959147 | Cnot4 |
| ENSMUSG( | 1885.983 | 0.008707 | 0.939942 | 0.993415 | Cnot6 |
| ENSMUSG( | 825.9561 | 0.049633 | 0.620552 | 0.911432 | Cnot6l |
| ENSMUSG( | 1878.854 | 0.001949 | 0.985025 | 0.997831 | Cnot7 |
| ENSMUSG( | 1276.987 | 0.098129 | 0.360667 | 0.79247 | Cnot8 |
| ENSMUSG( | 658.6529 | -0.03078 | 0.791845 | 0.959461 | Cnot9 |
| ENSMUSG( | 19610.45 | 0.031551 | 0.72419 | 0.944096 | Cnp |
| ENSMUSG( | 1082.258 | -0.10821 | 0.303873 | 0.752148 | Cnppd1 |
| ENSMUSG( | 1342.095 | -0.25829 | 0.038045 | 0.300991 | Cnpy2 |
| ENSMUSG( | 2909.019 | -0.33582 | 0.003428 | 0.073753 | Cnpy3 |
| ENSMUSG( | 490.0882 | -0.16511 | 0.245517 | 0.693175 | Cnpy4 |
| ENSMUSG( | 3459.649 | -0.599 | 3.23E-05 | 0.002865 | Cnr1 |
| ENSMUSG( | 9.584655 | -0.16015 | 0.813929 | NA | Cnr2 |
| ENSMUSG( | 1487.599 | 0.044192 | 0.740896 | 0.948028 | Cnrip1 |
| ENSMUSG( | 1098.814 | -0.07905 | 0.450246 | 0.835344 | Cnst |
| ENSMUSG( | 20.77359 | -0.37128 | 0.385632 | NA | Cntd1 |
| ENSMUSG( | 5696.551 | -0.30132 | 0.007411 | 0.118765 | Cntfr |
| ENSMUSG( | 590.6697 | -0.0517 | 0.655937 | 0.923929 | Cntln |
| ENSMUSG( | 12779.29 | -0.31118 | 0.02871 | 0.258426 | Cntn1 |
| ENSMUSG( | 4162.47 | -0.57737 | 0.00017 | 0.009401 | Cntn2 |
| ENSMUSG( | 685.5902 | -0.68783 | 6.37E-06 | 0.000807 | Cntn3 |
| ENSMUSG( | 816.4248 | -0.58511 | 0.0001 | 0.006438 | Cntn4 |
| ENSMUSG( | 151.3585 | -0.159 | 0.434406 | 0.828978 | Cntn5 |
| ENSMUSG( | 672.9356 | -0.11837 | 0.247585 | 0.696522 | Cntn6 |
| ENSMUSG( | 9024.274 | -0.21382 | 0.07266 | 0.416299 | Cntnap1 |
| ENSMUSG( | 2789.368 | -0.44476 | 0.000514 | 0.020832 | Cntnap2 |
| ENSMUSG( | 221.7294 | -0.47483 | 0.007789 | 0.12245 | Cntnap3 |
| ENSMUSG( | 587.5882 | -0.47106 | 0.005354 | 0.097684 | Cntnap4 |
| ENSMUSG( | 877.8855 | -0.49203 | 0.003832 | 0.079288 | Cntnap5a |
| ENSMUSG( | 98.14938 | -0.35571 | 0.163359 | 0.594631 | Cntnap5b |
| ENSMUSG( | 35.20901 | 0.008407 | 0.978841 | 0.997428 | Cntnap5c |
| ENSMUSG( | 642.0709 | 0.01612 | 0.885553 | 0.983008 | Cntrl |
| ENSMUSG( | 502.3708 | 0.188565 | 0.170722 | 0.605271 | Cntrob |
| ENSMUSG( | 1318.895 | -0.07492 | 0.434639 | 0.828978 | Coa3 |
| ENSMUSG( | 97.7054 | 0.104087 | 0.600969 | 0.904492 | Coa4 |
| ENSMUSG( | 5962.729 | 0.047603 | 0.6706 | 0.928683 | Coa5 |
| ENSMUSG( | 311.3609 | -0.12798 | 0.398702 | 0.812878 | Coa6 |
| ENSMUSG( | 299.9002 | 0.137306 | 0.358723 | 0.790019 | Coa7 |
| ENSMUSG( | 914.2864 | 0.087188 | 0.375753 | 0.800976 | Coa8 |
| ENSMUSG( | 915.1483 | 0.178636 | 0.069865 | 0.408574 | Coasy |
| ENSMUSG( | 1744.966 | 0.23351 | 0.029961 | 0.262543 | Cobl |
| ENSMUSG( | 308.3513 | -0.33865 | 0.016649 | 0.192198 | Cobll1 |
| ENSMUSG( | 1267.2 | -0.43539 | 0.001376 | 0.040585 | Coch |
| ENSMUSG( | 1258.001 | 0.138179 | 0.216457 | 0.661421 | Cog1 |
| ENSMUSG( | 809.425 | -0.01443 | 0.90194 | 0.985633 | Cog2 |
| ENSMUSG( | 1351.219 | 0.081795 | 0.390476 | 0.808053 | Cog3 |
| ENSMUSG( | 1300.89 | 0.169838 | 0.111946 | 0.50438 | Cog4 |
| ENSMUSG( | 843.0985 | 0.033453 | 0.737801 | 0.947329 | Cog5 |
| ENSMUSG( | 1310.222 | 0.079289 | 0.38287 | 0.803237 | Cog6 |
| ENSMUSG( | 1588.128 | 0.119717 | 0.230028 | 0.678005 | Cog7 |
| ENSMUSG( | 722.8955 | 0.087388 | 0.429517 | 0.826446 | Cog8 |
| ENSMUSG( | 363.7846 | 0.1183 | 0.358571 | 0.789791 | Coil |
| ENSMUSG( | 9.10504 | -1.22433 | 0.133525 | NA | Col10a1 |
| ENSMUSG( | 455.7759 | 0.339175 | 0.105225 | 0.492417 | Col11a1 |
| ENSMUSG( | 1725.555 | -0.1024 | 0.36855 | 0.796202 | Col11a2 |
| ENSMUSG( | 850.8485 | -0.00769 | 0.950006 | 0.993821 | Col12a1 |
| ENSMUSG( | 61.05371 | 0.157209 | 0.519808 | 0.874979 | Col13a1 |
| ENSMUSG( | 76.26932 | -0.45722 | 0.059062 | 0.375388 | Col14a1 |
| ENSMUSG( | 273.8456 | 0.085168 | 0.551075 | 0.886038 | Col15a1 |
| ENSMUSG( | 866.4556 | 0.136637 | 0.330255 | 0.774247 | Col16a1 |
| ENSMUSG( | 21.52258 | -0.79974 | 0.057607 | 0.370247 | Col17a1 |
| ENSMUSG( | 175.3533 | 0.08435 | 0.622288 | 0.911432 | Col18a1 |
| ENSMUSG( | 341.1475 | 0.059362 | 0.711321 | 0.940028 | Col19a1 |
| ENSMUSG( | 1340.683 | -0.16627 | 0.094283 | 0.471647 | Col1a1 |
| ENSMUSG( | 1784.31 | 0.007168 | 0.95144 | 0.993821 | Col1a2 |
| ENSMUSG( | 206.4079 | 0.58056 | 0.005035 | 0.094254 | Col20a1 |
| ENSMUSG( | 86.22791 | -0.06726 | 0.783006 | 0.957558 | Col22a1 |
| ENSMUSG( | 994.3072 | 0.115171 | 0.242996 | 0.690307 | Col23a1 |
| ENSMUSG( | 220.7661 | 0.594271 | 0.257307 | 0.708457 | Col24a1 |
| ENSMUSG( | 1107.331 | -0.47106 | 0.000329 | 0.015441 | Col25a1 |
| ENSMUSG( | 354.0344 | 0.101166 | 0.436638 | 0.829402 | Col26a1 |
| ENSMUSG( | 434.485 | 0.397009 | 0.008584 | 0.129874 | Col27a1 |
| ENSMUSG( | 31.06609 | 0.642686 | 0.133932 | 0.547197 | Col28a1 |
| ENSMUSG( | 40.22893 | 0.082427 | 0.784476 | 0.958003 | Col2a1 |
| ENSMUSG( | 501.4583 | -0.22933 | 0.172958 | 0.608361 | Col3a1 |
| ENSMUSG( | 2894.654 | 0.105872 | 0.520665 | 0.875186 | Col4a1 |
| ENSMUSG( | 2721.529 | 0.33265 | 0.034189 | 0.283507 | Col4a2 |
| ENSMUSG( | 33.30745 | 0.109629 | 0.788592 | 0.959226 | Col4a3 |
| ENSMUSG( | 63.34289 | 0.014516 | 0.958822 | 0.993874 | Col4a4 |
| ENSMUSG( | 343.9591 | 0.203211 | 0.168335 | 0.601667 | Col4a5 |
| ENSMUSG( | 79.68278 | 0.012102 | 0.95853 | 0.993874 | Col4a6 |
| ENSMUSG( | 1144.404 | 0.563888 | 0.16397 | 0.594869 | Col5a1 |
| ENSMUSG( | 312.2899 | 0.339933 | 0.014588 | 0.177737 | Col5a2 |
| ENSMUSG( | 175.9666 | 0.480646 | 0.030828 | 0.266969 | Col5a3 |
| ENSMUSG( | 2301.512 | -0.04851 | 0.795922 | 0.960062 | Col6a1 |
| ENSMUSG( | 1216.814 | -0.28249 | 0.002183 | 0.054296 | Col6a2 |
| ENSMUSG( | 153.0799 | -0.32184 | 0.099929 | 0.482395 | Col6a3 |
| ENSMUSG( | 114.3576 | 0.037893 | 0.899258 | 0.985633 | Col6a4 |
| ENSMUSG( | 49.909 | 0.010005 | 0.972126 | 0.996585 | Col6a5 |
| ENSMUSG( | 118.8501 | 0.932858 | 0.001194 | 0.037142 | Col7a1 |
| ENSMUSG( | 166.8552 | -0.29679 | 0.136064 | 0.550226 | Col8a1 |
| ENSMUSG( | 257.8295 | -0.67902 | 5.55E-05 | 0.004288 | Col8a2 |
| ENSMUSG( | 38.77505 | -0.49701 | 0.125315 | 0.532314 | Col9a1 |
| ENSMUSG( | 401.9496 | -0.01963 | 0.867948 | 0.976631 | Col9a2 |
| ENSMUSG( | 1156.9 | -0.10134 | 0.425854 | 0.824898 | Col9a3 |
| ENSMUSG( | 19.56849 | -1.0881 | 0.015325 | NA | Colec11 |
| ENSMUSG( | 448.4237 | 0.038877 | 0.828983 | 0.969228 | Colec12 |
| ENSMUSG( | 1941.212 | -0.20791 | 0.093745 | 0.470609 | Colgalt1 |
| ENSMUSG( | 344.2296 | -0.19834 | 0.269951 | 0.719883 | Colgalt2 |
| ENSMUSG( | 79.81732 | -0.50806 | 0.048621 | 0.337278 | Colq |
| ENSMUSG( | 325.5323 | 0.055465 | 0.687143 | 0.933063 | Commd1 |
| ENSMUSG( | 716.4378 | 0.08182 | 0.474515 | 0.849425 | Commd10 |
| ENSMUSG( | 258.2956 | 0.008119 | 0.956154 | 0.993874 | Commd2 |
| ENSMUSG( | 1215.494 | -0.00732 | 0.935733 | 0.993363 | Commd3 |
| ENSMUSG( | 1142.516 | 0.133496 | 0.308958 | 0.757108 | Commd4 |
| ENSMUSG( | 1020.947 | 0.006134 | 0.954588 | 0.993874 | Commd6 |
| ENSMUSG( | 740.1564 | 0.018607 | 0.863109 | 0.975055 | Commd7 |
| ENSMUSG( | 1416.366 | 0.068915 | 0.526835 | 0.877267 | Commd8 |
| ENSMUSG( | 676.4124 | 0.043482 | 0.67323 | 0.929007 | Commd9 |
| ENSMUSG( | 19.0935 | 0.171003 | 0.704201 | NA | Comp |
| ENSMUSG( | 3391.65 | -0.36518 | 0.00077 | 0.027266 | Comt |
| ENSMUSG( | 361.867 | -0.17822 | 0.150468 | 0.57507 | Comtd1 |
| ENSMUSG( | 2729.325 | 0.155188 | 0.060012 | 0.378891 | Cop1 |
| ENSMUSG( | 7411.934 | 0.05931 | 0.450372 | 0.835344 | Copa |
| ENSMUSG( | 1550.245 | -0.05255 | 0.676206 | 0.930542 | Copb1 |
| ENSMUSG( | 2359.186 | 0.139091 | 0.122561 | 0.528427 | Copb2 |
| ENSMUSG( | 2487.822 | 0.000779 | 0.993104 | 0.999327 | Cope |
| ENSMUSG( | 6812.726 | 0.139564 | 0.196982 | 0.63659 | Copg1 |
| ENSMUSG( | 1974.505 | 0.033932 | 0.707999 | 0.939519 | Copg2 |
| ENSMUSG( | 551.4205 | 0.077558 | 0.479174 | 0.854046 | Coprs |
| ENSMUSG( | 1381.487 | -0.0807 | 0.49298 | 0.860552 | Cops2 |
| ENSMUSG( | 1137.99 | -0.03701 | 0.697421 | 0.935662 | Cops3 |
| ENSMUSG( | 1476.495 | 0.067376 | 0.476751 | 0.851736 | Cops4 |
| ENSMUSG( | 1634.082 | 0.034733 | 0.716455 | 0.941697 | Cops5 |
| ENSMUSG( | 2615.159 | 0.03394 | 0.719624 | 0.941986 | Cops6 |
| ENSMUSG( | 3248.298 | 0.010109 | 0.911777 | 0.988483 | Cops7a |
| ENSMUSG( | 586.0518 | 0.076437 | 0.543792 | 0.884287 | Cops7b |
| ENSMUSG( | 2023.321 | 0.020555 | 0.818359 | 0.965837 | Cops8 |
| ENSMUSG( | 2677.25 | 0.061223 | 0.519596 | 0.874979 | Cops9 |
| ENSMUSG( | 2469.28 | 0.02798 | 0.772161 | 0.954994 | Copz1 |
| ENSMUSG( | 380.0329 | 0.102645 | 0.422709 | 0.823527 | Copz2 |
| ENSMUSG( | 926.0566 | -0.05246 | 0.677407 | 0.930542 | Coq10a |
| ENSMUSG( | 435.5422 | -0.05489 | 0.701701 | 0.937441 | Coq10b |
| ENSMUSG( | 3181.489 | -0.01255 | 0.880222 | 0.981469 | Coq2 |
| ENSMUSG( | 295.4757 | 0.170086 | 0.318516 | 0.764234 | Coq3 |
| ENSMUSG( | 831.4601 | 0.042508 | 0.669581 | 0.928147 | Coq4 |
| ENSMUSG( | 896.5994 | 0.075666 | 0.487166 | 0.858812 | Coq5 |
| ENSMUSG( | 503.9695 | -0.02675 | 0.805915 | 0.962659 | Coq6 |
| ENSMUSG( | 681.8154 | -0.01513 | 0.906405 | 0.986998 | Coq7 |

| | | | | | |
|---|---|---|---|---|---|
| ENSMUSG( | 3853.054 | 0.076213 | 0.555107 | 0.939565 | Cnnm1 |
| ENSMUSG( | 952.7644 | 0.235765 | 0.172441 | 0.843608 | Cnnm2 |
| ENSMUSG( | 1500.008 | 0.055303 | 0.712124 | 0.964363 | Cnnm3 |
| ENSMUSG( | 530.9946 | 0.253998 | 0.142604 | 0.831221 | Cnnm4 |
| ENSMUSG( | 2195.477 | 0.049144 | 0.631864 | 0.953251 | Cnot1 |
| ENSMUSG( | 1337.956 | 0.154059 | 0.262547 | 0.885357 | Cnot10 |
| ENSMUSG( | 607.3188 | 0.009506 | 0.942729 | 0.994774 | Cnot11 |
| ENSMUSG( | 1079.679 | 0.151097 | 0.221604 | 0.866104 | Cnot2 |
| ENSMUSG( | 914.1597 | 0.185312 | 0.227032 | 0.868507 | Cnot3 |
| ENSMUSG( | 599.5417 | -0.03744 | 0.788552 | 0.976395 | Cnot4 |
| ENSMUSG( | 1743.42 | 0.048075 | 0.712352 | 0.964363 | Cnot6 |
| ENSMUSG( | 790.9838 | 0.188006 | 0.141562 | 0.830678 | Cnot6l |
| ENSMUSG( | 1745.134 | 0.052743 | 0.700159 | 0.961335 | Cnot7 |
| ENSMUSG( | 1122.216 | -0.0067 | 0.95939 | 0.994798 | Cnot8 |
| ENSMUSG( | 606.8436 | 0.002003 | 0.987329 | 0.99751 | Cnot9 |
| ENSMUSG( | 16472.1 | -0.2127 | 0.083071 | 0.77681 | Cnp |
| ENSMUSG( | 1065.024 | 0.118411 | 0.351405 | 0.912093 | Cnppd1 |
| ENSMUSG( | 1309.698 | -0.04676 | 0.733326 | 0.967228 | Cnpy2 |
| ENSMUSG( | 2938.535 | -0.01653 | 0.89896 | 0.990558 | Cnpy3 |
| ENSMUSG( | 498.5628 | 0.154731 | 0.320594 | 0.903887 | Cnpy4 |
| ENSMUSG( | 4191.5 | 0.275786 | 0.108512 | 0.804969 | Cnr1 |
| ENSMUSG( | 10.28948 | 0.346584 | 0.628484 | NA | Cnr2 |
| ENSMUSG( | 1293.548 | -0.09718 | 0.576733 | 0.942912 | Cnrip1 |
| ENSMUSG( | 1043.813 | 0.042644 | 0.71182 | 0.964363 | Cnst |
| ENSMUSG( | 24.0044 | 0.298645 | 0.471012 | 0.923329 | Cntd1 |
| ENSMUSG( | 5838.872 | 0.056035 | 0.673672 | 0.95694 | Cntfr |
| ENSMUSG( | 524.2503 | -0.13542 | 0.320718 | 0.904019 | Cntln |
| ENSMUSG( | 13351.62 | 0.100037 | 0.550297 | 0.938699 | Cntn1 |
| ENSMUSG( | 4555.322 | 0.01467 | 0.94149 | 0.994578 | Cntn2 |
| ENSMUSG( | 766.7496 | -0.01104 | 0.9495 | 0.994798 | Cntn3 |
| ENSMUSG( | 976.5765 | 0.252404 | 0.122086 | 0.813384 | Cntn4 |
| ENSMUSG( | 153.0446 | 0.156738 | 0.478372 | 0.923329 | Cntn5 |
| ENSMUSG( | 675.2001 | 0.158222 | 0.182254 | 0.850579 | Cntn6 |
| ENSMUSG( | 8920.807 | 0.032543 | 0.821389 | 0.979198 | Cntnap1 |
| ENSMUSG( | 3114.117 | 0.173386 | 0.249028 | 0.877904 | Cntnap2 |
| ENSMUSG( | 266.7992 | 0.350915 | 0.071553 | 0.761942 | Cntnap3 |
| ENSMUSG( | 653.6435 | 0.145751 | 0.422862 | 0.917257 | Cntnap4 |
| ENSMUSG( | 986.4174 | 0.153584 | 0.422139 | 0.917257 | Cntnap5a |
| ENSMUSG( | 121.7752 | 0.528503 | 0.042619 | 0.690304 | Cntnap5b |
| ENSMUSG( | 39.12341 | 0.555502 | 0.096182 | 0.796133 | Cntnap5c |
| ENSMUSG( | 605.8187 | 0.120373 | 0.419397 | 0.917257 | Cntrl |
| ENSMUSG( | 419.5235 | -0.06487 | 0.696858 | 0.960867 | Cntrob |
| ENSMUSG( | 1120.555 | -0.29529 | 0.029389 | 0.618981 | Coa3 |
| ENSMUSG( | 76.98477 | -0.33387 | 0.162974 | 0.837925 | Coa4 |
| ENSMUSG( | 5075.114 | -0.15424 | 0.244412 | 0.877904 | Coa5 |
| ENSMUSG( | 280.926 | -0.17227 | 0.348836 | 0.911652 | Coa6 |
| ENSMUSG( | 262.2937 | 0.022569 | 0.892536 | 0.990558 | Coa7 |
| ENSMUSG( | 772.6466 | -0.13467 | 0.287789 | 0.894622 | Coa8 |
| ENSMUSG( | 770.5281 | -0.04191 | 0.734027 | 0.967228 | Coasy |
| ENSMUSG( | 1477.649 | 0.030887 | 0.793312 | 0.97681 | Cobl |
| ENSMUSG( | 378.115 | 0.504406 | 0.0024 | 0.192605 | Cobll1 |
| ENSMUSG( | 1400.628 | 0.153704 | 0.29123 | 0.897515 | Coch |
| ENSMUSG( | 1128.336 | 0.09764 | 0.452684 | 0.922316 | Cog1 |
| ENSMUSG( | 706.9165 | -0.13956 | 0.280664 | 0.892356 | Cog2 |
| ENSMUSG( | 1191.18 | -0.01287 | 0.910432 | 0.992124 | Cog3 |
| ENSMUSG( | 1083.686 | -0.08903 | 0.508455 | 0.929089 | Cog4 |
| ENSMUSG( | 750.8258 | -0.03255 | 0.785918 | 0.976285 | Cog5 |
| ENSMUSG( | 1145.457 | -0.03876 | 0.752225 | 0.970461 | Cog6 |
| ENSMUSG( | 1299.779 | -0.19806 | 0.138821 | 0.829437 | Cog7 |
| ENSMUSG( | 624.7861 | -0.06142 | 0.627613 | 0.953234 | Cog8 |
| ENSMUSG( | 331.9641 | 0.118867 | 0.421964 | 0.917257 | Coil |
| ENSMUSG( | 10.98255 | -0.18897 | 0.779673 | NA | Col10a1 |
| ENSMUSG( | 354.5365 | -0.10691 | 0.654707 | 0.955015 | Col11a1 |
| ENSMUSG( | 1703.368 | 0.128606 | 0.419991 | 0.917257 | Col11a2 |
| ENSMUSG( | 813.4735 | 0.130168 | 0.421376 | 0.917257 | Col12a1 |
| ENSMUSG( | 60.62692 | 0.39525 | 0.140731 | 0.830678 | Col13a1 |
| ENSMUSG( | 82.56615 | 0.077457 | 0.76727 | 0.974236 | Col14a1 |
| ENSMUSG( | 226.4814 | -0.18242 | 0.390367 | 0.917257 | Col15a1 |
| ENSMUSG( | 739.0279 | -0.05163 | 0.769787 | 0.974236 | Col16a1 |
| ENSMUSG( | 20.79292 | -0.53673 | 0.254711 | 0.882296 | Col17a1 |
| ENSMUSG( | 162.2437 | 0.142061 | 0.480924 | 0.924548 | Col18a1 |
| ENSMUSG( | 312.4439 | 0.081617 | 0.70037 | 0.961472 | Col19a1 |
| ENSMUSG( | 1289.813 | 0.000167 | 0.99905 | 0.999503 | Col1a1 |
| ENSMUSG( | 1804.919 | 0.298699 | 0.036163 | 0.668395 | Col1a2 |
| ENSMUSG( | 168.0578 | 0.301452 | 0.20653 | 0.864412 | Col20a1 |
| ENSMUSG( | 73.24191 | -0.2869 | 0.282806 | 0.892412 | Col22a1 |
| ENSMUSG( | 928.7338 | 0.180552 | 0.14339 | 0.831311 | Col23a1 |
| ENSMUSG( | 163.875 | 0.053176 | 0.925641 | 0.99306 | Col24a1 |
| ENSMUSG( | 1220.776 | 0.11411 | 0.431666 | 0.917504 | Col25a1 |
| ENSMUSG( | 288.191 | -0.22348 | 0.162457 | 0.837724 | Col26a1 |
| ENSMUSG( | 401.7739 | 0.438083 | 0.041996 | 0.690304 | Col27a1 |
| ENSMUSG( | 27.67886 | 0.611549 | 0.172572 | 0.843702 | Col28a1 |
| ENSMUSG( | 36.13335 | 0.02048 | 0.954229 | 0.994798 | Col2a1 |
| ENSMUSG( | 470.2006 | -0.13194 | 0.446586 | 0.921734 | Col3a1 |
| ENSMUSG( | 2682.392 | 0.159541 | 0.426986 | 0.917257 | Col4a1 |
| ENSMUSG( | 2380.592 | 0.231139 | 0.197945 | 0.862161 | Col4a2 |
| ENSMUSG( | 34.55451 | 0.470458 | 0.212618 | 0.864412 | Col4a3 |
| ENSMUSG( | 56.81735 | -0.02821 | 0.92392 | 0.992841 | Col4a4 |
| ENSMUSG( | 320.5629 | 0.267383 | 0.117304 | 0.813384 | Col4a5 |
| ENSMUSG( | 78.06121 | 0.229638 | 0.381645 | 0.917257 | Col4a6 |
| ENSMUSG( | 874.8103 | 0.104489 | 0.797642 | 0.97681 | Col5a1 |
| ENSMUSG( | 268.0025 | 0.187579 | 0.249809 | 0.877904 | Col5a2 |
| ENSMUSG( | 140.7809 | 0.154176 | 0.573443 | 0.942912 | Col5a3 |
| ENSMUSG( | 2151.026 | 0.02201 | 0.922043 | 0.992644 | Col6a1 |
| ENSMUSG( | 1214.131 | -0.00252 | 0.985483 | 0.997084 | Col6a2 |
| ENSMUSG( | 141.8766 | -0.24994 | 0.259514 | 0.883971 | Col6a3 |
| ENSMUSG( | 97.7162 | -0.16447 | 0.642042 | 0.953903 | Col6a4 |
| ENSMUSG( | 54.02576 | 0.470217 | 0.101293 | 0.801456 | Col6a5 |
| ENSMUSG( | 92.05681 | 0.559276 | 0.055529 | 0.723362 | Col7a1 |
| ENSMUSG( | 191.9044 | 0.366106 | 0.036336 | 0.669285 | Col8a1 |
| ENSMUSG( | 305.3969 | 0.160095 | 0.346386 | 0.910966 | Col8a2 |
| ENSMUSG( | 40.00376 | -0.09228 | 0.768548 | 0.974236 | Col9a1 |
| ENSMUSG( | 395.0038 | 0.190911 | 0.168678 | 0.840222 | Col9a2 |
| ENSMUSG( | 1225.631 | 0.320102 | 0.009538 | 0.406458 | Col9a3 |
| ENSMUSG( | 23.67609 | -0.09216 | 0.827058 | 0.980159 | Colec11 |
| ENSMUSG( | 497.3563 | 0.562994 | 0.004967 | 0.28972 | Colec12 |
| ENSMUSG( | 2057.425 | 0.231712 | 0.12116 | 0.813384 | Colgalt1 |
| ENSMUSG( | 350.9494 | 0.138609 | 0.460534 | 0.923329 | Colgalt2 |
| ENSMUSG( | 68.95003 | -0.69591 | 0.00855 | 0.391682 | Colq |
| ENSMUSG( | 265.4442 | -0.27403 | 0.082895 | 0.77681 | Commd1 |
| ENSMUSG( | 591.7713 | -0.21509 | 0.13406 | 0.822912 | Commd10 |
| ENSMUSG( | 233.7975 | -0.00631 | 0.969065 | 0.99594 | Commd2 |
| ENSMUSG( | 1124.511 | 0.033089 | 0.789178 | 0.976593 | Commd3 |
| ENSMUSG( | 970.4875 | -0.06622 | 0.693586 | 0.960767 | Commd4 |
| ENSMUSG( | 879.1597 | -0.16889 | 0.273667 | 0.889848 | Commd6 |
| ENSMUSG( | 653.2864 | -0.07742 | 0.540709 | 0.937411 | Commd7 |
| ENSMUSG( | 1233.367 | -0.06653 | 0.648294 | 0.95422 | Commd8 |
| ENSMUSG( | 562.5994 | -0.23117 | 0.074934 | 0.76762 | Commd9 |
| ENSMUSG( | 14.10496 | -0.49317 | 0.360247 | 0.914223 | Comp |
| ENSMUSG( | 3416.648 | -0.05121 | 0.673544 | 0.95694 | Comt |
| ENSMUSG( | 350.5668 | 0.00444 | 0.974624 | 0.996416 | Comtd1 |
| ENSMUSG( | 2276.278 | -0.09845 | 0.346411 | 0.910966 | Cop1 |
| ENSMUSG( | 6337.441 | -0.12647 | 0.209172 | 0.864412 | Copa |
| ENSMUSG( | 1501.479 | 0.125419 | 0.352076 | 0.912093 | Copb1 |
| ENSMUSG( | 2008.848 | -0.05532 | 0.618952 | 0.952598 | Copb2 |
| ENSMUSG( | 2154.71 | -0.15183 | 0.212744 | 0.864412 | Cope |
| ENSMUSG( | 5649.666 | -0.13346 | 0.305514 | 0.898538 | Copg1 |
| ENSMUSG( | 1707.212 | -0.12016 | 0.256245 | 0.883082 | Copg2 |
| ENSMUSG( | 482.2566 | -0.04467 | 0.741869 | 0.969293 | Coprs |
| ENSMUSG( | 1272.737 | -0.05444 | 0.749326 | 0.96998 | Cops2 |
| ENSMUSG( | 1020.575 | -0.0811 | 0.494986 | 0.927372 | Cops3 |
| ENSMUSG( | 1257.03 | -0.13217 | 0.23826 | 0.87259 | Cops4 |
| ENSMUSG( | 1436.871 | -0.0685 | 0.542484 | 0.938034 | Cops5 |
| ENSMUSG( | 2188.543 | -0.22212 | 0.055472 | 0.723362 | Cops6 |
| ENSMUSG( | 2740.57 | -0.2215 | 0.055427 | 0.723362 | Cops7a |
| ENSMUSG( | 512.8948 | -0.03541 | 0.799391 | 0.97681 | Cops7b |
| ENSMUSG( | 1765.5 | -0.10807 | 0.352476 | 0.912093 | Cops8 |
| ENSMUSG( | 2185.084 | -0.27125 | 0.032174 | 0.634374 | Cops9 |
| ENSMUSG( | 2113.437 | -0.15983 | 0.200043 | 0.863016 | Copz1 |
| ENSMUSG( | 312.1696 | -0.21148 | 0.177848 | 0.847901 | Copz2 |
| ENSMUSG( | 825.6422 | -0.11529 | 0.39739 | 0.917257 | Coq10a |
| ENSMUSG( | 467.5403 | 0.386509 | 0.01256 | 0.451781 | Coq10b |
| ENSMUSG( | 2843.376 | -0.0698 | 0.5061 | 0.927825 | Coq2 |
| ENSMUSG( | 246.5182 | -0.08771 | 0.654496 | 0.955015 | Coq3 |
| ENSMUSG( | 731.7745 | -0.05723 | 0.633701 | 0.953889 | Coq4 |
| ENSMUSG( | 768.7871 | -0.10446 | 0.432629 | 0.918256 | Coq5 |
| ENSMUSG( | 452.5421 | -0.0638 | 0.665988 | 0.956766 | Coq6 |
| ENSMUSG( | 594.9188 | -0.14927 | 0.31816 | 0.902849 | Coq7 |
| ENSMUSG( | 886.9767 | -0.03421 | 0.771631 | 0.974236 | Coq8a |
| ENSMUSG( | 624.1401 | 0.043582 | 0.751932 | 0.970461 | Coq8b |
| ENSMUSG( | 1571.686 | -0.07198 | 0.497935 | 0.927825 | Coq9 |
| ENSMUSG( | 1687 | -0.19494 | 0.203978 | 0.864412 | Coro1a |
| ENSMUSG( | 1502.288 | 0.020832 | 0.871908 | 0.988779 | Coro1b |
| ENSMUSG( | 2081.483 | -0.06706 | 0.536883 | 0.937169 | Coro1c |
| ENSMUSG( | 993.7057 | 0.086184 | 0.543587 | 0.938063 | Coro2a |
| ENSMUSG( | 5025.34 | -0.15523 | 0.199963 | 0.863016 | Coro2b |
| ENSMUSG( | 1058.558 | 0.00854 | 0.985973 | 0.997188 | Coro6 |
| ENSMUSG( | 1924.696 | -0.08552 | 0.53336 | 0.935162 | Coro7 |

| | | | | | | | | | | | |
|---|---|---|---|---|---|---|---|---|---|---|---|
| ENSMUSG( | Cnst | 1098.814 | -0.07905 | 0.450246 | 0.835344 | ENSMUSG( | Cnst | 1043.813 | 0.042644 | 0.71182 | 0.964363 |
| ENSMUSG( | Cntd1 | 20.77359 | -0.37128 | 0.385632 | NA | ENSMUSG( | Cntd1 | 24.0044 | 0.298645 | 0.471012 | 0.923329 |
| ENSMUSG( | Cntfr | 5696.551 | -0.30132 | 0.007411 | 0.118765 | ENSMUSG( | Cntfr | 5838.872 | 0.056035 | 0.673672 | 0.95694 |
| ENSMUSG( | Cntln | 590.6697 | -0.0517 | 0.655937 | 0.923929 | ENSMUSG( | Cntln | 524.2503 | -0.13542 | 0.320718 | 0.904019 |
| ENSMUSG( | Cntn1 | 12779.29 | -0.31118 | 0.02871 | 0.258426 | ENSMUSG( | Cntn1 | 13351.62 | 0.100037 | 0.550297 | 0.938699 |
| ENSMUSG( | Cntn2 | 4162.47 | -0.57737 | 0.00017 | 0.009401 | ENSMUSG( | Cntn2 | 4555.322 | 0.01467 | 0.94149 | 0.994578 |
| ENSMUSG( | Cntn3 | 685.5902 | -0.68783 | 6.37E-06 | 0.000807 | ENSMUSG( | Cntn3 | 766.7496 | -0.01104 | 0.9495 | 0.994798 |
| ENSMUSG( | Cntn4 | 816.4248 | -0.58511 | 0.0001 | 0.006438 | ENSMUSG( | Cntn4 | 976.5765 | 0.252404 | 0.122086 | 0.813384 |
| ENSMUSG( | Cntn5 | 151.3585 | -0.159 | 0.434406 | 0.828978 | ENSMUSG( | Cntn5 | 153.0446 | 0.156738 | 0.478372 | 0.923329 |
| ENSMUSG( | Cntn6 | 672.9356 | -0.11837 | 0.247585 | 0.696522 | ENSMUSG( | Cntn6 | 675.2001 | 0.158222 | 0.182254 | 0.850579 |
| ENSMUSG( | Cntnap1 | 9024.274 | -0.21382 | 0.07266 | 0.416299 | ENSMUSG( | Cntnap1 | 8920.807 | 0.032543 | 0.821389 | 0.979198 |
| ENSMUSG( | Cntnap2 | 2789.368 | -0.44476 | 0.000514 | 0.020832 | ENSMUSG( | Cntnap2 | 3114.117 | 0.173386 | 0.249028 | 0.877904 |
| ENSMUSG( | Cntnap3 | 221.7294 | -0.47483 | 0.007789 | 0.12245 | ENSMUSG( | Cntnap3 | 266.7992 | 0.350915 | 0.071553 | 0.761942 |
| ENSMUSG( | Cntnap4 | 587.5882 | -0.47106 | 0.005354 | 0.097684 | ENSMUSG( | Cntnap4 | 653.6435 | 0.145751 | 0.422862 | 0.917257 |
| ENSMUSG( | Cntnap5a | 877.8855 | -0.49203 | 0.003832 | 0.079288 | ENSMUSG( | Cntnap5a | 986.4174 | 0.153584 | 0.422139 | 0.917257 |
| ENSMUSG( | Cntnap5b | 98.14938 | -0.35571 | 0.163359 | 0.594631 | ENSMUSG( | Cntnap5b | 121.7752 | 0.528503 | 0.042619 | 0.690304 |
| ENSMUSG( | Cntnap5c | 35.20901 | 0.008407 | 0.978841 | 0.997428 | ENSMUSG( | Cntnap5c | 39.12341 | 0.555502 | 0.096182 | 0.796133 |
| ENSMUSG( | Cntrl | 642.0709 | 0.01612 | 0.885553 | 0.983008 | ENSMUSG( | Cntrl | 605.8187 | 0.120373 | 0.419397 | 0.917257 |
| ENSMUSG( | Cntrob | 502.3708 | 0.188565 | 0.170722 | 0.605271 | ENSMUSG( | Cntrob | 419.5235 | -0.06487 | 0.696858 | 0.960867 |
| ENSMUSG( | Coa3 | 1318.895 | -0.07492 | 0.434639 | 0.828978 | ENSMUSG( | Coa3 | 1120.555 | -0.29529 | 0.029389 | 0.618981 |
| ENSMUSG( | Coa4 | 97.7054 | 0.104087 | 0.600969 | 0.904492 | ENSMUSG( | Coa4 | 76.98477 | -0.33387 | 0.162974 | 0.837925 |
| ENSMUSG( | Coa5 | 5962.729 | 0.047603 | 0.6706 | 0.928683 | ENSMUSG( | Coa5 | 5075.114 | -0.15424 | 0.244412 | 0.877904 |
| ENSMUSG( | Coa6 | 311.3609 | -0.12798 | 0.398702 | 0.812878 | ENSMUSG( | Coa6 | 280.926 | -0.17227 | 0.348836 | 0.911652 |
| ENSMUSG( | Coa7 | 299.9002 | 0.137306 | 0.358723 | 0.790019 | ENSMUSG( | Coa7 | 262.2937 | 0.022569 | 0.892536 | 0.990558 |
| ENSMUSG( | Coa8 | 914.2864 | 0.087188 | 0.375753 | 0.800976 | ENSMUSG( | Coa8 | 772.6466 | -0.13467 | 0.287789 | 0.894622 |
| ENSMUSG( | Coasy | 915.1483 | 0.178636 | 0.069865 | 0.408574 | ENSMUSG( | Coasy | 770.5281 | -0.04191 | 0.734027 | 0.967228 |
| ENSMUSG( | Cobl | 1744.966 | 0.23351 | 0.029961 | 0.262543 | ENSMUSG( | Cobl | 1477.649 | 0.030887 | 0.793312 | 0.97681 |
| ENSMUSG( | Cobll1 | 308.3513 | -0.33865 | 0.016649 | 0.192198 | ENSMUSG( | Cobll1 | 378.115 | 0.504406 | 0.0024 | 0.192605 |
| ENSMUSG( | Coch | 1267.2 | -0.43539 | 0.001376 | 0.040585 | ENSMUSG( | Coch | 1400.628 | 0.153704 | 0.29123 | 0.897515 |
| ENSMUSG( | Cog1 | 1258.001 | 0.138179 | 0.216457 | 0.661421 | ENSMUSG( | Cog1 | 1128.336 | 0.09764 | 0.452684 | 0.922316 |
| ENSMUSG( | Cog2 | 809.425 | -0.01443 | 0.90194 | 0.985633 | ENSMUSG( | Cog2 | 706.9165 | -0.13956 | 0.280664 | 0.892356 |
| ENSMUSG( | Cog3 | 1351.219 | 0.081795 | 0.390476 | 0.808053 | ENSMUSG( | Cog3 | 1191.18 | -0.01287 | 0.910432 | 0.992124 |
| ENSMUSG( | Cog4 | 1300.89 | 0.169838 | 0.111946 | 0.50438 | ENSMUSG( | Cog4 | 1083.686 | -0.08903 | 0.508455 | 0.929089 |
| ENSMUSG( | Cog5 | 843.0985 | 0.033453 | 0.737801 | 0.947329 | ENSMUSG( | Cog5 | 750.8258 | -0.03255 | 0.785918 | 0.976285 |
| ENSMUSG( | Cog6 | 1310.222 | 0.079289 | 0.38287 | 0.803237 | ENSMUSG( | Cog6 | 1145.457 | -0.03876 | 0.752225 | 0.970461 |
| ENSMUSG( | Cog7 | 1588.128 | 0.119717 | 0.230028 | 0.678005 | ENSMUSG( | Cog7 | 1299.779 | -0.19806 | 0.138821 | 0.829437 |
| ENSMUSG( | Cog8 | 722.8955 | 0.087388 | 0.429517 | 0.826446 | ENSMUSG( | Cog8 | 624.7861 | -0.06142 | 0.627613 | 0.953234 |
| ENSMUSG( | Coil | 363.7846 | 0.1183 | 0.358571 | 0.789791 | ENSMUSG( | Coil | 331.9641 | 0.118867 | 0.421964 | 0.917257 |
| ENSMUSG( | Col10a1 | 9.10504 | -1.22433 | 0.133525 | NA | ENSMUSG( | Col10a1 | 10.98255 | -0.18897 | 0.779673 | NA |
| ENSMUSG( | Col11a1 | 455.7759 | 0.339175 | 0.105225 | 0.492417 | ENSMUSG( | Col11a1 | 354.5365 | -0.10691 | 0.654707 | 0.955015 |
| ENSMUSG( | Col11a2 | 1725.555 | -0.1024 | 0.36855 | 0.796202 | ENSMUSG( | Col11a2 | 1703.368 | 0.128606 | 0.419991 | 0.917257 |
| ENSMUSG( | Col12a1 | 850.8485 | -0.00769 | 0.950006 | 0.993821 | ENSMUSG( | Col12a1 | 813.4735 | 0.130168 | 0.421376 | 0.917257 |
| ENSMUSG( | Col13a1 | 61.05371 | 0.157209 | 0.519808 | 0.874979 | ENSMUSG( | Col13a1 | 60.62692 | 0.39525 | 0.140731 | 0.830678 |
| ENSMUSG( | Col14a1 | 76.26932 | -0.45722 | 0.059062 | 0.375388 | ENSMUSG( | Col14a1 | 82.56615 | 0.077457 | 0.76727 | 0.974236 |
| ENSMUSG( | Col15a1 | 273.8456 | 0.085168 | 0.551075 | 0.886038 | ENSMUSG( | Col15a1 | 226.4814 | -0.18242 | 0.390367 | 0.917257 |
| ENSMUSG( | Col16a1 | 866.4556 | 0.136637 | 0.330255 | 0.774247 | ENSMUSG( | Col16a1 | 739.0279 | -0.05163 | 0.769787 | 0.974236 |
| ENSMUSG( | Col17a1 | 21.52258 | -0.79974 | 0.057607 | 0.370247 | ENSMUSG( | Col17a1 | 20.79292 | -0.53673 | 0.254711 | 0.882296 |
| ENSMUSG( | Col18a1 | 175.3533 | 0.08435 | 0.622288 | 0.911432 | ENSMUSG( | Col18a1 | 162.2437 | 0.142061 | 0.480924 | 0.924548 |
| ENSMUSG( | Col19a1 | 341.1475 | 0.059362 | 0.711321 | 0.940028 | ENSMUSG( | Col19a1 | 312.4439 | 0.081617 | 0.70037 | 0.961472 |
| ENSMUSG( | Col1a1 | 1340.683 | -0.16627 | 0.094283 | 0.471647 | ENSMUSG( | Col1a1 | 1289.813 | 0.000167 | 0.99905 | 0.999503 |
| ENSMUSG( | Col1a2 | 1784.31 | 0.007168 | 0.95144 | 0.993821 | ENSMUSG( | Col1a2 | 1804.919 | 0.298699 | 0.036163 | 0.668395 |
| ENSMUSG( | Col20a1 | 206.4079 | 0.58056 | 0.005035 | 0.094254 | ENSMUSG( | Col20a1 | 168.0578 | 0.301452 | 0.20653 | 0.864412 |
| ENSMUSG( | Col22a1 | 86.22791 | -0.06726 | 0.783006 | 0.957558 | ENSMUSG( | Col22a1 | 73.24191 | -0.2869 | 0.282806 | 0.892412 |
| ENSMUSG( | Col23a1 | 994.3072 | 0.115171 | 0.242996 | 0.690307 | ENSMUSG( | Col23a1 | 928.7338 | 0.180552 | 0.14339 | 0.831311 |
| ENSMUSG( | Col24a1 | 220.7661 | 0.594271 | 0.257307 | 0.708457 | ENSMUSG( | Col24a1 | 163.875 | 0.053176 | 0.925641 | 0.99306 |
| ENSMUSG( | Col25a1 | 1107.331 | -0.47106 | 0.000329 | 0.015441 | ENSMUSG( | Col25a1 | 1220.776 | 0.11411 | 0.431666 | 0.917504 |
| ENSMUSG( | Col26a1 | 354.0344 | 0.101166 | 0.436638 | 0.829402 | ENSMUSG( | Col26a1 | 288.191 | -0.22348 | 0.162457 | 0.837724 |
| ENSMUSG( | Col27a1 | 434.485 | 0.397009 | 0.008584 | 0.129874 | ENSMUSG( | Col27a1 | 401.7739 | 0.438083 | 0.041996 | 0.690304 |
| ENSMUSG( | Col28a1 | 31.06609 | 0.642686 | 0.133932 | 0.547197 | ENSMUSG( | Col28a1 | 27.67886 | 0.611549 | 0.172572 | 0.843702 |
| ENSMUSG( | Col2a1 | 40.22893 | 0.082427 | 0.784476 | 0.958003 | ENSMUSG( | Col2a1 | 36.13335 | 0.02048 | 0.954229 | 0.994798 |
| ENSMUSG( | Col3a1 | 501.4583 | -0.22933 | 0.172958 | 0.608361 | ENSMUSG( | Col3a1 | 470.2006 | -0.13194 | 0.446586 | 0.921734 |
| ENSMUSG( | Col4a1 | 2894.654 | 0.105872 | 0.520665 | 0.875186 | ENSMUSG( | Col4a1 | 2682.392 | 0.159541 | 0.426986 | 0.917257 |
| ENSMUSG( | Col4a2 | 2721.529 | 0.33265 | 0.034189 | 0.283507 | ENSMUSG( | Col4a2 | 2380.592 | 0.231139 | 0.197945 | 0.862161 |
| ENSMUSG( | Col4a3 | 33.30745 | 0.109629 | 0.788592 | 0.959226 | ENSMUSG( | Col4a3 | 34.55451 | 0.470458 | 0.212618 | 0.864412 |
| ENSMUSG( | Col4a4 | 63.34289 | 0.014516 | 0.958822 | 0.993874 | ENSMUSG( | Col4a4 | 56.81735 | -0.02821 | 0.92392 | 0.992841 |
| ENSMUSG( | Col4a5 | 343.9591 | 0.203211 | 0.168335 | 0.601667 | ENSMUSG( | Col4a5 | 320.5629 | 0.267383 | 0.117304 | 0.813384 |
| ENSMUSG( | Col4a6 | 79.68278 | 0.012102 | 0.95853 | 0.993874 | ENSMUSG( | Col4a6 | 78.06121 | 0.229638 | 0.381645 | 0.917257 |
| ENSMUSG( | Col5a1 | 1144.404 | 0.563888 | 0.16397 | 0.594869 | ENSMUSG( | Col5a1 | 874.8103 | 0.104489 | 0.797642 | 0.97681 |
| ENSMUSG( | Col5a2 | 312.2899 | 0.339933 | 0.014588 | 0.177737 | ENSMUSG( | Col5a2 | 268.0025 | 0.187579 | 0.249809 | 0.877904 |
| ENSMUSG( | Col5a3 | 175.9666 | 0.480646 | 0.030828 | 0.266969 | ENSMUSG( | Col5a3 | 140.7809 | 0.154176 | 0.573443 | 0.942912 |
| ENSMUSG( | Col6a1 | 2301.512 | -0.04851 | 0.795922 | 0.960062 | ENSMUSG( | Col6a1 | 2151.026 | 0.02201 | 0.922043 | 0.992644 |
| ENSMUSG( | Col6a2 | 1216.814 | -0.28249 | 0.002183 | 0.054296 | ENSMUSG( | Col6a2 | 1214.131 | -0.00252 | 0.985483 | 0.997084 |
| ENSMUSG( | Col6a3 | 153.0799 | -0.32184 | 0.099929 | 0.482395 | ENSMUSG( | Col6a3 | 141.8766 | -0.24994 | 0.259514 | 0.883971 |
| ENSMUSG( | Col6a4 | 114.3576 | 0.037893 | 0.899258 | 0.985633 | ENSMUSG( | Col6a4 | 97.7162 | -0.16447 | 0.642042 | 0.953903 |
| ENSMUSG( | Col6a5 | 49.909 | 0.010005 | 0.972126 | 0.996585 | ENSMUSG( | Col6a5 | 54.02576 | 0.470217 | 0.101293 | 0.801456 |
| ENSMUSG( | Col7a1 | 118.8501 | 0.932858 | 0.001194 | 0.037142 | ENSMUSG( | Col7a1 | 92.05681 | 0.559276 | 0.055529 | 0.723362 |
| ENSMUSG( | Col8a1 | 166.8552 | -0.29679 | 0.136064 | 0.550226 | ENSMUSG( | Col8a1 | 191.9044 | 0.366106 | 0.036336 | 0.669285 |
| ENSMUSG( | Col8a2 | 257.8295 | -0.67902 | 5.55E-05 | 0.004288 | ENSMUSG( | Col8a2 | 305.3969 | 0.160095 | 0.346386 | 0.910966 |
| ENSMUSG( | Col9a1 | 38.77505 | -0.49701 | 0.125315 | 0.532314 | ENSMUSG( | Col9a1 | 40.00376 | -0.09228 | 0.768548 | 0.974236 |
| ENSMUSG( | Col9a2 | 401.9496 | -0.01963 | 0.867948 | 0.976631 | ENSMUSG( | Col9a2 | 395.0038 | 0.190911 | 0.168678 | 0.840222 |
| ENSMUSG( | Col9a3 | 1156.9 | -0.10134 | 0.425854 | 0.824898 | ENSMUSG( | Col9a3 | 1225.631 | 0.320102 | 0.009538 | 0.406458 |
| ENSMUSG( | Colec11 | 19.56849 | -1.0881 | 0.015325 | NA | ENSMUSG( | Colec11 | 23.67609 | -0.09216 | 0.827058 | 0.980159 |
| ENSMUSG( | Colec12 | 448.4237 | 0.038877 | 0.828983 | 0.969228 | ENSMUSG( | Colec12 | 497.3563 | 0.562994 | 0.004967 | 0.28972 |
| ENSMUSG( | Colgalt1 | 1941.212 | -0.20791 | 0.093745 | 0.470609 | ENSMUSG( | Colgalt1 | 2057.425 | 0.231712 | 0.12116 | 0.813384 |
| ENSMUSG( | Colgalt2 | 344.2296 | -0.19834 | 0.269951 | 0.719883 | ENSMUSG( | Colgalt2 | 350.9494 | 0.138609 | 0.460534 | 0.923329 |
| ENSMUSG( | Colq | 79.81732 | -0.50806 | 0.048621 | 0.337278 | ENSMUSG( | Colq | 68.95003 | -0.69591 | 0.00855 | 0.391682 |
| ENSMUSG( | Commd1 | 325.5323 | 0.055465 | 0.687143 | 0.933063 | ENSMUSG( | Commd1 | 265.4442 | -0.27403 | 0.082895 | 0.77681 |
| ENSMUSG( | Commd10 | 716.4378 | 0.08182 | 0.474515 | 0.849425 | ENSMUSG( | Commd10 | 591.7713 | -0.21509 | 0.13406 | 0.822912 |
| ENSMUSG( | Commd2 | 258.2956 | 0.008119 | 0.956154 | 0.993874 | ENSMUSG( | Commd2 | 233.7975 | -0.00631 | 0.969065 | 0.99594 |
| ENSMUSG( | Commd3 | 1215.494 | -0.00732 | 0.935733 | 0.993363 | ENSMUSG( | Commd3 | 1124.511 | 0.033089 | 0.789178 | 0.976593 |
| ENSMUSG( | Commd4 | 1142.516 | 0.133496 | 0.308958 | 0.757108 | ENSMUSG( | Commd4 | 970.4875 | -0.06622 | 0.693586 | 0.960767 |
| ENSMUSG( | Commd6 | 1020.947 | 0.006134 | 0.954588 | 0.993874 | ENSMUSG( | Commd6 | 879.1597 | -0.16889 | 0.273667 | 0.889848 |
| ENSMUSG( | Commd7 | 740.1564 | 0.018607 | 0.863109 | 0.975055 | ENSMUSG( | Commd7 | 653.2864 | -0.07742 | 0.540709 | 0.937411 |
| ENSMUSG( | Commd8 | 1416.366 | 0.068915 | 0.526835 | 0.877267 | ENSMUSG( | Commd8 | 1233.367 | -0.06653 | 0.648294 | 0.95422 |
| ENSMUSG( | Commd9 | 676.4124 | 0.043482 | 0.67323 | 0.929007 | ENSMUSG( | Commd9 | 562.5994 | -0.23117 | 0.074934 | 0.76762 |
| ENSMUSG( | Comp | 19.0935 | 0.171003 | 0.704201 | NA | ENSMUSG( | Comp | 14.10496 | -0.49317 | 0.360247 | 0.914223 |
| ENSMUSG( | Comt | 3391.65 | -0.36518 | 0.00077 | 0.027266 | ENSMUSG( | Comt | 3416.648 | -0.05121 | 0.673544 | 0.95694 |
| ENSMUSG( | Comtd1 | 361.867 | -0.17822 | 0.150468 | 0.57507 | ENSMUSG( | Comtd1 | 350.5668 | 0.00444 | 0.974624 | 0.996416 |
| ENSMUSG( | Cop1 | 2729.325 | 0.155188 | 0.060012 | 0.378891 | ENSMUSG( | Cop1 | 2276.278 | -0.09845 | 0.346411 | 0.910966 |
| ENSMUSG( | Copa | 7411.934 | 0.05931 | 0.450372 | 0.835344 | ENSMUSG( | Copa | 6337.441 | -0.12647 | 0.209172 | 0.864412 |
| ENSMUSG( | Copb1 | 1550.245 | -0.05255 | 0.676206 | 0.930542 | ENSMUSG( | Copb1 | 1501.479 | 0.125419 | 0.352076 | 0.912093 |
| ENSMUSG( | Copb2 | 2359.186 | 0.139091 | 0.122561 | 0.528427 | ENSMUSG( | Copb2 | 2008.848 | -0.05532 | 0.618952 | 0.952598 |
| ENSMUSG( | Cope | 2487.822 | 0.000779 | 0.993104 | 0.999327 | ENSMUSG( | Cope | 2154.71 | -0.15183 | 0.212744 | 0.864412 |
| ENSMUSG( | Copg1 | 6812.726 | 0.139564 | 0.196982 | 0.63659 | ENSMUSG( | Copg1 | 5649.666 | -0.13346 | 0.305514 | 0.898538 |
| ENSMUSG( | Copg2 | 1974.505 | 0.033932 | 0.707999 | 0.939519 | ENSMUSG( | Copg2 | 1707.212 | -0.12016 | 0.256245 | 0.883082 |
| ENSMUSG( | Coprs | 551.4205 | 0.077558 | 0.479174 | 0.854046 | ENSMUSG( | Coprs | 482.2566 | -0.04467 | 0.741869 | 0.969293 |
| ENSMUSG( | Cops2 | 1381.487 | -0.0807 | 0.49298 | 0.860552 | ENSMUSG( | Cops2 | 1272.737 | -0.05444 | 0.749326 | 0.96998 |
| ENSMUSG( | Cops3 | 1137.99 | -0.03701 | 0.697421 | 0.935662 | ENSMUSG( | Cops3 | 1020.575 | -0.0811 | 0.494986 | 0.927372 |
| ENSMUSG( | Cops4 | 1476.495 | 0.067376 | 0.476751 | 0.851736 | ENSMUSG( | Cops4 | 1257.03 | -0.13217 | 0.23826 | 0.87259 |
| ENSMUSG( | Cops5 | 1634.082 | 0.034733 | 0.716455 | 0.941697 | ENSMUSG( | Cops5 | 1436.871 | -0.0685 | 0.542484 | 0.938034 |
| ENSMUSG( | Cops6 | 2615.159 | 0.03394 | 0.719624 | 0.941986 | ENSMUSG( | Cops6 | 2188.543 | -0.22212 | 0.055472 | 0.723362 |
| ENSMUSG( | Cops7a | 3248.298 | 0.010109 | 0.911777 | 0.988483 | ENSMUSG( | Cops7a | 2740.57 | -0.2215 | 0.055427 | 0.723362 |
| ENSMUSG( | Cops7b | 586.0518 | 0.076437 | 0.543792 | 0.884287 | ENSMUSG( | Cops7b | 512.8948 | -0.03541 | 0.799391 | 0.97681 |
| ENSMUSG( | Cops8 | 2023.321 | 0.020555 | 0.818359 | 0.965837 | ENSMUSG( | Cops8 | 1765.5 | -0.10807 | 0.352476 | 0.912093 |
| ENSMUSG( | Cops9 | 2677.25 | 0.061223 | 0.519596 | 0.874979 | ENSMUSG( | Cops9 | 2185.084 | -0.27125 | 0.032174 | 0.634374 |
| ENSMUSG( | Copz1 | 2469.28 | 0.02798 | 0.772161 | 0.954994 | ENSMUSG( | Copz1 | 2113.437 | -0.15983 | 0.200043 | 0.863016 |
| ENSMUSG( | Copz2 | 380.0329 | 0.102645 | 0.422709 | 0.823527 | ENSMUSG( | Copz2 | 312.1696 | -0.21148 | 0.177848 | 0.847901 |
| ENSMUSG( | Coq10a | 926.0566 | -0.05246 | 0.677407 | 0.930542 | ENSMUSG( | Coq10a | 825.6422 | -0.11529 | 0.39739 | 0.917257 |
| ENSMUSG( | Coq10b | 435.5422 | -0.05489 | 0.701701 | 0.937441 | ENSMUSG( | Coq10b | 467.5403 | 0.386509 | 0.01256 | 0.451781 |
| ENSMUSG( | Coq2 | 3181.489 | -0.01255 | 0.880222 | 0.981469 | ENSMUSG( | Coq2 | 2843.376 | -0.0698 | 0.5061 | 0.927825 |
| ENSMUSG( | Coq3 | 295.4757 | 0.170086 | 0.318516 | 0.764234 | ENSMUSG( | Coq3 | 246.5182 | -0.08771 | 0.654496 | 0.955015 |
| ENSMUSG( | Coq4 | 831.4601 | 0.042508 | 0.669581 | 0.928147 | ENSMUSG( | Coq4 | 731.7745 | -0.05723 | 0.633701 | 0.953889 |
| ENSMUSG( | Coq5 | 896.5994 | 0.075666 | 0.487166 | 0.858812 | ENSMUSG( | Coq5 | 768.7871 | -0.10446 | 0.432629 | 0.918256 |
| ENSMUSG( | Coq6 | 503.9695 | -0.02675 | 0.805915 | 0.962659 | ENSMUSG( | Coq6 | 452.5421 | -0.0638 | 0.665988 | 0.956766 |
| ENSMUSG( | Coq7 | 681.8154 | -0.01513 | 0.906405 | 0.986998 | ENSMUSG( | Coq7 | 594.9188 | -0.14927 | 0.31816 | 0.902849 |
| ENSMUSG( | Coq8a | 1018.679 | 0.095231 | 0.370224 | 0.797542 | ENSMUSG( | Coq8a | 886.9767 | -0.03421 | 0.771631 | 0.974236 |
| ENSMUSG( | Coq8b | 688.2992 | 0.057887 | 0.644523 | 0.917773 | ENSMUSG( | Coq8b | 624.1401 | 0.043582 | 0.751932 | 0.970461 |
| ENSMUSG( | Coq9 | 1825.853 | 0.09138 | 0.292156 | 0.74239 | ENSMUSG( | Coq9 | 1571.686 | -0.07198 | 0.497935 | 0.927825 |
| ENSMUSG( | Coro1a | 1974.361 | -0.00014 | 0.999243 | 0.99989 | ENSMUSG( | Coro1a | 1687 | -0.19494 | 0.203978 | 0.864412 |
| ENSMUSG( | Coro1b | 1657.063 | 0.031615 | 0.730768 | 0.944956 | ENSMUSG( | Coro1b | 1502.288 | 0.020832 | 0.871908 | 0.988779 |
| ENSMUSG( | Coro1c | 2394.681 | 0.068254 | 0.458995 | 0.840244 | ENSMUSG( | Coro1c | 2081.483 | -0.06706 | 0.536883 | 0.937169 |
| ENSMUSG( | Coro2a | 1111.39 | 0.134683 | 0.233527 | 0.681508 | ENSMUSG( | Coro2a | 993.7057 | 0.086184 | 0.543587 | 0.938063 |
| ENSMUSG( | Coro2b | 5894.47 | 0.040537 | 0.705144 | 0.938303 | ENSMUSG( | Coro2b | 5025.34 | -0.15523 | 0.199963 | 0.863016 |
| ENSMUSG( | Coro6 | 1344.793 | 0.408632 | 0.369866 | 0.797346 | ENSMUSG( | Coro6 | 1058.558 | 0.00854 | 0.985973 | 0.997188 |
| ENSMUSG( | Coro7 | 2163.838 | -0.01208 | 0.915728 | 0.988634 | ENSMUSG( | Coro7 | 1924.696 | -0.08552 | 0.53336 | 0.935162 |
| ENSMUSG( | Cort | 302.1706 | -0.74021 | 1.89E-06 | 0.000324 | ENSMUSG( | Cort | 343.0412 | -0.00772 | 0.958988 | 0.994798 |
| ENSMUSG( | Cotl1 | 2952.381 | 0.124205 | 0.177496 | 0.614774 | ENSMUSG( | Cotl1 | 2457.863 | -0.13974 | 0.287911 | 0.894644 |
| ENSMUSG( | COX1 | 2063017 | -0.02863 | 0.832616 | 0.970333 | ENSMUSG( | COX1 | 1867827 | -0.0487 | 0.761435 | 0.9728 |
| ENSMUSG( | Cox10 | 857.6407 | -0.06405 | 0.571338 | 0.893319 | ENSMUSG( | Cox10 | 812.9812 | 0.054285 | 0.654003 | 0.955015 |
| ENSMUSG( | Cox11 | 305.1223 | -0.0964 | 0.468279 | 0.845336 | ENSMUSG( | Cox11 | 295.8617 | 0.086501 | 0.57186 | 0.942347 |
| ENSMUSG( | Cox14 | 1580.797 | 0.014635 | 0.890703 | 0.983764 | ENSMUSG( | Cox14 | 1341.883 | -0.20086 | 0.158507 | 0.837724 |
| ENSMUSG( | Cox15 | 610.6457 | -0.119 | 0.261907 | 0.712129 | ENSMUSG( | Cox15 | 551.5493 | -0.14449 | 0.257567 | 0.883401 |
| ENSMUSG( | Cox16 | 107.5324 | -0.08525 | 0.666419 | 0.927243 | ENSMUSG( | Cox16 | 94.4926 | -0.17742 | 0.450927 | 0.922202 |
| ENSMUSG( | Cox17 | 1358.457 | -0.03798 | 0.712909 | 0.94066 | ENSMUSG( | Cox17 | 1174.31 | -0.20017 | 0.103762 | 0.804969 |
| ENSMUSG( | Cox18 | 298.9286 | -0.03513 | 0.786978 | 0.959037 | ENSMUSG( | Cox18 | 290.3825 | 0.149569 | 0.310084 | 0.8987 |
| ENSMUSG( | Cox19 | 857.2658 | 0.089652 | 0.439121 | 0.830714 | ENSMUSG( | Cox19 | 751.9827 | -0.01859 | 0.890609 | 0.990558 |
| ENSMUSG( | COX2 | 7476.991 | 0.064371 | 0.656223 | 0.923929 | ENSMUSG( | COX2 | 5877.104 | -0.39063 | 0.056689 | 0.723362 |
| ENSMUSG( | Cox20 | 550.0803 | -0.05866 | 0.677814 | 0.930542 | ENSMUSG( | Cox20 | 485.3316 | -0.16041 | 0.290694 | 0.897019 |
| ENSMUSG( | COX3 | 5300.78 | -0.00551 | 0.977435 | 0.997428 | ENSMUSG( | COX3 | 4494.649 | -0.22528 | 0.587462 | 0.944975 |
| ENSMUSG( | Cox4i1 | 30417.22 | 0.017894 | 0.807517 | 0.963257 | ENSMUSG( | Cox4i1 | 26247.08 | -0.14309 | 0.178094 | 0.848262 |
| ENSMUSG( | Cox4i2 | 100.1005 | -0.41001 | 0.084409 | 0.447861 | ENSMUSG( | Cox4i2 | 100.4357 | -0.09092 | 0.711273 | 0.964363 |
| ENSMUSG( | Cox5a | 3659.847 | -0.05501 | 0.592628 | 0.902094 | ENSMUSG( | Cox5a | 3198.171 | -0.18397 | 0.148487 | 0.836241 |
| ENSMUSG( | Cox5b | 10377.42 | -0.05587 | 0.444972 | 0.833239 | ENSMUSG( | Cox5b | 8858.734 | -0.2555 | 0.018066 | 0.540045 |
| ENSMUSG( | Cox5b-ps | 126.3629 | -0.23756 | 0.243916 | 0.691355 | ENSMUSG( | Cox5b-ps | 104.1941 | -0.5699 | 0.032989 | 0.63923 |
| ENSMUSG( | Cox6a1 | 25613.79 | 0.009014 | 0.903902 | 0.98623 | ENSMUSG( | Cox6a1 | 21756.31 | -0.20094 | 0.046979 | 0.700664 |
| ENSMUSG( | Cox6a2 | 292.3987 | -0.05063 | 0.696182 | 0.935007 | ENSMUSG( | Cox6a2 | 262.2809 | -0.09702 | 0.521584 | 0.932571 |
| ENSMUSG( | Cox6b1 | 10012.78 | -0.01689 | 0.83019 | 0.96974 | ENSMUSG( | Cox6b1 | 8354.894 | -0.28625 | 0.009584 | 0.406458 |
| ENSMUSG( | Cox6b2 | 85.58447 | -0.12161 | 0.58261 | 0.897918 | ENSMUSG( | Cox6b2 | 76.52621 | -0.18264 | 0.450568 | 0.922202 |

| | | | | | |
|---|---|---|---|---|---|
| ENSMUSG( | 1018.679 | 0.095231 | 0.370224 | 0.797542 | Coq8a |
| ENSMUSG( | 688.2992 | 0.057887 | 0.644523 | 0.917773 | Coq8b |
| ENSMUSG( | 1825.853 | 0.09138 | 0.292156 | 0.74239 | Coq9 |
| ENSMUSG( | 1974.361 | -0.00014 | 0.999243 | 0.99989 | Coro1a |
| ENSMUSG( | 1657.063 | 0.031615 | 0.730768 | 0.944956 | Coro1b |
| ENSMUSG( | 2394.681 | 0.068254 | 0.458995 | 0.840244 | Coro1c |
| ENSMUSG( | 1111.39 | 0.134683 | 0.233527 | 0.681508 | Coro2a |
| ENSMUSG( | 5894.47 | 0.040537 | 0.705144 | 0.938303 | Coro2b |
| ENSMUSG( | 1344.793 | 0.408632 | 0.369866 | 0.797346 | Coro6 |
| ENSMUSG( | 2163.838 | -0.01208 | 0.915728 | 0.988634 | Coro7 |
| ENSMUSG( | 302.1706 | -0.74021 | 1.89E-06 | 0.000324 | Cort |
| ENSMUSG( | 2952.381 | 0.124205 | 0.177496 | 0.614774 | Cotl1 |
| ENSMUSG( | 2063017 | -0.02863 | 0.832616 | 0.970333 | COX1 |
| ENSMUSG( | 857.6407 | -0.06405 | 0.571338 | 0.893319 | Cox10 |
| ENSMUSG( | 305.1223 | -0.0964 | 0.468279 | 0.845336 | Cox11 |
| ENSMUSG( | 1580.797 | 0.014635 | 0.890703 | 0.983764 | Cox14 |
| ENSMUSG( | 610.6457 | -0.119 | 0.261907 | 0.712129 | Cox15 |
| ENSMUSG( | 107.5324 | -0.08525 | 0.666419 | 0.927243 | Cox16 |
| ENSMUSG( | 1358.457 | -0.03798 | 0.712909 | 0.94066 | Cox17 |
| ENSMUSG( | 298.9286 | -0.03513 | 0.786978 | 0.959037 | Cox18 |
| ENSMUSG( | 857.2658 | 0.089652 | 0.439121 | 0.830714 | Cox19 |
| ENSMUSG( | 7476.991 | 0.064371 | 0.656223 | 0.923929 | COX2 |
| ENSMUSG( | 550.0803 | -0.05866 | 0.677814 | 0.930542 | Cox20 |
| ENSMUSG( | 5300.78 | -0.00551 | 0.977435 | 0.997428 | COX3 |
| ENSMUSG( | 30417.22 | 0.017894 | 0.807517 | 0.963257 | Cox4i1 |
| ENSMUSG( | 100.1005 | -0.41001 | 0.084409 | 0.447861 | Cox4i2 |
| ENSMUSG( | 3659.847 | -0.05501 | 0.592628 | 0.902094 | Cox5a |
| ENSMUSG( | 10377.42 | -0.05587 | 0.444972 | 0.833239 | Cox5b |
| ENSMUSG( | 126.3629 | -0.23756 | 0.243916 | 0.691355 | Cox5b-ps |
| ENSMUSG( | 25613.79 | 0.009014 | 0.903902 | 0.98623 | Cox6a1 |
| ENSMUSG( | 292.3987 | -0.05063 | 0.696182 | 0.935007 | Cox6a2 |
| ENSMUSG( | 10012.78 | -0.01689 | 0.83019 | 0.96974 | Cox6b1 |
| ENSMUSG( | 85.58447 | -0.12161 | 0.58261 | 0.897918 | Cox6b2 |
| ENSMUSG( | 11813.68 | 0.054503 | 0.514537 | 0.872055 | Cox6c |
| ENSMUSG( | 62.1596 | 0.123649 | 0.615764 | 0.910647 | Cox7a1 |
| ENSMUSG( | 5539.858 | 0.043051 | 0.600643 | 0.90443 | Cox7a2 |
| ENSMUSG( | 6080.98 | 0.047565 | 0.573686 | 0.893584 | Cox7a2l |
| ENSMUSG( | 4848.425 | -0.05726 | 0.559142 | 0.888638 | Cox7b |
| ENSMUSG( | 6242.983 | 0.026727 | 0.781133 | 0.957558 | Cox7c |
| ENSMUSG( | 34552.59 | 0.000482 | 0.995467 | 0.99943 | Cox8a |
| ENSMUSG( | 15.89983 | -0.41562 | 0.420222 | NA | Cox8b |
| ENSMUSG( | 405.4205 | -0.20736 | 0.215512 | 0.66017 | Cp |
| ENSMUSG( | 29.6986 | -0.22061 | 0.581405 | 0.897705 | Cpa2 |
| ENSMUSG( | 2599.406 | -0.42213 | 0.008308 | 0.127252 | Cpd |
| ENSMUSG( | 321407.1 | 0.09524 | 0.370922 | 0.798261 | Cpe |
| ENSMUSG( | 1344.347 | -0.0062 | 0.955284 | 0.993874 | Cpeb1 |
| ENSMUSG( | 1538.889 | -0.01122 | 0.907989 | 0.9878 | Cpeb2 |
| ENSMUSG( | 1183.312 | -0.07554 | 0.46759 | 0.844819 | Cpeb3 |
| ENSMUSG( | 2518.282 | -0.30499 | 0.021026 | 0.220525 | Cpeb4 |
| ENSMUSG( | 171.0893 | 0.14799 | 0.35813 | 0.789704 | Cped1 |
| ENSMUSG( | 1861.229 | -0.0624 | 0.539955 | 0.88315 | Cplane1 |
| ENSMUSG( | 22.8785 | 0.083579 | 0.827984 | 0.968966 | Cplane2 |
| ENSMUSG( | 21595.5 | 0.080432 | 0.340025 | 0.777959 | Cplx1 |
| ENSMUSG( | 54831.73 | 0.041054 | 0.633743 | 0.914073 | Cplx2 |
| ENSMUSG( | 520.2242 | 0.020981 | 0.864406 | 0.975398 | Cplx2 |
| ENSMUSG( | 1119.798 | 0.195468 | 0.068646 | 0.405903 | Cplx3 |
| ENSMUSG( | 352.7308 | -0.20402 | 0.314965 | 0.762155 | Cpm |
| ENSMUSG( | 1484.519 | 0.062136 | 0.516116 | 0.872701 | Cpne2 |
| ENSMUSG( | 913.0259 | 0.102777 | 0.417976 | 0.821964 | Cpne3 |
| ENSMUSG( | 1016.202 | -0.07531 | 0.508559 | 0.868389 | Cpne4 |
| ENSMUSG( | 2354.981 | -0.25021 | 0.066512 | 0.39868 | Cpne5 |
| ENSMUSG( | 4492.791 | 0.469464 | 0.004386 | 0.086463 | Cpne6 |
| ENSMUSG( | 2375.884 | 0.615097 | 0.000926 | 0.031085 | Cpne7 |
| ENSMUSG( | 426.4924 | -0.04679 | 0.70057 | 0.93743 | Cpne8 |
| ENSMUSG( | 1638.371 | 0.188874 | 0.169179 | 0.603271 | Cpne9 |
| ENSMUSG( | 1021.761 | -0.35677 | 0.00044 | 0.019063 | Cpox |
| ENSMUSG( | 1710.456 | -0.03889 | 0.714916 | 0.941144 | Cpped1 |
| ENSMUSG( | 1113.305 | -0.20746 | 0.085959 | 0.451492 | Cpq |
| ENSMUSG( | 9.08372 | 0.491411 | 0.438836 | NA | Cps1 |
| ENSMUSG( | 1841.014 | 0.132799 | 0.172477 | 0.607771 | Cpsf1 |
| ENSMUSG( | 1305.711 | 0.134804 | 0.263563 | 0.714284 | Cpsf2 |
| ENSMUSG( | 1247.694 | 0.024758 | 0.828811 | 0.969097 | Cpsf3 |
| ENSMUSG( | 567.7388 | -0.06983 | 0.653491 | 0.922819 | Cpsf4 |
| ENSMUSG( | 1249.556 | -0.02651 | 0.797452 | 0.960507 | Cpsf6 |
| ENSMUSG( | 2304.901 | 0.011292 | 0.90289 | 0.986055 | Cpsf7 |
| ENSMUSG( | 2642.803 | -0.04144 | 0.731276 | 0.944956 | Cpt1a |
| ENSMUSG( | 3558.048 | -0.06989 | 0.530824 | 0.879289 | Cpt1c |
| ENSMUSG( | 337.0553 | 0.071467 | 0.576226 | 0.895164 | Cpt2 |
| ENSMUSG( | 461.3432 | 0.025565 | 0.831185 | 0.969889 | Cptp |
| ENSMUSG( | 366.7393 | -0.2627 | 0.039275 | 0.305097 | Cpxm1 |
| ENSMUSG( | 126.9374 | -0.30783 | 0.141564 | 0.558606 | Cpxm2 |
| ENSMUSG( | 58.50503 | -0.22478 | 0.402717 | 0.81552 | Cpz |
| ENSMUSG( | 747.6013 | -0.04701 | 0.778531 | 0.956215 | Cr1l |
| ENSMUSG( | 21.33141 | -0.30189 | 0.549302 | 0.885741 | Crabp1 |
| ENSMUSG( | 73.74432 | -0.40512 | 0.107271 | 0.495579 | Crabp2 |
| ENSMUSG( | 1433.045 | 0.115827 | 0.178868 | 0.616298 | Cracd |
| ENSMUSG( | 6411.913 | 0.058115 | 0.64267 | 0.916865 | Cracdl |
| ENSMUSG( | 81.22131 | -0.37566 | 0.100257 | 0.48338 | Cracr2a |
| ENSMUSG( | 114.5666 | 0.113102 | 0.580364 | 0.896952 | Cracr2b |
| ENSMUSG( | 149.9534 | -0.00861 | 0.960259 | 0.993991 | Cradd |
| ENSMUSG( | 1465.831 | 0.079576 | 0.380376 | 0.801757 | Cramp1 |
| ENSMUSG( | 2598.32 | 0.164883 | 0.047825 | 0.333933 | Crat |
| ENSMUSG( | 23.89836 | -0.40402 | 0.312936 | 0.76033 | Crb1 |
| ENSMUSG( | 278.0581 | 0.093717 | 0.538752 | 0.882772 | Crb2 |
| ENSMUSG( | 18.63984 | -0.08026 | 0.869404 | NA | Crb3 |
| ENSMUSG( | 966.5307 | 0.011365 | 0.910814 | 0.988386 | Crbn |
| ENSMUSG( | 768.1508 | 0.034138 | 0.763799 | 0.954013 | Crcp |
| ENSMUSG( | 1112.853 | 0.214358 | 0.071555 | 0.413178 | Creb1 |
| ENSMUSG( | 1351.33 | -0.20682 | 0.070352 | 0.40908 | Creb3 |
| ENSMUSG( | 270.4077 | -0.16099 | 0.321572 | 0.766705 | Creb3l1 |
| ENSMUSG( | 645.2099 | -0.03707 | 0.821418 | 0.966248 | Creb3l2 |
| ENSMUSG( | 14.52274 | -0.1738 | 0.779997 | NA | Creb3l3 |
| ENSMUSG( | 17.75931 | -0.69507 | 0.128811 | NA | Creb3l4 |
| ENSMUSG( | 123.3768 | -0.51801 | 0.010027 | 0.142701 | Creb5 |
| ENSMUSG( | 2055.405 | 0.073439 | 0.450671 | 0.835475 | Crebbp |
| ENSMUSG( | 1174.211 | 0.147383 | 0.159184 | 0.588305 | Crebl2 |
| ENSMUSG( | 1350.154 | 0.14277 | 0.159364 | 0.58868 | Crebrf |
| ENSMUSG( | 1326.885 | -0.1447 | 0.160839 | 0.590628 | Crebzf |
| ENSMUSG( | 1393.227 | -0.33692 | 0.015722 | 0.185786 | Creg1 |
| ENSMUSG( | 3045.003 | -0.29026 | 0.08325 | 0.445247 | Creg2 |
| ENSMUSG( | 2985.514 | -0.27914 | 0.018199 | 0.203238 | Creld1 |
| ENSMUSG( | 640.4631 | -0.00126 | 0.992922 | 0.999327 | Creld2 |
| ENSMUSG( | 460.2017 | 0.054547 | 0.658887 | 0.924506 | Crem |
| ENSMUSG( | 257.9942 | -0.73036 | 0.000355 | 0.016296 | Crh |
| ENSMUSG( | 531.2404 | -0.39883 | 0.001184 | 0.037077 | Crhbp |
| ENSMUSG( | 1037.193 | -0.0062 | 0.959419 | 0.993991 | Crhr1 |
| ENSMUSG( | 58.1888 | -0.04154 | 0.866823 | 0.976541 | Crhr2 |
| ENSMUSG( | 1872.054 | -0.08113 | 0.532459 | 0.879905 | Crim1 |
| ENSMUSG( | 91.12728 | -0.13067 | 0.557354 | 0.888279 | Crip1 |
| ENSMUSG( | 4460.588 | 0.015236 | 0.846974 | 0.972374 | Crip2 |
| ENSMUSG( | 68.95793 | 0.435804 | 0.08211 | 0.442642 | Crip3 |
| ENSMUSG( | 1087.233 | 0.033031 | 0.789681 | 0.959461 | Cript |
| ENSMUSG( | 89.41266 | -0.10139 | 0.648292 | 0.919949 | Cripto |
| ENSMUSG( | 122.0513 | -0.00686 | 0.974169 | 0.99691 | Crispld1 |
| ENSMUSG( | 96.56861 | -0.02156 | 0.922322 | 0.99045 | Crispld2 |
| ENSMUSG( | 4016.785 | 0.095991 | 0.310266 | 0.758431 | Crk |
| ENSMUSG( | 1706.342 | 0.186806 | 0.034271 | 0.284043 | Crkl |
| ENSMUSG( | 212.951 | 0.085317 | 0.594194 | 0.902869 | Crlf1 |
| ENSMUSG( | 830.4585 | -0.10051 | 0.36872 | 0.796356 | Crlf2 |
| ENSMUSG( | 508.56 | 0.16099 | 0.171439 | 0.605943 | Crlf3 |
| ENSMUSG( | 708.4283 | 0.046436 | 0.70094 | 0.93743 | Crls1 |
| ENSMUSG( | 4285.913 | 0.086549 | 0.516016 | 0.872701 | Crmp1 |
| ENSMUSG( | 17.6807 | 0.304188 | 0.504308 | NA | Crnde |
| ENSMUSG( | 443.4003 | 0.015661 | 0.898296 | 0.985633 | Crnkl1 |
| ENSMUSG( | 2000.076 | 0.300242 | 0.039979 | 0.3082 | Crocc |
| ENSMUSG( | 52.69693 | 0.63943 | 0.097145 | 0.47759 | Crocc2 |
| ENSMUSG( | 2836.392 | 0.151539 | 0.282437 | 0.73186 | Crot |
| ENSMUSG( | 155.7725 | -0.05846 | 0.73763 | 0.947329 | Crppa |
| ENSMUSG( | 2219.207 | -0.26933 | 0.070188 | 0.40908 | Crtac1 |
| ENSMUSG( | 274.9696 | -0.18044 | 0.336677 | 0.777 | Crtap |
| ENSMUSG( | 9194.981 | 0.126267 | 0.165629 | 0.59714 | Crtc1 |
| ENSMUSG( | 980.7799 | -0.04656 | 0.685953 | 0.932981 | Crtc2 |
| ENSMUSG( | 850.3839 | 0.041161 | 0.679737 | 0.930983 | Crtc3 |
| ENSMUSG( | 551.9353 | -0.0302 | 0.77833 | 0.956215 | Cry1 |
| ENSMUSG( | 3290.911 | 0.138232 | 0.145945 | 0.567409 | Cry2 |
| ENSMUSG( | 5220.011 | -0.00144 | 0.988416 | 0.999109 | Cryab |
| ENSMUSG( | 17.14335 | 0.454539 | 0.358862 | NA | Cryba2 |
| ENSMUSG( | 43.79235 | 0.447117 | 0.120045 | 0.523983 | Cryba4 |
| ENSMUSG( | 408.977 | 0.325903 | 0.018283 | 0.203658 | Crybb1 |
| ENSMUSG( | 47.42893 | 0.104936 | 0.724808 | 0.944096 | Crybb3 |
| ENSMUSG( | 54.28505 | -0.25072 | 0.346227 | 0.779153 | Crybg1 |
| ENSMUSG( | 22.1694 | -0.21778 | 0.60363 | 0.904887 | Crybg2 |
| ENSMUSG( | 360.4368 | -0.08292 | 0.561069 | 0.888934 | Crybg3 |
| ENSMUSG( | 15.97997 | -0.68942 | 0.156148 | NA | Crygn |
| ENSMUSG( | 378.0326 | -0.02148 | 0.881439 | 0.981798 | Cryl1 |
| ENSMUSG( | 1965.422 | -0.27809 | 0.121463 | 0.525713 | Crym |
| ENSMUSG( | 744.0903 | 0.006493 | 0.962771 | 0.994824 | Cryz |
| ENSMUSG( | 1430.742 | 0.021703 | 0.816392 | 0.965455 | Cryzl1 |

| | | | | | |
|---|---|---|---|---|---|
| ENSMUSG( | 343.0412 | -0.00772 | 0.958988 | 0.994798 | Cort |
| ENSMUSG( | 2457.863 | -0.13974 | 0.287911 | 0.894644 | Cotl1 |
| ENSMUSG( | 1867827 | -0.0487 | 0.761435 | 0.9728 | COX1 |
| ENSMUSG( | 812.9812 | 0.054285 | 0.654003 | 0.955015 | Cox10 |
| ENSMUSG( | 295.8617 | 0.086501 | 0.57186 | 0.942347 | Cox11 |
| ENSMUSG( | 1341.883 | -0.20086 | 0.158507 | 0.837724 | Cox14 |
| ENSMUSG( | 551.5493 | -0.14449 | 0.257567 | 0.883401 | Cox15 |
| ENSMUSG( | 94.4926 | -0.17742 | 0.450927 | 0.922202 | Cox16 |
| ENSMUSG( | 1174.31 | -0.20017 | 0.103762 | 0.804969 | Cox17 |
| ENSMUSG( | 290.3825 | 0.149569 | 0.310084 | 0.8987 | Cox18 |
| ENSMUSG( | 751.9827 | -0.01859 | 0.890609 | 0.990558 | Cox19 |
| ENSMUSG( | 5877.104 | -0.39063 | 0.056689 | 0.723362 | COX2 |
| ENSMUSG( | 485.3316 | -0.16041 | 0.290694 | 0.897019 | Cox20 |
| ENSMUSG( | 4494.649 | -0.22528 | 0.587462 | 0.944975 | COX3 |
| ENSMUSG( | 26247.08 | -0.14309 | 0.178094 | 0.848262 | Cox4i1 |
| ENSMUSG( | 100.4357 | -0.09092 | 0.711273 | 0.964363 | Cox4i2 |
| ENSMUSG( | 3198.171 | -0.18397 | 0.148487 | 0.836241 | Cox5a |
| ENSMUSG( | 8858.734 | -0.2555 | 0.018066 | 0.540045 | Cox5b |
| ENSMUSG( | 104.1941 | -0.5699 | 0.032989 | 0.63923 | Cox5b-ps |
| ENSMUSG( | 21756.31 | -0.20094 | 0.046979 | 0.700664 | Cox6a1 |
| ENSMUSG( | 262.2809 | -0.09702 | 0.521584 | 0.932571 | Cox6a2 |
| ENSMUSG( | 8354.894 | -0.28625 | 0.009584 | 0.406458 | Cox6b1 |
| ENSMUSG( | 76.52621 | -0.18264 | 0.450568 | 0.922202 | Cox6b2 |
| ENSMUSG( | 9966.111 | -0.17411 | 0.111025 | 0.807885 | Cox6c |
| ENSMUSG( | 47.56051 | -0.39792 | 0.198016 | 0.862161 | Cox7a1 |
| ENSMUSG( | 4610.507 | -0.22893 | 0.040592 | 0.683847 | Cox7a2 |
| ENSMUSG( | 5227.122 | -0.12387 | 0.287175 | 0.894091 | Cox7a2l |
| ENSMUSG( | 4042.431 | -0.33552 | 0.008274 | 0.388416 | Cox7b |
| ENSMUSG( | 5351.066 | -0.15545 | 0.18853 | 0.85698 | Cox7c |
| ENSMUSG( | 29012.19 | -0.24717 | 0.02486 | 0.592568 | Cox8a |
| ENSMUSG( | 14.48473 | -0.42583 | 0.438638 | 0.920721 | Cox8b |
| ENSMUSG( | 438.0296 | 0.288296 | 0.152425 | 0.837724 | Cp |
| ENSMUSG( | 25.9995 | -0.37266 | 0.398026 | 0.917257 | Cpa2 |
| ENSMUSG( | 2841.143 | 0.136393 | 0.483354 | 0.92499 | Cpd |
| ENSMUSG( | 281210.4 | -0.02112 | 0.865995 | 0.987156 | Cpe |
| ENSMUSG( | 1326.239 | 0.216778 | 0.075897 | 0.768524 | Cpeb1 |
| ENSMUSG( | 1488.376 | 0.158543 | 0.161953 | 0.837724 | Cpeb2 |
| ENSMUSG( | 1087.34 | -0.04691 | 0.711645 | 0.964363 | Cpeb3 |
| ENSMUSG( | 2488.257 | -0.05555 | 0.688364 | 0.959742 | Cpeb4 |
| ENSMUSG( | 167.9171 | 0.344292 | 0.074466 | 0.766854 | Cped1 |
| ENSMUSG( | 1735.558 | 0.005449 | 0.962688 | 0.995284 | Cplane1 |
| ENSMUSG( | 24.11593 | 0.432728 | 0.298718 | 0.898538 | Cplane2 |
| ENSMUSG( | 18178.41 | -0.15132 | 0.148183 | 0.836241 | Cplx1 |
| ENSMUSG( | 48462.28 | -0.04558 | 0.665107 | 0.956766 | Cplx2 |
| ENSMUSG( | 449.3242 | -0.13742 | 0.343078 | 0.909646 | Cplx2 |
| ENSMUSG( | 913.1286 | -0.12314 | 0.317578 | 0.902602 | Cplx3 |
| ENSMUSG( | 462.9226 | 0.768048 | 0.004408 | 0.269982 | Cpm |
| ENSMUSG( | 11.19797 | 0.162477 | 0.805804 | NA | Cpn1 |
| ENSMUSG( | 1232.659 | -0.2167 | 0.095701 | 0.796133 | Cpne2 |
| ENSMUSG( | 793.9782 | -0.03417 | 0.823132 | 0.979198 | Cpne3 |
| ENSMUSG( | 962.1459 | 0.032417 | 0.829429 | 0.980224 | Cpne4 |
| ENSMUSG( | 2145.888 | -0.25218 | 0.110787 | 0.807582 | Cpne5 |
| ENSMUSG( | 3491.128 | 0.04343 | 0.831065 | 0.98051 | Cpne6 |
| ENSMUSG( | 1874.483 | 0.249193 | 0.208397 | 0.864412 | Cpne7 |
| ENSMUSG( | 388.4419 | -0.05148 | 0.728938 | 0.966913 | Cpne8 |
| ENSMUSG( | 1348.079 | -0.1031 | 0.481899 | 0.924578 | Cpne9 |
| ENSMUSG( | 1079.709 | 0.092723 | 0.474963 | 0.923329 | Cpox |
| ENSMUSG( | 1474.952 | -0.20978 | 0.137156 | 0.828492 | Cpped1 |
| ENSMUSG( | 1071.533 | -0.04212 | 0.742601 | 0.969415 | Cpq |
| ENSMUSG( | 8.060035 | 0.411646 | 0.55947 | NA | Cps1 |
| ENSMUSG( | 1584.418 | -0.02803 | 0.818653 | 0.979198 | Cpsf1 |
| ENSMUSG( | 1152.213 | 0.041697 | 0.773245 | 0.974525 | Cpsf2 |
| ENSMUSG( | 1097.661 | -0.07758 | 0.537118 | 0.937216 | Cpsf3 |
| ENSMUSG( | 549.172 | 0.105613 | 0.493663 | 0.927372 | Cpsf4 |
| ENSMUSG( | 1316.814 | 0.370724 | 0.004427 | 0.270226 | Cpsf6 |
| ENSMUSG( | 2161.998 | 0.093339 | 0.437214 | 0.920144 | Cpsf7 |
| ENSMUSG( | 2403.83 | -0.04314 | 0.740137 | 0.968811 | Cpt1a |
| ENSMUSG( | 3489.226 | 0.143052 | 0.269733 | 0.887508 | Cpt1c |
| ENSMUSG( | 283.131 | -0.16349 | 0.281055 | 0.892356 | Cpt2 |
| ENSMUSG( | 429.1273 | 0.089745 | 0.529038 | 0.934795 | Cptp |
| ENSMUSG( | 376.1627 | 0.093814 | 0.511089 | 0.929571 | Cpxm1 |
| ENSMUSG( | 116.5365 | -0.26513 | 0.252065 | 0.879902 | Cpxm2 |
| ENSMUSG( | 49.96632 | -0.40794 | 0.179938 | 0.850372 | Cpz |
| ENSMUSG( | 711.3775 | 0.084837 | 0.645144 | 0.953988 | Cr1l |
| ENSMUSG( | 25.3662 | 0.389226 | 0.469765 | 0.923329 | Crabp1 |
| ENSMUSG( | 77.13562 | 0.003746 | 0.988646 | 0.997862 | Crabp2 |
| ENSMUSG( | 1252.827 | -0.00287 | 0.979359 | 0.996468 | Cracd |
| ENSMUSG( | 5827.874 | 0.052262 | 0.706296 | 0.963467 | Cracdl |
| ENSMUSG( | 71.5321 | -0.47115 | 0.061357 | 0.736111 | Cracr2a |
| ENSMUSG( | 132.8357 | 0.736972 | 0.001902 | 0.163326 | Cracr2b |
| ENSMUSG( | 131.3571 | -0.14138 | 0.506099 | 0.927825 | Cradd |
| ENSMUSG( | 1331.368 | 0.072649 | 0.559182 | 0.939587 | Cramp1 |
| ENSMUSG( | 2175.101 | -0.07708 | 0.474967 | 0.923329 | Crat |
| ENSMUSG( | 20.38584 | -0.67499 | 0.1528 | 0.837724 | Crb1 |
| ENSMUSG( | 273.5564 | 0.313859 | 0.09932 | 0.800296 | Crb2 |
| ENSMUSG( | 16.64786 | -0.12824 | 0.804148 | 0.977191 | Crb3 |
| ENSMUSG( | 865.4839 | -0.04052 | 0.736932 | 0.967389 | Crbn |
| ENSMUSG( | 660.3252 | -0.14234 | 0.307851 | 0.898538 | Crcp |
| ENSMUSG( | 997.0361 | 0.167251 | 0.198574 | 0.862705 | Creb1 |
| ENSMUSG( | 1326.448 | 0.018272 | 0.895075 | 0.990558 | Creb3 |
| ENSMUSG( | 264.6706 | 0.040866 | 0.817918 | 0.979198 | Creb3l1 |
| ENSMUSG( | 639.106 | 0.206566 | 0.284111 | 0.892412 | Creb3l2 |
| ENSMUSG( | 14.9092 | 0.227425 | 0.702604 | 0.962215 | Creb3l3 |
| ENSMUSG( | 14.71397 | -1.03364 | 0.049966 | 0.708648 | Creb3l4 |
| ENSMUSG( | 132.7829 | 0.02552 | 0.909534 | 0.991958 | Creb5 |
| ENSMUSG( | 1912.89 | 0.131459 | 0.243833 | 0.877904 | Crebbp |
| ENSMUSG( | 1038.699 | 0.064377 | 0.575004 | 0.942912 | Crebl2 |
| ENSMUSG( | 1209.533 | 0.096971 | 0.400092 | 0.917257 | Crebrf |
| ENSMUSG( | 1432.584 | 0.333318 | 0.048295 | 0.706142 | Crebzf |
| ENSMUSG( | 1495.542 | 0.154667 | 0.301224 | 0.898538 | Creg1 |
| ENSMUSG( | 3343.922 | 0.258076 | 0.178302 | 0.848538 | Creg2 |
| ENSMUSG( | 3099.844 | 0.112131 | 0.403499 | 0.917257 | Creld1 |
| ENSMUSG( | 682.9275 | 0.426234 | 0.004839 | 0.287322 | Creld2 |
| ENSMUSG( | 382.5738 | -0.22798 | 0.151773 | 0.837724 | Crem |
| ENSMUSG( | 261.1473 | -0.36328 | 0.110595 | 0.807582 | Crh |
| ENSMUSG( | 551.7271 | 0.010132 | 0.943369 | 0.994798 | Crhbp |
| ENSMUSG( | 950.7906 | 0.016124 | 0.90219 | 0.991012 | Crhr1 |
| ENSMUSG( | 57.70393 | 0.178691 | 0.512629 | 0.930164 | Crhr2 |
| ENSMUSG( | 1902.066 | 0.228901 | 0.127197 | 0.817278 | Crim1 |
| ENSMUSG( | 86.85423 | 0.024113 | 0.919943 | 0.992644 | Crip1 |
| ENSMUSG( | 3904.755 | -0.10281 | 0.32113 | 0.904019 | Crip2 |
| ENSMUSG( | 68.70501 | 0.644907 | 0.031529 | 0.631192 | Crip3 |
| ENSMUSG( | 944.7051 | -0.11292 | 0.483386 | 0.92499 | Cript |
| ENSMUSG( | 78.42798 | -0.21393 | 0.405241 | 0.917257 | Cripto |
| ENSMUSG( | 122.5508 | 0.277736 | 0.221272 | 0.866104 | Crispld1 |
| ENSMUSG( | 109.4195 | 0.569721 | 0.01291 | 0.458131 | Crispld2 |
| ENSMUSG( | 3502.503 | -0.03175 | 0.791822 | 0.97681 | Crk |
| ENSMUSG( | 1461.481 | 0.014199 | 0.893501 | 0.990558 | Crkl |
| ENSMUSG( | 206.8831 | 0.262423 | 0.105849 | 0.804969 | Crlf1 |
| ENSMUSG( | 781.8641 | -0.00451 | 0.972735 | 0.996416 | Crlf2 |
| ENSMUSG( | 476.5866 | 0.241592 | 0.083601 | 0.77681 | Crlf3 |
| ENSMUSG( | 650.4159 | 0.066096 | 0.601301 | 0.947749 | Crls1 |
| ENSMUSG( | 3608.097 | -0.14567 | 0.33941 | 0.907972 | Crmp1 |
| ENSMUSG( | 14.92397 | 0.125361 | 0.809054 | 0.97873 | Crnde |
| ENSMUSG( | 406.4781 | 0.032655 | 0.81823 | 0.979198 | Crnkl1 |
| ENSMUSG( | 1687.228 | 0.090213 | 0.554126 | 0.939565 | Crocc |
| ENSMUSG( | 37.84385 | 0.047208 | 0.898373 | 0.990558 | Crocc2 |
| ENSMUSG( | 2345.379 | -0.1286 | 0.379698 | 0.916995 | Crot |
| ENSMUSG( | 132.7312 | -0.2808 | 0.178107 | 0.848262 | Crppa |
| ENSMUSG( | 2300.213 | 0.118286 | 0.465812 | 0.923329 | Crtac1 |
| ENSMUSG( | 286.437 | 0.216846 | 0.299589 | 0.898538 | Crtap |
| ENSMUSG( | 8127.716 | 0.043156 | 0.690189 | 0.960071 | Crtc1 |
| ENSMUSG( | 901.1144 | -0.01801 | 0.898301 | 0.990558 | Crtc2 |
| ENSMUSG( | 765.1755 | 0.004244 | 0.971406 | 0.996236 | Crtc3 |
| ENSMUSG( | 599.0289 | 0.434757 | 0.003485 | 0.235685 | Cry1 |
| ENSMUSG( | 3017.971 | 0.15765 | 0.183333 | 0.850984 | Cry2 |
| ENSMUSG( | 4342.652 | -0.27974 | 0.020434 | 0.565891 | Cryab |
| ENSMUSG( | 10.74465 | -0.6263 | 0.316484 | NA | Cryba2 |
| ENSMUSG( | 31.45685 | -0.2027 | 0.560046 | 0.940119 | Cryba4 |
| ENSMUSG( | 336.5324 | 0.05294 | 0.726943 | 0.966887 | Crybb1 |
| ENSMUSG( | 43.42713 | 0.09433 | 0.779286 | 0.976106 | Crybb3 |
| ENSMUSG( | 69.37276 | 0.643955 | 0.017149 | 0.52741 | Crybg1 |
| ENSMUSG( | 24.9007 | 0.337878 | 0.450348 | 0.922202 | Crybg2 |
| ENSMUSG( | 352.1501 | 0.127649 | 0.450443 | 0.922202 | Crybg3 |
| ENSMUSG( | 16.31435 | -0.28123 | 0.572925 | 0.942783 | Crygn |
| ENSMUSG( | 326.6375 | -0.17903 | 0.274129 | 0.889874 | Cryl1 |
| ENSMUSG( | 1777.131 | -0.30529 | 0.153355 | 0.837724 | Crym |
| ENSMUSG( | 630.3913 | -0.21922 | 0.172017 | 0.843003 | Cryz |
| ENSMUSG( | 1246.036 | -0.11408 | 0.331425 | 0.907861 | Cryzl1 |
| ENSMUSG( | 597.3865 | -0.33007 | 0.020865 | 0.571913 | Cryzl2 |
| ENSMUSG( | 11383.54 | -0.22896 | 0.064117 | 0.736482 | Cs |
| ENSMUSG( | 903.4428 | -0.00428 | 0.971853 | 0.996416 | Csad |
| ENSMUSG( | 6351.218 | -0.11258 | 0.353081 | 0.912263 | Csdc2 |
| ENSMUSG( | 4624.228 | -0.07488 | 0.493652 | 0.927372 | Csde1 |
| ENSMUSG( | 940.2755 | 0.114635 | 0.332875 | 0.907907 | Cse1l |
| ENSMUSG( | 950.6832 | 0.215083 | 0.265028 | 0.886134 | Csf1 |
| ENSMUSG( | 8472.486 | 0.223612 | 0.213308 | 0.864412 | Csf1r |
| ENSMUSG( | 734.0554 | -0.01339 | 0.953294 | 0.994798 | Csf2ra |

| | | | | | | | | | | | |
|---|---|---|---|---|---|---|---|---|---|---|---|
| ENSMUSG( | Cox6c | 11813.68 | 0.054503 | 0.514537 | 0.872055 | ENSMUSG( | Cox6c | 9966.111 | -0.17411 | 0.111025 | 0.807885 |
| ENSMUSG( | Cox7a1 | 62.1596 | 0.123649 | 0.615764 | 0.910647 | ENSMUSG( | Cox7a1 | 47.56051 | -0.39792 | 0.198016 | 0.862161 |
| ENSMUSG( | Cox7a2 | 5539.858 | 0.043051 | 0.600643 | 0.90443 | ENSMUSG( | Cox7a2 | 4610.507 | -0.22893 | 0.040592 | 0.683847 |
| ENSMUSG( | Cox7a2l | 6080.98 | 0.047565 | 0.573686 | 0.893584 | ENSMUSG( | Cox7a2l | 5227.122 | -0.12387 | 0.287175 | 0.894091 |
| ENSMUSG( | Cox7b | 4848.425 | -0.05726 | 0.559142 | 0.888638 | ENSMUSG( | Cox7b | 4042.431 | -0.33552 | 0.008274 | 0.388416 |
| ENSMUSG( | Cox7c | 6242.983 | 0.026727 | 0.781133 | 0.957558 | ENSMUSG( | Cox7c | 5351.066 | -0.15545 | 0.18853 | 0.85698 |
| ENSMUSG( | Cox8a | 34552.59 | 0.000482 | 0.995467 | 0.99943 | ENSMUSG( | Cox8a | 29012.19 | -0.24717 | 0.02486 | 0.592568 |
| ENSMUSG( | Cox8b | 15.89983 | -0.41562 | 0.420222 | NA | ENSMUSG( | Cox8b | 14.48473 | -0.42583 | 0.438638 | 0.920721 |
| ENSMUSG( | Cp | 405.4205 | -0.20736 | 0.215512 | 0.66017 | ENSMUSG( | Cp | 438.0296 | 0.288296 | 0.152425 | 0.837724 |
| ENSMUSG( | Cpa2 | 29.6986 | -0.22061 | 0.581405 | 0.897705 | ENSMUSG( | Cpa2 | 25.9995 | -0.37266 | 0.398026 | 0.917257 |
| ENSMUSG( | Cpd | 2599.406 | -0.42213 | 0.008308 | 0.127252 | ENSMUSG( | Cpd | 2841.143 | 0.136393 | 0.483354 | 0.92499 |
| ENSMUSG( | Cpe | 321407.1 | 0.09524 | 0.370922 | 0.798261 | ENSMUSG( | Cpe | 281210.4 | -0.02112 | 0.865995 | 0.987156 |
| ENSMUSG( | Cpeb1 | 1344.347 | -0.0062 | 0.955284 | 0.993874 | ENSMUSG( | Cpeb1 | 1326.239 | 0.216778 | 0.075897 | 0.768524 |
| ENSMUSG( | Cpeb2 | 1538.889 | -0.01122 | 0.907989 | 0.9878 | ENSMUSG( | Cpeb2 | 1488.376 | 0.158543 | 0.161953 | 0.837724 |
| ENSMUSG( | Cpeb3 | 1183.312 | -0.07554 | 0.46759 | 0.844819 | ENSMUSG( | Cpeb3 | 1087.34 | -0.04691 | 0.711645 | 0.964363 |
| ENSMUSG( | Cpeb4 | 2518.282 | -0.30499 | 0.021026 | 0.220525 | ENSMUSG( | Cpeb4 | 2488.257 | -0.05555 | 0.688364 | 0.959742 |
| ENSMUSG( | Cped1 | 171.0893 | 0.14799 | 0.35813 | 0.789704 | ENSMUSG( | Cped1 | 167.9171 | 0.344292 | 0.074466 | 0.766854 |
| ENSMUSG( | Cplane1 | 1861.229 | -0.0624 | 0.539955 | 0.88315 | ENSMUSG( | Cplane1 | 1735.558 | 0.005449 | 0.962688 | 0.995284 |
| ENSMUSG( | Cplane2 | 22.8785 | 0.083579 | 0.827984 | 0.968966 | ENSMUSG( | Cplane2 | 24.11593 | 0.432728 | 0.298718 | 0.898538 |
| ENSMUSG( | Cplx1 | 21595.5 | 0.080432 | 0.340025 | 0.777959 | ENSMUSG( | Cplx1 | 18178.41 | -0.15132 | 0.148183 | 0.836241 |
| ENSMUSG( | Cplx2 | 54831.73 | 0.041054 | 0.633743 | 0.914073 | ENSMUSG( | Cplx2 | 48462.28 | -0.04558 | 0.665107 | 0.956766 |
| ENSMUSG( | Cplx2 | 520.2242 | 0.020981 | 0.864406 | 0.975398 | ENSMUSG( | Cplx2 | 449.3242 | -0.13742 | 0.343078 | 0.909646 |
| ENSMUSG( | Cplx3 | 1119.798 | 0.195468 | 0.068646 | 0.405903 | ENSMUSG( | Cplx3 | 913.1286 | -0.12314 | 0.317578 | 0.902602 |
| ENSMUSG( | Cpm | 352.7308 | -0.20402 | 0.314965 | 0.762155 | ENSMUSG( | Cpm | 462.9226 | 0.768048 | 0.004408 | 0.269982 |
| ENSMUSG( | Cpne2 | 1484.519 | 0.062136 | 0.516116 | 0.872701 | ENSMUSG( | Cpne2 | 1232.659 | -0.2167 | 0.095701 | 0.796133 |
| ENSMUSG( | Cpne3 | 913.0259 | 0.102777 | 0.417976 | 0.821964 | ENSMUSG( | Cpne3 | 793.9782 | -0.03417 | 0.823132 | 0.979198 |
| ENSMUSG( | Cpne4 | 1016.202 | -0.07531 | 0.508559 | 0.868389 | ENSMUSG( | Cpne4 | 962.1459 | 0.032417 | 0.829429 | 0.980224 |
| ENSMUSG( | Cpne5 | 2354.981 | -0.25021 | 0.066512 | 0.39868 | ENSMUSG( | Cpne5 | 2145.888 | -0.25218 | 0.110787 | 0.807582 |
| ENSMUSG( | Cpne6 | 4492.791 | 0.469464 | 0.004386 | 0.086463 | ENSMUSG( | Cpne6 | 3491.128 | 0.04343 | 0.831065 | 0.98051 |
| ENSMUSG( | Cpne7 | 2375.884 | 0.615097 | 0.000926 | 0.031085 | ENSMUSG( | Cpne7 | 1874.483 | 0.249193 | 0.208397 | 0.864412 |
| ENSMUSG( | Cpne8 | 426.4924 | -0.04679 | 0.70057 | 0.93743 | ENSMUSG( | Cpne8 | 388.4419 | -0.05148 | 0.728938 | 0.966913 |
| ENSMUSG( | Cpne9 | 1638.371 | 0.188874 | 0.169179 | 0.603271 | ENSMUSG( | Cpne9 | 1348.079 | -0.1031 | 0.481899 | 0.924578 |
| ENSMUSG( | Cpox | 1021.761 | -0.35677 | 0.00044 | 0.019063 | ENSMUSG( | Cpox | 1079.709 | 0.092723 | 0.474963 | 0.923329 |
| ENSMUSG( | Cpped1 | 1710.456 | -0.03889 | 0.714916 | 0.941144 | ENSMUSG( | Cpped1 | 1474.952 | -0.20978 | 0.137156 | 0.828492 |
| ENSMUSG( | Cpq | 1113.305 | -0.20746 | 0.085959 | 0.451492 | ENSMUSG( | Cpq | 1071.533 | -0.04212 | 0.742601 | 0.969415 |
| ENSMUSG( | Cps1 | 9.08372 | 0.491411 | 0.438836 | NA | ENSMUSG( | Cps1 | 8.060035 | 0.411646 | 0.55947 | NA |
| ENSMUSG( | Cpsf1 | 1841.014 | 0.132799 | 0.172477 | 0.607771 | ENSMUSG( | Cpsf1 | 1584.418 | -0.02803 | 0.818653 | 0.979198 |
| ENSMUSG( | Cpsf2 | 1305.711 | 0.134804 | 0.263563 | 0.714284 | ENSMUSG( | Cpsf2 | 1152.213 | 0.041697 | 0.773245 | 0.974525 |
| ENSMUSG( | Cpsf3 | 1247.694 | 0.024758 | 0.828811 | 0.969097 | ENSMUSG( | Cpsf3 | 1097.661 | -0.07758 | 0.537118 | 0.937216 |
| ENSMUSG( | Cpsf4 | 567.7388 | -0.06983 | 0.653491 | 0.922819 | ENSMUSG( | Cpsf4 | 549.172 | 0.105613 | 0.493663 | 0.927372 |
| ENSMUSG( | Cpsf6 | 1249.556 | -0.02651 | 0.797452 | 0.960507 | ENSMUSG( | Cpsf6 | 1316.814 | 0.370724 | 0.004427 | 0.270226 |
| ENSMUSG( | Cpsf7 | 2304.901 | 0.011292 | 0.90289 | 0.986055 | ENSMUSG( | Cpsf7 | 2161.998 | 0.093339 | 0.437214 | 0.920144 |
| ENSMUSG( | Cpt1a | 2642.803 | -0.04144 | 0.731276 | 0.944956 | ENSMUSG( | Cpt1a | 2403.83 | -0.04314 | 0.740137 | 0.968811 |
| ENSMUSG( | Cpt1c | 3558.048 | -0.06989 | 0.530824 | 0.879289 | ENSMUSG( | Cpt1c | 3489.226 | 0.143052 | 0.269733 | 0.887508 |
| ENSMUSG( | Cpt2 | 337.0553 | 0.071467 | 0.576226 | 0.895164 | ENSMUSG( | Cpt2 | 283.131 | -0.16349 | 0.281055 | 0.892356 |
| ENSMUSG( | Cptp | 461.3432 | 0.025565 | 0.831185 | 0.969889 | ENSMUSG( | Cptp | 429.1273 | 0.089745 | 0.529038 | 0.934795 |
| ENSMUSG( | Cpxm1 | 366.7393 | -0.2627 | 0.039275 | 0.305097 | ENSMUSG( | Cpxm1 | 376.1627 | 0.093814 | 0.511089 | 0.929571 |
| ENSMUSG( | Cpxm2 | 126.9374 | -0.30783 | 0.141564 | 0.558606 | ENSMUSG( | Cpxm2 | 116.5365 | -0.26513 | 0.252065 | 0.879902 |
| ENSMUSG( | Cpz | 58.50503 | -0.22478 | 0.402717 | 0.81552 | ENSMUSG( | Cpz | 49.96632 | -0.40794 | 0.179938 | 0.850372 |
| ENSMUSG( | Cr1l | 747.6013 | -0.04701 | 0.778531 | 0.956215 | ENSMUSG( | Cr1l | 711.3775 | 0.084837 | 0.645144 | 0.953988 |
| ENSMUSG( | Crabp1 | 21.33141 | -0.30189 | 0.549302 | 0.885741 | ENSMUSG( | Crabp1 | 25.3662 | 0.389226 | 0.469765 | 0.923329 |
| ENSMUSG( | Crabp2 | 73.74432 | -0.40512 | 0.107271 | 0.495579 | ENSMUSG( | Crabp2 | 77.13562 | 0.003746 | 0.988646 | 0.997862 |
| ENSMUSG( | Cracd | 1433.045 | 0.115827 | 0.178868 | 0.616298 | ENSMUSG( | Cracd | 1252.827 | -0.00287 | 0.979359 | 0.996468 |
| ENSMUSG( | Cracdl | 6411.913 | 0.058115 | 0.64267 | 0.916865 | ENSMUSG( | Cracdl | 5827.874 | 0.052262 | 0.706296 | 0.963467 |
| ENSMUSG( | Cracr2a | 81.22131 | -0.37566 | 0.100257 | 0.48338 | ENSMUSG( | Cracr2a | 71.5321 | -0.47115 | 0.061357 | 0.736111 |
| ENSMUSG( | Cracr2b | 114.5666 | 0.113102 | 0.580364 | 0.896952 | ENSMUSG( | Cracr2b | 132.8357 | 0.736972 | 0.001902 | 0.163326 |
| ENSMUSG( | Cradd | 149.9534 | -0.00861 | 0.960259 | 0.993991 | ENSMUSG( | Cradd | 131.3571 | -0.14138 | 0.506099 | 0.927825 |
| ENSMUSG( | Cramp1 | 1465.831 | 0.079576 | 0.380376 | 0.801757 | ENSMUSG( | Cramp1 | 1331.368 | 0.072649 | 0.559182 | 0.939587 |
| ENSMUSG( | Crat | 2598.32 | 0.164883 | 0.047825 | 0.333933 | ENSMUSG( | Crat | 2175.101 | -0.07708 | 0.474967 | 0.923329 |
| ENSMUSG( | Crb1 | 23.89836 | -0.40402 | 0.312936 | 0.76033 | ENSMUSG( | Crb1 | 20.38584 | -0.67499 | 0.1528 | 0.837724 |
| ENSMUSG( | Crb2 | 278.0581 | 0.093717 | 0.538752 | 0.882772 | ENSMUSG( | Crb2 | 273.5564 | 0.313859 | 0.09932 | 0.800296 |
| ENSMUSG( | Crb3 | 18.63984 | -0.08026 | 0.869404 | NA | ENSMUSG( | Crb3 | 16.64786 | -0.12824 | 0.804148 | 0.977191 |
| ENSMUSG( | Crbn | 966.5307 | 0.011365 | 0.910814 | 0.988386 | ENSMUSG( | Crbn | 865.4839 | -0.04052 | 0.736932 | 0.967389 |
| ENSMUSG( | Crcp | 768.1508 | 0.034138 | 0.763799 | 0.954013 | ENSMUSG( | Crcp | 660.3252 | -0.14234 | 0.307851 | 0.898538 |
| ENSMUSG( | Creb1 | 1112.853 | 0.214358 | 0.071555 | 0.413178 | ENSMUSG( | Creb1 | 997.0361 | 0.167251 | 0.198574 | 0.862705 |
| ENSMUSG( | Creb3 | 1351.33 | -0.20682 | 0.070352 | 0.40908 | ENSMUSG( | Creb3 | 1326.448 | 0.018272 | 0.895075 | 0.990558 |
| ENSMUSG( | Creb3l1 | 270.4077 | -0.16099 | 0.321572 | 0.766705 | ENSMUSG( | Creb3l1 | 264.6706 | 0.040866 | 0.817918 | 0.979198 |
| ENSMUSG( | Creb3l2 | 645.2099 | -0.03707 | 0.821418 | 0.966248 | ENSMUSG( | Creb3l2 | 639.106 | 0.206566 | 0.284111 | 0.892412 |
| ENSMUSG( | Creb3l3 | 14.52274 | -0.1738 | 0.779997 | NA | ENSMUSG( | Creb3l3 | 14.9092 | 0.227425 | 0.702604 | 0.962215 |
| ENSMUSG( | Creb3l4 | 17.75931 | -0.69507 | 0.128811 | NA | ENSMUSG( | Creb3l4 | 14.71397 | -1.03364 | 0.049966 | 0.708648 |
| ENSMUSG( | Creb5 | 123.3768 | -0.51801 | 0.010027 | 0.142701 | ENSMUSG( | Creb5 | 132.7829 | 0.02552 | 0.909534 | 0.991958 |
| ENSMUSG( | Crebbp | 2055.405 | 0.073439 | 0.450671 | 0.835475 | ENSMUSG( | Crebbp | 1912.89 | 0.131459 | 0.243833 | 0.877904 |
| ENSMUSG( | Crebl2 | 1174.211 | 0.147383 | 0.159184 | 0.588305 | ENSMUSG( | Crebl2 | 1038.699 | 0.064377 | 0.575004 | 0.942912 |
| ENSMUSG( | Crebrf | 1350.154 | 0.14277 | 0.159364 | 0.58868 | ENSMUSG( | Crebrf | 1209.533 | 0.096971 | 0.400092 | 0.917257 |
| ENSMUSG( | Crebzf | 1326.885 | -0.1447 | 0.160839 | 0.590628 | ENSMUSG( | Crebzf | 1432.584 | 0.333318 | 0.048295 | 0.706142 |
| ENSMUSG( | Creg1 | 1393.227 | -0.33692 | 0.015722 | 0.185786 | ENSMUSG( | Creg1 | 1495.542 | 0.154667 | 0.301224 | 0.898538 |
| ENSMUSG( | Creg2 | 3045.003 | -0.29026 | 0.08325 | 0.445247 | ENSMUSG( | Creg2 | 3343.922 | 0.258076 | 0.178302 | 0.848538 |
| ENSMUSG( | Creld1 | 2985.514 | -0.27914 | 0.018199 | 0.203238 | ENSMUSG( | Creld1 | 3099.844 | 0.112131 | 0.403499 | 0.917257 |
| ENSMUSG( | Creld2 | 640.4631 | -0.00126 | 0.992922 | 0.999327 | ENSMUSG( | Creld2 | 682.9275 | 0.426234 | 0.004839 | 0.287322 |
| ENSMUSG( | Crem | 460.2017 | 0.054547 | 0.658887 | 0.924506 | ENSMUSG( | Crem | 382.5738 | -0.22798 | 0.151773 | 0.837724 |
| ENSMUSG( | Crh | 257.9942 | -0.73036 | 0.000355 | 0.016296 | ENSMUSG( | Crh | 261.1473 | -0.36328 | 0.110595 | 0.807582 |
| ENSMUSG( | Crhbp | 531.2404 | -0.39883 | 0.001184 | 0.037077 | ENSMUSG( | Crhbp | 551.7271 | 0.010132 | 0.943369 | 0.994798 |
| ENSMUSG( | Crhr1 | 1037.193 | -0.0062 | 0.959419 | 0.993991 | ENSMUSG( | Crhr1 | 950.7906 | 0.016124 | 0.90219 | 0.991012 |
| ENSMUSG( | Crhr2 | 58.1888 | -0.04154 | 0.866823 | 0.976541 | ENSMUSG( | Crhr2 | 57.70393 | 0.178691 | 0.512629 | 0.930164 |
| ENSMUSG( | Crim1 | 1872.054 | -0.08113 | 0.532459 | 0.879905 | ENSMUSG( | Crim1 | 1902.066 | 0.228901 | 0.127197 | 0.817278 |
| ENSMUSG( | Crip1 | 91.12728 | -0.13067 | 0.557354 | 0.888279 | ENSMUSG( | Crip1 | 86.85423 | 0.024113 | 0.919943 | 0.992644 |
| ENSMUSG( | Crip2 | 4460.588 | 0.015236 | 0.846974 | 0.972374 | ENSMUSG( | Crip2 | 3904.755 | -0.10281 | 0.32113 | 0.904019 |
| ENSMUSG( | Crip3 | 68.95793 | 0.435804 | 0.08211 | 0.442642 | ENSMUSG( | Crip3 | 68.70501 | 0.644907 | 0.031529 | 0.631192 |
| ENSMUSG( | Cript | 1087.233 | 0.033031 | 0.789681 | 0.959461 | ENSMUSG( | Cript | 944.7051 | -0.11292 | 0.483386 | 0.92499 |
| ENSMUSG( | Cripto | 89.41266 | -0.10139 | 0.648292 | 0.919949 | ENSMUSG( | Cripto | 78.42798 | -0.21393 | 0.405241 | 0.917257 |
| ENSMUSG( | Crispld1 | 122.0513 | -0.00686 | 0.974169 | 0.99691 | ENSMUSG( | Crispld1 | 122.5508 | 0.277736 | 0.221272 | 0.866104 |
| ENSMUSG( | Crispld2 | 96.56861 | -0.02156 | 0.922322 | 0.99045 | ENSMUSG( | Crispld2 | 109.4195 | 0.569721 | 0.01291 | 0.458131 |
| ENSMUSG( | Crk | 4016.785 | 0.095991 | 0.310266 | 0.758431 | ENSMUSG( | Crk | 3502.503 | -0.03175 | 0.791822 | 0.97681 |
| ENSMUSG( | Crkl | 1706.342 | 0.186806 | 0.034271 | 0.284043 | ENSMUSG( | Crkl | 1461.481 | 0.014199 | 0.893501 | 0.990558 |
| ENSMUSG( | Crlf1 | 212.951 | 0.085317 | 0.594194 | 0.902869 | ENSMUSG( | Crlf1 | 206.8831 | 0.262423 | 0.105849 | 0.804969 |
| ENSMUSG( | Crlf2 | 830.4585 | -0.10051 | 0.36872 | 0.796356 | ENSMUSG( | Crlf2 | 781.8641 | -0.00451 | 0.972735 | 0.996416 |
| ENSMUSG( | Crlf3 | 508.56 | 0.16099 | 0.171439 | 0.605943 | ENSMUSG( | Crlf3 | 476.5866 | 0.241592 | 0.083601 | 0.77681 |
| ENSMUSG( | Crls1 | 708.4283 | 0.046436 | 0.70094 | 0.93743 | ENSMUSG( | Crls1 | 650.4159 | 0.066096 | 0.601301 | 0.947749 |
| ENSMUSG( | Crmp1 | 4285.913 | 0.086549 | 0.516016 | 0.872701 | ENSMUSG( | Crmp1 | 3608.097 | -0.14567 | 0.33941 | 0.907972 |
| ENSMUSG( | Crnde | 17.6807 | 0.304188 | 0.504308 | NA | ENSMUSG( | Crnde | 14.92397 | 0.125361 | 0.809054 | 0.97873 |
| ENSMUSG( | Crnkl1 | 443.4003 | 0.015661 | 0.898296 | 0.985633 | ENSMUSG( | Crnkl1 | 406.4781 | 0.032655 | 0.81823 | 0.979198 |
| ENSMUSG( | Crocc | 2000.076 | 0.300242 | 0.039979 | 0.3082 | ENSMUSG( | Crocc | 1687.228 | 0.090213 | 0.554126 | 0.939565 |
| ENSMUSG( | Crocc2 | 52.69693 | 0.63943 | 0.097145 | 0.47759 | ENSMUSG( | Crocc2 | 37.84385 | 0.047208 | 0.898373 | 0.990558 |
| ENSMUSG( | Crot | 2836.392 | 0.151539 | 0.282437 | 0.73186 | ENSMUSG( | Crot | 2345.379 | -0.1286 | 0.379698 | 0.916995 |
| ENSMUSG( | Crppa | 155.7725 | -0.05846 | 0.73763 | 0.947329 | ENSMUSG( | Crppa | 132.7312 | -0.2808 | 0.178107 | 0.848262 |
| ENSMUSG( | Crtac1 | 2219.207 | -0.26933 | 0.070188 | 0.40908 | ENSMUSG( | Crtac1 | 2300.213 | 0.118286 | 0.465812 | 0.923329 |
| ENSMUSG( | Crtap | 274.9696 | -0.18044 | 0.336677 | 0.777 | ENSMUSG( | Crtap | 286.437 | 0.216846 | 0.299589 | 0.898538 |
| ENSMUSG( | Crtc1 | 9194.981 | 0.126267 | 0.165629 | 0.59714 | ENSMUSG( | Crtc1 | 8127.716 | 0.043156 | 0.690189 | 0.960071 |
| ENSMUSG( | Crtc2 | 980.7799 | -0.04656 | 0.685953 | 0.932981 | ENSMUSG( | Crtc2 | 901.1144 | -0.01801 | 0.898301 | 0.990558 |
| ENSMUSG( | Crtc3 | 850.3839 | 0.041161 | 0.679737 | 0.930983 | ENSMUSG( | Crtc3 | 765.1755 | 0.004244 | 0.971406 | 0.996236 |
| ENSMUSG( | Cry1 | 551.9353 | -0.0302 | 0.77833 | 0.956215 | ENSMUSG( | Cry1 | 599.0289 | 0.434757 | 0.003485 | 0.235685 |
| ENSMUSG( | Cry2 | 3290.911 | 0.138232 | 0.145945 | 0.567409 | ENSMUSG( | Cry2 | 3017.971 | 0.15765 | 0.183333 | 0.850984 |
| ENSMUSG( | Cryab | 5220.011 | -0.00144 | 0.988416 | 0.999109 | ENSMUSG( | Cryab | 4342.652 | -0.27974 | 0.020434 | 0.565891 |
| ENSMUSG( | Cryba2 | 17.14335 | 0.454539 | 0.358862 | NA | ENSMUSG( | Cryba2 | 10.74465 | -0.6263 | 0.316484 | NA |
| ENSMUSG( | Cryba4 | 43.79235 | 0.447117 | 0.120045 | 0.523983 | ENSMUSG( | Cryba4 | 31.45685 | -0.2027 | 0.560046 | 0.940119 |
| ENSMUSG( | Crybb1 | 408.977 | 0.325903 | 0.018283 | 0.203658 | ENSMUSG( | Crybb1 | 336.5324 | 0.05294 | 0.726943 | 0.966887 |
| ENSMUSG( | Crybb3 | 47.42893 | 0.104936 | 0.724808 | 0.944096 | ENSMUSG( | Crybb3 | 43.42713 | 0.09433 | 0.779286 | 0.976106 |
| ENSMUSG( | Crybg1 | 54.28505 | -0.25072 | 0.346227 | 0.779153 | ENSMUSG( | Crybg1 | 69.37276 | 0.643955 | 0.017149 | 0.52741 |
| ENSMUSG( | Crybg2 | 22.1694 | -0.21778 | 0.60363 | 0.904887 | ENSMUSG( | Crybg2 | 24.9007 | 0.337878 | 0.450348 | 0.922202 |
| ENSMUSG( | Crybg3 | 360.4368 | -0.08292 | 0.561069 | 0.888934 | ENSMUSG( | Crybg3 | 352.1501 | 0.127649 | 0.450443 | 0.922202 |
| ENSMUSG( | Crygn | 15.97997 | -0.68942 | 0.156148 | NA | ENSMUSG( | Crygn | 16.31435 | -0.28123 | 0.572925 | 0.942783 |
| ENSMUSG( | Cryl1 | 378.0326 | -0.02148 | 0.881439 | 0.981798 | ENSMUSG( | Cryl1 | 326.6375 | -0.17903 | 0.274129 | 0.889874 |
| ENSMUSG( | Crym | 1965.422 | -0.27809 | 0.121463 | 0.525713 | ENSMUSG( | Crym | 1777.131 | -0.30529 | 0.153355 | 0.837724 |
| ENSMUSG( | Cryz | 744.0903 | 0.006493 | 0.962771 | 0.994824 | ENSMUSG( | Cryz | 630.3913 | -0.21922 | 0.172017 | 0.843003 |
| ENSMUSG( | Cryzl1 | 1430.742 | 0.021703 | 0.816392 | 0.965455 | ENSMUSG( | Cryzl1 | 1246.036 | -0.11408 | 0.331425 | 0.907861 |
| ENSMUSG( | Cryzl2 | 739.2448 | 0.041015 | 0.757815 | 0.951582 | ENSMUSG( | Cryzl2 | 597.3865 | -0.33007 | 0.020865 | 0.571913 |
| ENSMUSG( | Cs | 13784.08 | 0.064617 | 0.493861 | 0.860611 | ENSMUSG( | Cs | 11383.54 | -0.22896 | 0.064117 | 0.736482 |
| ENSMUSG( | Csad | 1059.474 | 0.181578 | 0.060861 | 0.381147 | ENSMUSG( | Csad | 903.4428 | -0.00428 | 0.971853 | 0.996416 |
| ENSMUSG( | Csdc2 | 7683.649 | 0.167247 | 0.042514 | 0.316636 | ENSMUSG( | Csdc2 | 6351.218 | -0.11258 | 0.353081 | 0.912263 |
| ENSMUSG( | Csde1 | 5216.475 | 0.006859 | 0.934445 | 0.99323 | ENSMUSG( | Csde1 | 4624.228 | -0.07488 | 0.493652 | 0.927372 |
| ENSMUSG( | Cse1l | 1000.29 | 0.027092 | 0.769898 | 0.954689 | ENSMUSG( | Cse1l | 940.2755 | 0.114635 | 0.332875 | 0.907907 |
| ENSMUSG( | Csf1 | 825.9645 | -0.50317 | 0.001056 | 0.034188 | ENSMUSG( | Csf1 | 950.6832 | 0.215083 | 0.265028 | 0.886134 |
| ENSMUSG( | Csf1r | 8336.079 | -0.0891 | 0.526955 | 0.877267 | ENSMUSG( | Csf1r | 8472.486 | 0.223612 | 0.213308 | 0.864412 |
| ENSMUSG( | Csf2ra | 873.0895 | 0.215703 | 0.273558 | 0.724071 | ENSMUSG( | Csf2ra | 734.0554 | -0.01339 | 0.953294 | 0.994798 |
| ENSMUSG( | Csf2rb | 90.25594 | -0.02887 | 0.896657 | 0.985017 | ENSMUSG( | Csf2rb | 94.74726 | 0.382644 | 0.152183 | 0.837724 |
| ENSMUSG( | Csf2rb2 | 60.43703 | -0.2346 | 0.368049 | 0.795774 | ENSMUSG( | Csf2rb2 | 67.20066 | 0.347365 | 0.1977 | 0.862161 |
| ENSMUSG( | Csf3r | 435.9686 | -0.27128 | 0.123822 | 0.531039 | ENSMUSG( | Csf3r | 430.3223 | -0.01761 | 0.930827 | 0.99306 |
| ENSMUSG( | Csgalnact1 | 1026.907 | -0.10274 | 0.412047 | 0.819446 | ENSMUSG( | Csgalnact1 | 960.1921 | -0.02426 | 0.869257 | 0.988097 |
| ENSMUSG( | Csgalnact2 | 493.524 | -0.25493 | 0.099885 | 0.482328 | ENSMUSG( | Csgalnact2 | 497.8946 | 0.05374 | 0.746296 | 0.969654 |
| ENSMUSG( | Csk | 1071.534 | 0.067914 | 0.50084 | 0.863671 | ENSMUSG( | Csk | 957.4904 | 0.014803 | 0.897386 | 0.990558 |
| ENSMUSG( | Csmd1 | 1699.164 | -0.41151 | 0.000439 | 0.019063 | ENSMUSG( | Csmd1 | 2005.322 | 0.348186 | 0.023314 | 0.585577 |
| ENSMUSG( | Csmd2 | 1762.64 | -0.4411 | 0.000147 | 0.008459 | ENSMUSG( | Csmd2 | 1796.382 | -0.0813 | 0.545326 | 0.938063 |
| ENSMUSG( | Csmd3 | 430.3292 | -0.86905 | 2.61E-08 | 9.19E-06 | ENSMUSG( | Csmd3 | 524.3221 | 0.107074 | 0.576617 | 0.942912 |
| ENSMUSG( | Csnk1a1 | 5270.27 | 0.032148 | 0.681773 | 0.931619 | ENSMUSG( | Csnk1a1 | 4845.557 | 0.056881 | 0.569224 | 0.941777 |
| ENSMUSG( | Csnk1d | 3569.468 | 0.176802 | 0.055298 | 0.361957 | ENSMUSG( | Csnk1d | 3036.828 | -0.01566 | 0.887492 | 0.990266 |
| ENSMUSG( | Csnk1e | 3025.17 | 0.065495 | 0.553607 | 0.886216 | ENSMUSG( | Csnk1e | 2762.643 | 0.071496 | 0.554562 | 0.939565 |
| ENSMUSG( | Csnk1g1 | 732.3493 | -0.02084 | 0.871478 | 0.978114 | ENSMUSG( | Csnk1g1 | 663.0475 | -0.03676 | 0.798397 | 0.97681 |
| ENSMUSG( | Csnk1g2 | 12090.77 | 0.208645 | 0.008674 | 0.130875 | ENSMUSG( | Csnk1g2 | 10549.18 | 0.089576 | 0.375504 | 0.915491 |
| ENSMUSG( | Csnk1g3 | 1116.018 | 0.131319 | 0.175578 | 0.611004 | ENSMUSG( | Csnk1g3 | 1019.489 | 0.135141 | 0.284502 | 0.892412 |
| ENSMUSG( | Csnk2a1 | 2967.157 | 0.007676 | 0.941072 | 0.993415 | ENSMUSG( | Csnk2a1 | 2613.307 | -0.09545 | 0.469665 | 0.923329 |
| ENSMUSG( | Csnk2a2 | 1847.675 | 0.131736 | 0.173542 | 0.609013 | ENSMUSG( | Csnk2a2 | 1615.558 | 0.01513 | 0.896952 | 0.990558 |
| ENSMUSG( | Csnk2b | 1996.983 | -0.01201 | 0.905103 | 0.98645 | ENSMUSG( | Csnk2b | 1758.616 | -0.1162 | 0.42736 | 0.917257 |
| ENSMUSG( | Cspg4 | 881.2189 | -0.79342 | 0.000419 | 0.01851 | ENSMUSG( | Cspg4 | 1082.706 | 0.181497 | 0.691873 | 0.960344 |
| ENSMUSG( | Cspg4b | 187.1761 | -0.04095 | 0.827041 | 0.96865 | ENSMUSG( | Cspg4b | 181.6064 | 0.13222 | 0.450858 | 0.922202 |
| ENSMUSG( | Cspg5 | 25426.58 | -0.0699 | 0.576465 | 0.895276 | ENSMUSG( | Cspg5 | 24055.7 | 0.041977 | 0.765824 | 0.974236 |
| ENSMUSG( | Cspp1 | 936.8173 | 0.142088 | 0.177775 | 0.615225 | ENSMUSG( | Cspp1 | 816.9355 | 0.016397 | 0.902833 | 0.991089 |
| ENSMUSG( | Csrnp1 | 187.6521 | -0.33153 | 0.083928 | 0.446625 | ENSMUSG( | Csrnp1 | 176.7833 | -0.20752 | 0.397866 | 0.917257 |
| ENSMUSG( | Csrnp2 | 561.4946 | -0.07308 | 0.592876 | 0.902275 | ENSMUSG( | Csrnp2 | 518.6104 | -0.03533 | 0.814793 | 0.979198 |

| | | | | | |
|---|---|---|---|---|---|
| ENSMUSG( | 739.2448 | 0.041015 | 0.757815 | 0.951582 | Cryzl2 |
| ENSMUSG( | 13784.08 | 0.064617 | 0.493861 | 0.860611 | Cs |
| ENSMUSG( | 1059.474 | 0.181578 | 0.060861 | 0.381147 | Csad |
| ENSMUSG( | 7683.649 | 0.167247 | 0.042514 | 0.316636 | Csdc2 |
| ENSMUSG( | 5216.475 | 0.006859 | 0.934445 | 0.99323 | Csde1 |
| ENSMUSG( | 1000.29 | 0.027092 | 0.769898 | 0.954689 | Cse1l |
| ENSMUSG( | 825.9645 | -0.50317 | 0.001056 | 0.034188 | Csf1 |
| ENSMUSG( | 8336.079 | -0.0891 | 0.526955 | 0.877267 | Csf1r |
| ENSMUSG( | 873.0895 | 0.215703 | 0.273558 | 0.724071 | Csf2ra |
| ENSMUSG( | 90.25594 | -0.02887 | 0.896657 | 0.985017 | Csf2rb |
| ENSMUSG( | 60.43703 | -0.2346 | 0.368049 | 0.795774 | Csf2rb2 |
| ENSMUSG( | 435.9686 | -0.27128 | 0.123822 | 0.531039 | Csf3r |
| ENSMUSG( | 1026.907 | -0.10274 | 0.412047 | 0.819446 | Csgalnact1 |
| ENSMUSG( | 493.524 | -0.25493 | 0.099885 | 0.482328 | Csgalnact2 |
| ENSMUSG( | 1071.534 | 0.067914 | 0.50084 | 0.863671 | Csk |
| ENSMUSG( | 9.796736 | -0.14596 | 0.800388 | NA | Csl |
| ENSMUSG( | 1699.164 | -0.41151 | 0.000439 | 0.019063 | Csmd1 |
| ENSMUSG( | 1762.64 | -0.4411 | 0.000147 | 0.008459 | Csmd2 |
| ENSMUSG( | 430.3292 | -0.86905 | 2.61E-08 | 9.19E-06 | Csmd3 |
| ENSMUSG( | 5270.27 | 0.032148 | 0.681773 | 0.931619 | Csnk1a1 |
| ENSMUSG( | 3569.468 | 0.176802 | 0.055298 | 0.361957 | Csnk1d |
| ENSMUSG( | 3025.17 | 0.065495 | 0.553607 | 0.886216 | Csnk1e |
| ENSMUSG( | 732.3493 | -0.02084 | 0.871478 | 0.978114 | Csnk1g1 |
| ENSMUSG( | 12090.77 | 0.208645 | 0.008674 | 0.130875 | Csnk1g2 |
| ENSMUSG( | 1116.018 | 0.131319 | 0.175578 | 0.611004 | Csnk1g3 |
| ENSMUSG( | 2967.157 | 0.007676 | 0.941072 | 0.993415 | Csnk2a1 |
| ENSMUSG( | 1847.675 | 0.131736 | 0.173542 | 0.609013 | Csnk2a2 |
| ENSMUSG( | 1996.983 | -0.01201 | 0.905103 | 0.98645 | Csnk2b |
| ENSMUSG( | 881.2189 | -0.79342 | 0.000419 | 0.01851 | Cspg4 |
| ENSMUSG( | 187.1761 | -0.04095 | 0.827041 | 0.96865 | Cspg4b |
| ENSMUSG( | 25426.58 | -0.0699 | 0.576465 | 0.895276 | Cspg5 |
| ENSMUSG( | 936.8173 | 0.142088 | 0.177775 | 0.615225 | Cspp1 |
| ENSMUSG( | 187.6521 | -0.33153 | 0.083928 | 0.446625 | Csrnp1 |
| ENSMUSG( | 561.4946 | -0.07308 | 0.592876 | 0.902275 | Csrnp2 |
| ENSMUSG( | 868.7991 | -0.05036 | 0.625589 | 0.912249 | Csrnp3 |
| ENSMUSG( | 20843.69 | 0.149251 | 0.119879 | 0.523965 | Csrp1 |
| ENSMUSG( | 201.8687 | -0.00826 | 0.963317 | 0.994911 | Csrp2 |
| ENSMUSG( | 94894.08 | -0.04063 | 0.705099 | 0.938303 | Cst3 |
| ENSMUSG( | 206.7391 | -0.15973 | 0.326133 | 0.771128 | Cst6 |
| ENSMUSG( | 13.62662 | -0.50332 | 0.315968 | NA | Cst7 |
| ENSMUSG( | 39.09651 | -0.26532 | 0.410148 | 0.818258 | Cstad |
| ENSMUSG( | 514.7191 | -0.03292 | 0.82266 | 0.966352 | Cstb |
| ENSMUSG( | 576.1378 | -0.01906 | 0.867904 | 0.976631 | Cstf1 |
| ENSMUSG( | 2719.623 | 0.097372 | 0.34553 | 0.779153 | Cstf2 |
| ENSMUSG( | 1655.201 | -0.01954 | 0.860254 | 0.974842 | Cstf2t |
| ENSMUSG( | 546.6062 | -0.01498 | 0.898939 | 0.985633 | Cstf3 |
| ENSMUSG( | 1890.206 | 0.074412 | 0.576085 | 0.895114 | Cstpp1 |
| ENSMUSG( | 9.870085 | -0.51812 | 0.411346 | NA | Ct55 |
| ENSMUSG( | 10291.14 | 0.058347 | 0.446786 | 0.833933 | Ctbp1 |
| ENSMUSG( | 751.591 | 0.182371 | 0.079433 | 0.43547 | Ctbp2 |
| ENSMUSG( | 323.0052 | -0.39043 | 0.033942 | 0.282701 | Ctbs |
| ENSMUSG( | 1303.688 | 0.112413 | 0.311265 | 0.759247 | Ctc1 |
| ENSMUSG( | 1655.42 | 0.123309 | 0.165568 | 0.59714 | Ctcf |
| ENSMUSG( | 3690.692 | -0.06712 | 0.460773 | 0.840962 | Ctdnep1 |
| ENSMUSG( | 486.6646 | 0.04158 | 0.753957 | 0.950981 | Ctdp1 |
| ENSMUSG( | 3038.802 | 0.11082 | 0.2406 | 0.688247 | Ctdsp1 |
| ENSMUSG( | 4047.108 | 0.090373 | 0.298085 | 0.746708 | Ctdsp2 |
| ENSMUSG( | 1531.818 | 0.063712 | 0.525604 | 0.876728 | Ctdspl |
| ENSMUSG( | 338.4522 | 0.065133 | 0.59911 | 0.903801 | Ctdspl2 |
| ENSMUSG( | 95.21484 | -0.03014 | 0.879802 | 0.981436 | Ctf1 |
| ENSMUSG( | 233.6706 | 0.351371 | 0.025986 | 0.247245 | Cth |
| ENSMUSG( | 176.4848 | -0.39272 | 0.019455 | 0.210037 | Cthrc1 |
| ENSMUSG( | 2059.81 | 0.216771 | 0.014711 | 0.178407 | Ctif |
| ENSMUSG( | 221.5491 | 0.277306 | 0.1283 | 0.536641 | Ctla2a |
| ENSMUSG( | 4036.735 | 0.153232 | 0.145701 | 0.566867 | Ctnna1 |
| ENSMUSG( | 9011.947 | 0.19121 | 0.019979 | 0.213756 | Ctnna2 |
| ENSMUSG( | 51.18368 | -0.07163 | 0.796203 | 0.960088 | Ctnna3 |
| ENSMUSG( | 783.1745 | -0.08818 | 0.414255 | 0.819998 | Ctnnal1 |
| ENSMUSG( | 34957.41 | 0.116692 | 0.263285 | 0.714007 | Ctnnb1 |
| ENSMUSG( | 3992.973 | 0.047717 | 0.538051 | 0.882548 | Ctnnbip1 |
| ENSMUSG( | 826.8834 | 0.019809 | 0.849077 | 0.972601 | Ctnnbl1 |
| ENSMUSG( | 571.9725 | -0.05236 | 0.66394 | 0.926511 | Ctnnd1 |
| ENSMUSG( | 14530.5 | 0.153271 | 0.060128 | 0.37919 | Ctnnd2 |
| ENSMUSG( | 680.0659 | -0.25689 | 0.071522 | 0.413178 | Ctns |
| ENSMUSG( | 994.2529 | 0.000213 | 0.998563 | 0.99989 | Ctps1 |
| ENSMUSG( | 837.2624 | 0.089964 | 0.54057 | 0.88315 | Ctps2 |
| ENSMUSG( | 1447.066 | 0.054975 | 0.606081 | 0.906049 | Ctr9 |
| ENSMUSG( | 23.73202 | 0.341735 | 0.38786 | 0.805835 | Ctrl |
| ENSMUSG( | 5164.614 | -0.26691 | 0.013509 | 0.169727 | Ctsa |
| ENSMUSG( | 25581.38 | -0.32429 | 0.00895 | 0.132538 | Ctsb |
| ENSMUSG( | 414.681 | -0.2589 | 0.172155 | 0.607115 | Ctsc |
| ENSMUSG( | 11193.58 | -0.13954 | 0.271981 | 0.722255 | Ctsd |
| ENSMUSG( | 3008.88 | -0.21541 | 0.099669 | 0.482191 | Ctsf |
| ENSMUSG( | 688.4768 | -0.12129 | 0.504974 | 0.866883 | Ctsh |
| ENSMUSG( | 202.6728 | -0.22256 | 0.131366 | 0.543562 | Ctsk |
| ENSMUSG( | 4527.475 | -0.24848 | 0.059019 | 0.375291 | Ctsl |
| ENSMUSG( | 3357.75 | 0.011545 | 0.926986 | 0.991229 | Ctso |
| ENSMUSG( | 4702.742 | -0.1785 | 0.162854 | 0.593778 | Ctss |
| ENSMUSG( | 11.37515 | -0.24763 | 0.695021 | NA | Ctsw |
| ENSMUSG( | 2142.782 | -0.26458 | 0.046565 | 0.330304 | Ctsz |
| ENSMUSG( | 3080.06 | 0.12374 | 0.160269 | 0.589853 | Cttn |
| ENSMUSG( | 2834.803 | 0.030637 | 0.776218 | 0.955658 | Cttnbp2 |
| ENSMUSG( | 1074.585 | -0.07956 | 0.474218 | 0.849425 | Cttnbp2nl |
| ENSMUSG( | 180.5756 | -0.0494 | 0.769088 | 0.954674 | Ctu1 |
| ENSMUSG( | 554.9728 | 0.071442 | 0.579252 | 0.896692 | Ctu2 |
| ENSMUSG( | 22247.54 | 0.087073 | 0.297264 | 0.745805 | Ctxn1 |
| ENSMUSG( | 203.4018 | 0.07597 | 0.636347 | 0.914873 | Ctxn2 |
| ENSMUSG( | 102.7631 | -0.00698 | 0.972769 | 0.996585 | Ctxn3 |
| ENSMUSG( | 1706.584 | 0.252442 | 0.004717 | 0.090333 | Ctxnd1 |
| ENSMUSG( | 24.86982 | -0.33482 | 0.431977 | 0.82768 | Cubn |
| ENSMUSG( | 2674.999 | -0.05392 | 0.573537 | 0.893584 | Cuedc2 |
| ENSMUSG( | 4112.977 | 0.085136 | 0.365473 | 0.794085 | Cul1 |
| ENSMUSG( | 1151.436 | 0.005387 | 0.953086 | 0.993874 | Cul2 |
| ENSMUSG( | 3599.119 | 0.077964 | 0.429415 | 0.826446 | Cul3 |
| ENSMUSG( | 2292.204 | 0.127164 | 0.119466 | 0.523291 | Cul4a |
| ENSMUSG( | 775.5685 | 0.046493 | 0.695523 | 0.934979 | Cul4b |
| ENSMUSG( | 1499.329 | 0.1439 | 0.109987 | 0.500285 | Cul5 |
| ENSMUSG( | 1446.334 | 0.089536 | 0.320085 | 0.765523 | Cul7 |
| ENSMUSG( | 2567.763 | 0.104974 | 0.252897 | 0.703255 | Cul9 |
| ENSMUSG( | 1373.296 | -0.08968 | 0.325265 | 0.77061 | Cuta |
| ENSMUSG( | 14.42654 | -0.46332 | 0.36945 | NA | Cutal |
| ENSMUSG( | 143.4587 | 0.125308 | 0.544589 | 0.884649 | Cutc |
| ENSMUSG( | 5165.689 | -0.02573 | 0.757009 | 0.951513 | Cux1 |
| ENSMUSG( | 4129.7 | 0.184494 | 0.049061 | 0.338072 | Cux2 |
| ENSMUSG( | 14.64025 | 0.159702 | 0.741025 | NA | Cuzd1 |
| ENSMUSG( | 2087.611 | 0.05294 | 0.604424 | 0.905556 | Cwc15 |
| ENSMUSG( | 558.3834 | 0.108647 | 0.330307 | 0.774247 | Cwc22 |
| ENSMUSG( | 512.1368 | 0.032404 | 0.774248 | 0.955606 | Cwc25 |
| ENSMUSG( | 431.328 | 0.099123 | 0.441849 | 0.832669 | Cwc27 |
| ENSMUSG( | 363.2375 | -0.05317 | 0.679755 | 0.930983 | Cwf19l1 |
| ENSMUSG( | 280.0131 | 0.099153 | 0.553142 | 0.886207 | Cwf19l2 |
| ENSMUSG( | 145.0132 | 0.217394 | 0.20189 | 0.641612 | Cwh43 |
| ENSMUSG( | 33285.39 | -0.3773 | 0.007638 | 0.120942 | Cx3cl1 |
| ENSMUSG( | 3840.242 | -0.12669 | 0.461521 | 0.841487 | Cx3cr1 |
| ENSMUSG( | 877.7582 | -0.4769 | 0.00068 | 0.025176 | Cxadr |
| ENSMUSG( | 13.49426 | -1.09042 | 0.056884 | NA | Cxcl10 |
| ENSMUSG( | 3083.756 | 0.10666 | 0.750359 | 0.949806 | Cxcl12 |
| ENSMUSG( | 11528.7 | -0.12975 | 0.2414 | 0.689139 | Cxcl14 |
| ENSMUSG( | 93.97008 | -0.2452 | 0.233155 | 0.681401 | Cxcl16 |
| ENSMUSG( | 108.749 | -0.44085 | 0.030161 | 0.263854 | Cxcr4 |
| ENSMUSG( | 1407.743 | 0.242392 | 0.012251 | 0.160458 | Cxxc1 |
| ENSMUSG( | 778.0619 | -0.05911 | 0.625846 | 0.912305 | Cxxc4 |
| ENSMUSG( | 1931.29 | 0.000678 | 0.99485 | 0.999369 | Cxxc5 |
| ENSMUSG( | 2259.629 | -0.20463 | 0.095585 | 0.475062 | Cyb561 |
| ENSMUSG( | 537.1686 | -0.39329 | 0.011988 | 0.158212 | Cyb561a3 |
| ENSMUSG( | 767.1101 | -0.21397 | 0.097312 | 0.477629 | Cyb561d1 |
| ENSMUSG( | 413.9639 | -0.37564 | 0.004682 | 0.089972 | Cyb561d2 |
| ENSMUSG( | 2189.158 | 0.078382 | 0.381739 | 0.802879 | Cyb5a |
| ENSMUSG( | 7842.601 | 0.080997 | 0.378018 | 0.801561 | Cyb5b |
| ENSMUSG( | 608.8572 | 0.102488 | 0.429032 | 0.826446 | Cyb5d1 |
| ENSMUSG( | 626.3564 | -0.12798 | 0.382736 | 0.803227 | Cyb5d2 |
| ENSMUSG( | 555.826 | -0.15498 | 0.284777 | 0.733422 | Cyb5r1 |
| ENSMUSG( | 4361.908 | -0.07891 | 0.356297 | 0.788108 | Cyb5r3 |
| ENSMUSG( | 870.3801 | 0.176162 | 0.070402 | 0.40908 | Cyb5r4 |
| ENSMUSG( | 247.8224 | 0.241702 | 0.079238 | 0.435105 | Cyb5rl |
| ENSMUSG( | 414.9947 | -0.24918 | 0.114511 | 0.510543 | Cyba |
| ENSMUSG( | 92.21306 | -1.05638 | 0.00257 | 0.060158 | Cybb |
| ENSMUSG( | 689.2268 | -0.21068 | 0.108084 | 0.497059 | Cybc1 |
| ENSMUSG( | 588.8103 | 0.278781 | 0.065163 | 0.395697 | Cybrd1 |
| ENSMUSG( | 5008.368 | 0.031911 | 0.67729 | 0.930542 | Cyc1 |
| ENSMUSG( | 2082.787 | 0.024033 | 0.826352 | 0.968315 | Cycs |
| ENSMUSG( | 5113.487 | 0.156997 | 0.089569 | 0.459604 | Cyfip1 |
| ENSMUSG( | 24578.93 | 0.131491 | 0.178477 | 0.616198 | Cyfip2 |
| ENSMUSG( | 1105.973 | -0.11974 | 0.454538 | 0.837383 | Cygb |
| ENSMUSG( | 2852.369 | -0.05781 | 0.523906 | 0.87583 | Cyld |
| ENSMUSG( | 191.8334 | 0.143501 | 0.36194 | 0.792797 | Cyp11a1 |
| ENSMUSG( | 180.0593 | 0.239267 | 0.215966 | 0.660937 | Cyp1b1 |
| ENSMUSG( | 592.6962 | -0.14634 | 0.204639 | 0.645534 | Cyp20a1 |

| | | | | | |
|---|---|---|---|---|---|
| ENSMUSG( | 94.74726 | 0.382644 | 0.152183 | 0.837724 | Csf2rb |
| ENSMUSG( | 67.20066 | 0.347365 | 0.1977 | 0.862161 | Csf2rb2 |
| ENSMUSG( | 430.3223 | -0.01761 | 0.930827 | 0.99306 | Csf3r |
| ENSMUSG( | 960.1921 | -0.02426 | 0.869257 | 0.988097 | Csgalnact1 |
| ENSMUSG( | 497.8946 | 0.05374 | 0.746296 | 0.969654 | Csgalnact2 |
| ENSMUSG( | 957.4904 | 0.014803 | 0.897386 | 0.990558 | Csk |
| ENSMUSG( | 2005.322 | 0.348186 | 0.023314 | 0.585577 | Csmd1 |
| ENSMUSG( | 1796.382 | -0.0813 | 0.545326 | 0.938063 | Csmd2 |
| ENSMUSG( | 524.3221 | 0.107074 | 0.576617 | 0.942912 | Csmd3 |
| ENSMUSG( | 4845.557 | 0.056881 | 0.569224 | 0.941777 | Csnk1a1 |
| ENSMUSG( | 3036.828 | -0.01566 | 0.887492 | 0.990266 | Csnk1d |
| ENSMUSG( | 2762.643 | 0.071496 | 0.554562 | 0.939565 | Csnk1e |
| ENSMUSG( | 663.0475 | -0.03676 | 0.798397 | 0.97681 | Csnk1g1 |
| ENSMUSG( | 10549.18 | 0.089576 | 0.375504 | 0.915491 | Csnk1g2 |
| ENSMUSG( | 1019.489 | 0.135141 | 0.284502 | 0.892412 | Csnk1g3 |
| ENSMUSG( | 2613.307 | -0.09545 | 0.469665 | 0.923329 | Csnk2a1 |
| ENSMUSG( | 1615.558 | 0.01513 | 0.896952 | 0.990558 | Csnk2a2 |
| ENSMUSG( | 1758.616 | -0.1162 | 0.42736 | 0.917257 | Csnk2b |
| ENSMUSG( | 1082.706 | 0.181497 | 0.691873 | 0.960344 | Cspg4 |
| ENSMUSG( | 181.6064 | 0.13222 | 0.450858 | 0.922202 | Cspg4b |
| ENSMUSG( | 24055.7 | 0.041977 | 0.765824 | 0.974236 | Cspg5 |
| ENSMUSG( | 816.9355 | 0.016397 | 0.902833 | 0.991089 | Cspp1 |
| ENSMUSG( | 176.7833 | -0.20752 | 0.397866 | 0.917257 | Csrnp1 |
| ENSMUSG( | 518.6104 | -0.03533 | 0.814793 | 0.979198 | Csrnp2 |
| ENSMUSG( | 845.3629 | 0.137951 | 0.242874 | 0.877399 | Csrnp3 |
| ENSMUSG( | 17102.24 | -0.15458 | 0.216332 | 0.865167 | Csrp1 |
| ENSMUSG( | 202.1044 | 0.266765 | 0.22028 | 0.866104 | Csrp2 |
| ENSMUSG( | 85574.82 | -0.06986 | 0.551174 | 0.939377 | Cst3 |
| ENSMUSG( | 172.3075 | -0.44319 | 0.015056 | 0.492049 | Cst6 |
| ENSMUSG( | 11.36584 | -0.91036 | 0.128405 | NA | Cst7 |
| ENSMUSG( | 36.03669 | -0.22515 | 0.541472 | 0.937841 | Cstad |
| ENSMUSG( | 441.0047 | -0.225 | 0.141031 | 0.830678 | Cstb |
| ENSMUSG( | 515.9708 | -0.06354 | 0.646691 | 0.953988 | Cstf1 |
| ENSMUSG( | 2484.135 | 0.101286 | 0.468524 | 0.923329 | Cstf2 |
| ENSMUSG( | 1460.749 | -0.11935 | 0.419309 | 0.917257 | Cstf2t |
| ENSMUSG( | 517.7686 | 0.093835 | 0.505916 | 0.927825 | Cstf3 |
| ENSMUSG( | 1581.407 | -0.17855 | 0.239074 | 0.873744 | Cstpp1 |
| ENSMUSG( | 9.729288 | -0.2639 | 0.669797 | NA | Ct55 |
| ENSMUSG( | 8954.464 | -0.07604 | 0.473142 | 0.923329 | Ctbp1 |
| ENSMUSG( | 668.5513 | 0.114261 | 0.36993 | 0.915491 | Ctbp2 |
| ENSMUSG( | 337.8835 | 0.041665 | 0.834152 | 0.981237 | Ctbs |
| ENSMUSG( | 1219.47 | 0.188549 | 0.148096 | 0.836241 | Ctc1 |
| ENSMUSG( | 1488.463 | 0.084783 | 0.471643 | 0.923329 | Ctcf |
| ENSMUSG( | 3432.037 | -0.00734 | 0.950875 | 0.994798 | Ctdnep1 |
| ENSMUSG( | 435.287 | -0.01651 | 0.913983 | 0.992407 | Ctdp1 |
| ENSMUSG( | 2640.526 | -0.02672 | 0.829845 | 0.980224 | Ctdsp1 |
| ENSMUSG( | 3733.18 | 0.127001 | 0.278397 | 0.891628 | Ctdsp2 |
| ENSMUSG( | 1378.439 | 0.027891 | 0.811145 | 0.978974 | Ctdspl |
| ENSMUSG( | 327.0156 | 0.231303 | 0.132963 | 0.822912 | Ctdspl2 |
| ENSMUSG( | 82.45348 | -0.18833 | 0.421723 | 0.917257 | Ctf1 |
| ENSMUSG( | 195.782 | 0.122401 | 0.480342 | 0.92453 | Cth |
| ENSMUSG( | 211.2752 | 0.40474 | 0.033322 | 0.642852 | Cthrc1 |
| ENSMUSG( | 1834.622 | 0.157626 | 0.168979 | 0.840222 | Ctif |
| ENSMUSG( | 208.547 | 0.375685 | 0.089405 | 0.788575 | Ctla2a |
| ENSMUSG( | 3476.21 | -0.00552 | 0.95873 | 0.994798 | Ctnna1 |
| ENSMUSG( | 7367.166 | -0.11923 | 0.307469 | 0.898538 | Ctnna2 |
| ENSMUSG( | 42.43473 | -0.34213 | 0.286434 | 0.894091 | Ctnna3 |
| ENSMUSG( | 725.9282 | -0.03507 | 0.778753 | 0.976106 | Ctnnal1 |
| ENSMUSG( | 29443.98 | -0.11192 | 0.391407 | 0.917257 | Ctnnb1 |
| ENSMUSG( | 3523.197 | -0.04674 | 0.721904 | 0.966887 | Ctnnbip1 |
| ENSMUSG( | 701.2658 | -0.19361 | 0.120875 | 0.813384 | Ctnnbl1 |
| ENSMUSG( | 511.5032 | -0.11873 | 0.4851 | 0.92541 | Ctnnd1 |
| ENSMUSG( | 12180.12 | -0.08497 | 0.404937 | 0.917257 | Ctnnd2 |
| ENSMUSG( | 709.2681 | 0.147203 | 0.339745 | 0.907972 | Ctns |
| ENSMUSG( | 874.8137 | -0.10582 | 0.449653 | 0.922202 | Ctps1 |
| ENSMUSG( | 742.3634 | 0.011277 | 0.943319 | 0.994798 | Ctps2 |
| ENSMUSG( | 1294.254 | 0.003656 | 0.976297 | 0.996416 | Ctr9 |
| ENSMUSG( | 19.61552 | 0.101511 | 0.833546 | 0.981218 | Ctrl |
| ENSMUSG( | 5290.177 | 0.084624 | 0.520517 | 0.932079 | Ctsa |
| ENSMUSG( | 26423.88 | 0.058134 | 0.685017 | 0.958678 | Ctsb |
| ENSMUSG( | 484.9517 | 0.449112 | 0.027903 | 0.61023 | Ctsc |
| ENSMUSG( | 11164.1 | 0.125407 | 0.412228 | 0.917257 | Ctsd |
| ENSMUSG( | 11.10708 | 0.24905 | 0.738209 | NA | Ctse |
| ENSMUSG( | 3201.446 | 0.235054 | 0.112059 | 0.808673 | Ctsf |
| ENSMUSG( | 705.8088 | 0.22249 | 0.273743 | 0.889848 | Ctsh |
| ENSMUSG( | 210.6469 | 0.172112 | 0.322111 | 0.904023 | Ctsk |
| ENSMUSG( | 4801.241 | 0.197451 | 0.198121 | 0.862161 | Ctsl |
| ENSMUSG( | 3060.287 | 0.01248 | 0.92768 | 0.99306 | Ctso |
| ENSMUSG( | 4581.381 | 0.022767 | 0.877524 | 0.989644 | Ctss |
| ENSMUSG( | 11.18588 | -0.01341 | 0.981416 | NA | Ctsw |
| ENSMUSG( | 2128.737 | 0.001434 | 0.992862 | 0.998792 | Ctsz |
| ENSMUSG( | 2665.292 | -0.02294 | 0.836252 | 0.981335 | Cttn |
| ENSMUSG( | 2528.804 | -0.0315 | 0.819415 | 0.979198 | Cttnbp2 |
| ENSMUSG( | 955.8018 | -0.14769 | 0.247213 | 0.877904 | Cttnbp2nl |
| ENSMUSG( | 169.523 | 0.037816 | 0.833403 | 0.981196 | Ctu1 |
| ENSMUSG( | 505.5203 | 0.073165 | 0.574748 | 0.942912 | Ctu2 |
| ENSMUSG( | 19260.88 | -0.06155 | 0.637585 | 0.953903 | Ctxn1 |
| ENSMUSG( | 173.5718 | -0.12681 | 0.485271 | 0.92541 | Ctxn2 |
| ENSMUSG( | 88.59135 | -0.19674 | 0.474132 | 0.923329 | Ctxn3 |
| ENSMUSG( | 1336.194 | -0.18 | 0.108882 | 0.804969 | Ctxnd1 |
| ENSMUSG( | 22.98045 | -0.27989 | 0.550054 | 0.938699 | Cubn |
| ENSMUSG( | 2283.898 | -0.25428 | 0.033617 | 0.645688 | Cuedc2 |
| ENSMUSG( | 3755.596 | 0.092159 | 0.409224 | 0.917257 | Cul1 |
| ENSMUSG( | 1049.57 | 0.005111 | 0.964931 | 0.995433 | Cul2 |
| ENSMUSG( | 3201.718 | 0.008095 | 0.950955 | 0.994798 | Cul3 |
| ENSMUSG( | 1978.208 | -0.02729 | 0.793945 | 0.97681 | Cul4a |
| ENSMUSG( | 701.8535 | 0.021695 | 0.874614 | 0.989644 | Cul4b |
| ENSMUSG( | 1355.503 | 0.119431 | 0.325265 | 0.905339 | Cul5 |
| ENSMUSG( | 1294.526 | 0.041059 | 0.712444 | 0.964363 | Cul7 |
| ENSMUSG( | 2286.468 | 0.042735 | 0.710408 | 0.964363 | Cul9 |
| ENSMUSG( | 1235.588 | -0.1293 | 0.248869 | 0.877904 | Cuta |
| ENSMUSG( | 17.36747 | 0.372557 | 0.441096 | 0.921166 | Cutal |
| ENSMUSG( | 125.6397 | 0.012363 | 0.953834 | 0.994798 | Cutc |
| ENSMUSG( | 4542.202 | -0.13277 | 0.211813 | 0.864412 | Cux1 |
| ENSMUSG( | 3749.327 | 0.174494 | 0.101613 | 0.801601 | Cux2 |
| ENSMUSG( | 11.51474 | -0.27455 | 0.624559 | 0.953234 | Cuzd1 |
| ENSMUSG( | 1786.567 | -0.13432 | 0.325369 | 0.905339 | Cwc15 |
| ENSMUSG( | 505.421 | 0.083998 | 0.564438 | 0.941206 | Cwc22 |
| ENSMUSG( | 463.4174 | 0.010597 | 0.936621 | 0.993573 | Cwc25 |
| ENSMUSG( | 365.4892 | -0.11865 | 0.453606 | 0.922316 | Cwc27 |
| ENSMUSG( | 338.4025 | 0.018788 | 0.899742 | 0.990558 | Cwf19l1 |
| ENSMUSG( | 255.408 | 0.108049 | 0.549853 | 0.938699 | Cwf19l2 |
| ENSMUSG( | 130.3214 | 0.188427 | 0.358556 | 0.913759 | Cwh43 |
| ENSMUSG( | 34419.79 | 0.016831 | 0.915334 | 0.992525 | Cx3cl1 |
| ENSMUSG( | 4029.063 | 0.276519 | 0.167732 | 0.840222 | Cx3cr1 |
| ENSMUSG( | 980.8669 | 0.146786 | 0.352592 | 0.912093 | Cxadr |
| ENSMUSG( | 13.05838 | -0.83569 | 0.133792 | 0.822912 | Cxcl10 |
| ENSMUSG( | 3861.308 | 0.897796 | 0.027755 | 0.61023 | Cxcl12 |
| ENSMUSG( | 10717.88 | -0.06773 | 0.588581 | 0.944975 | Cxcl14 |
| ENSMUSG( | 89.59183 | -0.09881 | 0.666672 | 0.956766 | Cxcl16 |
| ENSMUSG( | 125.2483 | 0.263272 | 0.203839 | 0.864412 | Cxcr4 |
| ENSMUSG( | 1244.636 | 0.162296 | 0.198009 | 0.862161 | Cxxc1 |
| ENSMUSG( | 695.683 | -0.12445 | 0.518669 | 0.930939 | Cxxc4 |
| ENSMUSG( | 1738.564 | -0.03756 | 0.781429 | 0.976106 | Cxxc5 |
| ENSMUSG( | 2335.264 | 0.164946 | 0.269367 | 0.887103 | Cyb561 |
| ENSMUSG( | 552.3007 | -0.00836 | 0.96495 | 0.995433 | Cyb561a3 |
| ENSMUSG( | 761.7456 | 0.046752 | 0.737631 | 0.967686 | Cyb561d1 |
| ENSMUSG( | 420.0459 | -0.03848 | 0.808478 | 0.97873 | Cyb561d2 |
| ENSMUSG( | 1827.387 | -0.1824 | 0.14277 | 0.831221 | Cyb5a |
| ENSMUSG( | 6624.046 | -0.14151 | 0.259907 | 0.883971 | Cyb5b |
| ENSMUSG( | 501.3354 | -0.19805 | 0.166319 | 0.840082 | Cyb5d1 |
| ENSMUSG( | 595.9094 | 0.001637 | 0.991756 | 0.998712 | Cyb5d2 |
| ENSMUSG( | 562.923 | 0.155109 | 0.324119 | 0.905257 | Cyb5r1 |
| ENSMUSG( | 3904.18 | -0.13281 | 0.238449 | 0.872849 | Cyb5r3 |
| ENSMUSG( | 751.909 | 0.025956 | 0.828449 | 0.980224 | Cyb5r4 |
| ENSMUSG( | 196.4049 | -0.14827 | 0.396509 | 0.917257 | Cyb5rl |
| ENSMUSG( | 416.7724 | 0.044876 | 0.764297 | 0.974 | Cyba |
| ENSMUSG( | 95.34982 | -0.57492 | 0.102404 | 0.802599 | Cybb |
| ENSMUSG( | 699.2426 | 0.112772 | 0.460285 | 0.923329 | Cybc1 |
| ENSMUSG( | 528.9701 | 0.23958 | 0.097974 | 0.798438 | Cybrd1 |
| ENSMUSG( | 4258.27 | -0.17359 | 0.080964 | 0.77681 | Cyc1 |
| ENSMUSG( | 1762.094 | -0.20133 | 0.151722 | 0.837724 | Cycs |
| ENSMUSG( | 4259.482 | -0.10036 | 0.350959 | 0.911707 | Cyfip1 |
| ENSMUSG( | 21019.92 | -0.04759 | 0.665614 | 0.956766 | Cyfip2 |
| ENSMUSG( | 994.9424 | -0.16151 | 0.361899 | 0.914223 | Cygb |
| ENSMUSG( | 2676.684 | 0.027236 | 0.793196 | 0.97681 | Cyld |
| ENSMUSG( | 163.4375 | -0.04796 | 0.800783 | 0.97681 | Cyp11a1 |
| ENSMUSG( | 154.1055 | 0.066592 | 0.737382 | 0.967686 | Cyp1b1 |
| ENSMUSG( | 585.4425 | 0.092975 | 0.492669 | 0.927358 | Cyp20a1 |
| ENSMUSG( | 364.8251 | 0.106843 | 0.500226 | 0.927825 | Cyp26b1 |
| ENSMUSG( | 229.9639 | 0.10767 | 0.615602 | 0.951177 | Cyp27a1 |
| ENSMUSG( | 10.19227 | -0.23781 | 0.70203 | NA | Cyp27b1 |
| ENSMUSG( | 2620.013 | 0.188988 | 0.160617 | 0.837724 | Cyp2d22 |
| ENSMUSG( | 89.48018 | 0.500821 | 0.034387 | 0.653323 | Cyp2f2 |
| ENSMUSG( | 103.4217 | 0.011928 | 0.960978 | 0.99511 | Cyp2j12 |
| ENSMUSG( | 1580.99 | 0.122993 | 0.418845 | 0.917257 | Cyp2j6 |
| ENSMUSG( | 2954.465 | -0.04864 | 0.69939 | 0.961286 | Cyp2j9 |
| ENSMUSG( | 213.2853 | -0.02316 | 0.897679 | 0.990558 | Cyp2s1 |

| | | | | | | | | | | | |
|---|---|---|---|---|---|---|---|---|---|---|---|
| ENSMUSG( | Csrnp3 | 868.7991 | -0.05036 | 0.625589 | 0.912249 | ENSMUSG( | Csrnp3 | 845.3629 | 0.137951 | 0.242874 | 0.877399 |
| ENSMUSG( | Csrp1 | 20843.69 | 0.149251 | 0.119879 | 0.523965 | ENSMUSG( | Csrp1 | 17102.24 | -0.15458 | 0.216332 | 0.865167 |
| ENSMUSG( | Csrp2 | 201.8687 | -0.00826 | 0.963317 | 0.994911 | ENSMUSG( | Csrp2 | 202.1044 | 0.266765 | 0.22028 | 0.866104 |
| ENSMUSG( | Cst3 | 94894.08 | -0.04063 | 0.705099 | 0.938303 | ENSMUSG( | Cst3 | 85574.82 | -0.06986 | 0.551174 | 0.939377 |
| ENSMUSG( | Cst6 | 206.7391 | -0.15973 | 0.326133 | 0.771128 | ENSMUSG( | Cst6 | 172.3075 | -0.44319 | 0.015056 | 0.492049 |
| ENSMUSG( | Cst7 | 13.62662 | -0.50332 | 0.315968 | NA | ENSMUSG( | Cst7 | 11.36584 | -0.91036 | 0.128405 | NA |
| ENSMUSG( | Cstad | 39.09651 | -0.26532 | 0.410148 | 0.818258 | ENSMUSG( | Cstad | 36.03669 | -0.22515 | 0.541472 | 0.937841 |
| ENSMUSG( | Cstb | 514.7191 | -0.03292 | 0.82266 | 0.966352 | ENSMUSG( | Cstb | 441.0047 | -0.225 | 0.141031 | 0.830678 |
| ENSMUSG( | Cstf1 | 576.1378 | -0.01906 | 0.867904 | 0.976631 | ENSMUSG( | Cstf1 | 515.9708 | -0.06354 | 0.646691 | 0.953988 |
| ENSMUSG( | Cstf2 | 2719.623 | 0.097372 | 0.34553 | 0.779153 | ENSMUSG( | Cstf2 | 2484.135 | 0.101286 | 0.468524 | 0.923329 |
| ENSMUSG( | Cstf2t | 1655.201 | -0.01954 | 0.860254 | 0.974842 | ENSMUSG( | Cstf2t | 1460.749 | -0.11935 | 0.419309 | 0.917257 |
| ENSMUSG( | Cstf3 | 546.6062 | -0.01498 | 0.898939 | 0.985633 | ENSMUSG( | Cstf3 | 517.7686 | 0.093835 | 0.505916 | 0.927825 |
| ENSMUSG( | Cstpp1 | 1890.206 | 0.074412 | 0.576085 | 0.895114 | ENSMUSG( | Cstpp1 | 1581.407 | -0.17855 | 0.239074 | 0.873744 |
| ENSMUSG( | Ct55 | 9.870085 | -0.51812 | 0.411346 | NA | ENSMUSG( | Ct55 | 9.729288 | -0.2639 | 0.669797 | NA |
| ENSMUSG( | Ctbp1 | 10291.14 | 0.058347 | 0.446786 | 0.833933 | ENSMUSG( | Ctbp1 | 8954.464 | -0.07604 | 0.473142 | 0.923329 |
| ENSMUSG( | Ctbp2 | 751.591 | 0.182371 | 0.079433 | 0.43547 | ENSMUSG( | Ctbp2 | 668.5513 | 0.114261 | 0.36993 | 0.915491 |
| ENSMUSG( | Ctbs | 323.0052 | -0.39043 | 0.033942 | 0.282701 | ENSMUSG( | Ctbs | 337.8835 | 0.041665 | 0.834152 | 0.981237 |
| ENSMUSG( | Ctc1 | 1303.688 | 0.112413 | 0.311265 | 0.759247 | ENSMUSG( | Ctc1 | 1219.47 | 0.188549 | 0.148096 | 0.836241 |
| ENSMUSG( | Ctcf | 1655.42 | 0.123309 | 0.165568 | 0.59714 | ENSMUSG( | Ctcf | 1488.463 | 0.084783 | 0.471643 | 0.923329 |
| ENSMUSG( | Ctdnep1 | 3690.692 | -0.06712 | 0.460773 | 0.840962 | ENSMUSG( | Ctdnep1 | 3432.037 | -0.00734 | 0.950875 | 0.994798 |
| ENSMUSG( | Ctdp1 | 486.6646 | 0.04158 | 0.753957 | 0.950981 | ENSMUSG( | Ctdp1 | 435.287 | -0.01651 | 0.913983 | 0.992407 |
| ENSMUSG( | Ctdsp1 | 3038.802 | 0.11082 | 0.2406 | 0.688247 | ENSMUSG( | Ctdsp1 | 2640.526 | -0.02672 | 0.829845 | 0.980224 |
| ENSMUSG( | Ctdsp2 | 4047.108 | 0.090373 | 0.298085 | 0.746708 | ENSMUSG( | Ctdsp2 | 3733.18 | 0.127001 | 0.278397 | 0.891628 |
| ENSMUSG( | Ctdspl | 1531.818 | 0.063712 | 0.525604 | 0.876728 | ENSMUSG( | Ctdspl | 1378.439 | 0.027891 | 0.811145 | 0.978974 |
| ENSMUSG( | Ctdspl2 | 338.4522 | 0.065133 | 0.59911 | 0.903801 | ENSMUSG( | Ctdspl2 | 327.0156 | 0.231303 | 0.132963 | 0.822912 |
| ENSMUSG( | Ctf1 | 95.21484 | -0.03014 | 0.879802 | 0.981436 | ENSMUSG( | Ctf1 | 82.45348 | -0.18833 | 0.421723 | 0.917257 |
| ENSMUSG( | Cth | 233.6706 | 0.351371 | 0.025986 | 0.247245 | ENSMUSG( | Cth | 195.782 | 0.122401 | 0.480342 | 0.92453 |
| ENSMUSG( | Cthrc1 | 176.4848 | -0.39272 | 0.019455 | 0.210037 | ENSMUSG( | Cthrc1 | 211.2752 | 0.40474 | 0.033322 | 0.642852 |
| ENSMUSG( | Ctif | 2059.81 | 0.216771 | 0.014711 | 0.178407 | ENSMUSG( | Ctif | 1834.622 | 0.157626 | 0.168979 | 0.840222 |
| ENSMUSG( | Ctla2a | 221.5491 | 0.277306 | 0.1283 | 0.536641 | ENSMUSG( | Ctla2a | 208.547 | 0.375685 | 0.089405 | 0.788575 |
| ENSMUSG( | Ctnna1 | 4036.735 | 0.153232 | 0.145701 | 0.566867 | ENSMUSG( | Ctnna1 | 3476.21 | -0.00552 | 0.95873 | 0.994798 |
| ENSMUSG( | Ctnna2 | 9011.947 | 0.19121 | 0.019979 | 0.213756 | ENSMUSG( | Ctnna2 | 7367.166 | -0.11923 | 0.307469 | 0.898538 |
| ENSMUSG( | Ctnna3 | 51.18368 | -0.07163 | 0.796203 | 0.960088 | ENSMUSG( | Ctnna3 | 42.43473 | -0.34213 | 0.286434 | 0.894091 |
| ENSMUSG( | Ctnnal1 | 783.1745 | -0.08818 | 0.414255 | 0.819998 | ENSMUSG( | Ctnnal1 | 725.9282 | -0.03507 | 0.778753 | 0.976106 |
| ENSMUSG( | Ctnnb1 | 34957.41 | 0.116692 | 0.263285 | 0.714007 | ENSMUSG( | Ctnnb1 | 29443.98 | -0.11192 | 0.391407 | 0.917257 |
| ENSMUSG( | Ctnnbip1 | 3992.973 | 0.047717 | 0.538051 | 0.882548 | ENSMUSG( | Ctnnbip1 | 3523.197 | -0.04674 | 0.721904 | 0.966887 |
| ENSMUSG( | Ctnnbl1 | 826.8834 | 0.019809 | 0.849077 | 0.972601 | ENSMUSG( | Ctnnbl1 | 701.2658 | -0.19361 | 0.120875 | 0.813384 |
| ENSMUSG( | Ctnnd1 | 571.9725 | -0.05236 | 0.66394 | 0.926511 | ENSMUSG( | Ctnnd1 | 511.5032 | -0.11873 | 0.4851 | 0.92541 |
| ENSMUSG( | Ctnnd2 | 14530.5 | 0.153271 | 0.060128 | 0.37919 | ENSMUSG( | Ctnnd2 | 12180.12 | -0.08497 | 0.404937 | 0.917257 |
| ENSMUSG( | Ctns | 680.0659 | -0.25689 | 0.071522 | 0.413178 | ENSMUSG( | Ctns | 709.2681 | 0.147203 | 0.339745 | 0.907972 |
| ENSMUSG( | Ctps1 | 994.2529 | 0.000213 | 0.998563 | 0.99989 | ENSMUSG( | Ctps1 | 874.8137 | -0.10582 | 0.449653 | 0.922202 |
| ENSMUSG( | Ctps2 | 837.2624 | 0.089964 | 0.54057 | 0.88315 | ENSMUSG( | Ctps2 | 742.3634 | 0.011277 | 0.943319 | 0.994798 |
| ENSMUSG( | Ctr9 | 1447.066 | 0.054975 | 0.606081 | 0.906049 | ENSMUSG( | Ctr9 | 1294.254 | 0.003656 | 0.976297 | 0.996416 |
| ENSMUSG( | Ctrl | 23.73202 | 0.341735 | 0.38786 | 0.805835 | ENSMUSG( | Ctrl | 19.61552 | 0.101511 | 0.833546 | 0.981218 |
| ENSMUSG( | Ctsa | 5164.614 | -0.26691 | 0.013509 | 0.169727 | ENSMUSG( | Ctsa | 5290.177 | 0.084624 | 0.520517 | 0.932079 |
| ENSMUSG( | Ctsb | 25581.38 | -0.32429 | 0.00895 | 0.132538 | ENSMUSG( | Ctsb | 26423.88 | 0.058134 | 0.685017 | 0.958678 |
| ENSMUSG( | Ctsc | 414.681 | -0.2589 | 0.172155 | 0.607115 | ENSMUSG( | Ctsc | 484.9517 | 0.449112 | 0.027903 | 0.61023 |
| ENSMUSG( | Ctsd | 11193.58 | -0.13954 | 0.271981 | 0.722255 | ENSMUSG( | Ctsd | 11164.1 | 0.125407 | 0.412228 | 0.917257 |
| ENSMUSG( | Ctsf | 3008.88 | -0.21541 | 0.099669 | 0.482191 | ENSMUSG( | Ctsf | 3201.446 | 0.235054 | 0.112059 | 0.808673 |
| ENSMUSG( | Ctsh | 688.4768 | -0.12129 | 0.504974 | 0.866883 | ENSMUSG( | Ctsh | 705.8088 | 0.22249 | 0.273743 | 0.889848 |
| ENSMUSG( | Ctsk | 202.6728 | -0.22256 | 0.131366 | 0.543562 | ENSMUSG( | Ctsk | 210.6469 | 0.172112 | 0.322111 | 0.904023 |
| ENSMUSG( | Ctsl | 4527.475 | -0.24848 | 0.059019 | 0.375291 | ENSMUSG( | Ctsl | 4801.241 | 0.197451 | 0.198121 | 0.862161 |
| ENSMUSG( | Ctso | 3357.75 | 0.011545 | 0.926986 | 0.991229 | ENSMUSG( | Ctso | 3060.287 | 0.01248 | 0.92768 | 0.99306 |
| ENSMUSG( | Ctss | 4702.742 | -0.1785 | 0.162854 | 0.593778 | ENSMUSG( | Ctss | 4581.381 | 0.022767 | 0.877524 | 0.989644 |
| ENSMUSG( | Ctsw | 11.37515 | -0.24763 | 0.695021 | NA | ENSMUSG( | Ctsw | 11.18588 | -0.01341 | 0.981416 | NA |
| ENSMUSG( | Ctsz | 2142.782 | -0.26458 | 0.046565 | 0.330304 | ENSMUSG( | Ctsz | 2128.737 | 0.001434 | 0.992862 | 0.998792 |
| ENSMUSG( | Cttn | 3080.06 | 0.12374 | 0.160269 | 0.589853 | ENSMUSG( | Cttn | 2665.292 | -0.02294 | 0.836252 | 0.981335 |
| ENSMUSG( | Cttnbp2 | 2834.803 | 0.030637 | 0.776218 | 0.955658 | ENSMUSG( | Cttnbp2 | 2528.804 | -0.0315 | 0.819415 | 0.979198 |
| ENSMUSG( | Cttnbp2nl | 1074.585 | -0.07956 | 0.474218 | 0.849425 | ENSMUSG( | Cttnbp2nl | 955.8018 | -0.14769 | 0.247213 | 0.877904 |
| ENSMUSG( | Ctu1 | 180.5756 | -0.0494 | 0.769088 | 0.954674 | ENSMUSG( | Ctu1 | 169.523 | 0.037816 | 0.833403 | 0.981196 |
| ENSMUSG( | Ctu2 | 554.9728 | 0.071442 | 0.579252 | 0.896692 | ENSMUSG( | Ctu2 | 505.5203 | 0.073165 | 0.574748 | 0.942912 |
| ENSMUSG( | Ctxn1 | 22247.54 | 0.087073 | 0.297264 | 0.745805 | ENSMUSG( | Ctxn1 | 19260.88 | -0.06155 | 0.637585 | 0.953903 |
| ENSMUSG( | Ctxn2 | 203.4018 | 0.07597 | 0.636347 | 0.914873 | ENSMUSG( | Ctxn2 | 173.5718 | -0.12681 | 0.485271 | 0.92541 |
| ENSMUSG( | Ctxn3 | 102.7631 | -0.00698 | 0.972769 | 0.996585 | ENSMUSG( | Ctxn3 | 88.59135 | -0.19674 | 0.474132 | 0.923329 |
| ENSMUSG( | Ctxnd1 | 1706.584 | 0.252442 | 0.004717 | 0.090333 | ENSMUSG( | Ctxnd1 | 1336.194 | -0.18 | 0.108882 | 0.804969 |
| ENSMUSG( | Cubn | 24.86982 | -0.33482 | 0.431977 | 0.82768 | ENSMUSG( | Cubn | 22.98045 | -0.27989 | 0.550054 | 0.938699 |
| ENSMUSG( | Cuedc2 | 2674.999 | -0.05392 | 0.573537 | 0.893584 | ENSMUSG( | Cuedc2 | 2283.898 | -0.25428 | 0.033617 | 0.645688 |
| ENSMUSG( | Cul1 | 4112.977 | 0.085136 | 0.365473 | 0.794085 | ENSMUSG( | Cul1 | 3755.596 | 0.092159 | 0.409224 | 0.917257 |
| ENSMUSG( | Cul2 | 1151.436 | 0.005387 | 0.953086 | 0.993874 | ENSMUSG( | Cul2 | 1049.57 | 0.005111 | 0.964931 | 0.995433 |
| ENSMUSG( | Cul3 | 3599.119 | 0.077964 | 0.429415 | 0.826446 | ENSMUSG( | Cul3 | 3201.718 | 0.008095 | 0.950955 | 0.994798 |
| ENSMUSG( | Cul4a | 2292.204 | 0.127164 | 0.119466 | 0.523291 | ENSMUSG( | Cul4a | 1978.208 | -0.02729 | 0.793945 | 0.97681 |
| ENSMUSG( | Cul4b | 775.5685 | 0.046493 | 0.695523 | 0.934979 | ENSMUSG( | Cul4b | 701.8535 | 0.021695 | 0.874614 | 0.989644 |
| ENSMUSG( | Cul5 | 1499.329 | 0.1439 | 0.109987 | 0.500285 | ENSMUSG( | Cul5 | 1355.503 | 0.119431 | 0.325265 | 0.905339 |
| ENSMUSG( | Cul7 | 1446.334 | 0.089536 | 0.320085 | 0.765523 | ENSMUSG( | Cul7 | 1294.526 | 0.041059 | 0.712444 | 0.964363 |
| ENSMUSG( | Cul9 | 2567.763 | 0.104974 | 0.252897 | 0.703255 | ENSMUSG( | Cul9 | 2286.468 | 0.042735 | 0.710408 | 0.964363 |
| ENSMUSG( | Cuta | 1373.296 | -0.08968 | 0.325265 | 0.77061 | ENSMUSG( | Cuta | 1235.588 | -0.1293 | 0.248869 | 0.877904 |
| ENSMUSG( | Cutal | 14.42654 | -0.46332 | 0.36945 | NA | ENSMUSG( | Cutal | 17.36747 | 0.372557 | 0.441096 | 0.921166 |
| ENSMUSG( | Cutc | 143.4587 | 0.125308 | 0.544589 | 0.884649 | ENSMUSG( | Cutc | 125.6397 | 0.012363 | 0.953834 | 0.994798 |
| ENSMUSG( | Cux1 | 5165.689 | -0.02573 | 0.757009 | 0.951513 | ENSMUSG( | Cux1 | 4542.202 | -0.13277 | 0.211813 | 0.864412 |
| ENSMUSG( | Cux2 | 4129.7 | 0.184494 | 0.049061 | 0.338072 | ENSMUSG( | Cux2 | 3749.327 | 0.174494 | 0.101613 | 0.801601 |
| ENSMUSG( | Cuzd1 | 14.64025 | 0.159702 | 0.741025 | NA | ENSMUSG( | Cuzd1 | 11.51474 | -0.27455 | 0.624559 | 0.953234 |
| ENSMUSG( | Cwc15 | 2087.611 | 0.05294 | 0.604424 | 0.905556 | ENSMUSG( | Cwc15 | 1786.567 | -0.13432 | 0.325369 | 0.905339 |
| ENSMUSG( | Cwc22 | 558.3834 | 0.108647 | 0.330307 | 0.774247 | ENSMUSG( | Cwc22 | 505.421 | 0.083998 | 0.564438 | 0.941206 |
| ENSMUSG( | Cwc25 | 512.1368 | 0.032404 | 0.774248 | 0.955606 | ENSMUSG( | Cwc25 | 463.4174 | 0.010597 | 0.936621 | 0.993573 |
| ENSMUSG( | Cwc27 | 431.328 | 0.099123 | 0.441849 | 0.832669 | ENSMUSG( | Cwc27 | 365.4892 | -0.11865 | 0.453606 | 0.922316 |
| ENSMUSG( | Cwf19l1 | 363.2375 | -0.05317 | 0.679755 | 0.930983 | ENSMUSG( | Cwf19l1 | 338.4025 | 0.018788 | 0.899742 | 0.990558 |
| ENSMUSG( | Cwf19l2 | 280.0131 | 0.099153 | 0.553142 | 0.886207 | ENSMUSG( | Cwf19l2 | 255.408 | 0.108049 | 0.549853 | 0.938699 |
| ENSMUSG( | Cwh43 | 145.0132 | 0.217394 | 0.20189 | 0.641612 | ENSMUSG( | Cwh43 | 130.3214 | 0.188427 | 0.358556 | 0.913759 |
| ENSMUSG( | Cx3cl1 | 33285.39 | -0.3773 | 0.007638 | 0.120942 | ENSMUSG( | Cx3cl1 | 34419.79 | 0.016831 | 0.915334 | 0.992525 |
| ENSMUSG( | Cx3cr1 | 3840.242 | -0.12669 | 0.461521 | 0.841487 | ENSMUSG( | Cx3cr1 | 4029.063 | 0.276519 | 0.167732 | 0.840222 |
| ENSMUSG( | Cxadr | 877.7582 | -0.4769 | 0.00068 | 0.025176 | ENSMUSG( | Cxadr | 980.8669 | 0.146786 | 0.352592 | 0.912093 |
| ENSMUSG( | Cxcl10 | 13.49426 | -1.09042 | 0.056884 | NA | ENSMUSG( | Cxcl10 | 13.05838 | -0.83569 | 0.133792 | 0.822912 |
| ENSMUSG( | Cxcl12 | 3083.756 | 0.10666 | 0.750359 | 0.949806 | ENSMUSG( | Cxcl12 | 3861.308 | 0.897796 | 0.027755 | 0.61023 |
| ENSMUSG( | Cxcl14 | 11528.7 | -0.12975 | 0.2414 | 0.689139 | ENSMUSG( | Cxcl14 | 10717.88 | -0.06773 | 0.588581 | 0.944975 |
| ENSMUSG( | Cxcl16 | 93.97008 | -0.2452 | 0.233155 | 0.681401 | ENSMUSG( | Cxcl16 | 89.59183 | -0.09881 | 0.666672 | 0.956766 |
| ENSMUSG( | Cxcr4 | 108.749 | -0.44085 | 0.030161 | 0.263854 | ENSMUSG( | Cxcr4 | 125.2483 | 0.263272 | 0.203839 | 0.864412 |
| ENSMUSG( | Cxxc1 | 1407.743 | 0.242392 | 0.012251 | 0.160458 | ENSMUSG( | Cxxc1 | 1244.636 | 0.162296 | 0.198009 | 0.862161 |
| ENSMUSG( | Cxxc4 | 778.0619 | -0.05911 | 0.625846 | 0.912305 | ENSMUSG( | Cxxc4 | 695.683 | -0.12445 | 0.518669 | 0.930939 |
| ENSMUSG( | Cxxc5 | 1931.29 | 0.000678 | 0.99485 | 0.999369 | ENSMUSG( | Cxxc5 | 1738.564 | -0.03756 | 0.781429 | 0.976106 |
| ENSMUSG( | Cyb561 | 2259.629 | -0.20463 | 0.095585 | 0.475062 | ENSMUSG( | Cyb561 | 2335.264 | 0.164946 | 0.269367 | 0.887103 |
| ENSMUSG( | Cyb561a3 | 537.1686 | -0.39329 | 0.011988 | 0.158212 | ENSMUSG( | Cyb561a3 | 552.3007 | -0.00836 | 0.96495 | 0.995433 |
| ENSMUSG( | Cyb561d1 | 767.1101 | -0.21397 | 0.097312 | 0.477629 | ENSMUSG( | Cyb561d1 | 761.7456 | 0.046752 | 0.737631 | 0.967686 |
| ENSMUSG( | Cyb561d2 | 413.9639 | -0.37564 | 0.004682 | 0.089972 | ENSMUSG( | Cyb561d2 | 420.0459 | -0.03848 | 0.808478 | 0.97873 |
| ENSMUSG( | Cyb5a | 2189.158 | 0.078382 | 0.381739 | 0.802879 | ENSMUSG( | Cyb5a | 1827.387 | -0.1824 | 0.14277 | 0.831221 |
| ENSMUSG( | Cyb5b | 7842.601 | 0.080997 | 0.378018 | 0.801561 | ENSMUSG( | Cyb5b | 6624.046 | -0.14151 | 0.259907 | 0.883971 |
| ENSMUSG( | Cyb5d1 | 608.8572 | 0.102488 | 0.429032 | 0.826446 | ENSMUSG( | Cyb5d1 | 501.3354 | -0.19805 | 0.166319 | 0.840082 |
| ENSMUSG( | Cyb5d2 | 626.3564 | -0.12798 | 0.382736 | 0.803227 | ENSMUSG( | Cyb5d2 | 595.9094 | 0.001637 | 0.991756 | 0.998712 |
| ENSMUSG( | Cyb5r1 | 555.826 | -0.15498 | 0.284777 | 0.733422 | ENSMUSG( | Cyb5r1 | 562.923 | 0.155109 | 0.324119 | 0.905257 |
| ENSMUSG( | Cyb5r3 | 4361.908 | -0.07891 | 0.356297 | 0.788108 | ENSMUSG( | Cyb5r3 | 3904.18 | -0.13281 | 0.238449 | 0.872849 |
| ENSMUSG( | Cyb5r4 | 870.3801 | 0.176162 | 0.070402 | 0.40908 | ENSMUSG( | Cyb5r4 | 751.909 | 0.025956 | 0.828449 | 0.980224 |
| ENSMUSG( | Cyb5rl | 247.8224 | 0.241702 | 0.079238 | 0.435105 | ENSMUSG( | Cyb5rl | 196.4049 | -0.14827 | 0.396509 | 0.917257 |
| ENSMUSG( | Cyba | 414.9947 | -0.24918 | 0.114511 | 0.510543 | ENSMUSG( | Cyba | 416.7724 | 0.044876 | 0.764297 | 0.974 |
| ENSMUSG( | Cybb | 92.21306 | -1.05638 | 0.00257 | 0.060158 | ENSMUSG( | Cybb | 95.34982 | -0.57492 | 0.102404 | 0.802599 |
| ENSMUSG( | Cybc1 | 689.2268 | -0.21068 | 0.108084 | 0.497059 | ENSMUSG( | Cybc1 | 699.2426 | 0.112772 | 0.460285 | 0.923329 |
| ENSMUSG( | Cybrd1 | 588.8103 | 0.278781 | 0.065163 | 0.395697 | ENSMUSG( | Cybrd1 | 528.9701 | 0.23958 | 0.097974 | 0.798438 |
| ENSMUSG( | Cyc1 | 5008.368 | 0.031911 | 0.67729 | 0.930542 | ENSMUSG( | Cyc1 | 4258.27 | -0.17359 | 0.080964 | 0.77681 |
| ENSMUSG( | Cycs | 2082.787 | 0.024033 | 0.826352 | 0.968315 | ENSMUSG( | Cycs | 1762.094 | -0.20133 | 0.151722 | 0.837724 |
| ENSMUSG( | Cyfip1 | 5113.487 | 0.156997 | 0.089569 | 0.459604 | ENSMUSG( | Cyfip1 | 4259.482 | -0.10036 | 0.350959 | 0.911707 |
| ENSMUSG( | Cyfip2 | 24578.93 | 0.131491 | 0.178477 | 0.616198 | ENSMUSG( | Cyfip2 | 21019.92 | -0.04759 | 0.665614 | 0.956766 |
| ENSMUSG( | Cygb | 1105.973 | -0.11974 | 0.454538 | 0.837383 | ENSMUSG( | Cygb | 994.9424 | -0.16151 | 0.361899 | 0.914223 |
| ENSMUSG( | Cyld | 2852.369 | -0.05781 | 0.523906 | 0.87583 | ENSMUSG( | Cyld | 2676.684 | 0.027236 | 0.793196 | 0.97681 |
| ENSMUSG( | Cyp11a1 | 191.8334 | 0.143501 | 0.36194 | 0.792797 | ENSMUSG( | Cyp11a1 | 163.4375 | -0.04796 | 0.800783 | 0.97681 |
| ENSMUSG( | Cyp1b1 | 180.0593 | 0.239267 | 0.215966 | 0.660937 | ENSMUSG( | Cyp1b1 | 154.1055 | 0.066592 | 0.737382 | 0.967686 |
| ENSMUSG( | Cyp20a1 | 592.6962 | -0.14634 | 0.204639 | 0.645534 | ENSMUSG( | Cyp20a1 | 585.4425 | 0.092975 | 0.492669 | 0.927358 |
| ENSMUSG( | Cyp26b1 | 375.8293 | -0.08124 | 0.613432 | 0.909027 | ENSMUSG( | Cyp26b1 | 364.8251 | 0.106843 | 0.500226 | 0.927825 |
| ENSMUSG( | Cyp27a1 | 203.7744 | -0.5791 | 0.000493 | 0.020342 | ENSMUSG( | Cyp27a1 | 229.9639 | 0.10767 | 0.615602 | 0.951177 |
| ENSMUSG( | Cyp27b1 | 13.29367 | 0.288519 | 0.579306 | NA | ENSMUSG( | Cyp27b1 | 10.19227 | -0.23781 | 0.70203 | NA |
| ENSMUSG( | Cyp2d22 | 2811.134 | 0.12367 | 0.282808 | 0.73186 | ENSMUSG( | Cyp2d22 | 2620.013 | 0.188988 | 0.160617 | 0.837724 |
| ENSMUSG( | Cyp2f2 | 80.28948 | -0.04837 | 0.840806 | 0.971179 | ENSMUSG( | Cyp2f2 | 89.48018 | 0.500821 | 0.034387 | 0.653323 |
| ENSMUSG( | Cyp2j12 | 104.8959 | -0.23319 | 0.283166 | 0.73186 | ENSMUSG( | Cyp2j12 | 103.4217 | 0.011928 | 0.960978 | 0.99511 |
| ENSMUSG( | Cyp2j6 | 1526.639 | -0.25821 | 0.05359 | 0.35565 | ENSMUSG( | Cyp2j6 | 1580.99 | 0.122993 | 0.418845 | 0.917257 |
| ENSMUSG( | Cyp2j9 | 3513.776 | 0.177708 | 0.141714 | 0.558728 | ENSMUSG( | Cyp2j9 | 2954.465 | -0.04864 | 0.69939 | 0.961286 |
| ENSMUSG( | Cyp2s1 | 179.177 | -0.9543 | 2.33E-07 | 5.39E-05 | ENSMUSG( | Cyp2s1 | 213.2853 | -0.02316 | 0.897679 | 0.990558 |
| ENSMUSG( | Cyp2u1 | 339.5862 | -0.26101 | 0.049821 | 0.340731 | ENSMUSG( | Cyp2u1 | 355.6921 | 0.150246 | 0.323947 | 0.905257 |
| ENSMUSG( | Cyp39a1 | 80.61126 | -0.21486 | 0.374197 | 0.800445 | ENSMUSG( | Cyp39a1 | 91.07048 | 0.386454 | 0.097654 | 0.796743 |
| ENSMUSG( | Cyp3a13 | 66.59725 | -0.20477 | 0.496808 | 0.86175 | ENSMUSG( | Cyp3a13 | 81.51121 | 0.59054 | 0.021763 | 0.575341 |
| ENSMUSG( | Cyp46a1 | 9666.941 | -0.27216 | 0.003786 | 0.078636 | ENSMUSG( | Cyp46a1 | 9955.182 | 0.094044 | 0.391149 | 0.917257 |
| ENSMUSG( | Cyp4b1 | 16.75156 | 0.357131 | 0.542967 | NA | ENSMUSG( | Cyp4b1 | 12.54005 | -0.17117 | 0.758085 | 0.971858 |
| ENSMUSG( | Cyp4f13 | 496.322 | -0.20207 | 0.111139 | 0.502556 | ENSMUSG( | Cyp4f13 | 491.8041 | 0.054945 | 0.728465 | 0.966913 |
| ENSMUSG( | Cyp4f14 | 1123.114 | 0.034608 | 0.790675 | 0.959461 | ENSMUSG( | Cyp4f14 | 991.1397 | -0.05549 | 0.696705 | 0.960867 |
| ENSMUSG( | Cyp4f15 | 1322.667 | 0.073847 | 0.57433 | 0.894186 | ENSMUSG( | Cyp4f15 | 1364.315 | 0.408749 | 0.005462 | 0.303106 |
| ENSMUSG( | Cyp4f16 | 261.576 | -0.30514 | 0.036914 | 0.296508 | ENSMUSG( | Cyp4f16 | 254.0137 | -0.09636 | 0.579287 | 0.943942 |
| ENSMUSG( | Cyp4f17 | 205.3366 | -0.1515 | 0.36218 | 0.792943 | ENSMUSG( | Cyp4f17 | 201.0721 | 0.071851 | 0.705676 | 0.96315 |
| ENSMUSG( | Cyp4f18 | 13.30573 | -0.57984 | 0.394282 | NA | ENSMUSG( | Cyp4f18 | 12.35128 | -0.60154 | 0.421996 | 0.917257 |
| ENSMUSG( | Cyp4v3 | 559.7506 | -0.22962 | 0.099547 | 0.481986 | ENSMUSG( | Cyp4v3 | 552.9795 | 0.017543 | 0.910186 | 0.992081 |
| ENSMUSG( | Cyp4x1 | 528.6787 | -0.34365 | 0.01361 | 0.170359 | ENSMUSG( | Cyp4x1 | 518.7136 | -0.10502 | 0.518129 | 0.930939 |
| ENSMUSG( | Cyp4x1os | 24.14883 | 0.022189 | 0.956094 | 0.993874 | ENSMUSG( | Cyp4x1os | 21.76244 | -0.04233 | 0.924179 | 0.992937 |
| ENSMUSG( | Cyp51 | 8197.241 | -0.04635 | 0.742432 | 0.948478 | ENSMUSG( | Cyp51 | 7058.563 | -0.21629 | 0.134971 | 0.82437 |
| ENSMUSG( | Cyp7b1 | 528.5729 | -0.25038 | 0.163695 | 0.594869 | ENSMUSG( | Cyp7b1 | 547.2518 | 0.130088 | 0.501406 | 0.927825 |
| ENSMUSG( | Cyren | 156.3049 | -0.18109 | 0.278299 | 0.728302 | ENSMUSG( | Cyren | 149.273 | -0.02962 | 0.895947 | 0.990558 |
| ENSMUSG( | Cyria | 4485.941 | 0.15221 | 0.074269 | 0.422054 | ENSMUSG( | Cyria | 3816.118 | -0.04425 | 0.707681 | 0.964029 |
| ENSMUSG( | Cyrib | 3539.351 | 0.002179 | 0.980991 | 0.997512 | ENSMUSG( | Cyrib | 3054.616 | -0.16133 | 0.182686 | 0.850579 |
| ENSMUSG( | Cys1 | 198.6164 | 0.331814 | 0.031604 | 0.271081 | ENSMUSG( | Cys1 | 204.4053 | 0.638728 | 0.000719 | 0.08668 |
| ENSMUSG( | Cysltr1 | 40.38538 | -0.19586 | 0.526622 | 0.877267 | ENSMUSG( | Cysltr1 | 45.66802 | 0.4237 | 0.188481 | 0.85698 |
| ENSMUSG( | Cysltr2 | 23.4108 | -0.41732 | 0.398222 | 0.812473 | ENSMUSG( | Cysltr2 | 16.665 | -1.33596 | 0.023998 | 0.588226 |
| ENSMUSG( | Cystm1 | 1274.746 | 0.035464 | 0.719274 | 0.941986 | ENSMUSG( | Cystm1 | 1065.682 | -0.22527 | 0.075385 | 0.767994 |
| ENSMUSG( | CYTB | 1021482 | -0.03817 | 0.757863 | 0.951582 | ENSMUSG( | CYTB | 915546 | -0.08919 | 0.568162 | 0.941777 |
| ENSMUSG( | Cyth1 | 2487.751 | 0.068544 | 0.612062 | 0.908586 | ENSMUSG( | Cyth1 | 2281.687 | 0.088755 | 0.517214 | 0.930939 |

| | | | | | |
|---|---|---|---|---|---|
| ENSMUSG( | 375.8293 | -0.08124 | 0.613432 | 0.909027 | Cyp26b1 |
| ENSMUSG( | 203.7744 | -0.5791 | 0.000493 | 0.020342 | Cyp27a1 |
| ENSMUSG( | 13.29367 | 0.288519 | 0.579306 | NA | Cyp27b1 |
| ENSMUSG( | 2811.134 | 0.12367 | 0.282808 | 0.73186 | Cyp2d22 |
| ENSMUSG( | 80.28948 | -0.04837 | 0.840806 | 0.971179 | Cyp2f2 |
| ENSMUSG( | 104.8959 | -0.23319 | 0.283166 | 0.73186 | Cyp2j12 |
| ENSMUSG( | 1526.639 | -0.25821 | 0.05359 | 0.35565 | Cyp2j6 |
| ENSMUSG( | 3513.776 | 0.177708 | 0.141714 | 0.558728 | Cyp2j9 |
| ENSMUSG( | 179.177 | -0.9543 | 2.33E-07 | 5.39E-05 | Cyp2s1 |
| ENSMUSG( | 339.5862 | -0.26101 | 0.049821 | 0.340731 | Cyp2u1 |
| ENSMUSG( | 80.61126 | -0.21486 | 0.374197 | 0.800445 | Cyp39a1 |
| ENSMUSG( | 66.59725 | -0.20477 | 0.496808 | 0.86175 | Cyp3a13 |
| ENSMUSG( | 9666.941 | -0.27216 | 0.003786 | 0.078636 | Cyp46a1 |
| ENSMUSG( | 16.75156 | 0.357131 | 0.542967 | NA | Cyp4b1 |
| ENSMUSG( | 496.322 | -0.20207 | 0.111139 | 0.502556 | Cyp4f13 |
| ENSMUSG( | 1123.114 | 0.034608 | 0.790675 | 0.959461 | Cyp4f14 |
| ENSMUSG( | 1322.667 | 0.073847 | 0.57433 | 0.894186 | Cyp4f15 |
| ENSMUSG( | 261.576 | -0.30514 | 0.036914 | 0.296508 | Cyp4f16 |
| ENSMUSG( | 205.3366 | -0.1515 | 0.36218 | 0.792943 | Cyp4f17 |
| ENSMUSG( | 13.30573 | -0.57984 | 0.394282 | NA | Cyp4f18 |
| ENSMUSG( | 559.7506 | -0.22962 | 0.099547 | 0.481986 | Cyp4v3 |
| ENSMUSG( | 528.6787 | -0.34365 | 0.01361 | 0.170359 | Cyp4x1 |
| ENSMUSG( | 24.14883 | 0.022189 | 0.956094 | 0.993874 | Cyp4x1os |
| ENSMUSG( | 8197.241 | -0.04635 | 0.742432 | 0.948478 | Cyp51 |
| ENSMUSG( | 528.5729 | -0.25038 | 0.163695 | 0.594869 | Cyp7b1 |
| ENSMUSG( | 156.3049 | -0.18109 | 0.278299 | 0.728302 | Cyren |
| ENSMUSG( | 4485.941 | 0.15221 | 0.074269 | 0.422054 | Cyria |
| ENSMUSG( | 3539.351 | 0.002179 | 0.980991 | 0.997512 | Cyrib |
| ENSMUSG( | 198.6164 | 0.331814 | 0.031604 | 0.271081 | Cys1 |
| ENSMUSG( | 40.38538 | -0.19586 | 0.526622 | 0.877267 | Cysltr1 |
| ENSMUSG( | 23.4108 | -0.41732 | 0.398222 | 0.812473 | Cysltr2 |
| ENSMUSG( | 1274.746 | 0.035464 | 0.719274 | 0.941986 | Cystm1 |
| ENSMUSG( | 1021482 | -0.03817 | 0.757863 | 0.951582 | CYTB |
| ENSMUSG( | 2487.751 | 0.068544 | 0.612062 | 0.908586 | Cyth1 |
| ENSMUSG( | 2479.399 | 0.023067 | 0.817968 | 0.965795 | Cyth2 |
| ENSMUSG( | 1094.392 | 0.096985 | 0.325081 | 0.770608 | Cyth3 |
| ENSMUSG( | 727.5433 | -0.06978 | 0.579712 | 0.89687 | Cyth4 |
| ENSMUSG( | 27.06286 | -1.29892 | 0.030702 | 0.266671 | Cytip |
| ENSMUSG( | 132.8395 | -0.08934 | 0.764159 | 0.954055 | Cyyr1 |
| ENSMUSG( | 1190.856 | -0.12645 | 0.1708 | 0.605413 | Czib |
| ENSMUSG( | 219.0557 | 0.058319 | 0.687555 | 0.93324 | D030028A08Rik |
| ENSMUSG( | 36.7143 | 0.330914 | 0.304031 | 0.752269 | D030047H15Rik |
| ENSMUSG( | 598.0711 | 0.11123 | 0.41028 | 0.818258 | D030056L22Rik |
| ENSMUSG( | 154.4228 | 0.051095 | 0.767819 | 0.954621 | D130020L05Rik |
| ENSMUSG( | 88.73753 | -0.30369 | 0.203845 | 0.64414 | D130040H23Rik |
| ENSMUSG( | 796.5653 | -0.22388 | 0.198588 | 0.638599 | D130043K22Rik |
| ENSMUSG( | 297.8633 | -0.07869 | 0.603229 | 0.904887 | D16Ertd472e |
| ENSMUSG( | 383.6902 | -0.31263 | 0.016458 | 0.190546 | D17H6S53E |
| ENSMUSG( | 148.7076 | -0.61689 | 0.001899 | 0.049802 | D17H6S56E-5 |
| ENSMUSG( | 401.5626 | -0.0573 | 0.641439 | 0.916693 | D2hgdh |
| ENSMUSG( | 49.07918 | 0.118881 | 0.689619 | 0.933609 | D330041H03Rik |
| ENSMUSG( | 102.2476 | 0.167087 | 0.421319 | 0.823126 | D330050G23Rik |
| ENSMUSG( | 638.9386 | -0.03492 | 0.801464 | 0.961947 | D3Ertd751e |
| ENSMUSG( | 8918.29 | 0.165454 | 0.087611 | 0.454663 | D430019H16Rik |
| ENSMUSG( | 44.52845 | -0.46758 | 0.169654 | 0.603859 | D430020J02Rik |
| ENSMUSG( | 124.5317 | -0.11626 | 0.56976 | 0.893041 | D430036J16Rik |
| ENSMUSG( | 4677.793 | -0.25909 | 0.076138 | 0.42702 | D430041D05Rik |
| ENSMUSG( | 3917.009 | 0.006764 | 0.937419 | 0.993415 | D5Ertd579e |
| ENSMUSG( | 424.4315 | 0.129516 | 0.278965 | 0.728867 | D630003M21Rik |
| ENSMUSG( | 47.57637 | -0.08234 | 0.76198 | 0.95339 | D630023F18Rik |
| ENSMUSG( | 13.02939 | -0.02921 | 0.959397 | NA | D630024D03Rik |
| ENSMUSG( | 19.17041 | 0.663936 | 0.123537 | NA | D630033O11Rik |
| ENSMUSG( | 36.87793 | 0.374358 | 0.241286 | 0.689026 | D630039A03Rik |
| ENSMUSG( | 13.46014 | 0.130934 | 0.798745 | NA | D630044L22Rik |
| ENSMUSG( | 1630.31 | -0.13795 | 0.408901 | 0.818173 | D630045J12Rik |
| ENSMUSG( | 534.553 | 0.230439 | 0.038321 | 0.302021 | D6Wsu163e |
| ENSMUSG( | 160.4981 | 0.002072 | 0.990336 | 0.999232 | D730003I15Rik |
| ENSMUSG( | 95.23951 | -0.43606 | 0.03727 | 0.297943 | D7Ertd443e |
| ENSMUSG( | 32.45584 | -0.14653 | 0.658008 | 0.92425 | D830030K20Rik |
| ENSMUSG( | 32.71849 | -0.19565 | 0.552218 | 0.886141 | D830035M03Rik |
| ENSMUSG( | 2267.918 | 0.071784 | 0.453776 | 0.836798 | D8Ertd738e |
| ENSMUSG( | 19.28301 | 0.457935 | 0.290673 | NA | D930020B18Rik |
| ENSMUSG( | 8.790113 | -0.115 | 0.856621 | NA | D930028M14Rik |
| ENSMUSG( | 85.16046 | 0.054288 | 0.832458 | 0.970333 | D930048N14Rik |
| ENSMUSG( | 3324.019 | -0.12481 | 0.220849 | 0.666476 | Daam1 |
| ENSMUSG( | 12635.86 | 0.047553 | 0.668544 | 0.927864 | Daam2 |
| ENSMUSG( | 4963.488 | 0.078515 | 0.420747 | 0.822925 | Dab1 |
| ENSMUSG( | 692.4865 | 0.023463 | 0.852096 | 0.973168 | Dab2 |
| ENSMUSG( | 8248.576 | 0.208499 | 0.031565 | 0.270895 | Dab2ip |
| ENSMUSG( | 164.0831 | -0.70901 | 0.000244 | 0.01251 | Dach1 |
| ENSMUSG( | 13.67946 | 0.311459 | 0.527426 | NA | Dach2 |
| ENSMUSG( | 223.8056 | 0.065109 | 0.678458 | 0.930542 | Dact1 |
| ENSMUSG( | 1397.283 | 0.076352 | 0.579648 | 0.89687 | Dact2 |
| ENSMUSG( | 11666.89 | 0.231439 | 0.009831 | 0.141621 | Dact3 |
| ENSMUSG( | 2516.027 | -0.23952 | 0.029142 | 0.259574 | Dad1 |
| ENSMUSG( | 5687.377 | -0.26393 | 0.086882 | 0.453311 | Dag1 |
| ENSMUSG( | 4862.481 | -0.33656 | 0.014374 | 0.176495 | Dagla |
| ENSMUSG( | 1802.514 | -0.28914 | 0.022125 | 0.226843 | Daglb |
| ENSMUSG( | 88.6694 | -0.01545 | 0.950853 | 0.993821 | Dalir |
| ENSMUSG( | 2262.233 | 0.152912 | 0.158757 | 0.588112 | Dalrd3 |
| ENSMUSG( | 82.8246 | -0.14623 | 0.507874 | 0.86795 | Dancr |
| ENSMUSG( | 293.7984 | 0.030908 | 0.834355 | 0.970333 | Dand5 |
| ENSMUSG( | 374.0173 | -0.02929 | 0.834046 | 0.970333 | Dap |
| ENSMUSG( | 1952.679 | 0.073122 | 0.430431 | 0.826896 | Dap3 |
| ENSMUSG( | 2512.956 | 0.224788 | 0.040089 | 0.308906 | Dapk1 |
| ENSMUSG( | 81.391 | 0.206424 | 0.344596 | 0.779153 | Dapk2 |
| ENSMUSG( | 1038.468 | 0.174116 | 0.14268 | 0.561169 | Dapk3 |
| ENSMUSG( | 9.499647 | -0.04569 | 0.940081 | NA | Dapl1 |
| ENSMUSG( | 494.0905 | -0.12268 | 0.405468 | 0.81652 | Dapp1 |
| ENSMUSG( | 1710.909 | 0.046359 | 0.701925 | 0.937441 | Dars1 |
| ENSMUSG( | 481.7072 | 0.087618 | 0.438344 | 0.830406 | Dars2 |
| ENSMUSG( | 16.21725 | 0.254738 | 0.643572 | NA | Daw1 |
| ENSMUSG( | 691.8541 | 0.080515 | 0.443347 | 0.833108 | Daxx |
| ENSMUSG( | 1128.282 | 0.045054 | 0.678447 | 0.930542 | Dazap1 |
| ENSMUSG( | 8775.751 | 0.055016 | 0.632403 | 0.914073 | Dazap2 |
| ENSMUSG( | 70.74187 | 0.065856 | 0.783733 | 0.957645 | Dazl |
| ENSMUSG( | 50.61983 | -0.1912 | 0.528 | 0.878105 | Dbf4 |
| ENSMUSG( | 19.26032 | -0.73407 | 0.122773 | NA | Dbh |
| ENSMUSG( | 11428.05 | 0.10949 | 0.303068 | 0.751443 | Dbi |
| ENSMUSG( | 22.27614 | 0.026711 | 0.948159 | 0.993821 | Dbil5 |
| ENSMUSG( | 11555.2 | 0.085195 | 0.391964 | 0.809274 | Dbn1 |
| ENSMUSG( | 300.5618 | 0.169118 | 0.216166 | 0.661017 | Dbndd1 |
| ENSMUSG( | 5859.654 | 0.020579 | 0.859132 | 0.974707 | Dbndd2 |
| ENSMUSG( | 1983.123 | 0.114769 | 0.180149 | 0.617634 | Dbnl |
| ENSMUSG( | 3249.442 | 0.075519 | 0.572774 | 0.893584 | Dbp |
| ENSMUSG( | 1195.451 | -0.0349 | 0.800472 | 0.961745 | Dbpht2 |
| ENSMUSG( | 236.1246 | 0.001362 | 0.993291 | 0.999369 | Dbr1 |
| ENSMUSG( | 1745.333 | 0.112362 | 0.328193 | 0.772854 | Dbt |
| ENSMUSG( | 1418.228 | 0.169928 | 0.131928 | 0.544244 | Dbx2 |
| ENSMUSG( | 695.3945 | 0.100582 | 0.380577 | 0.801959 | Dcaf1 |
| ENSMUSG( | 558.8489 | -0.12125 | 0.317855 | 0.764145 | Dcaf10 |
| ENSMUSG( | 3166.377 | 0.150463 | 0.062591 | 0.38686 | Dcaf11 |
| ENSMUSG( | 2098.779 | 0.17569 | 0.06294 | 0.388168 | Dcaf12 |
| ENSMUSG( | 693.6699 | 0.038615 | 0.724582 | 0.944096 | Dcaf12l1 |
| ENSMUSG( | 19.19083 | 0.129217 | 0.764915 | NA | Dcaf12l2 |
| ENSMUSG( | 538.4669 | -0.06807 | 0.593275 | 0.902578 | Dcaf13 |
| ENSMUSG( | 581.6058 | -0.01968 | 0.887625 | 0.983411 | Dcaf15 |
| ENSMUSG( | 324.8933 | -0.02922 | 0.814454 | 0.964574 | Dcaf17 |
| ENSMUSG( | 512.5309 | 0.04959 | 0.702203 | 0.937536 | Dcaf4 |
| ENSMUSG( | 3146.047 | 0.09847 | 0.228551 | 0.675862 | Dcaf5 |
| ENSMUSG( | 2794.32 | 0.21011 | 0.020067 | 0.214281 | Dcaf6 |
| ENSMUSG( | 7187.57 | 0.049156 | 0.54212 | 0.883182 | Dcaf7 |
| ENSMUSG( | 4778.19 | 0.144563 | 0.077124 | 0.429655 | Dcaf8 |
| ENSMUSG( | 1154.272 | 0.185529 | 0.110376 | 0.50108 | Dcakd |
| ENSMUSG( | 354.5504 | -0.34974 | 0.040356 | 0.310226 | Dcbld1 |
| ENSMUSG( | 1193.686 | -0.13459 | 0.354184 | 0.786644 | Dcbld2 |
| ENSMUSG( | 664.5816 | -0.32278 | 0.097381 | 0.477629 | Dcc |
| ENSMUSG( | 341.7663 | -0.17455 | 0.228666 | 0.675954 | Dcdc2a |
| ENSMUSG( | 72.95788 | -1.52145 | 2.28E-09 | 1.00E-06 | Dcdc2b |
| ENSMUSG( | 11.4458 | 0.785357 | 0.177435 | NA | Dcdc2c |
| ENSMUSG( | 779.7352 | -0.13899 | 0.388512 | 0.806278 | Dchs1 |
| ENSMUSG( | 43.58722 | -0.88989 | 0.00895 | 0.132538 | Dchs2 |
| ENSMUSG( | 448.9174 | 0.151605 | 0.206143 | 0.647992 | Dck |
| ENSMUSG( | 21963.4 | 0.096061 | 0.192336 | 0.63139 | Dclk1 |
| ENSMUSG( | 3688.507 | 0.027055 | 0.727791 | 0.944355 | Dclk2 |
| ENSMUSG( | 685.3111 | 0.213063 | 0.217686 | 0.662824 | Dclk3 |
| ENSMUSG( | 280.1636 | -0.19875 | 0.154361 | 0.581452 | Dclre1a |
| ENSMUSG( | 247.0886 | 0.044844 | 0.778456 | 0.956215 | Dclre1b |
| ENSMUSG( | 267.0609 | 0.019645 | 0.9022 | 0.985633 | Dclre1c |
| ENSMUSG( | 801.5953 | -0.20863 | 0.139972 | 0.556732 | Dcn |
| ENSMUSG( | 513.0688 | -0.05124 | 0.654512 | 0.923227 | Dcp1a |
| ENSMUSG( | 556.9409 | 0.035582 | 0.761473 | 0.953106 | Dcp1b |
| ENSMUSG( | 1303.878 | 0.082489 | 0.38038 | 0.801757 | Dcp2 |
| ENSMUSG( | 308.2793 | 0.202775 | 0.141962 | 0.559166 | Dcps |
| ENSMUSG( | 42.99478 | 0.34135 | 0.307106 | 0.75508 | Dcst2 |
| ENSMUSG( | 29.47066 | -0.54185 | 0.120279 | 0.524495 | Dct |
| ENSMUSG( | 411.0718 | -0.01034 | 0.937034 | 0.993415 | Dctd |
| ENSMUSG( | 10971.5 | 0.169561 | 0.051694 | 0.34893 | Dctn1 |

| | | | | | |
|---|---|---|---|---|---|
| ENSMUSG( | 355.6921 | 0.150246 | 0.323947 | 0.905257 | Cyp2u1 |
| ENSMUSG( | 91.07048 | 0.386454 | 0.097654 | 0.796743 | Cyp39a1 |
| ENSMUSG( | 81.51121 | 0.59054 | 0.021763 | 0.575341 | Cyp3a13 |
| ENSMUSG( | 9955.182 | 0.094044 | 0.391149 | 0.917257 | Cyp46a1 |
| ENSMUSG( | 12.54005 | -0.17117 | 0.758085 | 0.971858 | Cyp4b1 |
| ENSMUSG( | 491.8041 | 0.054945 | 0.728465 | 0.966913 | Cyp4f13 |
| ENSMUSG( | 991.1397 | -0.05549 | 0.696705 | 0.960867 | Cyp4f14 |
| ENSMUSG( | 1364.315 | 0.408749 | 0.005462 | 0.303106 | Cyp4f15 |
| ENSMUSG( | 254.0137 | -0.09636 | 0.579287 | 0.943942 | Cyp4f16 |
| ENSMUSG( | 201.0721 | 0.071851 | 0.705676 | 0.96315 | Cyp4f17 |
| ENSMUSG( | 12.35128 | -0.60154 | 0.421996 | 0.917257 | Cyp4f18 |
| ENSMUSG( | 552.9795 | 0.017543 | 0.910186 | 0.992081 | Cyp4v3 |
| ENSMUSG( | 518.7136 | -0.10502 | 0.518129 | 0.930939 | Cyp4x1 |
| ENSMUSG( | 21.76244 | -0.04233 | 0.924179 | 0.992937 | Cyp4x1os |
| ENSMUSG( | 7058.563 | -0.21629 | 0.134971 | 0.82437 | Cyp51 |
| ENSMUSG( | 547.2518 | 0.130088 | 0.501406 | 0.927825 | Cyp7b1 |
| ENSMUSG( | 149.273 | -0.02962 | 0.895947 | 0.990558 | Cyren |
| ENSMUSG( | 3816.118 | -0.04425 | 0.707681 | 0.964029 | Cyria |
| ENSMUSG( | 3054.616 | -0.16133 | 0.182686 | 0.850579 | Cyrib |
| ENSMUSG( | 204.4053 | 0.638728 | 0.000719 | 0.08668 | Cys1 |
| ENSMUSG( | 45.66802 | 0.4237 | 0.188481 | 0.85698 | Cysltr1 |
| ENSMUSG( | 16.665 | -1.33596 | 0.023998 | 0.588226 | Cysltr2 |
| ENSMUSG( | 1065.682 | -0.22527 | 0.075385 | 0.767994 | Cystm1 |
| ENSMUSG( | 915546 | -0.08919 | 0.568162 | 0.941777 | CYTB |
| ENSMUSG( | 2281.687 | 0.088755 | 0.517214 | 0.930939 | Cyth1 |
| ENSMUSG( | 2207.51 | -0.04542 | 0.702879 | 0.962352 | Cyth2 |
| ENSMUSG( | 983.8322 | 0.061316 | 0.598701 | 0.947033 | Cyth3 |
| ENSMUSG( | 707.6285 | 0.123658 | 0.426221 | 0.917257 | Cyth4 |
| ENSMUSG( | 22.47522 | -1.90793 | 0.003017 | 0.219289 | Cytip |
| ENSMUSG( | 155.7965 | 0.597998 | 0.277905 | 0.891468 | Cyyr1 |
| ENSMUSG( | 1128.48 | -0.00833 | 0.95237 | 0.994798 | Czib |
| ENSMUSG( | 189.4975 | -0.08108 | 0.640052 | 0.953903 | D030028A08Rik |
| ENSMUSG( | 27.08153 | -0.21409 | 0.612841 | 0.951071 | D030047H15Rik |
| ENSMUSG( | 528.5912 | 0.016837 | 0.915997 | 0.992529 | D030056L22Rik |
| ENSMUSG( | 136.9377 | -0.03755 | 0.854612 | 0.985649 | D130020L05Rik |
| ENSMUSG( | 91.87744 | 0.100109 | 0.679934 | 0.957966 | D130040H23Rik |
| ENSMUSG( | 918.6273 | 0.440214 | 0.02584 | 0.601244 | D130043K22Rik |
| ENSMUSG( | 267.6619 | -0.1232 | 0.495824 | 0.927825 | D16Ertd472e |
| ENSMUSG( | 404.4249 | 0.123697 | 0.3932 | 0.917257 | D17H6S53E |
| ENSMUSG( | 146.7805 | -0.31718 | 0.163665 | 0.838291 | D17H6S56E-5 |
| ENSMUSG( | 358.9107 | -0.12464 | 0.425059 | 0.917257 | D2hgdh |
| ENSMUSG( | 46.57204 | 0.263359 | 0.433132 | 0.918434 | D330041H03Rik |
| ENSMUSG( | 85.52573 | -0.07908 | 0.736263 | 0.967344 | D330050G23Rik |
| ENSMUSG( | 554.7718 | -0.18481 | 0.212922 | 0.864412 | D3Ertd751e |
| ENSMUSG( | 8031.226 | 0.135205 | 0.211791 | 0.864412 | D430019H16Rik |
| ENSMUSG( | 46.29096 | -0.02292 | 0.945214 | 0.994798 | D430020J02Rik |
| ENSMUSG( | 115.8825 | -0.06225 | 0.784555 | 0.976207 | D430036J16Rik |
| ENSMUSG( | 4972.785 | 0.19617 | 0.264903 | 0.886134 | D430041D05Rik |
| ENSMUSG( | 3453.952 | -0.09033 | 0.388907 | 0.917257 | D5Ertd579e |
| ENSMUSG( | 361.0515 | -0.05842 | 0.710389 | 0.964363 | D630003M21Rik |
| ENSMUSG( | 44.48877 | -0.02306 | 0.939381 | 0.993968 | D630023F18Rik |
| ENSMUSG( | 14.71709 | 0.228524 | 0.646219 | 0.953988 | D630033O11Rik |
| ENSMUSG( | 26.071 | -0.34156 | 0.386369 | 0.917257 | D630039A03Rik |
| ENSMUSG( | 12.04415 | 0.132605 | 0.822446 | 0.979198 | D630044L22Rik |
| ENSMUSG( | 1813.245 | 0.41781 | 0.026342 | 0.603041 | D630045J12Rik |
| ENSMUSG( | 458.3667 | 0.068029 | 0.620271 | 0.952896 | D6Wsu163e |
| ENSMUSG( | 141.4675 | -0.11891 | 0.602618 | 0.94789 | D730003I15Rik |
| ENSMUSG( | 99.20369 | -0.00915 | 0.971139 | 0.996102 | D7Ertd443e |
| ENSMUSG( | 32.35357 | 0.075961 | 0.836963 | 0.981335 | D830030K20Rik |
| ENSMUSG( | 25.30942 | -0.73395 | 0.070203 | 0.757694 | D830035M03Rik |
| ENSMUSG( | 1833.289 | -0.29038 | 0.02326 | 0.585577 | D8Ertd738e |
| ENSMUSG( | 17.06559 | 0.363389 | 0.442251 | 0.921554 | D930020B18Rik |
| ENSMUSG( | 7.724298 | -0.1874 | 0.79621 | NA | D930028M14Rik |
| ENSMUSG( | 87.18779 | 0.387989 | 0.154941 | 0.837724 | D930048N14Rik |
| ENSMUSG( | 2932.738 | -0.22599 | 0.052192 | 0.712974 | Daam1 |
| ENSMUSG( | 10599.23 | -0.19736 | 0.077217 | 0.771155 | Daam2 |
| ENSMUSG( | 4261.383 | -0.09524 | 0.474048 | 0.923329 | Dab1 |
| ENSMUSG( | 622.3253 | -0.01783 | 0.90004 | 0.990558 | Dab2 |
| ENSMUSG( | 7211.994 | 0.097565 | 0.431218 | 0.917257 | Dab2ip |
| ENSMUSG( | 173.5491 | -0.20843 | 0.292288 | 0.897854 | Dach1 |
| ENSMUSG( | 10.73831 | -0.08369 | 0.886332 | NA | Dach2 |
| ENSMUSG( | 205.0117 | 0.071982 | 0.695465 | 0.960867 | Dact1 |
| ENSMUSG( | 1242.731 | 0.006809 | 0.962389 | 0.995237 | Dact2 |
| ENSMUSG( | 10139.2 | 0.101456 | 0.333455 | 0.907907 | Dact3 |
| ENSMUSG( | 2452.062 | -0.03324 | 0.794639 | 0.97681 | Dad1 |
| ENSMUSG( | 5982.832 | 0.16322 | 0.354623 | 0.912411 | Dag1 |
| ENSMUSG( | 5098.898 | 0.091454 | 0.569351 | 0.941777 | Dagla |
| ENSMUSG( | 1818.95 | 0.024355 | 0.870772 | 0.988549 | Daglb |
| ENSMUSG( | 79.47605 | -0.07039 | 0.780587 | 0.976106 | Dalir |
| ENSMUSG( | 2047.121 | 0.133319 | 0.404444 | 0.917257 | Dalrd3 |
| ENSMUSG( | 86.49613 | 0.256817 | 0.312516 | 0.900037 | Dancr |
| ENSMUSG( | 258.2448 | -0.08 | 0.640946 | 0.953903 | Dand5 |
| ENSMUSG( | 331.8394 | -0.10949 | 0.475451 | 0.923329 | Dap |
| ENSMUSG( | 1679.735 | -0.09585 | 0.422682 | 0.917257 | Dap3 |
| ENSMUSG( | 2246.111 | 0.173358 | 0.174607 | 0.844619 | Dapk1 |
| ENSMUSG( | 58.20028 | -0.54126 | 0.058838 | 0.728906 | Dapk2 |
| ENSMUSG( | 891.697 | 0.008534 | 0.94682 | 0.994798 | Dapk3 |
| ENSMUSG( | 9.244189 | 0.0961 | 0.885787 | NA | Dapl1 |
| ENSMUSG( | 416.7222 | -0.36754 | 0.009751 | 0.406458 | Dapp1 |
| ENSMUSG( | 1424.55 | -0.22556 | 0.083018 | 0.77681 | Dars1 |
| ENSMUSG( | 402.1261 | -0.17553 | 0.247438 | 0.877904 | Dars2 |
| ENSMUSG( | 14.52879 | 0.240321 | 0.682209 | 0.958678 | Daw1 |
| ENSMUSG( | 617.2695 | 0.015794 | 0.908248 | 0.991958 | Daxx |
| ENSMUSG( | 1020.072 | 0.021451 | 0.86312 | 0.986547 | Dazap1 |
| ENSMUSG( | 7538.654 | -0.11904 | 0.398936 | 0.917257 | Dazap2 |
| ENSMUSG( | 69.25073 | 0.241368 | 0.394666 | 0.917257 | Dazl |
| ENSMUSG( | 53.20271 | 0.233333 | 0.42267 | 0.917257 | Dbf4 |
| ENSMUSG( | 18.31053 | -0.56475 | 0.260454 | 0.884524 | Dbh |
| ENSMUSG( | 9252.409 | -0.24162 | 0.085209 | 0.779799 | Dbi |
| ENSMUSG( | 16.91889 | -0.52255 | 0.28597 | 0.893906 | Dbil5 |
| ENSMUSG( | 10179.14 | -0.00851 | 0.941348 | 0.994578 | Dbn1 |
| ENSMUSG( | 270.8573 | 0.139355 | 0.386883 | 0.917257 | Dbndd1 |
| ENSMUSG( | 5337.442 | 0.017747 | 0.896192 | 0.990558 | Dbndd2 |
| ENSMUSG( | 1671.982 | -0.10988 | 0.308068 | 0.898538 | Dbnl |
| ENSMUSG( | 4180.7 | 0.923962 | 3.27E-12 | 4.09E-09 | Dbp |
| ENSMUSG( | 1097.919 | -0.01774 | 0.922703 | 0.992644 | Dbpht2 |
| ENSMUSG( | 215.7052 | 0.012315 | 0.944411 | 0.994798 | Dbr1 |
| ENSMUSG( | 1512.34 | -0.03498 | 0.814185 | 0.979198 | Dbt |
| ENSMUSG( | 1191.749 | -0.06356 | 0.636345 | 0.953903 | Dbx2 |
| ENSMUSG( | 621.1908 | 0.042108 | 0.770465 | 0.974236 | Dcaf1 |
| ENSMUSG( | 542.1369 | 0.058884 | 0.677683 | 0.957106 | Dcaf10 |
| ENSMUSG( | 2689.802 | -0.04701 | 0.660552 | 0.956205 | Dcaf11 |
| ENSMUSG( | 1762.48 | -0.05812 | 0.63605 | 0.953903 | Dcaf12 |
| ENSMUSG( | 634.3343 | 0.042828 | 0.77365 | 0.974672 | Dcaf12l1 |
| ENSMUSG( | 15.68463 | -0.16818 | 0.748862 | 0.96984 | Dcaf12l2 |
| ENSMUSG( | 495.3842 | -0.03327 | 0.825801 | 0.979653 | Dcaf13 |
| ENSMUSG( | 557.0583 | 0.125539 | 0.425298 | 0.917257 | Dcaf15 |
| ENSMUSG( | 299.6154 | 0.016525 | 0.92289 | 0.992644 | Dcaf17 |
| ENSMUSG( | 442.4775 | -0.10369 | 0.465275 | 0.923329 | Dcaf4 |
| ENSMUSG( | 2776.339 | 0.008772 | 0.932301 | 0.99306 | Dcaf5 |
| ENSMUSG( | 2449.782 | 0.104133 | 0.346269 | 0.910966 | Dcaf6 |
| ENSMUSG( | 6287.999 | -0.06938 | 0.496715 | 0.927825 | Dcaf7 |
| ENSMUSG( | 4177.617 | 0.030207 | 0.760863 | 0.972695 | Dcaf8 |
| ENSMUSG( | 962.4657 | -0.06944 | 0.655279 | 0.955294 | Dcakd |
| ENSMUSG( | 391.5789 | 0.22691 | 0.244293 | 0.877904 | Dcbld1 |
| ENSMUSG( | 1240.601 | 0.243241 | 0.136435 | 0.827092 | Dcbld2 |
| ENSMUSG( | 715.4057 | 0.17621 | 0.348142 | 0.911652 | Dcc |
| ENSMUSG( | 293.0958 | -0.3676 | 0.020455 | 0.565891 | Dcdc2a |
| ENSMUSG( | 113.8938 | 0.39614 | 0.229522 | 0.868507 | Dcdc2b |
| ENSMUSG( | 802.7702 | 0.217518 | 0.237315 | 0.872173 | Dchs1 |
| ENSMUSG( | 50.9166 | -0.03158 | 0.918509 | 0.992529 | Dchs2 |
| ENSMUSG( | 391.9921 | 0.030007 | 0.838314 | 0.981601 | Dck |
| ENSMUSG( | 19094.92 | -0.03803 | 0.711485 | 0.964363 | Dclk1 |
| ENSMUSG( | 3264.678 | -0.05702 | 0.561473 | 0.940279 | Dclk2 |
| ENSMUSG( | 552.2321 | -0.14067 | 0.469283 | 0.923329 | Dclk3 |
| ENSMUSG( | 262.1858 | -0.11423 | 0.457058 | 0.923329 | Dclre1a |
| ENSMUSG( | 229.4261 | 0.108412 | 0.548357 | 0.938063 | Dclre1b |
| ENSMUSG( | 260.6426 | 0.21957 | 0.244629 | 0.877904 | Dclre1c |
| ENSMUSG( | 794.0917 | 0.040208 | 0.787199 | 0.97637 | Dcn |
| ENSMUSG( | 498.4063 | 0.135866 | 0.334086 | 0.907907 | Dcp1a |
| ENSMUSG( | 514.0353 | 0.078721 | 0.605511 | 0.94924 | Dcp1b |
| ENSMUSG( | 1153.503 | -0.00144 | 0.989766 | 0.997928 | Dcp2 |
| ENSMUSG( | 255.3168 | -0.06725 | 0.680526 | 0.958395 | Dcps |
| ENSMUSG( | 36.93157 | 0.184347 | 0.572221 | 0.94251 | Dcst2 |
| ENSMUSG( | 31.63053 | 0.007981 | 0.98487 | 0.997067 | Dct |
| ENSMUSG( | 361.1037 | -0.11631 | 0.411734 | 0.917257 | Dctd |
| ENSMUSG( | 9165.519 | -0.07714 | 0.455412 | 0.923316 | Dctn1 |
| ENSMUSG( | 4575.931 | -0.16846 | 0.17181 | 0.842999 | Dctn2 |
| ENSMUSG( | 1551.329 | -0.20585 | 0.077751 | 0.77321 | Dctn3 |
| ENSMUSG( | 2135.037 | 0.049298 | 0.647317 | 0.95422 | Dctn4 |
| ENSMUSG( | 1755.909 | -0.11424 | 0.368832 | 0.915491 | Dctn5 |
| ENSMUSG( | 1226.607 | -0.15181 | 0.206942 | 0.864412 | Dctn6 |
| ENSMUSG( | 310.8067 | -0.1776 | 0.275097 | 0.889874 | Dctpp1 |
| ENSMUSG( | 654.9339 | -0.00062 | 0.997226 | 0.999268 | Dcun1d1 |
| ENSMUSG( | 1019.199 | 0.039404 | 0.751417 | 0.970317 | Dcun1d2 |
| ENSMUSG( | 350.109 | 0.264974 | 0.075282 | 0.767994 | Dcun1d3 |
| ENSMUSG( | 1485.862 | 0.050789 | 0.681973 | 0.958678 | Dcun1d4 |
| ENSMUSG( | 790.401 | 0.086387 | 0.495853 | 0.927825 | Dcun1d5 |

| | | | | | | | | | | | |
|---|---|---|---|---|---|---|---|---|---|---|---|
| ENSMUSG( | Cyth2 | 2479.399 | 0.023067 | 0.817968 | 0.965795 | ENSMUSG( | Cyth2 | 2207.51 | -0.04542 | 0.702879 | 0.962352 |
| ENSMUSG( | Cyth3 | 1094.392 | 0.096985 | 0.325081 | 0.770608 | ENSMUSG( | Cyth3 | 983.8322 | 0.061316 | 0.598701 | 0.947033 |
| ENSMUSG( | Cyth4 | 727.5433 | -0.06978 | 0.579712 | 0.89687 | ENSMUSG( | Cyth4 | 707.6285 | 0.123658 | 0.426221 | 0.917257 |
| ENSMUSG( | Cytip | 27.06286 | -1.29892 | 0.030702 | 0.266671 | ENSMUSG( | Cytip | 22.47522 | -1.90793 | 0.003017 | 0.219289 |
| ENSMUSG( | Cyyr1 | 132.8395 | -0.08934 | 0.764159 | 0.954055 | ENSMUSG( | Cyyr1 | 155.7965 | 0.597998 | 0.277905 | 0.891468 |
| ENSMUSG( | Czib | 1190.856 | -0.12645 | 0.1708 | 0.605413 | ENSMUSG( | Czib | 1128.48 | -0.00833 | 0.95237 | 0.994798 |
| ENSMUSG( | D030028A( | 219.0557 | 0.058319 | 0.687555 | 0.93324 | ENSMUSG( | D030028A( | 189.4975 | -0.08108 | 0.640052 | 0.953903 |
| ENSMUSG( | D030047H: | 36.7143 | 0.330914 | 0.304031 | 0.752269 | ENSMUSG( | D030047H: | 27.08153 | -0.21409 | 0.612841 | 0.951071 |
| ENSMUSG( | D030056L2 | 598.0711 | 0.11123 | 0.41028 | 0.818258 | ENSMUSG( | D030056L2 | 528.5912 | 0.016837 | 0.915997 | 0.992529 |
| ENSMUSG( | D130020L( | 154.4228 | 0.051095 | 0.767819 | 0.954621 | ENSMUSG( | D130020L( | 136.9377 | -0.03755 | 0.854612 | 0.985649 |
| ENSMUSG( | D130040H: | 88.73753 | -0.30369 | 0.203845 | 0.64414 | ENSMUSG( | D130040H: | 91.87744 | 0.100109 | 0.679934 | 0.957966 |
| ENSMUSG( | D130043K: | 796.5653 | -0.22388 | 0.198588 | 0.638599 | ENSMUSG( | D130043K: | 918.6273 | 0.440214 | 0.02584 | 0.601244 |
| ENSMUSG( | D16Ertd47: | 297.8633 | -0.07869 | 0.603229 | 0.904887 | ENSMUSG( | D16Ertd47: | 267.6619 | -0.1232 | 0.495824 | 0.927825 |
| ENSMUSG( | D17H6S53I | 383.6902 | -0.31263 | 0.016458 | 0.190546 | ENSMUSG( | D17H6S53I | 404.4249 | 0.123697 | 0.3932 | 0.917257 |
| ENSMUSG( | D17H6S56I | 148.7076 | -0.61689 | 0.001899 | 0.049802 | ENSMUSG( | D17H6S56I | 146.7805 | -0.31718 | 0.163665 | 0.838291 |
| ENSMUSG( | D2hgdh | 401.5626 | -0.0573 | 0.641439 | 0.916693 | ENSMUSG( | D2hgdh | 358.9107 | -0.12464 | 0.425059 | 0.917257 |
| ENSMUSG( | D330041H( | 49.07918 | 0.118881 | 0.689619 | 0.933609 | ENSMUSG( | D330041H( | 46.57204 | 0.263359 | 0.433132 | 0.918434 |
| ENSMUSG( | D330050G: | 102.2476 | 0.167087 | 0.421319 | 0.823126 | ENSMUSG( | D330050G: | 85.52573 | -0.07908 | 0.736263 | 0.967344 |
| ENSMUSG( | D3Ertd751( | 638.9386 | -0.03492 | 0.801464 | 0.961947 | ENSMUSG( | D3Ertd751( | 554.7718 | -0.18481 | 0.212922 | 0.864412 |
| ENSMUSG( | D430019H: | 8918.29 | 0.165454 | 0.087611 | 0.454663 | ENSMUSG( | D430019H: | 8031.226 | 0.135205 | 0.211791 | 0.864412 |
| ENSMUSG( | D430020J0 | 44.52845 | -0.46758 | 0.169654 | 0.603859 | ENSMUSG( | D430020J0 | 46.29096 | -0.02292 | 0.945214 | 0.994798 |
| ENSMUSG( | D430036J1 | 124.5317 | -0.11626 | 0.56976 | 0.893041 | ENSMUSG( | D430036J1 | 115.8825 | -0.06225 | 0.784555 | 0.976207 |
| ENSMUSG( | D430041D( | 4677.793 | -0.25909 | 0.076138 | 0.42702 | ENSMUSG( | D430041D( | 4972.785 | 0.19617 | 0.264903 | 0.886134 |
| ENSMUSG( | D5Ertd579( | 3917.009 | 0.006764 | 0.937419 | 0.993415 | ENSMUSG( | D5Ertd579( | 3453.952 | -0.09033 | 0.388907 | 0.917257 |
| ENSMUSG( | D630003M | 424.4315 | 0.129516 | 0.278965 | 0.728867 | ENSMUSG( | D630003M | 361.0515 | -0.05842 | 0.710389 | 0.964363 |
| ENSMUSG( | D630023F1 | 47.57637 | -0.08234 | 0.76198 | 0.95339 | ENSMUSG( | D630023F1 | 44.48877 | -0.02306 | 0.939381 | 0.993968 |
| ENSMUSG( | D630033O: | 19.17041 | 0.663936 | 0.123537 | NA | ENSMUSG( | D630033O: | 14.71709 | 0.228524 | 0.646219 | 0.953988 |
| ENSMUSG( | D630039A( | 36.87793 | 0.374358 | 0.241286 | 0.689026 | ENSMUSG( | D630039A( | 26.071 | -0.34156 | 0.386369 | 0.917257 |
| ENSMUSG( | D630044L2 | 13.46014 | 0.130934 | 0.798745 | NA | ENSMUSG( | D630044L2 | 12.04415 | 0.132605 | 0.822446 | 0.979198 |
| ENSMUSG( | D630045J1 | 1630.31 | -0.13795 | 0.408901 | 0.818173 | ENSMUSG( | D630045J1 | 1813.245 | 0.41781 | 0.026342 | 0.603041 |
| ENSMUSG( | D6Wsu163 | 534.553 | 0.230439 | 0.038321 | 0.302021 | ENSMUSG( | D6Wsu163 | 458.3667 | 0.068029 | 0.620271 | 0.952896 |
| ENSMUSG( | D730003I1 | 160.4981 | 0.002072 | 0.990336 | 0.999232 | ENSMUSG( | D730003I1 | 141.4675 | -0.11891 | 0.602618 | 0.94789 |
| ENSMUSG( | D7Ertd443( | 95.23951 | -0.43606 | 0.03727 | 0.297943 | ENSMUSG( | D7Ertd443( | 99.20369 | -0.00915 | 0.971139 | 0.996102 |
| ENSMUSG( | D830030K: | 32.45584 | -0.14653 | 0.658008 | 0.92425 | ENSMUSG( | D830030K: | 32.35357 | 0.075961 | 0.836963 | 0.981335 |
| ENSMUSG( | D830035M | 32.71849 | -0.19565 | 0.552218 | 0.886141 | ENSMUSG( | D830035M | 25.30942 | -0.73395 | 0.070203 | 0.757694 |
| ENSMUSG( | D8Ertd738( | 2267.918 | 0.071784 | 0.453776 | 0.836798 | ENSMUSG( | D8Ertd738( | 1833.289 | -0.29038 | 0.02326 | 0.585577 |
| ENSMUSG( | D930020B1 | 19.28301 | 0.457935 | 0.290673 | NA | ENSMUSG( | D930020B1 | 17.06559 | 0.363389 | 0.442251 | 0.921554 |
| ENSMUSG( | D930028M | 8.790113 | -0.115 | 0.856621 | NA | ENSMUSG( | D930028M | 7.724298 | -0.1874 | 0.79621 | NA |
| ENSMUSG( | D930048N: | 85.16046 | 0.054288 | 0.832458 | 0.970333 | ENSMUSG( | D930048N: | 87.18779 | 0.387989 | 0.154941 | 0.837724 |
| ENSMUSG( | Daam1 | 3324.019 | -0.12481 | 0.220849 | 0.666476 | ENSMUSG( | Daam1 | 2932.738 | -0.22599 | 0.052192 | 0.712974 |
| ENSMUSG( | Daam2 | 12635.86 | 0.047553 | 0.668544 | 0.927864 | ENSMUSG( | Daam2 | 10599.23 | -0.19736 | 0.077217 | 0.771155 |
| ENSMUSG( | Dab1 | 4963.488 | 0.078515 | 0.420747 | 0.822925 | ENSMUSG( | Dab1 | 4261.383 | -0.09524 | 0.474048 | 0.923329 |
| ENSMUSG( | Dab2 | 692.4865 | 0.023463 | 0.852096 | 0.973168 | ENSMUSG( | Dab2 | 622.3253 | -0.01783 | 0.90004 | 0.990558 |
| ENSMUSG( | Dab2ip | 8248.576 | 0.208499 | 0.031565 | 0.270895 | ENSMUSG( | Dab2ip | 7211.994 | 0.097565 | 0.431218 | 0.917257 |
| ENSMUSG( | Dach1 | 164.0831 | -0.70901 | 0.000244 | 0.01251 | ENSMUSG( | Dach1 | 173.5491 | -0.20843 | 0.292288 | 0.897854 |
| ENSMUSG( | Dach2 | 13.67946 | 0.311459 | 0.527426 | NA | ENSMUSG( | Dach2 | 10.73831 | -0.08369 | 0.886332 | NA |
| ENSMUSG( | Dact1 | 223.8056 | 0.065109 | 0.678458 | 0.930542 | ENSMUSG( | Dact1 | 205.0117 | 0.071982 | 0.695465 | 0.960867 |
| ENSMUSG( | Dact2 | 1397.283 | 0.076352 | 0.579648 | 0.89687 | ENSMUSG( | Dact2 | 1242.731 | 0.006809 | 0.962389 | 0.995237 |
| ENSMUSG( | Dact3 | 11666.89 | 0.231439 | 0.009831 | 0.141621 | ENSMUSG( | Dact3 | 10139.2 | 0.101456 | 0.333455 | 0.907907 |
| ENSMUSG( | Dad1 | 2516.027 | -0.23952 | 0.029142 | 0.259574 | ENSMUSG( | Dad1 | 2452.062 | -0.03324 | 0.794639 | 0.97681 |
| ENSMUSG( | Dag1 | 5687.377 | -0.26393 | 0.086882 | 0.453311 | ENSMUSG( | Dag1 | 5982.832 | 0.16322 | 0.354623 | 0.912411 |
| ENSMUSG( | Dagla | 4862.481 | -0.33656 | 0.014374 | 0.176495 | ENSMUSG( | Dagla | 5098.898 | 0.091454 | 0.569351 | 0.941777 |
| ENSMUSG( | Daglb | 1802.514 | -0.28914 | 0.022125 | 0.226843 | ENSMUSG( | Daglb | 1818.95 | 0.024355 | 0.870772 | 0.988549 |
| ENSMUSG( | Dalir | 88.6694 | -0.01545 | 0.950853 | 0.993821 | ENSMUSG( | Dalir | 79.47605 | -0.07039 | 0.780587 | 0.976106 |
| ENSMUSG( | Dalrd3 | 2262.233 | 0.152912 | 0.158757 | 0.588112 | ENSMUSG( | Dalrd3 | 2047.121 | 0.133319 | 0.404444 | 0.917257 |
| ENSMUSG( | Dancr | 82.8246 | -0.14623 | 0.507874 | 0.86795 | ENSMUSG( | Dancr | 86.49613 | 0.256817 | 0.312516 | 0.900037 |
| ENSMUSG( | Dand5 | 293.7984 | 0.030908 | 0.834355 | 0.970333 | ENSMUSG( | Dand5 | 258.2448 | -0.08 | 0.640946 | 0.953903 |
| ENSMUSG( | Dap | 374.0173 | -0.02929 | 0.834046 | 0.970333 | ENSMUSG( | Dap | 331.8394 | -0.10949 | 0.475451 | 0.923329 |
| ENSMUSG( | Dap3 | 1952.679 | 0.073122 | 0.430431 | 0.826896 | ENSMUSG( | Dap3 | 1679.735 | -0.09585 | 0.422682 | 0.917257 |
| ENSMUSG( | Dapk1 | 2512.956 | 0.224788 | 0.040089 | 0.308906 | ENSMUSG( | Dapk1 | 2246.111 | 0.173358 | 0.174607 | 0.844619 |
| ENSMUSG( | Dapk2 | 81.391 | 0.206424 | 0.344596 | 0.779153 | ENSMUSG( | Dapk2 | 58.20028 | -0.54126 | 0.058838 | 0.728906 |
| ENSMUSG( | Dapk3 | 1038.468 | 0.174116 | 0.14268 | 0.561169 | ENSMUSG( | Dapk3 | 891.697 | 0.008534 | 0.94682 | 0.994798 |
| ENSMUSG( | Dapl1 | 9.499647 | -0.04569 | 0.940081 | NA | ENSMUSG( | Dapl1 | 9.244189 | 0.0961 | 0.885787 | NA |
| ENSMUSG( | Dapp1 | 494.0905 | -0.12268 | 0.405468 | 0.81652 | ENSMUSG( | Dapp1 | 416.7222 | -0.36754 | 0.009751 | 0.406458 |
| ENSMUSG( | Dars1 | 1710.909 | 0.046359 | 0.701925 | 0.937441 | ENSMUSG( | Dars1 | 1424.55 | -0.22556 | 0.083018 | 0.77681 |
| ENSMUSG( | Dars2 | 481.7072 | 0.087618 | 0.438344 | 0.830406 | ENSMUSG( | Dars2 | 402.1261 | -0.17553 | 0.247438 | 0.877904 |
| ENSMUSG( | Daw1 | 16.21725 | 0.254738 | 0.643572 | NA | ENSMUSG( | Daw1 | 14.52879 | 0.240321 | 0.682209 | 0.958678 |
| ENSMUSG( | Daxx | 691.8541 | 0.080515 | 0.443347 | 0.833108 | ENSMUSG( | Daxx | 617.2695 | 0.015794 | 0.908248 | 0.991958 |
| ENSMUSG( | Dazap1 | 1128.282 | 0.045054 | 0.678447 | 0.930542 | ENSMUSG( | Dazap1 | 1020.072 | 0.021451 | 0.86312 | 0.986547 |
| ENSMUSG( | Dazap2 | 8775.751 | 0.055016 | 0.632403 | 0.914073 | ENSMUSG( | Dazap2 | 7538.654 | -0.11904 | 0.398936 | 0.917257 |
| ENSMUSG( | Dazl | 70.74187 | 0.065856 | 0.783733 | 0.957645 | ENSMUSG( | Dazl | 69.25073 | 0.241368 | 0.394666 | 0.917257 |
| ENSMUSG( | Dbf4 | 50.61983 | -0.1912 | 0.528 | 0.878105 | ENSMUSG( | Dbf4 | 53.20271 | 0.233333 | 0.42267 | 0.917257 |
| ENSMUSG( | Dbh | 19.26032 | -0.73407 | 0.122773 | NA | ENSMUSG( | Dbh | 18.31053 | -0.56475 | 0.260454 | 0.884524 |
| ENSMUSG( | Dbi | 11428.05 | 0.10949 | 0.303068 | 0.751443 | ENSMUSG( | Dbi | 9252.409 | -0.24162 | 0.085209 | 0.779799 |
| ENSMUSG( | Dbil5 | 22.27614 | 0.026711 | 0.948159 | 0.993821 | ENSMUSG( | Dbil5 | 16.91889 | -0.52255 | 0.28597 | 0.893906 |
| ENSMUSG( | Dbn1 | 11555.2 | 0.085195 | 0.391964 | 0.809274 | ENSMUSG( | Dbn1 | 10179.14 | -0.00851 | 0.941348 | 0.994578 |
| ENSMUSG( | Dbndd1 | 300.5618 | 0.169118 | 0.216166 | 0.661017 | ENSMUSG( | Dbndd1 | 270.8573 | 0.139355 | 0.386883 | 0.917257 |
| ENSMUSG( | Dbndd2 | 5859.654 | 0.020579 | 0.859132 | 0.974707 | ENSMUSG( | Dbndd2 | 5337.442 | 0.017747 | 0.896192 | 0.990558 |
| ENSMUSG( | Dbnl | 1983.123 | 0.114769 | 0.180149 | 0.617634 | ENSMUSG( | Dbnl | 1671.982 | -0.10988 | 0.308068 | 0.898538 |
| ENSMUSG( | Dbp | 3249.442 | 0.075519 | 0.572774 | 0.893584 | ENSMUSG( | Dbp | 4180.7 | 0.923962 | 3.27E-12 | 4.09E-09 |
| ENSMUSG( | Dbpht2 | 1195.451 | -0.0349 | 0.800472 | 0.961745 | ENSMUSG( | Dbpht2 | 1097.919 | -0.01774 | 0.922703 | 0.992644 |
| ENSMUSG( | Dbr1 | 236.1246 | 0.001362 | 0.993291 | 0.999369 | ENSMUSG( | Dbr1 | 215.7052 | 0.012315 | 0.944411 | 0.994798 |
| ENSMUSG( | Dbt | 1745.333 | 0.112362 | 0.328193 | 0.772854 | ENSMUSG( | Dbt | 1512.34 | -0.03498 | 0.814185 | 0.979198 |
| ENSMUSG( | Dbx2 | 1418.228 | 0.169928 | 0.131928 | 0.544244 | ENSMUSG( | Dbx2 | 1191.749 | -0.06356 | 0.636345 | 0.953903 |
| ENSMUSG( | Dcaf1 | 695.3945 | 0.100582 | 0.380577 | 0.801959 | ENSMUSG( | Dcaf1 | 621.1908 | 0.042108 | 0.770465 | 0.974236 |
| ENSMUSG( | Dcaf10 | 558.8489 | -0.12125 | 0.317855 | 0.764145 | ENSMUSG( | Dcaf10 | 542.1369 | 0.058884 | 0.677683 | 0.957106 |
| ENSMUSG( | Dcaf11 | 3166.377 | 0.150463 | 0.062591 | 0.38686 | ENSMUSG( | Dcaf11 | 2689.802 | -0.04701 | 0.660552 | 0.956205 |
| ENSMUSG( | Dcaf12 | 2098.779 | 0.17569 | 0.06294 | 0.388168 | ENSMUSG( | Dcaf12 | 1762.48 | -0.05812 | 0.63605 | 0.953903 |
| ENSMUSG( | Dcaf12l1 | 693.6699 | 0.038615 | 0.724582 | 0.944096 | ENSMUSG( | Dcaf12l1 | 634.3343 | 0.042828 | 0.77365 | 0.974672 |
| ENSMUSG( | Dcaf12l2 | 19.19083 | 0.129217 | 0.764915 | NA | ENSMUSG( | Dcaf12l2 | 15.68463 | -0.16818 | 0.748862 | 0.96984 |
| ENSMUSG( | Dcaf13 | 538.4669 | -0.06807 | 0.593275 | 0.902578 | ENSMUSG( | Dcaf13 | 495.3842 | -0.03327 | 0.825801 | 0.979653 |
| ENSMUSG( | Dcaf15 | 581.6058 | -0.01968 | 0.887625 | 0.983411 | ENSMUSG( | Dcaf15 | 557.0583 | 0.125539 | 0.425298 | 0.917257 |
| ENSMUSG( | Dcaf17 | 324.8933 | -0.02922 | 0.814454 | 0.964574 | ENSMUSG( | Dcaf17 | 299.6154 | 0.016525 | 0.92289 | 0.992644 |
| ENSMUSG( | Dcaf4 | 512.5309 | 0.04959 | 0.702203 | 0.937536 | ENSMUSG( | Dcaf4 | 442.4775 | -0.10369 | 0.465275 | 0.923329 |
| ENSMUSG( | Dcaf5 | 3146.047 | 0.09847 | 0.228551 | 0.675862 | ENSMUSG( | Dcaf5 | 2776.339 | 0.008772 | 0.932301 | 0.99306 |
| ENSMUSG( | Dcaf6 | 2794.32 | 0.21011 | 0.020067 | 0.214281 | ENSMUSG( | Dcaf6 | 2449.782 | 0.104133 | 0.346269 | 0.910966 |
| ENSMUSG( | Dcaf7 | 7187.57 | 0.049156 | 0.54212 | 0.883182 | ENSMUSG( | Dcaf7 | 6287.999 | -0.06938 | 0.496715 | 0.927825 |
| ENSMUSG( | Dcaf8 | 4778.19 | 0.144563 | 0.077124 | 0.429655 | ENSMUSG( | Dcaf8 | 4177.617 | 0.030207 | 0.760863 | 0.972695 |
| ENSMUSG( | Dcakd | 1154.272 | 0.185529 | 0.110376 | 0.50108 | ENSMUSG( | Dcakd | 962.4657 | -0.06944 | 0.655279 | 0.955294 |
| ENSMUSG( | Dcbld1 | 354.5504 | -0.34974 | 0.040356 | 0.310226 | ENSMUSG( | Dcbld1 | 391.5789 | 0.22691 | 0.244293 | 0.877904 |
| ENSMUSG( | Dcbld2 | 1193.686 | -0.13459 | 0.354184 | 0.786644 | ENSMUSG( | Dcbld2 | 1240.601 | 0.243241 | 0.136435 | 0.827092 |
| ENSMUSG( | Dcc | 664.5816 | -0.32278 | 0.097381 | 0.477629 | ENSMUSG( | Dcc | 715.4057 | 0.17621 | 0.348142 | 0.911652 |
| ENSMUSG( | Dcdc2a | 341.7663 | -0.17455 | 0.228666 | 0.675954 | ENSMUSG( | Dcdc2a | 293.0958 | -0.3676 | 0.020455 | 0.565891 |
| ENSMUSG( | Dcdc2b | 72.95788 | -1.52145 | 2.28E-09 | 1.00E-06 | ENSMUSG( | Dcdc2b | 113.8938 | 0.39614 | 0.229522 | 0.868507 |
| ENSMUSG( | Dchs1 | 779.7352 | -0.13899 | 0.388512 | 0.806278 | ENSMUSG( | Dchs1 | 802.7702 | 0.217518 | 0.237315 | 0.872173 |
| ENSMUSG( | Dchs2 | 43.58722 | -0.88989 | 0.00895 | 0.132538 | ENSMUSG( | Dchs2 | 50.9166 | -0.03158 | 0.918509 | 0.992529 |
| ENSMUSG( | Dck | 448.9174 | 0.151605 | 0.206143 | 0.647992 | ENSMUSG( | Dck | 391.9921 | 0.030007 | 0.838314 | 0.981601 |
| ENSMUSG( | Dclk1 | 21963.4 | 0.096061 | 0.192336 | 0.63139 | ENSMUSG( | Dclk1 | 19094.92 | -0.03803 | 0.711485 | 0.964363 |
| ENSMUSG( | Dclk2 | 3688.507 | 0.027055 | 0.727791 | 0.944355 | ENSMUSG( | Dclk2 | 3264.678 | -0.05702 | 0.561473 | 0.940279 |
| ENSMUSG( | Dclk3 | 685.3111 | 0.213063 | 0.217686 | 0.662824 | ENSMUSG( | Dclk3 | 552.2321 | -0.14067 | 0.469283 | 0.923329 |
| ENSMUSG( | Dclre1a | 280.1636 | -0.19875 | 0.154361 | 0.581452 | ENSMUSG( | Dclre1a | 262.1858 | -0.11423 | 0.457058 | 0.923329 |
| ENSMUSG( | Dclre1b | 247.0886 | 0.044844 | 0.778456 | 0.956215 | ENSMUSG( | Dclre1b | 229.4261 | 0.108412 | 0.548357 | 0.938063 |
| ENSMUSG( | Dclre1c | 267.0609 | 0.019645 | 0.9022 | 0.985633 | ENSMUSG( | Dclre1c | 260.6426 | 0.21957 | 0.244629 | 0.877904 |
| ENSMUSG( | Dcn | 801.5953 | -0.20863 | 0.139972 | 0.556732 | ENSMUSG( | Dcn | 794.0917 | 0.040208 | 0.787199 | 0.97637 |
| ENSMUSG( | Dcp1a | 513.0688 | -0.05124 | 0.654512 | 0.923227 | ENSMUSG( | Dcp1a | 498.4063 | 0.135866 | 0.334086 | 0.907907 |
| ENSMUSG( | Dcp1b | 556.9409 | 0.035582 | 0.761473 | 0.953106 | ENSMUSG( | Dcp1b | 514.0353 | 0.078721 | 0.605511 | 0.94924 |
| ENSMUSG( | Dcp2 | 1303.878 | 0.082489 | 0.38038 | 0.801757 | ENSMUSG( | Dcp2 | 1153.503 | -0.00144 | 0.989766 | 0.997928 |
| ENSMUSG( | Dcps | 308.2793 | 0.202775 | 0.141962 | 0.559166 | ENSMUSG( | Dcps | 255.3168 | -0.06725 | 0.680526 | 0.958395 |
| ENSMUSG( | Dcst2 | 42.99478 | 0.34135 | 0.307106 | 0.75508 | ENSMUSG( | Dcst2 | 36.93157 | 0.184347 | 0.572221 | 0.94251 |
| ENSMUSG( | Dct | 29.47066 | -0.54185 | 0.120279 | 0.524495 | ENSMUSG( | Dct | 31.63053 | 0.007981 | 0.98487 | 0.997067 |
| ENSMUSG( | Dctd | 411.0718 | -0.01034 | 0.937034 | 0.993415 | ENSMUSG( | Dctd | 361.1037 | -0.11631 | 0.411734 | 0.917257 |
| ENSMUSG( | Dctn1 | 10971.5 | 0.169561 | 0.051694 | 0.34893 | ENSMUSG( | Dctn1 | 9165.519 | -0.07714 | 0.455412 | 0.923316 |
| ENSMUSG( | Dctn2 | 5438.943 | 0.067321 | 0.500789 | 0.863671 | ENSMUSG( | Dctn2 | 4575.931 | -0.16846 | 0.17181 | 0.842999 |
| ENSMUSG( | Dctn3 | 1838.027 | 0.023283 | 0.804552 | 0.962377 | ENSMUSG( | Dctn3 | 1551.329 | -0.20585 | 0.077751 | 0.77321 |
| ENSMUSG( | Dctn4 | 2422.029 | 0.14286 | 0.102634 | 0.488148 | ENSMUSG( | Dctn4 | 2135.037 | 0.049298 | 0.647317 | 0.95422 |
| ENSMUSG( | Dctn5 | 1989.4 | -0.01743 | 0.847483 | 0.972374 | ENSMUSG( | Dctn5 | 1755.909 | -0.11424 | 0.368832 | 0.915491 |
| ENSMUSG( | Dctn6 | 1398.761 | -0.03604 | 0.722479 | 0.943752 | ENSMUSG( | Dctn6 | 1226.607 | -0.15181 | 0.206942 | 0.864412 |
| ENSMUSG( | Dctpp1 | 363.8932 | 0.022221 | 0.864508 | 0.975398 | ENSMUSG( | Dctpp1 | 310.8067 | -0.1776 | 0.275097 | 0.889874 |
| ENSMUSG( | Dcun1d1 | 741.2795 | 0.096157 | 0.436322 | 0.829093 | ENSMUSG( | Dcun1d1 | 654.9339 | -0.00062 | 0.997226 | 0.999268 |
| ENSMUSG( | Dcun1d2 | 1174.825 | 0.177725 | 0.094116 | 0.471597 | ENSMUSG( | Dcun1d2 | 1019.199 | 0.039404 | 0.751417 | 0.970317 |
| ENSMUSG( | Dcun1d3 | 368.856 | 0.158817 | 0.193349 | 0.632794 | ENSMUSG( | Dcun1d3 | 350.109 | 0.264974 | 0.075282 | 0.767994 |
| ENSMUSG( | Dcun1d4 | 1651.926 | 0.090723 | 0.318333 | 0.764234 | ENSMUSG( | Dcun1d4 | 1485.862 | 0.050789 | 0.681973 | 0.958678 |
| ENSMUSG( | Dcun1d5 | 839.3192 | -0.00719 | 0.946774 | 0.993731 | ENSMUSG( | Dcun1d5 | 790.401 | 0.086387 | 0.495853 | 0.927825 |
| ENSMUSG( | Dcx | 878.827 | -0.17365 | 0.154008 | 0.580901 | ENSMUSG( | Dcx | 865.6218 | 0.050332 | 0.726538 | 0.966887 |
| ENSMUSG( | Dcxr | 397.848 | 0.112016 | 0.38372 | 0.803608 | ENSMUSG( | Dcxr | 332.4123 | -0.13449 | 0.351863 | 0.912093 |
| ENSMUSG( | Dda1 | 2876.057 | 0.101188 | 0.250423 | 0.700322 | ENSMUSG( | Dda1 | 2557.804 | 0.033637 | 0.73739 | 0.967686 |
| ENSMUSG( | Ddah1 | 15420.81 | 0.009393 | 0.94339 | 0.993415 | ENSMUSG( | Ddah1 | 12394.65 | -0.37854 | 0.015084 | 0.492049 |
| ENSMUSG( | Ddah2 | 805.2604 | -0.07896 | 0.485702 | 0.857713 | ENSMUSG( | Ddah2 | 731.2457 | -0.09296 | 0.500769 | 0.927825 |
| ENSMUSG( | Ddb1 | 5947.666 | 0.142097 | 0.070234 | 0.40908 | ENSMUSG( | Ddb1 | 5089.632 | -0.03521 | 0.717204 | 0.965964 |
| ENSMUSG( | Ddb2 | 408.1175 | -0.05482 | 0.73955 | 0.947768 | ENSMUSG( | Ddb2 | 388.1959 | 0.072764 | 0.728102 | 0.966913 |
| ENSMUSG( | Ddc | 314.6539 | -0.07711 | 0.60702 | 0.906685 | ENSMUSG( | Ddc | 330.3303 | 0.327859 | 0.088304 | 0.786756 |
| ENSMUSG( | Ddhd1 | 4220.198 | 0.026242 | 0.764597 | 0.954055 | ENSMUSG( | Ddhd1 | 3733.863 | -0.05886 | 0.580808 | 0.944266 |
| ENSMUSG( | Ddhd2 | 1541.116 | -0.01993 | 0.852661 | 0.973168 | ENSMUSG( | Ddhd2 | 1406.412 | -0.01303 | 0.921898 | 0.992644 |
| ENSMUSG( | Ddi2 | 1202.678 | 0.066067 | 0.494039 | 0.860611 | ENSMUSG( | Ddi2 | 1092.94 | 0.061909 | 0.60346 | 0.948137 |
| ENSMUSG( | Ddias | 13.4967 | -0.2928 | 0.591431 | NA | ENSMUSG( | Ddias | 13.96225 | 0.123665 | 0.817316 | 0.979198 |
| ENSMUSG( | Ddit3 | 693.5239 | 0.08313 | 0.417066 | 0.8209 | ENSMUSG( | Ddit3 | 633.3428 | 0.082463 | 0.619023 | 0.952598 |
| ENSMUSG( | Ddit4 | 1153.734 | -0.29974 | 0.034443 | 0.285182 | ENSMUSG( | Ddit4 | 1060.159 | -0.27129 | 0.077684 | 0.77321 |
| ENSMUSG( | Ddit4l | 594.437 | 0.13008 | 0.299259 | 0.748131 | ENSMUSG( | Ddit4l | 519.723 | 0.013577 | 0.918057 | 0.992529 |
| ENSMUSG( | Ddn | 91130.09 | 0.268447 | 0.000804 | 0.027991 | ENSMUSG( | Ddn | 80707.9 | 0.193054 | 0.059723 | 0.729747 |
| ENSMUSG( | Ddo | 619.6368 | -0.04851 | 0.765752 | 0.954173 | ENSMUSG( | Ddo | 498.1849 | -0.45183 | 0.002569 | 0.198937 |
| ENSMUSG( | Ddost | 3714.608 | -0.3426 | 0.012435 | 0.161344 | ENSMUSG( | Ddost | 3865.604 | 0.063681 | 0.6841 | 0.958678 |
| ENSMUSG( | Ddr1 | 4211.611 | -0.51742 | 0.000233 | 0.011976 | ENSMUSG( | Ddr1 | 4758.413 | 0.150331 | 0.384339 | 0.917257 |
| ENSMUSG( | Ddr2 | 1348.726 | 0.08096 | 0.535537 | 0.881043 | ENSMUSG( | Ddr2 | 1142.971 | -0.13215 | 0.308627 | 0.898538 |
| ENSMUSG( | Ddrgk1 | 3332.383 | -0.11995 | 0.167043 | 0.599655 | ENSMUSG( | Ddrgk1 | 3114.925 | -0.04363 | 0.690852 | 0.960071 |
| ENSMUSG( | Ddt | 502.8166 | -0.07227 | 0.602607 | 0.904887 | ENSMUSG( | Ddt | 429.2844 | -0.27962 | 0.063981 | 0.736482 |
| ENSMUSG( | Ddx1 | 3269.795 | 0.082673 | 0.400909 | 0.813897 | ENSMUSG( | Ddx1 | 2775.625 | -0.12547 | 0.297303 | 0.898538 |
| ENSMUSG( | Ddx10 | 357.9487 | 0.218271 | 0.11692 | 0.515759 | ENSMUSG( | Ddx10 | 306.1552 | 0.044101 | 0.779655 | 0.976106 |
| ENSMUSG( | Ddx11 | 63.09245 | 0.027447 | 0.912131 | 0.988483 | ENSMUSG( | Ddx11 | 57.08032 | 0.021629 | 0.938848 | 0.993898 |

| | | | | | |
|---|---|---|---|---|---|
| ENSMUSG( | 5438.943 | 0.067321 | 0.500789 | 0.863671 | Dctn2 |
| ENSMUSG( | 1838.027 | 0.023283 | 0.804552 | 0.962377 | Dctn3 |
| ENSMUSG( | 2422.029 | 0.14286 | 0.102634 | 0.488148 | Dctn4 |
| ENSMUSG( | 1989.4 | -0.01743 | 0.847483 | 0.972374 | Dctn5 |
| ENSMUSG( | 1398.761 | -0.03604 | 0.722479 | 0.943752 | Dctn6 |
| ENSMUSG( | 363.8932 | 0.022221 | 0.864508 | 0.975398 | Dctpp1 |
| ENSMUSG( | 741.2795 | 0.096157 | 0.436322 | 0.829093 | Dcun1d1 |
| ENSMUSG( | 1174.825 | 0.177725 | 0.094116 | 0.471597 | Dcun1d2 |
| ENSMUSG( | 368.856 | 0.158817 | 0.193349 | 0.632794 | Dcun1d3 |
| ENSMUSG( | 1651.926 | 0.090723 | 0.318333 | 0.764234 | Dcun1d4 |
| ENSMUSG( | 839.3192 | -0.00719 | 0.946774 | 0.993731 | Dcun1d5 |
| ENSMUSG( | 878.827 | -0.17365 | 0.154008 | 0.580901 | Dcx |
| ENSMUSG( | 397.848 | 0.112016 | 0.38372 | 0.803608 | Dcxr |
| ENSMUSG( | 2876.057 | 0.101188 | 0.250423 | 0.700322 | Dda1 |
| ENSMUSG( | 15420.81 | 0.009393 | 0.94339 | 0.993415 | Ddah1 |
| ENSMUSG( | 805.2604 | -0.07896 | 0.485702 | 0.857713 | Ddah2 |
| ENSMUSG( | 5947.666 | 0.142097 | 0.070234 | 0.40908 | Ddb1 |
| ENSMUSG( | 408.1175 | -0.05482 | 0.73955 | 0.947768 | Ddb2 |
| ENSMUSG( | 314.6539 | -0.07711 | 0.60702 | 0.906685 | Ddc |
| ENSMUSG( | 4220.198 | 0.026242 | 0.764597 | 0.954055 | Ddhd1 |
| ENSMUSG( | 1541.116 | -0.01993 | 0.852661 | 0.973168 | Ddhd2 |
| ENSMUSG( | 1202.678 | 0.066067 | 0.494039 | 0.860611 | Ddi2 |
| ENSMUSG( | 13.4967 | -0.2928 | 0.591431 | NA | Ddias |
| ENSMUSG( | 693.5239 | 0.08313 | 0.417066 | 0.8209 | Ddit3 |
| ENSMUSG( | 1153.734 | -0.29974 | 0.034443 | 0.285182 | Ddit4 |
| ENSMUSG( | 594.437 | 0.13008 | 0.299259 | 0.748131 | Ddit4l |
| ENSMUSG( | 91130.09 | 0.268447 | 0.000804 | 0.027991 | Ddn |
| ENSMUSG( | 619.6368 | -0.04851 | 0.765752 | 0.954173 | Ddo |
| ENSMUSG( | 3714.608 | -0.3426 | 0.012435 | 0.161344 | Ddost |
| ENSMUSG( | 4211.611 | -0.51742 | 0.000233 | 0.011976 | Ddr1 |
| ENSMUSG( | 1348.726 | 0.08096 | 0.535537 | 0.881043 | Ddr2 |
| ENSMUSG( | 3332.383 | -0.11995 | 0.167043 | 0.599655 | Ddrgk1 |
| ENSMUSG( | 502.8166 | -0.07227 | 0.602607 | 0.904887 | Ddt |
| ENSMUSG( | 3269.795 | 0.082673 | 0.400909 | 0.813897 | Ddx1 |
| ENSMUSG( | 357.9487 | 0.218271 | 0.11692 | 0.515759 | Ddx10 |
| ENSMUSG( | 63.09245 | 0.027447 | 0.912131 | 0.988483 | Ddx11 |
| ENSMUSG( | 9687.066 | 0.016755 | 0.858191 | 0.974672 | Ddx17 |
| ENSMUSG( | 389.0526 | 0.099303 | 0.404593 | 0.816132 | Ddx18 |
| ENSMUSG( | 650.3058 | 0.060165 | 0.564532 | 0.890434 | Ddx19a |
| ENSMUSG( | 814.5113 | 0.056324 | 0.557552 | 0.888279 | Ddx19b |
| ENSMUSG( | 328.1249 | 0.078828 | 0.541937 | 0.883182 | Ddx20 |
| ENSMUSG( | 1117.051 | 0.150742 | 0.163237 | 0.594466 | Ddx21 |
| ENSMUSG( | 2171.974 | 0.112498 | 0.219122 | 0.664602 | Ddx23 |
| ENSMUSG( | 8226.755 | -0.11812 | 0.426967 | 0.82532 | Ddx24 |
| ENSMUSG( | 999.3436 | 0.180487 | 0.070287 | 0.40908 | Ddx25 |
| ENSMUSG( | 787.2984 | 0.283946 | 0.03682 | 0.296095 | Ddx27 |
| ENSMUSG( | 236.5735 | 0.000696 | 0.996388 | 0.99943 | Ddx28 |
| ENSMUSG( | 180.4378 | -0.13091 | 0.413136 | 0.819541 | Ddx31 |
| ENSMUSG( | 469.6981 | 0.059947 | 0.642356 | 0.916865 | Ddx39a |
| ENSMUSG( | 4187.069 | 0.009396 | 0.921232 | 0.990436 | Ddx39b |
| ENSMUSG( | 3990.09 | -0.23237 | 0.016444 | 0.190546 | Ddx3x |
| ENSMUSG( | 1097.584 | -0.0697 | 0.57094 | 0.8933 | Ddx3y |
| ENSMUSG( | 63.03078 | 0.376143 | 0.146511 | 0.568435 | Ddx4 |
| ENSMUSG( | 1263.005 | 0.138201 | 0.15113 | 0.576105 | Ddx41 |
| ENSMUSG( | 2464.824 | 0.040833 | 0.650923 | 0.92145 | Ddx42 |
| ENSMUSG( | 10.79642 | -0.58629 | 0.33749 | NA | Ddx43 |
| ENSMUSG( | 2363.57 | 0.081158 | 0.333911 | 0.775602 | Ddx46 |
| ENSMUSG( | 995.6988 | 0.027045 | 0.800018 | 0.961634 | Ddx47 |
| ENSMUSG( | 879.6021 | 0.058273 | 0.64587 | 0.918224 | Ddx49 |
| ENSMUSG( | 13806.87 | -0.06164 | 0.435026 | 0.829093 | Ddx5 |
| ENSMUSG( | 1844.279 | 0.06612 | 0.499622 | 0.863218 | Ddx50 |
| ENSMUSG( | 466.2784 | 0.102541 | 0.379451 | 0.801757 | Ddx51 |
| ENSMUSG( | 664.1139 | 0.024032 | 0.81472 | 0.964668 | Ddx52 |
| ENSMUSG( | 1568.229 | 0.14217 | 0.148456 | 0.571935 | Ddx54 |
| ENSMUSG( | 714.2885 | 0.087188 | 0.474298 | 0.849425 | Ddx55 |
| ENSMUSG( | 841.193 | 0.038676 | 0.713591 | 0.940866 | Ddx56 |
| ENSMUSG( | 224.4159 | -0.1393 | 0.352925 | 0.784977 | Ddx59 |
| ENSMUSG( | 3719.629 | 0.083937 | 0.288082 | 0.738882 | Ddx6 |
| ENSMUSG( | 35.86065 | -0.91878 | 0.00533 | 0.097362 | Ddx60 |
| ENSMUSG( | 3541.668 | 0.191192 | 0.032021 | 0.273359 | Deaf1 |
| ENSMUSG( | 1503.801 | 0.051185 | 0.709086 | 0.939578 | Decr1 |
| ENSMUSG( | 810.4175 | 0.087334 | 0.430741 | 0.826896 | Decr2 |
| ENSMUSG( | 1089.902 | 0.032369 | 0.750532 | 0.949877 | Dedd |
| ENSMUSG( | 587.3586 | 0.011381 | 0.923072 | 0.990693 | Dedd2 |
| ENSMUSG( | 118.7148 | -0.16308 | 0.413916 | 0.819998 | Def6 |
| ENSMUSG( | 2040.683 | 0.111076 | 0.191474 | 0.630474 | Def8 |
| ENSMUSG( | 11.56735 | -0.28809 | 0.716679 | NA | Defb11 |
| ENSMUSG( | 8.777133 | 0.277996 | 0.656495 | NA | Defb42 |
| ENSMUSG( | 3561.456 | -0.35303 | 0.011464 | 0.154051 | Degs1 |
| ENSMUSG( | 82.10893 | 0.387658 | 0.223723 | 0.669845 | Degs2 |
| ENSMUSG( | 1312.464 | 0.029505 | 0.80824 | 0.96345 | Dek |
| ENSMUSG( | 1916.413 | -0.03199 | 0.763051 | 0.953667 | Dele1 |
| ENSMUSG( | 835.1879 | -0.10388 | 0.380287 | 0.801757 | Dennd10 |
| ENSMUSG( | 5479.343 | -0.02044 | 0.797686 | 0.960507 | Dennd11 |
| ENSMUSG( | 2360.282 | 0.100583 | 0.257485 | 0.708686 | Dennd1a |
| ENSMUSG( | 389.0317 | 0.158596 | 0.197623 | 0.637377 | Dennd1b |
| ENSMUSG( | 101.6101 | -0.08498 | 0.698785 | 0.936563 | Dennd1c |
| ENSMUSG( | 1528.398 | 0.109693 | 0.214142 | 0.658712 | Dennd2a |
| ENSMUSG( | 885.9262 | 0.009099 | 0.930242 | 0.991848 | Dennd2b |
| ENSMUSG( | 33.5806 | 0.112878 | 0.76442 | 0.954055 | Dennd2c |
| ENSMUSG( | 266.644 | 0.17663 | 0.280343 | 0.729291 | Dennd3 |
| ENSMUSG( | 2214.742 | 0.00437 | 0.962281 | 0.994543 | Dennd4a |
| ENSMUSG( | 2111.087 | 0.18521 | 0.058069 | 0.371677 | Dennd4b |
| ENSMUSG( | 743.5284 | 0.092978 | 0.547111 | 0.885118 | Dennd4c |
| ENSMUSG( | 24409.66 | 0.074516 | 0.399356 | 0.813048 | Dennd5a |
| ENSMUSG( | 1302.172 | 0.069351 | 0.439265 | 0.830889 | Dennd5b |
| ENSMUSG( | 877.9452 | 0.124507 | 0.229351 | 0.67735 | Dennd6a |
| ENSMUSG( | 2067.769 | 0.040985 | 0.769402 | 0.954674 | Dennd6b |
| ENSMUSG( | 1419.89 | -0.01966 | 0.846731 | 0.972371 | Denr |
| ENSMUSG( | 16.76262 | -0.62522 | 0.204077 | NA | Depdc1b |
| ENSMUSG( | 1573.157 | 0.142504 | 0.149701 | 0.57376 | Depdc5 |
| ENSMUSG( | 24.94513 | -0.55467 | 0.137953 | 0.552903 | Depdc7 |
| ENSMUSG( | 46.54795 | -0.77993 | 0.032686 | 0.27685 | Depp1 |
| ENSMUSG( | 595.2457 | 0.195683 | 0.100487 | 0.483976 | Deptor |
| ENSMUSG( | 168.3203 | 0.031128 | 0.87293 | 0.978753 | Dera |
| ENSMUSG( | 2001.579 | -0.29748 | 0.045065 | 0.325595 | Derl1 |
| ENSMUSG( | 846.2884 | -0.16884 | 0.205295 | 0.64654 | Derl2 |
| ENSMUSG( | 46.25702 | 0.164896 | 0.570461 | 0.893192 | Derl3 |
| ENSMUSG( | 45.52577 | -0.25485 | 0.392574 | 0.809274 | Des |
| ENSMUSG( | 10401.53 | -0.07525 | 0.329131 | 0.773233 | Desi1 |
| ENSMUSG( | 1104.477 | 0.077494 | 0.497318 | 0.861898 | Desi2 |
| ENSMUSG( | 129.5497 | 0.048514 | 0.804032 | 0.962169 | Det1 |
| ENSMUSG( | 1170.788 | 0.04884 | 0.633602 | 0.914073 | Dexi |
| ENSMUSG( | 894.5052 | 0.033912 | 0.783879 | 0.957645 | Dffa |
| ENSMUSG( | 263.0757 | 0.00037 | 0.99822 | 0.99989 | Dffb |
| ENSMUSG( | 714.4623 | -0.13531 | 0.31967 | 0.765305 | Dgat1 |
| ENSMUSG( | 2402.711 | -0.44172 | 0.000422 | 0.018541 | Dgat2 |
| ENSMUSG( | 22.27791 | 0.083096 | 0.845593 | 0.97218 | Dgat2l6 |
| ENSMUSG( | 2961.172 | -0.21645 | 0.072761 | 0.416435 | Dgcr2 |
| ENSMUSG( | 2535.902 | 0.003077 | 0.972713 | 0.996585 | Dgcr6 |
| ENSMUSG( | 910.4627 | 0.235791 | 0.039955 | 0.308163 | Dgcr8 |
| ENSMUSG( | 435.1279 | 0.1655 | 0.201375 | 0.641362 | Dgka |
| ENSMUSG( | 2986.132 | -0.06516 | 0.553003 | 0.886204 | Dgkb |
| ENSMUSG( | 3175.682 | 0.193715 | 0.046162 | 0.329267 | Dgkd |
| ENSMUSG( | 2312.818 | -0.4218 | 0.003009 | 0.067314 | Dgke |
| ENSMUSG( | 484.0764 | 0.45104 | 0.001484 | 0.042763 | Dgkg |
| ENSMUSG( | 1833.987 | -0.02142 | 0.830602 | 0.96974 | Dgkh |
| ENSMUSG( | 387.1107 | -0.10375 | 0.443064 | 0.833025 | Dgki |
| ENSMUSG( | 74.98626 | 0.258262 | 0.289748 | 0.739839 | Dgkk |
| ENSMUSG( | 1387.143 | 0.061845 | 0.576064 | 0.895114 | Dgkq |
| ENSMUSG( | 28437.44 | 0.270733 | 0.002886 | 0.065928 | Dgkz |
| ENSMUSG( | 362.3006 | -0.04387 | 0.741414 | 0.948208 | Dglucy |
| ENSMUSG( | 306.5675 | 0.000261 | 0.998605 | 0.99989 | Dguok |
| ENSMUSG( | 5678.419 | -0.39185 | 0.011919 | 0.157992 | Dhcr24 |
| ENSMUSG( | 2130.046 | -0.50538 | 0.003831 | 0.079288 | Dhcr7 |
| ENSMUSG( | 1732.671 | 0.104515 | 0.312728 | 0.760258 | Dhdds |
| ENSMUSG( | 581.0983 | 0.13591 | 0.226565 | 0.673549 | Dhdh |
| ENSMUSG( | 254.4476 | 0.122955 | 0.432599 | 0.827838 | Dhfr |
| ENSMUSG( | 425.9093 | 0.099795 | 0.436887 | 0.829501 | Dhodh |
| ENSMUSG( | 924.7106 | 0.011921 | 0.912879 | 0.988483 | Dhps |
| ENSMUSG( | 2655.093 | 0.02371 | 0.81322 | 0.964574 | Dhrs1 |
| ENSMUSG( | 291.1207 | 0.076766 | 0.570071 | 0.893041 | Dhrs11 |
| ENSMUSG( | 251.3254 | -0.25465 | 0.227205 | 0.674223 | Dhrs13 |
| ENSMUSG( | 12.61022 | -0.61718 | 0.234856 | NA | Dhrs13os |
| ENSMUSG( | 942.0321 | -0.37036 | 0.007045 | 0.115775 | Dhrs3 |
| ENSMUSG( | 846.1725 | 0.121089 | 0.333005 | 0.775602 | Dhrs4 |
| ENSMUSG( | 1238.061 | -0.34161 | 0.02297 | 0.232278 | Dhrs7 |
| ENSMUSG( | 870.4801 | -0.19259 | 0.122449 | 0.528223 | Dhrs7b |
| ENSMUSG( | 310.1054 | 0.056122 | 0.677751 | 0.930542 | Dhtkd1 |
| ENSMUSG( | 1918.598 | 0.005627 | 0.961029 | 0.993999 | Dhx15 |
| ENSMUSG( | 892.1137 | 0.166766 | 0.130863 | 0.54231 | Dhx16 |
| ENSMUSG( | 712.2802 | 0.286574 | 0.016385 | 0.190235 | Dhx29 |
| ENSMUSG( | 2836.661 | 0.148067 | 0.128186 | 0.536641 | Dhx30 |
| ENSMUSG( | 796.6592 | -0.03137 | 0.770319 | 0.954689 | Dhx32 |
| ENSMUSG( | 994.2007 | 0.164998 | 0.186382 | 0.623869 | Dhx33 |
| ENSMUSG( | 476.4318 | 0.041952 | 0.725185 | 0.944096 | Dhx34 |
| ENSMUSG( | 693.9763 | -0.01756 | 0.865774 | 0.975876 | Dhx35 |
| ENSMUSG( | 1745.476 | 0.098438 | 0.291853 | 0.741868 | Dhx36 |

| | | | | | |
|---|---|---|---|---|---|
| ENSMUSG( | 865.6218 | 0.050332 | 0.726538 | 0.966887 | Dcx |
| ENSMUSG( | 332.4123 | -0.13449 | 0.351863 | 0.912093 | Dcxr |
| ENSMUSG( | 2557.804 | 0.033637 | 0.73739 | 0.967686 | Dda1 |
| ENSMUSG( | 12394.65 | -0.37854 | 0.015084 | 0.492049 | Ddah1 |
| ENSMUSG( | 731.2457 | -0.09296 | 0.500769 | 0.927825 | Ddah2 |
| ENSMUSG( | 5089.632 | -0.03521 | 0.717204 | 0.965964 | Ddb1 |
| ENSMUSG( | 388.1959 | 0.072764 | 0.728102 | 0.966913 | Ddb2 |
| ENSMUSG( | 330.3303 | 0.327859 | 0.088304 | 0.786756 | Ddc |
| ENSMUSG( | 3733.863 | -0.05886 | 0.580808 | 0.944266 | Ddhd1 |
| ENSMUSG( | 1406.412 | -0.01303 | 0.921898 | 0.992644 | Ddhd2 |
| ENSMUSG( | 1092.94 | 0.061909 | 0.60346 | 0.948137 | Ddi2 |
| ENSMUSG( | 13.96225 | 0.123665 | 0.817316 | 0.979198 | Ddias |
| ENSMUSG( | 633.3428 | 0.082463 | 0.619023 | 0.952598 | Ddit3 |
| ENSMUSG( | 1060.159 | -0.27129 | 0.077684 | 0.77321 | Ddit4 |
| ENSMUSG( | 519.723 | 0.013577 | 0.918057 | 0.992529 | Ddit4l |
| ENSMUSG( | 80707.9 | 0.193054 | 0.059723 | 0.729747 | Ddn |
| ENSMUSG( | 498.1849 | -0.45183 | 0.002569 | 0.198937 | Ddo |
| ENSMUSG( | 3865.604 | 0.063681 | 0.6841 | 0.958678 | Ddost |
| ENSMUSG( | 4758.413 | 0.150331 | 0.384339 | 0.917257 | Ddr1 |
| ENSMUSG( | 1142.971 | -0.13215 | 0.308627 | 0.898538 | Ddr2 |
| ENSMUSG( | 3114.925 | -0.04363 | 0.690852 | 0.960071 | Ddrgk1 |
| ENSMUSG( | 429.2844 | -0.27962 | 0.063981 | 0.736482 | Ddt |
| ENSMUSG( | 2775.625 | -0.12547 | 0.297303 | 0.898538 | Ddx1 |
| ENSMUSG( | 306.1552 | 0.044101 | 0.779655 | 0.976106 | Ddx10 |
| ENSMUSG( | 57.08032 | 0.021629 | 0.938848 | 0.993898 | Ddx11 |
| ENSMUSG( | 9217.98 | 0.138447 | 0.372408 | 0.915491 | Ddx17 |
| ENSMUSG( | 346.3697 | 0.040009 | 0.79275 | 0.97681 | Ddx18 |
| ENSMUSG( | 573.846 | -0.03467 | 0.784102 | 0.976154 | Ddx19a |
| ENSMUSG( | 724.2392 | -0.01182 | 0.923771 | 0.99282 | Ddx19b |
| ENSMUSG( | 278.7667 | -0.12205 | 0.432992 | 0.918434 | Ddx20 |
| ENSMUSG( | 978.9397 | 0.038476 | 0.76356 | 0.974 | Ddx21 |
| ENSMUSG( | 1944.188 | 0.063862 | 0.544461 | 0.938063 | Ddx23 |
| ENSMUSG( | 7672.611 | -0.04506 | 0.758978 | 0.971858 | Ddx24 |
| ENSMUSG( | 840.8904 | -0.04388 | 0.707481 | 0.964029 | Ddx25 |
| ENSMUSG( | 667.5344 | 0.090222 | 0.567194 | 0.941777 | Ddx27 |
| ENSMUSG( | 213.6912 | -0.02941 | 0.868012 | 0.98781 | Ddx28 |
| ENSMUSG( | 161.1057 | -0.19085 | 0.28832 | 0.894794 | Ddx31 |
| ENSMUSG( | 413.113 | -0.03408 | 0.829742 | 0.980224 | Ddx39a |
| ENSMUSG( | 3798.569 | -0.00269 | 0.98273 | 0.996859 | Ddx39b |
| ENSMUSG( | 4010.628 | 0.058912 | 0.606901 | 0.949797 | Ddx3x |
| ENSMUSG( | 1095.99 | 0.19159 | 0.168357 | 0.840222 | Ddx3y |
| ENSMUSG( | 52.24853 | 0.122897 | 0.668851 | 0.956766 | Ddx4 |
| ENSMUSG( | 1068.249 | -0.07262 | 0.530914 | 0.934795 | Ddx41 |
| ENSMUSG( | 2146.739 | -0.09103 | 0.397781 | 0.917257 | Ddx42 |
| ENSMUSG( | 11.70267 | 0.035801 | 0.954148 | 0.994798 | Ddx43 |
| ENSMUSG( | 2070.112 | -0.03324 | 0.76521 | 0.974236 | Ddx46 |
| ENSMUSG( | 900.3406 | 0.006076 | 0.962398 | 0.995237 | Ddx47 |
| ENSMUSG( | 738.3736 | -0.18446 | 0.180424 | 0.850372 | Ddx49 |
| ENSMUSG( | 13626.34 | 0.16529 | 0.137707 | 0.828492 | Ddx5 |
| ENSMUSG( | 1751.601 | 0.178786 | 0.132106 | 0.822912 | Ddx50 |
| ENSMUSG( | 402.5629 | -0.04422 | 0.776104 | 0.975717 | Ddx51 |
| ENSMUSG( | 599.6793 | 0.000355 | 0.997784 | 0.999268 | Ddx52 |
| ENSMUSG( | 1333.291 | -0.05534 | 0.631705 | 0.953251 | Ddx54 |
| ENSMUSG( | 624.0459 | -0.033 | 0.820713 | 0.979198 | Ddx55 |
| ENSMUSG( | 730.4311 | -0.09719 | 0.439525 | 0.920721 | Ddx56 |
| ENSMUSG( | 199.125 | -0.22999 | 0.186888 | 0.856099 | Ddx59 |
| ENSMUSG( | 3302.504 | 0.008983 | 0.934433 | 0.99306 | Ddx6 |
| ENSMUSG( | 37.36 | -0.37586 | 0.320985 | 0.904019 | Ddx60 |
| ENSMUSG( | 3153.598 | 0.126598 | 0.264634 | 0.886134 | Deaf1 |
| ENSMUSG( | 1242.843 | -0.24394 | 0.102624 | 0.803503 | Decr1 |
| ENSMUSG( | 688.3255 | -0.11468 | 0.365659 | 0.914623 | Decr2 |
| ENSMUSG( | 956.1169 | -0.0811 | 0.525998 | 0.934374 | Dedd |
| ENSMUSG( | 562.3473 | 0.157018 | 0.278198 | 0.891468 | Dedd2 |
| ENSMUSG( | 116.252 | 0.071652 | 0.754849 | 0.971098 | Def6 |
| ENSMUSG( | 1715.554 | -0.12284 | 0.244949 | 0.877904 | Def8 |
| ENSMUSG( | 17.91532 | 1.032064 | 0.10224 | 0.80239 | Defb11 |
| ENSMUSG( | 3803.993 | 0.127706 | 0.422176 | 0.917257 | Degs1 |
| ENSMUSG( | 107.9375 | 1.241005 | 0.000435 | 0.061321 | Degs2 |
| ENSMUSG( | 1223.618 | 0.088938 | 0.559572 | 0.939807 | Dek |
| ENSMUSG( | 1767.557 | 0.007192 | 0.95316 | 0.994798 | Dele1 |
| ENSMUSG( | 754.774 | -0.1337 | 0.355033 | 0.912411 | Dennd10 |
| ENSMUSG( | 4847.185 | -0.10864 | 0.304491 | 0.898538 | Dennd11 |
| ENSMUSG( | 2126.549 | 0.070848 | 0.51906 | 0.931053 | Dennd1a |
| ENSMUSG( | 352.5725 | 0.142093 | 0.309579 | 0.898538 | Dennd1b |
| ENSMUSG( | 100.1452 | 0.151615 | 0.576384 | 0.942912 | Dennd1c |
| ENSMUSG( | 1316.978 | -0.05055 | 0.645878 | 0.953988 | Dennd2a |
| ENSMUSG( | 804.1907 | 0.000111 | 0.999288 | 0.999503 | Dennd2b |
| ENSMUSG( | 35.16467 | 0.508375 | 0.21014 | 0.864412 | Dennd2c |
| ENSMUSG( | 8.788251 | 0.203593 | 0.755643 | NA | Dennd2d |
| ENSMUSG( | 233.5734 | 0.084152 | 0.662434 | 0.956205 | Dennd3 |
| ENSMUSG( | 1996.842 | -0.02608 | 0.815968 | 0.979198 | Dennd4a |
| ENSMUSG( | 1875.936 | 0.11958 | 0.374274 | 0.915491 | Dennd4b |
| ENSMUSG( | 649.0787 | -0.02506 | 0.870624 | 0.988445 | Dennd4c |
| ENSMUSG( | 20507.46 | -0.16338 | 0.104014 | 0.804969 | Dennd5a |
| ENSMUSG( | 1188.453 | 0.071055 | 0.552926 | 0.939565 | Dennd5b |
| ENSMUSG( | 812.55 | 0.167689 | 0.170016 | 0.841056 | Dennd6a |
| ENSMUSG( | 1943.347 | 0.126781 | 0.459337 | 0.923329 | Dennd6b |
| ENSMUSG( | 1217.171 | -0.20664 | 0.099125 | 0.799802 | Denr |
| ENSMUSG( | 14.53413 | -0.7838 | 0.149515 | 0.836241 | Depdc1b |
| ENSMUSG( | 1398.449 | 0.076887 | 0.500721 | 0.927825 | Depdc5 |
| ENSMUSG( | 32.75133 | 0.487393 | 0.171655 | 0.842999 | Depdc7 |
| ENSMUSG( | 53.88309 | 0.049016 | 0.911473 | 0.992196 | Depp1 |
| ENSMUSG( | 528.0897 | 0.119719 | 0.394814 | 0.917257 | Deptor |
| ENSMUSG( | 144.0473 | -0.15061 | 0.420757 | 0.917257 | Dera |
| ENSMUSG( | 2155.633 | 0.19807 | 0.249832 | 0.877904 | Derl1 |
| ENSMUSG( | 917.8769 | 0.323752 | 0.028222 | 0.611882 | Derl2 |
| ENSMUSG( | 52.4149 | 0.698483 | 0.018591 | 0.545482 | Derl3 |
| ENSMUSG( | 47.67357 | 0.183207 | 0.579069 | 0.943849 | Des |
| ENSMUSG( | 8936.153 | -0.25615 | 0.019656 | 0.556192 | Desi1 |
| ENSMUSG( | 969.3778 | -0.03482 | 0.809483 | 0.978779 | Desi2 |
| ENSMUSG( | 115.0785 | -0.01497 | 0.944841 | 0.994798 | Det1 |
| ENSMUSG( | 1057.093 | 0.020788 | 0.86403 | 0.986552 | Dexi |
| ENSMUSG( | 795.1513 | -0.03627 | 0.776917 | 0.975717 | Dffa |
| ENSMUSG( | 234.3608 | -0.06616 | 0.719545 | 0.966876 | Dffb |
| ENSMUSG( | 674.9719 | -0.02004 | 0.895369 | 0.990558 | Dgat1 |
| ENSMUSG( | 2766.522 | 0.260275 | 0.095503 | 0.796133 | Dgat2 |
| ENSMUSG( | 16.68858 | -0.50903 | 0.335574 | 0.907907 | Dgat2l6 |
| ENSMUSG( | 3082.879 | 0.175202 | 0.219608 | 0.866104 | Dgcr2 |
| ENSMUSG( | 2111.986 | -0.27045 | 0.012934 | 0.458131 | Dgcr6 |
| ENSMUSG( | 794.7441 | 0.117007 | 0.415543 | 0.917257 | Dgcr8 |
| ENSMUSG( | 375.642 | 0.018949 | 0.917422 | 0.992529 | Dgka |
| ENSMUSG( | 2589.974 | -0.21828 | 0.127736 | 0.81845 | Dgkb |
| ENSMUSG( | 2786.027 | 0.09303 | 0.469181 | 0.923329 | Dgkd |
| ENSMUSG( | 2552.208 | 0.160301 | 0.296829 | 0.898538 | Dgke |
| ENSMUSG( | 434.4871 | 0.423005 | 0.01541 | 0.495791 | Dgkg |
| ENSMUSG( | 1790.179 | 0.175498 | 0.15468 | 0.837724 | Dgkh |
| ENSMUSG( | 339.1026 | -0.21792 | 0.144688 | 0.833302 | Dgki |
| ENSMUSG( | 64.28231 | 0.097257 | 0.721914 | 0.966887 | Dgkk |
| ENSMUSG( | 1302.949 | 0.149716 | 0.292699 | 0.898093 | Dgkq |
| ENSMUSG( | 23987.66 | 0.060835 | 0.552114 | 0.939565 | Dgkz |
| ENSMUSG( | 298.5131 | -0.35223 | 0.020002 | 0.5597 | Dglucy |
| ENSMUSG( | 281.5135 | 0.026296 | 0.876629 | 0.989644 | Dguok |
| ENSMUSG( | 6030.293 | 0.080535 | 0.640849 | 0.953903 | Dhcr24 |
| ENSMUSG( | 2245.854 | -0.03502 | 0.852042 | 0.985598 | Dhcr7 |
| ENSMUSG( | 1539.422 | 0.032985 | 0.7755 | 0.975354 | Dhdds |
| ENSMUSG( | 512.3549 | 0.036477 | 0.812986 | 0.979198 | Dhdh |
| ENSMUSG( | 218.7904 | -0.05064 | 0.763289 | 0.973945 | Dhfr |
| ENSMUSG( | 400.5893 | 0.188426 | 0.180005 | 0.850372 | Dhodh |
| ENSMUSG( | 843.5797 | 0.015471 | 0.900405 | 0.990558 | Dhps |
| ENSMUSG( | 2231.766 | -0.22097 | 0.096309 | 0.796133 | Dhrs1 |
| ENSMUSG( | 249.5385 | -0.09706 | 0.533239 | 0.935104 | Dhrs11 |
| ENSMUSG( | 256.9243 | 0.095633 | 0.661215 | 0.956205 | Dhrs13 |
| ENSMUSG( | 12.24297 | -0.40348 | 0.466214 | 0.923329 | Dhrs13os |
| ENSMUSG( | 900.8645 | -0.21218 | 0.139836 | 0.830678 | Dhrs3 |
| ENSMUSG( | 685.2442 | -0.23318 | 0.119649 | 0.813384 | Dhrs4 |
| ENSMUSG( | 1276.705 | 0.040917 | 0.796139 | 0.97681 | Dhrs7 |
| ENSMUSG( | 841.9605 | -0.01045 | 0.939116 | 0.993898 | Dhrs7b |
| ENSMUSG( | 296.0394 | 0.185016 | 0.214464 | 0.864421 | Dhtkd1 |
| ENSMUSG( | 1832.662 | 0.140362 | 0.272634 | 0.888825 | Dhx15 |
| ENSMUSG( | 778.3887 | 0.043793 | 0.751145 | 0.970317 | Dhx16 |
| ENSMUSG( | 592.9843 | 0.039615 | 0.75146 | 0.970317 | Dhx29 |
| ENSMUSG( | 2476.13 | 0.028777 | 0.800249 | 0.97681 | Dhx30 |
| ENSMUSG( | 720.3608 | -0.05586 | 0.662109 | 0.956205 | Dhx32 |
| ENSMUSG( | 898.0365 | 0.146223 | 0.277957 | 0.891468 | Dhx33 |
| ENSMUSG( | 420.551 | -0.04527 | 0.744988 | 0.96951 | Dhx34 |
| ENSMUSG( | 626.1358 | -0.04369 | 0.725415 | 0.966887 | Dhx35 |
| ENSMUSG( | 1593.852 | 0.101711 | 0.38745 | 0.917257 | Dhx36 |
| ENSMUSG( | 405.4264 | -0.11158 | 0.512015 | 0.930005 | Dhx37 |
| ENSMUSG( | 999.3525 | 0.058751 | 0.608725 | 0.950113 | Dhx38 |
| ENSMUSG( | 777.4749 | -0.03103 | 0.797938 | 0.97681 | Dhx40 |
| ENSMUSG( | 1057.746 | 0.061694 | 0.623556 | 0.953234 | Dhx57 |
| ENSMUSG( | 68.4834 | -0.0436 | 0.866137 | 0.987254 | Dhx58 |
| ENSMUSG( | 822.6814 | 0.01772 | 0.886432 | 0.990011 | Dhx8 |
| ENSMUSG( | 3057.005 | -0.0926 | 0.450578 | 0.922202 | Dhx9 |
| ENSMUSG( | 237.0092 | 0.17485 | 0.332745 | 0.907907 | Diablo |
| ENSMUSG( | 3894.858 | 0.034649 | 0.754261 | 0.970785 | Diaph1 |
| ENSMUSG( | 584.7466 | 0.075124 | 0.605198 | 0.949082 | Diaph2 |
| ENSMUSG( | 103.8157 | -0.30086 | 0.189957 | 0.857355 | Diaph3 |

| | | | | | | | | | | | |
|---|---|---|---|---|---|---|---|---|---|---|---|
| ENSMUSG( | Ddx17 | 9687.066 | 0.016755 | 0.858191 | 0.974672 | ENSMUSG( | Ddx17 | 9217.98 | 0.138447 | 0.372408 | 0.915491 |
| ENSMUSG( | Ddx18 | 389.0526 | 0.099303 | 0.404593 | 0.816132 | ENSMUSG( | Ddx18 | 346.3697 | 0.040009 | 0.79275 | 0.97681 |
| ENSMUSG( | Ddx19a | 650.3058 | 0.060165 | 0.564532 | 0.890434 | ENSMUSG( | Ddx19a | 573.846 | -0.03467 | 0.784102 | 0.976154 |
| ENSMUSG( | Ddx19b | 814.5113 | 0.056324 | 0.557552 | 0.888279 | ENSMUSG( | Ddx19b | 724.2392 | -0.01182 | 0.923771 | 0.99282 |
| ENSMUSG( | Ddx20 | 328.1249 | 0.078828 | 0.541937 | 0.883182 | ENSMUSG( | Ddx20 | 278.7667 | -0.12205 | 0.432992 | 0.918434 |
| ENSMUSG( | Ddx21 | 1117.051 | 0.150742 | 0.163237 | 0.594466 | ENSMUSG( | Ddx21 | 978.9397 | 0.038476 | 0.76356 | 0.974 |
| ENSMUSG( | Ddx23 | 2171.974 | 0.112498 | 0.219122 | 0.664602 | ENSMUSG( | Ddx23 | 1944.188 | 0.063862 | 0.544461 | 0.938063 |
| ENSMUSG( | Ddx24 | 8226.755 | -0.11812 | 0.426967 | 0.82532 | ENSMUSG( | Ddx24 | 7672.611 | -0.04506 | 0.758978 | 0.971858 |
| ENSMUSG( | Ddx25 | 999.3436 | 0.180487 | 0.070287 | 0.40908 | ENSMUSG( | Ddx25 | 840.8904 | -0.04388 | 0.707481 | 0.964029 |
| ENSMUSG( | Ddx27 | 787.2984 | 0.283946 | 0.03682 | 0.296095 | ENSMUSG( | Ddx27 | 667.5344 | 0.090222 | 0.567194 | 0.941777 |
| ENSMUSG( | Ddx28 | 236.5735 | 0.000696 | 0.996388 | 0.99943 | ENSMUSG( | Ddx28 | 213.6912 | -0.02941 | 0.868012 | 0.98781 |
| ENSMUSG( | Ddx31 | 180.4378 | -0.13091 | 0.413136 | 0.819541 | ENSMUSG( | Ddx31 | 161.1057 | -0.19085 | 0.28832 | 0.894794 |
| ENSMUSG( | Ddx39a | 469.6981 | 0.059947 | 0.642356 | 0.916865 | ENSMUSG( | Ddx39a | 413.113 | -0.03408 | 0.829742 | 0.980224 |
| ENSMUSG( | Ddx39b | 4187.069 | 0.009396 | 0.921232 | 0.990436 | ENSMUSG( | Ddx39b | 3798.569 | -0.00269 | 0.98273 | 0.996859 |
| ENSMUSG( | Ddx3x | 3990.09 | -0.23237 | 0.016444 | 0.190546 | ENSMUSG( | Ddx3x | 4010.628 | 0.058912 | 0.606901 | 0.949797 |
| ENSMUSG( | Ddx3y | 1097.584 | -0.0697 | 0.57094 | 0.8933 | ENSMUSG( | Ddx3y | 1095.99 | 0.19159 | 0.168357 | 0.840222 |
| ENSMUSG( | Ddx4 | 63.03078 | 0.376143 | 0.146511 | 0.568435 | ENSMUSG( | Ddx4 | 52.24853 | 0.122897 | 0.668851 | 0.956766 |
| ENSMUSG( | Ddx41 | 1263.005 | 0.138201 | 0.15113 | 0.576105 | ENSMUSG( | Ddx41 | 1068.249 | -0.07262 | 0.530914 | 0.934795 |
| ENSMUSG( | Ddx42 | 2464.824 | 0.040833 | 0.650923 | 0.92145 | ENSMUSG( | Ddx42 | 2146.739 | -0.09103 | 0.397781 | 0.917257 |
| ENSMUSG( | Ddx43 | 10.79642 | -0.58629 | 0.33749 | NA | ENSMUSG( | Ddx43 | 11.70267 | 0.035801 | 0.954148 | 0.994798 |
| ENSMUSG( | Ddx46 | 2363.57 | 0.081158 | 0.333911 | 0.775602 | ENSMUSG( | Ddx46 | 2070.112 | -0.03324 | 0.76521 | 0.974236 |
| ENSMUSG( | Ddx47 | 995.6988 | 0.027045 | 0.800018 | 0.961634 | ENSMUSG( | Ddx47 | 900.3406 | 0.006076 | 0.962398 | 0.995237 |
| ENSMUSG( | Ddx49 | 879.6021 | 0.058273 | 0.64587 | 0.918224 | ENSMUSG( | Ddx49 | 738.3736 | -0.18446 | 0.180424 | 0.850372 |
| ENSMUSG( | Ddx5 | 13806.87 | -0.06164 | 0.435026 | 0.829093 | ENSMUSG( | Ddx5 | 13626.34 | 0.16529 | 0.137707 | 0.828492 |
| ENSMUSG( | Ddx50 | 1844.279 | 0.06612 | 0.499622 | 0.863218 | ENSMUSG( | Ddx50 | 1751.601 | 0.178786 | 0.132106 | 0.822912 |
| ENSMUSG( | Ddx51 | 466.2784 | 0.102541 | 0.379451 | 0.801757 | ENSMUSG( | Ddx51 | 402.5629 | -0.04422 | 0.776104 | 0.975717 |
| ENSMUSG( | Ddx52 | 664.1139 | 0.024032 | 0.81472 | 0.964668 | ENSMUSG( | Ddx52 | 599.6793 | 0.000355 | 0.997784 | 0.999268 |
| ENSMUSG( | Ddx54 | 1568.229 | 0.14217 | 0.148456 | 0.571935 | ENSMUSG( | Ddx54 | 1333.291 | -0.05534 | 0.631705 | 0.953251 |
| ENSMUSG( | Ddx55 | 714.2885 | 0.087188 | 0.474298 | 0.849425 | ENSMUSG( | Ddx55 | 624.0459 | -0.033 | 0.820713 | 0.979198 |
| ENSMUSG( | Ddx56 | 841.193 | 0.038676 | 0.713591 | 0.940866 | ENSMUSG( | Ddx56 | 730.4311 | -0.09719 | 0.439525 | 0.920721 |
| ENSMUSG( | Ddx59 | 224.4159 | -0.1393 | 0.352925 | 0.784977 | ENSMUSG( | Ddx59 | 199.125 | -0.22999 | 0.186888 | 0.856099 |
| ENSMUSG( | Ddx6 | 3719.629 | 0.083937 | 0.288082 | 0.738882 | ENSMUSG( | Ddx6 | 3302.504 | 0.008983 | 0.934433 | 0.99306 |
| ENSMUSG( | Ddx60 | 35.86065 | -0.91878 | 0.00533 | 0.097362 | ENSMUSG( | Ddx60 | 37.36 | -0.37586 | 0.320985 | 0.904019 |
| ENSMUSG( | Deaf1 | 3541.668 | 0.191192 | 0.032021 | 0.273359 | ENSMUSG( | Deaf1 | 3153.598 | 0.126598 | 0.264634 | 0.886134 |
| ENSMUSG( | Decr1 | 1503.801 | 0.051185 | 0.709086 | 0.939578 | ENSMUSG( | Decr1 | 1242.843 | -0.24394 | 0.102624 | 0.803503 |
| ENSMUSG( | Decr2 | 810.4175 | 0.087334 | 0.430741 | 0.826896 | ENSMUSG( | Decr2 | 688.3255 | -0.11468 | 0.365659 | 0.914623 |
| ENSMUSG( | Dedd | 1089.902 | 0.032369 | 0.750532 | 0.949877 | ENSMUSG( | Dedd | 956.1169 | -0.0811 | 0.525998 | 0.934374 |
| ENSMUSG( | Dedd2 | 587.3586 | 0.011381 | 0.923072 | 0.990693 | ENSMUSG( | Dedd2 | 562.3473 | 0.157018 | 0.278198 | 0.891468 |
| ENSMUSG( | Def6 | 118.7148 | -0.16308 | 0.413916 | 0.819998 | ENSMUSG( | Def6 | 116.252 | 0.071652 | 0.754849 | 0.971098 |
| ENSMUSG( | Def8 | 2040.683 | 0.111076 | 0.191474 | 0.630474 | ENSMUSG( | Def8 | 1715.554 | -0.12284 | 0.244949 | 0.877904 |
| ENSMUSG( | Defb11 | 11.56735 | -0.28809 | 0.716679 | NA | ENSMUSG( | Defb11 | 17.91532 | 1.032064 | 0.10224 | 0.80239 |
| ENSMUSG( | Degs1 | 3561.456 | -0.35303 | 0.011464 | 0.154051 | ENSMUSG( | Degs1 | 3803.993 | 0.127706 | 0.422176 | 0.917257 |
| ENSMUSG( | Degs2 | 82.10893 | 0.387658 | 0.223723 | 0.669845 | ENSMUSG( | Degs2 | 107.9375 | 1.241005 | 0.000435 | 0.061321 |
| ENSMUSG( | Dek | 1312.464 | 0.029505 | 0.80824 | 0.96345 | ENSMUSG( | Dek | 1223.618 | 0.088938 | 0.559572 | 0.939807 |
| ENSMUSG( | Dele1 | 1916.413 | -0.03199 | 0.763051 | 0.953667 | ENSMUSG( | Dele1 | 1767.557 | 0.007192 | 0.95316 | 0.994798 |
| ENSMUSG( | Dennd10 | 835.1879 | -0.10388 | 0.380287 | 0.801757 | ENSMUSG( | Dennd10 | 754.774 | -0.1337 | 0.355033 | 0.912411 |
| ENSMUSG( | Dennd11 | 5479.343 | -0.02044 | 0.797686 | 0.960507 | ENSMUSG( | Dennd11 | 4847.185 | -0.10864 | 0.304491 | 0.898538 |
| ENSMUSG( | Dennd1a | 2360.282 | 0.100583 | 0.257485 | 0.708686 | ENSMUSG( | Dennd1a | 2126.549 | 0.070848 | 0.51906 | 0.931053 |
| ENSMUSG( | Dennd1b | 389.0317 | 0.158596 | 0.197623 | 0.637377 | ENSMUSG( | Dennd1b | 352.5725 | 0.142093 | 0.309579 | 0.898538 |
| ENSMUSG( | Dennd1c | 101.6101 | -0.08498 | 0.698785 | 0.936563 | ENSMUSG( | Dennd1c | 100.1452 | 0.151615 | 0.576384 | 0.942912 |
| ENSMUSG( | Dennd2a | 1528.398 | 0.109693 | 0.214142 | 0.658712 | ENSMUSG( | Dennd2a | 1316.978 | -0.05055 | 0.645878 | 0.953988 |
| ENSMUSG( | Dennd2b | 885.9262 | 0.009099 | 0.930242 | 0.991848 | ENSMUSG( | Dennd2b | 804.1907 | 0.000111 | 0.999288 | 0.999503 |
| ENSMUSG( | Dennd2c | 33.5806 | 0.112878 | 0.76442 | 0.954055 | ENSMUSG( | Dennd2c | 35.16467 | 0.508375 | 0.21014 | 0.864412 |
| ENSMUSG( | Dennd3 | 266.644 | 0.17663 | 0.280343 | 0.729291 | ENSMUSG( | Dennd3 | 233.5734 | 0.084152 | 0.662434 | 0.956205 |
| ENSMUSG( | Dennd4a | 2214.742 | 0.00437 | 0.962281 | 0.994543 | ENSMUSG( | Dennd4a | 1996.842 | -0.02608 | 0.815968 | 0.979198 |
| ENSMUSG( | Dennd4b | 2111.087 | 0.18521 | 0.058069 | 0.371677 | ENSMUSG( | Dennd4b | 1875.936 | 0.11958 | 0.374274 | 0.915491 |
| ENSMUSG( | Dennd4c | 743.5284 | 0.092978 | 0.547111 | 0.885118 | ENSMUSG( | Dennd4c | 649.0787 | -0.02506 | 0.870624 | 0.988445 |
| ENSMUSG( | Dennd5a | 24409.66 | 0.074516 | 0.399356 | 0.813048 | ENSMUSG( | Dennd5a | 20507.46 | -0.16338 | 0.104014 | 0.804969 |
| ENSMUSG( | Dennd5b | 1302.172 | 0.069351 | 0.439265 | 0.830889 | ENSMUSG( | Dennd5b | 1188.453 | 0.071055 | 0.552926 | 0.939565 |
| ENSMUSG( | Dennd6a | 877.9452 | 0.124507 | 0.229351 | 0.67735 | ENSMUSG( | Dennd6a | 812.55 | 0.167689 | 0.170016 | 0.841056 |
| ENSMUSG( | Dennd6b | 2067.769 | 0.040985 | 0.769402 | 0.954674 | ENSMUSG( | Dennd6b | 1943.347 | 0.126781 | 0.459337 | 0.923329 |
| ENSMUSG( | Denr | 1419.89 | -0.01966 | 0.846731 | 0.972371 | ENSMUSG( | Denr | 1217.171 | -0.20664 | 0.099125 | 0.799802 |
| ENSMUSG( | Depdc1b | 16.76262 | -0.62522 | 0.204077 | NA | ENSMUSG( | Depdc1b | 14.53413 | -0.7838 | 0.149515 | 0.836241 |
| ENSMUSG( | Depdc5 | 1573.157 | 0.142504 | 0.149701 | 0.57376 | ENSMUSG( | Depdc5 | 1398.449 | 0.076887 | 0.500721 | 0.927825 |
| ENSMUSG( | Depdc7 | 24.94513 | -0.55467 | 0.137953 | 0.552903 | ENSMUSG( | Depdc7 | 32.75133 | 0.487393 | 0.171655 | 0.842999 |
| ENSMUSG( | Depp1 | 46.54795 | -0.77993 | 0.032686 | 0.27685 | ENSMUSG( | Depp1 | 53.88309 | 0.049016 | 0.911473 | 0.992196 |
| ENSMUSG( | Deptor | 595.2457 | 0.195683 | 0.100487 | 0.483976 | ENSMUSG( | Deptor | 528.0897 | 0.119719 | 0.394814 | 0.917257 |
| ENSMUSG( | Dera | 168.3203 | 0.031128 | 0.87293 | 0.978753 | ENSMUSG( | Dera | 144.0473 | -0.15061 | 0.420757 | 0.917257 |
| ENSMUSG( | Derl1 | 2001.579 | -0.29748 | 0.045065 | 0.325595 | ENSMUSG( | Derl1 | 2155.633 | 0.19807 | 0.249832 | 0.877904 |
| ENSMUSG( | Derl2 | 846.2884 | -0.16884 | 0.205295 | 0.64654 | ENSMUSG( | Derl2 | 917.8769 | 0.323752 | 0.028222 | 0.611882 |
| ENSMUSG( | Derl3 | 46.25702 | 0.164896 | 0.570461 | 0.893192 | ENSMUSG( | Derl3 | 52.4149 | 0.698483 | 0.018591 | 0.545482 |
| ENSMUSG( | Des | 45.52577 | -0.25485 | 0.392574 | 0.809274 | ENSMUSG( | Des | 47.67357 | 0.183207 | 0.579069 | 0.943849 |
| ENSMUSG( | Desi1 | 10401.53 | -0.07525 | 0.329131 | 0.773233 | ENSMUSG( | Desi1 | 8936.153 | -0.25615 | 0.019656 | 0.556192 |
| ENSMUSG( | Desi2 | 1104.477 | 0.077494 | 0.497318 | 0.861898 | ENSMUSG( | Desi2 | 969.3778 | -0.03482 | 0.809483 | 0.978779 |
| ENSMUSG( | Det1 | 129.5497 | 0.048514 | 0.804032 | 0.962169 | ENSMUSG( | Det1 | 115.0785 | -0.01497 | 0.944841 | 0.994798 |
| ENSMUSG( | Dexi | 1170.788 | 0.04884 | 0.633602 | 0.914073 | ENSMUSG( | Dexi | 1057.093 | 0.020788 | 0.86403 | 0.986552 |
| ENSMUSG( | Dffa | 894.5052 | 0.033912 | 0.783879 | 0.957645 | ENSMUSG( | Dffa | 795.1513 | -0.03627 | 0.776917 | 0.975717 |
| ENSMUSG( | Dffb | 263.0757 | 0.00037 | 0.99822 | 0.99989 | ENSMUSG( | Dffb | 234.3608 | -0.06616 | 0.719545 | 0.966876 |
| ENSMUSG( | Dgat1 | 714.4623 | -0.13531 | 0.31967 | 0.765305 | ENSMUSG( | Dgat1 | 674.9719 | -0.02004 | 0.895369 | 0.990558 |
| ENSMUSG( | Dgat2 | 2402.711 | -0.44172 | 0.000422 | 0.018541 | ENSMUSG( | Dgat2 | 2766.522 | 0.260275 | 0.095503 | 0.796133 |
| ENSMUSG( | Dgat2l6 | 22.27791 | 0.083096 | 0.845593 | 0.97218 | ENSMUSG( | Dgat2l6 | 16.68858 | -0.50903 | 0.335574 | 0.907907 |
| ENSMUSG( | Dgcr2 | 2961.172 | -0.21645 | 0.072761 | 0.416435 | ENSMUSG( | Dgcr2 | 3082.879 | 0.175202 | 0.219608 | 0.866104 |
| ENSMUSG( | Dgcr6 | 2535.902 | 0.003077 | 0.972713 | 0.996585 | ENSMUSG( | Dgcr6 | 2111.986 | -0.27045 | 0.012934 | 0.458131 |
| ENSMUSG( | Dgcr8 | 910.4627 | 0.235791 | 0.039955 | 0.308163 | ENSMUSG( | Dgcr8 | 794.7441 | 0.117007 | 0.415543 | 0.917257 |
| ENSMUSG( | Dgka | 435.1279 | 0.1655 | 0.201375 | 0.641362 | ENSMUSG( | Dgka | 375.642 | 0.018949 | 0.917422 | 0.992529 |
| ENSMUSG( | Dgkb | 2986.132 | -0.06516 | 0.553003 | 0.886204 | ENSMUSG( | Dgkb | 2589.974 | -0.21828 | 0.127736 | 0.81845 |
| ENSMUSG( | Dgkd | 3175.682 | 0.193715 | 0.046162 | 0.329267 | ENSMUSG( | Dgkd | 2786.027 | 0.09303 | 0.469181 | 0.923329 |
| ENSMUSG( | Dgke | 2312.818 | -0.4218 | 0.003009 | 0.067314 | ENSMUSG( | Dgke | 2552.208 | 0.160301 | 0.296829 | 0.898538 |
| ENSMUSG( | Dgkg | 484.0764 | 0.45104 | 0.001484 | 0.042763 | ENSMUSG( | Dgkg | 434.4871 | 0.423005 | 0.01541 | 0.495791 |
| ENSMUSG( | Dgkh | 1833.987 | -0.02142 | 0.830602 | 0.96974 | ENSMUSG( | Dgkh | 1790.179 | 0.175498 | 0.15468 | 0.837724 |
| ENSMUSG( | Dgki | 387.1107 | -0.10375 | 0.443064 | 0.833025 | ENSMUSG( | Dgki | 339.1026 | -0.21792 | 0.144688 | 0.833302 |
| ENSMUSG( | Dgkk | 74.98626 | 0.258262 | 0.289748 | 0.739839 | ENSMUSG( | Dgkk | 64.28231 | 0.097257 | 0.721914 | 0.966887 |
| ENSMUSG( | Dgkq | 1387.143 | 0.061845 | 0.576064 | 0.895114 | ENSMUSG( | Dgkq | 1302.949 | 0.149716 | 0.292699 | 0.898093 |
| ENSMUSG( | Dgkz | 28437.44 | 0.270733 | 0.002886 | 0.065928 | ENSMUSG( | Dgkz | 23987.66 | 0.060835 | 0.552114 | 0.939565 |
| ENSMUSG( | Dglucy | 362.3006 | -0.04387 | 0.741414 | 0.948208 | ENSMUSG( | Dglucy | 298.5131 | -0.35223 | 0.020002 | 0.5597 |
| ENSMUSG( | Dguok | 306.5675 | 0.000261 | 0.998605 | 0.99989 | ENSMUSG( | Dguok | 281.5135 | 0.026296 | 0.876629 | 0.989644 |
| ENSMUSG( | Dhcr24 | 5678.419 | -0.39185 | 0.011919 | 0.157992 | ENSMUSG( | Dhcr24 | 6030.293 | 0.080535 | 0.640849 | 0.953903 |
| ENSMUSG( | Dhcr7 | 2130.046 | -0.50538 | 0.003831 | 0.079288 | ENSMUSG( | Dhcr7 | 2245.854 | -0.03502 | 0.852042 | 0.985598 |
| ENSMUSG( | Dhdds | 1732.671 | 0.104515 | 0.312728 | 0.760258 | ENSMUSG( | Dhdds | 1539.422 | 0.032985 | 0.7755 | 0.975354 |
| ENSMUSG( | Dhdh | 581.0983 | 0.13591 | 0.226565 | 0.673549 | ENSMUSG( | Dhdh | 512.3549 | 0.036477 | 0.812986 | 0.979198 |
| ENSMUSG( | Dhfr | 254.4476 | 0.122955 | 0.432599 | 0.827838 | ENSMUSG( | Dhfr | 218.7904 | -0.05064 | 0.763289 | 0.973945 |
| ENSMUSG( | Dhodh | 425.9093 | 0.099795 | 0.436887 | 0.829501 | ENSMUSG( | Dhodh | 400.5893 | 0.188426 | 0.180005 | 0.850372 |
| ENSMUSG( | Dhps | 924.7106 | 0.011921 | 0.912879 | 0.988483 | ENSMUSG( | Dhps | 843.5797 | 0.015471 | 0.900405 | 0.990558 |
| ENSMUSG( | Dhrs1 | 2655.093 | 0.02371 | 0.81322 | 0.964574 | ENSMUSG( | Dhrs1 | 2231.766 | -0.22097 | 0.096309 | 0.796133 |
| ENSMUSG( | Dhrs11 | 291.1207 | 0.076766 | 0.570071 | 0.893041 | ENSMUSG( | Dhrs11 | 249.5385 | -0.09706 | 0.533239 | 0.935104 |
| ENSMUSG( | Dhrs13 | 251.3254 | -0.25465 | 0.227205 | 0.674223 | ENSMUSG( | Dhrs13 | 256.9243 | 0.095633 | 0.661215 | 0.956205 |
| ENSMUSG( | Dhrs13os | 12.61022 | -0.61718 | 0.234856 | NA | ENSMUSG( | Dhrs13os | 12.24297 | -0.40348 | 0.466214 | 0.923329 |
| ENSMUSG( | Dhrs3 | 942.0321 | -0.37036 | 0.007045 | 0.115775 | ENSMUSG( | Dhrs3 | 900.8645 | -0.21218 | 0.139836 | 0.830678 |
| ENSMUSG( | Dhrs4 | 846.1725 | 0.121089 | 0.333005 | 0.775602 | ENSMUSG( | Dhrs4 | 685.2442 | -0.23318 | 0.119649 | 0.813384 |
| ENSMUSG( | Dhrs7 | 1238.061 | -0.34161 | 0.02297 | 0.232278 | ENSMUSG( | Dhrs7 | 1276.705 | 0.040917 | 0.796139 | 0.97681 |
| ENSMUSG( | Dhrs7b | 870.4801 | -0.19259 | 0.122449 | 0.528223 | ENSMUSG( | Dhrs7b | 841.9605 | -0.01045 | 0.939116 | 0.993898 |
| ENSMUSG( | Dhtkd1 | 310.1054 | 0.056122 | 0.677751 | 0.930542 | ENSMUSG( | Dhtkd1 | 296.0394 | 0.185016 | 0.214464 | 0.864421 |
| ENSMUSG( | Dhx15 | 1918.598 | 0.005627 | 0.961029 | 0.993999 | ENSMUSG( | Dhx15 | 1832.662 | 0.140362 | 0.272634 | 0.888825 |
| ENSMUSG( | Dhx16 | 892.1137 | 0.166766 | 0.130863 | 0.54231 | ENSMUSG( | Dhx16 | 778.3887 | 0.043793 | 0.751145 | 0.970317 |
| ENSMUSG( | Dhx29 | 712.2802 | 0.286574 | 0.016385 | 0.190235 | ENSMUSG( | Dhx29 | 592.9843 | 0.039615 | 0.75146 | 0.970317 |
| ENSMUSG( | Dhx30 | 2836.661 | 0.148067 | 0.128186 | 0.536641 | ENSMUSG( | Dhx30 | 2476.13 | 0.028777 | 0.800249 | 0.97681 |
| ENSMUSG( | Dhx32 | 796.6592 | -0.03137 | 0.770319 | 0.954689 | ENSMUSG( | Dhx32 | 720.3608 | -0.05586 | 0.662109 | 0.956205 |
| ENSMUSG( | Dhx33 | 994.2007 | 0.164998 | 0.186382 | 0.623869 | ENSMUSG( | Dhx33 | 898.0365 | 0.146223 | 0.277957 | 0.891468 |
| ENSMUSG( | Dhx34 | 476.4318 | 0.041952 | 0.725185 | 0.944096 | ENSMUSG( | Dhx34 | 420.551 | -0.04527 | 0.744988 | 0.96951 |
| ENSMUSG( | Dhx35 | 693.9763 | -0.01756 | 0.865774 | 0.975876 | ENSMUSG( | Dhx35 | 626.1358 | -0.04369 | 0.725415 | 0.966887 |
| ENSMUSG( | Dhx36 | 1745.476 | 0.098438 | 0.291853 | 0.741868 | ENSMUSG( | Dhx36 | 1593.852 | 0.101711 | 0.38745 | 0.917257 |
| ENSMUSG( | Dhx37 | 469.0726 | 0.040258 | 0.767024 | 0.954393 | ENSMUSG( | Dhx37 | 405.4264 | -0.11158 | 0.512015 | 0.930005 |
| ENSMUSG( | Dhx38 | 1134.556 | 0.152498 | 0.112442 | 0.505997 | ENSMUSG( | Dhx38 | 999.3525 | 0.058751 | 0.608725 | 0.950113 |
| ENSMUSG( | Dhx40 | 883.9419 | 0.07195 | 0.47766 | 0.852702 | ENSMUSG( | Dhx40 | 777.4749 | -0.03103 | 0.797938 | 0.97681 |
| ENSMUSG( | Dhx57 | 1187.227 | 0.123503 | 0.238174 | 0.686299 | ENSMUSG( | Dhx57 | 1057.746 | 0.061694 | 0.623556 | 0.953234 |
| ENSMUSG( | Dhx58 | 73.10209 | -0.11585 | 0.641779 | 0.916828 | ENSMUSG( | Dhx58 | 68.4834 | -0.0436 | 0.866137 | 0.987254 |
| ENSMUSG( | Dhx8 | 967.4259 | 0.210316 | 0.037158 | 0.297481 | ENSMUSG( | Dhx8 | 822.6814 | 0.01772 | 0.886432 | 0.990011 |
| ENSMUSG( | Dhx9 | 3648.984 | 0.149279 | 0.102244 | 0.487127 | ENSMUSG( | Dhx9 | 3057.005 | -0.0926 | 0.450578 | 0.922202 |
| ENSMUSG( | Diablo | 247.655 | 0.043827 | 0.7673 | 0.954491 | ENSMUSG( | Diablo | 237.0092 | 0.17485 | 0.332745 | 0.907907 |
| ENSMUSG( | Diaph1 | 4290.135 | 0.043134 | 0.65628 | 0.923929 | ENSMUSG( | Diaph1 | 3894.858 | 0.034649 | 0.754261 | 0.970785 |
| ENSMUSG( | Diaph2 | 620.0118 | -0.01682 | 0.878946 | 0.981436 | ENSMUSG( | Diaph2 | 584.7466 | 0.075124 | 0.605198 | 0.949082 |
| ENSMUSG( | Diaph3 | 125.0871 | -0.00262 | 0.989442 | 0.999111 | ENSMUSG( | Diaph3 | 103.8157 | -0.30086 | 0.189957 | 0.857355 |
| ENSMUSG( | Dicer1 | 1836.152 | 0.003317 | 0.97243 | 0.996585 | ENSMUSG( | Dicer1 | 1671.04 | 0.003152 | 0.978398 | 0.996459 |
| ENSMUSG( | Dido1 | 1674.455 | -0.14774 | 0.161667 | 0.591927 | ENSMUSG( | Dido1 | 1651.08 | 0.087466 | 0.533768 | 0.93547 |
| ENSMUSG( | Dimt1 | 320.6622 | 0.197029 | 0.231588 | 0.67949 | ENSMUSG( | Dimt1 | 310.3218 | 0.359665 | 0.027972 | 0.61023 |
| ENSMUSG( | Dio2 | 22398.22 | 0.225519 | 0.110472 | 0.50108 | ENSMUSG( | Dio2 | 17903.24 | -0.14762 | 0.356049 | 0.913331 |
| ENSMUSG( | Dio3 | 68.80113 | 0.267434 | 0.308588 | 0.756544 | ENSMUSG( | Dio3 | 57.79841 | 0.025967 | 0.932471 | 0.99306 |
| ENSMUSG( | Dio3os | 17.78785 | -0.15007 | 0.738258 | NA | ENSMUSG( | Dio3os | 19.01952 | 0.343646 | 0.476801 | 0.923329 |
| ENSMUSG( | Dip2a | 3516.571 | 0.006169 | 0.937749 | 0.993415 | ENSMUSG( | Dip2a | 3132.251 | -0.06034 | 0.548232 | 0.938063 |
| ENSMUSG( | Dip2b | 2291.496 | 0.117245 | 0.190425 | 0.62983 | ENSMUSG( | Dip2b | 2054.306 | 0.07195 | 0.507067 | 0.928624 |
| ENSMUSG( | Dip2c | 2271.366 | 0.096408 | 0.276743 | 0.726926 | ENSMUSG( | Dip2c | 1984.214 | -0.02543 | 0.831479 | 0.980611 |
| ENSMUSG( | Dipk1a | 707.8848 | -0.5725 | 8.79E-05 | 0.005891 | ENSMUSG( | Dipk1a | 765.5 | -0.01771 | 0.918956 | 0.992529 |
| ENSMUSG( | Dipk1b | 1360.217 | -0.299 | 0.012697 | 0.162421 | ENSMUSG( | Dipk1b | 1355.543 | -0.02166 | 0.874636 | 0.989644 |
| ENSMUSG( | Dipk1c | 444.9392 | -0.03451 | 0.810914 | 0.964318 | ENSMUSG( | Dipk1c | 409.6529 | 0.003176 | 0.984187 | 0.996963 |
| ENSMUSG( | Dipk2a | 919.7594 | -0.41454 | 0.021948 | 0.225738 | ENSMUSG( | Dipk2a | 1150.087 | 0.490839 | 0.024857 | 0.592568 |
| ENSMUSG( | Dipk2b | 30.28425 | -0.32872 | 0.43452 | 0.828978 | ENSMUSG( | Dipk2b | 37.54714 | 0.569355 | 0.26036 | 0.884524 |
| ENSMUSG( | Diras1 | 3747.722 | 0.129341 | 0.181717 | 0.618944 | ENSMUSG( | Diras1 | 3143.146 | -0.1106 | 0.319306 | 0.903453 |
| ENSMUSG( | Diras2 | 8783.229 | 0.015663 | 0.841223 | 0.971179 | ENSMUSG( | Diras2 | 7457 | -0.19384 | 0.058932 | 0.729034 |
| ENSMUSG( | Dis3 | 507.1463 | -0.06738 | 0.568831 | 0.892651 | ENSMUSG( | Dis3 | 456.3543 | -0.10309 | 0.447654 | 0.922202 |
| ENSMUSG( | Dis3l | 1074.49 | 0.15072 | 0.120223 | 0.524481 | ENSMUSG( | Dis3l | 909.2931 | -0.06102 | 0.62111 | 0.953199 |
| ENSMUSG( | Dis3l2 | 526.8129 | 0.055231 | 0.65643 | 0.923929 | ENSMUSG( | Dis3l2 | 469.7048 | -0.00666 | 0.961824 | 0.995237 |
| ENSMUSG( | Disc1 | 65.99485 | 0.314231 | 0.221986 | 0.667523 | ENSMUSG( | Disc1 | 57.27811 | 0.181413 | 0.524752 | 0.933907 |
| ENSMUSG( | Disp1 | 270.0067 | -0.15998 | 0.346175 | 0.779153 | ENSMUSG( | Disp1 | 247.3366 | -0.12652 | 0.556773 | 0.939565 |
| ENSMUSG( | Disp2 | 9787.301 | -0.26933 | 0.034164 | 0.283507 | ENSMUSG( | Disp2 | 10360.78 | 0.173875 | 0.23234 | 0.868507 |
| ENSMUSG( | Disp3 | 1503.771 | -0.1179 | 0.405762 | 0.81652 | ENSMUSG( | Disp3 | 1409.216 | -0.02899 | 0.859057 | 0.985884 |
| ENSMUSG( | Dixdc1 | 2551.503 | -0.13065 | 0.151201 | 0.576198 | ENSMUSG( | Dixdc1 | 2510.235 | 0.091323 | 0.447921 | 0.922202 |
| ENSMUSG( | Dkc1 | 571.1587 | 0.127298 | 0.23673 | 0.684086 | ENSMUSG( | Dkc1 | 503.3051 | 0.02702 | 0.848971 | 0.985321 |
| ENSMUSG( | Dkk2 | 18.85707 | -0.31745 | 0.48751 | NA | ENSMUSG( | Dkk2 | 25.13206 | 0.691451 | 0.116688 | 0.813384 |

| | | | | | |
|---|---|---|---|---|---|
| ENSMUSG( | 469.0726 | 0.040258 | 0.767024 | 0.954393 | Dhx37 |
| ENSMUSG( | 1134.556 | 0.152498 | 0.112442 | 0.505997 | Dhx38 |
| ENSMUSG( | 883.9419 | 0.07195 | 0.47766 | 0.852702 | Dhx40 |
| ENSMUSG( | 1187.227 | 0.123503 | 0.238174 | 0.686299 | Dhx57 |
| ENSMUSG( | 73.10209 | -0.11585 | 0.641779 | 0.916828 | Dhx58 |
| ENSMUSG( | 967.4259 | 0.210316 | 0.037158 | 0.297481 | Dhx8 |
| ENSMUSG( | 3648.984 | 0.149279 | 0.102244 | 0.487127 | Dhx9 |
| ENSMUSG( | 247.655 | 0.043827 | 0.7673 | 0.954491 | Diablo |
| ENSMUSG( | 4290.135 | 0.043134 | 0.65628 | 0.923929 | Diaph1 |
| ENSMUSG( | 620.0118 | -0.01682 | 0.878946 | 0.981436 | Diaph2 |
| ENSMUSG( | 125.0871 | -0.00262 | 0.989442 | 0.999111 | Diaph3 |
| ENSMUSG( | 1836.152 | 0.003317 | 0.97243 | 0.996585 | Dicer1 |
| ENSMUSG( | 1674.455 | -0.14774 | 0.161667 | 0.591927 | Dido1 |
| ENSMUSG( | 320.6622 | 0.197029 | 0.231588 | 0.67949 | Dimt1 |
| ENSMUSG( | 22398.22 | 0.225519 | 0.110472 | 0.50108 | Dio2 |
| ENSMUSG( | 68.80113 | 0.267434 | 0.308588 | 0.756544 | Dio3 |
| ENSMUSG( | 17.78785 | -0.15007 | 0.738258 | NA | Dio3os |
| ENSMUSG( | 3516.571 | 0.006169 | 0.937749 | 0.993415 | Dip2a |
| ENSMUSG( | 2291.496 | 0.117245 | 0.190425 | 0.62983 | Dip2b |
| ENSMUSG( | 2271.366 | 0.096408 | 0.276743 | 0.726926 | Dip2c |
| ENSMUSG( | 707.8848 | -0.5725 | 8.79E-05 | 0.005891 | Dipk1a |
| ENSMUSG( | 1360.217 | -0.299 | 0.012697 | 0.162421 | Dipk1b |
| ENSMUSG( | 444.9392 | -0.03451 | 0.810914 | 0.964318 | Dipk1c |
| ENSMUSG( | 919.7594 | -0.41454 | 0.021948 | 0.225738 | Dipk2a |
| ENSMUSG( | 30.28425 | -0.32872 | 0.43452 | 0.828978 | Dipk2b |
| ENSMUSG( | 3747.722 | 0.129341 | 0.181717 | 0.618944 | Diras1 |
| ENSMUSG( | 8783.229 | 0.015663 | 0.841223 | 0.971179 | Diras2 |
| ENSMUSG( | 507.1463 | -0.06738 | 0.568831 | 0.892651 | Dis3 |
| ENSMUSG( | 1074.49 | 0.15072 | 0.120223 | 0.524481 | Dis3l |
| ENSMUSG( | 526.8129 | 0.055231 | 0.65643 | 0.923929 | Dis3l2 |
| ENSMUSG( | 65.99485 | 0.314231 | 0.221986 | 0.667523 | Disc1 |
| ENSMUSG( | 270.0067 | -0.15998 | 0.346175 | 0.779153 | Disp1 |
| ENSMUSG( | 9787.301 | -0.26933 | 0.034164 | 0.283507 | Disp2 |
| ENSMUSG( | 1503.771 | -0.1179 | 0.405762 | 0.81652 | Disp3 |
| ENSMUSG( | 2551.503 | -0.13065 | 0.151201 | 0.576198 | Dixdc1 |
| ENSMUSG( | 571.1587 | 0.127298 | 0.23673 | 0.684086 | Dkc1 |
| ENSMUSG( | 18.85707 | -0.31745 | 0.48751 | NA | Dkk2 |
| ENSMUSG( | 11179.97 | -0.25124 | 0.045562 | 0.327592 | Dkk3 |
| ENSMUSG( | 660.3032 | -0.169 | 0.138022 | 0.552905 | Dkkl1 |
| ENSMUSG( | 2796.486 | 0.090732 | 0.295113 | 0.743798 | Dlat |
| ENSMUSG( | 2270.548 | 0.054815 | 0.562403 | 0.889448 | Dlc1 |
| ENSMUSG( | 2812.103 | 0.090813 | 0.379689 | 0.801757 | Dld |
| ENSMUSG( | 91.87679 | 0.11397 | 0.607934 | 0.907063 | Dlec1 |
| ENSMUSG( | 254.9098 | 0.074133 | 0.604556 | 0.905556 | Dleu2 |
| ENSMUSG( | 97.18024 | 0.130596 | 0.576285 | 0.895164 | Dleu7 |
| ENSMUSG( | 7179.763 | -0.01559 | 0.854023 | 0.973491 | Dlg1 |
| ENSMUSG( | 21791.38 | 0.112714 | 0.304388 | 0.752574 | Dlg2 |
| ENSMUSG( | 3364.999 | 0.13458 | 0.253208 | 0.70338 | Dlg3 |
| ENSMUSG( | 57368.56 | 0.113481 | 0.134184 | 0.547472 | Dlg4 |
| ENSMUSG( | 2993.803 | 0.272705 | 0.002016 | 0.051507 | Dlg5 |
| ENSMUSG( | 16001.04 | 0.083269 | 0.309639 | 0.757658 | Dlgap1 |
| ENSMUSG( | 2159.651 | 0.102796 | 0.245489 | 0.693175 | Dlgap2 |
| ENSMUSG( | 26460.41 | 0.236789 | 0.00445 | 0.086961 | Dlgap3 |
| ENSMUSG( | 13256.41 | 0.196882 | 0.011355 | 0.153509 | Dlgap4 |
| ENSMUSG( | 14.487 | -0.09028 | 0.866651 | NA | Dlgap5 |
| ENSMUSG( | 295.362 | -0.04269 | 0.833607 | 0.970333 | Dlk1 |
| ENSMUSG( | 946.2704 | 0.071369 | 0.561105 | 0.888934 | Dlk2 |
| ENSMUSG( | 167.3622 | -0.3327 | 0.044837 | 0.324745 | Dll1 |
| ENSMUSG( | 77.42804 | -0.30815 | 0.280623 | 0.729679 | Dll3 |
| ENSMUSG( | 133.525 | 0.25996 | 0.324118 | 0.769158 | Dll4 |
| ENSMUSG( | 4100.217 | 0.086946 | 0.294489 | 0.743674 | Dlst |
| ENSMUSG( | 660.6026 | -0.11616 | 0.424066 | 0.824452 | Dlx1 |
| ENSMUSG( | 444.532 | 0.090039 | 0.489884 | 0.859319 | Dlx1as |
| ENSMUSG( | 163.2649 | -0.33457 | 0.134334 | 0.547472 | Dlx2 |
| ENSMUSG( | 101.8905 | -0.59629 | 0.028109 | 0.25643 | Dlx5 |
| ENSMUSG( | 113.1671 | -0.41705 | 0.114697 | 0.510718 | Dlx6 |
| ENSMUSG( | 285.7178 | -0.13876 | 0.409752 | 0.818258 | Dlx6os1 |
| ENSMUSG( | 526.1276 | 0.05863 | 0.608585 | 0.907154 | Dmac1 |
| ENSMUSG( | 1067.828 | 0.08803 | 0.418523 | 0.821966 | Dmac2 |
| ENSMUSG( | 421.4965 | 0.026889 | 0.862384 | 0.975055 | Dmac2l |
| ENSMUSG( | 568.8976 | 0.055559 | 0.704759 | 0.938303 | Dmap1 |
| ENSMUSG( | 35.33619 | 0.731224 | 0.037886 | 0.30067 | Dmc1 |
| ENSMUSG( | 2293.579 | 0.02971 | 0.80529 | 0.962616 | Dmd |
| ENSMUSG( | 14.92776 | 0.148251 | 0.754428 | NA | Dmgdh |
| ENSMUSG( | 67.38936 | -0.53104 | 0.081275 | 0.440604 | Dmkn |
| ENSMUSG( | 751.5319 | -0.01064 | 0.936361 | 0.993415 | Dmp1 |
| ENSMUSG( | 1228.24 | 0.114199 | 0.321823 | 0.766739 | Dmpk |
| ENSMUSG( | 19.35193 | -0.21027 | 0.653747 | NA | Dmrt2 |
| ENSMUSG( | 15.72119 | 0.392301 | 0.402169 | NA | Dmrt3 |
| ENSMUSG( | 17.03579 | -0.11365 | 0.803313 | NA | Dmrta1 |
| ENSMUSG( | 196.0216 | 0.290755 | 0.096389 | 0.476986 | Dmrta2 |
| ENSMUSG( | 52.24031 | -0.31536 | 0.301916 | 0.750166 | Dmrtb1 |
| ENSMUSG( | 73.87677 | -0.03071 | 0.901077 | 0.985633 | Dmrtc1a |
| ENSMUSG( | 1448.295 | 0.079826 | 0.424423 | 0.824645 | Dmtf1 |
| ENSMUSG( | 8183.712 | 0.149894 | 0.098682 | 0.480598 | Dmtn |
| ENSMUSG( | 4829.692 | 0.116588 | 0.244891 | 0.692492 | Dmwd |
| ENSMUSG( | 1012.814 | -0.06285 | 0.507388 | 0.86795 | Dmxl1 |
| ENSMUSG( | 3385.968 | -0.10251 | 0.343809 | 0.779153 | Dmxl2 |
| ENSMUSG( | 47.66056 | -0.08066 | 0.785224 | 0.958473 | Dna2 |
| ENSMUSG( | 13.537 | -0.08069 | 0.889413 | NA | Dnaaf1 |
| ENSMUSG( | 150.2583 | 0.004678 | 0.979611 | 0.997428 | Dnaaf10 |
| ENSMUSG( | 57.61442 | -0.5192 | 0.131931 | 0.544244 | Dnaaf11 |
| ENSMUSG( | 29.7591 | 0.007864 | 0.983681 | 0.997768 | Dnaaf2 |
| ENSMUSG( | 38.90733 | -0.397 | 0.269978 | 0.719883 | Dnaaf3 |
| ENSMUSG( | 33.89178 | -0.19641 | 0.579964 | 0.896931 | Dnaaf4 |
| ENSMUSG( | 275.2795 | 0.077574 | 0.569605 | 0.893041 | Dnaaf5 |
| ENSMUSG( | 7777.765 | 0.043783 | 0.617832 | 0.910872 | Dnaaf9 |
| ENSMUSG( | 300.6297 | -0.06277 | 0.636132 | 0.914873 | Dnah1 |
| ENSMUSG( | 95.17167 | -0.01979 | 0.941968 | 0.993415 | Dnah10 |
| ENSMUSG( | 59.61797 | -0.13091 | 0.682557 | 0.93198 | Dnah11 |
| ENSMUSG( | 43.93105 | -0.09265 | 0.761912 | 0.95339 | Dnah12 |
| ENSMUSG( | 42.96341 | -0.15487 | 0.606537 | 0.906481 | Dnah14 |
| ENSMUSG( | 18.42374 | -0.51457 | 0.265626 | NA | Dnah17 |
| ENSMUSG( | 154.8652 | 0.111198 | 0.544404 | 0.884649 | Dnah2 |
| ENSMUSG( | 30.05308 | 0.195784 | 0.60292 | 0.904887 | Dnah3 |
| ENSMUSG( | 127.5129 | -0.33153 | 0.082186 | 0.442642 | Dnah5 |
| ENSMUSG( | 88.73038 | 0.081536 | 0.768042 | 0.954621 | Dnah6 |
| ENSMUSG( | 100.5281 | 0.180269 | 0.372952 | 0.799429 | Dnah7a |
| ENSMUSG( | 212.4318 | -0.02694 | 0.855623 | 0.973813 | Dnah7b |
| ENSMUSG( | 45.52099 | 0.097787 | 0.726212 | 0.944223 | Dnah7c |
| ENSMUSG( | 21.95142 | -0.24593 | 0.56378 | 0.890032 | Dnah8 |
| ENSMUSG( | 171.51 | -0.13588 | 0.541732 | 0.883182 | Dnah9 |
| ENSMUSG( | 411.4435 | 0.048524 | 0.709905 | 0.939669 | Dnai1 |
| ENSMUSG( | 137.749 | -0.29074 | 0.208573 | 0.650496 | Dnai2 |
| ENSMUSG( | 20.46355 | 0.160781 | 0.710702 | NA | Dnai3 |
| ENSMUSG( | 207.2679 | -0.0301 | 0.848592 | 0.972374 | Dnai4 |
| ENSMUSG( | 50.84946 | -0.1854 | 0.508901 | 0.868744 | Dnai7 |
| ENSMUSG( | 3240.309 | 0.012383 | 0.891411 | 0.983919 | Dnaja1 |
| ENSMUSG( | 2895.968 | 0.085322 | 0.319252 | 0.765223 | Dnaja2 |
| ENSMUSG( | 1619.97 | 0.053998 | 0.60133 | 0.904603 | Dnaja3 |
| ENSMUSG( | 1171.105 | 0.139815 | 0.166081 | 0.598098 | Dnaja4 |
| ENSMUSG( | 751.7078 | 0.150682 | 0.300655 | 0.749904 | Dnajb1 |
| ENSMUSG( | 2676.96 | -0.17679 | 0.196171 | 0.635538 | Dnajb11 |
| ENSMUSG( | 1925.368 | -0.35611 | 0.005961 | 0.104409 | Dnajb12 |
| ENSMUSG( | 26.62622 | -0.39611 | 0.290998 | 0.741182 | Dnajb13 |
| ENSMUSG( | 3455.882 | -0.13047 | 0.297654 | 0.746178 | Dnajb14 |
| ENSMUSG( | 3248.122 | 0.040178 | 0.627351 | 0.912941 | Dnajb2 |
| ENSMUSG( | 29.06232 | 0.065592 | 0.860492 | 0.9749 | Dnajb3 |
| ENSMUSG( | 1054.849 | 0.067943 | 0.521453 | 0.875186 | Dnajb4 |
| ENSMUSG( | 3992.07 | -0.00381 | 0.978905 | 0.997428 | Dnajb5 |
| ENSMUSG( | 1881.14 | 0.045501 | 0.602883 | 0.904887 | Dnajb6 |
| ENSMUSG( | 1282.83 | -0.44813 | 0.003967 | 0.080726 | Dnajb9 |
| ENSMUSG( | 1105.074 | 0.087877 | 0.423951 | 0.824419 | Dnajc1 |
| ENSMUSG( | 2183.218 | -0.25744 | 0.05418 | 0.357985 | Dnajc10 |
| ENSMUSG( | 1805.817 | 0.115192 | 0.262928 | 0.713397 | Dnajc11 |
| ENSMUSG( | 151.1097 | 0.21342 | 0.254365 | 0.704536 | Dnajc12 |
| ENSMUSG( | 1682.258 | 0.050133 | 0.587721 | 0.900598 | Dnajc13 |
| ENSMUSG( | 2234.962 | -0.144 | 0.126585 | 0.534155 | Dnajc14 |
| ENSMUSG( | 1362.314 | 0.034474 | 0.720079 | 0.941986 | Dnajc15 |
| ENSMUSG( | 1968.004 | -0.27397 | 0.076872 | 0.429306 | Dnajc16 |
| ENSMUSG( | 178.494 | 0.041754 | 0.824962 | 0.967386 | Dnajc17 |
| ENSMUSG( | 3336.478 | -0.1484 | 0.215199 | 0.659815 | Dnajc18 |
| ENSMUSG( | 834.6519 | -0.0751 | 0.530711 | 0.879289 | Dnajc19 |
| ENSMUSG( | 652.9448 | 0.13062 | 0.219702 | 0.665093 | Dnajc2 |
| ENSMUSG( | 556.9792 | 0.149011 | 0.216919 | 0.661816 | Dnajc21 |
| ENSMUSG( | 262.0954 | 0.146625 | 0.338737 | 0.777518 | Dnajc24 |
| ENSMUSG( | 165.107 | -0.30607 | 0.08508 | 0.449548 | Dnajc25 |
| ENSMUSG( | 6434.105 | 0.141762 | 0.061573 | 0.383386 | Dnajc27 |
| ENSMUSG( | 264.5512 | 0.16729 | 0.227354 | 0.67424 | Dnajc28 |
| ENSMUSG( | 3190.705 | 0.026203 | 0.837229 | 0.971111 | Dnajc3 |
| ENSMUSG( | 934.5928 | -0.02657 | 0.812377 | 0.964574 | Dnajc30 |
| ENSMUSG( | 475.2116 | -0.13289 | 0.232816 | 0.680928 | Dnajc4 |
| ENSMUSG( | 9651.095 | 0.037116 | 0.629679 | 0.913965 | Dnajc5 |
| ENSMUSG( | 5598.069 | 0.145919 | 0.180696 | 0.617841 | Dnajc6 |
| ENSMUSG( | 3480.351 | 0.014969 | 0.872375 | 0.978556 | Dnajc7 |
| ENSMUSG( | 2169.83 | 0.045859 | 0.63768 | 0.915811 | Dnajc8 |
| ENSMUSG( | 648.2204 | 0.005316 | 0.965761 | 0.995377 | Dnajc9 |
| ENSMUSG( | 4567.315 | 0.127489 | 0.181448 | 0.618684 | Dnal1 |

| | | | | | |
|---|---|---|---|---|---|
| ENSMUSG( | 1671.04 | 0.003152 | 0.978398 | 0.996459 | Dicer1 |
| ENSMUSG( | 1651.08 | 0.087466 | 0.533768 | 0.93547 | Dido1 |
| ENSMUSG( | 310.3218 | 0.359665 | 0.027972 | 0.61023 | Dimt1 |
| ENSMUSG( | 17903.24 | -0.14762 | 0.356049 | 0.913331 | Dio2 |
| ENSMUSG( | 57.79841 | 0.025967 | 0.932471 | 0.99306 | Dio3 |
| ENSMUSG( | 19.01952 | 0.343646 | 0.476801 | 0.923329 | Dio3os |
| ENSMUSG( | 3132.251 | -0.06034 | 0.548232 | 0.938063 | Dip2a |
| ENSMUSG( | 2054.306 | 0.07195 | 0.507067 | 0.928624 | Dip2b |
| ENSMUSG( | 1984.214 | -0.02543 | 0.831479 | 0.980611 | Dip2c |
| ENSMUSG( | 765.5 | -0.01771 | 0.918956 | 0.992529 | Dipk1a |
| ENSMUSG( | 1355.543 | -0.02166 | 0.874636 | 0.989644 | Dipk1b |
| ENSMUSG( | 409.6529 | 0.003176 | 0.984187 | 0.996963 | Dipk1c |
| ENSMUSG( | 1150.087 | 0.490839 | 0.024857 | 0.592568 | Dipk2a |
| ENSMUSG( | 37.54714 | 0.569355 | 0.26036 | 0.884524 | Dipk2b |
| ENSMUSG( | 3143.146 | -0.1106 | 0.319306 | 0.903453 | Diras1 |
| ENSMUSG( | 7457 | -0.19384 | 0.058932 | 0.729034 | Diras2 |
| ENSMUSG( | 456.3543 | -0.10309 | 0.447654 | 0.922202 | Dis3 |
| ENSMUSG( | 909.2931 | -0.06102 | 0.62111 | 0.953199 | Dis3l |
| ENSMUSG( | 469.7048 | -0.00666 | 0.961824 | 0.995237 | Dis3l2 |
| ENSMUSG( | 57.27811 | 0.181413 | 0.524752 | 0.933907 | Disc1 |
| ENSMUSG( | 247.3366 | -0.12652 | 0.556773 | 0.939565 | Disp1 |
| ENSMUSG( | 10360.78 | 0.173875 | 0.23234 | 0.868507 | Disp2 |
| ENSMUSG( | 1409.216 | -0.02899 | 0.859057 | 0.985884 | Disp3 |
| ENSMUSG( | 2510.235 | 0.091323 | 0.447921 | 0.922202 | Dixdc1 |
| ENSMUSG( | 503.3051 | 0.02702 | 0.848971 | 0.985321 | Dkc1 |
| ENSMUSG( | 25.13206 | 0.691451 | 0.116688 | 0.813384 | Dkk2 |
| ENSMUSG( | 11456.98 | 0.100498 | 0.483894 | 0.925289 | Dkk3 |
| ENSMUSG( | 615.8146 | -0.09407 | 0.491382 | 0.926825 | Dkkl1 |
| ENSMUSG( | 2364.861 | -0.12855 | 0.280185 | 0.892356 | Dlat |
| ENSMUSG( | 2015.847 | -0.02012 | 0.853236 | 0.985598 | Dlc1 |
| ENSMUSG( | 2427.518 | -0.06755 | 0.588723 | 0.944975 | Dld |
| ENSMUSG( | 85.08969 | 0.175079 | 0.471599 | 0.923329 | Dlec1 |
| ENSMUSG( | 247.1432 | 0.251171 | 0.139747 | 0.830678 | Dleu2 |
| ENSMUSG( | 83.40114 | -0.06149 | 0.820375 | 0.979198 | Dleu7 |
| ENSMUSG( | 6360.516 | -0.09935 | 0.360318 | 0.914223 | Dlg1 |
| ENSMUSG( | 19547.5 | 0.070863 | 0.566191 | 0.941665 | Dlg2 |
| ENSMUSG( | 2879.943 | -0.04415 | 0.733505 | 0.967228 | Dlg3 |
| ENSMUSG( | 50160.08 | -0.00272 | 0.976757 | 0.996416 | Dlg4 |
| ENSMUSG( | 2527.179 | 0.064178 | 0.576658 | 0.942912 | Dlg5 |
| ENSMUSG( | 14827.13 | 0.13057 | 0.201592 | 0.864412 | Dlgap1 |
| ENSMUSG( | 1784.321 | -0.18272 | 0.080958 | 0.77681 | Dlgap2 |
| ENSMUSG( | 23562.84 | 0.175256 | 0.142921 | 0.831221 | Dlgap3 |
| ENSMUSG( | 11674.5 | 0.104203 | 0.281159 | 0.892356 | Dlgap4 |
| ENSMUSG( | 12.81592 | -0.12776 | 0.829487 | 0.980224 | Dlgap5 |
| ENSMUSG( | 296.3065 | 0.213354 | 0.33732 | 0.907972 | Dlk1 |
| ENSMUSG( | 837.1084 | -0.01017 | 0.934316 | 0.99306 | Dlk2 |
| ENSMUSG( | 191.8724 | 0.342146 | 0.091925 | 0.791889 | Dll1 |
| ENSMUSG( | 79.44403 | 0.062006 | 0.819221 | 0.979198 | Dll3 |
| ENSMUSG( | 117.8352 | 0.198739 | 0.538943 | 0.937216 | Dll4 |
| ENSMUSG( | 3457.22 | -0.13977 | 0.189527 | 0.857355 | Dlst |
| ENSMUSG( | 671.453 | 0.189525 | 0.309027 | 0.898538 | Dlx1 |
| ENSMUSG( | 381.8065 | -0.08012 | 0.603838 | 0.948137 | Dlx1as |
| ENSMUSG( | 180.5457 | 0.221768 | 0.370369 | 0.915491 | Dlx2 |
| ENSMUSG( | 105.7695 | -0.1827 | 0.523715 | 0.932848 | Dlx5 |
| ENSMUSG( | 124.6198 | 0.137799 | 0.644418 | 0.953903 | Dlx6 |
| ENSMUSG( | 279.6242 | 0.059819 | 0.718832 | 0.966668 | Dlx6os1 |
| ENSMUSG( | 453.1253 | -0.10853 | 0.424049 | 0.917257 | Dmac1 |
| ENSMUSG( | 967.9337 | 0.070887 | 0.561956 | 0.940388 | Dmac2 |
| ENSMUSG( | 362.2255 | -0.1565 | 0.381743 | 0.917257 | Dmac2l |
| ENSMUSG( | 489.3295 | -0.1194 | 0.468235 | 0.923329 | Dmap1 |
| ENSMUSG( | 29.87365 | 0.53292 | 0.19636 | 0.862161 | Dmc1 |
| ENSMUSG( | 2115.906 | 0.06565 | 0.609746 | 0.950397 | Dmd |
| ENSMUSG( | 10.39411 | -0.62845 | 0.329384 | NA | Dmgdh |
| ENSMUSG( | 70.45418 | -0.06536 | 0.826152 | 0.979795 | Dmkn |
| ENSMUSG( | 673.0371 | -0.05988 | 0.675461 | 0.95694 | Dmp1 |
| ENSMUSG( | 1113.072 | 0.095521 | 0.500676 | 0.927825 | Dmpk |
| ENSMUSG( | 20.92579 | 0.263392 | 0.549777 | 0.938699 | Dmrt2 |
| ENSMUSG( | 15.06551 | 0.545299 | 0.281883 | 0.892412 | Dmrt3 |
| ENSMUSG( | 14.38882 | -0.31102 | 0.572025 | 0.942369 | Dmrta1 |
| ENSMUSG( | 180.5299 | 0.313633 | 0.100904 | 0.801456 | Dmrta2 |
| ENSMUSG( | 51.91495 | -0.04239 | 0.881419 | 0.989644 | Dmrtb1 |
| ENSMUSG( | 68.2992 | -0.00107 | 0.996846 | 0.999268 | Dmrtc1a |
| ENSMUSG( | 1305.787 | 0.049832 | 0.706856 | 0.963529 | Dmtf1 |
| ENSMUSG( | 7041.716 | -0.0115 | 0.911831 | 0.992196 | Dmtn |
| ENSMUSG( | 4203.452 | -0.01383 | 0.903436 | 0.991089 | Dmwd |
| ENSMUSG( | 1036.931 | 0.263936 | 0.021689 | 0.575341 | Dmxl1 |
| ENSMUSG( | 3398.057 | 0.176167 | 0.150195 | 0.837086 | Dmxl2 |
| ENSMUSG( | 52.35737 | 0.44971 | 0.16642 | 0.84011 | Dna2 |
| ENSMUSG( | 13.33073 | 0.194996 | 0.745185 | 0.96951 | Dnaaf1 |
| ENSMUSG( | 143.0483 | 0.124899 | 0.523315 | 0.932848 | Dnaaf10 |
| ENSMUSG( | 59.99132 | -0.07524 | 0.829625 | 0.980224 | Dnaaf11 |
| ENSMUSG( | 28.53502 | 0.138478 | 0.731791 | 0.967228 | Dnaaf2 |
| ENSMUSG( | 41.50478 | 0.115811 | 0.754073 | 0.970686 | Dnaaf3 |
| ENSMUSG( | 30.84798 | -0.23187 | 0.556453 | 0.939565 | Dnaaf4 |
| ENSMUSG( | 245.1328 | 0.010748 | 0.946531 | 0.994798 | Dnaaf5 |
| ENSMUSG( | 6449.305 | -0.23818 | 0.038336 | 0.679117 | Dnaaf9 |
| ENSMUSG( | 259.2828 | -0.21754 | 0.195584 | 0.861682 | Dnah1 |
| ENSMUSG( | 90.21389 | 0.098353 | 0.685196 | 0.958678 | Dnah10 |
| ENSMUSG( | 58.09731 | 0.070535 | 0.828613 | 0.980224 | Dnah11 |
| ENSMUSG( | 44.15704 | 0.193075 | 0.53954 | 0.937378 | Dnah12 |
| ENSMUSG( | 48.39699 | 0.444736 | 0.156877 | 0.837724 | Dnah14 |
| ENSMUSG( | 19.33793 | -0.05129 | 0.909463 | 0.991958 | Dnah17 |
| ENSMUSG( | 143.9489 | 0.169053 | 0.398393 | 0.917257 | Dnah2 |
| ENSMUSG( | 28.53476 | 0.347678 | 0.453699 | 0.922316 | Dnah3 |
| ENSMUSG( | 135.1157 | 0.137081 | 0.535294 | 0.936508 | Dnah5 |
| ENSMUSG( | 93.54354 | 0.483819 | 0.110831 | 0.807582 | Dnah6 |
| ENSMUSG( | 78.45713 | -0.27548 | 0.254929 | 0.882307 | Dnah7a |
| ENSMUSG( | 191.5602 | -0.04858 | 0.818892 | 0.979198 | Dnah7b |
| ENSMUSG( | 48.1506 | 0.485871 | 0.104685 | 0.804969 | Dnah7c |
| ENSMUSG( | 19.52168 | -0.32865 | 0.464109 | 0.923329 | Dnah8 |
| ENSMUSG( | 179.6159 | 0.273781 | 0.213327 | 0.864412 | Dnah9 |
| ENSMUSG( | 332.0579 | -0.33225 | 0.050407 | 0.708648 | Dnai1 |
| ENSMUSG( | 127.2234 | -0.25022 | 0.290839 | 0.897019 | Dnai2 |
| ENSMUSG( | 23.56027 | 0.778565 | 0.087893 | 0.785919 | Dnai3 |
| ENSMUSG( | 197.4239 | 0.099444 | 0.563684 | 0.941206 | Dnai4 |
| ENSMUSG( | 45.54032 | -0.20121 | 0.530501 | 0.934795 | Dnai7 |
| ENSMUSG( | 2923.82 | -0.01878 | 0.887252 | 0.990263 | Dnaja1 |
| ENSMUSG( | 2566.225 | 0.00401 | 0.972487 | 0.996416 | Dnaja2 |
| ENSMUSG( | 1399.166 | -0.1041 | 0.428685 | 0.917257 | Dnaja3 |
| ENSMUSG( | 1008.852 | -0.02186 | 0.861638 | 0.986298 | Dnaja4 |
| ENSMUSG( | 691.4193 | 0.176636 | 0.280003 | 0.892356 | Dnajb1 |
| ENSMUSG( | 2731.673 | 0.155078 | 0.321523 | 0.904019 | Dnajb11 |
| ENSMUSG( | 2052.36 | 0.121225 | 0.435383 | 0.919062 | Dnajb12 |
| ENSMUSG( | 27.42951 | -0.03483 | 0.928631 | 0.99306 | Dnajb13 |
| ENSMUSG( | 3464.933 | 0.145595 | 0.344769 | 0.91045 | Dnajb14 |
| ENSMUSG( | 2787.29 | -0.13666 | 0.184213 | 0.852665 | Dnajb2 |
| ENSMUSG( | 23.78365 | -0.27344 | 0.525618 | 0.934374 | Dnajb3 |
| ENSMUSG( | 993.023 | 0.154912 | 0.257923 | 0.883466 | Dnajb4 |
| ENSMUSG( | 3369.86 | -0.23653 | 0.167551 | 0.840222 | Dnajb5 |
| ENSMUSG( | 1620.384 | -0.12152 | 0.319756 | 0.903706 | Dnajb6 |
| ENSMUSG( | 1521.328 | 0.33058 | 0.068041 | 0.75212 | Dnajb9 |
| ENSMUSG( | 1048.672 | 0.200924 | 0.102853 | 0.803962 | Dnajc1 |
| ENSMUSG( | 2361.341 | 0.243062 | 0.134825 | 0.824049 | Dnajc10 |
| ENSMUSG( | 1548.209 | -0.05889 | 0.627626 | 0.953234 | Dnajc11 |
| ENSMUSG( | 131.7109 | 0.074498 | 0.730752 | 0.967228 | Dnajc12 |
| ENSMUSG( | 1554.858 | 0.093969 | 0.429088 | 0.917257 | Dnajc13 |
| ENSMUSG( | 2192.897 | 0.072806 | 0.529179 | 0.934795 | Dnajc14 |
| ENSMUSG( | 1211.596 | -0.03981 | 0.758851 | 0.971858 | Dnajc15 |
| ENSMUSG( | 2122.68 | 0.223764 | 0.208773 | 0.864412 | Dnajc16 |
| ENSMUSG( | 167.9088 | 0.127215 | 0.564966 | 0.941248 | Dnajc17 |
| ENSMUSG( | 3270.356 | 0.069173 | 0.627122 | 0.953234 | Dnajc18 |
| ENSMUSG( | 760.3792 | -0.07834 | 0.560531 | 0.940254 | Dnajc19 |
| ENSMUSG( | 555.5329 | -0.07076 | 0.611629 | 0.950771 | Dnajc2 |
| ENSMUSG( | 499.5593 | 0.104699 | 0.490116 | 0.926144 | Dnajc21 |
| ENSMUSG( | 204.4381 | -0.30789 | 0.070385 | 0.758336 | Dnajc24 |
| ENSMUSG( | 176.6376 | 0.171468 | 0.365538 | 0.914603 | Dnajc25 |
| ENSMUSG( | 5531.166 | -0.02262 | 0.814517 | 0.979198 | Dnajc27 |
| ENSMUSG( | 225.8377 | -0.01259 | 0.93763 | 0.993685 | Dnajc28 |
| ENSMUSG( | 3172.348 | 0.267898 | 0.051255 | 0.711023 | Dnajc3 |
| ENSMUSG( | 808.9203 | -0.18484 | 0.187339 | 0.856372 | Dnajc30 |
| ENSMUSG( | 425.6456 | -0.1841 | 0.174236 | 0.844317 | Dnajc4 |
| ENSMUSG( | 8509.537 | -0.05711 | 0.550231 | 0.938699 | Dnajc5 |
| ENSMUSG( | 5142.9 | 0.171634 | 0.160012 | 0.837724 | Dnajc6 |
| ENSMUSG( | 3096.059 | -0.05524 | 0.631335 | 0.953251 | Dnajc7 |
| ENSMUSG( | 1872.045 | -0.11669 | 0.384201 | 0.917257 | Dnajc8 |
| ENSMUSG( | 562.5066 | -0.1481 | 0.359668 | 0.914223 | Dnajc9 |
| ENSMUSG( | 3955.093 | -0.01686 | 0.881319 | 0.989644 | Dnal1 |
| ENSMUSG( | 952.556 | -0.06203 | 0.67301 | 0.95694 | Dnal4 |
| ENSMUSG( | 56.2693 | 0.573893 | 0.073257 | 0.763629 | Dnali1 |
| ENSMUSG( | 66.72337 | 0.23812 | 0.407682 | 0.917257 | Dnase1l1 |
| ENSMUSG( | 146.9766 | -0.00937 | 0.968808 | 0.99594 | Dnase1l2 |
| ENSMUSG( | 296.5036 | -0.04953 | 0.778554 | 0.976106 | Dnase2a |
| ENSMUSG( | 38.90413 | -0.17766 | 0.625843 | 0.953234 | Dnd1 |
| ENSMUSG( | 5941.917 | 0.119298 | 0.4853 | 0.92541 | Dner |
| ENSMUSG( | 233.0624 | 0.200496 | 0.278619 | 0.891628 | Dnhd1 |
| ENSMUSG( | 1799.394 | -0.05562 | 0.615305 | 0.951177 | Dnlz |
| ENSMUSG( | 20475.47 | -0.02408 | 0.868398 | 0.988031 | Dnm1 |
| ENSMUSG( | 3993.216 | -0.0799 | 0.535971 | 0.936952 | Dnm1l |

| | | | | | | | | | | | |
|---|---|---|---|---|---|---|---|---|---|---|---|
| ENSMUSG( | Dkk3 | 11179.97 | -0.25124 | 0.045562 | 0.327592 | ENSMUSG( | Dkk3 | 11456.98 | 0.100498 | 0.483894 | 0.925289 |
| ENSMUSG( | Dkkl1 | 660.3032 | -0.169 | 0.138022 | 0.552905 | ENSMUSG( | Dkkl1 | 615.8146 | -0.09407 | 0.491382 | 0.926825 |
| ENSMUSG( | Dlat | 2796.486 | 0.090732 | 0.295113 | 0.743798 | ENSMUSG( | Dlat | 2364.861 | -0.12855 | 0.280185 | 0.892356 |
| ENSMUSG( | Dlc1 | 2270.548 | 0.054815 | 0.562403 | 0.889448 | ENSMUSG( | Dlc1 | 2015.847 | -0.02012 | 0.853236 | 0.985598 |
| ENSMUSG( | Dld | 2812.103 | 0.090813 | 0.379689 | 0.801757 | ENSMUSG( | Dld | 2427.518 | -0.06755 | 0.588723 | 0.944975 |
| ENSMUSG( | Dlec1 | 91.87679 | 0.11397 | 0.607934 | 0.907063 | ENSMUSG( | Dlec1 | 85.08969 | 0.175079 | 0.471599 | 0.923329 |
| ENSMUSG( | Dleu2 | 254.9098 | 0.074133 | 0.604556 | 0.905556 | ENSMUSG( | Dleu2 | 247.1432 | 0.251171 | 0.139747 | 0.830678 |
| ENSMUSG( | Dleu7 | 97.18024 | 0.130596 | 0.576285 | 0.895164 | ENSMUSG( | Dleu7 | 83.40114 | -0.06149 | 0.820375 | 0.979198 |
| ENSMUSG( | Dlg1 | 7179.763 | -0.01559 | 0.854023 | 0.973491 | ENSMUSG( | Dlg1 | 6360.516 | -0.09935 | 0.360318 | 0.914223 |
| ENSMUSG( | Dlg2 | 21791.38 | 0.112714 | 0.304388 | 0.752574 | ENSMUSG( | Dlg2 | 19547.5 | 0.070863 | 0.566191 | 0.941665 |
| ENSMUSG( | Dlg3 | 3364.999 | 0.13458 | 0.253208 | 0.70338 | ENSMUSG( | Dlg3 | 2879.943 | -0.04415 | 0.733505 | 0.967228 |
| ENSMUSG( | Dlg4 | 57368.56 | 0.113481 | 0.134184 | 0.547472 | ENSMUSG( | Dlg4 | 50160.08 | -0.00272 | 0.976757 | 0.996416 |
| ENSMUSG( | Dlg5 | 2993.803 | 0.272705 | 0.002016 | 0.051507 | ENSMUSG( | Dlg5 | 2527.179 | 0.064178 | 0.576658 | 0.942912 |
| ENSMUSG( | Dlgap1 | 16001.04 | 0.083269 | 0.309639 | 0.757658 | ENSMUSG( | Dlgap1 | 14827.13 | 0.13057 | 0.201592 | 0.864412 |
| ENSMUSG( | Dlgap2 | 2159.651 | 0.102796 | 0.245489 | 0.693175 | ENSMUSG( | Dlgap2 | 1784.321 | -0.18272 | 0.080958 | 0.77681 |
| ENSMUSG( | Dlgap3 | 26460.41 | 0.236789 | 0.00445 | 0.086961 | ENSMUSG( | Dlgap3 | 23562.84 | 0.175256 | 0.142921 | 0.831221 |
| ENSMUSG( | Dlgap4 | 13256.41 | 0.196882 | 0.011355 | 0.153509 | ENSMUSG( | Dlgap4 | 11674.5 | 0.104203 | 0.281159 | 0.892356 |
| ENSMUSG( | Dlgap5 | 14.487 | -0.09028 | 0.866651 | NA | ENSMUSG( | Dlgap5 | 12.81592 | -0.12776 | 0.829487 | 0.980224 |
| ENSMUSG( | Dlk1 | 295.362 | -0.04269 | 0.833607 | 0.970333 | ENSMUSG( | Dlk1 | 296.3065 | 0.213354 | 0.33732 | 0.907972 |
| ENSMUSG( | Dlk2 | 946.2704 | 0.071369 | 0.561105 | 0.888934 | ENSMUSG( | Dlk2 | 837.1084 | -0.01017 | 0.934316 | 0.99306 |
| ENSMUSG( | Dll1 | 167.3622 | -0.3327 | 0.044837 | 0.324745 | ENSMUSG( | Dll1 | 191.8724 | 0.342146 | 0.091925 | 0.791889 |
| ENSMUSG( | Dll3 | 77.42804 | -0.30815 | 0.280623 | 0.729679 | ENSMUSG( | Dll3 | 79.44403 | 0.062006 | 0.819221 | 0.979198 |
| ENSMUSG( | Dll4 | 133.525 | 0.25996 | 0.324118 | 0.769158 | ENSMUSG( | Dll4 | 117.8352 | 0.198739 | 0.538943 | 0.937216 |
| ENSMUSG( | Dlst | 4100.217 | 0.086946 | 0.294489 | 0.743674 | ENSMUSG( | Dlst | 3457.22 | -0.13977 | 0.189527 | 0.857355 |
| ENSMUSG( | Dlx1 | 660.6026 | -0.11616 | 0.424066 | 0.824452 | ENSMUSG( | Dlx1 | 671.453 | 0.189525 | 0.309027 | 0.898538 |
| ENSMUSG( | Dlx1as | 444.532 | 0.090039 | 0.489884 | 0.859319 | ENSMUSG( | Dlx1as | 381.8065 | -0.08012 | 0.603838 | 0.948137 |
| ENSMUSG( | Dlx2 | 163.2649 | -0.33457 | 0.134334 | 0.547472 | ENSMUSG( | Dlx2 | 180.5457 | 0.221768 | 0.370369 | 0.915491 |
| ENSMUSG( | Dlx5 | 101.8905 | -0.59629 | 0.028109 | 0.25643 | ENSMUSG( | Dlx5 | 105.7695 | -0.1827 | 0.523715 | 0.932848 |
| ENSMUSG( | Dlx6 | 113.1671 | -0.41705 | 0.114697 | 0.510718 | ENSMUSG( | Dlx6 | 124.6198 | 0.137799 | 0.644418 | 0.953903 |
| ENSMUSG( | Dlx6os1 | 285.7178 | -0.13876 | 0.409752 | 0.818258 | ENSMUSG( | Dlx6os1 | 279.6242 | 0.059819 | 0.718832 | 0.966668 |
| ENSMUSG( | Dmac1 | 526.1276 | 0.05863 | 0.608585 | 0.907154 | ENSMUSG( | Dmac1 | 453.1253 | -0.10853 | 0.424049 | 0.917257 |
| ENSMUSG( | Dmac2 | 1067.828 | 0.08803 | 0.418523 | 0.821966 | ENSMUSG( | Dmac2 | 967.9337 | 0.070887 | 0.561956 | 0.940388 |
| ENSMUSG( | Dmac2l | 421.4965 | 0.026889 | 0.862384 | 0.975055 | ENSMUSG( | Dmac2l | 362.2255 | -0.1565 | 0.381743 | 0.917257 |
| ENSMUSG( | Dmap1 | 568.8976 | 0.055559 | 0.704759 | 0.938303 | ENSMUSG( | Dmap1 | 489.3295 | -0.1194 | 0.468235 | 0.923329 |
| ENSMUSG( | Dmc1 | 35.33619 | 0.731224 | 0.037886 | 0.30067 | ENSMUSG( | Dmc1 | 29.87365 | 0.53292 | 0.19636 | 0.862161 |
| ENSMUSG( | Dmd | 2293.579 | 0.02971 | 0.80529 | 0.962616 | ENSMUSG( | Dmd | 2115.906 | 0.06565 | 0.609746 | 0.950397 |
| ENSMUSG( | Dmgdh | 14.92776 | 0.148251 | 0.754428 | NA | ENSMUSG( | Dmgdh | 10.39411 | -0.62845 | 0.329384 | NA |
| ENSMUSG( | Dmkn | 67.38936 | -0.53104 | 0.081275 | 0.440604 | ENSMUSG( | Dmkn | 70.45418 | -0.06536 | 0.826152 | 0.979795 |
| ENSMUSG( | Dmp1 | 751.5319 | -0.01064 | 0.936361 | 0.993415 | ENSMUSG( | Dmp1 | 673.0371 | -0.05988 | 0.675461 | 0.95694 |
| ENSMUSG( | Dmpk | 1228.24 | 0.114199 | 0.321823 | 0.766739 | ENSMUSG( | Dmpk | 1113.072 | 0.095521 | 0.500676 | 0.927825 |
| ENSMUSG( | Dmrt2 | 19.35193 | -0.21027 | 0.653747 | NA | ENSMUSG( | Dmrt2 | 20.92579 | 0.263392 | 0.549777 | 0.938699 |
| ENSMUSG( | Dmrt3 | 15.72119 | 0.392301 | 0.402169 | NA | ENSMUSG( | Dmrt3 | 15.06551 | 0.545299 | 0.281883 | 0.892412 |
| ENSMUSG( | Dmrta1 | 17.03579 | -0.11365 | 0.803313 | NA | ENSMUSG( | Dmrta1 | 14.38882 | -0.31102 | 0.572025 | 0.942369 |
| ENSMUSG( | Dmrta2 | 196.0216 | 0.290755 | 0.096389 | 0.476986 | ENSMUSG( | Dmrta2 | 180.5299 | 0.313633 | 0.100904 | 0.801456 |
| ENSMUSG( | Dmrtb1 | 52.24031 | -0.31536 | 0.301916 | 0.750166 | ENSMUSG( | Dmrtb1 | 51.91495 | -0.04239 | 0.881419 | 0.989644 |
| ENSMUSG( | Dmrtc1a | 73.87677 | -0.03071 | 0.901077 | 0.985633 | ENSMUSG( | Dmrtc1a | 68.2992 | -0.00107 | 0.996846 | 0.999268 |
| ENSMUSG( | Dmtf1 | 1448.295 | 0.079826 | 0.424423 | 0.824645 | ENSMUSG( | Dmtf1 | 1305.787 | 0.049832 | 0.706856 | 0.963529 |
| ENSMUSG( | Dmtn | 8183.712 | 0.149894 | 0.098682 | 0.480598 | ENSMUSG( | Dmtn | 7041.716 | -0.0115 | 0.911831 | 0.992196 |
| ENSMUSG( | Dmwd | 4829.692 | 0.116588 | 0.244891 | 0.692492 | ENSMUSG( | Dmwd | 4203.452 | -0.01383 | 0.903436 | 0.991089 |
| ENSMUSG( | Dmxl1 | 1012.814 | -0.06285 | 0.507388 | 0.86795 | ENSMUSG( | Dmxl1 | 1036.931 | 0.263936 | 0.021689 | 0.575341 |
| ENSMUSG( | Dmxl2 | 3385.968 | -0.10251 | 0.343809 | 0.779153 | ENSMUSG( | Dmxl2 | 3398.057 | 0.176167 | 0.150195 | 0.837086 |
| ENSMUSG( | Dna2 | 47.66056 | -0.08066 | 0.785224 | 0.958473 | ENSMUSG( | Dna2 | 52.35737 | 0.44971 | 0.16642 | 0.84011 |
| ENSMUSG( | Dnaaf1 | 13.537 | -0.08069 | 0.889413 | NA | ENSMUSG( | Dnaaf1 | 13.33073 | 0.194996 | 0.745185 | 0.96951 |
| ENSMUSG( | Dnaaf10 | 150.2583 | 0.004678 | 0.979611 | 0.997428 | ENSMUSG( | Dnaaf10 | 143.0483 | 0.124899 | 0.523315 | 0.932848 |
| ENSMUSG( | Dnaaf11 | 57.61442 | -0.5192 | 0.131931 | 0.544244 | ENSMUSG( | Dnaaf11 | 59.99132 | -0.07524 | 0.829625 | 0.980224 |
| ENSMUSG( | Dnaaf2 | 29.7591 | 0.007864 | 0.983681 | 0.997768 | ENSMUSG( | Dnaaf2 | 28.53502 | 0.138478 | 0.731791 | 0.967228 |
| ENSMUSG( | Dnaaf3 | 38.90733 | -0.397 | 0.269978 | 0.719883 | ENSMUSG( | Dnaaf3 | 41.50478 | 0.115811 | 0.754073 | 0.970686 |
| ENSMUSG( | Dnaaf4 | 33.89178 | -0.19641 | 0.579964 | 0.896931 | ENSMUSG( | Dnaaf4 | 30.84798 | -0.23187 | 0.556453 | 0.939565 |
| ENSMUSG( | Dnaaf5 | 275.2795 | 0.077574 | 0.569605 | 0.893041 | ENSMUSG( | Dnaaf5 | 245.1328 | 0.010748 | 0.946531 | 0.994798 |
| ENSMUSG( | Dnaaf9 | 7777.765 | 0.043783 | 0.617832 | 0.910872 | ENSMUSG( | Dnaaf9 | 6449.305 | -0.23818 | 0.038336 | 0.679117 |
| ENSMUSG( | Dnah1 | 300.6297 | -0.06277 | 0.636132 | 0.914873 | ENSMUSG( | Dnah1 | 259.2828 | -0.21754 | 0.195584 | 0.861682 |
| ENSMUSG( | Dnah10 | 95.17167 | -0.01979 | 0.941968 | 0.993415 | ENSMUSG( | Dnah10 | 90.21389 | 0.098353 | 0.685196 | 0.958678 |
| ENSMUSG( | Dnah11 | 59.61797 | -0.13091 | 0.682557 | 0.93198 | ENSMUSG( | Dnah11 | 58.09731 | 0.070535 | 0.828613 | 0.980224 |
| ENSMUSG( | Dnah12 | 43.93105 | -0.09265 | 0.761912 | 0.95339 | ENSMUSG( | Dnah12 | 44.15704 | 0.193075 | 0.53954 | 0.937378 |
| ENSMUSG( | Dnah14 | 42.96341 | -0.15487 | 0.606537 | 0.906481 | ENSMUSG( | Dnah14 | 48.39699 | 0.444736 | 0.156877 | 0.837724 |
| ENSMUSG( | Dnah17 | 18.42374 | -0.51457 | 0.265626 | NA | ENSMUSG( | Dnah17 | 19.33793 | -0.05129 | 0.909463 | 0.991958 |
| ENSMUSG( | Dnah2 | 154.8652 | 0.111198 | 0.544404 | 0.884649 | ENSMUSG( | Dnah2 | 143.9489 | 0.169053 | 0.398393 | 0.917257 |
| ENSMUSG( | Dnah3 | 30.05308 | 0.195784 | 0.60292 | 0.904887 | ENSMUSG( | Dnah3 | 28.53476 | 0.347678 | 0.453699 | 0.922316 |
| ENSMUSG( | Dnah5 | 127.5129 | -0.33153 | 0.082186 | 0.442642 | ENSMUSG( | Dnah5 | 135.1157 | 0.137081 | 0.535294 | 0.936508 |
| ENSMUSG( | Dnah6 | 88.73038 | 0.081536 | 0.768042 | 0.954621 | ENSMUSG( | Dnah6 | 93.54354 | 0.483819 | 0.110831 | 0.807582 |
| ENSMUSG( | Dnah7a | 100.5281 | 0.180269 | 0.372952 | 0.799429 | ENSMUSG( | Dnah7a | 78.45713 | -0.27548 | 0.254929 | 0.882307 |
| ENSMUSG( | Dnah7b | 212.4318 | -0.02694 | 0.855623 | 0.973813 | ENSMUSG( | Dnah7b | 191.5602 | -0.04858 | 0.818892 | 0.979198 |
| ENSMUSG( | Dnah7c | 45.52099 | 0.097787 | 0.726212 | 0.944223 | ENSMUSG( | Dnah7c | 48.1506 | 0.485871 | 0.104685 | 0.804969 |
| ENSMUSG( | Dnah8 | 21.95142 | -0.24593 | 0.56378 | 0.890032 | ENSMUSG( | Dnah8 | 19.52168 | -0.32865 | 0.464109 | 0.923329 |
| ENSMUSG( | Dnah9 | 171.51 | -0.13588 | 0.541732 | 0.883182 | ENSMUSG( | Dnah9 | 179.6159 | 0.273781 | 0.213327 | 0.864412 |
| ENSMUSG( | Dnai1 | 411.4435 | 0.048524 | 0.709905 | 0.939669 | ENSMUSG( | Dnai1 | 332.0579 | -0.33225 | 0.050407 | 0.708648 |
| ENSMUSG( | Dnai2 | 137.749 | -0.29074 | 0.208573 | 0.650496 | ENSMUSG( | Dnai2 | 127.2234 | -0.25022 | 0.290839 | 0.897019 |
| ENSMUSG( | Dnai3 | 20.46355 | 0.160781 | 0.710702 | NA | ENSMUSG( | Dnai3 | 23.56027 | 0.778565 | 0.087893 | 0.785919 |
| ENSMUSG( | Dnai4 | 207.2679 | -0.0301 | 0.848592 | 0.972374 | ENSMUSG( | Dnai4 | 197.4239 | 0.099444 | 0.563684 | 0.941206 |
| ENSMUSG( | Dnai7 | 50.84946 | -0.1854 | 0.508901 | 0.868744 | ENSMUSG( | Dnai7 | 45.54032 | -0.20121 | 0.530501 | 0.934795 |
| ENSMUSG( | Dnaja1 | 3240.309 | 0.012383 | 0.891411 | 0.983919 | ENSMUSG( | Dnaja1 | 2923.82 | -0.01878 | 0.887252 | 0.990263 |
| ENSMUSG( | Dnaja2 | 2895.968 | 0.085322 | 0.319252 | 0.765223 | ENSMUSG( | Dnaja2 | 2566.225 | 0.00401 | 0.972487 | 0.996416 |
| ENSMUSG( | Dnaja3 | 1619.97 | 0.053998 | 0.60133 | 0.904603 | ENSMUSG( | Dnaja3 | 1399.166 | -0.1041 | 0.428685 | 0.917257 |
| ENSMUSG( | Dnaja4 | 1171.105 | 0.139815 | 0.166081 | 0.598098 | ENSMUSG( | Dnaja4 | 1008.852 | -0.02186 | 0.861638 | 0.986298 |
| ENSMUSG( | Dnajb1 | 751.7078 | 0.150682 | 0.300655 | 0.749904 | ENSMUSG( | Dnajb1 | 691.4193 | 0.176636 | 0.280003 | 0.892356 |
| ENSMUSG( | Dnajb11 | 2676.96 | -0.17679 | 0.196171 | 0.635538 | ENSMUSG( | Dnajb11 | 2731.673 | 0.155078 | 0.321523 | 0.904019 |
| ENSMUSG( | Dnajb12 | 1925.368 | -0.35611 | 0.005961 | 0.104409 | ENSMUSG( | Dnajb12 | 2052.36 | 0.121225 | 0.435383 | 0.919062 |
| ENSMUSG( | Dnajb13 | 26.62622 | -0.39611 | 0.290998 | 0.741182 | ENSMUSG( | Dnajb13 | 27.42951 | -0.03483 | 0.928631 | 0.99306 |
| ENSMUSG( | Dnajb14 | 3455.882 | -0.13047 | 0.297654 | 0.746178 | ENSMUSG( | Dnajb14 | 3464.933 | 0.145595 | 0.344769 | 0.91045 |
| ENSMUSG( | Dnajb2 | 3248.122 | 0.040178 | 0.627351 | 0.912941 | ENSMUSG( | Dnajb2 | 2787.29 | -0.13666 | 0.184213 | 0.852665 |
| ENSMUSG( | Dnajb3 | 29.06232 | 0.065592 | 0.860492 | 0.9749 | ENSMUSG( | Dnajb3 | 23.78365 | -0.27344 | 0.525618 | 0.934374 |
| ENSMUSG( | Dnajb4 | 1054.849 | 0.067943 | 0.521453 | 0.875186 | ENSMUSG( | Dnajb4 | 993.023 | 0.154912 | 0.257923 | 0.883466 |
| ENSMUSG( | Dnajb5 | 3992.07 | -0.00381 | 0.978905 | 0.997428 | ENSMUSG( | Dnajb5 | 3369.86 | -0.23653 | 0.167551 | 0.840222 |
| ENSMUSG( | Dnajb6 | 1881.14 | 0.045501 | 0.602883 | 0.904887 | ENSMUSG( | Dnajb6 | 1620.384 | -0.12152 | 0.319756 | 0.903706 |
| ENSMUSG( | Dnajb9 | 1282.83 | -0.44813 | 0.003967 | 0.080726 | ENSMUSG( | Dnajb9 | 1521.328 | 0.33058 | 0.068041 | 0.75212 |
| ENSMUSG( | Dnajc1 | 1105.074 | 0.087877 | 0.423951 | 0.824419 | ENSMUSG( | Dnajc1 | 1048.672 | 0.200924 | 0.102853 | 0.803962 |
| ENSMUSG( | Dnajc10 | 2183.218 | -0.25744 | 0.05418 | 0.357985 | ENSMUSG( | Dnajc10 | 2361.341 | 0.243062 | 0.134825 | 0.824049 |
| ENSMUSG( | Dnajc11 | 1805.817 | 0.115192 | 0.262928 | 0.713397 | ENSMUSG( | Dnajc11 | 1548.209 | -0.05889 | 0.627626 | 0.953234 |
| ENSMUSG( | Dnajc12 | 151.1097 | 0.21342 | 0.254365 | 0.704536 | ENSMUSG( | Dnajc12 | 131.7109 | 0.074498 | 0.730752 | 0.967228 |
| ENSMUSG( | Dnajc13 | 1682.258 | 0.050133 | 0.587721 | 0.900598 | ENSMUSG( | Dnajc13 | 1554.858 | 0.093969 | 0.429088 | 0.917257 |
| ENSMUSG( | Dnajc14 | 2234.962 | -0.144 | 0.126585 | 0.534155 | ENSMUSG( | Dnajc14 | 2192.897 | 0.072806 | 0.529179 | 0.934795 |
| ENSMUSG( | Dnajc15 | 1362.314 | 0.034474 | 0.720079 | 0.941986 | ENSMUSG( | Dnajc15 | 1211.596 | -0.03981 | 0.758851 | 0.971858 |
| ENSMUSG( | Dnajc16 | 1968.004 | -0.27397 | 0.076872 | 0.429306 | ENSMUSG( | Dnajc16 | 2122.68 | 0.223764 | 0.208773 | 0.864412 |
| ENSMUSG( | Dnajc17 | 178.494 | 0.041754 | 0.824962 | 0.967386 | ENSMUSG( | Dnajc17 | 167.9088 | 0.127215 | 0.564966 | 0.941248 |
| ENSMUSG( | Dnajc18 | 3336.478 | -0.1484 | 0.215199 | 0.659815 | ENSMUSG( | Dnajc18 | 3270.356 | 0.069173 | 0.627122 | 0.953234 |
| ENSMUSG( | Dnajc19 | 834.6519 | -0.0751 | 0.530711 | 0.879289 | ENSMUSG( | Dnajc19 | 760.3792 | -0.07834 | 0.560531 | 0.940254 |
| ENSMUSG( | Dnajc2 | 652.9448 | 0.13062 | 0.219702 | 0.665093 | ENSMUSG( | Dnajc2 | 555.5329 | -0.07076 | 0.611629 | 0.950771 |
| ENSMUSG( | Dnajc21 | 556.9792 | 0.149011 | 0.216919 | 0.661816 | ENSMUSG( | Dnajc21 | 499.5593 | 0.104699 | 0.490116 | 0.926144 |
| ENSMUSG( | Dnajc24 | 262.0954 | 0.146625 | 0.338737 | 0.777518 | ENSMUSG( | Dnajc24 | 204.4381 | -0.30789 | 0.070385 | 0.758336 |
| ENSMUSG( | Dnajc25 | 165.107 | -0.30607 | 0.08508 | 0.449548 | ENSMUSG( | Dnajc25 | 176.6376 | 0.171468 | 0.365538 | 0.914603 |
| ENSMUSG( | Dnajc27 | 6434.105 | 0.141762 | 0.061573 | 0.383386 | ENSMUSG( | Dnajc27 | 5531.166 | -0.02262 | 0.814517 | 0.979198 |
| ENSMUSG( | Dnajc28 | 264.5512 | 0.16729 | 0.227354 | 0.67424 | ENSMUSG( | Dnajc28 | 225.8377 | -0.01259 | 0.93763 | 0.993685 |
| ENSMUSG( | Dnajc3 | 3190.705 | 0.026203 | 0.837229 | 0.971111 | ENSMUSG( | Dnajc3 | 3172.348 | 0.267898 | 0.051255 | 0.711023 |
| ENSMUSG( | Dnajc30 | 934.5928 | -0.02657 | 0.812377 | 0.964574 | ENSMUSG( | Dnajc30 | 808.9203 | -0.18484 | 0.187339 | 0.856372 |
| ENSMUSG( | Dnajc4 | 475.2116 | -0.13289 | 0.232816 | 0.680928 | ENSMUSG( | Dnajc4 | 425.6456 | -0.1841 | 0.174236 | 0.844317 |
| ENSMUSG( | Dnajc5 | 9651.095 | 0.037116 | 0.629679 | 0.913965 | ENSMUSG( | Dnajc5 | 8509.537 | -0.05711 | 0.550231 | 0.938699 |
| ENSMUSG( | Dnajc6 | 5598.069 | 0.145919 | 0.180696 | 0.617841 | ENSMUSG( | Dnajc6 | 5142.9 | 0.171634 | 0.160012 | 0.837724 |
| ENSMUSG( | Dnajc7 | 3480.351 | 0.014969 | 0.872375 | 0.978556 | ENSMUSG( | Dnajc7 | 3096.059 | -0.05524 | 0.631335 | 0.953251 |
| ENSMUSG( | Dnajc8 | 2169.83 | 0.045859 | 0.63768 | 0.915811 | ENSMUSG( | Dnajc8 | 1872.045 | -0.11669 | 0.384201 | 0.917257 |
| ENSMUSG( | Dnajc9 | 648.2204 | 0.005316 | 0.965761 | 0.995377 | ENSMUSG( | Dnajc9 | 562.5066 | -0.1481 | 0.359668 | 0.914223 |
| ENSMUSG( | Dnal1 | 4567.315 | 0.127489 | 0.181448 | 0.618684 | ENSMUSG( | Dnal1 | 3955.093 | -0.01686 | 0.881319 | 0.989644 |
| ENSMUSG( | Dnal4 | 1055.838 | -0.02884 | 0.783186 | 0.957558 | ENSMUSG( | Dnal4 | 952.556 | -0.06203 | 0.67301 | 0.95694 |
| ENSMUSG( | Dnali1 | 45.26098 | -0.2865 | 0.36547 | 0.794085 | ENSMUSG( | Dnali1 | 56.2693 | 0.573893 | 0.073257 | 0.763629 |
| ENSMUSG( | Dnase1l1 | 66.89007 | -0.03576 | 0.890344 | 0.983687 | ENSMUSG( | Dnase1l1 | 66.72337 | 0.23812 | 0.407682 | 0.917257 |
| ENSMUSG( | Dnase1l2 | 177.9163 | 0.261358 | 0.131899 | 0.544244 | ENSMUSG( | Dnase1l2 | 146.9766 | -0.00937 | 0.968808 | 0.99594 |
| ENSMUSG( | Dnase2a | 319.6105 | -0.11335 | 0.424361 | 0.824645 | ENSMUSG( | Dnase2a | 296.5036 | -0.04953 | 0.778554 | 0.976106 |
| ENSMUSG( | Dnd1 | 45.40128 | -0.0174 | 0.956435 | 0.993874 | ENSMUSG( | Dnd1 | 38.90413 | -0.17766 | 0.625843 | 0.953234 |
| ENSMUSG( | Dner | 5336.644 | -0.50308 | 0.001402 | 0.040991 | ENSMUSG( | Dner | 5941.917 | 0.119298 | 0.4853 | 0.92541 |
| ENSMUSG( | Dnhd1 | 254.991 | 0.191908 | 0.271492 | 0.722082 | ENSMUSG( | Dnhd1 | 233.0624 | 0.200496 | 0.278619 | 0.891628 |
| ENSMUSG( | Dnlz | 2073.187 | 0.085771 | 0.315466 | 0.762458 | ENSMUSG( | Dnlz | 1799.394 | -0.05562 | 0.615305 | 0.951177 |
| ENSMUSG( | Dnm1 | 24221.33 | 0.187435 | 0.181173 | 0.618139 | ENSMUSG( | Dnm1 | 20475.47 | -0.02408 | 0.868398 | 0.988031 |
| ENSMUSG( | Dnm1l | 4459.883 | -0.02584 | 0.765193 | 0.954127 | ENSMUSG( | Dnm1l | 3993.216 | -0.0799 | 0.535971 | 0.936952 |
| ENSMUSG( | Dnm2 | 3162.655 | 0.037936 | 0.630672 | 0.914073 | ENSMUSG( | Dnm2 | 2784.832 | -0.0607 | 0.54999 | 0.938699 |
| ENSMUSG( | Dnm3 | 1797.212 | 0.080585 | 0.346889 | 0.779684 | ENSMUSG( | Dnm3 | 1600.117 | 0.01479 | 0.890345 | 0.990558 |
| ENSMUSG( | Dnmbp | 475.7384 | -0.02463 | 0.850201 | 0.972638 | ENSMUSG( | Dnmbp | 434.4246 | -0.0147 | 0.917887 | 0.992529 |
| ENSMUSG( | Dnmt1 | 1385.862 | 0.187504 | 0.072115 | 0.414057 | ENSMUSG( | Dnmt1 | 1223.928 | 0.103373 | 0.423227 | 0.917257 |
| ENSMUSG( | Dnmt3a | 4687.174 | -0.00485 | 0.948972 | 0.993821 | ENSMUSG( | Dnmt3a | 4184.913 | -0.06474 | 0.529619 | 0.934795 |
| ENSMUSG( | Dnmt3b | 39.95129 | 0.39829 | 0.238588 | 0.686308 | ENSMUSG( | Dnmt3b | 32.59562 | 0.11912 | 0.744545 | 0.96951 |
| ENSMUSG( | Dnpep | 1019.174 | 0.03027 | 0.752495 | 0.950536 | ENSMUSG( | Dnpep | 906.8306 | -0.03829 | 0.754684 | 0.971098 |
| ENSMUSG( | Dnph1 | 66.28179 | 0.08608 | 0.717614 | 0.941697 | ENSMUSG( | Dnph1 | 58.28149 | -0.02982 | 0.917102 | 0.992529 |
| ENSMUSG( | Dnttip1 | 594.5673 | 0.045211 | 0.707181 | 0.93949 | ENSMUSG( | Dnttip1 | 535.1497 | 0.005136 | 0.972619 | 0.996416 |
| ENSMUSG( | Dnttip2 | 659.9171 | -0.2909 | 0.040722 | 0.311854 | ENSMUSG( | Dnttip2 | 686.3677 | 0.10338 | 0.509291 | 0.929089 |
| ENSMUSG( | Doc2a | 1227.934 | -0.00623 | 0.957821 | 0.993874 | ENSMUSG( | Doc2a | 1073.301 | -0.1308 | 0.326226 | 0.905339 |
| ENSMUSG( | Doc2b | 1457.612 | 0.143354 | 0.199976 | 0.64027 | ENSMUSG( | Doc2b | 1280.225 | 0.040422 | 0.722353 | 0.966887 |
| ENSMUSG( | Doc2g | 120.8296 | -0.19093 | 0.555867 | 0.887457 | ENSMUSG( | Doc2g | 124.9654 | 0.162412 | 0.813705 | 0.979198 |
| ENSMUSG( | Dock1 | 3464.527 | 0.17625 | 0.124443 | 0.531714 | ENSMUSG( | Dock1 | 2943.892 | -0.02083 | 0.873666 | 0.989644 |
| ENSMUSG( | Dock10 | 1447.838 | -0.18685 | 0.117418 | 0.517109 | ENSMUSG( | Dock10 | 1483.699 | 0.158716 | 0.278817 | 0.891737 |
| ENSMUSG( | Dock11 | 470.6551 | 0.439254 | 0.001344 | 0.039921 | ENSMUSG( | Dock11 | 404.6163 | 0.285506 | 0.078255 | 0.77321 |
| ENSMUSG( | Dock2 | 348.6061 | -0.00732 | 0.953607 | 0.993874 | ENSMUSG( | Dock2 | 307.1976 | -0.11255 | 0.482766 | 0.924615 |
| ENSMUSG( | Dock3 | 6305.095 | -0.03106 | 0.745364 | 0.949158 | ENSMUSG( | Dock3 | 5804.072 | -0.00078 | 0.994253 | 0.998813 |
| ENSMUSG( | Dock4 | 2412.232 | 0.007952 | 0.933871 | 0.993206 | ENSMUSG( | Dock4 | 2297.933 | 0.13551 | 0.221952 | 0.866104 |
| ENSMUSG( | Dock5 | 407.6218 | -0.15387 | 0.234453 | 0.681922 | ENSMUSG( | Dock5 | 408.3096 | 0.126755 | 0.37948 | 0.916995 |
| ENSMUSG( | Dock6 | 911.7257 | 0.189462 | 0.096924 | 0.477375 | ENSMUSG( | Dock6 | 842.5084 | 0.229391 | 0.177067 | 0.846973 |
| ENSMUSG( | Dock7 | 1656.367 | 0.038868 | 0.699818 | 0.937251 | ENSMUSG( | Dock7 | 1479.345 | -0.0183 | 0.863414 | 0.986547 |
| ENSMUSG( | Dock8 | 289.1777 | -0.07085 | 0.609762 | 0.907755 | ENSMUSG( | Dock8 | 271.1064 | 0.020562 | 0.905305 | 0.991089 |
| ENSMUSG( | Dock9 | 3660.122 | 0.168101 | 0.120562 | 0.524506 | ENSMUSG( | Dock9 | 3547.606 | 0.334376 | 0.015964 | 0.507531 |
| ENSMUSG( | Dohh | 587.947 | -0.02606 | 0.81424 | 0.964574 | ENSMUSG( | Dohh | 509.5491 | -0.18179 | 0.204256 | 0.864412 |
| ENSMUSG( | Dok1 | 77.94181 | -0.01835 | 0.937347 | 0.993415 | ENSMUSG( | Dok1 | 78.74469 | 0.287707 | 0.276587 | 0.890889 |

| | | | | | |
|---|---|---|---|---|---|
| ENSMUSG( | 1055.838 | -0.02884 | 0.783186 | 0.957558 | Dnal4 |
| ENSMUSG( | 45.26098 | -0.2865 | 0.36547 | 0.794085 | Dnali1 |
| ENSMUSG( | 66.89007 | -0.03576 | 0.890344 | 0.983687 | Dnase1l1 |
| ENSMUSG( | 177.9163 | 0.261358 | 0.131899 | 0.544244 | Dnase1l2 |
| ENSMUSG( | 319.6105 | -0.11335 | 0.424361 | 0.824645 | Dnase2a |
| ENSMUSG( | 45.40128 | -0.0174 | 0.956435 | 0.993874 | Dnd1 |
| ENSMUSG( | 5336.644 | -0.50308 | 0.001402 | 0.040991 | Dner |
| ENSMUSG( | 254.991 | 0.191908 | 0.271492 | 0.722082 | Dnhd1 |
| ENSMUSG( | 2073.187 | 0.085771 | 0.315466 | 0.762458 | Dnlz |
| ENSMUSG( | 24221.33 | 0.187435 | 0.181173 | 0.618139 | Dnm1 |
| ENSMUSG( | 4459.883 | -0.02584 | 0.765193 | 0.954127 | Dnm1l |
| ENSMUSG( | 3162.655 | 0.037936 | 0.630672 | 0.914073 | Dnm2 |
| ENSMUSG( | 1797.212 | 0.080585 | 0.346889 | 0.779684 | Dnm3 |
| ENSMUSG( | 475.7384 | -0.02463 | 0.850201 | 0.972638 | Dnmbp |
| ENSMUSG( | 1385.862 | 0.187504 | 0.072115 | 0.414057 | Dnmt1 |
| ENSMUSG( | 4687.174 | -0.00485 | 0.948972 | 0.993821 | Dnmt3a |
| ENSMUSG( | 39.95129 | 0.39829 | 0.238588 | 0.686308 | Dnmt3b |
| ENSMUSG( | 1019.174 | 0.03027 | 0.752495 | 0.950536 | Dnpep |
| ENSMUSG( | 66.28179 | 0.08608 | 0.717614 | 0.941697 | Dnph1 |
| ENSMUSG( | 594.5673 | 0.045211 | 0.707181 | 0.93949 | Dnttip1 |
| ENSMUSG( | 659.9171 | -0.2909 | 0.040722 | 0.311854 | Dnttip2 |
| ENSMUSG( | 1227.934 | -0.00623 | 0.957821 | 0.993874 | Doc2a |
| ENSMUSG( | 1457.612 | 0.143354 | 0.199976 | 0.64027 | Doc2b |
| ENSMUSG( | 120.8296 | -0.19093 | 0.555867 | 0.887457 | Doc2g |
| ENSMUSG( | 3464.527 | 0.17625 | 0.124443 | 0.531714 | Dock1 |
| ENSMUSG( | 1447.838 | -0.18685 | 0.117418 | 0.517109 | Dock10 |
| ENSMUSG( | 470.6551 | 0.439254 | 0.001344 | 0.039921 | Dock11 |
| ENSMUSG( | 348.6061 | -0.00732 | 0.953607 | 0.993874 | Dock2 |
| ENSMUSG( | 6305.095 | -0.03106 | 0.745364 | 0.949158 | Dock3 |
| ENSMUSG( | 2412.232 | 0.007952 | 0.933871 | 0.993206 | Dock4 |
| ENSMUSG( | 407.6218 | -0.15387 | 0.234453 | 0.681922 | Dock5 |
| ENSMUSG( | 911.7257 | 0.189462 | 0.096924 | 0.477375 | Dock6 |
| ENSMUSG( | 1656.367 | 0.038868 | 0.699818 | 0.937251 | Dock7 |
| ENSMUSG( | 289.1777 | -0.07085 | 0.609762 | 0.907755 | Dock8 |
| ENSMUSG( | 3660.122 | 0.168101 | 0.120562 | 0.524506 | Dock9 |
| ENSMUSG( | 587.947 | -0.02606 | 0.81424 | 0.964574 | Dohh |
| ENSMUSG( | 77.94181 | -0.01835 | 0.937347 | 0.993415 | Dok1 |
| ENSMUSG( | 37.1061 | 0.587799 | 0.068868 | 0.406642 | Dok2 |
| ENSMUSG( | 330.096 | 0.656398 | 7.41E-05 | 0.005296 | Dok3 |
| ENSMUSG( | 541.9 | 0.036462 | 0.754201 | 0.950981 | Dok4 |
| ENSMUSG( | 603.1762 | -0.00374 | 0.976243 | 0.997428 | Dok5 |
| ENSMUSG( | 4062.797 | 0.100024 | 0.482491 | 0.856002 | Dok6 |
| ENSMUSG( | 730.7947 | -0.13362 | 0.191338 | 0.630245 | Dok7 |
| ENSMUSG( | 748.226 | -0.20696 | 0.190602 | 0.629951 | Dolk |
| ENSMUSG( | 653.8578 | -0.324 | 0.012669 | 0.162184 | Dolpp1 |
| ENSMUSG( | 469.0575 | 0.070224 | 0.574816 | 0.894572 | Donson |
| ENSMUSG( | 1250.184 | -0.00284 | 0.980323 | 0.997512 | Dop1a |
| ENSMUSG( | 3322.176 | 0.048246 | 0.650704 | 0.921301 | Dop1b |
| ENSMUSG( | 2195.822 | 0.113896 | 0.374144 | 0.800445 | Dot1l |
| ENSMUSG( | 482.1939 | -0.18848 | 0.19418 | 0.63334 | Dpagt1 |
| ENSMUSG( | 1617.034 | -0.01898 | 0.843247 | 0.971847 | Dpcd |
| ENSMUSG( | 1429.375 | 0.054202 | 0.659299 | 0.924506 | Dpf1 |
| ENSMUSG( | 1519.358 | 0.182805 | 0.050832 | 0.345845 | Dpf2 |
| ENSMUSG( | 252.9769 | -0.06973 | 0.628482 | 0.913637 | Dpf3 |
| ENSMUSG( | 232.0787 | 0.04105 | 0.776023 | 0.955606 | Dph1 |
| ENSMUSG( | 231.0278 | 0.003516 | 0.980817 | 0.997512 | Dph2 |
| ENSMUSG( | 1247.223 | 0.025931 | 0.796305 | 0.960088 | Dph3 |
| ENSMUSG( | 253.0256 | 0.041168 | 0.765524 | 0.954173 | Dph5 |
| ENSMUSG( | 365.6959 | -0.09936 | 0.524376 | 0.87583 | Dph6 |
| ENSMUSG( | 490.7591 | -0.06471 | 0.618915 | 0.911432 | Dph7 |
| ENSMUSG( | 228.6894 | -0.21845 | 0.279135 | 0.729078 | Dpm1 |
| ENSMUSG( | 1219.963 | -0.13963 | 0.165186 | 0.596597 | Dpm2 |
| ENSMUSG( | 424.6422 | -0.25524 | 0.052874 | 0.352925 | Dpm3 |
| ENSMUSG( | 3568.836 | -0.2738 | 0.041466 | 0.314241 | Dpp10 |
| ENSMUSG( | 1028.4 | -0.02079 | 0.859264 | 0.974707 | Dpp3 |
| ENSMUSG( | 180.9154 | 0.158524 | 0.348594 | 0.78132 | Dpp4 |
| ENSMUSG( | 8368.642 | -0.31905 | 0.014128 | 0.174128 | Dpp6 |
| ENSMUSG( | 800.1286 | 0.083777 | 0.463791 | 0.842812 | Dpp7 |
| ENSMUSG( | 8881.666 | 0.135171 | 0.219058 | 0.664602 | Dpp8 |
| ENSMUSG( | 1779.001 | 0.162746 | 0.116596 | 0.514987 | Dpp9 |
| ENSMUSG( | 3563.907 | -0.26992 | 0.043727 | 0.320351 | Dpy19l1 |
| ENSMUSG( | 9.745387 | -0.49561 | 0.408665 | NA | Dpy19l2 |
| ENSMUSG( | 3247.279 | -0.08861 | 0.482241 | 0.855884 | Dpy19l3 |
| ENSMUSG( | 1224.144 | -0.19031 | 0.161505 | 0.59148 | Dpy19l4 |
| ENSMUSG( | 453.5252 | 0.073061 | 0.575129 | 0.894572 | Dpy30 |
| ENSMUSG( | 143.2882 | -0.13392 | 0.467946 | 0.84521 | Dpyd |
| ENSMUSG( | 4020.391 | -0.0842 | 0.520952 | 0.875186 | Dpysl2 |
| ENSMUSG( | 2195.252 | 0.112205 | 0.24338 | 0.690747 | Dpysl3 |
| ENSMUSG( | 1597.612 | 0.259199 | 0.078921 | 0.43467 | Dpysl4 |
| ENSMUSG( | 1116.104 | 0.14674 | 0.247846 | 0.696522 | Dpysl5 |
| ENSMUSG( | 19.8624 | -0.06743 | 0.881354 | NA | Dqx1 |
| ENSMUSG( | 529.8924 | -0.05693 | 0.671962 | 0.9288 | Dr1 |
| ENSMUSG( | 46.53445 | 0.366642 | 0.213013 | 0.656982 | Dram1 |
| ENSMUSG( | 650.209 | -0.338 | 0.034348 | 0.284534 | Dram2 |
| ENSMUSG( | 7710.195 | 0.016368 | 0.847222 | 0.972374 | Drap1 |
| ENSMUSG( | 74.6292 | -0.57106 | 0.055858 | 0.364301 | Draxin |
| ENSMUSG( | 432.5276 | 0.062356 | 0.588612 | 0.900921 | Drc1 |
| ENSMUSG( | 329.3702 | -0.04864 | 0.702857 | 0.937735 | Drc3 |
| ENSMUSG( | 87.10579 | -0.36504 | 0.355926 | 0.788009 | Drc7 |
| ENSMUSG( | 701.7886 | -1.29254 | 5.62E-12 | 4.34E-09 | Drd1 |
| ENSMUSG( | 611.3404 | -1.19423 | 0.006177 | 0.107367 | Drd2 |
| ENSMUSG( | 10.65344 | -0.82798 | 0.15852 | NA | Drd4 |
| ENSMUSG( | 107.2825 | -0.37359 | 0.079346 | 0.435382 | Drd5 |
| ENSMUSG( | 1307.098 | 0.00437 | 0.965163 | 0.995303 | Drg1 |
| ENSMUSG( | 1187.563 | 0.017893 | 0.87871 | 0.981436 | Drg2 |
| ENSMUSG( | 3559.785 | 0.184303 | 0.061087 | 0.381679 | Drosha |
| ENSMUSG( | 3900.312 | 0.120028 | 0.148396 | 0.571935 | Drp2 |
| ENSMUSG( | 22.25014 | 0.264753 | 0.542858 | 0.883579 | Dsc2 |
| ENSMUSG( | 39.67065 | 0.117214 | 0.748248 | 0.949604 | Dsc3 |
| ENSMUSG( | 2112.452 | -0.24358 | 0.102321 | 0.487127 | Dscam |
| ENSMUSG( | 1752.04 | 0.083866 | 0.555723 | 0.887457 | Dscaml1 |
| ENSMUSG( | 211.4265 | -0.09285 | 0.571402 | 0.893319 | Dse |
| ENSMUSG( | 325.465 | -0.43491 | 0.026436 | 0.248861 | Dsel |
| ENSMUSG( | 82.80422 | -0.31817 | 0.237785 | 0.685809 | Dsg2 |
| ENSMUSG( | 304.387 | 0.055964 | 0.70071 | 0.93743 | Dsn1 |
| ENSMUSG( | 94.0462 | 0.433962 | 0.069123 | 0.407256 | Dsp |
| ENSMUSG( | 30.66988 | -0.13145 | 0.713303 | 0.940785 | Dspp |
| ENSMUSG( | 21008.77 | -0.14167 | 0.240292 | 0.687899 | Dst |
| ENSMUSG( | 3532.452 | 0.059129 | 0.531009 | 0.879289 | Dstn |
| ENSMUSG( | 1582.194 | 0.122188 | 0.174636 | 0.610304 | Dstyk |
| ENSMUSG( | 732.8312 | -0.10347 | 0.436987 | 0.829501 | Dtd1 |
| ENSMUSG( | 305.7586 | 0.063918 | 0.686982 | 0.933041 | Dtd2 |
| ENSMUSG( | 123.3625 | 0.150926 | 0.43064 | 0.826896 | Dtl |
| ENSMUSG( | 11506.63 | 0.091858 | 0.418965 | 0.821966 | Dtna |
| ENSMUSG( | 1618.674 | 0.096396 | 0.412421 | 0.819452 | Dtnb |
| ENSMUSG( | 2386.585 | 0.044664 | 0.587315 | 0.900401 | Dtnbp1 |
| ENSMUSG( | 133.1684 | 0.182997 | 0.381133 | 0.802629 | Dtwd1 |
| ENSMUSG( | 118.8578 | 0.008363 | 0.965463 | 0.995377 | Dtwd2 |
| ENSMUSG( | 2878.191 | 0.071447 | 0.422971 | 0.823662 | Dtx1 |
| ENSMUSG( | 255.0612 | -0.20097 | 0.19576 | 0.635109 | Dtx2 |
| ENSMUSG( | 7405.894 | 0.223805 | 0.030874 | 0.267227 | Dtx3 |
| ENSMUSG( | 158.1423 | 0.093896 | 0.58129 | 0.897612 | Dtx3l |
| ENSMUSG( | 1868.716 | 0.149566 | 0.097529 | 0.477842 | Dtx4 |
| ENSMUSG( | 593.5193 | -0.04201 | 0.685942 | 0.932981 | Dtymk |
| ENSMUSG( | 754.1615 | 0.164327 | 0.1374 | 0.552453 | Dus1l |
| ENSMUSG( | 386.3615 | 0.006267 | 0.961024 | 0.993999 | Dus2 |
| ENSMUSG( | 1814.217 | 0.221717 | 0.119299 | 0.523036 | Dus3l |
| ENSMUSG( | 134.0947 | 0.120137 | 0.503772 | 0.866417 | Dus4l |
| ENSMUSG( | 620.3799 | -0.8622 | 0.000157 | 0.00887 | Dusp1 |
| ENSMUSG( | 124.2536 | 0.057914 | 0.770461 | 0.954689 | Dusp10 |
| ENSMUSG( | 2424.201 | 0.074478 | 0.452225 | 0.836145 | Dusp11 |
| ENSMUSG( | 364.8686 | -0.0135 | 0.912603 | 0.988483 | Dusp12 |
| ENSMUSG( | 10.43858 | 0.401292 | 0.511984 | NA | Dusp13b |
| ENSMUSG( | 885.1921 | -0.06685 | 0.699013 | 0.936791 | Dusp14 |
| ENSMUSG( | 695.084 | 0.010432 | 0.933821 | 0.993206 | Dusp15 |
| ENSMUSG( | 583.5374 | -0.12584 | 0.299135 | 0.748131 | Dusp16 |
| ENSMUSG( | 1587.303 | 0.079316 | 0.435778 | 0.829093 | Dusp18 |
| ENSMUSG( | 781.6576 | 0.027924 | 0.83619 | 0.971074 | Dusp19 |
| ENSMUSG( | 28.86747 | -0.10236 | 0.77558 | 0.955606 | Dusp2 |
| ENSMUSG( | 1093.762 | 0.04833 | 0.600471 | 0.90443 | Dusp22 |
| ENSMUSG( | 154.9109 | 0.168704 | 0.334647 | 0.776386 | Dusp23 |
| ENSMUSG( | 3165.379 | 0.026583 | 0.770304 | 0.954689 | Dusp26 |
| ENSMUSG( | 978.1204 | 0.082878 | 0.412499 | 0.819452 | Dusp28 |
| ENSMUSG( | 9410.247 | -0.00717 | 0.922447 | 0.99045 | Dusp3 |
| ENSMUSG( | 144.9819 | 0.0885 | 0.605562 | 0.905824 | Dusp4 |
| ENSMUSG( | 249.0746 | -0.49495 | 0.005642 | 0.100551 | Dusp5 |
| ENSMUSG( | 1155.115 | -0.3062 | 0.012875 | 0.163922 | Dusp6 |
| ENSMUSG( | 2396.39 | 0.016076 | 0.882662 | 0.982283 | Dusp7 |
| ENSMUSG( | 5942.36 | 0.158223 | 0.070107 | 0.40908 | Dusp8 |
| ENSMUSG( | 18.58289 | -0.12581 | 0.772402 | NA | Dusp9 |
| ENSMUSG( | 625.4253 | -0.00068 | 0.995948 | 0.99943 | Dut |
| ENSMUSG( | 122.0503 | -0.25658 | 0.154985 | 0.582587 | Duxbl1 |
| ENSMUSG( | 4701.441 | 0.141606 | 0.148973 | 0.572729 | Dvl1 |
| ENSMUSG( | 501.6234 | -0.00285 | 0.984625 | 0.997815 | Dvl2 |
| ENSMUSG( | 2478.512 | 0.091905 | 0.321794 | 0.766739 | Dvl3 |
| ENSMUSG( | 409.9858 | -0.03548 | 0.791578 | 0.959461 | Dxo |
| ENSMUSG( | 1052.397 | 0.048158 | 0.655667 | 0.923929 | Dym |
| ENSMUSG( | 15733.31 | 0.017288 | 0.843919 | 0.971962 | Dync1h1 |

| | | | | | |
|---|---|---|---|---|---|
| ENSMUSG( | 2784.832 | -0.0607 | 0.54999 | 0.938699 | Dnm2 |
| ENSMUSG( | 1600.117 | 0.01479 | 0.890345 | 0.990558 | Dnm3 |
| ENSMUSG( | 434.4246 | -0.0147 | 0.917887 | 0.992529 | Dnmbp |
| ENSMUSG( | 1223.928 | 0.103373 | 0.423227 | 0.917257 | Dnmt1 |
| ENSMUSG( | 4184.913 | -0.06474 | 0.529619 | 0.934795 | Dnmt3a |
| ENSMUSG( | 32.59562 | 0.11912 | 0.744545 | 0.96951 | Dnmt3b |
| ENSMUSG( | 906.8306 | -0.03829 | 0.754684 | 0.971098 | Dnpep |
| ENSMUSG( | 58.28149 | -0.02982 | 0.917102 | 0.992529 | Dnph1 |
| ENSMUSG( | 535.1497 | 0.005136 | 0.972619 | 0.996416 | Dnttip1 |
| ENSMUSG( | 686.3677 | 0.10338 | 0.509291 | 0.929089 | Dnttip2 |
| ENSMUSG( | 1073.301 | -0.1308 | 0.326226 | 0.905339 | Doc2a |
| ENSMUSG( | 1280.225 | 0.040422 | 0.722353 | 0.966887 | Doc2b |
| ENSMUSG( | 124.9654 | 0.162412 | 0.813705 | 0.979198 | Doc2g |
| ENSMUSG( | 2943.892 | -0.02083 | 0.873666 | 0.989644 | Dock1 |
| ENSMUSG( | 1483.699 | 0.158716 | 0.278817 | 0.891737 | Dock10 |
| ENSMUSG( | 404.6163 | 0.285506 | 0.078255 | 0.77321 | Dock11 |
| ENSMUSG( | 307.1976 | -0.11255 | 0.482766 | 0.924615 | Dock2 |
| ENSMUSG( | 5804.072 | -0.00078 | 0.994253 | 0.998813 | Dock3 |
| ENSMUSG( | 2297.933 | 0.13551 | 0.221952 | 0.866104 | Dock4 |
| ENSMUSG( | 408.3096 | 0.126755 | 0.37948 | 0.916995 | Dock5 |
| ENSMUSG( | 842.5084 | 0.229391 | 0.177067 | 0.846973 | Dock6 |
| ENSMUSG( | 1479.345 | -0.0183 | 0.863414 | 0.986547 | Dock7 |
| ENSMUSG( | 271.1064 | 0.020562 | 0.905305 | 0.991089 | Dock8 |
| ENSMUSG( | 3547.606 | 0.334376 | 0.015964 | 0.507531 | Dock9 |
| ENSMUSG( | 509.5491 | -0.18179 | 0.204256 | 0.864412 | Dohh |
| ENSMUSG( | 78.74469 | 0.287707 | 0.276587 | 0.890889 | Dok1 |
| ENSMUSG( | 25.34245 | -0.23038 | 0.571533 | 0.942116 | Dok2 |
| ENSMUSG( | 252.4804 | 0.217713 | 0.375826 | 0.915491 | Dok3 |
| ENSMUSG( | 499.789 | 0.067424 | 0.636856 | 0.953903 | Dok4 |
| ENSMUSG( | 497.134 | -0.31981 | 0.075284 | 0.767994 | Dok5 |
| ENSMUSG( | 3767.989 | 0.148572 | 0.335335 | 0.907907 | Dok6 |
| ENSMUSG( | 691.5378 | -0.01741 | 0.895889 | 0.990558 | Dok7 |
| ENSMUSG( | 747.2005 | 0.069319 | 0.715329 | 0.965063 | Dolk |
| ENSMUSG( | 722.3299 | 0.244423 | 0.087788 | 0.785783 | Dolpp1 |
| ENSMUSG( | 431.0786 | 0.091708 | 0.599642 | 0.947071 | Donson |
| ENSMUSG( | 1188.232 | 0.12161 | 0.35834 | 0.913759 | Dop1a |
| ENSMUSG( | 2994.547 | 0.021284 | 0.868726 | 0.988097 | Dop1b |
| ENSMUSG( | 2070.627 | 0.210466 | 0.189608 | 0.857355 | Dot1l |
| ENSMUSG( | 520.5309 | 0.299038 | 0.067659 | 0.75212 | Dpagt1 |
| ENSMUSG( | 1343.459 | -0.30492 | 0.012972 | 0.458131 | Dpcd |
| ENSMUSG( | 1284.039 | 0.009271 | 0.953032 | 0.994798 | Dpf1 |
| ENSMUSG( | 1333.979 | 0.081689 | 0.499244 | 0.927825 | Dpf2 |
| ENSMUSG( | 209.0203 | -0.37512 | 0.032143 | 0.634374 | Dpf3 |
| ENSMUSG( | 209.4699 | 0.028034 | 0.876506 | 0.989644 | Dph1 |
| ENSMUSG( | 200.2825 | -0.13895 | 0.420512 | 0.917257 | Dph2 |
| ENSMUSG( | 1062.45 | -0.17648 | 0.143063 | 0.831221 | Dph3 |
| ENSMUSG( | 222.5777 | -0.06596 | 0.685062 | 0.958678 | Dph5 |
| ENSMUSG( | 323.2475 | -0.1985 | 0.270848 | 0.888062 | Dph6 |
| ENSMUSG( | 458.8808 | 0.012338 | 0.931849 | 0.99306 | Dph7 |
| ENSMUSG( | 241.2838 | 0.22057 | 0.367032 | 0.915058 | Dpm1 |
| ENSMUSG( | 1143.527 | -0.05238 | 0.656283 | 0.955533 | Dpm2 |
| ENSMUSG( | 410.2731 | -0.08146 | 0.582626 | 0.944266 | Dpm3 |
| ENSMUSG( | 3966.636 | 0.300175 | 0.044968 | 0.693959 | Dpp10 |
| ENSMUSG( | 949.5385 | 0.018711 | 0.878458 | 0.989644 | Dpp3 |
| ENSMUSG( | 165.777 | 0.183482 | 0.343221 | 0.909646 | Dpp4 |
| ENSMUSG( | 8783.522 | 0.108768 | 0.469868 | 0.923329 | Dpp6 |
| ENSMUSG( | 738.9652 | 0.125094 | 0.381194 | 0.917202 | Dpp7 |
| ENSMUSG( | 7541.903 | -0.06608 | 0.558639 | 0.939574 | Dpp8 |
| ENSMUSG( | 1547.662 | 0.03403 | 0.779816 | 0.976106 | Dpp9 |
| ENSMUSG( | 3720.748 | 0.135489 | 0.375977 | 0.915491 | Dpy19l1 |
| ENSMUSG( | 13.92939 | 0.766031 | 0.143429 | 0.831311 | Dpy19l2 |
| ENSMUSG( | 3069.666 | 0.020353 | 0.880391 | 0.989644 | Dpy19l3 |
| ENSMUSG( | 1204.526 | 0.041081 | 0.784127 | 0.976154 | Dpy19l4 |
| ENSMUSG( | 389.6485 | -0.10759 | 0.538285 | 0.937216 | Dpy30 |
| ENSMUSG( | 136.2368 | -0.02782 | 0.898944 | 0.990558 | Dpyd |
| ENSMUSG( | 7.929887 | -0.11748 | 0.86364 | NA | Dpys |
| ENSMUSG( | 3533.742 | -0.19815 | 0.242906 | 0.877399 | Dpysl2 |
| ENSMUSG( | 1947.406 | 0.034524 | 0.799971 | 0.97681 | Dpysl3 |
| ENSMUSG( | 1348.022 | 0.047421 | 0.752107 | 0.970461 | Dpysl4 |
| ENSMUSG( | 948.6899 | -0.0523 | 0.717668 | 0.965964 | Dpysl5 |
| ENSMUSG( | 16.29255 | -0.38697 | 0.479645 | 0.924021 | Dqx1 |
| ENSMUSG( | 513.9381 | 0.111748 | 0.542939 | 0.938063 | Dr1 |
| ENSMUSG( | 41.71427 | 0.30563 | 0.36054 | 0.914223 | Dram1 |
| ENSMUSG( | 709.2383 | 0.197023 | 0.27364 | 0.889848 | Dram2 |
| ENSMUSG( | 6576.536 | -0.182 | 0.176943 | 0.846973 | Drap1 |
| ENSMUSG( | 91.07719 | 0.337076 | 0.330311 | 0.907251 | Draxin |
| ENSMUSG( | 372.0812 | -0.09645 | 0.518879 | 0.930939 | Drc1 |
| ENSMUSG( | 308.0277 | 0.036143 | 0.822667 | 0.979198 | Drc3 |
| ENSMUSG( | 92.50354 | 0.100356 | 0.766794 | 0.974236 | Drc7 |
| ENSMUSG( | 916.7616 | 0.029749 | 0.844268 | 0.983576 | Drd1 |
| ENSMUSG( | 777.9256 | 0.008484 | 0.95781 | 0.994798 | Drd2 |
| ENSMUSG( | 12.72832 | 0.053015 | 0.921752 | 0.992644 | Drd4 |
| ENSMUSG( | 126.1148 | 0.370811 | 0.08428 | 0.778656 | Drd5 |
| ENSMUSG( | 1155.078 | -0.08765 | 0.5105 | 0.929374 | Drg1 |
| ENSMUSG( | 1041.116 | -0.09666 | 0.474501 | 0.923329 | Drg2 |
| ENSMUSG( | 2988.834 | -0.04734 | 0.709055 | 0.964042 | Drosha |
| ENSMUSG( | 3257.646 | -0.1331 | 0.227493 | 0.868507 | Drp2 |
| ENSMUSG( | 19.47627 | 0.190552 | 0.690139 | 0.960071 | Dsc2 |
| ENSMUSG( | 42.04015 | 0.534507 | 0.157193 | 0.837724 | Dsc3 |
| ENSMUSG( | 2241.434 | 0.205178 | 0.256726 | 0.883082 | Dscam |
| ENSMUSG( | 1881.114 | 0.518679 | 0.002705 | 0.205145 | Dscaml1 |
| ENSMUSG( | 210.7735 | 0.16641 | 0.340429 | 0.907972 | Dse |
| ENSMUSG( | 392.4851 | 0.382734 | 0.100328 | 0.801311 | Dsel |
| ENSMUSG( | 96.06693 | 0.380748 | 0.14735 | 0.836241 | Dsg2 |
| ENSMUSG( | 269.0468 | -0.04096 | 0.813547 | 0.979198 | Dsn1 |
| ENSMUSG( | 114.7818 | 1.119403 | 2.35E-05 | 0.006966 | Dsp |
| ENSMUSG( | 24.59817 | -0.5359 | 0.195475 | 0.861579 | Dspp |
| ENSMUSG( | 20464.79 | 0.057029 | 0.696804 | 0.960867 | Dst |
| ENSMUSG( | 3088.916 | -0.06248 | 0.625672 | 0.953234 | Dstn |
| ENSMUSG( | 1399.369 | 0.039076 | 0.721447 | 0.966887 | Dstyk |
| ENSMUSG( | 648.0359 | -0.19893 | 0.209564 | 0.864412 | Dtd1 |
| ENSMUSG( | 262.0599 | -0.12211 | 0.491205 | 0.926604 | Dtd2 |
| ENSMUSG( | 98.72713 | -0.22923 | 0.303568 | 0.898538 | Dtl |
| ENSMUSG( | 9601.659 | -0.16721 | 0.21224 | 0.864412 | Dtna |
| ENSMUSG( | 1427.464 | 0.003935 | 0.976389 | 0.996416 | Dtnb |
| ENSMUSG( | 2002.077 | -0.20219 | 0.066892 | 0.751729 | Dtnbp1 |
| ENSMUSG( | 116.5852 | 0.07324 | 0.742503 | 0.969415 | Dtwd1 |
| ENSMUSG( | 102.3276 | -0.18457 | 0.457874 | 0.923329 | Dtwd2 |
| ENSMUSG( | 2542.638 | -0.01712 | 0.869251 | 0.988097 | Dtx1 |
| ENSMUSG( | 246.3918 | -0.01813 | 0.918379 | 0.992529 | Dtx2 |
| ENSMUSG( | 6175.3 | -0.02506 | 0.847916 | 0.985242 | Dtx3 |
| ENSMUSG( | 151.8131 | 0.242801 | 0.181142 | 0.850372 | Dtx3l |
| ENSMUSG( | 1533.877 | -0.15298 | 0.164927 | 0.838743 | Dtx4 |
| ENSMUSG( | 548.0864 | 0.00236 | 0.987163 | 0.997487 | Dtymk |
| ENSMUSG( | 673.1871 | 0.111958 | 0.423165 | 0.917257 | Dus1l |
| ENSMUSG( | 322.9855 | -0.25305 | 0.105551 | 0.804969 | Dus2 |
| ENSMUSG( | 1603.02 | 0.137022 | 0.439453 | 0.920721 | Dus3l |
| ENSMUSG( | 119.4553 | 0.048504 | 0.814666 | 0.979198 | Dus4l |
| ENSMUSG( | 625.1603 | -0.48079 | 0.249513 | 0.877904 | Dusp1 |
| ENSMUSG( | 110.9639 | 0.008923 | 0.967266 | 0.995606 | Dusp10 |
| ENSMUSG( | 2340.55 | 0.234824 | 0.072542 | 0.763629 | Dusp11 |
| ENSMUSG( | 348.3983 | 0.120207 | 0.419288 | 0.917257 | Dusp12 |
| ENSMUSG( | 7.137435 | -0.33031 | 0.672245 | NA | Dusp13b |
| ENSMUSG( | 802.1305 | -0.08972 | 0.670544 | 0.95694 | Dusp14 |
| ENSMUSG( | 627.0333 | -0.01548 | 0.923492 | 0.992815 | Dusp15 |
| ENSMUSG( | 522.5086 | -0.17549 | 0.221339 | 0.866104 | Dusp16 |
| ENSMUSG( | 1305.937 | -0.22686 | 0.088188 | 0.786756 | Dusp18 |
| ENSMUSG( | 673.4341 | -0.14297 | 0.352551 | 0.912093 | Dusp19 |
| ENSMUSG( | 26.65414 | -0.08968 | 0.816782 | 0.979198 | Dusp2 |
| ENSMUSG( | 950.8212 | -0.09051 | 0.439881 | 0.920926 | Dusp22 |
| ENSMUSG( | 138.214 | 0.103399 | 0.594369 | 0.945408 | Dusp23 |
| ENSMUSG( | 2814.342 | -0.04457 | 0.689557 | 0.960071 | Dusp26 |
| ENSMUSG( | 862.9635 | -0.0133 | 0.918259 | 0.992529 | Dusp28 |
| ENSMUSG( | 8358.583 | -0.08211 | 0.394263 | 0.917257 | Dusp3 |
| ENSMUSG( | 142.3355 | 0.278915 | 0.164513 | 0.838699 | Dusp4 |
| ENSMUSG( | 219.5125 | -0.6191 | 0.005606 | 0.303106 | Dusp5 |
| ENSMUSG( | 1070.871 | -0.2513 | 0.097431 | 0.796721 | Dusp6 |
| ENSMUSG( | 2236.501 | 0.082909 | 0.477521 | 0.923329 | Dusp7 |
| ENSMUSG( | 5326.285 | 0.114766 | 0.300721 | 0.898538 | Dusp8 |
| ENSMUSG( | 15.2344 | -0.43923 | 0.399903 | 0.917257 | Dusp9 |
| ENSMUSG( | 528.6029 | -0.23345 | 0.159767 | 0.837724 | Dut |
| ENSMUSG( | 110.4563 | -0.25937 | 0.261265 | 0.884926 | Duxbl1 |
| ENSMUSG( | 4214.165 | 0.096772 | 0.396568 | 0.917257 | Dvl1 |
| ENSMUSG( | 490.4599 | 0.198186 | 0.23606 | 0.872173 | Dvl2 |
| ENSMUSG( | 2174.998 | -0.01825 | 0.891093 | 0.990558 | Dvl3 |
| ENSMUSG( | 382.9718 | 0.03738 | 0.817948 | 0.979198 | Dxo |
| ENSMUSG( | 959.0373 | 0.046635 | 0.725954 | 0.966887 | Dym |
| ENSMUSG( | 14045.64 | -0.04027 | 0.708039 | 0.964042 | Dync1h1 |
| ENSMUSG( | 1436.389 | -0.1645 | 0.410018 | 0.917257 | Dync1i1 |
| ENSMUSG( | 3718.512 | -0.04656 | 0.77735 | 0.975838 | Dync1i2 |
| ENSMUSG( | 1340.898 | -0.06651 | 0.643996 | 0.953903 | Dync1li1 |
| ENSMUSG( | 11041.95 | -0.15321 | 0.234825 | 0.871042 | Dync1li2 |
| ENSMUSG( | 757.8229 | 0.113205 | 0.367928 | 0.915491 | Dync2h1 |
| ENSMUSG( | 517.829 | 0.050453 | 0.716748 | 0.965859 | Dync2i1 |
| ENSMUSG( | 955.8926 | -0.02789 | 0.824764 | 0.979198 | Dync2i2 |
| ENSMUSG( | 368.0549 | -0.13373 | 0.387063 | 0.917257 | Dync2li1 |
| ENSMUSG( | 4058.437 | -0.11761 | 0.406727 | 0.917257 | Dynll1 |
| ENSMUSG( | 20908.87 | -0.14034 | 0.29627 | 0.898538 | Dynll2 |

| | | | | | |
|---|---|---|---|---|---|
| ENSMUSG( | Dok2 | 37.1061 | 0.587799 | 0.068868 | 0.406642 |
| ENSMUSG( | Dok3 | 330.096 | 0.656398 | 7.41E-05 | 0.005296 |
| ENSMUSG( | Dok4 | 541.9 | 0.036462 | 0.754201 | 0.950981 |
| ENSMUSG( | Dok5 | 603.1762 | -0.00374 | 0.976243 | 0.997428 |
| ENSMUSG( | Dok6 | 4062.797 | 0.100024 | 0.482491 | 0.856002 |
| ENSMUSG( | Dok7 | 730.7947 | -0.13362 | 0.191338 | 0.630245 |
| ENSMUSG( | Dolk | 748.226 | -0.20696 | 0.190602 | 0.629951 |
| ENSMUSG( | Dolpp1 | 653.8578 | -0.324 | 0.012669 | 0.162184 |
| ENSMUSG( | Donson | 469.0575 | 0.070224 | 0.574816 | 0.894572 |
| ENSMUSG( | Dop1a | 1250.184 | -0.00284 | 0.980323 | 0.997512 |
| ENSMUSG( | Dop1b | 3322.176 | 0.048246 | 0.650704 | 0.921301 |
| ENSMUSG( | Dot1l | 2195.822 | 0.113896 | 0.374144 | 0.800445 |
| ENSMUSG( | Dpagt1 | 482.1939 | -0.18848 | 0.19418 | 0.63334 |
| ENSMUSG( | Dpcd | 1617.034 | -0.01898 | 0.843247 | 0.971847 |
| ENSMUSG( | Dpf1 | 1429.375 | 0.054202 | 0.659299 | 0.924506 |
| ENSMUSG( | Dpf2 | 1519.358 | 0.182805 | 0.050832 | 0.345845 |
| ENSMUSG( | Dpf3 | 252.9769 | -0.06973 | 0.628482 | 0.913637 |
| ENSMUSG( | Dph1 | 232.0787 | 0.04105 | 0.776023 | 0.955606 |
| ENSMUSG( | Dph2 | 231.0278 | 0.003516 | 0.980817 | 0.997512 |
| ENSMUSG( | Dph3 | 1247.223 | 0.025931 | 0.796305 | 0.960088 |
| ENSMUSG( | Dph5 | 253.0256 | 0.041168 | 0.765524 | 0.954173 |
| ENSMUSG( | Dph6 | 365.6959 | -0.09936 | 0.524376 | 0.87583 |
| ENSMUSG( | Dph7 | 490.7591 | -0.06471 | 0.618915 | 0.911432 |
| ENSMUSG( | Dpm1 | 228.6894 | -0.21845 | 0.279135 | 0.729078 |
| ENSMUSG( | Dpm2 | 1219.963 | -0.13963 | 0.165186 | 0.596597 |
| ENSMUSG( | Dpm3 | 424.6422 | -0.25524 | 0.052874 | 0.352925 |
| ENSMUSG( | Dpp10 | 3568.836 | -0.2738 | 0.041466 | 0.314241 |
| ENSMUSG( | Dpp3 | 1028.4 | -0.02079 | 0.859264 | 0.974707 |
| ENSMUSG( | Dpp4 | 180.9154 | 0.158524 | 0.348594 | 0.78132 |
| ENSMUSG( | Dpp6 | 8368.642 | -0.31905 | 0.014128 | 0.174128 |
| ENSMUSG( | Dpp7 | 800.1286 | 0.083777 | 0.463791 | 0.842812 |
| ENSMUSG( | Dpp8 | 8881.666 | 0.135171 | 0.219058 | 0.664602 |
| ENSMUSG( | Dpp9 | 1779.001 | 0.162746 | 0.116596 | 0.514987 |
| ENSMUSG( | Dpy19l1 | 3563.907 | -0.26992 | 0.043727 | 0.320351 |
| ENSMUSG( | Dpy19l2 | 9.745387 | -0.49561 | 0.408665 | NA |
| ENSMUSG( | Dpy19l3 | 3247.279 | -0.08861 | 0.482241 | 0.855884 |
| ENSMUSG( | Dpy19l4 | 1224.144 | -0.19031 | 0.161505 | 0.59148 |
| ENSMUSG( | Dpy30 | 453.5252 | 0.073061 | 0.575129 | 0.894572 |
| ENSMUSG( | Dpyd | 143.2882 | -0.13392 | 0.467946 | 0.84521 |
| ENSMUSG( | Dpysl2 | 4020.391 | -0.0842 | 0.520952 | 0.875186 |
| ENSMUSG( | Dpysl3 | 2195.252 | 0.112205 | 0.24338 | 0.690747 |
| ENSMUSG( | Dpysl4 | 1597.612 | 0.259199 | 0.078921 | 0.43467 |
| ENSMUSG( | Dpysl5 | 1116.104 | 0.14674 | 0.247846 | 0.696522 |
| ENSMUSG( | Dqx1 | 19.8624 | -0.06743 | 0.881354 | NA |
| ENSMUSG( | Dr1 | 529.8924 | -0.05693 | 0.671962 | 0.9288 |
| ENSMUSG( | Dram1 | 46.53445 | 0.366642 | 0.213013 | 0.656982 |
| ENSMUSG( | Dram2 | 650.209 | -0.338 | 0.034348 | 0.284534 |
| ENSMUSG( | Drap1 | 7710.195 | 0.016368 | 0.847222 | 0.972374 |
| ENSMUSG( | Draxin | 74.6292 | -0.57106 | 0.055858 | 0.364301 |
| ENSMUSG( | Drc1 | 432.5276 | 0.062356 | 0.588612 | 0.900921 |
| ENSMUSG( | Drc3 | 329.3702 | -0.04864 | 0.702857 | 0.937735 |
| ENSMUSG( | Drc7 | 87.10579 | -0.36504 | 0.355926 | 0.788009 |
| ENSMUSG( | Drd1 | 701.7886 | -1.29254 | 5.62E-12 | 4.34E-09 |
| ENSMUSG( | Drd2 | 611.3404 | -1.19423 | 0.006177 | 0.107367 |
| ENSMUSG( | Drd4 | 10.65344 | -0.82798 | 0.15852 | NA |
| ENSMUSG( | Drd5 | 107.2825 | -0.37359 | 0.079346 | 0.435382 |
| ENSMUSG( | Drg1 | 1307.098 | 0.00437 | 0.965163 | 0.995303 |
| ENSMUSG( | Drg2 | 1187.563 | 0.017893 | 0.87871 | 0.981436 |
| ENSMUSG( | Drosha | 3559.785 | 0.184303 | 0.061087 | 0.381679 |
| ENSMUSG( | Drp2 | 3900.312 | 0.120028 | 0.148396 | 0.571935 |
| ENSMUSG( | Dsc2 | 22.25014 | 0.264753 | 0.542858 | 0.883579 |
| ENSMUSG( | Dsc3 | 39.67065 | 0.117214 | 0.748248 | 0.949604 |
| ENSMUSG( | Dscam | 2112.452 | -0.24358 | 0.102321 | 0.487127 |
| ENSMUSG( | Dscaml1 | 1752.04 | 0.083866 | 0.555723 | 0.887457 |
| ENSMUSG( | Dse | 211.4265 | -0.09285 | 0.571402 | 0.893319 |
| ENSMUSG( | Dsel | 325.465 | -0.43491 | 0.026436 | 0.248861 |
| ENSMUSG( | Dsg2 | 82.80422 | -0.31817 | 0.237785 | 0.685809 |
| ENSMUSG( | Dsn1 | 304.387 | 0.055964 | 0.70071 | 0.93743 |
| ENSMUSG( | Dsp | 94.0462 | 0.433962 | 0.069123 | 0.407256 |
| ENSMUSG( | Dspp | 30.66988 | -0.13145 | 0.713303 | 0.940785 |
| ENSMUSG( | Dst | 21008.77 | -0.14167 | 0.240292 | 0.687899 |
| ENSMUSG( | Dstn | 3532.452 | 0.059129 | 0.531009 | 0.879289 |
| ENSMUSG( | Dstyk | 1582.194 | 0.122188 | 0.174636 | 0.610304 |
| ENSMUSG( | Dtd1 | 732.8312 | -0.10347 | 0.436987 | 0.829501 |
| ENSMUSG( | Dtd2 | 305.7586 | 0.063918 | 0.686982 | 0.933041 |
| ENSMUSG( | Dtl | 123.3625 | 0.150926 | 0.43064 | 0.826896 |
| ENSMUSG( | Dtna | 11506.63 | 0.091858 | 0.418965 | 0.821966 |
| ENSMUSG( | Dtnb | 1618.674 | 0.096396 | 0.412421 | 0.819452 |
| ENSMUSG( | Dtnbp1 | 2386.585 | 0.044664 | 0.587315 | 0.900401 |
| ENSMUSG( | Dtwd1 | 133.1684 | 0.182997 | 0.381133 | 0.802629 |
| ENSMUSG( | Dtwd2 | 118.8578 | 0.008363 | 0.965463 | 0.995377 |
| ENSMUSG( | Dtx1 | 2878.191 | 0.071447 | 0.422971 | 0.823662 |
| ENSMUSG( | Dtx2 | 255.0612 | -0.20097 | 0.19576 | 0.635109 |
| ENSMUSG( | Dtx3 | 7405.894 | 0.223805 | 0.030874 | 0.267227 |
| ENSMUSG( | Dtx3l | 158.1423 | 0.093896 | 0.58129 | 0.897612 |
| ENSMUSG( | Dtx4 | 1868.716 | 0.149566 | 0.097529 | 0.477842 |
| ENSMUSG( | Dtymk | 593.5193 | -0.04201 | 0.685942 | 0.932981 |
| ENSMUSG( | Dus1l | 754.1615 | 0.164327 | 0.1374 | 0.552453 |
| ENSMUSG( | Dus2 | 386.3615 | 0.006267 | 0.961024 | 0.993999 |
| ENSMUSG( | Dus3l | 1814.217 | 0.221717 | 0.119299 | 0.523036 |
| ENSMUSG( | Dus4l | 134.0947 | 0.120137 | 0.503772 | 0.866417 |
| ENSMUSG( | Dusp1 | 620.3799 | -0.8622 | 0.000157 | 0.00887 |
| ENSMUSG( | Dusp10 | 124.2536 | 0.057914 | 0.770461 | 0.954689 |
| ENSMUSG( | Dusp11 | 2424.201 | 0.074478 | 0.452225 | 0.836145 |
| ENSMUSG( | Dusp12 | 364.8686 | -0.0135 | 0.912603 | 0.988483 |
| ENSMUSG( | Dusp13b | 10.43858 | 0.401292 | 0.511984 | NA |
| ENSMUSG( | Dusp14 | 885.1921 | -0.06685 | 0.699013 | 0.936791 |
| ENSMUSG( | Dusp15 | 695.084 | 0.010432 | 0.933821 | 0.993206 |
| ENSMUSG( | Dusp16 | 583.5374 | -0.12584 | 0.299135 | 0.748131 |
| ENSMUSG( | Dusp18 | 1587.303 | 0.079316 | 0.435778 | 0.829093 |
| ENSMUSG( | Dusp19 | 781.6576 | 0.027924 | 0.83619 | 0.971074 |
| ENSMUSG( | Dusp2 | 28.86747 | -0.10236 | 0.77558 | 0.955606 |
| ENSMUSG( | Dusp22 | 1093.762 | 0.04833 | 0.600471 | 0.90443 |
| ENSMUSG( | Dusp23 | 154.9109 | 0.168704 | 0.334647 | 0.776386 |
| ENSMUSG( | Dusp26 | 3165.379 | 0.026583 | 0.770304 | 0.954689 |
| ENSMUSG( | Dusp28 | 978.1204 | 0.082878 | 0.412499 | 0.819452 |
| ENSMUSG( | Dusp3 | 9410.247 | -0.00717 | 0.922447 | 0.99045 |
| ENSMUSG( | Dusp4 | 144.9819 | 0.0885 | 0.605562 | 0.905824 |
| ENSMUSG( | Dusp5 | 249.0746 | -0.49495 | 0.005642 | 0.100551 |
| ENSMUSG( | Dusp6 | 1155.115 | -0.3062 | 0.012875 | 0.163922 |
| ENSMUSG( | Dusp7 | 2396.39 | 0.016076 | 0.882662 | 0.982283 |
| ENSMUSG( | Dusp8 | 5942.36 | 0.158223 | 0.070107 | 0.40908 |
| ENSMUSG( | Dusp9 | 18.58289 | -0.12581 | 0.772402 | NA |
| ENSMUSG( | Dut | 625.4253 | -0.00068 | 0.995948 | 0.99943 |
| ENSMUSG( | Duxbl1 | 122.0503 | -0.25658 | 0.154985 | 0.582587 |
| ENSMUSG( | Dvl1 | 4701.441 | 0.141606 | 0.148973 | 0.572729 |
| ENSMUSG( | Dvl2 | 501.6234 | -0.00285 | 0.984625 | 0.997815 |
| ENSMUSG( | Dvl3 | 2478.512 | 0.091905 | 0.321794 | 0.766739 |
| ENSMUSG( | Dxo | 409.9858 | -0.03548 | 0.791578 | 0.959461 |
| ENSMUSG( | Dym | 1052.397 | 0.048158 | 0.655667 | 0.923929 |
| ENSMUSG( | Dync1h1 | 15733.31 | 0.017288 | 0.843919 | 0.971962 |
| ENSMUSG( | Dync1i1 | 1715.822 | 0.087632 | 0.624479 | 0.911887 |
| ENSMUSG( | Dync1i2 | 4423.196 | 0.184417 | 0.185482 | 0.623185 |
| ENSMUSG( | Dync1li1 | 1542.123 | 0.070929 | 0.578647 | 0.896692 |
| ENSMUSG( | Dync1li2 | 12626.09 | -0.02883 | 0.739185 | 0.947768 |
| ENSMUSG( | Dync2h1 | 771.4431 | -0.1054 | 0.351701 | 0.783381 |
| ENSMUSG( | Dync2i1 | 563.5182 | 0.0282 | 0.802735 | 0.962169 |
| ENSMUSG( | Dync2i2 | 1115.578 | 0.148821 | 0.153321 | 0.57989 |
| ENSMUSG( | Dync2li1 | 424.5006 | 0.01242 | 0.928798 | 0.991747 |
| ENSMUSG( | Dynll1 | 4749.053 | 0.072325 | 0.506989 | 0.867833 |
| ENSMUSG( | Dynll2 | 24527.18 | 0.055926 | 0.514543 | 0.872055 |
| ENSMUSG( | Dynlrb1 | 3786.652 | -0.01305 | 0.883694 | 0.982622 |
| ENSMUSG( | Dynlrb2 | 30.02354 | -0.61528 | 0.081322 | 0.440708 |
| ENSMUSG( | Dynlt1a | 147.8984 | 0.065423 | 0.736032 | 0.946267 |
| ENSMUSG( | Dynlt1b | 39.48097 | -0.42793 | 0.153421 | 0.579959 |
| ENSMUSG( | Dynlt1c | 27.56916 | 0.128786 | 0.757237 | 0.951513 |
| ENSMUSG( | Dynlt1f | 69.03915 | 0.116611 | 0.655447 | 0.923929 |
| ENSMUSG( | Dynlt2b | 152.3698 | -0.13513 | 0.457359 | 0.839558 |
| ENSMUSG( | Dynlt3 | 3527.586 | 0.085516 | 0.503893 | 0.866426 |
| ENSMUSG( | Dynlt4 | 11.22475 | -0.61397 | 0.286398 | NA |
| ENSMUSG( | Dyrk1a | 2024.35 | 0.1164 | 0.200825 | 0.640988 |
| ENSMUSG( | Dyrk1b | 675.5893 | -0.02759 | 0.795868 | 0.960062 |
| ENSMUSG( | Dyrk2 | 1144.967 | 0.009223 | 0.927305 | 0.991229 |
| ENSMUSG( | Dyrk3 | 148.53 | -0.15817 | 0.47194 | 0.847528 |
| ENSMUSG( | Dyrk4 | 11.33562 | -0.18107 | 0.740805 | NA |
| ENSMUSG( | Dysf | 142.4436 | 0.150519 | 0.441507 | 0.832509 |
| ENSMUSG( | Dzank1 | 5100.419 | 0.073211 | 0.352115 | 0.783738 |
| ENSMUSG( | Dzip1 | 4313.229 | 0.213538 | 0.018377 | 0.203658 |
| ENSMUSG( | Dzip1l | 809.6422 | 0.264956 | 0.081747 | 0.441786 |
| ENSMUSG( | Dzip3 | 1347.573 | -0.04727 | 0.616516 | 0.910866 |
| ENSMUSG( | E030013I1 | 36.27995 | -0.23015 | 0.463962 | 0.842812 |
| ENSMUSG( | E130006D0 | 33.74267 | 0.11353 | 0.723281 | 0.943761 |
| ENSMUSG( | E130008D0 | 72.59868 | -0.3575 | 0.247872 | 0.696522 |
| ENSMUSG( | E130102H2 | 65.81485 | 0.076513 | 0.782146 | 0.957558 |
| ENSMUSG( | E130215H2 | 43.10047 | -0.29291 | 0.305458 | 0.753541 |
| ENSMUSG( | E130307A1 | 1453.368 | 0.018052 | 0.850507 | 0.972638 |
| ENSMUSG( | E130308A1 | 478.9404 | -0.09567 | 0.443751 | 0.833108 |
| ENSMUSG( | E130311K1 | 310.6902 | -0.33409 | 0.016243 | 0.189127 |

| | | | | | |
|---|---|---|---|---|---|
| ENSMUSG( | Dok2 | 25.34245 | -0.23038 | 0.571533 | 0.942116 |
| ENSMUSG( | Dok3 | 252.4804 | 0.217713 | 0.375826 | 0.915491 |
| ENSMUSG( | Dok4 | 499.789 | 0.067424 | 0.636856 | 0.953903 |
| ENSMUSG( | Dok5 | 497.134 | -0.31981 | 0.075284 | 0.767994 |
| ENSMUSG( | Dok6 | 3767.989 | 0.148572 | 0.335335 | 0.907907 |
| ENSMUSG( | Dok7 | 691.5378 | -0.01741 | 0.895889 | 0.990558 |
| ENSMUSG( | Dolk | 747.2005 | 0.069319 | 0.715329 | 0.965063 |
| ENSMUSG( | Dolpp1 | 722.3299 | 0.244423 | 0.087788 | 0.785783 |
| ENSMUSG( | Donson | 431.0786 | 0.091708 | 0.599642 | 0.947071 |
| ENSMUSG( | Dop1a | 1188.232 | 0.12161 | 0.35834 | 0.913759 |
| ENSMUSG( | Dop1b | 2994.547 | 0.021284 | 0.868726 | 0.988097 |
| ENSMUSG( | Dot1l | 2070.627 | 0.210466 | 0.189608 | 0.857355 |
| ENSMUSG( | Dpagt1 | 520.5309 | 0.299038 | 0.067659 | 0.75212 |
| ENSMUSG( | Dpcd | 1343.459 | -0.30492 | 0.012972 | 0.458131 |
| ENSMUSG( | Dpf1 | 1284.039 | 0.009271 | 0.953032 | 0.994798 |
| ENSMUSG( | Dpf2 | 1333.979 | 0.081689 | 0.499244 | 0.927825 |
| ENSMUSG( | Dpf3 | 209.0203 | -0.37512 | 0.032143 | 0.634374 |
| ENSMUSG( | Dph1 | 209.4699 | 0.028034 | 0.876506 | 0.989644 |
| ENSMUSG( | Dph2 | 200.2825 | -0.13895 | 0.420512 | 0.917257 |
| ENSMUSG( | Dph3 | 1062.45 | -0.17648 | 0.143063 | 0.831221 |
| ENSMUSG( | Dph5 | 222.5777 | -0.06596 | 0.685062 | 0.958678 |
| ENSMUSG( | Dph6 | 323.2475 | -0.1985 | 0.270848 | 0.888062 |
| ENSMUSG( | Dph7 | 458.8808 | 0.012338 | 0.931849 | 0.99306 |
| ENSMUSG( | Dpm1 | 241.2838 | 0.22057 | 0.367032 | 0.915058 |
| ENSMUSG( | Dpm2 | 1143.527 | -0.05238 | 0.656283 | 0.955533 |
| ENSMUSG( | Dpm3 | 410.2731 | -0.08146 | 0.582626 | 0.944266 |
| ENSMUSG( | Dpp10 | 3966.636 | 0.300175 | 0.044968 | 0.693959 |
| ENSMUSG( | Dpp3 | 949.5385 | 0.018711 | 0.878458 | 0.989644 |
| ENSMUSG( | Dpp4 | 165.777 | 0.183482 | 0.343221 | 0.909646 |
| ENSMUSG( | Dpp6 | 8783.522 | 0.108768 | 0.469868 | 0.923329 |
| ENSMUSG( | Dpp7 | 738.9652 | 0.125094 | 0.381194 | 0.917202 |
| ENSMUSG( | Dpp8 | 7541.903 | -0.06608 | 0.558639 | 0.939574 |
| ENSMUSG( | Dpp9 | 1547.662 | 0.03403 | 0.779816 | 0.976106 |
| ENSMUSG( | Dpy19l1 | 3720.748 | 0.135489 | 0.375977 | 0.915491 |
| ENSMUSG( | Dpy19l2 | 13.92939 | 0.766031 | 0.143429 | 0.831311 |
| ENSMUSG( | Dpy19l3 | 3069.666 | 0.020353 | 0.880391 | 0.989644 |
| ENSMUSG( | Dpy19l4 | 1204.526 | 0.041081 | 0.784127 | 0.976154 |
| ENSMUSG( | Dpy30 | 389.6485 | -0.10759 | 0.538285 | 0.937216 |
| ENSMUSG( | Dpyd | 136.2368 | -0.02782 | 0.898944 | 0.990558 |
| ENSMUSG( | Dpysl2 | 3533.742 | -0.19815 | 0.242906 | 0.877399 |
| ENSMUSG( | Dpysl3 | 1947.406 | 0.034524 | 0.799971 | 0.97681 |
| ENSMUSG( | Dpysl4 | 1348.022 | 0.047421 | 0.752107 | 0.970461 |
| ENSMUSG( | Dpysl5 | 948.6899 | -0.0523 | 0.717668 | 0.965964 |
| ENSMUSG( | Dqx1 | 16.29255 | -0.38697 | 0.479645 | 0.924021 |
| ENSMUSG( | Dr1 | 513.9381 | 0.111748 | 0.542939 | 0.938063 |
| ENSMUSG( | Dram1 | 41.71427 | 0.30563 | 0.36054 | 0.914223 |
| ENSMUSG( | Dram2 | 709.2383 | 0.197023 | 0.27364 | 0.889848 |
| ENSMUSG( | Drap1 | 6576.536 | -0.182 | 0.176943 | 0.846973 |
| ENSMUSG( | Draxin | 91.07719 | 0.337076 | 0.330311 | 0.907251 |
| ENSMUSG( | Drc1 | 372.0812 | -0.09645 | 0.518879 | 0.930939 |
| ENSMUSG( | Drc3 | 308.0277 | 0.036143 | 0.822667 | 0.979198 |
| ENSMUSG( | Drc7 | 92.50354 | 0.100356 | 0.766794 | 0.974236 |
| ENSMUSG( | Drd1 | 916.7616 | 0.029749 | 0.844268 | 0.983576 |
| ENSMUSG( | Drd2 | 777.9256 | 0.008484 | 0.95781 | 0.994798 |
| ENSMUSG( | Drd4 | 12.72832 | 0.053015 | 0.921752 | 0.992644 |
| ENSMUSG( | Drd5 | 126.1148 | 0.370811 | 0.08428 | 0.778656 |
| ENSMUSG( | Drg1 | 1155.078 | -0.08765 | 0.5105 | 0.929374 |
| ENSMUSG( | Drg2 | 1041.116 | -0.09666 | 0.474501 | 0.923329 |
| ENSMUSG( | Drosha | 2988.834 | -0.04734 | 0.709055 | 0.964042 |
| ENSMUSG( | Drp2 | 3257.646 | -0.1331 | 0.227493 | 0.868507 |
| ENSMUSG( | Dsc2 | 19.47627 | 0.190552 | 0.690139 | 0.960071 |
| ENSMUSG( | Dsc3 | 42.04015 | 0.534507 | 0.157193 | 0.837724 |
| ENSMUSG( | Dscam | 2241.434 | 0.205178 | 0.256726 | 0.883082 |
| ENSMUSG( | Dscaml1 | 1881.114 | 0.518679 | 0.002705 | 0.205145 |
| ENSMUSG( | Dse | 210.7735 | 0.16641 | 0.340429 | 0.907972 |
| ENSMUSG( | Dsel | 392.4851 | 0.382734 | 0.100328 | 0.801311 |
| ENSMUSG( | Dsg2 | 96.06693 | 0.380748 | 0.14735 | 0.836241 |
| ENSMUSG( | Dsn1 | 269.0468 | -0.04096 | 0.813547 | 0.979198 |
| ENSMUSG( | Dsp | 114.7818 | 1.119403 | 2.35E-05 | 0.006966 |
| ENSMUSG( | Dspp | 24.59817 | -0.5359 | 0.195475 | 0.861579 |
| ENSMUSG( | Dst | 20464.79 | 0.057029 | 0.696804 | 0.960867 |
| ENSMUSG( | Dstn | 3088.916 | -0.06248 | 0.625672 | 0.953234 |
| ENSMUSG( | Dstyk | 1399.369 | 0.039076 | 0.721447 | 0.966887 |
| ENSMUSG( | Dtd1 | 648.0359 | -0.19893 | 0.209564 | 0.864412 |
| ENSMUSG( | Dtd2 | 262.0599 | -0.12211 | 0.491205 | 0.926604 |
| ENSMUSG( | Dtl | 98.72713 | -0.22923 | 0.303568 | 0.898538 |
| ENSMUSG( | Dtna | 9601.659 | -0.16721 | 0.21224 | 0.864412 |
| ENSMUSG( | Dtnb | 1427.464 | 0.003935 | 0.976389 | 0.996416 |
| ENSMUSG( | Dtnbp1 | 2002.077 | -0.20219 | 0.066892 | 0.751729 |
| ENSMUSG( | Dtwd1 | 116.5852 | 0.07324 | 0.742503 | 0.969415 |
| ENSMUSG( | Dtwd2 | 102.3276 | -0.18457 | 0.457874 | 0.923329 |
| ENSMUSG( | Dtx1 | 2542.638 | -0.01712 | 0.869251 | 0.988097 |
| ENSMUSG( | Dtx2 | 246.3918 | -0.01813 | 0.918379 | 0.992529 |
| ENSMUSG( | Dtx3 | 6175.3 | -0.02506 | 0.847916 | 0.985242 |
| ENSMUSG( | Dtx3l | 151.8131 | 0.242801 | 0.181142 | 0.850372 |
| ENSMUSG( | Dtx4 | 1533.877 | -0.15298 | 0.164927 | 0.838743 |
| ENSMUSG( | Dtymk | 548.0864 | 0.00236 | 0.987163 | 0.997487 |
| ENSMUSG( | Dus1l | 673.1871 | 0.111958 | 0.423165 | 0.917257 |
| ENSMUSG( | Dus2 | 322.9855 | -0.25305 | 0.105551 | 0.804969 |
| ENSMUSG( | Dus3l | 1603.02 | 0.137022 | 0.439453 | 0.920721 |
| ENSMUSG( | Dus4l | 119.4553 | 0.048504 | 0.814666 | 0.979198 |
| ENSMUSG( | Dusp1 | 625.1603 | -0.48079 | 0.249513 | 0.877904 |
| ENSMUSG( | Dusp10 | 110.9639 | 0.008923 | 0.967266 | 0.995606 |
| ENSMUSG( | Dusp11 | 2340.55 | 0.234824 | 0.072542 | 0.763629 |
| ENSMUSG( | Dusp12 | 348.3983 | 0.120207 | 0.419288 | 0.917257 |
| ENSMUSG( | Dusp13b | 7.137435 | -0.33031 | 0.672245 | NA |
| ENSMUSG( | Dusp14 | 802.1305 | -0.08972 | 0.670544 | 0.95694 |
| ENSMUSG( | Dusp15 | 627.0333 | -0.01548 | 0.923492 | 0.992815 |
| ENSMUSG( | Dusp16 | 522.5086 | -0.17549 | 0.221339 | 0.866104 |
| ENSMUSG( | Dusp18 | 1305.937 | -0.22686 | 0.088188 | 0.786756 |
| ENSMUSG( | Dusp19 | 673.4341 | -0.14297 | 0.352551 | 0.912093 |
| ENSMUSG( | Dusp2 | 26.65414 | -0.08968 | 0.816782 | 0.979198 |
| ENSMUSG( | Dusp22 | 950.8212 | -0.09051 | 0.439881 | 0.920926 |
| ENSMUSG( | Dusp23 | 138.214 | 0.103399 | 0.594369 | 0.945408 |
| ENSMUSG( | Dusp26 | 2814.342 | -0.04457 | 0.689557 | 0.960071 |
| ENSMUSG( | Dusp28 | 862.9635 | -0.0133 | 0.918259 | 0.992529 |
| ENSMUSG( | Dusp3 | 8358.583 | -0.08211 | 0.394263 | 0.917257 |
| ENSMUSG( | Dusp4 | 142.3355 | 0.278915 | 0.164513 | 0.838699 |
| ENSMUSG( | Dusp5 | 219.5125 | -0.6191 | 0.005606 | 0.303106 |
| ENSMUSG( | Dusp6 | 1070.871 | -0.2513 | 0.097431 | 0.796721 |
| ENSMUSG( | Dusp7 | 2236.501 | 0.082909 | 0.477521 | 0.923329 |
| ENSMUSG( | Dusp8 | 5326.285 | 0.114766 | 0.300721 | 0.898538 |
| ENSMUSG( | Dusp9 | 15.2344 | -0.43923 | 0.399903 | 0.917257 |
| ENSMUSG( | Dut | 528.6029 | -0.23345 | 0.159767 | 0.837724 |
| ENSMUSG( | Duxbl1 | 110.4563 | -0.25937 | 0.261265 | 0.884926 |
| ENSMUSG( | Dvl1 | 4214.165 | 0.096772 | 0.396568 | 0.917257 |
| ENSMUSG( | Dvl2 | 490.4599 | 0.198186 | 0.23606 | 0.872173 |
| ENSMUSG( | Dvl3 | 2174.998 | -0.01825 | 0.891093 | 0.990558 |
| ENSMUSG( | Dxo | 382.9718 | 0.03738 | 0.817948 | 0.979198 |
| ENSMUSG( | Dym | 959.0373 | 0.046635 | 0.725954 | 0.966887 |
| ENSMUSG( | Dync1h1 | 14045.64 | -0.04027 | 0.708039 | 0.964042 |
| ENSMUSG( | Dync1i1 | 1436.389 | -0.1645 | 0.410018 | 0.917257 |
| ENSMUSG( | Dync1i2 | 3718.512 | -0.04656 | 0.77735 | 0.975838 |
| ENSMUSG( | Dync1li1 | 1340.898 | -0.06651 | 0.643996 | 0.953903 |
| ENSMUSG( | Dync1li2 | 11041.95 | -0.15321 | 0.234825 | 0.871042 |
| ENSMUSG( | Dync2h1 | 757.8229 | 0.113205 | 0.367928 | 0.915491 |
| ENSMUSG( | Dync2i1 | 517.829 | 0.050453 | 0.716748 | 0.965859 |
| ENSMUSG( | Dync2i2 | 955.8926 | -0.02789 | 0.824764 | 0.979198 |
| ENSMUSG( | Dync2li1 | 368.0549 | -0.13373 | 0.387063 | 0.917257 |
| ENSMUSG( | Dynll1 | 4058.437 | -0.11761 | 0.406727 | 0.917257 |
| ENSMUSG( | Dynll2 | 20908.87 | -0.14034 | 0.29627 | 0.898538 |
| ENSMUSG( | Dynlrb1 | 3048.417 | -0.40001 | 0.001061 | 0.111298 |
| ENSMUSG( | Dynlrb2 | 34.73511 | 0.126391 | 0.709 | 0.964042 |
| ENSMUSG( | Dynlt1a | 134.6835 | 0.068191 | 0.768164 | 0.974236 |
| ENSMUSG( | Dynlt1b | 34.43338 | -0.52849 | 0.155613 | 0.837724 |
| ENSMUSG( | Dynlt1c | 27.21429 | 0.356456 | 0.381238 | 0.917202 |
| ENSMUSG( | Dynlt1f | 65.4059 | 0.220429 | 0.384885 | 0.917257 |
| ENSMUSG( | Dynlt2b | 149.1042 | 0.066037 | 0.744051 | 0.96951 |
| ENSMUSG( | Dynlt3 | 3093.468 | -0.02779 | 0.875133 | 0.989644 |
| ENSMUSG( | Dynlt4 | 18.89779 | 1.061831 | 0.042842 | 0.690304 |
| ENSMUSG( | Dyrk1a | 1783.634 | 0.019971 | 0.860904 | 0.986226 |
| ENSMUSG( | Dyrk1b | 613.0924 | -0.03812 | 0.764142 | 0.974 |
| ENSMUSG( | Dyrk2 | 1074.76 | 0.094146 | 0.431699 | 0.917504 |
| ENSMUSG( | Dyrk3 | 125.4181 | -0.39721 | 0.109048 | 0.804969 |
| ENSMUSG( | Dyrk4 | 11.22962 | 0.08827 | 0.881996 | NA |
| ENSMUSG( | Dysf | 129.2878 | 0.157687 | 0.472935 | 0.923329 |
| ENSMUSG( | Dzank1 | 4432.275 | -0.06251 | 0.526193 | 0.934374 |
| ENSMUSG( | Dzip1 | 3687.547 | 0.036125 | 0.742144 | 0.969415 |
| ENSMUSG( | Dzip1l | 743.2364 | 0.290781 | 0.046892 | 0.700664 |
| ENSMUSG( | Dzip3 | 1260.423 | 0.028624 | 0.799217 | 0.97681 |
| ENSMUSG( | E030013I1 | 34.1084 | -0.14882 | 0.662819 | 0.956205 |
| ENSMUSG( | E130006D0 | 27.80086 | -0.21294 | 0.599932 | 0.947322 |
| ENSMUSG( | E130008D0 | 75.228 | 0.020094 | 0.950342 | 0.994798 |
| ENSMUSG( | E130102H2 | 69.00534 | 0.475174 | 0.116436 | 0.813384 |
| ENSMUSG( | E130215H2 | 39.35264 | -0.27566 | 0.387625 | 0.917257 |
| ENSMUSG( | E130307A1 | 1280.687 | -0.08269 | 0.4936 | 0.927372 |
| ENSMUSG( | E130308A1 | 443.1813 | -0.04858 | 0.737987 | 0.967709 |
| ENSMUSG( | E130311K1 | 319.1403 | 0.037971 | 0.818863 | 0.979198 |

| | | | | | |
|---|---|---|---|---|---|
| ENSMUSG( | 1715.822 | 0.087632 | 0.624479 | 0.91187 | Dync1i1 |
| ENSMUSG( | 4423.196 | 0.184417 | 0.185482 | 0.623185 | Dync1i2 |
| ENSMUSG( | 1542.123 | 0.070929 | 0.578647 | 0.896692 | Dync1li1 |
| ENSMUSG( | 12626.09 | -0.02883 | 0.739185 | 0.947768 | Dync1li2 |
| ENSMUSG( | 771.4431 | -0.1054 | 0.351701 | 0.783381 | Dync2h1 |
| ENSMUSG( | 563.5182 | 0.0282 | 0.802735 | 0.962169 | Dync2i1 |
| ENSMUSG( | 1115.578 | 0.148821 | 0.153321 | 0.57989 | Dync2i2 |
| ENSMUSG( | 424.5006 | 0.01242 | 0.928798 | 0.991747 | Dync2li1 |
| ENSMUSG( | 4749.053 | 0.072325 | 0.506989 | 0.867833 | Dynll1 |
| ENSMUSG( | 24527.18 | 0.055926 | 0.514543 | 0.872055 | Dynll2 |
| ENSMUSG( | 3786.652 | -0.01305 | 0.883694 | 0.982622 | Dynlrb1 |
| ENSMUSG( | 30.02354 | -0.61528 | 0.081322 | 0.440708 | Dynlrb2 |
| ENSMUSG( | 147.8984 | 0.065423 | 0.736032 | 0.946267 | Dynlt1a |
| ENSMUSG( | 39.48097 | -0.42793 | 0.153421 | 0.579959 | Dynlt1b |
| ENSMUSG( | 27.56916 | 0.128786 | 0.757237 | 0.951513 | Dynlt1c |
| ENSMUSG( | 69.03915 | 0.116611 | 0.655447 | 0.923929 | Dynlt1f |
| ENSMUSG( | 152.3698 | -0.13513 | 0.457359 | 0.839558 | Dynlt2b |
| ENSMUSG( | 3527.586 | 0.085516 | 0.503893 | 0.866426 | Dynlt3 |
| ENSMUSG( | 11.22475 | -0.61397 | 0.286398 | NA | Dynlt4 |
| ENSMUSG( | 9.860648 | 0.194356 | 0.753592 | NA | Dynlt5 |
| ENSMUSG( | 2024.35 | 0.1164 | 0.200825 | 0.640988 | Dyrk1a |
| ENSMUSG( | 675.5893 | -0.02759 | 0.795868 | 0.960062 | Dyrk1b |
| ENSMUSG( | 1144.967 | 0.009223 | 0.927305 | 0.991229 | Dyrk2 |
| ENSMUSG( | 148.53 | -0.15817 | 0.47194 | 0.847528 | Dyrk3 |
| ENSMUSG( | 11.33562 | -0.18107 | 0.740805 | NA | Dyrk4 |
| ENSMUSG( | 142.4436 | 0.150519 | 0.441507 | 0.832509 | Dysf |
| ENSMUSG( | 5100.419 | 0.073211 | 0.352115 | 0.783738 | Dzank1 |
| ENSMUSG( | 4313.229 | 0.213538 | 0.018377 | 0.203658 | Dzip1 |
| ENSMUSG( | 809.6422 | 0.264956 | 0.081747 | 0.441786 | Dzip1l |
| ENSMUSG( | 1347.573 | -0.04727 | 0.616516 | 0.910866 | Dzip3 |
| ENSMUSG( | 36.27995 | -0.23015 | 0.463962 | 0.842812 | E030013I19Rik |
| ENSMUSG( | 33.74952 | 0.11353 | 0.723281 | 0.943761 | E130006D01Rik |
| ENSMUSG( | 72.59868 | -0.3575 | 0.247872 | 0.696522 | E130008D07Rik |
| ENSMUSG( | 65.81485 | 0.076513 | 0.782146 | 0.957558 | E130102H24Rik |
| ENSMUSG( | 9.868994 | -0.09319 | 0.873163 | NA | E130201H02Rik |
| ENSMUSG( | 43.10047 | -0.29291 | 0.305458 | 0.753541 | E130215H24Rik |
| ENSMUSG( | 1453.368 | 0.018052 | 0.850507 | 0.972638 | E130307A14Rik |
| ENSMUSG( | 478.9404 | -0.09567 | 0.443571 | 0.833108 | E130308A19Rik |
| ENSMUSG( | 310.6902 | -0.33409 | 0.016243 | 0.189127 | E130311K13Rik |
| ENSMUSG( | 92.13965 | -0.09528 | 0.672895 | 0.929007 | E130317F20Rik |
| ENSMUSG( | 9.468681 | -0.26124 | 0.682481 | NA | E230015B07Rik |
| ENSMUSG( | 109.4871 | 0.156555 | 0.572093 | 0.893584 | E230016M11Rik |
| ENSMUSG( | 103.0623 | -0.04575 | 0.825495 | 0.967871 | E230029C05Rik |
| ENSMUSG( | 610.5737 | -0.07972 | 0.482964 | 0.856216 | E2f1 |
| ENSMUSG( | 61.82077 | -0.01669 | 0.950373 | 0.993821 | E2f2 |
| ENSMUSG( | 777.7729 | -0.02082 | 0.832755 | 0.970333 | E2f3 |
| ENSMUSG( | 861.2017 | 0.109926 | 0.355992 | 0.788009 | E2f4 |
| ENSMUSG( | 520.5395 | 0.039402 | 0.789329 | 0.959379 | E2f5 |
| ENSMUSG( | 1605.831 | 0.100819 | 0.282683 | 0.73186 | E2f6 |
| ENSMUSG( | 16.67978 | -0.42878 | 0.388396 | NA | E2f7 |
| ENSMUSG( | 31.87104 | -0.43956 | 0.237204 | 0.684843 | E2f8 |
| ENSMUSG( | 12.01558 | 0.155996 | 0.79411 | NA | E330011O21Rik |
| ENSMUSG( | 29.53039 | -0.06101 | 0.87721 | 0.981265 | E330013P04Rik |
| ENSMUSG( | 961.7353 | 0.23839 | 0.03895 | 0.304318 | E4f1 |
| ENSMUSG( | 602.5369 | -0.02736 | 0.803253 | 0.962169 | Eaf1 |
| ENSMUSG( | 34.5934 | -0.30756 | 0.445747 | 0.833239 | Eaf2 |
| ENSMUSG( | 1436.136 | 0.126122 | 0.223226 | 0.669382 | Eapp |
| ENSMUSG( | 22.40747 | -1.2511 | 0.062615 | 0.38686 | Ear2 |
| ENSMUSG( | 153.3052 | 0.115246 | 0.493324 | 0.860611 | Ears2 |
| ENSMUSG( | 604.5859 | -0.28203 | 0.027706 | 0.255176 | Ebag9 |
| ENSMUSG( | 88.25986 | -0.38604 | 0.136121 | 0.550232 | Ebf1 |
| ENSMUSG( | 12.05612 | 0.061399 | 0.908491 | NA | Ebf2 |
| ENSMUSG( | 30.31615 | -0.09108 | 0.789177 | 0.959299 | Ebf3 |
| ENSMUSG( | 401.5218 | 0.156836 | 0.24086 | 0.68836 | Ebf4 |
| ENSMUSG( | 57.81899 | -0.41902 | 0.102282 | 0.487127 | Ebi3 |
| ENSMUSG( | 986.353 | 0.096369 | 0.370931 | 0.798261 | Ebna1bp2 |
| ENSMUSG( | 803.7774 | -0.38325 | 0.004973 | 0.093585 | Ebp |
| ENSMUSG( | 504.4377 | -0.10381 | 0.351222 | 0.783067 | Ebpl |
| ENSMUSG( | 930.7168 | 0.156748 | 0.14378 | 0.562581 | Ecd |
| ENSMUSG( | 2596.886 | -0.27957 | 0.058217 | 0.37191 | Ece1 |
| ENSMUSG( | 93.90901 | 0.215818 | 0.411473 | 0.819309 | Ece2 |
| ENSMUSG( | 250.2152 | -0.7604 | 0.000621 | 0.02374 | Ecel1 |
| ENSMUSG( | 1829.642 | 0.036953 | 0.686285 | 0.932993 | Ech1 |
| ENSMUSG( | 1952.559 | 0.116441 | 0.421229 | 0.823126 | Echdc1 |
| ENSMUSG( | 481.6571 | -0.08835 | 0.622371 | 0.911432 | Echdc2 |
| ENSMUSG( | 82.34846 | 0.046742 | 0.837046 | 0.971111 | Echdc3 |
| ENSMUSG( | 2647.989 | 0.056234 | 0.568467 | 0.892252 | Echs1 |
| ENSMUSG( | 1253.671 | 0.138175 | 0.167461 | 0.600006 | Eci1 |
| ENSMUSG( | 1918.707 | 0.066773 | 0.554449 | 0.886657 | Eci2 |
| ENSMUSG( | 324.977 | -0.11202 | 0.416074 | 0.82084 | Ecm1 |
| ENSMUSG( | 154.1515 | -0.33341 | 0.118092 | 0.519373 | Ecm2 |
| ENSMUSG( | 2805.427 | 0.074211 | 0.35502 | 0.787423 | Ecpas |
| ENSMUSG( | 302.0217 | -0.34243 | 0.451047 | 0.835567 | Ecrg4 |
| ENSMUSG( | 186.0318 | -0.31747 | 0.082178 | 0.442642 | Ecscr |
| ENSMUSG( | 1145.28 | 0.122231 | 0.270962 | 0.72132 | Ecsit |
| ENSMUSG( | 28.88148 | -0.26007 | 0.457106 | 0.839312 | Ect2 |
| ENSMUSG( | 12.41904 | 0.189849 | 0.719377 | NA | Ect2l |
| ENSMUSG( | 294.7544 | -0.1716 | 0.367526 | 0.795314 | Eda |
| ENSMUSG( | 11.48671 | -0.38986 | 0.516031 | NA | Eda2r |
| ENSMUSG( | 16.33154 | -0.0924 | 0.84709 | NA | Edar |
| ENSMUSG( | 463.591 | 0.099464 | 0.430352 | 0.826896 | Edc3 |
| ENSMUSG( | 1804.72 | 0.271576 | 0.014572 | 0.177737 | Edc4 |
| ENSMUSG( | 866.6864 | -0.14668 | 0.340299 | 0.777962 | Edem1 |
| ENSMUSG( | 896.8515 | -0.13678 | 0.301026 | 0.749979 | Edem2 |
| ENSMUSG( | 1591.36 | -0.27378 | 0.062439 | 0.386467 | Edem3 |
| ENSMUSG( | 3401.134 | -0.04499 | 0.598554 | 0.903801 | Edf1 |
| ENSMUSG( | 2079.157 | -0.55531 | 0.001509 | 0.043083 | Edil3 |
| ENSMUSG( | 83.96001 | 0.006469 | 0.984018 | 0.99781 | Edn1 |
| ENSMUSG( | 182.5857 | -0.08146 | 0.697861 | 0.935828 | Edn3 |
| ENSMUSG( | 302.3017 | -0.34576 | 0.095463 | 0.474828 | Ednra |
| ENSMUSG( | 4234.64 | -0.19998 | 0.174124 | 0.60991 | Ednrb |
| ENSMUSG( | 573.0115 | 0.183776 | 0.133873 | 0.547197 | Edrf1 |
| ENSMUSG( | 1417.809 | 0.211522 | 0.055988 | 0.364712 | Eea1 |
| ENSMUSG( | 337.6961 | -0.13757 | 0.314403 | 0.761558 | Eed |
| ENSMUSG( | 90916.3 | 0.029921 | 0.712513 | 0.940507 | Eef1a1 |
| ENSMUSG( | 29929.43 | 0.127035 | 0.109957 | 0.500285 | Eef1a2 |
| ENSMUSG( | 444.9826 | 0.159558 | 0.20057 | 0.640661 | Eef1akmt1 |
| ENSMUSG( | 432.4297 | 0.016714 | 0.901486 | 0.985633 | Eef1akmt2 |
| ENSMUSG( | 23.33968 | 0.826524 | 0.04898 | 0.337919 | Eef1akmt4 |
| ENSMUSG( | 6864.289 | 0.065083 | 0.443157 | 0.833103 | Eef1b2 |
| ENSMUSG( | 2832.052 | 0.149816 | 0.103716 | 0.489428 | Eef1d |
| ENSMUSG( | 315.1169 | -0.002 | 0.987496 | 0.998796 | Eef1e1 |
| ENSMUSG( | 10445.86 | 0.09777 | 0.293274 | 0.742763 | Eef1g |
| ENSMUSG( | 75126.01 | 0.097227 | 0.192014 | 0.630811 | Eef2 |
| ENSMUSG( | 1555.599 | 0.159574 | 0.104387 | 0.490485 | Eef2k |
| ENSMUSG( | 204.0576 | 0.074346 | 0.621701 | 0.911432 | Eef2kmt |
| ENSMUSG( | 93.55146 | 0.279526 | 0.184278 | 0.621284 | Eef2-ps2 |
| ENSMUSG( | 454.005 | 0.050547 | 0.692023 | 0.934101 | Eefsec |
| ENSMUSG( | 4709.349 | -0.06515 | 0.398162 | 0.812452 | Eeig1 |
| ENSMUSG( | 1693.516 | 0.055575 | 0.641748 | 0.916828 | Eeig2 |
| ENSMUSG( | 405.7036 | -0.28958 | 0.052395 | 0.35132 | Eepd1 |
| ENSMUSG( | 30.09917 | -0.53808 | 0.120595 | 0.524506 | Efcab10 |
| ENSMUSG( | 11.20207 | -0.30018 | 0.601678 | NA | Efcab11 |
| ENSMUSG( | 136.7993 | -0.02816 | 0.889072 | 0.983434 | Efcab12 |
| ENSMUSG( | 3151.134 | -0.1109 | 0.367863 | 0.795671 | Efcab14 |
| ENSMUSG( | 85.857 | 0.215481 | 0.400602 | 0.813717 | Efcab15 |
| ENSMUSG( | 1261.419 | 0.057362 | 0.624377 | 0.91187 | Efcab2 |
| ENSMUSG( | 91.53432 | -0.08668 | 0.697491 | 0.935662 | Efcab5 |
| ENSMUSG( | 283.4123 | 0.161438 | 0.269832 | 0.719729 | Efcab6 |
| ENSMUSG( | 49.77817 | -0.04873 | 0.865231 | 0.975565 | Efcab7 |
| ENSMUSG( | 15.62786 | -0.33717 | 0.466085 | NA | Efcab9 |
| ENSMUSG( | 133.6857 | 0.043084 | 0.834909 | 0.970426 | Efcc1 |
| ENSMUSG( | 431.3899 | -0.28245 | 0.033402 | 0.279991 | Efemp1 |
| ENSMUSG( | 718.5956 | -0.00374 | 0.973924 | 0.99675 | Efemp2 |
| ENSMUSG( | 15.67321 | 0.011134 | 0.983656 | NA | Efhb |
| ENSMUSG( | 68.85366 | 0.040881 | 0.874545 | 0.9795 | Efhc1 |
| ENSMUSG( | 18.13563 | -0.62146 | 0.194098 | NA | Efhc2 |
| ENSMUSG( | 1586.911 | 0.089091 | 0.444021 | 0.833239 | Efhd1 |
| ENSMUSG( | 10181.1 | 0.134149 | 0.107841 | 0.49674 | Efhd2 |
| ENSMUSG( | 387.3967 | 0.164498 | 0.219596 | 0.664984 | Efl1 |
| ENSMUSG( | 175.1123 | -0.21169 | 0.320919 | 0.765984 | Efna1 |
| ENSMUSG( | 409.9233 | -0.41187 | 0.005271 | 0.09683 | Efna2 |
| ENSMUSG( | 2013.815 | -0.353 | 0.002904 | 0.066108 | Efna3 |
| ENSMUSG( | 26.6608 | -0.39341 | 0.320464 | 0.765921 | Efna4 |
| ENSMUSG( | 463.3962 | -0.14188 | 0.406243 | 0.816525 | Efna5 |
| ENSMUSG( | 558.7306 | -0.51321 | 0.003676 | 0.077033 | Efnb1 |
| ENSMUSG( | 1519.018 | -0.34511 | 0.014655 | 0.178054 | Efnb2 |
| ENSMUSG( | 4066.289 | -0.53083 | 2.35E-06 | 0.000367 | Efnb3 |
| ENSMUSG( | 3461.648 | 0.164595 | 0.06307 | 0.388234 | Efr3a |
| ENSMUSG( | 8040.363 | 0.18707 | 0.011235 | 0.152627 | Efr3b |
| ENSMUSG( | 498.097 | -0.17765 | 0.136133 | 0.550232 | Efs |
| ENSMUSG( | 2557.626 | 0.131292 | 0.189858 | 0.628983 | Eftud2 |
| ENSMUSG( | 38.30867 | -0.07363 | 0.818401 | 0.965837 | Egf |
| ENSMUSG( | 246.5737 | -0.01216 | 0.940769 | 0.993415 | Egfem1 |
| ENSMUSG( | 277.0103 | 0.006117 | 0.972144 | 0.996585 | Egfl6 |
| ENSMUSG( | 1097.084 | -0.05373 | 0.72459 | 0.944096 | Egfl7 |
| ENSMUSG( | 43.10297 | -0.44952 | 0.156336 | 0.584143 | Egfl8 |
| ENSMUSG( | 248.6952 | 0.046694 | 0.809814 | 0.96372 | Egflam |

| | | | | | |
|---|---|---|---|---|---|
| ENSMUSG( | 3048.417 | -0.40001 | 0.001061 | 0.111298 | Dynlrb1 |
| ENSMUSG( | 34.73511 | 0.126391 | 0.709 | 0.964042 | Dynlrb2 |
| ENSMUSG( | 134.6835 | 0.068191 | 0.768164 | 0.974236 | Dynlt1a |
| ENSMUSG( | 34.43338 | -0.52849 | 0.155613 | 0.837724 | Dynlt1b |
| ENSMUSG( | 27.21429 | 0.356456 | 0.381238 | 0.917202 | Dynlt1c |
| ENSMUSG( | 65.4059 | 0.220429 | 0.384885 | 0.917257 | Dynlt1f |
| ENSMUSG( | 149.1042 | 0.066037 | 0.744051 | 0.96951 | Dynlt2b |
| ENSMUSG( | 3093.468 | -0.02779 | 0.875133 | 0.989644 | Dynlt3 |
| ENSMUSG( | 18.89779 | 1.061831 | 0.042842 | 0.690304 | Dynlt4 |
| ENSMUSG( | 1783.634 | 0.019971 | 0.860904 | 0.986226 | Dyrk1a |
| ENSMUSG( | 613.0924 | -0.03812 | 0.764142 | 0.974 | Dyrk1b |
| ENSMUSG( | 1074.76 | 0.094146 | 0.431699 | 0.917504 | Dyrk2 |
| ENSMUSG( | 125.4181 | -0.39721 | 0.109048 | 0.804969 | Dyrk3 |
| ENSMUSG( | 11.22962 | 0.08827 | 0.881996 | NA | Dyrk4 |
| ENSMUSG( | 129.2878 | 0.157687 | 0.472935 | 0.923329 | Dysf |
| ENSMUSG( | 4432.275 | -0.06251 | 0.526193 | 0.934374 | Dzank1 |
| ENSMUSG( | 3687.547 | 0.036125 | 0.742144 | 0.969415 | Dzip1 |
| ENSMUSG( | 743.2364 | 0.290781 | 0.046892 | 0.700664 | Dzip1l |
| ENSMUSG( | 1260.423 | 0.028624 | 0.799217 | 0.97681 | Dzip3 |
| ENSMUSG( | 34.1084 | -0.14882 | 0.662819 | 0.956205 | E030013I19Rik |
| ENSMUSG( | 27.80086 | -0.21294 | 0.599932 | 0.947322 | E130006D01Rik |
| ENSMUSG( | 75.228 | 0.020094 | 0.950342 | 0.994798 | E130008D07Rik |
| ENSMUSG( | 69.00534 | 0.475174 | 0.116436 | 0.813384 | E130102H24Rik |
| ENSMUSG( | 39.35264 | -0.27566 | 0.387625 | 0.917257 | E130215H24Rik |
| ENSMUSG( | 1280.687 | -0.08269 | 0.4936 | 0.927372 | E130307A14Rik |
| ENSMUSG( | 443.1813 | -0.04858 | 0.737987 | 0.967709 | E130308A19Rik |
| ENSMUSG( | 319.1403 | 0.037971 | 0.818863 | 0.979198 | E130311K13Rik |
| ENSMUSG( | 87.20384 | 0.025689 | 0.911373 | 0.992196 | E130317F20Rik |
| ENSMUSG( | 99.63068 | 0.150927 | 0.602823 | 0.94789 | E230016M11Rik |
| ENSMUSG( | 99.35023 | 0.134426 | 0.566852 | 0.941777 | E230029C05Rik |
| ENSMUSG( | 592.8553 | 0.103131 | 0.424652 | 0.917257 | E2f1 |
| ENSMUSG( | 59.60781 | 0.155156 | 0.564607 | 0.941206 | E2f2 |
| ENSMUSG( | 726.6211 | 0.045638 | 0.733505 | 0.967228 | E2f3 |
| ENSMUSG( | 751.8581 | -0.01327 | 0.924484 | 0.99306 | E2f4 |
| ENSMUSG( | 449.0081 | -0.12776 | 0.44803 | 0.922202 | E2f5 |
| ENSMUSG( | 1392.87 | -0.04353 | 0.737656 | 0.967686 | E2f6 |
| ENSMUSG( | 19.13841 | 0.243876 | 0.602331 | 0.94789 | E2f7 |
| ENSMUSG( | 31.85944 | -0.10339 | 0.792177 | 0.97681 | E2f8 |
| ENSMUSG( | 9.572671 | -0.26812 | 0.682957 | NA | E330011O21Rik |
| ENSMUSG( | 22.25704 | -0.70285 | 0.120506 | 0.813384 | E330013P04Rik |
| ENSMUSG( | 824.0005 | 0.07029 | 0.643563 | 0.953903 | E4f1 |
| ENSMUSG( | 566.0463 | 0.06015 | 0.647691 | 0.95422 | Eaf1 |
| ENSMUSG( | 32.71866 | -0.19031 | 0.656046 | 0.955533 | Eaf2 |
| ENSMUSG( | 1206.836 | -0.11119 | 0.416679 | 0.917257 | Eapp |
| ENSMUSG( | 19.36484 | -1.57809 | 0.022228 | 0.577034 | Ear2 |
| ENSMUSG( | 132.506 | -0.04368 | 0.824167 | 0.979198 | Ears2 |
| ENSMUSG( | 659.1976 | 0.241689 | 0.096324 | 0.796133 | Ebag9 |
| ENSMUSG( | 112.7403 | 0.573175 | 0.023164 | 0.585577 | Ebf1 |
| ENSMUSG( | 10.18107 | -0.14856 | 0.80535 | NA | Ebf2 |
| ENSMUSG( | 25.2945 | -0.33036 | 0.409628 | 0.917257 | Ebf3 |
| ENSMUSG( | 384.6952 | 0.28895 | 0.039462 | 0.680377 | Ebf4 |
| ENSMUSG( | 56.41774 | -0.20264 | 0.458212 | 0.923329 | Ebi3 |
| ENSMUSG( | 852.7975 | -0.0551 | 0.674672 | 0.95694 | Ebna1bp2 |
| ENSMUSG( | 831.6528 | 0.013761 | 0.926414 | 0.99306 | Ebp |
| ENSMUSG( | 454.7625 | -0.14053 | 0.318921 | 0.903344 | Ebpl |
| ENSMUSG( | 790.6218 | -0.04133 | 0.742723 | 0.969415 | Ecd |
| ENSMUSG( | 2734.036 | 0.153082 | 0.35765 | 0.913759 | Ece1 |
| ENSMUSG( | 81.91443 | 0.093015 | 0.744625 | 0.96951 | Ece2 |
| ENSMUSG( | 277.0995 | -0.1047 | 0.592478 | 0.945041 | Ecel1 |
| ENSMUSG( | 1552.404 | -0.1755 | 0.118294 | 0.813384 | Ech1 |
| ENSMUSG( | 1577.097 | -0.24378 | 0.134416 | 0.823094 | Echdc1 |
| ENSMUSG( | 484.5531 | 0.192992 | 0.203414 | 0.864412 | Echdc2 |
| ENSMUSG( | 63.26823 | -0.48412 | 0.064311 | 0.737747 | Echdc3 |
| ENSMUSG( | 2234.703 | -0.17184 | 0.18278 | 0.850628 | Echs1 |
| ENSMUSG( | 1029.312 | -0.16851 | 0.192842 | 0.859764 | Eci1 |
| ENSMUSG( | 1632.874 | -0.1344 | 0.25039 | 0.87792 | Eci2 |
| ENSMUSG( | 325.2426 | 0.164878 | 0.291119 | 0.897421 | Ecm1 |
| ENSMUSG( | 182.001 | 0.418898 | 0.078117 | 0.77321 | Ecm2 |
| ENSMUSG( | 2546.822 | 0.064141 | 0.528337 | 0.934795 | Ecpas |
| ENSMUSG( | 329.8821 | 0.189311 | 0.426176 | 0.917257 | Ecrg4 |
| ENSMUSG( | 186.0581 | -0.01603 | 0.9375 | 0.993685 | Ecscr |
| ENSMUSG( | 997.9382 | -0.00716 | 0.957236 | 0.994798 | Ecsit |
| ENSMUSG( | 30.98027 | 0.236246 | 0.518205 | 0.930939 | Ect2 |
| ENSMUSG( | 12.00848 | 0.377264 | 0.517108 | 0.930939 | Ect2l |
| ENSMUSG( | 309.4406 | 0.243751 | 0.23858 | 0.872849 | Eda |
| ENSMUSG( | 14.74057 | 0.5619 | 0.29743 | 0.898538 | Eda2r |
| ENSMUSG( | 18.82682 | 0.556167 | 0.239188 | 0.873981 | Edar |
| ENSMUSG( | 411.6919 | 0.032929 | 0.818747 | 0.979198 | Edc3 |
| ENSMUSG( | 1548.893 | 0.111833 | 0.378688 | 0.916995 | Edc4 |
| ENSMUSG( | 906.2193 | 0.249694 | 0.166156 | 0.840082 | Edem1 |
| ENSMUSG( | 892.4029 | 0.124951 | 0.429655 | 0.917257 | Edem2 |
| ENSMUSG( | 1721.419 | 0.229712 | 0.168531 | 0.840222 | Edem3 |
| ENSMUSG( | 2933.329 | -0.21331 | 0.06769 | 0.75212 | Edf1 |
| ENSMUSG( | 2501.346 | 0.289096 | 0.164023 | 0.838291 | Edil3 |
| ENSMUSG( | 109.7477 | 0.927041 | 0.006635 | 0.338858 | Edn1 |
| ENSMUSG( | 218.6796 | 0.653316 | 0.008824 | 0.394786 | Edn3 |
| ENSMUSG( | 292.4293 | -0.14365 | 0.532267 | 0.934795 | Ednra |
| ENSMUSG( | 4263.408 | 0.094856 | 0.552866 | 0.939565 | Ednrb |
| ENSMUSG( | 505.4084 | 0.098271 | 0.474666 | 0.923329 | Edrf1 |
| ENSMUSG( | 1271.436 | 0.166431 | 0.201745 | 0.864412 | Eea1 |
| ENSMUSG( | 332.0403 | 0.085673 | 0.58742 | 0.944975 | Eed |
| ENSMUSG( | 77558.93 | -0.16481 | 0.144179 | 0.83148 | Eef1a1 |
| ENSMUSG( | 25233.63 | -0.09608 | 0.377203 | 0.915912 | Eef1a2 |
| ENSMUSG( | 346.1095 | -0.31383 | 0.039214 | 0.679972 | Eef1akmt1 |
| ENSMUSG( | 385.4609 | -0.05397 | 0.724756 | 0.966887 | Eef1akmt2 |
| ENSMUSG( | 18.69681 | 0.498499 | 0.315116 | 0.901306 | Eef1akmt4 |
| ENSMUSG( | 5950.455 | -0.0802 | 0.481163 | 0.924548 | Eef1b2 |
| ENSMUSG( | 2403.042 | -0.05529 | 0.644687 | 0.953903 | Eef1d |
| ENSMUSG( | 276.4936 | -0.11875 | 0.445253 | 0.921734 | Eef1e1 |
| ENSMUSG( | 8887.767 | -0.10134 | 0.367612 | 0.915217 | Eef1g |
| ENSMUSG( | 63632.81 | -0.11403 | 0.240671 | 0.875927 | Eef2 |
| ENSMUSG( | 1471.423 | 0.263974 | 0.033979 | 0.64909 | Eef2k |
| ENSMUSG( | 183.9283 | 0.044049 | 0.807871 | 0.978622 | Eef2kmt |
| ENSMUSG( | 75.44126 | -0.07412 | 0.762142 | 0.973255 | Eef2-ps2 |
| ENSMUSG( | 395.8996 | -0.08323 | 0.586861 | 0.944975 | Eefsec |
| ENSMUSG( | 4184.841 | -0.14009 | 0.154307 | 0.837724 | Eeig1 |
| ENSMUSG( | 1541.054 | 0.048494 | 0.735551 | 0.967344 | Eeig2 |
| ENSMUSG( | 362.8304 | -0.3505 | 0.033965 | 0.64909 | Eepd1 |
| ENSMUSG( | 29.62216 | -0.30622 | 0.406598 | 0.917257 | Efcab10 |
| ENSMUSG( | 10.19686 | -0.25326 | 0.676473 | NA | Efcab11 |
| ENSMUSG( | 132.695 | 0.168964 | 0.513501 | 0.930482 | Efcab12 |
| ENSMUSG( | 2978.713 | -0.00096 | 0.994389 | 0.998813 | Efcab14 |
| ENSMUSG( | 75.89488 | 0.12788 | 0.673906 | 0.95694 | Efcab15 |
| ENSMUSG( | 1051.053 | -0.21314 | 0.146679 | 0.836044 | Efcab2 |
| ENSMUSG( | 87.53039 | 0.057661 | 0.81228 | 0.979148 | Efcab5 |
| ENSMUSG( | 253.5048 | 0.109153 | 0.497668 | 0.927825 | Efcab6 |
| ENSMUSG( | 47.9106 | 0.082174 | 0.800256 | 0.97681 | Efcab7 |
| ENSMUSG( | 14.09854 | -0.36212 | 0.47853 | 0.923329 | Efcab9 |
| ENSMUSG( | 132.5926 | 0.267993 | 0.217224 | 0.865589 | Efcc1 |
| ENSMUSG( | 442.1943 | 0.076171 | 0.592806 | 0.945305 | Efemp1 |
| ENSMUSG( | 655.5727 | 0.003848 | 0.975931 | 0.996416 | Efemp2 |
| ENSMUSG( | 18.04052 | 0.647882 | 0.201114 | 0.86439 | Efhb |
| ENSMUSG( | 74.07475 | 0.493729 | 0.073482 | 0.763629 | Efhc1 |
| ENSMUSG( | 24.6803 | 0.569385 | 0.212891 | 0.864412 | Efhc2 |
| ENSMUSG( | 1275.068 | -0.29226 | 0.03619 | 0.668395 | Efhd1 |
| ENSMUSG( | 8505.911 | -0.11705 | 0.286078 | 0.893906 | Efhd2 |
| ENSMUSG( | 333.9776 | 0.007706 | 0.960073 | 0.994798 | Efl1 |
| ENSMUSG( | 183.8823 | 0.215794 | 0.374534 | 0.915491 | Efna1 |
| ENSMUSG( | 458.1114 | 0.199892 | 0.186645 | 0.856099 | Efna2 |
| ENSMUSG( | 2197.932 | 0.186538 | 0.168023 | 0.840222 | Efna3 |
| ENSMUSG( | 28.50587 | 0.118351 | 0.784402 | 0.976196 | Efna4 |
| ENSMUSG( | 504.3437 | 0.357875 | 0.067518 | 0.75212 | Efna5 |
| ENSMUSG( | 565.5111 | -0.16219 | 0.366458 | 0.914703 | Efnb1 |
| ENSMUSG( | 1762.154 | 0.354908 | 0.017885 | 0.536449 | Efnb2 |
| ENSMUSG( | 4321.362 | -0.0362 | 0.783549 | 0.976106 | Efnb3 |
| ENSMUSG( | 2965.723 | -0.00957 | 0.930152 | 0.99306 | Efr3a |
| ENSMUSG( | 6848.774 | -0.00111 | 0.990687 | 0.998287 | Efr3b |
| ENSMUSG( | 483.5782 | 0.018896 | 0.895319 | 0.990558 | Efs |
| ENSMUSG( | 2215.65 | -0.01131 | 0.923593 | 0.992815 | Eftud2 |
| ENSMUSG( | 42.0511 | 0.433278 | 0.169846 | 0.840806 | Egf |
| ENSMUSG( | 222.4182 | -0.04169 | 0.815951 | 0.979198 | Egfem1 |
| ENSMUSG( | 244.2213 | -0.1002 | 0.640762 | 0.953903 | Egfl6 |
| ENSMUSG( | 1094.764 | 0.207994 | 0.206456 | 0.864412 | Egfl7 |
| ENSMUSG( | 44.22822 | -0.08975 | 0.794117 | 0.97681 | Egfl8 |
| ENSMUSG( | 252.8085 | 0.35533 | 0.09747 | 0.796721 | Egflam |
| ENSMUSG( | 1218.845 | -0.06038 | 0.712348 | 0.964363 | Egfr |
| ENSMUSG( | 5286.696 | -0.01319 | 0.905828 | 0.991089 | Egln1 |
| ENSMUSG( | 3313.838 | 0.114082 | 0.243727 | 0.877904 | Egln2 |
| ENSMUSG( | 230.0282 | 0.005001 | 0.979107 | 0.996468 | Egln3 |
| ENSMUSG( | 4377.879 | -0.35177 | 0.350392 | 0.911652 | Egr1 |
| ENSMUSG( | 154.7261 | -1.06182 | 0.068501 | 0.752776 | Egr2 |
| ENSMUSG( | 2226.023 | -0.08162 | 0.485224 | 0.92541 | Egr3 |
| ENSMUSG( | 922.4199 | -0.50967 | 0.021967 | 0.575341 | Egr4 |
| ENSMUSG( | 1263.825 | 0.100962 | 0.466676 | 0.923329 | Ehbp1 |
| ENSMUSG( | 730.7124 | -0.0061 | 0.960005 | 0.994798 | Ehbp1l1 |
| ENSMUSG( | 1459.245 | -0.03215 | 0.771374 | 0.974236 | Ehd1 |
| ENSMUSG( | 301.1441 | 0.638005 | 0.001584 | 0.148357 | Ehd2 |
| ENSMUSG( | 2093.277 | -0.02443 | 0.861154 | 0.986226 | Ehd3 |

| | | | | | | | | | | | |
|---|---|---|---|---|---|---|---|---|---|---|---|
| ENSMUSG( | E130317F2 | 92.13965 | -0.09528 | 0.672895 | 0.929007 | ENSMUSG( | E130317F2 | 87.20384 | 0.025689 | 0.911373 | 0.992196 |
| ENSMUSG( | E230016M | 109.4871 | 0.156555 | 0.572093 | 0.893584 | ENSMUSG( | E230016M | 99.63068 | 0.150927 | 0.602823 | 0.94789 |
| ENSMUSG( | E230029C0 | 103.0623 | -0.04575 | 0.825495 | 0.967871 | ENSMUSG( | E230029C0 | 99.35023 | 0.134426 | 0.566852 | 0.941777 |
| ENSMUSG( | E2f1 | 610.5737 | -0.07972 | 0.482964 | 0.856216 | ENSMUSG( | E2f1 | 592.8553 | 0.103131 | 0.424652 | 0.917257 |
| ENSMUSG( | E2f2 | 61.82077 | -0.01669 | 0.950373 | 0.993821 | ENSMUSG( | E2f2 | 59.60781 | 0.155156 | 0.564607 | 0.941206 |
| ENSMUSG( | E2f3 | 777.7729 | -0.02082 | 0.832755 | 0.970333 | ENSMUSG( | E2f3 | 726.6211 | 0.045638 | 0.733505 | 0.967228 |
| ENSMUSG( | E2f4 | 861.2017 | 0.109926 | 0.355992 | 0.788009 | ENSMUSG( | E2f4 | 751.8581 | -0.01327 | 0.924484 | 0.99306 |
| ENSMUSG( | E2f5 | 520.5395 | 0.039402 | 0.789329 | 0.959379 | ENSMUSG( | E2f5 | 449.0081 | -0.12776 | 0.44803 | 0.922202 |
| ENSMUSG( | E2f6 | 1605.831 | 0.100819 | 0.282683 | 0.73186 | ENSMUSG( | E2f6 | 1392.87 | -0.04353 | 0.737656 | 0.967686 |
| ENSMUSG( | E2f7 | 16.67978 | -0.42878 | 0.388396 | NA | ENSMUSG( | E2f7 | 19.13841 | 0.243876 | 0.602331 | 0.94789 |
| ENSMUSG( | E2f8 | 31.87104 | -0.43956 | 0.237204 | 0.684843 | ENSMUSG( | E2f8 | 31.85944 | -0.10339 | 0.792177 | 0.97681 |
| ENSMUSG( | E330011O2 | 12.01558 | 0.155996 | 0.79411 | NA | ENSMUSG( | E330011O2 | 9.572671 | -0.26812 | 0.682957 | NA |
| ENSMUSG( | E330013P0 | 29.53039 | -0.06101 | 0.87721 | 0.981265 | ENSMUSG( | E330013P0 | 22.25704 | -0.70285 | 0.120506 | 0.813384 |
| ENSMUSG( | E4f1 | 961.7353 | 0.23839 | 0.03895 | 0.304318 | ENSMUSG( | E4f1 | 824.0005 | 0.07029 | 0.643563 | 0.953903 |
| ENSMUSG( | Eaf1 | 602.5369 | -0.02736 | 0.803253 | 0.962169 | ENSMUSG( | Eaf1 | 566.0463 | 0.06015 | 0.647691 | 0.95422 |
| ENSMUSG( | Eaf2 | 34.5934 | -0.30756 | 0.445747 | 0.833239 | ENSMUSG( | Eaf2 | 32.71866 | -0.19031 | 0.656046 | 0.955533 |
| ENSMUSG( | Eapp | 1436.136 | 0.126122 | 0.223226 | 0.669382 | ENSMUSG( | Eapp | 1206.836 | -0.11119 | 0.416679 | 0.917257 |
| ENSMUSG( | Ear2 | 22.40747 | -1.2511 | 0.062615 | 0.38686 | ENSMUSG( | Ear2 | 19.36484 | -1.57809 | 0.022228 | 0.577034 |
| ENSMUSG( | Ears2 | 153.3052 | 0.115246 | 0.493324 | 0.860611 | ENSMUSG( | Ears2 | 132.506 | -0.04368 | 0.824167 | 0.979198 |
| ENSMUSG( | Ebag9 | 604.5859 | -0.28203 | 0.027706 | 0.255176 | ENSMUSG( | Ebag9 | 659.1976 | 0.241689 | 0.096324 | 0.796133 |
| ENSMUSG( | Ebf1 | 88.25986 | -0.38604 | 0.136121 | 0.550232 | ENSMUSG( | Ebf1 | 112.7403 | 0.573175 | 0.023164 | 0.585577 |
| ENSMUSG( | Ebf2 | 12.05612 | 0.061399 | 0.908491 | NA | ENSMUSG( | Ebf2 | 10.18107 | -0.14856 | 0.80535 | NA |
| ENSMUSG( | Ebf3 | 30.31615 | -0.09108 | 0.789177 | 0.959299 | ENSMUSG( | Ebf3 | 25.2945 | -0.33036 | 0.409628 | 0.917257 |
| ENSMUSG( | Ebf4 | 401.5218 | 0.156836 | 0.24086 | 0.68836 | ENSMUSG( | Ebf4 | 384.6952 | 0.28895 | 0.039462 | 0.680377 |
| ENSMUSG( | Ebi3 | 57.81899 | -0.41902 | 0.102282 | 0.487127 | ENSMUSG( | Ebi3 | 56.41774 | -0.20264 | 0.458212 | 0.923329 |
| ENSMUSG( | Ebna1bp2 | 986.353 | 0.096369 | 0.370931 | 0.798261 | ENSMUSG( | Ebna1bp2 | 852.7975 | -0.0551 | 0.674672 | 0.95694 |
| ENSMUSG( | Ebp | 803.7774 | -0.38325 | 0.004973 | 0.093585 | ENSMUSG( | Ebp | 831.6528 | 0.013761 | 0.926414 | 0.99306 |
| ENSMUSG( | Ebpl | 504.4377 | -0.10381 | 0.351222 | 0.783067 | ENSMUSG( | Ebpl | 454.7625 | -0.14053 | 0.318921 | 0.903344 |
| ENSMUSG( | Ecd | 930.7168 | 0.156748 | 0.14378 | 0.562581 | ENSMUSG( | Ecd | 790.6218 | -0.04133 | 0.742723 | 0.969415 |
| ENSMUSG( | Ece1 | 2596.886 | -0.27957 | 0.058217 | 0.37191 | ENSMUSG( | Ece1 | 2734.036 | 0.153082 | 0.35765 | 0.913759 |
| ENSMUSG( | Ece2 | 93.90901 | 0.215818 | 0.411473 | 0.819309 | ENSMUSG( | Ece2 | 81.91443 | 0.093015 | 0.744625 | 0.96951 |
| ENSMUSG( | Ecel1 | 250.2152 | -0.7604 | 0.000621 | 0.02374 | ENSMUSG( | Ecel1 | 277.0995 | -0.1047 | 0.592478 | 0.945041 |
| ENSMUSG( | Ech1 | 1829.642 | 0.036953 | 0.686285 | 0.932993 | ENSMUSG( | Ech1 | 1552.404 | -0.1755 | 0.118294 | 0.813384 |
| ENSMUSG( | Echdc1 | 1952.559 | 0.116441 | 0.421229 | 0.823126 | ENSMUSG( | Echdc1 | 1577.097 | -0.24378 | 0.134416 | 0.823094 |
| ENSMUSG( | Echdc2 | 481.6571 | -0.08835 | 0.622371 | 0.911432 | ENSMUSG( | Echdc2 | 484.5531 | 0.192992 | 0.203414 | 0.864412 |
| ENSMUSG( | Echdc3 | 82.34846 | 0.046742 | 0.837046 | 0.971111 | ENSMUSG( | Echdc3 | 63.26823 | -0.48412 | 0.064311 | 0.737747 |
| ENSMUSG( | Echs1 | 2647.989 | 0.056234 | 0.568467 | 0.892252 | ENSMUSG( | Echs1 | 2234.703 | -0.17184 | 0.18278 | 0.850628 |
| ENSMUSG( | Eci1 | 1253.671 | 0.138175 | 0.167461 | 0.600006 | ENSMUSG( | Eci1 | 1029.312 | -0.16851 | 0.192842 | 0.859764 |
| ENSMUSG( | Eci2 | 1918.707 | 0.066773 | 0.554449 | 0.886657 | ENSMUSG( | Eci2 | 1632.874 | -0.1344 | 0.25039 | 0.87792 |
| ENSMUSG( | Ecm1 | 324.977 | -0.11202 | 0.416074 | 0.82084 | ENSMUSG( | Ecm1 | 325.2426 | 0.164878 | 0.291119 | 0.897421 |
| ENSMUSG( | Ecm2 | 154.1515 | -0.33341 | 0.118092 | 0.519373 | ENSMUSG( | Ecm2 | 182.001 | 0.418898 | 0.078117 | 0.77321 |
| ENSMUSG( | Ecpas | 2805.427 | 0.074211 | 0.35502 | 0.787423 | ENSMUSG( | Ecpas | 2546.822 | 0.064141 | 0.528337 | 0.934795 |
| ENSMUSG( | Ecrg4 | 302.0217 | -0.34243 | 0.451047 | 0.835567 | ENSMUSG( | Ecrg4 | 329.8821 | 0.189311 | 0.426176 | 0.917257 |
| ENSMUSG( | Ecscr | 186.0318 | -0.31747 | 0.082178 | 0.442642 | ENSMUSG( | Ecscr | 186.0581 | -0.01603 | 0.9375 | 0.993685 |
| ENSMUSG( | Ecsit | 1145.28 | 0.122231 | 0.270962 | 0.72132 | ENSMUSG( | Ecsit | 997.9382 | -0.00716 | 0.957236 | 0.994798 |
| ENSMUSG( | Ect2 | 28.88148 | -0.26007 | 0.457106 | 0.839312 | ENSMUSG( | Ect2 | 30.98027 | 0.236246 | 0.518205 | 0.930939 |
| ENSMUSG( | Ect2l | 12.41904 | 0.189849 | 0.719377 | NA | ENSMUSG( | Ect2l | 12.00848 | 0.377264 | 0.517108 | 0.930939 |
| ENSMUSG( | Eda | 294.7544 | -0.1716 | 0.367526 | 0.795314 | ENSMUSG( | Eda | 309.4406 | 0.243751 | 0.23858 | 0.872849 |
| ENSMUSG( | Eda2r | 11.48671 | -0.38986 | 0.516031 | NA | ENSMUSG( | Eda2r | 14.74057 | 0.5619 | 0.29743 | 0.898538 |
| ENSMUSG( | Edar | 16.33154 | -0.0924 | 0.84709 | NA | ENSMUSG( | Edar | 18.82682 | 0.556167 | 0.239188 | 0.873981 |
| ENSMUSG( | Edc3 | 463.591 | 0.099464 | 0.430352 | 0.826896 | ENSMUSG( | Edc3 | 411.6919 | 0.032929 | 0.818747 | 0.979198 |
| ENSMUSG( | Edc4 | 1804.72 | 0.271576 | 0.014572 | 0.177737 | ENSMUSG( | Edc4 | 1548.893 | 0.111833 | 0.378688 | 0.916995 |
| ENSMUSG( | Edem1 | 866.6864 | -0.14668 | 0.340299 | 0.777962 | ENSMUSG( | Edem1 | 906.2193 | 0.249694 | 0.166156 | 0.840082 |
| ENSMUSG( | Edem2 | 896.8515 | -0.13678 | 0.301026 | 0.749979 | ENSMUSG( | Edem2 | 892.4029 | 0.124951 | 0.429655 | 0.917257 |
| ENSMUSG( | Edem3 | 1591.36 | -0.27378 | 0.062439 | 0.386467 | ENSMUSG( | Edem3 | 1721.419 | 0.229712 | 0.168531 | 0.840222 |
| ENSMUSG( | Edf1 | 3401.134 | -0.04499 | 0.598554 | 0.903801 | ENSMUSG( | Edf1 | 2933.329 | -0.21331 | 0.06769 | 0.75212 |
| ENSMUSG( | Edil3 | 2079.157 | -0.55531 | 0.001509 | 0.043083 | ENSMUSG( | Edil3 | 2501.346 | 0.289096 | 0.164023 | 0.838291 |
| ENSMUSG( | Edn1 | 83.96001 | 0.006469 | 0.984018 | 0.99781 | ENSMUSG( | Edn1 | 109.7477 | 0.927041 | 0.006635 | 0.338858 |
| ENSMUSG( | Edn3 | 182.5857 | -0.08146 | 0.697861 | 0.935828 | ENSMUSG( | Edn3 | 218.6796 | 0.653316 | 0.008824 | 0.394786 |
| ENSMUSG( | Ednra | 302.3017 | -0.34576 | 0.095463 | 0.474828 | ENSMUSG( | Ednra | 292.4293 | -0.14365 | 0.532267 | 0.934795 |
| ENSMUSG( | Ednrb | 4234.64 | -0.19998 | 0.174124 | 0.60991 | ENSMUSG( | Ednrb | 4263.408 | 0.094856 | 0.552866 | 0.939565 |
| ENSMUSG( | Edrf1 | 573.0115 | 0.183776 | 0.133873 | 0.547197 | ENSMUSG( | Edrf1 | 505.4084 | 0.098271 | 0.474666 | 0.923329 |
| ENSMUSG( | Eea1 | 1417.809 | 0.211522 | 0.055988 | 0.364712 | ENSMUSG( | Eea1 | 1271.436 | 0.166431 | 0.201745 | 0.864412 |
| ENSMUSG( | Eed | 337.6961 | -0.13757 | 0.314403 | 0.761558 | ENSMUSG( | Eed | 332.0403 | 0.085673 | 0.58742 | 0.944975 |
| ENSMUSG( | Eef1a1 | 90916.3 | 0.029921 | 0.712513 | 0.940507 | ENSMUSG( | Eef1a1 | 77558.93 | -0.16481 | 0.144179 | 0.83148 |
| ENSMUSG( | Eef1a2 | 29929.43 | 0.127035 | 0.109957 | 0.500285 | ENSMUSG( | Eef1a2 | 25233.63 | -0.09608 | 0.377203 | 0.915912 |
| ENSMUSG( | Eef1akmt1 | 444.9826 | 0.159558 | 0.20057 | 0.640661 | ENSMUSG( | Eef1akmt1 | 346.1095 | -0.31383 | 0.039214 | 0.679972 |
| ENSMUSG( | Eef1akmt2 | 432.4297 | 0.016714 | 0.901486 | 0.985633 | ENSMUSG( | Eef1akmt2 | 385.4609 | -0.05397 | 0.724756 | 0.966887 |
| ENSMUSG( | Eef1akmt4 | 23.33968 | 0.826524 | 0.04898 | 0.337919 | ENSMUSG( | Eef1akmt4 | 18.69681 | 0.498499 | 0.315116 | 0.901306 |
| ENSMUSG( | Eef1b2 | 6864.289 | 0.065083 | 0.443157 | 0.833103 | ENSMUSG( | Eef1b2 | 5950.455 | -0.0802 | 0.481163 | 0.924548 |
| ENSMUSG( | Eef1d | 2832.052 | 0.149816 | 0.103716 | 0.489428 | ENSMUSG( | Eef1d | 2403.042 | -0.05529 | 0.644687 | 0.953903 |
| ENSMUSG( | Eef1e1 | 315.1169 | -0.002 | 0.987496 | 0.998796 | ENSMUSG( | Eef1e1 | 276.4936 | -0.11875 | 0.445253 | 0.921734 |
| ENSMUSG( | Eef1g | 10445.86 | 0.09777 | 0.293274 | 0.742763 | ENSMUSG( | Eef1g | 8887.767 | -0.10134 | 0.367612 | 0.915217 |
| ENSMUSG( | Eef2 | 75126.01 | 0.097227 | 0.192014 | 0.630811 | ENSMUSG( | Eef2 | 63632.81 | -0.11403 | 0.240671 | 0.875927 |
| ENSMUSG( | Eef2k | 1555.599 | 0.159574 | 0.104387 | 0.490485 | ENSMUSG( | Eef2k | 1471.423 | 0.263974 | 0.033979 | 0.64909 |
| ENSMUSG( | Eef2kmt | 204.0576 | 0.074346 | 0.621701 | 0.911432 | ENSMUSG( | Eef2kmt | 183.9283 | 0.044049 | 0.807871 | 0.978622 |
| ENSMUSG( | Eef2-ps2 | 93.55146 | 0.279526 | 0.184278 | 0.621284 | ENSMUSG( | Eef2-ps2 | 75.44126 | -0.07412 | 0.762142 | 0.973255 |
| ENSMUSG( | Eefsec | 454.005 | 0.050547 | 0.692023 | 0.934101 | ENSMUSG( | Eefsec | 395.8996 | -0.08323 | 0.586861 | 0.944975 |
| ENSMUSG( | Eeig1 | 4709.349 | -0.06515 | 0.398162 | 0.812452 | ENSMUSG( | Eeig1 | 4184.841 | -0.14009 | 0.154307 | 0.837724 |
| ENSMUSG( | Eeig2 | 1693.516 | 0.055575 | 0.641748 | 0.916828 | ENSMUSG( | Eeig2 | 1541.054 | 0.048494 | 0.735551 | 0.967344 |
| ENSMUSG( | Eepd1 | 405.7036 | -0.28958 | 0.052395 | 0.35132 | ENSMUSG( | Eepd1 | 362.8304 | -0.3505 | 0.033965 | 0.64909 |
| ENSMUSG( | Efcab10 | 30.09917 | -0.53808 | 0.120595 | 0.524506 | ENSMUSG( | Efcab10 | 29.62216 | -0.30622 | 0.406598 | 0.917257 |
| ENSMUSG( | Efcab11 | 11.20207 | -0.30018 | 0.601678 | NA | ENSMUSG( | Efcab11 | 10.19686 | -0.25326 | 0.676473 | NA |
| ENSMUSG( | Efcab12 | 136.7993 | -0.02816 | 0.889072 | 0.983434 | ENSMUSG( | Efcab12 | 132.695 | 0.168964 | 0.513501 | 0.930482 |
| ENSMUSG( | Efcab14 | 3151.134 | -0.1109 | 0.367863 | 0.795671 | ENSMUSG( | Efcab14 | 2978.713 | -0.00096 | 0.994389 | 0.998813 |
| ENSMUSG( | Efcab15 | 85.857 | 0.215481 | 0.400602 | 0.813717 | ENSMUSG( | Efcab15 | 75.89488 | 0.12788 | 0.673906 | 0.95694 |
| ENSMUSG( | Efcab2 | 1261.419 | 0.057362 | 0.624377 | 0.91187 | ENSMUSG( | Efcab2 | 1051.053 | -0.21314 | 0.146679 | 0.836044 |
| ENSMUSG( | Efcab5 | 91.53432 | -0.08668 | 0.697491 | 0.935662 | ENSMUSG( | Efcab5 | 87.53039 | 0.057661 | 0.81228 | 0.979148 |
| ENSMUSG( | Efcab6 | 283.4123 | 0.161438 | 0.269832 | 0.719729 | ENSMUSG( | Efcab6 | 253.5048 | 0.109153 | 0.497668 | 0.927825 |
| ENSMUSG( | Efcab7 | 49.77817 | -0.04873 | 0.865231 | 0.975565 | ENSMUSG( | Efcab7 | 47.9106 | 0.082174 | 0.800256 | 0.97681 |
| ENSMUSG( | Efcab9 | 15.62786 | -0.33717 | 0.466085 | NA | ENSMUSG( | Efcab9 | 14.09854 | -0.36212 | 0.47853 | 0.923329 |
| ENSMUSG( | Efcc1 | 133.6857 | 0.043084 | 0.834909 | 0.970426 | ENSMUSG( | Efcc1 | 132.5926 | 0.267993 | 0.217224 | 0.865589 |
| ENSMUSG( | Efemp1 | 431.3899 | -0.28245 | 0.033402 | 0.279991 | ENSMUSG( | Efemp1 | 442.1943 | 0.076171 | 0.592806 | 0.945305 |
| ENSMUSG( | Efemp2 | 718.5956 | -0.00374 | 0.973924 | 0.99675 | ENSMUSG( | Efemp2 | 655.5727 | 0.003848 | 0.975931 | 0.996416 |
| ENSMUSG( | Efhb | 15.67321 | 0.011134 | 0.983656 | NA | ENSMUSG( | Efhb | 18.04052 | 0.647882 | 0.201114 | 0.86439 |
| ENSMUSG( | Efhc1 | 68.85366 | 0.040881 | 0.874545 | 0.9795 | ENSMUSG( | Efhc1 | 74.07475 | 0.493729 | 0.073482 | 0.763629 |
| ENSMUSG( | Efhc2 | 18.13563 | -0.62146 | 0.194098 | NA | ENSMUSG( | Efhc2 | 24.6803 | 0.569385 | 0.212891 | 0.864412 |
| ENSMUSG( | Efhd1 | 1586.911 | 0.089091 | 0.444021 | 0.833239 | ENSMUSG( | Efhd1 | 1275.068 | -0.29226 | 0.03619 | 0.668395 |
| ENSMUSG( | Efhd2 | 10181.1 | 0.134149 | 0.107841 | 0.49674 | ENSMUSG( | Efhd2 | 8505.911 | -0.11705 | 0.286078 | 0.893906 |
| ENSMUSG( | Efl1 | 387.3967 | 0.164498 | 0.219596 | 0.664984 | ENSMUSG( | Efl1 | 333.9776 | 0.007706 | 0.960073 | 0.994798 |
| ENSMUSG( | Efna1 | 175.1123 | -0.21169 | 0.320919 | 0.765984 | ENSMUSG( | Efna1 | 183.8823 | 0.215794 | 0.374534 | 0.915491 |
| ENSMUSG( | Efna2 | 409.9233 | -0.41187 | 0.005271 | 0.09683 | ENSMUSG( | Efna2 | 458.1114 | 0.199892 | 0.186645 | 0.856099 |
| ENSMUSG( | Efna3 | 2013.815 | -0.353 | 0.002904 | 0.066108 | ENSMUSG( | Efna3 | 2197.932 | 0.186538 | 0.168023 | 0.840222 |
| ENSMUSG( | Efna4 | 26.6608 | -0.39341 | 0.320464 | 0.765921 | ENSMUSG( | Efna4 | 28.50587 | 0.118351 | 0.784402 | 0.976196 |
| ENSMUSG( | Efna5 | 463.3962 | -0.14188 | 0.406243 | 0.816525 | ENSMUSG( | Efna5 | 504.3437 | 0.357875 | 0.067518 | 0.75212 |
| ENSMUSG( | Efnb1 | 558.7306 | -0.51321 | 0.003676 | 0.077033 | ENSMUSG( | Efnb1 | 565.5111 | -0.16219 | 0.366458 | 0.914703 |
| ENSMUSG( | Efnb2 | 1519.018 | -0.34511 | 0.014655 | 0.178054 | ENSMUSG( | Efnb2 | 1762.154 | 0.354908 | 0.017885 | 0.536449 |
| ENSMUSG( | Efnb3 | 4066.289 | -0.53083 | 2.35E-06 | 0.000367 | ENSMUSG( | Efnb3 | 4321.362 | -0.0362 | 0.783549 | 0.976106 |
| ENSMUSG( | Efr3a | 3461.648 | 0.164595 | 0.06307 | 0.388234 | ENSMUSG( | Efr3a | 2965.723 | -0.00957 | 0.930152 | 0.99306 |
| ENSMUSG( | Efr3b | 8040.363 | 0.18707 | 0.011235 | 0.152627 | ENSMUSG( | Efr3b | 6848.774 | -0.00111 | 0.990687 | 0.998287 |
| ENSMUSG( | Efs | 498.097 | -0.17765 | 0.136133 | 0.550232 | ENSMUSG( | Efs | 483.5782 | 0.018896 | 0.895319 | 0.990558 |
| ENSMUSG( | Eftud2 | 2557.626 | 0.131292 | 0.189858 | 0.628983 | ENSMUSG( | Eftud2 | 2215.65 | -0.01131 | 0.923593 | 0.992815 |
| ENSMUSG( | Egf | 38.30867 | -0.07363 | 0.818401 | 0.965837 | ENSMUSG( | Egf | 42.0511 | 0.433278 | 0.169846 | 0.840806 |
| ENSMUSG( | Egfem1 | 246.5737 | -0.01216 | 0.940769 | 0.993415 | ENSMUSG( | Egfem1 | 222.4182 | -0.04169 | 0.815951 | 0.979198 |
| ENSMUSG( | Egfl6 | 277.0103 | 0.006117 | 0.972144 | 0.996585 | ENSMUSG( | Egfl6 | 244.2213 | -0.1002 | 0.640762 | 0.953903 |
| ENSMUSG( | Egfl7 | 1097.084 | -0.05373 | 0.72459 | 0.944096 | ENSMUSG( | Egfl7 | 1094.764 | 0.207994 | 0.206456 | 0.864412 |
| ENSMUSG( | Egfl8 | 43.10297 | -0.44952 | 0.156336 | 0.584143 | ENSMUSG( | Egfl8 | 44.22822 | -0.08975 | 0.794117 | 0.97681 |
| ENSMUSG( | Egflam | 248.6952 | 0.046694 | 0.809814 | 0.96372 | ENSMUSG( | Egflam | 252.8085 | 0.35533 | 0.09747 | 0.796721 |
| ENSMUSG( | Egfr | 1276.017 | -0.20745 | 0.149545 | 0.573617 | ENSMUSG( | Egfr | 1218.845 | -0.06038 | 0.712348 | 0.964363 |
| ENSMUSG( | Egln1 | 6030.561 | 0.098221 | 0.232603 | 0.680892 | ENSMUSG( | Egln1 | 5286.696 | -0.01319 | 0.905828 | 0.991089 |
| ENSMUSG( | Egln2 | 3773.229 | 0.214242 | 0.006466 | 0.110506 | ENSMUSG( | Egln2 | 3313.838 | 0.114082 | 0.243727 | 0.877904 |
| ENSMUSG( | Egln3 | 267.9863 | 0.18642 | 0.174489 | 0.610304 | ENSMUSG( | Egln3 | 230.0282 | 0.005001 | 0.979107 | 0.996468 |
| ENSMUSG( | Egr1 | 4855.967 | -0.3154 | 0.32229 | 0.767291 | ENSMUSG( | Egr1 | 4377.879 | -0.35177 | 0.350392 | 0.911652 |
| ENSMUSG( | Egr2 | 151.4157 | -1.64299 | 5.74E-08 | 1.79E-05 | ENSMUSG( | Egr2 | 154.7261 | -1.06182 | 0.068501 | 0.752776 |
| ENSMUSG( | Egr3 | 2576.174 | 0.07072 | 0.501283 | 0.864244 | ENSMUSG( | Egr3 | 2226.023 | -0.08162 | 0.485224 | 0.92541 |
| ENSMUSG( | Egr4 | 983.1013 | -0.61021 | 0.002807 | 0.06458 | ENSMUSG( | Egr4 | 922.4199 | -0.50967 | 0.021967 | 0.575341 |
| ENSMUSG( | Ehbp1 | 1433.832 | 0.195353 | 0.051065 | 0.346852 | ENSMUSG( | Ehbp1 | 1263.825 | 0.100962 | 0.466676 | 0.923329 |
| ENSMUSG( | Ehbp1l1 | 779.2829 | -0.08825 | 0.374604 | 0.800445 | ENSMUSG( | Ehbp1l1 | 730.7124 | -0.0061 | 0.960005 | 0.994798 |
| ENSMUSG( | Ehd1 | 1673.609 | 0.094546 | 0.293597 | 0.743041 | ENSMUSG( | Ehd1 | 1459.245 | -0.03215 | 0.771374 | 0.974236 |
| ENSMUSG( | Ehd2 | 291.3102 | 0.312711 | 0.051299 | 0.347567 | ENSMUSG( | Ehd2 | 301.1441 | 0.638005 | 0.001584 | 0.148357 |
| ENSMUSG( | Ehd3 | 2416.344 | 0.120224 | 0.334822 | 0.776386 | ENSMUSG( | Ehd3 | 2093.277 | -0.02443 | 0.861154 | 0.986226 |
| ENSMUSG( | Ehd4 | 350.3034 | -0.00613 | 0.961628 | 0.994286 | ENSMUSG( | Ehd4 | 328.7276 | 0.079649 | 0.583737 | 0.94434 |
| ENSMUSG( | Ehhadh | 112.3732 | -0.04547 | 0.807599 | 0.963257 | ENSMUSG( | Ehhadh | 105.4345 | 0.047868 | 0.820458 | 0.979198 |
| ENSMUSG( | Ehmt1 | 980.0234 | 0.121704 | 0.282837 | 0.73186 | ENSMUSG( | Ehmt1 | 873.7059 | 0.056034 | 0.678817 | 0.957617 |
| ENSMUSG( | Ehmt2 | 9297.658 | 0.164043 | 0.029475 | 0.260274 | ENSMUSG( | Ehmt2 | 8118.32 | 0.045074 | 0.639817 | 0.953903 |
| ENSMUSG( | Ei24 | 5554.517 | -0.04915 | 0.642713 | 0.916865 | ENSMUSG( | Ei24 | 5065.609 | -0.04633 | 0.70852 | 0.964042 |
| ENSMUSG( | Eid1 | 5042.188 | -0.09067 | 0.402224 | 0.815103 | ENSMUSG( | Eid1 | 4656.786 | -0.05428 | 0.725546 | 0.966887 |
| ENSMUSG( | Eid2 | 705.9347 | -0.09387 | 0.515574 | 0.872701 | ENSMUSG( | Eid2 | 602.501 | -0.30262 | 0.081487 | 0.77681 |
| ENSMUSG( | Eid2b | 466.1288 | 0.124223 | 0.326095 | 0.771128 | ENSMUSG( | Eid2b | 424.5849 | 0.117539 | 0.428184 | 0.917257 |
| ENSMUSG( | Eif1 | 9090.393 | 0.048245 | 0.586894 | 0.900235 | ENSMUSG( | Eif1 | 7775.222 | -0.1386 | 0.230283 | 0.868507 |
| ENSMUSG( | Eif1a | 509.611 | -0.01849 | 0.901159 | 0.985633 | ENSMUSG( | Eif1a | 463.1413 | -0.03336 | 0.851738 | 0.985591 |
| ENSMUSG( | Eif1ad | 734.7278 | 0.046464 | 0.679675 | 0.930983 | ENSMUSG( | Eif1ad | 668.2316 | 0.038748 | 0.773139 | 0.974463 |
| ENSMUSG( | Eif1ax | 4470.122 | 0.031449 | 0.797184 | 0.960507 | ENSMUSG( | Eif1ax | 3928.511 | -0.07722 | 0.635379 | 0.953903 |
| ENSMUSG( | Eif1b | 1849.979 | 0.12306 | 0.252053 | 0.701908 | ENSMUSG( | Eif1b | 1601.471 | -0.0261 | 0.856003 | 0.985884 |
| ENSMUSG( | Eif2a | 1096.066 | -0.05945 | 0.559108 | 0.888638 | ENSMUSG( | Eif2a | 983.3126 | -0.1086 | 0.375757 | 0.915491 |
| ENSMUSG( | Eif2ak1 | 2605.305 | -0.06692 | 0.499034 | 0.862973 | ENSMUSG( | Eif2ak1 | 2385.032 | -0.05021 | 0.661022 | 0.956205 |
| ENSMUSG( | Eif2ak2 | 627.5247 | 0.053923 | 0.679736 | 0.930983 | ENSMUSG( | Eif2ak2 | 536.6879 | -0.13676 | 0.40442 | 0.917257 |
| ENSMUSG( | Eif2ak3 | 973.6417 | -0.11824 | 0.42558 | 0.824898 | ENSMUSG( | Eif2ak3 | 1029.404 | 0.304901 | 0.083021 | 0.77681 |
| ENSMUSG( | Eif2ak4 | 685.4852 | 0.080436 | 0.435103 | 0.829093 | ENSMUSG( | Eif2ak4 | 635.9193 | 0.134921 | 0.327581 | 0.906255 |
| ENSMUSG( | Eif2b1 | 546.1351 | 0.101814 | 0.438969 | 0.830714 | ENSMUSG( | Eif2b1 | 468.568 | -0.07545 | 0.614815 | 0.951177 |
| ENSMUSG( | Eif2b2 | 675.0168 | 0.013726 | 0.901389 | 0.985633 | ENSMUSG( | Eif2b2 | 592.1867 | -0.09659 | 0.457406 | 0.923329 |
| ENSMUSG( | Eif2b3 | 255.2393 | 0.039504 | 0.807306 | 0.963257 | ENSMUSG( | Eif2b3 | 223.3171 | -0.07852 | 0.667114 | 0.956766 |
| ENSMUSG( | Eif2b4 | 953.0783 | 0.116518 | 0.241612 | 0.689179 | ENSMUSG( | Eif2b4 | 825.9944 | -0.02949 | 0.819883 | 0.979198 |
| ENSMUSG( | Eif2b5 | 1811.26 | 0.082523 | 0.340292 | 0.777962 | ENSMUSG( | Eif2b5 | 1549.826 | -0.09888 | 0.349005 | 0.911652 |
| ENSMUSG( | Eif2d | 831.4357 | 0.110618 | 0.393245 | 0.809274 | ENSMUSG( | Eif2d | 743.9493 | 0.063761 | 0.68112 | 0.958395 |
| ENSMUSG( | Eif2s1 | 759.0236 | 0.084787 | 0.444943 | 0.833239 | ENSMUSG( | Eif2s1 | 675.818 | 0.018232 | 0.884348 | 0.989688 |
| ENSMUSG( | Eif2s2 | 1667.407 | 0.07241 | 0.425969 | 0.824898 | ENSMUSG( | Eif2s2 | 1448.553 | -0.0676 | 0.608435 | 0.950113 |
| ENSMUSG( | Eif2s3x | 1131.333 | 0.082439 | 0.369282 | 0.796723 | ENSMUSG( | Eif2s3x | 956.3837 | -0.13673 | 0.229929 | 0.868507 |

| | | | | | |
|---|---|---|---|---|---|
| ENSMUSG( | 1276.017 | -0.20745 | 0.149545 | 0.573617 | Egfr |
| ENSMUSG( | 6030.561 | 0.098221 | 0.232603 | 0.680892 | Egln1 |
| ENSMUSG( | 3773.229 | 0.214242 | 0.006466 | 0.110506 | Egln2 |
| ENSMUSG( | 267.9863 | 0.18642 | 0.174489 | 0.610304 | Egln3 |
| ENSMUSG( | 4855.967 | -0.3154 | 0.32229 | 0.767291 | Egr1 |
| ENSMUSG( | 151.4157 | -1.64299 | 5.74E-08 | 1.79E-05 | Egr2 |
| ENSMUSG( | 2576.174 | 0.07072 | 0.501283 | 0.864244 | Egr3 |
| ENSMUSG( | 983.1013 | -0.61021 | 0.002807 | 0.06458 | Egr4 |
| ENSMUSG( | 1433.832 | 0.195353 | 0.051065 | 0.346852 | Ehbp1 |
| ENSMUSG( | 779.2829 | -0.08825 | 0.374604 | 0.800445 | Ehbp1l1 |
| ENSMUSG( | 1673.609 | 0.094546 | 0.293597 | 0.743041 | Ehd1 |
| ENSMUSG( | 291.3102 | 0.312711 | 0.051299 | 0.347567 | Ehd2 |
| ENSMUSG( | 2416.344 | 0.120224 | 0.334822 | 0.776386 | Ehd3 |
| ENSMUSG( | 350.3034 | -0.00613 | 0.961628 | 0.994286 | Ehd4 |
| ENSMUSG( | 112.3732 | -0.04547 | 0.807599 | 0.963257 | Ehhadh |
| ENSMUSG( | 980.0234 | 0.121704 | 0.282837 | 0.73186 | Ehmt1 |
| ENSMUSG( | 9297.658 | 0.164043 | 0.029475 | 0.260274 | Ehmt2 |
| ENSMUSG( | 5554.517 | -0.04915 | 0.642713 | 0.916865 | Ei24 |
| ENSMUSG( | 5042.188 | -0.09067 | 0.402224 | 0.815103 | Eid1 |
| ENSMUSG( | 705.9347 | -0.09387 | 0.515574 | 0.872701 | Eid2 |
| ENSMUSG( | 466.1288 | 0.124223 | 0.326095 | 0.771128 | Eid2b |
| ENSMUSG( | 9090.393 | 0.048245 | 0.586894 | 0.900235 | Eif1 |
| ENSMUSG( | 509.611 | -0.01849 | 0.901159 | 0.985633 | Eif1a |
| ENSMUSG( | 734.7278 | 0.046464 | 0.679675 | 0.930983 | Eif1ad |
| ENSMUSG( | 4470.122 | 0.031449 | 0.797184 | 0.960507 | Eif1ax |
| ENSMUSG( | 1849.979 | 0.12306 | 0.252053 | 0.701908 | Eif1b |
| ENSMUSG( | 1096.066 | -0.05945 | 0.559108 | 0.888638 | Eif2a |
| ENSMUSG( | 2605.305 | -0.06692 | 0.499034 | 0.862973 | Eif2ak1 |
| ENSMUSG( | 627.5247 | 0.053923 | 0.679736 | 0.930983 | Eif2ak2 |
| ENSMUSG( | 973.6417 | -0.11824 | 0.42558 | 0.824898 | Eif2ak3 |
| ENSMUSG( | 685.4852 | 0.080436 | 0.435103 | 0.829093 | Eif2ak4 |
| ENSMUSG( | 546.1351 | 0.101814 | 0.438969 | 0.830714 | Eif2b1 |
| ENSMUSG( | 675.0168 | 0.013726 | 0.901389 | 0.985633 | Eif2b2 |
| ENSMUSG( | 255.2393 | 0.039504 | 0.807306 | 0.963257 | Eif2b3 |
| ENSMUSG( | 953.0783 | 0.116518 | 0.241612 | 0.689179 | Eif2b4 |
| ENSMUSG( | 1811.26 | 0.082523 | 0.340292 | 0.777962 | Eif2b5 |
| ENSMUSG( | 831.4357 | 0.110618 | 0.393245 | 0.809274 | Eif2d |
| ENSMUSG( | 759.0236 | 0.084787 | 0.444943 | 0.833239 | Eif2s1 |
| ENSMUSG( | 1667.407 | 0.07241 | 0.425969 | 0.824898 | Eif2s2 |
| ENSMUSG( | 1131.333 | 0.082439 | 0.369282 | 0.796723 | Eif2s3x |
| ENSMUSG( | 1272.578 | 0.060367 | 0.602176 | 0.904887 | Eif2s3y |
| ENSMUSG( | 5967.754 | 0.145428 | 0.146856 | 0.56887 | Eif3a |
| ENSMUSG( | 2926.73 | 0.151845 | 0.082767 | 0.444372 | Eif3b |
| ENSMUSG( | 4619.426 | 0.110032 | 0.22165 | 0.667258 | Eif3c |
| ENSMUSG( | 1789.144 | 0.0204 | 0.844884 | 0.972023 | Eif3d |
| ENSMUSG( | 2177.215 | 0.020696 | 0.8295 | 0.969343 | Eif3e |
| ENSMUSG( | 7373.382 | 0.088743 | 0.249862 | 0.699512 | Eif3f |
| ENSMUSG( | 1526.276 | -0.01608 | 0.858467 | 0.974683 | Eif3g |
| ENSMUSG( | 5132.38 | 0.05331 | 0.504632 | 0.866883 | Eif3h |
| ENSMUSG( | 2501.769 | 0.089444 | 0.308548 | 0.756544 | Eif3i |
| ENSMUSG( | 1344.661 | 0.12467 | 0.315251 | 0.762288 | Eif3j1 |
| ENSMUSG( | 22.02044 | 0.330901 | 0.413942 | 0.819998 | Eif3j2 |
| ENSMUSG( | 2812.271 | 0.001048 | 0.989639 | 0.999144 | Eif3k |
| ENSMUSG( | 2010.349 | -0.00059 | 0.99536 | 0.99943 | Eif3l |
| ENSMUSG( | 1358.485 | 0.041348 | 0.697886 | 0.935828 | Eif3m |
| ENSMUSG( | 5186.251 | 0.006814 | 0.951721 | 0.993821 | Eif4a1 |
| ENSMUSG( | 8508.288 | 0.030377 | 0.718059 | 0.941697 | Eif4a2 |
| ENSMUSG( | 1204.546 | 0.063344 | 0.526914 | 0.877267 | Eif4a3 |
| ENSMUSG( | 5926.241 | 1.94E-05 | 0.999814 | 0.99989 | Eif4b |
| ENSMUSG( | 2257.114 | -0.00384 | 0.968389 | 0.996369 | Eif4e |
| ENSMUSG( | 1439.43 | 0.091779 | 0.295992 | 0.744386 | Eif4e2 |
| ENSMUSG( | 385.2931 | 0.097798 | 0.453987 | 0.836798 | Eif4e3 |
| ENSMUSG( | 153.4976 | 0.253194 | 0.128534 | 0.536774 | Eif4ebp1 |
| ENSMUSG( | 3693.505 | 0.177827 | 0.065905 | 0.397536 | Eif4ebp2 |
| ENSMUSG( | 12.09173 | 0.253979 | 0.633629 | NA | Eif4ebp3 |
| ENSMUSG( | 1362.548 | 0.099059 | 0.301114 | 0.749979 | Eif4enif1 |
| ENSMUSG( | 8556.198 | 0.136778 | 0.087797 | 0.454663 | Eif4g1 |
| ENSMUSG( | 26762.93 | 0.095198 | 0.338694 | 0.777518 | Eif4g2 |
| ENSMUSG( | 8712.006 | 0.083808 | 0.300916 | 0.749979 | Eif4g3 |
| ENSMUSG( | 11381.8 | 0.150883 | 0.042337 | 0.316636 | Eif4h |
| ENSMUSG( | 7909.787 | 0.037885 | 0.728109 | 0.944355 | Eif5 |
| ENSMUSG( | 9475.7 | 0.004078 | 0.960388 | 0.993991 | Eif5a |
| ENSMUSG( | 1152.689 | 0.040047 | 0.691483 | 0.933881 | Eif5a2 |
| ENSMUSG( | 2373.742 | 0.197317 | 0.069345 | 0.40782 | Eif5b |
| ENSMUSG( | 990.2693 | 0.1045 | 0.346358 | 0.779153 | Eif6 |
| ENSMUSG( | 888.1946 | 0.076405 | 0.53521 | 0.880733 | Eipr1 |
| ENSMUSG( | 728.7117 | -0.06348 | 0.565793 | 0.89104 | Elac1 |
| ENSMUSG( | 1161.724 | 0.194151 | 0.043343 | 0.319117 | Elac2 |
| ENSMUSG( | 1305.769 | -0.21765 | 0.052781 | 0.352597 | Elapor1 |
| ENSMUSG( | 813.2278 | -0.32085 | 0.010924 | 0.150598 | Elapor2 |
| ENSMUSG( | 1330.788 | 0.06093 | 0.505119 | 0.866883 | Elavl1 |
| ENSMUSG( | 658.3427 | -0.07303 | 0.564663 | 0.890434 | Elavl2 |
| ENSMUSG( | 7745.789 | 0.099917 | 0.28476 | 0.733422 | Elavl3 |
| ENSMUSG( | 959.6119 | -0.03916 | 0.749307 | 0.949664 | Elavl4 |
| ENSMUSG( | 10.31499 | -0.03606 | 0.959182 | NA | Eldr |
| ENSMUSG( | 272.4718 | -0.1646 | 0.346231 | 0.779153 | Elf1 |
| ENSMUSG( | 1085.434 | 0.071139 | 0.459193 | 0.840244 | Elf2 |
| ENSMUSG( | 90.78302 | 0.009384 | 0.966052 | 0.995377 | Elf4 |
| ENSMUSG( | 83.28809 | 0.163547 | 0.550302 | 0.886038 | Elf5 |
| ENSMUSG( | 1239.085 | -0.08064 | 0.53673 | 0.882043 | Elfn1 |
| ENSMUSG( | 5549.921 | 0.111153 | 0.361794 | 0.792797 | Elfn2 |
| ENSMUSG( | 2399.237 | 0.069796 | 0.442179 | 0.832774 | Elk1 |
| ENSMUSG( | 424.4374 | -0.13626 | 0.355692 | 0.787945 | Elk3 |
| ENSMUSG( | 834.7484 | 0.012207 | 0.912064 | 0.988483 | Elk4 |
| ENSMUSG( | 656.7659 | 0.025693 | 0.837629 | 0.971111 | Ell |
| ENSMUSG( | 582.7017 | 0.080324 | 0.476506 | 0.851672 | Ell2 |
| ENSMUSG( | 2440.392 | -0.08701 | 0.495109 | 0.860798 | Elmo1 |
| ENSMUSG( | 5977.398 | 0.11592 | 0.14521 | 0.565771 | Elmo2 |
| ENSMUSG( | 70.93688 | 0.095036 | 0.686218 | 0.932981 | Elmo3 |
| ENSMUSG( | 6353.671 | -0.7027 | 3.52E-08 | 1.19E-05 | Elmod1 |
| ENSMUSG( | 499.5116 | -0.28619 | 0.063677 | 0.390618 | Elmod2 |
| ENSMUSG( | 997.5887 | 0.094224 | 0.374427 | 0.800445 | Elmod3 |
| ENSMUSG( | 389.1607 | 0.061093 | 0.680135 | 0.930983 | Eln |
| ENSMUSG( | 1102.89 | 0.122181 | 0.21689 | 0.661816 | Eloa |
| ENSMUSG( | 4329.889 | -0.05982 | 0.517907 | 0.873754 | Elob |
| ENSMUSG( | 3289.856 | -0.02154 | 0.820046 | 0.965888 | Eloc |
| ENSMUSG( | 1409.56 | 0.033455 | 0.71476 | 0.941144 | Elof1 |
| ENSMUSG( | 1094.675 | -0.46325 | 0.003622 | 0.076313 | Elovl1 |
| ENSMUSG( | 1634.56 | -0.10527 | 0.5017 | 0.864596 | Elovl2 |
| ENSMUSG( | 1960.317 | -0.53731 | 0.00018 | 0.009729 | Elovl4 |
| ENSMUSG( | 5010.204 | -0.35264 | 0.024626 | 0.240618 | Elovl5 |
| ENSMUSG( | 2150.469 | -0.06348 | 0.659474 | 0.924506 | Elovl6 |
| ENSMUSG( | 520.1298 | -0.60874 | 0.001292 | 0.039144 | Elovl7 |
| ENSMUSG( | 1579.993 | 0.099215 | 0.302307 | 0.750677 | Elp1 |
| ENSMUSG( | 2034.469 | 0.070338 | 0.516943 | 0.873055 | Elp2 |
| ENSMUSG( | 1887.536 | 0.066755 | 0.517409 | 0.873295 | Elp3 |
| ENSMUSG( | 526.2767 | 0.012655 | 0.919401 | 0.990247 | Elp4 |
| ENSMUSG( | 2488.298 | 0.068105 | 0.395037 | 0.810154 | Elp5 |
| ENSMUSG( | 298.0772 | -0.12359 | 0.384796 | 0.804031 | Elp6 |
| ENSMUSG( | 802.0126 | -0.25639 | 0.019533 | 0.210473 | Emb |
| ENSMUSG( | 3347.967 | -0.26552 | 0.021936 | 0.225738 | Emc1 |
| ENSMUSG( | 7748.965 | -0.2669 | 0.035117 | 0.288256 | Emc10 |
| ENSMUSG( | 1543.999 | 0.079884 | 0.456492 | 0.838832 | Emc2 |
| ENSMUSG( | 4500.154 | -0.22601 | 0.068979 | 0.406851 | Emc3 |
| ENSMUSG( | 2985.363 | -0.57024 | 3.49E-07 | 7.75E-05 | Emc4 |
| ENSMUSG( | 1424.639 | 0.04299 | 0.651264 | 0.921451 | Emc6 |
| ENSMUSG( | 5119.837 | -0.23357 | 0.06893 | 0.406851 | Emc7 |
| ENSMUSG( | 4031.533 | 0.023703 | 0.800265 | 0.961745 | Emc8 |
| ENSMUSG( | 464.3434 | -0.08308 | 0.49811 | 0.862463 | Emc9 |
| ENSMUSG( | 320.1885 | -0.12779 | 0.516357 | 0.872701 | Emcn |
| ENSMUSG( | 569.8848 | -0.06779 | 0.622293 | 0.911432 | Emd |
| ENSMUSG( | 10.4435 | -0.32747 | 0.608613 | NA | Eme1 |
| ENSMUSG( | 432.1219 | 0.187715 | 0.116472 | 0.514944 | Eme2 |
| ENSMUSG( | 711.6941 | 0.087558 | 0.401684 | 0.814923 | Emg1 |
| ENSMUSG( | 451.0747 | -0.45093 | 0.006504 | 0.110626 | Emid1 |
| ENSMUSG( | 177.8458 | -0.47928 | 0.006216 | 0.107827 | Emilin1 |
| ENSMUSG( | 175.2495 | -0.34613 | 0.046354 | 0.329764 | Emilin2 |
| ENSMUSG( | 2260.933 | 0.007372 | 0.935632 | 0.993363 | Eml1 |
| ENSMUSG( | 6545.792 | 0.036323 | 0.703797 | 0.937933 | Eml2 |
| ENSMUSG( | 588.6062 | 0.087634 | 0.42622 | 0.824898 | Eml3 |
| ENSMUSG( | 1186.1 | 0.226448 | 0.03454 | 0.285252 | Eml4 |
| ENSMUSG( | 1796.326 | -0.09069 | 0.391348 | 0.808825 | Eml5 |
| ENSMUSG( | 1117.655 | 0.198008 | 0.039926 | 0.308163 | Eml6 |
| ENSMUSG( | 60.47382 | -0.45918 | 0.123456 | 0.530552 | Emp1 |
| ENSMUSG( | 4438.645 | 0.07108 | 0.583485 | 0.898521 | Emp2 |
| ENSMUSG( | 111.1351 | -0.35413 | 0.075896 | 0.426699 | Emp3 |
| ENSMUSG( | 1510.966 | 0.183104 | 0.085795 | 0.451465 | Emsy |
| ENSMUSG( | 507.5977 | 0.117447 | 0.37739 | 0.801213 | Emx1 |
| ENSMUSG( | 1556.131 | 0.330475 | 0.003874 | 0.079851 | Emx2 |
| ENSMUSG( | 303.7717 | 0.127188 | 0.400249 | 0.813641 | Emx2os |
| ENSMUSG( | 16.96939 | 0.244522 | 0.649033 | NA | En2 |
| ENSMUSG( | 8950.365 | 0.180307 | 0.103477 | 0.489288 | Enah |
| ENSMUSG( | 12957.2 | 0.168748 | 0.163843 | 0.594869 | Enc1 |
| ENSMUSG( | 2924.573 | -0.47058 | 0.003532 | 0.075287 | Endod1 |
| ENSMUSG( | 393.0407 | -0.00172 | 0.989706 | 0.999144 | Endog |
| ENSMUSG( | 178.3421 | -0.2867 | 0.189386 | 0.627932 | Endou |
| ENSMUSG( | 1597.942 | 0.032931 | 0.722593 | 0.943752 | Endov |
| ENSMUSG( | 1363.793 | 0.063741 | 0.751641 | 0.950317 | Eng |

| | | | | | |
|---|---|---|---|---|---|
| ENSMUSG( | 328.7276 | 0.079649 | 0.583737 | 0.94434 | Ehd4 |
| ENSMUSG( | 105.4345 | 0.047868 | 0.820458 | 0.979198 | Ehhadh |
| ENSMUSG( | 873.7059 | 0.056034 | 0.678817 | 0.957617 | Ehmt1 |
| ENSMUSG( | 8118.32 | 0.045074 | 0.639817 | 0.953903 | Ehmt2 |
| ENSMUSG( | 5065.609 | -0.04633 | 0.70852 | 0.964042 | Ei24 |
| ENSMUSG( | 4656.786 | -0.05428 | 0.725546 | 0.966887 | Eid1 |
| ENSMUSG( | 602.501 | -0.30262 | 0.081487 | 0.77681 | Eid2 |
| ENSMUSG( | 424.5849 | 0.117539 | 0.428184 | 0.917257 | Eid2b |
| ENSMUSG( | 7775.222 | -0.1386 | 0.230283 | 0.868507 | Eif1 |
| ENSMUSG( | 463.1413 | -0.03336 | 0.851738 | 0.985591 | Eif1a |
| ENSMUSG( | 668.2316 | 0.038748 | 0.773139 | 0.974463 | Eif1ad |
| ENSMUSG( | 3928.511 | -0.07722 | 0.635379 | 0.953903 | Eif1ax |
| ENSMUSG( | 1601.471 | -0.0261 | 0.856003 | 0.985884 | Eif1b |
| ENSMUSG( | 983.3126 | -0.1086 | 0.375757 | 0.915491 | Eif2a |
| ENSMUSG( | 2385.032 | -0.05021 | 0.661022 | 0.956205 | Eif2ak1 |
| ENSMUSG( | 536.6879 | -0.13676 | 0.40442 | 0.917257 | Eif2ak2 |
| ENSMUSG( | 1029.404 | 0.304901 | 0.083021 | 0.77681 | Eif2ak3 |
| ENSMUSG( | 635.9193 | 0.134921 | 0.327581 | 0.906255 | Eif2ak4 |
| ENSMUSG( | 468.568 | -0.07545 | 0.614815 | 0.951177 | Eif2b1 |
| ENSMUSG( | 592.1867 | -0.09659 | 0.457406 | 0.923329 | Eif2b2 |
| ENSMUSG( | 223.3171 | -0.07852 | 0.667114 | 0.956766 | Eif2b3 |
| ENSMUSG( | 825.9944 | -0.02949 | 0.819883 | 0.979198 | Eif2b4 |
| ENSMUSG( | 1549.826 | -0.09888 | 0.349005 | 0.911652 | Eif2b5 |
| ENSMUSG( | 743.9493 | 0.063761 | 0.68112 | 0.958395 | Eif2d |
| ENSMUSG( | 675.818 | 0.018232 | 0.884348 | 0.989688 | Eif2s1 |
| ENSMUSG( | 1448.553 | -0.0676 | 0.608435 | 0.950113 | Eif2s2 |
| ENSMUSG( | 956.3837 | -0.13673 | 0.229929 | 0.868507 | Eif2s3x |
| ENSMUSG( | 1135.109 | 0.000642 | 0.997161 | 0.999268 | Eif2s3y |
| ENSMUSG( | 5155.03 | -0.00648 | 0.956497 | 0.994798 | Eif3a |
| ENSMUSG( | 2511.594 | -0.0166 | 0.875047 | 0.989644 | Eif3b |
| ENSMUSG( | 3807.729 | -0.18389 | 0.096884 | 0.79645 | Eif3c |
| ENSMUSG( | 1561.753 | -0.10509 | 0.38832 | 0.917257 | Eif3d |
| ENSMUSG( | 1938.827 | -0.04893 | 0.69875 | 0.960908 | Eif3e |
| ENSMUSG( | 6294.866 | -0.10082 | 0.362701 | 0.914223 | Eif3f |
| ENSMUSG( | 1345.59 | -0.1158 | 0.329267 | 0.907083 | Eif3g |
| ENSMUSG( | 4414.031 | -0.11677 | 0.34636 | 0.910966 | Eif3h |
| ENSMUSG( | 2074.721 | -0.18785 | 0.104392 | 0.804969 | Eif3i |
| ENSMUSG( | 1158.693 | -0.03854 | 0.802742 | 0.97681 | Eif3j1 |
| ENSMUSG( | 19.56635 | 0.216379 | 0.643029 | 0.953903 | Eif3j2 |
| ENSMUSG( | 2391.437 | -0.20787 | 0.078459 | 0.77321 | Eif3k |
| ENSMUSG( | 1762.418 | -0.11479 | 0.339017 | 0.907972 | Eif3l |
| ENSMUSG( | 1186.919 | -0.08431 | 0.540621 | 0.937383 | Eif3m |
| ENSMUSG( | 4655.086 | -0.03921 | 0.770654 | 0.974236 | Eif4a1 |
| ENSMUSG( | 7595.863 | -0.02849 | 0.784695 | 0.976231 | Eif4a2 |
| ENSMUSG( | 1041.262 | -0.09083 | 0.45493 | 0.922891 | Eif4a3 |
| ENSMUSG( | 5244.325 | -0.08619 | 0.452169 | 0.922316 | Eif4b |
| ENSMUSG( | 2001.214 | -0.08815 | 0.541807 | 0.937841 | Eif4e |
| ENSMUSG( | 1283.199 | 0.029159 | 0.791451 | 0.97681 | Eif4e2 |
| ENSMUSG( | 342.9862 | 0.026456 | 0.862138 | 0.986355 | Eif4e3 |
| ENSMUSG( | 129.8923 | 0.038355 | 0.847955 | 0.985242 | Eif4ebp1 |
| ENSMUSG( | 3263.856 | 0.095252 | 0.394823 | 0.917257 | Eif4ebp2 |
| ENSMUSG( | 11.37598 | 0.365616 | 0.530633 | NA | Eif4ebp3 |
| ENSMUSG( | 1235.316 | 0.082884 | 0.486208 | 0.92541 | Eif4enif1 |
| ENSMUSG( | 7538.721 | 0.044148 | 0.659764 | 0.956205 | Eif4g1 |
| ENSMUSG( | 23165.59 | -0.05261 | 0.674045 | 0.95694 | Eif4g2 |
| ENSMUSG( | 7591.72 | -0.04414 | 0.668827 | 0.956766 | Eif4g3 |
| ENSMUSG( | 9703.365 | -0.03744 | 0.696279 | 0.960867 | Eif4h |
| ENSMUSG( | 6906.775 | -0.08796 | 0.549956 | 0.938699 | Eif5 |
| ENSMUSG( | 7949.876 | -0.24662 | 0.057171 | 0.723362 | Eif5a |
| ENSMUSG( | 1019.05 | -0.05215 | 0.713399 | 0.964766 | Eif5a2 |
| ENSMUSG( | 2210.788 | 0.252167 | 0.203609 | 0.864412 | Eif5b |
| ENSMUSG( | 855.3759 | -0.0475 | 0.713882 | 0.964986 | Eif6 |
| ENSMUSG( | 784.5251 | -0.01388 | 0.918707 | 0.992529 | Eipr1 |
| ENSMUSG( | 676.2712 | -0.01334 | 0.918621 | 0.992529 | Elac1 |
| ENSMUSG( | 985.4997 | -0.00324 | 0.977501 | 0.996416 | Elac2 |
| ENSMUSG( | 1331.501 | 0.116874 | 0.349287 | 0.911652 | Elapor1 |
| ENSMUSG( | 848.1933 | 0.088557 | 0.555072 | 0.939565 | Elapor2 |
| ENSMUSG( | 1188.625 | 0.00258 | 0.982014 | 0.996756 | Elavl1 |
| ENSMUSG( | 604.899 | -0.05215 | 0.734846 | 0.967344 | Elavl2 |
| ENSMUSG( | 7071.612 | 0.104636 | 0.44914 | 0.922202 | Elavl3 |
| ENSMUSG( | 882.7927 | -0.0169 | 0.911994 | 0.992196 | Elavl4 |
| ENSMUSG( | 9.970288 | 0.14689 | 0.827462 | NA | Eldr |
| ENSMUSG( | 259.6374 | -0.03043 | 0.858477 | 0.985884 | Elf1 |
| ENSMUSG( | 993.276 | 0.080323 | 0.51275 | 0.930164 | Elf2 |
| ENSMUSG( | 92.65233 | 0.336663 | 0.180521 | 0.850372 | Elf4 |
| ENSMUSG( | 64.12825 | -0.38206 | 0.237947 | 0.87259 | Elf5 |
| ENSMUSG( | 1218.388 | 0.140158 | 0.338649 | 0.907972 | Elfn1 |
| ENSMUSG( | 5397.447 | 0.290601 | 0.038665 | 0.679972 | Elfn2 |
| ENSMUSG( | 2158.805 | 0.036744 | 0.763332 | 0.973945 | Elk1 |
| ENSMUSG( | 406.6357 | 0.019257 | 0.906302 | 0.991302 | Elk3 |
| ENSMUSG( | 758.5793 | 0.001364 | 0.991491 | 0.998617 | Elk4 |
| ENSMUSG( | 593.2136 | 0.004109 | 0.975938 | 0.996416 | Ell |
| ENSMUSG( | 529.5239 | 0.07245 | 0.588213 | 0.944975 | Ell2 |
| ENSMUSG( | 2221.011 | -0.09317 | 0.539366 | 0.937216 | Elmo1 |
| ENSMUSG( | 4959.068 | -0.15705 | 0.122681 | 0.814249 | Elmo2 |
| ENSMUSG( | 61.21662 | -0.05027 | 0.856687 | 0.985884 | Elmo3 |
| ENSMUSG( | 7328.296 | 0.064726 | 0.665257 | 0.956766 | Elmod1 |
| ENSMUSG( | 559.3126 | 0.312363 | 0.061491 | 0.736111 | Elmod2 |
| ENSMUSG( | 942.9292 | 0.198767 | 0.101702 | 0.801719 | Elmod3 |
| ENSMUSG( | 380.6576 | 0.256956 | 0.117287 | 0.813384 | Eln |
| ENSMUSG( | 971.2321 | 0.027104 | 0.816095 | 0.979198 | Eloa |
| ENSMUSG( | 3719 | -0.24298 | 0.085434 | 0.779799 | Elob |
| ENSMUSG( | 2843.049 | -0.18311 | 0.158003 | 0.837724 | Eloc |
| ENSMUSG( | 1197.453 | -0.17679 | 0.138222 | 0.828492 | Elof1 |
| ENSMUSG( | 1236.403 | 0.19447 | 0.286118 | 0.893906 | Elovl1 |
| ENSMUSG( | 1533.494 | -0.01491 | 0.932362 | 0.99306 | Elovl2 |
| ENSMUSG( | 2224.526 | 0.14288 | 0.412611 | 0.917257 | Elovl4 |
| ENSMUSG( | 5064.031 | -0.02978 | 0.858624 | 0.985884 | Elovl5 |
| ENSMUSG( | 2167.315 | 0.223344 | 0.162538 | 0.837724 | Elovl6 |
| ENSMUSG( | 609.3442 | 0.183299 | 0.390119 | 0.917257 | Elovl7 |
| ENSMUSG( | 1397.582 | 0.019297 | 0.875144 | 0.989644 | Elp1 |
| ENSMUSG( | 1775.452 | -0.05282 | 0.662962 | 0.956205 | Elp2 |
| ENSMUSG( | 1575.57 | -0.19183 | 0.111679 | 0.808384 | Elp3 |
| ENSMUSG( | 439.9368 | -0.25787 | 0.113127 | 0.809623 | Elp4 |
| ENSMUSG( | 2133.264 | -0.11068 | 0.329141 | 0.907083 | Elp5 |
| ENSMUSG( | 250.6532 | -0.37957 | 0.022705 | 0.582319 | Elp6 |
| ENSMUSG( | 826.2793 | 0.110507 | 0.447872 | 0.922202 | Emb |
| ENSMUSG( | 3440.166 | 0.095222 | 0.463739 | 0.923329 | Emc1 |
| ENSMUSG( | 7842.084 | 0.051313 | 0.733354 | 0.967228 | Emc10 |
| ENSMUSG( | 1332.537 | -0.08069 | 0.526331 | 0.934374 | Emc2 |
| ENSMUSG( | 4413.916 | -0.00203 | 0.98813 | 0.997862 | Emc3 |
| ENSMUSG( | 3102.512 | -0.13635 | 0.276501 | 0.890889 | Emc4 |
| ENSMUSG( | 1270.735 | -0.0224 | 0.863006 | 0.986547 | Emc6 |
| ENSMUSG( | 5123.08 | 0.04818 | 0.737109 | 0.967476 | Emc7 |
| ENSMUSG( | 3420.192 | -0.19007 | 0.121377 | 0.813384 | Emc8 |
| ENSMUSG( | 417.054 | -0.13129 | 0.375053 | 0.915491 | Emc9 |
| ENSMUSG( | 348.0703 | 0.374997 | 0.140804 | 0.830678 | Emcn |
| ENSMUSG( | 504.8293 | -0.15515 | 0.31918 | 0.903431 | Emd |
| ENSMUSG( | 10.37305 | -0.0092 | 0.988362 | NA | Eme1 |
| ENSMUSG( | 389.4784 | 0.165349 | 0.25709 | 0.883082 | Eme2 |
| ENSMUSG( | 644.068 | 0.064253 | 0.614371 | 0.951177 | Emg1 |
| ENSMUSG( | 500.4611 | 0.156721 | 0.361372 | 0.914223 | Emid1 |
| ENSMUSG( | 181.4029 | -0.09934 | 0.633802 | 0.953889 | Emilin1 |
| ENSMUSG( | 187.2753 | 0.144712 | 0.533635 | 0.93533 | Emilin2 |
| ENSMUSG( | 1937.77 | -0.1738 | 0.113305 | 0.809623 | Eml1 |
| ENSMUSG( | 6091.063 | 0.095152 | 0.371199 | 0.915491 | Eml2 |
| ENSMUSG( | 560.6084 | 0.212033 | 0.117594 | 0.813384 | Eml3 |
| ENSMUSG( | 999.3264 | 0.009502 | 0.932904 | 0.99306 | Eml4 |
| ENSMUSG( | 1707.618 | 0.031329 | 0.804841 | 0.977232 | Eml5 |
| ENSMUSG( | 979.0199 | 0.087339 | 0.472029 | 0.923329 | Eml6 |
| ENSMUSG( | 71.32803 | 0.319666 | 0.259142 | 0.883971 | Emp1 |
| ENSMUSG( | 3511.721 | -0.35784 | 0.003522 | 0.237302 | Emp2 |
| ENSMUSG( | 106.9227 | -0.19381 | 0.37018 | 0.915491 | Emp3 |
| ENSMUSG( | 1441.327 | 0.303035 | 0.010101 | 0.40957 | Emsy |
| ENSMUSG( | 408.9165 | -0.24658 | 0.103695 | 0.804969 | Emx1 |
| ENSMUSG( | 1325.385 | 0.146253 | 0.334438 | 0.907907 | Emx2 |
| ENSMUSG( | 250.6316 | -0.16601 | 0.322665 | 0.904023 | Emx2os |
| ENSMUSG( | 19.4926 | 0.857964 | 0.170994 | 0.842562 | En2 |
| ENSMUSG( | 7690.811 | 0.015435 | 0.909938 | 0.991958 | Enah |
| ENSMUSG( | 10743.32 | -0.10164 | 0.463918 | 0.923329 | Enc1 |
| ENSMUSG( | 3373.454 | 0.243801 | 0.214458 | 0.864421 | Endod1 |
| ENSMUSG( | 328.3218 | -0.26544 | 0.089292 | 0.788502 | Endog |
| ENSMUSG( | 170.9534 | -0.13327 | 0.502885 | 0.927825 | Endou |
| ENSMUSG( | 1437.9 | -0.00045 | 0.996761 | 0.999268 | Endov |
| ENSMUSG( | 1615.328 | 0.737671 | 0.001814 | 0.15837 | Eng |
| ENSMUSG( | 329.5644 | 0.012062 | 0.937438 | 0.993685 | Engase |
| ENSMUSG( | 3560.864 | -0.21438 | 0.079844 | 0.776154 | Enho |
| ENSMUSG( | 192.1747 | -0.09474 | 0.593953 | 0.945408 | Enkd1 |
| ENSMUSG( | 56.92078 | 0.214429 | 0.484222 | 0.92541 | Enkur |
| ENSMUSG( | 16773.44 | -0.20804 | 0.109973 | 0.806572 | Eno1 |
| ENSMUSG( | 8.947096 | 0.236236 | 0.711624 | NA | Eno1b |
| ENSMUSG( | 12108.57 | -0.043 | 0.773067 | 0.974463 | Eno2 |
| ENSMUSG( | 288.1615 | -0.0175 | 0.93311 | 0.99306 | Eno3 |
| ENSMUSG( | 68.83826 | 0.1005 | 0.697441 | 0.960867 | Eno4 |
| ENSMUSG( | 965.6635 | -0.04087 | 0.74589 | 0.96951 | Enoph1 |
| ENSMUSG( | 492.1482 | -0.03747 | 0.778363 | 0.976106 | Enox1 |
| ENSMUSG( | 227.347 | -0.21072 | 0.21975 | 0.866104 | Enox2 |
| ENSMUSG( | 104.9915 | 0.554563 | 0.085621 | 0.779799 | Enpep |

| | | | | | | | | | | |
|---|---|---|---|---|---|---|---|---|---|---|
| ENSMUSG( | Eif2s3y | 1272.578 | 0.060367 | 0.602176 | 0.904887 | ENSMUSG( | Eif2s3y | 1135.109 | 0.000642 | 0.997161 | 0.999268 |
| ENSMUSG( | Eif3a | 5967.754 | 0.145428 | 0.146856 | 0.56887 | ENSMUSG( | Eif3a | 5155.03 | -0.00648 | 0.956497 | 0.994798 |
| ENSMUSG( | Eif3b | 2926.73 | 0.151845 | 0.082767 | 0.444372 | ENSMUSG( | Eif3b | 2511.594 | -0.0166 | 0.875047 | 0.989644 |
| ENSMUSG( | Eif3c | 4619.426 | 0.110032 | 0.22165 | 0.667258 | ENSMUSG( | Eif3c | 3807.729 | -0.18389 | 0.096884 | 0.79645 |
| ENSMUSG( | Eif3d | 1789.144 | 0.0204 | 0.844884 | 0.972023 | ENSMUSG( | Eif3d | 1561.753 | -0.10509 | 0.38832 | 0.917257 |
| ENSMUSG( | Eif3e | 2177.215 | 0.020696 | 0.8295 | 0.969343 | ENSMUSG( | Eif3e | 1938.827 | -0.04893 | 0.69875 | 0.960908 |
| ENSMUSG( | Eif3f | 7373.382 | 0.088743 | 0.249862 | 0.699512 | ENSMUSG( | Eif3f | 6294.866 | -0.10082 | 0.362701 | 0.914223 |
| ENSMUSG( | Eif3g | 1526.276 | -0.01608 | 0.858467 | 0.974683 | ENSMUSG( | Eif3g | 1345.59 | -0.1158 | 0.329267 | 0.907083 |
| ENSMUSG( | Eif3h | 5132.38 | 0.05331 | 0.504632 | 0.866883 | ENSMUSG( | Eif3h | 4414.031 | -0.11677 | 0.34636 | 0.910966 |
| ENSMUSG( | Eif3i | 2501.769 | 0.089444 | 0.308548 | 0.756544 | ENSMUSG( | Eif3i | 2074.721 | -0.18785 | 0.104392 | 0.804969 |
| ENSMUSG( | Eif3j1 | 1344.661 | 0.12467 | 0.315251 | 0.762288 | ENSMUSG( | Eif3j1 | 1158.693 | -0.03854 | 0.802742 | 0.97681 |
| ENSMUSG( | Eif3j2 | 22.02044 | 0.330901 | 0.413942 | 0.819998 | ENSMUSG( | Eif3j2 | 19.56635 | 0.216379 | 0.643029 | 0.953903 |
| ENSMUSG( | Eif3k | 2812.271 | 0.001048 | 0.989639 | 0.999144 | ENSMUSG( | Eif3k | 2391.437 | -0.20787 | 0.078459 | 0.77321 |
| ENSMUSG( | Eif3l | 2010.349 | -0.00059 | 0.99536 | 0.99943 | ENSMUSG( | Eif3l | 1762.418 | -0.11479 | 0.339017 | 0.907972 |
| ENSMUSG( | Eif3m | 1358.485 | 0.041348 | 0.697886 | 0.935828 | ENSMUSG( | Eif3m | 1186.919 | -0.08431 | 0.540621 | 0.937383 |
| ENSMUSG( | Eif4a1 | 5186.251 | 0.006814 | 0.951721 | 0.993821 | ENSMUSG( | Eif4a1 | 4655.086 | -0.03921 | 0.770654 | 0.974236 |
| ENSMUSG( | Eif4a2 | 8508.288 | 0.030377 | 0.718059 | 0.941697 | ENSMUSG( | Eif4a2 | 7595.863 | -0.02849 | 0.784695 | 0.976231 |
| ENSMUSG( | Eif4a3 | 1204.546 | 0.063344 | 0.526914 | 0.877267 | ENSMUSG( | Eif4a3 | 1041.262 | -0.09083 | 0.45493 | 0.922891 |
| ENSMUSG( | Eif4b | 5926.241 | 1.94E-05 | 0.999814 | 0.99989 | ENSMUSG( | Eif4b | 5244.325 | -0.08619 | 0.452169 | 0.922316 |
| ENSMUSG( | Eif4e | 2257.114 | -0.00384 | 0.968389 | 0.996369 | ENSMUSG( | Eif4e | 2001.214 | -0.08815 | 0.541807 | 0.937841 |
| ENSMUSG( | Eif4e2 | 1439.43 | 0.091779 | 0.295992 | 0.744386 | ENSMUSG( | Eif4e2 | 1283.199 | 0.029159 | 0.791451 | 0.97681 |
| ENSMUSG( | Eif4e3 | 385.2931 | 0.097798 | 0.453987 | 0.836798 | ENSMUSG( | Eif4e3 | 342.9862 | 0.026456 | 0.862138 | 0.986355 |
| ENSMUSG( | Eif4ebp1 | 153.4976 | 0.253194 | 0.128534 | 0.536774 | ENSMUSG( | Eif4ebp1 | 129.8923 | 0.038355 | 0.847955 | 0.985242 |
| ENSMUSG( | Eif4ebp2 | 3693.505 | 0.177827 | 0.065905 | 0.397536 | ENSMUSG( | Eif4ebp2 | 3263.856 | 0.095252 | 0.394823 | 0.917257 |
| ENSMUSG( | Eif4ebp3 | 12.09173 | 0.253979 | 0.633629 | NA | ENSMUSG( | Eif4ebp3 | 11.37598 | 0.365616 | 0.530633 | NA |
| ENSMUSG( | Eif4enif1 | 1362.548 | 0.099059 | 0.301114 | 0.749979 | ENSMUSG( | Eif4enif1 | 1235.316 | 0.082884 | 0.486208 | 0.92541 |
| ENSMUSG( | Eif4g1 | 8556.198 | 0.136778 | 0.087797 | 0.454663 | ENSMUSG( | Eif4g1 | 7538.721 | 0.044148 | 0.659764 | 0.956205 |
| ENSMUSG( | Eif4g2 | 26762.93 | 0.095198 | 0.338694 | 0.777518 | ENSMUSG( | Eif4g2 | 23165.59 | -0.05261 | 0.674045 | 0.95694 |
| ENSMUSG( | Eif4g3 | 8712.006 | 0.083808 | 0.300916 | 0.749979 | ENSMUSG( | Eif4g3 | 7591.72 | -0.04414 | 0.668827 | 0.956766 |
| ENSMUSG( | Eif4h | 11381.8 | 0.150883 | 0.042337 | 0.316636 | ENSMUSG( | Eif4h | 9703.365 | -0.03744 | 0.696279 | 0.960867 |
| ENSMUSG( | Eif5 | 7909.787 | 0.037885 | 0.728109 | 0.944355 | ENSMUSG( | Eif5 | 6906.775 | -0.08796 | 0.549956 | 0.938699 |
| ENSMUSG( | Eif5a | 9475.7 | 0.004078 | 0.960388 | 0.993991 | ENSMUSG( | Eif5a | 7949.876 | -0.24662 | 0.057171 | 0.723362 |
| ENSMUSG( | Eif5a2 | 1152.689 | 0.040047 | 0.691483 | 0.933881 | ENSMUSG( | Eif5a2 | 1019.05 | -0.05215 | 0.713399 | 0.964766 |
| ENSMUSG( | Eif5b | 2373.742 | 0.197317 | 0.069345 | 0.40782 | ENSMUSG( | Eif5b | 2210.788 | 0.252167 | 0.203609 | 0.864412 |
| ENSMUSG( | Eif6 | 990.2693 | 0.1045 | 0.346358 | 0.779153 | ENSMUSG( | Eif6 | 855.3759 | -0.0475 | 0.713882 | 0.964986 |
| ENSMUSG( | Eipr1 | 888.1946 | 0.076405 | 0.53521 | 0.880733 | ENSMUSG( | Eipr1 | 784.5251 | -0.01388 | 0.918707 | 0.992529 |
| ENSMUSG( | Elac1 | 728.7117 | -0.06348 | 0.565793 | 0.89104 | ENSMUSG( | Elac1 | 676.2712 | -0.01334 | 0.918621 | 0.992529 |
| ENSMUSG( | Elac2 | 1161.724 | 0.194151 | 0.043343 | 0.319117 | ENSMUSG( | Elac2 | 985.4997 | -0.00324 | 0.977501 | 0.996416 |
| ENSMUSG( | Elapor1 | 1305.769 | -0.21765 | 0.052781 | 0.352597 | ENSMUSG( | Elapor1 | 1331.501 | 0.116874 | 0.349287 | 0.911652 |
| ENSMUSG( | Elapor2 | 813.2278 | -0.32085 | 0.010924 | 0.150598 | ENSMUSG( | Elapor2 | 848.1933 | 0.088557 | 0.555072 | 0.939565 |
| ENSMUSG( | Elavl1 | 1330.788 | 0.06093 | 0.505119 | 0.866883 | ENSMUSG( | Elavl1 | 1188.625 | 0.00258 | 0.982014 | 0.996756 |
| ENSMUSG( | Elavl2 | 658.3427 | -0.07303 | 0.564663 | 0.890434 | ENSMUSG( | Elavl2 | 604.899 | -0.05215 | 0.734846 | 0.967344 |
| ENSMUSG( | Elavl3 | 7745.789 | 0.099917 | 0.28476 | 0.733422 | ENSMUSG( | Elavl3 | 7071.612 | 0.104636 | 0.44914 | 0.922202 |
| ENSMUSG( | Elavl4 | 959.6119 | -0.03916 | 0.749307 | 0.949664 | ENSMUSG( | Elavl4 | 882.7927 | -0.0169 | 0.911994 | 0.992196 |
| ENSMUSG( | Eldr | 10.31499 | -0.03606 | 0.959182 | NA | ENSMUSG( | Eldr | 9.970288 | 0.14689 | 0.827462 | NA |
| ENSMUSG( | Elf1 | 272.4718 | -0.1646 | 0.346231 | 0.779153 | ENSMUSG( | Elf1 | 259.6374 | -0.03043 | 0.858477 | 0.985884 |
| ENSMUSG( | Elf2 | 1085.434 | 0.071139 | 0.459193 | 0.840244 | ENSMUSG( | Elf2 | 993.276 | 0.080323 | 0.51275 | 0.930164 |
| ENSMUSG( | Elf4 | 90.78302 | 0.009384 | 0.966052 | 0.995377 | ENSMUSG( | Elf4 | 92.65233 | 0.336663 | 0.180521 | 0.850372 |
| ENSMUSG( | Elf5 | 83.28809 | 0.163547 | 0.550302 | 0.886038 | ENSMUSG( | Elf5 | 64.12825 | -0.38206 | 0.237947 | 0.87259 |
| ENSMUSG( | Elfn1 | 1239.085 | -0.08064 | 0.53673 | 0.882043 | ENSMUSG( | Elfn1 | 1218.388 | 0.140158 | 0.338649 | 0.907972 |
| ENSMUSG( | Elfn2 | 5549.921 | 0.111153 | 0.361794 | 0.792797 | ENSMUSG( | Elfn2 | 5397.447 | 0.290601 | 0.038665 | 0.679972 |
| ENSMUSG( | Elk1 | 2399.237 | 0.069796 | 0.442179 | 0.832774 | ENSMUSG( | Elk1 | 2158.805 | 0.036744 | 0.763332 | 0.973945 |
| ENSMUSG( | Elk3 | 424.4374 | -0.13626 | 0.355692 | 0.787945 | ENSMUSG( | Elk3 | 406.6357 | 0.019257 | 0.906302 | 0.991302 |
| ENSMUSG( | Elk4 | 834.7484 | 0.012207 | 0.912064 | 0.988483 | ENSMUSG( | Elk4 | 758.5793 | 0.001364 | 0.991491 | 0.998617 |
| ENSMUSG( | Ell | 656.7659 | 0.025693 | 0.837629 | 0.971111 | ENSMUSG( | Ell | 593.2136 | 0.004109 | 0.975938 | 0.996416 |
| ENSMUSG( | Ell2 | 582.7017 | 0.080324 | 0.476506 | 0.851672 | ENSMUSG( | Ell2 | 529.5239 | 0.07245 | 0.588213 | 0.944975 |
| ENSMUSG( | Elmo1 | 2440.392 | -0.08701 | 0.495109 | 0.860798 | ENSMUSG( | Elmo1 | 2221.011 | -0.09317 | 0.539366 | 0.937216 |
| ENSMUSG( | Elmo2 | 5977.398 | 0.11592 | 0.14521 | 0.565771 | ENSMUSG( | Elmo2 | 4959.068 | -0.15705 | 0.122681 | 0.814249 |
| ENSMUSG( | Elmo3 | 70.93688 | 0.095036 | 0.686218 | 0.932981 | ENSMUSG( | Elmo3 | 61.21662 | -0.05027 | 0.856687 | 0.985884 |
| ENSMUSG( | Elmod1 | 6353.671 | -0.7027 | 3.52E-08 | 1.19E-05 | ENSMUSG( | Elmod1 | 7328.296 | 0.064726 | 0.665257 | 0.956766 |
| ENSMUSG( | Elmod2 | 499.5116 | -0.28619 | 0.063677 | 0.390618 | ENSMUSG( | Elmod2 | 559.3126 | 0.312363 | 0.061491 | 0.736111 |
| ENSMUSG( | Elmod3 | 997.5887 | 0.094224 | 0.374427 | 0.800445 | ENSMUSG( | Elmod3 | 942.9292 | 0.198767 | 0.101702 | 0.801719 |
| ENSMUSG( | Eln | 389.1607 | 0.061093 | 0.680135 | 0.930983 | ENSMUSG( | Eln | 380.6576 | 0.256956 | 0.117287 | 0.813384 |
| ENSMUSG( | Eloa | 1102.89 | 0.122181 | 0.21689 | 0.661816 | ENSMUSG( | Eloa | 971.2321 | 0.027104 | 0.816095 | 0.979198 |
| ENSMUSG( | Elob | 4329.889 | -0.05982 | 0.517907 | 0.873754 | ENSMUSG( | Elob | 3719 | -0.24298 | 0.085434 | 0.779799 |
| ENSMUSG( | Eloc | 3289.856 | -0.02154 | 0.820046 | 0.965888 | ENSMUSG( | Eloc | 2843.049 | -0.18311 | 0.158003 | 0.837724 |
| ENSMUSG( | Elof1 | 1409.56 | 0.033455 | 0.71476 | 0.941144 | ENSMUSG( | Elof1 | 1197.453 | -0.17679 | 0.138222 | 0.828492 |
| ENSMUSG( | Elovl1 | 1094.675 | -0.46325 | 0.003622 | 0.076313 | ENSMUSG( | Elovl1 | 1236.403 | 0.19447 | 0.286118 | 0.893906 |
| ENSMUSG( | Elovl2 | 1634.56 | -0.10527 | 0.5017 | 0.864596 | ENSMUSG( | Elovl2 | 1533.494 | -0.01491 | 0.932362 | 0.99306 |
| ENSMUSG( | Elovl4 | 1960.317 | -0.53731 | 0.00018 | 0.009729 | ENSMUSG( | Elovl4 | 2224.526 | 0.14288 | 0.412611 | 0.917257 |
| ENSMUSG( | Elovl5 | 5010.204 | -0.35264 | 0.024626 | 0.240618 | ENSMUSG( | Elovl5 | 5064.031 | -0.02978 | 0.858624 | 0.985884 |
| ENSMUSG( | Elovl6 | 2150.469 | -0.06348 | 0.659474 | 0.924506 | ENSMUSG( | Elovl6 | 2167.315 | 0.223344 | 0.162538 | 0.837724 |
| ENSMUSG( | Elovl7 | 520.1298 | -0.60874 | 0.001292 | 0.039144 | ENSMUSG( | Elovl7 | 609.3442 | 0.183299 | 0.390119 | 0.917257 |
| ENSMUSG( | Elp1 | 1579.993 | 0.099215 | 0.302307 | 0.750677 | ENSMUSG( | Elp1 | 1397.582 | 0.019297 | 0.875144 | 0.989644 |
| ENSMUSG( | Elp2 | 2034.469 | 0.070338 | 0.516943 | 0.873055 | ENSMUSG( | Elp2 | 1775.452 | -0.05282 | 0.662962 | 0.956205 |
| ENSMUSG( | Elp3 | 1887.536 | 0.066755 | 0.517409 | 0.873295 | ENSMUSG( | Elp3 | 1575.57 | -0.19183 | 0.111679 | 0.808384 |
| ENSMUSG( | Elp4 | 526.2767 | 0.012655 | 0.919401 | 0.990247 | ENSMUSG( | Elp4 | 439.9368 | -0.25787 | 0.113127 | 0.809623 |
| ENSMUSG( | Elp5 | 2488.298 | 0.068105 | 0.395037 | 0.810154 | ENSMUSG( | Elp5 | 2133.264 | -0.11068 | 0.329141 | 0.907083 |
| ENSMUSG( | Elp6 | 298.0772 | -0.12359 | 0.384796 | 0.804031 | ENSMUSG( | Elp6 | 250.6532 | -0.37957 | 0.022705 | 0.582319 |
| ENSMUSG( | Emb | 802.0126 | -0.25639 | 0.019533 | 0.210473 | ENSMUSG( | Emb | 826.2793 | 0.110507 | 0.447872 | 0.922202 |
| ENSMUSG( | Emc1 | 3347.967 | -0.26552 | 0.021936 | 0.225738 | ENSMUSG( | Emc1 | 3440.166 | 0.095222 | 0.463739 | 0.923329 |
| ENSMUSG( | Emc10 | 7748.965 | -0.2669 | 0.035117 | 0.288256 | ENSMUSG( | Emc10 | 7842.084 | 0.051313 | 0.733354 | 0.967228 |
| ENSMUSG( | Emc2 | 1543.999 | 0.079884 | 0.456492 | 0.838832 | ENSMUSG( | Emc2 | 1332.537 | -0.08069 | 0.526331 | 0.934374 |
| ENSMUSG( | Emc3 | 4500.154 | -0.22601 | 0.068979 | 0.406851 | ENSMUSG( | Emc3 | 4413.916 | -0.00203 | 0.98813 | 0.997862 |
| ENSMUSG( | Emc4 | 2985.363 | -0.57024 | 3.49E-07 | 7.75E-05 | ENSMUSG( | Emc4 | 3102.512 | -0.13635 | 0.276501 | 0.890889 |
| ENSMUSG( | Emc6 | 1424.639 | 0.04299 | 0.651264 | 0.921451 | ENSMUSG( | Emc6 | 1270.735 | -0.0224 | 0.863006 | 0.986547 |
| ENSMUSG( | Emc7 | 5119.837 | -0.23357 | 0.06893 | 0.406851 | ENSMUSG( | Emc7 | 5123.08 | 0.04818 | 0.737109 | 0.967476 |
| ENSMUSG( | Emc8 | 4031.533 | 0.023703 | 0.800265 | 0.961745 | ENSMUSG( | Emc8 | 3420.192 | -0.19007 | 0.121377 | 0.813384 |
| ENSMUSG( | Emc9 | 464.3434 | -0.08308 | 0.49811 | 0.862463 | ENSMUSG( | Emc9 | 417.054 | -0.13129 | 0.375053 | 0.915491 |
| ENSMUSG( | Emcn | 320.1885 | -0.12779 | 0.516357 | 0.872701 | ENSMUSG( | Emcn | 348.0703 | 0.374997 | 0.140804 | 0.830678 |
| ENSMUSG( | Emd | 569.8848 | -0.06779 | 0.622293 | 0.911432 | ENSMUSG( | Emd | 504.8293 | -0.15515 | 0.31918 | 0.903431 |
| ENSMUSG( | Eme1 | 10.4435 | -0.32747 | 0.608613 | NA | ENSMUSG( | Eme1 | 10.37305 | -0.0092 | 0.988362 | NA |
| ENSMUSG( | Eme2 | 432.1219 | 0.187715 | 0.116472 | 0.514944 | ENSMUSG( | Eme2 | 389.4784 | 0.165349 | 0.25709 | 0.883082 |
| ENSMUSG( | Emg1 | 711.6941 | 0.087558 | 0.401684 | 0.814923 | ENSMUSG( | Emg1 | 644.068 | 0.064253 | 0.614371 | 0.951177 |
| ENSMUSG( | Emid1 | 451.0747 | -0.45093 | 0.006504 | 0.110626 | ENSMUSG( | Emid1 | 500.4611 | 0.156721 | 0.361372 | 0.914223 |
| ENSMUSG( | Emilin1 | 177.8458 | -0.47928 | 0.006216 | 0.107827 | ENSMUSG( | Emilin1 | 181.4029 | -0.09934 | 0.633802 | 0.953889 |
| ENSMUSG( | Emilin2 | 175.2495 | -0.34613 | 0.046354 | 0.329764 | ENSMUSG( | Emilin2 | 187.2753 | 0.144712 | 0.533635 | 0.93533 |
| ENSMUSG( | Eml1 | 2260.933 | 0.007372 | 0.935632 | 0.993363 | ENSMUSG( | Eml1 | 1937.77 | -0.1738 | 0.113305 | 0.809623 |
| ENSMUSG( | Eml2 | 6545.792 | 0.036323 | 0.703797 | 0.937933 | ENSMUSG( | Eml2 | 6091.063 | 0.095152 | 0.371199 | 0.915491 |
| ENSMUSG( | Eml3 | 588.6062 | 0.087634 | 0.42622 | 0.824898 | ENSMUSG( | Eml3 | 560.6084 | 0.212033 | 0.117594 | 0.813384 |
| ENSMUSG( | Eml4 | 1186.1 | 0.226448 | 0.03454 | 0.285252 | ENSMUSG( | Eml4 | 999.3264 | 0.009502 | 0.932904 | 0.99306 |
| ENSMUSG( | Eml5 | 1796.326 | -0.09069 | 0.391348 | 0.808825 | ENSMUSG( | Eml5 | 1707.618 | 0.031329 | 0.804841 | 0.977232 |
| ENSMUSG( | Eml6 | 1117.655 | 0.198008 | 0.039926 | 0.308163 | ENSMUSG( | Eml6 | 979.0199 | 0.087339 | 0.472029 | 0.923329 |
| ENSMUSG( | Emp1 | 60.47382 | -0.45918 | 0.123456 | 0.530552 | ENSMUSG( | Emp1 | 71.32803 | 0.319666 | 0.259142 | 0.883971 |
| ENSMUSG( | Emp2 | 4438.645 | 0.07108 | 0.583485 | 0.898521 | ENSMUSG( | Emp2 | 3511.721 | -0.35784 | 0.003522 | 0.237302 |
| ENSMUSG( | Emp3 | 111.1351 | -0.35413 | 0.075896 | 0.426699 | ENSMUSG( | Emp3 | 106.9227 | -0.19381 | 0.37018 | 0.915491 |
| ENSMUSG( | Emsy | 1510.966 | 0.183104 | 0.085795 | 0.451465 | ENSMUSG( | Emsy | 1441.327 | 0.303035 | 0.010101 | 0.40957 |
| ENSMUSG( | Emx1 | 507.5977 | 0.117447 | 0.37739 | 0.801213 | ENSMUSG( | Emx1 | 408.9165 | -0.24658 | 0.103695 | 0.804969 |
| ENSMUSG( | Emx2 | 1556.131 | 0.330475 | 0.003874 | 0.079851 | ENSMUSG( | Emx2 | 1325.385 | 0.146253 | 0.334438 | 0.907907 |
| ENSMUSG( | Emx2os | 303.7717 | 0.127188 | 0.400249 | 0.813641 | ENSMUSG( | Emx2os | 250.6316 | -0.16601 | 0.322665 | 0.904023 |
| ENSMUSG( | En2 | 16.96939 | 0.244522 | 0.649033 | NA | ENSMUSG( | En2 | 19.4926 | 0.857964 | 0.170994 | 0.842562 |
| ENSMUSG( | Enah | 8950.365 | 0.180307 | 0.103477 | 0.489288 | ENSMUSG( | Enah | 7690.811 | 0.015435 | 0.909938 | 0.991958 |
| ENSMUSG( | Enc1 | 12957.2 | 0.168748 | 0.163843 | 0.594869 | ENSMUSG( | Enc1 | 10743.32 | -0.10164 | 0.463918 | 0.923329 |
| ENSMUSG( | Endod1 | 2924.573 | -0.47058 | 0.003532 | 0.075287 | ENSMUSG( | Endod1 | 3373.454 | 0.243801 | 0.214458 | 0.864421 |
| ENSMUSG( | Endog | 393.0407 | -0.00172 | 0.989706 | 0.999144 | ENSMUSG( | Endog | 328.3218 | -0.26544 | 0.089292 | 0.788502 |
| ENSMUSG( | Endou | 178.3421 | -0.2867 | 0.189386 | 0.627932 | ENSMUSG( | Endou | 170.9534 | -0.13327 | 0.502885 | 0.927825 |
| ENSMUSG( | Endov | 1597.942 | 0.032931 | 0.722593 | 0.943752 | ENSMUSG( | Endov | 1437.9 | -0.00045 | 0.996761 | 0.999268 |
| ENSMUSG( | Eng | 1363.793 | 0.063741 | 0.751641 | 0.950317 | ENSMUSG( | Eng | 1615.328 | 0.737671 | 0.001814 | 0.15837 |
| ENSMUSG( | Engase | 356.5083 | -0.03682 | 0.777258 | 0.955886 | ENSMUSG( | Engase | 329.5644 | 0.012062 | 0.937438 | 0.993685 |
| ENSMUSG( | Enho | 4293.337 | 0.065893 | 0.499103 | 0.862973 | ENSMUSG( | Enho | 3560.864 | -0.21438 | 0.079844 | 0.776154 |
| ENSMUSG( | Enkd1 | 212.8635 | -0.074 | 0.634644 | 0.914073 | ENSMUSG( | Enkd1 | 192.1747 | -0.09474 | 0.593953 | 0.945408 |
| ENSMUSG( | Enkur | 51.60302 | -0.36989 | 0.190058 | 0.629259 | ENSMUSG( | Enkur | 56.92078 | 0.214429 | 0.484222 | 0.92541 |
| ENSMUSG( | Eno1 | 19841.91 | 0.017524 | 0.841236 | 0.971179 | ENSMUSG( | Eno1 | 16773.44 | -0.20804 | 0.109973 | 0.806572 |
| ENSMUSG( | Eno2 | 13727.5 | 0.051482 | 0.71466 | 0.941144 | ENSMUSG( | Eno2 | 12108.57 | -0.043 | 0.773067 | 0.974463 |
| ENSMUSG( | Eno3 | 343.9003 | 0.222547 | 0.231239 | 0.679229 | ENSMUSG( | Eno3 | 288.1615 | -0.0175 | 0.93311 | 0.99306 |
| ENSMUSG( | Eno4 | 73.13649 | -0.00103 | 0.996622 | 0.99943 | ENSMUSG( | Eno4 | 68.83826 | 0.1005 | 0.697441 | 0.960867 |
| ENSMUSG( | Enoph1 | 1105.322 | 0.082434 | 0.414216 | 0.819998 | ENSMUSG( | Enoph1 | 965.6635 | -0.04087 | 0.74589 | 0.96951 |
| ENSMUSG( | Enox1 | 523.7554 | -0.12401 | 0.250591 | 0.700403 | ENSMUSG( | Enox1 | 492.1482 | -0.03747 | 0.778363 | 0.976106 |
| ENSMUSG( | Enox2 | 260.5331 | -0.06928 | 0.614255 | 0.909419 | ENSMUSG( | Enox2 | 227.347 | -0.21072 | 0.21975 | 0.866104 |
| ENSMUSG( | Enpep | 86.94557 | -0.24253 | 0.387963 | 0.805835 | ENSMUSG( | Enpep | 104.9915 | 0.554563 | 0.085621 | 0.779799 |
| ENSMUSG( | Enpp1 | 184.1619 | -0.22363 | 0.19611 | 0.635538 | ENSMUSG( | Enpp1 | 171.7357 | -0.14538 | 0.437735 | 0.920339 |
| ENSMUSG( | Enpp2 | 10438.94 | -0.26344 | 0.148048 | 0.571232 | ENSMUSG( | Enpp2 | 12336.47 | 0.460792 | 0.001582 | 0.148357 |
| ENSMUSG( | Enpp3 | 18.24244 | -0.35097 | 0.419959 | NA | ENSMUSG( | Enpp3 | 17.79163 | -0.13348 | 0.771506 | 0.974236 |
| ENSMUSG( | Enpp4 | 812.3156 | -0.45203 | 0.013988 | 0.172788 | ENSMUSG( | Enpp4 | 922.9448 | 0.218259 | 0.282829 | 0.892412 |
| ENSMUSG( | Enpp5 | 7708.671 | -0.44667 | 0.00026 | 0.013103 | ENSMUSG( | Enpp5 | 8273.387 | 0.062432 | 0.652593 | 0.954799 |
| ENSMUSG( | Enpp6 | 676.4794 | -0.81072 | 2.27E-05 | 0.002131 | ENSMUSG( | Enpp6 | 859.5926 | 0.257414 | 0.268952 | 0.886755 |
| ENSMUSG( | Ensa | 8074.309 | 0.095678 | 0.258499 | 0.709726 | ENSMUSG( | Ensa | 6852.168 | -0.11144 | 0.330957 | 0.907513 |
| ENSMUSG( | Entpd1 | 697.5907 | 0.075517 | 0.608921 | 0.907377 | ENSMUSG( | Entpd1 | 745.2433 | 0.501333 | 0.009933 | 0.407479 |
| ENSMUSG( | Entpd2 | 505.1787 | -0.23139 | 0.185396 | 0.623185 | ENSMUSG( | Entpd2 | 510.9506 | 0.08649 | 0.648021 | 0.95422 |
| ENSMUSG( | Entpd3 | 339.3034 | -0.51527 | 0.000102 | 0.006477 | ENSMUSG( | Entpd3 | 355.8722 | -0.06649 | 0.668751 | 0.956766 |
| ENSMUSG( | Entpd5 | 2479.372 | -0.07351 | 0.56593 | 0.89104 | ENSMUSG( | Entpd5 | 2369.878 | 0.065753 | 0.634615 | 0.953903 |
| ENSMUSG( | Entpd6 | 1909.993 | -0.27761 | 0.02818 | 0.256543 | ENSMUSG( | Entpd6 | 1985.953 | 0.118852 | 0.398089 | 0.917257 |
| ENSMUSG( | Entpd7 | 247.2063 | -0.34186 | 0.103133 | 0.48855 | ENSMUSG( | Entpd7 | 279.0249 | 0.292987 | 0.188032 | 0.85698 |
| ENSMUSG( | Entr1 | 3372.945 | 0.097544 | 0.263764 | 0.714352 | ENSMUSG( | Entr1 | 3037.706 | 0.066197 | 0.579701 | 0.944209 |
| ENSMUSG( | Entrep1 | 883.0914 | -0.143 | 0.295506 | 0.743814 | ENSMUSG( | Entrep1 | 844.1038 | 0.0033 | 0.982365 | 0.996756 |
| ENSMUSG( | Entrep2 | 2513.953 | -0.11209 | 0.436003 | 0.829093 | ENSMUSG( | Entrep2 | 2572.938 | 0.222826 | 0.213156 | 0.864412 |
| ENSMUSG( | Entrep3 | 1719.326 | -0.11221 | 0.331654 | 0.775246 | ENSMUSG( | Entrep3 | 1702.896 | 0.132135 | 0.318585 | 0.903164 |
| ENSMUSG( | Eny2 | 1210.636 | -0.04007 | 0.668404 | 0.927864 | ENSMUSG( | Eny2 | 1135.658 | 0.039938 | 0.751108 | 0.970317 |
| ENSMUSG( | Eogt | 1584.426 | -0.01077 | 0.928955 | 0.991749 | ENSMUSG( | Eogt | 1609.143 | 0.291192 | 0.056704 | 0.723362 |
| ENSMUSG( | Eola1 | 205.8259 | 0.27112 | 0.088654 | 0.45666 | ENSMUSG( | Eola1 | 167.987 | -0.04027 | 0.839022 | 0.981774 |
| ENSMUSG( | Eomes | 24.31064 | -0.3808 | 0.329 | 0.773233 | ENSMUSG( | Eomes | 28.04994 | 0.250806 | 0.581867 | 0.944266 |
| ENSMUSG( | Ep300 | 1855.942 | -0.04031 | 0.726449 | 0.944223 | ENSMUSG( | Ep300 | 1866.941 | 0.239133 | 0.057341 | 0.723362 |
| ENSMUSG( | Ep400 | 2696 | 0.093834 | 0.294776 | 0.743798 | ENSMUSG( | Ep400 | 2448.141 | 0.087315 | 0.465779 | 0.923329 |
| ENSMUSG( | Epas1 | 17157.16 | 0.036333 | 0.769743 | 0.954674 | ENSMUSG( | Epas1 | 15435.18 | -2.85E-05 | 0.999816 | 0.99993 |
| ENSMUSG( | Epb41 | 1807.415 | -0.03929 | 0.754655 | 0.951007 | ENSMUSG( | Epb41 | 1695.836 | 0.043863 | 0.755212 | 0.971098 |
| ENSMUSG( | Epb41l1 | 31726.08 | 0.191078 | 0.034513 | 0.285239 | ENSMUSG( | Epb41l1 | 27168.1 | 0.019358 | 0.859296 | 0.985884 |
| ENSMUSG( | Epb41l2 | 10560.98 | 0.092561 | 0.376506 | 0.801213 | ENSMUSG( | Epb41l2 | 8771.065 | -0.18035 | 0.138435 | 0.828492 |
| ENSMUSG( | Epb41l3 | 4299.995 | -0.28306 | 0.000289 | 0.014061 | ENSMUSG( | Epb41l3 | 4208.734 | -0.06342 | 0.548019 | 0.938063 |

| | | | | | |
|---|---|---|---|---|---|
| ENSMUSG( | 356.5083 | -0.03682 | 0.777258 | 0.955886 | Engase |
| ENSMUSG( | 4293.337 | 0.065893 | 0.499103 | 0.862973 | Enho |
| ENSMUSG( | 212.8635 | -0.074 | 0.634644 | 0.914073 | Enkd1 |
| ENSMUSG( | 51.60302 | -0.36989 | 0.190058 | 0.629259 | Enkur |
| ENSMUSG( | 19841.91 | 0.017524 | 0.841236 | 0.971179 | Eno1 |
| ENSMUSG( | 13727.5 | 0.051482 | 0.71466 | 0.941144 | Eno2 |
| ENSMUSG( | 343.9003 | 0.222547 | 0.231239 | 0.679229 | Eno3 |
| ENSMUSG( | 73.13649 | -0.00103 | 0.996622 | 0.99943 | Eno4 |
| ENSMUSG( | 1105.322 | 0.082434 | 0.414216 | 0.819998 | Enoph1 |
| ENSMUSG( | 523.7554 | -0.12401 | 0.250591 | 0.700403 | Enox1 |
| ENSMUSG( | 260.5331 | -0.06928 | 0.614255 | 0.909419 | Enox2 |
| ENSMUSG( | 86.94557 | -0.24253 | 0.387963 | 0.805835 | Enpep |
| ENSMUSG( | 184.1619 | -0.22363 | 0.19611 | 0.635538 | Enpp1 |
| ENSMUSG( | 10438.94 | -0.26344 | 0.148048 | 0.571232 | Enpp2 |
| ENSMUSG( | 18.24244 | -0.35097 | 0.419959 | NA | Enpp3 |
| ENSMUSG( | 812.3156 | -0.45203 | 0.013988 | 0.172788 | Enpp4 |
| ENSMUSG( | 7708.671 | -0.44667 | 0.00026 | 0.013103 | Enpp5 |
| ENSMUSG( | 676.4794 | -0.81072 | 2.27E-05 | 0.002131 | Enpp6 |
| ENSMUSG( | 8074.309 | 0.095678 | 0.258499 | 0.709726 | Ensa |
| ENSMUSG( | 697.5907 | 0.075517 | 0.608921 | 0.907377 | Entpd1 |
| ENSMUSG( | 505.1787 | -0.23139 | 0.185396 | 0.623185 | Entpd2 |
| ENSMUSG( | 339.3034 | -0.51527 | 0.000102 | 0.006477 | Entpd3 |
| ENSMUSG( | 2479.372 | -0.07351 | 0.56593 | 0.89104 | Entpd5 |
| ENSMUSG( | 1909.993 | -0.27761 | 0.02818 | 0.256543 | Entpd6 |
| ENSMUSG( | 247.2063 | -0.34186 | 0.103133 | 0.48855 | Entpd7 |
| ENSMUSG( | 3372.945 | 0.097544 | 0.263764 | 0.714352 | Entr1 |
| ENSMUSG( | 883.0914 | -0.143 | 0.295506 | 0.743814 | Entrep1 |
| ENSMUSG( | 2513.953 | -0.11209 | 0.436003 | 0.829093 | Entrep2 |
| ENSMUSG( | 1719.326 | -0.11221 | 0.331654 | 0.775246 | Entrep3 |
| ENSMUSG( | 1210.636 | -0.04007 | 0.668404 | 0.927864 | Eny2 |
| ENSMUSG( | 1584.426 | -0.01077 | 0.928955 | 0.991749 | Eogt |
| ENSMUSG( | 205.8259 | 0.27112 | 0.088654 | 0.45666 | Eola1 |
| ENSMUSG( | 24.31064 | -0.3808 | 0.329 | 0.773233 | Eomes |
| ENSMUSG( | 1855.942 | -0.04031 | 0.726449 | 0.944223 | Ep300 |
| ENSMUSG( | 2696 | 0.093834 | 0.294776 | 0.743798 | Ep400 |
| ENSMUSG( | 17157.16 | 0.036333 | 0.769743 | 0.954674 | Epas1 |
| ENSMUSG( | 1807.415 | -0.03929 | 0.754655 | 0.951007 | Epb41 |
| ENSMUSG( | 31726.08 | 0.191078 | 0.034513 | 0.285239 | Epb41l1 |
| ENSMUSG( | 10560.98 | 0.092561 | 0.376506 | 0.801213 | Epb41l2 |
| ENSMUSG( | 4299.995 | -0.28306 | 0.000289 | 0.014061 | Epb41l3 |
| ENSMUSG( | 131.0411 | -0.14794 | 0.422094 | 0.823495 | Epb41l4a |
| ENSMUSG( | 240.8688 | 0.049539 | 0.750119 | 0.949664 | Epb41l4aos |
| ENSMUSG( | 1545.728 | 0.209589 | 0.016931 | 0.194376 | Epb41l4b |
| ENSMUSG( | 704.4173 | 0.139571 | 0.301456 | 0.749979 | Epb41l5 |
| ENSMUSG( | 967.7467 | 0.065925 | 0.52717 | 0.877445 | Epc1 |
| ENSMUSG( | 913.4709 | 0.078595 | 0.43246 | 0.82768 | Epc2 |
| ENSMUSG( | 24.32248 | -0.12937 | 0.731288 | 0.944956 | Epcam |
| ENSMUSG( | 92.09826 | -0.28248 | 0.172534 | 0.607843 | Epcip |
| ENSMUSG( | 3425.819 | -0.29292 | 0.029173 | 0.259574 | Epdr1 |
| ENSMUSG( | 1231.707 | 0.081989 | 0.417353 | 0.821339 | Epg5 |
| ENSMUSG( | 10.46295 | -0.36149 | 0.546587 | NA | Epha1 |
| ENSMUSG( | 1748.249 | -0.1481 | 0.193222 | 0.632632 | Epha10 |
| ENSMUSG( | 72.97512 | -0.01595 | 0.953233 | 0.993874 | Epha2 |
| ENSMUSG( | 159.7212 | -0.5571 | 0.006937 | 0.114527 | Epha3 |
| ENSMUSG( | 4282.619 | -0.46537 | 0.004113 | 0.082763 | Epha4 |
| ENSMUSG( | 2623.72 | -0.2728 | 0.047569 | 0.333288 | Epha5 |
| ENSMUSG( | 230.6702 | -0.1578 | 0.430639 | 0.826896 | Epha6 |
| ENSMUSG( | 1594.16 | -0.45926 | 0.001586 | 0.04465 | Epha7 |
| ENSMUSG( | 255.2471 | -0.24854 | 0.108028 | 0.497059 | Epha8 |
| ENSMUSG( | 4543.9 | -0.0021 | 0.984455 | 0.99781 | Ephb1 |
| ENSMUSG( | 724.8621 | -0.07944 | 0.655854 | 0.923929 | Ephb2 |
| ENSMUSG( | 1133.419 | -0.33725 | 0.021795 | 0.225679 | Ephb3 |
| ENSMUSG( | 331.2779 | -0.12943 | 0.49891 | 0.862973 | Ephb4 |
| ENSMUSG( | 3988.587 | -0.09411 | 0.56419 | 0.890139 | Ephb6 |
| ENSMUSG( | 1043.313 | -0.34725 | 0.006492 | 0.110626 | Ephx1 |
| ENSMUSG( | 1581.736 | -0.03577 | 0.785378 | 0.958528 | Ephx2 |
| ENSMUSG( | 1234.476 | -0.22865 | 0.041224 | 0.313778 | Ephx4 |
| ENSMUSG( | 1351.541 | 0.024049 | 0.80225 | 0.962119 | Epm2a |
| ENSMUSG( | 2173.309 | 0.026689 | 0.775877 | 0.955606 | Epm2aip1 |
| ENSMUSG( | 30139.12 | 0.190371 | 0.008573 | 0.129829 | Epn1 |
| ENSMUSG( | 3935.334 | 0.011434 | 0.901133 | 0.985633 | Epn2 |
| ENSMUSG( | 60.60453 | -0.79104 | 0.050325 | 0.342969 | Epn3 |
| ENSMUSG( | 4071.795 | 0.18402 | 0.036338 | 0.293685 | Epop |
| ENSMUSG( | 163.0181 | -0.33173 | 0.069335 | 0.40782 | Epor |
| ENSMUSG( | 3405.073 | 0.03456 | 0.720071 | 0.941986 | Eprs1 |
| ENSMUSG( | 11598.25 | -0.0422 | 0.680247 | 0.931028 | Eps15 |
| ENSMUSG( | 1427.153 | 0.164692 | 0.16691 | 0.599655 | Eps15l1 |
| ENSMUSG( | 3104.088 | 0.086867 | 0.424223 | 0.824554 | Eps8 |
| ENSMUSG( | 368.7706 | 0.568831 | 0.019405 | 0.209852 | Eps8l1 |
| ENSMUSG( | 77.51944 | -0.27796 | 0.242678 | 0.690216 | Eps8l2 |
| ENSMUSG( | 24.97392 | -0.40166 | 0.29408 | 0.743459 | Eps8l3 |
| ENSMUSG( | 51.3996 | -0.35601 | 0.225874 | 0.672685 | Epsti1 |
| ENSMUSG( | 50.18287 | 0.324189 | 0.266128 | 0.716441 | Epx |
| ENSMUSG( | 31.80899 | -0.85481 | 0.014899 | 0.179275 | Eqtn |
| ENSMUSG( | 458.5951 | 0.104123 | 0.418982 | 0.821966 | Eral1 |
| ENSMUSG( | 544.7043 | -0.2289 | 0.099434 | 0.481872 | Erap1 |
| ENSMUSG( | 190.7418 | -0.34296 | 0.046013 | 0.328532 | Erbb2 |
| ENSMUSG( | 564.4971 | -0.66376 | 0.000306 | 0.014613 | Erbb3 |
| ENSMUSG( | 1289.136 | -0.28268 | 0.077821 | 0.431216 | Erbb4 |
| ENSMUSG( | 7230.01 | -0.08669 | 0.422392 | 0.823508 | Erbin |
| ENSMUSG( | 2123.397 | 0.168979 | 0.076823 | 0.429306 | Erc1 |
| ENSMUSG( | 3286.214 | -0.02706 | 0.757793 | 0.951582 | Erc2 |
| ENSMUSG( | 477.166 | 0.064216 | 0.6897 | 0.933609 | Ercc1 |
| ENSMUSG( | 1089.756 | 0.167375 | 0.167041 | 0.599655 | Ercc2 |
| ENSMUSG( | 732.0908 | 0.231134 | 0.071007 | 0.411334 | Ercc3 |
| ENSMUSG( | 619.3449 | 0.143589 | 0.178962 | 0.616298 | Ercc4 |
| ENSMUSG( | 997.4532 | 0.035452 | 0.71209 | 0.940256 | Ercc5 |
| ENSMUSG( | 789.2081 | 0.109949 | 0.296286 | 0.744386 | Ercc6 |
| ENSMUSG( | 667.8007 | -0.09122 | 0.412791 | 0.819518 | Ercc6l2 |
| ENSMUSG( | 304.068 | 0.01495 | 0.909075 | 0.98794 | Ercc8 |
| ENSMUSG( | 1245.826 | -0.12729 | 0.256062 | 0.706828 | Erf |
| ENSMUSG( | 30.91398 | -1.01721 | 0.009098 | 0.134276 | Erfe |
| ENSMUSG( | 188.8728 | 0.045014 | 0.857644 | 0.974667 | Erg |
| ENSMUSG( | 1350.992 | -0.50701 | 0.00025 | 0.012746 | Erg28 |
| ENSMUSG( | 4909.383 | -0.2885 | 0.025473 | 0.246501 | Ergic1 |
| ENSMUSG( | 1655.797 | -0.17723 | 0.161463 | 0.59148 | Ergic2 |
| ENSMUSG( | 4421.83 | -0.24453 | 0.015169 | 0.181362 | Ergic3 |
| ENSMUSG( | 731.7561 | -0.04644 | 0.681091 | 0.931301 | Erh |
| ENSMUSG( | 290.8129 | 0.087982 | 0.531517 | 0.879536 | Eri1 |
| ENSMUSG( | 278.3008 | -0.05599 | 0.746568 | 0.949289 | Eri2 |
| ENSMUSG( | 3882.487 | 0.087521 | 0.397132 | 0.812088 | Eri3 |
| ENSMUSG( | 88.83528 | 0.104415 | 0.646426 | 0.918612 | Erich1 |
| ENSMUSG( | 34.14199 | 0.123881 | 0.727705 | 0.944355 | Erich2 |
| ENSMUSG( | 458.9915 | -0.19036 | 0.149919 | 0.574325 | Erich3 |
| ENSMUSG( | 83.21753 | -0.01382 | 0.955689 | 0.993874 | Erich6 |
| ENSMUSG( | 1576.743 | -0.23068 | 0.087001 | 0.453311 | Erlec1 |
| ENSMUSG( | 772.5795 | -0.34032 | 0.006859 | 0.11446 | Erlin1 |
| ENSMUSG( | 2536.341 | -0.17193 | 0.159156 | 0.588305 | Erlin2 |
| ENSMUSG( | 113.6828 | -0.16922 | 0.415037 | 0.820184 | Ermap |
| ENSMUSG( | 238.6538 | -0.02165 | 0.886038 | 0.983008 | Ermard |
| ENSMUSG( | 15.26323 | 0.438468 | 0.38582 | NA | Ermardl1 |
| ENSMUSG( | 33.3487 | 0.156724 | 0.661097 | 0.925033 | Ermardl2 |
| ENSMUSG( | 1437.81 | -0.11377 | 0.314469 | 0.761558 | Ermn |
| ENSMUSG( | 2275.538 | -0.30551 | 0.038176 | 0.301526 | Ermp1 |
| ENSMUSG( | 400.9293 | -0.17737 | 0.182232 | 0.619591 | Ern1 |
| ENSMUSG( | 809.2534 | -0.26076 | 0.108119 | 0.497078 | Ero1a |
| ENSMUSG( | 539.6075 | -0.15285 | 0.349353 | 0.78176 | Ero1b |
| ENSMUSG( | 3341.076 | -0.20994 | 0.041001 | 0.313031 | Erp29 |
| ENSMUSG( | 1230.336 | -0.26382 | 0.077689 | 0.430958 | Erp44 |
| ENSMUSG( | 492.5248 | -0.0416 | 0.800991 | 0.96188 | Errfi1 |
| ENSMUSG( | 44.32571 | 0.216656 | 0.519952 | 0.874979 | Erv3 |
| ENSMUSG( | 815.0865 | 0.224084 | 0.214915 | 0.659462 | Esam |
| ENSMUSG( | 680.0316 | -0.0141 | 0.909744 | 0.988084 | Esco1 |
| ENSMUSG( | 1495.475 | 0.007173 | 0.941362 | 0.993415 | Esd |
| ENSMUSG( | 584.7963 | 0.294451 | 0.006928 | 0.114527 | Esf1 |
| ENSMUSG( | 64.72017 | -0.38732 | 0.113759 | 0.508873 | Espl1 |
| ENSMUSG( | 68.16057 | 0.281472 | 0.278653 | 0.728705 | Espn |
| ENSMUSG( | 15.3419 | -0.1799 | 0.704563 | NA | Espnl |
| ENSMUSG( | 45.4806 | 0.382026 | 0.192731 | 0.631789 | Esr1 |
| ENSMUSG( | 1501.697 | 0.078596 | 0.49965 | 0.863218 | Esrra |
| ENSMUSG( | 57.85954 | -0.09784 | 0.717027 | 0.941697 | Esrrb |
| ENSMUSG( | 306.6514 | 0.067071 | 0.618492 | 0.911166 | Esrrg |
| ENSMUSG( | 628.42 | -0.00758 | 0.952037 | 0.993821 | Ess2 |
| ENSMUSG( | 407.5948 | -0.17406 | 0.361833 | 0.792797 | Esyt1 |
| ENSMUSG( | 1459.694 | -0.24406 | 0.101359 | 0.484996 | Esyt2 |
| ENSMUSG( | 43.75806 | -0.29825 | 0.370558 | 0.797778 | Esyt3 |
| ENSMUSG( | 185.4689 | -0.08699 | 0.584741 | 0.89892 | Etaa1 |
| ENSMUSG( | 14.26691 | 0.237878 | 0.659239 | NA | Etaa1os |
| ENSMUSG( | 2195.603 | 0.058852 | 0.558236 | 0.888427 | Etf1 |
| ENSMUSG( | 1627.006 | 0.010922 | 0.923889 | 0.991046 | Etfa |
| ENSMUSG( | 247.6222 | 0.08078 | 0.582535 | 0.897918 | Etfb |
| ENSMUSG( | 77.35028 | -0.25872 | 0.244188 | 0.691435 | Etfbkmt |
| ENSMUSG( | 1599.981 | 0.094287 | 0.390851 | 0.80826 | Etfdh |
| ENSMUSG( | 719.2486 | 0.06649 | 0.585381 | 0.899171 | Etfrf1 |
| ENSMUSG( | 280.7556 | -0.04164 | 0.756041 | 0.951479 | Ethe1 |
| ENSMUSG( | 2519.869 | 0.068612 | 0.536206 | 0.881449 | Etl4 |
| ENSMUSG( | 7473.938 | 0.005488 | 0.942782 | 0.993415 | Etnk1 |
| ENSMUSG( | 247.4997 | 0.149878 | 0.400138 | 0.813641 | Etnk2 |

| | | | | | |
|---|---|---|---|---|---|
| ENSMUSG( | 171.7357 | -0.14538 | 0.437735 | 0.920339 | Enpp1 |
| ENSMUSG( | 12336.47 | 0.460792 | 0.001582 | 0.148357 | Enpp2 |
| ENSMUSG( | 17.79163 | -0.13348 | 0.771506 | 0.974236 | Enpp3 |
| ENSMUSG( | 922.9448 | 0.218259 | 0.282829 | 0.892412 | Enpp4 |
| ENSMUSG( | 8273.387 | 0.062432 | 0.652593 | 0.954799 | Enpp5 |
| ENSMUSG( | 859.5926 | 0.257414 | 0.268952 | 0.886755 | Enpp6 |
| ENSMUSG( | 6852.168 | -0.11144 | 0.330957 | 0.907513 | Ensa |
| ENSMUSG( | 745.2433 | 0.501333 | 0.009933 | 0.407479 | Entpd1 |
| ENSMUSG( | 510.9506 | 0.08649 | 0.648021 | 0.95422 | Entpd2 |
| ENSMUSG( | 355.8722 | -0.06649 | 0.668751 | 0.956766 | Entpd3 |
| ENSMUSG( | 2369.878 | 0.065753 | 0.634615 | 0.953903 | Entpd5 |
| ENSMUSG( | 1985.953 | 0.118852 | 0.398089 | 0.917257 | Entpd6 |
| ENSMUSG( | 279.0249 | 0.292987 | 0.188032 | 0.85698 | Entpd7 |
| ENSMUSG( | 3037.706 | 0.066197 | 0.579701 | 0.944209 | Entr1 |
| ENSMUSG( | 844.1038 | 0.0033 | 0.982365 | 0.996756 | Entrep1 |
| ENSMUSG( | 2572.938 | 0.222826 | 0.213156 | 0.864412 | Entrep2 |
| ENSMUSG( | 1702.896 | 0.132135 | 0.318585 | 0.903164 | Entrep3 |
| ENSMUSG( | 1135.658 | 0.039938 | 0.751108 | 0.970317 | Eny2 |
| ENSMUSG( | 1609.143 | 0.291192 | 0.056704 | 0.723362 | Eogt |
| ENSMUSG( | 167.987 | -0.04027 | 0.839022 | 0.981774 | Eola1 |
| ENSMUSG( | 28.04994 | 0.250806 | 0.581867 | 0.944266 | Eomes |
| ENSMUSG( | 1866.941 | 0.239133 | 0.057341 | 0.723362 | Ep300 |
| ENSMUSG( | 2448.141 | 0.087315 | 0.465779 | 0.923329 | Ep400 |
| ENSMUSG( | 15435.18 | -2.85E-05 | 0.999816 | 0.99993 | Epas1 |
| ENSMUSG( | 1695.836 | 0.043863 | 0.755212 | 0.971098 | Epb41 |
| ENSMUSG( | 27168.1 | 0.019358 | 0.859296 | 0.985884 | Epb41l1 |
| ENSMUSG( | 8771.065 | -0.18035 | 0.138435 | 0.828492 | Epb41l2 |
| ENSMUSG( | 4208.734 | -0.06342 | 0.548019 | 0.938063 | Epb41l3 |
| ENSMUSG( | 124.9267 | -0.02646 | 0.896185 | 0.990558 | Epb41l4a |
| ENSMUSG( | 214.9012 | -0.02496 | 0.899032 | 0.990558 | Epb41l4aos |
| ENSMUSG( | 1387.613 | 0.170434 | 0.117542 | 0.813384 | Epb41l4b |
| ENSMUSG( | 612.6924 | 0.004136 | 0.977704 | 0.996416 | Epb41l5 |
| ENSMUSG( | 890.2243 | 0.09524 | 0.4249 | 0.917257 | Epc1 |
| ENSMUSG( | 846.0466 | 0.123651 | 0.305062 | 0.898538 | Epc2 |
| ENSMUSG( | 25.57283 | 0.302003 | 0.481069 | 0.924548 | Epcam |
| ENSMUSG( | 96.75102 | 0.13482 | 0.563196 | 0.941158 | Epcip |
| ENSMUSG( | 3577.042 | 0.116712 | 0.451796 | 0.922202 | Epdr1 |
| ENSMUSG( | 1114.182 | 0.064447 | 0.585538 | 0.944975 | Epg5 |
| ENSMUSG( | 12.13741 | 0.34916 | 0.529608 | 0.934795 | Epha1 |
| ENSMUSG( | 1775.142 | 0.163858 | 0.16859 | 0.840222 | Epha10 |
| ENSMUSG( | 78.37892 | 0.45272 | 0.14075 | 0.830678 | Epha2 |
| ENSMUSG( | 199.0963 | 0.382469 | 0.072797 | 0.763629 | Epha3 |
| ENSMUSG( | 5018.062 | 0.288832 | 0.133851 | 0.822912 | Epha4 |
| ENSMUSG( | 2880.12 | 0.267063 | 0.105611 | 0.804969 | Epha5 |
| ENSMUSG( | 233.7919 | 0.158042 | 0.462959 | 0.923329 | Epha6 |
| ENSMUSG( | 1849.544 | 0.267028 | 0.111103 | 0.807885 | Epha7 |
| ENSMUSG( | 344.6219 | 0.797368 | 1.21E-06 | 0.000604 | Epha8 |
| ENSMUSG( | 4157.706 | 0.012024 | 0.922991 | 0.992644 | Ephb1 |
| ENSMUSG( | 794.7435 | 0.432694 | 0.036722 | 0.671468 | Ephb2 |
| ENSMUSG( | 1211.965 | 0.145218 | 0.367746 | 0.915289 | Ephb3 |
| ENSMUSG( | 398.0408 | 0.61924 | 0.01046 | 0.41614 | Ephb4 |
| ENSMUSG( | 3953.215 | 0.152203 | 0.42086 | 0.917257 | Ephb6 |
| ENSMUSG( | 1186.89 | 0.303462 | 0.034246 | 0.652059 | Ephx1 |
| ENSMUSG( | 1291.485 | -0.38137 | 0.007247 | 0.357969 | Ephx2 |
| ENSMUSG( | 8.32667 | 0.753371 | 0.303988 | NA | Ephx3 |
| ENSMUSG( | 1225.282 | 0.028078 | 0.810168 | 0.978779 | Ephx4 |
| ENSMUSG( | 1202.947 | -0.04784 | 0.733944 | 0.967228 | Epm2a |
| ENSMUSG( | 1984.99 | 0.031669 | 0.783455 | 0.976106 | Epm2aip1 |
| ENSMUSG( | 25804.55 | 0.016458 | 0.865188 | 0.98675 | Epn1 |
| ENSMUSG( | 3542.072 | -0.02195 | 0.835375 | 0.981237 | Epn2 |
| ENSMUSG( | 56.06641 | -0.74454 | 0.051887 | 0.712864 | Epn3 |
| ENSMUSG( | 3745.498 | 0.210337 | 0.037175 | 0.673514 | Epop |
| ENSMUSG( | 163.5587 | -0.03145 | 0.873278 | 0.98947 | Epor |
| ENSMUSG( | 2917.982 | -0.1463 | 0.18373 | 0.85191 | Eprs1 |
| ENSMUSG( | 10328.83 | -0.11132 | 0.395427 | 0.917257 | Eps15 |
| ENSMUSG( | 1257.177 | 0.070198 | 0.63003 | 0.953251 | Eps15l1 |
| ENSMUSG( | 2650.853 | -0.10242 | 0.439085 | 0.920721 | Eps8 |
| ENSMUSG( | 349.9516 | 0.659608 | 0.171914 | 0.842999 | Eps8l1 |
| ENSMUSG( | 75.2016 | -0.0678 | 0.799388 | 0.97681 | Eps8l2 |
| ENSMUSG( | 20.93011 | -0.64824 | 0.149453 | 0.836241 | Eps8l3 |
| ENSMUSG( | 44.78218 | -0.54321 | 0.112745 | 0.809623 | Epsti1 |
| ENSMUSG( | 43.42915 | 0.187616 | 0.55359 | 0.939565 | Epx |
| ENSMUSG( | 34.4311 | -0.27542 | 0.456184 | 0.923329 | Eqtn |
| ENSMUSG( | 404.7071 | 0.014926 | 0.917118 | 0.992529 | Eral1 |
| ENSMUSG( | 552.3138 | 0.094549 | 0.565594 | 0.941404 | Erap1 |
| ENSMUSG( | 203.4657 | 0.141698 | 0.462353 | 0.923329 | Erbb2 |
| ENSMUSG( | 681.0238 | 0.223041 | 0.323147 | 0.904023 | Erbb3 |
| ENSMUSG( | 1417.949 | 0.266192 | 0.168073 | 0.840222 | Erbb4 |
| ENSMUSG( | 6518.122 | -0.11932 | 0.358691 | 0.913759 | Erbin |
| ENSMUSG( | 1841.81 | 0.031447 | 0.771376 | 0.974236 | Erc1 |
| ENSMUSG( | 3079.69 | 0.051302 | 0.664857 | 0.956766 | Erc2 |
| ENSMUSG( | 415.7851 | -0.06903 | 0.690991 | 0.960071 | Ercc1 |
| ENSMUSG( | 939.9796 | 0.017147 | 0.897728 | 0.990558 | Ercc2 |
| ENSMUSG( | 613.1423 | -0.00033 | 0.998262 | 0.999313 | Ercc3 |
| ENSMUSG( | 544.1439 | 0.047775 | 0.717518 | 0.965964 | Ercc4 |
| ENSMUSG( | 898.5409 | 0.001453 | 0.990489 | 0.998182 | Ercc5 |
| ENSMUSG( | 716.6979 | 0.098545 | 0.435174 | 0.919062 | Ercc6 |
| ENSMUSG( | 629.759 | 0.011299 | 0.929062 | 0.99306 | Ercc6l2 |
| ENSMUSG( | 271.8298 | -0.03535 | 0.818356 | 0.979198 | Ercc8 |
| ENSMUSG( | 1195.268 | 0.024401 | 0.848009 | 0.985242 | Erf |
| ENSMUSG( | 27.3636 | -1.07318 | 0.025484 | 0.598732 | Erfe |
| ENSMUSG( | 214.503 | 0.632797 | 0.039971 | 0.681706 | Erg |
| ENSMUSG( | 1426.028 | -0.03569 | 0.816891 | 0.979198 | Erg28 |
| ENSMUSG( | 5032.335 | 0.068646 | 0.648423 | 0.95422 | Ergic1 |
| ENSMUSG( | 1652.799 | 0.093345 | 0.501153 | 0.927825 | Ergic2 |
| ENSMUSG( | 4319.184 | -0.03216 | 0.780555 | 0.976106 | Ergic3 |
| ENSMUSG( | 637.3616 | -0.18999 | 0.261636 | 0.884926 | Erh |
| ENSMUSG( | 278.218 | 0.212232 | 0.21537 | 0.865167 | Eri1 |
| ENSMUSG( | 277.7735 | 0.211626 | 0.255341 | 0.882761 | Eri2 |
| ENSMUSG( | 3274.896 | -0.13931 | 0.300919 | 0.898538 | Eri3 |
| ENSMUSG( | 83.22525 | 0.172336 | 0.500873 | 0.927825 | Erich1 |
| ENSMUSG( | 30.13543 | 0.042351 | 0.910898 | 0.992196 | Erich2 |
| ENSMUSG( | 420.346 | -0.17258 | 0.249195 | 0.877904 | Erich3 |
| ENSMUSG( | 79.61131 | 0.116605 | 0.647919 | 0.95422 | Erich6 |
| ENSMUSG( | 1599.473 | 0.091334 | 0.574959 | 0.942912 | Erlec1 |
| ENSMUSG( | 823.08 | 0.131829 | 0.333362 | 0.907907 | Erlin1 |
| ENSMUSG( | 2486.723 | 0.045876 | 0.732602 | 0.967228 | Erlin2 |
| ENSMUSG( | 112.838 | 0.069371 | 0.751968 | 0.970461 | Ermap |
| ENSMUSG( | 217.8274 | -0.01325 | 0.940042 | 0.994334 | Ermard |
| ENSMUSG( | 14.47645 | 0.536511 | 0.32288 | 0.904023 | Ermardl1 |
| ENSMUSG( | 26.88162 | -0.19202 | 0.626336 | 0.953234 | Ermardl2 |
| ENSMUSG( | 1276.339 | -0.19764 | 0.218385 | 0.866104 | Ermn |
| ENSMUSG( | 2437.098 | 0.176808 | 0.308589 | 0.898538 | Ermp1 |
| ENSMUSG( | 417.0749 | 0.209923 | 0.173634 | 0.844317 | Ern1 |
| ENSMUSG( | 869.8714 | 0.225886 | 0.23101 | 0.868507 | Ero1a |
| ENSMUSG( | 564.3179 | 0.243693 | 0.164977 | 0.838743 | Ero1b |
| ENSMUSG( | 3241.563 | -0.01976 | 0.869067 | 0.988097 | Erp29 |
| ENSMUSG( | 1334.73 | 0.245241 | 0.13817 | 0.828492 | Erp44 |
| ENSMUSG( | 459.6201 | 0.029639 | 0.858815 | 0.985884 | Errfi1 |
| ENSMUSG( | 36.9528 | -0.03285 | 0.929832 | 0.99306 | Erv3 |
| ENSMUSG( | 812.6482 | 0.465608 | 0.031605 | 0.631941 | Esam |
| ENSMUSG( | 604.6061 | -0.08918 | 0.523743 | 0.932848 | Esco1 |
| ENSMUSG( | 11.82858 | 0.387433 | 0.53551 | 0.936648 | Esco2 |
| ENSMUSG( | 1255.647 | -0.24185 | 0.039968 | 0.681706 | Esd |
| ENSMUSG( | 498.0714 | 0.111994 | 0.389622 | 0.917257 | Esf1 |
| ENSMUSG( | 72.85883 | 0.247773 | 0.311465 | 0.900022 | Espl1 |
| ENSMUSG( | 57.85072 | 0.080476 | 0.781026 | 0.976106 | Espn |
| ENSMUSG( | 19.32932 | 0.701555 | 0.129418 | 0.820358 | Espnl |
| ENSMUSG( | 35.84959 | -0.00347 | 0.992151 | 0.99874 | Esr1 |
| ENSMUSG( | 1352.263 | 0.043057 | 0.733435 | 0.967228 | Esrra |
| ENSMUSG( | 52.95016 | -0.09045 | 0.760623 | 0.972688 | Esrrb |
| ENSMUSG( | 288.6808 | 0.146727 | 0.438483 | 0.920721 | Esrrg |
| ENSMUSG( | 594.2367 | 0.103781 | 0.494436 | 0.927372 | Ess2 |
| ENSMUSG( | 452.3833 | 0.386357 | 0.072898 | 0.763629 | Esyt1 |
| ENSMUSG( | 1544.472 | 0.196384 | 0.221824 | 0.866104 | Esyt2 |
| ENSMUSG( | 42.49316 | -0.14155 | 0.69574 | 0.960867 | Esyt3 |
| ENSMUSG( | 192.9857 | 0.27589 | 0.119939 | 0.813384 | Etaa1 |
| ENSMUSG( | 11.37858 | -0.16488 | 0.794077 | NA | Etaa1os |
| ENSMUSG( | 1971.52 | 0.01369 | 0.918582 | 0.992529 | Etf1 |
| ENSMUSG( | 1351.173 | -0.27115 | 0.026294 | 0.603041 | Etfa |
| ENSMUSG( | 203.9835 | -0.22361 | 0.182463 | 0.850579 | Etfb |
| ENSMUSG( | 70.03127 | -0.27942 | 0.266361 | 0.886134 | Etfbkmt |
| ENSMUSG( | 1290.928 | -0.27006 | 0.022465 | 0.580801 | Etfdh |
| ENSMUSG( | 628.5091 | -0.06163 | 0.710229 | 0.964363 | Etfrf1 |
| ENSMUSG( | 251.6165 | -0.09382 | 0.546234 | 0.938063 | Ethe1 |
| ENSMUSG( | 2344.019 | 0.126516 | 0.280629 | 0.892356 | Etl4 |
| ENSMUSG( | 6927.991 | 0.054157 | 0.587662 | 0.944975 | Etnk1 |
| ENSMUSG( | 193.2254 | -0.29951 | 0.125811 | 0.81516 | Etnk2 |
| ENSMUSG( | 2408.441 | 0.176096 | 0.243803 | 0.877904 | Etnppl |
| ENSMUSG( | 587.8279 | 0.511103 | 0.013992 | 0.472248 | Ets1 |
| ENSMUSG( | 1496.88 | 0.271322 | 0.090677 | 0.791074 | Ets2 |
| ENSMUSG( | 1607.719 | 0.023746 | 0.874469 | 0.989644 | Etv1 |
| ENSMUSG( | 662.4741 | 0.112585 | 0.386125 | 0.917257 | Etv3 |
| ENSMUSG( | 768.4066 | -0.1321 | 0.276219 | 0.890749 | Etv4 |
| ENSMUSG( | 2723.435 | -0.09499 | 0.389041 | 0.917257 | Etv5 |
| ENSMUSG( | 690.7367 | 0.192697 | 0.116206 | 0.813384 | Etv6 |
| ENSMUSG( | 966.8399 | 0.116685 | 0.506363 | 0.927825 | Eva1a |
| ENSMUSG( | 149.9434 | 0.377467 | 0.096881 | 0.79645 | Eva1b |

| | | | | | | | | | | | |
|---|---|---|---|---|---|---|---|---|---|---|---|
| ENSMUSG( | Epb41l4a | 131.0411 | -0.14794 | 0.422094 | 0.823495 | ENSMUSG( | Epb41l4a | 124.9267 | -0.02646 | 0.896185 | 0.990558 |
| ENSMUSG( | Epb41l4ao | 240.8688 | 0.049539 | 0.750119 | 0.949664 | ENSMUSG( | Epb41l4ao | 214.9012 | -0.02496 | 0.899032 | 0.990558 |
| ENSMUSG( | Epb41l4b | 1545.728 | 0.209589 | 0.016931 | 0.194376 | ENSMUSG( | Epb41l4b | 1387.613 | 0.170434 | 0.117542 | 0.813384 |
| ENSMUSG( | Epb41l5 | 704.4173 | 0.139571 | 0.301456 | 0.749979 | ENSMUSG( | Epb41l5 | 612.6924 | 0.004136 | 0.977704 | 0.996416 |
| ENSMUSG( | Epc1 | 967.7467 | 0.065925 | 0.52717 | 0.877445 | ENSMUSG( | Epc1 | 890.2243 | 0.09524 | 0.4249 | 0.917257 |
| ENSMUSG( | Epc2 | 913.4709 | 0.078595 | 0.43246 | 0.82768 | ENSMUSG( | Epc2 | 846.0466 | 0.123651 | 0.305062 | 0.898538 |
| ENSMUSG( | Epcam | 24.32248 | -0.12937 | 0.731288 | 0.944956 | ENSMUSG( | Epcam | 25.57283 | 0.302003 | 0.481069 | 0.924548 |
| ENSMUSG( | Epcip | 92.09826 | -0.28248 | 0.172534 | 0.607843 | ENSMUSG( | Epcip | 96.75102 | 0.13482 | 0.563196 | 0.941158 |
| ENSMUSG( | Epdr1 | 3425.819 | -0.29292 | 0.029173 | 0.259574 | ENSMUSG( | Epdr1 | 3577.042 | 0.116712 | 0.451796 | 0.922202 |
| ENSMUSG( | Epg5 | 1231.707 | 0.081989 | 0.417353 | 0.821339 | ENSMUSG( | Epg5 | 1114.182 | 0.064447 | 0.585538 | 0.944975 |
| ENSMUSG( | Epha1 | 10.46295 | -0.36149 | 0.546587 | NA | ENSMUSG( | Epha1 | 12.13741 | 0.34916 | 0.529608 | 0.934795 |
| ENSMUSG( | Epha10 | 1748.249 | -0.1481 | 0.193222 | 0.632632 | ENSMUSG( | Epha10 | 1775.142 | 0.163858 | 0.16859 | 0.840222 |
| ENSMUSG( | Epha2 | 72.97512 | -0.01595 | 0.953233 | 0.993874 | ENSMUSG( | Epha2 | 78.37892 | 0.45272 | 0.14075 | 0.830678 |
| ENSMUSG( | Epha3 | 159.7212 | -0.5571 | 0.006937 | 0.114527 | ENSMUSG( | Epha3 | 199.0963 | 0.382469 | 0.072797 | 0.763629 |
| ENSMUSG( | Epha4 | 4282.619 | -0.46537 | 0.004113 | 0.082763 | ENSMUSG( | Epha4 | 5018.062 | 0.288832 | 0.133851 | 0.822912 |
| ENSMUSG( | Epha5 | 2623.72 | -0.2728 | 0.047569 | 0.333288 | ENSMUSG( | Epha5 | 2880.12 | 0.267063 | 0.105611 | 0.804969 |
| ENSMUSG( | Epha6 | 230.6702 | -0.1578 | 0.430639 | 0.826896 | ENSMUSG( | Epha6 | 233.7919 | 0.158042 | 0.462959 | 0.923329 |
| ENSMUSG( | Epha7 | 1594.16 | -0.45926 | 0.001586 | 0.04465 | ENSMUSG( | Epha7 | 1849.544 | 0.267028 | 0.111103 | 0.807885 |
| ENSMUSG( | Epha8 | 255.2471 | -0.24854 | 0.108028 | 0.497059 | ENSMUSG( | Epha8 | 344.6219 | 0.797368 | 1.21E-06 | 0.000604 |
| ENSMUSG( | Ephb1 | 4543.9 | -0.0021 | 0.984455 | 0.99781 | ENSMUSG( | Ephb1 | 4157.706 | 0.012024 | 0.922991 | 0.992644 |
| ENSMUSG( | Ephb2 | 724.8621 | -0.07944 | 0.655854 | 0.923929 | ENSMUSG( | Ephb2 | 794.7435 | 0.432694 | 0.036722 | 0.671468 |
| ENSMUSG( | Ephb3 | 1133.419 | -0.33725 | 0.021795 | 0.225679 | ENSMUSG( | Ephb3 | 1211.965 | 0.145218 | 0.367746 | 0.915289 |
| ENSMUSG( | Ephb4 | 331.2779 | -0.12943 | 0.49891 | 0.862973 | ENSMUSG( | Ephb4 | 398.0408 | 0.61924 | 0.01046 | 0.41614 |
| ENSMUSG( | Ephb6 | 3988.587 | -0.09411 | 0.56419 | 0.890139 | ENSMUSG( | Ephb6 | 3953.215 | 0.152203 | 0.42086 | 0.917257 |
| ENSMUSG( | Ephx1 | 1043.313 | -0.34725 | 0.006492 | 0.110626 | ENSMUSG( | Ephx1 | 1186.89 | 0.303462 | 0.034246 | 0.652059 |
| ENSMUSG( | Ephx2 | 1581.736 | -0.03577 | 0.785378 | 0.958528 | ENSMUSG( | Ephx2 | 1291.485 | -0.38137 | 0.007247 | 0.357969 |
| ENSMUSG( | Ephx4 | 1234.476 | -0.22865 | 0.041224 | 0.313778 | ENSMUSG( | Ephx4 | 1225.282 | 0.028078 | 0.810168 | 0.978779 |
| ENSMUSG( | Epm2a | 1351.541 | 0.024049 | 0.80225 | 0.962119 | ENSMUSG( | Epm2a | 1202.947 | -0.04784 | 0.733944 | 0.967228 |
| ENSMUSG( | Epm2aip1 | 2173.309 | 0.026689 | 0.775877 | 0.955606 | ENSMUSG( | Epm2aip1 | 1984.99 | 0.031669 | 0.783455 | 0.976106 |
| ENSMUSG( | Epn1 | 30139.12 | 0.190371 | 0.008573 | 0.129829 | ENSMUSG( | Epn1 | 25804.55 | 0.016458 | 0.865188 | 0.98675 |
| ENSMUSG( | Epn2 | 3935.334 | 0.011434 | 0.901133 | 0.985633 | ENSMUSG( | Epn2 | 3542.072 | -0.02195 | 0.835375 | 0.981237 |
| ENSMUSG( | Epn3 | 60.60453 | -0.79104 | 0.050325 | 0.342969 | ENSMUSG( | Epn3 | 56.06641 | -0.74454 | 0.051887 | 0.712864 |
| ENSMUSG( | Epop | 4071.795 | 0.18402 | 0.036338 | 0.293685 | ENSMUSG( | Epop | 3745.498 | 0.210337 | 0.037175 | 0.673514 |
| ENSMUSG( | Epor | 163.0181 | -0.33173 | 0.069335 | 0.40782 | ENSMUSG( | Epor | 163.5587 | -0.03145 | 0.873278 | 0.98947 |
| ENSMUSG( | Eprs1 | 3405.073 | 0.03456 | 0.720071 | 0.941986 | ENSMUSG( | Eprs1 | 2917.982 | -0.1463 | 0.18373 | 0.85191 |
| ENSMUSG( | Eps15 | 11598.25 | -0.0422 | 0.680247 | 0.931028 | ENSMUSG( | Eps15 | 10328.83 | -0.11132 | 0.395427 | 0.917257 |
| ENSMUSG( | Eps15l1 | 1427.153 | 0.164692 | 0.16691 | 0.599655 | ENSMUSG( | Eps15l1 | 1257.177 | 0.070198 | 0.63003 | 0.953251 |
| ENSMUSG( | Eps8 | 3104.088 | 0.086867 | 0.424223 | 0.824554 | ENSMUSG( | Eps8 | 2650.853 | -0.10242 | 0.439085 | 0.920721 |
| ENSMUSG( | Eps8l1 | 368.7706 | 0.568831 | 0.019405 | 0.209852 | ENSMUSG( | Eps8l1 | 349.9516 | 0.659608 | 0.171914 | 0.842999 |
| ENSMUSG( | Eps8l2 | 77.51944 | -0.27796 | 0.242678 | 0.690216 | ENSMUSG( | Eps8l2 | 75.2016 | -0.0678 | 0.799388 | 0.97681 |
| ENSMUSG( | Eps8l3 | 24.97392 | -0.40166 | 0.29408 | 0.743459 | ENSMUSG( | Eps8l3 | 20.93011 | -0.64824 | 0.149453 | 0.836241 |
| ENSMUSG( | Epsti1 | 51.3996 | -0.35601 | 0.225874 | 0.672685 | ENSMUSG( | Epsti1 | 44.78218 | -0.54321 | 0.112745 | 0.809623 |
| ENSMUSG( | Epx | 50.18287 | 0.324189 | 0.266128 | 0.716441 | ENSMUSG( | Epx | 43.42915 | 0.187616 | 0.55359 | 0.939565 |
| ENSMUSG( | Eqtn | 31.80899 | -0.85481 | 0.014899 | 0.179275 | ENSMUSG( | Eqtn | 34.4311 | -0.27542 | 0.456184 | 0.923329 |
| ENSMUSG( | Eral1 | 458.5951 | 0.104123 | 0.418982 | 0.821966 | ENSMUSG( | Eral1 | 404.7071 | 0.014926 | 0.917118 | 0.992529 |
| ENSMUSG( | Erap1 | 544.7043 | -0.2289 | 0.099434 | 0.481872 | ENSMUSG( | Erap1 | 552.3138 | 0.094549 | 0.565594 | 0.941404 |
| ENSMUSG( | Erbb2 | 190.7418 | -0.34296 | 0.046013 | 0.328532 | ENSMUSG( | Erbb2 | 203.4657 | 0.141698 | 0.462353 | 0.923329 |
| ENSMUSG( | Erbb3 | 564.4971 | -0.66376 | 0.000306 | 0.014613 | ENSMUSG( | Erbb3 | 681.0238 | 0.223041 | 0.323147 | 0.904023 |
| ENSMUSG( | Erbb4 | 1289.136 | -0.28268 | 0.077821 | 0.431216 | ENSMUSG( | Erbb4 | 1417.949 | 0.266192 | 0.168073 | 0.840222 |
| ENSMUSG( | Erbin | 7230.01 | -0.08669 | 0.422392 | 0.823508 | ENSMUSG( | Erbin | 6518.122 | -0.11932 | 0.358691 | 0.913759 |
| ENSMUSG( | Erc1 | 2123.397 | 0.168979 | 0.076823 | 0.429306 | ENSMUSG( | Erc1 | 1841.81 | 0.031447 | 0.771376 | 0.974236 |
| ENSMUSG( | Erc2 | 3286.214 | -0.02706 | 0.757793 | 0.951582 | ENSMUSG( | Erc2 | 3079.69 | 0.051302 | 0.664857 | 0.956766 |
| ENSMUSG( | Ercc1 | 477.166 | 0.064216 | 0.6897 | 0.933609 | ENSMUSG( | Ercc1 | 415.7851 | -0.06903 | 0.690991 | 0.960071 |
| ENSMUSG( | Ercc2 | 1089.756 | 0.167375 | 0.167041 | 0.599655 | ENSMUSG( | Ercc2 | 939.9796 | 0.017147 | 0.897728 | 0.990558 |
| ENSMUSG( | Ercc3 | 732.0908 | 0.231134 | 0.071007 | 0.411334 | ENSMUSG( | Ercc3 | 613.1423 | -0.00033 | 0.998262 | 0.999313 |
| ENSMUSG( | Ercc4 | 619.3449 | 0.143589 | 0.178962 | 0.616298 | ENSMUSG( | Ercc4 | 544.1439 | 0.047775 | 0.717518 | 0.965964 |
| ENSMUSG( | Ercc5 | 997.4532 | 0.035452 | 0.71209 | 0.940256 | ENSMUSG( | Ercc5 | 898.5409 | 0.001453 | 0.990489 | 0.998182 |
| ENSMUSG( | Ercc6 | 789.2081 | 0.109949 | 0.296286 | 0.744386 | ENSMUSG( | Ercc6 | 716.6979 | 0.098545 | 0.435174 | 0.919062 |
| ENSMUSG( | Ercc6l2 | 667.8007 | -0.09122 | 0.412791 | 0.819518 | ENSMUSG( | Ercc6l2 | 629.759 | 0.011299 | 0.929062 | 0.99306 |
| ENSMUSG( | Ercc8 | 304.068 | 0.01495 | 0.909075 | 0.98794 | ENSMUSG( | Ercc8 | 271.8298 | -0.03535 | 0.818356 | 0.979198 |
| ENSMUSG( | Erf | 1245.826 | -0.12729 | 0.256062 | 0.706828 | ENSMUSG( | Erf | 1195.268 | 0.024401 | 0.848009 | 0.985242 |
| ENSMUSG( | Erfe | 30.91398 | -1.01721 | 0.009098 | 0.134276 | ENSMUSG( | Erfe | 27.3636 | -1.07318 | 0.025484 | 0.598732 |
| ENSMUSG( | Erg | 188.8728 | 0.045014 | 0.857644 | 0.974667 | ENSMUSG( | Erg | 214.503 | 0.632797 | 0.039971 | 0.681706 |
| ENSMUSG( | Erg28 | 1350.992 | -0.50701 | 0.00025 | 0.012746 | ENSMUSG( | Erg28 | 1426.028 | -0.03569 | 0.816891 | 0.979198 |
| ENSMUSG( | Ergic1 | 4909.383 | -0.2885 | 0.025473 | 0.246501 | ENSMUSG( | Ergic1 | 5032.335 | 0.068646 | 0.648423 | 0.95422 |
| ENSMUSG( | Ergic2 | 1655.797 | -0.17723 | 0.161463 | 0.59148 | ENSMUSG( | Ergic2 | 1652.799 | 0.093345 | 0.501153 | 0.927825 |
| ENSMUSG( | Ergic3 | 4421.83 | -0.24453 | 0.015169 | 0.181362 | ENSMUSG( | Ergic3 | 4319.184 | -0.03216 | 0.780555 | 0.976106 |
| ENSMUSG( | Erh | 731.7561 | -0.04644 | 0.681091 | 0.931301 | ENSMUSG( | Erh | 637.3616 | -0.18999 | 0.261636 | 0.884926 |
| ENSMUSG( | Eri1 | 290.8129 | 0.087982 | 0.531517 | 0.879536 | ENSMUSG( | Eri1 | 278.218 | 0.212232 | 0.21537 | 0.865167 |
| ENSMUSG( | Eri2 | 278.3008 | -0.05599 | 0.746568 | 0.949289 | ENSMUSG( | Eri2 | 277.7735 | 0.211626 | 0.255341 | 0.882761 |
| ENSMUSG( | Eri3 | 3882.487 | 0.087521 | 0.397132 | 0.812088 | ENSMUSG( | Eri3 | 3274.896 | -0.13931 | 0.300919 | 0.898538 |
| ENSMUSG( | Erich1 | 88.83528 | 0.104415 | 0.646426 | 0.918612 | ENSMUSG( | Erich1 | 83.22525 | 0.172336 | 0.500873 | 0.927825 |
| ENSMUSG( | Erich2 | 34.14199 | 0.123881 | 0.727705 | 0.944355 | ENSMUSG( | Erich2 | 30.13543 | 0.042351 | 0.910898 | 0.992196 |
| ENSMUSG( | Erich3 | 458.9915 | -0.19036 | 0.149919 | 0.574325 | ENSMUSG( | Erich3 | 420.346 | -0.17258 | 0.249195 | 0.877904 |
| ENSMUSG( | Erich6 | 83.21753 | -0.01382 | 0.955689 | 0.993874 | ENSMUSG( | Erich6 | 79.61131 | 0.116605 | 0.647919 | 0.95422 |
| ENSMUSG( | Erlec1 | 1576.743 | -0.23068 | 0.087001 | 0.453311 | ENSMUSG( | Erlec1 | 1599.473 | 0.091334 | 0.574959 | 0.942912 |
| ENSMUSG( | Erlin1 | 772.5795 | -0.34032 | 0.006859 | 0.11446 | ENSMUSG( | Erlin1 | 823.08 | 0.131829 | 0.333362 | 0.907907 |
| ENSMUSG( | Erlin2 | 2536.341 | -0.17193 | 0.159156 | 0.588305 | ENSMUSG( | Erlin2 | 2486.723 | 0.045876 | 0.732602 | 0.967228 |
| ENSMUSG( | Ermap | 113.6828 | -0.16922 | 0.415037 | 0.820184 | ENSMUSG( | Ermap | 112.838 | 0.069371 | 0.751968 | 0.970461 |
| ENSMUSG( | Ermard | 238.6538 | -0.02165 | 0.886038 | 0.983008 | ENSMUSG( | Ermard | 217.8274 | -0.01325 | 0.940042 | 0.994334 |
| ENSMUSG( | Ermardl1 | 15.26323 | 0.438468 | 0.38582 | NA | ENSMUSG( | Ermardl1 | 14.47645 | 0.536511 | 0.32288 | 0.904023 |
| ENSMUSG( | Ermardl2 | 33.3487 | 0.156724 | 0.661097 | 0.925033 | ENSMUSG( | Ermardl2 | 26.88162 | -0.19202 | 0.626336 | 0.953234 |
| ENSMUSG( | Ermn | 1437.81 | -0.11377 | 0.314469 | 0.761558 | ENSMUSG( | Ermn | 1276.339 | -0.19764 | 0.218385 | 0.866104 |
| ENSMUSG( | Ermp1 | 2275.538 | -0.30551 | 0.038176 | 0.301526 | ENSMUSG( | Ermp1 | 2437.098 | 0.176808 | 0.308589 | 0.898538 |
| ENSMUSG( | Ern1 | 400.9293 | -0.17737 | 0.182232 | 0.619591 | ENSMUSG( | Ern1 | 417.0749 | 0.209923 | 0.173634 | 0.844317 |
| ENSMUSG( | Ero1a | 809.2534 | -0.26076 | 0.108119 | 0.497078 | ENSMUSG( | Ero1a | 869.8714 | 0.225886 | 0.23101 | 0.868507 |
| ENSMUSG( | Ero1b | 539.6075 | -0.15285 | 0.349353 | 0.78176 | ENSMUSG( | Ero1b | 564.3179 | 0.243693 | 0.164977 | 0.838743 |
| ENSMUSG( | Erp29 | 3341.076 | -0.20994 | 0.041001 | 0.313031 | ENSMUSG( | Erp29 | 3241.563 | -0.01976 | 0.869067 | 0.988097 |
| ENSMUSG( | Erp44 | 1230.336 | -0.26382 | 0.077689 | 0.430958 | ENSMUSG( | Erp44 | 1334.73 | 0.245241 | 0.13817 | 0.828492 |
| ENSMUSG( | Errfi1 | 492.5248 | -0.0416 | 0.800991 | 0.96188 | ENSMUSG( | Errfi1 | 459.6201 | 0.029639 | 0.858815 | 0.985884 |
| ENSMUSG( | Erv3 | 44.32571 | 0.216656 | 0.519952 | 0.874979 | ENSMUSG( | Erv3 | 36.9528 | -0.03285 | 0.929832 | 0.99306 |
| ENSMUSG( | Esam | 815.0865 | 0.224084 | 0.214915 | 0.659462 | ENSMUSG( | Esam | 812.6482 | 0.465608 | 0.031605 | 0.631941 |
| ENSMUSG( | Esco1 | 680.0316 | -0.0141 | 0.909744 | 0.988084 | ENSMUSG( | Esco1 | 604.6061 | -0.08918 | 0.523743 | 0.932848 |
| ENSMUSG( | Esd | 1495.475 | 0.007173 | 0.941362 | 0.993415 | ENSMUSG( | Esd | 1255.647 | -0.24185 | 0.039968 | 0.681706 |
| ENSMUSG( | Esf1 | 584.7963 | 0.294451 | 0.006928 | 0.114527 | ENSMUSG( | Esf1 | 498.0714 | 0.111994 | 0.389622 | 0.917257 |
| ENSMUSG( | Espl1 | 64.72017 | -0.38732 | 0.113759 | 0.508873 | ENSMUSG( | Espl1 | 72.85883 | 0.247773 | 0.311465 | 0.900022 |
| ENSMUSG( | Espn | 68.16057 | 0.281472 | 0.278653 | 0.728705 | ENSMUSG( | Espn | 57.85072 | 0.080476 | 0.781026 | 0.976106 |
| ENSMUSG( | Espnl | 15.3419 | -0.1799 | 0.704563 | NA | ENSMUSG( | Espnl | 19.32932 | 0.701555 | 0.129418 | 0.820358 |
| ENSMUSG( | Esr1 | 45.4806 | 0.382026 | 0.192731 | 0.631789 | ENSMUSG( | Esr1 | 35.84959 | -0.00347 | 0.992151 | 0.99874 |
| ENSMUSG( | Esrra | 1501.697 | 0.078596 | 0.49965 | 0.863218 | ENSMUSG( | Esrra | 1352.263 | 0.043057 | 0.733435 | 0.967228 |
| ENSMUSG( | Esrrb | 57.85954 | -0.09784 | 0.717027 | 0.941697 | ENSMUSG( | Esrrb | 52.95016 | -0.09045 | 0.760623 | 0.972688 |
| ENSMUSG( | Esrrg | 306.6514 | 0.067071 | 0.618492 | 0.911166 | ENSMUSG( | Esrrg | 288.6808 | 0.146727 | 0.438483 | 0.920721 |
| ENSMUSG( | Ess2 | 628.42 | -0.00758 | 0.952037 | 0.993821 | ENSMUSG( | Ess2 | 594.2367 | 0.103781 | 0.494436 | 0.927372 |
| ENSMUSG( | Esyt1 | 407.5948 | -0.17406 | 0.361833 | 0.792797 | ENSMUSG( | Esyt1 | 452.3833 | 0.386357 | 0.072898 | 0.763629 |
| ENSMUSG( | Esyt2 | 1459.694 | -0.24406 | 0.101359 | 0.484996 | ENSMUSG( | Esyt2 | 1544.472 | 0.196384 | 0.221824 | 0.866104 |
| ENSMUSG( | Esyt3 | 43.75806 | -0.29825 | 0.370558 | 0.797778 | ENSMUSG( | Esyt3 | 42.49316 | -0.14155 | 0.69574 | 0.960867 |
| ENSMUSG( | Etaa1 | 185.4689 | -0.08699 | 0.584741 | 0.89892 | ENSMUSG( | Etaa1 | 192.9857 | 0.27589 | 0.119939 | 0.813384 |
| ENSMUSG( | Etaa1os | 14.26691 | 0.237878 | 0.659239 | NA | ENSMUSG( | Etaa1os | 11.37858 | -0.16488 | 0.794077 | NA |
| ENSMUSG( | Etf1 | 2195.603 | 0.058852 | 0.558236 | 0.888427 | ENSMUSG( | Etf1 | 1971.52 | 0.01369 | 0.918582 | 0.992529 |
| ENSMUSG( | Etfa | 1627.006 | 0.010922 | 0.923889 | 0.991046 | ENSMUSG( | Etfa | 1351.173 | -0.27115 | 0.026294 | 0.603041 |
| ENSMUSG( | Etfb | 247.6222 | 0.08078 | 0.582535 | 0.897918 | ENSMUSG( | Etfb | 203.9835 | -0.22361 | 0.182463 | 0.850579 |
| ENSMUSG( | Etfbkmt | 77.35028 | -0.25872 | 0.244188 | 0.691435 | ENSMUSG( | Etfbkmt | 70.03127 | -0.27942 | 0.266361 | 0.886134 |
| ENSMUSG( | Etfdh | 1599.981 | 0.094287 | 0.390851 | 0.80826 | ENSMUSG( | Etfdh | 1290.928 | -0.27006 | 0.022465 | 0.580801 |
| ENSMUSG( | Etfrf1 | 719.2486 | 0.06649 | 0.585381 | 0.899171 | ENSMUSG( | Etfrf1 | 628.5091 | -0.06163 | 0.710229 | 0.964363 |
| ENSMUSG( | Ethe1 | 280.7556 | -0.04164 | 0.756041 | 0.951479 | ENSMUSG( | Ethe1 | 251.6165 | -0.09382 | 0.546234 | 0.938063 |
| ENSMUSG( | Etl4 | 2519.869 | 0.068612 | 0.536206 | 0.881449 | ENSMUSG( | Etl4 | 2344.019 | 0.126516 | 0.280629 | 0.892356 |
| ENSMUSG( | Etnk1 | 7473.938 | 0.005488 | 0.942782 | 0.993415 | ENSMUSG( | Etnk1 | 6927.991 | 0.054157 | 0.587662 | 0.944975 |
| ENSMUSG( | Etnk2 | 247.4997 | 0.149878 | 0.400138 | 0.813641 | ENSMUSG( | Etnk2 | 193.2254 | -0.29951 | 0.125811 | 0.81516 |
| ENSMUSG( | Etnppl | 3109.255 | 0.594214 | 1.24E-05 | 0.001322 | ENSMUSG( | Etnppl | 2408.441 | 0.176096 | 0.243803 | 0.877904 |
| ENSMUSG( | Ets1 | 530.0419 | -0.02178 | 0.894149 | 0.984512 | ENSMUSG( | Ets1 | 587.8279 | 0.511103 | 0.013992 | 0.472248 |
| ENSMUSG( | Ets2 | 1469.747 | -0.04012 | 0.782071 | 0.957558 | ENSMUSG( | Ets2 | 1496.88 | 0.271322 | 0.090677 | 0.791074 |
| ENSMUSG( | Etv1 | 1775.494 | 0.046637 | 0.665758 | 0.927054 | ENSMUSG( | Etv1 | 1607.719 | 0.023746 | 0.874469 | 0.989644 |
| ENSMUSG( | Etv3 | 703.5978 | 0.015397 | 0.888724 | 0.983434 | ENSMUSG( | Etv3 | 662.4741 | 0.112585 | 0.386125 | 0.917257 |
| ENSMUSG( | Etv4 | 897.6712 | 0.05061 | 0.633117 | 0.914073 | ENSMUSG( | Etv4 | 768.4066 | -0.1321 | 0.276219 | 0.890749 |
| ENSMUSG( | Etv5 | 3235.093 | 0.133945 | 0.156587 | 0.584667 | ENSMUSG( | Etv5 | 2723.435 | -0.09499 | 0.389041 | 0.917257 |
| ENSMUSG( | Etv6 | 753.0248 | 0.17421 | 0.115298 | 0.512226 | ENSMUSG( | Etv6 | 690.7367 | 0.192697 | 0.116206 | 0.813384 |
| ENSMUSG( | Eva1a | 994.4577 | -0.07252 | 0.612532 | 0.908747 | ENSMUSG( | Eva1a | 966.8399 | 0.116685 | 0.506363 | 0.927825 |
| ENSMUSG( | Eva1b | 151.4383 | 0.138605 | 0.490736 | 0.859582 | ENSMUSG( | Eva1b | 149.9434 | 0.377467 | 0.096881 | 0.79645 |
| ENSMUSG( | Eva1c | 68.35871 | -0.57558 | 0.026168 | 0.247488 | ENSMUSG( | Eva1c | 65.88337 | -0.38735 | 0.137286 | 0.828492 |
| ENSMUSG( | Evc | 691.7264 | 0.032595 | 0.748934 | 0.949604 | ENSMUSG( | Evc | 614.1753 | -0.04139 | 0.735629 | 0.967344 |
| ENSMUSG( | Evc2 | 724.686 | -0.06612 | 0.641515 | 0.916693 | ENSMUSG( | Evc2 | 683.9685 | 0.042984 | 0.785128 | 0.976285 |
| ENSMUSG( | Evi2a | 1086.779 | -0.38595 | 0.001982 | 0.051182 | ENSMUSG( | Evi2a | 1092.677 | -0.07631 | 0.634893 | 0.953903 |
| ENSMUSG( | Evi5 | 3313.651 | 0.030354 | 0.764542 | 0.954055 | ENSMUSG( | Evi5 | 2995.152 | 0.004553 | 0.97364 | 0.996416 |
| ENSMUSG( | Evi5l | 4554.71 | 0.080669 | 0.357103 | 0.788702 | ENSMUSG( | Evi5l | 3992.648 | -0.02947 | 0.770965 | 0.974236 |
| ENSMUSG( | Evl | 6162.138 | 0.051718 | 0.528652 | 0.878365 | ENSMUSG( | Evl | 5479.549 | -0.01948 | 0.869566 | 0.988097 |
| ENSMUSG( | Evpl | 87.42844 | -0.16439 | 0.447014 | 0.834051 | ENSMUSG( | Evpl | 74.86 | -0.3462 | 0.171374 | 0.842999 |
| ENSMUSG( | Ewsr1 | 5957.347 | -0.05701 | 0.544672 | 0.884649 | ENSMUSG( | Ewsr1 | 5447.916 | -0.0474 | 0.699195 | 0.961139 |
| ENSMUSG( | Exd1 | 98.56783 | 0.429344 | 0.047691 | 0.33359 | ENSMUSG( | Exd1 | 75.85554 | -0.03842 | 0.881122 | 0.989644 |
| ENSMUSG( | Exd2 | 3393.033 | 0.122046 | 0.295554 | 0.743814 | ENSMUSG( | Exd2 | 3105.908 | 0.136859 | 0.295232 | 0.898538 |
| ENSMUSG( | Exo1 | 13.25869 | 0.079187 | 0.883381 | NA | ENSMUSG( | Exo1 | 15.7737 | 0.741939 | 0.154508 | 0.837724 |
| ENSMUSG( | Exo5 | 166.1436 | -0.10381 | 0.602633 | 0.904887 | ENSMUSG( | Exo5 | 149.1134 | -0.15972 | 0.470468 | 0.923329 |
| ENSMUSG( | Exoc1 | 1218.536 | 0.120058 | 0.285215 | 0.733952 | ENSMUSG( | Exoc1 | 1069.965 | 0.01939 | 0.884855 | 0.989688 |
| ENSMUSG( | Exoc2 | 1106.321 | -0.01596 | 0.876 | 0.980452 | ENSMUSG( | Exoc2 | 966.2289 | -0.14531 | 0.252664 | 0.88018 |
| ENSMUSG( | Exoc3 | 2986.149 | 0.194251 | 0.037245 | 0.29789 | ENSMUSG( | Exoc3 | 2439.183 | -0.11916 | 0.298867 | 0.898538 |
| ENSMUSG( | Exoc3l | 29.01886 | -0.45005 | 0.212344 | 0.656417 | ENSMUSG( | Exoc3l | 27.32599 | -0.30475 | 0.443026 | 0.921554 |
| ENSMUSG( | Exoc3l4 | 56.6972 | 0.064516 | 0.806512 | 0.962804 | ENSMUSG( | Exoc3l4 | 42.81194 | -0.56568 | 0.083192 | 0.77681 |
| ENSMUSG( | Exoc4 | 2122.018 | 0.054529 | 0.530327 | 0.87918 | ENSMUSG( | Exoc4 | 1750.987 | -0.24204 | 0.024047 | 0.588226 |
| ENSMUSG( | Exoc5 | 1571.372 | 0.056649 | 0.557557 | 0.888279 | ENSMUSG( | Exoc5 | 1449.128 | 0.086202 | 0.542265 | 0.938034 |
| ENSMUSG( | Exoc6 | 888.3853 | 0.073287 | 0.470823 | 0.846741 | ENSMUSG( | Exoc6 | 786.1467 | -0.01285 | 0.917667 | 0.992529 |
| ENSMUSG( | Exoc6b | 2734.093 | -0.00871 | 0.935662 | 0.993363 | ENSMUSG( | Exoc6b | 2432.309 | -0.07432 | 0.581 | 0.944266 |
| ENSMUSG( | Exoc7 | 1666.198 | 0.13584 | 0.284269 | 0.73274 | ENSMUSG( | Exoc7 | 1421.01 | -0.05268 | 0.698471 | 0.960908 |
| ENSMUSG( | Exoc8 | 661.3827 | 0.290063 | 0.008175 | 0.125925 | ENSMUSG( | Exoc8 | 547.9264 | 0.033818 | 0.796295 | 0.97681 |
| ENSMUSG( | Exog | 528.389 | -0.08345 | 0.529985 | 0.878883 | ENSMUSG( | Exog | 480.8478 | -0.08019 | 0.581575 | 0.944266 |
| ENSMUSG( | Exosc1 | 323.7399 | -0.01552 | 0.918136 | 0.989645 | ENSMUSG( | Exosc1 | 283.3111 | -0.14199 | 0.389058 | 0.917257 |
| ENSMUSG( | Exosc10 | 1257.164 | 0.045679 | 0.645157 | 0.917934 | ENSMUSG( | Exosc10 | 1147.759 | 0.052864 | 0.675572 | 0.95694 |
| ENSMUSG( | Exosc2 | 362.6947 | 0.031656 | 0.807841 | 0.963257 | ENSMUSG( | Exosc2 | 312.2478 | -0.13859 | 0.356488 | 0.913358 |
| ENSMUSG( | Exosc3 | 177.6436 | 0.204186 | 0.208224 | 0.650319 | ENSMUSG( | Exosc3 | 151.5641 | 0.016441 | 0.93017 | 0.99306 |
| ENSMUSG( | Exosc4 | 510.7076 | 0.050084 | 0.723298 | 0.943761 | ENSMUSG( | Exosc4 | 443.257 | -0.09657 | 0.528737 | 0.934795 |

| | | | | | |
|---|---|---|---|---|---|
| ENSMUSG( | 3109.255 | 0.594214 | 1.24E-05 | 0.001322 | Etnppl |
| ENSMUSG( | 530.0419 | -0.02178 | 0.894149 | 0.984512 | Ets1 |
| ENSMUSG( | 1469.747 | -0.04012 | 0.782071 | 0.957558 | Ets2 |
| ENSMUSG( | 1775.494 | 0.046637 | 0.665758 | 0.927054 | Etv1 |
| ENSMUSG( | 703.5978 | 0.015397 | 0.888724 | 0.983434 | Etv3 |
| ENSMUSG( | 897.6712 | 0.05061 | 0.633117 | 0.914073 | Etv4 |
| ENSMUSG( | 3235.093 | 0.133945 | 0.156587 | 0.584667 | Etv5 |
| ENSMUSG( | 753.0248 | 0.17421 | 0.115298 | 0.512226 | Etv6 |
| ENSMUSG( | 994.4577 | -0.07252 | 0.612532 | 0.908747 | Eva1a |
| ENSMUSG( | 151.4383 | 0.138605 | 0.490736 | 0.859582 | Eva1b |
| ENSMUSG( | 68.35871 | -0.57558 | 0.026168 | 0.247488 | Eva1c |
| ENSMUSG( | 691.7264 | 0.032595 | 0.748934 | 0.949604 | Evc |
| ENSMUSG( | 724.686 | -0.06612 | 0.641515 | 0.916693 | Evc2 |
| ENSMUSG( | 1086.779 | -0.38595 | 0.001982 | 0.051182 | Evi2a |
| ENSMUSG( | 3313.651 | 0.030354 | 0.764542 | 0.954055 | Evi5 |
| ENSMUSG( | 4554.71 | 0.080669 | 0.357103 | 0.788702 | Evi5l |
| ENSMUSG( | 6162.138 | 0.051718 | 0.528652 | 0.878365 | Evl |
| ENSMUSG( | 87.42844 | -0.16439 | 0.447014 | 0.834051 | Evpl |
| ENSMUSG( | 5957.347 | -0.05701 | 0.544672 | 0.884649 | Ewsr1 |
| ENSMUSG( | 98.56783 | 0.429344 | 0.047691 | 0.33359 | Exd1 |
| ENSMUSG( | 3393.033 | 0.122046 | 0.295554 | 0.743814 | Exd2 |
| ENSMUSG( | 13.25869 | 0.079187 | 0.883381 | NA | Exo1 |
| ENSMUSG( | 166.1436 | -0.10381 | 0.602633 | 0.904887 | Exo5 |
| ENSMUSG( | 1218.536 | 0.120058 | 0.285215 | 0.733952 | Exoc1 |
| ENSMUSG( | 1106.321 | -0.01596 | 0.876 | 0.980452 | Exoc2 |
| ENSMUSG( | 2986.149 | 0.194251 | 0.037245 | 0.29789 | Exoc3 |
| ENSMUSG( | 29.01886 | -0.45005 | 0.212344 | 0.656417 | Exoc3l |
| ENSMUSG( | 56.6972 | 0.064516 | 0.806512 | 0.962804 | Exoc3l4 |
| ENSMUSG( | 2122.018 | 0.054529 | 0.530327 | 0.87918 | Exoc4 |
| ENSMUSG( | 1571.372 | 0.056649 | 0.557557 | 0.888279 | Exoc5 |
| ENSMUSG( | 888.3853 | 0.073287 | 0.470823 | 0.846741 | Exoc6 |
| ENSMUSG( | 2734.093 | -0.00871 | 0.935662 | 0.993363 | Exoc6b |
| ENSMUSG( | 1666.198 | 0.13584 | 0.284269 | 0.73274 | Exoc7 |
| ENSMUSG( | 661.3827 | 0.290063 | 0.008175 | 0.125925 | Exoc8 |
| ENSMUSG( | 528.389 | -0.08345 | 0.529985 | 0.878883 | Exog |
| ENSMUSG( | 323.7399 | -0.01552 | 0.918136 | 0.989645 | Exosc1 |
| ENSMUSG( | 1257.164 | 0.045679 | 0.645157 | 0.917934 | Exosc10 |
| ENSMUSG( | 362.6947 | 0.031656 | 0.807841 | 0.963257 | Exosc2 |
| ENSMUSG( | 177.6436 | 0.204186 | 0.208224 | 0.650319 | Exosc3 |
| ENSMUSG( | 510.7076 | 0.050084 | 0.723298 | 0.943761 | Exosc4 |
| ENSMUSG( | 597.0085 | 0.139681 | 0.252018 | 0.701908 | Exosc5 |
| ENSMUSG( | 337.2601 | 0.040544 | 0.79037 | 0.959461 | Exosc7 |
| ENSMUSG( | 326.8308 | 0.000798 | 0.994857 | 0.999369 | Exosc8 |
| ENSMUSG( | 707.8124 | 0.084015 | 0.491316 | 0.859873 | Exosc9 |
| ENSMUSG( | 295.6341 | -0.02542 | 0.858214 | 0.974672 | Exph5 |
| ENSMUSG( | 1388.837 | -0.32217 | 0.030176 | 0.263854 | Ext1 |
| ENSMUSG( | 1286.616 | -0.30994 | 0.047183 | 0.33245 | Ext2 |
| ENSMUSG( | 2504.055 | 0.084602 | 0.52886 | 0.878365 | Extl1 |
| ENSMUSG( | 2569.285 | -0.31951 | 0.023345 | 0.233882 | Extl2 |
| ENSMUSG( | 7002.927 | -0.08928 | 0.480982 | 0.855135 | Extl3 |
| ENSMUSG( | 760.2782 | 0.455595 | 9.07E-05 | 0.006031 | Eya1 |
| ENSMUSG( | 163.7026 | 0.337961 | 0.081766 | 0.441786 | Eya2 |
| ENSMUSG( | 990.8101 | 0.105873 | 0.327958 | 0.772842 | Eya3 |
| ENSMUSG( | 197.9435 | -0.03286 | 0.826089 | 0.968159 | Eya4 |
| ENSMUSG( | 1428.928 | 0.163969 | 0.159962 | 0.589679 | Ezh1 |
| ENSMUSG( | 286.6818 | 0.169455 | 0.213478 | 0.65774 | Ezh2 |
| ENSMUSG( | 12499.07 | 0.110999 | 0.301297 | 0.749979 | Ezr |
| ENSMUSG( | 509.8063 | -0.20563 | 0.181867 | 0.618944 | F11r |
| ENSMUSG( | 16.47223 | -0.37868 | 0.431327 | NA | F12 |
| ENSMUSG( | 198.6716 | 0.452484 | 0.009195 | 0.135112 | F13a1 |
| ENSMUSG( | 251.377 | -0.38554 | 0.043538 | 0.31954 | F2r |
| ENSMUSG( | 38.1706 | -0.30855 | 0.419608 | 0.822309 | F2rl2 |
| ENSMUSG( | 2596.042 | -0.54611 | 0.000173 | 0.00947 | F3 |
| ENSMUSG( | 16.11061 | 0.133927 | 0.784507 | NA | F420014N23Rik |
| ENSMUSG( | 118.9563 | 0.076864 | 0.904711 | 0.986316 | F5 |
| ENSMUSG( | 128.5699 | 0.156236 | 0.415978 | 0.82084 | F630040K05Rik |
| ENSMUSG( | 755.8668 | 0.078709 | 0.452124 | 0.836145 | F730043M19Rik |
| ENSMUSG( | 63.32001 | -0.50401 | 0.04715 | 0.33245 | F8 |
| ENSMUSG( | 310.6168 | -0.0988 | 0.573735 | 0.893584 | F8a |
| ENSMUSG( | 14.49749 | -0.0593 | 0.902977 | NA | F9 |
| ENSMUSG( | 2099.456 | -0.71851 | 4.34E-05 | 0.003572 | Fa2h |
| ENSMUSG( | 4390.335 | -0.21686 | 0.068634 | 0.405903 | Faah |
| ENSMUSG( | 540.1447 | 0.245331 | 0.067012 | 0.400593 | Faap100 |
| ENSMUSG( | 681.0862 | 0.065752 | 0.547358 | 0.885128 | Faap20 |
| ENSMUSG( | 133.6858 | 0.055915 | 0.781229 | 0.957558 | Faap24 |
| ENSMUSG( | 992.1882 | -0.05261 | 0.740332 | 0.947823 | Fabp3 |
| ENSMUSG( | 5491.776 | 0.119248 | 0.284403 | 0.732908 | Fabp5 |
| ENSMUSG( | 3507.824 | 0.209024 | 0.210274 | 0.652595 | Fabp7 |
| ENSMUSG( | 127.8795 | 0.276955 | 0.123447 | 0.530552 | Fadd |
| ENSMUSG( | 41028.89 | 0.041815 | 0.685243 | 0.932981 | Fads1 |
| ENSMUSG( | 29564.86 | 0.120728 | 0.283808 | 0.732203 | Fads2 |
| ENSMUSG( | 1252.975 | -0.26215 | 0.054554 | 0.35941 | Fads3 |
| ENSMUSG( | 1145.94 | -0.19236 | 0.176894 | 0.613937 | Fads6 |
| ENSMUSG( | 958.6222 | -0.087 | 0.360285 | 0.791947 | Faf1 |
| ENSMUSG( | 1910.783 | -0.26315 | 0.034878 | 0.287263 | Faf2 |
| ENSMUSG( | 621.8611 | 0.058007 | 0.636242 | 0.914873 | Fah |
| ENSMUSG( | 336.0609 | -0.10831 | 0.45137 | 0.835567 | Fahd1 |
| ENSMUSG( | 494.0826 | -0.06918 | 0.583598 | 0.89861 | Fahd2a |
| ENSMUSG( | 702.8847 | -0.02851 | 0.780554 | 0.957491 | Faim |
| ENSMUSG( | 16462.67 | -0.23969 | 0.060034 | 0.378891 | Faim2 |
| ENSMUSG( | 90232.22 | 0.396138 | 8.54E-06 | 0.001033 | Fam107a |
| ENSMUSG( | 784.779 | 0.050701 | 0.707524 | 0.93949 | Fam107b |
| ENSMUSG( | 130.0487 | 0.010717 | 0.955258 | 0.993874 | Fam110a |
| ENSMUSG( | 615.581 | 0.242982 | 0.148863 | 0.572605 | Fam110b |
| ENSMUSG( | 117.7984 | 0.427402 | 0.02486 | 0.242612 | Fam110d |
| ENSMUSG( | 88.86241 | 0.144251 | 0.5337 | 0.880095 | Fam111a |
| ENSMUSG( | 383.7418 | 0.096272 | 0.459891 | 0.840338 | Fam114a1 |
| ENSMUSG( | 1032.399 | 0.114134 | 0.257961 | 0.709319 | Fam114a2 |
| ENSMUSG( | 203.4392 | -0.19013 | 0.264431 | 0.715323 | Fam117a |
| ENSMUSG( | 1178.932 | 0.035162 | 0.71382 | 0.940903 | Fam117b |
| ENSMUSG( | 369.1909 | -0.01953 | 0.883058 | 0.982422 | Fam118a |
| ENSMUSG( | 431.4877 | 0.227497 | 0.069876 | 0.408574 | Fam118b |
| ENSMUSG( | 6656.032 | 0.048581 | 0.637268 | 0.915436 | Fam120a |
| ENSMUSG( | 1036.083 | -0.04494 | 0.722047 | 0.943535 | Fam120aos |
| ENSMUSG( | 1352.863 | 0.073417 | 0.459434 | 0.840328 | Fam120b |
| ENSMUSG( | 1887.773 | 0.055377 | 0.671508 | 0.928781 | Fam120c |
| ENSMUSG( | 1106.388 | 0.061548 | 0.559969 | 0.888917 | Fam124a |
| ENSMUSG( | 7620.748 | 0.055044 | 0.723431 | 0.943859 | Fam131a |
| ENSMUSG( | 7767.327 | 0.04329 | 0.574854 | 0.894572 | Fam131b |
| ENSMUSG( | 60.07442 | 0.027992 | 0.912619 | 0.988483 | Fam131c |
| ENSMUSG( | 549.8681 | -0.05578 | 0.625516 | 0.912249 | Fam133b |
| ENSMUSG( | 1018.98 | -0.15938 | 0.186357 | 0.623869 | Fam135a |
| ENSMUSG( | 933.3289 | -0.01524 | 0.887844 | 0.983434 | Fam135b |
| ENSMUSG( | 450.6913 | -0.01758 | 0.89078 | 0.983764 | Fam136a |
| ENSMUSG( | 355.0875 | 0.278395 | 0.03489 | 0.287263 | Fam13a |
| ENSMUSG( | 1457.416 | 0.109335 | 0.274354 | 0.724547 | Fam13b |
| ENSMUSG( | 2231.804 | -0.07364 | 0.460582 | 0.840787 | Fam13c |
| ENSMUSG( | 1917.725 | 0.078503 | 0.435962 | 0.829093 | Fam149a |
| ENSMUSG( | 1972.476 | 0.075599 | 0.422155 | 0.823495 | Fam149b |
| ENSMUSG( | 27.24997 | -0.03606 | 0.922193 | 0.99045 | Fam151a |
| ENSMUSG( | 185.5891 | -0.16361 | 0.30901 | 0.757121 | Fam151b |
| ENSMUSG( | 124.642 | -0.12203 | 0.489153 | 0.859092 | Fam161a |
| ENSMUSG( | 316.2418 | 0.044595 | 0.739684 | 0.947823 | Fam161b |
| ENSMUSG( | 944.1851 | 0.014101 | 0.88816 | 0.983434 | Fam162a |
| ENSMUSG( | 1253.612 | 0.038677 | 0.77545 | 0.955606 | Fam163a |
| ENSMUSG( | 5401.315 | -0.10296 | 0.306016 | 0.753712 | Fam163b |
| ENSMUSG( | 118.8852 | 0.475387 | 0.029207 | 0.259574 | Fam167a |
| ENSMUSG( | 34.7535 | 1.385002 | 5.76E-05 | 0.004407 | Fam167b |
| ENSMUSG( | 11627.15 | 0.006048 | 0.944022 | 0.993503 | Fam168a |
| ENSMUSG( | 15766.66 | 0.087437 | 0.307523 | 0.755462 | Fam168b |
| ENSMUSG( | 902.4217 | -0.18936 | 0.062374 | 0.386294 | Fam169a |
| ENSMUSG( | 9.204231 | 0.007554 | 0.990704 | NA | Fam169b |
| ENSMUSG( | 2876.997 | -0.19623 | 0.112805 | 0.506698 | Fam171a1 |
| ENSMUSG( | 3757.813 | -0.18612 | 0.159227 | 0.588305 | Fam171a2 |
| ENSMUSG( | 17264.83 | -0.03187 | 0.818768 | 0.965888 | Fam171b |
| ENSMUSG( | 1589.162 | -0.21699 | 0.070771 | 0.410404 | Fam174a |
| ENSMUSG( | 962.0256 | -0.13729 | 0.299597 | 0.748647 | Fam174b |
| ENSMUSG( | 99.17152 | 0.083376 | 0.704826 | 0.938303 | Fam174c |
| ENSMUSG( | 13.95601 | 0.026847 | 0.960167 | NA | Fam178b |
| ENSMUSG( | 117.1552 | -0.23864 | 0.328025 | 0.772854 | Fam180a |
| ENSMUSG( | 68.65734 | -0.51272 | 0.03692 | 0.296508 | Fam181a |
| ENSMUSG( | 1105.512 | 0.134109 | 0.191836 | 0.630811 | Fam181b |
| ENSMUSG( | 874.3291 | 0.174944 | 0.103132 | 0.48855 | Fam184a |
| ENSMUSG( | 284.665 | -0.29797 | 0.03757 | 0.299456 | Fam184b |
| ENSMUSG( | 331.3661 | 0.098568 | 0.479519 | 0.854046 | Fam185a |
| ENSMUSG( | 11.22421 | -0.20901 | 0.719283 | NA | Fam186a |
| ENSMUSG( | 32.28512 | -0.14966 | 0.664902 | 0.926824 | Fam186b |
| ENSMUSG( | 1306.771 | 0.029775 | 0.749499 | 0.949664 | Fam193a |
| ENSMUSG( | 3343.442 | 0.265775 | 0.075062 | 0.42422 | Fam193b |
| ENSMUSG( | 292.6608 | -0.05986 | 0.701717 | 0.937441 | Fam199x |
| ENSMUSG( | 627.5895 | -0.03716 | 0.759507 | 0.952498 | Fam204a |
| ENSMUSG( | 1076.849 | -0.04775 | 0.766288 | 0.954173 | Fam20a |
| ENSMUSG( | 4228.919 | -0.04771 | 0.669711 | 0.928147 | Fam20b |
| ENSMUSG( | 1431.966 | -0.16292 | 0.233654 | 0.681508 | Fam20c |
| ENSMUSG( | 1184.577 | 0.101036 | 0.349215 | 0.78176 | Fam210a |
| ENSMUSG( | 1200.661 | -0.04808 | 0.648904 | 0.920025 | Fam210b |
| ENSMUSG( | 886.1685 | 0.036505 | 0.793993 | 0.959799 | Fam216a |
| ENSMUSG( | 26.12061 | -0.34535 | 0.347019 | 0.7797 | Fam216b |

| | | | | | |
|---|---|---|---|---|---|
| ENSMUSG( | 65.88337 | -0.38735 | 0.137286 | 0.828492 | Eva1c |
| ENSMUSG( | 614.1753 | -0.04139 | 0.735629 | 0.967344 | Evc |
| ENSMUSG( | 683.9685 | 0.042984 | 0.785128 | 0.976285 | Evc2 |
| ENSMUSG( | 1092.677 | -0.07631 | 0.634893 | 0.953903 | Evi2a |
| ENSMUSG( | 2995.152 | 0.004553 | 0.97364 | 0.996416 | Evi5 |
| ENSMUSG( | 3992.648 | -0.02947 | 0.770965 | 0.974236 | Evi5l |
| ENSMUSG( | 5479.549 | -0.01948 | 0.869566 | 0.988097 | Evl |
| ENSMUSG( | 74.86 | -0.3462 | 0.171374 | 0.842999 | Evpl |
| ENSMUSG( | 5447.916 | -0.0474 | 0.699195 | 0.961139 | Ewsr1 |
| ENSMUSG( | 75.85554 | -0.03842 | 0.881122 | 0.989644 | Exd1 |
| ENSMUSG( | 3105.908 | 0.136859 | 0.295232 | 0.898538 | Exd2 |
| ENSMUSG( | 15.7737 | 0.741939 | 0.154508 | 0.837724 | Exo1 |
| ENSMUSG( | 149.1134 | -0.15972 | 0.470468 | 0.923329 | Exo5 |
| ENSMUSG( | 1069.965 | 0.01939 | 0.884855 | 0.989688 | Exoc1 |
| ENSMUSG( | 966.2289 | -0.14531 | 0.252664 | 0.88018 | Exoc2 |
| ENSMUSG( | 2439.183 | -0.11916 | 0.298867 | 0.898538 | Exoc3 |
| ENSMUSG( | 27.32599 | -0.30475 | 0.443026 | 0.921554 | Exoc3l |
| ENSMUSG( | 9.70964 | 0.940239 | 0.131251 | NA | Exoc3l2 |
| ENSMUSG( | 42.81194 | -0.56568 | 0.083192 | 0.77681 | Exoc3l4 |
| ENSMUSG( | 1750.987 | -0.24204 | 0.024047 | 0.588226 | Exoc4 |
| ENSMUSG( | 1449.128 | 0.086202 | 0.542265 | 0.938034 | Exoc5 |
| ENSMUSG( | 786.1467 | -0.01285 | 0.917667 | 0.992529 | Exoc6 |
| ENSMUSG( | 2432.309 | -0.07432 | 0.581 | 0.944266 | Exoc6b |
| ENSMUSG( | 1421.01 | -0.05268 | 0.698471 | 0.960908 | Exoc7 |
| ENSMUSG( | 547.9264 | 0.033818 | 0.796295 | 0.97681 | Exoc8 |
| ENSMUSG( | 480.8478 | -0.08019 | 0.581575 | 0.944266 | Exog |
| ENSMUSG( | 283.3111 | -0.14199 | 0.389058 | 0.917257 | Exosc1 |
| ENSMUSG( | 1147.759 | 0.052864 | 0.675572 | 0.95694 | Exosc10 |
| ENSMUSG( | 312.2478 | -0.13859 | 0.356488 | 0.913358 | Exosc2 |
| ENSMUSG( | 151.5641 | 0.016441 | 0.93017 | 0.99306 | Exosc3 |
| ENSMUSG( | 443.257 | -0.09657 | 0.528737 | 0.934795 | Exosc4 |
| ENSMUSG( | 498.9634 | -0.10995 | 0.433156 | 0.918434 | Exosc5 |
| ENSMUSG( | 287.9534 | -0.15793 | 0.378166 | 0.916962 | Exosc7 |
| ENSMUSG( | 296.0506 | -0.01605 | 0.911937 | 0.992196 | Exosc8 |
| ENSMUSG( | 593.0143 | -0.16748 | 0.266672 | 0.886134 | Exosc9 |
| ENSMUSG( | 260.7232 | -0.10806 | 0.571907 | 0.942347 | Exph5 |
| ENSMUSG( | 1487.685 | 0.163535 | 0.304768 | 0.898538 | Ext1 |
| ENSMUSG( | 1433.844 | 0.281123 | 0.122637 | 0.814249 | Ext2 |
| ENSMUSG( | 2433.936 | 0.265753 | 0.111558 | 0.808371 | Extl1 |
| ENSMUSG( | 2784.594 | 0.196743 | 0.244999 | 0.877904 | Extl2 |
| ENSMUSG( | 6811.575 | 0.102633 | 0.500696 | 0.927825 | Extl3 |
| ENSMUSG( | 661.7369 | 0.337611 | 0.012201 | 0.44712 | Eya1 |
| ENSMUSG( | 135.3205 | 0.068064 | 0.758327 | 0.971858 | Eya2 |
| ENSMUSG( | 868.9223 | 0.00057 | 0.996462 | 0.999178 | Eya3 |
| ENSMUSG( | 191.0829 | 0.129579 | 0.441152 | 0.921166 | Eya4 |
| ENSMUSG( | 1274.146 | 0.108365 | 0.428431 | 0.917257 | Ezh1 |
| ENSMUSG( | 272.0161 | 0.288169 | 0.079094 | 0.77458 | Ezh2 |
| ENSMUSG( | 10563.11 | -0.10751 | 0.419127 | 0.917257 | Ezr |
| ENSMUSG( | 578.1234 | 0.413946 | 0.032989 | 0.63923 | F11r |
| ENSMUSG( | 16.10486 | -0.15738 | 0.750711 | 0.970209 | F12 |
| ENSMUSG( | 169.4897 | 0.272125 | 0.175602 | 0.845305 | F13a1 |
| ENSMUSG( | 292.5938 | 0.338311 | 0.122608 | 0.814249 | F2r |
| ENSMUSG( | 45.33656 | 0.458825 | 0.223338 | 0.867404 | F2rl2 |
| ENSMUSG( | 2710.276 | -0.09818 | 0.614188 | 0.951177 | F3 |
| ENSMUSG( | 17.01016 | 0.526041 | 0.274072 | 0.889874 | F420014N23Rik |
| ENSMUSG( | 145.6714 | 0.808956 | 0.013536 | 0.464932 | F5 |
| ENSMUSG( | 111.1058 | -0.01682 | 0.9453 | 0.994798 | F630040K05Rik |
| ENSMUSG( | 644.6988 | -0.11705 | 0.417663 | 0.917257 | F730043M19Rik |
| ENSMUSG( | 59.12262 | -0.40574 | 0.135276 | 0.825081 | F8 |
| ENSMUSG( | 7.745806 | -0.67431 | 0.379706 | NA | F830016B08Rik |
| ENSMUSG( | 271.4463 | -0.23281 | 0.20489 | 0.864412 | F8a |
| ENSMUSG( | 15.04254 | 0.313302 | 0.545119 | 0.938063 | F9 |
| ENSMUSG( | 2558.294 | 0.207964 | 0.322322 | 0.904023 | Fa2h |
| ENSMUSG( | 4268.896 | -0.01841 | 0.887079 | 0.990263 | Faah |
| ENSMUSG( | 454.7202 | 0.03419 | 0.836039 | 0.981237 | Faap100 |
| ENSMUSG( | 585.3824 | -0.11004 | 0.454461 | 0.922444 | Faap20 |
| ENSMUSG( | 113.111 | -0.16038 | 0.48754 | 0.925427 | Faap24 |
| ENSMUSG( | 869.1724 | -0.17586 | 0.334221 | 0.907907 | Fabp3 |
| ENSMUSG( | 4603.761 | -0.12407 | 0.350957 | 0.911707 | Fabp5 |
| ENSMUSG( | 2545.857 | -0.48582 | 0.019075 | 0.549166 | Fabp7 |
| ENSMUSG( | 109.7742 | 0.104153 | 0.626676 | 0.953234 | Fadd |
| ENSMUSG( | 35232.63 | -0.13204 | 0.268681 | 0.886755 | Fads1 |
| ENSMUSG( | 25221.29 | -0.06805 | 0.577902 | 0.943131 | Fads2 |
| ENSMUSG( | 1294.087 | 0.114047 | 0.462818 | 0.923329 | Fads3 |
| ENSMUSG( | 1133.124 | 0.055206 | 0.735081 | 0.967344 | Fads6 |
| ENSMUSG( | 892.6898 | -0.02272 | 0.846985 | 0.985027 | Faf1 |
| ENSMUSG( | 1952.791 | 0.08337 | 0.567327 | 0.941777 | Faf2 |
| ENSMUSG( | 524.4636 | -0.16852 | 0.259217 | 0.883971 | Fah |
| ENSMUSG( | 290.9205 | -0.2716 | 0.08377 | 0.77681 | Fahd1 |
| ENSMUSG( | 427.2129 | -0.23464 | 0.157806 | 0.837724 | Fahd2a |
| ENSMUSG( | 621.4761 | -0.12281 | 0.34667 | 0.91112 | Faim |
| ENSMUSG( | 16629.36 | 0.070596 | 0.633192 | 0.953844 | Faim2 |
| ENSMUSG( | 72338.35 | 0.053085 | 0.669347 | 0.956766 | Fam107a |
| ENSMUSG( | 672.6244 | -0.13583 | 0.425627 | 0.917257 | Fam107b |
| ENSMUSG( | 124.0079 | 0.153191 | 0.460288 | 0.923329 | Fam110a |
| ENSMUSG( | 537.7398 | 0.126838 | 0.497596 | 0.927825 | Fam110b |
| ENSMUSG( | 115.803 | 0.631336 | 0.008382 | 0.388416 | Fam110d |
| ENSMUSG( | 84.82567 | 0.26661 | 0.280513 | 0.892356 | Fam111a |
| ENSMUSG( | 327.01 | -0.10428 | 0.503468 | 0.927825 | Fam114a1 |
| ENSMUSG( | 927.8665 | 0.073548 | 0.57078 | 0.941973 | Fam114a2 |
| ENSMUSG( | 195.4591 | -0.03362 | 0.853986 | 0.98563 | Fam117a |
| ENSMUSG( | 1067.127 | 0.015553 | 0.899689 | 0.990558 | Fam117b |
| ENSMUSG( | 348.3729 | 0.072696 | 0.65589 | 0.955533 | Fam118a |
| ENSMUSG( | 383.0747 | 0.151995 | 0.305734 | 0.898538 | Fam118b |
| ENSMUSG( | 5821.168 | -0.07135 | 0.543078 | 0.938063 | Fam120a |
| ENSMUSG( | 913.0247 | -0.15238 | 0.378207 | 0.916962 | Fam120aos |
| ENSMUSG( | 1261.062 | 0.139978 | 0.257743 | 0.883401 | Fam120b |
| ENSMUSG( | 1592.275 | -0.1718 | 0.146715 | 0.836044 | Fam120c |
| ENSMUSG( | 962.4525 | -0.07673 | 0.594381 | 0.945408 | Fam124a |
| ENSMUSG( | 6337.18 | -0.21889 | 0.180764 | 0.850372 | Fam131a |
| ENSMUSG( | 6776.449 | -0.08258 | 0.386425 | 0.917257 | Fam131b |
| ENSMUSG( | 50.63365 | -0.22894 | 0.452421 | 0.922316 | Fam131c |
| ENSMUSG( | 498.046 | -0.07662 | 0.576393 | 0.942912 | Fam133b |
| ENSMUSG( | 1009.439 | 0.087625 | 0.503375 | 0.927825 | Fam135a |
| ENSMUSG( | 935.8535 | 0.251995 | 0.029728 | 0.61992 | Fam135b |
| ENSMUSG( | 390.4473 | -0.16977 | 0.247496 | 0.877904 | Fam136a |
| ENSMUSG( | 323.8937 | 0.289158 | 0.06618 | 0.74792 | Fam13a |
| ENSMUSG( | 1236.424 | -0.09485 | 0.407678 | 0.917257 | Fam13b |
| ENSMUSG( | 2087.074 | 0.00212 | 0.984615 | 0.996963 | Fam13c |
| ENSMUSG( | 1630.536 | -0.12359 | 0.299725 | 0.898538 | Fam149a |
| ENSMUSG( | 1689.456 | -0.10565 | 0.346453 | 0.910966 | Fam149b |
| ENSMUSG( | 18.94936 | -0.91744 | 0.059726 | 0.729747 | Fam151a |
| ENSMUSG( | 157.7837 | -0.40373 | 0.045298 | 0.693959 | Fam151b |
| ENSMUSG( | 116.2158 | -0.0568 | 0.782563 | 0.976106 | Fam161a |
| ENSMUSG( | 300.6487 | 0.170132 | 0.297107 | 0.898538 | Fam161b |
| ENSMUSG( | 814.4747 | -0.1522 | 0.242896 | 0.877399 | Fam162a |
| ENSMUSG( | 958.2656 | -0.51651 | 0.000522 | 0.068787 | Fam163a |
| ENSMUSG( | 5115.612 | 0.01206 | 0.921671 | 0.992644 | Fam163b |
| ENSMUSG( | 134.0097 | 0.969094 | 1.31E-05 | 0.004603 | Fam167a |
| ENSMUSG( | 18.91414 | 0.1567 | 0.749562 | 0.96998 | Fam167b |
| ENSMUSG( | 9949.072 | -0.18081 | 0.063937 | 0.736482 | Fam168a |
| ENSMUSG( | 13178.74 | -0.1652 | 0.121853 | 0.813384 | Fam168b |
| ENSMUSG( | 888.981 | 0.039992 | 0.75023 | 0.970113 | Fam169a |
| ENSMUSG( | 9.815301 | 0.385341 | 0.571583 | NA | Fam169b |
| ENSMUSG( | 3037.703 | 0.230157 | 0.097188 | 0.796721 | Fam171a1 |
| ENSMUSG( | 3901.902 | 0.193895 | 0.176279 | 0.846547 | Fam171a2 |
| ENSMUSG( | 16798.57 | 0.155826 | 0.297822 | 0.898538 | Fam171b |
| ENSMUSG( | 1537.096 | -0.03705 | 0.794391 | 0.97681 | Fam174a |
| ENSMUSG( | 986.7683 | 0.20433 | 0.157048 | 0.837724 | Fam174b |
| ENSMUSG( | 90.58874 | 0.10395 | 0.677464 | 0.957106 | Fam174c |
| ENSMUSG( | 103.5374 | -0.32681 | 0.196672 | 0.862161 | Fam180a |
| ENSMUSG( | 68.54817 | -0.19461 | 0.47647 | 0.923329 | Fam181a |
| ENSMUSG( | 956.2635 | -0.01792 | 0.89817 | 0.990558 | Fam181b |
| ENSMUSG( | 822.4819 | 0.258243 | 0.046554 | 0.698795 | Fam184a |
| ENSMUSG( | 267.688 | -0.20487 | 0.21153 | 0.864412 | Fam184b |
| ENSMUSG( | 270.2568 | -0.22456 | 0.16092 | 0.837724 | Fam185a |
| ENSMUSG( | 9.891574 | -0.35454 | 0.597306 | NA | Fam186a |
| ENSMUSG( | 33.07291 | 0.200223 | 0.589856 | 0.944975 | Fam186b |
| ENSMUSG( | 1171.329 | -0.01775 | 0.875487 | 0.989644 | Fam193a |
| ENSMUSG( | 2936.032 | 0.16506 | 0.340472 | 0.907972 | Fam193b |
| ENSMUSG( | 284.3967 | 0.118097 | 0.548253 | 0.938063 | Fam199x |
| ENSMUSG( | 562.3128 | -0.09374 | 0.522103 | 0.932661 | Fam204a |
| ENSMUSG( | 1048.416 | 0.147667 | 0.415622 | 0.917257 | Fam20a |
| ENSMUSG( | 4037.258 | 0.088292 | 0.491152 | 0.926604 | Fam20b |
| ENSMUSG( | 1525.538 | 0.28455 | 0.086657 | 0.782073 | Fam20c |
| ENSMUSG( | 1066.266 | 0.061236 | 0.682002 | 0.958678 | Fam210a |
| ENSMUSG( | 996.9425 | -0.33824 | 0.008756 | 0.393969 | Fam210b |
| ENSMUSG( | 791.6378 | -0.0234 | 0.881272 | 0.989644 | Fam216a |
| ENSMUSG( | 31.20875 | 0.458349 | 0.376659 | 0.915491 | Fam216b |
| ENSMUSG( | 67.09862 | -0.32847 | 0.24073 | 0.87596 | Fam217a |
| ENSMUSG( | 1715.9 | -0.00019 | 0.998652 | 0.999406 | Fam217b |
| ENSMUSG( | 7562.901 | -0.15234 | 0.166283 | 0.840082 | Fam219a |
| ENSMUSG( | 19.11502 | 0.110046 | 0.81527 | 0.979198 | Fam219aos |
| ENSMUSG( | 676.0038 | 0.023383 | 0.853847 | 0.98563 | Fam219b |
| ENSMUSG( | 12.3888 | 0.074544 | 0.893059 | 0.990558 | Fam221a |
| ENSMUSG( | 23.06119 | -0.31587 | 0.475021 | 0.923329 | Fam221b |
| ENSMUSG( | 260.3628 | 0.048532 | 0.762992 | 0.973795 | Fam222a |
| ENSMUSG( | 667.7719 | -0.01754 | 0.89326 | 0.990558 | Fam222b |

| | | | | | | | | | | | |
|---|---|---|---|---|---|---|---|---|---|---|---|
| ENSMUSG( | Exosc5 | 597.0085 | 0.139681 | 0.252018 | 0.701908 | ENSMUSG( | Exosc5 | 498.9634 | -0.10995 | 0.433156 | 0.918434 |
| ENSMUSG( | Exosc7 | 337.2601 | 0.040544 | 0.79037 | 0.959461 | ENSMUSG( | Exosc7 | 287.9534 | -0.15793 | 0.378166 | 0.916962 |
| ENSMUSG( | Exosc8 | 326.8308 | 0.000798 | 0.994857 | 0.999369 | ENSMUSG( | Exosc8 | 296.0506 | -0.01605 | 0.911937 | 0.992196 |
| ENSMUSG( | Exosc9 | 707.8124 | 0.084015 | 0.491316 | 0.859873 | ENSMUSG( | Exosc9 | 593.0143 | -0.16748 | 0.266672 | 0.886134 |
| ENSMUSG( | Exph5 | 295.6341 | -0.02542 | 0.858214 | 0.974672 | ENSMUSG( | Exph5 | 260.7232 | -0.10806 | 0.571907 | 0.942347 |
| ENSMUSG( | Ext1 | 1388.837 | -0.32217 | 0.030176 | 0.263854 | ENSMUSG( | Ext1 | 1487.685 | 0.163535 | 0.304768 | 0.898538 |
| ENSMUSG( | Ext2 | 1286.616 | -0.30994 | 0.047183 | 0.33245 | ENSMUSG( | Ext2 | 1433.844 | 0.281123 | 0.122637 | 0.814249 |
| ENSMUSG( | Extl1 | 2504.055 | 0.084602 | 0.52886 | 0.878365 | ENSMUSG( | Extl1 | 2433.936 | 0.265753 | 0.111558 | 0.808371 |
| ENSMUSG( | Extl2 | 2569.285 | -0.31951 | 0.023345 | 0.233882 | ENSMUSG( | Extl2 | 2784.594 | 0.196743 | 0.244999 | 0.877904 |
| ENSMUSG( | Extl3 | 7002.927 | -0.08928 | 0.480982 | 0.855135 | ENSMUSG( | Extl3 | 6811.575 | 0.102633 | 0.500696 | 0.927825 |
| ENSMUSG( | Eya1 | 760.2782 | 0.455595 | 9.07E-05 | 0.006031 | ENSMUSG( | Eya1 | 661.7369 | 0.337611 | 0.012201 | 0.44712 |
| ENSMUSG( | Eya2 | 163.7026 | 0.337961 | 0.081766 | 0.441786 | ENSMUSG( | Eya2 | 135.3205 | 0.068064 | 0.758327 | 0.971858 |
| ENSMUSG( | Eya3 | 990.8101 | 0.105873 | 0.327958 | 0.772842 | ENSMUSG( | Eya3 | 868.9223 | 0.00057 | 0.996462 | 0.999178 |
| ENSMUSG( | Eya4 | 197.9435 | -0.03286 | 0.826089 | 0.968159 | ENSMUSG( | Eya4 | 191.0829 | 0.129579 | 0.441152 | 0.921166 |
| ENSMUSG( | Ezh1 | 1428.928 | 0.163969 | 0.159962 | 0.589679 | ENSMUSG( | Ezh1 | 1274.146 | 0.108365 | 0.428431 | 0.917257 |
| ENSMUSG( | Ezh2 | 286.6818 | 0.169455 | 0.213478 | 0.65774 | ENSMUSG( | Ezh2 | 272.0161 | 0.288169 | 0.079094 | 0.77458 |
| ENSMUSG( | Ezr | 12499.07 | 0.110999 | 0.301297 | 0.749979 | ENSMUSG( | Ezr | 10563.11 | -0.10751 | 0.419127 | 0.917257 |
| ENSMUSG( | F11r | 509.8063 | -0.20563 | 0.181867 | 0.618944 | ENSMUSG( | F11r | 578.1234 | 0.413946 | 0.032989 | 0.63923 |
| ENSMUSG( | F12 | 16.47223 | -0.37868 | 0.431327 | NA | ENSMUSG( | F12 | 16.10486 | -0.15738 | 0.750711 | 0.970209 |
| ENSMUSG( | F13a1 | 198.6716 | 0.452484 | 0.009195 | 0.135112 | ENSMUSG( | F13a1 | 169.4897 | 0.272125 | 0.175602 | 0.845305 |
| ENSMUSG( | F2r | 251.377 | -0.38554 | 0.043538 | 0.31954 | ENSMUSG( | F2r | 292.5938 | 0.338311 | 0.122608 | 0.814249 |
| ENSMUSG( | F2rl2 | 38.1706 | -0.30855 | 0.419608 | 0.822309 | ENSMUSG( | F2rl2 | 45.33656 | 0.458825 | 0.223338 | 0.867404 |
| ENSMUSG( | F3 | 2596.042 | -0.54611 | 0.000173 | 0.00947 | ENSMUSG( | F3 | 2710.276 | -0.09818 | 0.614188 | 0.951177 |
| ENSMUSG( | F420014N2 | 16.11061 | 0.133927 | 0.784507 | NA | ENSMUSG( | F420014N2 | 17.01016 | 0.526041 | 0.274072 | 0.889874 |
| ENSMUSG( | F5 | 118.9563 | 0.076864 | 0.904711 | 0.986316 | ENSMUSG( | F5 | 145.6714 | 0.808956 | 0.013536 | 0.464932 |
| ENSMUSG( | F630040K0 | 128.5699 | 0.156236 | 0.415978 | 0.82084 | ENSMUSG( | F630040K0 | 111.1058 | -0.01682 | 0.9453 | 0.994798 |
| ENSMUSG( | F730043M | 755.8668 | 0.078709 | 0.452124 | 0.836145 | ENSMUSG( | F730043M | 644.6988 | -0.11705 | 0.417663 | 0.917257 |
| ENSMUSG( | F8 | 63.32001 | -0.50401 | 0.04715 | 0.33245 | ENSMUSG( | F8 | 59.12262 | -0.40574 | 0.135276 | 0.825081 |
| ENSMUSG( | F8a | 310.6168 | -0.0988 | 0.573735 | 0.893584 | ENSMUSG( | F8a | 271.4463 | -0.23281 | 0.20489 | 0.864412 |
| ENSMUSG( | F9 | 14.49749 | -0.0593 | 0.902977 | NA | ENSMUSG( | F9 | 15.04254 | 0.313302 | 0.545119 | 0.938063 |
| ENSMUSG( | Fa2h | 2099.456 | -0.71851 | 4.34E-05 | 0.003572 | ENSMUSG( | Fa2h | 2558.294 | 0.207964 | 0.322322 | 0.904023 |
| ENSMUSG( | Faah | 4390.335 | -0.21686 | 0.068634 | 0.405903 | ENSMUSG( | Faah | 4268.896 | -0.01841 | 0.887079 | 0.990263 |
| ENSMUSG( | Faap100 | 540.1447 | 0.245331 | 0.067012 | 0.400593 | ENSMUSG( | Faap100 | 454.7202 | 0.03419 | 0.836039 | 0.981237 |
| ENSMUSG( | Faap20 | 681.0862 | 0.065752 | 0.547358 | 0.885128 | ENSMUSG( | Faap20 | 585.3824 | -0.11004 | 0.454461 | 0.922444 |
| ENSMUSG( | Faap24 | 133.6858 | 0.055915 | 0.781229 | 0.957558 | ENSMUSG( | Faap24 | 113.111 | -0.16038 | 0.48754 | 0.925427 |
| ENSMUSG( | Fabp3 | 992.1882 | -0.05261 | 0.740332 | 0.947823 | ENSMUSG( | Fabp3 | 869.1724 | -0.17586 | 0.334221 | 0.907907 |
| ENSMUSG( | Fabp5 | 5491.776 | 0.119248 | 0.284403 | 0.732908 | ENSMUSG( | Fabp5 | 4603.761 | -0.12407 | 0.350957 | 0.911707 |
| ENSMUSG( | Fabp7 | 3507.824 | 0.209024 | 0.210274 | 0.652595 | ENSMUSG( | Fabp7 | 2545.857 | -0.48582 | 0.019075 | 0.549166 |
| ENSMUSG( | Fadd | 127.8795 | 0.276955 | 0.123447 | 0.530552 | ENSMUSG( | Fadd | 109.7742 | 0.104153 | 0.626676 | 0.953234 |
| ENSMUSG( | Fads1 | 41028.89 | 0.041815 | 0.685243 | 0.932981 | ENSMUSG( | Fads1 | 35232.63 | -0.13204 | 0.268681 | 0.886755 |
| ENSMUSG( | Fads2 | 29564.86 | 0.120728 | 0.283808 | 0.732203 | ENSMUSG( | Fads2 | 25221.29 | -0.06805 | 0.577902 | 0.943131 |
| ENSMUSG( | Fads3 | 1252.975 | -0.26215 | 0.054554 | 0.35941 | ENSMUSG( | Fads3 | 1294.087 | 0.114047 | 0.462818 | 0.923329 |
| ENSMUSG( | Fads6 | 1145.94 | -0.19236 | 0.176894 | 0.613937 | ENSMUSG( | Fads6 | 1133.124 | 0.055206 | 0.735081 | 0.967344 |
| ENSMUSG( | Faf1 | 958.6222 | -0.087 | 0.360285 | 0.791947 | ENSMUSG( | Faf1 | 892.6898 | -0.02272 | 0.846985 | 0.985027 |
| ENSMUSG( | Faf2 | 1910.783 | -0.26315 | 0.034878 | 0.287263 | ENSMUSG( | Faf2 | 1952.791 | 0.08337 | 0.567327 | 0.941777 |
| ENSMUSG( | Fah | 621.8611 | 0.058007 | 0.636242 | 0.914873 | ENSMUSG( | Fah | 524.4636 | -0.16852 | 0.259217 | 0.883971 |
| ENSMUSG( | Fahd1 | 336.0609 | -0.10831 | 0.45137 | 0.835567 | ENSMUSG( | Fahd1 | 290.9205 | -0.2716 | 0.08377 | 0.77681 |
| ENSMUSG( | Fahd2a | 494.0826 | -0.06918 | 0.583598 | 0.89861 | ENSMUSG( | Fahd2a | 427.2129 | -0.23464 | 0.157806 | 0.837724 |
| ENSMUSG( | Faim | 702.8847 | -0.02851 | 0.780554 | 0.957491 | ENSMUSG( | Faim | 621.4761 | -0.12281 | 0.34667 | 0.91112 |
| ENSMUSG( | Faim2 | 16462.67 | -0.23969 | 0.060034 | 0.378891 | ENSMUSG( | Faim2 | 16629.36 | 0.070596 | 0.633192 | 0.953844 |
| ENSMUSG( | Fam107a | 90232.22 | 0.396138 | 8.54E-06 | 0.001033 | ENSMUSG( | Fam107a | 72338.35 | 0.053085 | 0.669347 | 0.956766 |
| ENSMUSG( | Fam107b | 784.779 | 0.050701 | 0.707524 | 0.93949 | ENSMUSG( | Fam107b | 672.6244 | -0.13583 | 0.425627 | 0.917257 |
| ENSMUSG( | Fam110a | 130.0487 | 0.010717 | 0.955258 | 0.993874 | ENSMUSG( | Fam110a | 124.0079 | 0.153191 | 0.460288 | 0.923329 |
| ENSMUSG( | Fam110b | 615.581 | 0.242982 | 0.148863 | 0.572605 | ENSMUSG( | Fam110b | 537.7398 | 0.126838 | 0.497596 | 0.927825 |
| ENSMUSG( | Fam110d | 117.7984 | 0.427402 | 0.02486 | 0.242612 | ENSMUSG( | Fam110d | 115.803 | 0.631336 | 0.008382 | 0.388416 |
| ENSMUSG( | Fam111a | 88.86241 | 0.144251 | 0.5337 | 0.880095 | ENSMUSG( | Fam111a | 84.82567 | 0.26661 | 0.280513 | 0.892356 |
| ENSMUSG( | Fam114a1 | 383.7418 | 0.096272 | 0.459891 | 0.840338 | ENSMUSG( | Fam114a1 | 327.01 | -0.10428 | 0.503468 | 0.927825 |
| ENSMUSG( | Fam114a2 | 1032.399 | 0.114134 | 0.257961 | 0.709319 | ENSMUSG( | Fam114a2 | 927.8665 | 0.073548 | 0.57078 | 0.941973 |
| ENSMUSG( | Fam117a | 203.4392 | -0.19013 | 0.264431 | 0.715323 | ENSMUSG( | Fam117a | 195.4591 | -0.03362 | 0.853986 | 0.98563 |
| ENSMUSG( | Fam117b | 1178.932 | 0.035162 | 0.71382 | 0.940903 | ENSMUSG( | Fam117b | 1067.127 | 0.015553 | 0.899689 | 0.990558 |
| ENSMUSG( | Fam118a | 369.1909 | -0.01953 | 0.883058 | 0.982422 | ENSMUSG( | Fam118a | 348.3729 | 0.072696 | 0.65589 | 0.955533 |
| ENSMUSG( | Fam118b | 431.4877 | 0.227497 | 0.069876 | 0.408574 | ENSMUSG( | Fam118b | 383.0747 | 0.151995 | 0.305734 | 0.898538 |
| ENSMUSG( | Fam120a | 6656.032 | 0.048581 | 0.637268 | 0.915436 | ENSMUSG( | Fam120a | 5821.168 | -0.07135 | 0.543078 | 0.938063 |
| ENSMUSG( | Fam120aos | 1036.083 | -0.04494 | 0.722047 | 0.943535 | ENSMUSG( | Fam120aos | 913.0247 | -0.15238 | 0.378207 | 0.916962 |
| ENSMUSG( | Fam120b | 1352.863 | 0.073417 | 0.459434 | 0.840328 | ENSMUSG( | Fam120b | 1261.062 | 0.139978 | 0.257743 | 0.883401 |
| ENSMUSG( | Fam120c | 1887.773 | 0.055377 | 0.671508 | 0.928781 | ENSMUSG( | Fam120c | 1592.275 | -0.1718 | 0.146715 | 0.836044 |
| ENSMUSG( | Fam124a | 1106.388 | 0.061548 | 0.559969 | 0.888917 | ENSMUSG( | Fam124a | 962.4525 | -0.07673 | 0.594381 | 0.945408 |
| ENSMUSG( | Fam131a | 7620.748 | 0.055044 | 0.723431 | 0.943859 | ENSMUSG( | Fam131a | 6337.18 | -0.21889 | 0.180764 | 0.850372 |
| ENSMUSG( | Fam131b | 7767.327 | 0.04329 | 0.574854 | 0.894572 | ENSMUSG( | Fam131b | 6776.449 | -0.08258 | 0.386425 | 0.917257 |
| ENSMUSG( | Fam131c | 60.07442 | 0.027992 | 0.912619 | 0.988483 | ENSMUSG( | Fam131c | 50.63365 | -0.22894 | 0.452421 | 0.922316 |
| ENSMUSG( | Fam133b | 549.8681 | -0.05578 | 0.625516 | 0.912249 | ENSMUSG( | Fam133b | 498.046 | -0.07662 | 0.576393 | 0.942912 |
| ENSMUSG( | Fam135a | 1018.98 | -0.15938 | 0.186357 | 0.623869 | ENSMUSG( | Fam135a | 1009.439 | 0.087625 | 0.503375 | 0.927825 |
| ENSMUSG( | Fam135b | 933.3289 | -0.01524 | 0.887844 | 0.983434 | ENSMUSG( | Fam135b | 935.8535 | 0.251995 | 0.029728 | 0.61992 |
| ENSMUSG( | Fam136a | 450.6913 | -0.01758 | 0.89078 | 0.983764 | ENSMUSG( | Fam136a | 390.4473 | -0.16977 | 0.247496 | 0.877904 |
| ENSMUSG( | Fam13a | 355.0875 | 0.278395 | 0.03489 | 0.287263 | ENSMUSG( | Fam13a | 323.8937 | 0.289158 | 0.06618 | 0.74792 |
| ENSMUSG( | Fam13b | 1457.416 | 0.109335 | 0.274354 | 0.724547 | ENSMUSG( | Fam13b | 1236.424 | -0.09485 | 0.407678 | 0.917257 |
| ENSMUSG( | Fam13c | 2231.804 | -0.07364 | 0.460582 | 0.840787 | ENSMUSG( | Fam13c | 2087.074 | 0.00212 | 0.984615 | 0.996963 |
| ENSMUSG( | Fam149a | 1917.725 | 0.078503 | 0.435962 | 0.829093 | ENSMUSG( | Fam149a | 1630.536 | -0.12359 | 0.299725 | 0.898538 |
| ENSMUSG( | Fam149b | 1972.476 | 0.075599 | 0.422155 | 0.823495 | ENSMUSG( | Fam149b | 1689.456 | -0.10565 | 0.346453 | 0.910966 |
| ENSMUSG( | Fam151a | 27.24997 | -0.03606 | 0.922193 | 0.99045 | ENSMUSG( | Fam151a | 18.94936 | -0.91744 | 0.059726 | 0.729747 |
| ENSMUSG( | Fam151b | 185.5891 | -0.16361 | 0.30901 | 0.757121 | ENSMUSG( | Fam151b | 157.7837 | -0.40373 | 0.045298 | 0.693959 |
| ENSMUSG( | Fam161a | 124.642 | -0.12203 | 0.489153 | 0.859092 | ENSMUSG( | Fam161a | 116.2158 | -0.0568 | 0.782563 | 0.976106 |
| ENSMUSG( | Fam161b | 316.2418 | 0.044595 | 0.739684 | 0.947823 | ENSMUSG( | Fam161b | 300.6487 | 0.170132 | 0.297107 | 0.898538 |
| ENSMUSG( | Fam162a | 944.1851 | 0.014101 | 0.88816 | 0.983434 | ENSMUSG( | Fam162a | 814.4747 | -0.1522 | 0.242896 | 0.877399 |
| ENSMUSG( | Fam163a | 1253.612 | 0.038677 | 0.77545 | 0.955606 | ENSMUSG( | Fam163a | 958.2656 | -0.51651 | 0.000522 | 0.068787 |
| ENSMUSG( | Fam163b | 5401.315 | -0.10296 | 0.306016 | 0.753712 | ENSMUSG( | Fam163b | 5115.612 | 0.01206 | 0.921671 | 0.992644 |
| ENSMUSG( | Fam167a | 118.8852 | 0.475387 | 0.029207 | 0.259574 | ENSMUSG( | Fam167a | 134.0097 | 0.969094 | 1.31E-05 | 0.004603 |
| ENSMUSG( | Fam167b | 34.7535 | 1.385002 | 5.76E-05 | 0.004407 | ENSMUSG( | Fam167b | 18.91414 | 0.1567 | 0.749562 | 0.96998 |
| ENSMUSG( | Fam168a | 11627.15 | 0.006048 | 0.944022 | 0.993503 | ENSMUSG( | Fam168a | 9949.072 | -0.18081 | 0.063937 | 0.736482 |
| ENSMUSG( | Fam168b | 15766.66 | 0.087437 | 0.307523 | 0.755462 | ENSMUSG( | Fam168b | 13178.74 | -0.1652 | 0.121853 | 0.813384 |
| ENSMUSG( | Fam169a | 902.4217 | -0.18936 | 0.062374 | 0.386294 | ENSMUSG( | Fam169a | 888.981 | 0.039992 | 0.75023 | 0.970113 |
| ENSMUSG( | Fam169b | 9.204231 | 0.007554 | 0.990704 | NA | ENSMUSG( | Fam169b | 9.815301 | 0.385341 | 0.571583 | NA |
| ENSMUSG( | Fam171a1 | 2876.997 | -0.19623 | 0.112805 | 0.506698 | ENSMUSG( | Fam171a1 | 3037.703 | 0.230157 | 0.097188 | 0.796721 |
| ENSMUSG( | Fam171a2 | 3757.813 | -0.18612 | 0.159227 | 0.588305 | ENSMUSG( | Fam171a2 | 3901.902 | 0.193895 | 0.176279 | 0.846547 |
| ENSMUSG( | Fam171b | 17264.83 | -0.03187 | 0.818768 | 0.965888 | ENSMUSG( | Fam171b | 16798.57 | 0.155826 | 0.297822 | 0.898538 |
| ENSMUSG( | Fam174a | 1589.162 | -0.21699 | 0.070771 | 0.410404 | ENSMUSG( | Fam174a | 1537.096 | -0.03705 | 0.794391 | 0.97681 |
| ENSMUSG( | Fam174b | 962.0256 | -0.13729 | 0.299597 | 0.748647 | ENSMUSG( | Fam174b | 986.7683 | 0.20433 | 0.157048 | 0.837724 |
| ENSMUSG( | Fam174c | 99.17152 | 0.083376 | 0.704826 | 0.938303 | ENSMUSG( | Fam174c | 90.58874 | 0.10395 | 0.677464 | 0.957106 |
| ENSMUSG( | Fam180a | 117.1552 | -0.23864 | 0.328025 | 0.772854 | ENSMUSG( | Fam180a | 103.5374 | -0.32681 | 0.196672 | 0.862161 |
| ENSMUSG( | Fam181a | 68.65734 | -0.51272 | 0.03692 | 0.296508 | ENSMUSG( | Fam181a | 68.54817 | -0.19461 | 0.47647 | 0.923329 |
| ENSMUSG( | Fam181b | 1105.512 | 0.134109 | 0.191836 | 0.630811 | ENSMUSG( | Fam181b | 956.2635 | -0.01792 | 0.89817 | 0.990558 |
| ENSMUSG( | Fam184a | 874.3291 | 0.174944 | 0.103132 | 0.48855 | ENSMUSG( | Fam184a | 822.4819 | 0.258243 | 0.046554 | 0.698795 |
| ENSMUSG( | Fam184b | 284.665 | -0.29797 | 0.03757 | 0.299456 | ENSMUSG( | Fam184b | 267.688 | -0.20487 | 0.21153 | 0.864412 |
| ENSMUSG( | Fam185a | 331.3661 | 0.098568 | 0.479519 | 0.854046 | ENSMUSG( | Fam185a | 270.2568 | -0.22456 | 0.16092 | 0.837724 |
| ENSMUSG( | Fam186a | 11.22421 | -0.20901 | 0.719283 | NA | ENSMUSG( | Fam186a | 9.891574 | -0.35454 | 0.597306 | NA |
| ENSMUSG( | Fam186b | 32.28512 | -0.14966 | 0.664902 | 0.926824 | ENSMUSG( | Fam186b | 33.07291 | 0.200223 | 0.589856 | 0.944975 |
| ENSMUSG( | Fam193a | 1306.771 | 0.029775 | 0.749499 | 0.949664 | ENSMUSG( | Fam193a | 1171.329 | -0.01775 | 0.875487 | 0.989644 |
| ENSMUSG( | Fam193b | 3343.442 | 0.265775 | 0.075062 | 0.42422 | ENSMUSG( | Fam193b | 2936.032 | 0.16506 | 0.340472 | 0.907972 |
| ENSMUSG( | Fam199x | 292.6608 | -0.05986 | 0.701717 | 0.937441 | ENSMUSG( | Fam199x | 284.3967 | 0.118097 | 0.548253 | 0.938063 |
| ENSMUSG( | Fam204a | 627.5895 | -0.03716 | 0.759507 | 0.952498 | ENSMUSG( | Fam204a | 562.3128 | -0.09374 | 0.522103 | 0.932661 |
| ENSMUSG( | Fam20a | 1076.849 | -0.04775 | 0.766288 | 0.954173 | ENSMUSG( | Fam20a | 1048.416 | 0.147667 | 0.415622 | 0.917257 |
| ENSMUSG( | Fam20b | 4228.919 | -0.04771 | 0.669711 | 0.928147 | ENSMUSG( | Fam20b | 4037.258 | 0.088292 | 0.491152 | 0.926604 |
| ENSMUSG( | Fam20c | 1431.966 | -0.16292 | 0.233654 | 0.681508 | ENSMUSG( | Fam20c | 1525.538 | 0.28455 | 0.086657 | 0.782073 |
| ENSMUSG( | Fam210a | 1184.577 | 0.101036 | 0.349215 | 0.78176 | ENSMUSG( | Fam210a | 1066.266 | 0.061236 | 0.682002 | 0.958678 |
| ENSMUSG( | Fam210b | 1200.661 | -0.04808 | 0.648904 | 0.920025 | ENSMUSG( | Fam210b | 996.9425 | -0.33824 | 0.008756 | 0.393969 |
| ENSMUSG( | Fam216a | 886.1685 | 0.036505 | 0.793993 | 0.959799 | ENSMUSG( | Fam216a | 791.6378 | -0.0234 | 0.881272 | 0.989644 |
| ENSMUSG( | Fam216b | 26.12061 | -0.34535 | 0.347019 | 0.7797 | ENSMUSG( | Fam216b | 31.20875 | 0.458349 | 0.376659 | 0.915491 |
| ENSMUSG( | Fam217a | 77.52869 | -0.1395 | 0.552937 | 0.886204 | ENSMUSG( | Fam217a | 67.09862 | -0.32847 | 0.24073 | 0.87596 |
| ENSMUSG( | Fam217b | 1942.92 | 0.090549 | 0.306144 | 0.753712 | ENSMUSG( | Fam217b | 1715.9 | -0.00019 | 0.998652 | 0.999406 |
| ENSMUSG( | Fam219a | 8819.956 | 0.027583 | 0.740428 | 0.947823 | ENSMUSG( | Fam219a | 7562.901 | -0.15234 | 0.166283 | 0.840082 |
| ENSMUSG( | Fam219aos | 22.8541 | 0.338624 | 0.404594 | 0.816132 | ENSMUSG( | Fam219aos | 19.11502 | 0.110046 | 0.81527 | 0.979198 |
| ENSMUSG( | Fam219b | 754.9324 | 0.074312 | 0.481906 | 0.855573 | ENSMUSG( | Fam219b | 676.0038 | 0.023383 | 0.853847 | 0.98563 |
| ENSMUSG( | Fam221a | 12.32884 | -0.24223 | 0.6442 | NA | ENSMUSG( | Fam221a | 12.3888 | 0.074544 | 0.893059 | 0.990558 |
| ENSMUSG( | Fam221b | 25.57888 | -0.3129 | 0.414823 | 0.820184 | ENSMUSG( | Fam221b | 23.06119 | -0.31587 | 0.475021 | 0.923329 |
| ENSMUSG( | Fam222a | 249.1494 | -0.36169 | 0.013781 | 0.171544 | ENSMUSG( | Fam222a | 260.3628 | 0.048532 | 0.762992 | 0.973795 |
| ENSMUSG( | Fam222b | 757.618 | 0.0809 | 0.435551 | 0.829093 | ENSMUSG( | Fam222b | 667.7719 | -0.01754 | 0.89326 | 0.990558 |
| ENSMUSG( | Fam227a | 236.0191 | 0.13997 | 0.377063 | 0.801213 | ENSMUSG( | Fam227a | 218.2869 | 0.191116 | 0.294889 | 0.898538 |
| ENSMUSG( | Fam227b | 24.16641 | 0.114405 | 0.769315 | 0.954674 | ENSMUSG( | Fam227b | 20.25865 | -0.17774 | 0.701123 | 0.961756 |
| ENSMUSG( | Fam228a | 81.85626 | 0.251467 | 0.346142 | 0.779153 | ENSMUSG( | Fam228a | 68.5619 | 0.04023 | 0.891015 | 0.990558 |
| ENSMUSG( | Fam228b | 177.458 | -0.30109 | 0.06042 | 0.3797 | ENSMUSG( | Fam228b | 179.7277 | 0.033072 | 0.883214 | 0.989644 |
| ENSMUSG( | Fam229b | 291.8301 | 0.011514 | 0.932661 | 0.992701 | ENSMUSG( | Fam229b | 267.5007 | 0.027403 | 0.876053 | 0.989644 |
| ENSMUSG( | Fam234a | 911.7961 | -0.34496 | 0.017925 | 0.200989 | ENSMUSG( | Fam234a | 933.5532 | 0.017562 | 0.918193 | 0.992529 |
| ENSMUSG( | Fam234b | 10708.58 | -0.14759 | 0.18126 | 0.618305 | ENSMUSG( | Fam234b | 10471.5 | 0.061748 | 0.631886 | 0.953251 |
| ENSMUSG( | Fam241a | 116.6273 | -0.50928 | 0.011363 | 0.153509 | ENSMUSG( | Fam241a | 119.4058 | -0.13348 | 0.517985 | 0.930939 |
| ENSMUSG( | Fam241b | 454.3803 | -0.04409 | 0.725586 | 0.944223 | ENSMUSG( | Fam241b | 407.7115 | -0.0955 | 0.521991 | 0.932571 |
| ENSMUSG( | Fam32a | 2731.265 | -0.01181 | 0.899096 | 0.985633 | ENSMUSG( | Fam32a | 2347.097 | -0.18708 | 0.079431 | 0.77458 |
| ENSMUSG( | Fam3a | 710.1896 | -0.25474 | 0.046229 | 0.329472 | ENSMUSG( | Fam3a | 736.4887 | 0.131847 | 0.355118 | 0.912411 |
| ENSMUSG( | Fam3c | 2517.24 | -0.29586 | 0.015315 | 0.18239 | ENSMUSG( | Fam3c | 2589.134 | 0.068894 | 0.627898 | 0.953234 |
| ENSMUSG( | Fam43a | 423.3451 | 0.025906 | 0.862956 | 0.975055 | ENSMUSG( | Fam43a | 367.1996 | -0.11622 | 0.466424 | 0.923329 |
| ENSMUSG( | Fam43b | 502.6787 | 0.036895 | 0.811806 | 0.964426 | ENSMUSG( | Fam43b | 410.2644 | -0.30252 | 0.090483 | 0.791074 |
| ENSMUSG( | Fam47e | 13.08227 | -0.27414 | 0.608912 | NA | ENSMUSG( | Fam47e | 14.45923 | 0.299233 | 0.60762 | 0.950113 |
| ENSMUSG( | Fam50a | 670.199 | 0.181031 | 0.16775 | 0.60061 | ENSMUSG( | Fam50a | 591.0515 | 0.087271 | 0.582424 | 0.944266 |
| ENSMUSG( | Fam53a | 794.5031 | 0.036266 | 0.742904 | 0.948478 | ENSMUSG( | Fam53a | 786.5377 | 0.264593 | 0.025591 | 0.599234 |
| ENSMUSG( | Fam53b | 1130.574 | -0.05289 | 0.617334 | 0.910866 | ENSMUSG( | Fam53b | 1045.868 | -0.00813 | 0.94314 | 0.994798 |
| ENSMUSG( | Fam53c | 2919.771 | 0.136128 | 0.108827 | 0.497931 | ENSMUSG( | Fam53c | 2623.367 | 0.098121 | 0.339498 | 0.907972 |
| ENSMUSG( | Fam72a | 42.48039 | -0.61413 | 0.035379 | 0.289531 | ENSMUSG( | Fam72a | 37.92484 | -0.6698 | 0.05204 | 0.712974 |
| ENSMUSG( | Fam76a | 1337.234 | -0.04426 | 0.660187 | 0.924627 | ENSMUSG( | Fam76a | 1299.149 | 0.136454 | 0.256668 | 0.883082 |
| ENSMUSG( | Fam76b | 383.6749 | -0.23049 | 0.060513 | 0.379845 | ENSMUSG( | Fam76b | 418.1385 | 0.286295 | 0.050014 | 0.708648 |
| ENSMUSG( | Fam78a | 148.812 | -0.24007 | 0.202186 | 0.642173 | ENSMUSG( | Fam78a | 131.8151 | -0.31764 | 0.13537 | 0.825081 |
| ENSMUSG( | Fam78b | 994.7625 | 0.04808 | 0.636312 | 0.914873 | ENSMUSG( | Fam78b | 879.9112 | -0.04232 | 0.74737 | 0.969763 |
| ENSMUSG( | Fam81a | 10907.26 | 0.171753 | 0.030307 | 0.264512 | ENSMUSG( | Fam81a | 9273.396 | -0.0236 | 0.83019 | 0.980224 |
| ENSMUSG( | Fam83d | 71.9267 | -0.04089 | 0.884672 | 0.982647 | ENSMUSG( | Fam83d | 73.33689 | 0.299821 | 0.349132 | 0.911652 |
| ENSMUSG( | Fam83f | 15.00395 | -0.26154 | 0.601269 | NA | ENSMUSG( | Fam83f | 11.78648 | -0.72793 | 0.211371 | 0.864412 |
| ENSMUSG( | Fam83h | 108.1525 | -0.37077 | 0.055497 | 0.36248 | ENSMUSG( | Fam83h | 99.58396 | -0.32076 | 0.147142 | 0.836241 |
| ENSMUSG( | Fam89a | 33.88696 | -0.24135 | 0.473446 | 0.848919 | ENSMUSG( | Fam89a | 33.73669 | -0.00428 | 0.990764 | 0.998287 |
| ENSMUSG( | Fam8a1 | 2683.85 | -0.32801 | 0.009797 | 0.141256 | ENSMUSG( | Fam8a1 | 2803.808 | 0.085805 | 0.577257 | 0.942992 |
| ENSMUSG( | Fam91a1 | 2142.073 | 0.094238 | 0.296936 | 0.745676 | ENSMUSG( | Fam91a1 | 1927.615 | 0.058643 | 0.610592 | 0.950771 |
| ENSMUSG( | Fam98a | 1087.358 | 0.135608 | 0.195657 | 0.635109 | ENSMUSG( | Fam98a | 955.5807 | 0.030891 | 0.812128 | 0.979148 |

| | | | | | | | | | | | |
|---|---|---|---|---|---|---|---|---|---|---|---|
| ENSMUSG( | 77.52869 | -0.1395 | 0.552937 | 0.886204 | Fam217a | ENSMUSG( | 218.2869 | 0.191116 | 0.294889 | 0.898538 | Fam227a |
| ENSMUSG( | 1942.92 | 0.090549 | 0.306144 | 0.753712 | Fam217b | ENSMUSG( | 20.25865 | -0.17774 | 0.701123 | 0.961756 | Fam227b |
| ENSMUSG( | 8819.956 | 0.027583 | 0.740428 | 0.947823 | Fam219a | ENSMUSG( | 68.5619 | 0.04023 | 0.891015 | 0.990558 | Fam228a |
| ENSMUSG( | 22.8541 | 0.338624 | 0.404594 | 0.816132 | Fam219aos | ENSMUSG( | 179.7277 | 0.033072 | 0.883214 | 0.989644 | Fam228b |
| ENSMUSG( | 754.9324 | 0.074312 | 0.481906 | 0.855573 | Fam219b | ENSMUSG( | 267.5007 | 0.027403 | 0.876053 | 0.989644 | Fam229b |
| ENSMUSG( | 12.32884 | -0.24223 | 0.6442 | NA | Fam221a | ENSMUSG( | 933.5532 | 0.017562 | 0.918193 | 0.992529 | Fam234a |
| ENSMUSG( | 25.57888 | -0.3129 | 0.414823 | 0.820184 | Fam221b | ENSMUSG( | 10471.5 | 0.061748 | 0.631886 | 0.953251 | Fam234b |
| ENSMUSG( | 249.1494 | -0.36169 | 0.013781 | 0.171544 | Fam222a | ENSMUSG( | 119.4058 | -0.13348 | 0.517985 | 0.930939 | Fam241a |
| ENSMUSG( | 757.618 | 0.0809 | 0.435551 | 0.829093 | Fam222b | ENSMUSG( | 407.7115 | -0.0955 | 0.521991 | 0.932571 | Fam241b |
| ENSMUSG( | 236.0191 | 0.13997 | 0.377063 | 0.801213 | Fam227a | ENSMUSG( | 2347.097 | -0.18708 | 0.079431 | 0.77458 | Fam32a |
| ENSMUSG( | 24.16641 | 0.114405 | 0.769315 | 0.954674 | Fam227b | ENSMUSG( | 736.4887 | 0.131847 | 0.355118 | 0.912411 | Fam3a |
| ENSMUSG( | 81.85626 | 0.251467 | 0.346142 | 0.779153 | Fam228a | ENSMUSG( | 2589.134 | 0.068894 | 0.627898 | 0.953234 | Fam3c |
| ENSMUSG( | 177.458 | -0.30109 | 0.06042 | 0.3797 | Fam228b | ENSMUSG( | 367.1996 | -0.11622 | 0.466424 | 0.923329 | Fam43a |
| ENSMUSG( | 291.8301 | 0.011514 | 0.932661 | 0.992701 | Fam229b | ENSMUSG( | 410.2644 | -0.30252 | 0.090483 | 0.791074 | Fam43b |
| ENSMUSG( | 911.7961 | -0.34496 | 0.017925 | 0.200989 | Fam234a | ENSMUSG( | 14.45923 | 0.299233 | 0.60762 | 0.950113 | Fam47e |
| ENSMUSG( | 10708.58 | -0.14759 | 0.18126 | 0.618305 | Fam234b | ENSMUSG( | 591.0515 | 0.087271 | 0.582424 | 0.944266 | Fam50a |
| ENSMUSG( | 116.6273 | -0.50928 | 0.011363 | 0.153509 | Fam241a | ENSMUSG( | 786.5377 | 0.264593 | 0.025591 | 0.599234 | Fam53a |
| ENSMUSG( | 454.3803 | -0.04409 | 0.725586 | 0.944223 | Fam241b | ENSMUSG( | 1045.868 | -0.00813 | 0.94314 | 0.994798 | Fam53b |
| ENSMUSG( | 2731.265 | -0.01181 | 0.899096 | 0.985633 | Fam32a | ENSMUSG( | 2623.367 | 0.098121 | 0.339498 | 0.907972 | Fam53c |
| ENSMUSG( | 710.1896 | -0.25474 | 0.046229 | 0.329472 | Fam3a | ENSMUSG( | 37.92484 | -0.6698 | 0.05204 | 0.712974 | Fam72a |
| ENSMUSG( | 2517.24 | -0.29586 | 0.015315 | 0.18239 | Fam3c | ENSMUSG( | 1299.149 | 0.136454 | 0.256668 | 0.883082 | Fam76a |
| ENSMUSG( | 423.3451 | 0.025906 | 0.862956 | 0.975055 | Fam43a | ENSMUSG( | 418.1385 | 0.286295 | 0.050014 | 0.708648 | Fam76b |
| ENSMUSG( | 502.6787 | 0.036895 | 0.811806 | 0.964426 | Fam43b | ENSMUSG( | 131.8151 | -0.31764 | 0.13537 | 0.825081 | Fam78a |
| ENSMUSG( | 13.08227 | -0.27414 | 0.608912 | NA | Fam47e | ENSMUSG( | 879.9112 | -0.04232 | 0.74737 | 0.969763 | Fam78b |
| ENSMUSG( | 670.199 | 0.181031 | 0.16775 | 0.60061 | Fam50a | ENSMUSG( | 9273.396 | -0.0236 | 0.83019 | 0.980224 | Fam81a |
| ENSMUSG( | 794.5031 | 0.036266 | 0.742904 | 0.948478 | Fam53a | ENSMUSG( | 73.33689 | 0.299821 | 0.349132 | 0.911652 | Fam83d |
| ENSMUSG( | 1130.574 | -0.05289 | 0.617334 | 0.910866 | Fam53b | ENSMUSG( | 11.78648 | -0.72793 | 0.211371 | 0.864412 | Fam83f |
| ENSMUSG( | 2919.771 | 0.136128 | 0.108827 | 0.497931 | Fam53c | ENSMUSG( | 99.58396 | -0.32076 | 0.147142 | 0.836241 | Fam83h |
| ENSMUSG( | 42.48039 | -0.61413 | 0.035379 | 0.289531 | Fam72a | ENSMUSG( | 33.73669 | -0.00428 | 0.990764 | 0.998287 | Fam89a |
| ENSMUSG( | 1337.234 | -0.04426 | 0.660187 | 0.924627 | Fam76a | ENSMUSG( | 2803.808 | 0.085805 | 0.577257 | 0.942992 | Fam8a1 |
| ENSMUSG( | 383.6749 | -0.23049 | 0.060513 | 0.379845 | Fam76b | ENSMUSG( | 1927.615 | 0.058643 | 0.610592 | 0.950771 | Fam91a1 |
| ENSMUSG( | 148.812 | -0.24007 | 0.202186 | 0.642173 | Fam78a | ENSMUSG( | 955.5807 | 0.030891 | 0.812128 | 0.979148 | Fam98a |
| ENSMUSG( | 994.7625 | 0.04808 | 0.636312 | 0.914873 | Fam78b | ENSMUSG( | 796.4005 | -0.06879 | 0.651194 | 0.954477 | Fam98b |
| ENSMUSG( | 10907.26 | 0.171753 | 0.030307 | 0.264512 | Fam81a | ENSMUSG( | 1181.93 | -0.06942 | 0.53341 | 0.935162 | Fam98c |
| ENSMUSG( | 71.9267 | -0.04089 | 0.884672 | 0.982647 | Fam83d | ENSMUSG( | 306.1851 | -0.03316 | 0.83521 | 0.981237 | Fan1 |
| ENSMUSG( | 11.52652 | 0.455934 | 0.416519 | NA | Fam83e | ENSMUSG( | 50.65133 | 0.322889 | 0.315234 | 0.901306 | Fanca |
| ENSMUSG( | 15.00395 | -0.26154 | 0.601269 | NA | Fam83f | ENSMUSG( | 57.42768 | 0.399834 | 0.165099 | 0.838743 | Fancb |
| ENSMUSG( | 108.1525 | -0.37077 | 0.055497 | 0.36248 | Fam83h | ENSMUSG( | 236.4742 | -0.00695 | 0.966684 | 0.995475 | Fancc |
| ENSMUSG( | 33.88696 | -0.24135 | 0.473446 | 0.848919 | Fam89a | ENSMUSG( | 132.2617 | 0.171863 | 0.400836 | 0.917257 | Fancd2 |
| ENSMUSG( | 2683.85 | -0.32801 | 0.009797 | 0.141256 | Fam8a1 | ENSMUSG( | 373.3651 | -0.05376 | 0.722444 | 0.966887 | Fance |
| ENSMUSG( | 2142.073 | 0.094238 | 0.296936 | 0.745676 | Fam91a1 | ENSMUSG( | 52.60193 | -0.43128 | 0.126045 | 0.81516 | Fancf |
| ENSMUSG( | 1087.358 | 0.135608 | 0.195657 | 0.635109 | Fam98a | ENSMUSG( | 260.1025 | 0.083161 | 0.625541 | 0.953234 | Fancg |
| ENSMUSG( | 889.9974 | -0.00902 | 0.934435 | 0.99323 | Fam98b | ENSMUSG( | 22.03364 | -0.16754 | 0.694206 | 0.960867 | Fanci |
| ENSMUSG( | 1324.618 | -0.00796 | 0.936673 | 0.993415 | Fam98c | ENSMUSG( | 140.2886 | -0.03513 | 0.860823 | 0.986226 | Fancl |
| ENSMUSG( | 349.7976 | 0.077825 | 0.560346 | 0.888934 | Fan1 | ENSMUSG( | 419.6238 | 0.040056 | 0.792584 | 0.97681 | Fancm |
| ENSMUSG( | 52.35232 | 0.145129 | 0.598644 | 0.903801 | Fanca | ENSMUSG( | 90.54018 | -0.11746 | 0.631781 | 0.953251 | Fank1 |
| ENSMUSG( | 51.25677 | -0.18197 | 0.507123 | 0.867946 | Fancb | ENSMUSG( | 75.73333 | 0.120115 | 0.624074 | 0.953234 | Fap |
| ENSMUSG( | 254.8835 | -0.06772 | 0.623453 | 0.911646 | Fancc | ENSMUSG( | 1683.084 | 0.059287 | 0.647401 | 0.95422 | Far1 |
| ENSMUSG( | 137.8182 | 0.011694 | 0.945364 | 0.99365 | Fancd2 | ENSMUSG( | 415.0797 | 0.009915 | 0.949118 | 0.994798 | Far2 |
| ENSMUSG( | 427.5414 | 0.062028 | 0.626388 | 0.912685 | Fance | ENSMUSG( | 5460.643 | -0.0228 | 0.845918 | 0.98432 | Farp1 |
| ENSMUSG( | 59.35667 | -0.35292 | 0.158894 | 0.588112 | Fancf | ENSMUSG( | 299.3636 | 0.171672 | 0.272565 | 0.888825 | Farp2 |
| ENSMUSG( | 290.2359 | 0.120898 | 0.440251 | 0.831851 | Fancg | ENSMUSG( | 620.5168 | -0.17575 | 0.205387 | 0.864412 | Fars2 |
| ENSMUSG( | 26.52288 | 0.0935 | 0.808657 | 0.963455 | Fanci | ENSMUSG( | 711.9324 | 0.140287 | 0.313215 | 0.90077 | Farsa |
| ENSMUSG( | 162.968 | 0.136939 | 0.456443 | 0.838832 | Fancl | ENSMUSG( | 971.1343 | -0.03125 | 0.795492 | 0.97681 | Farsb |
| ENSMUSG( | 438.6224 | -0.10941 | 0.412263 | 0.819452 | Fancm | ENSMUSG( | 72.29221 | -0.03739 | 0.89873 | 0.990558 | Fas |
| ENSMUSG( | 97.0226 | -0.19764 | 0.345602 | 0.779153 | Fank1 | ENSMUSG( | 9979.109 | -0.00398 | 0.970503 | 0.99594 | Fasn |
| ENSMUSG( | 76.24562 | -0.1351 | 0.555652 | 0.887457 | Fap | ENSMUSG( | 3171.593 | 0.004754 | 0.964467 | 0.995433 | Fastk |
| ENSMUSG( | 1789.544 | -0.02849 | 0.792106 | 0.959461 | Far1 | ENSMUSG( | 315.6028 | 0.238917 | 0.093262 | 0.795363 | Fastkd1 |
| ENSMUSG( | 418.2516 | -0.25372 | 0.05265 | 0.352304 | Far2 | ENSMUSG( | 389.8063 | 0.028815 | 0.847424 | 0.985161 | Fastkd2 |
| ENSMUSG( | 6352.926 | 0.143755 | 0.117415 | 0.517109 | Farp1 | ENSMUSG( | 224.0605 | 0.155304 | 0.391877 | 0.917257 | Fastkd3 |
| ENSMUSG( | 320.0176 | 0.100721 | 0.441749 | 0.832577 | Farp2 | ENSMUSG( | 88.72142 | -0.45108 | 0.048063 | 0.704961 | Fastkd5 |
| ENSMUSG( | 736.4748 | 0.057023 | 0.607327 | 0.906826 | Fars2 | ENSMUSG( | 3364.646 | -0.03256 | 0.795968 | 0.97681 | Fat1 |
| ENSMUSG( | 742.5877 | -0.00617 | 0.955822 | 0.993874 | Farsa | ENSMUSG( | 41.46463 | 0.779317 | 0.160724 | 0.837724 | Fat2 |
| ENSMUSG( | 1121.756 | 0.116396 | 0.257254 | 0.708432 | Farsb | ENSMUSG( | 2736.073 | 0.106622 | 0.511784 | 0.930005 | Fat3 |
| ENSMUSG( | 85.19555 | 0.137787 | 0.574537 | 0.894423 | Fas | ENSMUSG( | 768.9455 | 0.161758 | 0.30137 | 0.898538 | Fat4 |
| ENSMUSG( | 11817.16 | 0.205692 | 0.046266 | 0.329472 | Fasn | ENSMUSG( | 6981.253 | -0.19899 | 0.103606 | 0.804969 | Fau |
| ENSMUSG( | 3649.355 | 0.137137 | 0.094152 | 0.471632 | Fastk | ENSMUSG( | 803.5324 | 0.11461 | 0.425736 | 0.917257 | Faxc |
| ENSMUSG( | 327.6798 | 0.087067 | 0.550423 | 0.886038 | Fastkd1 | ENSMUSG( | 2989.846 | 0.103001 | 0.335375 | 0.907907 | Fbf1 |
| ENSMUSG( | 435.946 | 0.075705 | 0.58824 | 0.900712 | Fastkd2 | ENSMUSG( | 2028.475 | 0.147875 | 0.1877 | 0.85698 | Fbh1 |
| ENSMUSG( | 221.3304 | -0.14864 | 0.381281 | 0.802629 | Fastkd3 | ENSMUSG( | 438.9336 | -0.05555 | 0.724604 | 0.966887 | Fbl |
| ENSMUSG( | 102.5158 | -0.27798 | 0.165359 | 0.596691 | Fastkd5 | ENSMUSG( | 34.85878 | -0.43119 | 0.21731 | 0.865589 | Fblim1 |
| ENSMUSG( | 3517.226 | -0.18085 | 0.199178 | 0.639621 | Fat1 | ENSMUSG( | 409.4386 | -0.24153 | 0.196346 | 0.862161 | Fbll1 |
| ENSMUSG( | 28.04085 | -0.62073 | 0.124362 | 0.531714 | Fat2 | ENSMUSG( | 816.6634 | 0.217309 | 0.114852 | 0.811703 | Fbln1 |
| ENSMUSG( | 2316.375 | -0.73894 | 8.14E-06 | 0.001 | Fat3 | ENSMUSG( | 290.0687 | 0.135011 | 0.411016 | 0.917257 | Fbln2 |
| ENSMUSG( | 662.1521 | -0.6021 | 6.19E-05 | 0.00467 | Fat4 | ENSMUSG( | 156.0504 | 0.005827 | 0.975199 | 0.996416 | Fbln5 |
| ENSMUSG( | 8469.025 | 0.096657 | 0.201938 | 0.641612 | Fau | ENSMUSG( | 124.5446 | -0.23978 | 0.355358 | 0.912555 | Fbln7 |
| ENSMUSG( | 887.2567 | 0.127159 | 0.355118 | 0.787533 | Faxc | ENSMUSG( | 277.9187 | -0.20673 | 0.277881 | 0.891468 | Fbn1 |
| ENSMUSG( | 3554.153 | 0.317574 | 0.000763 | 0.027082 | Fbf1 | ENSMUSG( | 150.5579 | 0.346989 | 0.076116 | 0.769376 | Fbn2 |
| ENSMUSG( | 2259.327 | 0.186565 | 0.023103 | 0.233188 | Fbh1 | ENSMUSG( | 10.32379 | 0.488563 | 0.440672 | NA | Fbp1 |
| ENSMUSG( | 517.184 | 0.153815 | 0.214492 | 0.659193 | Fbl | ENSMUSG( | 1589.246 | 0.074723 | 0.497547 | 0.927825 | Fbrs |
| ENSMUSG( | 43.53047 | -0.03004 | 0.92615 | 0.991229 | Fblim1 | ENSMUSG( | 2904.364 | 0.018928 | 0.854508 | 0.985649 | Fbrsl1 |
| ENSMUSG( | 477.8299 | -0.0481 | 0.768876 | 0.954674 | Fbll1 | ENSMUSG( | 306.6908 | 0.139011 | 0.42518 | 0.917257 | Fbxl12 |
| ENSMUSG( | 795.2834 | -0.12692 | 0.324049 | 0.769107 | Fbln1 | ENSMUSG( | 33.33882 | -0.01074 | 0.976462 | 0.996416 | Fbxl12os |
| ENSMUSG( | 309.1606 | 0.047795 | 0.757449 | 0.951513 | Fbln2 | ENSMUSG( | 10.25436 | 0.810143 | 0.209679 | NA | Fbxl13 |
| ENSMUSG( | 150.1941 | -0.40664 | 0.026036 | 0.247245 | Fbln5 | ENSMUSG( | 1000.622 | 0.105486 | 0.366877 | 0.914852 | Fbxl14 |
| ENSMUSG( | 128.8412 | -0.44863 | 0.05916 | 0.375449 | Fbln7 | ENSMUSG( | 469.3474 | -0.05923 | 0.699149 | 0.961139 | Fbxl15 |
| ENSMUSG( | 9.712274 | -0.49269 | 0.401328 | NA | Fbl-ps2 | ENSMUSG( | 62784.56 | 0.077303 | 0.419202 | 0.917257 | Fbxl16 |
| ENSMUSG( | 294.9296 | -0.32881 | 0.022577 | 0.229429 | Fbn1 | ENSMUSG( | 1478.667 | -0.00042 | 0.997281 | 0.999268 | Fbxl17 |
| ENSMUSG( | 141.9123 | -0.08141 | 0.670485 | 0.928683 | Fbn2 | ENSMUSG( | 1734.701 | 0.184644 | 0.147665 | 0.836241 | Fbxl18 |
| ENSMUSG( | 9.474578 | -0.04981 | 0.931966 | NA | Fbp1 | ENSMUSG( | 2589.66 | 0.196271 | 0.056808 | 0.723362 | Fbxl19 |
| ENSMUSG( | 1795.699 | 0.154949 | 0.096536 | 0.476986 | Fbrs | ENSMUSG( | 1439.494 | 0.020113 | 0.87062 | 0.988445 | Fbxl2 |
| ENSMUSG( | 3168.324 | -0.00019 | 0.998293 | 0.99989 | Fbrsl1 | ENSMUSG( | 2027.838 | 0.089378 | 0.509948 | 0.929089 | Fbxl20 |
| ENSMUSG( | 320.3203 | -0.01082 | 0.941346 | 0.993415 | Fbxl12 | ENSMUSG( | 80.66346 | -0.12826 | 0.61485 | 0.951177 | Fbxl21 |
| ENSMUSG( | 35.63018 | -0.09512 | 0.773535 | 0.955447 | Fbxl12os | ENSMUSG( | 2342.661 | -0.05614 | 0.599226 | 0.947045 | Fbxl3 |
| ENSMUSG( | 11.48538 | 0.829153 | 0.140112 | NA | Fbxl13 | ENSMUSG( | 477.5964 | 0.109132 | 0.4796 | 0.924021 | Fbxl4 |
| ENSMUSG( | 1011.894 | -0.13233 | 0.190609 | 0.629951 | Fbxl14 | ENSMUSG( | 1935.193 | -0.0145 | 0.890099 | 0.990558 | Fbxl5 |
| ENSMUSG( | 526.2397 | 0.005019 | 0.969621 | 0.996369 | Fbxl15 | ENSMUSG( | 945.4148 | 0.107826 | 0.444808 | 0.921734 | Fbxl6 |
| ENSMUSG( | 68940.47 | 0.077781 | 0.327016 | 0.771858 | Fbxl16 | ENSMUSG( | 57.49067 | 0.322851 | 0.254081 | 0.881335 | Fbxl7 |
| ENSMUSG( | 1691.748 | 0.119468 | 0.223347 | 0.669382 | Fbxl17 | ENSMUSG( | 18.71421 | 0.280037 | 0.545192 | 0.938063 | Fbxl8 |
| ENSMUSG( | 1932.204 | 0.22123 | 0.017632 | 0.199023 | Fbxl18 | ENSMUSG( | 41.74918 | 0.090475 | 0.810553 | 0.978779 | Fbxl9 |
| ENSMUSG( | 2974.557 | 0.31575 | 0.000353 | 0.016287 | Fbxl19 | ENSMUSG( | 1270.267 | 0.139671 | 0.426011 | 0.917257 | Fbxo10 |
| ENSMUSG( | 1456.228 | -0.2269 | 0.036945 | 0.296508 | Fbxl2 | ENSMUSG( | 2939.463 | 0.099288 | 0.381092 | 0.917202 | Fbxo11 |
| ENSMUSG( | 2060.637 | -0.13804 | 0.2386 | 0.686308 | Fbxl20 | ENSMUSG( | 143.2635 | -0.32839 | 0.144845 | 0.83339 | Fbxo16 |
| ENSMUSG( | 93.19335 | 0.047071 | 0.83724 | 0.971111 | Fbxl21 | ENSMUSG( | 77.542 | -0.1001 | 0.683456 | 0.958678 | Fbxo17 |
| ENSMUSG( | 13.23416 | 0.406917 | 0.452713 | NA | Fbxl22 | ENSMUSG( | 4804.231 | -0.14549 | 0.279426 | 0.892356 | Fbxo2 |
| ENSMUSG( | 2585.251 | -0.03942 | 0.657596 | 0.92425 | Fbxl3 | ENSMUSG( | 2433.802 | -0.03863 | 0.773604 | 0.974672 | Fbxo21 |
| ENSMUSG( | 504.3132 | -0.00621 | 0.965298 | 0.995368 | Fbxl4 | ENSMUSG( | 853.356 | -0.06713 | 0.608132 | 0.950113 | Fbxo22 |
| ENSMUSG( | 2099.581 | -0.04917 | 0.588365 | 0.90082 | Fbxl5 | ENSMUSG( | 10.53737 | -0.27848 | 0.639426 | NA | Fbxo24 |
| ENSMUSG( | 946.0285 | -0.16719 | 0.148679 | 0.572413 | Fbxl6 | ENSMUSG( | 888.6593 | 0.027796 | 0.840252 | 0.98236 | Fbxo25 |
| ENSMUSG( | 63.25213 | 0.329301 | 0.203595 | 0.643764 | Fbxl7 | ENSMUSG( | 229.7459 | -0.08156 | 0.645422 | 0.953988 | Fbxo27 |
| ENSMUSG( | 18.11642 | -0.11762 | 0.784037 | NA | Fbxl8 | ENSMUSG( | 1073.148 | 0.000599 | 0.996026 | 0.999163 | Fbxo28 |
| ENSMUSG( | 45.67983 | 0.056451 | 0.868525 | 0.976897 | Fbxl9 | ENSMUSG( | 1860.921 | 0.020568 | 0.857725 | 0.985884 | Fbxo3 |
| ENSMUSG( | 1309.99 | -0.04458 | 0.762007 | 0.95339 | Fbxo10 | ENSMUSG( | 240.5984 | 0.005256 | 0.978106 | 0.996416 | Fbxo30 |
| ENSMUSG( | 3209.843 | 0.086671 | 0.345824 | 0.779153 | Fbxo11 | ENSMUSG( | 5972.87 | 0.128821 | 0.248957 | 0.877904 | Fbxo31 |
| ENSMUSG( | 160.6107 | -0.26519 | 0.180682 | 0.617841 | Fbxo16 | ENSMUSG( | 1044.071 | -0.06419 | 0.593583 | 0.945408 | Fbxo32 |
| ENSMUSG( | 83.8989 | -0.1399 | 0.520827 | 0.875186 | Fbxo17 | ENSMUSG( | 749.7907 | 0.12594 | 0.317021 | 0.902088 | Fbxo33 |
| ENSMUSG( | 5884.878 | 0.172875 | 0.073832 | 0.42034 | Fbxo2 | ENSMUSG( | 1987.607 | 0.015096 | 0.919703 | 0.992644 | Fbxo34 |
| ENSMUSG( | 2814.186 | 0.111183 | 0.351124 | 0.783067 | Fbxo21 | ENSMUSG( | 104.315 | -0.14901 | 0.589605 | 0.944975 | Fbxo36 |
| ENSMUSG( | 988.1268 | 0.090956 | 0.392694 | 0.809274 | Fbxo22 | ENSMUSG( | 1014.823 | 0.129826 | 0.271671 | 0.888062 | Fbxo38 |
| ENSMUSG( | 11.59485 | -0.25152 | 0.646429 | NA | Fbxo24 | ENSMUSG( | 188.1116 | 0.234021 | 0.220029 | 0.866104 | Fbxo4 |
| ENSMUSG( | 977.1139 | 0.033157 | 0.782933 | 0.957558 | Fbxo25 | ENSMUSG( | 5098.65 | 0.209527 | 0.14254 | 0.831221 | Fbxo41 |
| ENSMUSG( | 259.2912 | -0.01076 | 0.947885 | 0.993821 | Fbxo27 | ENSMUSG( | 892.2621 | -0.03478 | 0.789757 | 0.976792 | Fbxo42 |
| ENSMUSG( | 1189.901 | 0.032672 | 0.730507 | 0.944956 | Fbxo28 | ENSMUSG( | 3503.904 | -0.07388 | 0.451809 | 0.922202 | Fbxo44 |
| ENSMUSG( | 2130.789 | 0.141766 | 0.128668 | 0.537025 | Fbxo3 | ENSMUSG( | 1678.24 | 0.033806 | 0.793853 | 0.97681 | Fbxo45 |
| ENSMUSG( | 271.1548 | 0.087203 | 0.625336 | 0.912249 | Fbxo30 | ENSMUSG( | 287.9609 | 0.181058 | 0.285078 | 0.892533 | Fbxo46 |
| ENSMUSG( | 6713.878 | 0.192347 | 0.02626 | 0.247758 | Fbxo31 | ENSMUSG( | 12.3721 | 0.225845 | 0.686582 | 0.95899 | Fbxo47 |
| ENSMUSG( | 1114.599 | -0.14594 | 0.197838 | 0.637707 | Fbxo32 | ENSMUSG( | 32.55671 | 0.112066 | 0.778311 | 0.976106 | Fbxo5 |
| ENSMUSG( | 752.5084 | -0.13126 | 0.20902 | 0.650854 | Fbxo33 | ENSMUSG( | 438.9216 | -0.05836 | 0.676825 | 0.957106 | Fbxo6 |
| ENSMUSG( | 2179.133 | 0.007045 | 0.95331 | 0.993874 | Fbxo34 | ENSMUSG( | 895.8607 | 0.074719 | 0.569838 | 0.941777 | Fbxo7 |
| ENSMUSG( | 109.0761 | -0.27002 | 0.243635 | 0.690987 | Fbxo36 | ENSMUSG( | 795.9569 | -0.04345 | 0.722716 | 0.966887 | Fbxo8 |
| ENSMUSG( | 1128.261 | 0.162801 | 0.099488 | 0.481972 | Fbxo38 | ENSMUSG( | 2603.65 | -0.09773 | 0.355625 | 0.912879 | Fbxo9 |
| ENSMUSG( | 207.5784 | 0.257875 | 0.115463 | 0.512397 | Fbxo4 | ENSMUSG( | 28.89827 | 0.126747 | 0.735799 | 0.967344 | Fbxw10 |
| ENSMUSG( | 5498.257 | 0.158713 | 0.128922 | 0.537838 | Fbxo41 | ENSMUSG( | 3941.34 | -0.0951 | 0.388501 | 0.917257 | Fbxw11 |
| ENSMUSG( | 1009.372 | 0.050464 | 0.627647 | 0.912961 | Fbxo42 | ENSMUSG( | 206.1058 | 0.047317 | 0.801368 | 0.97681 | Fbxw17 |
| ENSMUSG( | 4072.696 | 0.091218 | 0.234504 | 0.681922 | Fbxo44 | ENSMUSG( | 1991.409 | -0.09428 | 0.418904 | 0.917257 | Fbxw2 |
| ENSMUSG( | 1954.491 | 0.201025 | 0.026666 | 0.249727 | Fbxo45 | ENSMUSG( | 864.0739 | -0.07849 | 0.503934 | 0.927825 | Fbxw4 |
| ENSMUSG( | 304.995 | 0.072456 | 0.618394 | 0.911104 | Fbxo46 | ENSMUSG( | 3431.739 | -0.08824 | 0.38477 | 0.917257 | Fbxw5 |
| ENSMUSG( | 13.71343 | 0.216054 | 0.675036 | NA | Fbxo47 | ENSMUSG( | 4525.625 | 0.177192 | 0.127236 | 0.817278 | Fbxw7 |
| ENSMUSG( | 31.85627 | -0.27035 | 0.425076 | 0.82478 | Fbxo5 | ENSMUSG( | 2072.688 | -0.13324 | 0.202539 | 0.864412 | Fbxw8 |
| ENSMUSG( | 473.1522 | -0.10653 | 0.359424 | 0.791011 | Fbxo6 | ENSMUSG( | 490.356 | -0.01944 | 0.899835 | 0.990558 | Fbxw9 |
| ENSMUSG( | 869.4258 | -0.29645 | 0.006937 | 0.114527 | Fbxo7 | ENSMUSG( | 654.6294 | -0.03051 | 0.832296 | 0.98075 | Fcer1g |
| ENSMUSG( | 883.4104 | -0.00805 | 0.942033 | 0.993415 | Fbxo8 | ENSMUSG( | 313.9743 | -0.05159 | 0.72867 | 0.966913 | Fcf1 |
| ENSMUSG( | 3031.501 | 0.074478 | 0.364306 | 0.793796 | Fbxo9 | ENSMUSG( | 266.136 | 0.251687 | 0.284486 | 0.892412 | Fcgr1 |
| ENSMUSG( | 30.90105 | 0.054757 | 0.873122 | 0.978753 | Fbxw10 | ENSMUSG( | 303.3006 | 0.145225 | 0.485223 | 0.92541 | Fcgr2b |
| ENSMUSG( | 4624.221 | 0.098962 | 0.232814 | 0.680928 | Fbxw11 | ENSMUSG( | 977.3312 | 0.339648 | 0.014504 | 0.48395 | Fcgr3 |
| ENSMUSG( | 208.6973 | -0.20872 | 0.192394 | 0.631451 | Fbxw17 | ENSMUSG( | 25.36518 | -0.83677 | 0.115748 | 0.813384 | Fcgr4 |
| ENSMUSG( | 2294.012 | 0.04775 | 0.562841 | 0.88962 | Fbxw2 | ENSMUSG( | 1344.53 | 0.288012 | 0.074416 | 0.766789 | Fcgrt |
| ENSMUSG( | 999.9825 | 0.076718 | 0.418713 | 0.821966 | Fbxw4 | ENSMUSG( | 2711.245 | -0.03818 | 0.780041 | 0.976106 | Fcho1 |
| ENSMUSG( | 4046.007 | 0.116668 | 0.151225 | 0.576198 | Fbxw5 | ENSMUSG( | 1127.442 | 0.26259 | 0.045431 | 0.693959 | Fcho2 |
| ENSMUSG( | 4853.389 | 0.112169 | 0.258911 | 0.709814 | Fbxw7 | ENSMUSG( | 255.3169 | -0.09718 | 0.600725 | 0.947749 | Fchsd1 |
| ENSMUSG( | 2544.887 | 0.187846 | 0.020416 | 0.215588 | Fbxw8 | ENSMUSG( | 1528.831 | -0.1009 | 0.360502 | 0.914223 | Fchsd2 |
| ENSMUSG( | 562.3467 | 0.100116 | 0.456858 | 0.839183 | Fbxw9 | ENSMUSG( | 20.90068 | -0.20046 | 0.639758 | 0.953903 | Fcna |
| ENSMUSG( | 690.6619 | -0.15126 | 0.220541 | 0.666079 | Fcer1g | ENSMUSG( | 84.48997 | -0.4866 | 0.051652 | 0.711848 | Fcor |
| ENSMUSG( | 330.0342 | -0.17792 | 0.164039 | 0.594869 | Fcf1 | ENSMUSG( | 41.24244 | 0.538205 | 0.123938 | 0.814691 | Fcrl1 |
| ENSMUSG( | 12.69669 | 0.398798 | 0.436779 | NA | Fcgbp | ENSMUSG( | 2391.541 | 0.126217 | 0.487726 | 0.925427 | Fcrl2 |
| ENSMUSG( | 235.9747 | -0.39482 | 0.054447 | 0.359033 | Fcgr1 | ENSMUSG( | 11.15483 | -0.1376 | 0.809271 | NA | Fcrlb |
| ENSMUSG( | 274.979 | -0.44782 | 0.027518 | 0.254761 | Fcgr2b | ENSMUSG( | 497.6919 | 0.183353 | 0.2344 | 0.870096 | Fcsk |
| ENSMUSG( | 977.4467 | 0.086103 | 0.508052 | 0.867981 | Fcgr3 | ENSMUSG( | 3022.171 | -0.26108 | 0.019718 | 0.556192 | Fdft1 |

| | | | | | | | | | | | |
|---|---|---|---|---|---|---|---|---|---|---|---|
| ENSMUSG( | Fam98b | 889.9974 | -0.00902 | 0.934435 | 0.99323 | ENSMUSG( | Fam98b | 796.4005 | -0.06879 | 0.651194 | 0.954477 |
| ENSMUSG( | Fam98c | 1324.618 | -0.00796 | 0.936673 | 0.993415 | ENSMUSG( | Fam98c | 1181.93 | -0.06942 | 0.53341 | 0.935162 |
| ENSMUSG( | Fan1 | 349.7976 | 0.077825 | 0.560346 | 0.888934 | ENSMUSG( | Fan1 | 306.1851 | -0.03316 | 0.83521 | 0.981237 |
| ENSMUSG( | Fanca | 52.35232 | 0.145129 | 0.598644 | 0.903801 | ENSMUSG( | Fanca | 50.65133 | 0.322889 | 0.315234 | 0.901306 |
| ENSMUSG( | Fancb | 51.25677 | -0.18197 | 0.507123 | 0.867946 | ENSMUSG( | Fancb | 57.42768 | 0.399834 | 0.165099 | 0.838743 |
| ENSMUSG( | Fancc | 254.8835 | -0.06772 | 0.623453 | 0.911646 | ENSMUSG( | Fancc | 236.4742 | -0.00695 | 0.966684 | 0.995475 |
| ENSMUSG( | Fancd2 | 137.8182 | 0.011694 | 0.945364 | 0.99365 | ENSMUSG( | Fancd2 | 132.2617 | 0.171863 | 0.400836 | 0.917257 |
| ENSMUSG( | Fance | 427.5414 | 0.062028 | 0.626388 | 0.912685 | ENSMUSG( | Fance | 373.3651 | -0.05376 | 0.722444 | 0.966887 |
| ENSMUSG( | Fancf | 59.35667 | -0.35292 | 0.158894 | 0.588112 | ENSMUSG( | Fancf | 52.60193 | -0.43128 | 0.126045 | 0.81516 |
| ENSMUSG( | Fancg | 290.2359 | 0.120898 | 0.440251 | 0.831851 | ENSMUSG( | Fancg | 260.1025 | 0.083161 | 0.625541 | 0.953234 |
| ENSMUSG( | Fanci | 26.52288 | 0.0935 | 0.808657 | 0.963455 | ENSMUSG( | Fanci | 22.03364 | -0.16754 | 0.694206 | 0.960867 |
| ENSMUSG( | Fancl | 162.968 | 0.136939 | 0.456443 | 0.838832 | ENSMUSG( | Fancl | 140.2886 | -0.03513 | 0.860823 | 0.986226 |
| ENSMUSG( | Fancm | 438.6224 | -0.10941 | 0.412263 | 0.819452 | ENSMUSG( | Fancm | 419.6238 | 0.040056 | 0.792584 | 0.97681 |
| ENSMUSG( | Fank1 | 97.0226 | -0.19764 | 0.345602 | 0.779153 | ENSMUSG( | Fank1 | 90.54018 | -0.11746 | 0.631781 | 0.953251 |
| ENSMUSG( | Fap | 76.24562 | -0.1351 | 0.555652 | 0.887457 | ENSMUSG( | Fap | 75.73333 | 0.120115 | 0.624074 | 0.953234 |
| ENSMUSG( | Far1 | 1789.544 | -0.02849 | 0.792106 | 0.959461 | ENSMUSG( | Far1 | 1683.084 | 0.059287 | 0.647401 | 0.95422 |
| ENSMUSG( | Far2 | 418.2516 | -0.25372 | 0.05265 | 0.352304 | ENSMUSG( | Far2 | 415.0797 | 0.009915 | 0.949118 | 0.994798 |
| ENSMUSG( | Farp1 | 6352.926 | 0.143755 | 0.117415 | 0.517109 | ENSMUSG( | Farp1 | 5460.643 | -0.0228 | 0.845918 | 0.98432 |
| ENSMUSG( | Farp2 | 320.0176 | 0.100721 | 0.441749 | 0.832577 | ENSMUSG( | Farp2 | 299.3636 | 0.171672 | 0.272565 | 0.888825 |
| ENSMUSG( | Fars2 | 736.4748 | 0.057023 | 0.607327 | 0.906826 | ENSMUSG( | Fars2 | 620.5168 | -0.17575 | 0.205387 | 0.864412 |
| ENSMUSG( | Farsa | 742.5877 | -0.00617 | 0.955822 | 0.993874 | ENSMUSG( | Farsa | 711.9324 | 0.140287 | 0.313215 | 0.90077 |
| ENSMUSG( | Farsb | 1121.756 | 0.116396 | 0.257254 | 0.708432 | ENSMUSG( | Farsb | 971.1343 | -0.03125 | 0.795492 | 0.97681 |
| ENSMUSG( | Fas | 85.19555 | 0.137787 | 0.574537 | 0.894423 | ENSMUSG( | Fas | 72.29221 | -0.03739 | 0.89873 | 0.990558 |
| ENSMUSG( | Fasn | 11817.16 | 0.205692 | 0.046266 | 0.329472 | ENSMUSG( | Fasn | 9979.109 | -0.00398 | 0.970503 | 0.99594 |
| ENSMUSG( | Fastk | 3649.355 | 0.137137 | 0.094152 | 0.471632 | ENSMUSG( | Fastk | 3171.593 | 0.004754 | 0.964467 | 0.995433 |
| ENSMUSG( | Fastkd1 | 327.6798 | 0.087067 | 0.550423 | 0.886038 | ENSMUSG( | Fastkd1 | 315.6028 | 0.238917 | 0.093262 | 0.795363 |
| ENSMUSG( | Fastkd2 | 435.946 | 0.075705 | 0.58824 | 0.900712 | ENSMUSG( | Fastkd2 | 389.8063 | 0.028815 | 0.847424 | 0.985161 |
| ENSMUSG( | Fastkd3 | 221.3304 | -0.14864 | 0.381281 | 0.802629 | ENSMUSG( | Fastkd3 | 224.0605 | 0.155304 | 0.391877 | 0.917257 |
| ENSMUSG( | Fastkd5 | 102.5158 | -0.27798 | 0.165359 | 0.596691 | ENSMUSG( | Fastkd5 | 88.72142 | -0.45108 | 0.048063 | 0.704961 |
| ENSMUSG( | Fat1 | 3517.226 | -0.18085 | 0.199178 | 0.639621 | ENSMUSG( | Fat1 | 3364.646 | -0.03256 | 0.795968 | 0.97681 |
| ENSMUSG( | Fat2 | 28.04085 | -0.62073 | 0.124362 | 0.531714 | ENSMUSG( | Fat2 | 41.46463 | 0.779317 | 0.160724 | 0.837724 |
| ENSMUSG( | Fat3 | 2316.375 | -0.73894 | 8.14E-06 | 0.001 | ENSMUSG( | Fat3 | 2736.073 | 0.106622 | 0.511784 | 0.930005 |
| ENSMUSG( | Fat4 | 662.1521 | -0.6021 | 6.19E-05 | 0.00467 | ENSMUSG( | Fat4 | 768.9455 | 0.161758 | 0.30137 | 0.898538 |
| ENSMUSG( | Fau | 8469.025 | 0.096657 | 0.201938 | 0.641612 | ENSMUSG( | Fau | 6981.253 | -0.19899 | 0.103606 | 0.804969 |
| ENSMUSG( | Faxc | 887.2567 | 0.127159 | 0.355118 | 0.787533 | ENSMUSG( | Faxc | 803.5324 | 0.11461 | 0.425736 | 0.917257 |
| ENSMUSG( | Fbf1 | 3554.153 | 0.317574 | 0.000763 | 0.027082 | ENSMUSG( | Fbf1 | 2989.846 | 0.103001 | 0.335375 | 0.907907 |
| ENSMUSG( | Fbh1 | 2259.327 | 0.186565 | 0.023103 | 0.233188 | ENSMUSG( | Fbh1 | 2028.475 | 0.147875 | 0.1877 | 0.85698 |
| ENSMUSG( | Fbl | 517.184 | 0.153815 | 0.214492 | 0.659193 | ENSMUSG( | Fbl | 438.9336 | -0.05555 | 0.724604 | 0.966887 |
| ENSMUSG( | Fblim1 | 43.53047 | -0.03004 | 0.92615 | 0.991229 | ENSMUSG( | Fblim1 | 34.85878 | -0.43119 | 0.21731 | 0.865589 |
| ENSMUSG( | Fbll1 | 477.8299 | -0.0481 | 0.768876 | 0.954674 | ENSMUSG( | Fbll1 | 409.4386 | -0.24153 | 0.196346 | 0.862161 |
| ENSMUSG( | Fbln1 | 795.2834 | -0.12692 | 0.324049 | 0.769107 | ENSMUSG( | Fbln1 | 816.6634 | 0.217309 | 0.114852 | 0.811703 |
| ENSMUSG( | Fbln2 | 309.1606 | 0.047795 | 0.757449 | 0.951513 | ENSMUSG( | Fbln2 | 290.0687 | 0.135011 | 0.411016 | 0.917257 |
| ENSMUSG( | Fbln5 | 150.1941 | -0.40664 | 0.026036 | 0.247245 | ENSMUSG( | Fbln5 | 156.0504 | 0.005827 | 0.975199 | 0.996416 |
| ENSMUSG( | Fbln7 | 128.8412 | -0.44863 | 0.05916 | 0.375449 | ENSMUSG( | Fbln7 | 124.5446 | -0.23978 | 0.355358 | 0.912555 |
| ENSMUSG( | Fbn1 | 294.9296 | -0.32881 | 0.022577 | 0.229429 | ENSMUSG( | Fbn1 | 277.9187 | -0.20673 | 0.277881 | 0.891468 |
| ENSMUSG( | Fbn2 | 141.9123 | -0.08141 | 0.670485 | 0.928683 | ENSMUSG( | Fbn2 | 150.5579 | 0.346989 | 0.076116 | 0.769376 |
| ENSMUSG( | Fbp1 | 9.474578 | -0.04981 | 0.931966 | NA | ENSMUSG( | Fbp1 | 10.32379 | 0.488563 | 0.440672 | NA |
| ENSMUSG( | Fbrs | 1795.699 | 0.154949 | 0.096536 | 0.476986 | ENSMUSG( | Fbrs | 1589.246 | 0.074723 | 0.497547 | 0.927825 |
| ENSMUSG( | Fbrsl1 | 3168.324 | -0.00019 | 0.998293 | 0.99989 | ENSMUSG( | Fbrsl1 | 2904.364 | 0.018928 | 0.854508 | 0.985649 |
| ENSMUSG( | Fbxl12 | 320.3203 | -0.01082 | 0.941346 | 0.993415 | ENSMUSG( | Fbxl12 | 306.6908 | 0.139011 | 0.42518 | 0.917257 |
| ENSMUSG( | Fbxl12os | 35.63018 | -0.09512 | 0.773535 | 0.955447 | ENSMUSG( | Fbxl12os | 33.33882 | -0.01074 | 0.976462 | 0.996416 |
| ENSMUSG( | Fbxl13 | 11.48538 | 0.829153 | 0.140112 | NA | ENSMUSG( | Fbxl13 | 10.25436 | 0.810143 | 0.209679 | NA |
| ENSMUSG( | Fbxl14 | 1011.894 | -0.13233 | 0.190609 | 0.629951 | ENSMUSG( | Fbxl14 | 1000.622 | 0.105486 | 0.366877 | 0.914852 |
| ENSMUSG( | Fbxl15 | 526.2397 | 0.005019 | 0.969621 | 0.996369 | ENSMUSG( | Fbxl15 | 469.3474 | -0.05923 | 0.699149 | 0.961139 |
| ENSMUSG( | Fbxl16 | 68940.47 | 0.077781 | 0.327016 | 0.771858 | ENSMUSG( | Fbxl16 | 62784.56 | 0.077303 | 0.419202 | 0.917257 |
| ENSMUSG( | Fbxl17 | 1691.748 | 0.119468 | 0.223347 | 0.669382 | ENSMUSG( | Fbxl17 | 1478.667 | -0.00042 | 0.997281 | 0.999268 |
| ENSMUSG( | Fbxl18 | 1932.204 | 0.22123 | 0.017632 | 0.199023 | ENSMUSG( | Fbxl18 | 1734.701 | 0.184644 | 0.147665 | 0.836241 |
| ENSMUSG( | Fbxl19 | 2974.557 | 0.31575 | 0.000353 | 0.016287 | ENSMUSG( | Fbxl19 | 2589.66 | 0.196271 | 0.056808 | 0.723362 |
| ENSMUSG( | Fbxl2 | 1456.228 | -0.2269 | 0.036945 | 0.296508 | ENSMUSG( | Fbxl2 | 1439.494 | 0.020113 | 0.87062 | 0.988445 |
| ENSMUSG( | Fbxl20 | 2060.637 | -0.13804 | 0.2386 | 0.686308 | ENSMUSG( | Fbxl20 | 2027.838 | 0.089378 | 0.509948 | 0.929089 |
| ENSMUSG( | Fbxl21 | 93.19335 | 0.047071 | 0.83724 | 0.971111 | ENSMUSG( | Fbxl21 | 80.66346 | -0.12826 | 0.61485 | 0.951177 |
| ENSMUSG( | Fbxl3 | 2585.251 | -0.03942 | 0.657596 | 0.92425 | ENSMUSG( | Fbxl3 | 2342.661 | -0.05614 | 0.599226 | 0.947045 |
| ENSMUSG( | Fbxl4 | 504.3132 | -0.00621 | 0.965298 | 0.995368 | ENSMUSG( | Fbxl4 | 477.5964 | 0.109132 | 0.4796 | 0.924021 |
| ENSMUSG( | Fbxl5 | 2099.581 | -0.04917 | 0.588365 | 0.90082 | ENSMUSG( | Fbxl5 | 1935.193 | -0.0145 | 0.890099 | 0.990558 |
| ENSMUSG( | Fbxl6 | 946.0285 | -0.16719 | 0.148679 | 0.572413 | ENSMUSG( | Fbxl6 | 945.4148 | 0.107826 | 0.444808 | 0.921734 |
| ENSMUSG( | Fbxl7 | 63.25213 | 0.329301 | 0.203595 | 0.643764 | ENSMUSG( | Fbxl7 | 57.49067 | 0.322851 | 0.254081 | 0.881335 |
| ENSMUSG( | Fbxl8 | 18.11642 | -0.11762 | 0.784037 | NA | ENSMUSG( | Fbxl8 | 18.71421 | 0.280037 | 0.545192 | 0.938063 |
| ENSMUSG( | Fbxl9 | 45.67983 | 0.056451 | 0.868525 | 0.976897 | ENSMUSG( | Fbxl9 | 41.74918 | 0.090475 | 0.810553 | 0.978779 |
| ENSMUSG( | Fbxo10 | 1309.99 | -0.04458 | 0.762007 | 0.95339 | ENSMUSG( | Fbxo10 | 1270.267 | 0.139671 | 0.426011 | 0.917257 |
| ENSMUSG( | Fbxo11 | 3209.843 | 0.086671 | 0.345824 | 0.779153 | ENSMUSG( | Fbxo11 | 2939.463 | 0.099288 | 0.381092 | 0.917202 |
| ENSMUSG( | Fbxo16 | 160.6107 | -0.26519 | 0.180682 | 0.617841 | ENSMUSG( | Fbxo16 | 143.2635 | -0.32839 | 0.144845 | 0.83339 |
| ENSMUSG( | Fbxo17 | 83.8989 | -0.1399 | 0.520827 | 0.875186 | ENSMUSG( | Fbxo17 | 77.542 | -0.1001 | 0.683456 | 0.958678 |
| ENSMUSG( | Fbxo2 | 5884.878 | 0.172875 | 0.073832 | 0.42034 | ENSMUSG( | Fbxo2 | 4804.231 | -0.14549 | 0.279426 | 0.892356 |
| ENSMUSG( | Fbxo21 | 2814.186 | 0.111183 | 0.351124 | 0.783067 | ENSMUSG( | Fbxo21 | 2433.802 | -0.03863 | 0.773604 | 0.974672 |
| ENSMUSG( | Fbxo22 | 988.1268 | 0.090956 | 0.392694 | 0.809274 | ENSMUSG( | Fbxo22 | 853.356 | -0.06713 | 0.608132 | 0.950113 |
| ENSMUSG( | Fbxo24 | 11.59485 | -0.25152 | 0.646429 | NA | ENSMUSG( | Fbxo24 | 10.53737 | -0.27848 | 0.639426 | NA |
| ENSMUSG( | Fbxo25 | 977.1139 | 0.033157 | 0.782933 | 0.957558 | ENSMUSG( | Fbxo25 | 888.6593 | 0.027796 | 0.840252 | 0.98236 |
| ENSMUSG( | Fbxo27 | 259.2912 | -0.01076 | 0.947885 | 0.993821 | ENSMUSG( | Fbxo27 | 229.7459 | -0.08156 | 0.645422 | 0.953988 |
| ENSMUSG( | Fbxo28 | 1189.901 | 0.032672 | 0.730507 | 0.944956 | ENSMUSG( | Fbxo28 | 1073.148 | 0.000599 | 0.996026 | 0.999163 |
| ENSMUSG( | Fbxo3 | 2130.789 | 0.141766 | 0.128668 | 0.537025 | ENSMUSG( | Fbxo3 | 1860.921 | 0.020568 | 0.857725 | 0.985884 |
| ENSMUSG( | Fbxo30 | 271.1548 | 0.087203 | 0.625336 | 0.912249 | ENSMUSG( | Fbxo30 | 240.5984 | 0.005256 | 0.978106 | 0.996416 |
| ENSMUSG( | Fbxo31 | 6713.878 | 0.192347 | 0.02626 | 0.247758 | ENSMUSG( | Fbxo31 | 5972.87 | 0.128821 | 0.248957 | 0.877904 |
| ENSMUSG( | Fbxo32 | 1114.599 | -0.14594 | 0.197838 | 0.637707 | ENSMUSG( | Fbxo32 | 1044.071 | -0.06419 | 0.593583 | 0.945408 |
| ENSMUSG( | Fbxo33 | 752.5084 | -0.13126 | 0.20902 | 0.650854 | ENSMUSG( | Fbxo33 | 749.7907 | 0.12594 | 0.317021 | 0.902088 |
| ENSMUSG( | Fbxo34 | 2179.133 | 0.007045 | 0.95331 | 0.993874 | ENSMUSG( | Fbxo34 | 1987.607 | 0.015096 | 0.919703 | 0.992644 |
| ENSMUSG( | Fbxo36 | 109.0761 | -0.27002 | 0.243635 | 0.690987 | ENSMUSG( | Fbxo36 | 104.315 | -0.14901 | 0.589605 | 0.944975 |
| ENSMUSG( | Fbxo38 | 1128.261 | 0.162801 | 0.099488 | 0.481972 | ENSMUSG( | Fbxo38 | 1014.823 | 0.129826 | 0.271671 | 0.888062 |
| ENSMUSG( | Fbxo4 | 207.5784 | 0.257875 | 0.115463 | 0.512397 | ENSMUSG( | Fbxo4 | 188.1116 | 0.234021 | 0.220029 | 0.866104 |
| ENSMUSG( | Fbxo41 | 5498.257 | 0.158713 | 0.128922 | 0.537838 | ENSMUSG( | Fbxo41 | 5098.65 | 0.209527 | 0.14254 | 0.831221 |
| ENSMUSG( | Fbxo42 | 1009.372 | 0.050464 | 0.627647 | 0.912961 | ENSMUSG( | Fbxo42 | 892.2621 | -0.03478 | 0.789757 | 0.976792 |
| ENSMUSG( | Fbxo44 | 4072.696 | 0.091218 | 0.234504 | 0.681922 | ENSMUSG( | Fbxo44 | 3503.904 | -0.07388 | 0.451809 | 0.922202 |
| ENSMUSG( | Fbxo45 | 1954.491 | 0.201025 | 0.026666 | 0.249727 | ENSMUSG( | Fbxo45 | 1678.24 | 0.033806 | 0.793853 | 0.97681 |
| ENSMUSG( | Fbxo46 | 304.995 | 0.072456 | 0.618394 | 0.911104 | ENSMUSG( | Fbxo46 | 287.9609 | 0.181058 | 0.285078 | 0.892533 |
| ENSMUSG( | Fbxo47 | 13.71343 | 0.216054 | 0.675036 | NA | ENSMUSG( | Fbxo47 | 12.3721 | 0.225845 | 0.686582 | 0.95899 |
| ENSMUSG( | Fbxo5 | 31.85627 | -0.27035 | 0.425076 | 0.82478 | ENSMUSG( | Fbxo5 | 32.55671 | 0.112066 | 0.778311 | 0.976106 |
| ENSMUSG( | Fbxo6 | 473.1522 | -0.10653 | 0.359424 | 0.791011 | ENSMUSG( | Fbxo6 | 438.9216 | -0.05836 | 0.676825 | 0.957106 |
| ENSMUSG( | Fbxo7 | 869.4258 | -0.29645 | 0.006937 | 0.114527 | ENSMUSG( | Fbxo7 | 895.8607 | 0.074719 | 0.569838 | 0.941777 |
| ENSMUSG( | Fbxo8 | 883.4104 | -0.00805 | 0.942033 | 0.993415 | ENSMUSG( | Fbxo8 | 795.9569 | -0.04345 | 0.722716 | 0.966887 |
| ENSMUSG( | Fbxo9 | 3031.501 | 0.074478 | 0.364306 | 0.793796 | ENSMUSG( | Fbxo9 | 2603.65 | -0.09773 | 0.355625 | 0.912879 |
| ENSMUSG( | Fbxw10 | 30.90105 | 0.054757 | 0.873122 | 0.978753 | ENSMUSG( | Fbxw10 | 28.89827 | 0.126747 | 0.735799 | 0.967344 |
| ENSMUSG( | Fbxw11 | 4624.221 | 0.098962 | 0.232814 | 0.680928 | ENSMUSG( | Fbxw11 | 3941.34 | -0.0951 | 0.388501 | 0.917257 |
| ENSMUSG( | Fbxw17 | 208.6973 | -0.20872 | 0.192394 | 0.631451 | ENSMUSG( | Fbxw17 | 206.1058 | 0.047317 | 0.801368 | 0.97681 |
| ENSMUSG( | Fbxw2 | 2294.012 | 0.04775 | 0.562841 | 0.88962 | ENSMUSG( | Fbxw2 | 1991.409 | -0.09428 | 0.418904 | 0.917257 |
| ENSMUSG( | Fbxw4 | 999.9825 | 0.076718 | 0.418713 | 0.821966 | ENSMUSG( | Fbxw4 | 864.0739 | -0.07849 | 0.503934 | 0.927825 |
| ENSMUSG( | Fbxw5 | 4046.007 | 0.116668 | 0.151225 | 0.576198 | ENSMUSG( | Fbxw5 | 3431.739 | -0.08824 | 0.38477 | 0.917257 |
| ENSMUSG( | Fbxw7 | 4853.389 | 0.112169 | 0.258911 | 0.709814 | ENSMUSG( | Fbxw7 | 4525.625 | 0.177192 | 0.127236 | 0.817278 |
| ENSMUSG( | Fbxw8 | 2544.887 | 0.187846 | 0.020416 | 0.215588 | ENSMUSG( | Fbxw8 | 2072.688 | -0.13324 | 0.202539 | 0.864412 |
| ENSMUSG( | Fbxw9 | 562.3467 | 0.100116 | 0.456858 | 0.839183 | ENSMUSG( | Fbxw9 | 490.356 | -0.01944 | 0.899835 | 0.990558 |
| ENSMUSG( | Fcer1g | 690.6619 | -0.15126 | 0.220541 | 0.666079 | ENSMUSG( | Fcer1g | 654.6294 | -0.03051 | 0.832296 | 0.98075 |
| ENSMUSG( | Fcf1 | 330.0342 | -0.17792 | 0.164039 | 0.594869 | ENSMUSG( | Fcf1 | 313.9743 | -0.05159 | 0.72867 | 0.966913 |
| ENSMUSG( | Fcgr1 | 235.9747 | -0.39482 | 0.054447 | 0.359033 | ENSMUSG( | Fcgr1 | 266.136 | 0.251687 | 0.284486 | 0.892412 |
| ENSMUSG( | Fcgr2b | 274.979 | -0.44782 | 0.027518 | 0.254761 | ENSMUSG( | Fcgr2b | 303.3006 | 0.145225 | 0.485223 | 0.92541 |
| ENSMUSG( | Fcgr3 | 977.4467 | 0.086103 | 0.508052 | 0.867981 | ENSMUSG( | Fcgr3 | 977.3312 | 0.339648 | 0.014504 | 0.48395 |
| ENSMUSG( | Fcgr4 | 25.09842 | -1.25301 | 0.017866 | 0.200827 | ENSMUSG( | Fcgr4 | 25.36518 | -0.83677 | 0.115748 | 0.813384 |
| ENSMUSG( | Fcgrt | 1297.109 | -0.07567 | 0.547195 | 0.885118 | ENSMUSG( | Fcgrt | 1344.53 | 0.288012 | 0.074416 | 0.766789 |
| ENSMUSG( | Fcho1 | 3240.607 | 0.20277 | 0.079491 | 0.435569 | ENSMUSG( | Fcho1 | 2711.245 | -0.03818 | 0.780041 | 0.976106 |
| ENSMUSG( | Fcho2 | 1155.11 | 0.075734 | 0.54785 | 0.885153 | ENSMUSG( | Fcho2 | 1127.442 | 0.26259 | 0.045431 | 0.693959 |
| ENSMUSG( | Fchsd1 | 310.3503 | 0.179579 | 0.254027 | 0.704086 | ENSMUSG( | Fchsd1 | 255.3169 | -0.09718 | 0.600725 | 0.947749 |
| ENSMUSG( | Fchsd2 | 1765.399 | 0.047004 | 0.642498 | 0.916865 | ENSMUSG( | Fchsd2 | 1528.831 | -0.1009 | 0.360502 | 0.914223 |
| ENSMUSG( | Fcna | 21.83716 | -0.34311 | 0.379129 | 0.801757 | ENSMUSG( | Fcna | 20.90068 | -0.20046 | 0.639758 | 0.953903 |
| ENSMUSG( | Fcor | 98.65859 | -0.27646 | 0.196526 | 0.635942 | ENSMUSG( | Fcor | 84.48997 | -0.4866 | 0.051652 | 0.711848 |
| ENSMUSG( | Fcrl1 | 33.58055 | -0.3265 | 0.319385 | 0.765296 | ENSMUSG( | Fcrl1 | 41.24244 | 0.538205 | 0.123938 | 0.814691 |
| ENSMUSG( | Fcrl2 | 2461.18 | -0.06288 | 0.666894 | 0.92729 | ENSMUSG( | Fcrl2 | 2391.541 | 0.126217 | 0.487726 | 0.925427 |
| ENSMUSG( | Fcrlb | 13.10871 | 0.082293 | 0.868336 | NA | ENSMUSG( | Fcrlb | 11.15483 | -0.1376 | 0.809271 | NA |
| ENSMUSG( | Fcsk | 566.1425 | 0.277276 | 0.024922 | 0.242787 | ENSMUSG( | Fcsk | 497.6919 | 0.183353 | 0.2344 | 0.870096 |
| ENSMUSG( | Fdft1 | 3433.484 | -0.15535 | 0.108469 | 0.497337 | ENSMUSG( | Fdft1 | 3022.171 | -0.26108 | 0.019718 | 0.556192 |
| ENSMUSG( | Fdps | 2051.283 | -0.08284 | 0.402849 | 0.815656 | ENSMUSG( | Fdps | 1728.361 | -0.32791 | 0.009159 | 0.401114 |
| ENSMUSG( | Fdx1 | 290.195 | 0.036823 | 0.778139 | 0.956168 | ENSMUSG( | Fdx1 | 260.8635 | -0.00988 | 0.94991 | 0.994798 |
| ENSMUSG( | Fdx2 | 15.7812 | 0.562105 | 0.272205 | NA | ENSMUSG( | Fdx2 | 12.45305 | 0.214795 | 0.703326 | 0.962481 |
| ENSMUSG( | Fdxacb1 | 102.0316 | -0.36721 | 0.079065 | 0.43502 | ENSMUSG( | Fdxacb1 | 108.9536 | 0.123151 | 0.570059 | 0.941777 |
| ENSMUSG( | Fdxr | 623.1139 | 0.036218 | 0.733416 | 0.944956 | ENSMUSG( | Fdxr | 548.145 | -0.0708 | 0.596543 | 0.946541 |
| ENSMUSG( | Fech | 2521.44 | 0.016968 | 0.847713 | 0.972374 | ENSMUSG( | Fech | 2186.826 | -0.13055 | 0.267614 | 0.886134 |
| ENSMUSG( | Fem1a | 1772.492 | 0.06658 | 0.517263 | 0.873295 | ENSMUSG( | Fem1a | 1595.803 | 0.035101 | 0.781369 | 0.976106 |
| ENSMUSG( | Fem1al | 16.05903 | 0.075052 | 0.881322 | NA | ENSMUSG( | Fem1al | 13.63747 | -0.09886 | 0.860395 | 0.986137 |
| ENSMUSG( | Fem1b | 2719.201 | 0.256942 | 0.001583 | 0.04465 | ENSMUSG( | Fem1b | 2353.829 | 0.118253 | 0.246655 | 0.877904 |
| ENSMUSG( | Fem1c | 501.9913 | -0.06082 | 0.632654 | 0.914073 | ENSMUSG( | Fem1c | 459.2892 | -0.05108 | 0.70401 | 0.962625 |
| ENSMUSG( | Fen1 | 200.5056 | -0.07429 | 0.674761 | 0.929638 | ENSMUSG( | Fen1 | 177.5562 | -0.15544 | 0.417774 | 0.917257 |
| ENSMUSG( | Fendrr | 21.40405 | 0.430363 | 0.418999 | 0.821966 | ENSMUSG( | Fendrr | 16.99361 | 0.105117 | 0.839552 | 0.982077 |
| ENSMUSG( | Fer | 488.9383 | 0.158827 | 0.243564 | 0.690987 | ENSMUSG( | Fer | 403.6391 | -0.13008 | 0.350473 | 0.911652 |
| ENSMUSG( | Fermt1 | 22.85172 | 0.393673 | 0.432255 | 0.82768 | ENSMUSG( | Fermt1 | 16.63344 | -0.2033 | 0.685195 | 0.958678 |
| ENSMUSG( | Fermt2 | 7214.154 | 0.063468 | 0.602755 | 0.904887 | ENSMUSG( | Fermt2 | 5972.042 | -0.22179 | 0.080906 | 0.77681 |
| ENSMUSG( | Fermt3 | 297.9955 | -0.01999 | 0.887237 | 0.983326 | ENSMUSG( | Fermt3 | 284.3691 | 0.120766 | 0.464661 | 0.923329 |
| ENSMUSG( | Fes | 217.8209 | 0.06644 | 0.657732 | 0.92425 | ENSMUSG( | Fes | 217.2532 | 0.323579 | 0.153851 | 0.837724 |
| ENSMUSG( | Fez1 | 6276.096 | 0.032215 | 0.764951 | 0.954127 | ENSMUSG( | Fez1 | 5505.287 | -0.07978 | 0.518003 | 0.930939 |
| ENSMUSG( | Fez2 | 2321.639 | 0.061565 | 0.556607 | 0.887713 | ENSMUSG( | Fez2 | 1937.912 | -0.20064 | 0.108551 | 0.804969 |
| ENSMUSG( | Fezf2 | 752.4224 | 0.117642 | 0.334077 | 0.775728 | ENSMUSG( | Fezf2 | 677.6493 | 0.084209 | 0.488914 | 0.925573 |
| ENSMUSG( | Fgd1 | 818.2423 | 0.071755 | 0.540016 | 0.88315 | ENSMUSG( | Fgd1 | 709.3745 | -0.07143 | 0.599234 | 0.947045 |
| ENSMUSG( | Fgd2 | 476.6456 | 0.021818 | 0.855678 | 0.973813 | ENSMUSG( | Fgd2 | 470.7282 | 0.249603 | 0.153183 | 0.837724 |
| ENSMUSG( | Fgd3 | 867.5488 | -0.19655 | 0.092799 | 0.468183 | ENSMUSG( | Fgd3 | 727.8885 | -0.46763 | 0.000411 | 0.058872 |
| ENSMUSG( | Fgd4 | 963.795 | 0.043047 | 0.687976 | 0.93324 | ENSMUSG( | Fgd4 | 855.7766 | -0.03663 | 0.797263 | 0.97681 |
| ENSMUSG( | Fgd5 | 200.9203 | 0.510396 | 0.004882 | 0.092505 | ENSMUSG( | Fgd5 | 208.0922 | 0.825643 | 0.000869 | 0.099327 |
| ENSMUSG( | Fgd6 | 2353.82 | 0.242562 | 0.019861 | 0.212919 | ENSMUSG( | Fgd6 | 1869.9 | -0.14593 | 0.219531 | 0.866104 |
| ENSMUSG( | Fgf1 | 8822.348 | 0.037105 | 0.745261 | 0.949158 | ENSMUSG( | Fgf1 | 7245.04 | -0.27742 | 0.043727 | 0.690304 |
| ENSMUSG( | Fgf10 | 94.15238 | -0.18745 | 0.445682 | 0.833239 | ENSMUSG( | Fgf10 | 93.66645 | 0.069043 | 0.794227 | 0.97681 |
| ENSMUSG( | Fgf11 | 770.5436 | -0.04738 | 0.629513 | 0.913965 | ENSMUSG( | Fgf11 | 704.638 | -0.04069 | 0.748133 | 0.96984 |
| ENSMUSG( | Fgf12 | 2294.933 | -0.01948 | 0.846274 | 0.972226 | ENSMUSG( | Fgf12 | 2057.063 | -0.07139 | 0.624858 | 0.953234 |
| ENSMUSG( | Fgf13 | 6598.424 | 0.100483 | 0.333623 | 0.775602 | ENSMUSG( | Fgf13 | 5691.74 | -0.0569 | 0.659784 | 0.956205 |
| ENSMUSG( | Fgf14 | 444.7533 | -0.23771 | 0.047647 | 0.33359 | ENSMUSG( | Fgf14 | 433.2802 | -0.0416 | 0.784954 | 0.976285 |

| | | | | | |
|---|---|---|---|---|---|
| ENSMUSG( | 25.09842 | -1.25301 | 0.017866 | 0.200827 | Fcgr4 |
| ENSMUSG( | 1297.109 | -0.07567 | 0.547195 | 0.885118 | Fcgrt |
| ENSMUSG( | 3240.607 | 0.20277 | 0.079491 | 0.435569 | Fcho1 |
| ENSMUSG( | 1155.11 | 0.075734 | 0.54785 | 0.885153 | Fcho2 |
| ENSMUSG( | 310.3503 | 0.179579 | 0.254027 | 0.704086 | Fchsd1 |
| ENSMUSG( | 1765.399 | 0.047004 | 0.642498 | 0.916865 | Fchsd2 |
| ENSMUSG( | 21.83716 | -0.34311 | 0.379129 | 0.801757 | Fcna |
| ENSMUSG( | 98.65859 | -0.27646 | 0.196526 | 0.635942 | Fcor |
| ENSMUSG( | 33.58055 | -0.3265 | 0.319385 | 0.765296 | Fcrl1 |
| ENSMUSG( | 2461.18 | -0.06288 | 0.666894 | 0.92729 | Fcrl2 |
| ENSMUSG( | 13.10871 | 0.082293 | 0.868336 | NA | Fcrlb |
| ENSMUSG( | 566.1425 | 0.277276 | 0.024922 | 0.242787 | Fcsk |
| ENSMUSG( | 3433.484 | -0.15535 | 0.108469 | 0.497337 | Fdft1 |
| ENSMUSG( | 2051.283 | -0.08284 | 0.402849 | 0.815656 | Fdps |
| ENSMUSG( | 290.195 | 0.036823 | 0.778139 | 0.956168 | Fdx1 |
| ENSMUSG( | 15.7812 | 0.562105 | 0.272205 | NA | Fdx2 |
| ENSMUSG( | 102.0316 | -0.36721 | 0.079065 | 0.43502 | Fdxacb1 |
| ENSMUSG( | 623.1139 | 0.036218 | 0.733416 | 0.944956 | Fdxr |
| ENSMUSG( | 2521.44 | 0.016968 | 0.847713 | 0.972374 | Fech |
| ENSMUSG( | 1772.492 | 0.06658 | 0.517263 | 0.873295 | Fem1a |
| ENSMUSG( | 16.05903 | 0.075052 | 0.881322 | NA | Fem1al |
| ENSMUSG( | 2719.201 | 0.256942 | 0.001583 | 0.04465 | Fem1b |
| ENSMUSG( | 501.9913 | -0.06082 | 0.632654 | 0.914073 | Fem1c |
| ENSMUSG( | 200.5056 | -0.07429 | 0.674761 | 0.929638 | Fen1 |
| ENSMUSG( | 21.40405 | 0.430363 | 0.418999 | 0.821966 | Fendrr |
| ENSMUSG( | 488.9383 | 0.158827 | 0.243564 | 0.690987 | Fer |
| ENSMUSG( | 22.85172 | 0.393673 | 0.432255 | 0.82768 | Fermt1 |
| ENSMUSG( | 7214.154 | 0.063468 | 0.602755 | 0.904887 | Fermt2 |
| ENSMUSG( | 297.9955 | -0.01999 | 0.887237 | 0.983326 | Fermt3 |
| ENSMUSG( | 217.8209 | 0.06644 | 0.657732 | 0.92425 | Fes |
| ENSMUSG( | 6276.096 | 0.032215 | 0.764951 | 0.954127 | Fez1 |
| ENSMUSG( | 2321.639 | 0.061565 | 0.556607 | 0.887713 | Fez2 |
| ENSMUSG( | 752.4224 | 0.117642 | 0.334077 | 0.775728 | Fezf2 |
| ENSMUSG( | 818.2423 | 0.071755 | 0.540016 | 0.88315 | Fgd1 |
| ENSMUSG( | 476.6456 | 0.021818 | 0.855678 | 0.973813 | Fgd2 |
| ENSMUSG( | 867.5488 | -0.19655 | 0.092799 | 0.468183 | Fgd3 |
| ENSMUSG( | 963.795 | 0.043047 | 0.687976 | 0.93324 | Fgd4 |
| ENSMUSG( | 200.9203 | 0.510396 | 0.004882 | 0.092505 | Fgd5 |
| ENSMUSG( | 2353.82 | 0.242562 | 0.019861 | 0.212919 | Fgd6 |
| ENSMUSG( | 8822.348 | 0.037105 | 0.745261 | 0.949158 | Fgf1 |
| ENSMUSG( | 94.15238 | -0.18745 | 0.445682 | 0.833239 | Fgf10 |
| ENSMUSG( | 770.5436 | -0.04738 | 0.629513 | 0.913965 | Fgf11 |
| ENSMUSG( | 2294.933 | -0.01948 | 0.846274 | 0.972226 | Fgf12 |
| ENSMUSG( | 6598.424 | 0.100483 | 0.333623 | 0.775602 | Fgf13 |
| ENSMUSG( | 444.7533 | -0.23771 | 0.047647 | 0.33359 | Fgf14 |
| ENSMUSG( | 16.15709 | -0.52869 | 0.262983 | NA | Fgf16 |
| ENSMUSG( | 19.11494 | 0.016858 | 0.96792 | NA | Fgf17 |
| ENSMUSG( | 36.35705 | -0.08102 | 0.816228 | 0.965455 | Fgf18 |
| ENSMUSG( | 795.5345 | 0.318406 | 0.027155 | 0.253245 | Fgf2 |
| ENSMUSG( | 45.54921 | 0.026806 | 0.926135 | 0.991229 | Fgf20 |
| ENSMUSG( | 63.40149 | -0.11914 | 0.617114 | 0.910866 | Fgf22 |
| ENSMUSG( | 21.65529 | -1.30091 | 0.005819 | 0.102695 | Fgf3 |
| ENSMUSG( | 49.18608 | -0.27301 | 0.332104 | 0.775373 | Fgf5 |
| ENSMUSG( | 32.85525 | -0.13369 | 0.696043 | 0.935007 | Fgf7 |
| ENSMUSG( | 404.0703 | -0.39961 | 0.000721 | 0.025975 | Fgf9 |
| ENSMUSG( | 85.58033 | -0.06857 | 0.765756 | 0.954173 | Fgfbp1 |
| ENSMUSG( | 303.4247 | -0.52562 | 0.000417 | 0.01851 | Fgfbp3 |
| ENSMUSG( | 4797.023 | 0.007727 | 0.950314 | 0.993821 | Fgfr1 |
| ENSMUSG( | 1802.13 | 0.028055 | 0.756163 | 0.951479 | Fgfr1op2 |
| ENSMUSG( | 2845.306 | -0.16672 | 0.193014 | 0.632208 | Fgfr2 |
| ENSMUSG( | 10170.13 | 0.040839 | 0.758776 | 0.952113 | Fgfr3 |
| ENSMUSG( | 2068.421 | -0.20284 | 0.099369 | 0.48179 | Fgfrl1 |
| ENSMUSG( | 339.6771 | -0.02311 | 0.862004 | 0.975055 | Fggy |
| ENSMUSG( | 67.9636 | -0.1851 | 0.450571 | 0.83542 | Fgl2 |
| ENSMUSG( | 32.55458 | -1.06889 | 0.041881 | 0.315367 | Fgr |
| ENSMUSG( | 2175.118 | 0.1004 | 0.333426 | 0.775602 | Fh1 |
| ENSMUSG( | 225.4445 | -0.16003 | 0.260191 | 0.710129 | Fhad1 |
| ENSMUSG( | 154.9942 | 0.023786 | 0.900918 | 0.985633 | Fhdc1 |
| ENSMUSG( | 108.0561 | 0.135823 | 0.532826 | 0.879905 | Fhip1a |
| ENSMUSG( | 2251.367 | 0.128678 | 0.153843 | 0.580901 | Fhip1b |
| ENSMUSG( | 887.6351 | -0.01965 | 0.84205 | 0.971273 | Fhip2a |
| ENSMUSG( | 1554.017 | 0.163944 | 0.125757 | 0.53257 | Fhip2b |
| ENSMUSG( | 120.8387 | 0.120047 | 0.526613 | 0.877267 | Fhit |
| ENSMUSG( | 2884.004 | 0.056701 | 0.581262 | 0.897612 | Fhl1 |
| ENSMUSG( | 595.2126 | 0.044307 | 0.758126 | 0.951618 | Fhl2 |
| ENSMUSG( | 83.51982 | 0.224619 | 0.34287 | 0.779011 | Fhl3 |
| ENSMUSG( | 21.40291 | 0.286041 | 0.531103 | 0.8793 | Fhl4 |
| ENSMUSG( | 192.6188 | 0.040947 | 0.7867 | 0.958856 | Fhod1 |
| ENSMUSG( | 1284.179 | 0.124816 | 0.385389 | 0.804048 | Fhod3 |
| ENSMUSG( | 914.619 | 0.209475 | 0.227724 | 0.67477 | Fibcd1 |
| ENSMUSG( | 100.3128 | -0.1458 | 0.542068 | 0.883182 | Fibin |
| ENSMUSG( | 1181.256 | 0.030393 | 0.78052 | 0.957491 | Fibp |
| ENSMUSG( | 666.3818 | -0.30797 | 0.02047 | 0.21574 | Ficd |
| ENSMUSG( | 849.2328 | 0.090171 | 0.413197 | 0.819541 | Fig4 |
| ENSMUSG( | 47.93651 | -0.79238 | 0.007188 | 0.117052 | Fign |
| ENSMUSG( | 32.22937 | 0.123891 | 0.732435 | 0.944956 | Fignl1 |
| ENSMUSG( | 76.92494 | -0.06729 | 0.76072 | 0.952907 | Fignl2 |
| ENSMUSG( | 191.6222 | -0.60062 | 0.006579 | 0.111324 | Filip1 |
| ENSMUSG( | 101.13 | -0.29513 | 0.234037 | 0.681906 | Filip1l |
| ENSMUSG( | 1890.97 | 0.101722 | 0.24783 | 0.696522 | Fip1l1 |
| ENSMUSG( | 1993.103 | -0.07693 | 0.677752 | 0.930542 | Firre |
| ENSMUSG( | 28.50491 | -0.20213 | 0.569683 | 0.893041 | Firrm |
| ENSMUSG( | 5813.841 | -0.0107 | 0.90485 | 0.986401 | Fis1 |
| ENSMUSG( | 1043.02 | -0.24356 | 0.081253 | 0.440604 | Fitm2 |
| ENSMUSG( | 529.9866 | -0.00406 | 0.977392 | 0.997428 | Fiz1 |
| ENSMUSG( | 5426.05 | -0.2915 | 0.026288 | 0.247758 | Fjx1 |
| ENSMUSG( | 726.8229 | -0.35968 | 0.009988 | 0.142653 | Fkbp10 |
| ENSMUSG( | 30.12288 | -0.4899 | 0.186912 | 0.62471 | Fkbp11 |
| ENSMUSG( | 365.6093 | -0.1787 | 0.259771 | 0.709951 | Fkbp14 |
| ENSMUSG( | 1825.954 | 0.06436 | 0.45242 | 0.836145 | Fkbp15 |
| ENSMUSG( | 11324.44 | -0.00444 | 0.967453 | 0.99607 | Fkbp1a |
| ENSMUSG( | 946.3668 | 0.174087 | 0.148528 | 0.571966 | Fkbp1b |
| ENSMUSG( | 2697.673 | -0.19693 | 0.042771 | 0.316989 | Fkbp2 |
| ENSMUSG( | 2081.478 | 0.047966 | 0.644461 | 0.917773 | Fkbp3 |
| ENSMUSG( | 2421.402 | 0.203033 | 0.019463 | 0.210037 | Fkbp4 |
| ENSMUSG( | 1346.752 | 0.83506 | 2.54E-07 | 5.79E-05 | Fkbp5 |
| ENSMUSG( | 128.2165 | -0.32068 | 0.123676 | 0.530834 | Fkbp7 |
| ENSMUSG( | 11940.67 | 0.025982 | 0.751635 | 0.950317 | Fkbp8 |
| ENSMUSG( | 3192.378 | -0.14162 | 0.219625 | 0.664984 | Fkbp9 |
| ENSMUSG( | 113.3895 | 0.09308 | 0.665892 | 0.927054 | Fkbpl |
| ENSMUSG( | 1711.814 | -0.37112 | 0.001712 | 0.046884 | Fkrp |
| ENSMUSG( | 2312.986 | 0.004956 | 0.970225 | 0.996369 | Fktn |
| ENSMUSG( | 696.466 | 0.088469 | 0.445581 | 0.833239 | Flad1 |
| ENSMUSG( | 1390.767 | 0.017291 | 0.878423 | 0.981431 | Flcn |
| ENSMUSG( | 133.6172 | 0.122479 | 0.565937 | 0.89104 | Fli1 |
| ENSMUSG( | 1946.199 | 0.193558 | 0.023991 | 0.23699 | Flii |
| ENSMUSG( | 3441.944 | 0.112213 | 0.241496 | 0.689139 | Flna |
| ENSMUSG( | 2037.732 | 0.183759 | 0.11592 | 0.514015 | Flnb |
| ENSMUSG( | 141.5028 | 0.358943 | 0.045785 | 0.328083 | Flnc |
| ENSMUSG( | 3676.192 | 0.130371 | 0.337034 | 0.777008 | Flot1 |
| ENSMUSG( | 5414.808 | 0.047924 | 0.551775 | 0.886141 | Flot2 |
| ENSMUSG( | 719.0908 | -0.10097 | 0.603269 | 0.904887 | Flrt1 |
| ENSMUSG( | 1375.949 | -0.41781 | 0.028368 | 0.257483 | Flrt2 |
| ENSMUSG( | 604.4835 | -0.55266 | 0.000791 | 0.027736 | Flrt3 |
| ENSMUSG( | 2384.431 | 0.282356 | 0.441962 | 0.832762 | Flt1 |
| ENSMUSG( | 222.5589 | 0.184343 | 0.229903 | 0.677881 | Flt3 |
| ENSMUSG( | 56.82794 | -0.14511 | 0.585434 | 0.899171 | Flt3l |
| ENSMUSG( | 155.7481 | 0.423238 | 0.080762 | 0.439438 | Flt4 |
| ENSMUSG( | 295.6637 | -0.24619 | 0.138155 | 0.553165 | Flvcr1 |
| ENSMUSG( | 117.6702 | 0.256111 | 0.278778 | 0.728705 | Flvcr2 |
| ENSMUSG( | 6554.432 | 0.117044 | 0.319779 | 0.765305 | Flywch1 |
| ENSMUSG( | 237.2389 | -0.26806 | 0.103115 | 0.48855 | Flywch2 |
| ENSMUSG( | 741.5803 | 0.068649 | 0.537307 | 0.882504 | Fmc1 |
| ENSMUSG( | 379.6001 | 0.077866 | 0.613284 | 0.908943 | Fmn1 |
| ENSMUSG( | 6390.51 | -0.00304 | 0.983191 | 0.997554 | Fmn2 |
| ENSMUSG( | 4974.982 | 0.18452 | 0.055982 | 0.364712 | Fmnl1 |
| ENSMUSG( | 4977.958 | 0.027107 | 0.841689 | 0.971179 | Fmnl2 |
| ENSMUSG( | 709.6306 | -0.00118 | 0.992862 | 0.999327 | Fmnl3 |
| ENSMUSG( | 762.9219 | 0.223724 | 0.124442 | 0.531714 | Fmo1 |
| ENSMUSG( | 26.04124 | 0.803524 | 0.028977 | 0.259144 | Fmo2 |
| ENSMUSG( | 9.405733 | 0.85532 | 0.165886 | NA | Fmo4 |
| ENSMUSG( | 279.8101 | -0.04879 | 0.788787 | 0.959226 | Fmo5 |
| ENSMUSG( | 996.064 | -0.01043 | 0.937031 | 0.993415 | Fmod |
| ENSMUSG( | 1467.966 | -0.0069 | 0.957102 | 0.993874 | Fmr1 |
| ENSMUSG( | 3191.217 | 0.202467 | 0.183357 | 0.62047 | Fn1 |
| ENSMUSG( | 495.6081 | -0.10256 | 0.497129 | 0.861754 | Fn3k |
| ENSMUSG( | 913.2152 | 0.060961 | 0.544813 | 0.884712 | Fn3krp |
| ENSMUSG( | 6806.646 | -0.02582 | 0.737073 | 0.946839 | Fnbp1 |
| ENSMUSG( | 2598.772 | 0.132533 | 0.142996 | 0.561716 | Fnbp1l |
| ENSMUSG( | 1694.997 | -0.12546 | 0.218733 | 0.664559 | Fnbp4 |
| ENSMUSG( | 134.2431 | -0.11818 | 0.491859 | 0.860249 | Fndc1 |
| ENSMUSG( | 812.733 | -0.28386 | 0.018459 | 0.204231 | Fndc10 |
| ENSMUSG( | 1307.293 | 0.112884 | 0.232169 | 0.67996 | Fndc3a |
| ENSMUSG( | 561.0905 | 0.259378 | 0.017108 | 0.195465 | Fndc3b |
| ENSMUSG( | 4617.764 | -0.0089 | 0.920247 | 0.990247 | Fndc4 |
| ENSMUSG( | 4766.728 | -0.25758 | 0.009272 | 0.135856 | Fndc5 |
| ENSMUSG( | 10.7274 | -0.72238 | 0.252649 | NA | Fndc7 |
| ENSMUSG( | 102.9129 | -0.51228 | 0.018479 | 0.204273 | Fndc9 |

| | | | | | |
|---|---|---|---|---|---|
| ENSMUSG( | 1728.361 | -0.32791 | 0.009159 | 0.401114 | Fdps |
| ENSMUSG( | 260.8635 | -0.00988 | 0.94991 | 0.994798 | Fdx1 |
| ENSMUSG( | 12.45305 | 0.214795 | 0.703326 | 0.962481 | Fdx2 |
| ENSMUSG( | 108.9536 | 0.123151 | 0.570059 | 0.941777 | Fdxacb1 |
| ENSMUSG( | 548.145 | -0.0708 | 0.596543 | 0.946541 | Fdxr |
| ENSMUSG( | 2186.826 | -0.13055 | 0.267614 | 0.886134 | Fech |
| ENSMUSG( | 1595.803 | 0.035101 | 0.781369 | 0.976106 | Fem1a |
| ENSMUSG( | 13.63747 | -0.09886 | 0.860395 | 0.986137 | Fem1al |
| ENSMUSG( | 2353.829 | 0.118253 | 0.246655 | 0.877904 | Fem1b |
| ENSMUSG( | 459.2892 | -0.05108 | 0.70401 | 0.962625 | Fem1c |
| ENSMUSG( | 177.5562 | -0.15544 | 0.417774 | 0.917257 | Fen1 |
| ENSMUSG( | 16.99361 | 0.105117 | 0.839552 | 0.982077 | Fendrr |
| ENSMUSG( | 403.6391 | -0.13008 | 0.350473 | 0.911652 | Fer |
| ENSMUSG( | 16.63344 | -0.2033 | 0.685195 | 0.958678 | Fermt1 |
| ENSMUSG( | 5972.042 | -0.22179 | 0.080906 | 0.77681 | Fermt2 |
| ENSMUSG( | 284.3691 | 0.120766 | 0.464661 | 0.923329 | Fermt3 |
| ENSMUSG( | 217.2532 | 0.323579 | 0.153851 | 0.837724 | Fes |
| ENSMUSG( | 5505.287 | -0.07978 | 0.518003 | 0.930939 | Fez1 |
| ENSMUSG( | 1937.912 | -0.20064 | 0.108551 | 0.804969 | Fez2 |
| ENSMUSG( | 677.6493 | 0.084209 | 0.488914 | 0.925573 | Fezf2 |
| ENSMUSG( | 709.3745 | -0.07143 | 0.599234 | 0.947045 | Fgd1 |
| ENSMUSG( | 470.7282 | 0.249603 | 0.153183 | 0.837724 | Fgd2 |
| ENSMUSG( | 727.8885 | -0.46763 | 0.000411 | 0.058872 | Fgd3 |
| ENSMUSG( | 855.7766 | -0.03663 | 0.797263 | 0.97681 | Fgd4 |
| ENSMUSG( | 208.0922 | 0.825643 | 0.000869 | 0.099327 | Fgd5 |
| ENSMUSG( | 1869.9 | -0.14593 | 0.219531 | 0.866104 | Fgd6 |
| ENSMUSG( | 7245.04 | -0.27742 | 0.043727 | 0.690304 | Fgf1 |
| ENSMUSG( | 93.66645 | 0.069043 | 0.794227 | 0.97681 | Fgf10 |
| ENSMUSG( | 704.638 | -0.04069 | 0.748133 | 0.96984 | Fgf11 |
| ENSMUSG( | 2057.063 | -0.07139 | 0.624858 | 0.953234 | Fgf12 |
| ENSMUSG( | 5691.74 | -0.0569 | 0.659784 | 0.956205 | Fgf13 |
| ENSMUSG( | 433.2802 | -0.0416 | 0.784954 | 0.976285 | Fgf14 |
| ENSMUSG( | 14.4325 | -0.5699 | 0.292365 | 0.897854 | Fgf16 |
| ENSMUSG( | 16.95164 | -0.0742 | 0.875262 | 0.989644 | Fgf17 |
| ENSMUSG( | 40.30117 | 0.45328 | 0.190837 | 0.857813 | Fgf18 |
| ENSMUSG( | 677.2926 | 0.129753 | 0.465399 | 0.923329 | Fgf2 |
| ENSMUSG( | 42.77743 | 0.11045 | 0.720198 | 0.966887 | Fgf20 |
| ENSMUSG( | 66.51155 | 0.285592 | 0.2636 | 0.886134 | Fgf22 |
| ENSMUSG( | 21.26335 | -0.96938 | 0.031015 | 0.629564 | Fgf3 |
| ENSMUSG( | 53.42032 | 0.260079 | 0.406779 | 0.917257 | Fgf5 |
| ENSMUSG( | 32.60713 | 0.105368 | 0.784746 | 0.976231 | Fgf7 |
| ENSMUSG( | 396.2453 | -0.17294 | 0.259843 | 0.883971 | Fgf9 |
| ENSMUSG( | 74.59315 | -0.19166 | 0.428292 | 0.917257 | Fgfbp1 |
| ENSMUSG( | 349.4185 | 0.186521 | 0.205589 | 0.864412 | Fgfbp3 |
| ENSMUSG( | 4796.058 | 0.267063 | 0.069229 | 0.755467 | Fgfr1 |
| ENSMUSG( | 1655.12 | 0.049428 | 0.654312 | 0.955015 | Fgfr1op2 |
| ENSMUSG( | 2844.341 | 0.107274 | 0.45874 | 0.923329 | Fgfr2 |
| ENSMUSG( | 10141.5 | 0.289952 | 0.047782 | 0.703953 | Fgfr3 |
| ENSMUSG( | 1876.433 | -0.21475 | 0.118316 | 0.813384 | Fgfrl1 |
| ENSMUSG( | 302.7088 | -0.09285 | 0.54626 | 0.938063 | Fggy |
| ENSMUSG( | 70.34554 | 0.191382 | 0.461478 | 0.923329 | Fgl2 |
| ENSMUSG( | 26.61643 | -1.69168 | 0.003551 | 0.238323 | Fgr |
| ENSMUSG( | 1783.437 | -0.21205 | 0.094285 | 0.795735 | Fh1 |
| ENSMUSG( | 188.5012 | -0.44124 | 0.009913 | 0.407479 | Fhad1 |
| ENSMUSG( | 133.8603 | -0.12383 | 0.570732 | 0.941973 | Fhdc1 |
| ENSMUSG( | 96.42146 | 0.084906 | 0.728412 | 0.966913 | Fhip1a |
| ENSMUSG( | 2053.53 | 0.133426 | 0.212055 | 0.864412 | Fhip1b |
| ENSMUSG( | 842.4796 | 0.097398 | 0.402634 | 0.917257 | Fhip2a |
| ENSMUSG( | 1371.954 | 0.074524 | 0.555069 | 0.939565 | Fhip2b |
| ENSMUSG( | 102.5288 | -0.09925 | 0.662037 | 0.956205 | Fhit |
| ENSMUSG( | 2542.09 | -0.04266 | 0.763709 | 0.974 | Fhl1 |
| ENSMUSG( | 507.3471 | -0.15563 | 0.339929 | 0.907972 | Fhl2 |
| ENSMUSG( | 66.80474 | -0.12942 | 0.657304 | 0.955725 | Fhl3 |
| ENSMUSG( | 20.58358 | 0.4406 | 0.302194 | 0.898538 | Fhl4 |
| ENSMUSG( | 186.7922 | 0.222355 | 0.253308 | 0.88018 | Fhod1 |
| ENSMUSG( | 1199.115 | 0.197805 | 0.23378 | 0.868899 | Fhod3 |
| ENSMUSG( | 871.0092 | 0.333438 | 0.063081 | 0.736482 | Fibcd1 |
| ENSMUSG( | 113.6121 | 0.460822 | 0.053164 | 0.715818 | Fibin |
| ENSMUSG( | 1076.672 | 0.03153 | 0.822656 | 0.979198 | Fibp |
| ENSMUSG( | 697.6997 | 0.111532 | 0.463871 | 0.923329 | Ficd |
| ENSMUSG( | 754.8793 | 0.019449 | 0.883503 | 0.989644 | Fig4 |
| ENSMUSG( | 56.09716 | 0.033297 | 0.906128 | 0.991235 | Fign |
| ENSMUSG( | 27.36007 | -0.09108 | 0.815646 | 0.979198 | Fignl1 |
| ENSMUSG( | 82.15629 | 0.378204 | 0.105869 | 0.804969 | Fignl2 |
| ENSMUSG( | 199.9459 | -0.15318 | 0.476533 | 0.923329 | Filip1 |
| ENSMUSG( | 111.2024 | 0.251645 | 0.290658 | 0.897019 | Filip1l |
| ENSMUSG( | 1663.793 | -0.00065 | 0.996015 | 0.999163 | Fip1l1 |
| ENSMUSG( | 1997.36 | 0.192424 | 0.3828 | 0.917257 | Firre |
| ENSMUSG( | 27.32965 | -0.02702 | 0.944613 | 0.994798 | Firrm |
| ENSMUSG( | 4967.562 | -0.20621 | 0.115374 | 0.813375 | Fis1 |
| ENSMUSG( | 1065.138 | 0.099471 | 0.526503 | 0.934374 | Fitm2 |
| ENSMUSG( | 466.8797 | -0.10517 | 0.514366 | 0.930939 | Fiz1 |
| ENSMUSG( | 5444.206 | 0.00472 | 0.974563 | 0.996416 | Fjx1 |
| ENSMUSG( | 739.5965 | -0.01252 | 0.933454 | 0.99306 | Fkbp10 |
| ENSMUSG( | 29.67391 | -0.24475 | 0.530012 | 0.934795 | Fkbp11 |
| ENSMUSG( | 381.8261 | 0.218178 | 0.196349 | 0.862161 | Fkbp14 |
| ENSMUSG( | 1577.178 | -0.08897 | 0.415446 | 0.917257 | Fkbp15 |
| ENSMUSG( | 9724.525 | -0.18365 | 0.219241 | 0.866104 | Fkbp1a |
| ENSMUSG( | 816.2806 | 0.016366 | 0.910836 | 0.992196 | Fkbp1b |
| ENSMUSG( | 2540.726 | -0.09875 | 0.476244 | 0.923329 | Fkbp2 |
| ENSMUSG( | 1851.839 | -0.02331 | 0.857415 | 0.985884 | Fkbp3 |
| ENSMUSG( | 2082.19 | 0.041127 | 0.711317 | 0.964363 | Fkbp4 |
| ENSMUSG( | 1225.972 | 0.833462 | 8.07E-08 | 5.44E-05 | Fkbp5 |
| ENSMUSG( | 122.526 | -0.15613 | 0.489237 | 0.925673 | Fkbp7 |
| ENSMUSG( | 10278.4 | -0.1425 | 0.189118 | 0.85698 | Fkbp8 |
| ENSMUSG( | 3041.127 | -0.00743 | 0.954786 | 0.994798 | Fkbp9 |
| ENSMUSG( | 93.23475 | -0.19688 | 0.422222 | 0.917257 | Fkbpl |
| ENSMUSG( | 1743.476 | -0.02224 | 0.871505 | 0.988779 | Fkrp |
| ENSMUSG( | 2135.441 | 0.044873 | 0.752236 | 0.970461 | Fktn |
| ENSMUSG( | 611.981 | -0.00977 | 0.948067 | 0.994798 | Flad1 |
| ENSMUSG( | 1338.298 | 0.171347 | 0.257729 | 0.883401 | Flcn |
| ENSMUSG( | 133.4262 | 0.377897 | 0.098483 | 0.798438 | Fli1 |
| ENSMUSG( | 1691.373 | 0.064774 | 0.552725 | 0.939565 | Flii |
| ENSMUSG( | 3233.455 | 0.196597 | 0.204644 | 0.864412 | Flna |
| ENSMUSG( | 1927.242 | 0.28616 | 0.061432 | 0.736111 | Flnb |
| ENSMUSG( | 133.9463 | 0.468098 | 0.021965 | 0.575341 | Flnc |
| ENSMUSG( | 3047.283 | -0.14615 | 0.347485 | 0.911652 | Flot1 |
| ENSMUSG( | 4617.965 | -0.14659 | 0.162091 | 0.837724 | Flot2 |
| ENSMUSG( | 737.5864 | 0.24248 | 0.253148 | 0.88018 | Flrt1 |
| ENSMUSG( | 1685.034 | 0.438507 | 0.040205 | 0.681706 | Flrt2 |
| ENSMUSG( | 812.8552 | 0.570986 | 0.004401 | 0.269982 | Flrt3 |
| ENSMUSG( | 2638.279 | 0.766777 | 0.102615 | 0.803503 | Flt1 |
| ENSMUSG( | 199.7739 | 0.151272 | 0.380548 | 0.917202 | Flt3 |
| ENSMUSG( | 60.64671 | 0.30386 | 0.375344 | 0.915491 | Flt3l |
| ENSMUSG( | 171.2667 | 0.886221 | 0.003846 | 0.247715 | Flt4 |
| ENSMUSG( | 323.9156 | 0.29125 | 0.105854 | 0.804969 | Flvcr1 |
| ENSMUSG( | 143.0813 | 0.962759 | 0.001177 | 0.12275 | Flvcr2 |
| ENSMUSG( | 5692.009 | -0.01969 | 0.872306 | 0.988944 | Flywch1 |
| ENSMUSG( | 216.1085 | -0.28209 | 0.154262 | 0.837724 | Flywch2 |
| ENSMUSG( | 584.6787 | -0.38016 | 0.008602 | 0.391682 | Fmc1 |
| ENSMUSG( | 358.512 | 0.185056 | 0.293338 | 0.898538 | Fmn1 |
| ENSMUSG( | 5800.123 | -0.01328 | 0.922807 | 0.992644 | Fmn2 |
| ENSMUSG( | 4213.409 | -0.02128 | 0.85053 | 0.985541 | Fmnl1 |
| ENSMUSG( | 4378.289 | -0.07803 | 0.613064 | 0.951154 | Fmnl2 |
| ENSMUSG( | 723.5761 | 0.314931 | 0.07884 | 0.774307 | Fmnl3 |
| ENSMUSG( | 668.191 | 0.112311 | 0.446363 | 0.921734 | Fmo1 |
| ENSMUSG( | 32.60664 | 1.491412 | 6.97E-05 | 0.016495 | Fmo2 |
| ENSMUSG( | 245.3857 | -0.16387 | 0.321916 | 0.904023 | Fmo5 |
| ENSMUSG( | 936.3198 | 0.082607 | 0.537185 | 0.937216 | Fmod |
| ENSMUSG( | 1400.958 | 0.120315 | 0.436242 | 0.919879 | Fmr1 |
| ENSMUSG( | 3448.968 | 0.638796 | 0.001013 | 0.109511 | Fn1 |
| ENSMUSG( | 445.0932 | -0.1495 | 0.41276 | 0.917257 | Fn3k |
| ENSMUSG( | 773.7831 | -0.15737 | 0.231491 | 0.868507 | Fn3krp |
| ENSMUSG( | 5816.663 | -0.21846 | 0.023197 | 0.585577 | Fnbp1 |
| ENSMUSG( | 2424.112 | 0.194182 | 0.121915 | 0.813384 | Fnbp1l |
| ENSMUSG( | 1625.792 | 0.027962 | 0.826475 | 0.979922 | Fnbp4 |
| ENSMUSG( | 149.9047 | 0.42222 | 0.044405 | 0.693697 | Fndc1 |
| ENSMUSG( | 831.8101 | 0.069203 | 0.620187 | 0.952896 | Fndc10 |
| ENSMUSG( | 1206.627 | 0.14644 | 0.217554 | 0.865589 | Fndc3a |
| ENSMUSG( | 487.5785 | 0.133306 | 0.300764 | 0.898538 | Fndc3b |
| ENSMUSG( | 4076.603 | -0.10125 | 0.320566 | 0.903887 | Fndc4 |
| ENSMUSG( | 4423.187 | -0.20036 | 0.100086 | 0.801311 | Fndc5 |
| ENSMUSG( | 9.133814 | -1.08792 | 0.112525 | NA | Fndc7 |
| ENSMUSG( | 104.7772 | -0.16901 | 0.487181 | 0.92541 | Fndc9 |
| ENSMUSG( | 915.0943 | 0.12005 | 0.31993 | 0.903771 | Fnip1 |
| ENSMUSG( | 487.999 | 0.011643 | 0.928689 | 0.99306 | Fnip2 |
| ENSMUSG( | 1749.17 | -0.13022 | 0.298465 | 0.898538 | Fnta |
| ENSMUSG( | 700.757 | -0.10569 | 0.404495 | 0.917257 | Fntb |
| ENSMUSG( | 1099.961 | 0.018091 | 0.91 | 0.991958 | Focad |
| ENSMUSG( | 410.6144 | 0.125377 | 0.426371 | 0.917257 | Folh1 |
| ENSMUSG( | 138.5538 | 0.546117 | 0.332152 | 0.907907 | Folr1 |
| ENSMUSG( | 28.30159 | -0.53092 | 0.199802 | 0.863016 | Folr2 |
| ENSMUSG( | 461.9455 | -0.96707 | 0.052869 | 0.71525 | Fos |
| ENSMUSG( | 225.7317 | -0.26382 | 0.317418 | 0.902486 | Fosb |
| ENSMUSG( | 768.9868 | -0.06324 | 0.63256 | 0.953698 | Fosl2 |
| ENSMUSG( | 223.9209 | 0.313497 | 0.101127 | 0.801456 | Foxc1 |
| ENSMUSG( | 123.8643 | 0.396902 | 0.047212 | 0.700856 | Foxc2 |
| ENSMUSG( | 105.7541 | 0.425682 | 0.041773 | 0.690304 | Foxd1 |

| | | | | | | | | | | | |
|---|---|---|---|---|---|---|---|---|---|---|---|
| ENSMUSG( | Fgf16 | 16.15709 | -0.52869 | 0.262983 | NA | ENSMUSG( | Fgf16 | 14.4325 | -0.5699 | 0.292365 | 0.897854 |
| ENSMUSG( | Fgf17 | 19.11494 | 0.016858 | 0.96792 | NA | ENSMUSG( | Fgf17 | 16.95164 | -0.0742 | 0.875262 | 0.989644 |
| ENSMUSG( | Fgf18 | 36.35705 | -0.08102 | 0.816228 | 0.965455 | ENSMUSG( | Fgf18 | 40.30117 | 0.45328 | 0.190837 | 0.857813 |
| ENSMUSG( | Fgf2 | 795.5345 | 0.318406 | 0.027155 | 0.253245 | ENSMUSG( | Fgf2 | 677.2926 | 0.129753 | 0.465399 | 0.923329 |
| ENSMUSG( | Fgf20 | 45.54921 | 0.026806 | 0.926135 | 0.991229 | ENSMUSG( | Fgf20 | 42.77743 | 0.11045 | 0.720198 | 0.966887 |
| ENSMUSG( | Fgf22 | 63.40149 | -0.11914 | 0.617114 | 0.910866 | ENSMUSG( | Fgf22 | 66.51155 | 0.285592 | 0.2636 | 0.886134 |
| ENSMUSG( | Fgf3 | 21.65529 | -1.30091 | 0.005819 | 0.102695 | ENSMUSG( | Fgf3 | 21.26335 | -0.96938 | 0.031015 | 0.629564 |
| ENSMUSG( | Fgf5 | 49.18608 | -0.27301 | 0.332104 | 0.775373 | ENSMUSG( | Fgf5 | 53.42032 | 0.260079 | 0.406779 | 0.917257 |
| ENSMUSG( | Fgf7 | 32.85525 | -0.13369 | 0.696043 | 0.935007 | ENSMUSG( | Fgf7 | 32.60713 | 0.105368 | 0.784746 | 0.976231 |
| ENSMUSG( | Fgf9 | 404.0703 | -0.39961 | 0.000721 | 0.025975 | ENSMUSG( | Fgf9 | 396.2453 | -0.17294 | 0.259843 | 0.883971 |
| ENSMUSG( | Fgfbp1 | 85.58033 | -0.06857 | 0.765756 | 0.954173 | ENSMUSG( | Fgfbp1 | 74.59315 | -0.19166 | 0.428292 | 0.917257 |
| ENSMUSG( | Fgfbp3 | 303.4247 | -0.52562 | 0.000417 | 0.01851 | ENSMUSG( | Fgfbp3 | 349.4185 | 0.186521 | 0.205589 | 0.864412 |
| ENSMUSG( | Fgfr1 | 4797.023 | 0.007727 | 0.950314 | 0.993821 | ENSMUSG( | Fgfr1 | 4796.058 | 0.267063 | 0.069229 | 0.755467 |
| ENSMUSG( | Fgfr1op2 | 1802.13 | 0.028055 | 0.756163 | 0.951479 | ENSMUSG( | Fgfr1op2 | 1655.12 | 0.049428 | 0.654312 | 0.955015 |
| ENSMUSG( | Fgfr2 | 2845.306 | -0.16672 | 0.193014 | 0.632208 | ENSMUSG( | Fgfr2 | 2844.341 | 0.107274 | 0.45874 | 0.923329 |
| ENSMUSG( | Fgfr3 | 10170.13 | 0.040839 | 0.758776 | 0.952113 | ENSMUSG( | Fgfr3 | 10141.5 | 0.289952 | 0.047782 | 0.703953 |
| ENSMUSG( | Fgfrl1 | 2068.421 | -0.20284 | 0.099369 | 0.48179 | ENSMUSG( | Fgfrl1 | 1876.433 | -0.21475 | 0.118316 | 0.813384 |
| ENSMUSG( | Fggy | 339.6771 | -0.02311 | 0.862004 | 0.975055 | ENSMUSG( | Fggy | 302.7088 | -0.09285 | 0.54626 | 0.938063 |
| ENSMUSG( | Fgl2 | 67.9636 | -0.1851 | 0.450571 | 0.83542 | ENSMUSG( | Fgl2 | 70.34554 | 0.191382 | 0.461478 | 0.923329 |
| ENSMUSG( | Fgr | 32.55458 | -1.06889 | 0.041881 | 0.315367 | ENSMUSG( | Fgr | 26.61643 | -1.69168 | 0.003551 | 0.238323 |
| ENSMUSG( | Fh1 | 2175.118 | 0.1004 | 0.333426 | 0.775602 | ENSMUSG( | Fh1 | 1783.437 | -0.21205 | 0.094285 | 0.795735 |
| ENSMUSG( | Fhad1 | 225.4445 | -0.16003 | 0.260191 | 0.710129 | ENSMUSG( | Fhad1 | 188.5012 | -0.44124 | 0.009913 | 0.407479 |
| ENSMUSG( | Fhdc1 | 154.9942 | 0.023786 | 0.900918 | 0.985633 | ENSMUSG( | Fhdc1 | 133.8603 | -0.12383 | 0.570732 | 0.941973 |
| ENSMUSG( | Fhip1a | 108.0561 | 0.135823 | 0.532826 | 0.879905 | ENSMUSG( | Fhip1a | 96.42146 | 0.084906 | 0.728412 | 0.966913 |
| ENSMUSG( | Fhip1b | 2251.367 | 0.128678 | 0.153843 | 0.580901 | ENSMUSG( | Fhip1b | 2053.53 | 0.133426 | 0.212055 | 0.864412 |
| ENSMUSG( | Fhip2a | 887.6351 | -0.01965 | 0.84205 | 0.971273 | ENSMUSG( | Fhip2a | 842.4796 | 0.097398 | 0.402634 | 0.917257 |
| ENSMUSG( | Fhip2b | 1554.017 | 0.163944 | 0.125757 | 0.53257 | ENSMUSG( | Fhip2b | 1371.954 | 0.074524 | 0.555069 | 0.939565 |
| ENSMUSG( | Fhit | 120.8387 | 0.120047 | 0.526613 | 0.877267 | ENSMUSG( | Fhit | 102.5288 | -0.09925 | 0.662037 | 0.956205 |
| ENSMUSG( | Fhl1 | 2884.004 | 0.056701 | 0.581262 | 0.897612 | ENSMUSG( | Fhl1 | 2542.09 | -0.04266 | 0.763709 | 0.974 |
| ENSMUSG( | Fhl2 | 595.2126 | 0.044307 | 0.758126 | 0.951618 | ENSMUSG( | Fhl2 | 507.3471 | -0.15563 | 0.339929 | 0.907972 |
| ENSMUSG( | Fhl3 | 83.51982 | 0.224619 | 0.34287 | 0.779011 | ENSMUSG( | Fhl3 | 66.80474 | -0.12942 | 0.657304 | 0.955725 |
| ENSMUSG( | Fhl4 | 21.40291 | 0.286041 | 0.531103 | 0.8793 | ENSMUSG( | Fhl4 | 20.58358 | 0.4406 | 0.302194 | 0.898538 |
| ENSMUSG( | Fhod1 | 192.6188 | 0.040947 | 0.7867 | 0.958856 | ENSMUSG( | Fhod1 | 186.7922 | 0.222355 | 0.253308 | 0.88018 |
| ENSMUSG( | Fhod3 | 1284.179 | 0.124816 | 0.385389 | 0.804048 | ENSMUSG( | Fhod3 | 1199.115 | 0.197805 | 0.23378 | 0.868899 |
| ENSMUSG( | Fibcd1 | 914.619 | 0.209475 | 0.227724 | 0.67477 | ENSMUSG( | Fibcd1 | 871.0092 | 0.333438 | 0.063081 | 0.736482 |
| ENSMUSG( | Fibin | 100.3128 | -0.1458 | 0.542068 | 0.883182 | ENSMUSG( | Fibin | 113.6121 | 0.460822 | 0.053164 | 0.715818 |
| ENSMUSG( | Fibp | 1181.256 | 0.030393 | 0.78052 | 0.957491 | ENSMUSG( | Fibp | 1076.672 | 0.03153 | 0.822656 | 0.979198 |
| ENSMUSG( | Ficd | 666.3818 | -0.30797 | 0.02047 | 0.21574 | ENSMUSG( | Ficd | 697.6997 | 0.111532 | 0.463871 | 0.923329 |
| ENSMUSG( | Fig4 | 849.2328 | 0.090171 | 0.413197 | 0.819541 | ENSMUSG( | Fig4 | 754.8793 | 0.019449 | 0.883503 | 0.989644 |
| ENSMUSG( | Fign | 47.93651 | -0.79238 | 0.007188 | 0.117052 | ENSMUSG( | Fign | 56.09716 | 0.033297 | 0.906128 | 0.991235 |
| ENSMUSG( | Fignl1 | 32.22937 | 0.123891 | 0.732435 | 0.944956 | ENSMUSG( | Fignl1 | 27.36007 | -0.09108 | 0.815646 | 0.979198 |
| ENSMUSG( | Fignl2 | 76.92494 | -0.06729 | 0.76072 | 0.952907 | ENSMUSG( | Fignl2 | 82.15629 | 0.378204 | 0.105869 | 0.804969 |
| ENSMUSG( | Filip1 | 191.6222 | -0.60062 | 0.006579 | 0.111324 | ENSMUSG( | Filip1 | 199.9459 | -0.15318 | 0.476533 | 0.923329 |
| ENSMUSG( | Filip1l | 101.13 | -0.29513 | 0.234037 | 0.681906 | ENSMUSG( | Filip1l | 111.2024 | 0.251645 | 0.290658 | 0.897019 |
| ENSMUSG( | Fip1l1 | 1890.97 | 0.101722 | 0.24783 | 0.696522 | ENSMUSG( | Fip1l1 | 1663.793 | -0.00065 | 0.996015 | 0.999163 |
| ENSMUSG( | Firre | 1993.103 | -0.07693 | 0.677752 | 0.930542 | ENSMUSG( | Firre | 1997.36 | 0.192424 | 0.3828 | 0.917257 |
| ENSMUSG( | Firrm | 28.50491 | -0.20213 | 0.569683 | 0.893041 | ENSMUSG( | Firrm | 27.32965 | -0.02702 | 0.944613 | 0.994798 |
| ENSMUSG( | Fis1 | 5813.841 | -0.0107 | 0.90485 | 0.986401 | ENSMUSG( | Fis1 | 4967.562 | -0.20621 | 0.115374 | 0.813375 |
| ENSMUSG( | Fitm2 | 1043.02 | -0.24356 | 0.081253 | 0.440604 | ENSMUSG( | Fitm2 | 1065.138 | 0.099471 | 0.526503 | 0.934374 |
| ENSMUSG( | Fiz1 | 529.9866 | -0.00406 | 0.977392 | 0.997428 | ENSMUSG( | Fiz1 | 466.8797 | -0.10517 | 0.514366 | 0.930939 |
| ENSMUSG( | Fjx1 | 5426.05 | -0.2915 | 0.026288 | 0.247758 | ENSMUSG( | Fjx1 | 5444.206 | 0.00472 | 0.974563 | 0.996416 |
| ENSMUSG( | Fkbp10 | 726.8229 | -0.35968 | 0.009988 | 0.142653 | ENSMUSG( | Fkbp10 | 739.5965 | -0.01252 | 0.933454 | 0.99306 |
| ENSMUSG( | Fkbp11 | 30.12288 | -0.4899 | 0.186912 | 0.62471 | ENSMUSG( | Fkbp11 | 29.67391 | -0.24475 | 0.530012 | 0.934795 |
| ENSMUSG( | Fkbp14 | 365.6093 | -0.1787 | 0.259771 | 0.709951 | ENSMUSG( | Fkbp14 | 381.8261 | 0.218178 | 0.196349 | 0.862161 |
| ENSMUSG( | Fkbp15 | 1825.954 | 0.06436 | 0.45242 | 0.836145 | ENSMUSG( | Fkbp15 | 1577.178 | -0.08897 | 0.415446 | 0.917257 |
| ENSMUSG( | Fkbp1a | 11324.44 | -0.00444 | 0.967453 | 0.99607 | ENSMUSG( | Fkbp1a | 9724.525 | -0.18365 | 0.219241 | 0.866104 |
| ENSMUSG( | Fkbp1b | 946.3668 | 0.174087 | 0.148528 | 0.571966 | ENSMUSG( | Fkbp1b | 816.2806 | 0.016366 | 0.910836 | 0.992196 |
| ENSMUSG( | Fkbp2 | 2697.673 | -0.19693 | 0.042771 | 0.316989 | ENSMUSG( | Fkbp2 | 2540.726 | -0.09875 | 0.476244 | 0.923329 |
| ENSMUSG( | Fkbp3 | 2081.478 | 0.047966 | 0.644461 | 0.917773 | ENSMUSG( | Fkbp3 | 1851.839 | -0.02331 | 0.857415 | 0.985884 |
| ENSMUSG( | Fkbp4 | 2421.402 | 0.203033 | 0.019463 | 0.210037 | ENSMUSG( | Fkbp4 | 2082.19 | 0.041127 | 0.711317 | 0.964363 |
| ENSMUSG( | Fkbp5 | 1346.752 | 0.83506 | 2.54E-07 | 5.79E-05 | ENSMUSG( | Fkbp5 | 1225.972 | 0.833462 | 8.07E-08 | 5.44E-05 |
| ENSMUSG( | Fkbp7 | 128.2165 | -0.32068 | 0.123676 | 0.530834 | ENSMUSG( | Fkbp7 | 122.526 | -0.15613 | 0.489237 | 0.925673 |
| ENSMUSG( | Fkbp8 | 11940.67 | 0.025982 | 0.751635 | 0.950317 | ENSMUSG( | Fkbp8 | 10278.4 | -0.1425 | 0.189118 | 0.85698 |
| ENSMUSG( | Fkbp9 | 3192.378 | -0.14162 | 0.219625 | 0.664984 | ENSMUSG( | Fkbp9 | 3041.127 | -0.00743 | 0.954786 | 0.994798 |
| ENSMUSG( | Fkbpl | 113.3895 | 0.09308 | 0.665892 | 0.927054 | ENSMUSG( | Fkbpl | 93.23475 | -0.19688 | 0.422222 | 0.917257 |
| ENSMUSG( | Fkrp | 1711.814 | -0.37112 | 0.001712 | 0.046884 | ENSMUSG( | Fkrp | 1743.476 | -0.02224 | 0.871505 | 0.988779 |
| ENSMUSG( | Fktn | 2312.986 | 0.004956 | 0.970225 | 0.996369 | ENSMUSG( | Fktn | 2135.441 | 0.044873 | 0.752236 | 0.970461 |
| ENSMUSG( | Flad1 | 696.466 | 0.088469 | 0.445581 | 0.833239 | ENSMUSG( | Flad1 | 611.981 | -0.00977 | 0.948067 | 0.994798 |
| ENSMUSG( | Flcn | 1390.767 | 0.017291 | 0.878423 | 0.981431 | ENSMUSG( | Flcn | 1338.298 | 0.171347 | 0.257729 | 0.883401 |
| ENSMUSG( | Fli1 | 133.6172 | 0.122479 | 0.565937 | 0.89104 | ENSMUSG( | Fli1 | 133.4262 | 0.377897 | 0.098483 | 0.798438 |
| ENSMUSG( | Flii | 1946.199 | 0.193558 | 0.023991 | 0.23699 | ENSMUSG( | Flii | 1691.373 | 0.064774 | 0.552725 | 0.939565 |
| ENSMUSG( | Flna | 3441.944 | 0.112213 | 0.241496 | 0.689139 | ENSMUSG( | Flna | 3233.455 | 0.196597 | 0.204644 | 0.864412 |
| ENSMUSG( | Flnb | 2037.732 | 0.183759 | 0.11592 | 0.514015 | ENSMUSG( | Flnb | 1927.242 | 0.28616 | 0.061432 | 0.736111 |
| ENSMUSG( | Flnc | 141.5028 | 0.358943 | 0.045785 | 0.328083 | ENSMUSG( | Flnc | 133.9463 | 0.468098 | 0.021965 | 0.575341 |
| ENSMUSG( | Flot1 | 3676.192 | 0.130371 | 0.337034 | 0.777008 | ENSMUSG( | Flot1 | 3047.283 | -0.14615 | 0.347485 | 0.911652 |
| ENSMUSG( | Flot2 | 5414.808 | 0.047924 | 0.551775 | 0.886141 | ENSMUSG( | Flot2 | 4617.965 | -0.14659 | 0.162091 | 0.837724 |
| ENSMUSG( | Flrt1 | 719.0908 | -0.10097 | 0.603269 | 0.904887 | ENSMUSG( | Flrt1 | 737.5864 | 0.24248 | 0.253148 | 0.88018 |
| ENSMUSG( | Flrt2 | 1375.949 | -0.41781 | 0.028368 | 0.257483 | ENSMUSG( | Flrt2 | 1685.034 | 0.438507 | 0.040205 | 0.681706 |
| ENSMUSG( | Flrt3 | 604.4835 | -0.55266 | 0.000791 | 0.027736 | ENSMUSG( | Flrt3 | 812.8552 | 0.570986 | 0.004401 | 0.269982 |
| ENSMUSG( | Flt1 | 2384.431 | 0.282356 | 0.441962 | 0.832762 | ENSMUSG( | Flt1 | 2638.279 | 0.766777 | 0.102615 | 0.803503 |
| ENSMUSG( | Flt3 | 222.5589 | 0.184343 | 0.229903 | 0.677881 | ENSMUSG( | Flt3 | 199.7739 | 0.151272 | 0.380548 | 0.917202 |
| ENSMUSG( | Flt3l | 56.82794 | -0.14511 | 0.585434 | 0.899171 | ENSMUSG( | Flt3l | 60.64671 | 0.30386 | 0.375344 | 0.915491 |
| ENSMUSG( | Flt4 | 155.7481 | 0.423238 | 0.080762 | 0.439438 | ENSMUSG( | Flt4 | 171.2667 | 0.886221 | 0.003846 | 0.247715 |
| ENSMUSG( | Flvcr1 | 295.6637 | -0.24619 | 0.138155 | 0.553165 | ENSMUSG( | Flvcr1 | 323.9156 | 0.29125 | 0.105854 | 0.804969 |
| ENSMUSG( | Flvcr2 | 117.6702 | 0.256111 | 0.278778 | 0.728705 | ENSMUSG( | Flvcr2 | 143.0813 | 0.962759 | 0.001177 | 0.12275 |
| ENSMUSG( | Flywch1 | 6554.432 | 0.117044 | 0.319779 | 0.765305 | ENSMUSG( | Flywch1 | 5692.009 | -0.01969 | 0.872306 | 0.988944 |
| ENSMUSG( | Flywch2 | 237.2389 | -0.26806 | 0.103115 | 0.48855 | ENSMUSG( | Flywch2 | 216.1085 | -0.28209 | 0.154262 | 0.837724 |
| ENSMUSG( | Fmc1 | 741.5803 | 0.068649 | 0.537307 | 0.882504 | ENSMUSG( | Fmc1 | 584.6787 | -0.38016 | 0.008602 | 0.391682 |
| ENSMUSG( | Fmn1 | 379.6001 | 0.077866 | 0.613284 | 0.908943 | ENSMUSG( | Fmn1 | 358.512 | 0.185056 | 0.293338 | 0.898538 |
| ENSMUSG( | Fmn2 | 6390.51 | -0.00304 | 0.983191 | 0.997554 | ENSMUSG( | Fmn2 | 5800.123 | -0.01328 | 0.922807 | 0.992644 |
| ENSMUSG( | Fmnl1 | 4974.982 | 0.18452 | 0.055982 | 0.364712 | ENSMUSG( | Fmnl1 | 4213.409 | -0.02128 | 0.85053 | 0.985541 |
| ENSMUSG( | Fmnl2 | 4977.958 | 0.027107 | 0.841689 | 0.971179 | ENSMUSG( | Fmnl2 | 4378.289 | -0.07803 | 0.613064 | 0.951154 |
| ENSMUSG( | Fmnl3 | 709.6306 | -0.00118 | 0.992862 | 0.999327 | ENSMUSG( | Fmnl3 | 723.5761 | 0.314931 | 0.07884 | 0.774307 |
| ENSMUSG( | Fmo1 | 762.9219 | 0.223724 | 0.124442 | 0.531714 | ENSMUSG( | Fmo1 | 668.191 | 0.112311 | 0.446363 | 0.921734 |
| ENSMUSG( | Fmo2 | 26.04124 | 0.803524 | 0.028977 | 0.259144 | ENSMUSG( | Fmo2 | 32.60664 | 1.491412 | 6.97E-05 | 0.016495 |
| ENSMUSG( | Fmo5 | 279.8101 | -0.04879 | 0.788787 | 0.959226 | ENSMUSG( | Fmo5 | 245.3857 | -0.16387 | 0.321916 | 0.904023 |
| ENSMUSG( | Fmod | 996.064 | -0.01043 | 0.937031 | 0.993415 | ENSMUSG( | Fmod | 936.3198 | 0.082607 | 0.537185 | 0.937216 |
| ENSMUSG( | Fmr1 | 1467.966 | -0.0069 | 0.957102 | 0.993874 | ENSMUSG( | Fmr1 | 1400.958 | 0.120315 | 0.436242 | 0.919879 |
| ENSMUSG( | Fn1 | 3191.217 | 0.202467 | 0.183357 | 0.62047 | ENSMUSG( | Fn1 | 3448.968 | 0.638796 | 0.001013 | 0.109511 |
| ENSMUSG( | Fn3k | 495.6081 | -0.10256 | 0.497129 | 0.861754 | ENSMUSG( | Fn3k | 445.0932 | -0.1495 | 0.41276 | 0.917257 |
| ENSMUSG( | Fn3krp | 913.2152 | 0.060961 | 0.544813 | 0.884712 | ENSMUSG( | Fn3krp | 773.7831 | -0.15737 | 0.231491 | 0.868507 |
| ENSMUSG( | Fnbp1 | 6806.646 | -0.02582 | 0.737073 | 0.946839 | ENSMUSG( | Fnbp1 | 5816.663 | -0.21846 | 0.023197 | 0.585577 |
| ENSMUSG( | Fnbp1l | 2598.772 | 0.132533 | 0.142996 | 0.561716 | ENSMUSG( | Fnbp1l | 2424.112 | 0.194182 | 0.121915 | 0.813384 |
| ENSMUSG( | Fnbp4 | 1694.997 | -0.12546 | 0.218733 | 0.664559 | ENSMUSG( | Fnbp4 | 1625.792 | 0.027962 | 0.826475 | 0.979922 |
| ENSMUSG( | Fndc1 | 134.2431 | -0.11818 | 0.491859 | 0.860249 | ENSMUSG( | Fndc1 | 149.9047 | 0.42222 | 0.044405 | 0.693697 |
| ENSMUSG( | Fndc10 | 812.733 | -0.28386 | 0.018459 | 0.204231 | ENSMUSG( | Fndc10 | 831.8101 | 0.069203 | 0.620187 | 0.952896 |
| ENSMUSG( | Fndc3a | 1307.293 | 0.112884 | 0.232169 | 0.67996 | ENSMUSG( | Fndc3a | 1206.627 | 0.14644 | 0.217554 | 0.865589 |
| ENSMUSG( | Fndc3b | 561.0905 | 0.259378 | 0.017108 | 0.195465 | ENSMUSG( | Fndc3b | 487.5785 | 0.133306 | 0.300764 | 0.898538 |
| ENSMUSG( | Fndc4 | 4617.764 | -0.0089 | 0.920247 | 0.990247 | ENSMUSG( | Fndc4 | 4076.603 | -0.10125 | 0.320566 | 0.903887 |
| ENSMUSG( | Fndc5 | 4766.728 | -0.25758 | 0.009272 | 0.135856 | ENSMUSG( | Fndc5 | 4423.187 | -0.20036 | 0.100086 | 0.801311 |
| ENSMUSG( | Fndc7 | 10.7274 | -0.72238 | 0.252649 | NA | ENSMUSG( | Fndc7 | 9.133814 | -1.08792 | 0.112525 | NA |
| ENSMUSG( | Fndc9 | 102.9129 | -0.51228 | 0.018479 | 0.204273 | ENSMUSG( | Fndc9 | 104.7772 | -0.16901 | 0.487181 | 0.92541 |
| ENSMUSG( | Fnip1 | 892.8068 | -0.22554 | 0.034463 | 0.285197 | ENSMUSG( | Fnip1 | 915.0943 | 0.12005 | 0.31993 | 0.903771 |
| ENSMUSG( | Fnip2 | 504.2133 | -0.17038 | 0.128971 | 0.537903 | ENSMUSG( | Fnip2 | 487.999 | 0.011643 | 0.928689 | 0.99306 |
| ENSMUSG( | Fnta | 2107.875 | 0.142396 | 0.154435 | 0.581463 | ENSMUSG( | Fnta | 1749.17 | -0.13022 | 0.298465 | 0.898538 |
| ENSMUSG( | Fntb | 808.65 | 0.041417 | 0.698002 | 0.935901 | ENSMUSG( | Fntb | 700.757 | -0.10569 | 0.404495 | 0.917257 |
| ENSMUSG( | Focad | 1325.26 | 0.274995 | 0.041166 | 0.313607 | ENSMUSG( | Focad | 1099.961 | 0.018091 | 0.91 | 0.991958 |
| ENSMUSG( | Folh1 | 447.3512 | 0.100907 | 0.488795 | 0.859092 | ENSMUSG( | Folh1 | 410.6144 | 0.125377 | 0.426371 | 0.917257 |
| ENSMUSG( | Folr1 | 121.1354 | -0.04895 | 0.9447 | 0.993583 | ENSMUSG( | Folr1 | 138.5538 | 0.546117 | 0.332152 | 0.907907 |
| ENSMUSG( | Folr2 | 30.06405 | -0.65031 | 0.089629 | 0.45961 | ENSMUSG( | Folr2 | 28.30159 | -0.53092 | 0.199802 | 0.863016 |
| ENSMUSG( | Fos | 476.538 | -1.24915 | 0.001778 | 0.04746 | ENSMUSG( | Fos | 461.9455 | -0.96707 | 0.052869 | 0.71525 |
| ENSMUSG( | Fosb | 252.3502 | -0.20679 | 0.385358 | 0.804048 | ENSMUSG( | Fosb | 225.7317 | -0.26382 | 0.317418 | 0.902486 |
| ENSMUSG( | Fosl2 | 855.7834 | -0.02809 | 0.798984 | 0.960922 | ENSMUSG( | Fosl2 | 768.9868 | -0.06324 | 0.63256 | 0.953698 |
| ENSMUSG( | Foxc1 | 207.854 | -0.16931 | 0.336362 | 0.777 | ENSMUSG( | Foxc1 | 223.9209 | 0.313497 | 0.101127 | 0.801456 |
| ENSMUSG( | Foxc2 | 114.2686 | -0.0768 | 0.678681 | 0.930611 | ENSMUSG( | Foxc2 | 123.8643 | 0.396902 | 0.047212 | 0.700856 |
| ENSMUSG( | Foxd1 | 89.17411 | -0.32417 | 0.13411 | 0.547472 | ENSMUSG( | Foxd1 | 105.7541 | 0.425682 | 0.041773 | 0.690304 |
| ENSMUSG( | Foxd2 | 17.56086 | -0.32306 | 0.506481 | NA | ENSMUSG( | Foxd2 | 16.3127 | -0.28004 | 0.576202 | 0.942912 |
| ENSMUSG( | Foxf1 | 39.6908 | -0.15671 | 0.633483 | 0.914073 | ENSMUSG( | Foxf1 | 43.6223 | 0.402882 | 0.289855 | 0.896726 |
| ENSMUSG( | Foxf2 | 167.5467 | 0.301586 | 0.207622 | 0.64962 | ENSMUSG( | Foxf2 | 182.6626 | 0.756378 | 0.005036 | 0.292089 |
| ENSMUSG( | Foxg1 | 1911.897 | 0.04416 | 0.632625 | 0.914073 | ENSMUSG( | Foxg1 | 1840.197 | 0.192482 | 0.159261 | 0.837724 |
| ENSMUSG( | Foxj1 | 345.1018 | -0.28526 | 0.059024 | 0.375291 | ENSMUSG( | Foxj1 | 378.1404 | 0.256147 | 0.215956 | 0.865167 |
| ENSMUSG( | Foxj2 | 927.7492 | 0.083962 | 0.416308 | 0.82084 | ENSMUSG( | Foxj2 | 852.3723 | 0.109486 | 0.342575 | 0.909646 |
| ENSMUSG( | Foxj3 | 1341.725 | -0.03191 | 0.743928 | 0.948497 | ENSMUSG( | Foxj3 | 1210.063 | -0.0647 | 0.596737 | 0.946541 |
| ENSMUSG( | Foxk1 | 1520.561 | 0.098874 | 0.292196 | 0.74239 | ENSMUSG( | Foxk1 | 1371.84 | 0.0753 | 0.546069 | 0.938063 |
| ENSMUSG( | Foxk2 | 4693.518 | 0.130353 | 0.125082 | 0.532139 | ENSMUSG( | Foxk2 | 4135.204 | 0.037535 | 0.709633 | 0.96416 |
| ENSMUSG( | Foxl2 | 30.47442 | 0.120606 | 0.762297 | 0.953459 | ENSMUSG( | Foxl2 | 35.02344 | 0.740808 | 0.082834 | 0.77681 |
| ENSMUSG( | Foxl2os | 16.00396 | 0.633504 | 0.241905 | NA | ENSMUSG( | Foxl2os | 15.39678 | 0.79071 | 0.149028 | 0.836241 |
| ENSMUSG( | Foxm1 | 96.0075 | -0.28509 | 0.17977 | 0.617507 | ENSMUSG( | Foxm1 | 90.24856 | -0.17194 | 0.47056 | 0.923329 |
| ENSMUSG( | Foxn2 | 577.4231 | -0.19891 | 0.064779 | 0.394562 | ENSMUSG( | Foxn2 | 562.4054 | -0.0012 | 0.992439 | 0.99874 |
| ENSMUSG( | Foxn3 | 2031.616 | -0.05395 | 0.562579 | 0.889465 | ENSMUSG( | Foxn3 | 1819.08 | -0.10838 | 0.417364 | 0.917257 |
| ENSMUSG( | Foxo1 | 1044.839 | 0.020689 | 0.842993 | 0.971739 | ENSMUSG( | Foxo1 | 903.2306 | -0.13461 | 0.302859 | 0.898538 |
| ENSMUSG( | Foxo3 | 3438.349 | 0.371561 | 9.56E-06 | 0.001107 | ENSMUSG( | Foxo3 | 2825.611 | 0.097628 | 0.376234 | 0.915491 |
| ENSMUSG( | Foxo4 | 2816.366 | 0.174137 | 0.075057 | 0.42422 | ENSMUSG( | Foxo4 | 2362.338 | -0.06237 | 0.597436 | 0.946609 |
| ENSMUSG( | Foxo6 | 518.8004 | 0.178844 | 0.24376 | 0.691156 | ENSMUSG( | Foxo6 | 433.7539 | -0.07319 | 0.713829 | 0.964986 |
| ENSMUSG( | Foxo6os | 23.56292 | 0.310178 | 0.468453 | 0.845415 | ENSMUSG( | Foxo6os | 15.98802 | -0.57148 | 0.28205 | 0.892412 |
| ENSMUSG( | Foxp1 | 1744.562 | -0.0948 | 0.30294 | 0.751443 | ENSMUSG( | Foxp1 | 1673.935 | 0.052916 | 0.670912 | 0.95694 |
| ENSMUSG( | Foxp2 | 219.3227 | -0.11396 | 0.512961 | 0.871762 | ENSMUSG( | Foxp2 | 206.8478 | -0.02598 | 0.91282 | 0.992298 |
| ENSMUSG( | Foxp4 | 1423.826 | 0.054065 | 0.666935 | 0.92729 | ENSMUSG( | Foxp4 | 1251.425 | -0.05203 | 0.714609 | 0.964986 |
| ENSMUSG( | Foxq1 | 170.6574 | 0.596594 | 0.007501 | 0.119875 | ENSMUSG( | Foxq1 | 159.8852 | 0.681766 | 0.018523 | 0.545482 |
| ENSMUSG( | Foxr2 | 24.54467 | 0.077681 | 0.840013 | 0.971179 | ENSMUSG( | Foxr2 | 24.70038 | 0.327138 | 0.424245 | 0.917257 |
| ENSMUSG( | Foxred1 | 1278.862 | 0.121768 | 0.213563 | 0.657797 | ENSMUSG( | Foxred1 | 1085.718 | -0.08195 | 0.508676 | 0.929089 |
| ENSMUSG( | Foxred2 | 747.2109 | -0.38531 | 0.008999 | 0.132931 | ENSMUSG( | Foxred2 | 797.912 | 0.102949 | 0.525648 | 0.934374 |
| ENSMUSG( | Foxs1 | 20.01321 | -0.35748 | 0.432765 | NA | ENSMUSG( | Foxs1 | 18.20585 | -0.32906 | 0.499886 | 0.927825 |
| ENSMUSG( | Fpgs | 234.1501 | -0.10164 | 0.521441 | 0.875186 | ENSMUSG( | Fpgs | 203.815 | -0.23261 | 0.194141 | 0.860096 |
| ENSMUSG( | Fpgt | 371.2443 | 0.052974 | 0.669731 | 0.928147 | ENSMUSG( | Fpgt | 357.0552 | 0.211982 | 0.194202 | 0.860096 |
| ENSMUSG( | Fra10ac1 | 340.3977 | -0.03807 | 0.800363 | 0.961745 | ENSMUSG( | Fra10ac1 | 317.0371 | 0.018418 | 0.911267 | 0.992196 |
| ENSMUSG( | Fras1 | 800.6609 | -0.64754 | 1.71E-07 | 4.07E-05 | ENSMUSG( | Fras1 | 912.7067 | 0.073586 | 0.574557 | 0.942912 |
| ENSMUSG( | Frat1 | 281.751 | 0.068002 | 0.687763 | 0.93324 | ENSMUSG( | Frat1 | 255.7967 | 0.063431 | 0.734935 | 0.967344 |

| | | | | | |
|---|---|---|---|---|---|
| ENSMUSG( | 892.8068 | -0.22554 | 0.034463 | 0.285197 | Fnip1 |
| ENSMUSG( | 504.2133 | -0.17038 | 0.128971 | 0.537903 | Fnip2 |
| ENSMUSG( | 2107.875 | 0.142396 | 0.154435 | 0.581463 | Fnta |
| ENSMUSG( | 808.65 | 0.041417 | 0.698002 | 0.935901 | Fntb |
| ENSMUSG( | 1325.26 | 0.274995 | 0.041166 | 0.313607 | Focad |
| ENSMUSG( | 447.3512 | 0.100907 | 0.488795 | 0.859092 | Folh1 |
| ENSMUSG( | 121.1354 | -0.04895 | 0.9447 | 0.993583 | Folr1 |
| ENSMUSG( | 30.06405 | -0.65031 | 0.089629 | 0.45961 | Folr2 |
| ENSMUSG( | 476.538 | -1.24915 | 0.001778 | 0.04746 | Fos |
| ENSMUSG( | 252.3502 | -0.20679 | 0.385358 | 0.804048 | Fosb |
| ENSMUSG( | 855.7834 | -0.02809 | 0.798984 | 0.960922 | Fosl2 |
| ENSMUSG( | 207.854 | -0.16931 | 0.336362 | 0.777 | Foxc1 |
| ENSMUSG( | 114.2686 | -0.0768 | 0.678681 | 0.930611 | Foxc2 |
| ENSMUSG( | 89.17411 | -0.32417 | 0.13411 | 0.547472 | Foxd1 |
| ENSMUSG( | 17.56086 | -0.32306 | 0.506481 | NA | Foxd2 |
| ENSMUSG( | 39.6908 | -0.15671 | 0.633483 | 0.914073 | Foxf1 |
| ENSMUSG( | 167.5467 | 0.301586 | 0.207622 | 0.64962 | Foxf2 |
| ENSMUSG( | 1911.897 | 0.04416 | 0.632625 | 0.914073 | Foxg1 |
| ENSMUSG( | 345.1018 | -0.28526 | 0.059024 | 0.375291 | Foxj1 |
| ENSMUSG( | 927.7492 | 0.083962 | 0.416308 | 0.82084 | Foxj2 |
| ENSMUSG( | 1341.725 | -0.03191 | 0.743928 | 0.948497 | Foxj3 |
| ENSMUSG( | 1520.561 | 0.098874 | 0.292196 | 0.74239 | Foxk1 |
| ENSMUSG( | 4693.518 | 0.130353 | 0.125082 | 0.532139 | Foxk2 |
| ENSMUSG( | 30.47442 | 0.120606 | 0.762297 | 0.953459 | Foxl2 |
| ENSMUSG( | 16.00396 | 0.633504 | 0.241905 | NA | Foxl2os |
| ENSMUSG( | 96.0075 | -0.28509 | 0.17977 | 0.617507 | Foxm1 |
| ENSMUSG( | 577.4231 | -0.19891 | 0.064779 | 0.394562 | Foxn2 |
| ENSMUSG( | 2031.616 | -0.05395 | 0.562579 | 0.889465 | Foxn3 |
| ENSMUSG( | 1044.839 | 0.020689 | 0.842993 | 0.971739 | Foxo1 |
| ENSMUSG( | 3438.349 | 0.371561 | 9.56E-06 | 0.001107 | Foxo3 |
| ENSMUSG( | 2816.366 | 0.174137 | 0.075057 | 0.42422 | Foxo4 |
| ENSMUSG( | 518.8004 | 0.178844 | 0.24376 | 0.691156 | Foxo6 |
| ENSMUSG( | 23.56292 | 0.310178 | 0.468453 | 0.845415 | Foxo6os |
| ENSMUSG( | 1744.562 | -0.0948 | 0.30294 | 0.751443 | Foxp1 |
| ENSMUSG( | 219.3227 | -0.11396 | 0.512961 | 0.871762 | Foxp2 |
| ENSMUSG( | 1423.826 | 0.054065 | 0.666935 | 0.92729 | Foxp4 |
| ENSMUSG( | 170.6574 | 0.596594 | 0.007501 | 0.119875 | Foxq1 |
| ENSMUSG( | 24.54467 | 0.077681 | 0.840013 | 0.971179 | Foxr2 |
| ENSMUSG( | 1278.862 | 0.121768 | 0.213563 | 0.657797 | Foxred1 |
| ENSMUSG( | 747.2109 | -0.38531 | 0.008999 | 0.132931 | Foxred2 |
| ENSMUSG( | 20.01321 | -0.35748 | 0.432765 | NA | Foxs1 |
| ENSMUSG( | 234.1501 | -0.10164 | 0.521441 | 0.875186 | Fpgs |
| ENSMUSG( | 371.2443 | 0.052974 | 0.669731 | 0.928147 | Fpgt |
| ENSMUSG( | 340.3977 | -0.03807 | 0.800363 | 0.961745 | Fra10ac1 |
| ENSMUSG( | 800.6609 | -0.64754 | 1.71E-07 | 4.07E-05 | Fras1 |
| ENSMUSG( | 281.751 | 0.068002 | 0.687763 | 0.93324 | Frat1 |
| ENSMUSG( | 143.4618 | 0.02093 | 0.915979 | 0.988634 | Frat2 |
| ENSMUSG( | 184.8745 | -0.29941 | 0.055665 | 0.363338 | Frem1 |
| ENSMUSG( | 234.4926 | -0.66156 | 0.00022 | 0.011467 | Frem2 |
| ENSMUSG( | 15.2904 | -1.25537 | 0.018345 | NA | Frem3 |
| ENSMUSG( | 95.90846 | -0.0633 | 0.774638 | 0.955606 | Frey1 |
| ENSMUSG( | 424.2139 | 0.034296 | 0.785473 | 0.958571 | Frg1 |
| ENSMUSG( | 374.6631 | 0.007896 | 0.951251 | 0.993821 | Frg2f1 |
| ENSMUSG( | 14.5687 | 0.250226 | 0.637952 | NA | Frk |
| ENSMUSG( | 132.3019 | -0.246 | 0.235108 | 0.682364 | Frmd3 |
| ENSMUSG( | 2706.074 | -0.08669 | 0.31732 | 0.763823 | Frmd4a |
| ENSMUSG( | 797.9075 | -0.15077 | 0.361018 | 0.792797 | Frmd4b |
| ENSMUSG( | 991.7037 | -0.04975 | 0.687565 | 0.93324 | Frmd5 |
| ENSMUSG( | 1009.415 | -0.22815 | 0.068668 | 0.405903 | Frmd6 |
| ENSMUSG( | 41.68904 | -0.32636 | 0.476047 | 0.851133 | Frmd7 |
| ENSMUSG( | 861.9591 | -0.09332 | 0.363166 | 0.793211 | Frmd8 |
| ENSMUSG( | 1389.333 | 0.254956 | 0.018167 | 0.20312 | Frmpd1 |
| ENSMUSG( | 59.87243 | -0.29926 | 0.272481 | 0.722478 | Frmpd2 |
| ENSMUSG( | 2468.044 | 0.408113 | 6.45E-05 | 0.004822 | Frmpd3 |
| ENSMUSG( | 1032.539 | 0.074215 | 0.432668 | 0.827873 | Frmpd4 |
| ENSMUSG( | 132.9866 | -0.25818 | 0.175805 | 0.611006 | Frrs1 |
| ENSMUSG( | 6608.045 | -0.35838 | 0.015146 | 0.181303 | Frrs1l |
| ENSMUSG( | 3496.801 | 0.045455 | 0.586569 | 0.900023 | Frs2 |
| ENSMUSG( | 844.5979 | 0.038523 | 0.766112 | 0.954173 | Frs3 |
| ENSMUSG( | 3156.068 | 0.023831 | 0.821489 | 0.966248 | Fry |
| ENSMUSG( | 2540.33 | -0.00871 | 0.921648 | 0.990436 | Fryl |
| ENSMUSG( | 711.9483 | -0.54971 | 1.90E-05 | 0.001887 | Frzb |
| ENSMUSG( | 37.66952 | -0.21295 | 0.495038 | 0.860798 | Fsbp |
| ENSMUSG( | 6353.271 | 0.069416 | 0.528264 | 0.878236 | Fscn1 |
| ENSMUSG( | 1288.102 | 0.06107 | 0.566777 | 0.891644 | Fsd1 |
| ENSMUSG( | 516.0658 | -0.17933 | 0.138755 | 0.554336 | Fsd1l |
| ENSMUSG( | 55.94578 | 0.323689 | 0.304161 | 0.752287 | Fsd2 |
| ENSMUSG( | 101.2002 | 0.062001 | 0.772022 | 0.954994 | Fst |
| ENSMUSG( | 1724.486 | -0.15401 | 0.208664 | 0.650496 | Fstl1 |
| ENSMUSG( | 58.3381 | 0.108431 | 0.682812 | 0.932022 | Fstl3 |
| ENSMUSG( | 402.6135 | -0.54341 | 0.001121 | 0.035587 | Fstl4 |
| ENSMUSG( | 400.1026 | -0.63183 | 0.000458 | 0.019492 | Fstl5 |
| ENSMUSG( | 148310.2 | 0.125715 | 0.103335 | 0.488943 | Fth1 |
| ENSMUSG( | 15753.81 | 0.093725 | 0.377224 | 0.801213 | Ftl1 |
| ENSMUSG( | 43.38596 | 0.254614 | 0.40148 | 0.814815 | Ftl1-ps2 |
| ENSMUSG( | 5352.569 | 0.061708 | 0.496451 | 0.86175 | Fto |
| ENSMUSG( | 485.4482 | 0.046151 | 0.689179 | 0.933485 | Ftsj1 |
| ENSMUSG( | 944.9474 | 0.176891 | 0.0862 | 0.451605 | Ftsj3 |
| ENSMUSG( | 854.6939 | 0.357734 | 0.008768 | 0.131926 | Ftx |
| ENSMUSG( | 1569.368 | -0.05019 | 0.567221 | 0.891666 | Fubp1 |
| ENSMUSG( | 1575.054 | 0.023473 | 0.792408 | 0.959461 | Fubp3 |
| ENSMUSG( | 1858.139 | -0.08155 | 0.453741 | 0.836798 | Fuca1 |
| ENSMUSG( | 1862.474 | -0.18538 | 0.12431 | 0.531714 | Fuca2 |
| ENSMUSG( | 872.7305 | 0.017333 | 0.874364 | 0.979432 | Fundc1 |
| ENSMUSG( | 989.5846 | -0.05305 | 0.637023 | 0.915191 | Fundc2 |
| ENSMUSG( | 574.2064 | -0.02219 | 0.844718 | 0.972023 | Fuom |
| ENSMUSG( | 1456.305 | -0.138 | 0.294877 | 0.743798 | Furin |
| ENSMUSG( | 5457.147 | -0.07477 | 0.38035 | 0.801757 | Fus |
| ENSMUSG( | 643.3067 | 0.07895 | 0.56553 | 0.890871 | Fut10 |
| ENSMUSG( | 599.8212 | -0.24225 | 0.066599 | 0.398819 | Fut11 |
| ENSMUSG( | 15.34862 | -0.66276 | 0.180959 | NA | Fut2 |
| ENSMUSG( | 11.52629 | 0.091713 | 0.864997 | NA | Fut4 |
| ENSMUSG( | 9.939858 | -0.17858 | 0.783335 | NA | Fut7 |
| ENSMUSG( | 2722.678 | -0.27947 | 0.029845 | 0.261951 | Fut8 |
| ENSMUSG( | 2222.959 | -0.18873 | 0.208149 | 0.650319 | Fut9 |
| ENSMUSG( | 694.0421 | 0.069614 | 0.527385 | 0.877599 | Fuz |
| ENSMUSG( | 69.98066 | 0.289279 | 0.251807 | 0.701908 | Fv1 |
| ENSMUSG( | 175.1075 | 0.113999 | 0.522612 | 0.875389 | Fxn |
| ENSMUSG( | 1861.514 | 0.0824 | 0.360209 | 0.791947 | Fxr1 |
| ENSMUSG( | 3436.739 | 0.166793 | 0.109743 | 0.500285 | Fxr2 |
| ENSMUSG( | 2396.502 | -0.14102 | 0.175219 | 0.610405 | Fxyd1 |
| ENSMUSG( | 153.1077 | -0.08031 | 0.629808 | 0.913965 | Fxyd2 |
| ENSMUSG( | 33.81343 | 0.270029 | 0.472995 | 0.84867 | Fxyd3 |
| ENSMUSG( | 903.6491 | 0.125066 | 0.321311 | 0.766497 | Fxyd5 |
| ENSMUSG( | 4974.735 | -0.23238 | 0.088236 | 0.455783 | Fxyd6 |
| ENSMUSG( | 2327.371 | -0.16423 | 0.152884 | 0.579254 | Fxyd7 |
| ENSMUSG( | 289.1641 | 0.19129 | 0.245184 | 0.692812 | Fyb1 |
| ENSMUSG( | 54.42489 | -0.009 | 0.975129 | 0.997302 | Fyb2 |
| ENSMUSG( | 4032.012 | 0.111884 | 0.184396 | 0.621442 | Fyco1 |
| ENSMUSG( | 8633.232 | -0.00288 | 0.972986 | 0.996705 | Fyn |
| ENSMUSG( | 1275.417 | -0.00285 | 0.975325 | 0.997302 | Fyttd1 |
| ENSMUSG( | 615.7126 | 0.022122 | 0.867397 | 0.976559 | Fzd1 |
| ENSMUSG( | 178.7447 | -0.65843 | 0.000445 | 0.019233 | Fzd10 |
| ENSMUSG( | 216.0402 | 0.137257 | 0.336737 | 0.777 | Fzd10os |
| ENSMUSG( | 860.7951 | -0.01865 | 0.911377 | 0.988483 | Fzd2 |
| ENSMUSG( | 3107.422 | -0.28993 | 0.054347 | 0.358773 | Fzd3 |
| ENSMUSG( | 947.5206 | 0.038405 | 0.769713 | 0.954674 | Fzd4 |
| ENSMUSG( | 71.88599 | -0.39331 | 0.104565 | 0.490752 | Fzd5 |
| ENSMUSG( | 464.2056 | -0.09787 | 0.587483 | 0.900403 | Fzd6 |
| ENSMUSG( | 375.6484 | -0.11613 | 0.397759 | 0.812161 | Fzd7 |
| ENSMUSG( | 152.5075 | -0.33982 | 0.0914 | 0.464738 | Fzd8 |
| ENSMUSG( | 269.4078 | -0.50687 | 0.007004 | 0.115211 | Fzd9 |
| ENSMUSG( | 1402.444 | 0.112572 | 0.361808 | 0.792797 | Fzr1 |
| ENSMUSG( | 145.0137 | -0.58293 | 0.007724 | 0.122031 | G0s2 |
| ENSMUSG( | 355.8786 | -0.05229 | 0.70916 | 0.939599 | G2e3 |
| ENSMUSG( | 2260.046 | 0.102516 | 0.269645 | 0.719729 | G3bp1 |
| ENSMUSG( | 4305.866 | 0.031405 | 0.724103 | 0.944096 | G3bp2 |
| ENSMUSG( | 14.23521 | 0.158009 | 0.767533 | NA | G430095P16Rik |
| ENSMUSG( | 64.91386 | 1.04823 | 0.000495 | 0.020378 | G530011O06Rik |
| ENSMUSG( | 104.3817 | -0.07551 | 0.74866 | 0.949604 | G630016G05Rik |
| ENSMUSG( | 12.05665 | -0.37519 | 0.4975 | NA | G630093K05Rik |
| ENSMUSG( | 1911.448 | -0.18423 | 0.083855 | 0.446475 | G6pc3 |
| ENSMUSG( | 1501.196 | 0.09407 | 0.288971 | 0.739705 | G6pdx |
| ENSMUSG( | 9634.06 | -0.12212 | 0.37461 | 0.800445 | Gaa |
| ENSMUSG( | 1322.567 | -0.15824 | 0.186011 | 0.623496 | Gab1 |
| ENSMUSG( | 1243.796 | 0.001159 | 0.990151 | 0.999144 | Gab2 |
| ENSMUSG( | 34.44627 | 0.537627 | 0.100603 | 0.48402 | Gab3 |
| ENSMUSG( | 5164.718 | 0.079723 | 0.381414 | 0.802629 | Gabarap |
| ENSMUSG( | 11163.34 | 0.055973 | 0.621684 | 0.911432 | Gabarapl1 |
| ENSMUSG( | 3045.355 | 0.016377 | 0.882788 | 0.982356 | Gabarapl2 |
| ENSMUSG( | 41276.73 | -0.17158 | 0.138459 | 0.553773 | Gabbr1 |
| ENSMUSG( | 14494.25 | -0.25207 | 0.063201 | 0.388743 | Gabbr2 |
| ENSMUSG( | 1038.25 | 0.107363 | 0.402324 | 0.815103 | Gabpa |
| ENSMUSG( | 448.4059 | 0.131938 | 0.303271 | 0.75169 | Gabpb1 |
| ENSMUSG( | 697.9805 | 0.012607 | 0.92037 | 0.990276 | Gabpb2 |
| ENSMUSG( | 6452.521 | -0.71279 | 4.21E-05 | 0.003506 | Gabra1 |
| ENSMUSG( | 4490.366 | 1.549417 | 4.17E-23 | 6.15E-20 | Gabra2 |
| ENSMUSG( | 4357.62 | -0.13964 | 0.24018 | 0.687899 | Gabra3 |

| | | | | | |
|---|---|---|---|---|---|
| ENSMUSG( | 16.3127 | -0.28004 | 0.576202 | 0.942912 | Foxd2 |
| ENSMUSG( | 9.349705 | 0.093207 | 0.888501 | NA | Foxd2os |
| ENSMUSG( | 43.6223 | 0.402882 | 0.289855 | 0.896726 | Foxf1 |
| ENSMUSG( | 182.6626 | 0.756378 | 0.005036 | 0.292089 | Foxf2 |
| ENSMUSG( | 1840.197 | 0.192482 | 0.159261 | 0.837724 | Foxg1 |
| ENSMUSG( | 378.1404 | 0.256147 | 0.215956 | 0.865167 | Foxj1 |
| ENSMUSG( | 852.3723 | 0.109486 | 0.342575 | 0.909646 | Foxj2 |
| ENSMUSG( | 1210.063 | -0.0647 | 0.596737 | 0.946541 | Foxj3 |
| ENSMUSG( | 1371.84 | 0.0753 | 0.546069 | 0.938063 | Foxk1 |
| ENSMUSG( | 4135.204 | 0.037535 | 0.709633 | 0.96416 | Foxk2 |
| ENSMUSG( | 35.02344 | 0.740808 | 0.082834 | 0.77681 | Foxl2 |
| ENSMUSG( | 15.39678 | 0.79071 | 0.149028 | 0.836241 | Foxl2os |
| ENSMUSG( | 90.24856 | -0.17194 | 0.47056 | 0.923329 | Foxm1 |
| ENSMUSG( | 562.4054 | -0.0012 | 0.992439 | 0.99874 | Foxn2 |
| ENSMUSG( | 1819.08 | -0.10838 | 0.417364 | 0.917257 | Foxn3 |
| ENSMUSG( | 903.2306 | -0.13461 | 0.302859 | 0.898538 | Foxo1 |
| ENSMUSG( | 2825.611 | 0.097628 | 0.376234 | 0.915491 | Foxo3 |
| ENSMUSG( | 2362.338 | -0.06237 | 0.597436 | 0.946609 | Foxo4 |
| ENSMUSG( | 433.7539 | -0.07319 | 0.713829 | 0.964986 | Foxo6 |
| ENSMUSG( | 15.98802 | -0.57148 | 0.28205 | 0.892412 | Foxo6os |
| ENSMUSG( | 1673.935 | 0.052916 | 0.670912 | 0.95694 | Foxp1 |
| ENSMUSG( | 206.8478 | -0.02598 | 0.91282 | 0.992298 | Foxp2 |
| ENSMUSG( | 1251.425 | -0.05203 | 0.714609 | 0.964986 | Foxp4 |
| ENSMUSG( | 159.8852 | 0.681766 | 0.018523 | 0.545482 | Foxq1 |
| ENSMUSG( | 24.70038 | 0.327138 | 0.424245 | 0.917257 | Foxr2 |
| ENSMUSG( | 1085.718 | -0.08195 | 0.508676 | 0.929089 | Foxred1 |
| ENSMUSG( | 797.912 | 0.102949 | 0.525648 | 0.934374 | Foxred2 |
| ENSMUSG( | 18.20585 | -0.32906 | 0.499886 | 0.927825 | Foxs1 |
| ENSMUSG( | 203.815 | -0.23261 | 0.194141 | 0.860096 | Fpgs |
| ENSMUSG( | 357.0552 | 0.211982 | 0.194202 | 0.860096 | Fpgt |
| ENSMUSG( | 317.0371 | 0.018418 | 0.911267 | 0.992196 | Fra10ac1 |
| ENSMUSG( | 912.7067 | 0.073586 | 0.574557 | 0.942912 | Fras1 |
| ENSMUSG( | 255.7967 | 0.063431 | 0.734935 | 0.967344 | Frat1 |
| ENSMUSG( | 130.8039 | 0.018045 | 0.934569 | 0.99306 | Frat2 |
| ENSMUSG( | 179.7982 | -0.09918 | 0.565666 | 0.941404 | Frem1 |
| ENSMUSG( | 238.3205 | -0.27481 | 0.128188 | 0.818934 | Frem2 |
| ENSMUSG( | 17.08061 | -0.4553 | 0.359269 | 0.914061 | Frem3 |
| ENSMUSG( | 78.53116 | -0.38742 | 0.116474 | 0.813384 | Frey1 |
| ENSMUSG( | 388.5468 | 0.046807 | 0.741123 | 0.968811 | Frg1 |
| ENSMUSG( | 328.027 | -0.10352 | 0.510753 | 0.929545 | Frg2f1 |
| ENSMUSG( | 17.17717 | 0.8886 | 0.062817 | 0.736482 | Frk |
| ENSMUSG( | 120.6972 | -0.2443 | 0.262974 | 0.885357 | Frmd3 |
| ENSMUSG( | 2421.731 | -0.13935 | 0.18161 | 0.850579 | Frmd4a |
| ENSMUSG( | 798.2795 | 0.129559 | 0.497105 | 0.927825 | Frmd4b |
| ENSMUSG( | 905.6462 | -0.03999 | 0.766345 | 0.974236 | Frmd5 |
| ENSMUSG( | 928.7165 | -0.19214 | 0.194064 | 0.860096 | Frmd6 |
| ENSMUSG( | 51.39175 | 0.503049 | 0.561967 | 0.940388 | Frmd7 |
| ENSMUSG( | 774.8367 | -0.13214 | 0.296645 | 0.898538 | Frmd8 |
| ENSMUSG( | 1034.06 | -0.33868 | 0.007631 | 0.37235 | Frmpd1 |
| ENSMUSG( | 60.46124 | 0.035393 | 0.904085 | 0.991089 | Frmpd2 |
| ENSMUSG( | 2048.63 | 0.165958 | 0.170864 | 0.842157 | Frmpd3 |
| ENSMUSG( | 961.6596 | 0.135496 | 0.225239 | 0.868468 | Frmpd4 |
| ENSMUSG( | 129.7536 | -0.033 | 0.884449 | 0.989688 | Frrs1 |
| ENSMUSG( | 7328.311 | 0.226079 | 0.189828 | 0.857355 | Frrs1l |
| ENSMUSG( | 3048.488 | -0.08452 | 0.483315 | 0.92499 | Frs2 |
| ENSMUSG( | 750.7571 | -0.03315 | 0.794596 | 0.97681 | Frs3 |
| ENSMUSG( | 3017.938 | 0.162425 | 0.213006 | 0.864412 | Fry |
| ENSMUSG( | 2394.903 | 0.089357 | 0.403986 | 0.917257 | Fryl |
| ENSMUSG( | 798.615 | 0.097117 | 0.563101 | 0.941158 | Frzb |
| ENSMUSG( | 27.56386 | -1.09169 | 0.010897 | 0.422321 | Fsbp |
| ENSMUSG( | 5236.34 | -0.23013 | 0.079142 | 0.77458 | Fscn1 |
| ENSMUSG( | 1080.753 | -0.18084 | 0.123368 | 0.814691 | Fsd1 |
| ENSMUSG( | 514.5801 | 0.085378 | 0.527938 | 0.934616 | Fsd1l |
| ENSMUSG( | 38.94087 | -0.46884 | 0.160034 | 0.837724 | Fsd2 |
| ENSMUSG( | 9.20264 | 0.096825 | 0.881323 | NA | Fsip1 |
| ENSMUSG( | 116.8161 | 0.673089 | 0.003372 | 0.232514 | Fst |
| ENSMUSG( | 1779.06 | 0.205008 | 0.143865 | 0.831311 | Fstl1 |
| ENSMUSG( | 51.06085 | 0.000238 | 0.999332 | 0.999503 | Fstl3 |
| ENSMUSG( | 455.1264 | 0.13732 | 0.507906 | 0.929089 | Fstl4 |
| ENSMUSG( | 492.145 | 0.289389 | 0.133665 | 0.822912 | Fstl5 |
| ENSMUSG( | 123059.9 | -0.14595 | 0.145587 | 0.83423 | Fth1 |
| ENSMUSG( | 13994.14 | 0.0195 | 0.878639 | 0.989644 | Ftl1 |
| ENSMUSG( | 35.51581 | -0.05548 | 0.873789 | 0.989644 | Ftl1-ps2 |
| ENSMUSG( | 4531.552 | -0.15412 | 0.165369 | 0.838866 | Fto |
| ENSMUSG( | 417.2175 | -0.12097 | 0.393777 | 0.917257 | Ftsj1 |
| ENSMUSG( | 823.7333 | 0.054218 | 0.646617 | 0.953988 | Ftsj3 |
| ENSMUSG( | 749.8883 | 0.256083 | 0.130512 | 0.822225 | Ftx |
| ENSMUSG( | 1505.649 | 0.099481 | 0.497461 | 0.927825 | Fubp1 |
| ENSMUSG( | 1412.209 | -0.02367 | 0.828159 | 0.980224 | Fubp3 |
| ENSMUSG( | 1805.196 | 0.104415 | 0.415443 | 0.917257 | Fuca1 |
| ENSMUSG( | 1780.398 | -0.03864 | 0.766615 | 0.974236 | Fuca2 |
| ENSMUSG( | 759.3818 | -0.12301 | 0.423343 | 0.917257 | Fundc1 |
| ENSMUSG( | 858.1657 | -0.20581 | 0.125257 | 0.81516 | Fundc2 |
| ENSMUSG( | 510.1556 | -0.09863 | 0.461349 | 0.923329 | Fuom |
| ENSMUSG( | 1508.431 | 0.230965 | 0.111523 | 0.808371 | Furin |
| ENSMUSG( | 4971.226 | -0.07726 | 0.495103 | 0.927372 | Fus |
| ENSMUSG( | 593.8625 | 0.11572 | 0.403086 | 0.917257 | Fut10 |
| ENSMUSG( | 621.2976 | 0.140225 | 0.365264 | 0.914449 | Fut11 |
| ENSMUSG( | 12.05595 | -1.2978 | 0.029522 | 0.619333 | Fut2 |
| ENSMUSG( | 10.76255 | 0.128707 | 0.831253 | NA | Fut4 |
| ENSMUSG( | 9.525205 | -0.0749 | 0.910013 | NA | Fut7 |
| ENSMUSG( | 2732.808 | 0.016487 | 0.905487 | 0.991089 | Fut8 |
| ENSMUSG( | 2444.045 | 0.341144 | 0.06266 | 0.736482 | Fut9 |
| ENSMUSG( | 587.9325 | -0.14199 | 0.30487 | 0.898538 | Fuz |
| ENSMUSG( | 64.58092 | 0.356114 | 0.282391 | 0.892412 | Fv1 |
| ENSMUSG( | 146.1616 | -0.14757 | 0.436802 | 0.919879 | Fxn |
| ENSMUSG( | 1671.855 | 0.039206 | 0.734365 | 0.967228 | Fxr1 |
| ENSMUSG( | 2919.273 | -0.03292 | 0.789359 | 0.976593 | Fxr2 |
| ENSMUSG( | 2224.743 | -0.08696 | 0.470845 | 0.923329 | Fxyd1 |
| ENSMUSG( | 143.1194 | -0.02589 | 0.905749 | 0.991089 | Fxyd2 |
| ENSMUSG( | 27.40386 | -0.03837 | 0.928265 | 0.99306 | Fxyd3 |
| ENSMUSG( | 798.152 | 0.036514 | 0.786156 | 0.976285 | Fxyd5 |
| ENSMUSG( | 5158.917 | 0.15043 | 0.334225 | 0.907907 | Fxyd6 |
| ENSMUSG( | 2061.949 | -0.25482 | 0.063286 | 0.736482 | Fxyd7 |
| ENSMUSG( | 257.826 | 0.133835 | 0.42827 | 0.917257 | Fyb1 |
| ENSMUSG( | 50.20574 | 0.001545 | 0.996409 | 0.999178 | Fyb2 |
| ENSMUSG( | 3383.41 | -0.12899 | 0.31969 | 0.903664 | Fyco1 |
| ENSMUSG( | 7517.153 | -0.13909 | 0.255583 | 0.883043 | Fyn |
| ENSMUSG( | 1169.138 | 0.011712 | 0.923786 | 0.99282 | Fyttd1 |
| ENSMUSG( | 676.4249 | 0.522336 | 0.000901 | 0.101185 | Fzd1 |
| ENSMUSG( | 193.1868 | -0.08008 | 0.705527 | 0.96315 | Fzd10 |
| ENSMUSG( | 178.8624 | -0.12886 | 0.481045 | 0.924548 | Fzd10os |
| ENSMUSG( | 1063.87 | 0.768071 | 8.67E-05 | 0.018748 | Fzd2 |
| ENSMUSG( | 3239.697 | 0.112202 | 0.521402 | 0.932489 | Fzd3 |
| ENSMUSG( | 912.0084 | 0.197048 | 0.309723 | 0.898538 | Fzd4 |
| ENSMUSG( | 84.90401 | 0.378188 | 0.200706 | 0.86439 | Fzd5 |
| ENSMUSG( | 479.3003 | 0.264864 | 0.246806 | 0.877904 | Fzd6 |
| ENSMUSG( | 406.1376 | 0.365608 | 0.015409 | 0.495791 | Fzd7 |
| ENSMUSG( | 203.1549 | 0.702602 | 0.00105 | 0.111298 | Fzd8 |
| ENSMUSG( | 291.2361 | 0.043066 | 0.837838 | 0.981376 | Fzd9 |
| ENSMUSG( | 1195.865 | -0.07813 | 0.556282 | 0.939565 | Fzr1 |
| ENSMUSG( | 186.8868 | 0.455153 | 0.064007 | 0.736482 | G0s2 |
| ENSMUSG( | 373.0901 | 0.338521 | 0.026977 | 0.603041 | G2e3 |
| ENSMUSG( | 1994.865 | 0.011237 | 0.920782 | 0.992644 | G3bp1 |
| ENSMUSG( | 3874.915 | -0.00643 | 0.959964 | 0.994798 | G3bp2 |
| ENSMUSG( | 11.39353 | -0.16788 | 0.78534 | NA | G430095P16Rik |
| ENSMUSG( | 33.71582 | -0.43038 | 0.291988 | 0.897854 | G530011O06Rik |
| ENSMUSG( | 100.1539 | 0.062991 | 0.814626 | 0.979198 | G630016G05Rik |
| ENSMUSG( | 12.70898 | 0.01705 | 0.976627 | 0.996416 | G630093K05Rik |
| ENSMUSG( | 1924.164 | 0.110433 | 0.383924 | 0.917257 | G6pc3 |
| ENSMUSG( | 1272.363 | -0.1188 | 0.346358 | 0.910966 | G6pdx |
| ENSMUSG( | 10046.13 | 0.262576 | 0.111585 | 0.808371 | Gaa |
| ENSMUSG( | 1265.917 | -0.01263 | 0.916229 | 0.992529 | Gab1 |
| ENSMUSG( | 1075.156 | -0.15702 | 0.179571 | 0.850372 | Gab2 |
| ENSMUSG( | 27.89661 | 0.237263 | 0.558582 | 0.93957 | Gab3 |
| ENSMUSG( | 4353.774 | -0.14977 | 0.245023 | 0.877904 | Gabarap |
| ENSMUSG( | 9463.606 | -0.15812 | 0.276901 | 0.89114 | Gabarapl1 |
| ENSMUSG( | 2571.958 | -0.21437 | 0.115715 | 0.813384 | Gabarapl2 |
| ENSMUSG( | 39684.77 | -0.00999 | 0.939195 | 0.993898 | Gabbr1 |
| ENSMUSG( | 15021.63 | 0.131437 | 0.399553 | 0.917257 | Gabbr2 |
| ENSMUSG( | 902.5461 | -0.02874 | 0.828042 | 0.980224 | Gabpa |
| ENSMUSG( | 404.7829 | 0.102149 | 0.480849 | 0.924548 | Gabpb1 |
| ENSMUSG( | 655.224 | 0.103268 | 0.500631 | 0.927825 | Gabpb2 |
| ENSMUSG( | 8059.466 | 0.274656 | 0.169171 | 0.840432 | Gabra1 |
| ENSMUSG( | 4437.829 | 1.703753 | 2.92E-21 | 6.39E-18 | Gabra2 |
| ENSMUSG( | 4242.539 | 0.056353 | 0.671581 | 0.95694 | Gabra3 |
| ENSMUSG( | 2884.023 | 0.150443 | 0.371314 | 0.915491 | Gabra4 |
| ENSMUSG( | 2286.118 | 0.280285 | 0.052173 | 0.712974 | Gabra5 |
| ENSMUSG( | 4592.629 | 0.164833 | 0.275136 | 0.889874 | Gabrb1 |
| ENSMUSG( | 1226.574 | 0.459131 | 0.371665 | 0.915491 | Gabrb2 |
| ENSMUSG( | 7602.723 | 0.191179 | 0.245411 | 0.877904 | Gabrb3 |
| ENSMUSG( | 2492.826 | -0.09633 | 0.523473 | 0.932848 | Gabrd |
| ENSMUSG( | 17.01224 | -0.28357 | 0.562501 | 0.940761 | Gabre |
| ENSMUSG( | 1340.957 | -0.03356 | 0.843341 | 0.983281 | Gabrg1 |
| ENSMUSG( | 4412.633 | 0.242416 | 0.218787 | 0.866104 | Gabrg2 |
| ENSMUSG( | 880.3182 | 0.288743 | 0.065979 | 0.746371 | Gabrg3 |
| ENSMUSG( | 40.58847 | -0.24525 | 0.46727 | 0.923329 | Gabrq |
| ENSMUSG( | 13.99897 | 0.596565 | 0.247791 | 0.877904 | Gabrr2 |

| | | | | | | | | | | |
|---|---|---|---|---|---|---|---|---|---|---|
| ENSMUSG( | Frat2 | 143.4618 | 0.02093 | 0.915979 | 0.988634 | ENSMUSG( | Frat2 | 130.8039 | 0.018045 | 0.934569 | 0.99306 |
| ENSMUSG( | Frem1 | 184.8745 | -0.29941 | 0.055665 | 0.363338 | ENSMUSG( | Frem1 | 179.7982 | -0.09918 | 0.565666 | 0.941404 |
| ENSMUSG( | Frem2 | 234.4926 | -0.66156 | 0.00022 | 0.011467 | ENSMUSG( | Frem2 | 238.3205 | -0.27481 | 0.128188 | 0.818934 |
| ENSMUSG( | Frem3 | 15.2904 | -1.25537 | 0.018345 | NA | ENSMUSG( | Frem3 | 17.08061 | -0.4553 | 0.359269 | 0.914061 |
| ENSMUSG( | Frey1 | 95.90846 | -0.0633 | 0.774638 | 0.955606 | ENSMUSG( | Frey1 | 78.53116 | -0.38742 | 0.116474 | 0.813384 |
| ENSMUSG( | Frg1 | 424.2139 | 0.034296 | 0.785473 | 0.958571 | ENSMUSG( | Frg1 | 388.5468 | 0.046807 | 0.741123 | 0.968811 |
| ENSMUSG( | Frg2f1 | 374.6631 | 0.007896 | 0.951251 | 0.993821 | ENSMUSG( | Frg2f1 | 328.027 | -0.10352 | 0.510753 | 0.929545 |
| ENSMUSG( | Frk | 14.5687 | 0.250226 | 0.637952 | NA | ENSMUSG( | Frk | 17.17717 | 0.8886 | 0.062817 | 0.736482 |
| ENSMUSG( | Frmd3 | 132.3019 | -0.246 | 0.235108 | 0.682364 | ENSMUSG( | Frmd3 | 120.6972 | -0.2443 | 0.262974 | 0.885357 |
| ENSMUSG( | Frmd4a | 2706.074 | -0.08669 | 0.31732 | 0.763823 | ENSMUSG( | Frmd4a | 2421.731 | -0.13935 | 0.18161 | 0.850579 |
| ENSMUSG( | Frmd4b | 797.9075 | -0.15077 | 0.361018 | 0.792797 | ENSMUSG( | Frmd4b | 798.2795 | 0.129559 | 0.497105 | 0.927825 |
| ENSMUSG( | Frmd5 | 991.7037 | -0.04975 | 0.687565 | 0.93324 | ENSMUSG( | Frmd5 | 905.6462 | -0.03999 | 0.766345 | 0.974236 |
| ENSMUSG( | Frmd6 | 1009.415 | -0.22815 | 0.068668 | 0.405903 | ENSMUSG( | Frmd6 | 928.7165 | -0.19214 | 0.194064 | 0.860096 |
| ENSMUSG( | Frmd7 | 41.68904 | -0.32636 | 0.476047 | 0.851133 | ENSMUSG( | Frmd7 | 51.39175 | 0.503049 | 0.561967 | 0.940388 |
| ENSMUSG( | Frmd8 | 861.9591 | -0.09332 | 0.363166 | 0.793211 | ENSMUSG( | Frmd8 | 774.8367 | -0.13214 | 0.296645 | 0.898538 |
| ENSMUSG( | Frmpd1 | 1389.333 | 0.254956 | 0.018167 | 0.20312 | ENSMUSG( | Frmpd1 | 1034.06 | -0.33868 | 0.007631 | 0.37235 |
| ENSMUSG( | Frmpd2 | 59.87243 | -0.29926 | 0.272481 | 0.722478 | ENSMUSG( | Frmpd2 | 60.46124 | 0.035393 | 0.904085 | 0.991089 |
| ENSMUSG( | Frmpd3 | 2468.044 | 0.408113 | 6.45E-05 | 0.004822 | ENSMUSG( | Frmpd3 | 2048.63 | 0.165958 | 0.170864 | 0.842157 |
| ENSMUSG( | Frmpd4 | 1032.539 | 0.074215 | 0.432668 | 0.827873 | ENSMUSG( | Frmpd4 | 961.6596 | 0.135496 | 0.225239 | 0.868468 |
| ENSMUSG( | Frrs1 | 132.9866 | -0.25818 | 0.175805 | 0.611006 | ENSMUSG( | Frrs1 | 129.7536 | -0.033 | 0.884449 | 0.989688 |
| ENSMUSG( | Frrs1l | 6608.045 | -0.35838 | 0.015146 | 0.181303 | ENSMUSG( | Frrs1l | 7328.311 | 0.226079 | 0.189828 | 0.857355 |
| ENSMUSG( | Frs2 | 3496.801 | 0.045455 | 0.586569 | 0.900023 | ENSMUSG( | Frs2 | 3048.488 | -0.08452 | 0.483315 | 0.92499 |
| ENSMUSG( | Frs3 | 844.5979 | 0.038523 | 0.766112 | 0.954173 | ENSMUSG( | Frs3 | 750.7571 | -0.03315 | 0.794596 | 0.97681 |
| ENSMUSG( | Fry | 3156.068 | 0.023831 | 0.821489 | 0.966248 | ENSMUSG( | Fry | 3017.938 | 0.162425 | 0.213006 | 0.864412 |
| ENSMUSG( | Fryl | 2540.33 | -0.00871 | 0.921648 | 0.990436 | ENSMUSG( | Fryl | 2394.903 | 0.089357 | 0.403986 | 0.917257 |
| ENSMUSG( | Frzb | 711.9483 | -0.54971 | 1.90E-05 | 0.001887 | ENSMUSG( | Frzb | 798.615 | 0.097117 | 0.563101 | 0.941158 |
| ENSMUSG( | Fsbp | 37.66952 | -0.21295 | 0.495038 | 0.860798 | ENSMUSG( | Fsbp | 27.56386 | -1.09169 | 0.010897 | 0.422321 |
| ENSMUSG( | Fscn1 | 6353.271 | 0.069416 | 0.528264 | 0.878236 | ENSMUSG( | Fscn1 | 5236.34 | -0.23013 | 0.079142 | 0.77458 |
| ENSMUSG( | Fsd1 | 1288.102 | 0.06107 | 0.566777 | 0.891644 | ENSMUSG( | Fsd1 | 1080.753 | -0.18084 | 0.123368 | 0.814691 |
| ENSMUSG( | Fsd1l | 516.0658 | -0.17933 | 0.138755 | 0.554336 | ENSMUSG( | Fsd1l | 514.5801 | 0.085378 | 0.527938 | 0.934616 |
| ENSMUSG( | Fsd2 | 55.94578 | 0.323689 | 0.304161 | 0.752287 | ENSMUSG( | Fsd2 | 38.94087 | -0.46884 | 0.160034 | 0.837724 |
| ENSMUSG( | Fst | 101.2002 | 0.062001 | 0.772022 | 0.954994 | ENSMUSG( | Fst | 116.8161 | 0.673089 | 0.003372 | 0.232514 |
| ENSMUSG( | Fstl1 | 1724.486 | -0.15401 | 0.208664 | 0.650496 | ENSMUSG( | Fstl1 | 1779.06 | 0.205008 | 0.143865 | 0.831311 |
| ENSMUSG( | Fstl3 | 58.3381 | 0.108431 | 0.682812 | 0.932022 | ENSMUSG( | Fstl3 | 51.06085 | 0.000238 | 0.999332 | 0.999503 |
| ENSMUSG( | Fstl4 | 402.6135 | -0.54341 | 0.001121 | 0.035587 | ENSMUSG( | Fstl4 | 455.1264 | 0.13732 | 0.507906 | 0.929089 |
| ENSMUSG( | Fstl5 | 400.1026 | -0.63183 | 0.000458 | 0.019492 | ENSMUSG( | Fstl5 | 492.145 | 0.289389 | 0.133665 | 0.822912 |
| ENSMUSG( | Fth1 | 148310.2 | 0.125715 | 0.103335 | 0.488943 | ENSMUSG( | Fth1 | 123059.9 | -0.14595 | 0.145587 | 0.83423 |
| ENSMUSG( | Ftl1 | 15753.81 | 0.093725 | 0.377224 | 0.801213 | ENSMUSG( | Ftl1 | 13994.14 | 0.0195 | 0.878639 | 0.989644 |
| ENSMUSG( | Ftl1-ps2 | 43.38596 | 0.254614 | 0.40148 | 0.814815 | ENSMUSG( | Ftl1-ps2 | 35.51581 | -0.05548 | 0.873789 | 0.989644 |
| ENSMUSG( | Fto | 5352.569 | 0.061708 | 0.496451 | 0.86175 | ENSMUSG( | Fto | 4531.552 | -0.15412 | 0.165369 | 0.838866 |
| ENSMUSG( | Ftsj1 | 485.4482 | 0.046151 | 0.689179 | 0.933485 | ENSMUSG( | Ftsj1 | 417.2175 | -0.12097 | 0.393777 | 0.917257 |
| ENSMUSG( | Ftsj3 | 944.9474 | 0.176891 | 0.0862 | 0.451605 | ENSMUSG( | Ftsj3 | 823.7333 | 0.054218 | 0.646617 | 0.953988 |
| ENSMUSG( | Ftx | 854.6939 | 0.357734 | 0.008768 | 0.131926 | ENSMUSG( | Ftx | 749.8883 | 0.256083 | 0.130512 | 0.822225 |
| ENSMUSG( | Fubp1 | 1569.368 | -0.05019 | 0.567221 | 0.891666 | ENSMUSG( | Fubp1 | 1505.649 | 0.099481 | 0.497461 | 0.927825 |
| ENSMUSG( | Fubp3 | 1575.054 | 0.023473 | 0.792408 | 0.959461 | ENSMUSG( | Fubp3 | 1412.209 | -0.02367 | 0.828159 | 0.980224 |
| ENSMUSG( | Fuca1 | 1858.139 | -0.08155 | 0.453741 | 0.836798 | ENSMUSG( | Fuca1 | 1805.196 | 0.104415 | 0.415443 | 0.917257 |
| ENSMUSG( | Fuca2 | 1862.474 | -0.18538 | 0.12431 | 0.531714 | ENSMUSG( | Fuca2 | 1780.398 | -0.03864 | 0.766615 | 0.974236 |
| ENSMUSG( | Fundc1 | 872.7305 | 0.017333 | 0.874364 | 0.979432 | ENSMUSG( | Fundc1 | 759.3818 | -0.12301 | 0.423343 | 0.917257 |
| ENSMUSG( | Fundc2 | 989.5846 | -0.05305 | 0.637023 | 0.915191 | ENSMUSG( | Fundc2 | 858.1657 | -0.20581 | 0.125257 | 0.81516 |
| ENSMUSG( | Fuom | 574.2064 | -0.02219 | 0.844718 | 0.972023 | ENSMUSG( | Fuom | 510.1556 | -0.09863 | 0.461349 | 0.923329 |
| ENSMUSG( | Furin | 1456.305 | -0.138 | 0.294877 | 0.743798 | ENSMUSG( | Furin | 1508.431 | 0.230965 | 0.111523 | 0.808371 |
| ENSMUSG( | Fus | 5457.147 | -0.07477 | 0.38035 | 0.801757 | ENSMUSG( | Fus | 4971.226 | -0.07726 | 0.495103 | 0.927372 |
| ENSMUSG( | Fut10 | 643.3067 | 0.07895 | 0.56553 | 0.890871 | ENSMUSG( | Fut10 | 593.8625 | 0.11572 | 0.403086 | 0.917257 |
| ENSMUSG( | Fut11 | 599.8212 | -0.24225 | 0.066599 | 0.398819 | ENSMUSG( | Fut11 | 621.2976 | 0.140225 | 0.365264 | 0.914449 |
| ENSMUSG( | Fut2 | 15.34862 | -0.66276 | 0.180959 | NA | ENSMUSG( | Fut2 | 12.05595 | -1.2978 | 0.029522 | 0.619333 |
| ENSMUSG( | Fut4 | 11.52629 | 0.091713 | 0.864997 | NA | ENSMUSG( | Fut4 | 10.76255 | 0.128707 | 0.831253 | NA |
| ENSMUSG( | Fut7 | 9.939858 | -0.17858 | 0.783335 | NA | ENSMUSG( | Fut7 | 9.525205 | -0.0749 | 0.910013 | NA |
| ENSMUSG( | Fut8 | 2722.678 | -0.27947 | 0.029845 | 0.261951 | ENSMUSG( | Fut8 | 2732.808 | 0.016487 | 0.905487 | 0.991089 |
| ENSMUSG( | Fut9 | 2222.959 | -0.18873 | 0.208149 | 0.650319 | ENSMUSG( | Fut9 | 2444.045 | 0.341144 | 0.06266 | 0.736482 |
| ENSMUSG( | Fuz | 694.0421 | 0.069614 | 0.527385 | 0.877599 | ENSMUSG( | Fuz | 587.9325 | -0.14199 | 0.30487 | 0.898538 |
| ENSMUSG( | Fv1 | 69.98066 | 0.289279 | 0.251807 | 0.701908 | ENSMUSG( | Fv1 | 64.58092 | 0.356114 | 0.282391 | 0.892412 |
| ENSMUSG( | Fxn | 175.1075 | 0.113999 | 0.522612 | 0.875389 | ENSMUSG( | Fxn | 146.1616 | -0.14757 | 0.436802 | 0.919879 |
| ENSMUSG( | Fxr1 | 1861.514 | 0.0824 | 0.360209 | 0.791947 | ENSMUSG( | Fxr1 | 1671.855 | 0.039206 | 0.734365 | 0.967228 |
| ENSMUSG( | Fxr2 | 3436.739 | 0.166793 | 0.109743 | 0.500285 | ENSMUSG( | Fxr2 | 2919.273 | -0.03292 | 0.789359 | 0.976593 |
| ENSMUSG( | Fxyd1 | 2396.502 | -0.14102 | 0.175219 | 0.610405 | ENSMUSG( | Fxyd1 | 2224.743 | -0.08696 | 0.470845 | 0.923329 |
| ENSMUSG( | Fxyd2 | 153.1077 | -0.08031 | 0.629808 | 0.913965 | ENSMUSG( | Fxyd2 | 143.1194 | -0.02589 | 0.905749 | 0.991089 |
| ENSMUSG( | Fxyd3 | 33.81343 | 0.270029 | 0.472995 | 0.84867 | ENSMUSG( | Fxyd3 | 27.40386 | -0.03837 | 0.928265 | 0.99306 |
| ENSMUSG( | Fxyd5 | 903.6491 | 0.125066 | 0.321311 | 0.766497 | ENSMUSG( | Fxyd5 | 798.152 | 0.036514 | 0.786156 | 0.976285 |
| ENSMUSG( | Fxyd6 | 4974.735 | -0.23238 | 0.088236 | 0.455783 | ENSMUSG( | Fxyd6 | 5158.917 | 0.15043 | 0.334225 | 0.907907 |
| ENSMUSG( | Fxyd7 | 2327.371 | -0.16423 | 0.152884 | 0.579254 | ENSMUSG( | Fxyd7 | 2061.949 | -0.25482 | 0.063286 | 0.736482 |
| ENSMUSG( | Fyb1 | 289.1641 | 0.19129 | 0.245184 | 0.692812 | ENSMUSG( | Fyb1 | 257.826 | 0.133835 | 0.42827 | 0.917257 |
| ENSMUSG( | Fyb2 | 54.42489 | -0.009 | 0.975129 | 0.997302 | ENSMUSG( | Fyb2 | 50.20574 | 0.001545 | 0.996409 | 0.999178 |
| ENSMUSG( | Fyco1 | 4032.012 | 0.111884 | 0.184396 | 0.621442 | ENSMUSG( | Fyco1 | 3383.41 | -0.12899 | 0.31969 | 0.903664 |
| ENSMUSG( | Fyn | 8633.232 | -0.00288 | 0.972986 | 0.996705 | ENSMUSG( | Fyn | 7517.153 | -0.13909 | 0.255583 | 0.883043 |
| ENSMUSG( | Fyttd1 | 1275.417 | -0.00285 | 0.975325 | 0.997302 | ENSMUSG( | Fyttd1 | 1169.138 | 0.011712 | 0.923786 | 0.99282 |
| ENSMUSG( | Fzd1 | 615.7126 | 0.022122 | 0.867397 | 0.976559 | ENSMUSG( | Fzd1 | 676.4249 | 0.522336 | 0.000901 | 0.101185 |
| ENSMUSG( | Fzd10 | 178.7447 | -0.65843 | 0.000445 | 0.019233 | ENSMUSG( | Fzd10 | 193.1868 | -0.08008 | 0.705527 | 0.96315 |
| ENSMUSG( | Fzd10os | 216.0402 | 0.137257 | 0.336737 | 0.777 | ENSMUSG( | Fzd10os | 178.8624 | -0.12886 | 0.481045 | 0.924548 |
| ENSMUSG( | Fzd2 | 860.7951 | -0.01865 | 0.911377 | 0.988483 | ENSMUSG( | Fzd2 | 1063.87 | 0.768071 | 8.67E-05 | 0.018748 |
| ENSMUSG( | Fzd3 | 3107.422 | -0.28993 | 0.054347 | 0.358773 | ENSMUSG( | Fzd3 | 3239.697 | 0.112202 | 0.521402 | 0.932489 |
| ENSMUSG( | Fzd4 | 947.5206 | 0.038405 | 0.769713 | 0.954674 | ENSMUSG( | Fzd4 | 912.0084 | 0.197048 | 0.309723 | 0.898538 |
| ENSMUSG( | Fzd5 | 71.88599 | -0.39331 | 0.104565 | 0.490752 | ENSMUSG( | Fzd5 | 84.90401 | 0.378188 | 0.200706 | 0.86439 |
| ENSMUSG( | Fzd6 | 464.2056 | -0.09787 | 0.587483 | 0.900403 | ENSMUSG( | Fzd6 | 479.3003 | 0.264864 | 0.246806 | 0.877904 |
| ENSMUSG( | Fzd7 | 375.6484 | -0.11613 | 0.397759 | 0.812161 | ENSMUSG( | Fzd7 | 406.1376 | 0.365608 | 0.015409 | 0.495791 |
| ENSMUSG( | Fzd8 | 152.5075 | -0.33982 | 0.0914 | 0.464738 | ENSMUSG( | Fzd8 | 203.1549 | 0.702602 | 0.00105 | 0.111298 |
| ENSMUSG( | Fzd9 | 269.4078 | -0.50687 | 0.007004 | 0.115211 | ENSMUSG( | Fzd9 | 291.2361 | 0.043066 | 0.837838 | 0.981376 |
| ENSMUSG( | Fzr1 | 1402.444 | 0.112572 | 0.361808 | 0.792797 | ENSMUSG( | Fzr1 | 1195.865 | -0.07813 | 0.556282 | 0.939565 |
| ENSMUSG( | G0s2 | 145.0137 | -0.58293 | 0.007724 | 0.122031 | ENSMUSG( | G0s2 | 186.8868 | 0.455153 | 0.064007 | 0.736482 |
| ENSMUSG( | G2e3 | 355.8786 | -0.05229 | 0.70916 | 0.939599 | ENSMUSG( | G2e3 | 373.0901 | 0.338521 | 0.026977 | 0.603041 |
| ENSMUSG( | G3bp1 | 2260.046 | 0.102516 | 0.269645 | 0.719729 | ENSMUSG( | G3bp1 | 1994.865 | 0.011237 | 0.920782 | 0.992644 |
| ENSMUSG( | G3bp2 | 4305.866 | 0.031405 | 0.724103 | 0.944096 | ENSMUSG( | G3bp2 | 3874.915 | -0.00643 | 0.959964 | 0.994798 |
| ENSMUSG( | G430095P1 | 14.23521 | 0.158009 | 0.767533 | NA | ENSMUSG( | G430095P1 | 11.39353 | -0.16788 | 0.78534 | NA |
| ENSMUSG( | G530011O( | 64.91386 | 1.04823 | 0.000495 | 0.020378 | ENSMUSG( | G530011O( | 33.71582 | -0.43038 | 0.291988 | 0.897854 |
| ENSMUSG( | G630016G( | 104.3817 | -0.07551 | 0.74866 | 0.949604 | ENSMUSG( | G630016G( | 100.1539 | 0.062991 | 0.814626 | 0.979198 |
| ENSMUSG( | G630093K( | 12.05665 | -0.37519 | 0.4975 | NA | ENSMUSG( | G630093K( | 12.70898 | 0.01705 | 0.976627 | 0.996416 |
| ENSMUSG( | G6pc3 | 1911.448 | -0.18423 | 0.083855 | 0.446475 | ENSMUSG( | G6pc3 | 1924.164 | 0.110433 | 0.383924 | 0.917257 |
| ENSMUSG( | G6pdx | 1501.196 | 0.09407 | 0.288971 | 0.739705 | ENSMUSG( | G6pdx | 1272.363 | -0.1188 | 0.346358 | 0.910966 |
| ENSMUSG( | Gaa | 9634.06 | -0.12212 | 0.37461 | 0.800445 | ENSMUSG( | Gaa | 10046.13 | 0.262576 | 0.111585 | 0.808371 |
| ENSMUSG( | Gab1 | 1322.567 | -0.15824 | 0.186011 | 0.623496 | ENSMUSG( | Gab1 | 1265.917 | -0.01263 | 0.916229 | 0.992529 |
| ENSMUSG( | Gab2 | 1243.796 | 0.001159 | 0.990151 | 0.999144 | ENSMUSG( | Gab2 | 1075.156 | -0.15702 | 0.179571 | 0.850372 |
| ENSMUSG( | Gab3 | 34.44627 | 0.537627 | 0.100603 | 0.48402 | ENSMUSG( | Gab3 | 27.89661 | 0.237263 | 0.558582 | 0.93957 |
| ENSMUSG( | Gabarap | 5164.718 | 0.079723 | 0.381414 | 0.802629 | ENSMUSG( | Gabarap | 4353.774 | -0.14977 | 0.245023 | 0.877904 |
| ENSMUSG( | Gabarapl1 | 11163.34 | 0.055973 | 0.621684 | 0.911432 | ENSMUSG( | Gabarapl1 | 9463.606 | -0.15812 | 0.276901 | 0.89114 |
| ENSMUSG( | Gabarapl2 | 3045.355 | 0.016377 | 0.882788 | 0.982356 | ENSMUSG( | Gabarapl2 | 2571.958 | -0.21437 | 0.115715 | 0.813384 |
| ENSMUSG( | Gabbr1 | 41276.73 | -0.17158 | 0.138459 | 0.553773 | ENSMUSG( | Gabbr1 | 39684.77 | -0.00999 | 0.939195 | 0.993898 |
| ENSMUSG( | Gabbr2 | 14494.25 | -0.25207 | 0.063201 | 0.388743 | ENSMUSG( | Gabbr2 | 15021.63 | 0.131437 | 0.399553 | 0.917257 |
| ENSMUSG( | Gabpa | 1038.25 | 0.107363 | 0.402324 | 0.815103 | ENSMUSG( | Gabpa | 902.5461 | -0.02874 | 0.828042 | 0.980224 |
| ENSMUSG( | Gabpb1 | 448.4059 | 0.131938 | 0.303271 | 0.75169 | ENSMUSG( | Gabpb1 | 404.7829 | 0.102149 | 0.480849 | 0.924548 |
| ENSMUSG( | Gabpb2 | 697.9805 | 0.012607 | 0.92037 | 0.990276 | ENSMUSG( | Gabpb2 | 655.224 | 0.103268 | 0.500631 | 0.927825 |
| ENSMUSG( | Gabra1 | 6452.521 | -0.71279 | 4.21E-05 | 0.003506 | ENSMUSG( | Gabra1 | 8059.466 | 0.274656 | 0.169171 | 0.840432 |
| ENSMUSG( | Gabra2 | 4490.366 | 1.549417 | 4.17E-23 | 6.15E-20 | ENSMUSG( | Gabra2 | 4437.829 | 1.703753 | 2.92E-21 | 6.39E-18 |
| ENSMUSG( | Gabra3 | 4357.62 | -0.13964 | 0.24018 | 0.687899 | ENSMUSG( | Gabra3 | 4242.539 | 0.056353 | 0.671581 | 0.95694 |
| ENSMUSG( | Gabra4 | 2483.879 | -0.61185 | 3.58E-06 | 0.000492 | ENSMUSG( | Gabra4 | 2884.023 | 0.150443 | 0.371314 | 0.915491 |
| ENSMUSG( | Gabra5 | 2079.784 | -0.26198 | 0.037533 | 0.299308 | ENSMUSG( | Gabra5 | 2286.118 | 0.280285 | 0.052173 | 0.712974 |
| ENSMUSG( | Gabrb1 | 4586.255 | -0.10729 | 0.427793 | 0.825894 | ENSMUSG( | Gabrb1 | 4592.629 | 0.164833 | 0.275136 | 0.889874 |
| ENSMUSG( | Gabrb2 | 904.2318 | -0.76433 | 0.072007 | 0.414057 | ENSMUSG( | Gabrb2 | 1226.574 | 0.459131 | 0.371665 | 0.915491 |
| ENSMUSG( | Gabrb3 | 6755.067 | -0.45105 | 0.000947 | 0.031617 | ENSMUSG( | Gabrb3 | 7602.723 | 0.191179 | 0.245411 | 0.877904 |
| ENSMUSG( | Gabrd | 2545.588 | -0.32572 | 0.008112 | 0.125423 | ENSMUSG( | Gabrd | 2492.826 | -0.09633 | 0.523473 | 0.932848 |
| ENSMUSG( | Gabre | 17.78583 | -0.41846 | 0.402012 | NA | ENSMUSG( | Gabre | 17.01224 | -0.28357 | 0.562501 | 0.940761 |
| ENSMUSG( | Gabrg1 | 1384.391 | -0.21932 | 0.163502 | 0.594869 | ENSMUSG( | Gabrg1 | 1340.957 | -0.03356 | 0.843341 | 0.983281 |
| ENSMUSG( | Gabrg2 | 3712 | -0.57572 | 0.00074 | 0.026499 | ENSMUSG( | Gabrg2 | 4412.633 | 0.242416 | 0.218787 | 0.866104 |
| ENSMUSG( | Gabrg3 | 772.1637 | -0.37008 | 0.007094 | 0.116231 | ENSMUSG( | Gabrg3 | 880.3182 | 0.288743 | 0.065979 | 0.746371 |
| ENSMUSG( | Gabrq | 40.41064 | -0.53461 | 0.165303 | 0.596691 | ENSMUSG( | Gabrq | 40.58847 | -0.24525 | 0.46727 | 0.923329 |
| ENSMUSG( | Gabrr2 | 13.23969 | 0.194706 | 0.69545 | NA | ENSMUSG( | Gabrr2 | 13.99897 | 0.596565 | 0.247791 | 0.877904 |
| ENSMUSG( | Gad1 | 4623.29 | -0.10818 | 0.456946 | 0.839183 | ENSMUSG( | Gad1 | 4194.55 | -0.1246 | 0.478163 | 0.923329 |
| ENSMUSG( | Gad2 | 2486.532 | -0.11803 | 0.423571 | 0.823976 | ENSMUSG( | Gad2 | 2366.134 | 0.005805 | 0.97603 | 0.996416 |
| ENSMUSG( | Gadd45a | 149.5137 | -0.49207 | 0.003786 | 0.078636 | ENSMUSG( | Gadd45a | 138.5912 | -0.43032 | 0.023181 | 0.585577 |
| ENSMUSG( | Gadd45b | 250.5064 | -0.62962 | 0.005398 | 0.098038 | ENSMUSG( | Gadd45b | 250.9461 | -0.30193 | 0.181172 | 0.850372 |
| ENSMUSG( | Gadd45g | 345.548 | -0.31749 | 0.139776 | 0.55636 | ENSMUSG( | Gadd45g | 315.4379 | -0.30504 | 0.16494 | 0.838743 |
| ENSMUSG( | Gadd45gip | 746.6256 | 0.032876 | 0.759532 | 0.952498 | ENSMUSG( | Gadd45gip | 630.9188 | -0.19284 | 0.133928 | 0.822912 |
| ENSMUSG( | Gadl1 | 55.11573 | -0.1852 | 0.487487 | 0.858812 | ENSMUSG( | Gadl1 | 40.592 | -0.94733 | 0.004687 | 0.282119 |
| ENSMUSG( | Gak | 3462.305 | 0.122167 | 0.164775 | 0.595869 | ENSMUSG( | Gak | 3111.482 | 0.085141 | 0.414212 | 0.917257 |
| ENSMUSG( | Gal3st1 | 741.1894 | -0.2063 | 0.063822 | 0.390932 | ENSMUSG( | Gal3st1 | 728.5306 | 0.023972 | 0.876702 | 0.989644 |
| ENSMUSG( | Gal3st3 | 1914.891 | -0.17759 | 0.251039 | 0.700713 | ENSMUSG( | Gal3st3 | 2052.667 | 0.28813 | 0.076948 | 0.770558 |
| ENSMUSG( | Galc | 1831.617 | -0.00868 | 0.93075 | 0.992033 | ENSMUSG( | Galc | 1729.527 | 0.093928 | 0.440337 | 0.921166 |
| ENSMUSG( | Gale | 406.3915 | 0.034912 | 0.807802 | 0.963257 | ENSMUSG( | Gale | 381.3891 | 0.126568 | 0.428873 | 0.917257 |
| ENSMUSG( | Galk1 | 320.3151 | 0.036223 | 0.808991 | 0.963548 | ENSMUSG( | Galk1 | 300.7821 | 0.113998 | 0.495921 | 0.927825 |
| ENSMUSG( | Galk2 | 407.0704 | -0.091 | 0.481711 | 0.855573 | ENSMUSG( | Galk2 | 373.0542 | -0.08312 | 0.616365 | 0.951413 |
| ENSMUSG( | Galm | 99.22356 | 0.084203 | 0.682393 | 0.93198 | ENSMUSG( | Galm | 92.75347 | 0.147858 | 0.509636 | 0.929089 |
| ENSMUSG( | Galns | 314.2357 | -0.29073 | 0.07862 | 0.433892 | ENSMUSG( | Galns | 328.9076 | 0.13127 | 0.476743 | 0.923329 |
| ENSMUSG( | Galnt1 | 3757.491 | -0.03148 | 0.791336 | 0.959461 | ENSMUSG( | Galnt1 | 3392.062 | -0.05755 | 0.663203 | 0.956359 |
| ENSMUSG( | Galnt10 | 525.362 | -0.27384 | 0.054146 | 0.357985 | ENSMUSG( | Galnt10 | 560.164 | 0.194006 | 0.261177 | 0.884926 |
| ENSMUSG( | Galnt11 | 1617.914 | -0.13149 | 0.240419 | 0.688002 | ENSMUSG( | Galnt11 | 1634.522 | 0.168463 | 0.190097 | 0.857355 |
| ENSMUSG( | Galnt12 | 78.9852 | 0.202329 | 0.358459 | 0.789759 | ENSMUSG( | Galnt12 | 74.1326 | 0.293245 | 0.231103 | 0.868507 |
| ENSMUSG( | Galnt13 | 492.4258 | -0.65805 | 2.46E-06 | 0.00038 | ENSMUSG( | Galnt13 | 597.7835 | 0.236249 | 0.145497 | 0.833991 |
| ENSMUSG( | Galnt14 | 596.4124 | -0.275 | 0.046272 | 0.329472 | ENSMUSG( | Galnt14 | 607.0239 | 0.06245 | 0.695932 | 0.960867 |
| ENSMUSG( | Galnt15 | 218.5333 | 0.448678 | 0.022795 | 0.230795 | ENSMUSG( | Galnt15 | 186.6672 | 0.300957 | 0.237227 | 0.872173 |
| ENSMUSG( | Galnt16 | 4748.306 | -0.28245 | 0.014845 | 0.179046 | ENSMUSG( | Galnt16 | 4967.15 | 0.130301 | 0.354094 | 0.912411 |
| ENSMUSG( | Galnt17 | 2079.643 | -0.23729 | 0.128203 | 0.536641 | ENSMUSG( | Galnt17 | 2199.712 | 0.202371 | 0.279542 | 0.892356 |
| ENSMUSG( | Galnt18 | 1335.325 | -0.12364 | 0.385418 | 0.804048 | ENSMUSG( | Galnt18 | 1401.611 | 0.279504 | 0.090263 | 0.790963 |
| ENSMUSG( | Galnt2 | 2873.212 | -0.22485 | 0.098962 | 0.480953 | ENSMUSG( | Galnt2 | 2971.799 | 0.150702 | 0.354803 | 0.912411 |
| ENSMUSG( | Galnt3 | 53.21095 | -0.28761 | 0.296983 | 0.745676 | ENSMUSG( | Galnt3 | 51.17734 | -0.08124 | 0.802196 | 0.97681 |
| ENSMUSG( | Galnt4 | 282.3358 | -0.06579 | 0.68506 | 0.932981 | ENSMUSG( | Galnt4 | 277.4809 | 0.155951 | 0.406481 | 0.917257 |
| ENSMUSG( | Galnt6 | 493.8434 | -0.90618 | 9.49E-08 | 2.52E-05 | ENSMUSG( | Galnt6 | 557.8534 | -0.13902 | 0.530642 | 0.934795 |
| ENSMUSG( | Galnt7 | 269.3846 | -0.06534 | 0.680509 | 0.931072 | ENSMUSG( | Galnt7 | 291.0243 | 0.407006 | 0.021246 | 0.575231 |
| ENSMUSG( | Galnt9 | 3029.034 | 0.023785 | 0.879842 | 0.981436 | ENSMUSG( | Galnt9 | 3042.743 | 0.295703 | 0.070145 | 0.757694 |
| ENSMUSG( | Galntl6 | 563.8239 | -0.42008 | 0.006658 | 0.11214 | ENSMUSG( | Galntl6 | 632.8858 | 0.208762 | 0.23035 | 0.868507 |
| ENSMUSG( | Galt | 2383.503 | 0.17091 | 0.062457 | 0.386467 | ENSMUSG( | Galt | 2078.838 | 0.048851 | 0.654279 | 0.955015 |

| | | | | | |
|---|---|---|---|---|---|
| ENSMUSG( | 2483.879 | -0.61185 | 3.58E-06 | 0.000492 | Gabra4 |
| ENSMUSG( | 2079.784 | -0.26198 | 0.037533 | 0.299308 | Gabra5 |
| ENSMUSG( | 4586.255 | -0.10729 | 0.427793 | 0.825894 | Gabrb1 |
| ENSMUSG( | 904.2318 | -0.76433 | 0.072007 | 0.414057 | Gabrb2 |
| ENSMUSG( | 6755.067 | -0.45105 | 0.000947 | 0.031617 | Gabrb3 |
| ENSMUSG( | 2545.588 | -0.32572 | 0.008112 | 0.125423 | Gabrd |
| ENSMUSG( | 17.78583 | -0.41846 | 0.402012 | NA | Gabre |
| ENSMUSG( | 1384.391 | -0.21932 | 0.163502 | 0.594869 | Gabrg1 |
| ENSMUSG( | 3712 | -0.57572 | 0.00074 | 0.026499 | Gabrg2 |
| ENSMUSG( | 772.1637 | -0.37008 | 0.007094 | 0.116231 | Gabrg3 |
| ENSMUSG( | 40.41064 | -0.53461 | 0.165303 | 0.596691 | Gabrq |
| ENSMUSG( | 13.23969 | 0.194706 | 0.69545 | NA | Gabrr2 |
| ENSMUSG( | 4623.29 | -0.10818 | 0.456946 | 0.839183 | Gad1 |
| ENSMUSG( | 2486.532 | -0.11803 | 0.423571 | 0.823976 | Gad2 |
| ENSMUSG( | 149.5137 | -0.49207 | 0.003786 | 0.078636 | Gadd45a |
| ENSMUSG( | 250.5064 | -0.62962 | 0.005398 | 0.098038 | Gadd45b |
| ENSMUSG( | 345.548 | -0.31749 | 0.139776 | 0.55636 | Gadd45g |
| ENSMUSG( | 746.6256 | 0.032876 | 0.759532 | 0.952498 | Gadd45gip1 |
| ENSMUSG( | 55.11573 | -0.1852 | 0.487487 | 0.858812 | Gadl1 |
| ENSMUSG( | 3462.305 | 0.122167 | 0.164775 | 0.595869 | Gak |
| ENSMUSG( | 741.1894 | -0.2063 | 0.063822 | 0.390932 | Gal3st1 |
| ENSMUSG( | 1914.891 | -0.17759 | 0.251039 | 0.700713 | Gal3st3 |
| ENSMUSG( | 1831.617 | -0.00868 | 0.93075 | 0.992033 | Galc |
| ENSMUSG( | 406.3915 | 0.034912 | 0.807802 | 0.963257 | Gale |
| ENSMUSG( | 320.3151 | 0.036223 | 0.808991 | 0.963548 | Galk1 |
| ENSMUSG( | 407.0704 | -0.091 | 0.481711 | 0.855573 | Galk2 |
| ENSMUSG( | 99.22356 | 0.084203 | 0.682393 | 0.93198 | Galm |
| ENSMUSG( | 314.2357 | -0.29073 | 0.07862 | 0.433892 | Galns |
| ENSMUSG( | 3757.491 | -0.03148 | 0.791336 | 0.959461 | Galnt1 |
| ENSMUSG( | 525.362 | -0.27384 | 0.054146 | 0.357985 | Galnt10 |
| ENSMUSG( | 1617.914 | -0.13149 | 0.240419 | 0.688002 | Galnt11 |
| ENSMUSG( | 78.9852 | 0.202329 | 0.358459 | 0.789759 | Galnt12 |
| ENSMUSG( | 492.4258 | -0.65805 | 2.46E-06 | 0.00038 | Galnt13 |
| ENSMUSG( | 596.4124 | -0.275 | 0.046272 | 0.329472 | Galnt14 |
| ENSMUSG( | 218.5333 | 0.448678 | 0.022795 | 0.230795 | Galnt15 |
| ENSMUSG( | 4748.306 | -0.28245 | 0.014845 | 0.179046 | Galnt16 |
| ENSMUSG( | 2079.643 | -0.23729 | 0.128203 | 0.536641 | Galnt17 |
| ENSMUSG( | 1335.325 | -0.12364 | 0.385418 | 0.804048 | Galnt18 |
| ENSMUSG( | 2873.212 | -0.22485 | 0.098962 | 0.480953 | Galnt2 |
| ENSMUSG( | 53.21095 | -0.28761 | 0.296983 | 0.745676 | Galnt3 |
| ENSMUSG( | 282.3358 | -0.06579 | 0.68506 | 0.932981 | Galnt4 |
| ENSMUSG( | 493.8434 | -0.90618 | 9.49E-08 | 2.52E-05 | Galnt6 |
| ENSMUSG( | 269.3846 | -0.06534 | 0.680509 | 0.931072 | Galnt7 |
| ENSMUSG( | 3029.034 | 0.023785 | 0.879842 | 0.981436 | Galnt9 |
| ENSMUSG( | 563.8239 | -0.42008 | 0.006658 | 0.11214 | Galntl6 |
| ENSMUSG( | 2383.503 | 0.17091 | 0.062457 | 0.386467 | Galt |
| ENSMUSG( | 921.7539 | -0.08366 | 0.424764 | 0.824675 | Gamt |
| ENSMUSG( | 964.4753 | 0.028141 | 0.80055 | 0.961745 | Gan |
| ENSMUSG( | 4633.227 | -0.2096 | 0.066346 | 0.398419 | Ganab |
| ENSMUSG( | 505.845 | -0.03117 | 0.788567 | 0.959226 | Ganc |
| ENSMUSG( | 5402.387 | -0.03384 | 0.689788 | 0.933609 | Gap43 |
| ENSMUSG( | 44129.55 | 0.083048 | 0.352064 | 0.783738 | Gapdh |
| ENSMUSG( | 2786.714 | 0.115429 | 0.162034 | 0.592429 | Gapvd1 |
| ENSMUSG( | 253.5735 | -0.06085 | 0.667325 | 0.92729 | Gar1 |
| ENSMUSG( | 831.6519 | 0.077302 | 0.473577 | 0.848919 | Garem1 |
| ENSMUSG( | 1229.987 | 0.029639 | 0.759677 | 0.952532 | Garem2 |
| ENSMUSG( | 15.29723 | -0.00312 | 0.995156 | NA | Garin1a |
| ENSMUSG( | 184.0036 | -0.17996 | 0.286226 | 0.735517 | Garin5a |
| ENSMUSG( | 4005.931 | 0.213378 | 0.027332 | 0.253619 | Garnl3 |
| ENSMUSG( | 1505.506 | -0.0065 | 0.949943 | 0.993821 | Garre1 |
| ENSMUSG( | 2140.605 | 0.092858 | 0.482919 | 0.856216 | Gars1 |
| ENSMUSG( | 966.9893 | 0.055235 | 0.59362 | 0.90281 | Gart |
| ENSMUSG( | 553.8633 | -0.35453 | 0.023766 | 0.235641 | Gas1 |
| ENSMUSG( | 326.1626 | -0.16996 | 0.219186 | 0.664648 | Gas2 |
| ENSMUSG( | 3861.297 | 0.269205 | 0.000974 | 0.032367 | Gas2l1 |
| ENSMUSG( | 41.50648 | -0.18586 | 0.530472 | 0.879241 | Gas2l3 |
| ENSMUSG( | 4480.099 | 0.053251 | 0.505233 | 0.866883 | Gas5 |
| ENSMUSG( | 5223.892 | -0.08338 | 0.423179 | 0.823662 | Gas6 |
| ENSMUSG( | 3848.492 | 0.092635 | 0.270119 | 0.719904 | Gas7 |
| ENSMUSG( | 526.392 | 0.115228 | 0.379244 | 0.801757 | Gas8 |
| ENSMUSG( | 322.9973 | -0.09152 | 0.656198 | 0.923929 | Gask1a |
| ENSMUSG( | 122.6504 | -0.71461 | 0.000153 | 0.008659 | Gask1b |
| ENSMUSG( | 153.3269 | 0.072843 | 0.767703 | 0.954621 | Gata2 |
| ENSMUSG( | 3803.166 | 0.073012 | 0.463393 | 0.842789 | Gatad1 |
| ENSMUSG( | 1277.265 | -0.0116 | 0.896157 | 0.984747 | Gatad2a |
| ENSMUSG( | 2198.204 | 0.018005 | 0.855041 | 0.973665 | Gatad2b |
| ENSMUSG( | 631.4832 | 0.099825 | 0.3811 | 0.802629 | Gatb |
| ENSMUSG( | 765.5898 | 0.113212 | 0.272334 | 0.722365 | Gatc |
| ENSMUSG( | 666.5939 | -0.1694 | 0.21615 | 0.661017 | Gatd1 |
| ENSMUSG( | 2860.471 | 0.008151 | 0.93322 | 0.99297 | Gatd3a |
| ENSMUSG( | 3890.494 | -0.05912 | 0.55988 | 0.888917 | Gatm |
| ENSMUSG( | 1267.804 | -0.13319 | 0.330885 | 0.775123 | Gba1 |
| ENSMUSG( | 2387.027 | 0.110067 | 0.309177 | 0.757302 | Gba2 |
| ENSMUSG( | 420.5579 | 0.131959 | 0.283857 | 0.732215 | Gbe1 |
| ENSMUSG( | 2364.811 | 0.163813 | 0.089715 | 0.45977 | Gbf1 |
| ENSMUSG( | 36.62743 | -0.31744 | 0.316017 | 0.763108 | Gbgt1 |
| ENSMUSG( | 14.37411 | -0.69018 | 0.158359 | NA | Gbp11 |
| ENSMUSG( | 131.4116 | -0.15903 | 0.41178 | 0.81942 | Gbp2 |
| ENSMUSG( | 365.7918 | -0.26543 | 0.170433 | 0.605271 | Gbp3 |
| ENSMUSG( | 43.99457 | -0.12918 | 0.66922 | 0.928136 | Gbp4 |
| ENSMUSG( | 88.36551 | -0.08337 | 0.774025 | 0.955606 | Gbp5 |
| ENSMUSG( | 28.56291 | -0.68523 | 0.060689 | 0.380656 | Gbp6 |
| ENSMUSG( | 249.7847 | -0.12105 | 0.484634 | 0.856665 | Gbp7 |
| ENSMUSG( | 157.5136 | -0.43881 | 0.042394 | 0.316636 | Gbp9 |
| ENSMUSG( | 468.9805 | 0.177307 | 0.199182 | 0.639621 | Gca |
| ENSMUSG( | 86.66792 | 0.160234 | 0.475844 | 0.850919 | Gcat |
| ENSMUSG( | 666.6937 | 0.241098 | 0.025877 | 0.247245 | Gcc1 |
| ENSMUSG( | 1237.715 | 0.100829 | 0.271773 | 0.722082 | Gcc2 |
| ENSMUSG( | 1281.778 | 0.163768 | 0.131111 | 0.543058 | Gcdh |
| ENSMUSG( | 227.8775 | -0.0655 | 0.642779 | 0.916865 | Gcfc2 |
| ENSMUSG( | 48.77313 | 0.123263 | 0.708848 | 0.939519 | Gch1 |
| ENSMUSG( | 20.23246 | -0.46253 | 0.31558 | NA | Gck |
| ENSMUSG( | 1668.124 | 0.104072 | 0.249917 | 0.699512 | Gclc |
| ENSMUSG( | 2293.75 | -0.01005 | 0.936925 | 0.993415 | Gclm |
| ENSMUSG( | 1938.316 | 0.135941 | 0.198526 | 0.638526 | Gcn1 |
| ENSMUSG( | 109.8173 | -0.58995 | 0.007952 | 0.124138 | Gcnt1 |
| ENSMUSG( | 804.1681 | -0.36559 | 0.032279 | 0.274548 | Gcnt2 |
| ENSMUSG( | 499.1939 | -0.21297 | 0.143733 | 0.562581 | Gcnt4 |
| ENSMUSG( | 3096.268 | 0.013541 | 0.899109 | 0.985633 | Gcsh |
| ENSMUSG( | 4244.87 | 0.036913 | 0.747017 | 0.949504 | Gda |
| ENSMUSG( | 1556.389 | 0.005191 | 0.960258 | 0.993991 | Gdap1 |
| ENSMUSG( | 1849.079 | 0.041185 | 0.725142 | 0.944096 | Gdap1l1 |
| ENSMUSG( | 534.7396 | -0.04991 | 0.692467 | 0.934101 | Gdap2 |
| ENSMUSG( | 10753.31 | -0.28026 | 0.007131 | 0.116294 | Gde1 |
| ENSMUSG( | 1325.118 | -0.03745 | 0.854024 | 0.973491 | Gdf1 |
| ENSMUSG( | 597.5884 | -0.31977 | 0.023308 | 0.233882 | Gdf10 |
| ENSMUSG( | 2005.455 | -0.09373 | 0.423759 | 0.824145 | Gdf11 |
| ENSMUSG( | 31.34115 | 0.005155 | 0.988663 | 0.999109 | Gdf9 |
| ENSMUSG( | 13555.17 | 0.043235 | 0.698311 | 0.936084 | Gdi1 |
| ENSMUSG( | 6585.591 | 0.073702 | 0.427479 | 0.825539 | Gdi2 |
| ENSMUSG( | 16.02264 | -1.36973 | 0.007659 | NA | Gdnf |
| ENSMUSG( | 1494.875 | -0.42924 | 0.004173 | 0.083766 | Gdpd1 |
| ENSMUSG( | 671.3851 | 0.286077 | 0.027455 | 0.254456 | Gdpd2 |
| ENSMUSG( | 24.0462 | -0.39286 | 0.317786 | 0.764145 | Gdpd3 |
| ENSMUSG( | 1341.572 | -0.31525 | 0.002362 | 0.057092 | Gdpd5 |
| ENSMUSG( | 143.0511 | 0.115796 | 0.588141 | 0.900712 | Gdpgp1 |
| ENSMUSG( | 101.2816 | -0.2924 | 0.139022 | 0.555099 | Gem |
| ENSMUSG( | 116.7869 | 0.141656 | 0.487202 | 0.858812 | Gemin2 |
| ENSMUSG( | 12.79914 | -0.40007 | 0.458673 | NA | Gemin4 |
| ENSMUSG( | 585.005 | 0.034012 | 0.760536 | 0.952907 | Gemin5 |
| ENSMUSG( | 66.62118 | -0.14787 | 0.55049 | 0.886038 | Gemin6 |
| ENSMUSG( | 679.5999 | 0.099674 | 0.322827 | 0.76823 | Gemin7 |
| ENSMUSG( | 234.9916 | -0.0764 | 0.599359 | 0.903801 | Gemin8 |
| ENSMUSG( | 36.91309 | 0.338891 | 0.298226 | 0.746947 | Gen1 |
| ENSMUSG( | 1586.226 | -0.19649 | 0.101442 | 0.485079 | Get1 |
| ENSMUSG( | 2780.89 | -0.02073 | 0.825886 | 0.968061 | Get3 |
| ENSMUSG( | 975.5925 | -0.06868 | 0.497025 | 0.861754 | Get4 |
| ENSMUSG( | 31570.25 | 0.231114 | 0.066439 | 0.39868 | Gfap |
| ENSMUSG( | 455.559 | 0.131179 | 0.269515 | 0.719729 | Gfer |
| ENSMUSG( | 1765.018 | 0.114505 | 0.217889 | 0.663176 | Gfm1 |
| ENSMUSG( | 1138.112 | 0.12962 | 0.204828 | 0.645861 | Gfm2 |
| ENSMUSG( | 4599.553 | 0.465169 | 8.64E-06 | 0.001038 | Gfod1 |
| ENSMUSG( | 625.7108 | 0.239284 | 0.028489 | 0.257525 | Gfod2 |
| ENSMUSG( | 2447.503 | 0.112986 | 0.255622 | 0.706195 | Gfpt1 |
| ENSMUSG( | 250.5698 | -0.01029 | 0.946801 | 0.993731 | Gfpt2 |
| ENSMUSG( | 4460.168 | -0.37148 | 0.006941 | 0.114527 | Gfra1 |
| ENSMUSG( | 2539.595 | -0.14003 | 0.323158 | 0.768415 | Gfra2 |
| ENSMUSG( | 2148.865 | 0.052483 | 0.573228 | 0.893584 | Gfra4 |
| ENSMUSG( | 729.7374 | 0.038395 | 0.724334 | 0.944096 | Gfus |
| ENSMUSG( | 2446.855 | 0.215759 | 0.011219 | 0.152581 | Gga1 |
| ENSMUSG( | 747.3149 | -0.00911 | 0.939727 | 0.993415 | Gga2 |
| ENSMUSG( | 1747.011 | 0.248366 | 0.034937 | 0.287507 | Gga3 |
| ENSMUSG( | 822.6607 | -0.03841 | 0.743696 | 0.948478 | Ggact |
| ENSMUSG( | 346.809 | -0.11633 | 0.372409 | 0.798768 | Ggct |
| ENSMUSG( | 528.4594 | 0.030332 | 0.812301 | 0.964574 | Ggcx |
| ENSMUSG( | 580.4374 | -0.31692 | 0.0474 | 0.332824 | Ggh |
| ENSMUSG( | 68.26149 | -0.33941 | 0.177807 | 0.615225 | Ggn |
| ENSMUSG( | 58.21423 | 0.266447 | 0.308353 | 0.756439 | Ggnbp1 |

| | | | | | |
|---|---|---|---|---|---|
| ENSMUSG( | 4194.55 | -0.1246 | 0.478163 | 0.923329 | Gad1 |
| ENSMUSG( | 2366.134 | 0.005805 | 0.97603 | 0.996416 | Gad2 |
| ENSMUSG( | 138.5912 | -0.43032 | 0.023181 | 0.585577 | Gadd45a |
| ENSMUSG( | 250.9461 | -0.30193 | 0.181172 | 0.850372 | Gadd45b |
| ENSMUSG( | 315.4379 | -0.30504 | 0.16494 | 0.838743 | Gadd45g |
| ENSMUSG( | 630.9188 | -0.19284 | 0.133928 | 0.822912 | Gadd45gip1 |
| ENSMUSG( | 40.592 | -0.94733 | 0.004687 | 0.282119 | Gadl1 |
| ENSMUSG( | 3111.482 | 0.085141 | 0.414212 | 0.917257 | Gak |
| ENSMUSG( | 10.49909 | -0.65592 | 0.304866 | NA | Gal |
| ENSMUSG( | 728.5306 | 0.023972 | 0.876702 | 0.989644 | Gal3st1 |
| ENSMUSG( | 2052.667 | 0.28813 | 0.076948 | 0.770558 | Gal3st3 |
| ENSMUSG( | 1729.527 | 0.093928 | 0.440337 | 0.921166 | Galc |
| ENSMUSG( | 381.3891 | 0.126568 | 0.428873 | 0.917257 | Gale |
| ENSMUSG( | 300.7821 | 0.113998 | 0.495921 | 0.927825 | Galk1 |
| ENSMUSG( | 373.0542 | -0.08312 | 0.616365 | 0.951413 | Galk2 |
| ENSMUSG( | 92.75347 | 0.147858 | 0.509636 | 0.929089 | Galm |
| ENSMUSG( | 328.9076 | 0.13127 | 0.476743 | 0.923329 | Galns |
| ENSMUSG( | 3392.062 | -0.05755 | 0.663203 | 0.956359 | Galnt1 |
| ENSMUSG( | 560.164 | 0.194006 | 0.261177 | 0.884926 | Galnt10 |
| ENSMUSG( | 1634.522 | 0.168463 | 0.190097 | 0.857355 | Galnt11 |
| ENSMUSG( | 74.1326 | 0.293245 | 0.231103 | 0.868507 | Galnt12 |
| ENSMUSG( | 597.7835 | 0.236249 | 0.145497 | 0.833991 | Galnt13 |
| ENSMUSG( | 607.0239 | 0.06245 | 0.695932 | 0.960867 | Galnt14 |
| ENSMUSG( | 186.6672 | 0.300957 | 0.237227 | 0.872173 | Galnt15 |
| ENSMUSG( | 4967.15 | 0.130301 | 0.354094 | 0.912411 | Galnt16 |
| ENSMUSG( | 2199.712 | 0.202371 | 0.279542 | 0.892356 | Galnt17 |
| ENSMUSG( | 1401.611 | 0.279504 | 0.090263 | 0.790963 | Galnt18 |
| ENSMUSG( | 2971.799 | 0.150702 | 0.354803 | 0.912411 | Galnt2 |
| ENSMUSG( | 51.17734 | -0.08124 | 0.802196 | 0.97681 | Galnt3 |
| ENSMUSG( | 277.4809 | 0.155951 | 0.406481 | 0.917257 | Galnt4 |
| ENSMUSG( | 557.8534 | -0.13902 | 0.530642 | 0.934795 | Galnt6 |
| ENSMUSG( | 291.0243 | 0.407006 | 0.021246 | 0.575231 | Galnt7 |
| ENSMUSG( | 3042.743 | 0.295703 | 0.070145 | 0.757694 | Galnt9 |
| ENSMUSG( | 632.8858 | 0.208762 | 0.23035 | 0.868507 | Galntl6 |
| ENSMUSG( | 2078.838 | 0.048851 | 0.654279 | 0.955015 | Galt |
| ENSMUSG( | 772.5753 | -0.34313 | 0.004093 | 0.25867 | Gamt |
| ENSMUSG( | 881.2442 | 0.0397 | 0.75396 | 0.970671 | Gan |
| ENSMUSG( | 4707.024 | 0.112493 | 0.387063 | 0.917257 | Ganab |
| ENSMUSG( | 460.6237 | -0.03463 | 0.804368 | 0.977232 | Ganc |
| ENSMUSG( | 4690.064 | -0.18062 | 0.198447 | 0.862367 | Gap43 |
| ENSMUSG( | 36561.02 | -0.19859 | 0.139207 | 0.830678 | Gapdh |
| ENSMUSG( | 2488.06 | 0.056694 | 0.619297 | 0.952814 | Gapvd1 |
| ENSMUSG( | 222.1717 | -0.18234 | 0.262763 | 0.885357 | Gar1 |
| ENSMUSG( | 778.1858 | 0.151105 | 0.191173 | 0.858412 | Garem1 |
| ENSMUSG( | 1202.725 | 0.223792 | 0.071835 | 0.763548 | Garem2 |
| ENSMUSG( | 13.10782 | -0.15225 | 0.79406 | 0.97681 | Garin1a |
| ENSMUSG( | 168.1968 | -0.16831 | 0.36602 | 0.914623 | Garin5a |
| ENSMUSG( | 3511.943 | 0.108426 | 0.304048 | 0.898538 | Garnl3 |
| ENSMUSG( | 1388.619 | 0.033179 | 0.799952 | 0.97681 | Garre1 |
| ENSMUSG( | 1809.858 | -0.12524 | 0.390318 | 0.917257 | Gars1 |
| ENSMUSG( | 829.2247 | -0.12548 | 0.351276 | 0.912093 | Gart |
| ENSMUSG( | 604.2882 | 0.187519 | 0.292795 | 0.898229 | Gas1 |
| ENSMUSG( | 301.3656 | -0.13989 | 0.420951 | 0.917257 | Gas2 |
| ENSMUSG( | 3460.76 | 0.22695 | 0.034946 | 0.659157 | Gas2l1 |
| ENSMUSG( | 14.77795 | 1.257325 | 0.022797 | 0.582823 | Gas2l2 |
| ENSMUSG( | 45.94043 | 0.36561 | 0.228931 | 0.868507 | Gas2l3 |
| ENSMUSG( | 3891.13 | -0.08778 | 0.438404 | 0.920721 | Gas5 |
| ENSMUSG( | 5147.381 | 0.143045 | 0.279804 | 0.892356 | Gas6 |
| ENSMUSG( | 3323.828 | -0.05915 | 0.606178 | 0.949497 | Gas7 |
| ENSMUSG( | 464.7267 | 0.026824 | 0.855664 | 0.985738 | Gas8 |
| ENSMUSG( | 303.0377 | 0.008339 | 0.970323 | 0.99594 | Gask1a |
| ENSMUSG( | 119.7828 | -0.45258 | 0.031731 | 0.631941 | Gask1b |
| ENSMUSG( | 163.3652 | 0.501035 | 0.058379 | 0.728256 | Gata2 |
| ENSMUSG( | 3245.91 | -0.11983 | 0.350454 | 0.911652 | Gatad1 |
| ENSMUSG( | 1155.716 | -0.03092 | 0.780745 | 0.976106 | Gatad2a |
| ENSMUSG( | 2051.49 | 0.088264 | 0.470124 | 0.923329 | Gatad2b |
| ENSMUSG( | 522.4817 | -0.18188 | 0.182126 | 0.850579 | Gatb |
| ENSMUSG( | 682.2494 | 0.045021 | 0.741333 | 0.968811 | Gatc |
| ENSMUSG( | 602.1482 | -0.19688 | 0.157581 | 0.837724 | Gatd1 |
| ENSMUSG( | 2332.006 | -0.33613 | 0.014888 | 0.489529 | Gatd3a |
| ENSMUSG( | 3573.059 | -0.03774 | 0.756334 | 0.971598 | Gatm |
| ENSMUSG( | 1297.635 | 0.203859 | 0.211638 | 0.864412 | Gba1 |
| ENSMUSG( | 2218.365 | 0.166809 | 0.181105 | 0.850372 | Gba2 |
| ENSMUSG( | 371.9152 | 0.048186 | 0.722528 | 0.966887 | Gbe1 |
| ENSMUSG( | 2026.511 | -0.00859 | 0.940342 | 0.994334 | Gbf1 |
| ENSMUSG( | 44.3513 | 0.500943 | 0.125013 | 0.81516 | Gbgt1 |
| ENSMUSG( | 14.34653 | -0.38789 | 0.448839 | 0.922202 | Gbp11 |
| ENSMUSG( | 145.7878 | 0.402227 | 0.055069 | 0.723362 | Gbp2 |
| ENSMUSG( | 351.7873 | -0.09001 | 0.606771 | 0.949772 | Gbp3 |
| ENSMUSG( | 53.69245 | 0.661883 | 0.031097 | 0.629763 | Gbp4 |
| ENSMUSG( | 100.2897 | 0.529192 | 0.112011 | 0.808673 | Gbp5 |
| ENSMUSG( | 25.39304 | -0.80602 | 0.051266 | 0.711023 | Gbp6 |
| ENSMUSG( | 231.747 | -0.05884 | 0.756879 | 0.971858 | Gbp7 |
| ENSMUSG( | 168.0949 | 0.067957 | 0.774435 | 0.974931 | Gbp9 |
| ENSMUSG( | 9.265011 | 0.336494 | 0.656775 | NA | Gbx1 |
| ENSMUSG( | 391.3591 | -0.07944 | 0.613031 | 0.951154 | Gca |
| ENSMUSG( | 71.60986 | -0.12629 | 0.612661 | 0.951071 | Gcat |
| ENSMUSG( | 573.9721 | 0.083913 | 0.507848 | 0.929089 | Gcc1 |
| ENSMUSG( | 1108.417 | 0.051412 | 0.640209 | 0.953903 | Gcc2 |
| ENSMUSG( | 1088.698 | -0.03736 | 0.765647 | 0.974236 | Gcdh |
| ENSMUSG( | 219.834 | 0.104431 | 0.517639 | 0.930939 | Gcfc2 |
| ENSMUSG( | 51.24604 | 0.52006 | 0.13933 | 0.830678 | Gch1 |
| ENSMUSG( | 20.66686 | -0.1019 | 0.831194 | 0.980539 | Gck |
| ENSMUSG( | 1494.274 | 0.055746 | 0.614752 | 0.951177 | Gclc |
| ENSMUSG( | 1976.144 | -0.18049 | 0.198784 | 0.862972 | Gclm |
| ENSMUSG( | 1732.162 | 0.086453 | 0.534248 | 0.935936 | Gcn1 |
| ENSMUSG( | 129.6995 | 0.225726 | 0.322922 | 0.904023 | Gcnt1 |
| ENSMUSG( | 895.7083 | 0.235738 | 0.230601 | 0.868507 | Gcnt2 |
| ENSMUSG( | 530.5142 | 0.234093 | 0.172143 | 0.84324 | Gcnt4 |
| ENSMUSG( | 2594.756 | -0.24079 | 0.072367 | 0.763629 | Gcsh |
| ENSMUSG( | 3785.586 | -0.02744 | 0.841884 | 0.983044 | Gda |
| ENSMUSG( | 1440.652 | 0.046574 | 0.720868 | 0.966887 | Gdap1 |
| ENSMUSG( | 1616.948 | -0.08055 | 0.552839 | 0.939565 | Gdap1l1 |
| ENSMUSG( | 493.8073 | -0.0195 | 0.905768 | 0.991089 | Gdap2 |
| ENSMUSG( | 10678.33 | -0.01652 | 0.89294 | 0.990558 | Gde1 |
| ENSMUSG( | 1455.477 | 0.473041 | 0.22515 | 0.868468 | Gdf1 |
| ENSMUSG( | 712.9849 | 0.448061 | 0.004978 | 0.28972 | Gdf10 |
| ENSMUSG( | 1843.075 | -0.07074 | 0.648301 | 0.95422 | Gdf11 |
| ENSMUSG( | 10.66205 | 0.442362 | 0.475055 | NA | Gdf5 |
| ENSMUSG( | 28.59172 | 0.006996 | 0.98557 | 0.997084 | Gdf9 |
| ENSMUSG( | 11607.4 | -0.14075 | 0.295085 | 0.898538 | Gdi1 |
| ENSMUSG( | 5707.26 | -0.07317 | 0.548201 | 0.938063 | Gdi2 |
| ENSMUSG( | 20.59007 | -0.08542 | 0.852396 | 0.985598 | Gdnf |
| ENSMUSG( | 1619.154 | 0.103122 | 0.544627 | 0.938063 | Gdpd1 |
| ENSMUSG( | 559.7768 | 0.042137 | 0.775778 | 0.975472 | Gdpd2 |
| ENSMUSG( | 29.95502 | 0.514298 | 0.176298 | 0.846547 | Gdpd3 |
| ENSMUSG( | 1380.026 | 0.054987 | 0.658935 | 0.956205 | Gdpd5 |
| ENSMUSG( | 121.3237 | -0.0829 | 0.726969 | 0.966887 | Gdpgp1 |
| ENSMUSG( | 101.742 | 0.010104 | 0.962111 | 0.995237 | Gem |
| ENSMUSG( | 108.9856 | 0.21737 | 0.375918 | 0.915491 | Gemin2 |
| ENSMUSG( | 13.16288 | -0.03884 | 0.944322 | 0.994798 | Gemin4 |
| ENSMUSG( | 538.9014 | 0.067068 | 0.599346 | 0.947071 | Gemin5 |
| ENSMUSG( | 58.69457 | -0.27839 | 0.316566 | 0.901868 | Gemin6 |
| ENSMUSG( | 594.6227 | -0.02205 | 0.873143 | 0.989445 | Gemin7 |
| ENSMUSG( | 207.0601 | -0.18403 | 0.279421 | 0.892356 | Gemin8 |
| ENSMUSG( | 37.44318 | 0.587042 | 0.090122 | 0.790741 | Gen1 |
| ENSMUSG( | 1543.852 | 0.001925 | 0.988454 | 0.997862 | Get1 |
| ENSMUSG( | 2345.532 | -0.25677 | 0.028415 | 0.613486 | Get3 |
| ENSMUSG( | 903.0143 | -0.02201 | 0.849808 | 0.985522 | Get4 |
| ENSMUSG( | 24095.04 | -0.28752 | 0.030514 | 0.627354 | Gfap |
| ENSMUSG( | 388.7748 | -0.05356 | 0.69857 | 0.960908 | Gfer |
| ENSMUSG( | 1461.443 | -0.16701 | 0.154052 | 0.837724 | Gfm1 |
| ENSMUSG( | 979.4694 | -0.03275 | 0.779117 | 0.976106 | Gfm2 |
| ENSMUSG( | 3823.584 | 0.2316 | 0.078063 | 0.77321 | Gfod1 |
| ENSMUSG( | 523.4303 | -0.00346 | 0.980557 | 0.996479 | Gfod2 |
| ENSMUSG( | 2139.766 | -0.00574 | 0.961541 | 0.99511 | Gfpt1 |
| ENSMUSG( | 219.5626 | -0.12587 | 0.477534 | 0.923329 | Gfpt2 |
| ENSMUSG( | 4211.677 | -0.25427 | 0.055419 | 0.723362 | Gfra1 |
| ENSMUSG( | 2434.814 | 0.014724 | 0.92879 | 0.99306 | Gfra2 |
| ENSMUSG( | 1869.796 | -0.08316 | 0.463499 | 0.923329 | Gfra4 |
| ENSMUSG( | 644.2121 | -0.04943 | 0.697017 | 0.960867 | Gfus |
| ENSMUSG( | 9.777646 | 1.197613 | 0.119584 | NA | Gfy |
| ENSMUSG( | 2085.571 | 0.030913 | 0.774636 | 0.974931 | Gga1 |
| ENSMUSG( | 671.8221 | -0.04594 | 0.76625 | 0.974236 | Gga2 |
| ENSMUSG( | 1494.006 | 0.07405 | 0.607911 | 0.950113 | Gga3 |
| ENSMUSG( | 700.8094 | -0.24957 | 0.090015 | 0.790741 | Ggact |
| ENSMUSG( | 309.6237 | -0.18255 | 0.240141 | 0.875205 | Ggct |
| ENSMUSG( | 516.3886 | 0.231354 | 0.141504 | 0.830678 | Ggcx |
| ENSMUSG( | 606.31 | 0.096666 | 0.564366 | 0.941206 | Ggh |
| ENSMUSG( | 60.82751 | -0.40845 | 0.126297 | 0.81516 | Ggn |
| ENSMUSG( | 48.35153 | 0.021002 | 0.9477 | 0.994798 | Ggnbp1 |
| ENSMUSG( | 1954.134 | 0.033199 | 0.787194 | 0.97637 | Ggnbp2 |
| ENSMUSG( | 1095.526 | 0.093514 | 0.474693 | 0.923329 | Ggps1 |
| ENSMUSG( | 128.1384 | 0.464444 | 0.086416 | 0.781091 | Ggt1 |
| ENSMUSG( | 132.7184 | 0.311029 | 0.25607 | 0.883082 | Ggt5 |
| ENSMUSG( | 4633.6 | 0.100854 | 0.426143 | 0.917257 | Ggt7 |
| ENSMUSG( | 225.5042 | 0.40969 | 0.097607 | 0.796743 | Ggta1 |
| ENSMUSG( | 466.3757 | 0.075443 | 0.63782 | 0.953903 | Ghdc |

| | | | | | | | | | | | |
|---|---|---|---|---|---|---|---|---|---|---|---|
| ENSMUSG( | Gamt | 921.7539 | -0.08366 | 0.424764 | 0.824675 | ENSMUSG( | Gamt | 772.5753 | -0.34313 | 0.004093 | 0.25867 |
| ENSMUSG( | Gan | 964.4753 | 0.028141 | 0.80055 | 0.961745 | ENSMUSG( | Gan | 881.2442 | 0.0397 | 0.75396 | 0.970671 |
| ENSMUSG( | Ganab | 4633.227 | -0.2096 | 0.066346 | 0.398419 | ENSMUSG( | Ganab | 4707.024 | 0.112493 | 0.387063 | 0.917257 |
| ENSMUSG( | Ganc | 505.845 | -0.03117 | 0.788567 | 0.959226 | ENSMUSG( | Ganc | 460.6237 | -0.03463 | 0.804368 | 0.977232 |
| ENSMUSG( | Gap43 | 5402.387 | -0.03384 | 0.689788 | 0.933609 | ENSMUSG( | Gap43 | 4690.064 | -0.18062 | 0.198447 | 0.862367 |
| ENSMUSG( | Gapdh | 44129.55 | 0.083048 | 0.352064 | 0.783738 | ENSMUSG( | Gapdh | 36561.02 | -0.19859 | 0.139207 | 0.830678 |
| ENSMUSG( | Gapvd1 | 2786.714 | 0.115429 | 0.162034 | 0.592429 | ENSMUSG( | Gapvd1 | 2488.06 | 0.056694 | 0.619297 | 0.952814 |
| ENSMUSG( | Gar1 | 253.5735 | -0.06085 | 0.667325 | 0.92729 | ENSMUSG( | Gar1 | 222.1717 | -0.18234 | 0.262763 | 0.885357 |
| ENSMUSG( | Garem1 | 831.6519 | 0.077302 | 0.473577 | 0.848919 | ENSMUSG( | Garem1 | 778.1858 | 0.151105 | 0.191173 | 0.858412 |
| ENSMUSG( | Garem2 | 1229.987 | 0.029639 | 0.759677 | 0.952532 | ENSMUSG( | Garem2 | 1202.725 | 0.223792 | 0.071835 | 0.763548 |
| ENSMUSG( | Garin1a | 15.29723 | -0.00312 | 0.995156 | NA | ENSMUSG( | Garin1a | 13.10782 | -0.15225 | 0.79406 | 0.97681 |
| ENSMUSG( | Garin5a | 184.0036 | -0.17996 | 0.286226 | 0.735517 | ENSMUSG( | Garin5a | 168.1968 | -0.16831 | 0.36602 | 0.914623 |
| ENSMUSG( | Garnl3 | 4005.931 | 0.213378 | 0.027332 | 0.253619 | ENSMUSG( | Garnl3 | 3511.943 | 0.108426 | 0.304048 | 0.898538 |
| ENSMUSG( | Garre1 | 1505.506 | -0.0065 | 0.949943 | 0.993821 | ENSMUSG( | Garre1 | 1388.619 | 0.033179 | 0.799952 | 0.97681 |
| ENSMUSG( | Gars1 | 2140.605 | 0.092858 | 0.482919 | 0.856216 | ENSMUSG( | Gars1 | 1809.858 | -0.12524 | 0.390318 | 0.917257 |
| ENSMUSG( | Gart | 966.9893 | 0.055235 | 0.59362 | 0.90281 | ENSMUSG( | Gart | 829.2247 | -0.12548 | 0.351276 | 0.912093 |
| ENSMUSG( | Gas1 | 553.8633 | -0.35453 | 0.023766 | 0.235641 | ENSMUSG( | Gas1 | 604.2882 | 0.187519 | 0.292795 | 0.898229 |
| ENSMUSG( | Gas2 | 326.1626 | -0.16996 | 0.219186 | 0.664648 | ENSMUSG( | Gas2 | 301.3656 | -0.13989 | 0.420951 | 0.917257 |
| ENSMUSG( | Gas2l1 | 3861.297 | 0.269205 | 0.000974 | 0.032367 | ENSMUSG( | Gas2l1 | 3460.76 | 0.22695 | 0.034946 | 0.659157 |
| ENSMUSG( | Gas2l3 | 41.50648 | -0.18586 | 0.530472 | 0.879241 | ENSMUSG( | Gas2l3 | 45.94043 | 0.36561 | 0.228931 | 0.868507 |
| ENSMUSG( | Gas5 | 4480.099 | 0.053251 | 0.505233 | 0.866883 | ENSMUSG( | Gas5 | 3891.13 | -0.08778 | 0.438404 | 0.920721 |
| ENSMUSG( | Gas6 | 5223.892 | -0.08338 | 0.423179 | 0.823662 | ENSMUSG( | Gas6 | 5147.381 | 0.143045 | 0.279804 | 0.892356 |
| ENSMUSG( | Gas7 | 3848.492 | 0.092635 | 0.270119 | 0.719904 | ENSMUSG( | Gas7 | 3323.828 | -0.05915 | 0.606178 | 0.949497 |
| ENSMUSG( | Gas8 | 526.392 | 0.115228 | 0.379244 | 0.801757 | ENSMUSG( | Gas8 | 464.7267 | 0.026824 | 0.855664 | 0.985738 |
| ENSMUSG( | Gask1a | 322.9973 | -0.09152 | 0.656198 | 0.923929 | ENSMUSG( | Gask1a | 303.0377 | 0.008339 | 0.970323 | 0.99594 |
| ENSMUSG( | Gask1b | 122.6504 | -0.71461 | 0.000153 | 0.008659 | ENSMUSG( | Gask1b | 119.7828 | -0.45258 | 0.031731 | 0.631941 |
| ENSMUSG( | Gata2 | 153.3269 | 0.072843 | 0.767703 | 0.954621 | ENSMUSG( | Gata2 | 163.3652 | 0.501035 | 0.058379 | 0.728256 |
| ENSMUSG( | Gatad1 | 3803.166 | 0.073012 | 0.463393 | 0.842789 | ENSMUSG( | Gatad1 | 3245.91 | -0.11983 | 0.350454 | 0.911652 |
| ENSMUSG( | Gatad2a | 1277.265 | -0.0116 | 0.896157 | 0.984747 | ENSMUSG( | Gatad2a | 1155.716 | -0.03092 | 0.780745 | 0.976106 |
| ENSMUSG( | Gatad2b | 2198.204 | 0.018005 | 0.855041 | 0.973665 | ENSMUSG( | Gatad2b | 2051.49 | 0.088264 | 0.470124 | 0.923329 |
| ENSMUSG( | Gatb | 631.4832 | 0.099825 | 0.3811 | 0.802629 | ENSMUSG( | Gatb | 522.4817 | -0.18188 | 0.182126 | 0.850579 |
| ENSMUSG( | Gatc | 765.5898 | 0.113212 | 0.272334 | 0.722365 | ENSMUSG( | Gatc | 682.2494 | 0.045021 | 0.741333 | 0.968811 |
| ENSMUSG( | Gatd1 | 666.5939 | -0.1694 | 0.21615 | 0.661017 | ENSMUSG( | Gatd1 | 602.1482 | -0.19688 | 0.157581 | 0.837724 |
| ENSMUSG( | Gatd3a | 2860.471 | 0.008151 | 0.93322 | 0.99297 | ENSMUSG( | Gatd3a | 2332.006 | -0.33613 | 0.014888 | 0.489529 |
| ENSMUSG( | Gatm | 3890.494 | -0.05912 | 0.55988 | 0.888917 | ENSMUSG( | Gatm | 3573.059 | -0.03774 | 0.756334 | 0.971598 |
| ENSMUSG( | Gba1 | 1267.804 | -0.13319 | 0.330885 | 0.775123 | ENSMUSG( | Gba1 | 1297.635 | 0.203859 | 0.211638 | 0.864412 |
| ENSMUSG( | Gba2 | 2387.027 | 0.110067 | 0.309177 | 0.757302 | ENSMUSG( | Gba2 | 2218.365 | 0.166809 | 0.181105 | 0.850372 |
| ENSMUSG( | Gbe1 | 420.5579 | 0.131959 | 0.283857 | 0.732215 | ENSMUSG( | Gbe1 | 371.9152 | 0.048186 | 0.722528 | 0.966887 |
| ENSMUSG( | Gbf1 | 2364.811 | 0.163813 | 0.089715 | 0.45977 | ENSMUSG( | Gbf1 | 2026.511 | -0.00859 | 0.940342 | 0.994334 |
| ENSMUSG( | Gbgt1 | 36.62743 | -0.31744 | 0.316017 | 0.763108 | ENSMUSG( | Gbgt1 | 44.3513 | 0.500943 | 0.125013 | 0.81516 |
| ENSMUSG( | Gbp11 | 14.37411 | -0.69018 | 0.158359 | NA | ENSMUSG( | Gbp11 | 14.34653 | -0.38789 | 0.448839 | 0.922202 |
| ENSMUSG( | Gbp2 | 131.4116 | -0.15903 | 0.41178 | 0.81942 | ENSMUSG( | Gbp2 | 145.7878 | 0.402227 | 0.055069 | 0.723362 |
| ENSMUSG( | Gbp3 | 365.7918 | -0.26543 | 0.170433 | 0.605271 | ENSMUSG( | Gbp3 | 351.7873 | -0.09001 | 0.606771 | 0.949772 |
| ENSMUSG( | Gbp4 | 43.99457 | -0.12918 | 0.66922 | 0.928136 | ENSMUSG( | Gbp4 | 53.69245 | 0.661883 | 0.031097 | 0.629763 |
| ENSMUSG( | Gbp5 | 88.36551 | -0.08337 | 0.774025 | 0.955606 | ENSMUSG( | Gbp5 | 100.2897 | 0.529192 | 0.112011 | 0.808673 |
| ENSMUSG( | Gbp6 | 28.56291 | -0.68523 | 0.060689 | 0.380656 | ENSMUSG( | Gbp6 | 25.39304 | -0.80602 | 0.051266 | 0.711023 |
| ENSMUSG( | Gbp7 | 249.7847 | -0.12105 | 0.484634 | 0.856665 | ENSMUSG( | Gbp7 | 231.747 | -0.05884 | 0.756879 | 0.971858 |
| ENSMUSG( | Gbp9 | 157.5136 | -0.43881 | 0.042394 | 0.316636 | ENSMUSG( | Gbp9 | 168.0949 | 0.067957 | 0.774435 | 0.974931 |
| ENSMUSG( | Gca | 468.9805 | 0.177307 | 0.199182 | 0.639621 | ENSMUSG( | Gca | 391.3591 | -0.07944 | 0.613031 | 0.951154 |
| ENSMUSG( | Gcat | 86.66792 | 0.160234 | 0.475844 | 0.850919 | ENSMUSG( | Gcat | 71.60986 | -0.12629 | 0.612661 | 0.951071 |
| ENSMUSG( | Gcc1 | 666.6937 | 0.241098 | 0.025877 | 0.247245 | ENSMUSG( | Gcc1 | 573.9721 | 0.083913 | 0.507848 | 0.929089 |
| ENSMUSG( | Gcc2 | 1237.715 | 0.100829 | 0.271773 | 0.722082 | ENSMUSG( | Gcc2 | 1108.417 | 0.051412 | 0.640209 | 0.953903 |
| ENSMUSG( | Gcdh | 1281.778 | 0.163768 | 0.131111 | 0.543058 | ENSMUSG( | Gcdh | 1088.698 | -0.03736 | 0.765647 | 0.974236 |
| ENSMUSG( | Gcfc2 | 227.8775 | -0.0655 | 0.642779 | 0.916865 | ENSMUSG( | Gcfc2 | 219.834 | 0.104431 | 0.517639 | 0.930939 |
| ENSMUSG( | Gch1 | 48.77313 | 0.123263 | 0.708848 | 0.939519 | ENSMUSG( | Gch1 | 51.24604 | 0.52006 | 0.13933 | 0.830678 |
| ENSMUSG( | Gck | 20.23246 | -0.46253 | 0.31558 | NA | ENSMUSG( | Gck | 20.66686 | -0.1019 | 0.831194 | 0.980539 |
| ENSMUSG( | Gclc | 1668.124 | 0.104072 | 0.249917 | 0.699512 | ENSMUSG( | Gclc | 1494.274 | 0.055746 | 0.614752 | 0.951177 |
| ENSMUSG( | Gclm | 2293.75 | -0.01005 | 0.936925 | 0.993415 | ENSMUSG( | Gclm | 1976.144 | -0.18049 | 0.198784 | 0.862972 |
| ENSMUSG( | Gcn1 | 1938.316 | 0.135941 | 0.198526 | 0.638526 | ENSMUSG( | Gcn1 | 1732.162 | 0.086453 | 0.534248 | 0.935936 |
| ENSMUSG( | Gcnt1 | 109.8173 | -0.58995 | 0.007952 | 0.124138 | ENSMUSG( | Gcnt1 | 129.6995 | 0.225726 | 0.322922 | 0.904023 |
| ENSMUSG( | Gcnt2 | 804.1681 | -0.36559 | 0.032279 | 0.274548 | ENSMUSG( | Gcnt2 | 895.7083 | 0.235738 | 0.230601 | 0.868507 |
| ENSMUSG( | Gcnt4 | 499.1939 | -0.21297 | 0.143733 | 0.562581 | ENSMUSG( | Gcnt4 | 530.5142 | 0.234093 | 0.172143 | 0.84324 |
| ENSMUSG( | Gcsh | 3096.268 | 0.013541 | 0.899109 | 0.985633 | ENSMUSG( | Gcsh | 2594.756 | -0.24079 | 0.072367 | 0.763629 |
| ENSMUSG( | Gda | 4244.87 | 0.036913 | 0.747017 | 0.949504 | ENSMUSG( | Gda | 3785.586 | -0.02744 | 0.841884 | 0.983044 |
| ENSMUSG( | Gdap1 | 1556.389 | 0.005191 | 0.960258 | 0.993991 | ENSMUSG( | Gdap1 | 1440.652 | 0.046574 | 0.720868 | 0.966887 |
| ENSMUSG( | Gdap1l1 | 1849.079 | 0.041185 | 0.725142 | 0.944096 | ENSMUSG( | Gdap1l1 | 1616.948 | -0.08055 | 0.552839 | 0.939565 |
| ENSMUSG( | Gdap2 | 534.7396 | -0.04991 | 0.692467 | 0.934101 | ENSMUSG( | Gdap2 | 493.8073 | -0.0195 | 0.905768 | 0.991089 |
| ENSMUSG( | Gde1 | 10753.31 | -0.28026 | 0.007131 | 0.116294 | ENSMUSG( | Gde1 | 10678.33 | -0.01652 | 0.89294 | 0.990558 |
| ENSMUSG( | Gdf1 | 1325.118 | -0.03745 | 0.854024 | 0.973491 | ENSMUSG( | Gdf1 | 1455.477 | 0.473041 | 0.22515 | 0.868468 |
| ENSMUSG( | Gdf10 | 597.5884 | -0.31977 | 0.023308 | 0.233882 | ENSMUSG( | Gdf10 | 712.9849 | 0.448061 | 0.004978 | 0.28972 |
| ENSMUSG( | Gdf11 | 2005.455 | -0.09373 | 0.423759 | 0.824145 | ENSMUSG( | Gdf11 | 1843.075 | -0.07074 | 0.648301 | 0.95422 |
| ENSMUSG( | Gdf9 | 31.34115 | 0.005155 | 0.988663 | 0.999109 | ENSMUSG( | Gdf9 | 28.59172 | 0.006996 | 0.98557 | 0.997084 |
| ENSMUSG( | Gdi1 | 13555.17 | 0.043235 | 0.698311 | 0.936084 | ENSMUSG( | Gdi1 | 11607.4 | -0.14075 | 0.295085 | 0.898538 |
| ENSMUSG( | Gdi2 | 6585.591 | 0.073702 | 0.427479 | 0.825539 | ENSMUSG( | Gdi2 | 5707.26 | -0.07317 | 0.548201 | 0.938063 |
| ENSMUSG( | Gdnf | 16.02264 | -1.36973 | 0.007659 | NA | ENSMUSG( | Gdnf | 20.59007 | -0.08542 | 0.852396 | 0.985598 |
| ENSMUSG( | Gdpd1 | 1494.875 | -0.42924 | 0.004173 | 0.083766 | ENSMUSG( | Gdpd1 | 1619.154 | 0.103122 | 0.544627 | 0.938063 |
| ENSMUSG( | Gdpd2 | 671.3851 | 0.286077 | 0.027455 | 0.254456 | ENSMUSG( | Gdpd2 | 559.7768 | 0.042137 | 0.775778 | 0.975472 |
| ENSMUSG( | Gdpd3 | 24.0462 | -0.39286 | 0.317786 | 0.764145 | ENSMUSG( | Gdpd3 | 29.95502 | 0.514298 | 0.176298 | 0.846547 |
| ENSMUSG( | Gdpd5 | 1341.572 | -0.31525 | 0.002362 | 0.057092 | ENSMUSG( | Gdpd5 | 1380.026 | 0.054987 | 0.658935 | 0.956205 |
| ENSMUSG( | Gdpgp1 | 143.0511 | 0.115796 | 0.588141 | 0.900712 | ENSMUSG( | Gdpgp1 | 121.3237 | -0.0829 | 0.726969 | 0.966887 |
| ENSMUSG( | Gem | 101.2816 | -0.2924 | 0.139022 | 0.555099 | ENSMUSG( | Gem | 101.742 | 0.010104 | 0.962111 | 0.995237 |
| ENSMUSG( | Gemin2 | 116.7869 | 0.141656 | 0.487202 | 0.858812 | ENSMUSG( | Gemin2 | 108.9856 | 0.21737 | 0.375918 | 0.915491 |
| ENSMUSG( | Gemin4 | 12.79914 | -0.40007 | 0.458673 | NA | ENSMUSG( | Gemin4 | 13.16288 | -0.03884 | 0.944322 | 0.994798 |
| ENSMUSG( | Gemin5 | 585.005 | 0.034012 | 0.760536 | 0.952907 | ENSMUSG( | Gemin5 | 538.9014 | 0.067068 | 0.599346 | 0.947071 |
| ENSMUSG( | Gemin6 | 66.62118 | -0.14787 | 0.55049 | 0.886038 | ENSMUSG( | Gemin6 | 58.69457 | -0.27839 | 0.316566 | 0.901868 |
| ENSMUSG( | Gemin7 | 679.5999 | 0.099674 | 0.322827 | 0.76823 | ENSMUSG( | Gemin7 | 594.6227 | -0.02205 | 0.873143 | 0.989445 |
| ENSMUSG( | Gemin8 | 234.9916 | -0.0764 | 0.599359 | 0.903801 | ENSMUSG( | Gemin8 | 207.0601 | -0.18403 | 0.279421 | 0.892356 |
| ENSMUSG( | Gen1 | 36.91309 | 0.338891 | 0.298226 | 0.746947 | ENSMUSG( | Gen1 | 37.44318 | 0.587042 | 0.090122 | 0.790741 |
| ENSMUSG( | Get1 | 1586.226 | -0.19649 | 0.101442 | 0.485079 | ENSMUSG( | Get1 | 1543.852 | 0.001925 | 0.988454 | 0.997862 |
| ENSMUSG( | Get3 | 2780.89 | -0.02073 | 0.825886 | 0.968061 | ENSMUSG( | Get3 | 2345.532 | -0.25677 | 0.028415 | 0.613486 |
| ENSMUSG( | Get4 | 975.5925 | -0.06868 | 0.497025 | 0.861754 | ENSMUSG( | Get4 | 903.0143 | -0.02201 | 0.849808 | 0.985522 |
| ENSMUSG( | Gfap | 31570.25 | 0.231114 | 0.066439 | 0.39868 | ENSMUSG( | Gfap | 24095.04 | -0.28752 | 0.030514 | 0.627354 |
| ENSMUSG( | Gfer | 455.559 | 0.131179 | 0.269515 | 0.719729 | ENSMUSG( | Gfer | 388.7748 | -0.05356 | 0.69857 | 0.960908 |
| ENSMUSG( | Gfm1 | 1765.018 | 0.114505 | 0.217889 | 0.663176 | ENSMUSG( | Gfm1 | 1461.443 | -0.16701 | 0.154052 | 0.837724 |
| ENSMUSG( | Gfm2 | 1138.112 | 0.12962 | 0.204828 | 0.645861 | ENSMUSG( | Gfm2 | 979.4694 | -0.03275 | 0.779117 | 0.976106 |
| ENSMUSG( | Gfod1 | 4599.553 | 0.465169 | 8.64E-06 | 0.001038 | ENSMUSG( | Gfod1 | 3823.584 | 0.2316 | 0.078063 | 0.77321 |
| ENSMUSG( | Gfod2 | 625.7108 | 0.239284 | 0.028489 | 0.257525 | ENSMUSG( | Gfod2 | 523.4303 | -0.00346 | 0.980557 | 0.996479 |
| ENSMUSG( | Gfpt1 | 2447.503 | 0.112986 | 0.255622 | 0.706195 | ENSMUSG( | Gfpt1 | 2139.766 | -0.00574 | 0.961541 | 0.99511 |
| ENSMUSG( | Gfpt2 | 250.5698 | -0.01029 | 0.946801 | 0.993731 | ENSMUSG( | Gfpt2 | 219.5626 | -0.12587 | 0.477534 | 0.923329 |
| ENSMUSG( | Gfra1 | 4460.168 | -0.37148 | 0.006941 | 0.114527 | ENSMUSG( | Gfra1 | 4211.677 | -0.25427 | 0.055419 | 0.723362 |
| ENSMUSG( | Gfra2 | 2539.595 | -0.14003 | 0.323158 | 0.768415 | ENSMUSG( | Gfra2 | 2434.814 | 0.014724 | 0.92879 | 0.99306 |
| ENSMUSG( | Gfra4 | 2148.865 | 0.052483 | 0.573228 | 0.893584 | ENSMUSG( | Gfra4 | 1869.796 | -0.08316 | 0.463499 | 0.923329 |
| ENSMUSG( | Gfus | 729.7374 | 0.038395 | 0.724334 | 0.944096 | ENSMUSG( | Gfus | 644.2121 | -0.04943 | 0.697017 | 0.960867 |
| ENSMUSG( | Gga1 | 2446.855 | 0.215759 | 0.011219 | 0.152581 | ENSMUSG( | Gga1 | 2085.571 | 0.030913 | 0.774636 | 0.974931 |
| ENSMUSG( | Gga2 | 747.3149 | -0.00911 | 0.939727 | 0.993415 | ENSMUSG( | Gga2 | 671.8221 | -0.04594 | 0.76625 | 0.974236 |
| ENSMUSG( | Gga3 | 1747.011 | 0.248366 | 0.034937 | 0.287507 | ENSMUSG( | Gga3 | 1494.006 | 0.07405 | 0.607911 | 0.950113 |
| ENSMUSG( | Ggact | 822.6607 | -0.03841 | 0.743696 | 0.948478 | ENSMUSG( | Ggact | 700.8094 | -0.24957 | 0.090015 | 0.790741 |
| ENSMUSG( | Ggct | 346.809 | -0.11633 | 0.372409 | 0.798768 | ENSMUSG( | Ggct | 309.6237 | -0.18255 | 0.240141 | 0.875205 |
| ENSMUSG( | Ggcx | 528.4594 | 0.030332 | 0.812301 | 0.964574 | ENSMUSG( | Ggcx | 516.3886 | 0.231354 | 0.141504 | 0.830678 |
| ENSMUSG( | Ggh | 580.4374 | -0.31692 | 0.0474 | 0.332824 | ENSMUSG( | Ggh | 606.31 | 0.096666 | 0.564366 | 0.941206 |
| ENSMUSG( | Ggn | 68.26149 | -0.33941 | 0.177807 | 0.615225 | ENSMUSG( | Ggn | 60.82751 | -0.40845 | 0.126297 | 0.81516 |
| ENSMUSG( | Ggnbp1 | 58.21423 | 0.266447 | 0.308353 | 0.756439 | ENSMUSG( | Ggnbp1 | 48.35153 | 0.021002 | 0.9477 | 0.994798 |
| ENSMUSG( | Ggnbp2 | 2114.68 | -0.00489 | 0.960148 | 0.993991 | ENSMUSG( | Ggnbp2 | 1954.134 | 0.033199 | 0.787194 | 0.97637 |
| ENSMUSG( | Ggps1 | 1182.114 | 0.049487 | 0.590481 | 0.90113 | ENSMUSG( | Ggps1 | 1095.526 | 0.093514 | 0.474693 | 0.923329 |
| ENSMUSG( | Ggt1 | 117.9027 | -0.03124 | 0.879713 | 0.981436 | ENSMUSG( | Ggt1 | 128.1384 | 0.464444 | 0.086416 | 0.781091 |
| ENSMUSG( | Ggt5 | 141.3953 | 0.210028 | 0.335774 | 0.777 | ENSMUSG( | Ggt5 | 132.7184 | 0.311029 | 0.25607 | 0.883082 |
| ENSMUSG( | Ggt7 | 4661.896 | -0.15527 | 0.179286 | 0.616869 | ENSMUSG( | Ggt7 | 4633.6 | 0.100854 | 0.426143 | 0.917257 |
| ENSMUSG( | Ggta1 | 201.958 | -0.17027 | 0.383883 | 0.803646 | ENSMUSG( | Ggta1 | 225.5042 | 0.40969 | 0.097607 | 0.796743 |
| ENSMUSG( | Ghdc | 491.6673 | -0.04318 | 0.764794 | 0.954127 | ENSMUSG( | Ghdc | 466.3757 | 0.075443 | 0.63782 | 0.953903 |
| ENSMUSG( | Ghitm | 11997.73 | 0.062025 | 0.526417 | 0.877267 | ENSMUSG( | Ghitm | 10139.12 | -0.1609 | 0.210048 | 0.864412 |
| ENSMUSG( | Ghr | 484.3061 | -0.15158 | 0.430134 | 0.826789 | ENSMUSG( | Ghr | 465.1581 | 0.005093 | 0.974994 | 0.996416 |
| ENSMUSG( | Gid4 | 6654.749 | 0.174521 | 0.051495 | 0.34814 | ENSMUSG( | Gid4 | 5609.86 | -0.04579 | 0.674788 | 0.95694 |
| ENSMUSG( | Gid8 | 1180.035 | 0.049997 | 0.61299 | 0.908902 | ENSMUSG( | Gid8 | 1013.392 | -0.12575 | 0.29519 | 0.898538 |
| ENSMUSG( | Gigyf1 | 2178.042 | 0.112061 | 0.325876 | 0.771128 | ENSMUSG( | Gigyf1 | 1924.109 | 0.024642 | 0.85924 | 0.985884 |
| ENSMUSG( | Gigyf2 | 1785.316 | 0.265757 | 0.001632 | 0.045399 | ENSMUSG( | Gigyf2 | 1510.502 | 0.061696 | 0.583817 | 0.944379 |
| ENSMUSG( | Gimap1 | 52.4688 | -0.02337 | 0.937861 | 0.993415 | ENSMUSG( | Gimap1 | 50.60683 | 0.167929 | 0.629001 | 0.953251 |
| ENSMUSG( | Gimap4 | 19.44122 | -0.22933 | 0.614594 | NA | ENSMUSG( | Gimap4 | 19.52339 | 0.05017 | 0.914547 | 0.992407 |
| ENSMUSG( | Gimap5 | 25.75303 | -0.36555 | 0.398737 | 0.812878 | ENSMUSG( | Gimap5 | 22.06355 | -0.5294 | 0.3018 | 0.898538 |
| ENSMUSG( | Gimap6 | 227.591 | -0.03155 | 0.884215 | 0.982647 | ENSMUSG( | Gimap6 | 205.1568 | -0.043 | 0.869235 | 0.988097 |
| ENSMUSG( | Gimap8 | 45.35327 | 0.084109 | 0.787496 | 0.959163 | ENSMUSG( | Gimap8 | 48.54991 | 0.538041 | 0.135572 | 0.825163 |
| ENSMUSG( | Gimap9 | 14.62491 | -0.47707 | 0.316387 | NA | ENSMUSG( | Gimap9 | 17.5097 | 0.359944 | 0.442284 | 0.921554 |
| ENSMUSG( | Gin1 | 150.5275 | 0.082595 | 0.623395 | 0.911646 | ENSMUSG( | Gin1 | 142.7776 | 0.201413 | 0.300468 | 0.898538 |
| ENSMUSG( | Ginm1 | 763.1432 | -0.36358 | 0.014228 | 0.175234 | ENSMUSG( | Ginm1 | 810.0701 | 0.103768 | 0.538679 | 0.937216 |
| ENSMUSG( | Gins1 | 26.71086 | -0.04807 | 0.895448 | 0.984512 | ENSMUSG( | Gins1 | 24.38042 | -0.04268 | 0.917729 | 0.992529 |
| ENSMUSG( | Gins2 | 76.65482 | 0.02485 | 0.916614 | 0.988856 | ENSMUSG( | Gins2 | 61.03366 | -0.38485 | 0.14119 | 0.830678 |
| ENSMUSG( | Gins3 | 57.84734 | 0.159851 | 0.535025 | 0.880733 | ENSMUSG( | Gins3 | 52.02604 | 0.1086 | 0.714284 | 0.964986 |
| ENSMUSG( | Gins4 | 427.8749 | -0.00143 | 0.990059 | 0.999144 | ENSMUSG( | Gins4 | 372.513 | -0.13973 | 0.319516 | 0.903464 |
| ENSMUSG( | Gipc1 | 3134.49 | 0.063922 | 0.463942 | 0.842812 | ENSMUSG( | Gipc1 | 2679.526 | -0.123 | 0.232152 | 0.868507 |
| ENSMUSG( | Gipc2 | 23.87614 | -1.16694 | 0.003685 | 0.077133 | ENSMUSG( | Gipc2 | 23.88039 | -0.79015 | 0.055333 | 0.723362 |
| ENSMUSG( | Gipc3 | 74.16698 | -0.10208 | 0.673294 | 0.929007 | ENSMUSG( | Gipc3 | 82.43736 | 0.462717 | 0.079111 | 0.77458 |
| ENSMUSG( | Gipr | 103.7534 | -0.32532 | 0.214331 | 0.659043 | ENSMUSG( | Gipr | 95.88529 | -0.27118 | 0.422767 | 0.917257 |
| ENSMUSG( | Git1 | 23646.99 | 0.298082 | 0.00049 | 0.020296 | ENSMUSG( | Git1 | 20265.45 | 0.134166 | 0.229685 | 0.868507 |
| ENSMUSG( | Git2 | 1232.076 | 0.037982 | 0.709592 | 0.939669 | ENSMUSG( | Git2 | 1183.577 | 0.1898 | 0.16397 | 0.838291 |
| ENSMUSG( | Gja1 | 16265.94 | -0.33728 | 0.045022 | 0.325572 | ENSMUSG( | Gja1 | 16145.86 | -0.06805 | 0.698339 | 0.960908 |
| ENSMUSG( | Gja4 | 79.22011 | -0.30241 | 0.297739 | 0.746188 | ENSMUSG( | Gja4 | 110.5993 | 0.848716 | 0.0156 | 0.499582 |
| ENSMUSG( | Gjb1 | 514.5127 | -0.79139 | 2.00E-05 | 0.001952 | ENSMUSG( | Gjb1 | 583.9221 | -0.0409 | 0.84982 | 0.985522 |
| ENSMUSG( | Gjb2 | 785.8233 | -0.29348 | 0.080733 | 0.439425 | ENSMUSG( | Gjb2 | 812.4443 | 0.089578 | 0.578295 | 0.943377 |
| ENSMUSG( | Gjb6 | 5199.753 | 0.184189 | 0.15063 | 0.575407 | ENSMUSG( | Gjb6 | 4873.254 | 0.263266 | 0.082092 | 0.77681 |
| ENSMUSG( | Gjc1 | 223.5707 | -0.04019 | 0.819348 | 0.965888 | ENSMUSG( | Gjc1 | 219.9547 | 0.193463 | 0.442352 | 0.921554 |
| ENSMUSG( | Gjc2 | 979.6957 | -0.88786 | 1.45E-07 | 3.74E-05 | ENSMUSG( | Gjc2 | 1127.953 | -0.07377 | 0.684132 | 0.958678 |
| ENSMUSG( | Gjc3 | 3040.332 | -0.22864 | 0.011086 | 0.151746 | ENSMUSG( | Gjc3 | 2972.264 | -0.01544 | 0.908033 | 0.991958 |
| ENSMUSG( | Gjd2 | 200.2219 | -0.80441 | 0.000112 | 0.006972 | ENSMUSG( | Gjd2 | 223.4807 | -0.10889 | 0.627111 | 0.953234 |
| ENSMUSG( | Gjd2os | 81.32224 | 0.265608 | 0.264283 | 0.715323 | ENSMUSG( | Gjd2os | 73.03056 | 0.20292 | 0.478413 | 0.923329 |
| ENSMUSG( | Gjd3 | 25.80475 | -0.37733 | 0.329399 | 0.773311 | ENSMUSG( | Gjd3 | 26.05761 | -0.06434 | 0.871323 | 0.988779 |
| ENSMUSG( | Gk | 1021.173 | 0.106366 | 0.411933 | 0.81942 | ENSMUSG( | Gk | 885.4183 | -0.03908 | 0.796676 | 0.97681 |
| ENSMUSG( | Gk5 | 91.45392 | 0.01556 | 0.942727 | 0.993415 | ENSMUSG( | Gk5 | 85.92423 | 0.112251 | 0.632587 | 0.953698 |
| ENSMUSG( | Gkap1 | 666.1253 | 0.073992 | 0.509099 | 0.868944 | ENSMUSG( | Gkap1 | 612.2166 | 0.094065 | 0.488134 | 0.925453 |
| ENSMUSG( | Gkn3 | 15.21542 | 0.085174 | 0.876036 | NA | ENSMUSG( | Gkn3 | 30.09982 | 1.815403 | 5.11E-05 | 0.01279 |

| | | | | | |
|---|---|---|---|---|---|
| ENSMUSG( | 2114.68 | -0.00489 | 0.960148 | 0.993991 | Ggnbp2 |
| ENSMUSG( | 1182.114 | 0.049487 | 0.590481 | 0.90113 | Ggps1 |
| ENSMUSG( | 117.9027 | -0.03124 | 0.879713 | 0.981436 | Ggt1 |
| ENSMUSG( | 141.3953 | 0.210028 | 0.335774 | 0.777 | Ggt5 |
| ENSMUSG( | 4661.896 | -0.15527 | 0.179286 | 0.616869 | Ggt7 |
| ENSMUSG( | 201.958 | -0.17027 | 0.383883 | 0.803646 | Ggta1 |
| ENSMUSG( | 491.6673 | -0.04318 | 0.764794 | 0.954127 | Ghdc |
| ENSMUSG( | 11997.73 | 0.062025 | 0.526417 | 0.877267 | Ghitm |
| ENSMUSG( | 484.3061 | -0.15158 | 0.430134 | 0.826789 | Ghr |
| ENSMUSG( | 6654.749 | 0.174521 | 0.051495 | 0.34814 | Gid4 |
| ENSMUSG( | 1180.035 | 0.049997 | 0.61299 | 0.908902 | Gid8 |
| ENSMUSG( | 2178.042 | 0.112061 | 0.325876 | 0.771128 | Gigyf1 |
| ENSMUSG( | 1785.316 | 0.265757 | 0.001632 | 0.045399 | Gigyf2 |
| ENSMUSG( | 52.4688 | -0.02337 | 0.937861 | 0.993415 | Gimap1 |
| ENSMUSG( | 19.44122 | -0.22933 | 0.614594 | NA | Gimap4 |
| ENSMUSG( | 25.75303 | -0.36555 | 0.398737 | 0.812878 | Gimap5 |
| ENSMUSG( | 227.591 | -0.03155 | 0.884215 | 0.982647 | Gimap6 |
| ENSMUSG( | 45.35327 | 0.084109 | 0.787496 | 0.959163 | Gimap8 |
| ENSMUSG( | 14.62491 | -0.47707 | 0.316387 | NA | Gimap9 |
| ENSMUSG( | 150.5275 | 0.082595 | 0.623395 | 0.911646 | Gin1 |
| ENSMUSG( | 763.1432 | -0.36358 | 0.014228 | 0.175234 | Ginm1 |
| ENSMUSG( | 26.71086 | -0.04807 | 0.895448 | 0.984512 | Gins1 |
| ENSMUSG( | 76.65482 | 0.02485 | 0.916614 | 0.988856 | Gins2 |
| ENSMUSG( | 57.84734 | 0.159851 | 0.535025 | 0.880733 | Gins3 |
| ENSMUSG( | 427.8749 | -0.00143 | 0.990059 | 0.999144 | Gins4 |
| ENSMUSG( | 3134.49 | 0.063922 | 0.463942 | 0.842812 | Gipc1 |
| ENSMUSG( | 23.87614 | -1.16694 | 0.003685 | 0.077133 | Gipc2 |
| ENSMUSG( | 74.16698 | -0.10208 | 0.673294 | 0.929007 | Gipc3 |
| ENSMUSG( | 103.7534 | -0.32532 | 0.214331 | 0.659043 | Gipr |
| ENSMUSG( | 23646.99 | 0.298082 | 0.00049 | 0.020296 | Git1 |
| ENSMUSG( | 1232.076 | 0.037982 | 0.709592 | 0.939669 | Git2 |
| ENSMUSG( | 16265.94 | -0.33728 | 0.045022 | 0.325572 | Gja1 |
| ENSMUSG( | 79.22011 | -0.30241 | 0.297739 | 0.746188 | Gja4 |
| ENSMUSG( | 514.5127 | -0.79139 | 2.00E-05 | 0.001952 | Gjb1 |
| ENSMUSG( | 785.8233 | -0.29348 | 0.080733 | 0.439425 | Gjb2 |
| ENSMUSG( | 5199.753 | 0.184189 | 0.15063 | 0.575407 | Gjb6 |
| ENSMUSG( | 223.5707 | -0.04019 | 0.819348 | 0.965888 | Gjc1 |
| ENSMUSG( | 979.6957 | -0.88786 | 1.45E-07 | 3.74E-05 | Gjc2 |
| ENSMUSG( | 3040.332 | -0.22864 | 0.011086 | 0.151746 | Gjc3 |
| ENSMUSG( | 200.2219 | -0.80441 | 0.000112 | 0.006972 | Gjd2 |
| ENSMUSG( | 81.32224 | 0.265608 | 0.264283 | 0.715323 | Gjd2os |
| ENSMUSG( | 25.80475 | -0.37733 | 0.329399 | 0.773311 | Gjd3 |
| ENSMUSG( | 1021.173 | 0.106366 | 0.411933 | 0.81942 | Gk |
| ENSMUSG( | 91.45392 | 0.01556 | 0.942727 | 0.993415 | Gk5 |
| ENSMUSG( | 666.1253 | 0.073992 | 0.509099 | 0.868944 | Gkap1 |
| ENSMUSG( | 15.21542 | 0.085174 | 0.876036 | NA | Gkn3 |
| ENSMUSG( | 403.6016 | -0.58956 | 0.0024 | 0.057587 | Gla |
| ENSMUSG( | 784.8243 | -0.16978 | 0.224149 | 0.669929 | Glb1 |
| ENSMUSG( | 468.0066 | -0.00686 | 0.955206 | 0.993874 | Glb1l |
| ENSMUSG( | 29.64973 | -0.10619 | 0.755727 | 0.95146 | Glb1l2 |
| ENSMUSG( | 49.08833 | -0.03171 | 0.913504 | 0.988483 | Glcci1 |
| ENSMUSG( | 755.072 | -0.43216 | 0.018357 | 0.203658 | Glce |
| ENSMUSG( | 3822.878 | 0.060767 | 0.598062 | 0.903724 | Gldc |
| ENSMUSG( | 39.60428 | -0.20201 | 0.513867 | 0.872055 | Gldn |
| ENSMUSG( | 1693.37 | 0.083688 | 0.331916 | 0.775288 | Gle1 |
| ENSMUSG( | 6371.732 | -0.16438 | 0.325523 | 0.770741 | Glg1 |
| ENSMUSG( | 169.6058 | -0.37258 | 0.027205 | 0.253245 | Gli1 |
| ENSMUSG( | 343.9648 | -0.03457 | 0.788455 | 0.959226 | Gli2 |
| ENSMUSG( | 849.7661 | 0.024899 | 0.843475 | 0.971847 | Gli3 |
| ENSMUSG( | 30.68051 | -0.18289 | 0.588653 | 0.900921 | Glipr1 |
| ENSMUSG( | 271.6868 | 0.257882 | 0.119987 | 0.523983 | Glipr2 |
| ENSMUSG( | 16.8597 | 0.161286 | 0.749159 | NA | Glis1 |
| ENSMUSG( | 681.9314 | 0.002173 | 0.983211 | 0.997554 | Glis2 |
| ENSMUSG( | 434.7548 | 0.110253 | 0.440675 | 0.832102 | Glis3 |
| ENSMUSG( | 236.6175 | -0.14719 | 0.35502 | 0.787423 | Glmn |
| ENSMUSG( | 1793.321 | -0.25975 | 0.03824 | 0.301606 | Glmp |
| ENSMUSG( | 139.3371 | 0.221481 | 0.249496 | 0.699055 | Glns-ps1 |
| ENSMUSG( | 4837.498 | 0.052478 | 0.599694 | 0.903906 | Glo1 |
| ENSMUSG( | 24.29116 | 0.330389 | 0.409814 | 0.818258 | Glo1-ps |
| ENSMUSG( | 1633.297 | 0.078079 | 0.48792 | 0.858824 | Glod4 |
| ENSMUSG( | 43.00691 | -0.1528 | 0.671849 | 0.928781 | Glp1r |
| ENSMUSG( | 82.8616 | 0.047881 | 0.849827 | 0.972638 | Glp2r |
| ENSMUSG( | 320.4865 | -0.66116 | 2.35E-05 | 0.00217 | Glra2 |
| ENSMUSG( | 93.90564 | -0.66902 | 0.005505 | 0.099427 | Glra3 |
| ENSMUSG( | 4889.488 | -0.48734 | 0.002336 | 0.056722 | Glrb |
| ENSMUSG( | 395.4415 | -0.06432 | 0.619456 | 0.911432 | Glrx |
| ENSMUSG( | 2202.365 | 0.046935 | 0.600939 | 0.904492 | Glrx2 |
| ENSMUSG( | 2334.84 | 0.050538 | 0.575092 | 0.894572 | Glrx3 |
| ENSMUSG( | 1521.695 | 0.013038 | 0.89065 | 0.983764 | Glrx5 |
| ENSMUSG( | 7105.921 | 0.164274 | 0.067641 | 0.402445 | Gls |
| ENSMUSG( | 750.1318 | -0.00976 | 0.928634 | 0.991668 | Gls2 |
| ENSMUSG( | 639.5505 | 0.116865 | 0.356192 | 0.788108 | Glt1d1 |
| ENSMUSG( | 9.972145 | -0.49448 | 0.464055 | NA | Glt28d2 |
| ENSMUSG( | 1094.771 | 0.051824 | 0.665887 | 0.927054 | Glt8d1 |
| ENSMUSG( | 367.8089 | -0.45984 | 0.005133 | 0.095477 | Glt8d2 |
| ENSMUSG( | 4154.637 | -0.00157 | 0.987188 | 0.998764 | Gltp |
| ENSMUSG( | 29130.61 | 0.111585 | 0.279666 | 0.729119 | Glud1 |
| ENSMUSG( | 204394.1 | 0.248037 | 0.023448 | 0.234187 | Glul |
| ENSMUSG( | 11.27743 | 0.532862 | 0.402076 | NA | Glycam1 |
| ENSMUSG( | 174.7374 | -0.09244 | 0.576285 | 0.895164 | Glyctk |
| ENSMUSG( | 5216.481 | 0.158581 | 0.066826 | 0.399821 | Glyr1 |
| ENSMUSG( | 25.71027 | 0.489239 | 0.223451 | 0.669469 | Gm10030 |
| ENSMUSG( | 214.0457 | -0.01107 | 0.943791 | 0.993503 | Gm10033 |
| ENSMUSG( | 54.45915 | 0.062568 | 0.830493 | 0.96974 | Gm10036 |
| ENSMUSG( | 9.740169 | -0.35062 | 0.621845 | NA | Gm10051 |
| ENSMUSG( | 14.92236 | 0.427363 | 0.391985 | NA | Gm10060 |
| ENSMUSG( | 26.78863 | 0.403783 | 0.303528 | 0.751968 | Gm10069 |
| ENSMUSG( | 911.3909 | 0.158118 | 0.110737 | 0.501952 | Gm10076 |
| ENSMUSG( | 74.96418 | -0.0353 | 0.879417 | 0.981436 | Gm10136 |
| ENSMUSG( | 89.78757 | 0.180439 | 0.444555 | 0.833239 | Gm10138 |
| ENSMUSG( | 24.55834 | 0.084244 | 0.827179 | 0.968725 | Gm10167 |
| ENSMUSG( | 887.6815 | 0.047122 | 0.750166 | 0.949664 | Gm10222 |
| ENSMUSG( | 42.15211 | -0.04979 | 0.862711 | 0.975055 | Gm10244 |
| ENSMUSG( | 71.46504 | 0.008977 | 0.968861 | 0.996369 | Gm10250 |
| ENSMUSG( | 51.7335 | 0.184816 | 0.495033 | 0.860798 | Gm10269 |
| ENSMUSG( | 8.759954 | -0.44312 | 0.499122 | NA | Gm10284 |
| ENSMUSG( | 54.67973 | 0.030904 | 0.904432 | 0.986246 | Gm10288 |
| ENSMUSG( | 11.47291 | -0.05519 | 0.919153 | NA | Gm10293 |
| ENSMUSG( | 23.72253 | 0.13912 | 0.726879 | 0.944223 | Gm10327 |
| ENSMUSG( | 20.16479 | 0.033033 | 0.940376 | NA | Gm10335 |
| ENSMUSG( | 11.93483 | 0.851054 | 0.126776 | NA | Gm10384 |
| ENSMUSG( | 19.17315 | 0.09556 | 0.835345 | NA | Gm10406 |
| ENSMUSG( | 8.731535 | 1.503817 | 0.022761 | NA | Gm10416 |
| ENSMUSG( | 25.55279 | -1.20957 | 0.002916 | 0.066108 | Gm10421 |
| ENSMUSG( | 281.0018 | -0.04212 | 0.802085 | 0.962119 | Gm1043 |
| ENSMUSG( | 137.9463 | 0.036491 | 0.838314 | 0.971111 | Gm10432 |
| ENSMUSG( | 9.583031 | -0.29466 | 0.643079 | NA | Gm10433 |
| ENSMUSG( | 122.7498 | -0.14749 | 0.462549 | 0.842321 | Gm10509 |
| ENSMUSG( | 205.0081 | 0.069587 | 0.638102 | 0.916107 | Gm10516 |
| ENSMUSG( | 95.78769 | 0.121261 | 0.559475 | 0.888669 | Gm10575 |
| ENSMUSG( | 54.17278 | -0.0081 | 0.975804 | 0.997428 | Gm10635 |
| ENSMUSG( | 11.39844 | -0.39142 | 0.543212 | NA | Gm10640 |
| ENSMUSG( | 89.88138 | 0.336657 | 0.123421 | 0.530552 | Gm10644 |
| ENSMUSG( | 81.00351 | 0.065608 | 0.793992 | 0.959799 | Gm10654 |
| ENSMUSG( | 17.27407 | -0.13511 | 0.76912 | NA | Gm10710 |
| ENSMUSG( | 15.36361 | -0.67264 | 0.188465 | NA | Gm10754 |
| ENSMUSG( | 8.995593 | -0.04043 | 0.950956 | NA | Gm10778 |
| ENSMUSG( | 14.5173 | -0.40738 | 0.401821 | NA | Gm10785 |
| ENSMUSG( | 116.194 | -0.40383 | 0.052493 | 0.35151 | Gm10790 |
| ENSMUSG( | 11.56752 | -0.01646 | 0.975468 | NA | Gm10814 |
| ENSMUSG( | 67.87957 | -0.47167 | 0.066334 | 0.398419 | Gm10863 |
| ENSMUSG( | 14.81219 | 0.069784 | 0.892304 | NA | Gm10865 |
| ENSMUSG( | 16.2061 | 0.349713 | 0.446739 | NA | Gm10941 |
| ENSMUSG( | 21.78564 | 0.034436 | 0.939093 | 0.993415 | Gm11110 |
| ENSMUSG( | 31.95295 | -0.60988 | 0.135928 | 0.549949 | Gm11149 |
| ENSMUSG( | 11.59372 | 0.273993 | 0.644236 | NA | Gm11185 |
| ENSMUSG( | 14.98709 | 0.110159 | 0.822298 | NA | Gm11249 |
| ENSMUSG( | 19.01465 | 0.033447 | 0.941855 | NA | Gm11263 |
| ENSMUSG( | 158.6699 | 0.257915 | 0.117672 | 0.517948 | Gm11266 |
| ENSMUSG( | 21.00258 | 0.200423 | 0.654448 | NA | Gm11353 |
| ENSMUSG( | 26.43305 | -0.27909 | 0.476921 | 0.851851 | Gm11423 |
| ENSMUSG( | 32.25069 | -0.17974 | 0.584005 | 0.898666 | Gm11451 |
| ENSMUSG( | 193.634 | -0.00983 | 0.94974 | 0.993821 | Gm11478 |
| ENSMUSG( | 8.127113 | 0.877028 | 0.176459 | NA | Gm11598 |
| ENSMUSG( | 286.1481 | 0.36461 | 0.008832 | 0.132321 | Gm11627 |
| ENSMUSG( | 40.36781 | -0.45874 | 0.125341 | 0.532314 | Gm11650 |
| ENSMUSG( | 54.3566 | 0.768837 | 0.005612 | 0.100463 | Gm11681 |
| ENSMUSG( | 138.6398 | -0.06438 | 0.716961 | 0.941697 | Gm11696 |
| ENSMUSG( | 8.356743 | -0.69419 | 0.305704 | NA | Gm11732 |
| ENSMUSG( | 35.89553 | 0.347331 | 0.26647 | 0.716441 | Gm11762 |
| ENSMUSG( | 275.1477 | 0.209347 | 0.153886 | 0.580901 | Gm11767 |
| ENSMUSG( | 14.74136 | 0.106598 | 0.831779 | NA | Gm11826 |
| ENSMUSG( | 14.09424 | 0.687334 | 0.19804 | NA | Gm11827 |
| ENSMUSG( | 62.04995 | -0.06354 | 0.792375 | 0.959461 | Gm11837 |
| ENSMUSG( | 16.96891 | 0.480507 | 0.31326 | NA | Gm11839 |
| ENSMUSG( | 9.593547 | 0.473029 | 0.428765 | NA | Gm11841 |
| ENSMUSG( | 30.61753 | 0.0581 | 0.868311 | 0.976834 | Gm11914 |
| ENSMUSG( | 18.71053 | -0.1356 | 0.770083 | NA | Gm11944 |

| | | | | | |
|---|---|---|---|---|---|
| ENSMUSG( | 10139.12 | -0.1609 | 0.210048 | 0.864412 | Ghitm |
| ENSMUSG( | 465.1581 | 0.005093 | 0.974994 | 0.996416 | Ghr |
| ENSMUSG( | 5609.86 | -0.04579 | 0.674788 | 0.95694 | Gid4 |
| ENSMUSG( | 1013.392 | -0.12575 | 0.29519 | 0.898538 | Gid8 |
| ENSMUSG( | 1924.109 | 0.024642 | 0.85924 | 0.985884 | Gigyf1 |
| ENSMUSG( | 1510.502 | 0.061696 | 0.583817 | 0.944379 | Gigyf2 |
| ENSMUSG( | 50.60683 | 0.167929 | 0.629001 | 0.953251 | Gimap1 |
| ENSMUSG( | 19.52339 | 0.05017 | 0.914547 | 0.992407 | Gimap4 |
| ENSMUSG( | 22.06355 | -0.5294 | 0.3018 | 0.898538 | Gimap5 |
| ENSMUSG( | 205.1568 | -0.043 | 0.869235 | 0.988097 | Gimap6 |
| ENSMUSG( | 48.54991 | 0.538041 | 0.135572 | 0.825163 | Gimap8 |
| ENSMUSG( | 17.5097 | 0.359944 | 0.442284 | 0.921554 | Gimap9 |
| ENSMUSG( | 142.7776 | 0.201413 | 0.300468 | 0.898538 | Gin1 |
| ENSMUSG( | 810.0701 | 0.103768 | 0.538679 | 0.937216 | Ginm1 |
| ENSMUSG( | 24.38042 | -0.04268 | 0.917729 | 0.992529 | Gins1 |
| ENSMUSG( | 61.03366 | -0.38485 | 0.14119 | 0.830678 | Gins2 |
| ENSMUSG( | 52.02604 | 0.1086 | 0.714284 | 0.964986 | Gins3 |
| ENSMUSG( | 372.513 | -0.13973 | 0.319516 | 0.903464 | Gins4 |
| ENSMUSG( | 2679.526 | -0.123 | 0.232152 | 0.868507 | Gipc1 |
| ENSMUSG( | 23.88039 | -0.79015 | 0.055333 | 0.723362 | Gipc2 |
| ENSMUSG( | 82.43736 | 0.462717 | 0.079111 | 0.77458 | Gipc3 |
| ENSMUSG( | 95.88529 | -0.27118 | 0.422767 | 0.917257 | Gipr |
| ENSMUSG( | 20265.45 | 0.134166 | 0.229685 | 0.868507 | Git1 |
| ENSMUSG( | 1183.577 | 0.1898 | 0.16397 | 0.838291 | Git2 |
| ENSMUSG( | 16145.86 | -0.06805 | 0.698339 | 0.960908 | Gja1 |
| ENSMUSG( | 110.5993 | 0.848716 | 0.0156 | 0.499582 | Gja4 |
| ENSMUSG( | 583.9221 | -0.0409 | 0.84982 | 0.985522 | Gjb1 |
| ENSMUSG( | 812.4443 | 0.089578 | 0.578295 | 0.943377 | Gjb2 |
| ENSMUSG( | 4873.254 | 0.263266 | 0.082092 | 0.77681 | Gjb6 |
| ENSMUSG( | 219.9547 | 0.193463 | 0.442352 | 0.921554 | Gjc1 |
| ENSMUSG( | 1127.953 | -0.07377 | 0.684132 | 0.958678 | Gjc2 |
| ENSMUSG( | 2972.264 | -0.01544 | 0.908033 | 0.991958 | Gjc3 |
| ENSMUSG( | 223.4807 | -0.10889 | 0.627111 | 0.953234 | Gjd2 |
| ENSMUSG( | 73.03056 | 0.20292 | 0.478413 | 0.923329 | Gjd2os |
| ENSMUSG( | 26.05761 | -0.06434 | 0.871323 | 0.988779 | Gjd3 |
| ENSMUSG( | 885.4183 | -0.03908 | 0.796676 | 0.97681 | Gk |
| ENSMUSG( | 85.92423 | 0.112251 | 0.632587 | 0.953698 | Gk5 |
| ENSMUSG( | 612.2166 | 0.094065 | 0.488134 | 0.925453 | Gkap1 |
| ENSMUSG( | 30.09982 | 1.815403 | 5.11E-05 | 0.01279 | Gkn3 |
| ENSMUSG( | 468.4869 | 0.172259 | 0.381419 | 0.917257 | Gla |
| ENSMUSG( | 849.6876 | 0.322039 | 0.073347 | 0.763629 | Glb1 |
| ENSMUSG( | 426.8198 | 0.000934 | 0.995467 | 0.999116 | Glb1l |
| ENSMUSG( | 33.4926 | 0.49882 | 0.157653 | 0.837724 | Glb1l2 |
| ENSMUSG( | 49.45673 | 0.274269 | 0.421216 | 0.917257 | Glcci1 |
| ENSMUSG( | 857.6767 | 0.234409 | 0.25645 | 0.883082 | Glce |
| ENSMUSG( | 3282.782 | -0.11176 | 0.338821 | 0.907972 | Gldc |
| ENSMUSG( | 38.60839 | -0.00596 | 0.98533 | 0.997084 | Gldn |
| ENSMUSG( | 1486.906 | -0.02115 | 0.841945 | 0.983044 | Gle1 |
| ENSMUSG( | 6678.989 | 0.241226 | 0.22475 | 0.868316 | Glg1 |
| ENSMUSG( | 188.7471 | 0.231301 | 0.199855 | 0.863016 | Gli1 |
| ENSMUSG( | 322.0008 | 0.049644 | 0.739151 | 0.968416 | Gli2 |
| ENSMUSG( | 752.1825 | -0.06172 | 0.683251 | 0.958678 | Gli3 |
| ENSMUSG( | 32.77711 | 0.262471 | 0.449396 | 0.922202 | Glipr1 |
| ENSMUSG( | 199.3539 | -0.39585 | 0.038303 | 0.679117 | Glipr2 |
| ENSMUSG( | 17.50896 | 0.47616 | 0.336993 | 0.907972 | Glis1 |
| ENSMUSG( | 621.1025 | 0.000628 | 0.995865 | 0.999116 | Glis2 |
| ENSMUSG( | 403.6465 | 0.165364 | 0.27092 | 0.888062 | Glis3 |
| ENSMUSG( | 227.66 | 0.010201 | 0.954285 | 0.994798 | Glmn |
| ENSMUSG( | 1869.717 | 0.141869 | 0.379954 | 0.916995 | Glmp |
| ENSMUSG( | 107.1725 | -0.28809 | 0.210939 | 0.864412 | Glns-ps1 |
| ENSMUSG( | 4036.697 | -0.2109 | 0.108036 | 0.804969 | Glo1 |
| ENSMUSG( | 19.37036 | -0.02136 | 0.961346 | 0.99511 | Glo1-ps |
| ENSMUSG( | 1403.62 | -0.09355 | 0.427504 | 0.917257 | Glod4 |
| ENSMUSG( | 46.50018 | 0.318612 | 0.404823 | 0.917257 | Glp1r |
| ENSMUSG( | 77.16853 | 0.125342 | 0.671372 | 0.95694 | Glp2r |
| ENSMUSG( | 376.3867 | 0.140053 | 0.46113 | 0.923329 | Glra2 |
| ENSMUSG( | 114.1071 | 0.2246 | 0.341478 | 0.908653 | Glra3 |
| ENSMUSG( | 5582.46 | 0.19897 | 0.28453 | 0.892412 | Glrb |
| ENSMUSG( | 346.9505 | -0.1827 | 0.222485 | 0.866791 | Glrx |
| ENSMUSG( | 1974.302 | -0.00101 | 0.992753 | 0.99874 | Glrx2 |
| ENSMUSG( | 1944.105 | -0.21965 | 0.054612 | 0.723362 | Glrx3 |
| ENSMUSG( | 1284.67 | -0.21978 | 0.112682 | 0.809623 | Glrx5 |
| ENSMUSG( | 6215.109 | 0.04981 | 0.622114 | 0.953234 | Gls |
| ENSMUSG( | 665.7929 | -0.08366 | 0.511989 | 0.930005 | Gls2 |
| ENSMUSG( | 568.4851 | 0.045769 | 0.759324 | 0.972002 | Glt1d1 |
| ENSMUSG( | 9.911417 | -0.24733 | 0.69394 | NA | Glt28d2 |
| ENSMUSG( | 1045.454 | 0.184983 | 0.265927 | 0.886134 | Glt8d1 |
| ENSMUSG( | 435.0283 | 0.317225 | 0.059433 | 0.729747 | Glt8d2 |
| ENSMUSG( | 3544.897 | -0.20093 | 0.12848 | 0.81925 | Gltp |
| ENSMUSG( | 24260.4 | -0.15079 | 0.261886 | 0.884926 | Glud1 |
| ENSMUSG( | 166095.6 | -0.0737 | 0.597553 | 0.946609 | Glul |
| ENSMUSG( | 146.2927 | -0.35208 | 0.067836 | 0.75212 | Glyctk |
| ENSMUSG( | 4505.187 | 0.008343 | 0.93683 | 0.993669 | Glyr1 |
| ENSMUSG( | 20.34734 | 0.1181 | 0.796307 | 0.97681 | Gm10030 |
| ENSMUSG( | 215.2572 | 0.265541 | 0.12656 | 0.815653 | Gm10033 |
| ENSMUSG( | 44.83096 | -0.26958 | 0.393808 | 0.917257 | Gm10036 |
| ENSMUSG( | 11.80697 | 0.021824 | 0.970414 | 0.99594 | Gm10060 |
| ENSMUSG( | 19.86872 | -0.17851 | 0.711947 | 0.964363 | Gm10069 |
| ENSMUSG( | 761.1108 | -0.09637 | 0.517334 | 0.930939 | Gm10076 |
| ENSMUSG( | 57.11834 | -0.6206 | 0.030939 | 0.629448 | Gm10136 |
| ENSMUSG( | 74.93441 | -0.0531 | 0.842513 | 0.983044 | Gm10138 |
| ENSMUSG( | 21.08613 | -0.12649 | 0.773016 | 0.974463 | Gm10167 |
| ENSMUSG( | 709.2869 | -0.36253 | 0.088107 | 0.786635 | Gm10222 |
| ENSMUSG( | 34.26583 | -0.39216 | 0.252618 | 0.88018 | Gm10244 |
| ENSMUSG( | 57.93612 | -0.37463 | 0.188023 | 0.85698 | Gm10250 |
| ENSMUSG( | 44.84087 | 0.03061 | 0.921597 | 0.992644 | Gm10269 |
| ENSMUSG( | 41.07646 | -0.56332 | 0.073414 | 0.763629 | Gm10288 |
| ENSMUSG( | 8.168416 | -0.85966 | 0.209214 | NA | Gm10293 |
| ENSMUSG( | 18.03777 | -0.40904 | 0.369242 | 0.915491 | Gm10327 |
| ENSMUSG( | 20.02991 | 0.22791 | 0.643935 | 0.953903 | Gm10335 |
| ENSMUSG( | 12.54095 | 1.188294 | 0.043587 | 0.690304 | Gm10384 |
| ENSMUSG( | 15.10595 | -0.3289 | 0.540158 | 0.937383 | Gm10406 |
| ENSMUSG( | 30.38133 | -0.19847 | 0.623271 | 0.953234 | Gm10421 |
| ENSMUSG( | 291.8454 | 0.323646 | 0.065463 | 0.744625 | Gm1043 |
| ENSMUSG( | 118.8058 | -0.13237 | 0.513502 | 0.930482 | Gm10432 |
| ENSMUSG( | 125.5951 | 0.192302 | 0.380612 | 0.917202 | Gm10509 |
| ENSMUSG( | 191.6655 | 0.126253 | 0.515281 | 0.930939 | Gm10516 |
| ENSMUSG( | 83.12989 | 0.000865 | 0.997192 | 0.999268 | Gm10575 |
| ENSMUSG( | 44.14771 | -0.35382 | 0.259885 | 0.883971 | Gm10635 |
| ENSMUSG( | 67.83185 | -0.16334 | 0.581901 | 0.944266 | Gm10644 |
| ENSMUSG( | 71.99396 | -0.02407 | 0.935731 | 0.993406 | Gm10654 |
| ENSMUSG( | 14.35343 | -0.43996 | 0.391473 | 0.917257 | Gm10710 |
| ENSMUSG( | 15.47194 | -0.37855 | 0.478103 | 0.923329 | Gm10754 |
| ENSMUSG( | 12.97789 | -0.47436 | 0.385994 | 0.917257 | Gm10785 |
| ENSMUSG( | 115.0485 | -0.11263 | 0.676473 | 0.95694 | Gm10790 |
| ENSMUSG( | 10.72648 | 0.070967 | 0.906395 | NA | Gm10814 |
| ENSMUSG( | 68.04176 | -0.13161 | 0.655959 | 0.955533 | Gm10863 |
| ENSMUSG( | 10.05044 | -0.85139 | 0.180391 | NA | Gm10865 |
| ENSMUSG( | 15.36111 | 0.439091 | 0.374346 | 0.915491 | Gm10941 |
| ENSMUSG( | 18.59949 | -0.18845 | 0.694819 | 0.960867 | Gm11110 |
| ENSMUSG( | 29.36087 | -0.52933 | 0.257654 | 0.883401 | Gm11149 |
| ENSMUSG( | 9.390034 | -0.11251 | 0.865829 | NA | Gm11185 |
| ENSMUSG( | 8.609883 | -1.63873 | 0.022888 | NA | Gm11249 |
| ENSMUSG( | 13.87451 | -0.74549 | 0.196526 | 0.862161 | Gm11263 |
| ENSMUSG( | 145.5361 | 0.258946 | 0.22053 | 0.866104 | Gm11266 |
| ENSMUSG( | 16.16179 | -0.34939 | 0.500589 | 0.927825 | Gm11353 |
| ENSMUSG( | 26.81728 | 0.041968 | 0.913388 | 0.992298 | Gm11423 |
| ENSMUSG( | 27.79003 | -0.36346 | 0.338328 | 0.907972 | Gm11451 |
| ENSMUSG( | 139.6111 | -0.79111 | 6.47E-05 | 0.015522 | Gm11478 |
| ENSMUSG( | 252.7954 | 0.280713 | 0.081918 | 0.77681 | Gm11627 |
| ENSMUSG( | 38.79409 | -0.31412 | 0.358837 | 0.913759 | Gm11650 |
| ENSMUSG( | 32.84198 | -0.32373 | 0.35491 | 0.912411 | Gm11681 |
| ENSMUSG( | 124.7423 | -0.09838 | 0.638517 | 0.953903 | Gm11696 |
| ENSMUSG( | 6.68442 | -1.2115 | 0.156399 | NA | Gm11732 |
| ENSMUSG( | 28.00109 | -0.04852 | 0.898626 | 0.990558 | Gm11762 |
| ENSMUSG( | 242.1321 | 0.121663 | 0.552895 | 0.939565 | Gm11767 |
| ENSMUSG( | 11.00598 | -0.56457 | 0.351439 | NA | Gm11826 |
| ENSMUSG( | 14.08483 | 0.935003 | 0.101636 | 0.801601 | Gm11827 |
| ENSMUSG( | 57.59677 | -0.02419 | 0.930558 | 0.99306 | Gm11837 |
| ENSMUSG( | 16.9118 | 0.719312 | 0.188651 | 0.85698 | Gm11839 |
| ENSMUSG( | 26.83842 | -0.05691 | 0.881389 | 0.989644 | Gm11914 |
| ENSMUSG( | 22.30054 | 0.593348 | 0.155762 | 0.837724 | Gm11944 |
| ENSMUSG( | 56.51614 | 0.222537 | 0.462687 | 0.923329 | Gm11992 |
| ENSMUSG( | 22.33388 | -0.7479 | 0.083758 | 0.77681 | Gm12034 |
| ENSMUSG( | 11.59508 | -0.31716 | 0.578056 | 0.943163 | Gm12165 |
| ENSMUSG( | 41.77474 | -0.1706 | 0.641764 | 0.953903 | Gm12174 |
| ENSMUSG( | 33.61207 | -0.17809 | 0.617544 | 0.951952 | Gm12183 |
| ENSMUSG( | 28.14148 | 0.014203 | 0.970249 | 0.99594 | Gm12216 |
| ENSMUSG( | 31.27017 | -0.47339 | 0.275135 | 0.889874 | Gm12224 |
| ENSMUSG( | 18.2312 | -0.50804 | 0.312332 | 0.900037 | Gm12251 |
| ENSMUSG( | 351.9456 | 0.246482 | 0.274662 | 0.889874 | Gm12258 |
| ENSMUSG( | 20.03415 | -0.00771 | 0.985929 | 0.997188 | Gm12326 |
| ENSMUSG( | 92.55577 | -0.44322 | 0.08019 | 0.77681 | Gm12346 |
| ENSMUSG( | 25.79371 | -0.17707 | 0.678471 | 0.957506 | Gm12359 |
| ENSMUSG( | 84.79944 | -0.30388 | 0.222833 | 0.867222 | Gm12371 |
| ENSMUSG( | 15.29515 | -0.25605 | 0.610596 | 0.950771 | Gm12416 |
| ENSMUSG( | 35.42531 | -0.21645 | 0.514772 | 0.930939 | Gm12500 |
| ENSMUSG( | 26.03185 | 0.131571 | 0.780861 | 0.976106 | Gm12522 |

| | | | | | | | | | | | |
|---|---|---|---|---|---|---|---|---|---|---|---|
| ENSMUSG( | Gla | 403.6016 | -0.58956 | 0.0024 | 0.057587 | ENSMUSG( | Gla | 468.4869 | 0.172259 | 0.381419 | 0.917257 |
| ENSMUSG( | Glb1 | 784.8243 | -0.16978 | 0.224149 | 0.669929 | ENSMUSG( | Glb1 | 849.6876 | 0.322039 | 0.073347 | 0.763629 |
| ENSMUSG( | Glb1l | 468.0066 | -0.00686 | 0.955206 | 0.993874 | ENSMUSG( | Glb1l | 426.8198 | 0.000934 | 0.995467 | 0.999116 |
| ENSMUSG( | Glb1l2 | 29.64973 | -0.10619 | 0.755727 | 0.95146 | ENSMUSG( | Glb1l2 | 33.4926 | 0.49882 | 0.157653 | 0.837724 |
| ENSMUSG( | Glcci1 | 49.08833 | -0.03171 | 0.913504 | 0.988483 | ENSMUSG( | Glcci1 | 49.45673 | 0.274269 | 0.421216 | 0.917257 |
| ENSMUSG( | Glce | 755.072 | -0.43216 | 0.018357 | 0.203658 | ENSMUSG( | Glce | 857.6767 | 0.234409 | 0.25645 | 0.883082 |
| ENSMUSG( | Gldc | 3822.878 | 0.060767 | 0.598062 | 0.903724 | ENSMUSG( | Gldc | 3282.782 | -0.11176 | 0.338821 | 0.907972 |
| ENSMUSG( | Gldn | 39.60428 | -0.20201 | 0.513867 | 0.872055 | ENSMUSG( | Gldn | 38.60839 | -0.00596 | 0.98533 | 0.997084 |
| ENSMUSG( | Gle1 | 1693.37 | 0.083688 | 0.331916 | 0.775288 | ENSMUSG( | Gle1 | 1486.906 | -0.02115 | 0.841945 | 0.983044 |
| ENSMUSG( | Glg1 | 6371.732 | -0.16438 | 0.325523 | 0.770741 | ENSMUSG( | Glg1 | 6678.989 | 0.241226 | 0.22475 | 0.868316 |
| ENSMUSG( | Gli1 | 169.6058 | -0.37258 | 0.027205 | 0.253245 | ENSMUSG( | Gli1 | 188.7471 | 0.231301 | 0.199855 | 0.863016 |
| ENSMUSG( | Gli2 | 343.9648 | -0.03457 | 0.788455 | 0.959226 | ENSMUSG( | Gli2 | 322.0008 | 0.049644 | 0.739151 | 0.968416 |
| ENSMUSG( | Gli3 | 849.7661 | 0.024899 | 0.843475 | 0.971847 | ENSMUSG( | Gli3 | 752.1825 | -0.06172 | 0.683251 | 0.958678 |
| ENSMUSG( | Glipr1 | 30.68051 | -0.18289 | 0.588653 | 0.900921 | ENSMUSG( | Glipr1 | 32.77711 | 0.262471 | 0.449396 | 0.922202 |
| ENSMUSG( | Glipr2 | 271.6868 | 0.257882 | 0.119987 | 0.523983 | ENSMUSG( | Glipr2 | 199.3539 | -0.39585 | 0.038303 | 0.679117 |
| ENSMUSG( | Glis1 | 16.8597 | 0.161286 | 0.749159 | NA | ENSMUSG( | Glis1 | 17.50896 | 0.47616 | 0.336993 | 0.907972 |
| ENSMUSG( | Glis2 | 681.9314 | 0.002173 | 0.983211 | 0.997554 | ENSMUSG( | Glis2 | 621.1025 | 0.000628 | 0.995865 | 0.999116 |
| ENSMUSG( | Glis3 | 434.7548 | 0.110253 | 0.440675 | 0.832102 | ENSMUSG( | Glis3 | 403.6465 | 0.165364 | 0.27092 | 0.888062 |
| ENSMUSG( | Glmn | 236.6175 | -0.14719 | 0.35502 | 0.787423 | ENSMUSG( | Glmn | 227.66 | 0.010201 | 0.954285 | 0.994798 |
| ENSMUSG( | Glmp | 1793.321 | -0.25975 | 0.03824 | 0.301606 | ENSMUSG( | Glmp | 1869.717 | 0.141869 | 0.379954 | 0.916995 |
| ENSMUSG( | Glns-ps1 | 139.3371 | 0.221481 | 0.249496 | 0.699055 | ENSMUSG( | Glns-ps1 | 107.1725 | -0.28809 | 0.210939 | 0.864412 |
| ENSMUSG( | Glo1 | 4837.498 | 0.052478 | 0.599694 | 0.903906 | ENSMUSG( | Glo1 | 4036.697 | -0.2109 | 0.108036 | 0.804969 |
| ENSMUSG( | Glo1-ps | 24.29116 | 0.330389 | 0.409814 | 0.818258 | ENSMUSG( | Glo1-ps | 19.37036 | -0.02136 | 0.961346 | 0.99511 |
| ENSMUSG( | Glod4 | 1633.297 | 0.078079 | 0.48792 | 0.858824 | ENSMUSG( | Glod4 | 1403.62 | -0.09355 | 0.427504 | 0.917257 |
| ENSMUSG( | Glp1r | 43.00691 | -0.1528 | 0.671849 | 0.928781 | ENSMUSG( | Glp1r | 46.50018 | 0.318612 | 0.404823 | 0.917257 |
| ENSMUSG( | Glp2r | 82.8616 | 0.047881 | 0.849827 | 0.972638 | ENSMUSG( | Glp2r | 77.16853 | 0.125342 | 0.671372 | 0.95694 |
| ENSMUSG( | Glra2 | 320.4865 | -0.66116 | 2.35E-05 | 0.00217 | ENSMUSG( | Glra2 | 376.3867 | 0.140053 | 0.46113 | 0.923329 |
| ENSMUSG( | Glra3 | 93.90564 | -0.66902 | 0.005505 | 0.099427 | ENSMUSG( | Glra3 | 114.1071 | 0.2246 | 0.341478 | 0.908653 |
| ENSMUSG( | Glrb | 4889.488 | -0.48734 | 0.002336 | 0.056722 | ENSMUSG( | Glrb | 5582.46 | 0.19897 | 0.28453 | 0.892412 |
| ENSMUSG( | Glrx | 395.4415 | -0.06432 | 0.619456 | 0.911432 | ENSMUSG( | Glrx | 346.9505 | -0.1827 | 0.222485 | 0.866791 |
| ENSMUSG( | Glrx2 | 2202.365 | 0.046935 | 0.600939 | 0.904492 | ENSMUSG( | Glrx2 | 1974.302 | -0.00101 | 0.992753 | 0.99874 |
| ENSMUSG( | Glrx3 | 2334.84 | 0.050538 | 0.575092 | 0.894572 | ENSMUSG( | Glrx3 | 1944.105 | -0.21965 | 0.054612 | 0.723362 |
| ENSMUSG( | Glrx5 | 1521.695 | 0.013038 | 0.89065 | 0.983764 | ENSMUSG( | Glrx5 | 1284.67 | -0.21978 | 0.112682 | 0.809623 |
| ENSMUSG( | Gls | 7105.921 | 0.164274 | 0.067641 | 0.402445 | ENSMUSG( | Gls | 6215.109 | 0.04981 | 0.622114 | 0.953234 |
| ENSMUSG( | Gls2 | 750.1318 | -0.00976 | 0.928634 | 0.991668 | ENSMUSG( | Gls2 | 665.7929 | -0.08366 | 0.511989 | 0.930005 |
| ENSMUSG( | Glt1d1 | 639.5505 | 0.116865 | 0.356192 | 0.788108 | ENSMUSG( | Glt1d1 | 568.4851 | 0.045769 | 0.759324 | 0.972002 |
| ENSMUSG( | Glt28d2 | 9.972145 | -0.49448 | 0.464055 | NA | ENSMUSG( | Glt28d2 | 9.911417 | -0.24733 | 0.69394 | NA |
| ENSMUSG( | Glt8d1 | 1094.771 | 0.051824 | 0.665887 | 0.927054 | ENSMUSG( | Glt8d1 | 1045.454 | 0.184983 | 0.265927 | 0.886134 |
| ENSMUSG( | Glt8d2 | 367.8089 | -0.45984 | 0.005133 | 0.095477 | ENSMUSG( | Glt8d2 | 435.0283 | 0.317225 | 0.059433 | 0.729747 |
| ENSMUSG( | Gltp | 4154.637 | -0.00157 | 0.987188 | 0.998764 | ENSMUSG( | Gltp | 3544.897 | -0.20093 | 0.12848 | 0.81925 |
| ENSMUSG( | Glud1 | 29130.61 | 0.111585 | 0.279666 | 0.729119 | ENSMUSG( | Glud1 | 24260.4 | -0.15079 | 0.261886 | 0.884926 |
| ENSMUSG( | Glul | 204394.1 | 0.248037 | 0.023448 | 0.234187 | ENSMUSG( | Glul | 166095.6 | -0.0737 | 0.597553 | 0.946609 |
| ENSMUSG( | Glyctk | 174.7374 | -0.09244 | 0.576285 | 0.895164 | ENSMUSG( | Glyctk | 146.2927 | -0.35208 | 0.067836 | 0.75212 |
| ENSMUSG( | Glyr1 | 5216.481 | 0.158581 | 0.066826 | 0.399821 | ENSMUSG( | Glyr1 | 4505.187 | 0.008343 | 0.93683 | 0.993669 |
| ENSMUSG( | Gm10030 | 25.71027 | 0.489239 | 0.223451 | 0.669469 | ENSMUSG( | Gm10030 | 20.34734 | 0.1181 | 0.796307 | 0.97681 |
| ENSMUSG( | Gm10033 | 214.0457 | -0.01107 | 0.943791 | 0.993503 | ENSMUSG( | Gm10033 | 215.2572 | 0.265541 | 0.12656 | 0.815653 |
| ENSMUSG( | Gm10036 | 54.45915 | 0.062568 | 0.830493 | 0.96974 | ENSMUSG( | Gm10036 | 44.83096 | -0.26958 | 0.393808 | 0.917257 |
| ENSMUSG( | Gm10060 | 14.92236 | 0.427363 | 0.391985 | NA | ENSMUSG( | Gm10060 | 11.80697 | 0.021824 | 0.970414 | 0.99594 |
| ENSMUSG( | Gm10069 | 26.78863 | 0.403783 | 0.303528 | 0.751968 | ENSMUSG( | Gm10069 | 19.86872 | -0.17851 | 0.711947 | 0.964363 |
| ENSMUSG( | Gm10076 | 911.3909 | 0.158118 | 0.110737 | 0.501952 | ENSMUSG( | Gm10076 | 761.1108 | -0.09637 | 0.517334 | 0.930939 |
| ENSMUSG( | Gm10136 | 74.96418 | -0.0353 | 0.879417 | 0.981436 | ENSMUSG( | Gm10136 | 57.11834 | -0.6206 | 0.030939 | 0.629448 |
| ENSMUSG( | Gm10138 | 89.78757 | 0.180439 | 0.444555 | 0.833239 | ENSMUSG( | Gm10138 | 74.93441 | -0.0531 | 0.842513 | 0.983044 |
| ENSMUSG( | Gm10167 | 24.55834 | 0.084244 | 0.827179 | 0.968725 | ENSMUSG( | Gm10167 | 21.08613 | -0.12649 | 0.773016 | 0.974463 |
| ENSMUSG( | Gm10222 | 887.6815 | 0.047122 | 0.750166 | 0.949664 | ENSMUSG( | Gm10222 | 709.2869 | -0.36253 | 0.088107 | 0.786635 |
| ENSMUSG( | Gm10244 | 42.15211 | -0.04979 | 0.862711 | 0.975055 | ENSMUSG( | Gm10244 | 34.26583 | -0.39216 | 0.252618 | 0.88018 |
| ENSMUSG( | Gm10250 | 71.46504 | 0.008977 | 0.968861 | 0.996369 | ENSMUSG( | Gm10250 | 57.93612 | -0.37463 | 0.188023 | 0.85698 |
| ENSMUSG( | Gm10269 | 51.7335 | 0.184816 | 0.495033 | 0.860798 | ENSMUSG( | Gm10269 | 44.84087 | 0.03061 | 0.921597 | 0.992644 |
| ENSMUSG( | Gm10288 | 54.67973 | 0.030904 | 0.904432 | 0.986246 | ENSMUSG( | Gm10288 | 41.07646 | -0.56332 | 0.073414 | 0.763629 |
| ENSMUSG( | Gm10293 | 11.47291 | -0.05519 | 0.919153 | NA | ENSMUSG( | Gm10293 | 8.168416 | -0.85966 | 0.209214 | NA |
| ENSMUSG( | Gm10327 | 23.72253 | 0.13912 | 0.726879 | 0.944223 | ENSMUSG( | Gm10327 | 18.03777 | -0.40904 | 0.369242 | 0.915491 |
| ENSMUSG( | Gm10335 | 20.16479 | 0.033033 | 0.940376 | NA | ENSMUSG( | Gm10335 | 20.02991 | 0.22791 | 0.643935 | 0.953903 |
| ENSMUSG( | Gm10384 | 11.93483 | 0.851054 | 0.126776 | NA | ENSMUSG( | Gm10384 | 12.54095 | 1.188294 | 0.043587 | 0.690304 |
| ENSMUSG( | Gm10406 | 19.17315 | 0.09556 | 0.835345 | NA | ENSMUSG( | Gm10406 | 15.10595 | -0.3289 | 0.540158 | 0.937383 |
| ENSMUSG( | Gm10421 | 25.55279 | -1.20957 | 0.002916 | 0.066108 | ENSMUSG( | Gm10421 | 30.38133 | -0.19847 | 0.623271 | 0.953234 |
| ENSMUSG( | Gm1043 | 281.0018 | -0.04212 | 0.802085 | 0.962119 | ENSMUSG( | Gm1043 | 291.8454 | 0.323646 | 0.065463 | 0.744625 |
| ENSMUSG( | Gm10432 | 137.9463 | 0.036491 | 0.838314 | 0.971111 | ENSMUSG( | Gm10432 | 118.8058 | -0.13237 | 0.513502 | 0.930482 |
| ENSMUSG( | Gm10509 | 122.7498 | -0.14749 | 0.462549 | 0.842321 | ENSMUSG( | Gm10509 | 125.5951 | 0.192302 | 0.380612 | 0.917202 |
| ENSMUSG( | Gm10516 | 205.0081 | 0.069587 | 0.638102 | 0.916107 | ENSMUSG( | Gm10516 | 191.6655 | 0.126253 | 0.515281 | 0.930939 |
| ENSMUSG( | Gm10575 | 95.78769 | 0.121261 | 0.559475 | 0.888669 | ENSMUSG( | Gm10575 | 83.12989 | 0.000865 | 0.997192 | 0.999268 |
| ENSMUSG( | Gm10635 | 54.17278 | -0.0081 | 0.975804 | 0.997428 | ENSMUSG( | Gm10635 | 44.14771 | -0.35382 | 0.259885 | 0.883971 |
| ENSMUSG( | Gm10644 | 89.88138 | 0.336657 | 0.123421 | 0.530552 | ENSMUSG( | Gm10644 | 67.83185 | -0.16334 | 0.581901 | 0.944266 |
| ENSMUSG( | Gm10654 | 81.00351 | 0.065608 | 0.793992 | 0.959799 | ENSMUSG( | Gm10654 | 71.99396 | -0.02407 | 0.935731 | 0.993406 |
| ENSMUSG( | Gm10710 | 17.27407 | -0.13511 | 0.76912 | NA | ENSMUSG( | Gm10710 | 14.35343 | -0.43996 | 0.391473 | 0.917257 |
| ENSMUSG( | Gm10754 | 15.36361 | -0.67264 | 0.188465 | NA | ENSMUSG( | Gm10754 | 15.47194 | -0.37855 | 0.478103 | 0.923329 |
| ENSMUSG( | Gm10785 | 14.5173 | -0.40738 | 0.401821 | NA | ENSMUSG( | Gm10785 | 12.97789 | -0.47436 | 0.385994 | 0.917257 |
| ENSMUSG( | Gm10790 | 116.194 | -0.40383 | 0.052493 | 0.35151 | ENSMUSG( | Gm10790 | 115.0485 | -0.11263 | 0.676473 | 0.95694 |
| ENSMUSG( | Gm10814 | 11.56752 | -0.01646 | 0.975468 | NA | ENSMUSG( | Gm10814 | 10.72648 | 0.070967 | 0.906395 | NA |
| ENSMUSG( | Gm10863 | 67.87957 | -0.47167 | 0.066334 | 0.398419 | ENSMUSG( | Gm10863 | 68.04176 | -0.13161 | 0.655959 | 0.955533 |
| ENSMUSG( | Gm10865 | 14.81219 | 0.069784 | 0.892304 | NA | ENSMUSG( | Gm10865 | 10.05044 | -0.85139 | 0.180391 | NA |
| ENSMUSG( | Gm10941 | 16.2061 | 0.349713 | 0.446739 | NA | ENSMUSG( | Gm10941 | 15.36111 | 0.439091 | 0.374346 | 0.915491 |
| ENSMUSG( | Gm11110 | 21.78564 | 0.034436 | 0.939093 | 0.993415 | ENSMUSG( | Gm11110 | 18.59949 | -0.18845 | 0.694819 | 0.960867 |
| ENSMUSG( | Gm11149 | 31.95295 | -0.60988 | 0.135928 | 0.549949 | ENSMUSG( | Gm11149 | 29.36087 | -0.52933 | 0.257654 | 0.883401 |
| ENSMUSG( | Gm11185 | 11.59372 | 0.273993 | 0.644236 | NA | ENSMUSG( | Gm11185 | 9.390034 | -0.11251 | 0.865829 | NA |
| ENSMUSG( | Gm11249 | 14.98709 | 0.110159 | 0.822298 | NA | ENSMUSG( | Gm11249 | 8.609883 | -1.63873 | 0.022888 | NA |
| ENSMUSG( | Gm11263 | 19.01465 | 0.033447 | 0.941855 | NA | ENSMUSG( | Gm11263 | 13.87451 | -0.74549 | 0.196526 | 0.862161 |
| ENSMUSG( | Gm11266 | 158.6699 | 0.257915 | 0.117672 | 0.517948 | ENSMUSG( | Gm11266 | 145.5361 | 0.258946 | 0.22053 | 0.866104 |
| ENSMUSG( | Gm11353 | 21.00258 | 0.200423 | 0.654448 | NA | ENSMUSG( | Gm11353 | 16.16179 | -0.34939 | 0.500589 | 0.927825 |
| ENSMUSG( | Gm11423 | 26.43305 | -0.27909 | 0.476921 | 0.851851 | ENSMUSG( | Gm11423 | 26.81728 | 0.041968 | 0.913388 | 0.992298 |
| ENSMUSG( | Gm11451 | 32.25069 | -0.17974 | 0.584005 | 0.898666 | ENSMUSG( | Gm11451 | 27.79003 | -0.36346 | 0.338328 | 0.907972 |
| ENSMUSG( | Gm11478 | 193.634 | -0.00983 | 0.94974 | 0.993821 | ENSMUSG( | Gm11478 | 139.6111 | -0.79111 | 6.47E-05 | 0.015522 |
| ENSMUSG( | Gm11627 | 286.1481 | 0.36461 | 0.008832 | 0.132321 | ENSMUSG( | Gm11627 | 252.7954 | 0.280713 | 0.081918 | 0.77681 |
| ENSMUSG( | Gm11650 | 40.36781 | -0.45874 | 0.125341 | 0.532314 | ENSMUSG( | Gm11650 | 38.79409 | -0.31412 | 0.358837 | 0.913759 |
| ENSMUSG( | Gm11681 | 54.3566 | 0.768837 | 0.005612 | 0.100463 | ENSMUSG( | Gm11681 | 32.84198 | -0.32373 | 0.35491 | 0.912411 |
| ENSMUSG( | Gm11696 | 138.6398 | -0.06438 | 0.716961 | 0.941697 | ENSMUSG( | Gm11696 | 124.7423 | -0.09838 | 0.638517 | 0.953903 |
| ENSMUSG( | Gm11732 | 8.356743 | -0.69419 | 0.305704 | NA | ENSMUSG( | Gm11732 | 6.68442 | -1.2115 | 0.156399 | NA |
| ENSMUSG( | Gm11762 | 35.89553 | 0.347331 | 0.26647 | 0.716441 | ENSMUSG( | Gm11762 | 28.00109 | -0.04852 | 0.898626 | 0.990558 |
| ENSMUSG( | Gm11767 | 275.1477 | 0.209347 | 0.153886 | 0.580901 | ENSMUSG( | Gm11767 | 242.1321 | 0.121663 | 0.552895 | 0.939565 |
| ENSMUSG( | Gm11826 | 14.74136 | 0.106598 | 0.831779 | NA | ENSMUSG( | Gm11826 | 11.00598 | -0.56457 | 0.351439 | NA |
| ENSMUSG( | Gm11827 | 14.09424 | 0.687334 | 0.19804 | NA | ENSMUSG( | Gm11827 | 14.08483 | 0.935003 | 0.101636 | 0.801601 |
| ENSMUSG( | Gm11837 | 62.04995 | -0.06354 | 0.792375 | 0.959461 | ENSMUSG( | Gm11837 | 57.59677 | -0.02419 | 0.930558 | 0.99306 |
| ENSMUSG( | Gm11839 | 16.96891 | 0.480507 | 0.31326 | NA | ENSMUSG( | Gm11839 | 16.9118 | 0.719312 | 0.188651 | 0.85698 |
| ENSMUSG( | Gm11914 | 30.61753 | 0.0581 | 0.868311 | 0.976834 | ENSMUSG( | Gm11914 | 26.83842 | -0.05691 | 0.881389 | 0.989644 |
| ENSMUSG( | Gm11944 | 18.71053 | -0.1356 | 0.770083 | NA | ENSMUSG( | Gm11944 | 22.30054 | 0.593348 | 0.155762 | 0.837724 |
| ENSMUSG( | Gm11992 | 56.36818 | -0.05566 | 0.847301 | 0.972374 | ENSMUSG( | Gm11992 | 56.51614 | 0.222537 | 0.462687 | 0.923329 |
| ENSMUSG( | Gm12034 | 31.6577 | 0.105984 | 0.753426 | 0.950753 | ENSMUSG( | Gm12034 | 22.33388 | -0.7479 | 0.083758 | 0.77681 |
| ENSMUSG( | Gm12165 | 13.44981 | -0.11976 | 0.81131 | NA | ENSMUSG( | Gm12165 | 11.59508 | -0.31716 | 0.578056 | 0.943163 |
| ENSMUSG( | Gm12174 | 53.93137 | 0.314563 | 0.309743 | 0.757658 | ENSMUSG( | Gm12174 | 41.77474 | -0.1706 | 0.641764 | 0.953903 |
| ENSMUSG( | Gm12183 | 34.08493 | -0.44802 | 0.188335 | 0.627268 | ENSMUSG( | Gm12183 | 33.61207 | -0.17809 | 0.617544 | 0.951952 |
| ENSMUSG( | Gm12216 | 34.32395 | 0.311682 | 0.351499 | 0.783381 | ENSMUSG( | Gm12216 | 28.14148 | 0.014203 | 0.970249 | 0.99594 |
| ENSMUSG( | Gm12224 | 39.21253 | -0.07441 | 0.84494 | 0.972023 | ENSMUSG( | Gm12224 | 31.27017 | -0.47339 | 0.275135 | 0.889874 |
| ENSMUSG( | Gm12251 | 21.83449 | -0.1538 | 0.74721 | 0.949526 | ENSMUSG( | Gm12251 | 18.2312 | -0.50804 | 0.312332 | 0.900037 |
| ENSMUSG( | Gm12258 | 330.0527 | -0.21724 | 0.205625 | 0.647118 | ENSMUSG( | Gm12258 | 351.9456 | 0.246482 | 0.274662 | 0.889874 |
| ENSMUSG( | Gm12326 | 20.12598 | -0.28504 | 0.505196 | NA | ENSMUSG( | Gm12326 | 20.03415 | -0.00771 | 0.985929 | 0.997188 |
| ENSMUSG( | Gm12346 | 120.7812 | 0.112461 | 0.545881 | 0.884974 | ENSMUSG( | Gm12346 | 92.55577 | -0.44322 | 0.08019 | 0.77681 |
| ENSMUSG( | Gm12359 | 32.28319 | 0.164386 | 0.651651 | 0.921676 | ENSMUSG( | Gm12359 | 25.79371 | -0.17707 | 0.678471 | 0.957506 |
| ENSMUSG( | Gm12371 | 95.73454 | -0.20277 | 0.377964 | 0.801561 | ENSMUSG( | Gm12371 | 84.79944 | -0.30388 | 0.222833 | 0.867222 |
| ENSMUSG( | Gm12416 | 19.04324 | 0.147216 | 0.728588 | NA | ENSMUSG( | Gm12416 | 15.29515 | -0.25605 | 0.610596 | 0.950771 |
| ENSMUSG( | Gm12500 | 42.17231 | 0.023559 | 0.937868 | 0.993415 | ENSMUSG( | Gm12500 | 35.42531 | -0.21645 | 0.514772 | 0.930939 |
| ENSMUSG( | Gm12522 | 25.12464 | -0.21355 | 0.636536 | 0.914969 | ENSMUSG( | Gm12522 | 26.03185 | 0.131571 | 0.780861 | 0.976106 |
| ENSMUSG( | Gm12532 | 30.51986 | 0.246933 | 0.490174 | 0.859319 | ENSMUSG( | Gm12532 | 23.09132 | -0.29116 | 0.486245 | 0.92541 |
| ENSMUSG( | Gm12592 | 56.60389 | 0.035387 | 0.901748 | 0.985633 | ENSMUSG( | Gm12592 | 51.39261 | 0.027972 | 0.928125 | 0.99306 |
| ENSMUSG( | Gm12630 | 10.11233 | 0.056834 | 0.921538 | NA | ENSMUSG( | Gm12630 | 10.25138 | 0.359634 | 0.543932 | NA |
| ENSMUSG( | Gm12655 | 60.47539 | 0.434706 | 0.079269 | 0.435105 | ENSMUSG( | Gm12655 | 50.92814 | 0.229417 | 0.420422 | 0.917257 |
| ENSMUSG( | Gm12669 | 27.956 | 0.173838 | 0.671835 | 0.928781 | ENSMUSG( | Gm12669 | 22.50532 | -0.25055 | 0.614205 | 0.951177 |
| ENSMUSG( | Gm12758 | 38.84792 | -0.24539 | 0.41599 | 0.82084 | ENSMUSG( | Gm12758 | 34.19213 | -0.3217 | 0.365925 | 0.914623 |
| ENSMUSG( | Gm12915 | 10.47273 | -0.66291 | 0.271994 | NA | ENSMUSG( | Gm12915 | 9.647096 | -0.60838 | 0.341596 | NA |
| ENSMUSG( | Gm12918 | 42.53149 | -0.04645 | 0.878827 | 0.981436 | ENSMUSG( | Gm12918 | 32.9878 | -0.57765 | 0.094714 | 0.796133 |
| ENSMUSG( | Gm12992 | 323.823 | -0.11012 | 0.405781 | 0.81652 | ENSMUSG( | Gm12992 | 296.598 | -0.09262 | 0.545496 | 0.938063 |
| ENSMUSG( | Gm13031 | 101.7384 | 0.155812 | 0.444572 | 0.833239 | ENSMUSG( | Gm13031 | 83.19722 | -0.17302 | 0.476091 | 0.923329 |
| ENSMUSG( | Gm13033 | 17.85292 | -0.39934 | 0.386701 | NA | ENSMUSG( | Gm13033 | 17.45455 | -0.13655 | 0.788338 | 0.97637 |
| ENSMUSG( | Gm13199 | 29.10023 | 0.411434 | 0.275765 | 0.725669 | ENSMUSG( | Gm13199 | 28.57847 | 0.602759 | 0.122211 | 0.813384 |
| ENSMUSG( | Gm13270 | 16.85809 | 0.42253 | 0.353101 | NA | ENSMUSG( | Gm13270 | 14.70124 | 0.260847 | 0.628563 | 0.953251 |
| ENSMUSG( | Gm13292 | 28.87678 | -0.15018 | 0.660311 | 0.924627 | ENSMUSG( | Gm13292 | 22.79576 | -0.61798 | 0.133792 | 0.822912 |
| ENSMUSG( | Gm13293 | 46.4604 | -0.58857 | 0.035915 | 0.291124 | ENSMUSG( | Gm13293 | 46.51312 | -0.22691 | 0.510277 | 0.929089 |
| ENSMUSG( | Gm13375 | 537.8721 | -0.02618 | 0.817609 | 0.965785 | ENSMUSG( | Gm13375 | 508.2003 | 0.072937 | 0.632178 | 0.953452 |
| ENSMUSG( | Gm13387 | 29.65506 | 0.040784 | 0.910251 | 0.9881 | ENSMUSG( | Gm13387 | 28.5323 | 0.17554 | 0.65456 | 0.955015 |
| ENSMUSG( | Gm13408 | 34.21079 | -0.06806 | 0.840337 | 0.971179 | ENSMUSG( | Gm13408 | 26.83276 | -0.61494 | 0.168985 | 0.840222 |
| ENSMUSG( | Gm13416 | 30.1386 | 0.15456 | 0.656963 | 0.924069 | ENSMUSG( | Gm13416 | 30.93674 | 0.473082 | 0.192248 | 0.859726 |
| ENSMUSG( | Gm13421 | 57.82513 | 0.297605 | 0.234998 | 0.682364 | ENSMUSG( | Gm13421 | 47.87901 | 0.045723 | 0.874557 | 0.989644 |
| ENSMUSG( | Gm13436 | 97.52382 | 0.043736 | 0.850848 | 0.972638 | ENSMUSG( | Gm13436 | 73.50732 | -0.57436 | 0.020482 | 0.565891 |
| ENSMUSG( | Gm13446 | 125.0648 | -0.13329 | 0.554102 | 0.886397 | ENSMUSG( | Gm13446 | 112.4949 | -0.17368 | 0.435978 | 0.919569 |
| ENSMUSG( | Gm13464 | 28.37399 | 0.115101 | 0.763918 | 0.954015 | ENSMUSG( | Gm13464 | 20.71776 | -0.6558 | 0.181018 | 0.850372 |
| ENSMUSG( | Gm13498 | 34.3813 | 0.48283 | 0.151051 | 0.576088 | ENSMUSG( | Gm13498 | 27.36085 | 0.13009 | 0.736769 | 0.967344 |
| ENSMUSG( | Gm13680 | 26.27344 | 0.164744 | 0.683426 | 0.932153 | ENSMUSG( | Gm13680 | 21.89312 | -0.12198 | 0.783201 | 0.976106 |
| ENSMUSG( | Gm13754 | 12.82499 | 0.053708 | 0.920347 | NA | ENSMUSG( | Gm13754 | 10.33513 | -0.28859 | 0.637708 | NA |
| ENSMUSG( | Gm13872 | 117.2117 | -0.03289 | 0.881423 | 0.981798 | ENSMUSG( | Gm13872 | 87.55713 | -0.67401 | 0.007952 | 0.382597 |
| ENSMUSG( | Gm13889 | 621.5381 | -0.3443 | 0.020004 | 0.213886 | ENSMUSG( | Gm13889 | 569.312 | -0.33446 | 0.072529 | 0.763629 |
| ENSMUSG( | Gm14023 | 25.32955 | 0.161251 | 0.676505 | 0.930542 | ENSMUSG( | Gm14023 | 18.86196 | -0.47292 | 0.292482 | 0.897982 |
| ENSMUSG( | Gm14113 | 39.29963 | 0.239722 | 0.436302 | 0.829093 | ENSMUSG( | Gm14113 | 37.33845 | 0.323163 | 0.357782 | 0.913759 |
| ENSMUSG( | Gm14121 | 22.47894 | -0.16132 | 0.719855 | 0.941986 | ENSMUSG( | Gm14121 | 16.43801 | -0.99765 | 0.063487 | 0.736482 |
| ENSMUSG( | Gm14150 | 18.03498 | 0.119893 | 0.789865 | NA | ENSMUSG( | Gm14150 | 14.6847 | -0.28112 | 0.659305 | 0.956205 |
| ENSMUSG( | Gm14165 | 20.67616 | -0.23049 | 0.602102 | NA | ENSMUSG( | Gm14165 | 12.98543 | -1.95615 | 0.001308 | 0.130907 |
| ENSMUSG( | Gm14169 | 229.3515 | 0.102529 | 0.540198 | 0.88315 | ENSMUSG( | Gm14169 | 189.3169 | -0.19334 | 0.313473 | 0.900883 |
| ENSMUSG( | Gm14204 | 202.6754 | -0.10876 | 0.493817 | 0.860611 | ENSMUSG( | Gm14204 | 192 | -0.00354 | 0.984629 | 0.996963 |
| ENSMUSG( | Gm14230 | 34.27083 | -0.09737 | 0.759244 | 0.952284 | ENSMUSG( | Gm14230 | 32.97108 | 0.087711 | 0.802328 | 0.97681 |
| ENSMUSG( | Gm14276 | 85.11145 | -0.11563 | 0.59525 | 0.903007 | ENSMUSG( | Gm14276 | 73.4417 | -0.28757 | 0.239695 | 0.874882 |
| ENSMUSG( | Gm14295 | 245.0153 | 0.140232 | 0.395798 | 0.811098 | ENSMUSG( | Gm14295 | 224.5428 | 0.146731 | 0.430139 | 0.917257 |
| ENSMUSG( | Gm14296 | 28.47573 | 0.010139 | 0.979574 | 0.997428 | ENSMUSG( | Gm14296 | 26.11648 | 0.041378 | 0.918955 | 0.992529 |

| | | | | | |
|---|---|---|---|---|---|
| ENSMUSG( | 56.36818 | -0.05566 | 0.847301 | 0.972374 | Gm11992 |
| ENSMUSG( | 31.6577 | 0.105984 | 0.753426 | 0.950753 | Gm12034 |
| ENSMUSG( | 13.44981 | -0.11976 | 0.81131 | NA | Gm12165 |
| ENSMUSG( | 53.93137 | 0.314563 | 0.309743 | 0.757658 | Gm12174 |
| ENSMUSG( | 34.08493 | -0.44802 | 0.188335 | 0.627268 | Gm12183 |
| ENSMUSG( | 34.32395 | 0.311682 | 0.351499 | 0.783381 | Gm12216 |
| ENSMUSG( | 39.21253 | -0.07441 | 0.84494 | 0.972023 | Gm12224 |
| ENSMUSG( | 21.83449 | -0.1538 | 0.74721 | 0.949526 | Gm12251 |
| ENSMUSG( | 330.0527 | -0.21724 | 0.205625 | 0.647118 | Gm12258 |
| ENSMUSG( | 15.64871 | 0.254249 | 0.598162 | NA | Gm12286 |
| ENSMUSG( | 20.12598 | -0.28504 | 0.505196 | NA | Gm12326 |
| ENSMUSG( | 120.7812 | 0.112461 | 0.545881 | 0.884974 | Gm12346 |
| ENSMUSG( | 32.28319 | 0.164386 | 0.651651 | 0.921676 | Gm12359 |
| ENSMUSG( | 95.73454 | -0.20277 | 0.377964 | 0.801561 | Gm12371 |
| ENSMUSG( | 8.871626 | -0.16604 | 0.790247 | NA | Gm12411 |
| ENSMUSG( | 19.04324 | 0.147216 | 0.728588 | NA | Gm12416 |
| ENSMUSG( | 42.17231 | 0.023559 | 0.937868 | 0.993415 | Gm12500 |
| ENSMUSG( | 25.12464 | -0.21355 | 0.636536 | 0.914969 | Gm12522 |
| ENSMUSG( | 30.51986 | 0.246933 | 0.490174 | 0.859319 | Gm12532 |
| ENSMUSG( | 56.60389 | 0.035387 | 0.901748 | 0.985633 | Gm12592 |
| ENSMUSG( | 10.11233 | 0.056834 | 0.921538 | NA | Gm12630 |
| ENSMUSG( | 60.47539 | 0.434706 | 0.079269 | 0.435105 | Gm12655 |
| ENSMUSG( | 27.956 | 0.173838 | 0.671835 | 0.928781 | Gm12669 |
| ENSMUSG( | 38.84792 | -0.24539 | 0.41599 | 0.82084 | Gm12758 |
| ENSMUSG( | 8.465314 | 0.614436 | 0.352486 | NA | Gm12791 |
| ENSMUSG( | 10.47273 | -0.66291 | 0.271994 | NA | Gm12915 |
| ENSMUSG( | 42.53149 | -0.04645 | 0.878827 | 0.981436 | Gm12918 |
| ENSMUSG( | 323.823 | -0.11012 | 0.405781 | 0.81652 | Gm12992 |
| ENSMUSG( | 101.7384 | 0.155812 | 0.444572 | 0.833239 | Gm13031 |
| ENSMUSG( | 17.85292 | -0.39934 | 0.386701 | NA | Gm13033 |
| ENSMUSG( | 29.10023 | 0.411434 | 0.275765 | 0.725669 | Gm13199 |
| ENSMUSG( | 10.5681 | -0.09871 | 0.86225 | NA | Gm13226 |
| ENSMUSG( | 16.85809 | 0.42253 | 0.353101 | NA | Gm13270 |
| ENSMUSG( | 28.87678 | -0.15018 | 0.660311 | 0.924627 | Gm13292 |
| ENSMUSG( | 46.4604 | -0.58857 | 0.035915 | 0.291124 | Gm13293 |
| ENSMUSG( | 537.8721 | -0.02618 | 0.817609 | 0.965785 | Gm13375 |
| ENSMUSG( | 29.65506 | 0.040784 | 0.910251 | 0.9881 | Gm13387 |
| ENSMUSG( | 34.21079 | -0.06806 | 0.840337 | 0.971179 | Gm13408 |
| ENSMUSG( | 30.1386 | 0.15456 | 0.656963 | 0.924069 | Gm13416 |
| ENSMUSG( | 57.82513 | 0.297605 | 0.234998 | 0.682364 | Gm13421 |
| ENSMUSG( | 97.52382 | 0.043736 | 0.850848 | 0.972638 | Gm13436 |
| ENSMUSG( | 125.0648 | -0.13329 | 0.554102 | 0.886397 | Gm13446 |
| ENSMUSG( | 28.37399 | 0.115101 | 0.763918 | 0.954015 | Gm13464 |
| ENSMUSG( | 11.02172 | 0.115741 | 0.837103 | NA | Gm13483 |
| ENSMUSG( | 34.3813 | 0.48283 | 0.151051 | 0.576088 | Gm13498 |
| ENSMUSG( | 26.27344 | 0.164744 | 0.683426 | 0.932153 | Gm13680 |
| ENSMUSG( | 12.82499 | 0.053708 | 0.920347 | NA | Gm13754 |
| ENSMUSG( | 117.2117 | -0.03289 | 0.881423 | 0.981798 | Gm13872 |
| ENSMUSG( | 621.5381 | -0.3443 | 0.020004 | 0.213886 | Gm13889 |
| ENSMUSG( | 25.32955 | 0.161251 | 0.676505 | 0.930542 | Gm14023 |
| ENSMUSG( | 39.29963 | 0.239722 | 0.436302 | 0.829093 | Gm14113 |
| ENSMUSG( | 22.47894 | -0.16132 | 0.719855 | 0.941986 | Gm14121 |
| ENSMUSG( | 18.03498 | 0.119893 | 0.789865 | NA | Gm14150 |
| ENSMUSG( | 20.67616 | -0.23049 | 0.602102 | NA | Gm14165 |
| ENSMUSG( | 229.3515 | 0.102529 | 0.540198 | 0.88315 | Gm14169 |
| ENSMUSG( | 202.6754 | -0.10876 | 0.493817 | 0.860611 | Gm14204 |
| ENSMUSG( | 34.27083 | -0.09737 | 0.759244 | 0.952284 | Gm14230 |
| ENSMUSG( | 85.11145 | -0.11563 | 0.59525 | 0.903007 | Gm14276 |
| ENSMUSG( | 245.0153 | 0.140232 | 0.395798 | 0.811098 | Gm14295 |
| ENSMUSG( | 28.47573 | 0.010139 | 0.979574 | 0.997428 | Gm14296 |
| ENSMUSG( | 318.1563 | -0.01246 | 0.935116 | 0.993293 | Gm14305 |
| ENSMUSG( | 27.63819 | 0.31505 | 0.392209 | 0.809274 | Gm14317 |
| ENSMUSG( | 65.18139 | 0.12445 | 0.610509 | 0.908189 | Gm14322 |
| ENSMUSG( | 514.2771 | 0.184048 | 0.163871 | 0.594869 | Gm14325 |
| ENSMUSG( | 300.7356 | 0.132367 | 0.371953 | 0.798768 | Gm14326 |
| ENSMUSG( | 132.9003 | 0.293663 | 0.109051 | 0.498536 | Gm14399 |
| ENSMUSG( | 170.4929 | 0.320771 | 0.077761 | 0.431059 | Gm14410 |
| ENSMUSG( | 42.23512 | 0.051997 | 0.862807 | 0.975055 | Gm14418 |
| ENSMUSG( | 14.39096 | -0.16969 | 0.724812 | NA | Gm14419 |
| ENSMUSG( | 113.9549 | 0.554977 | 0.005167 | 0.096 | Gm14569 |
| ENSMUSG( | 12.42658 | -0.08475 | 0.885685 | NA | Gm14584 |
| ENSMUSG( | 67.06629 | -0.12964 | 0.603513 | 0.904887 | Gm14586 |
| ENSMUSG( | 9.68388 | 0.343549 | 0.564649 | NA | Gm14633 |
| ENSMUSG( | 36.75289 | 0.205616 | 0.53293 | 0.879905 | Gm14634 |
| ENSMUSG( | 12.96806 | 0.289666 | 0.597702 | NA | Gm14662 |
| ENSMUSG( | 12.74699 | -0.42241 | 0.409679 | NA | Gm14824 |
| ENSMUSG( | 248.1151 | 0.418747 | 0.05697 | 0.368322 | Gm14827 |
| ENSMUSG( | 8.643473 | 0.205595 | 0.74366 | NA | Gm14963 |
| ENSMUSG( | 21.11345 | 0.293274 | 0.47706 | NA | Gm14966 |
| ENSMUSG( | 23.16579 | 0.028429 | 0.942868 | 0.993415 | Gm15201 |
| ENSMUSG( | 40.24762 | -0.19066 | 0.557823 | 0.888427 | Gm15246 |
| ENSMUSG( | 79.32978 | 0.276955 | 0.208444 | 0.650319 | Gm15328 |
| ENSMUSG( | 9.134635 | -0.24308 | 0.698329 | NA | Gm15382 |
| ENSMUSG( | 19.90067 | -0.24734 | 0.553273 | NA | Gm15408 |
| ENSMUSG( | 219.1334 | -0.12339 | 0.40189 | 0.814965 | Gm15417 |
| ENSMUSG( | 176.5185 | 0.113942 | 0.500188 | 0.863455 | Gm15427 |
| ENSMUSG( | 547.3198 | -0.06739 | 0.627619 | 0.912961 | Gm15446 |
| ENSMUSG( | 162.6502 | 0.139265 | 0.427219 | 0.825333 | Gm15459 |
| ENSMUSG( | 9.601903 | 1.450237 | 0.019363 | NA | Gm15471 |
| ENSMUSG( | 134.2175 | -0.30488 | 0.101383 | 0.484996 | Gm15500 |
| ENSMUSG( | 59.51812 | 0.181882 | 0.479402 | 0.854046 | Gm15501 |
| ENSMUSG( | 47.32327 | 0.071626 | 0.792306 | 0.959461 | Gm15510 |
| ENSMUSG( | 17.96705 | 0.596737 | 0.185525 | NA | Gm15542 |
| ENSMUSG( | 36.43105 | -0.27191 | 0.408472 | 0.818167 | Gm15545 |
| ENSMUSG( | 51.19858 | -0.16346 | 0.550839 | 0.886038 | Gm15559 |
| ENSMUSG( | 81.81602 | -0.19651 | 0.39385 | 0.809643 | Gm15612 |
| ENSMUSG( | 234.5 | -0.05691 | 0.707128 | 0.93949 | Gm15663 |
| ENSMUSG( | 15.96989 | 0.135864 | 0.830385 | NA | Gm15697 |
| ENSMUSG( | 216.6287 | 0.101011 | 0.567269 | 0.891666 | Gm15706 |
| ENSMUSG( | 18.74753 | -0.37425 | 0.395127 | NA | Gm15708 |
| ENSMUSG( | 107.5539 | 0.049973 | 0.805726 | 0.962616 | Gm15710 |
| ENSMUSG( | 36.60117 | -0.18365 | 0.547954 | 0.885153 | Gm15743 |
| ENSMUSG( | 68.48981 | -0.06866 | 0.774339 | 0.955606 | Gm15772 |
| ENSMUSG( | 69.49204 | 0.291124 | 0.23023 | 0.678139 | Gm15787 |
| ENSMUSG( | 46.50439 | 0.520647 | 0.076203 | 0.427093 | Gm15816 |
| ENSMUSG( | 9.526304 | -0.46001 | 0.434311 | NA | Gm15850 |
| ENSMUSG( | 27.32345 | -0.00308 | 0.993394 | 0.999369 | Gm15867 |
| ENSMUSG( | 10.93752 | 0.653832 | 0.249294 | NA | Gm15939 |
| ENSMUSG( | 9.88357 | -0.52624 | 0.404088 | NA | Gm15972 |
| ENSMUSG( | 20.35494 | -0.23981 | 0.599669 | NA | Gm15997 |
| ENSMUSG( | 13.78275 | 0.356087 | 0.484184 | NA | Gm16023 |
| ENSMUSG( | 75.10199 | -0.07595 | 0.762552 | 0.953519 | Gm1604a |
| ENSMUSG( | 16.27413 | 0.193771 | 0.675392 | NA | Gm16062 |
| ENSMUSG( | 47.63386 | 0.011679 | 0.968562 | 0.996369 | Gm16124 |
| ENSMUSG( | 29.11545 | 0.359412 | 0.312674 | 0.760241 | Gm16201 |
| ENSMUSG( | 51.99795 | -0.08918 | 0.738899 | 0.947768 | Gm16238 |
| ENSMUSG( | 18.32551 | -0.0803 | 0.875369 | NA | Gm16244 |
| ENSMUSG( | 14.2431 | -0.13274 | 0.79004 | NA | Gm16364 |
| ENSMUSG( | 13.14516 | 0.316411 | 0.564599 | NA | Gm16537 |
| ENSMUSG( | 72.96137 | -0.33317 | 0.221725 | 0.667354 | Gm16576 |
| ENSMUSG( | 37.76422 | 0.05455 | 0.873086 | 0.978753 | Gm16580 |
| ENSMUSG( | 9.90638 | -0.3571 | 0.536336 | NA | Gm16630 |
| ENSMUSG( | 228.8099 | -0.03109 | 0.849906 | 0.972638 | Gm16638 |
| ENSMUSG( | 29.98106 | -0.31363 | 0.376157 | 0.801127 | Gm16675 |
| ENSMUSG( | 24.55739 | 0.150973 | 0.692642 | 0.934101 | Gm16701 |
| ENSMUSG( | 72.24215 | -0.38279 | 0.088891 | 0.457425 | Gm16740 |
| ENSMUSG( | 18.5982 | 0.52814 | 0.282778 | NA | Gm16793 |
| ENSMUSG( | 159.6806 | 0.007004 | 0.970501 | 0.996369 | Gm16835 |
| ENSMUSG( | 25.2684 | 0.568031 | 0.19917 | 0.639621 | Gm16845 |
| ENSMUSG( | 37.36666 | 0.025952 | 0.935542 | 0.993363 | Gm16861 |
| ENSMUSG( | 59.81021 | -1.58049 | 2.32E-08 | 8.35E-06 | Gm16867 |
| ENSMUSG( | 15.64719 | -0.11692 | 0.816834 | NA | Gm16894 |
| ENSMUSG( | 19.70336 | -0.22287 | 0.610384 | NA | Gm16897 |
| ENSMUSG( | 18.14021 | -0.01698 | 0.970929 | NA | Gm16958 |
| ENSMUSG( | 157.8643 | -0.11162 | 0.495802 | 0.861294 | Gm16973 |
| ENSMUSG( | 18.47162 | -0.21133 | 0.643415 | NA | Gm17102 |
| ENSMUSG( | 52.37146 | 0.131133 | 0.621818 | 0.911432 | Gm17244 |
| ENSMUSG( | 17.28107 | -0.2609 | 0.579305 | NA | Gm17315 |
| ENSMUSG( | 13.8875 | 0.417835 | 0.444371 | NA | Gm17344 |
| ENSMUSG( | 12.71965 | 0.070227 | 0.893104 | NA | Gm17382 |
| ENSMUSG( | 20.31602 | -0.14933 | 0.731161 | NA | Gm17383 |
| ENSMUSG( | 52.60658 | -0.06389 | 0.819021 | 0.965888 | Gm17396 |
| ENSMUSG( | 34.15422 | -0.75143 | 0.023371 | 0.233882 | Gm17455 |
| ENSMUSG( | 49.0606 | -0.28876 | 0.367038 | 0.795164 | Gm17484 |
| ENSMUSG( | 22.70918 | 0.190182 | 0.667306 | 0.92729 | Gm17597 |
| ENSMUSG( | 16.53489 | -0.19686 | 0.659739 | NA | Gm17634 |
| ENSMUSG( | 20.04757 | -0.26312 | 0.57596 | NA | Gm17733 |
| ENSMUSG( | 209.6217 | 0.206561 | 0.206454 | 0.648468 | Gm17750 |
| ENSMUSG( | 8.650192 | -0.62603 | 0.327258 | NA | Gm17767 |
| ENSMUSG( | 11.64056 | 0.427928 | 0.464175 | NA | Gm17823 |
| ENSMUSG( | 40.2687 | -0.09397 | 0.773547 | 0.955447 | Gm17891 |
| ENSMUSG( | 151.0968 | 0.630424 | 0.02628 | 0.247758 | Gm17922 |
| ENSMUSG( | 14.37381 | -1.37294 | 0.020074 | NA | Gm17971 |
| ENSMUSG( | 10.59131 | -0.18485 | 0.765762 | NA | Gm18101 |
| ENSMUSG( | 17.33329 | 0.273632 | 0.548821 | NA | Gm18180 |
| ENSMUSG( | 13.31175 | 0.362302 | 0.488294 | NA | Gm1848 |
| ENSMUSG( | 26.47045 | -0.53135 | 0.236143 | 0.683188 | Gm18755 |
| ENSMUSG( | 50.1294 | 0.260248 | 0.327612 | 0.772692 | Gm19026 |

| | | | | | |
|---|---|---|---|---|---|
| ENSMUSG( | 23.09132 | -0.29116 | 0.486245 | 0.92541 | Gm12532 |
| ENSMUSG( | 51.39261 | 0.027972 | 0.928125 | 0.99306 | Gm12592 |
| ENSMUSG( | 10.25138 | 0.359634 | 0.543932 | NA | Gm12630 |
| ENSMUSG( | 50.92814 | 0.229417 | 0.420422 | 0.917257 | Gm12655 |
| ENSMUSG( | 22.50532 | -0.25055 | 0.614205 | 0.951177 | Gm12669 |
| ENSMUSG( | 34.19213 | -0.3217 | 0.365925 | 0.914623 | Gm12758 |
| ENSMUSG( | 9.647096 | -0.60838 | 0.341596 | NA | Gm12915 |
| ENSMUSG( | 32.9878 | -0.57765 | 0.094714 | 0.796133 | Gm12918 |
| ENSMUSG( | 296.598 | -0.09262 | 0.545496 | 0.938063 | Gm12992 |
| ENSMUSG( | 83.19722 | -0.17302 | 0.476091 | 0.923329 | Gm13031 |
| ENSMUSG( | 17.45455 | -0.13655 | 0.788338 | 0.97637 | Gm13033 |
| ENSMUSG( | 28.57847 | 0.602759 | 0.122211 | 0.813384 | Gm13199 |
| ENSMUSG( | 14.70124 | 0.260847 | 0.628563 | 0.953251 | Gm13270 |
| ENSMUSG( | 22.79576 | -0.61798 | 0.133792 | 0.822912 | Gm13292 |
| ENSMUSG( | 46.51312 | -0.22691 | 0.510277 | 0.929089 | Gm13293 |
| ENSMUSG( | 508.2003 | 0.072937 | 0.632178 | 0.953452 | Gm13375 |
| ENSMUSG( | 28.5323 | 0.17554 | 0.65456 | 0.955015 | Gm13387 |
| ENSMUSG( | 26.83276 | -0.61494 | 0.168985 | 0.840222 | Gm13408 |
| ENSMUSG( | 30.93674 | 0.473082 | 0.192248 | 0.859726 | Gm13416 |
| ENSMUSG( | 47.87901 | 0.045723 | 0.874557 | 0.989644 | Gm13421 |
| ENSMUSG( | 73.50732 | -0.57436 | 0.020482 | 0.565891 | Gm13436 |
| ENSMUSG( | 112.4949 | -0.17368 | 0.435978 | 0.919569 | Gm13446 |
| ENSMUSG( | 20.71776 | -0.6558 | 0.181018 | 0.850372 | Gm13464 |
| ENSMUSG( | 27.36085 | 0.13009 | 0.736769 | 0.967344 | Gm13498 |
| ENSMUSG( | 21.89312 | -0.12198 | 0.783201 | 0.976106 | Gm13680 |
| ENSMUSG( | 10.33513 | -0.28859 | 0.637708 | NA | Gm13754 |
| ENSMUSG( | 87.55713 | -0.67401 | 0.007952 | 0.382597 | Gm13872 |
| ENSMUSG( | 569.312 | -0.33446 | 0.072529 | 0.763629 | Gm13889 |
| ENSMUSG( | 18.86196 | -0.47292 | 0.292482 | 0.897982 | Gm14023 |
| ENSMUSG( | 37.33845 | 0.323163 | 0.357782 | 0.913759 | Gm14113 |
| ENSMUSG( | 16.43801 | -0.99765 | 0.063487 | 0.736482 | Gm14121 |
| ENSMUSG( | 14.6847 | -0.28112 | 0.659305 | 0.956205 | Gm14150 |
| ENSMUSG( | 12.98543 | -1.95615 | 0.001308 | 0.130907 | Gm14165 |
| ENSMUSG( | 189.3169 | -0.19334 | 0.313473 | 0.900883 | Gm14169 |
| ENSMUSG( | 192 | -0.00354 | 0.984629 | 0.996963 | Gm14204 |
| ENSMUSG( | 32.97108 | 0.087711 | 0.802328 | 0.97681 | Gm14230 |
| ENSMUSG( | 73.4417 | -0.28757 | 0.239695 | 0.874882 | Gm14276 |
| ENSMUSG( | 224.5428 | 0.146731 | 0.430139 | 0.917257 | Gm14295 |
| ENSMUSG( | 26.11648 | 0.041378 | 0.918955 | 0.992529 | Gm14296 |
| ENSMUSG( | 275.9918 | -0.16022 | 0.327125 | 0.905768 | Gm14305 |
| ENSMUSG( | 21.50918 | -0.14895 | 0.746887 | 0.969763 | Gm14317 |
| ENSMUSG( | 56.94466 | 0.028764 | 0.91868 | 0.992529 | Gm14322 |
| ENSMUSG( | 424.955 | -0.10296 | 0.509782 | 0.929089 | Gm14325 |
| ENSMUSG( | 270.7287 | 0.102148 | 0.500761 | 0.927825 | Gm14326 |
| ENSMUSG( | 111.1522 | 0.041247 | 0.853891 | 0.98563 | Gm14399 |
| ENSMUSG( | 133.6886 | -0.08524 | 0.664744 | 0.956766 | Gm14410 |
| ENSMUSG( | 40.75686 | 0.181068 | 0.595635 | 0.945941 | Gm14418 |
| ENSMUSG( | 11.25029 | -0.70797 | 0.227195 | NA | Gm14419 |
| ENSMUSG( | 83.93554 | -0.01516 | 0.949553 | 0.994798 | Gm14569 |
| ENSMUSG( | 9.323102 | -0.65724 | 0.374633 | NA | Gm14584 |
| ENSMUSG( | 51.09941 | -0.74766 | 0.014773 | 0.487333 | Gm14586 |
| ENSMUSG( | 33.09849 | 0.162798 | 0.644812 | 0.953903 | Gm14634 |
| ENSMUSG( | 10.20396 | -0.08798 | 0.889604 | NA | Gm14662 |
| ENSMUSG( | 11.68521 | -0.34095 | 0.557462 | 0.939565 | Gm14824 |
| ENSMUSG( | 195.7765 | 0.015347 | 0.955479 | 0.994798 | Gm14827 |
| ENSMUSG( | 21.13654 | 0.541446 | 0.217769 | 0.865589 | Gm14966 |
| ENSMUSG( | 21.86141 | 0.11838 | 0.782142 | 0.976106 | Gm15201 |
| ENSMUSG( | 11.40899 | 0.385601 | 0.498681 | NA | Gm15222 |
| ENSMUSG( | 44.3464 | 0.347898 | 0.333918 | 0.907907 | Gm15246 |
| ENSMUSG( | 72.12167 | 0.25767 | 0.351985 | 0.912093 | Gm15328 |
| ENSMUSG( | 10.63551 | 0.434032 | 0.462591 | NA | Gm15382 |
| ENSMUSG( | 17.7481 | -0.30764 | 0.509954 | 0.929089 | Gm15408 |
| ENSMUSG( | 203.4712 | -0.06656 | 0.68381 | 0.958678 | Gm15417 |
| ENSMUSG( | 130.7566 | -0.55595 | 0.012037 | 0.445877 | Gm15427 |
| ENSMUSG( | 530.9942 | 0.111046 | 0.423103 | 0.917257 | Gm15446 |
| ENSMUSG( | 140.2994 | -0.02129 | 0.91683 | 0.992529 | Gm15459 |
| ENSMUSG( | 120.7051 | -0.36782 | 0.148542 | 0.836241 | Gm15500 |
| ENSMUSG( | 49.25288 | -0.09532 | 0.744102 | 0.96951 | Gm15501 |
| ENSMUSG( | 41.26683 | -0.05899 | 0.849155 | 0.985321 | Gm15510 |
| ENSMUSG( | 14.25526 | 0.250965 | 0.633901 | 0.953891 | Gm15542 |
| ENSMUSG( | 33.78606 | -0.20881 | 0.565964 | 0.941499 | Gm15545 |
| ENSMUSG( | 46.43067 | -0.16026 | 0.601408 | 0.947749 | Gm15559 |
| ENSMUSG( | 72.6144 | -0.24759 | 0.3953 | 0.917257 | Gm15612 |
| ENSMUSG( | 212.1334 | -0.09129 | 0.627838 | 0.953234 | Gm15663 |
| ENSMUSG( | 13.66475 | -0.04754 | 0.945735 | 0.994798 | Gm15697 |
| ENSMUSG( | 194.0798 | 0.039932 | 0.843014 | 0.983211 | Gm15706 |
| ENSMUSG( | 18.08719 | -0.16872 | 0.74194 | 0.969314 | Gm15708 |
| ENSMUSG( | 84.46916 | -0.43288 | 0.101392 | 0.801456 | Gm15710 |
| ENSMUSG( | 33.01057 | -0.20307 | 0.553237 | 0.939565 | Gm15743 |
| ENSMUSG( | 52.34151 | -0.66554 | 0.021077 | 0.573311 | Gm15772 |
| ENSMUSG( | 60.50082 | 0.16205 | 0.5486 | 0.938063 | Gm15787 |
| ENSMUSG( | 30.72081 | -0.38858 | 0.310886 | 0.899422 | Gm15816 |
| ENSMUSG( | 9.307133 | -0.22004 | 0.721912 | NA | Gm15850 |
| ENSMUSG( | 21.88107 | -0.41595 | 0.346868 | 0.911434 | Gm15867 |
| ENSMUSG( | 8.244405 | 0.194888 | 0.772149 | NA | Gm15939 |
| ENSMUSG( | 10.74165 | 0.054393 | 0.928451 | NA | Gm15972 |
| ENSMUSG( | 21.50977 | 0.169539 | 0.703319 | 0.962481 | Gm15997 |
| ENSMUSG( | 16.46329 | 1.01465 | 0.057191 | 0.723362 | Gm16023 |
| ENSMUSG( | 64.58901 | -0.26855 | 0.337532 | 0.907972 | Gm1604a |
| ENSMUSG( | 10.73482 | -0.89613 | 0.179782 | NA | Gm16062 |
| ENSMUSG( | 49.05705 | 0.333146 | 0.282158 | 0.892412 | Gm16124 |
| ENSMUSG( | 25.0175 | 0.188439 | 0.66939 | 0.956766 | Gm16201 |
| ENSMUSG( | 41.84028 | -0.52347 | 0.112876 | 0.809623 | Gm16238 |
| ENSMUSG( | 16.53179 | -0.11255 | 0.823675 | 0.979198 | Gm16244 |
| ENSMUSG( | 13.86835 | 0.048093 | 0.929281 | 0.99306 | Gm16364 |
| ENSMUSG( | 12.10119 | 0.41223 | 0.52238 | 0.932681 | Gm16537 |
| ENSMUSG( | 65.10267 | -0.3941 | 0.173431 | 0.844317 | Gm16576 |
| ENSMUSG( | 26.7661 | -0.78379 | 0.04641 | 0.697814 | Gm16580 |
| ENSMUSG( | 9.295474 | -0.29374 | 0.642967 | NA | Gm16630 |
| ENSMUSG( | 214.92 | 0.06564 | 0.72659 | 0.966887 | Gm16638 |
| ENSMUSG( | 25.04182 | -0.60272 | 0.1677 | 0.840222 | Gm16675 |
| ENSMUSG( | 22.32829 | 0.117351 | 0.791371 | 0.97681 | Gm16701 |
| ENSMUSG( | 71.69373 | -0.10715 | 0.660775 | 0.956205 | Gm16740 |
| ENSMUSG( | 15.1447 | 0.245176 | 0.636275 | 0.953903 | Gm16793 |
| ENSMUSG( | 140.7525 | -0.08556 | 0.694629 | 0.960867 | Gm16835 |
| ENSMUSG( | 21.90509 | 0.461247 | 0.28165 | 0.892412 | Gm16845 |
| ENSMUSG( | 34.07014 | 0.028031 | 0.938308 | 0.993793 | Gm16861 |
| ENSMUSG( | 59.76073 | -1.10292 | 0.00013 | 0.02529 | Gm16867 |
| ENSMUSG( | 16.66714 | 0.333886 | 0.521652 | 0.932571 | Gm16894 |
| ENSMUSG( | 17.33012 | -0.34139 | 0.485123 | 0.92541 | Gm16897 |
| ENSMUSG( | 14.81521 | -0.37269 | 0.48827 | 0.925453 | Gm16958 |
| ENSMUSG( | 154.5006 | 0.101581 | 0.575595 | 0.942912 | Gm16973 |
| ENSMUSG( | 16.22578 | -0.32006 | 0.517099 | 0.930939 | Gm17102 |
| ENSMUSG( | 43.25034 | -0.14506 | 0.639521 | 0.953903 | Gm17244 |
| ENSMUSG( | 16.10537 | -0.23399 | 0.641231 | 0.953903 | Gm17315 |
| ENSMUSG( | 11.92764 | 0.257533 | 0.646832 | 0.953988 | Gm17344 |
| ENSMUSG( | 12.02537 | 0.150595 | 0.798878 | 0.97681 | Gm17382 |
| ENSMUSG( | 17.59629 | -0.35552 | 0.486062 | 0.92541 | Gm17383 |
| ENSMUSG( | 49.67436 | 0.018512 | 0.951799 | 0.994798 | Gm17396 |
| ENSMUSG( | 36.10803 | -0.20548 | 0.568713 | 0.941777 | Gm17455 |
| ENSMUSG( | 51.55313 | 0.143995 | 0.636613 | 0.953903 | Gm17484 |
| ENSMUSG( | 21.13554 | 0.271465 | 0.568892 | 0.941777 | Gm17597 |
| ENSMUSG( | 12.20537 | -1.04094 | 0.076962 | 0.770558 | Gm17634 |
| ENSMUSG( | 18.24376 | -0.26495 | 0.563854 | 0.941206 | Gm17733 |
| ENSMUSG( | 171.6287 | -0.11162 | 0.598076 | 0.946676 | Gm17750 |
| ENSMUSG( | 9.863172 | -0.40172 | 0.50977 | NA | Gm17751 |
| ENSMUSG( | 9.177814 | -0.09836 | 0.87794 | NA | Gm17767 |
| ENSMUSG( | 13.97881 | 1.008043 | 0.095945 | 0.796133 | Gm17823 |
| ENSMUSG( | 38.7388 | 0.076901 | 0.815834 | 0.979198 | Gm17891 |
| ENSMUSG( | 106.3113 | -0.08597 | 0.819067 | 0.979198 | Gm17922 |
| ENSMUSG( | 21.30584 | 0.357568 | 0.445849 | 0.921734 | Gm17971 |
| ENSMUSG( | 13.1748 | -0.216 | 0.693318 | 0.96076 | Gm18180 |
| ENSMUSG( | 9.875353 | -0.1958 | 0.755256 | NA | Gm1848 |
| ENSMUSG( | 9.263684 | 0.618164 | 0.337758 | NA | Gm18706 |
| ENSMUSG( | 9.3816 | -0.17828 | 0.803713 | NA | Gm18726 |
| ENSMUSG( | 22.81333 | -0.72153 | 0.11231 | 0.809018 | Gm18755 |
| ENSMUSG( | 40.08756 | -0.1047 | 0.743626 | 0.96951 | Gm19026 |
| ENSMUSG( | 42.03002 | -0.20062 | 0.534576 | 0.936107 | Gm19345 |
| ENSMUSG( | 978.6629 | 0.043177 | 0.74367 | 0.96951 | Gm19410 |
| ENSMUSG( | 93.81438 | -0.28627 | 0.21531 | 0.865167 | Gm19412 |
| ENSMUSG( | 96.76423 | -0.08472 | 0.745122 | 0.96951 | Gm19500 |
| ENSMUSG( | 168.3812 | -0.49073 | 0.024946 | 0.593727 | Gm19522 |
| ENSMUSG( | 103.1039 | -0.54788 | 0.013689 | 0.468333 | Gm19554 |
| ENSMUSG( | 65.9748 | 0.057119 | 0.843311 | 0.983281 | Gm19605 |
| ENSMUSG( | 37.02487 | 0.126785 | 0.731498 | 0.967228 | Gm19705 |
| ENSMUSG( | 761.8182 | -0.00771 | 0.954724 | 0.994798 | Gm1976 |
| ENSMUSG( | 158.4401 | -0.2959 | 0.104478 | 0.804969 | Gm19935 |
| ENSMUSG( | 260.4156 | -0.04197 | 0.820055 | 0.979198 | Gm20008 |
| ENSMUSG( | 36.58933 | 0.339403 | 0.344666 | 0.91045 | Gm20063 |
| ENSMUSG( | 111.5449 | -0.36888 | 0.130985 | 0.822884 | Gm2011 |
| ENSMUSG( | 27.30373 | 0.010816 | 0.97884 | 0.996468 | Gm20257 |
| ENSMUSG( | 1585.093 | -0.09099 | 0.546741 | 0.938063 | Gm20300 |
| ENSMUSG( | 67.8098 | -0.36924 | 0.2196 | 0.866104 | Gm20383 |
| ENSMUSG( | 10.35493 | 0.154969 | 0.794293 | NA | Gm20517 |
| ENSMUSG( | 17.26716 | -0.09445 | 0.851565 | 0.985541 | Gm20522 |
| ENSMUSG( | 107.6888 | -0.22172 | 0.289675 | 0.896726 | Gm20544 |
| ENSMUSG( | 34.88292 | -0.4906 | 0.147528 | 0.836241 | Gm20597 |
| ENSMUSG( | 194.3303 | 0.049835 | 0.793846 | 0.97681 | Gm20604 |

| | | | | | | | | | |
|---|---|---|---|---|---|---|---|---|---|
| ENSMUSG( | Gm14305 | 318.1563 | -0.01246 | 0.935116 | 0.993293 | ENSMUSG( | Gm14305 | 275.9918 | -0.16022 | 0.327125 | 0.905768 |
| ENSMUSG( | Gm14317 | 27.63819 | 0.31505 | 0.392209 | 0.809274 | ENSMUSG( | Gm14317 | 21.50918 | -0.14895 | 0.746887 | 0.969763 |
| ENSMUSG( | Gm14322 | 65.18139 | 0.12445 | 0.610509 | 0.908189 | ENSMUSG( | Gm14322 | 56.94466 | 0.028764 | 0.91868 | 0.992529 |
| ENSMUSG( | Gm14325 | 514.2771 | 0.184048 | 0.163871 | 0.594869 | ENSMUSG( | Gm14325 | 424.955 | -0.10296 | 0.509782 | 0.929089 |
| ENSMUSG( | Gm14326 | 300.7356 | 0.132367 | 0.371953 | 0.798768 | ENSMUSG( | Gm14326 | 270.7287 | 0.102148 | 0.500761 | 0.927825 |
| ENSMUSG( | Gm14399 | 132.9003 | 0.293663 | 0.109051 | 0.498536 | ENSMUSG( | Gm14399 | 111.1522 | 0.041247 | 0.853891 | 0.98563 |
| ENSMUSG( | Gm14410 | 170.4929 | 0.320771 | 0.077761 | 0.431059 | ENSMUSG( | Gm14410 | 133.6886 | -0.08524 | 0.664744 | 0.956766 |
| ENSMUSG( | Gm14418 | 42.23512 | 0.051997 | 0.862807 | 0.975055 | ENSMUSG( | Gm14418 | 40.75686 | 0.181068 | 0.595635 | 0.945941 |
| ENSMUSG( | Gm14419 | 14.39096 | -0.16969 | 0.724812 | NA | ENSMUSG( | Gm14419 | 11.25029 | -0.70797 | 0.227195 | NA |
| ENSMUSG( | Gm14569 | 113.9549 | 0.554977 | 0.005167 | 0.096 | ENSMUSG( | Gm14569 | 83.93554 | -0.01516 | 0.949553 | 0.994798 |
| ENSMUSG( | Gm14584 | 12.42658 | -0.08475 | 0.885685 | NA | ENSMUSG( | Gm14584 | 9.323102 | -0.65724 | 0.374633 | NA |
| ENSMUSG( | Gm14586 | 67.06629 | -0.12964 | 0.603513 | 0.904887 | ENSMUSG( | Gm14586 | 51.09941 | -0.74766 | 0.014773 | 0.487333 |
| ENSMUSG( | Gm14634 | 36.75289 | 0.205616 | 0.53293 | 0.879905 | ENSMUSG( | Gm14634 | 33.09849 | 0.162798 | 0.644812 | 0.953903 |
| ENSMUSG( | Gm14662 | 12.96806 | 0.289666 | 0.597702 | NA | ENSMUSG( | Gm14662 | 10.20396 | -0.08798 | 0.889604 | NA |
| ENSMUSG( | Gm14824 | 12.74699 | -0.42241 | 0.409679 | NA | ENSMUSG( | Gm14824 | 11.68521 | -0.34095 | 0.557462 | 0.939565 |
| ENSMUSG( | Gm14827 | 248.1151 | 0.418747 | 0.05697 | 0.368322 | ENSMUSG( | Gm14827 | 195.7765 | 0.015347 | 0.955479 | 0.994798 |
| ENSMUSG( | Gm14966 | 21.11345 | 0.293274 | 0.47706 | NA | ENSMUSG( | Gm14966 | 21.13654 | 0.541446 | 0.217769 | 0.865589 |
| ENSMUSG( | Gm15201 | 23.16579 | 0.028429 | 0.942868 | 0.993415 | ENSMUSG( | Gm15201 | 21.86141 | 0.11838 | 0.782142 | 0.976106 |
| ENSMUSG( | Gm15246 | 40.24762 | -0.19066 | 0.557823 | 0.888427 | ENSMUSG( | Gm15246 | 44.3464 | 0.347898 | 0.333918 | 0.907907 |
| ENSMUSG( | Gm15328 | 79.32978 | 0.276955 | 0.208444 | 0.650319 | ENSMUSG( | Gm15328 | 72.12167 | 0.25767 | 0.351985 | 0.912093 |
| ENSMUSG( | Gm15382 | 9.134635 | -0.24308 | 0.698329 | NA | ENSMUSG( | Gm15382 | 10.63551 | 0.434032 | 0.462591 | NA |
| ENSMUSG( | Gm15408 | 19.90067 | -0.24734 | 0.553273 | NA | ENSMUSG( | Gm15408 | 17.7481 | -0.30764 | 0.509954 | 0.929089 |
| ENSMUSG( | Gm15417 | 219.1334 | -0.12339 | 0.40189 | 0.814965 | ENSMUSG( | Gm15417 | 203.4712 | -0.06656 | 0.68381 | 0.958678 |
| ENSMUSG( | Gm15427 | 176.5185 | 0.113942 | 0.500188 | 0.863455 | ENSMUSG( | Gm15427 | 130.7566 | -0.55595 | 0.012037 | 0.445877 |
| ENSMUSG( | Gm15446 | 547.3198 | -0.06739 | 0.627619 | 0.912961 | ENSMUSG( | Gm15446 | 530.9942 | 0.111046 | 0.423103 | 0.917257 |
| ENSMUSG( | Gm15459 | 162.6502 | 0.139265 | 0.427219 | 0.825333 | ENSMUSG( | Gm15459 | 140.2994 | -0.02129 | 0.91683 | 0.992529 |
| ENSMUSG( | Gm15500 | 134.2175 | -0.30488 | 0.101383 | 0.484996 | ENSMUSG( | Gm15500 | 120.7051 | -0.36782 | 0.148542 | 0.836241 |
| ENSMUSG( | Gm15501 | 59.51812 | 0.181882 | 0.479402 | 0.854046 | ENSMUSG( | Gm15501 | 49.25288 | -0.09532 | 0.744102 | 0.96951 |
| ENSMUSG( | Gm15510 | 47.32327 | 0.071626 | 0.792306 | 0.959461 | ENSMUSG( | Gm15510 | 41.26683 | -0.05899 | 0.849155 | 0.985321 |
| ENSMUSG( | Gm15542 | 17.96705 | 0.596737 | 0.185525 | NA | ENSMUSG( | Gm15542 | 14.25526 | 0.250965 | 0.633901 | 0.953891 |
| ENSMUSG( | Gm15545 | 36.43105 | -0.27191 | 0.408472 | 0.818167 | ENSMUSG( | Gm15545 | 33.78606 | -0.20881 | 0.565964 | 0.941499 |
| ENSMUSG( | Gm15559 | 51.19858 | -0.16346 | 0.550839 | 0.886038 | ENSMUSG( | Gm15559 | 46.43067 | -0.16026 | 0.601408 | 0.947749 |
| ENSMUSG( | Gm15612 | 81.81602 | -0.19651 | 0.39385 | 0.809643 | ENSMUSG( | Gm15612 | 72.6144 | -0.24759 | 0.3953 | 0.917257 |
| ENSMUSG( | Gm15663 | 234.5 | -0.05691 | 0.707128 | 0.93949 | ENSMUSG( | Gm15663 | 212.1334 | -0.09129 | 0.627838 | 0.953234 |
| ENSMUSG( | Gm15697 | 15.96989 | 0.135864 | 0.830385 | NA | ENSMUSG( | Gm15697 | 13.66475 | -0.04754 | 0.945735 | 0.994798 |
| ENSMUSG( | Gm15706 | 216.6287 | 0.101011 | 0.567269 | 0.891666 | ENSMUSG( | Gm15706 | 194.0798 | 0.039932 | 0.843014 | 0.983211 |
| ENSMUSG( | Gm15708 | 18.74753 | -0.37425 | 0.395127 | NA | ENSMUSG( | Gm15708 | 18.08719 | -0.16872 | 0.74194 | 0.969314 |
| ENSMUSG( | Gm15710 | 107.5539 | 0.049973 | 0.805726 | 0.962616 | ENSMUSG( | Gm15710 | 84.46916 | -0.43288 | 0.101392 | 0.801456 |
| ENSMUSG( | Gm15743 | 36.60117 | -0.18365 | 0.547954 | 0.885153 | ENSMUSG( | Gm15743 | 33.01057 | -0.20307 | 0.553237 | 0.939565 |
| ENSMUSG( | Gm15772 | 68.48981 | -0.06866 | 0.774339 | 0.955606 | ENSMUSG( | Gm15772 | 52.34151 | -0.66554 | 0.021077 | 0.573311 |
| ENSMUSG( | Gm15787 | 69.49204 | 0.291124 | 0.23023 | 0.678139 | ENSMUSG( | Gm15787 | 60.50082 | 0.16205 | 0.5486 | 0.938063 |
| ENSMUSG( | Gm15816 | 46.50439 | 0.520647 | 0.076203 | 0.427093 | ENSMUSG( | Gm15816 | 30.72081 | -0.38858 | 0.310886 | 0.899422 |
| ENSMUSG( | Gm15850 | 9.526304 | -0.46001 | 0.434311 | NA | ENSMUSG( | Gm15850 | 9.307133 | -0.22004 | 0.721912 | NA |
| ENSMUSG( | Gm15867 | 27.32345 | -0.00308 | 0.993394 | 0.999369 | ENSMUSG( | Gm15867 | 21.88107 | -0.41595 | 0.346868 | 0.911434 |
| ENSMUSG( | Gm15939 | 10.93752 | 0.653832 | 0.249294 | NA | ENSMUSG( | Gm15939 | 8.244405 | 0.194888 | 0.772149 | NA |
| ENSMUSG( | Gm15972 | 9.88357 | -0.52624 | 0.404088 | NA | ENSMUSG( | Gm15972 | 10.74165 | 0.054393 | 0.928451 | NA |
| ENSMUSG( | Gm15997 | 20.35494 | -0.23981 | 0.599669 | NA | ENSMUSG( | Gm15997 | 21.50977 | 0.169539 | 0.703319 | 0.962481 |
| ENSMUSG( | Gm16023 | 13.78275 | 0.356087 | 0.484184 | NA | ENSMUSG( | Gm16023 | 16.46329 | 1.01465 | 0.057191 | 0.723362 |
| ENSMUSG( | Gm1604a | 75.10199 | -0.07595 | 0.762552 | 0.953519 | ENSMUSG( | Gm1604a | 64.58901 | -0.26855 | 0.337532 | 0.907972 |
| ENSMUSG( | Gm16062 | 16.27413 | 0.193771 | 0.675392 | NA | ENSMUSG( | Gm16062 | 10.73482 | -0.89613 | 0.179782 | NA |
| ENSMUSG( | Gm16124 | 47.63386 | 0.011679 | 0.968562 | 0.996369 | ENSMUSG( | Gm16124 | 49.05705 | 0.333146 | 0.282158 | 0.892412 |
| ENSMUSG( | Gm16201 | 29.11545 | 0.359412 | 0.312674 | 0.760241 | ENSMUSG( | Gm16201 | 25.0175 | 0.188439 | 0.66939 | 0.956766 |
| ENSMUSG( | Gm16238 | 51.99795 | -0.08918 | 0.738899 | 0.947768 | ENSMUSG( | Gm16238 | 41.84028 | -0.52347 | 0.112876 | 0.809623 |
| ENSMUSG( | Gm16244 | 18.32551 | -0.0803 | 0.875369 | NA | ENSMUSG( | Gm16244 | 16.53179 | -0.11255 | 0.823675 | 0.979198 |
| ENSMUSG( | Gm16364 | 14.2431 | -0.13274 | 0.79004 | NA | ENSMUSG( | Gm16364 | 13.86835 | 0.048093 | 0.929281 | 0.99306 |
| ENSMUSG( | Gm16537 | 13.14516 | 0.316411 | 0.564599 | NA | ENSMUSG( | Gm16537 | 12.10119 | 0.41223 | 0.52238 | 0.932681 |
| ENSMUSG( | Gm16576 | 72.96137 | -0.33317 | 0.221725 | 0.667354 | ENSMUSG( | Gm16576 | 65.10267 | -0.3941 | 0.173431 | 0.844317 |
| ENSMUSG( | Gm16580 | 37.76422 | 0.05455 | 0.873086 | 0.978753 | ENSMUSG( | Gm16580 | 26.7661 | -0.78379 | 0.04641 | 0.697814 |
| ENSMUSG( | Gm16630 | 9.90638 | -0.3571 | 0.536336 | NA | ENSMUSG( | Gm16630 | 9.295474 | -0.29374 | 0.642967 | NA |
| ENSMUSG( | Gm16638 | 228.8099 | -0.03109 | 0.849906 | 0.972638 | ENSMUSG( | Gm16638 | 214.92 | 0.06564 | 0.72659 | 0.966887 |
| ENSMUSG( | Gm16675 | 29.98106 | -0.31363 | 0.376157 | 0.801127 | ENSMUSG( | Gm16675 | 25.04182 | -0.60272 | 0.1677 | 0.840222 |
| ENSMUSG( | Gm16701 | 24.55739 | 0.150973 | 0.692642 | 0.934101 | ENSMUSG( | Gm16701 | 22.32829 | 0.117351 | 0.791371 | 0.97681 |
| ENSMUSG( | Gm16740 | 72.24215 | -0.38279 | 0.088891 | 0.457425 | ENSMUSG( | Gm16740 | 71.69373 | -0.10715 | 0.660775 | 0.956205 |
| ENSMUSG( | Gm16793 | 18.5982 | 0.52814 | 0.282778 | NA | ENSMUSG( | Gm16793 | 15.1447 | 0.245176 | 0.636275 | 0.953903 |
| ENSMUSG( | Gm16835 | 159.6806 | 0.007004 | 0.970501 | 0.996369 | ENSMUSG( | Gm16835 | 140.7525 | -0.08556 | 0.694629 | 0.960867 |
| ENSMUSG( | Gm16845 | 25.2684 | 0.568031 | 0.19917 | 0.639621 | ENSMUSG( | Gm16845 | 21.90509 | 0.461247 | 0.28165 | 0.892412 |
| ENSMUSG( | Gm16861 | 37.36666 | 0.025952 | 0.935542 | 0.993363 | ENSMUSG( | Gm16861 | 34.07014 | 0.028031 | 0.938308 | 0.993793 |
| ENSMUSG( | Gm16867 | 59.81021 | -1.58049 | 2.32E-08 | 8.35E-06 | ENSMUSG( | Gm16867 | 59.76073 | -1.10292 | 0.00013 | 0.02529 |
| ENSMUSG( | Gm16894 | 15.64719 | -0.11692 | 0.816834 | NA | ENSMUSG( | Gm16894 | 16.66714 | 0.333886 | 0.521652 | 0.932571 |
| ENSMUSG( | Gm16897 | 19.70336 | -0.22287 | 0.610384 | NA | ENSMUSG( | Gm16897 | 17.33012 | -0.34139 | 0.485123 | 0.92541 |
| ENSMUSG( | Gm16958 | 18.14021 | -0.01698 | 0.970929 | NA | ENSMUSG( | Gm16958 | 14.81521 | -0.37269 | 0.48827 | 0.925453 |
| ENSMUSG( | Gm16973 | 157.8643 | -0.11162 | 0.495802 | 0.861294 | ENSMUSG( | Gm16973 | 154.5006 | 0.101581 | 0.575595 | 0.942912 |
| ENSMUSG( | Gm17102 | 18.47162 | -0.21133 | 0.643415 | NA | ENSMUSG( | Gm17102 | 16.22578 | -0.32006 | 0.517099 | 0.930939 |
| ENSMUSG( | Gm17244 | 52.37146 | 0.131133 | 0.621818 | 0.911432 | ENSMUSG( | Gm17244 | 43.25034 | -0.14506 | 0.639521 | 0.953903 |
| ENSMUSG( | Gm17315 | 17.28107 | -0.2609 | 0.579305 | NA | ENSMUSG( | Gm17315 | 16.10537 | -0.23399 | 0.641231 | 0.953903 |
| ENSMUSG( | Gm17344 | 13.8875 | 0.417835 | 0.444371 | NA | ENSMUSG( | Gm17344 | 11.92764 | 0.257533 | 0.646832 | 0.953988 |
| ENSMUSG( | Gm17382 | 12.71965 | 0.070227 | 0.893104 | NA | ENSMUSG( | Gm17382 | 12.02537 | 0.150595 | 0.798878 | 0.97681 |
| ENSMUSG( | Gm17383 | 20.31602 | -0.14933 | 0.731161 | NA | ENSMUSG( | Gm17383 | 17.59629 | -0.35552 | 0.486062 | 0.92541 |
| ENSMUSG( | Gm17396 | 52.60658 | -0.06389 | 0.819021 | 0.965888 | ENSMUSG( | Gm17396 | 49.67436 | 0.018512 | 0.951799 | 0.994798 |
| ENSMUSG( | Gm17455 | 34.15422 | -0.75143 | 0.023371 | 0.233882 | ENSMUSG( | Gm17455 | 36.10803 | -0.20548 | 0.568713 | 0.941777 |
| ENSMUSG( | Gm17484 | 49.0606 | -0.28876 | 0.367038 | 0.795164 | ENSMUSG( | Gm17484 | 51.55313 | 0.143995 | 0.636613 | 0.953903 |
| ENSMUSG( | Gm17597 | 22.70918 | 0.190182 | 0.667306 | 0.92729 | ENSMUSG( | Gm17597 | 21.13554 | 0.271465 | 0.568892 | 0.941777 |
| ENSMUSG( | Gm17634 | 16.53489 | -0.19686 | 0.659739 | NA | ENSMUSG( | Gm17634 | 12.20537 | -1.04094 | 0.076962 | 0.770558 |
| ENSMUSG( | Gm17733 | 20.04757 | -0.26312 | 0.57596 | NA | ENSMUSG( | Gm17733 | 18.24376 | -0.26495 | 0.563854 | 0.941206 |
| ENSMUSG( | Gm17750 | 209.6217 | 0.206561 | 0.206454 | 0.648468 | ENSMUSG( | Gm17750 | 171.6287 | -0.11162 | 0.598076 | 0.946676 |
| ENSMUSG( | Gm17767 | 8.650192 | -0.62603 | 0.327258 | NA | ENSMUSG( | Gm17767 | 9.177814 | -0.09836 | 0.87794 | NA |
| ENSMUSG( | Gm17823 | 11.64056 | 0.427928 | 0.464175 | NA | ENSMUSG( | Gm17823 | 13.97881 | 1.008043 | 0.095945 | 0.796133 |
| ENSMUSG( | Gm17891 | 40.2687 | -0.09397 | 0.773547 | 0.955447 | ENSMUSG( | Gm17891 | 38.7388 | 0.076901 | 0.815834 | 0.979198 |
| ENSMUSG( | Gm17922 | 151.0968 | 0.630424 | 0.02628 | 0.247758 | ENSMUSG( | Gm17922 | 106.3113 | -0.08597 | 0.819067 | 0.979198 |
| ENSMUSG( | Gm17971 | 14.37381 | -1.37294 | 0.020074 | NA | ENSMUSG( | Gm17971 | 21.30584 | 0.357568 | 0.445849 | 0.921734 |
| ENSMUSG( | Gm18180 | 17.33329 | 0.273632 | 0.548821 | NA | ENSMUSG( | Gm18180 | 13.1748 | -0.216 | 0.693318 | 0.96076 |
| ENSMUSG( | Gm1848 | 13.31175 | 0.362302 | 0.488294 | NA | ENSMUSG( | Gm1848 | 9.875353 | -0.1958 | 0.755256 | NA |
| ENSMUSG( | Gm18755 | 26.47045 | -0.53135 | 0.236143 | 0.683188 | ENSMUSG( | Gm18755 | 22.81333 | -0.72153 | 0.11231 | 0.809018 |
| ENSMUSG( | Gm19026 | 50.1294 | 0.260248 | 0.327612 | 0.772692 | ENSMUSG( | Gm19026 | 40.08756 | -0.1047 | 0.743626 | 0.96951 |
| ENSMUSG( | Gm19345 | 47.5545 | -0.14352 | 0.633492 | 0.914073 | ENSMUSG( | Gm19345 | 42.03002 | -0.20062 | 0.534576 | 0.936107 |
| ENSMUSG( | Gm19410 | 993.5278 | -0.1922 | 0.080528 | 0.438901 | ENSMUSG( | Gm19410 | 978.6629 | 0.043177 | 0.74367 | 0.96951 |
| ENSMUSG( | Gm19412 | 111.6889 | -0.03572 | 0.862129 | 0.975055 | ENSMUSG( | Gm19412 | 93.81438 | -0.28627 | 0.21531 | 0.865167 |
| ENSMUSG( | Gm19500 | 100.0641 | -0.28701 | 0.203647 | 0.643764 | ENSMUSG( | Gm19500 | 96.76423 | -0.08472 | 0.745122 | 0.96951 |
| ENSMUSG( | Gm19522 | 208.0519 | -0.098 | 0.603329 | 0.904887 | ENSMUSG( | Gm19522 | 168.3812 | -0.49073 | 0.024946 | 0.593727 |
| ENSMUSG( | Gm19554 | 127.8805 | -0.14518 | 0.440517 | 0.831996 | ENSMUSG( | Gm19554 | 103.1039 | -0.54788 | 0.013689 | 0.468333 |
| ENSMUSG( | Gm19605 | 73.94552 | 0.099026 | 0.701632 | 0.937441 | ENSMUSG( | Gm19605 | 65.9748 | 0.057119 | 0.843311 | 0.983281 |
| ENSMUSG( | Gm19705 | 40.14733 | 0.054762 | 0.859151 | 0.974707 | ENSMUSG( | Gm19705 | 37.02487 | 0.126785 | 0.731498 | 0.967228 |
| ENSMUSG( | Gm1976 | 864.8116 | 0.093693 | 0.386333 | 0.804623 | ENSMUSG( | Gm1976 | 761.8182 | -0.00771 | 0.954724 | 0.994798 |
| ENSMUSG( | Gm19935 | 185.3933 | -0.09887 | 0.554461 | 0.886657 | ENSMUSG( | Gm19935 | 158.4401 | -0.2959 | 0.104478 | 0.804969 |
| ENSMUSG( | Gm20008 | 284.5482 | -0.04466 | 0.760998 | 0.952907 | ENSMUSG( | Gm20008 | 260.4156 | -0.04197 | 0.820055 | 0.979198 |
| ENSMUSG( | Gm20063 | 36.85417 | 0.100621 | 0.764506 | 0.954055 | ENSMUSG( | Gm20063 | 36.58933 | 0.339403 | 0.344666 | 0.91045 |
| ENSMUSG( | Gm2011 | 132.8826 | -0.10919 | 0.603275 | 0.904887 | ENSMUSG( | Gm2011 | 111.5449 | -0.36888 | 0.130985 | 0.822884 |
| ENSMUSG( | Gm20257 | 30.11979 | 0.060353 | 0.863811 | 0.975244 | ENSMUSG( | Gm20257 | 27.30373 | 0.010816 | 0.97884 | 0.996468 |
| ENSMUSG( | Gm20300 | 1805.236 | 0.021723 | 0.853659 | 0.973491 | ENSMUSG( | Gm20300 | 1585.093 | -0.09099 | 0.546741 | 0.938063 |
| ENSMUSG( | Gm20383 | 84.09144 | 0.041182 | 0.858158 | 0.974672 | ENSMUSG( | Gm20383 | 67.8098 | -0.36924 | 0.2196 | 0.866104 |
| ENSMUSG( | Gm20517 | 10.04639 | -0.19013 | 0.744412 | NA | ENSMUSG( | Gm20517 | 10.35493 | 0.154969 | 0.794293 | NA |
| ENSMUSG( | Gm20522 | 18.93305 | -0.0465 | 0.911793 | NA | ENSMUSG( | Gm20522 | 17.26716 | -0.09445 | 0.851565 | 0.985541 |
| ENSMUSG( | Gm20544 | 126.449 | -0.02548 | 0.885974 | 0.983008 | ENSMUSG( | Gm20544 | 107.6888 | -0.22172 | 0.289675 | 0.896726 |
| ENSMUSG( | Gm20597 | 40.30937 | -0.31304 | 0.291468 | 0.741366 | ENSMUSG( | Gm20597 | 34.88292 | -0.4906 | 0.147528 | 0.836241 |
| ENSMUSG( | Gm20604 | 204.0757 | -0.06239 | 0.686765 | 0.933041 | ENSMUSG( | Gm20604 | 194.3303 | 0.049835 | 0.793846 | 0.97681 |
| ENSMUSG( | Gm2061 | 223.5251 | -0.12272 | 0.451455 | 0.835567 | ENSMUSG( | Gm2061 | 210.1194 | -0.03753 | 0.838236 | 0.981575 |
| ENSMUSG( | Gm20646 | 107.0453 | -0.04264 | 0.840669 | 0.971179 | ENSMUSG( | Gm20646 | 95.89062 | -0.10716 | 0.650296 | 0.95422 |
| ENSMUSG( | Gm20743 | 123.5608 | 0.022741 | 0.930317 | 0.991848 | ENSMUSG( | Gm20743 | 101.4686 | -0.29366 | 0.232162 | 0.868507 |
| ENSMUSG( | Gm20751 | 27.18147 | -0.28426 | 0.456865 | 0.839183 | ENSMUSG( | Gm20751 | 30.32659 | 0.330002 | 0.472732 | 0.923329 |
| ENSMUSG( | Gm20752 | 20.40174 | 0.002237 | 0.995668 | NA | ENSMUSG( | Gm20752 | 15.1193 | -0.68873 | 0.171577 | 0.842999 |
| ENSMUSG( | Gm20753 | 19.1088 | -0.26799 | 0.557668 | NA | ENSMUSG( | Gm20753 | 20.35592 | 0.228851 | 0.64052 | 0.953903 |
| ENSMUSG( | Gm20754 | 40.39169 | 0.131964 | 0.686829 | 0.933041 | ENSMUSG( | Gm20754 | 35.77353 | 0.058008 | 0.872145 | 0.988907 |
| ENSMUSG( | Gm20939 | 33.70173 | -0.30464 | 0.368032 | 0.795774 | ENSMUSG( | Gm20939 | 34.70756 | 0.092952 | 0.803127 | 0.976937 |
| ENSMUSG( | Gm21092 | 43.23337 | 0.183851 | 0.576338 | 0.895164 | ENSMUSG( | Gm21092 | 36.10013 | -0.04121 | 0.910006 | 0.991958 |
| ENSMUSG( | Gm21811 | 140.3235 | 0.011333 | 0.957023 | 0.993874 | ENSMUSG( | Gm21811 | 123.3461 | -0.09619 | 0.640109 | 0.953903 |
| ENSMUSG( | Gm21814 | 16.6992 | 0.003252 | 0.994702 | NA | ENSMUSG( | Gm21814 | 14.7711 | -0.08865 | 0.867789 | 0.98781 |
| ENSMUSG( | Gm21926 | 618.3855 | 0.127317 | 0.274064 | 0.724254 | ENSMUSG( | Gm21926 | 528.0414 | -0.06178 | 0.653202 | 0.955015 |
| ENSMUSG( | Gm21992 | 128.8542 | 0.140459 | 0.50746 | 0.86795 | ENSMUSG( | Gm21992 | 115.5686 | 0.11602 | 0.645377 | 0.953988 |
| ENSMUSG( | Gm22009 | 31.26475 | -1.2627 | 0.000668 | 0.024972 | ENSMUSG( | Gm22009 | 44.52627 | 0.253653 | 0.488025 | 0.925453 |
| ENSMUSG( | Gm2237 | 32.87096 | -0.19215 | 0.591268 | 0.90113 | ENSMUSG( | Gm2237 | 33.8214 | 0.170286 | 0.648768 | 0.95422 |
| ENSMUSG( | Gm22442 | 15.49626 | -0.25223 | 0.609329 | NA | ENSMUSG( | Gm22442 | 12.529 | -0.70898 | 0.230764 | 0.868507 |
| ENSMUSG( | Gm22540 | 11.91464 | -0.00939 | 0.985896 | NA | ENSMUSG( | Gm22540 | 11.32029 | 0.107539 | 0.848644 | NA |
| ENSMUSG( | Gm22753 | 11.11711 | 0.251857 | 0.6498 | NA | ENSMUSG( | Gm22753 | 10.22325 | 0.196023 | 0.774131 | NA |
| ENSMUSG( | Gm23217 | 9.40916 | -0.6251 | 0.325184 | NA | ENSMUSG( | Gm23217 | 8.548474 | -0.58042 | 0.406776 | NA |
| ENSMUSG( | Gm23301 | 9.297795 | -0.50144 | 0.403113 | NA | ENSMUSG( | Gm23301 | 10.44509 | 0.148444 | 0.800499 | NA |
| ENSMUSG( | Gm23318 | 10.37557 | -0.40083 | 0.488338 | NA | ENSMUSG( | Gm23318 | 11.64053 | 0.256526 | 0.661274 | 0.956205 |
| ENSMUSG( | Gm23458 | 14.93222 | -0.36231 | 0.458459 | NA | ENSMUSG( | Gm23458 | 11.44934 | -0.97333 | 0.10461 | NA |
| ENSMUSG( | Gm24119 | 16.08115 | -0.11937 | 0.80396 | NA | ENSMUSG( | Gm24119 | 13.39912 | -0.32586 | 0.565746 | 0.941404 |
| ENSMUSG( | Gm2464 | 70.55169 | 0.166231 | 0.466148 | 0.844453 | ENSMUSG( | Gm2464 | 57.1029 | -0.16619 | 0.539699 | 0.937378 |
| ENSMUSG( | Gm25047 | 48.62653 | -0.01536 | 0.958629 | 0.993874 | ENSMUSG( | Gm25047 | 46.83342 | 0.122975 | 0.708836 | 0.964042 |
| ENSMUSG( | Gm25117 | 25.35419 | -0.31079 | 0.413136 | 0.819541 | ENSMUSG( | Gm25117 | 21.15018 | -0.57907 | 0.192796 | 0.859764 |
| ENSMUSG( | Gm25129 | 9.643839 | -0.40591 | 0.530063 | NA | ENSMUSG( | Gm25129 | 13.92105 | 0.862589 | 0.156614 | 0.837724 |
| ENSMUSG( | Gm25492 | 12.879 | 0.137701 | 0.789613 | NA | ENSMUSG( | Gm25492 | 11.45713 | 0.085332 | 0.880895 | 0.989644 |
| ENSMUSG( | Gm25596 | 32.93605 | 0.130536 | 0.693337 | 0.934202 | ENSMUSG( | Gm25596 | 26.43129 | -0.26357 | 0.504888 | 0.927825 |
| ENSMUSG( | Gm26397 | 13.26223 | -0.1538 | 0.777378 | NA | ENSMUSG( | Gm26397 | 16.06458 | 0.578921 | 0.297722 | 0.898538 |
| ENSMUSG( | Gm266 | 225.5157 | 0.074038 | 0.62488 | 0.912167 | ENSMUSG( | Gm266 | 192.1416 | -0.12598 | 0.455537 | 0.923329 |
| ENSMUSG( | Gm26644 | 47.99851 | 0.416887 | 0.142821 | 0.561453 | ENSMUSG( | Gm26644 | 38.80237 | 0.127709 | 0.70842 | 0.964042 |
| ENSMUSG( | Gm26671 | 18.55958 | 0.100917 | 0.811908 | NA | ENSMUSG( | Gm26671 | 15.25711 | -0.20764 | 0.671144 | 0.95694 |
| ENSMUSG( | Gm26684 | 47.46216 | 0.088863 | 0.772937 | 0.955272 | ENSMUSG( | Gm26684 | 36.96378 | -0.37995 | 0.302385 | 0.898538 |
| ENSMUSG( | Gm26760 | 39.22954 | 0.144537 | 0.644498 | 0.917773 | ENSMUSG( | Gm26760 | 29.323 | -0.47043 | 0.198065 | 0.862161 |
| ENSMUSG( | Gm26782 | 695.8144 | -0.01913 | 0.888734 | 0.983434 | ENSMUSG( | Gm26782 | 574.6292 | -0.3136 | 0.041918 | 0.690304 |
| ENSMUSG( | Gm26788 | 11.34782 | -0.03906 | 0.944368 | NA | ENSMUSG( | Gm26788 | 13.56917 | 0.715406 | 0.208016 | 0.864412 |
| ENSMUSG( | Gm26797 | 22.06677 | -0.07846 | 0.849644 | 0.972638 | ENSMUSG( | Gm26797 | 22.28381 | 0.256138 | 0.599469 | 0.947071 |
| ENSMUSG( | Gm2694 | 113.5135 | -0.04182 | 0.836777 | 0.971111 | ENSMUSG( | Gm2694 | 102.4449 | -0.08693 | 0.712813 | 0.96439 |
| ENSMUSG( | Gm27239 | 30.94247 | -0.46875 | 0.164801 | 0.595869 | ENSMUSG( | Gm27239 | 34.00031 | 0.134446 | 0.715148 | 0.965063 |
| ENSMUSG( | Gm27343 | 20.83221 | 0.036395 | 0.93295 | NA | ENSMUSG( | Gm27343 | 17.76323 | -0.18984 | 0.683275 | 0.958678 |
| ENSMUSG( | Gm2762 | 77.4547 | 0.339701 | 0.139501 | 0.555676 | ENSMUSG( | Gm2762 | 62.28292 | 0.015548 | 0.954641 | 0.994798 |
| ENSMUSG( | Gm28154 | 11.07055 | 0.37619 | 0.503916 | NA | ENSMUSG( | Gm28154 | 11.36757 | 0.650677 | 0.266472 | NA |

| | | | | | |
|---|---|---|---|---|---|
| ENSMUSG( | 9.611906 | 0.466268 | 0.440495 | NA | Gm19243 |
| ENSMUSG( | 47.5545 | -0.14352 | 0.633492 | 0.914073 | Gm19345 |
| ENSMUSG( | 993.5278 | -0.1922 | 0.080528 | 0.438901 | Gm19410 |
| ENSMUSG( | 111.6889 | -0.03572 | 0.862129 | 0.975055 | Gm19412 |
| ENSMUSG( | 100.0641 | -0.28701 | 0.203647 | 0.643764 | Gm19500 |
| ENSMUSG( | 208.0519 | -0.098 | 0.603329 | 0.904887 | Gm19522 |
| ENSMUSG( | 127.8805 | -0.14518 | 0.440517 | 0.831996 | Gm19554 |
| ENSMUSG( | 73.94552 | 0.099026 | 0.701632 | 0.937441 | Gm19605 |
| ENSMUSG( | 40.14733 | 0.054762 | 0.859151 | 0.974707 | Gm19705 |
| ENSMUSG( | 864.8116 | 0.093693 | 0.386333 | 0.804623 | Gm1976 |
| ENSMUSG( | 185.3933 | -0.09887 | 0.554461 | 0.886657 | Gm19935 |
| ENSMUSG( | 9.601142 | 0.025688 | 0.966154 | NA | Gm19951 |
| ENSMUSG( | 284.5482 | -0.04466 | 0.760998 | 0.952907 | Gm20008 |
| ENSMUSG( | 36.85417 | 0.100621 | 0.764506 | 0.954055 | Gm20063 |
| ENSMUSG( | 132.8826 | -0.10919 | 0.603275 | 0.904887 | Gm2011 |
| ENSMUSG( | 30.11979 | 0.060353 | 0.863811 | 0.975244 | Gm20257 |
| ENSMUSG( | 1805.236 | 0.021723 | 0.853659 | 0.973491 | Gm20300 |
| ENSMUSG( | 84.09144 | 0.041182 | 0.858158 | 0.974672 | Gm20383 |
| ENSMUSG( | 10.04639 | -0.19013 | 0.744412 | NA | Gm20517 |
| ENSMUSG( | 18.93305 | -0.0465 | 0.911793 | NA | Gm20522 |
| ENSMUSG( | 126.449 | -0.02548 | 0.885974 | 0.983008 | Gm20544 |
| ENSMUSG( | 40.30937 | -0.31304 | 0.291468 | 0.741366 | Gm20597 |
| ENSMUSG( | 204.0757 | -0.06239 | 0.686765 | 0.933041 | Gm20604 |
| ENSMUSG( | 223.5251 | -0.12272 | 0.451455 | 0.835567 | Gm2061 |
| ENSMUSG( | 107.0453 | -0.04264 | 0.840669 | 0.971179 | Gm20646 |
| ENSMUSG( | 123.5608 | 0.022741 | 0.930317 | 0.991848 | Gm20743 |
| ENSMUSG( | 27.18147 | -0.28426 | 0.456865 | 0.839183 | Gm20751 |
| ENSMUSG( | 20.40174 | 0.002237 | 0.995668 | NA | Gm20752 |
| ENSMUSG( | 19.1088 | -0.26799 | 0.557668 | NA | Gm20753 |
| ENSMUSG( | 40.39169 | 0.131964 | 0.686829 | 0.933041 | Gm20754 |
| ENSMUSG( | 33.70173 | -0.30464 | 0.368032 | 0.795774 | Gm20939 |
| ENSMUSG( | 43.23337 | 0.183851 | 0.576338 | 0.895164 | Gm21092 |
| ENSMUSG( | 13.14239 | -0.43528 | 0.411689 | NA | Gm21388 |
| ENSMUSG( | 140.3235 | 0.011333 | 0.957023 | 0.993874 | Gm21811 |
| ENSMUSG( | 16.6992 | 0.003252 | 0.994702 | NA | Gm21814 |
| ENSMUSG( | 618.3855 | 0.127317 | 0.274064 | 0.724254 | Gm21926 |
| ENSMUSG( | 128.8542 | 0.140459 | 0.50746 | 0.86795 | Gm21992 |
| ENSMUSG( | 31.26475 | -1.2627 | 0.000668 | 0.024972 | Gm22009 |
| ENSMUSG( | 32.87096 | -0.19215 | 0.591268 | 0.90113 | Gm2237 |
| ENSMUSG( | 15.49626 | -0.25223 | 0.609329 | NA | Gm22442 |
| ENSMUSG( | 11.91464 | -0.00939 | 0.985896 | NA | Gm22540 |
| ENSMUSG( | 11.11711 | 0.251857 | 0.6498 | NA | Gm22753 |
| ENSMUSG( | 9.40916 | -0.6251 | 0.325184 | NA | Gm23217 |
| ENSMUSG( | 9.297795 | -0.50144 | 0.403113 | NA | Gm23301 |
| ENSMUSG( | 10.37557 | -0.40083 | 0.488338 | NA | Gm23318 |
| ENSMUSG( | 14.93222 | -0.36231 | 0.458459 | NA | Gm23458 |
| ENSMUSG( | 16.08115 | -0.11937 | 0.80396 | NA | Gm24119 |
| ENSMUSG( | 70.55169 | 0.166231 | 0.466148 | 0.844453 | Gm2464 |
| ENSMUSG( | 11.11751 | -0.0387 | 0.948652 | NA | Gm24873 |
| ENSMUSG( | 48.62653 | -0.01536 | 0.958629 | 0.993874 | Gm25047 |
| ENSMUSG( | 25.35419 | -0.31079 | 0.413136 | 0.819541 | Gm25117 |
| ENSMUSG( | 9.643839 | -0.40591 | 0.530063 | NA | Gm25129 |
| ENSMUSG( | 12.879 | 0.137701 | 0.789613 | NA | Gm25492 |
| ENSMUSG( | 32.93605 | 0.130536 | 0.693337 | 0.934202 | Gm25596 |
| ENSMUSG( | 13.26223 | -0.1538 | 0.777378 | NA | Gm26397 |
| ENSMUSG( | 9.393764 | 0.833009 | 0.196135 | NA | Gm26496 |
| ENSMUSG( | 225.5157 | 0.074038 | 0.62488 | 0.912167 | Gm266 |
| ENSMUSG( | 47.99851 | 0.416887 | 0.142821 | 0.561453 | Gm26644 |
| ENSMUSG( | 18.55958 | 0.100917 | 0.811908 | NA | Gm26671 |
| ENSMUSG( | 8.084089 | -0.39206 | 0.558687 | NA | Gm26682 |
| ENSMUSG( | 47.46216 | 0.088863 | 0.772937 | 0.955272 | Gm26684 |
| ENSMUSG( | 39.22954 | 0.144537 | 0.644498 | 0.917773 | Gm26760 |
| ENSMUSG( | 695.8144 | -0.01913 | 0.888734 | 0.983434 | Gm26782 |
| ENSMUSG( | 11.34782 | -0.03906 | 0.944368 | NA | Gm26788 |
| ENSMUSG( | 22.06677 | -0.07846 | 0.849644 | 0.972638 | Gm26797 |
| ENSMUSG( | 113.5135 | -0.04182 | 0.836777 | 0.971111 | Gm2694 |
| ENSMUSG( | 30.94247 | -0.46875 | 0.164801 | 0.595869 | Gm27239 |
| ENSMUSG( | 20.83221 | 0.036395 | 0.93295 | NA | Gm27343 |
| ENSMUSG( | 77.4547 | 0.339701 | 0.139501 | 0.555676 | Gm2762 |
| ENSMUSG( | 11.07055 | 0.37619 | 0.503916 | NA | Gm28154 |
| ENSMUSG( | 11.34886 | 0.21573 | 0.700096 | NA | Gm28729 |
| ENSMUSG( | 42.38777 | -0.25569 | 0.376008 | 0.801127 | Gm2885 |
| ENSMUSG( | 11.46949 | -0.3557 | 0.535295 | NA | Gm2897 |
| ENSMUSG( | 13.54382 | -0.15177 | 0.771006 | NA | Gm29257 |
| ENSMUSG( | 12.2403 | -0.01383 | 0.978812 | NA | Gm29443 |
| ENSMUSG( | 18.63752 | 0.478541 | 0.26649 | NA | Gm29483 |
| ENSMUSG( | 71.45719 | 0.095128 | 0.691069 | 0.933881 | Gm29508 |
| ENSMUSG( | 96.54558 | 0.237576 | 0.358212 | 0.78975 | Gm29514 |
| ENSMUSG( | 10.58649 | 0.196873 | 0.742949 | NA | Gm29543 |
| ENSMUSG( | 129.5274 | -0.0419 | 0.848685 | 0.972374 | Gm29595 |
| ENSMUSG( | 23.01597 | -0.09207 | 0.830059 | 0.96974 | Gm29642 |
| ENSMUSG( | 80.89339 | 0.132951 | 0.573263 | 0.893584 | Gm29674 |
| ENSMUSG( | 21.71278 | 0.435147 | 0.400239 | 0.813641 | Gm29683 |
| ENSMUSG( | 11.69299 | 0.634314 | 0.313404 | NA | Gm2974 |
| ENSMUSG( | 15.08693 | -0.34502 | 0.491043 | NA | Gm29773 |
| ENSMUSG( | 8186.258 | -0.17953 | 0.112792 | 0.506698 | Gm2a |
| ENSMUSG( | 57.35977 | 0.092177 | 0.729382 | 0.944738 | Gm30082 |
| ENSMUSG( | 14.95931 | -0.10815 | 0.829408 | NA | Gm30092 |
| ENSMUSG( | 22.01229 | 0.182147 | 0.651956 | 0.921946 | Gm30108 |
| ENSMUSG( | 9.365774 | 0.100328 | 0.872333 | NA | Gm30400 |
| ENSMUSG( | 39.99044 | -0.08999 | 0.762247 | 0.953459 | Gm30539 |
| ENSMUSG( | 38.05929 | -0.19305 | 0.561276 | 0.889052 | Gm3055 |
| ENSMUSG( | 29.41084 | 0.10852 | 0.75728 | 0.951513 | Gm30571 |
| ENSMUSG( | 13.7734 | -0.17241 | 0.730837 | NA | Gm30606 |
| ENSMUSG( | 12.9379 | 0.087189 | 0.871289 | NA | Gm30648 |
| ENSMUSG( | 60.43515 | -0.15421 | 0.601488 | 0.904603 | Gm30731 |
| ENSMUSG( | 17.01626 | 0.989765 | 0.038273 | NA | Gm30875 |
| ENSMUSG( | 8.311038 | 0.210449 | 0.769198 | NA | Gm31160 |
| ENSMUSG( | 38.04391 | -0.28564 | 0.349803 | 0.781943 | Gm31308 |
| ENSMUSG( | 55.66298 | -0.37368 | 0.152682 | 0.579011 | Gm31518 |
| ENSMUSG( | 9.032055 | -0.01369 | 0.983322 | NA | Gm31592 |
| ENSMUSG( | 127.6429 | 0.016501 | 0.936446 | 0.993415 | Gm31593 |
| ENSMUSG( | 7.991513 | -1.25138 | 0.101225 | NA | Gm31616 |
| ENSMUSG( | 12.99229 | -0.42995 | 0.459411 | NA | Gm31645 |
| ENSMUSG( | 27.35701 | 0.255798 | 0.49182 | 0.860249 | Gm31728 |
| ENSMUSG( | 23.83856 | -0.3561 | 0.388117 | 0.805915 | Gm3200 |
| ENSMUSG( | 18.26495 | -0.31455 | 0.540889 | NA | Gm32014 |
| ENSMUSG( | 76.86486 | 0.012754 | 0.954754 | 0.993874 | Gm32200 |
| ENSMUSG( | 12.73149 | 0.140111 | 0.797838 | NA | Gm3222 |
| ENSMUSG( | 14.34894 | 0.147962 | 0.762367 | NA | Gm3226 |
| ENSMUSG( | 191.0385 | 0.043826 | 0.787736 | 0.959226 | Gm32261 |
| ENSMUSG( | 70.41411 | -0.00317 | 0.989151 | 0.999111 | Gm32282 |
| ENSMUSG( | 123.1805 | -0.05165 | 0.777019 | 0.955828 | Gm32391 |
| ENSMUSG( | 121.1738 | 0.050351 | 0.805599 | 0.962616 | Gm32618 |
| ENSMUSG( | 56.97157 | 0.408351 | 0.14873 | 0.572474 | Gm32633 |
| ENSMUSG( | 28.1245 | -0.07156 | 0.843488 | 0.971847 | Gm32687 |
| ENSMUSG( | 29.71134 | 0.137045 | 0.703353 | 0.937735 | Gm32834 |
| ENSMUSG( | 44.65768 | 0.122473 | 0.681787 | 0.931619 | Gm32856 |
| ENSMUSG( | 32.18318 | 0.263046 | 0.52515 | 0.87651 | Gm32926 |
| ENSMUSG( | 24.42904 | 0.158759 | 0.68701 | 0.933041 | Gm33045 |
| ENSMUSG( | 14.6367 | -0.06527 | 0.895525 | NA | Gm33302 |
| ENSMUSG( | 72.09386 | -0.07389 | 0.774272 | 0.955606 | Gm33680 |
| ENSMUSG( | 12.8962 | -0.80462 | 0.122541 | NA | Gm33727 |
| ENSMUSG( | 23.27787 | -0.17123 | 0.668554 | 0.927864 | Gm3375 |
| ENSMUSG( | 808.4672 | -0.26144 | 0.044727 | 0.324745 | Gm33989 |
| ENSMUSG( | 9.71253 | 0.568235 | 0.355584 | NA | Gm34121 |
| ENSMUSG( | 82.92585 | -0.07687 | 0.740498 | 0.947823 | Gm34267 |
| ENSMUSG( | 58.33668 | 0.107809 | 0.666006 | 0.927057 | Gm34304 |
| ENSMUSG( | 49.79036 | -0.1026 | 0.703909 | 0.938005 | Gm34354 |
| ENSMUSG( | 24.86706 | -0.30713 | 0.426281 | 0.824898 | Gm34466 |
| ENSMUSG( | 42.15128 | 0.002737 | 0.993067 | 0.999327 | Gm34567 |
| ENSMUSG( | 16.63411 | -0.04284 | 0.923297 | NA | Gm34574 |
| ENSMUSG( | 23.58627 | -0.14775 | 0.705526 | 0.938511 | Gm34655 |
| ENSMUSG( | 21.47315 | 0.100182 | 0.801048 | 0.96188 | Gm35021 |
| ENSMUSG( | 29.62504 | 0.160641 | 0.656682 | 0.923929 | Gm35315 |
| ENSMUSG( | 15.02938 | 0.228787 | 0.641249 | NA | Gm35340 |
| ENSMUSG( | 19.40457 | -0.31994 | 0.513955 | NA | Gm35394 |
| ENSMUSG( | 12.99077 | 0.80452 | 0.134223 | NA | Gm3558 |
| ENSMUSG( | 47.95254 | 0.434 | 0.1245 | 0.531714 | Gm35612 |
| ENSMUSG( | 22.91552 | -0.26588 | 0.526552 | 0.877267 | Gm3571 |
| ENSMUSG( | 40.56099 | -0.49421 | 0.094498 | 0.471713 | Gm35853 |
| ENSMUSG( | 15.6984 | 0.644587 | 0.212485 | NA | Gm36001 |
| ENSMUSG( | 13.11639 | -0.56412 | 0.266274 | NA | Gm36012 |
| ENSMUSG( | 16.17321 | -0.30204 | 0.523409 | NA | Gm3604 |
| ENSMUSG( | 73.24566 | -0.32557 | 0.216171 | 0.661017 | Gm3625 |
| ENSMUSG( | 115.8693 | -0.0422 | 0.823236 | 0.966603 | Gm36251 |
| ENSMUSG( | 23.16071 | 0.229782 | 0.553085 | 0.886204 | Gm36266 |
| ENSMUSG( | 24.5213 | 0.275376 | 0.462011 | 0.841812 | Gm36283 |
| ENSMUSG( | 55.19678 | 0.315427 | 0.24924 | 0.698698 | Gm3636 |
| ENSMUSG( | 18.94434 | 0.26331 | 0.636146 | NA | Gm36457 |
| ENSMUSG( | 13.99403 | 0.104873 | 0.832002 | NA | Gm36529 |
| ENSMUSG( | 36.06691 | -0.09971 | 0.764492 | 0.954055 | Gm3667 |
| ENSMUSG( | 83.32746 | 0.11408 | 0.624473 | 0.91187 | Gm36742 |
| ENSMUSG( | 116.6946 | 0.087453 | 0.656046 | 0.923929 | Gm36908 |
| ENSMUSG( | 74.57623 | -0.02925 | 0.899272 | 0.985633 | Gm3693 |
| ENSMUSG( | 24.08779 | 0.368383 | 0.344206 | 0.779153 | Gm3696 |
| ENSMUSG( | 44.79296 | 0.051086 | 0.878021 | 0.981431 | Gm3739 |
| ENSMUSG( | 23.87859 | 0.038978 | 0.921158 | 0.990436 | Gm3756 |

| | | | | | |
|---|---|---|---|---|---|
| ENSMUSG( | 210.1194 | -0.03753 | 0.838236 | 0.981575 | Gm2061 |
| ENSMUSG( | 95.89062 | -0.10716 | 0.650296 | 0.95422 | Gm20646 |
| ENSMUSG( | 101.4686 | -0.29366 | 0.232162 | 0.868507 | Gm20743 |
| ENSMUSG( | 30.32659 | 0.330002 | 0.472732 | 0.923329 | Gm20751 |
| ENSMUSG( | 15.1193 | -0.68873 | 0.171577 | 0.842999 | Gm20752 |
| ENSMUSG( | 20.35592 | 0.228851 | 0.64052 | 0.953903 | Gm20753 |
| ENSMUSG( | 35.77353 | 0.058008 | 0.872145 | 0.988907 | Gm20754 |
| ENSMUSG( | 11.38445 | 0.425337 | 0.483953 | NA | Gm20878 |
| ENSMUSG( | 34.70756 | 0.092952 | 0.803127 | 0.976937 | Gm20939 |
| ENSMUSG( | 36.10013 | -0.04121 | 0.910006 | 0.991958 | Gm21092 |
| ENSMUSG( | 123.3461 | -0.09619 | 0.640109 | 0.953903 | Gm21811 |
| ENSMUSG( | 14.7711 | -0.08865 | 0.867789 | 0.98781 | Gm21814 |
| ENSMUSG( | 528.0414 | -0.06178 | 0.653202 | 0.955015 | Gm21926 |
| ENSMUSG( | 115.5686 | 0.11602 | 0.645377 | 0.953988 | Gm21992 |
| ENSMUSG( | 44.52627 | 0.253653 | 0.488025 | 0.925453 | Gm22009 |
| ENSMUSG( | 33.8214 | 0.170286 | 0.648768 | 0.95422 | Gm2237 |
| ENSMUSG( | 12.529 | -0.70898 | 0.230764 | 0.868507 | Gm22442 |
| ENSMUSG( | 74.19208 | -4.17488 | 0.000445 | 0.061321 | Gm22513 |
| ENSMUSG( | 11.32029 | 0.107539 | 0.848644 | NA | Gm22540 |
| ENSMUSG( | 10.22325 | 0.196023 | 0.774131 | NA | Gm22753 |
| ENSMUSG( | 8.548474 | -0.58042 | 0.406776 | NA | Gm23217 |
| ENSMUSG( | 10.44509 | 0.148444 | 0.800499 | NA | Gm23301 |
| ENSMUSG( | 11.64053 | 0.256526 | 0.661274 | 0.956205 | Gm23318 |
| ENSMUSG( | 11.44934 | -0.97333 | 0.10461 | NA | Gm23458 |
| ENSMUSG( | 13.39912 | -0.32586 | 0.565746 | 0.941404 | Gm24119 |
| ENSMUSG( | 7.598295 | 1.111917 | 0.18234 | NA | Gm24616 |
| ENSMUSG( | 57.1029 | -0.16619 | 0.539699 | 0.937378 | Gm2464 |
| ENSMUSG( | 46.83342 | 0.122975 | 0.708836 | 0.964042 | Gm25047 |
| ENSMUSG( | 21.15018 | -0.57907 | 0.192796 | 0.859764 | Gm25117 |
| ENSMUSG( | 13.92105 | 0.862589 | 0.156614 | 0.837724 | Gm25129 |
| ENSMUSG( | 7.612793 | 0.609968 | 0.395123 | NA | Gm25328 |
| ENSMUSG( | 11.45713 | 0.085332 | 0.880895 | 0.989644 | Gm25492 |
| ENSMUSG( | 26.43129 | -0.26357 | 0.504888 | 0.927825 | Gm25596 |
| ENSMUSG( | 16.06458 | 0.578921 | 0.297722 | 0.898538 | Gm26397 |
| ENSMUSG( | 192.1416 | -0.12598 | 0.455537 | 0.923329 | Gm266 |
| ENSMUSG( | 38.80237 | 0.127709 | 0.70842 | 0.964042 | Gm26644 |
| ENSMUSG( | 15.25711 | -0.20764 | 0.671144 | 0.95694 | Gm26671 |
| ENSMUSG( | 36.96378 | -0.37995 | 0.302385 | 0.898538 | Gm26684 |
| ENSMUSG( | 29.323 | -0.47043 | 0.198065 | 0.862161 | Gm26760 |
| ENSMUSG( | 574.6292 | -0.3136 | 0.041918 | 0.690304 | Gm26782 |
| ENSMUSG( | 13.56917 | 0.715406 | 0.208016 | 0.864412 | Gm26788 |
| ENSMUSG( | 22.28381 | 0.256138 | 0.599469 | 0.947071 | Gm26797 |
| ENSMUSG( | 102.4449 | -0.08693 | 0.712813 | 0.96439 | Gm2694 |
| ENSMUSG( | 34.00031 | 0.134446 | 0.715148 | 0.965063 | Gm27239 |
| ENSMUSG( | 17.76323 | -0.18984 | 0.683275 | 0.958678 | Gm27343 |
| ENSMUSG( | 62.28292 | 0.015548 | 0.954641 | 0.994798 | Gm2762 |
| ENSMUSG( | 11.36757 | 0.650677 | 0.266472 | NA | Gm28154 |
| ENSMUSG( | 14.42284 | 1.02369 | 0.059255 | 0.729429 | Gm28729 |
| ENSMUSG( | 34.66557 | -0.65164 | 0.08221 | 0.77681 | Gm2885 |
| ENSMUSG( | 10.12465 | -0.5016 | 0.438986 | NA | Gm2897 |
| ENSMUSG( | 13.12563 | -0.04648 | 0.940352 | 0.994334 | Gm29257 |
| ENSMUSG( | 9.846084 | -0.38001 | 0.538579 | NA | Gm29443 |
| ENSMUSG( | 12.94931 | -0.29594 | 0.596742 | 0.946541 | Gm29483 |
| ENSMUSG( | 55.85587 | -0.36966 | 0.179857 | 0.850372 | Gm29508 |
| ENSMUSG( | 83.20556 | 0.080955 | 0.755304 | 0.971129 | Gm29514 |
| ENSMUSG( | 107.731 | -0.34727 | 0.226934 | 0.868507 | Gm29595 |
| ENSMUSG( | 23.96703 | 0.264832 | 0.514696 | 0.930939 | Gm29642 |
| ENSMUSG( | 68.433 | -0.06423 | 0.827586 | 0.980224 | Gm29674 |
| ENSMUSG( | 16.4086 | -0.12424 | 0.812473 | 0.979148 | Gm29683 |
| ENSMUSG( | 14.61484 | -0.10748 | 0.841082 | 0.9826 | Gm29773 |
| ENSMUSG( | 7729.393 | -0.0711 | 0.581979 | 0.944266 | Gm2a |
| ENSMUSG( | 44.61526 | -0.40283 | 0.180113 | 0.850372 | Gm30082 |
| ENSMUSG( | 14.81576 | 0.101774 | 0.848749 | 0.985321 | Gm30092 |
| ENSMUSG( | 19.28913 | 0.100018 | 0.830015 | 0.980224 | Gm30108 |
| ENSMUSG( | 8.836523 | 0.20995 | 0.749475 | NA | Gm30400 |
| ENSMUSG( | 32.60073 | -0.4373 | 0.217942 | 0.866082 | Gm30539 |
| ENSMUSG( | 20.31358 | -3.30936 | 2.61E-08 | 2.08E-05 | Gm3055 |
| ENSMUSG( | 30.19044 | 0.459686 | 0.23738 | 0.872173 | Gm30571 |
| ENSMUSG( | 9.684103 | -1.21723 | 0.066768 | NA | Gm30606 |
| ENSMUSG( | 12.55062 | 0.247097 | 0.66515 | 0.956766 | Gm30648 |
| ENSMUSG( | 51.47885 | -0.3647 | 0.252846 | 0.88018 | Gm30731 |
| ENSMUSG( | 13.17855 | 0.638348 | 0.240071 | 0.875205 | Gm30875 |
| ENSMUSG( | 32.03351 | -0.53801 | 0.127747 | 0.81845 | Gm31308 |
| ENSMUSG( | 52.44767 | -0.25322 | 0.421884 | 0.917257 | Gm31518 |
| ENSMUSG( | 7.831854 | -0.17165 | 0.803739 | NA | Gm31592 |
| ENSMUSG( | 127.0296 | 0.255944 | 0.259762 | 0.883971 | Gm31593 |
| ENSMUSG( | 10.50505 | 0.034411 | 0.955088 | NA | Gm31616 |
| ENSMUSG( | 12.98937 | -0.10115 | 0.860482 | 0.986137 | Gm31645 |
| ENSMUSG( | 19.85577 | -0.44335 | 0.314239 | 0.900949 | Gm31728 |
| ENSMUSG( | 22.9617 | -0.2105 | 0.626287 | 0.953234 | Gm3200 |
| ENSMUSG( | 17.7877 | -0.1198 | 0.795466 | 0.97681 | Gm32014 |
| ENSMUSG( | 61.82046 | -0.34137 | 0.209149 | 0.864412 | Gm32200 |
| ENSMUSG( | 13.48403 | 0.482428 | 0.400308 | 0.917257 | Gm3222 |
| ENSMUSG( | 10.95862 | -0.38557 | 0.509238 | NA | Gm3226 |
| ENSMUSG( | 165.8447 | -0.09145 | 0.651963 | 0.9547 | Gm32261 |
| ENSMUSG( | 63.20341 | -0.05889 | 0.822295 | 0.979198 | Gm32282 |
| ENSMUSG( | 109.6177 | -0.14084 | 0.574412 | 0.942912 | Gm32391 |
| ENSMUSG( | 108.3409 | -0.00236 | 0.991555 | 0.998625 | Gm32618 |
| ENSMUSG( | 47.3033 | 0.157705 | 0.614783 | 0.951177 | Gm32633 |
| ENSMUSG( | 13.21494 | -7.09086 | 2.85E-08 | 2.17E-05 | Gm32687 |
| ENSMUSG( | 21.92727 | -0.52525 | 0.231691 | 0.868507 | Gm32834 |
| ENSMUSG( | 36.29662 | -0.18069 | 0.621129 | 0.953199 | Gm32856 |
| ENSMUSG( | 27.98836 | 0.133534 | 0.765541 | 0.974236 | Gm32926 |
| ENSMUSG( | 19.97979 | -0.18504 | 0.687936 | 0.959742 | Gm33045 |
| ENSMUSG( | 11.23042 | -0.61512 | 0.300741 | NA | Gm33302 |
| ENSMUSG( | 57.49676 | -0.51345 | 0.071536 | 0.761942 | Gm33680 |
| ENSMUSG( | 14.46868 | -0.07148 | 0.888466 | 0.990558 | Gm33727 |
| ENSMUSG( | 20.58644 | -0.31432 | 0.493926 | 0.927372 | Gm3375 |
| ENSMUSG( | 856.5087 | 0.184694 | 0.236501 | 0.872173 | Gm33989 |
| ENSMUSG( | 68.50019 | -0.39141 | 0.15654 | 0.837724 | Gm34267 |
| ENSMUSG( | 53.07925 | 0.083115 | 0.77212 | 0.974236 | Gm34304 |
| ENSMUSG( | 47.05172 | 0.012759 | 0.965298 | 0.995433 | Gm34354 |
| ENSMUSG( | 22.98183 | -0.27863 | 0.51279 | 0.930164 | Gm34466 |
| ENSMUSG( | 37.29877 | -0.08185 | 0.816483 | 0.979198 | Gm34567 |
| ENSMUSG( | 11.59248 | -0.9436 | 0.148111 | 0.836241 | Gm34574 |
| ENSMUSG( | 21.04026 | -0.22861 | 0.599249 | 0.947045 | Gm34655 |
| ENSMUSG( | 18.04254 | -0.16479 | 0.728598 | 0.966913 | Gm35021 |
| ENSMUSG( | 26.29553 | 0.128891 | 0.753787 | 0.970671 | Gm35315 |
| ENSMUSG( | 11.59712 | -0.31013 | 0.601081 | 0.947749 | Gm35340 |
| ENSMUSG( | 19.00394 | -0.07242 | 0.888011 | 0.990442 | Gm35394 |
| ENSMUSG( | 39.48432 | 0.14047 | 0.669235 | 0.956766 | Gm35612 |
| ENSMUSG( | 18.75955 | -0.5939 | 0.23661 | 0.872173 | Gm3571 |
| ENSMUSG( | 40.3835 | -0.19171 | 0.548156 | 0.938063 | Gm35853 |
| ENSMUSG( | 12.04516 | 0.152545 | 0.804847 | 0.977232 | Gm36001 |
| ENSMUSG( | 12.88957 | -0.25767 | 0.64076 | 0.953903 | Gm36012 |
| ENSMUSG( | 17.21022 | 0.140718 | 0.771348 | 0.974236 | Gm3604 |
| ENSMUSG( | 67.22976 | -0.30001 | 0.292336 | 0.897854 | Gm3625 |
| ENSMUSG( | 82.42734 | -0.92659 | 0.000241 | 0.041026 | Gm36251 |
| ENSMUSG( | 19.20121 | -0.01172 | 0.978915 | 0.996468 | Gm36266 |
| ENSMUSG( | 15.7381 | -0.83182 | 0.091728 | 0.791889 | Gm36283 |
| ENSMUSG( | 45.03483 | 0.026612 | 0.933423 | 0.99306 | Gm3636 |
| ENSMUSG( | 18.08849 | 0.38536 | 0.499295 | 0.927825 | Gm36457 |
| ENSMUSG( | 11.27951 | -0.25048 | 0.664429 | NA | Gm36529 |
| ENSMUSG( | 28.16053 | -0.58069 | 0.122064 | 0.813384 | Gm3667 |
| ENSMUSG( | 62.47698 | -0.47218 | 0.106241 | 0.804969 | Gm36742 |
| ENSMUSG( | 99.7172 | -0.12126 | 0.613824 | 0.951177 | Gm36908 |
| ENSMUSG( | 69.20478 | 0.030159 | 0.904781 | 0.991089 | Gm3693 |
| ENSMUSG( | 18.7679 | -0.08134 | 0.86013 | 0.986137 | Gm3696 |
| ENSMUSG( | 44.48667 | 0.306539 | 0.370362 | 0.915491 | Gm3739 |
| ENSMUSG( | 18.60675 | -0.44556 | 0.33592 | 0.907913 | Gm3756 |
| ENSMUSG( | 3719.303 | 0.503954 | 0.001363 | 0.134176 | Gm3764 |
| ENSMUSG( | 90.61385 | 0.126575 | 0.659885 | 0.956205 | Gm38413 |
| ENSMUSG( | 33.88033 | 0.251136 | 0.464898 | 0.923329 | Gm38576 |
| ENSMUSG( | 16.05299 | -0.60169 | 0.269996 | 0.887508 | Gm39043 |
| ENSMUSG( | 14.74787 | 0.633485 | 0.212913 | 0.864412 | Gm39090 |
| ENSMUSG( | 9.89126 | 0.167099 | 0.781168 | NA | Gm39213 |
| ENSMUSG( | 54.76021 | -0.1488 | 0.627916 | 0.953234 | Gm39468 |
| ENSMUSG( | 167.782 | -0.23977 | 0.20657 | 0.864412 | Gm39822 |
| ENSMUSG( | 13.01628 | 0.573301 | 0.305644 | 0.898538 | Gm4013 |
| ENSMUSG( | 43.67758 | -0.69856 | 0.026427 | 0.603041 | Gm40518 |
| ENSMUSG( | 17.54914 | -0.6523 | 0.183004 | 0.850741 | Gm40893 |
| ENSMUSG( | 9.385516 | -0.46768 | 0.483759 | NA | Gm41414 |
| ENSMUSG( | 8.694769 | 0.05261 | 0.937579 | NA | Gm41609 |
| ENSMUSG( | 87.61149 | 0.000752 | 0.997688 | 0.999268 | Gm4202 |
| ENSMUSG( | 159.9214 | -0.1071 | 0.601339 | 0.947749 | Gm4204 |
| ENSMUSG( | 10.66855 | -0.36203 | 0.540207 | NA | Gm42064 |
| ENSMUSG( | 97.73164 | -0.32117 | 0.166662 | 0.840183 | Gm42067 |
| ENSMUSG( | 32.12982 | 0.087011 | 0.827906 | 0.980224 | Gm4211 |
| ENSMUSG( | 114.3869 | -0.3189 | 0.115584 | 0.813384 | Gm4221 |
| ENSMUSG( | 79.30116 | -0.45996 | 0.056753 | 0.723362 | Gm4332 |
| ENSMUSG( | 11.23626 | -0.30221 | 0.614387 | NA | Gm4335 |
| ENSMUSG( | 103.48 | -0.38641 | 0.079895 | 0.776218 | Gm4366 |
| ENSMUSG( | 90.44475 | 0.190662 | 0.424561 | 0.917257 | Gm4419 |
| ENSMUSG( | 42.74707 | -0.14993 | 0.638732 | 0.953903 | Gm4544 |
| ENSMUSG( | 14.87837 | -0.37461 | 0.464935 | 0.923329 | Gm45650 |
| ENSMUSG( | 144.7573 | 0.213387 | 0.302103 | 0.898538 | Gm45871 |
| ENSMUSG( | 291.5212 | -0.05017 | 0.754877 | 0.971098 | Gm46430 |
| ENSMUSG( | 11.99636 | -0.44865 | 0.422106 | 0.917257 | Gm4654 |

| | | | | | | | | | | | |
|---|---|---|---|---|---|---|---|---|---|---|---|
| ENSMUSG( | Gm28729 | 11.34886 | 0.21573 | 0.700096 | NA | ENSMUSG( | Gm28729 | 14.42284 | 1.02369 | 0.059255 | 0.729429 |
| ENSMUSG( | Gm2885 | 42.38777 | -0.25569 | 0.376008 | 0.801127 | ENSMUSG( | Gm2885 | 34.66557 | -0.65164 | 0.08221 | 0.77681 |
| ENSMUSG( | Gm2897 | 11.46949 | -0.3557 | 0.535295 | NA | ENSMUSG( | Gm2897 | 10.12465 | -0.5016 | 0.438986 | NA |
| ENSMUSG( | Gm29257 | 13.54382 | -0.15177 | 0.771006 | NA | ENSMUSG( | Gm29257 | 13.12563 | -0.04648 | 0.940352 | 0.994334 |
| ENSMUSG( | Gm29443 | 12.2403 | -0.01383 | 0.978812 | NA | ENSMUSG( | Gm29443 | 9.846084 | -0.38001 | 0.538579 | NA |
| ENSMUSG( | Gm29483 | 18.63752 | 0.478541 | 0.26649 | NA | ENSMUSG( | Gm29483 | 12.94931 | -0.29594 | 0.596742 | 0.946541 |
| ENSMUSG( | Gm29508 | 71.45719 | 0.095128 | 0.691069 | 0.933881 | ENSMUSG( | Gm29508 | 55.85587 | -0.36966 | 0.179857 | 0.850372 |
| ENSMUSG( | Gm29514 | 96.54558 | 0.237576 | 0.358212 | 0.78975 | ENSMUSG( | Gm29514 | 83.20556 | 0.080955 | 0.755304 | 0.971129 |
| ENSMUSG( | Gm29595 | 129.5274 | -0.0419 | 0.848685 | 0.972374 | ENSMUSG( | Gm29595 | 107.731 | -0.34727 | 0.226934 | 0.868507 |
| ENSMUSG( | Gm29642 | 23.01597 | -0.09207 | 0.830059 | 0.96974 | ENSMUSG( | Gm29642 | 23.96703 | 0.264832 | 0.514696 | 0.930939 |
| ENSMUSG( | Gm29674 | 80.89339 | 0.132951 | 0.573263 | 0.893584 | ENSMUSG( | Gm29674 | 68.433 | -0.06423 | 0.827586 | 0.980224 |
| ENSMUSG( | Gm29683 | 21.71278 | 0.435147 | 0.400239 | 0.813641 | ENSMUSG( | Gm29683 | 16.4086 | -0.12424 | 0.812473 | 0.979148 |
| ENSMUSG( | Gm29773 | 15.08693 | -0.34502 | 0.491043 | NA | ENSMUSG( | Gm29773 | 14.61484 | -0.10748 | 0.841082 | 0.9826 |
| ENSMUSG( | Gm2a | 8186.258 | -0.17953 | 0.112792 | 0.506698 | ENSMUSG( | Gm2a | 7729.393 | -0.0711 | 0.581979 | 0.944266 |
| ENSMUSG( | Gm30082 | 57.35977 | 0.092177 | 0.729382 | 0.944738 | ENSMUSG( | Gm30082 | 44.61526 | -0.40283 | 0.180113 | 0.850372 |
| ENSMUSG( | Gm30092 | 14.95931 | -0.10815 | 0.829408 | NA | ENSMUSG( | Gm30092 | 14.81576 | 0.101774 | 0.848749 | 0.985321 |
| ENSMUSG( | Gm30108 | 22.01229 | 0.182147 | 0.651956 | 0.921946 | ENSMUSG( | Gm30108 | 19.28913 | 0.100018 | 0.830015 | 0.980224 |
| ENSMUSG( | Gm30400 | 9.365774 | 0.100328 | 0.872333 | NA | ENSMUSG( | Gm30400 | 8.836523 | 0.20995 | 0.749475 | NA |
| ENSMUSG( | Gm30539 | 39.99044 | -0.08999 | 0.762247 | 0.953459 | ENSMUSG( | Gm30539 | 32.60073 | -0.4373 | 0.217942 | 0.866082 |
| ENSMUSG( | Gm3055 | 38.05929 | -0.19305 | 0.561276 | 0.889052 | ENSMUSG( | Gm3055 | 20.31358 | -3.30936 | 2.61E-08 | 2.08E-05 |
| ENSMUSG( | Gm30571 | 29.41084 | 0.10852 | 0.75728 | 0.951513 | ENSMUSG( | Gm30571 | 30.19044 | 0.459686 | 0.23738 | 0.872173 |
| ENSMUSG( | Gm30606 | 13.7734 | -0.17241 | 0.730837 | NA | ENSMUSG( | Gm30606 | 9.684103 | -1.21723 | 0.066768 | NA |
| ENSMUSG( | Gm30648 | 12.9379 | 0.087189 | 0.871289 | NA | ENSMUSG( | Gm30648 | 12.55062 | 0.247097 | 0.66515 | 0.956766 |
| ENSMUSG( | Gm30731 | 60.43515 | -0.15421 | 0.601488 | 0.904603 | ENSMUSG( | Gm30731 | 51.47885 | -0.3647 | 0.252846 | 0.88018 |
| ENSMUSG( | Gm30875 | 17.01626 | 0.989765 | 0.038273 | NA | ENSMUSG( | Gm30875 | 13.17855 | 0.638348 | 0.240071 | 0.875205 |
| ENSMUSG( | Gm31308 | 38.04391 | -0.28564 | 0.349803 | 0.781943 | ENSMUSG( | Gm31308 | 32.03351 | -0.53801 | 0.127747 | 0.81845 |
| ENSMUSG( | Gm31518 | 55.66298 | -0.37368 | 0.152682 | 0.579011 | ENSMUSG( | Gm31518 | 52.44767 | -0.25322 | 0.421884 | 0.917257 |
| ENSMUSG( | Gm31592 | 9.032055 | -0.01369 | 0.983322 | NA | ENSMUSG( | Gm31592 | 7.831854 | -0.17165 | 0.803739 | NA |
| ENSMUSG( | Gm31593 | 127.6429 | 0.016501 | 0.936446 | 0.993415 | ENSMUSG( | Gm31593 | 127.0296 | 0.255944 | 0.259762 | 0.883971 |
| ENSMUSG( | Gm31616 | 7.991513 | -1.25138 | 0.101225 | NA | ENSMUSG( | Gm31616 | 10.50505 | 0.034411 | 0.955088 | NA |
| ENSMUSG( | Gm31645 | 12.99229 | -0.42995 | 0.459411 | NA | ENSMUSG( | Gm31645 | 12.98937 | -0.10115 | 0.860482 | 0.986137 |
| ENSMUSG( | Gm31728 | 27.35701 | 0.255798 | 0.49182 | 0.860249 | ENSMUSG( | Gm31728 | 19.85577 | -0.44335 | 0.314239 | 0.900949 |
| ENSMUSG( | Gm3200 | 23.83856 | -0.3561 | 0.388117 | 0.805915 | ENSMUSG( | Gm3200 | 22.9617 | -0.2105 | 0.626287 | 0.953234 |
| ENSMUSG( | Gm32014 | 18.26495 | -0.31455 | 0.540889 | NA | ENSMUSG( | Gm32014 | 17.7877 | -0.1198 | 0.795466 | 0.97681 |
| ENSMUSG( | Gm32200 | 76.86486 | 0.012754 | 0.954754 | 0.993874 | ENSMUSG( | Gm32200 | 61.82046 | -0.34137 | 0.209149 | 0.864412 |
| ENSMUSG( | Gm3222 | 12.73149 | 0.140111 | 0.797838 | NA | ENSMUSG( | Gm3222 | 13.48403 | 0.482428 | 0.400308 | 0.917257 |
| ENSMUSG( | Gm3226 | 14.34894 | 0.147962 | 0.762367 | NA | ENSMUSG( | Gm3226 | 10.95862 | -0.38557 | 0.509238 | NA |
| ENSMUSG( | Gm32261 | 191.0385 | 0.043826 | 0.787736 | 0.959226 | ENSMUSG( | Gm32261 | 165.8447 | -0.09145 | 0.651963 | 0.9547 |
| ENSMUSG( | Gm32282 | 70.41411 | -0.00317 | 0.989151 | 0.999111 | ENSMUSG( | Gm32282 | 63.20341 | -0.05889 | 0.822295 | 0.979198 |
| ENSMUSG( | Gm32391 | 123.1805 | -0.05165 | 0.777019 | 0.955828 | ENSMUSG( | Gm32391 | 109.6177 | -0.14084 | 0.574412 | 0.942912 |
| ENSMUSG( | Gm32618 | 121.1738 | 0.050351 | 0.805599 | 0.962616 | ENSMUSG( | Gm32618 | 108.3409 | -0.00236 | 0.991555 | 0.998625 |
| ENSMUSG( | Gm32633 | 56.97157 | 0.408351 | 0.14873 | 0.572474 | ENSMUSG( | Gm32633 | 47.3033 | 0.157705 | 0.614783 | 0.951177 |
| ENSMUSG( | Gm32687 | 28.1245 | -0.07156 | 0.843488 | 0.971847 | ENSMUSG( | Gm32687 | 13.21494 | -7.09086 | 2.85E-08 | 2.17E-05 |
| ENSMUSG( | Gm32834 | 29.71134 | 0.137045 | 0.703353 | 0.937735 | ENSMUSG( | Gm32834 | 21.92727 | -0.52525 | 0.231691 | 0.868507 |
| ENSMUSG( | Gm32856 | 44.65768 | 0.122473 | 0.681787 | 0.931619 | ENSMUSG( | Gm32856 | 36.29662 | -0.18069 | 0.621129 | 0.953199 |
| ENSMUSG( | Gm32926 | 32.18318 | 0.263046 | 0.52515 | 0.87651 | ENSMUSG( | Gm32926 | 27.98836 | 0.133534 | 0.765541 | 0.974236 |
| ENSMUSG( | Gm33045 | 24.42904 | 0.158759 | 0.68701 | 0.933041 | ENSMUSG( | Gm33045 | 19.97979 | -0.18504 | 0.687936 | 0.959742 |
| ENSMUSG( | Gm33302 | 14.6367 | -0.06527 | 0.895525 | NA | ENSMUSG( | Gm33302 | 11.23042 | -0.61512 | 0.300741 | NA |
| ENSMUSG( | Gm33680 | 72.09386 | -0.07389 | 0.774272 | 0.955606 | ENSMUSG( | Gm33680 | 57.49676 | -0.51345 | 0.071536 | 0.761942 |
| ENSMUSG( | Gm33727 | 12.8962 | -0.80462 | 0.122541 | NA | ENSMUSG( | Gm33727 | 14.46868 | -0.07148 | 0.888466 | 0.990558 |
| ENSMUSG( | Gm3375 | 23.27787 | -0.17123 | 0.668554 | 0.927864 | ENSMUSG( | Gm3375 | 20.58644 | -0.31432 | 0.493926 | 0.927372 |
| ENSMUSG( | Gm33989 | 808.4672 | -0.26144 | 0.044727 | 0.324745 | ENSMUSG( | Gm33989 | 856.5087 | 0.184694 | 0.236501 | 0.872173 |
| ENSMUSG( | Gm34267 | 82.92585 | -0.07687 | 0.740498 | 0.947823 | ENSMUSG( | Gm34267 | 68.50019 | -0.39141 | 0.15654 | 0.837724 |
| ENSMUSG( | Gm34304 | 58.33668 | 0.107809 | 0.666006 | 0.927057 | ENSMUSG( | Gm34304 | 53.07925 | 0.083115 | 0.77212 | 0.974236 |
| ENSMUSG( | Gm34354 | 49.79036 | -0.1026 | 0.703909 | 0.938005 | ENSMUSG( | Gm34354 | 47.05172 | 0.012759 | 0.965298 | 0.995433 |
| ENSMUSG( | Gm34466 | 24.86706 | -0.30713 | 0.426281 | 0.824898 | ENSMUSG( | Gm34466 | 22.98183 | -0.27863 | 0.51279 | 0.930164 |
| ENSMUSG( | Gm34567 | 42.15128 | 0.002737 | 0.993067 | 0.999327 | ENSMUSG( | Gm34567 | 37.29877 | -0.08185 | 0.816483 | 0.979198 |
| ENSMUSG( | Gm34574 | 16.63411 | -0.04284 | 0.923297 | NA | ENSMUSG( | Gm34574 | 11.59248 | -0.9436 | 0.148111 | 0.836241 |
| ENSMUSG( | Gm34655 | 23.58627 | -0.14775 | 0.705526 | 0.938511 | ENSMUSG( | Gm34655 | 21.04026 | -0.22861 | 0.599249 | 0.947045 |
| ENSMUSG( | Gm35021 | 21.47315 | 0.100182 | 0.801048 | 0.96188 | ENSMUSG( | Gm35021 | 18.04254 | -0.16479 | 0.728598 | 0.966913 |
| ENSMUSG( | Gm35315 | 29.62504 | 0.160641 | 0.656682 | 0.923929 | ENSMUSG( | Gm35315 | 26.29553 | 0.128891 | 0.753787 | 0.970671 |
| ENSMUSG( | Gm35340 | 15.02938 | 0.228787 | 0.641249 | NA | ENSMUSG( | Gm35340 | 11.59712 | -0.31013 | 0.601081 | 0.947749 |
| ENSMUSG( | Gm35394 | 19.40457 | -0.31994 | 0.513955 | NA | ENSMUSG( | Gm35394 | 19.00394 | -0.07242 | 0.888011 | 0.990442 |
| ENSMUSG( | Gm35612 | 47.95254 | 0.434 | 0.1245 | 0.531714 | ENSMUSG( | Gm35612 | 39.48432 | 0.14047 | 0.669235 | 0.956766 |
| ENSMUSG( | Gm3571 | 22.91552 | -0.26588 | 0.526552 | 0.877267 | ENSMUSG( | Gm3571 | 18.75955 | -0.5939 | 0.23661 | 0.872173 |
| ENSMUSG( | Gm35853 | 40.56099 | -0.49421 | 0.094498 | 0.471713 | ENSMUSG( | Gm35853 | 40.3835 | -0.19171 | 0.548156 | 0.938063 |
| ENSMUSG( | Gm36001 | 15.6984 | 0.644587 | 0.212485 | NA | ENSMUSG( | Gm36001 | 12.04516 | 0.152545 | 0.804847 | 0.977232 |
| ENSMUSG( | Gm36012 | 13.11639 | -0.56412 | 0.266274 | NA | ENSMUSG( | Gm36012 | 12.88957 | -0.25767 | 0.64076 | 0.953903 |
| ENSMUSG( | Gm3604 | 16.17321 | -0.30204 | 0.523409 | NA | ENSMUSG( | Gm3604 | 17.21022 | 0.140718 | 0.771348 | 0.974236 |
| ENSMUSG( | Gm3625 | 73.24566 | -0.32557 | 0.216171 | 0.661017 | ENSMUSG( | Gm3625 | 67.22976 | -0.30001 | 0.292336 | 0.897854 |
| ENSMUSG( | Gm36251 | 115.8693 | -0.0422 | 0.823236 | 0.966603 | ENSMUSG( | Gm36251 | 82.42734 | -0.92659 | 0.000241 | 0.041026 |
| ENSMUSG( | Gm36266 | 23.16071 | 0.229782 | 0.553085 | 0.886204 | ENSMUSG( | Gm36266 | 19.20121 | -0.01172 | 0.978915 | 0.996468 |
| ENSMUSG( | Gm36283 | 24.5213 | 0.275376 | 0.462011 | 0.841812 | ENSMUSG( | Gm36283 | 15.7381 | -0.83182 | 0.091728 | 0.791889 |
| ENSMUSG( | Gm3636 | 55.19678 | 0.315427 | 0.24924 | 0.698698 | ENSMUSG( | Gm3636 | 45.03483 | 0.026612 | 0.933423 | 0.99306 |
| ENSMUSG( | Gm36457 | 18.94434 | 0.26331 | 0.636146 | NA | ENSMUSG( | Gm36457 | 18.08849 | 0.38536 | 0.499295 | 0.927825 |
| ENSMUSG( | Gm36529 | 13.99403 | 0.104873 | 0.832002 | NA | ENSMUSG( | Gm36529 | 11.27951 | -0.25048 | 0.664429 | NA |
| ENSMUSG( | Gm3667 | 36.06691 | -0.09971 | 0.764492 | 0.954055 | ENSMUSG( | Gm3667 | 28.16053 | -0.58069 | 0.122064 | 0.813384 |
| ENSMUSG( | Gm36742 | 83.32746 | 0.11408 | 0.624473 | 0.91187 | ENSMUSG( | Gm36742 | 62.47698 | -0.47218 | 0.106241 | 0.804969 |
| ENSMUSG( | Gm36908 | 116.6946 | 0.087453 | 0.656046 | 0.923929 | ENSMUSG( | Gm36908 | 99.7172 | -0.12126 | 0.613824 | 0.951177 |
| ENSMUSG( | Gm3693 | 74.57623 | -0.02925 | 0.899272 | 0.985633 | ENSMUSG( | Gm3693 | 69.20478 | 0.030159 | 0.904781 | 0.991089 |
| ENSMUSG( | Gm3696 | 24.08779 | 0.368383 | 0.344206 | 0.779153 | ENSMUSG( | Gm3696 | 18.7679 | -0.08134 | 0.86013 | 0.986137 |
| ENSMUSG( | Gm3739 | 44.79296 | 0.051086 | 0.878021 | 0.981431 | ENSMUSG( | Gm3739 | 44.48667 | 0.306539 | 0.370362 | 0.915491 |
| ENSMUSG( | Gm3756 | 23.87859 | 0.038978 | 0.921158 | 0.990436 | ENSMUSG( | Gm3756 | 18.60675 | -0.44556 | 0.33592 | 0.907913 |
| ENSMUSG( | Gm3764 | 3646.236 | 0.210278 | 0.088356 | 0.455976 | ENSMUSG( | Gm3764 | 3719.303 | 0.503954 | 0.001363 | 0.134176 |
| ENSMUSG( | Gm38413 | 95.05765 | 0.007995 | 0.975881 | 0.997428 | ENSMUSG( | Gm38413 | 90.61385 | 0.126575 | 0.659885 | 0.956205 |
| ENSMUSG( | Gm38576 | 29.35746 | -0.44584 | 0.199396 | 0.640054 | ENSMUSG( | Gm38576 | 33.88033 | 0.251136 | 0.464898 | 0.923329 |
| ENSMUSG( | Gm39043 | 14.45359 | -1.39382 | 0.016446 | NA | ENSMUSG( | Gm39043 | 16.05299 | -0.60169 | 0.269996 | 0.887508 |
| ENSMUSG( | Gm39090 | 16.84586 | 0.710702 | 0.11797 | NA | ENSMUSG( | Gm39090 | 14.74787 | 0.633485 | 0.212913 | 0.864412 |
| ENSMUSG( | Gm39213 | 10.43368 | 0.039648 | 0.94289 | NA | ENSMUSG( | Gm39213 | 9.89126 | 0.167099 | 0.781168 | NA |
| ENSMUSG( | Gm39468 | 66.00264 | 0.116954 | 0.669997 | 0.928357 | ENSMUSG( | Gm39468 | 54.76021 | -0.1488 | 0.627916 | 0.953234 |
| ENSMUSG( | Gm39822 | 200.3385 | 0.027051 | 0.861224 | 0.975026 | ENSMUSG( | Gm39822 | 167.782 | -0.23977 | 0.20657 | 0.864412 |
| ENSMUSG( | Gm4013 | 11.68524 | 0.080191 | 0.885831 | NA | ENSMUSG( | Gm4013 | 13.01628 | 0.573301 | 0.305644 | 0.898538 |
| ENSMUSG( | Gm40518 | 55.85976 | -0.18293 | 0.49973 | 0.863218 | ENSMUSG( | Gm40518 | 43.67758 | -0.69856 | 0.026427 | 0.603041 |
| ENSMUSG( | Gm40893 | 27.66877 | 0.425148 | 0.2788 | 0.728705 | ENSMUSG( | Gm40893 | 17.54914 | -0.6523 | 0.183004 | 0.850741 |
| ENSMUSG( | Gm41414 | 10.17062 | -0.54458 | 0.386206 | NA | ENSMUSG( | Gm41414 | 9.385516 | -0.46768 | 0.483759 | NA |
| ENSMUSG( | Gm41609 | 9.367258 | -0.03917 | 0.948435 | NA | ENSMUSG( | Gm41609 | 8.694769 | 0.05261 | 0.937579 | NA |
| ENSMUSG( | Gm4202 | 100.9915 | 0.154435 | 0.550148 | 0.886038 | ENSMUSG( | Gm4202 | 87.61149 | 0.000752 | 0.997688 | 0.999268 |
| ENSMUSG( | Gm4204 | 180.1001 | -0.01484 | 0.925617 | 0.991204 | ENSMUSG( | Gm4204 | 159.9214 | -0.1071 | 0.601339 | 0.947749 |
| ENSMUSG( | Gm42064 | 13.55603 | 0.053032 | 0.916397 | NA | ENSMUSG( | Gm42064 | 10.66855 | -0.36203 | 0.540207 | NA |
| ENSMUSG( | Gm42067 | 116.9517 | -0.04286 | 0.820647 | 0.965888 | ENSMUSG( | Gm42067 | 97.73164 | -0.32117 | 0.166662 | 0.840183 |
| ENSMUSG( | Gm4211 | 36.22259 | 0.126329 | 0.704877 | 0.938303 | ENSMUSG( | Gm4211 | 32.12982 | 0.087011 | 0.827906 | 0.980224 |
| ENSMUSG( | Gm4221 | 131.0387 | -0.1806 | 0.315032 | 0.762205 | ENSMUSG( | Gm4221 | 114.3869 | -0.3189 | 0.115584 | 0.813384 |
| ENSMUSG( | Gm4332 | 108.5126 | 0.215533 | 0.269416 | 0.719729 | ENSMUSG( | Gm4332 | 79.30116 | -0.45996 | 0.056753 | 0.723362 |
| ENSMUSG( | Gm4335 | 14.94535 | 0.307457 | 0.527701 | NA | ENSMUSG( | Gm4335 | 11.23626 | -0.30221 | 0.614387 | NA |
| ENSMUSG( | Gm4366 | 139.6555 | 0.240178 | 0.162506 | 0.593257 | ENSMUSG( | Gm4366 | 103.48 | -0.38641 | 0.079895 | 0.776218 |
| ENSMUSG( | Gm4419 | 90.64528 | -0.0463 | 0.822573 | 0.966352 | ENSMUSG( | Gm4419 | 90.44475 | 0.190662 | 0.424561 | 0.917257 |
| ENSMUSG( | Gm4544 | 52.79474 | 0.189352 | 0.478166 | 0.853453 | ENSMUSG( | Gm4544 | 42.74707 | -0.14993 | 0.638732 | 0.953903 |
| ENSMUSG( | Gm45650 | 19.69 | 0.222714 | 0.595977 | NA | ENSMUSG( | Gm45650 | 14.87837 | -0.37461 | 0.464935 | 0.923329 |
| ENSMUSG( | Gm45871 | 155.181 | 0.131858 | 0.468322 | 0.845336 | ENSMUSG( | Gm45871 | 144.7573 | 0.213387 | 0.302103 | 0.898538 |
| ENSMUSG( | Gm46430 | 298.2005 | -0.2701 | 0.075051 | 0.42422 | ENSMUSG( | Gm46430 | 291.5212 | -0.05017 | 0.754877 | 0.971098 |
| ENSMUSG( | Gm4654 | 15.12923 | -0.0025 | 0.995768 | NA | ENSMUSG( | Gm4654 | 11.99636 | -0.44865 | 0.422106 | 0.917257 |
| ENSMUSG( | Gm46637 | 46.40848 | 0.100393 | 0.748498 | 0.949604 | ENSMUSG( | Gm46637 | 36.69105 | -0.33143 | 0.311034 | 0.899422 |
| ENSMUSG( | Gm4691 | 12.47243 | 0.189341 | 0.718636 | NA | ENSMUSG( | Gm4691 | 7.038678 | -1.74665 | 0.030371 | NA |
| ENSMUSG( | Gm47113 | 31.26713 | 0.393329 | 0.302017 | 0.750166 | ENSMUSG( | Gm47113 | 20.30662 | -0.63383 | 0.174642 | 0.844619 |
| ENSMUSG( | Gm4724 | 16.12258 | 0.154515 | 0.751864 | NA | ENSMUSG( | Gm4724 | 13.41201 | -0.10056 | 0.849877 | 0.985523 |
| ENSMUSG( | Gm47415 | 18.09875 | 0.011495 | 0.980072 | NA | ENSMUSG( | Gm47415 | 18.73754 | 0.345231 | 0.474095 | 0.923329 |
| ENSMUSG( | Gm4787 | 27.41446 | -0.02772 | 0.944066 | 0.993503 | ENSMUSG( | Gm4787 | 24.21248 | -0.10993 | 0.784178 | 0.976154 |
| ENSMUSG( | Gm4804 | 14.97446 | -0.29785 | 0.543521 | NA | ENSMUSG( | Gm4804 | 12.73516 | -0.51287 | 0.34036 | 0.907972 |
| ENSMUSG( | Gm4829 | 13.14261 | -0.72945 | 0.18074 | NA | ENSMUSG( | Gm4829 | 13.71439 | -0.22924 | 0.683739 | 0.958678 |
| ENSMUSG( | Gm4833 | 23.34977 | 0.327928 | 0.38614 | 0.804623 | ENSMUSG( | Gm4833 | 15.56507 | -0.63251 | 0.197803 | 0.862161 |
| ENSMUSG( | Gm48551 | 23.02305 | -0.22262 | 0.570799 | 0.8933 | ENSMUSG( | Gm48551 | 22.00975 | -0.06989 | 0.869407 | 0.988097 |
| ENSMUSG( | Gm48552 | 10.45888 | -0.28929 | 0.635722 | NA | ENSMUSG( | Gm48552 | 11.05073 | 0.176647 | 0.781267 | NA |
| ENSMUSG( | Gm4876 | 153.8166 | -0.05396 | 0.753335 | 0.950753 | ENSMUSG( | Gm4876 | 144.5936 | 0.02438 | 0.901212 | 0.990558 |
| ENSMUSG( | Gm4924 | 108.1223 | -0.14193 | 0.468105 | 0.845319 | ENSMUSG( | Gm4924 | 126.067 | 0.537424 | 0.010618 | 0.419324 |
| ENSMUSG( | Gm4949 | 210.2662 | 0.360951 | 0.018666 | 0.205542 | ENSMUSG( | Gm4949 | 147.0417 | -0.41705 | 0.043359 | 0.690304 |
| ENSMUSG( | Gm50340 | 17.64874 | 0.447239 | 0.403393 | NA | ENSMUSG( | Gm50340 | 12.5658 | -0.23734 | 0.671355 | 0.95694 |
| ENSMUSG( | Gm5083 | 77.23006 | 0.202269 | 0.429184 | 0.826446 | ENSMUSG( | Gm5083 | 62.81304 | -0.15004 | 0.626699 | 0.953234 |
| ENSMUSG( | Gm5086 | 38.76636 | 0.034042 | 0.919554 | 0.990247 | ENSMUSG( | Gm5086 | 33.00078 | -0.15131 | 0.674785 | 0.95694 |
| ENSMUSG( | Gm5093 | 22.56469 | 0.654358 | 0.093056 | 0.468892 | ENSMUSG( | Gm5093 | 18.54798 | 0.377301 | 0.408167 | 0.917257 |
| ENSMUSG( | Gm5105 | 11.37513 | -0.4304 | 0.431489 | NA | ENSMUSG( | Gm5105 | 9.431492 | -0.73195 | 0.251454 | NA |
| ENSMUSG( | Gm5113 | 873.1834 | 0.019501 | 0.852696 | 0.973168 | ENSMUSG( | Gm5113 | 809.694 | 0.069284 | 0.573385 | 0.942912 |
| ENSMUSG( | Gm5141 | 55.56545 | -0.03863 | 0.892111 | 0.984156 | ENSMUSG( | Gm5141 | 50.65755 | -0.02263 | 0.940812 | 0.994558 |
| ENSMUSG( | Gm51425 | 60.45072 | -0.18413 | 0.471621 | 0.847331 | ENSMUSG( | Gm51425 | 66.89158 | 0.362314 | 0.152397 | 0.837724 |
| ENSMUSG( | Gm5148 | 74.97071 | 0.146445 | 0.531835 | 0.879628 | ENSMUSG( | Gm5148 | 68.46639 | 0.158186 | 0.555034 | 0.939565 |
| ENSMUSG( | Gm5165 | 73.55672 | 0.102199 | 0.683398 | 0.932153 | ENSMUSG( | Gm5165 | 75.34368 | 0.390368 | 0.156974 | 0.837724 |
| ENSMUSG( | Gm5174 | 13.69345 | -0.30753 | 0.568426 | NA | ENSMUSG( | Gm5174 | 15.94977 | 0.361254 | 0.481362 | 0.924564 |
| ENSMUSG( | Gm5176 | 16.00481 | -0.59061 | 0.220887 | NA | ENSMUSG( | Gm5176 | 13.79775 | -0.785 | 0.149559 | 0.836241 |
| ENSMUSG( | Gm5242 | 50.58107 | 0.26397 | 0.353558 | 0.785899 | ENSMUSG( | Gm5242 | 38.83664 | -0.19586 | 0.56127 | 0.940254 |
| ENSMUSG( | Gm527 | 88.0947 | -0.19565 | 0.346262 | 0.779153 | ENSMUSG( | Gm527 | 82.00904 | -0.1287 | 0.586882 | 0.944975 |
| ENSMUSG( | Gm53058 | 34.84099 | -0.48382 | 0.134992 | 0.548629 | ENSMUSG( | Gm53058 | 36.53728 | -0.04739 | 0.887476 | 0.990266 |
| ENSMUSG( | Gm5321 | 13.57086 | 0.12047 | 0.806668 | NA | ENSMUSG( | Gm5321 | 10.54373 | -0.36405 | 0.541286 | NA |
| ENSMUSG( | Gm5345 | 44.66882 | -0.24501 | 0.388159 | 0.805915 | ENSMUSG( | Gm5345 | 38.76842 | -0.45319 | 0.203567 | 0.864412 |
| ENSMUSG( | Gm536 | 9.736482 | -0.24807 | 0.697214 | NA | ENSMUSG( | Gm536 | 9.017462 | -0.17932 | 0.778652 | NA |
| ENSMUSG( | Gm5421 | 26.30108 | 0.228445 | 0.553329 | 0.886216 | ENSMUSG( | Gm5421 | 20.43913 | -0.23267 | 0.598231 | 0.946676 |
| ENSMUSG( | Gm5422 | 12.54502 | 0.772127 | 0.15096 | NA | ENSMUSG( | Gm5422 | 7.814647 | -0.16087 | 0.821951 | NA |
| ENSMUSG( | Gm5425 | 10.14439 | -0.52216 | 0.390255 | NA | ENSMUSG( | Gm5425 | 8.720149 | -0.79624 | 0.241018 | NA |
| ENSMUSG( | Gm5431 | 12.37878 | -0.50424 | 0.358526 | NA | ENSMUSG( | Gm5431 | 12.07097 | -0.24708 | 0.674212 | 0.95694 |
| ENSMUSG( | Gm5436 | 13.77092 | 0.069932 | 0.890352 | NA | ENSMUSG( | Gm5436 | 10.64274 | -0.45792 | 0.454465 | NA |
| ENSMUSG( | Gm5451 | 51.20867 | 0.076121 | 0.813166 | 0.964574 | ENSMUSG( | Gm5451 | 39.09433 | -0.51635 | 0.171576 | 0.842999 |
| ENSMUSG( | Gm5525 | 14.22259 | -0.2556 | 0.613956 | NA | ENSMUSG( | Gm5525 | 11.24127 | -0.80563 | 0.180249 | NA |
| ENSMUSG( | Gm5535 | 29.48924 | -0.33886 | 0.328517 | 0.773036 | ENSMUSG( | Gm5535 | 29.1196 | -0.06541 | 0.861613 | 0.986298 |
| ENSMUSG( | Gm5577 | 154.4191 | 0.057834 | 0.762865 | 0.953524 | ENSMUSG( | Gm5577 | 138.0918 | 0.014689 | 0.952601 | 0.994798 |
| ENSMUSG( | Gm5586 | 25.57975 | -0.235 | 0.603581 | 0.904887 | ENSMUSG( | Gm5586 | 18.12208 | -1.27763 | 0.025676 | 0.599234 |
| ENSMUSG( | Gm5617 | 106.8822 | 0.163787 | 0.414314 | 0.819998 | ENSMUSG( | Gm5617 | 88.75815 | -0.123 | 0.611368 | 0.950771 |
| ENSMUSG( | Gm5620 | 16.05085 | 0.27526 | 0.556228 | NA | ENSMUSG( | Gm5620 | 12.53356 | -0.19201 | 0.735111 | 0.967344 |
| ENSMUSG( | Gm5628 | 66.03969 | -0.29373 | 0.24421 | 0.691435 | ENSMUSG( | Gm5628 | 56.94379 | -0.48819 | 0.090147 | 0.790741 |
| ENSMUSG( | Gm5778 | 15.15062 | 0.609629 | 0.26502 | NA | ENSMUSG( | Gm5778 | 11.53501 | 0.11276 | 0.845821 | 0.984272 |
| ENSMUSG( | Gm5781 | 20.5084 | 0.07326 | 0.863191 | NA | ENSMUSG( | Gm5781 | 12.79972 | -1.26434 | 0.024173 | 0.588924 |

| | | | | | |
|---|---|---|---|---|---|
| ENSMUSG( | 3646.236 | 0.210278 | 0.088356 | 0.455976 | Gm3764 |
| ENSMUSG( | 95.05765 | 0.007995 | 0.975881 | 0.997428 | Gm38413 |
| ENSMUSG( | 29.35746 | -0.44584 | 0.199396 | 0.640054 | Gm38576 |
| ENSMUSG( | 14.45359 | -1.39382 | 0.016446 | NA | Gm39043 |
| ENSMUSG( | 16.84586 | 0.710702 | 0.11797 | NA | Gm39090 |
| ENSMUSG( | 10.43368 | 0.039648 | 0.94289 | NA | Gm39213 |
| ENSMUSG( | 66.00264 | 0.116954 | 0.669997 | 0.928357 | Gm39468 |
| ENSMUSG( | 200.3385 | 0.027051 | 0.861224 | 0.975026 | Gm39822 |
| ENSMUSG( | 11.68524 | 0.080191 | 0.885831 | NA | Gm4013 |
| ENSMUSG( | 55.85976 | -0.18293 | 0.49973 | 0.863218 | Gm40518 |
| ENSMUSG( | 27.66877 | 0.425148 | 0.2788 | 0.728705 | Gm40893 |
| ENSMUSG( | 10.17062 | -0.54458 | 0.386206 | NA | Gm41414 |
| ENSMUSG( | 9.367258 | -0.03917 | 0.948435 | NA | Gm41609 |
| ENSMUSG( | 100.9915 | 0.154435 | 0.550148 | 0.886038 | Gm4202 |
| ENSMUSG( | 180.1001 | -0.01484 | 0.925617 | 0.991204 | Gm4204 |
| ENSMUSG( | 13.55603 | 0.053032 | 0.916397 | NA | Gm42064 |
| ENSMUSG( | 116.9517 | -0.04286 | 0.820647 | 0.965888 | Gm42067 |
| ENSMUSG( | 36.22259 | 0.126329 | 0.704877 | 0.938303 | Gm4211 |
| ENSMUSG( | 131.0387 | -0.1806 | 0.315032 | 0.762205 | Gm4221 |
| ENSMUSG( | 108.5126 | 0.215533 | 0.269416 | 0.719729 | Gm4332 |
| ENSMUSG( | 14.94535 | 0.307457 | 0.527701 | NA | Gm4335 |
| ENSMUSG( | 139.6555 | 0.240178 | 0.162506 | 0.593257 | Gm4366 |
| ENSMUSG( | 90.64528 | -0.0463 | 0.822573 | 0.966352 | Gm4419 |
| ENSMUSG( | 8.668074 | 0.813362 | 0.225681 | NA | Gm4521 |
| ENSMUSG( | 52.79474 | 0.189352 | 0.478166 | 0.853453 | Gm4544 |
| ENSMUSG( | 19.69 | 0.222714 | 0.595977 | NA | Gm45650 |
| ENSMUSG( | 155.181 | 0.131858 | 0.468322 | 0.845336 | Gm45871 |
| ENSMUSG( | 298.2005 | -0.2701 | 0.075051 | 0.42422 | Gm46430 |
| ENSMUSG( | 15.12923 | -0.0025 | 0.995768 | NA | Gm4654 |
| ENSMUSG( | 46.40848 | 0.100393 | 0.748498 | 0.949604 | Gm46637 |
| ENSMUSG( | 12.47243 | 0.189341 | 0.718636 | NA | Gm4691 |
| ENSMUSG( | 10.3564 | -0.20202 | 0.746655 | NA | Gm4707 |
| ENSMUSG( | 31.26713 | 0.393329 | 0.302017 | 0.750166 | Gm47113 |
| ENSMUSG( | 16.12258 | 0.154515 | 0.751864 | NA | Gm4724 |
| ENSMUSG( | 18.09875 | 0.011495 | 0.980072 | NA | Gm47415 |
| ENSMUSG( | 13.92948 | -1.06552 | 0.241501 | NA | Gm4779 |
| ENSMUSG( | 27.41446 | -0.02772 | 0.944066 | 0.993503 | Gm4787 |
| ENSMUSG( | 8.828101 | -0.17443 | 0.785093 | NA | Gm4799 |
| ENSMUSG( | 6.07992 | 4.990985 | 0.000196 | NA | Gm4800 |
| ENSMUSG( | 14.97446 | -0.29785 | 0.543521 | NA | Gm4804 |
| ENSMUSG( | 13.14261 | -0.72945 | 0.18074 | NA | Gm4829 |
| ENSMUSG( | 23.34977 | 0.327928 | 0.38614 | 0.804623 | Gm4833 |
| ENSMUSG( | 23.02305 | -0.22262 | 0.570799 | 0.8933 | Gm48551 |
| ENSMUSG( | 10.45888 | -0.28929 | 0.635722 | NA | Gm48552 |
| ENSMUSG( | 153.8166 | -0.05396 | 0.753335 | 0.950753 | Gm4876 |
| ENSMUSG( | 108.1223 | -0.14193 | 0.468105 | 0.845319 | Gm4924 |
| ENSMUSG( | 8.646824 | 0.146525 | 0.818687 | NA | Gm4948 |
| ENSMUSG( | 210.2662 | 0.360951 | 0.018666 | 0.205542 | Gm4949 |
| ENSMUSG( | 17.64874 | 0.447239 | 0.403393 | NA | Gm50340 |
| ENSMUSG( | 9.076933 | 0.378557 | 0.557343 | NA | Gm5064 |
| ENSMUSG( | 77.23006 | 0.202269 | 0.429184 | 0.826446 | Gm5083 |
| ENSMUSG( | 38.76636 | 0.034042 | 0.919554 | 0.990247 | Gm5086 |
| ENSMUSG( | 22.56469 | 0.654358 | 0.093056 | 0.468892 | Gm5093 |
| ENSMUSG( | 11.37513 | -0.4304 | 0.431489 | NA | Gm5105 |
| ENSMUSG( | 873.1834 | 0.019501 | 0.852696 | 0.973168 | Gm5113 |
| ENSMUSG( | 55.56545 | -0.03863 | 0.892111 | 0.984156 | Gm5141 |
| ENSMUSG( | 60.45072 | -0.18413 | 0.471621 | 0.847331 | Gm51425 |
| ENSMUSG( | 74.97071 | 0.146445 | 0.531835 | 0.879628 | Gm5148 |
| ENSMUSG( | 73.55672 | 0.102199 | 0.683398 | 0.932153 | Gm5165 |
| ENSMUSG( | 13.69345 | -0.30753 | 0.568426 | NA | Gm5174 |
| ENSMUSG( | 16.00481 | -0.59061 | 0.220887 | NA | Gm5176 |
| ENSMUSG( | 50.58107 | 0.26397 | 0.353558 | 0.785899 | Gm5242 |
| ENSMUSG( | 88.0947 | -0.19565 | 0.346262 | 0.779153 | Gm527 |
| ENSMUSG( | 34.84099 | -0.48382 | 0.134992 | 0.548629 | Gm53058 |
| ENSMUSG( | 13.57086 | 0.12047 | 0.806668 | NA | Gm5321 |
| ENSMUSG( | 44.66882 | -0.24501 | 0.388159 | 0.805915 | Gm5345 |
| ENSMUSG( | 9.736482 | -0.24807 | 0.697214 | NA | Gm536 |
| ENSMUSG( | 26.30108 | 0.228445 | 0.553329 | 0.886216 | Gm5421 |
| ENSMUSG( | 12.54502 | 0.772127 | 0.15096 | NA | Gm5422 |
| ENSMUSG( | 10.14439 | -0.52216 | 0.390255 | NA | Gm5425 |
| ENSMUSG( | 12.37878 | -0.50424 | 0.358526 | NA | Gm5431 |
| ENSMUSG( | 13.77092 | 0.069932 | 0.890352 | NA | Gm5436 |
| ENSMUSG( | 51.20867 | 0.076121 | 0.813166 | 0.964574 | Gm5451 |
| ENSMUSG( | 14.22259 | -0.2556 | 0.613956 | NA | Gm5525 |
| ENSMUSG( | 29.48924 | -0.33886 | 0.328517 | 0.773036 | Gm5535 |
| ENSMUSG( | 154.4191 | 0.057834 | 0.762865 | 0.953524 | Gm5577 |
| ENSMUSG( | 25.57975 | -0.235 | 0.603581 | 0.904887 | Gm5586 |
| ENSMUSG( | 106.8822 | 0.163787 | 0.414314 | 0.819998 | Gm5617 |
| ENSMUSG( | 16.05085 | 0.27526 | 0.556228 | NA | Gm5620 |
| ENSMUSG( | 10.91205 | -0.33444 | 0.55342 | NA | Gm5621 |
| ENSMUSG( | 66.03969 | -0.29373 | 0.24421 | 0.691435 | Gm5628 |
| ENSMUSG( | 15.15062 | 0.609629 | 0.26502 | NA | Gm5778 |
| ENSMUSG( | 20.5084 | 0.07326 | 0.863191 | NA | Gm5781 |
| ENSMUSG( | 35.43005 | 0.368854 | 0.252997 | 0.703275 | Gm57854 |
| ENSMUSG( | 12.08266 | -0.25806 | 0.635185 | NA | Gm5786 |
| ENSMUSG( | 37.23985 | -0.36687 | 0.228999 | 0.676568 | Gm5805 |
| ENSMUSG( | 35.82992 | 0.064523 | 0.841751 | 0.971179 | Gm5820 |
| ENSMUSG( | 95.35488 | 0.196315 | 0.350101 | 0.781943 | Gm5835 |
| ENSMUSG( | 19.00311 | -0.16328 | 0.713927 | NA | Gm5846 |
| ENSMUSG( | 12.10369 | -0.23517 | 0.674738 | NA | Gm5860 |
| ENSMUSG( | 43.529 | 0.122981 | 0.675165 | 0.929878 | Gm5869 |
| ENSMUSG( | 11.1161 | -0.31378 | 0.584041 | NA | Gm5879 |
| ENSMUSG( | 36.88769 | -0.08703 | 0.793792 | 0.959799 | Gm5881 |
| ENSMUSG( | 26.62021 | -0.09322 | 0.809753 | 0.96372 | Gm5904 |
| ENSMUSG( | 97.55516 | -0.04768 | 0.811544 | 0.964426 | Gm5905 |
| ENSMUSG( | 38.31784 | 0.078063 | 0.815525 | 0.965133 | Gm5914 |
| ENSMUSG( | 23.91137 | -0.25955 | 0.503527 | 0.866217 | Gm6013 |
| ENSMUSG( | 51.80819 | -0.03252 | 0.90317 | 0.986055 | Gm6136 |
| ENSMUSG( | 2249.185 | 0.142835 | 0.169546 | 0.603608 | Gm6145 |
| ENSMUSG( | 18.18222 | -0.10501 | 0.814832 | NA | Gm6192 |
| ENSMUSG( | 35.45482 | 0.239499 | 0.448684 | 0.834689 | Gm6204 |
| ENSMUSG( | 42.48422 | 0.169451 | 0.572993 | 0.893584 | Gm6206 |
| ENSMUSG( | 16.87695 | -0.21292 | 0.683283 | NA | Gm6210 |
| ENSMUSG( | 12.80717 | -0.37426 | 0.50432 | NA | Gm6225 |
| ENSMUSG( | 29.82971 | 0.145187 | 0.668969 | 0.927963 | Gm6252 |
| ENSMUSG( | 9.32655 | -0.05718 | 0.931873 | NA | Gm6254 |
| ENSMUSG( | 12.01008 | -0.1685 | 0.753739 | NA | Gm6345 |
| ENSMUSG( | 20.83816 | 0.077236 | 0.859173 | NA | Gm6352 |
| ENSMUSG( | 30.52749 | -0.12633 | 0.719171 | 0.941986 | Gm6368 |
| ENSMUSG( | 10.77816 | -0.5698 | 0.325585 | NA | Gm6392 |
| ENSMUSG( | 15.74804 | -0.14496 | 0.762702 | NA | Gm6450 |
| ENSMUSG( | 22.4185 | -0.04802 | 0.910035 | 0.9881 | Gm6451 |
| ENSMUSG( | 56.62855 | -0.24856 | 0.363456 | 0.793339 | Gm6483 |
| ENSMUSG( | 11.00985 | -0.47007 | 0.408284 | NA | Gm6501 |
| ENSMUSG( | 26.84772 | 0.365202 | 0.304158 | 0.752287 | Gm6533 |
| ENSMUSG( | 32.64801 | -0.27041 | 0.421346 | 0.823126 | Gm6552 |
| ENSMUSG( | 12.06259 | -0.1432 | 0.796138 | NA | Gm6569 |
| ENSMUSG( | 16.13595 | -0.38427 | 0.416718 | NA | Gm6627 |
| ENSMUSG( | 20.73762 | -0.0397 | 0.932152 | NA | Gm6682 |
| ENSMUSG( | 85.58031 | -0.10488 | 0.656627 | 0.923929 | Gm6712 |
| ENSMUSG( | 19.2512 | -0.54913 | 0.225044 | NA | Gm6807 |
| ENSMUSG( | 53.55287 | 0.167998 | 0.531402 | 0.879512 | Gm6863 |
| ENSMUSG( | 12.07433 | -0.34153 | 0.518728 | NA | Gm6883 |
| ENSMUSG( | 43.58739 | 0.03156 | 0.912935 | 0.988483 | Gm7008 |
| ENSMUSG( | 12.78234 | -0.03186 | 0.956877 | NA | Gm7067 |
| ENSMUSG( | 300.0646 | -0.28348 | 0.122257 | 0.527707 | Gm7072 |
| ENSMUSG( | 11.19697 | -0.13284 | 0.811619 | NA | Gm7114 |
| ENSMUSG( | 13.07933 | -0.61993 | 0.219561 | NA | Gm7143 |
| ENSMUSG( | 201.7 | -0.16444 | 0.275291 | 0.725109 | Gm715 |
| ENSMUSG( | 52.03 | -0.04023 | 0.883418 | 0.982465 | Gm7160 |
| ENSMUSG( | 10.37942 | -0.34076 | 0.595865 | NA | Gm7299 |
| ENSMUSG( | 17.99495 | 0.789668 | 0.19425 | NA | Gm7353 |
| ENSMUSG( | 42.64607 | 0.504517 | 0.092471 | 0.467108 | Gm7361 |
| ENSMUSG( | 12.6054 | 0.389089 | 0.465098 | NA | Gm7363 |
| ENSMUSG( | 11.26282 | 0.283729 | 0.665081 | NA | Gm7389 |
| ENSMUSG( | 49.05328 | 0.186983 | 0.502304 | 0.864924 | Gm7392 |
| ENSMUSG( | 10.45036 | -0.05171 | 0.928543 | NA | Gm7434 |
| ENSMUSG( | 17.31323 | 0.045373 | 0.91834 | NA | Gm7435 |
| ENSMUSG( | 28.07386 | -0.25184 | 0.487852 | 0.858824 | Gm7467 |
| ENSMUSG( | 10.04821 | 0.088117 | 0.884701 | NA | Gm7517 |
| ENSMUSG( | 141.2915 | 0.076536 | 0.692348 | 0.934101 | Gm7536 |
| ENSMUSG( | 11.9462 | -0.16081 | 0.760077 | NA | Gm7600 |
| ENSMUSG( | 20.83686 | -0.14708 | 0.724904 | NA | Gm7628 |
| ENSMUSG( | 8.299087 | 0.635143 | 0.361294 | NA | Gm7658 |
| ENSMUSG( | 29.03813 | -0.22231 | 0.53353 | 0.880095 | Gm7666 |
| ENSMUSG( | 374.8563 | 0.244162 | 0.062497 | 0.386467 | Gm7694 |
| ENSMUSG( | 8.296511 | 0.113105 | 0.869104 | NA | Gm7854 |
| ENSMUSG( | 22.63941 | 0.293489 | 0.465734 | 0.844222 | Gm7895 |
| ENSMUSG( | 34.63904 | 0.126431 | 0.705066 | 0.938303 | Gm7901 |
| ENSMUSG( | 23.96631 | -0.48967 | 0.210634 | 0.653372 | Gm7908 |
| ENSMUSG( | 49.05867 | -0.22558 | 0.437133 | 0.829501 | Gm7932 |
| ENSMUSG( | 12.54482 | 0.129069 | 0.822545 | NA | Gm807 |
| ENSMUSG( | 41.72 | -0.50687 | 0.109395 | 0.499391 | Gm8098 |
| ENSMUSG( | 23.24328 | -0.12776 | 0.766175 | 0.954173 | Gm8130 |
| ENSMUSG( | 10.98289 | 0.231342 | 0.676253 | NA | Gm8188 |
| ENSMUSG( | 13.44146 | 0.091651 | 0.855847 | NA | Gm8276 |
| ENSMUSG( | 25.73745 | 0.308977 | 0.408882 | 0.818173 | Gm8281 |
| ENSMUSG( | 76.22796 | -0.01295 | 0.954262 | 0.993874 | Gm8292 |

| | | | | | |
|---|---|---|---|---|---|
| ENSMUSG( | 36.69105 | -0.33143 | 0.311034 | 0.899422 | Gm46637 |
| ENSMUSG( | 7.038678 | -1.74665 | 0.030371 | NA | Gm4691 |
| ENSMUSG( | 20.30662 | -0.63383 | 0.174642 | 0.844619 | Gm47113 |
| ENSMUSG( | 13.41201 | -0.10056 | 0.849877 | 0.985523 | Gm4724 |
| ENSMUSG( | 18.73754 | 0.345231 | 0.474095 | 0.923329 | Gm47415 |
| ENSMUSG( | 24.21248 | -0.10993 | 0.784178 | 0.976154 | Gm4787 |
| ENSMUSG( | 12.73516 | -0.51287 | 0.34036 | 0.907972 | Gm4804 |
| ENSMUSG( | 13.71439 | -0.22924 | 0.683739 | 0.958678 | Gm4829 |
| ENSMUSG( | 15.56507 | -0.63251 | 0.197803 | 0.862161 | Gm4833 |
| ENSMUSG( | 22.00975 | -0.06989 | 0.869407 | 0.988097 | Gm48551 |
| ENSMUSG( | 11.05073 | 0.176647 | 0.781267 | NA | Gm48552 |
| ENSMUSG( | 144.5936 | 0.02438 | 0.901212 | 0.990558 | Gm4876 |
| ENSMUSG( | 126.067 | 0.537424 | 0.010618 | 0.419324 | Gm4924 |
| ENSMUSG( | 147.0417 | -0.41705 | 0.043359 | 0.690304 | Gm4949 |
| ENSMUSG( | 12.5658 | -0.23734 | 0.671355 | 0.95694 | Gm50340 |
| ENSMUSG( | 62.81304 | -0.15004 | 0.626699 | 0.953234 | Gm5083 |
| ENSMUSG( | 33.00078 | -0.15131 | 0.674785 | 0.95694 | Gm5086 |
| ENSMUSG( | 18.54798 | 0.377301 | 0.408167 | 0.917257 | Gm5093 |
| ENSMUSG( | 9.431492 | -0.73195 | 0.251454 | NA | Gm5105 |
| ENSMUSG( | 809.694 | 0.069284 | 0.573385 | 0.942912 | Gm5113 |
| ENSMUSG( | 50.65755 | -0.02263 | 0.940812 | 0.994558 | Gm5141 |
| ENSMUSG( | 66.89158 | 0.362314 | 0.152397 | 0.837724 | Gm51425 |
| ENSMUSG( | 68.46639 | 0.158186 | 0.555034 | 0.939565 | Gm5148 |
| ENSMUSG( | 75.34368 | 0.390368 | 0.156974 | 0.837724 | Gm5165 |
| ENSMUSG( | 15.94977 | 0.361254 | 0.481362 | 0.924564 | Gm5174 |
| ENSMUSG( | 13.79775 | -0.785 | 0.149559 | 0.836241 | Gm5176 |
| ENSMUSG( | 38.83664 | -0.19586 | 0.56127 | 0.940254 | Gm5242 |
| ENSMUSG( | 82.00904 | -0.1287 | 0.586882 | 0.944975 | Gm527 |
| ENSMUSG( | 36.53728 | -0.04739 | 0.887476 | 0.990266 | Gm53058 |
| ENSMUSG( | 10.54373 | -0.36405 | 0.541286 | NA | Gm5321 |
| ENSMUSG( | 38.76842 | -0.45319 | 0.203567 | 0.864412 | Gm5345 |
| ENSMUSG( | 9.017462 | -0.17932 | 0.778652 | NA | Gm536 |
| ENSMUSG( | 20.43913 | -0.23267 | 0.598231 | 0.946676 | Gm5421 |
| ENSMUSG( | 7.814647 | -0.16087 | 0.821951 | NA | Gm5422 |
| ENSMUSG( | 8.720149 | -0.79624 | 0.241018 | NA | Gm5425 |
| ENSMUSG( | 12.07097 | -0.24708 | 0.674212 | 0.95694 | Gm5431 |
| ENSMUSG( | 10.64274 | -0.45792 | 0.454465 | NA | Gm5436 |
| ENSMUSG( | 39.09433 | -0.51635 | 0.171576 | 0.842999 | Gm5451 |
| ENSMUSG( | 11.24127 | -0.80563 | 0.180249 | NA | Gm5525 |
| ENSMUSG( | 29.1196 | -0.06541 | 0.861613 | 0.986298 | Gm5535 |
| ENSMUSG( | 138.0918 | 0.014689 | 0.952601 | 0.994798 | Gm5577 |
| ENSMUSG( | 18.12208 | -1.27763 | 0.025676 | 0.599234 | Gm5586 |
| ENSMUSG( | 88.75815 | -0.123 | 0.611368 | 0.950771 | Gm5617 |
| ENSMUSG( | 12.53356 | -0.19201 | 0.735111 | 0.967344 | Gm5620 |
| ENSMUSG( | 56.94379 | -0.48819 | 0.090147 | 0.790741 | Gm5628 |
| ENSMUSG( | 11.53501 | 0.11276 | 0.845821 | 0.984272 | Gm5778 |
| ENSMUSG( | 12.79972 | -1.26434 | 0.024173 | 0.588924 | Gm5781 |
| ENSMUSG( | 36.2054 | 0.638134 | 0.063285 | 0.736482 | Gm57854 |
| ENSMUSG( | 10.08153 | -0.54036 | 0.407301 | NA | Gm5786 |
| ENSMUSG( | 28.77585 | -1.02914 | 0.00786 | 0.379307 | Gm5805 |
| ENSMUSG( | 28.75765 | -0.32096 | 0.376406 | 0.915491 | Gm5820 |
| ENSMUSG( | 82.46441 | 0.022817 | 0.932312 | 0.99306 | Gm5835 |
| ENSMUSG( | 13.70596 | -0.99391 | 0.076044 | 0.769092 | Gm5846 |
| ENSMUSG( | 11.90315 | -0.01893 | 0.973223 | 0.996416 | Gm5860 |
| ENSMUSG( | 31.68478 | -0.61394 | 0.089452 | 0.788594 | Gm5869 |
| ENSMUSG( | 35.29854 | 0.026113 | 0.943438 | 0.994798 | Gm5881 |
| ENSMUSG( | 24.01741 | -0.15855 | 0.707548 | 0.964029 | Gm5904 |
| ENSMUSG( | 73.80937 | -0.68881 | 0.008885 | 0.394945 | Gm5905 |
| ENSMUSG( | 35.91139 | 0.122975 | 0.752781 | 0.970521 | Gm5914 |
| ENSMUSG( | 22.36201 | -0.15357 | 0.727693 | 0.966887 | Gm6013 |
| ENSMUSG( | 7.578827 | 0.262606 | 0.751748 | NA | Gm6089 |
| ENSMUSG( | 43.97771 | -0.28178 | 0.396568 | 0.917257 | Gm6136 |
| ENSMUSG( | 1995.852 | 0.067795 | 0.582726 | 0.944266 | Gm6145 |
| ENSMUSG( | 16.16484 | -0.19243 | 0.688974 | 0.959742 | Gm6192 |
| ENSMUSG( | 29.47665 | -0.02478 | 0.946473 | 0.994798 | Gm6204 |
| ENSMUSG( | 40.83805 | 0.298326 | 0.35828 | 0.913759 | Gm6206 |
| ENSMUSG( | 17.68259 | 0.182839 | 0.699412 | 0.961286 | Gm6210 |
| ENSMUSG( | 15.54496 | 0.481297 | 0.390931 | 0.917257 | Gm6225 |
| ENSMUSG( | 25.4616 | -0.08009 | 0.847462 | 0.985161 | Gm6252 |
| ENSMUSG( | 11.54197 | -0.04564 | 0.93659 | 0.993573 | Gm6345 |
| ENSMUSG( | 14.80017 | -0.7602 | 0.143043 | 0.831221 | Gm6352 |
| ENSMUSG( | 22.85434 | -0.83833 | 0.06137 | 0.736111 | Gm6368 |
| ENSMUSG( | 11.2238 | -1.10245 | 0.084329 | NA | Gm6450 |
| ENSMUSG( | 20.99273 | -0.02196 | 0.963603 | 0.995433 | Gm6451 |
| ENSMUSG( | 51.58857 | -0.24206 | 0.416191 | 0.917257 | Gm6483 |
| ENSMUSG( | 10.32589 | -0.39282 | 0.523876 | NA | Gm6501 |
| ENSMUSG( | 20.00147 | -0.20374 | 0.636766 | 0.953903 | Gm6533 |
| ENSMUSG( | 28.67665 | -0.41773 | 0.282477 | 0.892412 | Gm6552 |
| ENSMUSG( | 9.51558 | 0.163135 | 0.79173 | NA | Gm6566 |
| ENSMUSG( | 9.347124 | -0.69579 | 0.350984 | NA | Gm6569 |
| ENSMUSG( | 16.44063 | -0.01704 | 0.972694 | 0.996416 | Gm6627 |
| ENSMUSG( | 16.46487 | -0.46199 | 0.346004 | 0.910966 | Gm6682 |
| ENSMUSG( | 78.48723 | -0.0878 | 0.723597 | 0.966887 | Gm6712 |
| ENSMUSG( | 14.72767 | -1.34915 | 0.016754 | 0.521279 | Gm6807 |
| ENSMUSG( | 43.59383 | -0.17129 | 0.602477 | 0.94789 | Gm6863 |
| ENSMUSG( | 10.52224 | -0.54142 | 0.385273 | NA | Gm6883 |
| ENSMUSG( | 35.84454 | -0.24679 | 0.465276 | 0.923329 | Gm7008 |
| ENSMUSG( | 15.1788 | 0.633347 | 0.212497 | 0.864412 | Gm7067 |
| ENSMUSG( | 325.4868 | 0.221805 | 0.284448 | 0.892412 | Gm7072 |
| ENSMUSG( | 9.410188 | -0.33881 | 0.591411 | NA | Gm7114 |
| ENSMUSG( | 11.96071 | -0.62546 | 0.29529 | 0.898538 | Gm7143 |
| ENSMUSG( | 180.9892 | -0.22667 | 0.226539 | 0.868507 | Gm715 |
| ENSMUSG( | 49.9302 | 0.102853 | 0.727718 | 0.966887 | Gm7160 |
| ENSMUSG( | 7.376535 | -1.21758 | 0.139492 | NA | Gm7299 |
| ENSMUSG( | 13.20435 | 0.267328 | 0.676538 | 0.95694 | Gm7353 |
| ENSMUSG( | 36.90195 | 0.389195 | 0.255829 | 0.883082 | Gm7361 |
| ENSMUSG( | 10.63001 | 0.198018 | 0.749014 | NA | Gm7363 |
| ENSMUSG( | 8.054543 | -0.3741 | 0.635861 | NA | Gm7389 |
| ENSMUSG( | 37.50569 | -0.3687 | 0.282849 | 0.892412 | Gm7392 |
| ENSMUSG( | 16.40925 | 0.182141 | 0.704813 | 0.962677 | Gm7435 |
| ENSMUSG( | 27.52807 | -0.03941 | 0.920731 | 0.992644 | Gm7467 |
| ENSMUSG( | 112.8505 | -0.33267 | 0.12213 | 0.813384 | Gm7536 |
| ENSMUSG( | 9.873867 | -0.51826 | 0.416212 | NA | Gm7600 |
| ENSMUSG( | 19.97458 | -0.0036 | 0.994287 | 0.998813 | Gm7628 |
| ENSMUSG( | 28.56819 | -0.02431 | 0.948449 | 0.994798 | Gm7666 |
| ENSMUSG( | 305.5384 | -0.08138 | 0.645915 | 0.953988 | Gm7694 |
| ENSMUSG( | 11.26169 | 1.044156 | 0.082529 | NA | Gm7854 |
| ENSMUSG( | 17.64396 | -0.19874 | 0.676064 | 0.95694 | Gm7895 |
| ENSMUSG( | 27.37182 | -0.27869 | 0.471394 | 0.923329 | Gm7901 |
| ENSMUSG( | 18.60947 | -1.07437 | 0.02256 | 0.581149 | Gm7908 |
| ENSMUSG( | 52.528 | 0.226963 | 0.447797 | 0.922202 | Gm7932 |
| ENSMUSG( | 12.14828 | 0.303117 | 0.581813 | 0.944266 | Gm807 |
| ENSMUSG( | 42.25714 | -0.17744 | 0.587124 | 0.944975 | Gm8098 |
| ENSMUSG( | 16.02263 | -1.14988 | 0.038894 | 0.679972 | Gm8130 |
| ENSMUSG( | 12.20607 | 0.094686 | 0.861934 | 0.98632 | Gm8276 |
| ENSMUSG( | 18.99216 | -0.33016 | 0.479761 | 0.924021 | Gm8281 |
| ENSMUSG( | 66.10585 | -0.16452 | 0.559044 | 0.939587 | Gm8292 |
| ENSMUSG( | 28.22864 | -0.6383 | 0.094606 | 0.796133 | Gm8355 |
| ENSMUSG( | 54.08695 | -0.61068 | 0.050241 | 0.708648 | Gm8451 |
| ENSMUSG( | 10.41002 | 0.163795 | 0.783599 | NA | Gm8494 |
| ENSMUSG( | 10.37749 | -0.74257 | 0.226191 | NA | Gm8546 |
| ENSMUSG( | 29.21898 | 0.050658 | 0.889811 | 0.990558 | Gm867 |
| ENSMUSG( | 18.09097 | -0.33133 | 0.473658 | 0.923329 | Gm8692 |
| ENSMUSG( | 14.41344 | 0.012092 | 0.982306 | 0.996756 | Gm8717 |
| ENSMUSG( | 74.93188 | -0.25258 | 0.304958 | 0.898538 | Gm8730 |
| ENSMUSG( | 24.94995 | 0.23568 | 0.644786 | 0.953903 | Gm8738 |
| ENSMUSG( | 9.621478 | -0.293 | 0.65315 | NA | Gm8797 |
| ENSMUSG( | 11.96292 | -0.58443 | 0.308333 | 0.898538 | Gm8971 |
| ENSMUSG( | 24.6359 | -0.2137 | 0.620625 | 0.952971 | Gm8983 |
| ENSMUSG( | 16.0149 | -0.03584 | 0.949014 | 0.994798 | Gm9294 |
| ENSMUSG( | 14.31506 | 0.425017 | 0.441743 | 0.921554 | Gm9320 |
| ENSMUSG( | 10.98719 | -0.17172 | 0.765838 | NA | Gm9333 |
| ENSMUSG( | 11.80359 | -0.32201 | 0.574095 | 0.942912 | Gm9347 |
| ENSMUSG( | 14.72766 | -0.0536 | 0.919416 | 0.992644 | Gm9403 |
| ENSMUSG( | 49.46911 | -0.4266 | 0.206637 | 0.864412 | Gm9493 |
| ENSMUSG( | 81.59905 | -0.21665 | 0.423964 | 0.917257 | Gm9616 |
| ENSMUSG( | 31.35812 | -0.05242 | 0.891719 | 0.990558 | Gm9625 |
| ENSMUSG( | 637.0888 | -0.26576 | 0.046328 | 0.697814 | Gm973 |
| ENSMUSG( | 12.48059 | -0.36545 | 0.512953 | 0.930358 | Gm9755 |
| ENSMUSG( | 103.188 | -0.09552 | 0.654864 | 0.955015 | Gm9776 |
| ENSMUSG( | 108.7028 | -0.41337 | 0.131166 | 0.822884 | Gm9794 |
| ENSMUSG( | 125.5476 | -0.10375 | 0.630018 | 0.953251 | Gm9828 |
| ENSMUSG( | 379.4243 | -0.02247 | 0.89145 | 0.990558 | Gm9899 |
| ENSMUSG( | 62.41868 | 0.475899 | 0.14136 | 0.830678 | Gm9917 |
| ENSMUSG( | 13.61331 | 0.009449 | 0.985489 | 0.997084 | Gm9936 |
| ENSMUSG( | 11.34711 | 0.224129 | 0.712454 | NA | Gm9946 |
| ENSMUSG( | 21.42349 | 0.141493 | 0.748849 | 0.96984 | Gm9958 |
| ENSMUSG( | 48.68653 | -0.06597 | 0.833111 | 0.981149 | Gm9962 |
| ENSMUSG( | 818.5811 | -0.02858 | 0.820594 | 0.979198 | Gmcl1 |
| ENSMUSG( | 165.8035 | -0.29677 | 0.094396 | 0.795735 | Gmds |
| ENSMUSG( | 634.5673 | -0.00263 | 0.984227 | 0.996963 | Gmeb1 |
| ENSMUSG( | 638.7548 | -0.0557 | 0.662466 | 0.956205 | Gmeb2 |
| ENSMUSG( | 7897.501 | -0.09044 | 0.620038 | 0.952896 | Gmfb |
| ENSMUSG( | 62.85358 | -0.24898 | 0.396447 | 0.917257 | Gmfg |
| ENSMUSG( | 347.5191 | 0.287177 | 0.104882 | 0.804969 | Gmip |
| ENSMUSG( | 18.60716 | 0.589375 | 0.200025 | 0.863016 | Gmnc |
| ENSMUSG( | 47.29847 | -0.19461 | 0.552213 | 0.939565 | Gmnn |
| ENSMUSG( | 709.4877 | 0.055037 | 0.729476 | 0.967022 | Gmppa |

| | | | | | | | | | | | |
|---|---|---|---|---|---|---|---|---|---|---|---|
| ENSMUSG( | Gm57854 | 35.43005 | 0.368854 | 0.252997 | 0.703275 | ENSMUSG( | Gm57854 | 36.2054 | 0.638134 | 0.063285 | 0.736482 |
| ENSMUSG( | Gm5786 | 12.08266 | -0.25806 | 0.635185 | NA | ENSMUSG( | Gm5786 | 10.08153 | -0.54036 | 0.407301 | NA |
| ENSMUSG( | Gm5805 | 37.23985 | -0.36687 | 0.228999 | 0.676568 | ENSMUSG( | Gm5805 | 28.77585 | -1.02914 | 0.00786 | 0.379307 |
| ENSMUSG( | Gm5820 | 35.82992 | 0.064523 | 0.841751 | 0.971179 | ENSMUSG( | Gm5820 | 28.75765 | -0.32096 | 0.376406 | 0.915491 |
| ENSMUSG( | Gm5835 | 95.35488 | 0.196315 | 0.350101 | 0.781943 | ENSMUSG( | Gm5835 | 82.46441 | 0.022817 | 0.932312 | 0.99306 |
| ENSMUSG( | Gm5846 | 19.00311 | -0.16328 | 0.713927 | NA | ENSMUSG( | Gm5846 | 13.70596 | -0.99391 | 0.076044 | 0.769092 |
| ENSMUSG( | Gm5860 | 12.10369 | -0.23517 | 0.674738 | NA | ENSMUSG( | Gm5860 | 11.90315 | -0.01893 | 0.973223 | 0.996416 |
| ENSMUSG( | Gm5869 | 43.529 | 0.122981 | 0.675165 | 0.929878 | ENSMUSG( | Gm5869 | 31.68478 | -0.61394 | 0.089452 | 0.788594 |
| ENSMUSG( | Gm5881 | 36.88769 | -0.08703 | 0.793792 | 0.959799 | ENSMUSG( | Gm5881 | 35.29854 | 0.026113 | 0.943438 | 0.994798 |
| ENSMUSG( | Gm5904 | 26.62021 | -0.09322 | 0.809753 | 0.96372 | ENSMUSG( | Gm5904 | 24.01741 | -0.15855 | 0.707548 | 0.964029 |
| ENSMUSG( | Gm5905 | 97.55516 | -0.04768 | 0.811544 | 0.964426 | ENSMUSG( | Gm5905 | 73.80937 | -0.68881 | 0.008885 | 0.394945 |
| ENSMUSG( | Gm5914 | 38.31784 | 0.078063 | 0.815525 | 0.965133 | ENSMUSG( | Gm5914 | 35.91139 | 0.122975 | 0.752781 | 0.970521 |
| ENSMUSG( | Gm6013 | 23.91137 | -0.25955 | 0.503527 | 0.866217 | ENSMUSG( | Gm6013 | 22.36201 | -0.15357 | 0.727693 | 0.966887 |
| ENSMUSG( | Gm6136 | 51.80819 | -0.03252 | 0.90317 | 0.986055 | ENSMUSG( | Gm6136 | 43.97771 | -0.28178 | 0.396568 | 0.917257 |
| ENSMUSG( | Gm6145 | 2249.185 | 0.142835 | 0.169546 | 0.603608 | ENSMUSG( | Gm6145 | 1995.852 | 0.067795 | 0.582726 | 0.944266 |
| ENSMUSG( | Gm6192 | 18.18222 | -0.10501 | 0.814832 | NA | ENSMUSG( | Gm6192 | 16.16484 | -0.19243 | 0.688974 | 0.959742 |
| ENSMUSG( | Gm6204 | 35.45482 | 0.239499 | 0.448684 | 0.834689 | ENSMUSG( | Gm6204 | 29.47665 | -0.02478 | 0.946473 | 0.994798 |
| ENSMUSG( | Gm6206 | 42.48422 | 0.169451 | 0.572993 | 0.893584 | ENSMUSG( | Gm6206 | 40.83805 | 0.298326 | 0.35828 | 0.913759 |
| ENSMUSG( | Gm6210 | 16.87695 | -0.21292 | 0.683283 | NA | ENSMUSG( | Gm6210 | 17.68259 | 0.182839 | 0.699412 | 0.961286 |
| ENSMUSG( | Gm6225 | 12.80717 | -0.37426 | 0.50432 | NA | ENSMUSG( | Gm6225 | 15.54496 | 0.481297 | 0.390931 | 0.917257 |
| ENSMUSG( | Gm6252 | 29.82971 | 0.145187 | 0.668969 | 0.927963 | ENSMUSG( | Gm6252 | 25.4616 | -0.08009 | 0.847462 | 0.985161 |
| ENSMUSG( | Gm6345 | 12.01008 | -0.1685 | 0.753739 | NA | ENSMUSG( | Gm6345 | 11.54197 | -0.04564 | 0.93659 | 0.993573 |
| ENSMUSG( | Gm6352 | 20.83816 | 0.077236 | 0.859173 | NA | ENSMUSG( | Gm6352 | 14.80017 | -0.7602 | 0.143043 | 0.831221 |
| ENSMUSG( | Gm6368 | 30.52749 | -0.12633 | 0.719171 | 0.941986 | ENSMUSG( | Gm6368 | 22.85434 | -0.83833 | 0.06137 | 0.736111 |
| ENSMUSG( | Gm6450 | 15.74804 | -0.14496 | 0.762702 | NA | ENSMUSG( | Gm6450 | 11.2238 | -1.10245 | 0.084329 | NA |
| ENSMUSG( | Gm6451 | 22.4185 | -0.04802 | 0.910035 | 0.9881 | ENSMUSG( | Gm6451 | 20.99273 | -0.02196 | 0.963603 | 0.995433 |
| ENSMUSG( | Gm6483 | 56.62855 | -0.24856 | 0.363456 | 0.793339 | ENSMUSG( | Gm6483 | 51.58857 | -0.24206 | 0.416191 | 0.917257 |
| ENSMUSG( | Gm6501 | 11.00985 | -0.47007 | 0.408284 | NA | ENSMUSG( | Gm6501 | 10.32589 | -0.39282 | 0.523876 | NA |
| ENSMUSG( | Gm6533 | 26.84772 | 0.365202 | 0.304158 | 0.752287 | ENSMUSG( | Gm6533 | 20.00147 | -0.20374 | 0.636766 | 0.953903 |
| ENSMUSG( | Gm6552 | 32.64801 | -0.27041 | 0.421346 | 0.823126 | ENSMUSG( | Gm6552 | 28.67665 | -0.41773 | 0.282477 | 0.892412 |
| ENSMUSG( | Gm6569 | 12.06259 | -0.1432 | 0.796138 | NA | ENSMUSG( | Gm6569 | 9.347124 | -0.69579 | 0.350984 | NA |
| ENSMUSG( | Gm6627 | 16.13595 | -0.38427 | 0.416718 | NA | ENSMUSG( | Gm6627 | 16.44063 | -0.01704 | 0.972694 | 0.996416 |
| ENSMUSG( | Gm6682 | 20.73762 | -0.0397 | 0.932152 | NA | ENSMUSG( | Gm6682 | 16.46487 | -0.46199 | 0.346004 | 0.910966 |
| ENSMUSG( | Gm6712 | 85.58031 | -0.10488 | 0.656627 | 0.923929 | ENSMUSG( | Gm6712 | 78.48723 | -0.0878 | 0.723597 | 0.966887 |
| ENSMUSG( | Gm6807 | 19.2512 | -0.54913 | 0.225044 | NA | ENSMUSG( | Gm6807 | 14.72767 | -1.34915 | 0.016754 | 0.521279 |
| ENSMUSG( | Gm6863 | 53.55287 | 0.167998 | 0.531402 | 0.879512 | ENSMUSG( | Gm6863 | 43.59383 | -0.17129 | 0.602477 | 0.94789 |
| ENSMUSG( | Gm6883 | 12.07433 | -0.34153 | 0.518728 | NA | ENSMUSG( | Gm6883 | 10.52224 | -0.54142 | 0.385273 | NA |
| ENSMUSG( | Gm7008 | 43.58739 | 0.03156 | 0.912935 | 0.988483 | ENSMUSG( | Gm7008 | 35.84454 | -0.24679 | 0.465276 | 0.923329 |
| ENSMUSG( | Gm7067 | 12.78234 | -0.03186 | 0.956877 | NA | ENSMUSG( | Gm7067 | 15.1788 | 0.633347 | 0.212497 | 0.864412 |
| ENSMUSG( | Gm7072 | 300.0646 | -0.28348 | 0.122257 | 0.527707 | ENSMUSG( | Gm7072 | 325.4868 | 0.221805 | 0.284448 | 0.892412 |
| ENSMUSG( | Gm7114 | 11.19697 | -0.13284 | 0.811619 | NA | ENSMUSG( | Gm7114 | 9.410188 | -0.33881 | 0.591411 | NA |
| ENSMUSG( | Gm7143 | 13.07933 | -0.61993 | 0.219561 | NA | ENSMUSG( | Gm7143 | 11.96071 | -0.62546 | 0.29529 | 0.898538 |
| ENSMUSG( | Gm715 | 201.7 | -0.16444 | 0.275291 | 0.725109 | ENSMUSG( | Gm715 | 180.9892 | -0.22667 | 0.226539 | 0.868507 |
| ENSMUSG( | Gm7160 | 52.03 | -0.04023 | 0.883418 | 0.982465 | ENSMUSG( | Gm7160 | 49.9302 | 0.102853 | 0.727718 | 0.966887 |
| ENSMUSG( | Gm7299 | 10.37942 | -0.34076 | 0.595865 | NA | ENSMUSG( | Gm7299 | 7.376535 | -1.21758 | 0.139492 | NA |
| ENSMUSG( | Gm7353 | 17.99495 | 0.789668 | 0.19425 | NA | ENSMUSG( | Gm7353 | 13.20435 | 0.267328 | 0.676538 | 0.95694 |
| ENSMUSG( | Gm7361 | 42.64607 | 0.504517 | 0.092471 | 0.467108 | ENSMUSG( | Gm7361 | 36.90195 | 0.389195 | 0.255829 | 0.883082 |
| ENSMUSG( | Gm7363 | 12.6054 | 0.389089 | 0.465098 | NA | ENSMUSG( | Gm7363 | 10.63001 | 0.198018 | 0.749014 | NA |
| ENSMUSG( | Gm7389 | 11.26282 | 0.283729 | 0.665081 | NA | ENSMUSG( | Gm7389 | 8.054543 | -0.3741 | 0.635861 | NA |
| ENSMUSG( | Gm7392 | 49.05328 | 0.186983 | 0.502304 | 0.864924 | ENSMUSG( | Gm7392 | 37.50569 | -0.3687 | 0.282849 | 0.892412 |
| ENSMUSG( | Gm7435 | 17.31323 | 0.045373 | 0.91834 | NA | ENSMUSG( | Gm7435 | 16.40925 | 0.182141 | 0.704813 | 0.962677 |
| ENSMUSG( | Gm7467 | 28.07386 | -0.25184 | 0.487852 | 0.858824 | ENSMUSG( | Gm7467 | 27.52807 | -0.03941 | 0.920731 | 0.992644 |
| ENSMUSG( | Gm7536 | 141.2915 | 0.076536 | 0.692348 | 0.934101 | ENSMUSG( | Gm7536 | 112.8505 | -0.33267 | 0.12213 | 0.813384 |
| ENSMUSG( | Gm7600 | 11.9462 | -0.16081 | 0.760077 | NA | ENSMUSG( | Gm7600 | 9.873867 | -0.51826 | 0.416212 | NA |
| ENSMUSG( | Gm7628 | 20.83686 | -0.14708 | 0.724904 | NA | ENSMUSG( | Gm7628 | 19.97458 | -0.0036 | 0.994287 | 0.998813 |
| ENSMUSG( | Gm7666 | 29.03813 | -0.22231 | 0.53353 | 0.880095 | ENSMUSG( | Gm7666 | 28.56819 | -0.02431 | 0.948449 | 0.994798 |
| ENSMUSG( | Gm7694 | 374.8563 | 0.244162 | 0.062497 | 0.386467 | ENSMUSG( | Gm7694 | 305.5384 | -0.08138 | 0.645915 | 0.953988 |
| ENSMUSG( | Gm7854 | 8.296511 | 0.113105 | 0.869104 | NA | ENSMUSG( | Gm7854 | 11.26169 | 1.044156 | 0.082529 | NA |
| ENSMUSG( | Gm7895 | 22.63941 | 0.293489 | 0.465734 | 0.844222 | ENSMUSG( | Gm7895 | 17.64396 | -0.19874 | 0.676064 | 0.95694 |
| ENSMUSG( | Gm7901 | 34.63904 | 0.126431 | 0.705066 | 0.938303 | ENSMUSG( | Gm7901 | 27.37182 | -0.27869 | 0.471394 | 0.923329 |
| ENSMUSG( | Gm7908 | 23.96631 | -0.48967 | 0.210634 | 0.653372 | ENSMUSG( | Gm7908 | 18.60947 | -1.07437 | 0.02256 | 0.581149 |
| ENSMUSG( | Gm7932 | 49.05867 | -0.22558 | 0.437133 | 0.829501 | ENSMUSG( | Gm7932 | 52.528 | 0.226963 | 0.447797 | 0.922202 |
| ENSMUSG( | Gm807 | 12.54482 | 0.129069 | 0.822545 | NA | ENSMUSG( | Gm807 | 12.14828 | 0.303117 | 0.581813 | 0.944266 |
| ENSMUSG( | Gm8098 | 41.72 | -0.50687 | 0.109395 | 0.499391 | ENSMUSG( | Gm8098 | 42.25714 | -0.17744 | 0.587124 | 0.944975 |
| ENSMUSG( | Gm8130 | 23.24328 | -0.12776 | 0.766175 | 0.954173 | ENSMUSG( | Gm8130 | 16.02263 | -1.14988 | 0.038894 | 0.679972 |
| ENSMUSG( | Gm8276 | 13.44146 | 0.091651 | 0.855847 | NA | ENSMUSG( | Gm8276 | 12.20607 | 0.094686 | 0.861934 | 0.98632 |
| ENSMUSG( | Gm8281 | 25.73745 | 0.308977 | 0.408882 | 0.818173 | ENSMUSG( | Gm8281 | 18.99216 | -0.33016 | 0.479761 | 0.924021 |
| ENSMUSG( | Gm8292 | 76.22796 | -0.01295 | 0.954262 | 0.993874 | ENSMUSG( | Gm8292 | 66.10585 | -0.16452 | 0.559044 | 0.939587 |
| ENSMUSG( | Gm8355 | 36.4122 | -0.12107 | 0.713893 | 0.940903 | ENSMUSG( | Gm8355 | 28.22864 | -0.6383 | 0.094606 | 0.796133 |
| ENSMUSG( | Gm8451 | 68.88453 | -0.08884 | 0.723074 | 0.943761 | ENSMUSG( | Gm8451 | 54.08695 | -0.61068 | 0.050241 | 0.708648 |
| ENSMUSG( | Gm8494 | 14.47734 | 0.723816 | 0.168905 | NA | ENSMUSG( | Gm8494 | 10.41002 | 0.163795 | 0.783599 | NA |
| ENSMUSG( | Gm8546 | 13.67153 | -0.12081 | 0.81251 | NA | ENSMUSG( | Gm8546 | 10.37749 | -0.74257 | 0.226191 | NA |
| ENSMUSG( | Gm867 | 33.38852 | 0.149176 | 0.683557 | 0.932154 | ENSMUSG( | Gm867 | 29.21898 | 0.050658 | 0.889811 | 0.990558 |
| ENSMUSG( | Gm8692 | 24.83499 | 0.341818 | 0.361879 | 0.792797 | ENSMUSG( | Gm8692 | 18.09097 | -0.33133 | 0.473658 | 0.923329 |
| ENSMUSG( | Gm8717 | 17.49074 | 0.317881 | 0.483781 | NA | ENSMUSG( | Gm8717 | 14.41344 | 0.012092 | 0.982306 | 0.996756 |
| ENSMUSG( | Gm8730 | 98.19825 | 0.245417 | 0.25384 | 0.704011 | ENSMUSG( | Gm8730 | 74.93188 | -0.25258 | 0.304958 | 0.898538 |
| ENSMUSG( | Gm8738 | 30.75495 | 0.594355 | 0.112677 | 0.506698 | ENSMUSG( | Gm8738 | 24.94995 | 0.23568 | 0.644786 | 0.953903 |
| ENSMUSG( | Gm8797 | 11.03286 | -0.09661 | 0.861829 | NA | ENSMUSG( | Gm8797 | 9.621478 | -0.293 | 0.65315 | NA |
| ENSMUSG( | Gm8971 | 13.11116 | -0.54421 | 0.288858 | NA | ENSMUSG( | Gm8971 | 11.96292 | -0.58443 | 0.308333 | 0.898538 |
| ENSMUSG( | Gm8983 | 28.71499 | -0.00811 | 0.983101 | 0.997554 | ENSMUSG( | Gm8983 | 24.6359 | -0.2137 | 0.620625 | 0.952971 |
| ENSMUSG( | Gm9294 | 18.34954 | 0.133759 | 0.777191 | NA | ENSMUSG( | Gm9294 | 16.0149 | -0.03584 | 0.949014 | 0.994798 |
| ENSMUSG( | Gm9320 | 13.09063 | -0.04851 | 0.929347 | NA | ENSMUSG( | Gm9320 | 14.31506 | 0.425017 | 0.441743 | 0.921554 |
| ENSMUSG( | Gm9333 | 14.50569 | 0.36387 | 0.447921 | NA | ENSMUSG( | Gm9333 | 10.98719 | -0.17172 | 0.765838 | NA |
| ENSMUSG( | Gm9347 | 14.12161 | -0.06478 | 0.896478 | NA | ENSMUSG( | Gm9347 | 11.80359 | -0.32201 | 0.574095 | 0.942912 |
| ENSMUSG( | Gm9403 | 15.87729 | -0.1482 | 0.756591 | NA | ENSMUSG( | Gm9403 | 14.72766 | -0.0536 | 0.919416 | 0.992644 |
| ENSMUSG( | Gm9493 | 56.6912 | -0.25326 | 0.359765 | 0.791346 | ENSMUSG( | Gm9493 | 49.46911 | -0.4266 | 0.206637 | 0.864412 |
| ENSMUSG( | Gm9616 | 104.4119 | 0.247454 | 0.194232 | 0.633383 | ENSMUSG( | Gm9616 | 81.59905 | -0.21665 | 0.423964 | 0.917257 |
| ENSMUSG( | Gm9625 | 39.97549 | 0.388516 | 0.195685 | 0.635109 | ENSMUSG( | Gm9625 | 31.35812 | -0.05242 | 0.891719 | 0.990558 |
| ENSMUSG( | Gm973 | 736.3007 | -0.10257 | 0.419735 | 0.822433 | ENSMUSG( | Gm973 | 637.0888 | -0.26576 | 0.046328 | 0.697814 |
| ENSMUSG( | Gm9755 | 16.33254 | 0.193972 | 0.677065 | NA | ENSMUSG( | Gm9755 | 12.48059 | -0.36545 | 0.512953 | 0.930358 |
| ENSMUSG( | Gm9776 | 107.2528 | -0.27109 | 0.164623 | 0.595625 | ENSMUSG( | Gm9776 | 103.188 | -0.09552 | 0.654864 | 0.955015 |
| ENSMUSG( | Gm9794 | 149.2459 | 0.261875 | 0.192711 | 0.631789 | ENSMUSG( | Gm9794 | 108.7028 | -0.41337 | 0.131166 | 0.822884 |
| ENSMUSG( | Gm9828 | 134.2469 | -0.16936 | 0.382432 | 0.803227 | ENSMUSG( | Gm9828 | 125.5476 | -0.10375 | 0.630018 | 0.953251 |
| ENSMUSG( | Gm9899 | 446.7619 | 0.170121 | 0.277051 | 0.727148 | ENSMUSG( | Gm9899 | 379.4243 | -0.02247 | 0.89145 | 0.990558 |
| ENSMUSG( | Gm9917 | 59.58827 | 0.077709 | 0.768455 | 0.954674 | ENSMUSG( | Gm9917 | 62.41868 | 0.475899 | 0.14136 | 0.830678 |
| ENSMUSG( | Gm9936 | 14.64277 | -0.05137 | 0.916003 | NA | ENSMUSG( | Gm9936 | 13.61331 | 0.009449 | 0.985489 | 0.997084 |
| ENSMUSG( | Gm9946 | 11.62759 | 0.018596 | 0.978934 | NA | ENSMUSG( | Gm9946 | 11.34711 | 0.224129 | 0.712454 | NA |
| ENSMUSG( | Gm9958 | 21.71841 | -0.11103 | 0.787983 | 0.959226 | ENSMUSG( | Gm9958 | 21.42349 | 0.141493 | 0.748849 | 0.96984 |
| ENSMUSG( | Gm9962 | 48.28587 | -0.35011 | 0.230241 | 0.678139 | ENSMUSG( | Gm9962 | 48.68653 | -0.06597 | 0.833111 | 0.981149 |
| ENSMUSG( | Gmcl1 | 850.7324 | -0.19139 | 0.086535 | 0.452045 | ENSMUSG( | Gmcl1 | 818.5811 | -0.02858 | 0.820594 | 0.979198 |
| ENSMUSG( | Gmds | 199.3321 | -0.01688 | 0.914718 | 0.988522 | ENSMUSG( | Gmds | 165.8035 | -0.29677 | 0.094396 | 0.795735 |
| ENSMUSG( | Gmeb1 | 691.7946 | -0.02231 | 0.846078 | 0.972226 | ENSMUSG( | Gmeb1 | 634.5673 | -0.00263 | 0.984227 | 0.996963 |
| ENSMUSG( | Gmeb2 | 706.1354 | -0.03773 | 0.733037 | 0.944956 | ENSMUSG( | Gmeb2 | 638.7548 | -0.0557 | 0.662466 | 0.956205 |
| ENSMUSG( | Gmfb | 8939.394 | 0.00321 | 0.982823 | 0.997522 | ENSMUSG( | Gmfb | 7897.501 | -0.09044 | 0.620038 | 0.952896 |
| ENSMUSG( | Gmfg | 71.62583 | -0.10589 | 0.672819 | 0.929007 | ENSMUSG( | Gmfg | 62.85358 | -0.24898 | 0.396447 | 0.917257 |
| ENSMUSG( | Gmip | 375.7112 | 0.250501 | 0.104089 | 0.489938 | ENSMUSG( | Gmip | 347.5191 | 0.287177 | 0.104882 | 0.804969 |
| ENSMUSG( | Gmnc | 17.45655 | 0.208181 | 0.64812 | NA | ENSMUSG( | Gmnc | 18.60716 | 0.589375 | 0.200025 | 0.863016 |
| ENSMUSG( | Gmnn | 56.14945 | 0.062876 | 0.813374 | 0.964574 | ENSMUSG( | Gmnn | 47.29847 | -0.19461 | 0.552213 | 0.939565 |
| ENSMUSG( | Gmppa | 834.2568 | 0.245611 | 0.042672 | 0.316769 | ENSMUSG( | Gmppa | 709.4877 | 0.055037 | 0.729476 | 0.967022 |
| ENSMUSG( | Gmppb | 272.1709 | 0.054798 | 0.736827 | 0.94664 | ENSMUSG( | Gmppb | 237.5411 | -0.07466 | 0.69143 | 0.960071 |
| ENSMUSG( | Gmpr | 685.4732 | -0.10581 | 0.505422 | 0.86689 | ENSMUSG( | Gmpr | 583.1721 | -0.32412 | 0.065782 | 0.745828 |
| ENSMUSG( | Gmpr2 | 538.9672 | 0.022139 | 0.83518 | 0.970426 | ENSMUSG( | Gmpr2 | 491.3948 | 0.025562 | 0.84278 | 0.98313 |
| ENSMUSG( | Gmps | 1215.144 | 0.116022 | 0.280378 | 0.729291 | ENSMUSG( | Gmps | 1091.768 | 0.072645 | 0.57739 | 0.943044 |
| ENSMUSG( | Gna11 | 6218.343 | 0.113074 | 0.191625 | 0.630674 | ENSMUSG( | Gna11 | 5494.541 | 0.028153 | 0.779574 | 0.976106 |
| ENSMUSG( | Gna12 | 8997.072 | 0.097371 | 0.266201 | 0.716441 | ENSMUSG( | Gna12 | 7548.726 | -0.14375 | 0.188904 | 0.85698 |
| ENSMUSG( | Gna13 | 5104.527 | 0.172031 | 0.08626 | 0.451772 | ENSMUSG( | Gna13 | 4313.834 | -0.04227 | 0.736125 | 0.967344 |
| ENSMUSG( | Gna15 | 93.7873 | 0.014776 | 0.945494 | 0.99365 | ENSMUSG( | Gna15 | 98.46954 | 0.412056 | 0.084714 | 0.779799 |
| ENSMUSG( | Gnai1 | 2675.438 | -0.07201 | 0.394873 | 0.810072 | ENSMUSG( | Gnai1 | 2470.833 | -0.0349 | 0.813311 | 0.979198 |
| ENSMUSG( | Gnai2 | 17231.37 | -0.06023 | 0.536993 | 0.882123 | ENSMUSG( | Gnai2 | 15205.78 | -0.1566 | 0.178982 | 0.849041 |
| ENSMUSG( | Gnai3 | 1390.837 | -0.00251 | 0.982406 | 0.997512 | ENSMUSG( | Gnai3 | 1252.229 | -0.03937 | 0.771944 | 0.974236 |
| ENSMUSG( | Gnal | 6732.773 | -0.23075 | 0.022545 | 0.229407 | ENSMUSG( | Gnal | 6418.77 | -0.09148 | 0.374955 | 0.915491 |
| ENSMUSG( | Gnao1 | 41443.38 | 0.013821 | 0.884212 | 0.982647 | ENSMUSG( | Gnao1 | 34990.64 | -0.21346 | 0.063681 | 0.736482 |
| ENSMUSG( | Gnaq | 5910.783 | 0.172766 | 0.028742 | 0.258426 | ENSMUSG( | Gnaq | 5073.427 | 0.005152 | 0.965883 | 0.995433 |
| ENSMUSG( | Gnas | 62267.32 | 0.044878 | 0.553376 | 0.886216 | ENSMUSG( | Gnas | 53410.84 | -0.13292 | 0.217317 | 0.865589 |
| ENSMUSG( | Gnaz | 2172.456 | 0.142572 | 0.179761 | 0.617507 | ENSMUSG( | Gnaz | 1923.219 | 0.060825 | 0.610044 | 0.950639 |
| ENSMUSG( | Gnb1 | 28451.12 | 0.097916 | 0.199927 | 0.640239 | ENSMUSG( | Gnb1 | 24214.06 | -0.09975 | 0.393543 | 0.917257 |
| ENSMUSG( | Gnb1l | 102.1054 | 0.085834 | 0.67757 | 0.930542 | ENSMUSG( | Gnb1l | 99.46891 | 0.263434 | 0.237883 | 0.87259 |
| ENSMUSG( | Gnb2 | 8551.015 | 0.01444 | 0.882506 | 0.982244 | ENSMUSG( | Gnb2 | 7352.726 | -0.1589 | 0.208864 | 0.864412 |
| ENSMUSG( | Gnb4 | 1253.769 | 0.006719 | 0.944356 | 0.993543 | ENSMUSG( | Gnb4 | 1117.231 | -0.05626 | 0.637334 | 0.953903 |
| ENSMUSG( | Gnb5 | 2155.108 | -0.06287 | 0.599906 | 0.903978 | ENSMUSG( | Gnb5 | 1914.413 | -0.14169 | 0.330173 | 0.907251 |
| ENSMUSG( | Gne | 729.2545 | 0.134262 | 0.305859 | 0.753712 | ENSMUSG( | Gne | 638.1181 | 0.021762 | 0.881922 | 0.989644 |
| ENSMUSG( | Gng10 | 1476.958 | 0.028626 | 0.797639 | 0.960507 | ENSMUSG( | Gng10 | 1271.472 | -0.14324 | 0.331331 | 0.907861 |
| ENSMUSG( | Gng11 | 269.2676 | -0.49434 | 0.001994 | 0.051341 | ENSMUSG( | Gng11 | 278.8389 | -0.0734 | 0.681768 | 0.958678 |
| ENSMUSG( | Gng12 | 7465.765 | 0.128625 | 0.288764 | 0.739578 | ENSMUSG( | Gng12 | 6252.057 | -0.11611 | 0.432306 | 0.918158 |
| ENSMUSG( | Gng13 | 1879.941 | 0.034869 | 0.677051 | 0.930542 | ENSMUSG( | Gng13 | 1534.682 | -0.29976 | 0.008762 | 0.393969 |
| ENSMUSG( | Gng2 | 3269.129 | 0.020225 | 0.824951 | 0.967386 | ENSMUSG( | Gng2 | 2875.118 | -0.08637 | 0.503628 | 0.927825 |
| ENSMUSG( | Gng3 | 5481.286 | 0.066953 | 0.619754 | 0.911432 | ENSMUSG( | Gng3 | 4611.501 | -0.17056 | 0.325986 | 0.905339 |
| ENSMUSG( | Gng4 | 2144.46 | 0.022081 | 0.851088 | 0.972638 | ENSMUSG( | Gng4 | 1900.096 | -0.06374 | 0.74131 | 0.968811 |
| ENSMUSG( | Gng5 | 1326.177 | 0.111364 | 0.457195 | 0.839356 | ENSMUSG( | Gng5 | 1115.027 | -0.12566 | 0.456313 | 0.923329 |
| ENSMUSG( | Gng7 | 15142.03 | -0.44928 | 0.00483 | 0.091852 | ENSMUSG( | Gng7 | 15446.46 | -0.08942 | 0.450032 | 0.922202 |
| ENSMUSG( | Gng8 | 67.07596 | -0.12164 | 0.621296 | 0.911432 | ENSMUSG( | Gng8 | 57.30979 | -0.31764 | 0.259169 | 0.883971 |
| ENSMUSG( | Gngt2 | 109.1628 | 0.062613 | 0.792273 | 0.959461 | ENSMUSG( | Gngt2 | 84.79069 | -0.44053 | 0.100186 | 0.801311 |
| ENSMUSG( | Gnl1 | 10100.22 | 0.116186 | 0.116798 | 0.515501 | ENSMUSG( | Gnl1 | 8694.586 | -0.04557 | 0.62925 | 0.953251 |
| ENSMUSG( | Gnl2 | 1536.187 | -0.00662 | 0.955149 | 0.993874 | ENSMUSG( | Gnl2 | 1411.682 | 0.016515 | 0.910837 | 0.992196 |
| ENSMUSG( | Gnl3 | 568.2545 | 0.072987 | 0.523518 | 0.875628 | ENSMUSG( | Gnl3 | 523.1311 | 0.10905 | 0.463287 | 0.923329 |
| ENSMUSG( | Gnl3l | 3751.815 | 0.027429 | 0.762878 | 0.953524 | ENSMUSG( | Gnl3l | 3227.049 | -0.14426 | 0.221218 | 0.866104 |
| ENSMUSG( | Gnmt | 19.32537 | 0.239712 | 0.593258 | NA | ENSMUSG( | Gnmt | 21.37761 | 0.697594 | 0.1255 | 0.81516 |
| ENSMUSG( | Gnpat | 2066.904 | 0.093715 | 0.261034 | 0.710889 | ENSMUSG( | Gnpat | 1841.154 | 0.031132 | 0.772967 | 0.974463 |
| ENSMUSG( | Gnpda1 | 624.0778 | -0.02049 | 0.86026 | 0.974842 | ENSMUSG( | Gnpda1 | 541.8699 | -0.16599 | 0.217464 | 0.865589 |
| ENSMUSG( | Gnpda2 | 589.0245 | -0.01549 | 0.894441 | 0.984512 | ENSMUSG( | Gnpda2 | 524.7684 | -0.08737 | 0.525195 | 0.934374 |
| ENSMUSG( | Gnpnat1 | 170.6952 | -0.11752 | 0.476689 | 0.851717 | ENSMUSG( | Gnpnat1 | 160.4017 | -0.03405 | 0.854153 | 0.98563 |
| ENSMUSG( | Gnptab | 2575.407 | -0.10706 | 0.417642 | 0.821707 | ENSMUSG( | Gnptab | 2634.545 | 0.22494 | 0.138243 | 0.828492 |
| ENSMUSG( | Gnptg | 3555.483 | -0.13231 | 0.167929 | 0.600753 | ENSMUSG( | Gnptg | 3341.377 | -0.03961 | 0.695225 | 0.960867 |
| ENSMUSG( | Gnrh1 | 11.18292 | -0.28973 | 0.655102 | NA | ENSMUSG( | Gnrh1 | 13.91694 | 0.581716 | 0.302065 | 0.898538 |
| ENSMUSG( | Gns | 3907.583 | -0.30275 | 0.019164 | 0.208619 | ENSMUSG( | Gns | 4197.835 | 0.186776 | 0.249363 | 0.877904 |
| ENSMUSG( | Golga1 | 1059.221 | 0.112331 | 0.230173 | 0.678139 | ENSMUSG( | Golga1 | 954.9766 | 0.08307 | 0.498009 | 0.927825 |
| ENSMUSG( | Golga2 | 2943.317 | 0.145887 | 0.107809 | 0.49674 | ENSMUSG( | Golga2 | 2538.451 | -0.0088 | 0.933465 | 0.99306 |
| ENSMUSG( | Golga3 | 1364.025 | 0.165296 | 0.089237 | 0.458482 | ENSMUSG( | Golga3 | 1190.038 | 0.04525 | 0.684 | 0.958678 |

| | | | | | |
|---|---|---|---|---|---|
| ENSMUSG( | 36.4122 | -0.12107 | 0.713893 | 0.940903 | Gm8355 |
| ENSMUSG( | 68.88453 | -0.08884 | 0.723074 | 0.943761 | Gm8451 |
| ENSMUSG( | 14.47734 | 0.723816 | 0.168905 | NA | Gm8494 |
| ENSMUSG( | 13.67153 | -0.12081 | 0.81251 | NA | Gm8546 |
| ENSMUSG( | 33.38852 | 0.149176 | 0.683557 | 0.932154 | Gm867 |
| ENSMUSG( | 24.83499 | 0.341818 | 0.361879 | 0.792797 | Gm8692 |
| ENSMUSG( | 17.49074 | 0.317881 | 0.483781 | NA | Gm8717 |
| ENSMUSG( | 98.19825 | 0.245417 | 0.25384 | 0.704011 | Gm8730 |
| ENSMUSG( | 30.75495 | 0.594355 | 0.112677 | 0.506698 | Gm8738 |
| ENSMUSG( | 11.03286 | -0.09661 | 0.861829 | NA | Gm8797 |
| ENSMUSG( | 13.11116 | -0.54421 | 0.288858 | NA | Gm8971 |
| ENSMUSG( | 28.71499 | -0.00811 | 0.983101 | 0.997554 | Gm8983 |
| ENSMUSG( | 18.34954 | 0.133759 | 0.777191 | NA | Gm9294 |
| ENSMUSG( | 13.09063 | -0.04851 | 0.929347 | NA | Gm9320 |
| ENSMUSG( | 14.50569 | 0.36387 | 0.447921 | NA | Gm9333 |
| ENSMUSG( | 14.12161 | -0.06478 | 0.896478 | NA | Gm9347 |
| ENSMUSG( | 14.44482 | 0.081572 | 0.88273 | NA | Gm9396 |
| ENSMUSG( | 15.87729 | -0.1482 | 0.756591 | NA | Gm9403 |
| ENSMUSG( | 56.6912 | -0.25326 | 0.359765 | 0.791346 | Gm9493 |
| ENSMUSG( | 9.330316 | -0.11626 | 0.861958 | NA | Gm9575 |
| ENSMUSG( | 104.4119 | 0.247454 | 0.194232 | 0.633383 | Gm9616 |
| ENSMUSG( | 39.97549 | 0.388516 | 0.195685 | 0.635109 | Gm9625 |
| ENSMUSG( | 736.3007 | -0.10257 | 0.419735 | 0.822433 | Gm973 |
| ENSMUSG( | 16.33254 | 0.193972 | 0.677065 | NA | Gm9755 |
| ENSMUSG( | 10.52693 | 0.006688 | 0.990762 | NA | Gm9767 |
| ENSMUSG( | 107.2528 | -0.27109 | 0.164623 | 0.595625 | Gm9776 |
| ENSMUSG( | 149.2459 | 0.261875 | 0.192711 | 0.631789 | Gm9794 |
| ENSMUSG( | 134.2469 | -0.16936 | 0.382432 | 0.803227 | Gm9828 |
| ENSMUSG( | 446.7619 | 0.170121 | 0.277051 | 0.727148 | Gm9899 |
| ENSMUSG( | 59.58827 | 0.077709 | 0.768455 | 0.954674 | Gm9917 |
| ENSMUSG( | 14.64277 | -0.05137 | 0.916003 | NA | Gm9936 |
| ENSMUSG( | 11.62759 | 0.018596 | 0.978934 | NA | Gm9946 |
| ENSMUSG( | 21.71841 | -0.11103 | 0.787983 | 0.959226 | Gm9958 |
| ENSMUSG( | 48.28587 | -0.35011 | 0.230241 | 0.678139 | Gm9962 |
| ENSMUSG( | 850.7324 | -0.19139 | 0.086535 | 0.452045 | Gmcl1 |
| ENSMUSG( | 199.3321 | -0.01688 | 0.914718 | 0.988522 | Gmds |
| ENSMUSG( | 691.7946 | -0.02231 | 0.846078 | 0.972226 | Gmeb1 |
| ENSMUSG( | 706.1354 | -0.03773 | 0.733037 | 0.944956 | Gmeb2 |
| ENSMUSG( | 8939.394 | 0.00321 | 0.982823 | 0.997522 | Gmfb |
| ENSMUSG( | 71.62583 | -0.10589 | 0.672819 | 0.929007 | Gmfg |
| ENSMUSG( | 375.7112 | 0.250501 | 0.104089 | 0.489938 | Gmip |
| ENSMUSG( | 17.45655 | 0.208181 | 0.64812 | NA | Gmnc |
| ENSMUSG( | 56.14945 | 0.062876 | 0.813374 | 0.964574 | Gmnn |
| ENSMUSG( | 834.2568 | 0.245611 | 0.042672 | 0.316769 | Gmppa |
| ENSMUSG( | 272.1709 | 0.054798 | 0.736827 | 0.94664 | Gmppb |
| ENSMUSG( | 685.4732 | -0.10581 | 0.505422 | 0.86689 | Gmpr |
| ENSMUSG( | 538.9672 | 0.022139 | 0.83518 | 0.970426 | Gmpr2 |
| ENSMUSG( | 1215.144 | 0.116022 | 0.280378 | 0.729291 | Gmps |
| ENSMUSG( | 6218.343 | 0.113074 | 0.191625 | 0.630674 | Gna11 |
| ENSMUSG( | 8997.072 | 0.097371 | 0.266201 | 0.716441 | Gna12 |
| ENSMUSG( | 5104.527 | 0.172031 | 0.08626 | 0.451772 | Gna13 |
| ENSMUSG( | 93.7873 | 0.014776 | 0.945494 | 0.99365 | Gna15 |
| ENSMUSG( | 2675.438 | -0.07201 | 0.394873 | 0.810072 | Gnai1 |
| ENSMUSG( | 17231.37 | -0.06023 | 0.536993 | 0.882123 | Gnai2 |
| ENSMUSG( | 1390.837 | -0.00251 | 0.982406 | 0.997512 | Gnai3 |
| ENSMUSG( | 6732.773 | -0.23075 | 0.022545 | 0.229407 | Gnal |
| ENSMUSG( | 41443.38 | 0.013821 | 0.884212 | 0.982647 | Gnao1 |
| ENSMUSG( | 5910.783 | 0.172766 | 0.028742 | 0.258426 | Gnaq |
| ENSMUSG( | 62267.32 | 0.044878 | 0.553376 | 0.886216 | Gnas |
| ENSMUSG( | 2172.456 | 0.142572 | 0.179761 | 0.617507 | Gnaz |
| ENSMUSG( | 28451.12 | 0.097916 | 0.199927 | 0.640239 | Gnb1 |
| ENSMUSG( | 102.1054 | 0.085834 | 0.67757 | 0.930542 | Gnb1l |
| ENSMUSG( | 8551.015 | 0.01444 | 0.882506 | 0.982244 | Gnb2 |
| ENSMUSG( | 1253.769 | 0.006719 | 0.944356 | 0.993543 | Gnb4 |
| ENSMUSG( | 2155.108 | -0.06287 | 0.599906 | 0.903978 | Gnb5 |
| ENSMUSG( | 729.2545 | 0.134262 | 0.305859 | 0.753712 | Gne |
| ENSMUSG( | 1476.958 | 0.028626 | 0.797639 | 0.960507 | Gng10 |
| ENSMUSG( | 269.2676 | -0.49434 | 0.001994 | 0.051341 | Gng11 |
| ENSMUSG( | 7465.765 | 0.128625 | 0.288764 | 0.739578 | Gng12 |
| ENSMUSG( | 1879.941 | 0.034869 | 0.677051 | 0.930542 | Gng13 |
| ENSMUSG( | 3269.129 | 0.020225 | 0.824951 | 0.967386 | Gng2 |
| ENSMUSG( | 5481.286 | 0.066953 | 0.619754 | 0.911432 | Gng3 |
| ENSMUSG( | 2144.46 | 0.022081 | 0.851088 | 0.972638 | Gng4 |
| ENSMUSG( | 1326.177 | 0.111364 | 0.457195 | 0.839356 | Gng5 |
| ENSMUSG( | 15142.03 | -0.44928 | 0.00483 | 0.091852 | Gng7 |
| ENSMUSG( | 67.07596 | -0.12164 | 0.621296 | 0.911432 | Gng8 |
| ENSMUSG( | 109.1628 | 0.062613 | 0.792273 | 0.959461 | Gngt2 |
| ENSMUSG( | 10100.22 | 0.116186 | 0.116798 | 0.515501 | Gnl1 |
| ENSMUSG( | 1536.187 | -0.00662 | 0.955149 | 0.993874 | Gnl2 |
| ENSMUSG( | 568.2545 | 0.072987 | 0.523518 | 0.875628 | Gnl3 |
| ENSMUSG( | 3751.815 | 0.027429 | 0.762878 | 0.953524 | Gnl3l |
| ENSMUSG( | 19.32537 | 0.239712 | 0.593258 | NA | Gnmt |
| ENSMUSG( | 2066.904 | 0.093715 | 0.261034 | 0.710889 | Gnpat |
| ENSMUSG( | 624.0778 | -0.02049 | 0.86026 | 0.974842 | Gnpda1 |
| ENSMUSG( | 589.0245 | -0.01549 | 0.894441 | 0.984512 | Gnpda2 |
| ENSMUSG( | 170.6952 | -0.11752 | 0.476689 | 0.851717 | Gnpnat1 |
| ENSMUSG( | 2575.407 | -0.10706 | 0.417642 | 0.821707 | Gnptab |
| ENSMUSG( | 3555.483 | -0.13231 | 0.167929 | 0.600753 | Gnptg |
| ENSMUSG( | 11.18292 | -0.28973 | 0.655102 | NA | Gnrh1 |
| ENSMUSG( | 3907.583 | -0.30275 | 0.019164 | 0.208619 | Gns |
| ENSMUSG( | 1059.221 | 0.112331 | 0.230173 | 0.678139 | Golga1 |
| ENSMUSG( | 2943.317 | 0.145887 | 0.107809 | 0.49674 | Golga2 |
| ENSMUSG( | 1364.025 | 0.165296 | 0.089237 | 0.458482 | Golga3 |
| ENSMUSG( | 4019.528 | -0.00178 | 0.983221 | 0.997554 | Golga4 |
| ENSMUSG( | 1109.662 | 0.133528 | 0.185619 | 0.623185 | Golga5 |
| ENSMUSG( | 6786.516 | -0.01781 | 0.873853 | 0.979333 | Golga7 |
| ENSMUSG( | 6536.246 | 0.169876 | 0.051992 | 0.350064 | Golga7b |
| ENSMUSG( | 3883.272 | -0.22425 | 0.005724 | 0.101308 | Golgb1 |
| ENSMUSG( | 1864.974 | -0.03083 | 0.788789 | 0.959226 | Golim4 |
| ENSMUSG( | 1770.999 | 0.003882 | 0.976103 | 0.997428 | Golm1 |
| ENSMUSG( | 4239.142 | -0.25171 | 0.05718 | 0.369067 | Golm2 |
| ENSMUSG( | 2635.202 | 0.071351 | 0.445209 | 0.833239 | Golph3 |
| ENSMUSG( | 486.6169 | 0.107206 | 0.365025 | 0.79407 | Golph3l |
| ENSMUSG( | 1024.95 | -0.41881 | 0.004531 | 0.088122 | Golt1b |
| ENSMUSG( | 2035.01 | 0.101831 | 0.261023 | 0.710889 | Gon4l |
| ENSMUSG( | 102.7163 | -0.19287 | 0.346121 | 0.779153 | Gon7 |
| ENSMUSG( | 1088.484 | -0.05064 | 0.61201 | 0.908586 | Gopc |
| ENSMUSG( | 160.3083 | -0.18722 | 0.264846 | 0.715848 | Gorab |
| ENSMUSG( | 915.9324 | -0.02166 | 0.83748 | 0.971111 | Gorasp1 |
| ENSMUSG( | 2287.758 | -0.19972 | 0.024491 | 0.23959 | Gorasp2 |
| ENSMUSG( | 1444.474 | 0.055698 | 0.541059 | 0.883182 | Gosr1 |
| ENSMUSG( | 4347.32 | 0.033854 | 0.715854 | 0.941484 | Gosr2 |
| ENSMUSG( | 4068.094 | 0.042486 | 0.770018 | 0.954689 | Got1 |
| ENSMUSG( | 20.47531 | 0.188452 | 0.674585 | NA | Got1l1 |
| ENSMUSG( | 4165.683 | 0.021954 | 0.86312 | 0.975055 | Got2 |
| ENSMUSG( | 28.89394 | 0.218383 | 0.534592 | 0.880719 | Gp1bb |
| ENSMUSG( | 21.40059 | -0.03028 | 0.942432 | 0.993415 | Gp5 |
| ENSMUSG( | 30.82696 | -0.2278 | 0.558765 | 0.888638 | Gp9 |
| ENSMUSG( | 1361.697 | -0.0893 | 0.492684 | 0.860462 | Gpaa1 |
| ENSMUSG( | 796.8766 | 0.079388 | 0.445014 | 0.833239 | Gpalpp1 |
| ENSMUSG( | 6467.747 | 0.188369 | 0.106973 | 0.494682 | Gpam |
| ENSMUSG( | 226.7054 | 0.011536 | 0.948971 | 0.993821 | Gpank1 |
| ENSMUSG( | 85.947 | -0.76056 | 0.000513 | 0.020832 | Gpat3 |
| ENSMUSG( | 3324.189 | -0.07286 | 0.565211 | 0.890779 | Gpat4 |
| ENSMUSG( | 820.1895 | 0.124384 | 0.234725 | 0.682178 | Gpatch1 |
| ENSMUSG( | 754.5933 | 0.044577 | 0.683924 | 0.93244 | Gpatch11 |
| ENSMUSG( | 673.1768 | 0.08793 | 0.435207 | 0.829093 | Gpatch2 |
| ENSMUSG( | 1126.653 | 0.013573 | 0.883254 | 0.982422 | Gpatch2l |
| ENSMUSG( | 124.5145 | -0.12118 | 0.512162 | 0.871563 | Gpatch3 |
| ENSMUSG( | 560.1924 | 0.097392 | 0.449345 | 0.835057 | Gpatch4 |
| ENSMUSG( | 3727.808 | 0.125551 | 0.178708 | 0.616298 | Gpatch8 |
| ENSMUSG( | 1515.605 | 0.103188 | 0.365062 | 0.79407 | Gpbp1 |
| ENSMUSG( | 987.5973 | 0.039473 | 0.684396 | 0.93282 | Gpbp1l1 |
| ENSMUSG( | 3096.961 | -0.19376 | 0.1377 | 0.552903 | Gpc1 |
| ENSMUSG( | 35.68712 | 0.599919 | 0.061025 | 0.381679 | Gpc2 |
| ENSMUSG( | 204.2825 | -0.1685 | 0.340051 | 0.777959 | Gpc3 |
| ENSMUSG( | 548.6062 | -0.46353 | 0.0013 | 0.039202 | Gpc4 |
| ENSMUSG( | 4506.972 | 0.02665 | 0.823533 | 0.966603 | Gpc5 |
| ENSMUSG( | 404.0536 | 0.192024 | 0.248097 | 0.69682 | Gpc6 |
| ENSMUSG( | 2757.395 | 0.05553 | 0.504724 | 0.866883 | Gpcpd1 |
| ENSMUSG( | 3751.952 | 0.204213 | 0.037802 | 0.300541 | Gpd1 |
| ENSMUSG( | 3934.15 | 0.120621 | 0.120774 | 0.524807 | Gpd1l |
| ENSMUSG( | 5569.414 | 0.136088 | 0.221614 | 0.667258 | Gpd2 |
| ENSMUSG( | 103.087 | 0.008181 | 0.975407 | 0.997302 | Gper1 |
| ENSMUSG( | 1618.244 | 0.048253 | 0.646633 | 0.918796 | Gphn |
| ENSMUSG( | 20733.81 | 0.092746 | 0.323533 | 0.768536 | Gpi1 |
| ENSMUSG( | 115.2102 | 0.229537 | 0.240755 | 0.688247 | Gpi-ps |
| ENSMUSG( | 1776.312 | 0.053937 | 0.586211 | 0.899709 | Gpkow |
| ENSMUSG( | 3288.088 | -0.11472 | 0.362457 | 0.792943 | Gpld1 |
| ENSMUSG( | 40349.75 | -0.28281 | 0.056562 | 0.366826 | Gpm6a |
| ENSMUSG( | 95233.3 | -0.03286 | 0.803345 | 0.962169 | Gpm6b |
| ENSMUSG( | 777.8492 | 0.094056 | 0.390389 | 0.807976 | Gpn1 |
| ENSMUSG( | 204.5415 | 0.208106 | 0.208327 | 0.650319 | Gpn2 |
| ENSMUSG( | 360.4441 | 0.037698 | 0.801417 | 0.961947 | Gpn3 |
| ENSMUSG( | 96.30105 | -0.59876 | 0.029105 | 0.259574 | Gpnmb |
| ENSMUSG( | 120.2802 | -0.11134 | 0.61095 | 0.908472 | Gpr101 |
| ENSMUSG( | 2327.431 | -0.11737 | 0.426153 | 0.824898 | Gpr107 |
| ENSMUSG( | 963.2336 | -0.06734 | 0.644805 | 0.917836 | Gpr108 |

| | | | | | |
|---|---|---|---|---|---|
| ENSMUSG( | 237.5411 | -0.07466 | 0.69143 | 0.960071 | Gmppb |
| ENSMUSG( | 583.1721 | -0.32412 | 0.065782 | 0.745828 | Gmpr |
| ENSMUSG( | 491.3948 | 0.025562 | 0.84278 | 0.98313 | Gmpr2 |
| ENSMUSG( | 1091.768 | 0.072645 | 0.57739 | 0.943044 | Gmps |
| ENSMUSG( | 5494.541 | 0.028153 | 0.779574 | 0.976106 | Gna11 |
| ENSMUSG( | 7548.726 | -0.14375 | 0.188904 | 0.85698 | Gna12 |
| ENSMUSG( | 4313.834 | -0.04227 | 0.736125 | 0.967344 | Gna13 |
| ENSMUSG( | 98.46954 | 0.412056 | 0.084714 | 0.779799 | Gna15 |
| ENSMUSG( | 2470.833 | -0.0349 | 0.813311 | 0.979198 | Gnai1 |
| ENSMUSG( | 15205.78 | -0.1566 | 0.178982 | 0.849041 | Gnai2 |
| ENSMUSG( | 1252.229 | -0.03937 | 0.771944 | 0.974236 | Gnai3 |
| ENSMUSG( | 6418.77 | -0.09148 | 0.374955 | 0.915491 | Gnal |
| ENSMUSG( | 34990.64 | -0.21346 | 0.063681 | 0.736482 | Gnao1 |
| ENSMUSG( | 5073.427 | 0.005152 | 0.965883 | 0.995433 | Gnaq |
| ENSMUSG( | 53410.84 | -0.13292 | 0.217317 | 0.865589 | Gnas |
| ENSMUSG( | 1923.219 | 0.060825 | 0.610044 | 0.950639 | Gnaz |
| ENSMUSG( | 24214.06 | -0.09975 | 0.393543 | 0.917257 | Gnb1 |
| ENSMUSG( | 99.46891 | 0.263434 | 0.237883 | 0.87259 | Gnb1l |
| ENSMUSG( | 7352.726 | -0.1589 | 0.208864 | 0.864412 | Gnb2 |
| ENSMUSG( | 1117.231 | -0.05626 | 0.637334 | 0.953903 | Gnb4 |
| ENSMUSG( | 1914.413 | -0.14169 | 0.330173 | 0.907251 | Gnb5 |
| ENSMUSG( | 638.1181 | 0.021762 | 0.881922 | 0.989644 | Gne |
| ENSMUSG( | 1271.472 | -0.14324 | 0.331331 | 0.907861 | Gng10 |
| ENSMUSG( | 278.8389 | -0.0734 | 0.681768 | 0.958678 | Gng11 |
| ENSMUSG( | 6252.057 | -0.11611 | 0.432306 | 0.918158 | Gng12 |
| ENSMUSG( | 1534.682 | -0.29976 | 0.008762 | 0.393969 | Gng13 |
| ENSMUSG( | 2875.118 | -0.08637 | 0.503628 | 0.927825 | Gng2 |
| ENSMUSG( | 4611.501 | -0.17056 | 0.325986 | 0.905339 | Gng3 |
| ENSMUSG( | 1900.096 | -0.06374 | 0.74131 | 0.968811 | Gng4 |
| ENSMUSG( | 1115.027 | -0.12566 | 0.456313 | 0.923329 | Gng5 |
| ENSMUSG( | 15446.46 | -0.08942 | 0.450032 | 0.922202 | Gng7 |
| ENSMUSG( | 57.30979 | -0.31764 | 0.259169 | 0.883971 | Gng8 |
| ENSMUSG( | 84.79069 | -0.44053 | 0.100186 | 0.801311 | Gngt2 |
| ENSMUSG( | 8694.586 | -0.04557 | 0.62925 | 0.953251 | Gnl1 |
| ENSMUSG( | 1411.682 | 0.016515 | 0.910837 | 0.992196 | Gnl2 |
| ENSMUSG( | 523.1311 | 0.10905 | 0.463287 | 0.923329 | Gnl3 |
| ENSMUSG( | 3227.049 | -0.14426 | 0.221218 | 0.866104 | Gnl3l |
| ENSMUSG( | 21.37761 | 0.697594 | 0.1255 | 0.81516 | Gnmt |
| ENSMUSG( | 1841.154 | 0.031132 | 0.772967 | 0.974463 | Gnpat |
| ENSMUSG( | 541.8699 | -0.16599 | 0.217464 | 0.865589 | Gnpda1 |
| ENSMUSG( | 524.7684 | -0.08737 | 0.525195 | 0.934374 | Gnpda2 |
| ENSMUSG( | 160.4017 | -0.03405 | 0.854153 | 0.98563 | Gnpnat1 |
| ENSMUSG( | 2634.545 | 0.22494 | 0.138243 | 0.828492 | Gnptab |
| ENSMUSG( | 3341.377 | -0.03961 | 0.695225 | 0.960867 | Gnptg |
| ENSMUSG( | 13.91694 | 0.581716 | 0.302065 | 0.898538 | Gnrh1 |
| ENSMUSG( | 4197.835 | 0.186776 | 0.249363 | 0.877904 | Gns |
| ENSMUSG( | 954.9766 | 0.08307 | 0.498009 | 0.927825 | Golga1 |
| ENSMUSG( | 2538.451 | -0.0088 | 0.933465 | 0.99306 | Golga2 |
| ENSMUSG( | 1190.038 | 0.04525 | 0.684 | 0.958678 | Golga3 |
| ENSMUSG( | 3567.475 | -0.07998 | 0.439583 | 0.920721 | Golga4 |
| ENSMUSG( | 955.2429 | -0.02892 | 0.802802 | 0.97681 | Golga5 |
| ENSMUSG( | 5901.893 | -0.15951 | 0.265202 | 0.886134 | Golga7 |
| ENSMUSG( | 5500.525 | -0.05508 | 0.569672 | 0.941777 | Golga7b |
| ENSMUSG( | 3809.239 | -0.00189 | 0.984592 | 0.996963 | Golgb1 |
| ENSMUSG( | 1730.428 | 0.023471 | 0.842198 | 0.983044 | Golim4 |
| ENSMUSG( | 1812.78 | 0.326598 | 0.051395 | 0.711116 | Golm1 |
| ENSMUSG( | 4305.092 | 0.073346 | 0.619191 | 0.952774 | Golm2 |
| ENSMUSG( | 2251.624 | -0.11826 | 0.412152 | 0.917257 | Golph3 |
| ENSMUSG( | 427.4297 | -0.00303 | 0.983867 | 0.996963 | Golph3l |
| ENSMUSG( | 1104.152 | 0.098255 | 0.544627 | 0.938063 | Golt1b |
| ENSMUSG( | 1819.563 | 0.048563 | 0.674424 | 0.95694 | Gon4l |
| ENSMUSG( | 98.31492 | -0.07172 | 0.777273 | 0.975838 | Gon7 |
| ENSMUSG( | 1020.697 | 0.029585 | 0.810674 | 0.978779 | Gopc |
| ENSMUSG( | 153.9975 | -0.03206 | 0.867249 | 0.98775 | Gorab |
| ENSMUSG( | 807.3084 | -0.11666 | 0.333065 | 0.907907 | Gorasp1 |
| ENSMUSG( | 2184.364 | -0.05733 | 0.597954 | 0.946676 | Gorasp2 |
| ENSMUSG( | 1257.965 | -0.07657 | 0.495247 | 0.927372 | Gosr1 |
| ENSMUSG( | 3685.885 | -0.18058 | 0.127833 | 0.81845 | Gosr2 |
| ENSMUSG( | 3494.934 | -0.13192 | 0.404537 | 0.917257 | Got1 |
| ENSMUSG( | 18.34562 | 0.17603 | 0.70896 | 0.964042 | Got1l1 |
| ENSMUSG( | 3610.536 | -0.12495 | 0.335609 | 0.907907 | Got2 |
| ENSMUSG( | 22.57523 | -0.20911 | 0.615667 | 0.951194 | Gp1bb |
| ENSMUSG( | 18.93153 | -0.11567 | 0.797544 | 0.97681 | Gp5 |
| ENSMUSG( | 33.24886 | 0.270195 | 0.491742 | 0.927018 | Gp9 |
| ENSMUSG( | 1356.744 | 0.170184 | 0.231826 | 0.868507 | Gpaa1 |
| ENSMUSG( | 713.2565 | 0.030207 | 0.801068 | 0.97681 | Gpalpp1 |
| ENSMUSG( | 5493.268 | -0.00544 | 0.967163 | 0.995581 | Gpam |
| ENSMUSG( | 218.588 | 0.166346 | 0.383949 | 0.917257 | Gpank1 |
| ENSMUSG( | 92.95418 | -0.17658 | 0.435384 | 0.919062 | Gpat3 |
| ENSMUSG( | 3278.548 | 0.156948 | 0.298834 | 0.898538 | Gpat4 |
| ENSMUSG( | 718.9039 | 0.012559 | 0.922547 | 0.992644 | Gpatch1 |
| ENSMUSG( | 668.6361 | -0.03423 | 0.798436 | 0.97681 | Gpatch11 |
| ENSMUSG( | 589.9835 | -0.02354 | 0.849366 | 0.985321 | Gpatch2 |
| ENSMUSG( | 1016.036 | -0.01917 | 0.874885 | 0.989644 | Gpatch2l |
| ENSMUSG( | 122.0026 | 0.093199 | 0.658823 | 0.956205 | Gpatch3 |
| ENSMUSG( | 462.8615 | -0.19353 | 0.210618 | 0.864412 | Gpatch4 |
| ENSMUSG( | 3304.455 | 0.046188 | 0.756435 | 0.971656 | Gpatch8 |
| ENSMUSG( | 1390.522 | 0.116442 | 0.49797 | 0.927825 | Gpbp1 |
| ENSMUSG( | 961.1258 | 0.223484 | 0.060795 | 0.734852 | Gpbp1l1 |
| ENSMUSG( | 3229.118 | 0.199845 | 0.19017 | 0.857355 | Gpc1 |
| ENSMUSG( | 33.80676 | 0.716097 | 0.080009 | 0.776379 | Gpc2 |
| ENSMUSG( | 207.4422 | 0.151289 | 0.410592 | 0.917257 | Gpc3 |
| ENSMUSG( | 598.5838 | 0.09417 | 0.543559 | 0.938063 | Gpc4 |
| ENSMUSG( | 4205.126 | 0.093617 | 0.517105 | 0.930939 | Gpc5 |
| ENSMUSG( | 365.1999 | 0.179893 | 0.378888 | 0.916995 | Gpc6 |
| ENSMUSG( | 2760.466 | 0.31242 | 0.008255 | 0.388416 | Gpcpd1 |
| ENSMUSG( | 3213.621 | 0.03255 | 0.752463 | 0.970521 | Gpd1 |
| ENSMUSG( | 3383.296 | -0.04573 | 0.690906 | 0.960071 | Gpd1l |
| ENSMUSG( | 4844.616 | 0.006679 | 0.957309 | 0.994798 | Gpd2 |
| ENSMUSG( | 138.9512 | 0.992797 | 0.001989 | 0.169168 | Gper1 |
| ENSMUSG( | 1412.722 | -0.08027 | 0.561939 | 0.940388 | Gphn |
| ENSMUSG( | 17597.56 | -0.11396 | 0.347453 | 0.911652 | Gpi1 |
| ENSMUSG( | 86.681 | -0.32758 | 0.160856 | 0.837724 | Gpi-ps |
| ENSMUSG( | 1531.782 | -0.11017 | 0.382243 | 0.917257 | Gpkow |
| ENSMUSG( | 3018.108 | -0.09048 | 0.507921 | 0.929089 | Gpld1 |
| ENSMUSG( | 43201.09 | 0.191415 | 0.271886 | 0.888383 | Gpm6a |
| ENSMUSG( | 84625.11 | -0.10804 | 0.483539 | 0.924993 | Gpm6b |
| ENSMUSG( | 651.9279 | -0.1488 | 0.228852 | 0.868507 | Gpn1 |
| ENSMUSG( | 175.9295 | 0.051172 | 0.777349 | 0.975838 | Gpn2 |
| ENSMUSG( | 325.8233 | 0.011487 | 0.946032 | 0.994798 | Gpn3 |
| ENSMUSG( | 101.7865 | -0.10297 | 0.684307 | 0.958678 | Gpnmb |
| ENSMUSG( | 122.6442 | 0.191194 | 0.391345 | 0.917257 | Gpr101 |
| ENSMUSG( | 2306.343 | 0.129699 | 0.430123 | 0.917257 | Gpr107 |
| ENSMUSG( | 925.7888 | 0.093907 | 0.602598 | 0.94789 | Gpr108 |
| ENSMUSG( | 432.6856 | 0.046741 | 0.786369 | 0.976303 | Gpr12 |
| ENSMUSG( | 447.085 | 0.104397 | 0.556123 | 0.939565 | Gpr135 |
| ENSMUSG( | 3751.119 | 0.058456 | 0.675794 | 0.95694 | Gpr137 |
| ENSMUSG( | 426.6863 | 0.116892 | 0.459471 | 0.923329 | Gpr137b |
| ENSMUSG( | 1003.329 | 0.234163 | 0.12596 | 0.81516 | Gpr137c |
| ENSMUSG( | 70.08526 | -0.16651 | 0.529754 | 0.934795 | Gpr139 |
| ENSMUSG( | 1187.667 | 0.185017 | 0.349713 | 0.911652 | Gpr146 |
| ENSMUSG( | 52.78422 | 0.399106 | 0.212859 | 0.864412 | Gpr149 |
| ENSMUSG( | 119.7808 | 0.098943 | 0.63119 | 0.953251 | Gpr150 |
| ENSMUSG( | 22.67336 | 0.822138 | 0.057429 | 0.723727 | Gpr151 |
| ENSMUSG( | 1290.935 | -0.01199 | 0.913392 | 0.992298 | Gpr153 |
| ENSMUSG( | 2199.586 | 0.194075 | 0.229989 | 0.868507 | Gpr155 |
| ENSMUSG( | 73.3712 | 0.133629 | 0.643651 | 0.953903 | Gpr156 |
| ENSMUSG( | 78.67233 | 0.290614 | 0.259218 | 0.883971 | Gpr157 |
| ENSMUSG( | 4571.359 | 0.123492 | 0.450771 | 0.922202 | Gpr158 |
| ENSMUSG( | 14.06384 | -0.07364 | 0.887731 | 0.990399 | Gpr15lg |
| ENSMUSG( | 46.87109 | 0.147679 | 0.671093 | 0.95694 | Gpr160 |
| ENSMUSG( | 663.0574 | 0.304452 | 0.040679 | 0.683847 | Gpr161 |
| ENSMUSG( | 7089.651 | 0.112325 | 0.382522 | 0.917257 | Gpr162 |
| ENSMUSG( | 126.7336 | 0.45982 | 0.074108 | 0.766789 | Gpr165 |
| ENSMUSG( | 4922.131 | 0.489407 | 0.232851 | 0.868507 | Gpr17 |
| ENSMUSG( | 582.8057 | 0.170651 | 0.276349 | 0.890889 | Gpr173 |
| ENSMUSG( | 460.1443 | 0.150811 | 0.370365 | 0.915491 | Gpr176 |
| ENSMUSG( | 46.41798 | -0.99729 | 0.00165 | 0.150169 | Gpr179 |
| ENSMUSG( | 890.2572 | 0.123564 | 0.401993 | 0.917257 | Gpr180 |
| ENSMUSG( | 95.64431 | 0.920953 | 0.000293 | 0.046205 | Gpr182 |
| ENSMUSG( | 88.04604 | 0.818785 | 0.009625 | 0.406458 | Gpr183 |
| ENSMUSG( | 496.0011 | -0.11425 | 0.524532 | 0.933764 | Gpr19 |
| ENSMUSG( | 89.60406 | 0.079539 | 0.78771 | 0.97637 | Gpr21 |
| ENSMUSG( | 1018.702 | 0.516235 | 0.002936 | 0.217424 | Gpr22 |
| ENSMUSG( | 56.43744 | 0.025737 | 0.924969 | 0.99306 | Gpr25 |
| ENSMUSG( | 5421.969 | 0.028296 | 0.875574 | 0.989644 | Gpr26 |
| ENSMUSG( | 1584.384 | 0.251995 | 0.068183 | 0.75212 | Gpr27 |
| ENSMUSG( | 285.5065 | -0.36986 | 0.05049 | 0.708889 | Gpr3 |
| ENSMUSG( | 500.509 | 0.485302 | 0.052053 | 0.712974 | Gpr34 |
| ENSMUSG( | 34.40511 | 0.113524 | 0.76552 | 0.974236 | Gpr35 |
| ENSMUSG( | 1808.485 | 0.136543 | 0.731748 | 0.967228 | Gpr37 |
| ENSMUSG( | 17890.23 | -0.01144 | 0.945518 | 0.994798 | Gpr37l1 |
| ENSMUSG( | 17.40047 | -0.57507 | 0.238495 | 0.872849 | Gpr39 |
| ENSMUSG( | 225.7357 | 0.79617 | 6.96E-06 | 0.002837 | Gpr4 |
| ENSMUSG( | 514.6665 | -0.02203 | 0.873039 | 0.989445 | Gpr45 |
| ENSMUSG( | 124.8417 | 0.367138 | 0.102185 | 0.802316 | Gpr52 |
| ENSMUSG( | 9.51746 | 0.005244 | 0.993282 | NA | Gpr55 |
| ENSMUSG( | 317.7689 | 0.228093 | 0.267523 | 0.886134 | Gpr6 |

| | | | | | | | | | | | |
|---|---|---|---|---|---|---|---|---|---|---|---|
| ENSMUSG( | Golga4 | 4019.528 | -0.00178 | 0.983221 | 0.997554 | ENSMUSG( | Golga4 | 3567.475 | -0.07998 | 0.439583 | 0.920721 |
| ENSMUSG( | Golga5 | 1109.662 | 0.133528 | 0.185619 | 0.623185 | ENSMUSG( | Golga5 | 955.2429 | -0.02892 | 0.802802 | 0.97681 |
| ENSMUSG( | Golga7 | 6786.516 | -0.01781 | 0.873853 | 0.979333 | ENSMUSG( | Golga7 | 5901.893 | -0.15951 | 0.265202 | 0.886134 |
| ENSMUSG( | Golga7b | 6536.246 | 0.169876 | 0.051992 | 0.350064 | ENSMUSG( | Golga7b | 5500.525 | -0.05508 | 0.569672 | 0.941777 |
| ENSMUSG( | Golgb1 | 3883.272 | -0.22425 | 0.005724 | 0.101308 | ENSMUSG( | Golgb1 | 3809.239 | -0.00189 | 0.984592 | 0.996963 |
| ENSMUSG( | Golim4 | 1864.974 | -0.03083 | 0.788789 | 0.959226 | ENSMUSG( | Golim4 | 1730.428 | 0.023471 | 0.842198 | 0.983044 |
| ENSMUSG( | Golm1 | 1770.999 | 0.003882 | 0.976103 | 0.997428 | ENSMUSG( | Golm1 | 1812.78 | 0.326598 | 0.051395 | 0.711116 |
| ENSMUSG( | Golm2 | 4239.142 | -0.25171 | 0.05718 | 0.369067 | ENSMUSG( | Golm2 | 4305.092 | 0.073346 | 0.619191 | 0.952774 |
| ENSMUSG( | Golph3 | 2635.202 | 0.071351 | 0.445209 | 0.833239 | ENSMUSG( | Golph3 | 2251.624 | -0.11826 | 0.412152 | 0.917257 |
| ENSMUSG( | Golph3l | 486.6169 | 0.107206 | 0.365025 | 0.79407 | ENSMUSG( | Golph3l | 427.4297 | -0.00303 | 0.983867 | 0.996963 |
| ENSMUSG( | Golt1b | 1024.95 | -0.41881 | 0.004531 | 0.088122 | ENSMUSG( | Golt1b | 1104.152 | 0.098255 | 0.544627 | 0.938063 |
| ENSMUSG( | Gon4l | 2035.01 | 0.101831 | 0.261023 | 0.710889 | ENSMUSG( | Gon4l | 1819.563 | 0.048563 | 0.674424 | 0.95694 |
| ENSMUSG( | Gon7 | 102.7163 | -0.19287 | 0.346121 | 0.779153 | ENSMUSG( | Gon7 | 98.31492 | -0.07172 | 0.777273 | 0.975838 |
| ENSMUSG( | Gopc | 1088.484 | -0.05064 | 0.61201 | 0.908586 | ENSMUSG( | Gopc | 1020.697 | 0.029585 | 0.810674 | 0.978779 |
| ENSMUSG( | Gorab | 160.3083 | -0.18722 | 0.264846 | 0.715848 | ENSMUSG( | Gorab | 153.9975 | -0.03206 | 0.867249 | 0.98775 |
| ENSMUSG( | Gorasp1 | 915.9324 | -0.02166 | 0.83748 | 0.971111 | ENSMUSG( | Gorasp1 | 807.3084 | -0.11666 | 0.333065 | 0.907907 |
| ENSMUSG( | Gorasp2 | 2287.758 | -0.19972 | 0.024491 | 0.23959 | ENSMUSG( | Gorasp2 | 2184.364 | -0.05733 | 0.597954 | 0.946676 |
| ENSMUSG( | Gosr1 | 1444.474 | 0.055698 | 0.541059 | 0.883182 | ENSMUSG( | Gosr1 | 1257.965 | -0.07657 | 0.495247 | 0.927372 |
| ENSMUSG( | Gosr2 | 4347.32 | 0.033854 | 0.715854 | 0.941484 | ENSMUSG( | Gosr2 | 3685.885 | -0.18058 | 0.127833 | 0.81845 |
| ENSMUSG( | Got1 | 4068.094 | 0.042486 | 0.770018 | 0.954689 | ENSMUSG( | Got1 | 3494.934 | -0.13192 | 0.404537 | 0.917257 |
| ENSMUSG( | Got1l1 | 20.47531 | 0.188452 | 0.674585 | NA | ENSMUSG( | Got1l1 | 18.34562 | 0.17603 | 0.70896 | 0.964042 |
| ENSMUSG( | Got2 | 4165.683 | 0.021954 | 0.86312 | 0.975055 | ENSMUSG( | Got2 | 3610.536 | -0.12495 | 0.335609 | 0.907907 |
| ENSMUSG( | Gp1bb | 28.89394 | 0.218383 | 0.534592 | 0.880719 | ENSMUSG( | Gp1bb | 22.57523 | -0.20911 | 0.615667 | 0.951194 |
| ENSMUSG( | Gp5 | 21.40059 | -0.03028 | 0.942432 | 0.993415 | ENSMUSG( | Gp5 | 18.93153 | -0.11567 | 0.797544 | 0.97681 |
| ENSMUSG( | Gp9 | 30.82696 | -0.2278 | 0.558765 | 0.888638 | ENSMUSG( | Gp9 | 33.24886 | 0.270195 | 0.491742 | 0.927018 |
| ENSMUSG( | Gpaa1 | 1361.697 | -0.0893 | 0.492684 | 0.860462 | ENSMUSG( | Gpaa1 | 1356.744 | 0.170184 | 0.231826 | 0.868507 |
| ENSMUSG( | Gpalpp1 | 796.8766 | 0.079388 | 0.445014 | 0.833239 | ENSMUSG( | Gpalpp1 | 713.2565 | 0.030207 | 0.801068 | 0.97681 |
| ENSMUSG( | Gpam | 6467.747 | 0.188369 | 0.106973 | 0.494682 | ENSMUSG( | Gpam | 5493.268 | -0.00544 | 0.967163 | 0.995581 |
| ENSMUSG( | Gpank1 | 226.7054 | 0.011536 | 0.948971 | 0.993821 | ENSMUSG( | Gpank1 | 218.588 | 0.166346 | 0.383949 | 0.917257 |
| ENSMUSG( | Gpat3 | 85.947 | -0.76056 | 0.000513 | 0.020832 | ENSMUSG( | Gpat3 | 92.95418 | -0.17658 | 0.435384 | 0.919062 |
| ENSMUSG( | Gpat4 | 3324.189 | -0.07286 | 0.565211 | 0.890779 | ENSMUSG( | Gpat4 | 3278.548 | 0.156948 | 0.298834 | 0.898538 |
| ENSMUSG( | Gpatch1 | 820.1895 | 0.124384 | 0.234725 | 0.682178 | ENSMUSG( | Gpatch1 | 718.9039 | 0.012559 | 0.922547 | 0.992644 |
| ENSMUSG( | Gpatch11 | 754.5933 | 0.044577 | 0.683924 | 0.93244 | ENSMUSG( | Gpatch11 | 668.6361 | -0.03423 | 0.798436 | 0.97681 |
| ENSMUSG( | Gpatch2 | 673.1768 | 0.08793 | 0.435207 | 0.829093 | ENSMUSG( | Gpatch2 | 589.9835 | -0.02354 | 0.849366 | 0.985321 |
| ENSMUSG( | Gpatch2l | 1126.653 | 0.013573 | 0.883254 | 0.982422 | ENSMUSG( | Gpatch2l | 1016.036 | -0.01917 | 0.874885 | 0.989644 |
| ENSMUSG( | Gpatch3 | 124.5145 | -0.12118 | 0.512162 | 0.871563 | ENSMUSG( | Gpatch3 | 122.0026 | 0.093199 | 0.658823 | 0.956205 |
| ENSMUSG( | Gpatch4 | 560.1924 | 0.097392 | 0.449345 | 0.835057 | ENSMUSG( | Gpatch4 | 462.8615 | -0.19353 | 0.210618 | 0.864412 |
| ENSMUSG( | Gpatch8 | 3727.808 | 0.125551 | 0.178708 | 0.616298 | ENSMUSG( | Gpatch8 | 3304.455 | 0.046188 | 0.756435 | 0.971656 |
| ENSMUSG( | Gpbp1 | 1515.605 | 0.103188 | 0.365062 | 0.79407 | ENSMUSG( | Gpbp1 | 1390.522 | 0.116442 | 0.49797 | 0.927825 |
| ENSMUSG( | Gpbp1l1 | 987.5973 | 0.039473 | 0.684396 | 0.93282 | ENSMUSG( | Gpbp1l1 | 961.1258 | 0.223484 | 0.060795 | 0.734852 |
| ENSMUSG( | Gpc1 | 3096.961 | -0.19376 | 0.1377 | 0.552903 | ENSMUSG( | Gpc1 | 3229.118 | 0.199845 | 0.19017 | 0.857355 |
| ENSMUSG( | Gpc2 | 35.68712 | 0.599919 | 0.061025 | 0.381679 | ENSMUSG( | Gpc2 | 33.80676 | 0.716097 | 0.080009 | 0.776379 |
| ENSMUSG( | Gpc3 | 204.2825 | -0.1685 | 0.340051 | 0.777959 | ENSMUSG( | Gpc3 | 207.4422 | 0.151289 | 0.410592 | 0.917257 |
| ENSMUSG( | Gpc4 | 548.6062 | -0.46353 | 0.0013 | 0.039202 | ENSMUSG( | Gpc4 | 598.5838 | 0.09417 | 0.543559 | 0.938063 |
| ENSMUSG( | Gpc5 | 4506.972 | 0.02665 | 0.823533 | 0.966603 | ENSMUSG( | Gpc5 | 4205.126 | 0.093617 | 0.517105 | 0.930939 |
| ENSMUSG( | Gpc6 | 404.0536 | 0.192024 | 0.248097 | 0.69682 | ENSMUSG( | Gpc6 | 365.1999 | 0.179893 | 0.378888 | 0.916995 |
| ENSMUSG( | Gpcpd1 | 2757.395 | 0.05553 | 0.504724 | 0.866883 | ENSMUSG( | Gpcpd1 | 2760.466 | 0.31242 | 0.008255 | 0.388416 |
| ENSMUSG( | Gpd1 | 3751.952 | 0.204213 | 0.037802 | 0.300541 | ENSMUSG( | Gpd1 | 3213.621 | 0.03255 | 0.752463 | 0.970521 |
| ENSMUSG( | Gpd1l | 3934.15 | 0.120621 | 0.120774 | 0.524807 | ENSMUSG( | Gpd1l | 3383.296 | -0.04573 | 0.690906 | 0.960071 |
| ENSMUSG( | Gpd2 | 5569.414 | 0.136088 | 0.221614 | 0.667258 | ENSMUSG( | Gpd2 | 4844.616 | 0.006679 | 0.957309 | 0.994798 |
| ENSMUSG( | Gper1 | 103.087 | 0.008181 | 0.975407 | 0.997302 | ENSMUSG( | Gper1 | 138.9512 | 0.992797 | 0.001989 | 0.169168 |
| ENSMUSG( | Gphn | 1618.244 | 0.048253 | 0.646633 | 0.918796 | ENSMUSG( | Gphn | 1412.722 | -0.08027 | 0.561939 | 0.940388 |
| ENSMUSG( | Gpi1 | 20733.81 | 0.092746 | 0.323533 | 0.768536 | ENSMUSG( | Gpi1 | 17597.56 | -0.11396 | 0.347453 | 0.911652 |
| ENSMUSG( | Gpi-ps | 115.2102 | 0.229537 | 0.240755 | 0.688247 | ENSMUSG( | Gpi-ps | 86.681 | -0.32758 | 0.160856 | 0.837724 |
| ENSMUSG( | Gpkow | 1776.312 | 0.053937 | 0.586211 | 0.899709 | ENSMUSG( | Gpkow | 1531.782 | -0.11017 | 0.382243 | 0.917257 |
| ENSMUSG( | Gpld1 | 3288.088 | -0.11472 | 0.362457 | 0.792943 | ENSMUSG( | Gpld1 | 3018.108 | -0.09048 | 0.507921 | 0.929089 |
| ENSMUSG( | Gpm6a | 40349.75 | -0.28281 | 0.056562 | 0.366826 | ENSMUSG( | Gpm6a | 43201.09 | 0.191415 | 0.271886 | 0.888383 |
| ENSMUSG( | Gpm6b | 95233.3 | -0.03286 | 0.803345 | 0.962169 | ENSMUSG( | Gpm6b | 84625.11 | -0.10804 | 0.483539 | 0.924993 |
| ENSMUSG( | Gpn1 | 777.8492 | 0.094056 | 0.390389 | 0.807976 | ENSMUSG( | Gpn1 | 651.9279 | -0.1488 | 0.228852 | 0.868507 |
| ENSMUSG( | Gpn2 | 204.5415 | 0.208106 | 0.208327 | 0.650319 | ENSMUSG( | Gpn2 | 175.9295 | 0.051172 | 0.777349 | 0.975838 |
| ENSMUSG( | Gpn3 | 360.4441 | 0.037698 | 0.801417 | 0.961947 | ENSMUSG( | Gpn3 | 325.8233 | 0.011487 | 0.946032 | 0.994798 |
| ENSMUSG( | Gpnmb | 96.30105 | -0.59876 | 0.029105 | 0.259574 | ENSMUSG( | Gpnmb | 101.7865 | -0.10297 | 0.684307 | 0.958678 |
| ENSMUSG( | Gpr101 | 120.2802 | -0.11134 | 0.61095 | 0.908472 | ENSMUSG( | Gpr101 | 122.6442 | 0.191194 | 0.391345 | 0.917257 |
| ENSMUSG( | Gpr107 | 2327.431 | -0.11737 | 0.426153 | 0.824898 | ENSMUSG( | Gpr107 | 2306.343 | 0.129699 | 0.430123 | 0.917257 |
| ENSMUSG( | Gpr108 | 963.2336 | -0.06734 | 0.644805 | 0.917836 | ENSMUSG( | Gpr108 | 925.7888 | 0.093907 | 0.602598 | 0.94789 |
| ENSMUSG( | Gpr12 | 401.3443 | -0.48114 | 0.003252 | 0.071142 | ENSMUSG( | Gpr12 | 432.6856 | 0.046741 | 0.786369 | 0.976303 |
| ENSMUSG( | Gpr135 | 397.9599 | -0.55468 | 0.000473 | 0.019871 | ENSMUSG( | Gpr135 | 447.085 | 0.104397 | 0.556123 | 0.939565 |
| ENSMUSG( | Gpr137 | 4043.224 | 0.003412 | 0.975363 | 0.997302 | ENSMUSG( | Gpr137 | 3751.119 | 0.058456 | 0.675794 | 0.95694 |
| ENSMUSG( | Gpr137b | 431.6405 | -0.12474 | 0.388761 | 0.806561 | ENSMUSG( | Gpr137b | 426.6863 | 0.116892 | 0.459471 | 0.923329 |
| ENSMUSG( | Gpr137c | 870.7794 | -0.47746 | 0.000537 | 0.021456 | ENSMUSG( | Gpr137c | 1003.329 | 0.234163 | 0.12596 | 0.81516 |
| ENSMUSG( | Gpr139 | 53.18372 | -1.71644 | 1.82E-08 | 6.71E-06 | ENSMUSG( | Gpr139 | 70.08526 | -0.16651 | 0.529754 | 0.934795 |
| ENSMUSG( | Gpr146 | 1131.004 | -0.23615 | 0.147683 | 0.57034 | ENSMUSG( | Gpr146 | 1187.667 | 0.185017 | 0.349713 | 0.911652 |
| ENSMUSG( | Gpr149 | 34.82727 | -1.31298 | 0.000261 | 0.013121 | ENSMUSG( | Gpr149 | 52.78422 | 0.399106 | 0.212859 | 0.864412 |
| ENSMUSG( | Gpr150 | 131.4292 | 0.104212 | 0.557385 | 0.888279 | ENSMUSG( | Gpr150 | 119.7808 | 0.098943 | 0.63119 | 0.953251 |
| ENSMUSG( | Gpr151 | 18.10663 | 0.006939 | 0.987975 | NA | ENSMUSG( | Gpr151 | 22.67336 | 0.822138 | 0.057429 | 0.723727 |
| ENSMUSG( | Gpr153 | 1288.898 | -0.3024 | 0.001052 | 0.034136 | ENSMUSG( | Gpr153 | 1290.935 | -0.01199 | 0.913392 | 0.992298 |
| ENSMUSG( | Gpr155 | 1894.265 | -0.55615 | 4.32E-05 | 0.003572 | ENSMUSG( | Gpr155 | 2199.586 | 0.194075 | 0.229989 | 0.868507 |
| ENSMUSG( | Gpr156 | 82.15294 | 0.15878 | 0.514071 | 0.872055 | ENSMUSG( | Gpr156 | 73.3712 | 0.133629 | 0.643651 | 0.953903 |
| ENSMUSG( | Gpr157 | 68.95059 | -0.39294 | 0.113175 | 0.50752 | ENSMUSG( | Gpr157 | 78.67233 | 0.290614 | 0.259218 | 0.883971 |
| ENSMUSG( | Gpr158 | 4023.34 | -0.57039 | 3.35E-05 | 0.002933 | ENSMUSG( | Gpr158 | 4571.359 | 0.123492 | 0.450771 | 0.922202 |
| ENSMUSG( | Gpr15lg | 15.67532 | 0.012032 | 0.979143 | NA | ENSMUSG( | Gpr15lg | 14.06384 | -0.07364 | 0.887731 | 0.990399 |
| ENSMUSG( | Gpr160 | 42.58814 | -0.45264 | 0.185726 | 0.623185 | ENSMUSG( | Gpr160 | 46.87109 | 0.147679 | 0.671093 | 0.95694 |
| ENSMUSG( | Gpr161 | 702.0181 | 0.20464 | 0.148889 | 0.572605 | ENSMUSG( | Gpr161 | 663.0574 | 0.304452 | 0.040679 | 0.683847 |
| ENSMUSG( | Gpr162 | 7306.125 | -0.07051 | 0.518148 | 0.873754 | ENSMUSG( | Gpr162 | 7089.651 | 0.112325 | 0.382522 | 0.917257 |
| ENSMUSG( | Gpr165 | 117.8744 | 0.001789 | 0.993945 | 0.999369 | ENSMUSG( | Gpr165 | 126.7336 | 0.45982 | 0.074108 | 0.766789 |
| ENSMUSG( | Gpr17 | 4049.217 | -0.32795 | 0.315912 | 0.762967 | ENSMUSG( | Gpr17 | 4922.131 | 0.489407 | 0.232851 | 0.868507 |
| ENSMUSG( | Gpr173 | 585.5306 | -0.08601 | 0.556612 | 0.887713 | ENSMUSG( | Gpr173 | 582.8057 | 0.170651 | 0.276349 | 0.890889 |
| ENSMUSG( | Gpr176 | 456.7033 | -0.14071 | 0.377049 | 0.801213 | ENSMUSG( | Gpr176 | 460.1443 | 0.150811 | 0.370365 | 0.915491 |
| ENSMUSG( | Gpr179 | 60.33593 | -0.36882 | 0.130583 | 0.541841 | ENSMUSG( | Gpr179 | 46.41798 | -0.99729 | 0.00165 | 0.150169 |
| ENSMUSG( | Gpr180 | 901.923 | -0.11134 | 0.408073 | 0.818167 | ENSMUSG( | Gpr180 | 890.2572 | 0.123564 | 0.401993 | 0.917257 |
| ENSMUSG( | Gpr182 | 81.73067 | 0.302606 | 0.182451 | 0.619759 | ENSMUSG( | Gpr182 | 95.64431 | 0.920953 | 0.000293 | 0.046205 |
| ENSMUSG( | Gpr183 | 69.38169 | -0.04697 | 0.881335 | 0.981798 | ENSMUSG( | Gpr183 | 88.04604 | 0.818785 | 0.009625 | 0.406458 |
| ENSMUSG( | Gpr19 | 484.7596 | -0.49652 | 0.001779 | 0.04746 | ENSMUSG( | Gpr19 | 496.0011 | -0.11425 | 0.524532 | 0.933764 |
| ENSMUSG( | Gpr21 | 81.9034 | -0.51616 | 0.073112 | 0.417421 | ENSMUSG( | Gpr21 | 89.60406 | 0.079539 | 0.78771 | 0.97637 |
| ENSMUSG( | Gpr22 | 833.7899 | -0.30514 | 0.028015 | 0.255998 | ENSMUSG( | Gpr22 | 1018.702 | 0.516235 | 0.002936 | 0.217424 |
| ENSMUSG( | Gpr25 | 62.57059 | 0.070582 | 0.772161 | 0.954994 | ENSMUSG( | Gpr25 | 56.43744 | 0.025737 | 0.924969 | 0.99306 |
| ENSMUSG( | Gpr26 | 5433.792 | -0.25017 | 0.099785 | 0.482257 | ENSMUSG( | Gpr26 | 5421.969 | 0.028296 | 0.875574 | 0.989644 |
| ENSMUSG( | Gpr27 | 1651.678 | 0.110426 | 0.329848 | 0.773811 | ENSMUSG( | Gpr27 | 1584.384 | 0.251995 | 0.068183 | 0.75212 |
| ENSMUSG( | Gpr3 | 281.3369 | -0.75637 | 6.42E-05 | 0.004822 | ENSMUSG( | Gpr3 | 285.5065 | -0.36986 | 0.05049 | 0.708889 |
| ENSMUSG( | Gpr34 | 443.3318 | -0.1042 | 0.625136 | 0.912249 | ENSMUSG( | Gpr34 | 500.509 | 0.485302 | 0.052053 | 0.712974 |
| ENSMUSG( | Gpr35 | 33.06743 | -0.25465 | 0.513826 | 0.872055 | ENSMUSG( | Gpr35 | 34.40511 | 0.113524 | 0.76552 | 0.974236 |
| ENSMUSG( | Gpr37 | 1579.167 | -0.58706 | 0.003724 | 0.077639 | ENSMUSG( | Gpr37 | 1808.485 | 0.136543 | 0.731748 | 0.967228 |
| ENSMUSG( | Gpr37l1 | 17875.71 | -0.30162 | 0.042513 | 0.316636 | ENSMUSG( | Gpr37l1 | 17890.23 | -0.01144 | 0.945518 | 0.994798 |
| ENSMUSG( | Gpr39 | 20.12474 | -0.3556 | 0.380987 | NA | ENSMUSG( | Gpr39 | 17.40047 | -0.57507 | 0.238495 | 0.872849 |
| ENSMUSG( | Gpr4 | 196.4753 | 0.219182 | 0.169077 | 0.603241 | ENSMUSG( | Gpr4 | 225.7357 | 0.79617 | 6.96E-06 | 0.002837 |
| ENSMUSG( | Gpr45 | 570.1024 | 0.006491 | 0.95729 | 0.993874 | ENSMUSG( | Gpr45 | 514.6665 | -0.02203 | 0.873039 | 0.989445 |
| ENSMUSG( | Gpr52 | 91.99373 | -0.91717 | 0.000105 | 0.006643 | ENSMUSG( | Gpr52 | 124.8417 | 0.367138 | 0.102185 | 0.802316 |
| ENSMUSG( | Gpr55 | 12.1999 | 0.396849 | 0.466822 | NA | ENSMUSG( | Gpr55 | 9.51746 | 0.005244 | 0.993282 | NA |
| ENSMUSG( | Gpr6 | 229.5874 | -1.22405 | 4.26E-09 | 1.77E-06 | ENSMUSG( | Gpr6 | 317.7689 | 0.228093 | 0.267523 | 0.886134 |
| ENSMUSG( | Gpr61 | 433.7158 | -0.22528 | 0.124638 | 0.531714 | ENSMUSG( | Gpr61 | 465.822 | 0.255623 | 0.122809 | 0.814249 |
| ENSMUSG( | Gpr62 | 548.24 | -0.5506 | 0.000596 | 0.022994 | ENSMUSG( | Gpr62 | 610.0563 | 0.08504 | 0.65705 | 0.955625 |
| ENSMUSG( | Gpr63 | 67.13107 | -0.15365 | 0.554532 | 0.886683 | ENSMUSG( | Gpr63 | 72.81215 | 0.344925 | 0.21105 | 0.864412 |
| ENSMUSG( | Gpr68 | 380.907 | -0.46141 | 0.002355 | 0.057092 | ENSMUSG( | Gpr68 | 373.1898 | -0.21745 | 0.181685 | 0.850579 |
| ENSMUSG( | Gpr75 | 214.2439 | -0.31976 | 0.084688 | 0.448558 | ENSMUSG( | Gpr75 | 223.2561 | 0.091049 | 0.627681 | 0.953234 |
| ENSMUSG( | Gpr83 | 837.7055 | -0.73085 | 7.18E-08 | 2.12E-05 | ENSMUSG( | Gpr83 | 968.8081 | 0.048993 | 0.752718 | 0.970521 |
| ENSMUSG( | Gpr84 | 45.07752 | -0.3565 | 0.291172 | 0.741182 | ENSMUSG( | Gpr84 | 50.75761 | 0.28106 | 0.398403 | 0.917257 |
| ENSMUSG( | Gpr85 | 875.2301 | -0.27545 | 0.0651 | 0.395697 | ENSMUSG( | Gpr85 | 989.6339 | 0.344609 | 0.02947 | 0.618981 |
| ENSMUSG( | Gpr88 | 3895.811 | -1.26958 | 2.24E-10 | 1.21E-07 | ENSMUSG( | Gpr88 | 5048.776 | 0.019214 | 0.901132 | 0.990558 |
| ENSMUSG( | Gpr89 | 1129.822 | -0.17932 | 0.196664 | 0.635942 | ENSMUSG( | Gpr89 | 1087.312 | -0.01073 | 0.946641 | 0.994798 |
| ENSMUSG( | Gprasp1 | 15385.63 | 0.093515 | 0.513967 | 0.872055 | ENSMUSG( | Gprasp1 | 13903.09 | 0.072422 | 0.590154 | 0.944975 |
| ENSMUSG( | Gprasp2 | 2393.197 | 0.081789 | 0.408372 | 0.818167 | ENSMUSG( | Gprasp2 | 2180.054 | 0.077637 | 0.549801 | 0.938699 |
| ENSMUSG( | Gprc5a | 12.5044 | -0.17011 | 0.764272 | NA | ENSMUSG( | Gprc5a | 11.69712 | -0.05388 | 0.92797 | 0.99306 |
| ENSMUSG( | Gprc5b | 18826.19 | -0.06332 | 0.584389 | 0.898802 | ENSMUSG( | Gprc5b | 16430.35 | -0.19237 | 0.106932 | 0.804969 |
| ENSMUSG( | Gprc5c | 141.6255 | -0.05482 | 0.833902 | 0.970333 | ENSMUSG( | Gprc5c | 168.8021 | 0.666096 | 0.029705 | 0.61992 |
| ENSMUSG( | Gprin1 | 3202.45 | 0.22096 | 0.019924 | 0.213312 | ENSMUSG( | Gprin1 | 2681.679 | -0.01477 | 0.900019 | 0.990558 |
| ENSMUSG( | Gprin2 | 143.4221 | -0.25864 | 0.179409 | 0.61692 | ENSMUSG( | Gprin2 | 119.0436 | -0.57369 | 0.006109 | 0.322135 |
| ENSMUSG( | Gprin3 | 87.12965 | -0.74309 | 0.001516 | 0.043205 | ENSMUSG( | Gprin3 | 93.12714 | -0.18014 | 0.462785 | 0.923329 |
| ENSMUSG( | Gps1 | 3117.625 | 0.047549 | 0.55022 | 0.886038 | ENSMUSG( | Gps1 | 2708.88 | -0.08956 | 0.394012 | 0.917257 |
| ENSMUSG( | Gps2 | 650.3145 | 0.051044 | 0.675398 | 0.930049 | ENSMUSG( | Gps2 | 593.201 | 0.051493 | 0.739466 | 0.968466 |
| ENSMUSG( | Gpsm1 | 1489.523 | 0.041726 | 0.725572 | 0.944223 | ENSMUSG( | Gpsm1 | 1319.788 | -0.04154 | 0.765879 | 0.974236 |
| ENSMUSG( | Gpsm2 | 576.6257 | 0.165336 | 0.142734 | 0.561249 | ENSMUSG( | Gpsm2 | 504.8212 | 0.046624 | 0.747044 | 0.969763 |
| ENSMUSG( | Gpsm3 | 117.223 | -0.05769 | 0.757043 | 0.951513 | ENSMUSG( | Gpsm3 | 112.943 | 0.09593 | 0.64046 | 0.953903 |
| ENSMUSG( | Gpt | 958.3168 | 0.162398 | 0.128441 | 0.536774 | ENSMUSG( | Gpt | 893.5758 | 0.226126 | 0.082506 | 0.77681 |
| ENSMUSG( | Gpt2 | 4433.198 | 0.348539 | 3.73E-05 | 0.00318 | ENSMUSG( | Gpt2 | 3702.324 | 0.113647 | 0.333729 | 0.907907 |
| ENSMUSG( | Gpx1 | 2785.693 | -0.10893 | 0.243935 | 0.691355 | ENSMUSG( | Gpx1 | 2565.145 | -0.07967 | 0.486668 | 0.92541 |
| ENSMUSG( | Gpx3 | 314.5916 | -0.102 | 0.534969 | 0.880733 | ENSMUSG( | Gpx3 | 325.1962 | 0.249969 | 0.098802 | 0.798438 |
| ENSMUSG( | Gpx4 | 13112.05 | 0.037759 | 0.608216 | 0.907154 | ENSMUSG( | Gpx4 | 11137.64 | -0.17051 | 0.131412 | 0.822912 |
| ENSMUSG( | Gpx6 | 12.70087 | -1.47297 | 0.009705 | NA | ENSMUSG( | Gpx6 | 19.50003 | 0.365332 | 0.444804 | 0.921734 |
| ENSMUSG( | Gpx7 | 199.8426 | -0.03559 | 0.872144 | 0.978459 | ENSMUSG( | Gpx7 | 195.7925 | 0.177725 | 0.441595 | 0.921554 |
| ENSMUSG( | Gpx8 | 417.4719 | -0.00802 | 0.94984 | 0.993821 | ENSMUSG( | Gpx8 | 405.4945 | 0.176131 | 0.20868 | 0.864412 |
| ENSMUSG( | Gramd1a | 3552.885 | -0.1469 | 0.180191 | 0.617634 | ENSMUSG( | Gramd1a | 3471.787 | 0.060893 | 0.66981 | 0.95694 |
| ENSMUSG( | Gramd1b | 5362.434 | 0.041499 | 0.700124 | 0.937391 | ENSMUSG( | Gramd1b | 5079.862 | 0.153314 | 0.227624 | 0.868507 |
| ENSMUSG( | Gramd1c | 108.618 | 0.162887 | 0.410053 | 0.818258 | ENSMUSG( | Gramd1c | 100.9728 | 0.234855 | 0.37265 | 0.915491 |
| ENSMUSG( | Gramd2a | 271.4032 | 0.18427 | 0.225302 | 0.672044 | ENSMUSG( | Gramd2a | 228.7512 | -0.03939 | 0.825866 | 0.979664 |
| ENSMUSG( | Gramd2b | 1839.345 | -0.09719 | 0.440921 | 0.832371 | ENSMUSG( | Gramd2b | 1517.579 | -0.41235 | 0.002418 | 0.192605 |
| ENSMUSG( | Gramd4 | 2428.051 | -0.04807 | 0.628632 | 0.913658 | ENSMUSG( | Gramd4 | 2195.385 | -0.06784 | 0.567811 | 0.941777 |
| ENSMUSG( | Grap | 295.229 | -0.08148 | 0.609657 | 0.907714 | ENSMUSG( | Grap | 274.03 | -0.01214 | 0.951877 | 0.994798 |
| ENSMUSG( | Grap2 | 16.47258 | 0.312308 | 0.486465 | NA | ENSMUSG( | Grap2 | 12.8083 | -0.10557 | 0.844601 | 0.983642 |
| ENSMUSG( | Grb10 | 920.8734 | 0.104753 | 0.414207 | 0.819998 | ENSMUSG( | Grb10 | 851.0011 | 0.141524 | 0.326698 | 0.905371 |
| ENSMUSG( | Grb14 | 1029.618 | 0.077407 | 0.471338 | 0.847103 | ENSMUSG( | Grb14 | 927.1426 | 0.040919 | 0.745019 | 0.96951 |
| ENSMUSG( | Grb2 | 4127.331 | 0.022173 | 0.804796 | 0.962473 | ENSMUSG( | Grb2 | 3689.327 | -0.03485 | 0.780097 | 0.976106 |
| ENSMUSG( | Grb7 | 44.6239 | -0.07071 | 0.813908 | 0.964574 | ENSMUSG( | Grb7 | 43.78578 | 0.166727 | 0.629454 | 0.953251 |
| ENSMUSG( | Grcc10 | 638.773 | -0.07023 | 0.496712 | 0.86175 | ENSMUSG( | Grcc10 | 570.0586 | -0.1367 | 0.287457 | 0.894091 |
| ENSMUSG( | Greb1 | 56.48882 | -0.62659 | 0.028893 | 0.258922 | ENSMUSG( | Greb1 | 64.48855 | 0.092268 | 0.744567 | 0.96951 |
| ENSMUSG( | Greb1l | 250.202 | -0.07333 | 0.625327 | 0.912249 | ENSMUSG( | Greb1l | 237.9994 | 0.057576 | 0.727686 | 0.966887 |
| ENSMUSG( | Grem1 | 58.3399 | -0.19902 | 0.434185 | 0.828931 | ENSMUSG( | Grem1 | 59.9395 | 0.133382 | 0.627924 | 0.953234 |
| ENSMUSG( | Grem2 | 653.0257 | -0.42787 | 0.021403 | 0.223539 | ENSMUSG( | Grem2 | 705.4353 | 0.098843 | 0.583537 | 0.944266 |
| ENSMUSG( | Grhl1 | 365.3906 | 0.198383 | 0.106453 | 0.494174 | ENSMUSG( | Grhl1 | 341.9066 | 0.270028 | 0.052415 | 0.714514 |

| | | | | | |
|---|---|---|---|---|---|
| ENSMUSG( | 401.3443 | -0.48114 | 0.003252 | 0.071142 | Gpr12 |
| ENSMUSG( | 397.9599 | -0.55468 | 0.000473 | 0.019871 | Gpr135 |
| ENSMUSG( | 4043.224 | 0.003412 | 0.975363 | 0.997302 | Gpr137 |
| ENSMUSG( | 431.6405 | -0.12474 | 0.388761 | 0.806561 | Gpr137b |
| ENSMUSG( | 870.7794 | -0.47746 | 0.000537 | 0.021456 | Gpr137c |
| ENSMUSG( | 53.18372 | -1.71644 | 1.82E-08 | 6.71E-06 | Gpr139 |
| ENSMUSG( | 1131.004 | -0.23615 | 0.147683 | 0.57034 | Gpr146 |
| ENSMUSG( | 34.82727 | -1.31298 | 0.000261 | 0.013121 | Gpr149 |
| ENSMUSG( | 131.4292 | 0.104212 | 0.557385 | 0.888279 | Gpr150 |
| ENSMUSG( | 18.10663 | 0.006939 | 0.987975 | NA | Gpr151 |
| ENSMUSG( | 1288.898 | -0.3024 | 0.001052 | 0.034136 | Gpr153 |
| ENSMUSG( | 1894.265 | -0.55615 | 4.32E-05 | 0.003572 | Gpr155 |
| ENSMUSG( | 82.15294 | 0.15878 | 0.514071 | 0.872055 | Gpr156 |
| ENSMUSG( | 68.95059 | -0.39294 | 0.113175 | 0.50752 | Gpr157 |
| ENSMUSG( | 4023.34 | -0.57039 | 3.35E-05 | 0.002933 | Gpr158 |
| ENSMUSG( | 15.67532 | 0.012032 | 0.979143 | NA | Gpr15lg |
| ENSMUSG( | 42.58814 | -0.45264 | 0.185726 | 0.623185 | Gpr160 |
| ENSMUSG( | 702.0181 | 0.20464 | 0.148889 | 0.572605 | Gpr161 |
| ENSMUSG( | 7306.125 | -0.07051 | 0.518148 | 0.873754 | Gpr162 |
| ENSMUSG( | 117.8744 | 0.001789 | 0.993945 | 0.999369 | Gpr165 |
| ENSMUSG( | 4049.217 | -0.32795 | 0.315912 | 0.762967 | Gpr17 |
| ENSMUSG( | 585.5306 | -0.08601 | 0.556612 | 0.887713 | Gpr173 |
| ENSMUSG( | 456.7033 | -0.14071 | 0.377049 | 0.801213 | Gpr176 |
| ENSMUSG( | 60.33593 | -0.36882 | 0.130583 | 0.541841 | Gpr179 |
| ENSMUSG( | 901.923 | -0.11134 | 0.408073 | 0.818167 | Gpr180 |
| ENSMUSG( | 81.73067 | 0.302606 | 0.182451 | 0.619759 | Gpr182 |
| ENSMUSG( | 69.38169 | -0.04697 | 0.881335 | 0.981798 | Gpr183 |
| ENSMUSG( | 484.7596 | -0.49652 | 0.001779 | 0.04746 | Gpr19 |
| ENSMUSG( | 81.9034 | -0.51616 | 0.073112 | 0.417421 | Gpr21 |
| ENSMUSG( | 833.7899 | -0.30514 | 0.028015 | 0.255998 | Gpr22 |
| ENSMUSG( | 62.57059 | 0.070582 | 0.772161 | 0.954994 | Gpr25 |
| ENSMUSG( | 5433.792 | -0.25017 | 0.099785 | 0.482257 | Gpr26 |
| ENSMUSG( | 1651.678 | 0.110426 | 0.329848 | 0.773811 | Gpr27 |
| ENSMUSG( | 281.3369 | -0.75637 | 6.42E-05 | 0.004822 | Gpr3 |
| ENSMUSG( | 443.3318 | -0.1042 | 0.625136 | 0.912249 | Gpr34 |
| ENSMUSG( | 33.06743 | -0.25465 | 0.513826 | 0.872055 | Gpr35 |
| ENSMUSG( | 1579.167 | -0.58706 | 0.003724 | 0.077639 | Gpr37 |
| ENSMUSG( | 17875.71 | -0.30162 | 0.042513 | 0.316636 | Gpr37l1 |
| ENSMUSG( | 20.12474 | -0.3556 | 0.380987 | NA | Gpr39 |
| ENSMUSG( | 196.4753 | 0.219182 | 0.169077 | 0.603241 | Gpr4 |
| ENSMUSG( | 570.1024 | 0.006491 | 0.95729 | 0.993874 | Gpr45 |
| ENSMUSG( | 9.101448 | -0.40951 | 0.517306 | NA | Gpr50 |
| ENSMUSG( | 91.99373 | -0.91717 | 0.000105 | 0.006643 | Gpr52 |
| ENSMUSG( | 12.1999 | 0.396849 | 0.466822 | NA | Gpr55 |
| ENSMUSG( | 229.5874 | -1.22405 | 4.26E-09 | 1.77E-06 | Gpr6 |
| ENSMUSG( | 433.7158 | -0.22528 | 0.124638 | 0.531714 | Gpr61 |
| ENSMUSG( | 548.24 | -0.5506 | 0.000596 | 0.022994 | Gpr62 |
| ENSMUSG( | 67.13107 | -0.15365 | 0.554532 | 0.886683 | Gpr63 |
| ENSMUSG( | 380.907 | -0.46141 | 0.002355 | 0.057092 | Gpr68 |
| ENSMUSG( | 214.2439 | -0.31976 | 0.084688 | 0.448558 | Gpr75 |
| ENSMUSG( | 837.7055 | -0.73085 | 7.18E-08 | 2.12E-05 | Gpr83 |
| ENSMUSG( | 45.07752 | -0.3565 | 0.291172 | 0.741182 | Gpr84 |
| ENSMUSG( | 875.2301 | -0.27545 | 0.0651 | 0.395697 | Gpr85 |
| ENSMUSG( | 3895.811 | -1.26958 | 2.24E-10 | 1.21E-07 | Gpr88 |
| ENSMUSG( | 1129.822 | -0.17932 | 0.196664 | 0.635942 | Gpr89 |
| ENSMUSG( | 15385.63 | 0.093515 | 0.513967 | 0.872055 | Gprasp1 |
| ENSMUSG( | 2393.197 | 0.081789 | 0.408372 | 0.818167 | Gprasp2 |
| ENSMUSG( | 12.5044 | -0.17011 | 0.764272 | NA | Gprc5a |
| ENSMUSG( | 18826.19 | -0.06332 | 0.584389 | 0.898802 | Gprc5b |
| ENSMUSG( | 141.6255 | -0.05482 | 0.833902 | 0.970333 | Gprc5c |
| ENSMUSG( | 3202.45 | 0.22096 | 0.019924 | 0.213312 | Gprin1 |
| ENSMUSG( | 143.4221 | -0.25864 | 0.179409 | 0.61692 | Gprin2 |
| ENSMUSG( | 87.12965 | -0.74309 | 0.001516 | 0.043205 | Gprin3 |
| ENSMUSG( | 3117.625 | 0.047549 | 0.55022 | 0.886038 | Gps1 |
| ENSMUSG( | 650.3145 | 0.051044 | 0.675398 | 0.930049 | Gps2 |
| ENSMUSG( | 1489.523 | 0.041726 | 0.725572 | 0.944223 | Gpsm1 |
| ENSMUSG( | 576.6257 | 0.165336 | 0.142734 | 0.561249 | Gpsm2 |
| ENSMUSG( | 117.223 | -0.05769 | 0.757043 | 0.951513 | Gpsm3 |
| ENSMUSG( | 958.3168 | 0.162398 | 0.128441 | 0.536774 | Gpt |
| ENSMUSG( | 4433.198 | 0.348539 | 3.73E-05 | 0.00318 | Gpt2 |
| ENSMUSG( | 2785.693 | -0.10893 | 0.243935 | 0.691355 | Gpx1 |
| ENSMUSG( | 314.5916 | -0.102 | 0.534969 | 0.880733 | Gpx3 |
| ENSMUSG( | 13112.05 | 0.037759 | 0.608216 | 0.907154 | Gpx4 |
| ENSMUSG( | 12.70087 | -1.47297 | 0.009705 | NA | Gpx6 |
| ENSMUSG( | 199.8426 | -0.03559 | 0.872144 | 0.978459 | Gpx7 |
| ENSMUSG( | 417.4719 | -0.00802 | 0.94984 | 0.993821 | Gpx8 |
| ENSMUSG( | 3552.885 | -0.1469 | 0.180191 | 0.617634 | Gramd1a |
| ENSMUSG( | 5362.434 | 0.041499 | 0.700124 | 0.937391 | Gramd1b |
| ENSMUSG( | 108.618 | 0.162887 | 0.410053 | 0.818258 | Gramd1c |
| ENSMUSG( | 271.4032 | 0.18427 | 0.225302 | 0.672044 | Gramd2a |
| ENSMUSG( | 1839.345 | -0.09719 | 0.440921 | 0.832371 | Gramd2b |
| ENSMUSG( | 2428.051 | -0.04807 | 0.628632 | 0.913658 | Gramd4 |
| ENSMUSG( | 295.229 | -0.08148 | 0.609657 | 0.907714 | Grap |
| ENSMUSG( | 16.47258 | 0.312308 | 0.486465 | NA | Grap2 |
| ENSMUSG( | 920.8734 | 0.104753 | 0.414207 | 0.819998 | Grb10 |
| ENSMUSG( | 1029.618 | 0.077407 | 0.471338 | 0.847103 | Grb14 |
| ENSMUSG( | 4127.331 | 0.022173 | 0.804796 | 0.962473 | Grb2 |
| ENSMUSG( | 44.6239 | -0.07071 | 0.813908 | 0.964574 | Grb7 |
| ENSMUSG( | 638.773 | -0.07023 | 0.496712 | 0.86175 | Grcc10 |
| ENSMUSG( | 56.48882 | -0.62659 | 0.028893 | 0.258922 | Greb1 |
| ENSMUSG( | 250.202 | -0.07333 | 0.625327 | 0.912249 | Greb1l |
| ENSMUSG( | 58.3399 | -0.19902 | 0.434185 | 0.828931 | Grem1 |
| ENSMUSG( | 653.0257 | -0.42787 | 0.021403 | 0.223539 | Grem2 |
| ENSMUSG( | 365.3906 | 0.198383 | 0.106453 | 0.494174 | Grhl1 |
| ENSMUSG( | 21.03614 | 0.178409 | 0.66515 | NA | Grhl3 |
| ENSMUSG( | 629.1669 | 0.087654 | 0.448282 | 0.834051 | Grhpr |
| ENSMUSG( | 12445.19 | -0.16071 | 0.114178 | 0.509818 | Gria1 |
| ENSMUSG( | 9315.647 | -0.3086 | 0.014623 | 0.177929 | Gria2 |
| ENSMUSG( | 6417.452 | -0.19723 | 0.074671 | 0.423665 | Gria3 |
| ENSMUSG( | 2929.574 | -0.39824 | 0.000421 | 0.018541 | Gria4 |
| ENSMUSG( | 2456.373 | -0.26748 | 0.064398 | 0.393421 | Grid1 |
| ENSMUSG( | 969.1876 | -0.15497 | 0.308937 | 0.757108 | Grid2 |
| ENSMUSG( | 75.05367 | -0.63146 | 0.005795 | 0.102397 | Grid2ip |
| ENSMUSG( | 17.06093 | -0.21664 | 0.638441 | NA | Grifin |
| ENSMUSG( | 468.602 | 0.01915 | 0.888647 | 0.983434 | Grik1 |
| ENSMUSG( | 1876.377 | -0.25536 | 0.025431 | 0.246258 | Grik2 |
| ENSMUSG( | 2426.95 | -0.06094 | 0.717184 | 0.941697 | Grik3 |
| ENSMUSG( | 901.2053 | -0.20574 | 0.146138 | 0.567605 | Grik4 |
| ENSMUSG( | 12167.89 | 0.11068 | 0.227572 | 0.674441 | Grik5 |
| ENSMUSG( | 19857.73 | -0.3065 | 0.008929 | 0.132538 | Grin1 |
| ENSMUSG( | 131.1553 | -0.35994 | 0.131517 | 0.543959 | Grin1os |
| ENSMUSG( | 3709.53 | -0.3787 | 0.011245 | 0.15263 | Grin2a |
| ENSMUSG( | 6395.813 | -0.37881 | 0.02046 | 0.21574 | Grin2b |
| ENSMUSG( | 6193.911 | 0.057779 | 0.600672 | 0.90443 | Grin2c |
| ENSMUSG( | 668.6037 | -0.04269 | 0.732485 | 0.944956 | Grin2d |
| ENSMUSG( | 1536.022 | -0.24691 | 0.093365 | 0.469793 | Grin3a |
| ENSMUSG( | 26.37469 | -0.20828 | 0.579271 | 0.896692 | Grin3b |
| ENSMUSG( | 19335.83 | -0.30437 | 0.002954 | 0.066464 | Grina |
| ENSMUSG( | 466.7449 | -0.17976 | 0.116293 | 0.514678 | Grip1 |
| ENSMUSG( | 910.373 | 0.201139 | 0.097593 | 0.477947 | Grip2 |
| ENSMUSG( | 2246.627 | 0.186703 | 0.111842 | 0.504332 | Gripap1 |
| ENSMUSG( | 15451.2 | 0.180905 | 0.033969 | 0.282701 | Grk2 |
| ENSMUSG( | 5724.047 | 0.063725 | 0.459077 | 0.840244 | Grk3 |
| ENSMUSG( | 109.7334 | 0.174232 | 0.355398 | 0.78783 | Grk4 |
| ENSMUSG( | 974.3159 | 0.044162 | 0.720024 | 0.941986 | Grk5 |
| ENSMUSG( | 2784.308 | 0.17863 | 0.04657 | 0.330304 | Grk6 |
| ENSMUSG( | 847.9675 | -0.62891 | 9.42E-05 | 0.006211 | Grm1 |
| ENSMUSG( | 2344.934 | -0.29985 | 0.024913 | 0.242787 | Grm2 |
| ENSMUSG( | 4955.109 | -0.10101 | 0.470457 | 0.84674 | Grm3 |
| ENSMUSG( | 1142.133 | -0.3963 | 0.0011 | 0.035114 | Grm4 |
| ENSMUSG( | 3366.514 | -0.48986 | 0.002944 | 0.066413 | Grm5 |
| ENSMUSG( | 1410.1 | -0.47758 | 0.000833 | 0.028872 | Grm7 |
| ENSMUSG( | 373.1663 | -0.56647 | 0.001668 | 0.046175 | Grm8 |
| ENSMUSG( | 3512.443 | -0.21633 | 0.116599 | 0.514987 | Grn |
| ENSMUSG( | 319.8303 | -0.11368 | 0.493422 | 0.860611 | Grp |
| ENSMUSG( | 982.1526 | 0.066891 | 0.523375 | 0.875628 | Grpel1 |
| ENSMUSG( | 477.3387 | -0.16271 | 0.259545 | 0.709948 | Grpel2 |
| ENSMUSG( | 1973.939 | -0.02293 | 0.804245 | 0.962169 | Grsf1 |
| ENSMUSG( | 356.9862 | 0.016952 | 0.918546 | 0.989954 | Grtp1 |
| ENSMUSG( | 270.596 | 0.174668 | 0.289547 | 0.739839 | Grwd1 |
| ENSMUSG( | 578.3832 | 0.069914 | 0.573515 | 0.893584 | Gsap |
| ENSMUSG( | 150.0988 | -0.09981 | 0.604642 | 0.905556 | Gsdmd |
| ENSMUSG( | 168.9846 | -0.03953 | 0.819242 | 0.965888 | Gsdme |
| ENSMUSG( | 2321.269 | 0.126268 | 0.238506 | 0.686308 | Gse1 |
| ENSMUSG( | 3595.913 | -0.2371 | 0.031903 | 0.27264 | Gsg1l |
| ENSMUSG( | 10796.95 | 0.115891 | 0.12761 | 0.535393 | Gsk3a |
| ENSMUSG( | 4985.401 | 0.058595 | 0.502275 | 0.864924 | Gsk3b |
| ENSMUSG( | 579.0426 | 0.151104 | 0.217224 | 0.662164 | Gskip |
| ENSMUSG( | 5957.475 | -0.47806 | 0.001769 | 0.047422 | Gsn |
| ENSMUSG( | 2485.029 | 0.086524 | 0.346376 | 0.779153 | Gspt1 |
| ENSMUSG( | 485.9373 | -0.09857 | 0.452739 | 0.836191 | Gspt2 |
| ENSMUSG( | 692.1195 | -0.05593 | 0.697275 | 0.935662 | Gsr |
| ENSMUSG( | 513.2972 | -0.03484 | 0.803023 | 0.962169 | Gss |
| ENSMUSG( | 29.94744 | -0.70209 | 0.0876 | 0.454663 | Gsta3 |
| ENSMUSG( | 1245.956 | -0.08914 | 0.330954 | 0.775174 | Gsta4 |
| ENSMUSG( | 197.5595 | -0.05715 | 0.718028 | 0.941697 | Gstcd |
| ENSMUSG( | 799.02 | 0.141235 | 0.272586 | 0.722478 | Gstk1 |
| ENSMUSG( | 23471.34 | 0.288048 | 0.00312 | 0.069321 | Gstm1 |

| | | | | | |
|---|---|---|---|---|---|
| ENSMUSG( | 465.822 | 0.255623 | 0.122809 | 0.814249 | Gpr61 |
| ENSMUSG( | 610.0563 | 0.08504 | 0.65705 | 0.955625 | Gpr62 |
| ENSMUSG( | 72.81215 | 0.344925 | 0.21105 | 0.864412 | Gpr63 |
| ENSMUSG( | 373.1898 | -0.21745 | 0.181685 | 0.850579 | Gpr68 |
| ENSMUSG( | 223.2561 | 0.091049 | 0.627681 | 0.953234 | Gpr75 |
| ENSMUSG( | 968.8081 | 0.048993 | 0.752718 | 0.970521 | Gpr83 |
| ENSMUSG( | 50.75761 | 0.28106 | 0.398403 | 0.917257 | Gpr84 |
| ENSMUSG( | 989.6339 | 0.344609 | 0.02947 | 0.618981 | Gpr85 |
| ENSMUSG( | 5048.776 | 0.019214 | 0.901132 | 0.990558 | Gpr88 |
| ENSMUSG( | 1087.312 | -0.01073 | 0.946641 | 0.994798 | Gpr89 |
| ENSMUSG( | 13903.09 | 0.072422 | 0.590154 | 0.944975 | Gprasp1 |
| ENSMUSG( | 2180.054 | 0.077637 | 0.549801 | 0.938699 | Gprasp2 |
| ENSMUSG( | 11.69712 | -0.05388 | 0.92797 | 0.99306 | Gprc5a |
| ENSMUSG( | 16430.35 | -0.19237 | 0.106932 | 0.804969 | Gprc5b |
| ENSMUSG( | 168.8021 | 0.666096 | 0.029705 | 0.61992 | Gprc5c |
| ENSMUSG( | 2681.679 | -0.01477 | 0.900019 | 0.990558 | Gprin1 |
| ENSMUSG( | 119.0436 | -0.57369 | 0.006109 | 0.322135 | Gprin2 |
| ENSMUSG( | 93.12714 | -0.18014 | 0.462785 | 0.923329 | Gprin3 |
| ENSMUSG( | 2708.88 | -0.08956 | 0.394012 | 0.917257 | Gps1 |
| ENSMUSG( | 593.201 | 0.051493 | 0.739466 | 0.968466 | Gps2 |
| ENSMUSG( | 1319.788 | -0.04154 | 0.765879 | 0.974236 | Gpsm1 |
| ENSMUSG( | 504.8212 | 0.046624 | 0.747044 | 0.969763 | Gpsm2 |
| ENSMUSG( | 112.943 | 0.09593 | 0.64046 | 0.953903 | Gpsm3 |
| ENSMUSG( | 893.5758 | 0.226126 | 0.082506 | 0.77681 | Gpt |
| ENSMUSG( | 3702.324 | 0.113647 | 0.333729 | 0.907907 | Gpt2 |
| ENSMUSG( | 2565.145 | -0.07967 | 0.486668 | 0.92541 | Gpx1 |
| ENSMUSG( | 325.1962 | 0.249969 | 0.098802 | 0.798438 | Gpx3 |
| ENSMUSG( | 11137.64 | -0.17051 | 0.131412 | 0.822912 | Gpx4 |
| ENSMUSG( | 19.50003 | 0.365332 | 0.444804 | 0.921734 | Gpx6 |
| ENSMUSG( | 195.7925 | 0.177725 | 0.441595 | 0.921554 | Gpx7 |
| ENSMUSG( | 405.4945 | 0.176131 | 0.20868 | 0.864412 | Gpx8 |
| ENSMUSG( | 3471.787 | 0.060893 | 0.66981 | 0.95694 | Gramd1a |
| ENSMUSG( | 5079.862 | 0.153314 | 0.227624 | 0.868507 | Gramd1b |
| ENSMUSG( | 100.9728 | 0.234855 | 0.37265 | 0.915491 | Gramd1c |
| ENSMUSG( | 228.7512 | -0.03939 | 0.825866 | 0.979664 | Gramd2a |
| ENSMUSG( | 1517.579 | -0.41235 | 0.002418 | 0.192605 | Gramd2b |
| ENSMUSG( | 2195.385 | -0.06784 | 0.567811 | 0.941777 | Gramd4 |
| ENSMUSG( | 274.03 | -0.01214 | 0.951877 | 0.994798 | Grap |
| ENSMUSG( | 12.8083 | -0.10557 | 0.844601 | 0.983642 | Grap2 |
| ENSMUSG( | 851.0011 | 0.141524 | 0.326698 | 0.905371 | Grb10 |
| ENSMUSG( | 927.1426 | 0.040919 | 0.745019 | 0.96951 | Grb14 |
| ENSMUSG( | 3689.327 | -0.03485 | 0.780097 | 0.976106 | Grb2 |
| ENSMUSG( | 43.78578 | 0.166727 | 0.629454 | 0.953251 | Grb7 |
| ENSMUSG( | 570.0586 | -0.1367 | 0.287457 | 0.894091 | Grcc10 |
| ENSMUSG( | 64.48855 | 0.092268 | 0.744567 | 0.96951 | Greb1 |
| ENSMUSG( | 237.9994 | 0.057576 | 0.727686 | 0.966887 | Greb1l |
| ENSMUSG( | 59.9395 | 0.133382 | 0.627924 | 0.953234 | Grem1 |
| ENSMUSG( | 705.4353 | 0.098843 | 0.583537 | 0.944266 | Grem2 |
| ENSMUSG( | 341.9066 | 0.270028 | 0.052415 | 0.714514 | Grhl1 |
| ENSMUSG( | 13.23332 | -0.98685 | 0.096489 | 0.796133 | Grhl3 |
| ENSMUSG( | 511.7245 | -0.24963 | 0.067803 | 0.75212 | Grhpr |
| ENSMUSG( | 12652.42 | 0.157908 | 0.188218 | 0.85698 | Gria1 |
| ENSMUSG( | 10817.11 | 0.385303 | 0.009202 | 0.401972 | Gria2 |
| ENSMUSG( | 6690.667 | 0.192065 | 0.124214 | 0.814691 | Gria3 |
| ENSMUSG( | 3028.104 | -0.00551 | 0.966594 | 0.995475 | Gria4 |
| ENSMUSG( | 2589.238 | 0.16633 | 0.332575 | 0.907907 | Grid1 |
| ENSMUSG( | 983.6884 | 0.160402 | 0.350607 | 0.911652 | Grid2 |
| ENSMUSG( | 72.11701 | -0.43369 | 0.078167 | 0.77321 | Grid2ip |
| ENSMUSG( | 12.83961 | -0.95427 | 0.092086 | 0.791889 | Grifin |
| ENSMUSG( | 459.8434 | 0.231756 | 0.160616 | 0.837724 | Grik1 |
| ENSMUSG( | 1977.182 | 0.171751 | 0.172046 | 0.843003 | Grik2 |
| ENSMUSG( | 2613.704 | 0.401452 | 0.040234 | 0.681706 | Grik3 |
| ENSMUSG( | 902.7239 | 0.079881 | 0.610938 | 0.950771 | Grik4 |
| ENSMUSG( | 12311.09 | 0.390476 | 0.001654 | 0.150169 | Grik5 |
| ENSMUSG( | 21151.07 | 0.158847 | 0.233065 | 0.868507 | Grin1 |
| ENSMUSG( | 117.4048 | -0.38955 | 0.168964 | 0.840222 | Grin1os |
| ENSMUSG( | 4093.489 | 0.195349 | 0.236761 | 0.872173 | Grin2a |
| ENSMUSG( | 7086.131 | 0.208893 | 0.277031 | 0.891231 | Grin2b |
| ENSMUSG( | 5528.324 | 0.000744 | 0.9953 | 0.999116 | Grin2c |
| ENSMUSG( | 651.2861 | 0.150967 | 0.256228 | 0.883082 | Grin2d |
| ENSMUSG( | 1596.414 | 0.144028 | 0.38754 | 0.917257 | Grin3a |
| ENSMUSG( | 26.3842 | 0.068324 | 0.862807 | 0.986547 | Grin3b |
| ENSMUSG( | 19821.98 | 0.05334 | 0.653811 | 0.955015 | Grina |
| ENSMUSG( | 435.3408 | -0.11522 | 0.430818 | 0.917257 | Grip1 |
| ENSMUSG( | 793.3825 | 0.074648 | 0.615942 | 0.951413 | Grip2 |
| ENSMUSG( | 1903.682 | -0.01999 | 0.889478 | 0.990558 | Gripap1 |
| ENSMUSG( | 12829.66 | -0.08174 | 0.397244 | 0.917257 | Grk2 |
| ENSMUSG( | 5078.251 | -0.01211 | 0.904128 | 0.991089 | Grk3 |
| ENSMUSG( | 104.2938 | 0.294406 | 0.165238 | 0.838743 | Grk4 |
| ENSMUSG( | 858.5603 | -0.05299 | 0.652897 | 0.954897 | Grk5 |
| ENSMUSG( | 2400.423 | 0.025687 | 0.807805 | 0.978622 | Grk6 |
| ENSMUSG( | 1026.357 | 0.249395 | 0.17356 | 0.844317 | Grm1 |
| ENSMUSG( | 2361.375 | 0.006965 | 0.963989 | 0.995433 | Grm2 |
| ENSMUSG( | 4298.309 | -0.2485 | 0.095809 | 0.796133 | Grm3 |
| ENSMUSG( | 1186.603 | 0.013989 | 0.922914 | 0.992644 | Grm4 |
| ENSMUSG( | 3966.143 | 0.281266 | 0.138042 | 0.828492 | Grm5 |
| ENSMUSG( | 1509.584 | 0.030256 | 0.853382 | 0.985598 | Grm7 |
| ENSMUSG( | 432.0559 | 0.180716 | 0.358552 | 0.913759 | Grm8 |
| ENSMUSG( | 3656.566 | 0.176253 | 0.266493 | 0.886134 | Grn |
| ENSMUSG( | 323.9636 | 0.189969 | 0.307069 | 0.898538 | Grp |
| ENSMUSG( | 801.3651 | -0.2676 | 0.041201 | 0.687358 | Grpel1 |
| ENSMUSG( | 435.0941 | -0.15916 | 0.289497 | 0.896591 | Grpel2 |
| ENSMUSG( | 1766.721 | -0.07841 | 0.5179 | 0.930939 | Grsf1 |
| ENSMUSG( | 301.149 | -0.20706 | 0.217251 | 0.865589 | Grtp1 |
| ENSMUSG( | 233.9532 | 0.023367 | 0.896237 | 0.990558 | Grwd1 |
| ENSMUSG( | 497.2235 | -0.0928 | 0.521179 | 0.932489 | Gsap |
| ENSMUSG( | 151.5207 | 0.204011 | 0.400171 | 0.917257 | Gsdmd |
| ENSMUSG( | 145.8637 | -0.21144 | 0.28705 | 0.894091 | Gsdme |
| ENSMUSG( | 2190.078 | 0.224069 | 0.066727 | 0.751678 | Gse1 |
| ENSMUSG( | 3579.216 | 0.031037 | 0.810531 | 0.978779 | Gsg1l |
| ENSMUSG( | 9364.548 | -0.02404 | 0.821881 | 0.979198 | Gsk3a |
| ENSMUSG( | 4476.562 | 0.015313 | 0.910017 | 0.991958 | Gsk3b |
| ENSMUSG( | 474.6904 | -0.16035 | 0.338369 | 0.907972 | Gskip |
| ENSMUSG( | 6683.773 | 0.162665 | 0.379532 | 0.916995 | Gsn |
| ENSMUSG( | 2174.428 | -0.03148 | 0.799552 | 0.97681 | Gspt1 |
| ENSMUSG( | 465.3784 | 0.048199 | 0.767583 | 0.974236 | Gspt2 |
| ENSMUSG( | 622.9361 | -0.10039 | 0.564064 | 0.941206 | Gsr |
| ENSMUSG( | 454.4973 | -0.12186 | 0.443945 | 0.921734 | Gss |
| ENSMUSG( | 34.44423 | 0.075662 | 0.829748 | 0.980224 | Gsta3 |
| ENSMUSG( | 1141.735 | -0.0746 | 0.520058 | 0.931904 | Gsta4 |
| ENSMUSG( | 182.7294 | -0.02105 | 0.907273 | 0.991683 | Gstcd |
| ENSMUSG( | 646.4824 | -0.21015 | 0.125194 | 0.81516 | Gstk1 |
| ENSMUSG( | 19054.3 | -0.03251 | 0.803858 | 0.977 | Gstm1 |
| ENSMUSG( | 80.29255 | -0.29278 | 0.243129 | 0.877399 | Gstm2 |
| ENSMUSG( | 266.8346 | -0.21909 | 0.182134 | 0.850579 | Gstm4 |
| ENSMUSG( | 9529.482 | -0.11366 | 0.377928 | 0.916784 | Gstm5 |
| ENSMUSG( | 163.7019 | -0.04119 | 0.82424 | 0.979198 | Gstm6 |
| ENSMUSG( | 548.7313 | -0.13617 | 0.320594 | 0.903887 | Gstm7 |
| ENSMUSG( | 638.6662 | -0.05516 | 0.67071 | 0.95694 | Gsto1 |
| ENSMUSG( | 34.75531 | -0.74376 | 0.054285 | 0.723362 | Gsto2 |
| ENSMUSG( | 1598.507 | -0.20009 | 0.10794 | 0.804969 | Gstp1 |
| ENSMUSG( | 317.1844 | -0.34129 | 0.049674 | 0.708648 | Gstt1 |
| ENSMUSG( | 91.12355 | 0.5586 | 0.022112 | 0.576679 | Gstt2 |
| ENSMUSG( | 80.50833 | -0.31123 | 0.213557 | 0.864421 | Gstt3 |
| ENSMUSG( | 1243.101 | -0.06099 | 0.623642 | 0.953234 | Gstz1 |
| ENSMUSG( | 16.64863 | -0.2302 | 0.640667 | 0.953903 | Gsx1 |
| ENSMUSG( | 171.8929 | 0.168192 | 0.396079 | 0.917257 | Gt(ROSA)26Sor |
| ENSMUSG( | 1342.86 | 0.092187 | 0.408238 | 0.917257 | Gtdc1 |
| ENSMUSG( | 952.8577 | 0.028786 | 0.848925 | 0.985321 | Gtf2a1 |
| ENSMUSG( | 560.8816 | 0.063969 | 0.64646 | 0.953988 | Gtf2a2 |
| ENSMUSG( | 464.2353 | -0.01198 | 0.937915 | 0.993731 | Gtf2b |
| ENSMUSG( | 442.9085 | -0.18673 | 0.187084 | 0.856099 | Gtf2e1 |
| ENSMUSG( | 233.8908 | 0.122127 | 0.445575 | 0.921734 | Gtf2e2 |
| ENSMUSG( | 1339.642 | -0.11319 | 0.409694 | 0.917257 | Gtf2f1 |
| ENSMUSG( | 254.8855 | -0.07696 | 0.676314 | 0.95694 | Gtf2f2 |
| ENSMUSG( | 1010.194 | -0.02562 | 0.840012 | 0.98221 | Gtf2h1 |
| ENSMUSG( | 425.6974 | 0.126477 | 0.396781 | 0.917257 | Gtf2h2 |
| ENSMUSG( | 408.4373 | 0.005876 | 0.967793 | 0.995833 | Gtf2h3 |
| ENSMUSG( | 268.9204 | 0.022883 | 0.895205 | 0.990558 | Gtf2h4 |
| ENSMUSG( | 1859.731 | -0.15206 | 0.284937 | 0.892412 | Gtf2h5 |
| ENSMUSG( | 8576.94 | 0.00872 | 0.928089 | 0.99306 | Gtf2i |
| ENSMUSG( | 697.3875 | -0.02798 | 0.820116 | 0.979198 | Gtf2ird1 |
| ENSMUSG( | 488.326 | 0.082508 | 0.534333 | 0.93599 | Gtf2ird2 |
| ENSMUSG( | 788.2894 | -0.07652 | 0.535579 | 0.936675 | Gtf3a |
| ENSMUSG( | 3680.035 | 0.072795 | 0.55247 | 0.939565 | Gtf3c1 |
| ENSMUSG( | 841.1402 | 0.024853 | 0.869416 | 0.988097 | Gtf3c2 |
| ENSMUSG( | 505.1693 | 0.048685 | 0.730076 | 0.967228 | Gtf3c3 |
| ENSMUSG( | 1133.32 | 0.048602 | 0.672255 | 0.95694 | Gtf3c4 |
| ENSMUSG( | 523.348 | 0.044031 | 0.744228 | 0.96951 | Gtf3c5 |
| ENSMUSG( | 1010.759 | -0.1601 | 0.262311 | 0.885336 | Gtf3c6 |
| ENSMUSG( | 1272.893 | -0.01722 | 0.882815 | 0.989644 | Gtpbp1 |
| ENSMUSG( | 336.1466 | -0.01732 | 0.911762 | 0.992196 | Gtpbp10 |
| ENSMUSG( | 1626.119 | 0.079777 | 0.597568 | 0.946609 | Gtpbp2 |
| ENSMUSG( | 586.8581 | 0.163501 | 0.325394 | 0.905339 | Gtpbp3 |
| ENSMUSG( | 655.4829 | 0.028476 | 0.827972 | 0.980224 | Gtpbp4 |
| ENSMUSG( | 1311.562 | 0.02086 | 0.851445 | 0.985541 | Gtpbp6 |
| ENSMUSG( | 559.5658 | 0.086931 | 0.497116 | 0.927825 | Gtpbp8 |
| ENSMUSG( | 17.41957 | -0.37444 | 0.419605 | 0.917257 | Gtse1 |

| | | | | | | | | | | | |
|---|---|---|---|---|---|---|---|---|---|---|---|
| ENSMUSG( | Grhl3 | 21.03614 | 0.178409 | 0.66515 | NA | ENSMUSG( | Grhl3 | 13.23332 | -0.98685 | 0.096489 | 0.796133 |
| ENSMUSG( | Grhpr | 629.1669 | 0.087654 | 0.448282 | 0.834051 | ENSMUSG( | Grhpr | 511.7245 | -0.24963 | 0.067803 | 0.75212 |
| ENSMUSG( | Gria1 | 12445.19 | -0.16071 | 0.114178 | 0.509818 | ENSMUSG( | Gria1 | 12652.42 | 0.157908 | 0.188218 | 0.85698 |
| ENSMUSG( | Gria2 | 9315.647 | -0.3086 | 0.014623 | 0.177929 | ENSMUSG( | Gria2 | 10817.11 | 0.385303 | 0.009202 | 0.401972 |
| ENSMUSG( | Gria3 | 6417.452 | -0.19723 | 0.074671 | 0.423665 | ENSMUSG( | Gria3 | 6690.667 | 0.192065 | 0.124214 | 0.814691 |
| ENSMUSG( | Gria4 | 2929.574 | -0.39824 | 0.000421 | 0.018541 | ENSMUSG( | Gria4 | 3028.104 | -0.00551 | 0.966594 | 0.995475 |
| ENSMUSG( | Grid1 | 2456.373 | -0.26748 | 0.064398 | 0.393421 | ENSMUSG( | Grid1 | 2589.238 | 0.16633 | 0.332575 | 0.907907 |
| ENSMUSG( | Grid2 | 969.1876 | -0.15497 | 0.308937 | 0.757108 | ENSMUSG( | Grid2 | 983.6884 | 0.160402 | 0.350607 | 0.911652 |
| ENSMUSG( | Grid2ip | 75.05367 | -0.63146 | 0.005795 | 0.102397 | ENSMUSG( | Grid2ip | 72.11701 | -0.43369 | 0.078167 | 0.77321 |
| ENSMUSG( | Grifin | 17.06093 | -0.21664 | 0.638441 | NA | ENSMUSG( | Grifin | 12.83961 | -0.95427 | 0.092086 | 0.791889 |
| ENSMUSG( | Grik1 | 468.602 | 0.01915 | 0.888647 | 0.983434 | ENSMUSG( | Grik1 | 459.8434 | 0.231756 | 0.160616 | 0.837724 |
| ENSMUSG( | Grik2 | 1876.377 | -0.25536 | 0.025431 | 0.246258 | ENSMUSG( | Grik2 | 1977.182 | 0.171751 | 0.172046 | 0.843003 |
| ENSMUSG( | Grik3 | 2426.95 | -0.06094 | 0.717184 | 0.941697 | ENSMUSG( | Grik3 | 2613.704 | 0.401452 | 0.040234 | 0.681706 |
| ENSMUSG( | Grik4 | 901.2053 | -0.20574 | 0.146138 | 0.567605 | ENSMUSG( | Grik4 | 902.7239 | 0.079881 | 0.610938 | 0.950771 |
| ENSMUSG( | Grik5 | 12167.89 | 0.11068 | 0.227572 | 0.674441 | ENSMUSG( | Grik5 | 12311.09 | 0.390476 | 0.001654 | 0.150169 |
| ENSMUSG( | Grin1 | 19857.73 | -0.3065 | 0.008929 | 0.132538 | ENSMUSG( | Grin1 | 21151.07 | 0.158847 | 0.233065 | 0.868507 |
| ENSMUSG( | Grin1os | 131.1553 | -0.35994 | 0.131517 | 0.543959 | ENSMUSG( | Grin1os | 117.4048 | -0.38955 | 0.168964 | 0.840222 |
| ENSMUSG( | Grin2a | 3709.53 | -0.3787 | 0.011245 | 0.15263 | ENSMUSG( | Grin2a | 4093.489 | 0.195349 | 0.236761 | 0.872173 |
| ENSMUSG( | Grin2b | 6395.813 | -0.37881 | 0.02046 | 0.21574 | ENSMUSG( | Grin2b | 7086.131 | 0.208893 | 0.277031 | 0.891231 |
| ENSMUSG( | Grin2c | 6193.911 | 0.057779 | 0.600672 | 0.90443 | ENSMUSG( | Grin2c | 5528.324 | 0.000744 | 0.9953 | 0.999116 |
| ENSMUSG( | Grin2d | 668.6037 | -0.04269 | 0.732485 | 0.944956 | ENSMUSG( | Grin2d | 651.2861 | 0.150967 | 0.256228 | 0.883082 |
| ENSMUSG( | Grin3a | 1536.022 | -0.24691 | 0.093365 | 0.469793 | ENSMUSG( | Grin3a | 1596.414 | 0.144028 | 0.38754 | 0.917257 |
| ENSMUSG( | Grin3b | 26.37469 | -0.20828 | 0.579271 | 0.896692 | ENSMUSG( | Grin3b | 26.3842 | 0.068324 | 0.862807 | 0.986547 |
| ENSMUSG( | Grina | 19335.83 | -0.30437 | 0.002954 | 0.066464 | ENSMUSG( | Grina | 19821.98 | 0.05334 | 0.653811 | 0.955015 |
| ENSMUSG( | Grip1 | 466.7449 | -0.17976 | 0.116293 | 0.514678 | ENSMUSG( | Grip1 | 435.3408 | -0.11522 | 0.430818 | 0.917257 |
| ENSMUSG( | Grip2 | 910.373 | 0.201139 | 0.097593 | 0.477947 | ENSMUSG( | Grip2 | 793.3825 | 0.074648 | 0.615942 | 0.951413 |
| ENSMUSG( | Gripap1 | 2246.627 | 0.186703 | 0.111842 | 0.504332 | ENSMUSG( | Gripap1 | 1903.682 | -0.01999 | 0.889478 | 0.990558 |
| ENSMUSG( | Grk2 | 15451.2 | 0.180905 | 0.033969 | 0.282701 | ENSMUSG( | Grk2 | 12829.66 | -0.08174 | 0.397244 | 0.917257 |
| ENSMUSG( | Grk3 | 5724.047 | 0.063725 | 0.459077 | 0.840244 | ENSMUSG( | Grk3 | 5078.251 | -0.01211 | 0.904128 | 0.991089 |
| ENSMUSG( | Grk4 | 109.7334 | 0.174232 | 0.355398 | 0.78783 | ENSMUSG( | Grk4 | 104.2938 | 0.294406 | 0.165238 | 0.838743 |
| ENSMUSG( | Grk5 | 974.3159 | 0.044162 | 0.720024 | 0.941986 | ENSMUSG( | Grk5 | 858.5603 | -0.05299 | 0.652897 | 0.954897 |
| ENSMUSG( | Grk6 | 2784.308 | 0.17863 | 0.04657 | 0.330304 | ENSMUSG( | Grk6 | 2400.423 | 0.025687 | 0.807805 | 0.978622 |
| ENSMUSG( | Grm1 | 847.9675 | -0.62891 | 9.42E-05 | 0.006211 | ENSMUSG( | Grm1 | 1026.357 | 0.249395 | 0.17356 | 0.844317 |
| ENSMUSG( | Grm2 | 2344.934 | -0.29985 | 0.024913 | 0.242787 | ENSMUSG( | Grm2 | 2361.375 | 0.006965 | 0.963989 | 0.995433 |
| ENSMUSG( | Grm3 | 4955.109 | -0.10101 | 0.470457 | 0.84674 | ENSMUSG( | Grm3 | 4298.309 | -0.2485 | 0.095809 | 0.796133 |
| ENSMUSG( | Grm4 | 1142.133 | -0.3963 | 0.0011 | 0.035114 | ENSMUSG( | Grm4 | 1186.603 | 0.013989 | 0.922914 | 0.992644 |
| ENSMUSG( | Grm5 | 3366.514 | -0.48986 | 0.002944 | 0.066413 | ENSMUSG( | Grm5 | 3966.143 | 0.281266 | 0.138042 | 0.828492 |
| ENSMUSG( | Grm7 | 1410.1 | -0.47758 | 0.000833 | 0.028872 | ENSMUSG( | Grm7 | 1509.584 | 0.030256 | 0.853382 | 0.985598 |
| ENSMUSG( | Grm8 | 373.1663 | -0.56647 | 0.001668 | 0.046175 | ENSMUSG( | Grm8 | 432.0559 | 0.180716 | 0.358552 | 0.913759 |
| ENSMUSG( | Grn | 3512.443 | -0.21633 | 0.116599 | 0.514987 | ENSMUSG( | Grn | 3656.566 | 0.176253 | 0.266493 | 0.886134 |
| ENSMUSG( | Grp | 319.8303 | -0.11368 | 0.493422 | 0.860611 | ENSMUSG( | Grp | 323.9636 | 0.189969 | 0.307069 | 0.898538 |
| ENSMUSG( | Grpel1 | 982.1526 | 0.066891 | 0.523375 | 0.875628 | ENSMUSG( | Grpel1 | 801.3651 | -0.2676 | 0.041201 | 0.687358 |
| ENSMUSG( | Grpel2 | 477.3387 | -0.16271 | 0.259545 | 0.709948 | ENSMUSG( | Grpel2 | 435.0941 | -0.15916 | 0.289497 | 0.896591 |
| ENSMUSG( | Grsf1 | 1973.939 | -0.02293 | 0.804245 | 0.962169 | ENSMUSG( | Grsf1 | 1766.721 | -0.07841 | 0.5179 | 0.930939 |
| ENSMUSG( | Grtp1 | 356.9862 | 0.016952 | 0.918546 | 0.989954 | ENSMUSG( | Grtp1 | 301.149 | -0.20706 | 0.217251 | 0.865589 |
| ENSMUSG( | Grwd1 | 270.596 | 0.174668 | 0.289547 | 0.739839 | ENSMUSG( | Grwd1 | 233.9532 | 0.023367 | 0.896237 | 0.990558 |
| ENSMUSG( | Gsap | 578.3832 | 0.069914 | 0.573515 | 0.893584 | ENSMUSG( | Gsap | 497.2235 | -0.0928 | 0.521179 | 0.932489 |
| ENSMUSG( | Gsdmd | 150.0988 | -0.09981 | 0.604642 | 0.905556 | ENSMUSG( | Gsdmd | 151.5207 | 0.204011 | 0.400171 | 0.917257 |
| ENSMUSG( | Gsdme | 168.9846 | -0.03953 | 0.819242 | 0.965888 | ENSMUSG( | Gsdme | 145.8637 | -0.21144 | 0.28705 | 0.894091 |
| ENSMUSG( | Gse1 | 2321.269 | 0.126268 | 0.238506 | 0.686308 | ENSMUSG( | Gse1 | 2190.078 | 0.224069 | 0.066727 | 0.751678 |
| ENSMUSG( | Gsg1l | 3595.913 | -0.2371 | 0.031903 | 0.27264 | ENSMUSG( | Gsg1l | 3579.216 | 0.031037 | 0.810531 | 0.978779 |
| ENSMUSG( | Gsk3a | 10796.95 | 0.115891 | 0.12761 | 0.535393 | ENSMUSG( | Gsk3a | 9364.548 | -0.02404 | 0.821881 | 0.979198 |
| ENSMUSG( | Gsk3b | 4985.401 | 0.058595 | 0.502275 | 0.864924 | ENSMUSG( | Gsk3b | 4476.562 | 0.015313 | 0.910017 | 0.991958 |
| ENSMUSG( | Gskip | 579.0426 | 0.151104 | 0.217224 | 0.662164 | ENSMUSG( | Gskip | 474.6904 | -0.16035 | 0.338369 | 0.907972 |
| ENSMUSG( | Gsn | 5957.475 | -0.47806 | 0.001769 | 0.047422 | ENSMUSG( | Gsn | 6683.773 | 0.162665 | 0.379532 | 0.916995 |
| ENSMUSG( | Gspt1 | 2485.029 | 0.086524 | 0.346376 | 0.779153 | ENSMUSG( | Gspt1 | 2174.428 | -0.03148 | 0.799552 | 0.97681 |
| ENSMUSG( | Gspt2 | 485.9373 | -0.09857 | 0.452739 | 0.836191 | ENSMUSG( | Gspt2 | 465.3784 | 0.048199 | 0.767583 | 0.974236 |
| ENSMUSG( | Gsr | 692.1195 | -0.05593 | 0.697275 | 0.935662 | ENSMUSG( | Gsr | 622.9361 | -0.10039 | 0.564064 | 0.941206 |
| ENSMUSG( | Gss | 513.2972 | -0.03484 | 0.803023 | 0.962169 | ENSMUSG( | Gss | 454.4973 | -0.12186 | 0.443945 | 0.921734 |
| ENSMUSG( | Gsta3 | 29.94744 | -0.70209 | 0.0876 | 0.454663 | ENSMUSG( | Gsta3 | 34.44423 | 0.075662 | 0.829748 | 0.980224 |
| ENSMUSG( | Gsta4 | 1245.956 | -0.08914 | 0.330954 | 0.775174 | ENSMUSG( | Gsta4 | 1141.735 | -0.0746 | 0.520058 | 0.931904 |
| ENSMUSG( | Gstcd | 197.5595 | -0.05715 | 0.718028 | 0.941697 | ENSMUSG( | Gstcd | 182.7294 | -0.02105 | 0.907273 | 0.991683 |
| ENSMUSG( | Gstk1 | 799.02 | 0.141235 | 0.272586 | 0.722478 | ENSMUSG( | Gstk1 | 646.4824 | -0.21015 | 0.125194 | 0.81516 |
| ENSMUSG( | Gstm1 | 23471.34 | 0.288048 | 0.00312 | 0.069321 | ENSMUSG( | Gstm1 | 19054.3 | -0.03251 | 0.803858 | 0.977 |
| ENSMUSG( | Gstm2 | 87.5837 | -0.31219 | 0.174563 | 0.610304 | ENSMUSG( | Gstm2 | 80.29255 | -0.29278 | 0.243129 | 0.877399 |
| ENSMUSG( | Gstm4 | 298.9909 | -0.14534 | 0.273107 | 0.723468 | ENSMUSG( | Gstm4 | 266.8346 | -0.21909 | 0.182134 | 0.850579 |
| ENSMUSG( | Gstm5 | 11807.37 | 0.230929 | 0.022138 | 0.226843 | ENSMUSG( | Gstm5 | 9529.482 | -0.11366 | 0.377928 | 0.916784 |
| ENSMUSG( | Gstm6 | 188.4367 | 0.103328 | 0.557304 | 0.888279 | ENSMUSG( | Gstm6 | 163.7019 | -0.04119 | 0.82424 | 0.979198 |
| ENSMUSG( | Gstm7 | 608.6917 | -0.10466 | 0.368878 | 0.796486 | ENSMUSG( | Gstm7 | 548.7313 | -0.13617 | 0.320594 | 0.903887 |
| ENSMUSG( | Gsto1 | 738.6012 | 0.094983 | 0.375741 | 0.800976 | ENSMUSG( | Gsto1 | 638.6662 | -0.05516 | 0.67071 | 0.95694 |
| ENSMUSG( | Gsto2 | 42.01282 | -0.43353 | 0.178913 | 0.616298 | ENSMUSG( | Gsto2 | 34.75531 | -0.74376 | 0.054285 | 0.723362 |
| ENSMUSG( | Gstp1 | 1848.825 | -0.0391 | 0.674862 | 0.929651 | ENSMUSG( | Gstp1 | 1598.507 | -0.20009 | 0.10794 | 0.804969 |
| ENSMUSG( | Gstt1 | 351.347 | -0.29836 | 0.042979 | 0.318178 | ENSMUSG( | Gstt1 | 317.1844 | -0.34129 | 0.049674 | 0.708648 |
| ENSMUSG( | Gstt2 | 80.48637 | -0.02482 | 0.914313 | 0.988483 | ENSMUSG( | Gstt2 | 91.12355 | 0.5586 | 0.022112 | 0.576679 |
| ENSMUSG( | Gstt3 | 84.22251 | -0.45579 | 0.059109 | 0.375389 | ENSMUSG( | Gstt3 | 80.50833 | -0.31123 | 0.213557 | 0.864421 |
| ENSMUSG( | Gstz1 | 1484.56 | 0.179466 | 0.067054 | 0.400593 | ENSMUSG( | Gstz1 | 1243.101 | -0.06099 | 0.623642 | 0.953234 |
| ENSMUSG( | Gsx1 | 16.14641 | -0.66392 | 0.168599 | NA | ENSMUSG( | Gsx1 | 16.64863 | -0.2302 | 0.640667 | 0.953903 |
| ENSMUSG( | Gt(ROSA)2( | 160.0852 | -0.30933 | 0.081528 | 0.441102 | ENSMUSG( | Gt(ROSA)2( | 171.8929 | 0.168192 | 0.396079 | 0.917257 |
| ENSMUSG( | Gtdc1 | 1501.083 | 0.142464 | 0.132335 | 0.544251 | ENSMUSG( | Gtdc1 | 1342.86 | 0.092187 | 0.408238 | 0.917257 |
| ENSMUSG( | Gtf2a1 | 1008.237 | -0.07152 | 0.467343 | 0.844819 | ENSMUSG( | Gtf2a1 | 952.8577 | 0.028786 | 0.848925 | 0.985321 |
| ENSMUSG( | Gtf2a2 | 593.878 | -0.04137 | 0.742109 | 0.948478 | ENSMUSG( | Gtf2a2 | 560.8816 | 0.063969 | 0.64646 | 0.953988 |
| ENSMUSG( | Gtf2b | 526.6442 | 0.088501 | 0.494272 | 0.860611 | ENSMUSG( | Gtf2b | 464.2353 | -0.01198 | 0.937915 | 0.993731 |
| ENSMUSG( | Gtf2e1 | 532.7302 | 0.088888 | 0.436308 | 0.829093 | ENSMUSG( | Gtf2e1 | 442.9085 | -0.18673 | 0.187084 | 0.856099 |
| ENSMUSG( | Gtf2e2 | 250.3649 | 0.055537 | 0.703249 | 0.937735 | ENSMUSG( | Gtf2e2 | 233.8908 | 0.122127 | 0.445575 | 0.921734 |
| ENSMUSG( | Gtf2f1 | 1566.479 | 0.071673 | 0.553925 | 0.886397 | ENSMUSG( | Gtf2f1 | 1339.642 | -0.11319 | 0.409694 | 0.917257 |
| ENSMUSG( | Gtf2f2 | 292.0488 | 0.053515 | 0.756774 | 0.951513 | ENSMUSG( | Gtf2f2 | 254.8855 | -0.07696 | 0.676314 | 0.95694 |
| ENSMUSG( | Gtf2h1 | 1139.945 | 0.057613 | 0.58392 | 0.898666 | ENSMUSG( | Gtf2h1 | 1010.194 | -0.02562 | 0.840012 | 0.98221 |
| ENSMUSG( | Gtf2h2 | 474.5584 | 0.165364 | 0.189325 | 0.627932 | ENSMUSG( | Gtf2h2 | 425.6974 | 0.126477 | 0.396781 | 0.917257 |
| ENSMUSG( | Gtf2h3 | 448.4988 | 0.006362 | 0.960825 | 0.993991 | ENSMUSG( | Gtf2h3 | 408.4373 | 0.005876 | 0.967793 | 0.995833 |
| ENSMUSG( | Gtf2h4 | 308.3712 | 0.148612 | 0.320939 | 0.765984 | ENSMUSG( | Gtf2h4 | 268.9204 | 0.022883 | 0.895205 | 0.990558 |
| ENSMUSG( | Gtf2h5 | 2143.777 | -0.00209 | 0.982562 | 0.997512 | ENSMUSG( | Gtf2h5 | 1859.731 | -0.15206 | 0.284937 | 0.892412 |
| ENSMUSG( | Gtf2i | 9867.51 | 0.140811 | 0.076764 | 0.429306 | ENSMUSG( | Gtf2i | 8576.94 | 0.00872 | 0.928089 | 0.99306 |
| ENSMUSG( | Gtf2ird1 | 700.511 | -0.30049 | 0.006495 | 0.110626 | ENSMUSG( | Gtf2ird1 | 697.3875 | -0.02798 | 0.820116 | 0.979198 |
| ENSMUSG( | Gtf2ird2 | 533.5983 | 0.066672 | 0.550776 | 0.886038 | ENSMUSG( | Gtf2ird2 | 488.326 | 0.082508 | 0.534333 | 0.93599 |
| ENSMUSG( | Gtf3a | 882.7097 | -0.01696 | 0.86618 | 0.97613 | ENSMUSG( | Gtf3a | 788.2894 | -0.07652 | 0.535579 | 0.936675 |
| ENSMUSG( | Gtf3c1 | 3887.15 | -0.04022 | 0.699679 | 0.937143 | ENSMUSG( | Gtf3c1 | 3680.035 | 0.072795 | 0.55247 | 0.939565 |
| ENSMUSG( | Gtf3c2 | 950.6053 | 0.110704 | 0.386302 | 0.804623 | ENSMUSG( | Gtf3c2 | 841.1402 | 0.024853 | 0.869416 | 0.988097 |
| ENSMUSG( | Gtf3c3 | 554.2017 | 0.043056 | 0.727842 | 0.944355 | ENSMUSG( | Gtf3c3 | 505.1693 | 0.048685 | 0.730076 | 0.967228 |
| ENSMUSG( | Gtf3c4 | 1330.256 | 0.230656 | 0.015033 | 0.180217 | ENSMUSG( | Gtf3c4 | 1133.32 | 0.048602 | 0.672255 | 0.95694 |
| ENSMUSG( | Gtf3c5 | 596.477 | 0.144942 | 0.187587 | 0.626062 | ENSMUSG( | Gtf3c5 | 523.348 | 0.044031 | 0.744228 | 0.96951 |
| ENSMUSG( | Gtf3c6 | 1186.956 | 0.044128 | 0.647957 | 0.919759 | ENSMUSG( | Gtf3c6 | 1010.759 | -0.1601 | 0.262311 | 0.885336 |
| ENSMUSG( | Gtpbp1 | 1479.382 | 0.145089 | 0.135423 | 0.548884 | ENSMUSG( | Gtpbp1 | 1272.893 | -0.01722 | 0.882815 | 0.989644 |
| ENSMUSG( | Gtpbp10 | 383.3446 | 0.095096 | 0.516154 | 0.872701 | ENSMUSG( | Gtpbp10 | 336.1466 | -0.01732 | 0.911762 | 0.992196 |
| ENSMUSG( | Gtpbp2 | 1893.383 | 0.241925 | 0.043455 | 0.319227 | ENSMUSG( | Gtpbp2 | 1626.119 | 0.079777 | 0.597568 | 0.946609 |
| ENSMUSG( | Gtpbp3 | 628.5075 | 0.089778 | 0.484705 | 0.856698 | ENSMUSG( | Gtpbp3 | 586.8581 | 0.163501 | 0.325394 | 0.905339 |
| ENSMUSG( | Gtpbp4 | 760.7736 | 0.186354 | 0.081123 | 0.440221 | ENSMUSG( | Gtpbp4 | 655.4829 | 0.028476 | 0.827972 | 0.980224 |
| ENSMUSG( | Gtpbp6 | 1489.748 | 0.116061 | 0.301528 | 0.749979 | ENSMUSG( | Gtpbp6 | 1311.562 | 0.02086 | 0.851445 | 0.985541 |
| ENSMUSG( | Gtpbp8 | 598.8046 | 0.011554 | 0.915226 | 0.988634 | ENSMUSG( | Gtpbp8 | 559.5658 | 0.086931 | 0.497116 | 0.927825 |
| ENSMUSG( | Gtse1 | 20.79101 | -0.09951 | 0.810497 | NA | ENSMUSG( | Gtse1 | 17.41957 | -0.37444 | 0.419605 | 0.917257 |
| ENSMUSG( | Guca1b | 23.80093 | 0.471906 | 0.244572 | 0.69183 | ENSMUSG( | Guca1b | 19.8692 | 0.242326 | 0.602827 | 0.94789 |
| ENSMUSG( | Gucd1 | 491.3478 | -0.09049 | 0.460818 | 0.840962 | ENSMUSG( | Gucd1 | 448.8686 | -0.07471 | 0.638754 | 0.953903 |
| ENSMUSG( | Gucy1a1 | 1544.765 | 0.002547 | 0.986319 | 0.998326 | ENSMUSG( | Gucy1a1 | 1321.846 | -0.18952 | 0.306221 | 0.898538 |
| ENSMUSG( | Gucy1a2 | 96.66083 | -0.09205 | 0.654403 | 0.923227 | ENSMUSG( | Gucy1a2 | 86.80921 | -0.1326 | 0.552784 | 0.939565 |
| ENSMUSG( | Gucy1b1 | 2392.045 | 0.012814 | 0.922785 | 0.990631 | ENSMUSG( | Gucy1b1 | 2191.542 | 0.026049 | 0.86554 | 0.987023 |
| ENSMUSG( | Gucy1b2 | 10.21182 | -0.55125 | 0.335651 | NA | ENSMUSG( | Gucy1b2 | 10.08356 | -0.28705 | 0.633887 | NA |
| ENSMUSG( | Gucy2e | 327.8112 | -0.0992 | 0.534607 | 0.880719 | ENSMUSG( | Gucy2e | 297.0625 | -0.11723 | 0.48913 | 0.925673 |
| ENSMUSG( | Gucy2f | 39.55666 | -0.31288 | 0.384019 | 0.803646 | ENSMUSG( | Gucy2f | 36.4056 | -0.25458 | 0.530121 | 0.934795 |
| ENSMUSG( | Gucy2g | 150.5609 | -0.1658 | 0.401262 | 0.814475 | ENSMUSG( | Gucy2g | 137.2957 | -0.16856 | 0.430048 | 0.917257 |
| ENSMUSG( | Guf1 | 1273.937 | -0.12094 | 0.275831 | 0.725669 | ENSMUSG( | Guf1 | 1260.131 | 0.120764 | 0.349757 | 0.911652 |
| ENSMUSG( | Guk1 | 4824.927 | 0.005644 | 0.941827 | 0.993415 | ENSMUSG( | Guk1 | 4022.154 | -0.26352 | 0.011511 | 0.436431 |
| ENSMUSG( | Gulp1 | 128.4119 | 0.171717 | 0.466107 | 0.844453 | ENSMUSG( | Gulp1 | 114.684 | 0.100216 | 0.725597 | 0.966887 |
| ENSMUSG( | Gusb | 904.422 | -0.13369 | 0.25134 | 0.700869 | ENSMUSG( | Gusb | 903.0169 | 0.134958 | 0.372781 | 0.915491 |
| ENSMUSG( | Gxylt1 | 819.7184 | -0.0244 | 0.854689 | 0.973665 | ENSMUSG( | Gxylt1 | 773.3492 | 0.076319 | 0.59545 | 0.945853 |
| ENSMUSG( | Gxylt2 | 114.6702 | 0.023257 | 0.910307 | 0.9881 | ENSMUSG( | Gxylt2 | 114.3403 | 0.286905 | 0.205801 | 0.864412 |
| ENSMUSG( | Gyg1 | 1005.604 | 0.048729 | 0.602607 | 0.904887 | ENSMUSG( | Gyg1 | 851.5964 | -0.16812 | 0.142413 | 0.831221 |
| ENSMUSG( | Gypa | 21.43372 | -0.90821 | 0.119171 | 0.522984 | ENSMUSG( | Gypa | 26.29635 | 0.045917 | 0.936497 | 0.993573 |
| ENSMUSG( | Gypc | 87.54419 | -0.37684 | 0.125366 | 0.532314 | ENSMUSG( | Gypc | 88.93449 | -0.02875 | 0.909368 | 0.991958 |
| ENSMUSG( | Gys1 | 1079.834 | 0.09713 | 0.334006 | 0.775712 | ENSMUSG( | Gys1 | 956.1042 | 0.014577 | 0.901113 | 0.990558 |
| ENSMUSG( | Gzf1 | 1186.935 | 0.148494 | 0.111532 | 0.503511 | ENSMUSG( | Gzf1 | 1015.248 | -0.03299 | 0.778101 | 0.976106 |
| ENSMUSG( | Gzma | 15.94399 | -1.10305 | 0.151679 | NA | ENSMUSG( | Gzma | 15.14582 | -1.01365 | 0.194439 | 0.860096 |
| ENSMUSG( | Gzmm | 33.93537 | 0.051223 | 0.875936 | 0.980448 | ENSMUSG( | Gzmm | 29.51611 | -0.0406 | 0.918057 | 0.992529 |
| ENSMUSG( | H13 | 2736.344 | -0.25215 | 0.035184 | 0.288514 | ENSMUSG( | H13 | 2773.4 | 0.0689 | 0.611247 | 0.950771 |
| ENSMUSG( | H19 | 67.5177 | -0.05227 | 0.852758 | 0.973168 | ENSMUSG( | H19 | 45.17948 | -1.20328 | 0.000451 | 0.061774 |
| ENSMUSG( | H1f0 | 4891.383 | 0.097266 | 0.402325 | 0.815103 | ENSMUSG( | H1f0 | 3811.048 | -0.37942 | 0.000837 | 0.097764 |
| ENSMUSG( | H1f10 | 1417.626 | 0.078815 | 0.469732 | 0.846186 | ENSMUSG( | H1f10 | 1211.722 | -0.11006 | 0.416242 | 0.917257 |
| ENSMUSG( | H1f11-ps | 18.11383 | -0.00688 | 0.987239 | NA | ENSMUSG( | H1f11-ps | 15.63305 | -0.13094 | 0.795126 | 0.97681 |
| ENSMUSG( | H1f2 | 1884.991 | 0.067804 | 0.708628 | 0.939519 | ENSMUSG( | H1f2 | 1407.695 | -0.56524 | 0.000229 | 0.039667 |
| ENSMUSG( | H1f4 | 206.2619 | 0.04646 | 0.860597 | 0.9749 | ENSMUSG( | H1f4 | 141.5349 | -0.91596 | 0.000118 | 0.02332 |
| ENSMUSG( | H2-Aa | 125.5785 | -0.44004 | 0.017298 | 0.197176 | ENSMUSG( | H2-Aa | 117.6668 | -0.33512 | 0.118695 | 0.813384 |
| ENSMUSG( | H2-Ab1 | 128.8923 | -0.47334 | 0.012294 | 0.160458 | ENSMUSG( | H2-Ab1 | 121.5258 | -0.36603 | 0.078351 | 0.77321 |
| ENSMUSG( | H2ac19 | 30.72353 | 0.13138 | 0.735415 | 0.946109 | ENSMUSG( | H2ac19 | 22.7796 | -0.52625 | 0.267854 | 0.886134 |
| ENSMUSG( | H2ac25 | 304.8384 | 0.144513 | 0.429681 | 0.826446 | ENSMUSG( | H2ac25 | 274.0794 | 0.095043 | 0.646399 | 0.953988 |
| ENSMUSG( | H2ac6 | 109.4226 | -0.23451 | 0.31827 | 0.764234 | ENSMUSG( | H2ac6 | 90.34149 | -0.58623 | 0.020847 | 0.571913 |
| ENSMUSG( | H2aj | 1467.615 | -0.10744 | 0.282608 | 0.73186 | ENSMUSG( | H2aj | 1299.343 | -0.19893 | 0.107712 | 0.804969 |
| ENSMUSG( | H2ax | 872.5428 | -0.13439 | 0.269753 | 0.719729 | ENSMUSG( | H2ax | 850.1816 | 0.061648 | 0.624077 | 0.953234 |
| ENSMUSG( | H2az1 | 1328.366 | -0.0877 | 0.358335 | 0.789759 | ENSMUSG( | H2az1 | 1244.114 | -0.00849 | 0.939234 | 0.993898 |
| ENSMUSG( | H2az2 | 1717.403 | -0.02121 | 0.863462 | 0.975167 | ENSMUSG( | H2az2 | 1491.932 | -0.16661 | 0.214717 | 0.864841 |
| ENSMUSG( | H2bc15 | 17.69677 | -0.24526 | 0.605349 | NA | ENSMUSG( | H2bc15 | 14.29007 | -0.6636 | 0.213892 | 0.864421 |
| ENSMUSG( | H2bc21 | 593.6685 | -0.11195 | 0.495103 | 0.860798 | ENSMUSG( | H2bc21 | 510.1487 | -0.30243 | 0.087837 | 0.78582 |
| ENSMUSG( | H2bc27 | 387.2897 | -0.03791 | 0.828037 | 0.968966 | ENSMUSG( | H2bc27 | 322.7615 | -0.32185 | 0.138306 | 0.828492 |
| ENSMUSG( | H2bc4 | 1909.279 | 0.123844 | 0.394616 | 0.810072 | ENSMUSG( | H2bc4 | 1512.305 | -0.29831 | 0.055074 | 0.723362 |
| ENSMUSG( | H2bc6 | 332.5427 | 0.075557 | 0.719717 | 0.941986 | ENSMUSG( | H2bc6 | 242.24 | -0.64789 | 0.001706 | 0.152157 |
| ENSMUSG( | H2bc8 | 25.66713 | -0.00935 | 0.980966 | 0.997512 | ENSMUSG( | H2bc8 | 23.48207 | 0.02335 | 0.957956 | 0.994798 |
| ENSMUSG( | H2-D1 | 2210.765 | -0.28815 | 0.008962 | 0.132538 | ENSMUSG( | H2-D1 | 2267.341 | 0.069251 | 0.628021 | 0.953234 |
| ENSMUSG( | H2-DMa | 546.3346 | -0.37065 | 0.001271 | 0.038891 | ENSMUSG( | H2-DMa | 542.0095 | -0.10289 | 0.429068 | 0.917257 |
| ENSMUSG( | H2-DMb1 | 84.43255 | -0.07039 | 0.743041 | 0.948478 | ENSMUSG( | H2-DMb1 | 88.14803 | 0.328721 | 0.186762 | 0.856099 |
| ENSMUSG( | H2-DMb2 | 20.80512 | 0.039542 | 0.924492 | NA | ENSMUSG( | H2-DMb2 | 22.32767 | 0.489749 | 0.276121 | 0.890749 |
| ENSMUSG( | H2-Eb1 | 110.9092 | -0.55488 | 0.008089 | 0.125198 | ENSMUSG( | H2-Eb1 | 113.8003 | -0.16983 | 0.421118 | 0.917257 |

| | | | | | |
|---|---|---|---|---|---|
| ENSMUSG( | 87.5837 | -0.31219 | 0.174563 | 0.610304 | Gstm2 |
| ENSMUSG( | 298.9909 | -0.14534 | 0.273107 | 0.723468 | Gstm4 |
| ENSMUSG( | 11807.37 | 0.230929 | 0.022138 | 0.226843 | Gstm5 |
| ENSMUSG( | 188.4367 | 0.103328 | 0.557304 | 0.888279 | Gstm6 |
| ENSMUSG( | 608.6917 | -0.10466 | 0.368878 | 0.796486 | Gstm7 |
| ENSMUSG( | 738.6012 | 0.094983 | 0.375741 | 0.800976 | Gsto1 |
| ENSMUSG( | 42.01282 | -0.43353 | 0.178913 | 0.616298 | Gsto2 |
| ENSMUSG( | 1848.825 | -0.0391 | 0.674862 | 0.929651 | Gstp1 |
| ENSMUSG( | 351.347 | -0.29836 | 0.042979 | 0.318178 | Gstt1 |
| ENSMUSG( | 80.48637 | -0.02482 | 0.914313 | 0.988483 | Gstt2 |
| ENSMUSG( | 84.22251 | -0.45579 | 0.059109 | 0.375389 | Gstt3 |
| ENSMUSG( | 1484.56 | 0.179466 | 0.067054 | 0.400593 | Gstz1 |
| ENSMUSG( | 16.14641 | -0.66392 | 0.168599 | NA | Gsx1 |
| ENSMUSG( | 160.0852 | -0.30933 | 0.081528 | 0.441102 | Gt(ROSA)26Sor |
| ENSMUSG( | 1501.083 | 0.142464 | 0.132335 | 0.544251 | Gtdc1 |
| ENSMUSG( | 1008.237 | -0.07152 | 0.467343 | 0.844819 | Gtf2a1 |
| ENSMUSG( | 593.878 | -0.04137 | 0.742109 | 0.948478 | Gtf2a2 |
| ENSMUSG( | 526.6442 | 0.088501 | 0.494272 | 0.860611 | Gtf2b |
| ENSMUSG( | 532.7302 | 0.088888 | 0.436308 | 0.829093 | Gtf2e1 |
| ENSMUSG( | 250.3649 | 0.055537 | 0.703249 | 0.937735 | Gtf2e2 |
| ENSMUSG( | 1566.479 | 0.071673 | 0.553925 | 0.886397 | Gtf2f1 |
| ENSMUSG( | 292.0488 | 0.053515 | 0.756774 | 0.951513 | Gtf2f2 |
| ENSMUSG( | 1139.945 | 0.057613 | 0.58392 | 0.898666 | Gtf2h1 |
| ENSMUSG( | 474.5584 | 0.165364 | 0.189325 | 0.627932 | Gtf2h2 |
| ENSMUSG( | 448.4988 | 0.006362 | 0.960825 | 0.993991 | Gtf2h3 |
| ENSMUSG( | 308.3712 | 0.148612 | 0.320939 | 0.765984 | Gtf2h4 |
| ENSMUSG( | 2143.777 | -0.00209 | 0.982562 | 0.997512 | Gtf2h5 |
| ENSMUSG( | 9867.51 | 0.140811 | 0.076764 | 0.429306 | Gtf2i |
| ENSMUSG( | 700.511 | -0.30049 | 0.006495 | 0.110626 | Gtf2ird1 |
| ENSMUSG( | 533.5983 | 0.066672 | 0.550776 | 0.886038 | Gtf2ird2 |
| ENSMUSG( | 882.7097 | -0.01696 | 0.86618 | 0.97613 | Gtf3a |
| ENSMUSG( | 3887.15 | -0.04022 | 0.699679 | 0.937143 | Gtf3c1 |
| ENSMUSG( | 950.6053 | 0.110704 | 0.386302 | 0.804623 | Gtf3c2 |
| ENSMUSG( | 554.2017 | 0.043056 | 0.727842 | 0.944355 | Gtf3c3 |
| ENSMUSG( | 1330.256 | 0.230656 | 0.015033 | 0.180217 | Gtf3c4 |
| ENSMUSG( | 596.477 | 0.144942 | 0.187587 | 0.626062 | Gtf3c5 |
| ENSMUSG( | 1186.956 | 0.044128 | 0.647957 | 0.919759 | Gtf3c6 |
| ENSMUSG( | 1479.382 | 0.145089 | 0.135423 | 0.548884 | Gtpbp1 |
| ENSMUSG( | 383.3446 | 0.095096 | 0.516154 | 0.872701 | Gtpbp10 |
| ENSMUSG( | 1893.383 | 0.241925 | 0.043455 | 0.319227 | Gtpbp2 |
| ENSMUSG( | 628.5075 | 0.089778 | 0.484705 | 0.856698 | Gtpbp3 |
| ENSMUSG( | 760.7736 | 0.186354 | 0.081123 | 0.440221 | Gtpbp4 |
| ENSMUSG( | 1489.748 | 0.116061 | 0.301528 | 0.749979 | Gtpbp6 |
| ENSMUSG( | 598.8046 | 0.011554 | 0.915226 | 0.988634 | Gtpbp8 |
| ENSMUSG( | 20.79101 | -0.09951 | 0.810497 | NA | Gtse1 |
| ENSMUSG( | 23.80093 | 0.471906 | 0.244572 | 0.69183 | Guca1b |
| ENSMUSG( | 491.3478 | -0.09049 | 0.460818 | 0.840962 | Gucd1 |
| ENSMUSG( | 1544.765 | 0.002547 | 0.986319 | 0.998326 | Gucy1a1 |
| ENSMUSG( | 96.66083 | -0.09205 | 0.654403 | 0.923227 | Gucy1a2 |
| ENSMUSG( | 2392.045 | 0.012814 | 0.922785 | 0.990631 | Gucy1b1 |
| ENSMUSG( | 10.21182 | -0.55125 | 0.335651 | NA | Gucy1b2 |
| ENSMUSG( | 327.8112 | -0.0992 | 0.534607 | 0.880719 | Gucy2e |
| ENSMUSG( | 39.55666 | -0.31288 | 0.384019 | 0.803646 | Gucy2f |
| ENSMUSG( | 150.5609 | -0.1658 | 0.401262 | 0.814475 | Gucy2g |
| ENSMUSG( | 1273.937 | -0.12094 | 0.275831 | 0.725669 | Guf1 |
| ENSMUSG( | 4824.927 | 0.005644 | 0.941827 | 0.993415 | Guk1 |
| ENSMUSG( | 128.4119 | 0.171717 | 0.466107 | 0.844453 | Gulp1 |
| ENSMUSG( | 904.422 | -0.13369 | 0.25134 | 0.700869 | Gusb |
| ENSMUSG( | 9.801035 | 0.315337 | 0.653299 | NA | Gvin2 |
| ENSMUSG( | 819.7184 | -0.0244 | 0.854689 | 0.973665 | Gxylt1 |
| ENSMUSG( | 114.6702 | 0.023257 | 0.910307 | 0.9881 | Gxylt2 |
| ENSMUSG( | 1005.604 | 0.048729 | 0.602607 | 0.904887 | Gyg1 |
| ENSMUSG( | 21.43372 | -0.90821 | 0.119171 | 0.522984 | Gypa |
| ENSMUSG( | 87.54419 | -0.37684 | 0.125366 | 0.532314 | Gypc |
| ENSMUSG( | 1079.834 | 0.09713 | 0.334006 | 0.775712 | Gys1 |
| ENSMUSG( | 1186.935 | 0.148494 | 0.111532 | 0.503511 | Gzf1 |
| ENSMUSG( | 15.94399 | -1.10305 | 0.151679 | NA | Gzma |
| ENSMUSG( | 33.93537 | 0.051223 | 0.875936 | 0.980448 | Gzmm |
| ENSMUSG( | 2736.344 | -0.25215 | 0.035184 | 0.288514 | H13 |
| ENSMUSG( | 67.5177 | -0.05227 | 0.852758 | 0.973168 | H19 |
| ENSMUSG( | 4891.383 | 0.097266 | 0.402325 | 0.815103 | H1f0 |
| ENSMUSG( | 1417.626 | 0.078815 | 0.469732 | 0.846186 | H1f10 |
| ENSMUSG( | 18.11383 | -0.00688 | 0.987239 | NA | H1f11-ps |
| ENSMUSG( | 1884.991 | 0.067804 | 0.708628 | 0.939519 | H1f2 |
| ENSMUSG( | 206.2619 | 0.04646 | 0.860597 | 0.9749 | H1f4 |
| ENSMUSG( | 125.5785 | -0.44004 | 0.017298 | 0.197176 | H2-Aa |
| ENSMUSG( | 128.8923 | -0.47334 | 0.012294 | 0.160458 | H2-Ab1 |
| ENSMUSG( | 30.72353 | 0.13138 | 0.735415 | 0.946109 | H2ac19 |
| ENSMUSG( | 304.8384 | 0.144513 | 0.429681 | 0.826446 | H2ac25 |
| ENSMUSG( | 109.4226 | -0.23451 | 0.31827 | 0.764234 | H2ac6 |
| ENSMUSG( | 1467.615 | -0.10744 | 0.282608 | 0.73186 | H2aj |
| ENSMUSG( | 872.5428 | -0.13439 | 0.269753 | 0.719729 | H2ax |
| ENSMUSG( | 1328.366 | -0.0877 | 0.358335 | 0.789759 | H2az1 |
| ENSMUSG( | 1717.403 | -0.02121 | 0.863462 | 0.975167 | H2az2 |
| ENSMUSG( | 17.69677 | -0.24526 | 0.605349 | NA | H2bc15 |
| ENSMUSG( | 593.6685 | -0.11195 | 0.495103 | 0.860798 | H2bc21 |
| ENSMUSG( | 387.2897 | -0.03791 | 0.828037 | 0.968966 | H2bc27 |
| ENSMUSG( | 1909.279 | 0.123844 | 0.394616 | 0.810072 | H2bc4 |
| ENSMUSG( | 332.5427 | 0.075557 | 0.719717 | 0.941986 | H2bc6 |
| ENSMUSG( | 25.66713 | -0.00935 | 0.980966 | 0.997512 | H2bc8 |
| ENSMUSG( | 8.460762 | 0.588982 | 0.358889 | NA | H2bc9 |
| ENSMUSG( | 2210.765 | -0.28815 | 0.008962 | 0.132538 | H2-D1 |
| ENSMUSG( | 546.3346 | -0.37065 | 0.001271 | 0.038891 | H2-DMa |
| ENSMUSG( | 84.43255 | -0.07039 | 0.743041 | 0.948478 | H2-DMb1 |
| ENSMUSG( | 20.80512 | 0.039542 | 0.924492 | NA | H2-DMb2 |
| ENSMUSG( | 110.9092 | -0.55488 | 0.008089 | 0.125198 | H2-Eb1 |
| ENSMUSG( | 1449.869 | -0.3068 | 0.01619 | 0.189046 | H2-K1 |
| ENSMUSG( | 14.45285 | -0.3203 | 0.510689 | NA | H2-K2 |
| ENSMUSG( | 85.83899 | -0.37026 | 0.101235 | 0.484996 | H2-M3 |
| ENSMUSG( | 60.57632 | 0.220904 | 0.41029 | 0.818258 | H2-M5 |
| ENSMUSG( | 24.72813 | 0.020857 | 0.959163 | 0.99396 | H2-M6-ps |
| ENSMUSG( | 37.43102 | -0.48941 | 0.126353 | 0.533885 | H2-Oa |
| ENSMUSG( | 32.75258 | -0.3126 | 0.336045 | 0.777 | H2-Ob |
| ENSMUSG( | 26.72327 | -0.4943 | 0.199909 | 0.640239 | H2-Q1 |
| ENSMUSG( | 18.09748 | 0.532138 | 0.239974 | NA | H2-Q2 |
| ENSMUSG( | 209.2781 | -0.20773 | 0.181776 | 0.618944 | H2-Q4 |
| ENSMUSG( | 22.41095 | -0.4847 | 0.226265 | 0.67315 | H2-T10 |
| ENSMUSG( | 711.8216 | -0.08898 | 0.454572 | 0.837383 | H2-T22 |
| ENSMUSG( | 578.5268 | -0.17366 | 0.278826 | 0.728705 | H2-T23 |
| ENSMUSG( | 95.92905 | -0.27328 | 0.24125 | 0.689026 | H2-T24 |
| ENSMUSG( | 18.22183 | -0.6929 | 0.129792 | NA | H2-T-ps |
| ENSMUSG( | 115.2899 | 0.123205 | 0.607464 | 0.906876 | H3c15 |
| ENSMUSG( | 27.60241 | 0.462624 | 0.209129 | 0.651071 | H3c4 |
| ENSMUSG( | 3292.967 | 0.006838 | 0.944881 | 0.993612 | H3f3a |
| ENSMUSG( | 3947.352 | 0.009143 | 0.934803 | 0.993231 | H3f3b |
| ENSMUSG( | 29.07209 | -0.32893 | 0.385692 | 0.804413 | H4c12 |
| ENSMUSG( | 15.8271 | -0.06589 | 0.888429 | NA | H4c14 |
| ENSMUSG( | 10.23941 | -0.15319 | 0.828242 | NA | H4c17 |
| ENSMUSG( | 15.92613 | 0.311302 | 0.530154 | NA | H4c3 |
| ENSMUSG( | 60.0763 | 0.050955 | 0.869944 | 0.977655 | H4c8 |
| ENSMUSG( | 89.45236 | 0.107163 | 0.641885 | 0.916865 | H4c9 |
| ENSMUSG( | 15.78587 | -0.80696 | 0.123376 | NA | H60b |
| ENSMUSG( | 278.0529 | -0.34064 | 0.112932 | 0.506708 | H6pd |
| ENSMUSG( | 13.32059 | -0.50111 | 0.386979 | NA | Haao |
| ENSMUSG( | 6860.362 | 0.107028 | 0.210043 | 0.652413 | Habp4 |
| ENSMUSG( | 424.9409 | -0.12801 | 0.351212 | 0.783067 | Hacd1 |
| ENSMUSG( | 1765.663 | -0.16432 | 0.256768 | 0.707935 | Hacd2 |
| ENSMUSG( | 7467.262 | -0.31396 | 0.026101 | 0.247245 | Hacd3 |
| ENSMUSG( | 278.7641 | -0.0086 | 0.962041 | 0.994538 | Hacd4 |
| ENSMUSG( | 958.505 | 0.051875 | 0.613142 | 0.908902 | Hace1 |
| ENSMUSG( | 55.33233 | -0.01465 | 0.955903 | 0.993874 | Hacl1 |
| ENSMUSG( | 2233.115 | 0.036 | 0.755445 | 0.951395 | Hadh |
| ENSMUSG( | 5157.04 | 0.09286 | 0.335211 | 0.776511 | Hadha |
| ENSMUSG( | 3875.214 | 0.095048 | 0.410478 | 0.818434 | Hadhb |
| ENSMUSG( | 1706.588 | -0.04898 | 0.62191 | 0.911432 | Hagh |
| ENSMUSG( | 1953.97 | 0.019526 | 0.891878 | 0.983979 | Haghl |
| ENSMUSG( | 2581.504 | 0.20299 | 0.219455 | 0.664984 | Hap1 |
| ENSMUSG( | 1584.985 | -0.4097 | 0.029075 | 0.259547 | Hapln1 |
| ENSMUSG( | 828.9488 | -0.52095 | 7.27E-05 | 0.005238 | Hapln2 |
| ENSMUSG( | 93.61519 | -0.25136 | 0.277701 | 0.727559 | Hapln3 |
| ENSMUSG( | 3093.035 | -0.34402 | 0.008139 | 0.125721 | Hapln4 |
| ENSMUSG( | 1539.33 | -0.02557 | 0.801419 | 0.961947 | Hapstr1 |
| ENSMUSG( | 90.17768 | -0.17402 | 0.431857 | 0.82768 | Harbi1 |
| ENSMUSG( | 1969.553 | 0.0365 | 0.748196 | 0.949604 | Hars1 |
| ENSMUSG( | 812.1345 | 0.109935 | 0.284995 | 0.733568 | Hars2 |
| ENSMUSG( | 25.66937 | -0.18339 | 0.639599 | 0.916434 | Has1 |
| ENSMUSG( | 9.280827 | -0.15983 | 0.795191 | NA | Has2 |
| ENSMUSG( | 12.8339 | -0.43261 | 0.390914 | NA | Has2os |
| ENSMUSG( | 63.78528 | -0.34366 | 0.181524 | 0.618685 | Has3 |
| ENSMUSG( | 303.7414 | 0.078193 | 0.54406 | 0.884411 | Hat1 |
| ENSMUSG( | 261.728 | -0.09949 | 0.472351 | 0.848048 | Haus1 |
| ENSMUSG( | 765.4884 | 0.06325 | 0.575621 | 0.894836 | Haus2 |
| ENSMUSG( | 147.3899 | 0.13074 | 0.460351 | 0.840676 | Haus3 |
| ENSMUSG( | 74.2019 | -0.2578 | 0.269646 | 0.719729 | Haus4 |
| ENSMUSG( | 32.61902 | 0.13325 | 0.705232 | 0.938303 | Haus5 |
| ENSMUSG( | 300.9711 | 0.000275 | 0.998385 | 0.99989 | Haus6 |
| ENSMUSG( | 218.7272 | -0.0486 | 0.760718 | 0.952907 | Haus7 |
| ENSMUSG( | 227.3231 | 0.049793 | 0.731798 | 0.944956 | Haus8 |

| | | | | | |
|---|---|---|---|---|---|
| ENSMUSG( | 19.8692 | 0.242326 | 0.602827 | 0.94789 | Guca1b |
| ENSMUSG( | 448.8686 | -0.07471 | 0.638754 | 0.953903 | Gucd1 |
| ENSMUSG( | 1321.846 | -0.18952 | 0.306221 | 0.898538 | Gucy1a1 |
| ENSMUSG( | 86.80921 | -0.1326 | 0.552784 | 0.939565 | Gucy1a2 |
| ENSMUSG( | 2191.542 | 0.026049 | 0.86554 | 0.987023 | Gucy1b1 |
| ENSMUSG( | 10.08356 | -0.28705 | 0.633887 | NA | Gucy1b2 |
| ENSMUSG( | 297.0625 | -0.11723 | 0.48913 | 0.925673 | Gucy2e |
| ENSMUSG( | 36.4056 | -0.25458 | 0.530121 | 0.934795 | Gucy2f |
| ENSMUSG( | 137.2957 | -0.16856 | 0.430048 | 0.917257 | Gucy2g |
| ENSMUSG( | 1260.131 | 0.120764 | 0.349757 | 0.911652 | Guf1 |
| ENSMUSG( | 4022.154 | -0.26352 | 0.011511 | 0.436431 | Guk1 |
| ENSMUSG( | 114.684 | 0.100216 | 0.725597 | 0.966887 | Gulp1 |
| ENSMUSG( | 903.0169 | 0.134958 | 0.372781 | 0.915491 | Gusb |
| ENSMUSG( | 9.592795 | 0.13497 | 0.839741 | NA | Gvin1 |
| ENSMUSG( | 773.3492 | 0.076319 | 0.59545 | 0.945853 | Gxylt1 |
| ENSMUSG( | 114.3403 | 0.286905 | 0.205801 | 0.864412 | Gxylt2 |
| ENSMUSG( | 851.5964 | -0.16812 | 0.142413 | 0.831221 | Gyg1 |
| ENSMUSG( | 26.29635 | 0.045917 | 0.936497 | 0.993573 | Gypa |
| ENSMUSG( | 88.93449 | -0.02875 | 0.909368 | 0.991958 | Gypc |
| ENSMUSG( | 956.1042 | 0.014577 | 0.901113 | 0.990558 | Gys1 |
| ENSMUSG( | 1015.248 | -0.03299 | 0.778101 | 0.976106 | Gzf1 |
| ENSMUSG( | 15.14582 | -1.01365 | 0.194439 | 0.860096 | Gzma |
| ENSMUSG( | 29.51611 | -0.0406 | 0.918057 | 0.992529 | Gzmm |
| ENSMUSG( | 2773.4 | 0.0689 | 0.611247 | 0.950771 | H13 |
| ENSMUSG( | 45.17948 | -1.20328 | 0.000451 | 0.061774 | H19 |
| ENSMUSG( | 3811.048 | -0.37942 | 0.000837 | 0.097764 | H1f0 |
| ENSMUSG( | 1211.722 | -0.11006 | 0.416242 | 0.917257 | H1f10 |
| ENSMUSG( | 15.63305 | -0.13094 | 0.795126 | 0.97681 | H1f11-ps |
| ENSMUSG( | 1407.695 | -0.56524 | 0.000229 | 0.039667 | H1f2 |
| ENSMUSG( | 141.5349 | -0.91596 | 0.000118 | 0.02332 | H1f4 |
| ENSMUSG( | 117.6668 | -0.33512 | 0.118695 | 0.813384 | H2-Aa |
| ENSMUSG( | 121.5258 | -0.36603 | 0.078351 | 0.77321 | H2-Ab1 |
| ENSMUSG( | 22.7796 | -0.52625 | 0.267854 | 0.886134 | H2ac19 |
| ENSMUSG( | 274.0794 | 0.095043 | 0.646399 | 0.953988 | H2ac25 |
| ENSMUSG( | 90.34149 | -0.58623 | 0.020847 | 0.571913 | H2ac6 |
| ENSMUSG( | 1299.343 | -0.19893 | 0.107712 | 0.804969 | H2aj |
| ENSMUSG( | 850.1816 | 0.061648 | 0.624077 | 0.953234 | H2ax |
| ENSMUSG( | 1244.114 | -0.00849 | 0.939234 | 0.993898 | H2az1 |
| ENSMUSG( | 1491.932 | -0.16661 | 0.214717 | 0.864841 | H2az2 |
| ENSMUSG( | 14.29007 | -0.6636 | 0.213892 | 0.864421 | H2bc15 |
| ENSMUSG( | 510.1487 | -0.30243 | 0.087837 | 0.78582 | H2bc21 |
| ENSMUSG( | 322.7615 | -0.32185 | 0.138306 | 0.828492 | H2bc27 |
| ENSMUSG( | 1512.305 | -0.29831 | 0.055074 | 0.723362 | H2bc4 |
| ENSMUSG( | 242.24 | -0.64789 | 0.001706 | 0.152157 | H2bc6 |
| ENSMUSG( | 23.48207 | 0.02335 | 0.957956 | 0.994798 | H2bc8 |
| ENSMUSG( | 2267.341 | 0.069251 | 0.628021 | 0.953234 | H2-D1 |
| ENSMUSG( | 542.0095 | -0.10289 | 0.429068 | 0.917257 | H2-DMa |
| ENSMUSG( | 88.14803 | 0.328721 | 0.186762 | 0.856099 | H2-DMb1 |
| ENSMUSG( | 22.32767 | 0.489749 | 0.276121 | 0.890749 | H2-DMb2 |
| ENSMUSG( | 113.8003 | -0.16983 | 0.421118 | 0.917257 | H2-Eb1 |
| ENSMUSG( | 1546.488 | 0.163875 | 0.294444 | 0.898538 | H2-K1 |
| ENSMUSG( | 13.27984 | -0.27079 | 0.628593 | 0.953251 | H2-K2 |
| ENSMUSG( | 93.04153 | 0.151945 | 0.505877 | 0.927825 | H2-M3 |
| ENSMUSG( | 47.52509 | -0.20598 | 0.483897 | 0.925289 | H2-M5 |
| ENSMUSG( | 27.74784 | 0.588254 | 0.128577 | 0.81925 | H2-M6-ps |
| ENSMUSG( | 39.42997 | -0.01825 | 0.955183 | 0.994798 | H2-Oa |
| ENSMUSG( | 38.00246 | 0.40416 | 0.231281 | 0.868507 | H2-Ob |
| ENSMUSG( | 27.06209 | -0.17367 | 0.667147 | 0.956766 | H2-Q1 |
| ENSMUSG( | 15.52785 | 0.387466 | 0.453387 | 0.922316 | H2-Q2 |
| ENSMUSG( | 216.3588 | 0.173121 | 0.382095 | 0.917257 | H2-Q4 |
| ENSMUSG( | 23.09152 | -0.07177 | 0.864767 | 0.986591 | H2-T10 |
| ENSMUSG( | 639.4262 | -0.12652 | 0.337869 | 0.907972 | H2-T22 |
| ENSMUSG( | 571.7866 | 0.075167 | 0.683855 | 0.958678 | H2-T23 |
| ENSMUSG( | 94.78561 | 0.00344 | 0.990893 | 0.998302 | H2-T24 |
| ENSMUSG( | 18.37699 | -0.35734 | 0.442459 | 0.921554 | H2-T-ps |
| ENSMUSG( | 90.18884 | -0.3386 | 0.19249 | 0.859726 | H3c15 |
| ENSMUSG( | 23.79235 | 0.318585 | 0.434925 | 0.918876 | H3c4 |
| ENSMUSG( | 2861.032 | -0.13687 | 0.299171 | 0.898538 | H3f3a |
| ENSMUSG( | 3473.585 | -0.09642 | 0.494288 | 0.927372 | H3f3b |
| ENSMUSG( | 25.41841 | -0.51242 | 0.215501 | 0.865167 | H4c12 |
| ENSMUSG( | 15.41204 | 0.164506 | 0.753562 | 0.970671 | H4c14 |
| ENSMUSG( | 11.38697 | -0.41563 | 0.490189 | NA | H4c3 |
| ENSMUSG( | 46.42508 | -0.46686 | 0.120464 | 0.813384 | H4c8 |
| ENSMUSG( | 63.4161 | -0.71717 | 0.011265 | 0.432745 | H4c9 |
| ENSMUSG( | 16.92924 | -0.2224 | 0.656741 | 0.955567 | H60b |
| ENSMUSG( | 309.7262 | 0.261115 | 0.277698 | 0.891468 | H6pd |
| ENSMUSG( | 11.55443 | -0.63914 | 0.267316 | 0.886134 | Haao |
| ENSMUSG( | 5792.958 | -0.1128 | 0.276389 | 0.890889 | Habp4 |
| ENSMUSG( | 401.4293 | -0.02647 | 0.868801 | 0.988097 | Hacd1 |
| ENSMUSG( | 1797.933 | 0.161284 | 0.312016 | 0.900037 | Hacd2 |
| ENSMUSG( | 7639.782 | 0.040387 | 0.800786 | 0.97681 | Hacd3 |
| ENSMUSG( | 245.1409 | -0.10986 | 0.536706 | 0.937108 | Hacd4 |
| ENSMUSG( | 886.0151 | 0.093639 | 0.490234 | 0.926267 | Hace1 |
| ENSMUSG( | 43.15453 | -0.52081 | 0.092502 | 0.793518 | Hacl1 |
| ENSMUSG( | 1869.557 | -0.21989 | 0.114176 | 0.811037 | Hadh |
| ENSMUSG( | 4256.005 | -0.19959 | 0.078503 | 0.77321 | Hadha |
| ENSMUSG( | 3203.523 | -0.19288 | 0.131217 | 0.822884 | Hadhb |
| ENSMUSG( | 1474.19 | -0.21322 | 0.124656 | 0.81516 | Hagh |
| ENSMUSG( | 1758.26 | -0.02006 | 0.899956 | 0.990558 | Haghl |
| ENSMUSG( | 2231.505 | 0.052864 | 0.782713 | 0.976106 | Hap1 |
| ENSMUSG( | 1673.444 | 0.046176 | 0.822999 | 0.979198 | Hapln1 |
| ENSMUSG( | 980.0916 | 0.269661 | 0.197728 | 0.862161 | Hapln2 |
| ENSMUSG( | 102.9837 | 0.287132 | 0.226705 | 0.868507 | Hapln3 |
| ENSMUSG( | 3160.83 | 0.011094 | 0.942512 | 0.994774 | Hapln4 |
| ENSMUSG( | 1390.31 | -0.05536 | 0.671569 | 0.95694 | Hapstr1 |
| ENSMUSG( | 89.24231 | 0.076604 | 0.74377 | 0.96951 | Harbi1 |
| ENSMUSG( | 1723.129 | -0.08381 | 0.554268 | 0.939565 | Hars1 |
| ENSMUSG( | 725.7448 | 0.058634 | 0.681124 | 0.958395 | Hars2 |
| ENSMUSG( | 24.76079 | 2.27E-05 | 0.999955 | 0.999955 | Has1 |
| ENSMUSG( | 12.24383 | 0.81659 | 0.146236 | 0.835973 | Has2 |
| ENSMUSG( | 13.30261 | -0.03231 | 0.950504 | 0.994798 | Has2os |
| ENSMUSG( | 69.25003 | 0.18414 | 0.493173 | 0.927372 | Has3 |
| ENSMUSG( | 9.359672 | 0.261893 | 0.695803 | NA | Haspin |
| ENSMUSG( | 266.7344 | -0.03702 | 0.821121 | 0.979198 | Hat1 |
| ENSMUSG( | 231.3967 | -0.1843 | 0.257015 | 0.883082 | Haus1 |
| ENSMUSG( | 649.862 | -0.14967 | 0.338261 | 0.907972 | Haus2 |
| ENSMUSG( | 145.2648 | 0.342753 | 0.073435 | 0.763629 | Haus3 |
| ENSMUSG( | 66.88893 | -0.2767 | 0.315242 | 0.901306 | Haus4 |
| ENSMUSG( | 26.45184 | -0.19889 | 0.633297 | 0.953844 | Haus5 |
| ENSMUSG( | 282.0512 | 0.074671 | 0.637441 | 0.953903 | Haus6 |
| ENSMUSG( | 201.7186 | -0.02894 | 0.878512 | 0.989644 | Haus7 |
| ENSMUSG( | 199.9276 | -0.05177 | 0.758769 | 0.971858 | Haus8 |
| ENSMUSG( | 276.441 | 0.380002 | 0.052862 | 0.71525 | Havcr2 |
| ENSMUSG( | 683.086 | 0.009099 | 0.948447 | 0.994798 | Hax1 |
| ENSMUSG( | 7698.084 | -0.43441 | 0.317273 | 0.902322 | Hba-a1 |
| ENSMUSG( | 2973.887 | -0.6507 | 0.143739 | 0.831311 | Hba-a2 |
| ENSMUSG( | 14766.36 | -0.56505 | 0.159865 | 0.837724 | Hbb-b1 |
| ENSMUSG( | 2138.005 | -0.64584 | 0.143308 | 0.831311 | Hbb-bt |
| ENSMUSG( | 1146.971 | -0.0868 | 0.527462 | 0.934466 | Hbegf |
| ENSMUSG( | 788.2857 | 0.326933 | 0.012826 | 0.457834 | Hbp1 |
| ENSMUSG( | 1384.364 | -0.05174 | 0.629874 | 0.953251 | Hbs1l |
| ENSMUSG( | 14.98662 | -0.41951 | 0.414056 | 0.917257 | Hc |
| ENSMUSG( | 68.29843 | -0.16439 | 0.549749 | 0.938699 | Hcar1 |
| ENSMUSG( | 627.4059 | 0.08745 | 0.561404 | 0.940254 | Hccs |
| ENSMUSG( | 2904.812 | 0.049539 | 0.677046 | 0.957106 | Hcfc1 |
| ENSMUSG( | 2017.676 | -0.15342 | 0.263738 | 0.886134 | Hcfc1r1 |
| ENSMUSG( | 741.8288 | 0.10437 | 0.408541 | 0.917257 | Hcfc2 |
| ENSMUSG( | 146.8343 | 0.100229 | 0.616365 | 0.951413 | Hck |
| ENSMUSG( | 312.221 | -0.15747 | 0.370919 | 0.915491 | Hcls1 |
| ENSMUSG( | 1555.011 | 0.193244 | 0.289853 | 0.896726 | Hcn1 |
| ENSMUSG( | 6714.525 | 0.155385 | 0.249154 | 0.877904 | Hcn2 |
| ENSMUSG( | 586.0447 | -0.04618 | 0.72761 | 0.966887 | Hcn3 |
| ENSMUSG( | 205.3857 | 0.016941 | 0.921281 | 0.992644 | Hcn4 |
| ENSMUSG( | 107.8733 | 0.083887 | 0.696915 | 0.960867 | Hcrtr1 |
| ENSMUSG( | 50.69411 | 0.70056 | 0.050377 | 0.708648 | Hcrtr2 |
| ENSMUSG( | 19.16366 | -0.34016 | 0.443978 | 0.921734 | Hcst |
| ENSMUSG( | 1019.857 | -0.0317 | 0.77915 | 0.976106 | Hdac1 |
| ENSMUSG( | 405.5142 | 0.277949 | 0.132418 | 0.822912 | Hdac10 |
| ENSMUSG( | 4506.721 | -0.05687 | 0.646219 | 0.953988 | Hdac11 |
| ENSMUSG( | 1482.393 | 0.004499 | 0.974221 | 0.996416 | Hdac2 |
| ENSMUSG( | 2167.846 | -0.05216 | 0.677664 | 0.957106 | Hdac3 |
| ENSMUSG( | 2280.83 | 0.311676 | 0.007483 | 0.367166 | Hdac4 |
| ENSMUSG( | 8472.607 | 0.19214 | 0.082928 | 0.77681 | Hdac5 |
| ENSMUSG( | 1471.552 | -0.02834 | 0.845774 | 0.984272 | Hdac6 |
| ENSMUSG( | 1061.916 | 0.078844 | 0.582285 | 0.944266 | Hdac7 |
| ENSMUSG( | 631.1512 | -0.25541 | 0.078789 | 0.774307 | Hdac8 |
| ENSMUSG( | 639.7379 | -0.21599 | 0.094333 | 0.795735 | Hdac9 |
| ENSMUSG( | 87.71939 | -0.22752 | 0.373385 | 0.915491 | Hdc |
| ENSMUSG( | 386.9253 | -0.11719 | 0.436208 | 0.919879 | Hddc2 |
| ENSMUSG( | 508.3338 | -0.17006 | 0.214372 | 0.864421 | Hddc3 |
| ENSMUSG( | 6001.361 | -0.07652 | 0.547675 | 0.938063 | Hdgf |
| ENSMUSG( | 1863.134 | 0.109775 | 0.423779 | 0.917257 | Hdgfl2 |
| ENSMUSG( | 2253.756 | 0.0266 | 0.853053 | 0.985598 | Hdgfl3 |
| ENSMUSG( | 4685.271 | -0.19786 | 0.124364 | 0.814691 | Hdhd2 |
| ENSMUSG( | 74.12396 | -0.15495 | 0.598164 | 0.946676 | Hdhd3 |
| ENSMUSG( | 1070.074 | 0.094184 | 0.43665 | 0.919879 | Hdhd5 |
| ENSMUSG( | 4208.791 | 0.018903 | 0.851227 | 0.985541 | Hdlbp |
| ENSMUSG( | 238.636 | 0.240158 | 0.133952 | 0.822912 | Hdx |

| | | | | | |
|---|---|---|---|---|---|
| ENSMUSG( | H2-K1 | 1449.869 | -0.3068 | 0.01619 | 0.189046 |
| ENSMUSG( | H2-K2 | 14.45285 | -0.3203 | 0.510689 | NA |
| ENSMUSG( | H2-M3 | 85.83899 | -0.37026 | 0.101235 | 0.484996 |
| ENSMUSG( | H2-M5 | 60.57632 | 0.220904 | 0.41029 | 0.818258 |
| ENSMUSG( | H2-M6-ps | 24.72813 | 0.020857 | 0.959163 | 0.99396 |
| ENSMUSG( | H2-Oa | 37.43102 | -0.48941 | 0.126353 | 0.533885 |
| ENSMUSG( | H2-Ob | 32.75258 | -0.3126 | 0.336045 | 0.777 |
| ENSMUSG( | H2-Q1 | 26.72327 | -0.4943 | 0.199909 | 0.640239 |
| ENSMUSG( | H2-Q2 | 18.09748 | 0.532138 | 0.239974 | NA |
| ENSMUSG( | H2-Q4 | 209.2781 | -0.20773 | 0.181776 | 0.618944 |
| ENSMUSG( | H2-T10 | 22.41095 | -0.4847 | 0.226265 | 0.67315 |
| ENSMUSG( | H2-T22 | 711.8216 | -0.08898 | 0.454572 | 0.837383 |
| ENSMUSG( | H2-T23 | 578.5268 | -0.17366 | 0.278826 | 0.728705 |
| ENSMUSG( | H2-T24 | 95.92905 | -0.27328 | 0.24125 | 0.689026 |
| ENSMUSG( | H2-T-ps | 18.22183 | -0.6929 | 0.129792 | NA |
| ENSMUSG( | H3c15 | 115.2899 | 0.123205 | 0.607464 | 0.906876 |
| ENSMUSG( | H3c4 | 27.60241 | 0.462624 | 0.209129 | 0.651071 |
| ENSMUSG( | H3f3a | 3292.967 | 0.006838 | 0.944881 | 0.993612 |
| ENSMUSG( | H3f3b | 3947.352 | 0.009143 | 0.934803 | 0.993231 |
| ENSMUSG( | H4c12 | 29.07209 | -0.32893 | 0.385692 | 0.804413 |
| ENSMUSG( | H4c14 | 15.8271 | -0.06589 | 0.888429 | NA |
| ENSMUSG( | H4c3 | 15.92613 | 0.311302 | 0.530154 | NA |
| ENSMUSG( | H4c8 | 60.0763 | 0.050955 | 0.869944 | 0.977655 |
| ENSMUSG( | H4c9 | 89.45236 | 0.107163 | 0.641885 | 0.916865 |
| ENSMUSG( | H60b | 15.78587 | -0.80696 | 0.123376 | NA |
| ENSMUSG( | H6pd | 278.0529 | -0.34064 | 0.112932 | 0.506708 |
| ENSMUSG( | Haao | 13.32059 | -0.50111 | 0.386979 | NA |
| ENSMUSG( | Habp4 | 6860.362 | 0.107028 | 0.210043 | 0.652413 |
| ENSMUSG( | Hacd1 | 424.9409 | -0.12801 | 0.351212 | 0.783067 |
| ENSMUSG( | Hacd2 | 1765.663 | -0.16432 | 0.256768 | 0.707935 |
| ENSMUSG( | Hacd3 | 7467.262 | -0.31396 | 0.026101 | 0.247245 |
| ENSMUSG( | Hacd4 | 278.7641 | -0.0086 | 0.962041 | 0.994538 |
| ENSMUSG( | Hace1 | 958.505 | 0.051875 | 0.613142 | 0.908902 |
| ENSMUSG( | Hacl1 | 55.33233 | -0.01465 | 0.955903 | 0.993874 |
| ENSMUSG( | Hadh | 2233.115 | 0.036 | 0.755445 | 0.951395 |
| ENSMUSG( | Hadha | 5157.04 | 0.09286 | 0.335211 | 0.776511 |
| ENSMUSG( | Hadhb | 3875.214 | 0.095048 | 0.410478 | 0.818434 |
| ENSMUSG( | Hagh | 1706.588 | -0.04898 | 0.62191 | 0.911432 |
| ENSMUSG( | Haghl | 1953.97 | 0.019526 | 0.891878 | 0.983979 |
| ENSMUSG( | Hap1 | 2581.504 | 0.20299 | 0.219455 | 0.664984 |
| ENSMUSG( | Hapln1 | 1584.985 | -0.4097 | 0.029075 | 0.259547 |
| ENSMUSG( | Hapln2 | 828.9488 | -0.52095 | 7.27E-05 | 0.005238 |
| ENSMUSG( | Hapln3 | 93.61519 | -0.25136 | 0.277701 | 0.727559 |
| ENSMUSG( | Hapln4 | 3093.035 | -0.34402 | 0.008139 | 0.125721 |
| ENSMUSG( | Hapstr1 | 1539.33 | -0.02557 | 0.801419 | 0.961947 |
| ENSMUSG( | Harbi1 | 90.17768 | -0.17402 | 0.431857 | 0.82768 |
| ENSMUSG( | Hars1 | 1969.553 | 0.0365 | 0.748196 | 0.949604 |
| ENSMUSG( | Hars2 | 812.1345 | 0.109935 | 0.284995 | 0.733568 |
| ENSMUSG( | Has1 | 25.66937 | -0.18339 | 0.639599 | 0.916434 |
| ENSMUSG( | Has2 | 9.280827 | -0.15983 | 0.795191 | NA |
| ENSMUSG( | Has2os | 12.8339 | -0.43261 | 0.390914 | NA |
| ENSMUSG( | Has3 | 63.78528 | -0.34366 | 0.181524 | 0.618685 |
| ENSMUSG( | Hat1 | 303.7414 | 0.078193 | 0.54406 | 0.884411 |
| ENSMUSG( | Haus1 | 261.728 | -0.09949 | 0.472351 | 0.848048 |
| ENSMUSG( | Haus2 | 765.4884 | 0.06325 | 0.575621 | 0.894836 |
| ENSMUSG( | Haus3 | 147.3899 | 0.13074 | 0.460351 | 0.840676 |
| ENSMUSG( | Haus4 | 74.2019 | -0.2578 | 0.269646 | 0.719729 |
| ENSMUSG( | Haus5 | 32.61902 | 0.13325 | 0.705232 | 0.938303 |
| ENSMUSG( | Haus6 | 300.9711 | 0.000275 | 0.998385 | 0.99989 |
| ENSMUSG( | Haus7 | 218.7272 | -0.0486 | 0.760718 | 0.952907 |
| ENSMUSG( | Haus8 | 227.3231 | 0.049793 | 0.731798 | 0.944956 |
| ENSMUSG( | Havcr2 | 259.9468 | -0.05413 | 0.738076 | 0.947329 |
| ENSMUSG( | Hax1 | 762.9455 | 0.058637 | 0.619559 | 0.911432 |
| ENSMUSG( | Hba-a1 | 7900.489 | -0.66631 | 0.086902 | 0.453311 |
| ENSMUSG( | Hba-a2 | 3132.996 | -0.79968 | 0.045895 | 0.328083 |
| ENSMUSG( | Hbb-b1 | 15540.84 | -0.71281 | 0.048463 | 0.336526 |
| ENSMUSG( | Hbb-bt | 2266.671 | -0.77076 | 0.060173 | 0.379324 |
| ENSMUSG( | Hbegf | 1280.262 | -0.03895 | 0.74824 | 0.949604 |
| ENSMUSG( | Hbp1 | 785.4345 | 0.060309 | 0.572628 | 0.893584 |
| ENSMUSG( | Hbs1l | 1604.879 | 0.105055 | 0.231385 | 0.679437 |
| ENSMUSG( | Hc | 20.404 | 0.193252 | 0.643528 | NA |
| ENSMUSG( | Hcar1 | 70.04779 | -0.40592 | 0.084794 | 0.448764 |
| ENSMUSG( | Hccs | 660.061 | -0.02646 | 0.831779 | 0.97005 |
| ENSMUSG( | Hcfc1 | 3204.539 | 0.060904 | 0.570039 | 0.893041 |
| ENSMUSG( | Hcfc1r1 | 2342.415 | 0.016064 | 0.855889 | 0.973813 |
| ENSMUSG( | Hcfc2 | 808.1942 | 0.087685 | 0.40938 | 0.818258 |
| ENSMUSG( | Hck | 163.2729 | 0.125087 | 0.485002 | 0.857036 |
| ENSMUSG( | Hcls1 | 369.2335 | 0.066484 | 0.664695 | 0.926824 |
| ENSMUSG( | Hcn1 | 1401.3 | -0.40068 | 0.011223 | 0.152581 |
| ENSMUSG( | Hcn2 | 6998.174 | 0.008144 | 0.936522 | 0.993415 |
| ENSMUSG( | Hcn3 | 617.8162 | -0.17092 | 0.158644 | 0.588112 |
| ENSMUSG( | Hcn4 | 185.9864 | -0.59378 | 0.000772 | 0.027292 |
| ENSMUSG( | Hcrtr1 | 119.1067 | 0.087417 | 0.679339 | 0.930983 |
| ENSMUSG( | Hcrtr2 | 38.95208 | -0.27544 | 0.437521 | 0.829914 |
| ENSMUSG( | Hcst | 20.1206 | -0.49626 | 0.226253 | NA |
| ENSMUSG( | Hdac1 | 959.2832 | -0.52317 | 8.95E-08 | 2.42E-05 |
| ENSMUSG( | Hdac10 | 439.6091 | 0.237199 | 0.083126 | 0.444839 |
| ENSMUSG( | Hdac11 | 5231.343 | 0.103933 | 0.34137 | 0.778828 |
| ENSMUSG( | Hdac2 | 1674.047 | 0.088469 | 0.395262 | 0.81041 |
| ENSMUSG( | Hdac3 | 2470.617 | 0.056145 | 0.559005 | 0.888638 |
| ENSMUSG( | Hdac4 | 2627.868 | 0.433453 | 3.74E-07 | 8.21E-05 |
| ENSMUSG( | Hdac5 | 9348.703 | 0.205304 | 0.018331 | 0.203658 |
| ENSMUSG( | Hdac6 | 1756.84 | 0.209459 | 0.095285 | 0.47423 |
| ENSMUSG( | Hdac7 | 1151.566 | 0.04173 | 0.726716 | 0.944223 |
| ENSMUSG( | Hdac8 | 753.7592 | 0.006355 | 0.959043 | 0.993937 |
| ENSMUSG( | Hdac9 | 739.2606 | -0.05584 | 0.589421 | 0.90113 |
| ENSMUSG( | Hdc | 102.5058 | -0.06074 | 0.768289 | 0.954641 |
| ENSMUSG( | Hddc2 | 459.9048 | 0.120497 | 0.366513 | 0.794765 |
| ENSMUSG( | Hddc3 | 600.948 | 0.045721 | 0.682529 | 0.93198 |
| ENSMUSG( | Hdgf | 6864.685 | 0.044686 | 0.562772 | 0.889597 |
| ENSMUSG( | Hdgfl2 | 2092.665 | 0.175658 | 0.163866 | 0.594869 |
| ENSMUSG( | Hdgfl3 | 2494.569 | 0.05419 | 0.546862 | 0.884974 |
| ENSMUSG( | Hdhd2 | 5588.118 | 0.050794 | 0.621882 | 0.911432 |
| ENSMUSG( | Hdhd3 | 87.95372 | 0.086528 | 0.743198 | 0.948478 |
| ENSMUSG( | Hdhd5 | 1173.579 | 0.089287 | 0.404134 | 0.816132 |
| ENSMUSG( | Hdlbp | 4912.805 | 0.190214 | 0.019438 | 0.210037 |
| ENSMUSG( | Hdx | 236.7415 | -0.04066 | 0.772177 | 0.954994 |
| ENSMUSG( | Heatr1 | 685.9094 | 0.243323 | 0.036044 | 0.291881 |
| ENSMUSG( | Heatr3 | 742.2251 | -0.08089 | 0.44672 | 0.833905 |
| ENSMUSG( | Heatr5a | 776.9695 | 0.137313 | 0.20061 | 0.640661 |
| ENSMUSG( | Heatr5b | 1177.404 | 0.064218 | 0.560613 | 0.888934 |
| ENSMUSG( | Heatr6 | 706.4011 | 0.062687 | 0.559245 | 0.888638 |
| ENSMUSG( | Hebp1 | 509.4297 | -0.0745 | 0.571564 | 0.893319 |
| ENSMUSG( | Hebp2 | 105.8565 | -0.08911 | 0.649297 | 0.920273 |
| ENSMUSG( | Heca | 1045.029 | 0.141596 | 0.203638 | 0.643764 |
| ENSMUSG( | Hectd1 | 3150.699 | 0.055436 | 0.570197 | 0.893152 |
| ENSMUSG( | Hectd2 | 476.4647 | 0.068465 | 0.540241 | 0.88315 |
| ENSMUSG( | Hectd2os | 35.59754 | -0.25276 | 0.508277 | 0.868 |
| ENSMUSG( | Hectd3 | 1500.632 | 0.115876 | 0.239511 | 0.687308 |
| ENSMUSG( | Hectd4 | 9647.972 | 0.073586 | 0.396413 | 0.811742 |
| ENSMUSG( | Hecw1 | 1320.981 | -0.11725 | 0.399318 | 0.813048 |
| ENSMUSG( | Hecw2 | 993.068 | -0.43979 | 2.20E-05 | 0.002114 |
| ENSMUSG( | Heg1 | 1065.63 | 0.096158 | 0.581839 | 0.897862 |
| ENSMUSG( | Helb | 175.7495 | -0.08387 | 0.641442 | 0.916693 |
| ENSMUSG( | Hells | 85.05281 | -0.3559 | 0.115 | 0.51132 |
| ENSMUSG( | Helq | 347.057 | -0.04093 | 0.760807 | 0.952907 |
| ENSMUSG( | Helz | 1519.28 | 0.082856 | 0.436196 | 0.829093 |
| ENSMUSG( | Helz2 | 240.3546 | 0.117655 | 0.56663 | 0.891644 |
| ENSMUSG( | Hemk1 | 439.9587 | -0.21604 | 0.115134 | 0.511776 |
| ENSMUSG( | Henmt1 | 24.7933 | -0.26807 | 0.469188 | 0.845876 |
| ENSMUSG( | Hepacam | 11113.05 | -0.05585 | 0.628331 | 0.913637 |
| ENSMUSG( | Heph | 823.2518 | 0.300251 | 0.017508 | 0.198444 |
| ENSMUSG( | Herc1 | 4612.019 | -0.07349 | 0.513273 | 0.871762 |
| ENSMUSG( | Herc2 | 6884.119 | -0.14683 | 0.233538 | 0.681508 |
| ENSMUSG( | Herc3 | 2389.645 | 0.149889 | 0.167418 | 0.599984 |
| ENSMUSG( | Herc4 | 728.4796 | 0.159667 | 0.114191 | 0.509818 |
| ENSMUSG( | Herc6 | 241.821 | -0.19645 | 0.185713 | 0.623185 |
| ENSMUSG( | Herpud1 | 1517.088 | -0.27114 | 0.056375 | 0.366006 |
| ENSMUSG( | Herpud2 | 1454.506 | -0.05903 | 0.582974 | 0.898094 |
| ENSMUSG( | Hes1 | 310.8298 | -0.02874 | 0.837299 | 0.971111 |
| ENSMUSG( | Hes3 | 22.72177 | -0.7706 | 0.082712 | 0.444235 |
| ENSMUSG( | Hes5 | 601.8803 | 0.186728 | 0.187802 | 0.626399 |
| ENSMUSG( | Hes6 | 418.5671 | -0.04976 | 0.684583 | 0.93282 |
| ENSMUSG( | Hes7 | 24.64784 | -0.39421 | 0.302447 | 0.750794 |
| ENSMUSG( | Hexa | 2751.441 | -0.10265 | 0.385062 | 0.804048 |
| ENSMUSG( | Hexb | 8111.561 | -0.05351 | 0.74028 | 0.947823 |
| ENSMUSG( | Hexd | 908.0476 | 0.065678 | 0.660102 | 0.924627 |
| ENSMUSG( | Hexim1 | 1377.311 | 0.041129 | 0.72422 | 0.944096 |
| ENSMUSG( | Hexim2 | 177.885 | -0.05083 | 0.751809 | 0.950381 |
| ENSMUSG( | Hey1 | 1704.348 | 0.072641 | 0.477602 | 0.852702 |
| ENSMUSG( | Hey2 | 182.7584 | 0.230314 | 0.173792 | 0.60923 |
| ENSMUSG( | Heyl | 322.9145 | -0.35506 | 0.007568 | 0.120352 |
| ENSMUSG( | Hfe | 309.5383 | -0.02781 | 0.831391 | 0.969889 |
| ENSMUSG( | Hfm1 | 69.36003 | 0.048928 | 0.846778 | 0.972371 |
| ENSMUSG( | Hgf | 287.0815 | -0.23405 | 0.169331 | 0.603554 |
| ENSMUSG( | Hgh1 | 248.5157 | -0.26382 | 0.076951 | 0.429361 |
| ENSMUSG( | Hgs | 1996.06 | 0.192986 | 0.05613 | 0.36528 |
| ENSMUSG( | Hgsnat | 1565.066 | -0.30595 | 0.028406 | 0.257525 |

| | | | | | |
|---|---|---|---|---|---|
| ENSMUSG( | H2-K1 | 1546.488 | 0.163875 | 0.294444 | 0.898538 |
| ENSMUSG( | H2-K2 | 13.27984 | -0.27079 | 0.628593 | 0.953251 |
| ENSMUSG( | H2-M3 | 93.04153 | 0.151945 | 0.505877 | 0.927825 |
| ENSMUSG( | H2-M5 | 47.52509 | -0.20598 | 0.483897 | 0.925289 |
| ENSMUSG( | H2-M6-ps | 27.74784 | 0.588254 | 0.128577 | 0.81925 |
| ENSMUSG( | H2-Oa | 39.42997 | -0.01825 | 0.955183 | 0.994798 |
| ENSMUSG( | H2-Ob | 38.00246 | 0.40416 | 0.231281 | 0.868507 |
| ENSMUSG( | H2-Q1 | 27.06209 | -0.17367 | 0.667147 | 0.956766 |
| ENSMUSG( | H2-Q2 | 15.52785 | 0.387466 | 0.453387 | 0.922316 |
| ENSMUSG( | H2-Q4 | 216.3588 | 0.173121 | 0.382095 | 0.917257 |
| ENSMUSG( | H2-T10 | 23.09152 | -0.07177 | 0.864767 | 0.986591 |
| ENSMUSG( | H2-T22 | 639.4262 | -0.12652 | 0.337869 | 0.907972 |
| ENSMUSG( | H2-T23 | 571.7866 | 0.075167 | 0.683855 | 0.958678 |
| ENSMUSG( | H2-T24 | 94.78561 | 0.00344 | 0.990893 | 0.998302 |
| ENSMUSG( | H2-T-ps | 18.37699 | -0.35734 | 0.442459 | 0.921554 |
| ENSMUSG( | H3c15 | 90.18884 | -0.3386 | 0.19249 | 0.859726 |
| ENSMUSG( | H3c4 | 23.79235 | 0.318585 | 0.434925 | 0.918876 |
| ENSMUSG( | H3f3a | 2861.032 | -0.13687 | 0.299171 | 0.898538 |
| ENSMUSG( | H3f3b | 3473.585 | -0.09642 | 0.494288 | 0.927372 |
| ENSMUSG( | H4c12 | 25.41841 | -0.51242 | 0.215501 | 0.865167 |
| ENSMUSG( | H4c14 | 15.41204 | 0.164506 | 0.753562 | 0.970671 |
| ENSMUSG( | H4c3 | 11.38697 | -0.41563 | 0.490189 | NA |
| ENSMUSG( | H4c8 | 46.42508 | -0.46686 | 0.120464 | 0.813384 |
| ENSMUSG( | H4c9 | 63.4161 | -0.71717 | 0.011265 | 0.432745 |
| ENSMUSG( | H60b | 16.92924 | -0.2224 | 0.656741 | 0.955567 |
| ENSMUSG( | H6pd | 309.7262 | 0.261115 | 0.277698 | 0.891468 |
| ENSMUSG( | Haao | 11.55443 | -0.63914 | 0.267316 | 0.886134 |
| ENSMUSG( | Habp4 | 5792.958 | -0.1128 | 0.276389 | 0.890889 |
| ENSMUSG( | Hacd1 | 401.4293 | -0.02647 | 0.868801 | 0.988097 |
| ENSMUSG( | Hacd2 | 1797.933 | 0.161284 | 0.312016 | 0.900037 |
| ENSMUSG( | Hacd3 | 7639.782 | 0.040387 | 0.800786 | 0.97681 |
| ENSMUSG( | Hacd4 | 245.1409 | -0.10986 | 0.536706 | 0.937108 |
| ENSMUSG( | Hace1 | 886.0151 | 0.093639 | 0.490234 | 0.926267 |
| ENSMUSG( | Hacl1 | 43.15453 | -0.52081 | 0.092502 | 0.793518 |
| ENSMUSG( | Hadh | 1869.557 | -0.21989 | 0.114176 | 0.811037 |
| ENSMUSG( | Hadha | 4256.005 | -0.19959 | 0.078503 | 0.77321 |
| ENSMUSG( | Hadhb | 3203.523 | -0.19288 | 0.131217 | 0.822884 |
| ENSMUSG( | Hagh | 1474.19 | -0.21322 | 0.124656 | 0.81516 |
| ENSMUSG( | Haghl | 1758.26 | -0.02006 | 0.899956 | 0.990558 |
| ENSMUSG( | Hap1 | 2231.505 | 0.052864 | 0.782713 | 0.976106 |
| ENSMUSG( | Hapln1 | 1673.444 | 0.046176 | 0.822999 | 0.979198 |
| ENSMUSG( | Hapln2 | 980.0916 | 0.269661 | 0.197728 | 0.862161 |
| ENSMUSG( | Hapln3 | 102.9837 | 0.287132 | 0.226705 | 0.868507 |
| ENSMUSG( | Hapln4 | 3160.83 | 0.011094 | 0.942512 | 0.994774 |
| ENSMUSG( | Hapstr1 | 1390.31 | -0.05536 | 0.671569 | 0.95694 |
| ENSMUSG( | Harbi1 | 89.24231 | 0.076604 | 0.74377 | 0.96951 |
| ENSMUSG( | Hars1 | 1723.129 | -0.08381 | 0.554268 | 0.939565 |
| ENSMUSG( | Hars2 | 725.7448 | 0.058634 | 0.681124 | 0.958395 |
| ENSMUSG( | Has1 | 24.76079 | 2.27E-05 | 0.999955 | 0.999955 |
| ENSMUSG( | Has2 | 12.24383 | 0.81659 | 0.146236 | 0.835973 |
| ENSMUSG( | Has2os | 13.30261 | -0.03231 | 0.950504 | 0.994798 |
| ENSMUSG( | Has3 | 69.25003 | 0.18414 | 0.493173 | 0.927372 |
| ENSMUSG( | Hat1 | 266.7344 | -0.03702 | 0.821121 | 0.979198 |
| ENSMUSG( | Haus1 | 231.3967 | -0.1843 | 0.257015 | 0.883082 |
| ENSMUSG( | Haus2 | 649.862 | -0.14967 | 0.338261 | 0.907972 |
| ENSMUSG( | Haus3 | 145.2648 | 0.342753 | 0.073435 | 0.763629 |
| ENSMUSG( | Haus4 | 66.88893 | -0.2767 | 0.315242 | 0.901306 |
| ENSMUSG( | Haus5 | 26.45184 | -0.19889 | 0.633297 | 0.953844 |
| ENSMUSG( | Haus6 | 282.0512 | 0.074671 | 0.637441 | 0.953903 |
| ENSMUSG( | Haus7 | 201.7186 | -0.02894 | 0.878512 | 0.989644 |
| ENSMUSG( | Haus8 | 199.9276 | -0.05177 | 0.758769 | 0.971858 |
| ENSMUSG( | Havcr2 | 276.441 | 0.380002 | 0.052862 | 0.71525 |
| ENSMUSG( | Hax1 | 683.086 | 0.009099 | 0.948447 | 0.994798 |
| ENSMUSG( | Hba-a1 | 7698.084 | -0.43441 | 0.317273 | 0.902322 |
| ENSMUSG( | Hba-a2 | 2973.887 | -0.6507 | 0.143739 | 0.831311 |
| ENSMUSG( | Hbb-b1 | 14766.36 | -0.56505 | 0.159865 | 0.837724 |
| ENSMUSG( | Hbb-bt | 2138.005 | -0.64584 | 0.143308 | 0.831311 |
| ENSMUSG( | Hbegf | 1146.971 | -0.0868 | 0.527462 | 0.934466 |
| ENSMUSG( | Hbp1 | 788.2857 | 0.326933 | 0.012826 | 0.457834 |
| ENSMUSG( | Hbs1l | 1384.364 | -0.05174 | 0.629874 | 0.953251 |
| ENSMUSG( | Hc | 14.98662 | -0.41951 | 0.414056 | 0.917257 |
| ENSMUSG( | Hcar1 | 68.29843 | -0.16439 | 0.549749 | 0.938699 |
| ENSMUSG( | Hccs | 627.4059 | 0.08745 | 0.561404 | 0.940254 |
| ENSMUSG( | Hcfc1 | 2904.812 | 0.049539 | 0.677046 | 0.957106 |
| ENSMUSG( | Hcfc1r1 | 2017.676 | -0.15342 | 0.263738 | 0.886134 |
| ENSMUSG( | Hcfc2 | 741.8288 | 0.10437 | 0.408541 | 0.917257 |
| ENSMUSG( | Hck | 146.8343 | 0.100229 | 0.616365 | 0.951413 |
| ENSMUSG( | Hcls1 | 312.221 | -0.15747 | 0.370919 | 0.915491 |
| ENSMUSG( | Hcn1 | 1555.011 | 0.193244 | 0.289853 | 0.896726 |
| ENSMUSG( | Hcn2 | 6714.525 | 0.155385 | 0.249154 | 0.877904 |
| ENSMUSG( | Hcn3 | 586.0447 | -0.04618 | 0.72761 | 0.966887 |
| ENSMUSG( | Hcn4 | 205.3857 | 0.016941 | 0.921281 | 0.992644 |
| ENSMUSG( | Hcrtr1 | 107.8733 | 0.083887 | 0.696915 | 0.960867 |
| ENSMUSG( | Hcrtr2 | 50.69411 | 0.70056 | 0.050377 | 0.708648 |
| ENSMUSG( | Hcst | 19.16366 | -0.34016 | 0.443978 | 0.921734 |
| ENSMUSG( | Hdac1 | 1019.857 | -0.0317 | 0.77915 | 0.976106 |
| ENSMUSG( | Hdac10 | 405.5142 | 0.277949 | 0.132418 | 0.822912 |
| ENSMUSG( | Hdac11 | 4506.721 | -0.05687 | 0.646219 | 0.953988 |
| ENSMUSG( | Hdac2 | 1482.393 | 0.004499 | 0.974221 | 0.996416 |
| ENSMUSG( | Hdac3 | 2167.846 | -0.05216 | 0.677664 | 0.957106 |
| ENSMUSG( | Hdac4 | 2280.83 | 0.311676 | 0.007483 | 0.367166 |
| ENSMUSG( | Hdac5 | 8472.607 | 0.19214 | 0.082928 | 0.77681 |
| ENSMUSG( | Hdac6 | 1471.552 | -0.02834 | 0.845774 | 0.984272 |
| ENSMUSG( | Hdac7 | 1061.916 | 0.078844 | 0.582285 | 0.944266 |
| ENSMUSG( | Hdac8 | 631.1512 | -0.25541 | 0.078789 | 0.774307 |
| ENSMUSG( | Hdac9 | 639.7379 | -0.21599 | 0.094333 | 0.795735 |
| ENSMUSG( | Hdc | 87.71939 | -0.22752 | 0.373385 | 0.915491 |
| ENSMUSG( | Hddc2 | 386.9253 | -0.11719 | 0.436208 | 0.919879 |
| ENSMUSG( | Hddc3 | 508.3338 | -0.17006 | 0.214372 | 0.864421 |
| ENSMUSG( | Hdgf | 6001.361 | -0.07652 | 0.547675 | 0.938063 |
| ENSMUSG( | Hdgfl2 | 1863.134 | 0.109775 | 0.423779 | 0.917257 |
| ENSMUSG( | Hdgfl3 | 2253.756 | 0.0266 | 0.853053 | 0.985598 |
| ENSMUSG( | Hdhd2 | 4685.271 | -0.19786 | 0.124364 | 0.814691 |
| ENSMUSG( | Hdhd3 | 74.12396 | -0.15495 | 0.598164 | 0.946676 |
| ENSMUSG( | Hdhd5 | 1070.074 | 0.094184 | 0.43665 | 0.919879 |
| ENSMUSG( | Hdlbp | 4208.791 | 0.018903 | 0.851227 | 0.985541 |
| ENSMUSG( | Hdx | 238.636 | 0.240158 | 0.133952 | 0.822912 |
| ENSMUSG( | Heatr1 | 598.4957 | 0.132686 | 0.350079 | 0.911652 |
| ENSMUSG( | Heatr3 | 717.7477 | 0.094527 | 0.456826 | 0.923329 |
| ENSMUSG( | Heatr5a | 695.5292 | 0.089151 | 0.472908 | 0.923329 |
| ENSMUSG( | Heatr5b | 1061.532 | 0.03774 | 0.764432 | 0.974 |
| ENSMUSG( | Heatr6 | 631.9386 | 0.011769 | 0.92971 | 0.99306 |
| ENSMUSG( | Hebp1 | 449.8295 | -0.1801 | 0.299666 | 0.898538 |
| ENSMUSG( | Hebp2 | 83.53516 | -0.57149 | 0.018858 | 0.548832 |
| ENSMUSG( | Heca | 971.2244 | 0.189833 | 0.238023 | 0.87259 |
| ENSMUSG( | Hectd1 | 2914.987 | 0.100514 | 0.35407 | 0.912411 |
| ENSMUSG( | Hectd2 | 435.0832 | 0.067638 | 0.644593 | 0.953903 |
| ENSMUSG( | Hectd2os | 32.49963 | -0.2777 | 0.458514 | 0.923329 |
| ENSMUSG( | Hectd3 | 1332.97 | 0.044917 | 0.720473 | 0.966887 |
| ENSMUSG( | Hectd4 | 8892.918 | 0.107391 | 0.33356 | 0.907907 |
| ENSMUSG( | Hecw1 | 1359.318 | 0.230227 | 0.152642 | 0.837724 |
| ENSMUSG( | Hecw2 | 929.9001 | -0.35056 | 0.005056 | 0.292317 |
| ENSMUSG( | Heg1 | 1141.634 | 0.525609 | 0.009533 | 0.406458 |
| ENSMUSG( | Helb | 174.3502 | 0.168068 | 0.422065 | 0.917257 |
| ENSMUSG( | Hells | 88.54843 | 0.052096 | 0.821536 | 0.979198 |
| ENSMUSG( | Helq | 316.0933 | -0.03659 | 0.826257 | 0.979795 |
| ENSMUSG( | Helz | 1388.594 | 0.091725 | 0.420293 | 0.917257 |
| ENSMUSG( | Helz2 | 237.1678 | 0.344146 | 0.10476 | 0.804969 |
| ENSMUSG( | Hemk1 | 410.9457 | -0.13771 | 0.409252 | 0.917257 |
| ENSMUSG( | Henmt1 | 25.7311 | 0.1363 | 0.725563 | 0.966887 |
| ENSMUSG( | Hepacam | 10384.35 | 0.019186 | 0.882808 | 0.989644 |
| ENSMUSG( | Heph | 738.157 | 0.25596 | 0.092352 | 0.793008 |
| ENSMUSG( | Herc1 | 4480.948 | 0.113743 | 0.393811 | 0.917257 |
| ENSMUSG( | Herc2 | 6956.522 | 0.155923 | 0.295928 | 0.898538 |
| ENSMUSG( | Herc3 | 2139.963 | 0.102362 | 0.375144 | 0.915491 |
| ENSMUSG( | Herc4 | 674.2182 | 0.205156 | 0.093522 | 0.795507 |
| ENSMUSG( | Herc6 | 234.8503 | 0.000245 | 0.998827 | 0.999503 |
| ENSMUSG( | Herpud1 | 1787.394 | 0.453377 | 0.004679 | 0.282119 |
| ENSMUSG( | Herpud2 | 1390.482 | 0.079732 | 0.509885 | 0.929089 |
| ENSMUSG( | Hes1 | 272.34 | -0.13299 | 0.459254 | 0.923329 |
| ENSMUSG( | Hes3 | 19.08003 | -1.10912 | 0.027743 | 0.61023 |
| ENSMUSG( | Hes5 | 584.7955 | 0.351596 | 0.028686 | 0.614171 |
| ENSMUSG( | Hes6 | 389.1262 | 0.00357 | 0.980398 | 0.996479 |
| ENSMUSG( | Hes7 | 22.38522 | -0.42025 | 0.307045 | 0.898538 |
| ENSMUSG( | Hexa | 2601.78 | 0.009567 | 0.945763 | 0.994798 |
| ENSMUSG( | Hexb | 8217.034 | 0.246483 | 0.177104 | 0.846973 |
| ENSMUSG( | Hexd | 824.0274 | 0.053823 | 0.77163 | 0.974236 |
| ENSMUSG( | Hexim1 | 1237.726 | -0.00257 | 0.98661 | 0.997372 |
| ENSMUSG( | Hexim2 | 160.3841 | -0.09117 | 0.626218 | 0.953234 |
| ENSMUSG( | Hey1 | 1559.477 | 0.080966 | 0.530278 | 0.934795 |
| ENSMUSG( | Hey2 | 189.058 | 0.542306 | 0.009838 | 0.406458 |
| ENSMUSG( | Heyl | 327.1752 | -0.03326 | 0.821265 | 0.979198 |
| ENSMUSG( | Hfe | 322.6831 | 0.350686 | 0.034573 | 0.655419 |
| ENSMUSG( | Hfm1 | 64.54403 | 0.093903 | 0.7326 | 0.967228 |
| ENSMUSG( | Hgf | 275.9744 | -0.05942 | 0.757611 | 0.971858 |
| ENSMUSG( | Hgh1 | 240.4571 | -0.07742 | 0.635669 | 0.953903 |
| ENSMUSG( | Hgs | 1654.78 | -0.07458 | 0.555469 | 0.939565 |
| ENSMUSG( | Hgsnat | 1707.999 | 0.226759 | 0.175606 | 0.845305 |

| | | | | | |
|---|---|---|---|---|---|
| ENSMUSG( | 259.9468 | -0.05413 | 0.738076 | 0.947329 | Havcr2 |
| ENSMUSG( | 762.9455 | 0.058637 | 0.619559 | 0.911432 | Hax1 |
| ENSMUSG( | 7900.489 | -0.66631 | 0.086902 | 0.453311 | Hba-a1 |
| ENSMUSG( | 3132.996 | -0.79968 | 0.045895 | 0.328083 | Hba-a2 |
| ENSMUSG( | 15540.84 | -0.71281 | 0.048463 | 0.336526 | Hbb-b1 |
| ENSMUSG( | 2266.671 | -0.77076 | 0.060173 | 0.379324 | Hbb-bt |
| ENSMUSG( | 1280.262 | -0.03895 | 0.74824 | 0.949604 | Hbegf |
| ENSMUSG( | 785.4345 | 0.060309 | 0.572628 | 0.893584 | Hbp1 |
| ENSMUSG( | 12.33055 | 0.408867 | 0.487956 | NA | Hbq1b |
| ENSMUSG( | 1604.879 | 0.105055 | 0.231385 | 0.679437 | Hbs1l |
| ENSMUSG( | 20.404 | 0.193252 | 0.643528 | NA | Hc |
| ENSMUSG( | 70.04779 | -0.40592 | 0.084794 | 0.448764 | Hcar1 |
| ENSMUSG( | 660.061 | -0.02646 | 0.831779 | 0.97005 | Hccs |
| ENSMUSG( | 3204.539 | 0.060904 | 0.570039 | 0.893041 | Hcfc1 |
| ENSMUSG( | 2342.415 | 0.016064 | 0.855889 | 0.973813 | Hcfc1r1 |
| ENSMUSG( | 808.1942 | 0.087685 | 0.40938 | 0.818258 | Hcfc2 |
| ENSMUSG( | 163.2729 | 0.125087 | 0.485002 | 0.857036 | Hck |
| ENSMUSG( | 369.2335 | 0.066484 | 0.664695 | 0.926824 | Hcls1 |
| ENSMUSG( | 1401.3 | -0.40068 | 0.011223 | 0.152581 | Hcn1 |
| ENSMUSG( | 6998.174 | 0.008144 | 0.936522 | 0.993415 | Hcn2 |
| ENSMUSG( | 617.8162 | -0.17092 | 0.158644 | 0.588112 | Hcn3 |
| ENSMUSG( | 185.9864 | -0.59378 | 0.000772 | 0.027292 | Hcn4 |
| ENSMUSG( | 119.1067 | 0.087417 | 0.679339 | 0.930983 | Hcrtr1 |
| ENSMUSG( | 38.95208 | -0.27544 | 0.437521 | 0.829914 | Hcrtr2 |
| ENSMUSG( | 20.1206 | -0.49626 | 0.226253 | NA | Hcst |
| ENSMUSG( | 959.2832 | -0.52317 | 8.95E-08 | 2.42E-05 | Hdac1 |
| ENSMUSG( | 439.6091 | 0.237199 | 0.083126 | 0.444839 | Hdac10 |
| ENSMUSG( | 5231.343 | 0.103933 | 0.34137 | 0.778828 | Hdac11 |
| ENSMUSG( | 1674.047 | 0.088469 | 0.395262 | 0.81041 | Hdac2 |
| ENSMUSG( | 2470.617 | 0.056145 | 0.559005 | 0.888638 | Hdac3 |
| ENSMUSG( | 2627.868 | 0.433453 | 3.74E-07 | 8.21E-05 | Hdac4 |
| ENSMUSG( | 9348.703 | 0.205304 | 0.018331 | 0.203658 | Hdac5 |
| ENSMUSG( | 1756.84 | 0.209459 | 0.095285 | 0.47423 | Hdac6 |
| ENSMUSG( | 1151.566 | 0.04173 | 0.726716 | 0.944223 | Hdac7 |
| ENSMUSG( | 753.7592 | 0.006355 | 0.959043 | 0.993937 | Hdac8 |
| ENSMUSG( | 739.2606 | -0.05584 | 0.589421 | 0.90113 | Hdac9 |
| ENSMUSG( | 102.5058 | -0.06074 | 0.768289 | 0.954641 | Hdc |
| ENSMUSG( | 459.9048 | 0.120497 | 0.366513 | 0.794765 | Hddc2 |
| ENSMUSG( | 600.948 | 0.045721 | 0.682529 | 0.93198 | Hddc3 |
| ENSMUSG( | 6864.685 | 0.044686 | 0.562772 | 0.889597 | Hdgf |
| ENSMUSG( | 2092.665 | 0.175658 | 0.163866 | 0.594869 | Hdgfl2 |
| ENSMUSG( | 2494.569 | 0.05419 | 0.546862 | 0.884974 | Hdgfl3 |
| ENSMUSG( | 5588.118 | 0.050794 | 0.621882 | 0.911432 | Hdhd2 |
| ENSMUSG( | 87.95372 | 0.086528 | 0.743198 | 0.948478 | Hdhd3 |
| ENSMUSG( | 1173.579 | 0.089287 | 0.404134 | 0.816132 | Hdhd5 |
| ENSMUSG( | 4912.805 | 0.190214 | 0.019438 | 0.210037 | Hdlbp |
| ENSMUSG( | 236.7415 | -0.04066 | 0.772177 | 0.954994 | Hdx |
| ENSMUSG( | 685.9094 | 0.243323 | 0.036044 | 0.291881 | Heatr1 |
| ENSMUSG( | 742.2251 | -0.08089 | 0.44672 | 0.833905 | Heatr3 |
| ENSMUSG( | 776.9695 | 0.137313 | 0.20061 | 0.640661 | Heatr5a |
| ENSMUSG( | 1177.404 | 0.064218 | 0.560613 | 0.888934 | Heatr5b |
| ENSMUSG( | 706.4011 | 0.062687 | 0.559245 | 0.888638 | Heatr6 |
| ENSMUSG( | 509.4297 | -0.0745 | 0.571564 | 0.893319 | Hebp1 |
| ENSMUSG( | 105.8565 | -0.08911 | 0.649297 | 0.920273 | Hebp2 |
| ENSMUSG( | 1045.029 | 0.141596 | 0.203638 | 0.643764 | Heca |
| ENSMUSG( | 3150.699 | 0.055436 | 0.570197 | 0.893152 | Hectd1 |
| ENSMUSG( | 476.4647 | 0.068465 | 0.540241 | 0.88315 | Hectd2 |
| ENSMUSG( | 35.59754 | -0.25276 | 0.508277 | 0.868 | Hectd2os |
| ENSMUSG( | 1500.632 | 0.115876 | 0.239511 | 0.687308 | Hectd3 |
| ENSMUSG( | 9647.972 | 0.073586 | 0.396413 | 0.811742 | Hectd4 |
| ENSMUSG( | 1320.981 | -0.11725 | 0.399318 | 0.813048 | Hecw1 |
| ENSMUSG( | 993.068 | -0.43979 | 2.20E-05 | 0.002114 | Hecw2 |
| ENSMUSG( | 1065.63 | 0.096158 | 0.581839 | 0.897862 | Heg1 |
| ENSMUSG( | 175.7495 | -0.08387 | 0.641442 | 0.916693 | Helb |
| ENSMUSG( | 85.05281 | -0.3559 | 0.115 | 0.51132 | Hells |
| ENSMUSG( | 347.057 | -0.04093 | 0.760807 | 0.952907 | Helq |
| ENSMUSG( | 1519.28 | 0.082856 | 0.436196 | 0.829093 | Helz |
| ENSMUSG( | 240.3546 | 0.117655 | 0.56663 | 0.891644 | Helz2 |
| ENSMUSG( | 439.9587 | -0.21604 | 0.115134 | 0.511776 | Hemk1 |
| ENSMUSG( | 24.7933 | -0.26807 | 0.469188 | 0.845876 | Henmt1 |
| ENSMUSG( | 11113.05 | -0.05585 | 0.628331 | 0.913637 | Hepacam |
| ENSMUSG( | 823.2518 | 0.300251 | 0.017508 | 0.198444 | Heph |
| ENSMUSG( | 4612.019 | -0.07349 | 0.513273 | 0.871762 | Herc1 |
| ENSMUSG( | 6884.119 | -0.14683 | 0.233538 | 0.681508 | Herc2 |
| ENSMUSG( | 2389.645 | 0.149889 | 0.167418 | 0.599984 | Herc3 |
| ENSMUSG( | 728.4796 | 0.159667 | 0.114191 | 0.509818 | Herc4 |
| ENSMUSG( | 241.821 | -0.19645 | 0.185713 | 0.623185 | Herc6 |
| ENSMUSG( | 1517.088 | -0.27114 | 0.056375 | 0.366006 | Herpud1 |
| ENSMUSG( | 1454.506 | -0.05903 | 0.582974 | 0.898094 | Herpud2 |
| ENSMUSG( | 310.8298 | -0.02874 | 0.837299 | 0.971111 | Hes1 |
| ENSMUSG( | 22.72177 | -0.7706 | 0.082712 | 0.444235 | Hes3 |
| ENSMUSG( | 601.8803 | 0.186728 | 0.187802 | 0.626399 | Hes5 |
| ENSMUSG( | 418.5671 | -0.04976 | 0.684583 | 0.93282 | Hes6 |
| ENSMUSG( | 24.64784 | -0.39421 | 0.302447 | 0.750794 | Hes7 |
| ENSMUSG( | 2751.441 | -0.10265 | 0.385062 | 0.804048 | Hexa |
| ENSMUSG( | 8111.561 | -0.05351 | 0.74028 | 0.947823 | Hexb |
| ENSMUSG( | 908.0476 | 0.065678 | 0.660102 | 0.924627 | Hexd |
| ENSMUSG( | 1377.311 | 0.041129 | 0.72422 | 0.944096 | Hexim1 |
| ENSMUSG( | 177.885 | -0.05083 | 0.751809 | 0.950381 | Hexim2 |
| ENSMUSG( | 1704.348 | 0.072641 | 0.477602 | 0.852702 | Hey1 |
| ENSMUSG( | 182.7584 | 0.230314 | 0.173792 | 0.60923 | Hey2 |
| ENSMUSG( | 322.9145 | -0.35506 | 0.007568 | 0.120352 | Heyl |
| ENSMUSG( | 309.5383 | -0.02781 | 0.831391 | 0.969889 | Hfe |
| ENSMUSG( | 69.36003 | 0.048928 | 0.846778 | 0.972371 | Hfm1 |
| ENSMUSG( | 287.0815 | -0.23405 | 0.169331 | 0.603554 | Hgf |
| ENSMUSG( | 248.5157 | -0.26382 | 0.076951 | 0.429361 | Hgh1 |
| ENSMUSG( | 1996.06 | 0.192986 | 0.05613 | 0.36528 | Hgs |
| ENSMUSG( | 1565.066 | -0.30595 | 0.028406 | 0.257525 | Hgsnat |
| ENSMUSG( | 31.21757 | -0.36455 | 0.28307 | 0.73186 | Hhat |
| ENSMUSG( | 598.2261 | -0.39555 | 0.002433 | 0.058029 | Hhatl |
| ENSMUSG( | 76.2502 | 0.296006 | 0.242233 | 0.689699 | Hhex |
| ENSMUSG( | 364.1867 | -0.51302 | 0.002135 | 0.053442 | Hhip |
| ENSMUSG( | 506.9318 | -0.13673 | 0.32656 | 0.771368 | Hhipl1 |
| ENSMUSG( | 17.70427 | 0.741923 | 0.140791 | NA | Hhipl2 |
| ENSMUSG( | 3193.826 | 0.104596 | 0.416011 | 0.82084 | Hibadh |
| ENSMUSG( | 640.8107 | 0.10628 | 0.374152 | 0.800445 | Hibch |
| ENSMUSG( | 91.78721 | -0.27746 | 0.207149 | 0.649517 | Hic1 |
| ENSMUSG( | 173.7807 | 0.047937 | 0.781731 | 0.957558 | Hic2 |
| ENSMUSG( | 2538.366 | -0.02202 | 0.820164 | 0.965888 | Hid1 |
| ENSMUSG( | 1508.248 | 0.022622 | 0.80262 | 0.962169 | Hif1a |
| ENSMUSG( | 1753.721 | 0.031144 | 0.726259 | 0.944223 | Hif1an |
| ENSMUSG( | 335.3507 | 1.152566 | 1.53E-07 | 3.77E-05 | Hif3a |
| ENSMUSG( | 2065.787 | 0.010946 | 0.911652 | 0.988483 | Higd1a |
| ENSMUSG( | 74.72946 | -0.13303 | 0.59774 | 0.90372 | Higd1b |
| ENSMUSG( | 2326.102 | 0.041192 | 0.677752 | 0.930542 | Higd2a |
| ENSMUSG( | 587.4736 | -0.07866 | 0.542768 | 0.883563 | Hikeshi |
| ENSMUSG( | 110.7341 | -0.45733 | 0.025399 | 0.246101 | Hilpda |
| ENSMUSG( | 528.2595 | -0.01218 | 0.915938 | 0.988634 | Hinfp |
| ENSMUSG( | 3521.041 | 0.025993 | 0.789144 | 0.959299 | Hint1 |
| ENSMUSG( | 485.4769 | -0.05438 | 0.634768 | 0.914073 | Hint2 |
| ENSMUSG( | 394.5164 | 0.024527 | 0.86837 | 0.976834 | Hint3 |
| ENSMUSG( | 4051.322 | 0.256538 | 0.002668 | 0.062087 | Hip1 |
| ENSMUSG( | 4618.349 | 0.107699 | 0.224134 | 0.669929 | Hip1r |
| ENSMUSG( | 5344.258 | 0.113231 | 0.20859 | 0.650496 | Hipk1 |
| ENSMUSG( | 10848.7 | 0.031547 | 0.791547 | 0.959461 | Hipk2 |
| ENSMUSG( | 1797.632 | -0.00814 | 0.937886 | 0.993415 | Hipk3 |
| ENSMUSG( | 40.30891 | -0.49067 | 0.164368 | 0.59535 | Hipk4 |
| ENSMUSG( | 795.471 | 0.008236 | 0.941018 | 0.993415 | Hira |
| ENSMUSG( | 488.395 | -0.01088 | 0.937955 | 0.993415 | Hirip3 |
| ENSMUSG( | 4399.87 | 0.080333 | 0.443312 | 0.833108 | Hivep1 |
| ENSMUSG( | 11029.44 | 0.095432 | 0.28268 | 0.73186 | Hivep2 |
| ENSMUSG( | 3200.412 | 0.044263 | 0.654415 | 0.923227 | Hivep3 |
| ENSMUSG( | 23.07374 | -0.86053 | 0.027026 | 0.252658 | Hjv |
| ENSMUSG( | 13059.97 | 0.054578 | 0.506324 | 0.867425 | Hk1 |
| ENSMUSG( | 29.98049 | 0.09996 | 0.770567 | 0.954722 | Hk1os |
| ENSMUSG( | 240.9206 | 0.215369 | 0.162475 | 0.593257 | Hk2 |
| ENSMUSG( | 120.3562 | -0.0369 | 0.852583 | 0.973168 | Hk3 |
| ENSMUSG( | 504.8202 | 0.149719 | 0.389154 | 0.806967 | Hkdc1 |
| ENSMUSG( | 565.4348 | -0.22322 | 0.066497 | 0.39868 | Hlcs |
| ENSMUSG( | 8096.431 | 0.087562 | 0.352919 | 0.784977 | Hlf |
| ENSMUSG( | 590.8561 | 0.077134 | 0.483354 | 0.856304 | Hltf |
| ENSMUSG( | 36.99105 | -0.34545 | 0.288351 | 0.739103 | Hlx |
| ENSMUSG( | 3153.221 | 0.098415 | 0.291853 | 0.741868 | Hmbox1 |
| ENSMUSG( | 471.4661 | 0.005157 | 0.968007 | 0.996284 | Hmbs |
| ENSMUSG( | 354.4796 | 0.021322 | 0.880609 | 0.981542 | Hmces |
| ENSMUSG( | 247.8752 | -0.02477 | 0.890276 | 0.983687 | Hmcn1 |
| ENSMUSG( | 39.40864 | -0.09533 | 0.755544 | 0.951395 | Hmcn2 |
| ENSMUSG( | 1704.523 | 0.107218 | 0.322114 | 0.767096 | Hmg20a |
| ENSMUSG( | 812.9324 | 0.074081 | 0.521947 | 0.875186 | Hmg20b |
| ENSMUSG( | 1412.621 | 0.029875 | 0.752117 | 0.950417 | Hmga1 |
| ENSMUSG( | 8.170938 | 0.389703 | 0.542243 | NA | Hmga1b |
| ENSMUSG( | 50.11299 | -0.04752 | 0.877392 | 0.981333 | Hmga2-ps1 |
| ENSMUSG( | 3965.77 | 0.068061 | 0.465831 | 0.844222 | Hmgb1 |
| ENSMUSG( | 304.9723 | 0.006388 | 0.96399 | 0.995094 | Hmgb2 |
| ENSMUSG( | 673.2076 | -0.16372 | 0.178118 | 0.615613 | Hmgb3 |
| ENSMUSG( | 545.1285 | 0.101623 | 0.402234 | 0.815103 | Hmgcl |
| ENSMUSG( | 364.1936 | 0.112618 | 0.38173 | 0.802879 | Hmgcll1 |
| ENSMUSG( | 5406.023 | -0.3536 | 0.030332 | 0.264512 | Hmgcr |

| | | | | | |
|---|---|---|---|---|---|
| ENSMUSG( | 598.4957 | 0.132686 | 0.350079 | 0.911652 | Heatr1 |
| ENSMUSG( | 717.7477 | 0.094527 | 0.456826 | 0.923329 | Heatr3 |
| ENSMUSG( | 695.5292 | 0.089151 | 0.472908 | 0.923329 | Heatr5a |
| ENSMUSG( | 1061.532 | 0.03774 | 0.764432 | 0.974 | Heatr5b |
| ENSMUSG( | 631.9386 | 0.011769 | 0.92971 | 0.99306 | Heatr6 |
| ENSMUSG( | 449.8295 | -0.1801 | 0.299666 | 0.898538 | Hebp1 |
| ENSMUSG( | 83.53516 | -0.57149 | 0.018858 | 0.548832 | Hebp2 |
| ENSMUSG( | 971.2244 | 0.189833 | 0.238023 | 0.87259 | Heca |
| ENSMUSG( | 2914.987 | 0.100514 | 0.35407 | 0.912411 | Hectd1 |
| ENSMUSG( | 435.0832 | 0.067638 | 0.644593 | 0.953903 | Hectd2 |
| ENSMUSG( | 32.49963 | -0.2777 | 0.458514 | 0.923329 | Hectd2os |
| ENSMUSG( | 1332.97 | 0.044917 | 0.720473 | 0.966887 | Hectd3 |
| ENSMUSG( | 8892.918 | 0.107391 | 0.33356 | 0.907907 | Hectd4 |
| ENSMUSG( | 1359.318 | 0.230227 | 0.152642 | 0.837724 | Hecw1 |
| ENSMUSG( | 929.9001 | -0.35056 | 0.005056 | 0.292317 | Hecw2 |
| ENSMUSG( | 1141.634 | 0.525609 | 0.009533 | 0.406458 | Heg1 |
| ENSMUSG( | 174.3502 | 0.168068 | 0.422065 | 0.917257 | Helb |
| ENSMUSG( | 88.54843 | 0.052096 | 0.821536 | 0.979198 | Hells |
| ENSMUSG( | 316.0933 | -0.03659 | 0.826257 | 0.979795 | Helq |
| ENSMUSG( | 1388.594 | 0.091725 | 0.420293 | 0.917257 | Helz |
| ENSMUSG( | 237.1678 | 0.344146 | 0.10476 | 0.804969 | Helz2 |
| ENSMUSG( | 410.9457 | -0.13771 | 0.409252 | 0.917257 | Hemk1 |
| ENSMUSG( | 25.7311 | 0.1363 | 0.725563 | 0.966887 | Henmt1 |
| ENSMUSG( | 10384.35 | 0.019186 | 0.882808 | 0.989644 | Hepacam |
| ENSMUSG( | 738.157 | 0.25596 | 0.092352 | 0.793008 | Heph |
| ENSMUSG( | 4480.948 | 0.113743 | 0.393811 | 0.917257 | Herc1 |
| ENSMUSG( | 6956.522 | 0.155923 | 0.295928 | 0.898538 | Herc2 |
| ENSMUSG( | 2139.963 | 0.102362 | 0.375144 | 0.915491 | Herc3 |
| ENSMUSG( | 674.2182 | 0.205156 | 0.093522 | 0.795507 | Herc4 |
| ENSMUSG( | 234.8503 | 0.000245 | 0.998827 | 0.999503 | Herc6 |
| ENSMUSG( | 1787.394 | 0.453377 | 0.004679 | 0.282119 | Herpud1 |
| ENSMUSG( | 1390.482 | 0.079732 | 0.509885 | 0.929089 | Herpud2 |
| ENSMUSG( | 272.34 | -0.13299 | 0.459254 | 0.923329 | Hes1 |
| ENSMUSG( | 19.08003 | -1.10912 | 0.027743 | 0.61023 | Hes3 |
| ENSMUSG( | 584.7955 | 0.351596 | 0.028686 | 0.614171 | Hes5 |
| ENSMUSG( | 389.1262 | 0.00357 | 0.980398 | 0.996479 | Hes6 |
| ENSMUSG( | 22.38522 | -0.42025 | 0.307045 | 0.898538 | Hes7 |
| ENSMUSG( | 2601.78 | 0.009567 | 0.945763 | 0.994798 | Hexa |
| ENSMUSG( | 8217.034 | 0.246483 | 0.177104 | 0.846973 | Hexb |
| ENSMUSG( | 824.0274 | 0.053823 | 0.77163 | 0.974236 | Hexd |
| ENSMUSG( | 1237.726 | -0.00257 | 0.98661 | 0.997372 | Hexim1 |
| ENSMUSG( | 160.3841 | -0.09117 | 0.626218 | 0.953234 | Hexim2 |
| ENSMUSG( | 1559.477 | 0.080966 | 0.530278 | 0.934795 | Hey1 |
| ENSMUSG( | 189.058 | 0.542306 | 0.009838 | 0.406458 | Hey2 |
| ENSMUSG( | 327.1752 | -0.03326 | 0.821265 | 0.979198 | Heyl |
| ENSMUSG( | 322.6831 | 0.350686 | 0.034573 | 0.655419 | Hfe |
| ENSMUSG( | 64.54403 | 0.093903 | 0.7326 | 0.967228 | Hfm1 |
| ENSMUSG( | 275.9744 | -0.05942 | 0.757611 | 0.971858 | Hgf |
| ENSMUSG( | 240.4571 | -0.07742 | 0.635669 | 0.953903 | Hgh1 |
| ENSMUSG( | 1654.78 | -0.07458 | 0.555469 | 0.939565 | Hgs |
| ENSMUSG( | 1707.999 | 0.226759 | 0.175606 | 0.845305 | Hgsnat |
| ENSMUSG( | 29.66632 | -0.19395 | 0.604217 | 0.948139 | Hhat |
| ENSMUSG( | 618.628 | 0.000433 | 0.997782 | 0.999268 | Hhatl |
| ENSMUSG( | 79.51692 | 0.646835 | 0.012448 | 0.451463 | Hhex |
| ENSMUSG( | 412.3537 | 0.162913 | 0.403472 | 0.917257 | Hhip |
| ENSMUSG( | 503.2296 | 0.117082 | 0.448499 | 0.922202 | Hhipl1 |
| ENSMUSG( | 16.33677 | 0.76014 | 0.158541 | 0.837724 | Hhipl2 |
| ENSMUSG( | 2706.017 | -0.1086 | 0.490062 | 0.926143 | Hibadh |
| ENSMUSG( | 538.2946 | -0.13498 | 0.349187 | 0.911652 | Hibch |
| ENSMUSG( | 109.311 | 0.490525 | 0.045638 | 0.693959 | Hic1 |
| ENSMUSG( | 165.0643 | 0.176347 | 0.372606 | 0.915491 | Hic2 |
| ENSMUSG( | 2281.33 | -0.05938 | 0.618743 | 0.952502 | Hid1 |
| ENSMUSG( | 1403.137 | 0.079471 | 0.499961 | 0.927825 | Hif1a |
| ENSMUSG( | 1602.53 | 0.040185 | 0.720751 | 0.966887 | Hif1an |
| ENSMUSG( | 294.051 | 1.083353 | 7.90E-05 | 0.017291 | Hif3a |
| ENSMUSG( | 1755.063 | -0.20163 | 0.125725 | 0.81516 | Higd1a |
| ENSMUSG( | 77.79725 | 0.264331 | 0.405824 | 0.917257 | Higd1b |
| ENSMUSG( | 1982.719 | -0.15889 | 0.216498 | 0.865167 | Higd2a |
| ENSMUSG( | 531.0568 | -0.10977 | 0.453108 | 0.922316 | Hikeshi |
| ENSMUSG( | 119.2698 | 0.068392 | 0.731424 | 0.967228 | Hilpda |
| ENSMUSG( | 484.8163 | 0.017465 | 0.918135 | 0.992529 | Hinfp |
| ENSMUSG( | 3013.267 | -0.16185 | 0.201829 | 0.864412 | Hint1 |
| ENSMUSG( | 430.4543 | -0.14347 | 0.344536 | 0.91045 | Hint2 |
| ENSMUSG( | 334.1774 | -0.19113 | 0.207181 | 0.864412 | Hint3 |
| ENSMUSG( | 3679.544 | 0.248828 | 0.013445 | 0.463609 | Hip1 |
| ENSMUSG( | 3963.38 | -0.06408 | 0.526031 | 0.934374 | Hip1r |
| ENSMUSG( | 4763.29 | 0.051942 | 0.62897 | 0.953251 | Hipk1 |
| ENSMUSG( | 8806.546 | -0.31921 | 0.022139 | 0.576679 | Hipk2 |
| ENSMUSG( | 1737.275 | 0.158049 | 0.221702 | 0.866104 | Hipk3 |
| ENSMUSG( | 35.4526 | -0.56964 | 0.156178 | 0.837724 | Hipk4 |
| ENSMUSG( | 693.6667 | -0.1271 | 0.377577 | 0.91645 | Hira |
| ENSMUSG( | 474.2915 | 0.161115 | 0.349195 | 0.911652 | Hirip3 |
| ENSMUSG( | 3692.891 | -0.15925 | 0.149265 | 0.836241 | Hivep1 |
| ENSMUSG( | 10298.22 | 0.164209 | 0.160166 | 0.837724 | Hivep2 |
| ENSMUSG( | 3215.608 | 0.312259 | 0.004742 | 0.284457 | Hivep3 |
| ENSMUSG( | 24.36427 | -0.34545 | 0.401952 | 0.917257 | Hjv |
| ENSMUSG( | 11423.67 | -0.06329 | 0.545695 | 0.938063 | Hk1 |
| ENSMUSG( | 26.37976 | -0.04474 | 0.91377 | 0.992405 | Hk1os |
| ENSMUSG( | 216.1356 | 0.183702 | 0.323052 | 0.904023 | Hk2 |
| ENSMUSG( | 109.5945 | -0.0288 | 0.896748 | 0.990558 | Hk3 |
| ENSMUSG( | 442.806 | 0.045066 | 0.787372 | 0.97637 | Hkdc1 |
| ENSMUSG( | 500.11 | -0.31477 | 0.013427 | 0.463609 | Hlcs |
| ENSMUSG( | 7362.688 | 0.080572 | 0.552624 | 0.939565 | Hlf |
| ENSMUSG( | 550.1728 | 0.137992 | 0.283746 | 0.892412 | Hltf |
| ENSMUSG( | 39.41776 | 0.120824 | 0.708262 | 0.964042 | Hlx |
| ENSMUSG( | 2720.587 | -0.05924 | 0.586065 | 0.944975 | Hmbox1 |
| ENSMUSG( | 416.3368 | -0.08838 | 0.557158 | 0.939565 | Hmbs |
| ENSMUSG( | 318.3217 | -0.02574 | 0.87552 | 0.989644 | Hmces |
| ENSMUSG( | 245.4092 | 0.2231 | 0.354704 | 0.912411 | Hmcn1 |
| ENSMUSG( | 38.60954 | 0.138329 | 0.694924 | 0.960867 | Hmcn2 |
| ENSMUSG( | 1486.879 | -0.01375 | 0.911923 | 0.992196 | Hmg20a |
| ENSMUSG( | 747.3702 | 0.098287 | 0.469336 | 0.923329 | Hmg20b |
| ENSMUSG( | 1244.254 | -0.07145 | 0.575747 | 0.942912 | Hmga1 |
| ENSMUSG( | 52.63416 | 0.367258 | 0.246252 | 0.877904 | Hmga2-ps1 |
| ENSMUSG( | 3482.04 | -0.04078 | 0.748504 | 0.96984 | Hmgb1 |
| ENSMUSG( | 269.9569 | -0.07447 | 0.63694 | 0.953903 | Hmgb2 |
| ENSMUSG( | 617.4465 | -0.15153 | 0.321127 | 0.904019 | Hmgb3 |
| ENSMUSG( | 478.4757 | -0.00311 | 0.982779 | 0.996859 | Hmgcl |
| ENSMUSG( | 331.5554 | 0.101111 | 0.533209 | 0.935104 | Hmgcll1 |
| ENSMUSG( | 5495.489 | -0.01097 | 0.950851 | 0.994798 | Hmgcr |
| ENSMUSG( | 15734.79 | -0.45175 | 0.003033 | 0.219421 | Hmgcs1 |
| ENSMUSG( | 651.3115 | -0.37026 | 0.010143 | 0.410151 | Hmgcs2 |
| ENSMUSG( | 2127.544 | -0.14596 | 0.327693 | 0.906255 | Hmgn1 |
| ENSMUSG( | 1690.309 | -0.08101 | 0.518612 | 0.930939 | Hmgn2 |
| ENSMUSG( | 10.24945 | -0.00558 | 0.99256 | NA | Hmgn2-ps |
| ENSMUSG( | 1698.133 | 0.042707 | 0.735861 | 0.967344 | Hmgn3 |
| ENSMUSG( | 654.8802 | 0.135933 | 0.392937 | 0.917257 | Hmgn5 |
| ENSMUSG( | 1095.205 | 0.071382 | 0.560257 | 0.940131 | Hmgxb3 |
| ENSMUSG( | 466.7957 | 0.046701 | 0.72134 | 0.966887 | Hmgxb4 |
| ENSMUSG( | 20.73914 | 0.200112 | 0.650451 | 0.95422 | Hmmr |
| ENSMUSG( | 155.2028 | -0.12492 | 0.52119 | 0.932489 | Hmox1 |
| ENSMUSG( | 1276.726 | -0.17313 | 0.198036 | 0.862161 | Hmox2 |
| ENSMUSG( | 16.62732 | -1.29948 | 0.010477 | 0.41614 | Hnf1b |
| ENSMUSG( | 562.7262 | 0.068429 | 0.654837 | 0.955015 | Hnmt |
| ENSMUSG( | 2732.979 | -0.07511 | 0.619709 | 0.952896 | Hnrnpa0 |
| ENSMUSG( | 4140.97 | -0.07321 | 0.490619 | 0.92646 | Hnrnpa1 |
| ENSMUSG( | 8826.384 | 0.049124 | 0.670899 | 0.95694 | Hnrnpa2b1 |
| ENSMUSG( | 4407.802 | 0.014528 | 0.905868 | 0.991089 | Hnrnpa3 |
| ENSMUSG( | 2710.772 | 0.011077 | 0.937667 | 0.993685 | Hnrnpab |
| ENSMUSG( | 3995.639 | -0.10016 | 0.470172 | 0.923329 | Hnrnpc |
| ENSMUSG( | 3024.641 | -0.07882 | 0.546575 | 0.938063 | Hnrnpd |
| ENSMUSG( | 4084.771 | 0.151786 | 0.211562 | 0.864412 | Hnrnpdl |
| ENSMUSG( | 1191.138 | 0.076586 | 0.508233 | 0.929089 | Hnrnpf |
| ENSMUSG( | 4119.342 | 0.091039 | 0.428982 | 0.917257 | Hnrnph1 |
| ENSMUSG( | 1869.317 | -0.0744 | 0.508853 | 0.929089 | Hnrnph2 |
| ENSMUSG( | 1302.229 | -0.11176 | 0.412314 | 0.917257 | Hnrnph3 |
| ENSMUSG( | 6838.117 | -0.10689 | 0.354342 | 0.912411 | Hnrnpk |
| ENSMUSG( | 5831.362 | -0.14665 | 0.228979 | 0.868507 | Hnrnpl |
| ENSMUSG( | 1216.225 | -0.11331 | 0.333747 | 0.907907 | Hnrnpll |
| ENSMUSG( | 2642.659 | 0.036156 | 0.769208 | 0.974236 | Hnrnpm |
| ENSMUSG( | 1818.321 | 0.052658 | 0.650055 | 0.95422 | Hnrnpr |
| ENSMUSG( | 9941.538 | 0.050423 | 0.632004 | 0.953312 | Hnrnpu |
| ENSMUSG( | 5166.594 | -0.07887 | 0.464632 | 0.923329 | Hnrnpul1 |
| ENSMUSG( | 5097.153 | -0.07399 | 0.54799 | 0.938063 | Hnrnpul2 |
| ENSMUSG( | 90.0986 | -0.11541 | 0.651512 | 0.95449 | Hoga1 |
| ENSMUSG( | 2377.983 | -0.00664 | 0.956923 | 0.994798 | Homer1 |
| ENSMUSG( | 3378.655 | 0.177985 | 0.158155 | 0.837724 | Homer2 |
| ENSMUSG( | 909.4023 | 0.023907 | 0.839394 | 0.982012 | Homer3 |
| ENSMUSG( | 306.0422 | 0.069749 | 0.666496 | 0.956766 | Homez |
| ENSMUSG( | 1220.205 | 0.275833 | 0.031528 | 0.631192 | Hook1 |
| ENSMUSG( | 365.6268 | 0.018265 | 0.925889 | 0.99306 | Hook2 |
| ENSMUSG( | 1584.298 | 0.259389 | 0.035033 | 0.659157 | Hook3 |
| ENSMUSG( | 3886.818 | -0.357 | 0.031724 | 0.631941 | Hopx |
| ENSMUSG( | 27.27289 | -0.32364 | 0.546005 | 0.938063 | Hp |
| ENSMUSG( | 5713.884 | -0.05478 | 0.59717 | 0.946609 | Hp1bp3 |
| ENSMUSG( | 12974.63 | -0.1519 | 0.312409 | 0.900037 | Hpca |
| ENSMUSG( | 748.7125 | -0.18788 | 0.24608 | 0.877904 | Hpcal1 |
| ENSMUSG( | 54731.81 | 0.087063 | 0.406497 | 0.917257 | Hpcal4 |

| | | | | | | | | | | | |
|---|---|---|---|---|---|---|---|---|---|---|---|
| ENSMUSG( | Hhat | 31.21757 | -0.36455 | 0.28307 | 0.73186 | ENSMUSG( | Hhat | 29.66632 | -0.19395 | 0.604217 | 0.948139 |
| ENSMUSG( | Hhatl | 598.2261 | -0.39555 | 0.002433 | 0.058029 | ENSMUSG( | Hhatl | 618.628 | 0.000433 | 0.997782 | 0.999268 |
| ENSMUSG( | Hhex | 76.2502 | 0.296006 | 0.242233 | 0.689699 | ENSMUSG( | Hhex | 79.51692 | 0.646835 | 0.012448 | 0.451463 |
| ENSMUSG( | Hhip | 364.1867 | -0.51302 | 0.002135 | 0.053442 | ENSMUSG( | Hhip | 412.3537 | 0.162913 | 0.403472 | 0.917257 |
| ENSMUSG( | Hhipl1 | 506.9318 | -0.13673 | 0.32656 | 0.771368 | ENSMUSG( | Hhipl1 | 503.2296 | 0.117082 | 0.448499 | 0.922202 |
| ENSMUSG( | Hhipl2 | 17.70427 | 0.741923 | 0.140791 | NA | ENSMUSG( | Hhipl2 | 16.33677 | 0.76014 | 0.158541 | 0.837724 |
| ENSMUSG( | Hibadh | 3193.826 | 0.104596 | 0.416011 | 0.82084 | ENSMUSG( | Hibadh | 2706.017 | -0.1086 | 0.490062 | 0.926143 |
| ENSMUSG( | Hibch | 640.8107 | 0.10628 | 0.374152 | 0.800445 | ENSMUSG( | Hibch | 538.2946 | -0.13498 | 0.349187 | 0.911652 |
| ENSMUSG( | Hic1 | 91.78721 | -0.27746 | 0.207149 | 0.649517 | ENSMUSG( | Hic1 | 109.311 | 0.490525 | 0.045638 | 0.693959 |
| ENSMUSG( | Hic2 | 173.7807 | 0.047937 | 0.781731 | 0.957558 | ENSMUSG( | Hic2 | 165.0643 | 0.176347 | 0.372606 | 0.915491 |
| ENSMUSG( | Hid1 | 2538.366 | -0.02202 | 0.820164 | 0.965888 | ENSMUSG( | Hid1 | 2281.33 | -0.05938 | 0.618743 | 0.952502 |
| ENSMUSG( | Hif1a | 1508.248 | 0.022622 | 0.80262 | 0.962169 | ENSMUSG( | Hif1a | 1403.137 | 0.079471 | 0.499961 | 0.927825 |
| ENSMUSG( | Hif1an | 1753.721 | 0.031144 | 0.726259 | 0.944223 | ENSMUSG( | Hif1an | 1602.53 | 0.040185 | 0.720751 | 0.966887 |
| ENSMUSG( | Hif3a | 335.3507 | 1.152566 | 1.53E-07 | 3.77E-05 | ENSMUSG( | Hif3a | 294.051 | 1.083353 | 7.90E-05 | 0.017291 |
| ENSMUSG( | Higd1a | 2065.787 | 0.010946 | 0.911652 | 0.988483 | ENSMUSG( | Higd1a | 1755.063 | -0.20163 | 0.125725 | 0.81516 |
| ENSMUSG( | Higd1b | 74.72946 | -0.13303 | 0.59774 | 0.90372 | ENSMUSG( | Higd1b | 77.79725 | 0.264331 | 0.405824 | 0.917257 |
| ENSMUSG( | Higd2a | 2326.102 | 0.041192 | 0.677752 | 0.930542 | ENSMUSG( | Higd2a | 1982.719 | -0.15889 | 0.216498 | 0.865167 |
| ENSMUSG( | Hikeshi | 587.4736 | -0.07866 | 0.542768 | 0.883563 | ENSMUSG( | Hikeshi | 531.0568 | -0.10977 | 0.453108 | 0.922316 |
| ENSMUSG( | Hilpda | 110.7341 | -0.45733 | 0.025399 | 0.246101 | ENSMUSG( | Hilpda | 119.2698 | 0.068392 | 0.731424 | 0.967228 |
| ENSMUSG( | Hinfp | 528.2595 | -0.01218 | 0.915938 | 0.988634 | ENSMUSG( | Hinfp | 484.8163 | 0.017465 | 0.918135 | 0.992529 |
| ENSMUSG( | Hint1 | 3521.041 | 0.025993 | 0.789144 | 0.959299 | ENSMUSG( | Hint1 | 3013.267 | -0.16185 | 0.201829 | 0.864412 |
| ENSMUSG( | Hint2 | 485.4769 | -0.05438 | 0.634768 | 0.914073 | ENSMUSG( | Hint2 | 430.4543 | -0.14347 | 0.344536 | 0.91045 |
| ENSMUSG( | Hint3 | 394.5164 | 0.024527 | 0.86837 | 0.976834 | ENSMUSG( | Hint3 | 334.1774 | -0.19113 | 0.207181 | 0.864412 |
| ENSMUSG( | Hip1 | 4051.322 | 0.256538 | 0.002668 | 0.062087 | ENSMUSG( | Hip1 | 3679.544 | 0.248828 | 0.013445 | 0.463609 |
| ENSMUSG( | Hip1r | 4618.349 | 0.107699 | 0.224134 | 0.669929 | ENSMUSG( | Hip1r | 3963.38 | -0.06408 | 0.526031 | 0.934374 |
| ENSMUSG( | Hipk1 | 5344.258 | 0.113231 | 0.20859 | 0.650496 | ENSMUSG( | Hipk1 | 4763.29 | 0.051942 | 0.62897 | 0.953251 |
| ENSMUSG( | Hipk2 | 10848.7 | 0.031547 | 0.791547 | 0.959461 | ENSMUSG( | Hipk2 | 8806.546 | -0.31921 | 0.022139 | 0.576679 |
| ENSMUSG( | Hipk3 | 1797.632 | -0.00814 | 0.937886 | 0.993415 | ENSMUSG( | Hipk3 | 1737.275 | 0.158049 | 0.221702 | 0.866104 |
| ENSMUSG( | Hipk4 | 40.30891 | -0.49067 | 0.164368 | 0.59535 | ENSMUSG( | Hipk4 | 35.4526 | -0.56964 | 0.156178 | 0.837724 |
| ENSMUSG( | Hira | 795.471 | 0.008236 | 0.941018 | 0.993415 | ENSMUSG( | Hira | 693.6667 | -0.1271 | 0.377577 | 0.91645 |
| ENSMUSG( | Hirip3 | 488.395 | -0.01088 | 0.937955 | 0.993415 | ENSMUSG( | Hirip3 | 474.2915 | 0.161115 | 0.349195 | 0.911652 |
| ENSMUSG( | Hivep1 | 4399.87 | 0.080333 | 0.443312 | 0.833108 | ENSMUSG( | Hivep1 | 3692.891 | -0.15925 | 0.149265 | 0.836241 |
| ENSMUSG( | Hivep2 | 11029.44 | 0.095432 | 0.28268 | 0.73186 | ENSMUSG( | Hivep2 | 10298.22 | 0.164209 | 0.160166 | 0.837724 |
| ENSMUSG( | Hivep3 | 3200.412 | 0.044263 | 0.654415 | 0.923227 | ENSMUSG( | Hivep3 | 3215.608 | 0.312259 | 0.004742 | 0.284457 |
| ENSMUSG( | Hjv | 23.07374 | -0.86053 | 0.027026 | 0.252658 | ENSMUSG( | Hjv | 24.36427 | -0.34545 | 0.401952 | 0.917257 |
| ENSMUSG( | Hk1 | 13059.97 | 0.054578 | 0.506324 | 0.867425 | ENSMUSG( | Hk1 | 11423.67 | -0.06329 | 0.545695 | 0.938063 |
| ENSMUSG( | Hk1os | 29.98049 | 0.09996 | 0.770567 | 0.954722 | ENSMUSG( | Hk1os | 26.37976 | -0.04474 | 0.91377 | 0.992405 |
| ENSMUSG( | Hk2 | 240.9206 | 0.215369 | 0.162475 | 0.593257 | ENSMUSG( | Hk2 | 216.1356 | 0.183702 | 0.323052 | 0.904023 |
| ENSMUSG( | Hk3 | 120.3562 | -0.0369 | 0.852583 | 0.973168 | ENSMUSG( | Hk3 | 109.5945 | -0.0288 | 0.896748 | 0.990558 |
| ENSMUSG( | Hkdc1 | 504.8202 | 0.149719 | 0.389154 | 0.806967 | ENSMUSG( | Hkdc1 | 442.806 | 0.045066 | 0.787372 | 0.97637 |
| ENSMUSG( | Hlcs | 565.4348 | -0.22322 | 0.066497 | 0.39868 | ENSMUSG( | Hlcs | 500.11 | -0.31477 | 0.013427 | 0.463609 |
| ENSMUSG( | Hlf | 8096.431 | 0.087562 | 0.352919 | 0.784977 | ENSMUSG( | Hlf | 7362.688 | 0.080572 | 0.552624 | 0.939565 |
| ENSMUSG( | Hltf | 590.8561 | 0.077134 | 0.483354 | 0.856304 | ENSMUSG( | Hltf | 550.1728 | 0.137992 | 0.283746 | 0.892412 |
| ENSMUSG( | Hlx | 36.99105 | -0.34545 | 0.288351 | 0.739103 | ENSMUSG( | Hlx | 39.41776 | 0.120824 | 0.708262 | 0.964042 |
| ENSMUSG( | Hmbox1 | 3153.221 | 0.098415 | 0.291853 | 0.741868 | ENSMUSG( | Hmbox1 | 2720.587 | -0.05924 | 0.586065 | 0.944975 |
| ENSMUSG( | Hmbs | 471.4661 | 0.005157 | 0.968007 | 0.996284 | ENSMUSG( | Hmbs | 416.3368 | -0.08838 | 0.557158 | 0.939565 |
| ENSMUSG( | Hmces | 354.4796 | 0.021322 | 0.880609 | 0.981542 | ENSMUSG( | Hmces | 318.3217 | -0.02574 | 0.87552 | 0.989644 |
| ENSMUSG( | Hmcn1 | 247.8752 | -0.02477 | 0.890276 | 0.983687 | ENSMUSG( | Hmcn1 | 245.4092 | 0.2231 | 0.354704 | 0.912411 |
| ENSMUSG( | Hmcn2 | 39.40864 | -0.09533 | 0.755544 | 0.951395 | ENSMUSG( | Hmcn2 | 38.60954 | 0.138329 | 0.694924 | 0.960867 |
| ENSMUSG( | Hmg20a | 1704.523 | 0.107218 | 0.322114 | 0.767096 | ENSMUSG( | Hmg20a | 1486.879 | -0.01375 | 0.911923 | 0.992196 |
| ENSMUSG( | Hmg20b | 812.9324 | 0.074081 | 0.521947 | 0.875186 | ENSMUSG( | Hmg20b | 747.3702 | 0.098287 | 0.469336 | 0.923329 |
| ENSMUSG( | Hmga1 | 1412.621 | 0.029875 | 0.752117 | 0.950417 | ENSMUSG( | Hmga1 | 1244.254 | -0.07145 | 0.575747 | 0.942912 |
| ENSMUSG( | Hmga2-ps1 | 50.11299 | -0.04752 | 0.877392 | 0.981333 | ENSMUSG( | Hmga2-ps1 | 52.63416 | 0.367258 | 0.246252 | 0.877904 |
| ENSMUSG( | Hmgb1 | 3965.77 | 0.068061 | 0.465831 | 0.844222 | ENSMUSG( | Hmgb1 | 3482.04 | -0.04078 | 0.748504 | 0.96984 |
| ENSMUSG( | Hmgb2 | 304.9723 | 0.006388 | 0.96399 | 0.995094 | ENSMUSG( | Hmgb2 | 269.9569 | -0.07447 | 0.63694 | 0.953903 |
| ENSMUSG( | Hmgb3 | 673.2076 | -0.16372 | 0.178118 | 0.615613 | ENSMUSG( | Hmgb3 | 617.4465 | -0.15153 | 0.321127 | 0.904019 |
| ENSMUSG( | Hmgcl | 545.1285 | 0.101623 | 0.402234 | 0.815103 | ENSMUSG( | Hmgcl | 478.4757 | -0.00311 | 0.982779 | 0.996859 |
| ENSMUSG( | Hmgcll1 | 364.1936 | 0.112618 | 0.38173 | 0.802879 | ENSMUSG( | Hmgcll1 | 331.5554 | 0.101111 | 0.533209 | 0.935104 |
| ENSMUSG( | Hmgcr | 5406.023 | -0.3536 | 0.030332 | 0.264512 | ENSMUSG( | Hmgcr | 5495.489 | -0.01097 | 0.950851 | 0.994798 |
| ENSMUSG( | Hmgcs1 | 19873.59 | -0.01 | 0.946233 | 0.99369 | ENSMUSG( | Hmgcs1 | 15734.79 | -0.45175 | 0.003033 | 0.219421 |
| ENSMUSG( | Hmgcs2 | 850.3169 | 0.155234 | 0.136542 | 0.550651 | ENSMUSG( | Hmgcs2 | 651.3115 | -0.37026 | 0.010143 | 0.410151 |
| ENSMUSG( | Hmgn1 | 2493.179 | 0.050889 | 0.668844 | 0.927963 | ENSMUSG( | Hmgn1 | 2127.544 | -0.14596 | 0.327693 | 0.906255 |
| ENSMUSG( | Hmgn2 | 1842.523 | -0.09792 | 0.342715 | 0.778971 | ENSMUSG( | Hmgn2 | 1690.309 | -0.08101 | 0.518612 | 0.930939 |
| ENSMUSG( | Hmgn2-ps | 11.59704 | 0.065016 | 0.904396 | NA | ENSMUSG( | Hmgn2-ps | 10.24945 | -0.00558 | 0.99256 | NA |
| ENSMUSG( | Hmgn3 | 1861.18 | 0.041606 | 0.68902 | 0.933441 | ENSMUSG( | Hmgn3 | 1698.133 | 0.042707 | 0.735861 | 0.967344 |
| ENSMUSG( | Hmgn5 | 745.2409 | 0.240543 | 0.051973 | 0.350064 | ENSMUSG( | Hmgn5 | 654.8802 | 0.135933 | 0.392937 | 0.917257 |
| ENSMUSG( | Hmgxb3 | 1213.946 | 0.097258 | 0.364581 | 0.793796 | ENSMUSG( | Hmgxb3 | 1095.205 | 0.071382 | 0.560257 | 0.940131 |
| ENSMUSG( | Hmgxb4 | 512.9449 | 0.04771 | 0.677999 | 0.930542 | ENSMUSG( | Hmgxb4 | 466.7957 | 0.046701 | 0.72134 | 0.966887 |
| ENSMUSG( | Hmmr | 20.59443 | -0.12046 | 0.772986 | NA | ENSMUSG( | Hmmr | 20.73914 | 0.200112 | 0.650451 | 0.95422 |
| ENSMUSG( | Hmox1 | 180.3578 | 0.050498 | 0.756564 | 0.951479 | ENSMUSG( | Hmox1 | 155.2028 | -0.12492 | 0.52119 | 0.932489 |
| ENSMUSG( | Hmox2 | 1471.452 | -0.0236 | 0.818525 | 0.965843 | ENSMUSG( | Hmox2 | 1276.726 | -0.17313 | 0.198036 | 0.862161 |
| ENSMUSG( | Hnf1b | 15.48249 | -2.33524 | 2.19E-05 | NA | ENSMUSG( | Hnf1b | 16.62732 | -1.29948 | 0.010477 | 0.41614 |
| ENSMUSG( | Hnmt | 550.0863 | -0.27375 | 0.059165 | 0.375449 | ENSMUSG( | Hnmt | 562.7262 | 0.068429 | 0.654837 | 0.955015 |
| ENSMUSG( | Hnrnpa0 | 2995.976 | -0.07574 | 0.461439 | 0.841432 | ENSMUSG( | Hnrnpa0 | 2732.979 | -0.07511 | 0.619709 | 0.952896 |
| ENSMUSG( | Hnrnpa1 | 4471.687 | -0.12022 | 0.157265 | 0.585553 | ENSMUSG( | Hnrnpa1 | 4140.97 | -0.07321 | 0.490619 | 0.92646 |
| ENSMUSG( | Hnrnpa2b1 | 9692.577 | 0.052645 | 0.56344 | 0.889914 | ENSMUSG( | Hnrnpa2b1 | 8826.384 | 0.049124 | 0.670899 | 0.95694 |
| ENSMUSG( | Hnrnpa3 | 4844.925 | 0.020999 | 0.833221 | 0.970333 | ENSMUSG( | Hnrnpa3 | 4407.802 | 0.014528 | 0.905868 | 0.991089 |
| ENSMUSG( | Hnrnpab | 2994.977 | 0.032944 | 0.739087 | 0.947768 | ENSMUSG( | Hnrnpab | 2710.772 | 0.011077 | 0.937667 | 0.993685 |
| ENSMUSG( | Hnrnpc | 4669.332 | 0.084854 | 0.419602 | 0.822309 | ENSMUSG( | Hnrnpc | 3995.639 | -0.10016 | 0.470172 | 0.923329 |
| ENSMUSG( | Hnrnpd | 3369.077 | -0.03281 | 0.736082 | 0.946267 | ENSMUSG( | Hnrnpd | 3024.641 | -0.07882 | 0.546575 | 0.938063 |
| ENSMUSG( | Hnrnpdl | 4019.575 | -0.16599 | 0.058204 | 0.37191 | ENSMUSG( | Hnrnpdl | 4084.771 | 0.151786 | 0.211562 | 0.864412 |
| ENSMUSG( | Hnrnpf | 1317.516 | 0.099642 | 0.303853 | 0.752148 | ENSMUSG( | Hnrnpf | 1191.138 | 0.076586 | 0.508233 | 0.929089 |
| ENSMUSG( | Hnrnph1 | 4381.329 | 0.003337 | 0.967668 | 0.996228 | ENSMUSG( | Hnrnph1 | 4119.342 | 0.091039 | 0.428982 | 0.917257 |
| ENSMUSG( | Hnrnph2 | 2057.664 | -0.06374 | 0.550175 | 0.886038 | ENSMUSG( | Hnrnph2 | 1869.317 | -0.0744 | 0.508853 | 0.929089 |
| ENSMUSG( | Hnrnph3 | 1537.78 | 0.104496 | 0.352149 | 0.783738 | ENSMUSG( | Hnrnph3 | 1302.229 | -0.11176 | 0.412314 | 0.917257 |
| ENSMUSG( | Hnrnpk | 7740.021 | -0.01446 | 0.868649 | 0.976897 | ENSMUSG( | Hnrnpk | 6838.117 | -0.10689 | 0.354342 | 0.912411 |
| ENSMUSG( | Hnrnpl | 6323.542 | -0.1853 | 0.033327 | 0.279795 | ENSMUSG( | Hnrnpl | 5831.362 | -0.14665 | 0.228979 | 0.868507 |
| ENSMUSG( | Hnrnpll | 1329.35 | -0.12505 | 0.207077 | 0.649503 | ENSMUSG( | Hnrnpll | 1216.225 | -0.11331 | 0.333747 | 0.907907 |
| ENSMUSG( | Hnrnpm | 2992.264 | 0.126477 | 0.199553 | 0.640054 | ENSMUSG( | Hnrnpm | 2642.659 | 0.036156 | 0.769208 | 0.974236 |
| ENSMUSG( | Hnrnpr | 1988.36 | 0.044857 | 0.636741 | 0.91503 | ENSMUSG( | Hnrnpr | 1818.321 | 0.052658 | 0.650055 | 0.95422 |
| ENSMUSG( | Hnrnpu | 10531.93 | -0.05096 | 0.534381 | 0.880719 | ENSMUSG( | Hnrnpu | 9941.538 | 0.050423 | 0.632004 | 0.953312 |
| ENSMUSG( | Hnrnpul1 | 5935.06 | 0.054712 | 0.487716 | 0.858824 | ENSMUSG( | Hnrnpul1 | 5166.594 | -0.07887 | 0.464632 | 0.923329 |
| ENSMUSG( | Hnrnpul2 | 5925.759 | 0.09359 | 0.304703 | 0.752574 | ENSMUSG( | Hnrnpul2 | 5097.153 | -0.07399 | 0.54799 | 0.938063 |
| ENSMUSG( | Hoga1 | 107.5009 | 0.135425 | 0.578922 | 0.896692 | ENSMUSG( | Hoga1 | 90.0986 | -0.11541 | 0.651512 | 0.95449 |
| ENSMUSG( | Homer1 | 2649.001 | 0.038956 | 0.67211 | 0.9288 | ENSMUSG( | Homer1 | 2377.983 | -0.00664 | 0.956923 | 0.994798 |
| ENSMUSG( | Homer2 | 3790.19 | 0.236067 | 0.038473 | 0.302696 | ENSMUSG( | Homer2 | 3378.655 | 0.177985 | 0.158155 | 0.837724 |
| ENSMUSG( | Homer3 | 1005.779 | 0.047145 | 0.624098 | 0.91187 | ENSMUSG( | Homer3 | 909.4023 | 0.023907 | 0.839394 | 0.982012 |
| ENSMUSG( | Homez | 336.1848 | 0.077721 | 0.565541 | 0.890871 | ENSMUSG( | Homez | 306.0422 | 0.069749 | 0.666496 | 0.956766 |
| ENSMUSG( | Hook1 | 1188.075 | -0.05915 | 0.581931 | 0.897918 | ENSMUSG( | Hook1 | 1220.205 | 0.275833 | 0.031528 | 0.631192 |
| ENSMUSG( | Hook2 | 412.0115 | 0.092659 | 0.605314 | 0.905737 | ENSMUSG( | Hook2 | 365.6268 | 0.018265 | 0.925889 | 0.99306 |
| ENSMUSG( | Hook3 | 1583.957 | -0.0009 | 0.993037 | 0.999327 | ENSMUSG( | Hook3 | 1584.298 | 0.259389 | 0.035033 | 0.659157 |
| ENSMUSG( | Hopx | 5058.067 | 0.155256 | 0.233497 | 0.681508 | ENSMUSG( | Hopx | 3886.818 | -0.357 | 0.031724 | 0.631941 |
| ENSMUSG( | Hp | 23.53597 | -1.20244 | 0.029574 | 0.260563 | ENSMUSG( | Hp | 27.27289 | -0.32364 | 0.546005 | 0.938063 |
| ENSMUSG( | Hp1bp3 | 6341.812 | -0.02125 | 0.782355 | 0.957558 | ENSMUSG( | Hp1bp3 | 5713.884 | -0.05478 | 0.59717 | 0.946609 |
| ENSMUSG( | Hpca | 14697.07 | -0.05564 | 0.653869 | 0.922914 | ENSMUSG( | Hpca | 12974.63 | -0.1519 | 0.312409 | 0.900037 |
| ENSMUSG( | Hpcal1 | 913.1574 | 0.123876 | 0.361763 | 0.792797 | ENSMUSG( | Hpcal1 | 748.7125 | -0.18788 | 0.24608 | 0.877904 |
| ENSMUSG( | Hpcal4 | 64342.34 | 0.272406 | 0.003537 | 0.075287 | ENSMUSG( | Hpcal4 | 54731.81 | 0.087063 | 0.406497 | 0.917257 |
| ENSMUSG( | Hpdl | 41.27794 | -0.23029 | 0.500593 | 0.863514 | ENSMUSG( | Hpdl | 37.20992 | -0.28445 | 0.437079 | 0.920144 |
| ENSMUSG( | Hpf1 | 1215.398 | 0.036839 | 0.739894 | 0.947823 | ENSMUSG( | Hpf1 | 1040.32 | -0.15159 | 0.239716 | 0.874882 |
| ENSMUSG( | Hpgd | 286.8051 | 0.22519 | 0.180729 | 0.617841 | ENSMUSG( | Hpgd | 277.674 | 0.374463 | 0.050873 | 0.709741 |
| ENSMUSG( | Hpgds | 462.8031 | -0.32431 | 0.027314 | 0.253597 | ENSMUSG( | Hpgds | 489.935 | 0.134057 | 0.475174 | 0.923329 |
| ENSMUSG( | Hpn | 35.97376 | 0.196447 | 0.546683 | 0.884974 | ENSMUSG( | Hpn | 32.92952 | 0.224559 | 0.540213 | 0.937383 |
| ENSMUSG( | Hprt1 | 1547.692 | -0.09875 | 0.416745 | 0.8209 | ENSMUSG( | Hprt1 | 1369.146 | -0.19276 | 0.190238 | 0.857355 |
| ENSMUSG( | Hps1 | 529.9705 | 0.149231 | 0.207919 | 0.650048 | ENSMUSG( | Hps1 | 464.758 | 0.044687 | 0.794966 | 0.97681 |
| ENSMUSG( | Hps3 | 475.19 | 0.020479 | 0.860641 | 0.9749 | ENSMUSG( | Hps3 | 453.7639 | 0.157247 | 0.268308 | 0.886493 |
| ENSMUSG( | Hps4 | 604.2049 | 0.287694 | 0.036443 | 0.294207 | ENSMUSG( | Hps4 | 537.2437 | 0.228452 | 0.111226 | 0.807885 |
| ENSMUSG( | Hps5 | 894.1828 | 0.028115 | 0.79573 | 0.960062 | ENSMUSG( | Hps5 | 794.7819 | -0.04518 | 0.727413 | 0.966887 |
| ENSMUSG( | Hps6 | 124.2363 | 0.073944 | 0.72518 | 0.944096 | ENSMUSG( | Hps6 | 104.8249 | -0.13429 | 0.554598 | 0.939565 |
| ENSMUSG( | Hpse | 49.87481 | -0.18953 | 0.496602 | 0.86175 | ENSMUSG( | Hpse | 50.02762 | 0.119658 | 0.702724 | 0.962215 |
| ENSMUSG( | Hpx | 39.78439 | 0.008918 | 0.977008 | 0.997428 | ENSMUSG( | Hpx | 26.80472 | -1.03545 | 0.011481 | 0.436431 |
| ENSMUSG( | Hr | 2254.271 | 0.088546 | 0.329491 | 0.773311 | ENSMUSG( | Hr | 1928.116 | -0.09103 | 0.457795 | 0.923329 |
| ENSMUSG( | Hras | 4582.494 | -0.04233 | 0.617619 | 0.910872 | ENSMUSG( | Hras | 3977.079 | -0.19038 | 0.112115 | 0.808673 |
| ENSMUSG( | Hrc | 10.56496 | -0.50696 | 0.390637 | NA | ENSMUSG( | Hrc | 10.05143 | -0.27436 | 0.6859 | NA |
| ENSMUSG( | Hrct1 | 24.90314 | -0.02139 | 0.960148 | 0.993991 | ENSMUSG( | Hrct1 | 28.118 | 0.570228 | 0.175835 | 0.845715 |
| ENSMUSG( | Hrh1 | 659.1933 | -0.36028 | 0.02557 | 0.246694 | ENSMUSG( | Hrh1 | 710.8776 | 0.149767 | 0.375022 | 0.915491 |
| ENSMUSG( | Hrh2 | 365.5616 | 0.043666 | 0.795866 | 0.960062 | ENSMUSG( | Hrh2 | 336.9606 | 0.087217 | 0.638403 | 0.953903 |
| ENSMUSG( | Hrh3 | 3463.722 | -0.15504 | 0.108459 | 0.497337 | ENSMUSG( | Hrh3 | 4051.994 | 0.525473 | 2.53E-05 | 0.007381 |
| ENSMUSG( | Hrk | 3133.461 | 0.117072 | 0.181148 | 0.618139 | ENSMUSG( | Hrk | 2675.188 | -0.07016 | 0.532387 | 0.934795 |
| ENSMUSG( | Hrob | 16.59087 | -0.19685 | 0.667595 | NA | ENSMUSG( | Hrob | 17.91943 | 0.300907 | 0.510134 | 0.929089 |
| ENSMUSG( | Hs1bp3 | 431.8433 | 0.226258 | 0.098822 | 0.480628 | ENSMUSG( | Hs1bp3 | 360.6103 | -0.01028 | 0.947565 | 0.994798 |
| ENSMUSG( | Hs2st1 | 2440.437 | -0.16903 | 0.17806 | 0.615613 | ENSMUSG( | Hs2st1 | 2450.725 | 0.116541 | 0.430696 | 0.917257 |
| ENSMUSG( | Hs3st1 | 761.1972 | -0.46026 | 0.009852 | 0.141792 | ENSMUSG( | Hs3st1 | 872.6517 | 0.237174 | 0.229964 | 0.868507 |
| ENSMUSG( | Hs3st2 | 1593.874 | -0.26818 | 0.042628 | 0.316734 | ENSMUSG( | Hs3st2 | 1643.517 | 0.103344 | 0.445895 | 0.921734 |
| ENSMUSG( | Hs3st3a1 | 54.37909 | -0.77526 | 0.010306 | 0.144599 | ENSMUSG( | Hs3st3a1 | 64.70351 | 0.070507 | 0.805119 | 0.977358 |
| ENSMUSG( | Hs3st3b1 | 109.0912 | -0.09391 | 0.669472 | 0.928147 | ENSMUSG( | Hs3st3b1 | 98.98254 | -0.11375 | 0.643097 | 0.953903 |
| ENSMUSG( | Hs3st4 | 2912.007 | 0.080553 | 0.385052 | 0.804048 | ENSMUSG( | Hs3st4 | 2648.748 | 0.077508 | 0.48735 | 0.92541 |
| ENSMUSG( | Hs3st5 | 113.5999 | -0.73323 | 0.000168 | 0.009317 | ENSMUSG( | Hs3st5 | 128.3147 | -0.02024 | 0.920262 | 0.992644 |
| ENSMUSG( | Hs3st6 | 9.11815 | 0.0365 | 0.956914 | NA | ENSMUSG( | Hs3st6 | 8.896783 | 0.210961 | 0.746693 | NA |
| ENSMUSG( | Hs6st1 | 2745.958 | -0.13149 | 0.294296 | 0.743534 | ENSMUSG( | Hs6st1 | 2873.434 | 0.263143 | 0.069369 | 0.755467 |
| ENSMUSG( | Hs6st2 | 728.372 | -0.3568 | 0.007277 | 0.117913 | ENSMUSG( | Hs6st2 | 816.0352 | 0.253517 | 0.112044 | 0.808673 |
| ENSMUSG( | Hs6st3 | 130.9556 | 0.086013 | 0.688845 | 0.933441 | ENSMUSG( | Hs6st3 | 134.2259 | 0.42147 | 0.131095 | 0.822884 |
| ENSMUSG( | Hsbp1 | 8431.305 | 0.061811 | 0.510928 | 0.870755 | ENSMUSG( | Hsbp1 | 7129.106 | -0.15974 | 0.220068 | 0.866104 |
| ENSMUSG( | Hsbp1l1 | 22.3723 | -0.16178 | 0.711308 | 0.940028 | ENSMUSG( | Hsbp1l1 | 21.00859 | -0.06793 | 0.887161 | 0.990263 |
| ENSMUSG( | Hscb | 146.2761 | -0.03538 | 0.837513 | 0.971111 | ENSMUSG( | Hscb | 127.8655 | -0.1762 | 0.402215 | 0.917257 |
| ENSMUSG( | Hsd11b1 | 2381.036 | 0.00721 | 0.957608 | 0.993874 | ENSMUSG( | Hsd11b1 | 2089.372 | -0.10095 | 0.505074 | 0.927825 |
| ENSMUSG( | Hsd17b10 | 1165.392 | 0.103329 | 0.2724 | 0.722422 | ENSMUSG( | Hsd17b10 | 959.2789 | -0.19807 | 0.120245 | 0.813384 |
| ENSMUSG( | Hsd17b11 | 2176.335 | -0.13205 | 0.311337 | 0.759267 | ENSMUSG( | Hsd17b11 | 2201.26 | 0.170029 | 0.235949 | 0.87215 |
| ENSMUSG( | Hsd17b12 | 4285.179 | -0.23992 | 0.076749 | 0.429306 | ENSMUSG( | Hsd17b12 | 4170.41 | -0.03763 | 0.803175 | 0.976937 |
| ENSMUSG( | Hsd17b4 | 3205.576 | 0.158028 | 0.145442 | 0.566269 | ENSMUSG( | Hsd17b4 | 2659.428 | -0.11284 | 0.349227 | 0.911652 |
| ENSMUSG( | Hsd17b7 | 723.8279 | -0.22619 | 0.204713 | 0.645625 | ENSMUSG( | Hsd17b7 | 710.5345 | 0.004076 | 0.983025 | 0.996891 |
| ENSMUSG( | Hsd17b8 | 21.37351 | -0.25105 | 0.52448 | 0.875844 | ENSMUSG( | Hsd17b8 | 17.93209 | -0.49439 | 0.284538 | 0.892412 |
| ENSMUSG( | Hsd3b7 | 15.63043 | 0.062514 | 0.895956 | NA | ENSMUSG( | Hsd3b7 | 15.58613 | 0.352385 | 0.50215 | 0.927825 |
| ENSMUSG( | Hsdl1 | 1345.702 | 0.025464 | 0.778073 | 0.956168 | ENSMUSG( | Hsdl1 | 1216.166 | 0.002732 | 0.979872 | 0.996479 |
| ENSMUSG( | Hsdl2 | 2476.409 | 0.055151 | 0.666188 | 0.927057 | ENSMUSG( | Hsdl2 | 1993.796 | -0.32244 | 0.025204 | 0.597422 |
| ENSMUSG( | Hsf1 | 1161.939 | 0.195413 | 0.067661 | 0.402445 | ENSMUSG( | Hsf1 | 1021.896 | 0.099292 | 0.461653 | 0.923329 |
| ENSMUSG( | Hsf2 | 513.0014 | -0.06248 | 0.593592 | 0.90281 | ENSMUSG( | Hsf2 | 492.4269 | 0.095637 | 0.514884 | 0.930939 |
| ENSMUSG( | Hsf4 | 245.05 | -0.06229 | 0.663802 | 0.926511 | ENSMUSG( | Hsf4 | 216.5521 | -0.15722 | 0.392253 | 0.917257 |
| ENSMUSG( | Hsf5 | 16.05344 | -0.76826 | 0.099399 | NA | ENSMUSG( | Hsf5 | 18.8339 | 0.088845 | 0.843728 | 0.983462 |

| | | | | | |
|---|---|---|---|---|---|
| ENSMUSG( | 19873.59 | -0.01 | 0.946233 | 0.99369 | Hmgcs1 |
| ENSMUSG( | 850.3169 | 0.155234 | 0.136542 | 0.550651 | Hmgcs2 |
| ENSMUSG( | 2493.179 | 0.050889 | 0.668844 | 0.927963 | Hmgn1 |
| ENSMUSG( | 1842.523 | -0.09792 | 0.342715 | 0.778971 | Hmgn2 |
| ENSMUSG( | 11.59704 | 0.065016 | 0.904396 | NA | Hmgn2-ps |
| ENSMUSG( | 1861.18 | 0.041606 | 0.68902 | 0.933441 | Hmgn3 |
| ENSMUSG( | 745.2409 | 0.240543 | 0.051973 | 0.350064 | Hmgn5 |
| ENSMUSG( | 1213.946 | 0.097258 | 0.364581 | 0.793796 | Hmgxb3 |
| ENSMUSG( | 512.9449 | 0.04771 | 0.677999 | 0.930542 | Hmgxb4 |
| ENSMUSG( | 20.59443 | -0.12046 | 0.772986 | NA | Hmmr |
| ENSMUSG( | 180.3578 | 0.050498 | 0.756564 | 0.951479 | Hmox1 |
| ENSMUSG( | 1471.452 | -0.0236 | 0.818525 | 0.965843 | Hmox2 |
| ENSMUSG( | 15.48249 | -2.33524 | 2.19E-05 | NA | Hnf1b |
| ENSMUSG( | 550.0863 | -0.27375 | 0.059165 | 0.375449 | Hnmt |
| ENSMUSG( | 2995.976 | -0.07574 | 0.461439 | 0.841432 | Hnrnpa0 |
| ENSMUSG( | 4471.687 | -0.12022 | 0.157265 | 0.585553 | Hnrnpa1 |
| ENSMUSG( | 9692.577 | 0.052645 | 0.56344 | 0.889914 | Hnrnpa2b1 |
| ENSMUSG( | 4844.925 | 0.020999 | 0.833221 | 0.970333 | Hnrnpa3 |
| ENSMUSG( | 2994.977 | 0.032944 | 0.739087 | 0.947768 | Hnrnpab |
| ENSMUSG( | 4669.332 | 0.084854 | 0.419602 | 0.822309 | Hnrnpc |
| ENSMUSG( | 3369.077 | -0.03281 | 0.736082 | 0.946267 | Hnrnpd |
| ENSMUSG( | 4019.575 | -0.16599 | 0.058204 | 0.37191 | Hnrnpdl |
| ENSMUSG( | 1317.516 | 0.099642 | 0.303853 | 0.752148 | Hnrnpf |
| ENSMUSG( | 4381.329 | 0.003337 | 0.967668 | 0.996228 | Hnrnph1 |
| ENSMUSG( | 2057.664 | -0.06374 | 0.550175 | 0.886038 | Hnrnph2 |
| ENSMUSG( | 1537.78 | 0.104496 | 0.352149 | 0.783738 | Hnrnph3 |
| ENSMUSG( | 7740.021 | -0.01446 | 0.868649 | 0.976897 | Hnrnpk |
| ENSMUSG( | 6323.542 | -0.1853 | 0.033327 | 0.279795 | Hnrnpl |
| ENSMUSG( | 1329.35 | -0.12505 | 0.207077 | 0.649503 | Hnrnpll |
| ENSMUSG( | 2992.264 | 0.126477 | 0.199553 | 0.640054 | Hnrnpm |
| ENSMUSG( | 1988.36 | 0.044857 | 0.636741 | 0.91503 | Hnrnpr |
| ENSMUSG( | 10531.93 | -0.05096 | 0.534381 | 0.880719 | Hnrnpu |
| ENSMUSG( | 5935.06 | 0.054712 | 0.487716 | 0.858824 | Hnrnpul1 |
| ENSMUSG( | 5925.759 | 0.09359 | 0.304703 | 0.752574 | Hnrnpul2 |
| ENSMUSG( | 107.5009 | 0.135425 | 0.578922 | 0.896692 | Hoga1 |
| ENSMUSG( | 2649.001 | 0.038956 | 0.67211 | 0.9288 | Homer1 |
| ENSMUSG( | 3790.19 | 0.236067 | 0.038473 | 0.302696 | Homer2 |
| ENSMUSG( | 1005.779 | 0.047145 | 0.624098 | 0.91187 | Homer3 |
| ENSMUSG( | 336.1848 | 0.077721 | 0.565541 | 0.890871 | Homez |
| ENSMUSG( | 1188.075 | -0.05915 | 0.581931 | 0.897918 | Hook1 |
| ENSMUSG( | 412.0115 | 0.092659 | 0.605314 | 0.905737 | Hook2 |
| ENSMUSG( | 1583.957 | -0.0009 | 0.993037 | 0.999327 | Hook3 |
| ENSMUSG( | 5058.067 | 0.155256 | 0.233497 | 0.681508 | Hopx |
| ENSMUSG( | 23.53597 | -1.20244 | 0.029574 | 0.260563 | Hp |
| ENSMUSG( | 6341.812 | -0.02125 | 0.782355 | 0.957558 | Hp1bp3 |
| ENSMUSG( | 14697.07 | -0.05564 | 0.653869 | 0.922914 | Hpca |
| ENSMUSG( | 913.1574 | 0.123876 | 0.361763 | 0.792797 | Hpcal1 |
| ENSMUSG( | 64342.34 | 0.272406 | 0.003537 | 0.075287 | Hpcal4 |
| ENSMUSG( | 41.27794 | -0.23029 | 0.500593 | 0.863514 | Hpdl |
| ENSMUSG( | 1215.398 | 0.036839 | 0.739894 | 0.947823 | Hpf1 |
| ENSMUSG( | 286.8051 | 0.22519 | 0.180729 | 0.617841 | Hpgd |
| ENSMUSG( | 462.8031 | -0.32431 | 0.027314 | 0.253597 | Hpgds |
| ENSMUSG( | 35.97376 | 0.196447 | 0.546683 | 0.884974 | Hpn |
| ENSMUSG( | 1547.692 | -0.09875 | 0.416745 | 0.8209 | Hprt1 |
| ENSMUSG( | 529.9705 | 0.149231 | 0.207919 | 0.650048 | Hps1 |
| ENSMUSG( | 475.19 | 0.020479 | 0.860641 | 0.9749 | Hps3 |
| ENSMUSG( | 604.2049 | 0.287694 | 0.036443 | 0.294207 | Hps4 |
| ENSMUSG( | 894.1828 | 0.028115 | 0.79573 | 0.960062 | Hps5 |
| ENSMUSG( | 124.2363 | 0.073944 | 0.72518 | 0.944096 | Hps6 |
| ENSMUSG( | 49.87481 | -0.18953 | 0.496602 | 0.86175 | Hpse |
| ENSMUSG( | 39.78439 | 0.008918 | 0.977008 | 0.997428 | Hpx |
| ENSMUSG( | 2254.271 | 0.088546 | 0.329491 | 0.773311 | Hr |
| ENSMUSG( | 4582.494 | -0.04233 | 0.617619 | 0.910872 | Hras |
| ENSMUSG( | 10.56496 | -0.50696 | 0.390637 | NA | Hrc |
| ENSMUSG( | 24.90314 | -0.02139 | 0.960148 | 0.993991 | Hrct1 |
| ENSMUSG( | 659.1933 | -0.36028 | 0.02557 | 0.246694 | Hrh1 |
| ENSMUSG( | 365.5616 | 0.043666 | 0.795866 | 0.960062 | Hrh2 |
| ENSMUSG( | 3463.722 | -0.15504 | 0.108459 | 0.497337 | Hrh3 |
| ENSMUSG( | 3133.461 | 0.117072 | 0.181148 | 0.618139 | Hrk |
| ENSMUSG( | 16.59087 | -0.19685 | 0.667595 | NA | Hrob |
| ENSMUSG( | 431.8433 | 0.226258 | 0.098822 | 0.480628 | Hs1bp3 |
| ENSMUSG( | 2440.437 | -0.16903 | 0.17806 | 0.615613 | Hs2st1 |
| ENSMUSG( | 761.1972 | -0.46026 | 0.009852 | 0.141792 | Hs3st1 |
| ENSMUSG( | 1593.874 | -0.26818 | 0.042628 | 0.316734 | Hs3st2 |
| ENSMUSG( | 54.37909 | -0.77526 | 0.010306 | 0.144599 | Hs3st3a1 |
| ENSMUSG( | 109.0912 | -0.09391 | 0.669472 | 0.928147 | Hs3st3b1 |
| ENSMUSG( | 2912.007 | 0.080553 | 0.385052 | 0.804048 | Hs3st4 |
| ENSMUSG( | 113.5999 | -0.73323 | 0.000168 | 0.009317 | Hs3st5 |
| ENSMUSG( | 9.11815 | 0.0365 | 0.956914 | NA | Hs3st6 |
| ENSMUSG( | 2745.958 | -0.13149 | 0.294296 | 0.743534 | Hs6st1 |
| ENSMUSG( | 728.372 | -0.3568 | 0.007277 | 0.117913 | Hs6st2 |
| ENSMUSG( | 130.9556 | 0.086013 | 0.688845 | 0.933441 | Hs6st3 |
| ENSMUSG( | 8431.305 | 0.061811 | 0.510928 | 0.870755 | Hsbp1 |
| ENSMUSG( | 22.3723 | -0.16178 | 0.711308 | 0.940028 | Hsbp1l1 |
| ENSMUSG( | 146.2761 | -0.03538 | 0.837513 | 0.971111 | Hscb |
| ENSMUSG( | 2381.036 | 0.00721 | 0.957608 | 0.993874 | Hsd11b1 |
| ENSMUSG( | 1165.392 | 0.103329 | 0.2724 | 0.722422 | Hsd17b10 |
| ENSMUSG( | 2176.335 | -0.13205 | 0.311337 | 0.759267 | Hsd17b11 |
| ENSMUSG( | 4285.179 | -0.23992 | 0.076749 | 0.429306 | Hsd17b12 |
| ENSMUSG( | 3205.576 | 0.158028 | 0.145442 | 0.566269 | Hsd17b4 |
| ENSMUSG( | 723.8279 | -0.22619 | 0.204713 | 0.645625 | Hsd17b7 |
| ENSMUSG( | 21.37351 | -0.25105 | 0.52448 | 0.875844 | Hsd17b8 |
| ENSMUSG( | 15.63043 | 0.062514 | 0.895956 | NA | Hsd3b7 |
| ENSMUSG( | 1345.702 | 0.025464 | 0.778073 | 0.956168 | Hsdl1 |
| ENSMUSG( | 2476.409 | 0.055151 | 0.666188 | 0.927057 | Hsdl2 |
| ENSMUSG( | 1161.939 | 0.195413 | 0.067661 | 0.402445 | Hsf1 |
| ENSMUSG( | 513.0014 | -0.06248 | 0.593592 | 0.90281 | Hsf2 |
| ENSMUSG( | 245.05 | -0.06229 | 0.663802 | 0.926511 | Hsf4 |
| ENSMUSG( | 16.05344 | -0.76826 | 0.099399 | NA | Hsf5 |
| ENSMUSG( | 12449.45 | 0.118203 | 0.19112 | 0.630245 | Hsp90aa1 |
| ENSMUSG( | 37959.74 | 0.105973 | 0.250533 | 0.700403 | Hsp90ab1 |
| ENSMUSG( | 11211.78 | -0.06737 | 0.627103 | 0.912825 | Hsp90b1 |
| ENSMUSG( | 3736.053 | 0.105329 | 0.373617 | 0.800375 | Hspa12a |
| ENSMUSG( | 292.463 | 0.342269 | 0.07892 | 0.43467 | Hspa12b |
| ENSMUSG( | 993.841 | -0.37655 | 0.045447 | 0.327333 | Hspa13 |
| ENSMUSG( | 672.5843 | 0.073398 | 0.521203 | 0.875186 | Hspa14 |
| ENSMUSG( | 79.56711 | -0.10627 | 0.670636 | 0.928683 | Hspa1a |
| ENSMUSG( | 57.64428 | 0.134748 | 0.62353 | 0.911646 | Hspa1b |
| ENSMUSG( | 11.90852 | 0.24447 | 0.649281 | NA | Hspa1l |
| ENSMUSG( | 1200.843 | 0.230833 | 0.048905 | 0.337919 | Hspa2 |
| ENSMUSG( | 5756.622 | 0.219337 | 0.009773 | 0.141149 | Hspa4 |
| ENSMUSG( | 3323.129 | 0.058421 | 0.505274 | 0.866883 | Hspa4l |
| ENSMUSG( | 10017.11 | 0.055216 | 0.715074 | 0.941207 | Hspa5 |
| ENSMUSG( | 27792.59 | 0.179233 | 0.065758 | 0.397094 | Hspa8 |
| ENSMUSG( | 6618.433 | 0.118011 | 0.197169 | 0.636662 | Hspa9 |
| ENSMUSG( | 150.3811 | 0.44596 | 0.069654 | 0.408352 | Hspb1 |
| ENSMUSG( | 18.44705 | 0.175809 | 0.683999 | NA | Hspb2 |
| ENSMUSG( | 35.57813 | 0.039933 | 0.899802 | 0.985633 | Hspb3 |
| ENSMUSG( | 476.7153 | 0.023063 | 0.851625 | 0.973046 | Hspb6 |
| ENSMUSG( | 9.184417 | 0.342666 | 0.586015 | NA | Hspb7 |
| ENSMUSG( | 1377.017 | 0.108931 | 0.341905 | 0.778828 | Hspb8 |
| ENSMUSG( | 117.2449 | -0.23563 | 0.196617 | 0.635942 | Hspbap1 |
| ENSMUSG( | 1757.347 | 0.041704 | 0.716423 | 0.941697 | Hspbp1 |
| ENSMUSG( | 4455.403 | 0.110729 | 0.211049 | 0.654035 | Hspd1 |
| ENSMUSG( | 1384.439 | -0.01688 | 0.866801 | 0.976541 | Hspe1 |
| ENSMUSG( | 15.56225 | 0.174039 | 0.723776 | NA | Hspe1-rs1 |
| ENSMUSG( | 796.7612 | -0.00551 | 0.97348 | 0.99675 | Hspg2 |
| ENSMUSG( | 4700.971 | 0.392933 | 0.000586 | 0.022791 | Hsph1 |
| ENSMUSG( | 273.778 | -0.03269 | 0.814128 | 0.964574 | Htatip2 |
| ENSMUSG( | 2448.832 | 0.141162 | 0.134246 | 0.547472 | Htatsf1 |
| ENSMUSG( | 515.7279 | -0.15811 | 0.301316 | 0.749979 | Htr1a |
| ENSMUSG( | 203.0148 | -0.65585 | 0.000563 | 0.02226 | Htr1b |
| ENSMUSG( | 53.56212 | -1.45454 | 1.39E-05 | 0.001454 | Htr1d |
| ENSMUSG( | 33.94511 | -1.04242 | 0.004305 | 0.085156 | Htr1f |
| ENSMUSG( | 485.1255 | -0.55839 | 0.002098 | 0.053006 | Htr2a |
| ENSMUSG( | 716.279 | -0.48343 | 0.002265 | 0.055678 | Htr2c |
| ENSMUSG( | 265.0423 | -0.04217 | 0.787423 | 0.959147 | Htr3a |
| ENSMUSG( | 49.62342 | -0.15035 | 0.57374 | 0.893584 | Htr4 |
| ENSMUSG( | 481.6781 | -0.34942 | 0.049108 | 0.338229 | Htr5a |
| ENSMUSG( | 93.52504 | 0.224199 | 0.368866 | 0.796486 | Htr5b |
| ENSMUSG( | 164.093 | -0.45492 | 0.014847 | 0.179046 | Htr6 |
| ENSMUSG( | 318.0533 | 0.270729 | 0.071608 | 0.413189 | Htr7 |
| ENSMUSG( | 11560.34 | -0.28053 | 0.072759 | 0.416435 | Htra1 |
| ENSMUSG( | 897.733 | 0.015697 | 0.887837 | 0.983434 | Htra2 |
| ENSMUSG( | 606.3647 | -0.27837 | 0.030661 | 0.266555 | Htra3 |
| ENSMUSG( | 35.70555 | -0.2783 | 0.382746 | 0.803227 | Htra4 |
| ENSMUSG( | 4093.77 | 0.108115 | 0.312083 | 0.759715 | Htt |
| ENSMUSG( | 551.5866 | -0.11241 | 0.348418 | 0.78132 | Hunk |
| ENSMUSG( | 1340.875 | 0.083169 | 0.543058 | 0.883579 | Hus1 |
| ENSMUSG( | 12660.87 | 0.040066 | 0.665477 | 0.927054 | Huwe1 |
| ENSMUSG( | 202.6801 | -0.54493 | 0.006092 | 0.10614 | Hvcn1 |
| ENSMUSG( | 177.8665 | 0.151498 | 0.397919 | 0.812161 | Hyal1 |
| ENSMUSG( | 321.4198 | -0.27153 | 0.18864 | 0.627639 | Hyal2 |
| ENSMUSG( | 13.45486 | 0.18896 | 0.734799 | NA | Hyal3 |
| ENSMUSG( | 938.6535 | 0.105325 | 0.404444 | 0.816132 | Hycc1 |
| ENSMUSG( | 1127.62 | -0.13816 | 0.157776 | 0.585927 | Hycc2 |
| ENSMUSG( | 50.9966 | -0.37374 | 0.282143 | 0.73186 | Hydin |
| ENSMUSG( | 313.5954 | 0.516118 | 0.000252 | 0.012746 | Hyi |

| | | | | | |
|---|---|---|---|---|---|
| ENSMUSG( | 37.20992 | -0.28445 | 0.437079 | 0.920144 | Hpdl |
| ENSMUSG( | 1040.32 | -0.15159 | 0.239716 | 0.874882 | Hpf1 |
| ENSMUSG( | 277.674 | 0.374463 | 0.050873 | 0.709741 | Hpgd |
| ENSMUSG( | 489.935 | 0.134057 | 0.475174 | 0.923329 | Hpgds |
| ENSMUSG( | 32.92952 | 0.224559 | 0.540213 | 0.937383 | Hpn |
| ENSMUSG( | 1369.146 | -0.19276 | 0.190238 | 0.857355 | Hprt1 |
| ENSMUSG( | 464.758 | 0.044687 | 0.794966 | 0.97681 | Hps1 |
| ENSMUSG( | 453.7639 | 0.157247 | 0.268308 | 0.886493 | Hps3 |
| ENSMUSG( | 537.2437 | 0.228452 | 0.111226 | 0.807885 | Hps4 |
| ENSMUSG( | 794.7819 | -0.04518 | 0.727413 | 0.966887 | Hps5 |
| ENSMUSG( | 104.8249 | -0.13429 | 0.554598 | 0.939565 | Hps6 |
| ENSMUSG( | 50.02762 | 0.119658 | 0.702724 | 0.962215 | Hpse |
| ENSMUSG( | 26.80472 | -1.03545 | 0.011481 | 0.436431 | Hpx |
| ENSMUSG( | 1928.116 | -0.09103 | 0.457795 | 0.923329 | Hr |
| ENSMUSG( | 3977.079 | -0.19038 | 0.112115 | 0.808673 | Hras |
| ENSMUSG( | 10.05143 | -0.27436 | 0.6859 | NA | Hrc |
| ENSMUSG( | 28.118 | 0.570228 | 0.175835 | 0.845715 | Hrct1 |
| ENSMUSG( | 710.8776 | 0.149767 | 0.375022 | 0.915491 | Hrh1 |
| ENSMUSG( | 336.9606 | 0.087217 | 0.638403 | 0.953903 | Hrh2 |
| ENSMUSG( | 4051.994 | 0.525473 | 2.53E-05 | 0.007381 | Hrh3 |
| ENSMUSG( | 2675.188 | -0.07016 | 0.532387 | 0.934795 | Hrk |
| ENSMUSG( | 17.91943 | 0.300907 | 0.510134 | 0.929089 | Hrob |
| ENSMUSG( | 360.6103 | -0.01028 | 0.947565 | 0.994798 | Hs1bp3 |
| ENSMUSG( | 2450.725 | 0.116541 | 0.430696 | 0.917257 | Hs2st1 |
| ENSMUSG( | 872.6517 | 0.237174 | 0.229964 | 0.868507 | Hs3st1 |
| ENSMUSG( | 1643.517 | 0.103344 | 0.445895 | 0.921734 | Hs3st2 |
| ENSMUSG( | 64.70351 | 0.070507 | 0.805119 | 0.977358 | Hs3st3a1 |
| ENSMUSG( | 98.98254 | -0.11375 | 0.643097 | 0.953903 | Hs3st3b1 |
| ENSMUSG( | 2648.748 | 0.077508 | 0.48735 | 0.92541 | Hs3st4 |
| ENSMUSG( | 128.3147 | -0.02024 | 0.920262 | 0.992644 | Hs3st5 |
| ENSMUSG( | 8.896783 | 0.210961 | 0.746693 | NA | Hs3st6 |
| ENSMUSG( | 2873.434 | 0.263143 | 0.069369 | 0.755467 | Hs6st1 |
| ENSMUSG( | 816.0352 | 0.253517 | 0.112044 | 0.808673 | Hs6st2 |
| ENSMUSG( | 134.2259 | 0.42147 | 0.131095 | 0.822884 | Hs6st3 |
| ENSMUSG( | 7129.106 | -0.15974 | 0.220068 | 0.866104 | Hsbp1 |
| ENSMUSG( | 21.00859 | -0.06793 | 0.887161 | 0.990263 | Hsbp1l1 |
| ENSMUSG( | 127.8655 | -0.1762 | 0.402215 | 0.917257 | Hscb |
| ENSMUSG( | 2089.372 | -0.10095 | 0.505074 | 0.927825 | Hsd11b1 |
| ENSMUSG( | 959.2789 | -0.19807 | 0.120245 | 0.813384 | Hsd17b10 |
| ENSMUSG( | 2201.26 | 0.170029 | 0.235949 | 0.87215 | Hsd17b11 |
| ENSMUSG( | 4170.41 | -0.03763 | 0.803175 | 0.976937 | Hsd17b12 |
| ENSMUSG( | 2659.428 | -0.11284 | 0.349227 | 0.911652 | Hsd17b4 |
| ENSMUSG( | 710.5345 | 0.004076 | 0.983025 | 0.996891 | Hsd17b7 |
| ENSMUSG( | 17.93209 | -0.49439 | 0.284538 | 0.892412 | Hsd17b8 |
| ENSMUSG( | 15.58613 | 0.352385 | 0.50215 | 0.927825 | Hsd3b7 |
| ENSMUSG( | 1216.166 | 0.002732 | 0.979872 | 0.996479 | Hsdl1 |
| ENSMUSG( | 1993.796 | -0.32244 | 0.025204 | 0.597422 | Hsdl2 |
| ENSMUSG( | 1021.896 | 0.099292 | 0.461653 | 0.923329 | Hsf1 |
| ENSMUSG( | 492.4269 | 0.095637 | 0.514884 | 0.930939 | Hsf2 |
| ENSMUSG( | 10.42664 | 0.667398 | 0.273239 | NA | Hsf2bp |
| ENSMUSG( | 216.5521 | -0.15722 | 0.392253 | 0.917257 | Hsf4 |
| ENSMUSG( | 18.8339 | 0.088845 | 0.843728 | 0.983462 | Hsf5 |
| ENSMUSG( | 10361.1 | -0.14614 | 0.235729 | 0.871888 | Hsp90aa1 |
| ENSMUSG( | 32028.6 | -0.1172 | 0.337924 | 0.907972 | Hsp90ab1 |
| ENSMUSG( | 11119.14 | 0.176291 | 0.256062 | 0.883082 | Hsp90b1 |
| ENSMUSG( | 3216.847 | -0.05768 | 0.657553 | 0.955772 | Hspa12a |
| ENSMUSG( | 346.7641 | 0.975107 | 3.34E-05 | 0.009128 | Hspa12b |
| ENSMUSG( | 1198.379 | 0.429747 | 0.038683 | 0.679972 | Hspa13 |
| ENSMUSG( | 580.5849 | -0.08633 | 0.526057 | 0.934374 | Hspa14 |
| ENSMUSG( | 85.32079 | 0.32553 | 0.233528 | 0.868507 | Hspa1a |
| ENSMUSG( | 63.42745 | 0.599249 | 0.043279 | 0.690304 | Hspa1b |
| ENSMUSG( | 11.43919 | 0.343321 | 0.569948 | NA | Hspa1l |
| ENSMUSG( | 1120.108 | 0.29302 | 0.021836 | 0.575341 | Hspa2 |
| ENSMUSG( | 4729.977 | -0.07344 | 0.501554 | 0.927825 | Hspa4 |
| ENSMUSG( | 2846.239 | -0.12385 | 0.243073 | 0.877399 | Hspa4l |
| ENSMUSG( | 10629.58 | 0.462381 | 0.003403 | 0.233777 | Hspa5 |
| ENSMUSG( | 23727.6 | -0.00338 | 0.977581 | 0.996416 | Hspa8 |
| ENSMUSG( | 5484.681 | -0.1587 | 0.176835 | 0.846973 | Hspa9 |
| ENSMUSG( | 168.0519 | 0.938511 | 0.000442 | 0.061321 | Hspb1 |
| ENSMUSG( | 14.48677 | -0.26134 | 0.605146 | 0.949082 | Hspb2 |
| ENSMUSG( | 28.76995 | -0.29409 | 0.453325 | 0.922316 | Hspb3 |
| ENSMUSG( | 396.077 | -0.26308 | 0.082957 | 0.77681 | Hspb6 |
| ENSMUSG( | 1126.421 | -0.20984 | 0.091217 | 0.791889 | Hspb8 |
| ENSMUSG( | 103.6161 | -0.31154 | 0.189993 | 0.857355 | Hspbap1 |
| ENSMUSG( | 1485.889 | -0.18238 | 0.171342 | 0.842999 | Hspbp1 |
| ENSMUSG( | 3791.631 | -0.08734 | 0.470047 | 0.923329 | Hspd1 |
| ENSMUSG( | 1186.681 | -0.20487 | 0.120401 | 0.813384 | Hspe1 |
| ENSMUSG( | 13.58306 | 0.050109 | 0.926411 | 0.99306 | Hspe1-rs1 |
| ENSMUSG( | 815.1301 | 0.320527 | 0.10497 | 0.804969 | Hspg2 |
| ENSMUSG( | 3723.821 | 0.015019 | 0.908182 | 0.991958 | Hsph1 |
| ENSMUSG( | 241.8257 | -0.11935 | 0.451968 | 0.922316 | Htatip2 |
| ENSMUSG( | 2054.907 | -0.09737 | 0.412324 | 0.917257 | Htatsf1 |
| ENSMUSG( | 547.8279 | 0.282202 | 0.104426 | 0.804969 | Htr1a |
| ENSMUSG( | 260.6188 | 0.380786 | 0.031819 | 0.631941 | Htr1b |
| ENSMUSG( | 58.70286 | -0.65824 | 0.021892 | 0.575341 | Htr1d |
| ENSMUSG( | 43.18523 | 0.106467 | 0.764079 | 0.974 | Htr1f |
| ENSMUSG( | 607.346 | 0.392368 | 0.050084 | 0.708648 | Htr2a |
| ENSMUSG( | 847.9666 | 0.296276 | 0.045824 | 0.695584 | Htr2c |
| ENSMUSG( | 274.1245 | 0.313397 | 0.076986 | 0.770558 | Htr3a |
| ENSMUSG( | 54.72685 | 0.409292 | 0.1851 | 0.855288 | Htr4 |
| ENSMUSG( | 542.1245 | 0.277023 | 0.164975 | 0.838743 | Htr5a |
| ENSMUSG( | 100.4462 | 0.646324 | 0.011203 | 0.431317 | Htr5b |
| ENSMUSG( | 179.108 | 0.10928 | 0.588456 | 0.944975 | Htr6 |
| ENSMUSG( | 292.5935 | 0.297968 | 0.062178 | 0.736482 | Htr7 |
| ENSMUSG( | 11839.35 | 0.074403 | 0.672959 | 0.95694 | Htra1 |
| ENSMUSG( | 808.8947 | -0.01294 | 0.925828 | 0.99306 | Htra2 |
| ENSMUSG( | 532.8777 | -0.38956 | 0.008857 | 0.394786 | Htra3 |
| ENSMUSG( | 41.64174 | 0.400579 | 0.212719 | 0.864412 | Htra4 |
| ENSMUSG( | 3667.54 | 0.063718 | 0.634566 | 0.953903 | Htt |
| ENSMUSG( | 467.2525 | -0.33317 | 0.014559 | 0.484847 | Hunk |
| ENSMUSG( | 1115.499 | -0.1859 | 0.183927 | 0.852278 | Hus1 |
| ENSMUSG( | 11679.39 | 0.076898 | 0.485576 | 0.92541 | Huwe1 |
| ENSMUSG( | 212.6693 | -0.07712 | 0.657837 | 0.955976 | Hvcn1 |
| ENSMUSG( | 160.8017 | 0.148376 | 0.499367 | 0.927825 | Hyal1 |
| ENSMUSG( | 343.4586 | 0.206656 | 0.393949 | 0.917257 | Hyal2 |
| ENSMUSG( | 13.30793 | 0.40085 | 0.494908 | 0.927372 | Hyal3 |
| ENSMUSG( | 856.1418 | 0.107864 | 0.416468 | 0.917257 | Hycc1 |
| ENSMUSG( | 1156.507 | 0.19576 | 0.117737 | 0.813384 | Hycc2 |
| ENSMUSG( | 65.06676 | 0.587438 | 0.072995 | 0.763629 | Hydin |
| ENSMUSG( | 228.5377 | -0.08654 | 0.612316 | 0.951071 | Hyi |
| ENSMUSG( | 95.07399 | -0.05015 | 0.827069 | 0.980159 | Hykk |
| ENSMUSG( | 90.94001 | -0.1379 | 0.539294 | 0.937216 | Hyls1 |
| ENSMUSG( | 3689.329 | 0.246143 | 0.121549 | 0.813384 | Hyou1 |
| ENSMUSG( | 94.09008 | 0.091513 | 0.69529 | 0.960867 | Hypk |
| ENSMUSG( | 9.664522 | 0.038239 | 0.952086 | NA | I730030J21Rik |
| ENSMUSG( | 84.21877 | 0.046504 | 0.869281 | 0.988097 | I830077J02Rik |
| ENSMUSG( | 302.7038 | -0.02595 | 0.869943 | 0.988097 | Iah1 |
| ENSMUSG( | 1289.737 | 0.065407 | 0.583402 | 0.944266 | Iars1 |
| ENSMUSG( | 2161.733 | 0.003971 | 0.972698 | 0.996416 | Iars2 |
| ENSMUSG( | 208.7561 | -0.27079 | 0.098487 | 0.798438 | Iba57 |
| ENSMUSG( | 16.28083 | 1.777281 | 0.003663 | 0.242109 | Ibsp |
| ENSMUSG( | 640.4967 | 0.142688 | 0.245976 | 0.877904 | Ibtk |
| ENSMUSG( | 947.2563 | -0.06059 | 0.712714 | 0.96439 | Ica1 |
| ENSMUSG( | 871.0549 | -0.0306 | 0.79854 | 0.97681 | Ica1l |
| ENSMUSG( | 215.8225 | 0.639647 | 0.001032 | 0.110223 | Icam1 |
| ENSMUSG( | 123.0051 | 0.220254 | 0.390306 | 0.917257 | Icam2 |
| ENSMUSG( | 13.41181 | -0.78596 | 0.181961 | 0.850579 | Icam4 |
| ENSMUSG( | 7710.77 | 0.252709 | 0.051114 | 0.711023 | Icam5 |
| ENSMUSG( | 4277.686 | -0.0607 | 0.637742 | 0.953903 | Ice1 |
| ENSMUSG( | 274.6797 | 0.135663 | 0.430586 | 0.917257 | Ice2 |
| ENSMUSG( | 2881.968 | 0.129358 | 0.327714 | 0.906255 | Icmt |
| ENSMUSG( | 297.2891 | 0.14033 | 0.577045 | 0.942912 | Icosl |
| ENSMUSG( | 691.5228 | 0.670981 | 0.000675 | 0.083211 | Id1 |
| ENSMUSG( | 3244.65 | -0.37704 | 0.001655 | 0.150169 | Id2 |
| ENSMUSG( | 1489.199 | 0.578607 | 9.66E-06 | 0.003679 | Id3 |
| ENSMUSG( | 4785.968 | 0.297333 | 0.056386 | 0.723362 | Id4 |
| ENSMUSG( | 2265.302 | -0.09738 | 0.439323 | 0.920721 | Ide |
| ENSMUSG( | 2117.364 | -0.21756 | 0.100975 | 0.801456 | Idh1 |
| ENSMUSG( | 3068.303 | 0.153042 | 0.149094 | 0.836241 | Idh2 |
| ENSMUSG( | 3739.253 | -0.20266 | 0.118979 | 0.813384 | Idh3a |
| ENSMUSG( | 5045.377 | -0.18351 | 0.109003 | 0.804969 | Idh3b |
| ENSMUSG( | 3188.705 | -0.08177 | 0.505174 | 0.927825 | Idh3g |
| ENSMUSG( | 713.1833 | -0.54169 | 0.006515 | 0.337628 | Idi1 |
| ENSMUSG( | 422.7752 | -0.21632 | 0.129308 | 0.820358 | Idnk |
| ENSMUSG( | 58.14041 | -0.57109 | 0.069661 | 0.756038 | Ido1 |
| ENSMUSG( | 40.47589 | -0.40085 | 0.276523 | 0.890889 | Ido2 |
| ENSMUSG( | 21793.93 | 0.134049 | 0.311103 | 0.899422 | Ids |
| ENSMUSG( | 566.8387 | -0.01622 | 0.910279 | 0.99212 | Idua |
| ENSMUSG( | 242.4833 | -0.47839 | 0.081432 | 0.77681 | Ier2 |
| ENSMUSG( | 243.4224 | 0.678519 | 0.000655 | 0.081977 | Ier3 |
| ENSMUSG( | 997.624 | 0.133017 | 0.362544 | 0.914223 | Ier3ip1 |
| ENSMUSG( | 1746.351 | -0.1021 | 0.478199 | 0.923329 | Ier5 |
| ENSMUSG( | 468.7513 | -0.23064 | 0.133551 | 0.822912 | Ier5l |
| ENSMUSG( | 1997.186 | 0.026585 | 0.854668 | 0.985649 | Iffo1 |
| ENSMUSG( | 514.4187 | -0.23795 | 0.079724 | 0.77542 | Iffo2 |
| ENSMUSG( | 43.15615 | -0.16774 | 0.620747 | 0.953074 | Ifi203 |
| ENSMUSG( | 21.26232 | -0.52183 | 0.255747 | 0.883082 | Ifi204 |
| ENSMUSG( | 24.26055 | -0.88546 | 0.103685 | 0.804969 | Ifi207 |

| | | | | | | | | | |
|---|---|---|---|---|---|---|---|---|---|
| ENSMUSG( | Hsp90aa1 | 12449.45 | 0.118203 | 0.19112 | 0.630245 | ENSMUSG( | Hsp90aa1 | 10361.1 | -0.14614 | 0.235729 | 0.871888 |
| ENSMUSG( | Hsp90ab1 | 37959.74 | 0.105973 | 0.250533 | 0.700403 | ENSMUSG( | Hsp90ab1 | 32028.6 | -0.1172 | 0.337924 | 0.907972 |
| ENSMUSG( | Hsp90b1 | 11211.78 | -0.06737 | 0.627103 | 0.912825 | ENSMUSG( | Hsp90b1 | 11119.14 | 0.176291 | 0.256062 | 0.883082 |
| ENSMUSG( | Hspa12a | 3736.053 | 0.105329 | 0.373617 | 0.800375 | ENSMUSG( | Hspa12a | 3216.847 | -0.05768 | 0.657553 | 0.955772 |
| ENSMUSG( | Hspa12b | 292.463 | 0.342269 | 0.07892 | 0.43467 | ENSMUSG( | Hspa12b | 346.7641 | 0.975107 | 3.34E-05 | 0.009128 |
| ENSMUSG( | Hspa13 | 993.841 | -0.37655 | 0.045447 | 0.327333 | ENSMUSG( | Hspa13 | 1198.379 | 0.429747 | 0.038683 | 0.679972 |
| ENSMUSG( | Hspa14 | 672.5843 | 0.073398 | 0.521203 | 0.875186 | ENSMUSG( | Hspa14 | 580.5849 | -0.08633 | 0.526057 | 0.934374 |
| ENSMUSG( | Hspa1a | 79.56711 | -0.10627 | 0.670636 | 0.928683 | ENSMUSG( | Hspa1a | 85.32079 | 0.32553 | 0.233528 | 0.868507 |
| ENSMUSG( | Hspa1b | 57.64428 | 0.134748 | 0.62353 | 0.911646 | ENSMUSG( | Hspa1b | 63.42745 | 0.599249 | 0.043279 | 0.690304 |
| ENSMUSG( | Hspa1l | 11.90852 | 0.24447 | 0.649281 | NA | ENSMUSG( | Hspa1l | 11.43919 | 0.343321 | 0.569948 | NA |
| ENSMUSG( | Hspa2 | 1200.843 | 0.230833 | 0.048905 | 0.337919 | ENSMUSG( | Hspa2 | 1120.108 | 0.29302 | 0.021836 | 0.575341 |
| ENSMUSG( | Hspa4 | 5756.622 | 0.219337 | 0.009773 | 0.141149 | ENSMUSG( | Hspa4 | 4729.977 | -0.07344 | 0.501554 | 0.927825 |
| ENSMUSG( | Hspa4l | 3323.129 | 0.058421 | 0.505274 | 0.866883 | ENSMUSG( | Hspa4l | 2846.239 | -0.12385 | 0.243073 | 0.877399 |
| ENSMUSG( | Hspa5 | 10017.11 | 0.055216 | 0.715074 | 0.941207 | ENSMUSG( | Hspa5 | 10629.58 | 0.462381 | 0.003403 | 0.233777 |
| ENSMUSG( | Hspa8 | 27792.59 | 0.179233 | 0.065758 | 0.397094 | ENSMUSG( | Hspa8 | 23727.6 | -0.00338 | 0.977581 | 0.996416 |
| ENSMUSG( | Hspa9 | 6618.433 | 0.118011 | 0.197169 | 0.636662 | ENSMUSG( | Hspa9 | 5484.681 | -0.1587 | 0.176835 | 0.846973 |
| ENSMUSG( | Hspb1 | 150.3811 | 0.44596 | 0.069654 | 0.408352 | ENSMUSG( | Hspb1 | 168.0519 | 0.938511 | 0.000442 | 0.061321 |
| ENSMUSG( | Hspb2 | 18.44705 | 0.175809 | 0.683999 | NA | ENSMUSG( | Hspb2 | 14.48677 | -0.26134 | 0.605146 | 0.949082 |
| ENSMUSG( | Hspb3 | 35.57813 | 0.039933 | 0.899802 | 0.985633 | ENSMUSG( | Hspb3 | 28.76995 | -0.29409 | 0.453325 | 0.922316 |
| ENSMUSG( | Hspb6 | 476.7153 | 0.023063 | 0.851625 | 0.973046 | ENSMUSG( | Hspb6 | 396.077 | -0.26308 | 0.082957 | 0.77681 |
| ENSMUSG( | Hspb8 | 1377.017 | 0.108931 | 0.341905 | 0.778828 | ENSMUSG( | Hspb8 | 1126.421 | -0.20984 | 0.091217 | 0.791889 |
| ENSMUSG( | Hspbap1 | 117.2449 | -0.23563 | 0.196617 | 0.635942 | ENSMUSG( | Hspbap1 | 103.6161 | -0.31154 | 0.189993 | 0.857355 |
| ENSMUSG( | Hspbp1 | 1757.347 | 0.041704 | 0.716423 | 0.941697 | ENSMUSG( | Hspbp1 | 1485.889 | -0.18238 | 0.171342 | 0.842999 |
| ENSMUSG( | Hspd1 | 4455.403 | 0.110729 | 0.211049 | 0.654035 | ENSMUSG( | Hspd1 | 3791.631 | -0.08734 | 0.470047 | 0.923329 |
| ENSMUSG( | Hspe1 | 1384.439 | -0.01688 | 0.866801 | 0.976541 | ENSMUSG( | Hspe1 | 1186.681 | -0.20487 | 0.120401 | 0.813384 |
| ENSMUSG( | Hspe1-rs1 | 15.56225 | 0.174039 | 0.723776 | NA | ENSMUSG( | Hspe1-rs1 | 13.58306 | 0.050109 | 0.926411 | 0.99306 |
| ENSMUSG( | Hspg2 | 796.7612 | -0.00551 | 0.97348 | 0.99675 | ENSMUSG( | Hspg2 | 815.1301 | 0.320527 | 0.10497 | 0.804969 |
| ENSMUSG( | Hsph1 | 4700.971 | 0.392933 | 0.000586 | 0.022791 | ENSMUSG( | Hsph1 | 3723.821 | 0.015019 | 0.908182 | 0.991958 |
| ENSMUSG( | Htatip2 | 273.778 | -0.03269 | 0.814128 | 0.964574 | ENSMUSG( | Htatip2 | 241.8257 | -0.11935 | 0.451968 | 0.922316 |
| ENSMUSG( | Htatsf1 | 2448.832 | 0.141162 | 0.134246 | 0.547472 | ENSMUSG( | Htatsf1 | 2054.907 | -0.09737 | 0.412324 | 0.917257 |
| ENSMUSG( | Htr1a | 515.7279 | -0.15811 | 0.301316 | 0.749979 | ENSMUSG( | Htr1a | 547.8279 | 0.282202 | 0.104426 | 0.804969 |
| ENSMUSG( | Htr1b | 203.0148 | -0.65585 | 0.000563 | 0.02226 | ENSMUSG( | Htr1b | 260.6188 | 0.380786 | 0.031819 | 0.631941 |
| ENSMUSG( | Htr1d | 53.56212 | -1.45454 | 1.39E-05 | 0.001454 | ENSMUSG( | Htr1d | 58.70286 | -0.65824 | 0.021892 | 0.575341 |
| ENSMUSG( | Htr1f | 33.94511 | -1.04242 | 0.004305 | 0.085156 | ENSMUSG( | Htr1f | 43.18523 | 0.106467 | 0.764079 | 0.974 |
| ENSMUSG( | Htr2a | 485.1255 | -0.55839 | 0.002098 | 0.053006 | ENSMUSG( | Htr2a | 607.346 | 0.392368 | 0.050084 | 0.708648 |
| ENSMUSG( | Htr2c | 716.279 | -0.48343 | 0.002265 | 0.055678 | ENSMUSG( | Htr2c | 847.9666 | 0.296276 | 0.045824 | 0.695584 |
| ENSMUSG( | Htr3a | 265.0423 | -0.04217 | 0.787423 | 0.959147 | ENSMUSG( | Htr3a | 274.1245 | 0.313397 | 0.076986 | 0.770558 |
| ENSMUSG( | Htr4 | 49.62342 | -0.15035 | 0.57374 | 0.893584 | ENSMUSG( | Htr4 | 54.72685 | 0.409292 | 0.1851 | 0.855288 |
| ENSMUSG( | Htr5a | 481.6781 | -0.34942 | 0.049108 | 0.338229 | ENSMUSG( | Htr5a | 542.1245 | 0.277023 | 0.164975 | 0.838743 |
| ENSMUSG( | Htr5b | 93.52504 | 0.224199 | 0.368866 | 0.796486 | ENSMUSG( | Htr5b | 100.4462 | 0.646324 | 0.011203 | 0.431317 |
| ENSMUSG( | Htr6 | 164.093 | -0.45492 | 0.014847 | 0.179046 | ENSMUSG( | Htr6 | 179.108 | 0.10928 | 0.588456 | 0.944975 |
| ENSMUSG( | Htr7 | 318.0533 | 0.270729 | 0.071608 | 0.413189 | ENSMUSG( | Htr7 | 292.5935 | 0.297968 | 0.062178 | 0.736482 |
| ENSMUSG( | Htra1 | 11560.34 | -0.28053 | 0.072759 | 0.416435 | ENSMUSG( | Htra1 | 11839.35 | 0.074403 | 0.672959 | 0.95694 |
| ENSMUSG( | Htra2 | 897.733 | 0.015697 | 0.887837 | 0.983434 | ENSMUSG( | Htra2 | 808.8947 | -0.01294 | 0.925828 | 0.99306 |
| ENSMUSG( | Htra3 | 606.3647 | -0.27837 | 0.030661 | 0.266555 | ENSMUSG( | Htra3 | 532.8777 | -0.38956 | 0.008857 | 0.394786 |
| ENSMUSG( | Htra4 | 35.70555 | -0.2783 | 0.382746 | 0.803227 | ENSMUSG( | Htra4 | 41.64174 | 0.400579 | 0.212719 | 0.864412 |
| ENSMUSG( | Htt | 4093.77 | 0.108115 | 0.312083 | 0.759715 | ENSMUSG( | Htt | 3667.54 | 0.063718 | 0.634566 | 0.953903 |
| ENSMUSG( | Hunk | 551.5866 | -0.11241 | 0.348418 | 0.78132 | ENSMUSG( | Hunk | 467.2525 | -0.33317 | 0.014559 | 0.484847 |
| ENSMUSG( | Hus1 | 1340.875 | 0.083169 | 0.543058 | 0.883579 | ENSMUSG( | Hus1 | 1115.499 | -0.1859 | 0.183927 | 0.852278 |
| ENSMUSG( | Huwe1 | 12660.87 | 0.040066 | 0.665477 | 0.927054 | ENSMUSG( | Huwe1 | 11679.39 | 0.076898 | 0.485576 | 0.92541 |
| ENSMUSG( | Hvcn1 | 202.6801 | -0.54493 | 0.006092 | 0.10614 | ENSMUSG( | Hvcn1 | 212.6693 | -0.07712 | 0.657837 | 0.955976 |
| ENSMUSG( | Hyal1 | 177.8665 | 0.151498 | 0.397919 | 0.812161 | ENSMUSG( | Hyal1 | 160.8017 | 0.148376 | 0.499367 | 0.927825 |
| ENSMUSG( | Hyal2 | 321.4198 | -0.27153 | 0.18864 | 0.627639 | ENSMUSG( | Hyal2 | 343.4586 | 0.206656 | 0.393949 | 0.917257 |
| ENSMUSG( | Hyal3 | 13.45486 | 0.18896 | 0.734799 | NA | ENSMUSG( | Hyal3 | 13.30793 | 0.40085 | 0.494908 | 0.927372 |
| ENSMUSG( | Hycc1 | 938.6535 | 0.105325 | 0.404444 | 0.816132 | ENSMUSG( | Hycc1 | 856.1418 | 0.107864 | 0.416468 | 0.917257 |
| ENSMUSG( | Hycc2 | 1127.62 | -0.13816 | 0.157776 | 0.585927 | ENSMUSG( | Hycc2 | 1156.507 | 0.19576 | 0.117737 | 0.813384 |
| ENSMUSG( | Hydin | 50.9966 | -0.37374 | 0.282143 | 0.73186 | ENSMUSG( | Hydin | 65.06676 | 0.587438 | 0.072995 | 0.763629 |
| ENSMUSG( | Hyi | 313.5954 | 0.516118 | 0.000252 | 0.012746 | ENSMUSG( | Hyi | 228.5377 | -0.08654 | 0.612316 | 0.951071 |
| ENSMUSG( | Hykk | 109.2391 | 0.081219 | 0.699549 | 0.937045 | ENSMUSG( | Hykk | 95.07399 | -0.05015 | 0.827069 | 0.980159 |
| ENSMUSG( | Hyls1 | 102.9013 | -0.04849 | 0.81693 | 0.96566 | ENSMUSG( | Hyls1 | 90.94001 | -0.1379 | 0.539294 | 0.937216 |
| ENSMUSG( | Hyou1 | 3461.146 | -0.20885 | 0.153906 | 0.580901 | ENSMUSG( | Hyou1 | 3689.329 | 0.246143 | 0.121549 | 0.813384 |
| ENSMUSG( | Hypk | 91.72155 | -0.278 | 0.196708 | 0.63596 | ENSMUSG( | Hypk | 94.09008 | 0.091513 | 0.69529 | 0.960867 |
| ENSMUSG( | I730030J21 | 9.808822 | -0.22411 | 0.703162 | NA | ENSMUSG( | I730030J21 | 9.664522 | 0.038239 | 0.952086 | NA |
| ENSMUSG( | I830077J02 | 77.1182 | -0.55453 | 0.023403 | 0.233882 | ENSMUSG( | I830077J02 | 84.21877 | 0.046504 | 0.869281 | 0.988097 |
| ENSMUSG( | Iah1 | 337.3966 | 0.013222 | 0.927718 | 0.991229 | ENSMUSG( | Iah1 | 302.7038 | -0.02595 | 0.869943 | 0.988097 |
| ENSMUSG( | Iars1 | 1410.015 | 0.050864 | 0.608933 | 0.907377 | ENSMUSG( | Iars1 | 1289.737 | 0.065407 | 0.583402 | 0.944266 |
| ENSMUSG( | Iars2 | 2427.779 | 0.066825 | 0.453876 | 0.836798 | ENSMUSG( | Iars2 | 2161.733 | 0.003971 | 0.972698 | 0.996416 |
| ENSMUSG( | Iba57 | 251.7483 | 0.016668 | 0.905849 | 0.986695 | ENSMUSG( | Iba57 | 208.7561 | -0.27079 | 0.098487 | 0.798438 |
| ENSMUSG( | Ibtk | 682.7056 | 0.05946 | 0.578025 | 0.89624 | ENSMUSG( | Ibtk | 640.4967 | 0.142688 | 0.245976 | 0.877904 |
| ENSMUSG( | Ica1 | 1096.06 | 0.09432 | 0.550408 | 0.886038 | ENSMUSG( | Ica1 | 947.2563 | -0.06059 | 0.712714 | 0.96439 |
| ENSMUSG( | Ica1l | 1037.463 | 0.199878 | 0.03869 | 0.303018 | ENSMUSG( | Ica1l | 871.0549 | -0.0306 | 0.79854 | 0.97681 |
| ENSMUSG( | Icam1 | 164.322 | -0.37856 | 0.071472 | 0.413144 | ENSMUSG( | Icam1 | 215.8225 | 0.639647 | 0.001032 | 0.110223 |
| ENSMUSG( | Icam2 | 114.2274 | -0.28735 | 0.248212 | 0.696902 | ENSMUSG( | Icam2 | 123.0051 | 0.220254 | 0.390306 | 0.917257 |
| ENSMUSG( | Icam4 | 14.5299 | -0.8893 | 0.09011 | NA | ENSMUSG( | Icam4 | 13.41181 | -0.78596 | 0.181961 | 0.850579 |
| ENSMUSG( | Icam5 | 7600.129 | -0.0501 | 0.657542 | 0.92425 | ENSMUSG( | Icam5 | 7710.77 | 0.252709 | 0.051114 | 0.711023 |
| ENSMUSG( | Ice1 | 4555.327 | -0.14974 | 0.131162 | 0.543132 | ENSMUSG( | Ice1 | 4277.686 | -0.0607 | 0.637742 | 0.953903 |
| ENSMUSG( | Ice2 | 290.8511 | 0.025189 | 0.845006 | 0.972023 | ENSMUSG( | Ice2 | 274.6797 | 0.135663 | 0.430586 | 0.917257 |
| ENSMUSG( | Icmt | 2900.806 | -0.12343 | 0.254979 | 0.705501 | ENSMUSG( | Icmt | 2881.968 | 0.129358 | 0.327714 | 0.906255 |
| ENSMUSG( | Icosl | 274.1364 | -0.40053 | 0.083612 | 0.445968 | ENSMUSG( | Icosl | 297.2891 | 0.14033 | 0.577045 | 0.942912 |
| ENSMUSG( | Id1 | 624.9141 | 0.173014 | 0.310562 | 0.758631 | ENSMUSG( | Id1 | 691.5228 | 0.670981 | 0.000675 | 0.083211 |
| ENSMUSG( | Id2 | 3858.612 | -0.12155 | 0.183314 | 0.62047 | ENSMUSG( | Id2 | 3244.65 | -0.37704 | 0.001655 | 0.150169 |
| ENSMUSG( | Id3 | 1342.091 | 0.063215 | 0.484424 | 0.856665 | ENSMUSG( | Id3 | 1489.199 | 0.578607 | 9.66E-06 | 0.003679 |
| ENSMUSG( | Id4 | 4977.48 | 0.155713 | 0.132202 | 0.544244 | ENSMUSG( | Id4 | 4785.968 | 0.297333 | 0.056386 | 0.723362 |
| ENSMUSG( | Ide | 2704.91 | 0.145474 | 0.154609 | 0.581712 | ENSMUSG( | Ide | 2265.302 | -0.09738 | 0.439323 | 0.920721 |
| ENSMUSG( | Idh1 | 2584.45 | 0.098173 | 0.382171 | 0.803163 | ENSMUSG( | Idh1 | 2117.364 | -0.21756 | 0.100975 | 0.801456 |
| ENSMUSG( | Idh2 | 3505.223 | 0.261783 | 0.001816 | 0.04829 | ENSMUSG( | Idh2 | 3068.303 | 0.153042 | 0.149094 | 0.836241 |
| ENSMUSG( | Idh3a | 4394.262 | 0.004112 | 0.964723 | 0.995155 | ENSMUSG( | Idh3a | 3739.253 | -0.20266 | 0.118979 | 0.813384 |
| ENSMUSG( | Idh3b | 6012.45 | 0.060914 | 0.483691 | 0.85636 | ENSMUSG( | Idh3b | 5045.377 | -0.18351 | 0.109003 | 0.804969 |
| ENSMUSG( | Idh3g | 3786.381 | 0.144884 | 0.124041 | 0.531559 | ENSMUSG( | Idh3g | 3188.705 | -0.08177 | 0.505174 | 0.927825 |
| ENSMUSG( | Idi1 | 935.8689 | 0.029498 | 0.868822 | 0.976897 | ENSMUSG( | Idi1 | 713.1833 | -0.54169 | 0.006515 | 0.337628 |
| ENSMUSG( | Idnk | 474.6998 | -0.15346 | 0.210303 | 0.652595 | ENSMUSG( | Idnk | 422.7752 | -0.21632 | 0.129308 | 0.820358 |
| ENSMUSG( | Ido1 | 54.56765 | -1.20003 | 0.003253 | 0.071142 | ENSMUSG( | Ido1 | 58.14041 | -0.57109 | 0.069661 | 0.756038 |
| ENSMUSG( | Ido2 | 51.54087 | 0.086387 | 0.747358 | 0.949604 | ENSMUSG( | Ido2 | 40.47589 | -0.40085 | 0.276523 | 0.890889 |
| ENSMUSG( | Ids | 22160.48 | -0.08583 | 0.451647 | 0.835696 | ENSMUSG( | Ids | 21793.93 | 0.134049 | 0.311103 | 0.899422 |
| ENSMUSG( | Idua | 571.618 | -0.27985 | 0.015749 | 0.185786 | ENSMUSG( | Idua | 566.8387 | -0.01622 | 0.910279 | 0.99212 |
| ENSMUSG( | Ier2 | 266.2002 | -0.48538 | 0.031006 | 0.268075 | ENSMUSG( | Ier2 | 242.4833 | -0.47839 | 0.081432 | 0.77681 |
| ENSMUSG( | Ier3 | 184.7654 | -0.33677 | 0.067154 | 0.400611 | ENSMUSG( | Ier3 | 243.4224 | 0.678519 | 0.000655 | 0.081977 |
| ENSMUSG( | Ier3ip1 | 932.1596 | -0.34952 | 0.003623 | 0.076313 | ENSMUSG( | Ier3ip1 | 997.624 | 0.133017 | 0.362544 | 0.914223 |
| ENSMUSG( | Ier5 | 1901.449 | -0.12434 | 0.337089 | 0.777008 | ENSMUSG( | Ier5 | 1746.351 | -0.1021 | 0.478199 | 0.923329 |
| ENSMUSG( | Ier5l | 506.9423 | -0.27356 | 0.030762 | 0.266819 | ENSMUSG( | Ier5l | 468.7513 | -0.23064 | 0.133551 | 0.822912 |
| ENSMUSG( | Iffo1 | 2038.574 | -0.19118 | 0.080492 | 0.438901 | ENSMUSG( | Iffo1 | 1997.186 | 0.026585 | 0.854668 | 0.985649 |
| ENSMUSG( | Iffo2 | 575.6435 | -0.17551 | 0.139006 | 0.555099 | ENSMUSG( | Iffo2 | 514.4187 | -0.23795 | 0.079724 | 0.77542 |
| ENSMUSG( | Ifi203 | 47.89032 | -0.15206 | 0.62966 | 0.913965 | ENSMUSG( | Ifi203 | 43.15615 | -0.16774 | 0.620747 | 0.953074 |
| ENSMUSG( | Ifi204 | 19.89672 | -1.11031 | 0.009928 | NA | ENSMUSG( | Ifi204 | 21.26232 | -0.52183 | 0.255747 | 0.883082 |
| ENSMUSG( | Ifi207 | 26.32455 | -0.87035 | 0.083096 | 0.444826 | ENSMUSG( | Ifi207 | 24.26055 | -0.88546 | 0.103685 | 0.804969 |
| ENSMUSG( | Ifi209 | 25.59806 | -0.59255 | 0.13714 | 0.551825 | ENSMUSG( | Ifi209 | 20.06278 | -1.19311 | 0.014167 | 0.477228 |
| ENSMUSG( | Ifi213 | 11.49345 | -0.72711 | 0.252591 | NA | ENSMUSG( | Ifi213 | 11.37711 | -0.41179 | 0.496311 | NA |
| ENSMUSG( | Ifi27 | 836.5467 | -0.31206 | 0.005188 | 0.096057 | ENSMUSG( | Ifi27 | 764.0111 | -0.30182 | 0.023986 | 0.588226 |
| ENSMUSG( | Ifi27l2a | 36.87281 | -0.50903 | 0.123558 | 0.530552 | ENSMUSG( | Ifi27l2a | 28.25555 | -1.26271 | 0.002675 | 0.204614 |
| ENSMUSG( | Ifi30 | 179.2454 | -0.33355 | 0.094664 | 0.472011 | ENSMUSG( | Ifi30 | 200.1189 | 0.271585 | 0.158884 | 0.837724 |
| ENSMUSG( | Ifi35 | 120.2342 | -0.19084 | 0.355911 | 0.788009 | ENSMUSG( | Ifi35 | 126.0612 | 0.228643 | 0.306778 | 0.898538 |
| ENSMUSG( | Ifi44 | 41.89906 | -0.41446 | 0.216943 | 0.661816 | ENSMUSG( | Ifi44 | 38.64648 | -0.33159 | 0.370674 | 0.915491 |
| ENSMUSG( | Ifi47 | 38.42779 | -0.57243 | 0.073113 | 0.417421 | ENSMUSG( | Ifi47 | 42.58546 | 0.06907 | 0.83375 | 0.981218 |
| ENSMUSG( | Ifih1 | 193.4088 | -0.1934 | 0.280989 | 0.73004 | ENSMUSG( | Ifih1 | 183.4598 | -0.06891 | 0.697843 | 0.960867 |
| ENSMUSG( | Ifit1 | 97.2935 | -0.28798 | 0.181745 | 0.618944 | ENSMUSG( | Ifit1 | 102.9931 | 0.172037 | 0.47424 | 0.923329 |
| ENSMUSG( | Ifit1bl1 | 43.29113 | 0.101185 | 0.734506 | 0.945584 | ENSMUSG( | Ifit1bl1 | 38.9518 | 0.060684 | 0.849073 | 0.985321 |
| ENSMUSG( | Ifit1bl2 | 11.17191 | 0.112339 | 0.83622 | NA | ENSMUSG( | Ifit1bl2 | 10.31403 | 0.142995 | 0.808811 | NA |
| ENSMUSG( | Ifit2 | 298.1492 | -0.43845 | 0.004643 | 0.08986 | ENSMUSG( | Ifit2 | 280.7684 | -0.33817 | 0.043176 | 0.690304 |
| ENSMUSG( | Ifit3 | 298.5508 | -0.31192 | 0.058702 | 0.373994 | ENSMUSG( | Ifit3 | 321.8039 | 0.182734 | 0.264057 | 0.886134 |
| ENSMUSG( | Ifit3b | 178.4486 | -0.70653 | 0.001219 | 0.037795 | ENSMUSG( | Ifit3b | 197.2642 | -0.07084 | 0.686692 | 0.959003 |
| ENSMUSG( | Ifitm1 | 46.82074 | -0.08077 | 0.775644 | 0.955606 | ENSMUSG( | Ifitm1 | 64.59151 | 0.973012 | 0.000245 | 0.041307 |
| ENSMUSG( | Ifitm10 | 399.3063 | 0.030888 | 0.826965 | 0.96865 | ENSMUSG( | Ifitm10 | 355.1203 | -0.03386 | 0.833898 | 0.981218 |
| ENSMUSG( | Ifitm2 | 208.2171 | -0.28829 | 0.097854 | 0.478264 | ENSMUSG( | Ifitm2 | 206.5729 | -0.02441 | 0.899786 | 0.990558 |
| ENSMUSG( | Ifitm3 | 495.1093 | -0.42432 | 0.002038 | 0.051756 | ENSMUSG( | Ifitm3 | 514.5865 | -0.00978 | 0.940445 | 0.994373 |
| ENSMUSG( | Ifitm6 | 16.00331 | -0.77585 | 0.236137 | NA | ENSMUSG( | Ifitm6 | 15.89226 | -0.47363 | 0.460205 | 0.923329 |
| ENSMUSG( | Ifitm7 | 27.47864 | -0.07076 | 0.853688 | 0.973491 | ENSMUSG( | Ifitm7 | 23.518 | -0.22931 | 0.599191 | 0.947045 |
| ENSMUSG( | Ifnar1 | 1719.816 | -0.00266 | 0.985035 | 0.997831 | ENSMUSG( | Ifnar1 | 1764.58 | 0.327773 | 0.062659 | 0.736482 |
| ENSMUSG( | Ifnar2 | 353.3023 | -0.16682 | 0.338996 | 0.777518 | ENSMUSG( | Ifnar2 | 372.7684 | 0.259898 | 0.177312 | 0.847005 |
| ENSMUSG( | Ifngr1 | 896.0289 | -0.29407 | 0.045564 | 0.327592 | ENSMUSG( | Ifngr1 | 1036.414 | 0.389235 | 0.043606 | 0.690304 |
| ENSMUSG( | Ifngr2 | 1811.699 | -0.16649 | 0.182822 | 0.620372 | ENSMUSG( | Ifngr2 | 1783.607 | 0.064354 | 0.646324 | 0.953988 |
| ENSMUSG( | Ifrd1 | 1272.001 | 0.178507 | 0.054785 | 0.360194 | ENSMUSG( | Ifrd1 | 1175.937 | 0.214864 | 0.083885 | 0.777053 |
| ENSMUSG( | Ifrd2 | 272.1281 | 0.182165 | 0.210214 | 0.65257 | ENSMUSG( | Ifrd2 | 259.5987 | 0.305316 | 0.070392 | 0.758336 |
| ENSMUSG( | Ift122 | 1014.614 | 0.047493 | 0.639746 | 0.916434 | ENSMUSG( | Ift122 | 864.9377 | -0.14895 | 0.205934 | 0.864412 |
| ENSMUSG( | Ift140 | 906.9355 | 0.010312 | 0.924301 | 0.99114 | ENSMUSG( | Ift140 | 839.1004 | 0.058108 | 0.670761 | 0.95694 |
| ENSMUSG( | Ift172 | 1831.647 | 0.216237 | 0.020034 | 0.214064 | ENSMUSG( | Ift172 | 1531.009 | -0.02364 | 0.853364 | 0.985598 |
| ENSMUSG( | Ift20 | 1262.377 | 0.049472 | 0.660938 | 0.92497 | ENSMUSG( | Ift20 | 1130.128 | -0.00488 | 0.970686 | 0.99594 |
| ENSMUSG( | Ift22 | 1521.38 | 0.106685 | 0.258512 | 0.709726 | ENSMUSG( | Ift22 | 1250.772 | -0.19494 | 0.088564 | 0.786756 |
| ENSMUSG( | Ift25 | 171.4863 | 0.009423 | 0.953222 | 0.993874 | ENSMUSG( | Ift25 | 142.8603 | -0.27211 | 0.157651 | 0.837724 |
| ENSMUSG( | Ift27 | 1092.545 | 0.09296 | 0.328257 | 0.772854 | ENSMUSG( | Ift27 | 916.5733 | -0.15119 | 0.213473 | 0.864421 |
| ENSMUSG( | Ift43 | 262.5734 | -0.16068 | 0.304488 | 0.752574 | ENSMUSG( | Ift43 | 231.0557 | -0.2694 | 0.141631 | 0.830678 |
| ENSMUSG( | Ift46 | 874.7407 | -0.0355 | 0.726927 | 0.944223 | ENSMUSG( | Ift46 | 758.3538 | -0.18697 | 0.131032 | 0.822884 |
| ENSMUSG( | Ift52 | 674.6783 | -0.01984 | 0.856985 | 0.974366 | ENSMUSG( | Ift52 | 576.7339 | -0.20982 | 0.11256 | 0.809623 |
| ENSMUSG( | Ift56 | 216.1449 | 0.191545 | 0.239541 | 0.687308 | ENSMUSG( | Ift56 | 208.3781 | 0.341371 | 0.059935 | 0.7301 |
| ENSMUSG( | Ift57 | 1255.732 | 0.060226 | 0.53233 | 0.879905 | ENSMUSG( | Ift57 | 1075.236 | -0.12386 | 0.303795 | 0.898538 |
| ENSMUSG( | Ift70a1 | 57.82565 | 0.255866 | 0.346255 | 0.779153 | ENSMUSG( | Ift70a1 | 49.26848 | 0.082953 | 0.794062 | 0.97681 |
| ENSMUSG( | Ift70b | 225.9605 | 0.189803 | 0.180491 | 0.617841 | ENSMUSG( | Ift70b | 205.7671 | 0.189 | 0.259005 | 0.883971 |
| ENSMUSG( | Ift74 | 567.3969 | 0.015911 | 0.897194 | 0.985474 | ENSMUSG( | Ift74 | 495.0686 | -0.119 | 0.415561 | 0.917257 |
| ENSMUSG( | Ift80 | 826.3613 | 0.098501 | 0.390669 | 0.808144 | ENSMUSG( | Ift80 | 693.7132 | -0.14209 | 0.285861 | 0.893906 |
| ENSMUSG( | Ift81 | 612.2493 | 0.109642 | 0.314085 | 0.761319 | ENSMUSG( | Ift81 | 520.291 | -0.09419 | 0.477569 | 0.923329 |
| ENSMUSG( | Ift88 | 359.3083 | 0.059962 | 0.689588 | 0.933609 | ENSMUSG( | Ift88 | 336.893 | 0.141962 | 0.38118 | 0.917202 |
| ENSMUSG( | Iftap | 594.9544 | 0.050238 | 0.704485 | 0.938303 | ENSMUSG( | Iftap | 507.5948 | -0.15101 | 0.339309 | 0.907972 |
| ENSMUSG( | Igbp1 | 1244.395 | 0.144919 | 0.113065 | 0.507167 | ENSMUSG( | Igbp1 | 1081.709 | 0.011536 | 0.922205 | 0.992644 |
| ENSMUSG( | Igdcc3 | 71.87763 | 0.10108 | 0.732121 | 0.944956 | ENSMUSG( | Igdcc3 | 77.43484 | 0.546855 | 0.062128 | 0.736482 |
| ENSMUSG( | Igdcc4 | 4154.558 | -0.02438 | 0.83417 | 0.970333 | ENSMUSG( | Igdcc4 | 3847.559 | 0.023947 | 0.855092 | 0.985649 |
| ENSMUSG( | Igf1 | 225.634 | -0.52229 | 0.002657 | 0.06191 | ENSMUSG( | Igf1 | 268.8416 | 0.277366 | 0.168707 | 0.840222 |
| ENSMUSG( | Igf1r | 2248.333 | 0.232491 | 0.12003 | 0.523983 | ENSMUSG( | Igf1r | 2291.865 | 0.524948 | 0.003681 | 0.242404 |
| ENSMUSG( | Igf2 | 125.5348 | 0.219432 | 0.276248 | 0.72598 | ENSMUSG( | Igf2 | 91.64242 | -0.43875 | 0.063789 | 0.736482 |
| ENSMUSG( | Igf2bp2 | 53.30595 | -0.72165 | 0.014995 | 0.180023 | ENSMUSG( | Igf2bp2 | 46.82913 | -0.88616 | 0.004345 | 0.268942 |

| | | | | | |
|---|---|---|---|---|---|
| ENSMUSG( | 109.2391 | 0.081219 | 0.699549 | 0.937045 | Hykk |
| ENSMUSG( | 102.9013 | -0.04849 | 0.81693 | 0.96566 | Hyls1 |
| ENSMUSG( | 3461.146 | -0.20885 | 0.153906 | 0.580901 | Hyou1 |
| ENSMUSG( | 91.72155 | -0.278 | 0.196708 | 0.63596 | Hypk |
| ENSMUSG( | 9.808822 | -0.22411 | 0.703162 | NA | I730030J21Rik |
| ENSMUSG( | 77.1182 | -0.55453 | 0.023403 | 0.233882 | I830077J02Rik |
| ENSMUSG( | 337.3966 | 0.013222 | 0.927718 | 0.991229 | Iah1 |
| ENSMUSG( | 1410.015 | 0.050864 | 0.608933 | 0.907377 | Iars1 |
| ENSMUSG( | 2427.779 | 0.066825 | 0.453876 | 0.836798 | Iars2 |
| ENSMUSG( | 251.7483 | 0.016668 | 0.905849 | 0.986695 | Iba57 |
| ENSMUSG( | 682.7056 | 0.05946 | 0.578025 | 0.89624 | Ibtk |
| ENSMUSG( | 1096.06 | 0.09432 | 0.550408 | 0.886038 | Ica1 |
| ENSMUSG( | 1037.463 | 0.199878 | 0.03869 | 0.303018 | Ica1l |
| ENSMUSG( | 164.322 | -0.37856 | 0.071472 | 0.413144 | Icam1 |
| ENSMUSG( | 114.2274 | -0.28735 | 0.248212 | 0.696902 | Icam2 |
| ENSMUSG( | 14.5299 | -0.8893 | 0.09011 | NA | Icam4 |
| ENSMUSG( | 7600.129 | -0.0501 | 0.657542 | 0.92425 | Icam5 |
| ENSMUSG( | 4555.327 | -0.14974 | 0.131162 | 0.543132 | Ice1 |
| ENSMUSG( | 290.8511 | 0.025189 | 0.845006 | 0.972023 | Ice2 |
| ENSMUSG( | 2900.806 | -0.12343 | 0.254979 | 0.705501 | Icmt |
| ENSMUSG( | 8.485042 | -0.05992 | 0.931155 | NA | Icos |
| ENSMUSG( | 274.1364 | -0.40053 | 0.083612 | 0.445968 | Icosl |
| ENSMUSG( | 624.9141 | 0.173014 | 0.310562 | 0.758631 | Id1 |
| ENSMUSG( | 3858.612 | -0.12155 | 0.183314 | 0.62047 | Id2 |
| ENSMUSG( | 1342.091 | 0.063215 | 0.484424 | 0.856665 | Id3 |
| ENSMUSG( | 4977.48 | 0.155713 | 0.132202 | 0.544244 | Id4 |
| ENSMUSG( | 2704.91 | 0.145474 | 0.154609 | 0.581712 | Ide |
| ENSMUSG( | 2584.45 | 0.098173 | 0.382171 | 0.803163 | Idh1 |
| ENSMUSG( | 3505.223 | 0.261783 | 0.001816 | 0.04829 | Idh2 |
| ENSMUSG( | 4394.262 | 0.004112 | 0.964723 | 0.995155 | Idh3a |
| ENSMUSG( | 6012.45 | 0.060914 | 0.483691 | 0.85636 | Idh3b |
| ENSMUSG( | 3786.381 | 0.144884 | 0.124041 | 0.531559 | Idh3g |
| ENSMUSG( | 935.8689 | 0.029498 | 0.868822 | 0.976897 | Idi1 |
| ENSMUSG( | 474.6998 | -0.15346 | 0.210303 | 0.652595 | Idnk |
| ENSMUSG( | 54.56765 | -1.20003 | 0.003253 | 0.071142 | Ido1 |
| ENSMUSG( | 51.54087 | 0.086387 | 0.747358 | 0.949604 | Ido2 |
| ENSMUSG( | 22160.48 | -0.08583 | 0.451647 | 0.835696 | Ids |
| ENSMUSG( | 571.618 | -0.27985 | 0.015749 | 0.185786 | Idua |
| ENSMUSG( | 266.2002 | -0.48538 | 0.031006 | 0.268075 | Ier2 |
| ENSMUSG( | 184.7654 | -0.33677 | 0.067154 | 0.400611 | Ier3 |
| ENSMUSG( | 932.1596 | -0.34952 | 0.003623 | 0.076313 | Ier3ip1 |
| ENSMUSG( | 1901.449 | -0.12434 | 0.337089 | 0.777008 | Ier5 |
| ENSMUSG( | 506.9423 | -0.27356 | 0.030762 | 0.266819 | Ier5l |
| ENSMUSG( | 2038.574 | -0.19118 | 0.080492 | 0.438901 | Iffo1 |
| ENSMUSG( | 575.6435 | -0.17551 | 0.139006 | 0.555099 | Iffo2 |
| ENSMUSG( | 47.89032 | -0.15206 | 0.62966 | 0.913965 | Ifi203 |
| ENSMUSG( | 19.89672 | -1.11031 | 0.009928 | NA | Ifi204 |
| ENSMUSG( | 26.32455 | -0.87035 | 0.083096 | 0.444826 | Ifi207 |
| ENSMUSG( | 25.59806 | -0.59255 | 0.13714 | 0.551825 | Ifi209 |
| ENSMUSG( | 11.49345 | -0.72711 | 0.252591 | NA | Ifi213 |
| ENSMUSG( | 836.5467 | -0.31206 | 0.005188 | 0.096057 | Ifi27 |
| ENSMUSG( | 36.87281 | -0.50903 | 0.123558 | 0.530552 | Ifi27l2a |
| ENSMUSG( | 179.2454 | -0.33355 | 0.094664 | 0.472011 | Ifi30 |
| ENSMUSG( | 120.2342 | -0.19084 | 0.355911 | 0.788009 | Ifi35 |
| ENSMUSG( | 41.89906 | -0.41446 | 0.216943 | 0.661816 | Ifi44 |
| ENSMUSG( | 38.42779 | -0.57243 | 0.073113 | 0.417421 | Ifi47 |
| ENSMUSG( | 193.4088 | -0.1934 | 0.280989 | 0.73004 | Ifih1 |
| ENSMUSG( | 97.2935 | -0.28798 | 0.181745 | 0.618944 | Ifit1 |
| ENSMUSG( | 43.29113 | 0.101185 | 0.734506 | 0.945584 | Ifit1bl1 |
| ENSMUSG( | 11.17191 | 0.112339 | 0.83622 | NA | Ifit1bl2 |
| ENSMUSG( | 298.1492 | -0.43845 | 0.004643 | 0.08986 | Ifit2 |
| ENSMUSG( | 298.5508 | -0.31192 | 0.058702 | 0.373994 | Ifit3 |
| ENSMUSG( | 178.4486 | -0.70653 | 0.001219 | 0.037795 | Ifit3b |
| ENSMUSG( | 46.82074 | -0.08077 | 0.775644 | 0.955606 | Ifitm1 |
| ENSMUSG( | 399.3063 | 0.030888 | 0.826965 | 0.96865 | Ifitm10 |
| ENSMUSG( | 208.2171 | -0.28829 | 0.097854 | 0.478264 | Ifitm2 |
| ENSMUSG( | 495.1093 | -0.42432 | 0.002038 | 0.051756 | Ifitm3 |
| ENSMUSG( | 16.00331 | -0.77585 | 0.236137 | NA | Ifitm6 |
| ENSMUSG( | 27.47864 | -0.07076 | 0.853688 | 0.973491 | Ifitm7 |
| ENSMUSG( | 1719.816 | -0.00266 | 0.985035 | 0.997831 | Ifnar1 |
| ENSMUSG( | 353.3023 | -0.16682 | 0.338996 | 0.777518 | Ifnar2 |
| ENSMUSG( | 896.0289 | -0.29407 | 0.045564 | 0.327592 | Ifngr1 |
| ENSMUSG( | 1811.699 | -0.16649 | 0.182822 | 0.620372 | Ifngr2 |
| ENSMUSG( | 1272.001 | 0.178507 | 0.054785 | 0.360194 | Ifrd1 |
| ENSMUSG( | 272.1281 | 0.182165 | 0.210214 | 0.65257 | Ifrd2 |
| ENSMUSG( | 1014.614 | 0.047493 | 0.639746 | 0.916434 | Ift122 |
| ENSMUSG( | 906.9355 | 0.010312 | 0.924301 | 0.99114 | Ift140 |
| ENSMUSG( | 1831.647 | 0.216237 | 0.020034 | 0.214064 | Ift172 |
| ENSMUSG( | 1262.377 | 0.049472 | 0.660938 | 0.92497 | Ift20 |
| ENSMUSG( | 1521.38 | 0.106685 | 0.258512 | 0.709726 | Ift22 |
| ENSMUSG( | 171.4863 | 0.009423 | 0.953222 | 0.993874 | Ift25 |
| ENSMUSG( | 1092.545 | 0.09296 | 0.328257 | 0.772854 | Ift27 |
| ENSMUSG( | 262.5734 | -0.16068 | 0.304488 | 0.752574 | Ift43 |
| ENSMUSG( | 874.7407 | -0.0355 | 0.726927 | 0.944223 | Ift46 |
| ENSMUSG( | 674.6783 | -0.01984 | 0.856985 | 0.974366 | Ift52 |
| ENSMUSG( | 216.1449 | 0.191545 | 0.239541 | 0.687308 | Ift56 |
| ENSMUSG( | 1255.732 | 0.060226 | 0.53233 | 0.879905 | Ift57 |
| ENSMUSG( | 57.82565 | 0.255866 | 0.346255 | 0.779153 | Ift70a1 |
| ENSMUSG( | 225.9605 | 0.189803 | 0.180491 | 0.617841 | Ift70b |
| ENSMUSG( | 567.3969 | 0.015911 | 0.897194 | 0.985474 | Ift74 |
| ENSMUSG( | 826.3613 | 0.098501 | 0.390669 | 0.808144 | Ift80 |
| ENSMUSG( | 612.2493 | 0.109642 | 0.314085 | 0.761319 | Ift81 |
| ENSMUSG( | 359.3083 | 0.059962 | 0.689588 | 0.933609 | Ift88 |
| ENSMUSG( | 594.9544 | 0.050238 | 0.704485 | 0.938303 | Iftap |
| ENSMUSG( | 1244.395 | 0.144919 | 0.113065 | 0.507167 | Igbp1 |
| ENSMUSG( | 71.87763 | 0.10108 | 0.732121 | 0.944956 | Igdcc3 |
| ENSMUSG( | 4154.558 | -0.02438 | 0.83417 | 0.970333 | Igdcc4 |
| ENSMUSG( | 225.634 | -0.52229 | 0.002657 | 0.06191 | Igf1 |
| ENSMUSG( | 2248.333 | 0.232491 | 0.12003 | 0.523983 | Igf1r |
| ENSMUSG( | 125.5348 | 0.219432 | 0.276248 | 0.72598 | Igf2 |
| ENSMUSG( | 53.30595 | -0.72165 | 0.014995 | 0.180023 | Igf2bp2 |
| ENSMUSG( | 12.79128 | -0.11937 | 0.819827 | NA | Igf2os |
| ENSMUSG( | 1255.151 | -0.11362 | 0.413533 | 0.819888 | Igf2r |
| ENSMUSG( | 3642.976 | -0.27944 | 0.008856 | 0.132506 | Igfbp2 |
| ENSMUSG( | 437.788 | 0.311932 | 0.00946 | 0.137493 | Igfbp3 |
| ENSMUSG( | 3105.38 | -0.14316 | 0.209529 | 0.65169 | Igfbp4 |
| ENSMUSG( | 16625.8 | 0.060509 | 0.611484 | 0.908586 | Igfbp5 |
| ENSMUSG( | 1101.598 | -0.25931 | 0.010049 | 0.142735 | Igfbp6 |
| ENSMUSG( | 1416.291 | -0.19588 | 0.253761 | 0.704011 | Igfbp7 |
| ENSMUSG( | 220.7487 | -0.89116 | 0.000638 | 0.024356 | Igfbpl1 |
| ENSMUSG( | 596.783 | -0.08197 | 0.611279 | 0.908586 | Igfn1 |
| ENSMUSG( | 9.27238 | 1.619718 | 0.071274 | NA | Igha |
| ENSMUSG( | 30.25175 | 0.285143 | 0.450298 | 0.835344 | Ighd |
| ENSMUSG( | 105.4026 | 0.605041 | 0.002113 | 0.053164 | Ighg2c |
| ENSMUSG( | 1242.238 | 0.39306 | 0.000196 | 0.01044 | Ighm |
| ENSMUSG( | 297.8598 | -0.03053 | 0.844474 | 0.972023 | Ighmbp2 |
| ENSMUSG( | 2533.635 | 0.029032 | 0.775484 | 0.955606 | Igip |
| ENSMUSG( | 37.34076 | -0.03641 | 0.919202 | 0.990247 | Igkc |
| ENSMUSG( | 1897.61 | -0.18896 | 0.047323 | 0.332697 | Iglon5 |
| ENSMUSG( | 358.0666 | -0.40909 | 0.411764 | 0.81942 | Igsf1 |
| ENSMUSG( | 101.4461 | -0.44101 | 0.028466 | 0.257525 | Igsf10 |
| ENSMUSG( | 1739.152 | -0.11536 | 0.336668 | 0.777 | Igsf11 |
| ENSMUSG( | 3320.257 | -0.05843 | 0.607604 | 0.906878 | Igsf21 |
| ENSMUSG( | 682.8374 | -0.28089 | 0.084096 | 0.447085 | Igsf3 |
| ENSMUSG( | 15.14175 | -0.01354 | 0.976938 | NA | Igsf5 |
| ENSMUSG( | 56.75012 | -0.59251 | 0.047714 | 0.33359 | Igsf6 |
| ENSMUSG( | 8112.033 | -0.38543 | 0.003159 | 0.06974 | Igsf8 |
| ENSMUSG( | 180.7563 | -0.11785 | 0.516529 | 0.872788 | Igsf9 |
| ENSMUSG( | 2063.222 | -0.02368 | 0.835491 | 0.970557 | Igsf9b |
| ENSMUSG( | 212.0091 | -0.27525 | 0.094244 | 0.471647 | Igtp |
| ENSMUSG( | 9.699878 | 0.267806 | 0.666933 | NA | Ihh |
| ENSMUSG( | 13.4514 | 0.020638 | 0.968408 | NA | Iho1 |
| ENSMUSG( | 50.53299 | -0.59234 | 0.073213 | 0.417696 | Iigp1 |
| ENSMUSG( | 9.980008 | -0.8783 | 0.159919 | NA | Iigp1c |
| ENSMUSG( | 2614.259 | 0.103105 | 0.247288 | 0.696236 | Ik |
| ENSMUSG( | 557.6348 | -0.24841 | 0.098597 | 0.480543 | Ikbip |
| ENSMUSG( | 1579.539 | 0.142642 | 0.104234 | 0.49005 | Ikbkb |
| ENSMUSG( | 87.67239 | 0.103887 | 0.618046 | 0.910929 | Ikbke |
| ENSMUSG( | 1041.937 | 0.10654 | 0.275064 | 0.72509 | Ikbkg |
| ENSMUSG( | 254.539 | -0.05748 | 0.721481 | 0.943059 | Ikzf1 |
| ENSMUSG( | 74.1601 | 0.138861 | 0.570333 | 0.893192 | Ikzf2 |
| ENSMUSG( | 423.8883 | 0.15279 | 0.196628 | 0.635942 | Ikzf4 |
| ENSMUSG( | 543.9337 | 0.044142 | 0.702277 | 0.937536 | Ikzf5 |
| ENSMUSG( | 301.4276 | -0.7783 | 0.000126 | 0.007476 | Il10ra |
| ENSMUSG( | 8.950107 | 0.004651 | 0.993953 | NA | Il10rb |
| ENSMUSG( | 11.56175 | 0.484417 | 0.371381 | NA | Il11 |
| ENSMUSG( | 1753.914 | 0.032588 | 0.742629 | 0.948478 | Il11ra1 |
| ENSMUSG( | 40.41078 | -0.15501 | 0.597794 | 0.90372 | Il12a |
| ENSMUSG( | 73.23139 | -0.54959 | 0.037206 | 0.297718 | Il12rb1 |
| ENSMUSG( | 269.0741 | -0.18667 | 0.321785 | 0.766739 | Il13ra1 |
| ENSMUSG( | 11.96383 | -0.00059 | 0.999137 | NA | Il13ra2 |
| ENSMUSG( | 72.88303 | -0.08617 | 0.717912 | 0.941697 | Il15 |
| ENSMUSG( | 121.5908 | -0.07909 | 0.695739 | 0.934979 | Il15ra |
| ENSMUSG( | 198.301 | 0.077933 | 0.621066 | 0.911432 | Il16 |
| ENSMUSG( | 11.99948 | -0.23542 | 0.679765 | NA | Il17b |
| ENSMUSG( | 361.006 | -0.01405 | 0.912604 | 0.988483 | Il17d |
| ENSMUSG( | 944.1485 | -0.23683 | 0.111591 | 0.50362 | Il17ra |
| ENSMUSG( | 39.12014 | -0.46749 | 0.168487 | 0.601952 | Il17rb |

| | | | | | |
|---|---|---|---|---|---|
| ENSMUSG( | 20.06278 | -1.19311 | 0.014167 | 0.477228 | Ifi209 |
| ENSMUSG( | 11.37711 | -0.41179 | 0.496311 | NA | Ifi213 |
| ENSMUSG( | 764.0111 | -0.30182 | 0.023986 | 0.588226 | Ifi27 |
| ENSMUSG( | 28.25555 | -1.26271 | 0.002675 | 0.204614 | Ifi27l2a |
| ENSMUSG( | 200.1189 | 0.271585 | 0.158884 | 0.837724 | Ifi30 |
| ENSMUSG( | 126.0612 | 0.228643 | 0.306778 | 0.898538 | Ifi35 |
| ENSMUSG( | 38.64648 | -0.33159 | 0.370674 | 0.915491 | Ifi44 |
| ENSMUSG( | 42.58546 | 0.06907 | 0.83375 | 0.981218 | Ifi47 |
| ENSMUSG( | 183.4598 | -0.06891 | 0.697843 | 0.960867 | Ifih1 |
| ENSMUSG( | 102.9931 | 0.172037 | 0.47424 | 0.923329 | Ifit1 |
| ENSMUSG( | 38.9518 | 0.060684 | 0.849073 | 0.985321 | Ifit1bl1 |
| ENSMUSG( | 10.31403 | 0.142995 | 0.808811 | NA | Ifit1bl2 |
| ENSMUSG( | 280.7684 | -0.33817 | 0.043176 | 0.690304 | Ifit2 |
| ENSMUSG( | 321.8039 | 0.182734 | 0.264057 | 0.886134 | Ifit3 |
| ENSMUSG( | 197.2642 | -0.07084 | 0.686692 | 0.959003 | Ifit3b |
| ENSMUSG( | 64.59151 | 0.973012 | 0.000245 | 0.041307 | Ifitm1 |
| ENSMUSG( | 355.1203 | -0.03386 | 0.833898 | 0.981218 | Ifitm10 |
| ENSMUSG( | 206.5729 | -0.02441 | 0.899786 | 0.990558 | Ifitm2 |
| ENSMUSG( | 514.5865 | -0.00978 | 0.940445 | 0.994373 | Ifitm3 |
| ENSMUSG( | 15.89226 | -0.47363 | 0.460205 | 0.923329 | Ifitm6 |
| ENSMUSG( | 23.518 | -0.22931 | 0.599191 | 0.947045 | Ifitm7 |
| ENSMUSG( | 1764.58 | 0.327773 | 0.062659 | 0.736482 | Ifnar1 |
| ENSMUSG( | 372.7684 | 0.259898 | 0.177312 | 0.847005 | Ifnar2 |
| ENSMUSG( | 1036.414 | 0.389235 | 0.043606 | 0.690304 | Ifngr1 |
| ENSMUSG( | 1783.607 | 0.064354 | 0.646324 | 0.953988 | Ifngr2 |
| ENSMUSG( | 1175.937 | 0.214864 | 0.083885 | 0.777053 | Ifrd1 |
| ENSMUSG( | 259.5987 | 0.305316 | 0.070392 | 0.758336 | Ifrd2 |
| ENSMUSG( | 864.9377 | -0.14895 | 0.205934 | 0.864412 | Ift122 |
| ENSMUSG( | 839.1004 | 0.058108 | 0.670761 | 0.95694 | Ift140 |
| ENSMUSG( | 1531.009 | -0.02364 | 0.853364 | 0.985598 | Ift172 |
| ENSMUSG( | 1130.128 | -0.00488 | 0.970686 | 0.99594 | Ift20 |
| ENSMUSG( | 1250.772 | -0.19494 | 0.088564 | 0.786756 | Ift22 |
| ENSMUSG( | 142.8603 | -0.27211 | 0.157651 | 0.837724 | Ift25 |
| ENSMUSG( | 916.5733 | -0.15119 | 0.213473 | 0.864421 | Ift27 |
| ENSMUSG( | 231.0557 | -0.2694 | 0.141631 | 0.830678 | Ift43 |
| ENSMUSG( | 758.3538 | -0.18697 | 0.131032 | 0.822884 | Ift46 |
| ENSMUSG( | 576.7339 | -0.20982 | 0.11256 | 0.809623 | Ift52 |
| ENSMUSG( | 208.3781 | 0.341371 | 0.059935 | 0.7301 | Ift56 |
| ENSMUSG( | 1075.236 | -0.12386 | 0.303795 | 0.898538 | Ift57 |
| ENSMUSG( | 49.26848 | 0.082953 | 0.794062 | 0.97681 | Ift70a1 |
| ENSMUSG( | 205.7671 | 0.189 | 0.259005 | 0.883971 | Ift70b |
| ENSMUSG( | 495.0686 | -0.119 | 0.415561 | 0.917257 | Ift74 |
| ENSMUSG( | 693.7132 | -0.14209 | 0.285861 | 0.893906 | Ift80 |
| ENSMUSG( | 520.291 | -0.09419 | 0.477569 | 0.923329 | Ift81 |
| ENSMUSG( | 336.893 | 0.141962 | 0.38118 | 0.917202 | Ift88 |
| ENSMUSG( | 507.5948 | -0.15101 | 0.339309 | 0.907972 | Iftap |
| ENSMUSG( | 1081.709 | 0.011536 | 0.922205 | 0.992644 | Igbp1 |
| ENSMUSG( | 77.43484 | 0.546855 | 0.062128 | 0.736482 | Igdcc3 |
| ENSMUSG( | 3847.559 | 0.023947 | 0.855092 | 0.985649 | Igdcc4 |
| ENSMUSG( | 268.8416 | 0.277366 | 0.168707 | 0.840222 | Igf1 |
| ENSMUSG( | 2291.865 | 0.524948 | 0.003681 | 0.242404 | Igf1r |
| ENSMUSG( | 91.64242 | -0.43875 | 0.063789 | 0.736482 | Igf2 |
| ENSMUSG( | 46.82913 | -0.88616 | 0.004345 | 0.268942 | Igf2bp2 |
| ENSMUSG( | 10.45259 | -0.45309 | 0.465434 | NA | Igf2os |
| ENSMUSG( | 1256.016 | 0.160908 | 0.312767 | 0.900219 | Igf2r |
| ENSMUSG( | 3454.866 | -0.15383 | 0.194049 | 0.860096 | Igfbp2 |
| ENSMUSG( | 425.6901 | 0.479635 | 0.000493 | 0.06693 | Igfbp3 |
| ENSMUSG( | 3076.655 | 0.102529 | 0.427777 | 0.917257 | Igfbp4 |
| ENSMUSG( | 15192.75 | 0.070642 | 0.577838 | 0.943131 | Igfbp5 |
| ENSMUSG( | 1018.747 | -0.21279 | 0.057359 | 0.723362 | Igfbp6 |
| ENSMUSG( | 1595.797 | 0.403047 | 0.040669 | 0.683847 | Igfbp7 |
| ENSMUSG( | 269.7312 | 0.102619 | 0.66295 | 0.956205 | Igfbpl1 |
| ENSMUSG( | 487.96 | -0.42824 | 0.028013 | 0.61023 | Igfn1 |
| ENSMUSG( | 11.98478 | 2.269698 | 0.002942 | 0.217424 | Igha |
| ENSMUSG( | 27.92658 | 0.298503 | 0.44447 | 0.921734 | Ighd |
| ENSMUSG( | 95.25697 | 0.592595 | 0.008631 | 0.391682 | Ighg2c |
| ENSMUSG( | 995.4542 | 0.051322 | 0.687653 | 0.959673 | Ighm |
| ENSMUSG( | 272.6882 | -0.01823 | 0.908681 | 0.991958 | Ighmbp2 |
| ENSMUSG( | 2310.946 | 0.027798 | 0.853048 | 0.985598 | Igip |
| ENSMUSG( | 29.52108 | -0.47451 | 0.25003 | 0.877904 | Igkc |
| ENSMUSG( | 1862.588 | 0.031791 | 0.770846 | 0.974236 | Iglon5 |
| ENSMUSG( | 315.1485 | -0.54136 | 0.016727 | 0.521279 | Igsf1 |
| ENSMUSG( | 119.9272 | 0.327551 | 0.107247 | 0.804969 | Igsf10 |
| ENSMUSG( | 1864.459 | 0.338713 | 0.015129 | 0.492251 | Igsf11 |
| ENSMUSG( | 3200.648 | 0.106282 | 0.440552 | 0.921166 | Igsf21 |
| ENSMUSG( | 762.4465 | 0.309793 | 0.08496 | 0.779799 | Igsf3 |
| ENSMUSG( | 15.21206 | 0.291821 | 0.593563 | 0.945408 | Igsf5 |
| ENSMUSG( | 68.05212 | 0.251088 | 0.346345 | 0.910966 | Igsf6 |
| ENSMUSG( | 8798.627 | 0.143224 | 0.353451 | 0.912263 | Igsf8 |
| ENSMUSG( | 164.6204 | -0.1032 | 0.656605 | 0.955533 | Igsf9 |
| ENSMUSG( | 1952.573 | 0.088387 | 0.542222 | 0.938034 | Igsf9b |
| ENSMUSG( | 210.3733 | -0.00088 | 0.996412 | 0.999178 | Igtp |
| ENSMUSG( | 11.81219 | -0.11828 | 0.84249 | 0.983044 | Iho1 |
| ENSMUSG( | 50.54666 | -0.24674 | 0.447692 | 0.922202 | Iigp1 |
| ENSMUSG( | 9.706206 | -0.58697 | 0.348665 | NA | Iigp1c |
| ENSMUSG( | 2292.023 | -0.0088 | 0.942324 | 0.994774 | Ik |
| ENSMUSG( | 562.7905 | 0.061335 | 0.714824 | 0.964986 | Ikbip |
| ENSMUSG( | 1384.403 | 0.033223 | 0.787661 | 0.97637 | Ikbkb |
| ENSMUSG( | 67.19615 | -0.40863 | 0.13351 | 0.822912 | Ikbke |
| ENSMUSG( | 923.5434 | 0.028395 | 0.807201 | 0.978393 | Ikbkg |
| ENSMUSG( | 240.5114 | 0.051263 | 0.773916 | 0.97478 | Ikzf1 |
| ENSMUSG( | 69.45129 | 0.223825 | 0.394512 | 0.917257 | Ikzf2 |
| ENSMUSG( | 11.08259 | -0.46233 | 0.519041 | NA | Ikzf3 |
| ENSMUSG( | 393.5522 | 0.198713 | 0.15413 | 0.837724 | Ikzf4 |
| ENSMUSG( | 504.322 | 0.096124 | 0.464011 | 0.923329 | Ikzf5 |
| ENSMUSG( | 338.941 | -0.05511 | 0.834966 | 0.981237 | Il10ra |
| ENSMUSG( | 12.96383 | 1.140684 | 0.058511 | 0.728256 | Il10rb |
| ENSMUSG( | 10.37987 | 0.471301 | 0.439683 | NA | Il11 |
| ENSMUSG( | 1560.698 | -0.03571 | 0.750581 | 0.970184 | Il11ra1 |
| ENSMUSG( | 35.93992 | -0.2363 | 0.478514 | 0.923329 | Il12a |
| ENSMUSG( | 75.60216 | -0.12934 | 0.629066 | 0.953251 | Il12rb1 |
| ENSMUSG( | 284.9123 | 0.252978 | 0.235341 | 0.871741 | Il13ra1 |
| ENSMUSG( | 12.43343 | 0.311342 | 0.589263 | 0.944975 | Il13ra2 |
| ENSMUSG( | 63.85799 | -0.20767 | 0.43065 | 0.917257 | Il15 |
| ENSMUSG( | 120.0579 | 0.16985 | 0.468052 | 0.923329 | Il15ra |
| ENSMUSG( | 195.246 | 0.301847 | 0.129784 | 0.820802 | Il16 |
| ENSMUSG( | 325.7912 | -0.04673 | 0.76025 | 0.972575 | Il17d |
| ENSMUSG( | 922.9035 | -0.01926 | 0.903859 | 0.991089 | Il17ra |
| ENSMUSG( | 46.76466 | 0.368321 | 0.300382 | 0.898538 | Il17rb |
| ENSMUSG( | 227.5032 | -0.23577 | 0.15309 | 0.837724 | Il17rc |
| ENSMUSG( | 300.6622 | -0.07106 | 0.733589 | 0.967228 | Il17rd |
| ENSMUSG( | 957.6943 | -0.13253 | 0.271936 | 0.888383 | Il18 |
| ENSMUSG( | 170.1012 | -0.20041 | 0.29463 | 0.898538 | Il18bp |
| ENSMUSG( | 27.6987 | 0.764686 | 0.063402 | 0.736482 | Il1a |
| ENSMUSG( | 165.2149 | 0.21771 | 0.372167 | 0.915491 | Il1r1 |
| ENSMUSG( | 9.808425 | -0.28072 | 0.66444 | NA | Il1r2 |
| ENSMUSG( | 1315.576 | 0.386956 | 0.064309 | 0.737747 | Il1rap |
| ENSMUSG( | 331.6463 | 0.231017 | 0.218609 | 0.866104 | Il1rapl1 |
| ENSMUSG( | 94.43487 | 0.733468 | 0.004045 | 0.257635 | Il1rapl2 |
| ENSMUSG( | 19.45617 | -0.33169 | 0.453806 | 0.922316 | Il1rl1 |
| ENSMUSG( | 50.14702 | 0.333891 | 0.25844 | 0.883971 | Il1rl2 |
| ENSMUSG( | 13.30596 | -0.10906 | 0.838724 | 0.981622 | Il20ra |
| ENSMUSG( | 26.12688 | 0.149448 | 0.723495 | 0.966887 | Il20rb |
| ENSMUSG( | 102.8764 | 0.509143 | 0.066536 | 0.750492 | Il21r |
| ENSMUSG( | 43.05723 | -0.32236 | 0.296128 | 0.898538 | Il23a |
| ENSMUSG( | 16.35901 | 0.874157 | 0.118635 | 0.813384 | Il27ra |
| ENSMUSG( | 8.837254 | 0.948528 | 0.163096 | NA | Il2ra |
| ENSMUSG( | 22.62541 | -0.30441 | 0.562972 | 0.941158 | Il2rb |
| ENSMUSG( | 82.77885 | 0.129417 | 0.6261 | 0.953234 | Il2rg |
| ENSMUSG( | 14.92279 | -0.12045 | 0.82161 | 0.979198 | Il31ra |
| ENSMUSG( | 1491.602 | -0.43453 | 0.004893 | 0.289578 | Il33 |
| ENSMUSG( | 1727.579 | 0.061064 | 0.68541 | 0.958678 | Il34 |
| ENSMUSG( | 76.7099 | -0.13193 | 0.583243 | 0.944266 | Il3ra |
| ENSMUSG( | 29.29844 | -0.32331 | 0.446152 | 0.921734 | Il4i1 |
| ENSMUSG( | 297.7437 | -0.00368 | 0.987187 | 0.997487 | Il4ra |
| ENSMUSG( | 381.1687 | 0.615573 | 0.00673 | 0.339232 | Il6ra |
| ENSMUSG( | 4824.886 | 0.086889 | 0.539144 | 0.937216 | Il6st |
| ENSMUSG( | 37.01329 | 0.157007 | 0.723348 | 0.966887 | Il7r |
| ENSMUSG( | 18.76556 | -0.42785 | 0.37634 | 0.915491 | Ildr1 |
| ENSMUSG( | 5714.385 | 0.075947 | 0.550347 | 0.938699 | Ildr2 |
| ENSMUSG( | 1377.582 | -0.07579 | 0.573264 | 0.942912 | Ilf2 |
| ENSMUSG( | 1798.334 | 0.057199 | 0.608729 | 0.950113 | Ilf3 |
| ENSMUSG( | 3957.114 | -0.0782 | 0.477693 | 0.923329 | Ilk |
| ENSMUSG( | 1079.751 | -0.06619 | 0.582289 | 0.944266 | Ilkap |
| ENSMUSG( | 15778.42 | -0.03095 | 0.758521 | 0.971858 | Ilrun |
| ENSMUSG( | 828.996 | 0.083087 | 0.615416 | 0.951177 | Ilvbl |
| ENSMUSG( | 408.6352 | -0.13614 | 0.342532 | 0.909646 | Immp1l |
| ENSMUSG( | 73.61109 | -0.24258 | 0.316557 | 0.901868 | Immp2l |
| ENSMUSG( | 3600.322 | -0.13316 | 0.218537 | 0.866104 | Immt |
| ENSMUSG( | 512.8529 | -0.09012 | 0.515958 | 0.930939 | Imp3 |
| ENSMUSG( | 787.2859 | -0.12562 | 0.353304 | 0.912263 | Imp4 |
| ENSMUSG( | 1131.021 | -0.09797 | 0.451737 | 0.922202 | Impa1 |
| ENSMUSG( | 221.6607 | -0.3944 | 0.02187 | 0.575341 | Impa2 |
| ENSMUSG( | 2945.47 | 0.026941 | 0.825214 | 0.979354 | Impact |
| ENSMUSG( | 1324.73 | 0.029121 | 0.82756 | 0.980224 | Impdh1 |
| ENSMUSG( | 854.5486 | 0.070343 | 0.604041 | 0.948139 | Impdh2 |
| ENSMUSG( | 13.37665 | 0.42135 | 0.433288 | 0.918434 | Impdh2-ps |
| ENSMUSG( | 18.95846 | 0.684777 | 0.165173 | 0.838743 | Impg1 |

| | | | | | | | | | | | |
|---|---|---|---|---|---|---|---|---|---|---|---|
| ENSMUSG( | Igf2os | 12.79128 | -0.11937 | 0.819827 | NA | ENSMUSG( | Igf2os | 10.45259 | -0.45309 | 0.465434 | NA |
| ENSMUSG( | Igf2r | 1255.151 | -0.11362 | 0.413533 | 0.819888 | ENSMUSG( | Igf2r | 1256.016 | 0.160908 | 0.312767 | 0.900219 |
| ENSMUSG( | Igfbp2 | 3642.976 | -0.27944 | 0.008856 | 0.132506 | ENSMUSG( | Igfbp2 | 3454.866 | -0.15383 | 0.194049 | 0.860096 |
| ENSMUSG( | Igfbp3 | 437.788 | 0.311932 | 0.00946 | 0.137493 | ENSMUSG( | Igfbp3 | 425.6901 | 0.479635 | 0.000493 | 0.06693 |
| ENSMUSG( | Igfbp4 | 3105.38 | -0.14316 | 0.209529 | 0.65169 | ENSMUSG( | Igfbp4 | 3076.655 | 0.102529 | 0.427777 | 0.917257 |
| ENSMUSG( | Igfbp5 | 16625.8 | 0.060509 | 0.611484 | 0.908586 | ENSMUSG( | Igfbp5 | 15192.75 | 0.070642 | 0.577838 | 0.943131 |
| ENSMUSG( | Igfbp6 | 1101.598 | -0.25931 | 0.010049 | 0.142735 | ENSMUSG( | Igfbp6 | 1018.747 | -0.21279 | 0.057359 | 0.723362 |
| ENSMUSG( | Igfbp7 | 1416.291 | -0.19588 | 0.253761 | 0.704011 | ENSMUSG( | Igfbp7 | 1595.797 | 0.403047 | 0.040669 | 0.683847 |
| ENSMUSG( | Igfbpl1 | 220.7487 | -0.89116 | 0.000638 | 0.024356 | ENSMUSG( | Igfbpl1 | 269.7312 | 0.102619 | 0.66295 | 0.956205 |
| ENSMUSG( | Igfn1 | 596.783 | -0.08197 | 0.611279 | 0.908586 | ENSMUSG( | Igfn1 | 487.96 | -0.42824 | 0.028013 | 0.61023 |
| ENSMUSG( | Igha | 9.27238 | 1.619718 | 0.071274 | NA | ENSMUSG( | Igha | 11.98478 | 2.269698 | 0.002942 | 0.217424 |
| ENSMUSG( | Ighd | 30.25175 | 0.285143 | 0.450298 | 0.835344 | ENSMUSG( | Ighd | 27.92658 | 0.298503 | 0.44447 | 0.921734 |
| ENSMUSG( | Ighg2c | 105.4026 | 0.605041 | 0.002113 | 0.053164 | ENSMUSG( | Ighg2c | 95.25697 | 0.592595 | 0.008631 | 0.391682 |
| ENSMUSG( | Ighm | 1242.238 | 0.39306 | 0.000196 | 0.01044 | ENSMUSG( | Ighm | 995.4542 | 0.051322 | 0.687653 | 0.959673 |
| ENSMUSG( | Ighmbp2 | 297.8598 | -0.03053 | 0.844474 | 0.972023 | ENSMUSG( | Ighmbp2 | 272.6882 | -0.01823 | 0.908681 | 0.991958 |
| ENSMUSG( | Igip | 2533.635 | 0.029032 | 0.775484 | 0.955606 | ENSMUSG( | Igip | 2310.946 | 0.027798 | 0.853048 | 0.985598 |
| ENSMUSG( | Igkc | 37.34076 | -0.03641 | 0.919202 | 0.990247 | ENSMUSG( | Igkc | 29.52108 | -0.47451 | 0.25003 | 0.877904 |
| ENSMUSG( | Iglon5 | 1897.61 | -0.18896 | 0.047323 | 0.332697 | ENSMUSG( | Iglon5 | 1862.588 | 0.031791 | 0.770846 | 0.974236 |
| ENSMUSG( | Igsf1 | 358.0666 | -0.40909 | 0.411764 | 0.81942 | ENSMUSG( | Igsf1 | 315.1485 | -0.54136 | 0.016727 | 0.521279 |
| ENSMUSG( | Igsf10 | 101.4461 | -0.44101 | 0.028466 | 0.257525 | ENSMUSG( | Igsf10 | 119.9272 | 0.327551 | 0.107247 | 0.804969 |
| ENSMUSG( | Igsf11 | 1739.152 | -0.11536 | 0.336668 | 0.777 | ENSMUSG( | Igsf11 | 1864.459 | 0.338713 | 0.015129 | 0.492251 |
| ENSMUSG( | Igsf21 | 3320.257 | -0.05843 | 0.607604 | 0.906878 | ENSMUSG( | Igsf21 | 3200.648 | 0.106282 | 0.440552 | 0.921166 |
| ENSMUSG( | Igsf3 | 682.8374 | -0.28089 | 0.084096 | 0.447085 | ENSMUSG( | Igsf3 | 762.4465 | 0.309793 | 0.08496 | 0.779799 |
| ENSMUSG( | Igsf5 | 15.14175 | -0.01354 | 0.976938 | NA | ENSMUSG( | Igsf5 | 15.21206 | 0.291821 | 0.593563 | 0.945408 |
| ENSMUSG( | Igsf6 | 56.75012 | -0.59251 | 0.047714 | 0.33359 | ENSMUSG( | Igsf6 | 68.05212 | 0.251088 | 0.346345 | 0.910966 |
| ENSMUSG( | Igsf8 | 8112.033 | -0.38543 | 0.003159 | 0.06974 | ENSMUSG( | Igsf8 | 8798.627 | 0.143224 | 0.353451 | 0.912263 |
| ENSMUSG( | Igsf9 | 180.7563 | -0.11785 | 0.516529 | 0.872788 | ENSMUSG( | Igsf9 | 164.6204 | -0.1032 | 0.656605 | 0.955533 |
| ENSMUSG( | Igsf9b | 2063.222 | -0.02368 | 0.835491 | 0.970557 | ENSMUSG( | Igsf9b | 1952.573 | 0.088387 | 0.542222 | 0.938034 |
| ENSMUSG( | Igtp | 212.0091 | -0.27525 | 0.094244 | 0.471647 | ENSMUSG( | Igtp | 210.3733 | -0.00088 | 0.996412 | 0.999178 |
| ENSMUSG( | Iho1 | 13.4514 | 0.020638 | 0.968408 | NA | ENSMUSG( | Iho1 | 11.81219 | -0.11828 | 0.84249 | 0.983044 |
| ENSMUSG( | Iigp1 | 50.53299 | -0.59234 | 0.073213 | 0.417696 | ENSMUSG( | Iigp1 | 50.54666 | -0.24674 | 0.447692 | 0.922202 |
| ENSMUSG( | Iigp1c | 9.980008 | -0.8783 | 0.159919 | NA | ENSMUSG( | Iigp1c | 9.706206 | -0.58697 | 0.348665 | NA |
| ENSMUSG( | Ik | 2614.259 | 0.103105 | 0.247288 | 0.696236 | ENSMUSG( | Ik | 2292.023 | -0.0088 | 0.942324 | 0.994774 |
| ENSMUSG( | Ikbip | 557.6348 | -0.24841 | 0.098597 | 0.480543 | ENSMUSG( | Ikbip | 562.7905 | 0.061335 | 0.714824 | 0.964986 |
| ENSMUSG( | Ikbkb | 1579.539 | 0.142642 | 0.104234 | 0.49005 | ENSMUSG( | Ikbkb | 1384.403 | 0.033223 | 0.787661 | 0.97637 |
| ENSMUSG( | Ikbke | 87.67239 | 0.103887 | 0.618046 | 0.910929 | ENSMUSG( | Ikbke | 67.19615 | -0.40863 | 0.13351 | 0.822912 |
| ENSMUSG( | Ikbkg | 1041.937 | 0.10654 | 0.275064 | 0.72509 | ENSMUSG( | Ikbkg | 923.5434 | 0.028395 | 0.807201 | 0.978393 |
| ENSMUSG( | Ikzf1 | 254.539 | -0.05748 | 0.721481 | 0.943059 | ENSMUSG( | Ikzf1 | 240.5114 | 0.051263 | 0.773916 | 0.97478 |
| ENSMUSG( | Ikzf2 | 74.1601 | 0.138861 | 0.570333 | 0.893192 | ENSMUSG( | Ikzf2 | 69.45129 | 0.223825 | 0.394512 | 0.917257 |
| ENSMUSG( | Ikzf4 | 423.8883 | 0.15279 | 0.196628 | 0.635942 | ENSMUSG( | Ikzf4 | 393.5522 | 0.198713 | 0.15413 | 0.837724 |
| ENSMUSG( | Ikzf5 | 543.9337 | 0.044142 | 0.702277 | 0.937536 | ENSMUSG( | Ikzf5 | 504.322 | 0.096124 | 0.464011 | 0.923329 |
| ENSMUSG( | Il10ra | 301.4276 | -0.7783 | 0.000126 | 0.007476 | ENSMUSG( | Il10ra | 338.941 | -0.05511 | 0.834966 | 0.981237 |
| ENSMUSG( | Il10rb | 8.950107 | 0.004651 | 0.993953 | NA | ENSMUSG( | Il10rb | 12.96383 | 1.140684 | 0.058511 | 0.728256 |
| ENSMUSG( | Il11 | 11.56175 | 0.484417 | 0.371381 | NA | ENSMUSG( | Il11 | 10.37987 | 0.471301 | 0.439683 | NA |
| ENSMUSG( | Il11ra1 | 1753.914 | 0.032588 | 0.742629 | 0.948478 | ENSMUSG( | Il11ra1 | 1560.698 | -0.03571 | 0.750581 | 0.970184 |
| ENSMUSG( | Il12a | 40.41078 | -0.15501 | 0.597794 | 0.90372 | ENSMUSG( | Il12a | 35.93992 | -0.2363 | 0.478514 | 0.923329 |
| ENSMUSG( | Il12rb1 | 73.23139 | -0.54959 | 0.037206 | 0.297718 | ENSMUSG( | Il12rb1 | 75.60216 | -0.12934 | 0.629066 | 0.953251 |
| ENSMUSG( | Il13ra1 | 269.0741 | -0.18667 | 0.321785 | 0.766739 | ENSMUSG( | Il13ra1 | 284.9123 | 0.252978 | 0.235341 | 0.871741 |
| ENSMUSG( | Il13ra2 | 11.96383 | -0.00059 | 0.999137 | NA | ENSMUSG( | Il13ra2 | 12.43343 | 0.311342 | 0.589263 | 0.944975 |
| ENSMUSG( | Il15 | 72.88303 | -0.08617 | 0.717912 | 0.941697 | ENSMUSG( | Il15 | 63.85799 | -0.20767 | 0.43065 | 0.917257 |
| ENSMUSG( | Il15ra | 121.5908 | -0.07909 | 0.695739 | 0.934979 | ENSMUSG( | Il15ra | 120.0579 | 0.16985 | 0.468052 | 0.923329 |
| ENSMUSG( | Il16 | 198.301 | 0.077933 | 0.621066 | 0.911432 | ENSMUSG( | Il16 | 195.246 | 0.301847 | 0.129784 | 0.820802 |
| ENSMUSG( | Il17d | 361.006 | -0.01405 | 0.912604 | 0.988483 | ENSMUSG( | Il17d | 325.7912 | -0.04673 | 0.76025 | 0.972575 |
| ENSMUSG( | Il17ra | 944.1485 | -0.23683 | 0.111591 | 0.50362 | ENSMUSG( | Il17ra | 922.9035 | -0.01926 | 0.903859 | 0.991089 |
| ENSMUSG( | Il17rb | 39.12014 | -0.46749 | 0.168487 | 0.601952 | ENSMUSG( | Il17rb | 46.76466 | 0.368321 | 0.300382 | 0.898538 |
| ENSMUSG( | Il17rc | 259.9988 | -0.11569 | 0.418443 | 0.821966 | ENSMUSG( | Il17rc | 227.5032 | -0.23577 | 0.15309 | 0.837724 |
| ENSMUSG( | Il17rd | 321.3497 | -0.16183 | 0.392447 | 0.809274 | ENSMUSG( | Il17rd | 300.6622 | -0.07106 | 0.733589 | 0.967228 |
| ENSMUSG( | Il18 | 1146.169 | 0.116894 | 0.338804 | 0.777518 | ENSMUSG( | Il18 | 957.6943 | -0.13253 | 0.271936 | 0.888383 |
| ENSMUSG( | Il18bp | 203.9318 | 0.062499 | 0.717569 | 0.941697 | ENSMUSG( | Il18bp | 170.1012 | -0.20041 | 0.29463 | 0.898538 |
| ENSMUSG( | Il1a | 22.80658 | 0.009389 | 0.982548 | 0.997512 | ENSMUSG( | Il1a | 27.6987 | 0.764686 | 0.063402 | 0.736482 |
| ENSMUSG( | Il1r1 | 186.8745 | 0.278421 | 0.16404 | 0.594869 | ENSMUSG( | Il1r1 | 165.2149 | 0.21771 | 0.372167 | 0.915491 |
| ENSMUSG( | Il1rap | 1087.749 | -0.44565 | 0.010137 | 0.143118 | ENSMUSG( | Il1rap | 1315.576 | 0.386956 | 0.064309 | 0.737747 |
| ENSMUSG( | Il1rapl1 | 276.0691 | -0.62629 | 0.000215 | 0.011305 | ENSMUSG( | Il1rapl1 | 331.6463 | 0.231017 | 0.218609 | 0.866104 |
| ENSMUSG( | Il1rapl2 | 75.58565 | -0.10189 | 0.680366 | 0.931044 | ENSMUSG( | Il1rapl2 | 94.43487 | 0.733468 | 0.004045 | 0.257635 |
| ENSMUSG( | Il1rl1 | 20.95386 | -0.36715 | 0.382693 | NA | ENSMUSG( | Il1rl1 | 19.45617 | -0.33169 | 0.453806 | 0.922316 |
| ENSMUSG( | Il1rl2 | 43.77193 | -0.34736 | 0.219567 | 0.664984 | ENSMUSG( | Il1rl2 | 50.14702 | 0.333891 | 0.25844 | 0.883971 |
| ENSMUSG( | Il20ra | 12.35647 | -0.66111 | 0.213327 | NA | ENSMUSG( | Il20ra | 13.30596 | -0.10906 | 0.838724 | 0.981622 |
| ENSMUSG( | Il20rb | 27.32906 | -0.01132 | 0.976776 | 0.997428 | ENSMUSG( | Il20rb | 26.12688 | 0.149448 | 0.723495 | 0.966887 |
| ENSMUSG( | Il21r | 94.87616 | 0.040648 | 0.881951 | 0.982069 | ENSMUSG( | Il21r | 102.8764 | 0.509143 | 0.066536 | 0.750492 |
| ENSMUSG( | Il23a | 60.86326 | 0.405818 | 0.10206 | 0.486742 | ENSMUSG( | Il23a | 43.05723 | -0.32236 | 0.296128 | 0.898538 |
| ENSMUSG( | Il27ra | 13.35512 | 0.119776 | 0.836378 | NA | ENSMUSG( | Il27ra | 16.35901 | 0.874157 | 0.118635 | 0.813384 |
| ENSMUSG( | Il2rb | 21.88567 | -0.75064 | 0.183257 | 0.62047 | ENSMUSG( | Il2rb | 22.62541 | -0.30441 | 0.562972 | 0.941158 |
| ENSMUSG( | Il2rg | 74.53134 | -0.50113 | 0.052942 | 0.35324 | ENSMUSG( | Il2rg | 82.77885 | 0.129417 | 0.6261 | 0.953234 |
| ENSMUSG( | Il31ra | 16.28624 | -0.1605 | 0.743078 | NA | ENSMUSG( | Il31ra | 14.92279 | -0.12045 | 0.82161 | 0.979198 |
| ENSMUSG( | Il33 | 1914.645 | 0.054296 | 0.729406 | 0.944738 | ENSMUSG( | Il33 | 1491.602 | -0.43453 | 0.004893 | 0.289578 |
| ENSMUSG( | Il34 | 1692.01 | -0.28555 | 0.036599 | 0.295048 | ENSMUSG( | Il34 | 1727.579 | 0.061064 | 0.68541 | 0.958678 |
| ENSMUSG( | Il3ra | 96.11111 | 0.237276 | 0.238585 | 0.686308 | ENSMUSG( | Il3ra | 76.7099 | -0.13193 | 0.583243 | 0.944266 |
| ENSMUSG( | Il4i1 | 34.75667 | -0.12697 | 0.696869 | 0.935464 | ENSMUSG( | Il4i1 | 29.29844 | -0.32331 | 0.446152 | 0.921734 |
| ENSMUSG( | Il4ra | 307.0869 | -0.19897 | 0.29302 | 0.742763 | ENSMUSG( | Il4ra | 297.7437 | -0.00368 | 0.987187 | 0.997487 |
| ENSMUSG( | Il6ra | 325.8001 | -0.05305 | 0.766312 | 0.954173 | ENSMUSG( | Il6ra | 381.1687 | 0.615573 | 0.00673 | 0.339232 |
| ENSMUSG( | Il6st | 4545.86 | -0.37948 | 0.026106 | 0.247245 | ENSMUSG( | Il6st | 4824.886 | 0.086889 | 0.539144 | 0.937216 |
| ENSMUSG( | Il7r | 34.36422 | -0.37272 | 0.259926 | 0.709951 | ENSMUSG( | Il7r | 37.01329 | 0.157007 | 0.723348 | 0.966887 |
| ENSMUSG( | Ildr1 | 21.85253 | -0.21802 | 0.594224 | 0.902869 | ENSMUSG( | Ildr1 | 18.76556 | -0.42785 | 0.37634 | 0.915491 |
| ENSMUSG( | Ildr2 | 5815.422 | -0.14713 | 0.192205 | 0.631194 | ENSMUSG( | Ildr2 | 5714.385 | 0.075947 | 0.550347 | 0.938699 |
| ENSMUSG( | Ilf2 | 1545.296 | -0.00983 | 0.932159 | 0.992515 | ENSMUSG( | Ilf2 | 1377.582 | -0.07579 | 0.573264 | 0.942912 |
| ENSMUSG( | Ilf3 | 2061.646 | 0.17654 | 0.062053 | 0.385335 | ENSMUSG( | Ilf3 | 1798.334 | 0.057199 | 0.608729 | 0.950113 |
| ENSMUSG( | Ilk | 4572.427 | 0.072227 | 0.39014 | 0.80794 | ENSMUSG( | Ilk | 3957.114 | -0.0782 | 0.477693 | 0.923329 |
| ENSMUSG( | Ilkap | 1230.876 | 0.044368 | 0.6591 | 0.924506 | ENSMUSG( | Ilkap | 1079.751 | -0.06619 | 0.582289 | 0.944266 |
| ENSMUSG( | Ilrun | 18568.55 | 0.165925 | 0.03693 | 0.296508 | ENSMUSG( | Ilrun | 15778.42 | -0.03095 | 0.758521 | 0.971858 |
| ENSMUSG( | Ilvbl | 795.688 | -0.32774 | 0.016548 | 0.191177 | ENSMUSG( | Ilvbl | 828.996 | 0.083087 | 0.615416 | 0.951177 |
| ENSMUSG( | Immp1l | 489.1058 | 0.121693 | 0.323374 | 0.768488 | ENSMUSG( | Immp1l | 408.6352 | -0.13614 | 0.342532 | 0.909646 |
| ENSMUSG( | Immp2l | 77.83229 | -0.35748 | 0.103693 | 0.489428 | ENSMUSG( | Immp2l | 73.61109 | -0.24258 | 0.316557 | 0.901868 |
| ENSMUSG( | Immt | 4382.704 | 0.16555 | 0.034521 | 0.285239 | ENSMUSG( | Immt | 3600.322 | -0.13316 | 0.218537 | 0.866104 |
| ENSMUSG( | Imp3 | 581.0275 | 0.005176 | 0.964093 | 0.995137 | ENSMUSG( | Imp3 | 512.8529 | -0.09012 | 0.515958 | 0.930939 |
| ENSMUSG( | Imp4 | 935.1967 | 0.107471 | 0.337437 | 0.777412 | ENSMUSG( | Imp4 | 787.2859 | -0.12562 | 0.353304 | 0.912263 |
| ENSMUSG( | Impa1 | 1266.422 | -0.03463 | 0.751166 | 0.950161 | ENSMUSG( | Impa1 | 1131.021 | -0.09797 | 0.451737 | 0.922202 |
| ENSMUSG( | Impa2 | 254.3927 | -0.25387 | 0.08824 | 0.455783 | ENSMUSG( | Impa2 | 221.6607 | -0.3944 | 0.02187 | 0.575341 |
| ENSMUSG( | Impact | 3249.047 | 0.043 | 0.656918 | 0.924069 | ENSMUSG( | Impact | 2945.47 | 0.026941 | 0.825214 | 0.979354 |
| ENSMUSG( | Impdh1 | 1464.636 | 0.048627 | 0.698155 | 0.936028 | ENSMUSG( | Impdh1 | 1324.73 | 0.029121 | 0.82756 | 0.980224 |
| ENSMUSG( | Impdh2 | 954.2539 | 0.120233 | 0.31153 | 0.759296 | ENSMUSG( | Impdh2 | 854.5486 | 0.070343 | 0.604041 | 0.948139 |
| ENSMUSG( | Impdh2-ps | 14.10579 | 0.336152 | 0.49918 | NA | ENSMUSG( | Impdh2-ps | 13.37665 | 0.42135 | 0.433288 | 0.918434 |
| ENSMUSG( | Impg1 | 11.92927 | -1.08437 | 0.054797 | NA | ENSMUSG( | Impg1 | 18.95846 | 0.684777 | 0.165173 | 0.838743 |
| ENSMUSG( | Impg2 | 27.88484 | -0.262 | 0.469618 | 0.846167 | ENSMUSG( | Impg2 | 31.22062 | 0.306853 | 0.396957 | 0.917257 |
| ENSMUSG( | Ina | 5186.733 | 0.048492 | 0.670719 | 0.928683 | ENSMUSG( | Ina | 4362.167 | -0.18991 | 0.13719 | 0.828492 |
| ENSMUSG( | Inafm1 | 979.1531 | -0.00828 | 0.93475 | 0.993231 | ENSMUSG( | Inafm1 | 895.8433 | 0.00432 | 0.970216 | 0.99594 |
| ENSMUSG( | Inafm2 | 1533.417 | -0.14196 | 0.125206 | 0.532314 | ENSMUSG( | Inafm2 | 1506.985 | 0.078164 | 0.50657 | 0.928006 |
| ENSMUSG( | Inava | 194.1708 | -0.22611 | 0.144473 | 0.564381 | ENSMUSG( | Inava | 184.167 | -0.09738 | 0.595785 | 0.946047 |
| ENSMUSG( | Inca1 | 76.9264 | 0.012145 | 0.956007 | 0.993874 | ENSMUSG( | Inca1 | 65.6928 | -0.15127 | 0.583333 | 0.944266 |
| ENSMUSG( | Incenp | 462.9267 | 0.057652 | 0.617889 | 0.910872 | ENSMUSG( | Incenp | 400.3027 | -0.10106 | 0.485689 | 0.92541 |
| ENSMUSG( | Inf2 | 5312.783 | -0.06764 | 0.577344 | 0.895776 | ENSMUSG( | Inf2 | 4633.824 | -0.20117 | 0.119441 | 0.813384 |
| ENSMUSG( | Ing1 | 1333.537 | 0.056929 | 0.547584 | 0.885153 | ENSMUSG( | Ing1 | 1180.532 | -0.02924 | 0.813008 | 0.979198 |
| ENSMUSG( | Ing2 | 490.8876 | -0.01671 | 0.90199 | 0.985633 | ENSMUSG( | Ing2 | 438.3699 | -0.07887 | 0.593283 | 0.945408 |
| ENSMUSG( | Ing3 | 328.8384 | 0.048983 | 0.714073 | 0.940959 | ENSMUSG( | Ing3 | 313.7674 | 0.174268 | 0.281187 | 0.892356 |
| ENSMUSG( | Ing4 | 21.71172 | 0.51794 | 0.243367 | 0.690747 | ENSMUSG( | Ing4 | 28.17808 | 1.321276 | 0.001817 | 0.15837 |
| ENSMUSG( | Ing5 | 331.8409 | -0.15406 | 0.324719 | 0.770182 | ENSMUSG( | Ing5 | 305.8433 | -0.10901 | 0.53911 | 0.937216 |
| ENSMUSG( | Inha | 716.4826 | 0.028407 | 0.859184 | 0.974707 | ENSMUSG( | Inha | 633.3092 | -0.06314 | 0.725999 | 0.966887 |
| ENSMUSG( | Inhba | 182.2833 | -0.5835 | 0.002959 | 0.066473 | ENSMUSG( | Inhba | 221.3123 | 0.295199 | 0.136621 | 0.827522 |
| ENSMUSG( | Inhbb | 330.4266 | -0.28443 | 0.133091 | 0.545688 | ENSMUSG( | Inhbb | 359.7202 | 0.237601 | 0.215406 | 0.865167 |
| ENSMUSG( | Inip | 630.9567 | 0.025789 | 0.805414 | 0.962616 | ENSMUSG( | Inip | 557.9039 | -0.06265 | 0.629758 | 0.953251 |
| ENSMUSG( | Inka1 | 61.60586 | -0.06938 | 0.780655 | 0.957491 | ENSMUSG( | Inka1 | 55.2499 | -0.09045 | 0.761505 | 0.972819 |
| ENSMUSG( | Inka2 | 2548.403 | 0.027209 | 0.795726 | 0.960062 | ENSMUSG( | Inka2 | 2162.935 | -0.18555 | 0.16437 | 0.83856 |
| ENSMUSG( | Inmt | 10.05538 | -0.58453 | 0.342781 | NA | ENSMUSG( | Inmt | 14.85162 | 0.735373 | 0.178569 | 0.849041 |
| ENSMUSG( | Ino80 | 987.6597 | 0.025356 | 0.798647 | 0.960896 | ENSMUSG( | Ino80 | 909.6137 | 0.057784 | 0.626272 | 0.953234 |
| ENSMUSG( | Ino80b | 375.6144 | 0.122334 | 0.393129 | 0.809274 | ENSMUSG( | Ino80b | 338.0597 | 0.086368 | 0.589223 | 0.944975 |
| ENSMUSG( | Ino80c | 989.6258 | -0.04551 | 0.670609 | 0.928683 | ENSMUSG( | Ino80c | 838.8835 | -0.27086 | 0.055258 | 0.723362 |
| ENSMUSG( | Ino80d | 778.6502 | 0.151774 | 0.170697 | 0.605271 | ENSMUSG( | Ino80d | 724.55 | 0.209533 | 0.094333 | 0.795735 |
| ENSMUSG( | Ino80dos | 129.8077 | -0.07157 | 0.714164 | 0.941003 | ENSMUSG( | Ino80dos | 120.0222 | -0.0312 | 0.890359 | 0.990558 |
| ENSMUSG( | Ino80e | 923.9834 | -0.06089 | 0.581613 | 0.897855 | ENSMUSG( | Ino80e | 817.7076 | -0.14907 | 0.251342 | 0.879497 |
| ENSMUSG( | Inpp1 | 780.9628 | -0.04753 | 0.671481 | 0.928781 | ENSMUSG( | Inpp1 | 717.4056 | -0.02938 | 0.84284 | 0.98313 |
| ENSMUSG( | Inpp4a | 2943.111 | 0.155844 | 0.125275 | 0.532314 | ENSMUSG( | Inpp4a | 2611.523 | 0.08286 | 0.465257 | 0.923329 |
| ENSMUSG( | Inpp4b | 140.0271 | -0.22867 | 0.185462 | 0.623185 | ENSMUSG( | Inpp4b | 149.8173 | 0.233367 | 0.205684 | 0.864412 |
| ENSMUSG( | Inpp5a | 1682.251 | -0.00539 | 0.952721 | 0.993874 | ENSMUSG( | Inpp5a | 1456.714 | -0.15918 | 0.166771 | 0.840183 |
| ENSMUSG( | Inpp5b | 1173.877 | 0.101435 | 0.294052 | 0.743459 | ENSMUSG( | Inpp5b | 1052.965 | 0.057649 | 0.659627 | 0.956205 |
| ENSMUSG( | Inpp5d | 897.6445 | 0.24991 | 0.03313 | 0.278602 | ENSMUSG( | Inpp5d | 825.9816 | 0.281262 | 0.038058 | 0.676422 |
| ENSMUSG( | Inpp5e | 728.4508 | 0.0706 | 0.596304 | 0.903254 | ENSMUSG( | Inpp5e | 694.4834 | 0.198763 | 0.207618 | 0.864412 |
| ENSMUSG( | Inpp5f | 2859.382 | -0.27695 | 0.004798 | 0.091446 | ENSMUSG( | Inpp5f | 2795.061 | -0.05942 | 0.60873 | 0.950113 |
| ENSMUSG( | Inpp5j | 1519.855 | 0.19381 | 0.164166 | 0.5949 | ENSMUSG( | Inpp5j | 1280.647 | -0.02761 | 0.857571 | 0.985884 |
| ENSMUSG( | Inpp5k | 1247.167 | 0.104438 | 0.396591 | 0.811742 | ENSMUSG( | Inpp5k | 1116.206 | 0.05382 | 0.698175 | 0.960868 |
| ENSMUSG( | Inppl1 | 2666.652 | 0.01347 | 0.867946 | 0.976631 | ENSMUSG( | Inppl1 | 2469.68 | 0.060646 | 0.556961 | 0.939565 |
| ENSMUSG( | Insc | 101.761 | -0.46081 | 0.02922 | 0.259574 | ENSMUSG( | Insc | 100.579 | -0.20573 | 0.376022 | 0.915491 |
| ENSMUSG( | Insig1 | 3813.774 | -0.41578 | 0.003932 | 0.080285 | ENSMUSG( | Insig1 | 3735.821 | -0.17921 | 0.261458 | 0.884926 |
| ENSMUSG( | Insig2 | 1248.798 | -0.17449 | 0.181808 | 0.618944 | ENSMUSG( | Insig2 | 1296.092 | 0.203495 | 0.16268 | 0.837724 |
| ENSMUSG( | Insl5 | 21.92546 | 0.279893 | 0.485942 | 0.858043 | ENSMUSG( | Insl5 | 26.00052 | 0.879403 | 0.05362 | 0.720293 |
| ENSMUSG( | Insm1 | 159.927 | -0.11273 | 0.527558 | 0.877641 | ENSMUSG( | Insm1 | 144.1423 | -0.13038 | 0.54239 | 0.938034 |
| ENSMUSG( | Insr | 1833.358 | -0.14754 | 0.247525 | 0.696522 | ENSMUSG( | Insr | 1932.192 | 0.26779 | 0.063799 | 0.736482 |
| ENSMUSG( | Insrr | 16.96861 | -0.02329 | 0.962455 | NA | ENSMUSG( | Insrr | 16.89278 | 0.221647 | 0.64556 | 0.953988 |
| ENSMUSG( | Insyn1 | 6985.019 | 0.053405 | 0.480459 | 0.854748 | ENSMUSG( | Insyn1 | 6333.316 | 0.039037 | 0.689199 | 0.959893 |
| ENSMUSG( | Insyn2a | 196.5973 | -0.14867 | 0.383942 | 0.803646 | ENSMUSG( | Insyn2a | 177.1375 | -0.1929 | 0.336952 | 0.907972 |
| ENSMUSG( | Insyn2b | 43.22805 | -0.08405 | 0.797258 | 0.960507 | ENSMUSG( | Insyn2b | 38.83734 | -0.15032 | 0.652277 | 0.954766 |
| ENSMUSG( | Ints1 | 2992.256 | 0.182996 | 0.054806 | 0.360194 | ENSMUSG( | Ints1 | 2616.373 | 0.071504 | 0.538868 | 0.937216 |
| ENSMUSG( | Ints10 | 1075.066 | 0.09511 | 0.347671 | 0.780695 | ENSMUSG( | Ints10 | 1008.725 | 0.179706 | 0.181628 | 0.850579 |
| ENSMUSG( | Ints11 | 758.1517 | 0.174519 | 0.096948 | 0.477375 | ENSMUSG( | Ints11 | 646.6731 | -0.00988 | 0.9399 | 0.994334 |
| ENSMUSG( | Ints12 | 204.6604 | 0.063399 | 0.686178 | 0.932981 | ENSMUSG( | Ints12 | 194.0763 | 0.166277 | 0.360384 | 0.914223 |
| ENSMUSG( | Ints13 | 903.2027 | 0.116063 | 0.249123 | 0.698492 | ENSMUSG( | Ints13 | 814.9668 | 0.091522 | 0.466494 | 0.923329 |
| ENSMUSG( | Ints14 | 624.1875 | 0.004461 | 0.971646 | 0.996528 | ENSMUSG( | Ints14 | 582.5127 | 0.077073 | 0.557666 | 0.939565 |
| ENSMUSG( | Ints15 | 727.5839 | 0.012415 | 0.903717 | 0.986094 | ENSMUSG( | Ints15 | 662.5453 | 0.01149 | 0.923874 | 0.992841 |
| ENSMUSG( | Ints2 | 549.8528 | -0.08676 | 0.469322 | 0.84601 | ENSMUSG( | Ints2 | 559.3198 | 0.226859 | 0.093552 | 0.795507 |
| ENSMUSG( | Ints3 | 1732.626 | 0.213647 | 0.018171 | 0.20312 | ENSMUSG( | Ints3 | 1509.887 | 0.093688 | 0.40898 | 0.917257 |

| | | | | | |
|---|---|---|---|---|---|
| ENSMUSG( | 259.9988 | -0.11569 | 0.418443 | 0.821966 | Il17rc |
| ENSMUSG( | 321.3497 | -0.16183 | 0.392447 | 0.809274 | Il17rd |
| ENSMUSG( | 1146.169 | 0.116894 | 0.338804 | 0.777518 | Il18 |
| ENSMUSG( | 203.9318 | 0.062499 | 0.717569 | 0.941697 | Il18bp |
| ENSMUSG( | 12.00961 | -0.14256 | 0.822588 | NA | Il18rap |
| ENSMUSG( | 22.80658 | 0.009389 | 0.982548 | 0.997512 | Il1a |
| ENSMUSG( | 186.8745 | 0.278421 | 0.16404 | 0.594869 | Il1r1 |
| ENSMUSG( | 1087.749 | -0.44565 | 0.010137 | 0.143118 | Il1rap |
| ENSMUSG( | 276.0691 | -0.62629 | 0.000215 | 0.011305 | Il1rapl1 |
| ENSMUSG( | 75.58565 | -0.10189 | 0.680366 | 0.931044 | Il1rapl2 |
| ENSMUSG( | 20.95386 | -0.36715 | 0.382693 | NA | Il1rl1 |
| ENSMUSG( | 43.77193 | -0.34736 | 0.219567 | 0.664984 | Il1rl2 |
| ENSMUSG( | 12.35647 | -0.66111 | 0.213327 | NA | Il20ra |
| ENSMUSG( | 27.32906 | -0.01132 | 0.976776 | 0.997428 | Il20rb |
| ENSMUSG( | 94.87616 | 0.040648 | 0.881951 | 0.982069 | Il21r |
| ENSMUSG( | 60.86326 | 0.405818 | 0.10206 | 0.486742 | Il23a |
| ENSMUSG( | 13.35512 | 0.119776 | 0.836378 | NA | Il27ra |
| ENSMUSG( | 21.88567 | -0.75064 | 0.183257 | 0.62047 | Il2rb |
| ENSMUSG( | 74.53134 | -0.50113 | 0.052942 | 0.35324 | Il2rg |
| ENSMUSG( | 16.28624 | -0.1605 | 0.743078 | NA | Il31ra |
| ENSMUSG( | 1914.645 | 0.054296 | 0.729406 | 0.944738 | Il33 |
| ENSMUSG( | 1692.01 | -0.28555 | 0.036599 | 0.295048 | Il34 |
| ENSMUSG( | 96.11111 | 0.237276 | 0.238585 | 0.686308 | Il3ra |
| ENSMUSG( | 34.75667 | -0.12697 | 0.696869 | 0.935464 | Il4i1 |
| ENSMUSG( | 307.0869 | -0.19897 | 0.29302 | 0.742763 | Il4ra |
| ENSMUSG( | 325.8001 | -0.05305 | 0.766312 | 0.954173 | Il6ra |
| ENSMUSG( | 4545.86 | -0.37948 | 0.026106 | 0.247245 | Il6st |
| ENSMUSG( | 34.36422 | -0.37272 | 0.259926 | 0.709951 | Il7r |
| ENSMUSG( | 21.85253 | -0.21802 | 0.594224 | 0.902869 | Ildr1 |
| ENSMUSG( | 5815.422 | -0.14713 | 0.192205 | 0.631194 | Ildr2 |
| ENSMUSG( | 1545.296 | -0.00983 | 0.932159 | 0.992515 | Ilf2 |
| ENSMUSG( | 2061.646 | 0.17654 | 0.062053 | 0.385335 | Ilf3 |
| ENSMUSG( | 4572.427 | 0.072227 | 0.39014 | 0.80794 | Ilk |
| ENSMUSG( | 1230.876 | 0.044368 | 0.6591 | 0.924506 | Ilkap |
| ENSMUSG( | 18568.55 | 0.165925 | 0.03693 | 0.296508 | Ilrun |
| ENSMUSG( | 795.688 | -0.32774 | 0.016548 | 0.191177 | Ilvbl |
| ENSMUSG( | 489.1058 | 0.121693 | 0.323374 | 0.768488 | Immp1l |
| ENSMUSG( | 77.83229 | -0.35748 | 0.103693 | 0.489428 | Immp2l |
| ENSMUSG( | 4382.704 | 0.16555 | 0.034521 | 0.285239 | Immt |
| ENSMUSG( | 581.0275 | 0.005176 | 0.964093 | 0.995137 | Imp3 |
| ENSMUSG( | 935.1967 | 0.107471 | 0.337437 | 0.777412 | Imp4 |
| ENSMUSG( | 1266.422 | -0.03463 | 0.751166 | 0.950161 | Impa1 |
| ENSMUSG( | 254.3927 | -0.25387 | 0.08824 | 0.455783 | Impa2 |
| ENSMUSG( | 3249.047 | 0.043 | 0.656918 | 0.924069 | Impact |
| ENSMUSG( | 1464.636 | 0.048627 | 0.698155 | 0.936028 | Impdh1 |
| ENSMUSG( | 954.2539 | 0.120233 | 0.31153 | 0.759296 | Impdh2 |
| ENSMUSG( | 14.10579 | 0.336152 | 0.49918 | NA | Impdh2-ps |
| ENSMUSG( | 11.92927 | -1.08437 | 0.054797 | NA | Impg1 |
| ENSMUSG( | 27.88484 | -0.262 | 0.469618 | 0.846167 | Impg2 |
| ENSMUSG( | 5186.733 | 0.048492 | 0.670719 | 0.928683 | Ina |
| ENSMUSG( | 979.1531 | -0.00828 | 0.93475 | 0.993231 | Inafm1 |
| ENSMUSG( | 1533.417 | -0.14196 | 0.125206 | 0.532314 | Inafm2 |
| ENSMUSG( | 194.1708 | -0.22611 | 0.144473 | 0.564381 | Inava |
| ENSMUSG( | 76.9264 | 0.012145 | 0.956007 | 0.993874 | Inca1 |
| ENSMUSG( | 462.9267 | 0.057652 | 0.617889 | 0.910872 | Incenp |
| ENSMUSG( | 5312.783 | -0.06764 | 0.577344 | 0.895776 | Inf2 |
| ENSMUSG( | 1333.537 | 0.056929 | 0.547584 | 0.885153 | Ing1 |
| ENSMUSG( | 490.8876 | -0.01671 | 0.90199 | 0.985633 | Ing2 |
| ENSMUSG( | 328.8384 | 0.048983 | 0.714073 | 0.940959 | Ing3 |
| ENSMUSG( | 21.71172 | 0.51794 | 0.243367 | 0.690747 | Ing4 |
| ENSMUSG( | 331.8409 | -0.15406 | 0.324719 | 0.770182 | Ing5 |
| ENSMUSG( | 716.4826 | 0.028407 | 0.859184 | 0.974707 | Inha |
| ENSMUSG( | 182.2833 | -0.5835 | 0.002959 | 0.066473 | Inhba |
| ENSMUSG( | 330.4266 | -0.28443 | 0.133091 | 0.545688 | Inhbb |
| ENSMUSG( | 7.938654 | -0.38729 | 0.580606 | NA | Inhbe |
| ENSMUSG( | 630.9567 | 0.025789 | 0.805414 | 0.962616 | Inip |
| ENSMUSG( | 61.60586 | -0.06938 | 0.780655 | 0.957491 | Inka1 |
| ENSMUSG( | 2548.403 | 0.027209 | 0.795726 | 0.960062 | Inka2 |
| ENSMUSG( | 10.05538 | -0.58453 | 0.342781 | NA | Inmt |
| ENSMUSG( | 987.6597 | 0.025356 | 0.798647 | 0.960896 | Ino80 |
| ENSMUSG( | 375.6144 | 0.122334 | 0.393129 | 0.809274 | Ino80b |
| ENSMUSG( | 989.6258 | -0.04551 | 0.670609 | 0.928683 | Ino80c |
| ENSMUSG( | 778.6502 | 0.151774 | 0.170697 | 0.605271 | Ino80d |
| ENSMUSG( | 129.8077 | -0.07157 | 0.714164 | 0.941003 | Ino80dos |
| ENSMUSG( | 923.9834 | -0.06089 | 0.581613 | 0.897855 | Ino80e |
| ENSMUSG( | 780.9628 | -0.04753 | 0.671481 | 0.928781 | Inpp1 |
| ENSMUSG( | 2943.111 | 0.155844 | 0.125275 | 0.532314 | Inpp4a |
| ENSMUSG( | 140.0271 | -0.22867 | 0.185462 | 0.623185 | Inpp4b |
| ENSMUSG( | 1682.251 | -0.00539 | 0.952721 | 0.993874 | Inpp5a |
| ENSMUSG( | 1173.877 | 0.101435 | 0.294052 | 0.743459 | Inpp5b |
| ENSMUSG( | 897.6445 | 0.24991 | 0.03313 | 0.278602 | Inpp5d |
| ENSMUSG( | 728.4508 | 0.0706 | 0.596304 | 0.903254 | Inpp5e |
| ENSMUSG( | 2859.382 | -0.27695 | 0.004798 | 0.091446 | Inpp5f |
| ENSMUSG( | 1519.855 | 0.19381 | 0.164166 | 0.5949 | Inpp5j |
| ENSMUSG( | 1247.167 | 0.104438 | 0.396591 | 0.811742 | Inpp5k |
| ENSMUSG( | 2666.652 | 0.01347 | 0.867946 | 0.976631 | Inppl1 |
| ENSMUSG( | 101.761 | -0.46081 | 0.02922 | 0.259574 | Insc |
| ENSMUSG( | 3813.774 | -0.41578 | 0.003932 | 0.080285 | Insig1 |
| ENSMUSG( | 1248.798 | -0.17449 | 0.181808 | 0.618944 | Insig2 |
| ENSMUSG( | 21.92546 | 0.279893 | 0.485942 | 0.858043 | Insl5 |
| ENSMUSG( | 10.25899 | -0.48764 | 0.47566 | NA | Insl6 |
| ENSMUSG( | 159.927 | -0.11273 | 0.527558 | 0.877641 | Insm1 |
| ENSMUSG( | 1833.358 | -0.14754 | 0.247525 | 0.696522 | Insr |
| ENSMUSG( | 16.96861 | -0.02329 | 0.962455 | NA | Insrr |
| ENSMUSG( | 6985.019 | 0.053405 | 0.480459 | 0.854748 | Insyn1 |
| ENSMUSG( | 196.5973 | -0.14867 | 0.383942 | 0.803646 | Insyn2a |
| ENSMUSG( | 43.22805 | -0.08405 | 0.797258 | 0.960507 | Insyn2b |
| ENSMUSG( | 2992.256 | 0.182996 | 0.054806 | 0.360194 | Ints1 |
| ENSMUSG( | 1075.066 | 0.09511 | 0.347671 | 0.780695 | Ints10 |
| ENSMUSG( | 758.1517 | 0.174519 | 0.096948 | 0.477375 | Ints11 |
| ENSMUSG( | 204.6604 | 0.063399 | 0.686178 | 0.932981 | Ints12 |
| ENSMUSG( | 903.2027 | 0.116063 | 0.249123 | 0.698492 | Ints13 |
| ENSMUSG( | 624.1875 | 0.004461 | 0.971646 | 0.996528 | Ints14 |
| ENSMUSG( | 727.5839 | 0.012415 | 0.903717 | 0.986094 | Ints15 |
| ENSMUSG( | 549.8528 | -0.08676 | 0.469322 | 0.84601 | Ints2 |
| ENSMUSG( | 1732.626 | 0.213647 | 0.018171 | 0.20312 | Ints3 |
| ENSMUSG( | 806.3945 | 0.151616 | 0.137632 | 0.552903 | Ints4 |
| ENSMUSG( | 433.8954 | 0.155213 | 0.304529 | 0.752574 | Ints5 |
| ENSMUSG( | 674.3342 | 0.002941 | 0.976908 | 0.997428 | Ints6 |
| ENSMUSG( | 1732.951 | -0.10257 | 0.42151 | 0.823126 | Ints6l |
| ENSMUSG( | 1261.459 | 0.040603 | 0.714968 | 0.941144 | Ints7 |
| ENSMUSG( | 648.5903 | 0.234855 | 0.057999 | 0.371396 | Ints8 |
| ENSMUSG( | 400.8054 | 0.198559 | 0.14345 | 0.562473 | Ints9 |
| ENSMUSG( | 388.4011 | -0.06546 | 0.578282 | 0.89641 | Intu |
| ENSMUSG( | 523.7596 | 0.002979 | 0.97857 | 0.997428 | Invs |
| ENSMUSG( | 6784.184 | 0.055427 | 0.553635 | 0.886216 | Ip6k1 |
| ENSMUSG( | 1587.086 | -0.08682 | 0.477416 | 0.852548 | Ip6k2 |
| ENSMUSG( | 43.03022 | -0.17147 | 0.549181 | 0.885723 | Ip6k3 |
| ENSMUSG( | 2015.756 | 0.167031 | 0.069052 | 0.407132 | Ipcef1 |
| ENSMUSG( | 1605.313 | 0.052077 | 0.666449 | 0.927243 | Ipmk |
| ENSMUSG( | 1253.202 | 0.061997 | 0.542972 | 0.883579 | Ipo11 |
| ENSMUSG( | 3220.382 | -0.04521 | 0.58311 | 0.898199 | Ipo13 |
| ENSMUSG( | 1129.002 | 0.105881 | 0.425182 | 0.82478 | Ipo4 |
| ENSMUSG( | 3745.395 | 0.140099 | 0.094546 | 0.471713 | Ipo5 |
| ENSMUSG( | 3207.487 | 0.129949 | 0.186324 | 0.623869 | Ipo7 |
| ENSMUSG( | 1976.5 | 0.133306 | 0.18347 | 0.620494 | Ipo8 |
| ENSMUSG( | 7263.257 | 0.023963 | 0.747462 | 0.949604 | Ipo9 |
| ENSMUSG( | 291.1477 | 0.143459 | 0.283673 | 0.732203 | Ipp |
| ENSMUSG( | 436.9566 | 0.219809 | 0.170538 | 0.605271 | Ippk |
| ENSMUSG( | 1206.85 | 0.010731 | 0.941486 | 0.993415 | Ipw |
| ENSMUSG( | 63.53277 | -0.05023 | 0.838568 | 0.971111 | Iqank1 |
| ENSMUSG( | 36.51704 | -0.96134 | 0.006183 | 0.107367 | Iqca1 |
| ENSMUSG( | 456.5683 | 0.084361 | 0.489276 | 0.859092 | Iqcb1 |
| ENSMUSG( | 473.0081 | 0.263404 | 0.033879 | 0.282701 | Iqcc |
| ENSMUSG( | 29.39358 | -0.26709 | 0.501168 | 0.864138 | Iqcd |
| ENSMUSG( | 1209.974 | 0.079806 | 0.399098 | 0.813048 | Iqce |
| ENSMUSG( | 128.3153 | -0.01331 | 0.946395 | 0.99369 | Iqcg |
| ENSMUSG( | 121.4729 | -0.15346 | 0.474134 | 0.849425 | Iqck |
| ENSMUSG( | 1070.678 | -0.01372 | 0.916312 | 0.988795 | Iqgap1 |
| ENSMUSG( | 576.7814 | 0.346521 | 0.023102 | 0.233188 | Iqgap2 |
| ENSMUSG( | 65.16678 | -0.16578 | 0.523142 | 0.875628 | Iqgap3 |
| ENSMUSG( | 10640.35 | 0.134252 | 0.146524 | 0.568435 | Iqsec1 |
| ENSMUSG( | 12833.8 | 0.146102 | 0.122183 | 0.527707 | Iqsec2 |
| ENSMUSG( | 11311.74 | 0.28829 | 0.000567 | 0.022339 | Iqsec3 |
| ENSMUSG( | 55.8348 | 0.110034 | 0.696125 | 0.935007 | Iqub |
| ENSMUSG( | 81.45182 | -0.07903 | 0.720752 | 0.942638 | Irag1 |
| ENSMUSG( | 89.57898 | -0.23563 | 0.421789 | 0.823375 | Irag2 |
| ENSMUSG( | 3624.035 | 0.111112 | 0.195524 | 0.63504 | Irak1 |
| ENSMUSG( | 323.6793 | 0.07603 | 0.548033 | 0.885153 | Irak1bp1 |
| ENSMUSG( | 1137.68 | 0.151464 | 0.164701 | 0.595775 | Irak2 |
| ENSMUSG( | 30.90527 | -0.11545 | 0.737832 | 0.947329 | Irak3 |
| ENSMUSG( | 120.2382 | 0.01718 | 0.929892 | 0.991848 | Irak4 |
| ENSMUSG( | 2003.156 | 0.081842 | 0.443542 | 0.833108 | Ireb2 |
| ENSMUSG( | 184.703 | -0.0587 | 0.718498 | 0.941911 | Irf1 |
| ENSMUSG( | 807.8508 | 0.292482 | 0.017387 | 0.197592 | Irf2 |
| ENSMUSG( | 930.5303 | 0.059486 | 0.67691 | 0.930542 | Irf2bp1 |
| ENSMUSG( | 833.9981 | -0.22112 | 0.034483 | 0.285222 | Irf2bp2 |
| ENSMUSG( | 1355.028 | -0.13109 | 0.217587 | 0.662824 | Irf2bpl |

| | | | | | |
|---|---|---|---|---|---|
| ENSMUSG( | 31.22062 | 0.306853 | 0.396957 | 0.917257 | Impg2 |
| ENSMUSG( | 4362.167 | -0.18991 | 0.13719 | 0.828492 | Ina |
| ENSMUSG( | 895.8433 | 0.00432 | 0.970216 | 0.99594 | Inafm1 |
| ENSMUSG( | 1506.985 | 0.078164 | 0.50657 | 0.928006 | Inafm2 |
| ENSMUSG( | 184.167 | -0.09738 | 0.595785 | 0.946047 | Inava |
| ENSMUSG( | 65.6928 | -0.15127 | 0.583333 | 0.944266 | Inca1 |
| ENSMUSG( | 400.3027 | -0.10106 | 0.485689 | 0.92541 | Incenp |
| ENSMUSG( | 4633.824 | -0.20117 | 0.119441 | 0.813384 | Inf2 |
| ENSMUSG( | 1180.532 | -0.02924 | 0.813008 | 0.979198 | Ing1 |
| ENSMUSG( | 438.3699 | -0.07887 | 0.593283 | 0.945408 | Ing2 |
| ENSMUSG( | 313.7674 | 0.174268 | 0.281187 | 0.892356 | Ing3 |
| ENSMUSG( | 28.17808 | 1.321276 | 0.001817 | 0.15837 | Ing4 |
| ENSMUSG( | 305.8433 | -0.10901 | 0.53911 | 0.937216 | Ing5 |
| ENSMUSG( | 633.3092 | -0.06314 | 0.725999 | 0.966887 | Inha |
| ENSMUSG( | 221.3123 | 0.295199 | 0.136621 | 0.827522 | Inhba |
| ENSMUSG( | 359.7202 | 0.237601 | 0.215406 | 0.865167 | Inhbb |
| ENSMUSG( | 557.9039 | -0.06265 | 0.629758 | 0.953251 | Inip |
| ENSMUSG( | 55.2499 | -0.09045 | 0.761505 | 0.972819 | Inka1 |
| ENSMUSG( | 2162.935 | -0.18555 | 0.16437 | 0.83856 | Inka2 |
| ENSMUSG( | 14.85162 | 0.735373 | 0.178569 | 0.849041 | Inmt |
| ENSMUSG( | 909.6137 | 0.057784 | 0.626272 | 0.953234 | Ino80 |
| ENSMUSG( | 338.0597 | 0.086368 | 0.589223 | 0.944975 | Ino80b |
| ENSMUSG( | 838.8835 | -0.27086 | 0.055258 | 0.723362 | Ino80c |
| ENSMUSG( | 724.55 | 0.209533 | 0.094333 | 0.795735 | Ino80d |
| ENSMUSG( | 120.0222 | -0.0312 | 0.890359 | 0.990558 | Ino80dos |
| ENSMUSG( | 817.7076 | -0.14907 | 0.251342 | 0.879497 | Ino80e |
| ENSMUSG( | 717.4056 | -0.02938 | 0.84284 | 0.98313 | Inpp1 |
| ENSMUSG( | 2611.523 | 0.08286 | 0.465257 | 0.923329 | Inpp4a |
| ENSMUSG( | 149.8173 | 0.233367 | 0.205684 | 0.864412 | Inpp4b |
| ENSMUSG( | 1456.714 | -0.15918 | 0.166771 | 0.840183 | Inpp5a |
| ENSMUSG( | 1052.965 | 0.057649 | 0.659627 | 0.956205 | Inpp5b |
| ENSMUSG( | 825.9816 | 0.281262 | 0.038058 | 0.676422 | Inpp5d |
| ENSMUSG( | 694.4834 | 0.198763 | 0.207618 | 0.864412 | Inpp5e |
| ENSMUSG( | 2795.061 | -0.05942 | 0.60873 | 0.950113 | Inpp5f |
| ENSMUSG( | 1280.647 | -0.02761 | 0.857571 | 0.985884 | Inpp5j |
| ENSMUSG( | 1116.206 | 0.05382 | 0.698175 | 0.960868 | Inpp5k |
| ENSMUSG( | 2469.68 | 0.060646 | 0.556961 | 0.939565 | Inppl1 |
| ENSMUSG( | 100.579 | -0.20573 | 0.376022 | 0.915491 | Insc |
| ENSMUSG( | 3735.821 | -0.17921 | 0.261458 | 0.884926 | Insig1 |
| ENSMUSG( | 1296.092 | 0.203495 | 0.16268 | 0.837724 | Insig2 |
| ENSMUSG( | 26.00052 | 0.879403 | 0.05362 | 0.720293 | Insl5 |
| ENSMUSG( | 144.1423 | -0.13038 | 0.54239 | 0.938034 | Insm1 |
| ENSMUSG( | 1932.192 | 0.26779 | 0.063799 | 0.736482 | Insr |
| ENSMUSG( | 16.89278 | 0.221647 | 0.64556 | 0.953988 | Insrr |
| ENSMUSG( | 6333.316 | 0.039037 | 0.689199 | 0.959893 | Insyn1 |
| ENSMUSG( | 177.1375 | -0.1929 | 0.336952 | 0.907972 | Insyn2a |
| ENSMUSG( | 38.83734 | -0.15032 | 0.652277 | 0.954766 | Insyn2b |
| ENSMUSG( | 2616.373 | 0.071504 | 0.538868 | 0.937216 | Ints1 |
| ENSMUSG( | 1008.725 | 0.179706 | 0.181628 | 0.850579 | Ints10 |
| ENSMUSG( | 646.6731 | -0.00988 | 0.9399 | 0.994334 | Ints11 |
| ENSMUSG( | 194.0763 | 0.166277 | 0.360384 | 0.914223 | Ints12 |
| ENSMUSG( | 814.9668 | 0.091522 | 0.466494 | 0.923329 | Ints13 |
| ENSMUSG( | 582.5127 | 0.077073 | 0.557666 | 0.939565 | Ints14 |
| ENSMUSG( | 662.5453 | 0.01149 | 0.923874 | 0.992841 | Ints15 |
| ENSMUSG( | 559.3198 | 0.226859 | 0.093552 | 0.795507 | Ints2 |
| ENSMUSG( | 1509.887 | 0.093688 | 0.40898 | 0.917257 | Ints3 |
| ENSMUSG( | 688.9995 | -0.03345 | 0.78606 | 0.976285 | Ints4 |
| ENSMUSG( | 384.5012 | 0.078651 | 0.634344 | 0.953903 | Ints5 |
| ENSMUSG( | 615.6108 | 0.002469 | 0.985606 | 0.997084 | Ints6 |
| ENSMUSG( | 1802.894 | 0.270044 | 0.150877 | 0.837173 | Ints6l |
| ENSMUSG( | 1154.721 | 0.056778 | 0.69065 | 0.960071 | Ints7 |
| ENSMUSG( | 583.0833 | 0.192733 | 0.191019 | 0.858187 | Ints8 |
| ENSMUSG( | 349.8689 | 0.085661 | 0.595206 | 0.945701 | Ints9 |
| ENSMUSG( | 385.0074 | 0.171829 | 0.212536 | 0.864412 | Intu |
| ENSMUSG( | 457.5041 | -0.1239 | 0.358748 | 0.913759 | Invs |
| ENSMUSG( | 6330.691 | 0.124043 | 0.243382 | 0.877758 | Ip6k1 |
| ENSMUSG( | 1495.45 | 0.01156 | 0.930745 | 0.99306 | Ip6k2 |
| ENSMUSG( | 32.7701 | -0.77864 | 0.026631 | 0.603041 | Ip6k3 |
| ENSMUSG( | 1656.326 | -0.13086 | 0.243375 | 0.877758 | Ipcef1 |
| ENSMUSG( | 1462.621 | 0.047669 | 0.724727 | 0.966887 | Ipmk |
| ENSMUSG( | 1083.533 | -0.09182 | 0.451387 | 0.922202 | Ipo11 |
| ENSMUSG( | 2968.247 | -0.01009 | 0.921384 | 0.992644 | Ipo13 |
| ENSMUSG( | 952.3745 | -0.12046 | 0.409338 | 0.917257 | Ipo4 |
| ENSMUSG( | 3209.829 | -0.03347 | 0.75001 | 0.970113 | Ipo5 |
| ENSMUSG( | 2805.923 | 0.012797 | 0.920869 | 0.992644 | Ipo7 |
| ENSMUSG( | 1687.386 | -0.05375 | 0.649715 | 0.95422 | Ipo8 |
| ENSMUSG( | 6300.654 | -0.12048 | 0.212297 | 0.864412 | Ipo9 |
| ENSMUSG( | 241.1549 | -0.13981 | 0.393906 | 0.917257 | Ipp |
| ENSMUSG( | 352.8531 | -0.12027 | 0.491055 | 0.926604 | Ippk |
| ENSMUSG( | 1165.648 | 0.172388 | 0.311777 | 0.900031 | Ipw |
| ENSMUSG( | 54.86841 | -0.22553 | 0.458511 | 0.923329 | Iqank1 |
| ENSMUSG( | 46.14498 | 0.165725 | 0.692181 | 0.960404 | Iqca1 |
| ENSMUSG( | 416.7093 | 0.086796 | 0.554236 | 0.939565 | Iqcb1 |
| ENSMUSG( | 378.8988 | -0.09625 | 0.547629 | 0.938063 | Iqcc |
| ENSMUSG( | 28.29527 | -0.09335 | 0.820299 | 0.979198 | Iqcd |
| ENSMUSG( | 1093.011 | 0.055469 | 0.672259 | 0.95694 | Iqce |
| ENSMUSG( | 120.0059 | 0.068446 | 0.735125 | 0.967344 | Iqcg |
| ENSMUSG( | 113.2979 | -0.08176 | 0.705974 | 0.96335 | Iqck |
| ENSMUSG( | 1034.716 | 0.156744 | 0.278454 | 0.891628 | Iqgap1 |
| ENSMUSG( | 572.4994 | 0.565424 | 0.000163 | 0.030394 | Iqgap2 |
| ENSMUSG( | 55.53642 | -0.38591 | 0.173857 | 0.844317 | Iqgap3 |
| ENSMUSG( | 9490.269 | 0.076804 | 0.547348 | 0.938063 | Iqsec1 |
| ENSMUSG( | 11736.49 | 0.158117 | 0.171559 | 0.842999 | Iqsec2 |
| ENSMUSG( | 9576.795 | 0.090474 | 0.384498 | 0.917257 | Iqsec3 |
| ENSMUSG( | 50.98929 | 0.119288 | 0.699957 | 0.961335 | Iqub |
| ENSMUSG( | 86.05172 | 0.344122 | 0.157037 | 0.837724 | Irag1 |
| ENSMUSG( | 81.49526 | -0.23624 | 0.428134 | 0.917257 | Irag2 |
| ENSMUSG( | 3123.94 | -0.04887 | 0.665128 | 0.956766 | Irak1 |
| ENSMUSG( | 295.175 | 0.07805 | 0.594686 | 0.945444 | Irak1bp1 |
| ENSMUSG( | 930.1205 | -0.15951 | 0.160847 | 0.837724 | Irak2 |
| ENSMUSG( | 27.73709 | -0.12311 | 0.761316 | 0.972718 | Irak3 |
| ENSMUSG( | 118.0111 | 0.220218 | 0.286039 | 0.893906 | Irak4 |
| ENSMUSG( | 1835.095 | 0.09544 | 0.458735 | 0.923329 | Ireb2 |
| ENSMUSG( | 192.6838 | 0.324218 | 0.077054 | 0.770558 | Irf1 |
| ENSMUSG( | 716.3714 | 0.216897 | 0.146545 | 0.835973 | Irf2 |
| ENSMUSG( | 819.4519 | -0.04274 | 0.791873 | 0.97681 | Irf2bp1 |
| ENSMUSG( | 815.5741 | -0.00684 | 0.953838 | 0.994798 | Irf2bp2 |
| ENSMUSG( | 1292.289 | 0.001982 | 0.988084 | 0.997862 | Irf2bpl |
| ENSMUSG( | 635.7698 | 0.321644 | 0.057904 | 0.728256 | Irf3 |
| ENSMUSG( | 201.2109 | 0.153311 | 0.379401 | 0.916995 | Irf5 |
| ENSMUSG( | 8.311901 | 0.812876 | 0.241353 | NA | Irf6 |
| ENSMUSG( | 157.2971 | 0.029219 | 0.893605 | 0.990558 | Irf7 |
| ENSMUSG( | 266.8938 | 0.310062 | 0.045565 | 0.693959 | Irf8 |
| ENSMUSG( | 435.7547 | 0.139879 | 0.39125 | 0.917257 | Irf9 |
| ENSMUSG( | 353.5054 | -0.15425 | 0.356004 | 0.913331 | Irgm1 |
| ENSMUSG( | 186.348 | 0.510849 | 0.014687 | 0.48727 | Irgm2 |
| ENSMUSG( | 3899.146 | 0.051588 | 0.627241 | 0.953234 | Irgq |
| ENSMUSG( | 412.7187 | 0.101867 | 0.480708 | 0.924548 | Irs1 |
| ENSMUSG( | 1136.572 | 0.124586 | 0.362322 | 0.914223 | Irs2 |
| ENSMUSG( | 17.46929 | -0.52837 | 0.266326 | 0.886134 | Irs4 |
| ENSMUSG( | 22.03516 | -0.88504 | 0.079505 | 0.77458 | Irx1 |
| ENSMUSG( | 3029.059 | -0.05001 | 0.657588 | 0.955772 | Isca1 |
| ENSMUSG( | 938.6858 | -0.0602 | 0.616151 | 0.951413 | Isca2 |
| ENSMUSG( | 1883.412 | -0.23526 | 0.042188 | 0.690304 | Iscu |
| ENSMUSG( | 84.4717 | 0.035453 | 0.884572 | 0.989688 | Isg15 |
| ENSMUSG( | 89.4144 | -0.19975 | 0.469127 | 0.923329 | Isg20 |
| ENSMUSG( | 342.2577 | 0.107845 | 0.447286 | 0.922202 | Isg20l2 |
| ENSMUSG( | 25.85771 | -0.68527 | 0.226653 | 0.868507 | Isl1 |
| ENSMUSG( | 998.4291 | 0.128 | 0.31599 | 0.901868 | Islr |
| ENSMUSG( | 3252.728 | 0.280993 | 0.090255 | 0.790963 | Islr2 |
| ENSMUSG( | 236.347 | 0.237183 | 0.197202 | 0.862161 | Ism1 |
| ENSMUSG( | 11.85223 | -0.0878 | 0.883852 | 0.989644 | Ism2 |
| ENSMUSG( | 1164.743 | 0.010483 | 0.94947 | 0.994798 | Isoc1 |
| ENSMUSG( | 1016.408 | -0.32893 | 0.016669 | 0.521279 | Isoc2a |
| ENSMUSG( | 141.468 | -0.03086 | 0.894039 | 0.990558 | Isoc2b |
| ENSMUSG( | 2125.068 | 0.135231 | 0.308992 | 0.898538 | Ist1 |
| ENSMUSG( | 500.8017 | -0.0565 | 0.668569 | 0.956766 | Isy1 |
| ENSMUSG( | 901.4106 | 0.306026 | 0.036613 | 0.670163 | Isyna1 |
| ENSMUSG( | 1416.541 | 0.14114 | 0.285004 | 0.89246 | Itch |
| ENSMUSG( | 7053.676 | 0.150274 | 0.362789 | 0.914223 | Itfg1 |
| ENSMUSG( | 431.9684 | -0.10679 | 0.431071 | 0.917257 | Itfg2 |
| ENSMUSG( | 534.6283 | 0.512791 | 0.256078 | 0.883082 | Itga1 |
| ENSMUSG( | 120.0993 | 0.534111 | 0.032988 | 0.63923 | Itga10 |
| ENSMUSG( | 96.81074 | 0.123954 | 0.603401 | 0.948137 | Itga11 |
| ENSMUSG( | 62.64904 | 0.085206 | 0.739819 | 0.968726 | Itga2b |
| ENSMUSG( | 663.9258 | 0.206951 | 0.152207 | 0.837724 | Itga3 |
| ENSMUSG( | 464.0873 | 0.115403 | 0.603597 | 0.948137 | Itga4 |
| ENSMUSG( | 242.105 | -0.16633 | 0.430975 | 0.917257 | Itga5 |
| ENSMUSG( | 1749.097 | 0.166262 | 0.374698 | 0.915491 | Itga6 |
| ENSMUSG( | 733.691 | 0.184119 | 0.259521 | 0.883971 | Itga7 |
| ENSMUSG( | 219.9989 | 0.226342 | 0.204428 | 0.864412 | Itga8 |
| ENSMUSG( | 392.5165 | 0.232499 | 0.260807 | 0.884853 | Itga9 |
| ENSMUSG( | 12.28166 | 0.059764 | 0.914149 | 0.992407 | Itgad |
| ENSMUSG( | 12.02496 | -0.03996 | 0.947428 | 0.994798 | Itgae |
| ENSMUSG( | 112.3111 | -1.24444 | 0.042484 | 0.690304 | Itgal |
| ENSMUSG( | 633.1556 | 0.069491 | 0.684088 | 0.958678 | Itgam |
| ENSMUSG( | 3294.023 | 0.157333 | 0.306019 | 0.898538 | Itgav |
| ENSMUSG( | 25.3663 | -0.37079 | 0.418577 | 0.917257 | Itgax |

| | | | | | |
|---|---|---|---|---|---|
| ENSMUSG( | Ints4 | 806.3945 | 0.151616 | 0.137632 | 0.552903 |
| ENSMUSG( | Ints5 | 433.8954 | 0.155213 | 0.304529 | 0.752574 |
| ENSMUSG( | Ints6 | 674.3342 | 0.002941 | 0.976908 | 0.997428 |
| ENSMUSG( | Ints6l | 1732.951 | -0.10257 | 0.42151 | 0.823126 |
| ENSMUSG( | Ints7 | 1261.459 | 0.040603 | 0.714968 | 0.941144 |
| ENSMUSG( | Ints8 | 648.5903 | 0.234855 | 0.057999 | 0.371396 |
| ENSMUSG( | Ints9 | 400.8054 | 0.198559 | 0.14345 | 0.562473 |
| ENSMUSG( | Intu | 388.4011 | -0.06546 | 0.578282 | 0.89641 |
| ENSMUSG( | Invs | 523.7596 | 0.002979 | 0.97857 | 0.997428 |
| ENSMUSG( | Ip6k1 | 6784.184 | 0.055427 | 0.553635 | 0.886216 |
| ENSMUSG( | Ip6k2 | 1587.086 | -0.08682 | 0.477416 | 0.852548 |
| ENSMUSG( | Ip6k3 | 43.03022 | -0.17147 | 0.549181 | 0.885723 |
| ENSMUSG( | Ipcef1 | 2015.756 | 0.167031 | 0.069052 | 0.407132 |
| ENSMUSG( | Ipmk | 1605.313 | 0.052077 | 0.666449 | 0.927243 |
| ENSMUSG( | Ipo11 | 1253.202 | 0.061997 | 0.542972 | 0.883579 |
| ENSMUSG( | Ipo13 | 3220.382 | -0.04521 | 0.58311 | 0.898199 |
| ENSMUSG( | Ipo4 | 1129.002 | 0.105881 | 0.425182 | 0.82478 |
| ENSMUSG( | Ipo5 | 3745.395 | 0.140099 | 0.094546 | 0.471713 |
| ENSMUSG( | Ipo7 | 3207.487 | 0.129949 | 0.186324 | 0.623869 |
| ENSMUSG( | Ipo8 | 1976.5 | 0.133306 | 0.18347 | 0.620494 |
| ENSMUSG( | Ipo9 | 7263.257 | 0.023963 | 0.747462 | 0.949604 |
| ENSMUSG( | Ipp | 291.1477 | 0.143459 | 0.283673 | 0.732203 |
| ENSMUSG( | Ippk | 436.9566 | 0.219809 | 0.170538 | 0.605271 |
| ENSMUSG( | Ipw | 1206.85 | 0.010731 | 0.941486 | 0.993415 |
| ENSMUSG( | Iqank1 | 63.53277 | -0.05023 | 0.838568 | 0.971111 |
| ENSMUSG( | Iqca1 | 36.51704 | -0.96134 | 0.006183 | 0.107367 |
| ENSMUSG( | Iqcb1 | 456.5683 | 0.084361 | 0.489276 | 0.859092 |
| ENSMUSG( | Iqcc | 473.0081 | 0.263404 | 0.033879 | 0.282701 |
| ENSMUSG( | Iqcd | 29.39358 | -0.26709 | 0.501168 | 0.864138 |
| ENSMUSG( | Iqce | 1209.974 | 0.079806 | 0.399098 | 0.813048 |
| ENSMUSG( | Iqcg | 128.3153 | -0.01331 | 0.946395 | 0.99369 |
| ENSMUSG( | Iqck | 121.4729 | -0.15346 | 0.474134 | 0.849425 |
| ENSMUSG( | Iqgap1 | 1070.678 | -0.01372 | 0.916312 | 0.988795 |
| ENSMUSG( | Iqgap2 | 576.7814 | 0.346521 | 0.023102 | 0.233188 |
| ENSMUSG( | Iqgap3 | 65.16678 | -0.16578 | 0.523142 | 0.875628 |
| ENSMUSG( | Iqsec1 | 10640.35 | 0.134252 | 0.146524 | 0.568435 |
| ENSMUSG( | Iqsec2 | 12833.8 | 0.146102 | 0.122183 | 0.527707 |
| ENSMUSG( | Iqsec3 | 11311.74 | 0.28829 | 0.000567 | 0.022339 |
| ENSMUSG( | Iqub | 55.8348 | 0.110034 | 0.696125 | 0.935007 |
| ENSMUSG( | Irag1 | 81.45182 | -0.07903 | 0.720752 | 0.942638 |
| ENSMUSG( | Irag2 | 89.57898 | -0.23563 | 0.421789 | 0.823375 |
| ENSMUSG( | Irak1 | 3624.035 | 0.111112 | 0.195524 | 0.63504 |
| ENSMUSG( | Irak1bp1 | 323.6793 | 0.07603 | 0.548033 | 0.885153 |
| ENSMUSG( | Irak2 | 1137.68 | 0.151464 | 0.164701 | 0.595775 |
| ENSMUSG( | Irak3 | 30.90527 | -0.11545 | 0.737832 | 0.947329 |
| ENSMUSG( | Irak4 | 120.2382 | 0.01718 | 0.929892 | 0.991848 |
| ENSMUSG( | Ireb2 | 2003.156 | 0.081842 | 0.443542 | 0.833108 |
| ENSMUSG( | Irf1 | 184.703 | -0.0587 | 0.718498 | 0.941911 |
| ENSMUSG( | Irf2 | 807.8508 | 0.292482 | 0.017387 | 0.197592 |
| ENSMUSG( | Irf2bp1 | 930.5303 | 0.059486 | 0.67691 | 0.930542 |
| ENSMUSG( | Irf2bp2 | 833.9981 | -0.22112 | 0.034483 | 0.285222 |
| ENSMUSG( | Irf2bpl | 1355.028 | -0.13109 | 0.217587 | 0.662824 |
| ENSMUSG( | Irf3 | 661.6655 | 0.176505 | 0.123208 | 0.530407 |
| ENSMUSG( | Irf5 | 212.8768 | 0.040099 | 0.822572 | 0.966352 |
| ENSMUSG( | Irf6 | 8.726886 | 0.702179 | 0.280873 | NA |
| ENSMUSG( | Irf7 | 158.373 | -0.24891 | 0.177611 | 0.61504 |
| ENSMUSG( | Irf8 | 265.6052 | 0.038398 | 0.787197 | 0.959147 |
| ENSMUSG( | Irf9 | 455.8537 | 0.000513 | 0.996917 | 0.999505 |
| ENSMUSG( | Irgm1 | 381.1176 | -0.22136 | 0.076094 | 0.42702 |
| ENSMUSG( | Irgm2 | 165.5415 | -0.07145 | 0.699103 | 0.936835 |
| ENSMUSG( | Irgq | 4510.523 | 0.195239 | 0.027208 | 0.253245 |
| ENSMUSG( | Irs1 | 445.8894 | 0.051612 | 0.681136 | 0.931301 |
| ENSMUSG( | Irs2 | 1278.378 | 0.192504 | 0.132926 | 0.545594 |
| ENSMUSG( | Irs4 | 20.87276 | -0.24102 | 0.628441 | NA |
| ENSMUSG( | Irx1 | 22.18933 | -1.21918 | 0.018137 | 0.20302 |
| ENSMUSG( | Isca1 | 3401 | 0.017794 | 0.8384 | 0.971111 |
| ENSMUSG( | Isca2 | 1049.738 | -0.00475 | 0.960722 | 0.993991 |
| ENSMUSG( | Iscu | 2189.048 | -0.05859 | 0.504392 | 0.866883 |
| ENSMUSG( | Isg15 | 78.54445 | -0.46393 | 0.082291 | 0.442706 |
| ENSMUSG( | Isg20 | 77.94774 | -1.01889 | 7.94E-05 | 0.005507 |
| ENSMUSG( | Isg20l2 | 346.4676 | -0.12383 | 0.31857 | 0.764234 |
| ENSMUSG( | Isl1 | 22.47027 | -1.70452 | 0.00315 | 0.06974 |
| ENSMUSG( | Islr | 941.966 | -0.32412 | 0.019022 | 0.207571 |
| ENSMUSG( | Islr2 | 3051.879 | -0.16869 | 0.278127 | 0.728086 |
| ENSMUSG( | Ism1 | 233.404 | -0.06917 | 0.640341 | 0.916434 |
| ENSMUSG( | Ism2 | 12.36446 | -0.27812 | 0.625119 | NA |
| ENSMUSG( | Isoc1 | 1276.78 | 0.011911 | 0.915691 | 0.988634 |
| ENSMUSG( | Isoc2a | 1248.917 | 0.020937 | 0.837087 | 0.971111 |
| ENSMUSG( | Isoc2b | 147.234 | -0.17412 | 0.367592 | 0.795314 |
| ENSMUSG( | Ist1 | 2201.925 | -0.03196 | 0.776489 | 0.955658 |
| ENSMUSG( | Isy1 | 568.806 | 0.045516 | 0.694559 | 0.9343 |
| ENSMUSG( | Isyna1 | 962.5803 | 0.231424 | 0.038183 | 0.301526 |
| ENSMUSG( | Itch | 1549.367 | 0.132202 | 0.230599 | 0.678947 |
| ENSMUSG( | Itfg1 | 6490.319 | -0.38292 | 0.008835 | 0.132321 |
| ENSMUSG( | Itfg2 | 505.4384 | 0.080355 | 0.464539 | 0.843387 |
| ENSMUSG( | Itga1 | 480.3269 | -0.03124 | 0.888689 | 0.983434 |
| ENSMUSG( | Itga10 | 141.79 | 0.7033 | 0.001594 | 0.044805 |
| ENSMUSG( | Itga11 | 84.36718 | -0.61959 | 0.008216 | 0.126198 |
| ENSMUSG( | Itga2b | 74.03265 | 0.286039 | 0.264975 | 0.715962 |
| ENSMUSG( | Itga3 | 601.1089 | -0.37072 | 0.005514 | 0.099488 |
| ENSMUSG( | Itga4 | 423.8221 | -0.44838 | 0.033316 | 0.279795 |
| ENSMUSG( | Itga5 | 237.6777 | -0.54874 | 0.0006 | 0.023008 |
| ENSMUSG( | Itga6 | 1725.605 | -0.14635 | 0.360247 | 0.791947 |
| ENSMUSG( | Itga7 | 795.926 | 0.144445 | 0.298285 | 0.74698 |
| ENSMUSG( | Itga8 | 243.3225 | 0.236582 | 0.19648 | 0.635942 |
| ENSMUSG( | Itga9 | 348.4569 | -0.40938 | 0.022179 | 0.227114 |
| ENSMUSG( | Itgad | 14.13357 | 0.204087 | 0.685553 | NA |
| ENSMUSG( | Itgae | 12.84717 | -0.13736 | 0.824979 | NA |
| ENSMUSG( | Itgal | 135.3491 | -0.80923 | 0.171439 | 0.605943 |
| ENSMUSG( | Itgam | 642.8969 | -0.15779 | 0.25131 | 0.700869 |
| ENSMUSG( | Itgav | 3272.776 | -0.13136 | 0.343868 | 0.779153 |
| ENSMUSG( | Itgax | 24.38763 | -0.84421 | 0.06049 | 0.379845 |
| ENSMUSG( | Itgb1 | 4851.432 | -0.09805 | 0.439506 | 0.831151 |
| ENSMUSG( | Itgb1bp1 | 738.4183 | -0.03104 | 0.78623 | 0.958774 |
| ENSMUSG( | Itgb1bp2 | 14.55751 | -0.57589 | 0.320299 | NA |
| ENSMUSG( | Itgb2 | 427.1769 | -0.36412 | 0.008337 | 0.127456 |
| ENSMUSG( | Itgb3 | 167.7877 | -0.146 | 0.426203 | 0.824898 |
| ENSMUSG( | Itgb3bp | 113.7773 | 0.131453 | 0.522655 | 0.875389 |
| ENSMUSG( | Itgb4 | 782.9553 | -0.51866 | 1.07E-07 | 2.80E-05 |
| ENSMUSG( | Itgb5 | 4296.732 | 0.009932 | 0.939324 | 0.993415 |
| ENSMUSG( | Itgb7 | 51.6608 | -0.23936 | 0.405575 | 0.81652 |
| ENSMUSG( | Itgb8 | 2991.897 | -0.06687 | 0.647076 | 0.919213 |
| ENSMUSG( | Itgbl1 | 155.5246 | -0.03414 | 0.847328 | 0.972374 |
| ENSMUSG( | Itih2 | 226.5848 | 0.247201 | 0.14067 | 0.557902 |
| ENSMUSG( | Itih3 | 1998.502 | -0.08264 | 0.385095 | 0.804048 |
| ENSMUSG( | Itih4 | 13.99539 | -0.20302 | 0.706476 | NA |
| ENSMUSG( | Itih5 | 3583.3 | 0.114625 | 0.346306 | 0.779153 |
| ENSMUSG( | Itk | 12.19832 | -0.6886 | 0.216139 | NA |
| ENSMUSG( | Itm2a | 2006.127 | -0.10361 | 0.621881 | 0.911432 |
| ENSMUSG( | Itm2b | 32078.24 | -0.21992 | 0.072623 | 0.416237 |
| ENSMUSG( | Itm2c | 31234.37 | -0.3037 | 0.011082 | 0.151746 |
| ENSMUSG( | Itpa | 859.875 | 0.016034 | 0.89452 | 0.984512 |
| ENSMUSG( | Itpk1 | 2254.673 | 0.167201 | 0.08758 | 0.454663 |
| ENSMUSG( | Itpka | 5864.732 | 0.116847 | 0.287363 | 0.737854 |
| ENSMUSG( | Itpkb | 2436.092 | -0.01101 | 0.923019 | 0.990693 |
| ENSMUSG( | Itpkc | 173.0113 | -0.193 | 0.275792 | 0.725669 |
| ENSMUSG( | Itpr1 | 13318.17 | -0.06359 | 0.62217 | 0.911432 |
| ENSMUSG( | Itpr2 | 2426.569 | -0.05046 | 0.674728 | 0.929638 |
| ENSMUSG( | Itpr3 | 160.8018 | 0.377578 | 0.053485 | 0.35564 |
| ENSMUSG( | Itprid1 | 36.4356 | -0.19537 | 0.576594 | 0.895391 |
| ENSMUSG( | Itprid2 | 4943.031 | 0.215448 | 0.030714 | 0.266671 |
| ENSMUSG( | Itprip | 44.7664 | -0.11334 | 0.726483 | 0.944223 |
| ENSMUSG( | Itpripl1 | 103.4072 | -0.47192 | 0.028533 | 0.257525 |
| ENSMUSG( | Itpripl2 | 492.1281 | 0.027382 | 0.857911 | 0.974667 |
| ENSMUSG( | Itsn1 | 8437.152 | 0.277228 | 0.009159 | 0.134809 |
| ENSMUSG( | Itsn2 | 1803.23 | 0.18753 | 0.038583 | 0.302696 |
| ENSMUSG( | Ivd | 4549.931 | 0.110565 | 0.190617 | 0.629951 |
| ENSMUSG( | Ivns1abp | 5502.368 | -0.06654 | 0.426467 | 0.825155 |
| ENSMUSG( | Iws1 | 778.1427 | 0.098307 | 0.361559 | 0.792797 |
| ENSMUSG( | Iyd | 32.13925 | -0.36298 | 0.301248 | 0.749979 |
| ENSMUSG( | Izumo4 | 744.4674 | 0.13787 | 0.446023 | 0.833467 |
| ENSMUSG( | Jade1 | 1312.183 | -0.0072 | 0.93741 | 0.993415 |
| ENSMUSG( | Jade2 | 1351.642 | 0.174085 | 0.163593 | 0.594869 |
| ENSMUSG( | Jade3 | 264.6501 | -0.09679 | 0.481666 | 0.855573 |
| ENSMUSG( | Jag1 | 219.1492 | -0.34076 | 0.081982 | 0.442365 |
| ENSMUSG( | Jag2 | 1399.439 | 0.144396 | 0.259931 | 0.709951 |
| ENSMUSG( | Jagn1 | 859.5961 | -0.35202 | 0.009529 | 0.138128 |
| ENSMUSG( | Jak1 | 4094.989 | 0.116882 | 0.137605 | 0.552903 |
| ENSMUSG( | Jak2 | 1615.102 | 0.130599 | 0.178354 | 0.616026 |
| ENSMUSG( | Jak3 | 306.6622 | 0.22263 | 0.111351 | 0.503375 |
| ENSMUSG( | Jakmip1 | 309.4895 | -0.01581 | 0.920937 | 0.990426 |
| ENSMUSG( | Jakmip2 | 822.3087 | 0.064058 | 0.605966 | 0.90596 |
| ENSMUSG( | Jakmip3 | 922.8495 | -0.00307 | 0.976741 | 0.997428 |
| ENSMUSG( | Jam2 | 4063.035 | -0.13479 | 0.312254 | 0.759791 |
| ENSMUSG( | Jam3 | 1306.812 | -0.34943 | 0.040561 | 0.311208 |
| ENSMUSG( | Jaml | 9.908053 | -0.35696 | 0.548266 | NA |
| ENSMUSG( | Jarid2 | 1083.293 | -0.23736 | 0.050656 | 0.344934 |
| ENSMUSG( | Jazf1 | 330.5096 | 0.305675 | 0.018664 | 0.205542 |

| | | | | | |
|---|---|---|---|---|---|
| ENSMUSG( | Ints4 | 688.9995 | -0.03345 | 0.78606 | 0.976285 |
| ENSMUSG( | Ints5 | 384.5012 | 0.078651 | 0.634344 | 0.953903 |
| ENSMUSG( | Ints6 | 615.6108 | 0.002469 | 0.985606 | 0.997084 |
| ENSMUSG( | Ints6l | 1802.894 | 0.270044 | 0.150877 | 0.837173 |
| ENSMUSG( | Ints7 | 1154.721 | 0.056778 | 0.69065 | 0.960071 |
| ENSMUSG( | Ints8 | 583.0833 | 0.192733 | 0.191019 | 0.858187 |
| ENSMUSG( | Ints9 | 349.8689 | 0.085661 | 0.595206 | 0.945701 |
| ENSMUSG( | Intu | 385.0074 | 0.171829 | 0.212536 | 0.864412 |
| ENSMUSG( | Invs | 457.5041 | -0.1239 | 0.358748 | 0.913759 |
| ENSMUSG( | Ip6k1 | 6330.691 | 0.124043 | 0.243382 | 0.877758 |
| ENSMUSG( | Ip6k2 | 1495.45 | 0.01156 | 0.930745 | 0.99306 |
| ENSMUSG( | Ip6k3 | 32.7701 | -0.77864 | 0.026631 | 0.603041 |
| ENSMUSG( | Ipcef1 | 1656.326 | -0.13086 | 0.243375 | 0.877758 |
| ENSMUSG( | Ipmk | 1462.621 | 0.047669 | 0.724727 | 0.966887 |
| ENSMUSG( | Ipo11 | 1083.533 | -0.09182 | 0.451387 | 0.922202 |
| ENSMUSG( | Ipo13 | 2968.247 | -0.01009 | 0.921384 | 0.992644 |
| ENSMUSG( | Ipo4 | 952.3745 | -0.12046 | 0.409338 | 0.917257 |
| ENSMUSG( | Ipo5 | 3209.829 | -0.03347 | 0.75001 | 0.970113 |
| ENSMUSG( | Ipo7 | 2805.923 | 0.012797 | 0.920869 | 0.992644 |
| ENSMUSG( | Ipo8 | 1687.386 | -0.05375 | 0.649715 | 0.95422 |
| ENSMUSG( | Ipo9 | 6300.654 | -0.12048 | 0.212297 | 0.864412 |
| ENSMUSG( | Ipp | 241.1549 | -0.13981 | 0.393906 | 0.917257 |
| ENSMUSG( | Ippk | 352.8531 | -0.12027 | 0.491055 | 0.926604 |
| ENSMUSG( | Ipw | 1165.648 | 0.172388 | 0.311777 | 0.900031 |
| ENSMUSG( | Iqank1 | 54.86841 | -0.22553 | 0.458511 | 0.923329 |
| ENSMUSG( | Iqca1 | 46.14498 | 0.165725 | 0.692181 | 0.960404 |
| ENSMUSG( | Iqcb1 | 416.7093 | 0.086796 | 0.554236 | 0.939565 |
| ENSMUSG( | Iqcc | 378.8988 | -0.09625 | 0.547629 | 0.938063 |
| ENSMUSG( | Iqcd | 28.29527 | -0.09335 | 0.820299 | 0.979198 |
| ENSMUSG( | Iqce | 1093.011 | 0.055469 | 0.672259 | 0.95694 |
| ENSMUSG( | Iqcg | 120.0059 | 0.068446 | 0.735125 | 0.967344 |
| ENSMUSG( | Iqck | 113.2979 | -0.08176 | 0.705974 | 0.96335 |
| ENSMUSG( | Iqgap1 | 1034.716 | 0.156744 | 0.278454 | 0.891628 |
| ENSMUSG( | Iqgap2 | 572.4994 | 0.565424 | 0.000163 | 0.030394 |
| ENSMUSG( | Iqgap3 | 55.53642 | -0.38591 | 0.173857 | 0.844317 |
| ENSMUSG( | Iqsec1 | 9490.269 | 0.076804 | 0.547348 | 0.938063 |
| ENSMUSG( | Iqsec2 | 11736.49 | 0.158117 | 0.171559 | 0.842999 |
| ENSMUSG( | Iqsec3 | 9576.795 | 0.090474 | 0.384498 | 0.917257 |
| ENSMUSG( | Iqub | 50.98929 | 0.119288 | 0.699957 | 0.961335 |
| ENSMUSG( | Irag1 | 86.05172 | 0.344122 | 0.157037 | 0.837724 |
| ENSMUSG( | Irag2 | 81.49526 | -0.23624 | 0.428134 | 0.917257 |
| ENSMUSG( | Irak1 | 3123.94 | -0.04887 | 0.665128 | 0.956766 |
| ENSMUSG( | Irak1bp1 | 295.175 | 0.07805 | 0.594686 | 0.945444 |
| ENSMUSG( | Irak2 | 930.1205 | -0.15951 | 0.160847 | 0.837724 |
| ENSMUSG( | Irak3 | 27.73709 | -0.12311 | 0.761316 | 0.972718 |
| ENSMUSG( | Irak4 | 118.0111 | 0.220218 | 0.286039 | 0.893906 |
| ENSMUSG( | Ireb2 | 1835.095 | 0.09544 | 0.458735 | 0.923329 |
| ENSMUSG( | Irf1 | 192.6838 | 0.324218 | 0.077054 | 0.770558 |
| ENSMUSG( | Irf2 | 716.3714 | 0.216897 | 0.146545 | 0.835973 |
| ENSMUSG( | Irf2bp1 | 819.4519 | -0.04274 | 0.791873 | 0.97681 |
| ENSMUSG( | Irf2bp2 | 815.5741 | -0.00684 | 0.953838 | 0.994798 |
| ENSMUSG( | Irf2bpl | 1292.289 | 0.001982 | 0.988084 | 0.997862 |
| ENSMUSG( | Irf3 | 635.7698 | 0.321644 | 0.057904 | 0.728256 |
| ENSMUSG( | Irf5 | 201.2109 | 0.153311 | 0.379401 | 0.916995 |
| ENSMUSG( | Irf6 | 8.311901 | 0.812876 | 0.241353 | NA |
| ENSMUSG( | Irf7 | 157.2971 | 0.029219 | 0.893605 | 0.990558 |
| ENSMUSG( | Irf8 | 266.8938 | 0.310062 | 0.045565 | 0.693959 |
| ENSMUSG( | Irf9 | 435.7547 | 0.139879 | 0.39125 | 0.917257 |
| ENSMUSG( | Irgm1 | 353.5054 | -0.15425 | 0.356004 | 0.913331 |
| ENSMUSG( | Irgm2 | 186.348 | 0.510849 | 0.014687 | 0.48727 |
| ENSMUSG( | Irgq | 3899.146 | 0.051588 | 0.627241 | 0.953234 |
| ENSMUSG( | Irs1 | 412.7187 | 0.101867 | 0.480708 | 0.924548 |
| ENSMUSG( | Irs2 | 1136.572 | 0.124586 | 0.362322 | 0.914223 |
| ENSMUSG( | Irs4 | 17.46929 | -0.52837 | 0.266326 | 0.886134 |
| ENSMUSG( | Irx1 | 22.03516 | -0.88504 | 0.079505 | 0.77458 |
| ENSMUSG( | Isca1 | 3029.059 | -0.05001 | 0.657588 | 0.955772 |
| ENSMUSG( | Isca2 | 938.6858 | -0.0602 | 0.616151 | 0.951413 |
| ENSMUSG( | Iscu | 1883.412 | -0.23526 | 0.042188 | 0.690304 |
| ENSMUSG( | Isg15 | 84.4717 | 0.035453 | 0.884572 | 0.989688 |
| ENSMUSG( | Isg20 | 89.4144 | -0.19975 | 0.469127 | 0.923329 |
| ENSMUSG( | Isg20l2 | 342.2577 | 0.107845 | 0.447286 | 0.922202 |
| ENSMUSG( | Isl1 | 25.85771 | -0.68527 | 0.226653 | 0.868507 |
| ENSMUSG( | Islr | 998.4291 | 0.128 | 0.31599 | 0.901868 |
| ENSMUSG( | Islr2 | 3252.728 | 0.280993 | 0.090255 | 0.790963 |
| ENSMUSG( | Ism1 | 236.347 | 0.237183 | 0.197202 | 0.862161 |
| ENSMUSG( | Ism2 | 11.85223 | -0.0878 | 0.883852 | 0.989644 |
| ENSMUSG( | Isoc1 | 1164.743 | 0.010483 | 0.94947 | 0.994798 |
| ENSMUSG( | Isoc2a | 1016.408 | -0.32893 | 0.016669 | 0.521279 |
| ENSMUSG( | Isoc2b | 141.468 | -0.03086 | 0.894039 | 0.990558 |
| ENSMUSG( | Ist1 | 2125.068 | 0.135231 | 0.308992 | 0.898538 |
| ENSMUSG( | Isy1 | 500.8017 | -0.0565 | 0.668569 | 0.956766 |
| ENSMUSG( | Isyna1 | 901.4106 | 0.306026 | 0.036613 | 0.670163 |
| ENSMUSG( | Itch | 1416.541 | 0.14114 | 0.285004 | 0.89246 |
| ENSMUSG( | Itfg1 | 7053.676 | 0.150274 | 0.362789 | 0.914223 |
| ENSMUSG( | Itfg2 | 431.9684 | -0.10679 | 0.431071 | 0.917257 |
| ENSMUSG( | Itga1 | 534.6283 | 0.512791 | 0.256078 | 0.883082 |
| ENSMUSG( | Itga10 | 120.0993 | 0.534111 | 0.032988 | 0.63923 |
| ENSMUSG( | Itga11 | 96.81074 | 0.123954 | 0.603401 | 0.948137 |
| ENSMUSG( | Itga2b | 62.64904 | 0.085206 | 0.739819 | 0.968726 |
| ENSMUSG( | Itga3 | 663.9258 | 0.206951 | 0.152207 | 0.837724 |
| ENSMUSG( | Itga4 | 464.0873 | 0.115403 | 0.603597 | 0.948137 |
| ENSMUSG( | Itga5 | 242.105 | -0.16633 | 0.430975 | 0.917257 |
| ENSMUSG( | Itga6 | 1749.097 | 0.166262 | 0.374698 | 0.915491 |
| ENSMUSG( | Itga7 | 733.691 | 0.184119 | 0.259521 | 0.883971 |
| ENSMUSG( | Itga8 | 219.9989 | 0.226342 | 0.204428 | 0.864412 |
| ENSMUSG( | Itga9 | 392.5165 | 0.232499 | 0.260807 | 0.884853 |
| ENSMUSG( | Itgad | 12.28166 | 0.059764 | 0.914149 | 0.992407 |
| ENSMUSG( | Itgae | 12.02496 | -0.03996 | 0.947428 | 0.994798 |
| ENSMUSG( | Itgal | 112.3111 | -1.24444 | 0.042484 | 0.690304 |
| ENSMUSG( | Itgam | 633.1556 | 0.069491 | 0.684088 | 0.958678 |
| ENSMUSG( | Itgav | 3294.023 | 0.157333 | 0.306019 | 0.898538 |
| ENSMUSG( | Itgax | 25.3663 | -0.37079 | 0.418577 | 0.917257 |
| ENSMUSG( | Itgb1 | 4813.803 | 0.148144 | 0.317857 | 0.902849 |
| ENSMUSG( | Itgb1bp1 | 642.5558 | -0.17341 | 0.17391 | 0.844317 |
| ENSMUSG( | Itgb1bp2 | 14.64324 | -0.26043 | 0.637656 | 0.953903 |
| ENSMUSG( | Itgb2 | 407.4976 | -0.21095 | 0.227569 | 0.868507 |
| ENSMUSG( | Itgb3 | 162.4612 | 0.044402 | 0.848678 | 0.985321 |
| ENSMUSG( | Itgb3bp | 90.21207 | -0.27274 | 0.279537 | 0.892356 |
| ENSMUSG( | Itgb4 | 770.2127 | -0.25843 | 0.058819 | 0.728906 |
| ENSMUSG( | Itgb5 | 3941.456 | 0.033419 | 0.831725 | 0.980656 |
| ENSMUSG( | Itgb7 | 51.86903 | 0.036666 | 0.90508 | 0.991089 |
| ENSMUSG( | Itgb8 | 2845.751 | 0.057219 | 0.726246 | 0.966887 |
| ENSMUSG( | Itgbl1 | 150.4319 | 0.139779 | 0.464524 | 0.923329 |
| ENSMUSG( | Itih2 | 213.1224 | 0.329884 | 0.050183 | 0.708648 |
| ENSMUSG( | Itih3 | 1871.367 | -0.00245 | 0.981883 | 0.996756 |
| ENSMUSG( | Itih4 | 12.40076 | -0.27099 | 0.642134 | 0.953903 |
| ENSMUSG( | Itih5 | 3347.005 | 0.183899 | 0.12301 | 0.814657 |
| ENSMUSG( | Itk | 12.45242 | -0.27105 | 0.633404 | 0.953864 |
| ENSMUSG( | Itm2a | 2466.548 | 0.690648 | 0.110279 | 0.807051 |
| ENSMUSG( | Itm2b | 32689.28 | 0.111041 | 0.436296 | 0.919879 |
| ENSMUSG( | Itm2c | 32098.22 | 0.062242 | 0.656691 | 0.955567 |
| ENSMUSG( | Itpa | 763.0684 | -0.06332 | 0.666899 | 0.956766 |
| ENSMUSG( | Itpk1 | 1932.822 | -0.00352 | 0.974975 | 0.996416 |
| ENSMUSG( | Itpka | 5221.047 | 0.052608 | 0.67458 | 0.95694 |
| ENSMUSG( | Itpkb | 2056.947 | -0.24141 | 0.02829 | 0.612554 |
| ENSMUSG( | Itpkc | 161.4232 | -0.10949 | 0.58653 | 0.944975 |
| ENSMUSG( | Itpr1 | 12944.65 | 0.124883 | 0.415361 | 0.917257 |
| ENSMUSG( | Itpr2 | 2240.184 | -0.00911 | 0.945245 | 0.994798 |
| ENSMUSG( | Itpr3 | 174.6032 | 0.814765 | 0.00019 | 0.034114 |
| ENSMUSG( | Itprid1 | 36.65095 | 0.115892 | 0.758785 | 0.971858 |
| ENSMUSG( | Itprid2 | 4318.548 | 0.100514 | 0.403606 | 0.917257 |
| ENSMUSG( | Itprip | 51.56882 | 0.543998 | 0.109333 | 0.805353 |
| ENSMUSG( | Itpripl1 | 110.0569 | 0.041489 | 0.873494 | 0.989644 |
| ENSMUSG( | Itpripl2 | 483.4475 | 0.241562 | 0.105352 | 0.804969 |
| ENSMUSG( | Itsn1 | 6703.157 | -0.10745 | 0.324852 | 0.905257 |
| ENSMUSG( | Itsn2 | 1662.261 | 0.220874 | 0.045126 | 0.693959 |
| ENSMUSG( | Ivd | 3727.695 | -0.20258 | 0.075799 | 0.768524 |
| ENSMUSG( | Ivns1abp | 5276.969 | 0.081473 | 0.470358 | 0.923329 |
| ENSMUSG( | Iws1 | 710.3806 | 0.107771 | 0.405509 | 0.917257 |
| ENSMUSG( | Iyd | 29.19266 | -0.35271 | 0.331628 | 0.907907 |
| ENSMUSG( | Izumo4 | 696.5182 | 0.208384 | 0.324645 | 0.905257 |
| ENSMUSG( | Jade1 | 1205.36 | 0.016418 | 0.882543 | 0.989644 |
| ENSMUSG( | Jade2 | 1195.547 | 0.095556 | 0.472511 | 0.923329 |
| ENSMUSG( | Jade3 | 243.318 | -0.07706 | 0.643294 | 0.953903 |
| ENSMUSG( | Jag1 | 262.0766 | 0.441994 | 0.030586 | 0.628105 |
| ENSMUSG( | Jag2 | 1307.148 | 0.215878 | 0.148473 | 0.836241 |
| ENSMUSG( | Jagn1 | 917.0329 | 0.1242 | 0.414559 | 0.917257 |
| ENSMUSG( | Jak1 | 3588.132 | 0.005913 | 0.95272 | 0.994798 |
| ENSMUSG( | Jak2 | 1433.21 | 0.055311 | 0.622286 | 0.953234 |
| ENSMUSG( | Jak3 | 273.7184 | 0.176368 | 0.392973 | 0.917257 |
| ENSMUSG( | Jakmip1 | 260.2686 | -0.26919 | 0.155341 | 0.837724 |
| ENSMUSG( | Jakmip2 | 733.7186 | 0.002996 | 0.98325 | 0.996963 |
| ENSMUSG( | Jakmip3 | 833.6754 | -0.02483 | 0.856346 | 0.985884 |
| ENSMUSG( | Jam2 | 3885.683 | 0.009926 | 0.946019 | 0.994798 |
| ENSMUSG( | Jam3 | 1476.768 | 0.286197 | 0.169606 | 0.840806 |
| ENSMUSG( | Jaml | 17.41415 | 1.311779 | 0.009694 | 0.406458 |
| ENSMUSG( | Jarid2 | 1049.767 | -0.04904 | 0.709047 | 0.964042 |
| ENSMUSG( | Jazf1 | 280.6843 | 0.109693 | 0.506166 | 0.927825 |

| | | | | | |
|---|---|---|---|---|---|
| ENSMUSG( | 661.6655 | 0.176505 | 0.123208 | 0.530407 | Irf3 |
| ENSMUSG( | 13.24507 | -0.2542 | 0.651652 | NA | Irf4 |
| ENSMUSG( | 212.8768 | 0.040099 | 0.822572 | 0.966352 | Irf5 |
| ENSMUSG( | 8.726886 | 0.702179 | 0.280873 | NA | Irf6 |
| ENSMUSG( | 158.373 | -0.24891 | 0.177611 | 0.61504 | Irf7 |
| ENSMUSG( | 265.6052 | 0.038398 | 0.787197 | 0.959147 | Irf8 |
| ENSMUSG( | 455.8537 | 0.000513 | 0.996917 | 0.999505 | Irf9 |
| ENSMUSG( | 381.1176 | -0.22136 | 0.076094 | 0.42702 | Irgm1 |
| ENSMUSG( | 165.5415 | -0.07145 | 0.699103 | 0.936835 | Irgm2 |
| ENSMUSG( | 4510.523 | 0.195239 | 0.027208 | 0.253245 | Irgq |
| ENSMUSG( | 445.8894 | 0.051612 | 0.681136 | 0.931301 | Irs1 |
| ENSMUSG( | 1278.378 | 0.192504 | 0.132926 | 0.545594 | Irs2 |
| ENSMUSG( | 20.87276 | -0.24102 | 0.628441 | NA | Irs4 |
| ENSMUSG( | 22.18933 | -1.21918 | 0.018137 | 0.20302 | Irx1 |
| ENSMUSG( | 9.818835 | -0.91137 | 0.187926 | NA | Irx3 |
| ENSMUSG( | 3401 | 0.017794 | 0.8384 | 0.971111 | Isca1 |
| ENSMUSG( | 1049.738 | -0.00475 | 0.960722 | 0.993991 | Isca2 |
| ENSMUSG( | 2189.048 | -0.05859 | 0.504392 | 0.866883 | Iscu |
| ENSMUSG( | 78.54445 | -0.46393 | 0.082291 | 0.442706 | Isg15 |
| ENSMUSG( | 77.94774 | -1.01889 | 7.94E-05 | 0.005507 | Isg20 |
| ENSMUSG( | 346.4676 | -0.12383 | 0.31857 | 0.764234 | Isg20l2 |
| ENSMUSG( | 22.47027 | -1.70452 | 0.00315 | 0.06974 | Isl1 |
| ENSMUSG( | 941.966 | -0.32412 | 0.019022 | 0.207571 | Islr |
| ENSMUSG( | 3051.879 | -0.16869 | 0.278127 | 0.728086 | Islr2 |
| ENSMUSG( | 233.404 | -0.06917 | 0.640341 | 0.916434 | Ism1 |
| ENSMUSG( | 12.36446 | -0.27812 | 0.625119 | NA | Ism2 |
| ENSMUSG( | 1276.78 | 0.011911 | 0.915691 | 0.988634 | Isoc1 |
| ENSMUSG( | 1248.917 | 0.020937 | 0.837087 | 0.971111 | Isoc2a |
| ENSMUSG( | 147.234 | -0.17412 | 0.367592 | 0.795314 | Isoc2b |
| ENSMUSG( | 2201.925 | -0.03196 | 0.776489 | 0.955658 | Ist1 |
| ENSMUSG( | 568.806 | 0.045516 | 0.694559 | 0.9343 | Isy1 |
| ENSMUSG( | 962.5803 | 0.231424 | 0.038183 | 0.301526 | Isyna1 |
| ENSMUSG( | 1549.367 | 0.132202 | 0.230599 | 0.678947 | Itch |
| ENSMUSG( | 6490.319 | -0.38292 | 0.008835 | 0.132321 | Itfg1 |
| ENSMUSG( | 505.4384 | 0.080355 | 0.464539 | 0.843387 | Itfg2 |
| ENSMUSG( | 480.3269 | -0.03124 | 0.888689 | 0.983434 | Itga1 |
| ENSMUSG( | 141.79 | 0.7033 | 0.001594 | 0.044805 | Itga10 |
| ENSMUSG( | 84.36718 | -0.61959 | 0.008216 | 0.126198 | Itga11 |
| ENSMUSG( | 74.03265 | 0.286039 | 0.264975 | 0.715962 | Itga2b |
| ENSMUSG( | 601.1089 | -0.37072 | 0.005514 | 0.099488 | Itga3 |
| ENSMUSG( | 423.8221 | -0.44838 | 0.033316 | 0.279795 | Itga4 |
| ENSMUSG( | 237.6777 | -0.54874 | 0.0006 | 0.023008 | Itga5 |
| ENSMUSG( | 1725.605 | -0.14635 | 0.360247 | 0.791947 | Itga6 |
| ENSMUSG( | 795.926 | 0.144445 | 0.298285 | 0.74698 | Itga7 |
| ENSMUSG( | 243.3225 | 0.236582 | 0.19648 | 0.635942 | Itga8 |
| ENSMUSG( | 348.4569 | -0.40938 | 0.022179 | 0.227114 | Itga9 |
| ENSMUSG( | 14.13357 | 0.204087 | 0.685553 | NA | Itgad |
| ENSMUSG( | 12.84717 | -0.13736 | 0.824979 | NA | Itgae |
| ENSMUSG( | 135.3491 | -0.80923 | 0.171439 | 0.605943 | Itgal |
| ENSMUSG( | 642.8969 | -0.15779 | 0.25131 | 0.700869 | Itgam |
| ENSMUSG( | 3272.776 | -0.13136 | 0.343868 | 0.779153 | Itgav |
| ENSMUSG( | 24.38763 | -0.84421 | 0.06049 | 0.379845 | Itgax |
| ENSMUSG( | 4851.432 | -0.09805 | 0.439506 | 0.831151 | Itgb1 |
| ENSMUSG( | 738.4183 | -0.03104 | 0.78623 | 0.958774 | Itgb1bp1 |
| ENSMUSG( | 14.55751 | -0.57589 | 0.320299 | NA | Itgb1bp2 |
| ENSMUSG( | 427.1769 | -0.36412 | 0.008337 | 0.127456 | Itgb2 |
| ENSMUSG( | 167.7877 | -0.146 | 0.426203 | 0.824898 | Itgb3 |
| ENSMUSG( | 113.7773 | 0.131453 | 0.522655 | 0.875389 | Itgb3bp |
| ENSMUSG( | 782.9553 | -0.51866 | 1.07E-07 | 2.80E-05 | Itgb4 |
| ENSMUSG( | 4296.732 | 0.009932 | 0.939324 | 0.993415 | Itgb5 |
| ENSMUSG( | 51.6608 | -0.23936 | 0.405575 | 0.81652 | Itgb7 |
| ENSMUSG( | 2991.897 | -0.06687 | 0.647076 | 0.919213 | Itgb8 |
| ENSMUSG( | 155.5246 | -0.03414 | 0.847328 | 0.972374 | Itgbl1 |
| ENSMUSG( | 226.5848 | 0.247201 | 0.14067 | 0.557902 | Itih2 |
| ENSMUSG( | 1998.502 | -0.08264 | 0.385095 | 0.804048 | Itih3 |
| ENSMUSG( | 13.99539 | -0.20302 | 0.706476 | NA | Itih4 |
| ENSMUSG( | 3583.3 | 0.114625 | 0.346306 | 0.779153 | Itih5 |
| ENSMUSG( | 12.19832 | -0.6886 | 0.216139 | NA | Itk |
| ENSMUSG( | 2006.127 | -0.10361 | 0.621881 | 0.911432 | Itm2a |
| ENSMUSG( | 32078.24 | -0.21992 | 0.072623 | 0.416237 | Itm2b |
| ENSMUSG( | 31234.37 | -0.3037 | 0.011082 | 0.151746 | Itm2c |
| ENSMUSG( | 859.875 | 0.016034 | 0.89452 | 0.984512 | Itpa |
| ENSMUSG( | 2254.673 | 0.167201 | 0.08758 | 0.454663 | Itpk1 |
| ENSMUSG( | 5864.732 | 0.116847 | 0.287363 | 0.737854 | Itpka |
| ENSMUSG( | 2436.092 | -0.01101 | 0.923019 | 0.990693 | Itpkb |
| ENSMUSG( | 173.0113 | -0.193 | 0.275792 | 0.725669 | Itpkc |
| ENSMUSG( | 13318.17 | -0.06359 | 0.62217 | 0.911432 | Itpr1 |
| ENSMUSG( | 2426.569 | -0.05046 | 0.674728 | 0.929638 | Itpr2 |
| ENSMUSG( | 160.8018 | 0.377578 | 0.053485 | 0.35564 | Itpr3 |
| ENSMUSG( | 36.4356 | -0.19537 | 0.576594 | 0.895391 | Itprid1 |
| ENSMUSG( | 4943.031 | 0.215448 | 0.030714 | 0.266671 | Itprid2 |
| ENSMUSG( | 44.7664 | -0.11334 | 0.726483 | 0.944223 | Itprip |
| ENSMUSG( | 103.4072 | -0.47192 | 0.028533 | 0.257525 | Itpripl1 |
| ENSMUSG( | 492.1281 | 0.027382 | 0.857911 | 0.974667 | Itpripl2 |
| ENSMUSG( | 8437.152 | 0.277228 | 0.009159 | 0.134809 | Itsn1 |
| ENSMUSG( | 1803.23 | 0.18753 | 0.038583 | 0.302696 | Itsn2 |
| ENSMUSG( | 4549.931 | 0.110565 | 0.190617 | 0.629951 | Ivd |
| ENSMUSG( | 5502.368 | -0.06654 | 0.426467 | 0.825155 | Ivns1abp |
| ENSMUSG( | 778.1427 | 0.098307 | 0.361559 | 0.792797 | Iws1 |
| ENSMUSG( | 32.13925 | -0.36298 | 0.301248 | 0.749979 | Iyd |
| ENSMUSG( | 744.4674 | 0.13787 | 0.405823 | 0.833467 | Izumo4 |
| ENSMUSG( | 1312.183 | -0.0072 | 0.93741 | 0.993415 | Jade1 |
| ENSMUSG( | 1351.642 | 0.174085 | 0.163593 | 0.594869 | Jade2 |
| ENSMUSG( | 264.6501 | -0.09679 | 0.481666 | 0.855573 | Jade3 |
| ENSMUSG( | 219.1492 | -0.34076 | 0.081982 | 0.442365 | Jag1 |
| ENSMUSG( | 1399.439 | 0.144396 | 0.259931 | 0.709951 | Jag2 |
| ENSMUSG( | 859.5961 | -0.35202 | 0.009529 | 0.138128 | Jagn1 |
| ENSMUSG( | 4094.989 | 0.116882 | 0.137605 | 0.552903 | Jak1 |
| ENSMUSG( | 1615.102 | 0.130599 | 0.178354 | 0.616026 | Jak2 |
| ENSMUSG( | 306.6622 | 0.22263 | 0.111351 | 0.503375 | Jak3 |
| ENSMUSG( | 309.4895 | -0.01581 | 0.920937 | 0.990426 | Jakmip1 |
| ENSMUSG( | 822.3087 | -0.064058 | 0.605966 | 0.90596 | Jakmip2 |
| ENSMUSG( | 922.8495 | -0.00307 | 0.976741 | 0.997428 | Jakmip3 |
| ENSMUSG( | 4063.035 | -0.13479 | 0.312254 | 0.759791 | Jam2 |
| ENSMUSG( | 1306.812 | -0.34943 | 0.040561 | 0.311208 | Jam3 |
| ENSMUSG( | 9.908053 | -0.35696 | 0.548266 | NA | Jaml |
| ENSMUSG( | 1083.293 | -0.23736 | 0.050656 | 0.344934 | Jarid2 |
| ENSMUSG( | 330.5096 | 0.305675 | 0.018664 | 0.205542 | Jazf1 |
| ENSMUSG( | 5051.157 | 0.182139 | 0.042263 | 0.316636 | Jcad |
| ENSMUSG( | 26.05449 | 0.324624 | 0.4621 | 0.841881 | Jchain |
| ENSMUSG( | 1147.639 | 0.131759 | 0.316868 | 0.763754 | Jdp2 |
| ENSMUSG( | 10.70325 | 0.15596 | 0.800125 | NA | Jhy |
| ENSMUSG( | 1411.078 | -0.26882 | 0.036222 | 0.293031 | Jkamp |
| ENSMUSG( | 2263.055 | -0.15355 | 0.170538 | 0.605271 | Jmjd1c |
| ENSMUSG( | 978.3021 | 0.06923 | 0.488793 | 0.859092 | Jmjd4 |
| ENSMUSG( | 638.8056 | -0.045 | 0.749087 | 0.949604 | Jmjd6 |
| ENSMUSG( | 1521.096 | -0.10216 | 0.423657 | 0.824045 | Jmjd8 |
| ENSMUSG( | 2087.354 | 0.075585 | 0.430173 | 0.826789 | Jmy |
| ENSMUSG( | 2539.435 | 0.149236 | 0.1365 | 0.550651 | Josd1 |
| ENSMUSG( | 2035.869 | -0.01944 | 0.859681 | 0.97481 | Josd2 |
| ENSMUSG( | 318.4166 | -0.22951 | 0.099234 | 0.481621 | Jph1 |
| ENSMUSG( | 112.1929 | 0.017038 | 0.940847 | 0.993415 | Jph2 |
| ENSMUSG( | 11025.21 | 0.154225 | 0.089597 | 0.459604 | Jph3 |
| ENSMUSG( | 19649.54 | 0.205497 | 0.011646 | 0.155574 | Jph4 |
| ENSMUSG( | 1658.656 | 0.050387 | 0.616932 | 0.910866 | Jpt1 |
| ENSMUSG( | 250.6342 | -0.05789 | 0.706029 | 0.938825 | Jpt2 |
| ENSMUSG( | 238.1567 | 0.015155 | 0.922838 | 0.990631 | Jpx |
| ENSMUSG( | 178.3365 | 0.122 | 0.494972 | 0.860798 | Jrk |
| ENSMUSG( | 117.9791 | -0.14921 | 0.419142 | 0.821966 | Jrkl |
| ENSMUSG( | 87.59548 | -0.26007 | 0.270461 | 0.72034 | Jsrp1 |
| ENSMUSG( | 1240.79 | -0.09826 | 0.377596 | 0.801546 | Jtb |
| ENSMUSG( | 1384.179 | 0.091295 | 0.429351 | 0.826446 | Jun |
| ENSMUSG( | 1063.501 | -1.02541 | 0.004609 | 0.089309 | Junb |
| ENSMUSG( | 5764.014 | -0.03127 | 0.744046 | 0.948497 | Jund |
| ENSMUSG( | 43.22354 | 0.279966 | 0.38468 | 0.804031 | Junos |
| ENSMUSG( | 1606.289 | 0.033461 | 0.716037 | 0.94152 | Jup |
| ENSMUSG( | 16919.03 | 0.174685 | 0.051843 | 0.349498 | Kalrn |
| ENSMUSG( | 1714.241 | 0.157712 | 0.083288 | 0.445262 | Kank1 |
| ENSMUSG( | 1015.811 | 0.056238 | 0.627535 | 0.912961 | Kank2 |
| ENSMUSG( | 712.6044 | 0.275228 | 0.039244 | 0.305097 | Kank3 |
| ENSMUSG( | 122.8351 | -0.3818 | 0.05112 | 0.346957 | Kank4 |
| ENSMUSG( | 2304.91 | 0.128551 | 0.157825 | 0.585927 | Kansl1 |
| ENSMUSG( | 417.1923 | -0.03857 | 0.757926 | 0.951587 | Kansl1l |
| ENSMUSG( | 1088.964 | 0.035323 | 0.733477 | 0.944956 | Kansl2 |
| ENSMUSG( | 2810.01 | 0.171696 | 0.052049 | 0.350227 | Kansl3 |
| ENSMUSG( | 452.7039 | -0.05578 | 0.684605 | 0.93282 | Kantr |
| ENSMUSG( | 1858.299 | 0.042601 | 0.671681 | 0.928781 | Kars1 |
| ENSMUSG( | 96.98046 | 0.151 | 0.48941 | 0.859092 | Kash5 |
| ENSMUSG( | 761.8117 | 0.173276 | 0.110288 | 0.500944 | Kat14 |
| ENSMUSG( | 1891.669 | 0.194144 | 0.18299 | 0.62047 | Kat2a |
| ENSMUSG( | 1233.629 | 0.062405 | 0.596986 | 0.903526 | Kat2b |
| ENSMUSG( | 1528.998 | 0.05805 | 0.551951 | 0.886141 | Kat5 |
| ENSMUSG( | 3431.792 | 0.104703 | 0.331467 | 0.77519 | Kat6a |
| ENSMUSG( | 1987.094 | -0.06499 | 0.45335 | 0.836798 | Kat6b |
| ENSMUSG( | 1491.874 | 0.074519 | 0.463584 | 0.842812 | Kat7 |
| ENSMUSG( | 472.5702 | 0.121224 | 0.358365 | 0.789759 | Kat8 |
| ENSMUSG( | 335.1734 | 0.035774 | 0.775608 | 0.955606 | Katna1 |
| ENSMUSG( | 2775.681 | 0.075743 | 0.362342 | 0.792943 | Katnal1 |

| | | | | | |
|---|---|---|---|---|---|
| ENSMUSG( | 4813.803 | 0.148144 | 0.317857 | 0.902849 | Itgb1 |
| ENSMUSG( | 642.5558 | -0.17341 | 0.17391 | 0.844317 | Itgb1bp1 |
| ENSMUSG( | 14.64324 | -0.26043 | 0.637656 | 0.953903 | Itgb1bp2 |
| ENSMUSG( | 407.4976 | -0.21095 | 0.227569 | 0.868507 | Itgb2 |
| ENSMUSG( | 162.4612 | 0.044402 | 0.848678 | 0.985321 | Itgb3 |
| ENSMUSG( | 90.21207 | -0.27274 | 0.279537 | 0.892356 | Itgb3bp |
| ENSMUSG( | 770.2127 | -0.25843 | 0.058819 | 0.728906 | Itgb4 |
| ENSMUSG( | 3941.456 | 0.033419 | 0.831725 | 0.980656 | Itgb5 |
| ENSMUSG( | 51.86903 | 0.036666 | 0.90508 | 0.991089 | Itgb7 |
| ENSMUSG( | 2845.751 | 0.057219 | 0.726246 | 0.966887 | Itgb8 |
| ENSMUSG( | 150.4319 | 0.139779 | 0.464524 | 0.923329 | Itgbl1 |
| ENSMUSG( | 213.1224 | 0.329884 | 0.050183 | 0.708648 | Itih2 |
| ENSMUSG( | 1871.367 | -0.00245 | 0.981883 | 0.996756 | Itih3 |
| ENSMUSG( | 12.40076 | -0.27099 | 0.642134 | 0.953903 | Itih4 |
| ENSMUSG( | 3347.005 | 0.183899 | 0.12301 | 0.814657 | Itih5 |
| ENSMUSG( | 12.45242 | -0.27105 | 0.633404 | 0.953864 | Itk |
| ENSMUSG( | 2466.548 | 0.690648 | 0.110279 | 0.807051 | Itm2a |
| ENSMUSG( | 32689.28 | 0.111041 | 0.436296 | 0.919879 | Itm2b |
| ENSMUSG( | 32098.22 | 0.062242 | 0.656691 | 0.955567 | Itm2c |
| ENSMUSG( | 763.0684 | -0.06332 | 0.666899 | 0.956766 | Itpa |
| ENSMUSG( | 1932.822 | -0.00352 | 0.974975 | 0.996416 | Itpk1 |
| ENSMUSG( | 5221.047 | 0.052608 | 0.67458 | 0.95694 | Itpka |
| ENSMUSG( | 2056.947 | -0.24141 | 0.02829 | 0.612554 | Itpkb |
| ENSMUSG( | 161.4232 | -0.10949 | 0.58653 | 0.944975 | Itpkc |
| ENSMUSG( | 12944.65 | 0.124883 | 0.415361 | 0.917257 | Itpr1 |
| ENSMUSG( | 2240.184 | -0.00911 | 0.945245 | 0.994798 | Itpr2 |
| ENSMUSG( | 174.6032 | 0.814765 | 0.00019 | 0.034114 | Itpr3 |
| ENSMUSG( | 36.65095 | 0.115892 | 0.758785 | 0.971858 | Itprid1 |
| ENSMUSG( | 4318.548 | 0.100514 | 0.403606 | 0.917257 | Itprid2 |
| ENSMUSG( | 51.56882 | 0.543998 | 0.109333 | 0.805353 | Itprip |
| ENSMUSG( | 110.0569 | 0.041489 | 0.873494 | 0.989644 | Itpripl1 |
| ENSMUSG( | 483.4475 | 0.241562 | 0.105352 | 0.804969 | Itpripl2 |
| ENSMUSG( | 6703.157 | -0.10745 | 0.324852 | 0.905257 | Itsn1 |
| ENSMUSG( | 1662.261 | 0.220874 | 0.045126 | 0.693959 | Itsn2 |
| ENSMUSG( | 3727.695 | -0.20258 | 0.075799 | 0.768524 | Ivd |
| ENSMUSG( | 5276.969 | 0.081473 | 0.470358 | 0.923329 | Ivns1abp |
| ENSMUSG( | 710.3806 | 0.107771 | 0.405509 | 0.917257 | Iws1 |
| ENSMUSG( | 29.19266 | -0.35271 | 0.331628 | 0.907907 | Iyd |
| ENSMUSG( | 696.5182 | 0.208384 | 0.324645 | 0.905257 | Izumo4 |
| ENSMUSG( | 1205.36 | 0.016418 | 0.882543 | 0.989644 | Jade1 |
| ENSMUSG( | 1195.547 | 0.095556 | 0.472511 | 0.923329 | Jade2 |
| ENSMUSG( | 243.318 | -0.07706 | 0.643294 | 0.953903 | Jade3 |
| ENSMUSG( | 262.0766 | 0.441994 | 0.030586 | 0.628105 | Jag1 |
| ENSMUSG( | 1307.148 | 0.215878 | 0.148473 | 0.836241 | Jag2 |
| ENSMUSG( | 917.0329 | 0.1242 | 0.414559 | 0.917257 | Jagn1 |
| ENSMUSG( | 3588.132 | 0.005913 | 0.95272 | 0.994798 | Jak1 |
| ENSMUSG( | 1433.21 | 0.055311 | 0.622286 | 0.953234 | Jak2 |
| ENSMUSG( | 273.7184 | 0.176368 | 0.392973 | 0.917257 | Jak3 |
| ENSMUSG( | 260.2686 | -0.26919 | 0.155341 | 0.837724 | Jakmip1 |
| ENSMUSG( | 733.7186 | 0.002996 | 0.98325 | 0.996963 | Jakmip2 |
| ENSMUSG( | 833.6754 | -0.02483 | 0.856346 | 0.985884 | Jakmip3 |
| ENSMUSG( | 3885.683 | 0.009926 | 0.946019 | 0.994798 | Jam2 |
| ENSMUSG( | 1476.768 | 0.286197 | 0.169606 | 0.840806 | Jam3 |
| ENSMUSG( | 17.41415 | 1.311779 | 0.009694 | 0.406458 | Jaml |
| ENSMUSG( | 1049.767 | -0.04904 | 0.709047 | 0.964042 | Jarid2 |
| ENSMUSG( | 280.6843 | 0.109693 | 0.506166 | 0.927825 | Jazf1 |
| ENSMUSG( | 4553.033 | 0.155259 | 0.226065 | 0.868507 | Jcad |
| ENSMUSG( | 23.46181 | 0.267206 | 0.585344 | 0.944975 | Jchain |
| ENSMUSG( | 980.5407 | -0.05337 | 0.709875 | 0.96416 | Jdp2 |
| ENSMUSG( | 1467.383 | 0.125845 | 0.397539 | 0.917257 | Jkamp |
| ENSMUSG( | 2227.564 | 0.077113 | 0.586021 | 0.944975 | Jmjd1c |
| ENSMUSG( | 865.7154 | -0.01112 | 0.927862 | 0.99306 | Jmjd4 |
| ENSMUSG( | 612.5685 | 0.105945 | 0.455059 | 0.923029 | Jmjd6 |
| ENSMUSG( | 1510.845 | 0.149236 | 0.325199 | 0.905339 | Jmjd8 |
| ENSMUSG( | 1921.555 | 0.103948 | 0.394877 | 0.917257 | Jmy |
| ENSMUSG( | 2222.1 | 0.03875 | 0.730925 | 0.967228 | Josd1 |
| ENSMUSG( | 1874.507 | 0.012035 | 0.934913 | 0.99306 | Josd2 |
| ENSMUSG( | 316.7354 | 0.041394 | 0.790397 | 0.97681 | Jph1 |
| ENSMUSG( | 87.369 | -0.50468 | 0.058631 | 0.728256 | Jph2 |
| ENSMUSG( | 9878.344 | 0.109637 | 0.345463 | 0.910581 | Jph3 |
| ENSMUSG( | 17381.81 | 0.125828 | 0.201304 | 0.864412 | Jph4 |
| ENSMUSG( | 1423.812 | -0.1247 | 0.302092 | 0.898538 | Jpt1 |
| ENSMUSG( | 228.5795 | -0.06054 | 0.721447 | 0.966887 | Jpt2 |
| ENSMUSG( | 207.1454 | -0.13108 | 0.442449 | 0.921554 | Jpx |
| ENSMUSG( | 146.914 | -0.15273 | 0.470371 | 0.923329 | Jrk |
| ENSMUSG( | 130.1636 | 0.39331 | 0.048576 | 0.706142 | Jrkl |
| ENSMUSG( | 89.31686 | 0.054542 | 0.840591 | 0.982495 | Jsrp1 |
| ENSMUSG( | 1200.185 | 0.077813 | 0.517287 | 0.930939 | Jtb |
| ENSMUSG( | 1280.524 | 0.134022 | 0.298089 | 0.898538 | Jun |
| ENSMUSG( | 1021.262 | -0.80749 | 0.039969 | 0.681706 | Junb |
| ENSMUSG( | 5011.956 | -0.17265 | 0.149965 | 0.837086 | Jund |
| ENSMUSG( | 35.24516 | -0.02828 | 0.938193 | 0.993731 | Junos |
| ENSMUSG( | 1398.443 | -0.10093 | 0.382246 | 0.917257 | Jup |
| ENSMUSG( | 14670.86 | 0.037678 | 0.743804 | 0.96951 | Kalrn |
| ENSMUSG( | 1445.06 | -0.06255 | 0.562642 | 0.940887 | Kank1 |
| ENSMUSG( | 862.1741 | -0.15329 | 0.22961 | 0.868507 | Kank2 |
| ENSMUSG( | 688.3874 | 0.432502 | 0.019894 | 0.558472 | Kank3 |
| ENSMUSG( | 130.0401 | 0.085775 | 0.683202 | 0.958678 | Kank4 |
| ENSMUSG( | 2101.569 | 0.131383 | 0.209569 | 0.864412 | Kansl1 |
| ENSMUSG( | 430.8111 | 0.303321 | 0.031098 | 0.629763 | Kansl1l |
| ENSMUSG( | 994.8977 | 0.041307 | 0.727878 | 0.966887 | Kansl2 |
| ENSMUSG( | 2506.337 | 0.114035 | 0.334077 | 0.907907 | Kansl3 |
| ENSMUSG( | 428.9377 | 0.050162 | 0.764399 | 0.974 | Kantr |
| ENSMUSG( | 1556.69 | -0.2083 | 0.090492 | 0.791074 | Kars1 |
| ENSMUSG( | 80.92124 | -0.08765 | 0.729196 | 0.966913 | Kash5 |
| ENSMUSG( | 673.1173 | 0.093247 | 0.484626 | 0.92541 | Kat14 |
| ENSMUSG( | 1598.735 | -0.0188 | 0.90006 | 0.990558 | Kat2a |
| ENSMUSG( | 1077.146 | -0.063 | 0.646874 | 0.953988 | Kat2b |
| ENSMUSG( | 1342.447 | -0.0479 | 0.676169 | 0.95694 | Kat5 |
| ENSMUSG( | 3078.247 | 0.060381 | 0.611166 | 0.950771 | Kat6a |
| ENSMUSG( | 1819.294 | -0.05206 | 0.659984 | 0.956205 | Kat6b |
| ENSMUSG( | 1343.053 | 0.040494 | 0.730863 | 0.967228 | Kat7 |
| ENSMUSG( | 422.1386 | 0.060333 | 0.688307 | 0.959742 | Kat8 |
| ENSMUSG( | 317.1379 | 0.145777 | 0.373007 | 0.915491 | Katna1 |
| ENSMUSG( | 2362.056 | -0.12484 | 0.257057 | 0.883082 | Katnal1 |
| ENSMUSG( | 151.1505 | -0.02079 | 0.913688 | 0.992378 | Katnal2 |
| ENSMUSG( | 933.623 | -0.04237 | 0.728118 | 0.966913 | Katnb1 |
| ENSMUSG( | 312.275 | 0.139329 | 0.510204 | 0.929089 | Katnbl1 |
| ENSMUSG( | 791.1644 | 0.137984 | 0.374312 | 0.915491 | Katnip |
| ENSMUSG( | 89.27144 | -0.31805 | 0.202752 | 0.864412 | Kazald1 |
| ENSMUSG( | 2587.829 | -0.00355 | 0.973309 | 0.996416 | Kazn |
| ENSMUSG( | 3849.206 | -0.07461 | 0.462328 | 0.923329 | Kbtbd11 |
| ENSMUSG( | 990.734 | 0.031638 | 0.80268 | 0.97681 | Kbtbd2 |
| ENSMUSG( | 121.9439 | -0.19584 | 0.326114 | 0.905339 | Kbtbd3 |
| ENSMUSG( | 478.2795 | -0.03871 | 0.785933 | 0.976285 | Kbtbd4 |
| ENSMUSG( | 14.2323 | -0.09422 | 0.861577 | 0.986298 | Kbtbd6 |
| ENSMUSG( | 432.5538 | 0.239608 | 0.151844 | 0.837724 | Kbtbd7 |
| ENSMUSG( | 107.7511 | 0.225977 | 0.307381 | 0.898538 | Kbtbd8 |
| ENSMUSG( | 1985.162 | -0.09552 | 0.409022 | 0.917257 | Kcmf1 |
| ENSMUSG( | 628.0542 | 0.582133 | 0.204748 | 0.864412 | Kcna1 |
| ENSMUSG( | 7692.143 | 0.015763 | 0.910743 | 0.992196 | Kcna2 |
| ENSMUSG( | 49.52904 | 0.438004 | 0.253486 | 0.88018 | Kcna3 |
| ENSMUSG( | 417.6431 | -0.00773 | 0.962271 | 0.995237 | Kcna4 |
| ENSMUSG( | 173.676 | 0.180155 | 0.350105 | 0.911652 | Kcna5 |
| ENSMUSG( | 3089.622 | 0.200555 | 0.256721 | 0.883082 | Kcna6 |
| ENSMUSG( | 1175.644 | -0.17024 | 0.348937 | 0.911652 | Kcnab1 |
| ENSMUSG( | 17494.45 | 0.09732 | 0.413646 | 0.917257 | Kcnab2 |
| ENSMUSG( | 1217.021 | -0.23692 | 0.158285 | 0.837724 | Kcnab3 |
| ENSMUSG( | 7318.681 | 0.178186 | 0.292034 | 0.897854 | Kcnb1 |
| ENSMUSG( | 452.1759 | 0.141721 | 0.468738 | 0.923329 | Kcnb2 |
| ENSMUSG( | 4915.257 | 0.066371 | 0.685353 | 0.958678 | Kcnc1 |
| ENSMUSG( | 2424.64 | 0.132499 | 0.451268 | 0.922202 | Kcnc2 |
| ENSMUSG( | 4100.361 | 0.211162 | 0.258323 | 0.883971 | Kcnc3 |
| ENSMUSG( | 2661.883 | 0.210041 | 0.174166 | 0.844317 | Kcnc4 |
| ENSMUSG( | 457.3367 | 0.135141 | 0.40987 | 0.917257 | Kcnd1 |
| ENSMUSG( | 2304.253 | 0.224173 | 0.147963 | 0.836241 | Kcnd2 |
| ENSMUSG( | 2100.499 | 0.378929 | 0.013211 | 0.460068 | Kcnd3 |
| ENSMUSG( | 30.25706 | 0.138262 | 0.719956 | 0.966887 | Kcnd3os |
| ENSMUSG( | 55.42487 | 0.424043 | 0.622973 | 0.953234 | Kcne2 |
| ENSMUSG( | 60.74795 | 0.182349 | 0.609429 | 0.950397 | Kcne4 |
| ENSMUSG( | 578.6243 | -0.30233 | 0.113567 | 0.809623 | Kcne5 |
| ENSMUSG( | 4131.755 | 0.244119 | 0.174976 | 0.845107 | Kcnf1 |
| ENSMUSG( | 1234.146 | 0.275667 | 0.057987 | 0.728256 | Kcng1 |
| ENSMUSG( | 290.9385 | 0.398838 | 0.010642 | 0.419324 | Kcng2 |
| ENSMUSG( | 7.31951 | 0.781701 | 0.284274 | NA | Kcng3 |
| ENSMUSG( | 499.2001 | -0.36703 | 0.018395 | 0.544291 | Kcng4 |
| ENSMUSG( | 2285.804 | 0.074989 | 0.69802 | 0.960867 | Kcnh1 |
| ENSMUSG( | 1000.079 | 0.092227 | 0.53067 | 0.934795 | Kcnh2 |
| ENSMUSG( | 7255.267 | 0.257842 | 0.063694 | 0.736482 | Kcnh3 |
| ENSMUSG( | 520.6521 | 0.195399 | 0.183286 | 0.850984 | Kcnh4 |
| ENSMUSG( | 158.8177 | 0.051036 | 0.835961 | 0.981237 | Kcnh5 |
| ENSMUSG( | 72.0183 | 0.039615 | 0.878867 | 0.989644 | Kcnh6 |
| ENSMUSG( | 2097.14 | 0.321868 | 0.069367 | 0.755467 | Kcnh7 |
| ENSMUSG( | 22.0941 | 0.446971 | 0.338391 | 0.907972 | Kcnh8 |
| ENSMUSG( | 513.8793 | -0.03136 | 0.844912 | 0.983863 | Kcnip1 |
| ENSMUSG( | 3597.219 | -0.12376 | 0.32799 | 0.906255 | Kcnip2 |
| ENSMUSG( | 6432.746 | -0.08237 | 0.468581 | 0.923329 | Kcnip3 |
| ENSMUSG( | 625.252 | -0.03779 | 0.843932 | 0.983571 | Kcnip4 |

| | | | | | | | | | | | |
|---|---|---|---|---|---|---|---|---|---|---|---|
| ENSMUSG( | Jcad | 5051.157 | 0.182139 | 0.042263 | 0.316636 | ENSMUSG( | Jcad | 4553.033 | 0.155259 | 0.226065 | 0.868507 |
| ENSMUSG( | Jchain | 26.05449 | 0.324624 | 0.4621 | 0.841881 | ENSMUSG( | Jchain | 23.46181 | 0.267206 | 0.585344 | 0.944975 |
| ENSMUSG( | Jdp2 | 1147.639 | 0.131759 | 0.316868 | 0.763754 | ENSMUSG( | Jdp2 | 980.5407 | -0.05337 | 0.709875 | 0.96416 |
| ENSMUSG( | Jkamp | 1411.078 | -0.26882 | 0.036222 | 0.293031 | ENSMUSG( | Jkamp | 1467.383 | 0.125845 | 0.397539 | 0.917257 |
| ENSMUSG( | Jmjd1c | 2263.055 | -0.15355 | 0.170538 | 0.605271 | ENSMUSG( | Jmjd1c | 2227.564 | 0.077113 | 0.586021 | 0.944975 |
| ENSMUSG( | Jmjd4 | 978.3021 | 0.06923 | 0.488793 | 0.859092 | ENSMUSG( | Jmjd4 | 865.7154 | -0.01112 | 0.927862 | 0.99306 |
| ENSMUSG( | Jmjd6 | 638.8056 | -0.045 | 0.749087 | 0.949604 | ENSMUSG( | Jmjd6 | 612.5685 | 0.105945 | 0.455059 | 0.923029 |
| ENSMUSG( | Jmjd8 | 1521.096 | -0.10216 | 0.423657 | 0.824045 | ENSMUSG( | Jmjd8 | 1510.845 | 0.149236 | 0.325199 | 0.905339 |
| ENSMUSG( | Jmy | 2087.354 | 0.075585 | 0.430173 | 0.826789 | ENSMUSG( | Jmy | 1921.555 | 0.103948 | 0.394877 | 0.917257 |
| ENSMUSG( | Josd1 | 2539.435 | 0.149236 | 0.1365 | 0.550651 | ENSMUSG( | Josd1 | 2222.1 | 0.03875 | 0.730925 | 0.967228 |
| ENSMUSG( | Josd2 | 2035.869 | -0.01944 | 0.859681 | 0.97481 | ENSMUSG( | Josd2 | 1874.507 | 0.012035 | 0.934913 | 0.99306 |
| ENSMUSG( | Jph1 | 318.4166 | -0.22951 | 0.099234 | 0.481621 | ENSMUSG( | Jph1 | 316.7354 | 0.041394 | 0.790397 | 0.97681 |
| ENSMUSG( | Jph2 | 112.1929 | 0.017038 | 0.940847 | 0.993415 | ENSMUSG( | Jph2 | 87.369 | -0.50468 | 0.058631 | 0.728256 |
| ENSMUSG( | Jph3 | 11025.21 | 0.154225 | 0.089597 | 0.459604 | ENSMUSG( | Jph3 | 9878.344 | 0.109637 | 0.345463 | 0.910581 |
| ENSMUSG( | Jph4 | 19649.54 | 0.205497 | 0.011646 | 0.155574 | ENSMUSG( | Jph4 | 17381.81 | 0.125828 | 0.201304 | 0.864412 |
| ENSMUSG( | Jpt1 | 1658.656 | 0.050387 | 0.616932 | 0.910866 | ENSMUSG( | Jpt1 | 1423.812 | -0.1247 | 0.302092 | 0.898538 |
| ENSMUSG( | Jpt2 | 250.6342 | -0.05789 | 0.706029 | 0.938825 | ENSMUSG( | Jpt2 | 228.5795 | -0.06054 | 0.721447 | 0.966887 |
| ENSMUSG( | Jpx | 238.1567 | 0.015155 | 0.922838 | 0.990631 | ENSMUSG( | Jpx | 207.1454 | -0.13108 | 0.442449 | 0.921554 |
| ENSMUSG( | Jrk | 178.3365 | 0.122 | 0.494972 | 0.860798 | ENSMUSG( | Jrk | 146.914 | -0.15273 | 0.470371 | 0.923329 |
| ENSMUSG( | Jrkl | 117.9791 | -0.14921 | 0.419142 | 0.821966 | ENSMUSG( | Jrkl | 130.1636 | 0.39331 | 0.048576 | 0.706142 |
| ENSMUSG( | Jsrp1 | 87.59548 | -0.26007 | 0.270461 | 0.72034 | ENSMUSG( | Jsrp1 | 89.31686 | 0.054542 | 0.840591 | 0.982495 |
| ENSMUSG( | Jtb | 1240.79 | -0.09826 | 0.377596 | 0.801546 | ENSMUSG( | Jtb | 1200.185 | 0.077813 | 0.517287 | 0.930939 |
| ENSMUSG( | Jun | 1384.179 | 0.091295 | 0.429351 | 0.826446 | ENSMUSG( | Jun | 1280.524 | 0.134022 | 0.298089 | 0.898538 |
| ENSMUSG( | Junb | 1063.501 | -1.02541 | 0.004609 | 0.089309 | ENSMUSG( | Junb | 1021.262 | -0.80749 | 0.039969 | 0.681706 |
| ENSMUSG( | Jund | 5764.014 | -0.03127 | 0.744046 | 0.948497 | ENSMUSG( | Jund | 5011.956 | -0.17265 | 0.149965 | 0.837086 |
| ENSMUSG( | Junos | 43.22354 | 0.279966 | 0.38468 | 0.804031 | ENSMUSG( | Junos | 35.24516 | -0.02828 | 0.938193 | 0.993731 |
| ENSMUSG( | Jup | 1606.289 | 0.033461 | 0.716037 | 0.94152 | ENSMUSG( | Jup | 1398.443 | -0.10093 | 0.382246 | 0.917257 |
| ENSMUSG( | Kalrn | 16919.03 | 0.174685 | 0.051843 | 0.349498 | ENSMUSG( | Kalrn | 14670.86 | 0.037678 | 0.743804 | 0.96951 |
| ENSMUSG( | Kank1 | 1714.241 | 0.157712 | 0.083288 | 0.445262 | ENSMUSG( | Kank1 | 1445.06 | -0.06255 | 0.562642 | 0.940887 |
| ENSMUSG( | Kank2 | 1015.811 | 0.056238 | 0.627535 | 0.912961 | ENSMUSG( | Kank2 | 862.1741 | -0.15329 | 0.22961 | 0.868507 |
| ENSMUSG( | Kank3 | 712.6044 | 0.275228 | 0.039244 | 0.305097 | ENSMUSG( | Kank3 | 688.3874 | 0.432502 | 0.019894 | 0.558472 |
| ENSMUSG( | Kank4 | 122.8351 | -0.3818 | 0.05112 | 0.346957 | ENSMUSG( | Kank4 | 130.0401 | 0.085775 | 0.683202 | 0.958678 |
| ENSMUSG( | Kansl1 | 2304.91 | 0.128551 | 0.157825 | 0.585927 | ENSMUSG( | Kansl1 | 2101.569 | 0.131383 | 0.209569 | 0.864412 |
| ENSMUSG( | Kansl1l | 417.1923 | -0.03857 | 0.757926 | 0.951587 | ENSMUSG( | Kansl1l | 430.8111 | 0.303321 | 0.031098 | 0.629763 |
| ENSMUSG( | Kansl2 | 1088.964 | 0.035323 | 0.733477 | 0.944956 | ENSMUSG( | Kansl2 | 994.8977 | 0.041307 | 0.727878 | 0.966887 |
| ENSMUSG( | Kansl3 | 2810.01 | 0.171696 | 0.052049 | 0.350227 | ENSMUSG( | Kansl3 | 2506.337 | 0.114035 | 0.334077 | 0.907907 |
| ENSMUSG( | Kantr | 452.7039 | -0.05578 | 0.684605 | 0.93282 | ENSMUSG( | Kantr | 428.9377 | 0.050162 | 0.764399 | 0.974 |
| ENSMUSG( | Kars1 | 1858.299 | 0.042601 | 0.671681 | 0.928781 | ENSMUSG( | Kars1 | 1556.69 | -0.2083 | 0.090492 | 0.791074 |
| ENSMUSG( | Kash5 | 96.98046 | 0.151 | 0.48941 | 0.859092 | ENSMUSG( | Kash5 | 80.92124 | -0.08765 | 0.729196 | 0.966913 |
| ENSMUSG( | Kat14 | 761.8117 | 0.173276 | 0.110288 | 0.500944 | ENSMUSG( | Kat14 | 673.1173 | 0.093247 | 0.484626 | 0.92541 |
| ENSMUSG( | Kat2a | 1891.669 | 0.194144 | 0.18299 | 0.62047 | ENSMUSG( | Kat2a | 1598.735 | -0.0188 | 0.90006 | 0.990558 |
| ENSMUSG( | Kat2b | 1233.629 | 0.062405 | 0.596986 | 0.903526 | ENSMUSG( | Kat2b | 1077.146 | -0.063 | 0.646874 | 0.953988 |
| ENSMUSG( | Kat5 | 1528.998 | 0.05805 | 0.551951 | 0.886141 | ENSMUSG( | Kat5 | 1342.447 | -0.0479 | 0.676169 | 0.95694 |
| ENSMUSG( | Kat6a | 3431.792 | 0.104703 | 0.331467 | 0.77519 | ENSMUSG( | Kat6a | 3078.247 | 0.060381 | 0.611166 | 0.950771 |
| ENSMUSG( | Kat6b | 1987.094 | -0.06499 | 0.45335 | 0.836798 | ENSMUSG( | Kat6b | 1819.294 | -0.05206 | 0.659984 | 0.956205 |
| ENSMUSG( | Kat7 | 1491.874 | 0.074519 | 0.463584 | 0.842812 | ENSMUSG( | Kat7 | 1343.053 | 0.040494 | 0.730863 | 0.967228 |
| ENSMUSG( | Kat8 | 472.5702 | 0.121224 | 0.358365 | 0.789759 | ENSMUSG( | Kat8 | 422.1386 | 0.060333 | 0.688307 | 0.959742 |
| ENSMUSG( | Katna1 | 335.1734 | 0.035774 | 0.775608 | 0.955606 | ENSMUSG( | Katna1 | 317.1379 | 0.145777 | 0.373007 | 0.915491 |
| ENSMUSG( | Katnal1 | 2775.681 | 0.075743 | 0.362342 | 0.792943 | ENSMUSG( | Katnal1 | 2362.056 | -0.12484 | 0.257057 | 0.883082 |
| ENSMUSG( | Katnal2 | 150.9098 | -0.31249 | 0.079646 | 0.435703 | ENSMUSG( | Katnal2 | 151.1505 | -0.02079 | 0.913688 | 0.992378 |
| ENSMUSG( | Katnb1 | 1095.898 | 0.146613 | 0.173623 | 0.609084 | ENSMUSG( | Katnb1 | 933.623 | -0.04237 | 0.728118 | 0.966913 |
| ENSMUSG( | Katnbl1 | 307.1282 | -0.16859 | 0.274062 | 0.722954 | ENSMUSG( | Katnbl1 | 312.275 | 0.139329 | 0.510204 | 0.929089 |
| ENSMUSG( | Katnip | 818.4567 | -0.03633 | 0.77706 | 0.955828 | ENSMUSG( | Katnip | 791.1644 | 0.137984 | 0.374312 | 0.915491 |
| ENSMUSG( | Kazald1 | 96.76352 | -0.37484 | 0.11416 | 0.509818 | ENSMUSG( | Kazald1 | 89.27144 | -0.31805 | 0.202752 | 0.864412 |
| ENSMUSG( | Kazn | 2773.562 | -0.07141 | 0.388183 | 0.805915 | ENSMUSG( | Kazn | 2587.829 | -0.00355 | 0.973309 | 0.996416 |
| ENSMUSG( | Kbtbd11 | 4562.439 | 0.145039 | 0.072076 | 0.414057 | ENSMUSG( | Kbtbd11 | 3849.206 | -0.07461 | 0.462328 | 0.923329 |
| ENSMUSG( | Kbtbd2 | 1084.41 | 0.02665 | 0.791152 | 0.959461 | ENSMUSG( | Kbtbd2 | 990.734 | 0.031638 | 0.80268 | 0.97681 |
| ENSMUSG( | Kbtbd3 | 135.8141 | -0.14277 | 0.425734 | 0.824898 | ENSMUSG( | Kbtbd3 | 121.9439 | -0.19584 | 0.326114 | 0.905339 |
| ENSMUSG( | Kbtbd4 | 528.3477 | -0.01819 | 0.894647 | 0.984512 | ENSMUSG( | Kbtbd4 | 478.2795 | -0.03871 | 0.785933 | 0.976285 |
| ENSMUSG( | Kbtbd6 | 15.5674 | -0.07585 | 0.870395 | NA | ENSMUSG( | Kbtbd6 | 14.2323 | -0.09422 | 0.861577 | 0.986298 |
| ENSMUSG( | Kbtbd7 | 432.7218 | -0.00931 | 0.937486 | 0.993415 | ENSMUSG( | Kbtbd7 | 432.5538 | 0.239608 | 0.151844 | 0.837724 |
| ENSMUSG( | Kbtbd8 | 104.5566 | -0.12748 | 0.529988 | 0.878883 | ENSMUSG( | Kbtbd8 | 107.7511 | 0.225977 | 0.307381 | 0.898538 |
| ENSMUSG( | Kcmf1 | 2244.964 | -0.00581 | 0.943325 | 0.993415 | ENSMUSG( | Kcmf1 | 1985.162 | -0.09552 | 0.409022 | 0.917257 |
| ENSMUSG( | Kcna1 | 497.6133 | -0.33023 | 0.158918 | 0.588112 | ENSMUSG( | Kcna1 | 628.0542 | 0.582133 | 0.204748 | 0.864412 |
| ENSMUSG( | Kcna2 | 8303.129 | -0.03565 | 0.772762 | 0.955272 | ENSMUSG( | Kcna2 | 7692.143 | 0.015763 | 0.910743 | 0.992196 |
| ENSMUSG( | Kcna3 | 41.90034 | -0.33427 | 0.318862 | 0.764629 | ENSMUSG( | Kcna3 | 49.52904 | 0.438004 | 0.253486 | 0.88018 |
| ENSMUSG( | Kcna4 | 376.953 | -0.65187 | 7.41E-05 | 0.005296 | ENSMUSG( | Kcna4 | 417.6431 | -0.00773 | 0.962271 | 0.995237 |
| ENSMUSG( | Kcna5 | 143.1092 | -0.72055 | 0.000218 | 0.011384 | ENSMUSG( | Kcna5 | 173.676 | 0.180155 | 0.350105 | 0.911652 |
| ENSMUSG( | Kcna6 | 2881.749 | -0.28117 | 0.049479 | 0.339723 | ENSMUSG( | Kcna6 | 3089.622 | 0.200555 | 0.256721 | 0.883082 |
| ENSMUSG( | Kcnab1 | 1239.678 | -0.29154 | 0.061192 | 0.381822 | ENSMUSG( | Kcnab1 | 1175.644 | -0.17024 | 0.348937 | 0.911652 |
| ENSMUSG( | Kcnab2 | 19646.93 | 0.158976 | 0.093542 | 0.469962 | ENSMUSG( | Kcnab2 | 17494.45 | 0.09732 | 0.413646 | 0.917257 |
| ENSMUSG( | Kcnab3 | 1463.721 | 0.009537 | 0.792396 | 0.959461 | ENSMUSG( | Kcnab3 | 1217.021 | -0.23692 | 0.158285 | 0.837724 |
| ENSMUSG( | Kcnb1 | 7161.145 | -0.15675 | 0.266785 | 0.716789 | ENSMUSG( | Kcnb1 | 7318.681 | 0.178186 | 0.292034 | 0.897854 |
| ENSMUSG( | Kcnb2 | 411.1567 | -0.43563 | 0.01 | 0.142653 | ENSMUSG( | Kcnb2 | 452.1759 | 0.141721 | 0.468738 | 0.923329 |
| ENSMUSG( | Kcnc1 | 4730.647 | -0.33472 | 0.013865 | 0.172062 | ENSMUSG( | Kcnc1 | 4915.257 | 0.066371 | 0.685353 | 0.958678 |
| ENSMUSG( | Kcnc2 | 2330.029 | -0.26404 | 0.067893 | 0.403231 | ENSMUSG( | Kcnc2 | 2424.64 | 0.132499 | 0.451268 | 0.922202 |
| ENSMUSG( | Kcnc3 | 3786.879 | -0.30116 | 0.057045 | 0.368342 | ENSMUSG( | Kcnc3 | 4100.361 | 0.211162 | 0.258323 | 0.883971 |
| ENSMUSG( | Kcnc4 | 2769.608 | 0.058658 | 0.654188 | 0.923091 | ENSMUSG( | Kcnc4 | 2661.883 | 0.210041 | 0.174166 | 0.844317 |
| ENSMUSG( | Kcnd1 | 448.6862 | -0.20122 | 0.163968 | 0.594869 | ENSMUSG( | Kcnd1 | 457.3367 | 0.135141 | 0.40987 | 0.917257 |
| ENSMUSG( | Kcnd2 | 2003.257 | -0.48375 | 0.000293 | 0.014095 | ENSMUSG( | Kcnd2 | 2304.253 | 0.224173 | 0.147963 | 0.836241 |
| ENSMUSG( | Kcnd3 | 1922.687 | -0.12599 | 0.376221 | 0.801127 | ENSMUSG( | Kcnd3 | 2100.499 | 0.378929 | 0.013211 | 0.460068 |
| ENSMUSG( | Kcnd3os | 29.89429 | -0.122 | 0.729098 | 0.944223 | ENSMUSG( | Kcnd3os | 30.25706 | 0.138262 | 0.719956 | 0.966887 |
| ENSMUSG( | Kcne2 | 47.0732 | -0.27961 | 0.776283 | 0.955658 | ENSMUSG( | Kcne2 | 55.42487 | 0.424043 | 0.622973 | 0.953234 |
| ENSMUSG( | Kcne4 | 50.42631 | -0.75767 | 0.016996 | 0.194647 | ENSMUSG( | Kcne4 | 60.74795 | 0.182349 | 0.609429 | 0.950397 |
| ENSMUSG( | Kcne5 | 634.746 | -0.2932 | 0.095657 | 0.475065 | ENSMUSG( | Kcne5 | 578.6243 | -0.30233 | 0.113567 | 0.809623 |
| ENSMUSG( | Kcnf1 | 3862.537 | -0.22362 | 0.170624 | 0.605271 | ENSMUSG( | Kcnf1 | 4131.755 | 0.244119 | 0.174976 | 0.845107 |
| ENSMUSG( | Kcng1 | 1229.475 | 0.005636 | 0.96724 | 0.995977 | ENSMUSG( | Kcng1 | 1234.146 | 0.275667 | 0.057987 | 0.728256 |
| ENSMUSG( | Kcng2 | 281.1929 | 0.050727 | 0.751248 | 0.950191 | ENSMUSG( | Kcng2 | 290.9385 | 0.398838 | 0.010642 | 0.419324 |
| ENSMUSG( | Kcng4 | 598.6193 | -0.09521 | 0.452188 | 0.836145 | ENSMUSG( | Kcng4 | 499.2001 | -0.36703 | 0.018395 | 0.544291 |
| ENSMUSG( | Kcnh1 | 2092.4 | -0.49743 | 0.002785 | 0.064345 | ENSMUSG( | Kcnh1 | 2285.804 | 0.074989 | 0.69802 | 0.960867 |
| ENSMUSG( | Kcnh2 | 979.3221 | -0.2516 | 0.021727 | 0.225476 | ENSMUSG( | Kcnh2 | 1000.079 | 0.092227 | 0.53067 | 0.934795 |
| ENSMUSG( | Kcnh3 | 7540.61 | 0.107025 | 0.326924 | 0.772858 | ENSMUSG( | Kcnh3 | 7255.267 | 0.257842 | 0.063694 | 0.736482 |
| ENSMUSG( | Kcnh4 | 502.7499 | -0.17957 | 0.198438 | 0.638387 | ENSMUSG( | Kcnh4 | 520.6521 | 0.195399 | 0.183286 | 0.850984 |
| ENSMUSG( | Kcnh5 | 148.5453 | -0.46414 | 0.035521 | 0.290108 | ENSMUSG( | Kcnh5 | 158.8177 | 0.051036 | 0.835961 | 0.981237 |
| ENSMUSG( | Kcnh6 | 77.12223 | -0.04188 | 0.863227 | 0.975055 | ENSMUSG( | Kcnh6 | 72.0183 | 0.039615 | 0.878867 | 0.989644 |
| ENSMUSG( | Kcnh7 | 1767.048 | -0.4643 | 0.004914 | 0.092785 | ENSMUSG( | Kcnh7 | 2097.14 | 0.321868 | 0.069367 | 0.755467 |
| ENSMUSG( | Kcnh8 | 18.82308 | -0.29407 | 0.549853 | NA | ENSMUSG( | Kcnh8 | 22.0941 | 0.446971 | 0.338391 | 0.907972 |
| ENSMUSG( | Kcnip1 | 572.0791 | 0.015468 | 0.908362 | 0.987809 | ENSMUSG( | Kcnip1 | 513.8793 | -0.03136 | 0.844912 | 0.983863 |
| ENSMUSG( | Kcnip2 | 4201.62 | 0.059088 | 0.546742 | 0.884974 | ENSMUSG( | Kcnip2 | 3597.219 | -0.12376 | 0.32799 | 0.906255 |
| ENSMUSG( | Kcnip3 | 6750.337 | -0.22012 | 0.014484 | 0.177206 | ENSMUSG( | Kcnip3 | 6432.746 | -0.08237 | 0.468581 | 0.923329 |
| ENSMUSG( | Kcnip4 | 674.7156 | -0.08 | 0.594 | 0.902869 | ENSMUSG( | Kcnip4 | 625.252 | -0.03779 | 0.843932 | 0.983571 |
| ENSMUSG( | Kcnj10 | 37914.17 | 0.073547 | 0.563628 | 0.88799 | ENSMUSG( | Kcnj10 | 31450.26 | -0.20453 | 0.129865 | 0.814691 |
| ENSMUSG( | Kcnj11 | 1467.822 | -0.15956 | 0.262995 | 0.713397 | ENSMUSG( | Kcnj11 | 1448.132 | 0.079319 | 0.624646 | 0.953234 |
| ENSMUSG( | Kcnj12 | 299.4161 | -0.04357 | 0.793009 | 0.95947 | ENSMUSG( | Kcnj12 | 293.6901 | 0.172738 | 0.356839 | 0.913358 |
| ENSMUSG( | Kcnj13 | 86.64341 | -0.1228 | 0.755771 | 0.95146 | ENSMUSG( | Kcnj13 | 101.6099 | 0.54033 | 0.149564 | 0.836241 |
| ENSMUSG( | Kcnj14 | 163.0951 | 0.020082 | 0.912322 | 0.988483 | ENSMUSG( | Kcnj14 | 144.3175 | -0.07893 | 0.711009 | 0.964363 |
| ENSMUSG( | Kcnj16 | 4417.855 | 0.081438 | 0.546452 | 0.884974 | ENSMUSG( | Kcnj16 | 3728.517 | -0.1428 | 0.319434 | 0.903461 |
| ENSMUSG( | Kcnj2 | 391.9178 | -0.25269 | 0.094265 | 0.471647 | ENSMUSG( | Kcnj2 | 511.1795 | 0.710573 | 0.000327 | 0.049389 |
| ENSMUSG( | Kcnj3 | 2987.592 | -0.5012 | 0.000292 | 0.014095 | ENSMUSG( | Kcnj3 | 3237.591 | 0.043912 | 0.771943 | 0.974236 |
| ENSMUSG( | Kcnj4 | 3935.419 | -0.40667 | 0.002151 | 0.053771 | ENSMUSG( | Kcnj4 | 4273.897 | 0.13055 | 0.399882 | 0.917257 |
| ENSMUSG( | Kcnj5 | 20.49057 | 0.260497 | 0.548151 | NA | ENSMUSG( | Kcnj5 | 21.32786 | 0.605596 | 0.175241 | 0.845305 |
| ENSMUSG( | Kcnj6 | 305.8444 | -0.10729 | 0.630475 | 0.914073 | ENSMUSG( | Kcnj6 | 344.202 | 0.479921 | 0.071222 | 0.761942 |
| ENSMUSG( | Kcnj8 | 184.8713 | -0.062 | 0.900621 | 0.985633 | ENSMUSG( | Kcnj8 | 208.079 | 0.51931 | 0.392971 | 0.917257 |
| ENSMUSG( | Kcnj9 | 4790.645 | -0.12331 | 0.396351 | 0.811742 | ENSMUSG( | Kcnj9 | 4884.137 | 0.201513 | 0.22425 | 0.867489 |
| ENSMUSG( | Kcnk1 | 4878.427 | -0.27549 | 0.048572 | 0.337113 | ENSMUSG( | Kcnk1 | 5226.889 | 0.202009 | 0.208131 | 0.864412 |
| ENSMUSG( | Kcnk10 | 345.1175 | -0.2064 | 0.257031 | 0.708297 | ENSMUSG( | Kcnk10 | 332.0106 | -0.04155 | 0.827123 | 0.980159 |
| ENSMUSG( | Kcnk12 | 319.8264 | 0.067122 | 0.707552 | 0.93949 | ENSMUSG( | Kcnk12 | 331.2907 | 0.419334 | 0.031199 | 0.631086 |
| ENSMUSG( | Kcnk13 | 163.4976 | -0.2957 | 0.172778 | 0.608361 | ENSMUSG( | Kcnk13 | 185.2119 | 0.344852 | 0.183048 | 0.850741 |
| ENSMUSG( | Kcnk2 | 3633.172 | -0.06767 | 0.558164 | 0.888427 | ENSMUSG( | Kcnk2 | 3618.311 | 0.184457 | 0.195014 | 0.861121 |
| ENSMUSG( | Kcnk3 | 928.1854 | -0.1717 | 0.173018 | 0.608361 | ENSMUSG( | Kcnk3 | 893.8172 | -0.00182 | 0.989865 | 0.997928 |
| ENSMUSG( | Kcnk4 | 527.8389 | 0.140631 | 0.292404 | 0.742521 | ENSMUSG( | Kcnk4 | 492.4233 | 0.213462 | 0.211323 | 0.864412 |
| ENSMUSG( | Kcnk5 | 68.20794 | -0.42264 | 0.106071 | 0.493403 | ENSMUSG( | Kcnk5 | 77.9305 | 0.250607 | 0.32417 | 0.905257 |
| ENSMUSG( | Kcnk6 | 222.6716 | -0.22053 | 0.259745 | 0.709951 | ENSMUSG( | Kcnk6 | 226.7189 | 0.123885 | 0.628381 | 0.953251 |
| ENSMUSG( | Kcnk9 | 2537.659 | -0.12891 | 0.230891 | 0.679066 | ENSMUSG( | Kcnk9 | 2525.834 | 0.12978 | 0.339544 | 0.907972 |
| ENSMUSG( | Kcnma1 | 7423.013 | -0.23929 | 0.072902 | 0.416802 | ENSMUSG( | Kcnma1 | 7787.935 | 0.176191 | 0.248997 | 0.877904 |
| ENSMUSG( | Kcnmb1 | 36.11095 | -0.59425 | 0.066022 | 0.397686 | ENSMUSG( | Kcnmb1 | 43.48848 | 0.295053 | 0.439099 | 0.920721 |
| ENSMUSG( | Kcnmb2 | 60.416 | -0.3609 | 0.19253 | 0.631514 | ENSMUSG( | Kcnmb2 | 66.54519 | 0.203922 | 0.45307 | 0.922316 |
| ENSMUSG( | Kcnmb4 | 2563.306 | 0.048155 | 0.617662 | 0.910872 | ENSMUSG( | Kcnmb4 | 2307.279 | 0.012986 | 0.909468 | 0.991958 |
| ENSMUSG( | Kcnmb4os1 | 31.15925 | -0.13088 | 0.717452 | 0.941697 | ENSMUSG( | Kcnmb4os1 | 28.88158 | -0.08025 | 0.837547 | 0.981335 |
| ENSMUSG( | Kcnmb4os2 | 172.6612 | 0.173492 | 0.323174 | 0.768415 | ENSMUSG( | Kcnmb4os2 | 146.4163 | -0.02182 | 0.912308 | 0.992225 |
| ENSMUSG( | Kcnn1 | 829.5082 | 0.021196 | 0.879125 | 0.981436 | ENSMUSG( | Kcnn1 | 794.2545 | 0.166039 | 0.275379 | 0.889984 |
| ENSMUSG( | Kcnn2 | 2165.793 | 0.400489 | 0.001666 | 0.046175 | ENSMUSG( | Kcnn2 | 2181.71 | 0.649355 | 8.89E-06 | 0.003541 |
| ENSMUSG( | Kcnn3 | 1419.456 | -0.21962 | 0.126866 | 0.534201 | ENSMUSG( | Kcnn3 | 1325.726 | -0.1403 | 0.307298 | 0.898538 |
| ENSMUSG( | Kcnn4 | 14.36553 | -0.68209 | 0.20421 | NA | ENSMUSG( | Kcnn4 | 15.48642 | -0.1041 | 0.835786 | 0.981237 |
| ENSMUSG( | Kcnq1 | 32.1937 | -0.05918 | 0.886056 | 0.983008 | ENSMUSG( | Kcnq1 | 26.36592 | -0.36225 | 0.416028 | 0.917257 |
| ENSMUSG( | Kcnq1ot1 | 783.0025 | 0.145005 | 0.374474 | 0.800445 | ENSMUSG( | Kcnq1ot1 | 802.9937 | 0.448208 | 0.004926 | 0.28966 |
| ENSMUSG( | Kcnq2 | 12592.89 | 0.045759 | 0.733833 | 0.945265 | ENSMUSG( | Kcnq2 | 12215.66 | 0.222622 | 0.165455 | 0.838866 |
| ENSMUSG( | Kcnq3 | 6115.769 | -0.27 | 0.014259 | 0.175484 | ENSMUSG( | Kcnq3 | 6319.064 | 0.10588 | 0.406492 | 0.917257 |
| ENSMUSG( | Kcnq4 | 555.2155 | 0.192182 | 0.210924 | 0.653771 | ENSMUSG( | Kcnq4 | 500.038 | 0.168359 | 0.30833 | 0.898538 |
| ENSMUSG( | Kcnq5 | 2407.76 | -0.3902 | 0.011376 | 0.153509 | ENSMUSG( | Kcnq5 | 2721.682 | 0.253277 | 0.162408 | 0.837724 |
| ENSMUSG( | Kcns1 | 686.7172 | -0.28177 | 0.073713 | 0.419813 | ENSMUSG( | Kcns1 | 683.0357 | -0.00807 | 0.96212 | 0.995237 |
| ENSMUSG( | Kcns2 | 683.6255 | -0.48893 | 0.004563 | 0.088525 | ENSMUSG( | Kcns2 | 712.426 | -0.05281 | 0.786175 | 0.976285 |
| ENSMUSG( | Kcns3 | 173.2731 | 0.030693 | 0.857694 | 0.974667 | ENSMUSG( | Kcns3 | 163.7989 | 0.137008 | 0.469594 | 0.923329 |
| ENSMUSG( | Kcnt1 | 4333.205 | 0.046317 | 0.692686 | 0.934101 | ENSMUSG( | Kcnt1 | 4058.773 | 0.122856 | 0.409688 | 0.917257 |
| ENSMUSG( | Kcnt2 | 415.1732 | -0.36373 | 0.027759 | 0.255176 | ENSMUSG( | Kcnt2 | 470.6287 | 0.285751 | 0.168736 | 0.840222 |
| ENSMUSG( | Kcnu1 | 75.24098 | 0.034947 | 0.882015 | 0.982069 | ENSMUSG( | Kcnu1 | 61.99013 | -0.24553 | 0.391499 | 0.917257 |
| ENSMUSG( | Kcnv1 | 1785.441 | -0.58008 | 0.000323 | 0.015194 | ENSMUSG( | Kcnv1 | 2043.965 | 0.137429 | 0.439275 | 0.920721 |
| ENSMUSG( | Kcp | 114.4659 | 0.494096 | 0.077114 | 0.429655 | ENSMUSG( | Kcp | 92.61458 | 0.196851 | 0.542658 | 0.938034 |
| ENSMUSG( | Kctd1 | 2049.559 | 0.086205 | 0.332693 | 0.775602 | ENSMUSG( | Kctd1 | 1891.119 | 0.120586 | 0.262193 | 0.885219 |
| ENSMUSG( | Kctd10 | 1752.598 | 0.026516 | 0.771838 | 0.954994 | ENSMUSG( | Kctd10 | 1551.684 | -0.05735 | 0.589406 | 0.944975 |
| ENSMUSG( | Kctd11 | 78.11911 | 0.251908 | 0.279462 | 0.729114 | ENSMUSG( | Kctd11 | 69.03974 | 0.172617 | 0.486814 | 0.92541 |
| ENSMUSG( | Kctd12 | 2235.49 | 0.062256 | 0.638542 | 0.916319 | ENSMUSG( | Kctd12 | 2229.745 | 0.303707 | 0.073464 | 0.763629 |
| ENSMUSG( | Kctd12b | 277.6524 | 0.127325 | 0.468528 | 0.845424 | ENSMUSG( | Kctd12b | 256.7737 | 0.162239 | 0.356839 | 0.913358 |
| ENSMUSG( | Kctd13 | 3851.158 | 0.106121 | 0.313448 | 0.760368 | ENSMUSG( | Kctd13 | 3237.233 | -0.12829 | 0.321972 | 0.904023 |
| ENSMUSG( | Kctd14 | 245.6718 | 0.101846 | 0.486485 | 0.858713 | ENSMUSG( | Kctd14 | 209.5763 | -0.10262 | 0.593032 | 0.945408 |
| ENSMUSG( | Kctd15 | 637.0288 | 0.128107 | 0.229745 | 0.677538 | ENSMUSG( | Kctd15 | 555.5123 | 0.004646 | 0.971095 | 0.996102 |
| ENSMUSG( | Kctd16 | 124.5018 | -0.12813 | 0.518273 | 0.873754 | ENSMUSG( | Kctd16 | 131.2781 | 0.272291 | 0.177238 | 0.846973 |
| ENSMUSG( | Kctd17 | 7091.14 | -0.02119 | 0.796379 | 0.960106 | ENSMUSG( | Kctd17 | 6346.077 | -0.07472 | 0.466058 | 0.923329 |

| | | | | | |
|---|---|---|---|---|---|
| ENSMUSG( | 150.9098 | -0.31249 | 0.079646 | 0.435703 | Katnal2 |
| ENSMUSG( | 1095.898 | 0.146613 | 0.173623 | 0.609084 | Katnb1 |
| ENSMUSG( | 307.1282 | -0.16859 | 0.274062 | 0.724254 | Katnbl1 |
| ENSMUSG( | 818.4567 | -0.03633 | 0.77706 | 0.955828 | Katnip |
| ENSMUSG( | 96.76352 | -0.37484 | 0.11416 | 0.509818 | Kazald1 |
| ENSMUSG( | 2773.562 | -0.07141 | 0.388183 | 0.805915 | Kazn |
| ENSMUSG( | 4562.439 | 0.145039 | 0.072076 | 0.414057 | Kbtbd11 |
| ENSMUSG( | 1084.41 | 0.02665 | 0.791152 | 0.959461 | Kbtbd2 |
| ENSMUSG( | 135.8141 | -0.14277 | 0.425734 | 0.824898 | Kbtbd3 |
| ENSMUSG( | 528.3477 | -0.01819 | 0.894647 | 0.984512 | Kbtbd4 |
| ENSMUSG( | 15.5674 | -0.07585 | 0.870395 | NA | Kbtbd6 |
| ENSMUSG( | 432.7218 | -0.00931 | 0.937486 | 0.993415 | Kbtbd7 |
| ENSMUSG( | 104.5566 | -0.12748 | 0.52988 | 0.878883 | Kbtbd8 |
| ENSMUSG( | 2244.964 | -0.00581 | 0.943325 | 0.993415 | Kcmf1 |
| ENSMUSG( | 497.6133 | -0.33023 | 0.158918 | 0.588112 | Kcna1 |
| ENSMUSG( | 8303.129 | -0.03565 | 0.772762 | 0.955272 | Kcna2 |
| ENSMUSG( | 41.90034 | -0.33427 | 0.318862 | 0.764629 | Kcna3 |
| ENSMUSG( | 376.953 | -0.65187 | 7.41E-05 | 0.005296 | Kcna4 |
| ENSMUSG( | 143.1092 | -0.72055 | 0.000218 | 0.011384 | Kcna5 |
| ENSMUSG( | 2881.749 | -0.28117 | 0.049479 | 0.339723 | Kcna6 |
| ENSMUSG( | 1239.678 | -0.29154 | 0.061192 | 0.381822 | Kcnab1 |
| ENSMUSG( | 19646.93 | 0.158976 | 0.093542 | 0.469962 | Kcnab2 |
| ENSMUSG( | 1463.721 | 0.039537 | 0.792396 | 0.959461 | Kcnab3 |
| ENSMUSG( | 7161.145 | -0.15675 | 0.266785 | 0.716789 | Kcnb1 |
| ENSMUSG( | 411.1567 | -0.43563 | 0.01 | 0.142653 | Kcnb2 |
| ENSMUSG( | 4730.647 | -0.33472 | 0.013865 | 0.172062 | Kcnc1 |
| ENSMUSG( | 2330.029 | -0.26404 | 0.067893 | 0.403231 | Kcnc2 |
| ENSMUSG( | 3786.879 | -0.30116 | 0.057045 | 0.368342 | Kcnc3 |
| ENSMUSG( | 2769.608 | 0.058658 | 0.654188 | 0.923091 | Kcnc4 |
| ENSMUSG( | 448.6862 | -0.20122 | 0.163968 | 0.594869 | Kcnd1 |
| ENSMUSG( | 2003.257 | -0.48375 | 0.000293 | 0.014095 | Kcnd2 |
| ENSMUSG( | 1922.687 | -0.12599 | 0.376221 | 0.801127 | Kcnd3 |
| ENSMUSG( | 29.89429 | -0.122 | 0.726092 | 0.944223 | Kcnd3os |
| ENSMUSG( | 47.0732 | -0.27961 | 0.776283 | 0.955658 | Kcne2 |
| ENSMUSG( | 50.42631 | -0.75767 | 0.016996 | 0.194647 | Kcne4 |
| ENSMUSG( | 634.746 | -0.2932 | 0.095657 | 0.475065 | Kcne5 |
| ENSMUSG( | 3862.537 | -0.22362 | 0.170624 | 0.605271 | Kcnf1 |
| ENSMUSG( | 1229.475 | 0.005636 | 0.96724 | 0.995977 | Kcng1 |
| ENSMUSG( | 281.1929 | 0.050727 | 0.751248 | 0.950191 | Kcng2 |
| ENSMUSG( | 598.6193 | -0.09521 | 0.452188 | 0.836145 | Kcng4 |
| ENSMUSG( | 2092.4 | -0.49743 | 0.002785 | 0.064345 | Kcnh1 |
| ENSMUSG( | 979.3221 | -0.2516 | 0.021727 | 0.225476 | Kcnh2 |
| ENSMUSG( | 7540.61 | 0.107025 | 0.326924 | 0.771858 | Kcnh3 |
| ENSMUSG( | 502.7499 | -0.17957 | 0.198438 | 0.638387 | Kcnh4 |
| ENSMUSG( | 148.5453 | -0.46414 | 0.035521 | 0.290108 | Kcnh5 |
| ENSMUSG( | 77.12223 | -0.04188 | 0.863227 | 0.975055 | Kcnh6 |
| ENSMUSG( | 1767.048 | -0.4643 | 0.004914 | 0.092785 | Kcnh7 |
| ENSMUSG( | 18.82308 | -0.29407 | 0.549853 | NA | Kcnh8 |
| ENSMUSG( | 572.0791 | 0.015468 | 0.908362 | 0.987809 | Kcnip1 |
| ENSMUSG( | 4201.62 | 0.059088 | 0.546742 | 0.884974 | Kcnip2 |
| ENSMUSG( | 6750.337 | -0.22012 | 0.014484 | 0.177206 | Kcnip3 |
| ENSMUSG( | 674.7156 | -0.08 | 0.594 | 0.902869 | Kcnip4 |
| ENSMUSG( | 37914.17 | 0.073547 | 0.563624 | 0.88999 | Kcnj10 |
| ENSMUSG( | 1467.822 | -0.15956 | 0.262995 | 0.713397 | Kcnj11 |
| ENSMUSG( | 299.4161 | -0.04357 | 0.793009 | 0.95947 | Kcnj12 |
| ENSMUSG( | 86.64341 | -0.1228 | 0.755771 | 0.95146 | Kcnj13 |
| ENSMUSG( | 163.0951 | 0.020082 | 0.912322 | 0.988483 | Kcnj14 |
| ENSMUSG( | 4417.855 | 0.081438 | 0.546452 | 0.884974 | Kcnj16 |
| ENSMUSG( | 391.9178 | -0.25269 | 0.094265 | 0.471647 | Kcnj2 |
| ENSMUSG( | 2987.592 | -0.5012 | 0.000292 | 0.014095 | Kcnj3 |
| ENSMUSG( | 3935.419 | -0.40667 | 0.002151 | 0.053771 | Kcnj4 |
| ENSMUSG( | 20.49057 | 0.260497 | 0.548151 | NA | Kcnj5 |
| ENSMUSG( | 305.8444 | -0.10729 | 0.630475 | 0.914073 | Kcnj6 |
| ENSMUSG( | 184.8713 | -0.062 | 0.900621 | 0.985633 | Kcnj8 |
| ENSMUSG( | 4790.645 | -0.12331 | 0.396351 | 0.811742 | Kcnj9 |
| ENSMUSG( | 4878.427 | -0.27549 | 0.048572 | 0.337113 | Kcnk1 |
| ENSMUSG( | 345.1175 | -0.2064 | 0.257031 | 0.708297 | Kcnk10 |
| ENSMUSG( | 319.8264 | 0.067122 | 0.707552 | 0.93949 | Kcnk12 |
| ENSMUSG( | 163.4976 | -0.2957 | 0.172778 | 0.608361 | Kcnk13 |
| ENSMUSG( | 3633.172 | -0.06767 | 0.558164 | 0.888427 | Kcnk2 |
| ENSMUSG( | 928.1854 | -0.1717 | 0.173018 | 0.608361 | Kcnk3 |
| ENSMUSG( | 527.8389 | 0.140631 | 0.292404 | 0.742521 | Kcnk4 |
| ENSMUSG( | 68.20794 | -0.42264 | 0.106071 | 0.493403 | Kcnk5 |
| ENSMUSG( | 222.6716 | -0.22053 | 0.259745 | 0.709951 | Kcnk6 |
| ENSMUSG( | 2537.659 | -0.12891 | 0.230891 | 0.679066 | Kcnk9 |
| ENSMUSG( | 7423.013 | -0.23929 | 0.072902 | 0.416802 | Kcnma1 |
| ENSMUSG( | 36.11095 | -0.59425 | 0.066022 | 0.397686 | Kcnmb1 |
| ENSMUSG( | 60.4116 | -0.3609 | 0.19253 | 0.631514 | Kcnmb2 |
| ENSMUSG( | 2563.306 | 0.048155 | 0.617662 | 0.910872 | Kcnmb4 |
| ENSMUSG( | 31.15925 | -0.13088 | 0.717452 | 0.941697 | Kcnmb4os1 |
| ENSMUSG( | 172.6612 | 0.173492 | 0.323174 | 0.768415 | Kcnmb4os2 |
| ENSMUSG( | 829.5082 | 0.021196 | 0.879125 | 0.981436 | Kcnn1 |
| ENSMUSG( | 2165.793 | 0.400489 | 0.001666 | 0.046175 | Kcnn2 |
| ENSMUSG( | 1419.456 | -0.21962 | 0.126866 | 0.534201 | Kcnn3 |
| ENSMUSG( | 14.36553 | -0.68209 | 0.20421 | NA | Kcnn4 |
| ENSMUSG( | 32.1937 | -0.05918 | 0.886056 | 0.983008 | Kcnq1 |
| ENSMUSG( | 783.0025 | 0.145005 | 0.374474 | 0.800445 | Kcnq1ot1 |
| ENSMUSG( | 12592.89 | 0.045759 | 0.733833 | 0.945265 | Kcnq2 |
| ENSMUSG( | 6115.769 | -0.27 | 0.014259 | 0.175484 | Kcnq3 |
| ENSMUSG( | 555.2155 | 0.192182 | 0.210924 | 0.653771 | Kcnq4 |
| ENSMUSG( | 2407.76 | -0.3902 | 0.011376 | 0.153509 | Kcnq5 |
| ENSMUSG( | 686.7172 | -0.28177 | 0.073713 | 0.419813 | Kcns1 |
| ENSMUSG( | 683.6255 | -0.48893 | 0.004563 | 0.088525 | Kcns2 |
| ENSMUSG( | 173.2731 | 0.030693 | 0.857694 | 0.974667 | Kcns3 |
| ENSMUSG( | 4333.205 | 0.046317 | 0.692686 | 0.934101 | Kcnt1 |
| ENSMUSG( | 415.1732 | -0.36373 | 0.027759 | 0.255176 | Kcnt2 |
| ENSMUSG( | 75.24098 | 0.034947 | 0.882015 | 0.982069 | Kcnu1 |
| ENSMUSG( | 1785.441 | -0.58008 | 0.000323 | 0.015194 | Kcnv1 |
| ENSMUSG( | 114.4659 | 0.494096 | 0.077114 | 0.429655 | Kcp |
| ENSMUSG( | 2049.559 | 0.086205 | 0.332693 | 0.775602 | Kctd1 |
| ENSMUSG( | 1752.598 | 0.026516 | 0.771838 | 0.954994 | Kctd10 |
| ENSMUSG( | 78.11911 | 0.251908 | 0.279462 | 0.729114 | Kctd11 |
| ENSMUSG( | 2235.49 | 0.062256 | 0.638542 | 0.916319 | Kctd12 |
| ENSMUSG( | 277.6524 | 0.127325 | 0.468528 | 0.845424 | Kctd12b |
| ENSMUSG( | 3851.158 | 0.106121 | 0.313448 | 0.760368 | Kctd13 |
| ENSMUSG( | 245.6718 | 0.101846 | 0.486485 | 0.858713 | Kctd14 |
| ENSMUSG( | 637.0288 | 0.128107 | 0.229745 | 0.677538 | Kctd15 |
| ENSMUSG( | 124.5018 | -0.12813 | 0.518273 | 0.873754 | Kctd16 |
| ENSMUSG( | 7091.14 | -0.02119 | 0.796379 | 0.960106 | Kctd17 |
| ENSMUSG( | 542.6966 | 0.034252 | 0.763202 | 0.953781 | Kctd18 |
| ENSMUSG( | 2435.746 | 0.115671 | 0.240341 | 0.687899 | Kctd2 |
| ENSMUSG( | 2419.09 | 0.114109 | 0.188782 | 0.627678 | Kctd20 |
| ENSMUSG( | 567.1589 | 0.234923 | 0.051145 | 0.346957 | Kctd21 |
| ENSMUSG( | 2726.985 | 0.152289 | 0.116618 | 0.514987 | Kctd3 |
| ENSMUSG( | 303.3585 | -0.04949 | 0.742553 | 0.948478 | Kctd4 |
| ENSMUSG( | 1728.686 | 0.078675 | 0.449463 | 0.835181 | Kctd5 |
| ENSMUSG( | 724.5052 | 0.01886 | 0.869172 | 0.977194 | Kctd6 |
| ENSMUSG( | 337.0354 | 0.004449 | 0.972752 | 0.996585 | Kctd7 |
| ENSMUSG( | 75.3021 | 0.068699 | 0.769069 | 0.954674 | Kctd8 |
| ENSMUSG( | 270.9528 | -0.02556 | 0.850012 | 0.972638 | Kctd9 |
| ENSMUSG( | 2681.914 | -0.42079 | 0.000463 | 0.019662 | Kdelr1 |
| ENSMUSG( | 934.5752 | -0.51181 | 0.000228 | 0.011828 | Kdelr2 |
| ENSMUSG( | 44.38488 | -0.48156 | 0.097035 | 0.477375 | Kdelr3 |
| ENSMUSG( | 1614.234 | 0.204862 | 0.048421 | 0.336526 | Kdm1a |
| ENSMUSG( | 521.8108 | 0.057546 | 0.642885 | 0.916865 | Kdm1b |
| ENSMUSG( | 2427.893 | 0.041552 | 0.617321 | 0.910866 | Kdm2a |
| ENSMUSG( | 689.9226 | 0.071222 | 0.505437 | 0.86689 | Kdm2b |
| ENSMUSG( | 1244.48 | 0.196806 | 0.052059 | 0.350227 | Kdm3a |
| ENSMUSG( | 2218.586 | 0.065688 | 0.44453 | 0.833239 | Kdm3b |
| ENSMUSG( | 1570.357 | 0.165308 | 0.094092 | 0.471597 | Kdm4a |
| ENSMUSG( | 1837.722 | 0.132279 | 0.236773 | 0.684086 | Kdm4b |
| ENSMUSG( | 711.117 | 0.091284 | 0.453432 | 0.836798 | Kdm4c |
| ENSMUSG( | 1288.691 | 0.010598 | 0.917912 | 0.989535 | Kdm5a |
| ENSMUSG( | 1878.274 | 0.223074 | 0.028151 | 0.256523 | Kdm5b |
| ENSMUSG( | 1896.022 | 0.151586 | 0.081051 | 0.440124 | Kdm5c |
| ENSMUSG( | 777.1048 | 0.038907 | 0.695079 | 0.93484 | Kdm5d |
| ENSMUSG( | 550.1877 | 0.066744 | 0.620284 | 0.911432 | Kdm6a |
| ENSMUSG( | 1284.139 | -0.15298 | 0.104839 | 0.491651 | Kdm6b |
| ENSMUSG( | 1056.053 | -0.09654 | 0.356301 | 0.788108 | Kdm7a |
| ENSMUSG( | 116.7803 | 0.231669 | 0.21913 | 0.664602 | Kdm8 |
| ENSMUSG( | 804.9155 | 0.08683 | 0.814298 | 0.964574 | Kdr |
| ENSMUSG( | 1293.839 | -0.20582 | 0.097519 | 0.477842 | Kdsr |
| ENSMUSG( | 3136.91 | 0.107044 | 0.190918 | 0.630245 | Keap1 |
| ENSMUSG( | 1268.633 | 0.003824 | 0.970803 | 0.996369 | Khdc4 |
| ENSMUSG( | 2182.344 | -0.03209 | 0.746638 | 0.949289 | Khdrbs1 |
| ENSMUSG( | 113.4259 | -0.14065 | 0.491326 | 0.859873 | Khdrbs2 |
| ENSMUSG( | 4114.884 | 0.146701 | 0.099844 | 0.482272 | Khdrbs3 |
| ENSMUSG( | 357.9828 | -0.0042 | 0.973826 | 0.99675 | Khk |
| ENSMUSG( | 653.8202 | 0.044676 | 0.681791 | 0.931619 | Khnyn |
| ENSMUSG( | 2328.147 | 0.168423 | 0.060341 | 0.3797 | Khsrp |
| ENSMUSG( | 199.7264 | -0.11518 | 0.498794 | 0.862973 | Kics2 |
| ENSMUSG( | 12203 | -0.10341 | 0.414774 | 0.820184 | Kidins220 |
| ENSMUSG( | 97.81969 | 0.086918 | 0.688878 | 0.933441 | Kif11 |
| ENSMUSG( | 2195.562 | -0.02682 | 0.79785 | 0.96053 | Kif13a |
| ENSMUSG( | 3175.312 | -0.23664 | 0.006384 | 0.109686 | Kif13b |
| ENSMUSG( | 21.18244 | -1.09607 | 0.010502 | NA | Kif14 |
| ENSMUSG( | 26.50624 | -0.41761 | 0.245839 | 0.693547 | Kif15 |
| ENSMUSG( | 552.3518 | 0.014611 | 0.904464 | 0.986246 | Kif16b |

| | | | | | |
|---|---|---|---|---|---|
| ENSMUSG( | 31450.26 | -0.20453 | 0.123685 | 0.814691 | Kcnj10 |
| ENSMUSG( | 1448.132 | 0.079319 | 0.624646 | 0.953234 | Kcnj11 |
| ENSMUSG( | 293.6901 | 0.172738 | 0.356839 | 0.913358 | Kcnj12 |
| ENSMUSG( | 101.6099 | 0.54033 | 0.149564 | 0.836241 | Kcnj13 |
| ENSMUSG( | 144.3175 | -0.07893 | 0.711009 | 0.964363 | Kcnj14 |
| ENSMUSG( | 3728.517 | -0.1428 | 0.319434 | 0.903461 | Kcnj16 |
| ENSMUSG( | 511.1795 | 0.710573 | 0.000327 | 0.049389 | Kcnj2 |
| ENSMUSG( | 3237.591 | 0.043912 | 0.771943 | 0.974236 | Kcnj3 |
| ENSMUSG( | 4273.897 | 0.13055 | 0.399882 | 0.917257 | Kcnj4 |
| ENSMUSG( | 21.32786 | 0.605596 | 0.175241 | 0.845305 | Kcnj5 |
| ENSMUSG( | 344.202 | 0.479921 | 0.071222 | 0.761942 | Kcnj6 |
| ENSMUSG( | 208.079 | 0.51931 | 0.392971 | 0.917257 | Kcnj8 |
| ENSMUSG( | 4884.137 | 0.201513 | 0.22425 | 0.867489 | Kcnj9 |
| ENSMUSG( | 5226.889 | 0.202009 | 0.208131 | 0.864412 | Kcnk1 |
| ENSMUSG( | 332.0106 | -0.04155 | 0.827123 | 0.980159 | Kcnk10 |
| ENSMUSG( | 331.2907 | 0.419334 | 0.031199 | 0.631086 | Kcnk12 |
| ENSMUSG( | 185.2119 | 0.344852 | 0.183048 | 0.850741 | Kcnk13 |
| ENSMUSG( | 3618.311 | 0.184457 | 0.195014 | 0.861121 | Kcnk2 |
| ENSMUSG( | 893.8172 | -0.00182 | 0.989865 | 0.997928 | Kcnk3 |
| ENSMUSG( | 492.4233 | 0.213462 | 0.211323 | 0.864412 | Kcnk4 |
| ENSMUSG( | 77.9305 | 0.250607 | 0.32417 | 0.905257 | Kcnk5 |
| ENSMUSG( | 226.7189 | 0.123885 | 0.628381 | 0.953251 | Kcnk6 |
| ENSMUSG( | 2525.834 | 0.12978 | 0.339544 | 0.907972 | Kcnk9 |
| ENSMUSG( | 7787.935 | 0.176191 | 0.248997 | 0.877904 | Kcnma1 |
| ENSMUSG( | 43.48848 | 0.295053 | 0.439099 | 0.920721 | Kcnmb1 |
| ENSMUSG( | 66.54519 | 0.203922 | 0.45307 | 0.922316 | Kcnmb2 |
| ENSMUSG( | 2307.279 | 0.012986 | 0.909468 | 0.991958 | Kcnmb4 |
| ENSMUSG( | 28.88158 | -0.08025 | 0.837547 | 0.981335 | Kcnmb4os1 |
| ENSMUSG( | 146.4163 | -0.02182 | 0.912308 | 0.992225 | Kcnmb4os2 |
| ENSMUSG( | 794.2545 | 0.166039 | 0.275379 | 0.889984 | Kcnn1 |
| ENSMUSG( | 2181.71 | 0.649355 | 8.89E-06 | 0.003541 | Kcnn2 |
| ENSMUSG( | 1325.726 | -0.1403 | 0.307298 | 0.898538 | Kcnn3 |
| ENSMUSG( | 15.48642 | -0.1041 | 0.835786 | 0.981237 | Kcnn4 |
| ENSMUSG( | 26.36592 | -0.36225 | 0.416028 | 0.917257 | Kcnq1 |
| ENSMUSG( | 802.9937 | 0.448208 | 0.004926 | 0.28966 | Kcnq1ot1 |
| ENSMUSG( | 12215.66 | 0.222622 | 0.165455 | 0.838866 | Kcnq2 |
| ENSMUSG( | 6319.064 | 0.10588 | 0.406492 | 0.917257 | Kcnq3 |
| ENSMUSG( | 500.038 | 0.168359 | 0.30833 | 0.898538 | Kcnq4 |
| ENSMUSG( | 2721.682 | 0.253277 | 0.162408 | 0.837724 | Kcnq5 |
| ENSMUSG( | 10.23109 | 0.916984 | 0.149738 | NA | Kcnrg |
| ENSMUSG( | 683.0357 | -0.00807 | 0.96212 | 0.995237 | Kcns1 |
| ENSMUSG( | 712.426 | -0.05281 | 0.786175 | 0.976285 | Kcns2 |
| ENSMUSG( | 163.7989 | 0.137008 | 0.469594 | 0.923329 | Kcns3 |
| ENSMUSG( | 4058.773 | 0.122856 | 0.409688 | 0.917257 | Kcnt1 |
| ENSMUSG( | 470.6287 | 0.285751 | 0.168736 | 0.840222 | Kcnt2 |
| ENSMUSG( | 61.99013 | -0.24553 | 0.391499 | 0.917257 | Kcnu1 |
| ENSMUSG( | 2043.965 | 0.137429 | 0.439275 | 0.920721 | Kcnv1 |
| ENSMUSG( | 92.61458 | 0.196851 | 0.542658 | 0.938034 | Kcp |
| ENSMUSG( | 1891.119 | 0.120586 | 0.262193 | 0.885219 | Kctd1 |
| ENSMUSG( | 1551.684 | -0.05735 | 0.589406 | 0.944975 | Kctd10 |
| ENSMUSG( | 69.03974 | 0.172617 | 0.486814 | 0.92541 | Kctd11 |
| ENSMUSG( | 2229.745 | 0.303707 | 0.073464 | 0.763629 | Kctd12 |
| ENSMUSG( | 256.7737 | 0.162239 | 0.356839 | 0.913358 | Kctd12b |
| ENSMUSG( | 3237.233 | -0.12829 | 0.321972 | 0.904023 | Kctd13 |
| ENSMUSG( | 209.5763 | -0.10262 | 0.593032 | 0.945408 | Kctd14 |
| ENSMUSG( | 555.5123 | 0.004646 | 0.971095 | 0.996102 | Kctd15 |
| ENSMUSG( | 131.2781 | 0.272291 | 0.177238 | 0.846973 | Kctd16 |
| ENSMUSG( | 6346.077 | -0.07472 | 0.466058 | 0.923329 | Kctd17 |
| ENSMUSG( | 503.964 | 0.090493 | 0.485817 | 0.92541 | Kctd18 |
| ENSMUSG( | 2103.953 | -0.03904 | 0.767362 | 0.974236 | Kctd2 |
| ENSMUSG( | 2093.337 | -0.03372 | 0.749433 | 0.96998 | Kctd20 |
| ENSMUSG( | 435.9493 | -0.25904 | 0.073786 | 0.765251 | Kctd21 |
| ENSMUSG( | 2336.544 | -0.02084 | 0.855266 | 0.985649 | Kctd3 |
| ENSMUSG( | 300.7387 | 0.188482 | 0.232028 | 0.868507 | Kctd4 |
| ENSMUSG( | 1459.086 | -0.14899 | 0.282155 | 0.892412 | Kctd5 |
| ENSMUSG( | 671.4193 | 0.065236 | 0.636857 | 0.953903 | Kctd6 |
| ENSMUSG( | 290.2826 | -0.15981 | 0.286936 | 0.894091 | Kctd7 |
| ENSMUSG( | 75.56345 | 0.323555 | 0.19764 | 0.862161 | Kctd8 |
| ENSMUSG( | 236.9092 | -0.15247 | 0.344629 | 0.91045 | Kctd9 |
| ENSMUSG( | 2902.393 | 0.106551 | 0.449857 | 0.922202 | Kdelr1 |
| ENSMUSG( | 1043.288 | 0.120846 | 0.440609 | 0.921166 | Kdelr2 |
| ENSMUSG( | 42.4135 | -0.28478 | 0.381947 | 0.917257 | Kdelr3 |
| ENSMUSG( | 1332.194 | -0.07412 | 0.543144 | 0.938063 | Kdm1a |
| ENSMUSG( | 465.2398 | -0.00474 | 0.973232 | 0.996416 | Kdm1b |
| ENSMUSG( | 2196.579 | 0.022691 | 0.832539 | 0.980805 | Kdm2a |
| ENSMUSG( | 602.7761 | -0.0448 | 0.725387 | 0.966887 | Kdm2b |
| ENSMUSG( | 1132.214 | 0.19733 | 0.126247 | 0.81516 | Kdm3a |
| ENSMUSG( | 1957.564 | -0.02662 | 0.795865 | 0.97681 | Kdm3b |
| ENSMUSG( | 1371.024 | 0.048012 | 0.6883 | 0.959742 | Kdm4a |
| ENSMUSG( | 1652.914 | 0.099615 | 0.422024 | 0.917257 | Kdm4b |
| ENSMUSG( | 624.8271 | -0.0084 | 0.950796 | 0.994798 | Kdm4c |
| ENSMUSG( | 14.16411 | 0.456343 | 0.417191 | 0.917257 | Kdm4d |
| ENSMUSG( | 1136.018 | -0.08302 | 0.469483 | 0.923329 | Kdm5a |
| ENSMUSG( | 1629.601 | 0.088198 | 0.44711 | 0.922066 | Kdm5b |
| ENSMUSG( | 1652.099 | 0.02672 | 0.796215 | 0.97681 | Kdm5c |
| ENSMUSG( | 706.7246 | 0.036952 | 0.787955 | 0.97637 | Kdm5d |
| ENSMUSG( | 542.7036 | 0.289706 | 0.08049 | 0.77681 | Kdm6a |
| ENSMUSG( | 1226.713 | -0.014 | 0.910432 | 0.992124 | Kdm6b |
| ENSMUSG( | 1003.593 | 0.026479 | 0.815768 | 0.979198 | Kdm7a |
| ENSMUSG( | 102.3301 | 0.126053 | 0.546496 | 0.938063 | Kdm8 |
| ENSMUSG( | 811.7531 | 0.368759 | 0.126397 | 0.815475 | Kdr |
| ENSMUSG( | 1373.546 | 0.235713 | 0.07582 | 0.768524 | Kdsr |
| ENSMUSG( | 2750.947 | -0.00094 | 0.992636 | 0.99874 | Keap1 |
| ENSMUSG( | 1301.528 | 0.327062 | 0.006927 | 0.343883 | Khdc4 |
| ENSMUSG( | 1948.636 | -0.09496 | 0.469223 | 0.923329 | Khdrbs1 |
| ENSMUSG( | 102.0415 | -0.17485 | 0.424446 | 0.917257 | Khdrbs2 |
| ENSMUSG( | 3411.269 | -0.12698 | 0.287567 | 0.894091 | Khdrbs3 |
| ENSMUSG( | 324.1651 | -0.01941 | 0.896701 | 0.990558 | Khk |
| ENSMUSG( | 610.5773 | 0.121822 | 0.496948 | 0.927825 | Khnyn |
| ENSMUSG( | 2022.004 | 0.033185 | 0.781966 | 0.976106 | Khsrp |
| ENSMUSG( | 184.846 | -0.0619 | 0.747338 | 0.969763 | Kics2 |
| ENSMUSG( | 11629.33 | 0.031191 | 0.823382 | 0.979198 | Kidins220 |
| ENSMUSG( | 94.26864 | 0.254705 | 0.292334 | 0.897854 | Kif11 |
| ENSMUSG( | 1935.205 | -0.12636 | 0.242547 | 0.877399 | Kif13a |
| ENSMUSG( | 2813.013 | -0.32741 | 0.003261 | 0.228473 | Kif13b |
| ENSMUSG( | 24.98207 | -0.13281 | 0.74257 | 0.969415 | Kif14 |
| ENSMUSG( | 26.65812 | -0.10946 | 0.772734 | 0.974463 | Kif15 |
| ENSMUSG( | 488.9795 | -0.07376 | 0.60683 | 0.949772 | Kif16b |
| ENSMUSG( | 720.9253 | 0.026463 | 0.848926 | 0.985321 | Kif17 |
| ENSMUSG( | 30.60552 | 0.096829 | 0.783892 | 0.976154 | Kif18a |
| ENSMUSG( | 15.69234 | -0.1155 | 0.820445 | 0.979198 | Kif18b |
| ENSMUSG( | 284.2725 | -0.07327 | 0.643616 | 0.953903 | Kif19a |
| ENSMUSG( | 65021.76 | -0.13517 | 0.173784 | 0.844317 | Kif1a |
| ENSMUSG( | 27148.09 | 0.049477 | 0.662555 | 0.956205 | Kif1b |
| ENSMUSG( | 16840.25 | -0.19708 | 0.103708 | 0.804969 | Kif1c |
| ENSMUSG( | 31.58069 | -0.17471 | 0.621386 | 0.953199 | Kif20a |
| ENSMUSG( | 33.696 | 0.322649 | 0.352596 | 0.912093 | Kif20b |
| ENSMUSG( | 3127.511 | 0.084763 | 0.433933 | 0.91869 | Kif21a |
| ENSMUSG( | 5361.755 | -0.17922 | 0.081382 | 0.77681 | Kif21b |
| ENSMUSG( | 67.39474 | -0.1 | 0.702193 | 0.962014 | Kif22 |
| ENSMUSG( | 50.25825 | 0.511206 | 0.122355 | 0.813384 | Kif23 |
| ENSMUSG( | 31.86106 | -0.04009 | 0.910973 | 0.992196 | Kif24 |
| ENSMUSG( | 689.6289 | -0.34398 | 0.019493 | 0.556192 | Kif26a |
| ENSMUSG( | 428.4589 | 0.174064 | 0.20405 | 0.864412 | Kif26b |
| ENSMUSG( | 175.6131 | -0.0208 | 0.917354 | 0.992529 | Kif27 |
| ENSMUSG( | 7.855677 | 0.431319 | 0.56177 | NA | Kif28 |
| ENSMUSG( | 1641.066 | 0.069402 | 0.581834 | 0.944266 | Kif2a |
| ENSMUSG( | 20.31368 | 0.379977 | 0.418424 | 0.917257 | Kif2c |
| ENSMUSG( | 1951.642 | 0.000653 | 0.99546 | 0.999116 | Kif3a |
| ENSMUSG( | 1805.973 | 0.016655 | 0.884993 | 0.989688 | Kif3b |
| ENSMUSG( | 5428.346 | 0.080587 | 0.443189 | 0.921575 | Kif3c |
| ENSMUSG( | 197.538 | -0.31921 | 0.084152 | 0.777886 | Kif4 |
| ENSMUSG( | 471255.1 | -0.12563 | 0.303853 | 0.898538 | Kif5a |
| ENSMUSG( | 25117.28 | -0.17824 | 0.171406 | 0.842999 | Kif5b |
| ENSMUSG( | 75794.62 | -0.11665 | 0.309874 | 0.898538 | Kif5c |
| ENSMUSG( | 51.37318 | 0.01675 | 0.959664 | 0.994798 | Kif6 |
| ENSMUSG( | 194.2517 | 0.091806 | 0.593459 | 0.945408 | Kif7 |
| ENSMUSG( | 159.2417 | 0.192854 | 0.336536 | 0.907972 | Kif9 |
| ENSMUSG( | 4676.554 | -0.11559 | 0.302093 | 0.898538 | Kifap3 |
| ENSMUSG( | 3501.577 | -0.16141 | 0.218835 | 0.866104 | Kifbp |
| ENSMUSG( | 42.93582 | -0.26044 | 0.465323 | 0.923329 | Kifc1 |
| ENSMUSG( | 34815.23 | 0.019694 | 0.834928 | 0.981237 | Kifc2 |
| ENSMUSG( | 3248.924 | 0.006981 | 0.944738 | 0.994798 | Kifc3 |
| ENSMUSG( | 71.28958 | -0.15464 | 0.572385 | 0.942515 | Kifc5b |
| ENSMUSG( | 247.7869 | 0.03987 | 0.816483 | 0.979198 | Kin |
| ENSMUSG( | 290.1853 | 0.13515 | 0.447597 | 0.922202 | Kirrel1 |
| ENSMUSG( | 1466.155 | 0.356413 | 0.005459 | 0.303106 | Kirrel2 |
| ENSMUSG( | 2102.8 | -0.07138 | 0.6373 | 0.953903 | Kirrel3 |
| ENSMUSG( | 298.8445 | 0.090966 | 0.560096 | 0.940119 | Kiss1r |
| ENSMUSG( | 1439.358 | 0.220732 | 0.267233 | 0.886134 | Kit |
| ENSMUSG( | 1062.758 | 0.577037 | 0.004397 | 0.269982 | Kitl |
| ENSMUSG( | 494.6037 | -0.15451 | 0.270947 | 0.888062 | Kiz |
| ENSMUSG( | 404.2238 | 0.538575 | 0.328292 | 0.906478 | Kl |
| ENSMUSG( | 19481.62 | -0.17919 | 0.169764 | 0.840806 | Klc1 |
| ENSMUSG( | 6069.215 | -0.15383 | 0.214954 | 0.865167 | Klc2 |
| ENSMUSG( | 95.6234 | -0.27955 | 0.248544 | 0.877904 | Klc3 |
| ENSMUSG( | 929.6984 | -0.08307 | 0.4772 | 0.923329 | Klc4 |
| ENSMUSG( | 465.5015 | -0.5711 | 0.003103 | 0.221768 | Klf10 |

| | | | | | | | | | | | |
|---|---|---|---|---|---|---|---|---|---|---|---|
| ENSMUSG( | Kctd18 | 542.6966 | 0.034252 | 0.763202 | 0.953781 | ENSMUSG( | Kctd18 | 503.964 | 0.090493 | 0.485817 | 0.92541 |
| ENSMUSG( | Kctd2 | 2435.746 | 0.115671 | 0.240341 | 0.687899 | ENSMUSG( | Kctd2 | 2103.953 | -0.03904 | 0.767362 | 0.974236 |
| ENSMUSG( | Kctd20 | 2419.09 | 0.114109 | 0.188782 | 0.627678 | ENSMUSG( | Kctd20 | 2093.337 | -0.03372 | 0.749433 | 0.96998 |
| ENSMUSG( | Kctd21 | 567.1589 | 0.234923 | 0.051145 | 0.346957 | ENSMUSG( | Kctd21 | 435.9493 | -0.25904 | 0.073786 | 0.765251 |
| ENSMUSG( | Kctd3 | 2726.985 | 0.152289 | 0.116618 | 0.514987 | ENSMUSG( | Kctd3 | 2336.544 | -0.02084 | 0.855266 | 0.985649 |
| ENSMUSG( | Kctd4 | 303.3585 | -0.04949 | 0.742553 | 0.948478 | ENSMUSG( | Kctd4 | 300.7387 | 0.188482 | 0.232028 | 0.868507 |
| ENSMUSG( | Kctd5 | 1728.686 | 0.078675 | 0.449463 | 0.835181 | ENSMUSG( | Kctd5 | 1459.086 | -0.14899 | 0.282155 | 0.892412 |
| ENSMUSG( | Kctd6 | 724.5052 | 0.01886 | 0.869172 | 0.977194 | ENSMUSG( | Kctd6 | 671.4193 | 0.065236 | 0.636857 | 0.953903 |
| ENSMUSG( | Kctd7 | 337.0354 | 0.004449 | 0.972752 | 0.996585 | ENSMUSG( | Kctd7 | 290.2826 | -0.15981 | 0.286936 | 0.894091 |
| ENSMUSG( | Kctd8 | 75.3021 | 0.068699 | 0.769069 | 0.954674 | ENSMUSG( | Kctd8 | 75.56345 | 0.323555 | 0.19764 | 0.862161 |
| ENSMUSG( | Kctd9 | 270.9528 | -0.02556 | 0.850012 | 0.972638 | ENSMUSG( | Kctd9 | 236.9092 | -0.15247 | 0.344629 | 0.91045 |
| ENSMUSG( | Kdelr1 | 2681.914 | -0.42079 | 0.000463 | 0.019662 | ENSMUSG( | Kdelr1 | 2902.393 | 0.106551 | 0.449857 | 0.922202 |
| ENSMUSG( | Kdelr2 | 934.5752 | -0.51181 | 0.000228 | 0.011828 | ENSMUSG( | Kdelr2 | 1043.288 | 0.120846 | 0.440609 | 0.921166 |
| ENSMUSG( | Kdelr3 | 44.38488 | -0.48156 | 0.097035 | 0.477375 | ENSMUSG( | Kdelr3 | 42.4135 | -0.28478 | 0.381947 | 0.917257 |
| ENSMUSG( | Kdm1a | 1614.234 | 0.204862 | 0.048421 | 0.336526 | ENSMUSG( | Kdm1a | 1332.194 | -0.07412 | 0.543144 | 0.938063 |
| ENSMUSG( | Kdm1b | 521.8108 | 0.057546 | 0.642885 | 0.916865 | ENSMUSG( | Kdm1b | 465.2398 | -0.00474 | 0.973232 | 0.996416 |
| ENSMUSG( | Kdm2a | 2427.893 | 0.041552 | 0.617321 | 0.910866 | ENSMUSG( | Kdm2a | 2196.579 | 0.022691 | 0.832539 | 0.980805 |
| ENSMUSG( | Kdm2b | 689.9226 | 0.071222 | 0.505437 | 0.86689 | ENSMUSG( | Kdm2b | 602.7761 | -0.0448 | 0.725387 | 0.966887 |
| ENSMUSG( | Kdm3a | 1244.48 | 0.196806 | 0.052059 | 0.350227 | ENSMUSG( | Kdm3a | 1132.214 | 0.19733 | 0.126247 | 0.81516 |
| ENSMUSG( | Kdm3b | 2218.586 | 0.065688 | 0.44453 | 0.833239 | ENSMUSG( | Kdm3b | 1957.564 | -0.02662 | 0.795865 | 0.97681 |
| ENSMUSG( | Kdm4a | 1570.357 | 0.165308 | 0.094092 | 0.471597 | ENSMUSG( | Kdm4a | 1371.024 | 0.048012 | 0.6883 | 0.959742 |
| ENSMUSG( | Kdm4b | 1837.722 | 0.132279 | 0.236773 | 0.684086 | ENSMUSG( | Kdm4b | 1652.914 | 0.099615 | 0.422024 | 0.917257 |
| ENSMUSG( | Kdm4c | 711.117 | 0.091284 | 0.453432 | 0.836798 | ENSMUSG( | Kdm4c | 624.8271 | -0.0084 | 0.950796 | 0.994798 |
| ENSMUSG( | Kdm5a | 1288.691 | 0.010598 | 0.917912 | 0.989535 | ENSMUSG( | Kdm5a | 1136.018 | -0.08302 | 0.469483 | 0.923329 |
| ENSMUSG( | Kdm5b | 1878.274 | 0.223074 | 0.028151 | 0.256523 | ENSMUSG( | Kdm5b | 1629.601 | 0.088198 | 0.44711 | 0.922066 |
| ENSMUSG( | Kdm5c | 1896.022 | 0.151586 | 0.081051 | 0.440124 | ENSMUSG( | Kdm5c | 1652.099 | 0.02672 | 0.796215 | 0.97681 |
| ENSMUSG( | Kdm5d | 777.1048 | 0.038907 | 0.695079 | 0.93484 | ENSMUSG( | Kdm5d | 706.7246 | 0.036952 | 0.787955 | 0.97637 |
| ENSMUSG( | Kdm6a | 550.1877 | 0.066744 | 0.620284 | 0.911432 | ENSMUSG( | Kdm6a | 542.7036 | 0.289706 | 0.08049 | 0.77681 |
| ENSMUSG( | Kdm6b | 1284.139 | -0.15298 | 0.104839 | 0.491651 | ENSMUSG( | Kdm6b | 1226.713 | -0.014 | 0.910432 | 0.992124 |
| ENSMUSG( | Kdm7a | 1056.053 | -0.09654 | 0.356301 | 0.788108 | ENSMUSG( | Kdm7a | 1003.593 | 0.026479 | 0.815768 | 0.979198 |
| ENSMUSG( | Kdm8 | 116.7803 | 0.231669 | 0.21913 | 0.664602 | ENSMUSG( | Kdm8 | 102.3301 | 0.126053 | 0.546496 | 0.938063 |
| ENSMUSG( | Kdr | 804.9155 | 0.08683 | 0.814298 | 0.964574 | ENSMUSG( | Kdr | 811.7531 | 0.368759 | 0.126397 | 0.815475 |
| ENSMUSG( | Kdsr | 1293.839 | -0.20582 | 0.097519 | 0.477842 | ENSMUSG( | Kdsr | 1373.546 | 0.235713 | 0.07582 | 0.768524 |
| ENSMUSG( | Keap1 | 3136.91 | 0.107044 | 0.190918 | 0.630245 | ENSMUSG( | Keap1 | 2750.947 | -0.00094 | 0.992636 | 0.99874 |
| ENSMUSG( | Khdc4 | 1268.633 | 0.003824 | 0.970803 | 0.996369 | ENSMUSG( | Khdc4 | 1301.528 | 0.327062 | 0.006927 | 0.343883 |
| ENSMUSG( | Khdrbs1 | 2182.344 | -0.03209 | 0.746638 | 0.949289 | ENSMUSG( | Khdrbs1 | 1948.636 | -0.09496 | 0.469223 | 0.923329 |
| ENSMUSG( | Khdrbs2 | 113.4259 | -0.14065 | 0.491326 | 0.859873 | ENSMUSG( | Khdrbs2 | 102.0415 | -0.17485 | 0.424446 | 0.917257 |
| ENSMUSG( | Khdrbs3 | 4114.884 | 0.146701 | 0.099844 | 0.482272 | ENSMUSG( | Khdrbs3 | 3411.269 | -0.12698 | 0.287567 | 0.894091 |
| ENSMUSG( | Khk | 357.9828 | -0.0042 | 0.973826 | 0.99675 | ENSMUSG( | Khk | 324.1651 | -0.01941 | 0.896701 | 0.990558 |
| ENSMUSG( | Khnyn | 653.8202 | 0.044676 | 0.681791 | 0.931619 | ENSMUSG( | Khnyn | 610.5773 | 0.121822 | 0.496948 | 0.927825 |
| ENSMUSG( | Khsrp | 2328.147 | 0.168423 | 0.060341 | 0.3797 | ENSMUSG( | Khsrp | 2022.004 | 0.033185 | 0.781966 | 0.976106 |
| ENSMUSG( | Kics2 | 199.7264 | -0.11518 | 0.498794 | 0.862973 | ENSMUSG( | Kics2 | 184.846 | -0.0619 | 0.747338 | 0.969763 |
| ENSMUSG( | Kidins220 | 12203 | -0.10341 | 0.414774 | 0.820184 | ENSMUSG( | Kidins220 | 11629.33 | 0.031191 | 0.823382 | 0.979198 |
| ENSMUSG( | Kif11 | 97.81969 | 0.086918 | 0.688878 | 0.933441 | ENSMUSG( | Kif11 | 94.26864 | 0.254705 | 0.292334 | 0.897854 |
| ENSMUSG( | Kif13a | 2195.562 | -0.02682 | 0.79785 | 0.96053 | ENSMUSG( | Kif13a | 1935.205 | -0.12636 | 0.242547 | 0.877399 |
| ENSMUSG( | Kif13b | 3175.312 | -0.23664 | 0.006384 | 0.109686 | ENSMUSG( | Kif13b | 2813.013 | -0.32741 | 0.003261 | 0.228473 |
| ENSMUSG( | Kif14 | 21.18244 | -1.09607 | 0.010502 | NA | ENSMUSG( | Kif14 | 24.98207 | -0.13281 | 0.74257 | 0.969415 |
| ENSMUSG( | Kif15 | 26.50624 | -0.41761 | 0.245839 | 0.693547 | ENSMUSG( | Kif15 | 26.65812 | -0.10946 | 0.772734 | 0.974463 |
| ENSMUSG( | Kif16b | 552.3518 | 0.014611 | 0.904464 | 0.986246 | ENSMUSG( | Kif16b | 488.9795 | -0.07376 | 0.60683 | 0.949772 |
| ENSMUSG( | Kif17 | 813.7654 | 0.106011 | 0.406673 | 0.816928 | ENSMUSG( | Kif17 | 720.9253 | 0.026463 | 0.848926 | 0.985321 |
| ENSMUSG( | Kif18a | 30.91783 | -0.15908 | 0.642936 | 0.916865 | ENSMUSG( | Kif18a | 30.60552 | 0.096829 | 0.783892 | 0.976154 |
| ENSMUSG( | Kif18b | 15.61929 | -0.47781 | 0.311104 | NA | ENSMUSG( | Kif18b | 15.69234 | -0.1155 | 0.820445 | 0.979198 |
| ENSMUSG( | Kif19a | 312.0478 | -0.07924 | 0.578778 | 0.896692 | ENSMUSG( | Kif19a | 284.2725 | -0.07327 | 0.643616 | 0.953903 |
| ENSMUSG( | Kif1a | 76489.67 | 0.066787 | 0.389068 | 0.806967 | ENSMUSG( | Kif1a | 65021.76 | -0.13517 | 0.173784 | 0.844317 |
| ENSMUSG( | Kif1b | 29863.1 | 0.054612 | 0.534677 | 0.880719 | ENSMUSG( | Kif1b | 27148.09 | 0.049477 | 0.662555 | 0.956205 |
| ENSMUSG( | Kif1c | 19712.63 | -0.00295 | 0.974724 | 0.997207 | ENSMUSG( | Kif1c | 16840.25 | -0.19708 | 0.103708 | 0.804969 |
| ENSMUSG( | Kif20a | 33.80028 | -0.25005 | 0.456347 | 0.838832 | ENSMUSG( | Kif20a | 31.58069 | -0.17471 | 0.621386 | 0.953199 |
| ENSMUSG( | Kif20b | 31.40033 | -0.12161 | 0.7235 | 0.943873 | ENSMUSG( | Kif20b | 33.696 | 0.322649 | 0.352596 | 0.912093 |
| ENSMUSG( | Kif21a | 3452.201 | 0.101351 | 0.252328 | 0.702378 | ENSMUSG( | Kif21a | 3127.511 | 0.084763 | 0.433933 | 0.91869 |
| ENSMUSG( | Kif21b | 6075.424 | -0.08233 | 0.323622 | 0.768544 | ENSMUSG( | Kif21b | 5361.755 | -0.17922 | 0.081382 | 0.77681 |
| ENSMUSG( | Kif22 | 76.0844 | -0.02519 | 0.922737 | 0.990631 | ENSMUSG( | Kif22 | 67.39474 | -0.1 | 0.702193 | 0.962014 |
| ENSMUSG( | Kif23 | 49.00638 | 0.179747 | 0.544697 | 0.884649 | ENSMUSG( | Kif23 | 50.25825 | 0.511206 | 0.122355 | 0.813384 |
| ENSMUSG( | Kif24 | 29.40248 | -0.60028 | 0.087138 | 0.453876 | ENSMUSG( | Kif24 | 31.86106 | -0.04009 | 0.910973 | 0.992196 |
| ENSMUSG( | Kif26a | 735.7204 | -0.4343 | 0.001008 | 0.03318 | ENSMUSG( | Kif26a | 689.6289 | -0.34398 | 0.019493 | 0.556192 |
| ENSMUSG( | Kif26b | 454.2538 | 0.074532 | 0.544623 | 0.884649 | ENSMUSG( | Kif26b | 428.4589 | 0.174064 | 0.20405 | 0.864412 |
| ENSMUSG( | Kif27 | 190.3334 | -0.04932 | 0.752316 | 0.950536 | ENSMUSG( | Kif27 | 175.6131 | -0.0208 | 0.917354 | 0.992529 |
| ENSMUSG( | Kif28 | 10.77412 | 0.968094 | 0.125399 | NA | ENSMUSG( | Kif28 | 7.855677 | 0.431319 | 0.56177 | NA |
| ENSMUSG( | Kif2a | 1731.519 | -0.04044 | 0.687803 | 0.93324 | ENSMUSG( | Kif2a | 1641.066 | 0.069402 | 0.581834 | 0.944266 |
| ENSMUSG( | Kif2c | 17.12682 | -0.37045 | 0.433321 | NA | ENSMUSG( | Kif2c | 20.31368 | 0.379977 | 0.418424 | 0.917257 |
| ENSMUSG( | Kif3a | 2257.393 | 0.148747 | 0.12747 | 0.535223 | ENSMUSG( | Kif3a | 1951.642 | 0.000653 | 0.99546 | 0.999116 |
| ENSMUSG( | Kif3b | 2141.731 | 0.231504 | 0.015184 | 0.181362 | ENSMUSG( | Kif3b | 1805.973 | 0.016655 | 0.884993 | 0.989688 |
| ENSMUSG( | Kif3c | 6303.362 | 0.233621 | 0.008878 | 0.132538 | ENSMUSG( | Kif3c | 5428.346 | 0.080587 | 0.443189 | 0.921575 |
| ENSMUSG( | Kif4 | 245.2776 | 0.05613 | 0.754574 | 0.951007 | ENSMUSG( | Kif4 | 197.538 | -0.31921 | 0.084152 | 0.777886 |
| ENSMUSG( | Kif5a | 573141.3 | 0.169926 | 0.07506 | 0.42422 | ENSMUSG( | Kif5a | 471255.1 | -0.12563 | 0.303853 | 0.898538 |
| ENSMUSG( | Kif5b | 30162.87 | 0.087981 | 0.375322 | 0.800806 | ENSMUSG( | Kif5b | 25117.28 | -0.17824 | 0.171406 | 0.842999 |
| ENSMUSG( | Kif5c | 88242.33 | 0.056468 | 0.537479 | 0.882536 | ENSMUSG( | Kif5c | 75794.62 | -0.11665 | 0.309874 | 0.898538 |
| ENSMUSG( | Kif6 | 51.51517 | -0.29229 | 0.292103 | 0.742388 | ENSMUSG( | Kif6 | 51.37318 | 0.01675 | 0.959664 | 0.994798 |
| ENSMUSG( | Kif7 | 197.3504 | -0.13725 | 0.391202 | 0.808628 | ENSMUSG( | Kif7 | 194.2517 | 0.091806 | 0.593459 | 0.945408 |
| ENSMUSG( | Kif9 | 170.1092 | 0.120783 | 0.520924 | 0.875186 | ENSMUSG( | Kif9 | 159.2417 | 0.192854 | 0.336536 | 0.907972 |
| ENSMUSG( | Kifap3 | 5362.529 | 0.014087 | 0.876876 | 0.980991 | ENSMUSG( | Kifap3 | 4676.554 | -0.11559 | 0.302093 | 0.898538 |
| ENSMUSG( | Kifbp | 4096.745 | 0.029946 | 0.769408 | 0.954674 | ENSMUSG( | Kifbp | 3501.577 | -0.16141 | 0.218835 | 0.866104 |
| ENSMUSG( | Kifc1 | 43.36423 | -0.57262 | 0.065757 | 0.397094 | ENSMUSG( | Kifc1 | 42.93582 | -0.26044 | 0.465323 | 0.923329 |
| ENSMUSG( | Kifc2 | 39951.96 | 0.144761 | 0.081046 | 0.440124 | ENSMUSG( | Kifc2 | 34815.23 | 0.019694 | 0.834928 | 0.981237 |
| ENSMUSG( | Kifc3 | 3650.492 | 0.074685 | 0.338185 | 0.777422 | ENSMUSG( | Kifc3 | 3248.924 | 0.006981 | 0.944738 | 0.994798 |
| ENSMUSG( | Kifc5b | 74.61974 | -0.33195 | 0.170951 | 0.605536 | ENSMUSG( | Kifc5b | 71.28958 | -0.15464 | 0.572385 | 0.942515 |
| ENSMUSG( | Kin | 274.5694 | 0.064263 | 0.66734 | 0.92729 | ENSMUSG( | Kin | 247.7869 | 0.03987 | 0.816483 | 0.979198 |
| ENSMUSG( | Kirrel1 | 277.1311 | -0.29067 | 0.069832 | 0.408574 | ENSMUSG( | Kirrel1 | 290.1853 | 0.13515 | 0.447597 | 0.922202 |
| ENSMUSG( | Kirrel2 | 1416.236 | 0.00625 | 0.958624 | 0.993874 | ENSMUSG( | Kirrel2 | 1466.155 | 0.356413 | 0.005459 | 0.303106 |
| ENSMUSG( | Kirrel3 | 2080.085 | -0.4037 | 0.001842 | 0.048742 | ENSMUSG( | Kirrel3 | 2102.8 | -0.07138 | 0.6373 | 0.953903 |
| ENSMUSG( | Kiss1r | 338.5583 | 0.170402 | 0.187226 | 0.625372 | ENSMUSG( | Kiss1r | 298.8445 | 0.090966 | 0.560096 | 0.940119 |
| ENSMUSG( | Kit | 1297.419 | -0.37081 | 0.037012 | 0.296902 | ENSMUSG( | Kit | 1439.358 | 0.220732 | 0.267233 | 0.886134 |
| ENSMUSG( | Kitl | 880.4962 | -0.19041 | 0.236415 | 0.683538 | ENSMUSG( | Kitl | 1062.758 | 0.577037 | 0.004397 | 0.269982 |
| ENSMUSG( | Kiz | 570.4923 | -0.00071 | 0.995723 | 0.99943 | ENSMUSG( | Kiz | 494.6037 | -0.15451 | 0.270947 | 0.888062 |
| ENSMUSG( | Kl | 370.7958 | 0.079624 | 0.903175 | 0.986055 | ENSMUSG( | Kl | 404.2238 | 0.538575 | 0.328292 | 0.906478 |
| ENSMUSG( | Klc1 | 23056.16 | 0.045801 | 0.66035 | 0.924627 | ENSMUSG( | Klc1 | 19481.62 | -0.17919 | 0.169764 | 0.840806 |
| ENSMUSG( | Klc2 | 7336.889 | 0.127953 | 0.264185 | 0.715323 | ENSMUSG( | Klc2 | 6069.215 | -0.15383 | 0.214954 | 0.865167 |
| ENSMUSG( | Klc3 | 108.9371 | -0.14949 | 0.523474 | 0.875628 | ENSMUSG( | Klc3 | 95.6234 | -0.27955 | 0.248544 | 0.877904 |
| ENSMUSG( | Klc4 | 1083.13 | 0.090399 | 0.341514 | 0.778828 | ENSMUSG( | Klc4 | 929.6984 | -0.08307 | 0.4772 | 0.923329 |
| ENSMUSG( | Klf10 | 511.4563 | -0.56303 | 1.66E-05 | 0.001679 | ENSMUSG( | Klf10 | 465.5015 | -0.5711 | 0.003103 | 0.221768 |
| ENSMUSG( | Klf11 | 165.7644 | -0.38145 | 0.027282 | 0.253589 | ENSMUSG( | Klf11 | 161.7035 | -0.14554 | 0.467149 | 0.923329 |
| ENSMUSG( | Klf12 | 1241.431 | 0.131931 | 0.379453 | 0.801757 | ENSMUSG( | Klf12 | 991.5326 | -0.25673 | 0.046073 | 0.696944 |
| ENSMUSG( | Klf13 | 6033.327 | 0.245676 | 0.004059 | 0.081988 | ENSMUSG( | Klf13 | 5434.881 | 0.217014 | 0.056465 | 0.723362 |
| ENSMUSG( | Klf15 | 1174.195 | 0.224658 | 0.047671 | 0.33359 | ENSMUSG( | Klf15 | 1019.889 | 0.096526 | 0.463197 | 0.923329 |
| ENSMUSG( | Klf16 | 1372.995 | -0.04755 | 0.656719 | 0.923929 | ENSMUSG( | Klf16 | 1217.284 | -0.12958 | 0.302592 | 0.898538 |
| ENSMUSG( | Klf2 | 418.1334 | -0.28472 | 0.154179 | 0.580901 | ENSMUSG( | Klf2 | 491.9402 | 0.445189 | 0.078526 | 0.77321 |
| ENSMUSG( | Klf3 | 1054.776 | -0.15366 | 0.193709 | 0.633085 | ENSMUSG( | Klf3 | 1024.322 | 0.031603 | 0.805995 | 0.977674 |
| ENSMUSG( | Klf4 | 223.5173 | -0.03006 | 0.885327 | 0.982905 | ENSMUSG( | Klf4 | 238.9078 | 0.41546 | 0.081249 | 0.77681 |
| ENSMUSG( | Klf5 | 191.6528 | 0.033903 | 0.837678 | 0.971111 | ENSMUSG( | Klf5 | 176.9719 | 0.063349 | 0.753497 | 0.970671 |
| ENSMUSG( | Klf6 | 813.3645 | 0.095634 | 0.403456 | 0.816132 | ENSMUSG( | Klf6 | 844.3258 | 0.442701 | 0.000191 | 0.034114 |
| ENSMUSG( | Klf7 | 1170.24 | -0.04857 | 0.653521 | 0.922819 | ENSMUSG( | Klf7 | 1146.71 | 0.155209 | 0.182 | 0.850579 |
| ENSMUSG( | Klf8 | 51.60862 | 0.080575 | 0.777591 | 0.956168 | ENSMUSG( | Klf8 | 48.92625 | 0.196378 | 0.51485 | 0.930939 |
| ENSMUSG( | Klf9 | 7111.453 | 0.225039 | 0.011421 | 0.153856 | ENSMUSG( | Klf9 | 6099.89 | 0.057233 | 0.630393 | 0.953251 |
| ENSMUSG( | Klhdc1 | 399.0564 | -0.0062 | 0.958101 | 0.993874 | ENSMUSG( | Klhdc1 | 356.0101 | -0.07435 | 0.609424 | 0.950397 |
| ENSMUSG( | Klhdc10 | 4350.859 | 0.049364 | 0.547393 | 0.885128 | ENSMUSG( | Klhdc10 | 3777.771 | -0.09204 | 0.3744 | 0.915491 |
| ENSMUSG( | Klhdc2 | 2648.951 | 0.040745 | 0.624014 | 0.91187 | ENSMUSG( | Klhdc2 | 2291.426 | -0.11167 | 0.29526 | 0.898538 |
| ENSMUSG( | Klhdc3 | 6152.025 | 0.183712 | 0.060416 | 0.3797 | ENSMUSG( | Klhdc3 | 5088.989 | -0.09107 | 0.414176 | 0.917257 |
| ENSMUSG( | Klhdc4 | 752.3082 | 0.056763 | 0.587715 | 0.900598 | ENSMUSG( | Klhdc4 | 657.0895 | -0.0642 | 0.615529 | 0.951177 |
| ENSMUSG( | Klhdc7a | 2366.747 | 0.05478 | 0.590649 | 0.90113 | ENSMUSG( | Klhdc7a | 1907.145 | -0.31346 | 0.026412 | 0.603041 |
| ENSMUSG( | Klhdc7b | 11.45341 | -0.0645 | 0.914598 | NA | ENSMUSG( | Klhdc7b | 8.766492 | -0.65177 | 0.337071 | NA |
| ENSMUSG( | Klhdc8a | 591.5986 | 0.072806 | 0.565596 | 0.890871 | ENSMUSG( | Klhdc8a | 516.584 | -0.05256 | 0.717758 | 0.965964 |
| ENSMUSG( | Klhdc8b | 1185.068 | 0.100432 | 0.308003 | 0.756252 | ENSMUSG( | Klhdc8b | 1078.036 | 0.093571 | 0.461229 | 0.923329 |
| ENSMUSG( | Klhdc9 | 111.0721 | 0.015122 | 0.943375 | 0.993415 | ENSMUSG( | Klhdc9 | 104.3375 | 0.09135 | 0.683728 | 0.958678 |
| ENSMUSG( | Klhl1 | 51.29206 | -0.30345 | 0.327017 | 0.771858 | ENSMUSG( | Klhl1 | 45.69226 | -0.39371 | 0.241865 | 0.876528 |
| ENSMUSG( | Klhl11 | 232.1127 | 0.001817 | 0.990949 | 0.999232 | ENSMUSG( | Klhl11 | 208.4732 | -0.03398 | 0.840937 | 0.9826 |
| ENSMUSG( | Klhl12 | 991.7816 | 0.210524 | 0.026831 | 0.250983 | ENSMUSG( | Klhl12 | 847.6408 | 0.031502 | 0.790239 | 0.97681 |
| ENSMUSG( | Klhl13 | 808.6029 | -0.25494 | 0.069737 | 0.408352 | ENSMUSG( | Klhl13 | 726.0552 | -0.30872 | 0.062988 | 0.736482 |
| ENSMUSG( | Klhl14 | 34.31805 | -0.15615 | 0.633067 | 0.914073 | ENSMUSG( | Klhl14 | 36.75255 | 0.2774 | 0.487328 | 0.92541 |
| ENSMUSG( | Klhl15 | 210.2548 | -0.42444 | 0.00816 | 0.125816 | ENSMUSG( | Klhl15 | 225.3877 | 0.063913 | 0.730582 | 0.967228 |
| ENSMUSG( | Klhl17 | 1100.848 | 0.231236 | 0.058498 | 0.373218 | ENSMUSG( | Klhl17 | 991.0441 | 0.203136 | 0.232476 | 0.868507 |
| ENSMUSG( | Klhl18 | 897.2671 | 0.168849 | 0.172467 | 0.607771 | ENSMUSG( | Klhl18 | 759.2946 | -0.03599 | 0.797498 | 0.97681 |
| ENSMUSG( | Klhl2 | 6721.368 | -0.03032 | 0.729482 | 0.944762 | ENSMUSG( | Klhl2 | 6129.393 | -0.02928 | 0.789937 | 0.976798 |
| ENSMUSG( | Klhl20 | 503.0831 | -0.0783 | 0.508928 | 0.868744 | ENSMUSG( | Klhl20 | 482.1589 | 0.070671 | 0.620539 | 0.952922 |
| ENSMUSG( | Klhl21 | 432.018 | -0.15675 | 0.350254 | 0.781943 | ENSMUSG( | Klhl21 | 406.1499 | -0.06217 | 0.684745 | 0.958678 |
| ENSMUSG( | Klhl22 | 2159.138 | 0.165579 | 0.193896 | 0.633085 | ENSMUSG( | Klhl22 | 1838.639 | -0.02542 | 0.85272 | 0.985598 |
| ENSMUSG( | Klhl23 | 2397.437 | 0.063393 | 0.459316 | 0.840244 | ENSMUSG( | Klhl23 | 2082.134 | -0.07781 | 0.501505 | 0.927825 |
| ENSMUSG( | Klhl24 | 2334.393 | 0.12248 | 0.23832 | 0.686308 | ENSMUSG( | Klhl24 | 2132.84 | 0.131509 | 0.241665 | 0.876528 |
| ENSMUSG( | Klhl25 | 312.7926 | 0.150042 | 0.285841 | 0.734989 | ENSMUSG( | Klhl25 | 272.241 | 0.028928 | 0.8627 | 0.986547 |
| ENSMUSG( | Klhl26 | 1440.837 | 0.150235 | 0.179383 | 0.61692 | ENSMUSG( | Klhl26 | 1263.727 | 0.043905 | 0.710461 | 0.964363 |
| ENSMUSG( | Klhl28 | 435.4397 | -0.15516 | 0.205084 | 0.646229 | ENSMUSG( | Klhl28 | 440.3422 | 0.140837 | 0.31719 | 0.902275 |
| ENSMUSG( | Klhl29 | 2088.494 | 0.195818 | 0.041066 | 0.313163 | ENSMUSG( | Klhl29 | 2010.252 | 0.342597 | 0.00253 | 0.197651 |
| ENSMUSG( | Klhl3 | 1787.978 | 0.319549 | 0.001372 | 0.040534 | ENSMUSG( | Klhl3 | 1496.696 | 0.087464 | 0.512413 | 0.930164 |
| ENSMUSG( | Klhl32 | 404.1077 | 0.001104 | 0.992833 | 0.999327 | ENSMUSG( | Klhl32 | 376.4942 | 0.062619 | 0.662108 | 0.956205 |
| ENSMUSG( | Klhl33 | 105.1121 | -0.01599 | 0.942768 | 0.993415 | ENSMUSG( | Klhl33 | 100.5442 | 0.130046 | 0.59065 | 0.944975 |
| ENSMUSG( | Klhl34 | 21.91835 | 0.728163 | 0.132595 | 0.544753 | ENSMUSG( | Klhl34 | 17.47556 | 0.451699 | 0.417945 | 0.917257 |
| ENSMUSG( | Klhl35 | 36.89066 | 0.434731 | 0.194931 | 0.63426 | ENSMUSG( | Klhl35 | 31.48869 | 0.286306 | 0.443074 | 0.921554 |
| ENSMUSG( | Klhl36 | 184.8812 | 0.015225 | 0.931861 | 0.9924 | ENSMUSG( | Klhl36 | 164.9459 | -0.04416 | 0.82864 | 0.980224 |
| ENSMUSG( | Klhl4 | 346.5475 | -0.02907 | 0.838398 | 0.971111 | ENSMUSG( | Klhl4 | 311.1584 | -0.07622 | 0.665456 | 0.956766 |
| ENSMUSG( | Klhl40 | 133.2099 | 0.026287 | 0.886161 | 0.983008 | ENSMUSG( | Klhl40 | 126.4765 | 0.13145 | 0.522638 | 0.932822 |
| ENSMUSG( | Klhl42 | 717.0753 | 0.067601 | 0.515738 | 0.872701 | ENSMUSG( | Klhl42 | 663.9816 | 0.110648 | 0.372656 | 0.915491 |
| ENSMUSG( | Klhl5 | 5341.948 | 0.060254 | 0.541023 | 0.883182 | ENSMUSG( | Klhl5 | 4510.09 | -0.16503 | 0.19143 | 0.858412 |
| ENSMUSG( | Klhl6 | 95.99928 | -0.09866 | 0.652164 | 0.92208 | ENSMUSG( | Klhl6 | 86.52437 | -0.10626 | 0.68585 | 0.958956 |
| ENSMUSG( | Klhl7 | 1728.876 | -0.20129 | 0.042513 | 0.316636 | ENSMUSG( | Klhl7 | 1686.921 | 0.00384 | 0.972865 | 0.996416 |
| ENSMUSG( | Klhl8 | 684.8883 | 0.16766 | 0.152713 | 0.579011 | ENSMUSG( | Klhl8 | 601.2493 | 0.061271 | 0.670346 | 0.95694 |
| ENSMUSG( | Klhl9 | 1556.165 | -0.06462 | 0.589779 | 0.90113 | ENSMUSG( | Klhl9 | 1454.579 | 0.006769 | 0.959139 | 0.994798 |
| ENSMUSG( | Klk6 | 107.2047 | -1.83585 | 2.36E-13 | 2.25E-10 | ENSMUSG( | Klk6 | 128.8042 | -0.53915 | 0.006472 | 0.336411 |
| ENSMUSG( | Klk8 | 123.8932 | 0.184203 | 0.3483 | 0.78132 | ENSMUSG( | Klk8 | 97.32526 | -0.234 | 0.410118 | 0.917257 |
| ENSMUSG( | Klra2 | 13.09282 | -0.37893 | 0.524227 | NA | ENSMUSG( | Klra2 | 12.16973 | -0.31099 | 0.626604 | 0.953234 |
| ENSMUSG( | Kmt2a | 4177.877 | -0.58765 | 8.04E-07 | 0.000161 | ENSMUSG( | Kmt2a | 4837.693 | 0.16283 | 0.226878 | 0.868507 |

| | | | | | |
|---|---|---|---|---|---|
| ENSMUSG( | 813.7654 | 0.106011 | 0.406673 | 0.816928 | Kif17 |
| ENSMUSG( | 30.91783 | -0.15908 | 0.642936 | 0.916865 | Kif18a |
| ENSMUSG( | 15.61929 | -0.47781 | 0.311104 | NA | Kif18b |
| ENSMUSG( | 312.0478 | -0.07924 | 0.578778 | 0.896692 | Kif19a |
| ENSMUSG( | 76489.67 | 0.066787 | 0.389068 | 0.806967 | Kif1a |
| ENSMUSG( | 29863.1 | 0.054612 | 0.534677 | 0.880719 | Kif1b |
| ENSMUSG( | 19712.63 | -0.00295 | 0.974724 | 0.997207 | Kif1c |
| ENSMUSG( | 33.80028 | -0.25005 | 0.456347 | 0.838832 | Kif20a |
| ENSMUSG( | 31.40033 | -0.12161 | 0.7235 | 0.943873 | Kif20b |
| ENSMUSG( | 3452.201 | 0.101351 | 0.252328 | 0.702378 | Kif21a |
| ENSMUSG( | 6075.424 | -0.08233 | 0.323622 | 0.768544 | Kif21b |
| ENSMUSG( | 76.0844 | -0.02519 | 0.922737 | 0.990631 | Kif22 |
| ENSMUSG( | 49.00638 | 0.179747 | 0.544697 | 0.884649 | Kif23 |
| ENSMUSG( | 29.40248 | -0.60028 | 0.087138 | 0.453876 | Kif24 |
| ENSMUSG( | 735.7204 | -0.4343 | 0.001008 | 0.03318 | Kif26a |
| ENSMUSG( | 454.2538 | 0.074532 | 0.544623 | 0.884649 | Kif26b |
| ENSMUSG( | 190.3334 | -0.04932 | 0.752316 | 0.950536 | Kif27 |
| ENSMUSG( | 10.77412 | 0.968094 | 0.125399 | NA | Kif28 |
| ENSMUSG( | 1731.519 | -0.04044 | 0.687803 | 0.93324 | Kif2a |
| ENSMUSG( | 17.12682 | -0.37045 | 0.433321 | NA | Kif2c |
| ENSMUSG( | 2257.393 | 0.148747 | 0.12747 | 0.535223 | Kif3a |
| ENSMUSG( | 2141.731 | 0.231504 | 0.015184 | 0.181362 | Kif3b |
| ENSMUSG( | 6303.362 | 0.233621 | 0.008878 | 0.132538 | Kif3c |
| ENSMUSG( | 245.2776 | 0.05613 | 0.754574 | 0.951007 | Kif4 |
| ENSMUSG( | 573141.3 | 0.169926 | 0.07506 | 0.42422 | Kif5a |
| ENSMUSG( | 30162.87 | 0.087981 | 0.375322 | 0.800806 | Kif5b |
| ENSMUSG( | 88242.33 | 0.056468 | 0.537479 | 0.882536 | Kif5c |
| ENSMUSG( | 51.51517 | -0.29229 | 0.292103 | 0.742388 | Kif6 |
| ENSMUSG( | 197.3504 | -0.13725 | 0.391202 | 0.808628 | Kif7 |
| ENSMUSG( | 170.1092 | 0.120783 | 0.520924 | 0.875186 | Kif9 |
| ENSMUSG( | 5362.529 | 0.014087 | 0.876876 | 0.980991 | Kifap3 |
| ENSMUSG( | 4096.745 | 0.029946 | 0.769408 | 0.954674 | Kifbp |
| ENSMUSG( | 43.36423 | -0.57262 | 0.065757 | 0.397094 | Kifc1 |
| ENSMUSG( | 39951.96 | 0.144761 | 0.081046 | 0.440124 | Kifc2 |
| ENSMUSG( | 3650.492 | 0.074685 | 0.338185 | 0.777422 | Kifc3 |
| ENSMUSG( | 74.61974 | -0.33195 | 0.170951 | 0.605536 | Kifc5b |
| ENSMUSG( | 274.5694 | 0.064263 | 0.66734 | 0.92729 | Kin |
| ENSMUSG( | 277.1311 | -0.29067 | 0.069832 | 0.408574 | Kirrel1 |
| ENSMUSG( | 1416.236 | 0.00625 | 0.958624 | 0.993874 | Kirrel2 |
| ENSMUSG( | 2080.085 | -0.4037 | 0.001842 | 0.048742 | Kirrel3 |
| ENSMUSG( | 338.5583 | 0.170402 | 0.187226 | 0.625372 | Kiss1r |
| ENSMUSG( | 1297.419 | -0.37081 | 0.037012 | 0.296902 | Kit |
| ENSMUSG( | 880.4962 | -0.19041 | 0.236415 | 0.683538 | Kitl |
| ENSMUSG( | 570.4923 | -0.00071 | 0.995723 | 0.99943 | Kiz |
| ENSMUSG( | 370.7958 | 0.079624 | 0.903175 | 0.986055 | Kl |
| ENSMUSG( | 23056.16 | 0.045801 | 0.66035 | 0.924627 | Klc1 |
| ENSMUSG( | 7336.889 | 0.127953 | 0.264185 | 0.715323 | Klc2 |
| ENSMUSG( | 108.9371 | -0.14949 | 0.523474 | 0.875628 | Klc3 |
| ENSMUSG( | 1083.13 | 0.090399 | 0.341514 | 0.778828 | Klc4 |
| ENSMUSG( | 8.348349 | -0.23365 | 0.721402 | NA | Klf1 |
| ENSMUSG( | 511.4563 | -0.56303 | 1.66E-05 | 0.001679 | Klf10 |
| ENSMUSG( | 165.7644 | -0.38145 | 0.027282 | 0.253589 | Klf11 |
| ENSMUSG( | 1241.431 | 0.131931 | 0.379453 | 0.801757 | Klf12 |
| ENSMUSG( | 6033.327 | 0.245676 | 0.004059 | 0.081988 | Klf13 |
| ENSMUSG( | 7.236839 | 0.935118 | 0.261023 | NA | Klf14 |
| ENSMUSG( | 1174.195 | 0.224658 | 0.047671 | 0.33359 | Klf15 |
| ENSMUSG( | 1372.995 | -0.04755 | 0.656719 | 0.923929 | Klf16 |
| ENSMUSG( | 418.1334 | -0.28472 | 0.154179 | 0.580901 | Klf2 |
| ENSMUSG( | 1054.776 | -0.15366 | 0.193709 | 0.633085 | Klf3 |
| ENSMUSG( | 223.5173 | -0.03006 | 0.885327 | 0.982905 | Klf4 |
| ENSMUSG( | 191.6528 | 0.033903 | 0.837678 | 0.971111 | Klf5 |
| ENSMUSG( | 813.3645 | 0.095634 | 0.403456 | 0.816132 | Klf6 |
| ENSMUSG( | 1170.24 | -0.04857 | 0.653521 | 0.922819 | Klf7 |
| ENSMUSG( | 51.60862 | 0.080575 | 0.777591 | 0.956168 | Klf8 |
| ENSMUSG( | 7111.453 | 0.225039 | 0.011421 | 0.153856 | Klf9 |
| ENSMUSG( | 399.0564 | -0.0062 | 0.958101 | 0.993874 | Klhdc1 |
| ENSMUSG( | 4350.859 | 0.049364 | 0.547393 | 0.885128 | Klhdc10 |
| ENSMUSG( | 2648.951 | 0.040745 | 0.624014 | 0.91187 | Klhdc2 |
| ENSMUSG( | 6152.025 | 0.183712 | 0.060416 | 0.3797 | Klhdc3 |
| ENSMUSG( | 752.3082 | 0.056763 | 0.587715 | 0.900598 | Klhdc4 |
| ENSMUSG( | 2366.747 | 0.05478 | 0.590649 | 0.90113 | Klhdc7a |
| ENSMUSG( | 11.45341 | -0.0645 | 0.914598 | NA | Klhdc7b |
| ENSMUSG( | 591.5986 | 0.072806 | 0.565596 | 0.890871 | Klhdc8a |
| ENSMUSG( | 1185.068 | 0.100432 | 0.308003 | 0.756252 | Klhdc8b |
| ENSMUSG( | 111.0721 | 0.015122 | 0.943375 | 0.993415 | Klhdc9 |
| ENSMUSG( | 51.29206 | -0.30345 | 0.327017 | 0.771858 | Klhl1 |
| ENSMUSG( | 9.249051 | 0.342739 | 0.591378 | NA | Klhl10 |
| ENSMUSG( | 232.1127 | 0.001817 | 0.990949 | 0.999232 | Klhl11 |
| ENSMUSG( | 991.7816 | 0.210524 | 0.026831 | 0.250983 | Klhl12 |
| ENSMUSG( | 808.6029 | -0.25494 | 0.069737 | 0.408352 | Klhl13 |
| ENSMUSG( | 34.31805 | -0.15615 | 0.633067 | 0.914073 | Klhl14 |
| ENSMUSG( | 210.2548 | -0.42444 | 0.00816 | 0.125816 | Klhl15 |
| ENSMUSG( | 1100.848 | 0.231236 | 0.058498 | 0.373218 | Klhl17 |
| ENSMUSG( | 897.2671 | 0.168849 | 0.172467 | 0.607771 | Klhl18 |
| ENSMUSG( | 6721.368 | -0.03032 | 0.729482 | 0.944762 | Klhl2 |
| ENSMUSG( | 503.0831 | -0.0783 | 0.508928 | 0.868744 | Klhl20 |
| ENSMUSG( | 432.018 | -0.15675 | 0.350254 | 0.781943 | Klhl21 |
| ENSMUSG( | 2159.138 | 0.165579 | 0.193896 | 0.633085 | Klhl22 |
| ENSMUSG( | 2397.437 | 0.063393 | 0.459316 | 0.840244 | Klhl23 |
| ENSMUSG( | 2334.393 | 0.12248 | 0.23832 | 0.686308 | Klhl24 |
| ENSMUSG( | 312.7926 | 0.150042 | 0.285841 | 0.734989 | Klhl25 |
| ENSMUSG( | 1440.837 | 0.150235 | 0.179383 | 0.61692 | Klhl26 |
| ENSMUSG( | 435.4397 | -0.15516 | 0.205084 | 0.646229 | Klhl28 |
| ENSMUSG( | 2088.494 | 0.195818 | 0.041066 | 0.313163 | Klhl29 |
| ENSMUSG( | 1787.978 | 0.319549 | 0.001372 | 0.040534 | Klhl3 |
| ENSMUSG( | 404.1077 | 0.001104 | 0.992833 | 0.999327 | Klhl32 |
| ENSMUSG( | 105.1121 | -0.01599 | 0.942768 | 0.993415 | Klhl33 |
| ENSMUSG( | 21.91835 | 0.728163 | 0.132595 | 0.544753 | Klhl34 |
| ENSMUSG( | 36.89066 | 0.434731 | 0.194931 | 0.63426 | Klhl35 |
| ENSMUSG( | 184.8812 | 0.015225 | 0.931861 | 0.9924 | Klhl36 |
| ENSMUSG( | 346.5475 | -0.02907 | 0.838398 | 0.971111 | Klhl4 |
| ENSMUSG( | 133.2099 | 0.026287 | 0.886161 | 0.983008 | Klhl40 |
| ENSMUSG( | 717.0753 | 0.067601 | 0.515738 | 0.872701 | Klhl42 |
| ENSMUSG( | 5341.948 | 0.060254 | 0.541023 | 0.883182 | Klhl5 |
| ENSMUSG( | 95.99928 | -0.09866 | 0.652164 | 0.92208 | Klhl6 |
| ENSMUSG( | 1728.876 | -0.20129 | 0.042513 | 0.316636 | Klhl7 |
| ENSMUSG( | 684.8883 | 0.16766 | 0.152713 | 0.579011 | Klhl8 |
| ENSMUSG( | 1556.165 | -0.06462 | 0.589779 | 0.90113 | Klhl9 |
| ENSMUSG( | 107.2047 | -1.83585 | 2.36E-13 | 2.25E-10 | Klk6 |
| ENSMUSG( | 123.8932 | 0.184203 | 0.3483 | 0.78132 | Klk8 |
| ENSMUSG( | 13.09282 | -0.37893 | 0.524227 | NA | Klra2 |
| ENSMUSG( | 8.837484 | 0.893791 | 0.15254 | NA | Klrg2 |
| ENSMUSG( | 4177.877 | -0.58765 | 8.04E-07 | 0.000161 | Kmt2a |
| ENSMUSG( | 1966.659 | 0.068619 | 0.505587 | 0.866932 | Kmt2b |
| ENSMUSG( | 3310.54 | -0.09961 | 0.370192 | 0.797542 | Kmt2c |
| ENSMUSG( | 3132.886 | -0.41206 | 1.08E-05 | 0.001199 | Kmt2d |
| ENSMUSG( | 3189.364 | 0.026579 | 0.753494 | 0.950753 | Kmt2e |
| ENSMUSG( | 2255.108 | 0.059896 | 0.519731 | 0.874979 | Kmt5a |
| ENSMUSG( | 1435.93 | 0.059791 | 0.519842 | 0.874979 | Kmt5b |
| ENSMUSG( | 667.6827 | 0.145608 | 0.21157 | 0.654835 | Kmt5c |
| ENSMUSG( | 4030.017 | 0.16733 | 0.176041 | 0.611564 | Kndc1 |
| ENSMUSG( | 15.50957 | -0.78979 | 0.164732 | NA | Knl1 |
| ENSMUSG( | 1066.333 | 0.046118 | 0.643268 | 0.917178 | Knop1 |
| ENSMUSG( | 76.59764 | -0.19725 | 0.435557 | 0.829093 | Knstrn |
| ENSMUSG( | 23.07859 | 0.116079 | 0.760888 | 0.952907 | Kntc1 |
| ENSMUSG( | 2343.172 | -0.07376 | 0.408063 | 0.818167 | Kpna1 |
| ENSMUSG( | 311.0302 | 0.106024 | 0.543034 | 0.883579 | Kpna2 |
| ENSMUSG( | 1570.95 | 0.107133 | 0.303734 | 0.752147 | Kpna3 |
| ENSMUSG( | 1650.147 | 0.059137 | 0.512741 | 0.871762 | Kpna4 |
| ENSMUSG( | 3298.993 | 0.087403 | 0.265729 | 0.716441 | Kpna6 |
| ENSMUSG( | 6036.699 | -0.02905 | 0.785274 | 0.958473 | Kpnb1 |
| ENSMUSG( | 661.6881 | -0.01366 | 0.900757 | 0.985633 | Kptn |
| ENSMUSG( | 2949.377 | -0.05428 | 0.617628 | 0.910872 | Kras |
| ENSMUSG( | 1599.14 | -0.03626 | 0.753031 | 0.950667 | Krba1 |
| ENSMUSG( | 109.3201 | 0.382285 | 0.101694 | 0.486141 | Krbox5 |
| ENSMUSG( | 1101.668 | -0.03583 | 0.773178 | 0.955282 | Krcc1 |
| ENSMUSG( | 1085.514 | -0.1275 | 0.339019 | 0.777518 | Kremen1 |
| ENSMUSG( | 15.26028 | -0.30621 | 0.579719 | NA | Kremen2 |
| ENSMUSG( | 781.8456 | 0.194941 | 0.147163 | 0.569203 | Kri1 |
| ENSMUSG( | 1153.972 | 0.101696 | 0.436972 | 0.829501 | Krit1 |
| ENSMUSG( | 816.0226 | 0.022926 | 0.823528 | 0.966603 | Krr1 |
| ENSMUSG( | 137.2283 | -0.06871 | 0.700653 | 0.93743 | Krt1 |
| ENSMUSG( | 228.3181 | -0.45271 | 0.0067 | 0.112611 | Krt10 |
| ENSMUSG( | 306.548 | -0.16731 | 0.233934 | 0.681906 | Krt12 |
| ENSMUSG( | 25.70243 | 0.106476 | 0.855118 | 0.973676 | Krt18 |
| ENSMUSG( | 16.15958 | -1.44936 | 0.007999 | NA | Krt19 |
| ENSMUSG( | 18.57384 | 0.007245 | 0.987687 | NA | Krt2 |
| ENSMUSG( | 760.6195 | -0.02586 | 0.82945 | 0.969343 | Krt222 |
| ENSMUSG( | 8.514238 | 0.266975 | 0.680329 | NA | Krt28 |
| ENSMUSG( | 40.80584 | 0.305371 | 0.38128 | 0.802629 | Krt73 |
| ENSMUSG( | 272.8206 | 0.105513 | 0.454656 | 0.837443 | Krt77 |
| ENSMUSG( | 14.83187 | -0.13147 | 0.824266 | NA | Krt8 |
| ENSMUSG( | 189.8592 | -0.09547 | 0.615124 | 0.910339 | Krt80 |
| ENSMUSG( | 173.817 | -0.39689 | 0.097775 | 0.478264 | Krt9 |
| ENSMUSG( | 1128.637 | -0.16867 | 0.09216 | 0.466065 | Krtcap2 |
| ENSMUSG( | 20.93189 | 0.003594 | 0.992917 | NA | Krtcap3 |
| ENSMUSG( | 1906.338 | 0.378484 | 0.001684 | 0.046443 | Ksr1 |
| ENSMUSG( | 1254.562 | 0.161948 | 0.124695 | 0.531714 | Ksr2 |

| | | | | | |
|---|---|---|---|---|---|
| ENSMUSG( | 161.7035 | -0.14554 | 0.467149 | 0.923329 | Klf11 |
| ENSMUSG( | 991.5326 | -0.25673 | 0.046073 | 0.696944 | Klf12 |
| ENSMUSG( | 5434.881 | 0.217014 | 0.056465 | 0.723362 | Klf13 |
| ENSMUSG( | 1019.889 | 0.096526 | 0.463197 | 0.923329 | Klf15 |
| ENSMUSG( | 1217.284 | -0.12958 | 0.302592 | 0.898538 | Klf16 |
| ENSMUSG( | 491.9402 | 0.445189 | 0.078526 | 0.77321 | Klf2 |
| ENSMUSG( | 1024.322 | 0.031603 | 0.805995 | 0.977674 | Klf3 |
| ENSMUSG( | 238.9078 | 0.41546 | 0.081249 | 0.77681 | Klf4 |
| ENSMUSG( | 176.9719 | 0.063349 | 0.753497 | 0.970671 | Klf5 |
| ENSMUSG( | 844.3258 | 0.442701 | 0.000191 | 0.034114 | Klf6 |
| ENSMUSG( | 1146.71 | 0.155209 | 0.182 | 0.850579 | Klf7 |
| ENSMUSG( | 48.92625 | 0.196378 | 0.51485 | 0.930939 | Klf8 |
| ENSMUSG( | 6099.89 | 0.057233 | 0.630393 | 0.953251 | Klf9 |
| ENSMUSG( | 356.0101 | -0.07435 | 0.609424 | 0.950397 | Klhdc1 |
| ENSMUSG( | 3777.771 | -0.09204 | 0.3744 | 0.915491 | Klhdc10 |
| ENSMUSG( | 2291.426 | -0.11167 | 0.29526 | 0.898538 | Klhdc2 |
| ENSMUSG( | 5088.989 | -0.09107 | 0.414176 | 0.917257 | Klhdc3 |
| ENSMUSG( | 657.0895 | -0.0642 | 0.615529 | 0.951177 | Klhdc4 |
| ENSMUSG( | 1907.145 | -0.31346 | 0.026412 | 0.603041 | Klhdc7a |
| ENSMUSG( | 8.766492 | -0.65177 | 0.337071 | NA | Klhdc7b |
| ENSMUSG( | 516.584 | -0.05256 | 0.717758 | 0.965964 | Klhdc8a |
| ENSMUSG( | 1078.036 | 0.093571 | 0.461229 | 0.923329 | Klhdc8b |
| ENSMUSG( | 104.3375 | 0.09135 | 0.683728 | 0.958678 | Klhdc9 |
| ENSMUSG( | 45.69226 | -0.39371 | 0.241865 | 0.876528 | Klhl1 |
| ENSMUSG( | 208.4732 | -0.03398 | 0.840937 | 0.9826 | Klhl11 |
| ENSMUSG( | 847.6408 | 0.031502 | 0.790239 | 0.97681 | Klhl12 |
| ENSMUSG( | 726.0552 | -0.30872 | 0.062988 | 0.736482 | Klhl13 |
| ENSMUSG( | 36.75255 | 0.2774 | 0.487328 | 0.92541 | Klhl14 |
| ENSMUSG( | 225.3877 | 0.063913 | 0.730582 | 0.967228 | Klhl15 |
| ENSMUSG( | 991.0441 | 0.203136 | 0.232476 | 0.868507 | Klhl17 |
| ENSMUSG( | 759.2946 | -0.03599 | 0.797498 | 0.97681 | Klhl18 |
| ENSMUSG( | 6129.393 | -0.02928 | 0.789937 | 0.976798 | Klhl2 |
| ENSMUSG( | 482.1589 | 0.070671 | 0.620539 | 0.952922 | Klhl20 |
| ENSMUSG( | 406.1499 | -0.06217 | 0.684745 | 0.958678 | Klhl21 |
| ENSMUSG( | 1838.639 | -0.02542 | 0.85272 | 0.985598 | Klhl22 |
| ENSMUSG( | 2082.134 | -0.07781 | 0.501505 | 0.927825 | Klhl23 |
| ENSMUSG( | 2132.84 | 0.131509 | 0.241665 | 0.876528 | Klhl24 |
| ENSMUSG( | 272.241 | 0.028928 | 0.8627 | 0.986547 | Klhl25 |
| ENSMUSG( | 1263.727 | 0.043905 | 0.710461 | 0.964363 | Klhl26 |
| ENSMUSG( | 440.3422 | 0.140837 | 0.31719 | 0.902275 | Klhl28 |
| ENSMUSG( | 2010.252 | 0.342597 | 0.00253 | 0.197651 | Klhl29 |
| ENSMUSG( | 1496.696 | 0.087464 | 0.512413 | 0.930164 | Klhl3 |
| ENSMUSG( | 376.4942 | 0.062619 | 0.662108 | 0.956205 | Klhl32 |
| ENSMUSG( | 100.5442 | 0.130046 | 0.59065 | 0.944975 | Klhl33 |
| ENSMUSG( | 17.47556 | 0.451699 | 0.417945 | 0.917257 | Klhl34 |
| ENSMUSG( | 31.48869 | 0.286306 | 0.443074 | 0.921554 | Klhl35 |
| ENSMUSG( | 164.9459 | -0.04416 | 0.82864 | 0.980224 | Klhl36 |
| ENSMUSG( | 311.1584 | -0.07622 | 0.665456 | 0.956766 | Klhl4 |
| ENSMUSG( | 126.4765 | 0.13145 | 0.522638 | 0.932822 | Klhl40 |
| ENSMUSG( | 663.9816 | 0.110648 | 0.372656 | 0.915491 | Klhl42 |
| ENSMUSG( | 4510.09 | -0.16503 | 0.19143 | 0.858412 | Klhl5 |
| ENSMUSG( | 86.52437 | -0.10626 | 0.68585 | 0.958956 | Klhl6 |
| ENSMUSG( | 1686.921 | 0.00384 | 0.972865 | 0.996416 | Klhl7 |
| ENSMUSG( | 601.2493 | 0.061271 | 0.670346 | 0.95694 | Klhl8 |
| ENSMUSG( | 1454.579 | 0.006769 | 0.959139 | 0.994798 | Klhl9 |
| ENSMUSG( | 128.8042 | -0.53915 | 0.006472 | 0.336411 | Klk6 |
| ENSMUSG( | 97.32526 | -0.234 | 0.410118 | 0.917257 | Klk8 |
| ENSMUSG( | 12.16973 | -0.31099 | 0.626604 | 0.953234 | Klra2 |
| ENSMUSG( | 4837.693 | 0.16283 | 0.226878 | 0.868507 | Kmt2a |
| ENSMUSG( | 1776.691 | 0.047307 | 0.707925 | 0.964042 | Kmt2b |
| ENSMUSG( | 3210.219 | 0.08258 | 0.44563 | 0.921734 | Kmt2c |
| ENSMUSG( | 3440.54 | 0.153998 | 0.193874 | 0.860096 | Kmt2d |
| ENSMUSG( | 2999.5 | 0.116503 | 0.266222 | 0.886134 | Kmt2e |
| ENSMUSG( | 1969.048 | -0.06589 | 0.602225 | 0.94789 | Kmt5a |
| ENSMUSG( | 1315.602 | 0.077623 | 0.493011 | 0.927358 | Kmt5b |
| ENSMUSG( | 573.5785 | -0.02209 | 0.892589 | 0.990558 | Kmt5c |
| ENSMUSG( | 3599.949 | 0.113116 | 0.438985 | 0.920721 | Kndc1 |
| ENSMUSG( | 17.11286 | -0.08944 | 0.857095 | 0.985884 | Knl1 |
| ENSMUSG( | 983.4821 | 0.076587 | 0.553269 | 0.939565 | Knop1 |
| ENSMUSG( | 73.09082 | -0.06006 | 0.825354 | 0.979388 | Knstrn |
| ENSMUSG( | 22.67284 | 0.319256 | 0.434694 | 0.918742 | Kntc1 |
| ENSMUSG( | 2303.834 | 0.144466 | 0.195512 | 0.861579 | Kpna1 |
| ENSMUSG( | 305.3107 | 0.302344 | 0.1153 | 0.813375 | Kpna2 |
| ENSMUSG( | 1374.309 | -0.01174 | 0.926453 | 0.99306 | Kpna3 |
| ENSMUSG( | 1502.822 | 0.055367 | 0.656521 | 0.955533 | Kpna4 |
| ENSMUSG( | 2899.697 | -0.01494 | 0.880783 | 0.989644 | Kpna6 |
| ENSMUSG( | 5644.019 | 0.047505 | 0.695806 | 0.960867 | Kpnb1 |
| ENSMUSG( | 583.5317 | -0.11433 | 0.404023 | 0.917257 | Kptn |
| ENSMUSG( | 2712.928 | -0.03099 | 0.850438 | 0.985541 | Kras |
| ENSMUSG( | 1481.105 | 0.014792 | 0.913944 | 0.992407 | Krba1 |
| ENSMUSG( | 86.20191 | -0.00569 | 0.982484 | 0.996767 | Krbox5 |
| ENSMUSG( | 998.171 | -0.0584 | 0.721256 | 0.966887 | Krcc1 |
| ENSMUSG( | 1055.771 | 0.06905 | 0.677498 | 0.957106 | Kremen1 |
| ENSMUSG( | 14.77006 | -0.11204 | 0.833277 | 0.981193 | Kremen2 |
| ENSMUSG( | 691.9818 | 0.111965 | 0.467537 | 0.923329 | Kri1 |
| ENSMUSG( | 1155.199 | 0.358216 | 0.02248 | 0.580801 | Krit1 |
| ENSMUSG( | 743.9981 | 0.024597 | 0.841357 | 0.982799 | Krr1 |
| ENSMUSG( | 115.0304 | -0.34572 | 0.149364 | 0.836241 | Krt1 |
| ENSMUSG( | 213.1894 | -0.38869 | 0.040465 | 0.682882 | Krt10 |
| ENSMUSG( | 270.5867 | -0.26814 | 0.085227 | 0.779799 | Krt12 |
| ENSMUSG( | 8.183984 | 0.223959 | 0.744931 | NA | Krt15 |
| ENSMUSG( | 20.94697 | -0.21841 | 0.730438 | 0.967228 | Krt18 |
| ENSMUSG( | 13.72931 | -1.86105 | 0.002778 | 0.209728 | Krt19 |
| ENSMUSG( | 15.7095 | -0.27122 | 0.62786 | 0.953234 | Krt2 |
| ENSMUSG( | 683.2166 | -0.07304 | 0.625855 | 0.953234 | Krt222 |
| ENSMUSG( | 28.68011 | -0.45026 | 0.248246 | 0.877904 | Krt73 |
| ENSMUSG( | 239.6432 | -0.00899 | 0.958762 | 0.994798 | Krt77 |
| ENSMUSG( | 17.7323 | 0.5848 | 0.24385 | 0.877904 | Krt8 |
| ENSMUSG( | 155.0907 | -0.43831 | 0.029 | 0.61427 | Krt80 |
| ENSMUSG( | 152.1959 | -0.5515 | 0.027074 | 0.603041 | Krt9 |
| ENSMUSG( | 1048.238 | -0.10994 | 0.369499 | 0.915491 | Krtcap2 |
| ENSMUSG( | 25.51255 | 0.776296 | 0.0637 | 0.736482 | Krtcap3 |
| ENSMUSG( | 1661.752 | 0.265774 | 0.038075 | 0.676422 | Ksr1 |
| ENSMUSG( | 1113.333 | 0.09127 | 0.459371 | 0.923329 | Ksr2 |
| ENSMUSG( | 200.5669 | 0.10709 | 0.567592 | 0.941777 | Kti12 |
| ENSMUSG( | 3803.214 | 0.090457 | 0.536293 | 0.93705 | Ktn1 |
| ENSMUSG( | 977.3646 | -0.18176 | 0.158662 | 0.837724 | Kxd1 |
| ENSMUSG( | 167.2979 | -0.09763 | 0.669334 | 0.956766 | Ky |
| ENSMUSG( | 385.5107 | -0.00833 | 0.955502 | 0.994798 | Kyat1 |
| ENSMUSG( | 255.8255 | -0.07475 | 0.637941 | 0.953903 | Kyat3 |
| ENSMUSG( | 4929.676 | 0.042995 | 0.715054 | 0.965063 | L1cam |
| ENSMUSG( | 523.8701 | 0.034482 | 0.802365 | 0.97681 | L2hgdh |
| ENSMUSG( | 74.42133 | 0.246955 | 0.318032 | 0.902849 | L3hypdh |
| ENSMUSG( | 248.8125 | 0.03707 | 0.880556 | 0.989644 | L3mbtl1 |
| ENSMUSG( | 827.7592 | 0.031984 | 0.814084 | 0.979198 | L3mbtl2 |
| ENSMUSG( | 853.4104 | 0.020919 | 0.874702 | 0.989644 | L3mbtl3 |
| ENSMUSG( | 21.125 | 0.202911 | 0.661001 | 0.956205 | L3mbtl4 |
| ENSMUSG( | 237.3576 | -0.07699 | 0.629947 | 0.953251 | Lacc1 |
| ENSMUSG( | 607.9237 | 0.006394 | 0.964633 | 0.995433 | Lactb |
| ENSMUSG( | 401.9593 | -0.00548 | 0.968748 | 0.99594 | Lactb2 |
| ENSMUSG( | 201.3483 | -0.14076 | 0.448264 | 0.922202 | Lad1 |
| ENSMUSG( | 674.7827 | 0.079957 | 0.609767 | 0.950397 | Lag3 |
| ENSMUSG( | 402.112 | -0.25407 | 0.078706 | 0.77411 | Lage3 |
| ENSMUSG( | 628.9527 | 0.050435 | 0.72682 | 0.966887 | Lair1 |
| ENSMUSG( | 162.3353 | 0.035629 | 0.888854 | 0.990558 | Lama1 |
| ENSMUSG( | 898.9361 | -0.01375 | 0.934813 | 0.99306 | Lama2 |
| ENSMUSG( | 218.9473 | -0.4495 | 0.011768 | 0.443295 | Lama3 |
| ENSMUSG( | 437.5923 | 0.032879 | 0.897488 | 0.990558 | Lama4 |
| ENSMUSG( | 527.5249 | 0.331686 | 0.102034 | 0.801851 | Lama5 |
| ENSMUSG( | 614.4428 | 0.303835 | 0.105321 | 0.804969 | Lamb1 |
| ENSMUSG( | 2002.828 | 0.307326 | 0.028847 | 0.614171 | Lamb2 |
| ENSMUSG( | 53.99471 | -0.08276 | 0.772089 | 0.974236 | Lamb3 |
| ENSMUSG( | 1847.276 | 0.317366 | 0.150634 | 0.837173 | Lamc1 |
| ENSMUSG( | 437.8324 | -0.15084 | 0.361467 | 0.914223 | Lamc2 |
| ENSMUSG( | 251.3043 | 0.512959 | 0.068928 | 0.754165 | Lamc3 |
| ENSMUSG( | 21585.08 | 0.089704 | 0.539355 | 0.937216 | Lamp1 |
| ENSMUSG( | 2995.22 | 0.239563 | 0.193292 | 0.860096 | Lamp2 |
| ENSMUSG( | 6280.368 | -0.07285 | 0.587381 | 0.944975 | Lamp5 |
| ENSMUSG( | 3152.141 | -0.21359 | 0.058364 | 0.728256 | Lamtor1 |
| ENSMUSG( | 1049.982 | -0.20667 | 0.140269 | 0.830678 | Lamtor2 |
| ENSMUSG( | 874.7813 | -0.01059 | 0.938855 | 0.993898 | Lamtor3 |
| ENSMUSG( | 996.1057 | -0.26515 | 0.043986 | 0.69104 | Lamtor4 |
| ENSMUSG( | 881.3191 | -0.20659 | 0.095494 | 0.796133 | Lamtor5 |
| ENSMUSG( | 3015.481 | -0.16251 | 0.176717 | 0.846973 | Lancl1 |
| ENSMUSG( | 1423.57 | -0.07488 | 0.616664 | 0.951597 | Lancl2 |
| ENSMUSG( | 35.47905 | 0.312283 | 0.395401 | 0.917257 | Lancl3 |
| ENSMUSG( | 14.40627 | 2.59284 | 1.70E-05 | 0.005604 | Lao1 |
| ENSMUSG( | 2864.071 | -0.36494 | 0.006714 | 0.339232 | Lap3 |
| ENSMUSG( | 10143.78 | -0.00932 | 0.948325 | 0.994798 | Laptm4a |
| ENSMUSG( | 3000.624 | 0.043567 | 0.796389 | 0.97681 | Laptm4b |
| ENSMUSG( | 3134.147 | 0.031057 | 0.858976 | 0.985884 | Laptm5 |
| ENSMUSG( | 4256.758 | 0.235473 | 0.246016 | 0.877904 | Large1 |
| ENSMUSG( | 15.50632 | 0.374194 | 0.473234 | 0.923329 | Large2 |
| ENSMUSG( | 3874.499 | 0.194187 | 0.062588 | 0.736482 | Larp1 |
| ENSMUSG( | 454.3768 | 0.004756 | 0.973874 | 0.996416 | Larp1b |
| ENSMUSG( | 798.9305 | 0.097707 | 0.411931 | 0.917257 | Larp4 |
| ENSMUSG( | 2239.258 | -0.00359 | 0.978559 | 0.996459 | Larp4b |
| ENSMUSG( | 519.7837 | -0.2735 | 0.097211 | 0.796721 | Larp6 |

| | | | | | | | | | | | |
|---|---|---|---|---|---|---|---|---|---|---|---|
| ENSMUSG( | Kmt2b | 1966.659 | 0.068619 | 0.505587 | 0.866932 | ENSMUSG( | Kmt2b | 1776.691 | 0.047307 | 0.707925 | 0.964042 |
| ENSMUSG( | Kmt2c | 3310.54 | -0.09961 | 0.370192 | 0.797542 | ENSMUSG( | Kmt2c | 3210.219 | 0.08258 | 0.44563 | 0.921734 |
| ENSMUSG( | Kmt2d | 3132.886 | -0.41206 | 1.08E-05 | 0.001199 | ENSMUSG( | Kmt2d | 3440.54 | 0.153998 | 0.193874 | 0.860096 |
| ENSMUSG( | Kmt2e | 3189.364 | 0.026579 | 0.753494 | 0.950753 | ENSMUSG( | Kmt2e | 2999.5 | 0.116503 | 0.266222 | 0.886134 |
| ENSMUSG( | Kmt5a | 2255.108 | 0.059896 | 0.519731 | 0.874979 | ENSMUSG( | Kmt5a | 1969.048 | -0.06589 | 0.602225 | 0.94789 |
| ENSMUSG( | Kmt5b | 1435.93 | 0.059791 | 0.519842 | 0.874979 | ENSMUSG( | Kmt5b | 1315.602 | 0.077623 | 0.493011 | 0.927358 |
| ENSMUSG( | Kmt5c | 667.6827 | 0.145608 | 0.21157 | 0.654835 | ENSMUSG( | Kmt5c | 573.5785 | -0.02209 | 0.892589 | 0.990558 |
| ENSMUSG( | Kndc1 | 4030.017 | 0.16733 | 0.176041 | 0.611564 | ENSMUSG( | Kndc1 | 3599.949 | 0.113116 | 0.438985 | 0.920721 |
| ENSMUSG( | Knl1 | 15.50957 | -0.78979 | 0.164732 | NA | ENSMUSG( | Knl1 | 17.11286 | -0.08944 | 0.857095 | 0.985884 |
| ENSMUSG( | Knop1 | 1066.333 | 0.046118 | 0.643268 | 0.917178 | ENSMUSG( | Knop1 | 983.4821 | 0.076587 | 0.553269 | 0.939565 |
| ENSMUSG( | Knstrn | 76.59764 | -0.19725 | 0.435557 | 0.829093 | ENSMUSG( | Knstrn | 73.09082 | -0.06006 | 0.825354 | 0.979388 |
| ENSMUSG( | Kntc1 | 23.07859 | 0.116079 | 0.760888 | 0.952907 | ENSMUSG( | Kntc1 | 22.67284 | 0.319256 | 0.434694 | 0.918742 |
| ENSMUSG( | Kpna1 | 2343.172 | -0.07376 | 0.408063 | 0.818167 | ENSMUSG( | Kpna1 | 2303.834 | 0.144466 | 0.195512 | 0.861579 |
| ENSMUSG( | Kpna2 | 311.0302 | 0.106024 | 0.543034 | 0.883579 | ENSMUSG( | Kpna2 | 305.3107 | 0.302344 | 0.1153 | 0.813375 |
| ENSMUSG( | Kpna3 | 1570.95 | 0.107133 | 0.303734 | 0.752147 | ENSMUSG( | Kpna3 | 1374.309 | -0.01174 | 0.926453 | 0.99306 |
| ENSMUSG( | Kpna4 | 1650.147 | 0.059137 | 0.512741 | 0.871762 | ENSMUSG( | Kpna4 | 1502.822 | 0.055367 | 0.656521 | 0.955533 |
| ENSMUSG( | Kpna6 | 3298.993 | 0.087403 | 0.265729 | 0.716441 | ENSMUSG( | Kpna6 | 2899.697 | -0.01494 | 0.880783 | 0.989644 |
| ENSMUSG( | Kpnb1 | 6036.699 | -0.02905 | 0.785274 | 0.958473 | ENSMUSG( | Kpnb1 | 5644.019 | 0.047505 | 0.695806 | 0.960867 |
| ENSMUSG( | Kptn | 661.6881 | -0.01366 | 0.900757 | 0.985633 | ENSMUSG( | Kptn | 583.5317 | -0.11433 | 0.404023 | 0.917257 |
| ENSMUSG( | Kras | 2949.377 | -0.05428 | 0.617628 | 0.910872 | ENSMUSG( | Kras | 2712.928 | -0.03099 | 0.850438 | 0.985541 |
| ENSMUSG( | Krba1 | 1599.14 | -0.03626 | 0.753031 | 0.950667 | ENSMUSG( | Krba1 | 1481.105 | 0.014792 | 0.913944 | 0.992407 |
| ENSMUSG( | Krbox5 | 109.3201 | 0.382285 | 0.101694 | 0.486141 | ENSMUSG( | Krbox5 | 86.20191 | -0.00569 | 0.982484 | 0.996767 |
| ENSMUSG( | Krcc1 | 1101.668 | -0.03583 | 0.773178 | 0.955282 | ENSMUSG( | Krcc1 | 998.171 | -0.0584 | 0.721256 | 0.966887 |
| ENSMUSG( | Kremen1 | 1085.514 | -0.1275 | 0.339019 | 0.777518 | ENSMUSG( | Kremen1 | 1055.771 | 0.06905 | 0.677498 | 0.957106 |
| ENSMUSG( | Kremen2 | 15.26028 | -0.30621 | 0.579719 | NA | ENSMUSG( | Kremen2 | 14.77006 | -0.11204 | 0.833277 | 0.981193 |
| ENSMUSG( | Kri1 | 781.8456 | 0.194941 | 0.147163 | 0.569203 | ENSMUSG( | Kri1 | 691.9818 | 0.111965 | 0.467537 | 0.923329 |
| ENSMUSG( | Krit1 | 1153.972 | 0.101696 | 0.436972 | 0.829501 | ENSMUSG( | Krit1 | 1155.199 | 0.358216 | 0.02248 | 0.580801 |
| ENSMUSG( | Krr1 | 816.0226 | 0.022926 | 0.823528 | 0.966603 | ENSMUSG( | Krr1 | 743.9981 | 0.024597 | 0.841357 | 0.982799 |
| ENSMUSG( | Krt1 | 137.2283 | -0.06871 | 0.700653 | 0.93743 | ENSMUSG( | Krt1 | 115.0304 | -0.34572 | 0.149364 | 0.836241 |
| ENSMUSG( | Krt10 | 228.3181 | -0.45271 | 0.0067 | 0.112611 | ENSMUSG( | Krt10 | 213.1894 | -0.38869 | 0.040465 | 0.682882 |
| ENSMUSG( | Krt12 | 306.548 | -0.16731 | 0.233934 | 0.681906 | ENSMUSG( | Krt12 | 270.5867 | -0.26814 | 0.085227 | 0.779799 |
| ENSMUSG( | Krt18 | 25.70243 | 0.106476 | 0.855118 | 0.973676 | ENSMUSG( | Krt18 | 20.94697 | -0.21841 | 0.730438 | 0.967228 |
| ENSMUSG( | Krt19 | 16.15958 | -1.44936 | 0.007999 | NA | ENSMUSG( | Krt19 | 13.72931 | -1.86105 | 0.002778 | 0.209728 |
| ENSMUSG( | Krt2 | 18.57384 | 0.007245 | 0.987687 | NA | ENSMUSG( | Krt2 | 15.7095 | -0.27122 | 0.62786 | 0.953234 |
| ENSMUSG( | Krt222 | 760.6195 | -0.02586 | 0.82945 | 0.969343 | ENSMUSG( | Krt222 | 683.2166 | -0.07304 | 0.625855 | 0.953234 |
| ENSMUSG( | Krt73 | 40.80584 | 0.305371 | 0.38128 | 0.802629 | ENSMUSG( | Krt73 | 28.68011 | -0.45026 | 0.248246 | 0.877904 |
| ENSMUSG( | Krt77 | 272.8206 | 0.105513 | 0.454656 | 0.837443 | ENSMUSG( | Krt77 | 239.6432 | -0.00899 | 0.958762 | 0.994798 |
| ENSMUSG( | Krt8 | 14.83187 | -0.13147 | 0.824266 | NA | ENSMUSG( | Krt8 | 17.7323 | 0.5848 | 0.24385 | 0.877904 |
| ENSMUSG( | Krt80 | 189.8592 | -0.09547 | 0.615124 | 0.910339 | ENSMUSG( | Krt80 | 155.0907 | -0.43831 | 0.029 | 0.61427 |
| ENSMUSG( | Krt9 | 173.817 | -0.39689 | 0.097775 | 0.478264 | ENSMUSG( | Krt9 | 152.1959 | -0.5515 | 0.027074 | 0.603041 |
| ENSMUSG( | Krtcap2 | 1128.637 | -0.16867 | 0.09216 | 0.466065 | ENSMUSG( | Krtcap2 | 1048.238 | -0.10994 | 0.369499 | 0.915491 |
| ENSMUSG( | Krtcap3 | 20.93189 | 0.003594 | 0.992917 | NA | ENSMUSG( | Krtcap3 | 25.51255 | 0.776296 | 0.0637 | 0.736482 |
| ENSMUSG( | Ksr1 | 1906.338 | 0.378484 | 0.001684 | 0.046443 | ENSMUSG( | Ksr1 | 1661.752 | 0.265774 | 0.038075 | 0.676422 |
| ENSMUSG( | Ksr2 | 1254.562 | 0.161948 | 0.124695 | 0.531714 | ENSMUSG( | Ksr2 | 1113.333 | 0.09127 | 0.459371 | 0.923329 |
| ENSMUSG( | Kti12 | 213.2485 | 0.025941 | 0.875561 | 0.980097 | ENSMUSG( | Kti12 | 200.5669 | 0.10709 | 0.567592 | 0.941777 |
| ENSMUSG( | Ktn1 | 3916.157 | -0.09648 | 0.435443 | 0.829093 | ENSMUSG( | Ktn1 | 3803.214 | 0.090457 | 0.536293 | 0.93705 |
| ENSMUSG( | Kxd1 | 1127.027 | -0.03036 | 0.753264 | 0.950753 | ENSMUSG( | Kxd1 | 977.3646 | -0.18176 | 0.158662 | 0.837724 |
| ENSMUSG( | Ky | 235.1655 | 0.545842 | 0.000439 | 0.019063 | ENSMUSG( | Ky | 167.2979 | -0.09763 | 0.669334 | 0.956766 |
| ENSMUSG( | Kyat1 | 440.0021 | 0.098551 | 0.455212 | 0.837897 | ENSMUSG( | Kyat1 | 385.5107 | -0.00833 | 0.955502 | 0.994798 |
| ENSMUSG( | Kyat3 | 299.6826 | 0.114613 | 0.44787 | 0.834051 | ENSMUSG( | Kyat3 | 255.8255 | -0.07475 | 0.637941 | 0.953903 |
| ENSMUSG( | L1cam | 4842.149 | -0.29299 | 0.005394 | 0.098038 | ENSMUSG( | L1cam | 4929.676 | 0.042995 | 0.715054 | 0.965063 |
| ENSMUSG( | L2hgdh | 612.3705 | 0.214253 | 0.041404 | 0.314189 | ENSMUSG( | L2hgdh | 523.8701 | 0.034482 | 0.802365 | 0.97681 |
| ENSMUSG( | L3hypdh | 72.97239 | -0.06048 | 0.792507 | 0.959461 | ENSMUSG( | L3hypdh | 74.42133 | 0.246955 | 0.318032 | 0.902849 |
| ENSMUSG( | L3mbtl1 | 317.2474 | 0.441134 | 0.037959 | 0.300779 | ENSMUSG( | L3mbtl1 | 248.8125 | 0.03707 | 0.880556 | 0.989644 |
| ENSMUSG( | L3mbtl2 | 999.0002 | 0.289586 | 0.011644 | 0.155574 | ENSMUSG( | L3mbtl2 | 827.7592 | 0.031984 | 0.814084 | 0.979198 |
| ENSMUSG( | L3mbtl3 | 995.6656 | 0.195937 | 0.057789 | 0.370784 | ENSMUSG( | L3mbtl3 | 853.4104 | 0.020919 | 0.874702 | 0.989644 |
| ENSMUSG( | L3mbtl4 | 25.50265 | 0.479512 | 0.222523 | 0.668434 | ENSMUSG( | L3mbtl4 | 21.125 | 0.202911 | 0.661001 | 0.956205 |
| ENSMUSG( | Lacc1 | 257.8619 | -0.1074 | 0.449139 | 0.834943 | ENSMUSG( | Lacc1 | 237.3576 | -0.07699 | 0.629947 | 0.953251 |
| ENSMUSG( | Lactb | 647.0459 | -0.07962 | 0.479746 | 0.854357 | ENSMUSG( | Lactb | 607.9237 | 0.006394 | 0.964633 | 0.995433 |
| ENSMUSG( | Lactb2 | 441.082 | -0.00931 | 0.942802 | 0.993415 | ENSMUSG( | Lactb2 | 401.9593 | -0.00548 | 0.968748 | 0.99594 |
| ENSMUSG( | Lad1 | 217.4368 | -0.18707 | 0.263016 | 0.713397 | ENSMUSG( | Lad1 | 201.3483 | -0.14076 | 0.448264 | 0.922202 |
| ENSMUSG( | Lag3 | 718.9118 | -0.01092 | 0.935687 | 0.993363 | ENSMUSG( | Lag3 | 674.7827 | 0.079957 | 0.609767 | 0.950397 |
| ENSMUSG( | Lage3 | 459.5893 | -0.12293 | 0.279967 | 0.729155 | ENSMUSG( | Lage3 | 402.112 | -0.25407 | 0.078706 | 0.77411 |
| ENSMUSG( | Lair1 | 638.0432 | -0.18716 | 0.111653 | 0.50376 | ENSMUSG( | Lair1 | 628.9527 | 0.050435 | 0.72682 | 0.966887 |
| ENSMUSG( | Lama1 | 174.209 | -0.05105 | 0.764558 | 0.954055 | ENSMUSG( | Lama1 | 162.3353 | 0.035629 | 0.888854 | 0.990558 |
| ENSMUSG( | Lama2 | 947.214 | -0.14074 | 0.268327 | 0.718663 | ENSMUSG( | Lama2 | 898.9361 | -0.01375 | 0.934813 | 0.99306 |
| ENSMUSG( | Lama3 | 267.0813 | -0.12009 | 0.439112 | 0.830714 | ENSMUSG( | Lama3 | 218.9473 | -0.4495 | 0.011768 | 0.443295 |
| ENSMUSG( | Lama4 | 426.2678 | -0.3457 | 0.078673 | 0.433892 | ENSMUSG( | Lama4 | 437.5923 | 0.032879 | 0.897488 | 0.990558 |
| ENSMUSG( | Lama5 | 506.2992 | -0.0443 | 0.740036 | 0.947823 | ENSMUSG( | Lama5 | 527.5249 | 0.331686 | 0.102034 | 0.801851 |
| ENSMUSG( | Lamb1 | 615.135 | 0.044534 | 0.787658 | 0.959218 | ENSMUSG( | Lamb1 | 614.4428 | 0.303835 | 0.105321 | 0.804969 |
| ENSMUSG( | Lamb2 | 1938.027 | -0.04379 | 0.713197 | 0.940734 | ENSMUSG( | Lamb2 | 2002.828 | 0.307326 | 0.028847 | 0.614171 |
| ENSMUSG( | Lamb3 | 59.90785 | -0.06463 | 0.80783 | 0.963257 | ENSMUSG( | Lamb3 | 53.99471 | -0.08276 | 0.772089 | 0.974236 |
| ENSMUSG( | Lamc1 | 1703.452 | -0.18209 | 0.299983 | 0.749035 | ENSMUSG( | Lamc1 | 1847.276 | 0.317366 | 0.150634 | 0.837173 |
| ENSMUSG( | Lamc2 | 433.4023 | -0.48921 | 0.001072 | 0.034583 | ENSMUSG( | Lamc2 | 437.8324 | -0.15084 | 0.361467 | 0.914223 |
| ENSMUSG( | Lamc3 | 241.1206 | 0.147317 | 0.492822 | 0.860541 | ENSMUSG( | Lamc3 | 251.3043 | 0.512959 | 0.068928 | 0.754165 |
| ENSMUSG( | Lamp1 | 22275.12 | -0.09048 | 0.470471 | 0.84674 | ENSMUSG( | Lamp1 | 21585.08 | 0.089704 | 0.539355 | 0.937216 |
| ENSMUSG( | Lamp2 | 2827.113 | -0.19673 | 0.235859 | 0.683028 | ENSMUSG( | Lamp2 | 2995.22 | 0.239563 | 0.193292 | 0.860096 |
| ENSMUSG( | Lamp5 | 6420.278 | -0.29459 | 0.016082 | 0.188199 | ENSMUSG( | Lamp5 | 6280.368 | -0.07285 | 0.587381 | 0.944975 |
| ENSMUSG( | Lamtor1 | 3780.474 | 0.051134 | 0.53757 | 0.882536 | ENSMUSG( | Lamtor1 | 3152.141 | -0.21359 | 0.058364 | 0.728256 |
| ENSMUSG( | Lamtor2 | 1247.203 | 0.033818 | 0.756479 | 0.951479 | ENSMUSG( | Lamtor2 | 1049.982 | -0.20667 | 0.140269 | 0.830678 |
| ENSMUSG( | Lamtor3 | 933.0042 | -0.09131 | 0.442052 | 0.832762 | ENSMUSG( | Lamtor3 | 874.7813 | -0.01059 | 0.938855 | 0.993898 |
| ENSMUSG( | Lamtor4 | 1203.202 | 0.028044 | 0.779323 | 0.95668 | ENSMUSG( | Lamtor4 | 996.1057 | -0.26515 | 0.043986 | 0.69104 |
| ENSMUSG( | Lamtor5 | 1062.275 | 0.073885 | 0.463079 | 0.842422 | ENSMUSG( | Lamtor5 | 881.3191 | -0.20659 | 0.095494 | 0.796133 |
| ENSMUSG( | Lancl1 | 3372.489 | -0.10528 | 0.334464 | 0.776332 | ENSMUSG( | Lancl1 | 3015.481 | -0.16251 | 0.176717 | 0.846973 |
| ENSMUSG( | Lancl2 | 1598.03 | -0.00526 | 0.965088 | 0.995303 | ENSMUSG( | Lancl2 | 1423.57 | -0.07488 | 0.616664 | 0.951597 |
| ENSMUSG( | Lancl3 | 38.4222 | 0.247226 | 0.438014 | 0.830072 | ENSMUSG( | Lancl3 | 35.47905 | 0.312283 | 0.395401 | 0.917257 |
| ENSMUSG( | Lap3 | 3682.308 | 0.114664 | 0.279532 | 0.729114 | ENSMUSG( | Lap3 | 2864.071 | -0.36494 | 0.006714 | 0.339232 |
| ENSMUSG( | Laptm4a | 10397.03 | -0.21545 | 0.093196 | 0.469454 | ENSMUSG( | Laptm4a | 10143.78 | -0.00932 | 0.948325 | 0.994798 |
| ENSMUSG( | Laptm4b | 2911.186 | -0.33515 | 0.024253 | 0.238311 | ENSMUSG( | Laptm4b | 3000.624 | 0.043567 | 0.796389 | 0.97681 |
| ENSMUSG( | Laptm5 | 2971.945 | -0.42746 | 0.00491 | 0.092785 | ENSMUSG( | Laptm5 | 3134.147 | 0.031057 | 0.858976 | 0.985884 |
| ENSMUSG( | Large1 | 4057.165 | -0.17403 | 0.302684 | 0.750923 | ENSMUSG( | Large1 | 4256.758 | 0.235473 | 0.246016 | 0.877904 |
| ENSMUSG( | Large2 | 15.18576 | 0.094026 | 0.844892 | NA | ENSMUSG( | Large2 | 15.50632 | 0.374194 | 0.473234 | 0.923329 |
| ENSMUSG( | Larp1 | 4268.033 | 0.202807 | 0.028202 | 0.256543 | ENSMUSG( | Larp1 | 3874.499 | 0.194187 | 0.062588 | 0.736482 |
| ENSMUSG( | Larp1b | 530.3117 | 0.183056 | 0.13207 | 0.544244 | ENSMUSG( | Larp1b | 454.3768 | 0.004756 | 0.973874 | 0.996416 |
| ENSMUSG( | Larp4 | 830.7066 | -0.05834 | 0.574875 | 0.894572 | ENSMUSG( | Larp4 | 798.9305 | 0.097707 | 0.411931 | 0.917257 |
| ENSMUSG( | Larp4b | 2540.325 | 0.092669 | 0.255392 | 0.706181 | ENSMUSG( | Larp4b | 2239.258 | -0.00359 | 0.978559 | 0.996459 |
| ENSMUSG( | Larp6 | 612.2484 | -0.05397 | 0.733287 | 0.944956 | ENSMUSG( | Larp6 | 519.7837 | -0.2735 | 0.097211 | 0.796721 |
| ENSMUSG( | Larp7 | 355.7479 | -0.15093 | 0.258353 | 0.709574 | ENSMUSG( | Larp7 | 326.5642 | -0.1312 | 0.405696 | 0.917257 |
| ENSMUSG( | Lars1 | 1276.909 | 0.059835 | 0.532375 | 0.879905 | ENSMUSG( | Lars1 | 1122.728 | -0.04307 | 0.707667 | 0.964029 |
| ENSMUSG( | Lars2 | 9845.432 | -1.43429 | 1.49E-36 | 4.84E-33 | ENSMUSG( | Lars2 | 7927.626 | -2.24551 | 8.00E-34 | 4.67E-30 |
| ENSMUSG( | Las1l | 1609.004 | 0.177186 | 0.16395 | 0.594869 | ENSMUSG( | Las1l | 1367.296 | -0.02219 | 0.888946 | 0.990558 |
| ENSMUSG( | Lasp1 | 9613.196 | 0.139329 | 0.127456 | 0.535223 | ENSMUSG( | Lasp1 | 8210.1 | -0.04305 | 0.688397 | 0.959742 |
| ENSMUSG( | Lat | 15.60273 | 0.47667 | 0.332861 | NA | ENSMUSG( | Lat | 15.67005 | 0.723077 | 0.187933 | 0.85698 |
| ENSMUSG( | Lat2 | 120.5506 | -0.28146 | 0.153968 | 0.580901 | ENSMUSG( | Lat2 | 122.153 | 0.053769 | 0.808687 | 0.97873 |
| ENSMUSG( | Lats1 | 1725.193 | 0.018775 | 0.854957 | 0.973665 | ENSMUSG( | Lats1 | 1611.219 | 0.087036 | 0.523527 | 0.932848 |
| ENSMUSG( | Lats2 | 507.4341 | 0.015751 | 0.908961 | 0.98794 | ENSMUSG( | Lats2 | 500.653 | 0.233384 | 0.138129 | 0.828492 |
| ENSMUSG( | Lax1 | 15.24484 | 0.181334 | 0.712789 | NA | ENSMUSG( | Lax1 | 15.09227 | 0.337782 | 0.554035 | 0.939565 |
| ENSMUSG( | Layn | 126.2747 | -0.4556 | 0.077298 | 0.429743 | ENSMUSG( | Layn | 153.0126 | 0.394725 | 0.141409 | 0.830678 |
| ENSMUSG( | Lbh | 2519.715 | -0.1151 | 0.182265 | 0.619591 | ENSMUSG( | Lbh | 2227.72 | -0.20996 | 0.100183 | 0.801311 |
| ENSMUSG( | Lbhd1 | 121.7847 | 0.03247 | 0.877752 | 0.981431 | ENSMUSG( | Lbhd1 | 109.0927 | -0.01912 | 0.928533 | 0.99306 |
| ENSMUSG( | Lbhd2 | 12.94324 | -0.6274 | 0.312926 | NA | ENSMUSG( | Lbhd2 | 14.82159 | 0.022738 | 0.96985 | 0.99594 |
| ENSMUSG( | Lbp | 339.2024 | -0.01189 | 0.945252 | 0.99365 | ENSMUSG( | Lbp | 397.7173 | 0.649439 | 0.001649 | 0.150169 |
| ENSMUSG( | Lbr | 437.176 | -0.40421 | 0.021443 | 0.22362 | ENSMUSG( | Lbr | 492.4687 | 0.238345 | 0.259344 | 0.883971 |
| ENSMUSG( | Lca5 | 369.7697 | 0.032257 | 0.815543 | 0.965133 | ENSMUSG( | Lca5 | 343.0025 | 0.080184 | 0.605515 | 0.94924 |
| ENSMUSG( | Lca5l | 70.97656 | -0.00472 | 0.984809 | 0.997815 | ENSMUSG( | Lca5l | 65.70109 | 0.059952 | 0.827461 | 0.980224 |
| ENSMUSG( | Lcat | 1940.605 | -0.15722 | 0.292278 | 0.742481 | ENSMUSG( | Lcat | 1965.478 | 0.152523 | 0.36773 | 0.915289 |
| ENSMUSG( | Lck | 50.76151 | 0.257543 | 0.349348 | 0.78176 | ENSMUSG( | Lck | 41.62128 | -0.06446 | 0.846307 | 0.984578 |
| ENSMUSG( | Lclat1 | 1367.79 | -0.37956 | 0.010882 | 0.150598 | ENSMUSG( | Lclat1 | 1465.924 | 0.11613 | 0.471574 | 0.923329 |
| ENSMUSG( | Lcmt1 | 1218.11 | 0.108603 | 0.321315 | 0.766497 | ENSMUSG( | Lcmt1 | 1020.727 | -0.1366 | 0.296017 | 0.898538 |
| ENSMUSG( | Lcmt2 | 133.4268 | 0.1288 | 0.573734 | 0.893584 | ENSMUSG( | Lcmt2 | 117.7937 | 0.043778 | 0.833589 | 0.981218 |
| ENSMUSG( | Lcn2 | 41.39427 | -1.14437 | 0.023325 | 0.233882 | ENSMUSG( | Lcn2 | 56.65577 | 0.210039 | 0.648494 | 0.95422 |
| ENSMUSG( | Lcor | 110.8873 | -0.28587 | 0.204645 | 0.645534 | ENSMUSG( | Lcor | 119.0294 | 0.214411 | 0.416103 | 0.917257 |
| ENSMUSG( | Lcorl | 578.1015 | -0.10902 | 0.345772 | 0.779153 | ENSMUSG( | Lcorl | 570.0284 | 0.116634 | 0.431219 | 0.917257 |
| ENSMUSG( | Lcp1 | 611.202 | -0.2542 | 0.032872 | 0.277278 | ENSMUSG( | Lcp1 | 545.5853 | -0.31723 | 0.033857 | 0.648168 |
| ENSMUSG( | Lcp2 | 127.7213 | 0.063575 | 0.728201 | 0.944355 | ENSMUSG( | Lcp2 | 121.1318 | 0.179848 | 0.373745 | 0.915491 |
| ENSMUSG( | Lct | 92.11568 | 0.226708 | 0.283802 | 0.732203 | ENSMUSG( | Lct | 86.93561 | 0.346298 | 0.276046 | 0.890679 |
| ENSMUSG( | Ldaf1 | 363.2078 | -0.04375 | 0.726786 | 0.944223 | ENSMUSG( | Ldaf1 | 351.913 | 0.129669 | 0.372778 | 0.915491 |
| ENSMUSG( | Ldah | 1286.506 | -0.01626 | 0.861129 | 0.975026 | ENSMUSG( | Ldah | 1105.809 | -0.19415 | 0.100145 | 0.801311 |
| ENSMUSG( | Ldb1 | 3121.238 | -0.03514 | 0.748628 | 0.949604 | ENSMUSG( | Ldb1 | 3040.648 | 0.155187 | 0.213006 | 0.864412 |
| ENSMUSG( | Ldb2 | 837.0759 | -0.02646 | 0.839347 | 0.971179 | ENSMUSG( | Ldb2 | 720.6465 | -0.20429 | 0.199568 | 0.863016 |
| ENSMUSG( | Ldb3 | 195.5843 | -0.08704 | 0.564673 | 0.890434 | ENSMUSG( | Ldb3 | 168.0399 | -0.26681 | 0.131204 | 0.822884 |
| ENSMUSG( | Ldha | 7527.635 | 0.114474 | 0.180766 | 0.617841 | ENSMUSG( | Ldha | 6173.866 | -0.19467 | 0.104477 | 0.804969 |
| ENSMUSG( | Ldhb | 17476.13 | 0.018575 | 0.839016 | 0.971179 | ENSMUSG( | Ldhb | 14551.47 | -0.25405 | 0.041377 | 0.688699 |
| ENSMUSG( | Ldhd | 588.0772 | 0.070239 | 0.572771 | 0.893584 | ENSMUSG( | Ldhd | 519.8574 | -0.01361 | 0.934667 | 0.99306 |
| ENSMUSG( | Ldlr | 1051.221 | -1.08262 | 1.48E-08 | 5.59E-06 | ENSMUSG( | Ldlr | 1268.657 | -0.06497 | 0.778312 | 0.976106 |
| ENSMUSG( | Ldlrad3 | 587.1554 | -0.48286 | 0.0043 | 0.085154 | ENSMUSG( | Ldlrad3 | 628.4067 | 0.029381 | 0.883769 | 0.989644 |
| ENSMUSG( | Ldlrad4 | 728.7751 | -0.13295 | 0.278437 | 0.728428 | ENSMUSG( | Ldlrad4 | 708.7169 | 0.063248 | 0.649284 | 0.95422 |
| ENSMUSG( | Ldlrap1 | 246.6872 | -0.41065 | 0.018559 | 0.20492 | ENSMUSG( | Ldlrap1 | 268.6901 | 0.140519 | 0.455178 | 0.923163 |
| ENSMUSG( | Ldoc1 | 25.25995 | 0.064879 | 0.864292 | 0.975358 | ENSMUSG( | Ldoc1 | 23.1039 | 0.063571 | 0.875497 | 0.989644 |
| ENSMUSG( | Lef1 | 237.6559 | 0.150172 | 0.307392 | 0.755324 | ENSMUSG( | Lef1 | 226.2453 | 0.278155 | 0.12033 | 0.813384 |
| ENSMUSG( | Lefty1 | 42.13897 | 0.195656 | 0.561988 | 0.889052 | ENSMUSG( | Lefty1 | 44.26344 | 0.549671 | 0.072811 | 0.763629 |
| ENSMUSG( | Lefty2 | 30.03084 | 0.585458 | 0.100718 | 0.484034 | ENSMUSG( | Lefty2 | 27.8112 | 0.612537 | 0.10834 | 0.804969 |
| ENSMUSG( | Lekr1 | 118.5806 | 0.103366 | 0.579805 | 0.89687 | ENSMUSG( | Lekr1 | 99.34012 | -0.14012 | 0.527051 | 0.934449 |
| ENSMUSG( | Lemd1 | 36.56774 | 0.049567 | 0.894182 | 0.984512 | ENSMUSG( | Lemd1 | 31.02267 | -0.19291 | 0.626846 | 0.953234 |
| ENSMUSG( | Lemd2 | 1450.361 | -0.05746 | 0.600158 | 0.904195 | ENSMUSG( | Lemd2 | 1422.898 | 0.156766 | 0.250699 | 0.878324 |
| ENSMUSG( | Lemd3 | 1094.81 | -0.35606 | 0.025837 | 0.247245 | ENSMUSG( | Lemd3 | 1190.527 | 0.175044 | 0.33137 | 0.907861 |
| ENSMUSG( | Leng1 | 305.7592 | 0.236965 | 0.06773 | 0.40256 | ENSMUSG( | Leng1 | 240.2197 | -0.18863 | 0.247435 | 0.877904 |
| ENSMUSG( | Leng8 | 5684.851 | 0.306961 | 0.122152 | 0.527707 | ENSMUSG( | Leng8 | 5243.837 | 0.336151 | 0.390656 | 0.917257 |
| ENSMUSG( | Leng9 | 174.3965 | 0.030031 | 0.850487 | 0.972638 | ENSMUSG( | Leng9 | 143.164 | -0.29377 | 0.127273 | 0.817278 |
| ENSMUSG( | Leo1 | 494.0921 | 0.159187 | 0.189036 | 0.627678 | ENSMUSG( | Leo1 | 383.5754 | -0.32033 | 0.033124 | 0.64093 |
| ENSMUSG( | Lepr | 140.032 | -0.20926 | 0.242226 | 0.689699 | ENSMUSG( | Lepr | 130.3827 | -0.12303 | 0.577143 | 0.942912 |
| ENSMUSG( | Leprot | 1295.485 | -0.33608 | 0.015454 | 0.183729 | ENSMUSG( | Leprot | 1381.517 | 0.139044 | 0.398703 | 0.917257 |
| ENSMUSG( | Leprotl1 | 2078.724 | -0.42215 | 0.003838 | 0.079297 | ENSMUSG( | Leprotl1 | 2276.136 | 0.139298 | 0.394796 | 0.917257 |
| ENSMUSG( | Letm1 | 4322.606 | 0.031969 | 0.711259 | 0.940028 | ENSMUSG( | Letm1 | 3837.443 | -0.04179 | 0.693761 | 0.960767 |
| ENSMUSG( | Letm2 | 340.0478 | -0.15137 | 0.27036 | 0.720226 | ENSMUSG( | Letm2 | 338.1916 | 0.109694 | 0.511343 | 0.92971 |
| ENSMUSG( | Letmd1 | 2844.451 | -0.01266 | 0.899905 | 0.985633 | ENSMUSG( | Letmd1 | 2610.724 | 0.01061 | 0.926047 | 0.99306 |
| ENSMUSG( | Lfng | 1149.187 | -0.44604 | 0.005905 | 0.103987 | ENSMUSG( | Lfng | 1450.073 | 0.492008 | 0.029814 | 0.61992 |
| ENSMUSG( | Lgals1 | 494.588 | -0.1554 | 0.209917 | 0.652297 | ENSMUSG( | Lgals1 | 429.3672 | -0.3121 | 0.029095 | 0.614779 |

| | | | | | |
|---|---|---|---|---|---|
| ENSMUSG( | 213.2485 | 0.025941 | 0.875561 | 0.980097 | Kti12 |
| ENSMUSG( | 3916.157 | -0.09648 | 0.435443 | 0.829093 | Ktn1 |
| ENSMUSG( | 1127.027 | -0.03036 | 0.753264 | 0.950753 | Kxd1 |
| ENSMUSG( | 235.1655 | 0.545842 | 0.000439 | 0.019063 | Ky |
| ENSMUSG( | 440.0021 | 0.098551 | 0.455212 | 0.837897 | Kyat1 |
| ENSMUSG( | 299.6826 | 0.114613 | 0.44787 | 0.834051 | Kyat3 |
| ENSMUSG( | 4842.149 | -0.29299 | 0.005394 | 0.098038 | L1cam |
| ENSMUSG( | 612.3705 | 0.214253 | 0.041404 | 0.314189 | L2hgdh |
| ENSMUSG( | 72.97239 | -0.06048 | 0.792507 | 0.959461 | L3hypdh |
| ENSMUSG( | 317.2474 | 0.441134 | 0.037959 | 0.300779 | L3mbtl1 |
| ENSMUSG( | 999.0002 | 0.289586 | 0.011644 | 0.155574 | L3mbtl2 |
| ENSMUSG( | 995.6656 | 0.195937 | 0.057789 | 0.370784 | L3mbtl3 |
| ENSMUSG( | 25.50265 | 0.479512 | 0.222523 | 0.668434 | L3mbtl4 |
| ENSMUSG( | 257.8619 | -0.1074 | 0.449139 | 0.834943 | Lacc1 |
| ENSMUSG( | 647.0459 | -0.07962 | 0.479746 | 0.854357 | Lactb |
| ENSMUSG( | 441.082 | -0.00931 | 0.942802 | 0.993415 | Lactb2 |
| ENSMUSG( | 217.4368 | -0.18707 | 0.263016 | 0.713397 | Lad1 |
| ENSMUSG( | 718.9118 | -0.01092 | 0.935687 | 0.993363 | Lag3 |
| ENSMUSG( | 459.5893 | -0.12293 | 0.279967 | 0.729155 | Lage3 |
| ENSMUSG( | 638.0432 | -0.18716 | 0.111653 | 0.50376 | Lair1 |
| ENSMUSG( | 174.209 | -0.05105 | 0.764558 | 0.954055 | Lama1 |
| ENSMUSG( | 947.214 | -0.14074 | 0.268327 | 0.718663 | Lama2 |
| ENSMUSG( | 267.0813 | -0.12009 | 0.439112 | 0.830714 | Lama3 |
| ENSMUSG( | 426.2678 | -0.3457 | 0.078673 | 0.433892 | Lama4 |
| ENSMUSG( | 506.2992 | -0.0443 | 0.740036 | 0.947823 | Lama5 |
| ENSMUSG( | 615.135 | 0.044534 | 0.787658 | 0.959218 | Lamb1 |
| ENSMUSG( | 1938.027 | -0.04379 | 0.713197 | 0.940734 | Lamb2 |
| ENSMUSG( | 59.90785 | -0.06463 | 0.80783 | 0.963257 | Lamb3 |
| ENSMUSG( | 1703.452 | -0.18209 | 0.299983 | 0.749035 | Lamc1 |
| ENSMUSG( | 433.4023 | -0.48921 | 0.001072 | 0.034583 | Lamc2 |
| ENSMUSG( | 241.1206 | 0.147317 | 0.492822 | 0.860541 | Lamc3 |
| ENSMUSG( | 22275.12 | -0.09048 | 0.470471 | 0.84674 | Lamp1 |
| ENSMUSG( | 2827.113 | -0.19673 | 0.235859 | 0.683028 | Lamp2 |
| ENSMUSG( | 6420.278 | -0.29459 | 0.016082 | 0.188199 | Lamp5 |
| ENSMUSG( | 3780.474 | 0.051134 | 0.53757 | 0.882536 | Lamtor1 |
| ENSMUSG( | 1247.203 | 0.033818 | 0.756479 | 0.951479 | Lamtor2 |
| ENSMUSG( | 933.0042 | -0.09131 | 0.442052 | 0.832762 | Lamtor3 |
| ENSMUSG( | 1203.202 | 0.028044 | 0.779323 | 0.95668 | Lamtor4 |
| ENSMUSG( | 1062.275 | 0.073885 | 0.463079 | 0.842422 | Lamtor5 |
| ENSMUSG( | 3372.489 | -0.10528 | 0.334464 | 0.776332 | Lancl1 |
| ENSMUSG( | 1598.03 | -0.00526 | 0.965088 | 0.995303 | Lancl2 |
| ENSMUSG( | 38.4222 | 0.247226 | 0.438014 | 0.830072 | Lancl3 |
| ENSMUSG( | 3682.308 | 0.114664 | 0.279532 | 0.729114 | Lap3 |
| ENSMUSG( | 10397.03 | -0.21545 | 0.093196 | 0.469454 | Laptm4a |
| ENSMUSG( | 2911.186 | -0.33515 | 0.024253 | 0.238311 | Laptm4b |
| ENSMUSG( | 2971.945 | -0.42746 | 0.00491 | 0.092785 | Laptm5 |
| ENSMUSG( | 4057.165 | -0.17403 | 0.302684 | 0.750923 | Large1 |
| ENSMUSG( | 15.18576 | 0.094026 | 0.844892 | NA | Large2 |
| ENSMUSG( | 4268.033 | 0.202807 | 0.028202 | 0.256543 | Larp1 |
| ENSMUSG( | 530.3117 | 0.183056 | 0.13207 | 0.544244 | Larp1b |
| ENSMUSG( | 830.7066 | -0.05834 | 0.574875 | 0.894572 | Larp4 |
| ENSMUSG( | 2540.325 | 0.092669 | 0.255392 | 0.706181 | Larp4b |
| ENSMUSG( | 612.2484 | -0.05397 | 0.733287 | 0.944956 | Larp6 |
| ENSMUSG( | 355.7479 | -0.15093 | 0.258353 | 0.709574 | Larp7 |
| ENSMUSG( | 1276.909 | 0.059835 | 0.532375 | 0.879905 | Lars1 |
| ENSMUSG( | 9845.432 | -1.43429 | 1.49E-36 | 4.84E-33 | Lars2 |
| ENSMUSG( | 1609.004 | 0.177186 | 0.16395 | 0.594869 | Las1l |
| ENSMUSG( | 9613.196 | 0.139329 | 0.127456 | 0.535223 | Lasp1 |
| ENSMUSG( | 15.60273 | 0.47667 | 0.332861 | NA | Lat |
| ENSMUSG( | 120.5506 | -0.28146 | 0.153968 | 0.580901 | Lat2 |
| ENSMUSG( | 1725.193 | 0.018775 | 0.854957 | 0.973665 | Lats1 |
| ENSMUSG( | 507.4341 | 0.015751 | 0.908961 | 0.98794 | Lats2 |
| ENSMUSG( | 15.24484 | 0.181334 | 0.712789 | NA | Lax1 |
| ENSMUSG( | 126.2747 | -0.4556 | 0.077298 | 0.429743 | Layn |
| ENSMUSG( | 2519.715 | -0.1151 | 0.182265 | 0.619591 | Lbh |
| ENSMUSG( | 121.7847 | 0.03247 | 0.877752 | 0.981431 | Lbhd1 |
| ENSMUSG( | 12.94324 | -0.6274 | 0.312926 | NA | Lbhd2 |
| ENSMUSG( | 339.2024 | -0.01189 | 0.945252 | 0.99365 | Lbp |
| ENSMUSG( | 437.176 | -0.40421 | 0.021443 | 0.22362 | Lbr |
| ENSMUSG( | 369.7697 | 0.032257 | 0.815543 | 0.965133 | Lca5 |
| ENSMUSG( | 70.97656 | -0.00472 | 0.984809 | 0.997815 | Lca5l |
| ENSMUSG( | 1940.605 | -0.15722 | 0.292278 | 0.742481 | Lcat |
| ENSMUSG( | 50.76151 | 0.257543 | 0.349348 | 0.78176 | Lck |
| ENSMUSG( | 1367.79 | -0.37956 | 0.010882 | 0.150598 | Lclat1 |
| ENSMUSG( | 1218.11 | 0.108603 | 0.321315 | 0.766497 | Lcmt1 |
| ENSMUSG( | 133.4268 | 0.1288 | 0.573734 | 0.893584 | Lcmt2 |
| ENSMUSG( | 41.39427 | -1.14437 | 0.023325 | 0.233882 | Lcn2 |
| ENSMUSG( | 110.8873 | -0.28587 | 0.204645 | 0.645534 | Lcor |
| ENSMUSG( | 578.1015 | -0.10902 | 0.345772 | 0.779153 | Lcorl |
| ENSMUSG( | 611.202 | -0.2542 | 0.032872 | 0.277278 | Lcp1 |
| ENSMUSG( | 127.7213 | 0.063575 | 0.728201 | 0.944355 | Lcp2 |
| ENSMUSG( | 92.11568 | 0.226708 | 0.283802 | 0.732203 | Lct |
| ENSMUSG( | 363.2078 | -0.04375 | 0.726786 | 0.944223 | Ldaf1 |
| ENSMUSG( | 1286.506 | -0.01626 | 0.861129 | 0.975026 | Ldah |
| ENSMUSG( | 3121.238 | -0.03514 | 0.748628 | 0.949604 | Ldb1 |
| ENSMUSG( | 837.0759 | -0.02646 | 0.839347 | 0.971179 | Ldb2 |
| ENSMUSG( | 195.5843 | -0.08704 | 0.564673 | 0.890434 | Ldb3 |
| ENSMUSG( | 7527.635 | 0.114474 | 0.180766 | 0.617841 | Ldha |
| ENSMUSG( | 17476.13 | 0.018575 | 0.839016 | 0.971179 | Ldhb |
| ENSMUSG( | 588.0772 | 0.070239 | 0.572771 | 0.893584 | Ldhd |
| ENSMUSG( | 1051.221 | -1.08262 | 1.48E-08 | 5.59E-06 | Ldlr |
| ENSMUSG( | 587.1554 | -0.48286 | 0.0043 | 0.085154 | Ldlrad3 |
| ENSMUSG( | 728.7751 | -0.13295 | 0.278437 | 0.728428 | Ldlrad4 |
| ENSMUSG( | 246.6872 | -0.41065 | 0.018559 | 0.20492 | Ldlrap1 |
| ENSMUSG( | 25.25995 | 0.064879 | 0.864292 | 0.975358 | Ldoc1 |
| ENSMUSG( | 237.6559 | 0.150172 | 0.307392 | 0.755324 | Lef1 |
| ENSMUSG( | 42.13897 | 0.195656 | 0.561988 | 0.889052 | Lefty1 |
| ENSMUSG( | 30.03084 | 0.585458 | 0.100718 | 0.484034 | Lefty2 |
| ENSMUSG( | 118.5806 | 0.103366 | 0.579805 | 0.89687 | Lekr1 |
| ENSMUSG( | 36.56774 | 0.049567 | 0.894182 | 0.984512 | Lemd1 |
| ENSMUSG( | 1450.361 | -0.05746 | 0.600158 | 0.904195 | Lemd2 |
| ENSMUSG( | 1094.81 | -0.35606 | 0.025837 | 0.247245 | Lemd3 |
| ENSMUSG( | 305.7592 | 0.236965 | 0.06773 | 0.40256 | Leng1 |
| ENSMUSG( | 5684.851 | 0.306961 | 0.122152 | 0.527707 | Leng8 |
| ENSMUSG( | 174.3965 | 0.030031 | 0.850487 | 0.972638 | Leng9 |
| ENSMUSG( | 494.0921 | 0.159187 | 0.189036 | 0.627678 | Leo1 |
| ENSMUSG( | 140.032 | -0.20926 | 0.242226 | 0.689699 | Lepr |
| ENSMUSG( | 1295.485 | -0.33608 | 0.015454 | 0.183729 | Leprot |
| ENSMUSG( | 2078.724 | -0.42215 | 0.003838 | 0.079297 | Leprotl1 |
| ENSMUSG( | 4322.606 | 0.031969 | 0.711259 | 0.940028 | Letm1 |
| ENSMUSG( | 340.0478 | -0.15137 | 0.27036 | 0.720226 | Letm2 |
| ENSMUSG( | 2844.451 | -0.01266 | 0.899905 | 0.985633 | Letmd1 |
| ENSMUSG( | 1149.187 | -0.44604 | 0.005905 | 0.103987 | Lfng |
| ENSMUSG( | 494.588 | -0.1554 | 0.209917 | 0.652297 | Lgals1 |
| ENSMUSG( | 30.44279 | -0.07326 | 0.835209 | 0.970426 | Lgals12 |
| ENSMUSG( | 45.70427 | -0.70052 | 0.045893 | 0.328083 | Lgals3 |
| ENSMUSG( | 1154.04 | -0.5805 | 0.00032 | 0.015127 | Lgals3bp |
| ENSMUSG( | 88.32561 | 0.02156 | 0.934897 | 0.993231 | Lgals4 |
| ENSMUSG( | 1128.48 | 0.031461 | 0.780455 | 0.957491 | Lgals8 |
| ENSMUSG( | 499.926 | -0.23155 | 0.112327 | 0.505954 | Lgals9 |
| ENSMUSG( | 2982.358 | 0.12845 | 0.175804 | 0.611006 | Lgalsl |
| ENSMUSG( | 3502.559 | -0.3708 | 0.00391 | 0.080172 | Lgi1 |
| ENSMUSG( | 2742.145 | -0.5131 | 0.000251 | 0.012746 | Lgi2 |
| ENSMUSG( | 4627.97 | -0.42386 | 0.004512 | 0.087865 | Lgi3 |
| ENSMUSG( | 3807.445 | -0.04732 | 0.668288 | 0.927864 | Lgi4 |
| ENSMUSG( | 6326.313 | -0.34885 | 0.010763 | 0.149466 | Lgmn |
| ENSMUSG( | 1634.784 | -0.30042 | 0.060235 | 0.379572 | Lgr4 |
| ENSMUSG( | 86.22164 | -0.53081 | 0.016079 | 0.188199 | Lgr5 |
| ENSMUSG( | 358.6004 | -0.34816 | 0.02883 | 0.258644 | Lgr6 |
| ENSMUSG( | 35.08952 | -0.24219 | 0.476987 | 0.851875 | Lhcgr |
| ENSMUSG( | 844.2326 | -0.37347 | 0.041011 | 0.313031 | Lhfpl2 |
| ENSMUSG( | 1195.23 | -0.43165 | 0.008941 | 0.132538 | Lhfpl3 |
| ENSMUSG( | 4702.124 | -0.14818 | 0.174745 | 0.610304 | Lhfpl4 |
| ENSMUSG( | 69.90288 | -0.35824 | 0.121468 | 0.525713 | Lhfpl5 |
| ENSMUSG( | 1526.283 | 0.053663 | 0.657919 | 0.92425 | Lhfpl6 |
| ENSMUSG( | 929.7227 | 0.066436 | 0.556025 | 0.887457 | Lhpp |
| ENSMUSG( | 6743.774 | 0.20475 | 0.041498 | 0.314241 | Lhx2 |
| ENSMUSG( | 460.8759 | 0.237536 | 0.166644 | 0.599201 | Lhx6 |
| ENSMUSG( | 36.25943 | 0.049611 | 0.901272 | 0.985633 | Lhx9 |
| ENSMUSG( | 860.7194 | -0.04581 | 0.650696 | 0.921301 | Lias |
| ENSMUSG( | 94.38932 | -0.1877 | 0.390585 | 0.808144 | Liat1 |
| ENSMUSG( | 41.21897 | -0.18421 | 0.529081 | 0.878551 | Lif |
| ENSMUSG( | 1401.184 | -0.09526 | 0.512952 | 0.871762 | Lifr |
| ENSMUSG( | 277.9035 | 0.060966 | 0.652691 | 0.922498 | Lig1 |
| ENSMUSG( | 985.8276 | 0.110084 | 0.277909 | 0.727904 | Lig3 |
| ENSMUSG( | 221.3351 | 0.222458 | 0.140984 | 0.557902 | Lig4 |
| ENSMUSG( | 16.40253 | -0.8577 | 0.085888 | NA | Lilra5 |
| ENSMUSG( | 8.297957 | -0.29626 | 0.674349 | NA | Lilrb4a |
| ENSMUSG( | 946.1561 | 0.092789 | 0.422441 | 0.823508 | Lima1 |
| ENSMUSG( | 8322.24 | -0.00238 | 0.981663 | 0.997512 | Limch1 |
| ENSMUSG( | 729.487 | -0.03031 | 0.780749 | 0.957491 | Limd1 |
| ENSMUSG( | 2099.751 | 0.035404 | 0.71182 | 0.940028 | Limd2 |
| ENSMUSG( | 2830.672 | 0.12096 | 0.23563 | 0.682729 | Limk1 |
| ENSMUSG( | 3797.191 | 0.152353 | 0.096113 | 0.476417 | Limk2 |
| ENSMUSG( | 1052.671 | 0.02895 | 0.78602 | 0.958774 | Lims1 |
| ENSMUSG( | 856.3813 | 0.014319 | 0.909213 | 0.98794 | Lims2 |
| ENSMUSG( | 142.1237 | -0.33828 | 0.116357 | 0.514816 | Lin28b |
| ENSMUSG( | 440.9465 | -0.10586 | 0.379739 | 0.801757 | Lin37 |

| | | | | | |
|---|---|---|---|---|---|
| ENSMUSG( | 326.5642 | -0.1312 | 0.405696 | 0.917257 | Larp7 |
| ENSMUSG( | 1122.728 | -0.04307 | 0.707667 | 0.964029 | Lars1 |
| ENSMUSG( | 7927.626 | -2.24551 | 8.00E-34 | 4.67E-30 | Lars2 |
| ENSMUSG( | 1367.296 | -0.02219 | 0.888946 | 0.990558 | Las1l |
| ENSMUSG( | 8210.1 | -0.04305 | 0.688397 | 0.959742 | Lasp1 |
| ENSMUSG( | 15.67005 | 0.723077 | 0.187933 | 0.85698 | Lat |
| ENSMUSG( | 122.153 | 0.053769 | 0.808687 | 0.97873 | Lat2 |
| ENSMUSG( | 1611.219 | 0.087036 | 0.523527 | 0.932848 | Lats1 |
| ENSMUSG( | 500.653 | 0.233384 | 0.138129 | 0.828492 | Lats2 |
| ENSMUSG( | 15.09227 | 0.337782 | 0.554035 | 0.939565 | Lax1 |
| ENSMUSG( | 153.0126 | 0.394725 | 0.141409 | 0.830678 | Layn |
| ENSMUSG( | 2227.72 | -0.20996 | 0.100183 | 0.801311 | Lbh |
| ENSMUSG( | 109.0927 | -0.01912 | 0.928533 | 0.99306 | Lbhd1 |
| ENSMUSG( | 14.82159 | 0.022738 | 0.96985 | 0.99594 | Lbhd2 |
| ENSMUSG( | 397.7173 | 0.649439 | 0.001649 | 0.150169 | Lbp |
| ENSMUSG( | 492.4687 | 0.238345 | 0.259344 | 0.883971 | Lbr |
| ENSMUSG( | 343.0025 | 0.080184 | 0.605515 | 0.94924 | Lca5 |
| ENSMUSG( | 65.70109 | 0.059952 | 0.827461 | 0.980224 | Lca5l |
| ENSMUSG( | 1965.478 | 0.152523 | 0.36773 | 0.915289 | Lcat |
| ENSMUSG( | 41.62128 | -0.06446 | 0.846307 | 0.984578 | Lck |
| ENSMUSG( | 1465.924 | 0.11613 | 0.471574 | 0.923329 | Lclat1 |
| ENSMUSG( | 1020.727 | -0.1366 | 0.296017 | 0.898538 | Lcmt1 |
| ENSMUSG( | 117.7937 | 0.043778 | 0.833589 | 0.981218 | Lcmt2 |
| ENSMUSG( | 56.65577 | 0.210039 | 0.648494 | 0.95422 | Lcn2 |
| ENSMUSG( | 119.0294 | 0.214411 | 0.416103 | 0.917257 | Lcor |
| ENSMUSG( | 570.0284 | 0.116634 | 0.431219 | 0.917257 | Lcorl |
| ENSMUSG( | 545.5853 | -0.31723 | 0.033857 | 0.648168 | Lcp1 |
| ENSMUSG( | 121.1318 | 0.179848 | 0.373745 | 0.915491 | Lcp2 |
| ENSMUSG( | 86.93561 | 0.346298 | 0.276046 | 0.890679 | Lct |
| ENSMUSG( | 7.744879 | 0.085026 | 0.902589 | NA | Lctl |
| ENSMUSG( | 351.913 | 0.129669 | 0.372778 | 0.915491 | Ldaf1 |
| ENSMUSG( | 1105.809 | -0.19415 | 0.100145 | 0.801311 | Ldah |
| ENSMUSG( | 3040.648 | 0.155187 | 0.213006 | 0.864412 | Ldb1 |
| ENSMUSG( | 720.6465 | -0.20429 | 0.199568 | 0.863016 | Ldb2 |
| ENSMUSG( | 168.0399 | -0.26681 | 0.131204 | 0.822884 | Ldb3 |
| ENSMUSG( | 6173.866 | -0.19467 | 0.104477 | 0.804969 | Ldha |
| ENSMUSG( | 14551.47 | -0.25405 | 0.041377 | 0.688699 | Ldhb |
| ENSMUSG( | 519.8574 | -0.01361 | 0.934667 | 0.99306 | Ldhd |
| ENSMUSG( | 1268.657 | -0.06497 | 0.778312 | 0.976106 | Ldlr |
| ENSMUSG( | 628.4067 | 0.029381 | 0.883769 | 0.989644 | Ldlrad3 |
| ENSMUSG( | 708.7169 | 0.063248 | 0.649284 | 0.95422 | Ldlrad4 |
| ENSMUSG( | 268.6901 | 0.140519 | 0.455178 | 0.923163 | Ldlrap1 |
| ENSMUSG( | 23.1039 | 0.063571 | 0.875497 | 0.989644 | Ldoc1 |
| ENSMUSG( | 226.2453 | 0.278155 | 0.12033 | 0.813384 | Lef1 |
| ENSMUSG( | 44.26344 | 0.549671 | 0.072811 | 0.763629 | Lefty1 |
| ENSMUSG( | 27.8112 | 0.612537 | 0.10834 | 0.804969 | Lefty2 |
| ENSMUSG( | 99.34012 | -0.14012 | 0.527051 | 0.934449 | Lekr1 |
| ENSMUSG( | 31.02267 | -0.19291 | 0.626846 | 0.953234 | Lemd1 |
| ENSMUSG( | 1422.898 | 0.156766 | 0.250699 | 0.878324 | Lemd2 |
| ENSMUSG( | 1190.527 | 0.175044 | 0.33137 | 0.907861 | Lemd3 |
| ENSMUSG( | 240.2197 | -0.18863 | 0.247435 | 0.877904 | Leng1 |
| ENSMUSG( | 5243.837 | 0.336151 | 0.390656 | 0.917257 | Leng8 |
| ENSMUSG( | 143.164 | -0.29377 | 0.127273 | 0.817278 | Leng9 |
| ENSMUSG( | 383.5754 | -0.32033 | 0.033124 | 0.64093 | Leo1 |
| ENSMUSG( | 130.3827 | -0.12303 | 0.577143 | 0.942912 | Lepr |
| ENSMUSG( | 1381.517 | 0.139044 | 0.398703 | 0.917257 | Leprot |
| ENSMUSG( | 2276.136 | 0.139298 | 0.394796 | 0.917257 | Leprotl1 |
| ENSMUSG( | 3837.443 | -0.04179 | 0.693761 | 0.960767 | Letm1 |
| ENSMUSG( | 338.1916 | 0.109694 | 0.511343 | 0.92971 | Letm2 |
| ENSMUSG( | 2610.724 | 0.01061 | 0.926047 | 0.99306 | Letmd1 |
| ENSMUSG( | 1450.073 | 0.492008 | 0.029814 | 0.61992 | Lfng |
| ENSMUSG( | 429.3672 | -0.3121 | 0.029095 | 0.614779 | Lgals1 |
| ENSMUSG( | 28.58868 | 0.03354 | 0.9314 | 0.99306 | Lgals12 |
| ENSMUSG( | 42.6234 | -0.60219 | 0.120105 | 0.813384 | Lgals3 |
| ENSMUSG( | 1155.218 | -0.25689 | 0.093741 | 0.795735 | Lgals3bp |
| ENSMUSG( | 82.0791 | 0.088711 | 0.774885 | 0.974931 | Lgals4 |
| ENSMUSG( | 991.8397 | -0.0749 | 0.522994 | 0.932848 | Lgals8 |
| ENSMUSG( | 454.5143 | -0.23097 | 0.122755 | 0.814249 | Lgals9 |
| ENSMUSG( | 2638.911 | 0.043059 | 0.766499 | 0.974236 | Lgalsl |
| ENSMUSG( | 3817.54 | 0.168732 | 0.241091 | 0.876181 | Lgi1 |
| ENSMUSG( | 2970.748 | 0.035958 | 0.836099 | 0.981237 | Lgi2 |
| ENSMUSG( | 5103.345 | 0.159018 | 0.398074 | 0.917257 | Lgi3 |
| ENSMUSG( | 3362.383 | -0.13903 | 0.274512 | 0.889874 | Lgi4 |
| ENSMUSG( | 6906.651 | 0.191667 | 0.264334 | 0.886134 | Lgmn |
| ENSMUSG( | 1572.668 | -0.12803 | 0.436517 | 0.919879 | Lgr4 |
| ENSMUSG( | 108.4547 | 0.396005 | 0.165729 | 0.839705 | Lgr5 |
| ENSMUSG( | 352.535 | -0.10895 | 0.561349 | 0.940254 | Lgr6 |
| ENSMUSG( | 34.46677 | 0.004615 | 0.989553 | 0.997928 | Lhcgr |
| ENSMUSG( | 834.6157 | -0.10831 | 0.606812 | 0.949772 | Lhfpl2 |
| ENSMUSG( | 1339.447 | 0.190943 | 0.264013 | 0.886134 | Lhfpl3 |
| ENSMUSG( | 4669.319 | 0.105021 | 0.424916 | 0.917257 | Lhfpl4 |
| ENSMUSG( | 74.20616 | 0.103512 | 0.66941 | 0.956766 | Lhfpl5 |
| ENSMUSG( | 1450.126 | 0.170723 | 0.210404 | 0.864412 | Lhfpl6 |
| ENSMUSG( | 770.6241 | -0.22105 | 0.134911 | 0.82429 | Lhpp |
| ENSMUSG( | 5560.143 | -0.07901 | 0.585902 | 0.944975 | Lhx2 |
| ENSMUSG( | 389.6276 | 0.026328 | 0.886382 | 0.990011 | Lhx6 |
| ENSMUSG( | 12.54995 | -0.70328 | 0.305201 | 0.898538 | Lhx8 |
| ENSMUSG( | 28.67832 | -0.40944 | 0.36987 | 0.915491 | Lhx9 |
| ENSMUSG( | 760.0729 | -0.1433 | 0.273783 | 0.889848 | Lias |
| ENSMUSG( | 85.04523 | -0.22802 | 0.333371 | 0.907907 | Liat1 |
| ENSMUSG( | 39.12747 | -0.06364 | 0.842875 | 0.98313 | Lif |
| ENSMUSG( | 1486.133 | 0.330244 | 0.057127 | 0.723362 | Lifr |
| ENSMUSG( | 256.7114 | 0.100429 | 0.517437 | 0.930939 | Lig1 |
| ENSMUSG( | 856.2803 | -0.0224 | 0.859191 | 0.985884 | Lig3 |
| ENSMUSG( | 186.1879 | -0.00148 | 0.993074 | 0.998813 | Lig4 |
| ENSMUSG( | 18.53185 | -0.07089 | 0.891658 | 0.990558 | Lilra5 |
| ENSMUSG( | 7.571629 | -0.26222 | 0.735106 | NA | Lilrb4a |
| ENSMUSG( | 892.8197 | 0.193481 | 0.150971 | 0.837173 | Lima1 |
| ENSMUSG( | 7088.021 | -0.2055 | 0.078113 | 0.77321 | Limch1 |
| ENSMUSG( | 657.1299 | -0.06421 | 0.620414 | 0.952896 | Limd1 |
| ENSMUSG( | 1836.822 | -0.08413 | 0.474228 | 0.923329 | Limd2 |
| ENSMUSG( | 2360.371 | -0.1359 | 0.206406 | 0.864412 | Limk1 |
| ENSMUSG( | 3313.826 | 0.032927 | 0.769385 | 0.974236 | Limk2 |
| ENSMUSG( | 976.0325 | 0.078638 | 0.51997 | 0.931892 | Lims1 |
| ENSMUSG( | 776.2339 | 0.006043 | 0.967782 | 0.995833 | Lims2 |
| ENSMUSG( | 141.3905 | -0.08972 | 0.72781 | 0.966887 | Lin28b |
| ENSMUSG( | 395.5311 | -0.15461 | 0.318563 | 0.903164 | Lin37 |
| ENSMUSG( | 1301.904 | -0.12843 | 0.391909 | 0.917257 | Lin52 |
| ENSMUSG( | 298.6324 | 0.138583 | 0.429289 | 0.917257 | Lin54 |
| ENSMUSG( | 344.0959 | 0.02963 | 0.849197 | 0.985321 | Lin7a |
| ENSMUSG( | 1029.654 | -0.08341 | 0.554988 | 0.939565 | Lin7b |
| ENSMUSG( | 2485.04 | -0.04681 | 0.763573 | 0.974 | Lin7c |
| ENSMUSG( | 86.34083 | 0.35605 | 0.164235 | 0.838505 | Lin9 |
| ENSMUSG( | 11336.09 | 0.273181 | 0.137738 | 0.828492 | Lingo1 |
| ENSMUSG( | 500.0572 | -0.08364 | 0.692691 | 0.96043 | Lingo2 |
| ENSMUSG( | 1431.127 | 0.062783 | 0.69039 | 0.960071 | Lingo3 |
| ENSMUSG( | 45.14086 | -0.50636 | 0.089554 | 0.789094 | Lingo4 |
| ENSMUSG( | 274.4696 | 0.13996 | 0.428205 | 0.917257 | Lins1 |
| ENSMUSG( | 825.6962 | 0.209286 | 0.286988 | 0.894091 | Lipa |
| ENSMUSG( | 11.28794 | -0.05126 | 0.933686 | NA | Lipc |
| ENSMUSG( | 307.7879 | -0.06184 | 0.676859 | 0.957106 | Lipe |
| ENSMUSG( | 124.3679 | 0.068203 | 0.764232 | 0.974 | Lipg |
| ENSMUSG( | 36.92443 | 0.221693 | 0.548761 | 0.938075 | Liph |
| ENSMUSG( | 125.4978 | -0.12323 | 0.587281 | 0.944975 | Lipo3 |
| ENSMUSG( | 72.00943 | 0.215861 | 0.383672 | 0.917257 | Lipt1 |
| ENSMUSG( | 78.51417 | -0.16222 | 0.52612 | 0.934374 | Lipt2 |
| ENSMUSG( | 690.5564 | -0.2169 | 0.085278 | 0.779799 | Litaf |
| ENSMUSG( | 3491.622 | -0.24319 | 0.142145 | 0.830709 | Lix1 |
| ENSMUSG( | 1552.992 | -0.07552 | 0.608235 | 0.950113 | Lix1l |
| ENSMUSG( | 3978.704 | -0.14935 | 0.148718 | 0.836241 | Llgl1 |
| ENSMUSG( | 42.07559 | 0.112639 | 0.72253 | 0.966887 | Llgl2 |
| ENSMUSG( | 273.5388 | -0.13294 | 0.47645 | 0.923329 | Llph |
| ENSMUSG( | 1619.103 | 0.103977 | 0.536931 | 0.937169 | Lman1 |
| ENSMUSG( | 3425.459 | 0.166309 | 0.244717 | 0.877904 | Lman2 |
| ENSMUSG( | 1448.807 | -0.04854 | 0.724496 | 0.966887 | Lman2l |
| ENSMUSG( | 981.1429 | 0.203636 | 0.1761 | 0.846252 | Lmbr1 |
| ENSMUSG( | 322.596 | 0.302482 | 0.094217 | 0.795735 | Lmbr1l |
| ENSMUSG( | 8035.549 | 0.124968 | 0.393626 | 0.917257 | Lmbrd1 |
| ENSMUSG( | 2861.077 | 0.149454 | 0.3998 | 0.917257 | Lmbrd2 |
| ENSMUSG( | 309.4018 | -0.72209 | 3.12E-06 | 0.001517 | Lmcd1 |
| ENSMUSG( | 767.1651 | 0.127239 | 0.415921 | 0.917257 | Lmf1 |
| ENSMUSG( | 992.8612 | 0.120094 | 0.501217 | 0.927825 | Lmf2 |
| ENSMUSG( | 488.7151 | 0.208025 | 0.269001 | 0.886755 | Lmln |
| ENSMUSG( | 768.0912 | -0.03347 | 0.796024 | 0.97681 | Lmna |
| ENSMUSG( | 250.389 | -0.18319 | 0.265569 | 0.886134 | Lmnb1 |
| ENSMUSG( | 466.9778 | 0.076523 | 0.617005 | 0.951666 | Lmnb2 |
| ENSMUSG( | 41.75009 | -0.16701 | 0.590831 | 0.944975 | Lmntd1 |
| ENSMUSG( | 377.9518 | -0.29242 | 0.06868 | 0.753052 | Lmo1 |
| ENSMUSG( | 580.1785 | 0.03226 | 0.795417 | 0.97681 | Lmo2 |
| ENSMUSG( | 2212.259 | 0.008595 | 0.959476 | 0.994798 | Lmo3 |
| ENSMUSG( | 4161.213 | -0.06383 | 0.697967 | 0.960867 | Lmo4 |
| ENSMUSG( | 2135.924 | 0.095638 | 0.494345 | 0.927372 | Lmo7 |
| ENSMUSG( | 22.19559 | -0.1395 | 0.742891 | 0.969415 | Lmod1 |
| ENSMUSG( | 6311.639 | 0.064408 | 0.71336 | 0.964766 | Lmtk2 |
| ENSMUSG( | 10538.4 | 0.244182 | 0.049424 | 0.708648 | Lmtk3 |
| ENSMUSG( | 9.302181 | 0.541433 | 0.411582 | NA | Lmx1a |
| ENSMUSG( | 21.80952 | 0.424482 | 0.310603 | 0.899422 | Lmx1b |
| ENSMUSG( | 15.83479 | -0.2093 | 0.67789 | 0.957166 | Lncpint |

| | | | | | | | | | | | |
|---|---|---|---|---|---|---|---|---|---|---|---|
| ENSMUSG( | Lgals12 | 30.44279 | -0.07326 | 0.835209 | 0.970426 | ENSMUSG( | Lgals12 | 28.58868 | 0.03354 | 0.9314 | 0.99306 |
| ENSMUSG( | Lgals3 | 45.70427 | -0.70052 | 0.045893 | 0.328083 | ENSMUSG( | Lgals3 | 42.6234 | -0.60219 | 0.120105 | 0.813384 |
| ENSMUSG( | Lgals3bp | 1154.04 | -0.5805 | 0.00032 | 0.015127 | ENSMUSG( | Lgals3bp | 1155.218 | -0.25689 | 0.093741 | 0.795735 |
| ENSMUSG( | Lgals4 | 88.32561 | 0.02156 | 0.934897 | 0.993231 | ENSMUSG( | Lgals4 | 82.0791 | 0.088711 | 0.774885 | 0.974931 |
| ENSMUSG( | Lgals8 | 1128.48 | 0.031461 | 0.780455 | 0.957491 | ENSMUSG( | Lgals8 | 991.8397 | -0.0749 | 0.522994 | 0.932848 |
| ENSMUSG( | Lgals9 | 499.926 | -0.23155 | 0.112327 | 0.505954 | ENSMUSG( | Lgals9 | 454.5143 | -0.23097 | 0.122755 | 0.814249 |
| ENSMUSG( | Lgalsl | 2982.358 | 0.12845 | 0.175804 | 0.611006 | ENSMUSG( | Lgalsl | 2638.911 | 0.043059 | 0.766499 | 0.974236 |
| ENSMUSG( | Lgi1 | 3502.559 | -0.3708 | 0.00391 | 0.080172 | ENSMUSG( | Lgi1 | 3817.54 | 0.168732 | 0.241091 | 0.876181 |
| ENSMUSG( | Lgi2 | 2742.145 | -0.5131 | 0.000251 | 0.012746 | ENSMUSG( | Lgi2 | 2970.748 | 0.035958 | 0.836099 | 0.981237 |
| ENSMUSG( | Lgi3 | 4627.97 | -0.42386 | 0.004512 | 0.087865 | ENSMUSG( | Lgi3 | 5103.345 | 0.159018 | 0.398074 | 0.917257 |
| ENSMUSG( | Lgi4 | 3807.445 | -0.04732 | 0.668288 | 0.927864 | ENSMUSG( | Lgi4 | 3362.383 | -0.13903 | 0.274512 | 0.889874 |
| ENSMUSG( | Lgmn | 6326.313 | -0.34885 | 0.010763 | 0.149466 | ENSMUSG( | Lgmn | 6906.651 | 0.191667 | 0.264334 | 0.886134 |
| ENSMUSG( | Lgr4 | 1634.784 | -0.30042 | 0.060235 | 0.379572 | ENSMUSG( | Lgr4 | 1572.668 | -0.12803 | 0.436517 | 0.919879 |
| ENSMUSG( | Lgr5 | 86.22164 | -0.53081 | 0.016079 | 0.188199 | ENSMUSG( | Lgr5 | 108.4547 | 0.396005 | 0.165729 | 0.839705 |
| ENSMUSG( | Lgr6 | 358.6004 | -0.34816 | 0.02883 | 0.258644 | ENSMUSG( | Lgr6 | 352.535 | -0.10895 | 0.561349 | 0.940254 |
| ENSMUSG( | Lhcgr | 35.08952 | -0.24219 | 0.476987 | 0.851875 | ENSMUSG( | Lhcgr | 34.46677 | 0.004615 | 0.989553 | 0.997928 |
| ENSMUSG( | Lhfpl2 | 844.2326 | -0.37347 | 0.041011 | 0.313031 | ENSMUSG( | Lhfpl2 | 834.6157 | -0.10831 | 0.606812 | 0.949772 |
| ENSMUSG( | Lhfpl3 | 1195.23 | -0.43165 | 0.008941 | 0.132538 | ENSMUSG( | Lhfpl3 | 1339.447 | 0.190943 | 0.264013 | 0.886134 |
| ENSMUSG( | Lhfpl4 | 4702.124 | -0.14818 | 0.174745 | 0.610304 | ENSMUSG( | Lhfpl4 | 4669.319 | 0.105021 | 0.424916 | 0.917257 |
| ENSMUSG( | Lhfpl5 | 69.90288 | -0.35824 | 0.121468 | 0.525713 | ENSMUSG( | Lhfpl5 | 74.20616 | 0.103512 | 0.66941 | 0.956766 |
| ENSMUSG( | Lhfpl6 | 1526.283 | 0.053663 | 0.657919 | 0.92425 | ENSMUSG( | Lhfpl6 | 1450.126 | 0.170723 | 0.210404 | 0.864412 |
| ENSMUSG( | Lhpp | 929.7227 | 0.066436 | 0.556025 | 0.887457 | ENSMUSG( | Lhpp | 770.6241 | -0.22105 | 0.134911 | 0.82429 |
| ENSMUSG( | Lhx2 | 6743.774 | 0.20475 | 0.041498 | 0.314241 | ENSMUSG( | Lhx2 | 5560.143 | -0.07901 | 0.585902 | 0.944975 |
| ENSMUSG( | Lhx6 | 460.8759 | 0.237536 | 0.166644 | 0.599201 | ENSMUSG( | Lhx6 | 389.6276 | 0.026328 | 0.886382 | 0.990011 |
| ENSMUSG( | Lhx9 | 36.25943 | 0.049611 | 0.901272 | 0.985633 | ENSMUSG( | Lhx9 | 28.67832 | -0.40944 | 0.36987 | 0.915491 |
| ENSMUSG( | Lias | 860.7194 | -0.04581 | 0.650696 | 0.921301 | ENSMUSG( | Lias | 760.0729 | -0.1433 | 0.273783 | 0.889848 |
| ENSMUSG( | Liat1 | 94.38932 | -0.1877 | 0.390585 | 0.808144 | ENSMUSG( | Liat1 | 85.04523 | -0.22802 | 0.333371 | 0.907907 |
| ENSMUSG( | Lif | 41.21897 | -0.18421 | 0.529081 | 0.878551 | ENSMUSG( | Lif | 39.12747 | -0.06364 | 0.842875 | 0.98313 |
| ENSMUSG( | Lifr | 1401.184 | -0.09526 | 0.512952 | 0.871762 | ENSMUSG( | Lifr | 1486.133 | 0.330244 | 0.057127 | 0.723362 |
| ENSMUSG( | Lig1 | 277.9035 | 0.060966 | 0.652691 | 0.922498 | ENSMUSG( | Lig1 | 256.7114 | 0.100429 | 0.517437 | 0.930939 |
| ENSMUSG( | Lig3 | 985.8276 | 0.110084 | 0.277909 | 0.727904 | ENSMUSG( | Lig3 | 856.2803 | -0.0224 | 0.859191 | 0.985884 |
| ENSMUSG( | Lig4 | 221.3351 | 0.222458 | 0.140984 | 0.557902 | ENSMUSG( | Lig4 | 186.1879 | -0.00148 | 0.993074 | 0.998813 |
| ENSMUSG( | Lilra5 | 16.40253 | -0.8577 | 0.085888 | NA | ENSMUSG( | Lilra5 | 18.53185 | -0.07089 | 0.891658 | 0.990558 |
| ENSMUSG( | Lilrb4a | 8.297957 | -0.29626 | 0.674349 | NA | ENSMUSG( | Lilrb4a | 7.571629 | -0.26222 | 0.735106 | NA |
| ENSMUSG( | Lima1 | 946.1561 | 0.092789 | 0.422441 | 0.823508 | ENSMUSG( | Lima1 | 892.8197 | 0.193481 | 0.150971 | 0.837173 |
| ENSMUSG( | Limch1 | 8322.24 | -0.00238 | 0.981663 | 0.997512 | ENSMUSG( | Limch1 | 7088.021 | -0.2055 | 0.078113 | 0.77321 |
| ENSMUSG( | Limd1 | 729.487 | -0.03031 | 0.780749 | 0.957491 | ENSMUSG( | Limd1 | 657.1299 | -0.06421 | 0.620414 | 0.952896 |
| ENSMUSG( | Limd2 | 2099.751 | 0.035404 | 0.71182 | 0.940028 | ENSMUSG( | Limd2 | 1836.822 | -0.08413 | 0.474228 | 0.923329 |
| ENSMUSG( | Limk1 | 2830.672 | 0.12096 | 0.23563 | 0.682729 | ENSMUSG( | Limk1 | 2360.371 | -0.1359 | 0.206406 | 0.864412 |
| ENSMUSG( | Limk2 | 3797.191 | 0.152353 | 0.096113 | 0.476417 | ENSMUSG( | Limk2 | 3313.826 | 0.032927 | 0.769385 | 0.974236 |
| ENSMUSG( | Lims1 | 1052.671 | 0.02895 | 0.78602 | 0.958774 | ENSMUSG( | Lims1 | 976.0325 | 0.078638 | 0.51997 | 0.931892 |
| ENSMUSG( | Lims2 | 856.3813 | 0.014319 | 0.909213 | 0.98794 | ENSMUSG( | Lims2 | 776.2339 | 0.006043 | 0.967782 | 0.995833 |
| ENSMUSG( | Lin28b | 142.1237 | -0.33828 | 0.116357 | 0.514816 | ENSMUSG( | Lin28b | 141.3905 | -0.08972 | 0.72781 | 0.966887 |
| ENSMUSG( | Lin37 | 440.9465 | -0.10586 | 0.379739 | 0.801757 | ENSMUSG( | Lin37 | 395.5311 | -0.15461 | 0.318563 | 0.903164 |
| ENSMUSG( | Lin52 | 1522.293 | 0.06139 | 0.635233 | 0.914296 | ENSMUSG( | Lin52 | 1301.904 | -0.12843 | 0.391909 | 0.917257 |
| ENSMUSG( | Lin54 | 311.0862 | 0.002648 | 0.98475 | 0.997815 | ENSMUSG( | Lin54 | 298.6324 | 0.138583 | 0.429289 | 0.917257 |
| ENSMUSG( | Lin7a | 404.74 | 0.22631 | 0.11371 | 0.508792 | ENSMUSG( | Lin7a | 344.0959 | 0.02963 | 0.849197 | 0.985321 |
| ENSMUSG( | Lin7b | 1125.607 | -0.09387 | 0.508 | 0.867981 | ENSMUSG( | Lin7b | 1029.654 | -0.08341 | 0.554988 | 0.939565 |
| ENSMUSG( | Lin7c | 2770.141 | 0.001454 | 0.990785 | 0.999232 | ENSMUSG( | Lin7c | 2485.04 | -0.04681 | 0.763573 | 0.974 |
| ENSMUSG( | Lin9 | 85.9858 | 0.094611 | 0.710192 | 0.9397 | ENSMUSG( | Lin9 | 86.34083 | 0.35605 | 0.164235 | 0.838505 |
| ENSMUSG( | Lingo1 | 10829.18 | -0.1229 | 0.43612 | 0.829093 | ENSMUSG( | Lingo1 | 11336.09 | 0.273181 | 0.137738 | 0.828492 |
| ENSMUSG( | Lingo2 | 465.4133 | -0.63811 | 0.000682 | 0.025185 | ENSMUSG( | Lingo2 | 500.0572 | -0.08364 | 0.692691 | 0.96043 |
| ENSMUSG( | Lingo3 | 1305.633 | -0.52266 | 0.000133 | 0.007813 | ENSMUSG( | Lingo3 | 1431.127 | 0.062783 | 0.69039 | 0.960071 |
| ENSMUSG( | Lingo4 | 64.1774 | 0.274642 | 0.282472 | 0.73186 | ENSMUSG( | Lingo4 | 45.14086 | -0.50636 | 0.089554 | 0.789094 |
| ENSMUSG( | Lins1 | 281.653 | -0.04178 | 0.78334 | 0.957558 | ENSMUSG( | Lins1 | 274.4696 | 0.13996 | 0.428205 | 0.917257 |
| ENSMUSG( | Lipa | 750.9055 | -0.35539 | 0.038025 | 0.300991 | ENSMUSG( | Lipa | 825.6962 | 0.209286 | 0.286988 | 0.894091 |
| ENSMUSG( | Lipc | 12.51727 | 0.041901 | 0.938744 | NA | ENSMUSG( | Lipc | 11.28794 | -0.05126 | 0.933686 | NA |
| ENSMUSG( | Lipe | 350.1032 | 0.03786 | 0.767967 | 0.954621 | ENSMUSG( | Lipe | 307.7879 | -0.06184 | 0.676859 | 0.957106 |
| ENSMUSG( | Lipg | 116.7411 | -0.41135 | 0.06009 | 0.379098 | ENSMUSG( | Lipg | 124.3679 | 0.068203 | 0.764232 | 0.974 |
| ENSMUSG( | Liph | 33.77942 | -0.3518 | 0.330083 | 0.774055 | ENSMUSG( | Liph | 36.92443 | 0.221693 | 0.548761 | 0.938075 |
| ENSMUSG( | Lipo3 | 127.0818 | -0.36349 | 0.102133 | 0.486947 | ENSMUSG( | Lipo3 | 125.4978 | -0.12323 | 0.587281 | 0.944975 |
| ENSMUSG( | Lipt1 | 75.2576 | 0.076169 | 0.741438 | 0.948208 | ENSMUSG( | Lipt1 | 72.00943 | 0.215861 | 0.383672 | 0.917257 |
| ENSMUSG( | Lipt2 | 90.79703 | 0.013673 | 0.947924 | 0.993821 | ENSMUSG( | Lipt2 | 78.51417 | -0.16222 | 0.52612 | 0.934374 |
| ENSMUSG( | Litaf | 763.9064 | -0.1971 | 0.080614 | 0.438981 | ENSMUSG( | Litaf | 690.5564 | -0.2169 | 0.085278 | 0.779799 |
| ENSMUSG( | Lix1 | 4289.444 | 0.093533 | 0.490955 | 0.859739 | ENSMUSG( | Lix1 | 3491.622 | -0.24319 | 0.142145 | 0.830709 |
| ENSMUSG( | Lix1l | 1758.937 | 0.021374 | 0.850982 | 0.972638 | ENSMUSG( | Lix1l | 1552.992 | -0.07552 | 0.608235 | 0.950113 |
| ENSMUSG( | Llgl1 | 4646.949 | 0.034719 | 0.676596 | 0.930542 | ENSMUSG( | Llgl1 | 3978.704 | -0.14935 | 0.148718 | 0.836241 |
| ENSMUSG( | Llgl2 | 46.87229 | 0.136494 | 0.638112 | 0.916107 | ENSMUSG( | Llgl2 | 42.07559 | 0.112639 | 0.72253 | 0.966887 |
| ENSMUSG( | Llph | 318.2285 | 0.047341 | 0.776716 | 0.955723 | ENSMUSG( | Llph | 273.5388 | -0.13294 | 0.47645 | 0.923329 |
| ENSMUSG( | Lman1 | 1542.057 | -0.32648 | 0.033117 | 0.278602 | ENSMUSG( | Lman1 | 1619.103 | 0.103977 | 0.536931 | 0.937169 |
| ENSMUSG( | Lman2 | 3438.099 | -0.09232 | 0.456896 | 0.839183 | ENSMUSG( | Lman2 | 3425.459 | 0.166309 | 0.244717 | 0.877904 |
| ENSMUSG( | Lman2l | 1619.39 | 0.001712 | 0.989255 | 0.999111 | ENSMUSG( | Lman2l | 1448.807 | -0.04854 | 0.724496 | 0.966887 |
| ENSMUSG( | Lmbr1 | 919.6806 | -0.26127 | 0.063672 | 0.390618 | ENSMUSG( | Lmbr1 | 981.1429 | 0.203636 | 0.1761 | 0.846252 |
| ENSMUSG( | Lmbr1l | 314.7334 | -0.03332 | 0.834335 | 0.970333 | ENSMUSG( | Lmbr1l | 322.596 | 0.302482 | 0.094217 | 0.795735 |
| ENSMUSG( | Lmbrd1 | 7963.235 | -0.17476 | 0.177395 | 0.614553 | ENSMUSG( | Lmbrd1 | 8035.549 | 0.124968 | 0.393626 | 0.917257 |
| ENSMUSG( | Lmbrd2 | 2589.302 | -0.43739 | 0.003896 | 0.080172 | ENSMUSG( | Lmbrd2 | 2861.077 | 0.149454 | 0.3998 | 0.917257 |
| ENSMUSG( | Lmcd1 | 370.2429 | -0.40679 | 0.001292 | 0.039144 | ENSMUSG( | Lmcd1 | 309.4018 | -0.72209 | 3.12E-06 | 0.001517 |
| ENSMUSG( | Lmf1 | 758.7076 | -0.18154 | 0.171211 | 0.605749 | ENSMUSG( | Lmf1 | 767.1651 | 0.127239 | 0.415921 | 0.917257 |
| ENSMUSG( | Lmf2 | 964.6201 | -0.246 | 0.101819 | 0.486216 | ENSMUSG( | Lmf2 | 992.8612 | 0.120094 | 0.501217 | 0.927825 |
| ENSMUSG( | Lmln | 472.0936 | -0.16624 | 0.297246 | 0.745805 | ENSMUSG( | Lmln | 488.7151 | 0.208025 | 0.269001 | 0.886755 |
| ENSMUSG( | Lmna | 797.5262 | -0.19701 | 0.086149 | 0.451605 | ENSMUSG( | Lmna | 768.0912 | -0.03347 | 0.796024 | 0.97681 |
| ENSMUSG( | Lmnb1 | 300.354 | 0.075628 | 0.562575 | 0.889465 | ENSMUSG( | Lmnb1 | 250.389 | -0.18319 | 0.265569 | 0.886134 |
| ENSMUSG( | Lmnb2 | 521.4737 | 0.123075 | 0.312038 | 0.759715 | ENSMUSG( | Lmnb2 | 466.9778 | 0.076523 | 0.617005 | 0.951666 |
| ENSMUSG( | Lmntd1 | 46.79773 | -0.11764 | 0.671814 | 0.928781 | ENSMUSG( | Lmntd1 | 41.75009 | -0.16701 | 0.590831 | 0.944975 |
| ENSMUSG( | Lmo1 | 431.5938 | -0.15961 | 0.188303 | 0.627268 | ENSMUSG( | Lmo1 | 377.9518 | -0.29242 | 0.06868 | 0.753052 |
| ENSMUSG( | Lmo2 | 566.7101 | -0.32453 | 0.002117 | 0.053164 | ENSMUSG( | Lmo2 | 580.1785 | 0.03226 | 0.795417 | 0.97681 |
| ENSMUSG( | Lmo3 | 2333.208 | -0.10516 | 0.419123 | 0.821966 | ENSMUSG( | Lmo3 | 2212.259 | 0.008595 | 0.959476 | 0.994798 |
| ENSMUSG( | Lmo4 | 4664.975 | 0.000472 | 0.996565 | 0.99943 | ENSMUSG( | Lmo4 | 4161.213 | -0.06383 | 0.697967 | 0.960867 |
| ENSMUSG( | Lmo7 | 2107.068 | -0.22342 | 0.035515 | 0.290108 | ENSMUSG( | Lmo7 | 2135.924 | 0.095638 | 0.494345 | 0.927372 |
| ENSMUSG( | Lmod1 | 28.04371 | 0.264362 | 0.476455 | 0.851672 | ENSMUSG( | Lmod1 | 22.19559 | -0.1395 | 0.742891 | 0.969415 |
| ENSMUSG( | Lmtk2 | 6179.323 | -0.28351 | 0.05246 | 0.35151 | ENSMUSG( | Lmtk2 | 6311.639 | 0.064408 | 0.71336 | 0.964766 |
| ENSMUSG( | Lmtk3 | 10188.99 | -0.11612 | 0.307254 | 0.755324 | ENSMUSG( | Lmtk3 | 10538.4 | 0.244182 | 0.049424 | 0.708648 |
| ENSMUSG( | Lmx1b | 22.28284 | 0.227078 | 0.590567 | 0.90113 | ENSMUSG( | Lmx1b | 21.80952 | 0.424482 | 0.310603 | 0.899422 |
| ENSMUSG( | Lncpint | 18.4458 | -0.05829 | 0.892775 | NA | ENSMUSG( | Lncpint | 15.83479 | -0.2093 | 0.67789 | 0.957166 |
| ENSMUSG( | Lnp1 | 12.81447 | -0.71078 | 0.25432 | NA | ENSMUSG( | Lnp1 | 13.99006 | -0.19758 | 0.76032 | 0.972575 |
| ENSMUSG( | Lnpep | 2719.266 | -0.24167 | 0.135103 | 0.548658 | ENSMUSG( | Lnpep | 2855.001 | 0.174569 | 0.324894 | 0.905257 |
| ENSMUSG( | Lnpk | 688.0998 | -0.60832 | 2.77E-05 | 0.002487 | ENSMUSG( | Lnpk | 830.3545 | 0.255375 | 0.129958 | 0.820944 |
| ENSMUSG( | Lnx1 | 761.6193 | -0.1048 | 0.437772 | 0.830072 | ENSMUSG( | Lnx1 | 767.7327 | 0.189586 | 0.229722 | 0.868507 |
| ENSMUSG( | Lnx2 | 183.546 | 0.123866 | 0.480868 | 0.855113 | ENSMUSG( | Lnx2 | 175.9658 | 0.276776 | 0.174899 | 0.845107 |
| ENSMUSG( | LOC115486 | 22.22154 | 0.373206 | 0.338558 | 0.777489 | ENSMUSG( | LOC115486 | 14.86554 | -0.55233 | 0.272379 | 0.888825 |
| ENSMUSG( | LOC118567 | 112.8012 | -0.19287 | 0.365467 | 0.794085 | ENSMUSG( | LOC118567 | 112.6727 | 0.092454 | 0.701866 | 0.962014 |
| ENSMUSG( | LOC118568 | 462.6456 | -0.00802 | 0.946779 | 0.993731 | ENSMUSG( | LOC118568 | 373.9657 | -0.39016 | 0.062 | 0.736482 |
| ENSMUSG( | LOC118568 | 33.60451 | -0.39767 | 0.258861 | 0.709814 | ENSMUSG( | LOC118568 | 29.66008 | -0.51583 | 0.233983 | 0.86917 |
| ENSMUSG( | Lockd | 16.83297 | -0.0873 | 0.856199 | NA | ENSMUSG( | Lockd | 13.39207 | -0.49447 | 0.384048 | 0.917257 |
| ENSMUSG( | Lonp1 | 3432.884 | 0.144906 | 0.066095 | 0.397686 | ENSMUSG( | Lonp1 | 2862.91 | -0.11058 | 0.306077 | 0.898538 |
| ENSMUSG( | Lonp2 | 2010.974 | 0.150757 | 0.10846 | 0.497337 | ENSMUSG( | Lonp2 | 1659.352 | -0.13663 | 0.240951 | 0.876146 |
| ENSMUSG( | Lonrf1 | 533.8551 | 0.045219 | 0.758605 | 0.952113 | ENSMUSG( | Lonrf1 | 487.9803 | 0.048238 | 0.781135 | 0.976106 |
| ENSMUSG( | Lonrf2 | 3578.161 | 0.219411 | 0.011654 | 0.155574 | ENSMUSG( | Lonrf2 | 2914.708 | -0.09889 | 0.37965 | 0.916995 |
| ENSMUSG( | Lonrf3 | 509.1665 | 0.397459 | 0.001607 | 0.044935 | ENSMUSG( | Lonrf3 | 429.7989 | 0.197046 | 0.157779 | 0.837724 |
| ENSMUSG( | Loricrin | 118.0175 | -0.00733 | 0.975757 | 0.997428 | ENSMUSG( | Loricrin | 103.604 | -0.13375 | 0.614536 | 0.951177 |
| ENSMUSG( | Lox | 72.21756 | -0.91259 | 0.00032 | 0.015127 | ENSMUSG( | Lox | 66.78377 | -0.84064 | 0.002193 | 0.183242 |
| ENSMUSG( | Loxl1 | 201.3899 | -0.59705 | 0.001403 | 0.040991 | ENSMUSG( | Loxl1 | 199.3402 | -0.30277 | 0.09666 | 0.796245 |
| ENSMUSG( | Loxl2 | 50.18462 | -0.03033 | 0.914162 | 0.988483 | ENSMUSG( | Loxl2 | 40.26238 | -0.40087 | 0.230438 | 0.868507 |
| ENSMUSG( | Loxl3 | 372.6351 | -0.24368 | 0.091142 | 0.463717 | ENSMUSG( | Loxl3 | 392.9719 | 0.18693 | 0.227902 | 0.868507 |
| ENSMUSG( | Lpar1 | 5124.578 | -0.17301 | 0.094543 | 0.471713 | ENSMUSG( | Lpar1 | 4524.121 | -0.27341 | 0.035889 | 0.668395 |
| ENSMUSG( | Lpar2 | 44.35525 | 0.619607 | 0.033026 | 0.278135 | ENSMUSG( | Lpar2 | 37.80766 | 0.452163 | 0.166482 | 0.840181 |
| ENSMUSG( | Lpar3 | 20.91404 | 0.181538 | 0.654987 | NA | ENSMUSG( | Lpar3 | 24.46282 | 0.794168 | 0.048704 | 0.706195 |
| ENSMUSG( | Lpar4 | 59.75995 | -0.4848 | 0.102551 | 0.48808 | ENSMUSG( | Lpar4 | 67.613 | 0.181041 | 0.535947 | 0.936952 |
| ENSMUSG( | Lpar5 | 15.2362 | -0.2049 | 0.673468 | NA | ENSMUSG( | Lpar5 | 16.17721 | 0.256887 | 0.594124 | 0.945408 |
| ENSMUSG( | Lpar6 | 155.3819 | -0.43446 | 0.011943 | 0.157992 | ENSMUSG( | Lpar6 | 214.0682 | 0.71883 | 0.001754 | 0.155216 |
| ENSMUSG( | Lpcat1 | 2147.1 | -0.15947 | 0.236675 | 0.684086 | ENSMUSG( | Lpcat1 | 2119.235 | 0.079087 | 0.608789 | 0.950121 |
| ENSMUSG( | Lpcat2 | 728.2302 | -0.36799 | 0.008036 | 0.124826 | ENSMUSG( | Lpcat2 | 763.2368 | 0.065634 | 0.714366 | 0.964986 |
| ENSMUSG( | Lpcat3 | 850.4179 | -0.25486 | 0.140593 | 0.557902 | ENSMUSG( | Lpcat3 | 869.2318 | 0.094909 | 0.641063 | 0.953903 |
| ENSMUSG( | Lpcat4 | 4186.761 | -0.25337 | 0.021745 | 0.225516 | ENSMUSG( | Lpcat4 | 4159.127 | 0.009327 | 0.938701 | 0.993898 |
| ENSMUSG( | Lpgat1 | 5524.85 | -0.16893 | 0.208966 | 0.650854 | ENSMUSG( | Lpgat1 | 5626.805 | 0.156355 | 0.313896 | 0.900949 |
| ENSMUSG( | Lpin1 | 857.4859 | -0.0427 | 0.695559 | 0.934979 | ENSMUSG( | Lpin1 | 800.1976 | 0.030671 | 0.81173 | 0.979148 |
| ENSMUSG( | Lpin2 | 2153.014 | 0.249509 | 0.008352 | 0.127563 | ENSMUSG( | Lpin2 | 1837.932 | 0.073048 | 0.515311 | 0.930939 |
| ENSMUSG( | Lpin3 | 58.617 | 0.230013 | 0.392575 | 0.809274 | ENSMUSG( | Lpin3 | 58.02914 | 0.451656 | 0.102117 | 0.802142 |
| ENSMUSG( | Lpl | 657.665 | -0.39162 | 0.037304 | 0.298061 | ENSMUSG( | Lpl | 712.2214 | 0.132117 | 0.532874 | 0.934795 |
| ENSMUSG( | Lpo | 52.72068 | 0.121201 | 0.656695 | 0.923929 | ENSMUSG( | Lpo | 45.71423 | -0.01131 | 0.970557 | 0.99594 |
| ENSMUSG( | Lpp | 479.3861 | 0.096896 | 0.453546 | 0.836798 | ENSMUSG( | Lpp | 427.8215 | 0.031148 | 0.832331 | 0.98075 |
| ENSMUSG( | Lppos | 465.4076 | 0.225311 | 0.126544 | 0.534155 | ENSMUSG( | Lppos | 366.4997 | -0.19999 | 0.187314 | 0.856372 |
| ENSMUSG( | Lpxn | 62.00851 | 0.301548 | 0.270087 | 0.719904 | ENSMUSG( | Lpxn | 61.50509 | 0.517364 | 0.058241 | 0.728256 |
| ENSMUSG( | Lrat | 15.65424 | -0.51603 | 0.267546 | NA | ENSMUSG( | Lrat | 14.92939 | -0.41391 | 0.422878 | 0.917257 |
| ENSMUSG( | Lratd1 | 898.3813 | 0.110433 | 0.376341 | 0.801127 | ENSMUSG( | Lratd1 | 818.7934 | 0.110184 | 0.417408 | 0.917257 |
| ENSMUSG( | Lratd2 | 282.3534 | -0.2332 | 0.081452 | 0.441102 | ENSMUSG( | Lratd2 | 291.9439 | 0.143854 | 0.356392 | 0.913358 |
| ENSMUSG( | Lrba | 1304.366 | 0.045318 | 0.67359 | 0.929191 | ENSMUSG( | Lrba | 1125.546 | -0.11357 | 0.324458 | 0.905257 |
| ENSMUSG( | Lrch1 | 587.3287 | 0.235634 | 0.033557 | 0.280855 | ENSMUSG( | Lrch1 | 503.5624 | 0.071891 | 0.578533 | 0.943548 |
| ENSMUSG( | Lrch2 | 428.6292 | -0.12896 | 0.304708 | 0.752574 | ENSMUSG( | Lrch2 | 404.0361 | -0.03711 | 0.819455 | 0.979198 |
| ENSMUSG( | Lrch3 | 1314.743 | 0.098941 | 0.257196 | 0.708432 | ENSMUSG( | Lrch3 | 1209.158 | 0.127662 | 0.300517 | 0.898538 |
| ENSMUSG( | Lrfn1 | 1980.921 | -0.15972 | 0.212982 | 0.656982 | ENSMUSG( | Lrfn1 | 2037.018 | 0.191391 | 0.206625 | 0.864412 |
| ENSMUSG( | Lrfn2 | 723.0316 | 0.386559 | 0.009244 | 0.135575 | ENSMUSG( | Lrfn2 | 738.0141 | 0.67071 | 0.000606 | 0.076367 |
| ENSMUSG( | Lrfn3 | 734.1605 | -0.12225 | 0.442842 | 0.83278 | ENSMUSG( | Lrfn3 | 765.9798 | 0.267938 | 0.167267 | 0.840222 |
| ENSMUSG( | Lrfn4 | 1416.376 | -0.17394 | 0.194313 | 0.633391 | ENSMUSG( | Lrfn4 | 1428.507 | 0.126563 | 0.358489 | 0.913759 |
| ENSMUSG( | Lrfn5 | 1266.373 | -0.2619 | 0.071054 | 0.411459 | ENSMUSG( | Lrfn5 | 1344.745 | 0.191845 | 0.263556 | 0.886134 |
| ENSMUSG( | Lrg1 | 42.11688 | 0.233845 | 0.427999 | 0.825894 | ENSMUSG( | Lrg1 | 35.18588 | 0.005539 | 0.987033 | 0.997487 |
| ENSMUSG( | Lrguk | 105.2576 | -0.39758 | 0.041826 | 0.315339 | ENSMUSG( | Lrguk | 102.8708 | -0.17794 | 0.403296 | 0.917257 |
| ENSMUSG( | Lrif1 | 241.9108 | -0.14859 | 0.37666 | 0.801213 | ENSMUSG( | Lrif1 | 251.4591 | 0.229615 | 0.201555 | 0.864412 |
| ENSMUSG( | Lrig1 | 2007.32 | -0.13816 | 0.230868 | 0.679066 | ENSMUSG( | Lrig1 | 2036.94 | 0.172384 | 0.206453 | 0.864412 |
| ENSMUSG( | Lrig2 | 1018.55 | -0.13137 | 0.363983 | 0.793415 | ENSMUSG( | Lrig2 | 1014.958 | 0.13256 | 0.403959 | 0.917257 |
| ENSMUSG( | Lrig3 | 92.50446 | -0.19464 | 0.386768 | 0.805046 | ENSMUSG( | Lrig3 | 107.4978 | 0.497172 | 0.085801 | 0.780123 |
| ENSMUSG( | Lrmda | 24.32224 | -0.52082 | 0.18636 | 0.623869 | ENSMUSG( | Lrmda | 21.64377 | -0.62114 | 0.177254 | 0.846973 |
| ENSMUSG( | Lrp1 | 24020.42 | -0.08067 | 0.470009 | 0.846591 | ENSMUSG( | Lrp1 | 23154.8 | 0.084435 | 0.498234 | 0.927825 |
| ENSMUSG( | Lrp10 | 1845.767 | -0.13533 | 0.310449 | 0.758585 | ENSMUSG( | Lrp10 | 1851.793 | 0.14652 | 0.345298 | 0.91052 |
| ENSMUSG( | Lrp11 | 4889.804 | -0.34911 | 0.037957 | 0.300779 | ENSMUSG( | Lrp11 | 5193.841 | 0.117759 | 0.517753 | 0.930939 |
| ENSMUSG( | Lrp12 | 1366.16 | -0.53349 | 0.001723 | 0.046889 | ENSMUSG( | Lrp12 | 1560.255 | 0.168955 | 0.387584 | 0.917257 |
| ENSMUSG( | Lrp1b | 846.9353 | -0.49248 | 0.001044 | 0.033922 | ENSMUSG( | Lrp1b | 997.4653 | 0.283085 | 0.095447 | 0.796133 |
| ENSMUSG( | Lrp2 | 20.01849 | 0.037491 | 0.933174 | NA | ENSMUSG( | Lrp2 | 22.50857 | 0.632841 | 0.194236 | 0.860096 |

| | | | | | |
|---|---|---|---|---|---|
| ENSMUSG( | 1522.293 | 0.06139 | 0.635233 | 0.914296 | Lin52 |
| ENSMUSG( | 311.0862 | 0.002648 | 0.98475 | 0.997815 | Lin54 |
| ENSMUSG( | 404.74 | 0.22631 | 0.11371 | 0.508792 | Lin7a |
| ENSMUSG( | 1125.607 | -0.09387 | 0.508 | 0.867981 | Lin7b |
| ENSMUSG( | 2770.141 | 0.001454 | 0.990785 | 0.999232 | Lin7c |
| ENSMUSG( | 85.9858 | 0.094611 | 0.710192 | 0.9397 | Lin9 |
| ENSMUSG( | 10829.18 | -0.1229 | 0.43612 | 0.829093 | Lingo1 |
| ENSMUSG( | 465.4133 | -0.63811 | 0.000682 | 0.025185 | Lingo2 |
| ENSMUSG( | 1305.633 | -0.52266 | 0.000133 | 0.007813 | Lingo3 |
| ENSMUSG( | 64.1774 | 0.274642 | 0.282472 | 0.73186 | Lingo4 |
| ENSMUSG( | 281.653 | -0.04178 | 0.78334 | 0.957558 | Lins1 |
| ENSMUSG( | 750.9055 | -0.35539 | 0.038025 | 0.300991 | Lipa |
| ENSMUSG( | 12.51727 | 0.041901 | 0.938744 | NA | Lipc |
| ENSMUSG( | 350.1032 | 0.03786 | 0.767967 | 0.954621 | Lipe |
| ENSMUSG( | 116.7411 | -0.41135 | 0.06009 | 0.379098 | Lipg |
| ENSMUSG( | 33.77942 | -0.3518 | 0.330083 | 0.774055 | Liph |
| ENSMUSG( | 127.0818 | -0.36349 | 0.102133 | 0.486947 | Lipo3 |
| ENSMUSG( | 75.2576 | 0.076169 | 0.741438 | 0.948208 | Lipt1 |
| ENSMUSG( | 90.79703 | 0.013673 | 0.947924 | 0.993821 | Lipt2 |
| ENSMUSG( | 763.9064 | -0.1971 | 0.080614 | 0.438981 | Litaf |
| ENSMUSG( | 4289.444 | 0.093533 | 0.490955 | 0.859739 | Lix1 |
| ENSMUSG( | 1758.937 | 0.021374 | 0.850982 | 0.972638 | Lix1l |
| ENSMUSG( | 4646.949 | 0.034719 | 0.676596 | 0.930542 | Llgl1 |
| ENSMUSG( | 46.87229 | 0.136494 | 0.638112 | 0.916107 | Llgl2 |
| ENSMUSG( | 318.2285 | 0.047341 | 0.776716 | 0.955723 | Llph |
| ENSMUSG( | 1542.057 | -0.32648 | 0.033117 | 0.278602 | Lman1 |
| ENSMUSG( | 3438.099 | -0.09232 | 0.456896 | 0.839183 | Lman2 |
| ENSMUSG( | 1619.39 | 0.001712 | 0.989255 | 0.999111 | Lman2l |
| ENSMUSG( | 919.6806 | -0.26127 | 0.063672 | 0.390618 | Lmbr1 |
| ENSMUSG( | 314.7334 | -0.03332 | 0.834335 | 0.970333 | Lmbr1l |
| ENSMUSG( | 7963.235 | -0.17476 | 0.177395 | 0.614553 | Lmbrd1 |
| ENSMUSG( | 2589.302 | -0.43739 | 0.003896 | 0.080172 | Lmbrd2 |
| ENSMUSG( | 370.2429 | -0.40679 | 0.001292 | 0.039144 | Lmcd1 |
| ENSMUSG( | 758.7076 | -0.18154 | 0.171211 | 0.605749 | Lmf1 |
| ENSMUSG( | 964.6201 | -0.246 | 0.101819 | 0.486216 | Lmf2 |
| ENSMUSG( | 472.0936 | -0.16624 | 0.297246 | 0.745805 | Lmln |
| ENSMUSG( | 797.5262 | -0.19701 | 0.086149 | 0.451605 | Lmna |
| ENSMUSG( | 300.354 | 0.075628 | 0.562575 | 0.889465 | Lmnb1 |
| ENSMUSG( | 521.4737 | 0.123075 | 0.312038 | 0.759715 | Lmnb2 |
| ENSMUSG( | 46.79773 | -0.11764 | 0.671814 | 0.928781 | Lmntd1 |
| ENSMUSG( | 431.5938 | -0.15961 | 0.188303 | 0.627268 | Lmo1 |
| ENSMUSG( | 566.7101 | -0.32453 | 0.002117 | 0.053164 | Lmo2 |
| ENSMUSG( | 2333.208 | -0.10516 | 0.419123 | 0.821966 | Lmo3 |
| ENSMUSG( | 4664.975 | 0.000472 | 0.996565 | 0.99943 | Lmo4 |
| ENSMUSG( | 2107.068 | -0.22342 | 0.035515 | 0.290108 | Lmo7 |
| ENSMUSG( | 28.04371 | 0.264362 | 0.476455 | 0.851672 | Lmod1 |
| ENSMUSG( | 6179.323 | -0.28351 | 0.05246 | 0.35151 | Lmtk2 |
| ENSMUSG( | 10188.99 | -0.11612 | 0.307254 | 0.755324 | Lmtk3 |
| ENSMUSG( | 22.28284 | 0.227078 | 0.590567 | 0.90113 | Lmx1b |
| ENSMUSG( | 18.4458 | -0.05829 | 0.892775 | NA | Lncpint |
| ENSMUSG( | 12.81447 | -0.71078 | 0.25432 | NA | Lnp1 |
| ENSMUSG( | 2719.266 | -0.24167 | 0.135103 | 0.548658 | Lnpep |
| ENSMUSG( | 688.0998 | -0.60832 | 2.77E-05 | 0.002487 | Lnpk |
| ENSMUSG( | 761.6193 | -0.1048 | 0.437772 | 0.830072 | Lnx1 |
| ENSMUSG( | 183.546 | 0.123866 | 0.480868 | 0.855113 | Lnx2 |
| ENSMUSG( | 8.268908 | -0.05953 | 0.930416 | NA | LOC108167922 |
| ENSMUSG( | 22.22154 | 0.373206 | 0.338558 | 0.777489 | LOC115486416 |
| ENSMUSG( | 112.8012 | -0.19287 | 0.365467 | 0.794085 | LOC118567373 |
| ENSMUSG( | 13.19825 | 0.038974 | 0.941561 | NA | LOC118567674 |
| ENSMUSG( | 462.6456 | -0.00802 | 0.946779 | 0.993731 | LOC118568041 |
| ENSMUSG( | 33.60451 | -0.39767 | 0.258861 | 0.709814 | LOC118568483 |
| ENSMUSG( | 16.83297 | -0.0873 | 0.856199 | NA | Lockd |
| ENSMUSG( | 3432.884 | 0.144906 | 0.066095 | 0.397686 | Lonp1 |
| ENSMUSG( | 2010.974 | 0.150757 | 0.10846 | 0.497337 | Lonp2 |
| ENSMUSG( | 533.8551 | 0.045219 | 0.758605 | 0.952113 | Lonrf1 |
| ENSMUSG( | 3578.161 | 0.219411 | 0.011654 | 0.155574 | Lonrf2 |
| ENSMUSG( | 509.1665 | 0.397459 | 0.001607 | 0.044935 | Lonrf3 |
| ENSMUSG( | 118.0175 | -0.00733 | 0.975757 | 0.997428 | Loricrin |
| ENSMUSG( | 72.21756 | -0.91259 | 0.00032 | 0.015127 | Lox |
| ENSMUSG( | 201.3899 | -0.59705 | 0.001403 | 0.040991 | Loxl1 |
| ENSMUSG( | 50.18462 | -0.03033 | 0.914162 | 0.988483 | Loxl2 |
| ENSMUSG( | 372.6351 | -0.24368 | 0.091142 | 0.463717 | Loxl3 |
| ENSMUSG( | 5124.578 | -0.17301 | 0.094543 | 0.471713 | Lpar1 |
| ENSMUSG( | 44.35525 | 0.619607 | 0.033026 | 0.278135 | Lpar2 |
| ENSMUSG( | 20.91404 | 0.181538 | 0.654987 | NA | Lpar3 |
| ENSMUSG( | 59.75995 | -0.4848 | 0.102551 | 0.48808 | Lpar4 |
| ENSMUSG( | 15.2362 | -0.2049 | 0.673468 | NA | Lpar5 |
| ENSMUSG( | 155.3819 | -0.43446 | 0.011943 | 0.157992 | Lpar6 |
| ENSMUSG( | 2147.1 | -0.15947 | 0.236675 | 0.684086 | Lpcat1 |
| ENSMUSG( | 728.2302 | -0.36799 | 0.008036 | 0.124826 | Lpcat2 |
| ENSMUSG( | 850.4179 | -0.25486 | 0.140593 | 0.557902 | Lpcat3 |
| ENSMUSG( | 4186.761 | -0.25337 | 0.021745 | 0.225516 | Lpcat4 |
| ENSMUSG( | 5524.85 | -0.16893 | 0.208966 | 0.650854 | Lpgat1 |
| ENSMUSG( | 857.4859 | -0.0427 | 0.695559 | 0.934979 | Lpin1 |
| ENSMUSG( | 2153.014 | 0.249509 | 0.008352 | 0.127563 | Lpin2 |
| ENSMUSG( | 58.617 | 0.230013 | 0.392575 | 0.809274 | Lpin3 |
| ENSMUSG( | 657.665 | -0.39162 | 0.037304 | 0.298061 | Lpl |
| ENSMUSG( | 52.72068 | 0.121201 | 0.656695 | 0.923929 | Lpo |
| ENSMUSG( | 479.3861 | 0.096896 | 0.453546 | 0.836798 | Lpp |
| ENSMUSG( | 465.4076 | 0.225311 | 0.126544 | 0.534155 | Lppos |
| ENSMUSG( | 62.00851 | 0.301548 | 0.270087 | 0.719904 | Lpxn |
| ENSMUSG( | 15.65424 | -0.51603 | 0.267546 | NA | Lrat |
| ENSMUSG( | 898.3813 | 0.110433 | 0.376341 | 0.801127 | Lratd1 |
| ENSMUSG( | 282.3534 | -0.2332 | 0.081452 | 0.441102 | Lratd2 |
| ENSMUSG( | 1304.366 | 0.045318 | 0.67359 | 0.929191 | Lrba |
| ENSMUSG( | 587.3287 | 0.235634 | 0.033557 | 0.280855 | Lrch1 |
| ENSMUSG( | 428.6292 | -0.12896 | 0.304708 | 0.752574 | Lrch2 |
| ENSMUSG( | 1314.743 | 0.098941 | 0.257196 | 0.708432 | Lrch3 |
| ENSMUSG( | 1980.921 | -0.15972 | 0.212982 | 0.656982 | Lrfn1 |
| ENSMUSG( | 723.0316 | 0.386559 | 0.009244 | 0.135575 | Lrfn2 |
| ENSMUSG( | 734.1605 | -0.12225 | 0.442842 | 0.83278 | Lrfn3 |
| ENSMUSG( | 1416.376 | -0.17394 | 0.194313 | 0.633391 | Lrfn4 |
| ENSMUSG( | 1266.373 | -0.2619 | 0.071054 | 0.411459 | Lrfn5 |
| ENSMUSG( | 42.11688 | 0.233845 | 0.427999 | 0.825894 | Lrg1 |
| ENSMUSG( | 105.2576 | -0.39758 | 0.041826 | 0.315339 | Lrguk |
| ENSMUSG( | 241.9108 | -0.14859 | 0.37666 | 0.801213 | Lrif1 |
| ENSMUSG( | 2007.32 | -0.13816 | 0.230868 | 0.679066 | Lrig1 |
| ENSMUSG( | 1018.55 | -0.13137 | 0.363983 | 0.793415 | Lrig2 |
| ENSMUSG( | 92.50446 | -0.19464 | 0.386768 | 0.805046 | Lrig3 |
| ENSMUSG( | 24.32224 | -0.52082 | 0.18636 | 0.623869 | Lrmda |
| ENSMUSG( | 24020.42 | -0.08067 | 0.470009 | 0.846591 | Lrp1 |
| ENSMUSG( | 1845.767 | -0.13533 | 0.310449 | 0.758585 | Lrp10 |
| ENSMUSG( | 4889.804 | -0.34911 | 0.037957 | 0.300779 | Lrp11 |
| ENSMUSG( | 1366.16 | -0.53349 | 0.001723 | 0.046889 | Lrp12 |
| ENSMUSG( | 846.9353 | -0.49248 | 0.001044 | 0.033922 | Lrp1b |
| ENSMUSG( | 20.01849 | 0.037491 | 0.933174 | NA | Lrp2 |
| ENSMUSG( | 10.63053 | -0.53026 | 0.375526 | NA | Lrp2bp |
| ENSMUSG( | 2782.933 | -0.02195 | 0.861965 | 0.975055 | Lrp3 |
| ENSMUSG( | 5776.597 | 0.053864 | 0.693585 | 0.934202 | Lrp4 |
| ENSMUSG( | 836.9917 | -0.08746 | 0.566608 | 0.891644 | Lrp5 |
| ENSMUSG( | 2270.372 | -0.02189 | 0.846262 | 0.972226 | Lrp6 |
| ENSMUSG( | 3810.014 | -0.02717 | 0.803611 | 0.962169 | Lrp8 |
| ENSMUSG( | 29.35044 | -0.75068 | 0.140182 | 0.557263 | Lrp8os2 |
| ENSMUSG( | 32.51542 | 0.320098 | 0.335011 | 0.776499 | Lrp8os3 |
| ENSMUSG( | 6971.033 | -0.30634 | 0.012292 | 0.160458 | Lrpap1 |
| ENSMUSG( | 1956.684 | 0.240221 | 0.025033 | 0.243138 | Lrpprc |
| ENSMUSG( | 332.3774 | 0.009557 | 0.943041 | 0.993415 | Lrrc1 |
| ENSMUSG( | 1702.683 | -0.58462 | 0.003953 | 0.080551 | Lrrc10b |
| ENSMUSG( | 614.6826 | 0.045226 | 0.709605 | 0.939669 | Lrrc14 |
| ENSMUSG( | 10.62036 | -0.82032 | 0.153786 | NA | Lrrc14b |
| ENSMUSG( | 12.40656 | -0.52786 | 0.356382 | NA | Lrrc18 |
| ENSMUSG( | 178.0261 | 0.072284 | 0.727725 | 0.944355 | Lrrc2 |
| ENSMUSG( | 782.5943 | 0.028282 | 0.796097 | 0.960088 | Lrrc20 |
| ENSMUSG( | 80.98117 | 0.314114 | 0.174529 | 0.610304 | Lrrc23 |
| ENSMUSG( | 1219.471 | -0.36744 | 0.002055 | 0.052087 | Lrrc24 |
| ENSMUSG( | 69.73372 | -0.17013 | 0.48108 | 0.855135 | Lrrc25 |
| ENSMUSG( | 217.0274 | 0.134821 | 0.403738 | 0.816132 | Lrrc27 |
| ENSMUSG( | 535.6603 | -0.08028 | 0.467227 | 0.844819 | Lrrc28 |
| ENSMUSG( | 516.0241 | -0.36528 | 0.025825 | 0.247245 | Lrrc3 |
| ENSMUSG( | 161.9058 | -0.43645 | 0.044363 | 0.323404 | Lrrc32 |
| ENSMUSG( | 30.77794 | -0.38803 | 0.267238 | 0.717525 | Lrrc36 |
| ENSMUSG( | 67.62933 | -0.27849 | 0.269797 | 0.719729 | Lrrc38 |
| ENSMUSG( | 10.26321 | -0.48154 | 0.423716 | NA | Lrrc39 |
| ENSMUSG( | 314.6161 | -0.32083 | 0.013376 | 0.168706 | Lrrc3b |
| ENSMUSG( | 2510.485 | -0.17997 | 0.200148 | 0.640302 | Lrrc4 |
| ENSMUSG( | 541.9678 | 0.085392 | 0.49046 | 0.859379 | Lrrc40 |
| ENSMUSG( | 1179.667 | 0.060376 | 0.541676 | 0.883182 | Lrrc41 |
| ENSMUSG( | 543.5561 | 0.03831 | 0.743598 | 0.948478 | Lrrc42 |
| ENSMUSG( | 15.35875 | 0.578532 | 0.29228 | NA | Lrrc43 |
| ENSMUSG( | 1643.936 | 0.358151 | 0.044841 | 0.324745 | Lrrc45 |
| ENSMUSG( | 21.50253 | -0.3495 | 0.461347 | 0.841432 | Lrrc46 |
| ENSMUSG( | 1189.788 | 0.073712 | 0.531448 | 0.879512 | Lrrc47 |
| ENSMUSG( | 1565.272 | 0.043219 | 0.717455 | 0.941697 | Lrrc49 |
| ENSMUSG( | 6315.761 | -0.29673 | 0.018382 | 0.203658 | Lrrc4b |
| ENSMUSG( | 2344.356 | -0.45525 | 0.001358 | 0.040233 | Lrrc4c |
| ENSMUSG( | 143.2098 | 0.145698 | 0.392164 | 0.809274 | Lrrc51 |
| ENSMUSG( | 725.3828 | -0.51894 | 0.001281 | 0.039067 | Lrrc55 |
| ENSMUSG( | 167.8644 | -0.09031 | 0.573121 | 0.893584 | Lrrc56 |

| | | | | | |
|---|---|---|---|---|---|
| ENSMUSG( | 13.99006 | -0.19758 | 0.76032 | 0.972575 | Lnp1 |
| ENSMUSG( | 2855.001 | 0.174569 | 0.324894 | 0.905257 | Lnpep |
| ENSMUSG( | 830.3545 | 0.255375 | 0.129958 | 0.820944 | Lnpk |
| ENSMUSG( | 767.7327 | 0.189586 | 0.229722 | 0.868507 | Lnx1 |
| ENSMUSG( | 175.9658 | 0.276776 | 0.174899 | 0.845107 | Lnx2 |
| ENSMUSG( | 14.86554 | -0.55233 | 0.272379 | 0.888825 | LOC115486416 |
| ENSMUSG( | 112.6727 | 0.092454 | 0.701866 | 0.962014 | LOC118567373 |
| ENSMUSG( | 373.9657 | -0.39016 | 0.062 | 0.736482 | LOC118568041 |
| ENSMUSG( | 29.66008 | -0.51583 | 0.233983 | 0.86917 | LOC118568483 |
| ENSMUSG( | 13.39207 | -0.49447 | 0.384048 | 0.917257 | Lockd |
| ENSMUSG( | 2862.91 | -0.11058 | 0.306077 | 0.898538 | Lonp1 |
| ENSMUSG( | 1659.352 | -0.13663 | 0.240951 | 0.876146 | Lonp2 |
| ENSMUSG( | 487.9803 | 0.048238 | 0.781135 | 0.976106 | Lonrf1 |
| ENSMUSG( | 2914.708 | -0.09889 | 0.37965 | 0.916995 | Lonrf2 |
| ENSMUSG( | 429.7989 | 0.197046 | 0.157779 | 0.837724 | Lonrf3 |
| ENSMUSG( | 103.604 | -0.13375 | 0.614536 | 0.951177 | Loricrin |
| ENSMUSG( | 66.78377 | -0.84064 | 0.002193 | 0.183242 | Lox |
| ENSMUSG( | 199.3402 | -0.30277 | 0.09666 | 0.796245 | Loxl1 |
| ENSMUSG( | 40.26238 | -0.40087 | 0.230438 | 0.868507 | Loxl2 |
| ENSMUSG( | 392.9719 | 0.18693 | 0.227902 | 0.868507 | Loxl3 |
| ENSMUSG( | 4524.121 | -0.27341 | 0.035889 | 0.668395 | Lpar1 |
| ENSMUSG( | 37.80766 | 0.452163 | 0.166482 | 0.840181 | Lpar2 |
| ENSMUSG( | 24.46282 | 0.794168 | 0.048704 | 0.706195 | Lpar3 |
| ENSMUSG( | 67.613 | 0.181041 | 0.535947 | 0.936952 | Lpar4 |
| ENSMUSG( | 16.17721 | 0.256887 | 0.594124 | 0.945408 | Lpar5 |
| ENSMUSG( | 214.0682 | 0.71883 | 0.001754 | 0.155216 | Lpar6 |
| ENSMUSG( | 2119.235 | 0.079087 | 0.608789 | 0.950121 | Lpcat1 |
| ENSMUSG( | 763.2368 | 0.065634 | 0.714366 | 0.964986 | Lpcat2 |
| ENSMUSG( | 869.2318 | 0.094909 | 0.641063 | 0.953903 | Lpcat3 |
| ENSMUSG( | 4159.127 | 0.009327 | 0.938701 | 0.993898 | Lpcat4 |
| ENSMUSG( | 5626.805 | 0.156355 | 0.313896 | 0.900949 | Lpgat1 |
| ENSMUSG( | 800.1976 | 0.030671 | 0.81173 | 0.979148 | Lpin1 |
| ENSMUSG( | 1837.932 | 0.073048 | 0.515311 | 0.930939 | Lpin2 |
| ENSMUSG( | 58.02914 | 0.451656 | 0.102117 | 0.802142 | Lpin3 |
| ENSMUSG( | 712.2214 | 0.132117 | 0.532874 | 0.934795 | Lpl |
| ENSMUSG( | 45.71423 | -0.01131 | 0.970557 | 0.99594 | Lpo |
| ENSMUSG( | 427.8215 | 0.031148 | 0.832331 | 0.98075 | Lpp |
| ENSMUSG( | 366.4997 | -0.19999 | 0.187314 | 0.856372 | Lppos |
| ENSMUSG( | 61.50509 | 0.517364 | 0.058241 | 0.728256 | Lpxn |
| ENSMUSG( | 14.92939 | -0.41391 | 0.422878 | 0.917257 | Lrat |
| ENSMUSG( | 818.7934 | 0.110184 | 0.417408 | 0.917257 | Lratd1 |
| ENSMUSG( | 291.9439 | 0.143854 | 0.356392 | 0.913358 | Lratd2 |
| ENSMUSG( | 1125.546 | -0.11357 | 0.324458 | 0.905257 | Lrba |
| ENSMUSG( | 503.5624 | 0.071891 | 0.578533 | 0.943548 | Lrch1 |
| ENSMUSG( | 404.0361 | -0.03711 | 0.819455 | 0.979198 | Lrch2 |
| ENSMUSG( | 1209.158 | 0.127662 | 0.300517 | 0.898538 | Lrch3 |
| ENSMUSG( | 2037.018 | 0.191391 | 0.206625 | 0.864412 | Lrfn1 |
| ENSMUSG( | 738.0141 | 0.67071 | 0.000606 | 0.076367 | Lrfn2 |
| ENSMUSG( | 765.9798 | 0.267938 | 0.167267 | 0.840222 | Lrfn3 |
| ENSMUSG( | 1428.507 | 0.126563 | 0.358489 | 0.913759 | Lrfn4 |
| ENSMUSG( | 1344.745 | 0.191845 | 0.263556 | 0.886134 | Lrfn5 |
| ENSMUSG( | 35.18588 | 0.005539 | 0.987033 | 0.997487 | Lrg1 |
| ENSMUSG( | 102.8708 | -0.17794 | 0.403296 | 0.917257 | Lrguk |
| ENSMUSG( | 251.4591 | 0.229615 | 0.201555 | 0.864412 | Lrif1 |
| ENSMUSG( | 2036.94 | 0.172384 | 0.206453 | 0.864412 | Lrig1 |
| ENSMUSG( | 1014.958 | 0.13256 | 0.403959 | 0.917257 | Lrig2 |
| ENSMUSG( | 107.4978 | 0.497172 | 0.085801 | 0.780123 | Lrig3 |
| ENSMUSG( | 21.64377 | -0.62114 | 0.177254 | 0.846973 | Lrmda |
| ENSMUSG( | 23154.8 | 0.084435 | 0.498234 | 0.927825 | Lrp1 |
| ENSMUSG( | 1851.793 | 0.14652 | 0.345298 | 0.91052 | Lrp10 |
| ENSMUSG( | 5193.841 | 0.117759 | 0.517753 | 0.930939 | Lrp11 |
| ENSMUSG( | 1560.255 | 0.168955 | 0.387584 | 0.917257 | Lrp12 |
| ENSMUSG( | 997.4653 | 0.283085 | 0.095447 | 0.796133 | Lrp1b |
| ENSMUSG( | 22.50857 | 0.632841 | 0.194236 | 0.860096 | Lrp2 |
| ENSMUSG( | 10.8448 | -0.19253 | 0.752199 | NA | Lrp2bp |
| ENSMUSG( | 2823.481 | 0.279452 | 0.06752 | 0.75212 | Lrp3 |
| ENSMUSG( | 4897.499 | -0.15685 | 0.219739 | 0.866104 | Lrp4 |
| ENSMUSG( | 832.7246 | 0.17092 | 0.357578 | 0.913759 | Lrp5 |
| ENSMUSG( | 2222.841 | 0.181928 | 0.150515 | 0.837173 | Lrp6 |
| ENSMUSG( | 3418.577 | -0.06904 | 0.584778 | 0.944975 | Lrp8 |
| ENSMUSG( | 33.70337 | -0.02562 | 0.946389 | 0.994798 | Lrp8os2 |
| ENSMUSG( | 20.77552 | -0.73835 | 0.132877 | 0.822912 | Lrp8os3 |
| ENSMUSG( | 7209.835 | 0.078205 | 0.583551 | 0.944266 | Lrpap1 |
| ENSMUSG( | 1609.543 | -0.04995 | 0.730747 | 0.967228 | Lrpprc |
| ENSMUSG( | 298.4364 | -0.04264 | 0.814227 | 0.979198 | Lrrc1 |
| ENSMUSG( | 1985.147 | 0.178335 | 0.207243 | 0.864412 | Lrrc10b |
| ENSMUSG( | 525.5872 | -0.13674 | 0.334617 | 0.907907 | Lrrc14 |
| ENSMUSG( | 10.0552 | -0.64922 | 0.285857 | NA | Lrrc14b |
| ENSMUSG( | 9.082865 | 0.322746 | 0.612176 | NA | Lrrc17 |
| ENSMUSG( | 13.79501 | 0.114324 | 0.832091 | 0.980738 | Lrrc18 |
| ENSMUSG( | 149.0585 | -0.19453 | 0.400816 | 0.917257 | Lrrc2 |
| ENSMUSG( | 706.5469 | 0.002996 | 0.980608 | 0.996479 | Lrrc20 |
| ENSMUSG( | 73.14891 | 0.300866 | 0.244686 | 0.877904 | Lrrc23 |
| ENSMUSG( | 1254.298 | 0.007907 | 0.955125 | 0.994798 | Lrrc24 |
| ENSMUSG( | 73.69919 | 0.269121 | 0.342656 | 0.909646 | Lrrc25 |
| ENSMUSG( | 182.7013 | -0.09265 | 0.619906 | 0.952896 | Lrrc27 |
| ENSMUSG( | 479.1931 | -0.13997 | 0.313416 | 0.900883 | Lrrc28 |
| ENSMUSG( | 553.7442 | 0.135702 | 0.466002 | 0.923329 | Lrrc3 |
| ENSMUSG( | 179.6571 | 0.175842 | 0.453471 | 0.922316 | Lrrc32 |
| ENSMUSG( | 31.7168 | 0.015337 | 0.967417 | 0.995667 | Lrrc36 |
| ENSMUSG( | 68.39417 | 0.046787 | 0.852648 | 0.985598 | Lrrc38 |
| ENSMUSG( | 10.07661 | -0.27858 | 0.649197 | NA | Lrrc39 |
| ENSMUSG( | 319.3592 | 0.004795 | 0.975359 | 0.996416 | Lrrc3b |
| ENSMUSG( | 2561.282 | 0.152736 | 0.340911 | 0.907972 | Lrrc4 |
| ENSMUSG( | 477.6878 | -0.01335 | 0.92231 | 0.992644 | Lrrc40 |
| ENSMUSG( | 1082.459 | 0.082179 | 0.472571 | 0.923329 | Lrrc41 |
| ENSMUSG( | 469.7446 | -0.1154 | 0.397275 | 0.917257 | Lrrc42 |
| ENSMUSG( | 14.95531 | 0.745031 | 0.156244 | 0.837724 | Lrrc43 |
| ENSMUSG( | 1440.677 | 0.253003 | 0.207823 | 0.864412 | Lrrc45 |
| ENSMUSG( | 24.14165 | 0.279136 | 0.529795 | 0.934795 | Lrrc46 |
| ENSMUSG( | 1049.839 | -0.02003 | 0.888598 | 0.990558 | Lrrc47 |
| ENSMUSG( | 1358.247 | -0.09891 | 0.436909 | 0.919983 | Lrrc49 |
| ENSMUSG( | 6942.403 | 0.252029 | 0.089294 | 0.788502 | Lrrc4b |
| ENSMUSG( | 2610.592 | 0.15944 | 0.319986 | 0.903771 | Lrrc4c |
| ENSMUSG( | 126.311 | 0.050833 | 0.795717 | 0.97681 | Lrrc51 |
| ENSMUSG( | 768.8459 | -0.03135 | 0.858134 | 0.985884 | Lrrc55 |
| ENSMUSG( | 151.4985 | -0.10855 | 0.569034 | 0.941777 | Lrrc56 |
| ENSMUSG( | 964.9726 | -0.13159 | 0.289041 | 0.895811 | Lrrc57 |
| ENSMUSG( | 5265.254 | 0.149145 | 0.336386 | 0.907972 | Lrrc58 |
| ENSMUSG( | 2191.678 | -0.07182 | 0.502644 | 0.927825 | Lrrc59 |
| ENSMUSG( | 697.898 | -0.18055 | 0.192197 | 0.859726 | Lrrc61 |
| ENSMUSG( | 1360.946 | 0.168016 | 0.171572 | 0.842999 | Lrrc7 |
| ENSMUSG( | 33.06585 | 0.318239 | 0.387717 | 0.917257 | Lrrc71 |
| ENSMUSG( | 1191.712 | 0.041065 | 0.745359 | 0.96951 | Lrrc73 |
| ENSMUSG( | 41.30481 | 0.136427 | 0.70218 | 0.962014 | Lrrc74b |
| ENSMUSG( | 732.6495 | -0.13583 | 0.284033 | 0.892412 | Lrrc75a |
| ENSMUSG( | 1528.868 | -0.09798 | 0.453615 | 0.922316 | Lrrc75b |
| ENSMUSG( | 1007.437 | 0.157303 | 0.274741 | 0.889874 | Lrrc8a |
| ENSMUSG( | 3700.452 | 0.19097 | 0.313971 | 0.900949 | Lrrc8b |
| ENSMUSG( | 1327.015 | 0.697904 | 0.0005 | 0.067381 | Lrrc8c |
| ENSMUSG( | 1225.754 | 0.189677 | 0.28787 | 0.894644 | Lrrc8d |
| ENSMUSG( | 96.76832 | -0.07518 | 0.744545 | 0.96951 | Lrrc9 |
| ENSMUSG( | 500.2797 | -0.07475 | 0.602016 | 0.94789 | Lrrcc1 |
| ENSMUSG( | 21.29758 | 0.083981 | 0.853317 | 0.985598 | Lrrd1 |
| ENSMUSG( | 1430.52 | -0.04082 | 0.815393 | 0.979198 | Lrrfip1 |
| ENSMUSG( | 930.4122 | -0.05499 | 0.672704 | 0.95694 | Lrrfip2 |
| ENSMUSG( | 25.06639 | 0.989772 | 0.026931 | 0.603041 | Lrriq1 |
| ENSMUSG( | 15.03592 | -0.15736 | 0.753575 | 0.970671 | Lrriq3 |
| ENSMUSG( | 212.2226 | 0.494621 | 0.011057 | 0.426625 | Lrrk1 |
| ENSMUSG( | 1019.473 | 0.053573 | 0.660596 | 0.956205 | Lrrk2 |
| ENSMUSG( | 2189.39 | 0.346799 | 0.354449 | 0.912411 | Lrrn1 |
| ENSMUSG( | 3657.598 | 0.296894 | 0.113235 | 0.809623 | Lrrn2 |
| ENSMUSG( | 1623.23 | 0.565886 | 0.001693 | 0.152125 | Lrrn3 |
| ENSMUSG( | 7.810645 | 0.934444 | 0.193681 | NA | Lrrn4 |
| ENSMUSG( | 53.22051 | -0.15398 | 0.586566 | 0.944975 | Lrrn4cl |
| ENSMUSG( | 1902.696 | 0.362838 | 0.00646 | 0.336411 | Lrrtm1 |
| ENSMUSG( | 1497.313 | 0.350337 | 0.045358 | 0.693959 | Lrrtm2 |
| ENSMUSG( | 1380.422 | 0.340035 | 0.076217 | 0.769674 | Lrrtm3 |
| ENSMUSG( | 1015.031 | 0.283985 | 0.137744 | 0.828492 | Lrrtm4 |
| ENSMUSG( | 810.9578 | 0.055297 | 0.666459 | 0.956766 | Lrsam1 |
| ENSMUSG( | 2184.842 | 0.311999 | 0.045976 | 0.69654 | Lrtm2 |
| ENSMUSG( | 331.0219 | 0.119568 | 0.477103 | 0.923329 | Lrwd1 |
| ENSMUSG( | 13291.01 | 0.049688 | 0.718467 | 0.966633 | Lsamp |
| ENSMUSG( | 362.6358 | -0.02422 | 0.864679 | 0.986591 | Lsg1 |
| ENSMUSG( | 305.1681 | 0.030209 | 0.861546 | 0.986298 | Lsm1 |
| ENSMUSG( | 146.8788 | 0.0235 | 0.903866 | 0.991089 | Lsm10 |
| ENSMUSG( | 1131.226 | -0.03228 | 0.787393 | 0.97637 | Lsm11 |
| ENSMUSG( | 778.6005 | -0.01507 | 0.914653 | 0.992407 | Lsm12 |
| ENSMUSG( | 2098.577 | -0.03174 | 0.799704 | 0.97681 | Lsm14a |
| ENSMUSG( | 1789.691 | 0.030969 | 0.810944 | 0.978799 | Lsm14b |
| ENSMUSG( | 176.4579 | -0.03242 | 0.874652 | 0.989644 | Lsm2 |
| ENSMUSG( | 211.303 | -0.0123 | 0.944892 | 0.994798 | Lsm3 |
| ENSMUSG( | 815.8315 | -0.19439 | 0.122064 | 0.813384 | Lsm4 |
| ENSMUSG( | 128.3026 | 0.039637 | 0.865167 | 0.98675 | Lsm5 |
| ENSMUSG( | 1456.048 | -0.19105 | 0.187208 | 0.856372 | Lsm6 |
| ENSMUSG( | 423.4601 | -0.24923 | 0.142117 | 0.830709 | Lsm7 |
| ENSMUSG( | 297.3748 | 0.07659 | 0.63663 | 0.953903 | Lsm8 |
| ENSMUSG( | 11.48245 | -0.44896 | 0.447818 | 0.922202 | Lsmem2 |

| | | | | | | | | | | | |
|---|---|---|---|---|---|---|---|---|---|---|---|
| ENSMUSG( | Lrp2bp | 10.63053 | -0.53026 | 0.375526 | NA | ENSMUSG( | Lrp2bp | 10.8448 | -0.19253 | 0.752199 | NA |
| ENSMUSG( | Lrp3 | 2782.933 | -0.02195 | 0.861965 | 0.975055 | ENSMUSG( | Lrp3 | 2823.481 | 0.279452 | 0.06752 | 0.75212 |
| ENSMUSG( | Lrp4 | 5776.597 | 0.053864 | 0.693585 | 0.934202 | ENSMUSG( | Lrp4 | 4897.499 | -0.15685 | 0.219739 | 0.866104 |
| ENSMUSG( | Lrp5 | 836.9917 | -0.08746 | 0.566608 | 0.891644 | ENSMUSG( | Lrp5 | 832.7246 | 0.17092 | 0.357578 | 0.913759 |
| ENSMUSG( | Lrp6 | 2270.372 | -0.02189 | 0.846262 | 0.972226 | ENSMUSG( | Lrp6 | 2222.841 | 0.181928 | 0.150515 | 0.837173 |
| ENSMUSG( | Lrp8 | 3810.014 | -0.02717 | 0.803611 | 0.962169 | ENSMUSG( | Lrp8 | 3418.577 | -0.06904 | 0.584778 | 0.944975 |
| ENSMUSG( | Lrp8os2 | 29.35044 | -0.75068 | 0.140182 | 0.557263 | ENSMUSG( | Lrp8os2 | 33.70337 | -0.02562 | 0.946389 | 0.994798 |
| ENSMUSG( | Lrp8os3 | 32.51542 | 0.320098 | 0.335011 | 0.776499 | ENSMUSG( | Lrp8os3 | 20.77552 | -0.73835 | 0.132877 | 0.822912 |
| ENSMUSG( | Lrpap1 | 6971.033 | -0.30634 | 0.012292 | 0.160458 | ENSMUSG( | Lrpap1 | 7209.835 | 0.078205 | 0.583551 | 0.944266 |
| ENSMUSG( | Lrpprc | 1956.684 | 0.240221 | 0.025033 | 0.243138 | ENSMUSG( | Lrpprc | 1609.543 | -0.04995 | 0.730747 | 0.967228 |
| ENSMUSG( | Lrrc1 | 332.3774 | 0.009557 | 0.943041 | 0.993415 | ENSMUSG( | Lrrc1 | 298.4364 | -0.04264 | 0.814227 | 0.979198 |
| ENSMUSG( | Lrrc10b | 1702.683 | -0.58462 | 0.003953 | 0.080551 | ENSMUSG( | Lrrc10b | 1985.147 | 0.178335 | 0.207243 | 0.864412 |
| ENSMUSG( | Lrrc14 | 614.6826 | 0.045226 | 0.709605 | 0.939669 | ENSMUSG( | Lrrc14 | 525.5872 | -0.13674 | 0.334617 | 0.907907 |
| ENSMUSG( | Lrrc14b | 10.62036 | -0.82032 | 0.153786 | NA | ENSMUSG( | Lrrc14b | 10.0552 | -0.64922 | 0.285857 | NA |
| ENSMUSG( | Lrrc18 | 12.40656 | -0.52786 | 0.356382 | NA | ENSMUSG( | Lrrc18 | 13.79501 | 0.114324 | 0.832091 | 0.980738 |
| ENSMUSG( | Lrrc2 | 178.0261 | 0.072284 | 0.727725 | 0.944355 | ENSMUSG( | Lrrc2 | 149.0585 | -0.19453 | 0.400816 | 0.917257 |
| ENSMUSG( | Lrrc20 | 782.5943 | 0.028282 | 0.796097 | 0.960088 | ENSMUSG( | Lrrc20 | 706.5469 | 0.002996 | 0.980608 | 0.996479 |
| ENSMUSG( | Lrrc23 | 80.98117 | 0.314114 | 0.174529 | 0.610304 | ENSMUSG( | Lrrc23 | 73.14891 | 0.300866 | 0.244686 | 0.877904 |
| ENSMUSG( | Lrrc24 | 1219.471 | -0.36744 | 0.002055 | 0.052087 | ENSMUSG( | Lrrc24 | 1254.298 | 0.007907 | 0.955125 | 0.994798 |
| ENSMUSG( | Lrrc25 | 69.73372 | -0.17013 | 0.48108 | 0.855135 | ENSMUSG( | Lrrc25 | 73.69919 | 0.269121 | 0.342656 | 0.909646 |
| ENSMUSG( | Lrrc27 | 217.0274 | 0.134821 | 0.403738 | 0.816132 | ENSMUSG( | Lrrc27 | 182.7013 | -0.09265 | 0.619906 | 0.952896 |
| ENSMUSG( | Lrrc28 | 535.6603 | -0.08028 | 0.467227 | 0.844819 | ENSMUSG( | Lrrc28 | 479.1931 | -0.13997 | 0.313416 | 0.900883 |
| ENSMUSG( | Lrrc3 | 516.0241 | -0.36528 | 0.025825 | 0.247245 | ENSMUSG( | Lrrc3 | 553.7442 | 0.135702 | 0.466002 | 0.923329 |
| ENSMUSG( | Lrrc32 | 161.9058 | -0.43645 | 0.044363 | 0.323404 | ENSMUSG( | Lrrc32 | 179.6571 | 0.175842 | 0.453471 | 0.922316 |
| ENSMUSG( | Lrrc36 | 30.77794 | -0.38803 | 0.267238 | 0.717525 | ENSMUSG( | Lrrc36 | 31.7168 | 0.015337 | 0.967417 | 0.995667 |
| ENSMUSG( | Lrrc38 | 67.62933 | -0.27849 | 0.269797 | 0.719729 | ENSMUSG( | Lrrc38 | 68.39417 | 0.046787 | 0.852648 | 0.985598 |
| ENSMUSG( | Lrrc39 | 10.26321 | -0.48154 | 0.423716 | NA | ENSMUSG( | Lrrc39 | 10.07661 | -0.27858 | 0.649197 | NA |
| ENSMUSG( | Lrrc3b | 314.6161 | -0.32083 | 0.013376 | 0.168706 | ENSMUSG( | Lrrc3b | 319.3592 | 0.004795 | 0.975359 | 0.996416 |
| ENSMUSG( | Lrrc4 | 2510.485 | -0.17997 | 0.200148 | 0.640302 | ENSMUSG( | Lrrc4 | 2561.282 | 0.152736 | 0.340911 | 0.907972 |
| ENSMUSG( | Lrrc40 | 541.9678 | 0.085392 | 0.49046 | 0.859379 | ENSMUSG( | Lrrc40 | 477.6878 | -0.01335 | 0.92231 | 0.992644 |
| ENSMUSG( | Lrrc41 | 1179.667 | 0.060376 | 0.541676 | 0.883182 | ENSMUSG( | Lrrc41 | 1082.459 | 0.082179 | 0.472571 | 0.923329 |
| ENSMUSG( | Lrrc42 | 543.5561 | 0.03831 | 0.743598 | 0.948478 | ENSMUSG( | Lrrc42 | 469.7446 | -0.1154 | 0.397275 | 0.917257 |
| ENSMUSG( | Lrrc43 | 15.35875 | 0.578532 | 0.29228 | NA | ENSMUSG( | Lrrc43 | 14.95531 | 0.745031 | 0.156244 | 0.837724 |
| ENSMUSG( | Lrrc45 | 1643.936 | 0.358151 | 0.044841 | 0.324745 | ENSMUSG( | Lrrc45 | 1440.677 | 0.253003 | 0.207823 | 0.864412 |
| ENSMUSG( | Lrrc46 | 21.50253 | -0.3495 | 0.461347 | 0.841432 | ENSMUSG( | Lrrc46 | 24.14165 | 0.279136 | 0.529795 | 0.934795 |
| ENSMUSG( | Lrrc47 | 1189.788 | 0.073712 | 0.531448 | 0.879512 | ENSMUSG( | Lrrc47 | 1049.839 | -0.02003 | 0.888598 | 0.990558 |
| ENSMUSG( | Lrrc49 | 1565.272 | 0.043219 | 0.717455 | 0.941697 | ENSMUSG( | Lrrc49 | 1358.247 | -0.09891 | 0.436909 | 0.919983 |
| ENSMUSG( | Lrrc4b | 6315.761 | -0.29673 | 0.018382 | 0.203658 | ENSMUSG( | Lrrc4b | 6942.403 | 0.252029 | 0.089294 | 0.788502 |
| ENSMUSG( | Lrrc4c | 2344.356 | -0.45525 | 0.001358 | 0.040233 | ENSMUSG( | Lrrc4c | 2610.592 | 0.15944 | 0.319986 | 0.903771 |
| ENSMUSG( | Lrrc51 | 143.2098 | 0.145698 | 0.392164 | 0.809274 | ENSMUSG( | Lrrc51 | 126.311 | 0.050833 | 0.795717 | 0.97681 |
| ENSMUSG( | Lrrc55 | 725.3828 | -0.51894 | 0.001281 | 0.039067 | ENSMUSG( | Lrrc55 | 768.8459 | -0.03135 | 0.858134 | 0.985884 |
| ENSMUSG( | Lrrc56 | 167.8644 | -0.09031 | 0.573121 | 0.893584 | ENSMUSG( | Lrrc56 | 151.4985 | -0.10855 | 0.569034 | 0.941777 |
| ENSMUSG( | Lrrc57 | 1115.407 | 0.02503 | 0.796766 | 0.960359 | ENSMUSG( | Lrrc57 | 964.9726 | -0.13159 | 0.289041 | 0.895811 |
| ENSMUSG( | Lrrc58 | 5872.332 | 0.194093 | 0.063268 | 0.388993 | ENSMUSG( | Lrrc58 | 5265.254 | 0.149145 | 0.336386 | 0.907972 |
| ENSMUSG( | Lrrc59 | 2485.888 | 0.024906 | 0.772617 | 0.955272 | ENSMUSG( | Lrrc59 | 2191.678 | -0.07182 | 0.502644 | 0.927825 |
| ENSMUSG( | Lrrc61 | 816.1145 | 0.010887 | 0.924392 | 0.99114 | ENSMUSG( | Lrrc61 | 697.898 | -0.18055 | 0.192197 | 0.859726 |
| ENSMUSG( | Lrrc7 | 1454.627 | 0.092888 | 0.347813 | 0.780835 | ENSMUSG( | Lrrc7 | 1360.946 | 0.168016 | 0.171572 | 0.842999 |
| ENSMUSG( | Lrrc71 | 39.57402 | 0.524329 | 0.134123 | 0.547472 | ENSMUSG( | Lrrc71 | 33.06585 | 0.318239 | 0.387717 | 0.917257 |
| ENSMUSG( | Lrrc73 | 1338.229 | 0.108277 | 0.363723 | 0.793343 | ENSMUSG( | Lrrc73 | 1191.712 | 0.041065 | 0.745359 | 0.96951 |
| ENSMUSG( | Lrrc74b | 40.08997 | -0.24435 | 0.43886 | 0.830714 | ENSMUSG( | Lrrc74b | 41.30481 | 0.136427 | 0.70218 | 0.962014 |
| ENSMUSG( | Lrrc75a | 846.8089 | 0.021178 | 0.831603 | 0.969984 | ENSMUSG( | Lrrc75a | 732.6495 | -0.13583 | 0.284033 | 0.892412 |
| ENSMUSG( | Lrrc75b | 1673.835 | -0.10209 | 0.342031 | 0.778828 | ENSMUSG( | Lrrc75b | 1528.868 | -0.09798 | 0.453615 | 0.922316 |
| ENSMUSG( | Lrrc8a | 1001.032 | -0.12798 | 0.288838 | 0.739652 | ENSMUSG( | Lrrc8a | 1007.437 | 0.157303 | 0.274741 | 0.889874 |
| ENSMUSG( | Lrrc8b | 3326.562 | -0.41377 | 0.012287 | 0.160458 | ENSMUSG( | Lrrc8b | 3700.452 | 0.19097 | 0.313971 | 0.900949 |
| ENSMUSG( | Lrrc8c | 1172.542 | 0.145102 | 0.412562 | 0.819466 | ENSMUSG( | Lrrc8c | 1327.015 | 0.697904 | 0.0005 | 0.067381 |
| ENSMUSG( | Lrrc8d | 1137.937 | -0.3107 | 0.044061 | 0.321777 | ENSMUSG( | Lrrc8d | 1225.754 | 0.189677 | 0.28787 | 0.894644 |
| ENSMUSG( | Lrrc9 | 108.4905 | -0.008 | 0.972695 | 0.996585 | ENSMUSG( | Lrrc9 | 96.76832 | -0.07518 | 0.744545 | 0.96951 |
| ENSMUSG( | Lrrcc1 | 532.9708 | -0.16932 | 0.15849 | 0.587906 | ENSMUSG( | Lrrcc1 | 500.2797 | -0.07475 | 0.602016 | 0.94789 |
| ENSMUSG( | Lrrd1 | 24.67062 | 0.241078 | 0.530738 | 0.879289 | ENSMUSG( | Lrrd1 | 21.29758 | 0.083981 | 0.853317 | 0.985598 |
| ENSMUSG( | Lrrfip1 | 1753.927 | 0.269845 | 0.057742 | 0.370624 | ENSMUSG( | Lrrfip1 | 1430.52 | -0.04082 | 0.815393 | 0.979198 |
| ENSMUSG( | Lrrfip2 | 1061.994 | 0.061793 | 0.524658 | 0.875871 | ENSMUSG( | Lrrfip2 | 930.4122 | -0.05499 | 0.672704 | 0.95694 |
| ENSMUSG( | Lrriq1 | 22.0543 | 0.466867 | 0.292614 | 0.742521 | ENSMUSG( | Lrriq1 | 25.06639 | 0.989772 | 0.026931 | 0.603041 |
| ENSMUSG( | Lrriq3 | 18.79814 | 0.227539 | 0.59229 | NA | ENSMUSG( | Lrriq3 | 15.03592 | -0.15736 | 0.753575 | 0.970671 |
| ENSMUSG( | Lrrk1 | 186.2053 | -0.12128 | 0.447526 | 0.834051 | ENSMUSG( | Lrrk1 | 212.2226 | 0.494621 | 0.011057 | 0.426625 |
| ENSMUSG( | Lrrk2 | 1100.701 | 0.004082 | 0.973405 | 0.99675 | ENSMUSG( | Lrrk2 | 1019.473 | 0.053573 | 0.660596 | 0.956205 |
| ENSMUSG( | Lrrn1 | 1762.92 | -0.5951 | 0.003935 | 0.080285 | ENSMUSG( | Lrrn1 | 2189.39 | 0.346799 | 0.354449 | 0.912411 |
| ENSMUSG( | Lrrn2 | 3336.141 | -0.23808 | 0.140813 | 0.557902 | ENSMUSG( | Lrrn2 | 3657.598 | 0.296894 | 0.113235 | 0.809623 |
| ENSMUSG( | Lrrn3 | 1322.322 | -0.25759 | 0.120555 | 0.524506 | ENSMUSG( | Lrrn3 | 1623.23 | 0.565886 | 0.001693 | 0.152125 |
| ENSMUSG( | Lrrn4 | 7.275076 | 0.536726 | 0.471463 | NA | ENSMUSG( | Lrrn4 | 7.810645 | 0.934444 | 0.193681 | NA |
| ENSMUSG( | Lrrn4cl | 56.76514 | -0.26034 | 0.339784 | 0.777939 | ENSMUSG( | Lrrn4cl | 53.22051 | -0.15398 | 0.586566 | 0.944975 |
| ENSMUSG( | Lrrtm1 | 1654.535 | -0.30591 | 0.008329 | 0.127443 | ENSMUSG( | Lrrtm1 | 1902.696 | 0.362838 | 0.00646 | 0.336411 |
| ENSMUSG( | Lrrtm2 | 1190.741 | -0.62997 | 8.72E-05 | 0.00587 | ENSMUSG( | Lrrtm2 | 1497.313 | 0.350337 | 0.045358 | 0.693959 |
| ENSMUSG( | Lrrtm3 | 1174.885 | -0.40892 | 0.014793 | 0.17866 | ENSMUSG( | Lrrtm3 | 1380.422 | 0.340035 | 0.076217 | 0.769674 |
| ENSMUSG( | Lrrtm4 | 891.8553 | -0.37537 | 0.02926 | 0.259574 | ENSMUSG( | Lrrtm4 | 1015.031 | 0.283985 | 0.137744 | 0.828492 |
| ENSMUSG( | Lrsam1 | 916.7011 | 0.134153 | 0.169011 | 0.603241 | ENSMUSG( | Lrsam1 | 810.9578 | 0.055297 | 0.666459 | 0.956766 |
| ENSMUSG( | Lrtm2 | 1861.307 | -0.44015 | 0.000595 | 0.022994 | ENSMUSG( | Lrtm2 | 2184.842 | 0.311999 | 0.045976 | 0.69654 |
| ENSMUSG( | Lrwd1 | 374.0567 | 0.197384 | 0.128509 | 0.536774 | ENSMUSG( | Lrwd1 | 331.0219 | 0.119568 | 0.477103 | 0.923329 |
| ENSMUSG( | Lsamp | 13729.57 | -0.12939 | 0.254419 | 0.704536 | ENSMUSG( | Lsamp | 13291.01 | 0.049688 | 0.718467 | 0.966633 |
| ENSMUSG( | Lsg1 | 414.9719 | 0.090573 | 0.471775 | 0.847504 | ENSMUSG( | Lsg1 | 362.6358 | -0.02422 | 0.864679 | 0.986591 |
| ENSMUSG( | Lsm1 | 330.8271 | 0.00063 | 0.996524 | 0.99943 | ENSMUSG( | Lsm1 | 305.1681 | 0.030209 | 0.861546 | 0.986298 |
| ENSMUSG( | Lsm10 | 163.653 | 0.069077 | 0.707494 | 0.93949 | ENSMUSG( | Lsm10 | 146.8788 | 0.0235 | 0.903866 | 0.991089 |
| ENSMUSG( | Lsm11 | 1417.395 | 0.328419 | 0.000685 | 0.025185 | ENSMUSG( | Lsm11 | 1131.226 | -0.03228 | 0.787393 | 0.97637 |
| ENSMUSG( | Lsm12 | 872.4431 | 0.048902 | 0.660199 | 0.924627 | ENSMUSG( | Lsm12 | 778.6005 | -0.01507 | 0.914653 | 0.992407 |
| ENSMUSG( | Lsm14a | 2354.359 | 0.034212 | 0.747833 | 0.949604 | ENSMUSG( | Lsm14a | 2098.577 | -0.03174 | 0.799704 | 0.97681 |
| ENSMUSG( | Lsm14b | 2031.151 | 0.127314 | 0.224006 | 0.669869 | ENSMUSG( | Lsm14b | 1789.691 | 0.030969 | 0.810944 | 0.978799 |
| ENSMUSG( | Lsm2 | 178.923 | -0.25727 | 0.115217 | 0.512007 | ENSMUSG( | Lsm2 | 176.4579 | -0.03242 | 0.874652 | 0.989644 |
| ENSMUSG( | Lsm3 | 231.213 | -0.01536 | 0.924596 | 0.99114 | ENSMUSG( | Lsm3 | 211.303 | -0.0123 | 0.944892 | 0.994798 |
| ENSMUSG( | Lsm4 | 962.6989 | 0.02452 | 0.816283 | 0.965455 | ENSMUSG( | Lsm4 | 815.8315 | -0.19439 | 0.122064 | 0.813384 |
| ENSMUSG( | Lsm5 | 134.1708 | -0.08569 | 0.640421 | 0.916434 | ENSMUSG( | Lsm5 | 128.3026 | 0.039637 | 0.865167 | 0.98675 |
| ENSMUSG( | Lsm6 | 1718.972 | 0.030455 | 0.812045 | 0.964574 | ENSMUSG( | Lsm6 | 1456.048 | -0.19105 | 0.187208 | 0.856372 |
| ENSMUSG( | Lsm7 | 500.0724 | -0.02419 | 0.862473 | 0.975055 | ENSMUSG( | Lsm7 | 423.4601 | -0.24923 | 0.142117 | 0.830709 |
| ENSMUSG( | Lsm8 | 317.2818 | 0.003582 | 0.979855 | 0.997428 | ENSMUSG( | Lsm8 | 297.3748 | 0.07659 | 0.63663 | 0.953903 |
| ENSMUSG( | Lsmem2 | 13.59202 | -0.20478 | 0.687961 | NA | ENSMUSG( | Lsmem2 | 11.48245 | -0.44896 | 0.447818 | 0.922202 |
| ENSMUSG( | Lsp1 | 289.8672 | -0.30752 | 0.078257 | 0.432625 | ENSMUSG( | Lsp1 | 261.0582 | -0.35736 | 0.06452 | 0.737935 |
| ENSMUSG( | Lsr | 411.3774 | 0.220873 | 0.299345 | 0.748131 | ENSMUSG( | Lsr | 401.7843 | 0.413964 | 0.082188 | 0.77681 |
| ENSMUSG( | Lss | 2915.143 | -0.0819 | 0.528706 | 0.878365 | ENSMUSG( | Lss | 2484.998 | -0.28604 | 0.019434 | 0.556192 |
| ENSMUSG( | Lst1 | 93.99672 | -0.20593 | 0.39931 | 0.813048 | ENSMUSG( | Lst1 | 72.97295 | -0.78999 | 0.004948 | 0.28972 |
| ENSMUSG( | Lta4h | 880.4498 | 0.019729 | 0.863924 | 0.975282 | ENSMUSG( | Lta4h | 771.9922 | -0.09026 | 0.509551 | 0.929089 |
| ENSMUSG( | Ltb | 18.31739 | -0.13692 | 0.764315 | NA | ENSMUSG( | Ltb | 14.89953 | -0.46084 | 0.403973 | 0.917257 |
| ENSMUSG( | Ltb4r1 | 12.66135 | -0.76472 | 0.166175 | NA | ENSMUSG( | Ltb4r1 | 12.47862 | -0.54636 | 0.355439 | 0.912555 |
| ENSMUSG( | Ltbp1 | 694.3441 | -0.28487 | 0.047793 | 0.333848 | ENSMUSG( | Ltbp1 | 675.9348 | -0.07769 | 0.625974 | 0.953234 |
| ENSMUSG( | Ltbp2 | 34.10339 | -0.17746 | 0.605104 | 0.905594 | ENSMUSG( | Ltbp2 | 36.11258 | 0.222582 | 0.548636 | 0.938063 |
| ENSMUSG( | Ltbp3 | 1712.862 | -0.05873 | 0.590081 | 0.90113 | ENSMUSG( | Ltbp3 | 1629.564 | 0.069589 | 0.59411 | 0.945408 |
| ENSMUSG( | Ltbp4 | 2431.408 | 0.171139 | 0.155701 | 0.583654 | ENSMUSG( | Ltbp4 | 2386.507 | 0.371864 | 0.027447 | 0.607058 |
| ENSMUSG( | Ltbr | 864.8408 | -0.024 | 0.844085 | 0.971962 | ENSMUSG( | Ltbr | 810.0118 | 0.059243 | 0.670182 | 0.95694 |
| ENSMUSG( | Ltc4s | 223.3573 | -0.22323 | 0.151941 | 0.577721 | ENSMUSG( | Ltc4s | 218.3095 | -0.00396 | 0.982057 | 0.996756 |
| ENSMUSG( | Ltf | 59.10704 | -1.50703 | 0.106891 | 0.494634 | ENSMUSG( | Ltf | 86.92226 | 0.223889 | 0.792925 | 0.97681 |
| ENSMUSG( | Ltk | 833.8403 | 0.373504 | 0.068575 | 0.405797 | ENSMUSG( | Ltk | 695.0306 | 0.137291 | 0.50094 | 0.927825 |
| ENSMUSG( | Ltn1 | 1302.435 | 0.108091 | 0.307064 | 0.75508 | ENSMUSG( | Ltn1 | 1213.797 | 0.170659 | 0.133162 | 0.822912 |
| ENSMUSG( | LTO1 | 429.2255 | -0.02471 | 0.86305 | 0.975055 | ENSMUSG( | LTO1 | 384.72 | -0.07433 | 0.656956 | 0.955625 |
| ENSMUSG( | Ltv1 | 366.87 | 0.208936 | 0.106082 | 0.493403 | ENSMUSG( | Ltv1 | 312.9337 | 0.023658 | 0.87122 | 0.988779 |
| ENSMUSG( | Luc7l | 2638.272 | -0.07291 | 0.429244 | 0.826446 | ENSMUSG( | Luc7l | 2564.397 | 0.113967 | 0.280885 | 0.892356 |
| ENSMUSG( | Luc7l2 | 6581.43 | 0.008448 | 0.937728 | 0.993415 | ENSMUSG( | Luc7l2 | 6038.896 | 0.02461 | 0.863604 | 0.986547 |
| ENSMUSG( | Luc7l3 | 3288.631 | -0.1203 | 0.23439 | 0.681922 | ENSMUSG( | Luc7l3 | 3445.715 | 0.272845 | 0.077584 | 0.77321 |
| ENSMUSG( | Lum | 65.33677 | -0.14297 | 0.615557 | 0.910481 | ENSMUSG( | Lum | 91.90367 | 0.966052 | 0.000698 | 0.085493 |
| ENSMUSG( | Lurap1 | 1482.494 | 0.047062 | 0.606752 | 0.906634 | ENSMUSG( | Lurap1 | 1290.251 | -0.08786 | 0.440327 | 0.921166 |
| ENSMUSG( | Lurap1l | 202.1532 | 0.265674 | 0.207742 | 0.64962 | ENSMUSG( | Lurap1l | 177.7096 | 0.164078 | 0.41965 | 0.917257 |
| ENSMUSG( | Luzp1 | 2071.04 | 0.30702 | 0.000367 | 0.016728 | ENSMUSG( | Luzp1 | 1788.703 | 0.166102 | 0.121883 | 0.813384 |
| ENSMUSG( | Luzp2 | 15882.89 | -0.22847 | 0.094454 | 0.471713 | ENSMUSG( | Luzp2 | 14880.59 | -0.14297 | 0.379047 | 0.916995 |
| ENSMUSG( | Lxn | 2377.409 | 0.075146 | 0.490089 | 0.859319 | ENSMUSG( | Lxn | 2075.638 | -0.05182 | 0.734448 | 0.967228 |
| ENSMUSG( | Ly6a | 813.8923 | 0.072687 | 0.716182 | 0.94152 | ENSMUSG( | Ly6a | 925.7571 | 0.651081 | 0.140651 | 0.830678 |
| ENSMUSG( | Ly6c1 | 1426.642 | 0.30694 | 0.118862 | 0.521914 | ENSMUSG( | Ly6c1 | 1645.879 | 0.881454 | 0.0386 | 0.679972 |
| ENSMUSG( | Ly6d | 14.91745 | -0.00033 | 0.999525 | NA | ENSMUSG( | Ly6d | 14.35861 | 0.133878 | 0.821939 | 0.979198 |
| ENSMUSG( | Ly6e | 8870.753 | -0.13147 | 0.305804 | 0.753712 | ENSMUSG( | Ly6e | 8590.101 | 0.050619 | 0.726886 | 0.966887 |
| ENSMUSG( | Ly6g6d | 18.79898 | 0.056366 | 0.894559 | NA | ENSMUSG( | Ly6g6d | 17.17398 | 0.056665 | 0.902379 | 0.991089 |
| ENSMUSG( | Ly6g6e | 75.27331 | -0.86823 | 0.000173 | 0.00947 | ENSMUSG( | Ly6g6e | 86.3295 | -0.07751 | 0.731572 | 0.967228 |
| ENSMUSG( | Ly6h | 6808.571 | -0.22819 | 0.013124 | 0.166048 | ENSMUSG( | Ly6h | 6790.227 | 0.041476 | 0.734084 | 0.967228 |
| ENSMUSG( | Ly6m | 17.48641 | -0.30777 | 0.518213 | NA | ENSMUSG( | Ly6m | 13.12513 | -0.97113 | 0.110696 | 0.807582 |
| ENSMUSG( | Ly75 | 91.72181 | 0.181646 | 0.502086 | 0.864904 | ENSMUSG( | Ly75 | 113.6655 | 0.948117 | 0.002197 | 0.183242 |
| ENSMUSG( | Ly86 | 817.0514 | -0.10133 | 0.495145 | 0.860798 | ENSMUSG( | Ly86 | 740.3618 | -0.11639 | 0.381293 | 0.917207 |
| ENSMUSG( | Ly9 | 18.03942 | -0.46561 | 0.326185 | NA | ENSMUSG( | Ly9 | 14.64728 | -0.82406 | 0.138831 | 0.829437 |
| ENSMUSG( | Ly96 | 38.76377 | -0.21306 | 0.519897 | 0.874979 | ENSMUSG( | Ly96 | 33.42792 | -0.39044 | 0.252811 | 0.88018 |
| ENSMUSG( | Lyar | 292.2313 | 0.19624 | 0.173235 | 0.608563 | ENSMUSG( | Lyar | 256.4816 | 0.083884 | 0.64621 | 0.953988 |
| ENSMUSG( | Lyl1 | 173.6139 | -0.2377 | 0.214226 | 0.658845 | ENSMUSG( | Lyl1 | 189.4972 | 0.288477 | 0.153262 | 0.837724 |
| ENSMUSG( | Lyn | 581.2945 | -0.03306 | 0.825891 | 0.968061 | ENSMUSG( | Lyn | 581.4924 | 0.231664 | 0.135682 | 0.825545 |
| ENSMUSG( | Lynx1 | 18009.82 | -0.06526 | 0.590036 | 0.90113 | ENSMUSG( | Lynx1 | 17314.56 | 0.092292 | 0.548265 | 0.938063 |
| ENSMUSG( | Lypd1 | 2906.689 | -0.07596 | 0.412046 | 0.819446 | ENSMUSG( | Lypd1 | 2766.549 | 0.051735 | 0.638383 | 0.953903 |
| ENSMUSG( | Lypd2 | 24.51585 | 0.061989 | 0.872372 | 0.978556 | ENSMUSG( | Lypd2 | 17.48173 | -0.76144 | 0.11713 | 0.813384 |
| ENSMUSG( | Lypd6 | 2322.518 | -0.14675 | 0.244303 | 0.691435 | ENSMUSG( | Lypd6 | 2048.838 | -0.24624 | 0.089943 | 0.790741 |
| ENSMUSG( | Lypd6b | 480.119 | -0.24405 | 0.095627 | 0.475062 | ENSMUSG( | Lypd6b | 492.533 | 0.11244 | 0.497215 | 0.927825 |
| ENSMUSG( | Lypla1 | 282.3134 | -0.02701 | 0.84469 | 0.972023 | ENSMUSG( | Lypla1 | 281.2221 | 0.209462 | 0.32032 | 0.903887 |
| ENSMUSG( | Lypla2 | 1967.105 | 0.039938 | 0.688945 | 0.933441 | ENSMUSG( | Lypla2 | 1689.483 | -0.13539 | 0.291337 | 0.897566 |
| ENSMUSG( | Lyplal1 | 133.8966 | 0.043495 | 0.818986 | 0.965888 | ENSMUSG( | Lyplal1 | 106.1208 | -0.37917 | 0.082987 | 0.77681 |
| ENSMUSG( | Lyrm1 | 204.9941 | 0.056775 | 0.749753 | 0.949664 | ENSMUSG( | Lyrm1 | 184.5851 | 0.019461 | 0.920026 | 0.992644 |
| ENSMUSG( | Lyrm2 | 387.1953 | -0.00567 | 0.965884 | 0.995377 | ENSMUSG( | Lyrm2 | 339.7804 | -0.12427 | 0.404939 | 0.917257 |
| ENSMUSG( | Lyrm4 | 615.5592 | 0.012506 | 0.913653 | 0.988483 | ENSMUSG( | Lyrm4 | 512.4744 | -0.26255 | 0.064623 | 0.738417 |
| ENSMUSG( | Lyrm7 | 300.1971 | 0.020005 | 0.893233 | 0.984512 | ENSMUSG( | Lyrm7 | 262.2223 | -0.10995 | 0.516269 | 0.930939 |
| ENSMUSG( | Lyrm9 | 1213.01 | 0.013707 | 0.886851 | 0.983255 | ENSMUSG( | Lyrm9 | 1120.349 | 0.053622 | 0.643583 | 0.953903 |
| ENSMUSG( | Lyset | 241.7068 | -0.25164 | 0.081623 | 0.441308 | ENSMUSG( | Lyset | 256.1053 | 0.188202 | 0.231242 | 0.868507 |
| ENSMUSG( | Lysmd1 | 171.6939 | 0.218295 | 0.195505 | 0.63504 | ENSMUSG( | Lysmd1 | 150.3077 | 0.095777 | 0.635317 | 0.953903 |
| ENSMUSG( | Lysmd2 | 1135.36 | 0.085494 | 0.416686 | 0.8209 | ENSMUSG( | Lysmd2 | 1002.855 | -0.00848 | 0.952324 | 0.994798 |
| ENSMUSG( | Lysmd3 | 137.7295 | -0.27947 | 0.143685 | 0.562581 | ENSMUSG( | Lysmd3 | 154.2037 | 0.32069 | 0.107867 | 0.804969 |
| ENSMUSG( | Lysmd4 | 817.1045 | 0.072619 | 0.512191 | 0.871563 | ENSMUSG( | Lysmd4 | 725.76 | 0.002547 | 0.983769 | 0.996963 |
| ENSMUSG( | Lyst | 1252.672 | 0.029473 | 0.809887 | 0.96372 | ENSMUSG( | Lyst | 1400.098 | 0.568771 | 0.000137 | 0.026415 |
| ENSMUSG( | Lyve1 | 103.6636 | 0.125189 | 0.628966 | 0.913816 | ENSMUSG( | Lyve1 | 108.6037 | 0.489704 | 0.071261 | 0.761942 |
| ENSMUSG( | Lyz2 | 683.3394 | -0.54464 | 0.019029 | 0.207571 | ENSMUSG( | Lyz2 | 666.2216 | -0.32814 | 0.174357 | 0.844317 |
| ENSMUSG( | Lyzl4 | 20.1369 | -0.25487 | 0.533455 | NA | ENSMUSG( | Lyzl4 | 20.14577 | 0.022952 | 0.957717 | 0.994798 |
| ENSMUSG( | Lzic | 393.7938 | 0.015365 | 0.904386 | 0.986246 | ENSMUSG( | Lzic | 334.4684 | -0.20354 | 0.202605 | 0.864412 |

| | | | | | |
|---|---|---|---|---|---|
| ENSMUSG( | 1115.407 | 0.02503 | 0.796766 | 0.960359 | Lrrc57 |
| ENSMUSG( | 5872.332 | 0.194093 | 0.063268 | 0.388993 | Lrrc58 |
| ENSMUSG( | 2485.888 | 0.024906 | 0.772617 | 0.955272 | Lrrc59 |
| ENSMUSG( | 816.1145 | 0.010887 | 0.924392 | 0.99114 | Lrrc61 |
| ENSMUSG( | 1454.627 | 0.092888 | 0.347813 | 0.780835 | Lrrc7 |
| ENSMUSG( | 39.57402 | 0.524329 | 0.134123 | 0.547472 | Lrrc71 |
| ENSMUSG( | 1338.229 | 0.108277 | 0.363723 | 0.793343 | Lrrc73 |
| ENSMUSG( | 40.08997 | -0.24435 | 0.43886 | 0.830714 | Lrrc74b |
| ENSMUSG( | 846.8089 | 0.021178 | 0.831603 | 0.969984 | Lrrc75a |
| ENSMUSG( | 1673.835 | -0.10209 | 0.342031 | 0.778828 | Lrrc75b |
| ENSMUSG( | 1001.032 | -0.12798 | 0.288838 | 0.739652 | Lrrc8a |
| ENSMUSG( | 3326.562 | -0.41377 | 0.012287 | 0.160458 | Lrrc8b |
| ENSMUSG( | 1172.542 | 0.145102 | 0.412562 | 0.819466 | Lrrc8c |
| ENSMUSG( | 1137.937 | -0.3107 | 0.044061 | 0.321777 | Lrrc8d |
| ENSMUSG( | 108.4905 | -0.008 | 0.972695 | 0.996585 | Lrrc9 |
| ENSMUSG( | 532.9708 | -0.16932 | 0.15849 | 0.587906 | Lrrcc1 |
| ENSMUSG( | 24.67062 | 0.241078 | 0.530738 | 0.879289 | Lrrd1 |
| ENSMUSG( | 1753.927 | 0.269845 | 0.057742 | 0.370624 | Lrrfip1 |
| ENSMUSG( | 1061.994 | 0.061793 | 0.524658 | 0.875871 | Lrrfip2 |
| ENSMUSG( | 22.0543 | 0.466867 | 0.292614 | 0.742521 | Lrriq1 |
| ENSMUSG( | 18.79814 | 0.227539 | 0.59229 | NA | Lrriq3 |
| ENSMUSG( | 186.2053 | -0.12128 | 0.447526 | 0.834051 | Lrrk1 |
| ENSMUSG( | 1100.701 | 0.004082 | 0.973405 | 0.99675 | Lrrk2 |
| ENSMUSG( | 1762.92 | -0.5951 | 0.003935 | 0.080285 | Lrrn1 |
| ENSMUSG( | 3336.141 | -0.23808 | 0.140813 | 0.557902 | Lrrn2 |
| ENSMUSG( | 1322.322 | -0.25759 | 0.120555 | 0.524506 | Lrrn3 |
| ENSMUSG( | 7.275076 | 0.536726 | 0.471463 | NA | Lrrn4 |
| ENSMUSG( | 56.76514 | -0.26034 | 0.339784 | 0.777939 | Lrrn4cl |
| ENSMUSG( | 1654.535 | -0.30591 | 0.008329 | 0.127443 | Lrrtm1 |
| ENSMUSG( | 1190.741 | -0.62997 | 8.72E-05 | 0.00587 | Lrrtm2 |
| ENSMUSG( | 1174.885 | -0.40892 | 0.014793 | 0.17866 | Lrrtm3 |
| ENSMUSG( | 891.8553 | -0.37537 | 0.02926 | 0.259574 | Lrrtm4 |
| ENSMUSG( | 916.7011 | 0.134153 | 0.169011 | 0.603241 | Lrsam1 |
| ENSMUSG( | 1861.307 | -0.44015 | 0.000595 | 0.022994 | Lrtm2 |
| ENSMUSG( | 374.0567 | 0.197384 | 0.128509 | 0.536774 | Lrwd1 |
| ENSMUSG( | 13729.57 | -0.12939 | 0.254419 | 0.704536 | Lsamp |
| ENSMUSG( | 414.9719 | 0.090573 | 0.471775 | 0.847504 | Lsg1 |
| ENSMUSG( | 330.8271 | 0.00063 | 0.996524 | 0.99943 | Lsm1 |
| ENSMUSG( | 163.653 | 0.069077 | 0.707494 | 0.93949 | Lsm10 |
| ENSMUSG( | 1417.395 | 0.328419 | 0.000685 | 0.025185 | Lsm11 |
| ENSMUSG( | 872.4431 | 0.048902 | 0.660199 | 0.924627 | Lsm12 |
| ENSMUSG( | 2354.359 | 0.034212 | 0.747833 | 0.949604 | Lsm14a |
| ENSMUSG( | 2031.151 | 0.127314 | 0.224006 | 0.669869 | Lsm14b |
| ENSMUSG( | 178.923 | -0.25727 | 0.115217 | 0.512007 | Lsm2 |
| ENSMUSG( | 231.213 | -0.01536 | 0.924596 | 0.99114 | Lsm3 |
| ENSMUSG( | 962.6989 | 0.02452 | 0.816283 | 0.965455 | Lsm4 |
| ENSMUSG( | 134.1708 | -0.08569 | 0.640421 | 0.916434 | Lsm5 |
| ENSMUSG( | 1718.972 | 0.030455 | 0.812045 | 0.964574 | Lsm6 |
| ENSMUSG( | 500.0724 | -0.02419 | 0.862473 | 0.975055 | Lsm7 |
| ENSMUSG( | 317.2818 | 0.003582 | 0.979855 | 0.997428 | Lsm8 |
| ENSMUSG( | 13.59202 | -0.20478 | 0.687961 | NA | Lsmem2 |
| ENSMUSG( | 289.8672 | -0.30752 | 0.078257 | 0.432625 | Lsp1 |
| ENSMUSG( | 411.3774 | 0.220873 | 0.299345 | 0.748131 | Lsr |
| ENSMUSG( | 2915.143 | -0.0819 | 0.528706 | 0.878365 | Lss |
| ENSMUSG( | 93.99672 | -0.20593 | 0.39931 | 0.813048 | Lst1 |
| ENSMUSG( | 880.4498 | 0.019729 | 0.863924 | 0.975282 | Lta4h |
| ENSMUSG( | 18.31739 | -0.13692 | 0.764315 | NA | Ltb |
| ENSMUSG( | 12.66135 | -0.76472 | 0.166175 | NA | Ltb4r1 |
| ENSMUSG( | 694.3441 | -0.28487 | 0.047793 | 0.333848 | Ltbp1 |
| ENSMUSG( | 34.10339 | -0.17746 | 0.605104 | 0.905594 | Ltbp2 |
| ENSMUSG( | 1712.862 | -0.05873 | 0.590081 | 0.90113 | Ltbp3 |
| ENSMUSG( | 2431.408 | 0.171139 | 0.155701 | 0.583654 | Ltbp4 |
| ENSMUSG( | 864.8408 | -0.024 | 0.844085 | 0.971962 | Ltbr |
| ENSMUSG( | 223.3573 | -0.22323 | 0.151941 | 0.577721 | Ltc4s |
| ENSMUSG( | 59.10704 | -1.50703 | 0.106891 | 0.494634 | Ltf |
| ENSMUSG( | 833.8403 | 0.373504 | 0.068575 | 0.405797 | Ltk |
| ENSMUSG( | 1302.435 | 0.108091 | 0.307064 | 0.75508 | Ltn1 |
| ENSMUSG( | 429.2255 | -0.02471 | 0.86305 | 0.975055 | LTO1 |
| ENSMUSG( | 366.87 | 0.208936 | 0.106082 | 0.493403 | Ltv1 |
| ENSMUSG( | 2638.272 | -0.07291 | 0.429244 | 0.826446 | Luc7l |
| ENSMUSG( | 6581.43 | 0.008448 | 0.937728 | 0.993415 | Luc7l2 |
| ENSMUSG( | 3288.631 | -0.1203 | 0.23439 | 0.681922 | Luc7l3 |
| ENSMUSG( | 65.33677 | -0.14297 | 0.615557 | 0.910481 | Lum |
| ENSMUSG( | 1482.494 | 0.047062 | 0.606752 | 0.906634 | Lurap1 |
| ENSMUSG( | 202.1532 | 0.265674 | 0.207742 | 0.64962 | Lurap1l |
| ENSMUSG( | 2071.04 | 0.30702 | 0.000367 | 0.016728 | Luzp1 |
| ENSMUSG( | 15882.89 | -0.22847 | 0.094454 | 0.471713 | Luzp2 |
| ENSMUSG( | 2377.409 | 0.075146 | 0.490089 | 0.859319 | Lxn |
| ENSMUSG( | 813.8923 | 0.072687 | 0.716182 | 0.94152 | Ly6a |
| ENSMUSG( | 1426.642 | 0.30694 | 0.118862 | 0.521914 | Ly6c1 |
| ENSMUSG( | 14.91745 | -0.00033 | 0.999525 | NA | Ly6d |
| ENSMUSG( | 8870.753 | -0.13147 | 0.305804 | 0.753712 | Ly6e |
| ENSMUSG( | 18.79898 | 0.056366 | 0.894559 | NA | Ly6g6d |
| ENSMUSG( | 75.27331 | -0.86823 | 0.000173 | 0.00947 | Ly6g6e |
| ENSMUSG( | 6808.571 | -0.22819 | 0.013124 | 0.166048 | Ly6h |
| ENSMUSG( | 17.48641 | -0.30777 | 0.518213 | NA | Ly6m |
| ENSMUSG( | 91.72181 | 0.181646 | 0.502086 | 0.864904 | Ly75 |
| ENSMUSG( | 817.0514 | -0.10133 | 0.495145 | 0.860798 | Ly86 |
| ENSMUSG( | 18.03942 | -0.46561 | 0.326185 | NA | Ly9 |
| ENSMUSG( | 38.76377 | -0.21306 | 0.519897 | 0.874979 | Ly96 |
| ENSMUSG( | 292.2313 | 0.19624 | 0.173235 | 0.608563 | Lyar |
| ENSMUSG( | 173.6139 | -0.2377 | 0.214226 | 0.658845 | Lyl1 |
| ENSMUSG( | 581.2945 | -0.03306 | 0.825891 | 0.968061 | Lyn |
| ENSMUSG( | 18009.82 | -0.06526 | 0.590036 | 0.90113 | Lynx1 |
| ENSMUSG( | 2906.689 | -0.07596 | 0.412046 | 0.819446 | Lypd1 |
| ENSMUSG( | 24.51585 | 0.061989 | 0.872372 | 0.978556 | Lypd2 |
| ENSMUSG( | 2322.518 | -0.14675 | 0.244303 | 0.691435 | Lypd6 |
| ENSMUSG( | 480.119 | -0.24405 | 0.095627 | 0.475062 | Lypd6b |
| ENSMUSG( | 282.3134 | -0.02701 | 0.84469 | 0.972023 | Lypla1 |
| ENSMUSG( | 1967.105 | 0.039938 | 0.688945 | 0.933441 | Lypla2 |
| ENSMUSG( | 133.8966 | 0.043495 | 0.818986 | 0.965888 | Lyplal1 |
| ENSMUSG( | 204.9941 | 0.056775 | 0.749753 | 0.949664 | Lyrm1 |
| ENSMUSG( | 387.1953 | -0.00567 | 0.965884 | 0.995377 | Lyrm2 |
| ENSMUSG( | 615.5592 | 0.012506 | 0.913653 | 0.988483 | Lyrm4 |
| ENSMUSG( | 300.1971 | 0.020005 | 0.893233 | 0.984512 | Lyrm7 |
| ENSMUSG( | 1213.01 | 0.013707 | 0.886851 | 0.983255 | Lyrm9 |
| ENSMUSG( | 241.7068 | -0.25164 | 0.081623 | 0.441308 | Lyset |
| ENSMUSG( | 171.6939 | 0.218295 | 0.195505 | 0.63504 | Lysmd1 |
| ENSMUSG( | 1135.36 | 0.085494 | 0.416686 | 0.8209 | Lysmd2 |
| ENSMUSG( | 137.7295 | -0.27947 | 0.143685 | 0.562581 | Lysmd3 |
| ENSMUSG( | 817.1045 | 0.072619 | 0.512191 | 0.871563 | Lysmd4 |
| ENSMUSG( | 1252.672 | 0.029473 | 0.809887 | 0.96372 | Lyst |
| ENSMUSG( | 103.6636 | 0.125189 | 0.628966 | 0.913816 | Lyve1 |
| ENSMUSG( | 683.3394 | -0.54464 | 0.019029 | 0.207571 | Lyz2 |
| ENSMUSG( | 20.1369 | -0.25487 | 0.533455 | NA | Lyzl4 |
| ENSMUSG( | 393.7938 | 0.015365 | 0.904386 | 0.986246 | Lzic |
| ENSMUSG( | 1058.13 | 0.121666 | 0.200096 | 0.640302 | Lztfl1 |
| ENSMUSG( | 5371.714 | 0.141891 | 0.10489 | 0.491708 | Lztr1 |
| ENSMUSG( | 2813.554 | 0.193418 | 0.051257 | 0.347426 | Lzts1 |
| ENSMUSG( | 1775.822 | 0.039183 | 0.73446 | 0.945584 | Lzts2 |
| ENSMUSG( | 6853.728 | 0.128308 | 0.199828 | 0.640239 | Lzts3 |
| ENSMUSG( | 152.5159 | 0.261418 | 0.114102 | 0.509818 | M5C1000I18Rik |
| ENSMUSG( | 3305.649 | -0.38796 | 0.003383 | 0.073288 | M6pr |
| ENSMUSG( | 12.94154 | -0.18255 | 0.729978 | NA | Mab21l1 |
| ENSMUSG( | 18394.05 | -0.25509 | 0.023881 | 0.236336 | Macf1 |
| ENSMUSG( | 1030.134 | -0.04401 | 0.708821 | 0.939519 | Macir |
| ENSMUSG( | 2338.912 | -0.29839 | 0.026124 | 0.247245 | Maco1 |
| ENSMUSG( | 493.4757 | 0.061952 | 0.59414 | 0.902869 | Macrod1 |
| ENSMUSG( | 894.4994 | 0.086828 | 0.410619 | 0.818493 | Macrod2 |
| ENSMUSG( | 2871.927 | -0.21592 | 0.010472 | 0.146428 | Macroh2a1 |
| ENSMUSG( | 670.2107 | -0.35138 | 0.004363 | 0.086207 | Macroh2a2 |
| ENSMUSG( | 524.8049 | -0.17554 | 0.119955 | 0.523983 | Mad1l1 |
| ENSMUSG( | 189.1801 | 0.123634 | 0.493995 | 0.860611 | Mad2l1 |
| ENSMUSG( | 135.4144 | 0.192689 | 0.393261 | 0.809274 | Mad2l1bp |
| ENSMUSG( | 349.7865 | -0.03341 | 0.816429 | 0.965455 | Mad2l2 |
| ENSMUSG( | 5468.947 | 0.084899 | 0.405856 | 0.81652 | Madd |
| ENSMUSG( | 1498.027 | 0.048502 | 0.691335 | 0.933881 | Maea |
| ENSMUSG( | 648.19 | 0.072011 | 0.519771 | 0.874979 | Maf |
| ENSMUSG( | 1811.367 | 0.123717 | 0.23744 | 0.685281 | Maf1 |
| ENSMUSG( | 56.03694 | 0.260395 | 0.334677 | 0.776386 | Mafa |
| ENSMUSG( | 554.4843 | 0.107934 | 0.362003 | 0.792829 | Mafb |
| ENSMUSG( | 63.69697 | 0.227136 | 0.37199 | 0.798768 | Maff |
| ENSMUSG( | 3957.974 | 0.165985 | 0.08796 | 0.454947 | Mafg |
| ENSMUSG( | 458.3803 | 0.066043 | 0.601357 | 0.904603 | Mafk |
| ENSMUSG( | 10458.26 | -0.62789 | 1.09E-05 | 0.001199 | Mag |
| ENSMUSG( | 11631.53 | -0.10011 | 0.214705 | 0.659193 | Maged1 |
| ENSMUSG( | 1263.14 | -0.04676 | 0.610288 | 0.908189 | Maged2 |
| ENSMUSG( | 2593.218 | -0.14559 | 0.08592 | 0.451489 | Magee1 |
| ENSMUSG( | 126.5022 | -0.3213 | 0.087382 | 0.454663 | Magee2 |
| ENSMUSG( | 1331.644 | -0.25921 | 0.052343 | 0.35132 | Mageh1 |
| ENSMUSG( | 53.61873 | 0.083632 | 0.767899 | 0.954621 | Magel2 |
| ENSMUSG( | 2413.251 | -0.03454 | 0.682622 | 0.93198 | Magi1 |
| ENSMUSG( | 3825.756 | -0.00585 | 0.945884 | 0.99369 | Magi2 |
| ENSMUSG( | 2056.276 | 0.15994 | 0.125831 | 0.532614 | Magi3 |
| ENSMUSG( | 31.47067 | -0.27061 | 0.451178 | 0.835567 | Magix |
| ENSMUSG( | 291.0403 | -0.13214 | 0.448156 | 0.834051 | Magoh |
| ENSMUSG( | 69.36033 | -0.19081 | 0.467182 | 0.844819 | Magohb |
| ENSMUSG( | 1151.508 | -0.08435 | 0.581108 | 0.897503 | Magt1 |

| | | | | | |
|---|---|---|---|---|---|
| ENSMUSG( | 261.0582 | -0.35736 | 0.06452 | 0.737935 | Lsp1 |
| ENSMUSG( | 401.7843 | 0.413964 | 0.082188 | 0.77681 | Lsr |
| ENSMUSG( | 2484.998 | -0.28604 | 0.019434 | 0.556192 | Lss |
| ENSMUSG( | 72.97295 | -0.78999 | 0.004948 | 0.28972 | Lst1 |
| ENSMUSG( | 771.9922 | -0.09026 | 0.509551 | 0.929089 | Lta4h |
| ENSMUSG( | 14.89953 | -0.46084 | 0.403973 | 0.917257 | Ltb |
| ENSMUSG( | 12.47862 | -0.54636 | 0.355439 | 0.912555 | Ltb4r1 |
| ENSMUSG( | 675.9348 | -0.07769 | 0.625974 | 0.953234 | Ltbp1 |
| ENSMUSG( | 36.11258 | 0.222582 | 0.548636 | 0.938063 | Ltbp2 |
| ENSMUSG( | 1629.564 | 0.069589 | 0.59411 | 0.945408 | Ltbp3 |
| ENSMUSG( | 2386.507 | 0.371864 | 0.027447 | 0.607058 | Ltbp4 |
| ENSMUSG( | 810.0118 | 0.059243 | 0.670182 | 0.95694 | Ltbr |
| ENSMUSG( | 218.3095 | -0.00396 | 0.982057 | 0.996756 | Ltc4s |
| ENSMUSG( | 86.92226 | 0.223889 | 0.792925 | 0.97681 | Ltf |
| ENSMUSG( | 695.0306 | 0.137291 | 0.50094 | 0.927825 | Ltk |
| ENSMUSG( | 1213.797 | 0.170659 | 0.133162 | 0.822912 | Ltn1 |
| ENSMUSG( | 384.72 | -0.07433 | 0.656956 | 0.955625 | LTO1 |
| ENSMUSG( | 312.9337 | 0.023658 | 0.87122 | 0.988779 | Ltv1 |
| ENSMUSG( | 2564.397 | 0.113967 | 0.280885 | 0.892356 | Luc7l |
| ENSMUSG( | 6038.896 | 0.02461 | 0.863604 | 0.986547 | Luc7l2 |
| ENSMUSG( | 3445.715 | 0.272845 | 0.077584 | 0.77321 | Luc7l3 |
| ENSMUSG( | 91.90367 | 0.966052 | 0.000698 | 0.085493 | Lum |
| ENSMUSG( | 1290.251 | -0.08786 | 0.440327 | 0.921166 | Lurap1 |
| ENSMUSG( | 177.7096 | 0.164078 | 0.41965 | 0.917257 | Lurap1l |
| ENSMUSG( | 1788.703 | 0.166102 | 0.121883 | 0.813384 | Luzp1 |
| ENSMUSG( | 14880.59 | -0.14297 | 0.379047 | 0.916995 | Luzp2 |
| ENSMUSG( | 2075.638 | -0.05182 | 0.734448 | 0.967228 | Lxn |
| ENSMUSG( | 925.7571 | 0.651081 | 0.140651 | 0.830678 | Ly6a |
| ENSMUSG( | 1645.879 | 0.881454 | 0.0386 | 0.679972 | Ly6c1 |
| ENSMUSG( | 12.62485 | -1.03228 | 0.157871 | 0.837724 | Ly6c2 |
| ENSMUSG( | 14.35861 | 0.133878 | 0.821939 | 0.979198 | Ly6d |
| ENSMUSG( | 8590.101 | 0.050619 | 0.726886 | 0.966887 | Ly6e |
| ENSMUSG( | 17.17398 | 0.056665 | 0.902379 | 0.991089 | Ly6g6d |
| ENSMUSG( | 86.3295 | -0.07751 | 0.731572 | 0.967228 | Ly6g6e |
| ENSMUSG( | 6790.227 | 0.041476 | 0.734084 | 0.967228 | Ly6h |
| ENSMUSG( | 13.12513 | -0.97113 | 0.110696 | 0.807582 | Ly6m |
| ENSMUSG( | 113.6655 | 0.948117 | 0.002197 | 0.183242 | Ly75 |
| ENSMUSG( | 740.3618 | -0.11639 | 0.381293 | 0.917207 | Ly86 |
| ENSMUSG( | 14.64728 | -0.82406 | 0.138831 | 0.829437 | Ly9 |
| ENSMUSG( | 33.42792 | -0.39044 | 0.252811 | 0.88018 | Ly96 |
| ENSMUSG( | 256.4816 | 0.083884 | 0.64621 | 0.953988 | Lyar |
| ENSMUSG( | 189.4972 | 0.288477 | 0.153262 | 0.837724 | Lyl1 |
| ENSMUSG( | 581.4924 | 0.231664 | 0.135682 | 0.825545 | Lyn |
| ENSMUSG( | 17314.56 | 0.092292 | 0.548265 | 0.938063 | Lynx1 |
| ENSMUSG( | 2766.549 | 0.051735 | 0.638383 | 0.953903 | Lypd1 |
| ENSMUSG( | 17.48173 | -0.76144 | 0.11713 | 0.813384 | Lypd2 |
| ENSMUSG( | 2048.838 | -0.24624 | 0.089943 | 0.790741 | Lypd6 |
| ENSMUSG( | 492.533 | 0.11244 | 0.497215 | 0.927825 | Lypd6b |
| ENSMUSG( | 281.2221 | 0.209462 | 0.32032 | 0.903887 | Lypla1 |
| ENSMUSG( | 1689.483 | -0.13539 | 0.291337 | 0.897566 | Lypla2 |
| ENSMUSG( | 106.1208 | -0.37917 | 0.082987 | 0.77681 | Lyplal1 |
| ENSMUSG( | 184.5851 | 0.019461 | 0.920026 | 0.992644 | Lyrm1 |
| ENSMUSG( | 339.7804 | -0.12427 | 0.404939 | 0.917257 | Lyrm2 |
| ENSMUSG( | 512.4744 | -0.26255 | 0.064623 | 0.738417 | Lyrm4 |
| ENSMUSG( | 262.2223 | -0.10995 | 0.516269 | 0.930939 | Lyrm7 |
| ENSMUSG( | 1120.349 | 0.053622 | 0.643583 | 0.953903 | Lyrm9 |
| ENSMUSG( | 256.1053 | 0.188202 | 0.231242 | 0.868507 | Lyset |
| ENSMUSG( | 150.3077 | 0.095777 | 0.635317 | 0.953903 | Lysmd1 |
| ENSMUSG( | 1002.855 | -0.00848 | 0.952324 | 0.994798 | Lysmd2 |
| ENSMUSG( | 154.2037 | 0.32069 | 0.107867 | 0.804969 | Lysmd3 |
| ENSMUSG( | 725.76 | 0.002547 | 0.983769 | 0.996963 | Lysmd4 |
| ENSMUSG( | 1400.098 | 0.568771 | 0.000137 | 0.026415 | Lyst |
| ENSMUSG( | 108.6037 | 0.489704 | 0.071261 | 0.761942 | Lyve1 |
| ENSMUSG( | 666.2216 | -0.32814 | 0.174357 | 0.844317 | Lyz2 |
| ENSMUSG( | 20.14577 | 0.022952 | 0.957717 | 0.994798 | Lyzl4 |
| ENSMUSG( | 334.4684 | -0.20354 | 0.202605 | 0.864412 | Lzic |
| ENSMUSG( | 984.6371 | 0.177233 | 0.13444 | 0.823094 | Lztfl1 |
| ENSMUSG( | 4613.083 | -0.02498 | 0.820524 | 0.979198 | Lztr1 |
| ENSMUSG( | 2407.133 | 0.019477 | 0.861398 | 0.986281 | Lzts1 |
| ENSMUSG( | 1705.803 | 0.188344 | 0.113947 | 0.810728 | Lzts2 |
| ENSMUSG( | 6142.456 | 0.083715 | 0.433155 | 0.918434 | Lzts3 |
| ENSMUSG( | 122.5457 | -0.07942 | 0.696821 | 0.960867 | M5C1000I18Rik |
| ENSMUSG( | 3483.336 | 0.060766 | 0.697611 | 0.960867 | M6pr |
| ENSMUSG( | 18245.92 | 0.003794 | 0.975741 | 0.996416 | Macf1 |
| ENSMUSG( | 946.5875 | -0.01913 | 0.88576 | 0.98977 | Macir |
| ENSMUSG( | 2563.648 | 0.243882 | 0.125521 | 0.81516 | Maco1 |
| ENSMUSG( | 406.4777 | -0.24721 | 0.108277 | 0.804969 | Macrod1 |
| ENSMUSG( | 788.0965 | -0.01563 | 0.916977 | 0.992529 | Macrod2 |
| ENSMUSG( | 2707.584 | -0.11318 | 0.316926 | 0.902088 | Macroh2a1 |
| ENSMUSG( | 680.8201 | -0.01496 | 0.915467 | 0.992529 | Macroh2a2 |
| ENSMUSG( | 472.7214 | -0.21458 | 0.120052 | 0.813384 | Mad1l1 |
| ENSMUSG( | 176.1242 | 0.173046 | 0.410084 | 0.917257 | Mad2l1 |
| ENSMUSG( | 118.9379 | 0.097096 | 0.694367 | 0.960867 | Mad2l1bp |
| ENSMUSG( | 312.5374 | -0.09305 | 0.523666 | 0.932848 | Mad2l2 |
| ENSMUSG( | 4982.749 | 0.086804 | 0.424114 | 0.917257 | Madd |
| ENSMUSG( | 1302.728 | -0.0886 | 0.514989 | 0.930939 | Maea |
| ENSMUSG( | 608.7405 | 0.156551 | 0.238232 | 0.87259 | Maf |
| ENSMUSG( | 1609.102 | 0.055547 | 0.639623 | 0.953903 | Maf1 |
| ENSMUSG( | 51.39563 | 0.2795 | 0.352132 | 0.912093 | Mafa |
| ENSMUSG( | 491.3491 | 0.034449 | 0.807892 | 0.978622 | Mafb |
| ENSMUSG( | 63.23449 | 0.450898 | 0.08347 | 0.77681 | Maff |
| ENSMUSG( | 3636.927 | 0.189738 | 0.090709 | 0.791074 | Mafg |
| ENSMUSG( | 413.2846 | 0.035615 | 0.800078 | 0.97681 | Mafk |
| ENSMUSG( | 11985.21 | 0.10399 | 0.579405 | 0.944046 | Mag |
| ENSMUSG( | 10947.66 | -0.00429 | 0.964913 | 0.995433 | Maged1 |
| ENSMUSG( | 1120.823 | -0.12663 | 0.253795 | 0.880692 | Maged2 |
| ENSMUSG( | 2433.871 | -0.05795 | 0.618246 | 0.952254 | Magee1 |
| ENSMUSG( | 116.6989 | -0.29133 | 0.166848 | 0.840183 | Magee2 |
| ENSMUSG( | 1358.163 | 0.079587 | 0.590961 | 0.944975 | Mageh1 |
| ENSMUSG( | 50.43609 | 0.141667 | 0.656196 | 0.955533 | Magel2 |
| ENSMUSG( | 2187.393 | -0.05022 | 0.628789 | 0.953251 | Magi1 |
| ENSMUSG( | 3446.021 | -0.04135 | 0.710966 | 0.964363 | Magi2 |
| ENSMUSG( | 1791.682 | 0.034322 | 0.788343 | 0.97637 | Magi3 |
| ENSMUSG( | 30.4057 | -0.05806 | 0.883842 | 0.989644 | Magix |
| ENSMUSG( | 267.9013 | -0.11486 | 0.576502 | 0.942912 | Magoh |
| ENSMUSG( | 66.2811 | -0.06103 | 0.832431 | 0.98075 | Magohb |
| ENSMUSG( | 1133.509 | 0.136388 | 0.401058 | 0.917257 | Magt1 |
| ENSMUSG( | 504.7634 | 0.07349 | 0.593717 | 0.945408 | Maip1 |
| ENSMUSG( | 61.54825 | 0.076135 | 0.79277 | 0.97681 | Mak |
| ENSMUSG( | 644.1829 | -0.02258 | 0.874002 | 0.989644 | Mak16 |
| ENSMUSG( | 19602.19 | 0.151521 | 0.388687 | 0.917257 | Mal |
| ENSMUSG( | 4657.191 | 0.13275 | 0.414222 | 0.917257 | Mal2 |
| ENSMUSG( | 7139.72 | 0.24594 | 0.169918 | 0.840806 | Malat1 |
| ENSMUSG( | 28.98951 | -0.04186 | 0.932333 | 0.99306 | Mall |
| ENSMUSG( | 223.3191 | -0.1193 | 0.489338 | 0.925673 | Malsu1 |
| ENSMUSG( | 255.8732 | 0.07657 | 0.63538 | 0.953903 | Malt1 |
| ENSMUSG( | 70.83589 | 0.028569 | 0.913177 | 0.992298 | Mamdc2 |
| ENSMUSG( | 59.21957 | 0.665555 | 0.025677 | 0.599234 | Mamdc4 |
| ENSMUSG( | 316.8836 | 0.085347 | 0.580293 | 0.944209 | Maml1 |
| ENSMUSG( | 486.2347 | 0.320634 | 0.015459 | 0.495973 | Maml2 |
| ENSMUSG( | 305.1336 | -0.02449 | 0.888224 | 0.990496 | Maml3 |
| ENSMUSG( | 735.8308 | 0.047835 | 0.711043 | 0.964363 | Mamld1 |
| ENSMUSG( | 43.47374 | -0.10405 | 0.758023 | 0.971858 | Mamstr |
| ENSMUSG( | 1347.22 | 0.382126 | 0.028686 | 0.614171 | Man1a |
| ENSMUSG( | 3548.371 | 0.188113 | 0.185906 | 0.855305 | Man1a2 |
| ENSMUSG( | 3107.517 | 0.078172 | 0.583415 | 0.944266 | Man1b1 |
| ENSMUSG( | 1821.648 | 0.265486 | 0.083439 | 0.77681 | Man1c1 |
| ENSMUSG( | 772.5161 | 0.434911 | 0.009811 | 0.406458 | Man2a1 |
| ENSMUSG( | 8902.172 | 0.118512 | 0.313713 | 0.900949 | Man2a2 |
| ENSMUSG( | 2646.608 | 0.30337 | 0.023881 | 0.588226 | Man2b1 |
| ENSMUSG( | 1538.269 | 0.098195 | 0.525766 | 0.934374 | Man2b2 |
| ENSMUSG( | 1576.507 | 0.049423 | 0.768811 | 0.974236 | Man2c1 |
| ENSMUSG( | 33.37117 | 0.161627 | 0.644291 | 0.953903 | Man2c1os |
| ENSMUSG( | 424.2393 | 0.196561 | 0.311161 | 0.899441 | Manba |
| ENSMUSG( | 1527.599 | -0.16235 | 0.166385 | 0.84011 | Manbal |
| ENSMUSG( | 1160.704 | 0.09642 | 0.612439 | 0.951071 | Manea |
| ENSMUSG( | 1981.186 | 0.061862 | 0.704557 | 0.962677 | Maneal |
| ENSMUSG( | 1927.62 | 0.25904 | 0.109468 | 0.805353 | Manf |
| ENSMUSG( | 333.7632 | -0.02182 | 0.91662 | 0.992529 | Mansc1 |
| ENSMUSG( | 1390.595 | -0.02761 | 0.824336 | 0.979198 | Maoa |
| ENSMUSG( | 2131.882 | -0.21066 | 0.13244 | 0.822912 | Maob |
| ENSMUSG( | 76.88389 | 0.341335 | 0.176948 | 0.846973 | Map10 |
| ENSMUSG( | 38742.9 | 0.094776 | 0.548242 | 0.938063 | Map1a |
| ENSMUSG( | 13825.65 | 0.061372 | 0.544701 | 0.938063 | Map1b |
| ENSMUSG( | 5507.162 | -0.23334 | 0.041304 | 0.688419 | Map1lc3a |
| ENSMUSG( | 8017.069 | -0.16554 | 0.135093 | 0.824827 | Map1lc3b |
| ENSMUSG( | 1444.248 | 0.119653 | 0.334861 | 0.907907 | Map1s |
| ENSMUSG( | 34821.64 | 0.005935 | 0.961572 | 0.99511 | Map2 |
| ENSMUSG( | 4001.575 | -0.11453 | 0.493551 | 0.927372 | Map2k1 |
| ENSMUSG( | 2713.474 | -0.0675 | 0.523805 | 0.932848 | Map2k2 |
| ENSMUSG( | 367.4799 | 0.113855 | 0.471494 | 0.923329 | Map2k3 |
| ENSMUSG( | 16.91078 | 0.169769 | 0.729444 | 0.967022 | Map2k3os |
| ENSMUSG( | 3176.694 | -0.06023 | 0.650411 | 0.95422 | Map2k4 |
| ENSMUSG( | 791.5703 | 0.015724 | 0.896317 | 0.990558 | Map2k5 |
| ENSMUSG( | 233.7749 | 0.322848 | 0.053901 | 0.722223 | Map2k6 |
| ENSMUSG( | 1604.756 | -0.07783 | 0.539334 | 0.937216 | Map2k7 |
| ENSMUSG( | 379.0845 | 0.430145 | 0.005509 | 0.303106 | Map3k1 |
| ENSMUSG( | 13320.26 | 0.125227 | 0.246249 | 0.877904 | Map3k10 |

| | | | | | | | | | | | |
|---|---|---|---|---|---|---|---|---|---|---|---|
| ENSMUSG( | Lztfl1 | 1058.13 | 0.121666 | 0.200096 | 0.640302 | ENSMUSG( | Lztfl1 | 984.6371 | 0.177233 | 0.13444 | 0.823094 |
| ENSMUSG( | Lztr1 | 5371.714 | 0.141891 | 0.10489 | 0.491708 | ENSMUSG( | Lztr1 | 4613.083 | -0.02498 | 0.820524 | 0.979198 |
| ENSMUSG( | Lzts1 | 2813.554 | 0.193418 | 0.051257 | 0.347426 | ENSMUSG( | Lzts1 | 2407.133 | 0.019477 | 0.861398 | 0.986281 |
| ENSMUSG( | Lzts2 | 1775.822 | 0.039183 | 0.73446 | 0.945584 | ENSMUSG( | Lzts2 | 1705.803 | 0.188344 | 0.113947 | 0.810728 |
| ENSMUSG( | Lzts3 | 6853.728 | 0.128308 | 0.199828 | 0.640239 | ENSMUSG( | Lzts3 | 6142.456 | 0.083715 | 0.433155 | 0.918434 |
| ENSMUSG( | M5C1000I1 | 152.5159 | 0.261418 | 0.114102 | 0.509818 | ENSMUSG( | M5C1000I1 | 122.5457 | -0.07942 | 0.696821 | 0.960867 |
| ENSMUSG( | M6pr | 3305.649 | -0.38796 | 0.003383 | 0.073288 | ENSMUSG( | M6pr | 3483.336 | 0.060766 | 0.697611 | 0.960867 |
| ENSMUSG( | Macf1 | 18394.05 | -0.25509 | 0.023881 | 0.236336 | ENSMUSG( | Macf1 | 18245.92 | 0.003794 | 0.975741 | 0.996416 |
| ENSMUSG( | Macir | 1030.134 | -0.04401 | 0.708821 | 0.939519 | ENSMUSG( | Macir | 946.5875 | -0.01913 | 0.88576 | 0.98977 |
| ENSMUSG( | Maco1 | 2338.912 | -0.29839 | 0.026124 | 0.247245 | ENSMUSG( | Maco1 | 2563.648 | 0.243882 | 0.125521 | 0.81516 |
| ENSMUSG( | Macrod1 | 493.4757 | 0.061952 | 0.59414 | 0.902869 | ENSMUSG( | Macrod1 | 406.4777 | -0.24721 | 0.108277 | 0.804969 |
| ENSMUSG( | Macrod2 | 894.4994 | 0.086828 | 0.410619 | 0.818493 | ENSMUSG( | Macrod2 | 788.0965 | -0.01563 | 0.916977 | 0.992529 |
| ENSMUSG( | Macroh2a1 | 2871.927 | -0.21592 | 0.010472 | 0.146428 | ENSMUSG( | Macroh2a1 | 2707.584 | -0.11318 | 0.316926 | 0.902088 |
| ENSMUSG( | Macroh2a2 | 670.2107 | -0.35138 | 0.004363 | 0.086207 | ENSMUSG( | Macroh2a2 | 680.8201 | -0.01496 | 0.915467 | 0.992529 |
| ENSMUSG( | Mad1l1 | 524.8049 | -0.17554 | 0.119955 | 0.523983 | ENSMUSG( | Mad1l1 | 472.7214 | -0.21458 | 0.120052 | 0.813384 |
| ENSMUSG( | Mad2l1 | 189.1801 | 0.123634 | 0.493995 | 0.860611 | ENSMUSG( | Mad2l1 | 176.1242 | 0.173046 | 0.410084 | 0.917257 |
| ENSMUSG( | Mad2l1bp | 135.4144 | 0.192689 | 0.393261 | 0.809274 | ENSMUSG( | Mad2l1bp | 118.9379 | 0.097096 | 0.694367 | 0.960867 |
| ENSMUSG( | Mad2l2 | 349.7865 | -0.03341 | 0.816429 | 0.965455 | ENSMUSG( | Mad2l2 | 312.5374 | -0.09305 | 0.523666 | 0.932848 |
| ENSMUSG( | Madd | 5468.947 | 0.084899 | 0.405856 | 0.81652 | ENSMUSG( | Madd | 4982.749 | 0.086804 | 0.424114 | 0.917257 |
| ENSMUSG( | Maea | 1498.027 | 0.048502 | 0.691335 | 0.933881 | ENSMUSG( | Maea | 1302.728 | -0.0886 | 0.514989 | 0.930939 |
| ENSMUSG( | Maf | 648.19 | 0.072011 | 0.519771 | 0.874979 | ENSMUSG( | Maf | 608.7405 | 0.156551 | 0.238232 | 0.87259 |
| ENSMUSG( | Maf1 | 1811.367 | 0.123717 | 0.23744 | 0.685281 | ENSMUSG( | Maf1 | 1609.102 | 0.055547 | 0.639623 | 0.953903 |
| ENSMUSG( | Mafa | 56.03694 | 0.260395 | 0.334677 | 0.776386 | ENSMUSG( | Mafa | 51.39563 | 0.2795 | 0.352132 | 0.912093 |
| ENSMUSG( | Mafb | 554.4843 | 0.107934 | 0.362003 | 0.792829 | ENSMUSG( | Mafb | 491.3491 | 0.034449 | 0.807892 | 0.978622 |
| ENSMUSG( | Maff | 63.69697 | 0.227136 | 0.37199 | 0.798768 | ENSMUSG( | Maff | 63.23449 | 0.450794 | 0.08347 | 0.77681 |
| ENSMUSG( | Mafg | 3957.974 | 0.165985 | 0.08796 | 0.454947 | ENSMUSG( | Mafg | 3636.927 | 0.189738 | 0.090709 | 0.791074 |
| ENSMUSG( | Mafk | 458.3803 | 0.066043 | 0.601357 | 0.904603 | ENSMUSG( | Mafk | 413.2846 | 0.035615 | 0.800078 | 0.97681 |
| ENSMUSG( | Mag | 10458.26 | -0.62789 | 1.09E-05 | 0.001199 | ENSMUSG( | Mag | 11985.21 | 0.10399 | 0.579405 | 0.944046 |
| ENSMUSG( | Maged1 | 11631.53 | -0.10011 | 0.214705 | 0.659193 | ENSMUSG( | Maged1 | 10947.66 | -0.00429 | 0.964913 | 0.995433 |
| ENSMUSG( | Maged2 | 1263.14 | -0.04676 | 0.610288 | 0.908189 | ENSMUSG( | Maged2 | 1120.823 | -0.12663 | 0.253795 | 0.880692 |
| ENSMUSG( | Magee1 | 2593.218 | -0.14559 | 0.08592 | 0.451489 | ENSMUSG( | Magee1 | 2433.871 | -0.05795 | 0.618246 | 0.952254 |
| ENSMUSG( | Magee2 | 126.5022 | -0.3213 | 0.087382 | 0.454663 | ENSMUSG( | Magee2 | 116.6989 | -0.29133 | 0.166848 | 0.840183 |
| ENSMUSG( | Mageh1 | 1331.644 | -0.25921 | 0.052343 | 0.35132 | ENSMUSG( | Mageh1 | 1358.163 | 0.079587 | 0.590961 | 0.944975 |
| ENSMUSG( | Magel2 | 53.61873 | 0.083632 | 0.767899 | 0.954621 | ENSMUSG( | Magel2 | 50.43609 | 0.141667 | 0.656196 | 0.955533 |
| ENSMUSG( | Magi1 | 2413.251 | -0.03454 | 0.682622 | 0.93198 | ENSMUSG( | Magi1 | 2187.393 | -0.05022 | 0.628789 | 0.953251 |
| ENSMUSG( | Magi2 | 3825.756 | -0.00585 | 0.945884 | 0.99369 | ENSMUSG( | Magi2 | 3446.021 | -0.04135 | 0.710966 | 0.964363 |
| ENSMUSG( | Magi3 | 2056.276 | 0.15994 | 0.125831 | 0.532614 | ENSMUSG( | Magi3 | 1791.682 | 0.034322 | 0.788343 | 0.97637 |
| ENSMUSG( | Magix | 31.47067 | -0.27061 | 0.451178 | 0.835567 | ENSMUSG( | Magix | 30.4057 | -0.05806 | 0.883842 | 0.989644 |
| ENSMUSG( | Magoh | 291.0403 | -0.13214 | 0.448156 | 0.834051 | ENSMUSG( | Magoh | 267.9013 | -0.11486 | 0.576502 | 0.942912 |
| ENSMUSG( | Magohb | 69.36033 | -0.19081 | 0.467182 | 0.844819 | ENSMUSG( | Magohb | 66.2811 | -0.06103 | 0.832431 | 0.98075 |
| ENSMUSG( | Magt1 | 1151.508 | -0.08435 | 0.581108 | 0.897503 | ENSMUSG( | Magt1 | 1133.509 | 0.136388 | 0.401058 | 0.917257 |
| ENSMUSG( | Maip1 | 539.9042 | -0.00316 | 0.97835 | 0.997428 | ENSMUSG( | Maip1 | 504.7634 | 0.07349 | 0.593717 | 0.945408 |
| ENSMUSG( | Mak | 65.52231 | -0.02964 | 0.909152 | 0.98794 | ENSMUSG( | Mak | 61.54825 | 0.076135 | 0.79277 | 0.97681 |
| ENSMUSG( | Mak16 | 730.0339 | 0.070143 | 0.551584 | 0.886141 | ENSMUSG( | Mak16 | 644.1829 | -0.02258 | 0.874002 | 0.989644 |
| ENSMUSG( | Mal | 17132.39 | -0.55935 | 5.60E-05 | 0.004302 | ENSMUSG( | Mal | 19602.19 | 0.151521 | 0.388687 | 0.917257 |
| ENSMUSG( | Mal2 | 4204.064 | -0.46817 | 0.00035 | 0.016176 | ENSMUSG( | Mal2 | 4657.191 | 0.13275 | 0.414222 | 0.917257 |
| ENSMUSG( | Malat1 | 6928.573 | -0.09967 | 0.479397 | 0.854046 | ENSMUSG( | Malat1 | 7139.72 | 0.24594 | 0.169918 | 0.840806 |
| ENSMUSG( | Mall | 30.17372 | -0.23691 | 0.60861 | 0.907154 | ENSMUSG( | Mall | 28.98951 | -0.04186 | 0.932333 | 0.99306 |
| ENSMUSG( | Malsu1 | 257.1797 | 0.028789 | 0.844057 | 0.971962 | ENSMUSG( | Malsu1 | 223.3191 | -0.1193 | 0.489338 | 0.925673 |
| ENSMUSG( | Malt1 | 269.2565 | -0.0528 | 0.711057 | 0.940028 | ENSMUSG( | Malt1 | 255.8732 | 0.07657 | 0.63538 | 0.953903 |
| ENSMUSG( | Mamdc2 | 67.65645 | -0.41499 | 0.118481 | 0.520662 | ENSMUSG( | Mamdc2 | 70.83589 | 0.028569 | 0.913177 | 0.992298 |
| ENSMUSG( | Mamdc4 | 59.57291 | 0.429866 | 0.137049 | 0.551733 | ENSMUSG( | Mamdc4 | 59.21957 | 0.665555 | 0.025677 | 0.599234 |
| ENSMUSG( | Maml1 | 319.5176 | -0.17 | 0.194475 | 0.63341 | ENSMUSG( | Maml1 | 316.8836 | 0.085347 | 0.580293 | 0.944209 |
| ENSMUSG( | Maml2 | 505.5083 | 0.176223 | 0.132302 | 0.544251 | ENSMUSG( | Maml2 | 486.2347 | 0.320634 | 0.015459 | 0.495973 |
| ENSMUSG( | Maml3 | 317.4961 | -0.19709 | 0.156196 | 0.583988 | ENSMUSG( | Maml3 | 305.1336 | -0.02449 | 0.888224 | 0.990496 |
| ENSMUSG( | Mamld1 | 843.2597 | 0.171186 | 0.087772 | 0.454663 | ENSMUSG( | Mamld1 | 735.8308 | 0.047835 | 0.711043 | 0.964363 |
| ENSMUSG( | Mamstr | 51.10512 | 0.07642 | 0.791252 | 0.959461 | ENSMUSG( | Mamstr | 43.47374 | -0.10405 | 0.758023 | 0.971858 |
| ENSMUSG( | Man1a | 1090.529 | -0.52194 | 0.00027 | 0.013409 | ENSMUSG( | Man1a | 1347.22 | 0.382126 | 0.028686 | 0.614171 |
| ENSMUSG( | Man1a2 | 3511.623 | -0.109 | 0.387497 | 0.805693 | ENSMUSG( | Man1a2 | 3548.371 | 0.188113 | 0.185906 | 0.855305 |
| ENSMUSG( | Man1b1 | 3145.619 | -0.16291 | 0.170521 | 0.605271 | ENSMUSG( | Man1b1 | 3107.517 | 0.078172 | 0.583415 | 0.944266 |
| ENSMUSG( | Man1c1 | 1776.307 | -0.06944 | 0.598246 | 0.903801 | ENSMUSG( | Man1c1 | 1821.648 | 0.265486 | 0.083439 | 0.77681 |
| ENSMUSG( | Man2a1 | 685.1701 | -0.15732 | 0.283557 | 0.732021 | ENSMUSG( | Man2a1 | 772.5161 | 0.434911 | 0.009811 | 0.406458 |
| ENSMUSG( | Man2a2 | 9181.355 | -0.06153 | 0.544891 | 0.884712 | ENSMUSG( | Man2a2 | 8902.172 | 0.118512 | 0.313713 | 0.900949 |
| ENSMUSG( | Man2b1 | 2602.001 | -0.00134 | 0.989501 | 0.999111 | ENSMUSG( | Man2b1 | 2646.608 | 0.30337 | 0.023881 | 0.588226 |
| ENSMUSG( | Man2b2 | 1544.052 | -0.1677 | 0.202392 | 0.642343 | ENSMUSG( | Man2b2 | 1538.269 | 0.098195 | 0.525766 | 0.934374 |
| ENSMUSG( | Man2c1 | 1864.84 | 0.257734 | 0.068801 | 0.406391 | ENSMUSG( | Man2c1 | 1576.507 | 0.049423 | 0.768811 | 0.974236 |
| ENSMUSG( | Man2c1os | 34.2956 | -0.01804 | 0.956152 | 0.993874 | ENSMUSG( | Man2c1os | 33.37117 | 0.161627 | 0.644291 | 0.953903 |
| ENSMUSG( | Manba | 412.0536 | -0.16223 | 0.333639 | 0.775602 | ENSMUSG( | Manba | 424.2393 | 0.196561 | 0.311161 | 0.899441 |
| ENSMUSG( | Manbal | 1640.12 | -0.22929 | 0.012188 | 0.160067 | ENSMUSG( | Manbal | 1527.599 | -0.16235 | 0.166385 | 0.84011 |
| ENSMUSG( | Manea | 1062.92 | -0.46709 | 0.004204 | 0.084295 | ENSMUSG( | Manea | 1160.704 | 0.09642 | 0.612439 | 0.951071 |
| ENSMUSG( | Maneal | 1939.576 | -0.28717 | 0.042487 | 0.316636 | ENSMUSG( | Maneal | 1981.186 | 0.061862 | 0.704557 | 0.962677 |
| ENSMUSG( | Manf | 1839.495 | -0.1393 | 0.357856 | 0.789405 | ENSMUSG( | Manf | 1927.62 | 0.25904 | 0.109468 | 0.805353 |
| ENSMUSG( | Mansc1 | 304.488 | -0.6288 | 0.000825 | 0.02866 | ENSMUSG( | Mansc1 | 333.7632 | -0.02182 | 0.91662 | 0.992529 |
| ENSMUSG( | Maoa | 1531.495 | -0.01759 | 0.884414 | 0.982647 | ENSMUSG( | Maoa | 1390.595 | -0.02761 | 0.824336 | 0.979198 |
| ENSMUSG( | Maob | 2747.994 | 0.251871 | 0.032762 | 0.277278 | ENSMUSG( | Maob | 2131.882 | -0.21066 | 0.13244 | 0.822912 |
| ENSMUSG( | Map10 | 79.99132 | 0.190806 | 0.453812 | 0.836798 | ENSMUSG( | Map10 | 76.88389 | 0.341335 | 0.176948 | 0.846973 |
| ENSMUSG( | Map1a | 41592.35 | 0.028215 | 0.821479 | 0.966248 | ENSMUSG( | Map1a | 38742.9 | 0.094776 | 0.548242 | 0.938063 |
| ENSMUSG( | Map1b | 15102.5 | 0.048232 | 0.514191 | 0.872055 | ENSMUSG( | Map1b | 13825.65 | 0.061372 | 0.544701 | 0.938063 |
| ENSMUSG( | Map1lc3a | 6561.908 | 0.014982 | 0.848064 | 0.972374 | ENSMUSG( | Map1lc3a | 5507.162 | -0.23334 | 0.041304 | 0.688419 |
| ENSMUSG( | Map1lc3b | 9658.126 | 0.107543 | 0.191153 | 0.630245 | ENSMUSG( | Map1lc3b | 8017.069 | -0.16554 | 0.135093 | 0.824827 |
| ENSMUSG( | Map1s | 1638.552 | 0.207263 | 0.053718 | 0.355948 | ENSMUSG( | Map1s | 1444.248 | 0.119653 | 0.334861 | 0.907907 |
| ENSMUSG( | Map2 | 39517.59 | 0.099185 | 0.379782 | 0.801757 | ENSMUSG( | Map2 | 34821.64 | 0.005935 | 0.961572 | 0.99511 |
| ENSMUSG( | Map2k1 | 4655.97 | 0.05882 | 0.648543 | 0.920007 | ENSMUSG( | Map2k1 | 4001.575 | -0.11453 | 0.493551 | 0.927372 |
| ENSMUSG( | Map2k2 | 3136.707 | 0.08205 | 0.341131 | 0.778828 | ENSMUSG( | Map2k2 | 2713.474 | -0.0675 | 0.523805 | 0.932848 |
| ENSMUSG( | Map2k3 | 389.9843 | 0.00897 | 0.943216 | 0.993415 | ENSMUSG( | Map2k3 | 367.4799 | 0.113855 | 0.471494 | 0.923329 |
| ENSMUSG( | Map2k3os | 18.41626 | 0.187551 | 0.673061 | NA | ENSMUSG( | Map2k3os | 16.91078 | 0.169769 | 0.729444 | 0.967022 |
| ENSMUSG( | Map2k4 | 3686.462 | 0.101968 | 0.380191 | 0.801757 | ENSMUSG( | Map2k4 | 3176.694 | -0.06023 | 0.650411 | 0.95422 |
| ENSMUSG( | Map2k5 | 889.6373 | 0.080903 | 0.452585 | 0.836145 | ENSMUSG( | Map2k5 | 791.5703 | 0.015724 | 0.896317 | 0.990558 |
| ENSMUSG( | Map2k6 | 248.0806 | 0.231947 | 0.101392 | 0.484996 | ENSMUSG( | Map2k6 | 233.7749 | 0.322848 | 0.053901 | 0.722223 |
| ENSMUSG( | Map2k7 | 1817.962 | 0.017655 | 0.8761 | 0.980497 | ENSMUSG( | Map2k7 | 1604.756 | -0.07783 | 0.539334 | 0.937216 |
| ENSMUSG( | Map3k1 | 347.5496 | -0.06372 | 0.686978 | 0.933041 | ENSMUSG( | Map3k1 | 379.0845 | 0.430145 | 0.005509 | 0.303106 |
| ENSMUSG( | Map3k10 | 15222.44 | 0.235018 | 0.002563 | 0.060076 | ENSMUSG( | Map3k10 | 13320.26 | 0.125227 | 0.246249 | 0.877904 |
| ENSMUSG( | Map3k11 | 792.0158 | -0.1004 | 0.355669 | 0.787945 | ENSMUSG( | Map3k11 | 760.497 | 0.059532 | 0.703437 | 0.962481 |
| ENSMUSG( | Map3k12 | 4473.529 | 0.197606 | 0.039952 | 0.308163 | ENSMUSG( | Map3k12 | 3847.159 | 0.036938 | 0.745342 | 0.96951 |
| ENSMUSG( | Map3k13 | 551.9376 | -0.00611 | 0.959215 | 0.99396 | ENSMUSG( | Map3k13 | 511.4601 | 0.040857 | 0.762861 | 0.973698 |
| ENSMUSG( | Map3k14 | 148.8323 | -0.19522 | 0.274162 | 0.724254 | ENSMUSG( | Map3k14 | 140.402 | -0.07543 | 0.700156 | 0.961335 |
| ENSMUSG( | Map3k15 | 96.64225 | 0.209501 | 0.332139 | 0.775373 | ENSMUSG( | Map3k15 | 83.53686 | 0.054864 | 0.834296 | 0.981237 |
| ENSMUSG( | Map3k19 | 406.7981 | 0.408654 | 0.00103 | 0.033612 | ENSMUSG( | Map3k19 | 329.9613 | 0.089852 | 0.576917 | 0.942912 |
| ENSMUSG( | Map3k2 | 602.0193 | -0.01014 | 0.940371 | 0.993415 | ENSMUSG( | Map3k2 | 583.0229 | 0.167849 | 0.266888 | 0.886134 |
| ENSMUSG( | Map3k20 | 215.6574 | 0.040407 | 0.822268 | 0.966352 | ENSMUSG( | Map3k20 | 228.0963 | 0.445714 | 0.017545 | 0.53154 |
| ENSMUSG( | Map3k21 | 75.69153 | 0.123875 | 0.613453 | 0.909027 | ENSMUSG( | Map3k21 | 71.21072 | 0.227049 | 0.365048 | 0.914449 |
| ENSMUSG( | Map3k3 | 732.2331 | 0.011205 | 0.921916 | 0.990436 | ENSMUSG( | Map3k3 | 698.0967 | 0.145623 | 0.35242 | 0.912093 |
| ENSMUSG( | Map3k4 | 1256.641 | 0.199408 | 0.086469 | 0.452045 | ENSMUSG( | Map3k4 | 1110.75 | 0.117877 | 0.361021 | 0.914223 |
| ENSMUSG( | Map3k5 | 1051.452 | 0.387379 | 0.000129 | 0.007629 | ENSMUSG( | Map3k5 | 842.0588 | 0.03882 | 0.744624 | 0.96951 |
| ENSMUSG( | Map3k6 | 148.2569 | 0.928766 | 1.04E-06 | 0.000197 | ENSMUSG( | Map3k6 | 117.0704 | 0.623597 | 0.019966 | 0.559604 |
| ENSMUSG( | Map3k7 | 2885.771 | 0.054494 | 0.544223 | 0.884587 | ENSMUSG( | Map3k7 | 2563.539 | -0.01845 | 0.866563 | 0.987419 |
| ENSMUSG( | Map3k8 | 70.2748 | -0.02096 | 0.92976 | 0.991848 | ENSMUSG( | Map3k8 | 70.73557 | 0.276173 | 0.336472 | 0.907972 |
| ENSMUSG( | Map3k9 | 1032.689 | 0.225537 | 0.051552 | 0.34826 | ENSMUSG( | Map3k9 | 922.8191 | 0.178227 | 0.194748 | 0.860433 |
| ENSMUSG( | Map4 | 40817.84 | -0.003 | 0.971899 | 0.996585 | ENSMUSG( | Map4 | 34346.43 | -0.24392 | 0.030111 | 0.621996 |
| ENSMUSG( | Map4k1 | 83.83385 | 0.022193 | 0.925104 | 0.991204 | ENSMUSG( | Map4k1 | 74.94817 | -0.0343 | 0.890439 | 0.990558 |
| ENSMUSG( | Map4k2 | 2692.147 | 0.333481 | 0.013546 | 0.170062 | ENSMUSG( | Map4k2 | 2398.547 | 0.27306 | 0.093217 | 0.795363 |
| ENSMUSG( | Map4k3 | 1605.875 | 0.26243 | 0.008077 | 0.125133 | ENSMUSG( | Map4k3 | 1443.673 | 0.227557 | 0.037357 | 0.673514 |
| ENSMUSG( | Map4k4 | 17967.92 | -0.03364 | 0.700203 | 0.937391 | ENSMUSG( | Map4k4 | 15289.75 | -0.24206 | 0.044193 | 0.693037 |
| ENSMUSG( | Map4k5 | 1497.072 | 0.079096 | 0.371816 | 0.798768 | ENSMUSG( | Map4k5 | 1369.134 | 0.088181 | 0.451443 | 0.922202 |
| ENSMUSG( | Map6 | 3686.107 | 0.104817 | 0.241245 | 0.689026 | ENSMUSG( | Map6 | 3133.348 | -0.09591 | 0.4532 | 0.922316 |
| ENSMUSG( | Map6d1 | 2396.091 | -0.11046 | 0.216532 | 0.661421 | ENSMUSG( | Map6d1 | 2073.087 | -0.27038 | 0.011814 | 0.444083 |
| ENSMUSG( | Map7 | 3370.551 | -0.02118 | 0.81646 | 0.965455 | ENSMUSG( | Map7 | 3215.983 | 0.11017 | 0.297834 | 0.898538 |
| ENSMUSG( | Map7d1 | 7380.108 | 0.090721 | 0.333645 | 0.775602 | ENSMUSG( | Map7d1 | 6619.929 | 0.046818 | 0.664447 | 0.956766 |
| ENSMUSG( | Map7d2 | 2266.197 | 0.168167 | 0.163537 | 0.594869 | ENSMUSG( | Map7d2 | 1919.353 | -0.0413 | 0.782161 | 0.976106 |
| ENSMUSG( | Map9 | 1892.892 | 0.154968 | 0.096938 | 0.477375 | ENSMUSG( | Map9 | 1653.761 | 0.037819 | 0.741177 | 0.968811 |
| ENSMUSG( | Mapk1 | 8579.399 | 0.096792 | 0.343946 | 0.779153 | ENSMUSG( | Mapk1 | 7508.325 | -0.01877 | 0.877798 | 0.989644 |
| ENSMUSG( | Mapk10 | 7302.086 | 0.14992 | 0.105427 | 0.492417 | ENSMUSG( | Mapk10 | 6179.802 | -0.06132 | 0.594281 | 0.945408 |
| ENSMUSG( | Mapk11 | 1350.386 | 0.199321 | 0.114373 | 0.51021 | ENSMUSG( | Mapk11 | 1128.945 | -0.04542 | 0.764331 | 0.974 |
| ENSMUSG( | Mapk12 | 194.3813 | -0.09503 | 0.576012 | 0.895114 | ENSMUSG( | Mapk12 | 191.1757 | 0.129812 | 0.514867 | 0.930939 |
| ENSMUSG( | Mapk13 | 26.89636 | -0.03666 | 0.919924 | 0.990247 | ENSMUSG( | Mapk13 | 23.44176 | -0.15584 | 0.702619 | 0.962215 |
| ENSMUSG( | Mapk14 | 2478.552 | 0.153701 | 0.09237 | 0.466744 | ENSMUSG( | Mapk14 | 2121.876 | -0.02283 | 0.839617 | 0.982077 |
| ENSMUSG( | Mapk15 | 56.49862 | 0.134227 | 0.733076 | 0.944956 | ENSMUSG( | Mapk15 | 50.34164 | 0.081414 | 0.837232 | 0.981335 |
| ENSMUSG( | Mapk1ip1 | 897.6389 | 0.125104 | 0.393847 | 0.809643 | ENSMUSG( | Mapk1ip1 | 783.4759 | 0.002188 | 0.989653 | 0.997928 |
| ENSMUSG( | Mapk1ip1l | 854.7385 | 0.104924 | 0.361825 | 0.792797 | ENSMUSG( | Mapk1ip1l | 745.3381 | -0.01904 | 0.879948 | 0.989644 |
| ENSMUSG( | Mapk3 | 8125.248 | 0.080851 | 0.319146 | 0.765081 | ENSMUSG( | Mapk3 | 7000.155 | -0.08172 | 0.453305 | 0.922316 |
| ENSMUSG( | Mapk4 | 2881.353 | 0.383596 | 0.000111 | 0.00695 | ENSMUSG( | Mapk4 | 2452.892 | 0.204703 | 0.081063 | 0.77681 |
| ENSMUSG( | Mapk6 | 1997.262 | 0.018439 | 0.838616 | 0.971111 | ENSMUSG( | Mapk6 | 1788.808 | -0.03254 | 0.765379 | 0.974236 |
| ENSMUSG( | Mapk7 | 757.4722 | 0.002183 | 0.983225 | 0.997554 | ENSMUSG( | Mapk7 | 671.9428 | -0.07236 | 0.583341 | 0.944266 |
| ENSMUSG( | Mapk8 | 1746.683 | 0.009202 | 0.93072 | 0.992033 | ENSMUSG( | Mapk8 | 1530.583 | -0.10618 | 0.383022 | 0.917257 |
| ENSMUSG( | Mapk8ip1 | 24623.4 | 0.134884 | 0.080096 | 0.437572 | ENSMUSG( | Mapk8ip1 | 21114.18 | -0.03703 | 0.702029 | 0.962014 |
| ENSMUSG( | Mapk8ip2 | 16711.45 | -0.02422 | 0.847977 | 0.972374 | ENSMUSG( | Mapk8ip2 | 16051.06 | 0.129648 | 0.420095 | 0.917257 |
| ENSMUSG( | Mapk8ip3 | 14284.48 | 0.196285 | 0.038589 | 0.302696 | ENSMUSG( | Mapk8ip3 | 11957.01 | -0.04161 | 0.722612 | 0.966887 |
| ENSMUSG( | Mapk9 | 6042.179 | 0.05297 | 0.480493 | 0.854748 | ENSMUSG( | Mapk9 | 5249.863 | -0.08506 | 0.382565 | 0.917257 |
| ENSMUSG( | Mapkap1 | 1887.199 | 0.071288 | 0.420182 | 0.822614 | ENSMUSG( | Mapkap1 | 1619.344 | -0.10536 | 0.361725 | 0.914223 |
| ENSMUSG( | Mapkapk2 | 1110.381 | -0.04193 | 0.668016 | 0.927753 | ENSMUSG( | Mapkapk2 | 986.3643 | -0.11782 | 0.324814 | 0.905257 |
| ENSMUSG( | Mapkapk3 | 232.5822 | 0.125707 | 0.415192 | 0.820369 | ENSMUSG( | Mapkapk3 | 217.5525 | 0.193894 | 0.254966 | 0.882307 |
| ENSMUSG( | Mapkapk5 | 54.55506 | -0.09615 | 0.721102 | 0.942868 | ENSMUSG( | Mapkapk5 | 51.50618 | 0.000343 | 0.999086 | 0.999503 |
| ENSMUSG( | Mapkbp1 | 2079.011 | 0.198491 | 0.042277 | 0.316636 | ENSMUSG( | Mapkbp1 | 1815.406 | 0.080466 | 0.495542 | 0.927692 |
| ENSMUSG( | Mapre1 | 6084.787 | 0.034286 | 0.732331 | 0.944956 | ENSMUSG( | Mapre1 | 5133.686 | -0.19598 | 0.164876 | 0.838743 |
| ENSMUSG( | Mapre2 | 6852.998 | 0.089332 | 0.273334 | 0.723833 | ENSMUSG( | Mapre2 | 5739.148 | -0.15796 | 0.156299 | 0.837724 |
| ENSMUSG( | Mapre3 | 4658.156 | -0.02378 | 0.84058 | 0.971179 | ENSMUSG( | Mapre3 | 4089.892 | -0.13547 | 0.324577 | 0.905257 |
| ENSMUSG( | Mapt | 13319.36 | 0.08852 | 0.264518 | 0.715437 | ENSMUSG( | Mapt | 11174.95 | -0.15222 | 0.151445 | 0.837724 |
| ENSMUSG( | Marchf1 | 570.69 | -0.27612 | 0.046414 | 0.330047 | ENSMUSG( | Marchf1 | 561.0485 | -0.04015 | 0.796083 | 0.97681 |
| ENSMUSG( | Marchf10 | 23.09789 | -0.11723 | 0.757415 | 0.951513 | ENSMUSG( | Marchf10 | 22.45405 | 0.09856 | 0.81256 | 0.979148 |
| ENSMUSG( | Marchf11 | 126.6433 | -0.08903 | 0.647518 | 0.919716 | ENSMUSG( | Marchf11 | 116.2871 | -0.07426 | 0.727598 | 0.966887 |
| ENSMUSG( | Marchf2 | 4724.869 | 0.099559 | 0.221935 | 0.667491 | ENSMUSG( | Marchf2 | 3920.673 | -0.17466 | 0.080884 | 0.77681 |
| ENSMUSG( | Marchf3 | 66.9561 | -0.001 | 0.996636 | 0.99943 | ENSMUSG( | Marchf3 | 60.54034 | -0.01011 | 0.969498 | 0.99594 |
| ENSMUSG( | Marchf4 | 582.342 | 0.094751 | 0.398921 | 0.812878 | ENSMUSG( | Marchf4 | 558.1947 | 0.23503 | 0.064516 | 0.737935 |
| ENSMUSG( | Marchf5 | 1346.293 | 0.029276 | 0.745789 | 0.949281 | ENSMUSG( | Marchf5 | 1226.394 | 0.025281 | 0.842045 | 0.983044 |
| ENSMUSG( | Marchf6 | 7311.803 | -0.46442 | 0.001901 | 0.049802 | ENSMUSG( | Marchf6 | 8142.034 | 0.150691 | 0.367281 | 0.915171 |
| ENSMUSG( | Marchf7 | 707.9511 | -0.2304 | 0.064609 | 0.393968 | ENSMUSG( | Marchf7 | 762.5611 | 0.25677 | 0.079324 | 0.77458 |
| ENSMUSG( | Marchf8 | 4608.481 | 0.008592 | 0.924696 | 0.99114 | ENSMUSG( | Marchf8 | 3969.998 | -0.15795 | 0.149686 | 0.836241 |
| ENSMUSG( | Marchf9 | 1066.242 | -0.13192 | 0.216711 | 0.661816 | ENSMUSG( | Marchf9 | 1046.063 | 0.085056 | 0.49337 | 0.927372 |

| | | | | | |
|---|---|---|---|---|---|
| ENSMUSG( | 539.9042 | -0.00316 | 0.97835 | 0.997428 | Maip1 |
| ENSMUSG( | 65.52231 | -0.02964 | 0.909152 | 0.98794 | Mak |
| ENSMUSG( | 730.0339 | 0.070143 | 0.551584 | 0.886141 | Mak16 |
| ENSMUSG( | 17132.39 | -0.55935 | 5.60E-05 | 0.004302 | Mal |
| ENSMUSG( | 4204.064 | -0.46817 | 0.00035 | 0.016176 | Mal2 |
| ENSMUSG( | 6928.573 | -0.09967 | 0.479397 | 0.854046 | Malat1 |
| ENSMUSG( | 30.17372 | -0.23691 | 0.60861 | 0.907154 | Mall |
| ENSMUSG( | 257.1797 | 0.028789 | 0.844057 | 0.971962 | Malsu1 |
| ENSMUSG( | 269.2565 | -0.0528 | 0.711057 | 0.940028 | Malt1 |
| ENSMUSG( | 67.65645 | -0.41499 | 0.118481 | 0.520662 | Mamdc2 |
| ENSMUSG( | 59.57291 | 0.429866 | 0.137049 | 0.551733 | Mamdc4 |
| ENSMUSG( | 319.5176 | -0.17 | 0.194475 | 0.63341 | Maml1 |
| ENSMUSG( | 505.5083 | 0.176223 | 0.132302 | 0.544251 | Maml2 |
| ENSMUSG( | 317.4961 | -0.19709 | 0.156196 | 0.583988 | Maml3 |
| ENSMUSG( | 843.2597 | 0.171186 | 0.087772 | 0.454663 | Mamld1 |
| ENSMUSG( | 51.10512 | 0.07642 | 0.791252 | 0.959461 | Mamstr |
| ENSMUSG( | 1090.529 | -0.52194 | 0.00027 | 0.013409 | Man1a |
| ENSMUSG( | 3511.623 | -0.109 | 0.387497 | 0.805693 | Man1a2 |
| ENSMUSG( | 3145.619 | -0.16291 | 0.170521 | 0.605271 | Man1b1 |
| ENSMUSG( | 1776.307 | -0.06944 | 0.598246 | 0.903801 | Man1c1 |
| ENSMUSG( | 685.1701 | -0.15732 | 0.283557 | 0.732021 | Man2a1 |
| ENSMUSG( | 9181.355 | -0.06153 | 0.544891 | 0.884712 | Man2a2 |
| ENSMUSG( | 2602.001 | -0.00134 | 0.989501 | 0.999111 | Man2b1 |
| ENSMUSG( | 1544.052 | -0.1677 | 0.202392 | 0.642343 | Man2b2 |
| ENSMUSG( | 1864.84 | 0.257734 | 0.068801 | 0.406391 | Man2c1 |
| ENSMUSG( | 34.2956 | -0.01804 | 0.956152 | 0.993874 | Man2c1os |
| ENSMUSG( | 412.0536 | -0.16223 | 0.333639 | 0.775602 | Manba |
| ENSMUSG( | 1640.12 | -0.22929 | 0.012188 | 0.160067 | Manbal |
| ENSMUSG( | 1062.92 | -0.46709 | 0.004204 | 0.084295 | Manea |
| ENSMUSG( | 1939.576 | -0.28717 | 0.042487 | 0.316636 | Maneal |
| ENSMUSG( | 1839.495 | -0.1393 | 0.357856 | 0.789405 | Manf |
| ENSMUSG( | 304.488 | -0.6288 | 0.000825 | 0.02866 | Mansc1 |
| ENSMUSG( | 11.26903 | 0.112889 | 0.848841 | NA | Mansc4 |
| ENSMUSG( | 1531.495 | -0.01759 | 0.884414 | 0.982647 | Maoa |
| ENSMUSG( | 2747.994 | 0.251871 | 0.032762 | 0.277278 | Maob |
| ENSMUSG( | 79.99132 | 0.190806 | 0.453812 | 0.836798 | Map10 |
| ENSMUSG( | 41592.35 | 0.028215 | 0.821479 | 0.966248 | Map1a |
| ENSMUSG( | 15102.5 | 0.048232 | 0.514191 | 0.872055 | Map1b |
| ENSMUSG( | 6561.908 | 0.014982 | 0.848064 | 0.972374 | Map1lc3a |
| ENSMUSG( | 9658.126 | 0.107543 | 0.191153 | 0.630245 | Map1lc3b |
| ENSMUSG( | 1638.552 | 0.207263 | 0.053718 | 0.355948 | Map1s |
| ENSMUSG( | 39517.59 | 0.099185 | 0.379782 | 0.801757 | Map2 |
| ENSMUSG( | 4655.97 | 0.05882 | 0.648543 | 0.920007 | Map2k1 |
| ENSMUSG( | 3136.707 | 0.08205 | 0.341131 | 0.778828 | Map2k2 |
| ENSMUSG( | 389.9843 | 0.00897 | 0.943216 | 0.993415 | Map2k3 |
| ENSMUSG( | 18.41626 | 0.187551 | 0.673061 | NA | Map2k3os |
| ENSMUSG( | 3686.462 | 0.101968 | 0.380191 | 0.801757 | Map2k4 |
| ENSMUSG( | 889.6373 | 0.080903 | 0.452585 | 0.836145 | Map2k5 |
| ENSMUSG( | 248.0806 | 0.231947 | 0.101392 | 0.484996 | Map2k6 |
| ENSMUSG( | 1817.962 | 0.017655 | 0.8761 | 0.980497 | Map2k7 |
| ENSMUSG( | 347.5496 | -0.06372 | 0.686978 | 0.933041 | Map3k1 |
| ENSMUSG( | 15222.44 | 0.235018 | 0.002563 | 0.060076 | Map3k10 |
| ENSMUSG( | 792.0158 | -0.1004 | 0.355669 | 0.787945 | Map3k11 |
| ENSMUSG( | 4473.529 | 0.197606 | 0.039952 | 0.308163 | Map3k12 |
| ENSMUSG( | 551.9376 | -0.00611 | 0.959215 | 0.99396 | Map3k13 |
| ENSMUSG( | 148.8323 | -0.19522 | 0.274162 | 0.724254 | Map3k14 |
| ENSMUSG( | 96.64225 | 0.209501 | 0.332139 | 0.775373 | Map3k15 |
| ENSMUSG( | 406.7981 | 0.408654 | 0.00103 | 0.033612 | Map3k19 |
| ENSMUSG( | 602.0193 | -0.01014 | 0.940371 | 0.993415 | Map3k2 |
| ENSMUSG( | 215.6574 | 0.040407 | 0.822268 | 0.966352 | Map3k20 |
| ENSMUSG( | 75.69153 | 0.123875 | 0.613453 | 0.909027 | Map3k21 |
| ENSMUSG( | 732.2331 | 0.011205 | 0.921916 | 0.990436 | Map3k3 |
| ENSMUSG( | 1256.641 | 0.199408 | 0.086469 | 0.452045 | Map3k4 |
| ENSMUSG( | 1051.452 | 0.387379 | 0.000129 | 0.007629 | Map3k5 |
| ENSMUSG( | 148.2569 | 0.928766 | 1.04E-06 | 0.000197 | Map3k6 |
| ENSMUSG( | 2885.771 | 0.054494 | 0.544223 | 0.884587 | Map3k7 |
| ENSMUSG( | 70.2748 | -0.02096 | 0.92976 | 0.991848 | Map3k8 |
| ENSMUSG( | 1032.689 | 0.225537 | 0.051552 | 0.34826 | Map3k9 |
| ENSMUSG( | 40817.84 | -0.003 | 0.971899 | 0.996585 | Map4 |
| ENSMUSG( | 83.83385 | 0.022193 | 0.925104 | 0.991204 | Map4k1 |
| ENSMUSG( | 2692.147 | 0.333481 | 0.013546 | 0.170062 | Map4k2 |
| ENSMUSG( | 1605.875 | 0.26243 | 0.008077 | 0.125133 | Map4k3 |
| ENSMUSG( | 17967.92 | -0.03364 | 0.700203 | 0.937391 | Map4k4 |
| ENSMUSG( | 1497.072 | 0.079096 | 0.371816 | 0.798768 | Map4k5 |
| ENSMUSG( | 3686.107 | 0.104817 | 0.241245 | 0.689026 | Map6 |
| ENSMUSG( | 2396.091 | -0.11046 | 0.216532 | 0.661421 | Map6d1 |
| ENSMUSG( | 3370.551 | -0.02118 | 0.81646 | 0.965455 | Map7 |
| ENSMUSG( | 7380.108 | 0.090721 | 0.333645 | 0.775602 | Map7d1 |
| ENSMUSG( | 2266.197 | 0.168167 | 0.163537 | 0.594869 | Map7d2 |
| ENSMUSG( | 1892.892 | 0.154968 | 0.096938 | 0.477375 | Map9 |
| ENSMUSG( | 8579.399 | 0.096792 | 0.343946 | 0.779153 | Mapk1 |
| ENSMUSG( | 7302.086 | 0.14992 | 0.105427 | 0.492417 | Mapk10 |
| ENSMUSG( | 1350.386 | 0.199321 | 0.114373 | 0.51021 | Mapk11 |
| ENSMUSG( | 194.3813 | -0.09503 | 0.576012 | 0.895114 | Mapk12 |
| ENSMUSG( | 26.89636 | -0.03666 | 0.919924 | 0.990247 | Mapk13 |
| ENSMUSG( | 2478.552 | 0.153701 | 0.09237 | 0.466744 | Mapk14 |
| ENSMUSG( | 56.49862 | 0.134227 | 0.733076 | 0.944956 | Mapk15 |
| ENSMUSG( | 897.6389 | 0.125104 | 0.393847 | 0.809643 | Mapk1ip1 |
| ENSMUSG( | 854.7385 | 0.104924 | 0.361825 | 0.792797 | Mapk1ip1l |
| ENSMUSG( | 8125.248 | 0.080851 | 0.319146 | 0.765081 | Mapk3 |
| ENSMUSG( | 2881.353 | 0.383596 | 0.000111 | 0.00695 | Mapk4 |
| ENSMUSG( | 1997.262 | 0.018439 | 0.838616 | 0.971111 | Mapk6 |
| ENSMUSG( | 757.4722 | 0.002183 | 0.983225 | 0.997554 | Mapk7 |
| ENSMUSG( | 1746.683 | 0.009202 | 0.93072 | 0.992033 | Mapk8 |
| ENSMUSG( | 24623.4 | 0.134884 | 0.080096 | 0.437572 | Mapk8ip1 |
| ENSMUSG( | 16711.45 | -0.02422 | 0.847977 | 0.972374 | Mapk8ip2 |
| ENSMUSG( | 14284.48 | 0.196285 | 0.038589 | 0.302696 | Mapk8ip3 |
| ENSMUSG( | 6042.179 | 0.05297 | 0.480493 | 0.854748 | Mapk9 |
| ENSMUSG( | 1887.199 | 0.071288 | 0.420182 | 0.822614 | Mapkap1 |
| ENSMUSG( | 1110.381 | -0.04193 | 0.668016 | 0.927753 | Mapkapk2 |
| ENSMUSG( | 232.5822 | 0.125707 | 0.415192 | 0.820369 | Mapkapk3 |
| ENSMUSG( | 54.55506 | -0.09615 | 0.721102 | 0.942868 | Mapkapk5 |
| ENSMUSG( | 2079.011 | 0.198491 | 0.042277 | 0.316636 | Mapkbp1 |
| ENSMUSG( | 6084.787 | 0.034286 | 0.732331 | 0.944956 | Mapre1 |
| ENSMUSG( | 6852.998 | 0.089332 | 0.273334 | 0.723833 | Mapre2 |
| ENSMUSG( | 4658.156 | -0.02378 | 0.84058 | 0.971179 | Mapre3 |
| ENSMUSG( | 13319.36 | 0.08852 | 0.264518 | 0.715437 | Mapt |
| ENSMUSG( | 570.69 | -0.27612 | 0.046414 | 0.330047 | Marchf1 |
| ENSMUSG( | 23.09789 | -0.11723 | 0.757415 | 0.951513 | Marchf10 |
| ENSMUSG( | 126.6433 | -0.08903 | 0.647518 | 0.919716 | Marchf11 |
| ENSMUSG( | 4724.869 | 0.099559 | 0.221935 | 0.667491 | Marchf2 |
| ENSMUSG( | 66.9561 | -0.001 | 0.996636 | 0.99943 | Marchf3 |
| ENSMUSG( | 582.342 | 0.094751 | 0.398921 | 0.812878 | Marchf4 |
| ENSMUSG( | 1346.293 | 0.029276 | 0.745789 | 0.949281 | Marchf5 |
| ENSMUSG( | 7311.803 | -0.46442 | 0.001901 | 0.049802 | Marchf6 |
| ENSMUSG( | 707.9511 | -0.2304 | 0.064609 | 0.393968 | Marchf7 |
| ENSMUSG( | 4608.481 | 0.008592 | 0.924696 | 0.99114 | Marchf8 |
| ENSMUSG( | 1066.242 | -0.13192 | 0.216711 | 0.661816 | Marchf9 |
| ENSMUSG( | 6500.518 | -0.07581 | 0.55348 | 0.886216 | Marcks |
| ENSMUSG( | 5566.837 | 0.154708 | 0.108482 | 0.497337 | Marcksl1 |
| ENSMUSG( | 6643.423 | 0.224296 | 0.014652 | 0.178054 | Marf1 |
| ENSMUSG( | 3772.22 | 0.177724 | 0.026456 | 0.24888 | Mark1 |
| ENSMUSG( | 3736.718 | 0.143817 | 0.128322 | 0.536641 | Mark2 |
| ENSMUSG( | 2457.494 | 0.202283 | 0.027969 | 0.255784 | Mark3 |
| ENSMUSG( | 3272.914 | 0.210596 | 0.02024 | 0.214698 | Mark4 |
| ENSMUSG( | 1274.442 | 0.081915 | 0.462163 | 0.841901 | Mars1 |
| ENSMUSG( | 509.9868 | -0.05016 | 0.684487 | 0.93282 | Mars2 |
| ENSMUSG( | 522.1323 | -0.03938 | 0.770117 | 0.954689 | Marveld1 |
| ENSMUSG( | 34.17567 | 0.322938 | 0.350029 | 0.781943 | Marveld2 |
| ENSMUSG( | 544.429 | -0.13197 | 0.394449 | 0.810072 | Mas1 |
| ENSMUSG( | 872.4631 | -0.25308 | 0.103887 | 0.489523 | Masp1 |
| ENSMUSG( | 20.43228 | -0.0048 | 0.99103 | NA | Masp2 |
| ENSMUSG( | 3234.461 | 0.335068 | 0.004774 | 0.091103 | Mast1 |
| ENSMUSG( | 6868.421 | 0.321152 | 0.000122 | 0.007362 | Mast2 |
| ENSMUSG( | 9327.947 | 0.185984 | 0.068971 | 0.406851 | Mast3 |
| ENSMUSG( | 1333.685 | 0.184381 | 0.145513 | 0.566408 | Mast4 |
| ENSMUSG( | 42.85505 | -0.27875 | 0.374832 | 0.800479 | Mastl |
| ENSMUSG( | 3623.741 | 0.164698 | 0.178093 | 0.615613 | Mat2a |
| ENSMUSG( | 1934.937 | 0.027555 | 0.786618 | 0.958856 | Mat2b |
| ENSMUSG( | 903.847 | 0.135188 | 0.260697 | 0.710468 | Matcap1 |
| ENSMUSG( | 148.2437 | 0.206794 | 0.218712 | 0.664559 | Matcap2 |
| ENSMUSG( | 6340.98 | 0.218546 | 0.082972 | 0.44459 | Matk |
| ENSMUSG( | 27.51158 | -1.27705 | 0.002794 | 0.064363 | Matn1 |
| ENSMUSG( | 643.1546 | -0.02358 | 0.886133 | 0.983008 | Matn2 |
| ENSMUSG( | 642.5483 | -0.54992 | 0.001082 | 0.034804 | Matn4 |
| ENSMUSG( | 2719.735 | -0.13975 | 0.250798 | 0.700403 | Matr3 |
| ENSMUSG( | 2765.571 | 0.056722 | 0.569988 | 0.893041 | Mau2 |
| ENSMUSG( | 395.7777 | 0.177045 | 0.146753 | 0.56887 | Mavs |
| ENSMUSG( | 1964.465 | 0.124751 | 0.177294 | 0.614553 | Max |
| ENSMUSG( | 11818.22 | 0.260949 | 0.001496 | 0.042956 | Maz |
| ENSMUSG( | 889.0499 | 0.215785 | 0.042465 | 0.316636 | Mb21d2 |
| ENSMUSG( | 1329.141 | 0.10099 | 0.273464 | 0.72394 | Mbd1 |
| ENSMUSG( | 2099.374 | -0.03512 | 0.724799 | 0.944096 | Mbd2 |
| ENSMUSG( | 5976.522 | 0.052008 | 0.503293 | 0.86596 | Mbd3 |
| ENSMUSG( | 203.4248 | -0.09836 | 0.549408 | 0.885825 | Mbd4 |
| ENSMUSG( | 1064.148 | 0.037356 | 0.730173 | 0.944956 | Mbd5 |
| ENSMUSG( | 1454.697 | 0.126503 | 0.422558 | 0.823508 | Mbd6 |
| ENSMUSG( | 404.0876 | -0.05858 | 0.693645 | 0.934202 | Mbip |

| | | | | | |
|---|---|---|---|---|---|
| ENSMUSG( | 760.497 | 0.059532 | 0.703437 | 0.962481 | Map3k11 |
| ENSMUSG( | 3847.159 | 0.036938 | 0.745342 | 0.96951 | Map3k12 |
| ENSMUSG( | 511.4601 | 0.040857 | 0.762861 | 0.973698 | Map3k13 |
| ENSMUSG( | 140.402 | -0.07543 | 0.700156 | 0.961335 | Map3k14 |
| ENSMUSG( | 83.53686 | 0.054864 | 0.834296 | 0.981237 | Map3k15 |
| ENSMUSG( | 329.9613 | 0.089852 | 0.576917 | 0.942912 | Map3k19 |
| ENSMUSG( | 583.0229 | 0.167849 | 0.266888 | 0.886134 | Map3k2 |
| ENSMUSG( | 228.0963 | 0.445714 | 0.017545 | 0.53154 | Map3k20 |
| ENSMUSG( | 71.21072 | 0.227049 | 0.365048 | 0.914449 | Map3k21 |
| ENSMUSG( | 698.0967 | 0.145623 | 0.35242 | 0.912093 | Map3k3 |
| ENSMUSG( | 1110.75 | 0.117877 | 0.361021 | 0.914223 | Map3k4 |
| ENSMUSG( | 842.0588 | 0.03882 | 0.744624 | 0.96951 | Map3k5 |
| ENSMUSG( | 117.0704 | 0.623597 | 0.019966 | 0.559604 | Map3k6 |
| ENSMUSG( | 2563.539 | -0.01845 | 0.866563 | 0.987419 | Map3k7 |
| ENSMUSG( | 70.73557 | 0.276173 | 0.336472 | 0.907972 | Map3k8 |
| ENSMUSG( | 922.8191 | 0.178227 | 0.194748 | 0.860433 | Map3k9 |
| ENSMUSG( | 34346.43 | -0.24392 | 0.030111 | 0.621996 | Map4 |
| ENSMUSG( | 74.94817 | -0.0343 | 0.890439 | 0.990558 | Map4k1 |
| ENSMUSG( | 2398.547 | 0.27306 | 0.093217 | 0.795363 | Map4k2 |
| ENSMUSG( | 1443.673 | 0.227557 | 0.037357 | 0.673514 | Map4k3 |
| ENSMUSG( | 15289.75 | -0.24206 | 0.044193 | 0.693037 | Map4k4 |
| ENSMUSG( | 1369.134 | 0.088181 | 0.451443 | 0.922202 | Map4k5 |
| ENSMUSG( | 3133.348 | -0.09591 | 0.4532 | 0.922316 | Map6 |
| ENSMUSG( | 2073.087 | -0.27038 | 0.011814 | 0.444083 | Map6d1 |
| ENSMUSG( | 3215.983 | 0.11017 | 0.297834 | 0.898538 | Map7 |
| ENSMUSG( | 6619.929 | 0.046818 | 0.664447 | 0.956766 | Map7d1 |
| ENSMUSG( | 1919.353 | -0.0413 | 0.782161 | 0.976106 | Map7d2 |
| ENSMUSG( | 1653.761 | 0.037819 | 0.741177 | 0.968811 | Map9 |
| ENSMUSG( | 7508.325 | -0.01877 | 0.877798 | 0.989644 | Mapk1 |
| ENSMUSG( | 6179.802 | -0.06132 | 0.594281 | 0.945408 | Mapk10 |
| ENSMUSG( | 1128.945 | -0.04542 | 0.764331 | 0.974 | Mapk11 |
| ENSMUSG( | 191.1757 | 0.129812 | 0.514867 | 0.930939 | Mapk12 |
| ENSMUSG( | 23.44176 | -0.15584 | 0.702619 | 0.962215 | Mapk13 |
| ENSMUSG( | 2121.876 | -0.02283 | 0.839617 | 0.982077 | Mapk14 |
| ENSMUSG( | 50.34164 | 0.081414 | 0.837232 | 0.981335 | Mapk15 |
| ENSMUSG( | 783.4759 | 0.002188 | 0.989653 | 0.997928 | Mapk1ip1 |
| ENSMUSG( | 745.3381 | -0.01904 | 0.879948 | 0.989644 | Mapk1ip1l |
| ENSMUSG( | 7000.155 | -0.08172 | 0.453305 | 0.922316 | Mapk3 |
| ENSMUSG( | 2452.892 | 0.204703 | 0.081063 | 0.77681 | Mapk4 |
| ENSMUSG( | 1788.808 | -0.03254 | 0.765379 | 0.974236 | Mapk6 |
| ENSMUSG( | 671.9428 | -0.07236 | 0.583341 | 0.944266 | Mapk7 |
| ENSMUSG( | 1530.583 | -0.10618 | 0.383022 | 0.917257 | Mapk8 |
| ENSMUSG( | 21114.18 | -0.03703 | 0.702029 | 0.962014 | Mapk8ip1 |
| ENSMUSG( | 16051.06 | 0.129648 | 0.420095 | 0.917257 | Mapk8ip2 |
| ENSMUSG( | 11957.01 | -0.04161 | 0.722612 | 0.966887 | Mapk8ip3 |
| ENSMUSG( | 5249.863 | -0.08506 | 0.382565 | 0.917257 | Mapk9 |
| ENSMUSG( | 1619.344 | -0.10536 | 0.361725 | 0.914223 | Mapkap1 |
| ENSMUSG( | 986.3643 | -0.11782 | 0.324814 | 0.905257 | Mapkapk2 |
| ENSMUSG( | 217.5525 | 0.193894 | 0.254966 | 0.882307 | Mapkapk3 |
| ENSMUSG( | 51.50618 | 0.000343 | 0.999086 | 0.999503 | Mapkapk5 |
| ENSMUSG( | 1815.406 | 0.080466 | 0.495542 | 0.927692 | Mapkbp1 |
| ENSMUSG( | 5133.686 | -0.19598 | 0.164876 | 0.838743 | Mapre1 |
| ENSMUSG( | 5739.148 | -0.15796 | 0.156299 | 0.837724 | Mapre2 |
| ENSMUSG( | 4089.892 | -0.13547 | 0.324577 | 0.905257 | Mapre3 |
| ENSMUSG( | 11174.95 | -0.15222 | 0.151445 | 0.837724 | Mapt |
| ENSMUSG( | 561.0485 | -0.04015 | 0.796083 | 0.97681 | Marchf1 |
| ENSMUSG( | 22.45405 | 0.09856 | 0.81256 | 0.979148 | Marchf10 |
| ENSMUSG( | 116.2871 | -0.07426 | 0.727598 | 0.966887 | Marchf11 |
| ENSMUSG( | 3920.673 | -0.17466 | 0.080884 | 0.77681 | Marchf2 |
| ENSMUSG( | 60.54034 | -0.01011 | 0.969498 | 0.99594 | Marchf3 |
| ENSMUSG( | 558.1947 | 0.23503 | 0.064516 | 0.737935 | Marchf4 |
| ENSMUSG( | 1226.394 | 0.025281 | 0.842045 | 0.983044 | Marchf5 |
| ENSMUSG( | 8142.034 | 0.150691 | 0.367281 | 0.915171 | Marchf6 |
| ENSMUSG( | 762.5611 | 0.25677 | 0.079324 | 0.77458 | Marchf7 |
| ENSMUSG( | 3969.998 | -0.15795 | 0.149686 | 0.836241 | Marchf8 |
| ENSMUSG( | 1046.063 | 0.085056 | 0.49337 | 0.927372 | Marchf9 |
| ENSMUSG( | 6277.636 | 0.089579 | 0.626152 | 0.953234 | Marcks |
| ENSMUSG( | 4696.188 | -0.06713 | 0.66394 | 0.956766 | Marcksl1 |
| ENSMUSG( | 5836.863 | 0.126596 | 0.235514 | 0.871807 | Marf1 |
| ENSMUSG( | 3221.838 | -0.00346 | 0.972514 | 0.996416 | Mark1 |
| ENSMUSG( | 3293.235 | 0.050758 | 0.623618 | 0.953234 | Mark2 |
| ENSMUSG( | 2127.282 | 0.060103 | 0.574264 | 0.942912 | Mark3 |
| ENSMUSG( | 2904.276 | 0.138821 | 0.193306 | 0.860096 | Mark4 |
| ENSMUSG( | 1164.863 | 0.094713 | 0.436719 | 0.919879 | Mars1 |
| ENSMUSG( | 481.6352 | 0.052092 | 0.699609 | 0.961335 | Mars2 |
| ENSMUSG( | 518.0372 | 0.209037 | 0.221921 | 0.866104 | Marveld1 |
| ENSMUSG( | 34.53613 | 0.594366 | 0.12609 | 0.81516 | Marveld2 |
| ENSMUSG( | 7.67993 | -0.35539 | 0.650917 | NA | Marveld3 |
| ENSMUSG( | 536.4007 | 0.101928 | 0.537715 | 0.937216 | Mas1 |
| ENSMUSG( | 951.427 | 0.269479 | 0.141981 | 0.830709 | Masp1 |
| ENSMUSG( | 19.56741 | 0.137148 | 0.768334 | 0.974236 | Masp2 |
| ENSMUSG( | 2662.677 | 0.059781 | 0.643386 | 0.953903 | Mast1 |
| ENSMUSG( | 5737.959 | 0.088588 | 0.383139 | 0.917257 | Mast2 |
| ENSMUSG( | 7830.872 | -0.04458 | 0.683865 | 0.958678 | Mast3 |
| ENSMUSG( | 1276.535 | 0.319988 | 0.024555 | 0.592201 | Mast4 |
| ENSMUSG( | 40.09103 | -0.21733 | 0.53147 | 0.934795 | Mastl |
| ENSMUSG( | 3402.456 | 0.247582 | 0.116447 | 0.813384 | Mat2a |
| ENSMUSG( | 1776.033 | 0.046475 | 0.709156 | 0.964042 | Mat2b |
| ENSMUSG( | 779.812 | -0.02518 | 0.869964 | 0.988097 | Matcap1 |
| ENSMUSG( | 128.4563 | 0.06869 | 0.730478 | 0.967228 | Matcap2 |
| ENSMUSG( | 5515.294 | 0.089365 | 0.533502 | 0.93519 | Matk |
| ENSMUSG( | 27.57178 | -0.80975 | 0.066871 | 0.751729 | Matn1 |
| ENSMUSG( | 706.806 | 0.485604 | 0.006659 | 0.339067 | Matn2 |
| ENSMUSG( | 752.618 | 0.226335 | 0.227155 | 0.868507 | Matn4 |
| ENSMUSG( | 2716.252 | 0.121577 | 0.457936 | 0.923329 | Matr3 |
| ENSMUSG( | 2562.844 | 0.107413 | 0.431425 | 0.917257 | Mau2 |
| ENSMUSG( | 355.5558 | 0.144792 | 0.334303 | 0.907907 | Mavs |
| ENSMUSG( | 1802.663 | 0.140481 | 0.298885 | 0.898538 | Max |
| ENSMUSG( | 10262.48 | 0.130315 | 0.194539 | 0.860277 | Maz |
| ENSMUSG( | 750.5131 | 0.004879 | 0.969821 | 0.99594 | Mb21d2 |
| ENSMUSG( | 1194.746 | 0.063799 | 0.591518 | 0.944975 | Mbd1 |
| ENSMUSG( | 1841.432 | -0.15252 | 0.24987 | 0.877904 | Mbd2 |
| ENSMUSG( | 5148.582 | -0.11279 | 0.301437 | 0.898538 | Mbd3 |
| ENSMUSG( | 190.3446 | -0.01035 | 0.960454 | 0.994931 | Mbd4 |
| ENSMUSG( | 999.3401 | 0.127154 | 0.340078 | 0.907972 | Mbd5 |
| ENSMUSG( | 1280.34 | 0.025373 | 0.893825 | 0.990558 | Mbd6 |
| ENSMUSG( | 392.673 | 0.126495 | 0.432922 | 0.918434 | Mbip |
| ENSMUSG( | 95.35107 | -0.04144 | 0.852798 | 0.985598 | Mblac1 |
| ENSMUSG( | 442.9456 | 0.044372 | 0.811356 | 0.978997 | Mblac2 |
| ENSMUSG( | 1448.686 | 0.10771 | 0.376589 | 0.915491 | Mbnl1 |
| ENSMUSG( | 2522.929 | 0.048973 | 0.689776 | 0.960071 | Mbnl2 |
| ENSMUSG( | 126.8418 | 0.037591 | 0.883595 | 0.989644 | Mboat1 |
| ENSMUSG( | 4368.213 | 0.026533 | 0.832336 | 0.98075 | Mboat2 |
| ENSMUSG( | 5976.096 | 0.153487 | 0.292592 | 0.897982 | Mboat7 |
| ENSMUSG( | 737983.4 | -0.13515 | 0.275475 | 0.889984 | Mbp |
| ENSMUSG( | 818.069 | 0.01268 | 0.923056 | 0.992644 | Mbtd1 |
| ENSMUSG( | 3055.482 | 0.126524 | 0.442976 | 0.921554 | Mbtps1 |
| ENSMUSG( | 659.6397 | 0.107545 | 0.558097 | 0.939565 | Mbtps2 |
| ENSMUSG( | 59.43308 | 0.63288 | 0.018983 | 0.549166 | Mc4r |
| ENSMUSG( | 66.40495 | 0.302503 | 0.358722 | 0.913759 | Mc5r |
| ENSMUSG( | 1400.244 | -0.04469 | 0.830072 | 0.980224 | Mcam |
| ENSMUSG( | 370.8359 | -0.08945 | 0.568439 | 0.941777 | Mcat |
| ENSMUSG( | 1027.799 | -0.02773 | 0.848696 | 0.985321 | Mcc |
| ENSMUSG( | 1093.253 | -0.14709 | 0.208517 | 0.864412 | Mccc1 |
| ENSMUSG( | 54.09891 | 0.210993 | 0.458902 | 0.923329 | Mccc1os |
| ENSMUSG( | 761.744 | -0.27653 | 0.01747 | 0.531273 | Mccc2 |
| ENSMUSG( | 577.241 | -0.19335 | 0.189331 | 0.85698 | Mcee |
| ENSMUSG( | 12.25281 | -0.08716 | 0.891662 | 0.990558 | Mcemp1 |
| ENSMUSG( | 200.793 | 0.120144 | 0.52237 | 0.932681 | Mcf2 |
| ENSMUSG( | 6660.821 | 0.193258 | 0.051398 | 0.711116 | Mcf2l |
| ENSMUSG( | 1857.446 | 0.169625 | 0.251984 | 0.879902 | Mcfd2 |
| ENSMUSG( | 770.6593 | 0.087182 | 0.592051 | 0.944975 | Mchr1 |
| ENSMUSG( | 2414.249 | 0.023996 | 0.816239 | 0.979198 | Mcl1 |
| ENSMUSG( | 36.90979 | -0.00244 | 0.994023 | 0.998813 | Mcm10 |
| ENSMUSG( | 138.3088 | 0.05897 | 0.770749 | 0.974236 | Mcm2 |
| ENSMUSG( | 154.1214 | 0.302077 | 0.100908 | 0.801456 | Mcm3 |
| ENSMUSG( | 1532.305 | 0.135484 | 0.295653 | 0.898538 | Mcm3ap |
| ENSMUSG( | 288.0412 | 0.077744 | 0.602144 | 0.94789 | Mcm4 |
| ENSMUSG( | 114.9151 | 0.059121 | 0.810709 | 0.978779 | Mcm5 |
| ENSMUSG( | 209.8531 | 0.093898 | 0.618079 | 0.952232 | Mcm6 |
| ENSMUSG( | 510.4698 | -0.12753 | 0.324471 | 0.905257 | Mcm7 |
| ENSMUSG( | 89.23355 | 0.062497 | 0.787562 | 0.97637 | Mcm8 |
| ENSMUSG( | 278.8158 | 0.070541 | 0.633536 | 0.953889 | Mcm9 |
| ENSMUSG( | 733.1078 | 0.145293 | 0.232572 | 0.868507 | Mcmbp |
| ENSMUSG( | 63.27939 | 0.023732 | 0.934156 | 0.99306 | Mcmdc2 |
| ENSMUSG( | 2389.748 | 0.162645 | 0.198252 | 0.862161 | Mcoln1 |
| ENSMUSG( | 192.7022 | -0.08233 | 0.632228 | 0.953452 | Mcph1 |
| ENSMUSG( | 2135.905 | -0.17238 | 0.107907 | 0.804969 | Mcrip1 |
| ENSMUSG( | 449.8196 | -0.11157 | 0.433659 | 0.918564 | Mcrip2 |
| ENSMUSG( | 2345.328 | -0.04317 | 0.674632 | 0.95694 | Mcrs1 |
| ENSMUSG( | 722.0634 | 0.364809 | 0.021804 | 0.575341 | Mctp1 |
| ENSMUSG( | 28.92071 | -0.1283 | 0.736291 | 0.967344 | Mctp2 |
| ENSMUSG( | 594.8962 | -0.11909 | 0.394341 | 0.917257 | Mcts1 |
| ENSMUSG( | 119.7163 | -0.23811 | 0.307696 | 0.898538 | Mcts2 |
| ENSMUSG( | 724.3708 | -0.02235 | 0.881524 | 0.989644 | Mcu |
| ENSMUSG( | 36.76495 | 0.379183 | 0.283866 | 0.892412 | Mcub |
| ENSMUSG( | 1941.795 | -0.24885 | 0.049023 | 0.707945 | Mcur1 |
| ENSMUSG( | 508.6683 | 0.20092 | 0.140708 | 0.830678 | Mdc1 |

| | | | | | | | | | | | |
|---|---|---|---|---|---|---|---|---|---|---|---|
| ENSMUSG( | Marcks | 6500.518 | -0.07581 | 0.55348 | 0.886216 | ENSMUSG( | Marcks | 6277.636 | 0.089579 | 0.626152 | 0.953234 |
| ENSMUSG( | Marcksl1 | 5566.837 | 0.154708 | 0.108482 | 0.497337 | ENSMUSG( | Marcksl1 | 4696.188 | -0.06713 | 0.66394 | 0.956766 |
| ENSMUSG( | Marf1 | 6643.423 | 0.224296 | 0.014652 | 0.178054 | ENSMUSG( | Marf1 | 5836.863 | 0.126596 | 0.235514 | 0.871807 |
| ENSMUSG( | Mark1 | 3772.22 | 0.177724 | 0.026456 | 0.24888 | ENSMUSG( | Mark1 | 3221.838 | -0.00346 | 0.972514 | 0.996416 |
| ENSMUSG( | Mark2 | 3736.718 | 0.143817 | 0.128322 | 0.536641 | ENSMUSG( | Mark2 | 3293.235 | 0.050758 | 0.623618 | 0.953234 |
| ENSMUSG( | Mark3 | 2457.494 | 0.202283 | 0.027969 | 0.255784 | ENSMUSG( | Mark3 | 2127.282 | 0.060103 | 0.574264 | 0.942912 |
| ENSMUSG( | Mark4 | 3272.914 | 0.210596 | 0.02024 | 0.214698 | ENSMUSG( | Mark4 | 2904.276 | 0.138821 | 0.193306 | 0.860096 |
| ENSMUSG( | Mars1 | 1274.442 | 0.081915 | 0.462163 | 0.841901 | ENSMUSG( | Mars1 | 1164.863 | 0.094713 | 0.436719 | 0.919879 |
| ENSMUSG( | Mars2 | 509.9868 | -0.05016 | 0.684487 | 0.93282 | ENSMUSG( | Mars2 | 481.6352 | 0.052092 | 0.699609 | 0.961335 |
| ENSMUSG( | Marveld1 | 522.1323 | -0.03938 | 0.770117 | 0.954689 | ENSMUSG( | Marveld1 | 518.0372 | 0.209037 | 0.221921 | 0.866104 |
| ENSMUSG( | Marveld2 | 34.17567 | 0.322938 | 0.350029 | 0.781943 | ENSMUSG( | Marveld2 | 34.53613 | 0.594366 | 0.12609 | 0.81516 |
| ENSMUSG( | Mas1 | 544.429 | -0.13197 | 0.394449 | 0.810072 | ENSMUSG( | Mas1 | 536.4007 | 0.101928 | 0.537715 | 0.937216 |
| ENSMUSG( | Masp1 | 872.4631 | -0.25308 | 0.103887 | 0.489523 | ENSMUSG( | Masp1 | 951.427 | 0.269479 | 0.141981 | 0.830709 |
| ENSMUSG( | Masp2 | 20.43228 | -0.0048 | 0.99103 | NA | ENSMUSG( | Masp2 | 19.56741 | 0.137148 | 0.768334 | 0.974236 |
| ENSMUSG( | Mast1 | 3234.461 | 0.335068 | 0.004774 | 0.091103 | ENSMUSG( | Mast1 | 2662.677 | 0.059781 | 0.643386 | 0.953903 |
| ENSMUSG( | Mast2 | 6868.421 | 0.321152 | 0.000122 | 0.007362 | ENSMUSG( | Mast2 | 5737.959 | 0.088588 | 0.383139 | 0.917257 |
| ENSMUSG( | Mast3 | 9327.947 | 0.185984 | 0.068971 | 0.406851 | ENSMUSG( | Mast3 | 7830.872 | -0.04458 | 0.683865 | 0.958678 |
| ENSMUSG( | Mast4 | 1333.685 | 0.184381 | 0.145513 | 0.566408 | ENSMUSG( | Mast4 | 1276.535 | 0.319988 | 0.024555 | 0.592201 |
| ENSMUSG( | Mastl | 42.85505 | -0.27875 | 0.374832 | 0.800479 | ENSMUSG( | Mastl | 40.09103 | -0.21733 | 0.53147 | 0.934795 |
| ENSMUSG( | Mat2a | 3623.741 | 0.164698 | 0.178093 | 0.615613 | ENSMUSG( | Mat2a | 3402.456 | 0.247582 | 0.116447 | 0.813384 |
| ENSMUSG( | Mat2b | 1934.937 | 0.027555 | 0.786618 | 0.958856 | ENSMUSG( | Mat2b | 1776.033 | 0.046475 | 0.709156 | 0.964042 |
| ENSMUSG( | Matcap1 | 903.847 | 0.135188 | 0.260697 | 0.710468 | ENSMUSG( | Matcap1 | 779.812 | -0.02518 | 0.869964 | 0.988097 |
| ENSMUSG( | Matcap2 | 148.2437 | 0.206794 | 0.218712 | 0.664559 | ENSMUSG( | Matcap2 | 128.4563 | 0.06869 | 0.730478 | 0.967228 |
| ENSMUSG( | Matk | 6340.98 | 0.218546 | 0.082972 | 0.44459 | ENSMUSG( | Matk | 5515.294 | 0.089365 | 0.533502 | 0.93519 |
| ENSMUSG( | Matn1 | 27.51158 | -1.27705 | 0.002794 | 0.064363 | ENSMUSG( | Matn1 | 27.57178 | -0.80975 | 0.066871 | 0.751729 |
| ENSMUSG( | Matn2 | 643.1546 | -0.02358 | 0.886133 | 0.983008 | ENSMUSG( | Matn2 | 706.806 | 0.485604 | 0.006659 | 0.339067 |
| ENSMUSG( | Matn4 | 642.5483 | -0.54992 | 0.001082 | 0.034804 | ENSMUSG( | Matn4 | 752.618 | 0.226335 | 0.227155 | 0.868507 |
| ENSMUSG( | Matr3 | 2719.735 | -0.13975 | 0.250798 | 0.700403 | ENSMUSG( | Matr3 | 2716.252 | 0.121577 | 0.457936 | 0.923329 |
| ENSMUSG( | Mau2 | 2765.571 | 0.056722 | 0.569988 | 0.893041 | ENSMUSG( | Mau2 | 2562.844 | 0.107413 | 0.431425 | 0.917257 |
| ENSMUSG( | Mavs | 395.7777 | 0.177045 | 0.146753 | 0.56887 | ENSMUSG( | Mavs | 355.5558 | 0.144792 | 0.334303 | 0.907907 |
| ENSMUSG( | Max | 1964.465 | 0.124751 | 0.177294 | 0.614553 | ENSMUSG( | Max | 1802.663 | 0.140481 | 0.298885 | 0.898538 |
| ENSMUSG( | Maz | 11818.22 | 0.260949 | 0.001496 | 0.042956 | ENSMUSG( | Maz | 10262.48 | 0.130315 | 0.194539 | 0.860277 |
| ENSMUSG( | Mb21d2 | 889.0499 | 0.215785 | 0.042465 | 0.316636 | ENSMUSG( | Mb21d2 | 750.5131 | 0.004879 | 0.969821 | 0.99594 |
| ENSMUSG( | Mbd1 | 1329.141 | 0.10099 | 0.273464 | 0.72394 | ENSMUSG( | Mbd1 | 1194.746 | 0.063799 | 0.591518 | 0.944975 |
| ENSMUSG( | Mbd2 | 2099.374 | -0.03512 | 0.724799 | 0.944096 | ENSMUSG( | Mbd2 | 1841.432 | -0.15252 | 0.24987 | 0.877904 |
| ENSMUSG( | Mbd3 | 5976.522 | 0.052008 | 0.503293 | 0.86596 | ENSMUSG( | Mbd3 | 5148.582 | -0.11279 | 0.301437 | 0.898538 |
| ENSMUSG( | Mbd4 | 203.4248 | -0.09836 | 0.549408 | 0.885825 | ENSMUSG( | Mbd4 | 190.3446 | -0.01035 | 0.960454 | 0.994931 |
| ENSMUSG( | Mbd5 | 1064.148 | 0.037356 | 0.730173 | 0.944956 | ENSMUSG( | Mbd5 | 999.3401 | 0.127154 | 0.340078 | 0.907972 |
| ENSMUSG( | Mbd6 | 1454.697 | 0.126503 | 0.422558 | 0.823508 | ENSMUSG( | Mbd6 | 1280.34 | 0.025373 | 0.893825 | 0.990558 |
| ENSMUSG( | Mbip | 404.0876 | -0.05858 | 0.693645 | 0.934202 | ENSMUSG( | Mbip | 392.673 | 0.126495 | 0.432922 | 0.918434 |
| ENSMUSG( | Mblac1 | 103.0859 | -0.08393 | 0.685594 | 0.932981 | ENSMUSG( | Mblac1 | 95.35107 | -0.04144 | 0.852798 | 0.985598 |
| ENSMUSG( | Mblac2 | 469.2521 | -0.048 | 0.684526 | 0.93282 | ENSMUSG( | Mblac2 | 442.9456 | 0.044372 | 0.811356 | 0.978997 |
| ENSMUSG( | Mbnl1 | 1475.179 | -0.10682 | 0.214746 | 0.659193 | ENSMUSG( | Mbnl1 | 1448.686 | 0.10771 | 0.376589 | 0.915491 |
| ENSMUSG( | Mbnl2 | 2718.334 | -0.00275 | 0.978677 | 0.997428 | ENSMUSG( | Mbnl2 | 2522.929 | 0.048973 | 0.689776 | 0.960071 |
| ENSMUSG( | Mboat1 | 119.6397 | -0.45729 | 0.043289 | 0.319014 | ENSMUSG( | Mboat1 | 126.8418 | 0.037591 | 0.883595 | 0.989644 |
| ENSMUSG( | Mboat2 | 4734.829 | -0.00966 | 0.930252 | 0.991848 | ENSMUSG( | Mboat2 | 4368.213 | 0.026533 | 0.832336 | 0.98075 |
| ENSMUSG( | Mboat7 | 5596.202 | -0.32163 | 0.011716 | 0.156231 | ENSMUSG( | Mboat7 | 5976.096 | 0.153487 | 0.292592 | 0.897982 |
| ENSMUSG( | Mbp | 866737.8 | 0.06492 | 0.48149 | 0.85549 | ENSMUSG( | Mbp | 737983.4 | -0.13515 | 0.275475 | 0.889984 |
| ENSMUSG( | Mbtd1 | 898.4458 | 0.019105 | 0.847074 | 0.972374 | ENSMUSG( | Mbtd1 | 818.069 | 0.01268 | 0.923056 | 0.992644 |
| ENSMUSG( | Mbtps1 | 3028.194 | -0.17543 | 0.141162 | 0.558043 | ENSMUSG( | Mbtps1 | 3055.482 | 0.126524 | 0.442976 | 0.921554 |
| ENSMUSG( | Mbtps2 | 611.8625 | -0.41025 | 0.012502 | 0.161575 | ENSMUSG( | Mbtps2 | 659.6397 | 0.107545 | 0.558097 | 0.939565 |
| ENSMUSG( | Mc4r | 54.95989 | 0.188345 | 0.520797 | 0.875186 | ENSMUSG( | Mc4r | 59.43308 | 0.63288 | 0.018983 | 0.549166 |
| ENSMUSG( | Mc5r | 58.38301 | -0.35627 | 0.289964 | 0.739839 | ENSMUSG( | Mc5r | 66.40495 | 0.302503 | 0.358722 | 0.913759 |
| ENSMUSG( | Mcam | 1271.449 | -0.67706 | 0.00012 | 0.007284 | ENSMUSG( | Mcam | 1400.244 | -0.04469 | 0.830072 | 0.980224 |
| ENSMUSG( | Mcat | 423.9436 | 0.033159 | 0.80582 | 0.962616 | ENSMUSG( | Mcat | 370.8359 | -0.08945 | 0.568439 | 0.941777 |
| ENSMUSG( | Mcc | 1195.396 | 0.138048 | 0.236844 | 0.684169 | ENSMUSG( | Mcc | 1027.799 | -0.02773 | 0.848696 | 0.985321 |
| ENSMUSG( | Mccc1 | 1297.693 | 0.080521 | 0.42573 | 0.824898 | ENSMUSG( | Mccc1 | 1093.253 | -0.14709 | 0.208517 | 0.864412 |
| ENSMUSG( | Mccc1os | 61.72152 | 0.322835 | 0.223824 | 0.669845 | ENSMUSG( | Mccc1os | 54.09891 | 0.210993 | 0.458902 | 0.923329 |
| ENSMUSG( | Mccc2 | 930.7986 | 0.045619 | 0.691367 | 0.933881 | ENSMUSG( | Mccc2 | 761.744 | -0.27653 | 0.01747 | 0.531273 |
| ENSMUSG( | Mcee | 703.7994 | 0.121066 | 0.273712 | 0.724244 | ENSMUSG( | Mcee | 577.241 | -0.19335 | 0.189331 | 0.85698 |
| ENSMUSG( | Mcemp1 | 13.24191 | -0.067 | 0.905162 | NA | ENSMUSG( | Mcemp1 | 12.25281 | -0.08716 | 0.891662 | 0.990558 |
| ENSMUSG( | Mcf2 | 196.6673 | -0.23079 | 0.143047 | 0.561716 | ENSMUSG( | Mcf2 | 200.793 | 0.120144 | 0.52237 | 0.932681 |
| ENSMUSG( | Mcf2l | 7347.83 | 0.206186 | 0.018291 | 0.203658 | ENSMUSG( | Mcf2l | 6660.821 | 0.193258 | 0.051398 | 0.711116 |
| ENSMUSG( | Mcfd2 | 1727.415 | -0.3263 | 0.015745 | 0.185786 | ENSMUSG( | Mcfd2 | 1857.446 | 0.169625 | 0.251984 | 0.879902 |
| ENSMUSG( | Mchr1 | 718.2139 | -0.41928 | 0.000936 | 0.031289 | ENSMUSG( | Mchr1 | 770.6593 | 0.087182 | 0.592051 | 0.944975 |
| ENSMUSG( | Mcl1 | 2613.645 | -0.01492 | 0.859563 | 0.97481 | ENSMUSG( | Mcl1 | 2414.249 | 0.023996 | 0.816239 | 0.979198 |
| ENSMUSG( | Mcm10 | 40.53282 | 0.005174 | 0.98673 | 0.998619 | ENSMUSG( | Mcm10 | 36.90979 | -0.00244 | 0.994023 | 0.998813 |
| ENSMUSG( | Mcm2 | 154.3972 | 0.098939 | 0.565299 | 0.890779 | ENSMUSG( | Mcm2 | 138.3088 | 0.05897 | 0.770749 | 0.974236 |
| ENSMUSG( | Mcm3 | 164.1916 | 0.230206 | 0.168828 | 0.602904 | ENSMUSG( | Mcm3 | 154.1214 | 0.302077 | 0.100908 | 0.801456 |
| ENSMUSG( | Mcm3ap | 1792.62 | 0.306701 | 0.005943 | 0.104409 | ENSMUSG( | Mcm3ap | 1532.305 | 0.135484 | 0.295653 | 0.898538 |
| ENSMUSG( | Mcm4 | 312.0013 | 0.043731 | 0.74055 | 0.947823 | ENSMUSG( | Mcm4 | 288.0412 | 0.077744 | 0.602144 | 0.94789 |
| ENSMUSG( | Mcm5 | 125.3026 | 0.021342 | 0.924289 | 0.99114 | ENSMUSG( | Mcm5 | 114.9151 | 0.059121 | 0.810709 | 0.978779 |
| ENSMUSG( | Mcm6 | 225.4625 | 0.021376 | 0.882908 | 0.982422 | ENSMUSG( | Mcm6 | 209.8531 | 0.093898 | 0.618079 | 0.952232 |
| ENSMUSG( | Mcm7 | 583.5944 | -0.00889 | 0.940725 | 0.993415 | ENSMUSG( | Mcm7 | 510.4698 | -0.12753 | 0.324471 | 0.905257 |
| ENSMUSG( | Mcm8 | 91.29483 | -0.15589 | 0.457768 | 0.839765 | ENSMUSG( | Mcm8 | 89.23355 | 0.062497 | 0.787562 | 0.97637 |
| ENSMUSG( | Mcm9 | 293.5814 | -0.04906 | 0.702861 | 0.937735 | ENSMUSG( | Mcm9 | 278.8158 | 0.070541 | 0.633536 | 0.953889 |
| ENSMUSG( | Mcmbp | 795.5519 | 0.115881 | 0.255909 | 0.706646 | ENSMUSG( | Mcmbp | 733.1078 | 0.145293 | 0.232572 | 0.868507 |
| ENSMUSG( | Mcmdc2 | 72.78252 | 0.151362 | 0.558298 | 0.888427 | ENSMUSG( | Mcmdc2 | 63.27939 | 0.023732 | 0.934156 | 0.99306 |
| ENSMUSG( | Mcoln1 | 2338.649 | -0.17235 | 0.126727 | 0.534201 | ENSMUSG( | Mcoln1 | 2389.748 | 0.162645 | 0.198252 | 0.862161 |
| ENSMUSG( | Mcph1 | 215.8545 | -0.01585 | 0.916174 | 0.988776 | ENSMUSG( | Mcph1 | 192.7022 | -0.08233 | 0.632228 | 0.953452 |
| ENSMUSG( | Mcrip1 | 2512.07 | 0.033065 | 0.678774 | 0.93066 | ENSMUSG( | Mcrip1 | 2135.905 | -0.17238 | 0.107907 | 0.804969 |
| ENSMUSG( | Mcrip2 | 517.1211 | 0.030196 | 0.798596 | 0.960896 | ENSMUSG( | Mcrip2 | 449.8196 | -0.11157 | 0.433659 | 0.918564 |
| ENSMUSG( | Mcrs1 | 2662.986 | 0.053911 | 0.523976 | 0.87583 | ENSMUSG( | Mcrs1 | 2345.328 | -0.04317 | 0.674632 | 0.95694 |
| ENSMUSG( | Mctp1 | 593.1358 | -0.49051 | 0.000455 | 0.019408 | ENSMUSG( | Mctp1 | 722.0634 | 0.364809 | 0.021804 | 0.575341 |
| ENSMUSG( | Mctp2 | 29.02587 | -0.39695 | 0.265118 | 0.715962 | ENSMUSG( | Mctp2 | 28.92071 | -0.1283 | 0.736291 | 0.967344 |
| ENSMUSG( | Mcts1 | 679.9247 | 0.005974 | 0.957191 | 0.993874 | ENSMUSG( | Mcts1 | 594.8962 | -0.11909 | 0.394341 | 0.917257 |
| ENSMUSG( | Mcts2 | 139.0338 | -0.07851 | 0.687874 | 0.93324 | ENSMUSG( | Mcts2 | 119.7163 | -0.23811 | 0.307696 | 0.898538 |
| ENSMUSG( | Mcu | 816.9391 | 0.059217 | 0.655311 | 0.923929 | ENSMUSG( | Mcu | 724.3708 | -0.02235 | 0.881524 | 0.989644 |
| ENSMUSG( | Mcub | 36.58788 | 0.111017 | 0.746443 | 0.949289 | ENSMUSG( | Mcub | 36.76495 | 0.379183 | 0.283866 | 0.892412 |
| ENSMUSG( | Mcur1 | 2297.455 | -0.0201 | 0.847789 | 0.972374 | ENSMUSG( | Mcur1 | 1941.795 | -0.24885 | 0.049023 | 0.707945 |
| ENSMUSG( | Mdc1 | 529.404 | 0.048198 | 0.659712 | 0.924524 | ENSMUSG( | Mdc1 | 508.6683 | 0.20092 | 0.140708 | 0.830678 |
| ENSMUSG( | Mdfi | 33.09184 | -0.03099 | 0.933344 | 0.993036 | ENSMUSG( | Mdfi | 36.26926 | 0.47522 | 0.167067 | 0.840222 |
| ENSMUSG( | Mdfic | 260.4441 | 0.075601 | 0.663832 | 0.926511 | ENSMUSG( | Mdfic | 221.2872 | -0.14481 | 0.515512 | 0.930939 |
| ENSMUSG( | Mdfic2 | 29.36883 | -0.33981 | 0.326938 | 0.771858 | ENSMUSG( | Mdfic2 | 25.64613 | -0.50374 | 0.202438 | 0.864412 |
| ENSMUSG( | Mdga1 | 1304.607 | 0.086549 | 0.430874 | 0.826896 | ENSMUSG( | Mdga1 | 1261.958 | 0.249516 | 0.045512 | 0.693959 |
| ENSMUSG( | Mdga2 | 1498.852 | -0.0425 | 0.76517 | 0.954127 | ENSMUSG( | Mdga2 | 1439.307 | 0.108642 | 0.476803 | 0.923329 |
| ENSMUSG( | Mdh1 | 12634.99 | 0.025131 | 0.841981 | 0.971263 | ENSMUSG( | Mdh1 | 10584.01 | -0.22934 | 0.145381 | 0.8336 |
| ENSMUSG( | Mdh1b | 19.62729 | -0.05142 | 0.902977 | NA | ENSMUSG( | Mdh1b | 17.60968 | -0.03111 | 0.95273 | 0.994798 |
| ENSMUSG( | Mdh2 | 7172.977 | 0.023107 | 0.830193 | 0.96974 | ENSMUSG( | Mdh2 | 6083.841 | -0.19241 | 0.16594 | 0.839855 |
| ENSMUSG( | Mdk | 1346.497 | -0.22745 | 0.039039 | 0.304527 | ENSMUSG( | Mdk | 1272.045 | -0.11502 | 0.364586 | 0.914223 |
| ENSMUSG( | Mdm1 | 217.5623 | 0.188275 | 0.235383 | 0.682423 | ENSMUSG( | Mdm1 | 198.3249 | 0.19317 | 0.30511 | 0.898538 |
| ENSMUSG( | Mdm2 | 1462.315 | -0.08065 | 0.405157 | 0.81652 | ENSMUSG( | Mdm2 | 1373.108 | 0.008134 | 0.942633 | 0.994774 |
| ENSMUSG( | Mdm4 | 2370.112 | -0.16734 | 0.079396 | 0.43547 | ENSMUSG( | Mdm4 | 2426.666 | 0.170957 | 0.157586 | 0.837724 |
| ENSMUSG( | Mdn1 | 1524.244 | -0.45722 | 2.71E-06 | 0.000405 | ENSMUSG( | Mdn1 | 1703.79 | 0.167739 | 0.231802 | 0.868507 |
| ENSMUSG( | Mdp1 | 978.6584 | -0.01328 | 0.896957 | 0.98528 | ENSMUSG( | Mdp1 | 885.4018 | -0.03743 | 0.78405 | 0.976154 |
| ENSMUSG( | Me1 | 3958.094 | 0.086493 | 0.461377 | 0.841432 | ENSMUSG( | Me1 | 3392.611 | -0.09319 | 0.518652 | 0.930939 |
| ENSMUSG( | Me2 | 595.5501 | -0.19427 | 0.100312 | 0.48338 | ENSMUSG( | Me2 | 527.0854 | -0.28936 | 0.037886 | 0.6763 |
| ENSMUSG( | Me3 | 1386.46 | -0.04446 | 0.697309 | 0.935662 | ENSMUSG( | Me3 | 1246.25 | -0.08712 | 0.522956 | 0.932848 |
| ENSMUSG( | Mea1 | 2269.145 | 0.072798 | 0.415753 | 0.82084 | ENSMUSG( | Mea1 | 1868.793 | -0.23005 | 0.059336 | 0.729747 |
| ENSMUSG( | Meaf6 | 2278.667 | 0.105452 | 0.234245 | 0.68192 | ENSMUSG( | Meaf6 | 2037.211 | 0.05129 | 0.646664 | 0.953988 |
| ENSMUSG( | Meak7 | 368.2434 | 0.230542 | 0.067143 | 0.400611 | ENSMUSG( | Meak7 | 310.3604 | 0.019287 | 0.894079 | 0.990558 |
| ENSMUSG( | Mecom | 50.03703 | -0.30405 | 0.350139 | 0.781943 | ENSMUSG( | Mecom | 59.81091 | 0.485652 | 0.172459 | 0.843608 |
| ENSMUSG( | Mecp2 | 4321.577 | 0.09323 | 0.338113 | 0.777422 | ENSMUSG( | Mecp2 | 3828.939 | 0.01556 | 0.88479 | 0.989688 |
| ENSMUSG( | Mecr | 576.3243 | 0.087787 | 0.4583 | 0.839958 | ENSMUSG( | Mecr | 480.6484 | -0.17383 | 0.237206 | 0.872173 |
| ENSMUSG( | Med1 | 1076.789 | 0.027807 | 0.775178 | 0.955606 | ENSMUSG( | Med1 | 1005.581 | 0.100384 | 0.425334 | 0.917257 |
| ENSMUSG( | Med10 | 605.4085 | -0.07807 | 0.489439 | 0.859092 | ENSMUSG( | Med10 | 546.5659 | -0.10178 | 0.441535 | 0.921554 |
| ENSMUSG( | Med11 | 179.1558 | -0.10357 | 0.523489 | 0.875628 | ENSMUSG( | Med11 | 169.0998 | -0.00041 | 0.998222 | 0.999313 |
| ENSMUSG( | Med12 | 1332.325 | 0.18243 | 0.076835 | 0.429306 | ENSMUSG( | Med12 | 1214.672 | 0.185543 | 0.185742 | 0.855305 |
| ENSMUSG( | Med12l | 1126.904 | -0.11946 | 0.198443 | 0.638387 | ENSMUSG( | Med12l | 1107.442 | 0.097433 | 0.400806 | 0.917257 |
| ENSMUSG( | Med13 | 2060.492 | 0.038819 | 0.739032 | 0.947768 | ENSMUSG( | Med13 | 1899.11 | 0.069904 | 0.625359 | 0.953234 |
| ENSMUSG( | Med13l | 3087.087 | 0.249302 | 0.007291 | 0.118022 | ENSMUSG( | Med13l | 2975.092 | 0.396787 | 0.000273 | 0.043871 |
| ENSMUSG( | Med14 | 1620.709 | 0.047154 | 0.593298 | 0.902578 | ENSMUSG( | Med14 | 1466.115 | 0.027427 | 0.798263 | 0.97681 |
| ENSMUSG( | Med15 | 1606.293 | 0.149221 | 0.167068 | 0.599655 | ENSMUSG( | Med15 | 1347.949 | -0.08642 | 0.492252 | 0.927315 |
| ENSMUSG( | Med16 | 2304.877 | 0.137647 | 0.105223 | 0.492417 | ENSMUSG( | Med16 | 2008.785 | 0.013231 | 0.899275 | 0.990558 |
| ENSMUSG( | Med17 | 610.3139 | 0.174869 | 0.15599 | 0.58375 | ENSMUSG( | Med17 | 527.0402 | 0.028899 | 0.83556 | 0.981237 |
| ENSMUSG( | Med18 | 52.09997 | 0.129416 | 0.703703 | 0.937885 | ENSMUSG( | Med18 | 45.84247 | 0.041894 | 0.899008 | 0.990558 |
| ENSMUSG( | Med19 | 653.1425 | 0.011316 | 0.921924 | 0.990436 | ENSMUSG( | Med19 | 563.447 | -0.15707 | 0.288575 | 0.894794 |
| ENSMUSG( | Med20 | 513.9425 | -0.05444 | 0.682382 | 0.93198 | ENSMUSG( | Med20 | 482.122 | 0.038936 | 0.817917 | 0.979198 |
| ENSMUSG( | Med21 | 360.9989 | -0.02774 | 0.821388 | 0.966248 | ENSMUSG( | Med21 | 322.1788 | -0.09248 | 0.522965 | 0.932848 |
| ENSMUSG( | Med22 | 1169.395 | 0.055416 | 0.548638 | 0.885686 | ENSMUSG( | Med22 | 991.3733 | -0.15634 | 0.177868 | 0.847901 |
| ENSMUSG( | Med23 | 982.2703 | -0.03521 | 0.785862 | 0.958774 | ENSMUSG( | Med23 | 942.4483 | 0.116735 | 0.40089 | 0.917257 |
| ENSMUSG( | Med24 | 3197.171 | 0.163401 | 0.173135 | 0.608373 | ENSMUSG( | Med24 | 2789.896 | 0.043262 | 0.738206 | 0.967839 |
| ENSMUSG( | Med25 | 4617.238 | 0.220011 | 0.008483 | 0.128832 | ENSMUSG( | Med25 | 4046.886 | 0.114667 | 0.243429 | 0.877758 |
| ENSMUSG( | Med26 | 187.1962 | -0.10874 | 0.547231 | 0.885118 | ENSMUSG( | Med26 | 172.3219 | -0.07365 | 0.711068 | 0.964363 |
| ENSMUSG( | Med27 | 382.8011 | -0.11588 | 0.538183 | 0.882553 | ENSMUSG( | Med27 | 334.9747 | -0.24673 | 0.243257 | 0.877679 |
| ENSMUSG( | Med28 | 1750.865 | 0.053461 | 0.581767 | 0.897862 | ENSMUSG( | Med28 | 1457.984 | -0.21547 | 0.050112 | 0.708648 |
| ENSMUSG( | Med29 | 222.8621 | 0.335745 | 0.037818 | 0.300541 | ENSMUSG( | Med29 | 185.0024 | 0.074378 | 0.688631 | 0.959742 |
| ENSMUSG( | Med30 | 364.9979 | -0.0603 | 0.620349 | 0.911432 | ENSMUSG( | Med30 | 326.8503 | -0.1169 | 0.44389 | 0.921734 |
| ENSMUSG( | Med31 | 202.8015 | -0.25816 | 0.090417 | 0.461669 | ENSMUSG( | Med31 | 196.9447 | -0.06211 | 0.718561 | 0.966633 |
| ENSMUSG( | Med4 | 492.0335 | -0.02583 | 0.855751 | 0.973813 | ENSMUSG( | Med4 | 450.3027 | -0.01971 | 0.907898 | 0.991958 |
| ENSMUSG( | Med6 | 248.9472 | -0.1489 | 0.377914 | 0.801561 | ENSMUSG( | Med6 | 230.9456 | -0.10366 | 0.603454 | 0.948137 |
| ENSMUSG( | Med7 | 310.6814 | -0.07549 | 0.581019 | 0.89745 | ENSMUSG( | Med7 | 306.8365 | 0.157437 | 0.297881 | 0.898538 |
| ENSMUSG( | Med8 | 526.9096 | -0.48052 | 0.000291 | 0.014095 | ENSMUSG( | Med8 | 538.3616 | -0.11149 | 0.439806 | 0.920879 |
| ENSMUSG( | Med9 | 1318.3 | 0.056605 | 0.579758 | 0.89687 | ENSMUSG( | Med9 | 1106.08 | -0.1896 | 0.110697 | 0.807582 |
| ENSMUSG( | Med9os | 33.59688 | 0.522569 | 0.135112 | 0.548658 | ENSMUSG( | Med9os | 23.8138 | -0.14588 | 0.725312 | 0.966887 |
| ENSMUSG( | Medag | 115.7119 | -0.13685 | 0.484147 | 0.856577 | ENSMUSG( | Medag | 100.4526 | -0.28016 | 0.208411 | 0.864412 |
| ENSMUSG( | Mef2a | 2853.069 | -0.11237 | 0.18556 | 0.623185 | ENSMUSG( | Mef2a | 2751.549 | 0.052111 | 0.638819 | 0.953903 |
| ENSMUSG( | Mef2c | 7907.642 | 0.104986 | 0.313476 | 0.760368 | ENSMUSG( | Mef2c | 6871.063 | -0.03289 | 0.799636 | 0.97681 |
| ENSMUSG( | Mef2d | 7137.486 | 0.121824 | 0.171843 | 0.606594 | ENSMUSG( | Mef2d | 6365.409 | 0.062536 | 0.536637 | 0.937108 |
| ENSMUSG( | Meg3 | 59910.6 | 0.39564 | 0.203112 | 0.643631 | ENSMUSG( | Meg3 | 48907.76 | 0.099368 | 0.659774 | 0.956205 |
| ENSMUSG( | Megf10 | 1643.517 | -0.2605 | 0.099718 | 0.482239 | ENSMUSG( | Megf10 | 1642.33 | 0.021661 | 0.9049 | 0.991089 |
| ENSMUSG( | Megf11 | 569.4479 | -0.45018 | 0.005928 | 0.104279 | ENSMUSG( | Megf11 | 609.5016 | 0.055132 | 0.748545 | 0.96984 |
| ENSMUSG( | Megf6 | 225.4569 | 0.528317 | 0.0047 | 0.090209 | ENSMUSG( | Megf6 | 232.7115 | 0.834543 | 0.000149 | 0.028037 |
| ENSMUSG( | Megf8 | 7042.984 | 0.002085 | 0.982947 | 0.997554 | ENSMUSG( | Megf8 | 6671.432 | 0.114249 | 0.361917 | 0.914223 |
| ENSMUSG( | Megf9 | 4005.953 | -0.40675 | 0.004821 | 0.091771 | ENSMUSG( | Megf9 | 4448.7 | 0.189944 | 0.265868 | 0.886134 |
| ENSMUSG( | Mei1 | 116.8569 | 0.089496 | 0.701835 | 0.937441 | ENSMUSG( | Mei1 | 99.71189 | -0.0808 | 0.751735 | 0.970386 |
| ENSMUSG( | Meig1 | 24.37786 | -0.34951 | 0.359436 | 0.791011 | ENSMUSG( | Meig1 | 26.56941 | 0.196967 | 0.61406 | 0.951177 |
| ENSMUSG( | Meiosin | 18.80672 | -0.72869 | 0.096255 | NA | ENSMUSG( | Meiosin | 17.40243 | -0.74761 | 0.152497 | 0.837724 |

| | | | | | |
|---|---|---|---|---|---|
| ENSMUSG( | 103.0859 | -0.08393 | 0.685594 | 0.932981 | Mblac1 |
| ENSMUSG( | 469.2521 | -0.048 | 0.684526 | 0.93282 | Mblac2 |
| ENSMUSG( | 1475.179 | -0.10682 | 0.214746 | 0.659193 | Mbnl1 |
| ENSMUSG( | 2718.334 | -0.00275 | 0.978677 | 0.997428 | Mbnl2 |
| ENSMUSG( | 119.6397 | -0.45729 | 0.043289 | 0.319014 | Mboat1 |
| ENSMUSG( | 4734.829 | -0.00966 | 0.930252 | 0.991848 | Mboat2 |
| ENSMUSG( | 5596.202 | -0.32163 | 0.011716 | 0.156231 | Mboat7 |
| ENSMUSG( | 866737.8 | 0.06492 | 0.48149 | 0.85549 | Mbp |
| ENSMUSG( | 898.4458 | 0.019105 | 0.847074 | 0.972374 | Mbtd1 |
| ENSMUSG( | 3028.194 | -0.17543 | 0.141162 | 0.558043 | Mbtps1 |
| ENSMUSG( | 611.8625 | -0.41025 | 0.012502 | 0.161575 | Mbtps2 |
| ENSMUSG( | 54.95989 | 0.188345 | 0.520797 | 0.875186 | Mc4r |
| ENSMUSG( | 58.38301 | -0.35627 | 0.289964 | 0.739839 | Mc5r |
| ENSMUSG( | 1271.449 | -0.67706 | 0.00012 | 0.007284 | Mcam |
| ENSMUSG( | 423.9436 | 0.033159 | 0.80582 | 0.962616 | Mcat |
| ENSMUSG( | 1195.396 | 0.138048 | 0.236844 | 0.684169 | Mcc |
| ENSMUSG( | 1297.693 | 0.080521 | 0.42573 | 0.824898 | Mccc1 |
| ENSMUSG( | 61.72152 | 0.322835 | 0.223824 | 0.669845 | Mccc1os |
| ENSMUSG( | 930.7986 | 0.045619 | 0.691367 | 0.933881 | Mccc2 |
| ENSMUSG( | 703.7994 | 0.121066 | 0.273712 | 0.724244 | Mcee |
| ENSMUSG( | 13.24191 | -0.067 | 0.905162 | NA | Mcemp1 |
| ENSMUSG( | 196.6673 | -0.23079 | 0.143047 | 0.561716 | Mcf2 |
| ENSMUSG( | 7347.83 | 0.206186 | 0.018291 | 0.203658 | Mcf2l |
| ENSMUSG( | 1727.415 | -0.3263 | 0.015745 | 0.185786 | Mcfd2 |
| ENSMUSG( | 718.2139 | -0.41928 | 0.000936 | 0.031289 | Mchr1 |
| ENSMUSG( | 2613.645 | -0.01492 | 0.859563 | 0.97481 | Mcl1 |
| ENSMUSG( | 40.53282 | 0.005174 | 0.98673 | 0.998619 | Mcm10 |
| ENSMUSG( | 154.3972 | 0.098939 | 0.565299 | 0.890779 | Mcm2 |
| ENSMUSG( | 164.1916 | 0.230206 | 0.168828 | 0.602904 | Mcm3 |
| ENSMUSG( | 1792.62 | 0.306701 | 0.005943 | 0.104409 | Mcm3ap |
| ENSMUSG( | 312.0013 | 0.043731 | 0.74055 | 0.947823 | Mcm4 |
| ENSMUSG( | 125.3026 | 0.021342 | 0.924289 | 0.99114 | Mcm5 |
| ENSMUSG( | 225.4625 | 0.021376 | 0.882908 | 0.982422 | Mcm6 |
| ENSMUSG( | 583.5944 | -0.00889 | 0.940725 | 0.993415 | Mcm7 |
| ENSMUSG( | 91.29483 | -0.15589 | 0.457768 | 0.839765 | Mcm8 |
| ENSMUSG( | 293.5814 | -0.04906 | 0.702861 | 0.937735 | Mcm9 |
| ENSMUSG( | 795.5519 | 0.115881 | 0.255909 | 0.706646 | Mcmbp |
| ENSMUSG( | 72.78252 | 0.151362 | 0.558298 | 0.888427 | Mcmdc2 |
| ENSMUSG( | 2338.649 | -0.17235 | 0.126727 | 0.534201 | Mcoln1 |
| ENSMUSG( | 215.8545 | -0.01585 | 0.916174 | 0.988776 | Mcph1 |
| ENSMUSG( | 2512.07 | 0.033065 | 0.678774 | 0.93066 | Mcrip1 |
| ENSMUSG( | 517.1211 | 0.030196 | 0.798596 | 0.960896 | Mcrip2 |
| ENSMUSG( | 2662.986 | 0.053911 | 0.523976 | 0.87583 | Mcrs1 |
| ENSMUSG( | 593.1358 | -0.49051 | 0.000455 | 0.019408 | Mctp1 |
| ENSMUSG( | 29.02587 | -0.39695 | 0.265118 | 0.715962 | Mctp2 |
| ENSMUSG( | 679.9247 | 0.005974 | 0.957191 | 0.993874 | Mcts1 |
| ENSMUSG( | 139.0338 | -0.07851 | 0.687874 | 0.93324 | Mcts2 |
| ENSMUSG( | 816.9391 | 0.059217 | 0.655311 | 0.923929 | Mcu |
| ENSMUSG( | 36.58788 | 0.111017 | 0.746443 | 0.949289 | Mcub |
| ENSMUSG( | 2297.455 | -0.0201 | 0.847789 | 0.972374 | Mcur1 |
| ENSMUSG( | 529.404 | 0.048198 | 0.659712 | 0.924524 | Mdc1 |
| ENSMUSG( | 33.09184 | -0.03099 | 0.933344 | 0.993036 | Mdfi |
| ENSMUSG( | 260.4441 | 0.075601 | 0.663832 | 0.926511 | Mdfic |
| ENSMUSG( | 29.36883 | -0.33981 | 0.326938 | 0.771858 | Mdfic2 |
| ENSMUSG( | 1304.607 | 0.086549 | 0.430874 | 0.826896 | Mdga1 |
| ENSMUSG( | 1498.852 | -0.0425 | 0.76517 | 0.954127 | Mdga2 |
| ENSMUSG( | 12634.99 | 0.025131 | 0.841981 | 0.971263 | Mdh1 |
| ENSMUSG( | 19.62729 | -0.05142 | 0.902977 | NA | Mdh1b |
| ENSMUSG( | 7172.977 | 0.023107 | 0.830193 | 0.96974 | Mdh2 |
| ENSMUSG( | 1346.497 | -0.22745 | 0.039039 | 0.304527 | Mdk |
| ENSMUSG( | 217.5623 | 0.188275 | 0.235383 | 0.682423 | Mdm1 |
| ENSMUSG( | 1462.315 | -0.08065 | 0.405157 | 0.81652 | Mdm2 |
| ENSMUSG( | 2370.112 | -0.16734 | 0.079396 | 0.43547 | Mdm4 |
| ENSMUSG( | 1524.244 | -0.45722 | 2.71E-06 | 0.000405 | Mdn1 |
| ENSMUSG( | 978.6584 | -0.01328 | 0.896957 | 0.98528 | Mdp1 |
| ENSMUSG( | 3958.094 | 0.086493 | 0.461377 | 0.841432 | Me1 |
| ENSMUSG( | 595.5501 | -0.19427 | 0.100312 | 0.48338 | Me2 |
| ENSMUSG( | 1386.46 | -0.04446 | 0.697309 | 0.935662 | Me3 |
| ENSMUSG( | 2269.145 | 0.072798 | 0.415753 | 0.82084 | Mea1 |
| ENSMUSG( | 2278.667 | 0.105452 | 0.234245 | 0.68192 | Meaf6 |
| ENSMUSG( | 368.2434 | 0.230542 | 0.067143 | 0.400611 | Meak7 |
| ENSMUSG( | 50.03703 | -0.30405 | 0.350139 | 0.781943 | Mecom |
| ENSMUSG( | 4321.577 | 0.09323 | 0.338113 | 0.777422 | Mecp2 |
| ENSMUSG( | 576.3243 | 0.087787 | 0.4583 | 0.839958 | Mecr |
| ENSMUSG( | 1076.789 | 0.027807 | 0.775178 | 0.955606 | Med1 |
| ENSMUSG( | 605.4085 | -0.07807 | 0.489439 | 0.859092 | Med10 |
| ENSMUSG( | 179.1558 | -0.10357 | 0.523489 | 0.875628 | Med11 |
| ENSMUSG( | 1332.325 | 0.18243 | 0.076835 | 0.429306 | Med12 |
| ENSMUSG( | 1126.904 | -0.11946 | 0.198443 | 0.638387 | Med12l |
| ENSMUSG( | 2060.492 | 0.038819 | 0.739032 | 0.947768 | Med13 |
| ENSMUSG( | 3087.087 | 0.249302 | 0.007291 | 0.118022 | Med13l |
| ENSMUSG( | 1620.709 | 0.047154 | 0.593298 | 0.902578 | Med14 |
| ENSMUSG( | 1606.293 | 0.149221 | 0.167068 | 0.599655 | Med15 |
| ENSMUSG( | 2304.877 | 0.137647 | 0.105223 | 0.492417 | Med16 |
| ENSMUSG( | 610.3139 | 0.174869 | 0.15599 | 0.58375 | Med17 |
| ENSMUSG( | 52.09997 | 0.129416 | 0.703703 | 0.937885 | Med18 |
| ENSMUSG( | 653.1425 | 0.011316 | 0.921924 | 0.990436 | Med19 |
| ENSMUSG( | 513.9425 | -0.05444 | 0.682382 | 0.93198 | Med20 |
| ENSMUSG( | 360.9989 | -0.02774 | 0.821388 | 0.966248 | Med21 |
| ENSMUSG( | 1169.395 | 0.055416 | 0.548638 | 0.885686 | Med22 |
| ENSMUSG( | 982.2703 | -0.03521 | 0.785862 | 0.958774 | Med23 |
| ENSMUSG( | 3197.171 | 0.163401 | 0.173135 | 0.608373 | Med24 |
| ENSMUSG( | 4617.238 | 0.220011 | 0.008483 | 0.128832 | Med25 |
| ENSMUSG( | 187.1962 | -0.10874 | 0.547231 | 0.885118 | Med26 |
| ENSMUSG( | 382.8011 | -0.11588 | 0.538183 | 0.882553 | Med27 |
| ENSMUSG( | 1750.865 | 0.053461 | 0.581767 | 0.897862 | Med28 |
| ENSMUSG( | 222.8621 | 0.335745 | 0.037818 | 0.300541 | Med29 |
| ENSMUSG( | 364.9979 | -0.0603 | 0.620349 | 0.911432 | Med30 |
| ENSMUSG( | 202.8015 | -0.25816 | 0.090417 | 0.461669 | Med31 |
| ENSMUSG( | 492.0335 | -0.02583 | 0.855751 | 0.973813 | Med4 |
| ENSMUSG( | 248.9472 | -0.1489 | 0.377914 | 0.801561 | Med6 |
| ENSMUSG( | 310.6814 | -0.07549 | 0.581019 | 0.89745 | Med7 |
| ENSMUSG( | 526.9096 | -0.48052 | 0.000291 | 0.014095 | Med8 |
| ENSMUSG( | 1318.3 | 0.056605 | 0.579758 | 0.89687 | Med9 |
| ENSMUSG( | 33.59688 | 0.522569 | 0.135112 | 0.548658 | Med9os |
| ENSMUSG( | 115.7119 | -0.13685 | 0.484147 | 0.856577 | Medag |
| ENSMUSG( | 2853.069 | -0.11237 | 0.18556 | 0.623185 | Mef2a |
| ENSMUSG( | 7907.642 | 0.104986 | 0.313476 | 0.760368 | Mef2c |
| ENSMUSG( | 7137.486 | 0.121824 | 0.171843 | 0.606594 | Mef2d |
| ENSMUSG( | 59910.6 | 0.39564 | 0.203112 | 0.643631 | Meg3 |
| ENSMUSG( | 1643.517 | -0.2605 | 0.099718 | 0.482239 | Megf10 |
| ENSMUSG( | 569.4479 | -0.45018 | 0.005928 | 0.104279 | Megf11 |
| ENSMUSG( | 225.4569 | 0.528317 | 0.0047 | 0.090209 | Megf6 |
| ENSMUSG( | 7042.984 | 0.002085 | 0.982947 | 0.997554 | Megf8 |
| ENSMUSG( | 4005.953 | -0.40675 | 0.004821 | 0.091771 | Megf9 |
| ENSMUSG( | 116.8569 | 0.089496 | 0.701835 | 0.937441 | Mei1 |
| ENSMUSG( | 24.37786 | -0.34951 | 0.359436 | 0.791011 | Meig1 |
| ENSMUSG( | 18.80672 | -0.72869 | 0.096255 | NA | Meiosin |
| ENSMUSG( | 157.6111 | -0.29839 | 0.115972 | 0.514015 | Meis1 |
| ENSMUSG( | 4105.95 | -0.32877 | 0.007103 | 0.11626 | Meis2 |
| ENSMUSG( | 1012.904 | 0.134517 | 0.278077 | 0.728086 | Meis3 |
| ENSMUSG( | 29.67621 | -0.8688 | 0.015792 | 0.18615 | Melk |
| ENSMUSG( | 34.11996 | -0.14863 | 0.647959 | 0.919759 | Meltf |
| ENSMUSG( | 650.5098 | 0.033431 | 0.787353 | 0.959147 | Memo1 |
| ENSMUSG( | 1990.877 | -0.24925 | 0.028501 | 0.257525 | Men1 |
| ENSMUSG( | 44.70563 | -0.36904 | 0.267457 | 0.717525 | Meox1 |
| ENSMUSG( | 858.9576 | 0.044409 | 0.76168 | 0.953201 | Mepce |
| ENSMUSG( | 4559.232 | 0.253698 | 0.041265 | 0.313941 | Mertk |
| ENSMUSG( | 1629.189 | -0.18146 | 0.158807 | 0.588112 | Mesd |
| ENSMUSG( | 15.27762 | -0.2201 | 0.635952 | NA | Mesp2 |
| ENSMUSG( | 812.9819 | -0.70129 | 1.49E-06 | 0.000263 | Mest |
| ENSMUSG( | 277.2778 | -0.31028 | 0.11749 | 0.517288 | Met |
| ENSMUSG( | 691.8834 | 0.101641 | 0.351446 | 0.783381 | Metap1 |
| ENSMUSG( | 584.1982 | -0.06588 | 0.67143 | 0.928781 | Metap1d |
| ENSMUSG( | 2903.905 | 0.089196 | 0.369944 | 0.797408 | Metap2 |
| ENSMUSG( | 6806.843 | -0.02278 | 0.840817 | 0.971179 | Metrn |
| ENSMUSG( | 278.7787 | -0.12075 | 0.395567 | 0.810727 | Metrnl |
| ENSMUSG( | 214.6944 | 0.099238 | 0.545374 | 0.884774 | Mettl1 |
| ENSMUSG( | 314.4014 | 0.022211 | 0.887153 | 0.983326 | Mettl13 |
| ENSMUSG( | 670.3091 | 0.089324 | 0.455027 | 0.837743 | Mettl14 |
| ENSMUSG( | 92.79394 | 0.027606 | 0.909737 | 0.988084 | Mettl15 |
| ENSMUSG( | 996.3188 | 0.298877 | 0.002538 | 0.059739 | Mettl16 |
| ENSMUSG( | 477.1164 | 0.173242 | 0.219541 | 0.664984 | Mettl17 |
| ENSMUSG( | 68.63289 | 0.159677 | 0.493104 | 0.860552 | Mettl18 |
| ENSMUSG( | 288.9946 | 0.158053 | 0.265195 | 0.715962 | Mettl2 |
| ENSMUSG( | 321.9641 | -0.35376 | 0.010138 | 0.143118 | Mettl21a |
| ENSMUSG( | 507.7541 | 0.042229 | 0.730527 | 0.944956 | Mettl22 |
| ENSMUSG( | 595.3393 | 0.08489 | 0.451382 | 0.835567 | Mettl23 |
| ENSMUSG( | 24.9844 | -0.67483 | 0.074127 | 0.421875 | Mettl24 |
| ENSMUSG( | 190.3781 | -0.05978 | 0.739169 | 0.947768 | Mettl25 |
| ENSMUSG( | 880.9468 | -0.01486 | 0.908449 | 0.987809 | Mettl25b |
| ENSMUSG( | 784.9062 | 0.057764 | 0.590658 | 0.90113 | Mettl26 |
| ENSMUSG( | 369.9216 | -0.00132 | 0.991898 | 0.999232 | Mettl27 |
| ENSMUSG( | 536.9013 | 0.176498 | 0.133797 | 0.547197 | Mettl3 |
| ENSMUSG( | 691.1057 | 0.190902 | 0.180803 | 0.617841 | Mettl4 |
| ENSMUSG( | 166.4573 | -0.13849 | 0.39482 | 0.810072 | Mettl5 |
| ENSMUSG( | 463.9896 | -0.06975 | 0.595506 | 0.903058 | Mettl6 |
| ENSMUSG( | 347.2684 | -0.04407 | 0.768578 | 0.954674 | Mettl8 |

| | | | | | |
|---|---|---|---|---|---|
| ENSMUSG( | 36.26926 | 0.47522 | 0.167067 | 0.840222 | Mdfi |
| ENSMUSG( | 221.2872 | -0.14481 | 0.515512 | 0.930939 | Mdfic |
| ENSMUSG( | 25.64613 | -0.50374 | 0.202438 | 0.864412 | Mdfic2 |
| ENSMUSG( | 1261.958 | 0.249516 | 0.045512 | 0.693959 | Mdga1 |
| ENSMUSG( | 1439.307 | 0.108642 | 0.476803 | 0.923329 | Mdga2 |
| ENSMUSG( | 10584.01 | -0.22934 | 0.145381 | 0.8336 | Mdh1 |
| ENSMUSG( | 17.60968 | -0.03111 | 0.95273 | 0.994798 | Mdh1b |
| ENSMUSG( | 6083.841 | -0.19241 | 0.16594 | 0.839855 | Mdh2 |
| ENSMUSG( | 1272.045 | -0.11502 | 0.364586 | 0.914223 | Mdk |
| ENSMUSG( | 198.3249 | 0.19317 | 0.30511 | 0.898538 | Mdm1 |
| ENSMUSG( | 1373.108 | 0.008134 | 0.942633 | 0.994774 | Mdm2 |
| ENSMUSG( | 2426.666 | 0.170957 | 0.157586 | 0.837724 | Mdm4 |
| ENSMUSG( | 1703.79 | 0.167739 | 0.231802 | 0.868507 | Mdn1 |
| ENSMUSG( | 885.4018 | -0.03743 | 0.78405 | 0.976154 | Mdp1 |
| ENSMUSG( | 3392.611 | -0.09319 | 0.518652 | 0.930939 | Me1 |
| ENSMUSG( | 527.0854 | -0.28936 | 0.037886 | 0.6763 | Me2 |
| ENSMUSG( | 1246.25 | -0.08712 | 0.522956 | 0.932848 | Me3 |
| ENSMUSG( | 1868.793 | -0.23005 | 0.059336 | 0.729747 | Mea1 |
| ENSMUSG( | 2037.211 | 0.05129 | 0.646664 | 0.953988 | Meaf6 |
| ENSMUSG( | 310.3604 | 0.019287 | 0.894079 | 0.990558 | Meak7 |
| ENSMUSG( | 59.81091 | 0.485652 | 0.172459 | 0.843608 | Mecom |
| ENSMUSG( | 3828.939 | 0.01556 | 0.88479 | 0.989688 | Mecp2 |
| ENSMUSG( | 480.6484 | -0.17383 | 0.237206 | 0.872173 | Mecr |
| ENSMUSG( | 1005.581 | 0.100384 | 0.425334 | 0.917257 | Med1 |
| ENSMUSG( | 546.5659 | -0.10178 | 0.441535 | 0.921554 | Med10 |
| ENSMUSG( | 169.0998 | -0.00041 | 0.998222 | 0.999313 | Med11 |
| ENSMUSG( | 1214.672 | 0.185543 | 0.185742 | 0.855305 | Med12 |
| ENSMUSG( | 1107.442 | 0.097433 | 0.400806 | 0.917257 | Med12l |
| ENSMUSG( | 1899.11 | 0.069904 | 0.625359 | 0.953234 | Med13 |
| ENSMUSG( | 2975.092 | 0.396787 | 0.000273 | 0.043871 | Med13l |
| ENSMUSG( | 1466.115 | 0.027427 | 0.798263 | 0.97681 | Med14 |
| ENSMUSG( | 1347.949 | -0.08642 | 0.492252 | 0.927315 | Med15 |
| ENSMUSG( | 2008.785 | 0.013231 | 0.899275 | 0.990558 | Med16 |
| ENSMUSG( | 527.0402 | 0.028899 | 0.83556 | 0.981237 | Med17 |
| ENSMUSG( | 45.84247 | 0.041894 | 0.899008 | 0.990558 | Med18 |
| ENSMUSG( | 563.447 | -0.15707 | 0.288575 | 0.894794 | Med19 |
| ENSMUSG( | 482.122 | 0.038936 | 0.817917 | 0.979198 | Med20 |
| ENSMUSG( | 322.1788 | -0.09248 | 0.522965 | 0.932848 | Med21 |
| ENSMUSG( | 991.3733 | -0.15634 | 0.177868 | 0.847901 | Med22 |
| ENSMUSG( | 942.4483 | 0.116735 | 0.40089 | 0.917257 | Med23 |
| ENSMUSG( | 2789.896 | 0.043262 | 0.738206 | 0.967839 | Med24 |
| ENSMUSG( | 4046.886 | 0.114667 | 0.243429 | 0.877758 | Med25 |
| ENSMUSG( | 172.3219 | -0.07365 | 0.711068 | 0.964363 | Med26 |
| ENSMUSG( | 334.9747 | -0.24673 | 0.243257 | 0.877679 | Med27 |
| ENSMUSG( | 1457.984 | -0.21547 | 0.050112 | 0.708648 | Med28 |
| ENSMUSG( | 185.0024 | 0.074378 | 0.688631 | 0.959742 | Med29 |
| ENSMUSG( | 326.8503 | -0.1169 | 0.44389 | 0.921734 | Med30 |
| ENSMUSG( | 196.9447 | -0.06211 | 0.718561 | 0.966633 | Med31 |
| ENSMUSG( | 450.3027 | -0.01971 | 0.907898 | 0.991958 | Med4 |
| ENSMUSG( | 230.9456 | -0.10366 | 0.603454 | 0.948137 | Med6 |
| ENSMUSG( | 306.8365 | 0.157437 | 0.297881 | 0.898538 | Med7 |
| ENSMUSG( | 538.3616 | -0.11149 | 0.439806 | 0.920879 | Med8 |
| ENSMUSG( | 1106.08 | -0.1896 | 0.110697 | 0.807582 | Med9 |
| ENSMUSG( | 23.8138 | -0.14588 | 0.725312 | 0.966887 | Med9os |
| ENSMUSG( | 100.4526 | -0.28016 | 0.208411 | 0.864412 | Medag |
| ENSMUSG( | 2751.549 | 0.052111 | 0.638819 | 0.953903 | Mef2a |
| ENSMUSG( | 6871.063 | -0.03289 | 0.799636 | 0.97681 | Mef2c |
| ENSMUSG( | 6365.409 | 0.062536 | 0.536637 | 0.937108 | Mef2d |
| ENSMUSG( | 48907.76 | 0.099368 | 0.659774 | 0.956205 | Meg3 |
| ENSMUSG( | 1642.33 | 0.021661 | 0.9049 | 0.991089 | Megf10 |
| ENSMUSG( | 609.5016 | 0.055132 | 0.748545 | 0.96984 | Megf11 |
| ENSMUSG( | 232.7115 | 0.834543 | 0.000149 | 0.028037 | Megf6 |
| ENSMUSG( | 6671.432 | 0.114249 | 0.361917 | 0.914223 | Megf8 |
| ENSMUSG( | 4448.7 | 0.189944 | 0.265868 | 0.886134 | Megf9 |
| ENSMUSG( | 99.71189 | -0.0808 | 0.751735 | 0.970386 | Mei1 |
| ENSMUSG( | 10.755 | 0.690561 | 0.249857 | NA | Mei4 |
| ENSMUSG( | 26.56941 | 0.196967 | 0.61406 | 0.951177 | Meig1 |
| ENSMUSG( | 17.40243 | -0.74761 | 0.152497 | 0.837724 | Meiosin |
| ENSMUSG( | 158.2797 | -0.00938 | 0.963979 | 0.995433 | Meis1 |
| ENSMUSG( | 4077.128 | -0.0649 | 0.669008 | 0.956766 | Meis2 |
| ENSMUSG( | 896.1235 | 0.050397 | 0.731043 | 0.967228 | Meis3 |
| ENSMUSG( | 32.87393 | -0.14504 | 0.689247 | 0.959893 | Melk |
| ENSMUSG( | 27.07944 | -0.58654 | 0.127428 | 0.817604 | Meltf |
| ENSMUSG( | 592.5585 | 0.02705 | 0.863466 | 0.986547 | Memo1 |
| ENSMUSG( | 2053.12 | 0.120351 | 0.363246 | 0.914223 | Men1 |
| ENSMUSG( | 43.03473 | -0.18955 | 0.585657 | 0.944975 | Meox1 |
| ENSMUSG( | 818.7852 | 0.171377 | 0.264751 | 0.886134 | Mepce |
| ENSMUSG( | 4297.072 | 0.34618 | 0.0196 | 0.556192 | Mertk |
| ENSMUSG( | 1590.487 | 0.025751 | 0.857026 | 0.985884 | Mesd |
| ENSMUSG( | 13.74181 | -0.29829 | 0.57366 | 0.942912 | Mesp2 |
| ENSMUSG( | 937.8841 | 0.065754 | 0.667837 | 0.956766 | Mest |
| ENSMUSG( | 319.1734 | 0.366569 | 0.071708 | 0.762854 | Met |
| ENSMUSG( | 638.2309 | 0.138549 | 0.277378 | 0.891468 | Metap1 |
| ENSMUSG( | 577.5044 | 0.168313 | 0.409082 | 0.917257 | Metap1d |
| ENSMUSG( | 2510.871 | -0.06494 | 0.638195 | 0.953903 | Metap2 |
| ENSMUSG( | 5785.172 | -0.23581 | 0.103842 | 0.804969 | Metrn |
| ENSMUSG( | 275.2604 | 0.116793 | 0.449502 | 0.922202 | Metrnl |
| ENSMUSG( | 185.933 | -0.04464 | 0.793965 | 0.97681 | Mettl1 |
| ENSMUSG( | 283.5182 | -0.00595 | 0.97093 | 0.996064 | Mettl13 |
| ENSMUSG( | 559.3918 | -0.16727 | 0.180267 | 0.850372 | Mettl14 |
| ENSMUSG( | 79.51159 | -0.16698 | 0.526489 | 0.934374 | Mettl15 |
| ENSMUSG( | 783.363 | -0.11729 | 0.349985 | 0.911652 | Mettl16 |
| ENSMUSG( | 457.1675 | 0.315044 | 0.071474 | 0.761942 | Mettl17 |
| ENSMUSG( | 68.55037 | 0.389908 | 0.144494 | 0.832876 | Mettl18 |
| ENSMUSG( | 272.5942 | 0.259064 | 0.106439 | 0.804969 | Mettl2 |
| ENSMUSG( | 351.7341 | 0.192747 | 0.208698 | 0.864412 | Mettl21a |
| ENSMUSG( | 461.2531 | 0.033431 | 0.808958 | 0.97873 | Mettl22 |
| ENSMUSG( | 521.8308 | -0.03135 | 0.817524 | 0.979198 | Mettl23 |
| ENSMUSG( | 22.16359 | -0.78138 | 0.067654 | 0.75212 | Mettl24 |
| ENSMUSG( | 175.4489 | -0.02758 | 0.882325 | 0.989644 | Mettl25 |
| ENSMUSG( | 833.6508 | 0.096917 | 0.538925 | 0.937216 | Mettl25b |
| ENSMUSG( | 665.9046 | -0.15161 | 0.249224 | 0.877904 | Mettl26 |
| ENSMUSG( | 331.2282 | -0.0534 | 0.719241 | 0.966844 | Mettl27 |
| ENSMUSG( | 494.5291 | 0.20711 | 0.193811 | 0.860096 | Mettl3 |
| ENSMUSG( | 581.952 | -0.032 | 0.828981 | 0.980224 | Mettl4 |
| ENSMUSG( | 148.9578 | -0.20506 | 0.303231 | 0.898538 | Mettl5 |
| ENSMUSG( | 448.1153 | 0.095571 | 0.53054 | 0.934795 | Mettl6 |
| ENSMUSG( | 324.5982 | 0.03801 | 0.809151 | 0.97873 | Mettl8 |
| ENSMUSG( | 2964.812 | -0.12909 | 0.370859 | 0.915491 | Mettl9 |
| ENSMUSG( | 155.2756 | 0.022657 | 0.899513 | 0.990558 | Mex3a |
| ENSMUSG( | 318.5921 | 0.174756 | 0.327557 | 0.906255 | Mex3b |
| ENSMUSG( | 1096.011 | 0.038206 | 0.768955 | 0.974236 | Mex3c |
| ENSMUSG( | 420.1071 | -0.02002 | 0.894321 | 0.990558 | Mex3d |
| ENSMUSG( | 470.7946 | 0.074797 | 0.649377 | 0.95422 | Mfap1a |
| ENSMUSG( | 337.9857 | -0.00658 | 0.968438 | 0.99594 | Mfap1b |
| ENSMUSG( | 176.1697 | 0.023225 | 0.89657 | 0.990558 | Mfap2 |
| ENSMUSG( | 880.1537 | 0.166985 | 0.38713 | 0.917257 | Mfap3 |
| ENSMUSG( | 5625.705 | 0.062277 | 0.681539 | 0.958678 | Mfap3l |
| ENSMUSG( | 98.71385 | 0.104216 | 0.651422 | 0.95449 | Mfap4 |
| ENSMUSG( | 25.59824 | -0.75136 | 0.070649 | 0.758714 | Mfap5 |
| ENSMUSG( | 4346.864 | -0.07638 | 0.470139 | 0.923329 | Mff |
| ENSMUSG( | 12555.42 | 0.143231 | 0.416328 | 0.917257 | Mfge8 |
| ENSMUSG( | 1502.563 | 0.150255 | 0.180508 | 0.850372 | Mfhas1 |
| ENSMUSG( | 4366.113 | 0.016345 | 0.881882 | 0.989644 | Mfn1 |
| ENSMUSG( | 6495.487 | -0.13484 | 0.216212 | 0.865167 | Mfn2 |
| ENSMUSG( | 192.7073 | 0.20436 | 0.350897 | 0.911707 | Mfng |
| ENSMUSG( | 67.77956 | 0.848234 | 0.380068 | 0.916995 | Mfrp |
| ENSMUSG( | 1850.282 | -0.01334 | 0.922349 | 0.992644 | Mfsd1 |
| ENSMUSG( | 635.928 | 0.131068 | 0.342026 | 0.909146 | Mfsd10 |
| ENSMUSG( | 1144.914 | 0.280646 | 0.0831 | 0.77681 | Mfsd11 |
| ENSMUSG( | 1524.608 | 0.173157 | 0.246298 | 0.877904 | Mfsd12 |
| ENSMUSG( | 1130.001 | -0.02285 | 0.866464 | 0.98737 | Mfsd13a |
| ENSMUSG( | 10.41144 | -0.34305 | 0.576513 | NA | Mfsd13b |
| ENSMUSG( | 1645.516 | 0.305805 | 0.087406 | 0.784337 | Mfsd14a |
| ENSMUSG( | 1267.777 | 0.313375 | 0.089998 | 0.790741 | Mfsd14b |
| ENSMUSG( | 2015.259 | 0.778862 | 7.68E-05 | 0.017018 | Mfsd2a |
| ENSMUSG( | 68.81874 | 0.468252 | 0.096928 | 0.79645 | Mfsd2b |
| ENSMUSG( | 222.1016 | 0.044952 | 0.807922 | 0.978622 | Mfsd3 |
| ENSMUSG( | 2121.173 | 0.141233 | 0.381168 | 0.917202 | Mfsd4a |
| ENSMUSG( | 16.33182 | -0.15318 | 0.753923 | 0.970671 | Mfsd4b1 |
| ENSMUSG( | 340.1284 | 0.06729 | 0.698044 | 0.960867 | Mfsd4b4 |
| ENSMUSG( | 1134.025 | 0.196324 | 0.261979 | 0.884926 | Mfsd5 |
| ENSMUSG( | 2703.282 | 0.133505 | 0.434244 | 0.91869 | Mfsd6 |
| ENSMUSG( | 695.1317 | 0.27644 | 0.073487 | 0.763629 | Mfsd8 |
| ENSMUSG( | 177.0249 | 0.191929 | 0.398373 | 0.917257 | Mfsd9 |
| ENSMUSG( | 1610.905 | 0.066432 | 0.564417 | 0.941206 | Mga |
| ENSMUSG( | 2037.783 | 0.09026 | 0.494921 | 0.927372 | Mgat1 |
| ENSMUSG( | 987.7778 | 0.188473 | 0.25254 | 0.88018 | Mgat2 |
| ENSMUSG( | 5344.574 | 0.139427 | 0.308584 | 0.898538 | Mgat3 |
| ENSMUSG( | 1103.353 | 0.185817 | 0.296628 | 0.898538 | Mgat4a |
| ENSMUSG( | 3376.293 | 0.151842 | 0.235029 | 0.871506 | Mgat4b |
| ENSMUSG( | 172.956 | 0.405832 | 0.092901 | 0.79479 | Mgat4c |
| ENSMUSG( | 4560.18 | -0.10902 | 0.51255 | 0.930164 | Mgat5 |
| ENSMUSG( | 3664.876 | 0.255131 | 0.075215 | 0.767994 | Mgat5b |
| ENSMUSG( | 35.41874 | -0.04606 | 0.891808 | 0.990558 | Mgl2 |
| ENSMUSG( | 20838.07 | -0.10454 | 0.34483 | 0.91045 | Mgll |
| ENSMUSG( | 248.0138 | -0.02369 | 0.883398 | 0.989644 | Mgme1 |
| ENSMUSG( | 40.81608 | -0.05315 | 0.867983 | 0.98781 | Mgmt |

| | | | | | | | | | | | |
|---|---|---|---|---|---|---|---|---|---|---|---|
| ENSMUSG( | Meis1 | 157.6111 | -0.29839 | 0.115972 | 0.514015 | ENSMUSG( | Meis1 | 158.2797 | -0.00938 | 0.963979 | 0.995433 |
| ENSMUSG( | Meis2 | 4105.95 | -0.32877 | 0.007103 | 0.11626 | ENSMUSG( | Meis2 | 4077.128 | -0.0649 | 0.669008 | 0.956766 |
| ENSMUSG( | Meis3 | 1012.904 | 0.134517 | 0.278077 | 0.728086 | ENSMUSG( | Meis3 | 896.1235 | 0.050397 | 0.731043 | 0.967228 |
| ENSMUSG( | Melk | 29.67621 | -0.8688 | 0.015792 | 0.18615 | ENSMUSG( | Melk | 32.87393 | -0.14504 | 0.689247 | 0.959893 |
| ENSMUSG( | Meltf | 34.11996 | -0.14863 | 0.647959 | 0.919759 | ENSMUSG( | Meltf | 27.07944 | -0.58654 | 0.127428 | 0.817604 |
| ENSMUSG( | Memo1 | 650.5098 | 0.033431 | 0.787353 | 0.959147 | ENSMUSG( | Memo1 | 592.5585 | 0.02705 | 0.863466 | 0.986547 |
| ENSMUSG( | Men1 | 1990.877 | -0.24925 | 0.028501 | 0.257525 | ENSMUSG( | Men1 | 2053.12 | 0.120351 | 0.363246 | 0.914223 |
| ENSMUSG( | Meox1 | 44.70563 | -0.36904 | 0.267457 | 0.717525 | ENSMUSG( | Meox1 | 43.03473 | -0.18955 | 0.585657 | 0.944975 |
| ENSMUSG( | Mepce | 858.9576 | 0.044409 | 0.76168 | 0.953201 | ENSMUSG( | Mepce | 818.7852 | 0.171377 | 0.264751 | 0.886134 |
| ENSMUSG( | Mertk | 4559.232 | 0.253698 | 0.041265 | 0.313941 | ENSMUSG( | Mertk | 4297.072 | 0.34618 | 0.0196 | 0.556192 |
| ENSMUSG( | Mesd | 1629.189 | -0.18146 | 0.158807 | 0.588112 | ENSMUSG( | Mesd | 1590.487 | 0.025751 | 0.857026 | 0.985884 |
| ENSMUSG( | Mesp2 | 15.27762 | -0.2201 | 0.635952 | NA | ENSMUSG( | Mesp2 | 13.74181 | -0.29829 | 0.57366 | 0.942912 |
| ENSMUSG( | Mest | 812.9819 | -0.70129 | 1.49E-06 | 0.000263 | ENSMUSG( | Mest | 937.8841 | 0.065754 | 0.667837 | 0.956766 |
| ENSMUSG( | Met | 277.2778 | -0.31028 | 0.11749 | 0.517288 | ENSMUSG( | Met | 319.1734 | 0.366569 | 0.071708 | 0.762854 |
| ENSMUSG( | Metap1 | 691.8834 | 0.101641 | 0.351446 | 0.783381 | ENSMUSG( | Metap1 | 638.2309 | 0.138549 | 0.277378 | 0.891468 |
| ENSMUSG( | Metap1d | 584.1982 | -0.06588 | 0.67143 | 0.928781 | ENSMUSG( | Metap1d | 577.5044 | 0.168313 | 0.409082 | 0.917257 |
| ENSMUSG( | Metap2 | 2903.905 | 0.089196 | 0.369944 | 0.797408 | ENSMUSG( | Metap2 | 2510.871 | -0.06494 | 0.638195 | 0.953903 |
| ENSMUSG( | Metrn | 6806.843 | -0.02278 | 0.840817 | 0.971179 | ENSMUSG( | Metrn | 5785.172 | -0.23581 | 0.103842 | 0.804969 |
| ENSMUSG( | Metrnl | 278.7787 | -0.12075 | 0.395567 | 0.810727 | ENSMUSG( | Metrnl | 275.2604 | 0.116793 | 0.449502 | 0.922202 |
| ENSMUSG( | Mettl1 | 214.6944 | 0.099238 | 0.545374 | 0.884774 | ENSMUSG( | Mettl1 | 185.933 | -0.04464 | 0.793965 | 0.97681 |
| ENSMUSG( | Mettl13 | 314.4014 | 0.022211 | 0.887153 | 0.983326 | ENSMUSG( | Mettl13 | 283.5182 | -0.00595 | 0.97093 | 0.996064 |
| ENSMUSG( | Mettl14 | 670.3091 | 0.089324 | 0.455027 | 0.837743 | ENSMUSG( | Mettl14 | 559.3918 | -0.16727 | 0.180267 | 0.850372 |
| ENSMUSG( | Mettl15 | 92.79394 | 0.027606 | 0.909737 | 0.988084 | ENSMUSG( | Mettl15 | 79.51159 | -0.16698 | 0.526489 | 0.934374 |
| ENSMUSG( | Mettl16 | 996.3188 | 0.298877 | 0.002538 | 0.059739 | ENSMUSG( | Mettl16 | 783.363 | -0.11729 | 0.349985 | 0.911652 |
| ENSMUSG( | Mettl17 | 477.1164 | 0.173242 | 0.219541 | 0.664984 | ENSMUSG( | Mettl17 | 457.1675 | 0.315044 | 0.071474 | 0.761942 |
| ENSMUSG( | Mettl18 | 68.63289 | 0.159677 | 0.493104 | 0.860552 | ENSMUSG( | Mettl18 | 68.55037 | 0.389908 | 0.144494 | 0.832876 |
| ENSMUSG( | Mettl2 | 288.9946 | 0.158053 | 0.265195 | 0.715962 | ENSMUSG( | Mettl2 | 272.5942 | 0.259064 | 0.106439 | 0.804969 |
| ENSMUSG( | Mettl21a | 321.9641 | -0.35376 | 0.010138 | 0.143118 | ENSMUSG( | Mettl21a | 351.7341 | 0.192747 | 0.208698 | 0.864412 |
| ENSMUSG( | Mettl22 | 507.7541 | 0.042229 | 0.730527 | 0.944956 | ENSMUSG( | Mettl22 | 461.2531 | 0.033431 | 0.808958 | 0.97873 |
| ENSMUSG( | Mettl23 | 595.3393 | 0.08489 | 0.451382 | 0.835567 | ENSMUSG( | Mettl23 | 521.8308 | -0.03135 | 0.817524 | 0.979198 |
| ENSMUSG( | Mettl24 | 24.9844 | -0.67483 | 0.074127 | 0.421875 | ENSMUSG( | Mettl24 | 22.16359 | -0.78138 | 0.067654 | 0.75212 |
| ENSMUSG( | Mettl25 | 190.3781 | -0.05978 | 0.739169 | 0.947768 | ENSMUSG( | Mettl25 | 175.4489 | -0.02758 | 0.882325 | 0.989644 |
| ENSMUSG( | Mettl25b | 880.9468 | -0.01486 | 0.908449 | 0.987809 | ENSMUSG( | Mettl25b | 833.6508 | 0.096917 | 0.538925 | 0.937216 |
| ENSMUSG( | Mettl26 | 784.9062 | 0.057764 | 0.590658 | 0.90113 | ENSMUSG( | Mettl26 | 665.9046 | -0.15161 | 0.249224 | 0.877904 |
| ENSMUSG( | Mettl27 | 369.9216 | -0.00132 | 0.991898 | 0.999232 | ENSMUSG( | Mettl27 | 331.2282 | -0.0534 | 0.719241 | 0.966844 |
| ENSMUSG( | Mettl3 | 536.9013 | 0.176498 | 0.133797 | 0.547197 | ENSMUSG( | Mettl3 | 494.5291 | 0.20711 | 0.193811 | 0.860096 |
| ENSMUSG( | Mettl4 | 691.1057 | 0.190902 | 0.180803 | 0.617841 | ENSMUSG( | Mettl4 | 581.952 | -0.032 | 0.828981 | 0.980224 |
| ENSMUSG( | Mettl5 | 166.4573 | -0.13849 | 0.39482 | 0.810072 | ENSMUSG( | Mettl5 | 148.9578 | -0.20506 | 0.303231 | 0.898538 |
| ENSMUSG( | Mettl6 | 463.9896 | -0.06975 | 0.595506 | 0.903058 | ENSMUSG( | Mettl6 | 448.1153 | 0.095571 | 0.53054 | 0.934795 |
| ENSMUSG( | Mettl8 | 347.2684 | -0.04407 | 0.768578 | 0.954674 | ENSMUSG( | Mettl8 | 324.5982 | 0.03801 | 0.809151 | 0.97873 |
| ENSMUSG( | Mettl9 | 3270.597 | -0.11074 | 0.30883 | 0.757024 | ENSMUSG( | Mettl9 | 2964.812 | -0.12909 | 0.370859 | 0.915491 |
| ENSMUSG( | Mex3a | 159.5805 | -0.17266 | 0.320927 | 0.765984 | ENSMUSG( | Mex3a | 155.2756 | 0.022657 | 0.899513 | 0.990558 |
| ENSMUSG( | Mex3b | 314.5563 | -0.13584 | 0.393203 | 0.809274 | ENSMUSG( | Mex3b | 318.5921 | 0.174756 | 0.327557 | 0.906255 |
| ENSMUSG( | Mex3c | 1186.572 | 0.002869 | 0.978521 | 0.997428 | ENSMUSG( | Mex3c | 1096.011 | 0.038206 | 0.768955 | 0.974236 |
| ENSMUSG( | Mex3d | 450.7472 | -0.09527 | 0.437622 | 0.830008 | ENSMUSG( | Mex3d | 420.1071 | -0.02002 | 0.894321 | 0.990558 |
| ENSMUSG( | Mfap1a | 523.3946 | 0.119922 | 0.338412 | 0.777489 | ENSMUSG( | Mfap1a | 470.7946 | 0.074797 | 0.649377 | 0.95422 |
| ENSMUSG( | Mfap1b | 390.3748 | 0.147572 | 0.289833 | 0.739839 | ENSMUSG( | Mfap1b | 337.9857 | -0.00658 | 0.968438 | 0.99594 |
| ENSMUSG( | Mfap2 | 175.8867 | -0.27446 | 0.141038 | 0.557961 | ENSMUSG( | Mfap2 | 176.1697 | 0.023225 | 0.89657 | 0.990558 |
| ENSMUSG( | Mfap3 | 814.3509 | -0.35021 | 0.035964 | 0.291377 | ENSMUSG( | Mfap3 | 880.1537 | 0.166985 | 0.38713 | 0.917257 |
| ENSMUSG( | Mfap3l | 5874.896 | -0.08225 | 0.515966 | 0.872701 | ENSMUSG( | Mfap3l | 5625.705 | 0.062277 | 0.681539 | 0.958678 |
| ENSMUSG( | Mfap4 | 105.979 | 0.040682 | 0.83599 | 0.970912 | ENSMUSG( | Mfap4 | 98.71385 | 0.104216 | 0.651422 | 0.95449 |
| ENSMUSG( | Mfap5 | 27.13855 | -0.83814 | 0.035276 | 0.288976 | ENSMUSG( | Mfap5 | 25.59824 | -0.75136 | 0.070649 | 0.758714 |
| ENSMUSG( | Mff | 5070.403 | 0.099794 | 0.266289 | 0.716441 | ENSMUSG( | Mff | 4346.864 | -0.07638 | 0.470139 | 0.923329 |
| ENSMUSG( | Mfge8 | 12253.1 | -0.2038 | 0.191366 | 0.630245 | ENSMUSG( | Mfge8 | 12555.42 | 0.143231 | 0.416328 | 0.917257 |
| ENSMUSG( | Mfhas1 | 1680.637 | 0.198964 | 0.039713 | 0.307318 | ENSMUSG( | Mfhas1 | 1502.563 | 0.150255 | 0.180508 | 0.850372 |
| ENSMUSG( | Mfn1 | 5170.436 | 0.228159 | 0.008708 | 0.131265 | ENSMUSG( | Mfn1 | 4366.113 | 0.016345 | 0.881882 | 0.989644 |
| ENSMUSG( | Mfn2 | 7618.642 | 0.060205 | 0.48915 | 0.859092 | ENSMUSG( | Mfn2 | 6495.487 | -0.13484 | 0.216212 | 0.865167 |
| ENSMUSG( | Mfng | 194.6723 | -0.04507 | 0.800309 | 0.961745 | ENSMUSG( | Mfng | 192.7073 | 0.20436 | 0.350897 | 0.911707 |
| ENSMUSG( | Mfrp | 56.86648 | 0.203305 | 0.851835 | 0.973149 | ENSMUSG( | Mfrp | 67.77956 | 0.848234 | 0.380068 | 0.916995 |
| ENSMUSG( | Mfsd1 | 2007.962 | -0.04709 | 0.714782 | 0.941144 | ENSMUSG( | Mfsd1 | 1850.282 | -0.01334 | 0.922349 | 0.992644 |
| ENSMUSG( | Mfsd10 | 624.1685 | -0.20126 | 0.125734 | 0.53257 | ENSMUSG( | Mfsd10 | 635.928 | 0.131068 | 0.342026 | 0.909146 |
| ENSMUSG( | Mfsd11 | 1070.986 | -0.17914 | 0.204231 | 0.64477 | ENSMUSG( | Mfsd11 | 1144.914 | 0.280646 | 0.0831 | 0.77681 |
| ENSMUSG( | Mfsd12 | 1499.659 | -0.14596 | 0.229514 | 0.67735 | ENSMUSG( | Mfsd12 | 1524.608 | 0.173157 | 0.246298 | 0.877904 |
| ENSMUSG( | Mfsd13a | 1137.883 | -0.29021 | 0.017112 | 0.195465 | ENSMUSG( | Mfsd13a | 1130.001 | -0.02285 | 0.866464 | 0.98737 |
| ENSMUSG( | Mfsd13b | 10.35609 | -0.68061 | 0.270519 | NA | ENSMUSG( | Mfsd13b | 10.41144 | -0.34305 | 0.576513 | NA |
| ENSMUSG( | Mfsd14a | 1425.306 | -0.39188 | 0.009981 | 0.142653 | ENSMUSG( | Mfsd14a | 1645.516 | 0.305805 | 0.087406 | 0.784337 |
| ENSMUSG( | Mfsd14b | 1140.676 | -0.25827 | 0.103113 | 0.48855 | ENSMUSG( | Mfsd14b | 1267.777 | 0.313375 | 0.089998 | 0.790741 |
| ENSMUSG( | Mfsd2a | 1775.728 | 0.232142 | 0.172178 | 0.607115 | ENSMUSG( | Mfsd2a | 2015.259 | 0.778862 | 7.68E-05 | 0.017018 |
| ENSMUSG( | Mfsd2b | 73.40139 | 0.409787 | 0.118681 | 0.521258 | ENSMUSG( | Mfsd2b | 68.81874 | 0.468252 | 0.096928 | 0.79645 |
| ENSMUSG( | Mfsd3 | 218.5473 | -0.29093 | 0.086119 | 0.451605 | ENSMUSG( | Mfsd3 | 222.1016 | 0.044952 | 0.807922 | 0.978622 |
| ENSMUSG( | Mfsd4a | 1906.024 | -0.47546 | 0.000988 | 0.032693 | ENSMUSG( | Mfsd4a | 2121.173 | 0.141233 | 0.381168 | 0.917202 |
| ENSMUSG( | Mfsd4b1 | 14.91096 | -0.83572 | 0.102295 | NA | ENSMUSG( | Mfsd4b1 | 16.33182 | -0.15318 | 0.753923 | 0.970671 |
| ENSMUSG( | Mfsd4b4 | 320.3187 | -0.41241 | 0.011404 | 0.153766 | ENSMUSG( | Mfsd4b4 | 340.1284 | 0.06729 | 0.698044 | 0.960867 |
| ENSMUSG( | Mfsd5 | 1073.733 | -0.23915 | 0.093974 | 0.47147 | ENSMUSG( | Mfsd5 | 1134.025 | 0.196324 | 0.261979 | 0.884926 |
| ENSMUSG( | Mfsd6 | 2472.095 | -0.42604 | 0.005125 | 0.095477 | ENSMUSG( | Mfsd6 | 2703.282 | 0.133505 | 0.434244 | 0.91869 |
| ENSMUSG( | Mfsd8 | 624.1676 | -0.3113 | 0.03196 | 0.272981 | ENSMUSG( | Mfsd8 | 695.1317 | 0.27644 | 0.073487 | 0.763629 |
| ENSMUSG( | Mfsd9 | 164.8505 | -0.3061 | 0.156015 | 0.58375 | ENSMUSG( | Mfsd9 | 177.0249 | 0.191929 | 0.398373 | 0.917257 |
| ENSMUSG( | Mga | 1634.538 | -0.16512 | 0.097489 | 0.477842 | ENSMUSG( | Mga | 1610.905 | 0.066432 | 0.564417 | 0.941206 |
| ENSMUSG( | Mgat1 | 2133.407 | -0.04824 | 0.645703 | 0.918146 | ENSMUSG( | Mgat1 | 2037.783 | 0.09026 | 0.494921 | 0.927372 |
| ENSMUSG( | Mgat2 | 865.577 | -0.50207 | 5.51E-05 | 0.004279 | ENSMUSG( | Mgat2 | 987.7778 | 0.188473 | 0.25254 | 0.88018 |
| ENSMUSG( | Mgat3 | 5447.535 | -0.075 | 0.505933 | 0.867008 | ENSMUSG( | Mgat3 | 5344.574 | 0.139427 | 0.308584 | 0.898538 |
| ENSMUSG( | Mgat4a | 1097.535 | -0.10122 | 0.504731 | 0.866883 | ENSMUSG( | Mgat4a | 1103.353 | 0.185817 | 0.296628 | 0.898538 |
| ENSMUSG( | Mgat4b | 3524.295 | 0.007747 | 0.939977 | 0.993415 | ENSMUSG( | Mgat4b | 3376.293 | 0.151842 | 0.235029 | 0.871506 |
| ENSMUSG( | Mgat4c | 141.7514 | -0.44812 | 0.061686 | 0.38394 | ENSMUSG( | Mgat4c | 172.956 | 0.405832 | 0.092901 | 0.79479 |
| ENSMUSG( | Mgat5 | 4644.162 | -0.3471 | 0.019207 | 0.208885 | ENSMUSG( | Mgat5 | 4560.18 | -0.10902 | 0.51255 | 0.930164 |
| ENSMUSG( | Mgat5b | 3574.192 | -0.07928 | 0.479847 | 0.854442 | ENSMUSG( | Mgat5b | 3664.876 | 0.255131 | 0.075215 | 0.767994 |
| ENSMUSG( | Mgl2 | 38.97956 | -0.06322 | 0.837852 | 0.971111 | ENSMUSG( | Mgl2 | 35.41874 | -0.04606 | 0.891808 | 0.990558 |
| ENSMUSG( | Mgll | 25158.54 | 0.167417 | 0.048156 | 0.335655 | ENSMUSG( | Mgll | 20838.07 | -0.10454 | 0.34483 | 0.91045 |
| ENSMUSG( | Mgme1 | 304.8849 | 0.296879 | 0.021023 | 0.220525 | ENSMUSG( | Mgme1 | 248.0138 | -0.02369 | 0.883398 | 0.989644 |
| ENSMUSG( | Mgmt | 48.85846 | 0.206283 | 0.466434 | 0.844661 | ENSMUSG( | Mgmt | 40.81608 | -0.05315 | 0.867983 | 0.98781 |
| ENSMUSG( | Mgp | 1046.988 | 0.068219 | 0.507697 | 0.86795 | ENSMUSG( | Mgp | 1148.439 | 0.554869 | 1.15E-05 | 0.004198 |
| ENSMUSG( | Mgrn1 | 8148.397 | 0.106331 | 0.216947 | 0.661816 | ENSMUSG( | Mgrn1 | 6858.069 | -0.12249 | 0.228366 | 0.868507 |
| ENSMUSG( | Mgst1 | 480.465 | -0.09362 | 0.564475 | 0.890434 | ENSMUSG( | Mgst1 | 458.1491 | 0.033583 | 0.82944 | 0.980224 |
| ENSMUSG( | Mgst3 | 2246.389 | -0.14852 | 0.085346 | 0.450474 | ENSMUSG( | Mgst3 | 2060.611 | -0.13016 | 0.264092 | 0.886134 |
| ENSMUSG( | Mia2 | 1796.637 | -0.25691 | 0.026118 | 0.247245 | ENSMUSG( | Mia2 | 1875.02 | 0.146067 | 0.27872 | 0.891628 |
| ENSMUSG( | Mia3 | 3392.814 | -0.20328 | 0.154823 | 0.582245 | ENSMUSG( | Mia3 | 3612.618 | 0.248214 | 0.143862 | 0.831311 |
| ENSMUSG( | Miat | 7978.533 | 0.366817 | 0.062012 | 0.385317 | ENSMUSG( | Miat | 7061.15 | 0.290391 | 0.197919 | 0.862161 |
| ENSMUSG( | Mib1 | 6816.537 | 0.054974 | 0.615455 | 0.910481 | ENSMUSG( | Mib1 | 6166.647 | 0.033415 | 0.820014 | 0.979198 |
| ENSMUSG( | Mib2 | 2827.788 | 0.168049 | 0.085986 | 0.451492 | ENSMUSG( | Mib2 | 2522.211 | 0.110613 | 0.359557 | 0.914223 |
| ENSMUSG( | Mical1 | 866.2088 | -0.1048 | 0.292596 | 0.742521 | ENSMUSG( | Mical1 | 745.2966 | -0.27814 | 0.022148 | 0.576679 |
| ENSMUSG( | Mical2 | 9812.037 | 0.424195 | 0.000642 | 0.024435 | ENSMUSG( | Mical2 | 7782.084 | 0.054036 | 0.674128 | 0.95694 |
| ENSMUSG( | Mical3 | 3980.914 | -0.04306 | 0.61919 | 0.911432 | ENSMUSG( | Mical3 | 3803.623 | 0.093912 | 0.362394 | 0.914223 |
| ENSMUSG( | Micall1 | 5765.866 | 0.044233 | 0.591286 | 0.90113 | ENSMUSG( | Micall1 | 4985.407 | -0.10942 | 0.292187 | 0.897854 |
| ENSMUSG( | Micall2 | 145.7785 | -0.40091 | 0.025372 | 0.245982 | ENSMUSG( | Micall2 | 136.6806 | -0.31363 | 0.106479 | 0.804969 |
| ENSMUSG( | Micos10 | 2345.339 | -0.01409 | 0.878235 | 0.981431 | ENSMUSG( | Micos10 | 2006.458 | -0.20691 | 0.096982 | 0.79645 |
| ENSMUSG( | Micos13 | 1502.527 | 0.039472 | 0.674471 | 0.929638 | ENSMUSG( | Micos13 | 1241.1 | -0.25868 | 0.034851 | 0.658559 |
| ENSMUSG( | Micu1 | 2617.527 | -0.06285 | 0.459668 | 0.840338 | ENSMUSG( | Micu1 | 2278.042 | -0.20247 | 0.060314 | 0.731659 |
| ENSMUSG( | Micu2 | 1091.595 | -0.0937 | 0.451646 | 0.835696 | ENSMUSG( | Micu2 | 953.3595 | -0.22779 | 0.095477 | 0.796133 |
| ENSMUSG( | Micu3 | 1097.903 | 0.035245 | 0.768975 | 0.954674 | ENSMUSG( | Micu3 | 1038.67 | 0.136723 | 0.340092 | 0.907972 |
| ENSMUSG( | Mid1 | 231.5396 | 0.179618 | 0.311233 | 0.759247 | ENSMUSG( | Mid1 | 133.2092 | -1.49295 | 3.01E-09 | 2.77E-06 |
| ENSMUSG( | Mid1ip1 | 7875.08 | 0.149705 | 0.157807 | 0.585927 | ENSMUSG( | Mid1ip1 | 6599.297 | -0.0905 | 0.495059 | 0.927372 |
| ENSMUSG( | Mid1-ps1 | 113.668 | -0.20491 | 0.363864 | 0.793343 | ENSMUSG( | Mid1-ps1 | 60.80147 | -3.24383 | 3.20E-19 | 6.22E-16 |
| ENSMUSG( | Mid2 | 866.9026 | 0.202112 | 0.062026 | 0.385317 | ENSMUSG( | Mid2 | 757.0979 | 0.079214 | 0.606087 | 0.949497 |
| ENSMUSG( | Mideas | 601.6706 | -0.12977 | 0.275165 | 0.72509 | ENSMUSG( | Mideas | 567.6131 | -0.02243 | 0.865031 | 0.98675 |
| ENSMUSG( | Midn | 1777.592 | -0.14756 | 0.120617 | 0.524506 | ENSMUSG( | Midn | 1666.876 | -0.061 | 0.611111 | 0.950771 |
| ENSMUSG( | Mief1 | 661.501 | 0.020516 | 0.854213 | 0.973491 | ENSMUSG( | Mief1 | 624.5719 | 0.122485 | 0.318306 | 0.902959 |
| ENSMUSG( | Mief2 | 495.9204 | -0.11929 | 0.377072 | 0.801213 | ENSMUSG( | Mief2 | 476.3617 | 0.045326 | 0.781041 | 0.976106 |
| ENSMUSG( | Mien1 | 2181.328 | 0.005982 | 0.948626 | 0.993821 | ENSMUSG( | Mien1 | 1785.647 | -0.32484 | 0.00859 | 0.391682 |
| ENSMUSG( | Mier1 | 1198.711 | -0.06322 | 0.590751 | 0.90113 | ENSMUSG( | Mier1 | 1163.389 | 0.117118 | 0.348985 | 0.911652 |
| ENSMUSG( | Mier2 | 892.7039 | 0.050327 | 0.663401 | 0.9265 | ENSMUSG( | Mier2 | 820.5528 | 0.076023 | 0.569842 | 0.941777 |
| ENSMUSG( | Mier3 | 413.0482 | -0.1572 | 0.187482 | 0.62584 | ENSMUSG( | Mier3 | 416.2348 | 0.134451 | 0.324865 | 0.905257 |
| ENSMUSG( | Mif | 3565.701 | -0.08328 | 0.416827 | 0.8209 | ENSMUSG( | Mif | 3122.54 | -0.20734 | 0.149077 | 0.836241 |
| ENSMUSG( | Mif4gd | 821.6313 | -0.10145 | 0.311624 | 0.759296 | ENSMUSG( | Mif4gd | 747.3442 | -0.10174 | 0.422749 | 0.917257 |
| ENSMUSG( | Miga1 | 3060.31 | -0.1317 | 0.23854 | 0.686308 | ENSMUSG( | Miga1 | 3033.596 | 0.114356 | 0.368635 | 0.915491 |
| ENSMUSG( | Miga2 | 3067.738 | -0.02157 | 0.861841 | 0.975055 | ENSMUSG( | Miga2 | 2926.659 | 0.113118 | 0.451389 | 0.922202 |
| ENSMUSG( | Miip | 560.6837 | 0.043183 | 0.781689 | 0.957558 | ENSMUSG( | Miip | 547.837 | 0.24152 | 0.204024 | 0.864412 |
| ENSMUSG( | Mill2 | 137.1799 | 0.214077 | 0.281706 | 0.731669 | ENSMUSG( | Mill2 | 127.4095 | 0.283352 | 0.231202 | 0.868507 |
| ENSMUSG( | Milr1 | 14.32942 | -0.49558 | 0.327414 | NA | ENSMUSG( | Milr1 | 15.88995 | 0.130917 | 0.797696 | 0.97681 |
| ENSMUSG( | Minar1 | 70.39249 | -0.00301 | 0.989927 | 0.999144 | ENSMUSG( | Minar1 | 68.12982 | 0.178927 | 0.505682 | 0.927825 |
| ENSMUSG( | Minar2 | 885.3146 | 0.031402 | 0.77604 | 0.955606 | ENSMUSG( | Minar2 | 764.1868 | -0.13052 | 0.337547 | 0.907972 |
| ENSMUSG( | Mindy1 | 857.3321 | -0.0043 | 0.969878 | 0.996369 | ENSMUSG( | Mindy1 | 851.5431 | 0.241006 | 0.085652 | 0.779799 |
| ENSMUSG( | Mindy2 | 4881.653 | -0.03489 | 0.783368 | 0.957558 | ENSMUSG( | Mindy2 | 4310.793 | -0.12961 | 0.373113 | 0.915491 |
| ENSMUSG( | Mindy3 | 925.6851 | 0.033064 | 0.759615 | 0.952528 | ENSMUSG( | Mindy3 | 836.631 | 0.004239 | 0.975227 | 0.996416 |
| ENSMUSG( | Mindy4 | 166.7302 | -0.02614 | 0.880384 | 0.981469 | ENSMUSG( | Mindy4 | 151.9186 | -0.01448 | 0.94026 | 0.994334 |
| ENSMUSG( | Mindy4b-p | 20.85869 | -0.0222 | 0.958621 | NA | ENSMUSG( | Mindy4b-p | 17.94643 | -0.19805 | 0.68085 | 0.958395 |
| ENSMUSG( | Mink1 | 13707.94 | 0.130517 | 0.129287 | 0.538528 | ENSMUSG( | Mink1 | 11838.86 | -0.02124 | 0.842325 | 0.983044 |
| ENSMUSG( | Minpp1 | 1337.234 | -0.18781 | 0.211292 | 0.654503 | ENSMUSG( | Minpp1 | 1442.514 | 0.296682 | 0.085651 | 0.779799 |
| ENSMUSG( | Mios | 481.5782 | 0.194314 | 0.109245 | 0.499001 | ENSMUSG( | Mios | 430.132 | 0.136557 | 0.331079 | 0.907593 |
| ENSMUSG( | Mipep | 854.2134 | 0.147383 | 0.162818 | 0.593778 | ENSMUSG( | Mipep | 679.5452 | -0.25751 | 0.054742 | 0.723362 |
| ENSMUSG( | Mipol1 | 160.9479 | 0.12807 | 0.445817 | 0.833275 | ENSMUSG( | Mipol1 | 138.272 | -0.0442 | 0.82117 | 0.979198 |
| ENSMUSG( | Mir100hg | 1809.35 | 0.08895 | 0.30356 | 0.751968 | ENSMUSG( | Mir100hg | 1700.429 | 0.174168 | 0.153104 | 0.837724 |
| ENSMUSG( | Mir124-2hg | 865.6771 | 0.117057 | 0.301392 | 0.749979 | ENSMUSG( | Mir124-2hg | 850.1765 | 0.314539 | 0.017066 | 0.52741 |
| ENSMUSG( | Mir124a-1l | 4017.567 | 0.182476 | 0.284874 | 0.733437 | ENSMUSG( | Mir124a-1l | 3539.42 | 0.088127 | 0.665185 | 0.956766 |
| ENSMUSG( | Mir132 | 16.83518 | -0.32472 | 0.553921 | NA | ENSMUSG( | Mir132 | 17.25731 | 0.080954 | 0.901935 | 0.99098 |
| ENSMUSG( | Mir212 | 22.57797 | -0.8319 | 0.107789 | 0.49674 | ENSMUSG( | Mir212 | 20.59232 | -0.8109 | 0.14816 | 0.836241 |
| ENSMUSG( | Mir219a-2 | 18.67594 | -0.40541 | 0.35433 | NA | ENSMUSG( | Mir219a-2 | 22.9106 | 0.493569 | 0.279359 | 0.892356 |
| ENSMUSG( | Mir223hg | 8.201904 | 0.194417 | 0.782865 | NA | ENSMUSG( | Mir223hg | 9.714302 | 0.811927 | 0.236183 | NA |
| ENSMUSG( | Mir22hg | 380.9221 | -0.27688 | 0.14313 | 0.561716 | ENSMUSG( | Mir22hg | 371.2801 | -0.06668 | 0.756334 | 0.971598 |
| ENSMUSG( | Mir23b | 9.184573 | 0.412468 | 0.545127 | NA | ENSMUSG( | Mir23b | 8.7816 | 0.53272 | 0.476557 | NA |
| ENSMUSG( | Mir3060 | 12.4253 | 0.567726 | 0.325372 | NA | ENSMUSG( | Mir3060 | 14.49197 | 1.103086 | 0.055721 | 0.723362 |
| ENSMUSG( | Mir3091 | 56.51939 | -0.0043 | 0.986901 | 0.998619 | ENSMUSG( | Mir3091 | 45.99341 | -0.37884 | 0.222558 | 0.866791 |
| ENSMUSG( | Mir374b | 14.41752 | 0.30495 | 0.531166 | NA | ENSMUSG( | Mir374b | 12.00033 | 0.104407 | 0.860452 | 0.986137 |
| ENSMUSG( | Mir421 | 16.97834 | 0.522542 | 0.264667 | NA | ENSMUSG( | Mir421 | 15.04793 | 0.506272 | 0.350688 | 0.911695 |
| ENSMUSG( | Mir433 | 17.89638 | 0.626128 | 0.160614 | NA | ENSMUSG( | Mir433 | 14.37836 | 0.276558 | 0.597391 | 0.946609 |
| ENSMUSG( | Mir5125 | 328.153 | 0.186912 | 0.405995 | 0.81652 | ENSMUSG( | Mir5125 | 310.3607 | 0.279845 | 0.290777 | 0.897019 |
| ENSMUSG( | Mir6236 | 2924.236 | -1.68395 | 4.82E-35 | 1.30E-31 | ENSMUSG( | Mir6236 | 2382.688 | -2.52618 | 3.88E-31 | 1.36E-27 |
| ENSMUSG( | Mir6955 | 14.13158 | 0.055235 | 0.914467 | NA | ENSMUSG( | Mir6955 | 12.49965 | -0.03194 | 0.953615 | 0.994798 |
| ENSMUSG( | Mir703 | 12.8599 | -1.19072 | 0.06948 | NA | ENSMUSG( | Mir703 | 17.50474 | 0.18462 | 0.776723 | 0.975717 |

| | | | | | |
|---|---|---|---|---|---|
| ENSMUSG( | 3270.597 | -0.11074 | 0.30883 | 0.757024 | Mettl9 |
| ENSMUSG( | 159.5805 | -0.17266 | 0.320927 | 0.765984 | Mex3a |
| ENSMUSG( | 314.5563 | -0.13584 | 0.393203 | 0.809274 | Mex3b |
| ENSMUSG( | 1186.572 | 0.002869 | 0.978521 | 0.997428 | Mex3c |
| ENSMUSG( | 450.7472 | -0.09527 | 0.437622 | 0.830008 | Mex3d |
| ENSMUSG( | 9.400138 | -0.8173 | 0.202403 | NA | Mexis |
| ENSMUSG( | 523.3946 | 0.119922 | 0.338412 | 0.777489 | Mfap1a |
| ENSMUSG( | 390.3748 | 0.147572 | 0.289833 | 0.739839 | Mfap1b |
| ENSMUSG( | 175.8867 | -0.27446 | 0.141038 | 0.557961 | Mfap2 |
| ENSMUSG( | 814.3509 | -0.35021 | 0.035964 | 0.291377 | Mfap3 |
| ENSMUSG( | 5874.896 | -0.08225 | 0.515966 | 0.872701 | Mfap3l |
| ENSMUSG( | 105.979 | 0.040682 | 0.83599 | 0.970912 | Mfap4 |
| ENSMUSG( | 27.13855 | -0.83814 | 0.035276 | 0.288976 | Mfap5 |
| ENSMUSG( | 5070.403 | 0.099794 | 0.266289 | 0.716441 | Mff |
| ENSMUSG( | 12253.1 | -0.2038 | 0.191366 | 0.630245 | Mfge8 |
| ENSMUSG( | 1680.637 | 0.198964 | 0.039713 | 0.307318 | Mfhas1 |
| ENSMUSG( | 5170.436 | 0.228159 | 0.008708 | 0.131265 | Mfn1 |
| ENSMUSG( | 7618.642 | 0.060205 | 0.48915 | 0.859092 | Mfn2 |
| ENSMUSG( | 194.6723 | -0.04507 | 0.800309 | 0.961745 | Mfng |
| ENSMUSG( | 56.86648 | 0.203305 | 0.851835 | 0.973149 | Mfrp |
| ENSMUSG( | 2007.962 | -0.04709 | 0.714782 | 0.941144 | Mfsd1 |
| ENSMUSG( | 624.1685 | -0.20126 | 0.125734 | 0.53257 | Mfsd10 |
| ENSMUSG( | 1070.986 | -0.17914 | 0.204231 | 0.64477 | Mfsd11 |
| ENSMUSG( | 1499.659 | -0.14596 | 0.229514 | 0.67735 | Mfsd12 |
| ENSMUSG( | 1137.883 | -0.29021 | 0.017112 | 0.195465 | Mfsd13a |
| ENSMUSG( | 10.35609 | -0.68061 | 0.270519 | NA | Mfsd13b |
| ENSMUSG( | 1425.306 | -0.39188 | 0.009981 | 0.142653 | Mfsd14a |
| ENSMUSG( | 1140.676 | -0.25827 | 0.103113 | 0.48855 | Mfsd14b |
| ENSMUSG( | 1775.728 | 0.232142 | 0.172178 | 0.607115 | Mfsd2a |
| ENSMUSG( | 73.40139 | 0.409787 | 0.118681 | 0.521258 | Mfsd2b |
| ENSMUSG( | 218.5473 | -0.29093 | 0.086119 | 0.451605 | Mfsd3 |
| ENSMUSG( | 1906.024 | -0.47546 | 0.000988 | 0.032693 | Mfsd4a |
| ENSMUSG( | 14.91096 | -0.83572 | 0.102295 | NA | Mfsd4b1 |
| ENSMUSG( | 320.3187 | -0.41241 | 0.011404 | 0.153766 | Mfsd4b4 |
| ENSMUSG( | 1073.733 | -0.23915 | 0.093974 | 0.47147 | Mfsd5 |
| ENSMUSG( | 2472.095 | -0.42604 | 0.005125 | 0.095477 | Mfsd6 |
| ENSMUSG( | 624.1676 | -0.3113 | 0.03196 | 0.272981 | Mfsd8 |
| ENSMUSG( | 164.8505 | -0.3061 | 0.156015 | 0.58375 | Mfsd9 |
| ENSMUSG( | 1634.538 | -0.16512 | 0.097489 | 0.477842 | Mga |
| ENSMUSG( | 2133.407 | -0.04824 | 0.645703 | 0.918146 | Mgat1 |
| ENSMUSG( | 865.577 | -0.50207 | 5.51E-05 | 0.004279 | Mgat2 |
| ENSMUSG( | 5447.535 | -0.075 | 0.505933 | 0.867008 | Mgat3 |
| ENSMUSG( | 1097.535 | -0.10122 | 0.504731 | 0.866883 | Mgat4a |
| ENSMUSG( | 3524.295 | 0.007747 | 0.939977 | 0.993415 | Mgat4b |
| ENSMUSG( | 141.7514 | -0.44812 | 0.061686 | 0.38394 | Mgat4c |
| ENSMUSG( | 4644.162 | -0.3471 | 0.019207 | 0.208885 | Mgat5 |
| ENSMUSG( | 3574.192 | -0.07928 | 0.479847 | 0.854442 | Mgat5b |
| ENSMUSG( | 38.97956 | -0.06322 | 0.837852 | 0.971111 | Mgl2 |
| ENSMUSG( | 25158.54 | 0.167417 | 0.048156 | 0.335655 | Mgll |
| ENSMUSG( | 304.8849 | 0.296879 | 0.021023 | 0.220525 | Mgme1 |
| ENSMUSG( | 48.85846 | 0.206283 | 0.466434 | 0.844661 | Mgmt |
| ENSMUSG( | 1046.988 | 0.068219 | 0.507697 | 0.86795 | Mgp |
| ENSMUSG( | 8148.397 | 0.106331 | 0.216947 | 0.661816 | Mgrn1 |
| ENSMUSG( | 480.465 | -0.09362 | 0.564475 | 0.890434 | Mgst1 |
| ENSMUSG( | 2246.389 | -0.14852 | 0.085346 | 0.450474 | Mgst3 |
| ENSMUSG( | 1796.637 | -0.25691 | 0.026118 | 0.247245 | Mia2 |
| ENSMUSG( | 3392.814 | -0.20328 | 0.154823 | 0.582245 | Mia3 |
| ENSMUSG( | 7978.533 | 0.366817 | 0.062012 | 0.385317 | Miat |
| ENSMUSG( | 6816.537 | 0.054974 | 0.615455 | 0.910481 | Mib1 |
| ENSMUSG( | 2827.788 | 0.168049 | 0.085986 | 0.451492 | Mib2 |
| ENSMUSG( | 866.2088 | -0.1048 | 0.292596 | 0.742521 | Mical1 |
| ENSMUSG( | 9812.037 | 0.424195 | 0.000642 | 0.024435 | Mical2 |
| ENSMUSG( | 3980.914 | -0.04306 | 0.61919 | 0.911432 | Mical3 |
| ENSMUSG( | 5765.866 | 0.044233 | 0.591286 | 0.90113 | Micall1 |
| ENSMUSG( | 145.7785 | -0.40091 | 0.025372 | 0.245982 | Micall2 |
| ENSMUSG( | 2345.339 | -0.01409 | 0.878235 | 0.981431 | Micos10 |
| ENSMUSG( | 1502.527 | 0.039472 | 0.674471 | 0.929638 | Micos13 |
| ENSMUSG( | 2617.527 | -0.06285 | 0.459668 | 0.840338 | Micu1 |
| ENSMUSG( | 1091.595 | -0.0937 | 0.451646 | 0.835696 | Micu2 |
| ENSMUSG( | 1097.903 | 0.035245 | 0.768975 | 0.954674 | Micu3 |
| ENSMUSG( | 231.5396 | 0.179618 | 0.311233 | 0.759247 | Mid1 |
| ENSMUSG( | 7875.08 | 0.149705 | 0.157807 | 0.585927 | Mid1ip1 |
| ENSMUSG( | 113.668 | -0.20491 | 0.363864 | 0.793343 | Mid1-ps1 |
| ENSMUSG( | 866.9026 | 0.202112 | 0.062026 | 0.385317 | Mid2 |
| ENSMUSG( | 601.6706 | -0.12977 | 0.275165 | 0.72509 | Mideas |
| ENSMUSG( | 1777.592 | -0.14756 | 0.120617 | 0.524506 | Midn |
| ENSMUSG( | 661.501 | 0.020516 | 0.854213 | 0.973491 | Mief1 |
| ENSMUSG( | 495.9204 | -0.11929 | 0.377072 | 0.801213 | Mief2 |
| ENSMUSG( | 2181.328 | 0.005982 | 0.948626 | 0.993821 | Mien1 |
| ENSMUSG( | 1198.711 | -0.06322 | 0.590751 | 0.90113 | Mier1 |
| ENSMUSG( | 892.7039 | 0.050327 | 0.663401 | 0.9265 | Mier2 |
| ENSMUSG( | 413.0482 | -0.1572 | 0.187482 | 0.62584 | Mier3 |
| ENSMUSG( | 3565.701 | -0.08328 | 0.416827 | 0.8209 | Mif |
| ENSMUSG( | 821.6313 | -0.10145 | 0.311624 | 0.759296 | Mif4gd |
| ENSMUSG( | 3060.31 | -0.1317 | 0.23854 | 0.686308 | Miga1 |
| ENSMUSG( | 3067.738 | -0.02157 | 0.861841 | 0.975055 | Miga2 |
| ENSMUSG( | 560.6837 | 0.043183 | 0.781689 | 0.957558 | Miip |
| ENSMUSG( | 137.1799 | 0.214077 | 0.281706 | 0.731669 | Mill2 |
| ENSMUSG( | 14.32942 | -0.49558 | 0.327414 | NA | Milr1 |
| ENSMUSG( | 70.39249 | -0.00301 | 0.989927 | 0.999144 | Minar1 |
| ENSMUSG( | 885.3146 | 0.031402 | 0.77604 | 0.955606 | Minar2 |
| ENSMUSG( | 857.3321 | -0.0043 | 0.969878 | 0.996369 | Mindy1 |
| ENSMUSG( | 4881.653 | -0.03489 | 0.783368 | 0.957558 | Mindy2 |
| ENSMUSG( | 925.6851 | 0.033064 | 0.759615 | 0.952528 | Mindy3 |
| ENSMUSG( | 166.7302 | -0.02614 | 0.880384 | 0.981469 | Mindy4 |
| ENSMUSG( | 20.85869 | -0.0222 | 0.958621 | NA | Mindy4b-ps |
| ENSMUSG( | 13707.94 | 0.130517 | 0.129287 | 0.538528 | Mink1 |
| ENSMUSG( | 1337.234 | -0.18781 | 0.211292 | 0.654503 | Minpp1 |
| ENSMUSG( | 481.5782 | 0.194314 | 0.109245 | 0.499001 | Mios |
| ENSMUSG( | 854.2134 | 0.147383 | 0.162818 | 0.593778 | Mipep |
| ENSMUSG( | 160.9479 | 0.12807 | 0.445817 | 0.833275 | Mipol1 |
| ENSMUSG( | 1809.35 | 0.08895 | 0.30356 | 0.751968 | Mir100hg |
| ENSMUSG( | 865.6771 | 0.117057 | 0.301392 | 0.749979 | Mir124-2hg |
| ENSMUSG( | 4017.567 | 0.182476 | 0.284874 | 0.733437 | Mir124a-1hg |
| ENSMUSG( | 16.83518 | -0.32472 | 0.553921 | NA | Mir132 |
| ENSMUSG( | 22.57797 | -0.8319 | 0.107789 | 0.49674 | Mir212 |
| ENSMUSG( | 18.67594 | -0.40541 | 0.35433 | NA | Mir219a-2 |
| ENSMUSG( | 8.201904 | 0.194417 | 0.782865 | NA | Mir223hg |
| ENSMUSG( | 380.9221 | -0.27688 | 0.14313 | 0.561716 | Mir22hg |
| ENSMUSG( | 9.184573 | 0.412468 | 0.545127 | NA | Mir23b |
| ENSMUSG( | 12.4253 | 0.567726 | 0.325372 | NA | Mir3060 |
| ENSMUSG( | 56.51939 | -0.0043 | 0.986901 | 0.998619 | Mir3091 |
| ENSMUSG( | 14.41752 | 0.30495 | 0.531166 | NA | Mir374b |
| ENSMUSG( | 16.97834 | 0.522542 | 0.264667 | NA | Mir421 |
| ENSMUSG( | 17.89638 | 0.626128 | 0.160614 | NA | Mir433 |
| ENSMUSG( | 328.153 | 0.186912 | 0.405995 | 0.81652 | Mir5125 |
| ENSMUSG( | 2924.236 | -1.68395 | 4.82E-35 | 1.30E-31 | Mir6236 |
| ENSMUSG( | 14.13158 | 0.055235 | 0.914467 | NA | Mir6955 |
| ENSMUSG( | 12.8599 | -1.19072 | 0.06948 | NA | Mir703 |
| ENSMUSG( | 39.72257 | 0.04555 | 0.890074 | 0.983687 | Mir7052 |
| ENSMUSG( | 11.66166 | -0.10344 | 0.856075 | NA | Mir7-1 |
| ENSMUSG( | 11.35464 | 0.402042 | 0.51368 | NA | Mir7220 |
| ENSMUSG( | 20.21794 | -0.18049 | 0.682858 | NA | Mir762 |
| ENSMUSG( | 26.71075 | -0.04212 | 0.920861 | 0.990409 | Mir7666 |
| ENSMUSG( | 9.468164 | -0.20868 | 0.733026 | NA | Mir8094 |
| ENSMUSG( | 1262.012 | 0.004742 | 0.961264 | 0.994061 | Mir9-2hg |
| ENSMUSG( | 1639.218 | -0.00151 | 0.988131 | 0.999109 | Mir9-3hg |
| ENSMUSG( | 10.94297 | 0.008666 | 0.987598 | NA | Mir99a |
| ENSMUSG( | 201.0127 | 0.301815 | 0.074289 | 0.422054 | Mir99ahg |
| ENSMUSG( | 817.3553 | 0.195476 | 0.141602 | 0.558606 | Mirg |
| ENSMUSG( | 632.0111 | 0.210325 | 0.058728 | 0.373994 | Mis12 |
| ENSMUSG( | 117.5401 | -0.10221 | 0.599958 | 0.903978 | Mis18a |
| ENSMUSG( | 20.05519 | -0.86614 | 0.048566 | NA | Mis18bp1 |
| ENSMUSG( | 10.76487 | 0.832388 | 0.162285 | NA | Misp |
| ENSMUSG( | 88.09677 | -0.24182 | 0.310062 | 0.758096 | Misp3 |
| ENSMUSG( | 137.3175 | -0.04366 | 0.799025 | 0.960922 | Mitd1 |
| ENSMUSG( | 244.6069 | -0.0102 | 0.947187 | 0.993821 | Mitf |
| ENSMUSG( | 202.5845 | 0.017441 | 0.916883 | 0.988885 | Mix23 |
| ENSMUSG( | 173.0391 | -0.77341 | 0.00023 | 0.011899 | Mki67 |
| ENSMUSG( | 272.993 | 0.375095 | 0.015314 | 0.18239 | Mkks |
| ENSMUSG( | 2305.48 | 0.156774 | 0.084097 | 0.447085 | Mkln1 |
| ENSMUSG( | 937.8119 | -0.14578 | 0.176184 | 0.611927 | Mknk1 |
| ENSMUSG( | 1708.476 | 0.080259 | 0.361936 | 0.792797 | Mknk2 |
| ENSMUSG( | 3380.67 | 0.009703 | 0.916795 | 0.988856 | Mkrn1 |
| ENSMUSG( | 1202.054 | 0.03766 | 0.673635 | 0.929191 | Mkrn2 |
| ENSMUSG( | 35.55992 | -0.18468 | 0.561103 | 0.888934 | Mkrn3 |
| ENSMUSG( | 222.681 | -0.06289 | 0.688458 | 0.933441 | Mks1 |
| ENSMUSG( | 337.4864 | 0.153019 | 0.343996 | 0.779153 | Mkx |
| ENSMUSG( | 22845.57 | 0.040227 | 0.715865 | 0.941484 | Mlc1 |
| ENSMUSG( | 6657.08 | 0.053084 | 0.665825 | 0.927054 | Mlec |
| ENSMUSG( | 26.52224 | -0.89649 | 0.015253 | 0.182053 | Mlf1 |
| ENSMUSG( | 18832.02 | 0.016005 | 0.862393 | 0.975055 | Mlf2 |
| ENSMUSG( | 357.8481 | -0.06636 | 0.620854 | 0.911432 | Mlh1 |
| ENSMUSG( | 675.0553 | 0.041404 | 0.691449 | 0.933881 | Mlh3 |
| ENSMUSG( | 1331.575 | -0.17842 | 0.060444 | 0.379704 | Mlip |
| ENSMUSG( | 10.92448 | -0.8738 | 0.123417 | NA | Mlkl |
| ENSMUSG( | 4279.12 | 0.254516 | 0.005548 | 0.09954 | Mllt1 |
| ENSMUSG( | 1035.326 | 0.040662 | 0.679959 | 0.930983 | Mllt10 |

| | | | | | |
|---|---|---|---|---|---|
| ENSMUSG( | 1148.439 | 0.554869 | 1.15E-05 | 0.004198 | Mgp |
| ENSMUSG( | 6858.069 | -0.12249 | 0.228366 | 0.868507 | Mgrn1 |
| ENSMUSG( | 458.1491 | 0.033583 | 0.82944 | 0.980224 | Mgst1 |
| ENSMUSG( | 2060.611 | -0.13016 | 0.264092 | 0.886134 | Mgst3 |
| ENSMUSG( | 1875.02 | 0.146067 | 0.27872 | 0.891628 | Mia2 |
| ENSMUSG( | 3612.618 | 0.248214 | 0.143862 | 0.831311 | Mia3 |
| ENSMUSG( | 7061.15 | 0.290391 | 0.197919 | 0.862161 | Miat |
| ENSMUSG( | 6166.647 | 0.033415 | 0.820014 | 0.979198 | Mib1 |
| ENSMUSG( | 2522.211 | 0.110613 | 0.359557 | 0.914223 | Mib2 |
| ENSMUSG( | 745.2966 | -0.27814 | 0.022148 | 0.576679 | Mical1 |
| ENSMUSG( | 7782.084 | 0.054036 | 0.674128 | 0.95694 | Mical2 |
| ENSMUSG( | 3803.623 | 0.093912 | 0.362394 | 0.914223 | Mical3 |
| ENSMUSG( | 4985.407 | -0.10942 | 0.292187 | 0.897854 | Micall1 |
| ENSMUSG( | 136.6806 | -0.31363 | 0.106479 | 0.804969 | Micall2 |
| ENSMUSG( | 2006.458 | -0.20691 | 0.096982 | 0.79645 | Micos10 |
| ENSMUSG( | 1241.1 | -0.25868 | 0.034851 | 0.658559 | Micos13 |
| ENSMUSG( | 2278.042 | -0.20247 | 0.060314 | 0.731659 | Micu1 |
| ENSMUSG( | 953.3595 | -0.22779 | 0.095477 | 0.796133 | Micu2 |
| ENSMUSG( | 1038.67 | 0.136723 | 0.340092 | 0.907972 | Micu3 |
| ENSMUSG( | 133.2092 | -1.49295 | 3.01E-09 | 2.77E-06 | Mid1 |
| ENSMUSG( | 6599.297 | -0.0905 | 0.495059 | 0.927372 | Mid1ip1 |
| ENSMUSG( | 60.80147 | -3.24383 | 3.20E-19 | 6.22E-16 | Mid1-ps1 |
| ENSMUSG( | 757.0979 | 0.079214 | 0.606087 | 0.949497 | Mid2 |
| ENSMUSG( | 567.6131 | -0.02243 | 0.865031 | 0.98675 | Mideas |
| ENSMUSG( | 1666.876 | -0.061 | 0.611111 | 0.950771 | Midn |
| ENSMUSG( | 624.5719 | 0.122485 | 0.318306 | 0.902959 | Mief1 |
| ENSMUSG( | 476.3617 | 0.045326 | 0.781041 | 0.976106 | Mief2 |
| ENSMUSG( | 1785.647 | -0.32484 | 0.00859 | 0.391682 | Mien1 |
| ENSMUSG( | 1163.389 | 0.117118 | 0.348985 | 0.911652 | Mier1 |
| ENSMUSG( | 820.5528 | 0.076023 | 0.569842 | 0.941777 | Mier2 |
| ENSMUSG( | 416.2348 | 0.134451 | 0.324865 | 0.905257 | Mier3 |
| ENSMUSG( | 3122.54 | -0.20734 | 0.149077 | 0.836241 | Mif |
| ENSMUSG( | 747.3442 | -0.10174 | 0.422749 | 0.917257 | Mif4gd |
| ENSMUSG( | 3033.596 | 0.114356 | 0.368635 | 0.915491 | Miga1 |
| ENSMUSG( | 2926.659 | 0.113118 | 0.451389 | 0.922202 | Miga2 |
| ENSMUSG( | 547.837 | 0.24152 | 0.204024 | 0.864412 | Miip |
| ENSMUSG( | 127.4095 | 0.283352 | 0.231202 | 0.868507 | Mill2 |
| ENSMUSG( | 15.88995 | 0.130917 | 0.797696 | 0.97681 | Milr1 |
| ENSMUSG( | 68.12982 | 0.178927 | 0.505682 | 0.927825 | Minar1 |
| ENSMUSG( | 764.1868 | -0.13052 | 0.337547 | 0.907972 | Minar2 |
| ENSMUSG( | 851.5431 | 0.241006 | 0.085652 | 0.779799 | Mindy1 |
| ENSMUSG( | 4310.793 | -0.12961 | 0.373113 | 0.915491 | Mindy2 |
| ENSMUSG( | 836.631 | 0.004239 | 0.975227 | 0.996416 | Mindy3 |
| ENSMUSG( | 151.9186 | -0.01448 | 0.94026 | 0.994334 | Mindy4 |
| ENSMUSG( | 17.94643 | -0.19805 | 0.68085 | 0.958395 | Mindy4b-ps |
| ENSMUSG( | 11838.86 | -0.02124 | 0.842325 | 0.983044 | Mink1 |
| ENSMUSG( | 1442.514 | 0.296682 | 0.085651 | 0.779799 | Minpp1 |
| ENSMUSG( | 430.132 | 0.136557 | 0.331079 | 0.907593 | Mios |
| ENSMUSG( | 679.5452 | -0.25751 | 0.054742 | 0.723362 | Mipep |
| ENSMUSG( | 138.272 | -0.0442 | 0.82117 | 0.979198 | Mipol1 |
| ENSMUSG( | 1700.429 | 0.174168 | 0.153104 | 0.837724 | Mir100hg |
| ENSMUSG( | 850.1765 | 0.314539 | 0.017066 | 0.52741 | Mir124-2hg |
| ENSMUSG( | 3539.42 | 0.088127 | 0.665185 | 0.956766 | Mir124a-1hg |
| ENSMUSG( | 17.25731 | 0.080954 | 0.901935 | 0.99098 | Mir132 |
| ENSMUSG( | 7.336732 | 1.302148 | 0.172045 | NA | Mir186 |
| ENSMUSG( | 20.59232 | -0.8109 | 0.14816 | 0.836241 | Mir212 |
| ENSMUSG( | 22.9106 | 0.493569 | 0.279359 | 0.892356 | Mir219a-2 |
| ENSMUSG( | 9.714302 | 0.811927 | 0.236183 | NA | Mir223hg |
| ENSMUSG( | 371.2801 | -0.06668 | 0.756334 | 0.971598 | Mir22hg |
| ENSMUSG( | 8.7816 | 0.53272 | 0.476557 | NA | Mir23b |
| ENSMUSG( | 14.49197 | 1.103086 | 0.055721 | 0.723362 | Mir3060 |
| ENSMUSG( | 45.99341 | -0.37884 | 0.222558 | 0.866791 | Mir3091 |
| ENSMUSG( | 12.00033 | 0.104407 | 0.860452 | 0.986137 | Mir374b |
| ENSMUSG( | 15.04793 | 0.506272 | 0.350688 | 0.911695 | Mir421 |
| ENSMUSG( | 14.37836 | 0.276558 | 0.597391 | 0.946609 | Mir433 |
| ENSMUSG( | 310.3607 | 0.279845 | 0.290777 | 0.897019 | Mir5125 |
| ENSMUSG( | 2382.688 | -2.52618 | 3.88E-31 | 1.36E-27 | Mir6236 |
| ENSMUSG( | 12.49965 | -0.03194 | 0.953615 | 0.994798 | Mir6955 |
| ENSMUSG( | 17.50474 | 0.18462 | 0.776723 | 0.975717 | Mir703 |
| ENSMUSG( | 31.4814 | -0.363 | 0.408624 | 0.917257 | Mir7052 |
| ENSMUSG( | 12.97473 | 0.47641 | 0.410317 | 0.917257 | Mir7-1 |
| ENSMUSG( | 17.22091 | -0.40802 | 0.403131 | 0.917257 | Mir762 |
| ENSMUSG( | 20.37718 | -0.65445 | 0.178807 | 0.849041 | Mir7666 |
| ENSMUSG( | 9.05749 | -0.01018 | 0.987817 | NA | Mir8094 |
| ENSMUSG( | 1244.552 | 0.222056 | 0.070896 | 0.760026 | Mir9-2hg |
| ENSMUSG( | 1567.347 | 0.134139 | 0.239508 | 0.874601 | Mir9-3hg |
| ENSMUSG( | 9.759759 | -0.10953 | 0.864634 | NA | Mir99a |
| ENSMUSG( | 197.5722 | 0.479409 | 0.022978 | 0.584403 | Mir99ahg |
| ENSMUSG( | 709.6107 | 0.059659 | 0.696953 | 0.960867 | Mirg |
| ENSMUSG( | 564.6325 | 0.149318 | 0.314771 | 0.90125 | Mis12 |
| ENSMUSG( | 115.2592 | 0.122694 | 0.570988 | 0.942052 | Mis18a |
| ENSMUSG( | 24.27978 | 0.111345 | 0.811367 | 0.978997 | Mis18bp1 |
| ENSMUSG( | 76.52328 | -0.41645 | 0.115598 | 0.813384 | Misp3 |
| ENSMUSG( | 136.5426 | 0.209174 | 0.284206 | 0.892412 | Mitd1 |
| ENSMUSG( | 230.622 | 0.074987 | 0.678553 | 0.957506 | Mitf |
| ENSMUSG( | 178.8722 | -0.08115 | 0.66924 | 0.956766 | Mix23 |
| ENSMUSG( | 197.9453 | 0.002597 | 0.990379 | 0.998129 | Mki67 |
| ENSMUSG( | 233.393 | 0.208435 | 0.217752 | 0.865589 | Mkks |
| ENSMUSG( | 2077.458 | 0.125964 | 0.257094 | 0.883082 | Mkln1 |
| ENSMUSG( | 921.2749 | 0.077277 | 0.513534 | 0.930482 | Mknk1 |
| ENSMUSG( | 1321.747 | -0.41699 | 0.001456 | 0.140378 | Mknk2 |
| ENSMUSG( | 3003.11 | -0.06496 | 0.571012 | 0.942052 | Mkrn1 |
| ENSMUSG( | 1062.804 | -0.05047 | 0.652295 | 0.954766 | Mkrn2 |
| ENSMUSG( | 36.53955 | 0.164007 | 0.616599 | 0.951597 | Mkrn3 |
| ENSMUSG( | 203.7454 | -0.03799 | 0.843549 | 0.983392 | Mks1 |
| ENSMUSG( | 275.7259 | -0.1679 | 0.308171 | 0.898538 | Mkx |
| ENSMUSG( | 19997.42 | -0.07523 | 0.527946 | 0.934616 | Mlc1 |
| ENSMUSG( | 6109.165 | 0.075478 | 0.574656 | 0.942912 | Mlec |
| ENSMUSG( | 30.72391 | -0.04042 | 0.914922 | 0.992407 | Mlf1 |
| ENSMUSG( | 16041.22 | -0.18616 | 0.161961 | 0.837724 | Mlf2 |
| ENSMUSG( | 323.109 | -0.08255 | 0.612873 | 0.951071 | Mlh1 |
| ENSMUSG( | 591.366 | -0.07657 | 0.558724 | 0.939574 | Mlh3 |
| ENSMUSG( | 1174.497 | -0.27859 | 0.013145 | 0.460068 | Mlip |
| ENSMUSG( | 13.88176 | 0.25348 | 0.631895 | 0.953251 | Mlkl |
| ENSMUSG( | 3786.265 | 0.177132 | 0.101299 | 0.801456 | Mllt1 |
| ENSMUSG( | 948.6378 | 0.055983 | 0.633758 | 0.953889 | Mllt10 |
| ENSMUSG( | 4197.391 | -0.09391 | 0.547164 | 0.938063 | Mllt11 |
| ENSMUSG( | 534.9246 | 0.308419 | 0.036867 | 0.672701 | Mllt3 |
| ENSMUSG( | 8292.116 | 0.079844 | 0.460064 | 0.923329 | Mllt6 |
| ENSMUSG( | 77.75244 | 0.516755 | 0.028961 | 0.614171 | Mlph |
| ENSMUSG( | 1048.673 | -0.03416 | 0.779832 | 0.976106 | Mlst8 |
| ENSMUSG( | 543.8318 | -0.07826 | 0.575224 | 0.942912 | Mlx |
| ENSMUSG( | 773.0079 | -0.09116 | 0.558373 | 0.93957 | Mlxip |
| ENSMUSG( | 189.7358 | 0.552123 | 0.021858 | 0.575341 | Mlxipl |
| ENSMUSG( | 531.8146 | -0.09045 | 0.537637 | 0.937216 | Mlycd |
| ENSMUSG( | 7.972499 | -0.37473 | 0.584681 | NA | Mm2pr |
| ENSMUSG( | 718.9017 | -0.06881 | 0.669414 | 0.956766 | Mmaa |
| ENSMUSG( | 1120.503 | -0.20312 | 0.083333 | 0.77681 | Mmab |
| ENSMUSG( | 363.8187 | -0.10172 | 0.579849 | 0.944209 | Mmachc |
| ENSMUSG( | 709.246 | -0.03865 | 0.765458 | 0.974236 | Mmadhc |
| ENSMUSG( | 5767.052 | 0.002884 | 0.984572 | 0.996963 | Mmd |
| ENSMUSG( | 13937.09 | -0.1023 | 0.394798 | 0.917257 | Mmd2 |
| ENSMUSG( | 231.0873 | 0.384121 | 0.031016 | 0.629564 | Mme |
| ENSMUSG( | 33.06856 | -0.03009 | 0.934774 | 0.99306 | Mmel1 |
| ENSMUSG( | 1830.703 | 0.007604 | 0.962692 | 0.995284 | Mmgt1 |
| ENSMUSG( | 709.5276 | -0.00642 | 0.964411 | 0.995433 | Mmgt2 |
| ENSMUSG( | 84.39989 | -0.55639 | 0.025392 | 0.598637 | Mmp11 |
| ENSMUSG( | 1050.512 | -0.70061 | 4.97E-06 | 0.002176 | Mmp14 |
| ENSMUSG( | 1602.065 | 0.074881 | 0.66429 | 0.956766 | Mmp15 |
| ENSMUSG( | 531.3693 | 0.06517 | 0.690628 | 0.960071 | Mmp16 |
| ENSMUSG( | 8425.111 | 0.128779 | 0.430034 | 0.917257 | Mmp17 |
| ENSMUSG( | 25.37752 | 0.304748 | 0.442885 | 0.921554 | Mmp19 |
| ENSMUSG( | 140.7132 | -0.03831 | 0.883596 | 0.989644 | Mmp2 |
| ENSMUSG( | 29.05654 | -0.12663 | 0.782899 | 0.976106 | Mmp23 |
| ENSMUSG( | 2017.594 | 0.218672 | 0.154729 | 0.837724 | Mmp24 |
| ENSMUSG( | 2482.642 | -0.23919 | 0.050257 | 0.708648 | Mmp24os1 |
| ENSMUSG( | 39.45577 | 0.031216 | 0.933606 | 0.99306 | Mmp25 |
| ENSMUSG( | 175.6372 | 0.43684 | 0.072152 | 0.763629 | Mmp28 |
| ENSMUSG( | 15.58644 | 0.689902 | 0.260388 | 0.884524 | Mmp8 |
| ENSMUSG( | 69.04521 | -0.04625 | 0.897834 | 0.990558 | Mmp9 |
| ENSMUSG( | 129.9169 | 0.496089 | 0.109416 | 0.805353 | Mmrn2 |
| ENSMUSG( | 1005.737 | -0.02002 | 0.87556 | 0.989644 | Mms19 |
| ENSMUSG( | 53.42039 | -0.02326 | 0.93925 | 0.993898 | Mms22l |
| ENSMUSG( | 1866.198 | -0.19465 | 0.094942 | 0.796133 | Mmut |
| ENSMUSG( | 815.8444 | 0.039473 | 0.736658 | 0.967344 | Mn1 |
| ENSMUSG( | 205.7813 | 0.078206 | 0.639778 | 0.953903 | Mnat1 |
| ENSMUSG( | 12.8802 | -0.51662 | 0.35465 | 0.912411 | Mnd1 |
| ENSMUSG( | 46.94471 | 0.058531 | 0.871654 | 0.988779 | Mndal |
| ENSMUSG( | 119.7531 | -0.07498 | 0.706795 | 0.963529 | Mns1 |
| ENSMUSG( | 1257.568 | 0.113354 | 0.339788 | 0.907972 | Mnt |
| ENSMUSG( | 1446.754 | -0.03421 | 0.785257 | 0.976285 | Mob1a |
| ENSMUSG( | 840.8271 | -0.02433 | 0.893091 | 0.990558 | Mob1b |
| ENSMUSG( | 724.8675 | 0.030822 | 0.812374 | 0.979148 | Mob2 |
| ENSMUSG( | 331.6356 | 0.008037 | 0.955117 | 0.994798 | Mob3a |
| ENSMUSG( | 188.8464 | -0.01784 | 0.927176 | 0.99306 | Mob3b |
| ENSMUSG( | 156.9547 | 0.061195 | 0.74353 | 0.96951 | Mob3c |
| ENSMUSG( | 857.9714 | -0.007 | 0.9643 | 0.995433 | Mob4 |
| ENSMUSG( | 213745.9 | -0.07844 | 0.531318 | 0.934795 | Mobp |

| | | | | | | | | | | | |
|---|---|---|---|---|---|---|---|---|---|---|---|
| ENSMUSG( | Mir7052 | 39.72257 | 0.04555 | 0.890074 | 0.983687 | ENSMUSG( | Mir7052 | 31.4814 | -0.363 | 0.408624 | 0.917257 |
| ENSMUSG( | Mir7-1 | 11.66166 | -0.10344 | 0.856075 | NA | ENSMUSG( | Mir7-1 | 12.97473 | 0.47641 | 0.410317 | 0.917257 |
| ENSMUSG( | Mir762 | 20.21794 | -0.18049 | 0.682858 | NA | ENSMUSG( | Mir762 | 17.22091 | -0.40802 | 0.403131 | 0.917257 |
| ENSMUSG( | Mir7666 | 26.71075 | -0.04212 | 0.920861 | 0.990409 | ENSMUSG( | Mir7666 | 20.37718 | -0.65445 | 0.178807 | 0.849041 |
| ENSMUSG( | Mir8094 | 9.468164 | -0.20868 | 0.733026 | NA | ENSMUSG( | Mir8094 | 9.05749 | -0.01018 | 0.987817 | NA |
| ENSMUSG( | Mir9-2hg | 1262.012 | 0.004742 | 0.961264 | 0.994061 | ENSMUSG( | Mir9-2hg | 1244.552 | 0.222056 | 0.070896 | 0.760026 |
| ENSMUSG( | Mir9-3hg | 1639.218 | -0.00151 | 0.988131 | 0.999109 | ENSMUSG( | Mir9-3hg | 1567.347 | 0.134139 | 0.239508 | 0.874601 |
| ENSMUSG( | Mir99a | 10.94297 | 0.008666 | 0.987598 | NA | ENSMUSG( | Mir99a | 9.759759 | -0.10953 | 0.864634 | NA |
| ENSMUSG( | Mir99ahg | 201.0127 | 0.301815 | 0.074289 | 0.422054 | ENSMUSG( | Mir99ahg | 197.5722 | 0.479409 | 0.022978 | 0.584403 |
| ENSMUSG( | Mirg | 817.3553 | 0.195476 | 0.141602 | 0.558606 | ENSMUSG( | Mirg | 709.6107 | 0.059659 | 0.696953 | 0.960867 |
| ENSMUSG( | Mis12 | 632.0111 | 0.210325 | 0.058728 | 0.373994 | ENSMUSG( | Mis12 | 564.6325 | 0.149318 | 0.314771 | 0.90125 |
| ENSMUSG( | Mis18a | 117.5401 | -0.10221 | 0.599958 | 0.903978 | ENSMUSG( | Mis18a | 115.2592 | 0.122694 | 0.570988 | 0.942052 |
| ENSMUSG( | Mis18bp1 | 20.05519 | -0.86614 | 0.048566 | NA | ENSMUSG( | Mis18bp1 | 24.27978 | 0.111345 | 0.811367 | 0.978997 |
| ENSMUSG( | Misp3 | 88.09677 | -0.24182 | 0.310062 | 0.758096 | ENSMUSG( | Misp3 | 76.52328 | -0.41645 | 0.115598 | 0.813384 |
| ENSMUSG( | Mitd1 | 137.3175 | -0.04366 | 0.799025 | 0.960922 | ENSMUSG( | Mitd1 | 136.5426 | 0.209174 | 0.284206 | 0.892412 |
| ENSMUSG( | Mitf | 244.6069 | -0.0102 | 0.947187 | 0.993821 | ENSMUSG( | Mitf | 230.622 | 0.074987 | 0.678553 | 0.957506 |
| ENSMUSG( | Mix23 | 202.5845 | 0.017441 | 0.916883 | 0.988885 | ENSMUSG( | Mix23 | 178.8722 | -0.08115 | 0.66924 | 0.956766 |
| ENSMUSG( | Mki67 | 173.0391 | -0.77341 | 0.00023 | 0.011899 | ENSMUSG( | Mki67 | 197.9453 | 0.002597 | 0.990379 | 0.998129 |
| ENSMUSG( | Mkks | 272.993 | 0.375095 | 0.015314 | 0.18239 | ENSMUSG( | Mkks | 233.393 | 0.208435 | 0.217752 | 0.865589 |
| ENSMUSG( | Mkln1 | 2305.48 | 0.156774 | 0.084097 | 0.447085 | ENSMUSG( | Mkln1 | 2077.458 | 0.125964 | 0.257094 | 0.883082 |
| ENSMUSG( | Mknk1 | 937.8119 | -0.14578 | 0.176184 | 0.611927 | ENSMUSG( | Mknk1 | 921.2749 | 0.077277 | 0.513534 | 0.930482 |
| ENSMUSG( | Mknk2 | 1708.476 | 0.080259 | 0.361936 | 0.792797 | ENSMUSG( | Mknk2 | 1321.747 | -0.41699 | 0.001456 | 0.140378 |
| ENSMUSG( | Mkrn1 | 3380.67 | 0.009703 | 0.916795 | 0.988856 | ENSMUSG( | Mkrn1 | 3003.11 | -0.06496 | 0.571012 | 0.942052 |
| ENSMUSG( | Mkrn2 | 1202.054 | 0.03766 | 0.673635 | 0.929191 | ENSMUSG( | Mkrn2 | 1062.804 | -0.05047 | 0.652295 | 0.954766 |
| ENSMUSG( | Mkrn3 | 35.55992 | -0.18468 | 0.561103 | 0.888934 | ENSMUSG( | Mkrn3 | 36.53955 | 0.164007 | 0.616599 | 0.951597 |
| ENSMUSG( | Mks1 | 222.681 | -0.06289 | 0.688458 | 0.933441 | ENSMUSG( | Mks1 | 203.7454 | -0.03799 | 0.843549 | 0.983392 |
| ENSMUSG( | Mkx | 337.4864 | 0.153019 | 0.343996 | 0.779153 | ENSMUSG( | Mkx | 275.7259 | -0.1679 | 0.308171 | 0.898538 |
| ENSMUSG( | Mlc1 | 22845.57 | 0.040227 | 0.715865 | 0.941484 | ENSMUSG( | Mlc1 | 19997.42 | -0.07523 | 0.527946 | 0.934616 |
| ENSMUSG( | Mlec | 6657.08 | 0.053084 | 0.665825 | 0.927054 | ENSMUSG( | Mlec | 6109.165 | 0.075478 | 0.574656 | 0.942912 |
| ENSMUSG( | Mlf1 | 26.52224 | -0.89649 | 0.015253 | 0.182053 | ENSMUSG( | Mlf1 | 30.72391 | -0.04042 | 0.914922 | 0.992407 |
| ENSMUSG( | Mlf2 | 18832.02 | 0.016005 | 0.862393 | 0.975055 | ENSMUSG( | Mlf2 | 16041.22 | -0.18616 | 0.161961 | 0.837724 |
| ENSMUSG( | Mlh1 | 357.8481 | -0.06636 | 0.620854 | 0.911432 | ENSMUSG( | Mlh1 | 323.109 | -0.08255 | 0.612873 | 0.951071 |
| ENSMUSG( | Mlh3 | 675.0553 | 0.041404 | 0.691449 | 0.933881 | ENSMUSG( | Mlh3 | 591.366 | -0.07657 | 0.558724 | 0.939574 |
| ENSMUSG( | Mlip | 1331.575 | -0.17842 | 0.060444 | 0.379704 | ENSMUSG( | Mlip | 1174.497 | -0.27859 | 0.013145 | 0.460068 |
| ENSMUSG( | Mlkl | 10.92448 | -0.8738 | 0.123417 | NA | ENSMUSG( | Mlkl | 13.88176 | 0.25348 | 0.631895 | 0.953251 |
| ENSMUSG( | Mllt1 | 4279.12 | 0.254516 | 0.005548 | 0.09954 | ENSMUSG( | Mllt1 | 3786.265 | 0.177132 | 0.101299 | 0.801456 |
| ENSMUSG( | Mllt10 | 1035.326 | 0.040662 | 0.679959 | 0.930983 | ENSMUSG( | Mllt10 | 948.6378 | 0.055983 | 0.633758 | 0.953889 |
| ENSMUSG( | Mllt11 | 4661.441 | -0.05597 | 0.639858 | 0.916434 | ENSMUSG( | Mllt11 | 4197.391 | -0.09391 | 0.547164 | 0.938063 |
| ENSMUSG( | Mllt3 | 543.8646 | 0.107253 | 0.386893 | 0.805056 | ENSMUSG( | Mllt3 | 534.9246 | 0.308419 | 0.036867 | 0.672701 |
| ENSMUSG( | Mllt6 | 9484.219 | 0.193444 | 0.02803 | 0.255998 | ENSMUSG( | Mllt6 | 8292.116 | 0.079844 | 0.460064 | 0.923329 |
| ENSMUSG( | Mlph | 80.77214 | 0.376667 | 0.083364 | 0.445503 | ENSMUSG( | Mlph | 77.75244 | 0.516755 | 0.028961 | 0.614171 |
| ENSMUSG( | Mlst8 | 1164.552 | -0.00406 | 0.968492 | 0.996369 | ENSMUSG( | Mlst8 | 1048.673 | -0.03416 | 0.779832 | 0.976106 |
| ENSMUSG( | Mlx | 594.1184 | -0.0939 | 0.44059 | 0.832037 | ENSMUSG( | Mlx | 543.8318 | -0.07826 | 0.575224 | 0.942912 |
| ENSMUSG( | Mlxip | 896.5606 | 0.062481 | 0.579884 | 0.896893 | ENSMUSG( | Mlxip | 773.0079 | -0.09116 | 0.558373 | 0.93957 |
| ENSMUSG( | Mlxipl | 182.9394 | 0.213917 | 0.221827 | 0.667414 | ENSMUSG( | Mlxipl | 189.7358 | 0.552123 | 0.021858 | 0.575341 |
| ENSMUSG( | Mlycd | 629.0168 | 0.124046 | 0.369425 | 0.796925 | ENSMUSG( | Mlycd | 531.8146 | -0.09045 | 0.537637 | 0.937216 |
| ENSMUSG( | Mm2pr | 10.16923 | 0.037964 | 0.946571 | NA | ENSMUSG( | Mm2pr | 7.972499 | -0.37473 | 0.584681 | NA |
| ENSMUSG( | Mmaa | 870.9558 | 0.212772 | 0.097778 | 0.478264 | ENSMUSG( | Mmaa | 718.9017 | -0.06881 | 0.669414 | 0.956766 |
| ENSMUSG( | Mmab | 1291.666 | -0.05358 | 0.585168 | 0.899171 | ENSMUSG( | Mmab | 1120.503 | -0.20312 | 0.083333 | 0.77681 |
| ENSMUSG( | Mmachc | 415.9467 | 0.031079 | 0.817482 | 0.965785 | ENSMUSG( | Mmachc | 363.8187 | -0.10172 | 0.579849 | 0.944209 |
| ENSMUSG( | Mmadhc | 812.3293 | 0.086153 | 0.408395 | 0.818167 | ENSMUSG( | Mmadhc | 709.246 | -0.03865 | 0.765458 | 0.974236 |
| ENSMUSG( | Mmd | 5926.491 | -0.19287 | 0.074233 | 0.422054 | ENSMUSG( | Mmd | 5767.052 | 0.002884 | 0.984572 | 0.996963 |
| ENSMUSG( | Mmd2 | 15788.75 | -0.00947 | 0.931928 | 0.9924 | ENSMUSG( | Mmd2 | 13937.09 | -0.1023 | 0.394798 | 0.917257 |
| ENSMUSG( | Mme | 178.1791 | -0.69713 | 0.000288 | 0.014061 | ENSMUSG( | Mme | 231.0873 | 0.384121 | 0.031016 | 0.629564 |
| ENSMUSG( | Mmel1 | 36.53931 | 0.008515 | 0.979497 | 0.997428 | ENSMUSG( | Mmel1 | 33.06856 | -0.03009 | 0.934774 | 0.99306 |
| ENSMUSG( | Mmgt1 | 1849.258 | -0.24361 | 0.100984 | 0.484745 | ENSMUSG( | Mmgt1 | 1830.703 | 0.007604 | 0.962692 | 0.995284 |
| ENSMUSG( | Mmgt2 | 723.5926 | -0.22955 | 0.086953 | 0.453311 | ENSMUSG( | Mmgt2 | 709.5276 | -0.00642 | 0.964411 | 0.995433 |
| ENSMUSG( | Mmp11 | 98.8594 | -0.35904 | 0.079779 | 0.436135 | ENSMUSG( | Mmp11 | 84.39989 | -0.55639 | 0.025392 | 0.598637 |
| ENSMUSG( | Mmp14 | 1309.02 | -0.26608 | 0.094319 | 0.471647 | ENSMUSG( | Mmp14 | 1050.512 | -0.70061 | 4.97E-06 | 0.002176 |
| ENSMUSG( | Mmp15 | 1548.456 | -0.31291 | 0.045256 | 0.326476 | ENSMUSG( | Mmp15 | 1602.065 | 0.074881 | 0.66429 | 0.956766 |
| ENSMUSG( | Mmp16 | 529.3766 | -0.2284 | 0.141234 | 0.558134 | ENSMUSG( | Mmp16 | 531.3693 | 0.06517 | 0.690628 | 0.960071 |
| ENSMUSG( | Mmp17 | 8158.476 | -0.24469 | 0.085895 | 0.451489 | ENSMUSG( | Mmp17 | 8425.111 | 0.128779 | 0.430034 | 0.917257 |
| ENSMUSG( | Mmp19 | 24.05722 | -0.08311 | 0.840193 | 0.971179 | ENSMUSG( | Mmp19 | 25.37752 | 0.304748 | 0.442885 | 0.921554 |
| ENSMUSG( | Mmp2 | 133.4361 | -0.52582 | 0.02203 | 0.226151 | ENSMUSG( | Mmp2 | 140.7132 | -0.03831 | 0.883596 | 0.989644 |
| ENSMUSG( | Mmp23 | 35.69649 | 0.153813 | 0.631041 | 0.914073 | ENSMUSG( | Mmp23 | 29.05654 | -0.12663 | 0.782899 | 0.976106 |
| ENSMUSG( | Mmp24 | 2085.654 | 0.048566 | 0.710986 | 0.940028 | ENSMUSG( | Mmp24 | 2017.594 | 0.218672 | 0.154729 | 0.837724 |
| ENSMUSG( | Mmp24os1 | 2986.195 | 0.037624 | 0.655548 | 0.923929 | ENSMUSG( | Mmp24os1 | 2482.642 | -0.23919 | 0.050257 | 0.708648 |
| ENSMUSG( | Mmp25 | 40.33707 | -0.22469 | 0.46683 | 0.844819 | ENSMUSG( | Mmp25 | 39.45577 | 0.031216 | 0.933606 | 0.99306 |
| ENSMUSG( | Mmp28 | 159.2432 | -0.10188 | 0.623786 | 0.91187 | ENSMUSG( | Mmp28 | 175.6372 | 0.43684 | 0.072152 | 0.763629 |
| ENSMUSG( | Mmp9 | 67.51147 | -0.36937 | 0.215818 | 0.660611 | ENSMUSG( | Mmp9 | 69.04521 | -0.04625 | 0.897834 | 0.990558 |
| ENSMUSG( | Mmrn2 | 111.7598 | -0.19569 | 0.450319 | 0.835344 | ENSMUSG( | Mmrn2 | 129.9169 | 0.496089 | 0.109416 | 0.805353 |
| ENSMUSG( | Mms19 | 1146.95 | 0.087364 | 0.333834 | 0.775602 | ENSMUSG( | Mms19 | 1005.737 | -0.02002 | 0.87556 | 0.989644 |
| ENSMUSG( | Mms22l | 50.92705 | -0.48568 | 0.095806 | 0.475512 | ENSMUSG( | Mms22l | 53.42039 | -0.02326 | 0.93925 | 0.993898 |
| ENSMUSG( | Mmut | 2272.673 | 0.111513 | 0.352981 | 0.784977 | ENSMUSG( | Mmut | 1866.198 | -0.19465 | 0.094942 | 0.796133 |
| ENSMUSG( | Mn1 | 870.6171 | -0.04096 | 0.69789 | 0.935828 | ENSMUSG( | Mn1 | 815.8444 | 0.039473 | 0.736658 | 0.967344 |
| ENSMUSG( | Mnat1 | 231.9461 | 0.156933 | 0.270534 | 0.720418 | ENSMUSG( | Mnat1 | 205.7813 | 0.078206 | 0.639778 | 0.953903 |
| ENSMUSG( | Mnd1 | 15.86522 | -0.13806 | 0.769292 | NA | ENSMUSG( | Mnd1 | 12.8802 | -0.51662 | 0.35465 | 0.912411 |
| ENSMUSG( | Mndal | 55.65463 | 0.248483 | 0.358142 | 0.789704 | ENSMUSG( | Mndal | 46.94471 | 0.058531 | 0.871654 | 0.988779 |
| ENSMUSG( | Mns1 | 123.5074 | -0.25897 | 0.14931 | 0.573248 | ENSMUSG( | Mns1 | 119.7531 | -0.07498 | 0.706795 | 0.963529 |
| ENSMUSG( | Mnt | 1298.234 | -0.06346 | 0.56109 | 0.888934 | ENSMUSG( | Mnt | 1257.568 | 0.113354 | 0.339788 | 0.907972 |
| ENSMUSG( | Mob1a | 1663.32 | 0.098073 | 0.378243 | 0.801688 | ENSMUSG( | Mob1a | 1446.754 | -0.03421 | 0.785257 | 0.976285 |
| ENSMUSG( | Mob1b | 909.933 | -0.05795 | 0.648764 | 0.920025 | ENSMUSG( | Mob1b | 840.8271 | -0.02433 | 0.893091 | 0.990558 |
| ENSMUSG( | Mob2 | 808.2842 | 0.072249 | 0.56902 | 0.892775 | ENSMUSG( | Mob2 | 724.8675 | 0.030822 | 0.812374 | 0.979148 |
| ENSMUSG( | Mob3a | 360.4985 | -0.01956 | 0.872168 | 0.978459 | ENSMUSG( | Mob3a | 331.6356 | 0.008037 | 0.955117 | 0.994798 |
| ENSMUSG( | Mob3b | 166.5292 | -0.74985 | 0.000296 | 0.014175 | ENSMUSG( | Mob3b | 188.8464 | -0.01784 | 0.927176 | 0.99306 |
| ENSMUSG( | Mob3c | 148.5333 | -0.40107 | 0.016218 | 0.189102 | ENSMUSG( | Mob3c | 156.9547 | 0.061195 | 0.74353 | 0.96951 |
| ENSMUSG( | Mob4 | 943.2261 | 0.004519 | 0.968892 | 0.996369 | ENSMUSG( | Mob4 | 857.9714 | -0.007 | 0.9643 | 0.995433 |
| ENSMUSG( | Mobp | 256950.4 | 0.181735 | 0.057478 | 0.370033 | ENSMUSG( | Mobp | 213745.9 | -0.07844 | 0.531318 | 0.934795 |
| ENSMUSG( | Mocos | 73.19207 | 0.279865 | 0.222591 | 0.668434 | ENSMUSG( | Mocos | 62.29585 | 0.085512 | 0.74596 | 0.96951 |
| ENSMUSG( | Mocs1 | 301.0388 | -0.01443 | 0.927085 | 0.991229 | ENSMUSG( | Mocs1 | 277.6407 | 0.024662 | 0.876834 | 0.989644 |
| ENSMUSG( | Mocs2 | 1452.8 | 0.088616 | 0.364534 | 0.793796 | ENSMUSG( | Mocs2 | 1243.021 | -0.09466 | 0.428986 | 0.917257 |
| ENSMUSG( | Mocs3 | 230.0902 | 0.089438 | 0.618371 | 0.911104 | ENSMUSG( | Mocs3 | 209.7175 | 0.084563 | 0.670935 | 0.95694 |
| ENSMUSG( | Mog | 5471.99 | -0.75257 | 1.52E-07 | 3.77E-05 | ENSMUSG( | Mog | 6347.675 | 0.045165 | 0.798353 | 0.97681 |
| ENSMUSG( | Mogs | 841.4888 | -0.27873 | 0.10272 | 0.488169 | ENSMUSG( | Mogs | 906.6271 | 0.218058 | 0.22309 | 0.867404 |
| ENSMUSG( | Mok | 257.9172 | 0.197426 | 0.188515 | 0.627572 | ENSMUSG( | Mok | 220.9592 | 0.031239 | 0.857908 | 0.985884 |
| ENSMUSG( | Mon1a | 744.1574 | 0.164967 | 0.108073 | 0.497059 | ENSMUSG( | Mon1a | 634.9896 | -0.01668 | 0.891999 | 0.990558 |
| ENSMUSG( | Mon1b | 1289.338 | 0.094151 | 0.325776 | 0.771061 | ENSMUSG( | Mon1b | 1122.864 | -0.03571 | 0.755528 | 0.971275 |
| ENSMUSG( | Mon2 | 2654.863 | 0.032947 | 0.708401 | 0.939519 | ENSMUSG( | Mon2 | 2525.37 | 0.154771 | 0.156534 | 0.837724 |
| ENSMUSG( | Morc2a | 1454.154 | 0.096641 | 0.271721 | 0.722082 | ENSMUSG( | Morc2a | 1289.214 | 0.021501 | 0.856265 | 0.985884 |
| ENSMUSG( | Morc2b | 59.48403 | -0.21962 | 0.426861 | 0.825231 | ENSMUSG( | Morc2b | 49.04605 | -0.59078 | 0.069243 | 0.755467 |
| ENSMUSG( | Morc3 | 593.608 | 0.063381 | 0.56984 | 0.893041 | ENSMUSG( | Morc3 | 565.4316 | 0.190192 | 0.171266 | 0.842999 |
| ENSMUSG( | Morc4 | 91.40183 | 0.170044 | 0.418727 | 0.821966 | ENSMUSG( | Morc4 | 81.588 | 0.125737 | 0.603857 | 0.948137 |
| ENSMUSG( | Morf4l1 | 6956.936 | 0.062053 | 0.463111 | 0.842422 | ENSMUSG( | Morf4l1 | 5897.975 | -0.15008 | 0.177536 | 0.847209 |
| ENSMUSG( | Morf4l2 | 3666.927 | 0.018858 | 0.864827 | 0.975565 | ENSMUSG( | Morf4l2 | 3217.811 | -0.09504 | 0.523067 | 0.932848 |
| ENSMUSG( | Morn1 | 165.4904 | -0.17245 | 0.318017 | 0.764184 | ENSMUSG( | Morn1 | 159.8143 | 0.011713 | 0.951622 | 0.994798 |
| ENSMUSG( | Morn2 | 424.9879 | 0.064809 | 0.615956 | 0.910822 | ENSMUSG( | Morn2 | 340.3688 | -0.33579 | 0.03315 | 0.64093 |
| ENSMUSG( | Morn4 | 3217.668 | 0.071134 | 0.395102 | 0.810183 | ENSMUSG( | Morn4 | 2711.674 | -0.15924 | 0.1795 | 0.850372 |
| ENSMUSG( | Morn5 | 21.56616 | -0.04455 | 0.91677 | 0.988856 | ENSMUSG( | Morn5 | 18.39698 | -0.2695 | 0.555399 | 0.939565 |
| ENSMUSG( | Morrbid | 14.40772 | -0.31309 | 0.533172 | NA | ENSMUSG( | Morrbid | 14.60064 | 0.06827 | 0.904193 | 0.991089 |
| ENSMUSG( | Mosmo | 1088.381 | -0.20977 | 0.08995 | 0.460542 | ENSMUSG( | Mosmo | 1161.731 | 0.245268 | 0.096621 | 0.796245 |
| ENSMUSG( | Mospd1 | 292.1307 | -0.01339 | 0.925523 | 0.991204 | ENSMUSG( | Mospd1 | 266.8911 | -0.00672 | 0.968199 | 0.995887 |
| ENSMUSG( | Mospd2 | 565.1245 | 0.153075 | 0.240167 | 0.687899 | ENSMUSG( | Mospd2 | 487.9968 | 0.001264 | 0.993147 | 0.998813 |
| ENSMUSG( | Mospd3 | 700.2832 | 0.041303 | 0.730458 | 0.944956 | ENSMUSG( | Mospd3 | 598.8908 | -0.15053 | 0.290413 | 0.897019 |
| ENSMUSG( | Mov10 | 553.9411 | 0.260611 | 0.122607 | 0.528484 | ENSMUSG( | Mov10 | 462.7052 | 0.017403 | 0.932268 | 0.99306 |
| ENSMUSG( | Moxd1 | 259.2248 | -0.33701 | 0.062197 | 0.386083 | ENSMUSG( | Moxd1 | 263.6572 | 0.001732 | 0.992615 | 0.99874 |
| ENSMUSG( | Mpc1 | 2730.619 | 0.11642 | 0.247034 | 0.69576 | ENSMUSG( | Mpc1 | 2296.764 | -0.11766 | 0.348554 | 0.911652 |
| ENSMUSG( | Mpc2 | 1862.697 | -0.12356 | 0.178834 | 0.616298 | ENSMUSG( | Mpc2 | 1628.252 | -0.25541 | 0.039283 | 0.679972 |
| ENSMUSG( | Mpdu1 | 1849.531 | -0.2311 | 0.022435 | 0.228899 | ENSMUSG( | Mpdu1 | 1817.055 | -0.00311 | 0.978659 | 0.996459 |
| ENSMUSG( | Mpdz | 2253.806 | 0.090825 | 0.462812 | 0.842421 | ENSMUSG( | Mpdz | 2004.915 | 0.020708 | 0.854249 | 0.98563 |
| ENSMUSG( | Mpeg1 | 2490.218 | -0.21083 | 0.104354 | 0.490474 | ENSMUSG( | Mpeg1 | 2449.667 | 0.021694 | 0.894245 | 0.990558 |
| ENSMUSG( | Mpg | 488.5874 | -0.16574 | 0.187813 | 0.626399 | ENSMUSG( | Mpg | 436.531 | -0.21678 | 0.153575 | 0.837724 |
| ENSMUSG( | Mphosph1( | 407.466 | 0.140315 | 0.255384 | 0.706181 | ENSMUSG( | Mphosph1( | 367.3455 | 0.104891 | 0.48047 | 0.924548 |
| ENSMUSG( | Mphosph6 | 297.6943 | 0.239339 | 0.093815 | 0.470818 | ENSMUSG( | Mphosph6 | 267.608 | 0.202927 | 0.208696 | 0.864412 |
| ENSMUSG( | Mphosph8 | 1708.7 | 0.35723 | 0.031643 | 0.271198 | ENSMUSG( | Mphosph8 | 1497.775 | 0.256384 | 0.074265 | 0.766789 |
| ENSMUSG( | Mphosph9 | 1072.785 | -0.03228 | 0.747917 | 0.949604 | ENSMUSG( | Mphosph9 | 990.9122 | 0.003253 | 0.980114 | 0.996479 |
| ENSMUSG( | Mpi | 1127.101 | 0.080613 | 0.452649 | 0.836145 | ENSMUSG( | Mpi | 979.7786 | -0.05399 | 0.650258 | 0.95422 |
| ENSMUSG( | Mpl | 83.21737 | 0.277589 | 0.197648 | 0.637377 | ENSMUSG( | Mpl | 66.95325 | -0.05529 | 0.836396 | 0.981335 |
| ENSMUSG( | Mplkip | 249.2146 | -0.10072 | 0.489602 | 0.859267 | ENSMUSG( | Mplkip | 234.4313 | -0.01141 | 0.945517 | 0.994798 |
| ENSMUSG( | Mpnd | 3797.776 | 0.11571 | 0.142968 | 0.561716 | ENSMUSG( | Mpnd | 3330.677 | 0.006324 | 0.953392 | 0.994798 |
| ENSMUSG( | Mpp1 | 523.7403 | -0.01966 | 0.879752 | 0.981436 | ENSMUSG( | Mpp1 | 502.2615 | 0.125093 | 0.38866 | 0.917257 |
| ENSMUSG( | Mpp2 | 5085.602 | 0.169062 | 0.09935 | 0.48179 | ENSMUSG( | Mpp2 | 4445.341 | 0.052532 | 0.659842 | 0.956205 |
| ENSMUSG( | Mpp3 | 1097.02 | 0.021149 | 0.865376 | 0.975565 | ENSMUSG( | Mpp3 | 948.3136 | -0.13439 | 0.324046 | 0.905257 |
| ENSMUSG( | Mpp4 | 33.33365 | 0.636912 | 0.086455 | 0.452045 | ENSMUSG( | Mpp4 | 23.93874 | -0.00928 | 0.986258 | 0.997188 |
| ENSMUSG( | Mpp7 | 132.3319 | -0.04537 | 0.795928 | 0.960062 | ENSMUSG( | Mpp7 | 118.3062 | -0.09324 | 0.654886 | 0.955015 |
| ENSMUSG( | Mppe1 | 548.7438 | -0.25883 | 0.039746 | 0.307429 | ENSMUSG( | Mppe1 | 565.1604 | 0.111008 | 0.453726 | 0.922316 |
| ENSMUSG( | Mpped1 | 8607.946 | 0.236148 | 0.007131 | 0.116294 | ENSMUSG( | Mpped1 | 7297.572 | 0.038786 | 0.692268 | 0.960404 |
| ENSMUSG( | Mpped2 | 587.6092 | -0.24283 | 0.083388 | 0.445503 | ENSMUSG( | Mpped2 | 528.5015 | -0.29003 | 0.078249 | 0.77321 |
| ENSMUSG( | Mprip | 12952.08 | 0.049573 | 0.521932 | 0.875186 | ENSMUSG( | Mprip | 11655.4 | 0.015078 | 0.889065 | 0.990558 |
| ENSMUSG( | Mpst | 1502.189 | 0.031499 | 0.727257 | 0.944223 | ENSMUSG( | Mpst | 1263.46 | -0.21176 | 0.125681 | 0.81516 |
| ENSMUSG( | Mpv17 | 1376.674 | 0.081617 | 0.389155 | 0.806967 | ENSMUSG( | Mpv17 | 1168.336 | -0.12788 | 0.293624 | 0.898538 |
| ENSMUSG( | Mpv17l | 2209.892 | -0.01114 | 0.914402 | 0.988483 | ENSMUSG( | Mpv17l | 1993.711 | -0.04394 | 0.776975 | 0.975717 |
| ENSMUSG( | Mpv17l2 | 1220.295 | -0.15353 | 0.212704 | 0.6567 | ENSMUSG( | Mpv17l2 | 1133.961 | -0.09523 | 0.501273 | 0.927825 |
| ENSMUSG( | Mpzl1 | 1423.073 | -0.34586 | 0.02577 | 0.247138 | ENSMUSG( | Mpzl1 | 1489.539 | 0.080812 | 0.662711 | 0.956205 |
| ENSMUSG( | Mpzl2 | 112.3363 | -0.26067 | 0.198274 | 0.638387 | ENSMUSG( | Mpzl2 | 123.4494 | 0.286137 | 0.159074 | 0.837724 |
| ENSMUSG( | Mpzl3 | 128.5136 | -0.32373 | 0.128221 | 0.536641 | ENSMUSG( | Mpzl3 | 125.2933 | -0.10522 | 0.639557 | 0.953903 |
| ENSMUSG( | Mr1 | 424.7537 | -0.17509 | 0.195628 | 0.635109 | ENSMUSG( | Mr1 | 394.3518 | -0.12135 | 0.410796 | 0.917257 |
| ENSMUSG( | Mrap | 17.36912 | -0.27689 | 0.556456 | NA | ENSMUSG( | Mrap | 19.14346 | 0.256772 | 0.574404 | 0.942912 |
| ENSMUSG( | Mrap2 | 127.0655 | -0.34616 | 0.074464 | 0.422899 | ENSMUSG( | Mrap2 | 138.304 | 0.181946 | 0.365755 | 0.914623 |
| ENSMUSG( | Mras | 15621.46 | 0.090274 | 0.24964 | 0.699258 | ENSMUSG( | Mras | 13201.32 | -0.12797 | 0.197119 | 0.862161 |
| ENSMUSG( | Mrc1 | 365.0113 | 0.022818 | 0.895455 | 0.984512 | ENSMUSG( | Mrc1 | 371.4406 | 0.326193 | 0.067284 | 0.75212 |
| ENSMUSG( | Mrc2 | 675.9576 | -0.36465 | 0.002448 | 0.058231 | ENSMUSG( | Mrc2 | 734.9807 | 0.170314 | 0.272142 | 0.888724 |
| ENSMUSG( | Mre11a | 355.854 | -0.07342 | 0.551 | 0.886038 | ENSMUSG( | Mre11a | 325.7754 | -0.06158 | 0.670807 | 0.95694 |
| ENSMUSG( | Mreg | 84.17341 | -0.00969 | 0.969966 | 0.996369 | ENSMUSG( | Mreg | 77.0437 | -0.02083 | 0.944764 | 0.994798 |
| ENSMUSG( | Mrfap1 | 7417.919 | 0.000149 | 0.998867 | 0.99989 | ENSMUSG( | Mrfap1 | 6560.059 | -0.09051 | 0.51015 | 0.929089 |
| ENSMUSG( | Mrgbp | 517.7027 | -0.04895 | 0.701836 | 0.937441 | ENSMUSG( | Mrgbp | 498.5316 | 0.115371 | 0.41268 | 0.917257 |
| ENSMUSG( | Mrgpre | 274.4234 | -0.26152 | 0.170848 | 0.605452 | ENSMUSG( | Mrgpre | 307.9902 | 0.345489 | 0.117161 | 0.813384 |
| ENSMUSG( | Mrgprf | 72.86058 | -0.27108 | 0.32039 | 0.765921 | ENSMUSG( | Mrgprf | 83.00162 | 0.388915 | 0.185439 | 0.855305 |
| ENSMUSG( | Mri1 | 365.1936 | -0.08918 | 0.556395 | 0.887681 | ENSMUSG( | Mri1 | 345.5501 | 0.028927 | 0.854952 | 0.985649 |

| | | | | | |
|---|---|---|---|---|---|
| ENSMUSG( | 4661.441 | -0.05597 | 0.639858 | 0.916434 | Mllt11 |
| ENSMUSG( | 543.8646 | 0.107253 | 0.386893 | 0.805056 | Mllt3 |
| ENSMUSG( | 9484.219 | 0.193444 | 0.02803 | 0.255998 | Mllt6 |
| ENSMUSG( | 80.77214 | 0.376667 | 0.083364 | 0.445503 | Mlph |
| ENSMUSG( | 1164.552 | -0.00406 | 0.968492 | 0.996369 | Mlst8 |
| ENSMUSG( | 594.1184 | -0.0939 | 0.44059 | 0.832037 | Mlx |
| ENSMUSG( | 896.5606 | 0.062481 | 0.579884 | 0.896893 | Mlxip |
| ENSMUSG( | 182.9394 | 0.213917 | 0.221827 | 0.667414 | Mlxipl |
| ENSMUSG( | 629.0168 | 0.124046 | 0.369425 | 0.796925 | Mlycd |
| ENSMUSG( | 10.16923 | 0.037964 | 0.946571 | NA | Mm2pr |
| ENSMUSG( | 870.9558 | 0.212772 | 0.097778 | 0.478264 | Mmaa |
| ENSMUSG( | 1291.666 | -0.05358 | 0.585168 | 0.899171 | Mmab |
| ENSMUSG( | 415.9467 | 0.031079 | 0.817482 | 0.965785 | Mmachc |
| ENSMUSG( | 812.3293 | 0.086153 | 0.408395 | 0.818167 | Mmadhc |
| ENSMUSG( | 5926.491 | -0.19287 | 0.074233 | 0.422054 | Mmd |
| ENSMUSG( | 15788.75 | -0.00947 | 0.931928 | 0.9924 | Mmd2 |
| ENSMUSG( | 178.1791 | -0.69713 | 0.000288 | 0.014061 | Mme |
| ENSMUSG( | 36.53931 | 0.008515 | 0.979497 | 0.997428 | Mmel1 |
| ENSMUSG( | 1849.258 | -0.24361 | 0.100984 | 0.484745 | Mmgt1 |
| ENSMUSG( | 723.5926 | -0.22955 | 0.086953 | 0.453311 | Mmgt2 |
| ENSMUSG( | 98.8594 | -0.35904 | 0.079779 | 0.436135 | Mmp11 |
| ENSMUSG( | 1309.02 | -0.26608 | 0.094319 | 0.471647 | Mmp14 |
| ENSMUSG( | 1548.456 | -0.31291 | 0.045256 | 0.326476 | Mmp15 |
| ENSMUSG( | 529.3766 | -0.2284 | 0.141234 | 0.558134 | Mmp16 |
| ENSMUSG( | 8158.476 | -0.24469 | 0.085895 | 0.451489 | Mmp17 |
| ENSMUSG( | 24.05722 | -0.08311 | 0.840193 | 0.971179 | Mmp19 |
| ENSMUSG( | 133.4361 | -0.52582 | 0.02203 | 0.226151 | Mmp2 |
| ENSMUSG( | 35.69649 | 0.153813 | 0.631041 | 0.914073 | Mmp23 |
| ENSMUSG( | 2085.654 | 0.048566 | 0.710986 | 0.940028 | Mmp24 |
| ENSMUSG( | 2986.195 | 0.037624 | 0.655548 | 0.923929 | Mmp24os1 |
| ENSMUSG( | 40.33707 | -0.22469 | 0.46683 | 0.844819 | Mmp25 |
| ENSMUSG( | 159.2432 | -0.10188 | 0.623786 | 0.91187 | Mmp28 |
| ENSMUSG( | 67.51147 | -0.36937 | 0.215818 | 0.660611 | Mmp9 |
| ENSMUSG( | 111.7598 | -0.19569 | 0.450319 | 0.835344 | Mmrn2 |
| ENSMUSG( | 1146.95 | 0.087364 | 0.333834 | 0.775602 | Mms19 |
| ENSMUSG( | 50.92705 | -0.48568 | 0.095806 | 0.475512 | Mms22l |
| ENSMUSG( | 2272.673 | 0.111513 | 0.352981 | 0.784977 | Mmut |
| ENSMUSG( | 870.6171 | -0.04096 | 0.69789 | 0.935828 | Mn1 |
| ENSMUSG( | 231.9461 | 0.156933 | 0.270534 | 0.720418 | Mnat1 |
| ENSMUSG( | 15.86522 | -0.13806 | 0.769292 | NA | Mnd1 |
| ENSMUSG( | 55.65463 | 0.248483 | 0.358142 | 0.789704 | Mndal |
| ENSMUSG( | 123.5074 | -0.25897 | 0.14931 | 0.573248 | Mns1 |
| ENSMUSG( | 1298.234 | -0.06346 | 0.56109 | 0.888934 | Mnt |
| ENSMUSG( | 1663.32 | 0.098073 | 0.378243 | 0.801688 | Mob1a |
| ENSMUSG( | 909.933 | -0.05795 | 0.648764 | 0.920025 | Mob1b |
| ENSMUSG( | 808.2842 | 0.072249 | 0.56902 | 0.892775 | Mob2 |
| ENSMUSG( | 360.4985 | -0.01956 | 0.872168 | 0.978459 | Mob3a |
| ENSMUSG( | 166.5292 | -0.74985 | 0.000296 | 0.014175 | Mob3b |
| ENSMUSG( | 148.5333 | -0.40107 | 0.016218 | 0.189102 | Mob3c |
| ENSMUSG( | 943.2261 | 0.004519 | 0.968892 | 0.996369 | Mob4 |
| ENSMUSG( | 256950.4 | 0.181735 | 0.057478 | 0.370033 | Mobp |
| ENSMUSG( | 73.19207 | 0.279865 | 0.222591 | 0.668434 | Mocos |
| ENSMUSG( | 301.0388 | -0.01443 | 0.927085 | 0.991229 | Mocs1 |
| ENSMUSG( | 1452.8 | 0.088616 | 0.364534 | 0.793796 | Mocs2 |
| ENSMUSG( | 230.0902 | 0.089438 | 0.618371 | 0.911104 | Mocs3 |
| ENSMUSG( | 5471.99 | -0.75257 | 1.52E-07 | 3.77E-05 | Mog |
| ENSMUSG( | 841.4888 | -0.27873 | 0.10272 | 0.488169 | Mogs |
| ENSMUSG( | 257.9172 | 0.197426 | 0.188515 | 0.627572 | Mok |
| ENSMUSG( | 744.1574 | 0.164967 | 0.108073 | 0.497059 | Mon1a |
| ENSMUSG( | 1289.338 | 0.094151 | 0.325776 | 0.771061 | Mon1b |
| ENSMUSG( | 2654.863 | 0.032947 | 0.708401 | 0.939519 | Mon2 |
| ENSMUSG( | 1454.154 | 0.096641 | 0.271721 | 0.722082 | Morc2a |
| ENSMUSG( | 59.48403 | -0.21962 | 0.426861 | 0.825231 | Morc2b |
| ENSMUSG( | 593.608 | 0.063381 | 0.56984 | 0.893041 | Morc3 |
| ENSMUSG( | 91.40183 | 0.170044 | 0.418727 | 0.821966 | Morc4 |
| ENSMUSG( | 6956.936 | 0.062053 | 0.463111 | 0.842422 | Morf4l1 |
| ENSMUSG( | 3666.927 | 0.018858 | 0.864827 | 0.975565 | Morf4l2 |
| ENSMUSG( | 165.4904 | -0.17245 | 0.318017 | 0.764184 | Morn1 |
| ENSMUSG( | 424.9879 | 0.064809 | 0.615956 | 0.910822 | Morn2 |
| ENSMUSG( | 3217.668 | 0.071134 | 0.395102 | 0.810183 | Morn4 |
| ENSMUSG( | 21.56616 | -0.04455 | 0.91677 | 0.988856 | Morn5 |
| ENSMUSG( | 14.40772 | -0.31309 | 0.533172 | NA | Morrbid |
| ENSMUSG( | 1088.381 | -0.20977 | 0.08995 | 0.460542 | Mosmo |
| ENSMUSG( | 292.1307 | -0.01339 | 0.925523 | 0.991204 | Mospd1 |
| ENSMUSG( | 565.1245 | 0.153075 | 0.240167 | 0.687899 | Mospd2 |
| ENSMUSG( | 700.2832 | 0.041303 | 0.730458 | 0.944956 | Mospd3 |
| ENSMUSG( | 553.9411 | 0.260611 | 0.122607 | 0.528484 | Mov10 |
| ENSMUSG( | 259.2248 | -0.33701 | 0.062197 | 0.386083 | Moxd1 |
| ENSMUSG( | 2730.619 | 0.11642 | 0.247034 | 0.69576 | Mpc1 |
| ENSMUSG( | 1862.697 | -0.12356 | 0.178834 | 0.616298 | Mpc2 |
| ENSMUSG( | 1849.531 | -0.2311 | 0.022435 | 0.228899 | Mpdu1 |
| ENSMUSG( | 2253.806 | 0.090825 | 0.462812 | 0.842421 | Mpdz |
| ENSMUSG( | 2490.218 | -0.21083 | 0.104354 | 0.490474 | Mpeg1 |
| ENSMUSG( | 488.5874 | -0.16574 | 0.187813 | 0.626399 | Mpg |
| ENSMUSG( | 407.466 | 0.140315 | 0.255384 | 0.706181 | Mphosph10 |
| ENSMUSG( | 297.6943 | 0.239339 | 0.093815 | 0.470818 | Mphosph6 |
| ENSMUSG( | 1708.7 | 0.35723 | 0.031643 | 0.271198 | Mphosph8 |
| ENSMUSG( | 1072.785 | -0.03228 | 0.747917 | 0.949604 | Mphosph9 |
| ENSMUSG( | 1127.101 | 0.080613 | 0.452649 | 0.836145 | Mpi |
| ENSMUSG( | 83.21737 | 0.277589 | 0.197648 | 0.637377 | Mpl |
| ENSMUSG( | 249.2146 | -0.10072 | 0.489602 | 0.859267 | Mplkip |
| ENSMUSG( | 3797.776 | 0.11571 | 0.142968 | 0.561716 | Mpnd |
| ENSMUSG( | 523.7403 | -0.01966 | 0.879752 | 0.981436 | Mpp1 |
| ENSMUSG( | 5085.602 | 0.169062 | 0.09935 | 0.48179 | Mpp2 |
| ENSMUSG( | 1097.02 | 0.021149 | 0.865376 | 0.975565 | Mpp3 |
| ENSMUSG( | 33.33365 | 0.636912 | 0.086455 | 0.452045 | Mpp4 |
| ENSMUSG( | 132.3319 | -0.04537 | 0.795928 | 0.960062 | Mpp7 |
| ENSMUSG( | 548.7438 | -0.25883 | 0.039746 | 0.307429 | Mppe1 |
| ENSMUSG( | 8607.946 | 0.236148 | 0.007131 | 0.116294 | Mpped1 |
| ENSMUSG( | 587.6092 | -0.24283 | 0.083388 | 0.445503 | Mpped2 |
| ENSMUSG( | 12952.08 | 0.049573 | 0.521932 | 0.875186 | Mprip |
| ENSMUSG( | 1502.189 | 0.031499 | 0.727257 | 0.944223 | Mpst |
| ENSMUSG( | 1376.674 | 0.081617 | 0.389155 | 0.806967 | Mpv17 |
| ENSMUSG( | 2209.892 | -0.01114 | 0.914402 | 0.988483 | Mpv17l |
| ENSMUSG( | 1220.295 | -0.15353 | 0.212704 | 0.6567 | Mpv17l2 |
| ENSMUSG( | 1423.073 | -0.34586 | 0.02577 | 0.247138 | Mpzl1 |
| ENSMUSG( | 112.3363 | -0.26067 | 0.198274 | 0.638387 | Mpzl2 |
| ENSMUSG( | 128.5136 | -0.32373 | 0.128221 | 0.536641 | Mpzl3 |
| ENSMUSG( | 424.7537 | -0.17509 | 0.195628 | 0.635109 | Mr1 |
| ENSMUSG( | 17.36912 | -0.27689 | 0.556456 | NA | Mrap |
| ENSMUSG( | 127.0655 | -0.34616 | 0.074464 | 0.422899 | Mrap2 |
| ENSMUSG( | 15621.46 | 0.090274 | 0.24964 | 0.699258 | Mras |
| ENSMUSG( | 365.0113 | 0.022818 | 0.895455 | 0.984512 | Mrc1 |
| ENSMUSG( | 675.9576 | -0.36465 | 0.002448 | 0.058231 | Mrc2 |
| ENSMUSG( | 355.854 | -0.07342 | 0.551 | 0.886038 | Mre11a |
| ENSMUSG( | 84.17341 | -0.00969 | 0.969966 | 0.996369 | Mreg |
| ENSMUSG( | 7417.919 | 0.000149 | 0.998867 | 0.99989 | Mrfap1 |
| ENSMUSG( | 517.7027 | -0.04895 | 0.701836 | 0.937441 | Mrgbp |
| ENSMUSG( | 274.4234 | -0.26152 | 0.170848 | 0.605452 | Mrgpre |
| ENSMUSG( | 72.86058 | -0.27108 | 0.32039 | 0.765921 | Mrgprf |
| ENSMUSG( | 365.1936 | -0.08918 | 0.556395 | 0.887681 | Mri1 |
| ENSMUSG( | 16.36204 | -0.14402 | 0.761811 | NA | Mrip-ps |
| ENSMUSG( | 8.861763 | 0.345574 | 0.608818 | NA | Mrln |
| ENSMUSG( | 335.9248 | -0.06506 | 0.631928 | 0.914073 | Mrm1 |
| ENSMUSG( | 187.6014 | 0.226213 | 0.147123 | 0.569203 | Mrm2 |
| ENSMUSG( | 184.6364 | -0.03715 | 0.817374 | 0.96575 | Mrm3 |
| ENSMUSG( | 186.4652 | 0.017873 | 0.923896 | 0.991046 | Mrnip |
| ENSMUSG( | 1070.265 | 0.042549 | 0.727105 | 0.944223 | Mro |
| ENSMUSG( | 2354.777 | 0.311975 | 0.004239 | 0.084545 | Mroh1 |
| ENSMUSG( | 29.70113 | 0.132833 | 0.722136 | 0.943535 | Mroh3 |
| ENSMUSG( | 65.8412 | -0.13264 | 0.650594 | 0.921301 | Mroh5 |
| ENSMUSG( | 290.7128 | 1.490885 | 9.42E-12 | 6.94E-09 | Mroh7 |
| ENSMUSG( | 28.97322 | 0.321894 | 0.361408 | 0.792797 | Mroh8 |
| ENSMUSG( | 378.9094 | 0.064205 | 0.667259 | 0.92729 | Mrpl1 |
| ENSMUSG( | 1415.811 | 0.044479 | 0.636407 | 0.914873 | Mrpl10 |
| ENSMUSG( | 907.5943 | 0.020874 | 0.838939 | 0.971179 | Mrpl11 |
| ENSMUSG( | 1537.82 | 0.01507 | 0.893126 | 0.984512 | Mrpl12 |
| ENSMUSG( | 944.2562 | 0.048991 | 0.653765 | 0.922914 | Mrpl13 |
| ENSMUSG( | 353.4705 | -0.00765 | 0.951338 | 0.993821 | Mrpl14 |
| ENSMUSG( | 623.5718 | 0.029632 | 0.821815 | 0.96635 | Mrpl15 |
| ENSMUSG( | 470.9251 | 0.051282 | 0.684985 | 0.932981 | Mrpl16 |
| ENSMUSG( | 1037.812 | 0.014749 | 0.878117 | 0.981431 | Mrpl17 |
| ENSMUSG( | 876.0201 | 0.11356 | 0.3277 | 0.772692 | Mrpl18 |
| ENSMUSG( | 552.5323 | 0.115161 | 0.297689 | 0.746178 | Mrpl19 |
| ENSMUSG( | 745.304 | -0.01065 | 0.92215 | 0.99045 | Mrpl2 |
| ENSMUSG( | 1251.905 | -0.02776 | 0.788443 | 0.959226 | Mrpl20 |
| ENSMUSG( | 518.0428 | 0.134415 | 0.264815 | 0.715848 | Mrpl21 |
| ENSMUSG( | 298.7707 | -0.06784 | 0.616578 | 0.910866 | Mrpl22 |
| ENSMUSG( | 552.5324 | 0.013878 | 0.90744 | 0.987566 | Mrpl23 |
| ENSMUSG( | 16.34573 | -0.10353 | 0.824655 | NA | Mrpl23-ps1 |
| ENSMUSG( | 1045.549 | -0.01192 | 0.899448 | 0.985633 | Mrpl24 |
| ENSMUSG( | 891.0852 | 0.082678 | 0.419289 | 0.821987 | Mrpl27 |
| ENSMUSG( | 1187.979 | 0.011972 | 0.897285 | 0.985507 | Mrpl28 |
| ENSMUSG( | 1366.511 | 0.08292 | 0.387374 | 0.805642 | Mrpl3 |
| ENSMUSG( | 1473.449 | 0.132523 | 0.188976 | 0.627678 | Mrpl30 |
| ENSMUSG( | 406.6613 | -0.14265 | 0.241862 | 0.689179 | Mrpl32 |
| ENSMUSG( | 664.5939 | -0.03654 | 0.746739 | 0.949289 | Mrpl33 |
| ENSMUSG( | 533.8229 | 0.09491 | 0.463588 | 0.842812 | Mrpl34 |

| | | | | | |
|---|---|---|---|---|---|
| ENSMUSG( | 62.29585 | 0.085512 | 0.74596 | 0.96951 | Mocos |
| ENSMUSG( | 277.6407 | 0.024662 | 0.876834 | 0.989644 | Mocs1 |
| ENSMUSG( | 1243.021 | -0.09466 | 0.428986 | 0.917257 | Mocs2 |
| ENSMUSG( | 209.7175 | 0.084563 | 0.670935 | 0.95694 | Mocs3 |
| ENSMUSG( | 6347.675 | 0.045165 | 0.798353 | 0.97681 | Mog |
| ENSMUSG( | 906.6271 | 0.218058 | 0.22309 | 0.867404 | Mogs |
| ENSMUSG( | 220.9592 | 0.031239 | 0.857908 | 0.985884 | Mok |
| ENSMUSG( | 634.9896 | -0.01668 | 0.891999 | 0.990558 | Mon1a |
| ENSMUSG( | 1122.864 | -0.03571 | 0.755528 | 0.971275 | Mon1b |
| ENSMUSG( | 2525.37 | 0.154771 | 0.156534 | 0.837724 | Mon2 |
| ENSMUSG( | 1289.214 | 0.021501 | 0.856265 | 0.985884 | Morc2a |
| ENSMUSG( | 49.04605 | -0.59078 | 0.069243 | 0.755467 | Morc2b |
| ENSMUSG( | 565.4316 | 0.190192 | 0.171266 | 0.842999 | Morc3 |
| ENSMUSG( | 81.588 | 0.125737 | 0.603857 | 0.948137 | Morc4 |
| ENSMUSG( | 5897.975 | -0.15008 | 0.177536 | 0.847209 | Morf4l1 |
| ENSMUSG( | 3217.811 | -0.09504 | 0.523067 | 0.932848 | Morf4l2 |
| ENSMUSG( | 159.8143 | 0.011713 | 0.951622 | 0.994798 | Morn1 |
| ENSMUSG( | 340.3688 | -0.33579 | 0.03315 | 0.64093 | Morn2 |
| ENSMUSG( | 7.864821 | 0.340435 | 0.6352 | NA | Morn3 |
| ENSMUSG( | 2711.674 | -0.15924 | 0.1795 | 0.850372 | Morn4 |
| ENSMUSG( | 18.39698 | -0.2695 | 0.555399 | 0.939565 | Morn5 |
| ENSMUSG( | 14.60064 | 0.06827 | 0.904193 | 0.991089 | Morrbid |
| ENSMUSG( | 1161.731 | 0.245268 | 0.096621 | 0.796245 | Mosmo |
| ENSMUSG( | 266.8911 | -0.00672 | 0.968199 | 0.995887 | Mospd1 |
| ENSMUSG( | 487.9968 | 0.001264 | 0.993147 | 0.998813 | Mospd2 |
| ENSMUSG( | 598.8908 | -0.15053 | 0.290413 | 0.897019 | Mospd3 |
| ENSMUSG( | 462.7052 | 0.017403 | 0.932268 | 0.99306 | Mov10 |
| ENSMUSG( | 263.6572 | 0.001732 | 0.992615 | 0.99874 | Moxd1 |
| ENSMUSG( | 2296.764 | -0.11766 | 0.348554 | 0.911652 | Mpc1 |
| ENSMUSG( | 1628.252 | -0.25541 | 0.039283 | 0.679972 | Mpc2 |
| ENSMUSG( | 1817.055 | -0.00311 | 0.978659 | 0.996459 | Mpdu1 |
| ENSMUSG( | 2004.915 | 0.020708 | 0.854249 | 0.98563 | Mpdz |
| ENSMUSG( | 2449.667 | 0.021694 | 0.894245 | 0.990558 | Mpeg1 |
| ENSMUSG( | 436.531 | -0.21678 | 0.153575 | 0.837724 | Mpg |
| ENSMUSG( | 367.3455 | 0.104891 | 0.48047 | 0.924548 | Mphosph10 |
| ENSMUSG( | 267.608 | 0.202927 | 0.208696 | 0.864412 | Mphosph6 |
| ENSMUSG( | 1497.775 | 0.256384 | 0.074265 | 0.766789 | Mphosph8 |
| ENSMUSG( | 990.9122 | 0.003253 | 0.980114 | 0.996479 | Mphosph9 |
| ENSMUSG( | 979.7786 | -0.05399 | 0.650258 | 0.95422 | Mpi |
| ENSMUSG( | 66.95325 | -0.05529 | 0.836396 | 0.981335 | Mpl |
| ENSMUSG( | 234.4313 | -0.01141 | 0.945517 | 0.994798 | Mplkip |
| ENSMUSG( | 3330.677 | 0.006324 | 0.953392 | 0.994798 | Mpnd |
| ENSMUSG( | 502.2615 | 0.125093 | 0.38866 | 0.917257 | Mpp1 |
| ENSMUSG( | 4445.341 | 0.052532 | 0.659842 | 0.956205 | Mpp2 |
| ENSMUSG( | 948.3136 | -0.13439 | 0.324046 | 0.905257 | Mpp3 |
| ENSMUSG( | 23.93874 | -0.00928 | 0.986258 | 0.997188 | Mpp4 |
| ENSMUSG( | 118.3062 | -0.09324 | 0.654886 | 0.955015 | Mpp7 |
| ENSMUSG( | 565.1604 | 0.111008 | 0.453726 | 0.922316 | Mppe1 |
| ENSMUSG( | 7297.572 | 0.038786 | 0.692268 | 0.960404 | Mpped1 |
| ENSMUSG( | 528.5015 | -0.29003 | 0.078249 | 0.77321 | Mpped2 |
| ENSMUSG( | 11655.4 | 0.015078 | 0.889065 | 0.990558 | Mprip |
| ENSMUSG( | 1263.46 | -0.21176 | 0.125681 | 0.81516 | Mpst |
| ENSMUSG( | 1168.336 | -0.12788 | 0.293624 | 0.898538 | Mpv17 |
| ENSMUSG( | 1993.711 | -0.04394 | 0.776975 | 0.975717 | Mpv17l |
| ENSMUSG( | 1133.961 | -0.09523 | 0.501273 | 0.927825 | Mpv17l2 |
| ENSMUSG( | 1489.539 | 0.080812 | 0.662711 | 0.956205 | Mpzl1 |
| ENSMUSG( | 123.4494 | 0.286137 | 0.159074 | 0.837724 | Mpzl2 |
| ENSMUSG( | 125.2933 | -0.10522 | 0.639557 | 0.953903 | Mpzl3 |
| ENSMUSG( | 394.3518 | -0.12135 | 0.410796 | 0.917257 | Mr1 |
| ENSMUSG( | 19.14346 | 0.256772 | 0.574404 | 0.942912 | Mrap |
| ENSMUSG( | 138.304 | 0.181946 | 0.365755 | 0.914623 | Mrap2 |
| ENSMUSG( | 13201.32 | -0.12797 | 0.197119 | 0.862161 | Mras |
| ENSMUSG( | 371.4406 | 0.326193 | 0.067284 | 0.75212 | Mrc1 |
| ENSMUSG( | 734.9807 | 0.170314 | 0.272142 | 0.888724 | Mrc2 |
| ENSMUSG( | 325.7754 | -0.06158 | 0.670807 | 0.95694 | Mre11a |
| ENSMUSG( | 77.0437 | -0.02083 | 0.944764 | 0.994798 | Mreg |
| ENSMUSG( | 6560.059 | -0.09051 | 0.51015 | 0.929089 | Mrfap1 |
| ENSMUSG( | 498.5316 | 0.115371 | 0.41268 | 0.917257 | Mrgbp |
| ENSMUSG( | 307.9902 | 0.345489 | 0.117161 | 0.813384 | Mrgpre |
| ENSMUSG( | 83.00162 | 0.388915 | 0.185439 | 0.855305 | Mrgprf |
| ENSMUSG( | 345.5501 | 0.028927 | 0.854952 | 0.985649 | Mri1 |
| ENSMUSG( | 15.02102 | -0.09138 | 0.856535 | 0.985884 | Mrip-ps |
| ENSMUSG( | 291.9118 | -0.21068 | 0.182983 | 0.850741 | Mrm1 |
| ENSMUSG( | 150.856 | -0.13576 | 0.471888 | 0.923329 | Mrm2 |
| ENSMUSG( | 145.5508 | -0.51054 | 0.010273 | 0.412737 | Mrm3 |
| ENSMUSG( | 167.5462 | -0.02621 | 0.901859 | 0.990969 | Mrnip |
| ENSMUSG( | 918.2636 | -0.13919 | 0.353525 | 0.912263 | Mro |
| ENSMUSG( | 1936.651 | 0.034366 | 0.802044 | 0.97681 | Mroh1 |
| ENSMUSG( | 26.96872 | 0.136734 | 0.73071 | 0.967228 | Mroh3 |
| ENSMUSG( | 68.20168 | 0.220759 | 0.393681 | 0.917257 | Mroh5 |
| ENSMUSG( | 152.5194 | 0.257632 | 0.37919 | 0.916995 | Mroh7 |
| ENSMUSG( | 24.61303 | 0.155882 | 0.696921 | 0.960867 | Mroh8 |
| ENSMUSG( | 325.9468 | -0.11242 | 0.529004 | 0.934795 | Mrpl1 |
| ENSMUSG( | 1230.34 | -0.09551 | 0.411057 | 0.917257 | Mrpl10 |
| ENSMUSG( | 778.7274 | -0.16225 | 0.243713 | 0.877904 | Mrpl11 |
| ENSMUSG( | 1344.748 | -0.1084 | 0.416911 | 0.917257 | Mrpl12 |
| ENSMUSG( | 798.6231 | -0.17657 | 0.216216 | 0.865167 | Mrpl13 |
| ENSMUSG( | 292.4345 | -0.30937 | 0.055105 | 0.723362 | Mrpl14 |
| ENSMUSG( | 536.412 | -0.14406 | 0.333552 | 0.907907 | Mrpl15 |
| ENSMUSG( | 393.3296 | -0.21089 | 0.162791 | 0.837724 | Mrpl16 |
| ENSMUSG( | 891.7316 | -0.16381 | 0.207689 | 0.864412 | Mrpl17 |
| ENSMUSG( | 783.1239 | 0.05864 | 0.650067 | 0.95422 | Mrpl18 |
| ENSMUSG( | 490.422 | 0.037807 | 0.786016 | 0.976285 | Mrpl19 |
| ENSMUSG( | 657.6124 | -0.10965 | 0.432861 | 0.918434 | Mrpl2 |
| ENSMUSG( | 1091.046 | -0.16245 | 0.191716 | 0.858764 | Mrpl20 |
| ENSMUSG( | 431.0495 | -0.13206 | 0.385439 | 0.917257 | Mrpl21 |
| ENSMUSG( | 255.3369 | -0.2718 | 0.094941 | 0.796133 | Mrpl22 |
| ENSMUSG( | 471.803 | -0.18378 | 0.177256 | 0.846973 | Mrpl23 |
| ENSMUSG( | 11.98033 | -0.82242 | 0.159783 | 0.837724 | Mrpl23-ps1 |
| ENSMUSG( | 923.5093 | -0.10255 | 0.361421 | 0.914223 | Mrpl24 |
| ENSMUSG( | 747.5829 | -0.1639 | 0.21414 | 0.864421 | Mrpl27 |
| ENSMUSG( | 1012.062 | -0.19025 | 0.098384 | 0.798438 | Mrpl28 |
| ENSMUSG( | 1170.837 | -0.09466 | 0.419004 | 0.917257 | Mrpl3 |
| ENSMUSG( | 1233.008 | -0.11595 | 0.368591 | 0.915491 | Mrpl30 |
| ENSMUSG( | 386.7632 | -0.0174 | 0.902935 | 0.991089 | Mrpl32 |
| ENSMUSG( | 570.5444 | -0.21643 | 0.087193 | 0.783264 | Mrpl33 |
| ENSMUSG( | 452.887 | -0.1155 | 0.453776 | 0.922316 | Mrpl34 |
| ENSMUSG( | 511.6098 | -0.17119 | 0.30553 | 0.898538 | Mrpl35 |
| ENSMUSG( | 449.4376 | -0.17735 | 0.214413 | 0.864421 | Mrpl36 |
| ENSMUSG( | 939.9739 | -0.02777 | 0.82947 | 0.980224 | Mrpl37 |
| ENSMUSG( | 780.9707 | -0.09844 | 0.486622 | 0.92541 | Mrpl38 |
| ENSMUSG( | 503.9416 | -0.07571 | 0.585416 | 0.944975 | Mrpl39 |
| ENSMUSG( | 1138.175 | -0.09392 | 0.425128 | 0.917257 | Mrpl4 |
| ENSMUSG( | 488.689 | -0.14314 | 0.361004 | 0.914223 | Mrpl40 |
| ENSMUSG( | 845.3467 | -0.05017 | 0.720966 | 0.966887 | Mrpl41 |
| ENSMUSG( | 1101.266 | -0.10149 | 0.441123 | 0.921166 | Mrpl42 |
| ENSMUSG( | 850.946 | -0.13035 | 0.329623 | 0.907083 | Mrpl43 |
| ENSMUSG( | 337.6532 | -0.03013 | 0.871823 | 0.988779 | Mrpl44 |
| ENSMUSG( | 734.178 | -0.09699 | 0.516868 | 0.930939 | Mrpl45 |
| ENSMUSG( | 350.3035 | -0.14611 | 0.343128 | 0.909646 | Mrpl46 |
| ENSMUSG( | 354.6647 | -0.13095 | 0.436675 | 0.919879 | Mrpl47 |
| ENSMUSG( | 2421.341 | -0.1067 | 0.325468 | 0.905339 | Mrpl48 |
| ENSMUSG( | 859.1954 | -0.14924 | 0.242209 | 0.876967 | Mrpl49 |
| ENSMUSG( | 778.176 | -0.14741 | 0.287201 | 0.894091 | Mrpl50 |
| ENSMUSG( | 1081.112 | -0.0873 | 0.509354 | 0.929089 | Mrpl51 |
| ENSMUSG( | 746.9255 | -0.19048 | 0.133113 | 0.822912 | Mrpl52 |
| ENSMUSG( | 609.6338 | -0.31607 | 0.04516 | 0.693959 | Mrpl54 |
| ENSMUSG( | 532.3476 | -0.21654 | 0.099689 | 0.801103 | Mrpl55 |
| ENSMUSG( | 595.873 | -0.26065 | 0.088812 | 0.788051 | Mrpl57 |
| ENSMUSG( | 796.5793 | -0.30278 | 0.039906 | 0.681706 | Mrpl58 |
| ENSMUSG( | 1084.682 | -0.22198 | 0.08267 | 0.77681 | Mrpl9 |
| ENSMUSG( | 568.2141 | -0.04168 | 0.745964 | 0.96951 | Mrps10 |
| ENSMUSG( | 397.1692 | -0.19218 | 0.167599 | 0.840222 | Mrps11 |
| ENSMUSG( | 715.8228 | -0.08058 | 0.571208 | 0.942052 | Mrps12 |
| ENSMUSG( | 603.6329 | -0.18289 | 0.220475 | 0.866104 | Mrps14 |
| ENSMUSG( | 706.8143 | -0.21894 | 0.084394 | 0.778891 | Mrps15 |
| ENSMUSG( | 488.2777 | -0.12663 | 0.423488 | 0.917257 | Mrps16 |
| ENSMUSG( | 1314.624 | -0.01868 | 0.865473 | 0.987011 | Mrps17 |
| ENSMUSG( | 1036.821 | -0.09114 | 0.432363 | 0.918158 | Mrps18a |
| ENSMUSG( | 280.2507 | -0.08878 | 0.600877 | 0.947749 | Mrps18b |
| ENSMUSG( | 276.2456 | -0.20673 | 0.210347 | 0.864412 | Mrps18c |
| ENSMUSG( | 664.7375 | -0.07591 | 0.601588 | 0.947749 | Mrps2 |
| ENSMUSG( | 954.7544 | -0.19433 | 0.121188 | 0.813384 | Mrps21 |
| ENSMUSG( | 263.8723 | -0.11986 | 0.451709 | 0.922202 | Mrps22 |
| ENSMUSG( | 727.8678 | -0.13169 | 0.34894 | 0.911652 | Mrps23 |
| ENSMUSG( | 885.809 | -0.23892 | 0.047027 | 0.700664 | Mrps24 |
| ENSMUSG( | 883.3914 | -0.23769 | 0.058471 | 0.728256 | Mrps25 |
| ENSMUSG( | 1229.261 | 0.023254 | 0.857108 | 0.985884 | Mrps26 |
| ENSMUSG( | 624.4699 | -0.20325 | 0.180636 | 0.850372 | Mrps27 |
| ENSMUSG( | 224.6374 | -0.00658 | 0.970588 | 0.99594 | Mrps28 |
| ENSMUSG( | 470.2282 | 0.048514 | 0.728599 | 0.966913 | Mrps30 |
| ENSMUSG( | 277.2638 | -0.0354 | 0.823995 | 0.979198 | Mrps31 |
| ENSMUSG( | 586.88 | -0.09654 | 0.494543 | 0.927372 | Mrps33 |
| ENSMUSG( | 858.265 | -0.14441 | 0.299706 | 0.898538 | Mrps34 |
| ENSMUSG( | 504.4759 | -0.05344 | 0.692181 | 0.960404 | Mrps35 |
| ENSMUSG( | 381.2079 | -0.15351 | 0.33084 | 0.907513 | Mrps36 |
| ENSMUSG( | 1004.668 | -0.01478 | 0.89653 | 0.990558 | Mrps5 |
| ENSMUSG( | 297.7377 | -0.00327 | 0.986135 | 0.997188 | Mrps6 |

| | | | | | | | | | | | |
|---|---|---|---|---|---|---|---|---|---|---|---|
| ENSMUSG( | Mrip-ps | 16.36204 | -0.14402 | 0.761811 | NA | ENSMUSG( | Mrip-ps | 15.02102 | -0.09138 | 0.856535 | 0.985884 |
| ENSMUSG( | Mrm1 | 335.9248 | -0.06506 | 0.631928 | 0.914073 | ENSMUSG( | Mrm1 | 291.9118 | -0.21068 | 0.182983 | 0.850741 |
| ENSMUSG( | Mrm2 | 187.6014 | 0.226213 | 0.147123 | 0.569203 | ENSMUSG( | Mrm2 | 150.856 | -0.13576 | 0.471888 | 0.923329 |
| ENSMUSG( | Mrm3 | 184.6364 | -0.03715 | 0.817374 | 0.96575 | ENSMUSG( | Mrm3 | 145.5508 | -0.51054 | 0.010273 | 0.412737 |
| ENSMUSG( | Mrnip | 186.4652 | 0.017873 | 0.923896 | 0.991046 | ENSMUSG( | Mrnip | 167.5462 | -0.02621 | 0.901859 | 0.990969 |
| ENSMUSG( | Mro | 1070.265 | 0.042549 | 0.727105 | 0.944223 | ENSMUSG( | Mro | 918.2636 | -0.13919 | 0.353525 | 0.912263 |
| ENSMUSG( | Mroh1 | 2354.777 | 0.311975 | 0.004239 | 0.084545 | ENSMUSG( | Mroh1 | 1936.651 | 0.034366 | 0.802044 | 0.97681 |
| ENSMUSG( | Mroh3 | 29.70113 | 0.132833 | 0.722136 | 0.943535 | ENSMUSG( | Mroh3 | 26.96872 | 0.136734 | 0.73071 | 0.967228 |
| ENSMUSG( | Mroh5 | 65.8412 | -0.13264 | 0.650594 | 0.921301 | ENSMUSG( | Mroh5 | 68.20168 | 0.220759 | 0.393681 | 0.917257 |
| ENSMUSG( | Mroh7 | 290.7128 | 1.490885 | 9.42E-12 | 6.94E-09 | ENSMUSG( | Mroh7 | 152.5194 | 0.257632 | 0.37919 | 0.916995 |
| ENSMUSG( | Mroh8 | 28.97322 | 0.321894 | 0.361408 | 0.792797 | ENSMUSG( | Mroh8 | 24.61303 | 0.155882 | 0.696921 | 0.960867 |
| ENSMUSG( | Mrpl1 | 378.9094 | 0.064205 | 0.667259 | 0.92729 | ENSMUSG( | Mrpl1 | 325.9468 | -0.11242 | 0.529004 | 0.934795 |
| ENSMUSG( | Mrpl10 | 1415.811 | 0.044479 | 0.636407 | 0.914873 | ENSMUSG( | Mrpl10 | 1230.34 | -0.09551 | 0.411057 | 0.917257 |
| ENSMUSG( | Mrpl11 | 907.5943 | 0.020874 | 0.838939 | 0.971179 | ENSMUSG( | Mrpl11 | 778.7274 | -0.16225 | 0.243713 | 0.877904 |
| ENSMUSG( | Mrpl12 | 1537.82 | 0.01507 | 0.893126 | 0.984512 | ENSMUSG( | Mrpl12 | 1344.748 | -0.1084 | 0.416911 | 0.917257 |
| ENSMUSG( | Mrpl13 | 944.2562 | 0.048991 | 0.653765 | 0.922914 | ENSMUSG( | Mrpl13 | 798.6231 | -0.17657 | 0.216216 | 0.865167 |
| ENSMUSG( | Mrpl14 | 353.4705 | -0.00765 | 0.951338 | 0.993821 | ENSMUSG( | Mrpl14 | 292.4345 | -0.30937 | 0.055105 | 0.723362 |
| ENSMUSG( | Mrpl15 | 623.5718 | 0.029632 | 0.821815 | 0.96635 | ENSMUSG( | Mrpl15 | 536.412 | -0.14406 | 0.333552 | 0.907907 |
| ENSMUSG( | Mrpl16 | 470.9251 | 0.051282 | 0.684985 | 0.932981 | ENSMUSG( | Mrpl16 | 393.3296 | -0.21089 | 0.162791 | 0.837724 |
| ENSMUSG( | Mrpl17 | 1037.812 | 0.014749 | 0.878117 | 0.981431 | ENSMUSG( | Mrpl17 | 891.7316 | -0.16381 | 0.207689 | 0.864412 |
| ENSMUSG( | Mrpl18 | 876.0201 | 0.11356 | 0.3277 | 0.772692 | ENSMUSG( | Mrpl18 | 783.1239 | 0.05864 | 0.650067 | 0.95422 |
| ENSMUSG( | Mrpl19 | 552.5323 | 0.115161 | 0.297689 | 0.746178 | ENSMUSG( | Mrpl19 | 490.422 | 0.037807 | 0.786016 | 0.976285 |
| ENSMUSG( | Mrpl2 | 745.304 | -0.01065 | 0.92215 | 0.99045 | ENSMUSG( | Mrpl2 | 657.6124 | -0.10965 | 0.432861 | 0.918434 |
| ENSMUSG( | Mrpl20 | 1251.905 | -0.02776 | 0.788443 | 0.959226 | ENSMUSG( | Mrpl20 | 1091.046 | -0.16245 | 0.191716 | 0.858764 |
| ENSMUSG( | Mrpl21 | 518.0428 | 0.134415 | 0.264815 | 0.715848 | ENSMUSG( | Mrpl21 | 431.0495 | -0.13206 | 0.385439 | 0.917257 |
| ENSMUSG( | Mrpl22 | 298.7707 | -0.06784 | 0.616578 | 0.910866 | ENSMUSG( | Mrpl22 | 255.3369 | -0.2718 | 0.094941 | 0.796133 |
| ENSMUSG( | Mrpl23 | 552.5324 | 0.013878 | 0.90744 | 0.987566 | ENSMUSG( | Mrpl23 | 471.803 | -0.18378 | 0.177256 | 0.846973 |
| ENSMUSG( | Mrpl23-ps1 | 16.34573 | -0.10353 | 0.824655 | NA | ENSMUSG( | Mrpl23-ps1 | 11.98033 | -0.82242 | 0.159783 | 0.837724 |
| ENSMUSG( | Mrpl24 | 1045.549 | -0.01192 | 0.899448 | 0.985633 | ENSMUSG( | Mrpl24 | 923.5093 | -0.10255 | 0.361421 | 0.914223 |
| ENSMUSG( | Mrpl27 | 891.0852 | 0.082678 | 0.419289 | 0.821987 | ENSMUSG( | Mrpl27 | 747.5829 | -0.1639 | 0.21414 | 0.864421 |
| ENSMUSG( | Mrpl28 | 1187.979 | 0.011972 | 0.897285 | 0.985507 | ENSMUSG( | Mrpl28 | 1012.062 | -0.19025 | 0.098384 | 0.798438 |
| ENSMUSG( | Mrpl3 | 1366.511 | 0.08292 | 0.387374 | 0.805642 | ENSMUSG( | Mrpl3 | 1170.837 | -0.09466 | 0.419004 | 0.917257 |
| ENSMUSG( | Mrpl30 | 1473.449 | 0.132523 | 0.188976 | 0.627678 | ENSMUSG( | Mrpl30 | 1233.008 | -0.11595 | 0.368591 | 0.915491 |
| ENSMUSG( | Mrpl32 | 406.6613 | -0.14265 | 0.241862 | 0.689179 | ENSMUSG( | Mrpl32 | 386.7632 | -0.0174 | 0.902935 | 0.991089 |
| ENSMUSG( | Mrpl33 | 664.5939 | -0.03654 | 0.746739 | 0.949289 | ENSMUSG( | Mrpl33 | 570.5444 | -0.21643 | 0.087193 | 0.783264 |
| ENSMUSG( | Mrpl34 | 533.8229 | 0.09491 | 0.463588 | 0.842812 | ENSMUSG( | Mrpl34 | 452.887 | -0.1155 | 0.453776 | 0.922316 |
| ENSMUSG( | Mrpl35 | 593.045 | 0.001607 | 0.989855 | 0.999144 | ENSMUSG( | Mrpl35 | 511.6098 | -0.17119 | 0.30553 | 0.898538 |
| ENSMUSG( | Mrpl36 | 523.1002 | 4.39E-05 | 0.99972 | 0.99989 | ENSMUSG( | Mrpl36 | 449.4376 | -0.17735 | 0.214413 | 0.864421 |
| ENSMUSG( | Mrpl37 | 1101.288 | 0.152434 | 0.150151 | 0.574666 | ENSMUSG( | Mrpl37 | 939.9739 | -0.02777 | 0.82947 | 0.980224 |
| ENSMUSG( | Mrpl38 | 889.6144 | 0.013745 | 0.912274 | 0.988483 | ENSMUSG( | Mrpl38 | 780.9707 | -0.09844 | 0.486622 | 0.92541 |
| ENSMUSG( | Mrpl39 | 592.3855 | 0.123974 | 0.309598 | 0.757658 | ENSMUSG( | Mrpl39 | 503.9416 | -0.07571 | 0.585416 | 0.944975 |
| ENSMUSG( | Mrpl4 | 1297.468 | 0.018453 | 0.845782 | 0.972191 | ENSMUSG( | Mrpl4 | 1138.175 | -0.09392 | 0.425128 | 0.917257 |
| ENSMUSG( | Mrpl40 | 543.5552 | -0.0938 | 0.429588 | 0.826446 | ENSMUSG( | Mrpl40 | 488.689 | -0.14314 | 0.361004 | 0.914223 |
| ENSMUSG( | Mrpl41 | 970.6717 | 0.082576 | 0.440894 | 0.832371 | ENSMUSG( | Mrpl41 | 845.3467 | -0.05017 | 0.720966 | 0.966887 |
| ENSMUSG( | Mrpl42 | 1283.086 | 0.075807 | 0.453996 | 0.836798 | ENSMUSG( | Mrpl42 | 1101.266 | -0.10149 | 0.441123 | 0.921166 |
| ENSMUSG( | Mrpl43 | 973.0806 | -0.0044 | 0.967792 | 0.996237 | ENSMUSG( | Mrpl43 | 850.946 | -0.13035 | 0.329623 | 0.907083 |
| ENSMUSG( | Mrpl44 | 391.1032 | 0.129385 | 0.432061 | 0.82768 | ENSMUSG( | Mrpl44 | 337.6532 | -0.03013 | 0.871823 | 0.988779 |
| ENSMUSG( | Mrpl45 | 858.1514 | 0.08985 | 0.460879 | 0.840979 | ENSMUSG( | Mrpl45 | 734.178 | -0.09699 | 0.516868 | 0.930939 |
| ENSMUSG( | Mrpl46 | 422.0494 | 0.13011 | 0.284442 | 0.732908 | ENSMUSG( | Mrpl46 | 350.3035 | -0.14611 | 0.343128 | 0.909646 |
| ENSMUSG( | Mrpl47 | 401.0847 | -0.03101 | 0.813103 | 0.964574 | ENSMUSG( | Mrpl47 | 354.6647 | -0.13095 | 0.436675 | 0.919879 |
| ENSMUSG( | Mrpl48 | 2883.031 | 0.12898 | 0.110055 | 0.500306 | ENSMUSG( | Mrpl48 | 2421.341 | -0.1067 | 0.325468 | 0.905339 |
| ENSMUSG( | Mrpl49 | 1011.225 | 0.05948 | 0.563525 | 0.88992 | ENSMUSG( | Mrpl49 | 859.1954 | -0.14924 | 0.242209 | 0.876967 |
| ENSMUSG( | Mrpl50 | 910.8897 | 0.047723 | 0.656263 | 0.923929 | ENSMUSG( | Mrpl50 | 778.176 | -0.14741 | 0.287201 | 0.894091 |
| ENSMUSG( | Mrpl51 | 1287.739 | 0.14944 | 0.114118 | 0.509818 | ENSMUSG( | Mrpl51 | 1081.112 | -0.0873 | 0.509354 | 0.929089 |
| ENSMUSG( | Mrpl52 | 884.0429 | 0.037007 | 0.741409 | 0.948208 | ENSMUSG( | Mrpl52 | 746.9255 | -0.19048 | 0.133113 | 0.822912 |
| ENSMUSG( | Mrpl54 | 705.4991 | -0.14198 | 0.215233 | 0.659815 | ENSMUSG( | Mrpl54 | 609.6338 | -0.31607 | 0.04516 | 0.693959 |
| ENSMUSG( | Mrpl55 | 617.6528 | -0.04986 | 0.669126 | 0.928101 | ENSMUSG( | Mrpl55 | 532.3476 | -0.21654 | 0.099689 | 0.801103 |
| ENSMUSG( | Mrpl57 | 683.3803 | -0.11757 | 0.309515 | 0.757658 | ENSMUSG( | Mrpl57 | 595.873 | -0.26065 | 0.088812 | 0.788051 |
| ENSMUSG( | Mrpl58 | 961.6932 | -0.00817 | 0.94464 | 0.993583 | ENSMUSG( | Mrpl58 | 796.5793 | -0.30278 | 0.039906 | 0.681706 |
| ENSMUSG( | Mrpl9 | 1297.512 | 0.038293 | 0.709319 | 0.939656 | ENSMUSG( | Mrpl9 | 1084.682 | -0.22198 | 0.08267 | 0.77681 |
| ENSMUSG( | Mrps10 | 624.1074 | -0.03705 | 0.723679 | 0.94403 | ENSMUSG( | Mrps10 | 568.2141 | -0.04168 | 0.745964 | 0.96951 |
| ENSMUSG( | Mrps11 | 474.0078 | 0.054714 | 0.703027 | 0.937735 | ENSMUSG( | Mrps11 | 397.1692 | -0.19218 | 0.167599 | 0.840222 |
| ENSMUSG( | Mrps12 | 841.9011 | 0.120135 | 0.339809 | 0.777939 | ENSMUSG( | Mrps12 | 715.8228 | -0.08058 | 0.571208 | 0.942052 |
| ENSMUSG( | Mrps14 | 719.4626 | 0.067533 | 0.592027 | 0.90183 | ENSMUSG( | Mrps14 | 603.6329 | -0.18289 | 0.220475 | 0.866104 |
| ENSMUSG( | Mrps15 | 872.9168 | 0.127878 | 0.249918 | 0.699512 | ENSMUSG( | Mrps15 | 706.8143 | -0.21894 | 0.084394 | 0.778891 |
| ENSMUSG( | Mrps16 | 558.4859 | 0.001673 | 0.988988 | 0.999111 | ENSMUSG( | Mrps16 | 488.2777 | -0.12663 | 0.423488 | 0.917257 |
| ENSMUSG( | Mrps17 | 1438.733 | -0.02538 | 0.76864 | 0.954674 | ENSMUSG( | Mrps17 | 1314.624 | -0.01868 | 0.865473 | 0.987011 |
| ENSMUSG( | Mrps18a | 1179.201 | 0.015715 | 0.861483 | 0.975026 | ENSMUSG( | Mrps18a | 1036.821 | -0.09114 | 0.432363 | 0.918158 |
| ENSMUSG( | Mrps18b | 316.4014 | -0.0172 | 0.9031 | 0.986055 | ENSMUSG( | Mrps18b | 280.2507 | -0.08878 | 0.600877 | 0.947749 |
| ENSMUSG( | Mrps18c | 316.0679 | -0.07218 | 0.579221 | 0.896692 | ENSMUSG( | Mrps18c | 276.2456 | -0.20673 | 0.210347 | 0.864412 |
| ENSMUSG( | Mrps2 | 774.8396 | 0.100678 | 0.408875 | 0.818173 | ENSMUSG( | Mrps2 | 664.7375 | -0.07591 | 0.601588 | 0.947749 |
| ENSMUSG( | Mrps21 | 1111.496 | -0.01355 | 0.893128 | 0.984512 | ENSMUSG( | Mrps21 | 954.7544 | -0.19433 | 0.121188 | 0.813384 |
| ENSMUSG( | Mrps22 | 312.5971 | 0.103585 | 0.457704 | 0.839765 | ENSMUSG( | Mrps22 | 263.8723 | -0.11986 | 0.451709 | 0.922202 |
| ENSMUSG( | Mrps23 | 841.9921 | 0.024815 | 0.841475 | 0.971179 | ENSMUSG( | Mrps23 | 727.8678 | -0.13169 | 0.34894 | 0.911652 |
| ENSMUSG( | Mrps24 | 1050.301 | -0.00376 | 0.969082 | 0.996369 | ENSMUSG( | Mrps24 | 885.809 | -0.23892 | 0.047027 | 0.700664 |
| ENSMUSG( | Mrps25 | 1067.989 | 0.053757 | 0.584537 | 0.898862 | ENSMUSG( | Mrps25 | 883.3914 | -0.23769 | 0.058471 | 0.728256 |
| ENSMUSG( | Mrps26 | 1324.375 | -0.02837 | 0.792384 | 0.959461 | ENSMUSG( | Mrps26 | 1229.261 | 0.023254 | 0.857108 | 0.985884 |
| ENSMUSG( | Mrps27 | 753.0088 | 0.076895 | 0.567462 | 0.891774 | ENSMUSG( | Mrps27 | 624.4699 | -0.20325 | 0.180636 | 0.850372 |
| ENSMUSG( | Mrps28 | 250.0318 | 0.043532 | 0.788114 | 0.959226 | ENSMUSG( | Mrps28 | 224.6374 | -0.00658 | 0.970588 | 0.99594 |
| ENSMUSG( | Mrps30 | 541.402 | 0.184528 | 0.119681 | 0.523665 | ENSMUSG( | Mrps30 | 470.2282 | 0.048514 | 0.728599 | 0.966913 |
| ENSMUSG( | Mrps31 | 294.5316 | -0.131 | 0.347454 | 0.78046 | ENSMUSG( | Mrps31 | 277.2638 | -0.0354 | 0.823995 | 0.979198 |
| ENSMUSG( | Mrps33 | 700.7226 | 0.150941 | 0.214995 | 0.659512 | ENSMUSG( | Mrps33 | 586.88 | -0.09654 | 0.494543 | 0.927372 |
| ENSMUSG( | Mrps34 | 992.3671 | 0.012553 | 0.912401 | 0.988483 | ENSMUSG( | Mrps34 | 858.265 | -0.14441 | 0.299706 | 0.898538 |
| ENSMUSG( | Mrps35 | 586.0436 | 0.1141 | 0.292431 | 0.742521 | ENSMUSG( | Mrps35 | 504.4759 | -0.05344 | 0.692181 | 0.960404 |
| ENSMUSG( | Mrps36 | 455.291 | 0.102066 | 0.405317 | 0.81652 | ENSMUSG( | Mrps36 | 381.2079 | -0.15351 | 0.33084 | 0.907513 |
| ENSMUSG( | Mrps5 | 1112.701 | 0.011424 | 0.908841 | 0.98794 | ENSMUSG( | Mrps5 | 1004.668 | -0.01478 | 0.89653 | 0.990558 |
| ENSMUSG( | Mrps6 | 349.4475 | 0.192982 | 0.216494 | 0.661421 | ENSMUSG( | Mrps6 | 297.7377 | -0.00327 | 0.986135 | 0.997188 |
| ENSMUSG( | Mrps7 | 1304.927 | 0.028304 | 0.765202 | 0.954127 | ENSMUSG( | Mrps7 | 1150.82 | -0.06754 | 0.553875 | 0.939565 |
| ENSMUSG( | Mrps9 | 752.5863 | -0.02446 | 0.848017 | 0.972374 | ENSMUSG( | Mrps9 | 700.1331 | 0.036006 | 0.796038 | 0.97681 |
| ENSMUSG( | Mrrf | 364.6609 | 0.12104 | 0.333066 | 0.775602 | ENSMUSG( | Mrrf | 315.6518 | -0.02966 | 0.852296 | 0.985598 |
| ENSMUSG( | Mrs2 | 3190.807 | 0.072782 | 0.412734 | 0.819518 | ENSMUSG( | Mrs2 | 2844.595 | 0.012006 | 0.915735 | 0.992529 |
| ENSMUSG( | Mrtfa | 1422.827 | 0.392671 | 0.011009 | 0.151333 | ENSMUSG( | Mrtfa | 1190.21 | 0.167717 | 0.286593 | 0.894091 |
| ENSMUSG( | Mrtfb | 4189.927 | 0.115586 | 0.264306 | 0.715323 | ENSMUSG( | Mrtfb | 3760.903 | 0.075356 | 0.521794 | 0.932571 |
| ENSMUSG( | Mrto4 | 693.2172 | 0.056256 | 0.594284 | 0.902869 | ENSMUSG( | Mrto4 | 594.3975 | -0.12645 | 0.341321 | 0.908511 |
| ENSMUSG( | Ms4a15 | 9.7428 | -1.70633 | 0.073372 | NA | ENSMUSG( | Ms4a15 | 19.62634 | 0.889473 | 0.515537 | 0.930939 |
| ENSMUSG( | Ms4a4a | 22.99388 | -0.0225 | 0.957294 | 0.993874 | ENSMUSG( | Ms4a4a | 18.40734 | -0.42394 | 0.442971 | 0.921554 |
| ENSMUSG( | Ms4a6b | 68.66952 | 0.044983 | 0.85859 | 0.974707 | ENSMUSG( | Ms4a6b | 76.49797 | 0.58417 | 0.042793 | 0.690304 |
| ENSMUSG( | Ms4a6c | 68.00212 | 0.058133 | 0.83396 | 0.970333 | ENSMUSG( | Ms4a6c | 62.17229 | 0.075848 | 0.801982 | 0.97681 |
| ENSMUSG( | Ms4a6d | 55.02253 | -0.19174 | 0.514317 | 0.872055 | ENSMUSG( | Ms4a6d | 59.58747 | 0.308642 | 0.314616 | 0.900952 |
| ENSMUSG( | Ms4a7 | 34.2597 | -0.2644 | 0.453889 | 0.836798 | ENSMUSG( | Ms4a7 | 30.4614 | -0.32123 | 0.401531 | 0.917257 |
| ENSMUSG( | Msantd1 | 70.03484 | -0.03617 | 0.875292 | 0.979998 | ENSMUSG( | Msantd1 | 64.97584 | 0.023996 | 0.931105 | 0.99306 |
| ENSMUSG( | Msantd2 | 446.2175 | 0.120648 | 0.381831 | 0.802968 | ENSMUSG( | Msantd2 | 401.479 | 0.086605 | 0.583637 | 0.944266 |
| ENSMUSG( | Msantd3 | 707.7996 | 0.230813 | 0.020842 | 0.218947 | ENSMUSG( | Msantd3 | 590.9207 | -0.01626 | 0.899123 | 0.990558 |
| ENSMUSG( | Msantd4 | 1435.59 | 0.091513 | 0.392594 | 0.809274 | ENSMUSG( | Msantd4 | 1287.194 | 0.041019 | 0.783582 | 0.976106 |
| ENSMUSG( | Msantd7 | 106.4475 | -0.36875 | 0.066037 | 0.397686 | ENSMUSG( | Msantd7 | 107.9063 | -0.03064 | 0.885723 | 0.98977 |
| ENSMUSG( | Msc | 28.758 | -0.2199 | 0.524634 | 0.875871 | ENSMUSG( | Msc | 26.4283 | -0.1609 | 0.683139 | 0.958678 |
| ENSMUSG( | Msh2 | 540.873 | 0.054509 | 0.651196 | 0.921451 | ENSMUSG( | Msh2 | 472.5644 | -0.06236 | 0.638836 | 0.953903 |
| ENSMUSG( | Msh3 | 533.2578 | 0.049691 | 0.688492 | 0.933441 | ENSMUSG( | Msh3 | 483.7828 | 0.038393 | 0.789049 | 0.976593 |
| ENSMUSG( | Msh5 | 17.35179 | -0.24757 | 0.587796 | NA | ENSMUSG( | Msh5 | 19.35978 | 0.3279 | 0.476789 | 0.923329 |
| ENSMUSG( | Msh6 | 441.1554 | 0.233441 | 0.053585 | 0.35565 | ENSMUSG( | Msh6 | 385.0107 | 0.113534 | 0.416139 | 0.917257 |
| ENSMUSG( | Msi1 | 2781.135 | -0.01525 | 0.865537 | 0.975676 | ENSMUSG( | Msi1 | 2398.973 | -0.18139 | 0.125298 | 0.81516 |
| ENSMUSG( | Msi2 | 6672.657 | 0.066279 | 0.541434 | 0.883182 | ENSMUSG( | Msi2 | 5657.973 | -0.14572 | 0.26711 | 0.886134 |
| ENSMUSG( | Msl1 | 8178.851 | 0.241754 | 0.005241 | 0.096494 | ENSMUSG( | Msl1 | 7282.894 | 0.179587 | 0.090544 | 0.791074 |
| ENSMUSG( | Msl2 | 1063.085 | -0.0065 | 0.94602 | 0.99369 | ENSMUSG( | Msl2 | 1036.513 | 0.181413 | 0.113583 | 0.809623 |
| ENSMUSG( | Msl3 | 569.5712 | 0.141785 | 0.230855 | 0.679066 | ENSMUSG( | Msl3 | 505.3697 | 0.06261 | 0.658413 | 0.956205 |
| ENSMUSG( | Msl3l2 | 283.4465 | 0.076821 | 0.596004 | 0.903152 | ENSMUSG( | Msl3l2 | 254.5669 | 0.033292 | 0.852585 | 0.985598 |
| ENSMUSG( | Msln | 90.21537 | 0.376895 | 0.075557 | 0.425975 | ENSMUSG( | Msln | 64.59813 | -0.31769 | 0.222574 | 0.866791 |
| ENSMUSG( | Msmo1 | 5326.89 | -0.50867 | 0.005392 | 0.098038 | ENSMUSG( | Msmo1 | 5493.418 | -0.10459 | 0.579804 | 0.944209 |
| ENSMUSG( | Msn | 2298.201 | -0.06955 | 0.595189 | 0.903007 | ENSMUSG( | Msn | 2103.686 | -0.05707 | 0.678354 | 0.957435 |
| ENSMUSG( | Msr1 | 19.92759 | 0.366917 | 0.390664 | NA | ENSMUSG( | Msr1 | 15.39941 | -0.10252 | 0.841055 | 0.9826 |
| ENSMUSG( | Msra | 860.698 | 0.107364 | 0.336956 | 0.777008 | ENSMUSG( | Msra | 721.7349 | -0.13904 | 0.368107 | 0.915491 |
| ENSMUSG( | Msrb1 | 597.085 | 0.052537 | 0.639742 | 0.916434 | ENSMUSG( | Msrb1 | 502.1268 | -0.18847 | 0.162874 | 0.837905 |
| ENSMUSG( | Msrb2 | 652.5005 | 0.050447 | 0.631326 | 0.914073 | ENSMUSG( | Msrb2 | 531.8487 | -0.28956 | 0.028789 | 0.614171 |
| ENSMUSG( | Msrb3 | 249.2846 | -0.11661 | 0.417521 | 0.821568 | ENSMUSG( | Msrb3 | 240.6363 | 0.063213 | 0.711879 | 0.964363 |
| ENSMUSG( | Mss51 | 16.34439 | -0.00692 | 0.987997 | NA | ENSMUSG( | Mss51 | 18.81774 | 0.614231 | 0.177394 | 0.847137 |
| ENSMUSG( | Mst1r | 51.44122 | 0.31195 | 0.276978 | 0.727074 | ENSMUSG( | Mst1r | 43.42274 | 0.128513 | 0.695623 | 0.960867 |
| ENSMUSG( | Msto1 | 635.3082 | -0.02668 | 0.812465 | 0.964574 | ENSMUSG( | Msto1 | 548.8028 | -0.18786 | 0.168359 | 0.840222 |
| ENSMUSG( | Msx1 | 183.356 | -0.2387 | 0.297561 | 0.746087 | ENSMUSG( | Msx1 | 197.0312 | 0.238016 | 0.294964 | 0.898538 |
| ENSMUSG( | Msx2 | 17.83357 | -0.02962 | 0.946192 | NA | ENSMUSG( | Msx2 | 18.43489 | 0.313663 | 0.499108 | 0.927825 |
| ENSMUSG( | Mt1 | 20177.6 | 0.121902 | 0.235423 | 0.682423 | ENSMUSG( | Mt1 | 16423.63 | -0.21077 | 0.104264 | 0.804969 |
| ENSMUSG( | Mt2 | 9541.749 | 0.091897 | 0.366136 | 0.794603 | ENSMUSG( | Mt2 | 7838.676 | -0.21551 | 0.071946 | 0.763629 |
| ENSMUSG( | Mt3 | 19688.66 | 0.091195 | 0.28344 | 0.73186 | ENSMUSG( | Mt3 | 15595.47 | -0.33201 | 0.008473 | 0.391601 |
| ENSMUSG( | Mta1 | 3236.717 | 0.044284 | 0.639195 | 0.916434 | ENSMUSG( | Mta1 | 2842.219 | -0.0634 | 0.568746 | 0.941777 |
| ENSMUSG( | Mta2 | 1409.938 | 0.051125 | 0.624181 | 0.91187 | ENSMUSG( | Mta2 | 1291.74 | 0.067667 | 0.583484 | 0.944266 |
| ENSMUSG( | Mta3 | 2333.537 | 0.231361 | 0.020283 | 0.214748 | ENSMUSG( | Mta3 | 2005.452 | 0.074202 | 0.56135 | 0.940254 |
| ENSMUSG( | Mtag2 | 31.49612 | -0.17664 | 0.603612 | 0.904887 | ENSMUSG( | Mtag2 | 28.59908 | -0.17479 | 0.643062 | 0.953903 |
| ENSMUSG( | Mtap | 277.8825 | -0.15319 | 0.318235 | 0.764234 | ENSMUSG( | Mtap | 253.7793 | -0.15546 | 0.380356 | 0.917202 |
| ENSMUSG( | Mtarc2 | 2804.136 | -0.1848 | 0.113323 | 0.507621 | ENSMUSG( | Mtarc2 | 2761.526 | 0.047077 | 0.709287 | 0.964042 |
| ENSMUSG( | Mtbp | 87.5457 | -0.18626 | 0.405903 | 0.81652 | ENSMUSG( | Mtbp | 88.24228 | 0.103932 | 0.655021 | 0.955053 |
| ENSMUSG( | Mtch1 | 12253.27 | 0.000764 | 0.993458 | 0.999369 | ENSMUSG( | Mtch1 | 10493.02 | -0.18547 | 0.123296 | 0.814691 |
| ENSMUSG( | Mtch2 | 2461.711 | -0.0574 | 0.474661 | 0.849485 | ENSMUSG( | Mtch2 | 2241.854 | -0.0592 | 0.561853 | 0.940388 |
| ENSMUSG( | Mtcl1 | 2145.969 | -0.19371 | 0.03734 | 0.298204 | ENSMUSG( | Mtcl1 | 1960.464 | -0.18628 | 0.091423 | 0.791889 |
| ENSMUSG( | Mtcl2 | 1000.019 | -0.00083 | 0.994324 | 0.999369 | ENSMUSG( | Mtcl2 | 850.3862 | -0.20229 | 0.169778 | 0.840806 |
| ENSMUSG( | Mtcl3 | 839.1222 | 0.129442 | 0.284082 | 0.732569 | ENSMUSG( | Mtcl3 | 738.952 | 0.033942 | 0.784708 | 0.976231 |
| ENSMUSG( | Mtcp1 | 51.44999 | -0.44338 | 0.137988 | 0.552905 | ENSMUSG( | Mtcp1 | 60.31204 | 0.335572 | 0.376433 | 0.915491 |
| ENSMUSG( | Mtdh | 2428.137 | -0.36118 | 0.017757 | 0.2 | ENSMUSG( | Mtdh | 2701.082 | 0.232672 | 0.210616 | 0.864412 |
| ENSMUSG( | Mterf1a | 51.68835 | 0.462069 | 0.087849 | 0.454663 | ENSMUSG( | Mterf1a | 46.77516 | 0.450395 | 0.140808 | 0.830678 |
| ENSMUSG( | Mterf1b | 33.39999 | 0.494482 | 0.159656 | 0.589221 | ENSMUSG( | Mterf1b | 32.76877 | 0.677003 | 0.060253 | 0.731553 |
| ENSMUSG( | Mterf2 | 239.4144 | 0.307479 | 0.039364 | 0.305596 | ENSMUSG( | Mterf2 | 186.6131 | -0.13522 | 0.437261 | 0.920144 |
| ENSMUSG( | Mterf3 | 313.2276 | -0.16225 | 0.256452 | 0.707303 | ENSMUSG( | Mterf3 | 318.5052 | 0.153103 | 0.333965 | 0.907907 |
| ENSMUSG( | Mterf4 | 252.39 | 0.108206 | 0.49406 | 0.860611 | ENSMUSG( | Mterf4 | 226.8933 | 0.072593 | 0.664238 | 0.956766 |
| ENSMUSG( | Mtf1 | 958.4213 | -0.01266 | 0.903287 | 0.986055 | ENSMUSG( | Mtf1 | 898.6474 | 0.066828 | 0.585375 | 0.944975 |
| ENSMUSG( | Mtf2 | 883.169 | -0.00822 | 0.944571 | 0.993583 | ENSMUSG( | Mtf2 | 831.9844 | 0.083765 | 0.531807 | 0.934795 |
| ENSMUSG( | Mtfmt | 302.6678 | -0.10328 | 0.465173 | 0.843971 | ENSMUSG( | Mtfmt | 284.9489 | 0.00918 | 0.964054 | 0.995433 |
| ENSMUSG( | Mtfp1 | 1281.728 | -0.02949 | 0.753845 | 0.950981 | ENSMUSG( | Mtfp1 | 1181.909 | 0.0024 | 0.985643 | 0.997084 |
| ENSMUSG( | Mtfr1 | 306.5784 | 0.026133 | 0.841639 | 0.971179 | ENSMUSG( | Mtfr1 | 279.1178 | 0.024 | 0.877744 | 0.989644 |
| ENSMUSG( | Mtfr1l | 2817.258 | -0.01912 | 0.814476 | 0.964574 | ENSMUSG( | Mtfr1l | 2450.353 | -0.15944 | 0.157101 | 0.837724 |
| ENSMUSG( | Mtfr2 | 18.52226 | -0.39001 | 0.365792 | NA | ENSMUSG( | Mtfr2 | 18.97422 | 0.01009 | 0.982569 | 0.996795 |

| | | | | | |
|---|---|---|---|---|---|
| ENSMUSG( | 593.045 | 0.001607 | 0.989855 | 0.999144 | Mrpl35 |
| ENSMUSG( | 523.1002 | 4.39E-05 | 0.99972 | 0.99989 | Mrpl36 |
| ENSMUSG( | 1101.288 | 0.152434 | 0.150151 | 0.574666 | Mrpl37 |
| ENSMUSG( | 889.6144 | 0.013745 | 0.912274 | 0.988483 | Mrpl38 |
| ENSMUSG( | 592.3855 | 0.123974 | 0.309598 | 0.757658 | Mrpl39 |
| ENSMUSG( | 1297.468 | 0.018453 | 0.845782 | 0.972191 | Mrpl4 |
| ENSMUSG( | 543.5552 | -0.0938 | 0.429588 | 0.826446 | Mrpl40 |
| ENSMUSG( | 970.6717 | 0.082576 | 0.440894 | 0.832371 | Mrpl41 |
| ENSMUSG( | 1283.086 | 0.075807 | 0.453996 | 0.836798 | Mrpl42 |
| ENSMUSG( | 973.0806 | -0.0044 | 0.967792 | 0.996237 | Mrpl43 |
| ENSMUSG( | 391.1032 | 0.129385 | 0.432061 | 0.82768 | Mrpl44 |
| ENSMUSG( | 858.1514 | 0.08985 | 0.460879 | 0.840979 | Mrpl45 |
| ENSMUSG( | 422.0494 | 0.13011 | 0.284442 | 0.732908 | Mrpl46 |
| ENSMUSG( | 401.0847 | -0.03101 | 0.813103 | 0.964574 | Mrpl47 |
| ENSMUSG( | 2883.031 | 0.12898 | 0.110055 | 0.500306 | Mrpl48 |
| ENSMUSG( | 1011.225 | 0.05948 | 0.563525 | 0.88992 | Mrpl49 |
| ENSMUSG( | 910.8897 | 0.047723 | 0.656263 | 0.923929 | Mrpl50 |
| ENSMUSG( | 1287.739 | 0.14944 | 0.114118 | 0.509818 | Mrpl51 |
| ENSMUSG( | 884.0429 | 0.037007 | 0.741409 | 0.948208 | Mrpl52 |
| ENSMUSG( | 705.4991 | -0.14198 | 0.215233 | 0.659815 | Mrpl54 |
| ENSMUSG( | 617.6528 | -0.04986 | 0.669126 | 0.928101 | Mrpl55 |
| ENSMUSG( | 683.3803 | -0.11757 | 0.309515 | 0.757658 | Mrpl57 |
| ENSMUSG( | 961.6932 | -0.00817 | 0.94464 | 0.993583 | Mrpl58 |
| ENSMUSG( | 1297.512 | 0.038293 | 0.709319 | 0.939656 | Mrpl9 |
| ENSMUSG( | 624.1074 | -0.03705 | 0.723679 | 0.94403 | Mrps10 |
| ENSMUSG( | 474.0078 | 0.054714 | 0.703027 | 0.937735 | Mrps11 |
| ENSMUSG( | 841.9011 | 0.120135 | 0.339809 | 0.777939 | Mrps12 |
| ENSMUSG( | 719.4626 | 0.067533 | 0.592027 | 0.90183 | Mrps14 |
| ENSMUSG( | 872.9168 | 0.127878 | 0.249918 | 0.699512 | Mrps15 |
| ENSMUSG( | 558.4859 | 0.001673 | 0.988988 | 0.999111 | Mrps16 |
| ENSMUSG( | 1438.733 | -0.02538 | 0.76864 | 0.954674 | Mrps17 |
| ENSMUSG( | 1179.201 | 0.015715 | 0.861483 | 0.975026 | Mrps18a |
| ENSMUSG( | 316.4014 | -0.0172 | 0.9031 | 0.986055 | Mrps18b |
| ENSMUSG( | 316.0679 | -0.07218 | 0.579221 | 0.896692 | Mrps18c |
| ENSMUSG( | 774.8396 | 0.100678 | 0.408875 | 0.818173 | Mrps2 |
| ENSMUSG( | 1111.496 | -0.01355 | 0.893128 | 0.984512 | Mrps21 |
| ENSMUSG( | 312.5971 | 0.103585 | 0.457704 | 0.839765 | Mrps22 |
| ENSMUSG( | 841.9921 | 0.024815 | 0.841475 | 0.971179 | Mrps23 |
| ENSMUSG( | 1050.301 | -0.00376 | 0.969082 | 0.996369 | Mrps24 |
| ENSMUSG( | 1067.989 | 0.053757 | 0.584537 | 0.898862 | Mrps25 |
| ENSMUSG( | 1324.375 | -0.02837 | 0.792384 | 0.959461 | Mrps26 |
| ENSMUSG( | 753.0088 | 0.076895 | 0.567462 | 0.891774 | Mrps27 |
| ENSMUSG( | 250.0318 | 0.043532 | 0.788114 | 0.959226 | Mrps28 |
| ENSMUSG( | 541.402 | 0.184528 | 0.119681 | 0.523665 | Mrps30 |
| ENSMUSG( | 294.5316 | -0.131 | 0.347454 | 0.78046 | Mrps31 |
| ENSMUSG( | 700.7226 | 0.150941 | 0.214995 | 0.659512 | Mrps33 |
| ENSMUSG( | 992.3671 | 0.012553 | 0.912401 | 0.988483 | Mrps34 |
| ENSMUSG( | 586.0436 | 0.1141 | 0.292431 | 0.742521 | Mrps35 |
| ENSMUSG( | 455.291 | 0.102066 | 0.405317 | 0.81652 | Mrps36 |
| ENSMUSG( | 1112.701 | 0.011424 | 0.908841 | 0.98794 | Mrps5 |
| ENSMUSG( | 349.4475 | 0.192982 | 0.216494 | 0.661421 | Mrps6 |
| ENSMUSG( | 1304.927 | 0.028304 | 0.765202 | 0.954127 | Mrps7 |
| ENSMUSG( | 752.5863 | -0.02446 | 0.848017 | 0.972374 | Mrps9 |
| ENSMUSG( | 364.6609 | 0.12104 | 0.333066 | 0.775602 | Mrrf |
| ENSMUSG( | 3190.807 | 0.072782 | 0.412734 | 0.819518 | Mrs2 |
| ENSMUSG( | 1422.827 | 0.392671 | 0.011009 | 0.151333 | Mrtfa |
| ENSMUSG( | 4189.927 | 0.115586 | 0.264306 | 0.715323 | Mrtfb |
| ENSMUSG( | 693.2172 | 0.056256 | 0.594284 | 0.902869 | Mrto4 |
| ENSMUSG( | 9.750412 | -0.11293 | 0.847512 | NA | Ms4a1 |
| ENSMUSG( | 9.7428 | -1.70633 | 0.073372 | NA | Ms4a15 |
| ENSMUSG( | 22.99388 | -0.0225 | 0.957294 | 0.993874 | Ms4a4a |
| ENSMUSG( | 68.66952 | 0.044983 | 0.85859 | 0.974707 | Ms4a6b |
| ENSMUSG( | 68.00212 | 0.058133 | 0.83396 | 0.970333 | Ms4a6c |
| ENSMUSG( | 55.02253 | -0.19174 | 0.514317 | 0.872055 | Ms4a6d |
| ENSMUSG( | 34.2597 | -0.2644 | 0.453889 | 0.836798 | Ms4a7 |
| ENSMUSG( | 70.03484 | -0.03617 | 0.875292 | 0.979998 | Msantd1 |
| ENSMUSG( | 446.2175 | 0.120648 | 0.381831 | 0.802968 | Msantd2 |
| ENSMUSG( | 707.7996 | 0.230813 | 0.020842 | 0.218947 | Msantd3 |
| ENSMUSG( | 1435.59 | 0.091513 | 0.392594 | 0.809274 | Msantd4 |
| ENSMUSG( | 106.4475 | -0.36875 | 0.066037 | 0.397686 | Msantd7 |
| ENSMUSG( | 28.758 | -0.2199 | 0.524634 | 0.875871 | Msc |
| ENSMUSG( | 540.873 | 0.054509 | 0.651196 | 0.921451 | Msh2 |
| ENSMUSG( | 533.2578 | 0.049691 | 0.688492 | 0.933441 | Msh3 |
| ENSMUSG( | 17.35179 | -0.24757 | 0.587796 | NA | Msh5 |
| ENSMUSG( | 441.1554 | 0.233441 | 0.053585 | 0.35565 | Msh6 |
| ENSMUSG( | 2781.135 | -0.01525 | 0.865537 | 0.975676 | Msi1 |
| ENSMUSG( | 6672.657 | 0.066279 | 0.541434 | 0.883182 | Msi2 |
| ENSMUSG( | 8178.851 | 0.241754 | 0.005241 | 0.096494 | Msl1 |
| ENSMUSG( | 1063.085 | -0.0065 | 0.94602 | 0.99369 | Msl2 |
| ENSMUSG( | 569.5712 | 0.141785 | 0.230855 | 0.679066 | Msl3 |
| ENSMUSG( | 283.4465 | 0.076821 | 0.596004 | 0.903152 | Msl3l2 |
| ENSMUSG( | 90.21537 | 0.376895 | 0.075557 | 0.425975 | Msln |
| ENSMUSG( | 11.42212 | 1.147803 | 0.052908 | NA | Mslnl |
| ENSMUSG( | 5326.89 | -0.50867 | 0.005392 | 0.098038 | Msmo1 |
| ENSMUSG( | 2298.201 | -0.06955 | 0.595189 | 0.903007 | Msn |
| ENSMUSG( | 19.92759 | 0.366917 | 0.390664 | NA | Msr1 |
| ENSMUSG( | 860.698 | 0.107364 | 0.336956 | 0.777008 | Msra |
| ENSMUSG( | 597.085 | 0.052537 | 0.639742 | 0.916434 | Msrb1 |
| ENSMUSG( | 652.5005 | 0.050447 | 0.631326 | 0.914073 | Msrb2 |
| ENSMUSG( | 249.2846 | -0.11661 | 0.417521 | 0.821568 | Msrb3 |
| ENSMUSG( | 16.34439 | -0.00692 | 0.987997 | NA | Mss51 |
| ENSMUSG( | 51.44122 | 0.31195 | 0.276978 | 0.727074 | Mst1r |
| ENSMUSG( | 635.3082 | -0.02668 | 0.812465 | 0.964574 | Msto1 |
| ENSMUSG( | 183.356 | -0.2387 | 0.297561 | 0.746087 | Msx1 |
| ENSMUSG( | 17.83357 | -0.02962 | 0.946192 | NA | Msx2 |
| ENSMUSG( | 20177.6 | 0.121902 | 0.235423 | 0.682423 | Mt1 |
| ENSMUSG( | 9541.749 | 0.091897 | 0.366136 | 0.794603 | Mt2 |
| ENSMUSG( | 19688.66 | 0.091195 | 0.28344 | 0.73186 | Mt3 |
| ENSMUSG( | 3236.717 | 0.044284 | 0.639195 | 0.916434 | Mta1 |
| ENSMUSG( | 1409.938 | 0.051125 | 0.624181 | 0.91187 | Mta2 |
| ENSMUSG( | 2333.537 | 0.231361 | 0.020283 | 0.214748 | Mta3 |
| ENSMUSG( | 31.49612 | -0.17664 | 0.603612 | 0.904887 | Mtag2 |
| ENSMUSG( | 277.8825 | -0.15319 | 0.318235 | 0.764234 | Mtap |
| ENSMUSG( | 2804.136 | -0.1848 | 0.113323 | 0.507621 | Mtarc2 |
| ENSMUSG( | 87.5457 | -0.18626 | 0.405903 | 0.81652 | Mtbp |
| ENSMUSG( | 12253.27 | 0.000764 | 0.993458 | 0.999369 | Mtch1 |
| ENSMUSG( | 2461.711 | -0.0574 | 0.474661 | 0.849485 | Mtch2 |
| ENSMUSG( | 2145.969 | -0.19371 | 0.03734 | 0.298204 | Mtcl1 |
| ENSMUSG( | 1000.019 | -0.00083 | 0.994324 | 0.999369 | Mtcl2 |
| ENSMUSG( | 839.1222 | 0.129442 | 0.284082 | 0.732569 | Mtcl3 |
| ENSMUSG( | 51.44999 | -0.44338 | 0.137988 | 0.552905 | Mtcp1 |
| ENSMUSG( | 2428.137 | -0.36118 | 0.017757 | 0.2 | Mtdh |
| ENSMUSG( | 51.68835 | 0.462069 | 0.087849 | 0.454663 | Mterf1a |
| ENSMUSG( | 33.39999 | 0.494482 | 0.159656 | 0.589221 | Mterf1b |
| ENSMUSG( | 239.4144 | 0.307479 | 0.039364 | 0.305596 | Mterf2 |
| ENSMUSG( | 313.2276 | -0.16225 | 0.256452 | 0.707303 | Mterf3 |
| ENSMUSG( | 252.39 | 0.108206 | 0.49406 | 0.860611 | Mterf4 |
| ENSMUSG( | 958.4213 | -0.01266 | 0.903287 | 0.986055 | Mtf1 |
| ENSMUSG( | 883.169 | -0.00822 | 0.944571 | 0.993583 | Mtf2 |
| ENSMUSG( | 302.6678 | -0.10328 | 0.465173 | 0.843971 | Mtfmt |
| ENSMUSG( | 1281.728 | -0.02949 | 0.753845 | 0.950981 | Mtfp1 |
| ENSMUSG( | 306.5784 | 0.026133 | 0.841639 | 0.971179 | Mtfr1 |
| ENSMUSG( | 2817.258 | -0.01912 | 0.814476 | 0.964574 | Mtfr1l |
| ENSMUSG( | 18.52226 | -0.39001 | 0.365792 | NA | Mtfr2 |
| ENSMUSG( | 372.6883 | 0.011706 | 0.923091 | 0.990693 | Mtg1 |
| ENSMUSG( | 436.0154 | 0.085638 | 0.542325 | 0.883182 | Mtg2 |
| ENSMUSG( | 1581.158 | 0.134283 | 0.172908 | 0.608361 | Mthfd1 |
| ENSMUSG( | 287.8829 | 0.169637 | 0.34186 | 0.778828 | Mthfd1l |
| ENSMUSG( | 170.3525 | -0.0806 | 0.681966 | 0.931729 | Mthfd2 |
| ENSMUSG( | 308.1418 | -0.00736 | 0.958812 | 0.993874 | Mthfd2l |
| ENSMUSG( | 992.8296 | -0.12106 | 0.320861 | 0.765984 | Mthfr |
| ENSMUSG( | 63.20004 | -0.19528 | 0.424069 | 0.824452 | Mthfs |
| ENSMUSG( | 1094.046 | 0.151761 | 0.231197 | 0.679229 | Mthfsd |
| ENSMUSG( | 179.2878 | -0.03267 | 0.841224 | 0.971179 | Mthfsl |
| ENSMUSG( | 694.8552 | 0.150119 | 0.163277 | 0.594466 | Mtif2 |
| ENSMUSG( | 425.6196 | 0.06851 | 0.588184 | 0.900712 | Mtif3 |
| ENSMUSG( | 634.5962 | 0.022751 | 0.839461 | 0.971179 | Mtln |
| ENSMUSG( | 233.5813 | 0.123466 | 0.415481 | 0.820741 | Mtm1 |
| ENSMUSG( | 1251.346 | 0.138276 | 0.152876 | 0.579254 | Mtmr1 |
| ENSMUSG( | 1153.573 | 0.11671 | 0.3015 | 0.749979 | Mtmr10 |
| ENSMUSG( | 535.456 | 0.092328 | 0.405546 | 0.81652 | Mtmr11 |
| ENSMUSG( | 1057.066 | 0.089953 | 0.420939 | 0.823101 | Mtmr12 |
| ENSMUSG( | 613.3582 | 0.138722 | 0.280154 | 0.729155 | Mtmr14 |
| ENSMUSG( | 1271.648 | 0.021161 | 0.842635 | 0.971663 | Mtmr2 |
| ENSMUSG( | 2999.997 | 0.106936 | 0.221603 | 0.667258 | Mtmr3 |
| ENSMUSG( | 1984.197 | 0.119684 | 0.214649 | 0.659193 | Mtmr4 |
| ENSMUSG( | 4257.901 | 0.02935 | 0.738088 | 0.947329 | Mtmr6 |
| ENSMUSG( | 875.7256 | 0.001013 | 0.993819 | 0.999369 | Mtmr7 |
| ENSMUSG( | 2897.196 | 0.107659 | 0.306122 | 0.753712 | Mtmr9 |
| ENSMUSG( | 142.4613 | -0.02668 | 0.888124 | 0.983434 | Mtnap1 |
| ENSMUSG( | 692.3478 | 0.104198 | 0.301107 | 0.749979 | Mto1 |
| ENSMUSG( | 2844.478 | 0.118633 | 0.17455 | 0.610304 | Mtor |
| ENSMUSG( | 439.9276 | 0.098474 | 0.409015 | 0.818231 | Mtpap |
| ENSMUSG( | 8201.609 | 0.004182 | 0.968068 | 0.996284 | Mtpn |
| ENSMUSG( | 669.6473 | 0.127544 | 0.280625 | 0.729679 | Mtr |
| ENSMUSG( | 468.3494 | 0.005356 | 0.967066 | 0.995924 | Mtres1 |
| ENSMUSG( | 1264.529 | 0.234713 | 0.023217 | 0.233609 | Mtrex |
| ENSMUSG( | 146.4257 | -0.02386 | 0.894303 | 0.984512 | Mtrf1 |

| | | | | | |
|---|---|---|---|---|---|
| ENSMUSG( | 1150.82 | -0.06754 | 0.553875 | 0.939565 | Mrps7 |
| ENSMUSG( | 700.1331 | 0.036006 | 0.796038 | 0.97681 | Mrps9 |
| ENSMUSG( | 315.6518 | -0.02966 | 0.852296 | 0.985598 | Mrrf |
| ENSMUSG( | 2844.595 | 0.012006 | 0.915735 | 0.992529 | Mrs2 |
| ENSMUSG( | 1190.21 | 0.167717 | 0.286593 | 0.894091 | Mrtfa |
| ENSMUSG( | 3760.903 | 0.075356 | 0.521794 | 0.932571 | Mrtfb |
| ENSMUSG( | 594.3975 | -0.12645 | 0.341321 | 0.908511 | Mrto4 |
| ENSMUSG( | 19.62634 | 0.889473 | 0.515537 | 0.930939 | Ms4a15 |
| ENSMUSG( | 18.40734 | -0.42394 | 0.442971 | 0.921554 | Ms4a4a |
| ENSMUSG( | 76.49797 | 0.58417 | 0.042793 | 0.690304 | Ms4a6b |
| ENSMUSG( | 62.17229 | 0.075848 | 0.801982 | 0.97681 | Ms4a6c |
| ENSMUSG( | 59.58747 | 0.308642 | 0.314616 | 0.900952 | Ms4a6d |
| ENSMUSG( | 30.4614 | -0.32123 | 0.401531 | 0.917257 | Ms4a7 |
| ENSMUSG( | 64.97584 | 0.023996 | 0.931105 | 0.99306 | Msantd1 |
| ENSMUSG( | 401.479 | 0.086605 | 0.583637 | 0.944266 | Msantd2 |
| ENSMUSG( | 590.9207 | -0.01626 | 0.899123 | 0.990558 | Msantd3 |
| ENSMUSG( | 1287.194 | 0.041019 | 0.783582 | 0.976106 | Msantd4 |
| ENSMUSG( | 107.9063 | -0.03064 | 0.885723 | 0.98977 | Msantd7 |
| ENSMUSG( | 26.4283 | -0.1609 | 0.683139 | 0.958678 | Msc |
| ENSMUSG( | 472.5644 | -0.06236 | 0.638836 | 0.953903 | Msh2 |
| ENSMUSG( | 483.7828 | 0.038393 | 0.789049 | 0.976593 | Msh3 |
| ENSMUSG( | 19.35978 | 0.3279 | 0.476789 | 0.923329 | Msh5 |
| ENSMUSG( | 385.0107 | 0.113534 | 0.416139 | 0.917257 | Msh6 |
| ENSMUSG( | 2398.973 | -0.18139 | 0.125298 | 0.81516 | Msi1 |
| ENSMUSG( | 5657.973 | -0.14572 | 0.26711 | 0.886134 | Msi2 |
| ENSMUSG( | 7282.894 | 0.179587 | 0.090544 | 0.791074 | Msl1 |
| ENSMUSG( | 1036.513 | 0.181413 | 0.113583 | 0.809623 | Msl2 |
| ENSMUSG( | 505.3697 | 0.06261 | 0.658413 | 0.956205 | Msl3 |
| ENSMUSG( | 254.5669 | 0.033292 | 0.852585 | 0.985598 | Msl3l2 |
| ENSMUSG( | 64.59813 | -0.31769 | 0.222574 | 0.866791 | Msln |
| ENSMUSG( | 5493.418 | -0.10459 | 0.579804 | 0.944209 | Msmo1 |
| ENSMUSG( | 2103.686 | -0.05707 | 0.678354 | 0.957435 | Msn |
| ENSMUSG( | 15.39941 | -0.10252 | 0.841055 | 0.9826 | Msr1 |
| ENSMUSG( | 721.7349 | -0.13904 | 0.368107 | 0.915491 | Msra |
| ENSMUSG( | 502.1268 | -0.18847 | 0.162874 | 0.837905 | Msrb1 |
| ENSMUSG( | 531.8487 | -0.28956 | 0.028789 | 0.614171 | Msrb2 |
| ENSMUSG( | 240.6363 | 0.063213 | 0.711879 | 0.964363 | Msrb3 |
| ENSMUSG( | 18.81774 | 0.614231 | 0.177394 | 0.847137 | Mss51 |
| ENSMUSG( | 43.42274 | 0.128513 | 0.695623 | 0.960867 | Mst1r |
| ENSMUSG( | 548.8028 | -0.18786 | 0.168359 | 0.840222 | Msto1 |
| ENSMUSG( | 197.0312 | 0.238016 | 0.294964 | 0.898538 | Msx1 |
| ENSMUSG( | 18.43489 | 0.313663 | 0.499108 | 0.927825 | Msx2 |
| ENSMUSG( | 16423.63 | -0.21077 | 0.104264 | 0.804969 | Mt1 |
| ENSMUSG( | 7838.676 | -0.21551 | 0.071946 | 0.763629 | Mt2 |
| ENSMUSG( | 15595.47 | -0.33201 | 0.008473 | 0.391601 | Mt3 |
| ENSMUSG( | 2842.219 | -0.0634 | 0.568746 | 0.941777 | Mta1 |
| ENSMUSG( | 1291.74 | 0.067667 | 0.583484 | 0.944266 | Mta2 |
| ENSMUSG( | 2005.452 | 0.074202 | 0.56135 | 0.940254 | Mta3 |
| ENSMUSG( | 28.59908 | -0.17479 | 0.643062 | 0.953903 | Mtag2 |
| ENSMUSG( | 253.7793 | -0.15546 | 0.380356 | 0.917202 | Mtap |
| ENSMUSG( | 2761.526 | 0.047077 | 0.709287 | 0.964042 | Mtarc2 |
| ENSMUSG( | 88.24228 | 0.103932 | 0.655021 | 0.955053 | Mtbp |
| ENSMUSG( | 10493.02 | -0.18547 | 0.123296 | 0.814691 | Mtch1 |
| ENSMUSG( | 2241.854 | -0.0592 | 0.561853 | 0.940388 | Mtch2 |
| ENSMUSG( | 1960.464 | -0.18628 | 0.091423 | 0.791889 | Mtcl1 |
| ENSMUSG( | 850.3862 | -0.20229 | 0.169778 | 0.840806 | Mtcl2 |
| ENSMUSG( | 738.952 | 0.033942 | 0.784708 | 0.976231 | Mtcl3 |
| ENSMUSG( | 60.31204 | 0.335572 | 0.376433 | 0.915491 | Mtcp1 |
| ENSMUSG( | 2701.082 | 0.232672 | 0.210616 | 0.864412 | Mtdh |
| ENSMUSG( | 46.77516 | 0.450395 | 0.140808 | 0.830678 | Mterf1a |
| ENSMUSG( | 32.76877 | 0.677003 | 0.060253 | 0.731553 | Mterf1b |
| ENSMUSG( | 186.6131 | -0.13522 | 0.437261 | 0.920144 | Mterf2 |
| ENSMUSG( | 318.5052 | 0.153103 | 0.333965 | 0.907907 | Mterf3 |
| ENSMUSG( | 226.8933 | 0.072593 | 0.664238 | 0.956766 | Mterf4 |
| ENSMUSG( | 898.6474 | 0.066828 | 0.585375 | 0.944975 | Mtf1 |
| ENSMUSG( | 831.9844 | 0.083765 | 0.531807 | 0.934795 | Mtf2 |
| ENSMUSG( | 284.9489 | 0.00918 | 0.964054 | 0.995433 | Mtfmt |
| ENSMUSG( | 1181.909 | 0.0024 | 0.985643 | 0.997084 | Mtfp1 |
| ENSMUSG( | 279.1178 | 0.024 | 0.877744 | 0.989644 | Mtfr1 |
| ENSMUSG( | 2450.353 | -0.15944 | 0.157101 | 0.837724 | Mtfr1l |
| ENSMUSG( | 18.97422 | 0.01009 | 0.982569 | 0.996795 | Mtfr2 |
| ENSMUSG( | 338.5739 | 0.010431 | 0.944487 | 0.994798 | Mtg1 |
| ENSMUSG( | 396.0681 | 0.075979 | 0.636772 | 0.953903 | Mtg2 |
| ENSMUSG( | 1284.675 | -0.20326 | 0.079608 | 0.774721 | Mthfd1 |
| ENSMUSG( | 289.4809 | 0.430469 | 0.017401 | 0.531019 | Mthfd1l |
| ENSMUSG( | 151.4938 | -0.15524 | 0.417217 | 0.917257 | Mthfd2 |
| ENSMUSG( | 280.2986 | -0.02367 | 0.893731 | 0.990558 | Mthfd2l |
| ENSMUSG( | 919.9785 | -0.066 | 0.639674 | 0.953903 | Mthfr |
| ENSMUSG( | 54.35085 | -0.41625 | 0.169863 | 0.840806 | Mthfs |
| ENSMUSG( | 960.6824 | 0.051583 | 0.758734 | 0.971858 | Mthfsd |
| ENSMUSG( | 153.6159 | -0.2279 | 0.216549 | 0.865167 | Mthfsl |
| ENSMUSG( | 615.3755 | 0.06735 | 0.603387 | 0.948137 | Mtif2 |
| ENSMUSG( | 356.9407 | -0.17026 | 0.247564 | 0.877904 | Mtif3 |
| ENSMUSG( | 519.614 | -0.30704 | 0.029904 | 0.61992 | Mtln |
| ENSMUSG( | 208.6536 | 0.072205 | 0.672489 | 0.95694 | Mtm1 |
| ENSMUSG( | 1087.021 | 0.000775 | 0.994819 | 0.998813 | Mtmr1 |
| ENSMUSG( | 1031.91 | 0.062994 | 0.652317 | 0.954766 | Mtmr10 |
| ENSMUSG( | 448.6878 | -0.15773 | 0.260298 | 0.884524 | Mtmr11 |
| ENSMUSG( | 917.8266 | -0.04635 | 0.728827 | 0.966913 | Mtmr12 |
| ENSMUSG( | 543.2515 | 0.060283 | 0.650002 | 0.95422 | Mtmr14 |
| ENSMUSG( | 1162.999 | 0.035066 | 0.768517 | 0.974236 | Mtmr2 |
| ENSMUSG( | 2681.529 | 0.055364 | 0.611446 | 0.950771 | Mtmr3 |
| ENSMUSG( | 1929.695 | 0.296385 | 0.006609 | 0.338858 | Mtmr4 |
| ENSMUSG( | 3859.531 | 0.013228 | 0.906984 | 0.991601 | Mtmr6 |
| ENSMUSG( | 776.2125 | -0.08068 | 0.572716 | 0.94262 | Mtmr7 |
| ENSMUSG( | 2463.338 | -0.09141 | 0.395348 | 0.917257 | Mtmr9 |
| ENSMUSG( | 134.8842 | 0.089297 | 0.659731 | 0.956205 | Mtnap1 |
| ENSMUSG( | 583.6094 | -0.12137 | 0.340043 | 0.907972 | Mto1 |
| ENSMUSG( | 2594.034 | 0.123788 | 0.294815 | 0.898538 | Mtor |
| ENSMUSG( | 411.3518 | 0.165281 | 0.244527 | 0.877904 | Mtpap |
| ENSMUSG( | 7167.567 | -0.12129 | 0.416136 | 0.917257 | Mtpn |
| ENSMUSG( | 586.7009 | 0.018693 | 0.878695 | 0.989644 | Mtr |
| ENSMUSG( | 417.8091 | -0.0638 | 0.695172 | 0.960867 | Mtres1 |
| ENSMUSG( | 1053.388 | -0.01353 | 0.913616 | 0.992378 | Mtrex |
| ENSMUSG( | 135.8366 | 0.0161 | 0.93759 | 0.993685 | Mtrf1 |
| ENSMUSG( | 162.512 | 0.384309 | 0.030311 | 0.623931 | Mtrf1l |
| ENSMUSG( | 280.1264 | -0.06452 | 0.694633 | 0.960867 | Mtrfr |
| ENSMUSG( | 34.79369 | -0.72487 | 0.041172 | 0.687358 | Mtrnr2l7 |
| ENSMUSG( | 274.2783 | 0.130423 | 0.378868 | 0.916995 | Mtrr |
| ENSMUSG( | 889.3536 | -0.17158 | 0.158727 | 0.837724 | Mtss1 |
| ENSMUSG( | 27063.21 | -0.29486 | 0.034959 | 0.659157 | Mtss2 |
| ENSMUSG( | 386.0362 | 0.195952 | 0.197371 | 0.862161 | Mttp |
| ENSMUSG( | 7464.647 | -0.04251 | 0.671256 | 0.95694 | Mturn |
| ENSMUSG( | 1399.349 | 0.019607 | 0.885167 | 0.989688 | Mtus1 |
| ENSMUSG( | 990.9277 | 0.146463 | 0.229252 | 0.868507 | Mtus2 |
| ENSMUSG( | 617.2676 | -0.0442 | 0.740638 | 0.968811 | Mtx1 |
| ENSMUSG( | 479.2067 | -0.02006 | 0.898654 | 0.990558 | Mtx2 |
| ENSMUSG( | 1478.595 | 0.028023 | 0.804486 | 0.977232 | Mtx3 |
| ENSMUSG( | 73.32591 | 0.313353 | 0.232555 | 0.868507 | Muc1 |
| ENSMUSG( | 7.929229 | 0.994907 | 0.226412 | NA | Muc15 |
| ENSMUSG( | 24.35372 | 1.255555 | 0.002223 | 0.183698 | Muc2 |
| ENSMUSG( | 213.3124 | -0.14189 | 0.618473 | 0.952254 | Muc3a |
| ENSMUSG( | 1266.206 | 0.018154 | 0.882232 | 0.989644 | Mul1 |
| ENSMUSG( | 523.161 | -0.02243 | 0.876888 | 0.989644 | Mus81 |
| ENSMUSG( | 28.98626 | 0.417893 | 0.360444 | 0.914223 | Musk |
| ENSMUSG( | 31.22855 | -0.26952 | 0.478955 | 0.923329 | Mustn1 |
| ENSMUSG( | 58.97036 | 0.328201 | 0.303506 | 0.898538 | Mutyh |
| ENSMUSG( | 632.8693 | 0.036398 | 0.769794 | 0.974236 | Mvb12a |
| ENSMUSG( | 6458.319 | 0.100164 | 0.357676 | 0.913759 | Mvb12b |
| ENSMUSG( | 809.1458 | -0.27362 | 0.020928 | 0.571913 | Mvd |
| ENSMUSG( | 1046.3 | -0.29676 | 0.015147 | 0.492251 | Mvk |
| ENSMUSG( | 312.2442 | 0.051021 | 0.731089 | 0.967228 | Mvp |
| ENSMUSG( | 18.36687 | -0.79774 | 0.088349 | 0.786756 | Mx1 |
| ENSMUSG( | 50.8292 | 0.026923 | 0.926034 | 0.99306 | Mx2 |
| ENSMUSG( | 749.066 | 0.17461 | 0.162342 | 0.837724 | Mxd1 |
| ENSMUSG( | 34.59309 | -0.46075 | 0.18424 | 0.852665 | Mxd3 |
| ENSMUSG( | 5555.724 | 0.24114 | 0.021564 | 0.575341 | Mxd4 |
| ENSMUSG( | 4910.623 | 0.162488 | 0.214545 | 0.864547 | Mxi1 |
| ENSMUSG( | 617.0357 | 0.126466 | 0.405589 | 0.917257 | Mxra7 |
| ENSMUSG( | 390.4347 | 0.03624 | 0.830458 | 0.980463 | Mxra8 |
| ENSMUSG( | 3124.395 | 0.222861 | 0.199002 | 0.863016 | Myadm |
| ENSMUSG( | 153.3138 | -0.03316 | 0.874795 | 0.989644 | Myadml2 |
| ENSMUSG( | 29.85817 | 0.938631 | 0.018924 | 0.548832 | Myb |
| ENSMUSG( | 1616.673 | -0.03867 | 0.736548 | 0.967344 | Mybbp1a |
| ENSMUSG( | 50.57538 | 0.38473 | 0.182086 | 0.850579 | Mybl1 |
| ENSMUSG( | 41.26144 | 0.495469 | 0.131045 | 0.822884 | Mybl2 |
| ENSMUSG( | 74.2738 | 0.682172 | 0.010382 | 0.41614 | Mybpc1 |
| ENSMUSG( | 10.66695 | -0.02436 | 0.968335 | NA | Mybpc2 |
| ENSMUSG( | 42.10724 | 0.046853 | 0.891029 | 0.990558 | Mybpc3 |
| ENSMUSG( | 39.92689 | 0.176225 | 0.591949 | 0.944975 | Mybph |
| ENSMUSG( | 92.72403 | -0.01557 | 0.958904 | 0.994798 | Myc |
| ENSMUSG( | 144.0338 | 0.067613 | 0.731384 | 0.967228 | Mycbp |
| ENSMUSG( | 7748.458 | 0.17066 | 0.184924 | 0.8547 | Mycbp2 |
| ENSMUSG( | 51.9012 | 0.233674 | 0.459756 | 0.923329 | Mycbpap |
| ENSMUSG( | 551.6669 | 0.128381 | 0.408634 | 0.917257 | Mycl |
| ENSMUSG( | 197.0082 | -0.08381 | 0.672497 | 0.95694 | Mycn |
| ENSMUSG( | 32.14348 | 0.492022 | 0.269054 | 0.886755 | Myct1 |
| ENSMUSG( | 200.6137 | -0.06756 | 0.715442 | 0.96507 | Myd88 |

| | | | | | | | | | | | |
|---|---|---|---|---|---|---|---|---|---|---|---|
| ENSMUSG( | Mtg1 | 372.6883 | 0.011706 | 0.923091 | 0.990693 | ENSMUSG( | Mtg1 | 338.5739 | 0.010431 | 0.944487 | 0.994798 |
| ENSMUSG( | Mtg2 | 436.0154 | 0.085638 | 0.542325 | 0.883182 | ENSMUSG( | Mtg2 | 396.0681 | 0.075979 | 0.636772 | 0.953903 |
| ENSMUSG( | Mthfd1 | 1581.158 | 0.134283 | 0.172908 | 0.608361 | ENSMUSG( | Mthfd1 | 1284.675 | -0.20326 | 0.079608 | 0.774721 |
| ENSMUSG( | Mthfd1l | 287.8829 | 0.169637 | 0.34186 | 0.778828 | ENSMUSG( | Mthfd1l | 289.4809 | 0.430469 | 0.017401 | 0.531019 |
| ENSMUSG( | Mthfd2 | 170.3525 | -0.0806 | 0.681966 | 0.931729 | ENSMUSG( | Mthfd2 | 151.4938 | -0.15524 | 0.417217 | 0.917257 |
| ENSMUSG( | Mthfd2l | 308.1418 | -0.00736 | 0.958812 | 0.993874 | ENSMUSG( | Mthfd2l | 280.2986 | -0.02367 | 0.893731 | 0.990558 |
| ENSMUSG( | Mthfr | 992.8296 | -0.12106 | 0.320861 | 0.765984 | ENSMUSG( | Mthfr | 919.9785 | -0.066 | 0.639674 | 0.953903 |
| ENSMUSG( | Mthfs | 63.20004 | -0.19528 | 0.424069 | 0.824452 | ENSMUSG( | Mthfs | 54.35085 | -0.41625 | 0.169863 | 0.840806 |
| ENSMUSG( | Mthfsd | 1094.046 | 0.151761 | 0.231197 | 0.679229 | ENSMUSG( | Mthfsd | 960.6824 | 0.051583 | 0.758734 | 0.971858 |
| ENSMUSG( | Mthfsl | 179.2878 | -0.03267 | 0.841224 | 0.971179 | ENSMUSG( | Mthfsl | 153.6159 | -0.2279 | 0.216549 | 0.865167 |
| ENSMUSG( | Mtif2 | 694.8552 | 0.150119 | 0.163277 | 0.594466 | ENSMUSG( | Mtif2 | 615.3755 | 0.06735 | 0.603387 | 0.948137 |
| ENSMUSG( | Mtif3 | 425.6196 | 0.06851 | 0.588184 | 0.900712 | ENSMUSG( | Mtif3 | 356.9407 | -0.17026 | 0.247564 | 0.877904 |
| ENSMUSG( | Mtln | 634.5962 | 0.022751 | 0.839461 | 0.971179 | ENSMUSG( | Mtln | 519.614 | -0.30704 | 0.029904 | 0.61992 |
| ENSMUSG( | Mtm1 | 233.5813 | 0.123466 | 0.415481 | 0.820741 | ENSMUSG( | Mtm1 | 208.6536 | 0.072205 | 0.672489 | 0.95694 |
| ENSMUSG( | Mtmr1 | 1251.346 | 0.138276 | 0.152876 | 0.579254 | ENSMUSG( | Mtmr1 | 1087.021 | 0.000775 | 0.994819 | 0.998813 |
| ENSMUSG( | Mtmr10 | 1153.573 | 0.11671 | 0.3015 | 0.749979 | ENSMUSG( | Mtmr10 | 1031.91 | 0.062994 | 0.652317 | 0.954766 |
| ENSMUSG( | Mtmr11 | 535.456 | 0.092328 | 0.405546 | 0.81652 | ENSMUSG( | Mtmr11 | 448.6878 | -0.15773 | 0.260298 | 0.884524 |
| ENSMUSG( | Mtmr12 | 1057.066 | 0.089953 | 0.420939 | 0.823101 | ENSMUSG( | Mtmr12 | 917.8266 | -0.04635 | 0.728827 | 0.966913 |
| ENSMUSG( | Mtmr14 | 613.3582 | 0.138722 | 0.280154 | 0.729155 | ENSMUSG( | Mtmr14 | 543.2515 | 0.060283 | 0.650002 | 0.95422 |
| ENSMUSG( | Mtmr2 | 1271.648 | 0.021161 | 0.842635 | 0.971663 | ENSMUSG( | Mtmr2 | 1162.999 | 0.035066 | 0.768517 | 0.974236 |
| ENSMUSG( | Mtmr3 | 2999.997 | 0.106936 | 0.221603 | 0.667258 | ENSMUSG( | Mtmr3 | 2681.529 | 0.055364 | 0.611446 | 0.950771 |
| ENSMUSG( | Mtmr4 | 1984.197 | 0.119684 | 0.214649 | 0.659193 | ENSMUSG( | Mtmr4 | 1929.695 | 0.296385 | 0.006609 | 0.338858 |
| ENSMUSG( | Mtmr6 | 4257.901 | 0.02935 | 0.738088 | 0.947329 | ENSMUSG( | Mtmr6 | 3859.531 | 0.013228 | 0.906984 | 0.991601 |
| ENSMUSG( | Mtmr7 | 875.7256 | 0.001013 | 0.993819 | 0.999369 | ENSMUSG( | Mtmr7 | 776.2125 | -0.08068 | 0.572716 | 0.94262 |
| ENSMUSG( | Mtmr9 | 2897.196 | 0.107659 | 0.306122 | 0.753712 | ENSMUSG( | Mtmr9 | 2463.338 | -0.09141 | 0.395348 | 0.917257 |
| ENSMUSG( | Mtnap1 | 142.4613 | -0.02668 | 0.888124 | 0.983434 | ENSMUSG( | Mtnap1 | 134.8842 | 0.089297 | 0.659731 | 0.956205 |
| ENSMUSG( | Mto1 | 692.3478 | 0.104198 | 0.301107 | 0.749979 | ENSMUSG( | Mto1 | 583.6094 | -0.12137 | 0.340043 | 0.907972 |
| ENSMUSG( | Mtor | 2844.478 | 0.118633 | 0.17455 | 0.610304 | ENSMUSG( | Mtor | 2594.034 | 0.123788 | 0.294815 | 0.898538 |
| ENSMUSG( | Mtpap | 439.9276 | 0.098474 | 0.409015 | 0.818231 | ENSMUSG( | Mtpap | 411.3518 | 0.165281 | 0.244527 | 0.877904 |
| ENSMUSG( | Mtpn | 8201.609 | 0.004182 | 0.968068 | 0.996284 | ENSMUSG( | Mtpn | 7167.567 | -0.12129 | 0.416136 | 0.917257 |
| ENSMUSG( | Mtr | 669.6473 | 0.127544 | 0.280625 | 0.729679 | ENSMUSG( | Mtr | 586.7009 | 0.018693 | 0.878695 | 0.989644 |
| ENSMUSG( | Mtres1 | 468.3494 | 0.005356 | 0.967066 | 0.995924 | ENSMUSG( | Mtres1 | 417.8091 | -0.0638 | 0.695172 | 0.960867 |
| ENSMUSG( | Mtrex | 1264.529 | 0.234713 | 0.023217 | 0.233609 | ENSMUSG( | Mtrex | 1053.388 | -0.01353 | 0.913616 | 0.992378 |
| ENSMUSG( | Mtrf1 | 146.4257 | -0.02386 | 0.894303 | 0.984512 | ENSMUSG( | Mtrf1 | 135.8366 | 0.0161 | 0.93759 | 0.993685 |
| ENSMUSG( | Mtrf1l | 170.349 | 0.262337 | 0.105346 | 0.492417 | ENSMUSG( | Mtrf1l | 162.512 | 0.384309 | 0.030311 | 0.623931 |
| ENSMUSG( | Mtrfr | 339.4689 | 0.222433 | 0.114578 | 0.510543 | ENSMUSG( | Mtrfr | 280.1264 | -0.06452 | 0.694633 | 0.960867 |
| ENSMUSG( | Mtrnr2l7 | 44.50001 | -0.16822 | 0.588619 | 0.900921 | ENSMUSG( | Mtrnr2l7 | 34.79369 | -0.72487 | 0.041172 | 0.687358 |
| ENSMUSG( | Mtrr | 313.4918 | 0.239237 | 0.063038 | 0.388234 | ENSMUSG( | Mtrr | 274.2783 | 0.130423 | 0.378868 | 0.916995 |
| ENSMUSG( | Mtss1 | 1001.136 | -0.09199 | 0.369646 | 0.797188 | ENSMUSG( | Mtss1 | 889.3536 | -0.17158 | 0.158727 | 0.837724 |
| ENSMUSG( | Mtss2 | 32260.96 | -0.04134 | 0.690041 | 0.933795 | ENSMUSG( | Mtss2 | 27063.21 | -0.29486 | 0.034959 | 0.659157 |
| ENSMUSG( | Mttp | 378.6066 | -0.12726 | 0.373781 | 0.80044 | ENSMUSG( | Mttp | 386.0362 | 0.195952 | 0.197371 | 0.862161 |
| ENSMUSG( | Mturn | 8832.501 | 0.169397 | 0.040651 | 0.311461 | ENSMUSG( | Mturn | 7464.647 | -0.04251 | 0.671256 | 0.95694 |
| ENSMUSG( | Mtus1 | 1519.501 | -0.01497 | 0.889009 | 0.983434 | ENSMUSG( | Mtus1 | 1399.349 | 0.019607 | 0.885167 | 0.989688 |
| ENSMUSG( | Mtus2 | 1119.196 | 0.221967 | 0.045207 | 0.326331 | ENSMUSG( | Mtus2 | 990.9277 | 0.146463 | 0.229252 | 0.868507 |
| ENSMUSG( | Mtx1 | 687.3685 | -0.00126 | 0.991146 | 0.999232 | ENSMUSG( | Mtx1 | 617.2676 | -0.0442 | 0.740638 | 0.968811 |
| ENSMUSG( | Mtx2 | 512.8697 | -0.09389 | 0.493935 | 0.860611 | ENSMUSG( | Mtx2 | 479.2067 | -0.02006 | 0.898654 | 0.990558 |
| ENSMUSG( | Mtx3 | 1660.706 | 0.093582 | 0.340893 | 0.778773 | ENSMUSG( | Mtx3 | 1478.595 | 0.028023 | 0.804486 | 0.977232 |
| ENSMUSG( | Muc1 | 72.71838 | 0.025364 | 0.912171 | 0.988483 | ENSMUSG( | Muc1 | 73.32591 | 0.313353 | 0.232555 | 0.868507 |
| ENSMUSG( | Muc2 | 15.25472 | -0.09946 | 0.834906 | NA | ENSMUSG( | Muc2 | 24.35372 | 1.255555 | 0.002223 | 0.183698 |
| ENSMUSG( | Muc3a | 268.632 | 0.257844 | 0.309737 | 0.757658 | ENSMUSG( | Muc3a | 213.3124 | -0.14189 | 0.618473 | 0.952254 |
| ENSMUSG( | Mul1 | 1347.173 | -0.07608 | 0.488894 | 0.859092 | ENSMUSG( | Mul1 | 1266.206 | 0.018154 | 0.882232 | 0.989644 |
| ENSMUSG( | Mus81 | 587.8447 | 0.038124 | 0.727471 | 0.944344 | ENSMUSG( | Mus81 | 523.161 | -0.02243 | 0.876888 | 0.989644 |
| ENSMUSG( | Musk | 24.77437 | -0.32234 | 0.488709 | 0.859092 | ENSMUSG( | Musk | 28.98626 | 0.417893 | 0.360444 | 0.914223 |
| ENSMUSG( | Mustn1 | 38.33177 | 0.037149 | 0.908516 | 0.987809 | ENSMUSG( | Mustn1 | 31.22855 | -0.26952 | 0.478955 | 0.923329 |
| ENSMUSG( | Mutyh | 59.93062 | 0.097084 | 0.703284 | 0.937735 | ENSMUSG( | Mutyh | 58.97036 | 0.328201 | 0.303506 | 0.898538 |
| ENSMUSG( | Mvb12a | 664.7666 | -0.09343 | 0.428935 | 0.826446 | ENSMUSG( | Mvb12a | 632.8693 | 0.036398 | 0.769794 | 0.974236 |
| ENSMUSG( | Mvb12b | 7198.989 | 0.141471 | 0.09405 | 0.471556 | ENSMUSG( | Mvb12b | 6458.319 | 0.100164 | 0.357676 | 0.913759 |
| ENSMUSG( | Mvd | 1020.227 | 0.138016 | 0.190207 | 0.629268 | ENSMUSG( | Mvd | 809.1458 | -0.27362 | 0.020928 | 0.571913 |
| ENSMUSG( | Mvk | 1253.552 | -0.0285 | 0.797562 | 0.960507 | ENSMUSG( | Mvk | 1046.3 | -0.29676 | 0.015147 | 0.492251 |
| ENSMUSG( | Mvp | 337.2113 | 0.003947 | 0.976589 | 0.997428 | ENSMUSG( | Mvp | 312.2442 | 0.051021 | 0.731089 | 0.967228 |
| ENSMUSG( | Mx1 | 20.57052 | -0.68876 | 0.095615 | NA | ENSMUSG( | Mx1 | 18.36687 | -0.79774 | 0.088349 | 0.786756 |
| ENSMUSG( | Mx2 | 54.21675 | -0.07943 | 0.759924 | 0.952768 | ENSMUSG( | Mx2 | 50.8292 | 0.026923 | 0.926034 | 0.99306 |
| ENSMUSG( | Mxd1 | 767.7603 | -0.02003 | 0.8546 | 0.973633 | ENSMUSG( | Mxd1 | 749.066 | 0.17461 | 0.162342 | 0.837724 |
| ENSMUSG( | Mxd3 | 41.09395 | -0.19976 | 0.499452 | 0.863218 | ENSMUSG( | Mxd3 | 34.59309 | -0.46075 | 0.18424 | 0.852665 |
| ENSMUSG( | Mxd4 | 6256.326 | 0.307252 | 0.000698 | 0.025426 | ENSMUSG( | Mxd4 | 5555.724 | 0.24114 | 0.021564 | 0.575341 |
| ENSMUSG( | Mxi1 | 5590.577 | 0.262916 | 0.002186 | 0.054296 | ENSMUSG( | Mxi1 | 4910.623 | 0.162488 | 0.214545 | 0.864547 |
| ENSMUSG( | Mxra7 | 612.6865 | -0.17066 | 0.199521 | 0.640054 | ENSMUSG( | Mxra7 | 617.0357 | 0.126466 | 0.405589 | 0.917257 |
| ENSMUSG( | Mxra8 | 381.1543 | -0.33026 | 0.018376 | 0.203658 | ENSMUSG( | Mxra8 | 390.4347 | 0.03624 | 0.830458 | 0.980463 |
| ENSMUSG( | Myadm | 2738.807 | -0.45692 | 0.003588 | 0.075902 | ENSMUSG( | Myadm | 3124.395 | 0.222861 | 0.199002 | 0.863016 |
| ENSMUSG( | Myadml2 | 145.4905 | -0.50546 | 0.015177 | 0.181362 | ENSMUSG( | Myadml2 | 153.3138 | -0.03316 | 0.874795 | 0.989644 |
| ENSMUSG( | Myb | 22.9266 | 0.02043 | 0.960259 | 0.993991 | ENSMUSG( | Myb | 29.85817 | 0.938631 | 0.018924 | 0.548832 |
| ENSMUSG( | Mybbp1a | 1865.336 | 0.105688 | 0.265456 | 0.716109 | ENSMUSG( | Mybbp1a | 1616.673 | -0.03867 | 0.736548 | 0.967344 |
| ENSMUSG( | Mybl1 | 42.5522 | -0.40083 | 0.172076 | 0.607115 | ENSMUSG( | Mybl1 | 50.57538 | 0.38473 | 0.182086 | 0.850579 |
| ENSMUSG( | Mybl2 | 44.95977 | 0.454633 | 0.110222 | 0.500785 | ENSMUSG( | Mybl2 | 41.26144 | 0.495469 | 0.131045 | 0.822884 |
| ENSMUSG( | Mybpc1 | 66.39989 | 0.18092 | 0.463124 | 0.842422 | ENSMUSG( | Mybpc1 | 74.2738 | 0.682172 | 0.010382 | 0.41614 |
| ENSMUSG( | Mybpc2 | 10.18896 | -0.51888 | 0.39741 | NA | ENSMUSG( | Mybpc2 | 10.66695 | -0.02436 | 0.968335 | NA |
| ENSMUSG( | Mybpc3 | 45.11729 | -0.04477 | 0.895064 | 0.984512 | ENSMUSG( | Mybpc3 | 42.10724 | 0.046853 | 0.891029 | 0.990558 |
| ENSMUSG( | Mybph | 40.09751 | -0.08645 | 0.803962 | 0.962169 | ENSMUSG( | Mybph | 39.92689 | 0.176225 | 0.591949 | 0.944975 |
| ENSMUSG( | Myc | 98.30413 | -0.14021 | 0.531844 | 0.879628 | ENSMUSG( | Myc | 92.72403 | -0.01557 | 0.958904 | 0.994798 |
| ENSMUSG( | Mycbp | 156.9144 | 0.054895 | 0.737762 | 0.947329 | ENSMUSG( | Mycbp | 144.0338 | 0.067613 | 0.731384 | 0.967228 |
| ENSMUSG( | Mycbp2 | 7879.573 | -0.04811 | 0.658699 | 0.924506 | ENSMUSG( | Mycbp2 | 7748.458 | 0.17066 | 0.184924 | 0.8547 |
| ENSMUSG( | Mycbpap | 53.50997 | 0.055721 | 0.862353 | 0.975055 | ENSMUSG( | Mycbpap | 51.9012 | 0.233674 | 0.459756 | 0.923329 |
| ENSMUSG( | Mycl | 591.9509 | 0.058618 | 0.634371 | 0.914073 | ENSMUSG( | Mycl | 551.6669 | 0.128381 | 0.408634 | 0.917257 |
| ENSMUSG( | Mycn | 204.0116 | -0.25952 | 0.127919 | 0.536163 | ENSMUSG( | Mycn | 197.0082 | -0.08381 | 0.672497 | 0.95694 |
| ENSMUSG( | Myct1 | 28.80068 | -0.08908 | 0.838494 | 0.971111 | ENSMUSG( | Myct1 | 32.14348 | 0.492022 | 0.269054 | 0.886755 |
| ENSMUSG( | Myd88 | 225.1068 | -0.01245 | 0.932357 | 0.992515 | ENSMUSG( | Myd88 | 200.6137 | -0.06756 | 0.715442 | 0.96507 |
| ENSMUSG( | Mydgf | 1581.615 | -0.36664 | 0.005486 | 0.0993 | ENSMUSG( | Mydgf | 1614.245 | -0.01226 | 0.932614 | 0.99306 |
| ENSMUSG( | Myef2 | 1723.905 | 0.008557 | 0.927857 | 0.991229 | ENSMUSG( | Myef2 | 1569.386 | 0.003317 | 0.977482 | 0.996416 |
| ENSMUSG( | Myef2l | 71.61028 | -0.03289 | 0.894487 | 0.984512 | ENSMUSG( | Myef2l | 63.17561 | -0.13946 | 0.602217 | 0.94789 |
| ENSMUSG( | Myg1 | 613.0842 | -0.03507 | 0.760402 | 0.952907 | ENSMUSG( | Myg1 | 555.4963 | -0.04788 | 0.728436 | 0.966913 |
| ENSMUSG( | Myh1 | 13.5193 | 0.210609 | 0.686974 | NA | ENSMUSG( | Myh1 | 12.26342 | 0.224497 | 0.699953 | 0.961335 |
| ENSMUSG( | Myh10 | 5773.079 | -0.03799 | 0.663489 | 0.9265 | ENSMUSG( | Myh10 | 5283.529 | -0.02408 | 0.830521 | 0.980472 |
| ENSMUSG( | Myh11 | 42.42265 | 0.207731 | 0.484013 | 0.856501 | ENSMUSG( | Myh11 | 47.28998 | 0.725436 | 0.01763 | 0.53154 |
| ENSMUSG( | Myh13 | 16.25858 | -0.11915 | 0.806595 | NA | ENSMUSG( | Myh13 | 13.41836 | -0.43475 | 0.415886 | 0.917257 |
| ENSMUSG( | Myh14 | 3591.742 | -0.02267 | 0.851258 | 0.972764 | ENSMUSG( | Myh14 | 3017.31 | -0.27083 | 0.012403 | 0.451463 |
| ENSMUSG( | Myh15 | 374.9717 | 0.252117 | 0.103272 | 0.488788 | ENSMUSG( | Myh15 | 273.4248 | -0.42306 | 0.014727 | 0.487333 |
| ENSMUSG( | Myh2 | 23.03713 | -0.03259 | 0.937684 | 0.993415 | ENSMUSG( | Myh2 | 17.78 | -0.55363 | 0.245047 | 0.877904 |
| ENSMUSG( | Myh3 | 88.68199 | 0.007362 | 0.976727 | 0.997428 | ENSMUSG( | Myh3 | 71.28273 | -0.38235 | 0.166254 | 0.840082 |
| ENSMUSG( | Myh4 | 9.389255 | -0.15698 | 0.816526 | NA | ENSMUSG( | Myh4 | 8.609076 | -0.11014 | 0.878795 | NA |
| ENSMUSG( | Myh7 | 658.3241 | 0.068243 | 0.571845 | 0.893506 | ENSMUSG( | Myh7 | 552.6989 | -0.17387 | 0.195481 | 0.861579 |
| ENSMUSG( | Myh7b | 241.7255 | 0.093856 | 0.597728 | 0.90372 | ENSMUSG( | Myh7b | 198.6922 | -0.21716 | 0.249807 | 0.877904 |
| ENSMUSG( | Myh8 | 23.22132 | 0.176612 | 0.650094 | 0.921 | ENSMUSG( | Myh8 | 19.66993 | -0.08687 | 0.85868 | 0.985884 |
| ENSMUSG( | Myh9 | 3729.737 | 0.10393 | 0.267876 | 0.718054 | ENSMUSG( | Myh9 | 3386.248 | 0.09612 | 0.40567 | 0.917257 |
| ENSMUSG( | Myl1 | 15.59598 | 0.031878 | 0.953286 | NA | ENSMUSG( | Myl1 | 19.31087 | 0.795781 | 0.095325 | 0.796133 |
| ENSMUSG( | Myl12a | 400.8588 | 0.016652 | 0.916233 | 0.988776 | ENSMUSG( | Myl12a | 393.1038 | 0.230543 | 0.201156 | 0.86439 |
| ENSMUSG( | Myl12b | 220.279 | 0.168602 | 0.287806 | 0.738524 | ENSMUSG( | Myl12b | 203.9117 | 0.195078 | 0.348456 | 0.911652 |
| ENSMUSG( | Myl2 | 35.75015 | 0.029606 | 0.94208 | 0.993415 | ENSMUSG( | Myl2 | 24.82789 | -0.91912 | 0.058633 | 0.728256 |
| ENSMUSG( | Myl4 | 638.373 | -0.13414 | 0.223996 | 0.669869 | ENSMUSG( | Myl4 | 529.5524 | -0.44195 | 0.001508 | 0.143602 |
| ENSMUSG( | Myl6 | 6642.117 | -0.03152 | 0.708102 | 0.939519 | ENSMUSG( | Myl6 | 5866.521 | -0.1245 | 0.253316 | 0.88018 |
| ENSMUSG( | Myl6b | 643.6299 | 0.074377 | 0.483845 | 0.856485 | ENSMUSG( | Myl6b | 525.6078 | -0.25447 | 0.048708 | 0.706195 |
| ENSMUSG( | Myl9 | 385.7028 | -0.22259 | 0.113505 | 0.508017 | ENSMUSG( | Myl9 | 395.2635 | 0.132434 | 0.488324 | 0.925453 |
| ENSMUSG( | Mylip | 473.2269 | 0.085761 | 0.44524 | 0.833239 | ENSMUSG( | Mylip | 445.9036 | 0.181003 | 0.183391 | 0.850984 |
| ENSMUSG( | Mylk | 725.5887 | -0.18392 | 0.161011 | 0.59072 | ENSMUSG( | Mylk | 687.1605 | -0.0694 | 0.583868 | 0.944379 |
| ENSMUSG( | Mylk3 | 43.7047 | -0.45722 | 0.144833 | 0.564979 | ENSMUSG( | Mylk3 | 39.81959 | -0.47505 | 0.160286 | 0.837724 |
| ENSMUSG( | Mylpf | 31.89739 | 0.17453 | 0.620947 | 0.911432 | ENSMUSG( | Mylpf | 27.02592 | -0.00834 | 0.983062 | 0.996891 |
| ENSMUSG( | Mymk | 15.49446 | -0.16395 | 0.758817 | NA | ENSMUSG( | Mymk | 14.98569 | -0.00559 | 0.991208 | 0.998389 |
| ENSMUSG( | Mynn | 416.7672 | 0.017059 | 0.886869 | 0.983255 | ENSMUSG( | Mynn | 385.0105 | 0.058597 | 0.665784 | 0.956766 |
| ENSMUSG( | Myo10 | 3899 | -0.07309 | 0.482668 | 0.856083 | ENSMUSG( | Myo10 | 3561.131 | -0.06707 | 0.570299 | 0.941823 |
| ENSMUSG( | Myo15a | 11.03824 | 0.306843 | 0.580762 | NA | ENSMUSG( | Myo15a | 10.52622 | 0.43762 | 0.461619 | NA |
| ENSMUSG( | Myo15b | 17.3625 | 0.113293 | 0.836311 | NA | ENSMUSG( | Myo15b | 14.12428 | -0.23423 | 0.704404 | 0.962677 |
| ENSMUSG( | Myo16 | 846.0986 | 0.039959 | 0.744087 | 0.948497 | ENSMUSG( | Myo16 | 791.3535 | 0.112593 | 0.429387 | 0.917257 |
| ENSMUSG( | Myo18a | 10149.86 | 0.132087 | 0.092128 | 0.466065 | ENSMUSG( | Myo18a | 8846.09 | 0.007433 | 0.943175 | 0.994798 |
| ENSMUSG( | Myo18b | 175.0599 | 0.171795 | 0.280153 | 0.729155 | ENSMUSG( | Myo18b | 133.4727 | -0.34437 | 0.113606 | 0.809623 |
| ENSMUSG( | Myo19 | 319.3044 | 0.264459 | 0.091536 | 0.465281 | ENSMUSG( | Myo19 | 290.3315 | 0.267015 | 0.164009 | 0.838291 |
| ENSMUSG( | Myo1a | 32.26701 | 0.26684 | 0.418096 | 0.821966 | ENSMUSG( | Myo1a | 31.46312 | 0.399648 | 0.364157 | 0.914223 |
| ENSMUSG( | Myo1b | 872.9851 | -0.19427 | 0.046995 | 0.331741 | ENSMUSG( | Myo1b | 845.3391 | -0.00718 | 0.955652 | 0.994798 |
| ENSMUSG( | Myo1c | 608.0296 | 0.001526 | 0.990673 | 0.999232 | ENSMUSG( | Myo1c | 620.765 | 0.321069 | 0.044625 | 0.693697 |
| ENSMUSG( | Myo1d | 382.3968 | -0.22718 | 0.062815 | 0.387645 | ENSMUSG( | Myo1d | 352.3213 | -0.18099 | 0.286091 | 0.893906 |
| ENSMUSG( | Myo1e | 751.5241 | 0.094543 | 0.350538 | 0.782297 | ENSMUSG( | Myo1e | 685.6193 | 0.099604 | 0.402401 | 0.917257 |
| ENSMUSG( | Myo1f | 235.7332 | -0.27049 | 0.112836 | 0.506698 | ENSMUSG( | Myo1f | 212.772 | -0.29743 | 0.147478 | 0.836241 |
| ENSMUSG( | Myo1g | 72.7606 | -0.76527 | 0.072814 | 0.416499 | ENSMUSG( | Myo1g | 65.95841 | -0.81906 | 0.068372 | 0.752306 |
| ENSMUSG( | Myo1h | 49.75884 | 0.006517 | 0.981432 | 0.997512 | ENSMUSG( | Myo1h | 51.679 | 0.376614 | 0.217556 | 0.865589 |
| ENSMUSG( | Myo3b | 47.42157 | -0.49426 | 0.084494 | 0.448139 | ENSMUSG( | Myo3b | 48.67359 | -0.11257 | 0.705083 | 0.96281 |
| ENSMUSG( | Myo5a | 15098.83 | 0.039541 | 0.683541 | 0.932154 | ENSMUSG( | Myo5a | 14059.64 | 0.102468 | 0.35619 | 0.913358 |
| ENSMUSG( | Myo5b | 905.2861 | 0.092682 | 0.405882 | 0.81652 | ENSMUSG( | Myo5b | 828.5518 | 0.10678 | 0.408403 | 0.917257 |
| ENSMUSG( | Myo5c | 93.4765 | -0.35721 | 0.125128 | 0.532139 | ENSMUSG( | Myo5c | 77.5817 | -0.66401 | 0.011454 | 0.436431 |
| ENSMUSG( | Myo6 | 17257.55 | 0.078512 | 0.55408 | 0.886397 | ENSMUSG( | Myo6 | 14263.87 | -0.21116 | 0.148387 | 0.836241 |
| ENSMUSG( | Myo7a | 954.9785 | 0.170885 | 0.108775 | 0.497919 | ENSMUSG( | Myo7a | 828.2654 | 0.03759 | 0.779399 | 0.976106 |
| ENSMUSG( | Myo9a | 2707.819 | -0.125 | 0.195759 | 0.635109 | ENSMUSG( | Myo9a | 2633.563 | 0.067227 | 0.543662 | 0.938063 |
| ENSMUSG( | Myo9b | 2656.803 | 0.17871 | 0.062339 | 0.386294 | ENSMUSG( | Myo9b | 2307.131 | 0.045101 | 0.704582 | 0.962677 |
| ENSMUSG( | Myoc | 1215.729 | -0.70948 | 0.034998 | 0.28772 | ENSMUSG( | Myoc | 1520.193 | 0.284868 | 0.23525 | 0.871741 |
| ENSMUSG( | Myof | 127.5198 | -0.24586 | 0.171231 | 0.605749 | ENSMUSG( | Myof | 126.3297 | 0.014835 | 0.941954 | 0.994708 |
| ENSMUSG( | Myom1 | 85.80991 | 0.190043 | 0.459797 | 0.840338 | ENSMUSG( | Myom1 | 75.89581 | 0.111947 | 0.647833 | 0.95422 |
| ENSMUSG( | Myom3 | 703.8964 | 0.063506 | 0.557466 | 0.888279 | ENSMUSG( | Myom3 | 592.2886 | -0.17734 | 0.220397 | 0.866104 |
| ENSMUSG( | Myorg | 3466.699 | -0.13524 | 0.368151 | 0.795774 | ENSMUSG( | Myorg | 3487.597 | 0.154632 | 0.374617 | 0.915491 |
| ENSMUSG( | Mypn | 84.83876 | -0.14934 | 0.550327 | 0.886038 | ENSMUSG( | Mypn | 76.43552 | -0.1859 | 0.47103 | 0.923329 |
| ENSMUSG( | Mypop | 1645.391 | 0.086379 | 0.4157 | 0.82084 | ENSMUSG( | Mypop | 1417.361 | -0.07718 | 0.532606 | 0.934795 |
| ENSMUSG( | Myrf | 1981.162 | -0.44825 | 0.002424 | 0.057903 | ENSMUSG( | Myrf | 2202.407 | 0.162059 | 0.428228 | 0.917257 |
| ENSMUSG( | Myrip | 3998.84 | 0.311196 | 8.26E-05 | 0.005675 | ENSMUSG( | Myrip | 3346.983 | 0.080938 | 0.423105 | 0.917257 |
| ENSMUSG( | Mysm1 | 762.6974 | -0.1625 | 0.172281 | 0.607345 | ENSMUSG( | Mysm1 | 760.6732 | 0.106529 | 0.464948 | 0.923329 |
| ENSMUSG( | Myt1 | 343.0612 | -0.30333 | 0.029072 | 0.259547 | ENSMUSG( | Myt1 | 354.05 | 0.073669 | 0.616389 | 0.951413 |
| ENSMUSG( | Myt1l | 4234.733 | -0.02063 | 0.823807 | 0.966603 | ENSMUSG( | Myt1l | 4238.052 | 0.241891 | 0.024062 | 0.588226 |
| ENSMUSG( | Myzap | 38.12431 | 0.474726 | 0.134451 | 0.547664 | ENSMUSG( | Myzap | 32.30721 | 0.301014 | 0.399488 | 0.917257 |
| ENSMUSG( | Mzf1 | 361.4612 | -0.01922 | 0.888855 | 0.983434 | ENSMUSG( | Mzf1 | 357.1441 | 0.213866 | 0.164378 | 0.83856 |
| ENSMUSG( | Mzt1 | 1602.635 | -0.0261 | 0.819533 | 0.965888 | ENSMUSG( | Mzt1 | 1435.854 | -0.07987 | 0.591275 | 0.944975 |
| ENSMUSG( | Mzt2 | 479.128 | -0.00746 | 0.952086 | 0.993821 | ENSMUSG( | Mzt2 | 398.9412 | -0.28981 | 0.072444 | 0.763629 |
| ENSMUSG( | N4bp1 | 1287.715 | 0.082433 | 0.361775 | 0.792797 | ENSMUSG( | N4bp1 | 1173.846 | 0.083798 | 0.454453 | 0.922444 |
| ENSMUSG( | N4bp2 | 571.6117 | 0.107548 | 0.406151 | 0.816525 | ENSMUSG( | N4bp2 | 534.0399 | 0.177279 | 0.174949 | 0.845107 |

| | | | | | |
|---|---|---|---|---|---|
| ENSMUSG( | 170.349 | 0.262337 | 0.105346 | 0.492417 | Mtrf1l |
| ENSMUSG( | 339.4689 | 0.222433 | 0.114578 | 0.510543 | Mtrfr |
| ENSMUSG( | 44.50001 | -0.16822 | 0.588619 | 0.900921 | Mtrnr2l7 |
| ENSMUSG( | 313.4918 | 0.239237 | 0.063038 | 0.388234 | Mtrr |
| ENSMUSG( | 1001.136 | -0.09199 | 0.369646 | 0.797188 | Mtss1 |
| ENSMUSG( | 32260.96 | -0.04134 | 0.690041 | 0.933795 | Mtss2 |
| ENSMUSG( | 378.6066 | -0.12726 | 0.373781 | 0.80044 | Mttp |
| ENSMUSG( | 8832.501 | 0.169397 | 0.040651 | 0.311461 | Mturn |
| ENSMUSG( | 1519.501 | -0.01497 | 0.889009 | 0.983434 | Mtus1 |
| ENSMUSG( | 1119.196 | 0.221967 | 0.045207 | 0.326331 | Mtus2 |
| ENSMUSG( | 687.3685 | -0.00126 | 0.991146 | 0.999232 | Mtx1 |
| ENSMUSG( | 512.8697 | -0.09389 | 0.493935 | 0.860611 | Mtx2 |
| ENSMUSG( | 1660.706 | 0.093582 | 0.340893 | 0.778773 | Mtx3 |
| ENSMUSG( | 72.71838 | 0.025364 | 0.912171 | 0.988483 | Muc1 |
| ENSMUSG( | 15.25472 | -0.09946 | 0.834906 | NA | Muc2 |
| ENSMUSG( | 268.632 | 0.257844 | 0.309737 | 0.757658 | Muc3a |
| ENSMUSG( | 1347.173 | -0.07608 | 0.488894 | 0.859092 | Mul1 |
| ENSMUSG( | 587.8447 | 0.038124 | 0.727471 | 0.944344 | Mus81 |
| ENSMUSG( | 24.77437 | -0.32234 | 0.488709 | 0.859092 | Musk |
| ENSMUSG( | 38.33177 | 0.037149 | 0.908516 | 0.987809 | Mustn1 |
| ENSMUSG( | 59.93062 | 0.097084 | 0.703284 | 0.937735 | Mutyh |
| ENSMUSG( | 664.7666 | -0.09343 | 0.428935 | 0.826446 | Mvb12a |
| ENSMUSG( | 7198.989 | 0.141471 | 0.09405 | 0.471556 | Mvb12b |
| ENSMUSG( | 1020.227 | 0.138016 | 0.190207 | 0.629268 | Mvd |
| ENSMUSG( | 1253.552 | -0.0285 | 0.797562 | 0.960507 | Mvk |
| ENSMUSG( | 337.2113 | 0.003947 | 0.976589 | 0.997428 | Mvp |
| ENSMUSG( | 20.57052 | -0.68876 | 0.095615 | NA | Mx1 |
| ENSMUSG( | 54.21675 | -0.07943 | 0.759924 | 0.952768 | Mx2 |
| ENSMUSG( | 767.7603 | -0.02003 | 0.8546 | 0.973633 | Mxd1 |
| ENSMUSG( | 41.09395 | -0.19976 | 0.499452 | 0.863218 | Mxd3 |
| ENSMUSG( | 6256.326 | 0.307252 | 0.000698 | 0.025426 | Mxd4 |
| ENSMUSG( | 5590.577 | 0.262916 | 0.002186 | 0.054296 | Mxi1 |
| ENSMUSG( | 612.6865 | -0.17066 | 0.199521 | 0.640054 | Mxra7 |
| ENSMUSG( | 381.1543 | -0.33026 | 0.018376 | 0.203658 | Mxra8 |
| ENSMUSG( | 2738.807 | -0.45692 | 0.003588 | 0.075902 | Myadm |
| ENSMUSG( | 145.4905 | -0.50546 | 0.015177 | 0.181362 | Myadml2 |
| ENSMUSG( | 22.9266 | 0.02043 | 0.960259 | 0.993991 | Myb |
| ENSMUSG( | 1865.336 | 0.105688 | 0.265456 | 0.716109 | Mybbp1a |
| ENSMUSG( | 42.5522 | -0.40083 | 0.172076 | 0.607115 | Mybl1 |
| ENSMUSG( | 44.95977 | 0.454633 | 0.110222 | 0.500785 | Mybl2 |
| ENSMUSG( | 66.39989 | 0.18092 | 0.463124 | 0.842422 | Mybpc1 |
| ENSMUSG( | 10.18896 | -0.51888 | 0.39741 | NA | Mybpc2 |
| ENSMUSG( | 45.11729 | -0.04477 | 0.895064 | 0.984512 | Mybpc3 |
| ENSMUSG( | 40.09751 | -0.08645 | 0.803962 | 0.962169 | Mybph |
| ENSMUSG( | 98.30413 | -0.14021 | 0.531844 | 0.879628 | Myc |
| ENSMUSG( | 156.9144 | 0.054895 | 0.737762 | 0.947329 | Mycbp |
| ENSMUSG( | 7879.573 | -0.04811 | 0.658699 | 0.924506 | Mycbp2 |
| ENSMUSG( | 53.50997 | 0.055721 | 0.862353 | 0.975055 | Mycbpap |
| ENSMUSG( | 591.9509 | 0.058618 | 0.634371 | 0.914073 | Mycl |
| ENSMUSG( | 204.0116 | -0.25952 | 0.127919 | 0.536163 | Mycn |
| ENSMUSG( | 28.80068 | -0.08908 | 0.838494 | 0.971111 | Myct1 |
| ENSMUSG( | 225.1068 | -0.01245 | 0.932357 | 0.992515 | Myd88 |
| ENSMUSG( | 1581.615 | -0.36664 | 0.005486 | 0.0993 | Mydgf |
| ENSMUSG( | 1723.905 | 0.008557 | 0.927857 | 0.991229 | Myef2 |
| ENSMUSG( | 71.61028 | -0.03289 | 0.894487 | 0.984512 | Myef2l |
| ENSMUSG( | 613.0842 | -0.03507 | 0.760402 | 0.952907 | Myg1 |
| ENSMUSG( | 13.5193 | 0.210609 | 0.686974 | NA | Myh1 |
| ENSMUSG( | 5773.079 | -0.03799 | 0.663489 | 0.9265 | Myh10 |
| ENSMUSG( | 42.42265 | 0.207731 | 0.484013 | 0.856501 | Myh11 |
| ENSMUSG( | 16.25858 | -0.11915 | 0.806595 | NA | Myh13 |
| ENSMUSG( | 3591.742 | -0.02267 | 0.851258 | 0.972764 | Myh14 |
| ENSMUSG( | 374.9717 | 0.252117 | 0.103272 | 0.488788 | Myh15 |
| ENSMUSG( | 23.03713 | -0.03259 | 0.937684 | 0.993415 | Myh2 |
| ENSMUSG( | 88.68199 | 0.007362 | 0.976727 | 0.997428 | Myh3 |
| ENSMUSG( | 9.389255 | -0.15698 | 0.816526 | NA | Myh4 |
| ENSMUSG( | 658.3241 | 0.068243 | 0.571845 | 0.893506 | Myh7 |
| ENSMUSG( | 241.7255 | 0.093856 | 0.597728 | 0.90372 | Myh7b |
| ENSMUSG( | 23.22132 | 0.176612 | 0.650094 | 0.921 | Myh8 |
| ENSMUSG( | 3729.737 | 0.10393 | 0.267876 | 0.718054 | Myh9 |
| ENSMUSG( | 8.917615 | 0.798303 | 0.254824 | NA | Myhas |
| ENSMUSG( | 15.59598 | 0.031878 | 0.953286 | NA | Myl1 |
| ENSMUSG( | 400.8588 | 0.016652 | 0.916233 | 0.988776 | Myl12a |
| ENSMUSG( | 220.279 | 0.168602 | 0.287806 | 0.738524 | Myl12b |
| ENSMUSG( | 35.75015 | 0.029606 | 0.94208 | 0.993415 | Myl2 |
| ENSMUSG( | 638.373 | -0.13414 | 0.223996 | 0.669869 | Myl4 |
| ENSMUSG( | 6642.117 | -0.03152 | 0.708102 | 0.939519 | Myl6 |
| ENSMUSG( | 643.6299 | 0.074377 | 0.483845 | 0.856485 | Myl6b |
| ENSMUSG( | 385.7028 | -0.22259 | 0.113505 | 0.508017 | Myl9 |
| ENSMUSG( | 473.2269 | 0.085761 | 0.44524 | 0.833239 | Mylip |
| ENSMUSG( | 725.5887 | -0.18392 | 0.161011 | 0.59072 | Mylk |
| ENSMUSG( | 43.7047 | -0.45722 | 0.144833 | 0.564979 | Mylk3 |
| ENSMUSG( | 31.89739 | 0.17453 | 0.620947 | 0.911432 | Mylpf |
| ENSMUSG( | 15.49446 | -0.16395 | 0.758817 | NA | Mymk |
| ENSMUSG( | 416.7672 | 0.017059 | 0.886869 | 0.983255 | Mynn |
| ENSMUSG( | 3899 | -0.07309 | 0.482668 | 0.856083 | Myo10 |
| ENSMUSG( | 11.03824 | 0.306843 | 0.580762 | NA | Myo15a |
| ENSMUSG( | 17.3625 | 0.113293 | 0.836311 | NA | Myo15b |
| ENSMUSG( | 846.0986 | 0.039959 | 0.744087 | 0.948497 | Myo16 |
| ENSMUSG( | 10149.86 | 0.132087 | 0.092128 | 0.466065 | Myo18a |
| ENSMUSG( | 175.0599 | 0.171795 | 0.280153 | 0.729155 | Myo18b |
| ENSMUSG( | 319.3044 | 0.264459 | 0.091536 | 0.465281 | Myo19 |
| ENSMUSG( | 32.26701 | 0.26684 | 0.418096 | 0.821966 | Myo1a |
| ENSMUSG( | 872.9851 | -0.19427 | 0.046995 | 0.331741 | Myo1b |
| ENSMUSG( | 608.0296 | 0.001526 | 0.990673 | 0.999232 | Myo1c |
| ENSMUSG( | 382.3968 | -0.22718 | 0.062815 | 0.387645 | Myo1d |
| ENSMUSG( | 751.5241 | 0.094543 | 0.350538 | 0.782297 | Myo1e |
| ENSMUSG( | 235.7332 | -0.27049 | 0.112836 | 0.506698 | Myo1f |
| ENSMUSG( | 72.7606 | -0.76527 | 0.072814 | 0.416499 | Myo1g |
| ENSMUSG( | 49.75884 | 0.006517 | 0.981432 | 0.997512 | Myo1h |
| ENSMUSG( | 47.42157 | -0.49426 | 0.084494 | 0.448139 | Myo3b |
| ENSMUSG( | 15098.83 | 0.039541 | 0.683541 | 0.932154 | Myo5a |
| ENSMUSG( | 905.2861 | 0.092682 | 0.405882 | 0.81652 | Myo5b |
| ENSMUSG( | 93.4765 | -0.35721 | 0.125128 | 0.532139 | Myo5c |
| ENSMUSG( | 17257.55 | 0.078512 | 0.55408 | 0.886397 | Myo6 |
| ENSMUSG( | 954.9785 | 0.170885 | 0.108775 | 0.497919 | Myo7a |
| ENSMUSG( | 2707.819 | -0.125 | 0.195759 | 0.635109 | Myo9a |
| ENSMUSG( | 2656.803 | 0.17871 | 0.062339 | 0.386294 | Myo9b |
| ENSMUSG( | 1215.729 | -0.70948 | 0.034998 | 0.28772 | Myoc |
| ENSMUSG( | 127.5198 | -0.24586 | 0.171231 | 0.605749 | Myof |
| ENSMUSG( | 85.80991 | 0.190043 | 0.459797 | 0.840338 | Myom1 |
| ENSMUSG( | 703.8964 | 0.063506 | 0.557466 | 0.888279 | Myom3 |
| ENSMUSG( | 3466.699 | -0.13524 | 0.368151 | 0.795774 | Myorg |
| ENSMUSG( | 84.83876 | -0.14934 | 0.550327 | 0.886038 | Mypn |
| ENSMUSG( | 1645.391 | 0.086379 | 0.4157 | 0.82084 | Mypop |
| ENSMUSG( | 1981.162 | -0.44825 | 0.002424 | 0.057903 | Myrf |
| ENSMUSG( | 3998.84 | 0.311196 | 8.26E-05 | 0.005675 | Myrip |
| ENSMUSG( | 762.6974 | -0.1625 | 0.172281 | 0.607345 | Mysm1 |
| ENSMUSG( | 343.0612 | -0.30333 | 0.029072 | 0.259547 | Myt1 |
| ENSMUSG( | 4234.733 | -0.02063 | 0.823807 | 0.966603 | Myt1l |
| ENSMUSG( | 38.12431 | 0.474726 | 0.134451 | 0.547664 | Myzap |
| ENSMUSG( | 361.4612 | -0.01922 | 0.888855 | 0.983434 | Mzf1 |
| ENSMUSG( | 1602.635 | -0.0261 | 0.819533 | 0.965888 | Mzt1 |
| ENSMUSG( | 479.128 | -0.00746 | 0.952086 | 0.993821 | Mzt2 |
| ENSMUSG( | 1287.715 | 0.082433 | 0.361775 | 0.792797 | N4bp1 |
| ENSMUSG( | 571.6117 | 0.107548 | 0.406151 | 0.816525 | N4bp2 |
| ENSMUSG( | 960.1676 | 0.029643 | 0.785613 | 0.95867 | N4bp2l1 |
| ENSMUSG( | 745.6135 | 0.016916 | 0.885145 | 0.982851 | N4bp2l2 |
| ENSMUSG( | 297.9273 | 0.05894 | 0.739487 | 0.947768 | N4bp3 |
| ENSMUSG( | 456.1469 | -0.22727 | 0.121804 | 0.526562 | N6amt1 |
| ENSMUSG( | 84.19223 | -0.1168 | 0.607065 | 0.906685 | NA |
| ENSMUSG( | 66.67283 | 0.171143 | 0.467619 | 0.844819 | NA |
| ENSMUSG( | 15.18749 | -0.38298 | 0.438301 | NA | NA |
| ENSMUSG( | 117.682 | 0.105242 | 0.572882 | 0.893584 | NA |
| ENSMUSG( | 381.086 | -0.00876 | 0.941913 | 0.993415 | NA |
| ENSMUSG( | 35.56141 | -0.12291 | 0.694619 | 0.9343 | NA |
| ENSMUSG( | 10.63855 | 0.165199 | 0.776223 | NA | NA |
| ENSMUSG( | 24.27333 | -0.17432 | 0.648227 | 0.919949 | NA |
| ENSMUSG( | 40.44742 | 0.152486 | 0.65632 | 0.923929 | NA |
| ENSMUSG( | 81.53609 | 0.152784 | 0.514746 | 0.872189 | NA |
| ENSMUSG( | 25.78109 | 0.231154 | 0.540665 | 0.88315 | NA |
| ENSMUSG( | 13.22371 | -0.14373 | 0.77824 | NA | NA |
| ENSMUSG( | 36.14783 | 0.743383 | 0.028463 | 0.257525 | NA |
| ENSMUSG( | 95.48916 | 0.006463 | 0.979146 | 0.997428 | NA |
| ENSMUSG( | 8.529076 | 0.141618 | 0.821496 | NA | NA |
| ENSMUSG( | 19.81136 | -0.4762 | 0.320379 | NA | NA |
| ENSMUSG( | 50.75926 | 0.503441 | 0.070441 | 0.40908 | NA |
| ENSMUSG( | 21.43278 | -0.12013 | 0.770079 | 0.954689 | NA |
| ENSMUSG( | 11.86057 | -0.06554 | 0.903141 | NA | NA |
| ENSMUSG( | 64.84578 | -0.15282 | 0.525842 | 0.876902 | NA |
| ENSMUSG( | 122.0201 | 0.331605 | 0.071841 | 0.414057 | NA |
| ENSMUSG( | 23.85162 | -0.70854 | 0.069685 | 0.408352 | NA |
| ENSMUSG( | 27.1535 | -0.37046 | 0.343584 | 0.779153 | NA |
| ENSMUSG( | 14.1576 | -0.237 | 0.641165 | NA | NA |
| ENSMUSG( | 21.15647 | 0.254 | 0.534884 | NA | NA |
| ENSMUSG( | 149.928 | -0.17008 | 0.329104 | 0.773233 | NA |
| ENSMUSG( | 15.92199 | -0.07691 | 0.876954 | NA | NA |
| ENSMUSG( | 53.48772 | -0.1366 | 0.599332 | 0.903801 | NA |
| ENSMUSG( | 110.4483 | 0.128896 | 0.500573 | 0.863514 | NA |

| | | | | | |
|---|---|---|---|---|---|
| ENSMUSG( | 1614.245 | -0.01226 | 0.932614 | 0.99306 | Mydgf |
| ENSMUSG( | 1569.386 | 0.003317 | 0.977482 | 0.996416 | Myef2 |
| ENSMUSG( | 63.17561 | -0.13946 | 0.602217 | 0.94789 | Myef2l |
| ENSMUSG( | 555.4963 | -0.04788 | 0.728436 | 0.966913 | Myg1 |
| ENSMUSG( | 12.26342 | 0.224497 | 0.699953 | 0.961335 | Myh1 |
| ENSMUSG( | 5283.529 | -0.02408 | 0.830521 | 0.980472 | Myh10 |
| ENSMUSG( | 47.28998 | 0.725436 | 0.01763 | 0.53154 | Myh11 |
| ENSMUSG( | 13.41836 | -0.43475 | 0.415886 | 0.917257 | Myh13 |
| ENSMUSG( | 3017.31 | -0.27083 | 0.012403 | 0.451463 | Myh14 |
| ENSMUSG( | 273.4248 | -0.42306 | 0.014727 | 0.487333 | Myh15 |
| ENSMUSG( | 17.78 | -0.55363 | 0.245047 | 0.877904 | Myh2 |
| ENSMUSG( | 71.28273 | -0.38235 | 0.166254 | 0.840082 | Myh3 |
| ENSMUSG( | 8.609076 | -0.11014 | 0.878795 | NA | Myh4 |
| ENSMUSG( | 552.6989 | -0.17387 | 0.195481 | 0.861579 | Myh7 |
| ENSMUSG( | 198.6922 | -0.21716 | 0.249807 | 0.877904 | Myh7b |
| ENSMUSG( | 19.66993 | -0.08687 | 0.85868 | 0.985884 | Myh8 |
| ENSMUSG( | 3386.248 | 0.09612 | 0.40567 | 0.917257 | Myh9 |
| ENSMUSG( | 19.31087 | 0.795781 | 0.095325 | 0.796133 | Myl1 |
| ENSMUSG( | 393.1038 | 0.230543 | 0.201156 | 0.86439 | Myl12a |
| ENSMUSG( | 203.9117 | 0.195078 | 0.348456 | 0.911652 | Myl12b |
| ENSMUSG( | 24.82789 | -0.91912 | 0.058633 | 0.728256 | Myl2 |
| ENSMUSG( | 529.5524 | -0.44195 | 0.001508 | 0.143602 | Myl4 |
| ENSMUSG( | 5866.521 | -0.1245 | 0.253316 | 0.88018 | Myl6 |
| ENSMUSG( | 525.6078 | -0.25447 | 0.048708 | 0.706195 | Myl6b |
| ENSMUSG( | 395.2635 | 0.132434 | 0.488324 | 0.925453 | Myl9 |
| ENSMUSG( | 445.9036 | 0.181003 | 0.183391 | 0.850984 | Mylip |
| ENSMUSG( | 687.1605 | -0.0694 | 0.583868 | 0.944379 | Mylk |
| ENSMUSG( | 39.81959 | -0.47505 | 0.160286 | 0.837724 | Mylk3 |
| ENSMUSG( | 11.05651 | -0.51315 | 0.396977 | NA | Mylk4 |
| ENSMUSG( | 27.02592 | -0.00834 | 0.983062 | 0.996891 | Mylpf |
| ENSMUSG( | 14.98569 | -0.00559 | 0.991208 | 0.998389 | Mymk |
| ENSMUSG( | 385.0105 | 0.058597 | 0.665784 | 0.956766 | Mynn |
| ENSMUSG( | 3561.131 | -0.06707 | 0.570299 | 0.941823 | Myo10 |
| ENSMUSG( | 10.52622 | 0.43762 | 0.461619 | NA | Myo15a |
| ENSMUSG( | 14.12428 | -0.23423 | 0.704404 | 0.962677 | Myo15b |
| ENSMUSG( | 791.3535 | 0.112593 | 0.429387 | 0.917257 | Myo16 |
| ENSMUSG( | 8846.09 | 0.007433 | 0.943175 | 0.994798 | Myo18a |
| ENSMUSG( | 133.4727 | -0.34437 | 0.113606 | 0.809623 | Myo18b |
| ENSMUSG( | 290.3315 | 0.267015 | 0.164009 | 0.838291 | Myo19 |
| ENSMUSG( | 31.46312 | 0.399648 | 0.364157 | 0.914223 | Myo1a |
| ENSMUSG( | 845.3391 | -0.00718 | 0.955652 | 0.994798 | Myo1b |
| ENSMUSG( | 620.765 | 0.321069 | 0.044625 | 0.693697 | Myo1c |
| ENSMUSG( | 352.3213 | -0.18099 | 0.286091 | 0.893906 | Myo1d |
| ENSMUSG( | 685.6193 | 0.099604 | 0.402401 | 0.917257 | Myo1e |
| ENSMUSG( | 212.772 | -0.29743 | 0.147478 | 0.836241 | Myo1f |
| ENSMUSG( | 65.95841 | -0.81906 | 0.068372 | 0.752306 | Myo1g |
| ENSMUSG( | 51.679 | 0.376614 | 0.217556 | 0.865589 | Myo1h |
| ENSMUSG( | 48.67359 | -0.11257 | 0.705083 | 0.96281 | Myo3b |
| ENSMUSG( | 14059.64 | 0.102468 | 0.35619 | 0.913358 | Myo5a |
| ENSMUSG( | 828.5518 | 0.10678 | 0.408403 | 0.917257 | Myo5b |
| ENSMUSG( | 77.5817 | -0.66401 | 0.011454 | 0.436431 | Myo5c |
| ENSMUSG( | 14263.87 | -0.21116 | 0.148387 | 0.836241 | Myo6 |
| ENSMUSG( | 828.2654 | 0.03759 | 0.779399 | 0.976106 | Myo7a |
| ENSMUSG( | 2633.563 | 0.067227 | 0.543662 | 0.938063 | Myo9a |
| ENSMUSG( | 2307.131 | 0.045101 | 0.704582 | 0.962677 | Myo9b |
| ENSMUSG( | 1520.193 | 0.284868 | 0.23525 | 0.871741 | Myoc |
| ENSMUSG( | 126.3297 | 0.014835 | 0.941954 | 0.994708 | Myof |
| ENSMUSG( | 75.89581 | 0.111947 | 0.647833 | 0.95422 | Myom1 |
| ENSMUSG( | 592.2886 | -0.17734 | 0.220397 | 0.866104 | Myom3 |
| ENSMUSG( | 3487.597 | 0.154632 | 0.374617 | 0.915491 | Myorg |
| ENSMUSG( | 76.43552 | -0.1859 | 0.47103 | 0.923329 | Mypn |
| ENSMUSG( | 1417.361 | -0.07718 | 0.532606 | 0.934795 | Mypop |
| ENSMUSG( | 2202.407 | 0.162059 | 0.428228 | 0.917257 | Myrf |
| ENSMUSG( | 6.892933 | 0.522514 | 0.484896 | NA | Myrfl |
| ENSMUSG( | 3346.983 | 0.080938 | 0.423105 | 0.917257 | Myrip |
| ENSMUSG( | 760.6732 | 0.106529 | 0.464948 | 0.923329 | Mysm1 |
| ENSMUSG( | 354.05 | 0.073669 | 0.616389 | 0.951413 | Myt1 |
| ENSMUSG( | 4238.052 | 0.241891 | 0.024062 | 0.588226 | Myt1l |
| ENSMUSG( | 32.30721 | 0.301014 | 0.399488 | 0.917257 | Myzap |
| ENSMUSG( | 357.1441 | 0.213866 | 0.164378 | 0.83856 | Mzf1 |
| ENSMUSG( | 1435.854 | -0.07987 | 0.591275 | 0.944975 | Mzt1 |
| ENSMUSG( | 398.9412 | -0.28981 | 0.072444 | 0.763629 | Mzt2 |
| ENSMUSG( | 1173.846 | 0.083798 | 0.454453 | 0.922444 | N4bp1 |
| ENSMUSG( | 534.0399 | 0.177279 | 0.174949 | 0.845107 | N4bp2 |
| ENSMUSG( | 885.9748 | 0.061582 | 0.644156 | 0.953903 | N4bp2l1 |
| ENSMUSG( | 709.113 | 0.142387 | 0.281443 | 0.892412 | N4bp2l2 |
| ENSMUSG( | 274.7237 | 0.09455 | 0.591235 | 0.944975 | N4bp3 |
| ENSMUSG( | 403.3378 | -0.32848 | 0.056175 | 0.723362 | N6amt1 |
| ENSMUSG( | 12.59506 | -0.71284 | 0.210459 | 0.864412 | NA |
| ENSMUSG( | 15.96299 | 0.513623 | 0.31515 | 0.901306 | NA |
| ENSMUSG( | 11.04532 | 0.149077 | 0.84693 | NA | NA |
| ENSMUSG( | 37.40635 | 0.09185 | 0.786805 | 0.976303 | NA |
| ENSMUSG( | 16.15763 | -0.45183 | 0.359196 | 0.914061 | NA |
| ENSMUSG( | 11.89616 | -0.60086 | 0.298029 | 0.898538 | NA |
| ENSMUSG( | 7.740628 | 0.276799 | 0.709407 | NA | NA |
| ENSMUSG( | 9.479064 | -0.13775 | 0.830663 | NA | NA |
| ENSMUSG( | 9.5911 | -0.56254 | 0.362595 | NA | NA |
| ENSMUSG( | 18.30025 | 0.337676 | 0.502816 | 0.927825 | NA |
| ENSMUSG( | 13.00278 | -0.49222 | 0.367421 | 0.915217 | NA |
| ENSMUSG( | 81.94945 | -0.23824 | 0.335872 | 0.907913 | NA |
| ENSMUSG( | 21.93833 | -1.40648 | 0.002486 | 0.195662 | NA |
| ENSMUSG( | 17.48813 | -0.34532 | 0.467004 | 0.923329 | NA |
| ENSMUSG( | 78.31497 | 0.170987 | 0.47542 | 0.923329 | NA |
| ENSMUSG( | 15.6943 | 0.024564 | 0.968096 | 0.995887 | NA |
| ENSMUSG( | 15.8212 | 0.06159 | 0.900315 | 0.990558 | NA |
| ENSMUSG( | 8.247402 | 0.811841 | 0.249125 | NA | NA |
| ENSMUSG( | 10.64853 | -0.13121 | 0.823484 | NA | NA |
| ENSMUSG( | 11.51821 | -0.38149 | 0.502739 | 0.927825 | NA |
| ENSMUSG( | 164.5119 | -0.21244 | 0.254226 | 0.881345 | NA |
| ENSMUSG( | 13.04141 | 0.550378 | 0.309709 | 0.898538 | NA |
| ENSMUSG( | 21.62652 | 0.220376 | 0.62519 | 0.953234 | NA |
| ENSMUSG( | 19.26822 | 0.019044 | 0.967073 | 0.995549 | NA |
| ENSMUSG( | 13.42743 | 0.127791 | 0.819393 | 0.979198 | NA |
| ENSMUSG( | 13.91015 | 0.044106 | 0.934538 | 0.99306 | NA |
| ENSMUSG( | 14.40278 | 0.586439 | 0.268745 | 0.886755 | NA |
| ENSMUSG( | 28.33789 | 0.206456 | 0.592194 | 0.944975 | NA |
| ENSMUSG( | 11.93086 | 0.371509 | 0.556564 | 0.939565 | NA |
| ENSMUSG( | 22.60692 | -0.15995 | 0.721174 | 0.966887 | NA |
| ENSMUSG( | 65.29784 | 0.330861 | 0.274873 | 0.889874 | NA |
| ENSMUSG( | 1385.476 | 0.224488 | 0.054268 | 0.723362 | NA |
| ENSMUSG( | 559.7511 | 0.469198 | 0.000403 | 0.058368 | NA |
| ENSMUSG( | 26.94154 | 0.548229 | 0.181098 | 0.850372 | NA |
| ENSMUSG( | 10.88443 | -0.53797 | 0.376058 | NA | NA |
| ENSMUSG( | 61.9148 | 0.011241 | 0.965386 | 0.995433 | NA |
| ENSMUSG( | 8.807009 | -0.40278 | 0.535122 | NA | NA |
| ENSMUSG( | 17.89905 | -0.15281 | 0.746237 | 0.969649 | NA |
| ENSMUSG( | 32.04586 | 0.205451 | 0.584894 | 0.944975 | NA |
| ENSMUSG( | 12.07427 | -0.03556 | 0.948942 | 0.994798 | NA |
| ENSMUSG( | 79.88162 | -0.23913 | 0.399093 | 0.917257 | NA |
| ENSMUSG( | 2865.546 | -0.14214 | 0.793526 | 0.97681 | NA |
| ENSMUSG( | 680.2407 | -0.2533 | 0.62506 | 0.953234 | NA |
| ENSMUSG( | 5714.118 | -0.21827 | 0.598256 | 0.946676 | NA |
| ENSMUSG( | 5955.481 | -0.36535 | 0.335507 | 0.907907 | NA |
| ENSMUSG( | 8261.567 | -0.3525 | 0.031809 | 0.631941 | NA |
| ENSMUSG( | 11.17887 | 0.851166 | 0.158221 | NA | NA |
| ENSMUSG( | 44.18254 | 0.249659 | 0.482194 | 0.924578 | NA |
| ENSMUSG( | 26.78699 | 2.223295 | 0.09731 | 0.796721 | NA |
| ENSMUSG( | 22.43273 | 0.243474 | 0.586621 | 0.944975 | NA |
| ENSMUSG( | 104.9404 | 0.070914 | 0.742885 | 0.969415 | NA |
| ENSMUSG( | 6.665955 | 0.365501 | 0.674028 | NA | NA |
| ENSMUSG( | 27.62972 | 0.081528 | 0.834633 | 0.981237 | NA |
| ENSMUSG( | 12.17541 | -0.00302 | 0.99585 | 0.999116 | NA |
| ENSMUSG( | 23.92921 | 0.254968 | 0.561826 | 0.940388 | NA |
| ENSMUSG( | 16.15627 | 0.911162 | 0.091583 | 0.791889 | NA |
| ENSMUSG( | 10.01923 | 0.378521 | 0.561562 | NA | NA |
| ENSMUSG( | 58.62265 | -0.2163 | 0.451391 | 0.922202 | NA |
| ENSMUSG( | 11.46363 | -1.36706 | 0.023478 | 0.585845 | NA |
| ENSMUSG( | 66.20855 | -0.27454 | 0.281201 | 0.892356 | NA |
| ENSMUSG( | 16.68646 | -0.45157 | 0.440523 | 0.921166 | NA |
| ENSMUSG( | 26.60456 | 0.760833 | 0.069973 | 0.757435 | NA |
| ENSMUSG( | 59.03953 | -0.2615 | 0.466018 | 0.923329 | NA |
| ENSMUSG( | 29.46864 | 0.083547 | 0.836966 | 0.981335 | NA |
| ENSMUSG( | 11.45719 | -0.16712 | 0.778871 | 0.976106 | NA |
| ENSMUSG( | 17.99576 | 0.483327 | 0.33265 | 0.907907 | NA |
| ENSMUSG( | 40.28923 | 0.383429 | 0.36 | 0.914223 | NA |
| ENSMUSG( | 38.04737 | -0.06577 | 0.840074 | 0.982217 | NA |
| ENSMUSG( | 13.49903 | -0.64604 | 0.294281 | 0.898538 | NA |
| ENSMUSG( | 42.43777 | -0.16556 | 0.589869 | 0.944975 | NA |
| ENSMUSG( | 14.13777 | -0.44063 | 0.407119 | 0.917257 | NA |
| ENSMUSG( | 15.74353 | 0.884146 | 0.106859 | 0.804969 | NA |
| ENSMUSG( | 10.41664 | -0.31032 | 0.609334 | NA | NA |
| ENSMUSG( | 46.5822 | 0.085516 | 0.790866 | 0.97681 | NA |
| ENSMUSG( | 18.88323 | -1.17243 | 0.012858 | 0.457834 | NA |
| ENSMUSG( | 331.527 | -0.12636 | 0.416286 | 0.917257 | NA |
| ENSMUSG( | 46.75187 | 0.084863 | 0.788243 | 0.97637 | NA |
| ENSMUSG( | 14.0876 | -3.68015 | 3.26E-06 | 0.001541 | NA |
| ENSMUSG( | 15.46146 | -0.54971 | 0.288385 | 0.894794 | NA |
| ENSMUSG( | 24.48176 | -0.01182 | 0.976003 | 0.996416 | NA |

| | | | | | |
|---|---|---|---|---|---|
| ENSMUSG( | N4bp2l1 | 960.1676 | 0.029643 | 0.785613 | 0.95867 |
| ENSMUSG( | N4bp2l2 | 745.6135 | 0.016916 | 0.885145 | 0.982851 |
| ENSMUSG( | N4bp3 | 297.9273 | 0.05894 | 0.739487 | 0.947768 |
| ENSMUSG( | N6amt1 | 456.1469 | -0.22727 | 0.121804 | 0.526562 |
| ENSMUSG( | NA | 84.19223 | -0.1168 | 0.607065 | 0.906685 |
| ENSMUSG( | NA | 66.67283 | 0.171143 | 0.467619 | 0.844819 |
| ENSMUSG( | NA | 15.18749 | -0.38298 | 0.438301 | NA |
| ENSMUSG( | NA | 117.682 | 0.105242 | 0.572882 | 0.893584 |
| ENSMUSG( | NA | 381.086 | -0.00876 | 0.941913 | 0.993415 |
| ENSMUSG( | NA | 35.56141 | -0.12291 | 0.694619 | 0.9343 |
| ENSMUSG( | NA | 24.27333 | -0.17432 | 0.648227 | 0.919949 |
| ENSMUSG( | NA | 40.44742 | 0.152486 | 0.65632 | 0.923929 |
| ENSMUSG( | NA | 81.53609 | 0.152784 | 0.514746 | 0.872189 |
| ENSMUSG( | NA | 25.78109 | 0.231154 | 0.540665 | 0.88315 |
| ENSMUSG( | NA | 13.22371 | -0.14373 | 0.77824 | NA |
| ENSMUSG( | NA | 36.14783 | 0.743383 | 0.028463 | 0.257525 |
| ENSMUSG( | NA | 95.48916 | 0.006463 | 0.979146 | 0.997428 |
| ENSMUSG( | NA | 19.81136 | -0.4762 | 0.320379 | NA |
| ENSMUSG( | NA | 50.75926 | 0.503441 | 0.070441 | 0.40908 |
| ENSMUSG( | NA | 21.43278 | -0.12013 | 0.770079 | 0.954689 |
| ENSMUSG( | NA | 11.86057 | -0.06554 | 0.903141 | NA |
| ENSMUSG( | NA | 64.84578 | -0.15282 | 0.525842 | 0.876902 |
| ENSMUSG( | NA | 122.0201 | 0.331605 | 0.071841 | 0.414057 |
| ENSMUSG( | NA | 23.85162 | -0.70854 | 0.069685 | 0.408352 |
| ENSMUSG( | NA | 27.1535 | -0.37046 | 0.343584 | 0.779153 |
| ENSMUSG( | NA | 14.1576 | -0.237 | 0.641165 | NA |
| ENSMUSG( | NA | 21.15647 | 0.254 | 0.534884 | NA |
| ENSMUSG( | NA | 149.928 | -0.17008 | 0.329104 | 0.773233 |
| ENSMUSG( | NA | 15.92199 | -0.07691 | 0.876954 | NA |
| ENSMUSG( | NA | 53.48772 | -0.1366 | 0.599332 | 0.903801 |
| ENSMUSG( | NA | 110.4483 | 0.128896 | 0.500573 | 0.863514 |
| ENSMUSG( | NA | 20.34754 | -0.34269 | 0.400694 | NA |
| ENSMUSG( | NA | 146.6646 | 0.177737 | 0.354483 | 0.786878 |
| ENSMUSG( | NA | 1003.19 | 0.030799 | 0.804066 | 0.962169 |
| ENSMUSG( | NA | 45.73217 | -0.29166 | 0.333867 | 0.775602 |
| ENSMUSG( | NA | 77.98993 | 0.211992 | 0.349305 | 0.78176 |
| ENSMUSG( | NA | 23.37217 | -0.32718 | 0.421655 | 0.823311 |
| ENSMUSG( | NA | 19.54734 | -0.13452 | 0.751105 | NA |
| ENSMUSG( | NA | 26.99167 | -0.09576 | 0.790326 | 0.959461 |
| ENSMUSG( | NA | 35.57335 | 0.049562 | 0.875487 | 0.980082 |
| ENSMUSG( | NA | 54.69076 | -0.17233 | 0.513357 | 0.871762 |
| ENSMUSG( | NA | 151.1995 | 0.077476 | 0.67466 | 0.929638 |
| ENSMUSG( | NA | 27.53518 | -0.05059 | 0.885922 | 0.983008 |
| ENSMUSG( | NA | 27.45088 | 0.043972 | 0.901501 | 0.985633 |
| ENSMUSG( | NA | 100.5908 | 0.124058 | 0.621654 | 0.911432 |
| ENSMUSG( | NA | 19.11156 | -0.08366 | 0.85624 | NA |
| ENSMUSG( | NA | 39.00653 | 0.059251 | 0.841211 | 0.971179 |
| ENSMUSG( | NA | 15.15949 | 0.340707 | 0.467885 | NA |
| ENSMUSG( | NA | 65.65015 | -0.08162 | 0.74156 | 0.948289 |
| ENSMUSG( | NA | 21.37703 | 0.262311 | 0.511533 | 0.870901 |
| ENSMUSG( | NA | 90.03016 | 0.11562 | 0.617089 | 0.910866 |
| ENSMUSG( | NA | 37.79747 | 0.186224 | 0.5783 | 0.89641 |
| ENSMUSG( | NA | 19.09839 | -0.01005 | 0.981274 | NA |
| ENSMUSG( | NA | 18.66889 | -0.39707 | 0.360993 | NA |
| ENSMUSG( | NA | 953.0742 | 0.00486 | 0.97051 | 0.996369 |
| ENSMUSG( | NA | 49.79394 | -0.1182 | 0.663548 | 0.9265 |
| ENSMUSG( | NA | 32.11252 | -0.11273 | 0.749349 | 0.949664 |
| ENSMUSG( | NA | 41.00528 | 0.216181 | 0.50283 | 0.865715 |
| ENSMUSG( | NA | 13.86655 | 0.357925 | 0.481091 | NA |
| ENSMUSG( | NA | 111.2099 | -0.11232 | 0.588499 | 0.900921 |
| ENSMUSG( | NA | 494.6208 | 0.19819 | 0.160229 | 0.589853 |
| ENSMUSG( | NA | 16.67936 | -0.15722 | 0.724256 | NA |
| ENSMUSG( | NA | 88.78723 | -0.05959 | 0.771356 | 0.954726 |
| ENSMUSG( | NA | 16.33138 | 0.178787 | 0.69393 | NA |
| ENSMUSG( | NA | 437.0647 | 0.010957 | 0.926533 | 0.991229 |
| ENSMUSG( | NA | 195.9693 | 0.100377 | 0.549672 | 0.885944 |
| ENSMUSG( | NA | 1602.652 | 0.099515 | 0.272241 | 0.7223 |
| ENSMUSG( | NA | 25.87262 | 0.245493 | 0.551402 | 0.886141 |
| ENSMUSG( | NA | 303.2443 | 0.293946 | 0.049689 | 0.340355 |
| ENSMUSG( | NA | 24.32698 | -0.3007 | 0.437985 | 0.830072 |
| ENSMUSG( | NA | 13.12932 | 0.416284 | 0.413424 | NA |
| ENSMUSG( | NA | 346.2748 | -0.07293 | 0.62471 | 0.912042 |
| ENSMUSG( | NA | 21.65521 | 0.118111 | 0.771102 | 0.954726 |
| ENSMUSG( | NA | 27.33897 | -0.17593 | 0.619747 | 0.911432 |
| ENSMUSG( | NA | 24.16938 | 0.046763 | 0.906614 | 0.98713 |
| ENSMUSG( | NA | 127.2508 | -0.18027 | 0.355355 | 0.78783 |
| ENSMUSG( | NA | 25.96521 | 0.544312 | 0.155078 | 0.5828 |
| ENSMUSG( | NA | 21.42822 | 0.236053 | 0.555632 | 0.887457 |
| ENSMUSG( | NA | 14.34971 | -0.07954 | 0.878447 | NA |
| ENSMUSG( | NA | 22.0474 | 0.808463 | 0.066495 | 0.39868 |
| ENSMUSG( | NA | 180.1973 | -0.02107 | 0.893192 | 0.984512 |
| ENSMUSG( | NA | 89.73384 | -0.0271 | 0.894098 | 0.984512 |
| ENSMUSG( | NA | 17.67093 | -0.42362 | 0.339794 | NA |
| ENSMUSG( | NA | 12.25662 | 1.134484 | 0.041361 | NA |
| ENSMUSG( | NA | 65.75911 | -0.03353 | 0.894633 | 0.984512 |
| ENSMUSG( | NA | 32.30008 | 0.195023 | 0.597577 | 0.903694 |
| ENSMUSG( | NA | 28.07008 | 0.406327 | 0.275079 | 0.72509 |
| ENSMUSG( | NA | 27.23009 | -0.17444 | 0.639663 | 0.916434 |
| ENSMUSG( | NA | 28.698 | -0.38687 | 0.293363 | 0.742763 |
| ENSMUSG( | NA | 24.60247 | 0.260495 | 0.491808 | 0.860249 |
| ENSMUSG( | NA | 51.19022 | 0.06354 | 0.820152 | 0.965888 |
| ENSMUSG( | NA | 12.16629 | 0.009466 | 0.986004 | NA |
| ENSMUSG( | NA | 28.75336 | 0.013911 | 0.969027 | 0.996369 |
| ENSMUSG( | NA | 16.24251 | 0.356756 | 0.464733 | NA |
| ENSMUSG( | NA | 14.84962 | 0.585908 | 0.265752 | NA |
| ENSMUSG( | NA | 13.49936 | 0.38392 | 0.488486 | NA |
| ENSMUSG( | NA | 35.75615 | -0.23281 | 0.465421 | 0.844064 |
| ENSMUSG( | NA | 15.762 | -0.43801 | 0.361851 | NA |
| ENSMUSG( | NA | 14.28481 | 0.545443 | 0.30469 | NA |
| ENSMUSG( | NA | 50.8244 | -0.10567 | 0.696405 | 0.935166 |
| ENSMUSG( | NA | 195.4782 | 0.026992 | 0.870704 | 0.977951 |
| ENSMUSG( | NA | 16.56244 | 0.468526 | 0.320531 | NA |
| ENSMUSG( | NA | 12.85775 | -0.14059 | 0.780291 | NA |
| ENSMUSG( | NA | 252.8672 | -0.08752 | 0.648934 | 0.920025 |
| ENSMUSG( | NA | 33.42039 | -0.13585 | 0.714947 | 0.941144 |
| ENSMUSG( | NA | 22.86203 | -0.66592 | 0.106665 | 0.494174 |
| ENSMUSG( | NA | 9.126465 | 0.131853 | 0.837514 | NA |
| ENSMUSG( | NA | 66.58867 | -0.08038 | 0.736696 | 0.946546 |
| ENSMUSG( | NA | 42.64685 | 0.011071 | 0.971236 | 0.996369 |
| ENSMUSG( | NA | 24.52871 | -0.76691 | 0.069258 | 0.407813 |
| ENSMUSG( | NA | 24.83779 | -0.4227 | 0.265489 | 0.716109 |
| ENSMUSG( | NA | 148.0169 | -0.17852 | 0.321541 | 0.766705 |
| ENSMUSG( | NA | 257.1961 | 0.171135 | 0.280878 | 0.729868 |
| ENSMUSG( | NA | 98.18784 | -0.2243 | 0.39085 | 0.80826 |
| ENSMUSG( | NA | 21.49071 | -0.38235 | 0.364893 | 0.79407 |
| ENSMUSG( | NA | 30.05902 | 0.201754 | 0.571978 | 0.893526 |
| ENSMUSG( | NA | 41.78521 | 0.509008 | 0.12659 | 0.534155 |
| ENSMUSG( | NA | 23.49512 | -0.15878 | 0.674603 | 0.929638 |
| ENSMUSG( | NA | 37.33397 | 0.129417 | 0.673329 | 0.929007 |
| ENSMUSG( | NA | 11.17383 | -0.22569 | 0.683636 | NA |
| ENSMUSG( | NA | 44.08166 | 0.137463 | 0.634737 | 0.914073 |
| ENSMUSG( | NA | 31.47698 | 0.249096 | 0.485318 | 0.857408 |
| ENSMUSG( | NA | 67.53653 | -0.08939 | 0.710183 | 0.9397 |
| ENSMUSG( | NA | 77.4573 | 0.047458 | 0.839556 | 0.971179 |
| ENSMUSG( | NA | 13.39409 | 0.319603 | 0.55166 | NA |
| ENSMUSG( | NA | 13.70824 | -0.05232 | 0.9145 | NA |
| ENSMUSG( | NA | 170.3186 | 0.150949 | 0.410178 | 0.818258 |
| ENSMUSG( | NA | 36.36136 | -0.43824 | 0.158997 | 0.588112 |
| ENSMUSG( | NA | 113.4041 | -0.19594 | 0.333744 | 0.775602 |
| ENSMUSG( | NA | 11.73129 | 0.099907 | 0.851378 | NA |
| ENSMUSG( | NA | 267.1909 | 0.038617 | 0.79426 | 0.959799 |
| ENSMUSG( | NA | 130.867 | -0.00671 | 0.972502 | 0.996585 |
| ENSMUSG( | NA | 4128.171 | 0.215383 | 0.030728 | 0.266671 |
| ENSMUSG( | NA | 39.25512 | -0.26165 | 0.420425 | 0.82289 |
| ENSMUSG( | NA | 18.81305 | -0.76014 | 0.132868 | NA |
| ENSMUSG( | NA | 55.42706 | 2.475976 | 2.62E-16 | 3.27E-13 |
| ENSMUSG( | NA | 24.05968 | 0.321842 | 0.425759 | 0.824898 |
| ENSMUSG( | NA | 81.27139 | 0.105712 | 0.631691 | 0.914073 |
| ENSMUSG( | NA | 29.8597 | 1.787302 | 7.59E-06 | 0.000947 |
| ENSMUSG( | NA | 58.71592 | 0.396531 | 0.127239 | 0.534725 |
| ENSMUSG( | NA | 88.83002 | -0.77638 | 0.003489 | 0.074758 |
| ENSMUSG( | NA | 10.63288 | 0.743067 | 0.209118 | NA |
| ENSMUSG( | NA | 144.4836 | -0.09689 | 0.643142 | 0.917078 |
| ENSMUSG( | NA | 306.3084 | -0.12781 | 0.339397 | 0.777656 |
| ENSMUSG( | NA | 407.0058 | 0.588316 | 3.57E-06 | 0.000492 |
| ENSMUSG( | NA | 39.63115 | -0.22673 | 0.505787 | 0.866936 |
| ENSMUSG( | NA | 12.91545 | 0.573753 | 0.282735 | NA |
| ENSMUSG( | NA | 81.98548 | -0.22642 | 0.31039 | 0.758554 |
| ENSMUSG( | NA | 37.16078 | -0.03548 | 0.917682 | 0.989377 |
| ENSMUSG( | NA | 306.1616 | -0.08763 | 0.546251 | 0.884974 |
| ENSMUSG( | NA | 18.4007 | 0.153154 | 0.744555 | NA |
| ENSMUSG( | NA | 489806.4 | 0.207701 | 0.107892 | 0.49674 |
| ENSMUSG( | NA | 636393 | 0.193178 | 0.21374 | 0.658097 |
| ENSMUSG( | NA | 765.5616 | 0.107054 | 0.536723 | 0.882043 |
| ENSMUSG( | NA | 218.1282 | -0.15732 | 0.806891 | 0.962973 |
| ENSMUSG( | NA | 105.8059 | -0.33365 | 0.219009 | 0.664602 |
| ENSMUSG( | NA | 1465.281 | 0.127751 | 0.299344 | 0.748131 |
| ENSMUSG( | NA | 886.1749 | 0.031352 | 0.950338 | 0.993821 |

| | | | | | |
|---|---|---|---|---|---|
| ENSMUSG( | N4bp2l1 | 885.9748 | 0.061582 | 0.644156 | 0.953903 |
| ENSMUSG( | N4bp2l2 | 709.113 | 0.142387 | 0.281443 | 0.892412 |
| ENSMUSG( | N4bp3 | 274.7237 | 0.09455 | 0.591235 | 0.944975 |
| ENSMUSG( | N6amt1 | 403.3378 | -0.32848 | 0.056175 | 0.723362 |
| ENSMUSG( | NA | 12.59506 | -0.71284 | 0.210459 | 0.864412 |
| ENSMUSG( | NA | 15.96299 | 0.513623 | 0.31515 | 0.901306 |
| ENSMUSG( | NA | 37.40635 | 0.09185 | 0.786805 | 0.976303 |
| ENSMUSG( | NA | 16.15763 | -0.45183 | 0.359196 | 0.914061 |
| ENSMUSG( | NA | 11.89616 | -0.60086 | 0.298029 | 0.898538 |
| ENSMUSG( | NA | 9.479064 | -0.13775 | 0.830663 | NA |
| ENSMUSG( | NA | 9.5911 | -0.56254 | 0.362595 | NA |
| ENSMUSG( | NA | 18.30025 | 0.337676 | 0.502816 | 0.927825 |
| ENSMUSG( | NA | 13.00278 | -0.49222 | 0.367421 | 0.915217 |
| ENSMUSG( | NA | 81.94945 | -0.23824 | 0.335872 | 0.907913 |
| ENSMUSG( | NA | 21.93833 | -1.40648 | 0.002486 | 0.195662 |
| ENSMUSG( | NA | 17.48813 | -0.34532 | 0.467004 | 0.923329 |
| ENSMUSG( | NA | 78.31497 | 0.170987 | 0.47542 | 0.923329 |
| ENSMUSG( | NA | 15.6943 | 0.024564 | 0.968096 | 0.995887 |
| ENSMUSG( | NA | 15.8212 | 0.06159 | 0.900315 | 0.990558 |
| ENSMUSG( | NA | 10.64853 | -0.13121 | 0.823484 | NA |
| ENSMUSG( | NA | 11.51821 | -0.38149 | 0.502739 | 0.927825 |
| ENSMUSG( | NA | 164.5119 | -0.21244 | 0.254226 | 0.881345 |
| ENSMUSG( | NA | 13.04141 | 0.550378 | 0.309709 | 0.898538 |
| ENSMUSG( | NA | 21.62652 | 0.220376 | 0.62519 | 0.953234 |
| ENSMUSG( | NA | 19.26822 | 0.019044 | 0.967073 | 0.995549 |
| ENSMUSG( | NA | 13.42743 | 0.127791 | 0.819393 | 0.979198 |
| ENSMUSG( | NA | 13.91015 | 0.044106 | 0.934538 | 0.99306 |
| ENSMUSG( | NA | 14.40278 | 0.586439 | 0.268745 | 0.886755 |
| ENSMUSG( | NA | 28.33789 | 0.206456 | 0.592194 | 0.944975 |
| ENSMUSG( | NA | 11.93086 | 0.371509 | 0.556564 | 0.939565 |
| ENSMUSG( | NA | 22.60692 | -0.15995 | 0.721174 | 0.966887 |
| ENSMUSG( | NA | 65.29784 | 0.330861 | 0.274873 | 0.889874 |
| ENSMUSG( | NA | 1385.476 | 0.224488 | 0.054268 | 0.723362 |
| ENSMUSG( | NA | 559.7511 | 0.469198 | 0.000403 | 0.058368 |
| ENSMUSG( | NA | 26.94154 | 0.548229 | 0.181098 | 0.850372 |
| ENSMUSG( | NA | 10.88443 | -0.53797 | 0.376058 | NA |
| ENSMUSG( | NA | 61.9148 | 0.011241 | 0.965386 | 0.995433 |
| ENSMUSG( | NA | 8.807009 | -0.40278 | 0.535122 | NA |
| ENSMUSG( | NA | 17.89905 | -0.15281 | 0.746237 | 0.969649 |
| ENSMUSG( | NA | 32.04586 | 0.205451 | 0.584894 | 0.944975 |
| ENSMUSG( | NA | 12.07427 | -0.03556 | 0.948942 | 0.994798 |
| ENSMUSG( | NA | 79.88162 | -0.23913 | 0.399093 | 0.917257 |
| ENSMUSG( | NA | 2865.546 | -0.14214 | 0.793526 | 0.97681 |
| ENSMUSG( | NA | 680.2407 | -0.2533 | 0.62506 | 0.953234 |
| ENSMUSG( | NA | 5714.118 | -0.21827 | 0.598256 | 0.946676 |
| ENSMUSG( | NA | 5955.481 | -0.36535 | 0.335507 | 0.907907 |
| ENSMUSG( | NA | 8261.567 | -0.3525 | 0.031809 | 0.631941 |
| ENSMUSG( | NA | 44.18254 | 0.249659 | 0.482194 | 0.924578 |
| ENSMUSG( | NA | 26.78699 | 2.223295 | 0.09731 | 0.796721 |
| ENSMUSG( | NA | 22.43273 | 0.243474 | 0.586621 | 0.944975 |
| ENSMUSG( | NA | 104.9404 | 0.070914 | 0.742885 | 0.969415 |
| ENSMUSG( | NA | 6.665955 | 0.365501 | 0.674028 | NA |
| ENSMUSG( | NA | 27.62972 | 0.081528 | 0.834633 | 0.981237 |
| ENSMUSG( | NA | 12.17541 | -0.00302 | 0.99585 | 0.999116 |
| ENSMUSG( | NA | 23.92921 | 0.254968 | 0.561826 | 0.940388 |
| ENSMUSG( | NA | 16.15627 | 0.911162 | 0.091583 | 0.791889 |
| ENSMUSG( | NA | 58.62265 | -0.2163 | 0.451391 | 0.922202 |
| ENSMUSG( | NA | 11.46363 | -1.36706 | 0.023478 | 0.585845 |
| ENSMUSG( | NA | 66.20855 | -0.27454 | 0.281201 | 0.892356 |
| ENSMUSG( | NA | 16.68646 | -0.45157 | 0.440523 | 0.921166 |
| ENSMUSG( | NA | 26.60456 | 0.760833 | 0.069973 | 0.757435 |
| ENSMUSG( | NA | 59.03953 | -0.2615 | 0.466018 | 0.923329 |
| ENSMUSG( | NA | 29.46864 | 0.083547 | 0.836966 | 0.981335 |
| ENSMUSG( | NA | 11.45719 | -0.16712 | 0.778871 | 0.976106 |
| ENSMUSG( | NA | 17.99576 | 0.483327 | 0.33265 | 0.907907 |
| ENSMUSG( | NA | 40.28923 | 0.383429 | 0.36 | 0.914223 |
| ENSMUSG( | NA | 38.04737 | -0.06577 | 0.840074 | 0.982217 |
| ENSMUSG( | NA | 13.49903 | -0.64604 | 0.294281 | 0.898538 |
| ENSMUSG( | NA | 42.43777 | -0.16556 | 0.589869 | 0.944975 |
| ENSMUSG( | NA | 14.13777 | -0.44063 | 0.407119 | 0.917257 |
| ENSMUSG( | NA | 15.74353 | 0.884146 | 0.106859 | 0.804969 |
| ENSMUSG( | NA | 10.41664 | -0.31032 | 0.609334 | NA |
| ENSMUSG( | NA | 46.5822 | 0.085516 | 0.790866 | 0.97681 |
| ENSMUSG( | NA | 18.88323 | -1.17243 | 0.012858 | 0.457834 |
| ENSMUSG( | NA | 331.527 | -0.12636 | 0.416286 | 0.917257 |
| ENSMUSG( | NA | 46.75187 | 0.084863 | 0.788243 | 0.97637 |
| ENSMUSG( | NA | 14.0876 | -3.68015 | 3.26E-06 | 0.001541 |
| ENSMUSG( | NA | 15.46146 | -0.54971 | 0.288385 | 0.894794 |
| ENSMUSG( | NA | 24.48176 | -0.01182 | 0.976003 | 0.996416 |
| ENSMUSG( | NA | 8.161166 | -0.40846 | 0.559256 | NA |
| ENSMUSG( | NA | 17.08392 | 0.136842 | 0.78746 | 0.97637 |
| ENSMUSG( | NA | 15.09506 | 0.05535 | 0.918938 | 0.992529 |
| ENSMUSG( | NA | 16.7805 | 0.288735 | 0.562052 | 0.94044 |
| ENSMUSG( | NA | 30.50783 | -0.67181 | 0.074978 | 0.76762 |
| ENSMUSG( | NA | 36.31751 | 0.575177 | 0.210772 | 0.864412 |
| ENSMUSG( | NA | 13.87565 | -0.29505 | 0.58899 | 0.944975 |
| ENSMUSG( | NA | 14.27386 | -0.33717 | 0.546929 | 0.938063 |
| ENSMUSG( | NA | 7.350145 | -1.42417 | 0.063019 | NA |
| ENSMUSG( | NA | 11.7858 | -0.84113 | 0.152889 | 0.837724 |
| ENSMUSG( | NA | 12.58695 | -0.526 | 0.354576 | 0.912411 |
| ENSMUSG( | NA | 14.34065 | -0.62818 | 0.252591 | 0.88018 |
| ENSMUSG( | NA | 21.98834 | 0.168909 | 0.717536 | 0.965964 |
| ENSMUSG( | NA | 8.558528 | 0.299425 | 0.661395 | NA |
| ENSMUSG( | NA | 31.18674 | 2.505986 | 6.46E-05 | 0.015522 |
| ENSMUSG( | NA | 17.56605 | -0.18246 | 0.71665 | 0.965807 |
| ENSMUSG( | NA | 274.5985 | -0.24305 | 0.215591 | 0.865167 |
| ENSMUSG( | NA | 13.49935 | -0.10166 | 0.855332 | 0.985649 |
| ENSMUSG( | NA | 32.34664 | -0.56297 | 0.112786 | 0.809623 |
| ENSMUSG( | NA | 138.5171 | -0.15104 | 0.446633 | 0.921734 |
| ENSMUSG( | NA | 10.05323 | 0.577801 | 0.358156 | NA |
| ENSMUSG( | NA | 26.51262 | -0.41225 | 0.440945 | 0.921166 |
| ENSMUSG( | NA | 71.66773 | 0.002524 | 0.993312 | 0.998813 |
| ENSMUSG( | NA | 17.82865 | 0.211765 | 0.645353 | 0.953988 |
| ENSMUSG( | NA | 15.44588 | 0.083134 | 0.879706 | 0.989644 |
| ENSMUSG( | NA | 15.44837 | -0.59057 | 0.276683 | 0.890889 |
| ENSMUSG( | NA | 52.13075 | 0.782783 | 0.015314 | 0.495791 |
| ENSMUSG( | NA | 83.28019 | 0.494404 | 0.087423 | 0.784337 |
| ENSMUSG( | NA | 9.248076 | 0.144783 | 0.821047 | NA |
| ENSMUSG( | NA | 15.37073 | -0.16495 | 0.783405 | 0.976106 |
| ENSMUSG( | NA | 230.096 | -0.16223 | 0.342276 | 0.909642 |
| ENSMUSG( | NA | 11.6722 | 0.193897 | 0.758051 | 0.971858 |
| ENSMUSG( | NA | 30.32973 | 0.45696 | 0.224209 | 0.867489 |
| ENSMUSG( | NA | 69.03297 | -0.0733 | 0.816288 | 0.979198 |
| ENSMUSG( | NA | 13.59654 | -0.97679 | 0.0786 | 0.773505 |
| ENSMUSG( | NA | 6.034431 | 0.812266 | 0.360948 | NA |
| ENSMUSG( | NA | 57.21583 | 0.362203 | 0.190686 | 0.857801 |
| ENSMUSG( | NA | 18.00253 | 0.2397 | 0.63972 | 0.953903 |
| ENSMUSG( | NA | 18.99581 | -0.35825 | 0.504821 | 0.927825 |
| ENSMUSG( | NA | 51.85251 | 0.129676 | 0.656806 | 0.955567 |
| ENSMUSG( | NA | 10.91928 | -0.29878 | 0.63801 | NA |
| ENSMUSG( | NA | 13.77059 | 0.22926 | 0.687125 | 0.959302 |
| ENSMUSG( | NA | 7.953109 | -0.36106 | 0.630265 | NA |
| ENSMUSG( | NA | 9.531496 | -0.21366 | 0.756812 | NA |
| ENSMUSG( | NA | 77.73879 | 0.189579 | 0.467971 | 0.923329 |
| ENSMUSG( | NA | 116.9467 | 0.317822 | 0.247318 | 0.877904 |
| ENSMUSG( | NA | 60.65015 | 0.474561 | 0.100007 | 0.801311 |
| ENSMUSG( | NA | 473.8937 | -0.4049 | 0.005185 | 0.296096 |
| ENSMUSG( | NA | 25.59305 | -0.47112 | 0.264408 | 0.886134 |
| ENSMUSG( | NA | 7.94032 | -0.69188 | 0.328518 | NA |
| ENSMUSG( | NA | 35.88362 | 0.049925 | 0.888871 | 0.990558 |
| ENSMUSG( | NA | 81.18496 | -0.09453 | 0.694123 | 0.960867 |
| ENSMUSG( | NA | 9.780293 | 0.08667 | 0.88698 | NA |
| ENSMUSG( | NA | 378.8975 | -0.13931 | 0.33005 | 0.907251 |
| ENSMUSG( | NA | 38.25326 | -0.41241 | 0.222471 | 0.866791 |
| ENSMUSG( | NA | 17.14846 | -0.71178 | 0.149128 | 0.836241 |
| ENSMUSG( | NA | 26.24069 | -0.36291 | 0.358834 | 0.913759 |
| ENSMUSG( | NA | 57.32067 | 0.488972 | 0.075926 | 0.768524 |
| ENSMUSG( | NA | 15.02222 | 0.394811 | 0.445584 | 0.921734 |
| ENSMUSG( | NA | 3256.382 | -0.20165 | 0.094182 | 0.795735 |
| ENSMUSG( | NA | 29.02338 | -0.42165 | 0.267787 | 0.886134 |
| ENSMUSG( | NA | 9.774681 | -0.2541 | 0.719829 | NA |
| ENSMUSG( | NA | 47.49955 | 0.271486 | 0.447621 | 0.922202 |
| ENSMUSG( | NA | 107.6517 | 0.003719 | 0.986052 | 0.997188 |
| ENSMUSG( | NA | 33.49378 | -0.02925 | 0.933564 | 0.99306 |
| ENSMUSG( | NA | 9.961631 | 0.513608 | 0.432767 | NA |
| ENSMUSG( | NA | 59.23543 | 0.412956 | 0.173761 | 0.844317 |
| ENSMUSG( | NA | 64.6843 | 0.120041 | 0.694858 | 0.960867 |
| ENSMUSG( | NA | 99.51405 | 0.011352 | 0.958136 | 0.994798 |
| ENSMUSG( | NA | 27.53495 | -0.04013 | 0.917008 | 0.992529 |
| ENSMUSG( | NA | 11.97855 | -0.5153 | 0.36292 | 0.914223 |
| ENSMUSG( | NA | 21.52187 | 0.058518 | 0.894801 | 0.990558 |
| ENSMUSG( | NA | 85.75639 | 0.180693 | 0.42233 | 0.917257 |
| ENSMUSG( | NA | 26.39568 | -0.52431 | 0.239367 | 0.87431 |
| ENSMUSG( | NA | 14.05243 | -0.0209 | 0.970433 | 0.99594 |
| ENSMUSG( | NA | 347.2521 | -0.09424 | 0.661523 | 0.956205 |
| ENSMUSG( | NA | 35.05591 | -0.30409 | 0.435939 | 0.919569 |
| ENSMUSG( | NA | 11.34769 | 0.826799 | 0.211853 | NA |
| ENSMUSG( | NA | 446.3991 | 0.074326 | 0.625786 | 0.953234 |

| | | | | | |
|---|---|---|---|---|---|
| ENSMUSG( | 20.34754 | -0.34269 | 0.400694 | NA | NA |
| ENSMUSG( | 146.6646 | 0.177737 | 0.354483 | 0.786878 | NA |
| ENSMUSG( | 1003.19 | 0.030799 | 0.804066 | 0.962169 | NA |
| ENSMUSG( | 45.73217 | -0.29166 | 0.333867 | 0.775602 | NA |
| ENSMUSG( | 13.03099 | -0.25235 | 0.639152 | NA | NA |
| ENSMUSG( | 12.32244 | 0.007804 | 0.98892 | NA | NA |
| ENSMUSG( | 77.98993 | 0.211992 | 0.349305 | 0.78176 | NA |
| ENSMUSG( | 10.2865 | -0.10804 | 0.851132 | NA | NA |
| ENSMUSG( | 23.37217 | -0.32718 | 0.421655 | 0.823311 | NA |
| ENSMUSG( | 19.54734 | -0.13452 | 0.751105 | NA | NA |
| ENSMUSG( | 26.99167 | -0.09576 | 0.790326 | 0.959461 | NA |
| ENSMUSG( | 35.57335 | 0.049562 | 0.875487 | 0.980082 | NA |
| ENSMUSG( | 54.69076 | -0.17233 | 0.513357 | 0.871762 | NA |
| ENSMUSG( | 151.1995 | 0.077476 | 0.67466 | 0.929638 | NA |
| ENSMUSG( | 27.53518 | -0.05059 | 0.885922 | 0.983008 | NA |
| ENSMUSG( | 27.45088 | 0.043972 | 0.901501 | 0.985633 | NA |
| ENSMUSG( | 8.601449 | 0.122404 | 0.851443 | NA | NA |
| ENSMUSG( | 100.5908 | 0.124058 | 0.621654 | 0.911432 | NA |
| ENSMUSG( | 19.11156 | -0.08366 | 0.85624 | NA | NA |
| ENSMUSG( | 39.00653 | 0.059251 | 0.841211 | 0.971179 | NA |
| ENSMUSG( | 15.15949 | 0.340707 | 0.467885 | NA | NA |
| ENSMUSG( | 65.65015 | -0.08162 | 0.74156 | 0.948289 | NA |
| ENSMUSG( | 21.37703 | 0.262311 | 0.511533 | 0.870901 | NA |
| ENSMUSG( | 90.03016 | 0.11562 | 0.617089 | 0.910866 | NA |
| ENSMUSG( | 37.79747 | 0.186224 | 0.5783 | 0.89641 | NA |
| ENSMUSG( | 10.42389 | 0.426171 | 0.488479 | NA | NA |
| ENSMUSG( | 19.09839 | -0.01005 | 0.981274 | NA | NA |
| ENSMUSG( | 18.66889 | -0.39707 | 0.360993 | NA | NA |
| ENSMUSG( | 953.0742 | 0.00486 | 0.97051 | 0.996369 | NA |
| ENSMUSG( | 49.79394 | -0.1182 | 0.663548 | 0.9265 | NA |
| ENSMUSG( | 32.11252 | -0.11273 | 0.749349 | 0.949664 | NA |
| ENSMUSG( | 41.00528 | 0.216181 | 0.50283 | 0.865715 | NA |
| ENSMUSG( | 11.79579 | 0.556703 | 0.302853 | NA | NA |
| ENSMUSG( | 13.86655 | 0.357925 | 0.481091 | NA | NA |
| ENSMUSG( | 111.2099 | -0.11232 | 0.588499 | 0.900921 | NA |
| ENSMUSG( | 494.6208 | 0.19819 | 0.160229 | 0.589853 | NA |
| ENSMUSG( | 16.67936 | -0.15722 | 0.724256 | NA | NA |
| ENSMUSG( | 88.78723 | -0.05959 | 0.771356 | 0.954726 | NA |
| ENSMUSG( | 10.8757 | -0.24997 | 0.655723 | NA | NA |
| ENSMUSG( | 16.33138 | 0.178787 | 0.69393 | NA | NA |
| ENSMUSG( | 437.0647 | 0.010957 | 0.926533 | 0.991229 | NA |
| ENSMUSG( | 9.389939 | 0.178377 | 0.781961 | NA | NA |
| ENSMUSG( | 195.9693 | 0.100377 | 0.549672 | 0.885944 | NA |
| ENSMUSG( | 1602.652 | 0.099515 | 0.272241 | 0.7223 | NA |
| ENSMUSG( | 8.237051 | -0.08067 | 0.911023 | NA | NA |
| ENSMUSG( | 8.553179 | 0.135047 | 0.830197 | NA | NA |
| ENSMUSG( | 25.87262 | 0.245493 | 0.551402 | 0.886141 | NA |
| ENSMUSG( | 303.2443 | 0.293946 | 0.049689 | 0.340355 | NA |
| ENSMUSG( | 24.32698 | -0.3007 | 0.437985 | 0.830072 | NA |
| ENSMUSG( | 13.12932 | 0.416284 | 0.413424 | NA | NA |
| ENSMUSG( | 346.2748 | -0.07293 | 0.62471 | 0.912042 | NA |
| ENSMUSG( | 21.65521 | 0.118111 | 0.771102 | 0.954726 | NA |
| ENSMUSG( | 27.33897 | -0.17593 | 0.619747 | 0.911432 | NA |
| ENSMUSG( | 24.16938 | 0.046763 | 0.906614 | 0.98713 | NA |
| ENSMUSG( | 127.2508 | -0.18027 | 0.355355 | 0.78783 | NA |
| ENSMUSG( | 25.96521 | 0.544312 | 0.155078 | 0.5828 | NA |
| ENSMUSG( | 21.42822 | 0.236053 | 0.555632 | 0.887457 | NA |
| ENSMUSG( | 14.34971 | -0.07954 | 0.878447 | NA | NA |
| ENSMUSG( | 22.0474 | 0.808463 | 0.066495 | 0.39868 | NA |
| ENSMUSG( | 180.1973 | -0.02107 | 0.893192 | 0.984512 | NA |
| ENSMUSG( | 89.73384 | -0.0271 | 0.894098 | 0.984512 | NA |
| ENSMUSG( | 17.67093 | -0.42362 | 0.339794 | NA | NA |
| ENSMUSG( | 12.25662 | 1.134484 | 0.041361 | NA | NA |
| ENSMUSG( | 65.75911 | -0.03353 | 0.894633 | 0.984512 | NA |
| ENSMUSG( | 10.99091 | 0.5758 | 0.303179 | NA | NA |
| ENSMUSG( | 32.30008 | 0.195023 | 0.597577 | 0.903694 | NA |
| ENSMUSG( | 28.07008 | 0.406327 | 0.275079 | 0.72509 | NA |
| ENSMUSG( | 27.23009 | -0.17444 | 0.639663 | 0.916434 | NA |
| ENSMUSG( | 28.698 | -0.38687 | 0.293363 | 0.742763 | NA |
| ENSMUSG( | 11.64977 | 0.313423 | 0.56362 | NA | NA |
| ENSMUSG( | 24.60247 | 0.260495 | 0.491808 | 0.860249 | NA |
| ENSMUSG( | 51.19022 | 0.06354 | 0.820152 | 0.965888 | NA |
| ENSMUSG( | 12.16629 | 0.009466 | 0.986004 | NA | NA |
| ENSMUSG( | 28.75336 | 0.013911 | 0.969027 | 0.996369 | NA |
| ENSMUSG( | 16.24251 | 0.356756 | 0.464733 | NA | NA |
| ENSMUSG( | 9.419069 | 0.361913 | 0.588652 | NA | NA |
| ENSMUSG( | 14.84962 | 0.585908 | 0.265752 | NA | NA |
| ENSMUSG( | 13.49936 | 0.38392 | 0.488486 | NA | NA |
| ENSMUSG( | 35.75615 | -0.23281 | 0.465421 | 0.844064 | NA |
| ENSMUSG( | 15.762 | -0.43801 | 0.361851 | NA | NA |
| ENSMUSG( | 14.28481 | 0.545443 | 0.30469 | NA | NA |
| ENSMUSG( | 50.8244 | -0.10567 | 0.696405 | 0.935166 | NA |
| ENSMUSG( | 195.4782 | 0.026992 | 0.870704 | 0.977951 | NA |
| ENSMUSG( | 16.56244 | 0.468526 | 0.320531 | NA | NA |
| ENSMUSG( | 12.85775 | -0.14059 | 0.780291 | NA | NA |
| ENSMUSG( | 252.8672 | -0.08752 | 0.648934 | 0.920025 | NA |
| ENSMUSG( | 33.42039 | -0.13585 | 0.714947 | 0.941144 | NA |
| ENSMUSG( | 22.86203 | -0.66592 | 0.106665 | 0.494174 | NA |
| ENSMUSG( | 9.126465 | 0.131853 | 0.837514 | NA | NA |
| ENSMUSG( | 66.58867 | -0.08038 | 0.736696 | 0.946546 | NA |
| ENSMUSG( | 42.64685 | 0.011071 | 0.971236 | 0.996369 | NA |
| ENSMUSG( | 12.51552 | -0.42845 | 0.555375 | NA | NA |
| ENSMUSG( | 24.52871 | -0.76691 | 0.069258 | 0.407813 | NA |
| ENSMUSG( | 24.83779 | -0.4227 | 0.265489 | 0.716109 | NA |
| ENSMUSG( | 148.0169 | -0.17852 | 0.321541 | 0.766705 | NA |
| ENSMUSG( | 257.1961 | 0.171135 | 0.280878 | 0.729868 | NA |
| ENSMUSG( | 98.18784 | -0.2243 | 0.39085 | 0.80826 | NA |
| ENSMUSG( | 21.49071 | -0.38235 | 0.364893 | 0.79407 | NA |
| ENSMUSG( | 30.05902 | 0.201754 | 0.571978 | 0.893526 | NA |
| ENSMUSG( | 8.529782 | 0.854714 | 0.260441 | NA | NA |
| ENSMUSG( | 10.36522 | 0.047091 | 0.934591 | NA | NA |
| ENSMUSG( | 41.78521 | 0.509008 | 0.12659 | 0.534155 | NA |
| ENSMUSG( | 23.49512 | -0.15878 | 0.674603 | 0.929638 | NA |
| ENSMUSG( | 37.33397 | 0.129417 | 0.673329 | 0.929007 | NA |
| ENSMUSG( | 11.17383 | -0.22569 | 0.683636 | NA | NA |
| ENSMUSG( | 44.08166 | 0.137463 | 0.634737 | 0.914073 | NA |
| ENSMUSG( | 31.47698 | 0.249096 | 0.485318 | 0.857408 | NA |
| ENSMUSG( | 67.53653 | -0.08939 | 0.710183 | 0.9397 | NA |
| ENSMUSG( | 77.4573 | 0.047458 | 0.839556 | 0.971179 | NA |
| ENSMUSG( | 13.39409 | 0.319603 | 0.55166 | NA | NA |
| ENSMUSG( | 13.70824 | -0.05232 | 0.9145 | NA | NA |
| ENSMUSG( | 170.3186 | 0.150949 | 0.410178 | 0.818258 | NA |
| ENSMUSG( | 36.36136 | -0.43824 | 0.158997 | 0.588112 | NA |
| ENSMUSG( | 113.4041 | -0.19594 | 0.333744 | 0.775602 | NA |
| ENSMUSG( | 10.23894 | 1.149499 | 0.064222 | NA | NA |
| ENSMUSG( | 11.73129 | 0.099907 | 0.851378 | NA | NA |
| ENSMUSG( | 267.1909 | 0.038617 | 0.79426 | 0.959799 | NA |
| ENSMUSG( | 130.867 | -0.00671 | 0.972502 | 0.996585 | NA |
| ENSMUSG( | 4128.171 | 0.215383 | 0.030728 | 0.266671 | NA |
| ENSMUSG( | 39.25512 | -0.26165 | 0.420425 | 0.82289 | NA |
| ENSMUSG( | 18.81305 | -0.76014 | 0.132868 | NA | NA |
| ENSMUSG( | 55.42706 | 2.475976 | 2.62E-16 | 3.27E-13 | NA |
| ENSMUSG( | 11.03213 | -0.20515 | 0.743275 | NA | NA |
| ENSMUSG( | 24.05968 | 0.321842 | 0.425759 | 0.824898 | NA |
| ENSMUSG( | 81.27139 | 0.105712 | 0.631691 | 0.914073 | NA |
| ENSMUSG( | 29.8597 | 1.787302 | 7.59E-06 | 0.000947 | NA |
| ENSMUSG( | 58.71592 | 0.396531 | 0.127239 | 0.534725 | NA |
| ENSMUSG( | 88.83002 | -0.77638 | 0.003489 | 0.074758 | NA |
| ENSMUSG( | 10.63288 | 0.743067 | 0.209118 | NA | NA |
| ENSMUSG( | 11.63717 | -0.01407 | 0.978973 | NA | NA |
| ENSMUSG( | 12.94434 | 0.724873 | 0.157558 | NA | NA |
| ENSMUSG( | 144.4836 | -0.09689 | 0.643142 | 0.917078 | NA |
| ENSMUSG( | 306.3084 | -0.12781 | 0.339397 | 0.777656 | NA |
| ENSMUSG( | 9.261557 | 0.135229 | 0.829042 | NA | NA |
| ENSMUSG( | 407.0058 | 0.588316 | 3.57E-06 | 0.000492 | NA |
| ENSMUSG( | 39.63115 | -0.22673 | 0.505787 | 0.866936 | NA |
| ENSMUSG( | 12.91545 | 0.573753 | 0.282735 | NA | NA |
| ENSMUSG( | 81.98548 | -0.22642 | 0.31039 | 0.758554 | NA |
| ENSMUSG( | 37.16078 | -0.03548 | 0.917682 | 0.989377 | NA |
| ENSMUSG( | 306.1616 | -0.08763 | 0.546251 | 0.884974 | NA |
| ENSMUSG( | 18.4007 | 0.153154 | 0.744555 | NA | NA |
| ENSMUSG( | 10.62653 | -0.14802 | 0.808076 | NA | NA |
| ENSMUSG( | 489806.4 | 0.207701 | 0.107892 | 0.49674 | NA |
| ENSMUSG( | 636393 | 0.193178 | 0.21374 | 0.658097 | NA |
| ENSMUSG( | 765.5616 | 0.107054 | 0.536723 | 0.882043 | NA |
| ENSMUSG( | 218.1282 | -0.15732 | 0.806891 | 0.962973 | NA |
| ENSMUSG( | 105.8059 | -0.33365 | 0.219009 | 0.664602 | NA |
| ENSMUSG( | 1465.281 | 0.127751 | 0.299344 | 0.748131 | NA |
| ENSMUSG( | 886.1749 | 0.031352 | 0.950338 | 0.993821 | NA |
| ENSMUSG( | 25.85554 | -0.29947 | 0.471094 | 0.846975 | NA |
| ENSMUSG( | 55.15565 | -0.11998 | 0.677366 | 0.930542 | NA |
| ENSMUSG( | 14.26742 | 0.061307 | 0.914975 | NA | NA |
| ENSMUSG( | 43.59095 | 0.131134 | 0.66377 | 0.926511 | NA |
| ENSMUSG( | 124.6798 | 0.002519 | 0.988699 | 0.999109 | NA |
| ENSMUSG( | 170.4916 | -0.08622 | 0.680577 | 0.931086 | NA |
| ENSMUSG( | 44.65534 | 0.252462 | 0.422107 | 0.823495 | NA |
| ENSMUSG( | 18.60359 | 0.010366 | 0.980639 | NA | NA |
| ENSMUSG( | 12.18315 | 0.377565 | 0.475311 | NA | NA |

| | | | | | |
|---|---|---|---|---|---|
| ENSMUSG( | 8.161166 | -0.40846 | 0.559256 | NA | NA |
| ENSMUSG( | 17.08392 | 0.136842 | 0.78746 | 0.97637 | NA |
| ENSMUSG( | 15.09506 | 0.05535 | 0.918938 | 0.992529 | NA |
| ENSMUSG( | 11.74281 | 0.765968 | 0.21276 | 0.864412 | NA |
| ENSMUSG( | 16.7805 | 0.288735 | 0.562052 | 0.94044 | NA |
| ENSMUSG( | 30.50783 | -0.67181 | 0.074978 | 0.76762 | NA |
| ENSMUSG( | 36.31751 | 0.575177 | 0.210772 | 0.864412 | NA |
| ENSMUSG( | 13.87565 | -0.29505 | 0.58899 | 0.944975 | NA |
| ENSMUSG( | 14.27386 | -0.33717 | 0.546929 | 0.938063 | NA |
| ENSMUSG( | 7.350145 | -1.42417 | 0.063019 | NA | NA |
| ENSMUSG( | 11.7858 | -0.84113 | 0.152889 | 0.837724 | NA |
| ENSMUSG( | 12.58695 | -0.526 | 0.354576 | 0.912411 | NA |
| ENSMUSG( | 14.34065 | -0.62818 | 0.252591 | 0.88018 | NA |
| ENSMUSG( | 21.98834 | 0.168909 | 0.717536 | 0.965964 | NA |
| ENSMUSG( | 8.558528 | 0.299425 | 0.661395 | NA | NA |
| ENSMUSG( | 31.18674 | 2.505986 | 6.46E-05 | 0.015522 | NA |
| ENSMUSG( | 17.56605 | -0.18246 | 0.71665 | 0.965807 | NA |
| ENSMUSG( | 274.5985 | -0.24305 | 0.215591 | 0.865167 | NA |
| ENSMUSG( | 13.49935 | -0.10166 | 0.855332 | 0.985649 | NA |
| ENSMUSG( | 32.34664 | -0.56297 | 0.112786 | 0.809623 | NA |
| ENSMUSG( | 138.5171 | -0.15104 | 0.446633 | 0.921734 | NA |
| ENSMUSG( | 10.05323 | 0.577801 | 0.358156 | NA | NA |
| ENSMUSG( | 26.51262 | -0.41225 | 0.440945 | 0.921166 | NA |
| ENSMUSG( | 71.66773 | 0.002524 | 0.993312 | 0.998813 | NA |
| ENSMUSG( | 17.82865 | 0.211765 | 0.645353 | 0.953988 | NA |
| ENSMUSG( | 15.44588 | 0.083134 | 0.879706 | 0.989644 | NA |
| ENSMUSG( | 15.44837 | -0.59057 | 0.276683 | 0.890889 | NA |
| ENSMUSG( | 52.13075 | 0.782783 | 0.015314 | 0.495791 | NA |
| ENSMUSG( | 83.28019 | 0.494404 | 0.087423 | 0.784337 | NA |
| ENSMUSG( | 9.248076 | 0.144783 | 0.821047 | NA | NA |
| ENSMUSG( | 15.37073 | -0.16495 | 0.783405 | 0.976106 | NA |
| ENSMUSG( | 230.096 | -0.16223 | 0.342276 | 0.909642 | NA |
| ENSMUSG( | 11.6722 | 0.193897 | 0.758051 | 0.971858 | NA |
| ENSMUSG( | 30.32973 | 0.45696 | 0.224209 | 0.867489 | NA |
| ENSMUSG( | 69.03297 | -0.0733 | 0.816288 | 0.979198 | NA |
| ENSMUSG( | 13.59654 | -0.97679 | 0.0786 | 0.773505 | NA |
| ENSMUSG( | 6.034431 | 0.812266 | 0.360948 | NA | NA |
| ENSMUSG( | 57.21583 | 0.362203 | 0.190686 | 0.857801 | NA |
| ENSMUSG( | 18.00253 | 0.2397 | 0.63972 | 0.953903 | NA |
| ENSMUSG( | 18.99581 | -0.35825 | 0.504821 | 0.927825 | NA |
| ENSMUSG( | 51.85251 | 0.129676 | 0.656806 | 0.955567 | NA |
| ENSMUSG( | 10.91928 | -0.29878 | 0.63801 | NA | NA |
| ENSMUSG( | 13.77059 | 0.22926 | 0.687125 | 0.959302 | NA |
| ENSMUSG( | 7.953109 | -0.36106 | 0.630265 | NA | NA |
| ENSMUSG( | 9.531496 | -0.21366 | 0.756812 | NA | NA |
| ENSMUSG( | 77.73879 | 0.189579 | 0.467971 | 0.923329 | NA |
| ENSMUSG( | 116.9467 | 0.317822 | 0.247318 | 0.877904 | NA |
| ENSMUSG( | 60.65015 | 0.474561 | 0.100007 | 0.801311 | NA |
| ENSMUSG( | 473.8937 | -0.4049 | 0.005185 | 0.296096 | NA |
| ENSMUSG( | 25.59305 | -0.47112 | 0.264408 | 0.886134 | NA |
| ENSMUSG( | 7.94032 | -0.69188 | 0.328518 | NA | NA |
| ENSMUSG( | 35.88362 | 0.049925 | 0.888871 | 0.990558 | NA |
| ENSMUSG( | 81.18496 | -0.09453 | 0.694123 | 0.960867 | NA |
| ENSMUSG( | 9.780293 | 0.08667 | 0.88698 | NA | NA |
| ENSMUSG( | 378.8975 | -0.13931 | 0.33005 | 0.907251 | NA |
| ENSMUSG( | 38.25326 | -0.41241 | 0.222471 | 0.866791 | NA |
| ENSMUSG( | 17.14846 | -0.71178 | 0.149128 | 0.836241 | NA |
| ENSMUSG( | 26.24069 | -0.36291 | 0.358834 | 0.913759 | NA |
| ENSMUSG( | 57.32067 | 0.488972 | 0.075926 | 0.768524 | NA |
| ENSMUSG( | 15.02222 | 0.394811 | 0.445584 | 0.921734 | NA |
| ENSMUSG( | 3256.382 | -0.20165 | 0.094182 | 0.795735 | NA |
| ENSMUSG( | 29.02338 | -0.42165 | 0.267787 | 0.886134 | NA |
| ENSMUSG( | 10.85814 | 0.792159 | 0.21439 | NA | NA |
| ENSMUSG( | 9.774681 | -0.2541 | 0.719829 | NA | NA |
| ENSMUSG( | 47.49955 | 0.271486 | 0.447621 | 0.922202 | NA |
| ENSMUSG( | 107.6517 | 0.003719 | 0.986052 | 0.997188 | NA |
| ENSMUSG( | 33.49378 | -0.02925 | 0.933564 | 0.99306 | NA |
| ENSMUSG( | 9.961631 | 0.513608 | 0.432767 | NA | NA |
| ENSMUSG( | 59.23543 | 0.412956 | 0.173761 | 0.844317 | NA |
| ENSMUSG( | 64.6843 | 0.120041 | 0.694858 | 0.960867 | NA |
| ENSMUSG( | 99.51405 | 0.011352 | 0.958136 | 0.994798 | NA |
| ENSMUSG( | 27.53495 | -0.04013 | 0.917008 | 0.992529 | NA |
| ENSMUSG( | 11.97855 | -0.5153 | 0.36292 | 0.914223 | NA |
| ENSMUSG( | 21.52187 | 0.058518 | 0.894801 | 0.990558 | NA |
| ENSMUSG( | 85.75639 | 0.180693 | 0.42233 | 0.917257 | NA |
| ENSMUSG( | 26.39568 | -0.52431 | 0.239367 | 0.87431 | NA |
| ENSMUSG( | 14.05243 | -0.0209 | 0.970433 | 0.99594 | NA |
| ENSMUSG( | 347.2521 | -0.09424 | 0.661523 | 0.956205 | NA |
| ENSMUSG( | 35.05591 | -0.30409 | 0.435939 | 0.919569 | NA |
| ENSMUSG( | 11.34769 | 0.826799 | 0.211853 | NA | NA |
| ENSMUSG( | 446.3991 | 0.074326 | 0.625786 | 0.953234 | NA |
| ENSMUSG( | 16.02758 | -0.71027 | 0.145005 | 0.83339 | NA |
| ENSMUSG( | 20.26847 | -0.48016 | 0.278214 | 0.891468 | NA |
| ENSMUSG( | 15.21035 | -0.45021 | 0.374823 | 0.915491 | NA |
| ENSMUSG( | 7.53809 | -0.29776 | 0.668594 | NA | NA |
| ENSMUSG( | 40.76565 | -0.48076 | 0.132991 | 0.822912 | NA |
| ENSMUSG( | 8.839189 | 0.629488 | 0.369363 | NA | NA |
| ENSMUSG( | 46.4083 | 0.253453 | 0.412455 | 0.917257 | NA |
| ENSMUSG( | 22.27681 | -0.11314 | 0.794831 | 0.97681 | NA |
| ENSMUSG( | 55.214 | -0.21194 | 0.453047 | 0.922316 | NA |
| ENSMUSG( | 54.63886 | -0.31299 | 0.311849 | 0.900037 | NA |
| ENSMUSG( | 25.41119 | -0.18929 | 0.657903 | 0.955993 | NA |
| ENSMUSG( | 83.03499 | -0.61126 | 0.02191 | 0.575341 | NA |
| ENSMUSG( | 26.85184 | 0.401764 | 0.284683 | 0.892412 | NA |
| ENSMUSG( | 46.44999 | -0.06654 | 0.836756 | 0.981335 | NA |
| ENSMUSG( | 27.90439 | -0.39857 | 0.283764 | 0.892412 | NA |
| ENSMUSG( | 10.45935 | -0.1738 | 0.785458 | NA | NA |
| ENSMUSG( | 18.9131 | 0.605776 | 0.253402 | 0.88018 | NA |
| ENSMUSG( | 21.81969 | 0.054847 | 0.909086 | 0.991958 | NA |
| ENSMUSG( | 8.857626 | 0.603214 | 0.351447 | NA | NA |
| ENSMUSG( | 388.128 | -0.38844 | 0.106775 | 0.804969 | NA |
| ENSMUSG( | 6654.488 | -0.28024 | 0.163296 | 0.838291 | NA |
| ENSMUSG( | 396.3731 | 0.152104 | 0.824485 | 0.979198 | NA |
| ENSMUSG( | 76.3322 | 0.327438 | 0.248749 | 0.877904 | NA |
| ENSMUSG( | 34.99633 | -0.40771 | 0.223733 | 0.867404 | NA |
| ENSMUSG( | 10.888 | 0.341444 | 0.580657 | NA | NA |
| ENSMUSG( | 88.79954 | -0.11508 | 0.633691 | 0.953889 | NA |
| ENSMUSG( | 12.49738 | -0.93195 | 0.118422 | 0.813384 | NA |
| ENSMUSG( | 13.31942 | 0.578493 | 0.322866 | 0.904023 | NA |
| ENSMUSG( | 8.474896 | -0.06967 | 0.917268 | NA | NA |
| ENSMUSG( | 10.49214 | -0.05431 | 0.927862 | NA | NA |
| ENSMUSG( | 52.39235 | -0.04308 | 0.885384 | 0.989688 | NA |
| ENSMUSG( | 16.9555 | 0.156866 | 0.74238 | 0.969415 | NA |
| ENSMUSG( | 227.0966 | -0.0078 | 0.967007 | 0.995549 | NA |
| ENSMUSG( | 23.54701 | -0.39144 | 0.349283 | 0.911652 | NA |
| ENSMUSG( | 36.61904 | -0.50648 | 0.155123 | 0.837724 | NA |
| ENSMUSG( | 13.08622 | -0.78633 | 0.186009 | 0.855305 | NA |
| ENSMUSG( | 21.14252 | -0.09951 | 0.826256 | 0.979795 | NA |
| ENSMUSG( | 9.906988 | -0.82604 | 0.203528 | NA | NA |
| ENSMUSG( | 13.52981 | 0.036354 | 0.946527 | 0.994798 | NA |
| ENSMUSG( | 184.7586 | -0.33139 | 0.096358 | 0.796133 | NA |
| ENSMUSG( | 9.406012 | -0.05793 | 0.932987 | NA | NA |
| ENSMUSG( | 37.80795 | 0.152169 | 0.657446 | 0.955725 | NA |
| ENSMUSG( | 27.20955 | -0.78386 | 0.054044 | 0.722658 | NA |
| ENSMUSG( | 38.37037 | 0.672005 | 0.050151 | 0.708648 | NA |
| ENSMUSG( | 21.17549 | -0.08072 | 0.856436 | 0.985884 | NA |
| ENSMUSG( | 16.10603 | 0.30743 | 0.547446 | 0.938063 | NA |
| ENSMUSG( | 82.0027 | -0.16968 | 0.461226 | 0.923329 | NA |
| ENSMUSG( | 12.6846 | 0.002667 | 0.99632 | 0.999178 | NA |
| ENSMUSG( | 10.57165 | 0.35083 | 0.566589 | NA | NA |
| ENSMUSG( | 255.5346 | -0.85295 | 7.50E-07 | 0.000398 | NA |
| ENSMUSG( | 13.24372 | -0.48148 | 0.371784 | 0.915491 | NA |
| ENSMUSG( | 12.80202 | 0.26695 | 0.616689 | 0.951597 | NA |
| ENSMUSG( | 16.16797 | 0.113793 | 0.819653 | 0.979198 | NA |
| ENSMUSG( | 23.37977 | 0.520962 | 0.356935 | 0.913358 | NA |
| ENSMUSG( | 104.7617 | -0.14188 | 0.512159 | 0.930074 | NA |
| ENSMUSG( | 119.1043 | 0.243275 | 0.322449 | 0.904023 | NA |
| ENSMUSG( | 9.411343 | 0.050686 | 0.934963 | NA | NA |
| ENSMUSG( | 1054.431 | -0.28335 | 0.027063 | 0.603041 | NA |
| ENSMUSG( | 14.63039 | -0.15326 | 0.77485 | 0.974931 | NA |
| ENSMUSG( | 296.6592 | -0.29849 | 0.121552 | 0.813384 | NA |
| ENSMUSG( | 14.63601 | -0.37945 | 0.466205 | 0.923329 | NA |
| ENSMUSG( | 600.934 | -0.05286 | 0.708158 | 0.964042 | NA |
| ENSMUSG( | 21.58057 | -0.56958 | 0.244842 | 0.877904 | NA |
| ENSMUSG( | 10.93116 | 0.099096 | 0.880389 | NA | NA |
| ENSMUSG( | 8.47474 | 0.555072 | 0.525801 | NA | NA |
| ENSMUSG( | 28.17509 | -0.11592 | 0.757891 | 0.971858 | NA |
| ENSMUSG( | 62.26946 | 0.086326 | 0.749759 | 0.96998 | NA |
| ENSMUSG( | 25.77972 | -0.11187 | 0.783544 | 0.976106 | NA |
| ENSMUSG( | 20.79406 | -0.07356 | 0.877396 | 0.989644 | NA |
| ENSMUSG( | 40.66502 | -0.23342 | 0.476218 | 0.923329 | NA |
| ENSMUSG( | 40.93599 | -0.02826 | 0.929817 | 0.99306 | NA |
| ENSMUSG( | 12.06589 | 0.019874 | 0.974608 | 0.996416 | NA |
| ENSMUSG( | 12.6781 | 0.073086 | 0.905148 | 0.991089 | NA |
| ENSMUSG( | 22.71732 | 0.198939 | 0.651021 | 0.954303 | NA |
| ENSMUSG( | 9.106528 | 0.506843 | 0.429455 | NA | NA |
| ENSMUSG( | 19.6888 | -0.34651 | 0.504685 | 0.927825 | NA |
| ENSMUSG( | 35.02733 | -0.33129 | 0.35679 | 0.913358 | NA |

| | | | | | | | | | | | |
|---|---|---|---|---|---|---|---|---|---|---|---|
| ENSMUSG( | NA | 25.85554 | -0.29947 | 0.471094 | 0.846975 | ENSMUSG( | NA | 16.02758 | -0.71027 | 0.145005 | 0.83339 |
| ENSMUSG( | NA | 55.15565 | -0.11998 | 0.677366 | 0.930542 | ENSMUSG( | NA | 20.26847 | -0.48016 | 0.278214 | 0.891468 |
| ENSMUSG( | NA | 14.26742 | 0.061307 | 0.914975 | NA | ENSMUSG( | NA | 15.21035 | -0.45021 | 0.374823 | 0.915491 |
| ENSMUSG( | NA | 43.59095 | 0.131134 | 0.66377 | 0.926511 | ENSMUSG( | NA | 40.76565 | -0.48076 | 0.132991 | 0.822912 |
| ENSMUSG( | NA | 124.6798 | 0.002519 | 0.988699 | 0.999109 | ENSMUSG( | NA | 8.839189 | 0.629488 | 0.369363 | NA |
| ENSMUSG( | NA | 170.4916 | -0.08622 | 0.680577 | 0.931086 | ENSMUSG( | NA | 46.4083 | 0.253453 | 0.412455 | 0.917257 |
| ENSMUSG( | NA | 44.65534 | 0.252462 | 0.422107 | 0.823495 | ENSMUSG( | NA | 22.27681 | -0.11314 | 0.794831 | 0.97681 |
| ENSMUSG( | NA | 18.60359 | 0.010366 | 0.980639 | NA | ENSMUSG( | NA | 55.214 | -0.21194 | 0.453047 | 0.922316 |
| ENSMUSG( | NA | 12.18315 | 0.377565 | 0.475311 | NA | ENSMUSG( | NA | 54.63886 | -0.31299 | 0.311849 | 0.900037 |
| ENSMUSG( | NA | 28.55621 | -0.12865 | 0.713732 | 0.940903 | ENSMUSG( | NA | 25.41119 | -0.18929 | 0.657903 | 0.955993 |
| ENSMUSG( | NA | 12.81816 | -0.05341 | 0.918261 | NA | ENSMUSG( | NA | 83.03499 | -0.61126 | 0.02191 | 0.575341 |
| ENSMUSG( | NA | 18.93676 | -0.09205 | 0.827614 | NA | ENSMUSG( | NA | 26.85184 | 0.401764 | 0.284683 | 0.892412 |
| ENSMUSG( | NA | 10.97004 | -0.69887 | 0.204773 | NA | ENSMUSG( | NA | 46.44999 | -0.06654 | 0.836756 | 0.981335 |
| ENSMUSG( | NA | 20.01162 | 0.359219 | 0.428786 | NA | ENSMUSG( | NA | 27.90439 | -0.39857 | 0.283764 | 0.892412 |
| ENSMUSG( | NA | 20.43468 | 0.2171 | 0.607488 | NA | ENSMUSG( | NA | 10.45935 | -0.1738 | 0.785458 | NA |
| ENSMUSG( | NA | 24.76337 | 0.407769 | 0.267838 | 0.718054 | ENSMUSG( | NA | 18.9131 | 0.605776 | 0.253402 | 0.88018 |
| ENSMUSG( | NA | 24.9184 | -0.45747 | 0.221911 | 0.667491 | ENSMUSG( | NA | 21.81969 | 0.054847 | 0.909086 | 0.991958 |
| ENSMUSG( | NA | 12.11318 | -0.02091 | 0.970919 | NA | ENSMUSG( | NA | 388.128 | -0.38844 | 0.106775 | 0.804969 |
| ENSMUSG( | NA | 30.81076 | -0.41905 | 0.214698 | 0.659193 | ENSMUSG( | NA | 6654.488 | -0.28024 | 0.163296 | 0.838291 |
| ENSMUSG( | NA | 13.14441 | 0.197725 | 0.698503 | NA | ENSMUSG( | NA | 396.3731 | 0.152104 | 0.824485 | 0.979198 |
| ENSMUSG( | NA | 87.02486 | -0.0282 | 0.913627 | 0.988483 | ENSMUSG( | NA | 76.3322 | 0.327438 | 0.248749 | 0.877904 |
| ENSMUSG( | NA | 31.66375 | 0.62271 | 0.123983 | 0.531451 | ENSMUSG( | NA | 34.99633 | -0.40771 | 0.223733 | 0.867404 |
| ENSMUSG( | NA | 101.0053 | 0.150435 | 0.467274 | 0.844819 | ENSMUSG( | NA | 10.888 | 0.341444 | 0.580657 | NA |
| ENSMUSG( | NA | 14.72636 | -0.13722 | 0.793908 | NA | ENSMUSG( | NA | 88.79954 | -0.11508 | 0.633691 | 0.953889 |
| ENSMUSG( | NA | 145.1001 | -0.01653 | 0.922093 | 0.99045 | ENSMUSG( | NA | 12.49738 | -0.93195 | 0.118422 | 0.813384 |
| ENSMUSG( | NA | 9.452881 | -0.33736 | 0.576077 | NA | ENSMUSG( | NA | 13.31942 | 0.578493 | 0.322866 | 0.904023 |
| ENSMUSG( | NA | 76.22826 | 0.099276 | 0.681597 | 0.931619 | ENSMUSG( | NA | 52.39235 | -0.04308 | 0.885384 | 0.989688 |
| ENSMUSG( | NA | 116.9455 | -0.45006 | 0.018777 | 0.206056 | ENSMUSG( | NA | 16.9555 | 0.156866 | 0.74238 | 0.969415 |
| ENSMUSG( | NA | 45.33972 | 0.341904 | 0.245863 | 0.693547 | ENSMUSG( | NA | 227.0966 | -0.0078 | 0.967007 | 0.995549 |
| ENSMUSG( | NA | 508.8893 | 0.173497 | 0.166646 | 0.599201 | ENSMUSG( | NA | 23.54701 | -0.39144 | 0.349283 | 0.911652 |
| ENSMUSG( | NA | 41.17927 | 0.165964 | 0.609301 | 0.907627 | ENSMUSG( | NA | 36.61904 | -0.50648 | 0.155123 | 0.837724 |
| ENSMUSG( | NA | 15.32174 | 0.154951 | 0.753082 | NA | ENSMUSG( | NA | 13.08622 | -0.78633 | 0.186009 | 0.855305 |
| ENSMUSG( | NA | 18.37007 | -0.16378 | 0.729747 | NA | ENSMUSG( | NA | 21.14252 | -0.09951 | 0.826256 | 0.979795 |
| ENSMUSG( | NA | 39.46393 | -0.40987 | 0.186395 | 0.623869 | ENSMUSG( | NA | 9.906988 | -0.82604 | 0.203528 | NA |
| ENSMUSG( | NA | 26.55616 | 0.002329 | 0.995153 | 0.99943 | ENSMUSG( | NA | 13.52981 | 0.036354 | 0.946527 | 0.994798 |
| ENSMUSG( | NA | 261.1064 | 0.12354 | 0.424845 | 0.824675 | ENSMUSG( | NA | 184.7586 | -0.33139 | 0.096358 | 0.796133 |
| ENSMUSG( | NA | 44.03045 | 0.642644 | 0.025648 | 0.246969 | ENSMUSG( | NA | 9.406012 | -0.05793 | 0.932987 | NA |
| ENSMUSG( | NA | 15.31573 | -0.16576 | 0.777036 | NA | ENSMUSG( | NA | 37.80795 | 0.152169 | 0.657446 | 0.955725 |
| ENSMUSG( | NA | 264.4583 | 0.13086 | 0.333144 | 0.775602 | ENSMUSG( | NA | 27.20955 | -0.78386 | 0.054044 | 0.722658 |
| ENSMUSG( | NA | 30.87672 | 0.124316 | 0.726586 | 0.944223 | ENSMUSG( | NA | 38.37037 | 0.672005 | 0.050151 | 0.708648 |
| ENSMUSG( | NA | 3090.628 | 0.142771 | 0.230231 | 0.678139 | ENSMUSG( | NA | 21.17549 | -0.08072 | 0.856436 | 0.985884 |
| ENSMUSG( | NA | 57.10743 | -0.15882 | 0.532647 | 0.879905 | ENSMUSG( | NA | 16.10603 | 0.30743 | 0.547446 | 0.938063 |
| ENSMUSG( | NA | 475.3699 | 0.02286 | 0.852175 | 0.973168 | ENSMUSG( | NA | 82.0027 | -0.16968 | 0.461226 | 0.923329 |
| ENSMUSG( | NA | 49.42544 | 0.066236 | 0.810421 | 0.964049 | ENSMUSG( | NA | 12.6846 | 0.002667 | 0.99632 | 0.999178 |
| ENSMUSG( | NA | 18.5141 | 0.44049 | 0.334046 | NA | ENSMUSG( | NA | 10.57165 | 0.35083 | 0.566589 | NA |
| ENSMUSG( | NA | 15.26201 | 0.198727 | 0.689208 | NA | ENSMUSG( | NA | 255.5346 | -0.85295 | 7.50E-07 | 0.000398 |
| ENSMUSG( | NA | 21.20706 | 0.224829 | 0.573693 | 0.893584 | ENSMUSG( | NA | 13.24372 | -0.48148 | 0.371784 | 0.915491 |
| ENSMUSG( | NA | 59.39771 | 0.613706 | 0.026414 | 0.248798 | ENSMUSG( | NA | 12.80202 | 0.26695 | 0.616689 | 0.951597 |
| ENSMUSG( | NA | 34.94553 | 0.074426 | 0.819714 | 0.965888 | ENSMUSG( | NA | 16.16797 | 0.113793 | 0.819653 | 0.979198 |
| ENSMUSG( | NA | 44.78684 | -0.26356 | 0.437843 | 0.830072 | ENSMUSG( | NA | 23.37977 | 0.520962 | 0.356935 | 0.913358 |
| ENSMUSG( | NA | 216.6634 | -0.01676 | 0.919451 | 0.990247 | ENSMUSG( | NA | 104.7617 | -0.14188 | 0.512159 | 0.930074 |
| ENSMUSG( | NA | 19.49547 | -0.23582 | 0.575299 | NA | ENSMUSG( | NA | 119.1043 | 0.243275 | 0.322449 | 0.904023 |
| ENSMUSG( | NA | 61.0789 | -0.17766 | 0.498947 | 0.862973 | ENSMUSG( | NA | 9.411343 | 0.050686 | 0.934963 | NA |
| ENSMUSG( | NA | 16.50138 | -0.3038 | 0.496813 | NA | ENSMUSG( | NA | 1054.431 | -0.28335 | 0.027063 | 0.603041 |
| ENSMUSG( | NA | 35.23777 | -0.2265 | 0.520032 | 0.874992 | ENSMUSG( | NA | 14.63039 | -0.15326 | 0.77485 | 0.974931 |
| ENSMUSG( | NA | 35.77642 | 0.076263 | 0.823577 | 0.966603 | ENSMUSG( | NA | 296.6592 | -0.29849 | 0.121552 | 0.813384 |
| ENSMUSG( | NA | 67.92379 | 0.208392 | 0.452213 | 0.836145 | ENSMUSG( | NA | 14.63601 | -0.37945 | 0.466205 | 0.923329 |
| ENSMUSG( | NA | 64.97808 | 0.060662 | 0.799789 | 0.961501 | ENSMUSG( | NA | 600.934 | -0.05286 | 0.708158 | 0.964042 |
| ENSMUSG( | NA | 12.7846 | -0.97418 | 0.117638 | NA | ENSMUSG( | NA | 21.58057 | -0.56958 | 0.244842 | 0.877904 |
| ENSMUSG( | NA | 36.04931 | 0.315375 | 0.319928 | 0.765375 | ENSMUSG( | NA | 10.93116 | 0.099096 | 0.880389 | NA |
| ENSMUSG( | NA | 23.36298 | -1.23301 | 0.02317 | 0.233568 | ENSMUSG( | NA | 28.17509 | -0.11592 | 0.757891 | 0.971858 |
| ENSMUSG( | NA | 79.63278 | -0.01475 | 0.949763 | 0.993821 | ENSMUSG( | NA | 62.26946 | 0.086326 | 0.749759 | 0.96998 |
| ENSMUSG( | NA | 66.66721 | -0.09335 | 0.717683 | 0.941697 | ENSMUSG( | NA | 25.77972 | -0.11187 | 0.783544 | 0.976106 |
| ENSMUSG( | NA | 30.82449 | -0.1371 | 0.708957 | 0.939519 | ENSMUSG( | NA | 20.79406 | -0.07356 | 0.877396 | 0.989644 |
| ENSMUSG( | NA | 77.69785 | -0.03445 | 0.881395 | 0.981798 | ENSMUSG( | NA | 40.66502 | -0.23342 | 0.476218 | 0.923329 |
| ENSMUSG( | NA | 19.55819 | -0.17451 | 0.684594 | NA | ENSMUSG( | NA | 40.93599 | -0.02826 | 0.929817 | 0.99306 |
| ENSMUSG( | NA | 50.53213 | 0.054251 | 0.844587 | 0.972023 | ENSMUSG( | NA | 12.06589 | 0.019874 | 0.974608 | 0.996416 |
| ENSMUSG( | NA | 197.8227 | -0.01489 | 0.920188 | 0.990247 | ENSMUSG( | NA | 12.6781 | 0.073086 | 0.905148 | 0.991089 |
| ENSMUSG( | NA | 24.43794 | 0.064375 | 0.868829 | 0.976897 | ENSMUSG( | NA | 22.71732 | 0.198939 | 0.651021 | 0.954303 |
| ENSMUSG( | NA | 41.94542 | -0.03403 | 0.915253 | 0.988634 | ENSMUSG( | NA | 9.106528 | 0.506843 | 0.429455 | NA |
| ENSMUSG( | NA | 14.76703 | 0.17859 | 0.738142 | NA | ENSMUSG( | NA | 19.6888 | -0.34651 | 0.504685 | 0.927825 |
| ENSMUSG( | NA | 18.6943 | -0.20069 | 0.644737 | NA | ENSMUSG( | NA | 35.02733 | -0.33129 | 0.35679 | 0.913358 |
| ENSMUSG( | NA | 34.54325 | 0.152217 | 0.653963 | 0.922934 | ENSMUSG( | NA | 10.95242 | 0.202502 | 0.738227 | NA |
| ENSMUSG( | NA | 212.7189 | -0.01826 | 0.91456 | 0.988483 | ENSMUSG( | NA | 12.50826 | 0.786112 | 0.18268 | 0.850579 |
| ENSMUSG( | NA | 80.43175 | -0.21542 | 0.312199 | 0.759769 | ENSMUSG( | NA | 8.808823 | -0.79242 | 0.238222 | NA |
| ENSMUSG( | NA | 88.31943 | -0.01573 | 0.941715 | 0.993415 | ENSMUSG( | NA | 51.10334 | -0.38753 | 0.188361 | 0.85698 |
| ENSMUSG( | NA | 26.90839 | 0.56771 | 0.124993 | 0.531982 | ENSMUSG( | NA | 86.06828 | -0.0657 | 0.801715 | 0.97681 |
| ENSMUSG( | NA | 42.75598 | -0.63609 | 0.056328 | 0.366006 | ENSMUSG( | NA | 168.404 | 0.089714 | 0.616406 | 0.951413 |
| ENSMUSG( | NA | 16.22873 | -0.10355 | 0.828894 | NA | ENSMUSG( | NA | 270.0788 | 0.274152 | 0.223783 | 0.867404 |
| ENSMUSG( | NA | 30.24077 | 0.257179 | 0.507377 | 0.86795 | ENSMUSG( | NA | 177.5594 | -0.17171 | 0.352349 | 0.912093 |
| ENSMUSG( | NA | 18.60829 | 0.39361 | 0.440381 | NA | ENSMUSG( | NA | 11.06248 | -0.23403 | 0.691486 | NA |
| ENSMUSG( | NA | 37.26012 | -0.12373 | 0.710604 | 0.939748 | ENSMUSG( | NA | 21.98294 | -0.37971 | 0.365428 | 0.914516 |
| ENSMUSG( | NA | 254.0003 | 0.055728 | 0.707895 | 0.939519 | ENSMUSG( | NA | 83.70643 | -0.1708 | 0.509173 | 0.929089 |
| ENSMUSG( | NA | 13.85136 | 0.002795 | 0.995712 | NA | ENSMUSG( | NA | 206.1982 | 0.051409 | 0.792075 | 0.97681 |
| ENSMUSG( | NA | 130.8625 | -0.14447 | 0.42138 | 0.823126 | ENSMUSG( | NA | 470.7023 | 0.148262 | 0.327934 | 0.906255 |
| ENSMUSG( | NA | 20.06449 | 0.019226 | 0.968102 | NA | ENSMUSG( | NA | 147.7778 | -0.17354 | 0.374537 | 0.915491 |
| ENSMUSG( | NA | 14.13901 | -0.07642 | 0.877227 | NA | ENSMUSG( | NA | 160.8182 | -0.08275 | 0.694958 | 0.960867 |
| ENSMUSG( | NA | 181.7077 | 0.033009 | 0.837873 | 0.971111 | ENSMUSG( | NA | 23.38411 | 0.442481 | 0.34031 | 0.907972 |
| ENSMUSG( | NA | 1472.025 | 0.025947 | 0.803852 | 0.962169 | ENSMUSG( | NA | 25.24132 | -0.10286 | 0.803406 | 0.976978 |
| ENSMUSG( | NA | 30.70606 | 0.206336 | 0.549049 | 0.885686 | ENSMUSG( | NA | 170.9869 | -0.28582 | 0.129791 | 0.820802 |
| ENSMUSG( | NA | 232.6372 | 0.029406 | 0.849948 | 0.972638 | ENSMUSG( | NA | 10.27664 | -0.27729 | 0.669681 | NA |
| ENSMUSG( | NA | 39.74545 | -0.43452 | 0.218585 | 0.66444 | ENSMUSG( | NA | 99.43578 | 0.10025 | 0.643626 | 0.953903 |
| ENSMUSG( | NA | 224.491 | 0.153919 | 0.274788 | 0.725078 | ENSMUSG( | NA | 14.99111 | -0.35753 | 0.515849 | 0.930939 |
| ENSMUSG( | NA | 37.60932 | 0.329272 | 0.313575 | 0.760494 | ENSMUSG( | NA | 72.25973 | -0.42297 | 0.130652 | 0.822659 |
| ENSMUSG( | NA | 61.04675 | -0.35952 | 0.176541 | 0.612904 | ENSMUSG( | NA | 60.8155 | 0.038418 | 0.898294 | 0.990558 |
| ENSMUSG( | NA | 58.345 | 0.10256 | 0.725492 | 0.944223 | ENSMUSG( | NA | 11.15414 | -0.53871 | 0.361755 | NA |
| ENSMUSG( | NA | 25.36572 | -0.51044 | 0.269712 | 0.719729 | ENSMUSG( | NA | 36.79059 | 0.477204 | 0.148519 | 0.836241 |
| ENSMUSG( | NA | 961.2496 | 0.180688 | 0.060733 | 0.380786 | ENSMUSG( | NA | 9.265101 | -0.16029 | 0.799371 | NA |
| ENSMUSG( | NA | 11.83674 | 0.354743 | 0.543089 | NA | ENSMUSG( | NA | 274.8702 | -0.01199 | 0.940249 | 0.994334 |
| ENSMUSG( | NA | 35.90286 | -0.40916 | 0.244098 | 0.691435 | ENSMUSG( | NA | 152.4746 | -0.15605 | 0.403675 | 0.917257 |
| ENSMUSG( | NA | 19.74054 | -0.06842 | 0.88145 | NA | ENSMUSG( | NA | 92.55244 | -0.15267 | 0.502983 | 0.927825 |
| ENSMUSG( | NA | 21.77046 | -0.10269 | 0.809093 | 0.963548 | ENSMUSG( | NA | 124.4557 | -0.03278 | 0.884629 | 0.989688 |
| ENSMUSG( | NA | 67.83143 | -0.25423 | 0.302055 | 0.750166 | ENSMUSG( | NA | 27.70854 | -0.64591 | 0.10134 | 0.801456 |
| ENSMUSG( | NA | 19.78156 | -0.29333 | 0.517145 | NA | ENSMUSG( | NA | 12.45187 | -0.11401 | 0.844177 | 0.983576 |
| ENSMUSG( | NA | 46.8264 | -0.06007 | 0.837494 | 0.971111 | ENSMUSG( | NA | 57.71717 | 0.005445 | 0.984485 | 0.996963 |
| ENSMUSG( | NA | 20.00947 | -0.50362 | 0.301952 | NA | ENSMUSG( | NA | 10.87046 | -0.19382 | 0.737555 | NA |
| ENSMUSG( | NA | 12.13314 | -0.38706 | 0.464742 | NA | ENSMUSG( | NA | 28.72991 | -0.36815 | 0.336438 | 0.907972 |
| ENSMUSG( | NA | 277.1556 | 0.125119 | 0.380609 | 0.801959 | ENSMUSG( | NA | 65.91055 | -0.2708 | 0.322593 | 0.904023 |
| ENSMUSG( | NA | 20.0606 | -0.03535 | 0.932978 | NA | ENSMUSG( | NA | 28.89386 | -0.21499 | 0.558913 | 0.939587 |
| ENSMUSG( | NA | 120.1835 | -0.60262 | 0.00731 | 0.118207 | ENSMUSG( | NA | 19.68654 | -0.24004 | 0.588098 | 0.944975 |
| ENSMUSG( | NA | 69.53051 | 0.096699 | 0.693657 | 0.934202 | ENSMUSG( | NA | 10.26027 | -0.09131 | 0.890353 | NA |
| ENSMUSG( | NA | 9.078544 | -0.22197 | 0.723623 | NA | ENSMUSG( | NA | 71.04554 | -0.17589 | 0.479411 | 0.924021 |
| ENSMUSG( | NA | 22.17385 | 0.312708 | 0.440258 | 0.831851 | ENSMUSG( | NA | 11.15324 | 0.062963 | 0.914262 | NA |
| ENSMUSG( | NA | 85.93294 | -0.07558 | 0.728162 | 0.944355 | ENSMUSG( | NA | 14.62704 | 0.073797 | 0.894926 | 0.990558 |
| ENSMUSG( | NA | 13.33879 | -0.42972 | 0.410583 | NA | ENSMUSG( | NA | 22.99301 | 0.404416 | 0.326687 | 0.905371 |
| ENSMUSG( | NA | 12.16296 | 0.329601 | 0.57472 | NA | ENSMUSG( | NA | 13.73435 | -0.19772 | 0.701976 | 0.962014 |
| ENSMUSG( | NA | 14.17021 | -0.53828 | 0.283202 | NA | ENSMUSG( | NA | 175.4605 | -0.68056 | 0.000662 | 0.082282 |
| ENSMUSG( | NA | 186.4813 | -0.02348 | 0.892673 | 0.984474 | ENSMUSG( | NA | 16.96177 | -0.76034 | 0.12185 | 0.813384 |
| ENSMUSG( | NA | 12.98063 | 0.179441 | 0.730254 | NA | ENSMUSG( | NA | 47.24925 | -0.43159 | 0.169633 | 0.840806 |
| ENSMUSG( | NA | 11.76804 | 0.125594 | 0.813719 | NA | ENSMUSG( | NA | 17.81868 | -0.47492 | 0.311569 | 0.900031 |
| ENSMUSG( | NA | 26.35714 | -0.3741 | 0.337696 | 0.777422 | ENSMUSG( | NA | 44.9734 | 0.643929 | 0.06391 | 0.736482 |
| ENSMUSG( | NA | 26.63401 | 0.492108 | 0.178811 | 0.616298 | ENSMUSG( | NA | 11.32974 | 0.855526 | 0.143569 | NA |
| ENSMUSG( | NA | 11.67093 | -0.25611 | 0.632276 | NA | ENSMUSG( | NA | 26.24538 | -0.41578 | 0.297723 | 0.898538 |
| ENSMUSG( | NA | 187.0386 | 0.006736 | 0.975304 | 0.997302 | ENSMUSG( | NA | 9.625461 | -0.14566 | 0.829617 | NA |
| ENSMUSG( | NA | 9.643548 | 0.196646 | 0.735666 | NA | ENSMUSG( | NA | 18.38442 | -0.26439 | 0.583493 | 0.944266 |
| ENSMUSG( | NA | 24.31061 | 0.234233 | 0.539854 | 0.88315 | ENSMUSG( | NA | 12.3293 | -0.81707 | 0.176091 | 0.846252 |
| ENSMUSG( | NA | 35.78769 | 0.084311 | 0.803071 | 0.962169 | ENSMUSG( | NA | 22.88118 | 0.52857 | 0.214884 | 0.865167 |
| ENSMUSG( | NA | 11.06557 | 0.014448 | 0.97985 | NA | ENSMUSG( | NA | 17.44207 | -0.42702 | 0.365715 | 0.914623 |
| ENSMUSG( | NA | 17.01076 | -0.47744 | 0.311709 | NA | ENSMUSG( | NA | 8.300776 | -1.27981 | 0.124 | NA |
| ENSMUSG( | NA | 16.1994 | 0.108285 | 0.814877 | NA | ENSMUSG( | NA | 14.96034 | 0.277293 | 0.60714 | 0.949917 |
| ENSMUSG( | NA | 714.8661 | 0.177687 | 0.127925 | 0.536163 | ENSMUSG( | NA | 276.7099 | -0.41639 | 0.05222 | 0.712974 |
| ENSMUSG( | NA | 344.5226 | -0.1275 | 0.392922 | 0.809274 | ENSMUSG( | NA | 13.4149 | 0.376382 | 0.501534 | 0.927825 |
| ENSMUSG( | NA | 13.88513 | 0.730085 | 0.173628 | NA | ENSMUSG( | NA | 131.3691 | -0.35516 | 0.075807 | 0.768524 |
| ENSMUSG( | NA | 15.73596 | 0.470485 | 0.318428 | NA | ENSMUSG( | NA | 8.760287 | -0.2926 | 0.662046 | NA |
| ENSMUSG( | NA | 64.52643 | -0.13828 | 0.583301 | 0.898408 | ENSMUSG( | NA | 19.44629 | -0.10333 | 0.814786 | 0.979198 |
| ENSMUSG( | NA | 10.15559 | -0.15018 | 0.796639 | NA | ENSMUSG( | NA | 15.88936 | 0.374542 | 0.539078 | 0.937216 |
| ENSMUSG( | NA | 11.62544 | 0.069313 | 0.896537 | NA | ENSMUSG( | NA | 218.2092 | -0.58413 | 0.000552 | 0.071583 |
| ENSMUSG( | NA | 11.74102 | 0.319196 | 0.586619 | NA | ENSMUSG( | NA | 22.78605 | -0.36149 | 0.392865 | 0.917257 |
| ENSMUSG( | NA | 14.09746 | -0.01215 | 0.980754 | NA | ENSMUSG( | NA | 66.08614 | 0.102316 | 0.719025 | 0.966777 |
| ENSMUSG( | NA | 11.92986 | 0.454867 | 0.405154 | NA | ENSMUSG( | NA | 12.8967 | 0.760298 | 0.167778 | 0.840222 |
| ENSMUSG( | NA | 25.21349 | 0.116715 | 0.752996 | 0.950667 | ENSMUSG( | NA | 179.1958 | 0.144098 | 0.569622 | 0.941777 |
| ENSMUSG( | NA | 765.4214 | 5.00268 | 8.23E-193 | 1.33E-188 | ENSMUSG( | NA | 53.77471 | -0.21047 | 0.532941 | 0.934795 |
| ENSMUSG( | NA | 81.94534 | 0.140665 | 0.571804 | 0.893506 | ENSMUSG( | NA | 49.67974 | -0.56935 | 0.062879 | 0.736482 |
| ENSMUSG( | NA | 21.13867 | 0.4095 | 0.341112 | NA | ENSMUSG( | NA | 11.98862 | -0.30289 | 0.611849 | 0.95083 |
| ENSMUSG( | NA | 16.93988 | -0.24076 | 0.625835 | NA | ENSMUSG( | NA | 21.09712 | 0.322178 | 0.4732 | 0.923329 |
| ENSMUSG( | NA | 21.86526 | 0.167881 | 0.678095 | 0.930542 | ENSMUSG( | NA | 15.29854 | -0.22122 | 0.662467 | 0.956205 |
| ENSMUSG( | NA | 49.70522 | 0.096055 | 0.735084 | 0.945899 | ENSMUSG( | NA | 48.70397 | 0.871462 | 0.040124 | 0.681706 |
| ENSMUSG( | NA | 57.38853 | -0.21401 | 0.449068 | 0.834943 | ENSMUSG( | NA | 203.6108 | -2.68443 | 8.09E-23 | 2.02E-19 |
| ENSMUSG( | NA | 21.96284 | 0.025094 | 0.957122 | 0.993874 | ENSMUSG( | NA | 80.44757 | 0.045944 | 0.855058 | 0.985649 |
| ENSMUSG( | NA | 31.09363 | 0.068483 | 0.854242 | 0.973491 | ENSMUSG( | NA | 18.2161 | -0.25986 | 0.594709 | 0.945444 |
| ENSMUSG( | NA | 17.16705 | -0.19216 | 0.666285 | NA | ENSMUSG( | NA | 67.98286 | 1.116513 | 4.91E-05 | 0.012613 |
| ENSMUSG( | NA | 72.81865 | 0.066595 | 0.771321 | 0.954726 | ENSMUSG( | NA | 233.6454 | -0.18919 | 0.309144 | 0.898538 |
| ENSMUSG( | NA | 31.21893 | 0.176875 | 0.6028 | 0.904887 | ENSMUSG( | NA | 232.5127 | 0.063387 | 0.7243 | 0.966887 |
| ENSMUSG( | NA | 17.92616 | -0.40014 | 0.369624 | NA | ENSMUSG( | NA | 62.59883 | -0.01955 | 0.946621 | 0.994798 |
| ENSMUSG( | NA | 18.10696 | -0.06257 | 0.885575 | NA | ENSMUSG( | NA | 267.3648 | -0.26611 | 0.09648 | 0.796133 |
| ENSMUSG( | NA | 17.44861 | 0.143699 | 0.757335 | NA | ENSMUSG( | NA | 31.86026 | -0.27491 | 0.470909 | 0.923329 |
| ENSMUSG( | NA | 34.48839 | 0.019055 | 0.958258 | 0.993874 | ENSMUSG( | NA | 29.02584 | 0.019882 | 0.957419 | 0.994798 |
| ENSMUSG( | NA | 20.98825 | 0.876074 | 0.070156 | NA | ENSMUSG( | NA | 18.35068 | 0.047109 | 0.920718 | 0.992644 |

| | | | | | |
|---|---|---|---|---|---|
| ENSMUSG( | 28.55621 | -0.12865 | 0.713732 | 0.940903 | NA |
| ENSMUSG( | 12.81816 | -0.05341 | 0.918261 | NA | NA |
| ENSMUSG( | 18.93676 | -0.09205 | 0.827614 | NA | NA |
| ENSMUSG( | 10.97004 | -0.69887 | 0.204773 | NA | NA |
| ENSMUSG( | 20.01162 | 0.359219 | 0.428786 | NA | NA |
| ENSMUSG( | 20.43468 | 0.2171 | 0.607488 | NA | NA |
| ENSMUSG( | 24.76337 | 0.407769 | 0.267838 | 0.718054 | NA |
| ENSMUSG( | 7.748591 | -0.75938 | 0.34841 | NA | NA |
| ENSMUSG( | 24.9184 | -0.45747 | 0.221911 | 0.667491 | NA |
| ENSMUSG( | 12.11318 | -0.02091 | 0.970919 | NA | NA |
| ENSMUSG( | 30.81076 | -0.41905 | 0.214698 | 0.659193 | NA |
| ENSMUSG( | 13.14441 | 0.197725 | 0.698503 | NA | NA |
| ENSMUSG( | 87.02486 | -0.0282 | 0.913627 | 0.988483 | NA |
| ENSMUSG( | 31.66375 | 0.62271 | 0.123983 | 0.531451 | NA |
| ENSMUSG( | 101.0053 | 0.150435 | 0.467274 | 0.844819 | NA |
| ENSMUSG( | 14.72636 | -0.13722 | 0.793908 | NA | NA |
| ENSMUSG( | 9.619425 | -0.15834 | 0.792786 | NA | NA |
| ENSMUSG( | 11.08914 | 0.452324 | 0.425736 | NA | NA |
| ENSMUSG( | 145.1001 | -0.01653 | 0.922093 | 0.99045 | NA |
| ENSMUSG( | 9.452881 | -0.33736 | 0.576077 | NA | NA |
| ENSMUSG( | 76.22826 | 0.099276 | 0.681597 | 0.931619 | NA |
| ENSMUSG( | 9.746852 | -0.02277 | 0.96994 | NA | NA |
| ENSMUSG( | 116.9455 | -0.45006 | 0.018777 | 0.206056 | NA |
| ENSMUSG( | 45.33972 | 0.341904 | 0.245863 | 0.693547 | NA |
| ENSMUSG( | 508.8893 | 0.173497 | 0.166646 | 0.599201 | NA |
| ENSMUSG( | 41.17927 | 0.165964 | 0.609301 | 0.907627 | NA |
| ENSMUSG( | 15.32174 | 0.154951 | 0.753082 | NA | NA |
| ENSMUSG( | 18.37007 | -0.16378 | 0.729747 | NA | NA |
| ENSMUSG( | 39.46393 | -0.40987 | 0.186395 | 0.623869 | NA |
| ENSMUSG( | 26.55616 | 0.002329 | 0.995153 | 0.99943 | NA |
| ENSMUSG( | 261.1064 | 0.12354 | 0.424845 | 0.824675 | NA |
| ENSMUSG( | 44.03045 | 0.642644 | 0.025648 | 0.246969 | NA |
| ENSMUSG( | 15.31573 | -0.16576 | 0.777036 | NA | NA |
| ENSMUSG( | 9.66936 | 0.121063 | 0.844293 | NA | NA |
| ENSMUSG( | 12.9552 | 0.324173 | 0.571802 | NA | NA |
| ENSMUSG( | 264.4583 | 0.13086 | 0.333144 | 0.775602 | NA |
| ENSMUSG( | 30.87672 | 0.124316 | 0.726586 | 0.944223 | NA |
| ENSMUSG( | 3090.628 | 0.142771 | 0.230231 | 0.678139 | NA |
| ENSMUSG( | 57.10743 | -0.15882 | 0.532647 | 0.879905 | NA |
| ENSMUSG( | 475.3699 | 0.02286 | 0.852175 | 0.973168 | NA |
| ENSMUSG( | 49.42544 | 0.066236 | 0.810421 | 0.964049 | NA |
| ENSMUSG( | 18.5141 | 0.44049 | 0.334046 | NA | NA |
| ENSMUSG( | 15.26201 | 0.198727 | 0.689208 | NA | NA |
| ENSMUSG( | 21.20706 | 0.224829 | 0.573693 | 0.893584 | NA |
| ENSMUSG( | 59.39771 | 0.613706 | 0.026414 | 0.248798 | NA |
| ENSMUSG( | 34.94553 | 0.074426 | 0.819714 | 0.965888 | NA |
| ENSMUSG( | 44.78684 | -0.26356 | 0.437843 | 0.830072 | NA |
| ENSMUSG( | 216.6634 | -0.01676 | 0.919451 | 0.990247 | NA |
| ENSMUSG( | 19.49547 | -0.23582 | 0.575299 | NA | NA |
| ENSMUSG( | 61.0789 | -0.17766 | 0.498947 | 0.862973 | NA |
| ENSMUSG( | 16.50138 | -0.3038 | 0.496813 | NA | NA |
| ENSMUSG( | 35.23777 | -0.2265 | 0.520032 | 0.874992 | NA |
| ENSMUSG( | 35.77642 | 0.076263 | 0.823577 | 0.966603 | NA |
| ENSMUSG( | 67.92379 | 0.208392 | 0.452213 | 0.836145 | NA |
| ENSMUSG( | 64.97808 | 0.060662 | 0.799789 | 0.961501 | NA |
| ENSMUSG( | 12.7846 | -0.97418 | 0.117638 | NA | NA |
| ENSMUSG( | 36.04931 | 0.315375 | 0.319928 | 0.765375 | NA |
| ENSMUSG( | 23.36298 | -1.23301 | 0.02317 | 0.233568 | NA |
| ENSMUSG( | 79.63278 | -0.01475 | 0.949763 | 0.993821 | NA |
| ENSMUSG( | 9.523797 | 0.478175 | 0.474799 | NA | NA |
| ENSMUSG( | 66.66721 | -0.09335 | 0.717683 | 0.941697 | NA |
| ENSMUSG( | 30.82449 | -0.1371 | 0.708957 | 0.939519 | NA |
| ENSMUSG( | 77.69785 | -0.03445 | 0.881395 | 0.981798 | NA |
| ENSMUSG( | 19.55819 | -0.17451 | 0.684594 | NA | NA |
| ENSMUSG( | 50.53213 | 0.054251 | 0.844587 | 0.972023 | NA |
| ENSMUSG( | 197.8227 | -0.01489 | 0.920188 | 0.990247 | NA |
| ENSMUSG( | 24.43794 | 0.064375 | 0.868829 | 0.976897 | NA |
| ENSMUSG( | 41.94542 | -0.03403 | 0.915253 | 0.988634 | NA |
| ENSMUSG( | 14.76703 | 0.17859 | 0.738142 | NA | NA |
| ENSMUSG( | 18.6943 | -0.20069 | 0.644737 | NA | NA |
| ENSMUSG( | 34.54325 | 0.152217 | 0.653963 | 0.922934 | NA |
| ENSMUSG( | 212.7189 | -0.01826 | 0.91456 | 0.988483 | NA |
| ENSMUSG( | 80.43175 | -0.21542 | 0.312199 | 0.759769 | NA |
| ENSMUSG( | 10.99551 | -0.31514 | 0.602723 | NA | NA |
| ENSMUSG( | 88.31943 | -0.01573 | 0.941715 | 0.993415 | NA |
| ENSMUSG( | 26.90839 | 0.56771 | 0.124993 | 0.531982 | NA |
| ENSMUSG( | 17.67236 | 0.391601 | 0.428866 | NA | NA |
| ENSMUSG( | 42.75598 | -0.63609 | 0.056328 | 0.366006 | NA |
| ENSMUSG( | 16.22873 | -0.10355 | 0.828894 | NA | NA |
| ENSMUSG( | 30.24077 | 0.257179 | 0.507377 | 0.86795 | NA |
| ENSMUSG( | 18.60829 | 0.39361 | 0.440381 | NA | NA |
| ENSMUSG( | 37.26012 | -0.12373 | 0.710604 | 0.939748 | NA |
| ENSMUSG( | 254.0003 | 0.055728 | 0.707895 | 0.939519 | NA |
| ENSMUSG( | 13.85136 | 0.002795 | 0.995712 | NA | NA |
| ENSMUSG( | 11.70266 | -0.11575 | 0.830367 | NA | NA |
| ENSMUSG( | 130.8625 | -0.14447 | 0.42138 | 0.823126 | NA |
| ENSMUSG( | 20.06449 | 0.019226 | 0.968102 | NA | NA |
| ENSMUSG( | 14.13901 | -0.07642 | 0.877227 | NA | NA |
| ENSMUSG( | 181.7077 | 0.033009 | 0.837873 | 0.971111 | NA |
| ENSMUSG( | 1472.025 | 0.025947 | 0.803852 | 0.962169 | NA |
| ENSMUSG( | 30.70606 | 0.206336 | 0.549049 | 0.885686 | NA |
| ENSMUSG( | 232.6372 | 0.029406 | 0.849948 | 0.972638 | NA |
| ENSMUSG( | 39.74545 | -0.43452 | 0.218585 | 0.66444 | NA |
| ENSMUSG( | 224.491 | 0.153919 | 0.274788 | 0.725078 | NA |
| ENSMUSG( | 37.60932 | 0.329272 | 0.313575 | 0.760494 | NA |
| ENSMUSG( | 61.04675 | -0.35952 | 0.176541 | 0.612904 | NA |
| ENSMUSG( | 58.345 | 0.10256 | 0.725492 | 0.944223 | NA |
| ENSMUSG( | 25.36572 | -0.51044 | 0.269712 | 0.719729 | NA |
| ENSMUSG( | 961.2496 | 0.180688 | 0.060733 | 0.380786 | NA |
| ENSMUSG( | 11.83674 | 0.354743 | 0.543089 | NA | NA |
| ENSMUSG( | 35.90286 | -0.40916 | 0.244098 | 0.691435 | NA |
| ENSMUSG( | 19.74054 | -0.06842 | 0.88145 | NA | NA |
| ENSMUSG( | 21.77046 | -0.10269 | 0.809093 | 0.963548 | NA |
| ENSMUSG( | 67.83143 | -0.25423 | 0.302055 | 0.750166 | NA |
| ENSMUSG( | 19.78156 | -0.29333 | 0.517145 | NA | NA |
| ENSMUSG( | 46.8264 | -0.06007 | 0.837494 | 0.971111 | NA |
| ENSMUSG( | 20.00947 | -0.50362 | 0.301952 | NA | NA |
| ENSMUSG( | 12.13314 | -0.38706 | 0.464742 | NA | NA |
| ENSMUSG( | 277.1556 | 0.125119 | 0.380609 | 0.801959 | NA |
| ENSMUSG( | 20.0606 | -0.03535 | 0.932978 | NA | NA |
| ENSMUSG( | 120.1835 | -0.60262 | 0.00731 | 0.118207 | NA |
| ENSMUSG( | 69.53051 | 0.096699 | 0.693657 | 0.934202 | NA |
| ENSMUSG( | 12.22176 | 0.507934 | 0.357699 | NA | NA |
| ENSMUSG( | 9.078544 | -0.22197 | 0.723623 | NA | NA |
| ENSMUSG( | 22.17385 | 0.312708 | 0.440258 | 0.831851 | NA |
| ENSMUSG( | 85.93294 | -0.07558 | 0.728162 | 0.944355 | NA |
| ENSMUSG( | 8.679516 | -0.33332 | 0.599299 | NA | NA |
| ENSMUSG( | 13.33879 | -0.42972 | 0.410583 | NA | NA |
| ENSMUSG( | 12.16296 | 0.329601 | 0.57472 | NA | NA |
| ENSMUSG( | 14.17021 | -0.53828 | 0.283202 | NA | NA |
| ENSMUSG( | 186.4813 | -0.02348 | 0.892673 | 0.984474 | NA |
| ENSMUSG( | 12.98063 | 0.179441 | 0.730254 | NA | NA |
| ENSMUSG( | 11.76804 | 0.125594 | 0.813719 | NA | NA |
| ENSMUSG( | 26.35714 | -0.3741 | 0.337696 | 0.777422 | NA |
| ENSMUSG( | 26.63401 | 0.492108 | 0.178811 | 0.616298 | NA |
| ENSMUSG( | 11.67093 | -0.25611 | 0.632276 | NA | NA |
| ENSMUSG( | 187.0386 | 0.006736 | 0.975304 | 0.997302 | NA |
| ENSMUSG( | 9.29024 | 0.274099 | 0.650423 | NA | NA |
| ENSMUSG( | 9.643548 | 0.196646 | 0.735666 | NA | NA |
| ENSMUSG( | 24.31061 | 0.234233 | 0.539854 | 0.88315 | NA |
| ENSMUSG( | 14.30073 | -0.98494 | 0.08829 | NA | NA |
| ENSMUSG( | 35.78769 | 0.084311 | 0.803071 | 0.962169 | NA |
| ENSMUSG( | 11.06557 | 0.014448 | 0.97985 | NA | NA |
| ENSMUSG( | 17.01076 | -0.47744 | 0.311709 | NA | NA |
| ENSMUSG( | 16.1994 | 0.108285 | 0.814877 | NA | NA |
| ENSMUSG( | 714.8661 | 0.177687 | 0.127925 | 0.536163 | NA |
| ENSMUSG( | 344.5226 | -0.1275 | 0.392922 | 0.809274 | NA |
| ENSMUSG( | 13.23982 | 0.296362 | 0.580343 | NA | NA |
| ENSMUSG( | 13.88513 | 0.730085 | 0.173628 | NA | NA |
| ENSMUSG( | 15.73596 | 0.470485 | 0.318428 | NA | NA |
| ENSMUSG( | 9.458096 | 0.192373 | 0.753701 | NA | NA |
| ENSMUSG( | 64.52643 | -0.13828 | 0.583301 | 0.898408 | NA |
| ENSMUSG( | 10.15559 | -0.15018 | 0.796639 | NA | NA |
| ENSMUSG( | 11.62544 | 0.069313 | 0.896537 | NA | NA |
| ENSMUSG( | 11.74102 | 0.319196 | 0.586619 | NA | NA |
| ENSMUSG( | 14.09746 | -0.01215 | 0.980754 | NA | NA |
| ENSMUSG( | 11.92986 | 0.454867 | 0.405154 | NA | NA |
| ENSMUSG( | 25.21349 | 0.116715 | 0.752996 | 0.950667 | NA |
| ENSMUSG( | 765.4214 | 5.00268 | 8.23E-193 | 1.33E-188 | NA |
| ENSMUSG( | 15.91842 | -0.46983 | 0.345426 | NA | NA |
| ENSMUSG( | 81.94534 | 0.140665 | 0.571804 | 0.893506 | NA |
| ENSMUSG( | 21.13867 | 0.4095 | 0.341112 | NA | NA |
| ENSMUSG( | 16.93988 | -0.24076 | 0.625835 | NA | NA |
| ENSMUSG( | 21.86526 | 0.167881 | 0.678095 | 0.930542 | NA |
| ENSMUSG( | 49.70522 | 0.096055 | 0.735084 | 0.945899 | NA |
| ENSMUSG( | 57.38853 | -0.21401 | 0.449068 | 0.834943 | NA |
| ENSMUSG( | 10.6817 | 0.079604 | 0.89261 | NA | NA |
| ENSMUSG( | 21.96284 | 0.025094 | 0.957122 | 0.993874 | NA |

| | | | | | |
|---|---|---|---|---|---|
| ENSMUSG( | 10.95242 | 0.202502 | 0.738227 | NA | NA |
| ENSMUSG( | 12.50826 | 0.786112 | 0.18268 | 0.850579 | NA |
| ENSMUSG( | 8.808823 | -0.79242 | 0.238222 | NA | NA |
| ENSMUSG( | 51.10334 | -0.38753 | 0.188361 | 0.85698 | NA |
| ENSMUSG( | 86.06828 | -0.0657 | 0.801715 | 0.97681 | NA |
| ENSMUSG( | 168.404 | 0.089714 | 0.616406 | 0.951413 | NA |
| ENSMUSG( | 270.0788 | 0.274152 | 0.223783 | 0.867404 | NA |
| ENSMUSG( | 177.5594 | -0.17171 | 0.352349 | 0.912093 | NA |
| ENSMUSG( | 11.06248 | -0.23403 | 0.691486 | NA | NA |
| ENSMUSG( | 21.98294 | -0.37971 | 0.365428 | 0.914516 | NA |
| ENSMUSG( | 83.70643 | -0.1708 | 0.509173 | 0.929089 | NA |
| ENSMUSG( | 206.1982 | 0.051409 | 0.792075 | 0.97681 | NA |
| ENSMUSG( | 470.7023 | 0.148262 | 0.327934 | 0.906255 | NA |
| ENSMUSG( | 147.7778 | -0.17354 | 0.374537 | 0.915491 | NA |
| ENSMUSG( | 160.8182 | -0.08275 | 0.694958 | 0.960867 | NA |
| ENSMUSG( | 23.38411 | 0.442481 | 0.34031 | 0.907972 | NA |
| ENSMUSG( | 25.24132 | -0.10286 | 0.803406 | 0.976978 | NA |
| ENSMUSG( | 170.9869 | -0.28582 | 0.129791 | 0.820802 | NA |
| ENSMUSG( | 10.27664 | -0.27729 | 0.669681 | NA | NA |
| ENSMUSG( | 99.43578 | 0.10025 | 0.643626 | 0.953903 | NA |
| ENSMUSG( | 14.99111 | -0.35753 | 0.515849 | 0.930939 | NA |
| ENSMUSG( | 72.25973 | -0.42297 | 0.130652 | 0.822659 | NA |
| ENSMUSG( | 60.8155 | 0.038418 | 0.898294 | 0.990558 | NA |
| ENSMUSG( | 11.15414 | -0.53871 | 0.361755 | NA | NA |
| ENSMUSG( | 36.79059 | 0.477204 | 0.148519 | 0.836241 | NA |
| ENSMUSG( | 9.265101 | -0.16029 | 0.799371 | NA | NA |
| ENSMUSG( | 274.8702 | -0.01199 | 0.940249 | 0.994334 | NA |
| ENSMUSG( | 152.4746 | -0.15605 | 0.403675 | 0.917257 | NA |
| ENSMUSG( | 92.55244 | -0.15267 | 0.502983 | 0.927825 | NA |
| ENSMUSG( | 124.4557 | -0.03278 | 0.884629 | 0.989688 | NA |
| ENSMUSG( | 11.69016 | 0.503944 | 0.457255 | 0.923329 | NA |
| ENSMUSG( | 27.70854 | -0.64591 | 0.10134 | 0.801456 | NA |
| ENSMUSG( | 12.45187 | -0.11401 | 0.844177 | 0.983576 | NA |
| ENSMUSG( | 57.71717 | 0.005445 | 0.984485 | 0.996963 | NA |
| ENSMUSG( | 10.87046 | -0.19382 | 0.737555 | NA | NA |
| ENSMUSG( | 28.72991 | -0.36815 | 0.336438 | 0.907972 | NA |
| ENSMUSG( | 65.91055 | -0.2708 | 0.322593 | 0.904023 | NA |
| ENSMUSG( | 28.89386 | -0.21499 | 0.558913 | 0.939587 | NA |
| ENSMUSG( | 19.68654 | -0.24004 | 0.588098 | 0.944975 | NA |
| ENSMUSG( | 10.26027 | -0.09131 | 0.890353 | NA | NA |
| ENSMUSG( | 71.04554 | -0.17589 | 0.479411 | 0.924021 | NA |
| ENSMUSG( | 11.15324 | 0.062963 | 0.914262 | NA | NA |
| ENSMUSG( | 14.62704 | 0.073797 | 0.894926 | 0.990558 | NA |
| ENSMUSG( | 22.99301 | 0.404416 | 0.326687 | 0.905371 | NA |
| ENSMUSG( | 13.73435 | -0.19772 | 0.701976 | 0.962014 | NA |
| ENSMUSG( | 175.4605 | -0.68056 | 0.000662 | 0.082282 | NA |
| ENSMUSG( | 16.96177 | -0.76034 | 0.12185 | 0.813384 | NA |
| ENSMUSG( | 47.24925 | -0.43159 | 0.169633 | 0.840806 | NA |
| ENSMUSG( | 17.81868 | -0.47492 | 0.311569 | 0.900031 | NA |
| ENSMUSG( | 44.9734 | 0.643929 | 0.06391 | 0.736482 | NA |
| ENSMUSG( | 11.32974 | 0.855526 | 0.143569 | NA | NA |
| ENSMUSG( | 26.24538 | -0.41578 | 0.297723 | 0.898538 | NA |
| ENSMUSG( | 9.625461 | -0.14566 | 0.829617 | NA | NA |
| ENSMUSG( | 18.38442 | -0.26439 | 0.583493 | 0.944266 | NA |
| ENSMUSG( | 12.3293 | -0.81707 | 0.176091 | 0.846252 | NA |
| ENSMUSG( | 22.88118 | 0.52857 | 0.214884 | 0.865167 | NA |
| ENSMUSG( | 17.44207 | -0.42702 | 0.365715 | 0.914623 | NA |
| ENSMUSG( | 8.300776 | -1.27981 | 0.124 | NA | NA |
| ENSMUSG( | 14.96034 | 0.277293 | 0.60714 | 0.949917 | NA |
| ENSMUSG( | 276.7099 | -0.41639 | 0.05222 | 0.712974 | NA |
| ENSMUSG( | 13.4149 | 0.376382 | 0.501534 | 0.927825 | NA |
| ENSMUSG( | 131.3691 | -0.35516 | 0.075807 | 0.768524 | NA |
| ENSMUSG( | 8.760287 | -0.2926 | 0.662046 | NA | NA |
| ENSMUSG( | 19.44629 | -0.10333 | 0.814786 | 0.979198 | NA |
| ENSMUSG( | 15.88936 | 0.374542 | 0.539078 | 0.937216 | NA |
| ENSMUSG( | 218.2092 | -0.58413 | 0.000552 | 0.071583 | NA |
| ENSMUSG( | 22.78605 | -0.36149 | 0.392865 | 0.917257 | NA |
| ENSMUSG( | 66.08614 | 0.102316 | 0.719025 | 0.966777 | NA |
| ENSMUSG( | 12.8967 | 0.760298 | 0.167778 | 0.840222 | NA |
| ENSMUSG( | 179.1958 | 0.144098 | 0.569622 | 0.941777 | NA |
| ENSMUSG( | 53.77471 | -0.21047 | 0.532941 | 0.934795 | NA |
| ENSMUSG( | 49.67974 | -0.56935 | 0.062879 | 0.736482 | NA |
| ENSMUSG( | 11.98862 | -0.30289 | 0.611849 | 0.95083 | NA |
| ENSMUSG( | 21.09712 | 0.322178 | 0.4732 | 0.923329 | NA |
| ENSMUSG( | 15.29854 | -0.22122 | 0.662467 | 0.956205 | NA |
| ENSMUSG( | 48.70397 | 0.871462 | 0.040124 | 0.681706 | NA |
| ENSMUSG( | 203.6108 | -2.68443 | 8.09E-23 | 2.02E-19 | NA |
| ENSMUSG( | 80.44757 | 0.045944 | 0.855058 | 0.985649 | NA |
| ENSMUSG( | 18.2161 | -0.25986 | 0.594709 | 0.945444 | NA |
| ENSMUSG( | 67.98286 | 1.116513 | 4.91E-05 | 0.012613 | NA |
| ENSMUSG( | 233.6454 | -0.18919 | 0.309144 | 0.898538 | NA |
| ENSMUSG( | 232.5127 | 0.063387 | 0.7243 | 0.966887 | NA |
| ENSMUSG( | 62.59883 | -0.01955 | 0.946621 | 0.994798 | NA |
| ENSMUSG( | 267.3648 | -0.26611 | 0.09648 | 0.796133 | NA |
| ENSMUSG( | 31.86026 | -0.27491 | 0.470909 | 0.923329 | NA |
| ENSMUSG( | 29.02584 | 0.019882 | 0.957419 | 0.994798 | NA |
| ENSMUSG( | 18.35068 | 0.047109 | 0.920718 | 0.992644 | NA |
| ENSMUSG( | 78.24299 | -0.36207 | 0.170818 | 0.842157 | NA |
| ENSMUSG( | 26.51356 | -0.23789 | 0.566972 | 0.941777 | NA |
| ENSMUSG( | 55.12446 | 0.776408 | 0.021119 | 0.573555 | NA |
| ENSMUSG( | 17.45964 | -0.4568 | 0.347702 | 0.911652 | NA |
| ENSMUSG( | 80.4407 | 0.289432 | 0.255921 | 0.883082 | NA |
| ENSMUSG( | 189.0169 | 0.259419 | 0.191409 | 0.858412 | NA |
| ENSMUSG( | 133.184 | 0.139096 | 0.536763 | 0.937108 | NA |
| ENSMUSG( | 24.6522 | 1.015199 | 0.02549 | 0.598732 | NA |
| ENSMUSG( | 24.89921 | 1.024467 | 0.049257 | 0.708275 | NA |
| ENSMUSG( | 164.9377 | -0.07241 | 0.741312 | 0.968811 | NA |
| ENSMUSG( | 9.359911 | 0.074401 | 0.904219 | NA | NA |
| ENSMUSG( | 95.71913 | 0.344952 | 0.137484 | 0.828492 | NA |
| ENSMUSG( | 8.786508 | 0.330068 | 0.628847 | NA | NA |
| ENSMUSG( | 79.51511 | -0.04376 | 0.878212 | 0.989644 | NA |
| ENSMUSG( | 26.06454 | -0.14217 | 0.725869 | 0.966887 | NA |
| ENSMUSG( | 14.95039 | -0.14543 | 0.774355 | 0.974931 | NA |
| ENSMUSG( | 36.2336 | 0.123341 | 0.754712 | 0.971098 | NA |
| ENSMUSG( | 29.28517 | 0.379155 | 0.37717 | 0.915912 | NA |
| ENSMUSG( | 47.05287 | -0.12626 | 0.696868 | 0.960867 | NA |
| ENSMUSG( | 134.9105 | 0.026972 | 0.905316 | 0.991089 | NA |
| ENSMUSG( | 20.63675 | 1.281563 | 0.022156 | 0.576679 | NA |
| ENSMUSG( | 51.80774 | 1.02891 | 0.014233 | 0.477949 | NA |
| ENSMUSG( | 19.44652 | 0.212154 | 0.653428 | 0.955015 | NA |
| ENSMUSG( | 30.15094 | -0.14048 | 0.730224 | 0.967228 | NA |
| ENSMUSG( | 43.55749 | -0.18134 | 0.600665 | 0.947749 | NA |
| ENSMUSG( | 91.44829 | -0.14749 | 0.621829 | 0.953234 | NA |
| ENSMUSG( | 11.04946 | -0.19011 | 0.756708 | NA | NA |
| ENSMUSG( | 26.79412 | 0.128479 | 0.747432 | 0.969763 | NA |
| ENSMUSG( | 18.8097 | -0.84159 | 0.087455 | 0.784337 | NA |
| ENSMUSG( | 24.99957 | -0.02941 | 0.941074 | 0.994558 | NA |
| ENSMUSG( | 25.59711 | 0.533495 | 0.303484 | 0.898538 | NA |
| ENSMUSG( | 47.93855 | 0.238292 | 0.451759 | 0.922202 | NA |
| ENSMUSG( | 19.5252 | -0.13437 | 0.779151 | 0.976106 | NA |
| ENSMUSG( | 13.80181 | 0.467147 | 0.396553 | 0.917257 | NA |
| ENSMUSG( | 22.07204 | 0.016705 | 0.970087 | 0.99594 | NA |
| ENSMUSG( | 38.91931 | 1.0051 | 0.037622 | 0.674819 | NA |
| ENSMUSG( | 37.15346 | -0.10377 | 0.748651 | 0.96984 | NA |
| ENSMUSG( | 54.20281 | -0.07117 | 0.800529 | 0.97681 | NA |
| ENSMUSG( | 44.89678 | -0.98483 | 0.001274 | 0.128998 | NA |
| ENSMUSG( | 52.47232 | -0.74459 | 0.013776 | 0.46927 | NA |
| ENSMUSG( | 32.9125 | -0.27135 | 0.45997 | 0.923329 | NA |
| ENSMUSG( | 11.37075 | -0.34336 | 0.562281 | NA | NA |
| ENSMUSG( | 248.9874 | 0.179523 | 0.320421 | 0.903887 | NA |
| ENSMUSG( | 50.69092 | 0.294237 | 0.340625 | 0.907972 | NA |
| ENSMUSG( | 13.00901 | 0.310095 | 0.565259 | 0.94131 | NA |
| ENSMUSG( | 17.67008 | -0.24822 | 0.622605 | 0.953234 | NA |
| ENSMUSG( | 11.12125 | 0.907461 | 0.131571 | NA | NA |
| ENSMUSG( | 15.87744 | -0.01872 | 0.969168 | 0.99594 | NA |
| ENSMUSG( | 15.0677 | -0.5492 | 0.27961 | 0.892356 | NA |
| ENSMUSG( | 19.96839 | -0.07139 | 0.890037 | 0.990558 | NA |
| ENSMUSG( | 109.5353 | -0.24169 | 0.296963 | 0.898538 | NA |
| ENSMUSG( | 2599.562 | -0.0885 | 0.508521 | 0.929089 | NA |
| ENSMUSG( | 12.3325 | -1.10834 | 0.06186 | 0.736482 | NA |
| ENSMUSG( | 69.786 | 0.021243 | 0.932887 | 0.99306 | NA |
| ENSMUSG( | 11.87318 | -0.3214 | 0.5734 | 0.942912 | NA |
| ENSMUSG( | 7.15343 | 0.375672 | 0.622778 | NA | NA |
| ENSMUSG( | 45.29017 | 0.030749 | 0.92122 | 0.992644 | NA |
| ENSMUSG( | 10.50874 | 0.39818 | 0.538977 | NA | NA |
| ENSMUSG( | 9.720584 | 0.668735 | 0.317588 | NA | NA |
| ENSMUSG( | 11.40586 | -0.49117 | 0.414298 | NA | NA |
| ENSMUSG( | 14.80832 | 0.253352 | 0.674878 | 0.95694 | NA |
| ENSMUSG( | 16.25794 | 0.989697 | 0.142726 | 0.831221 | NA |
| ENSMUSG( | 52.00501 | -0.20958 | 0.474938 | 0.923329 | NA |
| ENSMUSG( | 24.92487 | -0.72542 | 0.083742 | 0.77681 | NA |
| ENSMUSG( | 70.21034 | 0.676341 | 0.013053 | 0.458202 | NA |
| ENSMUSG( | 14.26724 | 0.377178 | 0.484841 | 0.92541 | NA |
| ENSMUSG( | 14.74524 | -0.34542 | 0.510291 | 0.929089 | NA |
| ENSMUSG( | 31.57372 | 0.252192 | 0.492985 | 0.927358 | NA |
| ENSMUSG( | 10.50516 | 0.593635 | 0.348132 | NA | NA |
| ENSMUSG( | 14.62441 | -0.46485 | 0.379108 | 0.916995 | NA |
| ENSMUSG( | 12.24059 | 0.091614 | 0.87406 | 0.989644 | NA |

| | | | | | | | | | | | |
|---|---|---|---|---|---|---|---|---|---|---|---|
| ENSMUSG( | NA | 380.811 | 0.176685 | 0.140887 | 0.557902 | ENSMUSG( | NA | 78.24299 | -0.36207 | 0.170818 | 0.842157 |
| ENSMUSG( | NA | 67.67039 | -0.07261 | 0.777913 | 0.956168 | ENSMUSG( | NA | 26.51356 | -0.23789 | 0.566972 | 0.941777 |
| ENSMUSG( | NA | 11.66084 | -0.1637 | 0.76509 | NA | ENSMUSG( | NA | 55.12446 | 0.776408 | 0.021119 | 0.573555 |
| ENSMUSG( | NA | 389.7218 | -0.15149 | 0.766877 | 0.954331 | ENSMUSG( | NA | 17.45964 | -0.4568 | 0.347702 | 0.911652 |
| ENSMUSG( | NA | 65.85481 | 0.311254 | 0.257229 | 0.708432 | ENSMUSG( | NA | 80.4407 | 0.289432 | 0.255921 | 0.883082 |
| ENSMUSG( | NA | 9.718569 | -0.09843 | 0.869747 | NA | ENSMUSG( | NA | 189.0169 | 0.259419 | 0.191409 | 0.858412 |
| ENSMUSG( | NA | 11.78177 | 0.072772 | 0.901165 | NA | ENSMUSG( | NA | 133.184 | 0.139096 | 0.536763 | 0.937108 |
| ENSMUSG( | NA | 43.45855 | 0.038003 | 0.893799 | 0.984512 | ENSMUSG( | NA | 24.6522 | 1.015199 | 0.02549 | 0.598732 |
| ENSMUSG( | NA | 20.08403 | 0.499465 | 0.276922 | NA | ENSMUSG( | NA | 24.89921 | 1.024467 | 0.049257 | 0.708275 |
| ENSMUSG( | NA | 23.00949 | -0.37648 | 0.347369 | 0.780377 | ENSMUSG( | NA | 164.9377 | -0.07241 | 0.741312 | 0.968811 |
| ENSMUSG( | NA | 13.34136 | -0.03853 | 0.941011 | NA | ENSMUSG( | NA | 9.359911 | 0.074401 | 0.904219 | NA |
| ENSMUSG( | NA | 214.1807 | 0.028389 | 0.917042 | 0.988991 | ENSMUSG( | NA | 95.71913 | 0.344952 | 0.137484 | 0.828492 |
| ENSMUSG( | NA | 59.67272 | 0.089534 | 0.731904 | 0.944956 | ENSMUSG( | NA | 8.786508 | 0.330068 | 0.628847 | NA |
| ENSMUSG( | NA | 13.15396 | 0.384024 | 0.54266 | NA | ENSMUSG( | NA | 79.51511 | -0.04376 | 0.878212 | 0.989644 |
| ENSMUSG( | NA | 53.54706 | -0.34347 | 0.23107 | 0.679229 | ENSMUSG( | NA | 26.06454 | -0.14217 | 0.725869 | 0.966887 |
| ENSMUSG( | NA | 8016.585 | 0.004091 | 0.977871 | 0.997428 | ENSMUSG( | NA | 14.95039 | -0.14543 | 0.774355 | 0.974931 |
| ENSMUSG( | NA | 11.02424 | -0.91872 | 0.118942 | NA | ENSMUSG( | NA | 36.2336 | 0.123341 | 0.754712 | 0.971098 |
| ENSMUSG( | NA | 13.16211 | 0.466221 | 0.399919 | NA | ENSMUSG( | NA | 29.28517 | 0.379155 | 0.37717 | 0.915912 |
| ENSMUSG( | NA | 23.7507 | -0.02953 | 0.942815 | 0.993415 | ENSMUSG( | NA | 47.05287 | -0.12626 | 0.696868 | 0.960867 |
| ENSMUSG( | NA | 69.47867 | 0.227434 | 0.335556 | 0.776986 | ENSMUSG( | NA | 134.9105 | 0.026972 | 0.905316 | 0.991089 |
| ENSMUSG( | NA | 10.15934 | -0.12948 | 0.828315 | NA | ENSMUSG( | NA | 20.63675 | 1.281563 | 0.022156 | 0.576679 |
| ENSMUSG( | NA | 133.5301 | 0.147923 | 0.404424 | 0.816132 | ENSMUSG( | NA | 51.80774 | 1.02891 | 0.014233 | 0.477949 |
| ENSMUSG( | NA | 20.51503 | -0.05472 | 0.900274 | NA | ENSMUSG( | NA | 19.44652 | 0.212154 | 0.653428 | 0.955015 |
| ENSMUSG( | NA | 466.1997 | -0.09075 | 0.568282 | 0.892252 | ENSMUSG( | NA | 30.15094 | -0.14048 | 0.730224 | 0.967228 |
| ENSMUSG( | NA | 54.01475 | 0.857502 | 0.003306 | 0.072126 | ENSMUSG( | NA | 43.55749 | -0.18134 | 0.600665 | 0.947749 |
| ENSMUSG( | NA | 15.18342 | -0.07557 | 0.881663 | NA | ENSMUSG( | NA | 91.44829 | -0.14749 | 0.621829 | 0.953234 |
| ENSMUSG( | NA | 10.29827 | 0.029987 | 0.960011 | NA | ENSMUSG( | NA | 11.04946 | -0.19011 | 0.756708 | NA |
| ENSMUSG( | NA | 26.66895 | 0.306876 | 0.403518 | 0.816132 | ENSMUSG( | NA | 26.79412 | 0.128479 | 0.747432 | 0.969763 |
| ENSMUSG( | NA | 23.58753 | 0.021388 | 0.958729 | 0.993874 | ENSMUSG( | NA | 18.8097 | -0.84159 | 0.087455 | 0.784337 |
| ENSMUSG( | NA | 9.256173 | -0.10947 | 0.865666 | NA | ENSMUSG( | NA | 24.99957 | -0.02941 | 0.941074 | 0.994558 |
| ENSMUSG( | NA | 30.98252 | 0.199137 | 0.567523 | 0.891774 | ENSMUSG( | NA | 25.59711 | 0.533495 | 0.303484 | 0.898538 |
| ENSMUSG( | NA | 18.64544 | 0.221792 | 0.608513 | NA | ENSMUSG( | NA | 47.93855 | 0.238292 | 0.451759 | 0.922202 |
| ENSMUSG( | NA | 10.9932 | 0.077614 | 0.887694 | NA | ENSMUSG( | NA | 19.5252 | -0.13437 | 0.779151 | 0.976106 |
| ENSMUSG( | NA | 17.83221 | -0.16775 | 0.706787 | NA | ENSMUSG( | NA | 13.80181 | 0.467147 | 0.396553 | 0.917257 |
| ENSMUSG( | NA | 51.98549 | 0.141283 | 0.642669 | 0.916865 | ENSMUSG( | NA | 22.07204 | 0.016705 | 0.970087 | 0.99594 |
| ENSMUSG( | NA | 12.04747 | -0.28681 | 0.584861 | NA | ENSMUSG( | NA | 38.91931 | 1.0051 | 0.037622 | 0.674819 |
| ENSMUSG( | NA | 296.3098 | 0.086249 | 0.537754 | 0.882548 | ENSMUSG( | NA | 37.15346 | -0.10377 | 0.748651 | 0.96984 |
| ENSMUSG( | NA | 502.8334 | 0.06842 | 0.55101 | 0.886038 | ENSMUSG( | NA | 54.20281 | -0.07117 | 0.800529 | 0.97681 |
| ENSMUSG( | NA | 41.77554 | 0.169749 | 0.5752 | 0.894597 | ENSMUSG( | NA | 44.89678 | -0.98483 | 0.001274 | 0.128998 |
| ENSMUSG( | NA | 16.71778 | 0.77417 | 0.089856 | NA | ENSMUSG( | NA | 52.47232 | -0.74459 | 0.013776 | 0.46927 |
| ENSMUSG( | NA | 9.566692 | -0.58886 | 0.368023 | NA | ENSMUSG( | NA | 32.9125 | -0.27135 | 0.45997 | 0.923329 |
| ENSMUSG( | NA | 85.02968 | 0.026727 | 0.898365 | 0.985633 | ENSMUSG( | NA | 11.37075 | -0.34336 | 0.562281 | NA |
| ENSMUSG( | NA | 47.41961 | 0.37107 | 0.189004 | 0.627678 | ENSMUSG( | NA | 248.9874 | 0.179523 | 0.320421 | 0.903887 |
| ENSMUSG( | NA | 21.24113 | -0.33804 | 0.444896 | 0.833239 | ENSMUSG( | NA | 50.69092 | 0.294237 | 0.340625 | 0.907972 |
| ENSMUSG( | NA | 34.36161 | 0.154241 | 0.624459 | 0.91187 | ENSMUSG( | NA | 13.00901 | 0.310095 | 0.565259 | 0.94131 |
| ENSMUSG( | NA | 22.66915 | 0.285046 | 0.499901 | 0.86333 | ENSMUSG( | NA | 17.67008 | -0.24822 | 0.622605 | 0.953234 |
| ENSMUSG( | NA | 8.987877 | 0.284047 | 0.645826 | NA | ENSMUSG( | NA | 15.87744 | -0.01872 | 0.969168 | 0.99594 |
| ENSMUSG( | NA | 32.65473 | 0.098011 | 0.774907 | 0.955606 | ENSMUSG( | NA | 15.0677 | -0.5492 | 0.27961 | 0.892356 |
| ENSMUSG( | NA | 16.9615 | 0.118464 | 0.807684 | NA | ENSMUSG( | NA | 19.96839 | -0.07139 | 0.890037 | 0.990558 |
| ENSMUSG( | NA | 38.15366 | 0.076894 | 0.815282 | 0.965035 | ENSMUSG( | NA | 109.5353 | -0.24169 | 0.296963 | 0.898538 |
| ENSMUSG( | NA | 39.30384 | 0.077177 | 0.818683 | 0.965888 | ENSMUSG( | NA | 2599.562 | -0.0885 | 0.508521 | 0.929089 |
| ENSMUSG( | NA | 13.00672 | -0.19579 | 0.708681 | NA | ENSMUSG( | NA | 12.3325 | -1.10834 | 0.06186 | 0.736482 |
| ENSMUSG( | NA | 32.15032 | -0.03119 | 0.925479 | 0.991204 | ENSMUSG( | NA | 69.786 | 0.021243 | 0.932887 | 0.99306 |
| ENSMUSG( | NA | 10.47335 | -0.06698 | 0.906661 | NA | ENSMUSG( | NA | 11.87318 | -0.3214 | 0.5734 | 0.942912 |
| ENSMUSG( | NA | 15.01806 | -0.01353 | 0.978468 | NA | ENSMUSG( | NA | 45.29017 | 0.030749 | 0.92122 | 0.992644 |
| ENSMUSG( | NA | 107.3428 | 0.006076 | 0.978763 | 0.997428 | ENSMUSG( | NA | 11.40586 | -0.49117 | 0.414298 | NA |
| ENSMUSG( | NA | 16.26662 | -0.28796 | 0.564705 | NA | ENSMUSG( | NA | 14.80832 | 0.253352 | 0.674878 | 0.95694 |
| ENSMUSG( | NA | 15.04823 | -2.17133 | 5.79E-05 | NA | ENSMUSG( | NA | 16.25794 | 0.989697 | 0.142726 | 0.831221 |
| ENSMUSG( | NA | 14.3565 | -0.237 | 0.625718 | NA | ENSMUSG( | NA | 52.00501 | -0.20958 | 0.474938 | 0.923329 |
| ENSMUSG( | NA | 105.744 | -0.143 | 0.533642 | 0.880095 | ENSMUSG( | NA | 24.92487 | -0.72542 | 0.083742 | 0.77681 |
| ENSMUSG( | NA | 325.6806 | 0.518042 | 0.002968 | 0.06659 | ENSMUSG( | NA | 70.21034 | 0.676341 | 0.013053 | 0.458202 |
| ENSMUSG( | NA | 188.0512 | -0.1484 | 0.487816 | 0.858824 | ENSMUSG( | NA | 14.26724 | 0.377178 | 0.484841 | 0.92541 |
| ENSMUSG( | NA | 17.93925 | -0.20153 | 0.67211 | NA | ENSMUSG( | NA | 14.74524 | -0.34542 | 0.510291 | 0.929089 |
| ENSMUSG( | NA | 32.36291 | -0.15072 | 0.672075 | 0.9288 | ENSMUSG( | NA | 31.57372 | 0.252192 | 0.492985 | 0.927358 |
| ENSMUSG( | NA | 76.44541 | -0.41631 | 0.077032 | 0.429516 | ENSMUSG( | NA | 10.50516 | 0.593635 | 0.348132 | NA |
| ENSMUSG( | NA | 12.3224 | 0.201035 | 0.697838 | NA | ENSMUSG( | NA | 14.62441 | -0.46485 | 0.379108 | 0.916995 |
| ENSMUSG( | NA | 193.9385 | 0.061095 | 0.717961 | 0.941697 | ENSMUSG( | NA | 12.24059 | 0.091614 | 0.87406 | 0.989644 |
| ENSMUSG( | NA | 113.6508 | -0.16883 | 0.383535 | 0.803428 | ENSMUSG( | NA | 22.89675 | 0.322297 | 0.444957 | 0.921734 |
| ENSMUSG( | NA | 4432.425 | -0.02854 | 0.792817 | 0.95947 | ENSMUSG( | NA | 206.2294 | 0.17721 | 0.281365 | 0.892412 |
| ENSMUSG( | NA | 178.5234 | -0.00647 | 0.969717 | 0.996369 | ENSMUSG( | NA | 42.3425 | -0.0229 | 0.941548 | 0.994578 |
| ENSMUSG( | NA | 22.9183 | 0.43683 | 0.259462 | 0.709948 | ENSMUSG( | NA | 60.69158 | 0.020242 | 0.943869 | 0.994798 |
| ENSMUSG( | NA | 15.85934 | 0.091335 | 0.840837 | NA | ENSMUSG( | NA | 93.52969 | 0.389652 | 0.098694 | 0.798438 |
| ENSMUSG( | NA | 21.27167 | 0.36259 | 0.433703 | 0.828485 | ENSMUSG( | NA | 28.58361 | 0.038754 | 0.92097 | 0.992644 |
| ENSMUSG( | NA | 32.15592 | 0.182075 | 0.646669 | 0.918796 | ENSMUSG( | NA | 37.99876 | 0.287946 | 0.384445 | 0.917257 |
| ENSMUSG( | NA | 53.77336 | 0.543218 | 0.04928 | 0.338836 | ENSMUSG( | NA | 25.20079 | -0.00434 | 0.991918 | 0.99872 |
| ENSMUSG( | NA | 13.99077 | -0.17232 | 0.720292 | NA | ENSMUSG( | NA | 25.08538 | 0.66475 | 0.103219 | 0.804311 |
| ENSMUSG( | NA | 30.10963 | -0.57537 | 0.105449 | 0.492417 | ENSMUSG( | NA | 34.74461 | 0.454168 | 0.219568 | 0.866104 |
| ENSMUSG( | NA | 23.51624 | 0.347549 | 0.382487 | 0.803227 | ENSMUSG( | NA | 24.47397 | 0.578417 | 0.201111 | 0.86439 |
| ENSMUSG( | NA | 41.4977 | -0.45004 | 0.151981 | 0.577721 | ENSMUSG( | NA | 14.79291 | -0.18841 | 0.720745 | 0.966887 |
| ENSMUSG( | NA | 27.54845 | -0.1871 | 0.635192 | 0.914296 | ENSMUSG( | NA | 40.71314 | -0.47329 | 0.162611 | 0.837724 |
| ENSMUSG( | NA | 14.67456 | 0.014424 | 0.976644 | NA | ENSMUSG( | NA | 178.8029 | -0.15429 | 0.376483 | 0.915491 |
| ENSMUSG( | NA | 8.074821 | 0.206623 | 0.770906 | NA | ENSMUSG( | NA | 70.94683 | -0.31636 | 0.193482 | 0.860096 |
| ENSMUSG( | NA | 29.81353 | -0.18686 | 0.657984 | 0.92425 | ENSMUSG( | NA | 33.736 | 0.057845 | 0.876258 | 0.989644 |
| ENSMUSG( | NA | 706.9915 | 0.092387 | 0.550238 | 0.886038 | ENSMUSG( | NA | 28.59772 | 0.552235 | 0.177143 | 0.846973 |
| ENSMUSG( | NA | 27.7975 | 0.084927 | 0.838559 | 0.971111 | ENSMUSG( | NA | 13.05634 | -0.11103 | 0.857608 | 0.985884 |
| ENSMUSG( | NA | 14.5229 | 0.19856 | 0.723505 | NA | ENSMUSG( | NA | 8.423624 | -0.62609 | 0.403183 | NA |
| ENSMUSG( | NA | 22.37114 | -0.36232 | 0.359539 | 0.791011 | ENSMUSG( | NA | 25.97897 | 0.048644 | 0.911592 | 0.992196 |
| ENSMUSG( | NA | 74.35359 | -0.07961 | 0.736175 | 0.946267 | ENSMUSG( | NA | 13.32692 | -1.72704 | 0.00604 | 0.320634 |
| ENSMUSG( | NA | 32.64654 | -0.15655 | 0.656989 | 0.924069 | ENSMUSG( | NA | 23.09737 | 0.079614 | 0.857136 | 0.985884 |
| ENSMUSG( | NA | 11.87998 | -0.23113 | 0.68277 | NA | ENSMUSG( | NA | 31.9913 | 0.410669 | 0.269793 | 0.887508 |
| ENSMUSG( | NA | 19.80802 | -0.43197 | 0.340731 | NA | ENSMUSG( | NA | 14.70594 | 0.556539 | 0.279556 | 0.892356 |
| ENSMUSG( | NA | 166.1089 | 0.009094 | 0.960038 | 0.993991 | ENSMUSG( | NA | 57.43551 | -0.26794 | 0.333543 | 0.907907 |
| ENSMUSG( | NA | 31.77027 | 0.035934 | 0.920441 | 0.990286 | ENSMUSG( | NA | 12.17985 | 0.219175 | 0.693481 | 0.960767 |
| ENSMUSG( | NA | 28.4661 | -0.96213 | 0.007134 | 0.116294 | ENSMUSG( | NA | 11.2633 | 0.380463 | 0.50648 | NA |
| ENSMUSG( | NA | 9.688407 | -0.30926 | 0.597429 | NA | ENSMUSG( | NA | 37.97472 | -0.28669 | 0.439944 | 0.920949 |
| ENSMUSG( | NA | 27.86814 | -0.01667 | 0.964317 | 0.995155 | ENSMUSG( | NA | 18.11099 | 0.511859 | 0.309412 | 0.898538 |
| ENSMUSG( | NA | 38.16458 | -0.10289 | 0.740611 | 0.947823 | ENSMUSG( | NA | 10.16046 | 0.74387 | 0.230603 | NA |
| ENSMUSG( | NA | 149.1711 | 0.131771 | 0.499233 | 0.863094 | ENSMUSG( | NA | 3526.118 | 0.207529 | 0.121342 | 0.813384 |
| ENSMUSG( | NA | 17.94177 | 0.164159 | 0.703362 | NA | ENSMUSG( | NA | 45.83089 | 0.053374 | 0.863658 | 0.986547 |
| ENSMUSG( | NA | 258.695 | 0.043792 | 0.770196 | 0.954689 | ENSMUSG( | NA | 16.08712 | -0.30614 | 0.538529 | 0.937216 |
| ENSMUSG( | NA | 232.6303 | 0.076139 | 0.630559 | 0.914073 | ENSMUSG( | NA | 31.97701 | -0.58514 | 0.155429 | 0.837724 |
| ENSMUSG( | NA | 549.938 | 0.216129 | 0.088799 | 0.457098 | ENSMUSG( | NA | 35.45391 | -0.58669 | 0.085494 | 0.779799 |
| ENSMUSG( | NA | 69.30739 | 0.164033 | 0.538126 | 0.882548 | ENSMUSG( | NA | 44.70558 | 0.324567 | 0.313982 | 0.900949 |
| ENSMUSG( | NA | 12.86802 | 0.388432 | 0.467224 | NA | ENSMUSG( | NA | 61.42023 | -0.43428 | 0.114121 | 0.811037 |
| ENSMUSG( | NA | 156.7959 | -1.28079 | 1.98E-05 | 0.001952 | ENSMUSG( | NA | 235.899 | 0.521814 | 0.01046 | 0.41614 |
| ENSMUSG( | NA | 11.53586 | 0.187421 | 0.745892 | NA | ENSMUSG( | NA | 20.49457 | -0.43801 | 0.349068 | 0.911652 |
| ENSMUSG( | NA | 65.41607 | -0.12384 | 0.614009 | 0.909352 | ENSMUSG( | NA | 12.01462 | -0.01147 | 0.985004 | 0.997071 |
| ENSMUSG( | NA | 49.22848 | 0.10057 | 0.742153 | 0.948478 | ENSMUSG( | NA | 68.87398 | 0.013943 | 0.955829 | 0.994798 |
| ENSMUSG( | NA | 179.7957 | -0.10769 | 0.522064 | 0.875186 | ENSMUSG( | NA | 51.74317 | -0.2844 | 0.340904 | 0.907972 |
| ENSMUSG( | NA | 38.74957 | 0.188439 | 0.547673 | 0.885153 | ENSMUSG( | NA | 36.47648 | -0.52658 | 0.134292 | 0.822912 |
| ENSMUSG( | NA | 57.29601 | 0.204261 | 0.434916 | 0.829093 | ENSMUSG( | NA | 44.49207 | -0.06312 | 0.850986 | 0.985541 |
| ENSMUSG( | NA | 14.12623 | 0.220195 | 0.679985 | NA | ENSMUSG( | NA | 19.57742 | 0.030236 | 0.947099 | 0.994798 |
| ENSMUSG( | NA | 27.44523 | -0.29238 | 0.417831 | 0.821964 | ENSMUSG( | NA | 14.18079 | 0.417349 | 0.505138 | 0.927825 |
| ENSMUSG( | NA | 21.60582 | 0.497993 | 0.265825 | 0.716441 | ENSMUSG( | NA | 9.490747 | -0.12719 | 0.84307 | NA |
| ENSMUSG( | NA | 13.60866 | 0.030219 | 0.954482 | NA | ENSMUSG( | NA | 159.6277 | -0.21314 | 0.253497 | 0.88018 |
| ENSMUSG( | NA | 17.39156 | 0.099011 | 0.834057 | NA | ENSMUSG( | NA | 11.53601 | -0.14042 | 0.812854 | 0.979148 |
| ENSMUSG( | NA | 11.69242 | -0.64294 | 0.27011 | NA | ENSMUSG( | NA | 40.03134 | 0.230055 | 0.482313 | 0.924578 |
| ENSMUSG( | NA | 16.56617 | -0.49068 | 0.302887 | NA | ENSMUSG( | NA | 21.1182 | -0.26074 | 0.565678 | 0.941404 |
| ENSMUSG( | NA | 9.653143 | -0.50413 | 0.434421 | NA | ENSMUSG( | NA | 57.59039 | 0.264402 | 0.357328 | 0.91373 |
| ENSMUSG( | NA | 132.9463 | 0.281704 | 0.194387 | 0.63341 | ENSMUSG( | NA | 12.25758 | -0.40789 | 0.48745 | 0.925427 |
| ENSMUSG( | NA | 22.34889 | -0.4107 | 0.337935 | 0.777422 | ENSMUSG( | NA | 52.09169 | 0.102499 | 0.716987 | 0.965864 |
| ENSMUSG( | NA | 45.04666 | 0.311661 | 0.291359 | 0.741366 | ENSMUSG( | NA | 9.871734 | -0.83299 | 0.24073 | NA |
| ENSMUSG( | NA | 20.86048 | 0.301128 | 0.479392 | NA | ENSMUSG( | NA | 247.1292 | 0.174714 | 0.448578 | 0.922202 |
| ENSMUSG( | NA | 9.589843 | 0.04984 | 0.942948 | NA | ENSMUSG( | NA | 517.9136 | -0.14283 | 0.335541 | 0.907907 |
| ENSMUSG( | NA | 122.1806 | -0.13149 | 0.485675 | 0.857713 | ENSMUSG( | NA | 26.07614 | -0.74171 | 0.072095 | 0.763629 |
| ENSMUSG( | NA | 23.59427 | -0.02385 | 0.953477 | 0.993874 | ENSMUSG( | NA | 46.61061 | -0.42671 | 0.199541 | 0.863016 |
| ENSMUSG( | NA | 28.81425 | -0.15776 | 0.645098 | 0.917934 | ENSMUSG( | NA | 13.61764 | 0.347728 | 0.536025 | 0.936952 |
| ENSMUSG( | NA | 149.4017 | -0.12574 | 0.463088 | 0.842422 | ENSMUSG( | NA | 9.377613 | -1.03528 | 0.113906 | NA |
| ENSMUSG( | NA | 15.91668 | -0.04695 | 0.925399 | NA | ENSMUSG( | NA | 197.1991 | 0.344455 | 0.083659 | 0.77681 |
| ENSMUSG( | NA | 48.87878 | 0.057405 | 0.836419 | 0.971111 | ENSMUSG( | NA | 8.131033 | -0.22627 | 0.766517 | NA |
| ENSMUSG( | NA | 20.16489 | -0.70004 | 0.090915 | NA | ENSMUSG( | NA | 24.94524 | 0.034151 | 0.93244 | 0.99306 |
| ENSMUSG( | NA | 19.59978 | 0.496118 | 0.28284 | NA | ENSMUSG( | NA | 98.23065 | 0.055421 | 0.827711 | 0.980224 |
| ENSMUSG( | NA | 21.67521 | -0.37364 | 0.370324 | 0.797542 | ENSMUSG( | NA | 39.77516 | 0.504653 | 0.193161 | 0.860096 |
| ENSMUSG( | NA | 14.51379 | 0.220461 | 0.681236 | NA | ENSMUSG( | NA | 17.91153 | 0.56264 | 0.294009 | 0.898538 |
| ENSMUSG( | NA | 51.60114 | 0.007011 | 0.979223 | 0.997428 | ENSMUSG( | NA | 59.37654 | -0.12749 | 0.644866 | 0.953903 |
| ENSMUSG( | NA | 159.1523 | 0.212986 | 0.225816 | 0.67268 | ENSMUSG( | NA | 23.35617 | -0.69776 | 0.114125 | 0.811037 |
| ENSMUSG( | NA | 44.66566 | 0.348458 | 0.253474 | 0.703636 | ENSMUSG( | NA | 26.08139 | -0.12288 | 0.752738 | 0.970521 |
| ENSMUSG( | NA | 68.58051 | -0.27686 | 0.309115 | 0.757263 | ENSMUSG( | NA | 808.6876 | 0.084375 | 0.597695 | 0.946676 |
| ENSMUSG( | NA | 1801.698 | -0.09056 | 0.328771 | 0.773233 | ENSMUSG( | NA | 29.44581 | -0.30629 | 0.414918 | 0.917257 |
| ENSMUSG( | NA | 16.52424 | -0.36507 | 0.465736 | NA | ENSMUSG( | NA | 15.58816 | 0.064524 | 0.896855 | 0.990558 |
| ENSMUSG( | NA | 29.92293 | 0.063517 | 0.871339 | 0.978114 | ENSMUSG( | NA | 454.0302 | 0.147037 | 0.305013 | 0.898538 |
| ENSMUSG( | NA | 32.23881 | 0.291667 | 0.456209 | 0.838832 | ENSMUSG( | NA | 9.799752 | 0.45032 | 0.483122 | NA |
| ENSMUSG( | NA | 14.80572 | -0.10318 | 0.855212 | NA | ENSMUSG( | NA | 7.480234 | -0.03466 | 0.965672 | NA |
| ENSMUSG( | NA | 8.05126 | -0.12022 | 0.866825 | NA | ENSMUSG( | NA | 22.59791 | 0.397661 | 0.356337 | 0.913358 |
| ENSMUSG( | NA | 56.17147 | 0.385089 | 0.160431 | 0.590122 | ENSMUSG( | NA | 15.2642 | 0.405435 | 0.448609 | 0.922202 |
| ENSMUSG( | NA | 15.28187 | 0.428709 | 0.429395 | NA | ENSMUSG( | NA | 30.48768 | -0.51528 | 0.163777 | 0.838291 |
| ENSMUSG( | NA | 85.1475 | -0.01654 | 0.943412 | 0.993415 | ENSMUSG( | NA | 31.22172 | -0.25764 | 0.482564 | 0.924615 |
| ENSMUSG( | NA | 124.6967 | 0.114834 | 0.520742 | 0.875186 | ENSMUSG( | NA | 205.5989 | -0.08109 | 0.638896 | 0.953903 |
| ENSMUSG( | NA | 13.32347 | -0.45093 | 0.447872 | NA | ENSMUSG( | NA | 20.22613 | -0.52825 | 0.280267 | 0.892356 |
| ENSMUSG( | NA | 28.90787 | 0.072018 | 0.837714 | 0.971111 | ENSMUSG( | NA | 30.75712 | 0.126427 | 0.728773 | 0.966913 |
| ENSMUSG( | NA | 8.867101 | -0.07564 | 0.909125 | NA | ENSMUSG( | NA | 19.12851 | 0.026685 | 0.952678 | 0.994798 |
| ENSMUSG( | NA | 47.98351 | -0.49514 | 0.067176 | 0.400611 | ENSMUSG( | NA | 14.01408 | -0.3263 | 0.531126 | 0.934795 |
| ENSMUSG( | NA | 69.0567 | 0.513327 | 0.053521 | 0.35564 | ENSMUSG( | NA | 58.20574 | 0.403787 | 0.292026 | 0.897854 |
| ENSMUSG( | NA | 497.8901 | -1.76663 | 3.12E-18 | 4.22E-15 | ENSMUSG( | NA | 16.85196 | -0.19165 | 0.689177 | 0.959893 |
| ENSMUSG( | NA | 24.42556 | 0.182627 | 0.629827 | 0.913965 | ENSMUSG( | NA | 19.44374 | -0.46259 | 0.293216 | 0.898538 |
| ENSMUSG( | NA | 87.85948 | -0.12332 | 0.563385 | 0.889914 | ENSMUSG( | NA | 49.78975 | 0.037802 | 0.914828 | 0.992407 |
| ENSMUSG( | NA | 28.15463 | 0.295186 | 0.402006 | 0.814965 | ENSMUSG( | NA | 68.60742 | 0.074367 | 0.769705 | 0.974236 |
| ENSMUSG( | NA | 21.60351 | -0.0459 | 0.91346 | 0.988483 | ENSMUSG( | NA | 23.56433 | 0.42144 | 0.330895 | 0.907513 |

| | | | | | |
|---|---|---|---|---|---|
| ENSMUSG( | 31.09363 | 0.068483 | 0.854242 | 0.973491 | NA |
| ENSMUSG( | 17.16705 | -0.19216 | 0.666285 | NA | NA |
| ENSMUSG( | 72.81865 | 0.066595 | 0.771321 | 0.954726 | NA |
| ENSMUSG( | 8.927096 | 0.64396 | 0.338601 | NA | NA |
| ENSMUSG( | 9.92879 | 0.16671 | 0.794696 | NA | NA |
| ENSMUSG( | 10.22466 | 0.01582 | 0.978469 | NA | NA |
| ENSMUSG( | 31.21893 | 0.176875 | 0.6028 | 0.904887 | NA |
| ENSMUSG( | 17.92616 | -0.40014 | 0.369624 | NA | NA |
| ENSMUSG( | 18.10696 | -0.06257 | 0.885575 | NA | NA |
| ENSMUSG( | 17.44861 | 0.143699 | 0.757335 | NA | NA |
| ENSMUSG( | 34.48839 | 0.019055 | 0.958258 | 0.993874 | NA |
| ENSMUSG( | 20.98825 | 0.876074 | 0.070156 | NA | NA |
| ENSMUSG( | 8.666844 | 0.522238 | 0.396715 | NA | NA |
| ENSMUSG( | 8.948045 | 0.626143 | 0.324742 | NA | NA |
| ENSMUSG( | 380.811 | 0.176685 | 0.140887 | 0.557902 | NA |
| ENSMUSG( | 67.67039 | -0.07261 | 0.777913 | 0.956168 | NA |
| ENSMUSG( | 11.66084 | -0.1637 | 0.76509 | NA | NA |
| ENSMUSG( | 389.7218 | -0.15149 | 0.766877 | 0.954331 | NA |
| ENSMUSG( | 65.85481 | 0.311254 | 0.257229 | 0.708432 | NA |
| ENSMUSG( | 9.718569 | -0.09843 | 0.869747 | NA | NA |
| ENSMUSG( | 11.78177 | 0.072772 | 0.901165 | NA | NA |
| ENSMUSG( | 43.45855 | 0.038003 | 0.893799 | 0.984512 | NA |
| ENSMUSG( | 20.08403 | 0.499465 | 0.276922 | NA | NA |
| ENSMUSG( | 23.00949 | -0.37648 | 0.347369 | 0.780377 | NA |
| ENSMUSG( | 13.34136 | -0.03853 | 0.941011 | NA | NA |
| ENSMUSG( | 214.1807 | 0.028389 | 0.917042 | 0.988991 | NA |
| ENSMUSG( | 11.19618 | 0.865591 | 0.119136 | NA | NA |
| ENSMUSG( | 59.67272 | 0.089534 | 0.731904 | 0.944956 | NA |
| ENSMUSG( | 13.15396 | 0.384024 | 0.54266 | NA | NA |
| ENSMUSG( | 53.54706 | -0.34347 | 0.23107 | 0.679229 | NA |
| ENSMUSG( | 9.610074 | 0.371812 | 0.53817 | NA | NA |
| ENSMUSG( | 8016.585 | 0.004091 | 0.977871 | 0.997428 | NA |
| ENSMUSG( | 11.02424 | -0.91872 | 0.118942 | NA | NA |
| ENSMUSG( | 13.16211 | 0.466221 | 0.399919 | NA | NA |
| ENSMUSG( | 23.7507 | -0.02953 | 0.942815 | 0.993415 | NA |
| ENSMUSG( | 69.47867 | 0.227434 | 0.335556 | 0.776986 | NA |
| ENSMUSG( | 10.15934 | -0.12948 | 0.828315 | NA | NA |
| ENSMUSG( | 133.5301 | 0.147923 | 0.404424 | 0.816132 | NA |
| ENSMUSG( | 10.17289 | 0.245162 | 0.689187 | NA | NA |
| ENSMUSG( | 20.51503 | -0.05472 | 0.900274 | NA | NA |
| ENSMUSG( | 466.1997 | -0.09075 | 0.568282 | 0.892252 | NA |
| ENSMUSG( | 54.01475 | 0.857502 | 0.003306 | 0.072126 | NA |
| ENSMUSG( | 15.18342 | -0.07557 | 0.881663 | NA | NA |
| ENSMUSG( | 10.29827 | 0.029987 | 0.960011 | NA | NA |
| ENSMUSG( | 26.66895 | 0.306876 | 0.403518 | 0.816132 | NA |
| ENSMUSG( | 23.58753 | 0.021388 | 0.958729 | 0.993874 | NA |
| ENSMUSG( | 9.256173 | -0.10947 | 0.865666 | NA | NA |
| ENSMUSG( | 30.98252 | 0.199137 | 0.567523 | 0.891774 | NA |
| ENSMUSG( | 18.64544 | 0.221792 | 0.608513 | NA | NA |
| ENSMUSG( | 10.9932 | 0.077614 | 0.887694 | NA | NA |
| ENSMUSG( | 17.83221 | -0.16775 | 0.706787 | NA | NA |
| ENSMUSG( | 11.47108 | 1.056006 | 0.069086 | NA | NA |
| ENSMUSG( | 51.98549 | 0.141283 | 0.642669 | 0.916865 | NA |
| ENSMUSG( | 12.04747 | -0.28681 | 0.584861 | NA | NA |
| ENSMUSG( | 296.3098 | 0.086249 | 0.537754 | 0.882548 | NA |
| ENSMUSG( | 502.8334 | 0.06842 | 0.55101 | 0.886038 | NA |
| ENSMUSG( | 41.77554 | 0.169749 | 0.5752 | 0.894597 | NA |
| ENSMUSG( | 16.71778 | 0.77417 | 0.089856 | NA | NA |
| ENSMUSG( | 9.566692 | -0.58886 | 0.368023 | NA | NA |
| ENSMUSG( | 10.54186 | 0.89068 | 0.136905 | NA | NA |
| ENSMUSG( | 85.02968 | 0.026727 | 0.898365 | 0.985633 | NA |
| ENSMUSG( | 47.41961 | 0.37107 | 0.189004 | 0.627678 | NA |
| ENSMUSG( | 21.24113 | -0.33804 | 0.444896 | 0.833239 | NA |
| ENSMUSG( | 34.36161 | 0.154241 | 0.624459 | 0.91187 | NA |
| ENSMUSG( | 22.66915 | 0.285046 | 0.499901 | 0.86333 | NA |
| ENSMUSG( | 8.987877 | 0.284047 | 0.645826 | NA | NA |
| ENSMUSG( | 32.65473 | 0.098011 | 0.774907 | 0.955606 | NA |
| ENSMUSG( | 16.9615 | 0.118464 | 0.807684 | NA | NA |
| ENSMUSG( | 38.15366 | 0.076894 | 0.815282 | 0.965035 | NA |
| ENSMUSG( | 39.30384 | 0.077177 | 0.818683 | 0.965888 | NA |
| ENSMUSG( | 13.00672 | -0.19579 | 0.708681 | NA | NA |
| ENSMUSG( | 32.15032 | -0.03119 | 0.925479 | 0.991204 | NA |
| ENSMUSG( | 10.47335 | -0.06698 | 0.906661 | NA | NA |
| ENSMUSG( | 8.268297 | -0.05431 | 0.94288 | NA | NA |
| ENSMUSG( | 15.01806 | -0.01353 | 0.978468 | NA | NA |
| ENSMUSG( | 107.3428 | 0.006076 | 0.978763 | 0.997428 | NA |
| ENSMUSG( | 7.915536 | 0.598895 | 0.368565 | NA | NA |
| ENSMUSG( | 16.26662 | -0.28796 | 0.564705 | NA | NA |
| ENSMUSG( | 15.04823 | -2.17133 | 5.79E-05 | NA | NA |
| ENSMUSG( | 10.79965 | 0.1319 | 0.820823 | NA | NA |
| ENSMUSG( | 14.3565 | -0.237 | 0.625718 | NA | NA |
| ENSMUSG( | 105.744 | -0.143 | 0.533642 | 0.880095 | NA |
| ENSMUSG( | 13.8455 | -0.62763 | 0.254725 | NA | NA |
| ENSMUSG( | 325.6806 | 0.518042 | 0.002968 | 0.06659 | NA |
| ENSMUSG( | 188.0512 | -0.1484 | 0.487816 | 0.858824 | NA |
| ENSMUSG( | 17.93925 | -0.20153 | 0.67211 | NA | NA |
| ENSMUSG( | 32.36291 | -0.15072 | 0.672075 | 0.9288 | NA |
| ENSMUSG( | 76.44541 | -0.41631 | 0.077032 | 0.429516 | NA |
| ENSMUSG( | 12.3224 | 0.201035 | 0.697838 | NA | NA |
| ENSMUSG( | 193.9385 | 0.061095 | 0.717961 | 0.941697 | NA |
| ENSMUSG( | 113.6508 | -0.16883 | 0.383535 | 0.803428 | NA |
| ENSMUSG( | 4432.425 | -0.02854 | 0.792817 | 0.95947 | NA |
| ENSMUSG( | 178.5234 | -0.00647 | 0.969717 | 0.996369 | NA |
| ENSMUSG( | 9.076651 | 0.661817 | 0.291172 | NA | NA |
| ENSMUSG( | 22.9183 | 0.43683 | 0.259462 | 0.709948 | NA |
| ENSMUSG( | 15.85934 | 0.091335 | 0.840837 | NA | NA |
| ENSMUSG( | 21.27167 | 0.36259 | 0.433703 | 0.828485 | NA |
| ENSMUSG( | 32.15592 | 0.182075 | 0.646669 | 0.918796 | NA |
| ENSMUSG( | 53.77336 | 0.543218 | 0.04928 | 0.338836 | NA |
| ENSMUSG( | 13.99077 | -0.17232 | 0.720292 | NA | NA |
| ENSMUSG( | 30.10963 | -0.57537 | 0.105449 | 0.492417 | NA |
| ENSMUSG( | 23.51624 | 0.347549 | 0.382487 | 0.803227 | NA |
| ENSMUSG( | 41.4977 | -0.45004 | 0.151981 | 0.577721 | NA |
| ENSMUSG( | 27.54845 | -0.1871 | 0.635192 | 0.914296 | NA |
| ENSMUSG( | 14.67456 | 0.014424 | 0.976644 | NA | NA |
| ENSMUSG( | 8.074821 | 0.206623 | 0.770906 | NA | NA |
| ENSMUSG( | 29.81353 | -0.18686 | 0.657984 | 0.92425 | NA |
| ENSMUSG( | 706.9915 | 0.092387 | 0.550238 | 0.886038 | NA |
| ENSMUSG( | 27.7975 | 0.084927 | 0.838559 | 0.971111 | NA |
| ENSMUSG( | 14.5229 | 0.19856 | 0.723505 | NA | NA |
| ENSMUSG( | 22.37114 | -0.36232 | 0.359539 | 0.791011 | NA |
| ENSMUSG( | 9.404352 | 0.241092 | 0.713587 | NA | NA |
| ENSMUSG( | 74.35359 | -0.07961 | 0.736175 | 0.946267 | NA |
| ENSMUSG( | 32.64654 | -0.15655 | 0.656989 | 0.924069 | NA |
| ENSMUSG( | 11.87998 | -0.23113 | 0.68277 | NA | NA |
| ENSMUSG( | 19.80802 | -0.43197 | 0.340731 | NA | NA |
| ENSMUSG( | 166.1089 | 0.009094 | 0.960038 | 0.993991 | NA |
| ENSMUSG( | 31.77027 | 0.035934 | 0.920441 | 0.990286 | NA |
| ENSMUSG( | 28.4661 | -0.96213 | 0.007134 | 0.116294 | NA |
| ENSMUSG( | 12.41288 | -0.1753 | 0.750481 | NA | NA |
| ENSMUSG( | 9.688407 | -0.30926 | 0.597429 | NA | NA |
| ENSMUSG( | 27.86814 | -0.01667 | 0.964317 | 0.995155 | NA |
| ENSMUSG( | 10.18731 | -0.32206 | 0.582592 | NA | NA |
| ENSMUSG( | 38.16458 | -0.10289 | 0.740611 | 0.947823 | NA |
| ENSMUSG( | 149.1711 | 0.131771 | 0.499233 | 0.863094 | NA |
| ENSMUSG( | 17.94177 | 0.164159 | 0.703362 | NA | NA |
| ENSMUSG( | 258.695 | 0.043792 | 0.770196 | 0.954689 | NA |
| ENSMUSG( | 232.6303 | 0.076139 | 0.630559 | 0.914073 | NA |
| ENSMUSG( | 549.938 | 0.216129 | 0.088799 | 0.457098 | NA |
| ENSMUSG( | 69.30739 | 0.164033 | 0.538126 | 0.882548 | NA |
| ENSMUSG( | 12.86802 | 0.388432 | 0.467224 | NA | NA |
| ENSMUSG( | 156.7959 | -1.28079 | 1.98E-05 | 0.001952 | NA |
| ENSMUSG( | 11.53586 | 0.187421 | 0.745892 | NA | NA |
| ENSMUSG( | 65.41607 | -0.12384 | 0.614009 | 0.909352 | NA |
| ENSMUSG( | 49.22848 | 0.10057 | 0.742153 | 0.948478 | NA |
| ENSMUSG( | 179.7957 | -0.10769 | 0.522064 | 0.875186 | NA |
| ENSMUSG( | 38.74957 | 0.188439 | 0.547673 | 0.885153 | NA |
| ENSMUSG( | 57.29601 | 0.204261 | 0.434916 | 0.829093 | NA |
| ENSMUSG( | 14.12623 | 0.220195 | 0.679985 | NA | NA |
| ENSMUSG( | 27.44523 | -0.29238 | 0.417831 | 0.821964 | NA |
| ENSMUSG( | 13.93861 | 0.223091 | 0.653442 | NA | NA |
| ENSMUSG( | 21.60582 | 0.497993 | 0.265825 | 0.716441 | NA |
| ENSMUSG( | 13.60866 | 0.030219 | 0.954482 | NA | NA |
| ENSMUSG( | 17.39156 | 0.099011 | 0.834057 | NA | NA |
| ENSMUSG( | 11.69242 | -0.64294 | 0.27011 | NA | NA |
| ENSMUSG( | 16.56617 | -0.49068 | 0.302887 | NA | NA |
| ENSMUSG( | 9.653143 | -0.50413 | 0.434421 | NA | NA |
| ENSMUSG( | 10.41304 | 0.44677 | 0.439877 | NA | NA |
| ENSMUSG( | 132.9463 | 0.281704 | 0.194387 | 0.63341 | NA |
| ENSMUSG( | 22.34889 | -0.4107 | 0.337935 | 0.777422 | NA |
| ENSMUSG( | 45.04666 | 0.311661 | 0.291359 | 0.741366 | NA |
| ENSMUSG( | 20.86048 | 0.301128 | 0.479392 | NA | NA |
| ENSMUSG( | 9.589843 | 0.04984 | 0.942948 | NA | NA |
| ENSMUSG( | 122.1806 | -0.13149 | 0.485675 | 0.857713 | NA |
| ENSMUSG( | 23.59427 | -0.02385 | 0.953477 | 0.993874 | NA |
| ENSMUSG( | 28.81425 | -0.15776 | 0.645098 | 0.917934 | NA |
| ENSMUSG( | 149.4017 | -0.12574 | 0.463088 | 0.842422 | NA |
| ENSMUSG( | 15.91668 | -0.04695 | 0.925399 | NA | NA |
| ENSMUSG( | 22.89675 | 0.322297 | 0.444957 | 0.921734 | NA |
| ENSMUSG( | 206.2294 | 0.17721 | 0.281365 | 0.892412 | NA |
| ENSMUSG( | 42.3425 | -0.0229 | 0.941548 | 0.994578 | NA |
| ENSMUSG( | 60.69158 | 0.020242 | 0.943869 | 0.994798 | NA |
| ENSMUSG( | 93.52969 | 0.389652 | 0.098694 | 0.798438 | NA |
| ENSMUSG( | 28.58361 | 0.038754 | 0.92097 | 0.992644 | NA |
| ENSMUSG( | 37.99876 | 0.287946 | 0.384445 | 0.917257 | NA |
| ENSMUSG( | 25.20079 | -0.00434 | 0.991918 | 0.99872 | NA |
| ENSMUSG( | 11.51096 | 0.143854 | 0.826682 | 0.980035 | NA |
| ENSMUSG( | 25.08538 | 0.66475 | 0.103219 | 0.804311 | NA |
| ENSMUSG( | 34.74461 | 0.454168 | 0.219568 | 0.866104 | NA |
| ENSMUSG( | 24.47397 | 0.578417 | 0.201111 | 0.86439 | NA |
| ENSMUSG( | 14.79291 | -0.18841 | 0.720745 | 0.966887 | NA |
| ENSMUSG( | 40.71314 | -0.47329 | 0.162611 | 0.837724 | NA |
| ENSMUSG( | 178.8029 | -0.15429 | 0.376483 | 0.915491 | NA |
| ENSMUSG( | 70.94683 | -0.31636 | 0.193482 | 0.860096 | NA |
| ENSMUSG( | 33.736 | 0.057845 | 0.876258 | 0.989644 | NA |
| ENSMUSG( | 28.59772 | 0.552235 | 0.177143 | 0.846973 | NA |
| ENSMUSG( | 13.05634 | -0.11103 | 0.857608 | 0.985884 | NA |
| ENSMUSG( | 8.423624 | -0.62609 | 0.403183 | NA | NA |
| ENSMUSG( | 25.97897 | 0.048644 | 0.911592 | 0.992196 | NA |
| ENSMUSG( | 13.32692 | -1.72704 | 0.00604 | 0.320634 | NA |
| ENSMUSG( | 23.09737 | 0.079614 | 0.857136 | 0.985884 | NA |
| ENSMUSG( | 31.9913 | 0.410669 | 0.269793 | 0.887508 | NA |
| ENSMUSG( | 14.70594 | 0.556539 | 0.279556 | 0.892356 | NA |
| ENSMUSG( | 57.43551 | -0.26794 | 0.333543 | 0.907907 | NA |
| ENSMUSG( | 12.17985 | 0.219175 | 0.693481 | 0.960767 | NA |
| ENSMUSG( | 11.2633 | 0.380463 | 0.50648 | NA | NA |
| ENSMUSG( | 37.97472 | -0.28669 | 0.439944 | 0.920949 | NA |
| ENSMUSG( | 18.11099 | 0.511859 | 0.309412 | 0.898538 | NA |
| ENSMUSG( | 10.16046 | 0.74387 | 0.230603 | NA | NA |
| ENSMUSG( | 3526.118 | 0.207529 | 0.121342 | 0.813384 | NA |
| ENSMUSG( | 45.83089 | 0.053374 | 0.863658 | 0.986547 | NA |
| ENSMUSG( | 16.08712 | -0.30614 | 0.538529 | 0.937216 | NA |
| ENSMUSG( | 31.97701 | -0.58514 | 0.155429 | 0.837724 | NA |
| ENSMUSG( | 35.45391 | -0.58669 | 0.085494 | 0.779799 | NA |
| ENSMUSG( | 44.70558 | 0.324567 | 0.313982 | 0.900949 | NA |
| ENSMUSG( | 61.42023 | -0.43428 | 0.114121 | 0.811037 | NA |
| ENSMUSG( | 235.899 | 0.521814 | 0.01046 | 0.41614 | NA |
| ENSMUSG( | 20.49457 | -0.43801 | 0.349068 | 0.911652 | NA |
| ENSMUSG( | 12.01462 | -0.01147 | 0.985004 | 0.997071 | NA |
| ENSMUSG( | 68.87398 | 0.013943 | 0.955829 | 0.994798 | NA |
| ENSMUSG( | 51.74317 | -0.2844 | 0.340904 | 0.907972 | NA |
| ENSMUSG( | 36.47648 | -0.52658 | 0.134292 | 0.822912 | NA |
| ENSMUSG( | 44.49207 | -0.06312 | 0.850986 | 0.985541 | NA |
| ENSMUSG( | 19.57742 | 0.030236 | 0.947099 | 0.994798 | NA |
| ENSMUSG( | 14.18079 | 0.417349 | 0.505138 | 0.927825 | NA |
| ENSMUSG( | 9.490747 | -0.12719 | 0.84307 | NA | NA |
| ENSMUSG( | 159.6277 | -0.21314 | 0.253497 | 0.88018 | NA |
| ENSMUSG( | 11.53601 | -0.14042 | 0.812854 | 0.979148 | NA |
| ENSMUSG( | 40.03134 | 0.230055 | 0.482313 | 0.924578 | NA |
| ENSMUSG( | 21.1182 | -0.26074 | 0.565678 | 0.941404 | NA |
| ENSMUSG( | 57.59039 | 0.264402 | 0.357328 | 0.91373 | NA |
| ENSMUSG( | 12.25758 | -0.40789 | 0.48745 | 0.925427 | NA |
| ENSMUSG( | 52.09169 | 0.102499 | 0.716987 | 0.965864 | NA |
| ENSMUSG( | 9.871734 | -0.83299 | 0.24073 | NA | NA |
| ENSMUSG( | 247.1292 | 0.174714 | 0.448578 | 0.922202 | NA |
| ENSMUSG( | 517.9136 | -0.14283 | 0.335541 | 0.907907 | NA |
| ENSMUSG( | 26.07614 | -0.74171 | 0.072095 | 0.763629 | NA |
| ENSMUSG( | 46.61061 | -0.42671 | 0.199541 | 0.863016 | NA |
| ENSMUSG( | 13.61764 | 0.347728 | 0.536025 | 0.936952 | NA |
| ENSMUSG( | 9.377613 | -1.03528 | 0.113906 | NA | NA |
| ENSMUSG( | 197.1991 | 0.344455 | 0.083659 | 0.77681 | NA |
| ENSMUSG( | 8.131033 | -0.22627 | 0.766517 | NA | NA |
| ENSMUSG( | 24.94524 | 0.034151 | 0.93244 | 0.99306 | NA |
| ENSMUSG( | 98.23065 | 0.055421 | 0.827711 | 0.980224 | NA |
| ENSMUSG( | 39.77516 | 0.504653 | 0.193161 | 0.860096 | NA |
| ENSMUSG( | 17.91153 | 0.56264 | 0.294009 | 0.898538 | NA |
| ENSMUSG( | 59.37654 | -0.12749 | 0.644866 | 0.953903 | NA |
| ENSMUSG( | 23.35617 | -0.69776 | 0.114125 | 0.811037 | NA |
| ENSMUSG( | 26.08139 | -0.12288 | 0.752738 | 0.970521 | NA |
| ENSMUSG( | 808.6876 | 0.084375 | 0.597695 | 0.946676 | NA |
| ENSMUSG( | 29.44581 | -0.30629 | 0.414918 | 0.917257 | NA |
| ENSMUSG( | 15.58816 | 0.064524 | 0.896855 | 0.990558 | NA |
| ENSMUSG( | 454.0302 | 0.147037 | 0.305013 | 0.898538 | NA |
| ENSMUSG( | 9.799752 | 0.45032 | 0.483122 | NA | NA |
| ENSMUSG( | 7.480234 | -0.03466 | 0.965672 | NA | NA |
| ENSMUSG( | 10.50607 | -0.55273 | 0.399995 | NA | NA |
| ENSMUSG( | 22.59791 | 0.397661 | 0.356337 | 0.913358 | NA |
| ENSMUSG( | 15.2642 | 0.405435 | 0.448609 | 0.922202 | NA |
| ENSMUSG( | 30.48768 | -0.51528 | 0.163777 | 0.838291 | NA |
| ENSMUSG( | 31.22172 | -0.25764 | 0.482564 | 0.924615 | NA |
| ENSMUSG( | 205.5989 | -0.08109 | 0.638896 | 0.953903 | NA |
| ENSMUSG( | 20.22613 | -0.52825 | 0.280267 | 0.892356 | NA |
| ENSMUSG( | 30.75712 | 0.126427 | 0.728773 | 0.966913 | NA |
| ENSMUSG( | 19.12851 | 0.026685 | 0.952678 | 0.994798 | NA |
| ENSMUSG( | 14.01408 | -0.3263 | 0.531126 | 0.934795 | NA |
| ENSMUSG( | 58.20574 | 0.403787 | 0.292026 | 0.897854 | NA |
| ENSMUSG( | 16.85196 | -0.19165 | 0.689177 | 0.959893 | NA |
| ENSMUSG( | 19.44374 | -0.46259 | 0.293216 | 0.898538 | NA |
| ENSMUSG( | 49.78975 | 0.037802 | 0.914828 | 0.992407 | NA |
| ENSMUSG( | 68.60742 | 0.074367 | 0.769705 | 0.974236 | NA |
| ENSMUSG( | 23.56433 | 0.42144 | 0.330895 | 0.907513 | NA |
| ENSMUSG( | 72.54957 | -0.27739 | 0.371847 | 0.915491 | NA |
| ENSMUSG( | 251.8332 | 0.115965 | 0.530317 | 0.934795 | NA |
| ENSMUSG( | 66.40871 | 0.003909 | 0.989044 | 0.997893 | NA |
| ENSMUSG( | 58.66294 | -0.3492 | 0.262489 | 0.885357 | NA |
| ENSMUSG( | 191.1311 | -0.04861 | 0.875969 | 0.989644 | NA |
| ENSMUSG( | 18.34442 | -0.49454 | 0.284063 | 0.892412 | NA |
| ENSMUSG( | 64.12372 | 0.514722 | 0.078342 | 0.77321 | NA |
| ENSMUSG( | 9.380983 | -0.41138 | 0.536407 | NA | NA |
| ENSMUSG( | 26.55456 | 0.203444 | 0.600441 | 0.947732 | NA |
| ENSMUSG( | 14.22837 | -0.19314 | 0.764848 | 0.974236 | NA |
| ENSMUSG( | 9.686808 | 0.048452 | 0.938155 | NA | NA |
| ENSMUSG( | 52.34855 | -0.21913 | 0.477081 | 0.923329 | NA |
| ENSMUSG( | 10.90201 | 0.401832 | 0.522484 | NA | NA |
| ENSMUSG( | 7.561088 | -0.70209 | 0.322729 | NA | NA |
| ENSMUSG( | 13.43512 | 0.180569 | 0.747724 | 0.96984 | NA |
| ENSMUSG( | 9.552482 | -0.08078 | 0.896074 | NA | NA |
| ENSMUSG( | 383.4198 | -0.28842 | 0.152296 | 0.837724 | NA |
| ENSMUSG( | 9.861103 | -0.50514 | 0.413538 | NA | NA |
| ENSMUSG( | 54.33542 | -0.37327 | 0.190745 | 0.857801 | NA |
| ENSMUSG( | 29.46384 | 0.202101 | 0.588436 | 0.944975 | NA |
| ENSMUSG( | 15.62015 | -0.37801 | 0.475085 | 0.923329 | NA |
| ENSMUSG( | 99.85308 | -0.47923 | 0.046612 | 0.699054 | NA |
| ENSMUSG( | 20.01034 | 0.402676 | 0.422381 | 0.917257 | NA |
| ENSMUSG( | 65.04248 | -0.18961 | 0.452315 | 0.922316 | NA |
| ENSMUSG( | 100.8075 | -0.12675 | 0.586522 | 0.944975 | NA |
| ENSMUSG( | 39.34949 | 0.302112 | 0.385282 | 0.917257 | NA |
| ENSMUSG( | 12.42745 | 0.257919 | 0.648312 | 0.95422 | NA |
| ENSMUSG( | 20.14666 | -0.04063 | 0.926304 | 0.99306 | NA |
| ENSMUSG( | 30.65531 | 0.286547 | 0.429685 | 0.917257 | NA |
| ENSMUSG( | 14.76344 | 0.625794 | 0.269955 | 0.887508 | NA |
| ENSMUSG( | 9.98414 | -0.31657 | 0.613053 | NA | NA |
| ENSMUSG( | 21.41713 | 0.516324 | 0.220726 | 0.866104 | NA |
| ENSMUSG( | 10.48887 | 0.251104 | 0.727533 | NA | NA |
| ENSMUSG( | 93.04702 | -0.41238 | 0.131521 | 0.822912 | NA |
| ENSMUSG( | 35.14416 | -0.18246 | 0.616236 | 0.951413 | NA |
| ENSMUSG( | 9.123455 | 0.561241 | 0.403047 | NA | NA |
| ENSMUSG( | 26.02347 | -0.23111 | 0.578568 | 0.943548 | NA |
| ENSMUSG( | 19.51763 | -0.60798 | 0.190526 | 0.857801 | NA |
| ENSMUSG( | 18.57202 | 0.055917 | 0.908546 | 0.991958 | NA |
| ENSMUSG( | 8.502177 | 0.500836 | 0.498207 | NA | NA |
| ENSMUSG( | 46.67642 | 0.988871 | 0.003428 | 0.234549 | NA |
| ENSMUSG( | 12.0454 | -0.02686 | 0.964167 | 0.995433 | NA |
| ENSMUSG( | 42.17307 | -0.63822 | 0.059233 | 0.729429 | NA |
| ENSMUSG( | 24.25176 | 0.222302 | 0.598064 | 0.946676 | NA |
| ENSMUSG( | 10.54461 | 0.43351 | 0.470297 | NA | NA |
| ENSMUSG( | 134.6043 | -0.05358 | 0.797461 | 0.97681 | NA |
| ENSMUSG( | 29.89892 | -0.07895 | 0.828787 | 0.980224 | NA |
| ENSMUSG( | 9.111511 | -0.27677 | 0.679908 | NA | NA |
| ENSMUSG( | 68.11736 | -0.07213 | 0.795439 | 0.97681 | NA |
| ENSMUSG( | 19.45365 | -0.2377 | 0.608235 | 0.950113 | NA |
| ENSMUSG( | 46.7802 | 0.072345 | 0.810816 | 0.978779 | NA |
| ENSMUSG( | 19.98017 | -0.65185 | 0.148068 | 0.836241 | NA |
| ENSMUSG( | 9.544236 | -0.40644 | 0.530953 | NA | NA |
| ENSMUSG( | 37.14746 | 0.78988 | 0.044845 | 0.693959 | NA |
| ENSMUSG( | 12.967 | 0.499194 | 0.413705 | 0.917257 | NA |
| ENSMUSG( | 20.76843 | 0.001091 | 0.998112 | 0.999313 | NA |
| ENSMUSG( | 39.08917 | -0.5647 | 0.108612 | 0.804969 | NA |
| ENSMUSG( | 117.8556 | -0.3085 | 0.18969 | 0.857355 | NA |
| ENSMUSG( | 28.26295 | -0.11436 | 0.767918 | 0.974236 | NA |
| ENSMUSG( | 49.08418 | 0.561965 | 0.060552 | 0.734037 | NA |
| ENSMUSG( | 26.17938 | 0.124491 | 0.75864 | 0.971858 | NA |
| ENSMUSG( | 11.59644 | -0.03435 | 0.953621 | 0.994798 | NA |
| ENSMUSG( | 8.133087 | -0.01231 | 0.985861 | NA | NA |
| ENSMUSG( | 16.69343 | -0.15403 | 0.748381 | 0.96984 | NA |
| ENSMUSG( | 106.6844 | 0.171896 | 0.415604 | 0.917257 | NA |

| | | | | | | | | | | | |
|---|---|---|---|---|---|---|---|---|---|---|---|
| ENSMUSG( | NA | 21.78772 | 0.453756 | 0.273671 | 0.724244 | ENSMUSG( | NA | 72.54957 | -0.27739 | 0.371847 | 0.915491 |
| ENSMUSG( | NA | 128.5485 | 0.17567 | 0.346358 | 0.779153 | ENSMUSG( | NA | 251.8332 | 0.115965 | 0.530317 | 0.934795 |
| ENSMUSG( | NA | 59.78923 | 0.095581 | 0.696541 | 0.935257 | ENSMUSG( | NA | 66.40871 | 0.003909 | 0.989044 | 0.997893 |
| ENSMUSG( | NA | 13.5225 | 1.021476 | 0.078497 | NA | ENSMUSG( | NA | 58.66294 | -0.3492 | 0.262489 | 0.885357 |
| ENSMUSG( | NA | 19.98348 | -0.77584 | 0.091489 | NA | ENSMUSG( | NA | 191.1311 | -0.04861 | 0.875969 | 0.989644 |
| ENSMUSG( | NA | 24.287 | 0.48659 | 0.215012 | 0.659512 | ENSMUSG( | NA | 18.34442 | -0.49454 | 0.284063 | 0.892412 |
| ENSMUSG( | NA | 25.19909 | 0.552898 | 0.184947 | 0.62251 | ENSMUSG( | NA | 64.12372 | 0.514722 | 0.078342 | 0.77321 |
| ENSMUSG( | NA | 257.539 | 0.024359 | 0.858995 | 0.974707 | ENSMUSG( | NA | 9.380983 | -0.41138 | 0.536407 | NA |
| ENSMUSG( | NA | 10.40332 | -0.01127 | 0.984453 | NA | ENSMUSG( | NA | 26.55456 | 0.203444 | 0.600441 | 0.947732 |
| ENSMUSG( | NA | 140.6865 | -0.03803 | 0.83344 | 0.970333 | ENSMUSG( | NA | 14.22837 | -0.19314 | 0.764848 | 0.974236 |
| ENSMUSG( | NA | 22.91934 | -0.54349 | 0.185789 | 0.623185 | ENSMUSG( | NA | 9.686808 | 0.048452 | 0.938155 | NA |
| ENSMUSG( | NA | 232.9491 | 0.01724 | 0.910429 | 0.9881 | ENSMUSG( | NA | 52.34855 | -0.21913 | 0.477081 | 0.923329 |
| ENSMUSG( | NA | 22.26155 | -0.08578 | 0.832942 | 0.970333 | ENSMUSG( | NA | 7.561088 | -0.70209 | 0.322729 | NA |
| ENSMUSG( | NA | 11.95708 | 0.071723 | 0.892688 | NA | ENSMUSG( | NA | 13.43512 | 0.180569 | 0.747724 | 0.96984 |
| ENSMUSG( | NA | 32.17049 | 0.076074 | 0.834247 | 0.970333 | ENSMUSG( | NA | 9.552482 | -0.08078 | 0.896074 | NA |
| ENSMUSG( | NA | 16.89194 | -0.1578 | 0.728618 | NA | ENSMUSG( | NA | 383.4198 | -0.28842 | 0.152296 | 0.837724 |
| ENSMUSG( | NA | 12.83773 | -0.47159 | 0.398609 | NA | ENSMUSG( | NA | 9.861103 | -0.50514 | 0.413538 | NA |
| ENSMUSG( | NA | 66.37326 | -0.15655 | 0.512499 | 0.871762 | ENSMUSG( | NA | 54.33542 | -0.37327 | 0.190745 | 0.857801 |
| ENSMUSG( | NA | 24.07003 | -0.30276 | 0.496654 | 0.86175 | ENSMUSG( | NA | 29.46384 | 0.202101 | 0.588436 | 0.944975 |
| ENSMUSG( | NA | 10.12229 | -0.87736 | 0.169032 | NA | ENSMUSG( | NA | 15.62015 | -0.37801 | 0.475085 | 0.923329 |
| ENSMUSG( | NA | 20.30699 | -0.00418 | 0.992191 | NA | ENSMUSG( | NA | 99.85308 | -0.47923 | 0.046612 | 0.699054 |
| ENSMUSG( | NA | 15.57516 | -0.52068 | 0.321089 | NA | ENSMUSG( | NA | 20.01034 | 0.402676 | 0.422381 | 0.917257 |
| ENSMUSG( | NA | 11.90697 | 0.29212 | 0.633599 | NA | ENSMUSG( | NA | 65.04248 | -0.18961 | 0.452315 | 0.922316 |
| ENSMUSG( | NA | 14.08115 | -0.60149 | 0.240271 | NA | ENSMUSG( | NA | 100.8075 | -0.12675 | 0.586522 | 0.944975 |
| ENSMUSG( | NA | 27.26855 | -0.20686 | 0.590251 | 0.90113 | ENSMUSG( | NA | 39.34949 | 0.302112 | 0.385282 | 0.917257 |
| ENSMUSG( | NA | 21.72961 | -0.08322 | 0.834854 | 0.970426 | ENSMUSG( | NA | 12.42745 | 0.257919 | 0.648312 | 0.95422 |
| ENSMUSG( | NA | 18.64604 | 0.091766 | 0.83693 | NA | ENSMUSG( | NA | 20.14666 | -0.04063 | 0.926304 | 0.99306 |
| ENSMUSG( | NA | 20.52804 | -0.10776 | 0.793755 | NA | ENSMUSG( | NA | 30.65531 | 0.286547 | 0.429685 | 0.917257 |
| ENSMUSG( | NA | 59.5729 | 0.049063 | 0.857852 | 0.974667 | ENSMUSG( | NA | 14.76344 | 0.625794 | 0.269955 | 0.887508 |
| ENSMUSG( | NA | 32.70178 | 0.017599 | 0.958439 | 0.993874 | ENSMUSG( | NA | 9.98414 | -0.31657 | 0.613053 | NA |
| ENSMUSG( | NA | 133.4646 | 0.224251 | 0.212702 | 0.6567 | ENSMUSG( | NA | 21.41713 | 0.516324 | 0.220726 | 0.866104 |
| ENSMUSG( | NA | 8.435563 | -0.5252 | 0.478986 | NA | ENSMUSG( | NA | 10.48887 | 0.251104 | 0.727533 | NA |
| ENSMUSG( | NA | 28.32453 | 0.42651 | 0.237601 | 0.685624 | ENSMUSG( | NA | 93.04702 | -0.41238 | 0.131521 | 0.822912 |
| ENSMUSG( | NA | 9.764407 | 0.450662 | 0.465596 | NA | ENSMUSG( | NA | 35.14416 | -0.18246 | 0.616236 | 0.951413 |
| ENSMUSG( | NA | 55.3464 | -0.10917 | 0.716567 | 0.941697 | ENSMUSG( | NA | 9.123455 | 0.561241 | 0.403047 | NA |
| ENSMUSG( | NA | 69.75408 | -0.25888 | 0.256555 | 0.707467 | ENSMUSG( | NA | 26.02347 | -0.23111 | 0.578568 | 0.943548 |
| ENSMUSG( | NA | 10.82519 | -0.13024 | 0.823095 | NA | ENSMUSG( | NA | 19.51763 | -0.60798 | 0.190526 | 0.857801 |
| ENSMUSG( | NA | 17.73704 | 0.09599 | 0.83119 | NA | ENSMUSG( | NA | 18.57202 | 0.055917 | 0.908546 | 0.991958 |
| ENSMUSG( | NA | 23.75116 | -0.57784 | 0.159412 | 0.58872 | ENSMUSG( | NA | 8.502177 | 0.500836 | 0.498207 | NA |
| ENSMUSG( | NA | 226.8688 | -0.14984 | 0.376989 | 0.801213 | ENSMUSG( | NA | 46.67642 | 0.988871 | 0.003428 | 0.234549 |
| ENSMUSG( | NA | 11.49119 | -0.10258 | 0.861876 | NA | ENSMUSG( | NA | 12.0454 | -0.02686 | 0.964167 | 0.995433 |
| ENSMUSG( | NA | 16.86562 | -0.2201 | 0.664191 | NA | ENSMUSG( | NA | 42.17307 | -0.63822 | 0.059233 | 0.729429 |
| ENSMUSG( | NA | 62.20295 | -0.03599 | 0.909189 | 0.98794 | ENSMUSG( | NA | 24.25176 | 0.222302 | 0.598064 | 0.946676 |
| ENSMUSG( | NA | 8.832897 | -0.80818 | 0.243815 | NA | ENSMUSG( | NA | 10.54461 | 0.43351 | 0.470297 | NA |
| ENSMUSG( | NA | 11.25993 | -0.20976 | 0.714437 | NA | ENSMUSG( | NA | 134.6043 | -0.05358 | 0.797461 | 0.97681 |
| ENSMUSG( | NA | 33.43533 | 0.146228 | 0.701897 | 0.937441 | ENSMUSG( | NA | 29.89892 | -0.07895 | 0.828787 | 0.980224 |
| ENSMUSG( | NA | 46.2013 | 0.142653 | 0.648836 | 0.920025 | ENSMUSG( | NA | 9.111511 | -0.27677 | 0.679908 | NA |
| ENSMUSG( | NA | 20.21507 | 0.116148 | 0.801593 | NA | ENSMUSG( | NA | 68.11736 | -0.07213 | 0.795439 | 0.97681 |
| ENSMUSG( | NA | 14.59763 | -0.04136 | 0.940644 | NA | ENSMUSG( | NA | 19.45365 | -0.2377 | 0.608235 | 0.950113 |
| ENSMUSG( | NA | 39.46987 | -0.01531 | 0.964467 | 0.995155 | ENSMUSG( | NA | 46.7802 | 0.072345 | 0.810816 | 0.978779 |
| ENSMUSG( | NA | 132.4298 | 0.188653 | 0.413307 | 0.819541 | ENSMUSG( | NA | 19.98017 | -0.65185 | 0.148068 | 0.836241 |
| ENSMUSG( | NA | 12.19329 | -0.04207 | 0.937904 | NA | ENSMUSG( | NA | 9.544236 | -0.40644 | 0.530953 | NA |
| ENSMUSG( | NA | 94.56494 | -0.27759 | 0.200872 | 0.640988 | ENSMUSG( | NA | 37.14746 | 0.78988 | 0.044845 | 0.693959 |
| ENSMUSG( | NA | 18.30172 | -0.08296 | 0.852351 | NA | ENSMUSG( | NA | 12.967 | 0.499194 | 0.413705 | 0.917257 |
| ENSMUSG( | NA | 11.27282 | 0.193968 | 0.753741 | NA | ENSMUSG( | NA | 20.76843 | 0.001091 | 0.998112 | 0.999313 |
| ENSMUSG( | NA | 13.74073 | -0.18392 | 0.724173 | NA | ENSMUSG( | NA | 39.08917 | -0.5647 | 0.108612 | 0.804969 |
| ENSMUSG( | NA | 174.0076 | 0.038838 | 0.818277 | 0.965837 | ENSMUSG( | NA | 117.8556 | -0.3085 | 0.18969 | 0.857355 |
| ENSMUSG( | NA | 12.46466 | -0.26648 | 0.639019 | NA | ENSMUSG( | NA | 28.26295 | -0.11436 | 0.767918 | 0.974236 |
| ENSMUSG( | NA | 65.6372 | -0.01343 | 0.961674 | 0.994286 | ENSMUSG( | NA | 49.08418 | 0.561965 | 0.060552 | 0.734037 |
| ENSMUSG( | NA | 107.6269 | -0.05617 | 0.777998 | 0.956168 | ENSMUSG( | NA | 26.17938 | 0.124491 | 0.75864 | 0.971858 |
| ENSMUSG( | NA | 44.84815 | -0.14187 | 0.632123 | 0.914073 | ENSMUSG( | NA | 11.59644 | -0.03435 | 0.953621 | 0.994798 |
| ENSMUSG( | NA | 34.82169 | -0.26547 | 0.421285 | 0.823126 | ENSMUSG( | NA | 16.69343 | -0.15403 | 0.748381 | 0.96984 |
| ENSMUSG( | NA | 18.89339 | 0.260107 | 0.576328 | NA | ENSMUSG( | NA | 106.6844 | 0.171896 | 0.415604 | 0.917257 |
| ENSMUSG( | NA | 11.49303 | -0.21576 | 0.722983 | NA | ENSMUSG( | NA | 53.16293 | -0.4614 | 0.16231 | 0.837724 |
| ENSMUSG( | NA | 93.32701 | 0.028419 | 0.90187 | 0.985633 | ENSMUSG( | NA | 23.2489 | 0.569236 | 0.199646 | 0.863016 |
| ENSMUSG( | NA | 27.98959 | -0.52368 | 0.148126 | 0.571232 | ENSMUSG( | NA | 22.21689 | -0.31028 | 0.471871 | 0.923329 |
| ENSMUSG( | NA | 37.7672 | 0.177246 | 0.566725 | 0.891644 | ENSMUSG( | NA | 19.20088 | 0.026054 | 0.956461 | 0.994798 |
| ENSMUSG( | NA | 74.56213 | -0.00482 | 0.983591 | 0.997738 | ENSMUSG( | NA | 33.98172 | -0.15044 | 0.662568 | 0.956205 |
| ENSMUSG( | NA | 10.79996 | 0.318786 | 0.599566 | NA | ENSMUSG( | NA | 37.2874 | -0.4087 | 0.228186 | 0.868507 |
| ENSMUSG( | NA | 83.73808 | 0.02061 | 0.924515 | 0.99114 | ENSMUSG( | NA | 721.2564 | 0.397192 | 0.013835 | 0.46927 |
| ENSMUSG( | NA | 10.40223 | -0.32258 | 0.608123 | NA | ENSMUSG( | NA | 9.217487 | 0.20538 | 0.780486 | NA |
| ENSMUSG( | NA | 82.5618 | 0.313176 | 0.175101 | 0.610399 | ENSMUSG( | NA | 197.0829 | -0.1335 | 0.54747 | 0.938063 |
| ENSMUSG( | NA | 42.55816 | 0.024434 | 0.935041 | 0.993279 | ENSMUSG( | NA | 11.25229 | 0.161763 | 0.79298 | NA |
| ENSMUSG( | NA | 49.31605 | -0.19548 | 0.527425 | 0.877599 | ENSMUSG( | NA | 31.62429 | 0.146536 | 0.692465 | 0.960404 |
| ENSMUSG( | NA | 34.28599 | -0.14545 | 0.656049 | 0.923929 | ENSMUSG( | NA | 60.22956 | 0.013289 | 0.962646 | 0.995284 |
| ENSMUSG( | NA | 14.77429 | -0.43148 | 0.394152 | NA | ENSMUSG( | NA | 214.4645 | -0.25129 | 0.218129 | 0.866104 |
| ENSMUSG( | NA | 50.85091 | 0.299081 | 0.268752 | 0.719042 | ENSMUSG( | NA | 23.88393 | 0.791625 | 0.057333 | 0.723362 |
| ENSMUSG( | NA | 13.95727 | -0.91277 | 0.084345 | NA | ENSMUSG( | NA | 26.74031 | -0.1376 | 0.746967 | 0.969763 |
| ENSMUSG( | NA | 51.79711 | 0.062766 | 0.81848 | 0.965843 | ENSMUSG( | NA | 36.67119 | -0.77135 | 0.026244 | 0.603041 |
| ENSMUSG( | NA | 10.24229 | -0.45116 | 0.501742 | NA | ENSMUSG( | NA | 179.0929 | 0.10695 | 0.674523 | 0.95694 |
| ENSMUSG( | NA | 74.81705 | 0.046558 | 0.860555 | 0.9749 | ENSMUSG( | NA | 97.55104 | 0.406229 | 0.141528 | 0.830678 |
| ENSMUSG( | NA | 13.20908 | -0.037 | 0.950787 | NA | ENSMUSG( | NA | 10.97532 | -0.27987 | 0.627425 | NA |
| ENSMUSG( | NA | 880.2653 | 0.008745 | 0.950708 | 0.993821 | ENSMUSG( | NA | 38.48496 | -0.35964 | 0.304217 | 0.898538 |
| ENSMUSG( | NA | 53.83667 | -0.13382 | 0.642576 | 0.916865 | ENSMUSG( | NA | 88.19318 | 0.045181 | 0.852217 | 0.985598 |
| ENSMUSG( | NA | 41.76572 | -0.52525 | 0.089166 | 0.458482 | ENSMUSG( | NA | 135.7058 | -0.1099 | 0.593925 | 0.945408 |
| ENSMUSG( | NA | 16.01959 | 0.592956 | 0.197457 | NA | ENSMUSG( | NA | 12.28768 | 0.401365 | 0.463654 | 0.923329 |
| ENSMUSG( | NA | 10.63903 | -0.54212 | 0.349268 | NA | ENSMUSG( | NA | 103.5621 | -0.07378 | 0.763721 | 0.974 |
| ENSMUSG( | NA | 93.04036 | -0.06739 | 0.738356 | 0.947329 | ENSMUSG( | NA | 12.88938 | -0.99649 | 0.068075 | 0.75212 |
| ENSMUSG( | NA | 26.28479 | 0.015541 | 0.966992 | 0.995911 | ENSMUSG( | NA | 46.96196 | -0.05638 | 0.858121 | 0.985884 |
| ENSMUSG( | NA | 9.705875 | 0.405131 | 0.535556 | NA | ENSMUSG( | NA | 20.03238 | 0.003054 | 0.994402 | 0.998813 |
| ENSMUSG( | NA | 298.663 | -0.48964 | 0.010128 | 0.143118 | ENSMUSG( | NA | 13.51031 | -0.01308 | 0.980145 | 0.996479 |
| ENSMUSG( | NA | 14.24005 | -0.4647 | 0.379383 | NA | ENSMUSG( | NA | 83.44397 | 0.380951 | 0.201585 | 0.864412 |
| ENSMUSG( | NA | 10.01271 | 0.012748 | 0.983079 | NA | ENSMUSG( | NA | 11.45138 | 0.279626 | 0.625777 | NA |
| ENSMUSG( | NA | 321.5994 | 0.171883 | 0.207221 | 0.649561 | ENSMUSG( | NA | 378.3557 | 0.091199 | 0.639606 | 0.953903 |
| ENSMUSG( | NA | 28.24259 | -0.01536 | 0.969418 | 0.996369 | ENSMUSG( | NA | 16.49083 | 0.90885 | 0.136511 | 0.827141 |
| ENSMUSG( | NA | 12.25437 | -3.56245 | 7.49E-06 | NA | ENSMUSG( | NA | 15.13387 | -0.57692 | 0.267041 | 0.886134 |
| ENSMUSG( | NA | 17.79592 | 0.533018 | 0.255381 | NA | ENSMUSG( | NA | 24.0858 | 0.364834 | 0.41903 | 0.917257 |
| ENSMUSG( | NA | 14.64675 | 1.197626 | 0.017722 | NA | ENSMUSG( | NA | 17.6997 | 0.434297 | 0.358068 | 0.913759 |
| ENSMUSG( | NA | 12.7636 | -0.39347 | 0.476838 | NA | ENSMUSG( | NA | 48.99542 | -0.09503 | 0.766468 | 0.974236 |
| ENSMUSG( | NA | 18.43132 | -0.44367 | 0.326552 | NA | ENSMUSG( | NA | 10.39523 | 0.237715 | 0.719749 | NA |
| ENSMUSG( | NA | 21.50864 | -0.08731 | 0.824778 | 0.967386 | ENSMUSG( | NA | 91.23397 | -0.27902 | 0.211784 | 0.864412 |
| ENSMUSG( | NA | 46.56157 | -0.0648 | 0.818818 | 0.965888 | ENSMUSG( | NA | 13.37458 | 0.529385 | 0.342949 | 0.909646 |
| ENSMUSG( | NA | 15.98712 | -0.17674 | 0.725472 | NA | ENSMUSG( | NA | 8.590902 | -0.18879 | 0.775259 | NA |
| ENSMUSG( | NA | 522.7214 | 0.014913 | 0.900743 | 0.985633 | ENSMUSG( | NA | 54.6459 | 0.430739 | 0.150475 | 0.837173 |
| ENSMUSG( | NA | 40.62877 | -0.00109 | 0.997162 | 0.999565 | ENSMUSG( | NA | 8.365078 | -1.16318 | 0.094649 | NA |
| ENSMUSG( | NA | 31.04999 | -0.33919 | 0.323183 | 0.768415 | ENSMUSG( | NA | 11.12982 | -0.21814 | 0.714605 | NA |
| ENSMUSG( | NA | 42.47425 | 0.157882 | 0.588876 | 0.90113 | ENSMUSG( | NA | 18.56797 | 0.154652 | 0.750919 | 0.970317 |
| ENSMUSG( | NA | 25.75633 | 0.153102 | 0.685518 | 0.932981 | ENSMUSG( | NA | 10.76572 | 0.179152 | 0.789672 | NA |
| ENSMUSG( | NA | 25.66964 | -0.02382 | 0.948973 | 0.993821 | ENSMUSG( | NA | 10.71883 | -0.41919 | 0.494949 | NA |
| ENSMUSG( | NA | 55.29254 | -0.05195 | 0.852847 | 0.973168 | ENSMUSG( | NA | 207.1108 | -0.34263 | 0.051055 | 0.711023 |
| ENSMUSG( | NA | 11.2082 | -0.52209 | 0.358093 | NA | ENSMUSG( | NA | 19.59184 | 0.132385 | 0.776793 | 0.975717 |
| ENSMUSG( | NA | 21.07243 | 0.140826 | 0.750831 | NA | ENSMUSG( | NA | 27.62953 | 0.073786 | 0.853829 | 0.98563 |
| ENSMUSG( | NA | 17.01354 | 0.300763 | 0.513692 | NA | ENSMUSG( | NA | 24.45288 | -0.45633 | 0.312316 | 0.900037 |
| ENSMUSG( | NA | 24.63935 | -0.01709 | 0.963427 | 0.994911 | ENSMUSG( | NA | 72.63174 | 0.186256 | 0.472379 | 0.923329 |
| ENSMUSG( | NA | 56.96792 | 0.068035 | 0.818387 | 0.965837 | ENSMUSG( | NA | 54.39871 | -0.6499 | 0.036474 | 0.66949 |
| ENSMUSG( | NA | 43.07177 | 0.034853 | 0.903715 | 0.986094 | ENSMUSG( | NA | 22.0062 | -0.57881 | 0.180689 | 0.850372 |
| ENSMUSG( | NA | 745.4922 | 0.303276 | 0.095558 | 0.475062 | ENSMUSG( | NA | 82.40986 | 0.061169 | 0.824131 | 0.979198 |
| ENSMUSG( | NA | 33.62152 | 0.088094 | 0.803496 | 0.962169 | ENSMUSG( | NA | 13.04824 | -0.19953 | 0.711931 | 0.964363 |
| ENSMUSG( | NA | 19.65809 | 0.120733 | 0.779516 | NA | ENSMUSG( | NA | 20.18202 | -0.21521 | 0.649249 | 0.95422 |
| ENSMUSG( | NA | 12.91125 | 0.911691 | 0.099368 | NA | ENSMUSG( | NA | 51.44839 | 0.171927 | 0.555054 | 0.939565 |
| ENSMUSG( | NA | 15.9724 | -0.31478 | 0.540682 | NA | ENSMUSG( | NA | 12.19062 | 0.365051 | 0.532826 | 0.934795 |
| ENSMUSG( | NA | 12.18463 | 0.458814 | 0.423527 | NA | ENSMUSG( | NA | 14.32584 | 0.745109 | 0.153043 | 0.837724 |
| ENSMUSG( | NA | 21.71641 | 0.334051 | 0.441268 | 0.832449 | ENSMUSG( | NA | 625.9694 | 0.267573 | 0.071163 | 0.761942 |
| ENSMUSG( | NA | 9.646024 | -0.2955 | 0.659954 | NA | ENSMUSG( | NA | 8.829397 | 0.563146 | 0.407433 | NA |
| ENSMUSG( | NA | 15.21487 | -0.13015 | 0.798828 | NA | ENSMUSG( | NA | 6.304426 | -0.15264 | 0.856381 | NA |
| ENSMUSG( | NA | 15.81319 | -0.12511 | 0.787956 | NA | ENSMUSG( | NA | 9.763164 | -0.64946 | 0.357568 | NA |
| ENSMUSG( | NA | 12.35204 | 0.205854 | 0.728044 | NA | ENSMUSG( | NA | 10.73616 | 0.600063 | 0.361679 | NA |
| ENSMUSG( | NA | 1259.826 | -0.02942 | 0.788311 | 0.959226 | ENSMUSG( | NA | 21.30004 | -0.22026 | 0.704396 | 0.962677 |
| ENSMUSG( | NA | 97.67909 | 0.385897 | 0.08408 | 0.447085 | ENSMUSG( | NA | 41.95168 | 0.495593 | 0.19347 | 0.860096 |
| ENSMUSG( | NA | 13.23876 | -0.63392 | 0.229994 | NA | ENSMUSG( | NA | 15.16203 | -0.60723 | 0.248706 | 0.877904 |
| ENSMUSG( | NA | 34.9713 | 0.510582 | 0.13924 | 0.555317 | ENSMUSG( | NA | 45.26477 | 0.026818 | 0.931146 | 0.99306 |
| ENSMUSG( | NA | 28.39002 | 0.418661 | 0.315302 | 0.762288 | ENSMUSG( | NA | 93.08279 | -0.27408 | 0.222183 | 0.866232 |
| ENSMUSG( | NA | 10.50714 | 0.171294 | 0.800363 | NA | ENSMUSG( | NA | 35.30091 | 0.443076 | 0.189083 | 0.85698 |
| ENSMUSG( | NA | 125.6532 | -0.11212 | 0.598276 | 0.903801 | ENSMUSG( | NA | 8.840646 | 0.087064 | 0.901939 | NA |
| ENSMUSG( | NA | 12.37359 | 0.07868 | 0.894911 | NA | ENSMUSG( | NA | 96.51863 | 0.298641 | 0.312997 | 0.900449 |
| ENSMUSG( | NA | 23.78226 | 0.03482 | 0.93302 | 0.992887 | ENSMUSG( | NA | 301.7859 | 0.425887 | 0.075734 | 0.768524 |
| ENSMUSG( | NA | 47.97269 | 0.391823 | 0.155531 | 0.583591 | ENSMUSG( | NA | 8.7143 | -0.28744 | 0.694965 | NA |
| ENSMUSG( | NA | 16.14031 | 0.287702 | 0.562308 | NA | ENSMUSG( | NA | 40.51125 | -0.34217 | 0.298332 | 0.898538 |
| ENSMUSG( | NA | 22.20705 | -0.21027 | 0.599451 | 0.903801 | ENSMUSG( | NA | 102.9856 | -0.47651 | 0.031764 | 0.631941 |
| ENSMUSG( | NA | 9.474376 | 0.280213 | 0.653548 | NA | ENSMUSG( | NA | 110.5684 | -0.10114 | 0.67472 | 0.95694 |
| ENSMUSG( | NA | 24.62469 | 0.143787 | 0.705027 | 0.938303 | ENSMUSG( | NA | 50.35214 | -0.13489 | 0.638029 | 0.953903 |
| ENSMUSG( | NA | 16.57183 | -0.01855 | 0.967332 | NA | ENSMUSG( | NA | 21.1338 | -0.71618 | 0.095393 | 0.796133 |
| ENSMUSG( | NA | 20.14294 | -0.50336 | 0.224021 | NA | ENSMUSG( | NA | 25.74411 | 0.824463 | 0.10148 | 0.801456 |
| ENSMUSG( | NA | 32.51497 | -0.09136 | 0.808772 | 0.963455 | ENSMUSG( | NA | 19.68946 | 0.22633 | 0.647055 | 0.954079 |
| ENSMUSG( | NA | 28.17827 | -0.11348 | 0.755531 | 0.951395 | ENSMUSG( | NA | 14.21599 | -0.03304 | 0.949307 | 0.994798 |
| ENSMUSG( | NA | 11.34986 | -0.36896 | 0.526445 | NA | ENSMUSG( | NA | 59.47339 | -0.18699 | 0.486301 | 0.92541 |
| ENSMUSG( | NA | 15.68287 | 0.288559 | 0.578117 | NA | ENSMUSG( | NA | 14.28925 | -0.21099 | 0.691288 | 0.960071 |
| ENSMUSG( | NA | 42.74012 | -0.23886 | 0.432455 | 0.82768 | ENSMUSG( | NA | 19.08421 | -1.11459 | 0.018768 | 0.547939 |
| ENSMUSG( | NA | 383.1434 | -0.05941 | 0.6417 | 0.916828 | ENSMUSG( | NA | 25.4713 | -0.79531 | 0.046007 | 0.69654 |
| ENSMUSG( | NA | 16.79621 | -0.47638 | 0.293839 | NA | ENSMUSG( | NA | 14.18697 | 0.560566 | 0.330306 | 0.907251 |
| ENSMUSG( | NA | 15.16597 | -0.24762 | 0.615724 | NA | ENSMUSG( | NA | 18.59147 | -0.65562 | 0.168341 | 0.840222 |
| ENSMUSG( | NA | 452.0022 | 0.079467 | 0.48905 | 0.859092 | ENSMUSG( | NA | 29.39196 | 2.206624 | 0.085841 | 0.780123 |
| ENSMUSG( | NA | 69.36007 | -0.0363 | 0.909371 | 0.98801 | ENSMUSG( | NA | 28.37334 | 0.170064 | 0.653344 | 0.955015 |
| ENSMUSG( | NA | 273.9417 | -0.0849 | 0.521681 | 0.875186 | ENSMUSG( | NA | 18.17811 | 0.009814 | 0.9862 | 0.997188 |
| ENSMUSG( | NA | 48.4574 | 0.407978 | 0.234857 | 0.682194 | ENSMUSG( | NA | 11.7331 | -0.01445 | 0.979308 | 0.996468 |
| ENSMUSG( | NA | 53.9151 | 0.13346 | 0.639229 | 0.916434 | ENSMUSG( | NA | 82.39615 | 0.173532 | 0.512793 | 0.930164 |
| ENSMUSG( | NA | 43.62062 | -0.53073 | 0.098352 | 0.479925 | ENSMUSG( | NA | 66.17206 | 0.160776 | 0.692754 | 0.96043 |
| ENSMUSG( | NA | 12.25421 | 0.115693 | 0.829874 | NA | ENSMUSG( | NA | 33.46438 | -0.19697 | 0.608927 | 0.950226 |

| | | | | | |
|---|---|---|---|---|---|
| ENSMUSG( | 48.87878 | 0.057405 | 0.836419 | 0.971111 | NA |
| ENSMUSG( | 13.15357 | 0.267504 | 0.615569 | NA | NA |
| ENSMUSG( | 20.16489 | -0.70004 | 0.090915 | NA | NA |
| ENSMUSG( | 19.59978 | 0.496118 | 0.28284 | NA | NA |
| ENSMUSG( | 21.67521 | -0.37364 | 0.370324 | 0.797542 | NA |
| ENSMUSG( | 14.51379 | 0.220461 | 0.681236 | NA | NA |
| ENSMUSG( | 51.60114 | 0.007011 | 0.979223 | 0.997428 | NA |
| ENSMUSG( | 159.1523 | 0.212986 | 0.225816 | 0.67268 | NA |
| ENSMUSG( | 44.66566 | 0.348458 | 0.253474 | 0.703636 | NA |
| ENSMUSG( | 68.58051 | -0.27686 | 0.309115 | 0.757263 | NA |
| ENSMUSG( | 1801.698 | -0.09056 | 0.328771 | 0.773233 | NA |
| ENSMUSG( | 16.52424 | -0.36507 | 0.465736 | NA | NA |
| ENSMUSG( | 29.92293 | 0.063517 | 0.871339 | 0.978114 | NA |
| ENSMUSG( | 10.33644 | 0.430027 | 0.51095 | NA | NA |
| ENSMUSG( | 32.23881 | 0.291667 | 0.456209 | 0.838832 | NA |
| ENSMUSG( | 14.80572 | -0.10318 | 0.855212 | NA | NA |
| ENSMUSG( | 8.05126 | -0.12022 | 0.866825 | NA | NA |
| ENSMUSG( | 56.17147 | 0.385089 | 0.160431 | 0.590122 | NA |
| ENSMUSG( | 15.28187 | 0.428709 | 0.429395 | NA | NA |
| ENSMUSG( | 85.1475 | -0.01654 | 0.943412 | 0.993415 | NA |
| ENSMUSG( | 124.6967 | 0.114834 | 0.520742 | 0.875186 | NA |
| ENSMUSG( | 13.17791 | 0.833242 | 0.143198 | NA | NA |
| ENSMUSG( | 13.32347 | -0.45093 | 0.447872 | NA | NA |
| ENSMUSG( | 28.90787 | 0.072018 | 0.837714 | 0.971111 | NA |
| ENSMUSG( | 8.867101 | -0.07564 | 0.909125 | NA | NA |
| ENSMUSG( | 47.98351 | -0.49514 | 0.067176 | 0.400611 | NA |
| ENSMUSG( | 69.0567 | 0.513327 | 0.053521 | 0.35564 | NA |
| ENSMUSG( | 497.8901 | -1.76663 | 3.12E-18 | 4.22E-15 | NA |
| ENSMUSG( | 24.42556 | 0.182627 | 0.629827 | 0.913965 | NA |
| ENSMUSG( | 87.85948 | -0.12332 | 0.563385 | 0.889914 | NA |
| ENSMUSG( | 28.15463 | 0.295186 | 0.402006 | 0.814965 | NA |
| ENSMUSG( | 21.60351 | -0.0459 | 0.91346 | 0.988483 | NA |
| ENSMUSG( | 21.78772 | 0.453756 | 0.273671 | 0.724244 | NA |
| ENSMUSG( | 128.5485 | 0.17567 | 0.346358 | 0.779153 | NA |
| ENSMUSG( | 59.78923 | 0.095581 | 0.696541 | 0.935257 | NA |
| ENSMUSG( | 13.5225 | 1.021476 | 0.078497 | NA | NA |
| ENSMUSG( | 19.98348 | -0.77584 | 0.091489 | NA | NA |
| ENSMUSG( | 24.287 | 0.48659 | 0.215012 | 0.659512 | NA |
| ENSMUSG( | 25.19909 | 0.552898 | 0.184947 | 0.62251 | NA |
| ENSMUSG( | 257.539 | 0.024359 | 0.858995 | 0.974707 | NA |
| ENSMUSG( | 10.40332 | -0.01127 | 0.984453 | NA | NA |
| ENSMUSG( | 140.6865 | -0.03803 | 0.83344 | 0.970333 | NA |
| ENSMUSG( | 22.91934 | -0.54349 | 0.185789 | 0.623185 | NA |
| ENSMUSG( | 232.9491 | 0.01724 | 0.910429 | 0.9881 | NA |
| ENSMUSG( | 22.26155 | -0.08578 | 0.832942 | 0.970333 | NA |
| ENSMUSG( | 11.95708 | 0.071723 | 0.892688 | NA | NA |
| ENSMUSG( | 32.17049 | 0.076074 | 0.834247 | 0.970333 | NA |
| ENSMUSG( | 16.89194 | -0.1578 | 0.728618 | NA | NA |
| ENSMUSG( | 12.83773 | -0.47159 | 0.398609 | NA | NA |
| ENSMUSG( | 12.29717 | 0.186876 | 0.751064 | NA | NA |
| ENSMUSG( | 66.37326 | -0.15655 | 0.512499 | 0.871762 | NA |
| ENSMUSG( | 24.07003 | -0.30276 | 0.496654 | 0.86175 | NA |
| ENSMUSG( | 10.12229 | -0.87736 | 0.169032 | NA | NA |
| ENSMUSG( | 20.30699 | -0.00418 | 0.992191 | NA | NA |
| ENSMUSG( | 15.57516 | -0.52068 | 0.321089 | NA | NA |
| ENSMUSG( | 11.90697 | 0.29212 | 0.633599 | NA | NA |
| ENSMUSG( | 14.08115 | -0.60149 | 0.240271 | NA | NA |
| ENSMUSG( | 27.26855 | -0.20686 | 0.590251 | 0.90113 | NA |
| ENSMUSG( | 21.72961 | -0.08322 | 0.834854 | 0.970426 | NA |
| ENSMUSG( | 18.64604 | 0.091766 | 0.83693 | NA | NA |
| ENSMUSG( | 20.52804 | -0.10776 | 0.793755 | NA | NA |
| ENSMUSG( | 59.5729 | 0.049063 | 0.857852 | 0.974667 | NA |
| ENSMUSG( | 32.70178 | 0.017599 | 0.958439 | 0.993874 | NA |
| ENSMUSG( | 133.4646 | 0.224251 | 0.212702 | 0.6567 | NA |
| ENSMUSG( | 8.435563 | -0.5252 | 0.478986 | NA | NA |
| ENSMUSG( | 28.32453 | 0.42651 | 0.237601 | 0.685624 | NA |
| ENSMUSG( | 9.764407 | 0.450662 | 0.465596 | NA | NA |
| ENSMUSG( | 55.3464 | -0.10917 | 0.716567 | 0.941697 | NA |
| ENSMUSG( | 69.75408 | -0.25888 | 0.256555 | 0.707467 | NA |
| ENSMUSG( | 10.82519 | -0.13024 | 0.823095 | NA | NA |
| ENSMUSG( | 9.789894 | 0.416227 | 0.482502 | NA | NA |
| ENSMUSG( | 17.73704 | 0.09599 | 0.83119 | NA | NA |
| ENSMUSG( | 23.75116 | -0.57784 | 0.159412 | 0.58872 | NA |
| ENSMUSG( | 226.8688 | -0.14984 | 0.376989 | 0.801213 | NA |
| ENSMUSG( | 11.49119 | -0.10258 | 0.861876 | NA | NA |
| ENSMUSG( | 16.86562 | -0.2201 | 0.664191 | NA | NA |
| ENSMUSG( | 62.20295 | -0.03599 | 0.909189 | 0.98794 | NA |
| ENSMUSG( | 8.832897 | -0.80818 | 0.243815 | NA | NA |
| ENSMUSG( | 10.39657 | 0.496304 | 0.381284 | NA | NA |
| ENSMUSG( | 11.25993 | -0.20976 | 0.714437 | NA | NA |
| ENSMUSG( | 33.43533 | 0.146228 | 0.701897 | 0.937441 | NA |
| ENSMUSG( | 8.548135 | 0.450162 | 0.497955 | NA | NA |
| ENSMUSG( | 46.2013 | 0.142653 | 0.648836 | 0.920025 | NA |
| ENSMUSG( | 20.21507 | 0.116148 | 0.801593 | NA | NA |
| ENSMUSG( | 14.59763 | -0.04136 | 0.940644 | NA | NA |
| ENSMUSG( | 9.148947 | 0.549646 | 0.364995 | NA | NA |
| ENSMUSG( | 39.46987 | -0.01531 | 0.964467 | 0.995155 | NA |
| ENSMUSG( | 132.4298 | 0.188653 | 0.413307 | 0.819541 | NA |
| ENSMUSG( | 12.19329 | -0.04207 | 0.937904 | NA | NA |
| ENSMUSG( | 94.56494 | -0.27759 | 0.200872 | 0.640988 | NA |
| ENSMUSG( | 18.30172 | -0.08296 | 0.852351 | NA | NA |
| ENSMUSG( | 11.27282 | 0.193968 | 0.753741 | NA | NA |
| ENSMUSG( | 13.74073 | -0.18392 | 0.724173 | NA | NA |
| ENSMUSG( | 174.0076 | 0.038838 | 0.818277 | 0.965837 | NA |
| ENSMUSG( | 12.46466 | -0.26648 | 0.639019 | NA | NA |
| ENSMUSG( | 65.6372 | -0.01343 | 0.961674 | 0.994286 | NA |
| ENSMUSG( | 107.6269 | -0.05617 | 0.777998 | 0.956168 | NA |
| ENSMUSG( | 12.58227 | 0.256685 | 0.619396 | NA | NA |
| ENSMUSG( | 44.84815 | -0.14187 | 0.632123 | 0.914073 | NA |
| ENSMUSG( | 34.82169 | -0.26547 | 0.421285 | 0.823126 | NA |
| ENSMUSG( | 18.89339 | 0.260107 | 0.576328 | NA | NA |
| ENSMUSG( | 11.49303 | -0.21576 | 0.722983 | NA | NA |
| ENSMUSG( | 93.32701 | 0.028419 | 0.90187 | 0.985633 | NA |
| ENSMUSG( | 27.98959 | -0.52368 | 0.148126 | 0.571232 | NA |
| ENSMUSG( | 37.7672 | 0.177246 | 0.566725 | 0.891644 | NA |
| ENSMUSG( | 74.56213 | -0.00482 | 0.983591 | 0.997738 | NA |
| ENSMUSG( | 10.26271 | -0.0158 | 0.978346 | NA | NA |
| ENSMUSG( | 10.79996 | 0.318786 | 0.599566 | NA | NA |
| ENSMUSG( | 83.73808 | 0.02061 | 0.924515 | 0.99114 | NA |
| ENSMUSG( | 10.40223 | -0.32258 | 0.608123 | NA | NA |
| ENSMUSG( | 82.5618 | 0.313176 | 0.175101 | 0.610399 | NA |
| ENSMUSG( | 42.55816 | 0.024434 | 0.935041 | 0.993279 | NA |
| ENSMUSG( | 49.31605 | -0.19548 | 0.527425 | 0.877599 | NA |
| ENSMUSG( | 34.28599 | -0.14545 | 0.656049 | 0.923929 | NA |
| ENSMUSG( | 14.77429 | -0.43148 | 0.394152 | NA | NA |
| ENSMUSG( | 10.79924 | 0.298665 | 0.589483 | NA | NA |
| ENSMUSG( | 50.85091 | 0.299081 | 0.268752 | 0.719042 | NA |
| ENSMUSG( | 13.95727 | -0.91277 | 0.084345 | NA | NA |
| ENSMUSG( | 51.79711 | 0.062766 | 0.81848 | 0.965843 | NA |
| ENSMUSG( | 10.24229 | -0.45116 | 0.501742 | NA | NA |
| ENSMUSG( | 74.81705 | 0.046558 | 0.860555 | 0.9749 | NA |
| ENSMUSG( | 13.20908 | -0.037 | 0.950708 | NA | NA |
| ENSMUSG( | 880.2653 | 0.008745 | 0.950708 | 0.993821 | NA |
| ENSMUSG( | 53.83667 | -0.13382 | 0.642576 | 0.916865 | NA |
| ENSMUSG( | 41.76572 | -0.52525 | 0.089166 | 0.458482 | NA |
| ENSMUSG( | 16.01959 | 0.592956 | 0.197457 | NA | NA |
| ENSMUSG( | 10.63903 | -0.54212 | 0.349268 | NA | NA |
| ENSMUSG( | 93.04036 | -0.06739 | 0.738356 | 0.947329 | NA |
| ENSMUSG( | 26.28479 | 0.015541 | 0.966992 | 0.995911 | NA |
| ENSMUSG( | 9.705875 | 0.405131 | 0.535556 | NA | NA |
| ENSMUSG( | 298.663 | -0.48964 | 0.010128 | 0.143118 | NA |
| ENSMUSG( | 14.24005 | -0.4647 | 0.379383 | NA | NA |
| ENSMUSG( | 10.01271 | 0.012748 | 0.983079 | NA | NA |
| ENSMUSG( | 321.5994 | 0.171883 | 0.207221 | 0.649561 | NA |
| ENSMUSG( | 28.24259 | -0.01536 | 0.969418 | 0.996369 | NA |
| ENSMUSG( | 12.25437 | -3.56245 | 7.49E-06 | NA | NA |
| ENSMUSG( | 17.79592 | 0.533018 | 0.255381 | NA | NA |
| ENSMUSG( | 14.64675 | 1.197626 | 0.017722 | NA | NA |
| ENSMUSG( | 12.7636 | -0.39347 | 0.476838 | NA | NA |
| ENSMUSG( | 18.43132 | -0.44367 | 0.326552 | NA | NA |
| ENSMUSG( | 21.50864 | -0.08731 | 0.824778 | 0.967386 | NA |
| ENSMUSG( | 46.56157 | -0.0648 | 0.818818 | 0.965888 | NA |
| ENSMUSG( | 15.98712 | -0.17674 | 0.725472 | NA | NA |
| ENSMUSG( | 522.7214 | 0.014913 | 0.900743 | 0.985633 | NA |
| ENSMUSG( | 40.62877 | -0.00109 | 0.997162 | 0.999565 | NA |
| ENSMUSG( | 31.04999 | -0.33919 | 0.323183 | 0.768415 | NA |
| ENSMUSG( | 42.47425 | 0.157882 | 0.588876 | 0.90113 | NA |
| ENSMUSG( | 25.75633 | 0.153102 | 0.685518 | 0.932981 | NA |
| ENSMUSG( | 25.66964 | -0.02382 | 0.948973 | 0.993821 | NA |
| ENSMUSG( | 55.29254 | -0.05195 | 0.852847 | 0.973168 | NA |
| ENSMUSG( | 11.2082 | -0.52209 | 0.358093 | NA | NA |
| ENSMUSG( | 9.45327 | 0.145583 | 0.80577 | NA | NA |
| ENSMUSG( | 21.07243 | 0.140826 | 0.750831 | NA | NA |
| ENSMUSG( | 17.01354 | 0.300763 | 0.513692 | NA | NA |
| ENSMUSG( | 24.63935 | -0.01709 | 0.963427 | 0.994911 | NA |
| ENSMUSG( | 56.96792 | 0.068035 | 0.818387 | 0.965837 | NA |
| ENSMUSG( | 43.07177 | 0.034853 | 0.903715 | 0.986094 | NA |
| ENSMUSG( | 745.4922 | 0.303276 | 0.095558 | 0.475062 | NA |
| ENSMUSG( | 53.16293 | -0.4614 | 0.16231 | 0.837724 | NA |
| ENSMUSG( | 23.2489 | 0.569236 | 0.199646 | 0.863016 | NA |
| ENSMUSG( | 22.21689 | -0.31028 | 0.471871 | 0.923329 | NA |
| ENSMUSG( | 19.20088 | 0.026054 | 0.956461 | 0.994798 | NA |
| ENSMUSG( | 9.917046 | 0.511008 | 0.417198 | NA | NA |
| ENSMUSG( | 33.98172 | -0.15044 | 0.662568 | 0.956205 | NA |
| ENSMUSG( | 37.2874 | -0.4087 | 0.228186 | 0.868507 | NA |
| ENSMUSG( | 721.2564 | 0.397192 | 0.013835 | 0.46927 | NA |
| ENSMUSG( | 9.217487 | 0.20538 | 0.780486 | NA | NA |
| ENSMUSG( | 197.0829 | -0.1335 | 0.54747 | 0.938063 | NA |
| ENSMUSG( | 11.25229 | 0.161763 | 0.79298 | NA | NA |
| ENSMUSG( | 31.62429 | 0.146536 | 0.692465 | 0.960404 | NA |
| ENSMUSG( | 60.22956 | 0.013289 | 0.962646 | 0.995284 | NA |
| ENSMUSG( | 214.4645 | -0.25129 | 0.218129 | 0.866104 | NA |
| ENSMUSG( | 23.88393 | 0.791625 | 0.057333 | 0.723362 | NA |
| ENSMUSG( | 26.74031 | -0.1376 | 0.746967 | 0.969763 | NA |
| ENSMUSG( | 36.67119 | -0.77135 | 0.026244 | 0.603041 | NA |
| ENSMUSG( | 179.0929 | 0.10695 | 0.674523 | 0.95694 | NA |
| ENSMUSG( | 97.55104 | 0.406229 | 0.141528 | 0.830678 | NA |
| ENSMUSG( | 10.97532 | -0.27987 | 0.627425 | NA | NA |
| ENSMUSG( | 38.48496 | -0.35964 | 0.304217 | 0.898538 | NA |
| ENSMUSG( | 88.19318 | 0.045181 | 0.852217 | 0.985598 | NA |
| ENSMUSG( | 135.7058 | -0.1099 | 0.593925 | 0.945408 | NA |
| ENSMUSG( | 10.3429 | 0.525428 | 0.385267 | NA | NA |
| ENSMUSG( | 12.28768 | 0.401365 | 0.463654 | 0.923329 | NA |
| ENSMUSG( | 6.300085 | 0.096692 | 0.911961 | NA | NA |
| ENSMUSG( | 103.5621 | -0.07378 | 0.763721 | 0.974 | NA |
| ENSMUSG( | 12.88938 | -0.99649 | 0.068075 | 0.75212 | NA |
| ENSMUSG( | 46.96196 | -0.05638 | 0.858121 | 0.985884 | NA |
| ENSMUSG( | 20.03238 | 0.003054 | 0.994402 | 0.998813 | NA |
| ENSMUSG( | 13.51031 | -0.01308 | 0.980145 | 0.996479 | NA |
| ENSMUSG( | 15.48023 | 1.420742 | 0.008844 | 0.394786 | NA |
| ENSMUSG( | 83.44397 | 0.380951 | 0.201585 | 0.864412 | NA |
| ENSMUSG( | 11.45138 | 0.279626 | 0.625777 | NA | NA |
| ENSMUSG( | 378.3557 | 0.091199 | 0.639606 | 0.953903 | NA |
| ENSMUSG( | 16.49083 | 0.90885 | 0.136511 | 0.827141 | NA |
| ENSMUSG( | 7.896701 | -0.00326 | 0.996498 | NA | NA |
| ENSMUSG( | 15.13387 | -0.57692 | 0.267041 | 0.886134 | NA |
| ENSMUSG( | 24.0858 | 0.364834 | 0.41903 | 0.917257 | NA |
| ENSMUSG( | 17.6997 | 0.434297 | 0.358068 | 0.913759 | NA |
| ENSMUSG( | 48.99542 | -0.09503 | 0.766468 | 0.974236 | NA |
| ENSMUSG( | 10.39523 | 0.237715 | 0.719749 | NA | NA |
| ENSMUSG( | 91.23397 | -0.27902 | 0.211784 | 0.864412 | NA |
| ENSMUSG( | 13.37458 | 0.529385 | 0.342949 | 0.909646 | NA |
| ENSMUSG( | 8.590902 | -0.18879 | 0.775259 | NA | NA |
| ENSMUSG( | 54.6459 | 0.430739 | 0.150475 | 0.837173 | NA |
| ENSMUSG( | 8.365078 | -1.16318 | 0.094649 | NA | NA |
| ENSMUSG( | 11.12982 | -0.21814 | 0.714605 | NA | NA |
| ENSMUSG( | 18.56797 | 0.154652 | 0.750919 | 0.970317 | NA |
| ENSMUSG( | 10.76572 | 0.179152 | 0.789672 | NA | NA |
| ENSMUSG( | 10.71883 | -0.41919 | 0.494949 | NA | NA |
| ENSMUSG( | 207.1108 | -0.34263 | 0.051055 | 0.711023 | NA |
| ENSMUSG( | 19.59184 | 0.132385 | 0.776793 | 0.975717 | NA |
| ENSMUSG( | 27.62953 | 0.073786 | 0.853829 | 0.98563 | NA |
| ENSMUSG( | 24.45288 | -0.45633 | 0.312316 | 0.900037 | NA |
| ENSMUSG( | 72.63174 | 0.186256 | 0.472379 | 0.923329 | NA |
| ENSMUSG( | 54.39871 | -0.6499 | 0.036474 | 0.66949 | NA |
| ENSMUSG( | 22.0062 | -0.57881 | 0.180689 | 0.850372 | NA |
| ENSMUSG( | 82.40986 | 0.061169 | 0.824131 | 0.979198 | NA |
| ENSMUSG( | 13.04824 | -0.19953 | 0.711931 | 0.964363 | NA |
| ENSMUSG( | 20.18202 | -0.21521 | 0.649249 | 0.95422 | NA |
| ENSMUSG( | 51.44839 | 0.171927 | 0.555054 | 0.939565 | NA |
| ENSMUSG( | 12.03584 | 0.175858 | 0.770986 | 0.974236 | NA |
| ENSMUSG( | 12.19062 | 0.365051 | 0.532826 | 0.934795 | NA |
| ENSMUSG( | 14.32584 | 0.745109 | 0.153043 | 0.837724 | NA |
| ENSMUSG( | 625.9694 | 0.267573 | 0.071163 | 0.761942 | NA |
| ENSMUSG( | 8.829397 | 0.563146 | 0.407433 | NA | NA |
| ENSMUSG( | 6.304426 | -0.15264 | 0.856381 | NA | NA |
| ENSMUSG( | 9.763164 | -0.64946 | 0.357568 | NA | NA |
| ENSMUSG( | 10.73616 | 0.600063 | 0.361679 | NA | NA |
| ENSMUSG( | 21.30004 | -0.22026 | 0.704396 | 0.962677 | NA |
| ENSMUSG( | 41.95168 | 0.495593 | 0.19347 | 0.860096 | NA |
| ENSMUSG( | 15.16203 | -0.60723 | 0.248706 | 0.877904 | NA |
| ENSMUSG( | 45.26477 | 0.026818 | 0.931146 | 0.99306 | NA |
| ENSMUSG( | 93.08279 | -0.27408 | 0.222183 | 0.866232 | NA |
| ENSMUSG( | 35.30091 | 0.443076 | 0.189083 | 0.85698 | NA |
| ENSMUSG( | 8.840646 | 0.087064 | 0.901939 | NA | NA |
| ENSMUSG( | 96.51863 | 0.298641 | 0.312997 | 0.900449 | NA |
| ENSMUSG( | 301.7859 | 0.425887 | 0.075734 | 0.768524 | NA |
| ENSMUSG( | 8.7143 | -0.28744 | 0.694965 | NA | NA |
| ENSMUSG( | 40.51125 | -0.34217 | 0.298332 | 0.898538 | NA |
| ENSMUSG( | 102.9856 | -0.47651 | 0.031764 | 0.631941 | NA |
| ENSMUSG( | 110.5684 | -0.10114 | 0.67472 | 0.95694 | NA |
| ENSMUSG( | 50.35214 | -0.13489 | 0.638029 | 0.953903 | NA |
| ENSMUSG( | 21.1338 | -0.71618 | 0.095393 | 0.796133 | NA |
| ENSMUSG( | 25.74411 | 0.824463 | 0.10148 | 0.801456 | NA |
| ENSMUSG( | 19.68946 | 0.22633 | 0.647055 | 0.954079 | NA |
| ENSMUSG( | 14.21599 | -0.03304 | 0.949307 | 0.994798 | NA |
| ENSMUSG( | 59.47339 | -0.18699 | 0.486301 | 0.92541 | NA |
| ENSMUSG( | 14.28925 | -0.21099 | 0.691288 | 0.960071 | NA |
| ENSMUSG( | 19.08421 | -1.11459 | 0.018768 | 0.547939 | NA |
| ENSMUSG( | 25.4713 | -0.79531 | 0.046007 | 0.69654 | NA |
| ENSMUSG( | 14.18697 | 0.560566 | 0.330306 | 0.907251 | NA |
| ENSMUSG( | 18.59147 | -0.65562 | 0.168341 | 0.840222 | NA |
| ENSMUSG( | 10.47781 | 1.104112 | 0.135374 | NA | NA |
| ENSMUSG( | 29.39196 | 2.206624 | 0.085841 | 0.780123 | NA |
| ENSMUSG( | 28.37334 | 0.170064 | 0.653344 | 0.955015 | NA |
| ENSMUSG( | 18.17811 | 0.009814 | 0.9862 | 0.997188 | NA |
| ENSMUSG( | 11.7331 | -0.01445 | 0.979308 | 0.996468 | NA |
| ENSMUSG( | 82.39615 | 0.173532 | 0.512793 | 0.930164 | NA |
| ENSMUSG( | 66.17206 | 0.160776 | 0.692754 | 0.96043 | NA |
| ENSMUSG( | 33.46438 | -0.19697 | 0.608927 | 0.950226 | NA |
| ENSMUSG( | 11.49222 | -0.15239 | 0.789301 | 0.976593 | NA |
| ENSMUSG( | 375.3597 | 0.045075 | 0.776933 | 0.975717 | NA |
| ENSMUSG( | 18.69081 | 0.090297 | 0.843272 | 0.983281 | NA |
| ENSMUSG( | 10.65946 | 0.065077 | 0.910817 | NA | NA |
| ENSMUSG( | 188.1575 | 0.039703 | 0.819569 | 0.979198 | NA |
| ENSMUSG( | 12.59875 | 0.259076 | 0.646884 | 0.953988 | NA |
| ENSMUSG( | 15.11397 | -0.54164 | 0.294015 | 0.898538 | NA |
| ENSMUSG( | 185.3587 | -0.11188 | 0.547506 | 0.938063 | NA |
| ENSMUSG( | 19.13746 | -0.33286 | 0.481196 | 0.924548 | NA |
| ENSMUSG( | 39.62872 | -0.04165 | 0.903828 | 0.991089 | NA |
| ENSMUSG( | 22.69305 | -0.44964 | 0.34845 | 0.911652 | NA |
| ENSMUSG( | 13.20404 | -0.79566 | 0.135553 | 0.825163 | NA |
| ENSMUSG( | 10.23928 | 0.297853 | 0.651748 | NA | NA |
| ENSMUSG( | 59.83198 | 0.190067 | 0.559581 | 0.939807 | NA |
| ENSMUSG( | 25.98685 | 0.095614 | 0.817081 | 0.979198 | NA |
| ENSMUSG( | 21.26476 | -0.4599 | 0.337478 | 0.907972 | NA |
| ENSMUSG( | 11.24375 | 0.22287 | 0.710805 | NA | NA |
| ENSMUSG( | 53.56872 | 0.341467 | 0.277031 | 0.891231 | NA |
| ENSMUSG( | 17.46556 | -0.26669 | 0.584654 | 0.944975 | NA |
| ENSMUSG( | 87.82864 | -0.07249 | 0.805927 | 0.977674 | NA |
| ENSMUSG( | 10.848 | -0.00232 | 0.997096 | NA | NA |
| ENSMUSG( | 8.943505 | 0.104015 | 0.876697 | NA | NA |
| ENSMUSG( | 11.74069 | 0.816129 | 0.211733 | 0.864412 | NA |
| ENSMUSG( | 30.99858 | -0.19523 | 0.59149 | 0.944975 | NA |
| ENSMUSG( | 11.58764 | 0.102263 | 0.859017 | 0.985884 | NA |
| ENSMUSG( | 12.74193 | 0.376876 | 0.487037 | 0.92541 | NA |
| ENSMUSG( | 1342.678 | 0.032546 | 0.785614 | 0.976285 | NA |
| ENSMUSG( | 18.04279 | -0.60097 | 0.232352 | 0.868507 | NA |
| ENSMUSG( | 9.69564 | 0.075666 | 0.904172 | NA | NA |
| ENSMUSG( | 36.84817 | 0.24546 | 0.488134 | 0.925453 | NA |
| ENSMUSG( | 10.92566 | 0.330901 | 0.588221 | NA | NA |
| ENSMUSG( | 75.46865 | -0.00776 | 0.976125 | 0.996416 | NA |
| ENSMUSG( | 131.4857 | 0.399998 | 0.125161 | 0.81516 | NA |
| ENSMUSG( | 29.86062 | -0.29482 | 0.433655 | 0.918564 | NA |
| ENSMUSG( | 109.0379 | 0.181486 | 0.522672 | 0.932822 | NA |
| ENSMUSG( | 16.446 | -0.24051 | 0.637767 | 0.953903 | NA |
| ENSMUSG( | 27.3105 | -0.04868 | 0.901868 | 0.990969 | NA |
| ENSMUSG( | 18.86038 | 0.478572 | 0.29999 | 0.898538 | NA |
| ENSMUSG( | 62.75021 | 0.105385 | 0.714786 | 0.964986 | NA |
| ENSMUSG( | 30.37716 | 0.738477 | 0.04467 | 0.693697 | NA |
| ENSMUSG( | 20.22711 | 0.291166 | 0.5015 | 0.927825 | NA |
| ENSMUSG( | 14.05543 | -0.80224 | 0.128005 | 0.81845 | NA |
| ENSMUSG( | 64.39566 | -0.54168 | 0.063261 | 0.736482 | NA |
| ENSMUSG( | 107.81 | -0.11152 | 0.606275 | 0.949497 | NA |
| ENSMUSG( | 79.37846 | -0.17188 | 0.492373 | 0.927315 | NA |
| ENSMUSG( | 9.302066 | -0.1213 | 0.848681 | NA | NA |
| ENSMUSG( | 8.338571 | -0.5532 | 0.447384 | NA | NA |
| ENSMUSG( | 9.000913 | -0.04155 | 0.948671 | NA | NA |
| ENSMUSG( | 12.43514 | 0.251323 | 0.640752 | 0.953903 | NA |
| ENSMUSG( | 10.52646 | -0.28126 | 0.639877 | NA | NA |
| ENSMUSG( | 25.25297 | -0.17484 | 0.655787 | 0.955533 | NA |
| ENSMUSG( | 164.3733 | 0.137637 | 0.529247 | 0.934795 | NA |
| ENSMUSG( | 8.149733 | -0.58123 | 0.418652 | NA | NA |
| ENSMUSG( | 18.61645 | -0.30701 | 0.546286 | 0.938063 | NA |
| ENSMUSG( | 31.84522 | -0.51821 | 0.151099 | 0.837173 | NA |
| ENSMUSG( | 2693.272 | 0.01526 | 0.912504 | 0.992261 | NA |

| | | | | | | | | | |
|---|---|---|---|---|---|---|---|---|---|
| ENSMUSG(NA | 76.12681 | 0.097164 | 0.675937 | 0.930542 | ENSMUSG(NA | 11.49222 | -0.15239 | 0.789301 | 0.976593 |
| ENSMUSG(NA | 9.129818 | -0.01655 | 0.979447 | NA | ENSMUSG(NA | 375.3597 | 0.045075 | 0.776933 | 0.975717 |
| ENSMUSG(NA | 20.87084 | 0.10751 | 0.809302 | NA | ENSMUSG(NA | 18.69081 | 0.090297 | 0.843272 | 0.983281 |
| ENSMUSG(NA | 106.4639 | 0.219527 | 0.253387 | 0.703636 | ENSMUSG(NA | 10.65946 | 0.065077 | 0.910817 | NA |
| ENSMUSG(NA | 23.23588 | 0.172953 | 0.66615 | 0.927057 | ENSMUSG(NA | 188.1575 | 0.039703 | 0.819569 | 0.979198 |
| ENSMUSG(NA | 76.48745 | -0.49234 | 0.051739 | 0.349085 | ENSMUSG(NA | 12.59875 | 0.259076 | 0.646884 | 0.953988 |
| ENSMUSG(NA | 21.79264 | -0.69218 | 0.105315 | 0.492417 | ENSMUSG(NA | 15.11397 | -0.54164 | 0.294015 | 0.898538 |
| ENSMUSG(NA | 13.18272 | 0.037436 | 0.94251 | NA | ENSMUSG(NA | 185.3587 | -0.11188 | 0.547506 | 0.938063 |
| ENSMUSG(NA | 11.26443 | -0.11394 | 0.843469 | NA | ENSMUSG(NA | 19.13746 | -0.33286 | 0.481196 | 0.924548 |
| ENSMUSG(NA | 86.28432 | 0.120994 | 0.568942 | 0.892739 | ENSMUSG(NA | 39.62872 | -0.04165 | 0.903828 | 0.991089 |
| ENSMUSG(NA | 10.0688 | 0.350799 | 0.565807 | NA | ENSMUSG(NA | 22.69305 | -0.44964 | 0.34845 | 0.911652 |
| ENSMUSG(NA | 83.22019 | -0.06817 | 0.750744 | 0.950042 | ENSMUSG(NA | 13.20404 | -0.79566 | 0.135553 | 0.825163 |
| ENSMUSG(NA | 9.707056 | -0.93631 | 0.175202 | NA | ENSMUSG(NA | 10.23928 | 0.297853 | 0.651748 | NA |
| ENSMUSG(NA | 15.74502 | 0.273482 | 0.580484 | NA | ENSMUSG(NA | 59.83198 | 0.190067 | 0.559581 | 0.939807 |
| ENSMUSG(NA | 10.89895 | -0.93469 | 0.117287 | NA | ENSMUSG(NA | 25.98685 | 0.095614 | 0.817081 | 0.979198 |
| ENSMUSG(NA | 42.82025 | -0.29672 | 0.384555 | 0.804007 | ENSMUSG(NA | 21.26476 | -0.4599 | 0.337478 | 0.907972 |
| ENSMUSG(NA | 23.36463 | -0.30136 | 0.429618 | 0.826446 | ENSMUSG(NA | 11.24375 | 0.22287 | 0.710805 | NA |
| ENSMUSG(NA | 29.47471 | 0.210707 | 0.571423 | 0.893319 | ENSMUSG(NA | 53.56872 | 0.341467 | 0.277031 | 0.891231 |
| ENSMUSG(NA | 105.5342 | -0.12117 | 0.547997 | 0.885153 | ENSMUSG(NA | 17.46556 | -0.26669 | 0.584654 | 0.944975 |
| ENSMUSG(NA | 93.44606 | -0.1321 | 0.528833 | 0.878365 | ENSMUSG(NA | 87.82864 | -0.07249 | 0.805927 | 0.977674 |
| ENSMUSG(NA | 10.04561 | -0.30111 | 0.632603 | NA | ENSMUSG(NA | 10.848 | -0.00232 | 0.997096 | NA |
| ENSMUSG(NA | 588.1459 | -0.04258 | 0.743554 | 0.948478 | ENSMUSG(NA | 8.943505 | 0.104015 | 0.876697 | NA |
| ENSMUSG(NA | 51.06596 | -0.52106 | 0.065391 | 0.396102 | ENSMUSG(NA | 30.99858 | -0.19523 | 0.59149 | 0.944975 |
| ENSMUSG(NA | 16.94287 | -0.22325 | 0.628614 | NA | ENSMUSG(NA | 11.58764 | 0.102263 | 0.859017 | 0.985884 |
| ENSMUSG(NA | 12.83312 | -0.42701 | 0.438366 | NA | ENSMUSG(NA | 12.74193 | 0.376876 | 0.487037 | 0.92541 |
| ENSMUSG(NA | 22.55415 | 0.171924 | 0.673097 | 0.929007 | ENSMUSG(NA | 1342.678 | 0.032546 | 0.785614 | 0.976285 |
| ENSMUSG(NA | 64.04169 | -0.11618 | 0.681635 | 0.931619 | ENSMUSG(NA | 18.04279 | -0.60097 | 0.232352 | 0.868507 |
| ENSMUSG(NA | 15.01744 | -0.43128 | 0.367194 | NA | ENSMUSG(NA | 9.69564 | 0.075666 | 0.904172 | NA |
| ENSMUSG(NA | 12.24063 | -0.09441 | 0.85854 | NA | ENSMUSG(NA | 36.84817 | 0.24546 | 0.488134 | 0.925453 |
| ENSMUSG(NA | 70.30702 | -0.07187 | 0.813435 | 0.964574 | ENSMUSG(NA | 10.92566 | 0.330901 | 0.588221 | NA |
| ENSMUSG(NA | 59.60509 | -0.2938 | 0.301669 | 0.749979 | ENSMUSG(NA | 75.46865 | -0.00776 | 0.976125 | 0.996416 |
| ENSMUSG(NA | 23.70466 | 0.074417 | 0.845936 | 0.972226 | ENSMUSG(NA | 131.4857 | 0.399998 | 0.125161 | 0.81516 |
| ENSMUSG(NA | 204.2323 | 0.293289 | 0.086849 | 0.453311 | ENSMUSG(NA | 29.86062 | -0.29482 | 0.433655 | 0.918564 |
| ENSMUSG(NA | 20.76936 | 0.115407 | 0.780938 | NA | ENSMUSG(NA | 109.0379 | 0.181486 | 0.522672 | 0.932822 |
| ENSMUSG(NA | 19.65113 | -0.24576 | 0.564166 | NA | ENSMUSG(NA | 16.446 | -0.24051 | 0.637767 | 0.953903 |
| ENSMUSG(NA | 142.0453 | 0.016104 | 0.929526 | 0.991848 | ENSMUSG(NA | 27.3105 | -0.04868 | 0.901868 | 0.990969 |
| ENSMUSG(NA | 17.56825 | -0.31943 | 0.477176 | NA | ENSMUSG(NA | 18.86038 | 0.478572 | 0.29999 | 0.898538 |
| ENSMUSG(NA | 9.591254 | 0.252454 | 0.724084 | NA | ENSMUSG(NA | 62.75021 | 0.105385 | 0.714786 | 0.964986 |
| ENSMUSG(NA | 15.01385 | -0.40772 | 0.388702 | NA | ENSMUSG(NA | 30.37716 | 0.738477 | 0.04467 | 0.693697 |
| ENSMUSG(NA | 21.46733 | 0.356375 | 0.383483 | 0.803428 | ENSMUSG(NA | 20.22711 | 0.291166 | 0.5015 | 0.927825 |
| ENSMUSG(NA | 16.56474 | 0.337995 | 0.519954 | NA | ENSMUSG(NA | 14.05543 | -0.80224 | 0.128005 | 0.81845 |
| ENSMUSG(NA | 118.1796 | -0.06174 | 0.781698 | 0.957558 | ENSMUSG(NA | 64.39566 | -0.54168 | 0.063261 | 0.736482 |
| ENSMUSG(NA | 23.84188 | 0.20696 | 0.612106 | 0.908586 | ENSMUSG(NA | 107.81 | -0.11152 | 0.606275 | 0.949497 |
| ENSMUSG(NA | 33.77622 | -0.06558 | 0.851971 | 0.973168 | ENSMUSG(NA | 79.37846 | -0.17188 | 0.492373 | 0.927315 |
| ENSMUSG(NA | 15.8741 | -0.06091 | 0.897802 | NA | ENSMUSG(NA | 9.302066 | -0.1213 | 0.848681 | NA |
| ENSMUSG(NA | 18.88822 | -0.26547 | 0.596613 | NA | ENSMUSG(NA | 8.338571 | -0.5532 | 0.447384 | NA |
| ENSMUSG(NA | 9.790521 | -0.53565 | 0.35874 | NA | ENSMUSG(NA | 12.43514 | 0.251323 | 0.640752 | 0.953903 |
| ENSMUSG(NA | 120.4825 | 0.278968 | 0.137812 | 0.552903 | ENSMUSG(NA | 10.52646 | -0.28126 | 0.639877 | NA |
| ENSMUSG(NA | 28.7182 | 0.262896 | 0.483096 | 0.856216 | ENSMUSG(NA | 25.25297 | -0.17484 | 0.655787 | 0.955533 |
| ENSMUSG(NA | 18.5202 | -0.52467 | 0.245766 | NA | ENSMUSG(NA | 164.3733 | 0.137637 | 0.529247 | 0.934795 |
| ENSMUSG(NA | 4115.45 | -1.52651 | 1.03E-05 | 0.00119 | ENSMUSG(NA | 18.61645 | -0.30701 | 0.546286 | 0.938063 |
| ENSMUSG(NA | 184.1498 | -0.00437 | 0.980636 | 0.997512 | ENSMUSG(NA | 31.84522 | -0.51821 | 0.151099 | 0.837173 |
| ENSMUSG(NA | 28.78782 | 0.062131 | 0.872779 | 0.978753 | ENSMUSG(NA | 2693.272 | 0.01526 | 0.912504 | 0.992261 |
| ENSMUSG(NA | 29.02044 | -0.29266 | 0.500334 | 0.863455 | ENSMUSG(NA | 49.91086 | 0.191287 | 0.552785 | 0.939565 |
| ENSMUSG(NA | 36.77531 | -1.05485 | 0.004957 | 0.093376 | ENSMUSG(NA | 11.07905 | 0.095081 | 0.86756 | NA |
| ENSMUSG(NA | 33.87152 | -0.20683 | 0.532448 | 0.879905 | ENSMUSG(NA | 389.9025 | -0.46785 | 0.002282 | 0.185073 |
| ENSMUSG(NA | 48.27139 | 0.15872 | 0.582331 | 0.897918 | ENSMUSG(NA | 35.08015 | -0.22801 | 0.502378 | 0.927825 |
| ENSMUSG(NA | 391.3566 | 0.03197 | 0.870505 | 0.977906 | ENSMUSG(NA | 22.05715 | -0.33435 | 0.446489 | 0.921734 |
| ENSMUSG(NA | 44.9842 | 0.287168 | 0.316223 | 0.763453 | ENSMUSG(NA | 17.32704 | -0.09638 | 0.838843 | 0.981696 |
| ENSMUSG(NA | 84.11652 | -0.07726 | 0.736447 | 0.946437 | ENSMUSG(NA | 724.2925 | 0.468304 | 0.054467 | 0.723362 |
| ENSMUSG(NA | 20.29068 | -0.37169 | 0.404788 | NA | ENSMUSG(NA | 14.80514 | -0.13839 | 0.786287 | 0.976303 |
| ENSMUSG(NA | 768.0985 | -0.09273 | 0.541055 | 0.883182 | ENSMUSG(NA | 25.24065 | -0.27334 | 0.505773 | 0.927825 |
| ENSMUSG(NA | 25.60042 | 0.514205 | 0.195224 | 0.6346 | ENSMUSG(NA | 13.89249 | -0.30347 | 0.56651 | 0.941777 |
| ENSMUSG(NA | 22.17236 | 0.343247 | 0.418399 | 0.821966 | ENSMUSG(NA | 25.68532 | 0.316102 | 0.433923 | 0.91869 |
| ENSMUSG(NA | 41.18168 | -0.68954 | 0.028543 | 0.257525 | ENSMUSG(NA | 85.18384 | 0.171013 | 0.509892 | 0.929089 |
| ENSMUSG(NA | 38.20074 | -0.15394 | 0.66161 | 0.925271 | ENSMUSG(NA | 7.791957 | 0.255323 | 0.711969 | NA |
| ENSMUSG(NA | 19.95066 | 0.303186 | 0.482292 | NA | ENSMUSG(NA | 21.09651 | 0.418523 | 0.330943 | 0.907513 |
| ENSMUSG(NA | 33.20643 | -0.14545 | 0.678024 | 0.930542 | ENSMUSG(NA | 17.1285 | 0.279065 | 0.586436 | 0.944975 |
| ENSMUSG(NA | 27.17117 | -0.28591 | 0.434529 | 0.828978 | ENSMUSG(NA | 628.3679 | -0.18232 | 0.158432 | 0.837724 |
| ENSMUSG(NA | 18.94563 | 0.488847 | 0.27917 | NA | ENSMUSG(NA | 11.82389 | -0.06938 | 0.905679 | 0.991089 |
| ENSMUSG(NA | 838.4521 | -0.18573 | 0.278858 | 0.728705 | ENSMUSG(NA | 187.2375 | -0.52906 | 0.019336 | 0.554338 |
| ENSMUSG(NA | 60.21968 | 0.011568 | 0.969404 | 0.996369 | ENSMUSG(NA | 7.014731 | -0.05105 | 0.949975 | NA |
| ENSMUSG(NA | 1458.021 | 0.038147 | 0.749728 | 0.949664 | ENSMUSG(NA | 73.44239 | -0.48344 | 0.117096 | 0.813384 |
| ENSMUSG(NA | 80.68308 | -0.3058 | 0.191348 | 0.630245 | ENSMUSG(NA | 11.21347 | -0.09689 | 0.867315 | NA |
| ENSMUSG(NA | 10.12304 | -0.23508 | 0.687372 | NA | ENSMUSG(NA | 140.755 | 0.14025 | 0.475061 | 0.923329 |
| ENSMUSG(NA | 9.185751 | -0.18973 | 0.757874 | NA | ENSMUSG(NA | 308.6161 | 0.283517 | 0.120545 | 0.813384 |
| ENSMUSG(NA | 250.2841 | -0.05028 | 0.794715 | 0.959799 | ENSMUSG(NA | 185.0548 | 0.248471 | 0.251826 | 0.879902 |
| ENSMUSG(NA | 22.90717 | 0.412682 | 0.298394 | 0.74713 | ENSMUSG(NA | 21.37235 | -0.60295 | 0.154806 | 0.837724 |
| ENSMUSG(NA | 18.15574 | -0.38571 | 0.397872 | NA | ENSMUSG(NA | 16.9236 | 0.598559 | 0.262882 | 0.885357 |
| ENSMUSG(NA | 17.67582 | 0.335657 | 0.485938 | NA | ENSMUSG(NA | 108.239 | 0.110577 | 0.629891 | 0.953251 |
| ENSMUSG(NA | 84.73409 | 0.008496 | 0.968152 | 0.996284 | ENSMUSG(NA | 9.963197 | 0.072482 | 0.913311 | NA |
| ENSMUSG(NA | 23.48176 | -0.371 | 0.338517 | 0.777489 | ENSMUSG(NA | 17.9452 | 0.712168 | 0.142129 | 0.830678 |
| ENSMUSG(NA | 106.1474 | -0.24548 | 0.275996 | 0.725669 | ENSMUSG(NA | 10.65716 | 1.305688 | 0.037237 | NA |
| ENSMUSG(NA | 11.75153 | 0.159468 | 0.777422 | NA | ENSMUSG(NA | 232.6494 | -0.17657 | 0.306217 | 0.898538 |
| ENSMUSG(NA | 112.9297 | -0.12751 | 0.503259 | 0.86596 | ENSMUSG(NA | 135.5472 | -0.07764 | 0.700609 | 0.961547 |
| ENSMUSG(NA | 46.46855 | -0.16381 | 0.565074 | 0.890735 | ENSMUSG(NA | 13.44356 | -0.26178 | 0.667328 | 0.956766 |
| ENSMUSG(NA | 20.18343 | -0.22279 | 0.611116 | NA | ENSMUSG(NA | 99.45421 | 0.153714 | 0.514464 | 0.930939 |
| ENSMUSG(NA | 32.64673 | 0.010398 | 0.97732 | 0.997428 | ENSMUSG(NA | 21.17686 | -0.71446 | 0.095029 | 0.796133 |
| ENSMUSG(NA | 24.40423 | -0.66745 | 0.090427 | 0.461669 | ENSMUSG(NA | 38.3805 | 0.644425 | 0.129444 | 0.820358 |
| ENSMUSG(NA | 512.8748 | 0.043568 | 0.755038 | 0.951276 | ENSMUSG(NA | 176.2725 | -0.07127 | 0.679712 | 0.957966 |
| ENSMUSG(NA | 15.16685 | -0.2999 | 0.551798 | NA | ENSMUSG(NA | 220.84 | 0.058676 | 0.749092 | 0.96998 |
| ENSMUSG(NA | 9.36326 | -0.52958 | 0.601286 | NA | ENSMUSG(NA | 196.5984 | 0.375003 | 0.052302 | 0.713538 |
| ENSMUSG(NA | 225.8838 | 0.095117 | 0.544616 | 0.884649 | ENSMUSG(NA | 10.04132 | -0.03856 | 0.950931 | NA |
| ENSMUSG(NA | 673.294 | -0.12298 | 0.265596 | 0.716205 | ENSMUSG(NA | 14.43531 | 0.295735 | 0.578516 | 0.943548 |
| ENSMUSG(NA | 28.63298 | -0.36478 | 0.325339 | 0.77061 | ENSMUSG(NA | 10.63806 | 0.155324 | 0.79636 | NA |
| ENSMUSG(NA | 37.87703 | -0.31661 | 0.321797 | 0.766739 | ENSMUSG(NA | 27.42512 | 0.303906 | 0.509716 | 0.929089 |
| ENSMUSG(NA | 13.4503 | -0.4296 | 0.406757 | NA | ENSMUSG(NA | 145.5312 | 0.098052 | 0.624334 | 0.953234 |
| ENSMUSG(NA | 36.62919 | 0.208491 | 0.507852 | 0.86795 | ENSMUSG(NA | 148.8631 | -0.02045 | 0.921471 | 0.992644 |
| ENSMUSG(NA | 45.16454 | -0.15756 | 0.580289 | 0.896952 | ENSMUSG(NA | 207.464 | -0.33253 | 0.092695 | 0.794394 |
| ENSMUSG(NA | 24.87478 | -0.23401 | 0.541745 | 0.883182 | ENSMUSG(NA | 14.8706 | 0.784499 | 0.163807 | 0.838291 |
| ENSMUSG(NA | 20.8056 | 0.181934 | 0.655086 | NA | ENSMUSG(NA | 11.18908 | -0.75583 | 0.204578 | NA |
| ENSMUSG(NA | 21.70951 | -0.23519 | 0.54573 | 0.884974 | ENSMUSG(NA | 36.87479 | -0.20255 | 0.566366 | 0.941745 |
| ENSMUSG(NA | 11.94112 | 0.144573 | 0.788692 | NA | ENSMUSG(NA | 48.45251 | 0.108548 | 0.721775 | 0.966887 |
| ENSMUSG(NA | 13.07226 | -0.12702 | 0.810042 | NA | ENSMUSG(NA | 32.93548 | -0.10633 | 0.757628 | 0.971858 |
| ENSMUSG(NA | 26.94514 | -0.00799 | 0.982107 | 0.997512 | ENSMUSG(NA | 12.40045 | 0.691409 | 0.245238 | 0.877904 |
| ENSMUSG(NA | 44.65329 | 0.397327 | 0.241682 | 0.689179 | ENSMUSG(NA | 12.97252 | -0.3231 | 0.588134 | 0.944975 |
| ENSMUSG(NA | 13.74593 | 0.268772 | 0.605764 | NA | ENSMUSG(NA | 30.23562 | -0.09985 | 0.791064 | 0.97681 |
| ENSMUSG(NA | 133.9444 | 0.085398 | 0.62808 | 0.91351 | ENSMUSG(NA | 14.0652 | -0.2393 | 0.650746 | 0.95422 |
| ENSMUSG(NA | 762.4847 | 0.112885 | 0.298453 | 0.74713 | ENSMUSG(NA | 248.2339 | -0.33693 | 0.046398 | 0.697814 |
| ENSMUSG(NA | 14.39215 | 0.365602 | 0.469484 | NA | ENSMUSG(NA | 12.46412 | -0.64908 | 0.267781 | 0.886134 |
| ENSMUSG(NA | 49.31965 | 0.121879 | 0.716523 | 0.94152 | ENSMUSG(NA | 18.98332 | 0.760449 | 0.115134 | 0.81257 |
| ENSMUSG(NA | 12.40577 | -0.05492 | 0.921719 | NA | ENSMUSG(NA | 26.57164 | 0.004829 | 0.913094 | 0.992298 |
| ENSMUSG(NA | 49.87928 | -0.15414 | 0.603177 | 0.904887 | ENSMUSG(NA | 62.54773 | 0.131071 | 0.618007 | 0.952232 |
| ENSMUSG(NA | 43.91465 | 0.00789 | 0.978117 | 0.997428 | ENSMUSG(NA | 10.07572 | -0.24595 | 0.695133 | NA |
| ENSMUSG(NA | 281.7488 | 0.044485 | 0.78797 | 0.959226 | ENSMUSG(NA | 190.1357 | 0.584287 | 0.003056 | 0.219421 |
| ENSMUSG(NA | 19.2657 | -0.63227 | 0.160275 | NA | ENSMUSG(NA | 9.369441 | -0.67991 | 0.333054 | NA |
| ENSMUSG(NA | 16.39583 | 0.658693 | 0.183783 | NA | ENSMUSG(NA | 15.44408 | 0.168691 | 0.740624 | 0.968811 |
| ENSMUSG(NA | 97.34036 | 0.183233 | 0.405256 | 0.81652 | ENSMUSG(NA | 24.38362 | -0.22413 | 0.570098 | 0.941777 |
| ENSMUSG(NA | 21.86555 | 0.00097 | 0.99806 | 0.99989 | ENSMUSG(NA | 59.74052 | 0.309745 | 0.249892 | 0.877904 |
| ENSMUSG(NA | 208.2415 | -0.1332 | 0.377156 | 0.801213 | ENSMUSG(NA | 14.41853 | 0.550004 | 0.326724 | 0.905371 |
| ENSMUSG(NA | 65.06646 | -0.09722 | 0.702891 | 0.937735 | ENSMUSG(NA | 27.57524 | 0.018564 | 0.961593 | 0.99511 |
| ENSMUSG(NA | 51.5842 | -0.2813 | 0.301591 | 0.749979 | ENSMUSG(NA | 18.67327 | -0.40394 | 0.373261 | 0.915491 |
| ENSMUSG(NA | 15.07008 | 0.473246 | 0.37773 | NA | ENSMUSG(NA | 17.43792 | 1.332473 | 0.077113 | 0.770558 |
| ENSMUSG(NA | 21.61837 | -0.14206 | 0.717116 | 0.941697 | ENSMUSG(NA | 33.00922 | 0.693799 | 0.243928 | 0.877904 |
| ENSMUSG(NA | 36.3196 | 0.046184 | 0.898358 | 0.985633 | ENSMUSG(NA | 42.92647 | 0.49877 | 0.119241 | 0.813384 |
| ENSMUSG(NA | 629.6854 | 0.331058 | 0.005183 | 0.096057 | ENSMUSG(NA | 219.9971 | 0.331069 | 0.067667 | 0.75212 |
| ENSMUSG(NA | 2891.822 | -0.04526 | 0.647797 | 0.919759 | ENSMUSG(NA | 31.28149 | 0.581233 | 0.115434 | 0.813375 |
| ENSMUSG(NA | 10.31817 | 0.166446 | 0.781052 | NA | ENSMUSG(NA | 132.3421 | -0.51224 | 0.014637 | 0.486524 |
| ENSMUSG(NA | 22.87362 | -0.0627 | 0.883526 | 0.982503 | ENSMUSG(NA | 50.8272 | -0.23005 | 0.416261 | 0.917257 |
| ENSMUSG(NA | 425.1888 | 0.399769 | 0.001304 | 0.039243 | ENSMUSG(NA | 37.99948 | -0.2033 | 0.595329 | 0.945798 |
| ENSMUSG(NA | 21488.63 | 0.217028 | 0.025539 | 0.246694 | ENSMUSG(NA | 24.0245 | -0.21871 | 0.61219 | 0.951071 |
| ENSMUSG(NA | 227.3637 | 0.141502 | 0.424156 | 0.824522 | ENSMUSG(NA | 50.31077 | 0.108943 | 0.712579 | 0.964363 |
| ENSMUSG(NA | 24.51194 | 0.296546 | 0.482127 | 0.855778 | ENSMUSG(NA | 40.37054 | 0.212786 | 0.496282 | 0.927825 |
| ENSMUSG(NA | 14.8819 | 0.538691 | 0.296356 | NA | ENSMUSG(NA | 56.67491 | 0.282767 | 0.362392 | 0.914223 |
| ENSMUSG(NA | 10.58122 | 0.477001 | 0.406848 | NA | ENSMUSG(NA | 16.18023 | -0.58664 | 0.252894 | 0.88018 |
| ENSMUSG(NA | 149.8918 | -0.09883 | 0.587081 | 0.900235 | ENSMUSG(NA | 24.41453 | 0.150801 | 0.723063 | 0.966887 |
| ENSMUSG(NA | 18.51291 | 0.769293 | 0.080441 | NA | ENSMUSG(NA | 402.4859 | -0.33355 | 0.036335 | 0.669285 |
| ENSMUSG(NA | 53.19073 | -0.15833 | 0.558129 | 0.888427 | ENSMUSG(NA | 10.33028 | 0.225763 | 0.732579 | NA |
| ENSMUSG(NA | 151.7496 | -0.19256 | 0.317009 | 0.763823 | ENSMUSG(NA | 43.77247 | 0.43092 | 0.227843 | 0.868507 |
| ENSMUSG(NA | 18.79112 | -0.32816 | 0.470102 | NA | ENSMUSG(NA | 179.3403 | 0.210634 | 0.247562 | 0.877904 |
| ENSMUSG(NA | 16.16834 | -0.60413 | 0.220738 | NA | ENSMUSG(NA | 42.43646 | -0.39357 | 0.211019 | 0.864412 |
| ENSMUSG(NA | 127.3342 | 0.160883 | 0.479515 | 0.854046 | ENSMUSG(NA | 29.732 | -0.16595 | 0.658729 | 0.956205 |
| ENSMUSG(NA | 11.64754 | 0.149532 | 0.777597 | NA | ENSMUSG(NA | 134.628 | 0.193922 | 0.364885 | 0.914449 |
| ENSMUSG(NA | 25.91518 | 0.230217 | 0.602986 | 0.904887 | ENSMUSG(NA | 13.99379 | -0.2124 | 0.700699 | 0.961547 |
| ENSMUSG(NA | 14.54148 | 0.643349 | 0.23501 | NA | ENSMUSG(NA | 71.34701 | 0.034327 | 0.897595 | 0.990558 |
| ENSMUSG(NA | 16.91491 | -0.0309 | 0.952268 | NA | ENSMUSG(NA | 11.35717 | -0.2784 | 0.625545 | NA |
| ENSMUSG(NA | 29.37052 | -0.30702 | 0.41993 | 0.822442 | ENSMUSG(NA | 23.26636 | -0.1575 | 0.741491 | 0.968944 |
| ENSMUSG(NA | 15.51585 | -0.14589 | 0.758401 | NA | ENSMUSG(NA | 68.94459 | -0.02432 | 0.930792 | 0.99306 |
| ENSMUSG(NA | 10.5105 | -0.02402 | 0.969744 | NA | ENSMUSG(NA | 45.40736 | 0.155574 | 0.665391 | 0.956766 |
| ENSMUSG(NA | 11.69109 | -0.12579 | 0.816332 | NA | ENSMUSG(NA | 10.69219 | -0.51207 | 0.392215 | NA |
| ENSMUSG(NA | 31.69967 | -0.1372 | 0.689524 | 0.933609 | ENSMUSG(NA | 11.39722 | -0.16117 | 0.78492 | NA |
| ENSMUSG(NA | 155.6622 | -0.29847 | 0.088663 | 0.45666 | ENSMUSG(NA | 42.33876 | -0.43752 | 0.232147 | 0.868507 |
| ENSMUSG(NA | 135.9729 | -0.11884 | 0.507003 | 0.867833 | ENSMUSG(NA | 10.06444 | 0.290264 | 0.682771 | NA |
| ENSMUSG(NA | 14.10681 | 1.0185 | 0.06718 | NA | ENSMUSG(NA | 96.99718 | -0.06281 | 0.783027 | 0.976106 |
| ENSMUSG(NA | 151.3633 | 0.19112 | 0.30833 | 0.756439 | ENSMUSG(NA | 22.1118 | 0.097004 | 0.817467 | 0.979198 |
| ENSMUSG(NA | 93.68033 | -0.24313 | 0.236762 | 0.684086 | ENSMUSG(NA | 8.685654 | 0.341651 | 0.603191 | NA |
| ENSMUSG(NA | 18.1113 | -0.01894 | 0.966552 | NA | ENSMUSG(NA | 20.78859 | -0.24588 | 0.576375 | 0.942912 |
| ENSMUSG(NA | 98.10108 | 0.487487 | 0.055349 | 0.361966 | ENSMUSG(NA | 14.95847 | -0.67919 | 0.187083 | 0.856099 |

| | | | | | |
|---|---|---|---|---|---|
| ENSMUSG( | 9.476674 | 0.844793 | 0.163598 | NA | NA |
| ENSMUSG( | 33.62152 | 0.088094 | 0.803496 | 0.962169 | NA |
| ENSMUSG( | 19.65809 | 0.120733 | 0.779516 | NA | NA |
| ENSMUSG( | 12.91125 | 0.911691 | 0.099368 | NA | NA |
| ENSMUSG( | 15.9724 | -0.31478 | 0.540682 | NA | NA |
| ENSMUSG( | 12.18463 | 0.458814 | 0.423527 | NA | NA |
| ENSMUSG( | 21.71641 | 0.334051 | 0.441268 | 0.832449 | NA |
| ENSMUSG( | 9.646024 | -0.2955 | 0.659954 | NA | NA |
| ENSMUSG( | 15.21487 | -0.13015 | 0.798828 | NA | NA |
| ENSMUSG( | 15.81319 | -0.12511 | 0.787956 | NA | NA |
| ENSMUSG( | 12.35204 | 0.205854 | 0.728044 | NA | NA |
| ENSMUSG( | 7.006028 | 0.914586 | 0.241316 | NA | NA |
| ENSMUSG( | 1259.826 | -0.02942 | 0.788311 | 0.959226 | NA |
| ENSMUSG( | 97.67909 | 0.385897 | 0.08408 | 0.447085 | NA |
| ENSMUSG( | 13.23876 | -0.63392 | 0.229994 | NA | NA |
| ENSMUSG( | 34.9713 | 0.510582 | 0.13924 | 0.555317 | NA |
| ENSMUSG( | 28.39002 | 0.418661 | 0.315302 | 0.762288 | NA |
| ENSMUSG( | 10.50714 | 0.171294 | 0.800363 | NA | NA |
| ENSMUSG( | 125.6532 | -0.11212 | 0.598276 | 0.903801 | NA |
| ENSMUSG( | 12.37359 | 0.07868 | 0.894911 | NA | NA |
| ENSMUSG( | 9.767332 | 0.029498 | 0.961577 | NA | NA |
| ENSMUSG( | 23.78226 | 0.03482 | 0.93302 | 0.992887 | NA |
| ENSMUSG( | 47.97269 | 0.391823 | 0.155531 | 0.583591 | NA |
| ENSMUSG( | 16.14031 | 0.287702 | 0.562308 | NA | NA |
| ENSMUSG( | 22.20705 | -0.21027 | 0.599451 | 0.903801 | NA |
| ENSMUSG( | 9.474376 | 0.280213 | 0.653548 | NA | NA |
| ENSMUSG( | 24.62469 | 0.143787 | 0.705027 | 0.938303 | NA |
| ENSMUSG( | 16.57183 | -0.01855 | 0.967332 | NA | NA |
| ENSMUSG( | 20.14294 | -0.50336 | 0.224021 | NA | NA |
| ENSMUSG( | 32.51497 | -0.09136 | 0.808772 | 0.963455 | NA |
| ENSMUSG( | 28.17827 | -0.11348 | 0.755531 | 0.951395 | NA |
| ENSMUSG( | 11.34986 | -0.36896 | 0.526445 | NA | NA |
| ENSMUSG( | 15.68287 | 0.288559 | 0.578117 | NA | NA |
| ENSMUSG( | 42.74012 | -0.23886 | 0.432455 | 0.82768 | NA |
| ENSMUSG( | 383.1434 | -0.05941 | 0.6417 | 0.916828 | NA |
| ENSMUSG( | 16.79621 | -0.47638 | 0.293839 | NA | NA |
| ENSMUSG( | 15.16597 | -0.24762 | 0.615724 | NA | NA |
| ENSMUSG( | 452.0022 | 0.079467 | 0.48905 | 0.859092 | NA |
| ENSMUSG( | 69.36007 | -0.0363 | 0.909371 | 0.98801 | NA |
| ENSMUSG( | 8.936028 | 1.250467 | 0.056271 | NA | NA |
| ENSMUSG( | 273.9417 | -0.0849 | 0.521681 | 0.875186 | NA |
| ENSMUSG( | 48.4574 | 0.407978 | 0.234857 | 0.682194 | NA |
| ENSMUSG( | 53.9151 | 0.13346 | 0.639229 | 0.916434 | NA |
| ENSMUSG( | 10.11617 | -0.05941 | 0.91738 | NA | NA |
| ENSMUSG( | 12.41412 | 0.561263 | 0.280208 | NA | NA |
| ENSMUSG( | 43.62062 | -0.53073 | 0.098352 | 0.479925 | NA |
| ENSMUSG( | 9.921122 | 0.462013 | 0.435283 | NA | NA |
| ENSMUSG( | 12.25421 | 0.115693 | 0.829874 | NA | NA |
| ENSMUSG( | 76.12681 | 0.097164 | 0.675937 | 0.930542 | NA |
| ENSMUSG( | 9.68213 | 0.320642 | 0.616121 | NA | NA |
| ENSMUSG( | 9.129818 | -0.01655 | 0.979447 | NA | NA |
| ENSMUSG( | 20.87084 | 0.10751 | 0.809302 | NA | NA |
| ENSMUSG( | 106.4639 | 0.219527 | 0.253387 | 0.703636 | NA |
| ENSMUSG( | 23.23588 | 0.172953 | 0.66615 | 0.927057 | NA |
| ENSMUSG( | 76.48745 | -0.49234 | 0.051739 | 0.349085 | NA |
| ENSMUSG( | 21.79264 | -0.69218 | 0.105315 | 0.492417 | NA |
| ENSMUSG( | 13.18272 | 0.037436 | 0.94251 | NA | NA |
| ENSMUSG( | 11.26443 | -0.11394 | 0.843469 | NA | NA |
| ENSMUSG( | 86.28432 | 0.120994 | 0.568942 | 0.892739 | NA |
| ENSMUSG( | 10.0688 | 0.350799 | 0.565807 | NA | NA |
| ENSMUSG( | 83.22019 | -0.06817 | 0.750744 | 0.950042 | NA |
| ENSMUSG( | 9.707056 | -0.93631 | 0.175202 | NA | NA |
| ENSMUSG( | 15.74502 | 0.273482 | 0.580484 | NA | NA |
| ENSMUSG( | 10.89895 | -0.93469 | 0.117287 | NA | NA |
| ENSMUSG( | 42.82025 | -0.29672 | 0.384555 | 0.804007 | NA |
| ENSMUSG( | 23.36463 | -0.30136 | 0.429618 | 0.826446 | NA |
| ENSMUSG( | 29.47471 | 0.210707 | 0.571423 | 0.893319 | NA |
| ENSMUSG( | 105.5342 | -0.12117 | 0.547997 | 0.885153 | NA |
| ENSMUSG( | 93.44606 | -0.1321 | 0.528833 | 0.878365 | NA |
| ENSMUSG( | 10.04561 | -0.30111 | 0.632603 | NA | NA |
| ENSMUSG( | 588.1459 | -0.04258 | 0.743554 | 0.948478 | NA |
| ENSMUSG( | 51.06596 | -0.52106 | 0.065391 | 0.396102 | NA |
| ENSMUSG( | 16.94287 | -0.22325 | 0.628614 | NA | NA |
| ENSMUSG( | 12.83312 | -0.42701 | 0.438366 | NA | NA |
| ENSMUSG( | 22.55415 | 0.171924 | 0.673097 | 0.929007 | NA |
| ENSMUSG( | 64.04169 | -0.11618 | 0.681635 | 0.931619 | NA |
| ENSMUSG( | 15.01744 | -0.43128 | 0.367194 | NA | NA |
| ENSMUSG( | 12.24063 | -0.09441 | 0.85854 | NA | NA |
| ENSMUSG( | 70.30702 | -0.07187 | 0.813435 | 0.964574 | NA |
| ENSMUSG( | 59.60509 | -0.2938 | 0.301669 | 0.749979 | NA |
| ENSMUSG( | 23.70466 | 0.074417 | 0.845936 | 0.972226 | NA |
| ENSMUSG( | 11.68057 | -0.19452 | 0.762182 | NA | NA |
| ENSMUSG( | 204.2323 | 0.293289 | 0.086849 | 0.453311 | NA |
| ENSMUSG( | 20.76936 | 0.115407 | 0.780938 | NA | NA |
| ENSMUSG( | 19.65113 | -0.24576 | 0.564166 | NA | NA |
| ENSMUSG( | 142.0453 | 0.016104 | 0.929526 | 0.991848 | NA |
| ENSMUSG( | 17.56825 | -0.31943 | 0.477176 | NA | NA |
| ENSMUSG( | 9.591254 | 0.252454 | 0.724084 | NA | NA |
| ENSMUSG( | 15.01385 | -0.40772 | 0.388702 | NA | NA |
| ENSMUSG( | 21.46733 | 0.356375 | 0.383483 | 0.803428 | NA |
| ENSMUSG( | 16.56474 | 0.337995 | 0.519954 | NA | NA |
| ENSMUSG( | 10.1049 | 0.442529 | 0.462499 | NA | NA |
| ENSMUSG( | 118.1796 | 0.06174 | 0.781698 | 0.957558 | NA |
| ENSMUSG( | 23.84188 | 0.20696 | 0.612106 | 0.908586 | NA |
| ENSMUSG( | 33.77622 | -0.06558 | 0.851971 | 0.973168 | NA |
| ENSMUSG( | 15.8741 | -0.06091 | 0.897802 | NA | NA |
| ENSMUSG( | 18.88822 | -0.26547 | 0.596613 | NA | NA |
| ENSMUSG( | 8.11496 | -0.08126 | 0.911725 | NA | NA |
| ENSMUSG( | 9.790521 | -0.53565 | 0.35874 | NA | NA |
| ENSMUSG( | 120.4825 | 0.278968 | 0.137812 | 0.552903 | NA |
| ENSMUSG( | 28.7182 | 0.262896 | 0.483096 | 0.856216 | NA |
| ENSMUSG( | 18.5202 | -0.52467 | 0.245766 | NA | NA |
| ENSMUSG( | 4115.45 | -1.52651 | 1.03E-05 | 0.00119 | NA |
| ENSMUSG( | 184.1498 | -0.00437 | 0.980636 | 0.997512 | NA |
| ENSMUSG( | 28.78782 | 0.062131 | 0.872779 | 0.978753 | NA |
| ENSMUSG( | 29.02044 | -0.29266 | 0.500334 | 0.863455 | NA |
| ENSMUSG( | 36.77531 | -1.05485 | 0.004957 | 0.093376 | NA |
| ENSMUSG( | 33.87152 | -0.20683 | 0.532448 | 0.879905 | NA |
| ENSMUSG( | 48.27139 | 0.15872 | 0.582331 | 0.897918 | NA |
| ENSMUSG( | 391.3566 | 0.03197 | 0.870505 | 0.977906 | NA |
| ENSMUSG( | 44.9842 | 0.287168 | 0.316223 | 0.763453 | NA |
| ENSMUSG( | 84.11652 | -0.07726 | 0.736447 | 0.946437 | NA |
| ENSMUSG( | 20.29068 | -0.37169 | 0.404788 | NA | NA |
| ENSMUSG( | 768.0985 | -0.09273 | 0.541056 | 0.883182 | NA |
| ENSMUSG( | 25.60042 | 0.514205 | 0.195224 | 0.6346 | NA |
| ENSMUSG( | 10.23087 | 0.11464 | 0.847686 | NA | NA |
| ENSMUSG( | 22.17236 | 0.343247 | 0.418399 | 0.821966 | NA |
| ENSMUSG( | 41.18168 | -0.68954 | 0.028543 | 0.257525 | NA |
| ENSMUSG( | 38.20074 | -0.15394 | 0.66161 | 0.925271 | NA |
| ENSMUSG( | 19.95066 | 0.303186 | 0.482292 | NA | NA |
| ENSMUSG( | 33.20643 | -0.14545 | 0.678024 | 0.930542 | NA |
| ENSMUSG( | 27.17117 | -0.28591 | 0.434529 | 0.828978 | NA |
| ENSMUSG( | 18.94563 | 0.488847 | 0.27917 | NA | NA |
| ENSMUSG( | 838.4521 | -0.18573 | 0.278858 | 0.728705 | NA |
| ENSMUSG( | 60.21968 | 0.011568 | 0.969404 | 0.996369 | NA |
| ENSMUSG( | 1458.021 | 0.038147 | 0.749728 | 0.949664 | NA |
| ENSMUSG( | 80.68308 | -0.3058 | 0.191348 | 0.630245 | NA |
| ENSMUSG( | 10.12304 | -0.23508 | 0.687372 | NA | NA |
| ENSMUSG( | 9.185751 | -0.18973 | 0.757874 | NA | NA |
| ENSMUSG( | 250.2841 | -0.05028 | 0.794715 | 0.959799 | NA |
| ENSMUSG( | 22.90717 | 0.412682 | 0.298394 | 0.74713 | NA |
| ENSMUSG( | 18.15574 | -0.38571 | 0.397872 | NA | NA |
| ENSMUSG( | 11.26573 | -0.41298 | 0.470294 | NA | NA |
| ENSMUSG( | 17.67582 | 0.335657 | 0.485938 | NA | NA |
| ENSMUSG( | 84.73409 | 0.008496 | 0.968152 | 0.996284 | NA |
| ENSMUSG( | 23.48176 | -0.371 | 0.338517 | 0.777489 | NA |
| ENSMUSG( | 106.1474 | -0.24548 | 0.275996 | 0.725669 | NA |
| ENSMUSG( | 11.75153 | 0.159468 | 0.777422 | NA | NA |
| ENSMUSG( | 112.9297 | -0.12751 | 0.503259 | 0.86596 | NA |
| ENSMUSG( | 46.46855 | -0.16381 | 0.565074 | 0.890735 | NA |
| ENSMUSG( | 20.18343 | -0.22279 | 0.611116 | NA | NA |
| ENSMUSG( | 32.64673 | 0.010398 | 0.97732 | 0.997428 | NA |
| ENSMUSG( | 24.40423 | -0.66745 | 0.090427 | 0.461669 | NA |
| ENSMUSG( | 512.8748 | 0.043568 | 0.755038 | 0.951276 | NA |
| ENSMUSG( | 15.16685 | -0.2999 | 0.551798 | NA | NA |
| ENSMUSG( | 9.36326 | -0.33258 | 0.601286 | NA | NA |
| ENSMUSG( | 225.8838 | 0.095117 | 0.544616 | 0.884649 | NA |
| ENSMUSG( | 673.294 | -0.12298 | 0.265596 | 0.716205 | NA |
| ENSMUSG( | 28.63298 | -0.36478 | 0.325339 | 0.77061 | NA |
| ENSMUSG( | 37.87703 | -0.31661 | 0.321797 | 0.766739 | NA |
| ENSMUSG( | 13.4503 | -0.4296 | 0.406757 | NA | NA |
| ENSMUSG( | 36.62919 | 0.208491 | 0.507852 | 0.86795 | NA |
| ENSMUSG( | 45.16454 | -0.15756 | 0.580289 | 0.896952 | NA |
| ENSMUSG( | 24.87478 | -0.23401 | 0.541745 | 0.883182 | NA |
| ENSMUSG( | 20.8056 | 0.181934 | 0.655086 | NA | NA |
| ENSMUSG( | 21.70951 | -0.23519 | 0.54573 | 0.884974 | NA |
| ENSMUSG( | 11.94112 | 0.144573 | 0.788692 | NA | NA |
| ENSMUSG( | 13.07226 | -0.12702 | 0.810042 | NA | NA |

| | | | | | |
|---|---|---|---|---|---|
| ENSMUSG( | 49.91086 | 0.191287 | 0.552785 | 0.939565 | NA |
| ENSMUSG( | 11.07905 | 0.095081 | 0.86756 | NA | NA |
| ENSMUSG( | 389.9025 | -0.46785 | 0.002282 | 0.185073 | NA |
| ENSMUSG( | 35.08015 | -0.22801 | 0.502378 | 0.927825 | NA |
| ENSMUSG( | 22.05715 | -0.33435 | 0.446489 | 0.921734 | NA |
| ENSMUSG( | 17.32704 | -0.09638 | 0.838843 | 0.981696 | NA |
| ENSMUSG( | 724.2925 | 0.468304 | 0.054467 | 0.723362 | NA |
| ENSMUSG( | 14.80514 | -0.13839 | 0.786287 | 0.976303 | NA |
| ENSMUSG( | 25.24065 | -0.27334 | 0.505773 | 0.927825 | NA |
| ENSMUSG( | 13.89249 | -0.30347 | 0.56651 | 0.941777 | NA |
| ENSMUSG( | 25.68532 | 0.316102 | 0.433923 | 0.91869 | NA |
| ENSMUSG( | 85.18384 | 0.171013 | 0.509892 | 0.929089 | NA |
| ENSMUSG( | 7.791957 | 0.255323 | 0.711969 | NA | NA |
| ENSMUSG( | 21.09651 | 0.418523 | 0.330943 | 0.907513 | NA |
| ENSMUSG( | 17.1285 | 0.279065 | 0.586436 | 0.944975 | NA |
| ENSMUSG( | 628.3679 | -0.18232 | 0.158432 | 0.837724 | NA |
| ENSMUSG( | 11.82389 | -0.06938 | 0.905679 | 0.991089 | NA |
| ENSMUSG( | 187.2735 | -0.52906 | 0.019336 | 0.554338 | NA |
| ENSMUSG( | 7.014731 | -0.05105 | 0.949975 | NA | NA |
| ENSMUSG( | 73.44239 | -0.48344 | 0.117096 | 0.813384 | NA |
| ENSMUSG( | 11.21347 | -0.09689 | 0.867315 | NA | NA |
| ENSMUSG( | 140.755 | 0.14025 | 0.475061 | 0.923329 | NA |
| ENSMUSG( | 308.6161 | 0.283517 | 0.120545 | 0.813384 | NA |
| ENSMUSG( | 185.0548 | 0.248471 | 0.251826 | 0.879902 | NA |
| ENSMUSG( | 21.37235 | -0.60295 | 0.154806 | 0.837724 | NA |
| ENSMUSG( | 16.9236 | 0.598559 | 0.262882 | 0.885357 | NA |
| ENSMUSG( | 108.239 | 0.110577 | 0.629891 | 0.953251 | NA |
| ENSMUSG( | 9.963197 | 0.072482 | 0.913311 | NA | NA |
| ENSMUSG( | 17.9452 | 0.712168 | 0.141219 | 0.830678 | NA |
| ENSMUSG( | 10.65716 | 1.305688 | 0.037237 | NA | NA |
| ENSMUSG( | 232.6494 | -0.17657 | 0.306217 | 0.898538 | NA |
| ENSMUSG( | 135.5472 | -0.07764 | 0.700609 | 0.961547 | NA |
| ENSMUSG( | 13.44356 | -0.26178 | 0.667328 | 0.956766 | NA |
| ENSMUSG( | 99.45421 | 0.153714 | 0.514464 | 0.930939 | NA |
| ENSMUSG( | 21.17686 | -0.71446 | 0.095029 | 0.796133 | NA |
| ENSMUSG( | 38.3805 | 0.644425 | 0.129444 | 0.820358 | NA |
| ENSMUSG( | 176.2725 | -0.07127 | 0.679712 | 0.957966 | NA |
| ENSMUSG( | 220.84 | 0.058676 | 0.749092 | 0.96998 | NA |
| ENSMUSG( | 196.5984 | 0.375003 | 0.052302 | 0.713538 | NA |
| ENSMUSG( | 10.04132 | -0.03856 | 0.950931 | NA | NA |
| ENSMUSG( | 14.43531 | 0.295735 | 0.578516 | 0.943548 | NA |
| ENSMUSG( | 10.63806 | 0.155324 | 0.79636 | NA | NA |
| ENSMUSG( | 27.42512 | 0.303906 | 0.509716 | 0.929089 | NA |
| ENSMUSG( | 145.5312 | 0.098052 | 0.624334 | 0.953234 | NA |
| ENSMUSG( | 148.8631 | -0.02045 | 0.921471 | 0.992644 | NA |
| ENSMUSG( | 207.464 | -0.33253 | 0.092695 | 0.794394 | NA |
| ENSMUSG( | 14.8706 | 0.784499 | 0.163807 | 0.838291 | NA |
| ENSMUSG( | 10.78646 | 0.122362 | 0.84256 | NA | NA |
| ENSMUSG( | 11.18908 | -0.75583 | 0.204578 | NA | NA |
| ENSMUSG( | 36.87479 | -0.20255 | 0.566366 | 0.941745 | NA |
| ENSMUSG( | 48.45251 | 0.108548 | 0.721775 | 0.966887 | NA |
| ENSMUSG( | 32.93548 | -0.10633 | 0.757628 | 0.971858 | NA |
| ENSMUSG( | 12.40045 | 0.691409 | 0.245238 | 0.877904 | NA |
| ENSMUSG( | 12.97252 | -0.3231 | 0.588134 | 0.944975 | NA |
| ENSMUSG( | 10.04677 | -0.67029 | 0.302594 | NA | NA |
| ENSMUSG( | 30.23562 | -0.09985 | 0.791064 | 0.97681 | NA |
| ENSMUSG( | 14.0652 | -0.2393 | 0.650746 | 0.95422 | NA |
| ENSMUSG( | 248.2339 | -0.33693 | 0.046398 | 0.697814 | NA |
| ENSMUSG( | 8.582791 | 1.124993 | 0.138952 | NA | NA |
| ENSMUSG( | 12.46412 | -0.64908 | 0.267781 | 0.886134 | NA |
| ENSMUSG( | 18.98332 | 0.760449 | 0.115134 | 0.81257 | NA |
| ENSMUSG( | 26.57164 | 0.044829 | 0.913094 | 0.992298 | NA |
| ENSMUSG( | 62.54773 | 0.131071 | 0.618007 | 0.952232 | NA |
| ENSMUSG( | 10.07572 | -0.24595 | 0.695133 | NA | NA |
| ENSMUSG( | 190.1357 | 0.584287 | 0.003056 | 0.219421 | NA |
| ENSMUSG( | 9.369441 | -0.67991 | 0.333054 | NA | NA |
| ENSMUSG( | 15.44408 | 0.168691 | 0.740624 | 0.968811 | NA |
| ENSMUSG( | 6.464863 | -1.32019 | 0.143392 | NA | NA |
| ENSMUSG( | 24.38362 | -0.22413 | 0.570098 | 0.941777 | NA |
| ENSMUSG( | 59.74052 | 0.309745 | 0.249892 | 0.877904 | NA |
| ENSMUSG( | 14.41853 | 0.550004 | 0.326724 | 0.905371 | NA |
| ENSMUSG( | 27.57524 | 0.018564 | 0.961593 | 0.99511 | NA |
| ENSMUSG( | 18.67327 | 0.40394 | 0.373261 | 0.915491 | NA |
| ENSMUSG( | 17.43792 | 1.332473 | 0.077113 | 0.770558 | NA |
| ENSMUSG( | 11.85655 | 0.628064 | 0.301428 | 0.898538 | NA |
| ENSMUSG( | 33.00922 | 0.693799 | 0.243928 | 0.877904 | NA |
| ENSMUSG( | 42.92647 | 0.49877 | 0.119241 | 0.813384 | NA |
| ENSMUSG( | 219.9971 | 0.331069 | 0.067667 | 0.75212 | NA |
| ENSMUSG( | 31.28149 | 0.581233 | 0.115434 | 0.813375 | NA |
| ENSMUSG( | 132.3421 | -0.51224 | 0.014637 | 0.486524 | NA |
| ENSMUSG( | 50.8272 | -0.23005 | 0.416261 | 0.917257 | NA |
| ENSMUSG( | 37.99948 | -0.2033 | 0.595329 | 0.945798 | NA |
| ENSMUSG( | 24.0245 | -0.21871 | 0.61219 | 0.951071 | NA |
| ENSMUSG( | 50.31077 | 0.108943 | 0.712579 | 0.964363 | NA |
| ENSMUSG( | 40.37054 | 0.212786 | 0.496282 | 0.927825 | NA |
| ENSMUSG( | 56.67491 | 0.282767 | 0.362392 | 0.914223 | NA |
| ENSMUSG( | 16.18023 | -0.58664 | 0.252894 | 0.88018 | NA |
| ENSMUSG( | 24.41453 | 0.150801 | 0.723063 | 0.966887 | NA |
| ENSMUSG( | 402.4859 | -0.33355 | 0.036335 | 0.669285 | NA |
| ENSMUSG( | 10.33028 | 0.225763 | 0.732579 | NA | NA |
| ENSMUSG( | 43.77247 | 0.43092 | 0.227843 | 0.868507 | NA |
| ENSMUSG( | 179.3403 | 0.210634 | 0.247562 | 0.877904 | NA |
| ENSMUSG( | 42.43646 | -0.39357 | 0.211019 | 0.864412 | NA |
| ENSMUSG( | 29.732 | -0.16595 | 0.658729 | 0.956205 | NA |
| ENSMUSG( | 134.628 | 0.193922 | 0.364885 | 0.914449 | NA |
| ENSMUSG( | 13.99379 | -0.2124 | 0.700699 | 0.961547 | NA |
| ENSMUSG( | 8.144613 | 0.023464 | 0.974029 | NA | NA |
| ENSMUSG( | 71.34701 | 0.034327 | 0.897595 | 0.990558 | NA |
| ENSMUSG( | 11.35717 | -0.2784 | 0.625545 | NA | NA |
| ENSMUSG( | 23.26636 | -0.1575 | 0.741491 | 0.968944 | NA |
| ENSMUSG( | 68.94459 | -0.02432 | 0.930792 | 0.99306 | NA |
| ENSMUSG( | 45.40736 | 0.155574 | 0.665391 | 0.956766 | NA |
| ENSMUSG( | 10.69219 | -0.51207 | 0.392215 | NA | NA |
| ENSMUSG( | 11.39722 | -0.16117 | 0.78492 | NA | NA |
| ENSMUSG( | 42.33876 | -0.43752 | 0.232147 | 0.868507 | NA |
| ENSMUSG( | 10.06444 | 0.290264 | 0.682771 | NA | NA |
| ENSMUSG( | 96.99718 | -0.06281 | 0.783027 | 0.976106 | NA |
| ENSMUSG( | 22.1118 | 0.097004 | 0.817467 | 0.979198 | NA |
| ENSMUSG( | 8.685654 | 0.341651 | 0.603191 | NA | NA |
| ENSMUSG( | 20.78859 | -0.24588 | 0.576375 | 0.942912 | NA |
| ENSMUSG( | 14.95847 | -0.67919 | 0.187083 | 0.856099 | NA |
| ENSMUSG( | 10.67291 | 0.634881 | 0.303946 | NA | NA |
| ENSMUSG( | 15.83238 | -0.70189 | 0.218208 | 0.866104 | NA |
| ENSMUSG( | 24.2312 | 0.183434 | 0.641562 | 0.953903 | NA |
| ENSMUSG( | 40.76981 | -0.24484 | 0.446507 | 0.921734 | NA |
| ENSMUSG( | 10.47308 | 0.773392 | 0.264996 | NA | NA |
| ENSMUSG( | 145.1532 | -0.53655 | 0.008511 | 0.391682 | NA |
| ENSMUSG( | 14.13122 | 0.246174 | 0.659307 | 0.956205 | NA |
| ENSMUSG( | 41.08817 | -0.08585 | 0.799542 | 0.97681 | NA |
| ENSMUSG( | 60.86801 | -0.39444 | 0.131252 | 0.822884 | NA |
| ENSMUSG( | 30.54684 | -0.11163 | 0.787604 | 0.97637 | NA |
| ENSMUSG( | 8.964938 | -0.01093 | 0.987351 | NA | NA |
| ENSMUSG( | 19.70914 | -0.08683 | 0.848386 | 0.985321 | NA |
| ENSMUSG( | 68.35556 | -0.24069 | 0.383073 | 0.917257 | NA |
| ENSMUSG( | 16.19071 | -0.25404 | 0.594612 | 0.945444 | NA |
| ENSMUSG( | 8.474423 | -0.42689 | 0.526927 | NA | NA |
| ENSMUSG( | 132.1782 | -0.00526 | 0.981798 | 0.996756 | NA |
| ENSMUSG( | 10.32627 | 0.265588 | 0.671795 | NA | NA |
| ENSMUSG( | 19.03363 | -0.26662 | 0.55845 | 0.93957 | NA |
| ENSMUSG( | 10.07488 | 0.592284 | 0.365358 | NA | NA |
| ENSMUSG( | 75.04074 | 0.316575 | 0.208827 | 0.864412 | NA |
| ENSMUSG( | 45.93044 | -0.32953 | 0.32157 | 0.904019 | NA |
| ENSMUSG( | 38.41949 | 0.147292 | 0.660573 | 0.956205 | NA |
| ENSMUSG( | 22.75812 | -0.04344 | 0.914492 | 0.992407 | NA |
| ENSMUSG( | 49.538 | -0.86034 | 0.005663 | 0.304277 | NA |
| ENSMUSG( | 10.97576 | 0.366055 | 0.549793 | NA | NA |
| ENSMUSG( | 11.06381 | -0.15588 | 0.802135 | NA | NA |
| ENSMUSG( | 18.37199 | -0.37023 | 0.539745 | 0.937378 | NA |
| ENSMUSG( | 217.8925 | 0.019848 | 0.908738 | 0.991958 | NA |
| ENSMUSG( | 16.25758 | -0.05511 | 0.915206 | 0.992488 | NA |
| ENSMUSG( | 25.02972 | -0.10853 | 0.792373 | 0.97681 | NA |
| ENSMUSG( | 138.2492 | 0.068852 | 0.717508 | 0.965964 | NA |
| ENSMUSG( | 47.47797 | 0.375455 | 0.211803 | 0.864412 | NA |
| ENSMUSG( | 17.46056 | 0.076838 | 0.896905 | 0.990558 | NA |
| ENSMUSG( | 19.60049 | 0.343172 | 0.438388 | 0.920721 | NA |
| ENSMUSG( | 10.38933 | -0.17431 | 0.778409 | NA | NA |
| ENSMUSG( | 539.3693 | 0.187481 | 0.174387 | 0.844317 | NA |
| ENSMUSG( | 3361.166 | -0.07963 | 0.607495 | 0.950113 | NA |
| ENSMUSG( | 13.47353 | -0.63637 | 0.250159 | 0.877904 | NA |
| ENSMUSG( | 20.34058 | -0.25856 | 0.587381 | 0.944975 | NA |
| ENSMUSG( | 209.8545 | -0.13499 | 0.502985 | 0.927825 | NA |
| ENSMUSG( | 32.37688 | 0.065316 | 0.853339 | 0.985598 | NA |
| ENSMUSG( | 2906.219 | 0.073073 | 0.662706 | 0.956205 | NA |
| ENSMUSG( | 8.573547 | 0.172736 | 0.790691 | NA | NA |
| ENSMUSG( | 10.19118 | 0.164627 | 0.795872 | NA | NA |
| ENSMUSG( | 13.57476 | -0.40394 | 0.463342 | 0.923329 | NA |
| ENSMUSG( | 100.67 | -0.24361 | 0.26635 | 0.886134 | NA |
| ENSMUSG( | 165.7062 | -0.13968 | 0.47221 | 0.923329 | NA |

| | | | | | | | | | | | |
|---|---|---|---|---|---|---|---|---|---|---|---|
| ENSMUSG( | NA | 170.7031 | -0.02214 | 0.908511 | 0.987809 | ENSMUSG( | NA | 10.67291 | 0.634881 | 0.303946 | NA |
| ENSMUSG( | NA | 622.5185 | 0.140614 | 0.279634 | 0.729119 | ENSMUSG( | NA | 15.83238 | -0.70189 | 0.218208 | 0.866104 |
| ENSMUSG( | NA | 11.57792 | -0.39267 | 0.490401 | NA | ENSMUSG( | NA | 24.2312 | 0.183434 | 0.641562 | 0.953903 |
| ENSMUSG( | NA | 14.79456 | -0.0333 | 0.947974 | NA | ENSMUSG( | NA | 40.76981 | -0.24484 | 0.446507 | 0.921734 |
| ENSMUSG( | NA | 124.2322 | -0.34645 | 0.102846 | 0.48834 | ENSMUSG( | NA | 10.47308 | 0.773392 | 0.264996 | NA |
| ENSMUSG( | NA | 14.12462 | 0.297041 | 0.56687 | NA | ENSMUSG( | NA | 145.1532 | -0.53655 | 0.008511 | 0.391682 |
| ENSMUSG( | NA | 22.70117 | -1.77931 | 0.0001 | 0.006438 | ENSMUSG( | NA | 14.13122 | 0.246174 | 0.659307 | 0.956205 |
| ENSMUSG( | NA | 486.1859 | -0.0651 | 0.570556 | 0.893192 | ENSMUSG( | NA | 41.08817 | -0.08585 | 0.799542 | 0.97681 |
| ENSMUSG( | NA | 320.7445 | 0.151686 | 0.33778 | 0.777422 | ENSMUSG( | NA | 60.86801 | -0.39444 | 0.131252 | 0.822884 |
| ENSMUSG( | NA | 84.12828 | -0.04273 | 0.86607 | 0.976074 | ENSMUSG( | NA | 30.54684 | -0.11163 | 0.787604 | 0.97637 |
| ENSMUSG( | NA | 14.67561 | 0.448181 | 0.378915 | NA | ENSMUSG( | NA | 8.964938 | -0.01093 | 0.987351 | NA |
| ENSMUSG( | NA | 17.77941 | -0.26378 | 0.564213 | NA | ENSMUSG( | NA | 19.70914 | -0.08683 | 0.848386 | 0.985321 |
| ENSMUSG( | NA | 25.89331 | -0.25775 | 0.518628 | 0.87417 | ENSMUSG( | NA | 68.35556 | -0.24069 | 0.383073 | 0.917257 |
| ENSMUSG( | NA | 33.6188 | -0.00527 | 0.988473 | 0.999109 | ENSMUSG( | NA | 16.19071 | -0.25404 | 0.594612 | 0.945444 |
| ENSMUSG( | NA | 12.26411 | -0.24745 | 0.682019 | NA | ENSMUSG( | NA | 132.1782 | -0.00526 | 0.981798 | 0.996756 |
| ENSMUSG( | NA | 15.52079 | 0.396516 | 0.431832 | NA | ENSMUSG( | NA | 10.32627 | 0.265588 | 0.671795 | NA |
| ENSMUSG( | NA | 25.20173 | 0.128858 | 0.731045 | 0.944956 | ENSMUSG( | NA | 19.03363 | -0.26662 | 0.55845 | 0.93957 |
| ENSMUSG( | NA | 42.03211 | 0.186415 | 0.540818 | 0.88315 | ENSMUSG( | NA | 10.07488 | 0.592284 | 0.365358 | NA |
| ENSMUSG( | NA | 85.50709 | 0.162351 | 0.449181 | 0.834943 | ENSMUSG( | NA | 75.04074 | 0.316575 | 0.208827 | 0.864412 |
| ENSMUSG( | NA | 202.6183 | -0.17175 | 0.358406 | 0.789759 | ENSMUSG( | NA | 45.93044 | -0.32953 | 0.32157 | 0.904019 |
| ENSMUSG( | NA | 315.5373 | 0.073627 | 0.605225 | 0.905688 | ENSMUSG( | NA | 38.41949 | 0.147292 | 0.660573 | 0.956205 |
| ENSMUSG( | NA | 390.7378 | 0.087828 | 0.491312 | 0.859873 | ENSMUSG( | NA | 22.75812 | -0.04344 | 0.914492 | 0.992407 |
| ENSMUSG( | NA | 22.40354 | 0.508042 | 0.195986 | 0.635396 | ENSMUSG( | NA | 49.538 | -0.86034 | 0.005663 | 0.304277 |
| ENSMUSG( | NA | 60.53471 | -0.15084 | 0.552675 | 0.886141 | ENSMUSG( | NA | 11.06381 | -0.15588 | 0.802135 | NA |
| ENSMUSG( | NA | 13.14219 | 0.83758 | 0.116099 | NA | ENSMUSG( | NA | 18.37199 | -0.37023 | 0.539745 | 0.937378 |
| ENSMUSG( | NA | 31.32453 | -0.13759 | 0.712222 | 0.94031 | ENSMUSG( | NA | 217.8925 | 0.019848 | 0.908738 | 0.991958 |
| ENSMUSG( | NA | 33.23006 | 0.2446 | 0.490393 | 0.859353 | ENSMUSG( | NA | 16.25758 | -0.05511 | 0.915206 | 0.992488 |
| ENSMUSG( | NA | 24.09021 | 0.075627 | 0.847125 | 0.972374 | ENSMUSG( | NA | 25.02972 | -0.10853 | 0.792373 | 0.97681 |
| ENSMUSG( | NA | 23.92851 | -0.5993 | 0.114344 | 0.51021 | ENSMUSG( | NA | 138.2492 | 0.068852 | 0.717508 | 0.965964 |
| ENSMUSG( | NA | 73.25115 | -0.27598 | 0.258727 | 0.709814 | ENSMUSG( | NA | 47.47797 | 0.375455 | 0.211803 | 0.864412 |
| ENSMUSG( | NA | 257.7397 | 0.021781 | 0.907524 | 0.987591 | ENSMUSG( | NA | 17.46056 | 0.076838 | 0.896905 | 0.990558 |
| ENSMUSG( | NA | 11.75887 | -0.22856 | 0.702496 | NA | ENSMUSG( | NA | 19.60049 | 0.343172 | 0.438388 | 0.920721 |
| ENSMUSG( | NA | 18.29469 | 0.08818 | 0.844322 | NA | ENSMUSG( | NA | 10.38933 | -0.17431 | 0.778409 | NA |
| ENSMUSG( | NA | 11.41726 | -0.28144 | 0.602657 | NA | ENSMUSG( | NA | 539.3693 | 0.187481 | 0.174387 | 0.844317 |
| ENSMUSG( | NA | 22.07331 | 0.351155 | 0.369069 | 0.796581 | ENSMUSG( | NA | 3361.166 | -0.07963 | 0.607495 | 0.950113 |
| ENSMUSG( | NA | 13.34571 | -0.29213 | 0.578325 | NA | ENSMUSG( | NA | 13.47353 | -0.63637 | 0.250159 | 0.877904 |
| ENSMUSG( | NA | 78.13739 | 0.034442 | 0.885342 | 0.982905 | ENSMUSG( | NA | 20.34058 | -0.25856 | 0.587381 | 0.944975 |
| ENSMUSG( | NA | 13.41922 | -0.37939 | 0.456284 | NA | ENSMUSG( | NA | 209.8545 | -0.13499 | 0.502985 | 0.927825 |
| ENSMUSG( | NA | 104.1373 | -0.09459 | 0.631543 | 0.914073 | ENSMUSG( | NA | 32.37688 | 0.065316 | 0.853339 | 0.985598 |
| ENSMUSG( | NA | 20.9574 | 0.836886 | 0.098317 | NA | ENSMUSG( | NA | 2906.219 | 0.073073 | 0.662706 | 0.956205 |
| ENSMUSG( | NA | 15.69242 | 0.484185 | 0.29153 | NA | ENSMUSG( | NA | 10.19118 | 0.164627 | 0.795872 | NA |
| ENSMUSG( | NA | 82.56223 | -0.54457 | 0.015925 | 0.18704 | ENSMUSG( | NA | 13.57476 | -0.40394 | 0.463342 | 0.923329 |
| ENSMUSG( | NA | 23.63155 | -0.24815 | 0.585529 | 0.899171 | ENSMUSG( | NA | 100.67 | -0.24361 | 0.26635 | 0.886134 |
| ENSMUSG( | NA | 155.836 | 0.14326 | 0.396863 | 0.811742 | ENSMUSG( | NA | 165.7062 | -0.13968 | 0.47221 | 0.923329 |
| ENSMUSG( | NA | 32.97601 | -0.06162 | 0.850933 | 0.972638 | ENSMUSG( | NA | 57.93944 | 0.022156 | 0.944142 | 0.994798 |
| ENSMUSG( | NA | 116.4171 | 0.102269 | 0.582089 | 0.897918 | ENSMUSG( | NA | 14.2562 | -0.0073 | 0.989177 | 0.997893 |
| ENSMUSG( | NA | 397.7465 | 0.029455 | 0.890967 | 0.983764 | ENSMUSG( | NA | 24.66651 | -0.09783 | 0.812839 | 0.979148 |
| ENSMUSG( | NA | 16.57392 | 0.018149 | 0.971753 | NA | ENSMUSG( | NA | 20.1329 | -1.03023 | 0.024154 | 0.588924 |
| ENSMUSG( | NA | 20.04598 | -0.58753 | 0.18843 | NA | ENSMUSG( | NA | 362.4564 | 0.521934 | 0.003456 | 0.23553 |
| ENSMUSG( | NA | 73.68964 | -0.04499 | 0.840847 | 0.971179 | ENSMUSG( | NA | 5188.124 | -0.008 | 0.965402 | 0.995433 |
| ENSMUSG( | NA | 33.49353 | 0.377041 | 0.271782 | 0.722082 | ENSMUSG( | NA | 55.18666 | -0.32104 | 0.277371 | 0.891468 |
| ENSMUSG( | NA | 25.65198 | -0.27968 | 0.471822 | 0.847504 | ENSMUSG( | NA | 573.2399 | -0.07127 | 0.640771 | 0.953903 |
| ENSMUSG( | NA | 44.38874 | 0.307305 | 0.395405 | 0.810601 | ENSMUSG( | NA | 17.47361 | 0.162719 | 0.736198 | 0.967344 |
| ENSMUSG( | NA | 20.32376 | 0.321122 | 0.479741 | NA | ENSMUSG( | NA | 281.5417 | -0.06471 | 0.709642 | 0.96416 |
| ENSMUSG( | NA | 34.58787 | 0.452205 | 0.18593 | 0.623352 | ENSMUSG( | NA | 14.72126 | 0.26929 | 0.641977 | 0.953903 |
| ENSMUSG( | NA | 29.27465 | -0.18657 | 0.585667 | 0.899171 | ENSMUSG( | NA | 109.1531 | 0.257405 | 0.325948 | 0.905339 |
| ENSMUSG( | NA | 8.966887 | 0.132249 | 0.838731 | NA | ENSMUSG( | NA | 148.6845 | -0.14841 | 0.472911 | 0.923329 |
| ENSMUSG( | NA | 24.30442 | 0.203132 | 0.647891 | 0.919759 | ENSMUSG( | NA | 12.48481 | 0.069827 | 0.905938 | 0.991089 |
| ENSMUSG( | NA | 39.75448 | 0.149718 | 0.614318 | 0.909419 | ENSMUSG( | NA | 10.4742 | 0.069653 | 0.911394 | NA |
| ENSMUSG( | NA | 38.87558 | -0.07682 | 0.820178 | 0.965888 | ENSMUSG( | NA | 13.35571 | 1.040861 | 0.083628 | 0.77681 |
| ENSMUSG( | NA | 12.28393 | -0.19988 | 0.734896 | NA | ENSMUSG( | NA | 396.5224 | 0.296526 | 0.044574 | 0.693697 |
| ENSMUSG( | NA | 14.58431 | -0.03054 | 0.949021 | NA | ENSMUSG( | NA | 23.91867 | -0.52357 | 0.217513 | 0.865589 |
| ENSMUSG( | NA | 13.74751 | 0.392639 | 0.491775 | NA | ENSMUSG( | NA | 12.21508 | 0.074455 | 0.891426 | 0.990558 |
| ENSMUSG( | NA | 16.55056 | -0.09941 | 0.827882 | NA | ENSMUSG( | NA | 81.93257 | 0.059613 | 0.80084 | 0.97681 |
| ENSMUSG( | NA | 13.14804 | 0.37023 | 0.462588 | NA | ENSMUSG( | NA | 20.67819 | -0.34751 | 0.421309 | 0.917257 |
| ENSMUSG( | NA | 18.14501 | -0.2793 | 0.517415 | NA | ENSMUSG( | NA | 33.26677 | 0.265455 | 0.549385 | 0.938519 |
| ENSMUSG( | NA | 18.61796 | -0.28938 | 0.535061 | NA | ENSMUSG( | NA | 24.1518 | 0.063963 | 0.877535 | 0.989644 |
| ENSMUSG( | NA | 16.2671 | 0.611486 | 0.229726 | NA | ENSMUSG( | NA | 42.78641 | 0.560318 | 0.13431 | 0.822912 |
| ENSMUSG( | NA | 51.5352 | -0.24636 | 0.381284 | 0.802629 | ENSMUSG( | NA | 18.97835 | -0.47177 | 0.296885 | 0.898538 |
| ENSMUSG( | NA | 9.774336 | -0.62156 | 0.340373 | NA | ENSMUSG( | NA | 18.94356 | 0.715369 | 0.143986 | 0.831311 |
| ENSMUSG( | NA | 19.73365 | -0.40042 | 0.335869 | NA | ENSMUSG( | NA | 264.2199 | -0.26312 | 0.126004 | 0.81516 |
| ENSMUSG( | NA | 12.73841 | -0.31418 | 0.563504 | NA | ENSMUSG( | NA | 14.781 | -0.11089 | 0.834573 | 0.981237 |
| ENSMUSG( | NA | 14.03775 | 0.15445 | 0.781921 | NA | ENSMUSG( | NA | 94.13846 | 0.122851 | 0.581751 | 0.944266 |
| ENSMUSG( | NA | 10.27217 | 0.212312 | 0.713256 | NA | ENSMUSG( | NA | 9.998116 | -0.33244 | 0.643535 | NA |
| ENSMUSG( | NA | 30.01053 | 0.013947 | 0.96973 | 0.996369 | ENSMUSG( | NA | 8.705801 | -0.13005 | 0.865698 | NA |
| ENSMUSG( | NA | 269.868 | 0.000129 | 0.99932 | 0.99989 | ENSMUSG( | NA | 9.565598 | -1.12055 | 0.122428 | NA |
| ENSMUSG( | NA | 11.17567 | -0.21665 | 0.695525 | NA | ENSMUSG( | NA | 11.48036 | 0.664522 | 0.259646 | 0.883971 |
| ENSMUSG( | NA | 78.20522 | 0.384916 | 0.077383 | 0.429743 | ENSMUSG( | NA | 52.98185 | -0.19976 | 0.490433 | 0.926343 |
| ENSMUSG( | NA | 20.68748 | -0.19297 | 0.658576 | NA | ENSMUSG( | NA | 37.28156 | 0.674095 | 0.120548 | 0.813384 |
| ENSMUSG( | NA | 107.7588 | 0.360254 | 0.107848 | 0.49674 | ENSMUSG( | NA | 17.12795 | 0.064446 | 0.904104 | 0.991089 |
| ENSMUSG( | NA | 169.1534 | 0.152697 | 0.370202 | 0.797542 | ENSMUSG( | NA | 17.62862 | 0.419187 | 0.402408 | 0.917257 |
| ENSMUSG( | NA | 16.73298 | -0.17566 | 0.71866 | NA | ENSMUSG( | NA | 21.96873 | 0.70262 | 0.103212 | 0.804311 |
| ENSMUSG( | NA | 10.7155 | -0.53027 | 0.397018 | NA | ENSMUSG( | NA | 21.70276 | -1.16563 | 0.043124 | 0.690304 |
| ENSMUSG( | NA | 43.92235 | -0.08488 | 0.783521 | 0.957558 | ENSMUSG( | NA | 7.635272 | -0.277 | 0.70535 | NA |
| ENSMUSG( | NA | 19.41747 | -0.20743 | 0.629073 | NA | ENSMUSG( | NA | 14.80473 | 1.38133 | 0.018382 | 0.544291 |
| ENSMUSG( | NA | 33.33179 | 0.456193 | 0.163146 | 0.59439 | ENSMUSG( | NA | 15.60231 | 0.382051 | 0.453693 | 0.922316 |
| ENSMUSG( | NA | 19.04421 | -0.11393 | 0.788439 | NA | ENSMUSG( | NA | 15.27881 | 0.136831 | 0.824803 | 0.979198 |
| ENSMUSG( | NA | 3207.778 | -0.07712 | 0.848532 | 0.972374 | ENSMUSG( | NA | 22.85119 | 0.430055 | 0.290199 | 0.896947 |
| ENSMUSG( | NA | 27.33295 | -0.45396 | 0.213868 | 0.658243 | ENSMUSG( | NA | 12.92956 | 0.138457 | 0.799604 | 0.97681 |
| ENSMUSG( | NA | 16.56705 | -0.16367 | 0.721466 | NA | ENSMUSG( | NA | 18.09312 | -0.1814 | 0.711141 | 0.964363 |
| ENSMUSG( | NA | 29.1755 | -0.29386 | 0.451174 | 0.835567 | ENSMUSG( | NA | 12.7319 | 0.737475 | 0.19843 | 0.862367 |
| ENSMUSG( | NA | 87.83919 | 0.223055 | 0.317282 | 0.763823 | ENSMUSG( | NA | 259.0339 | 0.447241 | 0.014393 | 0.48208 |
| ENSMUSG( | NA | 11.35408 | -0.05603 | 0.921328 | NA | ENSMUSG( | NA | 46.39184 | -0.00856 | 0.980035 | 0.996479 |
| ENSMUSG( | NA | 13.34689 | -0.22272 | 0.684042 | NA | ENSMUSG( | NA | 28.23133 | -0.11458 | 0.797976 | 0.97681 |
| ENSMUSG( | NA | 94.09243 | -0.09677 | 0.683352 | 0.932153 | ENSMUSG( | NA | 9.136393 | 0.240221 | 0.718782 | NA |
| ENSMUSG( | NA | 154.1923 | 0.086597 | 0.626239 | 0.912633 | ENSMUSG( | NA | 21.54846 | 0.420634 | 0.392078 | 0.917257 |
| ENSMUSG( | NA | 156.8959 | -0.05678 | 0.738287 | 0.947329 | ENSMUSG( | NA | 31.00981 | -0.07755 | 0.84253 | 0.983044 |
| ENSMUSG( | NA | 13.12324 | -0.33913 | 0.511231 | NA | ENSMUSG( | NA | 34.6968 | 0.092616 | 0.798598 | 0.97681 |
| ENSMUSG( | NA | 9.116823 | -0.76551 | 0.205059 | NA | ENSMUSG( | NA | 29.07289 | -0.16632 | 0.651394 | 0.95449 |
| ENSMUSG( | NA | 11.22621 | 0.082708 | 0.885227 | NA | ENSMUSG( | NA | 28.70382 | -0.01471 | 0.968009 | 0.995866 |
| ENSMUSG( | NA | 14.84113 | -0.43275 | 0.370959 | NA | ENSMUSG( | NA | 79.75886 | 0.273894 | 0.340075 | 0.907972 |
| ENSMUSG( | NA | 224.4657 | 0.10002 | 0.508222 | 0.867999 | ENSMUSG( | NA | 16.26996 | 0.187639 | 0.71375 | 0.964986 |
| ENSMUSG( | NA | 44.8776 | 0.332447 | 0.259555 | 0.709948 | ENSMUSG( | NA | 12.72981 | -0.48954 | 0.388555 | 0.917257 |
| ENSMUSG( | NA | 17.42207 | 0.570174 | 0.231125 | NA | ENSMUSG( | NA | 46.6436 | -0.25241 | 0.411225 | 0.917257 |
| ENSMUSG( | NA | 21.81153 | -0.17615 | 0.6592 | 0.924506 | ENSMUSG( | NA | 18.18496 | -0.57369 | 0.223394 | 0.867404 |
| ENSMUSG( | NA | 19.65244 | -0.34767 | 0.414844 | NA | ENSMUSG( | NA | 64.09131 | -0.07281 | 0.865837 | 0.987041 |
| ENSMUSG( | NA | 19.11628 | 0.314736 | 0.46126 | NA | ENSMUSG( | NA | 68.06644 | 0.243481 | 0.547722 | 0.938063 |
| ENSMUSG( | NA | 14.92128 | -0.21135 | 0.655704 | NA | ENSMUSG( | NA | 21.27058 | 2.531864 | 7.22E-05 | 0.01665 |
| ENSMUSG( | NA | 22.0339 | 0.20494 | 0.618101 | 0.910929 | ENSMUSG( | NA | 24.78347 | 0.384503 | 0.454418 | 0.922444 |
| ENSMUSG( | NA | 11.97164 | -0.54208 | 0.332706 | NA | ENSMUSG( | NA | 34.66294 | 0.798304 | 0.102286 | 0.80239 |
| ENSMUSG( | NA | 33.70467 | -0.39016 | 0.22726 | 0.674223 | ENSMUSG( | NA | 62.7026 | 0.003745 | 0.988784 | 0.997886 |
| ENSMUSG( | NA | 11.56565 | 0.309912 | 0.590657 | NA | ENSMUSG( | NA | 9.984398 | 0.014029 | 0.983303 | NA |
| ENSMUSG( | NA | 29.99621 | 0.601825 | 0.078163 | 0.432307 | ENSMUSG( | NA | 9.567653 | -0.61404 | 0.35267 | NA |
| ENSMUSG( | NA | 86.06314 | -0.12357 | 0.599743 | 0.903906 | ENSMUSG( | NA | 15.2714 | 0.758314 | 0.167809 | 0.840222 |
| ENSMUSG( | NA | 35.83026 | -0.41358 | 0.206337 | 0.648436 | ENSMUSG( | NA | 250.0467 | 0.025676 | 0.875842 | 0.989644 |
| ENSMUSG( | NA | 39.04376 | -0.28795 | 0.346542 | 0.779166 | ENSMUSG( | NA | 137.4559 | 0.401744 | 0.232983 | 0.868507 |
| ENSMUSG( | NA | 23.42657 | 0.498401 | 0.213454 | 0.65774 | ENSMUSG( | NA | 12.73565 | 0.026707 | 0.963562 | 0.995433 |
| ENSMUSG( | NA | 6794.406 | 0.024712 | 0.898335 | 0.985633 | ENSMUSG( | NA | 15.09301 | 0.067834 | 0.898982 | 0.990558 |
| ENSMUSG( | NA | 45.27922 | -0.36024 | 0.218187 | 0.663729 | ENSMUSG( | NA | 9.642799 | -0.74604 | 0.278336 | NA |
| ENSMUSG( | NA | 11.63299 | -0.48847 | 0.388819 | NA | ENSMUSG( | NA | 9.439097 | 0.524379 | 0.41361 | NA |
| ENSMUSG( | NA | 79.20657 | -0.01168 | 0.959875 | 0.993991 | ENSMUSG( | NA | 13.63079 | -0.59491 | 0.297455 | 0.898538 |
| ENSMUSG( | NA | 10217.33 | 0.014511 | 0.90432 | 0.986246 | ENSMUSG( | NA | 12.3138 | 0.06355 | 0.912071 | 0.992196 |
| ENSMUSG( | NA | 80.78156 | 0.028878 | 0.913694 | 0.988483 | ENSMUSG( | NA | 9.305442 | -1.30918 | 0.067794 | NA |
| ENSMUSG( | NA | 56.86008 | 0.063511 | 0.80981 | 0.96372 | ENSMUSG( | NA | 15.21588 | -0.54977 | 0.267557 | 0.886134 |
| ENSMUSG( | NA | 10.72797 | 0.590779 | 0.314819 | NA | ENSMUSG( | NA | 13.15765 | 0.014727 | 0.977998 | 0.996416 |
| ENSMUSG( | NA | 45.57091 | 0.2484 | 0.37767 | 0.801561 | ENSMUSG( | NA | 15.67023 | -1.23054 | 0.025846 | 0.601244 |
| ENSMUSG( | NA | 67.96493 | -0.06235 | 0.804031 | 0.962169 | ENSMUSG( | NA | 51.44601 | -0.4727 | 0.095417 | 0.796133 |
| ENSMUSG( | NA | 21.29732 | 0.08022 | 0.856667 | 0.974154 | ENSMUSG( | NA | 41.45052 | -0.10373 | 0.751279 | 0.970317 |
| ENSMUSG( | NA | 25.01828 | 0.396249 | 0.328964 | 0.773233 | ENSMUSG( | NA | 103.5053 | -0.09401 | 0.678232 | 0.957435 |
| ENSMUSG( | NA | 72.96072 | -0.13946 | 0.567133 | 0.891644 | ENSMUSG( | NA | 30.54021 | -0.37679 | 0.302296 | 0.898538 |
| ENSMUSG( | NA | 18.20392 | -0.15242 | 0.725399 | NA | ENSMUSG( | NA | 17.20083 | 0.547997 | 0.299228 | 0.898538 |
| ENSMUSG( | NA | 30.51103 | -0.2897 | 0.56561 | 0.890871 | ENSMUSG( | NA | 18.26561 | -0.06437 | 0.892931 | 0.990558 |
| ENSMUSG( | NA | 735.0519 | -0.29721 | 0.413587 | 0.819895 | ENSMUSG( | NA | 15.64885 | 0.204854 | 0.694287 | 0.960867 |
| ENSMUSG( | NA | 7358.256 | 0.003288 | 0.986157 | 0.998326 | ENSMUSG( | NA | 29.69818 | -0.38537 | 0.299488 | 0.898538 |
| ENSMUSG( | NA | 14.75215 | -0.46437 | 0.380826 | NA | ENSMUSG( | NA | 9.393108 | -1.40987 | 0.035886 | NA |
| ENSMUSG( | NA | 14.76508 | 0.576687 | 0.255182 | NA | ENSMUSG( | NA | 20.7194 | 0.65554 | 0.241909 | 0.876528 |
| ENSMUSG( | NA | 14.20962 | -0.74744 | 0.226229 | NA | ENSMUSG( | NA | 19.4924 | 0.008792 | 0.986706 | 0.997411 |
| ENSMUSG( | NA | 1821.191 | 0.677539 | 1.51E-13 | 1.53E-10 | ENSMUSG( | NA | 87.47786 | 0.08675 | 0.726163 | 0.966887 |
| ENSMUSG( | NA | 20.05733 | -0.06897 | 0.869721 | NA | ENSMUSG( | NA | 12.26379 | -0.90157 | 0.137556 | 0.828492 |
| ENSMUSG( | NA | 37.76337 | -0.0836 | 0.831421 | 0.969889 | ENSMUSG( | NA | 55.35063 | 0.59027 | 0.129266 | 0.820358 |
| ENSMUSG( | NA | 33.81982 | 0.285706 | 0.398781 | 0.812878 | ENSMUSG( | NA | 54.35489 | -0.79973 | 0.008188 | 0.388416 |
| ENSMUSG( | NA | 10.63275 | -0.42199 | 0.481279 | NA | ENSMUSG( | NA | 27.89872 | 0.20466 | 0.623647 | 0.953234 |
| ENSMUSG( | NA | 19.05228 | 1.481638 | 0.021218 | NA | ENSMUSG( | NA | 15.74766 | 0.390986 | 0.477884 | 0.923329 |
| ENSMUSG( | NA | 72.24839 | -0.23993 | 0.418879 | 0.821966 | ENSMUSG( | NA | 13.68746 | 0.928579 | 0.189303 | 0.85698 |
| ENSMUSG( | NA | 19.51734 | -0.82671 | 0.05021 | NA | ENSMUSG( | NA | 22.75194 | 0.343406 | 0.423187 | 0.917257 |
| ENSMUSG( | NA | 16.24887 | -0.03712 | 0.942393 | NA | ENSMUSG( | NA | 23.84865 | 0.593695 | 0.161021 | 0.837724 |
| ENSMUSG( | NA | 24.59505 | 1.957525 | 0.000851 | 0.029239 | ENSMUSG( | NA | 54.74408 | 0.279521 | 0.328874 | 0.906797 |
| ENSMUSG( | NA | 42.02908 | -0.23894 | 0.467789 | 0.845032 | ENSMUSG( | NA | 29.05436 | -0.13432 | 0.725027 | 0.966887 |
| ENSMUSG( | NA | 19.1835 | -0.23918 | 0.586713 | NA | ENSMUSG( | NA | 27.74671 | -0.09915 | 0.794093 | 0.97681 |
| ENSMUSG( | NA | 10.77107 | -0.19964 | 0.7535 | NA | ENSMUSG( | NA | 18.78543 | 0.215107 | 0.663431 | 0.956487 |
| ENSMUSG( | NA | 243.798 | 0.200793 | 0.308407 | 0.756439 | ENSMUSG( | NA | 83.4845 | 0.204521 | 0.420852 | 0.917257 |
| ENSMUSG( | NA | 15.26055 | 0.361105 | 0.469015 | NA | ENSMUSG( | NA | 17.58807 | -0.22138 | 0.686165 | 0.958956 |
| ENSMUSG( | NA | 31.55691 | -0.39032 | 0.314852 | 0.762059 | ENSMUSG( | NA | 8.595301 | 0.170429 | 0.800199 | NA |
| ENSMUSG( | NA | 23.82807 | -0.19164 | 0.622895 | 0.911589 | ENSMUSG( | NA | 20.38114 | 0.209056 | 0.68963 | 0.960071 |
| ENSMUSG( | NA | 31.75466 | 0.323414 | 0.37333 | 0.800133 | ENSMUSG( | NA | 187.7146 | -0.153 | 0.413554 | 0.917257 |
| ENSMUSG( | NA | 19.75427 | 0.244875 | 0.588574 | NA | ENSMUSG( | NA | 9.489104 | -0.02247 | 0.973062 | NA |
| ENSMUSG( | NA | 20.57171 | 0.315776 | 0.507801 | NA | ENSMUSG( | NA | 9.342837 | 0.169119 | 0.792044 | NA |
| ENSMUSG( | NA | 44.73151 | -0.72085 | 0.044753 | 0.324745 | ENSMUSG( | NA | 309.239 | -0.15703 | 0.321676 | 0.904023 |

| | | | | | |
|---|---|---|---|---|---|
| ENSMUSG( | 26.94514 | -0.00799 | 0.982107 | 0.997512 | NA |
| ENSMUSG( | 44.65329 | 0.397327 | 0.241682 | 0.689179 | NA |
| ENSMUSG( | 13.74593 | 0.268772 | 0.605764 | NA | NA |
| ENSMUSG( | 133.9444 | 0.085398 | 0.62808 | 0.91351 | NA |
| ENSMUSG( | 762.4847 | 0.112885 | 0.298453 | 0.74713 | NA |
| ENSMUSG( | 14.39215 | 0.365602 | 0.469484 | NA | NA |
| ENSMUSG( | 49.31965 | 0.121879 | 0.716153 | 0.94152 | NA |
| ENSMUSG( | 12.40577 | -0.05492 | 0.921719 | NA | NA |
| ENSMUSG( | 49.87928 | -0.15414 | 0.603177 | 0.904887 | NA |
| ENSMUSG( | 43.91465 | 0.00789 | 0.978117 | 0.997428 | NA |
| ENSMUSG( | 281.7488 | 0.044485 | 0.78797 | 0.959226 | NA |
| ENSMUSG( | 19.2657 | -0.63227 | 0.160275 | NA | NA |
| ENSMUSG( | 16.39583 | 0.658693 | 0.183783 | NA | NA |
| ENSMUSG( | 97.34036 | 0.183233 | 0.405256 | 0.81652 | NA |
| ENSMUSG( | 21.86555 | 0.00097 | 0.99806 | 0.99989 | NA |
| ENSMUSG( | 208.2415 | -0.1332 | 0.377156 | 0.801213 | NA |
| ENSMUSG( | 65.06646 | -0.09722 | 0.702891 | 0.937735 | NA |
| ENSMUSG( | 10.7681 | 0.387566 | 0.496069 | NA | NA |
| ENSMUSG( | 51.5842 | -0.2813 | 0.301591 | 0.749979 | NA |
| ENSMUSG( | 15.07008 | 0.473246 | 0.37773 | NA | NA |
| ENSMUSG( | 21.61837 | -0.14206 | 0.717116 | 0.941697 | NA |
| ENSMUSG( | 36.3196 | 0.046184 | 0.898358 | 0.985633 | NA |
| ENSMUSG( | 629.6854 | 0.331058 | 0.005183 | 0.096057 | NA |
| ENSMUSG( | 9.902648 | 0.601225 | 0.308576 | NA | NA |
| ENSMUSG( | 2891.822 | -0.04526 | 0.647797 | 0.919759 | NA |
| ENSMUSG( | 10.31817 | 0.166446 | 0.781052 | NA | NA |
| ENSMUSG( | 22.87362 | -0.0627 | 0.883526 | 0.982503 | NA |
| ENSMUSG( | 425.1888 | 0.399769 | 0.001304 | 0.039243 | NA |
| ENSMUSG( | 21488.63 | 0.217028 | 0.025539 | 0.246694 | NA |
| ENSMUSG( | 227.3637 | 0.141502 | 0.424156 | 0.824522 | NA |
| ENSMUSG( | 24.51194 | 0.296546 | 0.482127 | 0.855778 | NA |
| ENSMUSG( | 14.8819 | 0.538691 | 0.296356 | NA | NA |
| ENSMUSG( | 10.58122 | 0.477001 | 0.406848 | NA | NA |
| ENSMUSG( | 149.8918 | -0.09883 | 0.587081 | 0.900235 | NA |
| ENSMUSG( | 18.51291 | 0.769293 | 0.080441 | NA | NA |
| ENSMUSG( | 53.19073 | -0.15833 | 0.558129 | 0.888427 | NA |
| ENSMUSG( | 151.7496 | -0.19256 | 0.317009 | 0.763823 | NA |
| ENSMUSG( | 18.79112 | -0.32816 | 0.470102 | NA | NA |
| ENSMUSG( | 16.16834 | -0.60413 | 0.220738 | NA | NA |
| ENSMUSG( | 127.3342 | 0.160883 | 0.479515 | 0.854046 | NA |
| ENSMUSG( | 11.64754 | 0.149532 | 0.777597 | NA | NA |
| ENSMUSG( | 25.91518 | 0.230217 | 0.602986 | 0.904887 | NA |
| ENSMUSG( | 14.54148 | 0.643349 | 0.23501 | NA | NA |
| ENSMUSG( | 16.91491 | -0.0309 | 0.952268 | NA | NA |
| ENSMUSG( | 29.37052 | -0.30702 | 0.41993 | 0.822442 | NA |
| ENSMUSG( | 15.51585 | -0.14589 | 0.758401 | NA | NA |
| ENSMUSG( | 10.5105 | -0.02402 | 0.969744 | NA | NA |
| ENSMUSG( | 11.69109 | -0.12579 | 0.816332 | NA | NA |
| ENSMUSG( | 31.69967 | -0.1372 | 0.689524 | 0.933609 | NA |
| ENSMUSG( | 155.6622 | -0.29847 | 0.088663 | 0.45666 | NA |
| ENSMUSG( | 11.71862 | 0.315 | 0.6156 | NA | NA |
| ENSMUSG( | 135.9729 | -0.11884 | 0.507003 | 0.867833 | NA |
| ENSMUSG( | 14.10681 | 1.0185 | 0.06718 | NA | NA |
| ENSMUSG( | 151.3633 | 0.19112 | 0.30833 | 0.756439 | NA |
| ENSMUSG( | 93.68033 | -0.24313 | 0.236762 | 0.684086 | NA |
| ENSMUSG( | 18.1113 | -0.01894 | 0.966552 | NA | NA |
| ENSMUSG( | 98.10108 | 0.487487 | 0.055349 | 0.361966 | NA |
| ENSMUSG( | 170.7031 | -0.02214 | 0.908511 | 0.987809 | NA |
| ENSMUSG( | 622.5185 | 0.140614 | 0.279634 | 0.729119 | NA |
| ENSMUSG( | 11.57792 | -0.39267 | 0.490401 | NA | NA |
| ENSMUSG( | 14.79456 | -0.0333 | 0.947974 | NA | NA |
| ENSMUSG( | 124.2322 | -0.34645 | 0.102846 | 0.48834 | NA |
| ENSMUSG( | 12.26842 | 0.311906 | 0.559036 | NA | NA |
| ENSMUSG( | 14.12462 | 0.297041 | 0.56687 | NA | NA |
| ENSMUSG( | 22.70117 | -1.77931 | 0.0001 | 0.006438 | NA |
| ENSMUSG( | 486.1859 | -0.0651 | 0.570556 | 0.893192 | NA |
| ENSMUSG( | 320.7445 | 0.151686 | 0.33778 | 0.777422 | NA |
| ENSMUSG( | 8.510679 | 0.334601 | 0.628405 | NA | NA |
| ENSMUSG( | 84.12828 | -0.04273 | 0.86607 | 0.976074 | NA |
| ENSMUSG( | 14.67561 | 0.448181 | 0.378915 | NA | NA |
| ENSMUSG( | 17.77941 | -0.26378 | 0.564213 | NA | NA |
| ENSMUSG( | 25.89331 | -0.25775 | 0.518628 | 0.87417 | NA |
| ENSMUSG( | 33.6188 | -0.00527 | 0.988473 | 0.999109 | NA |
| ENSMUSG( | 12.26411 | -0.24745 | 0.682019 | NA | NA |
| ENSMUSG( | 15.52079 | 0.396516 | 0.431832 | NA | NA |
| ENSMUSG( | 25.20173 | 0.128858 | 0.731045 | 0.944956 | NA |
| ENSMUSG( | 42.03211 | 0.186415 | 0.540818 | 0.88315 | NA |
| ENSMUSG( | 85.50709 | 0.162351 | 0.449181 | 0.834943 | NA |
| ENSMUSG( | 202.6183 | -0.17175 | 0.358406 | 0.789759 | NA |
| ENSMUSG( | 315.5373 | 0.073627 | 0.605225 | 0.905688 | NA |
| ENSMUSG( | 390.7378 | 0.087828 | 0.491312 | 0.859873 | NA |
| ENSMUSG( | 22.40354 | 0.508042 | 0.195986 | 0.635396 | NA |
| ENSMUSG( | 60.53471 | -0.15084 | 0.552675 | 0.886141 | NA |
| ENSMUSG( | 13.14219 | 0.83758 | 0.116099 | NA | NA |
| ENSMUSG( | 31.32453 | -0.13759 | 0.712222 | 0.94031 | NA |
| ENSMUSG( | 33.23006 | 0.2446 | 0.490393 | 0.859353 | NA |
| ENSMUSG( | 24.09021 | 0.075627 | 0.847125 | 0.972374 | NA |
| ENSMUSG( | 23.92851 | -0.5993 | 0.114344 | 0.51021 | NA |
| ENSMUSG( | 73.25115 | -0.27598 | 0.258727 | 0.709814 | NA |
| ENSMUSG( | 257.7397 | 0.021781 | 0.907524 | 0.987591 | NA |
| ENSMUSG( | 11.75887 | -0.22856 | 0.702496 | NA | NA |
| ENSMUSG( | 18.29469 | 0.08818 | 0.844322 | NA | NA |
| ENSMUSG( | 11.41726 | -0.28144 | 0.602657 | NA | NA |
| ENSMUSG( | 9.47123 | 0.033225 | 0.956042 | NA | NA |
| ENSMUSG( | 22.07331 | 0.351155 | 0.369069 | 0.796581 | NA |
| ENSMUSG( | 13.34571 | -0.29213 | 0.578325 | NA | NA |
| ENSMUSG( | 78.13739 | 0.034442 | 0.885342 | 0.982905 | NA |
| ENSMUSG( | 13.41922 | -0.37939 | 0.456284 | NA | NA |
| ENSMUSG( | 104.1373 | -0.09459 | 0.631543 | 0.914073 | NA |
| ENSMUSG( | 20.9574 | 0.836886 | 0.098317 | NA | NA |
| ENSMUSG( | 15.69242 | 0.484185 | 0.29153 | NA | NA |
| ENSMUSG( | 82.56223 | -0.54457 | 0.015925 | 0.18704 | NA |
| ENSMUSG( | 23.63155 | -0.24815 | 0.585529 | 0.899171 | NA |
| ENSMUSG( | 155.836 | 0.14326 | 0.396863 | 0.811742 | NA |
| ENSMUSG( | 32.97601 | -0.06162 | 0.850933 | 0.972638 | NA |
| ENSMUSG( | 116.4171 | 0.102269 | 0.582089 | 0.897918 | NA |
| ENSMUSG( | 397.7465 | 0.029455 | 0.890967 | 0.983764 | NA |
| ENSMUSG( | 16.57392 | 0.018149 | 0.971753 | NA | NA |
| ENSMUSG( | 20.04598 | -0.58753 | 0.18843 | NA | NA |
| ENSMUSG( | 10.44953 | 0.200266 | 0.730396 | NA | NA |
| ENSMUSG( | 73.68964 | -0.04499 | 0.840847 | 0.971179 | NA |
| ENSMUSG( | 33.49353 | 0.377041 | 0.271782 | 0.722082 | NA |
| ENSMUSG( | 25.65198 | -0.27968 | 0.471822 | 0.847504 | NA |
| ENSMUSG( | 44.38874 | 0.307305 | 0.395405 | 0.810601 | NA |
| ENSMUSG( | 20.32376 | 0.321122 | 0.479741 | NA | NA |
| ENSMUSG( | 34.58787 | 0.452205 | 0.18593 | 0.623352 | NA |
| ENSMUSG( | 29.27465 | -0.18657 | 0.585667 | 0.899171 | NA |
| ENSMUSG( | 13.51884 | -0.65663 | 0.303023 | NA | NA |
| ENSMUSG( | 8.966887 | 0.132249 | 0.838731 | NA | NA |
| ENSMUSG( | 24.30442 | 0.203132 | 0.647891 | 0.919759 | NA |
| ENSMUSG( | 39.75448 | 0.149718 | 0.614318 | 0.909419 | NA |
| ENSMUSG( | 38.87558 | -0.07682 | 0.820178 | 0.965888 | NA |
| ENSMUSG( | 12.28393 | -0.19988 | 0.734896 | NA | NA |
| ENSMUSG( | 14.58431 | -0.03054 | 0.949021 | NA | NA |
| ENSMUSG( | 13.74751 | 0.392639 | 0.491775 | NA | NA |
| ENSMUSG( | 16.55056 | -0.09941 | 0.827882 | NA | NA |
| ENSMUSG( | 13.14804 | 0.37023 | 0.462588 | NA | NA |
| ENSMUSG( | 18.14501 | -0.2793 | 0.517415 | NA | NA |
| ENSMUSG( | 18.61796 | -0.28938 | 0.535061 | NA | NA |
| ENSMUSG( | 16.2671 | 0.611486 | 0.229726 | NA | NA |
| ENSMUSG( | 51.5352 | -0.24636 | 0.381284 | 0.802629 | NA |
| ENSMUSG( | 9.774336 | -0.62156 | 0.340373 | NA | NA |
| ENSMUSG( | 19.73365 | -0.40042 | 0.335869 | NA | NA |
| ENSMUSG( | 12.73841 | -0.31418 | 0.563504 | NA | NA |
| ENSMUSG( | 14.03775 | 0.15445 | 0.781921 | NA | NA |
| ENSMUSG( | 10.27217 | 0.212312 | 0.713256 | NA | NA |
| ENSMUSG( | 30.01053 | 0.013947 | 0.96973 | 0.996369 | NA |
| ENSMUSG( | 269.868 | 0.000129 | 0.99932 | 0.99989 | NA |
| ENSMUSG( | 11.17567 | -0.21665 | 0.695525 | NA | NA |
| ENSMUSG( | 78.20522 | 0.384916 | 0.077383 | 0.429743 | NA |
| ENSMUSG( | 20.68748 | -0.19297 | 0.658576 | NA | NA |
| ENSMUSG( | 107.7588 | 0.360254 | 0.107848 | 0.49674 | NA |
| ENSMUSG( | 169.1534 | 0.152697 | 0.370202 | 0.797542 | NA |
| ENSMUSG( | 16.73298 | -0.17566 | 0.71866 | NA | NA |
| ENSMUSG( | 10.7155 | -0.53027 | 0.397018 | NA | NA |
| ENSMUSG( | 43.92235 | -0.08488 | 0.783521 | 0.957558 | NA |
| ENSMUSG( | 19.41747 | -0.20743 | 0.629073 | NA | NA |
| ENSMUSG( | 33.33179 | 0.456193 | 0.163146 | 0.59439 | NA |
| ENSMUSG( | 19.04421 | -0.11393 | 0.788439 | NA | NA |
| ENSMUSG( | 3207.778 | -0.07712 | 0.848532 | 0.972374 | NA |
| ENSMUSG( | 27.33295 | -0.45396 | 0.213868 | 0.658243 | NA |
| ENSMUSG( | 11.74178 | 0.187352 | 0.751899 | NA | NA |
| ENSMUSG( | 16.56705 | -0.16367 | 0.721466 | NA | NA |
| ENSMUSG( | 29.1755 | -0.29386 | 0.451174 | 0.835567 | NA |
| ENSMUSG( | 87.83919 | 0.223055 | 0.317282 | 0.763823 | NA |
| ENSMUSG( | 11.35408 | -0.05603 | 0.921328 | NA | NA |
| ENSMUSG( | 13.34689 | -0.22272 | 0.684042 | NA | NA |
| ENSMUSG( | 94.09243 | -0.09677 | 0.683352 | 0.932153 | NA |

| | | | | | |
|---|---|---|---|---|---|
| ENSMUSG( | 57.93944 | 0.022156 | 0.944142 | 0.994798 | NA |
| ENSMUSG( | 14.2562 | -0.0073 | 0.989177 | 0.997893 | NA |
| ENSMUSG( | 24.66651 | -0.09783 | 0.812839 | 0.979148 | NA |
| ENSMUSG( | 20.1329 | -1.03023 | 0.024154 | 0.588924 | NA |
| ENSMUSG( | 362.4564 | 0.521934 | 0.003456 | 0.23553 | NA |
| ENSMUSG( | 5188.124 | -0.008 | 0.965402 | 0.995433 | NA |
| ENSMUSG( | 55.18666 | -0.32104 | 0.277371 | 0.891468 | NA |
| ENSMUSG( | 573.2399 | -0.07127 | 0.640771 | 0.953903 | NA |
| ENSMUSG( | 17.47361 | 0.162719 | 0.736198 | 0.967344 | NA |
| ENSMUSG( | 281.5417 | -0.06471 | 0.709642 | 0.96416 | NA |
| ENSMUSG( | 14.72126 | 0.26929 | 0.641977 | 0.953903 | NA |
| ENSMUSG( | 109.1531 | 0.257405 | 0.325948 | 0.905339 | NA |
| ENSMUSG( | 148.6845 | -0.14841 | 0.472911 | 0.923329 | NA |
| ENSMUSG( | 12.48481 | 0.069827 | 0.905938 | 0.991089 | NA |
| ENSMUSG( | 13.22461 | 0.367951 | 0.591642 | 0.944975 | NA |
| ENSMUSG( | 10.4742 | 0.069653 | 0.911394 | NA | NA |
| ENSMUSG( | 13.35571 | 1.040861 | 0.083628 | 0.77681 | NA |
| ENSMUSG( | 396.5224 | 0.296526 | 0.044574 | 0.693697 | NA |
| ENSMUSG( | 23.91867 | -0.52357 | 0.217513 | 0.865589 | NA |
| ENSMUSG( | 12.21508 | 0.074455 | 0.891426 | 0.990558 | NA |
| ENSMUSG( | 81.93257 | 0.059613 | 0.80084 | 0.97681 | NA |
| ENSMUSG( | 20.67819 | -0.34751 | 0.421309 | 0.917257 | NA |
| ENSMUSG( | 33.26677 | 0.265455 | 0.549385 | 0.938519 | NA |
| ENSMUSG( | 24.1518 | 0.063963 | 0.877535 | 0.989644 | NA |
| ENSMUSG( | 42.78641 | 0.560318 | 0.13431 | 0.822912 | NA |
| ENSMUSG( | 18.97835 | -0.47177 | 0.296885 | 0.898538 | NA |
| ENSMUSG( | 18.94356 | 0.715369 | 0.143986 | 0.831311 | NA |
| ENSMUSG( | 7.813056 | -0.03871 | 0.961122 | NA | NA |
| ENSMUSG( | 264.2199 | -0.26312 | 0.126004 | 0.81516 | NA |
| ENSMUSG( | 14.781 | -0.11089 | 0.834573 | 0.981237 | NA |
| ENSMUSG( | 94.13846 | 0.122851 | 0.581751 | 0.944266 | NA |
| ENSMUSG( | 9.998116 | -0.33244 | 0.643535 | NA | NA |
| ENSMUSG( | 8.705801 | -0.13005 | 0.865698 | NA | NA |
| ENSMUSG( | 9.565598 | -1.12055 | 0.122428 | NA | NA |
| ENSMUSG( | 11.48036 | 0.664522 | 0.259646 | 0.883971 | NA |
| ENSMUSG( | 8.141949 | 0.873557 | 0.250174 | NA | NA |
| ENSMUSG( | 52.98185 | -0.19976 | 0.490433 | 0.926343 | NA |
| ENSMUSG( | 8.304379 | 0.883962 | 0.258792 | NA | NA |
| ENSMUSG( | 9.033752 | 0.30252 | 0.653799 | NA | NA |
| ENSMUSG( | 37.28156 | 0.674095 | 0.120548 | 0.813384 | NA |
| ENSMUSG( | 17.12795 | 0.064446 | 0.904104 | 0.991089 | NA |
| ENSMUSG( | 17.62862 | 0.419187 | 0.402408 | 0.917257 | NA |
| ENSMUSG( | 21.96873 | 0.70262 | 0.103212 | 0.804311 | NA |
| ENSMUSG( | 21.70276 | -1.16563 | 0.043124 | 0.690304 | NA |
| ENSMUSG( | 7.635272 | -0.277 | 0.70535 | NA | NA |
| ENSMUSG( | 14.80473 | 1.38133 | 0.018382 | 0.544291 | NA |
| ENSMUSG( | 15.60231 | 0.382051 | 0.453693 | 0.922316 | NA |
| ENSMUSG( | 15.27881 | 0.136831 | 0.824803 | 0.979198 | NA |
| ENSMUSG( | 22.85119 | 0.430055 | 0.290199 | 0.896947 | NA |
| ENSMUSG( | 12.92956 | 0.138457 | 0.799604 | 0.97681 | NA |
| ENSMUSG( | 7.971094 | -0.68288 | 0.315591 | NA | NA |
| ENSMUSG( | 18.09312 | -0.1814 | 0.711141 | 0.964363 | NA |
| ENSMUSG( | 12.7319 | 0.737475 | 0.19843 | 0.862367 | NA |
| ENSMUSG( | 259.0339 | 0.447241 | 0.014393 | 0.48208 | NA |
| ENSMUSG( | 46.39184 | -0.00856 | 0.980035 | 0.996479 | NA |
| ENSMUSG( | 28.23133 | -0.11458 | 0.797976 | 0.97681 | NA |
| ENSMUSG( | 11.09008 | 1.513012 | 0.022158 | NA | NA |
| ENSMUSG( | 9.136393 | 0.240221 | 0.718782 | NA | NA |
| ENSMUSG( | 21.54846 | 0.420634 | 0.392078 | 0.917257 | NA |
| ENSMUSG( | 31.00981 | -0.07755 | 0.84253 | 0.983044 | NA |
| ENSMUSG( | 34.6968 | 0.092616 | 0.798598 | 0.97681 | NA |
| ENSMUSG( | 9.349734 | 0.354074 | 0.59932 | NA | NA |
| ENSMUSG( | 29.07289 | -0.16632 | 0.651394 | 0.95449 | NA |
| ENSMUSG( | 28.70382 | -0.01471 | 0.968009 | 0.995866 | NA |
| ENSMUSG( | 79.75886 | 0.273894 | 0.340075 | 0.907972 | NA |
| ENSMUSG( | 16.26996 | 0.187639 | 0.71375 | 0.964986 | NA |
| ENSMUSG( | 12.72981 | -0.48954 | 0.388555 | 0.917257 | NA |
| ENSMUSG( | 9.548803 | 1.094377 | 0.144952 | NA | NA |
| ENSMUSG( | 11.76247 | 1.845901 | 0.024295 | 0.589452 | NA |
| ENSMUSG( | 46.6436 | -0.25241 | 0.411225 | 0.917257 | NA |
| ENSMUSG( | 18.18496 | -0.57369 | 0.223394 | 0.867404 | NA |
| ENSMUSG( | 64.09131 | -0.07281 | 0.865837 | 0.987041 | NA |
| ENSMUSG( | 68.06644 | 0.243481 | 0.547722 | 0.938063 | NA |
| ENSMUSG( | 21.27058 | 2.531864 | 7.22E-05 | 0.01665 | NA |
| ENSMUSG( | 24.78347 | 0.384503 | 0.454418 | 0.922444 | NA |
| ENSMUSG( | 34.66294 | 0.798304 | 0.102286 | 0.80239 | NA |
| ENSMUSG( | 62.7026 | 0.003745 | 0.988784 | 0.997886 | NA |
| ENSMUSG( | 9.984398 | 0.014029 | 0.983303 | NA | NA |
| ENSMUSG( | 9.567653 | -0.61404 | 0.35267 | NA | NA |
| ENSMUSG( | 15.2714 | 0.758314 | 0.167809 | 0.840222 | NA |
| ENSMUSG( | 250.0467 | 0.025676 | 0.875842 | 0.989644 | NA |
| ENSMUSG( | 137.4559 | 0.401744 | 0.232983 | 0.868507 | NA |
| ENSMUSG( | 12.73565 | 0.026707 | 0.963562 | 0.995433 | NA |
| ENSMUSG( | 15.09301 | 0.067834 | 0.898982 | 0.990558 | NA |
| ENSMUSG( | 9.642799 | -0.74604 | 0.278336 | NA | NA |
| ENSMUSG( | 9.439097 | 0.524379 | 0.41361 | NA | NA |
| ENSMUSG( | 13.63079 | -0.59491 | 0.297455 | 0.898538 | NA |
| ENSMUSG( | 12.3138 | 0.06355 | 0.912071 | 0.992196 | NA |
| ENSMUSG( | 9.305442 | -1.30918 | 0.067794 | NA | NA |
| ENSMUSG( | 15.21588 | -0.54977 | 0.267557 | 0.886134 | NA |
| ENSMUSG( | 13.15765 | 0.014727 | 0.977998 | 0.996416 | NA |
| ENSMUSG( | 15.67023 | -1.23054 | 0.025846 | 0.601244 | NA |
| ENSMUSG( | 51.44601 | -0.4727 | 0.095417 | 0.796133 | NA |
| ENSMUSG( | 12.8823 | 0.646605 | 0.332407 | 0.907907 | NA |
| ENSMUSG( | 41.45052 | -0.10373 | 0.751279 | 0.970317 | NA |
| ENSMUSG( | 103.5053 | -0.09401 | 0.678232 | 0.957435 | NA |
| ENSMUSG( | 30.54021 | -0.37679 | 0.302296 | 0.898538 | NA |
| ENSMUSG( | 17.20083 | 0.547997 | 0.299228 | 0.898538 | NA |
| ENSMUSG( | 18.26561 | -0.06437 | 0.892931 | 0.990558 | NA |
| ENSMUSG( | 15.64885 | 0.204854 | 0.694287 | 0.960867 | NA |
| ENSMUSG( | 29.69818 | -0.38537 | 0.299488 | 0.898538 | NA |
| ENSMUSG( | 9.393108 | -1.40987 | 0.035886 | NA | NA |
| ENSMUSG( | 20.7194 | 0.65554 | 0.241909 | 0.876528 | NA |
| ENSMUSG( | 19.4924 | 0.008792 | 0.986706 | 0.997411 | NA |
| ENSMUSG( | 87.47786 | 0.08675 | 0.726163 | 0.966887 | NA |
| ENSMUSG( | 12.26379 | -0.90157 | 0.137556 | 0.828492 | NA |
| ENSMUSG( | 55.35063 | 0.59027 | 0.129266 | 0.820358 | NA |
| ENSMUSG( | 54.35489 | -0.79973 | 0.008188 | 0.388416 | NA |
| ENSMUSG( | 27.89872 | 0.20466 | 0.623647 | 0.953234 | NA |
| ENSMUSG( | 15.74766 | 0.390986 | 0.477884 | 0.923329 | NA |
| ENSMUSG( | 13.68746 | 0.928579 | 0.189303 | 0.85698 | NA |
| ENSMUSG( | 22.75194 | 0.343406 | 0.423187 | 0.917257 | NA |
| ENSMUSG( | 23.84865 | 0.593695 | 0.161021 | 0.837724 | NA |
| ENSMUSG( | 54.74408 | 0.279521 | 0.328874 | 0.906797 | NA |
| ENSMUSG( | 29.05436 | -0.13432 | 0.725027 | 0.966887 | NA |
| ENSMUSG( | 27.74671 | -0.09915 | 0.794093 | 0.97681 | NA |
| ENSMUSG( | 18.78543 | 0.215107 | 0.663431 | 0.956487 | NA |
| ENSMUSG( | 83.4845 | 0.204521 | 0.420852 | 0.917257 | NA |
| ENSMUSG( | 17.58807 | -0.22138 | 0.686165 | 0.958956 | NA |
| ENSMUSG( | 8.595301 | 0.170429 | 0.800199 | NA | NA |
| ENSMUSG( | 10.3392 | -0.28232 | 0.676246 | NA | NA |
| ENSMUSG( | 20.38114 | 0.209056 | 0.68963 | 0.960071 | NA |
| ENSMUSG( | 187.7146 | -0.153 | 0.413554 | 0.917257 | NA |
| ENSMUSG( | 9.489104 | -0.02247 | 0.973062 | NA | NA |
| ENSMUSG( | 9.342837 | 0.169119 | 0.792044 | NA | NA |
| ENSMUSG( | 309.239 | -0.15703 | 0.321676 | 0.904023 | NA |
| ENSMUSG( | 71.09068 | -0.4842 | 0.052176 | 0.712974 | NA |
| ENSMUSG( | 12.04866 | 0.248869 | 0.684293 | 0.958678 | NA |
| ENSMUSG( | 11.974 | 0.00561 | 0.992203 | 0.99874 | NA |
| ENSMUSG( | 11.36635 | 0.040051 | 0.949467 | NA | NA |
| ENSMUSG( | 31.61735 | -0.22959 | 0.513752 | 0.930586 | NA |
| ENSMUSG( | 1143.116 | 0.134849 | 0.374979 | 0.915491 | NA |
| ENSMUSG( | 11.52336 | 0.303231 | 0.608628 | 0.950113 | NA |
| ENSMUSG( | 122.3553 | -0.0476 | 0.820424 | 0.979198 | NA |
| ENSMUSG( | 49.99966 | -0.05719 | 0.849342 | 0.985321 | NA |
| ENSMUSG( | 658.657 | 0.338864 | 0.038343 | 0.679117 | NA |
| ENSMUSG( | 23.59078 | 0.022364 | 0.958473 | 0.994798 | NA |
| ENSMUSG( | 28.48328 | 0.187033 | 0.612515 | 0.951071 | NA |
| ENSMUSG( | 39.13752 | 0.214678 | 0.54064 | 0.937383 | NA |
| ENSMUSG( | 88.73819 | 0.686726 | 0.055841 | 0.723362 | NA |
| ENSMUSG( | 44.56828 | 1.058253 | 0.0086 | 0.391682 | NA |
| ENSMUSG( | 64.63213 | 0.391589 | 0.241279 | 0.876259 | NA |
| ENSMUSG( | 20.37656 | -0.18961 | 0.663669 | 0.956752 | NA |
| ENSMUSG( | 10.12476 | 0.220141 | 0.720081 | NA | NA |
| ENSMUSG( | 207.4734 | 1.132661 | 0.055032 | 0.723362 | NA |
| ENSMUSG( | 40.86627 | 0.258628 | 0.516277 | 0.930939 | NA |
| ENSMUSG( | 9.74644 | -0.1906 | 0.75632 | NA | NA |
| ENSMUSG( | 82.43646 | -0.23525 | 0.329543 | 0.907083 | NA |
| ENSMUSG( | 21.19915 | -0.11961 | 0.794696 | 0.97681 | NA |
| ENSMUSG( | 9.698497 | -0.08217 | 0.898312 | NA | NA |
| ENSMUSG( | 11.25099 | -0.28287 | 0.639028 | NA | NA |
| ENSMUSG( | 7.876436 | 0.329339 | 0.642464 | NA | NA |
| ENSMUSG( | 11.30392 | -0.15855 | 0.803076 | NA | NA |
| ENSMUSG( | 26.98596 | 0.132446 | 0.74122 | 0.968811 | NA |
| ENSMUSG( | 17.29211 | 0.0208 | 0.965539 | 0.995433 | NA |
| ENSMUSG( | 17.9073 | 0.027243 | 0.957252 | 0.994798 | NA |
| ENSMUSG( | 38.09309 | -0.03443 | 0.915172 | 0.992488 | NA |
| ENSMUSG( | 53.7296 | -0.57239 | 0.054876 | 0.723362 | NA |

| | | | | | | | | | | | |
|---|---|---|---|---|---|---|---|---|---|---|---|
| ENSMUSG( | NA | 12.20484 | -0.85856 | 0.128448 | NA | ENSMUSG( | NA | 71.09068 | -0.4842 | 0.052176 | 0.712974 |
| ENSMUSG( | NA | 52.3108 | 0.479295 | 0.084249 | 0.447434 | ENSMUSG( | NA | 12.04866 | 0.248869 | 0.684293 | 0.958678 |
| ENSMUSG( | NA | 9.668854 | -0.20002 | 0.735615 | NA | ENSMUSG( | NA | 11.974 | 0.00561 | 0.992203 | 0.99874 |
| ENSMUSG( | NA | 130.6097 | -0.19038 | 0.377954 | 0.801561 | ENSMUSG( | NA | 11.36635 | 0.040051 | 0.949467 | NA |
| ENSMUSG( | NA | 40.19096 | -0.30999 | 0.333643 | 0.775602 | ENSMUSG( | NA | 31.61735 | -0.22959 | 0.513752 | 0.930586 |
| ENSMUSG( | NA | 30.23584 | 0.243421 | 0.469568 | 0.846167 | ENSMUSG( | NA | 1143.116 | 0.134849 | 0.374979 | 0.915491 |
| ENSMUSG( | NA | 260.5226 | -0.0635 | 0.699483 | 0.937034 | ENSMUSG( | NA | 11.52336 | 0.303231 | 0.608628 | 0.950113 |
| ENSMUSG( | NA | 57.96506 | 0.185034 | 0.524183 | 0.87583 | ENSMUSG( | NA | 122.3553 | -0.0476 | 0.820424 | 0.979198 |
| ENSMUSG( | NA | 14.01709 | 0.430064 | 0.412527 | NA | ENSMUSG( | NA | 49.99966 | -0.05719 | 0.849342 | 0.985321 |
| ENSMUSG( | NA | 10.51297 | -0.18279 | 0.752727 | NA | ENSMUSG( | NA | 658.657 | 0.338864 | 0.038343 | 0.679117 |
| ENSMUSG( | NA | 48.15224 | 0.094782 | 0.741229 | 0.948165 | ENSMUSG( | NA | 23.59078 | 0.022364 | 0.958473 | 0.994798 |
| ENSMUSG( | NA | 14.65796 | -0.98675 | 0.057415 | NA | ENSMUSG( | NA | 28.48328 | 0.187033 | 0.612515 | 0.951071 |
| ENSMUSG( | NA | 36.27897 | -0.09349 | 0.770723 | 0.954722 | ENSMUSG( | NA | 39.13752 | 0.214678 | 0.54064 | 0.937383 |
| ENSMUSG( | NA | 207.2546 | 0.022407 | 0.883271 | 0.982422 | ENSMUSG( | NA | 88.73819 | 0.686726 | 0.055841 | 0.723362 |
| ENSMUSG( | NA | 13.93578 | 0.183012 | 0.712605 | NA | ENSMUSG( | NA | 44.56828 | 1.058253 | 0.0086 | 0.391682 |
| ENSMUSG( | NA | 200.8166 | -0.08573 | 0.577178 | 0.895659 | ENSMUSG( | NA | 64.63213 | 0.391589 | 0.241279 | 0.876259 |
| ENSMUSG( | NA | 30.85462 | -0.15151 | 0.720156 | 0.94201 | ENSMUSG( | NA | 20.37656 | -0.18961 | 0.663669 | 0.956752 |
| ENSMUSG( | NA | 20.40427 | 0.134035 | 0.776855 | NA | ENSMUSG( | NA | 10.12476 | 0.220141 | 0.720081 | NA |
| ENSMUSG( | NA | 22.05542 | -0.30984 | 0.513474 | 0.871762 | ENSMUSG( | NA | 207.4734 | 1.132661 | 0.055032 | 0.723362 |
| ENSMUSG( | NA | 36.61892 | -0.45116 | 0.151003 | 0.576088 | ENSMUSG( | NA | 40.86627 | 0.258628 | 0.516277 | 0.930939 |
| ENSMUSG( | NA | 122.184 | 0.04798 | 0.809865 | 0.96372 | ENSMUSG( | NA | 9.74644 | -0.1906 | 0.75632 | NA |
| ENSMUSG( | NA | 17.67093 | 0.489208 | 0.350125 | NA | ENSMUSG( | NA | 82.43646 | -0.23525 | 0.329543 | 0.907083 |
| ENSMUSG( | NA | 65.94393 | 0.480934 | 0.050189 | 0.342492 | ENSMUSG( | NA | 21.19915 | -0.11961 | 0.794696 | 0.97681 |
| ENSMUSG( | NA | 193.9815 | 0.077978 | 0.724026 | 0.944096 | ENSMUSG( | NA | 9.698497 | -0.08217 | 0.898312 | NA |
| ENSMUSG( | NA | 26.83157 | 0.286842 | 0.42233 | 0.823508 | ENSMUSG( | NA | 11.25099 | -0.28287 | 0.639028 | NA |
| ENSMUSG( | NA | 337.1665 | 0.103516 | 0.447261 | 0.834051 | ENSMUSG( | NA | 11.30392 | -0.15855 | 0.803076 | NA |
| ENSMUSG( | NA | 101.0255 | -0.07562 | 0.734283 | 0.945468 | ENSMUSG( | NA | 26.98596 | 0.132446 | 0.74122 | 0.968811 |
| ENSMUSG( | NA | 17.99367 | -0.15924 | 0.746826 | NA | ENSMUSG( | NA | 17.29211 | 0.0208 | 0.965539 | 0.995433 |
| ENSMUSG( | NA | 81.46233 | 0.451571 | 0.045335 | 0.326817 | ENSMUSG( | NA | 17.9073 | 0.027243 | 0.957252 | 0.994798 |
| ENSMUSG( | NA | 11.16202 | -1.42217 | 0.01514 | NA | ENSMUSG( | NA | 38.09309 | -0.03443 | 0.915172 | 0.992488 |
| ENSMUSG( | NA | 12.31128 | -0.21168 | 0.709221 | NA | ENSMUSG( | NA | 53.7296 | -0.57239 | 0.054876 | 0.723362 |
| ENSMUSG( | NA | 48.33372 | -0.09884 | 0.732455 | 0.944956 | ENSMUSG( | NA | 13.60583 | 0.181992 | 0.776619 | 0.975717 |
| ENSMUSG( | NA | 17.46735 | 0.447713 | 0.377796 | NA | ENSMUSG( | NA | 36.00081 | -0.07407 | 0.837954 | 0.981376 |
| ENSMUSG( | NA | 34.92991 | 0.358159 | 0.283048 | 0.73186 | ENSMUSG( | NA | 12.48012 | -0.3022 | 0.585841 | 0.944975 |
| ENSMUSG( | NA | 45.66925 | -0.05322 | 0.848252 | 0.972374 | ENSMUSG( | NA | 29.50765 | -0.18853 | 0.67289 | 0.95694 |
| ENSMUSG( | NA | 57.23953 | -0.19242 | 0.459034 | 0.840244 | ENSMUSG( | NA | 51.2063 | -0.17686 | 0.566024 | 0.94151 |
| ENSMUSG( | NA | 22.78073 | -0.23795 | 0.541738 | 0.883182 | ENSMUSG( | NA | 7.812057 | -0.95526 | 0.182966 | NA |
| ENSMUSG( | NA | 14.03383 | -0.21722 | 0.666148 | NA | ENSMUSG( | NA | 11.19749 | 0.186959 | 0.757132 | NA |
| ENSMUSG( | NA | 117.0315 | 0.707799 | 0.000115 | 0.007087 | ENSMUSG( | NA | 36.91534 | 0.063039 | 0.878668 | 0.989644 |
| ENSMUSG( | NA | 31.98245 | 0.104389 | 0.766398 | 0.954173 | ENSMUSG( | NA | 76.24283 | -0.48 | 0.058057 | 0.728256 |
| ENSMUSG( | NA | 18.57225 | -0.28289 | 0.503225 | NA | ENSMUSG( | NA | 49.20569 | -0.1862 | 0.536609 | 0.937108 |
| ENSMUSG( | NA | 46.09718 | -0.03076 | 0.913707 | 0.988483 | ENSMUSG( | NA | 1531.855 | -0.02571 | 0.82277 | 0.979198 |
| ENSMUSG( | NA | 14.85187 | -0.75113 | 0.165206 | NA | ENSMUSG( | NA | 50.70868 | -0.29916 | 0.319111 | 0.903431 |
| ENSMUSG( | NA | 16.69139 | -0.249 | 0.635477 | NA | ENSMUSG( | NA | 142.6918 | -0.33294 | 0.085012 | 0.779799 |
| ENSMUSG( | NA | 80.98966 | 0.057706 | 0.808097 | 0.963419 | ENSMUSG( | NA | 18.988 | -0.58771 | 0.231557 | 0.868507 |
| ENSMUSG( | NA | 16.70142 | 0.10531 | 0.819901 | NA | ENSMUSG( | NA | 8.799157 | -1.1457 | 0.102972 | NA |
| ENSMUSG( | NA | 16.64718 | 0.434216 | 0.374444 | NA | ENSMUSG( | NA | 33.89968 | -0.42293 | 0.301061 | 0.898538 |
| ENSMUSG( | NA | 14.72092 | 0.467797 | 0.396917 | NA | ENSMUSG( | NA | 211.1852 | -0.12189 | 0.505739 | 0.927825 |
| ENSMUSG( | NA | 39.64057 | 0.096583 | 0.786711 | 0.958856 | ENSMUSG( | NA | 23.76254 | -0.41494 | 0.300652 | 0.898538 |
| ENSMUSG( | NA | 39.14605 | 0.220116 | 0.51732 | 0.873295 | ENSMUSG( | NA | 58.67262 | 0.247147 | 0.392716 | 0.917257 |
| ENSMUSG( | NA | 243.4591 | -0.15422 | 0.365139 | 0.794085 | ENSMUSG( | NA | 60.61189 | 0.274409 | 0.44549 | 0.921734 |
| ENSMUSG( | NA | 46.42893 | -0.2999 | 0.285334 | 0.734038 | ENSMUSG( | NA | 9.071548 | -0.87951 | 0.18721 | NA |
| ENSMUSG( | NA | 12.56102 | -0.13582 | 0.798769 | NA | ENSMUSG( | NA | 107.061 | -0.44822 | 0.050209 | 0.708648 |
| ENSMUSG( | NA | 23.05417 | 0.114769 | 0.771583 | 0.954769 | ENSMUSG( | NA | 43.90004 | 0.238805 | 0.4971 | 0.927825 |
| ENSMUSG( | NA | 14.10127 | 0.354522 | 0.478475 | NA | ENSMUSG( | NA | 63.10346 | -0.03152 | 0.905389 | 0.991089 |
| ENSMUSG( | NA | 45.30503 | 0.207359 | 0.467577 | 0.844819 | ENSMUSG( | NA | 22.2941 | -0.18575 | 0.661771 | 0.956205 |
| ENSMUSG( | NA | 28.10474 | 0.067402 | 0.848655 | 0.972374 | ENSMUSG( | NA | 17.02557 | 0.780115 | 0.188023 | 0.85698 |
| ENSMUSG( | NA | 10.74624 | -0.26198 | 0.687007 | NA | ENSMUSG( | NA | 19.63278 | 0.311507 | 0.509457 | 0.929089 |
| ENSMUSG( | NA | 93.18414 | -0.0628 | 0.769013 | 0.954674 | ENSMUSG( | NA | 24.41769 | -0.14414 | 0.725502 | 0.966887 |
| ENSMUSG( | NA | 70.70774 | 0.169307 | 0.496873 | 0.861754 | ENSMUSG( | NA | 430.815 | -0.04787 | 0.746764 | 0.969763 |
| ENSMUSG( | NA | 8.898683 | -0.49948 | 0.425223 | NA | ENSMUSG( | NA | 23.31853 | 0.643273 | 0.121329 | 0.813384 |
| ENSMUSG( | NA | 31.53096 | 0.016586 | 0.960267 | 0.993991 | ENSMUSG( | NA | 18.25554 | 0.19206 | 0.677544 | 0.957106 |
| ENSMUSG( | NA | 31.00805 | 0.696488 | 0.041344 | 0.314189 | ENSMUSG( | NA | 30.59224 | 0.76954 | 0.064516 | 0.737935 |
| ENSMUSG( | NA | 62.2905 | -0.11041 | 0.689401 | 0.933609 | ENSMUSG( | NA | 103.6992 | -0.26489 | 0.282007 | 0.892412 |
| ENSMUSG( | NA | 39.05423 | -0.25349 | 0.442636 | 0.83278 | ENSMUSG( | NA | 359.5373 | 0.076144 | 0.586911 | 0.944975 |
| ENSMUSG( | NA | 41.87132 | -0.56154 | 0.068073 | 0.404152 | ENSMUSG( | NA | 35.90422 | 0.360013 | 0.328765 | 0.906727 |
| ENSMUSG( | NA | 25.35404 | 0.182737 | 0.673283 | 0.929007 | ENSMUSG( | NA | 46.40886 | 0.488066 | 0.223269 | 0.867404 |
| ENSMUSG( | NA | 15.14781 | -0.54316 | 0.341051 | NA | ENSMUSG( | NA | 10.10648 | -0.04006 | 0.94958 | NA |
| ENSMUSG( | NA | 76.86238 | -0.06853 | 0.803215 | 0.962169 | ENSMUSG( | NA | 16.28842 | -0.4447 | 0.372146 | 0.915491 |
| ENSMUSG( | NA | 78.67447 | 0.257529 | 0.315757 | 0.762707 | ENSMUSG( | NA | 358.0709 | 0.036979 | 0.823859 | 0.979198 |
| ENSMUSG( | NA | 15.4818 | -0.46006 | 0.366625 | NA | ENSMUSG( | NA | 261.1379 | 0.100212 | 0.582835 | 0.944266 |
| ENSMUSG( | NA | 77.75387 | -0.06543 | 0.786214 | 0.958774 | ENSMUSG( | NA | 17.407 | 0.209955 | 0.679736 | 0.957966 |
| ENSMUSG( | NA | 18.78135 | -0.24916 | 0.582411 | NA | ENSMUSG( | NA | 26.00289 | 0.255787 | 0.623173 | 0.953234 |
| ENSMUSG( | NA | 48.05075 | 0.251808 | 0.421894 | 0.823481 | ENSMUSG( | NA | 65.81461 | -0.07487 | 0.798801 | 0.97681 |
| ENSMUSG( | NA | 32.42198 | -0.33845 | 0.339332 | 0.777616 | ENSMUSG( | NA | 37.52039 | 0.599368 | 0.10843 | 0.804969 |
| ENSMUSG( | NA | 49.19992 | -0.21369 | 0.533447 | 0.880095 | ENSMUSG( | NA | 55.34167 | 0.194846 | 0.518835 | 0.930939 |
| ENSMUSG( | NA | 19.90778 | 0.257964 | 0.563547 | NA | ENSMUSG( | NA | 77.67433 | 0.175687 | 0.485344 | 0.92541 |
| ENSMUSG( | NA | 82.32924 | 0.050636 | 0.819028 | 0.965888 | ENSMUSG( | NA | 37.31813 | -0.57045 | 0.098218 | 0.798438 |
| ENSMUSG( | NA | 15.82525 | -0.42673 | 0.359956 | NA | ENSMUSG( | NA | 10.22884 | 0.030096 | 0.965117 | NA |
| ENSMUSG( | NA | 17.0724 | -0.06352 | 0.886176 | NA | ENSMUSG( | NA | 142.2206 | 0.111619 | 0.609485 | 0.950397 |
| ENSMUSG( | NA | 12.78636 | -0.00581 | 0.990919 | NA | ENSMUSG( | NA | 27.73471 | -0.04271 | 0.91416 | 0.992407 |
| ENSMUSG( | NA | 36.21843 | 0.306319 | 0.366242 | 0.794603 | ENSMUSG( | NA | 14.08441 | -0.47091 | 0.36532 | 0.914449 |
| ENSMUSG( | NA | 94.22397 | -0.08481 | 0.701281 | 0.93743 | ENSMUSG( | NA | 63.43908 | 0.068463 | 0.798896 | 0.97681 |
| ENSMUSG( | NA | 15.94446 | -0.31056 | 0.513168 | NA | ENSMUSG( | NA | 14.42832 | -0.10205 | 0.840984 | 0.9826 |
| ENSMUSG( | NA | 21.33344 | 0.041776 | 0.921253 | 0.990436 | ENSMUSG( | NA | 14.66248 | 0.769877 | 0.212326 | 0.864412 |
| ENSMUSG( | NA | 10.17275 | -0.25441 | 0.662904 | NA | ENSMUSG( | NA | 16.0134 | 0.445169 | 0.376911 | 0.915593 |
| ENSMUSG( | NA | 25.59289 | -0.38448 | 0.372546 | 0.798769 | ENSMUSG( | NA | 9.009568 | -0.1685 | 0.791323 | NA |
| ENSMUSG( | NA | 35.01655 | 0.326526 | 0.348248 | 0.78132 | ENSMUSG( | NA | 9.0183 | -0.14348 | 0.837089 | NA |
| ENSMUSG( | NA | 194.6721 | 0.077963 | 0.679397 | 0.930983 | ENSMUSG( | NA | 21.80934 | -0.0449 | 0.916224 | 0.992529 |
| ENSMUSG( | NA | 9.982853 | 0.207416 | 0.735769 | NA | ENSMUSG( | NA | 104.7334 | 0.06652 | 0.80043 | 0.97681 |
| ENSMUSG( | NA | 10.43138 | -0.44244 | 0.490465 | NA | ENSMUSG( | NA | 24.31011 | -0.62715 | 0.135518 | 0.825163 |
| ENSMUSG( | NA | 53.34677 | 0.528036 | 0.085502 | 0.450845 | ENSMUSG( | NA | 32.82453 | -0.16093 | 0.694132 | 0.960867 |
| ENSMUSG( | NA | 96.89703 | -0.21831 | 0.371955 | 0.798768 | ENSMUSG( | NA | 31.86647 | -0.05751 | 0.879663 | 0.989644 |
| ENSMUSG( | NA | 12.68135 | -0.04993 | 0.932988 | NA | ENSMUSG( | NA | 9.241831 | -0.07528 | 0.905306 | NA |
| ENSMUSG( | NA | 615.0416 | 0.475308 | 8.10E-05 | 0.005593 | ENSMUSG( | NA | 13.8115 | -0.10262 | 0.868365 | 0.988031 |
| ENSMUSG( | NA | 26.49891 | 0.129923 | 0.717978 | 0.941697 | ENSMUSG( | NA | 29.34019 | 0.472852 | 0.2237 | 0.867404 |
| ENSMUSG( | NA | 9.680707 | 0.966001 | 0.178788 | NA | ENSMUSG( | NA | 25.87644 | 0.438306 | 0.344159 | 0.910256 |
| ENSMUSG( | NA | 32.82738 | 0.19761 | 0.569326 | 0.893041 | ENSMUSG( | NA | 22.71431 | -0.15967 | 0.718436 | 0.966633 |
| ENSMUSG( | NA | 1151.308 | 0.379127 | 0.000858 | 0.029429 | ENSMUSG( | NA | 98.93245 | 0.201397 | 0.403347 | 0.917257 |
| ENSMUSG( | NA | 119.6142 | 0.16636 | 0.406097 | 0.816525 | ENSMUSG( | NA | 57.36331 | -0.02326 | 0.933471 | 0.99306 |
| ENSMUSG( | NA | 24.36688 | 0.249976 | 0.537781 | 0.882548 | ENSMUSG( | NA | 83.04991 | 0.189182 | 0.471119 | 0.923329 |
| ENSMUSG( | NA | 12.47714 | -0.26013 | 0.63068 | NA | ENSMUSG( | NA | 10.61445 | -0.40873 | 0.490329 | NA |
| ENSMUSG( | NA | 35.23194 | -0.21117 | 0.627009 | 0.912825 | ENSMUSG( | NA | 8.552823 | -0.2877 | 0.684862 | NA |
| ENSMUSG( | NA | 19.45997 | -0.09247 | 0.825755 | NA | ENSMUSG( | NA | 41.05876 | 0.074646 | 0.831475 | 0.980611 |
| ENSMUSG( | NA | 12.23751 | -0.19175 | 0.747189 | NA | ENSMUSG( | NA | 71.3462 | 0.094728 | 0.77248 | 0.974454 |
| ENSMUSG( | NA | 47.35393 | 0.018154 | 0.95102 | 0.993821 | ENSMUSG( | NA | 11.38654 | -0.21205 | 0.706194 | NA |
| ENSMUSG( | NA | 3595.657 | -0.00014 | 0.999007 | 0.99989 | ENSMUSG( | NA | 16.83634 | 0.581325 | 0.280153 | 0.892356 |
| ENSMUSG( | NA | 26.05562 | -0.18918 | 0.621868 | 0.911432 | ENSMUSG( | NA | 193.7757 | 0.393055 | 0.045139 | 0.693959 |
| ENSMUSG( | NA | 28.62058 | -0.26022 | 0.453978 | 0.836798 | ENSMUSG( | NA | 17.00633 | -0.14404 | 0.771138 | 0.974236 |
| ENSMUSG( | NA | 31.99057 | 0.228252 | 0.520394 | 0.875186 | ENSMUSG( | NA | 12.60314 | -0.36329 | 0.511853 | 0.930005 |
| ENSMUSG( | NA | 12.55955 | -0.17457 | 0.748619 | NA | ENSMUSG( | NA | 12.73277 | 0.138252 | 0.826341 | 0.979829 |
| ENSMUSG( | NA | 33.78757 | 0.204374 | 0.54027 | 0.88315 | ENSMUSG( | NA | 11.01533 | 0.18161 | 0.768943 | NA |
| ENSMUSG( | NA | 13.54113 | 0.238128 | 0.637096 | NA | ENSMUSG( | NA | 31.92241 | 0.092897 | 0.817859 | 0.979198 |
| ENSMUSG( | NA | 17.45058 | -0.24188 | 0.59774 | NA | ENSMUSG( | NA | 80.48113 | -0.24301 | 0.320178 | 0.903876 |
| ENSMUSG( | NA | 10.25014 | 0.120231 | 0.836407 | NA | ENSMUSG( | NA | 34.61374 | -0.62542 | 0.08152 | 0.77681 |
| ENSMUSG( | NA | 68.03879 | -0.07971 | 0.728351 | 0.944393 | ENSMUSG( | NA | 32.85344 | 0.359273 | 0.419562 | 0.917257 |
| ENSMUSG( | NA | 11.86201 | 0.094189 | 0.857244 | NA | ENSMUSG( | NA | 21.1249 | 0.055815 | 0.912414 | 0.992225 |
| ENSMUSG( | NA | 132.8003 | 0.159227 | 0.447885 | 0.834051 | ENSMUSG( | NA | 12.81547 | 0.040721 | 0.94132 | 0.994578 |
| ENSMUSG( | NA | 11.66659 | -0.13715 | 0.812085 | NA | ENSMUSG( | NA | 14.78468 | -0.71408 | 0.1627 | 0.837724 |
| ENSMUSG( | NA | 12.08747 | 0.635304 | 0.274923 | NA | ENSMUSG( | NA | 31.72547 | 0.605071 | 0.101939 | 0.801827 |
| ENSMUSG( | NA | 22.84353 | 0.00691 | 0.987478 | 0.998796 | ENSMUSG( | NA | 20.66721 | 1.202052 | 0.009018 | 0.397903 |
| ENSMUSG( | NA | 206.4501 | -0.07748 | 0.607704 | 0.906878 | ENSMUSG( | NA | 9.073139 | -1.1058 | 0.111081 | NA |
| ENSMUSG( | NA | 21.20829 | -0.73627 | 0.127561 | 0.535393 | ENSMUSG( | NA | 36.58748 | -0.39156 | 0.267652 | 0.886134 |
| ENSMUSG( | NA | 30.69562 | 0.075955 | 0.830741 | 0.969798 | ENSMUSG( | NA | 13.04237 | -0.41327 | 0.492193 | 0.927315 |
| ENSMUSG( | NA | 10.40982 | -0.77204 | 0.196297 | NA | ENSMUSG( | NA | 27.29517 | -0.40623 | 0.297841 | 0.898538 |
| ENSMUSG( | NA | 135.8441 | 0.239203 | 0.20101 | 0.641303 | ENSMUSG( | NA | 10.59247 | 0.787082 | 0.211209 | NA |
| ENSMUSG( | NA | 208.3395 | 0.291637 | 0.071993 | 0.414057 | ENSMUSG( | NA | 16.37179 | 0.561759 | 0.288245 | 0.894794 |
| ENSMUSG( | NA | 18.71752 | 0.33273 | 0.449623 | NA | ENSMUSG( | NA | 9.79098 | -1.44552 | 0.044978 | NA |
| ENSMUSG( | NA | 70.32668 | 0.45406 | 0.059584 | 0.377373 | ENSMUSG( | NA | 13.28365 | -0.50232 | 0.352456 | 0.912093 |
| ENSMUSG( | NA | 19.33647 | -0.31249 | 0.471477 | NA | ENSMUSG( | NA | 13.44667 | 0.164958 | 0.761742 | 0.973051 |
| ENSMUSG( | NA | 15.42733 | -0.4973 | 0.304812 | NA | ENSMUSG( | NA | 10.86852 | 0.339812 | 0.570939 | NA |
| ENSMUSG( | NA | 17.31999 | -0.1922 | 0.681193 | NA | ENSMUSG( | NA | 15.63915 | -0.08919 | 0.856213 | 0.985884 |
| ENSMUSG( | NA | 10.13384 | 0.303391 | 0.620332 | NA | ENSMUSG( | NA | 25.76506 | 0.188989 | 0.626987 | 0.953234 |
| ENSMUSG( | NA | 18.35663 | 0.478264 | 0.272559 | NA | ENSMUSG( | NA | 443.5589 | -0.51063 | 0.000111 | 0.022317 |
| ENSMUSG( | NA | 154.2056 | 0.148447 | 0.449776 | 0.835283 | ENSMUSG( | NA | 13.01293 | 0.051204 | 0.929621 | 0.99306 |
| ENSMUSG( | NA | 180.9294 | -0.05947 | 0.74348 | 0.948478 | ENSMUSG( | NA | 116.3242 | -0.4197 | 0.037402 | 0.673514 |
| ENSMUSG( | NA | 36.24625 | 0.466695 | 0.171126 | 0.605644 | ENSMUSG( | NA | 33.18363 | -0.48997 | 0.19574 | 0.861935 |
| ENSMUSG( | NA | 34.31262 | 0.123161 | 0.712899 | 0.94066 | ENSMUSG( | NA | 15.12509 | 0.080155 | 0.876765 | 0.989644 |
| ENSMUSG( | NA | 52.91958 | -0.03348 | 0.904366 | 0.986246 | ENSMUSG( | NA | 41.5933 | -0.97047 | 0.003161 | 0.224167 |
| ENSMUSG( | NA | 94.43667 | 0.077466 | 0.719685 | 0.941986 | ENSMUSG( | NA | 48.52299 | -0.41283 | 0.154732 | 0.837724 |
| ENSMUSG( | NA | 16.87953 | 0.088848 | 0.842309 | NA | ENSMUSG( | NA | 50.31908 | -0.08044 | 0.784428 | 0.976196 |
| ENSMUSG( | NA | 209.3189 | 0.143703 | 0.341115 | 0.778828 | ENSMUSG( | NA | 68.11709 | -0.37062 | 0.149374 | 0.836241 |
| ENSMUSG( | NA | 13.26196 | 0.083893 | 0.875889 | NA | ENSMUSG( | NA | 10.2728 | 0.01786 | 0.978055 | NA |
| ENSMUSG( | NA | 23.54898 | -0.6855 | 0.085119 | 0.44957 | ENSMUSG( | NA | 23.47814 | -1.16184 | 0.010247 | 0.412622 |
| ENSMUSG( | NA | 22.48052 | -0.42832 | 0.331582 | 0.77519 | ENSMUSG( | NA | 19.56452 | -0.42698 | 0.344083 | 0.910256 |
| ENSMUSG( | NA | 17.04579 | -0.00765 | 0.986788 | NA | ENSMUSG( | NA | 62.43027 | -0.28719 | 0.301039 | 0.898538 |
| ENSMUSG( | NA | 14.673 | 0.605615 | 0.329618 | NA | ENSMUSG( | NA | 608.6526 | -0.14761 | 0.258411 | 0.883971 |
| ENSMUSG( | NA | 86.64812 | -0.43238 | 0.07086 | 0.410627 | ENSMUSG( | NA | 97.51986 | -0.15714 | 0.482136 | 0.924578 |
| ENSMUSG( | NA | 110.0849 | -0.11162 | 0.626703 | 0.912825 | ENSMUSG( | NA | 22.87744 | -0.86301 | 0.067465 | 0.75212 |
| ENSMUSG( | NA | 10.30293 | 0.24933 | 0.693014 | NA | ENSMUSG( | NA | 937.0337 | -0.1594 | 0.225317 | 0.868468 |
| ENSMUSG( | NA | 30.40943 | -0.67052 | 0.061461 | 0.382832 | ENSMUSG( | NA | 21.70702 | 0.244553 | 0.559397 | 0.939768 |
| ENSMUSG( | NA | 41.87922 | 0.214583 | 0.468773 | 0.845636 | ENSMUSG( | NA | 13.35731 | -0.11344 | 0.859643 | 0.985947 |
| ENSMUSG( | NA | 11.27974 | -0.67689 | 0.211719 | NA | ENSMUSG( | NA | 21.14309 | -0.22377 | 0.657172 | 0.955644 |
| ENSMUSG( | NA | 11.1153 | -0.30031 | 0.603223 | NA | ENSMUSG( | NA | 45.80345 | 0.324875 | 0.359203 | 0.914061 |
| ENSMUSG( | NA | 50.21447 | -0.00258 | 0.992606 | 0.999327 | ENSMUSG( | NA | 9.409032 | -0.53298 | 0.439077 | NA |
| ENSMUSG( | NA | 17.02956 | 0.220353 | 0.672004 | NA | ENSMUSG( | NA | 149.9328 | -0.30994 | 0.111043 | 0.807885 |
| ENSMUSG( | NA | 11.73167 | -0.59061 | 0.296235 | NA | ENSMUSG( | NA | 14.63716 | -0.55162 | 0.296507 | 0.898538 |
| ENSMUSG( | NA | 48.80519 | -0.0458 | 0.887526 | 0.983411 | ENSMUSG( | NA | 820.7142 | -0.16831 | 0.228559 | 0.868507 |

| | | | | | |
|---|---|---|---|---|---|
| ENSMUSG( | 154.1923 | 0.086597 | 0.626239 | 0.912633 | NA |
| ENSMUSG( | 156.8959 | -0.05678 | 0.738287 | 0.947329 | NA |
| ENSMUSG( | 13.12324 | -0.33913 | 0.511231 | NA | NA |
| ENSMUSG( | 9.116823 | -0.76551 | 0.205059 | NA | NA |
| ENSMUSG( | 11.22621 | 0.082708 | 0.885227 | NA | NA |
| ENSMUSG( | 14.84113 | -0.43275 | 0.370959 | NA | NA |
| ENSMUSG( | 224.4657 | 0.10002 | 0.508222 | 0.867999 | NA |
| ENSMUSG( | 44.8776 | 0.332447 | 0.259555 | 0.709948 | NA |
| ENSMUSG( | 17.42207 | 0.570174 | 0.231125 | NA | NA |
| ENSMUSG( | 21.81153 | -0.17615 | 0.6592 | 0.924506 | NA |
| ENSMUSG( | 19.65244 | -0.34767 | 0.414844 | NA | NA |
| ENSMUSG( | 19.11628 | 0.314736 | 0.46126 | NA | NA |
| ENSMUSG( | 14.92128 | -0.21135 | 0.655704 | NA | NA |
| ENSMUSG( | 22.0339 | 0.20494 | 0.618101 | 0.910929 | NA |
| ENSMUSG( | 11.97164 | -0.54208 | 0.332706 | NA | NA |
| ENSMUSG( | 33.70467 | -0.39016 | 0.22726 | 0.674223 | NA |
| ENSMUSG( | 11.56565 | 0.309912 | 0.590657 | NA | NA |
| ENSMUSG( | 29.99621 | 0.601825 | 0.078163 | 0.432307 | NA |
| ENSMUSG( | 86.06314 | -0.12357 | 0.599743 | 0.903906 | NA |
| ENSMUSG( | 35.83026 | -0.41358 | 0.206337 | 0.648436 | NA |
| ENSMUSG( | 39.04376 | -0.28795 | 0.346542 | 0.779166 | NA |
| ENSMUSG( | 23.42657 | 0.498401 | 0.213454 | 0.65774 | NA |
| ENSMUSG( | 6794.406 | 0.024712 | 0.898335 | 0.985633 | NA |
| ENSMUSG( | 45.27922 | -0.36024 | 0.218187 | 0.663729 | NA |
| ENSMUSG( | 11.63299 | -0.48847 | 0.388819 | NA | NA |
| ENSMUSG( | 79.20657 | -0.01168 | 0.959875 | 0.993991 | NA |
| ENSMUSG( | 10217.33 | 0.014511 | 0.90432 | 0.986246 | NA |
| ENSMUSG( | 80.78156 | 0.028878 | 0.913694 | 0.988483 | NA |
| ENSMUSG( | 56.86008 | 0.063511 | 0.80981 | 0.96372 | NA |
| ENSMUSG( | 10.72797 | 0.590779 | 0.314819 | NA | NA |
| ENSMUSG( | 45.57091 | 0.2484 | 0.37767 | 0.801561 | NA |
| ENSMUSG( | 67.96493 | -0.06235 | 0.804031 | 0.962169 | NA |
| ENSMUSG( | 21.29732 | 0.08022 | 0.856667 | 0.974154 | NA |
| ENSMUSG( | 25.01828 | 0.396249 | 0.328964 | 0.773233 | NA |
| ENSMUSG( | 72.96072 | -0.13946 | 0.567133 | 0.891644 | NA |
| ENSMUSG( | 18.20392 | -0.15242 | 0.725399 | NA | NA |
| ENSMUSG( | 30.51103 | -0.2897 | 0.56561 | 0.890871 | NA |
| ENSMUSG( | 735.0519 | -0.29721 | 0.413587 | 0.819895 | NA |
| ENSMUSG( | 9.098038 | 0.552262 | 0.371649 | NA | NA |
| ENSMUSG( | 7358.256 | 0.003288 | 0.986157 | 0.998326 | NA |
| ENSMUSG( | 14.75215 | -0.46437 | 0.380826 | NA | NA |
| ENSMUSG( | 14.76508 | 0.576687 | 0.255182 | NA | NA |
| ENSMUSG( | 14.20962 | -0.74744 | 0.226229 | NA | NA |
| ENSMUSG( | 8.045937 | 0.322694 | 0.621092 | NA | NA |
| ENSMUSG( | 13.0515 | 1.046068 | 0.045247 | NA | NA |
| ENSMUSG( | 1821.191 | 0.677539 | 1.51E-13 | 1.53E-10 | NA |
| ENSMUSG( | 20.05733 | -0.06897 | 0.869721 | NA | NA |
| ENSMUSG( | 37.76337 | -0.0836 | 0.831421 | 0.969889 | NA |
| ENSMUSG( | 33.81982 | 0.285706 | 0.398781 | 0.812878 | NA |
| ENSMUSG( | 10.63275 | -0.42199 | 0.481279 | NA | NA |
| ENSMUSG( | 19.05228 | 1.481638 | 0.021218 | NA | NA |
| ENSMUSG( | 72.24839 | -0.23993 | 0.418879 | 0.821966 | NA |
| ENSMUSG( | 19.51734 | -0.82671 | 0.05021 | NA | NA |
| ENSMUSG( | 16.24887 | -0.03712 | 0.942393 | NA | NA |
| ENSMUSG( | 24.59505 | 1.957525 | 0.000851 | 0.029239 | NA |
| ENSMUSG( | 42.02908 | -0.23894 | 0.467789 | 0.845032 | NA |
| ENSMUSG( | 19.1835 | -0.23918 | 0.586713 | NA | NA |
| ENSMUSG( | 10.77107 | -0.19964 | 0.7535 | NA | NA |
| ENSMUSG( | 243.798 | 0.200793 | 0.308407 | 0.756439 | NA |
| ENSMUSG( | 15.26055 | 0.361105 | 0.469015 | NA | NA |
| ENSMUSG( | 31.55691 | -0.39032 | 0.314852 | 0.762059 | NA |
| ENSMUSG( | 23.82807 | -0.19164 | 0.622895 | 0.911589 | NA |
| ENSMUSG( | 31.75466 | 0.323414 | 0.37333 | 0.800133 | NA |
| ENSMUSG( | 19.75427 | 0.244875 | 0.588574 | NA | NA |
| ENSMUSG( | 20.57171 | 0.315776 | 0.507801 | NA | NA |
| ENSMUSG( | 44.73151 | -0.72085 | 0.044753 | 0.324745 | NA |
| ENSMUSG( | 12.20484 | -0.85856 | 0.128448 | NA | NA |
| ENSMUSG( | 52.3108 | 0.479295 | 0.084249 | 0.447434 | NA |
| ENSMUSG( | 9.668854 | -0.20002 | 0.735615 | NA | NA |
| ENSMUSG( | 130.6097 | -0.19038 | 0.377954 | 0.801561 | NA |
| ENSMUSG( | 40.19096 | -0.30999 | 0.333643 | 0.775602 | NA |
| ENSMUSG( | 30.23584 | 0.243421 | 0.469568 | 0.846167 | NA |
| ENSMUSG( | 260.5226 | -0.0635 | 0.699483 | 0.937034 | NA |
| ENSMUSG( | 57.96506 | 0.185034 | 0.524183 | 0.87583 | NA |
| ENSMUSG( | 11.29934 | 0.247775 | 0.660754 | NA | NA |
| ENSMUSG( | 14.01709 | 0.430064 | 0.412527 | NA | NA |
| ENSMUSG( | 10.51297 | -0.18279 | 0.752727 | NA | NA |
| ENSMUSG( | 9.478482 | 0.150151 | 0.797886 | NA | NA |
| ENSMUSG( | 48.15224 | 0.094782 | 0.741229 | 0.948165 | NA |
| ENSMUSG( | 14.65796 | -0.98675 | 0.057415 | NA | NA |
| ENSMUSG( | 36.27897 | -0.09349 | 0.770723 | 0.954722 | NA |
| ENSMUSG( | 207.2546 | 0.022407 | 0.883271 | 0.982422 | NA |
| ENSMUSG( | 13.93578 | 0.183012 | 0.712605 | NA | NA |
| ENSMUSG( | 200.8166 | -0.08573 | 0.577178 | 0.895659 | NA |
| ENSMUSG( | 30.85462 | -0.15151 | 0.720156 | 0.94201 | NA |
| ENSMUSG( | 20.40427 | 0.134035 | 0.776855 | NA | NA |
| ENSMUSG( | 22.05542 | -0.30984 | 0.513474 | 0.871762 | NA |
| ENSMUSG( | 36.61892 | -0.45116 | 0.151003 | 0.576088 | NA |
| ENSMUSG( | 122.184 | 0.04798 | 0.809865 | 0.96372 | NA |
| ENSMUSG( | 17.67093 | 0.489208 | 0.350125 | NA | NA |
| ENSMUSG( | 65.94393 | 0.480934 | 0.050189 | 0.342492 | NA |
| ENSMUSG( | 8.765277 | 0.685393 | 0.287131 | NA | NA |
| ENSMUSG( | 193.9815 | 0.077978 | 0.724026 | 0.944096 | NA |
| ENSMUSG( | 26.83157 | 0.286842 | 0.42233 | 0.823508 | NA |
| ENSMUSG( | 337.1665 | 0.103516 | 0.447261 | 0.834051 | NA |
| ENSMUSG( | 101.0255 | -0.07562 | 0.734283 | 0.945468 | NA |
| ENSMUSG( | 17.99367 | -0.15924 | 0.746826 | NA | NA |
| ENSMUSG( | 81.46233 | 0.451571 | 0.045335 | 0.326817 | NA |
| ENSMUSG( | 11.16202 | -1.42217 | 0.01514 | NA | NA |
| ENSMUSG( | 12.31128 | -0.21168 | 0.709221 | NA | NA |
| ENSMUSG( | 48.33372 | -0.09884 | 0.732455 | 0.944956 | NA |
| ENSMUSG( | 7.246341 | 0.514298 | 0.477634 | NA | NA |
| ENSMUSG( | 17.46735 | 0.447713 | 0.377796 | NA | NA |
| ENSMUSG( | 34.92991 | 0.358159 | 0.283048 | 0.73186 | NA |
| ENSMUSG( | 45.66925 | -0.05322 | 0.848252 | 0.972374 | NA |
| ENSMUSG( | 57.23953 | -0.19242 | 0.459034 | 0.840244 | NA |
| ENSMUSG( | 22.78073 | -0.23795 | 0.541738 | 0.883182 | NA |
| ENSMUSG( | 14.03383 | -0.21722 | 0.666148 | NA | NA |
| ENSMUSG( | 117.0315 | 0.707799 | 0.000115 | 0.007087 | NA |
| ENSMUSG( | 31.98245 | 0.104389 | 0.766398 | 0.954173 | NA |
| ENSMUSG( | 18.57225 | -0.28289 | 0.503225 | NA | NA |
| ENSMUSG( | 46.09718 | -0.03076 | 0.913707 | 0.988483 | NA |
| ENSMUSG( | 14.85187 | -0.75113 | 0.165206 | NA | NA |
| ENSMUSG( | 16.69139 | -0.249 | 0.635477 | NA | NA |
| ENSMUSG( | 80.98966 | 0.057706 | 0.808097 | 0.963419 | NA |
| ENSMUSG( | 16.70142 | 0.10531 | 0.819901 | NA | NA |
| ENSMUSG( | 16.64718 | 0.434216 | 0.374444 | NA | NA |
| ENSMUSG( | 14.72092 | 0.467797 | 0.396917 | NA | NA |
| ENSMUSG( | 39.64057 | 0.096583 | 0.786711 | 0.958856 | NA |
| ENSMUSG( | 39.14605 | 0.220116 | 0.51732 | 0.873295 | NA |
| ENSMUSG( | 243.4591 | -0.15422 | 0.365139 | 0.794085 | NA |
| ENSMUSG( | 46.42893 | -0.2999 | 0.285334 | 0.734038 | NA |
| ENSMUSG( | 12.56102 | -0.13582 | 0.798769 | NA | NA |
| ENSMUSG( | 23.05417 | 0.114769 | 0.771583 | 0.954769 | NA |
| ENSMUSG( | 14.10127 | 0.354522 | 0.478475 | NA | NA |
| ENSMUSG( | 45.30503 | 0.207359 | 0.467577 | 0.844819 | NA |
| ENSMUSG( | 28.10474 | 0.067402 | 0.848655 | 0.972374 | NA |
| ENSMUSG( | 10.74624 | -0.26198 | 0.687007 | NA | NA |
| ENSMUSG( | 93.18414 | -0.0628 | 0.769013 | 0.954674 | NA |
| ENSMUSG( | 70.70774 | 0.169307 | 0.496873 | 0.861754 | NA |
| ENSMUSG( | 8.898683 | -0.49948 | 0.425223 | NA | NA |
| ENSMUSG( | 31.53096 | 0.016586 | 0.960267 | 0.993991 | NA |
| ENSMUSG( | 31.00805 | 0.696488 | 0.041344 | 0.314189 | NA |
| ENSMUSG( | 62.2905 | -0.11041 | 0.689401 | 0.933609 | NA |
| ENSMUSG( | 39.05423 | -0.25349 | 0.442636 | 0.83278 | NA |
| ENSMUSG( | 41.87132 | -0.56154 | 0.068073 | 0.404152 | NA |
| ENSMUSG( | 25.35404 | 0.182737 | 0.673283 | 0.929007 | NA |
| ENSMUSG( | 15.14781 | -0.54316 | 0.341051 | NA | NA |
| ENSMUSG( | 76.86238 | -0.06853 | 0.803215 | 0.962169 | NA |
| ENSMUSG( | 78.67447 | 0.257529 | 0.315757 | 0.762707 | NA |
| ENSMUSG( | 15.4818 | -0.46006 | 0.366625 | NA | NA |
| ENSMUSG( | 77.75387 | -0.06543 | 0.786214 | 0.958774 | NA |
| ENSMUSG( | 18.78135 | -0.24916 | 0.582411 | NA | NA |
| ENSMUSG( | 48.05075 | 0.251808 | 0.421894 | 0.823481 | NA |
| ENSMUSG( | 8.707826 | -0.01801 | 0.977966 | NA | NA |
| ENSMUSG( | 32.42198 | -0.33845 | 0.339332 | 0.777616 | NA |
| ENSMUSG( | 49.19992 | -0.21369 | 0.533447 | 0.880095 | NA |
| ENSMUSG( | 19.90778 | 0.257964 | 0.563547 | NA | NA |
| ENSMUSG( | 82.32924 | 0.050636 | 0.819028 | 0.965888 | NA |
| ENSMUSG( | 15.82525 | -0.42673 | 0.359956 | NA | NA |
| ENSMUSG( | 17.0724 | -0.06352 | 0.886176 | NA | NA |
| ENSMUSG( | 12.78636 | -0.00581 | 0.990919 | NA | NA |
| ENSMUSG( | 36.21843 | 0.306319 | 0.366242 | 0.794603 | NA |
| ENSMUSG( | 94.22397 | -0.08481 | 0.701281 | 0.93743 | NA |
| ENSMUSG( | 15.94446 | -0.31056 | 0.513168 | NA | NA |
| ENSMUSG( | 21.33344 | 0.041776 | 0.921253 | 0.990436 | NA |
| ENSMUSG( | 10.17275 | -0.25441 | 0.662904 | NA | NA |
| ENSMUSG( | 25.59289 | -0.38448 | 0.372546 | 0.798769 | NA |

| | | | | | |
|---|---|---|---|---|---|
| ENSMUSG( | 13.60583 | 0.181992 | 0.776619 | 0.975717 | NA |
| ENSMUSG( | 36.00081 | -0.07407 | 0.837954 | 0.981376 | NA |
| ENSMUSG( | 12.48012 | -0.3022 | 0.585841 | 0.944975 | NA |
| ENSMUSG( | 29.50765 | -0.18853 | 0.67289 | 0.95694 | NA |
| ENSMUSG( | 51.2063 | -0.17686 | 0.566024 | 0.94151 | NA |
| ENSMUSG( | 7.812057 | -0.95526 | 0.182966 | NA | NA |
| ENSMUSG( | 11.19749 | 0.186959 | 0.757132 | NA | NA |
| ENSMUSG( | 36.91534 | 0.063039 | 0.878668 | 0.989644 | NA |
| ENSMUSG( | 76.24283 | -0.48 | 0.058057 | 0.728256 | NA |
| ENSMUSG( | 49.20569 | -0.1862 | 0.536609 | 0.937108 | NA |
| ENSMUSG( | 1531.855 | -0.02571 | 0.82277 | 0.979198 | NA |
| ENSMUSG( | 50.70868 | -0.29916 | 0.319111 | 0.903431 | NA |
| ENSMUSG( | 142.6918 | -0.33294 | 0.085012 | 0.779799 | NA |
| ENSMUSG( | 18.988 | -0.58771 | 0.231557 | 0.868507 | NA |
| ENSMUSG( | 8.799157 | -1.1457 | 0.102972 | NA | NA |
| ENSMUSG( | 33.89968 | -0.42293 | 0.301061 | 0.898538 | NA |
| ENSMUSG( | 211.1852 | -0.12189 | 0.505739 | 0.927825 | NA |
| ENSMUSG( | 23.76254 | -0.41494 | 0.300652 | 0.898538 | NA |
| ENSMUSG( | 58.67262 | 0.247147 | 0.392716 | 0.917257 | NA |
| ENSMUSG( | 60.61189 | 0.274409 | 0.44549 | 0.921734 | NA |
| ENSMUSG( | 9.071548 | -0.87951 | 0.18721 | NA | NA |
| ENSMUSG( | 107.061 | -0.44822 | 0.050209 | 0.708648 | NA |
| ENSMUSG( | 43.90004 | 0.238805 | 0.4971 | 0.927825 | NA |
| ENSMUSG( | 63.10346 | -0.03152 | 0.905389 | 0.991089 | NA |
| ENSMUSG( | 22.2941 | -0.18575 | 0.661771 | 0.956205 | NA |
| ENSMUSG( | 17.02557 | 0.780115 | 0.188023 | 0.85698 | NA |
| ENSMUSG( | 19.63278 | 0.311507 | 0.509457 | 0.929089 | NA |
| ENSMUSG( | 24.41769 | -0.14414 | 0.725502 | 0.966887 | NA |
| ENSMUSG( | 430.815 | -0.04787 | 0.746764 | 0.969763 | NA |
| ENSMUSG( | 23.31853 | 0.643273 | 0.121329 | 0.813384 | NA |
| ENSMUSG( | 18.25554 | 0.19206 | 0.677544 | 0.957106 | NA |
| ENSMUSG( | 30.59224 | 0.76954 | 0.064516 | 0.737935 | NA |
| ENSMUSG( | 103.6992 | -0.26489 | 0.282007 | 0.892412 | NA |
| ENSMUSG( | 359.5373 | 0.076144 | 0.586911 | 0.944975 | NA |
| ENSMUSG( | 35.90422 | 0.360013 | 0.328765 | 0.906727 | NA |
| ENSMUSG( | 10.89253 | 0.329702 | 0.598806 | NA | NA |
| ENSMUSG( | 46.40886 | 0.488066 | 0.223269 | 0.867404 | NA |
| ENSMUSG( | 10.10648 | -0.04006 | 0.94958 | NA | NA |
| ENSMUSG( | 16.28842 | -0.4447 | 0.372146 | 0.915491 | NA |
| ENSMUSG( | 358.0709 | 0.036979 | 0.823859 | 0.979198 | NA |
| ENSMUSG( | 261.1379 | 0.100212 | 0.582835 | 0.944266 | NA |
| ENSMUSG( | 17.407 | 0.209955 | 0.679736 | 0.957966 | NA |
| ENSMUSG( | 26.00289 | 0.255787 | 0.623173 | 0.953234 | NA |
| ENSMUSG( | 65.81461 | -0.07487 | 0.798801 | 0.97681 | NA |
| ENSMUSG( | 37.52039 | 0.599368 | 0.10843 | 0.804969 | NA |
| ENSMUSG( | 55.34167 | 0.194846 | 0.518835 | 0.930939 | NA |
| ENSMUSG( | 77.67433 | 0.175687 | 0.485344 | 0.92541 | NA |
| ENSMUSG( | 37.31813 | -0.57045 | 0.098218 | 0.798438 | NA |
| ENSMUSG( | 10.22884 | 0.030096 | 0.965117 | NA | NA |
| ENSMUSG( | 142.2206 | 0.111619 | 0.609485 | 0.950397 | NA |
| ENSMUSG( | 27.73471 | -0.04271 | 0.91416 | 0.992407 | NA |
| ENSMUSG( | 14.08441 | -0.47091 | 0.36532 | 0.914449 | NA |
| ENSMUSG( | 9.204935 | 0.103939 | 0.881973 | NA | NA |
| ENSMUSG( | 8.82797 | -0.58635 | 0.382039 | NA | NA |
| ENSMUSG( | 63.43908 | 0.068463 | 0.798896 | 0.97681 | NA |
| ENSMUSG( | 14.42832 | -0.10205 | 0.840984 | 0.9826 | NA |
| ENSMUSG( | 14.66248 | 0.769877 | 0.212326 | 0.864412 | NA |
| ENSMUSG( | 16.0134 | 0.445169 | 0.376911 | 0.915593 | NA |
| ENSMUSG( | 9.009568 | -0.1685 | 0.791323 | NA | NA |
| ENSMUSG( | 9.0183 | -0.14348 | 0.837089 | NA | NA |
| ENSMUSG( | 21.80934 | -0.0449 | 0.916224 | 0.992529 | NA |
| ENSMUSG( | 104.7334 | 0.06652 | 0.80043 | 0.97681 | NA |
| ENSMUSG( | 24.31011 | -0.62715 | 0.135518 | 0.825163 | NA |
| ENSMUSG( | 32.82453 | -0.16093 | 0.694132 | 0.960867 | NA |
| ENSMUSG( | 31.86647 | -0.05751 | 0.879663 | 0.989644 | NA |
| ENSMUSG( | 9.241831 | -0.07528 | 0.905306 | NA | NA |
| ENSMUSG( | 13.8115 | -0.10262 | 0.868365 | 0.988031 | NA |
| ENSMUSG( | 29.34019 | 0.472852 | 0.2237 | 0.867404 | NA |
| ENSMUSG( | 25.87644 | 0.438306 | 0.344159 | 0.910256 | NA |
| ENSMUSG( | 93.11214 | 10.05909 | 4.36E-17 | 7.63E-14 | NA |
| ENSMUSG( | 22.71431 | -0.15967 | 0.718436 | 0.966633 | NA |
| ENSMUSG( | 98.93245 | 0.201397 | 0.403347 | 0.917257 | NA |
| ENSMUSG( | 57.36331 | -0.02326 | 0.933471 | 0.99306 | NA |
| ENSMUSG( | 83.04991 | 0.189182 | 0.471119 | 0.923329 | NA |
| ENSMUSG( | 10.61445 | -0.40873 | 0.490329 | NA | NA |
| ENSMUSG( | 8.552823 | -0.2877 | 0.684862 | NA | NA |
| ENSMUSG( | 41.05876 | 0.074646 | 0.831475 | 0.980611 | NA |
| ENSMUSG( | 71.3462 | 0.094728 | 0.77248 | 0.974454 | NA |
| ENSMUSG( | 11.38654 | -0.21205 | 0.706194 | NA | NA |
| ENSMUSG( | 16.83634 | 0.581325 | 0.280153 | 0.892356 | NA |
| ENSMUSG( | 193.7757 | 0.393055 | 0.045139 | 0.693959 | NA |
| ENSMUSG( | 17.00633 | -0.14404 | 0.771138 | 0.974236 | NA |
| ENSMUSG( | 12.60314 | -0.36329 | 0.511853 | 0.930005 | NA |
| ENSMUSG( | 8.613873 | -0.21077 | 0.763281 | NA | NA |
| ENSMUSG( | 12.73277 | 0.138252 | 0.826341 | 0.979829 | NA |
| ENSMUSG( | 11.01533 | 0.18161 | 0.768943 | NA | NA |
| ENSMUSG( | 31.92241 | 0.092897 | 0.817859 | 0.979198 | NA |
| ENSMUSG( | 80.48113 | -0.24301 | 0.320178 | 0.903876 | NA |
| ENSMUSG( | 34.61374 | -0.62542 | 0.08152 | 0.77681 | NA |
| ENSMUSG( | 32.85344 | 0.359273 | 0.419562 | 0.917257 | NA |
| ENSMUSG( | 21.1249 | 0.055815 | 0.912414 | 0.992225 | NA |
| ENSMUSG( | 12.81547 | 0.040721 | 0.94132 | 0.994578 | NA |
| ENSMUSG( | 14.78468 | -0.71408 | 0.1627 | 0.837724 | NA |
| ENSMUSG( | 31.72547 | 0.605071 | 0.101939 | 0.801827 | NA |
| ENSMUSG( | 20.66721 | 1.202052 | 0.009018 | 0.397903 | NA |
| ENSMUSG( | 9.073139 | -1.1058 | 0.111081 | NA | NA |
| ENSMUSG( | 36.58748 | -0.39156 | 0.267652 | 0.886134 | NA |
| ENSMUSG( | 8.462924 | -0.36934 | 0.608876 | NA | NA |
| ENSMUSG( | 13.04237 | -0.41327 | 0.492193 | 0.927315 | NA |
| ENSMUSG( | 27.29517 | -0.40623 | 0.297841 | 0.898538 | NA |
| ENSMUSG( | 10.59247 | 0.787082 | 0.211209 | NA | NA |
| ENSMUSG( | 16.37179 | 0.561759 | 0.288245 | 0.894794 | NA |
| ENSMUSG( | 9.79098 | -1.44552 | 0.044978 | NA | NA |
| ENSMUSG( | 13.28365 | -0.50232 | 0.352456 | 0.912093 | NA |
| ENSMUSG( | 10.01365 | 0.181623 | 0.766143 | NA | NA |
| ENSMUSG( | 13.44667 | 0.164958 | 0.761742 | 0.973051 | NA |
| ENSMUSG( | 10.86852 | 0.339812 | 0.570939 | NA | NA |
| ENSMUSG( | 15.63915 | -0.08919 | 0.856213 | 0.985884 | NA |
| ENSMUSG( | 25.76506 | 0.188989 | 0.626987 | 0.953234 | NA |
| ENSMUSG( | 443.5589 | -0.51063 | 0.000111 | 0.022317 | NA |
| ENSMUSG( | 13.01293 | 0.051204 | 0.929621 | 0.99306 | NA |
| ENSMUSG( | 116.3242 | -0.4197 | 0.037402 | 0.673514 | NA |
| ENSMUSG( | 33.18363 | -0.48997 | 0.19574 | 0.861935 | NA |
| ENSMUSG( | 15.12509 | 0.080155 | 0.876765 | 0.989644 | NA |
| ENSMUSG( | 41.5933 | -0.97047 | 0.003161 | 0.224167 | NA |
| ENSMUSG( | 48.52299 | -0.41283 | 0.154732 | 0.837724 | NA |
| ENSMUSG( | 50.31908 | -0.08044 | 0.784428 | 0.976196 | NA |
| ENSMUSG( | 68.11709 | -0.37062 | 0.149374 | 0.836241 | NA |
| ENSMUSG( | 10.2728 | 0.01786 | 0.978055 | NA | NA |
| ENSMUSG( | 23.47814 | -1.16184 | 0.010247 | 0.412622 | NA |
| ENSMUSG( | 19.56452 | -0.42698 | 0.344083 | 0.910256 | NA |
| ENSMUSG( | 62.43027 | -0.28719 | 0.301039 | 0.898538 | NA |
| ENSMUSG( | 608.6526 | -0.14761 | 0.258411 | 0.883971 | NA |
| ENSMUSG( | 97.51986 | -0.15714 | 0.482136 | 0.924578 | NA |
| ENSMUSG( | 22.87744 | -0.86301 | 0.067465 | 0.75212 | NA |
| ENSMUSG( | 937.0337 | -0.1594 | 0.225317 | 0.868468 | NA |
| ENSMUSG( | 21.70702 | 0.244553 | 0.559397 | 0.939768 | NA |
| ENSMUSG( | 13.35731 | -0.11344 | 0.859643 | 0.985947 | NA |
| ENSMUSG( | 21.14309 | -0.22377 | 0.657172 | 0.955644 | NA |
| ENSMUSG( | 45.80345 | 0.324875 | 0.359203 | 0.914061 | NA |
| ENSMUSG( | 9.409032 | -0.53298 | 0.439077 | NA | NA |
| ENSMUSG( | 149.9328 | -0.30994 | 0.111043 | 0.807885 | NA |
| ENSMUSG( | 14.63716 | -0.55162 | 0.296507 | 0.898538 | NA |
| ENSMUSG( | 820.7142 | -0.16831 | 0.228559 | 0.868507 | NA |
| ENSMUSG( | 13.73934 | -0.77928 | 0.160413 | 0.837724 | NA |
| ENSMUSG( | 24.56649 | 0.080691 | 0.849652 | 0.985522 | NA |
| ENSMUSG( | 50.14318 | -0.77587 | 0.010421 | 0.41614 | NA |
| ENSMUSG( | 82.04419 | -1.06874 | 2.72E-05 | 0.007562 | NA |
| ENSMUSG( | 22.89299 | 0.175026 | 0.68315 | 0.958678 | NA |
| ENSMUSG( | 14.25615 | -0.44213 | 0.424447 | 0.917257 | NA |
| ENSMUSG( | 30.84733 | -0.7428 | 0.05304 | 0.71525 | NA |
| ENSMUSG( | 26.65058 | -0.11892 | 0.760364 | 0.972575 | NA |
| ENSMUSG( | 43.90757 | 0.306543 | 0.43052 | 0.917257 | NA |
| ENSMUSG( | 13.27873 | -0.09682 | 0.870864 | 0.98859 | NA |
| ENSMUSG( | 652.5631 | 0.304977 | 0.107692 | 0.804969 | NA |
| ENSMUSG( | 16.91529 | -0.67048 | 0.164083 | 0.838291 | NA |
| ENSMUSG( | 9.263401 | 0.728431 | 0.291344 | NA | NA |
| ENSMUSG( | 28.4069 | -0.11888 | 0.744493 | 0.96951 | NA |
| ENSMUSG( | 88.38825 | 0.381992 | 0.118995 | 0.813384 | NA |
| ENSMUSG( | 18.7882 | 0.479282 | 0.282625 | 0.892412 | NA |
| ENSMUSG( | 14.54567 | 0.243925 | 0.650603 | 0.95422 | NA |
| ENSMUSG( | 8.569284 | 0.520617 | 0.475794 | NA | NA |
| ENSMUSG( | 20.50153 | 0.018358 | 0.967053 | 0.995549 | NA |
| ENSMUSG( | 39.39731 | 0.301865 | 0.366582 | 0.914703 | NA |
| ENSMUSG( | 18.66855 | -0.00152 | 0.997397 | 0.999268 | NA |
| ENSMUSG( | 24.62916 | -0.5572 | 0.177886 | 0.847901 | NA |
| ENSMUSG( | 22.64807 | 0.267868 | 0.522618 | 0.932822 | NA |
| ENSMUSG( | 22.53623 | 0.340021 | 0.419247 | 0.917257 | NA |

| | | | | | | | | | | | |
|---|---|---|---|---|---|---|---|---|---|---|---|
| ENSMUSG( | NA | 31.33883 | -0.21817 | 0.529658 | 0.878883 | ENSMUSG( | NA | 13.73934 | -0.77928 | 0.160413 | 0.837724 |
| ENSMUSG( | NA | 24.52811 | 0.010686 | 0.978936 | 0.997428 | ENSMUSG( | NA | 24.56649 | 0.080691 | 0.849652 | 0.985522 |
| ENSMUSG( | NA | 26.7751 | -0.04028 | 0.911309 | 0.988483 | ENSMUSG( | NA | 50.14318 | -0.77587 | 0.010421 | 0.41614 |
| ENSMUSG( | NA | 23.49553 | -0.00573 | 0.988941 | 0.999111 | ENSMUSG( | NA | 82.04419 | -1.06874 | 2.72E-05 | 0.007562 |
| ENSMUSG( | NA | 128.1144 | 0.27452 | 0.191767 | 0.630811 | ENSMUSG( | NA | 22.89299 | 0.175026 | 0.68315 | 0.958678 |
| ENSMUSG( | NA | 65.0841 | -0.17898 | 0.456496 | 0.838832 | ENSMUSG( | NA | 14.25615 | -0.44213 | 0.424447 | 0.917257 |
| ENSMUSG( | NA | 10.37025 | -0.13247 | 0.815795 | NA | ENSMUSG( | NA | 30.84733 | -0.7428 | 0.05304 | 0.71525 |
| ENSMUSG( | NA | 25.92855 | 0.294741 | 0.465447 | 0.844064 | ENSMUSG( | NA | 26.65058 | -0.11892 | 0.760364 | 0.972575 |
| ENSMUSG( | NA | 27.28253 | -0.20243 | 0.605387 | 0.905764 | ENSMUSG( | NA | 43.90757 | 0.306543 | 0.43052 | 0.917257 |
| ENSMUSG( | NA | 14.45944 | 0.196153 | 0.713765 | NA | ENSMUSG( | NA | 13.27873 | -0.09682 | 0.870864 | 0.98859 |
| ENSMUSG( | NA | 21.91619 | 0.324662 | 0.408933 | 0.818173 | ENSMUSG( | NA | 652.5631 | 0.304977 | 0.107692 | 0.804969 |
| ENSMUSG( | NA | 54.87375 | -0.15827 | 0.546546 | 0.884974 | ENSMUSG( | NA | 16.91529 | -0.67048 | 0.164083 | 0.838291 |
| ENSMUSG( | NA | 14.93816 | -0.61938 | 0.226444 | NA | ENSMUSG( | NA | 9.263401 | 0.728431 | 0.291344 | NA |
| ENSMUSG( | NA | 9.247386 | 0.484929 | 0.470197 | NA | ENSMUSG( | NA | 28.4069 | -0.11888 | 0.744493 | 0.96951 |
| ENSMUSG( | NA | 13.26571 | -1.03849 | 0.041524 | NA | ENSMUSG( | NA | 88.38825 | 0.381992 | 0.118995 | 0.813384 |
| ENSMUSG( | NA | 73.55929 | -0.38996 | 0.149566 | 0.573617 | ENSMUSG( | NA | 18.7882 | 0.479282 | 0.282625 | 0.892412 |
| ENSMUSG( | NA | 18.55318 | 0.54196 | 0.288051 | NA | ENSMUSG( | NA | 14.54567 | 0.243925 | 0.650603 | 0.95422 |
| ENSMUSG( | NA | 89.73473 | -0.19611 | 0.378991 | 0.801757 | ENSMUSG( | NA | 8.569284 | 0.520617 | 0.475794 | NA |
| ENSMUSG( | NA | 31.14968 | -0.48004 | 0.23479 | 0.682188 | ENSMUSG( | NA | 20.50153 | 0.018358 | 0.967053 | 0.995549 |
| ENSMUSG( | NA | 420.1006 | -0.36915 | 0.0043 | 0.085154 | ENSMUSG( | NA | 39.39731 | 0.301865 | 0.366582 | 0.914703 |
| ENSMUSG( | NA | 8.783689 | -0.52348 | 0.439209 | NA | ENSMUSG( | NA | 18.66855 | -0.00152 | 0.997397 | 0.999268 |
| ENSMUSG( | NA | 26.0126 | 1.854406 | 0.001545 | 0.043953 | ENSMUSG( | NA | 24.62916 | -0.5572 | 0.177886 | 0.847901 |
| ENSMUSG( | NA | 147.3736 | -1.57145 | 0.084273 | 0.447434 | ENSMUSG( | NA | 22.64807 | 0.267868 | 0.522618 | 0.932822 |
| ENSMUSG( | NA | 11.92278 | 0.857727 | 0.11666 | NA | ENSMUSG( | NA | 22.53623 | 0.340021 | 0.419247 | 0.917257 |
| ENSMUSG( | NA | 30.9914 | 0.053171 | 0.878441 | 0.981431 | ENSMUSG( | NA | 26.5163 | -0.08853 | 0.81768 | 0.979198 |
| ENSMUSG( | NA | 20.94604 | 0.122612 | 0.757671 | NA | ENSMUSG( | NA | 8.327863 | -0.03889 | 0.959895 | NA |
| ENSMUSG( | NA | 37.36438 | 0.117184 | 0.705885 | 0.938711 | ENSMUSG( | NA | 16.80657 | 0.194187 | 0.703095 | 0.962428 |
| ENSMUSG( | NA | 218.3668 | 0.059751 | 0.741151 | 0.948141 | ENSMUSG( | NA | 35.51211 | -0.31504 | 0.354844 | 0.912411 |
| ENSMUSG( | NA | 17.78608 | 0.113437 | 0.800505 | NA | ENSMUSG( | NA | 9.37046 | 0.115817 | 0.863627 | NA |
| ENSMUSG( | NA | 14.83529 | -0.24929 | 0.601216 | NA | ENSMUSG( | NA | 98.9802 | -0.37045 | 0.123359 | 0.814691 |
| ENSMUSG( | NA | 14.25074 | -0.48433 | 0.320841 | NA | ENSMUSG( | NA | 656.4878 | 0.352056 | 0.046172 | 0.697814 |
| ENSMUSG( | NA | 9.79926 | -0.58912 | 0.356332 | NA | ENSMUSG( | NA | 14.03093 | -0.45869 | 0.3847 | 0.917257 |
| ENSMUSG( | NA | 154.4023 | -0.00816 | 0.970476 | 0.996369 | ENSMUSG( | NA | 148.5112 | 0.030292 | 0.884105 | 0.989644 |
| ENSMUSG( | NA | 17.48821 | -0.22644 | 0.605029 | NA | ENSMUSG( | NA | 14.90342 | -0.04736 | 0.929384 | 0.99306 |
| ENSMUSG( | NA | 8.977527 | -0.65248 | 0.344079 | NA | ENSMUSG( | NA | 8.871351 | 1.372711 | 0.056408 | NA |
| ENSMUSG( | NA | 70.01003 | -0.23605 | 0.37035 | 0.797542 | ENSMUSG( | NA | 33.98196 | -0.08468 | 0.808283 | 0.978622 |
| ENSMUSG( | NA | 289.0412 | -0.17468 | 0.295484 | 0.743814 | ENSMUSG( | NA | 17.75933 | 0.166945 | 0.723258 | 0.966887 |
| ENSMUSG( | NA | 191.341 | 0.044619 | 0.775872 | 0.955606 | ENSMUSG( | NA | 10.47681 | -0.43965 | 0.46322 | NA |
| ENSMUSG( | NA | 32.5651 | 0.139605 | 0.668384 | 0.927864 | ENSMUSG( | NA | 15.26474 | 0.134003 | 0.789023 | 0.976593 |
| ENSMUSG( | NA | 23.91523 | 0.232322 | 0.542207 | 0.883182 | ENSMUSG( | NA | 33.27255 | -0.5034 | 0.16047 | 0.837724 |
| ENSMUSG( | NA | 13.3732 | -1.13538 | 0.042905 | NA | ENSMUSG( | NA | 9.291286 | -0.2786 | 0.693249 | NA |
| ENSMUSG( | NA | 8.989355 | -1.30787 | 0.038703 | NA | ENSMUSG( | NA | 71.39673 | -0.22305 | 0.365226 | 0.914449 |
| ENSMUSG( | NA | 23.35333 | -0.67674 | 0.084205 | 0.447434 | ENSMUSG( | NA | 11.98404 | -0.72134 | 0.24533 | 0.877904 |
| ENSMUSG( | NA | 30.81038 | 0.243162 | 0.465636 | 0.84415 | ENSMUSG( | NA | 56.83411 | 0.530308 | 0.05545 | 0.723362 |
| ENSMUSG( | NA | 44.96967 | 0.062755 | 0.828799 | 0.969097 | ENSMUSG( | NA | 223.0361 | -0.23274 | 0.158856 | 0.837724 |
| ENSMUSG( | NA | 22.35244 | 0.312561 | 0.434143 | 0.828931 | ENSMUSG( | NA | 14.22411 | -0.26397 | 0.641192 | 0.953903 |
| ENSMUSG( | NA | 17.329 | 0.111284 | 0.805947 | NA | ENSMUSG( | NA | 73.59687 | 0.0639 | 0.801281 | 0.97681 |
| ENSMUSG( | NA | 68.50611 | 0.073033 | 0.771552 | 0.954769 | ENSMUSG( | NA | 14.71529 | -0.04235 | 0.934218 | 0.99306 |
| ENSMUSG( | NA | 9.813805 | -0.06117 | 0.91612 | NA | ENSMUSG( | NA | 36.96492 | -0.00755 | 0.983418 | 0.996963 |
| ENSMUSG( | NA | 11.63792 | 0.044638 | 0.936361 | NA | ENSMUSG( | NA | 33.6995 | 0.284396 | 0.463849 | 0.923329 |
| ENSMUSG( | NA | 21.99843 | 0.342236 | 0.452663 | 0.836145 | ENSMUSG( | NA | 36.64717 | 0.187835 | 0.570105 | 0.941777 |
| ENSMUSG( | NA | 94.62492 | -0.31705 | 0.141998 | 0.559166 | ENSMUSG( | NA | 285.5812 | 0.458871 | 0.01244 | 0.451463 |
| ENSMUSG( | NA | 31.56246 | -0.56459 | 0.106822 | 0.494622 | ENSMUSG( | NA | 30.12007 | -0.0337 | 0.92699 | 0.99306 |
| ENSMUSG( | NA | 199.8693 | 0.123985 | 0.495488 | 0.861026 | ENSMUSG( | NA | 28.19486 | 0.075611 | 0.848833 | 0.985321 |
| ENSMUSG( | NA | 124.2486 | 0.033669 | 0.861549 | 0.975026 | ENSMUSG( | NA | 21.25068 | 0.270662 | 0.573997 | 0.942912 |
| ENSMUSG( | NA | 33.32419 | 0.062922 | 0.858276 | 0.974672 | ENSMUSG( | NA | 176.3344 | -0.06814 | 0.756057 | 0.971384 |
| ENSMUSG( | NA | 14.79457 | 0.360874 | 0.448746 | NA | ENSMUSG( | NA | 13.0726 | -0.26157 | 0.652667 | 0.954799 |
| ENSMUSG( | NA | 39.1137 | 0.855752 | 0.035631 | 0.290384 | ENSMUSG( | NA | 9.610089 | -0.61475 | 0.366556 | NA |
| ENSMUSG( | NA | 44.02939 | -0.34702 | 0.238158 | 0.686299 | ENSMUSG( | NA | 86.42279 | 0.158572 | 0.538027 | 0.937216 |
| ENSMUSG( | NA | 21.18483 | -0.31412 | 0.481777 | NA | ENSMUSG( | NA | 131.6104 | -0.37341 | 0.106192 | 0.804969 |
| ENSMUSG( | NA | 61.15796 | -0.01104 | 0.966803 | 0.995784 | ENSMUSG( | NA | 33.14962 | 0.06597 | 0.85146 | 0.985541 |
| ENSMUSG( | NA | 84.8747 | 0.04647 | 0.837991 | 0.971111 | ENSMUSG( | NA | 31.89393 | -0.2901 | 0.420171 | 0.917257 |
| ENSMUSG( | NA | 18.53556 | -0.34094 | 0.506169 | NA | ENSMUSG( | NA | 15.56762 | -0.12138 | 0.80577 | 0.977674 |
| ENSMUSG( | NA | 11.00525 | 0.070524 | 0.902729 | NA | ENSMUSG( | NA | 42.88158 | 0.439402 | 0.15338 | 0.837724 |
| ENSMUSG( | NA | 10.11865 | -0.05912 | 0.918319 | NA | ENSMUSG( | NA | 41.08261 | 0.083905 | 0.803659 | 0.976978 |
| ENSMUSG( | NA | 12.65334 | -0.86165 | 0.10685 | NA | ENSMUSG( | NA | 10.48399 | -0.3747 | 0.536434 | NA |
| ENSMUSG( | NA | 37.59345 | 0.348298 | 0.280771 | 0.729868 | ENSMUSG( | NA | 8.313175 | -0.01162 | 0.986698 | NA |
| ENSMUSG( | NA | 7.945859 | 0.229113 | 0.736852 | NA | ENSMUSG( | NA | 12.6293 | -0.21432 | 0.693676 | 0.960767 |
| ENSMUSG( | NA | 59.63795 | -0.17838 | 0.514965 | 0.872348 | ENSMUSG( | NA | 73.51027 | 0.130529 | 0.59391 | 0.945408 |
| ENSMUSG( | NA | 202.5538 | 0.06263 | 0.68073 | 0.931139 | ENSMUSG( | NA | 26.15922 | -0.31304 | 0.4633 | 0.923329 |
| ENSMUSG( | NA | 79.69817 | -0.53971 | 0.026127 | 0.247245 | ENSMUSG( | NA | 80.36014 | -0.79284 | 0.000829 | 0.09749 |
| ENSMUSG( | NA | 53.96939 | -0.41619 | 0.212962 | 0.656982 | ENSMUSG( | NA | 99.32515 | 0.026412 | 0.90827 | 0.991958 |
| ENSMUSG( | NA | 13.1218 | -0.09279 | 0.873693 | NA | ENSMUSG( | NA | 51.13663 | -0.01785 | 0.949789 | 0.994798 |
| ENSMUSG( | NA | 18.11981 | -0.47493 | 0.306569 | NA | ENSMUSG( | NA | 17.44495 | 0.726318 | 0.149663 | 0.836241 |
| ENSMUSG( | NA | 142.7138 | -0.20144 | 0.292798 | 0.742763 | ENSMUSG( | NA | 22.32014 | -0.45183 | 0.298715 | 0.898538 |
| ENSMUSG( | NA | 649.3908 | 0.026546 | 0.826886 | 0.96865 | ENSMUSG( | NA | 71.98855 | 0.013501 | 0.959586 | 0.994798 |
| ENSMUSG( | NA | 18.82173 | -0.16127 | 0.703336 | NA | ENSMUSG( | NA | 78.05519 | -0.39889 | 0.122222 | 0.813384 |
| ENSMUSG( | NA | 125.3646 | 0.490195 | 0.012462 | 0.161344 | ENSMUSG( | NA | 9.407051 | 0.05218 | 0.938564 | NA |
| ENSMUSG( | NA | 7.912114 | -0.2413 | 0.754519 | NA | ENSMUSG( | NA | 197.3761 | 0.029322 | 0.873142 | 0.989445 |
| ENSMUSG( | NA | 25.92168 | 0.118224 | 0.744472 | 0.948797 | ENSMUSG( | NA | 36.12997 | 0.236757 | 0.475584 | 0.923329 |
| ENSMUSG( | NA | 13.45474 | 0.357853 | 0.494754 | NA | ENSMUSG( | NA | 865.0197 | 0.215017 | 0.092453 | 0.793483 |
| ENSMUSG( | NA | 79.82227 | 0.167501 | 0.459038 | 0.840244 | ENSMUSG( | NA | 61.32792 | 0.130371 | 0.652435 | 0.954799 |
| ENSMUSG( | NA | 47.96136 | -0.53293 | 0.060901 | 0.381246 | ENSMUSG( | NA | 189.5427 | -0.11921 | 0.489864 | 0.925987 |
| ENSMUSG( | NA | 14.6491 | -1.46907 | 0.016265 | NA | ENSMUSG( | NA | 4823.297 | -0.04075 | 0.812508 | 0.979148 |
| ENSMUSG( | NA | 154.2644 | -0.0056 | 0.974771 | 0.997207 | ENSMUSG( | NA | 48.96236 | -0.37392 | 0.237288 | 0.872173 |
| ENSMUSG( | NA | 15.01695 | 0.61632 | 0.237289 | NA | ENSMUSG( | NA | 12.3579 | -0.53447 | 0.366068 | 0.914623 |
| ENSMUSG( | NA | 413.704 | 0.063679 | 0.658757 | 0.924506 | ENSMUSG( | NA | 140.9084 | 0.077973 | 0.686927 | 0.959178 |
| ENSMUSG( | NA | 146.0288 | 0.168089 | 0.375797 | 0.800976 | ENSMUSG( | NA | 48.94096 | 0.32451 | 0.283432 | 0.892412 |
| ENSMUSG( | NA | 113.2868 | 0.004313 | 0.984875 | 0.997815 | ENSMUSG( | NA | 9.932551 | 0.596563 | 0.365036 | NA |
| ENSMUSG( | NA | 263.4217 | -0.08777 | 0.564569 | 0.890434 | ENSMUSG( | NA | 17.64745 | -0.19538 | 0.709608 | 0.96416 |
| ENSMUSG( | NA | 65.18268 | -0.10416 | 0.72449 | 0.944096 | ENSMUSG( | NA | 11.76328 | -0.06555 | 0.911735 | 0.992196 |
| ENSMUSG( | NA | 36.2355 | -0.04255 | 0.900283 | 0.985633 | ENSMUSG( | NA | 321.4355 | -0.04934 | 0.73899 | 0.968312 |
| ENSMUSG( | NA | 40.10066 | 0.25091 | 0.430216 | 0.826789 | ENSMUSG( | NA | 46.0789 | 0.463828 | 0.157448 | 0.837724 |
| ENSMUSG( | NA | 611.5774 | -0.05734 | 0.660339 | 0.924627 | ENSMUSG( | NA | 172.9018 | -0.14122 | 0.423112 | 0.917257 |
| ENSMUSG( | NA | 230.4404 | -0.68018 | 0.001911 | 0.049911 | ENSMUSG( | NA | 128.0818 | -0.35476 | 0.080242 | 0.77681 |
| ENSMUSG( | NA | 16.82833 | 0.305785 | 0.496834 | NA | ENSMUSG( | NA | 50.33504 | 0.444705 | 0.225025 | 0.868422 |
| ENSMUSG( | NA | 9.654279 | -0.81612 | 0.190537 | NA | ENSMUSG( | NA | 111.208 | 0.415697 | 0.157499 | 0.837724 |
| ENSMUSG( | NA | 198.4852 | -0.07337 | 0.628152 | 0.913532 | ENSMUSG( | NA | 25.64069 | -0.6757 | 0.099698 | 0.801103 |
| ENSMUSG( | NA | 26.82703 | -0.05588 | 0.890413 | 0.983687 | ENSMUSG( | NA | 16.28063 | 0.118084 | 0.819648 | 0.979198 |
| ENSMUSG( | NA | 30.83124 | 0.025043 | 0.942586 | 0.993415 | ENSMUSG( | NA | 11.60335 | 0.274042 | 0.639845 | 0.953903 |
| ENSMUSG( | NA | 18.48125 | -0.23806 | 0.669683 | NA | ENSMUSG( | NA | 47.96599 | 0.083614 | 0.795794 | 0.97681 |
| ENSMUSG( | NA | 29.01989 | 0.405044 | 0.244365 | 0.691487 | ENSMUSG( | NA | 121.7087 | -0.1577 | 0.464249 | 0.923329 |
| ENSMUSG( | NA | 8.372702 | 0.546886 | 0.441312 | NA | ENSMUSG( | NA | 12.26194 | -0.23263 | 0.670143 | 0.95694 |
| ENSMUSG( | NA | 44.44207 | -0.05123 | 0.860274 | 0.974842 | ENSMUSG( | NA | 13.84162 | -0.6372 | 0.220097 | 0.866104 |
| ENSMUSG( | NA | 15.19655 | -0.37606 | 0.428037 | NA | ENSMUSG( | NA | 124.7368 | 0.343747 | 0.145137 | 0.83339 |
| ENSMUSG( | NA | 228.9401 | -0.15784 | 0.352998 | 0.784977 | ENSMUSG( | NA | 69.05991 | 0.17825 | 0.648365 | 0.95422 |
| ENSMUSG( | NA | 15.77198 | -0.73442 | 0.151729 | NA | ENSMUSG( | NA | 33.84389 | 0.504733 | 0.177805 | 0.847901 |
| ENSMUSG( | NA | 38.16992 | 0.049594 | 0.869842 | 0.977607 | ENSMUSG( | NA | 30.64196 | 0.19444 | 0.642905 | 0.953903 |
| ENSMUSG( | NA | 12.43342 | -0.19496 | 0.715175 | NA | ENSMUSG( | NA | 20.94413 | 0.230175 | 0.599545 | 0.947071 |
| ENSMUSG( | NA | 12.13188 | -0.53555 | 0.325317 | NA | ENSMUSG( | NA | 12.66396 | -0.4006 | 0.482867 | 0.924615 |
| ENSMUSG( | NA | 14.59856 | -0.14336 | 0.766164 | NA | ENSMUSG( | NA | 13.09816 | -0.47645 | 0.382548 | 0.917257 |
| ENSMUSG( | NA | 12.16796 | 0.010304 | 0.98424 | NA | ENSMUSG( | NA | 11.72335 | -0.09445 | 0.871934 | 0.988779 |
| ENSMUSG( | NA | 373.1042 | -0.23592 | 0.139289 | 0.555376 | ENSMUSG( | NA | 11.26143 | -0.9888 | 0.100588 | NA |
| ENSMUSG( | NA | 42.10618 | 0.240136 | 0.434463 | 0.828978 | ENSMUSG( | NA | 29.30451 | 0.283845 | 0.474759 | 0.923329 |
| ENSMUSG( | NA | 18.24699 | -0.24696 | 0.57832 | NA | ENSMUSG( | NA | 55.91553 | -0.18105 | 0.54061 | 0.937383 |
| ENSMUSG( | NA | 26.32569 | -0.19103 | 0.620426 | 0.911432 | ENSMUSG( | NA | 18.36942 | -0.19247 | 0.675978 | 0.95694 |
| ENSMUSG( | NA | 194.2323 | 0.421962 | 0.012578 | 0.161972 | ENSMUSG( | NA | 148.8333 | -0.08938 | 0.647669 | 0.95422 |
| ENSMUSG( | NA | 575.5679 | 0.107011 | 0.386052 | 0.804623 | ENSMUSG( | NA | 17.33508 | -0.04728 | 0.921643 | 0.992644 |
| ENSMUSG( | NA | 59.86307 | -0.00903 | 0.970852 | 0.996369 | ENSMUSG( | NA | 3836.413 | -0.18312 | 0.116109 | 0.813384 |
| ENSMUSG( | NA | 16.07105 | 1.031178 | 0.112235 | NA | ENSMUSG( | NA | 447.0333 | -0.20318 | 0.202016 | 0.864412 |
| ENSMUSG( | NA | 14.46164 | 0.318049 | 0.530087 | NA | ENSMUSG( | NA | 11.47431 | 0.037701 | 0.949976 | 0.994798 |
| ENSMUSG( | NA | 11.71135 | -0.07877 | 0.88711 | NA | ENSMUSG( | NA | 48.90416 | 0.628072 | 0.048807 | 0.706228 |
| ENSMUSG( | NA | 22.99092 | 0.297523 | 0.442808 | 0.83278 | ENSMUSG( | NA | 15.22255 | -0.09146 | 0.861877 | 0.98632 |
| ENSMUSG( | NA | 46.57983 | 0.497648 | 0.071123 | 0.411712 | ENSMUSG( | NA | 16.50435 | 0.493688 | 0.301257 | 0.898538 |
| ENSMUSG( | NA | 14.18366 | -0.56971 | 0.283263 | NA | ENSMUSG( | NA | 47.17188 | 0.418541 | 0.292252 | 0.897854 |
| ENSMUSG( | NA | 25.15043 | -0.5282 | 0.173133 | 0.608373 | ENSMUSG( | NA | 8.799303 | -0.1084 | 0.864509 | NA |
| ENSMUSG( | NA | 23.5327 | 0.535701 | 0.177854 | 0.615225 | ENSMUSG( | NA | 10.49901 | -0.72333 | 0.234652 | NA |
| ENSMUSG( | NA | 157.0521 | -0.48142 | 0.004875 | 0.092488 | ENSMUSG( | NA | 11.75941 | -0.09279 | 0.869488 | 0.988097 |
| ENSMUSG( | NA | 10.75916 | 0.606418 | 0.322439 | NA | ENSMUSG( | NA | 16.55907 | -0.24729 | 0.629597 | 0.953251 |
| ENSMUSG( | NA | 29.66938 | -0.09152 | 0.796485 | 0.960163 | ENSMUSG( | NA | 24.07575 | 0.035637 | 0.934916 | 0.99306 |
| ENSMUSG( | NA | 34.34212 | -0.0758 | 0.828241 | 0.968966 | ENSMUSG( | NA | 7.593636 | 0.000513 | 0.999441 | NA |
| ENSMUSG( | NA | 49.77816 | 1.425747 | 2.29E-05 | 0.002131 | ENSMUSG( | NA | 19.36912 | -0.72287 | 0.147219 | 0.836241 |
| ENSMUSG( | NA | 41.64091 | 0.077722 | 0.808873 | 0.963495 | ENSMUSG( | NA | 20.75878 | -0.69939 | 0.119863 | 0.813384 |
| ENSMUSG( | NA | 29.59746 | -0.42958 | 0.230849 | 0.679066 | ENSMUSG( | NA | 211.0318 | 0.130287 | 0.590792 | 0.944975 |
| ENSMUSG( | NA | 226.4226 | -0.14054 | 0.376362 | 0.801127 | ENSMUSG( | NA | 50.94686 | -0.16331 | 0.565654 | 0.941404 |
| ENSMUSG( | NA | 29.73484 | 0.475484 | 0.171059 | 0.605644 | ENSMUSG( | NA | 22.24244 | 0.022651 | 0.959403 | 0.994798 |
| ENSMUSG( | NA | 44.81495 | -0.06802 | 0.812762 | 0.964574 | ENSMUSG( | NA | 12.30808 | -0.52563 | 0.352138 | 0.912093 |
| ENSMUSG( | NA | 22.47965 | -0.41537 | 0.328856 | 0.773233 | ENSMUSG( | NA | 20.77601 | 0.080211 | 0.861091 | 0.986226 |
| ENSMUSG( | NA | 14.38348 | 0.277424 | 0.582673 | NA | ENSMUSG( | NA | 18.97802 | -0.16416 | 0.722663 | 0.966887 |
| ENSMUSG( | NA | 20.74068 | 0.236714 | 0.559351 | NA | ENSMUSG( | NA | 64.97063 | 0.210678 | 0.519879 | 0.931824 |
| ENSMUSG( | NA | 115.3696 | 0.138918 | 0.581016 | 0.89745 | ENSMUSG( | NA | 18.56288 | 0.024244 | 0.957168 | 0.994798 |
| ENSMUSG( | NA | 14.31555 | -0.12664 | 0.820868 | NA | ENSMUSG( | NA | 21.49901 | 0.337913 | 0.570263 | 0.941823 |
| ENSMUSG( | NA | 295.276 | -0.19443 | 0.199904 | 0.640239 | ENSMUSG( | NA | 14.64787 | -0.48368 | 0.374767 | 0.915491 |
| ENSMUSG( | NA | 29.52239 | -0.06063 | 0.863042 | 0.975055 | ENSMUSG( | NA | 13.57792 | -0.01152 | 0.98345 | 0.996963 |
| ENSMUSG( | NA | 45.13071 | 0.008153 | 0.97688 | 0.997428 | ENSMUSG( | NA | 10.11286 | -0.251 | 0.678709 | NA |
| ENSMUSG( | NA | 16.10496 | -0.0125 | 0.979225 | NA | ENSMUSG( | NA | 19.86668 | -0.97556 | 0.029773 | 0.61992 |
| ENSMUSG( | NA | 59.04121 | 0.289949 | 0.280234 | 0.729246 | ENSMUSG( | NA | 9.760741 | 0.2776 | 0.655069 | NA |
| ENSMUSG( | NA | 68.25107 | 0.111001 | 0.630012 | 0.913965 | ENSMUSG( | NA | 28.57722 | -0.08086 | 0.851552 | 0.985541 |
| ENSMUSG( | NA | 14.14968 | -0.45824 | 0.359368 | NA | ENSMUSG( | NA | 30.3805 | 0.756456 | 0.124823 | 0.81516 |
| ENSMUSG( | NA | 28.61296 | 0.04667 | 0.900933 | 0.985633 | ENSMUSG( | NA | 18.96822 | 0.126792 | 0.795564 | 0.97681 |
| ENSMUSG( | NA | 40.19016 | -0.22119 | 0.492723 | 0.860462 | ENSMUSG( | NA | 18.751 | 0.032447 | 0.944936 | 0.994798 |
| ENSMUSG( | NA | 21.60329 | -0.38558 | 0.360477 | 0.792161 | ENSMUSG( | NA | 19.83744 | -0.22076 | 0.618979 | 0.952598 |
| ENSMUSG( | NA | 70.27625 | -0.25599 | 0.374559 | 0.800445 | ENSMUSG( | NA | 17.54829 | 0.43947 | 0.356904 | 0.913358 |
| ENSMUSG( | NA | 46.34876 | 0.319534 | 0.252021 | 0.701908 | ENSMUSG( | NA | 50.05397 | 0.083471 | 0.781022 | 0.976106 |
| ENSMUSG( | NA | 16.6184 | -0.43258 | 0.369352 | NA | ENSMUSG( | NA | 204.0776 | 0.104555 | 0.598159 | 0.946676 |
| ENSMUSG( | NA | 23.05804 | 0.21538 | 0.572346 | 0.893584 | ENSMUSG( | NA | 108.2904 | -0.22322 | 0.298237 | 0.898538 |
| ENSMUSG( | NA | 337.1343 | -0.16795 | 0.228511 | 0.675862 | ENSMUSG( | NA | 55.7295 | -0.53495 | 0.05861 | 0.728256 |
| ENSMUSG( | NA | 263.9818 | -0.07993 | 0.567866 | 0.892002 | ENSMUSG( | NA | 23.74118 | 0.833978 | 0.082351 | 0.77681 |

| | | | | | |
|---|---|---|---|---|---|
| ENSMUSG( | 35.01655 | 0.326526 | 0.348248 | 0.78132 | NA |
| ENSMUSG( | 194.6721 | 0.077963 | 0.679397 | 0.930983 | NA |
| ENSMUSG( | 9.982853 | 0.207416 | 0.735769 | NA | NA |
| ENSMUSG( | 10.43138 | -0.44244 | 0.490465 | NA | NA |
| ENSMUSG( | 53.34677 | 0.528036 | 0.085502 | 0.450845 | NA |
| ENSMUSG( | 96.89703 | -0.21831 | 0.371955 | 0.798768 | NA |
| ENSMUSG( | 12.68135 | -0.04993 | 0.932988 | NA | NA |
| ENSMUSG( | 615.0416 | 0.475308 | 8.10E-05 | 0.005593 | NA |
| ENSMUSG( | 26.49891 | 0.129923 | 0.717978 | 0.941697 | NA |
| ENSMUSG( | 9.680707 | 0.966001 | 0.178788 | NA | NA |
| ENSMUSG( | 32.82738 | 0.19761 | 0.569326 | 0.893041 | NA |
| ENSMUSG( | 1151.308 | 0.379127 | 0.000858 | 0.029429 | NA |
| ENSMUSG( | 119.6142 | 0.16636 | 0.406097 | 0.816525 | NA |
| ENSMUSG( | 24.36688 | 0.249976 | 0.537781 | 0.882548 | NA |
| ENSMUSG( | 12.47714 | -0.26013 | 0.63068 | NA | NA |
| ENSMUSG( | 35.23194 | -0.21117 | 0.627009 | 0.912825 | NA |
| ENSMUSG( | 19.45997 | -0.09247 | 0.825755 | NA | NA |
| ENSMUSG( | 12.23751 | -0.19175 | 0.747189 | NA | NA |
| ENSMUSG( | 47.35393 | 0.018154 | 0.95102 | 0.993821 | NA |
| ENSMUSG( | 3595.657 | -0.00014 | 0.999007 | 0.99989 | NA |
| ENSMUSG( | 26.05562 | -0.18918 | 0.621868 | 0.911432 | NA |
| ENSMUSG( | 28.62058 | -0.26022 | 0.453978 | 0.836798 | NA |
| ENSMUSG( | 31.99057 | 0.228252 | 0.520394 | 0.875186 | NA |
| ENSMUSG( | 12.55955 | -0.17457 | 0.748619 | NA | NA |
| ENSMUSG( | 33.78757 | 0.204374 | 0.54027 | 0.88315 | NA |
| ENSMUSG( | 13.54113 | 0.238128 | 0.637096 | NA | NA |
| ENSMUSG( | 17.45058 | -0.24188 | 0.59774 | NA | NA |
| ENSMUSG( | 10.25014 | 0.120231 | 0.836407 | NA | NA |
| ENSMUSG( | 68.03879 | -0.07971 | 0.728351 | 0.944393 | NA |
| ENSMUSG( | 11.86201 | 0.094189 | 0.857244 | NA | NA |
| ENSMUSG( | 12.46075 | 0.769561 | 0.141905 | NA | NA |
| ENSMUSG( | 132.8003 | 0.159227 | 0.447885 | 0.834051 | NA |
| ENSMUSG( | 11.66659 | -0.13715 | 0.812085 | NA | NA |
| ENSMUSG( | 12.08747 | 0.635304 | 0.274923 | NA | NA |
| ENSMUSG( | 22.84353 | 0.00691 | 0.987478 | 0.998796 | NA |
| ENSMUSG( | 206.4501 | -0.07748 | 0.607704 | 0.906878 | NA |
| ENSMUSG( | 21.20829 | -0.73627 | 0.127561 | 0.535393 | NA |
| ENSMUSG( | 30.69562 | 0.075955 | 0.830741 | 0.969798 | NA |
| ENSMUSG( | 10.40982 | -0.77204 | 0.196297 | NA | NA |
| ENSMUSG( | 135.8441 | 0.239203 | 0.20101 | 0.641303 | NA |
| ENSMUSG( | 208.3395 | 0.291637 | 0.071993 | 0.414057 | NA |
| ENSMUSG( | 18.71752 | -0.33273 | 0.449623 | NA | NA |
| ENSMUSG( | 70.32668 | 0.45406 | 0.059584 | 0.377373 | NA |
| ENSMUSG( | 19.33647 | -0.31249 | 0.471477 | NA | NA |
| ENSMUSG( | 15.42733 | -0.4973 | 0.304812 | NA | NA |
| ENSMUSG( | 9.310459 | -0.50559 | 0.442643 | NA | NA |
| ENSMUSG( | 17.31999 | -0.1922 | 0.681193 | NA | NA |
| ENSMUSG( | 10.13384 | 0.303391 | 0.620332 | NA | NA |
| ENSMUSG( | 18.35663 | 0.478264 | 0.272559 | NA | NA |
| ENSMUSG( | 154.2056 | 0.148447 | 0.449776 | 0.835283 | NA |
| ENSMUSG( | 180.9294 | -0.05947 | 0.74348 | 0.948478 | NA |
| ENSMUSG( | 36.24625 | 0.466695 | 0.171126 | 0.605644 | NA |
| ENSMUSG( | 34.31262 | 0.123161 | 0.712899 | 0.94066 | NA |
| ENSMUSG( | 52.91958 | -0.03348 | 0.904366 | 0.986246 | NA |
| ENSMUSG( | 94.43667 | 0.077466 | 0.719685 | 0.941986 | NA |
| ENSMUSG( | 16.87953 | 0.088848 | 0.842309 | NA | NA |
| ENSMUSG( | 209.3189 | 0.143703 | 0.341115 | 0.778828 | NA |
| ENSMUSG( | 13.26196 | 0.083893 | 0.875889 | NA | NA |
| ENSMUSG( | 23.54898 | -0.6855 | 0.085119 | 0.44957 | NA |
| ENSMUSG( | 22.48052 | -0.42832 | 0.331582 | 0.77519 | NA |
| ENSMUSG( | 17.04579 | -0.00765 | 0.986788 | NA | NA |
| ENSMUSG( | 14.673 | 0.605615 | 0.329618 | NA | NA |
| ENSMUSG( | 86.64812 | -0.43238 | 0.07086 | 0.410627 | NA |
| ENSMUSG( | 110.0849 | -0.11162 | 0.626703 | 0.912825 | NA |
| ENSMUSG( | 10.30293 | 0.24933 | 0.693014 | NA | NA |
| ENSMUSG( | 30.40943 | -0.67052 | 0.061461 | 0.382832 | NA |
| ENSMUSG( | 41.87922 | 0.214583 | 0.468773 | 0.845636 | NA |
| ENSMUSG( | 11.27974 | -0.67689 | 0.211719 | NA | NA |
| ENSMUSG( | 11.1153 | -0.30031 | 0.603223 | NA | NA |
| ENSMUSG( | 50.21447 | -0.00258 | 0.992606 | 0.999327 | NA |
| ENSMUSG( | 17.02956 | 0.220353 | 0.672004 | NA | NA |
| ENSMUSG( | 11.73167 | -0.59061 | 0.296235 | NA | NA |
| ENSMUSG( | 48.80519 | -0.0458 | 0.887526 | 0.983411 | NA |
| ENSMUSG( | 31.33883 | -0.21817 | 0.529658 | 0.878883 | NA |
| ENSMUSG( | 24.52811 | 0.010686 | 0.978936 | 0.997428 | NA |
| ENSMUSG( | 26.7751 | -0.04028 | 0.911309 | 0.988483 | NA |
| ENSMUSG( | 23.49553 | -0.00573 | 0.988941 | 0.999111 | NA |
| ENSMUSG( | 128.1144 | 0.27452 | 0.191767 | 0.630811 | NA |
| ENSMUSG( | 65.0841 | -0.17898 | 0.456496 | 0.838832 | NA |
| ENSMUSG( | 10.57897 | -0.2662 | 0.633321 | NA | NA |
| ENSMUSG( | 10.37025 | -0.13247 | 0.815795 | NA | NA |
| ENSMUSG( | 25.92855 | 0.294741 | 0.465447 | 0.844064 | NA |
| ENSMUSG( | 27.28253 | -0.20243 | 0.605387 | 0.905764 | NA |
| ENSMUSG( | 14.45944 | 0.196153 | 0.713765 | NA | NA |
| ENSMUSG( | 21.91619 | 0.324662 | 0.408933 | 0.818173 | NA |
| ENSMUSG( | 54.87375 | -0.15827 | 0.546546 | 0.884974 | NA |
| ENSMUSG( | 14.93816 | -0.61938 | 0.226444 | NA | NA |
| ENSMUSG( | 9.247386 | 0.484929 | 0.470197 | NA | NA |
| ENSMUSG( | 13.26571 | -1.03849 | 0.041524 | NA | NA |
| ENSMUSG( | 73.55929 | -0.38996 | 0.149566 | 0.573617 | NA |
| ENSMUSG( | 18.55318 | 0.54196 | 0.288051 | NA | NA |
| ENSMUSG( | 89.73473 | -0.19611 | 0.378991 | 0.801757 | NA |
| ENSMUSG( | 31.14968 | -0.48004 | 0.23479 | 0.682188 | NA |
| ENSMUSG( | 420.1006 | -0.36915 | 0.0043 | 0.085154 | NA |
| ENSMUSG( | 8.783689 | -0.52348 | 0.439209 | NA | NA |
| ENSMUSG( | 26.0126 | 1.854406 | 0.001545 | 0.043953 | NA |
| ENSMUSG( | 147.3736 | -1.57145 | 0.084273 | 0.447434 | NA |
| ENSMUSG( | 11.92278 | 0.857727 | 0.11666 | NA | NA |
| ENSMUSG( | 30.9914 | 0.053171 | 0.878441 | 0.981431 | NA |
| ENSMUSG( | 20.94604 | 0.122612 | 0.757671 | NA | NA |
| ENSMUSG( | 37.36438 | 0.117184 | 0.705885 | 0.938711 | NA |
| ENSMUSG( | 218.3668 | 0.059751 | 0.741151 | 0.948141 | NA |
| ENSMUSG( | 10.31509 | 0.487004 | 0.415304 | NA | NA |
| ENSMUSG( | 17.78608 | 0.113437 | 0.800505 | NA | NA |
| ENSMUSG( | 14.83529 | -0.24929 | 0.601216 | NA | NA |
| ENSMUSG( | 14.25074 | -0.48433 | 0.320841 | NA | NA |
| ENSMUSG( | 9.79926 | -0.58912 | 0.356332 | NA | NA |
| ENSMUSG( | 154.4023 | -0.00816 | 0.970476 | 0.996369 | NA |
| ENSMUSG( | 11.43949 | 0.389054 | 0.496388 | NA | NA |
| ENSMUSG( | 17.48821 | -0.22644 | 0.605029 | NA | NA |
| ENSMUSG( | 8.977527 | -0.65248 | 0.344079 | NA | NA |
| ENSMUSG( | 70.01003 | -0.23605 | 0.37035 | 0.797542 | NA |
| ENSMUSG( | 8.823926 | 0.186373 | 0.781065 | NA | NA |
| ENSMUSG( | 289.0412 | -0.17468 | 0.295484 | 0.743814 | NA |
| ENSMUSG( | 191.341 | 0.044619 | 0.775872 | 0.955606 | NA |
| ENSMUSG( | 32.5651 | 0.139605 | 0.668384 | 0.927864 | NA |
| ENSMUSG( | 23.91523 | 0.232322 | 0.542207 | 0.883182 | NA |
| ENSMUSG( | 13.3732 | -1.13538 | 0.042905 | NA | NA |
| ENSMUSG( | 8.989355 | -1.30787 | 0.038703 | NA | NA |
| ENSMUSG( | 23.35333 | -0.67674 | 0.084205 | 0.447434 | NA |
| ENSMUSG( | 30.81038 | 0.243162 | 0.465636 | 0.84415 | NA |
| ENSMUSG( | 44.96967 | 0.062755 | 0.828799 | 0.969097 | NA |
| ENSMUSG( | 22.35244 | 0.312561 | 0.434143 | 0.828931 | NA |
| ENSMUSG( | 17.329 | 0.111284 | 0.805947 | NA | NA |
| ENSMUSG( | 68.50611 | 0.073033 | 0.771552 | 0.954769 | NA |
| ENSMUSG( | 9.813805 | -0.06117 | 0.91612 | NA | NA |
| ENSMUSG( | 11.63792 | 0.044638 | 0.936361 | NA | NA |
| ENSMUSG( | 21.99843 | 0.342236 | 0.452663 | 0.836145 | NA |
| ENSMUSG( | 10.42657 | 0.500769 | 0.376562 | NA | NA |
| ENSMUSG( | 94.62492 | -0.31705 | 0.141998 | 0.559166 | NA |
| ENSMUSG( | 31.56246 | -0.56459 | 0.106822 | 0.494622 | NA |
| ENSMUSG( | 199.8693 | 0.123985 | 0.495488 | 0.861026 | NA |
| ENSMUSG( | 124.2486 | 0.033669 | 0.861549 | 0.975026 | NA |
| ENSMUSG( | 33.32419 | 0.062922 | 0.858276 | 0.974672 | NA |
| ENSMUSG( | 14.79457 | 0.360874 | 0.448746 | NA | NA |
| ENSMUSG( | 39.1137 | 0.855752 | 0.035631 | 0.290384 | NA |
| ENSMUSG( | 44.02939 | -0.34702 | 0.238158 | 0.686299 | NA |
| ENSMUSG( | 21.18483 | -0.31412 | 0.481777 | NA | NA |
| ENSMUSG( | 61.15796 | -0.01104 | 0.966803 | 0.995784 | NA |
| ENSMUSG( | 84.8747 | 0.04647 | 0.837991 | 0.971111 | NA |
| ENSMUSG( | 18.53556 | -0.34094 | 0.506169 | NA | NA |
| ENSMUSG( | 11.00525 | 0.070524 | 0.902729 | NA | NA |
| ENSMUSG( | 10.11865 | -0.05912 | 0.918319 | NA | NA |
| ENSMUSG( | 12.65334 | -0.86165 | 0.10685 | NA | NA |
| ENSMUSG( | 37.59345 | 0.348298 | 0.280771 | 0.729868 | NA |
| ENSMUSG( | 7.945859 | 0.229113 | 0.736852 | NA | NA |
| ENSMUSG( | 11.0995 | 1.013986 | 0.077932 | NA | NA |
| ENSMUSG( | 59.63795 | -0.17838 | 0.514965 | 0.872348 | NA |
| ENSMUSG( | 202.5538 | 0.06263 | 0.68073 | 0.931139 | NA |
| ENSMUSG( | 79.69817 | -0.53971 | 0.026127 | 0.247245 | NA |
| ENSMUSG( | 9.070272 | 0.359672 | 0.566101 | NA | NA |
| ENSMUSG( | 53.96939 | -0.41619 | 0.212962 | 0.656982 | NA |
| ENSMUSG( | 13.1218 | -0.09279 | 0.873693 | NA | NA |
| ENSMUSG( | 18.11981 | -0.47493 | 0.306569 | NA | NA |
| ENSMUSG( | 142.7138 | -0.20144 | 0.292798 | 0.742763 | NA |
| ENSMUSG( | 649.3908 | 0.026546 | 0.826886 | 0.96865 | NA |
| ENSMUSG( | 18.82173 | -0.16127 | 0.703336 | NA | NA |
| ENSMUSG( | 125.3646 | 0.490195 | 0.012462 | 0.161344 | NA |

| | | | | | |
|---|---|---|---|---|---|
| ENSMUSG( | 26.5163 | -0.08853 | 0.81768 | 0.979198 | NA |
| ENSMUSG( | 8.327863 | -0.03889 | 0.959895 | NA | NA |
| ENSMUSG( | 16.80657 | 0.194187 | 0.703095 | 0.962428 | NA |
| ENSMUSG( | 35.51211 | -0.31504 | 0.354844 | 0.912411 | NA |
| ENSMUSG( | 9.37046 | 0.115817 | 0.863627 | NA | NA |
| ENSMUSG( | 98.9802 | -0.37045 | 0.123359 | 0.814691 | NA |
| ENSMUSG( | 656.4878 | 0.352056 | 0.046172 | 0.697814 | NA |
| ENSMUSG( | 14.03093 | -0.45869 | 0.3847 | 0.917257 | NA |
| ENSMUSG( | 148.5112 | 0.030292 | 0.884105 | 0.989644 | NA |
| ENSMUSG( | 14.90342 | -0.04736 | 0.929384 | 0.99306 | NA |
| ENSMUSG( | 8.871351 | 1.372711 | 0.056408 | NA | NA |
| ENSMUSG( | 33.98196 | -0.08468 | 0.808283 | 0.978622 | NA |
| ENSMUSG( | 17.75933 | 0.166945 | 0.723258 | 0.966887 | NA |
| ENSMUSG( | 10.47681 | -0.43965 | 0.46322 | NA | NA |
| ENSMUSG( | 15.26474 | 0.134003 | 0.789023 | 0.976593 | NA |
| ENSMUSG( | 33.27255 | -0.5034 | 0.16047 | 0.837724 | NA |
| ENSMUSG( | 9.291286 | -0.2786 | 0.693249 | NA | NA |
| ENSMUSG( | 71.39673 | -0.22305 | 0.365226 | 0.914449 | NA |
| ENSMUSG( | 11.98404 | -0.72134 | 0.24533 | 0.877904 | NA |
| ENSMUSG( | 56.83411 | 0.530308 | 0.05545 | 0.723362 | NA |
| ENSMUSG( | 223.0361 | -0.23274 | 0.158856 | 0.837724 | NA |
| ENSMUSG( | 14.22411 | -0.26397 | 0.641192 | 0.953903 | NA |
| ENSMUSG( | 73.59687 | 0.0639 | 0.801281 | 0.97681 | NA |
| ENSMUSG( | 14.71529 | -0.04235 | 0.934218 | 0.99306 | NA |
| ENSMUSG( | 36.96492 | -0.00755 | 0.983418 | 0.996963 | NA |
| ENSMUSG( | 33.6995 | 0.284396 | 0.463849 | 0.923329 | NA |
| ENSMUSG( | 36.64717 | 0.187835 | 0.570105 | 0.941777 | NA |
| ENSMUSG( | 285.5812 | 0.458871 | 0.01244 | 0.451463 | NA |
| ENSMUSG( | 30.12007 | -0.0337 | 0.92699 | 0.99306 | NA |
| ENSMUSG( | 28.19486 | 0.075611 | 0.848833 | 0.985321 | NA |
| ENSMUSG( | 21.25068 | 0.270662 | 0.573997 | 0.942912 | NA |
| ENSMUSG( | 176.3344 | -0.06814 | 0.756057 | 0.971384 | NA |
| ENSMUSG( | 13.0726 | -0.26157 | 0.652667 | 0.954799 | NA |
| ENSMUSG( | 9.610089 | -0.61475 | 0.366556 | NA | NA |
| ENSMUSG( | 86.42279 | 0.158572 | 0.538027 | 0.937216 | NA |
| ENSMUSG( | 131.6104 | -0.37341 | 0.106192 | 0.804969 | NA |
| ENSMUSG( | 33.14962 | 0.06597 | 0.85146 | 0.985541 | NA |
| ENSMUSG( | 31.89393 | -0.2901 | 0.420171 | 0.917257 | NA |
| ENSMUSG( | 15.56762 | -0.12138 | 0.80577 | 0.977674 | NA |
| ENSMUSG( | 42.88158 | 0.439402 | 0.15338 | 0.837724 | NA |
| ENSMUSG( | 41.08261 | 0.083905 | 0.803659 | 0.976978 | NA |
| ENSMUSG( | 10.48399 | -0.3747 | 0.536434 | NA | NA |
| ENSMUSG( | 8.313175 | -0.01162 | 0.986698 | NA | NA |
| ENSMUSG( | 12.6293 | -0.21432 | 0.693676 | 0.960767 | NA |
| ENSMUSG( | 73.51027 | 0.130529 | 0.59391 | 0.945408 | NA |
| ENSMUSG( | 26.15922 | -0.31304 | 0.4633 | 0.923329 | NA |
| ENSMUSG( | 80.36014 | -0.79284 | 0.000829 | 0.09749 | NA |
| ENSMUSG( | 99.32515 | 0.026412 | 0.90827 | 0.991958 | NA |
| ENSMUSG( | 51.13663 | -0.01785 | 0.949789 | 0.994798 | NA |
| ENSMUSG( | 17.44495 | 0.726318 | 0.149663 | 0.836241 | NA |
| ENSMUSG( | 22.32014 | -0.45183 | 0.298715 | 0.898538 | NA |
| ENSMUSG( | 71.98855 | 0.013501 | 0.959586 | 0.994798 | NA |
| ENSMUSG( | 78.05519 | -0.39889 | 0.122222 | 0.813384 | NA |
| ENSMUSG( | 9.407051 | 0.05218 | 0.938564 | NA | NA |
| ENSMUSG( | 197.3761 | 0.029322 | 0.873142 | 0.989445 | NA |
| ENSMUSG( | 10.34788 | -0.27483 | 0.673339 | NA | NA |
| ENSMUSG( | 36.12997 | 0.236757 | 0.475584 | 0.923329 | NA |
| ENSMUSG( | 865.0197 | 0.215017 | 0.092453 | 0.793483 | NA |
| ENSMUSG( | 61.32792 | 0.130371 | 0.652435 | 0.954799 | NA |
| ENSMUSG( | 189.5427 | -0.11921 | 0.489864 | 0.925987 | NA |
| ENSMUSG( | 10.62071 | 1.140167 | 0.097623 | NA | NA |
| ENSMUSG( | 4823.297 | -0.04075 | 0.812508 | 0.979148 | NA |
| ENSMUSG( | 48.96236 | -0.37392 | 0.237288 | 0.872173 | NA |
| ENSMUSG( | 12.3579 | -0.53447 | 0.366068 | 0.914623 | NA |
| ENSMUSG( | 140.9084 | 0.077973 | 0.686927 | 0.959178 | NA |
| ENSMUSG( | 48.94096 | 0.32451 | 0.283432 | 0.892412 | NA |
| ENSMUSG( | 9.932551 | 0.596563 | 0.365036 | NA | NA |
| ENSMUSG( | 17.64745 | -0.19538 | 0.709608 | 0.96416 | NA |
| ENSMUSG( | 11.76328 | -0.06555 | 0.911735 | 0.992196 | NA |
| ENSMUSG( | 321.4355 | -0.04934 | 0.73899 | 0.968312 | NA |
| ENSMUSG( | 46.0789 | 0.463828 | 0.157448 | 0.837724 | NA |
| ENSMUSG( | 172.9018 | -0.14122 | 0.423112 | 0.917257 | NA |
| ENSMUSG( | 128.0818 | -0.35476 | 0.080242 | 0.77681 | NA |
| ENSMUSG( | 50.33504 | 0.444705 | 0.225025 | 0.868422 | NA |
| ENSMUSG( | 111.208 | 0.415697 | 0.157499 | 0.837724 | NA |
| ENSMUSG( | 25.64069 | -0.6757 | 0.099698 | 0.801103 | NA |
| ENSMUSG( | 16.28063 | 0.118084 | 0.819648 | 0.979198 | NA |
| ENSMUSG( | 11.60335 | 0.274042 | 0.639845 | 0.953903 | NA |
| ENSMUSG( | 47.96599 | 0.083614 | 0.795794 | 0.97681 | NA |
| ENSMUSG( | 121.7087 | -0.1577 | 0.464249 | 0.923329 | NA |
| ENSMUSG( | 12.26194 | -0.23263 | 0.670143 | 0.95694 | NA |
| ENSMUSG( | 13.84162 | -0.63729 | 0.220097 | 0.861004 | NA |
| ENSMUSG( | 124.7368 | 0.343747 | 0.145137 | 0.83339 | NA |
| ENSMUSG( | 69.05991 | 0.17825 | 0.648365 | 0.95422 | NA |
| ENSMUSG( | 33.84389 | 0.504733 | 0.177805 | 0.847901 | NA |
| ENSMUSG( | 30.64196 | 0.19444 | 0.642905 | 0.953903 | NA |
| ENSMUSG( | 20.94413 | 0.230175 | 0.599545 | 0.947071 | NA |
| ENSMUSG( | 12.66396 | -0.4006 | 0.482867 | 0.924615 | NA |
| ENSMUSG( | 13.09816 | -0.47645 | 0.382548 | 0.917257 | NA |
| ENSMUSG( | 11.72335 | -0.09445 | 0.871934 | 0.988779 | NA |
| ENSMUSG( | 11.26143 | -0.9888 | 0.100588 | NA | NA |
| ENSMUSG( | 29.30451 | 0.283845 | 0.474759 | 0.923329 | NA |
| ENSMUSG( | 55.91553 | -0.18105 | 0.54061 | 0.937383 | NA |
| ENSMUSG( | 18.36942 | -0.19247 | 0.675978 | 0.95694 | NA |
| ENSMUSG( | 148.8333 | -0.08938 | 0.647669 | 0.95422 | NA |
| ENSMUSG( | 17.33508 | -0.04728 | 0.921643 | 0.992644 | NA |
| ENSMUSG( | 3836.413 | -0.18312 | 0.116109 | 0.813384 | NA |
| ENSMUSG( | 447.0333 | -0.20318 | 0.202016 | 0.864412 | NA |
| ENSMUSG( | 8.685565 | 0.748048 | 0.334953 | NA | NA |
| ENSMUSG( | 11.47431 | 0.037701 | 0.949976 | 0.994798 | NA |
| ENSMUSG( | 48.90416 | 0.628072 | 0.048807 | 0.706228 | NA |
| ENSMUSG( | 15.22255 | -0.09146 | 0.861877 | 0.98632 | NA |
| ENSMUSG( | 16.50435 | 0.493688 | 0.301257 | 0.898538 | NA |
| ENSMUSG( | 47.17188 | 0.418541 | 0.292252 | 0.897854 | NA |
| ENSMUSG( | 8.799303 | -0.1084 | 0.864509 | NA | NA |
| ENSMUSG( | 10.49901 | -0.72333 | 0.234652 | NA | NA |
| ENSMUSG( | 11.75941 | -0.09279 | 0.869488 | 0.988097 | NA |
| ENSMUSG( | 16.55907 | -0.24729 | 0.629597 | 0.953251 | NA |
| ENSMUSG( | 24.07575 | 0.035637 | 0.934916 | 0.99306 | NA |
| ENSMUSG( | 7.593636 | 0.000513 | 0.999441 | NA | NA |
| ENSMUSG( | 19.36912 | -0.72287 | 0.147219 | 0.836241 | NA |
| ENSMUSG( | 20.75878 | -0.69939 | 0.119863 | 0.813384 | NA |
| ENSMUSG( | 211.0318 | 0.130287 | 0.590792 | 0.944975 | NA |
| ENSMUSG( | 50.94686 | -0.16331 | 0.565654 | 0.941404 | NA |
| ENSMUSG( | 22.24244 | 0.022651 | 0.959403 | 0.994798 | NA |
| ENSMUSG( | 12.30808 | -0.52563 | 0.352138 | 0.912093 | NA |
| ENSMUSG( | 20.77601 | 0.080211 | 0.861091 | 0.986226 | NA |
| ENSMUSG( | 18.97802 | -0.16416 | 0.722663 | 0.966887 | NA |
| ENSMUSG( | 64.97063 | 0.210678 | 0.519879 | 0.931824 | NA |
| ENSMUSG( | 18.56288 | 0.024244 | 0.957168 | 0.994798 | NA |
| ENSMUSG( | 21.49901 | 0.337913 | 0.570263 | 0.941823 | NA |
| ENSMUSG( | 14.64787 | -0.48368 | 0.374767 | 0.915491 | NA |
| ENSMUSG( | 13.57792 | -0.01152 | 0.98345 | 0.996963 | NA |
| ENSMUSG( | 10.11286 | -0.251 | 0.678709 | NA | NA |
| ENSMUSG( | 6.998319 | -0.43652 | 0.571502 | NA | NA |
| ENSMUSG( | 19.86668 | -0.97556 | 0.029773 | 0.61992 | NA |
| ENSMUSG( | 9.760741 | 0.2776 | 0.655069 | NA | NA |
| ENSMUSG( | 28.57722 | -0.08086 | 0.851552 | 0.985541 | NA |
| ENSMUSG( | 30.3805 | 0.756456 | 0.124823 | 0.81516 | NA |
| ENSMUSG( | 18.96822 | 0.126792 | 0.795564 | 0.97681 | NA |
| ENSMUSG( | 18.751 | 0.032447 | 0.944936 | 0.994798 | NA |
| ENSMUSG( | 19.83744 | -0.22076 | 0.618979 | 0.952598 | NA |
| ENSMUSG( | 17.54829 | 0.43947 | 0.356904 | 0.913358 | NA |
| ENSMUSG( | 50.05397 | 0.083471 | 0.781022 | 0.976106 | NA |
| ENSMUSG( | 204.0776 | 0.104555 | 0.598159 | 0.946676 | NA |
| ENSMUSG( | 108.2904 | -0.22322 | 0.298237 | 0.898538 | NA |
| ENSMUSG( | 55.7295 | -0.53495 | 0.05861 | 0.728256 | NA |
| ENSMUSG( | 23.74118 | 0.833978 | 0.082351 | 0.77681 | NA |
| ENSMUSG( | 42.83102 | -0.05087 | 0.882129 | 0.989644 | NA |
| ENSMUSG( | 46.64827 | 0.135848 | 0.653429 | 0.955015 | NA |
| ENSMUSG( | 66.57577 | 0.18847 | 0.486176 | 0.92541 | NA |
| ENSMUSG( | 12.69433 | -0.19783 | 0.714674 | 0.964986 | NA |
| ENSMUSG( | 141.5115 | -0.08278 | 0.675689 | 0.95694 | NA |
| ENSMUSG( | 42.38774 | -0.27731 | 0.388484 | 0.917257 | NA |
| ENSMUSG( | 65.01875 | 0.170775 | 0.576936 | 0.942912 | NA |
| ENSMUSG( | 10.49154 | -0.77152 | 0.222093 | NA | NA |
| ENSMUSG( | 9.774526 | -0.47063 | 0.45534 | NA | NA |
| ENSMUSG( | 62.374 | -0.42697 | 0.120731 | 0.813384 | NA |
| ENSMUSG( | 13.38455 | -0.45747 | 0.434535 | 0.91869 | NA |
| ENSMUSG( | 9.515216 | 0.604957 | 0.373759 | NA | NA |
| ENSMUSG( | 26.56599 | -0.40555 | 0.321322 | 0.904019 | NA |
| ENSMUSG( | 22.05453 | 0.435811 | 0.31108 | 0.899422 | NA |
| ENSMUSG( | 10.91086 | -0.22457 | 0.709344 | NA | NA |
| ENSMUSG( | 25.82291 | -0.6436 | 0.107147 | 0.804969 | NA |
| ENSMUSG( | 53.40481 | 0.066922 | 0.817861 | 0.979198 | NA |
| ENSMUSG( | 14.2977 | 0.960957 | 0.114826 | 0.811703 | NA |
| ENSMUSG( | 11.19714 | -0.48776 | 0.408675 | NA | NA |
| ENSMUSG( | 11.42804 | -0.03239 | 0.954894 | NA | NA |

| | | | | | | | | | | | |
|---|---|---|---|---|---|---|---|---|---|---|---|
| ENSMUSG( | NA | 1013.283 | -0.52437 | 7.25E-07 | 0.000149 | ENSMUSG( | NA | 42.83102 | -0.05087 | 0.882129 | 0.989644 |
| ENSMUSG( | NA | 55.73503 | -0.14768 | 0.573029 | 0.893584 | ENSMUSG( | NA | 46.64827 | 0.135848 | 0.653429 | 0.955015 |
| ENSMUSG( | NA | 12.10958 | 0.580528 | 0.375648 | NA | ENSMUSG( | NA | 66.57577 | 0.18847 | 0.486176 | 0.92541 |
| ENSMUSG( | NA | 66.41846 | -0.27045 | 0.287479 | 0.738036 | ENSMUSG( | NA | 12.69433 | -0.19783 | 0.714674 | 0.964986 |
| ENSMUSG( | NA | 81.23043 | 0.052265 | 0.83526 | 0.970426 | ENSMUSG( | NA | 141.5115 | -0.08278 | 0.675689 | 0.95694 |
| ENSMUSG( | NA | 205.3257 | -0.05435 | 0.746694 | 0.949289 | ENSMUSG( | NA | 42.38774 | -0.27731 | 0.388484 | 0.917257 |
| ENSMUSG( | NA | 19.25605 | 0.556867 | 0.221926 | NA | ENSMUSG( | NA | 65.01875 | 0.170775 | 0.576936 | 0.942912 |
| ENSMUSG( | NA | 21.47452 | -0.54801 | 0.190931 | 0.630245 | ENSMUSG( | NA | 10.49154 | -0.77152 | 0.222093 | NA |
| ENSMUSG( | NA | 86.16698 | -0.21125 | 0.361269 | 0.792797 | ENSMUSG( | NA | 9.774526 | -0.47063 | 0.45534 | NA |
| ENSMUSG( | NA | 64.81359 | 0.277705 | 0.280026 | 0.729155 | ENSMUSG( | NA | 62.374 | -0.42697 | 0.120731 | 0.813384 |
| ENSMUSG( | NA | 26.95192 | 0.069156 | 0.844857 | 0.972023 | ENSMUSG( | NA | 13.38455 | -0.45747 | 0.434535 | 0.91869 |
| ENSMUSG( | NA | 41.85044 | 0.508173 | 0.150412 | 0.574991 | ENSMUSG( | NA | 26.56599 | -0.40555 | 0.321322 | 0.904019 |
| ENSMUSG( | NA | 11.26791 | -0.06761 | 0.900746 | NA | ENSMUSG( | NA | 22.05453 | 0.435811 | 0.31108 | 0.899422 |
| ENSMUSG( | NA | 20.69342 | 0.431936 | 0.325994 | NA | ENSMUSG( | NA | 10.91086 | -0.22457 | 0.709344 | NA |
| ENSMUSG( | NA | 13.34331 | -0.61903 | 0.257671 | NA | ENSMUSG( | NA | 25.82291 | -0.6436 | 0.107147 | 0.804969 |
| ENSMUSG( | NA | 234.7045 | -0.06422 | 0.655317 | 0.923929 | ENSMUSG( | NA | 53.40481 | 0.066922 | 0.817861 | 0.979198 |
| ENSMUSG( | NA | 66.52422 | 0.191754 | 0.435672 | 0.829093 | ENSMUSG( | NA | 14.2977 | 0.960957 | 0.114826 | 0.811703 |
| ENSMUSG( | NA | 1729.637 | 0.055822 | 0.583373 | 0.898434 | ENSMUSG( | NA | 11.19714 | -0.48776 | 0.408675 | NA |
| ENSMUSG( | NA | 18.04549 | 0.508568 | 0.256536 | NA | ENSMUSG( | NA | 11.42804 | -0.03239 | 0.954894 | NA |
| ENSMUSG( | NA | 12.76613 | 0.139714 | 0.800774 | NA | ENSMUSG( | NA | 12.24207 | -0.04542 | 0.935179 | 0.993078 |
| ENSMUSG( | NA | 10.9084 | -0.87161 | 0.133881 | NA | ENSMUSG( | NA | 26.48545 | 0.134351 | 0.734066 | 0.967228 |
| ENSMUSG( | NA | 11.01291 | -0.11729 | 0.831897 | NA | ENSMUSG( | NA | 33.3218 | -0.37226 | 0.336194 | 0.907972 |
| ENSMUSG( | NA | 9.817443 | -0.17148 | 0.776919 | NA | ENSMUSG( | NA | 37.42062 | -0.73047 | 0.034534 | 0.655405 |
| ENSMUSG( | NA | 31.91604 | -0.5298 | 0.120304 | 0.524495 | ENSMUSG( | NA | 26.07328 | -0.4315 | 0.269211 | 0.886755 |
| ENSMUSG( | NA | 177.7524 | 0.056725 | 0.740946 | 0.948028 | ENSMUSG( | NA | 14.28218 | -0.32173 | 0.548458 | 0.938063 |
| ENSMUSG( | NA | 11.79359 | -0.28667 | 0.589422 | NA | ENSMUSG( | NA | 45.63077 | -0.09086 | 0.774577 | 0.974931 |
| ENSMUSG( | NA | 14.2923 | 0.390278 | 0.459267 | NA | ENSMUSG( | NA | 23.41759 | -0.25698 | 0.532688 | 0.934795 |
| ENSMUSG( | NA | 18.63065 | -0.49277 | 0.284147 | NA | ENSMUSG( | NA | 19.92366 | -0.0113 | 0.979382 | 0.996468 |
| ENSMUSG( | NA | 16.68986 | -0.2232 | 0.637706 | NA | ENSMUSG( | NA | 11.2359 | -0.33228 | 0.594185 | NA |
| ENSMUSG( | NA | 11.91903 | -0.37755 | 0.486532 | NA | ENSMUSG( | NA | 7.212321 | -2.29752 | 0.005811 | NA |
| ENSMUSG( | NA | 56.64289 | 0.047929 | 0.855795 | 0.973813 | ENSMUSG( | NA | 42.80983 | -0.12137 | 0.691535 | 0.960102 |
| ENSMUSG( | NA | 68.76629 | -0.07824 | 0.765117 | 0.954127 | ENSMUSG( | NA | 73.10768 | -0.30888 | 0.208115 | 0.864412 |
| ENSMUSG( | NA | 12.98667 | -0.29572 | 0.58675 | NA | ENSMUSG( | NA | 34.78528 | -0.30453 | 0.413975 | 0.917257 |
| ENSMUSG( | NA | 25.08784 | 0.163639 | 0.66269 | 0.926303 | ENSMUSG( | NA | 12.25733 | -0.46109 | 0.410094 | 0.917257 |
| ENSMUSG( | NA | 17.20561 | 0.271866 | 0.570451 | NA | ENSMUSG( | NA | 217.5113 | 0.106099 | 0.546214 | 0.938063 |
| ENSMUSG( | NA | 14.12135 | -0.60786 | 0.230971 | NA | ENSMUSG( | NA | 32.17211 | -0.00638 | 0.987238 | 0.997487 |
| ENSMUSG( | NA | 48.96121 | -0.77911 | 0.022452 | 0.228899 | ENSMUSG( | NA | 11.25429 | -0.34288 | 0.559025 | NA |
| ENSMUSG( | NA | 15.22351 | 0.110348 | 0.836285 | NA | ENSMUSG( | NA | 34.22178 | 0.009842 | 0.977629 | 0.996416 |
| ENSMUSG( | NA | 17.29482 | 0.165166 | 0.734987 | NA | ENSMUSG( | NA | 15.37158 | 0.457664 | 0.390837 | 0.917257 |
| ENSMUSG( | NA | 26.56448 | 0.101204 | 0.797587 | 0.960507 | ENSMUSG( | NA | 15.10468 | 0.303265 | 0.543932 | 0.938063 |
| ENSMUSG( | NA | 87.30191 | 0.550306 | 0.064371 | 0.393403 | ENSMUSG( | NA | 9.536745 | -0.42691 | 0.523398 | NA |
| ENSMUSG( | NA | 34.13216 | 0.246184 | 0.522575 | 0.875389 | ENSMUSG( | NA | 118.999 | -0.21976 | 0.315606 | 0.901725 |
| ENSMUSG( | NA | 22.63045 | -0.01763 | 0.96319 | 0.994911 | ENSMUSG( | NA | 15.82816 | 0.096176 | 0.841878 | 0.983044 |
| ENSMUSG( | NA | 237.8555 | -0.04135 | 0.794783 | 0.959799 | ENSMUSG( | NA | 17.68228 | -0.52981 | 0.262925 | 0.885357 |
| ENSMUSG( | NA | 47.17501 | 0.022797 | 0.938293 | 0.993415 | ENSMUSG( | NA | 35.85108 | -0.885 | 0.013397 | 0.463609 |
| ENSMUSG( | NA | 28.40897 | -0.28215 | 0.4267 | 0.825155 | ENSMUSG( | NA | 16.98518 | -0.08448 | 0.876897 | 0.989644 |
| ENSMUSG( | NA | 16.55796 | 0.548862 | 0.242377 | NA | ENSMUSG( | NA | 24.20399 | -0.11129 | 0.795439 | 0.97681 |
| ENSMUSG( | NA | 12.09114 | 1.363644 | 0.020985 | NA | ENSMUSG( | NA | 22.57768 | -0.25103 | 0.546712 | 0.938063 |
| ENSMUSG( | NA | 16.96755 | -0.77815 | 0.092948 | NA | ENSMUSG( | NA | 12.64995 | 0.355689 | 0.532314 | 0.934795 |
| ENSMUSG( | NA | 37.01982 | -0.06074 | 0.847603 | 0.972374 | ENSMUSG( | NA | 20.32035 | -0.33439 | 0.501132 | 0.927825 |
| ENSMUSG( | NA | 40.0066 | -0.03726 | 0.908203 | 0.9878 | ENSMUSG( | NA | 28.86819 | -1.44078 | 0.000231 | 0.039691 |
| ENSMUSG( | NA | 33.89792 | 0.109584 | 0.770644 | 0.954722 | ENSMUSG( | NA | 15.1532 | -0.04973 | 0.924732 | 0.99306 |
| ENSMUSG( | NA | 205.793 | 0.924115 | 9.75E-05 | 0.006367 | ENSMUSG( | NA | 140.8562 | 0.216477 | 0.29456 | 0.898538 |
| ENSMUSG( | NA | 23.90365 | -0.0202 | 0.956986 | 0.993874 | ENSMUSG( | NA | 31.64028 | -0.65073 | 0.070594 | 0.758714 |
| ENSMUSG( | NA | 20.27387 | -0.23417 | 0.594848 | NA | ENSMUSG( | NA | 375.6215 | 0.42745 | 0.350046 | 0.911652 |
| ENSMUSG( | NA | 18.95284 | 0.512023 | 0.305447 | NA | ENSMUSG( | NA | 135.0807 | -0.03506 | 0.883388 | 0.989644 |
| ENSMUSG( | NA | 59.79461 | -0.06527 | 0.802379 | 0.962119 | ENSMUSG( | NA | 65.83759 | 0.07744 | 0.758917 | 0.971858 |
| ENSMUSG( | NA | 9.230105 | 0.012263 | 0.984589 | NA | ENSMUSG( | NA | 109.0677 | 0.011866 | 0.964007 | 0.995433 |
| ENSMUSG( | NA | 10.34994 | 0.065508 | 0.91216 | NA | ENSMUSG( | NA | 9.020315 | -0.24648 | 0.735035 | NA |
| ENSMUSG( | NA | 17.88612 | -0.02326 | 0.957068 | NA | ENSMUSG( | NA | 231.794 | -0.11892 | 0.519077 | 0.931053 |
| ENSMUSG( | NA | 32.07299 | 0.039835 | 0.903257 | 0.986055 | ENSMUSG( | NA | 12.25744 | -0.17963 | 0.753905 | 0.970671 |
| ENSMUSG( | NA | 15.15265 | 0.148777 | 0.765139 | NA | ENSMUSG( | NA | 9.333652 | -0.20284 | 0.755852 | NA |
| ENSMUSG( | NA | 17.01822 | 0.00063 | 0.998894 | NA | ENSMUSG( | NA | 18.27404 | -0.37357 | 0.465524 | 0.923329 |
| ENSMUSG( | NA | 26.39934 | 0.113367 | 0.755159 | 0.951279 | ENSMUSG( | NA | 22.04455 | -0.47806 | 0.273807 | 0.889848 |
| ENSMUSG( | NA | 39.94222 | 0.512142 | 0.15662 | 0.584667 | ENSMUSG( | NA | 47.76307 | 0.016497 | 0.963192 | 0.995433 |
| ENSMUSG( | NA | 18.57369 | 0.042897 | 0.927461 | NA | ENSMUSG( | NA | 11.82908 | 0.194151 | 0.736485 | 0.967344 |
| ENSMUSG( | NA | 64.85647 | 0.215634 | 0.389665 | 0.807428 | ENSMUSG( | NA | 123.7957 | 0.037846 | 0.86267 | 0.986547 |
| ENSMUSG( | NA | 20.07816 | 0.11468 | 0.79542 | NA | ENSMUSG( | NA | 178.4514 | -0.06272 | 0.757635 | 0.971858 |
| ENSMUSG( | NA | 12.31115 | -0.05851 | 0.929726 | NA | ENSMUSG( | NA | 467.6754 | -0.04463 | 0.765828 | 0.974236 |
| ENSMUSG( | NA | 39.61697 | -0.72924 | 0.022478 | 0.229021 | ENSMUSG( | NA | 450.4104 | 0.101169 | 0.51508 | 0.930939 |
| ENSMUSG( | NA | 94.43734 | 0.641558 | 0.022001 | 0.226001 | ENSMUSG( | NA | 22.18394 | 0.489499 | 0.267418 | 0.886134 |
| ENSMUSG( | NA | 45.37999 | 0.205146 | 0.520351 | 0.875186 | ENSMUSG( | NA | 62.40043 | -0.23789 | 0.38654 | 0.917257 |
| ENSMUSG( | NA | 58.17057 | 0.467403 | 0.122226 | 0.527707 | ENSMUSG( | NA | 37.29866 | 0.90692 | 0.060931 | 0.73558 |
| ENSMUSG( | NA | 12.39207 | 0.315603 | 0.557321 | NA | ENSMUSG( | NA | 7.212457 | -0.68135 | 0.393077 | NA |
| ENSMUSG( | NA | 116.017 | -0.02877 | 0.884962 | 0.98282 | ENSMUSG( | NA | 12.61527 | -0.26919 | 0.669085 | 0.956766 |
| ENSMUSG( | NA | 12.99619 | 0.830927 | 0.144472 | NA | ENSMUSG( | NA | 15.11137 | 0.11017 | 0.836062 | 0.981237 |
| ENSMUSG( | NA | 15.66088 | -0.04044 | 0.931248 | NA | ENSMUSG( | NA | 13.91558 | -0.52268 | 0.322493 | 0.904023 |
| ENSMUSG( | NA | 101.0115 | 1.002476 | 0.000433 | 0.018963 | ENSMUSG( | NA | 10.32524 | -0.19327 | 0.753697 | NA |
| ENSMUSG( | NA | 13.86418 | -0.21824 | 0.666845 | NA | ENSMUSG( | NA | 12.15014 | 0.082569 | 0.88385 | 0.989644 |
| ENSMUSG( | NA | 145.8716 | -0.09053 | 0.642225 | 0.916865 | ENSMUSG( | NA | 11.33417 | -0.13897 | 0.806288 | NA |
| ENSMUSG( | NA | 662.5641 | 0.11261 | 0.47942 | 0.854046 | ENSMUSG( | NA | 1595.55 | -0.179 | 0.113639 | 0.809623 |
| ENSMUSG( | NA | 16.38792 | -0.05639 | 0.909088 | NA | ENSMUSG( | NA | 8.749257 | -0.73953 | 0.266088 | NA |
| ENSMUSG( | NA | 11.70264 | 0.371776 | 0.549753 | NA | ENSMUSG( | NA | 39.51012 | -0.40624 | 0.212834 | 0.864412 |
| ENSMUSG( | NA | 14.78194 | -0.16942 | 0.735328 | NA | ENSMUSG( | NA | 21.3732 | -0.00287 | 0.994821 | 0.998813 |
| ENSMUSG( | NA | 12.04996 | 0.641788 | 0.336268 | NA | ENSMUSG( | NA | 21.65837 | 0.77194 | 0.154921 | 0.837724 |
| ENSMUSG( | NA | 27.45145 | 0.150337 | 0.701459 | 0.937441 | ENSMUSG( | NA | 25.0448 | 0.818834 | 0.055769 | 0.723362 |
| ENSMUSG( | NA | 19.80264 | -0.1936 | 0.689389 | NA | ENSMUSG( | NA | 37.56007 | -0.36848 | 0.461995 | 0.923329 |
| ENSMUSG( | NA | 13.25773 | 0.204519 | 0.688499 | NA | ENSMUSG( | NA | 983.9592 | 0.20637 | 0.197618 | 0.862161 |
| ENSMUSG( | NA | 34.43466 | -0.10902 | 0.732425 | 0.944956 | ENSMUSG( | NA | 11.99207 | 0.423468 | 0.449814 | 0.922202 |
| ENSMUSG( | NA | 45.07382 | 0.414219 | 0.201661 | 0.641362 | ENSMUSG( | NA | 13.95523 | 0.432852 | 0.425487 | 0.917257 |
| ENSMUSG( | NA | 463.3774 | -0.07429 | 0.568438 | 0.892252 | ENSMUSG( | NA | 19.07844 | 0.180656 | 0.690493 | 0.960071 |
| ENSMUSG( | NA | 581.1441 | 0.169672 | 0.123536 | 0.530552 | ENSMUSG( | NA | 284.4994 | 0.050691 | 0.76529 | 0.974236 |
| ENSMUSG( | NA | 19.44691 | 0.0856 | 0.860402 | NA | ENSMUSG( | NA | 18.98802 | 0.294819 | 0.616318 | 0.951413 |
| ENSMUSG( | NA | 22.84251 | 0.099993 | 0.798164 | 0.96069 | ENSMUSG( | NA | 138.866 | 0.049523 | 0.818821 | 0.979198 |
| ENSMUSG( | NA | 27.078 | 0.169393 | 0.635334 | 0.914296 | ENSMUSG( | NA | 9.52931 | -0.09964 | 0.886022 | NA |
| ENSMUSG( | NA | 10.36833 | -0.21879 | 0.699512 | NA | ENSMUSG( | NA | 30.40916 | 0.218246 | 0.537688 | 0.937216 |
| ENSMUSG( | NA | 30.2951 | -0.27138 | 0.430897 | 0.826896 | ENSMUSG( | NA | 42.33024 | -0.08258 | 0.856584 | 0.985884 |
| ENSMUSG( | NA | 105.7471 | 0.116605 | 0.568549 | 0.892294 | ENSMUSG( | NA | 25.65405 | 0.369072 | 0.340634 | 0.907972 |
| ENSMUSG( | NA | 60.83049 | 0.407736 | 0.106018 | 0.493403 | ENSMUSG( | NA | 63.27542 | -0.80426 | 0.002254 | 0.184738 |
| ENSMUSG( | NA | 14.84987 | -1.27613 | 0.026927 | NA | ENSMUSG( | NA | 11.58846 | -0.75251 | 0.230101 | 0.868507 |
| ENSMUSG( | NA | 14.40947 | -0.30598 | 0.5454 | NA | ENSMUSG( | NA | 14.97943 | 1.009249 | 0.052804 | 0.71525 |
| ENSMUSG( | NA | 46.30595 | -0.26434 | 0.420032 | 0.824495 | ENSMUSG( | NA | 38.97534 | 0.048382 | 0.883332 | 0.989644 |
| ENSMUSG( | NA | 139.7563 | 0.081942 | 0.65958 | 0.924506 | ENSMUSG( | NA | 19.92238 | 0.32719 | 0.469182 | 0.923329 |
| ENSMUSG( | NA | 101.3457 | 0.239271 | 0.24453 | 0.69183 | ENSMUSG( | NA | 86.15414 | -0.20624 | 0.372837 | 0.915491 |
| ENSMUSG( | NA | 21.74232 | 0.175777 | 0.682778 | 0.932022 | ENSMUSG( | NA | 43.96823 | 0.150242 | 0.618599 | 0.952363 |
| ENSMUSG( | NA | 24.67154 | -0.40225 | 0.343635 | 0.779153 | ENSMUSG( | NA | 222.703 | -0.04056 | 0.833204 | 0.981192 |
| ENSMUSG( | NA | 22.76615 | -0.39956 | 0.356013 | 0.788009 | ENSMUSG( | NA | 8.663521 | -0.72977 | 0.318323 | NA |
| ENSMUSG( | NA | 11.3482 | 0.649892 | 0.256683 | NA | ENSMUSG( | NA | 59.96383 | 0.396242 | 0.147254 | 0.836241 |
| ENSMUSG( | NA | 34.47438 | 0.361771 | 0.258116 | 0.70936 | ENSMUSG( | NA | 12.96214 | 0.203777 | 0.708608 | 0.964042 |
| ENSMUSG( | NA | 39.11174 | 0.443409 | 0.165319 | 0.596691 | ENSMUSG( | NA | 18.9461 | -0.30169 | 0.545173 | 0.938063 |
| ENSMUSG( | NA | 36.22454 | -0.06569 | 0.833759 | 0.970333 | ENSMUSG( | NA | 16.95957 | -0.00055 | 0.999105 | 0.999503 |
| ENSMUSG( | NA | 60.17666 | 0.299827 | 0.235415 | 0.682423 | ENSMUSG( | NA | 21.1969 | 0.259832 | 0.612537 | 0.951177 |
| ENSMUSG( | NA | 408.4982 | 0.1747 | 0.137935 | 0.552903 | ENSMUSG( | NA | 13.94741 | 0.380959 | 0.494222 | 0.927372 |
| ENSMUSG( | NA | 20.12225 | 0.165079 | 0.723289 | NA | ENSMUSG( | NA | 855.8646 | -0.38745 | 0.004069 | 0.25824 |
| ENSMUSG( | NA | 20.18681 | 0.064798 | 0.882567 | NA | ENSMUSG( | NA | 28.91677 | 0.124733 | 0.734159 | 0.967228 |
| ENSMUSG( | NA | 30.82317 | 0.225471 | 0.521258 | 0.875186 | ENSMUSG( | NA | 17.78778 | -0.1122 | 0.81245 | 0.979148 |
| ENSMUSG( | NA | 36.20208 | -0.02294 | 0.944113 | 0.993503 | ENSMUSG( | NA | 16.36563 | 0.033838 | 0.948103 | 0.994798 |
| ENSMUSG( | NA | 23.80663 | 0.346073 | 0.385183 | 0.804048 | ENSMUSG( | NA | 9.444498 | 0.286313 | 0.650208 | NA |
| ENSMUSG( | NA | 11.91096 | -0.89239 | 0.101137 | NA | ENSMUSG( | NA | 18.06606 | 0.235916 | 0.602415 | 0.94789 |
| ENSMUSG( | NA | 17.15567 | 1.108584 | 0.04388 | NA | ENSMUSG( | NA | 11.68187 | -0.80984 | 0.156471 | 0.837724 |
| ENSMUSG( | NA | 62.68122 | -0.18439 | 0.480809 | 0.855113 | ENSMUSG( | NA | 62.99686 | 0.122672 | 0.639393 | 0.953903 |
| ENSMUSG( | NA | 15.64019 | -0.07469 | 0.878741 | NA | ENSMUSG( | NA | 12.44709 | -0.05604 | 0.920773 | 0.992644 |
| ENSMUSG( | NA | 580.0301 | -0.31146 | 0.070492 | 0.40908 | ENSMUSG( | NA | 20.96259 | -0.663 | 0.133241 | 0.822912 |
| ENSMUSG( | NA | 25.24371 | 0.34031 | 0.386129 | 0.804623 | ENSMUSG( | NA | 8.133185 | -0.14756 | 0.841159 | NA |
| ENSMUSG( | NA | 19.66507 | 0.162225 | 0.709177 | NA | ENSMUSG( | NA | 9.032614 | -0.27101 | 0.682327 | NA |
| ENSMUSG( | NA | 144.0272 | 0.25837 | 0.256271 | 0.706924 | ENSMUSG( | NA | 26.54696 | 0.122726 | 0.767644 | 0.974236 |
| ENSMUSG( | NA | 12.03011 | -0.17396 | 0.741676 | NA | ENSMUSG( | NA | 22.56368 | -0.55653 | 0.221115 | 0.866104 |
| ENSMUSG( | NA | 28.47364 | -0.17565 | 0.629876 | 0.913965 | ENSMUSG( | NA | 96.33083 | 0.067079 | 0.79313 | 0.97681 |
| ENSMUSG( | NA | 26.51093 | 0.489163 | 0.187169 | 0.625368 | ENSMUSG( | NA | 122.3446 | 0.308245 | 0.128021 | 0.81845 |
| ENSMUSG( | NA | 43.10863 | 0.009879 | 0.974273 | 0.996953 | ENSMUSG( | NA | 60.11288 | -0.47776 | 0.138441 | 0.828492 |
| ENSMUSG( | NA | 29.71924 | 0.082918 | 0.817724 | 0.965785 | ENSMUSG( | NA | 103.5443 | 0.084166 | 0.798184 | 0.97681 |
| ENSMUSG( | NA | 63.63252 | -0.09583 | 0.706896 | 0.93949 | ENSMUSG( | NA | 47.06341 | 0.006965 | 0.981418 | 0.996756 |
| ENSMUSG( | NA | 511.7439 | 0.182109 | 0.155367 | 0.583346 | ENSMUSG( | NA | 26.13693 | -0.13881 | 0.729713 | 0.96719 |
| ENSMUSG( | NA | 16.30154 | 0.942707 | 0.044696 | NA | ENSMUSG( | NA | 10.71641 | -0.28078 | 0.640579 | NA |
| ENSMUSG( | NA | 50.56412 | -0.35553 | 0.228742 | 0.675959 | ENSMUSG( | NA | 22.94363 | -0.22998 | 0.584571 | 0.944975 |
| ENSMUSG( | NA | 16.15375 | -0.55694 | 0.231564 | NA | ENSMUSG( | NA | 55.00645 | 0.747169 | 0.020232 | 0.563435 |
| ENSMUSG( | NA | 20.66388 | 0.484178 | 0.259899 | NA | ENSMUSG( | NA | 21.22069 | 0.336171 | 0.446503 | 0.921734 |
| ENSMUSG( | NA | 16.20848 | -0.4849 | 0.311371 | NA | ENSMUSG( | NA | 33.28157 | -0.48868 | 0.236456 | 0.872173 |
| ENSMUSG( | NA | 222.2279 | 0.117995 | 0.457945 | 0.839769 | ENSMUSG( | NA | 10.56373 | -0.19755 | 0.757539 | NA |
| ENSMUSG( | NA | 25.28463 | 0.459505 | 0.25799 | 0.709319 | ENSMUSG( | NA | 65.31069 | 0.231745 | 0.407649 | 0.917257 |
| ENSMUSG( | NA | 77.81787 | 0.018811 | 0.932387 | 0.992515 | ENSMUSG( | NA | 11.84561 | -1.03797 | 0.082584 | 0.77681 |
| ENSMUSG( | NA | 8.54217 | 0.248742 | 0.696912 | NA | ENSMUSG( | NA | 76.4696 | 0.033408 | 0.900121 | 0.990558 |
| ENSMUSG( | NA | 8.864524 | 1.152631 | 0.101628 | NA | ENSMUSG( | NA | 14.83786 | -0.49873 | 0.334772 | 0.907907 |
| ENSMUSG( | NA | 972.5091 | 0.280051 | 0.017396 | 0.197592 | ENSMUSG( | NA | 16.51509 | -0.41851 | 0.429457 | 0.917257 |
| ENSMUSG( | NA | 52.43066 | 0.255456 | 0.337611 | 0.777422 | ENSMUSG( | NA | 15.43301 | -0.14706 | 0.768322 | 0.974236 |
| ENSMUSG( | NA | 16.28673 | 0.530758 | 0.285999 | NA | ENSMUSG( | NA | 8.616208 | -0.25887 | 0.718035 | NA |
| ENSMUSG( | NA | 14.49861 | -0.53857 | 0.310519 | NA | ENSMUSG( | NA | 22.07807 | 0.114228 | 0.793384 | 0.97681 |
| ENSMUSG( | NA | 31.52423 | 0.098409 | 0.787608 | 0.959218 | ENSMUSG( | NA | 10.20755 | 0.486671 | 0.466283 | NA |
| ENSMUSG( | NA | 40.03989 | -0.22781 | 0.46383 | 0.842812 | ENSMUSG( | NA | 43.12577 | 0.432473 | 0.210952 | 0.864412 |
| ENSMUSG( | NA | 119.015 | -0.10423 | 0.604644 | 0.905556 | ENSMUSG( | NA | 37.70721 | 0.018432 | 0.958699 | 0.994798 |
| ENSMUSG( | NA | 18.46621 | 0.003336 | 0.994493 | NA | ENSMUSG( | NA | 206.3802 | 0.021793 | 0.92977 | 0.99306 |
| ENSMUSG( | NA | 66.84483 | 0.686158 | 0.003907 | 0.080172 | ENSMUSG( | NA | 14.7096 | -0.51657 | 0.312601 | 0.900037 |
| ENSMUSG( | NA | 13.65288 | -0.49241 | 0.335464 | NA | ENSMUSG( | NA | 1379.958 | -0.06567 | 0.572333 | 0.942515 |
| ENSMUSG( | NA | 78.12046 | -0.23419 | 0.285255 | 0.733952 | ENSMUSG( | NA | 10.67068 | -0.81497 | 0.196271 | NA |
| ENSMUSG( | NA | 304.4585 | 0.381373 | 0.083486 | 0.445586 | ENSMUSG( | NA | 54.52519 | -0.37816 | 0.187014 | 0.856099 |
| ENSMUSG( | NA | 19.09609 | -0.56735 | 0.189528 | NA | ENSMUSG( | NA | 10.22554 | -0.13634 | 0.827494 | NA |
| ENSMUSG( | NA | 9.824615 | 0.039956 | 0.949706 | NA | ENSMUSG( | NA | 84.25232 | -0.08816 | 0.738214 | 0.967839 |
| ENSMUSG( | NA | 10.45425 | -0.51021 | 0.434238 | NA | ENSMUSG( | NA | 14.32442 | -0.22485 | 0.671833 | 0.95694 |

| | | | | | |
|---|---|---|---|---|---|
| ENSMUSG( | 7.912114 | -0.2413 | 0.754519 | NA | NA |
| ENSMUSG( | 25.92168 | 0.118224 | 0.744472 | 0.948797 | NA |
| ENSMUSG( | 13.45474 | 0.357853 | 0.494754 | NA | NA |
| ENSMUSG( | 79.82227 | 0.167501 | 0.459038 | 0.840244 | NA |
| ENSMUSG( | 47.96136 | -0.53293 | 0.060901 | 0.381246 | NA |
| ENSMUSG( | 14.6491 | -1.46907 | 0.016265 | NA | NA |
| ENSMUSG( | 154.2644 | -0.0056 | 0.974771 | 0.997207 | NA |
| ENSMUSG( | 15.01695 | 0.61632 | 0.237289 | NA | NA |
| ENSMUSG( | 413.704 | 0.063679 | 0.658757 | 0.924506 | NA |
| ENSMUSG( | 146.0288 | 0.168089 | 0.375797 | 0.800976 | NA |
| ENSMUSG( | 113.2868 | 0.004313 | 0.984875 | 0.997815 | NA |
| ENSMUSG( | 263.4217 | -0.08777 | 0.564569 | 0.890434 | NA |
| ENSMUSG( | 13.08349 | -0.13229 | 0.801977 | NA | NA |
| ENSMUSG( | 65.18268 | -0.10416 | 0.72449 | 0.944096 | NA |
| ENSMUSG( | 36.2355 | -0.04255 | 0.900283 | 0.985633 | NA |
| ENSMUSG( | 40.10066 | 0.25091 | 0.430216 | 0.826789 | NA |
| ENSMUSG( | 611.5774 | -0.05734 | 0.660339 | 0.924627 | NA |
| ENSMUSG( | 230.4404 | -0.68018 | 0.001911 | 0.049911 | NA |
| ENSMUSG( | 16.82833 | 0.305785 | 0.496834 | NA | NA |
| ENSMUSG( | 9.654279 | -0.81612 | 0.190537 | NA | NA |
| ENSMUSG( | 198.4852 | -0.07337 | 0.628152 | 0.913532 | NA |
| ENSMUSG( | 26.82703 | -0.05588 | 0.890413 | 0.983687 | NA |
| ENSMUSG( | 30.83124 | 0.025043 | 0.942586 | 0.993415 | NA |
| ENSMUSG( | 18.48125 | -0.23806 | 0.669683 | NA | NA |
| ENSMUSG( | 29.01989 | 0.405044 | 0.244365 | 0.691487 | NA |
| ENSMUSG( | 8.372702 | 0.546886 | 0.441312 | NA | NA |
| ENSMUSG( | 44.44207 | -0.05123 | 0.860274 | 0.974842 | NA |
| ENSMUSG( | 15.19655 | -0.37606 | 0.428037 | NA | NA |
| ENSMUSG( | 228.9401 | -0.15784 | 0.352998 | 0.784977 | NA |
| ENSMUSG( | 15.77198 | -0.73442 | 0.151729 | NA | NA |
| ENSMUSG( | 38.16992 | 0.049594 | 0.869842 | 0.977607 | NA |
| ENSMUSG( | 12.43342 | -0.19496 | 0.715175 | NA | NA |
| ENSMUSG( | 12.13188 | -0.53555 | 0.325317 | NA | NA |
| ENSMUSG( | 14.59856 | -0.14336 | 0.766164 | NA | NA |
| ENSMUSG( | 12.8689 | 0.609488 | 0.256572 | NA | NA |
| ENSMUSG( | 12.16796 | 0.010304 | 0.98424 | NA | NA |
| ENSMUSG( | 373.1042 | -0.23592 | 0.139289 | 0.555376 | NA |
| ENSMUSG( | 42.10618 | 0.240136 | 0.434463 | 0.828978 | NA |
| ENSMUSG( | 18.24699 | -0.24696 | 0.57832 | NA | NA |
| ENSMUSG( | 26.32569 | -0.19103 | 0.620426 | 0.911432 | NA |
| ENSMUSG( | 194.2323 | 0.421962 | 0.012578 | 0.161972 | NA |
| ENSMUSG( | 575.5679 | 0.107011 | 0.386052 | 0.804623 | NA |
| ENSMUSG( | 59.86307 | -0.00903 | 0.970852 | 0.996369 | NA |
| ENSMUSG( | 16.07105 | 1.031178 | 0.112235 | NA | NA |
| ENSMUSG( | 14.46164 | 0.318049 | 0.530087 | NA | NA |
| ENSMUSG( | 11.71135 | -0.07877 | 0.88711 | NA | NA |
| ENSMUSG( | 22.99092 | 0.297523 | 0.442808 | 0.83278 | NA |
| ENSMUSG( | 46.57983 | 0.497648 | 0.071123 | 0.411712 | NA |
| ENSMUSG( | 14.18366 | -0.56971 | 0.283263 | NA | NA |
| ENSMUSG( | 25.15043 | -0.5282 | 0.173133 | 0.608373 | NA |
| ENSMUSG( | 23.5327 | 0.535701 | 0.177854 | 0.615225 | NA |
| ENSMUSG( | 157.0521 | -0.48142 | 0.004875 | 0.092488 | NA |
| ENSMUSG( | 10.75916 | 0.606418 | 0.322439 | NA | NA |
| ENSMUSG( | 29.66938 | -0.09152 | 0.796485 | 0.960163 | NA |
| ENSMUSG( | 10.24376 | 0.486019 | 0.498261 | NA | NA |
| ENSMUSG( | 34.34212 | -0.0758 | 0.828241 | 0.968966 | NA |
| ENSMUSG( | 49.77816 | 1.425747 | 2.29E-05 | 0.002131 | NA |
| ENSMUSG( | 41.64091 | 0.077722 | 0.808873 | 0.963495 | NA |
| ENSMUSG( | 29.59746 | -0.42958 | 0.230849 | 0.679066 | NA |
| ENSMUSG( | 226.4226 | -0.14054 | 0.376362 | 0.801127 | NA |
| ENSMUSG( | 29.73484 | 0.475484 | 0.171059 | 0.605644 | NA |
| ENSMUSG( | 44.81495 | -0.06802 | 0.812762 | 0.964574 | NA |
| ENSMUSG( | 22.47965 | -0.41537 | 0.328856 | 0.773233 | NA |
| ENSMUSG( | 14.38348 | 0.277424 | 0.582673 | NA | NA |
| ENSMUSG( | 20.74068 | 0.236714 | 0.559351 | NA | NA |
| ENSMUSG( | 115.3696 | 0.138918 | 0.581016 | 0.89745 | NA |
| ENSMUSG( | 14.31555 | -0.12664 | 0.820868 | NA | NA |
| ENSMUSG( | 295.276 | -0.19443 | 0.199904 | 0.640239 | NA |
| ENSMUSG( | 10.37211 | 0.474908 | 0.417542 | NA | NA |
| ENSMUSG( | 29.52239 | -0.06063 | 0.863042 | 0.975055 | NA |
| ENSMUSG( | 45.13071 | 0.008153 | 0.97688 | 0.997428 | NA |
| ENSMUSG( | 12.34274 | 0.103745 | 0.859566 | NA | NA |
| ENSMUSG( | 16.10496 | -0.0125 | 0.979225 | NA | NA |
| ENSMUSG( | 59.04121 | 0.289949 | 0.280234 | 0.729246 | NA |
| ENSMUSG( | 68.25107 | 0.111001 | 0.630012 | 0.913965 | NA |
| ENSMUSG( | 14.14968 | -0.45824 | 0.359368 | NA | NA |
| ENSMUSG( | 28.61296 | 0.04667 | 0.900933 | 0.985633 | NA |
| ENSMUSG( | 40.19016 | -0.22119 | 0.492723 | 0.860462 | NA |
| ENSMUSG( | 21.60329 | -0.38558 | 0.360477 | 0.792161 | NA |
| ENSMUSG( | 70.27625 | -0.25599 | 0.374559 | 0.800445 | NA |
| ENSMUSG( | 46.34876 | 0.319534 | 0.252021 | 0.701908 | NA |
| ENSMUSG( | 11.44892 | -0.17871 | 0.768167 | NA | NA |
| ENSMUSG( | 16.6184 | -0.43258 | 0.369352 | NA | NA |
| ENSMUSG( | 23.05804 | 0.21538 | 0.572346 | 0.893584 | NA |
| ENSMUSG( | 337.1343 | -0.16795 | 0.228511 | 0.675862 | NA |
| ENSMUSG( | 263.9818 | -0.07993 | 0.567866 | 0.892002 | NA |
| ENSMUSG( | 10.04921 | 0.055291 | 0.92222 | NA | NA |
| ENSMUSG( | 1013.283 | -0.52437 | 7.25E-07 | 0.000149 | NA |
| ENSMUSG( | 55.73503 | -0.14768 | 0.573029 | 0.893584 | NA |
| ENSMUSG( | 12.10958 | 0.580528 | 0.375648 | NA | NA |
| ENSMUSG( | 66.41846 | -0.27045 | 0.287479 | 0.738036 | NA |
| ENSMUSG( | 81.23043 | 0.052265 | 0.83526 | 0.970426 | NA |
| ENSMUSG( | 205.3257 | -0.05435 | 0.746694 | 0.949289 | NA |
| ENSMUSG( | 19.25605 | 0.556867 | 0.221926 | NA | NA |
| ENSMUSG( | 21.47452 | -0.54801 | 0.190931 | 0.630245 | NA |
| ENSMUSG( | 86.16698 | -0.21125 | 0.361269 | 0.792797 | NA |
| ENSMUSG( | 64.81359 | 0.277705 | 0.280026 | 0.729155 | NA |
| ENSMUSG( | 26.95192 | 0.069156 | 0.844857 | 0.972023 | NA |
| ENSMUSG( | 41.85044 | 0.508173 | 0.150412 | 0.574991 | NA |
| ENSMUSG( | 11.26791 | -0.06761 | 0.900746 | NA | NA |
| ENSMUSG( | 10.90268 | 0.255301 | 0.669816 | NA | NA |
| ENSMUSG( | 20.69342 | 0.431936 | 0.325994 | NA | NA |
| ENSMUSG( | 13.34331 | -0.61903 | 0.257671 | NA | NA |
| ENSMUSG( | 234.7045 | -0.06422 | 0.655317 | 0.923929 | NA |
| ENSMUSG( | 66.52422 | 0.191754 | 0.435672 | 0.829093 | NA |
| ENSMUSG( | 1729.637 | 0.055822 | 0.583373 | 0.898434 | NA |
| ENSMUSG( | 18.04549 | 0.508568 | 0.256536 | NA | NA |
| ENSMUSG( | 12.76613 | 0.139714 | 0.800774 | NA | NA |
| ENSMUSG( | 10.9084 | -0.87161 | 0.133881 | NA | NA |
| ENSMUSG( | 11.01291 | -0.11729 | 0.831897 | NA | NA |
| ENSMUSG( | 9.817443 | -0.17148 | 0.776919 | NA | NA |
| ENSMUSG( | 31.91604 | -0.5298 | 0.120304 | 0.524495 | NA |
| ENSMUSG( | 177.7524 | 0.056725 | 0.740946 | 0.948028 | NA |
| ENSMUSG( | 11.79359 | -0.28667 | 0.589422 | NA | NA |
| ENSMUSG( | 14.2923 | 0.390278 | 0.459267 | NA | NA |
| ENSMUSG( | 18.63065 | -0.49277 | 0.284147 | NA | NA |
| ENSMUSG( | 16.68986 | -0.2232 | 0.637706 | NA | NA |
| ENSMUSG( | 11.91903 | -0.37755 | 0.486532 | NA | NA |
| ENSMUSG( | 9.366902 | -0.0202 | 0.976677 | NA | NA |
| ENSMUSG( | 56.64289 | 0.047929 | 0.855795 | 0.973813 | NA |
| ENSMUSG( | 68.76629 | -0.07824 | 0.765117 | 0.954127 | NA |
| ENSMUSG( | 12.98667 | -0.29572 | 0.58675 | NA | NA |
| ENSMUSG( | 25.08784 | 0.163639 | 0.66269 | 0.926303 | NA |
| ENSMUSG( | 17.20561 | 0.271866 | 0.570451 | NA | NA |
| ENSMUSG( | 12.79984 | 1.348518 | 0.020019 | NA | NA |
| ENSMUSG( | 14.12135 | -0.60786 | 0.230971 | NA | NA |
| ENSMUSG( | 48.96121 | -0.77911 | 0.022452 | 0.228899 | NA |
| ENSMUSG( | 15.22351 | 0.110348 | 0.836285 | NA | NA |
| ENSMUSG( | 17.29482 | 0.165166 | 0.734987 | NA | NA |
| ENSMUSG( | 26.56448 | 0.101204 | 0.797587 | 0.960507 | NA |
| ENSMUSG( | 87.30191 | 0.550306 | 0.064371 | 0.393403 | NA |
| ENSMUSG( | 34.13216 | 0.246184 | 0.522575 | 0.875389 | NA |
| ENSMUSG( | 22.63045 | -0.01763 | 0.96319 | 0.994911 | NA |
| ENSMUSG( | 237.8555 | -0.04135 | 0.794783 | 0.959799 | NA |
| ENSMUSG( | 47.17501 | 0.022797 | 0.938293 | 0.993415 | NA |
| ENSMUSG( | 28.40897 | -0.28215 | 0.4267 | 0.825155 | NA |
| ENSMUSG( | 16.55796 | 0.548862 | 0.242377 | NA | NA |
| ENSMUSG( | 12.09114 | 1.363644 | 0.020985 | NA | NA |
| ENSMUSG( | 16.96755 | -0.77815 | 0.092948 | NA | NA |
| ENSMUSG( | 37.01982 | -0.06074 | 0.847603 | 0.972374 | NA |
| ENSMUSG( | 40.0066 | -0.03726 | 0.908203 | 0.9878 | NA |
| ENSMUSG( | 33.89792 | 0.109584 | 0.770644 | 0.954722 | NA |
| ENSMUSG( | 205.793 | 0.924115 | 9.75E-05 | 0.006367 | NA |
| ENSMUSG( | 23.90365 | -0.0202 | 0.956986 | 0.993874 | NA |
| ENSMUSG( | 20.27387 | -0.23417 | 0.594848 | NA | NA |
| ENSMUSG( | 18.95284 | 0.512023 | 0.305447 | NA | NA |
| ENSMUSG( | 59.79461 | -0.06527 | 0.802379 | 0.962119 | NA |
| ENSMUSG( | 9.230105 | 0.012263 | 0.984589 | NA | NA |
| ENSMUSG( | 10.34994 | 0.065508 | 0.91216 | NA | NA |
| ENSMUSG( | 17.88612 | -0.02326 | 0.957068 | NA | NA |
| ENSMUSG( | 32.07299 | 0.039835 | 0.903257 | 0.986055 | NA |
| ENSMUSG( | 15.15265 | 0.148777 | 0.765139 | NA | NA |
| ENSMUSG( | 17.01822 | 0.00063 | 0.998894 | NA | NA |
| ENSMUSG( | 26.39934 | 0.113367 | 0.755159 | 0.951279 | NA |
| ENSMUSG( | 39.94222 | 0.512142 | 0.15662 | 0.584667 | NA |
| ENSMUSG( | 18.57369 | 0.042897 | 0.927461 | NA | NA |
| ENSMUSG( | 64.85647 | 0.215634 | 0.389665 | 0.807428 | NA |
| ENSMUSG( | 9.401083 | 0.369038 | 0.545815 | NA | NA |

| | | | | | |
|---|---|---|---|---|---|
| ENSMUSG( | 12.24207 | -0.04542 | 0.935179 | 0.993078 | NA |
| ENSMUSG( | 26.48545 | 0.134351 | 0.734066 | 0.967228 | NA |
| ENSMUSG( | 33.3218 | -0.37226 | 0.336194 | 0.907972 | NA |
| ENSMUSG( | 37.42062 | -0.73047 | 0.034534 | 0.655405 | NA |
| ENSMUSG( | 26.07328 | -0.4315 | 0.269211 | 0.886755 | NA |
| ENSMUSG( | 14.28218 | -0.32173 | 0.548458 | 0.938063 | NA |
| ENSMUSG( | 45.63077 | -0.09086 | 0.774577 | 0.974931 | NA |
| ENSMUSG( | 23.41759 | -0.25698 | 0.532688 | 0.934795 | NA |
| ENSMUSG( | 19.92366 | -0.0113 | 0.979382 | 0.996468 | NA |
| ENSMUSG( | 11.2359 | -0.33228 | 0.594185 | NA | NA |
| ENSMUSG( | 7.212321 | -2.29752 | 0.005811 | NA | NA |
| ENSMUSG( | 42.80983 | -0.12137 | 0.691535 | 0.960102 | NA |
| ENSMUSG( | 73.10768 | -0.30888 | 0.208115 | 0.864412 | NA |
| ENSMUSG( | 34.78528 | -0.30453 | 0.413975 | 0.917257 | NA |
| ENSMUSG( | 12.25733 | -0.46109 | 0.410094 | 0.917257 | NA |
| ENSMUSG( | 217.5113 | 0.106099 | 0.546214 | 0.938063 | NA |
| ENSMUSG( | 12.48695 | 1.098836 | 0.055152 | 0.723362 | NA |
| ENSMUSG( | 32.17211 | -0.00638 | 0.987238 | 0.997487 | NA |
| ENSMUSG( | 11.25429 | -0.34288 | 0.559025 | NA | NA |
| ENSMUSG( | 7.558258 | -1.0104 | 0.166536 | NA | NA |
| ENSMUSG( | 34.22178 | 0.009842 | 0.977629 | 0.996416 | NA |
| ENSMUSG( | 15.37158 | 0.457664 | 0.390837 | 0.917257 | NA |
| ENSMUSG( | 15.10468 | 0.303265 | 0.543932 | 0.938063 | NA |
| ENSMUSG( | 9.536745 | -0.42691 | 0.523398 | NA | NA |
| ENSMUSG( | 118.999 | -0.21976 | 0.315606 | 0.901725 | NA |
| ENSMUSG( | 15.82816 | 0.096176 | 0.841878 | 0.983044 | NA |
| ENSMUSG( | 17.68228 | -0.52981 | 0.262925 | 0.885357 | NA |
| ENSMUSG( | 35.85108 | -0.885 | 0.013397 | 0.463609 | NA |
| ENSMUSG( | 16.98518 | -0.08448 | 0.876897 | 0.989644 | NA |
| ENSMUSG( | 24.20399 | -0.11129 | 0.795439 | 0.97681 | NA |
| ENSMUSG( | 22.57768 | -0.25103 | 0.546712 | 0.938063 | NA |
| ENSMUSG( | 12.64995 | 0.355689 | 0.532314 | 0.934795 | NA |
| ENSMUSG( | 20.32035 | -0.33439 | 0.501132 | 0.927825 | NA |
| ENSMUSG( | 28.86819 | -1.44078 | 0.000231 | 0.039691 | NA |
| ENSMUSG( | 15.15332 | -0.04973 | 0.924732 | 0.99306 | NA |
| ENSMUSG( | 140.8562 | 0.216477 | 0.29456 | 0.898538 | NA |
| ENSMUSG( | 31.64028 | -0.65073 | 0.070594 | 0.758714 | NA |
| ENSMUSG( | 375.6215 | 0.42745 | 0.350046 | 0.911652 | NA |
| ENSMUSG( | 135.0807 | -0.03506 | 0.883388 | 0.989644 | NA |
| ENSMUSG( | 65.83759 | 0.07744 | 0.758917 | 0.971858 | NA |
| ENSMUSG( | 109.0677 | 0.011866 | 0.964007 | 0.995433 | NA |
| ENSMUSG( | 9.020315 | -0.24648 | 0.735035 | NA | NA |
| ENSMUSG( | 231.794 | -0.11892 | 0.519077 | 0.931053 | NA |
| ENSMUSG( | 12.25744 | -0.17963 | 0.753905 | 0.970671 | NA |
| ENSMUSG( | 9.333652 | -0.20284 | 0.755852 | NA | NA |
| ENSMUSG( | 18.27404 | -0.37357 | 0.465524 | 0.923329 | NA |
| ENSMUSG( | 22.04455 | -0.47806 | 0.273807 | 0.889848 | NA |
| ENSMUSG( | 47.76307 | 0.016497 | 0.963192 | 0.995433 | NA |
| ENSMUSG( | 9.55512 | 0.902948 | 0.203388 | NA | NA |
| ENSMUSG( | 11.82908 | 0.194151 | 0.736485 | 0.967344 | NA |
| ENSMUSG( | 123.7957 | 0.037846 | 0.86267 | 0.986547 | NA |
| ENSMUSG( | 178.4514 | -0.06272 | 0.757635 | 0.971858 | NA |
| ENSMUSG( | 467.6754 | -0.04463 | 0.765828 | 0.974236 | NA |
| ENSMUSG( | 450.4104 | 0.101169 | 0.51508 | 0.930939 | NA |
| ENSMUSG( | 22.18394 | 0.489499 | 0.267418 | 0.886134 | NA |
| ENSMUSG( | 62.40043 | -0.23789 | 0.38654 | 0.917257 | NA |
| ENSMUSG( | 37.29866 | 0.90692 | 0.060931 | 0.73558 | NA |
| ENSMUSG( | 7.212457 | -0.68135 | 0.393077 | NA | NA |
| ENSMUSG( | 7.486296 | 0.582249 | 0.468476 | NA | NA |
| ENSMUSG( | 12.61527 | -0.26919 | 0.669085 | 0.956766 | NA |
| ENSMUSG( | 15.11137 | 0.11017 | 0.836062 | 0.981237 | NA |
| ENSMUSG( | 13.91558 | -0.52268 | 0.322493 | 0.904023 | NA |
| ENSMUSG( | 10.32524 | -0.19327 | 0.753697 | NA | NA |
| ENSMUSG( | 12.15014 | 0.082569 | 0.88385 | 0.989644 | NA |
| ENSMUSG( | 11.33417 | -0.13897 | 0.806288 | NA | NA |
| ENSMUSG( | 1595.55 | -0.179 | 0.113639 | 0.809623 | NA |
| ENSMUSG( | 8.749257 | -0.73953 | 0.266088 | NA | NA |
| ENSMUSG( | 39.51012 | -0.40624 | 0.212834 | 0.864412 | NA |
| ENSMUSG( | 21.3732 | -0.00287 | 0.994821 | 0.998813 | NA |
| ENSMUSG( | 21.65837 | 0.77194 | 0.154921 | 0.837724 | NA |
| ENSMUSG( | 25.0448 | 0.818834 | 0.055769 | 0.723362 | NA |
| ENSMUSG( | 37.56007 | -0.36848 | 0.461995 | 0.923329 | NA |
| ENSMUSG( | 983.9592 | 0.20637 | 0.197618 | 0.862161 | NA |
| ENSMUSG( | 11.99207 | 0.423468 | 0.449814 | 0.922202 | NA |
| ENSMUSG( | 12.17132 | 0.9079 | 0.152059 | 0.837724 | NA |
| ENSMUSG( | 13.95523 | 0.432852 | 0.425487 | 0.917257 | NA |
| ENSMUSG( | 19.07844 | 0.180656 | 0.690493 | 0.960071 | NA |
| ENSMUSG( | 284.4994 | 0.050691 | 0.76529 | 0.974236 | NA |
| ENSMUSG( | 18.98802 | 0.294819 | 0.616318 | 0.951413 | NA |
| ENSMUSG( | 138.866 | 0.049523 | 0.818821 | 0.979198 | NA |
| ENSMUSG( | 9.52931 | -0.09964 | 0.886022 | NA | NA |
| ENSMUSG( | 30.40916 | 0.218246 | 0.537688 | 0.937216 | NA |
| ENSMUSG( | 42.33024 | -0.08258 | 0.856584 | 0.985884 | NA |
| ENSMUSG( | 25.65405 | 0.369072 | 0.340634 | 0.907972 | NA |
| ENSMUSG( | 63.27542 | -0.80426 | 0.002254 | 0.184738 | NA |
| ENSMUSG( | 11.58846 | -0.75251 | 0.230101 | 0.868507 | NA |
| ENSMUSG( | 14.97943 | 1.009249 | 0.052804 | 0.71525 | NA |
| ENSMUSG( | 38.97534 | 0.048382 | 0.883332 | 0.989644 | NA |
| ENSMUSG( | 19.92238 | 0.32719 | 0.469182 | 0.923329 | NA |
| ENSMUSG( | 86.15414 | -0.20624 | 0.372837 | 0.915491 | NA |
| ENSMUSG( | 43.96823 | 0.150242 | 0.618599 | 0.952363 | NA |
| ENSMUSG( | 222.703 | -0.04056 | 0.833204 | 0.981192 | NA |
| ENSMUSG( | 8.663521 | -0.72977 | 0.318323 | NA | NA |
| ENSMUSG( | 9.681298 | 0.125101 | 0.849573 | NA | NA |
| ENSMUSG( | 59.96383 | 0.396242 | 0.147254 | 0.836241 | NA |
| ENSMUSG( | 12.96214 | 0.203777 | 0.708608 | 0.964042 | NA |
| ENSMUSG( | 18.9461 | -0.30169 | 0.545173 | 0.938063 | NA |
| ENSMUSG( | 16.95957 | -0.00055 | 0.999105 | 0.999503 | NA |
| ENSMUSG( | 21.1969 | 0.256032 | 0.615227 | 0.951177 | NA |
| ENSMUSG( | 13.94741 | 0.380959 | 0.494222 | 0.927372 | NA |
| ENSMUSG( | 855.8646 | -0.38745 | 0.004069 | 0.25824 | NA |
| ENSMUSG( | 28.91677 | 0.124733 | 0.734159 | 0.967228 | NA |
| ENSMUSG( | 17.78778 | -0.1122 | 0.81245 | 0.979148 | NA |
| ENSMUSG( | 16.36563 | 0.033838 | 0.948103 | 0.994798 | NA |
| ENSMUSG( | 9.444498 | 0.286313 | 0.650208 | NA | NA |
| ENSMUSG( | 18.06606 | 0.235916 | 0.602415 | 0.94789 | NA |
| ENSMUSG( | 11.68187 | -0.80984 | 0.156471 | 0.837724 | NA |
| ENSMUSG( | 62.99686 | 0.122672 | 0.639393 | 0.953903 | NA |
| ENSMUSG( | 12.44709 | -0.05604 | 0.920773 | 0.992644 | NA |
| ENSMUSG( | 20.96259 | -0.663 | 0.133241 | 0.822912 | NA |
| ENSMUSG( | 8.133185 | -0.14756 | 0.841159 | NA | NA |
| ENSMUSG( | 9.032614 | -0.27101 | 0.682327 | NA | NA |
| ENSMUSG( | 26.54696 | 0.122726 | 0.767644 | 0.974236 | NA |
| ENSMUSG( | 22.56368 | -0.55653 | 0.221115 | 0.866104 | NA |
| ENSMUSG( | 96.33083 | 0.067079 | 0.79313 | 0.97681 | NA |
| ENSMUSG( | 122.3446 | 0.308245 | 0.128021 | 0.81845 | NA |
| ENSMUSG( | 60.11288 | -0.47776 | 0.138441 | 0.828492 | NA |
| ENSMUSG( | 103.5443 | 0.084166 | 0.798184 | 0.97681 | NA |
| ENSMUSG( | 47.06341 | 0.006965 | 0.981418 | 0.996756 | NA |
| ENSMUSG( | 26.13693 | -0.13881 | 0.729713 | 0.96719 | NA |
| ENSMUSG( | 10.71641 | -0.28078 | 0.640579 | NA | NA |
| ENSMUSG( | 22.94363 | -0.22998 | 0.584571 | 0.944975 | NA |
| ENSMUSG( | 55.00645 | 0.747169 | 0.020232 | 0.563435 | NA |
| ENSMUSG( | 21.22069 | 0.336171 | 0.446503 | 0.921734 | NA |
| ENSMUSG( | 33.28157 | 0.44868 | 0.236456 | 0.872173 | NA |
| ENSMUSG( | 10.56373 | -0.19755 | 0.757539 | NA | NA |
| ENSMUSG( | 65.31069 | 0.231745 | 0.407649 | 0.917257 | NA |
| ENSMUSG( | 11.84561 | -1.03797 | 0.082584 | 0.77681 | NA |
| ENSMUSG( | 76.4696 | 0.033408 | 0.900121 | 0.990558 | NA |
| ENSMUSG( | 14.83786 | -0.49873 | 0.334772 | 0.907907 | NA |
| ENSMUSG( | 16.51509 | -0.41851 | 0.429457 | 0.917257 | NA |
| ENSMUSG( | 15.43301 | -0.14706 | 0.768322 | 0.974236 | NA |
| ENSMUSG( | 8.616208 | -0.25887 | 0.718035 | NA | NA |
| ENSMUSG( | 22.07807 | 0.114228 | 0.793384 | 0.97681 | NA |
| ENSMUSG( | 10.20755 | 0.486671 | 0.466283 | NA | NA |
| ENSMUSG( | 43.12577 | 0.432473 | 0.210952 | 0.864412 | NA |
| ENSMUSG( | 37.70721 | 0.018432 | 0.958699 | 0.994798 | NA |
| ENSMUSG( | 206.3802 | 0.021793 | 0.92977 | 0.99306 | NA |
| ENSMUSG( | 14.7096 | -0.51657 | 0.312601 | 0.900037 | NA |
| ENSMUSG( | 1379.958 | -0.06567 | 0.572333 | 0.942515 | NA |
| ENSMUSG( | 10.67068 | -0.81497 | 0.196271 | NA | NA |
| ENSMUSG( | 54.52519 | -0.37816 | 0.187014 | 0.856099 | NA |
| ENSMUSG( | 10.22554 | -0.13634 | 0.827494 | NA | NA |
| ENSMUSG( | 9.110002 | -0.13994 | 0.836874 | NA | NA |
| ENSMUSG( | 84.25232 | -0.08816 | 0.738214 | 0.967839 | NA |
| ENSMUSG( | 14.32442 | -0.22485 | 0.671833 | 0.95694 | NA |
| ENSMUSG( | 10.10019 | -1.40634 | 0.032847 | NA | NA |
| ENSMUSG( | 9.624657 | 0.46783 | 0.485159 | NA | NA |
| ENSMUSG( | 74.14413 | 0.428654 | 0.149452 | 0.836241 | NA |
| ENSMUSG( | 25.071 | -0.40794 | 0.356867 | 0.913358 | NA |
| ENSMUSG( | 16.11742 | -0.32966 | 0.510966 | 0.929546 | NA |
| ENSMUSG( | 11.84529 | 0.548934 | 0.403724 | 0.917257 | NA |
| ENSMUSG( | 27.57331 | -0.41345 | 0.299607 | 0.898538 | NA |
| ENSMUSG( | 10.84762 | -0.11394 | 0.853636 | NA | NA |
| ENSMUSG( | 89.04832 | 0.137665 | 0.563182 | 0.941158 | NA |
| ENSMUSG( | 12.76193 | 0.112522 | 0.83377 | 0.981218 | NA |
| ENSMUSG( | 22.74972 | 0.177718 | 0.698634 | 0.960908 | NA |
| ENSMUSG( | 435.828 | 0.037166 | 0.800408 | 0.97681 | NA |

| | | | | | |
|---|---|---|---|---|---|
| ENSMUSG( | NA | 13.54982 | -0.71693 | 0.175341 | NA |
| ENSMUSG( | NA | 158.7147 | -0.05131 | 0.791974 | 0.959461 |
| ENSMUSG( | NA | 687.2169 | 0.229715 | 0.087276 | 0.454452 |
| ENSMUSG( | NA | 12.19396 | 0.169513 | 0.754148 | NA |
| ENSMUSG( | NA | 14.57227 | 0.084761 | 0.877085 | NA |
| ENSMUSG( | NA | 10.84134 | 0.080616 | 0.885034 | NA |
| ENSMUSG( | NA | 68.59327 | 0.692629 | 0.029324 | 0.259647 |
| ENSMUSG( | NA | 74.88278 | 0.172901 | 0.440513 | 0.831996 |
| ENSMUSG( | NA | 9.087852 | -0.44383 | 0.482358 | NA |
| ENSMUSG( | NA | 15.92376 | 0.213569 | 0.659401 | NA |
| ENSMUSG( | NA | 67.76893 | 0.076721 | 0.745206 | 0.949158 |
| ENSMUSG( | NA | 549.1407 | -0.11927 | 0.297834 | 0.746309 |
| ENSMUSG( | NA | 18.23016 | 0.219787 | 0.606984 | NA |
| ENSMUSG( | NA | 14.3286 | -0.12258 | 0.804188 | NA |
| ENSMUSG( | NA | 17.09759 | 0.140785 | 0.76338 | NA |
| ENSMUSG( | NA | 12.01749 | 0.012985 | 0.982471 | NA |
| ENSMUSG( | NA | 38.81905 | -0.1496 | 0.633318 | 0.914073 |
| ENSMUSG( | NA | 238.1169 | -0.11945 | 0.546488 | 0.884974 |
| ENSMUSG( | NA | 25.33416 | 0.719914 | 0.124285 | 0.531714 |
| ENSMUSG( | NA | 11.41784 | 0.238652 | 0.696226 | NA |
| ENSMUSG( | NA | 12.86913 | -0.66038 | 0.212043 | NA |
| ENSMUSG( | NA | 14.04427 | 0.962768 | 0.058326 | NA |
| ENSMUSG( | NA | 13.21981 | -0.80709 | 0.110972 | NA |
| ENSMUSG( | NA | 498.4477 | 0.118637 | 0.393671 | 0.809643 |
| ENSMUSG( | NA | 28.3273 | -0.01761 | 0.960897 | 0.993991 |
| ENSMUSG( | NA | 14.26168 | 0.300799 | 0.559195 | NA |
| ENSMUSG( | NA | 34.227 | -1.15609 | 0.00506 | 0.094553 |
| ENSMUSG( | NA | 54.61727 | 0.183355 | 0.594747 | 0.903007 |
| ENSMUSG( | NA | 38.92664 | -0.17288 | 0.596035 | 0.903152 |
| ENSMUSG( | NA | 11.18557 | -0.41157 | 0.452437 | NA |
| ENSMUSG( | NA | 148.167 | 0.089571 | 0.628486 | 0.913637 |
| ENSMUSG( | NA | 37.89263 | -0.30501 | 0.351656 | 0.783381 |
| ENSMUSG( | NA | 15.31996 | 0.216077 | 0.679775 | NA |
| ENSMUSG( | NA | 29.04176 | -0.17404 | 0.612299 | 0.908728 |
| ENSMUSG( | NA | 22.5657 | -0.02386 | 0.951045 | 0.993821 |
| ENSMUSG( | NA | 39.93816 | -0.62055 | 0.048781 | 0.337919 |
| ENSMUSG( | NA | 153.9394 | 0.095816 | 0.584634 | 0.898884 |
| ENSMUSG( | NA | 11.97883 | 0.195607 | 0.709854 | NA |
| ENSMUSG( | NA | 99.38978 | -0.04717 | 0.82241 | 0.966352 |
| ENSMUSG( | NA | 52.14238 | -0.2919 | 0.289041 | 0.739705 |
| ENSMUSG( | NA | 306.1831 | -0.44723 | 0.063382 | 0.389418 |
| ENSMUSG( | NA | 9.324013 | -0.17516 | 0.785113 | NA |
| ENSMUSG( | NA | 12.47841 | -0.22066 | 0.721998 | NA |
| ENSMUSG( | NA | 38.86879 | 0.128542 | 0.671164 | 0.928781 |
| ENSMUSG( | NA | 11.025 | -0.27379 | 0.648345 | NA |
| ENSMUSG( | NA | 8.281318 | -0.92717 | 0.216433 | NA |
| ENSMUSG( | NA | 65.46421 | -0.39988 | 0.101388 | 0.484996 |
| ENSMUSG( | NA | 15.36976 | -0.28911 | 0.561486 | NA |
| ENSMUSG( | NA | 39.82111 | -0.09987 | 0.756604 | 0.951479 |
| ENSMUSG( | NA | 19.47905 | 0.091658 | 0.83353 | NA |
| ENSMUSG( | NA | 19.21303 | -0.52897 | 0.269107 | NA |
| ENSMUSG( | NA | 10.91531 | 0.113086 | 0.850223 | NA |
| ENSMUSG( | NA | 18.9308 | 0.23484 | 0.603553 | NA |
| ENSMUSG( | NA | 13.75892 | -0.30629 | 0.54431 | NA |
| ENSMUSG( | NA | 20.89645 | 0.094266 | 0.83771 | NA |
| ENSMUSG( | NA | 74.63205 | -0.06695 | 0.81265 | 0.964574 |
| ENSMUSG( | NA | 16.2213 | -0.61369 | 0.178255 | NA |
| ENSMUSG( | NA | 23.49595 | -0.10755 | 0.780901 | 0.957531 |
| ENSMUSG( | NA | 20.72372 | 0.315084 | 0.444701 | NA |
| ENSMUSG( | NA | 17.84055 | -0.49039 | 0.300741 | NA |
| ENSMUSG( | NA | 18.81003 | -0.17314 | 0.69161 | NA |
| ENSMUSG( | NA | 15.66715 | -0.04701 | 0.921593 | NA |
| ENSMUSG( | NA | 1055.424 | -0.08331 | 0.492483 | 0.860378 |
| ENSMUSG( | NA | 64.58557 | -0.48125 | 0.092259 | 0.466326 |
| ENSMUSG( | NA | 20.04506 | -0.23345 | 0.59154 | NA |
| ENSMUSG( | NA | 18.39503 | -0.53055 | 0.22302 | NA |
| ENSMUSG( | NA | 61.12209 | -0.01694 | 0.955574 | 0.993874 |
| ENSMUSG( | NA | 52.32137 | -0.43663 | 0.108325 | 0.497337 |
| ENSMUSG( | NA | 26.19851 | -0.72589 | 0.059684 | 0.377567 |
| ENSMUSG( | NA | 29.31465 | -0.76528 | 0.03568 | 0.290384 |
| ENSMUSG( | NA | 41.97587 | -0.13343 | 0.695931 | 0.934979 |
| ENSMUSG( | NA | 27.22792 | -0.01105 | 0.976819 | 0.997428 |
| ENSMUSG( | NA | 136.0242 | 0.090911 | 0.634761 | 0.914073 |
| ENSMUSG( | NA | 94.28762 | -0.03612 | 0.86373 | 0.975244 |
| ENSMUSG( | NA | 217.6673 | -0.09043 | 0.531258 | 0.879376 |
| ENSMUSG( | NA | 82.35619 | -0.0747 | 0.74512 | 0.949158 |
| ENSMUSG( | NA | 16.00978 | 0.335789 | 0.513457 | NA |
| ENSMUSG( | NA | 66.65285 | 0.140674 | 0.553421 | 0.886216 |
| ENSMUSG( | NA | 39.60003 | -0.50616 | 0.11729 | 0.516826 |
| ENSMUSG( | NA | 23.28813 | -0.23021 | 0.549035 | 0.885686 |
| ENSMUSG( | NA | 12.67975 | -0.20839 | 0.723781 | NA |
| ENSMUSG( | NA | 24.84899 | -0.46822 | 0.231089 | 0.679229 |
| ENSMUSG( | NA | 38.27664 | 0.344408 | 0.294212 | 0.743459 |
| ENSMUSG( | NA | 10.87411 | -0.36497 | 0.537956 | NA |
| ENSMUSG( | NA | 25.97245 | -0.14219 | 0.7004 | 0.93743 |
| ENSMUSG( | NA | 12.16494 | -0.94481 | 0.107177 | NA |
| ENSMUSG( | NA | 56.29603 | -0.00697 | 0.978607 | 0.997428 |
| ENSMUSG( | NA | 38.93497 | -0.56191 | 0.064995 | 0.395432 |
| ENSMUSG( | NA | 16.49978 | 0.569235 | 0.247881 | NA |
| ENSMUSG( | NA | 18.12405 | -0.33822 | 0.465612 | NA |
| ENSMUSG( | NA | 13.24024 | -0.61557 | 0.297571 | NA |
| ENSMUSG( | NA | 11.25765 | 0.175396 | 0.752066 | NA |
| ENSMUSG( | NA | 13.61342 | 0.025988 | 0.968806 | NA |
| ENSMUSG( | NA | 15.68177 | 0.343595 | 0.469488 | NA |
| ENSMUSG( | NA | 25.31852 | -0.51352 | 0.182178 | 0.619591 |
| ENSMUSG( | NA | 24.60934 | -0.37189 | 0.313883 | 0.761128 |
| ENSMUSG( | NA | 19.59915 | -0.30276 | 0.486076 | NA |
| ENSMUSG( | NA | 356.4439 | -0.1026 | 0.42273 | 0.823527 |
| ENSMUSG( | NA | 60.20759 | 0.130387 | 0.614141 | 0.909405 |
| ENSMUSG( | NA | 928.8983 | 0.322622 | 0.006514 | 0.110626 |
| ENSMUSG( | NA | 19.88955 | -0.03853 | 0.926169 | NA |
| ENSMUSG( | NA | 180.1752 | 0.147826 | 0.349547 | 0.781913 |
| ENSMUSG( | NA | 16.16534 | 0.15939 | 0.740261 | NA |
| ENSMUSG( | NA | 33.63469 | 0.602622 | 0.080361 | 0.438432 |
| ENSMUSG( | NA | 40.6497 | -0.01539 | 0.960096 | 0.993991 |
| ENSMUSG( | NA | 215.8525 | 0.08562 | 0.627254 | 0.912941 |
| ENSMUSG( | NA | 85.96166 | 0.030622 | 0.890911 | 0.983764 |
| ENSMUSG( | NA | 43.08333 | -0.01562 | 0.957669 | 0.993874 |
| ENSMUSG( | NA | 19.39496 | 0.064597 | 0.887725 | NA |
| ENSMUSG( | NA | 662.3096 | 0.23641 | 0.066584 | 0.398819 |
| ENSMUSG( | NA | 468.6425 | 0.16073 | 0.313222 | 0.76033 |
| ENSMUSG( | NA | 14.87609 | 0.184169 | 0.710701 | NA |
| ENSMUSG( | NA | 93.17276 | -0.3332 | 0.145824 | 0.567075 |
| ENSMUSG( | NA | 10.7397 | 0.383048 | 0.504337 | NA |
| ENSMUSG( | NA | 8.939059 | -0.15223 | 0.814272 | NA |
| ENSMUSG( | NA | 158.8911 | -0.02658 | 0.899344 | 0.985633 |
| ENSMUSG( | NA | 72.7677 | -0.46627 | 0.147251 | 0.569348 |
| ENSMUSG( | NA | 21.83963 | -0.09525 | 0.827055 | 0.96865 |
| ENSMUSG( | NA | 47.51293 | 0.055451 | 0.848229 | 0.972374 |
| ENSMUSG( | NA | 86.09343 | 0.095889 | 0.703357 | 0.937735 |
| ENSMUSG( | NA | 13.42733 | -0.46767 | 0.348341 | NA |
| ENSMUSG( | NA | 15.55959 | -0.34125 | 0.492911 | NA |
| ENSMUSG( | NA | 44.87021 | -0.37734 | 0.189077 | 0.627678 |
| ENSMUSG( | NA | 13.28558 | 0.253912 | 0.612741 | NA |
| ENSMUSG( | NA | 40.26675 | -0.36349 | 0.224552 | 0.670886 |
| ENSMUSG( | NA | 10.26129 | -0.38354 | 0.507655 | NA |
| ENSMUSG( | NA | 36.7863 | -0.58879 | 0.077385 | 0.429743 |
| ENSMUSG( | NA | 50.16408 | -0.07551 | 0.777096 | 0.955828 |
| ENSMUSG( | NA | 311.2135 | -0.00065 | 0.996186 | 0.99943 |
| ENSMUSG( | NA | 13.51558 | -0.37616 | 0.458237 | NA |
| ENSMUSG( | NA | 13.64576 | -0.93799 | 0.066934 | NA |
| ENSMUSG( | NA | 17.50486 | 0.488472 | 0.294947 | NA |
| ENSMUSG( | NA | 16.52544 | 0.584193 | 0.211077 | NA |
| ENSMUSG( | NA | 22.88292 | 0.181588 | 0.660994 | 0.92497 |
| ENSMUSG( | NA | 14.62785 | -0.0599 | 0.90478 | NA |
| ENSMUSG( | NA | 96.73028 | -0.19338 | 0.351557 | 0.783381 |
| ENSMUSG( | NA | 95.66822 | -0.10509 | 0.650765 | 0.921307 |
| ENSMUSG( | NA | 87.60581 | -0.01489 | 0.946793 | 0.993731 |
| ENSMUSG( | NA | 10.93108 | -0.08536 | 0.880382 | NA |
| ENSMUSG( | NA | 167.8401 | -0.05074 | 0.764109 | 0.954055 |
| ENSMUSG( | NA | 78.7764 | 0.178435 | 0.542051 | 0.883182 |
| ENSMUSG( | NA | 13.08691 | 0.293188 | 0.56121 | NA |
| ENSMUSG( | NA | 14.75276 | 0.107583 | 0.825576 | NA |
| ENSMUSG( | NA | 24.09415 | -0.56845 | 0.175998 | 0.611543 |
| ENSMUSG( | NA | 13.04002 | -0.27566 | 0.587675 | NA |
| ENSMUSG( | NA | 9.041704 | 0.047636 | 0.947374 | NA |
| ENSMUSG( | NA | 409.0307 | 0.009836 | 0.941892 | 0.993415 |
| ENSMUSG( | NA | 40.12892 | 0.06671 | 0.830846 | 0.969798 |
| ENSMUSG( | NA | 15.10021 | 0.267398 | 0.584284 | NA |
| ENSMUSG( | NA | 191.9898 | -0.20504 | 0.266852 | 0.71685 |
| ENSMUSG( | NA | 9.005604 | -0.10416 | 0.871851 | NA |
| ENSMUSG( | NA | 15.79362 | -0.30732 | 0.572628 | NA |
| ENSMUSG( | NA | 16.03519 | -0.2558 | 0.593987 | NA |
| ENSMUSG( | NA | 27.68952 | -0.18989 | 0.645344 | 0.917958 |
| ENSMUSG( | NA | 16.80375 | -0.02195 | 0.967509 | NA |
| ENSMUSG( | NA | 89.76476 | -0.06643 | 0.747946 | 0.949604 |
| ENSMUSG( | NA | 53.33181 | 0.168013 | 0.546068 | 0.884974 |
| ENSMUSG( | NA | 23.57377 | -0.3826 | 0.363253 | 0.793211 |

| | | | | | |
|---|---|---|---|---|---|
| ENSMUSG( | NA | 10.10019 | -1.40634 | 0.032847 | NA |
| ENSMUSG( | NA | 74.14413 | 0.428654 | 0.149452 | 0.836241 |
| ENSMUSG( | NA | 25.071 | -0.40794 | 0.356867 | 0.913358 |
| ENSMUSG( | NA | 16.11742 | -0.32966 | 0.510966 | 0.929546 |
| ENSMUSG( | NA | 11.84529 | 0.548934 | 0.403724 | 0.917257 |
| ENSMUSG( | NA | 27.57331 | -0.41345 | 0.299607 | 0.898538 |
| ENSMUSG( | NA | 10.84762 | -0.11394 | 0.853636 | NA |
| ENSMUSG( | NA | 89.04832 | 0.137665 | 0.563182 | 0.941158 |
| ENSMUSG( | NA | 12.76193 | 0.112522 | 0.83377 | 0.981218 |
| ENSMUSG( | NA | 22.74972 | 0.177718 | 0.698634 | 0.960908 |
| ENSMUSG( | NA | 435.828 | 0.037166 | 0.800408 | 0.97681 |
| ENSMUSG( | NA | 30.40819 | -0.82784 | 0.023462 | 0.585845 |
| ENSMUSG( | NA | 67.86446 | -0.13104 | 0.662343 | 0.956205 |
| ENSMUSG( | NA | 9.347427 | -0.42329 | 0.504044 | NA |
| ENSMUSG( | NA | 11.35703 | -1.15001 | 0.056721 | NA |
| ENSMUSG( | NA | 28.03863 | 0.193265 | 0.673364 | 0.95694 |
| ENSMUSG( | NA | 38.26519 | -0.17184 | 0.611589 | 0.950771 |
| ENSMUSG( | NA | 13.46285 | 1.553842 | 0.012556 | 0.451781 |
| ENSMUSG( | NA | 10.95213 | -0.36734 | 0.548701 | NA |
| ENSMUSG( | NA | 26.19671 | -0.01379 | 0.973127 | 0.996416 |
| ENSMUSG( | NA | 51.62893 | -0.19368 | 0.49202 | 0.927315 |
| ENSMUSG( | NA | 25.90879 | -0.08365 | 0.840371 | 0.982367 |
| ENSMUSG( | NA | 25.80545 | -0.24523 | 0.528017 | 0.934648 |
| ENSMUSG( | NA | 16.60587 | 0.081271 | 0.890124 | 0.990558 |
| ENSMUSG( | NA | 217.5348 | -0.3965 | 0.042505 | 0.690304 |
| ENSMUSG( | NA | 27.00807 | -0.49714 | 0.221374 | 0.866104 |
| ENSMUSG( | NA | 20.30008 | -0.02196 | 0.960785 | 0.995097 |
| ENSMUSG( | NA | 20.06475 | 0.29376 | 0.505365 | 0.927825 |
| ENSMUSG( | NA | 24.69954 | -0.00608 | 0.989316 | 0.997918 |
| ENSMUSG( | NA | 27.75201 | 0.073669 | 0.849361 | 0.985321 |
| ENSMUSG( | NA | 15.80491 | 0.013498 | 0.978654 | 0.996459 |
| ENSMUSG( | NA | 10.48818 | -0.06937 | 0.910362 | NA |
| ENSMUSG( | NA | 22.65791 | -0.62701 | 0.168545 | 0.840222 |
| ENSMUSG( | NA | 9.476946 | -2.05037 | 0.004183 | NA |
| ENSMUSG( | NA | 9.477223 | -0.3043 | 0.636989 | NA |
| ENSMUSG( | NA | 110.4355 | -0.04299 | 0.833707 | 0.981218 |
| ENSMUSG( | NA | 30.34451 | -0.5295 | 0.154452 | 0.837724 |
| ENSMUSG( | NA | 17.80935 | 0.032687 | 0.948171 | 0.994798 |
| ENSMUSG( | NA | 51.65687 | -0.07106 | 0.810942 | 0.978799 |
| ENSMUSG( | NA | 24.66746 | -0.34763 | 0.380542 | 0.917202 |
| ENSMUSG( | NA | 13.2868 | -0.21298 | 0.684494 | 0.958678 |
| ENSMUSG( | NA | 11.94403 | 0.974286 | 0.097629 | 0.796743 |
| ENSMUSG( | NA | 22.7648 | -0.04175 | 0.919708 | 0.992644 |
| ENSMUSG( | NA | 10.41622 | -0.7315 | 0.237901 | NA |
| ENSMUSG( | NA | 32.4001 | 0.320498 | 0.371253 | 0.915491 |
| ENSMUSG( | NA | 495.9298 | -0.00804 | 0.952735 | 0.994798 |
| ENSMUSG( | NA | 17.82601 | 0.228049 | 0.633077 | 0.953844 |
| ENSMUSG( | NA | 697.0745 | -0.43391 | 0.006919 | 0.343883 |
| ENSMUSG( | NA | 38.56907 | 0.272424 | 0.405374 | 0.917257 |
| ENSMUSG( | NA | 33.82114 | -0.29226 | 0.428698 | 0.917257 |
| ENSMUSG( | NA | 21.02071 | -0.3453 | 0.462872 | 0.923329 |
| ENSMUSG( | NA | 69.59796 | 0.062774 | 0.820394 | 0.979198 |
| ENSMUSG( | NA | 218.5923 | -0.18191 | 0.307837 | 0.898538 |
| ENSMUSG( | NA | 44.06308 | -0.19096 | 0.538284 | 0.937216 |
| ENSMUSG( | NA | 35.01476 | 0.586595 | 0.127278 | 0.817278 |
| ENSMUSG( | NA | 14.11505 | 0.002835 | 0.995731 | 0.999116 |
| ENSMUSG( | NA | 15.34825 | -0.3077 | 0.569003 | 0.941777 |
| ENSMUSG( | NA | 16.18605 | 0.34222 | 0.474892 | 0.923329 |
| ENSMUSG( | NA | 32.03094 | 0.090952 | 0.801185 | 0.97681 |
| ENSMUSG( | NA | 29.33234 | 0.023406 | 0.952178 | 0.994798 |
| ENSMUSG( | NA | 34.34319 | -0.07983 | 0.817603 | 0.979198 |
| ENSMUSG( | NA | 19.03291 | -0.05108 | 0.912105 | 0.992196 |
| ENSMUSG( | NA | 65.47919 | -0.13685 | 0.62779 | 0.953234 |
| ENSMUSG( | NA | 1675.147 | 0.099345 | 0.360742 | 0.914223 |
| ENSMUSG( | NA | 65.39072 | -0.16573 | 0.55502 | 0.939565 |
| ENSMUSG( | NA | 74.65892 | -0.12756 | 0.591474 | 0.944975 |
| ENSMUSG( | NA | 30.02796 | 0.500222 | 0.283448 | 0.892412 |
| ENSMUSG( | NA | 155.3515 | -0.89279 | 0.182276 | 0.850579 |
| ENSMUSG( | NA | 41.79899 | 0.354768 | 0.2825 | 0.892412 |
| ENSMUSG( | NA | 39.27088 | 0.227636 | 0.533433 | 0.935162 |
| ENSMUSG( | NA | 23.0131 | 0.150552 | 0.727705 | 0.966887 |
| ENSMUSG( | NA | 37.38181 | -0.55992 | 0.152421 | 0.837724 |
| ENSMUSG( | NA | 163.5105 | 0.697145 | 0.000546 | 0.071324 |
| ENSMUSG( | NA | 98.54775 | 0.100302 | 0.685251 | 0.958678 |
| ENSMUSG( | NA | 255.0964 | -0.19106 | 0.278275 | 0.891468 |
| ENSMUSG( | NA | 169.4696 | -0.13872 | 0.426078 | 0.917257 |
| ENSMUSG( | NA | 15.3076 | -0.05576 | 0.921976 | 0.992644 |
| ENSMUSG( | NA | 12.61791 | -0.20176 | 0.706932 | 0.963529 |
| ENSMUSG( | NA | 38.18168 | -0.27622 | 0.403376 | 0.917257 |
| ENSMUSG( | NA | 12.82717 | -0.45383 | 0.414163 | 0.917257 |
| ENSMUSG( | NA | 106.1338 | 0.041835 | 0.842577 | 0.983044 |
| ENSMUSG( | NA | 16.09828 | 0.733855 | 0.140107 | 0.830678 |
| ENSMUSG( | NA | 19.51168 | 0.624767 | 0.174039 | 0.844317 |
| ENSMUSG( | NA | 11.05761 | -0.10882 | 0.863624 | NA |
| ENSMUSG( | NA | 14.2904 | -0.27909 | 0.59734 | 0.946609 |
| ENSMUSG( | NA | 36.44851 | -0.55424 | 0.107733 | 0.804969 |
| ENSMUSG( | NA | 26.79785 | -0.63714 | 0.116216 | 0.813384 |
| ENSMUSG( | NA | 99.12006 | -0.15048 | 0.502517 | 0.927825 |
| ENSMUSG( | NA | 16.21768 | -0.11124 | 0.819775 | 0.979198 |
| ENSMUSG( | NA | 50.77324 | -0.23632 | 0.414564 | 0.917257 |
| ENSMUSG( | NA | 969.3482 | 0.18969 | 0.220766 | 0.866104 |
| ENSMUSG( | NA | 10.81113 | -0.83163 | 0.201943 | NA |
| ENSMUSG( | NA | 11.68548 | 0.311484 | 0.599102 | 0.947045 |
| ENSMUSG( | NA | 16.76446 | -0.70142 | 0.151057 | 0.837173 |
| ENSMUSG( | NA | 28.13004 | -0.1789 | 0.685528 | 0.958678 |
| ENSMUSG( | NA | 35.87811 | 0.602052 | 0.14963 | 0.836241 |
| ENSMUSG( | NA | 480.8118 | 0.102422 | 0.464267 | 0.923329 |
| ENSMUSG( | NA | 19.28479 | -0.29863 | 0.51089 | 0.929546 |
| ENSMUSG( | NA | 18.83169 | -0.49599 | 0.289708 | 0.896726 |
| ENSMUSG( | NA | 11.17692 | -0.3479 | 0.548205 | NA |
| ENSMUSG( | NA | 43.44408 | -0.18425 | 0.560603 | 0.940254 |
| ENSMUSG( | NA | 166.4064 | -0.42279 | 0.026313 | 0.603041 |
| ENSMUSG( | NA | 69.34762 | -0.48616 | 0.079286 | 0.77458 |
| ENSMUSG( | NA | 12.44819 | -0.38035 | 0.530858 | 0.934795 |
| ENSMUSG( | NA | 15.87939 | 0.359712 | 0.475849 | 0.923329 |
| ENSMUSG( | NA | 54.20127 | 0.116487 | 0.745738 | 0.96951 |
| ENSMUSG( | NA | 11.47573 | 0.283451 | 0.629575 | 0.953251 |
| ENSMUSG( | NA | 11.03031 | -1.27336 | 0.035058 | NA |
| ENSMUSG( | NA | 108.9962 | -0.02958 | 0.886574 | 0.990063 |
| ENSMUSG( | NA | 9.019895 | -0.35786 | 0.584948 | NA |
| ENSMUSG( | NA | 40.95606 | 0.018602 | 0.958092 | 0.994798 |
| ENSMUSG( | NA | 27.52213 | 0.591767 | 0.171949 | 0.842999 |
| ENSMUSG( | NA | 135.9496 | -0.19174 | 0.320971 | 0.904019 |
| ENSMUSG( | NA | 32.01463 | -0.36259 | 0.389756 | 0.917257 |
| ENSMUSG( | NA | 222.6291 | -0.14292 | 0.40961 | 0.917257 |
| ENSMUSG( | NA | 9.812582 | -0.58286 | 0.364961 | NA |
| ENSMUSG( | NA | 16.57301 | -0.09524 | 0.84527 | 0.983957 |
| ENSMUSG( | NA | 10.43343 | 0.69185 | 0.339872 | NA |
| ENSMUSG( | NA | 18.92112 | -0.86576 | 0.063624 | 0.736482 |
| ENSMUSG( | NA | 17.35549 | -0.57416 | 0.229365 | 0.868507 |
| ENSMUSG( | NA | 22.43596 | -0.23859 | 0.600162 | 0.947461 |
| ENSMUSG( | NA | 13.84978 | 0.183398 | 0.73269 | 0.967228 |
| ENSMUSG( | NA | 18.37621 | 0.193481 | 0.679877 | 0.957966 |
| ENSMUSG( | NA | 9.472301 | -0.34961 | 0.600567 | NA |
| ENSMUSG( | NA | 8.528729 | -0.38043 | 0.570225 | NA |
| ENSMUSG( | NA | 15.44838 | -0.08406 | 0.877771 | 0.989644 |
| ENSMUSG( | NA | 20.49819 | -0.48267 | 0.292234 | 0.897854 |
| ENSMUSG( | NA | 8.101016 | -0.56807 | 0.421482 | NA |
| ENSMUSG( | NA | 12.63366 | -0.04517 | 0.934796 | 0.99306 |
| ENSMUSG( | NA | 19.62705 | -0.6696 | 0.134292 | 0.822912 |
| ENSMUSG( | NA | 14.77324 | -0.81098 | 0.140475 | 0.830678 |
| ENSMUSG( | NA | 379.2834 | 0.059157 | 0.690471 | 0.960071 |
| ENSMUSG( | NA | 15.45464 | 0.368199 | 0.477557 | 0.923329 |
| ENSMUSG( | NA | 37.44488 | 0.01773 | 0.95807 | 0.994798 |
| ENSMUSG( | NA | 22.61712 | 0.368943 | 0.414689 | 0.917257 |
| ENSMUSG( | NA | 38.52588 | -0.32518 | 0.341025 | 0.908138 |
| ENSMUSG( | NA | 177.0336 | -0.31589 | 0.109933 | 0.806572 |
| ENSMUSG( | NA | 120.3465 | -0.16996 | 0.434457 | 0.91869 |
| ENSMUSG( | NA | 11.98347 | -0.83681 | 0.189325 | 0.85698 |
| ENSMUSG( | NA | 8.408328 | -0.93395 | 0.174168 | NA |
| ENSMUSG( | NA | 13.79664 | 0.447039 | 0.39214 | 0.917257 |
| ENSMUSG( | NA | 144.3582 | -0.04687 | 0.843622 | 0.983412 |
| ENSMUSG( | NA | 16.21535 | 0.048283 | 0.932324 | 0.99306 |
| ENSMUSG( | NA | 16.20765 | -0.32877 | 0.546412 | 0.938063 |
| ENSMUSG( | NA | 31.26828 | 0.221222 | 0.529468 | 0.934795 |
| ENSMUSG( | NA | 65.81986 | -0.08711 | 0.788832 | 0.976593 |
| ENSMUSG( | NA | 18.61003 | -0.16697 | 0.727581 | 0.966887 |
| ENSMUSG( | NA | 17.50012 | 0.082118 | 0.864657 | 0.986591 |
| ENSMUSG( | NA | 26.86186 | -0.21358 | 0.587393 | 0.944975 |
| ENSMUSG( | NA | 9.052265 | 0.277731 | 0.667041 | NA |
| ENSMUSG( | NA | 12.17676 | -0.13114 | 0.83503 | 0.981237 |
| ENSMUSG( | NA | 32.56523 | -0.68072 | 0.053038 | 0.71525 |
| ENSMUSG( | NA | 385.5498 | -0.13879 | 0.473903 | 0.923329 |
| ENSMUSG( | NA | 22.10662 | 0.338386 | 0.429759 | 0.917257 |
| ENSMUSG( | NA | 23.36485 | -0.0811 | 0.860274 | 0.986137 |
| ENSMUSG( | NA | 22.09194 | 0.012977 | 0.975266 | 0.996416 |
| ENSMUSG( | NA | 133.7114 | -0.16842 | 0.401866 | 0.917257 |
| ENSMUSG( | NA | 340.398 | 0.035989 | 0.804201 | 0.977191 |

| | | | | | |
|---|---|---|---|---|---|
| ENSMUSG( | 8.874604 | 0.404341 | 0.541679 | NA | NA |
| ENSMUSG( | 20.07816 | 0.11468 | 0.79542 | NA | NA |
| ENSMUSG( | 8.662231 | 1.589783 | 0.020804 | NA | NA |
| ENSMUSG( | 12.31115 | -0.05851 | 0.929726 | NA | NA |
| ENSMUSG( | 39.61697 | -0.72924 | 0.022478 | 0.229021 | NA |
| ENSMUSG( | 94.43734 | 0.641558 | 0.022001 | 0.226001 | NA |
| ENSMUSG( | 45.37999 | 0.205146 | 0.520351 | 0.875186 | NA |
| ENSMUSG( | 58.17057 | 0.467403 | 0.122226 | 0.527707 | NA |
| ENSMUSG( | 12.39207 | 0.315603 | 0.557321 | NA | NA |
| ENSMUSG( | 116.017 | -0.02877 | 0.884962 | 0.98282 | NA |
| ENSMUSG( | 12.99619 | 0.830927 | 0.144472 | NA | NA |
| ENSMUSG( | 15.66088 | -0.04044 | 0.931248 | NA | NA |
| ENSMUSG( | 101.0115 | 1.002476 | 0.000433 | 0.018963 | NA |
| ENSMUSG( | 13.86418 | -0.21824 | 0.666845 | NA | NA |
| ENSMUSG( | 145.8716 | -0.09053 | 0.642225 | 0.916865 | NA |
| ENSMUSG( | 662.5641 | 0.11261 | 0.47942 | 0.854046 | NA |
| ENSMUSG( | 16.38792 | -0.05639 | 0.909088 | NA | NA |
| ENSMUSG( | 11.70264 | 0.371776 | 0.549753 | NA | NA |
| ENSMUSG( | 14.78194 | -0.16942 | 0.735328 | NA | NA |
| ENSMUSG( | 12.04996 | 0.641788 | 0.336268 | NA | NA |
| ENSMUSG( | 27.45145 | 0.150337 | 0.701459 | 0.937441 | NA |
| ENSMUSG( | 19.80264 | -0.1936 | 0.689389 | NA | NA |
| ENSMUSG( | 13.25773 | 0.204519 | 0.688499 | NA | NA |
| ENSMUSG( | 34.43466 | -0.10902 | 0.732425 | 0.944956 | NA |
| ENSMUSG( | 45.07382 | 0.414219 | 0.201661 | 0.641362 | NA |
| ENSMUSG( | 463.3774 | -0.0729 | 0.568438 | 0.892252 | NA |
| ENSMUSG( | 581.1441 | 0.169672 | 0.123536 | 0.530552 | NA |
| ENSMUSG( | 19.44691 | 0.0856 | 0.860402 | NA | NA |
| ENSMUSG( | 22.84251 | 0.099993 | 0.798164 | 0.96069 | NA |
| ENSMUSG( | 27.078 | 0.169393 | 0.635334 | 0.914296 | NA |
| ENSMUSG( | 10.36833 | -0.21879 | 0.699512 | NA | NA |
| ENSMUSG( | 30.2951 | -0.27138 | 0.430897 | 0.826896 | NA |
| ENSMUSG( | 105.7471 | 0.116605 | 0.568549 | 0.892294 | NA |
| ENSMUSG( | 60.83049 | 0.407736 | 0.106018 | 0.493403 | NA |
| ENSMUSG( | 14.84987 | -1.27613 | 0.026927 | NA | NA |
| ENSMUSG( | 14.40947 | -0.30598 | 0.5454 | NA | NA |
| ENSMUSG( | 46.30595 | 0.464239 | 0.120442 | 0.524495 | NA |
| ENSMUSG( | 139.7563 | 0.081942 | 0.65958 | 0.924506 | NA |
| ENSMUSG( | 101.3457 | 0.239271 | 0.24453 | 0.69183 | NA |
| ENSMUSG( | 11.27703 | 0.371415 | 0.538514 | NA | NA |
| ENSMUSG( | 21.74232 | 0.175777 | 0.682778 | 0.932022 | NA |
| ENSMUSG( | 24.67154 | -0.40225 | 0.343635 | 0.779153 | NA |
| ENSMUSG( | 22.76615 | -0.39956 | 0.356013 | 0.788009 | NA |
| ENSMUSG( | 11.3482 | 0.649892 | 0.256683 | NA | NA |
| ENSMUSG( | 34.47438 | 0.361771 | 0.258116 | 0.70936 | NA |
| ENSMUSG( | 39.11174 | 0.443409 | 0.165319 | 0.596691 | NA |
| ENSMUSG( | 36.22454 | -0.06569 | 0.833759 | 0.970333 | NA |
| ENSMUSG( | 60.17666 | 0.299827 | 0.235415 | 0.682423 | NA |
| ENSMUSG( | 408.4982 | 0.1747 | 0.137935 | 0.552903 | NA |
| ENSMUSG( | 20.12225 | 0.165079 | 0.723289 | NA | NA |
| ENSMUSG( | 20.18681 | 0.064798 | 0.882567 | NA | NA |
| ENSMUSG( | 30.82317 | 0.225471 | 0.521258 | 0.875186 | NA |
| ENSMUSG( | 36.20208 | -0.02294 | 0.944113 | 0.993503 | NA |
| ENSMUSG( | 23.80663 | 0.346073 | 0.385183 | 0.804048 | NA |
| ENSMUSG( | 11.91096 | -0.89239 | 0.101137 | NA | NA |
| ENSMUSG( | 17.15567 | 1.108584 | 0.04388 | NA | NA |
| ENSMUSG( | 62.68122 | -0.18439 | 0.480809 | 0.855113 | NA |
| ENSMUSG( | 15.64019 | -0.07469 | 0.878741 | NA | NA |
| ENSMUSG( | 580.0301 | -0.31146 | 0.070492 | 0.40908 | NA |
| ENSMUSG( | 25.24371 | 0.34031 | 0.386129 | 0.804623 | NA |
| ENSMUSG( | 19.66507 | 0.162225 | 0.709177 | NA | NA |
| ENSMUSG( | 144.0272 | 0.25837 | 0.256271 | 0.706924 | NA |
| ENSMUSG( | 12.03011 | -0.17396 | 0.741676 | NA | NA |
| ENSMUSG( | 28.47364 | -0.17565 | 0.629876 | 0.913965 | NA |
| ENSMUSG( | 26.51093 | 0.489163 | 0.187169 | 0.625368 | NA |
| ENSMUSG( | 43.10863 | 0.009879 | 0.974273 | 0.996953 | NA |
| ENSMUSG( | 29.71924 | 0.082918 | 0.817724 | 0.965785 | NA |
| ENSMUSG( | 9.455081 | -0.127 | 0.835835 | NA | NA |
| ENSMUSG( | 63.63252 | -0.09583 | 0.706896 | 0.93949 | NA |
| ENSMUSG( | 511.7439 | 0.182109 | 0.155367 | 0.583346 | NA |
| ENSMUSG( | 16.30154 | 0.942707 | 0.044696 | NA | NA |
| ENSMUSG( | 50.56412 | -0.35553 | 0.228742 | 0.675959 | NA |
| ENSMUSG( | 16.15375 | -0.55694 | 0.231564 | NA | NA |
| ENSMUSG( | 20.66388 | 0.484178 | 0.259899 | NA | NA |
| ENSMUSG( | 16.20848 | -0.4849 | 0.311371 | NA | NA |
| ENSMUSG( | 222.2279 | 0.117995 | 0.457945 | 0.839769 | NA |
| ENSMUSG( | 25.28463 | 0.459505 | 0.25799 | 0.709319 | NA |
| ENSMUSG( | 77.81787 | 0.018811 | 0.932387 | 0.992515 | NA |
| ENSMUSG( | 10.94549 | 0.866866 | 0.133049 | NA | NA |
| ENSMUSG( | 8.54217 | 0.248742 | 0.696912 | NA | NA |
| ENSMUSG( | 8.864524 | 1.152631 | 0.101628 | NA | NA |
| ENSMUSG( | 972.5091 | 0.280051 | 0.017396 | 0.197592 | NA |
| ENSMUSG( | 52.43066 | 0.255456 | 0.337611 | 0.777422 | NA |
| ENSMUSG( | 16.28673 | 0.530758 | 0.285999 | NA | NA |
| ENSMUSG( | 14.49861 | -0.53857 | 0.310519 | NA | NA |
| ENSMUSG( | 31.52423 | 0.098409 | 0.787608 | 0.959218 | NA |
| ENSMUSG( | 40.03989 | -0.22781 | 0.46383 | 0.842812 | NA |
| ENSMUSG( | 119.015 | -0.10423 | 0.604644 | 0.905556 | NA |
| ENSMUSG( | 18.46621 | 0.003336 | 0.994493 | NA | NA |
| ENSMUSG( | 66.84483 | 0.686158 | 0.003907 | 0.080172 | NA |
| ENSMUSG( | 13.65288 | -0.49241 | 0.335464 | NA | NA |
| ENSMUSG( | 78.12046 | -0.23419 | 0.285255 | 0.733952 | NA |
| ENSMUSG( | 304.4585 | 0.381373 | 0.083486 | 0.445586 | NA |
| ENSMUSG( | 19.09609 | -0.56735 | 0.189528 | NA | NA |
| ENSMUSG( | 9.824615 | 0.039956 | 0.949706 | NA | NA |
| ENSMUSG( | 10.45425 | -0.51021 | 0.434238 | NA | NA |
| ENSMUSG( | 13.54982 | -0.71693 | 0.175341 | NA | NA |
| ENSMUSG( | 158.7147 | -0.05131 | 0.791974 | 0.959461 | NA |
| ENSMUSG( | 687.2169 | 0.229715 | 0.087276 | 0.454452 | NA |
| ENSMUSG( | 12.19396 | 0.169513 | 0.754148 | NA | NA |
| ENSMUSG( | 14.57227 | 0.084761 | 0.877085 | NA | NA |
| ENSMUSG( | 10.84134 | 0.080616 | 0.885034 | NA | NA |
| ENSMUSG( | 68.59327 | 0.692629 | 0.029324 | 0.259647 | NA |
| ENSMUSG( | 74.88278 | 0.172901 | 0.440513 | 0.831996 | NA |
| ENSMUSG( | 9.087852 | -0.44383 | 0.482358 | NA | NA |
| ENSMUSG( | 7.763325 | 0.745435 | 0.338742 | NA | NA |
| ENSMUSG( | 15.92376 | 0.213569 | 0.659401 | NA | NA |
| ENSMUSG( | 67.76893 | 0.076721 | 0.745206 | 0.949158 | NA |
| ENSMUSG( | 549.1407 | -0.11927 | 0.297834 | 0.746309 | NA |
| ENSMUSG( | 18.23016 | 0.219787 | 0.606984 | NA | NA |
| ENSMUSG( | 14.3286 | -0.12258 | 0.804188 | NA | NA |
| ENSMUSG( | 17.09759 | 0.140785 | 0.76338 | NA | NA |
| ENSMUSG( | 12.01749 | 0.012985 | 0.982471 | NA | NA |
| ENSMUSG( | 38.81905 | -0.1496 | 0.633318 | 0.914073 | NA |
| ENSMUSG( | 238.1169 | -0.11945 | 0.546488 | 0.884974 | NA |
| ENSMUSG( | 25.33416 | 0.719914 | 0.124285 | 0.531714 | NA |
| ENSMUSG( | 11.41784 | 0.238652 | 0.696226 | NA | NA |
| ENSMUSG( | 12.86913 | -0.66038 | 0.212043 | NA | NA |
| ENSMUSG( | 14.04427 | 0.962768 | 0.058326 | NA | NA |
| ENSMUSG( | 11.43309 | -0.06123 | 0.90989 | NA | NA |
| ENSMUSG( | 13.21981 | -0.80709 | 0.110972 | NA | NA |
| ENSMUSG( | 498.4477 | 0.118637 | 0.393671 | 0.809643 | NA |
| ENSMUSG( | 28.3273 | -0.01761 | 0.960897 | 0.993991 | NA |
| ENSMUSG( | 14.26168 | 0.300799 | 0.559195 | NA | NA |
| ENSMUSG( | 34.227 | -1.15609 | 0.00506 | 0.094553 | NA |
| ENSMUSG( | 54.61727 | 0.183355 | 0.594747 | 0.903007 | NA |
| ENSMUSG( | 38.92664 | -0.17288 | 0.596035 | 0.903152 | NA |
| ENSMUSG( | 11.18557 | -0.41157 | 0.452437 | NA | NA |
| ENSMUSG( | 148.167 | 0.089571 | 0.628486 | 0.913637 | NA |
| ENSMUSG( | 37.89263 | -0.30501 | 0.351656 | 0.783381 | NA |
| ENSMUSG( | 15.31996 | 0.216077 | 0.679775 | NA | NA |
| ENSMUSG( | 29.04176 | -0.17404 | 0.612299 | 0.908728 | NA |
| ENSMUSG( | 22.5657 | -0.02386 | 0.951045 | 0.993821 | NA |
| ENSMUSG( | 39.93816 | -0.62055 | 0.048781 | 0.337919 | NA |
| ENSMUSG( | 8.167239 | -0.24941 | 0.713666 | NA | NA |
| ENSMUSG( | 153.9394 | 0.095816 | 0.584634 | 0.898884 | NA |
| ENSMUSG( | 11.97883 | 0.195607 | 0.709854 | NA | NA |
| ENSMUSG( | 99.38978 | -0.04717 | 0.82241 | 0.966352 | NA |
| ENSMUSG( | 52.14238 | -0.2919 | 0.289041 | 0.739705 | NA |
| ENSMUSG( | 306.1831 | -0.44723 | 0.063382 | 0.389418 | NA |
| ENSMUSG( | 9.324013 | -0.17516 | 0.785113 | NA | NA |
| ENSMUSG( | 12.47841 | -0.22066 | 0.721998 | NA | NA |
| ENSMUSG( | 38.86879 | 0.128542 | 0.671164 | 0.928781 | NA |
| ENSMUSG( | 11.025 | -0.27379 | 0.648345 | NA | NA |
| ENSMUSG( | 8.281318 | -0.92717 | 0.216433 | NA | NA |
| ENSMUSG( | 65.46421 | -0.39988 | 0.101388 | 0.484996 | NA |
| ENSMUSG( | 15.36976 | -0.28911 | 0.561486 | NA | NA |
| ENSMUSG( | 12.36395 | 0.018501 | 0.972211 | NA | NA |
| ENSMUSG( | 39.82111 | -0.09987 | 0.756604 | 0.951479 | NA |
| ENSMUSG( | 19.47905 | 0.091658 | 0.83353 | NA | NA |
| ENSMUSG( | 19.21303 | -0.52897 | 0.269107 | NA | NA |
| ENSMUSG( | 10.91531 | 0.113086 | 0.850223 | NA | NA |
| ENSMUSG( | 18.9308 | 0.23484 | 0.603553 | NA | NA |
| ENSMUSG( | 13.75892 | -0.30629 | 0.54431 | NA | NA |
| ENSMUSG( | 20.89645 | 0.094266 | 0.83771 | NA | NA |
| ENSMUSG( | 74.63205 | -0.06695 | 0.81265 | 0.964574 | NA |
| ENSMUSG( | 16.2213 | -0.61369 | 0.178255 | NA | NA |
| ENSMUSG( | 11.55218 | 0.375713 | 0.541382 | NA | NA |

| | | | | | |
|---|---|---|---|---|---|
| ENSMUSG( | 10.51148 | -0.68524 | 0.300001 | NA | NA |
| ENSMUSG( | 30.40819 | -0.82784 | 0.023462 | 0.585845 | NA |
| ENSMUSG( | 67.86446 | -0.13104 | 0.662343 | 0.956205 | NA |
| ENSMUSG( | 9.347427 | -0.42329 | 0.504044 | NA | NA |
| ENSMUSG( | 11.35703 | -1.15001 | 0.056721 | NA | NA |
| ENSMUSG( | 28.03863 | 0.193265 | 0.673364 | 0.95694 | NA |
| ENSMUSG( | 38.26519 | -0.17184 | 0.611589 | 0.950771 | NA |
| ENSMUSG( | 13.46285 | 1.553842 | 0.012556 | 0.451781 | NA |
| ENSMUSG( | 10.95213 | -0.36734 | 0.548701 | NA | NA |
| ENSMUSG( | 26.19671 | -0.01379 | 0.973127 | 0.996416 | NA |
| ENSMUSG( | 51.62893 | -0.19368 | 0.49202 | 0.927315 | NA |
| ENSMUSG( | 25.90879 | -0.08365 | 0.840371 | 0.982367 | NA |
| ENSMUSG( | 25.80545 | -0.24523 | 0.528017 | 0.934648 | NA |
| ENSMUSG( | 16.60587 | 0.081271 | 0.890124 | 0.990558 | NA |
| ENSMUSG( | 217.5348 | -0.3965 | 0.042505 | 0.690304 | NA |
| ENSMUSG( | 27.00807 | -0.49714 | 0.221374 | 0.866104 | NA |
| ENSMUSG( | 20.30008 | -0.02196 | 0.960785 | 0.995097 | NA |
| ENSMUSG( | 20.06475 | 0.29376 | 0.505365 | 0.927825 | NA |
| ENSMUSG( | 24.69954 | -0.00608 | 0.989316 | 0.997918 | NA |
| ENSMUSG( | 27.75201 | 0.073669 | 0.849361 | 0.985321 | NA |
| ENSMUSG( | 15.80491 | 0.013498 | 0.978654 | 0.996459 | NA |
| ENSMUSG( | 10.48818 | -0.06937 | 0.910362 | NA | NA |
| ENSMUSG( | 22.65791 | -0.62701 | 0.168545 | 0.840222 | NA |
| ENSMUSG( | 9.476946 | -2.05037 | 0.004183 | NA | NA |
| ENSMUSG( | 9.477223 | -0.3043 | 0.636989 | NA | NA |
| ENSMUSG( | 110.4355 | -0.04299 | 0.833707 | 0.981218 | NA |
| ENSMUSG( | 30.34451 | -0.5295 | 0.154452 | 0.837724 | NA |
| ENSMUSG( | 17.80935 | 0.032687 | 0.948171 | 0.994798 | NA |
| ENSMUSG( | 51.65687 | -0.07106 | 0.810942 | 0.978799 | NA |
| ENSMUSG( | 24.66746 | -0.34763 | 0.380542 | 0.917202 | NA |
| ENSMUSG( | 13.2868 | -0.21298 | 0.684494 | 0.958678 | NA |
| ENSMUSG( | 9.267027 | -0.41166 | 0.534473 | NA | NA |
| ENSMUSG( | 11.94403 | 0.974286 | 0.097629 | 0.796743 | NA |
| ENSMUSG( | 22.7648 | -0.04175 | 0.919708 | 0.992644 | NA |
| ENSMUSG( | 10.41622 | -0.7315 | 0.237901 | NA | NA |
| ENSMUSG( | 32.4001 | 0.320498 | 0.371253 | 0.915491 | NA |
| ENSMUSG( | 495.9298 | -0.00804 | 0.952735 | 0.994798 | NA |
| ENSMUSG( | 17.82601 | 0.228049 | 0.633077 | 0.953844 | NA |
| ENSMUSG( | 697.0745 | -0.43391 | 0.006919 | 0.343883 | NA |
| ENSMUSG( | 38.56907 | 0.272424 | 0.405374 | 0.917257 | NA |
| ENSMUSG( | 33.82114 | -0.29226 | 0.428698 | 0.917257 | NA |
| ENSMUSG( | 21.02071 | -0.3453 | 0.462872 | 0.923329 | NA |
| ENSMUSG( | 69.59796 | 0.062774 | 0.820394 | 0.979198 | NA |
| ENSMUSG( | 218.5923 | -0.18191 | 0.307837 | 0.898538 | NA |
| ENSMUSG( | 44.06308 | -0.19096 | 0.538284 | 0.937216 | NA |
| ENSMUSG( | 35.01476 | 0.586595 | 0.127278 | 0.817278 | NA |
| ENSMUSG( | 14.11505 | 0.002835 | 0.995731 | 0.999116 | NA |
| ENSMUSG( | 15.34825 | -0.3077 | 0.569003 | 0.941777 | NA |
| ENSMUSG( | 16.18605 | 0.34222 | 0.474892 | 0.923329 | NA |
| ENSMUSG( | 32.03094 | 0.090952 | 0.801185 | 0.97681 | NA |
| ENSMUSG( | 29.33234 | 0.023406 | 0.952178 | 0.994798 | NA |
| ENSMUSG( | 34.34319 | -0.07983 | 0.817603 | 0.979198 | NA |
| ENSMUSG( | 19.03291 | -0.05108 | 0.912105 | 0.992196 | NA |
| ENSMUSG( | 65.47919 | -0.13685 | 0.62779 | 0.953234 | NA |
| ENSMUSG( | 1675.147 | 0.099345 | 0.360742 | 0.914223 | NA |
| ENSMUSG( | 65.39072 | -0.16573 | 0.55502 | 0.939565 | NA |
| ENSMUSG( | 74.65892 | -0.12756 | 0.591474 | 0.944975 | NA |
| ENSMUSG( | 30.02796 | 0.500222 | 0.283448 | 0.892412 | NA |
| ENSMUSG( | 155.3515 | -0.89279 | 0.182276 | 0.850579 | NA |
| ENSMUSG( | 41.79899 | 0.354768 | 0.2825 | 0.892412 | NA |
| ENSMUSG( | 39.27088 | 0.227636 | 0.533433 | 0.935162 | NA |
| ENSMUSG( | 23.0131 | 0.150552 | 0.727705 | 0.966887 | NA |
| ENSMUSG( | 37.38181 | -0.55992 | 0.152421 | 0.837724 | NA |
| ENSMUSG( | 163.5105 | 0.697145 | 0.000546 | 0.071324 | NA |
| ENSMUSG( | 98.54775 | 0.100302 | 0.685251 | 0.958678 | NA |
| ENSMUSG( | 255.0964 | -0.19106 | 0.278275 | 0.891468 | NA |
| ENSMUSG( | 169.4696 | -0.13872 | 0.426078 | 0.917257 | NA |
| ENSMUSG( | 15.3076 | -0.05576 | 0.921976 | 0.992644 | NA |
| ENSMUSG( | 12.61791 | -0.20176 | 0.706932 | 0.963529 | NA |
| ENSMUSG( | 38.18168 | -0.27622 | 0.403376 | 0.917257 | NA |
| ENSMUSG( | 12.82717 | -0.45383 | 0.414163 | 0.917257 | NA |
| ENSMUSG( | 106.1338 | 0.041835 | 0.842577 | 0.983044 | NA |
| ENSMUSG( | 16.09828 | 0.733855 | 0.140107 | 0.830678 | NA |
| ENSMUSG( | 19.51168 | 0.624767 | 0.174039 | 0.844317 | NA |
| ENSMUSG( | 11.05761 | -0.10882 | 0.863624 | NA | NA |
| ENSMUSG( | 14.2904 | -0.27909 | 0.59734 | 0.946609 | NA |
| ENSMUSG( | 36.44851 | -0.55424 | 0.107733 | 0.804969 | NA |
| ENSMUSG( | 7.864924 | -0.41455 | 0.589033 | NA | NA |
| ENSMUSG( | 26.79785 | -0.63714 | 0.116216 | 0.813384 | NA |
| ENSMUSG( | 99.12006 | -0.15048 | 0.502517 | 0.927825 | NA |
| ENSMUSG( | 16.21768 | -0.11124 | 0.819775 | 0.979198 | NA |
| ENSMUSG( | 50.77324 | -0.23632 | 0.414564 | 0.917257 | NA |
| ENSMUSG( | 969.3482 | 0.18969 | 0.220766 | 0.866104 | NA |
| ENSMUSG( | 10.81113 | -0.83163 | 0.201943 | NA | NA |
| ENSMUSG( | 11.68548 | 0.311484 | 0.599102 | 0.947045 | NA |
| ENSMUSG( | 16.76446 | -0.70142 | 0.151057 | 0.837173 | NA |
| ENSMUSG( | 28.13004 | -0.1789 | 0.685528 | 0.958678 | NA |
| ENSMUSG( | 35.87811 | 0.602052 | 0.14963 | 0.836241 | NA |
| ENSMUSG( | 480.8118 | 0.102422 | 0.464267 | 0.923329 | NA |
| ENSMUSG( | 19.28479 | -0.29863 | 0.51089 | 0.929546 | NA |
| ENSMUSG( | 18.83169 | -0.49599 | 0.289708 | 0.896726 | NA |
| ENSMUSG( | 11.17692 | -0.3479 | 0.548205 | NA | NA |
| ENSMUSG( | 43.44408 | -0.18425 | 0.560603 | 0.940254 | NA |
| ENSMUSG( | 166.4064 | -0.42279 | 0.026313 | 0.603041 | NA |
| ENSMUSG( | 69.34762 | -0.48616 | 0.079286 | 0.77458 | NA |
| ENSMUSG( | 12.44819 | -0.38035 | 0.530858 | 0.934795 | NA |
| ENSMUSG( | 15.87939 | 0.359712 | 0.475849 | 0.923329 | NA |
| ENSMUSG( | 54.20127 | 0.116487 | 0.745738 | 0.96951 | NA |
| ENSMUSG( | 11.47573 | 0.283451 | 0.629575 | 0.953251 | NA |
| ENSMUSG( | 11.03031 | -1.27336 | 0.035058 | NA | NA |
| ENSMUSG( | 108.9962 | -0.02958 | 0.886574 | 0.990063 | NA |
| ENSMUSG( | 9.019895 | -0.35786 | 0.584948 | NA | NA |
| ENSMUSG( | 40.95606 | 0.018602 | 0.958092 | 0.994798 | NA |
| ENSMUSG( | 27.52213 | 0.591767 | 0.171949 | 0.842999 | NA |
| ENSMUSG( | 135.9496 | -0.19174 | 0.320971 | 0.904019 | NA |
| ENSMUSG( | 32.01463 | -0.36259 | 0.389756 | 0.917257 | NA |
| ENSMUSG( | 222.6291 | -0.14292 | 0.40961 | 0.917257 | NA |
| ENSMUSG( | 9.812582 | -0.58286 | 0.364961 | NA | NA |
| ENSMUSG( | 9.772401 | 0.712587 | 0.269005 | NA | NA |
| ENSMUSG( | 16.57301 | -0.09524 | 0.84527 | 0.983957 | NA |
| ENSMUSG( | 10.43343 | 0.69185 | 0.339872 | NA | NA |
| ENSMUSG( | 18.92112 | -0.86576 | 0.063624 | 0.736482 | NA |
| ENSMUSG( | 17.35549 | -0.57416 | 0.229365 | 0.868507 | NA |
| ENSMUSG( | 22.43596 | -0.23859 | 0.600162 | 0.947461 | NA |
| ENSMUSG( | 13.84978 | 0.183398 | 0.73269 | 0.967228 | NA |
| ENSMUSG( | 18.37621 | 0.193481 | 0.679877 | 0.957966 | NA |
| ENSMUSG( | 9.472301 | -0.34961 | 0.600567 | NA | NA |
| ENSMUSG( | 8.528729 | -0.38043 | 0.570225 | NA | NA |
| ENSMUSG( | 15.44838 | -0.08406 | 0.877771 | 0.989644 | NA |
| ENSMUSG( | 20.49819 | -0.48267 | 0.292234 | 0.897854 | NA |
| ENSMUSG( | 8.101016 | -0.56807 | 0.421482 | NA | NA |
| ENSMUSG( | 12.63366 | -0.04517 | 0.934796 | 0.99306 | NA |
| ENSMUSG( | 19.62705 | -0.6696 | 0.134292 | 0.822912 | NA |
| ENSMUSG( | 9.368139 | 0.590994 | 0.349716 | NA | NA |
| ENSMUSG( | 14.77324 | -0.81098 | 0.140475 | 0.830678 | NA |
| ENSMUSG( | 379.2834 | 0.059157 | 0.690471 | 0.960071 | NA |
| ENSMUSG( | 15.45464 | 0.368199 | 0.477557 | 0.923329 | NA |
| ENSMUSG( | 8.230388 | 0.223857 | 0.746103 | NA | NA |
| ENSMUSG( | 37.44488 | 0.01773 | 0.95807 | 0.994798 | NA |
| ENSMUSG( | 22.61712 | 0.368943 | 0.414689 | 0.917257 | NA |
| ENSMUSG( | 38.52588 | -0.32518 | 0.341025 | 0.908138 | NA |
| ENSMUSG( | 177.0336 | -0.31589 | 0.109933 | 0.806572 | NA |
| ENSMUSG( | 120.3465 | -0.16996 | 0.434457 | 0.91869 | NA |
| ENSMUSG( | 11.98347 | -0.83681 | 0.189325 | 0.85698 | NA |
| ENSMUSG( | 8.408328 | -0.93395 | 0.174168 | NA | NA |
| ENSMUSG( | 13.79664 | 0.447039 | 0.39214 | 0.917257 | NA |
| ENSMUSG( | 144.3582 | -0.04687 | 0.843622 | 0.983412 | NA |
| ENSMUSG( | 16.21535 | 0.048283 | 0.932324 | 0.99306 | NA |
| ENSMUSG( | 16.20765 | -0.32877 | 0.546412 | 0.938063 | NA |
| ENSMUSG( | 31.26828 | 0.221222 | 0.529468 | 0.934795 | NA |
| ENSMUSG( | 65.81986 | -0.08711 | 0.788832 | 0.976593 | NA |
| ENSMUSG( | 18.61003 | -0.16697 | 0.727581 | 0.966887 | NA |
| ENSMUSG( | 17.50012 | 0.082118 | 0.864657 | 0.986591 | NA |
| ENSMUSG( | 26.86186 | -0.21358 | 0.587393 | 0.944975 | NA |
| ENSMUSG( | 9.052265 | 0.277731 | 0.667041 | NA | NA |
| ENSMUSG( | 12.17676 | -0.13114 | 0.83503 | 0.981237 | NA |
| ENSMUSG( | 32.56523 | -0.68072 | 0.053038 | 0.71525 | NA |
| ENSMUSG( | 385.5498 | -0.13879 | 0.473903 | 0.923329 | NA |
| ENSMUSG( | 22.10662 | 0.338386 | 0.429759 | 0.917257 | NA |
| ENSMUSG( | 23.36485 | -0.0811 | 0.860274 | 0.986137 | NA |
| ENSMUSG( | 22.09194 | 0.012977 | 0.975266 | 0.996416 | NA |
| ENSMUSG( | 133.7114 | -0.16842 | 0.401866 | 0.917257 | NA |
| ENSMUSG( | 340.398 | 0.035989 | 0.804201 | 0.977191 | NA |
| ENSMUSG( | 568.9999 | 0.195264 | 0.249176 | 0.877904 | NA |
| ENSMUSG( | 30.49398 | -0.41804 | 0.290226 | 0.896947 | NA |
| ENSMUSG( | 49.41306 | -0.17366 | 0.554418 | 0.939565 | NA |
| ENSMUSG( | 9.517996 | -0.21912 | 0.727834 | NA | NA |
| ENSMUSG( | 163.2527 | 0.138065 | 0.447012 | 0.921971 | NA |

| | | | | | |
|---|---|---|---|---|---|
| ENSMUSG( | NA | 21.15748 | -0.56085 | 0.25166 | NA |
| ENSMUSG( | NA | 13.71651 | 0.437198 | 0.409748 | NA |
| ENSMUSG( | NA | 55.30185 | 0.213478 | 0.44528 | 0.833239 |
| ENSMUSG( | NA | 29.78521 | -0.21105 | 0.567145 | 0.891644 |
| ENSMUSG( | NA | 13.25063 | -0.19296 | 0.714164 | NA |
| ENSMUSG( | NA | 8.732039 | -1.14494 | 0.069107 | NA |
| ENSMUSG( | NA | 11.82383 | -0.1548 | 0.780545 | NA |
| ENSMUSG( | NA | 39.48496 | 0.112641 | 0.734781 | 0.945584 |
| ENSMUSG( | NA | 11.72133 | -0.37462 | 0.487951 | NA |
| ENSMUSG( | NA | 36.96834 | 0.425618 | 0.168581 | 0.602155 |
| ENSMUSG( | NA | 14.06422 | 0.288359 | 0.570445 | NA |
| ENSMUSG( | NA | 16.7918 | 0.14262 | 0.760627 | NA |
| ENSMUSG( | NA | 25.79613 | -0.08156 | 0.823648 | 0.966603 |
| ENSMUSG( | NA | 17.74783 | 0.161453 | 0.731691 | NA |
| ENSMUSG( | NA | 64.38523 | 0.484086 | 0.06945 | 0.407995 |
| ENSMUSG( | NA | 10.66495 | -0.03805 | 0.945553 | NA |
| ENSMUSG( | NA | 114.9991 | -0.16792 | 0.426211 | 0.824898 |
| ENSMUSG( | NA | 8.026502 | -1.67258 | 0.018631 | NA |
| ENSMUSG( | NA | 9.956333 | -0.21577 | 0.71515 | NA |
| ENSMUSG( | NA | 26.36034 | -0.00641 | 0.986026 | 0.998326 |
| ENSMUSG( | NA | 11.88876 | -0.23024 | 0.679875 | NA |
| ENSMUSG( | NA | 20.08126 | -0.21568 | 0.627319 | NA |
| ENSMUSG( | NA | 14.42571 | -0.0075 | 0.987588 | NA |
| ENSMUSG( | NA | 15.09746 | 0.088368 | 0.865375 | NA |
| ENSMUSG( | NA | 53.48589 | -0.23749 | 0.372059 | 0.798768 |
| ENSMUSG( | NA | 13.59951 | 0.190493 | 0.714641 | NA |
| ENSMUSG( | NA | 28.28019 | -0.50049 | 0.150389 | 0.574991 |
| ENSMUSG( | NA | 20.51512 | 0.24689 | 0.571439 | NA |
| ENSMUSG( | NA | 18.18625 | 0.493057 | 0.261019 | NA |
| ENSMUSG( | NA | 10.66944 | 0.452134 | 0.453091 | NA |
| ENSMUSG( | NA | 44.51375 | -0.28246 | 0.381422 | 0.802629 |
| ENSMUSG( | NA | 62.69358 | 0.001215 | 0.996324 | 0.99943 |
| ENSMUSG( | NA | 149.7058 | -0.20629 | 0.221527 | 0.667258 |
| ENSMUSG( | NA | 11.98207 | -0.02416 | 0.96465 | NA |
| ENSMUSG( | NA | 62.78612 | -0.17113 | 0.52202 | 0.875186 |
| ENSMUSG( | NA | 206.8034 | -0.11739 | 0.494102 | 0.860611 |
| ENSMUSG( | NA | 25.33328 | -0.43271 | 0.246429 | 0.694958 |
| ENSMUSG( | NA | 24.4451 | 0.311353 | 0.40267 | 0.81552 |
| ENSMUSG( | NA | 51.13866 | -0.45176 | 0.091964 | 0.465937 |
| ENSMUSG( | NA | 53.25763 | 0.409063 | 0.250782 | 0.700403 |
| ENSMUSG( | NA | 17.00235 | -0.73181 | 0.211063 | NA |
| ENSMUSG( | NA | 46.35018 | 0.268022 | 0.40707 | 0.817163 |
| ENSMUSG( | NA | 167.318 | -0.01225 | 0.940862 | 0.993415 |
| ENSMUSG( | NA | 34.20248 | 0.14477 | 0.697482 | 0.935662 |
| ENSMUSG( | NA | 23.50467 | 0.329796 | 0.397549 | 0.812127 |
| ENSMUSG( | NA | 564.3165 | 0.207977 | 0.136955 | 0.55149 |
| ENSMUSG( | NA | 12.59574 | 0.529499 | 0.3798 | NA |
| ENSMUSG( | NA | 45.14816 | -0.37097 | 0.197099 | 0.636662 |
| ENSMUSG( | NA | 14.49841 | -0.12161 | 0.807485 | NA |
| ENSMUSG( | NA | 48.54665 | 0.152125 | 0.603018 | 0.904887 |
| ENSMUSG( | NA | 18.16707 | -0.34193 | 0.432961 | NA |
| ENSMUSG( | NA | 35.80285 | -0.18984 | 0.592941 | 0.90229 |
| ENSMUSG( | NA | 8.937982 | -0.20821 | 0.740475 | NA |
| ENSMUSG( | NA | 24.74481 | 0.098047 | 0.808156 | 0.963419 |
| ENSMUSG( | NA | 19.17065 | 0.458691 | 0.275932 | NA |
| ENSMUSG( | NA | 12.38842 | 0.565812 | 0.273471 | NA |
| ENSMUSG( | NA | 13.60923 | -0.22715 | 0.681454 | NA |
| ENSMUSG( | NA | 31.05786 | -0.3194 | 0.472527 | 0.848206 |
| ENSMUSG( | NA | 23.78676 | 0.17427 | 0.659376 | 0.924506 |
| ENSMUSG( | NA | 19.68258 | 0.374878 | 0.394726 | NA |
| ENSMUSG( | NA | 20.32904 | 0.590577 | 0.180939 | NA |
| ENSMUSG( | NA | 34.38633 | -0.89863 | 0.048196 | 0.335655 |
| ENSMUSG( | NA | 12.29094 | -0.47018 | 0.387527 | NA |
| ENSMUSG( | NA | 35.28601 | 0.343855 | 0.318604 | 0.764234 |
| ENSMUSG( | NA | 23.71913 | 0.25815 | 0.499796 | 0.86324 |
| ENSMUSG( | NA | 71.69468 | 0.277031 | 0.291528 | 0.741366 |
| ENSMUSG( | NA | 63.46334 | -0.29027 | 0.244304 | 0.691435 |
| ENSMUSG( | NA | 15.469 | -0.37593 | 0.4641 | NA |
| ENSMUSG( | NA | 37.7855 | 0.005056 | 0.987355 | 0.998796 |
| ENSMUSG( | NA | 28.25366 | 0.228936 | 0.560914 | 0.888934 |
| ENSMUSG( | NA | 14.54783 | -0.60075 | 0.206767 | NA |
| ENSMUSG( | NA | 27.62345 | 0.393423 | 0.314878 | 0.762059 |
| ENSMUSG( | NA | 13.42192 | 0.767364 | 0.164752 | NA |
| ENSMUSG( | NA | 78.77365 | 0.107246 | 0.663404 | 0.9265 |
| ENSMUSG( | NA | 21.4953 | 0.137287 | 0.733197 | 0.944956 |
| ENSMUSG( | NA | 47.63042 | -0.83558 | 0.005955 | 0.104409 |
| ENSMUSG( | NA | 20.34538 | 0.180414 | 0.680735 | NA |
| ENSMUSG( | NA | 694.2862 | -0.23492 | 0.077756 | 0.431059 |
| ENSMUSG( | NA | 16.70896 | 0.001022 | 0.998197 | NA |
| ENSMUSG( | NA | 19.58375 | -0.12949 | 0.787892 | NA |
| ENSMUSG( | NA | 18.78317 | 0.094638 | 0.830988 | NA |
| ENSMUSG( | NA | 426.9991 | 0.057836 | 0.619819 | 0.911432 |
| ENSMUSG( | NA | 31.74936 | -0.3334 | 0.328374 | 0.772854 |
| ENSMUSG( | NA | 10.80238 | 0.227534 | 0.701765 | NA |
| ENSMUSG( | NA | 19.89275 | -0.76452 | 0.069464 | NA |
| ENSMUSG( | NA | 57.04249 | 0.147782 | 0.621379 | 0.911432 |
| ENSMUSG( | NA | 37.21991 | 0.216398 | 0.501671 | 0.864596 |
| ENSMUSG( | NA | 25.38671 | -0.57089 | 0.126394 | 0.533885 |
| ENSMUSG( | NA | 166.2888 | -0.07743 | 0.660243 | 0.924627 |
| ENSMUSG( | NA | 18.34993 | -0.28545 | 0.52955 | NA |
| ENSMUSG( | NA | 15.2647 | 0.109551 | 0.850598 | NA |
| ENSMUSG( | NA | 98.65981 | -0.17136 | 0.401932 | 0.814965 |
| ENSMUSG( | NA | 44.55784 | -0.20017 | 0.514577 | 0.872055 |
| ENSMUSG( | NA | 9.815303 | 0.369166 | 0.590247 | NA |
| ENSMUSG( | NA | 68.5083 | 0.461466 | 0.05381 | 0.356147 |
| ENSMUSG( | NA | 19.74988 | -0.12107 | 0.778739 | NA |
| ENSMUSG( | NA | 13.24738 | -0.52552 | 0.293013 | NA |
| ENSMUSG( | NA | 196.2032 | -0.14101 | 0.397851 | 0.812161 |
| ENSMUSG( | NA | 126.7052 | -0.11708 | 0.516176 | 0.872701 |
| ENSMUSG( | NA | 35.12063 | 0.387119 | 0.259477 | 0.709948 |
| ENSMUSG( | NA | 17.12495 | -0.8023 | 0.099257 | NA |
| ENSMUSG( | NA | 17.52004 | 0.050459 | 0.910293 | NA |
| ENSMUSG( | NA | 30.52475 | -0.49819 | 0.167171 | 0.599823 |
| ENSMUSG( | NA | 12.32838 | -0.95984 | 0.070709 | NA |
| ENSMUSG( | NA | 22.26734 | -0.12511 | 0.752775 | 0.950621 |
| ENSMUSG( | NA | 20.94474 | 0.106431 | 0.79939 | NA |
| ENSMUSG( | NA | 295.5486 | 0.038978 | 0.797263 | 0.960507 |
| ENSMUSG( | NA | 46.5443 | 0.321021 | 0.254467 | 0.704536 |
| ENSMUSG( | NA | 16.14955 | -0.4897 | 0.301974 | NA |
| ENSMUSG( | NA | 16.17676 | -0.42406 | 0.351563 | NA |
| ENSMUSG( | NA | 15.4955 | -0.00141 | 0.997611 | NA |
| ENSMUSG( | NA | 17.71887 | 0.473011 | 0.335303 | NA |
| ENSMUSG( | NA | 14.45386 | 0.47457 | 0.331407 | NA |
| ENSMUSG( | NA | 85.97965 | -0.23837 | 0.323213 | 0.768415 |
| ENSMUSG( | NA | 59.31701 | -0.12538 | 0.620445 | 0.911432 |
| ENSMUSG( | NA | 9.698451 | 0.01985 | 0.976078 | NA |
| ENSMUSG( | NA | 36.48983 | 0.350473 | 0.268842 | 0.719099 |
| ENSMUSG( | NA | 9.845391 | -0.66401 | 0.294234 | NA |
| ENSMUSG( | NA | 26.8643 | -1.02804 | 0.005201 | 0.096074 |
| ENSMUSG( | NA | 10.1366 | 0.014607 | 0.981577 | NA |
| ENSMUSG( | NA | 21.95667 | -0.58382 | 0.16827 | 0.601572 |
| ENSMUSG( | NA | 16.1767 | -0.01818 | 0.968754 | NA |
| ENSMUSG( | NA | 135.1021 | -0.16005 | 0.41264 | 0.819518 |
| ENSMUSG( | NA | 98.10077 | -0.06735 | 0.73322 | 0.944956 |
| ENSMUSG( | NA | 32.58683 | 0.190907 | 0.590691 | 0.90113 |
| ENSMUSG( | NA | 80.24501 | -0.11049 | 0.620795 | 0.911432 |
| ENSMUSG( | NA | 11.26565 | -0.4752 | 0.430949 | NA |
| ENSMUSG( | NA | 41.70991 | 0.396577 | 0.227082 | 0.674223 |
| ENSMUSG( | NA | 24.15481 | -0.55932 | 0.149236 | 0.573248 |
| ENSMUSG( | NA | 12.06032 | 0.87941 | 0.116328 | NA |
| ENSMUSG( | NA | 26.30873 | 0.395369 | 0.337946 | 0.777422 |
| ENSMUSG( | NA | 9.797835 | -0.86609 | 0.167035 | NA |
| ENSMUSG( | NA | 26.29471 | -0.44411 | 0.242292 | 0.68971 |
| ENSMUSG( | NA | 11.99105 | 0.133052 | 0.810032 | NA |
| ENSMUSG( | NA | 328.7038 | 1.290531 | 0.013019 | 0.165126 |
| ENSMUSG( | NA | 13.72268 | 0.224926 | 0.665501 | NA |
| ENSMUSG( | NA | 149.0157 | 0.250895 | 0.156554 | 0.584667 |
| ENSMUSG( | NA | 17.92012 | -0.37224 | 0.44537 | NA |
| ENSMUSG( | NA | 25.57771 | 0.067174 | 0.857899 | 0.974667 |
| ENSMUSG( | NA | 11.23569 | 0.341573 | 0.54845 | NA |
| ENSMUSG( | NA | 30.40242 | 0.438935 | 0.234591 | 0.681922 |
| ENSMUSG( | NA | 197.3917 | -0.0856 | 0.634329 | 0.914073 |
| ENSMUSG( | NA | 31.58579 | -0.29721 | 0.43505 | 0.829093 |
| ENSMUSG( | NA | 35.21593 | 0.002483 | 0.993762 | 0.999369 |
| ENSMUSG( | NA | 10.87673 | 0.189348 | 0.744692 | NA |
| ENSMUSG( | NA | 46.65382 | -0.07913 | 0.792531 | 0.959461 |
| ENSMUSG( | NA | 229.4443 | 0.062104 | 0.663334 | 0.9265 |
| ENSMUSG( | NA | 11.92232 | -1.00046 | 0.081108 | NA |
| ENSMUSG( | NA | 26.19675 | -0.63426 | 0.085928 | 0.451489 |
| ENSMUSG( | NA | 14.9341 | -0.1644 | 0.757675 | NA |
| ENSMUSG( | NA | 98.98782 | 0.104092 | 0.601435 | 0.904603 |
| ENSMUSG( | NA | 14.63802 | -0.02343 | 0.962701 | NA |
| ENSMUSG( | NA | 316.1098 | -0.32889 | 0.046019 | 0.328532 |
| ENSMUSG( | NA | 14.85825 | 0.371952 | 0.50347 | NA |
| ENSMUSG( | NA | 230.4468 | 0.162762 | 0.246501 | 0.694958 |
| ENSMUSG( | NA | 22.293 | -0.34127 | 0.407038 | 0.817163 |
| ENSMUSG( | NA | 41.671 | -0.02461 | 0.932425 | 0.992515 |
| ENSMUSG( | NA | 9.963841 | -0.51222 | 0.409416 | NA |

| | | | | | |
|---|---|---|---|---|---|
| ENSMUSG( | NA | 568.9999 | 0.195264 | 0.249176 | 0.877904 |
| ENSMUSG( | NA | 30.49398 | -0.41804 | 0.290226 | 0.896947 |
| ENSMUSG( | NA | 49.41306 | -0.17366 | 0.554418 | 0.939565 |
| ENSMUSG( | NA | 9.517996 | -0.21912 | 0.727834 | NA |
| ENSMUSG( | NA | 163.2527 | 0.138065 | 0.447012 | 0.921971 |
| ENSMUSG( | NA | 20.70058 | 0.617038 | 0.21594 | 0.865167 |
| ENSMUSG( | NA | 37.09582 | 0.062502 | 0.851793 | 0.985591 |
| ENSMUSG( | NA | 91.55776 | 0.02604 | 0.914807 | 0.992407 |
| ENSMUSG( | NA | 24.51841 | -0.51831 | 0.231896 | 0.868507 |
| ENSMUSG( | NA | 152.3134 | 0.332836 | 0.075944 | 0.768524 |
| ENSMUSG( | NA | 1224.174 | -0.21053 | 0.130993 | 0.822884 |
| ENSMUSG( | NA | 12.07828 | 0.345927 | 0.546714 | 0.938063 |
| ENSMUSG( | NA | 51.22768 | -0.12496 | 0.699869 | 0.961335 |
| ENSMUSG( | NA | 30.54137 | -1.3968 | 0.000265 | 0.04337 |
| ENSMUSG( | NA | 11.62846 | 0.101214 | 0.871584 | 0.988779 |
| ENSMUSG( | NA | 45.66507 | 0.156173 | 0.649018 | 0.95422 |
| ENSMUSG( | NA | 69.26844 | 0.324718 | 0.203783 | 0.864412 |
| ENSMUSG( | NA | 40.13608 | -0.40157 | 0.221787 | 0.866104 |
| ENSMUSG( | NA | 15.75031 | 0.432507 | 0.403123 | 0.917257 |
| ENSMUSG( | NA | 34.87109 | -0.11608 | 0.739399 | 0.968458 |
| ENSMUSG( | NA | 27.87565 | -0.23741 | 0.527723 | 0.934603 |
| ENSMUSG( | NA | 9.516006 | -0.47319 | 0.44676 | NA |
| ENSMUSG( | NA | 14.92116 | -0.32654 | 0.532956 | 0.934795 |
| ENSMUSG( | NA | 9.860791 | -0.26526 | 0.68279 | NA |
| ENSMUSG( | NA | 29.29585 | 0.238462 | 0.542412 | 0.938034 |
| ENSMUSG( | NA | 11.66313 | 0.156419 | 0.780935 | 0.976106 |
| ENSMUSG( | NA | 7.337955 | -0.44282 | 0.568132 | NA |
| ENSMUSG( | NA | 39.18936 | -0.05095 | 0.889449 | 0.990558 |
| ENSMUSG( | NA | 14.50765 | -0.12928 | 0.810189 | 0.978779 |
| ENSMUSG( | NA | 23.63881 | 0.373021 | 0.393102 | 0.917257 |
| ENSMUSG( | NA | 10.67734 | 0.010822 | 0.987472 | NA |
| ENSMUSG( | NA | 85.53119 | -0.29068 | 0.207591 | 0.864412 |
| ENSMUSG( | NA | 22.18549 | 0.306806 | 0.47098 | 0.923329 |
| ENSMUSG( | NA | 9.971965 | 0.223667 | 0.726447 | NA |
| ENSMUSG( | NA | 165.1508 | -0.05212 | 0.803843 | 0.977 |
| ENSMUSG( | NA | 366.1288 | -0.28233 | 0.066988 | 0.75212 |
| ENSMUSG( | NA | 24.9882 | 0.202443 | 0.62877 | 0.953251 |
| ENSMUSG( | NA | 70.02962 | -0.31465 | 0.231934 | 0.868507 |
| ENSMUSG( | NA | 11.68626 | 0.819087 | 0.152293 | 0.837724 |
| ENSMUSG( | NA | 32.30619 | -0.26 | 0.501999 | 0.927825 |
| ENSMUSG( | NA | 18.1706 | 0.029862 | 0.949766 | 0.994798 |
| ENSMUSG( | NA | 18.13039 | 0.158513 | 0.732778 | 0.967228 |
| ENSMUSG( | NA | 14.54738 | -0.29808 | 0.596816 | 0.946541 |
| ENSMUSG( | NA | 17.17473 | 0.705661 | 0.138283 | 0.828492 |
| ENSMUSG( | NA | 15.02023 | -0.4906 | 0.339655 | 0.907972 |
| ENSMUSG( | NA | 19.60289 | -0.02003 | 0.965193 | 0.995433 |
| ENSMUSG( | NA | 270.738 | -0.14302 | 0.375985 | 0.915491 |
| ENSMUSG( | NA | 12.13046 | -0.44442 | 0.428038 | 0.917257 |
| ENSMUSG( | NA | 14.60842 | 0.488023 | 0.347601 | 0.911652 |
| ENSMUSG( | NA | 14.22349 | -0.76126 | 0.143856 | 0.831311 |
| ENSMUSG( | NA | 14.69581 | -0.19845 | 0.706035 | 0.96335 |
| ENSMUSG( | NA | 30.66946 | 0.545414 | 0.26375 | 0.886134 |
| ENSMUSG( | NA | 183.1372 | -0.25915 | 0.143485 | 0.831311 |
| ENSMUSG( | NA | 105.9158 | 0.299135 | 0.203 | 0.864412 |
| ENSMUSG( | NA | 21.78561 | 0.351556 | 0.404027 | 0.917257 |
| ENSMUSG( | NA | 27.83786 | 0.068045 | 0.855158 | 0.985649 |
| ENSMUSG( | NA | 9.01457 | -0.47119 | 0.482628 | NA |
| ENSMUSG( | NA | 28.22471 | 0.212784 | 0.602862 | 0.94789 |
| ENSMUSG( | NA | 160.0558 | 0.075788 | 0.6924 | 0.960404 |
| ENSMUSG( | NA | 17.70561 | -0.36454 | 0.474255 | 0.923329 |
| ENSMUSG( | NA | 9.842578 | 0.369568 | 0.566089 | NA |
| ENSMUSG( | NA | 20.10192 | -0.15798 | 0.717759 | 0.965964 |
| ENSMUSG( | NA | 12.87347 | 0.075647 | 0.891661 | 0.990558 |
| ENSMUSG( | NA | 43.26562 | 0.320407 | 0.303994 | 0.898538 |
| ENSMUSG( | NA | 71.80027 | -0.16167 | 0.574079 | 0.942912 |
| ENSMUSG( | NA | 13.4225 | -0.58535 | 0.334586 | 0.907907 |
| ENSMUSG( | NA | 75.39198 | -0.02802 | 0.934026 | 0.99306 |
| ENSMUSG( | NA | 9.414813 | -1.20677 | 0.070547 | NA |
| ENSMUSG( | NA | 49.69782 | 0.460697 | 0.171613 | 0.842999 |
| ENSMUSG( | NA | 25.40125 | 0.345897 | 0.372724 | 0.915491 |
| ENSMUSG( | NA | 53.63668 | -0.15019 | 0.672172 | 0.95694 |
| ENSMUSG( | NA | 12.04652 | 0.101264 | 0.858107 | 0.985884 |
| ENSMUSG( | NA | 34.39673 | -0.24434 | 0.482261 | 0.924578 |
| ENSMUSG( | NA | 26.59867 | -0.77725 | 0.05397 | 0.722223 |
| ENSMUSG( | NA | 196.8398 | -0.12701 | 0.479602 | 0.924021 |
| ENSMUSG( | NA | 15.31883 | -0.31696 | 0.534613 | 0.936107 |
| ENSMUSG( | NA | 14.52272 | 0.058127 | 0.908503 | 0.991958 |
| ENSMUSG( | NA | 15.13045 | 0.543657 | 0.399401 | 0.917257 |
| ENSMUSG( | NA | 13.631 | 0.215641 | 0.735786 | 0.967344 |
| ENSMUSG( | NA | 75.02526 | 0.071556 | 0.769126 | 0.974236 |
| ENSMUSG( | NA | 13.99327 | 0.412767 | 0.468492 | 0.923329 |
| ENSMUSG( | NA | 181.7445 | -0.0962 | 0.620192 | 0.952896 |
| ENSMUSG( | NA | 584.2 | 0.154497 | 0.297321 | 0.898538 |
| ENSMUSG( | NA | 7.965554 | -0.18673 | 0.787222 | NA |
| ENSMUSG( | NA | 15.36659 | 0.228218 | 0.67741 | 0.957106 |
| ENSMUSG( | NA | 41.13395 | -0.3835 | 0.226487 | 0.868507 |
| ENSMUSG( | NA | 21.19724 | 0.270078 | 0.581112 | 0.944266 |
| ENSMUSG( | NA | 13.593 | 0.527651 | 0.345035 | 0.91045 |
| ENSMUSG( | NA | 16.35109 | -0.29286 | 0.555505 | 0.939565 |
| ENSMUSG( | NA | 10.76515 | -0.56399 | 0.351124 | NA |
| ENSMUSG( | NA | 34.32679 | 0.509735 | 0.141993 | 0.830709 |
| ENSMUSG( | NA | 18.95793 | -0.07876 | 0.867763 | 0.98781 |
| ENSMUSG( | NA | 25.68259 | -0.41661 | 0.303466 | 0.898538 |
| ENSMUSG( | NA | 26.75966 | -0.36845 | 0.35818 | 0.913759 |
| ENSMUSG( | NA | 321.322 | 0.070048 | 0.741684 | 0.969125 |
| ENSMUSG( | NA | 35.74836 | 0.36997 | 0.338859 | 0.907972 |
| ENSMUSG( | NA | 225.1003 | -0.0325 | 0.850673 | 0.985541 |
| ENSMUSG( | NA | 11.49811 | -0.05725 | 0.923633 | 0.992815 |
| ENSMUSG( | NA | 20.97317 | -0.79751 | 0.093214 | 0.795363 |
| ENSMUSG( | NA | 38.76579 | 0.245593 | 0.442868 | 0.921554 |
| ENSMUSG( | NA | 9.568682 | -0.81101 | 0.205499 | NA |
| ENSMUSG( | NA | 114.5887 | -0.14079 | 0.523849 | 0.932848 |
| ENSMUSG( | NA | 10.63514 | 0.628973 | 0.303422 | NA |
| ENSMUSG( | NA | 29.15697 | 0.400454 | 0.289347 | 0.896442 |
| ENSMUSG( | NA | 7.698566 | -0.284 | 0.70521 | NA |
| ENSMUSG( | NA | 814.5748 | -0.02601 | 0.837464 | 0.981335 |
| ENSMUSG( | NA | 181.272 | 0.188671 | 0.371679 | 0.915491 |
| ENSMUSG( | NA | 29.0243 | 0.841139 | 0.037196 | 0.673514 |
| ENSMUSG( | NA | 406.5499 | -0.3285 | 0.015415 | 0.495791 |
| ENSMUSG( | NA | 18679.09 | 0.0858 | 0.506149 | 0.927825 |
| ENSMUSG( | NA | 98.26181 | -0.43798 | 0.06864 | 0.753052 |
| ENSMUSG( | NA | 22.50803 | -0.6223 | 0.139312 | 0.830678 |
| ENSMUSG( | NA | 133.7178 | -0.05872 | 0.767465 | 0.974236 |
| ENSMUSG( | NA | 32.88013 | -0.02823 | 0.934952 | 0.99306 |
| ENSMUSG( | NA | 29.71692 | 0.760762 | 0.05447 | 0.723362 |
| ENSMUSG( | NA | 64.82556 | -0.10362 | 0.696007 | 0.960867 |
| ENSMUSG( | NA | 11.52232 | -0.08016 | 0.889205 | 0.990558 |
| ENSMUSG( | NA | 19.37405 | 0.296224 | 0.602196 | 0.94789 |
| ENSMUSG( | NA | 9.784527 | -0.05341 | 0.931097 | NA |
| ENSMUSG( | NA | 16.64004 | -0.22807 | 0.654076 | 0.955015 |
| ENSMUSG( | NA | 736.3628 | -0.06883 | 0.632946 | 0.953844 |
| ENSMUSG( | NA | 65.85936 | -0.68891 | 0.007761 | 0.37555 |
| ENSMUSG( | NA | 10.18875 | -0.35214 | 0.572544 | NA |
| ENSMUSG( | NA | 47.29014 | -0.22286 | 0.51308 | 0.930358 |
| ENSMUSG( | NA | 11.92558 | 0.759046 | 0.275697 | 0.890209 |
| ENSMUSG( | NA | 22.12934 | 0.627639 | 0.150138 | 0.837086 |
| ENSMUSG( | NA | 127.8141 | -0.04 | 0.864273 | 0.986552 |
| ENSMUSG( | NA | 11.75465 | -1.09798 | 0.056177 | 0.723362 |
| ENSMUSG( | NA | 15.04587 | 0.104361 | 0.835689 | 0.981237 |
| ENSMUSG( | NA | 135.6324 | -0.13116 | 0.525716 | 0.934374 |
| ENSMUSG( | NA | 20.38823 | 0.806299 | 0.065923 | 0.746371 |
| ENSMUSG( | NA | 21.46928 | 0.55814 | 0.315344 | 0.901306 |
| ENSMUSG( | NA | 15.75351 | -0.12879 | 0.805072 | 0.977358 |
| ENSMUSG( | NA | 18.22633 | 0.040152 | 0.93258 | 0.99306 |
| ENSMUSG( | NA | 12.93019 | -0.39175 | 0.516263 | 0.930939 |
| ENSMUSG( | NA | 32.26841 | -0.40803 | 0.308912 | 0.898538 |
| ENSMUSG( | NA | 8.966632 | 0.055937 | 0.930954 | NA |
| ENSMUSG( | NA | 192.8873 | -0.06035 | 0.753127 | 0.970622 |
| ENSMUSG( | NA | 64.36051 | 0.040899 | 0.878142 | 0.989644 |
| ENSMUSG( | NA | 43.87724 | 0.121175 | 0.748082 | 0.96984 |
| ENSMUSG( | NA | 18.98854 | 0.101119 | 0.822036 | 0.979198 |
| ENSMUSG( | NA | 103.4457 | 0.181533 | 0.511579 | 0.929791 |
| ENSMUSG( | NA | 30.16187 | -0.45028 | 0.251846 | 0.879902 |
| ENSMUSG( | NA | 21.69955 | -0.41054 | 0.33685 | 0.907972 |
| ENSMUSG( | NA | 147.6893 | 0.040194 | 0.830953 | 0.98051 |
| ENSMUSG( | NA | 69.79293 | -0.45943 | 0.121119 | 0.813384 |
| ENSMUSG( | NA | 10.59916 | -0.14944 | 0.801047 | NA |
| ENSMUSG( | NA | 16.22505 | 1.284809 | 0.013201 | 0.460068 |
| ENSMUSG( | NA | 19.39648 | -0.03221 | 0.941624 | 0.994578 |
| ENSMUSG( | NA | 41.03478 | -0.04961 | 0.885114 | 0.989688 |
| ENSMUSG( | NA | 26.19911 | 0.443881 | 0.398439 | 0.917257 |
| ENSMUSG( | NA | 11.23709 | -0.56157 | 0.404488 | NA |
| ENSMUSG( | NA | 17.65424 | 1.357659 | 0.005587 | 0.303106 |
| ENSMUSG( | NA | 15.78559 | 0.059346 | 0.905101 | 0.991089 |
| ENSMUSG( | NA | 27.77203 | -0.49301 | 0.196731 | 0.862161 |
| ENSMUSG( | NA | 34.83446 | -0.29372 | 0.478626 | 0.923329 |
| ENSMUSG( | NA | 52.95129 | 0.197742 | 0.555709 | 0.939565 |
| ENSMUSG( | NA | 15.50323 | -0.60951 | 0.224622 | 0.868206 |

| | | | | | |
|---|---|---|---|---|---|
| ENSMUSG( | 23.49595 | -0.10755 | 0.780901 | 0.957531 | NA |
| ENSMUSG( | 20.72372 | 0.315084 | 0.444701 | NA | NA |
| ENSMUSG( | 17.84055 | -0.49039 | 0.300741 | NA | NA |
| ENSMUSG( | 18.81003 | -0.17314 | 0.69161 | NA | NA |
| ENSMUSG( | 15.66715 | -0.04701 | 0.921593 | NA | NA |
| ENSMUSG( | 1055.424 | -0.08331 | 0.492483 | 0.860378 | NA |
| ENSMUSG( | 64.58557 | -0.48125 | 0.092259 | 0.466326 | NA |
| ENSMUSG( | 20.04506 | -0.23345 | 0.59154 | NA | NA |
| ENSMUSG( | 10.04398 | 4.794355 | 3.05E-05 | NA | NA |
| ENSMUSG( | 18.39503 | -0.53055 | 0.22302 | NA | NA |
| ENSMUSG( | 61.12209 | -0.01694 | 0.955574 | 0.993874 | NA |
| ENSMUSG( | 11.63949 | -0.18395 | 0.728499 | NA | NA |
| ENSMUSG( | 52.32137 | -0.43663 | 0.108325 | 0.497337 | NA |
| ENSMUSG( | 26.19851 | -0.72589 | 0.059684 | 0.377567 | NA |
| ENSMUSG( | 29.31465 | -0.76528 | 0.03568 | 0.290384 | NA |
| ENSMUSG( | 8.104408 | -0.23077 | 0.744646 | NA | NA |
| ENSMUSG( | 41.97587 | -0.13343 | 0.695931 | 0.934979 | NA |
| ENSMUSG( | 27.22792 | -0.01105 | 0.976819 | 0.997428 | NA |
| ENSMUSG( | 136.0242 | 0.090911 | 0.634761 | 0.914073 | NA |
| ENSMUSG( | 94.28762 | -0.03612 | 0.86373 | 0.975244 | NA |
| ENSMUSG( | 217.6673 | -0.09043 | 0.531258 | 0.879376 | NA |
| ENSMUSG( | 82.35619 | -0.0747 | 0.74512 | 0.949158 | NA |
| ENSMUSG( | 16.00978 | 0.335789 | 0.513457 | NA | NA |
| ENSMUSG( | 66.65285 | 0.140674 | 0.553421 | 0.886216 | NA |
| ENSMUSG( | 39.60003 | -0.50616 | 0.11729 | 0.516826 | NA |
| ENSMUSG( | 23.28813 | -0.23021 | 0.549035 | 0.885686 | NA |
| ENSMUSG( | 12.67975 | -0.20839 | 0.723781 | NA | NA |
| ENSMUSG( | 24.84899 | -0.46822 | 0.231089 | 0.679229 | NA |
| ENSMUSG( | 10.56179 | 0.793751 | 0.244695 | NA | NA |
| ENSMUSG( | 38.27664 | 0.344408 | 0.294212 | 0.743459 | NA |
| ENSMUSG( | 10.87411 | -0.36497 | 0.537956 | NA | NA |
| ENSMUSG( | 8.518782 | 0.025155 | 0.969505 | NA | NA |
| ENSMUSG( | 25.97245 | -0.14219 | 0.7004 | 0.93743 | NA |
| ENSMUSG( | 12.16494 | -0.94481 | 0.107177 | NA | NA |
| ENSMUSG( | 56.29603 | -0.00697 | 0.978607 | 0.997428 | NA |
| ENSMUSG( | 38.93497 | -0.56191 | 0.064995 | 0.395432 | NA |
| ENSMUSG( | 16.49978 | 0.569235 | 0.247881 | NA | NA |
| ENSMUSG( | 18.12405 | -0.33822 | 0.465612 | NA | NA |
| ENSMUSG( | 10.33234 | -0.00549 | 0.992987 | NA | NA |
| ENSMUSG( | 13.24024 | -0.61557 | 0.297571 | NA | NA |
| ENSMUSG( | 11.25765 | 0.175396 | 0.752066 | NA | NA |
| ENSMUSG( | 13.61342 | 0.025988 | 0.968806 | NA | NA |
| ENSMUSG( | 15.68177 | 0.343595 | 0.469488 | NA | NA |
| ENSMUSG( | 25.31852 | -0.51352 | 0.182178 | 0.619591 | NA |
| ENSMUSG( | 24.60934 | -0.37189 | 0.313883 | 0.761128 | NA |
| ENSMUSG( | 19.59915 | -0.30276 | 0.486076 | NA | NA |
| ENSMUSG( | 356.4439 | -0.1026 | 0.42273 | 0.823527 | NA |
| ENSMUSG( | 60.20759 | 0.130387 | 0.614141 | 0.909405 | NA |
| ENSMUSG( | 928.8983 | 0.322622 | 0.006514 | 0.110626 | NA |
| ENSMUSG( | 19.88955 | -0.03853 | 0.926169 | NA | NA |
| ENSMUSG( | 180.1752 | 0.147826 | 0.349547 | 0.781913 | NA |
| ENSMUSG( | 16.16534 | 0.15939 | 0.740261 | NA | NA |
| ENSMUSG( | 33.63469 | 0.602622 | 0.080361 | 0.438432 | NA |
| ENSMUSG( | 40.6497 | -0.01539 | 0.960096 | 0.993991 | NA |
| ENSMUSG( | 9.726066 | 0.438665 | 0.55636 | NA | NA |
| ENSMUSG( | 215.8525 | 0.08562 | 0.627254 | 0.912941 | NA |
| ENSMUSG( | 85.96166 | 0.030622 | 0.890911 | 0.983764 | NA |
| ENSMUSG( | 43.08333 | -0.01562 | 0.957669 | 0.993874 | NA |
| ENSMUSG( | 19.39496 | 0.064597 | 0.887725 | NA | NA |
| ENSMUSG( | 662.3096 | 0.23641 | 0.066584 | 0.398819 | NA |
| ENSMUSG( | 468.6425 | 0.16073 | 0.313222 | 0.76033 | NA |
| ENSMUSG( | 14.87609 | 0.184169 | 0.710701 | NA | NA |
| ENSMUSG( | 93.17276 | -0.3332 | 0.145824 | 0.567075 | NA |
| ENSMUSG( | 10.7397 | 0.383048 | 0.504337 | NA | NA |
| ENSMUSG( | 8.939059 | -0.15223 | 0.814272 | NA | NA |
| ENSMUSG( | 158.8911 | -0.02658 | 0.899344 | 0.985633 | NA |
| ENSMUSG( | 72.7677 | -0.46627 | 0.147251 | 0.569348 | NA |
| ENSMUSG( | 21.83963 | -0.09525 | 0.827055 | 0.96865 | NA |
| ENSMUSG( | 47.51293 | 0.055451 | 0.848229 | 0.972374 | NA |
| ENSMUSG( | 9.738078 | 0.147439 | 0.799607 | NA | NA |
| ENSMUSG( | 86.09343 | 0.095889 | 0.703357 | 0.937735 | NA |
| ENSMUSG( | 13.42733 | -0.46767 | 0.348341 | NA | NA |
| ENSMUSG( | 15.55959 | -0.34125 | 0.492911 | NA | NA |
| ENSMUSG( | 44.87021 | -0.37734 | 0.189077 | 0.627678 | NA |
| ENSMUSG( | 13.28558 | 0.253912 | 0.612741 | NA | NA |
| ENSMUSG( | 40.26675 | -0.36349 | 0.224552 | 0.670886 | NA |
| ENSMUSG( | 10.26129 | -0.38354 | 0.507655 | NA | NA |
| ENSMUSG( | 36.7863 | -0.58879 | 0.077385 | 0.429743 | NA |
| ENSMUSG( | 50.16408 | -0.07551 | 0.777096 | 0.955828 | NA |
| ENSMUSG( | 311.2135 | -0.00065 | 0.996186 | 0.99943 | NA |
| ENSMUSG( | 13.51558 | -0.37616 | 0.458237 | NA | NA |
| ENSMUSG( | 13.64576 | -0.93799 | 0.066934 | NA | NA |
| ENSMUSG( | 17.50486 | 0.488472 | 0.294947 | NA | NA |
| ENSMUSG( | 16.52544 | 0.584193 | 0.211077 | NA | NA |
| ENSMUSG( | 22.88292 | 0.181588 | 0.660994 | 0.92497 | NA |
| ENSMUSG( | 14.62785 | -0.0599 | 0.90478 | NA | NA |
| ENSMUSG( | 96.73028 | -0.19338 | 0.351557 | 0.783381 | NA |
| ENSMUSG( | 95.66822 | -0.10509 | 0.650765 | 0.921307 | NA |
| ENSMUSG( | 87.60581 | -0.01489 | 0.946793 | 0.993731 | NA |
| ENSMUSG( | 10.93108 | -0.08536 | 0.880382 | NA | NA |
| ENSMUSG( | 167.8401 | -0.05074 | 0.764109 | 0.954055 | NA |
| ENSMUSG( | 78.7764 | 0.178435 | 0.542051 | 0.883182 | NA |
| ENSMUSG( | 13.08691 | 0.293188 | 0.56121 | NA | NA |
| ENSMUSG( | 14.75276 | 0.107583 | 0.825576 | NA | NA |
| ENSMUSG( | 24.09415 | -0.56845 | 0.175998 | 0.611543 | NA |
| ENSMUSG( | 13.04002 | -0.27566 | 0.587675 | NA | NA |
| ENSMUSG( | 9.041704 | 0.047636 | 0.947374 | NA | NA |
| ENSMUSG( | 409.0307 | 0.009836 | 0.941892 | 0.993415 | NA |
| ENSMUSG( | 40.12892 | 0.06671 | 0.830846 | 0.969798 | NA |
| ENSMUSG( | 15.10021 | 0.267398 | 0.584284 | NA | NA |
| ENSMUSG( | 14.59432 | 0.974759 | 0.078354 | NA | NA |
| ENSMUSG( | 191.9898 | -0.20504 | 0.266852 | 0.71685 | NA |
| ENSMUSG( | 9.005604 | -0.10416 | 0.871851 | NA | NA |
| ENSMUSG( | 15.79362 | -0.30732 | 0.572628 | NA | NA |
| ENSMUSG( | 16.03519 | -0.2558 | 0.593987 | NA | NA |
| ENSMUSG( | 9.784857 | 0.564438 | 0.388468 | NA | NA |
| ENSMUSG( | 27.68952 | -0.18989 | 0.645344 | 0.917958 | NA |
| ENSMUSG( | 16.80375 | -0.02195 | 0.967509 | NA | NA |
| ENSMUSG( | 89.76476 | -0.06643 | 0.747946 | 0.949604 | NA |
| ENSMUSG( | 53.33181 | 0.168013 | 0.546068 | 0.884974 | NA |
| ENSMUSG( | 23.57377 | -0.3826 | 0.363253 | 0.793211 | NA |
| ENSMUSG( | 21.15748 | -0.56085 | 0.25166 | NA | NA |
| ENSMUSG( | 13.71651 | 0.437198 | 0.409748 | NA | NA |
| ENSMUSG( | 55.30185 | 0.213478 | 0.44528 | 0.833239 | NA |
| ENSMUSG( | 29.78521 | -0.21105 | 0.567145 | 0.891644 | NA |
| ENSMUSG( | 13.25063 | -0.19296 | 0.714164 | NA | NA |
| ENSMUSG( | 10.40808 | -0.28198 | 0.667162 | NA | NA |
| ENSMUSG( | 8.732039 | -1.14494 | 0.069107 | NA | NA |
| ENSMUSG( | 11.82383 | -0.1548 | 0.780545 | NA | NA |
| ENSMUSG( | 39.48496 | 0.112641 | 0.734781 | 0.945584 | NA |
| ENSMUSG( | 11.72133 | -0.37462 | 0.487951 | NA | NA |
| ENSMUSG( | 36.96834 | 0.425618 | 0.168581 | 0.602155 | NA |
| ENSMUSG( | 14.06422 | 0.288359 | 0.570445 | NA | NA |
| ENSMUSG( | 16.7918 | 0.14262 | 0.760627 | NA | NA |
| ENSMUSG( | 25.79613 | -0.08156 | 0.823648 | 0.966603 | NA |
| ENSMUSG( | 17.74783 | 0.161453 | 0.731691 | NA | NA |
| ENSMUSG( | 64.38523 | 0.484086 | 0.06945 | 0.407995 | NA |
| ENSMUSG( | 10.66495 | -0.03805 | 0.945553 | NA | NA |
| ENSMUSG( | 114.9991 | -0.16792 | 0.426211 | 0.824898 | NA |
| ENSMUSG( | 8.026502 | -1.67258 | 0.018631 | NA | NA |
| ENSMUSG( | 9.956333 | -0.21577 | 0.71515 | NA | NA |
| ENSMUSG( | 26.36034 | -0.00641 | 0.986026 | 0.998326 | NA |
| ENSMUSG( | 11.88876 | -0.23024 | 0.679875 | NA | NA |
| ENSMUSG( | 20.08126 | -0.21568 | 0.627319 | NA | NA |
| ENSMUSG( | 14.42571 | -0.0075 | 0.987588 | NA | NA |
| ENSMUSG( | 15.09746 | 0.088368 | 0.865375 | NA | NA |
| ENSMUSG( | 53.48589 | -0.23749 | 0.372059 | 0.798768 | NA |
| ENSMUSG( | 13.59951 | 0.190493 | 0.714641 | NA | NA |
| ENSMUSG( | 28.28019 | -0.50049 | 0.150389 | 0.574991 | NA |
| ENSMUSG( | 20.51512 | 0.24689 | 0.571439 | NA | NA |
| ENSMUSG( | 18.18625 | 0.493057 | 0.261019 | NA | NA |
| ENSMUSG( | 11.80365 | 0.951825 | 0.088591 | NA | NA |
| ENSMUSG( | 10.66944 | 0.452134 | 0.453091 | NA | NA |
| ENSMUSG( | 44.51375 | -0.28246 | 0.381422 | 0.802629 | NA |
| ENSMUSG( | 62.69358 | 0.001215 | 0.996324 | 0.99943 | NA |
| ENSMUSG( | 149.7058 | -0.20629 | 0.221527 | 0.667258 | NA |
| ENSMUSG( | 11.98207 | -0.02416 | 0.96465 | NA | NA |
| ENSMUSG( | 62.78612 | -0.17113 | 0.52202 | 0.875186 | NA |
| ENSMUSG( | 11.03696 | 0.084512 | 0.888995 | NA | NA |
| ENSMUSG( | 206.8034 | -0.11739 | 0.494102 | 0.860611 | NA |
| ENSMUSG( | 25.33328 | -0.43271 | 0.246429 | 0.694958 | NA |
| ENSMUSG( | 24.4451 | 0.311353 | 0.40267 | 0.81552 | NA |
| ENSMUSG( | 51.13866 | -0.45176 | 0.091964 | 0.465937 | NA |
| ENSMUSG( | 53.25763 | 0.409063 | 0.250782 | 0.700403 | NA |
| ENSMUSG( | 17.00235 | -0.73181 | 0.211063 | NA | NA |
| ENSMUSG( | 46.35018 | 0.268022 | 0.40707 | 0.817163 | NA |
| ENSMUSG( | 167.318 | -0.01225 | 0.940862 | 0.993415 | NA |
| ENSMUSG( | 34.20248 | 0.14477 | 0.697482 | 0.935662 | NA |

| | | | | | |
|---|---|---|---|---|---|
| ENSMUSG( | 20.70058 | 0.617038 | 0.21594 | 0.865167 | NA |
| ENSMUSG( | 37.09582 | 0.062502 | 0.851793 | 0.985591 | NA |
| ENSMUSG( | 91.55776 | 0.02604 | 0.914807 | 0.992407 | NA |
| ENSMUSG( | 24.51841 | -0.51831 | 0.231896 | 0.868507 | NA |
| ENSMUSG( | 152.3134 | 0.332836 | 0.075944 | 0.768524 | NA |
| ENSMUSG( | 1224.174 | -0.21053 | 0.130993 | 0.822884 | NA |
| ENSMUSG( | 12.07828 | 0.345927 | 0.546714 | 0.938063 | NA |
| ENSMUSG( | 51.22768 | -0.12496 | 0.699869 | 0.961335 | NA |
| ENSMUSG( | 30.54137 | -1.3968 | 0.000265 | 0.04337 | NA |
| ENSMUSG( | 11.62846 | 0.101214 | 0.871584 | 0.988779 | NA |
| ENSMUSG( | 45.66507 | 0.156173 | 0.649018 | 0.95422 | NA |
| ENSMUSG( | 69.26844 | 0.324718 | 0.203783 | 0.864412 | NA |
| ENSMUSG( | 40.13608 | -0.40157 | 0.221787 | 0.866104 | NA |
| ENSMUSG( | 15.75031 | 0.432507 | 0.403123 | 0.917257 | NA |
| ENSMUSG( | 34.87109 | -0.11608 | 0.739399 | 0.968458 | NA |
| ENSMUSG( | 27.87565 | -0.23741 | 0.527723 | 0.934603 | NA |
| ENSMUSG( | 9.516006 | -0.47319 | 0.44676 | NA | NA |
| ENSMUSG( | 14.92116 | -0.32654 | 0.532956 | 0.934795 | NA |
| ENSMUSG( | 9.860791 | -0.26526 | 0.68279 | NA | NA |
| ENSMUSG( | 29.29585 | 0.238462 | 0.542412 | 0.938034 | NA |
| ENSMUSG( | 11.66313 | 0.156419 | 0.780935 | 0.976106 | NA |
| ENSMUSG( | 7.337955 | -0.44282 | 0.568132 | NA | NA |
| ENSMUSG( | 39.18936 | -0.05095 | 0.889449 | 0.990558 | NA |
| ENSMUSG( | 14.50765 | -0.12928 | 0.810189 | 0.978779 | NA |
| ENSMUSG( | 23.63881 | 0.373021 | 0.393102 | 0.917257 | NA |
| ENSMUSG( | 10.67734 | 0.010822 | 0.987472 | NA | NA |
| ENSMUSG( | 85.53119 | -0.29068 | 0.207591 | 0.864412 | NA |
| ENSMUSG( | 22.18549 | 0.306806 | 0.47098 | 0.923329 | NA |
| ENSMUSG( | 9.971965 | 0.223667 | 0.726447 | NA | NA |
| ENSMUSG( | 165.1508 | -0.05212 | 0.803843 | 0.977 | NA |
| ENSMUSG( | 366.1288 | -0.28233 | 0.066988 | 0.75212 | NA |
| ENSMUSG( | 24.9882 | 0.202443 | 0.62877 | 0.953251 | NA |
| ENSMUSG( | 70.02962 | -0.31465 | 0.231934 | 0.868507 | NA |
| ENSMUSG( | 11.68626 | 0.819087 | 0.152293 | 0.837724 | NA |
| ENSMUSG( | 32.30619 | -0.26 | 0.501999 | 0.927825 | NA |
| ENSMUSG( | 18.1706 | 0.029862 | 0.949766 | 0.994798 | NA |
| ENSMUSG( | 18.13039 | 0.158513 | 0.732778 | 0.967228 | NA |
| ENSMUSG( | 14.54738 | -0.29808 | 0.596816 | 0.946541 | NA |
| ENSMUSG( | 17.17473 | 0.705661 | 0.138283 | 0.828492 | NA |
| ENSMUSG( | 15.02023 | -0.4906 | 0.339655 | 0.907972 | NA |
| ENSMUSG( | 19.60289 | -0.02003 | 0.965193 | 0.995433 | NA |
| ENSMUSG( | 270.738 | -0.14302 | 0.375985 | 0.915491 | NA |
| ENSMUSG( | 12.13046 | -0.44442 | 0.428038 | 0.917257 | NA |
| ENSMUSG( | 14.60842 | 0.488023 | 0.347601 | 0.911652 | NA |
| ENSMUSG( | 14.22349 | -0.76126 | 0.143856 | 0.831311 | NA |
| ENSMUSG( | 14.69581 | -0.19845 | 0.706035 | 0.96335 | NA |
| ENSMUSG( | 30.66946 | 0.545414 | 0.26375 | 0.886134 | NA |
| ENSMUSG( | 183.1372 | -0.25915 | 0.143485 | 0.831311 | NA |
| ENSMUSG( | 105.9158 | 0.299135 | 0.203 | 0.864412 | NA |
| ENSMUSG( | 21.78561 | 0.351556 | 0.404027 | 0.917257 | NA |
| ENSMUSG( | 27.83786 | 0.068045 | 0.855158 | 0.985649 | NA |
| ENSMUSG( | 9.01457 | -0.47119 | 0.482628 | NA | NA |
| ENSMUSG( | 28.22471 | 0.212784 | 0.602862 | 0.94789 | NA |
| ENSMUSG( | 160.0558 | 0.075788 | 0.6924 | 0.960404 | NA |
| ENSMUSG( | 17.70561 | -0.36454 | 0.474255 | 0.923329 | NA |
| ENSMUSG( | 9.842578 | 0.369568 | 0.566089 | NA | NA |
| ENSMUSG( | 20.10192 | -0.15798 | 0.717759 | 0.965964 | NA |
| ENSMUSG( | 12.87347 | 0.075647 | 0.891661 | 0.990558 | NA |
| ENSMUSG( | 43.26562 | 0.320407 | 0.303994 | 0.898538 | NA |
| ENSMUSG( | 71.80027 | -0.16167 | 0.574079 | 0.942912 | NA |
| ENSMUSG( | 11.88221 | 0.352807 | 0.559017 | 0.939587 | NA |
| ENSMUSG( | 13.4225 | -0.58535 | 0.334586 | 0.907907 | NA |
| ENSMUSG( | 75.39198 | -0.02802 | 0.934026 | 0.99306 | NA |
| ENSMUSG( | 9.414813 | -1.20677 | 0.070547 | NA | NA |
| ENSMUSG( | 49.69782 | 0.460697 | 0.171613 | 0.842999 | NA |
| ENSMUSG( | 25.40125 | 0.345897 | 0.372724 | 0.915491 | NA |
| ENSMUSG( | 53.63668 | -0.15019 | 0.672172 | 0.95694 | NA |
| ENSMUSG( | 12.04652 | 0.101264 | 0.858107 | 0.985884 | NA |
| ENSMUSG( | 34.39673 | -0.24434 | 0.482261 | 0.924578 | NA |
| ENSMUSG( | 26.59867 | -0.77725 | 0.05397 | 0.722223 | NA |
| ENSMUSG( | 196.8398 | -0.12701 | 0.479602 | 0.924021 | NA |
| ENSMUSG( | 15.31883 | -0.31696 | 0.534613 | 0.936107 | NA |
| ENSMUSG( | 14.52272 | 0.058127 | 0.908503 | 0.991958 | NA |
| ENSMUSG( | 15.13045 | 0.543657 | 0.399401 | 0.917257 | NA |
| ENSMUSG( | 13.631 | 0.215641 | 0.735786 | 0.967344 | NA |
| ENSMUSG( | 75.02526 | 0.071556 | 0.769126 | 0.974236 | NA |
| ENSMUSG( | 13.99327 | 0.412767 | 0.468492 | 0.923329 | NA |
| ENSMUSG( | 181.7445 | -0.0962 | 0.620192 | 0.952896 | NA |
| ENSMUSG( | 584.2 | 0.154497 | 0.297321 | 0.898538 | NA |
| ENSMUSG( | 7.965554 | -0.18673 | 0.787222 | NA | NA |
| ENSMUSG( | 15.36659 | 0.228218 | 0.67741 | 0.957106 | NA |
| ENSMUSG( | 41.13395 | -0.3835 | 0.226487 | 0.868507 | NA |
| ENSMUSG( | 21.19724 | 0.270078 | 0.581112 | 0.944266 | NA |
| ENSMUSG( | 13.593 | 0.527651 | 0.345035 | 0.91045 | NA |
| ENSMUSG( | 16.35109 | -0.29286 | 0.555505 | 0.939565 | NA |
| ENSMUSG( | 10.76515 | -0.56399 | 0.351124 | NA | NA |
| ENSMUSG( | 34.32679 | 0.509735 | 0.141993 | 0.830709 | NA |
| ENSMUSG( | 18.95793 | -0.07876 | 0.867763 | 0.98781 | NA |
| ENSMUSG( | 12.28678 | 0.833671 | 0.142496 | 0.831221 | NA |
| ENSMUSG( | 25.68259 | -0.41661 | 0.303466 | 0.898538 | NA |
| ENSMUSG( | 26.75966 | -0.36845 | 0.35818 | 0.913759 | NA |
| ENSMUSG( | 321.322 | 0.070048 | 0.741684 | 0.969125 | NA |
| ENSMUSG( | 35.74836 | 0.36997 | 0.338859 | 0.907972 | NA |
| ENSMUSG( | 225.1003 | -0.0325 | 0.850673 | 0.985541 | NA |
| ENSMUSG( | 11.49811 | -0.05725 | 0.923633 | 0.992815 | NA |
| ENSMUSG( | 20.97317 | -0.79751 | 0.093214 | 0.795363 | NA |
| ENSMUSG( | 38.76579 | 0.245593 | 0.442868 | 0.921554 | NA |
| ENSMUSG( | 9.568682 | -0.81101 | 0.205499 | NA | NA |
| ENSMUSG( | 114.5887 | -0.14079 | 0.523849 | 0.932848 | NA |
| ENSMUSG( | 10.63514 | 0.628973 | 0.303422 | NA | NA |
| ENSMUSG( | 29.15697 | 0.400454 | 0.289347 | 0.896442 | NA |
| ENSMUSG( | 7.698566 | -0.284 | 0.70521 | NA | NA |
| ENSMUSG( | 814.5748 | -0.02601 | 0.837464 | 0.981335 | NA |
| ENSMUSG( | 181.272 | 0.188671 | 0.371679 | 0.915491 | NA |
| ENSMUSG( | 29.0243 | 0.841139 | 0.037196 | 0.673514 | NA |
| ENSMUSG( | 406.5499 | -0.3285 | 0.015415 | 0.495791 | NA |
| ENSMUSG( | 9.334858 | -0.32189 | 0.667413 | NA | NA |
| ENSMUSG( | 18679.09 | 0.0858 | 0.506149 | 0.927825 | NA |
| ENSMUSG( | 98.26181 | -0.43798 | 0.06864 | 0.753052 | NA |
| ENSMUSG( | 22.50803 | -0.6223 | 0.139312 | 0.830678 | NA |
| ENSMUSG( | 133.7178 | -0.05872 | 0.767465 | 0.974236 | NA |
| ENSMUSG( | 32.88013 | -0.02823 | 0.934952 | 0.99306 | NA |
| ENSMUSG( | 29.71692 | 0.760762 | 0.05447 | 0.723362 | NA |
| ENSMUSG( | 64.82556 | -0.10362 | 0.696007 | 0.960867 | NA |
| ENSMUSG( | 11.52232 | -0.08016 | 0.889205 | 0.990558 | NA |
| ENSMUSG( | 19.37405 | 0.296224 | 0.602196 | 0.94789 | NA |
| ENSMUSG( | 9.784527 | -0.05341 | 0.931097 | NA | NA |
| ENSMUSG( | 16.64004 | -0.22807 | 0.654076 | 0.955015 | NA |
| ENSMUSG( | 736.3628 | -0.06883 | 0.632946 | 0.953844 | NA |
| ENSMUSG( | 65.85936 | -0.68891 | 0.007761 | 0.37555 | NA |
| ENSMUSG( | 10.18875 | -0.35214 | 0.572544 | NA | NA |
| ENSMUSG( | 47.29014 | -0.22286 | 0.51308 | 0.930358 | NA |
| ENSMUSG( | 11.92558 | 0.759046 | 0.275697 | 0.890209 | NA |
| ENSMUSG( | 22.12934 | 0.627639 | 0.150138 | 0.837086 | NA |
| ENSMUSG( | 127.8141 | -0.04 | 0.864273 | 0.986552 | NA |
| ENSMUSG( | 11.75465 | -1.09798 | 0.056177 | 0.723362 | NA |
| ENSMUSG( | 15.04587 | 0.104361 | 0.835689 | 0.981237 | NA |
| ENSMUSG( | 135.6324 | -0.13116 | 0.525716 | 0.934374 | NA |
| ENSMUSG( | 20.38823 | 0.806299 | 0.065923 | 0.746371 | NA |
| ENSMUSG( | 21.46928 | 0.55814 | 0.315344 | 0.901306 | NA |
| ENSMUSG( | 15.75351 | -0.12879 | 0.805072 | 0.977358 | NA |
| ENSMUSG( | 18.22633 | 0.040152 | 0.93258 | 0.99306 | NA |
| ENSMUSG( | 12.93019 | -0.39175 | 0.516263 | 0.930939 | NA |
| ENSMUSG( | 32.26841 | -0.40803 | 0.308912 | 0.898538 | NA |
| ENSMUSG( | 8.966632 | 0.055937 | 0.930954 | NA | NA |
| ENSMUSG( | 192.8873 | -0.06035 | 0.753127 | 0.970622 | NA |
| ENSMUSG( | 64.36051 | 0.040899 | 0.878142 | 0.989644 | NA |
| ENSMUSG( | 43.87724 | 0.121175 | 0.748082 | 0.96984 | NA |
| ENSMUSG( | 18.98854 | 0.101119 | 0.822036 | 0.979198 | NA |
| ENSMUSG( | 103.4457 | 0.181533 | 0.511579 | 0.929791 | NA |
| ENSMUSG( | 30.16187 | -0.45028 | 0.251846 | 0.879902 | NA |
| ENSMUSG( | 21.69955 | -0.41054 | 0.33685 | 0.907972 | NA |
| ENSMUSG( | 147.6893 | 0.040194 | 0.830953 | 0.98051 | NA |
| ENSMUSG( | 69.79293 | -0.45943 | 0.121119 | 0.813384 | NA |
| ENSMUSG( | 10.59916 | -0.14944 | 0.801047 | NA | NA |
| ENSMUSG( | 16.22505 | 1.284809 | 0.013201 | 0.460068 | NA |
| ENSMUSG( | 19.39648 | -0.03221 | 0.941624 | 0.994578 | NA |
| ENSMUSG( | 41.03478 | -0.04961 | 0.885114 | 0.989688 | NA |
| ENSMUSG( | 26.19911 | 0.443881 | 0.398439 | 0.917257 | NA |
| ENSMUSG( | 8.542336 | 0.246104 | 0.737298 | NA | NA |
| ENSMUSG( | 11.23709 | -0.56157 | 0.404488 | NA | NA |
| ENSMUSG( | 17.65424 | 1.357659 | 0.005587 | 0.303106 | NA |
| ENSMUSG( | 15.78559 | 0.059346 | 0.905101 | 0.991089 | NA |
| ENSMUSG( | 11.2103 | 2.074682 | 0.004988 | NA | NA |
| ENSMUSG( | 27.77203 | -0.49301 | 0.196731 | 0.862161 | NA |
| ENSMUSG( | 34.83446 | -0.29372 | 0.478626 | 0.923329 | NA |
| ENSMUSG( | 52.95129 | 0.197742 | 0.555709 | 0.939565 | NA |
| ENSMUSG( | 15.50323 | -0.60951 | 0.224622 | 0.868206 | NA |

| | | | | | | | | | | | |
|---|---|---|---|---|---|---|---|---|---|---|---|
| ENSMUSG( | NA | 332.1371 | -0.38508 | 0.012121 | 0.159579 | ENSMUSG( | NA | 20.38929 | -0.47901 | 0.298148 | 0.898538 |
| ENSMUSG( | NA | 14.50274 | -0.79946 | 0.128136 | NA | ENSMUSG( | NA | 72.04741 | -0.21012 | 0.40835 | 0.917257 |
| ENSMUSG( | NA | 11.22347 | -0.12896 | 0.84004 | NA | ENSMUSG( | NA | 25.92897 | -0.58066 | 0.148789 | 0.836241 |
| ENSMUSG( | NA | 24.21292 | -0.44669 | 0.266034 | 0.716441 | ENSMUSG( | NA | 18.09724 | 0.433579 | 0.358835 | 0.913759 |
| ENSMUSG( | NA | 35.41381 | -0.36274 | 0.261944 | 0.712129 | ENSMUSG( | NA | 744.1166 | 0.236954 | 0.105441 | 0.804969 |
| ENSMUSG( | NA | 56.13046 | -0.18098 | 0.512888 | 0.871762 | ENSMUSG( | NA | 21.70793 | -0.16302 | 0.699653 | 0.961335 |
| ENSMUSG( | NA | 14.41078 | 0.252643 | 0.604374 | NA | ENSMUSG( | NA | 37.39255 | -0.07844 | 0.810661 | 0.978779 |
| ENSMUSG( | NA | 58.80551 | -0.23215 | 0.379914 | 0.801757 | ENSMUSG( | NA | 28.88314 | 0.081999 | 0.824588 | 0.979198 |
| ENSMUSG( | NA | 496.1366 | -0.01039 | 0.930071 | 0.991848 | ENSMUSG( | NA | 80.55504 | -0.48707 | 0.055923 | 0.723362 |
| ENSMUSG( | NA | 344.4894 | 0.182118 | 0.19343 | 0.632931 | ENSMUSG( | NA | 23.86501 | -0.43173 | 0.287531 | 0.894091 |
| ENSMUSG( | NA | 7.562286 | -0.16812 | 0.822657 | NA | ENSMUSG( | NA | 54.32637 | -0.15336 | 0.603139 | 0.948137 |
| ENSMUSG( | NA | 34.79113 | -0.63091 | 0.046484 | 0.330252 | ENSMUSG( | NA | 11.57187 | -0.04723 | 0.932824 | 0.99306 |
| ENSMUSG( | NA | 22.35792 | 0.09826 | 0.825896 | 0.968061 | ENSMUSG( | NA | 22.10799 | -0.44723 | 0.3156 | 0.901725 |
| ENSMUSG( | NA | 14.21958 | -0.12061 | 0.826423 | NA | ENSMUSG( | NA | 11.82459 | 0.018513 | 0.978684 | 0.996459 |
| ENSMUSG( | NA | 35.88591 | -0.26 | 0.430921 | 0.826896 | ENSMUSG( | NA | 34.34636 | 0.028211 | 0.937082 | 0.993685 |
| ENSMUSG( | NA | 25.33493 | 0.240769 | 0.549031 | 0.885686 | ENSMUSG( | NA | 8.11887 | 0.18135 | 0.794475 | NA |
| ENSMUSG( | NA | 46.15353 | 0.237355 | 0.393077 | 0.809274 | ENSMUSG( | NA | 9.065948 | 0.379785 | 0.571035 | NA |
| ENSMUSG( | NA | 86.52772 | 0.143511 | 0.560322 | 0.888934 | ENSMUSG( | NA | 161.1915 | 0.107623 | 0.58361 | 0.944266 |
| ENSMUSG( | NA | 13.67305 | -0.04214 | 0.932855 | NA | ENSMUSG( | NA | 24.26544 | -0.28111 | 0.514508 | 0.930939 |
| ENSMUSG( | NA | 39.69613 | -0.37013 | 0.234241 | 0.68192 | ENSMUSG( | NA | 20.04147 | 0.06517 | 0.882499 | 0.989644 |
| ENSMUSG( | NA | 14.71836 | 0.285129 | 0.558748 | NA | ENSMUSG( | NA | 26.6628 | -0.37609 | 0.330104 | 0.907251 |
| ENSMUSG( | NA | 179.8119 | 0.019334 | 0.903405 | 0.986055 | ENSMUSG( | NA | 22.22627 | 0.106331 | 0.823925 | 0.979198 |
| ENSMUSG( | NA | 21.61661 | 0.638107 | 0.107734 | 0.49674 | ENSMUSG( | NA | 14.31085 | 0.033789 | 0.947482 | 0.994798 |
| ENSMUSG( | NA | 22.18805 | -0.19439 | 0.627642 | 0.912961 | ENSMUSG( | NA | 40.84529 | -0.36751 | 0.305806 | 0.898538 |
| ENSMUSG( | NA | 21.19266 | -0.15143 | 0.733066 | NA | ENSMUSG( | NA | 61.03815 | -0.27575 | 0.322945 | 0.904023 |
| ENSMUSG( | NA | 538.7027 | -0.18218 | 0.149044 | 0.572864 | ENSMUSG( | NA | 38.48194 | -0.36053 | 0.274322 | 0.889874 |
| ENSMUSG( | NA | 274.3064 | 0.068468 | 0.623358 | 0.911646 | ENSMUSG( | NA | 75.94178 | 0.201869 | 0.409442 | 0.917257 |
| ENSMUSG( | NA | 20.64602 | 0.221703 | 0.608276 | NA | ENSMUSG( | NA | 14.45696 | 0.435873 | 0.413668 | 0.917257 |
| ENSMUSG( | NA | 154.2195 | 0.122125 | 0.537614 | 0.882536 | ENSMUSG( | NA | 46.96248 | -0.20984 | 0.550214 | 0.938699 |
| ENSMUSG( | NA | 15.46578 | -0.02521 | 0.957314 | NA | ENSMUSG( | NA | 17.19082 | -0.92235 | 0.055375 | 0.723362 |
| ENSMUSG( | NA | 46.71455 | 0.119869 | 0.672149 | 0.9288 | ENSMUSG( | NA | 44.59516 | -0.09709 | 0.772071 | 0.974236 |
| ENSMUSG( | NA | 23.28122 | -0.59212 | 0.13788 | 0.552903 | ENSMUSG( | NA | 16.85842 | -0.34707 | 0.491621 | 0.926889 |
| ENSMUSG( | NA | 25.25999 | 0.194369 | 0.641063 | 0.916692 | ENSMUSG( | NA | 38.32139 | 0.060953 | 0.859507 | 0.985947 |
| ENSMUSG( | NA | 18.32151 | -0.23207 | 0.630539 | NA | ENSMUSG( | NA | 149.6336 | -0.27151 | 0.151224 | 0.837494 |
| ENSMUSG( | NA | 33.84197 | 0.283677 | 0.408529 | 0.818167 | ENSMUSG( | NA | 134.444 | -0.14391 | 0.640401 | 0.953903 |
| ENSMUSG( | NA | 17.71267 | -0.21368 | 0.653144 | NA | ENSMUSG( | NA | 16.19668 | 0.272438 | 0.589517 | 0.944975 |
| ENSMUSG( | NA | 17.25965 | -0.42785 | 0.361333 | NA | ENSMUSG( | NA | 27.26783 | 0.413711 | 0.267384 | 0.886134 |
| ENSMUSG( | NA | 13.60317 | 1.279953 | 0.019848 | NA | ENSMUSG( | NA | 65.50397 | -0.14366 | 0.604997 | 0.949082 |
| ENSMUSG( | NA | 55.88965 | 1.165864 | 4.99E-05 | 0.00395 | ENSMUSG( | NA | 21.38197 | 0.405087 | 0.383184 | 0.917257 |
| ENSMUSG( | NA | 68.90958 | 0.012814 | 0.956429 | 0.993874 | ENSMUSG( | NA | 76.92051 | 0.382418 | 0.119162 | 0.813384 |
| ENSMUSG( | NA | 15.65189 | 0.149731 | 0.795533 | NA | ENSMUSG( | NA | 79.82759 | -0.06657 | 0.791267 | 0.97681 |
| ENSMUSG( | NA | 72.8735 | -1.41401 | 1.37E-06 | 0.000244 | ENSMUSG( | NA | 22.67091 | -0.30214 | 0.517786 | 0.930939 |
| ENSMUSG( | NA | 14.32994 | 0.177799 | 0.72912 | NA | ENSMUSG( | NA | 528.37 | 0.120642 | 0.4973 | 0.927825 |
| ENSMUSG( | NA | 14.34631 | -0.18214 | 0.712114 | NA | ENSMUSG( | NA | 89.89621 | -0.19466 | 0.42625 | 0.917257 |
| ENSMUSG( | NA | 350.7158 | -0.31649 | 0.045768 | 0.328083 | ENSMUSG( | NA | 15.75602 | -0.45585 | 0.363327 | 0.914223 |
| ENSMUSG( | NA | 29.01034 | 0.120774 | 0.731568 | 0.944956 | ENSMUSG( | NA | 26.96423 | -0.55572 | 0.154733 | 0.837724 |
| ENSMUSG( | NA | 31.43993 | -0.02768 | 0.936757 | 0.993415 | ENSMUSG( | NA | 15.13257 | 0.355384 | 0.532053 | 0.934795 |
| ENSMUSG( | NA | 30.36913 | -0.49099 | 0.182363 | 0.619591 | ENSMUSG( | NA | 19.70962 | 0.441735 | 0.320386 | 0.903887 |
| ENSMUSG( | NA | 159.5494 | 0.048533 | 0.786573 | 0.958856 | ENSMUSG( | NA | 384.0194 | 0.015279 | 0.916938 | 0.992529 |
| ENSMUSG( | NA | 34.58048 | 0.057532 | 0.869737 | 0.977607 | ENSMUSG( | NA | 14.11753 | -0.46376 | 0.376536 | 0.915491 |
| ENSMUSG( | NA | 33.59738 | -0.05904 | 0.86383 | 0.975244 | ENSMUSG( | NA | 14.50753 | -0.5912 | 0.254964 | 0.882307 |
| ENSMUSG( | NA | 21.40306 | -0.45652 | 0.256235 | 0.706924 | ENSMUSG( | NA | 37.39107 | 0.760251 | 0.029077 | 0.614779 |
| ENSMUSG( | NA | 25.05953 | -0.02511 | 0.946607 | 0.993731 | ENSMUSG( | NA | 37.17216 | -0.44104 | 0.195987 | 0.862161 |
| ENSMUSG( | NA | 20.05486 | -0.20799 | 0.623316 | NA | ENSMUSG( | NA | 51.57018 | 0.263973 | 0.457137 | 0.923329 |
| ENSMUSG( | NA | 16.35168 | 0.260226 | 0.650547 | NA | ENSMUSG( | NA | 40.61744 | 0.205632 | 0.5281 | 0.934677 |
| ENSMUSG( | NA | 53.37305 | 0.120446 | 0.688792 | 0.933441 | ENSMUSG( | NA | 30.08724 | -0.47299 | 0.224045 | 0.867404 |
| ENSMUSG( | NA | 63.3506 | 0.387293 | 0.135349 | 0.548884 | ENSMUSG( | NA | 113.049 | -0.17837 | 0.388713 | 0.917257 |
| ENSMUSG( | NA | 14.31515 | -0.16537 | 0.73765 | NA | ENSMUSG( | NA | 24.00252 | 0.094342 | 0.819912 | 0.979198 |
| ENSMUSG( | NA | 75.86832 | 0.035343 | 0.874157 | 0.979336 | ENSMUSG( | NA | 22.34399 | -0.55054 | 0.203816 | 0.864412 |
| ENSMUSG( | NA | 89.14652 | -0.00924 | 0.969031 | 0.996369 | ENSMUSG( | NA | 19.32644 | 0.344678 | 0.450162 | 0.922202 |
| ENSMUSG( | NA | 15.82824 | -0.38621 | 0.419433 | NA | ENSMUSG( | NA | 20.82131 | -0.07681 | 0.858965 | 0.985884 |
| ENSMUSG( | NA | 41.69278 | 0.099861 | 0.755997 | 0.951479 | ENSMUSG( | NA | 9.630261 | -0.37451 | 0.578517 | NA |
| ENSMUSG( | NA | 26.37245 | -0.34966 | 0.33488 | 0.776409 | ENSMUSG( | NA | 147.6692 | -0.50665 | 0.059162 | 0.729429 |
| ENSMUSG( | NA | 10.35677 | -0.22989 | 0.688552 | NA | ENSMUSG( | NA | 188.7796 | -0.77784 | 3.39E-05 | 0.009128 |
| ENSMUSG( | NA | 16.41246 | 0.114901 | 0.802384 | NA | ENSMUSG( | NA | 10.58947 | 0.057011 | 0.923176 | NA |
| ENSMUSG( | NA | 40.61638 | -0.12133 | 0.686206 | 0.932981 | ENSMUSG( | NA | 71.75579 | -0.19639 | 0.427924 | 0.917257 |
| ENSMUSG( | NA | 45.7351 | -0.30116 | 0.289017 | 0.739705 | ENSMUSG( | NA | 8.294161 | 0.065785 | 0.921801 | NA |
| ENSMUSG( | NA | 21.83024 | -0.2694 | 0.53378 | 0.880095 | ENSMUSG( | NA | 55.75473 | 0.146126 | 0.619665 | 0.952896 |
| ENSMUSG( | NA | 54.89178 | -1.16918 | 2.28E-05 | 0.002131 | ENSMUSG( | NA | 49.79263 | 0.115195 | 0.687765 | 0.959703 |
| ENSMUSG( | NA | 11.3788 | -0.44685 | 0.43043 | NA | ENSMUSG( | NA | 39.88199 | 0.136364 | 0.668624 | 0.956766 |
| ENSMUSG( | NA | 47.59336 | 0.118221 | 0.667662 | 0.927484 | ENSMUSG( | NA | 15.16581 | -0.15314 | 0.792398 | 0.97681 |
| ENSMUSG( | NA | 31.85257 | -0.13119 | 0.692543 | 0.934101 | ENSMUSG( | NA | 14.08547 | -0.02983 | 0.953323 | 0.994798 |
| ENSMUSG( | NA | 25.1329 | 0.087245 | 0.82045 | 0.965888 | ENSMUSG( | NA | 29.45396 | -0.33342 | 0.391912 | 0.917257 |
| ENSMUSG( | NA | 40.94011 | -0.02777 | 0.930835 | 0.992059 | ENSMUSG( | NA | 19.69489 | 0.219258 | 0.61996 | 0.952896 |
| ENSMUSG( | NA | 16.30587 | -0.25422 | 0.577406 | NA | ENSMUSG( | NA | 23.03446 | 0.415283 | 0.323246 | 0.904023 |
| ENSMUSG( | NA | 11.32042 | -0.41471 | 0.451 | NA | ENSMUSG( | NA | 17.31399 | -0.28874 | 0.546511 | 0.938063 |
| ENSMUSG( | NA | 24.50207 | 0.359764 | 0.340639 | 0.778521 | ENSMUSG( | NA | 18.70556 | 0.576039 | 0.216195 | 0.865167 |
| ENSMUSG( | NA | 17.86038 | 0.162125 | 0.73365 | NA | ENSMUSG( | NA | 12.69168 | 0.400762 | 0.467913 | 0.923329 |
| ENSMUSG( | NA | 10.34829 | -0.86661 | 0.154463 | NA | ENSMUSG( | NA | 47.18416 | -0.56617 | 0.085146 | 0.779799 |
| ENSMUSG( | NA | 237.1708 | -0.22553 | 0.116294 | 0.514678 | ENSMUSG( | NA | 21.09954 | 0.59161 | 0.212311 | 0.864412 |
| ENSMUSG( | NA | 92.87299 | 0.107626 | 0.601391 | 0.904603 | ENSMUSG( | NA | 98.21655 | -0.13906 | 0.520329 | 0.931964 |
| ENSMUSG( | NA | 9.689409 | -0.51132 | 0.427896 | NA | ENSMUSG( | NA | 9.590072 | -0.57896 | 0.42089 | NA |
| ENSMUSG( | NA | 8.030999 | -0.61462 | 0.390176 | NA | ENSMUSG( | NA | 13.45097 | 0.364233 | 0.506174 | 0.927825 |
| ENSMUSG( | NA | 19.26524 | 0.356015 | 0.415185 | NA | ENSMUSG( | NA | 22.42212 | -0.00225 | 0.995857 | 0.999116 |
| ENSMUSG( | NA | 13.79896 | 0.212396 | 0.661604 | NA | ENSMUSG( | NA | 190.0075 | 0.047331 | 0.809268 | 0.978779 |
| ENSMUSG( | NA | 146.6197 | -0.11492 | 0.556152 | 0.887523 | ENSMUSG( | NA | 9.782455 | -0.30195 | 0.619538 | NA |
| ENSMUSG( | NA | 14.69445 | 0.492746 | 0.348946 | NA | ENSMUSG( | NA | 14.36874 | 0.200918 | 0.710597 | 0.964363 |
| ENSMUSG( | NA | 19.25161 | 0.250619 | 0.585269 | NA | ENSMUSG( | NA | 14.08133 | -0.00748 | 0.988328 | 0.997862 |
| ENSMUSG( | NA | 28.92905 | 0.519581 | 0.141133 | 0.558043 | ENSMUSG( | NA | 31.76932 | -0.10994 | 0.755954 | 0.971371 |
| ENSMUSG( | NA | 10.07462 | 0.192747 | 0.758779 | NA | ENSMUSG( | NA | 69.58621 | 0.322826 | 0.229282 | 0.868507 |
| ENSMUSG( | NA | 31.15745 | -0.29381 | 0.443837 | 0.833239 | ENSMUSG( | NA | 342.2238 | -0.04587 | 0.748876 | 0.96984 |
| ENSMUSG( | NA | 8.7552 | -0.26951 | 0.699879 | NA | ENSMUSG( | NA | 393.601 | -2.98188 | 1.28E-30 | 3.73E-27 |
| ENSMUSG( | NA | 28.22545 | 0.02833 | 0.936173 | 0.993415 | ENSMUSG( | NA | 20.1226 | -0.17053 | 0.737581 | 0.967686 |
| ENSMUSG( | NA | 26.65197 | 0.091096 | 0.828258 | 0.968966 | ENSMUSG( | NA | 14.54822 | -0.46884 | 0.376659 | 0.915491 |
| ENSMUSG( | NA | 11.30136 | 0.680158 | 0.222932 | NA | ENSMUSG( | NA | 10.31825 | -0.00486 | 0.993898 | NA |
| ENSMUSG( | NA | 342.9353 | 0.004432 | 0.977294 | 0.997428 | ENSMUSG( | NA | 33.16975 | 0.190825 | 0.666003 | 0.956766 |
| ENSMUSG( | NA | 11.45732 | -0.64122 | 0.281767 | NA | ENSMUSG( | NA | 23.64291 | -0.28421 | 0.512137 | 0.930074 |
| ENSMUSG( | NA | 16.39976 | 0.493642 | 0.313708 | NA | ENSMUSG( | NA | 81.45392 | 0.206823 | 0.388962 | 0.917257 |
| ENSMUSG( | NA | 13.0238 | 0.042717 | 0.93727 | NA | ENSMUSG( | NA | 29.11216 | -0.29242 | 0.484463 | 0.92541 |
| ENSMUSG( | NA | 50.4974 | 0.08027 | 0.769514 | 0.954674 | ENSMUSG( | NA | 43.7518 | 0.213217 | 0.55434 | 0.939565 |
| ENSMUSG( | NA | 154.6557 | -0.07663 | 0.659142 | 0.924506 | ENSMUSG( | NA | 27.41034 | 0.742582 | 0.069788 | 0.756483 |
| ENSMUSG( | NA | 230.6057 | -0.09795 | 0.525871 | 0.876902 | ENSMUSG( | NA | 12.62374 | 0.468819 | 0.417138 | 0.917257 |
| ENSMUSG( | NA | 86.64182 | 0.108592 | 0.622424 | 0.911432 | ENSMUSG( | NA | 22.95795 | -0.31864 | 0.45582 | 0.923329 |
| ENSMUSG( | NA | 15.05936 | 0.83952 | 0.079483 | NA | ENSMUSG( | NA | 131.6856 | 0.167471 | 0.449927 | 0.922202 |
| ENSMUSG( | NA | 162.1033 | 0.171642 | 0.328252 | 0.772854 | ENSMUSG( | NA | 14.57389 | 0.672925 | 0.197704 | 0.862161 |
| ENSMUSG( | NA | 21.58302 | 0.489126 | 0.232194 | 0.67996 | ENSMUSG( | NA | 14.04488 | -0.63294 | 0.265144 | 0.886134 |
| ENSMUSG( | NA | 62.34033 | 0.051387 | 0.854296 | 0.973491 | ENSMUSG( | NA | 14.9423 | -0.36359 | 0.518867 | 0.930939 |
| ENSMUSG( | NA | 356.1105 | -0.02034 | 0.87334 | 0.978894 | ENSMUSG( | NA | 67.92746 | 0.236856 | 0.372056 | 0.915491 |
| ENSMUSG( | NA | 30.76227 | 0.341502 | 0.321932 | 0.766887 | ENSMUSG( | NA | 14.60286 | -0.06046 | 0.909919 | 0.991958 |
| ENSMUSG( | NA | 11.95977 | -0.2344 | 0.65667 | NA | ENSMUSG( | NA | 13.60541 | -0.08676 | 0.866661 | 0.987458 |
| ENSMUSG( | NA | 15.55657 | 0.245377 | 0.618542 | NA | ENSMUSG( | NA | 9.755664 | -1.02405 | 0.118935 | NA |
| ENSMUSG( | NA | 12.89366 | -0.15109 | 0.768431 | NA | ENSMUSG( | NA | 23.97141 | -0.29208 | 0.479759 | 0.924021 |
| ENSMUSG( | NA | 47.65213 | -0.2315 | 0.436985 | 0.829501 | ENSMUSG( | NA | 9.955674 | -0.44688 | 0.473646 | NA |
| ENSMUSG( | NA | 10.27697 | -0.44193 | 0.456468 | NA | ENSMUSG( | NA | 33.12493 | 0.454308 | 0.232945 | 0.868507 |
| ENSMUSG( | NA | 41.59516 | -0.43126 | 0.155609 | 0.583641 | ENSMUSG( | NA | 271.6714 | 0.053216 | 0.804798 | 0.977232 |
| ENSMUSG( | NA | 91.19284 | -0.20963 | 0.333493 | 0.775602 | ENSMUSG( | NA | 51.79139 | -0.26192 | 0.360287 | 0.914223 |
| ENSMUSG( | NA | 37.84461 | -0.07207 | 0.840396 | 0.971179 | ENSMUSG( | NA | 37.06513 | 0.303795 | 0.380663 | 0.917202 |
| ENSMUSG( | NA | 23.56253 | -0.2638 | 0.532773 | 0.879905 | ENSMUSG( | NA | 12.03132 | 0.266483 | 0.636716 | 0.953903 |
| ENSMUSG( | NA | 11.94276 | -0.20443 | 0.747686 | NA | ENSMUSG( | NA | 19.2132 | 0.161444 | 0.72579 | 0.966887 |
| ENSMUSG( | NA | 8.622283 | 0.253217 | 0.693967 | NA | ENSMUSG( | NA | 10.32882 | 0.313092 | 0.628509 | NA |
| ENSMUSG( | NA | 26.96121 | 0.108091 | 0.763329 | 0.953781 | ENSMUSG( | NA | 9.013218 | 0.375374 | 0.564893 | NA |
| ENSMUSG( | NA | 131.9938 | 0.179941 | 0.365659 | 0.794294 | ENSMUSG( | NA | 47.86545 | -0.18585 | 0.580565 | 0.944266 |
| ENSMUSG( | NA | 68.40518 | 0.366359 | 0.26351 | 0.714284 | ENSMUSG( | NA | 25.76273 | 0.215142 | 0.610447 | 0.950771 |
| ENSMUSG( | NA | 191.0927 | 0.070554 | 0.704723 | 0.938303 | ENSMUSG( | NA | 135.2855 | 0.108461 | 0.571628 | 0.942153 |
| ENSMUSG( | NA | 69.35427 | -0.1896 | 0.474314 | 0.849425 | ENSMUSG( | NA | 67.85119 | -1.24455 | 0.000323 | 0.049152 |
| ENSMUSG( | NA | 18.73784 | -0.24045 | 0.610718 | NA | ENSMUSG( | NA | 15.01213 | -0.09887 | 0.844503 | 0.983642 |
| ENSMUSG( | NA | 14.24581 | 0.033613 | 0.944266 | NA | ENSMUSG( | NA | 82.72489 | -0.4704 | 0.044127 | 0.692633 |
| ENSMUSG( | NA | 19.86337 | 0.198256 | 0.65858 | NA | ENSMUSG( | NA | 15.42429 | 0.693574 | 0.184446 | 0.853168 |
| ENSMUSG( | NA | 14.06182 | -0.00942 | 0.986112 | NA | ENSMUSG( | NA | 44.19442 | -0.02281 | 0.947385 | 0.994798 |
| ENSMUSG( | NA | 114.909 | 0.047454 | 0.813482 | 0.964574 | ENSMUSG( | NA | 28.70674 | -0.05398 | 0.882306 | 0.989644 |
| ENSMUSG( | NA | 26.30759 | -0.04855 | 0.893922 | 0.984512 | ENSMUSG( | NA | 15.67503 | -0.23527 | 0.632965 | 0.953844 |
| ENSMUSG( | NA | 74.76709 | 0.275823 | 0.243779 | 0.691156 | ENSMUSG( | NA | 17.57876 | -0.20479 | 0.667068 | 0.956766 |
| ENSMUSG( | NA | 42.85727 | -0.19437 | 0.532394 | 0.879905 | ENSMUSG( | NA | 21.25952 | -0.66888 | 0.119715 | 0.813384 |
| ENSMUSG( | NA | 25.56926 | -0.11246 | 0.766187 | 0.954173 | ENSMUSG( | NA | 83.92947 | -0.25299 | 0.269186 | 0.886755 |
| ENSMUSG( | NA | 19.19957 | -0.3983 | 0.380464 | NA | ENSMUSG( | NA | 38.95266 | 0.456059 | 0.192163 | 0.859726 |
| ENSMUSG( | NA | 38.86109 | 0.02701 | 0.934101 | 0.99323 | ENSMUSG( | NA | 9.348092 | 0.083657 | 0.895195 | NA |
| ENSMUSG( | NA | 118.471 | -0.13158 | 0.545222 | 0.884748 | ENSMUSG( | NA | 10.4125 | -0.06547 | 0.918024 | NA |
| ENSMUSG( | NA | 24.25864 | -0.1628 | 0.661666 | 0.925271 | ENSMUSG( | NA | 25.8669 | 0.853294 | 0.036057 | 0.668395 |
| ENSMUSG( | NA | 27.0525 | -0.06673 | 0.864038 | 0.975331 | ENSMUSG( | NA | 11.82863 | -0.0921 | 0.8719 | 0.988779 |
| ENSMUSG( | NA | 108.0181 | 0.111329 | 0.611402 | 0.908586 | ENSMUSG( | NA | 199.0576 | -0.08578 | 0.640206 | 0.953903 |
| ENSMUSG( | NA | 26.64175 | -0.21511 | 0.57283 | 0.893584 | ENSMUSG( | NA | 22.6271 | 0.170553 | 0.690762 | 0.960071 |
| ENSMUSG( | NA | 13.72068 | 0.275516 | 0.580901 | NA | ENSMUSG( | NA | 140.0486 | 0.109395 | 0.561224 | 0.940254 |
| ENSMUSG( | NA | 37.78482 | -0.19679 | 0.551823 | 0.886141 | ENSMUSG( | NA | 11.72107 | 0.196639 | 0.747392 | 0.969763 |
| ENSMUSG( | NA | 14.75461 | -0.03688 | 0.938907 | NA | ENSMUSG( | NA | 30.54602 | -0.62608 | 0.125969 | 0.81516 |
| ENSMUSG( | NA | 37.25806 | 0.105914 | 0.748983 | 0.949604 | ENSMUSG( | NA | 70.39189 | -0.47349 | 0.085524 | 0.779799 |
| ENSMUSG( | NA | 70.43116 | -0.23641 | 0.300405 | 0.749512 | ENSMUSG( | NA | 71.21871 | -0.16367 | 0.556122 | 0.939565 |
| ENSMUSG( | NA | 17.02719 | 0.008646 | 0.984948 | NA | ENSMUSG( | NA | 76.89125 | -0.29799 | 0.222028 | 0.866205 |
| ENSMUSG( | NA | 9.995776 | -0.8608 | 0.151924 | NA | ENSMUSG( | NA | 37.47958 | -0.08587 | 0.791699 | 0.97681 |
| ENSMUSG( | NA | 28.02315 | 0.3751 | 0.307352 | 0.755324 | ENSMUSG( | NA | 10.51638 | 0.880811 | 0.18263 | NA |
| ENSMUSG( | NA | 48.11856 | 0.246642 | 0.410381 | 0.818339 | ENSMUSG( | NA | 39.03803 | 0.617391 | 0.061605 | 0.736111 |
| ENSMUSG( | NA | 143.5591 | -0.19943 | 0.352151 | 0.783738 | ENSMUSG( | NA | 11.26177 | 0.391766 | 0.491935 | NA |
| ENSMUSG( | NA | 10.49364 | 0.369441 | 0.533025 | NA | ENSMUSG( | NA | 33.67244 | 0.083831 | 0.814449 | 0.979198 |
| ENSMUSG( | NA | 62.87149 | 0.303767 | 0.211983 | 0.655675 | ENSMUSG( | NA | 22.0211 | -0.16161 | 0.714755 | 0.964986 |
| ENSMUSG( | NA | 10.74838 | 0.750801 | 0.21172 | NA | ENSMUSG( | NA | 46.66096 | -0.07202 | 0.82006 | 0.979198 |
| ENSMUSG( | NA | 21.58537 | -0.18794 | 0.664745 | 0.926824 | ENSMUSG( | NA | 10.97064 | -0.18655 | 0.75013 | NA |
| ENSMUSG( | NA | 34.02041 | -0.05341 | 0.883139 | 0.982422 | ENSMUSG( | NA | 25.91826 | 0.1573 | 0.729689 | 0.96719 |
| ENSMUSG( | NA | 34.74375 | 0.026279 | 0.934033 | 0.99323 | ENSMUSG( | NA | 106.534 | -0.32981 | 0.137326 | 0.828492 |
| ENSMUSG( | NA | 95.15895 | 0.094215 | 0.701204 | 0.93743 | ENSMUSG( | NA | 17.94981 | 0.307324 | 0.546348 | 0.938063 |

| | | | | | |
|---|---|---|---|---|---|
| ENSMUSG( | 23.50467 | 0.329796 | 0.397549 | 0.812127 | NA |
| ENSMUSG( | 564.3165 | 0.207977 | 0.136955 | 0.55149 | NA |
| ENSMUSG( | 12.59574 | 0.529499 | 0.3798 | NA | NA |
| ENSMUSG( | 10.80415 | 0.238051 | 0.679978 | NA | NA |
| ENSMUSG( | 45.14816 | -0.37097 | 0.197099 | 0.636662 | NA |
| ENSMUSG( | 14.49841 | -0.12161 | 0.807485 | NA | NA |
| ENSMUSG( | 48.54665 | 0.152125 | 0.603018 | 0.904887 | NA |
| ENSMUSG( | 18.16707 | -0.34193 | 0.432961 | NA | NA |
| ENSMUSG( | 35.80285 | -0.18984 | 0.592941 | 0.90229 | NA |
| ENSMUSG( | 8.937982 | -0.20821 | 0.740475 | NA | NA |
| ENSMUSG( | 24.74481 | 0.098047 | 0.808156 | 0.963419 | NA |
| ENSMUSG( | 19.17065 | 0.458691 | 0.275932 | NA | NA |
| ENSMUSG( | 12.38842 | 0.565812 | 0.273471 | NA | NA |
| ENSMUSG( | 13.60923 | -0.22715 | 0.681454 | NA | NA |
| ENSMUSG( | 31.05786 | -0.3194 | 0.472527 | 0.848206 | NA |
| ENSMUSG( | 23.78676 | 0.17427 | 0.659376 | 0.924506 | NA |
| ENSMUSG( | 19.68258 | 0.374878 | 0.394726 | NA | NA |
| ENSMUSG( | 20.32904 | 0.590577 | 0.180939 | NA | NA |
| ENSMUSG( | 8.983019 | -1.44367 | 0.058152 | NA | NA |
| ENSMUSG( | 34.38633 | -0.89863 | 0.048196 | 0.335655 | NA |
| ENSMUSG( | 12.29094 | -0.47018 | 0.387527 | NA | NA |
| ENSMUSG( | 35.28601 | 0.343855 | 0.318604 | 0.764234 | NA |
| ENSMUSG( | 23.71913 | 0.25815 | 0.499796 | 0.86324 | NA |
| ENSMUSG( | 71.69468 | 0.277031 | 0.291528 | 0.741366 | NA |
| ENSMUSG( | 63.46334 | -0.29027 | 0.244304 | 0.691435 | NA |
| ENSMUSG( | 15.469 | -0.37593 | 0.4641 | NA | NA |
| ENSMUSG( | 37.7855 | 0.005056 | 0.987355 | 0.998796 | NA |
| ENSMUSG( | 28.25366 | 0.228936 | 0.560914 | 0.888934 | NA |
| ENSMUSG( | 14.54783 | -0.60075 | 0.206767 | NA | NA |
| ENSMUSG( | 27.62345 | 0.393423 | 0.314878 | 0.762059 | NA |
| ENSMUSG( | 13.42192 | 0.767364 | 0.164752 | NA | NA |
| ENSMUSG( | 78.77365 | 0.107246 | 0.663404 | 0.9265 | NA |
| ENSMUSG( | 21.4953 | 0.137287 | 0.733197 | 0.944956 | NA |
| ENSMUSG( | 47.63042 | -0.83558 | 0.005955 | 0.104409 | NA |
| ENSMUSG( | 20.34538 | 0.180414 | 0.680735 | NA | NA |
| ENSMUSG( | 694.2862 | -0.23492 | 0.077756 | 0.431059 | NA |
| ENSMUSG( | 16.70896 | 0.001022 | 0.998197 | NA | NA |
| ENSMUSG( | 19.58375 | -0.12949 | 0.787892 | NA | NA |
| ENSMUSG( | 18.78317 | 0.094638 | 0.830988 | NA | NA |
| ENSMUSG( | 426.9991 | 0.057836 | 0.619819 | 0.911432 | NA |
| ENSMUSG( | 31.74936 | -0.3334 | 0.328374 | 0.772854 | NA |
| ENSMUSG( | 10.80238 | 0.227534 | 0.701765 | NA | NA |
| ENSMUSG( | 19.89275 | -0.76452 | 0.069464 | NA | NA |
| ENSMUSG( | 57.04249 | 0.147782 | 0.621379 | 0.911432 | NA |
| ENSMUSG( | 37.21991 | 0.216398 | 0.501671 | 0.864596 | NA |
| ENSMUSG( | 25.38671 | -0.57089 | 0.126394 | 0.533885 | NA |
| ENSMUSG( | 166.2888 | -0.07743 | 0.660243 | 0.924627 | NA |
| ENSMUSG( | 18.34993 | -0.28545 | 0.52955 | NA | NA |
| ENSMUSG( | 15.2647 | 0.109551 | 0.850598 | NA | NA |
| ENSMUSG( | 98.65981 | -0.17136 | 0.401932 | 0.814965 | NA |
| ENSMUSG( | 44.55784 | -0.20017 | 0.514577 | 0.872055 | NA |
| ENSMUSG( | 9.815303 | 0.369166 | 0.590247 | NA | NA |
| ENSMUSG( | 68.5083 | 0.461466 | 0.05381 | 0.356147 | NA |
| ENSMUSG( | 19.74988 | -0.12107 | 0.778739 | NA | NA |
| ENSMUSG( | 13.24738 | -0.52552 | 0.293013 | NA | NA |
| ENSMUSG( | 10.55109 | 0.533063 | 0.343421 | NA | NA |
| ENSMUSG( | 196.2032 | -0.14101 | 0.397851 | 0.812161 | NA |
| ENSMUSG( | 126.7052 | -0.11708 | 0.516176 | 0.872701 | NA |
| ENSMUSG( | 35.12063 | 0.387119 | 0.259477 | 0.709948 | NA |
| ENSMUSG( | 12.77943 | 0.876239 | 0.204601 | NA | NA |
| ENSMUSG( | 17.12495 | -0.8023 | 0.099257 | NA | NA |
| ENSMUSG( | 11.24818 | 0.888908 | 0.166646 | NA | NA |
| ENSMUSG( | 17.52004 | 0.050459 | 0.910293 | NA | NA |
| ENSMUSG( | 30.52475 | -0.49819 | 0.167171 | 0.599823 | NA |
| ENSMUSG( | 12.32838 | -0.95984 | 0.070709 | NA | NA |
| ENSMUSG( | 22.26734 | -0.12511 | 0.752775 | 0.950621 | NA |
| ENSMUSG( | 20.94474 | 0.106431 | 0.79939 | NA | NA |
| ENSMUSG( | 295.5486 | 0.038978 | 0.797263 | 0.960507 | NA |
| ENSMUSG( | 46.5443 | 0.321021 | 0.254467 | 0.704536 | NA |
| ENSMUSG( | 16.14955 | -0.4897 | 0.301974 | NA | NA |
| ENSMUSG( | 9.574802 | 0.471307 | 0.44796 | NA | NA |
| ENSMUSG( | 16.17676 | -0.42406 | 0.351563 | NA | NA |
| ENSMUSG( | 15.4955 | -0.00141 | 0.997611 | NA | NA |
| ENSMUSG( | 17.71887 | 0.473011 | 0.335303 | NA | NA |
| ENSMUSG( | 14.45386 | 0.47457 | 0.331407 | NA | NA |
| ENSMUSG( | 9.028506 | 0.257538 | 0.706984 | NA | NA |
| ENSMUSG( | 85.97965 | -0.23837 | 0.323213 | 0.768415 | NA |
| ENSMUSG( | 59.31701 | -0.12538 | 0.620445 | 0.911432 | NA |
| ENSMUSG( | 9.698451 | 0.01985 | 0.976078 | NA | NA |
| ENSMUSG( | 36.48983 | 0.350473 | 0.268842 | 0.719099 | NA |
| ENSMUSG( | 9.845391 | -0.66401 | 0.294234 | NA | NA |
| ENSMUSG( | 26.8643 | -1.02804 | 0.005201 | 0.096074 | NA |
| ENSMUSG( | 10.1366 | 0.014607 | 0.981577 | NA | NA |
| ENSMUSG( | 21.95667 | -0.58382 | 0.16827 | 0.601572 | NA |
| ENSMUSG( | 16.1767 | -0.01818 | 0.968754 | NA | NA |
| ENSMUSG( | 135.1021 | -0.16005 | 0.41264 | 0.819518 | NA |
| ENSMUSG( | 98.10077 | -0.06735 | 0.73322 | 0.944956 | NA |
| ENSMUSG( | 32.58683 | 0.190907 | 0.590691 | 0.90113 | NA |
| ENSMUSG( | 80.24501 | -0.11049 | 0.620795 | 0.911432 | NA |
| ENSMUSG( | 11.26565 | -0.4752 | 0.430949 | NA | NA |
| ENSMUSG( | 10.92986 | -0.58655 | 0.336916 | NA | NA |
| ENSMUSG( | 41.70991 | 0.396577 | 0.227082 | 0.674223 | NA |
| ENSMUSG( | 24.15481 | -0.55932 | 0.149236 | 0.573248 | NA |
| ENSMUSG( | 12.06032 | 0.87941 | 0.116328 | NA | NA |
| ENSMUSG( | 26.30873 | 0.395369 | 0.337946 | 0.777422 | NA |
| ENSMUSG( | 9.797835 | -0.86609 | 0.167035 | NA | NA |
| ENSMUSG( | 9.712992 | -0.29495 | 0.621892 | NA | NA |
| ENSMUSG( | 26.29471 | -0.44411 | 0.242292 | 0.68971 | NA |
| ENSMUSG( | 11.99105 | 0.133052 | 0.810032 | NA | NA |
| ENSMUSG( | 9.359401 | 0.046263 | 0.945389 | NA | NA |
| ENSMUSG( | 10.93371 | -1.25528 | 0.056197 | NA | NA |
| ENSMUSG( | 328.7038 | 1.290531 | 0.013019 | 0.165126 | NA |
| ENSMUSG( | 13.72268 | 0.224926 | 0.665501 | NA | NA |
| ENSMUSG( | 149.0157 | 0.250895 | 0.156554 | 0.584667 | NA |
| ENSMUSG( | 9.286424 | -0.07392 | 0.906491 | NA | NA |
| ENSMUSG( | 17.92012 | -0.37224 | 0.44537 | NA | NA |
| ENSMUSG( | 25.57771 | 0.067174 | 0.857899 | 0.974667 | NA |
| ENSMUSG( | 11.23569 | 0.341573 | 0.54845 | NA | NA |
| ENSMUSG( | 30.40242 | 0.438935 | 0.234591 | 0.681922 | NA |
| ENSMUSG( | 197.3917 | -0.0856 | 0.634329 | 0.914073 | NA |
| ENSMUSG( | 31.58579 | -0.29721 | 0.43505 | 0.829093 | NA |
| ENSMUSG( | 35.21593 | 0.002483 | 0.993762 | 0.999369 | NA |
| ENSMUSG( | 9.256707 | 0.91407 | 0.148771 | NA | NA |
| ENSMUSG( | 10.87673 | 0.189348 | 0.744692 | NA | NA |
| ENSMUSG( | 46.65382 | -0.07913 | 0.792531 | 0.959461 | NA |
| ENSMUSG( | 229.4443 | 0.062104 | 0.663334 | 0.9265 | NA |
| ENSMUSG( | 11.92232 | -1.00046 | 0.081108 | NA | NA |
| ENSMUSG( | 26.19675 | -0.63426 | 0.085928 | 0.451489 | NA |
| ENSMUSG( | 14.9341 | -0.1644 | 0.757675 | NA | NA |
| ENSMUSG( | 98.98782 | 0.104092 | 0.601435 | 0.904603 | NA |
| ENSMUSG( | 14.63802 | -0.02343 | 0.962701 | NA | NA |
| ENSMUSG( | 316.1098 | -0.32889 | 0.046019 | 0.328532 | NA |
| ENSMUSG( | 14.85825 | 0.371952 | 0.50347 | NA | NA |
| ENSMUSG( | 230.4468 | 0.162762 | 0.246501 | 0.694958 | NA |
| ENSMUSG( | 22.293 | -0.34127 | 0.407038 | 0.817163 | NA |
| ENSMUSG( | 41.671 | -0.02461 | 0.932425 | 0.992515 | NA |
| ENSMUSG( | 9.963841 | -0.51222 | 0.409416 | NA | NA |
| ENSMUSG( | 332.1371 | -0.38508 | 0.012121 | 0.159579 | NA |
| ENSMUSG( | 14.50274 | -0.79946 | 0.128136 | NA | NA |
| ENSMUSG( | 11.22347 | -0.12896 | 0.84004 | NA | NA |
| ENSMUSG( | 24.21292 | -0.44669 | 0.266034 | 0.716441 | NA |
| ENSMUSG( | 35.41381 | -0.36274 | 0.261944 | 0.712129 | NA |
| ENSMUSG( | 56.13046 | -0.18098 | 0.512888 | 0.871762 | NA |
| ENSMUSG( | 14.41078 | 0.252643 | 0.604374 | NA | NA |
| ENSMUSG( | 58.80551 | -0.23215 | 0.379914 | 0.801757 | NA |
| ENSMUSG( | 496.1366 | -0.01039 | 0.930071 | 0.991848 | NA |
| ENSMUSG( | 344.4894 | 0.182118 | 0.19343 | 0.632931 | NA |
| ENSMUSG( | 7.562286 | -0.16812 | 0.822657 | NA | NA |
| ENSMUSG( | 34.79113 | -0.63091 | 0.046484 | 0.330252 | NA |
| ENSMUSG( | 22.35792 | 0.09826 | 0.825896 | 0.968061 | NA |
| ENSMUSG( | 14.21958 | -0.12061 | 0.826423 | NA | NA |
| ENSMUSG( | 35.88591 | -0.26 | 0.430921 | 0.826896 | NA |
| ENSMUSG( | 25.33493 | 0.240769 | 0.549031 | 0.885686 | NA |
| ENSMUSG( | 46.15353 | 0.237355 | 0.393077 | 0.809274 | NA |
| ENSMUSG( | 86.52772 | 0.143511 | 0.560322 | 0.888934 | NA |
| ENSMUSG( | 13.67305 | -0.04214 | 0.932855 | NA | NA |
| ENSMUSG( | 39.69613 | -0.37013 | 0.234241 | 0.68192 | NA |
| ENSMUSG( | 14.71836 | 0.285129 | 0.558748 | NA | NA |
| ENSMUSG( | 179.8119 | 0.019334 | 0.903405 | 0.986055 | NA |
| ENSMUSG( | 21.61661 | 0.638107 | 0.107734 | 0.49674 | NA |
| ENSMUSG( | 22.18805 | -0.19439 | 0.627642 | 0.912961 | NA |
| ENSMUSG( | 21.19266 | -0.15143 | 0.733066 | NA | NA |
| ENSMUSG( | 9.666273 | -0.52934 | 0.390106 | NA | NA |
| ENSMUSG( | 538.7027 | -0.18218 | 0.149044 | 0.572864 | NA |
| ENSMUSG( | 15.56348 | 0.315467 | 0.513019 | NA | NA |
| ENSMUSG( | 274.3064 | 0.068468 | 0.623358 | 0.911646 | NA |
| ENSMUSG( | 20.64602 | 0.221703 | 0.608276 | NA | NA |
| ENSMUSG( | 154.2195 | 0.122125 | 0.537614 | 0.882536 | NA |

| | | | | | |
|---|---|---|---|---|---|
| ENSMUSG( | 20.38929 | -0.47901 | 0.298148 | 0.898538 | NA |
| ENSMUSG( | 72.04741 | -0.21012 | 0.40835 | 0.917257 | NA |
| ENSMUSG( | 25.92897 | -0.58066 | 0.148789 | 0.836241 | NA |
| ENSMUSG( | 18.09724 | 0.433579 | 0.358835 | 0.913759 | NA |
| ENSMUSG( | 744.1166 | 0.236954 | 0.105441 | 0.804969 | NA |
| ENSMUSG( | 21.70793 | -0.16302 | 0.699653 | 0.961335 | NA |
| ENSMUSG( | 37.39255 | -0.07844 | 0.810661 | 0.978779 | NA |
| ENSMUSG( | 6.323227 | -0.26064 | 0.754673 | NA | NA |
| ENSMUSG( | 28.88314 | 0.081999 | 0.824588 | 0.979198 | NA |
| ENSMUSG( | 80.55504 | -0.48707 | 0.055923 | 0.723362 | NA |
| ENSMUSG( | 23.86501 | -0.43173 | 0.287531 | 0.894091 | NA |
| ENSMUSG( | 54.32637 | -0.15336 | 0.603139 | 0.948137 | NA |
| ENSMUSG( | 11.57187 | -0.04723 | 0.932824 | 0.99306 | NA |
| ENSMUSG( | 22.10799 | -0.44723 | 0.3156 | 0.901725 | NA |
| ENSMUSG( | 11.82459 | 0.018513 | 0.978684 | 0.996459 | NA |
| ENSMUSG( | 34.34636 | 0.028211 | 0.937082 | 0.993685 | NA |
| ENSMUSG( | 8.11887 | 0.18135 | 0.794475 | NA | NA |
| ENSMUSG( | 9.065948 | 0.379785 | 0.571035 | NA | NA |
| ENSMUSG( | 161.1915 | 0.107623 | 0.58361 | 0.944266 | NA |
| ENSMUSG( | 24.26544 | -0.28111 | 0.514508 | 0.930939 | NA |
| ENSMUSG( | 20.04147 | 0.06517 | 0.882499 | 0.989644 | NA |
| ENSMUSG( | 26.6628 | -0.37609 | 0.330104 | 0.907251 | NA |
| ENSMUSG( | 22.22627 | 0.106331 | 0.823925 | 0.979198 | NA |
| ENSMUSG( | 14.31085 | 0.033789 | 0.947482 | 0.994798 | NA |
| ENSMUSG( | 40.84529 | -0.36751 | 0.305806 | 0.898538 | NA |
| ENSMUSG( | 61.03815 | -0.27575 | 0.322945 | 0.904023 | NA |
| ENSMUSG( | 38.48194 | -0.36053 | 0.274322 | 0.889874 | NA |
| ENSMUSG( | 75.94178 | 0.201869 | 0.409442 | 0.917257 | NA |
| ENSMUSG( | 14.45696 | 0.435873 | 0.413668 | 0.917257 | NA |
| ENSMUSG( | 46.96248 | -0.20984 | 0.550214 | 0.938699 | NA |
| ENSMUSG( | 17.19082 | -0.92235 | 0.055375 | 0.723362 | NA |
| ENSMUSG( | 44.59516 | -0.09709 | 0.772071 | 0.974236 | NA |
| ENSMUSG( | 16.85842 | -0.34707 | 0.491621 | 0.926889 | NA |
| ENSMUSG( | 38.32139 | 0.060953 | 0.859507 | 0.985947 | NA |
| ENSMUSG( | 149.6336 | -0.27151 | 0.151224 | 0.837494 | NA |
| ENSMUSG( | 134.444 | -0.14391 | 0.640401 | 0.953903 | NA |
| ENSMUSG( | 16.19668 | 0.272438 | 0.589517 | 0.944975 | NA |
| ENSMUSG( | 27.26783 | 0.413711 | 0.267384 | 0.886134 | NA |
| ENSMUSG( | 65.50397 | -0.14366 | 0.604997 | 0.949082 | NA |
| ENSMUSG( | 21.38197 | 0.405087 | 0.383184 | 0.917257 | NA |
| ENSMUSG( | 76.92051 | 0.382418 | 0.119162 | 0.813384 | NA |
| ENSMUSG( | 79.82759 | -0.06657 | 0.791267 | 0.97681 | NA |
| ENSMUSG( | 22.67091 | -0.30214 | 0.517786 | 0.930939 | NA |
| ENSMUSG( | 528.37 | 0.120642 | 0.4973 | 0.927825 | NA |
| ENSMUSG( | 89.89621 | -0.19466 | 0.42625 | 0.917257 | NA |
| ENSMUSG( | 15.75602 | -0.45585 | 0.363327 | 0.914223 | NA |
| ENSMUSG( | 26.96423 | -0.55572 | 0.154733 | 0.837724 | NA |
| ENSMUSG( | 15.13257 | 0.355384 | 0.532053 | 0.934795 | NA |
| ENSMUSG( | 19.70962 | 0.441735 | 0.320386 | 0.903887 | NA |
| ENSMUSG( | 384.0194 | 0.015279 | 0.916938 | 0.992529 | NA |
| ENSMUSG( | 14.11753 | -0.46376 | 0.376536 | 0.915491 | NA |
| ENSMUSG( | 14.50753 | -0.5912 | 0.254964 | 0.882307 | NA |
| ENSMUSG( | 37.39107 | 0.760251 | 0.029077 | 0.614779 | NA |
| ENSMUSG( | 37.17216 | -0.44104 | 0.195987 | 0.862161 | NA |
| ENSMUSG( | 51.57018 | 0.263973 | 0.457137 | 0.923329 | NA |
| ENSMUSG( | 40.61744 | 0.205632 | 0.5281 | 0.934677 | NA |
| ENSMUSG( | 30.08724 | -0.47299 | 0.224045 | 0.867404 | NA |
| ENSMUSG( | 113.049 | -0.17837 | 0.388713 | 0.917257 | NA |
| ENSMUSG( | 24.00252 | 0.094342 | 0.819912 | 0.979198 | NA |
| ENSMUSG( | 22.34399 | -0.55054 | 0.203816 | 0.864412 | NA |
| ENSMUSG( | 19.32644 | 0.344678 | 0.450162 | 0.922202 | NA |
| ENSMUSG( | 20.82131 | -0.07681 | 0.858965 | 0.985884 | NA |
| ENSMUSG( | 9.630261 | -0.37451 | 0.578517 | NA | NA |
| ENSMUSG( | 147.6692 | -0.50665 | 0.059162 | 0.729429 | NA |
| ENSMUSG( | 11.54258 | 0.462996 | 0.468013 | 0.923329 | NA |
| ENSMUSG( | 8.439066 | -0.92332 | 0.20978 | NA | NA |
| ENSMUSG( | 188.7796 | -0.77784 | 3.39E-05 | 0.009128 | NA |
| ENSMUSG( | 10.58947 | 0.057011 | 0.923176 | NA | NA |
| ENSMUSG( | 71.75579 | -0.19639 | 0.427924 | 0.917257 | NA |
| ENSMUSG( | 8.294161 | 0.065785 | 0.921801 | NA | NA |
| ENSMUSG( | 55.75473 | 0.146126 | 0.619665 | 0.952896 | NA |
| ENSMUSG( | 49.79263 | 0.115195 | 0.687765 | 0.959703 | NA |
| ENSMUSG( | 39.88199 | 0.136364 | 0.668624 | 0.956766 | NA |
| ENSMUSG( | 15.16581 | -0.15314 | 0.792398 | 0.97681 | NA |
| ENSMUSG( | 14.08547 | -0.02983 | 0.953323 | 0.994798 | NA |
| ENSMUSG( | 29.45396 | -0.33342 | 0.391912 | 0.917257 | NA |
| ENSMUSG( | 19.69489 | 0.219258 | 0.61996 | 0.952896 | NA |
| ENSMUSG( | 23.03446 | 0.415283 | 0.323246 | 0.904023 | NA |
| ENSMUSG( | 8.812843 | 0.709085 | 0.340781 | NA | NA |
| ENSMUSG( | 17.31399 | -0.28874 | 0.546511 | 0.938063 | NA |
| ENSMUSG( | 13.8916 | 0.579082 | 0.374025 | 0.915491 | NA |
| ENSMUSG( | 18.70556 | 0.576039 | 0.216195 | 0.865167 | NA |
| ENSMUSG( | 12.69168 | 0.400762 | 0.467913 | 0.923329 | NA |
| ENSMUSG( | 16.09582 | 1.586547 | 0.029993 | 0.620326 | NA |
| ENSMUSG( | 47.18416 | -0.56617 | 0.085146 | 0.779799 | NA |
| ENSMUSG( | 21.09954 | 0.59161 | 0.212311 | 0.864412 | NA |
| ENSMUSG( | 98.21655 | -0.13906 | 0.520329 | 0.931964 | NA |
| ENSMUSG( | 9.590072 | -0.57896 | 0.42089 | NA | NA |
| ENSMUSG( | 13.45097 | 0.364233 | 0.506174 | 0.927825 | NA |
| ENSMUSG( | 22.42212 | -0.00225 | 0.995857 | 0.999116 | NA |
| ENSMUSG( | 190.0075 | 0.047331 | 0.809268 | 0.978779 | NA |
| ENSMUSG( | 9.782455 | -0.30195 | 0.619538 | NA | NA |
| ENSMUSG( | 14.36874 | 0.200918 | 0.710597 | 0.964363 | NA |
| ENSMUSG( | 14.08133 | -0.00748 | 0.988328 | 0.997862 | NA |
| ENSMUSG( | 31.76932 | -0.10994 | 0.755954 | 0.971371 | NA |
| ENSMUSG( | 69.58621 | 0.322826 | 0.229282 | 0.868507 | NA |
| ENSMUSG( | 342.2238 | -0.04587 | 0.748876 | 0.96984 | NA |
| ENSMUSG( | 393.601 | -2.98188 | 1.28E-30 | 3.73E-27 | NA |
| ENSMUSG( | 20.1226 | -0.17053 | 0.737581 | 0.967686 | NA |
| ENSMUSG( | 14.54822 | -0.46884 | 0.376659 | 0.915491 | NA |
| ENSMUSG( | 10.31825 | -0.00486 | 0.993898 | NA | NA |
| ENSMUSG( | 33.16975 | 0.190825 | 0.666003 | 0.956766 | NA |
| ENSMUSG( | 23.64291 | -0.28421 | 0.512137 | 0.930074 | NA |
| ENSMUSG( | 8.900753 | 1.146624 | 0.121481 | NA | NA |
| ENSMUSG( | 81.45392 | 0.206823 | 0.388962 | 0.917257 | NA |
| ENSMUSG( | 29.11216 | -0.29242 | 0.484463 | 0.92541 | NA |
| ENSMUSG( | 43.7518 | 0.213217 | 0.55434 | 0.939565 | NA |
| ENSMUSG( | 27.41034 | 0.742582 | 0.069788 | 0.756483 | NA |
| ENSMUSG( | 12.62374 | 0.468819 | 0.417138 | 0.917257 | NA |
| ENSMUSG( | 22.95795 | -0.31864 | 0.45582 | 0.923329 | NA |
| ENSMUSG( | 131.6856 | 0.167471 | 0.449927 | 0.922202 | NA |
| ENSMUSG( | 14.57389 | 0.672925 | 0.197704 | 0.862161 | NA |
| ENSMUSG( | 14.04488 | -0.63294 | 0.265144 | 0.886134 | NA |
| ENSMUSG( | 14.9423 | -0.36359 | 0.518867 | 0.930939 | NA |
| ENSMUSG( | 67.92746 | 0.236856 | 0.372056 | 0.915491 | NA |
| ENSMUSG( | 14.60286 | -0.06046 | 0.909919 | 0.991958 | NA |
| ENSMUSG( | 13.60541 | -0.08676 | 0.866661 | 0.987458 | NA |
| ENSMUSG( | 9.755664 | -1.02405 | 0.118935 | NA | NA |
| ENSMUSG( | 23.97141 | -0.29208 | 0.479759 | 0.924021 | NA |
| ENSMUSG( | 9.955674 | -0.44688 | 0.473646 | NA | NA |
| ENSMUSG( | 33.12493 | 0.454308 | 0.232945 | 0.868507 | NA |
| ENSMUSG( | 271.6714 | 0.053216 | 0.804798 | 0.977232 | NA |
| ENSMUSG( | 51.79139 | -0.26192 | 0.360287 | 0.914223 | NA |
| ENSMUSG( | 37.06513 | 0.303795 | 0.380663 | 0.917202 | NA |
| ENSMUSG( | 12.03132 | 0.266483 | 0.636716 | 0.953903 | NA |
| ENSMUSG( | 19.2132 | 0.161444 | 0.72579 | 0.966887 | NA |
| ENSMUSG( | 10.32882 | 0.313092 | 0.628509 | NA | NA |
| ENSMUSG( | 9.013218 | 0.375374 | 0.564893 | NA | NA |
| ENSMUSG( | 47.86545 | -0.18585 | 0.580565 | 0.944266 | NA |
| ENSMUSG( | 25.76273 | 0.215142 | 0.610447 | 0.950771 | NA |
| ENSMUSG( | 135.2855 | 0.108461 | 0.571628 | 0.942153 | NA |
| ENSMUSG( | 67.85119 | -1.24455 | 0.000323 | 0.049152 | NA |
| ENSMUSG( | 15.01213 | -0.09887 | 0.844503 | 0.983642 | NA |
| ENSMUSG( | 82.72489 | -0.4704 | 0.044127 | 0.692633 | NA |
| ENSMUSG( | 15.42429 | 0.693574 | 0.184446 | 0.853168 | NA |
| ENSMUSG( | 44.19442 | -0.02281 | 0.947385 | 0.994798 | NA |
| ENSMUSG( | 28.70674 | -0.05398 | 0.882306 | 0.989644 | NA |
| ENSMUSG( | 15.67503 | -0.23527 | 0.632965 | 0.953844 | NA |
| ENSMUSG( | 17.57876 | -0.20479 | 0.667068 | 0.956766 | NA |
| ENSMUSG( | 21.25952 | -0.66888 | 0.119715 | 0.813384 | NA |
| ENSMUSG( | 83.92947 | -0.25299 | 0.269186 | 0.886755 | NA |
| ENSMUSG( | 38.95266 | 0.456059 | 0.192163 | 0.859726 | NA |
| ENSMUSG( | 9.348092 | 0.083657 | 0.895195 | NA | NA |
| ENSMUSG( | 10.4125 | -0.06547 | 0.918024 | NA | NA |
| ENSMUSG( | 25.8669 | 0.853294 | 0.036057 | 0.668395 | NA |
| ENSMUSG( | 11.82863 | -0.0921 | 0.8719 | 0.988779 | NA |
| ENSMUSG( | 199.0576 | -0.08578 | 0.640206 | 0.953903 | NA |
| ENSMUSG( | 22.6271 | 0.170553 | 0.690762 | 0.960071 | NA |
| ENSMUSG( | 140.0486 | 0.109395 | 0.561224 | 0.940254 | NA |
| ENSMUSG( | 11.72107 | 0.196639 | 0.747392 | 0.969763 | NA |
| ENSMUSG( | 30.54602 | -0.62608 | 0.125969 | 0.81516 | NA |
| ENSMUSG( | 70.39189 | -0.47349 | 0.085524 | 0.779799 | NA |
| ENSMUSG( | 71.21871 | -0.16367 | 0.556122 | 0.939565 | NA |
| ENSMUSG( | 76.89125 | -0.29799 | 0.222028 | 0.866205 | NA |
| ENSMUSG( | 37.47958 | -0.08587 | 0.791699 | 0.97681 | NA |
| ENSMUSG( | 10.51638 | 0.880811 | 0.18263 | NA | NA |
| ENSMUSG( | 39.03803 | 0.617391 | 0.061605 | 0.736111 | NA |
| ENSMUSG( | 11.26177 | 0.391766 | 0.491935 | NA | NA |

| | | | | | |
|---|---|---|---|---|---|
| ENSMUSG( | NA | 12.49646 | -0.04106 | 0.938497 | NA |
| ENSMUSG( | NA | 66.30669 | 0.043766 | 0.868207 | 0.976786 |
| ENSMUSG( | NA | 9.644401 | -0.27862 | 0.645779 | NA |
| ENSMUSG( | NA | 55.73599 | 0.301462 | 0.247814 | 0.696522 |
| ENSMUSG( | NA | 10.4319 | -0.15934 | 0.786377 | NA |
| ENSMUSG( | NA | 9.254259 | -0.58022 | 0.384116 | NA |
| ENSMUSG( | NA | 51.58038 | 0.135116 | 0.616152 | 0.91085 |
| ENSMUSG( | NA | 178.3539 | -0.03735 | 0.839761 | 0.971179 |
| ENSMUSG( | NA | 427.6746 | -0.43087 | 0.004414 | 0.086676 |
| ENSMUSG( | NA | 16.08038 | 0.148704 | 0.748443 | NA |
| ENSMUSG( | NA | 23.13541 | 0.50925 | 0.217965 | 0.663176 |
| ENSMUSG( | NA | 23.80639 | -0.2918 | 0.487377 | 0.858812 |
| ENSMUSG( | NA | 19.63127 | 0.387048 | 0.351206 | NA |
| ENSMUSG( | NA | 17.93999 | -0.29581 | 0.504991 | NA |
| ENSMUSG( | NA | 47.71894 | -0.11654 | 0.692307 | 0.934101 |
| ENSMUSG( | NA | 12.03235 | 0.429483 | 0.437041 | NA |
| ENSMUSG( | NA | 38.34844 | -0.33366 | 0.280107 | 0.729155 |
| ENSMUSG( | NA | 58.80066 | 0.081609 | 0.7592 | 0.952284 |
| ENSMUSG( | NA | 514.509 | 0.101111 | 0.451078 | 0.835567 |
| ENSMUSG( | NA | 10.51119 | -0.33589 | 0.573005 | NA |
| ENSMUSG( | NA | 70.21228 | 0.071534 | 0.772358 | 0.955069 |
| ENSMUSG( | NA | 12.184 | -0.22369 | 0.673384 | NA |
| ENSMUSG( | NA | 61.686 | -0.23927 | 0.345044 | 0.779153 |
| ENSMUSG( | NA | 9.402414 | -0.01124 | 0.986231 | NA |
| ENSMUSG( | NA | 102.2354 | -0.14561 | 0.505325 | 0.866883 |
| ENSMUSG( | NA | 20.84002 | -0.2598 | 0.548805 | NA |
| ENSMUSG( | NA | 35.56092 | 0.466557 | 0.178602 | 0.616298 |
| ENSMUSG( | NA | 80.632 | -0.11875 | 0.584443 | 0.898802 |
| ENSMUSG( | NA | 23.49454 | 0.111871 | 0.773697 | 0.95556 |
| ENSMUSG( | NA | 56.56064 | -0.30617 | 0.247854 | 0.696522 |
| ENSMUSG( | NA | 40.5895 | 0.0073 | 0.982533 | 0.997512 |
| ENSMUSG( | NA | 3787.095 | -0.00126 | 0.991033 | 0.999232 |
| ENSMUSG( | NA | 31.72141 | -0.04592 | 0.900568 | 0.985633 |
| ENSMUSG( | NA | 117.2171 | 0.024757 | 0.911463 | 0.988483 |
| ENSMUSG( | NA | 12.67868 | -0.06379 | 0.909922 | NA |
| ENSMUSG( | NA | 38.83768 | -0.01862 | 0.950692 | 0.993821 |
| ENSMUSG( | NA | 42.89025 | 0.102636 | 0.732749 | 0.944956 |
| ENSMUSG( | NA | 12.37604 | -0.17103 | 0.746317 | NA |
| ENSMUSG( | NA | 10.19212 | 0.316814 | 0.602522 | NA |
| ENSMUSG( | NA | 16.28439 | -0.26901 | 0.570786 | NA |
| ENSMUSG( | NA | 27.46404 | 0.064424 | 0.860884 | 0.975025 |
| ENSMUSG( | NA | 38.56483 | -0.12729 | 0.693945 | 0.934202 |
| ENSMUSG( | NA | 83.47962 | 0.530257 | 0.027549 | 0.2548 |
| ENSMUSG( | NA | 264.4154 | 0.174547 | 0.259113 | 0.709814 |
| ENSMUSG( | NA | 15.76482 | -0.12423 | 0.786078 | NA |
| ENSMUSG( | NA | 37.57224 | 0.086035 | 0.790382 | 0.959461 |
| ENSMUSG( | NA | 178.1833 | -0.76285 | 0.000998 | 0.032953 |
| ENSMUSG( | NA | 10.73011 | -0.59698 | 0.320604 | NA |
| ENSMUSG( | NA | 21.91241 | -0.31926 | 0.442532 | 0.83278 |
| ENSMUSG( | NA | 16.30941 | -0.81551 | 0.110744 | NA |
| ENSMUSG( | NA | 15.69389 | -0.24522 | 0.609001 | NA |
| ENSMUSG( | NA | 18.4745 | -0.23911 | 0.584594 | NA |
| ENSMUSG( | NA | 8.806321 | 1.352293 | 0.056119 | NA |
| ENSMUSG( | NA | 29.59097 | -0.07995 | 0.819962 | 0.965888 |
| ENSMUSG( | NA | 44.96202 | -0.0025 | 0.994349 | 0.999369 |
| ENSMUSG( | NA | 13.02944 | -0.40029 | 0.437068 | NA |
| ENSMUSG( | NA | 14.65697 | -0.06309 | 0.896194 | NA |
| ENSMUSG( | NA | 29.60975 | -0.06155 | 0.861535 | 0.975026 |
| ENSMUSG( | NA | 511.7232 | -0.406 | 0.014048 | 0.173278 |
| ENSMUSG( | NA | 62.99359 | 0.110106 | 0.691127 | 0.933881 |
| ENSMUSG( | NA | 20.27636 | 0.59535 | 0.199663 | NA |
| ENSMUSG( | NA | 10.66514 | 0.422448 | 0.496092 | NA |
| ENSMUSG( | NA | 17.4061 | 0.528763 | 0.295023 | NA |
| ENSMUSG( | NA | 27.63074 | -0.3754 | 0.382662 | 0.803227 |
| ENSMUSG( | NA | 24.74657 | 0.248002 | 0.53938 | 0.883094 |
| ENSMUSG( | NA | 27.6965 | 0.479446 | 0.183954 | 0.620925 |
| ENSMUSG( | NA | 28.0791 | -0.04216 | 0.911708 | 0.988483 |
| ENSMUSG( | NA | 8.721198 | -0.14092 | 0.81904 | NA |
| ENSMUSG( | NA | 18.84827 | -0.02233 | 0.960282 | NA |
| ENSMUSG( | NA | 26.77851 | 0.375967 | 0.338166 | 0.777422 |
| ENSMUSG( | NA | 32.43474 | -0.31667 | 0.333759 | 0.775602 |
| ENSMUSG( | NA | 12.84978 | -0.13471 | 0.808414 | NA |
| ENSMUSG( | NA | 118.6228 | 0.163937 | 0.479306 | 0.854046 |
| ENSMUSG( | NA | 35.86796 | -0.28237 | 0.371165 | 0.798554 |
| ENSMUSG( | NA | 39.77314 | -0.04716 | 0.886805 | 0.983255 |
| ENSMUSG( | NA | 1081.475 | 0.043406 | 0.689539 | 0.933609 |
| ENSMUSG( | NA | 13.87737 | -0.81523 | 0.112541 | NA |
| ENSMUSG( | NA | 9.237475 | -0.57872 | 0.36633 | NA |
| ENSMUSG( | NA | 26.91355 | 0.056811 | 0.882496 | 0.982244 |
| ENSMUSG( | NA | 12.69998 | -0.11333 | 0.846306 | NA |
| ENSMUSG( | NA | 25.86309 | -0.40315 | 0.293795 | 0.743425 |
| ENSMUSG( | NA | 28.568 | 0.308979 | 0.440337 | 0.831851 |
| ENSMUSG( | NA | 62.6086 | -0.05509 | 0.834925 | 0.970426 |
| ENSMUSG( | NA | 38.08984 | -0.4 | 0.207704 | 0.64962 |
| ENSMUSG( | NA | 43.06804 | -0.24652 | 0.420828 | 0.822984 |
| ENSMUSG( | NA | 2676.946 | -0.47552 | 0.000474 | 0.01988 |
| ENSMUSG( | NA | 11.02616 | -0.10604 | 0.844467 | NA |
| ENSMUSG( | NA | 74.3217 | 0.039124 | 0.86693 | 0.976541 |
| ENSMUSG( | NA | 17.30452 | -0.30044 | 0.511943 | NA |
| ENSMUSG( | NA | 16.97146 | 0.164486 | 0.718363 | NA |
| ENSMUSG( | NA | 29.42793 | -0.27391 | 0.458778 | 0.840244 |
| ENSMUSG( | NA | 66.97598 | 0.516255 | 0.037691 | 0.299975 |
| ENSMUSG( | NA | 37.55639 | -0.11079 | 0.738481 | 0.947414 |
| ENSMUSG( | NA | 182.5503 | -0.20012 | 0.189298 | 0.627932 |
| ENSMUSG( | NA | 110.5413 | 0.14378 | 0.447446 | 0.834051 |
| ENSMUSG( | NA | 7.840173 | -0.08935 | 0.898042 | NA |
| ENSMUSG( | NA | 83.63013 | -0.33185 | 0.181856 | 0.618944 |
| ENSMUSG( | NA | 19.88148 | -0.58112 | 0.244295 | NA |
| ENSMUSG( | NA | 15.79677 | 0.27908 | 0.547154 | NA |
| ENSMUSG( | NA | 8.653621 | -0.72025 | 0.305823 | NA |
| ENSMUSG( | NA | 15.57715 | -0.50649 | 0.273284 | NA |
| ENSMUSG( | NA | 20.41993 | 0.319245 | 0.463786 | NA |
| ENSMUSG( | NA | 37.58266 | -0.14611 | 0.632444 | 0.914073 |
| ENSMUSG( | NA | 20.66055 | -0.88088 | 0.037415 | NA |
| ENSMUSG( | NA | 111.7646 | -0.05457 | 0.76933 | 0.954674 |
| ENSMUSG( | NA | 24.5917 | 0.064036 | 0.866856 | 0.976541 |
| ENSMUSG( | NA | 30.27701 | 0.032749 | 0.930521 | 0.991965 |
| ENSMUSG( | NA | 81.72892 | 0.140229 | 0.524372 | 0.87583 |
| ENSMUSG( | NA | 17.90454 | -0.51663 | 0.277287 | NA |
| ENSMUSG( | NA | 14.72852 | -0.17475 | 0.710326 | NA |
| ENSMUSG( | NA | 80.35265 | 0.166965 | 0.474391 | 0.849425 |
| ENSMUSG( | NA | 66.05073 | -0.16598 | 0.542023 | 0.883182 |
| ENSMUSG( | NA | 42.82259 | -0.54833 | 0.068779 | 0.406391 |
| ENSMUSG( | NA | 66.7256 | -0.27471 | 0.250735 | 0.700403 |
| ENSMUSG( | NA | 144.9608 | -0.10338 | 0.563141 | 0.889834 |
| ENSMUSG( | NA | 23.7776 | 0.007769 | 0.984578 | 0.997815 |
| ENSMUSG( | NA | 72.68179 | -0.23194 | 0.317408 | 0.763823 |
| ENSMUSG( | NA | 98.53444 | -0.17027 | 0.496223 | 0.861564 |
| ENSMUSG( | NA | 15.60698 | -0.90223 | 0.056748 | NA |
| ENSMUSG( | NA | 24.38665 | -0.32998 | 0.382923 | 0.803237 |
| ENSMUSG( | NA | 1881.198 | 0.164207 | 0.061869 | 0.384781 |
| ENSMUSG( | NA | 10.87586 | -0.14026 | 0.802868 | NA |
| ENSMUSG( | NA | 20.89623 | 0.03226 | 0.939678 | NA |
| ENSMUSG( | NA | 11.89638 | -0.55656 | 0.303208 | NA |
| ENSMUSG( | NA | 15.55229 | 0.371688 | 0.428959 | NA |
| ENSMUSG( | NA | 17.55542 | 0.47153 | 0.366852 | NA |
| ENSMUSG( | NA | 16.45465 | 0.420949 | 0.41382 | NA |
| ENSMUSG( | NA | 9.146223 | -0.35093 | 0.580085 | NA |
| ENSMUSG( | NA | 221.4211 | 0.001626 | 0.991963 | 0.999232 |
| ENSMUSG( | NA | 5577.446 | 0.11113 | 0.279518 | 0.729114 |
| ENSMUSG( | NA | 36.07487 | -0.04213 | 0.895403 | 0.984512 |
| ENSMUSG( | NA | 5447.289 | -0.13628 | 0.206672 | 0.648942 |
| ENSMUSG( | NA | 41.24381 | -0.65521 | 0.029261 | 0.259574 |
| ENSMUSG( | NA | 28.29225 | 0.046963 | 0.899405 | 0.985633 |
| ENSMUSG( | NA | 21.31253 | 0.114522 | 0.793763 | 0.959799 |
| ENSMUSG( | NA | 94.18158 | 0.135226 | 0.513554 | 0.871779 |
| ENSMUSG( | NA | 47.59662 | -0.08207 | 0.811168 | 0.964318 |
| ENSMUSG( | NA | 73.24337 | -0.24133 | 0.331468 | 0.77519 |
| ENSMUSG( | NA | 13.88852 | -0.55192 | 0.319839 | NA |
| ENSMUSG( | NA | 99.44611 | 0.024239 | 0.910726 | 0.988356 |
| ENSMUSG( | NA | 9.311005 | -0.4798 | 0.427312 | NA |
| ENSMUSG( | NA | 204.7302 | -0.02431 | 0.876866 | 0.980991 |
| ENSMUSG( | NA | 112.1326 | -0.07452 | 0.730883 | 0.944956 |
| ENSMUSG( | NA | 20.58735 | 0.275223 | 0.532679 | NA |
| ENSMUSG( | NA | 14.37038 | -0.01991 | 0.969049 | NA |
| ENSMUSG( | NA | 186.4292 | 0.153644 | 0.327843 | 0.772842 |
| ENSMUSG( | NA | 102.379 | -0.11481 | 0.556415 | 0.887681 |
| ENSMUSG( | NA | 284.7827 | -0.13296 | 0.418988 | 0.821966 |
| ENSMUSG( | NA | 11.13939 | 0.166066 | 0.761944 | NA |
| ENSMUSG( | NA | 32.29312 | 0.016573 | 0.962221 | 0.994543 |
| ENSMUSG( | NA | 60.54458 | 0.376633 | 0.126114 | 0.533394 |
| ENSMUSG( | NA | 148.9881 | -0.01135 | 0.947599 | 0.993821 |
| ENSMUSG( | NA | 26.60391 | 0.295944 | 0.433696 | 0.828485 |
| ENSMUSG( | NA | 41.22689 | 0.460236 | 0.160465 | 0.590122 |
| ENSMUSG( | NA | 13.16909 | 0.219653 | 0.716566 | NA |
| ENSMUSG( | NA | 18.38319 | 0.809768 | 0.061056 | NA |
| ENSMUSG( | NA | 1079.414 | -0.00897 | 0.964533 | 0.995155 |
| ENSMUSG( | NA | 120.6572 | 0.26973 | 0.318104 | 0.764234 |

| | | | | | |
|---|---|---|---|---|---|
| ENSMUSG( | NA | 15.9713 | -0.46985 | 0.33474 | 0.907907 |
| ENSMUSG( | NA | 16.53893 | 0.027976 | 0.95471 | 0.994798 |
| ENSMUSG( | NA | 138.8775 | -0.01217 | 0.954412 | 0.994798 |
| ENSMUSG( | NA | 12.85638 | -0.57741 | 0.342013 | 0.909146 |
| ENSMUSG( | NA | 92.71478 | -0.16093 | 0.506363 | 0.927825 |
| ENSMUSG( | NA | 85.37158 | -0.16656 | 0.50036 | 0.927825 |
| ENSMUSG( | NA | 13.58288 | -1.16985 | 0.037278 | 0.673514 |
| ENSMUSG( | NA | 12.45021 | 0.820506 | 0.154419 | 0.837724 |
| ENSMUSG( | NA | 34.41649 | 0.039641 | 0.90654 | 0.991494 |
| ENSMUSG( | NA | 12.54924 | 0.352111 | 0.578615 | 0.943548 |
| ENSMUSG( | NA | 60.13088 | 0.270232 | 0.46111 | 0.923329 |
| ENSMUSG( | NA | 1021.551 | 0.093531 | 0.659198 | 0.956205 |
| ENSMUSG( | NA | 230.0639 | -0.3461 | 0.042516 | 0.690304 |
| ENSMUSG( | NA | 16.67017 | -0.51639 | 0.298306 | 0.898538 |
| ENSMUSG( | NA | 12.61656 | -0.10543 | 0.85301 | 0.985598 |
| ENSMUSG( | NA | 206.7455 | 0.123333 | 0.487239 | 0.92541 |
| ENSMUSG( | NA | 21.19277 | -0.14972 | 0.738502 | 0.96799 |
| ENSMUSG( | NA | 9.691789 | 0.299565 | 0.663227 | NA |
| ENSMUSG( | NA | 138.0473 | -0.137 | 0.495158 | 0.927372 |
| ENSMUSG( | NA | 120.6822 | 0.201625 | 0.329806 | 0.907083 |
| ENSMUSG( | NA | 16.83179 | -0.90451 | 0.064088 | 0.736482 |
| ENSMUSG( | NA | 32.77858 | 0.327513 | 0.429063 | 0.917257 |
| ENSMUSG( | NA | 33.00478 | 0.001525 | 0.996839 | 0.999268 |
| ENSMUSG( | NA | 14.90314 | 0.168074 | 0.742597 | 0.969415 |
| ENSMUSG( | NA | 11.15433 | -0.73428 | 0.207268 | NA |
| ENSMUSG( | NA | 12.39811 | 0.152827 | 0.782292 | 0.976106 |
| ENSMUSG( | NA | 83.86028 | 0.297635 | 0.197862 | 0.862161 |
| ENSMUSG( | NA | 23.11415 | -5.08082 | 8.90E-10 | 8.66E-07 |
| ENSMUSG( | NA | 443.2544 | -3.46801 | 2.18E-07 | 0.000135 |
| ENSMUSG( | NA | 10.19466 | -5.74296 | 1.04E-05 | NA |
| ENSMUSG( | NA | 86.26316 | 0.032798 | 0.912862 | 0.992298 |
| ENSMUSG( | NA | 29.28437 | 0.032621 | 0.928631 | 0.99306 |
| ENSMUSG( | NA | 26.0493 | 0.27523 | 0.493579 | 0.927372 |
| ENSMUSG( | NA | 277.8132 | 1.145307 | 0.017597 | 0.53154 |
| ENSMUSG( | NA | 16.55708 | -0.31454 | 0.528193 | 0.934677 |
| ENSMUSG( | NA | 33.7588 | 0.08307 | 0.830812 | 0.98051 |
| ENSMUSG( | NA | 31.17766 | -0.07776 | 0.833977 | 0.981237 |
| ENSMUSG( | NA | 37.87468 | -0.07796 | 0.83416 | 0.981237 |
| ENSMUSG( | NA | 9.530536 | -0.24294 | 0.693292 | NA |
| ENSMUSG( | NA | 18.49345 | 0.116124 | 0.794729 | 0.97681 |
| ENSMUSG( | NA | 38.01082 | 0.124829 | 0.713107 | 0.964595 |
| ENSMUSG( | NA | 50.33396 | -0.4407 | 0.123991 | 0.814691 |
| ENSMUSG( | NA | 14.21802 | -0.59459 | 0.306649 | 0.898538 |
| ENSMUSG( | NA | 77.26593 | -0.35789 | 0.141198 | 0.830678 |
| ENSMUSG( | NA | 10.90045 | -0.0082 | 0.990909 | NA |
| ENSMUSG( | NA | 33.8404 | -0.02741 | 0.942621 | 0.994774 |
| ENSMUSG( | NA | 88.66583 | 0.173709 | 0.478786 | 0.923329 |
| ENSMUSG( | NA | 21.35451 | -0.11409 | 0.816488 | 0.979198 |
| ENSMUSG( | NA | 82.73292 | -0.38878 | 0.161604 | 0.837724 |
| ENSMUSG( | NA | 8.602864 | -0.25231 | 0.708363 | NA |
| ENSMUSG( | NA | 513.2726 | -0.04897 | 0.708317 | 0.964042 |
| ENSMUSG( | NA | 20.80966 | -0.66727 | 0.156754 | 0.837724 |
| ENSMUSG( | NA | 297.2576 | 0.021222 | 0.902883 | 0.991089 |
| ENSMUSG( | NA | 37.24948 | -0.13395 | 0.706984 | 0.963529 |
| ENSMUSG( | NA | 14.25635 | 0.133634 | 0.80064 | 0.97681 |
| ENSMUSG( | NA | 122.6149 | -0.19692 | 0.409272 | 0.917257 |
| ENSMUSG( | NA | 22.65783 | -0.03087 | 0.940625 | 0.99445 |
| ENSMUSG( | NA | 32.92503 | 0.524582 | 0.270823 | 0.888062 |
| ENSMUSG( | NA | 27.39668 | -0.6824 | 0.113733 | 0.809746 |
| ENSMUSG( | NA | 11.5688 | -0.70578 | 0.265042 | 0.886134 |
| ENSMUSG( | NA | 16.3607 | -0.75978 | 0.139878 | 0.830678 |
| ENSMUSG( | NA | 14.67506 | 0.012842 | 0.981363 | 0.996756 |
| ENSMUSG( | NA | 11.73582 | -0.47258 | 0.433696 | 0.918564 |
| ENSMUSG( | NA | 13.37833 | 0.457276 | 0.395863 | 0.917257 |
| ENSMUSG( | NA | 432.4103 | -0.19299 | 0.196188 | 0.862161 |
| ENSMUSG( | NA | 444.0515 | -0.06464 | 0.666942 | 0.956766 |
| ENSMUSG( | NA | 16.83111 | 0.695919 | 0.163987 | 0.838291 |
| ENSMUSG( | NA | 43.17955 | 0.002125 | 0.994575 | 0.998813 |
| ENSMUSG( | NA | 34.61798 | 0.197334 | 0.561218 | 0.940254 |
| ENSMUSG( | NA | 369.2445 | 0.277183 | 0.056802 | 0.723362 |
| ENSMUSG( | NA | 11.05817 | 0.207789 | 0.75784 | NA |
| ENSMUSG( | NA | 57.30165 | -0.05472 | 0.844491 | 0.983642 |
| ENSMUSG( | NA | 12.83706 | -0.17435 | 0.768398 | 0.974236 |
| ENSMUSG( | NA | 22.62474 | -0.03529 | 0.932301 | 0.99306 |
| ENSMUSG( | NA | 18.57391 | -0.50197 | 0.269766 | 0.887508 |
| ENSMUSG( | NA | 13.53278 | -1.16974 | 0.042388 | 0.690304 |
| ENSMUSG( | NA | 50.50381 | 0.006595 | 0.982059 | 0.996756 |
| ENSMUSG( | NA | 27.30274 | -0.0488 | 0.89962 | 0.990558 |
| ENSMUSG( | NA | 40.2037 | 0.099826 | 0.755059 | 0.971098 |
| ENSMUSG( | NA | 19.4475 | 0.007423 | 0.987466 | 0.99751 |
| ENSMUSG( | NA | 83.24644 | -0.04083 | 0.870428 | 0.988445 |
| ENSMUSG( | NA | 16.45551 | -0.8753 | 0.085695 | 0.779799 |
| ENSMUSG( | NA | 11.95289 | -0.36209 | 0.529033 | 0.934795 |
| ENSMUSG( | NA | 14.80687 | 0.193551 | 0.723621 | 0.966887 |
| ENSMUSG( | NA | 29.94897 | 0.338702 | 0.372449 | 0.915491 |
| ENSMUSG( | NA | 7.955201 | -0.2983 | 0.669143 | NA |
| ENSMUSG( | NA | 132.1701 | -0.23449 | 0.25682 | 0.883082 |
| ENSMUSG( | NA | 213.521 | 0.397386 | 0.108233 | 0.804969 |
| ENSMUSG( | NA | 58.9594 | 0.036372 | 0.897386 | 0.990558 |
| ENSMUSG( | NA | 41.28256 | 0.188726 | 0.617684 | 0.952043 |
| ENSMUSG( | NA | 106.749 | 0.005662 | 0.981967 | 0.996756 |
| ENSMUSG( | NA | 24.87412 | 0.136901 | 0.755369 | 0.971141 |
| ENSMUSG( | NA | 38.23988 | -0.19503 | 0.556178 | 0.939565 |
| ENSMUSG( | NA | 23.94525 | -0.34855 | 0.448008 | 0.922202 |
| ENSMUSG( | NA | 10.52877 | -0.44166 | 0.479675 | NA |
| ENSMUSG( | NA | 54.67084 | -0.21609 | 0.456247 | 0.923329 |
| ENSMUSG( | NA | 10.77688 | -0.69137 | 0.277768 | NA |
| ENSMUSG( | NA | 16.51817 | -0.34709 | 0.483485 | 0.92499 |
| ENSMUSG( | NA | 22.98159 | 0.176046 | 0.703603 | 0.962481 |
| ENSMUSG( | NA | 19.67361 | 0.243066 | 0.615318 | 0.951177 |
| ENSMUSG( | NA | 25.35692 | -0.83813 | 0.035006 | 0.659157 |
| ENSMUSG( | NA | 38.84409 | -0.31629 | 0.364502 | 0.914223 |
| ENSMUSG( | NA | 50.90176 | -0.18376 | 0.54632 | 0.938063 |
| ENSMUSG( | NA | 6.362387 | -0.38168 | 0.655149 | NA |
| ENSMUSG( | NA | 17.26143 | 0.540581 | 0.265624 | 0.886134 |
| ENSMUSG( | NA | 9.788286 | -0.15737 | 0.821081 | NA |
| ENSMUSG( | NA | 12.31617 | -0.45397 | 0.459009 | 0.923329 |
| ENSMUSG( | NA | 161.711 | -0.01529 | 0.932958 | 0.99306 |
| ENSMUSG( | NA | 13.32905 | -0.27153 | 0.644588 | 0.953903 |
| ENSMUSG( | NA | 15.06837 | 0.585605 | 0.246088 | 0.877904 |
| ENSMUSG( | NA | 20.60155 | -0.12527 | 0.796135 | 0.97681 |
| ENSMUSG( | NA | 30.21401 | -0.53223 | 0.161074 | 0.837724 |
| ENSMUSG( | NA | 337.9193 | 0.133531 | 0.435321 | 0.919062 |
| ENSMUSG( | NA | 343.1162 | -0.00321 | 0.984536 | 0.996963 |
| ENSMUSG( | NA | 20.00976 | -0.13005 | 0.769341 | 0.974236 |
| ENSMUSG( | NA | 11.91829 | -0.65111 | 0.26401 | 0.886134 |
| ENSMUSG( | NA | 26.00295 | 0.276348 | 0.471433 | 0.923329 |
| ENSMUSG( | NA | 9.324175 | -0.56652 | 0.371203 | NA |
| ENSMUSG( | NA | 19.10943 | 0.281841 | 0.539169 | 0.937216 |
| ENSMUSG( | NA | 18.73743 | -0.38528 | 0.420743 | 0.917257 |
| ENSMUSG( | NA | 13.63405 | 0.015834 | 0.977227 | 0.996416 |
| ENSMUSG( | NA | 24.85635 | -0.03628 | 0.930497 | 0.99306 |
| ENSMUSG( | NA | 10.31813 | -0.06605 | 0.915351 | NA |
| ENSMUSG( | NA | 11.76304 | -0.38146 | 0.497818 | 0.927825 |
| ENSMUSG( | NA | 25.26468 | -0.25042 | 0.561856 | 0.940388 |
| ENSMUSG( | NA | 180.1834 | -0.30691 | 0.106238 | 0.804969 |
| ENSMUSG( | NA | 331.8118 | 0.104358 | 0.564847 | 0.941206 |
| ENSMUSG( | NA | 23.09034 | 0.438447 | 0.30873 | 0.898538 |
| ENSMUSG( | NA | 115.9358 | -0.47799 | 0.039579 | 0.68111 |
| ENSMUSG( | NA | 14.85704 | -0.79777 | 0.120167 | 0.813384 |
| ENSMUSG( | NA | 21.1334 | -0.09107 | 0.853725 | 0.98563 |
| ENSMUSG( | NA | 39.15123 | -0.14547 | 0.660611 | 0.956205 |
| ENSMUSG( | NA | 15.79253 | 0.014772 | 0.976681 | 0.996416 |
| ENSMUSG( | NA | 35.21879 | 0.061316 | 0.887809 | 0.990399 |
| ENSMUSG( | NA | 19.07145 | 0.399614 | 0.457135 | 0.923329 |
| ENSMUSG( | NA | 9.51504 | -0.19316 | 0.764135 | NA |
| ENSMUSG( | NA | 20.33173 | -0.34487 | 0.425317 | 0.917257 |
| ENSMUSG( | NA | 15.69606 | 0.451169 | 0.35341 | 0.912263 |
| ENSMUSG( | NA | 86.68377 | 0.506795 | 0.110744 | 0.807582 |
| ENSMUSG( | NA | 24.37313 | 0.161373 | 0.691026 | 0.960071 |
| ENSMUSG( | NA | 121.495 | -0.42872 | 0.051342 | 0.711116 |
| ENSMUSG( | NA | 23.86687 | 0.20597 | 0.608216 | 0.950113 |
| ENSMUSG( | NA | 77.96955 | 0.132709 | 0.648131 | 0.95422 |
| ENSMUSG( | NA | 400.0778 | -0.2431 | 0.094523 | 0.796133 |
| ENSMUSG( | NA | 35.28379 | 0.023709 | 0.94981 | 0.994798 |
| ENSMUSG( | NA | 8.809161 | 0.152373 | 0.822734 | NA |
| ENSMUSG( | NA | 14.48743 | -0.16578 | 0.749228 | 0.96998 |
| ENSMUSG( | NA | 18.1851 | -0.10308 | 0.845706 | 0.984269 |
| ENSMUSG( | NA | 17.34952 | -0.56479 | 0.246699 | 0.877904 |
| ENSMUSG( | NA | 35.61776 | -0.13005 | 0.710285 | 0.964363 |
| ENSMUSG( | NA | 20.51514 | -0.27736 | 0.551145 | 0.939377 |
| ENSMUSG( | NA | 17.71974 | 0.793434 | 0.108875 | 0.804969 |
| ENSMUSG( | NA | 24.04074 | 0.435142 | 0.30769 | 0.898538 |
| ENSMUSG( | NA | 218.9856 | 0.377171 | 0.020976 | 0.572337 |
| ENSMUSG( | NA | 45.15293 | -0.01343 | 0.965872 | 0.995433 |
| ENSMUSG( | NA | 38.75156 | -0.20685 | 0.558033 | 0.939565 |
| ENSMUSG( | NA | 67.63674 | -0.09151 | 0.721513 | 0.966887 |
| ENSMUSG( | NA | 27.53936 | 0.000697 | 0.998664 | 0.999406 |

| | | | | | |
|---|---|---|---|---|---|
| ENSMUSG( | 15.46578 | -0.02521 | 0.957314 | NA | NA |
| ENSMUSG( | 46.71455 | 0.119869 | 0.672149 | 0.9288 | NA |
| ENSMUSG( | 23.28122 | -0.59212 | 0.13788 | 0.552903 | NA |
| ENSMUSG( | 12.87906 | 0.347946 | 0.535941 | NA | NA |
| ENSMUSG( | 25.25999 | 0.194369 | 0.641063 | 0.916692 | NA |
| ENSMUSG( | 18.32151 | -0.23207 | 0.630539 | NA | NA |
| ENSMUSG( | 33.84197 | 0.283677 | 0.408529 | 0.818167 | NA |
| ENSMUSG( | 17.71267 | -0.21368 | 0.653144 | NA | NA |
| ENSMUSG( | 17.25965 | -0.42785 | 0.361333 | NA | NA |
| ENSMUSG( | 13.60317 | 1.279953 | 0.019848 | NA | NA |
| ENSMUSG( | 55.88965 | 1.165864 | 4.99E-05 | 0.00395 | NA |
| ENSMUSG( | 68.90958 | 0.012814 | 0.956429 | 0.993874 | NA |
| ENSMUSG( | 15.65189 | 0.149731 | 0.795533 | NA | NA |
| ENSMUSG( | 72.8735 | -1.41401 | 1.37E-06 | 0.000244 | NA |
| ENSMUSG( | 14.32994 | 0.177799 | 0.72912 | NA | NA |
| ENSMUSG( | 14.34631 | -0.18214 | 0.712114 | NA | NA |
| ENSMUSG( | 350.7158 | -0.31649 | 0.045768 | 0.328083 | NA |
| ENSMUSG( | 29.01034 | 0.120774 | 0.731568 | 0.944956 | NA |
| ENSMUSG( | 31.43993 | -0.02768 | 0.936757 | 0.993415 | NA |
| ENSMUSG( | 30.36913 | -0.49099 | 0.182363 | 0.619591 | NA |
| ENSMUSG( | 159.5494 | 0.048533 | 0.786573 | 0.958856 | NA |
| ENSMUSG( | 34.58048 | 0.057532 | 0.869737 | 0.977607 | NA |
| ENSMUSG( | 33.59738 | -0.05904 | 0.86383 | 0.975244 | NA |
| ENSMUSG( | 21.40306 | -0.45652 | 0.256235 | 0.706924 | NA |
| ENSMUSG( | 25.05953 | -0.02511 | 0.946607 | 0.993731 | NA |
| ENSMUSG( | 20.05486 | -0.20799 | 0.623316 | NA | NA |
| ENSMUSG( | 16.35168 | 0.260226 | 0.650547 | NA | NA |
| ENSMUSG( | 53.37305 | 0.120446 | 0.688792 | 0.933441 | NA |
| ENSMUSG( | 63.3506 | 0.387293 | 0.135349 | 0.548884 | NA |
| ENSMUSG( | 14.31515 | -0.16537 | 0.73765 | NA | NA |
| ENSMUSG( | 75.86832 | 0.035343 | 0.874157 | 0.979336 | NA |
| ENSMUSG( | 89.14652 | -0.00924 | 0.969031 | 0.996369 | NA |
| ENSMUSG( | 15.82824 | -0.38621 | 0.419433 | NA | NA |
| ENSMUSG( | 41.69278 | 0.099861 | 0.755997 | 0.951479 | NA |
| ENSMUSG( | 26.37245 | -0.34966 | 0.33488 | 0.776409 | NA |
| ENSMUSG( | 10.35677 | -0.22989 | 0.688552 | NA | NA |
| ENSMUSG( | 16.41246 | 0.114901 | 0.802384 | NA | NA |
| ENSMUSG( | 40.61638 | -0.12133 | 0.686206 | 0.932981 | NA |
| ENSMUSG( | 45.7351 | -0.30116 | 0.289017 | 0.739705 | NA |
| ENSMUSG( | 21.83024 | -0.2694 | 0.53378 | 0.880095 | NA |
| ENSMUSG( | 54.89178 | -1.16918 | 2.28E-05 | 0.002131 | NA |
| ENSMUSG( | 11.3788 | -0.44685 | 0.43043 | NA | NA |
| ENSMUSG( | 47.59336 | 0.118221 | 0.667662 | 0.927484 | NA |
| ENSMUSG( | 9.435912 | 0.529282 | 0.391864 | NA | NA |
| ENSMUSG( | 31.85257 | -0.13119 | 0.692543 | 0.934101 | NA |
| ENSMUSG( | 25.1329 | 0.087245 | 0.82045 | 0.965888 | NA |
| ENSMUSG( | 40.94011 | -0.02777 | 0.930835 | 0.992059 | NA |
| ENSMUSG( | 16.30587 | -0.25422 | 0.577406 | NA | NA |
| ENSMUSG( | 11.32042 | -0.41471 | 0.451 | NA | NA |
| ENSMUSG( | 24.50207 | 0.359764 | 0.340639 | 0.778521 | NA |
| ENSMUSG( | 17.86038 | 0.162125 | 0.73365 | NA | NA |
| ENSMUSG( | 10.34829 | -0.86661 | 0.154463 | NA | NA |
| ENSMUSG( | 237.1708 | -0.22553 | 0.116294 | 0.514678 | NA |
| ENSMUSG( | 92.87299 | 0.107626 | 0.601391 | 0.904603 | NA |
| ENSMUSG( | 9.689409 | -0.51132 | 0.427896 | NA | NA |
| ENSMUSG( | 8.030999 | -0.61462 | 0.390176 | NA | NA |
| ENSMUSG( | 19.26524 | 0.356015 | 0.415185 | NA | NA |
| ENSMUSG( | 13.79896 | 0.212396 | 0.661604 | NA | NA |
| ENSMUSG( | 146.6197 | -0.11492 | 0.556152 | 0.887523 | NA |
| ENSMUSG( | 14.69445 | 0.492746 | 0.348946 | NA | NA |
| ENSMUSG( | 19.25161 | 0.250619 | 0.585269 | NA | NA |
| ENSMUSG( | 10.66924 | 0.40942 | 0.47657 | NA | NA |
| ENSMUSG( | 28.92905 | 0.519581 | 0.141133 | 0.558043 | NA |
| ENSMUSG( | 10.07462 | 0.192747 | 0.758779 | NA | NA |
| ENSMUSG( | 31.15745 | -0.29381 | 0.443837 | 0.833239 | NA |
| ENSMUSG( | 8.7552 | -0.26951 | 0.699879 | NA | NA |
| ENSMUSG( | 28.22545 | 0.02833 | 0.936173 | 0.993415 | NA |
| ENSMUSG( | 26.65197 | 0.091096 | 0.828258 | 0.968966 | NA |
| ENSMUSG( | 11.30136 | 0.680158 | 0.222932 | NA | NA |
| ENSMUSG( | 342.9353 | 0.004432 | 0.977294 | 0.997428 | NA |
| ENSMUSG( | 11.45732 | -0.64122 | 0.281767 | NA | NA |
| ENSMUSG( | 16.39976 | 0.493642 | 0.313708 | NA | NA |
| ENSMUSG( | 13.0238 | 0.042717 | 0.93727 | NA | NA |
| ENSMUSG( | 50.4974 | 0.08027 | 0.769514 | 0.954674 | NA |
| ENSMUSG( | 154.6557 | -0.07663 | 0.659142 | 0.924506 | NA |
| ENSMUSG( | 230.6057 | -0.09795 | 0.525871 | 0.876902 | NA |
| ENSMUSG( | 86.64182 | 0.108592 | 0.622424 | 0.911432 | NA |
| ENSMUSG( | 15.05936 | 0.83952 | 0.079483 | NA | NA |
| ENSMUSG( | 162.1033 | 0.171642 | 0.328252 | 0.772854 | NA |
| ENSMUSG( | 21.58302 | 0.489126 | 0.232194 | 0.67996 | NA |
| ENSMUSG( | 62.34033 | 0.051387 | 0.854296 | 0.973491 | NA |
| ENSMUSG( | 356.1105 | -0.02034 | 0.87334 | 0.978894 | NA |
| ENSMUSG( | 30.76227 | 0.341502 | 0.321932 | 0.766887 | NA |
| ENSMUSG( | 11.95977 | -0.2344 | 0.65667 | NA | NA |
| ENSMUSG( | 15.55657 | 0.245377 | 0.618542 | NA | NA |
| ENSMUSG( | 12.89366 | -0.15109 | 0.768431 | NA | NA |
| ENSMUSG( | 47.65213 | -0.2315 | 0.436985 | 0.829501 | NA |
| ENSMUSG( | 10.27697 | -0.44193 | 0.456468 | NA | NA |
| ENSMUSG( | 41.59516 | -0.43126 | 0.155609 | 0.583641 | NA |
| ENSMUSG( | 91.19284 | -0.20963 | 0.333493 | 0.775602 | NA |
| ENSMUSG( | 37.84461 | -0.07207 | 0.840396 | 0.971179 | NA |
| ENSMUSG( | 23.56253 | -0.2638 | 0.532773 | 0.879905 | NA |
| ENSMUSG( | 10.19423 | 0.65313 | 0.258888 | NA | NA |
| ENSMUSG( | 11.94276 | -0.20443 | 0.747686 | NA | NA |
| ENSMUSG( | 8.622283 | 0.253217 | 0.693967 | NA | NA |
| ENSMUSG( | 26.96121 | 0.108091 | 0.763329 | 0.953781 | NA |
| ENSMUSG( | 131.9938 | 0.179941 | 0.365659 | 0.794294 | NA |
| ENSMUSG( | 68.40518 | 0.366359 | 0.26351 | 0.714284 | NA |
| ENSMUSG( | 191.0927 | 0.070554 | 0.704723 | 0.938303 | NA |
| ENSMUSG( | 69.35427 | -0.1896 | 0.474314 | 0.849425 | NA |
| ENSMUSG( | 18.73784 | -0.24045 | 0.610718 | NA | NA |
| ENSMUSG( | 14.24581 | 0.033613 | 0.944266 | NA | NA |
| ENSMUSG( | 19.86337 | 0.198256 | 0.65858 | NA | NA |
| ENSMUSG( | 14.06182 | -0.00942 | 0.986112 | NA | NA |
| ENSMUSG( | 114.909 | 0.047454 | 0.813482 | 0.964574 | NA |
| ENSMUSG( | 26.30759 | -0.04855 | 0.893922 | 0.984512 | NA |
| ENSMUSG( | 74.76709 | 0.275823 | 0.243779 | 0.691156 | NA |
| ENSMUSG( | 42.85727 | -0.19437 | 0.532394 | 0.879905 | NA |
| ENSMUSG( | 25.56926 | -0.11246 | 0.766187 | 0.954173 | NA |
| ENSMUSG( | 19.19957 | -0.3983 | 0.380464 | NA | NA |
| ENSMUSG( | 38.86109 | 0.02701 | 0.934101 | 0.99323 | NA |
| ENSMUSG( | 118.471 | -0.13158 | 0.545222 | 0.884748 | NA |
| ENSMUSG( | 24.25864 | -0.1628 | 0.661666 | 0.925271 | NA |
| ENSMUSG( | 27.0525 | -0.06673 | 0.864038 | 0.975331 | NA |
| ENSMUSG( | 108.0181 | 0.111329 | 0.611402 | 0.908586 | NA |
| ENSMUSG( | 26.64175 | -0.21511 | 0.57283 | 0.893584 | NA |
| ENSMUSG( | 13.72068 | 0.275516 | 0.580901 | NA | NA |
| ENSMUSG( | 7.639078 | 0.020625 | 0.976875 | NA | NA |
| ENSMUSG( | 37.78482 | -0.19679 | 0.551823 | 0.886141 | NA |
| ENSMUSG( | 14.75461 | -0.03688 | 0.938907 | NA | NA |
| ENSMUSG( | 9.715822 | -0.18173 | 0.764397 | NA | NA |
| ENSMUSG( | 37.25806 | 0.105914 | 0.748983 | 0.949604 | NA |
| ENSMUSG( | 70.43116 | -0.23641 | 0.300405 | 0.749512 | NA |
| ENSMUSG( | 17.02719 | 0.008646 | 0.984948 | NA | NA |
| ENSMUSG( | 9.995776 | -0.8608 | 0.151924 | NA | NA |
| ENSMUSG( | 28.02315 | 0.3751 | 0.307352 | 0.755324 | NA |
| ENSMUSG( | 48.11856 | 0.246642 | 0.410381 | 0.818339 | NA |
| ENSMUSG( | 143.5591 | -0.19943 | 0.352151 | 0.783738 | NA |
| ENSMUSG( | 10.66449 | 0.540172 | 0.355667 | NA | NA |
| ENSMUSG( | 10.49364 | 0.369441 | 0.533025 | NA | NA |
| ENSMUSG( | 62.87149 | 0.303767 | 0.211983 | 0.655675 | NA |
| ENSMUSG( | 10.74838 | 0.750801 | 0.21172 | NA | NA |
| ENSMUSG( | 21.58537 | -0.18794 | 0.664745 | 0.926824 | NA |
| ENSMUSG( | 8.842697 | -0.15118 | 0.81629 | NA | NA |
| ENSMUSG( | 34.02041 | -0.05341 | 0.883139 | 0.982422 | NA |
| ENSMUSG( | 34.74375 | 0.026279 | 0.934033 | 0.99323 | NA |
| ENSMUSG( | 95.15895 | 0.094215 | 0.701204 | 0.93743 | NA |
| ENSMUSG( | 12.49646 | -0.04106 | 0.938497 | NA | NA |
| ENSMUSG( | 66.30669 | 0.043766 | 0.868207 | 0.976786 | NA |
| ENSMUSG( | 9.644401 | -0.27862 | 0.645779 | NA | NA |
| ENSMUSG( | 11.04699 | -0.11219 | 0.840631 | NA | NA |
| ENSMUSG( | 55.73599 | 0.301462 | 0.247814 | 0.696522 | NA |
| ENSMUSG( | 10.4319 | -0.15934 | 0.786377 | NA | NA |
| ENSMUSG( | 9.254259 | -0.58022 | 0.384116 | NA | NA |
| ENSMUSG( | 51.58038 | 0.135116 | 0.616152 | 0.91085 | NA |
| ENSMUSG( | 178.3539 | -0.03735 | 0.839761 | 0.971179 | NA |
| ENSMUSG( | 427.6746 | -0.43087 | 0.004414 | 0.086676 | NA |
| ENSMUSG( | 16.08038 | 0.148704 | 0.748443 | NA | NA |
| ENSMUSG( | 23.13541 | 0.50925 | 0.217965 | 0.663176 | NA |
| ENSMUSG( | 23.80639 | -0.2918 | 0.487377 | 0.858812 | NA |
| ENSMUSG( | 19.63127 | 0.387048 | 0.351206 | NA | NA |
| ENSMUSG( | 17.93999 | -0.29581 | 0.504991 | NA | NA |
| ENSMUSG( | 10.65903 | 0.284951 | 0.638292 | NA | NA |
| ENSMUSG( | 47.71894 | -0.11654 | 0.692307 | 0.934101 | NA |
| ENSMUSG( | 12.03235 | 0.429483 | 0.437041 | NA | NA |
| ENSMUSG( | 38.34844 | -0.33366 | 0.280107 | 0.729155 | NA |
| ENSMUSG( | 58.80066 | 0.081609 | 0.7592 | 0.952284 | NA |
| ENSMUSG( | 514.509 | 0.101111 | 0.451078 | 0.835567 | NA |

| | | | | | |
|---|---|---|---|---|---|
| ENSMUSG( | 33.67244 | 0.083831 | 0.814449 | 0.979198 | NA |
| ENSMUSG( | 22.0211 | -0.16161 | 0.714755 | 0.964986 | NA |
| ENSMUSG( | 46.66096 | -0.07202 | 0.82006 | 0.979198 | NA |
| ENSMUSG( | 10.97064 | -0.18655 | 0.75013 | NA | NA |
| ENSMUSG( | 25.91826 | 0.1573 | 0.729689 | 0.96719 | NA |
| ENSMUSG( | 106.534 | -0.32981 | 0.137326 | 0.828492 | NA |
| ENSMUSG( | 17.94981 | 0.307324 | 0.546348 | 0.938063 | NA |
| ENSMUSG( | 15.9713 | -0.46985 | 0.33474 | 0.907907 | NA |
| ENSMUSG( | 16.53893 | 0.027976 | 0.95471 | 0.994798 | NA |
| ENSMUSG( | 138.8775 | -0.01217 | 0.954412 | 0.994798 | NA |
| ENSMUSG( | 12.85638 | -0.57741 | 0.342013 | 0.909146 | NA |
| ENSMUSG( | 92.71478 | -0.16093 | 0.506363 | 0.927825 | NA |
| ENSMUSG( | 85.37158 | -0.16656 | 0.50036 | 0.927825 | NA |
| ENSMUSG( | 13.58288 | -1.16985 | 0.037278 | 0.673514 | NA |
| ENSMUSG( | 12.45021 | 0.820506 | 0.154419 | 0.837724 | NA |
| ENSMUSG( | 34.41649 | 0.039641 | 0.90654 | 0.991494 | NA |
| ENSMUSG( | 12.54924 | 0.352111 | 0.578615 | 0.943548 | NA |
| ENSMUSG( | 60.13088 | 0.270232 | 0.46111 | 0.923329 | NA |
| ENSMUSG( | 1021.551 | 0.093531 | 0.659198 | 0.956205 | NA |
| ENSMUSG( | 230.0639 | -0.3461 | 0.042516 | 0.690304 | NA |
| ENSMUSG( | 16.67017 | -0.51639 | 0.298306 | 0.898538 | NA |
| ENSMUSG( | 12.61656 | -0.10543 | 0.85301 | 0.985598 | NA |
| ENSMUSG( | 206.7455 | 0.123333 | 0.487239 | 0.92541 | NA |
| ENSMUSG( | 9.032315 | 0.054534 | 0.931346 | NA | NA |
| ENSMUSG( | 21.19277 | -0.14972 | 0.738502 | 0.96799 | NA |
| ENSMUSG( | 9.691789 | 0.299565 | 0.663227 | NA | NA |
| ENSMUSG( | 138.0473 | -0.137 | 0.495158 | 0.927372 | NA |
| ENSMUSG( | 120.6822 | 0.201625 | 0.329806 | 0.907083 | NA |
| ENSMUSG( | 16.83179 | -0.90451 | 0.064088 | 0.736482 | NA |
| ENSMUSG( | 32.77858 | 0.327513 | 0.429063 | 0.917257 | NA |
| ENSMUSG( | 33.00478 | 0.001525 | 0.996839 | 0.999268 | NA |
| ENSMUSG( | 14.90314 | 0.168074 | 0.742597 | 0.969415 | NA |
| ENSMUSG( | 11.15433 | -0.73428 | 0.207268 | NA | NA |
| ENSMUSG( | 12.39811 | 0.152827 | 0.782292 | 0.976106 | NA |
| ENSMUSG( | 83.86028 | 0.297635 | 0.197862 | 0.862161 | NA |
| ENSMUSG( | 23.11415 | -5.08082 | 8.90E-10 | 8.66E-07 | NA |
| ENSMUSG( | 443.2544 | -3.46801 | 2.18E-07 | 0.000135 | NA |
| ENSMUSG( | 10.19466 | -5.74296 | 1.04E-05 | NA | NA |
| ENSMUSG( | 86.26316 | 0.032798 | 0.912862 | 0.992298 | NA |
| ENSMUSG( | 29.28437 | 0.032621 | 0.928631 | 0.99306 | NA |
| ENSMUSG( | 26.0493 | 0.27523 | 0.493579 | 0.927372 | NA |
| ENSMUSG( | 277.8132 | 1.145307 | 0.017597 | 0.53154 | NA |
| ENSMUSG( | 16.55708 | -0.31454 | 0.528193 | 0.934677 | NA |
| ENSMUSG( | 33.7588 | 0.08307 | 0.830812 | 0.98051 | NA |
| ENSMUSG( | 31.17766 | -0.07776 | 0.833977 | 0.981237 | NA |
| ENSMUSG( | 37.87468 | -0.07796 | 0.83416 | 0.981237 | NA |
| ENSMUSG( | 9.530536 | -0.24294 | 0.693292 | NA | NA |
| ENSMUSG( | 18.49345 | 0.116124 | 0.794729 | 0.97681 | NA |
| ENSMUSG( | 38.01082 | 0.124829 | 0.713107 | 0.964595 | NA |
| ENSMUSG( | 50.33396 | -0.4407 | 0.123991 | 0.814691 | NA |
| ENSMUSG( | 14.21802 | -0.59459 | 0.306649 | 0.898538 | NA |
| ENSMUSG( | 77.26593 | -0.35789 | 0.141198 | 0.830678 | NA |
| ENSMUSG( | 10.90045 | -0.0082 | 0.990909 | NA | NA |
| ENSMUSG( | 33.8404 | -0.02741 | 0.942621 | 0.994774 | NA |
| ENSMUSG( | 88.66583 | 0.173709 | 0.478786 | 0.923329 | NA |
| ENSMUSG( | 21.35451 | -0.11409 | 0.816488 | 0.979198 | NA |
| ENSMUSG( | 82.73292 | -0.38878 | 0.161604 | 0.837724 | NA |
| ENSMUSG( | 8.602864 | -0.25231 | 0.708363 | NA | NA |
| ENSMUSG( | 513.2726 | -0.04897 | 0.708317 | 0.964042 | NA |
| ENSMUSG( | 20.80966 | -0.66727 | 0.156754 | 0.837724 | NA |
| ENSMUSG( | 297.2576 | 0.021222 | 0.902883 | 0.991089 | NA |
| ENSMUSG( | 37.24948 | -0.13395 | 0.706984 | 0.963529 | NA |
| ENSMUSG( | 14.25635 | 0.133634 | 0.80064 | 0.97681 | NA |
| ENSMUSG( | 122.6149 | -0.19692 | 0.409272 | 0.917257 | NA |
| ENSMUSG( | 22.65783 | -0.03087 | 0.940625 | 0.99445 | NA |
| ENSMUSG( | 32.92503 | 0.524582 | 0.270823 | 0.888062 | NA |
| ENSMUSG( | 27.39668 | -0.6824 | 0.113733 | 0.809746 | NA |
| ENSMUSG( | 11.5688 | -0.70578 | 0.265042 | 0.886134 | NA |
| ENSMUSG( | 16.3607 | -0.75978 | 0.139878 | 0.830678 | NA |
| ENSMUSG( | 14.67506 | 0.012842 | 0.981363 | 0.996756 | NA |
| ENSMUSG( | 11.73582 | -0.47258 | 0.433696 | 0.918564 | NA |
| ENSMUSG( | 13.37833 | 0.457276 | 0.395863 | 0.917257 | NA |
| ENSMUSG( | 432.4103 | -0.19299 | 0.196188 | 0.862161 | NA |
| ENSMUSG( | 444.0515 | -0.06464 | 0.666942 | 0.956766 | NA |
| ENSMUSG( | 16.83111 | 0.695919 | 0.163987 | 0.838291 | NA |
| ENSMUSG( | 43.17955 | 0.002125 | 0.994575 | 0.998813 | NA |
| ENSMUSG( | 34.61798 | 0.197334 | 0.561218 | 0.940254 | NA |
| ENSMUSG( | 369.2445 | 0.277183 | 0.056802 | 0.723362 | NA |
| ENSMUSG( | 11.05817 | 0.207789 | 0.75784 | NA | NA |
| ENSMUSG( | 57.30165 | -0.05472 | 0.844491 | 0.983642 | NA |
| ENSMUSG( | 12.83706 | -0.17435 | 0.768398 | 0.974236 | NA |
| ENSMUSG( | 22.62474 | -0.03529 | 0.932301 | 0.99306 | NA |
| ENSMUSG( | 18.57391 | -0.50197 | 0.269766 | 0.887508 | NA |
| ENSMUSG( | 13.53278 | -1.16974 | 0.042388 | 0.690304 | NA |
| ENSMUSG( | 50.50381 | 0.006595 | 0.982059 | 0.996756 | NA |
| ENSMUSG( | 27.30274 | -0.0488 | 0.89962 | 0.990558 | NA |
| ENSMUSG( | 40.2037 | 0.099826 | 0.755059 | 0.971098 | NA |
| ENSMUSG( | 19.4475 | 0.007423 | 0.987466 | 0.99751 | NA |
| ENSMUSG( | 83.24644 | -0.04083 | 0.870428 | 0.988445 | NA |
| ENSMUSG( | 16.45551 | -0.8753 | 0.085695 | 0.779799 | NA |
| ENSMUSG( | 11.95289 | -0.36209 | 0.529033 | 0.934795 | NA |
| ENSMUSG( | 14.80687 | 0.193551 | 0.723621 | 0.966887 | NA |
| ENSMUSG( | 29.94897 | 0.338702 | 0.372449 | 0.915491 | NA |
| ENSMUSG( | 7.955201 | -0.2983 | 0.669143 | NA | NA |
| ENSMUSG( | 132.1701 | -0.23449 | 0.25682 | 0.883082 | NA |
| ENSMUSG( | 213.521 | 0.397386 | 0.108233 | 0.804969 | NA |
| ENSMUSG( | 58.9594 | 0.036372 | 0.897386 | 0.990558 | NA |
| ENSMUSG( | 41.28256 | 0.188726 | 0.617684 | 0.952043 | NA |
| ENSMUSG( | 106.749 | 0.005662 | 0.981967 | 0.996756 | NA |
| ENSMUSG( | 24.87412 | 0.136901 | 0.755369 | 0.971141 | NA |
| ENSMUSG( | 38.23988 | -0.19503 | 0.556178 | 0.939565 | NA |
| ENSMUSG( | 23.94525 | -0.34855 | 0.448008 | 0.922202 | NA |
| ENSMUSG( | 10.52877 | -0.44166 | 0.479675 | NA | NA |
| ENSMUSG( | 54.67084 | -0.21609 | 0.456247 | 0.923329 | NA |
| ENSMUSG( | 12.26514 | 0.420948 | 0.445814 | 0.921734 | NA |
| ENSMUSG( | 10.77688 | -0.69137 | 0.277768 | NA | NA |
| ENSMUSG( | 16.51817 | -0.34709 | 0.483485 | 0.92499 | NA |
| ENSMUSG( | 22.98159 | 0.176046 | 0.703603 | 0.962481 | NA |
| ENSMUSG( | 19.67361 | 0.243066 | 0.615318 | 0.951177 | NA |
| ENSMUSG( | 25.35692 | -0.83813 | 0.035006 | 0.659157 | NA |
| ENSMUSG( | 38.84409 | -0.31629 | 0.364502 | 0.914223 | NA |
| ENSMUSG( | 50.90176 | -0.18376 | 0.54632 | 0.938063 | NA |
| ENSMUSG( | 6.362387 | -0.38168 | 0.655149 | NA | NA |
| ENSMUSG( | 9.20508 | 0.539239 | 0.467416 | NA | NA |
| ENSMUSG( | 17.26143 | 0.540581 | 0.265624 | 0.886134 | NA |
| ENSMUSG( | 9.788286 | -0.15737 | 0.821081 | NA | NA |
| ENSMUSG( | 12.31617 | -0.45397 | 0.459009 | 0.923329 | NA |
| ENSMUSG( | 161.711 | -0.01529 | 0.932958 | 0.99306 | NA |
| ENSMUSG( | 13.32905 | -0.27153 | 0.644588 | 0.953903 | NA |
| ENSMUSG( | 15.06837 | 0.585605 | 0.246088 | 0.877904 | NA |
| ENSMUSG( | 20.60155 | -0.12527 | 0.796135 | 0.97681 | NA |
| ENSMUSG( | 30.21401 | -0.53223 | 0.161074 | 0.837724 | NA |
| ENSMUSG( | 337.9193 | 0.133531 | 0.435321 | 0.919062 | NA |
| ENSMUSG( | 343.1162 | -0.00321 | 0.984536 | 0.996963 | NA |
| ENSMUSG( | 20.00976 | -0.13005 | 0.769341 | 0.974236 | NA |
| ENSMUSG( | 8.908866 | 0.132042 | 0.83831 | NA | NA |
| ENSMUSG( | 11.91829 | -0.65111 | 0.26401 | 0.886134 | NA |
| ENSMUSG( | 26.00295 | 0.276348 | 0.471433 | 0.923329 | NA |
| ENSMUSG( | 9.324175 | -0.56652 | 0.371203 | NA | NA |
| ENSMUSG( | 19.10943 | 0.281841 | 0.539169 | 0.937216 | NA |
| ENSMUSG( | 18.73743 | -0.38528 | 0.420743 | 0.917257 | NA |
| ENSMUSG( | 13.63405 | 0.015834 | 0.977227 | 0.996416 | NA |
| ENSMUSG( | 24.85635 | -0.03628 | 0.930497 | 0.99306 | NA |
| ENSMUSG( | 10.31813 | -0.06605 | 0.915351 | NA | NA |
| ENSMUSG( | 11.76304 | -0.38146 | 0.497818 | 0.927825 | NA |
| ENSMUSG( | 25.26468 | -0.25042 | 0.561856 | 0.940388 | NA |
| ENSMUSG( | 180.1834 | -0.30691 | 0.106238 | 0.804969 | NA |
| ENSMUSG( | 331.8118 | 0.104358 | 0.564847 | 0.941206 | NA |
| ENSMUSG( | 23.09034 | 0.438447 | 0.30873 | 0.898538 | NA |
| ENSMUSG( | 115.9358 | -0.47799 | 0.039579 | 0.68111 | NA |
| ENSMUSG( | 14.85704 | -0.79777 | 0.120167 | 0.813384 | NA |
| ENSMUSG( | 21.1334 | -0.09107 | 0.853725 | 0.98563 | NA |
| ENSMUSG( | 39.15123 | -0.14547 | 0.660611 | 0.956205 | NA |
| ENSMUSG( | 15.79253 | 0.014772 | 0.976681 | 0.996416 | NA |
| ENSMUSG( | 35.21879 | 0.061316 | 0.887809 | 0.990399 | NA |
| ENSMUSG( | 19.07145 | 0.399614 | 0.457135 | 0.923329 | NA |
| ENSMUSG( | 9.51504 | -0.19316 | 0.764135 | NA | NA |
| ENSMUSG( | 20.33173 | -0.34487 | 0.425317 | 0.917257 | NA |
| ENSMUSG( | 15.69606 | 0.451169 | 0.35341 | 0.912263 | NA |
| ENSMUSG( | 86.68377 | 0.506795 | 0.110744 | 0.807582 | NA |
| ENSMUSG( | 24.37313 | 0.161373 | 0.691026 | 0.960071 | NA |
| ENSMUSG( | 121.495 | -0.42872 | 0.051342 | 0.711116 | NA |
| ENSMUSG( | 23.86687 | 0.20597 | 0.608216 | 0.950113 | NA |
| ENSMUSG( | 77.96955 | 0.132709 | 0.648131 | 0.95422 | NA |
| ENSMUSG( | 400.0778 | -0.2431 | 0.094523 | 0.796133 | NA |
| ENSMUSG( | 35.28379 | 0.023709 | 0.94981 | 0.994798 | NA |
| ENSMUSG( | 8.809161 | 0.152373 | 0.822734 | NA | NA |
| ENSMUSG( | 14.48743 | -0.16578 | 0.749228 | 0.96998 | NA |

| | | | | | | | | | | |
|---|---|---|---|---|---|---|---|---|---|---|
| ENSMUSG( | NA | 51.16794 | -0.01315 | 0.962394 | 0.994586 | ENSMUSG( | NA | 32.76541 | 0.378856 | 0.340203 | 0.907972 |
| ENSMUSG( | NA | 11.88952 | 0.605775 | 0.270928 | NA | ENSMUSG( | NA | 15.32675 | 0.177621 | 0.722336 | 0.966887 |
| ENSMUSG( | NA | 29.62313 | -0.03319 | 0.93165 | 0.9924 | ENSMUSG( | NA | 401936.9 | -0.09091 | 0.601446 | 0.947749 |
| ENSMUSG( | NA | 887.9032 | 0.091622 | 0.420633 | 0.8229 | ENSMUSG( | NA | 540426.2 | -0.00734 | 0.972156 | 0.996416 |
| ENSMUSG( | NA | 167.3126 | -0.05684 | 0.762756 | 0.953519 | ENSMUSG( | NA | 702.2438 | 0.11919 | 0.599443 | 0.947071 |
| ENSMUSG( | NA | 149.6224 | 0.027765 | 0.877281 | 0.981277 | ENSMUSG( | NA | 256.5104 | 0.525109 | 0.52371 | 0.932848 |
| ENSMUSG( | NA | 12.16281 | -0.19509 | 0.727767 | NA | ENSMUSG( | NA | 108.7975 | 0.012629 | 0.967076 | 0.995549 |
| ENSMUSG( | NA | 23.79064 | -0.10214 | 0.806237 | 0.962804 | ENSMUSG( | NA | 1269.447 | -0.01717 | 0.900997 | 0.990558 |
| ENSMUSG( | NA | 28.80452 | 0.471377 | 0.253443 | 0.703636 | ENSMUSG( | NA | 827.1415 | 0.093466 | 0.881314 | 0.989644 |
| ENSMUSG( | NA | 21.90557 | 0.283034 | 0.478263 | 0.853496 | ENSMUSG( | NA | 26.80678 | 0.027236 | 0.960008 | 0.994798 |
| ENSMUSG( | NA | 14.75381 | 0.258811 | 0.618737 | NA | ENSMUSG( | NA | 44.57035 | -0.48258 | 0.162532 | 0.837724 |
| ENSMUSG( | NA | 26.68348 | 0.642839 | 0.092071 | 0.466032 | ENSMUSG( | NA | 17.51686 | 0.775596 | 0.216394 | 0.865167 |
| ENSMUSG( | NA | 17.68956 | -2.09143 | 0.000189 | NA | ENSMUSG( | NA | 38.07125 | -0.00544 | 0.987442 | 0.99751 |
| ENSMUSG( | NA | 8.289241 | -1.88119 | 0.007571 | NA | ENSMUSG( | NA | 116.3019 | 0.047839 | 0.842009 | 0.983044 |
| ENSMUSG( | NA | 48.79978 | -0.31798 | 0.268511 | 0.718807 | ENSMUSG( | NA | 162.301 | 0.023777 | 0.929615 | 0.99306 |
| ENSMUSG( | NA | 22.91177 | -0.80378 | 0.096845 | 0.477375 | ENSMUSG( | NA | 6900.762 | 0.563905 | 0.016706 | 0.521279 |
| ENSMUSG( | NA | 42.80247 | -0.13992 | 0.646088 | 0.918453 | ENSMUSG( | NA | 727.2601 | 0.016665 | 0.908676 | 0.991958 |
| ENSMUSG( | Naa10 | 365.2369 | -0.12156 | 0.333232 | 0.775602 | ENSMUSG( | Naa10 | 332.3922 | -0.12966 | 0.422572 | 0.917257 |
| ENSMUSG( | Naa15 | 1359.873 | 0.085106 | 0.356774 | 0.788403 | ENSMUSG( | Naa15 | 1237.926 | 0.079422 | 0.536599 | 0.937108 |
| ENSMUSG( | Naa16 | 198.6441 | 0.15641 | 0.331201 | 0.77519 | ENSMUSG( | Naa16 | 195.4067 | 0.36601 | 0.03281 | 0.637876 |
| ENSMUSG( | Naa20 | 696.9992 | -0.02553 | 0.845431 | 0.97218 | ENSMUSG( | Naa20 | 650.6714 | 0.042283 | 0.781988 | 0.976106 |
| ENSMUSG( | Naa25 | 1106.078 | -0.05234 | 0.631368 | 0.914073 | ENSMUSG( | Naa25 | 1066.228 | 0.112462 | 0.405522 | 0.917257 |
| ENSMUSG( | Naa30 | 772.0969 | 0.009858 | 0.940185 | 0.993415 | ENSMUSG( | Naa30 | 734.72 | 0.125032 | 0.488557 | 0.92554 |
| ENSMUSG( | Naa35 | 1327.942 | 0.122586 | 0.191552 | 0.630601 | ENSMUSG( | Naa35 | 1145.683 | -0.03087 | 0.792702 | 0.97681 |
| ENSMUSG( | Naa38 | 1060.16 | -0.0808 | 0.41998 | 0.822442 | ENSMUSG( | Naa38 | 926.7072 | -0.21118 | 0.112321 | 0.809018 |
| ENSMUSG( | Naa40 | 1064.522 | -0.00708 | 0.948068 | 0.993821 | ENSMUSG( | Naa40 | 1027.724 | 0.156346 | 0.229241 | 0.868507 |
| ENSMUSG( | Naa50 | 2001.21 | -0.00302 | 0.980805 | 0.997512 | ENSMUSG( | Naa50 | 1817.917 | -0.01655 | 0.926239 | 0.99306 |
| ENSMUSG( | Naa60 | 3491.668 | 0.039876 | 0.621615 | 0.911432 | ENSMUSG( | Naa60 | 3179.223 | 0.038116 | 0.704738 | 0.962677 |
| ENSMUSG( | Naa80 | 406.6976 | -0.07663 | 0.516273 | 0.872701 | ENSMUSG( | Naa80 | 382.2889 | 0.015224 | 0.911149 | 0.992196 |
| ENSMUSG( | Naaa | 3029.758 | -0.00064 | 0.99588 | 0.99943 | ENSMUSG( | Naaa | 2918.273 | 0.1582 | 0.277911 | 0.891468 |
| ENSMUSG( | Naalad2 | 49.93705 | -0.51429 | 0.090315 | 0.461669 | ENSMUSG( | Naalad2 | 62.63477 | 0.449034 | 0.140267 | 0.830678 |
| ENSMUSG( | Naaladl2 | 44.21782 | -0.96113 | 0.002864 | 0.065614 | ENSMUSG( | Naaladl2 | 52.58284 | -0.02015 | 0.948887 | 0.994798 |
| ENSMUSG( | Nab1 | 1810.767 | 0.009532 | 0.925886 | 0.991229 | ENSMUSG( | Nab1 | 1613.681 | -0.05731 | 0.698541 | 0.960908 |
| ENSMUSG( | Nab2 | 866.7868 | -0.26175 | 0.087823 | 0.454663 | ENSMUSG( | Nab2 | 769.7198 | -0.35131 | 0.063583 | 0.736482 |
| ENSMUSG( | Nabp1 | 175.4203 | -0.09312 | 0.659685 | 0.924524 | ENSMUSG( | Nabp1 | 172.6908 | 0.121758 | 0.569879 | 0.941777 |
| ENSMUSG( | Nabp2 | 2145.28 | -0.03729 | 0.682643 | 0.93198 | ENSMUSG( | Nabp2 | 1895.275 | -0.13172 | 0.307081 | 0.898538 |
| ENSMUSG( | Naca | 4562.253 | 0.096161 | 0.254708 | 0.705011 | ENSMUSG( | Naca | 3891.113 | -0.0965 | 0.435527 | 0.919062 |
| ENSMUSG( | Nacad | 1150.969 | -0.15694 | 0.132934 | 0.545594 | ENSMUSG( | Nacad | 1090.274 | -0.03571 | 0.781812 | 0.976106 |
| ENSMUSG( | Nacc1 | 6122.356 | 0.134244 | 0.111911 | 0.50438 | ENSMUSG( | Nacc1 | 5293.69 | -0.01318 | 0.896074 | 0.990558 |
| ENSMUSG( | Nacc2 | 16314.42 | 0.047059 | 0.682252 | 0.931962 | ENSMUSG( | Nacc2 | 13385.45 | -0.26861 | 0.049198 | 0.708275 |
| ENSMUSG( | Nadk | 4095.865 | 0.146665 | 0.128694 | 0.537025 | ENSMUSG( | Nadk | 3461.87 | -0.06919 | 0.559799 | 0.939812 |
| ENSMUSG( | Nadk2 | 3994.699 | 0.108356 | 0.427125 | 0.82532 | ENSMUSG( | Nadk2 | 3470.127 | -0.03008 | 0.862886 | 0.986547 |
| ENSMUSG( | Nadsyn1 | 176.5366 | -0.0535 | 0.741986 | 0.948478 | ENSMUSG( | Nadsyn1 | 163.6809 | 0.013228 | 0.946065 | 0.994798 |
| ENSMUSG( | Nae1 | 855.9598 | -0.03504 | 0.767928 | 0.954621 | ENSMUSG( | Nae1 | 791.492 | 0.007015 | 0.958131 | 0.994798 |
| ENSMUSG( | Naf1 | 528.1917 | 0.008016 | 0.954087 | 0.993874 | ENSMUSG( | Naf1 | 465.3688 | -0.09876 | 0.535377 | 0.936508 |
| ENSMUSG( | Naga | 948.8724 | -0.33985 | 0.009196 | 0.135112 | ENSMUSG( | Naga | 985.5315 | 0.063301 | 0.669945 | 0.95694 |
| ENSMUSG( | Nagk | 1162.654 | 0.069737 | 0.458168 | 0.839914 | ENSMUSG( | Nagk | 993.1444 | -0.11943 | 0.31775 | 0.902777 |
| ENSMUSG( | Naglu | 409.6716 | -0.42976 | 0.002683 | 0.062344 | ENSMUSG( | Naglu | 450.3445 | 0.148777 | 0.403549 | 0.917257 |
| ENSMUSG( | Nagpa | 346.5948 | -0.12357 | 0.396819 | 0.811742 | ENSMUSG( | Nagpa | 347.6167 | 0.161002 | 0.323204 | 0.904023 |
| ENSMUSG( | Naif1 | 143.61 | -0.19015 | 0.255702 | 0.706195 | ENSMUSG( | Naif1 | 138.4162 | -0.0259 | 0.890836 | 0.990558 |
| ENSMUSG( | Naip1 | 12.5672 | 0.131899 | 0.814068 | NA | ENSMUSG( | Naip1 | 10.37192 | -0.2127 | 0.736834 | NA |
| ENSMUSG( | Naip2 | 55.67633 | -0.15751 | 0.613738 | 0.909201 | ENSMUSG( | Naip2 | 53.41703 | 0.026529 | 0.93222 | 0.99306 |
| ENSMUSG( | Naip5 | 80.81009 | -0.49157 | 0.048973 | 0.337919 | ENSMUSG( | Naip5 | 80.86629 | -0.17651 | 0.557631 | 0.939565 |
| ENSMUSG( | Naip6 | 47.49215 | -0.4272 | 0.152016 | 0.577721 | ENSMUSG( | Naip6 | 47.07557 | -0.15196 | 0.636994 | 0.953903 |
| ENSMUSG( | Nalcn | 2173.799 | -0.4548 | 0.003208 | 0.070583 | ENSMUSG( | Nalcn | 2466.199 | 0.209144 | 0.220786 | 0.866104 |
| ENSMUSG( | Nalf1 | 2643.177 | -0.18911 | 0.042158 | 0.316282 | ENSMUSG( | Nalf1 | 2567.576 | 0.000961 | 0.993565 | 0.998813 |
| ENSMUSG( | Nalf2 | 970.3349 | 0.239572 | 0.012297 | 0.160458 | ENSMUSG( | Nalf2 | 861.6186 | 0.171457 | 0.138531 | 0.828492 |
| ENSMUSG( | Nampt | 1166.309 | -0.05934 | 0.540786 | 0.88315 | ENSMUSG( | Nampt | 1171.922 | 0.217098 | 0.048029 | 0.704961 |
| ENSMUSG( | Nanos1 | 1084.202 | 0.008186 | 0.934855 | 0.993231 | ENSMUSG( | Nanos1 | 999.8811 | 0.04013 | 0.744105 | 0.96951 |
| ENSMUSG( | Nanos2 | 40.06825 | -0.32542 | 0.376808 | 0.801213 | ENSMUSG( | Nanos2 | 38.01172 | -0.22407 | 0.594757 | 0.945444 |
| ENSMUSG( | Nanos3 | 22.05071 | -0.03939 | 0.924414 | 0.99114 | ENSMUSG( | Nanos3 | 20.10723 | -0.00314 | 0.994401 | 0.998813 |
| ENSMUSG( | Nans | 372.8805 | 0.039235 | 0.768225 | 0.954641 | ENSMUSG( | Nans | 310.0434 | -0.23792 | 0.135269 | 0.825081 |
| ENSMUSG( | Nap1l1 | 5082.426 | 0.04686 | 0.635559 | 0.914465 | ENSMUSG( | Nap1l1 | 4426.971 | -0.08629 | 0.454277 | 0.922444 |
| ENSMUSG( | Nap1l2 | 793.5891 | -0.23517 | 0.108327 | 0.497337 | ENSMUSG( | Nap1l2 | 794.82 | 0.041394 | 0.814421 | 0.979198 |
| ENSMUSG( | Nap1l3 | 479.2366 | -0.02842 | 0.863274 | 0.975055 | ENSMUSG( | Nap1l3 | 430.0776 | -0.0789 | 0.672125 | 0.95694 |
| ENSMUSG( | Nap1l4 | 3810.561 | 0.316357 | 0.003062 | 0.068418 | ENSMUSG( | Nap1l4 | 3092.241 | -0.00241 | 0.985257 | 0.997084 |
| ENSMUSG( | Nap1l5 | 4199.872 | 0.047165 | 0.729696 | 0.944813 | ENSMUSG( | Nap1l5 | 3601.732 | -0.13463 | 0.446734 | 0.921834 |
| ENSMUSG( | Napa | 5232.399 | -0.02685 | 0.82242 | 0.966352 | ENSMUSG( | Napa | 4578.877 | -0.14858 | 0.265969 | 0.886134 |
| ENSMUSG( | Napb | 7831.572 | 0.079717 | 0.436234 | 0.829093 | ENSMUSG( | Napb | 6886.917 | -0.02331 | 0.85778 | 0.985884 |
| ENSMUSG( | Napepld | 291.0775 | 0.266233 | 0.116659 | 0.515027 | ENSMUSG( | Napepld | 266.6188 | 0.278621 | 0.100364 | 0.801311 |
| ENSMUSG( | Napg | 2685.351 | 0.01502 | 0.874921 | 0.979746 | ENSMUSG( | Napg | 2439.526 | 0.005056 | 0.965444 | 0.995433 |
| ENSMUSG( | Naprt | 367.9891 | -0.00215 | 0.988256 | 0.999109 | ENSMUSG( | Naprt | 321.0411 | -0.12753 | 0.398654 | 0.917257 |
| ENSMUSG( | Napsa | 26.01754 | -1.22066 | 0.01488 | 0.179275 | ENSMUSG( | Napsa | 24.8046 | -1.05945 | 0.041439 | 0.688699 |
| ENSMUSG( | Narf | 1425.817 | 0.145534 | 0.132338 | 0.544251 | ENSMUSG( | Narf | 1310.354 | 0.168769 | 0.13959 | 0.830678 |
| ENSMUSG( | Nars1 | 3921.562 | 0.088454 | 0.344709 | 0.779153 | ENSMUSG( | Nars1 | 3399.23 | -0.05613 | 0.615522 | 0.951177 |
| ENSMUSG( | Nars2 | 328.7754 | -0.09086 | 0.46158 | 0.8415 | ENSMUSG( | Nars2 | 285.7222 | -0.24171 | 0.143604 | 0.831311 |
| ENSMUSG( | Nasp | 600.6874 | 0.125498 | 0.284236 | 0.73274 | ENSMUSG( | Nasp | 542.534 | 0.097584 | 0.471201 | 0.923329 |
| ENSMUSG( | Nat1 | 90.61949 | -0.1417 | 0.548777 | 0.885686 | ENSMUSG( | Nat1 | 79.09507 | -0.28156 | 0.255632 | 0.883043 |
| ENSMUSG( | Nat10 | 909.5075 | 0.158788 | 0.157473 | 0.585653 | ENSMUSG( | Nat10 | 808.8006 | 0.095573 | 0.497867 | 0.927825 |
| ENSMUSG( | Nat14 | 3014.825 | -0.20343 | 0.080804 | 0.439517 | ENSMUSG( | Nat14 | 2908.879 | -0.02962 | 0.829418 | 0.980224 |
| ENSMUSG( | Nat2 | 30.75329 | 0.180962 | 0.611656 | 0.908586 | ENSMUSG( | Nat2 | 26.57647 | 0.051812 | 0.895668 | 0.990558 |
| ENSMUSG( | Nat8 | 99.91793 | -0.08713 | 0.738951 | 0.947768 | ENSMUSG( | Nat8 | 88.88223 | -0.13532 | 0.59718 | 0.946609 |
| ENSMUSG( | Nat8f1 | 633.1842 | 0.049616 | 0.713956 | 0.940903 | ENSMUSG( | Nat8f1 | 569.631 | 0.018905 | 0.897778 | 0.990558 |
| ENSMUSG( | Nat8f2 | 23.47079 | 0.329809 | 0.40293 | 0.815716 | ENSMUSG( | Nat8f2 | 22.61365 | 0.479831 | 0.260422 | 0.884524 |
| ENSMUSG( | Nat8f3 | 154.8668 | 0.020715 | 0.920272 | 0.990247 | ENSMUSG( | Nat8f3 | 134.0635 | -0.12011 | 0.60722 | 0.949957 |
| ENSMUSG( | Nat8f4 | 1033.9 | 0.047005 | 0.760518 | 0.952907 | ENSMUSG( | Nat8f4 | 887.8126 | -0.1218 | 0.415176 | 0.917257 |
| ENSMUSG( | Nat8f5 | 95.77454 | 0.134518 | 0.603797 | 0.905053 | ENSMUSG( | Nat8f5 | 76.42045 | -0.23532 | 0.409145 | 0.917257 |
| ENSMUSG( | Nat8f6 | 25.04484 | 0.489424 | 0.203144 | 0.643631 | ENSMUSG( | Nat8f6 | 19.16387 | 0.009228 | 0.983957 | 0.996963 |
| ENSMUSG( | Nat8l | 13458.89 | -0.08718 | 0.467992 | 0.84521 | ENSMUSG( | Nat8l | 12727.56 | 0.024734 | 0.86449 | 0.986591 |
| ENSMUSG( | Nat9 | 321.2198 | 0.086189 | 0.511738 | 0.871106 | ENSMUSG( | Nat9 | 273.2002 | -0.10624 | 0.500654 | 0.927825 |
| ENSMUSG( | Natd1 | 663.0213 | 0.171421 | 0.106494 | 0.494174 | ENSMUSG( | Natd1 | 588.6569 | 0.102114 | 0.468906 | 0.923329 |
| ENSMUSG( | Nav1 | 4182.74 | -0.18527 | 0.028206 | 0.256543 | ENSMUSG( | Nav1 | 4222.979 | 0.115017 | 0.260089 | 0.884311 |
| ENSMUSG( | Nav2 | 2835.328 | 0.141054 | 0.112744 | 0.506698 | ENSMUSG( | Nav2 | 2614.374 | 0.173925 | 0.101878 | 0.801719 |
| ENSMUSG( | Nav3 | 5177.665 | 0.101038 | 0.225248 | 0.672044 | ENSMUSG( | Nav3 | 4573.795 | 0.013674 | 0.896639 | 0.990558 |
| ENSMUSG( | Naxd | 1704.974 | -0.01261 | 0.883265 | 0.982422 | ENSMUSG( | Naxd | 1491.866 | -0.1335 | 0.236902 | 0.872173 |
| ENSMUSG( | Naxe | 1650.221 | -0.1252 | 0.157135 | 0.58551 | ENSMUSG( | Naxe | 1487.887 | -0.15878 | 0.176975 | 0.846973 |
| ENSMUSG( | Nbas | 1086.883 | -0.21152 | 0.122793 | 0.529145 | ENSMUSG( | Nbas | 1071.924 | 0.031594 | 0.853384 | 0.985598 |
| ENSMUSG( | Nbdy | 567.5132 | 0.075494 | 0.546835 | 0.884974 | ENSMUSG( | Nbdy | 485.2741 | -0.11645 | 0.431916 | 0.917854 |
| ENSMUSG( | Nbea | 4216.01 | 0.052869 | 0.598542 | 0.903801 | ENSMUSG( | Nbea | 4073.158 | 0.217528 | 0.096729 | 0.796245 |
| ENSMUSG( | Nbeal1 | 654.3249 | -0.09138 | 0.558484 | 0.888521 | ENSMUSG( | Nbeal1 | 657.9186 | 0.191042 | 0.209303 | 0.864412 |
| ENSMUSG( | Nbeal2 | 475.6675 | 0.120996 | 0.31163 | 0.759296 | ENSMUSG( | Nbeal2 | 457.7792 | 0.273176 | 0.056186 | 0.723362 |
| ENSMUSG( | Nbl1 | 3212.344 | -0.0678 | 0.464439 | 0.843364 | ENSMUSG( | Nbl1 | 3116.069 | 0.114389 | 0.326212 | 0.905339 |
| ENSMUSG( | Nbn | 493.9583 | 0.24112 | 0.043799 | 0.320589 | ENSMUSG( | Nbn | 430.0014 | 0.116227 | 0.405607 | 0.917257 |
| ENSMUSG( | Nbr1 | 6141.272 | -0.02961 | 0.766119 | 0.954173 | ENSMUSG( | Nbr1 | 5525.832 | -0.06637 | 0.536058 | 0.936952 |
| ENSMUSG( | Ncald | 13831.09 | 0.065252 | 0.432929 | 0.828185 | ENSMUSG( | Ncald | 11704.22 | -0.15178 | 0.153742 | 0.837724 |
| ENSMUSG( | Ncam1 | 9986.927 | -0.30831 | 0.015544 | 0.184433 | ENSMUSG( | Ncam1 | 10442.08 | 0.105898 | 0.470351 | 0.923329 |
| ENSMUSG( | Ncam2 | 1137.295 | -0.43606 | 0.014697 | 0.178407 | ENSMUSG( | Ncam2 | 1307.461 | 0.262129 | 0.19542 | 0.861579 |
| ENSMUSG( | Ncan | 25322.95 | -0.19682 | 0.102053 | 0.486742 | ENSMUSG( | Ncan | 23921.23 | -0.0855 | 0.505139 | 0.927825 |
| ENSMUSG( | Ncapd2 | 788.3054 | 0.014446 | 0.897785 | 0.985633 | ENSMUSG( | Ncapd2 | 748.9053 | 0.135169 | 0.280238 | 0.892356 |
| ENSMUSG( | Ncapd3 | 628.6196 | 0.007535 | 0.961474 | 0.994205 | ENSMUSG( | Ncapd3 | 598.4757 | 0.133018 | 0.44981 | 0.922202 |
| ENSMUSG( | Ncapg | 20.30485 | -1.05574 | 0.039579 | NA | ENSMUSG( | Ncapg | 22.0529 | -0.36337 | 0.477285 | 0.923329 |
| ENSMUSG( | Ncapg2 | 194.7071 | -0.00707 | 0.969182 | 0.996369 | ENSMUSG( | Ncapg2 | 186.703 | 0.146743 | 0.426062 | 0.917257 |
| ENSMUSG( | Ncaph | 80.20079 | -0.16186 | 0.48979 | 0.859319 | ENSMUSG( | Ncaph | 75.74192 | -0.03191 | 0.903312 | 0.991089 |
| ENSMUSG( | Ncaph2 | 4042.454 | 0.100789 | 0.227167 | 0.674223 | ENSMUSG( | Ncaph2 | 3454.264 | -0.08356 | 0.415019 | 0.917257 |
| ENSMUSG( | Ncbp1 | 1540.589 | 0.09302 | 0.335716 | 0.777 | ENSMUSG( | Ncbp1 | 1380.978 | 0.046554 | 0.67455 | 0.95694 |
| ENSMUSG( | Ncbp2 | 1332.15 | 0.050808 | 0.613902 | 0.909277 | ENSMUSG( | Ncbp2 | 1164.831 | -0.07273 | 0.571457 | 0.942116 |
| ENSMUSG( | Ncbp2as2 | 1409.012 | 0.028503 | 0.798693 | 0.960896 | ENSMUSG( | Ncbp2as2 | 1186.206 | -0.21319 | 0.140964 | 0.830678 |
| ENSMUSG( | Ncbp3 | 590.1284 | 0.103313 | 0.353224 | 0.785373 | ENSMUSG( | Ncbp3 | 527.5448 | 0.051876 | 0.687662 | 0.959673 |
| ENSMUSG( | Nccrp1 | 18.88366 | -0.1506 | 0.723146 | NA | ENSMUSG( | Nccrp1 | 16.30846 | -0.29422 | 0.57707 | 0.942912 |
| ENSMUSG( | Ncdn | 33400.41 | 0.0626 | 0.519623 | 0.874979 | ENSMUSG( | Ncdn | 29192.41 | -0.05726 | 0.609026 | 0.950237 |
| ENSMUSG( | Nceh1 | 3685.805 | -0.44877 | 0.001763 | 0.047418 | ENSMUSG( | Nceh1 | 4194.989 | 0.223218 | 0.194314 | 0.860096 |
| ENSMUSG( | Ncf1 | 435.3406 | 0.033439 | 0.789824 | 0.959461 | ENSMUSG( | Ncf1 | 406.1233 | 0.10496 | 0.455964 | 0.923329 |
| ENSMUSG( | Ncf2 | 241.0065 | -0.0078 | 0.957669 | 0.993874 | ENSMUSG( | Ncf2 | 214.8037 | -0.06581 | 0.696464 | 0.960867 |
| ENSMUSG( | Ncf4 | 49.87119 | -0.11167 | 0.708735 | 0.939519 | ENSMUSG( | Ncf4 | 42.94044 | -0.28875 | 0.37913 | 0.916995 |
| ENSMUSG( | Nck1 | 336.5827 | 0.13777 | 0.297149 | 0.745805 | ENSMUSG( | Nck1 | 309.443 | 0.160357 | 0.286222 | 0.893906 |
| ENSMUSG( | Nck2 | 742.1888 | 0.050123 | 0.687635 | 0.93324 | ENSMUSG( | Nck2 | 652.1433 | -0.05994 | 0.676111 | 0.95694 |
| ENSMUSG( | Nckap1 | 14623.88 | 0.099012 | 0.228404 | 0.675849 | ENSMUSG( | Nckap1 | 12974.14 | 0.024125 | 0.805615 | 0.977674 |
| ENSMUSG( | Nckap1l | 876.2404 | 0.138412 | 0.219044 | 0.664602 | ENSMUSG( | Nckap1l | 783.1579 | 0.08783 | 0.529771 | 0.934795 |
| ENSMUSG( | Nckap5 | 339.2962 | -0.00665 | 0.958925 | 0.993914 | ENSMUSG( | Nckap5 | 289.7383 | -0.20652 | 0.169715 | 0.840806 |
| ENSMUSG( | Nckap5l | 884.6343 | 0.035789 | 0.72797 | 0.944355 | ENSMUSG( | Nckap5l | 812.9531 | 0.062043 | 0.600911 | 0.947749 |
| ENSMUSG( | Nckipsd | 1757.44 | 0.180516 | 0.087536 | 0.454663 | ENSMUSG( | Nckipsd | 1472.548 | -0.05712 | 0.630475 | 0.953251 |
| ENSMUSG( | Ncl | 7729.268 | -0.0009 | 0.992197 | 0.999327 | ENSMUSG( | Ncl | 6899.227 | -0.06232 | 0.626874 | 0.953234 |
| ENSMUSG( | Ncln | 2188.879 | -0.17674 | 0.224768 | 0.671058 | ENSMUSG( | Ncln | 2220.22 | 0.140942 | 0.430723 | 0.917257 |
| ENSMUSG( | Ncmap | 24.38786 | 0.376307 | 0.364621 | 0.793796 | ENSMUSG( | Ncmap | 22.68829 | 0.437542 | 0.338496 | 0.907972 |
| ENSMUSG( | Ncoa1 | 4045.52 | 0.157538 | 0.047369 | 0.332754 | ENSMUSG( | Ncoa1 | 3544.862 | 0.048003 | 0.651914 | 0.9547 |
| ENSMUSG( | Ncoa2 | 2150.781 | 0.100305 | 0.271699 | 0.722082 | ENSMUSG( | Ncoa2 | 1946.555 | 0.08205 | 0.437317 | 0.920144 |
| ENSMUSG( | Ncoa3 | 1444.696 | -0.12577 | 0.190175 | 0.629268 | ENSMUSG( | Ncoa3 | 1444.427 | 0.144164 | 0.264008 | 0.886134 |
| ENSMUSG( | Ncoa4 | 1056.468 | 0.003105 | 0.974451 | 0.997072 | ENSMUSG( | Ncoa4 | 920.3219 | -0.13239 | 0.260408 | 0.884524 |
| ENSMUSG( | Ncoa5 | 1324.78 | 0.167359 | 0.083841 | 0.446475 | ENSMUSG( | Ncoa5 | 1160.435 | 0.058704 | 0.588626 | 0.944975 |
| ENSMUSG( | Ncoa6 | 2070.292 | 0.028845 | 0.762173 | 0.953459 | ENSMUSG( | Ncoa6 | 1925.771 | 0.088024 | 0.423646 | 0.917257 |
| ENSMUSG( | Ncoa7 | 927.8252 | 0.05637 | 0.648734 | 0.920025 | ENSMUSG( | Ncoa7 | 881.4536 | 0.173716 | 0.223915 | 0.867404 |
| ENSMUSG( | Ncor1 | 7721.544 | 0.120463 | 0.188609 | 0.627639 | ENSMUSG( | Ncor1 | 6803.161 | 0.027428 | 0.790422 | 0.97681 |
| ENSMUSG( | Ncor2 | 6519.236 | -0.03181 | 0.751715 | 0.950337 | ENSMUSG( | Ncor2 | 6334.805 | 0.152681 | 0.202496 | 0.864412 |
| ENSMUSG( | Ncs1 | 18757.74 | 0.190733 | 0.020134 | 0.214568 | ENSMUSG( | Ncs1 | 16487.73 | 0.092649 | 0.358051 | 0.913759 |
| ENSMUSG( | Ncstn | 2737.881 | -0.31513 | 0.015728 | 0.185786 | ENSMUSG( | Ncstn | 2896.495 | 0.134439 | 0.373755 | 0.915491 |
| ENSMUSG( | ND1 | 684327.5 | -0.08464 | 0.608335 | 0.907154 | ENSMUSG( | ND1 | 626459.3 | -0.07378 | 0.722557 | 0.966887 |
| ENSMUSG( | ND2 | 407091.2 | -0.09528 | 0.5595 | 0.888669 | ENSMUSG( | ND2 | 378714.8 | -0.03837 | 0.860639 | 0.986137 |
| ENSMUSG( | ND3 | 835.36 | -0.22345 | 0.31332 | 0.76033 | ENSMUSG( | ND3 | 812.3942 | -0.03646 | 0.948278 | 0.994798 |
| ENSMUSG( | ND4 | 457567 | -0.02801 | 0.805201 | 0.962616 | ENSMUSG( | ND4 | 404536.8 | -0.11932 | 0.442257 | 0.921554 |
| ENSMUSG( | ND4L | 248.0795 | -0.08908 | 0.709907 | 0.939669 | ENSMUSG( | ND4L | 227.9615 | -0.07347 | 0.88073 | 0.989644 |
| ENSMUSG( | ND5 | 510938.7 | -0.05522 | 0.669833 | 0.928209 | ENSMUSG( | ND5 | 465776.9 | -0.05593 | 0.757278 | 0.971858 |
| ENSMUSG( | ND6 | 4694.734 | -0.26279 | 0.138224 | 0.553167 | ENSMUSG( | ND6 | 4786.228 | 0.069666 | 0.762661 | 0.973566 |
| ENSMUSG( | Ndc1 | 867.0912 | -0.11815 | 0.359429 | 0.791011 | ENSMUSG( | Ndc1 | 873.0906 | 0.169383 | 0.255313 | 0.882761 |
| ENSMUSG( | Ndc80 | 15.20264 | -0.16889 | 0.733576 | NA | ENSMUSG( | Ndc80 | 15.12097 | 0.109094 | 0.835344 | 0.981237 |
| ENSMUSG( | Nde1 | 2176.859 | 0.283984 | 0.01345 | 0.169431 | ENSMUSG( | Nde1 | 1755.656 | -0.05628 | 0.65287 | 0.954897 |
| ENSMUSG( | Ndel1 | 2003.23 | 0.143013 | 0.121581 | 0.52602 | ENSMUSG( | Ndel1 | 1704.338 | -0.05147 | 0.640065 | 0.953903 |
| ENSMUSG( | Ndfip1 | 11850.63 | -0.54522 | 0.000116 | 0.007099 | ENSMUSG( | Ndfip1 | 13577.71 | 0.165337 | 0.324633 | 0.905257 |
| ENSMUSG( | Ndfip2 | 2981.595 | -0.5201 | 7.18E-05 | 0.005196 | ENSMUSG( | Ndfip2 | 3246.803 | 0.044273 | 0.772081 | 0.974236 |

| | | | | | |
|---|---|---|---|---|---|
| ENSMUSG( | 10.51119 | -0.33589 | 0.573005 | NA | NA |
| ENSMUSG( | 70.21228 | 0.071534 | 0.772358 | 0.955069 | NA |
| ENSMUSG( | 12.184 | -0.22369 | 0.673384 | NA | NA |
| ENSMUSG( | 61.686 | -0.23927 | 0.345044 | 0.779153 | NA |
| ENSMUSG( | 9.402414 | -0.01124 | 0.986231 | NA | NA |
| ENSMUSG( | 102.2354 | -0.14561 | 0.505325 | 0.866883 | NA |
| ENSMUSG( | 20.84002 | -0.2598 | 0.548805 | NA | NA |
| ENSMUSG( | 35.56092 | 0.466557 | 0.178602 | 0.616298 | NA |
| ENSMUSG( | 80.632 | -0.11875 | 0.584443 | 0.898802 | NA |
| ENSMUSG( | 23.49454 | 0.111871 | 0.773697 | 0.95556 | NA |
| ENSMUSG( | 56.56064 | -0.30617 | 0.247854 | 0.696522 | NA |
| ENSMUSG( | 40.5895 | 0.0073 | 0.982533 | 0.997512 | NA |
| ENSMUSG( | 3787.095 | -0.00126 | 0.991033 | 0.999232 | NA |
| ENSMUSG( | 31.72141 | -0.04592 | 0.900568 | 0.985633 | NA |
| ENSMUSG( | 117.2171 | 0.024757 | 0.911463 | 0.988483 | NA |
| ENSMUSG( | 12.67868 | -0.06379 | 0.909922 | NA | NA |
| ENSMUSG( | 38.83768 | -0.01862 | 0.950692 | 0.993821 | NA |
| ENSMUSG( | 42.89025 | 0.102636 | 0.732749 | 0.944956 | NA |
| ENSMUSG( | 12.37604 | -0.17103 | 0.746317 | NA | NA |
| ENSMUSG( | 10.19212 | 0.316814 | 0.602522 | NA | NA |
| ENSMUSG( | 16.28439 | -0.26901 | 0.570786 | NA | NA |
| ENSMUSG( | 27.46404 | 0.064424 | 0.860884 | 0.975025 | NA |
| ENSMUSG( | 38.56483 | -0.12729 | 0.693945 | 0.934202 | NA |
| ENSMUSG( | 83.47962 | 0.530257 | 0.027549 | 0.2548 | NA |
| ENSMUSG( | 264.4154 | 0.174547 | 0.259113 | 0.709814 | NA |
| ENSMUSG( | 15.76482 | -0.12423 | 0.786078 | NA | NA |
| ENSMUSG( | 37.57224 | 0.086035 | 0.790382 | 0.959461 | NA |
| ENSMUSG( | 178.1833 | -0.76285 | 0.000998 | 0.032953 | NA |
| ENSMUSG( | 10.73011 | -0.59698 | 0.320604 | NA | NA |
| ENSMUSG( | 21.91241 | -0.31926 | 0.442532 | 0.83278 | NA |
| ENSMUSG( | 16.30941 | -0.81551 | 0.110744 | NA | NA |
| ENSMUSG( | 15.69389 | -0.24522 | 0.609001 | NA | NA |
| ENSMUSG( | 18.4745 | -0.23911 | 0.584594 | NA | NA |
| ENSMUSG( | 8.806321 | 1.352293 | 0.056119 | NA | NA |
| ENSMUSG( | 29.59097 | -0.07995 | 0.819962 | 0.965888 | NA |
| ENSMUSG( | 44.96202 | -0.0025 | 0.994349 | 0.999369 | NA |
| ENSMUSG( | 13.02944 | -0.40029 | 0.437068 | NA | NA |
| ENSMUSG( | 14.65697 | -0.06309 | 0.896194 | NA | NA |
| ENSMUSG( | 29.60975 | -0.06155 | 0.861535 | 0.975026 | NA |
| ENSMUSG( | 511.7232 | -0.406 | 0.014048 | 0.173278 | NA |
| ENSMUSG( | 62.99359 | 0.110106 | 0.691127 | 0.933881 | NA |
| ENSMUSG( | 20.27636 | 0.59535 | 0.199663 | NA | NA |
| ENSMUSG( | 10.66514 | 0.422448 | 0.496092 | NA | NA |
| ENSMUSG( | 17.4061 | 0.528763 | 0.295023 | NA | NA |
| ENSMUSG( | 27.63074 | -0.3754 | 0.382662 | 0.803227 | NA |
| ENSMUSG( | 24.74657 | 0.248002 | 0.53938 | 0.883094 | NA |
| ENSMUSG( | 27.6965 | 0.479446 | 0.183954 | 0.620925 | NA |
| ENSMUSG( | 28.0791 | -0.04216 | 0.911708 | 0.988483 | NA |
| ENSMUSG( | 8.721198 | -0.14092 | 0.81904 | NA | NA |
| ENSMUSG( | 18.84827 | -0.02233 | 0.960282 | NA | NA |
| ENSMUSG( | 26.77851 | 0.375967 | 0.338166 | 0.777422 | NA |
| ENSMUSG( | 32.43474 | -0.31667 | 0.333759 | 0.775602 | NA |
| ENSMUSG( | 12.84978 | -0.13471 | 0.808414 | NA | NA |
| ENSMUSG( | 118.6228 | 0.163937 | 0.479306 | 0.854046 | NA |
| ENSMUSG( | 35.86796 | -0.28237 | 0.371165 | 0.798554 | NA |
| ENSMUSG( | 39.77314 | -0.04716 | 0.886805 | 0.983255 | NA |
| ENSMUSG( | 1081.475 | 0.043406 | 0.689539 | 0.933609 | NA |
| ENSMUSG( | 13.87737 | -0.81523 | 0.112541 | NA | NA |
| ENSMUSG( | 9.938952 | 0.907245 | 0.148705 | NA | NA |
| ENSMUSG( | 9.237475 | -0.57872 | 0.36633 | NA | NA |
| ENSMUSG( | 26.91355 | 0.056811 | 0.882496 | 0.982244 | NA |
| ENSMUSG( | 12.69998 | -0.11333 | 0.846306 | NA | NA |
| ENSMUSG( | 25.86309 | -0.40315 | 0.293795 | 0.743425 | NA |
| ENSMUSG( | 28.568 | 0.308979 | 0.440337 | 0.831851 | NA |
| ENSMUSG( | 62.6086 | -0.05509 | 0.834925 | 0.970426 | NA |
| ENSMUSG( | 38.08984 | -0.4 | 0.207704 | 0.64962 | NA |
| ENSMUSG( | 43.06804 | -0.24652 | 0.420828 | 0.822984 | NA |
| ENSMUSG( | 2676.946 | -0.47552 | 0.000474 | 0.01988 | NA |
| ENSMUSG( | 11.02616 | -0.10604 | 0.844467 | NA | NA |
| ENSMUSG( | 74.3217 | 0.039124 | 0.86693 | 0.976541 | NA |
| ENSMUSG( | 17.30452 | -0.30044 | 0.511943 | NA | NA |
| ENSMUSG( | 16.97146 | 0.164486 | 0.718363 | NA | NA |
| ENSMUSG( | 29.42793 | -0.27391 | 0.458778 | 0.840244 | NA |
| ENSMUSG( | 66.97598 | 0.516255 | 0.037691 | 0.299975 | NA |
| ENSMUSG( | 37.55639 | -0.11079 | 0.738481 | 0.947414 | NA |
| ENSMUSG( | 182.5503 | -0.20012 | 0.189298 | 0.627932 | NA |
| ENSMUSG( | 110.5413 | 0.14378 | 0.447446 | 0.834051 | NA |
| ENSMUSG( | 7.840173 | -0.08935 | 0.898042 | NA | NA |
| ENSMUSG( | 83.63013 | -0.33185 | 0.181856 | 0.618944 | NA |
| ENSMUSG( | 19.88148 | -0.58112 | 0.244295 | NA | NA |
| ENSMUSG( | 15.79677 | 0.27908 | 0.547154 | NA | NA |
| ENSMUSG( | 13.8602 | 0.616335 | 0.249411 | NA | NA |
| ENSMUSG( | 8.653621 | -0.72025 | 0.305823 | NA | NA |
| ENSMUSG( | 15.57715 | -0.50649 | 0.273284 | NA | NA |
| ENSMUSG( | 20.41993 | 0.319245 | 0.463786 | NA | NA |
| ENSMUSG( | 37.58266 | -0.14611 | 0.632444 | 0.914073 | NA |
| ENSMUSG( | 20.66055 | -0.88088 | 0.037415 | NA | NA |
| ENSMUSG( | 111.7646 | -0.05457 | 0.76933 | 0.954674 | NA |
| ENSMUSG( | 24.5917 | 0.064036 | 0.866856 | 0.976541 | NA |
| ENSMUSG( | 30.27701 | 0.032749 | 0.930521 | 0.991965 | NA |
| ENSMUSG( | 81.72892 | 0.140229 | 0.524372 | 0.87583 | NA |
| ENSMUSG( | 17.90454 | -0.51663 | 0.277287 | NA | NA |
| ENSMUSG( | 14.72852 | -0.17475 | 0.710326 | NA | NA |
| ENSMUSG( | 80.35265 | 0.166965 | 0.474391 | 0.849425 | NA |
| ENSMUSG( | 66.05073 | -0.16598 | 0.542023 | 0.883182 | NA |
| ENSMUSG( | 42.82259 | -0.54833 | 0.068779 | 0.406391 | NA |
| ENSMUSG( | 66.7256 | -0.27471 | 0.250735 | 0.700403 | NA |
| ENSMUSG( | 144.9608 | -0.10338 | 0.563141 | 0.889834 | NA |
| ENSMUSG( | 23.7776 | 0.007769 | 0.984578 | 0.997815 | NA |
| ENSMUSG( | 72.68179 | -0.23194 | 0.317408 | 0.763823 | NA |
| ENSMUSG( | 98.53444 | -0.17027 | 0.496223 | 0.861564 | NA |
| ENSMUSG( | 15.60698 | -0.90223 | 0.056748 | NA | NA |
| ENSMUSG( | 24.38665 | -0.32998 | 0.382923 | 0.803237 | NA |
| ENSMUSG( | 1881.198 | 0.164207 | 0.061869 | 0.384781 | NA |
| ENSMUSG( | 10.87586 | -0.14026 | 0.802868 | NA | NA |
| ENSMUSG( | 20.89623 | 0.03226 | 0.939678 | NA | NA |
| ENSMUSG( | 11.89638 | -0.55656 | 0.303208 | NA | NA |
| ENSMUSG( | 15.55229 | 0.371688 | 0.428959 | NA | NA |
| ENSMUSG( | 17.55542 | 0.47153 | 0.366852 | NA | NA |
| ENSMUSG( | 16.45465 | 0.420949 | 0.41382 | NA | NA |
| ENSMUSG( | 9.146223 | -0.35093 | 0.580085 | NA | NA |
| ENSMUSG( | 221.4211 | 0.001626 | 0.991963 | 0.999232 | NA |
| ENSMUSG( | 5577.446 | 0.11113 | 0.279518 | 0.729114 | NA |
| ENSMUSG( | 36.07487 | -0.04213 | 0.895403 | 0.984512 | NA |
| ENSMUSG( | 5447.289 | -0.13628 | 0.206672 | 0.648942 | NA |
| ENSMUSG( | 41.24381 | -0.65521 | 0.029261 | 0.259574 | NA |
| ENSMUSG( | 28.29225 | 0.046963 | 0.899405 | 0.985633 | NA |
| ENSMUSG( | 21.31253 | 0.114522 | 0.793763 | 0.959799 | NA |
| ENSMUSG( | 94.18158 | 0.135226 | 0.513554 | 0.871779 | NA |
| ENSMUSG( | 47.59662 | -0.08207 | 0.811168 | 0.964318 | NA |
| ENSMUSG( | 73.24337 | -0.24133 | 0.331468 | 0.77519 | NA |
| ENSMUSG( | 13.88852 | -0.55192 | 0.319839 | NA | NA |
| ENSMUSG( | 99.44611 | 0.024239 | 0.910726 | 0.988356 | NA |
| ENSMUSG( | 9.311005 | -0.4798 | 0.427312 | NA | NA |
| ENSMUSG( | 204.7302 | -0.02431 | 0.876866 | 0.980991 | NA |
| ENSMUSG( | 112.1326 | -0.07452 | 0.730883 | 0.944956 | NA |
| ENSMUSG( | 20.58735 | 0.275223 | 0.532679 | NA | NA |
| ENSMUSG( | 14.37038 | -0.01991 | 0.969049 | NA | NA |
| ENSMUSG( | 186.4292 | 0.153644 | 0.327843 | 0.772842 | NA |
| ENSMUSG( | 102.379 | -0.11481 | 0.556415 | 0.887681 | NA |
| ENSMUSG( | 284.7827 | -0.13296 | 0.418988 | 0.821966 | NA |
| ENSMUSG( | 11.13939 | 0.166066 | 0.761944 | NA | NA |
| ENSMUSG( | 11.18062 | 0.28915 | 0.664352 | NA | NA |
| ENSMUSG( | 32.29312 | 0.016573 | 0.962221 | 0.994543 | NA |
| ENSMUSG( | 60.54458 | 0.376633 | 0.126114 | 0.533394 | NA |
| ENSMUSG( | 148.9881 | -0.01135 | 0.947599 | 0.993821 | NA |
| ENSMUSG( | 10.76023 | -0.35695 | 0.538037 | NA | NA |
| ENSMUSG( | 26.60391 | 0.295944 | 0.433696 | 0.828485 | NA |
| ENSMUSG( | 41.22689 | 0.460236 | 0.160465 | 0.590122 | NA |
| ENSMUSG( | 13.16909 | 0.219653 | 0.716566 | NA | NA |
| ENSMUSG( | 18.38319 | 0.809768 | 0.061056 | NA | NA |
| ENSMUSG( | 1079.414 | -0.00897 | 0.964533 | 0.995155 | NA |
| ENSMUSG( | 120.6572 | 0.26973 | 0.318104 | 0.764234 | NA |
| ENSMUSG( | 51.16794 | -0.01315 | 0.962394 | 0.994586 | NA |
| ENSMUSG( | 11.88952 | 0.605775 | 0.270928 | NA | NA |
| ENSMUSG( | 29.62313 | -0.03319 | 0.93165 | 0.9924 | NA |
| ENSMUSG( | 887.9032 | 0.091622 | 0.420633 | 0.8229 | NA |
| ENSMUSG( | 167.3126 | -0.05684 | 0.762756 | 0.953519 | NA |
| ENSMUSG( | 149.6224 | 0.027765 | 0.877281 | 0.981277 | NA |
| ENSMUSG( | 12.16281 | -0.19509 | 0.727767 | NA | NA |
| ENSMUSG( | 23.79064 | -0.10214 | 0.806237 | 0.962804 | NA |
| ENSMUSG( | 28.80452 | 0.471377 | 0.253443 | 0.703636 | NA |
| ENSMUSG( | 21.90557 | 0.283034 | 0.478263 | 0.853496 | NA |
| ENSMUSG( | 14.75381 | 0.258811 | 0.618737 | NA | NA |
| ENSMUSG( | 26.68348 | 0.642839 | 0.092071 | 0.466032 | NA |
| ENSMUSG( | 10.2435 | -0.42326 | 0.497024 | NA | NA |
| ENSMUSG( | 17.68956 | -2.09143 | 0.000189 | NA | NA |
| ENSMUSG( | 8.289241 | -1.88119 | 0.007571 | NA | NA |

| | | | | | |
|---|---|---|---|---|---|
| ENSMUSG( | 18.1851 | -0.10308 | 0.845706 | 0.984269 | NA |
| ENSMUSG( | 17.34952 | -0.56479 | 0.246699 | 0.877904 | NA |
| ENSMUSG( | 35.61776 | -0.13005 | 0.710285 | 0.964363 | NA |
| ENSMUSG( | 20.51514 | -0.27736 | 0.551145 | 0.939377 | NA |
| ENSMUSG( | 17.71974 | 0.793434 | 0.108875 | 0.804969 | NA |
| ENSMUSG( | 24.04074 | 0.435142 | 0.30769 | 0.898538 | NA |
| ENSMUSG( | 218.9856 | 0.377171 | 0.020976 | 0.572337 | NA |
| ENSMUSG( | 45.15293 | -0.01343 | 0.965872 | 0.995433 | NA |
| ENSMUSG( | 38.75156 | -0.20685 | 0.558033 | 0.939565 | NA |
| ENSMUSG( | 67.63674 | -0.09151 | 0.721513 | 0.966887 | NA |
| ENSMUSG( | 27.53936 | 0.000697 | 0.998664 | 0.999406 | NA |
| ENSMUSG( | 32.76541 | 0.378856 | 0.340203 | 0.907972 | NA |
| ENSMUSG( | 15.32675 | 0.177621 | 0.722336 | 0.966887 | NA |
| ENSMUSG( | 401936.9 | -0.09091 | 0.601446 | 0.947749 | NA |
| ENSMUSG( | 540426.2 | -0.00734 | 0.972156 | 0.996416 | NA |
| ENSMUSG( | 702.2438 | 0.11919 | 0.599443 | 0.947071 | NA |
| ENSMUSG( | 256.5104 | 0.525109 | 0.52371 | 0.932848 | NA |
| ENSMUSG( | 108.7975 | 0.012629 | 0.967076 | 0.995549 | NA |
| ENSMUSG( | 1269.447 | -0.01717 | 0.900997 | 0.990558 | NA |
| ENSMUSG( | 827.1415 | 0.093466 | 0.881314 | 0.989644 | NA |
| ENSMUSG( | 26.80678 | 0.027236 | 0.960008 | 0.994798 | NA |
| ENSMUSG( | 44.57035 | -0.48258 | 0.162532 | 0.837724 | NA |
| ENSMUSG( | 17.51686 | 0.775596 | 0.216394 | 0.865167 | NA |
| ENSMUSG( | 38.07125 | -0.00544 | 0.987442 | 0.99751 | NA |
| ENSMUSG( | 116.3019 | 0.047839 | 0.842009 | 0.983044 | NA |
| ENSMUSG( | 162.301 | 0.023777 | 0.929615 | 0.99306 | NA |
| ENSMUSG( | 6900.762 | 0.563905 | 0.016706 | 0.521279 | NA |
| ENSMUSG( | 727.2601 | 0.016665 | 0.908676 | 0.991958 | NA |
| ENSMUSG( | 332.3922 | -0.12966 | 0.422572 | 0.917257 | Naa10 |
| ENSMUSG( | 1237.926 | 0.079422 | 0.536599 | 0.937108 | Naa15 |
| ENSMUSG( | 195.4067 | 0.36601 | 0.03281 | 0.637876 | Naa16 |
| ENSMUSG( | 650.6714 | 0.042283 | 0.781988 | 0.976106 | Naa20 |
| ENSMUSG( | 1066.228 | 0.112462 | 0.405522 | 0.917257 | Naa25 |
| ENSMUSG( | 734.72 | 0.125032 | 0.488557 | 0.92554 | Naa30 |
| ENSMUSG( | 1145.683 | -0.03087 | 0.792702 | 0.97681 | Naa35 |
| ENSMUSG( | 926.7072 | -0.21118 | 0.112321 | 0.809018 | Naa38 |
| ENSMUSG( | 1027.724 | 0.156346 | 0.229241 | 0.868507 | Naa40 |
| ENSMUSG( | 1817.917 | -0.01655 | 0.926239 | 0.99306 | Naa50 |
| ENSMUSG( | 3179.223 | 0.038116 | 0.704738 | 0.962677 | Naa60 |
| ENSMUSG( | 382.2889 | 0.015224 | 0.911149 | 0.992196 | Naa80 |
| ENSMUSG( | 2918.273 | 0.1582 | 0.277911 | 0.891468 | Naaa |
| ENSMUSG( | 62.63477 | 0.449034 | 0.140267 | 0.830678 | Naalad2 |
| ENSMUSG( | 52.58284 | -0.02015 | 0.948887 | 0.994798 | Naaladl2 |
| ENSMUSG( | 1613.681 | -0.05731 | 0.698541 | 0.960908 | Nab1 |
| ENSMUSG( | 769.7198 | -0.35131 | 0.063583 | 0.736482 | Nab2 |
| ENSMUSG( | 172.6908 | 0.121758 | 0.569879 | 0.941777 | Nabp1 |
| ENSMUSG( | 1895.275 | -0.13172 | 0.307081 | 0.898538 | Nabp2 |
| ENSMUSG( | 3891.113 | -0.0965 | 0.435527 | 0.919062 | Naca |
| ENSMUSG( | 1090.274 | -0.03571 | 0.781812 | 0.976106 | Nacad |
| ENSMUSG( | 5293.69 | -0.01318 | 0.896074 | 0.990558 | Nacc1 |
| ENSMUSG( | 13385.45 | -0.26861 | 0.049198 | 0.708275 | Nacc2 |
| ENSMUSG( | 3461.87 | -0.06919 | 0.559799 | 0.939812 | Nadk |
| ENSMUSG( | 3470.127 | -0.03008 | 0.862886 | 0.986547 | Nadk2 |
| ENSMUSG( | 163.6809 | 0.013228 | 0.946065 | 0.994798 | Nadsyn1 |
| ENSMUSG( | 791.492 | 0.007015 | 0.958131 | 0.994798 | Nae1 |
| ENSMUSG( | 465.3688 | -0.09876 | 0.535377 | 0.936508 | Naf1 |
| ENSMUSG( | 985.5315 | 0.063301 | 0.669945 | 0.95694 | Naga |
| ENSMUSG( | 993.1444 | -0.11943 | 0.31775 | 0.902777 | Nagk |
| ENSMUSG( | 450.3445 | 0.148777 | 0.403549 | 0.917257 | Naglu |
| ENSMUSG( | 347.6167 | 0.161002 | 0.323204 | 0.904023 | Nagpa |
| ENSMUSG( | 138.4162 | -0.0259 | 0.890836 | 0.990558 | Naif1 |
| ENSMUSG( | 10.37192 | -0.2127 | 0.736834 | NA | Naip1 |
| ENSMUSG( | 53.41703 | 0.026529 | 0.93222 | 0.99306 | Naip2 |
| ENSMUSG( | 80.86629 | -0.17651 | 0.557631 | 0.939565 | Naip5 |
| ENSMUSG( | 47.07557 | -0.15196 | 0.636994 | 0.953903 | Naip6 |
| ENSMUSG( | 2466.199 | 0.209144 | 0.220786 | 0.866104 | Nalcn |
| ENSMUSG( | 2567.576 | 0.000961 | 0.993565 | 0.998813 | Nalf1 |
| ENSMUSG( | 861.6186 | 0.171457 | 0.138531 | 0.828492 | Nalf2 |
| ENSMUSG( | 1171.922 | 0.217098 | 0.048029 | 0.704961 | Nampt |
| ENSMUSG( | 999.8811 | 0.04013 | 0.744105 | 0.96951 | Nanos1 |
| ENSMUSG( | 38.01172 | -0.22407 | 0.594757 | 0.945444 | Nanos2 |
| ENSMUSG( | 20.10723 | -0.00314 | 0.994401 | 0.998813 | Nanos3 |
| ENSMUSG( | 310.0434 | -0.23792 | 0.135269 | 0.825081 | Nans |
| ENSMUSG( | 4426.971 | -0.08629 | 0.454277 | 0.922444 | Nap1l1 |
| ENSMUSG( | 794.82 | 0.041394 | 0.814421 | 0.979198 | Nap1l2 |
| ENSMUSG( | 430.0776 | -0.0789 | 0.672125 | 0.95694 | Nap1l3 |
| ENSMUSG( | 3092.241 | -0.00241 | 0.985257 | 0.997084 | Nap1l4 |
| ENSMUSG( | 3601.732 | -0.13463 | 0.446734 | 0.921834 | Nap1l5 |
| ENSMUSG( | 4578.877 | -0.14858 | 0.265969 | 0.886134 | Napa |
| ENSMUSG( | 6886.917 | -0.02331 | 0.85778 | 0.985884 | Napb |
| ENSMUSG( | 266.6188 | 0.278621 | 0.100364 | 0.801311 | Napepld |
| ENSMUSG( | 2439.526 | 0.005056 | 0.965444 | 0.995433 | Napg |
| ENSMUSG( | 321.0411 | -0.12753 | 0.398654 | 0.917257 | Naprt |
| ENSMUSG( | 24.8046 | -1.05945 | 0.041439 | 0.688699 | Napsa |
| ENSMUSG( | 1310.354 | 0.168769 | 0.13959 | 0.830678 | Narf |
| ENSMUSG( | 3399.23 | -0.05613 | 0.615522 | 0.951177 | Nars1 |
| ENSMUSG( | 285.7222 | -0.24171 | 0.143604 | 0.831311 | Nars2 |
| ENSMUSG( | 542.534 | 0.097584 | 0.471201 | 0.923329 | Nasp |
| ENSMUSG( | 79.09507 | -0.28156 | 0.255632 | 0.883043 | Nat1 |
| ENSMUSG( | 808.8006 | 0.095573 | 0.497867 | 0.927825 | Nat10 |
| ENSMUSG( | 2908.879 | -0.02962 | 0.829418 | 0.980224 | Nat14 |
| ENSMUSG( | 26.57647 | 0.051812 | 0.895668 | 0.990558 | Nat2 |
| ENSMUSG( | 88.88223 | -0.13532 | 0.59718 | 0.946609 | Nat8 |
| ENSMUSG( | 569.631 | 0.018905 | 0.897778 | 0.990558 | Nat8f1 |
| ENSMUSG( | 22.61365 | 0.479831 | 0.260422 | 0.884524 | Nat8f2 |
| ENSMUSG( | 134.0635 | -0.12011 | 0.60722 | 0.949957 | Nat8f3 |
| ENSMUSG( | 887.8126 | -0.1218 | 0.415176 | 0.917257 | Nat8f4 |
| ENSMUSG( | 76.42045 | -0.23532 | 0.409145 | 0.917257 | Nat8f5 |
| ENSMUSG( | 19.16387 | 0.009228 | 0.983957 | 0.996963 | Nat8f6 |
| ENSMUSG( | 12727.56 | 0.024734 | 0.86449 | 0.986591 | Nat8l |
| ENSMUSG( | 273.2002 | -0.10624 | 0.500654 | 0.927825 | Nat9 |
| ENSMUSG( | 588.6569 | 0.102114 | 0.468906 | 0.923329 | Natd1 |
| ENSMUSG( | 4222.979 | 0.115017 | 0.260089 | 0.884311 | Nav1 |
| ENSMUSG( | 2614.374 | 0.173925 | 0.101878 | 0.801719 | Nav2 |
| ENSMUSG( | 4573.795 | 0.013674 | 0.896639 | 0.990558 | Nav3 |
| ENSMUSG( | 1491.866 | -0.1335 | 0.236902 | 0.872173 | Naxd |
| ENSMUSG( | 1487.887 | -0.15878 | 0.176975 | 0.846973 | Naxe |
| ENSMUSG( | 1071.924 | 0.031594 | 0.853384 | 0.985598 | Nbas |
| ENSMUSG( | 485.2741 | -0.11645 | 0.431916 | 0.917854 | Nbdy |
| ENSMUSG( | 4073.158 | 0.217528 | 0.096729 | 0.796245 | Nbea |
| ENSMUSG( | 657.9186 | 0.191042 | 0.209303 | 0.864412 | Nbeal1 |
| ENSMUSG( | 457.7792 | 0.273176 | 0.056186 | 0.723362 | Nbeal2 |
| ENSMUSG( | 3116.069 | 0.114389 | 0.326212 | 0.905339 | Nbl1 |
| ENSMUSG( | 430.0014 | 0.116227 | 0.405607 | 0.917257 | Nbn |
| ENSMUSG( | 5525.832 | -0.06637 | 0.536058 | 0.936952 | Nbr1 |
| ENSMUSG( | 11704.22 | -0.15178 | 0.153742 | 0.837724 | Ncald |
| ENSMUSG( | 10442.08 | 0.105898 | 0.470351 | 0.923329 | Ncam1 |
| ENSMUSG( | 1307.461 | 0.262129 | 0.19542 | 0.861579 | Ncam2 |
| ENSMUSG( | 23921.23 | -0.0855 | 0.505139 | 0.927825 | Ncan |
| ENSMUSG( | 748.9053 | 0.135169 | 0.280238 | 0.892356 | Ncapd2 |
| ENSMUSG( | 598.4757 | 0.133018 | 0.44981 | 0.922202 | Ncapd3 |
| ENSMUSG( | 22.0529 | -0.36337 | 0.477285 | 0.923329 | Ncapg |
| ENSMUSG( | 186.703 | 0.146743 | 0.426062 | 0.917257 | Ncapg2 |
| ENSMUSG( | 75.74192 | -0.03191 | 0.903312 | 0.991089 | Ncaph |
| ENSMUSG( | 3454.264 | -0.08356 | 0.415019 | 0.917257 | Ncaph2 |
| ENSMUSG( | 1380.978 | 0.046554 | 0.67455 | 0.95694 | Ncbp1 |
| ENSMUSG( | 1164.831 | -0.07273 | 0.571457 | 0.942116 | Ncbp2 |
| ENSMUSG( | 1186.206 | -0.21319 | 0.140964 | 0.830678 | Ncbp2as2 |
| ENSMUSG( | 527.5448 | 0.051876 | 0.687662 | 0.959673 | Ncbp3 |
| ENSMUSG( | 16.30846 | -0.29422 | 0.57707 | 0.942912 | Nccrp1 |
| ENSMUSG( | 29192.41 | -0.05726 | 0.609026 | 0.950237 | Ncdn |
| ENSMUSG( | 4194.989 | 0.223218 | 0.194314 | 0.860096 | Nceh1 |
| ENSMUSG( | 406.1233 | 0.10496 | 0.455964 | 0.923329 | Ncf1 |
| ENSMUSG( | 214.8037 | -0.06581 | 0.696464 | 0.960867 | Ncf2 |
| ENSMUSG( | 42.94044 | -0.28875 | 0.37913 | 0.916995 | Ncf4 |
| ENSMUSG( | 309.443 | 0.160357 | 0.286222 | 0.893906 | Nck1 |
| ENSMUSG( | 652.1433 | -0.05994 | 0.676111 | 0.95694 | Nck2 |
| ENSMUSG( | 12974.14 | 0.024125 | 0.805615 | 0.977674 | Nckap1 |
| ENSMUSG( | 783.1579 | 0.08783 | 0.529771 | 0.934795 | Nckap1l |
| ENSMUSG( | 289.7383 | -0.20652 | 0.169715 | 0.840806 | Nckap5 |
| ENSMUSG( | 812.9531 | 0.062043 | 0.600911 | 0.947749 | Nckap5l |
| ENSMUSG( | 1472.548 | -0.05712 | 0.630475 | 0.953251 | Nckipsd |
| ENSMUSG( | 6899.227 | -0.06232 | 0.626874 | 0.953234 | Ncl |
| ENSMUSG( | 2220.22 | 0.140942 | 0.430723 | 0.917257 | Ncln |
| ENSMUSG( | 22.68829 | 0.437542 | 0.338496 | 0.907972 | Ncmap |
| ENSMUSG( | 3544.862 | 0.048003 | 0.651914 | 0.9547 | Ncoa1 |
| ENSMUSG( | 1946.555 | 0.08205 | 0.437317 | 0.920144 | Ncoa2 |
| ENSMUSG( | 1444.427 | 0.144164 | 0.264008 | 0.886134 | Ncoa3 |
| ENSMUSG( | 920.3219 | -0.13239 | 0.260408 | 0.884524 | Ncoa4 |
| ENSMUSG( | 1160.435 | 0.058704 | 0.588626 | 0.944975 | Ncoa5 |
| ENSMUSG( | 1925.771 | 0.088024 | 0.423646 | 0.917257 | Ncoa6 |
| ENSMUSG( | 881.4536 | 0.173716 | 0.223915 | 0.867404 | Ncoa7 |
| ENSMUSG( | 6803.161 | 0.027428 | 0.790422 | 0.97681 | Ncor1 |
| ENSMUSG( | 6334.805 | 0.152681 | 0.202496 | 0.864412 | Ncor2 |
| ENSMUSG( | 16487.73 | 0.092649 | 0.358051 | 0.913759 | Ncs1 |
| ENSMUSG( | 2896.495 | 0.134439 | 0.373755 | 0.915491 | Ncstn |
| ENSMUSG( | 626459.3 | -0.07378 | 0.722557 | 0.966887 | ND1 |
| ENSMUSG( | 378714.8 | -0.03837 | 0.860639 | 0.986137 | ND2 |

| | | | | | | | | | | | |
|---|---|---|---|---|---|---|---|---|---|---|---|
| ENSMUSG( | Ndn | 3138.617 | 0.175276 | 0.055364 | 0.361966 | ENSMUSG( | Ndn | 2757.425 | 0.071335 | 0.594782 | 0.945444 |
| ENSMUSG( | Ndnf | 471.2843 | -0.32492 | 0.055512 | 0.36248 | ENSMUSG( | Ndnf | 594.6424 | 0.582716 | 0.002419 | 0.192605 |
| ENSMUSG( | Ndor1 | 350.4582 | 0.035462 | 0.811092 | 0.964318 | ENSMUSG( | Ndor1 | 309.9123 | -0.04055 | 0.807822 | 0.978622 |
| ENSMUSG( | Ndp | 667.1414 | 0.027573 | 0.813829 | 0.964574 | ENSMUSG( | Ndp | 607.9638 | 0.028075 | 0.850004 | 0.985538 |
| ENSMUSG( | Ndrg1 | 3332.791 | -0.0492 | 0.574934 | 0.894572 | ENSMUSG( | Ndrg1 | 3040.651 | -0.04332 | 0.727409 | 0.966887 |
| ENSMUSG( | Ndrg2 | 125067.3 | 0.207084 | 0.068379 | 0.405144 | ENSMUSG( | Ndrg2 | 98271.41 | -0.22325 | 0.108749 | 0.804969 |
| ENSMUSG( | Ndrg3 | 13511.29 | 0.093057 | 0.345619 | 0.779153 | ENSMUSG( | Ndrg3 | 11310.82 | -0.15449 | 0.202252 | 0.864412 |
| ENSMUSG( | Ndrg4 | 31993.87 | -0.01962 | 0.873546 | 0.979057 | ENSMUSG( | Ndrg4 | 28071.2 | -0.13271 | 0.330288 | 0.907251 |
| ENSMUSG( | Ndst1 | 6197.817 | 0.014352 | 0.890053 | 0.983687 | ENSMUSG( | Ndst1 | 5905.05 | 0.142651 | 0.282886 | 0.892412 |
| ENSMUSG( | Ndst2 | 484.3612 | -0.11578 | 0.453378 | 0.836798 | ENSMUSG( | Ndst2 | 518.0026 | 0.33999 | 0.078276 | 0.77321 |
| ENSMUSG( | Ndst3 | 654.3215 | -0.08601 | 0.517399 | 0.873295 | ENSMUSG( | Ndst3 | 719.8676 | 0.431301 | 0.000784 | 0.093388 |
| ENSMUSG( | Ndst4 | 172.2822 | -1.05949 | 1.40E-05 | 0.00146 | ENSMUSG( | Ndst4 | 215.6412 | 0.039357 | 0.853431 | 0.985598 |
| ENSMUSG( | Ndufa1 | 2324.439 | -0.01294 | 0.903501 | 0.986055 | ENSMUSG( | Ndufa1 | 2024.119 | -0.15129 | 0.252189 | 0.879902 |
| ENSMUSG( | Ndufa10 | 4597.218 | 0.001667 | 0.986193 | 0.998326 | ENSMUSG( | Ndufa10 | 3841.265 | -0.26212 | 0.03199 | 0.633188 |
| ENSMUSG( | Ndufa11 | 1913.384 | -0.07764 | 0.497273 | 0.861898 | ENSMUSG( | Ndufa11 | 1636.252 | -0.27744 | 0.088438 | 0.786756 |
| ENSMUSG( | Ndufa12 | 2835.46 | 0.013023 | 0.879582 | 0.981436 | ENSMUSG( | Ndufa12 | 2307.15 | -0.33623 | 0.012489 | 0.451781 |
| ENSMUSG( | Ndufa13 | 5958.606 | -0.04239 | 0.586152 | 0.899709 | ENSMUSG( | Ndufa13 | 5106.516 | -0.22974 | 0.048471 | 0.706142 |
| ENSMUSG( | Ndufa2 | 2889.414 | 0.062395 | 0.524538 | 0.875849 | ENSMUSG( | Ndufa2 | 2373.967 | -0.25004 | 0.067057 | 0.75212 |
| ENSMUSG( | Ndufa3 | 3781.229 | -0.03957 | 0.611727 | 0.908586 | ENSMUSG( | Ndufa3 | 3102.71 | -0.36632 | 0.001518 | 0.143766 |
| ENSMUSG( | Ndufa4 | 7086.729 | -0.07806 | 0.320478 | 0.765921 | ENSMUSG( | Ndufa4 | 6114.304 | -0.24659 | 0.03258 | 0.636943 |
| ENSMUSG( | Ndufa4l2 | 92.47794 | -0.42343 | 0.169108 | 0.603241 | ENSMUSG( | Ndufa4l2 | 105.4871 | 0.271168 | 0.474177 | 0.923329 |
| ENSMUSG( | Ndufa5 | 2287.417 | 0.039525 | 0.686545 | 0.933041 | ENSMUSG( | Ndufa5 | 1898.458 | -0.24376 | 0.07071 | 0.758714 |
| ENSMUSG( | Ndufa6 | 2312.043 | -0.0451 | 0.631528 | 0.914073 | ENSMUSG( | Ndufa6 | 2021.664 | -0.17148 | 0.148804 | 0.836241 |
| ENSMUSG( | Ndufa7 | 2889.892 | -0.01947 | 0.814068 | 0.964574 | ENSMUSG( | Ndufa7 | 2384.455 | -0.32665 | 0.006026 | 0.320634 |
| ENSMUSG( | Ndufa8 | 3952.654 | 0.024125 | 0.771248 | 0.954726 | ENSMUSG( | Ndufa8 | 3238.745 | -0.30009 | 0.009617 | 0.406458 |
| ENSMUSG( | Ndufa9 | 3258.348 | 0.044673 | 0.634581 | 0.914073 | ENSMUSG( | Ndufa9 | 2747.639 | -0.18687 | 0.124071 | 0.814691 |
| ENSMUSG( | Ndufab1 | 1794.876 | 0.006196 | 0.949789 | 0.993821 | ENSMUSG( | Ndufab1 | 1554.935 | -0.14542 | 0.221345 | 0.866104 |
| ENSMUSG( | Ndufaf1 | 439.5051 | -0.01399 | 0.911449 | 0.988483 | ENSMUSG( | Ndufaf1 | 378.5086 | -0.1881 | 0.189997 | 0.857355 |
| ENSMUSG( | Ndufaf2 | 323.6231 | -0.03378 | 0.794352 | 0.959799 | ENSMUSG( | Ndufaf2 | 272.2255 | -0.27896 | 0.063972 | 0.736482 |
| ENSMUSG( | Ndufaf3 | 898.7031 | -0.08323 | 0.426005 | 0.824898 | ENSMUSG( | Ndufaf3 | 817.5995 | -0.0908 | 0.481778 | 0.924578 |
| ENSMUSG( | Ndufaf4 | 606.0955 | 0.016447 | 0.880777 | 0.981565 | ENSMUSG( | Ndufaf4 | 591.7593 | 0.203998 | 0.125859 | 0.81516 |
| ENSMUSG( | Ndufaf5 | 552.0546 | 0.085048 | 0.49012 | 0.859319 | ENSMUSG( | Ndufaf5 | 479.1184 | -0.06178 | 0.729839 | 0.967228 |
| ENSMUSG( | Ndufaf6 | 147.49 | -0.12349 | 0.505673 | 0.866932 | ENSMUSG( | Ndufaf6 | 134.0166 | -0.12976 | 0.526448 | 0.934374 |
| ENSMUSG( | Ndufaf7 | 996.0377 | 0.090282 | 0.343391 | 0.779153 | ENSMUSG( | Ndufaf7 | 865.8228 | -0.04536 | 0.716514 | 0.965773 |
| ENSMUSG( | Ndufaf8 | 554.9828 | -0.03076 | 0.77311 | 0.955272 | ENSMUSG( | Ndufaf8 | 444.4028 | -0.44607 | 0.002938 | 0.217424 |
| ENSMUSG( | Ndufb1 | 1696.227 | 0.015216 | 0.891566 | 0.983955 | ENSMUSG( | Ndufb1 | 1436.588 | -0.20729 | 0.139423 | 0.830678 |
| ENSMUSG( | Ndufb10 | 3356.922 | -0.00554 | 0.952156 | 0.993821 | ENSMUSG( | Ndufb10 | 2877.174 | -0.19056 | 0.113229 | 0.809623 |
| ENSMUSG( | Ndufb11 | 4205.914 | -0.02379 | 0.792463 | 0.959461 | ENSMUSG( | Ndufb11 | 3669.66 | -0.15504 | 0.182468 | 0.850579 |
| ENSMUSG( | Ndufb2 | 927.9536 | 0.015696 | 0.880806 | 0.981565 | ENSMUSG( | Ndufb2 | 776.3207 | -0.24461 | 0.078061 | 0.77321 |
| ENSMUSG( | Ndufb3 | 1690.035 | -0.08269 | 0.429876 | 0.826626 | ENSMUSG( | Ndufb3 | 1453.449 | -0.26453 | 0.045213 | 0.693959 |
| ENSMUSG( | Ndufb4 | 897.3947 | -0.11426 | 0.311679 | 0.759301 | ENSMUSG( | Ndufb4 | 787.2094 | -0.2354 | 0.074618 | 0.767423 |
| ENSMUSG( | Ndufb4b | 22.37961 | 0.100527 | 0.817564 | 0.965785 | ENSMUSG( | Ndufb4b | 17.46768 | -0.3949 | 0.446307 | 0.921734 |
| ENSMUSG( | Ndufb5 | 3523.627 | -0.06422 | 0.457877 | 0.839769 | ENSMUSG( | Ndufb5 | 3064.008 | -0.20795 | 0.068442 | 0.752603 |
| ENSMUSG( | Ndufb6 | 1438.895 | -0.05166 | 0.590204 | 0.90113 | ENSMUSG( | Ndufb6 | 1245.327 | -0.21276 | 0.186158 | 0.855305 |
| ENSMUSG( | Ndufb7 | 2291.032 | 0.021326 | 0.820829 | 0.965962 | ENSMUSG( | Ndufb7 | 1934.434 | -0.20917 | 0.105515 | 0.804969 |
| ENSMUSG( | Ndufb8 | 637.593 | -0.03933 | 0.710162 | 0.9397 | ENSMUSG( | Ndufb8 | 543.4895 | -0.24507 | 0.069507 | 0.75601 |
| ENSMUSG( | Ndufb9 | 7453.157 | 0.0595 | 0.43551 | 0.829093 | ENSMUSG( | Ndufb9 | 6319.899 | -0.15181 | 0.14246 | 0.831221 |
| ENSMUSG( | Ndufc1 | 1437.887 | 0.009188 | 0.93177 | 0.9924 | ENSMUSG( | Ndufc1 | 1211.324 | -0.23069 | 0.074828 | 0.767423 |
| ENSMUSG( | Ndufc2 | 2844.899 | -0.19996 | 0.044028 | 0.321685 | ENSMUSG( | Ndufc2 | 2611.536 | -0.17889 | 0.166858 | 0.840183 |
| ENSMUSG( | Ndufs1 | 4889.826 | 0.093302 | 0.320645 | 0.765984 | ENSMUSG( | Ndufs1 | 4064.733 | -0.17739 | 0.105696 | 0.804969 |
| ENSMUSG( | Ndufs2 | 7585.578 | 0.070645 | 0.402732 | 0.81552 | ENSMUSG( | Ndufs2 | 6339.478 | -0.18541 | 0.111235 | 0.807885 |
| ENSMUSG( | Ndufs3 | 2125.323 | -0.0524 | 0.585405 | 0.899171 | ENSMUSG( | Ndufs3 | 1843.726 | -0.20219 | 0.085642 | 0.779799 |
| ENSMUSG( | Ndufs4 | 1873.463 | -0.0585 | 0.530247 | 0.879138 | ENSMUSG( | Ndufs4 | 1620.901 | -0.21945 | 0.08201 | 0.77681 |
| ENSMUSG( | Ndufs5 | 1420.53 | -1.53272 | 1.25E-44 | 6.73E-41 | ENSMUSG( | Ndufs5 | 1796.998 | -0.2086 | 0.133553 | 0.822912 |
| ENSMUSG( | Ndufs6 | 1715.489 | -0.018 | 0.850642 | 0.972638 | ENSMUSG( | Ndufs6 | 1431.065 | -0.2906 | 0.017254 | 0.527468 |
| ENSMUSG( | Ndufs7 | 2385.886 | 0.038435 | 0.71949 | 0.941986 | ENSMUSG( | Ndufs7 | 2021.205 | -0.18108 | 0.198633 | 0.862745 |
| ENSMUSG( | Ndufs8 | 2567.572 | -0.11028 | 0.186584 | 0.623869 | ENSMUSG( | Ndufs8 | 2209.199 | -0.28971 | 0.012137 | 0.446654 |
| ENSMUSG( | Ndufv1 | 4673.234 | 0.088587 | 0.295533 | 0.743814 | ENSMUSG( | Ndufv1 | 3878.64 | -0.18683 | 0.10979 | 0.80637 |
| ENSMUSG( | Ndufv2 | 3274.112 | -0.01598 | 0.867527 | 0.97663 | ENSMUSG( | Ndufv2 | 2803.159 | -0.20482 | 0.117289 | 0.813384 |
| ENSMUSG( | Ndufv3 | 2280.054 | 0.006802 | 0.938955 | 0.993415 | ENSMUSG( | Ndufv3 | 1917.188 | -0.23756 | 0.061483 | 0.736111 |
| ENSMUSG( | Neat1 | 1379.008 | 0.08679 | 0.638689 | 0.91636 | ENSMUSG( | Neat1 | 1304.381 | 0.187974 | 0.387177 | 0.917257 |
| ENSMUSG( | Neb | 54.75968 | -0.02295 | 0.932951 | 0.992879 | ENSMUSG( | Neb | 62.64989 | 0.591791 | 0.031526 | 0.631192 |
| ENSMUSG( | Nebl | 3510.675 | 0.134285 | 0.357343 | 0.788775 | ENSMUSG( | Nebl | 3006.782 | -0.04483 | 0.776186 | 0.975717 |
| ENSMUSG( | Necab1 | 1323.441 | 0.003465 | 0.975853 | 0.997428 | ENSMUSG( | Necab1 | 1215.962 | 0.021914 | 0.891218 | 0.990558 |
| ENSMUSG( | Necab2 | 2306.411 | -0.11242 | 0.411699 | 0.81942 | ENSMUSG( | Necab2 | 2030.439 | -0.22156 | 0.153054 | 0.837724 |
| ENSMUSG( | Necab3 | 1660.598 | 0.053656 | 0.637306 | 0.915436 | ENSMUSG( | Necab3 | 1460.065 | -0.04924 | 0.676146 | 0.95694 |
| ENSMUSG( | Necap1 | 2586.638 | 0.024024 | 0.83973 | 0.971179 | ENSMUSG( | Necap1 | 2251.367 | -0.11228 | 0.42802 | 0.917257 |
| ENSMUSG( | Necap2 | 2285.402 | 0.076958 | 0.429569 | 0.826446 | ENSMUSG( | Necap2 | 1943.36 | -0.12501 | 0.267528 | 0.886134 |
| ENSMUSG( | Nectin1 | 2178.487 | -0.22399 | 0.144738 | 0.564884 | ENSMUSG( | Nectin1 | 2206.489 | 0.09393 | 0.557067 | 0.939565 |
| ENSMUSG( | Nectin2 | 351.0743 | -0.46933 | 0.002017 | 0.051507 | ENSMUSG( | Nectin2 | 376.1849 | 0.045134 | 0.820622 | 0.979198 |
| ENSMUSG( | Nectin3 | 1561.651 | -0.16402 | 0.265517 | 0.716109 | ENSMUSG( | Nectin3 | 1526.574 | 0.04654 | 0.77252 | 0.974454 |
| ENSMUSG( | Nectin4 | 150.7513 | -0.20037 | 0.230379 | 0.678421 | ENSMUSG( | Nectin4 | 148.8223 | 0.050709 | 0.789352 | 0.976593 |
| ENSMUSG( | Nedd1 | 159.4703 | 0.174352 | 0.299657 | 0.74868 | ENSMUSG( | Nedd1 | 152.3325 | 0.304312 | 0.107261 | 0.804969 |
| ENSMUSG( | Nedd4 | 16318.54 | -0.09297 | 0.305743 | 0.753712 | ENSMUSG( | Nedd4 | 15615.2 | 0.04964 | 0.598259 | 0.946676 |
| ENSMUSG( | Nedd4l | 3743.483 | 0.144074 | 0.116067 | 0.514235 | ENSMUSG( | Nedd4l | 3332.723 | 0.078694 | 0.487184 | 0.92541 |
| ENSMUSG( | Nedd8 | 2209.47 | -0.01497 | 0.886549 | 0.983204 | ENSMUSG( | Nedd8 | 1815.631 | -0.33674 | 0.024045 | 0.588226 |
| ENSMUSG( | Nedd9 | 308.48 | -0.21168 | 0.10989 | 0.500285 | ENSMUSG( | Nedd9 | 280.5329 | -0.20573 | 0.223407 | 0.867404 |
| ENSMUSG( | Nefh | 1695.492 | -0.03739 | 0.750996 | 0.950053 | ENSMUSG( | Nefh | 1504.492 | -0.11691 | 0.387515 | 0.917257 |
| ENSMUSG( | Nefl | 10680.01 | -0.0549 | 0.518414 | 0.873901 | ENSMUSG( | Nefl | 9213.239 | -0.22194 | 0.036034 | 0.668395 |
| ENSMUSG( | Nefm | 3925.113 | 0.115308 | 0.375442 | 0.800845 | ENSMUSG( | Nefm | 3349.098 | -0.07252 | 0.61112 | 0.950771 |
| ENSMUSG( | Negr1 | 6478.596 | -0.28107 | 0.05702 | 0.368322 | ENSMUSG( | Negr1 | 6817.592 | 0.147623 | 0.36985 | 0.915491 |
| ENSMUSG( | Neil1 | 312.8988 | 0.208171 | 0.14252 | 0.560815 | ENSMUSG( | Neil1 | 271.1569 | 0.078147 | 0.659611 | 0.956205 |
| ENSMUSG( | Neil2 | 73.24599 | 0.282951 | 0.235737 | 0.682914 | ENSMUSG( | Neil2 | 66.84101 | 0.28817 | 0.281082 | 0.892356 |
| ENSMUSG( | Nek1 | 1397.02 | 0.063218 | 0.589362 | 0.90113 | ENSMUSG( | Nek1 | 1216.109 | -0.07411 | 0.607085 | 0.949917 |
| ENSMUSG( | Nek10 | 80.64171 | -0.36217 | 0.11107 | 0.502387 | ENSMUSG( | Nek10 | 84.31083 | 0.039965 | 0.872431 | 0.989023 |
| ENSMUSG( | Nek11 | 39.54741 | -0.26491 | 0.380108 | 0.801757 | ENSMUSG( | Nek11 | 37.00525 | -0.187 | 0.589669 | 0.944975 |
| ENSMUSG( | Nek2 | 25.8449 | -0.33642 | 0.352619 | 0.784565 | ENSMUSG( | Nek2 | 23.99592 | -0.26286 | 0.505907 | 0.927825 |
| ENSMUSG( | Nek3 | 97.94979 | 0.284142 | 0.184182 | 0.621101 | ENSMUSG( | Nek3 | 76.5677 | -0.14666 | 0.557094 | 0.939565 |
| ENSMUSG( | Nek4 | 603.5321 | 0.114731 | 0.351967 | 0.783738 | ENSMUSG( | Nek4 | 521.9597 | -0.03779 | 0.786178 | 0.976285 |
| ENSMUSG( | Nek5 | 19.97921 | -0.85254 | 0.144601 | NA | ENSMUSG( | Nek5 | 20.94258 | -0.32923 | 0.514977 | 0.930939 |
| ENSMUSG( | Nek6 | 1463.507 | 0.081501 | 0.367261 | 0.795221 | ENSMUSG( | Nek6 | 1240.513 | -0.13214 | 0.280989 | 0.892356 |
| ENSMUSG( | Nek7 | 3798.242 | -0.00199 | 0.985968 | 0.998326 | ENSMUSG( | Nek7 | 3231.992 | -0.20937 | 0.143826 | 0.831311 |
| ENSMUSG( | Nek8 | 185.4321 | -0.0472 | 0.778691 | 0.956339 | ENSMUSG( | Nek8 | 162.2542 | -0.16399 | 0.445327 | 0.921734 |
| ENSMUSG( | Nek9 | 4026.629 | 0.087771 | 0.349497 | 0.781909 | ENSMUSG( | Nek9 | 3376.264 | -0.15626 | 0.163761 | 0.838291 |
| ENSMUSG( | Nelfa | 625.6925 | 0.158257 | 0.30302 | 0.751443 | ENSMUSG( | Nelfa | 526.9263 | -0.07128 | 0.670747 | 0.95694 |
| ENSMUSG( | Nelfb | 1467.861 | 0.129826 | 0.140276 | 0.557263 | ENSMUSG( | Nelfb | 1284.752 | 0.01631 | 0.894248 | 0.990558 |
| ENSMUSG( | Nelfcd | 1102.378 | 0.148572 | 0.22266 | 0.668434 | ENSMUSG( | Nelfcd | 927.7636 | -0.07714 | 0.595912 | 0.946125 |
| ENSMUSG( | Nelfe | 891.141 | 0.081846 | 0.444618 | 0.833239 | ENSMUSG( | Nelfe | 750.6623 | -0.15148 | 0.252325 | 0.880122 |
| ENSMUSG( | Nell1 | 1229.567 | -0.49409 | 0.000654 | 0.024667 | ENSMUSG( | Nell1 | 1325.786 | 0.038846 | 0.811901 | 0.979148 |
| ENSMUSG( | Nell2 | 15315.2 | -0.26061 | 0.076915 | 0.429306 | ENSMUSG( | Nell2 | 16508.2 | 0.231882 | 0.175154 | 0.845305 |
| ENSMUSG( | Nemf | 1097.128 | 0.018854 | 0.858249 | 0.974672 | ENSMUSG( | Nemf | 1013.92 | 0.062597 | 0.596805 | 0.946541 |
| ENSMUSG( | Nemp2 | 193.3475 | 0.069678 | 0.681263 | 0.931396 | ENSMUSG( | Nemp2 | 195.4301 | 0.359552 | 0.056998 | 0.723362 |
| ENSMUSG( | Nenf | 2113.227 | -0.27067 | 0.006426 | 0.110055 | ENSMUSG( | Nenf | 2005.315 | -0.14575 | 0.249775 | 0.877904 |
| ENSMUSG( | Neo1 | 5258.529 | -0.16467 | 0.205468 | 0.646873 | ENSMUSG( | Neo1 | 5225.771 | 0.092541 | 0.520736 | 0.932121 |
| ENSMUSG( | Nepro | 108.6934 | -0.00645 | 0.976529 | 0.997428 | ENSMUSG( | Nepro | 112.2767 | 0.339392 | 0.114486 | 0.811703 |
| ENSMUSG( | Nes | 606.8842 | 0.074346 | 0.680918 | 0.931244 | ENSMUSG( | Nes | 649.2046 | 0.505058 | 0.015781 | 0.502615 |
| ENSMUSG( | Net1 | 645.6868 | 0.088996 | 0.513251 | 0.871762 | ENSMUSG( | Net1 | 634.6736 | 0.301299 | 0.099961 | 0.801311 |
| ENSMUSG( | Neto1 | 2225.196 | -0.29547 | 0.007472 | 0.119529 | ENSMUSG( | Neto1 | 2404 | 0.205717 | 0.136306 | 0.827092 |
| ENSMUSG( | Neto2 | 1626.781 | -0.58389 | 0.000126 | 0.007476 | ENSMUSG( | Neto2 | 1926.561 | 0.223882 | 0.20543 | 0.864412 |
| ENSMUSG( | Neu1 | 1854.576 | -0.24329 | 0.069571 | 0.408118 | ENSMUSG( | Neu1 | 1981.185 | 0.22251 | 0.160135 | 0.837724 |
| ENSMUSG( | Neu2 | 161.4735 | -0.11434 | 0.539419 | 0.883094 | ENSMUSG( | Neu2 | 135.8905 | -0.34307 | 0.131394 | 0.822912 |
| ENSMUSG( | Neu3 | 65.03909 | -0.53047 | 0.039589 | 0.306803 | ENSMUSG( | Neu3 | 61.83453 | -0.3779 | 0.155837 | 0.837724 |
| ENSMUSG( | Neu4 | 531.1895 | -0.2836 | 0.058676 | 0.373994 | ENSMUSG( | Neu4 | 567.3743 | 0.188894 | 0.252802 | 0.88018 |
| ENSMUSG( | Neurl1a | 11144.24 | 0.253549 | 0.002912 | 0.066108 | ENSMUSG( | Neurl1a | 9913.718 | 0.190651 | 0.086812 | 0.782073 |
| ENSMUSG( | Neurl1b | 3290.344 | 0.237995 | 0.017056 | 0.195102 | ENSMUSG( | Neurl1b | 3040.19 | 0.277282 | 0.024448 | 0.590702 |
| ENSMUSG( | Neurl2 | 61.17785 | 0.422825 | 0.106657 | 0.494174 | ENSMUSG( | Neurl2 | 51.51781 | 0.204652 | 0.493273 | 0.927372 |
| ENSMUSG( | Neurl3 | 58.17691 | -0.91344 | 0.000875 | 0.029753 | ENSMUSG( | Neurl3 | 56.61765 | -0.66277 | 0.022825 | 0.582823 |
| ENSMUSG( | Neurl4 | 5006.935 | 0.133896 | 0.129343 | 0.538561 | ENSMUSG( | Neurl4 | 4414.233 | 0.043336 | 0.701974 | 0.962014 |
| ENSMUSG( | Neurod1 | 159.9451 | 0.092577 | 0.624117 | 0.91187 | ENSMUSG( | Neurod1 | 149.3987 | 0.153907 | 0.437451 | 0.920144 |
| ENSMUSG( | Neurod2 | 2165.395 | 0.179153 | 0.200434 | 0.640598 | ENSMUSG( | Neurod2 | 1858.16 | 0.009519 | 0.949822 | 0.994798 |
| ENSMUSG( | Neurod4 | 10.7139 | -0.1947 | 0.731241 | NA | ENSMUSG( | Neurod4 | 11.74769 | 0.307202 | 0.596368 | 0.946541 |
| ENSMUSG( | Neurod6 | 977.5383 | 0.017631 | 0.912941 | 0.988483 | ENSMUSG( | Neurod6 | 858.7844 | -0.09513 | 0.601483 | 0.947749 |
| ENSMUSG( | Neurog2 | 58.41749 | 0.064693 | 0.811854 | 0.964426 | ENSMUSG( | Neurog2 | 54.6835 | 0.157902 | 0.575845 | 0.942912 |
| ENSMUSG( | Nexmif | 458.9989 | 0.013747 | 0.919802 | 0.990247 | ENSMUSG( | Nexmif | 464.4861 | 0.30438 | 0.042001 | 0.690304 |
| ENSMUSG( | Nexn | 180.4167 | -0.78414 | 0.000884 | 0.02993 | ENSMUSG( | Nexn | 167.8605 | -0.72302 | 0.000879 | 0.099346 |
| ENSMUSG( | Nf1 | 3512.326 | 0.032377 | 0.72807 | 0.944355 | ENSMUSG( | Nf1 | 3177.873 | 0.013805 | 0.902298 | 0.991069 |
| ENSMUSG( | Nf2 | 8606.31 | 0.080343 | 0.342168 | 0.778828 | ENSMUSG( | Nf2 | 7409.957 | -0.08397 | 0.44579 | 0.921734 |
| ENSMUSG( | Nfam1 | 229.7625 | -0.15787 | 0.399874 | 0.813593 | ENSMUSG( | Nfam1 | 238.7789 | 0.229446 | 0.294351 | 0.898538 |
| ENSMUSG( | Nfasc | 12618.59 | -0.206 | 0.093558 | 0.469962 | ENSMUSG( | Nfasc | 13164.56 | 0.189643 | 0.20889 | 0.864412 |
| ENSMUSG( | Nfat5 | 2455.623 | -0.03029 | 0.752489 | 0.950536 | ENSMUSG( | Nfat5 | 2248.527 | -0.01489 | 0.892983 | 0.990558 |
| ENSMUSG( | Nfatc1 | 366.9787 | -0.14076 | 0.393012 | 0.809274 | ENSMUSG( | Nfatc1 | 338.1608 | -0.10356 | 0.498295 | 0.927825 |
| ENSMUSG( | Nfatc2 | 472.6968 | -0.09952 | 0.387808 | 0.805835 | ENSMUSG( | Nfatc2 | 422.5429 | -0.16419 | 0.280437 | 0.892356 |
| ENSMUSG( | Nfatc2ip | 167.7987 | 0.005891 | 0.97156 | 0.996503 | ENSMUSG( | Nfatc2ip | 155.8403 | 0.061358 | 0.736213 | 0.967344 |
| ENSMUSG( | Nfatc3 | 400.3722 | 0.179092 | 0.126871 | 0.534201 | ENSMUSG( | Nfatc3 | 380.1418 | 0.291602 | 0.033557 | 0.64525 |
| ENSMUSG( | Nfatc4 | 64.19232 | -0.01128 | 0.962869 | 0.994824 | ENSMUSG( | Nfatc4 | 70.2899 | 0.497978 | 0.092782 | 0.794753 |
| ENSMUSG( | Nfe2l1 | 14227.43 | -0.07792 | 0.400813 | 0.81387 | ENSMUSG( | Nfe2l1 | 13402.96 | 0.020879 | 0.854859 | 0.985649 |
| ENSMUSG( | Nfe2l2 | 851.891 | 0.071566 | 0.537629 | 0.882536 | ENSMUSG( | Nfe2l2 | 803.1454 | 0.167782 | 0.15182 | 0.837724 |
| ENSMUSG( | Nfe2l3 | 124.0338 | -0.88122 | 5.77E-06 | 0.000749 | ENSMUSG( | Nfe2l3 | 148.2524 | 0.049388 | 0.836938 | 0.981335 |
| ENSMUSG( | Nfia | 2889.624 | 0.120856 | 0.284085 | 0.732569 | ENSMUSG( | Nfia | 2517.774 | -0.00837 | 0.958581 | 0.994798 |
| ENSMUSG( | Nfib | 3864.509 | 0.157526 | 0.051455 | 0.348059 | ENSMUSG( | Nfib | 3539.529 | 0.170798 | 0.229043 | 0.868507 |
| ENSMUSG( | Nfic | 7533.281 | 0.169281 | 0.045632 | 0.327937 | ENSMUSG( | Nfic | 6636.965 | 0.077003 | 0.461878 | 0.923329 |
| ENSMUSG( | Nfil3 | 164.9377 | -0.06216 | 0.704678 | 0.938303 | ENSMUSG( | Nfil3 | 130.4276 | -0.52995 | 0.009795 | 0.406458 |
| ENSMUSG( | Nfix | 14453.36 | 0.052287 | 0.486944 | 0.858812 | ENSMUSG( | Nfix | 12380.26 | -0.12805 | 0.204475 | 0.864412 |
| ENSMUSG( | Nfkb1 | 688.8203 | -0.03866 | 0.725896 | 0.944223 | ENSMUSG( | Nfkb1 | 651.3283 | 0.07311 | 0.593413 | 0.945408 |
| ENSMUSG( | Nfkb2 | 241.3455 | 0.232676 | 0.124862 | 0.531843 | ENSMUSG( | Nfkb2 | 204.6614 | 0.035831 | 0.829851 | 0.980224 |
| ENSMUSG( | Nfkbia | 618.8492 | -0.26583 | 0.053293 | 0.354557 | ENSMUSG( | Nfkbia | 688.1309 | 0.311674 | 0.111954 | 0.808673 |
| ENSMUSG( | Nfkbib | 657.2655 | -0.01892 | 0.899576 | 0.985633 | ENSMUSG( | Nfkbib | 570.5692 | -0.16524 | 0.281619 | 0.892412 |
| ENSMUSG( | Nfkbid | 90.16783 | -0.0571 | 0.801391 | 0.961947 | ENSMUSG( | Nfkbid | 81.75432 | -0.05962 | 0.833317 | 0.981193 |
| ENSMUSG( | Nfkbie | 101.0194 | -0.39737 | 0.081607 | 0.441308 | ENSMUSG( | Nfkbie | 94.14396 | -0.32198 | 0.16439 | 0.83856 |
| ENSMUSG( | Nfkbil1 | 355.8038 | -0.07829 | 0.556541 | 0.887713 | ENSMUSG( | Nfkbil1 | 319.5972 | -0.13061 | 0.451429 | 0.922202 |
| ENSMUSG( | Nfkbiz | 177.2441 | -0.29516 | 0.062357 | 0.386294 | ENSMUSG( | Nfkbiz | 176.41 | -0.01594 | 0.932075 | 0.99306 |
| ENSMUSG( | Nfrkb | 893.6802 | -0.0418 | 0.71437 | 0.941045 | ENSMUSG( | Nfrkb | 829.2445 | 0.016602 | 0.909639 | 0.991958 |
| ENSMUSG( | Nfs1 | 1337.764 | 0.035519 | 0.748857 | 0.949604 | ENSMUSG( | Nfs1 | 1129.056 | -0.19679 | 0.13532 | 0.825081 |
| ENSMUSG( | Nfu1 | 922.8541 | 0.002726 | 0.979379 | 0.997428 | ENSMUSG( | Nfu1 | 797.436 | -0.16078 | 0.269186 | 0.886755 |
| ENSMUSG( | Nfx1 | 1903.005 | -0.04783 | 0.631806 | 0.914073 | ENSMUSG( | Nfx1 | 1797.308 | 0.058272 | 0.61828 | 0.952254 |
| ENSMUSG( | Nfxl1 | 285.6438 | -0.17603 | 0.194439 | 0.63341 | ENSMUSG( | Nfxl1 | 288.7873 | 0.132241 | 0.393331 | 0.917257 |
| ENSMUSG( | Nfya | 659.1457 | 0.171971 | 0.114371 | 0.51021 | ENSMUSG( | Nfya | 564.0699 | -0.00035 | 0.997842 | 0.999268 |
| ENSMUSG( | Nfyb | 1554.954 | 0.048528 | 0.63874 | 0.91636 | ENSMUSG( | Nfyb | 1445.045 | 0.10029 | 0.463014 | 0.923329 |

| | | | | | |
|---|---|---|---|---|---|
| ENSMUSG( | 48.79978 | -0.31798 | 0.268511 | 0.718807 | NA |
| ENSMUSG( | 22.91177 | -0.80378 | 0.096845 | 0.477375 | NA |
| ENSMUSG( | 42.80247 | -0.13992 | 0.646088 | 0.918453 | NA |
| ENSMUSG( | 365.23693 | -0.12156 | 0.333232 | 0.775602 | Naa10 |
| ENSMUSG( | 1359.873 | 0.085106 | 0.356774 | 0.788403 | Naa15 |
| ENSMUSG( | 198.6441 | 0.15641 | 0.331201 | 0.77519 | Naa16 |
| ENSMUSG( | 696.9992 | -0.02553 | 0.845431 | 0.97218 | Naa20 |
| ENSMUSG( | 1106.078 | -0.05234 | 0.631368 | 0.914073 | Naa25 |
| ENSMUSG( | 772.0969 | 0.009858 | 0.940185 | 0.993415 | Naa30 |
| ENSMUSG( | 1327.942 | 0.122586 | 0.191552 | 0.630601 | Naa35 |
| ENSMUSG( | 1060.16 | -0.0808 | 0.41998 | 0.822442 | Naa38 |
| ENSMUSG( | 1064.522 | -0.00708 | 0.948068 | 0.993821 | Naa40 |
| ENSMUSG( | 2001.21 | -0.00302 | 0.980805 | 0.997512 | Naa50 |
| ENSMUSG( | 3491.668 | 0.039876 | 0.621615 | 0.911432 | Naa60 |
| ENSMUSG( | 406.6976 | -0.07663 | 0.516273 | 0.872701 | Naa80 |
| ENSMUSG( | 3029.758 | -0.00064 | 0.99588 | 0.99943 | Naaa |
| ENSMUSG( | 49.93705 | -0.51429 | 0.090315 | 0.461669 | Naalad2 |
| ENSMUSG( | 44.21782 | -0.96113 | 0.002864 | 0.065614 | Naaladl2 |
| ENSMUSG( | 1810.767 | 0.009532 | 0.925886 | 0.991229 | Nab1 |
| ENSMUSG( | 866.7868 | -0.26175 | 0.087823 | 0.454663 | Nab2 |
| ENSMUSG( | 175.4203 | -0.09312 | 0.659685 | 0.924524 | Nabp1 |
| ENSMUSG( | 2145.28 | -0.03729 | 0.682643 | 0.93198 | Nabp2 |
| ENSMUSG( | 4562.253 | 0.096161 | 0.254708 | 0.705011 | Naca |
| ENSMUSG( | 1150.969 | -0.15694 | 0.132934 | 0.545594 | Nacad |
| ENSMUSG( | 6122.356 | 0.134244 | 0.111911 | 0.50438 | Nacc1 |
| ENSMUSG( | 16314.42 | 0.047059 | 0.682252 | 0.931962 | Nacc2 |
| ENSMUSG( | 4095.865 | 0.146665 | 0.128694 | 0.537025 | Nadk |
| ENSMUSG( | 3994.699 | 0.108356 | 0.427125 | 0.82532 | Nadk2 |
| ENSMUSG( | 176.5366 | -0.0535 | 0.741986 | 0.948478 | Nadsyn1 |
| ENSMUSG( | 855.9598 | -0.03504 | 0.767928 | 0.954621 | Nae1 |
| ENSMUSG( | 528.1917 | 0.008016 | 0.954087 | 0.993874 | Naf1 |
| ENSMUSG( | 948.8724 | -0.33985 | 0.009196 | 0.135112 | Naga |
| ENSMUSG( | 1162.654 | 0.069737 | 0.458168 | 0.839914 | Nagk |
| ENSMUSG( | 409.6716 | -0.42976 | 0.002683 | 0.062344 | Naglu |
| ENSMUSG( | 346.5948 | -0.12357 | 0.396819 | 0.811742 | Nagpa |
| ENSMUSG( | 143.61 | -0.19015 | 0.255702 | 0.706195 | Naif1 |
| ENSMUSG( | 12.5672 | 0.131899 | 0.814068 | NA | Naip1 |
| ENSMUSG( | 55.67633 | -0.15751 | 0.613738 | 0.909201 | Naip2 |
| ENSMUSG( | 80.81009 | -0.49157 | 0.048973 | 0.337919 | Naip5 |
| ENSMUSG( | 47.49215 | -0.4272 | 0.152016 | 0.577721 | Naip6 |
| ENSMUSG( | 2173.799 | -0.4548 | 0.003208 | 0.070583 | Nalcn |
| ENSMUSG( | 2643.177 | -0.18911 | 0.042158 | 0.316282 | Nalf1 |
| ENSMUSG( | 970.3349 | 0.239572 | 0.012297 | 0.160458 | Nalf2 |
| ENSMUSG( | 1166.309 | -0.05934 | 0.540786 | 0.88315 | Nampt |
| ENSMUSG( | 1084.202 | 0.008186 | 0.934855 | 0.993231 | Nanos1 |
| ENSMUSG( | 40.06825 | -0.32542 | 0.376808 | 0.801213 | Nanos2 |
| ENSMUSG( | 22.05071 | -0.03939 | 0.924414 | 0.99114 | Nanos3 |
| ENSMUSG( | 372.8805 | 0.039235 | 0.768225 | 0.954641 | Nans |
| ENSMUSG( | 5082.426 | 0.04686 | 0.635559 | 0.914465 | Nap1l1 |
| ENSMUSG( | 793.5891 | -0.23517 | 0.108327 | 0.497337 | Nap1l2 |
| ENSMUSG( | 479.2366 | -0.02842 | 0.863274 | 0.975055 | Nap1l3 |
| ENSMUSG( | 3810.561 | 0.316357 | 0.003062 | 0.068418 | Nap1l4 |
| ENSMUSG( | 4199.872 | 0.047165 | 0.729696 | 0.944813 | Nap1l5 |
| ENSMUSG( | 5232.399 | -0.02685 | 0.82242 | 0.966352 | Napa |
| ENSMUSG( | 7831.572 | 0.079717 | 0.436234 | 0.829093 | Napb |
| ENSMUSG( | 291.0775 | 0.266233 | 0.116659 | 0.515027 | Napepld |
| ENSMUSG( | 2685.351 | 0.01502 | 0.874921 | 0.979746 | Napg |
| ENSMUSG( | 367.9891 | -0.00215 | 0.988256 | 0.999109 | Naprt |
| ENSMUSG( | 26.01754 | -1.22066 | 0.01488 | 0.179275 | Napsa |
| ENSMUSG( | 1425.817 | 0.145534 | 0.132338 | 0.544251 | Narf |
| ENSMUSG( | 3921.562 | 0.088454 | 0.344709 | 0.779153 | Nars1 |
| ENSMUSG( | 328.7754 | -0.09086 | 0.46158 | 0.8415 | Nars2 |
| ENSMUSG( | 600.6874 | 0.125498 | 0.284236 | 0.73274 | Nasp |
| ENSMUSG( | 90.61949 | -0.1417 | 0.548777 | 0.885686 | Nat1 |
| ENSMUSG( | 909.5075 | 0.158788 | 0.157473 | 0.585653 | Nat10 |
| ENSMUSG( | 3014.825 | -0.20343 | 0.080804 | 0.439517 | Nat14 |
| ENSMUSG( | 30.75329 | 0.180962 | 0.611656 | 0.908586 | Nat2 |
| ENSMUSG( | 99.91793 | -0.08713 | 0.738951 | 0.947768 | Nat8 |
| ENSMUSG( | 633.1842 | 0.049616 | 0.713956 | 0.940903 | Nat8f1 |
| ENSMUSG( | 23.47079 | 0.329809 | 0.40293 | 0.815716 | Nat8f2 |
| ENSMUSG( | 154.8668 | 0.020715 | 0.920272 | 0.990247 | Nat8f3 |
| ENSMUSG( | 1033.9 | 0.047005 | 0.760518 | 0.952907 | Nat8f4 |
| ENSMUSG( | 95.77454 | 0.134518 | 0.603797 | 0.905053 | Nat8f5 |
| ENSMUSG( | 25.04484 | 0.489424 | 0.203144 | 0.643631 | Nat8f6 |
| ENSMUSG( | 11.99024 | -0.2473 | 0.660613 | NA | Nat8f7 |
| ENSMUSG( | 13458.89 | -0.08718 | 0.467992 | 0.84521 | Nat8l |
| ENSMUSG( | 321.2198 | 0.086189 | 0.511738 | 0.871106 | Nat9 |
| ENSMUSG( | 663.0213 | 0.171421 | 0.106494 | 0.494174 | Natd1 |
| ENSMUSG( | 4182.74 | -0.18527 | 0.028206 | 0.256543 | Nav1 |
| ENSMUSG( | 2835.328 | 0.141054 | 0.112744 | 0.506698 | Nav2 |
| ENSMUSG( | 5177.665 | 0.101038 | 0.225248 | 0.672044 | Nav3 |
| ENSMUSG( | 1704.974 | -0.01261 | 0.883265 | 0.982422 | Naxd |
| ENSMUSG( | 1650.221 | -0.1252 | 0.157135 | 0.58551 | Naxe |
| ENSMUSG( | 1086.883 | -0.21152 | 0.122793 | 0.529145 | Nbas |
| ENSMUSG( | 567.5132 | 0.075494 | 0.546835 | 0.884974 | Nbdy |
| ENSMUSG( | 4216.01 | 0.052869 | 0.598542 | 0.903801 | Nbea |
| ENSMUSG( | 654.3249 | -0.09138 | 0.558484 | 0.888521 | Nbeal1 |
| ENSMUSG( | 475.6675 | 0.120996 | 0.31163 | 0.759296 | Nbeal2 |
| ENSMUSG( | 3212.344 | -0.0678 | 0.464439 | 0.843364 | Nbl1 |
| ENSMUSG( | 493.9583 | 0.24112 | 0.043799 | 0.320589 | Nbn |
| ENSMUSG( | 6141.272 | -0.02961 | 0.766119 | 0.954173 | Nbr1 |
| ENSMUSG( | 13831.09 | 0.065252 | 0.432929 | 0.828185 | Ncald |
| ENSMUSG( | 9986.927 | -0.30831 | 0.015544 | 0.184433 | Ncam1 |
| ENSMUSG( | 1137.295 | -0.43606 | 0.014697 | 0.178407 | Ncam2 |
| ENSMUSG( | 25322.95 | -0.19682 | 0.102053 | 0.486742 | Ncan |
| ENSMUSG( | 788.3054 | 0.014446 | 0.897785 | 0.985633 | Ncapd2 |
| ENSMUSG( | 628.6196 | 0.007535 | 0.961474 | 0.994205 | Ncapd3 |
| ENSMUSG( | 20.30485 | -1.05574 | 0.039579 | NA | Ncapg |
| ENSMUSG( | 194.7071 | -0.00707 | 0.969182 | 0.996369 | Ncapg2 |
| ENSMUSG( | 80.20079 | -0.16186 | 0.48979 | 0.859319 | Ncaph |
| ENSMUSG( | 4042.454 | 0.100789 | 0.227167 | 0.674223 | Ncaph2 |
| ENSMUSG( | 1540.589 | 0.09302 | 0.335716 | 0.7777 | Ncbp1 |
| ENSMUSG( | 1332.15 | 0.050808 | 0.613902 | 0.909277 | Ncbp2 |
| ENSMUSG( | 1409.012 | 0.028503 | 0.798693 | 0.960896 | Ncbp2as2 |
| ENSMUSG( | 590.1284 | 0.103313 | 0.353224 | 0.785373 | Ncbp3 |
| ENSMUSG( | 18.88366 | -0.1506 | 0.723146 | NA | Nccrp1 |
| ENSMUSG( | 33400.41 | 0.0626 | 0.519623 | 0.874979 | Ncdn |
| ENSMUSG( | 3685.805 | -0.44877 | 0.001763 | 0.047418 | Nceh1 |
| ENSMUSG( | 435.3406 | 0.033439 | 0.789824 | 0.959461 | Ncf1 |
| ENSMUSG( | 241.0065 | -0.0078 | 0.957669 | 0.993874 | Ncf2 |
| ENSMUSG( | 49.87119 | -0.11167 | 0.708735 | 0.939519 | Ncf4 |
| ENSMUSG( | 336.5827 | 0.13777 | 0.297149 | 0.745805 | Nck1 |
| ENSMUSG( | 742.1888 | 0.050123 | 0.687635 | 0.93324 | Nck2 |
| ENSMUSG( | 14623.88 | 0.099012 | 0.228404 | 0.675849 | Nckap1 |
| ENSMUSG( | 876.2404 | 0.138412 | 0.219044 | 0.664602 | Nckap1l |
| ENSMUSG( | 339.2962 | -0.00665 | 0.958925 | 0.993914 | Nckap5 |
| ENSMUSG( | 884.6343 | 0.035789 | 0.72797 | 0.944355 | Nckap5l |
| ENSMUSG( | 1757.44 | 0.180516 | 0.087536 | 0.454663 | Nckipsd |
| ENSMUSG( | 7729.268 | -0.0009 | 0.992197 | 0.999327 | Ncl |
| ENSMUSG( | 2188.879 | -0.17674 | 0.224768 | 0.671058 | Ncln |
| ENSMUSG( | 24.38786 | 0.376307 | 0.364621 | 0.793796 | Ncmap |
| ENSMUSG( | 4045.52 | 0.157538 | 0.047369 | 0.332754 | Ncoa1 |
| ENSMUSG( | 2150.781 | 0.100305 | 0.271699 | 0.722082 | Ncoa2 |
| ENSMUSG( | 1444.696 | -0.12577 | 0.190175 | 0.629268 | Ncoa3 |
| ENSMUSG( | 1056.468 | 0.003105 | 0.974451 | 0.997072 | Ncoa4 |
| ENSMUSG( | 1324.78 | 0.167359 | 0.083841 | 0.446475 | Ncoa5 |
| ENSMUSG( | 2070.292 | 0.028845 | 0.762173 | 0.953459 | Ncoa6 |
| ENSMUSG( | 927.8252 | 0.05637 | 0.648734 | 0.920025 | Ncoa7 |
| ENSMUSG( | 7721.544 | 0.120463 | 0.188609 | 0.627639 | Ncor1 |
| ENSMUSG( | 6519.236 | -0.03181 | 0.751715 | 0.950337 | Ncor2 |
| ENSMUSG( | 18757.74 | 0.190733 | 0.020134 | 0.214568 | Ncs1 |
| ENSMUSG( | 2737.881 | -0.31513 | 0.015728 | 0.185786 | Ncstn |
| ENSMUSG( | 684327.5 | -0.08464 | 0.608335 | 0.907154 | ND1 |
| ENSMUSG( | 407091.2 | -0.09528 | 0.5595 | 0.888669 | ND2 |
| ENSMUSG( | 835.36 | -0.22345 | 0.31332 | 0.76033 | ND3 |
| ENSMUSG( | 457567 | -0.02801 | 0.805201 | 0.962616 | ND4 |
| ENSMUSG( | 248.0795 | -0.08908 | 0.709907 | 0.939669 | ND4L |
| ENSMUSG( | 510938.7 | -0.05522 | 0.669833 | 0.928209 | ND5 |
| ENSMUSG( | 4694.734 | -0.26279 | 0.138224 | 0.553167 | ND6 |
| ENSMUSG( | 867.0912 | -0.11815 | 0.359429 | 0.791011 | Ndc1 |
| ENSMUSG( | 15.20264 | -0.16889 | 0.733576 | NA | Ndc80 |
| ENSMUSG( | 2176.859 | 0.283984 | 0.01345 | 0.169431 | Nde1 |
| ENSMUSG( | 2003.23 | 0.143013 | 0.121581 | 0.52602 | Ndel1 |
| ENSMUSG( | 11850.63 | -0.54522 | 0.000116 | 0.007099 | Ndfip1 |
| ENSMUSG( | 2981.595 | -0.5201 | 7.18E-05 | 0.005196 | Ndfip2 |
| ENSMUSG( | 3138.617 | 0.175276 | 0.055364 | 0.361966 | Ndn |
| ENSMUSG( | 471.2843 | -0.32492 | 0.055512 | 0.36248 | Ndnf |
| ENSMUSG( | 350.4582 | 0.035462 | 0.811092 | 0.964318 | Ndor1 |
| ENSMUSG( | 667.1414 | 0.027573 | 0.813829 | 0.964574 | Ndp |
| ENSMUSG( | 3332.791 | -0.0492 | 0.574934 | 0.894572 | Ndrg1 |
| ENSMUSG( | 125067.3 | 0.207084 | 0.068379 | 0.405144 | Ndrg2 |
| ENSMUSG( | 13511.29 | 0.093057 | 0.345619 | 0.779153 | Ndrg3 |
| ENSMUSG( | 31993.87 | -0.01962 | 0.873546 | 0.979057 | Ndrg4 |
| ENSMUSG( | 6197.817 | 0.014352 | 0.890053 | 0.983687 | Ndst1 |
| ENSMUSG( | 484.3612 | -0.11578 | 0.453378 | 0.836798 | Ndst2 |
| ENSMUSG( | 654.3215 | -0.08601 | 0.517399 | 0.873295 | Ndst3 |
| ENSMUSG( | 172.2822 | -1.05949 | 1.40E-05 | 0.00146 | Ndst4 |
| ENSMUSG( | 2324.439 | -0.01294 | 0.903501 | 0.986055 | Ndufa1 |

| | | | | | |
|---|---|---|---|---|---|
| ENSMUSG( | 812.3942 | -0.03646 | 0.948278 | 0.994798 | ND3 |
| ENSMUSG( | 404536.8 | -0.11932 | 0.442257 | 0.921554 | ND4 |
| ENSMUSG( | 227.9615 | -0.07347 | 0.88073 | 0.989644 | ND4L |
| ENSMUSG( | 465776.9 | -0.05593 | 0.757278 | 0.971858 | ND5 |
| ENSMUSG( | 4786.228 | 0.069666 | 0.762661 | 0.973566 | ND6 |
| ENSMUSG( | 873.0906 | 0.169383 | 0.255313 | 0.882761 | Ndc1 |
| ENSMUSG( | 15.12097 | 0.109094 | 0.835344 | 0.981237 | Ndc80 |
| ENSMUSG( | 1755.656 | -0.05628 | 0.65287 | 0.954897 | Nde1 |
| ENSMUSG( | 1704.338 | -0.05147 | 0.640065 | 0.953903 | Ndel1 |
| ENSMUSG( | 13577.71 | 0.165337 | 0.324633 | 0.905257 | Ndfip1 |
| ENSMUSG( | 3246.803 | 0.044273 | 0.772081 | 0.974236 | Ndfip2 |
| ENSMUSG( | 2757.425 | 0.071335 | 0.594782 | 0.945444 | Ndn |
| ENSMUSG( | 594.6424 | 0.582716 | 0.002419 | 0.192605 | Ndnf |
| ENSMUSG( | 309.9123 | -0.04055 | 0.807822 | 0.978622 | Ndor1 |
| ENSMUSG( | 607.9638 | 0.028075 | 0.850004 | 0.985538 | Ndp |
| ENSMUSG( | 3040.651 | -0.04332 | 0.727409 | 0.966887 | Ndrg1 |
| ENSMUSG( | 98271.41 | -0.22325 | 0.108749 | 0.804969 | Ndrg2 |
| ENSMUSG( | 11310.82 | -0.15449 | 0.202252 | 0.864412 | Ndrg3 |
| ENSMUSG( | 28071.2 | -0.13271 | 0.330288 | 0.907251 | Ndrg4 |
| ENSMUSG( | 5905.05 | 0.142651 | 0.282886 | 0.892412 | Ndst1 |
| ENSMUSG( | 518.0026 | 0.33999 | 0.078276 | 0.77321 | Ndst2 |
| ENSMUSG( | 719.8676 | 0.431301 | 0.000784 | 0.093388 | Ndst3 |
| ENSMUSG( | 215.6412 | 0.039357 | 0.853431 | 0.985598 | Ndst4 |
| ENSMUSG( | 2024.119 | -0.15129 | 0.252189 | 0.879902 | Ndufa1 |
| ENSMUSG( | 3841.265 | -0.26212 | 0.03199 | 0.633188 | Ndufa10 |
| ENSMUSG( | 1636.252 | -0.27744 | 0.088438 | 0.786756 | Ndufa11 |
| ENSMUSG( | 2307.15 | -0.33623 | 0.012489 | 0.451781 | Ndufa12 |
| ENSMUSG( | 5106.516 | -0.22974 | 0.048471 | 0.706142 | Ndufa13 |
| ENSMUSG( | 2373.967 | -0.25004 | 0.067057 | 0.75212 | Ndufa2 |
| ENSMUSG( | 3102.71 | -0.36632 | 0.001518 | 0.143766 | Ndufa3 |
| ENSMUSG( | 6114.304 | -0.24659 | 0.03258 | 0.636943 | Ndufa4 |
| ENSMUSG( | 105.4871 | 0.271168 | 0.474177 | 0.923329 | Ndufa4l2 |
| ENSMUSG( | 1898.458 | -0.24376 | 0.07071 | 0.758714 | Ndufa5 |
| ENSMUSG( | 2021.664 | -0.17148 | 0.148804 | 0.836241 | Ndufa6 |
| ENSMUSG( | 2384.455 | -0.32665 | 0.006026 | 0.320634 | Ndufa7 |
| ENSMUSG( | 3238.745 | -0.30009 | 0.009617 | 0.406458 | Ndufa8 |
| ENSMUSG( | 2747.639 | -0.18687 | 0.124071 | 0.814691 | Ndufa9 |
| ENSMUSG( | 1554.935 | -0.14542 | 0.221345 | 0.866104 | Ndufab1 |
| ENSMUSG( | 378.5086 | -0.1881 | 0.189997 | 0.857355 | Ndufaf1 |
| ENSMUSG( | 272.2255 | -0.27896 | 0.063972 | 0.736482 | Ndufaf2 |
| ENSMUSG( | 817.5995 | -0.0908 | 0.481778 | 0.924578 | Ndufaf3 |
| ENSMUSG( | 591.7593 | 0.203998 | 0.125859 | 0.815516 | Ndufaf4 |
| ENSMUSG( | 479.1184 | -0.06178 | 0.729839 | 0.967228 | Ndufaf5 |
| ENSMUSG( | 134.0166 | -0.12976 | 0.526448 | 0.934374 | Ndufaf6 |
| ENSMUSG( | 865.8228 | -0.04536 | 0.716514 | 0.965773 | Ndufaf7 |
| ENSMUSG( | 444.4028 | -0.44607 | 0.002938 | 0.217424 | Ndufaf8 |
| ENSMUSG( | 1436.588 | -0.20729 | 0.139423 | 0.830678 | Ndufb1 |
| ENSMUSG( | 2877.174 | -0.19056 | 0.113229 | 0.809623 | Ndufb10 |
| ENSMUSG( | 3669.66 | -0.15504 | 0.182468 | 0.850579 | Ndufb11 |
| ENSMUSG( | 776.3207 | -0.24461 | 0.078061 | 0.77321 | Ndufb2 |
| ENSMUSG( | 1453.449 | -0.26453 | 0.045213 | 0.693959 | Ndufb3 |
| ENSMUSG( | 787.2094 | -0.2354 | 0.074618 | 0.767423 | Ndufb4 |
| ENSMUSG( | 17.46768 | -0.3949 | 0.446307 | 0.921734 | Ndufb4b |
| ENSMUSG( | 3064.008 | -0.20795 | 0.068442 | 0.752603 | Ndufb5 |
| ENSMUSG( | 1245.327 | -0.21276 | 0.186158 | 0.855305 | Ndufb6 |
| ENSMUSG( | 1934.434 | -0.20917 | 0.105515 | 0.804969 | Ndufb7 |
| ENSMUSG( | 543.4895 | -0.24507 | 0.069507 | 0.75601 | Ndufb8 |
| ENSMUSG( | 6319.899 | -0.15181 | 0.14246 | 0.831221 | Ndufb9 |
| ENSMUSG( | 1211.324 | -0.23069 | 0.074828 | 0.767423 | Ndufc1 |
| ENSMUSG( | 2611.536 | -0.17889 | 0.166858 | 0.840183 | Ndufc2 |
| ENSMUSG( | 4064.733 | -0.17739 | 0.105696 | 0.804969 | Ndufs1 |
| ENSMUSG( | 6339.478 | -0.18541 | 0.111235 | 0.807885 | Ndufs2 |
| ENSMUSG( | 1843.726 | -0.20219 | 0.085642 | 0.779799 | Ndufs3 |
| ENSMUSG( | 1620.901 | -0.21945 | 0.08201 | 0.77681 | Ndufs4 |
| ENSMUSG( | 1796.998 | -0.2086 | 0.133553 | 0.822912 | Ndufs5 |
| ENSMUSG( | 1431.065 | -0.2906 | 0.017254 | 0.527468 | Ndufs6 |
| ENSMUSG( | 2021.205 | -0.18108 | 0.198633 | 0.862745 | Ndufs7 |
| ENSMUSG( | 2209.199 | -0.28971 | 0.012137 | 0.446654 | Ndufs8 |
| ENSMUSG( | 3878.64 | -0.18683 | 0.10979 | 0.80637 | Ndufv1 |
| ENSMUSG( | 2803.159 | -0.20482 | 0.117289 | 0.813384 | Ndufv2 |
| ENSMUSG( | 1917.188 | -0.23756 | 0.061483 | 0.736111 | Ndufv3 |
| ENSMUSG( | 1304.381 | 0.187974 | 0.387177 | 0.917257 | Neat1 |
| ENSMUSG( | 62.64989 | 0.591791 | 0.031526 | 0.631192 | Neb |
| ENSMUSG( | 3006.782 | -0.04483 | 0.776186 | 0.975717 | Nebl |
| ENSMUSG( | 1215.962 | 0.021914 | 0.891218 | 0.990558 | Necab1 |
| ENSMUSG( | 2030.439 | -0.22156 | 0.153054 | 0.837724 | Necab2 |
| ENSMUSG( | 1460.065 | -0.04924 | 0.676146 | 0.95694 | Necab3 |
| ENSMUSG( | 2251.367 | -0.11228 | 0.42802 | 0.917257 | Necap1 |
| ENSMUSG( | 1943.36 | -0.12501 | 0.267528 | 0.886134 | Necap2 |
| ENSMUSG( | 2206.489 | 0.09393 | 0.557067 | 0.939565 | Nectin1 |
| ENSMUSG( | 376.1849 | 0.045134 | 0.820622 | 0.979198 | Nectin2 |
| ENSMUSG( | 1526.574 | 0.04654 | 0.77252 | 0.974454 | Nectin3 |
| ENSMUSG( | 148.8223 | 0.050709 | 0.789352 | 0.976593 | Nectin4 |
| ENSMUSG( | 152.3325 | 0.304312 | 0.107261 | 0.804969 | Nedd1 |
| ENSMUSG( | 15615.2 | 0.04964 | 0.598259 | 0.946676 | Nedd4 |
| ENSMUSG( | 3332.723 | 0.078694 | 0.487184 | 0.92541 | Nedd4l |
| ENSMUSG( | 1815.631 | -0.33674 | 0.024045 | 0.588226 | Nedd8 |
| ENSMUSG( | 280.5329 | -0.20573 | 0.223407 | 0.867404 | Nedd9 |
| ENSMUSG( | 1504.492 | -0.11691 | 0.387515 | 0.917257 | Nefh |
| ENSMUSG( | 9213.239 | -0.22194 | 0.036034 | 0.668395 | Nefl |
| ENSMUSG( | 3349.098 | -0.07252 | 0.61112 | 0.950071 | Nefm |
| ENSMUSG( | 6817.592 | 0.147623 | 0.36985 | 0.915491 | Negr1 |
| ENSMUSG( | 271.1569 | 0.078147 | 0.659611 | 0.956205 | Neil1 |
| ENSMUSG( | 66.84101 | 0.28817 | 0.281082 | 0.892356 | Neil2 |
| ENSMUSG( | 1216.109 | -0.07411 | 0.607085 | 0.949917 | Nek1 |
| ENSMUSG( | 84.31083 | 0.039965 | 0.872431 | 0.989023 | Nek10 |
| ENSMUSG( | 37.00525 | -0.187 | 0.589669 | 0.944975 | Nek11 |
| ENSMUSG( | 23.99592 | -0.26286 | 0.505907 | 0.927825 | Nek2 |
| ENSMUSG( | 76.5677 | -0.14666 | 0.557094 | 0.939565 | Nek3 |
| ENSMUSG( | 521.9597 | -0.03779 | 0.786178 | 0.976285 | Nek4 |
| ENSMUSG( | 20.94258 | -0.32923 | 0.514977 | 0.930093 | Nek5 |
| ENSMUSG( | 1240.513 | -0.13214 | 0.280989 | 0.892356 | Nek6 |
| ENSMUSG( | 3231.992 | -0.20937 | 0.143826 | 0.831311 | Nek7 |
| ENSMUSG( | 162.2542 | -0.16399 | 0.445327 | 0.921734 | Nek8 |
| ENSMUSG( | 3376.264 | -0.15626 | 0.163761 | 0.838291 | Nek9 |
| ENSMUSG( | 526.9263 | -0.07128 | 0.670747 | 0.95694 | Nelfa |
| ENSMUSG( | 1284.752 | 0.01631 | 0.894248 | 0.990558 | Nelfb |
| ENSMUSG( | 927.7636 | -0.07714 | 0.595912 | 0.946125 | Nelfcd |
| ENSMUSG( | 750.6623 | -0.15148 | 0.252325 | 0.880122 | Nelfe |
| ENSMUSG( | 1325.786 | 0.038846 | 0.811901 | 0.979148 | Nell1 |
| ENSMUSG( | 16508.2 | 0.231882 | 0.175154 | 0.845305 | Nell2 |
| ENSMUSG( | 1013.92 | 0.062597 | 0.596805 | 0.946541 | Nemf |
| ENSMUSG( | 195.4301 | 0.359552 | 0.056998 | 0.723362 | Nemp2 |
| ENSMUSG( | 2005.315 | -0.14575 | 0.249775 | 0.877904 | Nenf |
| ENSMUSG( | 5225.771 | 0.092541 | 0.520736 | 0.932121 | Neo1 |
| ENSMUSG( | 112.2767 | 0.339392 | 0.114486 | 0.811703 | Nepro |
| ENSMUSG( | 649.2046 | 0.505058 | 0.015781 | 0.502615 | Nes |
| ENSMUSG( | 634.6736 | 0.301299 | 0.099961 | 0.801311 | Net1 |
| ENSMUSG( | 2404 | 0.205717 | 0.136306 | 0.827092 | Neto1 |
| ENSMUSG( | 1926.561 | 0.223882 | 0.20543 | 0.864412 | Neto2 |
| ENSMUSG( | 1981.185 | 0.22251 | 0.160135 | 0.837724 | Neu1 |
| ENSMUSG( | 135.8905 | -0.34307 | 0.131394 | 0.822912 | Neu2 |
| ENSMUSG( | 61.83453 | -0.3779 | 0.155837 | 0.837724 | Neu3 |
| ENSMUSG( | 567.3743 | 0.188894 | 0.252802 | 0.88018 | Neu4 |
| ENSMUSG( | 9913.718 | 0.190651 | 0.086812 | 0.782073 | Neurl1a |
| ENSMUSG( | 3040.19 | 0.277282 | 0.024448 | 0.590702 | Neurl1b |
| ENSMUSG( | 51.51781 | 0.204652 | 0.493273 | 0.927372 | Neurl2 |
| ENSMUSG( | 56.61765 | -0.66277 | 0.022825 | 0.582823 | Neurl3 |
| ENSMUSG( | 4414.233 | 0.043336 | 0.701974 | 0.962014 | Neurl4 |
| ENSMUSG( | 149.3987 | 0.153907 | 0.437451 | 0.920144 | Neurod1 |
| ENSMUSG( | 1858.16 | 0.009519 | 0.949822 | 0.994798 | Neurod2 |
| ENSMUSG( | 11.74769 | 0.307202 | 0.596888 | 0.946541 | Neurod4 |
| ENSMUSG( | 858.7844 | -0.09513 | 0.601483 | 0.947749 | Neurod6 |
| ENSMUSG( | 54.6835 | 0.157902 | 0.575845 | 0.942912 | Neurog2 |
| ENSMUSG( | 464.4861 | 0.30438 | 0.042001 | 0.690304 | Nexmif |
| ENSMUSG( | 167.8605 | -0.72302 | 0.000879 | 0.099346 | Nexn |
| ENSMUSG( | 3177.873 | 0.013805 | 0.902298 | 0.991069 | Nf1 |
| ENSMUSG( | 7409.957 | -0.08397 | 0.44579 | 0.921734 | Nf2 |
| ENSMUSG( | 238.7789 | 0.229446 | 0.294351 | 0.898538 | Nfam1 |
| ENSMUSG( | 13164.56 | 0.189643 | 0.20889 | 0.864412 | Nfasc |
| ENSMUSG( | 2248.527 | -0.01489 | 0.892983 | 0.990558 | Nfat5 |
| ENSMUSG( | 338.1608 | -0.10356 | 0.498295 | 0.927825 | Nfatc1 |
| ENSMUSG( | 422.5429 | -0.16419 | 0.280437 | 0.892356 | Nfatc2 |
| ENSMUSG( | 155.8403 | 0.061358 | 0.736213 | 0.967344 | Nfatc2ip |
| ENSMUSG( | 380.1418 | 0.291602 | 0.033557 | 0.64525 | Nfatc3 |
| ENSMUSG( | 70.2899 | 0.497978 | 0.092782 | 0.794753 | Nfatc4 |
| ENSMUSG( | 13402.96 | 0.020879 | 0.854859 | 0.985649 | Nfe2l1 |
| ENSMUSG( | 803.1454 | 0.167782 | 0.15182 | 0.837724 | Nfe2l2 |
| ENSMUSG( | 148.2524 | 0.049388 | 0.836938 | 0.981335 | Nfe2l3 |
| ENSMUSG( | 2517.774 | -0.00837 | 0.958581 | 0.994798 | Nfia |
| ENSMUSG( | 3539.529 | 0.170798 | 0.229043 | 0.868507 | Nfib |
| ENSMUSG( | 6636.965 | 0.077003 | 0.461878 | 0.923329 | Nfic |
| ENSMUSG( | 130.4276 | -0.52995 | 0.009795 | 0.406458 | Nfil3 |
| ENSMUSG( | 12380.26 | -0.12805 | 0.204475 | 0.864412 | Nfix |
| ENSMUSG( | 651.3283 | 0.07311 | 0.593413 | 0.945408 | Nfkb1 |
| ENSMUSG( | 204.6614 | 0.035831 | 0.829851 | 0.980224 | Nfkb2 |
| ENSMUSG( | 688.1309 | 0.311674 | 0.111954 | 0.808673 | Nfkbia |
| ENSMUSG( | 570.5692 | -0.16524 | 0.281619 | 0.892412 | Nfkbib |

| | | | | | | | | | | | |
|---|---|---|---|---|---|---|---|---|---|---|---|
| ENSMUSG( | Nfyc | 838.2596 | 0.084838 | 0.519878 | 0.874979 | ENSMUSG( | Nfyc | 738.4512 | -0.01319 | 0.926443 | 0.99306 |
| ENSMUSG( | Ngb | 189.3546 | 0.239588 | 0.160933 | 0.590706 | ENSMUSG( | Ngb | 149.0093 | -0.19152 | 0.352263 | 0.912093 |
| ENSMUSG( | Ngdn | 413.0293 | 0.191882 | 0.105973 | 0.493403 | ENSMUSG( | Ngdn | 352.6297 | 0.007333 | 0.959147 | 0.994798 |
| ENSMUSG( | Ngef | 9723.852 | 0.004561 | 0.969697 | 0.996369 | ENSMUSG( | Ngef | 8059.315 | -0.2846 | 0.042603 | 0.690304 |
| ENSMUSG( | Ngf | 105.5186 | -0.60226 | 0.004705 | 0.090209 | ENSMUSG( | Ngf | 112.7416 | -0.07392 | 0.736252 | 0.967344 |
| ENSMUSG( | Ngfr | 49.88413 | -0.65149 | 0.02187 | 0.225738 | ENSMUSG( | Ngfr | 57.59068 | 0.093 | 0.739377 | 0.968458 |
| ENSMUSG( | Ngly1 | 682.4441 | 0.024153 | 0.833031 | 0.970333 | ENSMUSG( | Ngly1 | 593.5205 | -0.11359 | 0.381986 | 0.917257 |
| ENSMUSG( | Ngp | 94.70385 | -1.43909 | 0.182881 | 0.62044 | ENSMUSG( | Ngp | 131.568 | 0.1034 | 0.909777 | 0.991958 |
| ENSMUSG( | Ngrn | 1092.605 | 0.074763 | 0.481338 | 0.855314 | ENSMUSG( | Ngrn | 941.9754 | -0.08978 | 0.487301 | 0.92541 |
| ENSMUSG( | Nhej1 | 72.49774 | -0.01314 | 0.957253 | 0.993874 | ENSMUSG( | Nhej1 | 62.93745 | -0.17968 | 0.524198 | 0.933263 |
| ENSMUSG( | Nherf1 | 4623.136 | 0.164149 | 0.103808 | 0.489428 | ENSMUSG( | Nherf1 | 3924.861 | -0.03675 | 0.770349 | 0.974236 |
| ENSMUSG( | Nherf2 | 1388.79 | 0.28974 | 0.01074 | 0.149272 | ENSMUSG( | Nherf2 | 1483.123 | 0.687222 | 4.97E-05 | 0.012613 |
| ENSMUSG( | Nherf4 | 19.99701 | 0.031146 | 0.946234 | NA | ENSMUSG( | Nherf4 | 17.0451 | -0.13162 | 0.803578 | 0.976978 |
| ENSMUSG( | Nhlh1 | 14.32453 | 0.013953 | 0.977508 | NA | ENSMUSG( | Nhlh1 | 11.01917 | -0.50158 | 0.399579 | NA |
| ENSMUSG( | Nhlh2 | 8.716014 | -0.02798 | 0.965809 | NA | ENSMUSG( | Nhlh2 | 7.222672 | -0.26269 | 0.71813 | NA |
| ENSMUSG( | Nhlrc1 | 378.7571 | 0.071143 | 0.56755 | 0.891774 | ENSMUSG( | Nhlrc1 | 329.092 | -0.07379 | 0.626041 | 0.953234 |
| ENSMUSG( | Nhlrc2 | 904.941 | 0.083931 | 0.409836 | 0.818258 | ENSMUSG( | Nhlrc2 | 810.8156 | 0.036757 | 0.753391 | 0.970671 |
| ENSMUSG( | Nhlrc3 | 537.8439 | -0.37833 | 0.006057 | 0.105754 | ENSMUSG( | Nhlrc3 | 562.5192 | 0.049851 | 0.743853 | 0.96951 |
| ENSMUSG( | Nhp2 | 735.8608 | -0.09042 | 0.404576 | 0.816132 | ENSMUSG( | Nhp2 | 637.1128 | -0.25307 | 0.083766 | 0.77681 |
| ENSMUSG( | Nhs | 85.28882 | -0.36277 | 0.115986 | 0.514015 | ENSMUSG( | Nhs | 85.55564 | -0.07723 | 0.750299 | 0.970113 |
| ENSMUSG( | Nhsl1 | 3598.753 | 0.313302 | 0.038482 | 0.302696 | ENSMUSG( | Nhsl1 | 2747.605 | -0.18818 | 0.151949 | 0.837724 |
| ENSMUSG( | Nhsl2 | 1255.196 | -0.01403 | 0.895454 | 0.984512 | ENSMUSG( | Nhsl2 | 1255.402 | 0.245644 | 0.032496 | 0.63668 |
| ENSMUSG( | Nhsl3 | 1472.286 | 0.271478 | 0.031237 | 0.26893 | ENSMUSG( | Nhsl3 | 1229.23 | 0.03663 | 0.788469 | 0.97637 |
| ENSMUSG( | Niban1 | 214.2249 | -0.2874 | 0.164919 | 0.596162 | ENSMUSG( | Niban1 | 227.8184 | 0.176879 | 0.437657 | 0.920339 |
| ENSMUSG( | Niban2 | 2587.774 | -0.05841 | 0.47844 | 0.853567 | ENSMUSG( | Niban2 | 2354.716 | -0.06393 | 0.569005 | 0.941777 |
| ENSMUSG( | Niban3 | 56.46371 | 0.278186 | 0.32609 | 0.771128 | ENSMUSG( | Niban3 | 45.73644 | -0.06609 | 0.829502 | 0.980224 |
| ENSMUSG( | Nicn1 | 2272.965 | 0.069399 | 0.45035 | 0.835344 | ENSMUSG( | Nicn1 | 1930.047 | -0.13779 | 0.235333 | 0.871741 |
| ENSMUSG( | Nicol1 | 794.6425 | -0.3254 | 0.007945 | 0.124138 | ENSMUSG( | Nicol1 | 728.5482 | -0.31041 | 0.033455 | 0.643993 |
| ENSMUSG( | Nid1 | 777.8137 | -0.19583 | 0.29598 | 0.744386 | ENSMUSG( | Nid1 | 835.9382 | 0.282505 | 0.146336 | 0.835973 |
| ENSMUSG( | Nid2 | 335.3439 | -0.31627 | 0.06957 | 0.408118 | ENSMUSG( | Nid2 | 349.9066 | 0.100089 | 0.559775 | 0.939812 |
| ENSMUSG( | Nif3l1 | 430.4253 | 0.058081 | 0.64242 | 0.916865 | ENSMUSG( | Nif3l1 | 374.2535 | -0.07963 | 0.57663 | 0.942912 |
| ENSMUSG( | Nifk | 441.3258 | 0.108152 | 0.509992 | 0.869743 | ENSMUSG( | Nifk | 366.8071 | -0.18998 | 0.189256 | 0.85698 |
| ENSMUSG( | Nim1k | 1244.496 | 0.05265 | 0.639609 | 0.916434 | ENSMUSG( | Nim1k | 1172.326 | 0.143151 | 0.24756 | 0.877904 |
| ENSMUSG( | Nin | 1262.487 | -0.25816 | 0.005251 | 0.096573 | ENSMUSG( | Nin | 1278.781 | 0.058756 | 0.588362 | 0.944975 |
| ENSMUSG( | Ninj1 | 592.431 | -0.12167 | 0.295226 | 0.743798 | ENSMUSG( | Ninj1 | 601.8074 | 0.18681 | 0.167964 | 0.840222 |
| ENSMUSG( | Ninj2 | 77.98944 | -0.32675 | 0.178389 | 0.616026 | ENSMUSG( | Ninj2 | 86.3941 | 0.243551 | 0.331513 | 0.907907 |
| ENSMUSG( | Ninl | 259.139 | -0.08279 | 0.6161 | 0.91085 | ENSMUSG( | Ninl | 216.632 | -0.33451 | 0.085908 | 0.780123 |
| ENSMUSG( | Nip7 | 498.0745 | 0.04109 | 0.750184 | 0.949664 | ENSMUSG( | Nip7 | 437.1127 | -0.06834 | 0.64459 | 0.953903 |
| ENSMUSG( | Nipa1 | 1483.723 | -0.21905 | 0.140298 | 0.557263 | ENSMUSG( | Nipa1 | 1535.254 | 0.157192 | 0.364998 | 0.914449 |
| ENSMUSG( | Nipa2 | 1104.407 | -0.29226 | 0.043031 | 0.31819 | ENSMUSG( | Nipa2 | 1145.517 | 0.096925 | 0.558858 | 0.939587 |
| ENSMUSG( | Nipal2 | 237.4313 | -0.48729 | 0.003371 | 0.073204 | ENSMUSG( | Nipal2 | 244.3259 | -0.09907 | 0.578669 | 0.943548 |
| ENSMUSG( | Nipal3 | 3547.524 | -0.19392 | 0.198162 | 0.638387 | ENSMUSG( | Nipal3 | 3814.537 | 0.282599 | 0.096622 | 0.796245 |
| ENSMUSG( | Nipal4 | 249.924 | -0.57315 | 0.004453 | 0.086961 | ENSMUSG( | Nipal4 | 296.5936 | 0.247856 | 0.258193 | 0.883971 |
| ENSMUSG( | Nipbl | 1745.597 | 0.07527 | 0.550074 | 0.886038 | ENSMUSG( | Nipbl | 1661.562 | 0.199536 | 0.146559 | 0.835973 |
| ENSMUSG( | Nipsnap1 | 1067.565 | -0.03661 | 0.794287 | 0.959799 | ENSMUSG( | Nipsnap1 | 920.5126 | -0.20734 | 0.229048 | 0.868507 |
| ENSMUSG( | Nipsnap2 | 2501.114 | 0.001965 | 0.983804 | 0.99779 | ENSMUSG( | Nipsnap2 | 2137.348 | -0.19299 | 0.119413 | 0.813384 |
| ENSMUSG( | Nipsnap3b | 307.4761 | 0.012915 | 0.92688 | 0.991229 | ENSMUSG( | Nipsnap3b | 267.8698 | -0.12495 | 0.453336 | 0.922316 |
| ENSMUSG( | Nisch | 16103.81 | 0.154652 | 0.05766 | 0.370247 | ENSMUSG( | Nisch | 14275.86 | 0.079765 | 0.42107 | 0.917257 |
| ENSMUSG( | Nit1 | 979.0069 | 0.106366 | 0.335932 | 0.777 | ENSMUSG( | Nit1 | 838.7527 | -0.06962 | 0.589637 | 0.944975 |
| ENSMUSG( | Nit2 | 303.5218 | 0.031633 | 0.830331 | 0.96974 | ENSMUSG( | Nit2 | 259.8571 | -0.14547 | 0.363739 | 0.914223 |
| ENSMUSG( | Nkain1 | 806.5299 | -0.25384 | 0.042369 | 0.316636 | ENSMUSG( | Nkain1 | 814.8865 | 0.057624 | 0.709257 | 0.964042 |
| ENSMUSG( | Nkain2 | 1558.939 | -0.23658 | 0.032148 | 0.273917 | ENSMUSG( | Nkain2 | 1641.122 | 0.181386 | 0.222445 | 0.867923 |
| ENSMUSG( | Nkain3 | 40.95892 | -0.2647 | 0.380069 | 0.801757 | ENSMUSG( | Nkain3 | 42.41524 | 0.132737 | 0.67887 | 0.957617 |
| ENSMUSG( | Nkain4 | 2124.44 | 0.025159 | 0.809893 | 0.96372 | ENSMUSG( | Nkain4 | 1911.187 | -0.01029 | 0.934254 | 0.99306 |
| ENSMUSG( | Nkap | 349.5301 | 0.01128 | 0.930332 | 0.991848 | ENSMUSG( | Nkap | 311.8612 | -0.05439 | 0.725402 | 0.966887 |
| ENSMUSG( | Nkapd1 | 764.1973 | -0.00172 | 0.986921 | 0.998619 | ENSMUSG( | Nkapd1 | 705.474 | 0.030156 | 0.831113 | 0.98051 |
| ENSMUSG( | Nkd1 | 1512.83 | -0.07721 | 0.377305 | 0.801213 | ENSMUSG( | Nkd1 | 1403.431 | -0.02655 | 0.815361 | 0.979198 |
| ENSMUSG( | Nkd2 | 139.8734 | 0.287737 | 0.106273 | 0.49377 | ENSMUSG( | Nkd2 | 129.8454 | 0.337368 | 0.099654 | 0.801103 |
| ENSMUSG( | Nkiras1 | 1920.825 | -0.00735 | 0.940693 | 0.993415 | ENSMUSG( | Nkiras1 | 1653.421 | -0.17972 | 0.154211 | 0.837724 |
| ENSMUSG( | Nkiras2 | 571.7374 | 0.064982 | 0.538617 | 0.882729 | ENSMUSG( | Nkiras2 | 490.8987 | -0.10918 | 0.392613 | 0.917257 |
| ENSMUSG( | Nkpd1 | 35.95053 | -0.56789 | 0.075508 | 0.425931 | ENSMUSG( | Nkpd1 | 36.98199 | -0.16126 | 0.618983 | 0.952598 |
| ENSMUSG( | Nkrf | 423.5121 | -0.02653 | 0.850794 | 0.976538 | ENSMUSG( | Nkrf | 395.485 | 0.043611 | 0.787824 | 0.977637 |
| ENSMUSG( | Nktr | 4865.694 | 0.066763 | 0.485532 | 0.857637 | ENSMUSG( | Nktr | 4617.636 | 0.179246 | 0.121937 | 0.813384 |
| ENSMUSG( | Nkx2-1 | 19.07544 | -0.49158 | 0.306588 | NA | ENSMUSG( | Nkx2-1 | 19.76529 | -0.13971 | 0.783206 | 0.976106 |
| ENSMUSG( | Nkx2-2 | 280.7481 | -0.15754 | 0.248052 | 0.696814 | ENSMUSG( | Nkx2-2 | 280.4541 | 0.097406 | 0.611359 | 0.950771 |
| ENSMUSG( | Nkx2-2os | 120.7972 | 0.262401 | 0.177223 | 0.614553 | ENSMUSG( | Nkx2-2os | 109.9809 | 0.242209 | 0.318712 | 0.903297 |
| ENSMUSG( | Nkx3-1 | 24.66346 | 0.132882 | 0.745412 | 0.949158 | ENSMUSG( | Nkx3-1 | 21.72101 | 0.012755 | 0.97827 | 0.996416 |
| ENSMUSG( | Nkx6-2 | 314.8966 | -0.51268 | 0.001458 | 0.042158 | ENSMUSG( | Nkx6-2 | 321.0611 | -0.14957 | 0.388839 | 0.917257 |
| ENSMUSG( | Nle1 | 286.8218 | -0.07558 | 0.60324 | 0.904887 | ENSMUSG( | Nle1 | 275.7864 | 0.089709 | 0.620326 | 0.952896 |
| ENSMUSG( | Nlgn1 | 1476.482 | -0.35327 | 0.027689 | 0.255176 | ENSMUSG( | Nlgn1 | 1620.557 | 0.203726 | 0.271436 | 0.888062 |
| ENSMUSG( | Nlgn2 | 12047.25 | -0.17202 | 0.124671 | 0.531714 | ENSMUSG( | Nlgn2 | 12188.21 | 0.134789 | 0.288584 | 0.894794 |
| ENSMUSG( | Nlgn3 | 2244.169 | -0.32213 | 0.020324 | 0.214896 | ENSMUSG( | Nlgn3 | 2425.698 | 0.184856 | 0.241326 | 0.876259 |
| ENSMUSG( | Nlk | 2224.545 | 0.158694 | 0.069712 | 0.408352 | ENSMUSG( | Nlk | 1966.602 | 0.074933 | 0.48394 | 0.925289 |
| ENSMUSG( | Nln | 1404.514 | 0.058742 | 0.579311 | 0.896692 | ENSMUSG( | Nln | 1223.206 | -0.07567 | 0.553401 | 0.939565 |
| ENSMUSG( | Nlrc3 | 35.59956 | 0.071241 | 0.829381 | 0.969343 | ENSMUSG( | Nlrc3 | 33.47444 | 0.150825 | 0.703763 | 0.96251 |
| ENSMUSG( | Nlrc4 | 15.30372 | -0.20172 | 0.699529 | NA | ENSMUSG( | Nlrc4 | 12.42893 | -0.57236 | 0.350532 | 0.911652 |
| ENSMUSG( | Nlrc5 | 37.09013 | -0.30844 | 0.341352 | 0.778828 | ENSMUSG( | Nlrc5 | 31.8629 | -0.46858 | 0.227613 | 0.868507 |
| ENSMUSG( | Nlrp1b | 34.19006 | -0.17467 | 0.63126 | 0.914073 | ENSMUSG( | Nlrp1b | 34.79785 | 0.147343 | 0.719741 | 0.966887 |
| ENSMUSG( | Nlrp3 | 43.26631 | -0.04719 | 0.880046 | 0.981469 | ENSMUSG( | Nlrp3 | 47.34954 | 0.472855 | 0.155661 | 0.837724 |
| ENSMUSG( | Nlrp6 | 15.3571 | 0.80202 | 0.132785 | NA | ENSMUSG( | Nlrp6 | 12.11404 | 0.439458 | 0.478864 | 0.923329 |
| ENSMUSG( | Nlrx1 | 300.6471 | 0.011894 | 0.930563 | 0.991965 | ENSMUSG( | Nlrx1 | 276.3703 | 0.044861 | 0.781841 | 0.976106 |
| ENSMUSG( | Nmb | 111.3299 | -0.04275 | 0.868404 | 0.978464 | ENSMUSG( | Nmb | 110.8204 | 0.194239 | 0.36135 | 0.914223 |
| ENSMUSG( | Nmbr | 57.40223 | -0.6175 | 0.024202 | 0.238178 | ENSMUSG( | Nmbr | 71.73668 | 0.33889 | 0.193537 | 0.860096 |
| ENSMUSG( | Nmd3 | 513.0357 | -0.08711 | 0.455854 | 0.838601 | ENSMUSG( | Nmd3 | 455.297 | -0.16511 | 0.223832 | 0.867404 |
| ENSMUSG( | Nme1 | 1501.557 | 0.028133 | 0.805702 | 0.962616 | ENSMUSG( | Nme1 | 1280.353 | -0.17364 | 0.272308 | 0.888771 |
| ENSMUSG( | Nme2 | 43.25933 | 0.121197 | 0.680685 | 0.931139 | ENSMUSG( | Nme2 | 34.79232 | -0.24564 | 0.472664 | 0.923329 |
| ENSMUSG( | Nme4 | 72.39709 | 0.121836 | 0.587215 | 0.900334 | ENSMUSG( | Nme4 | 70.21076 | 0.292998 | 0.233113 | 0.868507 |
| ENSMUSG( | Nme5 | 177.1043 | -0.13788 | 0.421989 | 0.823495 | ENSMUSG( | Nme5 | 160.8303 | -0.159 | 0.40745 | 0.917257 |
| ENSMUSG( | Nme6 | 134.5178 | 0.278125 | 0.132158 | 0.544244 | ENSMUSG( | Nme6 | 115.2746 | 0.1224 | 0.564859 | 0.941206 |
| ENSMUSG( | Nme7 | 1025.307 | 0.061582 | 0.592324 | 0.902094 | ENSMUSG( | Nme7 | 859.7746 | -0.18722 | 0.157749 | 0.837724 |
| ENSMUSG( | Nmi | 201.8906 | 0.010318 | 0.949521 | 0.993821 | ENSMUSG( | Nmi | 175.0931 | -0.13203 | 0.462891 | 0.923329 |
| ENSMUSG( | Nmnat1 | 168.4385 | 0.105166 | 0.532164 | 0.886141 | ENSMUSG( | Nmnat1 | 158.5973 | 0.192846 | 0.296966 | 0.898538 |
| ENSMUSG( | Nmnat2 | 3655.273 | 0.060659 | 0.543935 | 0.884297 | ENSMUSG( | Nmnat2 | 3188.712 | -0.06583 | 0.567297 | 0.941777 |
| ENSMUSG( | Nmnat3 | 53.66938 | -0.26233 | 0.335645 | 0.777 | ENSMUSG( | Nmnat3 | 56.87174 | 0.162559 | 0.567751 | 0.941777 |
| ENSMUSG( | Nmral1 | 270.725 | 0.038829 | 0.824159 | 0.966864 | ENSMUSG( | Nmral1 | 242.9724 | 0.004803 | 0.979836 | 0.996479 |
| ENSMUSG( | Nmrk1 | 129.9189 | -0.06514 | 0.744764 | 0.94902 | ENSMUSG( | Nmrk1 | 115.3854 | -0.12435 | 0.599698 | 0.947071 |
| ENSMUSG( | Nmt1 | 3840.569 | 0.127371 | 0.199746 | 0.640239 | ENSMUSG( | Nmt1 | 3420.747 | 0.065162 | 0.571214 | 0.942052 |
| ENSMUSG( | Nmt2 | 2652.641 | 0.036931 | 0.66653 | 0.927247 | ENSMUSG( | Nmt2 | 2263.821 | -0.15919 | 0.207766 | 0.864412 |
| ENSMUSG( | Nnat | 9368.527 | -0.06361 | 0.62698 | 0.912825 | ENSMUSG( | Nnat | 8018.831 | -0.2572 | 0.081758 | 0.77681 |
| ENSMUSG( | Nnmt | 18.02908 | -0.04871 | 0.909953 | NA | ENSMUSG( | Nnmt | 16.18138 | -0.07878 | 0.869913 | 0.988097 |
| ENSMUSG( | Nnt | 10.65973 | -0.22646 | 0.714867 | NA | ENSMUSG( | Nnt | 9.331284 | -0.40515 | 0.523736 | NA |
| ENSMUSG( | Noa1 | 526.3188 | 0.151022 | 0.215712 | 0.660456 | ENSMUSG( | Noa1 | 457.287 | 0.017937 | 0.897211 | 0.990558 |
| ENSMUSG( | Nob1 | 552.9691 | -0.03997 | 0.71709 | 0.941697 | ENSMUSG( | Nob1 | 503.1595 | -0.04354 | 0.753156 | 0.970622 |
| ENSMUSG( | Noc2l | 2406.326 | 0.133278 | 0.165029 | 0.596428 | ENSMUSG( | Noc2l | 2089.814 | -0.00308 | 0.978165 | 0.996416 |
| ENSMUSG( | Noc3l | 433.2561 | 0.188407 | 0.130111 | 0.540593 | ENSMUSG( | Noc3l | 383.3094 | 0.105009 | 0.464013 | 0.923329 |
| ENSMUSG( | Noc4l | 419.4747 | 0.107596 | 0.557244 | 0.888279 | ENSMUSG( | Noc4l | 348.5681 | -0.15871 | 0.404517 | 0.917257 |
| ENSMUSG( | Noct | 903.1493 | -0.03484 | 0.793651 | 0.959799 | ENSMUSG( | Noct | 813.6279 | -0.07062 | 0.636867 | 0.953903 |
| ENSMUSG( | Nod1 | 243.5321 | -0.1164 | 0.513333 | 0.871762 | ENSMUSG( | Nod1 | 255.6461 | 0.294462 | 0.227918 | 0.868507 |
| ENSMUSG( | Nod2 | 15.95999 | -0.73539 | 0.122631 | NA | ENSMUSG( | Nod2 | 14.5553 | -0.69583 | 0.183962 | 0.852278 |
| ENSMUSG( | Nodal | 23.96119 | -0.59515 | 0.202819 | 0.643279 | ENSMUSG( | Nodal | 23.3354 | -0.29571 | 0.585484 | 0.944975 |
| ENSMUSG( | Nog | 365.94 | -0.47843 | 0.003308 | 0.072126 | ENSMUSG( | Nog | 420.9981 | 0.225509 | 0.213538 | 0.864421 |
| ENSMUSG( | Nol10 | 506.5975 | 0.147197 | 0.237781 | 0.685809 | ENSMUSG( | Nol10 | 432.6736 | -0.00297 | 0.987567 | 0.99767 |
| ENSMUSG( | Nol11 | 468.7763 | 0.001209 | 0.991942 | 0.999232 | ENSMUSG( | Nol11 | 443.6385 | 0.10669 | 0.4239 | 0.917257 |
| ENSMUSG( | Nol12 | 823.6854 | 0.101825 | 0.298592 | 0.747287 | ENSMUSG( | Nol12 | 676.1582 | -0.20905 | 0.098559 | 0.798438 |
| ENSMUSG( | Nol3 | 882.0782 | -0.02435 | 0.822964 | 0.96644 | ENSMUSG( | Nol3 | 753.0309 | -0.22531 | 0.114504 | 0.811703 |
| ENSMUSG( | Nol4 | 1388.3 | 0.033059 | 0.706529 | 0.939259 | ENSMUSG( | Nol4 | 1278.434 | 0.062913 | 0.569351 | 0.941777 |
| ENSMUSG( | Nol4l | 2270.664 | 0.181696 | 0.078079 | 0.432233 | ENSMUSG( | Nol4l | 1972.086 | 0.051587 | 0.682549 | 0.958678 |
| ENSMUSG( | Nol6 | 2952.013 | 0.075898 | 0.48074 | 0.855094 | ENSMUSG( | Nol6 | 2511.496 | -0.12441 | 0.317778 | 0.902777 |
| ENSMUSG( | Nol7 | 1127.811 | 0.064151 | 0.594132 | 0.902869 | ENSMUSG( | Nol7 | 981.6162 | -0.07313 | 0.586868 | 0.944975 |
| ENSMUSG( | Nol8 | 640.4942 | 0.115275 | 0.336338 | 0.777 | ENSMUSG( | Nol8 | 555.6073 | -0.02945 | 0.82926 | 0.980224 |
| ENSMUSG( | Nol9 | 607.2052 | 0.113272 | 0.328612 | 0.773148 | ENSMUSG( | Nol9 | 553.272 | 0.116474 | 0.379066 | 0.916995 |
| ENSMUSG( | Nolc1 | 1158.434 | 0.116021 | 0.240655 | 0.688247 | ENSMUSG( | Nolc1 | 999.7049 | -0.04161 | 0.74246 | 0.969415 |
| ENSMUSG( | Nom1 | 853.8447 | 0.04638 | 0.664209 | 0.926591 | ENSMUSG( | Nom1 | 741.1768 | -0.09866 | 0.476471 | 0.923329 |
| ENSMUSG( | Nomo1 | 5473.384 | -0.11558 | 0.386754 | 0.805046 | ENSMUSG( | Nomo1 | 5192.909 | 0.008175 | 0.959103 | 0.994798 |
| ENSMUSG( | Nono | 3468.17 | 0.013406 | 0.878484 | 0.981431 | ENSMUSG( | Nono | 3019.484 | -0.12135 | 0.292906 | 0.898413 |
| ENSMUSG( | Nop10 | 813.4089 | -0.07342 | 0.502259 | 0.864924 | ENSMUSG( | Nop10 | 716.7926 | -0.17874 | 0.186942 | 0.856099 |
| ENSMUSG( | Nop14 | 644.8658 | 0.169484 | 0.157173 | 0.585516 | ENSMUSG( | Nop14 | 551.1725 | -0.01326 | 0.922644 | 0.992644 |
| ENSMUSG( | Nop16 | 333.5234 | 0.192054 | 0.209386 | 0.65137 | ENSMUSG( | Nop16 | 286.7981 | 0.022733 | 0.899135 | 0.990558 |
| ENSMUSG( | Nop2 | 570.7333 | 0.150448 | 0.165352 | 0.596691 | ENSMUSG( | Nop2 | 486.5376 | -0.04016 | 0.763814 | 0.974 |
| ENSMUSG( | Nop53 | 3441.134 | 0.029284 | 0.727064 | 0.944223 | ENSMUSG( | Nop53 | 3051.268 | -0.05096 | 0.669006 | 0.956766 |
| ENSMUSG( | Nop56 | 1917.407 | 0.085472 | 0.42554 | 0.824898 | ENSMUSG( | Nop56 | 1709.022 | 0.024133 | 0.837592 | 0.981335 |
| ENSMUSG( | Nop58 | 797.2025 | 0.088192 | 0.401683 | 0.814923 | ENSMUSG( | Nop58 | 661.4826 | -0.1931 | 0.168268 | 0.840222 |
| ENSMUSG( | Nop9 | 338.1695 | 0.190975 | 0.154039 | 0.580901 | ENSMUSG( | Nop9 | 299.3792 | 0.117358 | 0.427357 | 0.917257 |
| ENSMUSG( | Nopchap1 | 1120.546 | 0.105476 | 0.283733 | 0.732203 | ENSMUSG( | Nopchap1 | 959.8556 | -0.07628 | 0.54588 | 0.938063 |
| ENSMUSG( | Norad | 34785.76 | 0.113516 | 0.15554 | 0.583591 | ENSMUSG( | Norad | 30047.14 | -0.0384 | 0.704379 | 0.962677 |
| ENSMUSG( | Nos1 | 888.6835 | 0.174972 | 0.106917 | 0.494634 | ENSMUSG( | Nos1 | 736.991 | -0.09425 | 0.488284 | 0.925453 |
| ENSMUSG( | Nos1ap | 1144.182 | 0.380086 | 0.006082 | 0.106072 | ENSMUSG( | Nos1ap | 942.1189 | 0.114965 | 0.446606 | 0.921734 |
| ENSMUSG( | Nos2 | 33.78285 | 0.168566 | 0.643613 | 0.917316 | ENSMUSG( | Nos2 | 36.00791 | 0.595989 | 0.106615 | 0.804969 |
| ENSMUSG( | Nos3 | 388.1407 | 0.398121 | 0.043456 | 0.319227 | ENSMUSG( | Nos3 | 418.5921 | 0.813121 | 0.000722 | 0.08668 |
| ENSMUSG( | Nosip | 1164.514 | -0.01885 | 0.867232 | 0.976559 | ENSMUSG( | Nosip | 1047.64 | -0.05995 | 0.675232 | 0.95694 |
| ENSMUSG( | Nostrin | 232.5353 | 0.276032 | 0.174504 | 0.610304 | ENSMUSG( | Nostrin | 221.9808 | 0.410742 | 0.06734 | 0.75212 |
| ENSMUSG( | Notch1 | 1542.44 | -0.12831 | 0.433783 | 0.828541 | ENSMUSG( | Notch1 | 1838.474 | 0.596044 | 0.002208 | 0.183318 |
| ENSMUSG( | Notch2 | 1388.972 | -0.30751 | 0.024144 | 0.237772 | ENSMUSG( | Notch2 | 1387.89 | -0.01971 | 0.899778 | 0.990558 |
| ENSMUSG( | Notch3 | 760.8927 | -0.15842 | 0.317914 | 0.764145 | ENSMUSG( | Notch3 | 866.565 | 0.46088 | 0.009724 | 0.406458 |
| ENSMUSG( | Notch4 | 398.4602 | 0.641142 | 0.000146 | 0.008459 | ENSMUSG( | Notch4 | 377.8752 | 0.745649 | 0.000973 | 0.106505 |
| ENSMUSG( | Notum | 159.3534 | 0.107977 | 0.561708 | 0.889052 | ENSMUSG( | Notum | 199.1889 | 0.90086 | 7.58E-05 | 0.017018 |
| ENSMUSG( | Nova1 | 1559.038 | 0.07405 | 0.434785 | 0.829025 | ENSMUSG( | Nova1 | 1375.658 | -0.02035 | 0.876421 | 0.989644 |
| ENSMUSG( | Nova2 | 3524.362 | 0.164433 | 0.066084 | 0.397686 | ENSMUSG( | Nova2 | 3039.873 | 0.011554 | 0.914919 | 0.992407 |
| ENSMUSG( | Noxo1 | 100.2271 | 0.152001 | 0.454746 | 0.837513 | ENSMUSG( | Noxo1 | 96.81048 | 0.3104 | 0.175085 | 0.845305 |
| ENSMUSG( | Noxred1 | 40.77795 | 0.291613 | 0.358539 | 0.789791 | ENSMUSG( | Noxred1 | 38.33074 | 0.380825 | 0.297161 | 0.898538 |
| ENSMUSG( | Npas1 | 296.6626 | 0.465292 | 0.006422 | 0.110055 | ENSMUSG( | Npas1 | 234.377 | 0.096251 | 0.586727 | 0.944975 |
| ENSMUSG( | Npas2 | 1920.613 | 0.200291 | 0.027555 | 0.2548 | ENSMUSG( | Npas2 | 1554.198 | -0.13887 | 0.188169 | 0.85698 |
| ENSMUSG( | Npas3 | 756.7756 | -0.00307 | 0.979578 | 0.997428 | ENSMUSG( | Npas3 | 632.4674 | -0.26857 | 0.037941 | 0.6763 |
| ENSMUSG( | Npas4 | 558.5984 | -0.59053 | 0.357181 | 0.788702 | ENSMUSG( | Npas4 | 519.1175 | -0.51754 | 0.418264 | 0.917257 |
| ENSMUSG( | Npat | 546.7853 | -0.03197 | 0.781289 | 0.957558 | ENSMUSG( | Npat | 521.6855 | 0.103534 | 0.463056 | 0.923329 |
| ENSMUSG( | Npb | 21.29479 | -0.94625 | 0.022587 | 0.229429 | ENSMUSG( | Npb | 27.73961 | 0.234807 | 0.539003 | 0.937216 |
| ENSMUSG( | Npbwr1 | 45.90949 | -0.48066 | 0.151504 | 0.576717 | ENSMUSG( | Npbwr1 | 48.82852 | 0.038287 | 0.918304 | 0.992529 |
| ENSMUSG( | Npc1 | 3336.281 | -0.33884 | 0.031166 | 0.268462 | ENSMUSG( | Npc1 | 3734.206 | 0.267832 | 0.12906 | 0.820358 |

| | | | | | |
|---|---|---|---|---|---|
| ENSMUSG( | 4597.218 | 0.001667 | 0.986193 | 0.998326 | Ndufa10 |
| ENSMUSG( | 1913.384 | -0.07764 | 0.497273 | 0.861898 | Ndufa11 |
| ENSMUSG( | 2835.46 | 0.013023 | 0.879582 | 0.981436 | Ndufa12 |
| ENSMUSG( | 5958.606 | -0.04239 | 0.586152 | 0.899709 | Ndufa13 |
| ENSMUSG( | 2889.414 | 0.062395 | 0.524538 | 0.875849 | Ndufa2 |
| ENSMUSG( | 3781.229 | -0.03957 | 0.611727 | 0.908586 | Ndufa3 |
| ENSMUSG( | 7086.729 | -0.07806 | 0.320478 | 0.765921 | Ndufa4 |
| ENSMUSG( | 92.47794 | -0.42343 | 0.169108 | 0.603241 | Ndufa4l2 |
| ENSMUSG( | 2287.417 | 0.039525 | 0.686545 | 0.933041 | Ndufa5 |
| ENSMUSG( | 2312.043 | -0.0451 | 0.631528 | 0.914073 | Ndufa6 |
| ENSMUSG( | 2889.892 | -0.01947 | 0.814068 | 0.964574 | Ndufa7 |
| ENSMUSG( | 3952.654 | 0.024125 | 0.771248 | 0.954726 | Ndufa8 |
| ENSMUSG( | 3258.348 | 0.044673 | 0.634581 | 0.914073 | Ndufa9 |
| ENSMUSG( | 1794.876 | 0.006196 | 0.949789 | 0.993821 | Ndufab1 |
| ENSMUSG( | 439.5051 | -0.01399 | 0.911449 | 0.988483 | Ndufaf1 |
| ENSMUSG( | 323.6231 | -0.03378 | 0.794352 | 0.959799 | Ndufaf2 |
| ENSMUSG( | 898.7031 | -0.08323 | 0.426005 | 0.824898 | Ndufaf3 |
| ENSMUSG( | 606.0955 | 0.016447 | 0.880777 | 0.981565 | Ndufaf4 |
| ENSMUSG( | 552.0546 | 0.085048 | 0.49012 | 0.859319 | Ndufaf5 |
| ENSMUSG( | 147.49 | -0.12349 | 0.505673 | 0.866932 | Ndufaf6 |
| ENSMUSG( | 996.0377 | 0.090282 | 0.343391 | 0.779153 | Ndufaf7 |
| ENSMUSG( | 554.9828 | -0.03076 | 0.77311 | 0.955272 | Ndufaf8 |
| ENSMUSG( | 1696.227 | 0.015216 | 0.891566 | 0.983955 | Ndufb1 |
| ENSMUSG( | 3356.922 | -0.00554 | 0.952156 | 0.993821 | Ndufb10 |
| ENSMUSG( | 4205.914 | -0.02379 | 0.792463 | 0.959461 | Ndufb11 |
| ENSMUSG( | 927.9536 | 0.015696 | 0.880806 | 0.981565 | Ndufb2 |
| ENSMUSG( | 1690.035 | -0.08269 | 0.429876 | 0.826626 | Ndufb3 |
| ENSMUSG( | 897.3947 | -0.11426 | 0.311679 | 0.759301 | Ndufb4 |
| ENSMUSG( | 22.37961 | 0.100527 | 0.817564 | 0.965785 | Ndufb4b |
| ENSMUSG( | 3523.627 | -0.06422 | 0.457877 | 0.839769 | Ndufb5 |
| ENSMUSG( | 1438.895 | -0.05166 | 0.590204 | 0.90113 | Ndufb6 |
| ENSMUSG( | 2291.032 | 0.021326 | 0.820829 | 0.965962 | Ndufb7 |
| ENSMUSG( | 637.593 | -0.03933 | 0.710162 | 0.9397 | Ndufb8 |
| ENSMUSG( | 7453.157 | 0.0595 | 0.43551 | 0.829093 | Ndufb9 |
| ENSMUSG( | 1437.887 | 0.009188 | 0.93177 | 0.9924 | Ndufc1 |
| ENSMUSG( | 2844.899 | -0.19996 | 0.044028 | 0.321685 | Ndufc2 |
| ENSMUSG( | 4889.826 | 0.093302 | 0.320645 | 0.765984 | Ndufs1 |
| ENSMUSG( | 7585.578 | 0.070645 | 0.402732 | 0.81552 | Ndufs2 |
| ENSMUSG( | 2125.323 | -0.0524 | 0.585405 | 0.899171 | Ndufs3 |
| ENSMUSG( | 1873.463 | -0.0585 | 0.530247 | 0.879138 | Ndufs4 |
| ENSMUSG( | 1420.53 | -1.53272 | 1.25E-44 | 6.73E-41 | Ndufs5 |
| ENSMUSG( | 1715.489 | -0.018 | 0.850642 | 0.972638 | Ndufs6 |
| ENSMUSG( | 2385.886 | 0.038435 | 0.71949 | 0.941986 | Ndufs7 |
| ENSMUSG( | 2567.572 | -0.11028 | 0.186584 | 0.623869 | Ndufs8 |
| ENSMUSG( | 4673.234 | 0.088587 | 0.295533 | 0.743814 | Ndufv1 |
| ENSMUSG( | 3274.112 | -0.01598 | 0.867527 | 0.97663 | Ndufv2 |
| ENSMUSG( | 2280.054 | 0.006802 | 0.938955 | 0.993415 | Ndufv3 |
| ENSMUSG( | 1379.008 | 0.08679 | 0.638689 | 0.91636 | Neat1 |
| ENSMUSG( | 54.75968 | -0.02295 | 0.932951 | 0.992879 | Neb |
| ENSMUSG( | 3510.675 | 0.134285 | 0.357343 | 0.788775 | Nebl |
| ENSMUSG( | 1323.441 | 0.003465 | 0.975853 | 0.997428 | Necab1 |
| ENSMUSG( | 2306.411 | -0.11242 | 0.411699 | 0.81942 | Necab2 |
| ENSMUSG( | 1660.598 | 0.053656 | 0.637306 | 0.915436 | Necab3 |
| ENSMUSG( | 2586.638 | 0.024024 | 0.83973 | 0.971179 | Necap1 |
| ENSMUSG( | 2285.402 | 0.076958 | 0.429569 | 0.826446 | Necap2 |
| ENSMUSG( | 2178.487 | -0.22399 | 0.144738 | 0.564884 | Nectin1 |
| ENSMUSG( | 351.0743 | -0.46933 | 0.002017 | 0.051507 | Nectin2 |
| ENSMUSG( | 1561.651 | -0.16402 | 0.265517 | 0.716109 | Nectin3 |
| ENSMUSG( | 150.7513 | -0.20037 | 0.230379 | 0.678421 | Nectin4 |
| ENSMUSG( | 159.4703 | 0.174352 | 0.299657 | 0.74868 | Nedd1 |
| ENSMUSG( | 16318.54 | -0.09297 | 0.305743 | 0.753712 | Nedd4 |
| ENSMUSG( | 3743.483 | 0.144074 | 0.116067 | 0.514235 | Nedd4l |
| ENSMUSG( | 2209.47 | -0.01497 | 0.886549 | 0.983204 | Nedd8 |
| ENSMUSG( | 308.48 | -0.21168 | 0.10989 | 0.500285 | Nedd9 |
| ENSMUSG( | 1695.492 | -0.03739 | 0.750996 | 0.950053 | Nefh |
| ENSMUSG( | 10680.01 | -0.0549 | 0.518414 | 0.873901 | Nefl |
| ENSMUSG( | 3925.113 | 0.115308 | 0.375442 | 0.800845 | Nefm |
| ENSMUSG( | 6478.596 | -0.28107 | 0.05702 | 0.368322 | Negr1 |
| ENSMUSG( | 312.8988 | 0.208171 | 0.14252 | 0.560815 | Neil1 |
| ENSMUSG( | 73.24599 | 0.282951 | 0.235737 | 0.682914 | Neil2 |
| ENSMUSG( | 12.18657 | -0.25411 | 0.645328 | NA | Neil3 |
| ENSMUSG( | 1397.02 | 0.063218 | 0.589362 | 0.90113 | Nek1 |
| ENSMUSG( | 80.64171 | -0.36217 | 0.11107 | 0.502387 | Nek10 |
| ENSMUSG( | 39.54741 | -0.26491 | 0.380108 | 0.801757 | Nek11 |
| ENSMUSG( | 25.8449 | -0.33642 | 0.352619 | 0.784565 | Nek2 |
| ENSMUSG( | 97.94979 | 0.284142 | 0.184182 | 0.621101 | Nek3 |
| ENSMUSG( | 603.5321 | 0.114731 | 0.351967 | 0.783738 | Nek4 |
| ENSMUSG( | 19.97921 | -0.85254 | 0.144601 | NA | Nek5 |
| ENSMUSG( | 1463.507 | 0.081501 | 0.367261 | 0.795221 | Nek6 |
| ENSMUSG( | 3798.242 | -0.00199 | 0.985968 | 0.998326 | Nek7 |
| ENSMUSG( | 185.4321 | -0.0472 | 0.778691 | 0.956339 | Nek8 |
| ENSMUSG( | 4026.629 | 0.087771 | 0.349497 | 0.781909 | Nek9 |
| ENSMUSG( | 625.6925 | 0.158257 | 0.30302 | 0.751443 | Nelfa |
| ENSMUSG( | 1467.861 | 0.129826 | 0.140276 | 0.557263 | Nelfb |
| ENSMUSG( | 1102.378 | 0.148572 | 0.22266 | 0.668434 | Nelfcd |
| ENSMUSG( | 891.141 | 0.081846 | 0.444618 | 0.833239 | Nelfe |
| ENSMUSG( | 1229.567 | -0.49409 | 0.000654 | 0.024667 | Nell1 |
| ENSMUSG( | 15315.2 | -0.26061 | 0.076915 | 0.429306 | Nell2 |
| ENSMUSG( | 1097.128 | 0.018854 | 0.858249 | 0.974672 | Nemf |
| ENSMUSG( | 193.3475 | 0.069678 | 0.681263 | 0.931396 | Nemp2 |
| ENSMUSG( | 2113.227 | -0.27067 | 0.006426 | 0.110055 | Nenf |
| ENSMUSG( | 5258.529 | -0.16467 | 0.205468 | 0.646873 | Neo1 |
| ENSMUSG( | 108.6934 | -0.00645 | 0.976529 | 0.997428 | Nepro |
| ENSMUSG( | 606.8842 | 0.074346 | 0.680918 | 0.931244 | Nes |
| ENSMUSG( | 645.6868 | 0.088996 | 0.513251 | 0.871762 | Net1 |
| ENSMUSG( | 2225.196 | -0.29547 | 0.007472 | 0.119529 | Neto1 |
| ENSMUSG( | 1626.781 | -0.58389 | 0.000126 | 0.007476 | Neto2 |
| ENSMUSG( | 1854.576 | -0.24329 | 0.069571 | 0.408118 | Neu1 |
| ENSMUSG( | 161.4735 | -0.11434 | 0.539419 | 0.883094 | Neu2 |
| ENSMUSG( | 65.03909 | -0.53047 | 0.039589 | 0.306803 | Neu3 |
| ENSMUSG( | 531.1895 | -0.2836 | 0.058676 | 0.373994 | Neu4 |
| ENSMUSG( | 11144.24 | 0.253549 | 0.002912 | 0.066108 | Neurl1a |
| ENSMUSG( | 3290.344 | 0.237995 | 0.017056 | 0.195102 | Neurl1b |
| ENSMUSG( | 61.17785 | 0.422825 | 0.106657 | 0.494174 | Neurl2 |
| ENSMUSG( | 58.17691 | -0.91344 | 0.000875 | 0.029753 | Neurl3 |
| ENSMUSG( | 5006.935 | 0.133896 | 0.129343 | 0.538561 | Neurl4 |
| ENSMUSG( | 159.9451 | 0.092577 | 0.624117 | 0.91187 | Neurod1 |
| ENSMUSG( | 2165.395 | 0.179153 | 0.200434 | 0.640598 | Neurod2 |
| ENSMUSG( | 10.7139 | -0.1947 | 0.731241 | NA | Neurod4 |
| ENSMUSG( | 977.5383 | 0.017631 | 0.912941 | 0.988483 | Neurod6 |
| ENSMUSG( | 58.41749 | 0.064693 | 0.811854 | 0.964426 | Neurog2 |
| ENSMUSG( | 458.9989 | 0.013747 | 0.919802 | 0.990247 | Nexmif |
| ENSMUSG( | 180.4167 | -0.78414 | 0.000884 | 0.02993 | Nexn |
| ENSMUSG( | 3512.326 | 0.032377 | 0.72807 | 0.944355 | Nf1 |
| ENSMUSG( | 8606.31 | 0.080343 | 0.342168 | 0.778828 | Nf2 |
| ENSMUSG( | 229.7625 | -0.15787 | 0.399874 | 0.813593 | Nfam1 |
| ENSMUSG( | 12618.59 | -0.206 | 0.093558 | 0.469962 | Nfasc |
| ENSMUSG( | 2455.623 | -0.03029 | 0.752489 | 0.950536 | Nfat5 |
| ENSMUSG( | 366.9787 | -0.14076 | 0.393012 | 0.809274 | Nfatc1 |
| ENSMUSG( | 472.6968 | -0.09952 | 0.387808 | 0.805835 | Nfatc2 |
| ENSMUSG( | 167.7987 | 0.005891 | 0.97156 | 0.996503 | Nfatc2ip |
| ENSMUSG( | 400.3722 | 0.179092 | 0.126871 | 0.534201 | Nfatc3 |
| ENSMUSG( | 64.19232 | -0.01128 | 0.962869 | 0.994824 | Nfatc4 |
| ENSMUSG( | 14227.43 | -0.07792 | 0.400813 | 0.81387 | Nfe2l1 |
| ENSMUSG( | 851.891 | 0.071566 | 0.537629 | 0.882536 | Nfe2l2 |
| ENSMUSG( | 124.0338 | -0.88122 | 5.77E-06 | 0.000749 | Nfe2l3 |
| ENSMUSG( | 2889.624 | 0.120856 | 0.284085 | 0.732569 | Nfia |
| ENSMUSG( | 3864.509 | 0.157526 | 0.051455 | 0.348059 | Nfib |
| ENSMUSG( | 7533.281 | 0.169281 | 0.045632 | 0.327937 | Nfic |
| ENSMUSG( | 164.9377 | -0.06216 | 0.704678 | 0.938303 | Nfil3 |
| ENSMUSG( | 14453.36 | 0.052287 | 0.486944 | 0.858812 | Nfix |
| ENSMUSG( | 688.8203 | -0.03866 | 0.725896 | 0.944223 | Nfkb1 |
| ENSMUSG( | 241.3455 | 0.232676 | 0.124862 | 0.531843 | Nfkb2 |
| ENSMUSG( | 618.8492 | -0.26583 | 0.053293 | 0.354557 | Nfkbia |
| ENSMUSG( | 657.2655 | -0.01892 | 0.899576 | 0.985633 | Nfkbib |
| ENSMUSG( | 90.16783 | -0.0571 | 0.801391 | 0.961947 | Nfkbid |
| ENSMUSG( | 101.0194 | -0.39737 | 0.081607 | 0.441308 | Nfkbie |
| ENSMUSG( | 355.8038 | -0.07829 | 0.556541 | 0.887713 | Nfkbil1 |
| ENSMUSG( | 177.2441 | -0.29516 | 0.062357 | 0.386294 | Nfkbiz |
| ENSMUSG( | 893.6802 | -0.0418 | 0.71437 | 0.941045 | Nfrkb |
| ENSMUSG( | 1337.764 | 0.035519 | 0.748857 | 0.949604 | Nfs1 |
| ENSMUSG( | 922.8541 | 0.002726 | 0.979379 | 0.997428 | Nfu1 |
| ENSMUSG( | 1903.005 | -0.04783 | 0.631806 | 0.914073 | Nfx1 |
| ENSMUSG( | 285.6438 | -0.17603 | 0.194439 | 0.63341 | Nfxl1 |
| ENSMUSG( | 659.1457 | 0.171971 | 0.114371 | 0.51021 | Nfya |
| ENSMUSG( | 1554.954 | 0.048528 | 0.63874 | 0.91636 | Nfyb |
| ENSMUSG( | 838.2596 | 0.084838 | 0.519878 | 0.874979 | Nfyc |
| ENSMUSG( | 189.3546 | 0.239588 | 0.160933 | 0.590706 | Ngb |
| ENSMUSG( | 413.0293 | 0.191882 | 0.105973 | 0.493403 | Ngdn |
| ENSMUSG( | 9723.852 | 0.004561 | 0.969697 | 0.996369 | Ngef |
| ENSMUSG( | 105.5186 | -0.60226 | 0.004705 | 0.090209 | Ngf |
| ENSMUSG( | 49.88413 | -0.65149 | 0.02187 | 0.225738 | Ngfr |
| ENSMUSG( | 682.4441 | 0.024153 | 0.833031 | 0.970333 | Ngly1 |
| ENSMUSG( | 94.70385 | -1.43909 | 0.182881 | 0.62044 | Ngp |
| ENSMUSG( | 1092.605 | 0.074763 | 0.481338 | 0.855314 | Ngrn |
| ENSMUSG( | 72.49774 | -0.01314 | 0.957253 | 0.993874 | Nhej1 |
| ENSMUSG( | 4623.136 | 0.164149 | 0.103808 | 0.489428 | Nherf1 |
| ENSMUSG( | 1388.79 | 0.28974 | 0.01074 | 0.149272 | Nherf2 |

| | | | | | |
|---|---|---|---|---|---|
| ENSMUSG( | 81.75432 | -0.05962 | 0.833317 | 0.981193 | Nfkbid |
| ENSMUSG( | 94.14396 | -0.32198 | 0.16439 | 0.83856 | Nfkbie |
| ENSMUSG( | 319.5972 | -0.13061 | 0.451429 | 0.922202 | Nfkbil1 |
| ENSMUSG( | 176.41 | -0.01594 | 0.932075 | 0.99306 | Nfkbiz |
| ENSMUSG( | 829.2445 | 0.016602 | 0.909639 | 0.991958 | Nfrkb |
| ENSMUSG( | 1129.056 | -0.19679 | 0.13532 | 0.825081 | Nfs1 |
| ENSMUSG( | 797.436 | -0.16078 | 0.269186 | 0.886755 | Nfu1 |
| ENSMUSG( | 1797.308 | 0.058272 | 0.61828 | 0.952254 | Nfx1 |
| ENSMUSG( | 288.7873 | 0.132241 | 0.393331 | 0.917257 | Nfxl1 |
| ENSMUSG( | 564.0699 | -0.00035 | 0.997842 | 0.999268 | Nfya |
| ENSMUSG( | 1445.045 | 0.10029 | 0.463014 | 0.923329 | Nfyb |
| ENSMUSG( | 738.4512 | -0.01319 | 0.926443 | 0.99306 | Nfyc |
| ENSMUSG( | 149.0093 | -0.19152 | 0.352263 | 0.912093 | Ngb |
| ENSMUSG( | 352.6297 | 0.007333 | 0.959147 | 0.994798 | Ngdn |
| ENSMUSG( | 8059.315 | -0.2846 | 0.042603 | 0.690304 | Ngef |
| ENSMUSG( | 112.7416 | -0.07392 | 0.736252 | 0.967344 | Ngf |
| ENSMUSG( | 57.59068 | 0.093 | 0.739377 | 0.968458 | Ngfr |
| ENSMUSG( | 593.5205 | -0.11359 | 0.381986 | 0.917257 | Ngly1 |
| ENSMUSG( | 131.568 | 0.1034 | 0.909777 | 0.991958 | Ngp |
| ENSMUSG( | 941.9754 | -0.08978 | 0.487301 | 0.92541 | Ngrn |
| ENSMUSG( | 62.93745 | -0.17968 | 0.524198 | 0.933263 | Nhej1 |
| ENSMUSG( | 3924.861 | -0.03675 | 0.770349 | 0.974236 | Nherf1 |
| ENSMUSG( | 1483.123 | 0.687222 | 4.97E-05 | 0.012613 | Nherf2 |
| ENSMUSG( | 17.0451 | -0.13162 | 0.803578 | 0.976978 | Nherf4 |
| ENSMUSG( | 11.01917 | -0.50158 | 0.399579 | NA | Nhlh1 |
| ENSMUSG( | 7.222672 | -0.26269 | 0.71813 | NA | Nhlh2 |
| ENSMUSG( | 329.092 | -0.07379 | 0.626041 | 0.953234 | Nhlrc1 |
| ENSMUSG( | 810.8156 | 0.036757 | 0.753391 | 0.970671 | Nhlrc2 |
| ENSMUSG( | 562.5192 | 0.049851 | 0.743853 | 0.96951 | Nhlrc3 |
| ENSMUSG( | 637.1128 | -0.25307 | 0.083766 | 0.77681 | Nhp2 |
| ENSMUSG( | 85.55564 | -0.07723 | 0.750299 | 0.970113 | Nhs |
| ENSMUSG( | 2747.605 | -0.18818 | 0.151949 | 0.837724 | Nhsl1 |
| ENSMUSG( | 1255.402 | 0.245644 | 0.032496 | 0.63668 | Nhsl2 |
| ENSMUSG( | 1229.23 | 0.03663 | 0.788469 | 0.97637 | Nhsl3 |
| ENSMUSG( | 227.8184 | 0.176879 | 0.437657 | 0.920339 | Niban1 |
| ENSMUSG( | 2354.716 | -0.06393 | 0.569005 | 0.941777 | Niban2 |
| ENSMUSG( | 45.73644 | -0.06609 | 0.829502 | 0.980224 | Niban3 |
| ENSMUSG( | 1930.047 | -0.13779 | 0.235333 | 0.871741 | Nicn1 |
| ENSMUSG( | 728.5482 | -0.31041 | 0.033455 | 0.643993 | Nicol1 |
| ENSMUSG( | 835.9382 | 0.282505 | 0.146336 | 0.835973 | Nid1 |
| ENSMUSG( | 349.9066 | 0.100089 | 0.559775 | 0.939812 | Nid2 |
| ENSMUSG( | 374.2535 | -0.07963 | 0.57663 | 0.942912 | Nif3l1 |
| ENSMUSG( | 366.8071 | -0.18998 | 0.189256 | 0.85698 | Nifk |
| ENSMUSG( | 1172.326 | 0.143151 | 0.24756 | 0.877904 | Nim1k |
| ENSMUSG( | 1278.781 | 0.058756 | 0.588362 | 0.944975 | Nin |
| ENSMUSG( | 601.8074 | 0.18681 | 0.167964 | 0.840222 | Ninj1 |
| ENSMUSG( | 86.3941 | 0.243551 | 0.331513 | 0.907907 | Ninj2 |
| ENSMUSG( | 216.632 | -0.33451 | 0.085908 | 0.780123 | Ninl |
| ENSMUSG( | 437.1127 | -0.06834 | 0.64459 | 0.953903 | Nip7 |
| ENSMUSG( | 1535.254 | 0.157192 | 0.364998 | 0.914449 | Nipa1 |
| ENSMUSG( | 1145.517 | 0.096925 | 0.558858 | 0.939587 | Nipa2 |
| ENSMUSG( | 244.3259 | -0.09907 | 0.578669 | 0.943548 | Nipal2 |
| ENSMUSG( | 3814.537 | 0.282599 | 0.096622 | 0.796245 | Nipal3 |
| ENSMUSG( | 296.5936 | 0.247856 | 0.258193 | 0.883971 | Nipal4 |
| ENSMUSG( | 1661.562 | 0.199536 | 0.146559 | 0.835973 | Nipbl |
| ENSMUSG( | 920.5126 | -0.20734 | 0.229048 | 0.868507 | Nipsnap1 |
| ENSMUSG( | 2137.348 | -0.19299 | 0.119413 | 0.813384 | Nipsnap2 |
| ENSMUSG( | 267.8698 | -0.12495 | 0.453336 | 0.922316 | Nipsnap3b |
| ENSMUSG( | 14275.86 | 0.079765 | 0.42107 | 0.917257 | Nisch |
| ENSMUSG( | 838.7527 | -0.06962 | 0.589637 | 0.944975 | Nit1 |
| ENSMUSG( | 259.8571 | -0.14547 | 0.363739 | 0.914223 | Nit2 |
| ENSMUSG( | 814.8865 | 0.057624 | 0.709257 | 0.964042 | Nkain1 |
| ENSMUSG( | 1641.122 | 0.181386 | 0.22445 | 0.867923 | Nkain2 |
| ENSMUSG( | 42.41524 | 0.132737 | 0.67887 | 0.957617 | Nkain3 |
| ENSMUSG( | 1911.187 | -0.01029 | 0.934254 | 0.99306 | Nkain4 |
| ENSMUSG( | 311.8612 | -0.05439 | 0.725402 | 0.966887 | Nkap |
| ENSMUSG( | 705.474 | 0.030156 | 0.831113 | 0.98051 | Nkapd1 |
| ENSMUSG( | 1403.431 | -0.02655 | 0.815361 | 0.979198 | Nkd1 |
| ENSMUSG( | 129.8454 | 0.337368 | 0.099654 | 0.801103 | Nkd2 |
| ENSMUSG( | 1653.421 | -0.17972 | 0.154211 | 0.837724 | Nkiras1 |
| ENSMUSG( | 490.8987 | -0.10918 | 0.392613 | 0.917257 | Nkiras2 |
| ENSMUSG( | 36.98199 | -0.16126 | 0.618983 | 0.952598 | Nkpd1 |
| ENSMUSG( | 395.485 | 0.043611 | 0.787824 | 0.97637 | Nkrf |
| ENSMUSG( | 4617.636 | 0.179246 | 0.121937 | 0.813384 | Nktr |
| ENSMUSG( | 19.76529 | -0.13971 | 0.783206 | 0.976106 | Nkx2-1 |
| ENSMUSG( | 280.4541 | 0.097406 | 0.611359 | 0.950771 | Nkx2-2 |
| ENSMUSG( | 109.9809 | 0.242209 | 0.318712 | 0.903297 | Nkx2-2os |
| ENSMUSG( | 21.72101 | 0.012755 | 0.97827 | 0.996416 | Nkx3-1 |
| ENSMUSG( | 321.0611 | -0.14957 | 0.388839 | 0.917257 | Nkx6-2 |
| ENSMUSG( | 275.7864 | 0.089709 | 0.620326 | 0.952896 | Nle1 |
| ENSMUSG( | 1620.557 | 0.203726 | 0.271436 | 0.888062 | Nlgn1 |
| ENSMUSG( | 12188.21 | 0.134789 | 0.288584 | 0.894794 | Nlgn2 |
| ENSMUSG( | 2425.698 | 0.184856 | 0.241326 | 0.876259 | Nlgn3 |
| ENSMUSG( | 1966.602 | 0.074933 | 0.48394 | 0.925289 | Nlk |
| ENSMUSG( | 1223.206 | -0.07567 | 0.553401 | 0.939565 | Nln |
| ENSMUSG( | 33.47444 | 0.150825 | 0.703763 | 0.96251 | Nlrc3 |
| ENSMUSG( | 12.42893 | -0.57236 | 0.350532 | 0.911652 | Nlrc4 |
| ENSMUSG( | 31.8629 | -0.46858 | 0.227613 | 0.868507 | Nlrc5 |
| ENSMUSG( | 34.79785 | 0.147343 | 0.719741 | 0.966887 | Nlrp1b |
| ENSMUSG( | 47.34954 | 0.472855 | 0.155661 | 0.837724 | Nlrp3 |
| ENSMUSG( | 12.11404 | 0.439863 | 0.478864 | 0.923329 | Nlrp6 |
| ENSMUSG( | 276.3703 | 0.044861 | 0.781841 | 0.976106 | Nlrx1 |
| ENSMUSG( | 110.8204 | 0.194239 | 0.36135 | 0.914223 | Nmb |
| ENSMUSG( | 71.73668 | 0.33889 | 0.193537 | 0.860096 | Nmbr |
| ENSMUSG( | 455.297 | -0.16511 | 0.223832 | 0.867404 | Nmd3 |
| ENSMUSG( | 1280.353 | -0.17364 | 0.272308 | 0.888771 | Nme1 |
| ENSMUSG( | 34.79232 | -0.24564 | 0.472664 | 0.923329 | Nme2 |
| ENSMUSG( | 70.21076 | 0.292998 | 0.233113 | 0.868507 | Nme4 |
| ENSMUSG( | 160.8303 | -0.159 | 0.40745 | 0.917257 | Nme5 |
| ENSMUSG( | 115.2746 | 0.1224 | 0.564859 | 0.941206 | Nme6 |
| ENSMUSG( | 859.7746 | -0.18722 | 0.157749 | 0.837724 | Nme7 |
| ENSMUSG( | 175.0931 | -0.13203 | 0.462891 | 0.923329 | Nmi |
| ENSMUSG( | 158.5973 | 0.192846 | 0.296966 | 0.898538 | Nmnat1 |
| ENSMUSG( | 3188.712 | -0.06583 | 0.567297 | 0.941777 | Nmnat2 |
| ENSMUSG( | 56.87174 | 0.162559 | 0.567751 | 0.941777 | Nmnat3 |
| ENSMUSG( | 242.9724 | 0.004803 | 0.979836 | 0.996479 | Nmral1 |
| ENSMUSG( | 115.3854 | -0.12435 | 0.599698 | 0.947071 | Nmrk1 |
| ENSMUSG( | 3420.747 | 0.065162 | 0.571214 | 0.942052 | Nmt1 |
| ENSMUSG( | 2263.821 | -0.15919 | 0.207766 | 0.864412 | Nmt2 |
| ENSMUSG( | 8018.831 | -0.2572 | 0.081758 | 0.77681 | Nnat |
| ENSMUSG( | 16.18138 | -0.07878 | 0.869913 | 0.988097 | Nnmt |
| ENSMUSG( | 9.331284 | -0.40515 | 0.523736 | NA | Nnt |
| ENSMUSG( | 457.287 | 0.017937 | 0.897211 | 0.990558 | Noa1 |
| ENSMUSG( | 503.1595 | -0.04354 | 0.753156 | 0.970622 | Nob1 |
| ENSMUSG( | 2089.814 | -0.00308 | 0.978165 | 0.996416 | Noc2l |
| ENSMUSG( | 383.3094 | 0.105009 | 0.464013 | 0.923329 | Noc3l |
| ENSMUSG( | 348.5681 | -0.15871 | 0.404517 | 0.917257 | Noc4l |
| ENSMUSG( | 813.6279 | -0.07062 | 0.636867 | 0.953903 | Noct |
| ENSMUSG( | 255.6461 | 0.294462 | 0.227918 | 0.868507 | Nod1 |
| ENSMUSG( | 14.5553 | -0.69583 | 0.183962 | 0.852278 | Nod2 |
| ENSMUSG( | 23.3354 | -0.29571 | 0.585484 | 0.944975 | Nodal |
| ENSMUSG( | 420.9981 | 0.225509 | 0.213538 | 0.864421 | Nog |
| ENSMUSG( | 432.6736 | -0.03977 | 0.787567 | 0.97637 | Nol10 |
| ENSMUSG( | 443.6385 | 0.10669 | 0.4239 | 0.917257 | Nol11 |
| ENSMUSG( | 676.1582 | -0.20905 | 0.098559 | 0.798438 | Nol12 |
| ENSMUSG( | 753.0309 | -0.22531 | 0.114504 | 0.811703 | Nol3 |
| ENSMUSG( | 1278.434 | 0.062913 | 0.569351 | 0.941777 | Nol4 |
| ENSMUSG( | 1972.086 | 0.051587 | 0.682549 | 0.958678 | Nol4l |
| ENSMUSG( | 2511.496 | -0.12441 | 0.317778 | 0.902777 | Nol6 |
| ENSMUSG( | 981.6162 | -0.07313 | 0.586868 | 0.944975 | Nol7 |
| ENSMUSG( | 555.6073 | -0.02945 | 0.82926 | 0.980224 | Nol8 |
| ENSMUSG( | 553.272 | 0.116474 | 0.379066 | 0.916995 | Nol9 |
| ENSMUSG( | 999.7049 | -0.04161 | 0.74246 | 0.969415 | Nolc1 |
| ENSMUSG( | 741.1768 | -0.09866 | 0.476471 | 0.923329 | Nom1 |
| ENSMUSG( | 5192.909 | 0.008175 | 0.959103 | 0.994798 | Nomo1 |
| ENSMUSG( | 3019.484 | -0.12135 | 0.292906 | 0.898413 | Nono |
| ENSMUSG( | 716.7926 | -0.17874 | 0.186942 | 0.856099 | Nop10 |
| ENSMUSG( | 551.1725 | -0.01326 | 0.922644 | 0.992644 | Nop14 |
| ENSMUSG( | 286.7981 | 0.022733 | 0.899135 | 0.990558 | Nop16 |
| ENSMUSG( | 486.5376 | -0.04016 | 0.763814 | 0.974 | Nop2 |
| ENSMUSG( | 3051.268 | -0.05096 | 0.669006 | 0.956766 | Nop53 |
| ENSMUSG( | 1709.022 | 0.024133 | 0.837592 | 0.981335 | Nop56 |
| ENSMUSG( | 661.4826 | -0.1931 | 0.168268 | 0.840222 | Nop58 |
| ENSMUSG( | 299.3792 | 0.117358 | 0.427357 | 0.917257 | Nop9 |
| ENSMUSG( | 959.8556 | -0.07628 | 0.54588 | 0.938063 | Nopchap1 |
| ENSMUSG( | 30047.14 | -0.0384 | 0.704379 | 0.962677 | Norad |
| ENSMUSG( | 736.991 | -0.09425 | 0.488284 | 0.925453 | Nos1 |
| ENSMUSG( | 942.1189 | 0.114965 | 0.446606 | 0.921734 | Nos1ap |
| ENSMUSG( | 36.00791 | 0.595989 | 0.106615 | 0.804969 | Nos2 |
| ENSMUSG( | 418.5921 | 0.813121 | 0.000722 | 0.08668 | Nos3 |
| ENSMUSG( | 1047.64 | -0.05995 | 0.675232 | 0.95694 | Nosip |
| ENSMUSG( | 221.9808 | 0.410742 | 0.06734 | 0.75212 | Nostrin |
| ENSMUSG( | 1838.474 | 0.596044 | 0.002208 | 0.183318 | Notch1 |
| ENSMUSG( | 1387.89 | -0.01971 | 0.899778 | 0.990558 | Notch2 |
| ENSMUSG( | 866.565 | 0.46088 | 0.009724 | 0.406458 | Notch3 |
| ENSMUSG( | 377.8752 | 0.745649 | 0.000973 | 0.106505 | Notch4 |
| ENSMUSG( | 199.1889 | 0.90086 | 7.58E-05 | 0.017018 | Notum |
| ENSMUSG( | 1375.658 | -0.02035 | 0.876421 | 0.989644 | Nova1 |

| | | | | | |
|---|---|---|---|---|---|
| ENSMUSG( | Npc1l1 | 86.5722 | 0.186516 | 0.423104 | 0.823662 |
| ENSMUSG( | Npc2 | 5610.883 | -0.14068 | 0.201161 | 0.641362 |
| ENSMUSG( | Npdc1 | 9068.627 | -0.1948 | 0.096979 | 0.477375 |
| ENSMUSG( | Npepl1 | 1042.382 | 0.106021 | 0.274163 | 0.724254 |
| ENSMUSG( | Npepps | 5337.181 | 0.169891 | 0.035115 | 0.288256 |
| ENSMUSG( | Npff | 39.96941 | 0.3854 | 0.25983 | 0.709951 |
| ENSMUSG( | Npffr1 | 35.47242 | 0.106568 | 0.742196 | 0.948478 |
| ENSMUSG( | Nphp1 | 644.7389 | 0.032443 | 0.75351 | 0.950753 |
| ENSMUSG( | Nphp3 | 418.9768 | -0.24322 | 0.05248 | 0.35151 |
| ENSMUSG( | Nphp4 | 575.9062 | 0.072902 | 0.553805 | 0.886394 |
| ENSMUSG( | Nphs1 | 301.6568 | 0.173741 | 0.235117 | 0.682364 |
| ENSMUSG( | Nphs1os | 36.65198 | -0.17233 | 0.639475 | 0.916434 |
| ENSMUSG( | Nphs2 | 15.7841 | -0.06931 | 0.897481 | NA |
| ENSMUSG( | Npl | 164.9555 | 0.308194 | 0.086355 | 0.452045 |
| ENSMUSG( | Nploc4 | 2287.747 | 0.125053 | 0.133772 | 0.547197 |
| ENSMUSG( | Npm1 | 3038.752 | -0.04927 | 0.624472 | 0.91187 |
| ENSMUSG( | Npm2 | 15.44514 | 0.347948 | 0.477253 | NA |
| ENSMUSG( | Npm3 | 184.1042 | 0.135754 | 0.384864 | 0.804031 |
| ENSMUSG( | Npnt | 1292.646 | -0.0731 | 0.664868 | 0.926824 |
| ENSMUSG( | Nppa | 10.40045 | -1.13139 | 0.147178 | NA |
| ENSMUSG( | Nppc | 178.9428 | -0.61631 | 0.002465 | 0.058361 |
| ENSMUSG( | Npr1 | 128.9001 | -0.31248 | 0.101789 | 0.486216 |
| ENSMUSG( | Npr2 | 2204.931 | 0.070424 | 0.414589 | 0.820184 |
| ENSMUSG( | Npr3 | 330.4608 | 0.025491 | 0.871228 | 0.978114 |
| ENSMUSG( | Nprl2 | 693.6934 | 0.046891 | 0.73214 | 0.944956 |
| ENSMUSG( | Nprl3 | 877.9907 | 0.008877 | 0.946189 | 0.99369 |
| ENSMUSG( | Npsr1 | 48.28876 | -0.68802 | 0.044126 | 0.322108 |
| ENSMUSG( | Nptn | 17488.89 | -0.61425 | 7.85E-07 | 0.000159 |
| ENSMUSG( | Nptx1 | 10092.01 | -0.31648 | 0.055853 | 0.364301 |
| ENSMUSG( | Nptx2 | 2298.464 | -0.57492 | 5.40E-05 | 0.004212 |
| ENSMUSG( | Nptxr | 40260.18 | -0.1028 | 0.401805 | 0.814965 |
| ENSMUSG( | Npy | 3812.252 | -0.77417 | 1.99E-09 | 8.98E-07 |
| ENSMUSG( | Npy1r | 701.8671 | -0.34521 | 0.040487 | 0.311082 |
| ENSMUSG( | Npy2r | 205.1565 | -0.31828 | 0.086163 | 0.451605 |
| ENSMUSG( | Npy5r | 105.1711 | -0.34576 | 0.136741 | 0.551176 |
| ENSMUSG( | Nqo1 | 259.6419 | 0.034172 | 0.820619 | 0.965888 |
| ENSMUSG( | Nqo2 | 678.3729 | 0.066393 | 0.667116 | 0.92729 |
| ENSMUSG( | Nr1d1 | 2903.091 | 0.195688 | 0.105485 | 0.492417 |
| ENSMUSG( | Nr1d2 | 2310.528 | 0.095682 | 0.331383 | 0.77519 |
| ENSMUSG( | Nr1h2 | 2068.322 | 0.176681 | 0.049003 | 0.337938 |
| ENSMUSG( | Nr1h3 | 139.5272 | 0.298335 | 0.093408 | 0.469793 |
| ENSMUSG( | Nr1i3 | 24.33574 | -0.34897 | 0.369175 | 0.796641 |
| ENSMUSG( | Nr2c1 | 256.8529 | -0.28714 | 0.13306 | 0.545688 |
| ENSMUSG( | Nr2c2 | 4317.523 | 0.059733 | 0.519729 | 0.874979 |
| ENSMUSG( | Nr2c2ap | 311.6272 | 0.019982 | 0.886709 | 0.983255 |
| ENSMUSG( | Nr2e1 | 1728.646 | -0.04265 | 0.724847 | 0.944096 |
| ENSMUSG( | Nr2f1 | 3485.297 | 0.195936 | 0.025656 | 0.246969 |
| ENSMUSG( | Nr2f2 | 458.8348 | 0.015147 | 0.928845 | 0.991747 |
| ENSMUSG( | Nr2f6 | 1259.673 | 0.075433 | 0.405514 | 0.81652 |
| ENSMUSG( | Nr3c1 | 2792.646 | -0.03425 | 0.747097 | 0.949526 |
| ENSMUSG( | Nr3c2 | 1148.873 | -0.08446 | 0.412461 | 0.819452 |
| ENSMUSG( | Nr4a1 | 2282.211 | -0.68222 | 7.14E-05 | 0.005192 |
| ENSMUSG( | Nr4a2 | 1312.134 | 0.11763 | 0.210725 | 0.653529 |
| ENSMUSG( | Nr4a3 | 751.8325 | 0.084712 | 0.399132 | 0.813048 |
| ENSMUSG( | Nr6a1 | 189.4328 | -0.2233 | 0.165109 | 0.596585 |
| ENSMUSG( | Nradd | 48.89396 | -0.40268 | 0.179995 | 0.617634 |
| ENSMUSG( | Nrap | 32.64834 | -0.01446 | 0.970467 | 0.996369 |
| ENSMUSG( | Nrarp | 2003.444 | 0.084378 | 0.436539 | 0.829373 |
| ENSMUSG( | Nras | 3423.33 | 0.006105 | 0.958785 | 0.993874 |
| ENSMUSG( | Nrbf2 | 205.7725 | 0.024712 | 0.900805 | 0.985633 |
| ENSMUSG( | Nrbp1 | 1828.127 | 0.120654 | 0.270632 | 0.720561 |
| ENSMUSG( | Nrbp2 | 5895.503 | 0.099089 | 0.590938 | 0.90113 |
| ENSMUSG( | Nrcam | 13306.94 | -0.04798 | 0.718618 | 0.941974 |
| ENSMUSG( | Nrdc | 5089.563 | -0.11072 | 0.140487 | 0.557685 |
| ENSMUSG( | Nrde2 | 369.218 | 0.075234 | 0.630276 | 0.914073 |
| ENSMUSG( | Nrep | 4019.184 | 0.106257 | 0.256176 | 0.706924 |
| ENSMUSG( | Nrf1 | 542.4099 | 0.197538 | 0.069395 | 0.40782 |
| ENSMUSG( | Nrg1 | 323.7025 | -0.1011 | 0.524414 | 0.87583 |
| ENSMUSG( | Nrg2 | 241.4858 | -0.07078 | 0.61245 | 0.908747 |
| ENSMUSG( | Nrg3 | 1048.247 | -0.49556 | 0.002722 | 0.063172 |
| ENSMUSG( | Nrg4 | 11.61529 | -0.41272 | 0.443418 | NA |
| ENSMUSG( | Nrgn | 60013.31 | 0.09104 | 0.348595 | 0.78132 |
| ENSMUSG( | Nrip1 | 687.2703 | -0.10514 | 0.305581 | 0.753616 |
| ENSMUSG( | Nrip2 | 177.5155 | 0.129982 | 0.507769 | 0.86795 |
| ENSMUSG( | Nrip3 | 3019.309 | 0.124373 | 0.190774 | 0.630215 |
| ENSMUSG( | Nrm | 139.9948 | -0.37331 | 0.038611 | 0.302696 |
| ENSMUSG( | Nrn1 | 5766.661 | -0.31146 | 0.003254 | 0.071142 |
| ENSMUSG( | Nrp1 | 800.3894 | -0.07418 | 0.68547 | 0.932981 |
| ENSMUSG( | Nrp2 | 1083.651 | -0.41883 | 0.020106 | 0.214413 |
| ENSMUSG( | Nrros | 291.4668 | -0.00458 | 0.978242 | 0.997428 |
| ENSMUSG( | Nrsn1 | 34606.39 | 0.222345 | 0.03289 | 0.277278 |
| ENSMUSG( | Nrsn2 | 2251.078 | -0.71633 | 1.85E-10 | 1.07E-07 |
| ENSMUSG( | Nrtn | 155.2788 | -0.07854 | 0.671496 | 0.928781 |
| ENSMUSG( | Nrxn1 | 15091.53 | -0.20345 | 0.151458 | 0.576717 |
| ENSMUSG( | Nrxn2 | 20810.85 | -0.11565 | 0.34414 | 0.779153 |
| ENSMUSG( | Nrxn3 | 3825.202 | -0.41876 | 0.001186 | 0.037079 |
| ENSMUSG( | Nsa2 | 1556.889 | 0.086607 | 0.383621 | 0.803504 |
| ENSMUSG( | Nscme3l | 10.82608 | -0.6796 | 0.246563 | NA |
| ENSMUSG( | Nsd1 | 4389.521 | -0.05975 | 0.521904 | 0.875186 |
| ENSMUSG( | Nsd2 | 2834.504 | 0.075904 | 0.363177 | 0.793211 |
| ENSMUSG( | Nsd3 | 2438.254 | -0.00936 | 0.926851 | 0.991229 |
| ENSMUSG( | Nsdhl | 1460.312 | -0.13766 | 0.332302 | 0.775474 |
| ENSMUSG( | Nsf | 12973.4 | 0.181818 | 0.155258 | 0.583155 |
| ENSMUSG( | Nsfl1c | 1535.27 | 0.053388 | 0.589146 | 0.90113 |
| ENSMUSG( | Nsg1 | 8031.302 | -0.43982 | 4.99E-05 | 0.00395 |
| ENSMUSG( | Nsg2 | 17230.65 | -0.42364 | 0.000252 | 0.012746 |
| ENSMUSG( | Nsl1 | 29.34348 | -0.01625 | 0.965404 | 0.995377 |
| ENSMUSG( | Nsmaf | 1217.346 | 0.126543 | 0.208447 | 0.650319 |
| ENSMUSG( | Nsmce1 | 375.8568 | 0.001554 | 0.990543 | 0.999232 |
| ENSMUSG( | Nsmce2 | 448.9543 | 0.160581 | 0.279435 | 0.729114 |
| ENSMUSG( | Nsmce3 | 755.6087 | -0.61655 | 1.91E-06 | 0.000324 |
| ENSMUSG( | Nsmce4a | 680.2453 | 0.069362 | 0.539935 | 0.88315 |
| ENSMUSG( | Nsmf | 46491.73 | 0.23259 | 0.006802 | 0.113745 |
| ENSMUSG( | Nsrp1 | 463.2242 | 0.079927 | 0.524174 | 0.87583 |
| ENSMUSG( | Nsun2 | 1673.443 | 0.104889 | 0.277619 | 0.72746 |
| ENSMUSG( | Nsun3 | 314.8393 | 0.008356 | 0.955908 | 0.993874 |
| ENSMUSG( | Nsun4 | 349.5936 | 0.13778 | 0.290759 | 0.740751 |
| ENSMUSG( | Nsun5 | 358.9904 | 0.071728 | 0.597564 | 0.903694 |
| ENSMUSG( | Nsun6 | 157.3169 | -0.04331 | 0.790539 | 0.959461 |
| ENSMUSG( | Nsun7 | 317.8613 | 0.056789 | 0.686027 | 0.932981 |
| ENSMUSG( | Nt5c | 716.925 | 0.066819 | 0.583872 | 0.898666 |
| ENSMUSG( | Nt5c1a | 92.11023 | 0.106273 | 0.623169 | 0.911599 |
| ENSMUSG( | Nt5c2 | 2925.488 | 0.068944 | 0.381952 | 0.802983 |
| ENSMUSG( | Nt5c3 | 1072.886 | -0.20536 | 0.044529 | 0.324173 |
| ENSMUSG( | Nt5c3b | 1318.843 | 0.088883 | 0.34231 | 0.778828 |
| ENSMUSG( | Nt5dc1 | 61.95924 | -0.24164 | 0.3423 | 0.778828 |
| ENSMUSG( | Nt5dc2 | 380.1357 | 0.160095 | 0.238234 | 0.686308 |
| ENSMUSG( | Nt5dc3 | 2626.471 | 0.210833 | 0.076083 | 0.42702 |
| ENSMUSG( | Nt5e | 237.5643 | -0.48362 | 0.01221 | 0.160227 |
| ENSMUSG( | Nt5m | 1334.804 | 0.085368 | 0.337744 | 0.777422 |
| ENSMUSG( | Ntan1 | 1346.862 | 0.006026 | 0.947499 | 0.993821 |
| ENSMUSG( | Ntaq1 | 463.328 | -0.06217 | 0.62675 | 0.912825 |
| ENSMUSG( | Ntf3 | 56.19566 | -0.30139 | 0.273385 | 0.72385 |
| ENSMUSG( | Nthl1 | 83.78553 | 0.252095 | 0.283449 | 0.73186 |
| ENSMUSG( | Ntm | 15260.7 | -0.21553 | 0.059232 | 0.375733 |
| ENSMUSG( | Ntmt1 | 505.7741 | -0.14947 | 0.348465 | 0.78132 |
| ENSMUSG( | Ntmt2 | 191.1062 | 0.582249 | 0.000381 | 0.017215 |
| ENSMUSG( | Ntn1 | 522.1964 | -0.45494 | 0.004738 | 0.09051 |
| ENSMUSG( | Ntn3 | 467.8534 | -0.06432 | 0.594952 | 0.903007 |
| ENSMUSG( | Ntn4 | 488.1515 | -0.17373 | 0.226546 | 0.673549 |
| ENSMUSG( | Ntn5 | 173.2394 | -0.15289 | 0.404414 | 0.816132 |
| ENSMUSG( | Ntng1 | 717.1214 | -0.47641 | 8.35E-05 | 0.005713 |
| ENSMUSG( | Ntng2 | 1549.418 | -0.09401 | 0.410539 | 0.818455 |
| ENSMUSG( | Ntpcr | 583.4327 | -0.1073 | 0.355619 | 0.787945 |
| ENSMUSG( | Ntrk1 | 23.32465 | -1.89014 | 9.15E-05 | 0.006056 |
| ENSMUSG( | Ntrk2 | 56407.18 | 0.08077 | 0.473566 | 0.848919 |
| ENSMUSG( | Ntrk3 | 3856.502 | -0.04767 | 0.679482 | 0.930983 |
| ENSMUSG( | Nts | 149.1341 | -0.99162 | 1.58E-06 | 0.000275 |
| ENSMUSG( | Ntsr1 | 332.5423 | -0.38225 | 0.027653 | 0.255168 |
| ENSMUSG( | Ntsr2 | 15135.74 | 0.015664 | 0.888674 | 0.983434 |
| ENSMUSG( | Nuak1 | 1956.273 | 0.21044 | 0.08669 | 0.452707 |
| ENSMUSG( | Nuak2 | 31.61192 | -0.42493 | 0.262699 | 0.713259 |
| ENSMUSG( | Nub1 | 2458.053 | 0.054239 | 0.536835 | 0.882123 |
| ENSMUSG( | Nubp1 | 506.4902 | -0.05994 | 0.62255 | 0.911432 |
| ENSMUSG( | Nubp2 | 933.008 | 0.11409 | 0.241731 | 0.689179 |
| ENSMUSG( | Nubpl | 149.9321 | -0.23087 | 0.207545 | 0.64962 |
| ENSMUSG( | Nucb1 | 3434.475 | -0.26737 | 0.027306 | 0.253597 |
| ENSMUSG( | Nucb2 | 509.8105 | -0.30825 | 0.014893 | 0.179275 |
| ENSMUSG( | Nucks1 | 2806.364 | 0.016897 | 0.851576 | 0.973046 |
| ENSMUSG( | Nudc | 3929.832 | -0.0019 | 0.981665 | 0.997512 |
| ENSMUSG( | Nudcd1 | 282.3328 | 0.143518 | 0.295416 | 0.743814 |
| ENSMUSG( | Nudcd2 | 557.6909 | 0.032037 | 0.832521 | 0.970333 |
| ENSMUSG( | Nudcd3 | 3963.35 | 0.005581 | 0.951048 | 0.993821 |
| ENSMUSG( | Nudt1 | 155.9462 | 0.078607 | 0.659438 | 0.924506 |
| ENSMUSG( | Nudt10 | 397.2792 | 0.039767 | 0.763741 | 0.954013 |
| ENSMUSG( | Nudt11 | 675.9869 | 0.010414 | 0.933161 | 0.99297 |
| ENSMUSG( | Nudt12 | 189.0995 | 0.217502 | 0.275922 | 0.725669 |
| ENSMUSG( | Nudt13 | 520.3273 | 0.024055 | 0.833926 | 0.970333 |

| | | | | | |
|---|---|---|---|---|---|
| ENSMUSG( | Npc1l1 | 72.45776 | -0.0499 | 0.853276 | 0.985598 |
| ENSMUSG( | Npc2 | 5169.517 | -0.10592 | 0.395473 | 0.917257 |
| ENSMUSG( | Npdc1 | 8905.842 | 0.031089 | 0.810606 | 0.978779 |
| ENSMUSG( | Npepl1 | 915.53 | 0.003531 | 0.977784 | 0.996416 |
| ENSMUSG( | Npepps | 4497.37 | -0.05227 | 0.614725 | 0.951177 |
| ENSMUSG( | Npff | 30.93809 | -0.0335 | 0.934714 | 0.99306 |
| ENSMUSG( | Npffr1 | 39.1097 | 0.610223 | 0.063873 | 0.736482 |
| ENSMUSG( | Nphp1 | 576.8633 | -0.02223 | 0.859889 | 0.9861 |
| ENSMUSG( | Nphp3 | 395.6777 | -0.12726 | 0.406277 | 0.917257 |
| ENSMUSG( | Nphp4 | 511.9495 | 0.004552 | 0.972783 | 0.996416 |
| ENSMUSG( | Nphs1 | 250.7223 | -0.09064 | 0.558455 | 0.93957 |
| ENSMUSG( | Nphs1os | 29.96449 | -0.5736 | 0.166308 | 0.840082 |
| ENSMUSG( | Nphs2 | 14.01011 | -0.18977 | 0.733169 | 0.967228 |
| ENSMUSG( | Npl | 134.6392 | 0.009893 | 0.959031 | 0.994798 |
| ENSMUSG( | Nploc4 | 1996.122 | 0.00342 | 0.973521 | 0.996416 |
| ENSMUSG( | Npm1 | 2689.069 | -0.1396 | 0.311052 | 0.899422 |
| ENSMUSG( | Npm2 | 12.64186 | 0.078254 | 0.889544 | 0.990558 |
| ENSMUSG( | Npm3 | 163.6645 | 0.055454 | 0.770808 | 0.974236 |
| ENSMUSG( | Npnt | 1375.979 | 0.360811 | 0.047586 | 0.701648 |
| ENSMUSG( | Nppa | 11.45606 | -0.50161 | 0.509983 | 0.929089 |
| ENSMUSG( | Nppc | 217.2619 | 0.268433 | 0.233384 | 0.868507 |
| ENSMUSG( | Npr1 | 137.8045 | 0.168984 | 0.399019 | 0.917257 |
| ENSMUSG( | Npr2 | 1979.013 | 0.02924 | 0.804459 | 0.977232 |
| ENSMUSG( | Npr3 | 344.6638 | 0.391803 | 0.018528 | 0.545482 |
| ENSMUSG( | Nprl2 | 619.9524 | -0.01009 | 0.948011 | 0.994798 |
| ENSMUSG( | Nprl3 | 852.0015 | 0.187149 | 0.168311 | 0.840222 |
| ENSMUSG( | Npsr1 | 52.50402 | -0.10841 | 0.708652 | 0.964042 |
| ENSMUSG( | Nptn | 19559.28 | 0.043745 | 0.768838 | 0.974236 |
| ENSMUSG( | Nptx1 | 11340.44 | 0.296768 | 0.111104 | 0.807885 |
| ENSMUSG( | Nptx2 | 2314.096 | -0.23483 | 0.117579 | 0.813384 |
| ENSMUSG( | Nptxr | 39431.59 | 0.109028 | 0.428076 | 0.917257 |
| ENSMUSG( | Npy | 4233.994 | -0.10066 | 0.506366 | 0.927825 |
| ENSMUSG( | Npy1r | 828.4475 | 0.400829 | 0.032195 | 0.634374 |
| ENSMUSG( | Npy2r | 235.6651 | 0.35546 | 0.081394 | 0.77681 |
| ENSMUSG( | Npy5r | 119.9308 | 0.312846 | 0.164234 | 0.838505 |
| ENSMUSG( | Nqo1 | 224.2981 | -0.13447 | 0.455334 | 0.923266 |
| ENSMUSG( | Nqo2 | 564.5062 | -0.20693 | 0.209564 | 0.864412 |
| ENSMUSG( | Nr1d1 | 2600.782 | 0.148744 | 0.258797 | 0.883971 |
| ENSMUSG( | Nr1d2 | 2551.79 | 0.59036 | 3.92E-07 | 0.000229 |
| ENSMUSG( | Nr1h2 | 1799.863 | 0.050319 | 0.661598 | 0.956205 |
| ENSMUSG( | Nr1h3 | 110.4397 | -0.08258 | 0.707909 | 0.964042 |
| ENSMUSG( | Nr1i3 | 16.93713 | -1.40402 | 0.005189 | 0.296096 |
| ENSMUSG( | Nr2c1 | 256.8575 | 0.003783 | 0.984943 | 0.997067 |
| ENSMUSG( | Nr2c2 | 3766.988 | -0.0659 | 0.538134 | 0.937216 |
| ENSMUSG( | Nr2c2ap | 269.1082 | -0.13911 | 0.400351 | 0.917257 |
| ENSMUSG( | Nr2e1 | 1489.771 | -0.21378 | 0.141312 | 0.830678 |
| ENSMUSG( | Nr2f1 | 3061.59 | 0.09485 | 0.405631 | 0.917257 |
| ENSMUSG( | Nr2f2 | 448.326 | 0.201825 | 0.228468 | 0.868507 |
| ENSMUSG( | Nr2f6 | 1166.084 | 0.122338 | 0.301947 | 0.898538 |
| ENSMUSG( | Nr3c1 | 2589.462 | 0.016155 | 0.891663 | 0.990558 |
| ENSMUSG( | Nr3c2 | 1052.171 | -0.0717 | 0.577939 | 0.943131 |
| ENSMUSG( | Nr4a1 | 2186.604 | -0.50112 | 0.025364 | 0.598637 |
| ENSMUSG( | Nr4a2 | 1175.14 | 0.067916 | 0.554898 | 0.939565 |
| ENSMUSG( | Nr4a3 | 689.0757 | 0.098196 | 0.436661 | 0.919879 |
| ENSMUSG( | Nr6a1 | 179.6878 | -0.09522 | 0.592415 | 0.945026 |
| ENSMUSG( | Nradd | 46.70624 | -0.22868 | 0.48754 | 0.925427 |
| ENSMUSG( | Nrap | 25.50821 | -0.49427 | 0.264545 | 0.886134 |
| ENSMUSG( | Nrarp | 1821.579 | 0.074406 | 0.585313 | 0.944975 |
| ENSMUSG( | Nras | 2941.082 | -0.17126 | 0.239378 | 0.87431 |
| ENSMUSG( | Nrbf2 | 187.092 | 0.021917 | 0.912049 | 0.992196 |
| ENSMUSG( | Nrbp1 | 1594.56 | -0.00395 | 0.975082 | 0.996416 |
| ENSMUSG( | Nrbp2 | 5276.952 | 0.044082 | 0.828906 | 0.980224 |
| ENSMUSG( | Nrcam | 12424.49 | 0.0263 | 0.862009 | 0.98632 |
| ENSMUSG( | Nrdc | 4668.303 | -0.09082 | 0.365451 | 0.914516 |
| ENSMUSG( | Nrde2 | 349.9127 | 0.191103 | 0.284877 | 0.892412 |
| ENSMUSG( | Nrep | 3362.136 | -0.145 | 0.250451 | 0.877932 |
| ENSMUSG( | Nrf1 | 483.8722 | 0.134464 | 0.350105 | 0.911652 |
| ENSMUSG( | Nrg1 | 310.7741 | 0.059619 | 0.736776 | 0.967344 |
| ENSMUSG( | Nrg2 | 231.3492 | 0.078506 | 0.617797 | 0.952134 |
| ENSMUSG( | Nrg3 | 1147.473 | 0.08101 | 0.650909 | 0.95422 |
| ENSMUSG( | Nrg4 | 11.09099 | -0.2677 | 0.644158 | NA |
| ENSMUSG( | Nrgn | 49555.42 | -0.20037 | 0.155162 | 0.837724 |
| ENSMUSG( | Nrip1 | 689.5887 | 0.17326 | 0.234849 | 0.871042 |
| ENSMUSG( | Nrip2 | 149.7503 | -0.09877 | 0.654801 | 0.955015 |
| ENSMUSG( | Nrip3 | 2613.675 | -0.02315 | 0.85202 | 0.985598 |
| ENSMUSG( | Nrm | 144.6835 | 0.029039 | 0.89477 | 0.990558 |
| ENSMUSG( | Nrn1 | 5672.454 | -0.07306 | 0.575596 | 0.942912 |
| ENSMUSG( | Nrp1 | 844.5595 | 0.339738 | 0.08695 | 0.782278 |
| ENSMUSG( | Nrp2 | 1146.845 | 0.048812 | 0.770554 | 0.974236 |
| ENSMUSG( | Nrros | 302.2 | 0.360549 | 0.09566 | 0.796133 |
| ENSMUSG( | Nrsn1 | 28941.98 | -0.01712 | 0.90131 | 0.990605 |
| ENSMUSG( | Nrsn2 | 2482.981 | -0.07937 | 0.59266 | 0.945158 |
| ENSMUSG( | Nrtn | 140.5582 | -0.10592 | 0.589844 | 0.944975 |
| ENSMUSG( | Nrxn1 | 15598.49 | 0.164297 | 0.327128 | 0.905768 |
| ENSMUSG( | Nrxn2 | 20628.95 | 0.130159 | 0.349195 | 0.911652 |
| ENSMUSG( | Nrxn3 | 4354.122 | 0.246182 | 0.098509 | 0.798438 |
| ENSMUSG( | Nsa2 | 1361.612 | -0.03466 | 0.80479 | 0.977232 |
| ENSMUSG( | Nscme3l | 15.00316 | 0.616915 | 0.34068 | 0.907972 |
| ENSMUSG( | Nsd1 | 4209.554 | 0.089353 | 0.407158 | 0.917257 |
| ENSMUSG( | Nsd2 | 2502.152 | -0.01341 | 0.894725 | 0.990558 |
| ENSMUSG( | Nsd3 | 2321.556 | 0.116441 | 0.315982 | 0.901868 |
| ENSMUSG( | Nsdhl | 1351.519 | -0.08885 | 0.555517 | 0.939565 |
| ENSMUSG( | Nsf | 10651.29 | -0.11647 | 0.421228 | 0.917257 |
| ENSMUSG( | Nsfl1c | 1360.256 | -0.02898 | 0.820692 | 0.979198 |
| ENSMUSG( | Nsg1 | 8673.535 | 0.084863 | 0.517993 | 0.930939 |
| ENSMUSG( | Nsg2 | 18563.74 | 0.092338 | 0.489629 | 0.925858 |
| ENSMUSG( | Nsl1 | 26.79494 | 0.024501 | 0.954621 | 0.994798 |
| ENSMUSG( | Nsmaf | 1077.055 | 0.04452 | 0.740633 | 0.968811 |
| ENSMUSG( | Nsmce1 | 317.6973 | -0.2326 | 0.131817 | 0.822912 |
| ENSMUSG( | Nsmce2 | 376.9368 | -0.08218 | 0.65659 | 0.955533 |
| ENSMUSG( | Nsmce3 | 877.175 | 0.147937 | 0.326428 | 0.905371 |
| ENSMUSG( | Nsmce4a | 621.241 | 0.072393 | 0.581976 | 0.944266 |
| ENSMUSG( | Nsmf | 40333.75 | 0.099704 | 0.364173 | 0.914223 |
| ENSMUSG( | Nsrp1 | 396.66 | -0.10886 | 0.493405 | 0.927372 |
| ENSMUSG( | Nsun2 | 1387.067 | -0.1707 | 0.1371 | 0.828492 |
| ENSMUSG( | Nsun3 | 284.7507 | -0.02377 | 0.887812 | 0.990399 |
| ENSMUSG( | Nsun4 | 294.3611 | -0.0947 | 0.534984 | 0.936475 |
| ENSMUSG( | Nsun5 | 339.0152 | 0.179421 | 0.290001 | 0.896726 |
| ENSMUSG( | Nsun6 | 145.3399 | -0.00733 | 0.969272 | 0.99594 |
| ENSMUSG( | Nsun7 | 263.5576 | -0.23546 | 0.238203 | 0.87259 |
| ENSMUSG( | Nt5c | 657.2621 | 0.082867 | 0.545084 | 0.938063 |
| ENSMUSG( | Nt5c1a | 77.09647 | -0.12009 | 0.639999 | 0.953903 |
| ENSMUSG( | Nt5c2 | 2664.87 | 0.067925 | 0.497383 | 0.927825 |
| ENSMUSG( | Nt5c3 | 1065.596 | 0.050444 | 0.698078 | 0.960867 |
| ENSMUSG( | Nt5c3b | 1105.256 | -0.15806 | 0.189684 | 0.857355 |
| ENSMUSG( | Nt5dc1 | 61.67725 | 0.051567 | 0.854111 | 0.98563 |
| ENSMUSG( | Nt5dc2 | 348.0142 | 0.171251 | 0.278251 | 0.891468 |
| ENSMUSG( | Nt5dc3 | 2133.703 | -0.11452 | 0.379175 | 0.916995 |
| ENSMUSG( | Nt5e | 258.7371 | 0.075057 | 0.662684 | 0.956205 |
| ENSMUSG( | Nt5m | 1106.569 | -0.19348 | 0.083211 | 0.77681 |
| ENSMUSG( | Ntan1 | 1182.998 | -0.10383 | 0.378706 | 0.916995 |
| ENSMUSG( | Ntaq1 | 416.6954 | -0.10524 | 0.456784 | 0.923329 |
| ENSMUSG( | Ntf3 | 54.97556 | -0.05775 | 0.851725 | 0.985591 |
| ENSMUSG( | Nthl1 | 79.87407 | 0.36643 | 0.124901 | 0.81516 |
| ENSMUSG( | Ntm | 14504.22 | -0.08623 | 0.546184 | 0.938063 |
| ENSMUSG( | Ntmt1 | 458.4811 | -0.16533 | 0.331974 | 0.907907 |
| ENSMUSG( | Ntmt2 | 143.9168 | 0.080266 | 0.683061 | 0.958678 |
| ENSMUSG( | Ntn1 | 535.4092 | -0.07147 | 0.675331 | 0.95694 |
| ENSMUSG( | Ntn3 | 446.6419 | 0.071953 | 0.619555 | 0.952896 |
| ENSMUSG( | Ntn4 | 487.9759 | 0.094219 | 0.568395 | 0.941777 |
| ENSMUSG( | Ntn5 | 157.9334 | -0.13756 | 0.503195 | 0.927825 |
| ENSMUSG( | Ntng1 | 726.12 | -0.13875 | 0.345199 | 0.91045 |
| ENSMUSG( | Ntng2 | 1533.667 | 0.146545 | 0.271594 | 0.888062 |
| ENSMUSG( | Ntpcr | 532.7932 | -0.1036 | 0.44619 | 0.921734 |
| ENSMUSG( | Ntrk1 | 31.10906 | -0.24266 | 0.508771 | 0.929089 |
| ENSMUSG( | Ntrk2 | 51270.89 | 0.075075 | 0.538316 | 0.937216 |
| ENSMUSG( | Ntrk3 | 3864.266 | 0.221465 | 0.099476 | 0.80042 |
| ENSMUSG( | Nts | 162.8857 | -0.33525 | 0.149709 | 0.836241 |
| ENSMUSG( | Ntsr1 | 342.949 | 0.00991 | 0.958232 | 0.994798 |
| ENSMUSG( | Ntsr2 | 13583.74 | -0.02669 | 0.831612 | 0.980656 |
| ENSMUSG( | Nuak1 | 1640.242 | -0.01937 | 0.883822 | 0.989644 |
| ENSMUSG( | Nuak2 | 34.90466 | 0.19063 | 0.599665 | 0.947071 |
| ENSMUSG( | Nub1 | 2130.964 | -0.09004 | 0.400287 | 0.917257 |
| ENSMUSG( | Nubp1 | 455.6017 | -0.09452 | 0.541126 | 0.93767 |
| ENSMUSG( | Nubp2 | 762.9991 | -0.20128 | 0.086045 | 0.780251 |
| ENSMUSG( | Nubpl | 137.32 | -0.23482 | 0.306175 | 0.898538 |
| ENSMUSG( | Nucb1 | 3656.271 | 0.191154 | 0.177202 | 0.846973 |
| ENSMUSG( | Nucb2 | 507.3759 | -0.03727 | 0.780792 | 0.976106 |
| ENSMUSG( | Nucks1 | 2491.768 | -0.06154 | 0.666707 | 0.956766 |
| ENSMUSG( | Nudc | 3240.066 | -0.30808 | 0.005271 | 0.297824 |
| ENSMUSG( | Nudcd1 | 257.2985 | 0.133728 | 0.429895 | 0.917257 |
| ENSMUSG( | Nudcd2 | 496.1043 | -0.04472 | 0.774553 | 0.974931 |
| ENSMUSG( | Nudcd3 | 3640.617 | 0.031153 | 0.776877 | 0.975717 |
| ENSMUSG( | Nudt1 | 124.9246 | -0.30986 | 0.119061 | 0.813384 |
| ENSMUSG( | Nudt10 | 352.1525 | -0.05207 | 0.758958 | 0.971858 |
| ENSMUSG( | Nudt11 | 619.155 | 0.016789 | 0.926764 | 0.99306 |
| ENSMUSG( | Nudt12 | 166.6036 | 0.129488 | 0.516823 | 0.930939 |
| ENSMUSG( | Nudt13 | 461.4396 | -0.05529 | 0.688157 | 0.959742 |

| | | | | | |
|---|---|---|---|---|---|
| ENSMUSG( | 19.99701 | 0.031146 | 0.946234 | NA | Nherf4 |
| ENSMUSG( | 14.32453 | 0.013953 | 0.977508 | NA | Nhlh1 |
| ENSMUSG( | 8.716014 | -0.02798 | 0.965809 | NA | Nhlh2 |
| ENSMUSG( | 378.7571 | 0.071143 | 0.56755 | 0.891774 | Nhlrc1 |
| ENSMUSG( | 904.941 | 0.083931 | 0.409836 | 0.818258 | Nhlrc2 |
| ENSMUSG( | 537.8439 | -0.37833 | 0.006057 | 0.105754 | Nhlrc3 |
| ENSMUSG( | 735.8608 | -0.09042 | 0.404576 | 0.816132 | Nhp2 |
| ENSMUSG( | 85.28882 | -0.36277 | 0.115986 | 0.514015 | Nhs |
| ENSMUSG( | 3598.753 | 0.313302 | 0.038482 | 0.302696 | Nhsl1 |
| ENSMUSG( | 1255.196 | -0.01403 | 0.895454 | 0.984512 | Nhsl2 |
| ENSMUSG( | 1472.286 | 0.271478 | 0.031237 | 0.26893 | Nhsl3 |
| ENSMUSG( | 214.2249 | -0.2874 | 0.164919 | 0.596162 | Niban1 |
| ENSMUSG( | 2587.774 | -0.05841 | 0.47844 | 0.853567 | Niban2 |
| ENSMUSG( | 56.46371 | 0.278186 | 0.32609 | 0.771128 | Niban3 |
| ENSMUSG( | 2272.965 | 0.069399 | 0.45035 | 0.835344 | Nicn1 |
| ENSMUSG( | 794.6425 | -0.3254 | 0.007945 | 0.124138 | Nicol1 |
| ENSMUSG( | 777.8137 | -0.19583 | 0.29598 | 0.744386 | Nid1 |
| ENSMUSG( | 335.3439 | -0.31627 | 0.06957 | 0.408118 | Nid2 |
| ENSMUSG( | 430.4253 | 0.058081 | 0.64242 | 0.916865 | Nif3l1 |
| ENSMUSG( | 441.3258 | 0.08152 | 0.509992 | 0.869743 | Nifk |
| ENSMUSG( | 1244.496 | 0.05265 | 0.639609 | 0.916434 | Nim1k |
| ENSMUSG( | 1262.487 | -0.25816 | 0.005251 | 0.096573 | Nin |
| ENSMUSG( | 592.431 | -0.12167 | 0.295226 | 0.743798 | Ninj1 |
| ENSMUSG( | 77.98944 | -0.32675 | 0.178389 | 0.616026 | Ninj2 |
| ENSMUSG( | 259.139 | -0.08279 | 0.6161 | 0.91085 | Ninl |
| ENSMUSG( | 498.0745 | 0.04109 | 0.750184 | 0.949664 | Nip7 |
| ENSMUSG( | 1483.723 | -0.21905 | 0.140298 | 0.557263 | Nipa1 |
| ENSMUSG( | 1104.407 | -0.29226 | 0.043031 | 0.31819 | Nipa2 |
| ENSMUSG( | 237.4313 | -0.48729 | 0.003371 | 0.073204 | Nipal2 |
| ENSMUSG( | 3547.524 | -0.19392 | 0.198162 | 0.638387 | Nipal3 |
| ENSMUSG( | 249.924 | -0.57315 | 0.004453 | 0.086961 | Nipal4 |
| ENSMUSG( | 1745.597 | 0.07527 | 0.550074 | 0.886038 | Nipbl |
| ENSMUSG( | 1067.565 | -0.03661 | 0.794287 | 0.959799 | Nipsnap1 |
| ENSMUSG( | 2501.114 | 0.001965 | 0.983804 | 0.99779 | Nipsnap2 |
| ENSMUSG( | 307.4761 | 0.012915 | 0.92688 | 0.991229 | Nipsnap3b |
| ENSMUSG( | 16103.81 | 0.154652 | 0.05766 | 0.370247 | Nisch |
| ENSMUSG( | 979.0069 | 0.106366 | 0.335932 | 0.777 | Nit1 |
| ENSMUSG( | 303.5218 | 0.031633 | 0.830331 | 0.96974 | Nit2 |
| ENSMUSG( | 806.5299 | -0.25384 | 0.042369 | 0.316636 | Nkain1 |
| ENSMUSG( | 1558.939 | -0.23658 | 0.032148 | 0.273917 | Nkain2 |
| ENSMUSG( | 40.95892 | -0.2647 | 0.380069 | 0.801757 | Nkain3 |
| ENSMUSG( | 2124.44 | 0.025159 | 0.809893 | 0.96372 | Nkain4 |
| ENSMUSG( | 349.5301 | 0.01128 | 0.930332 | 0.991848 | Nkap |
| ENSMUSG( | 764.1973 | -0.00172 | 0.986921 | 0.998619 | Nkapd1 |
| ENSMUSG( | 1512.83 | -0.07721 | 0.377305 | 0.801213 | Nkd1 |
| ENSMUSG( | 139.8734 | 0.287737 | 0.106273 | 0.49377 | Nkd2 |
| ENSMUSG( | 1920.825 | -0.00735 | 0.940693 | 0.993415 | Nkiras1 |
| ENSMUSG( | 571.7374 | 0.064982 | 0.538617 | 0.882729 | Nkiras2 |
| ENSMUSG( | 35.95053 | -0.56789 | 0.075508 | 0.425931 | Nkpd1 |
| ENSMUSG( | 423.5121 | -0.02653 | 0.850794 | 0.972638 | Nkrf |
| ENSMUSG( | 4865.694 | 0.066763 | 0.485532 | 0.857637 | Nktr |
| ENSMUSG( | 19.07544 | -0.49158 | 0.306588 | NA | Nkx2-1 |
| ENSMUSG( | 280.7481 | -0.15754 | 0.248052 | 0.696814 | Nkx2-2 |
| ENSMUSG( | 120.7972 | 0.262401 | 0.177223 | 0.614553 | Nkx2-2os |
| ENSMUSG( | 24.66346 | 0.132882 | 0.745412 | 0.949158 | Nkx3-1 |
| ENSMUSG( | 314.8966 | -0.51268 | 0.001458 | 0.042158 | Nkx6-2 |
| ENSMUSG( | 286.8218 | -0.07558 | 0.60324 | 0.904887 | Nle1 |
| ENSMUSG( | 1476.482 | -0.35327 | 0.027689 | 0.255176 | Nlgn1 |
| ENSMUSG( | 12047.25 | -0.17202 | 0.124671 | 0.531714 | Nlgn2 |
| ENSMUSG( | 2244.169 | -0.32213 | 0.020324 | 0.214896 | Nlgn3 |
| ENSMUSG( | 2224.545 | 0.158694 | 0.069712 | 0.408352 | Nlk |
| ENSMUSG( | 1404.514 | 0.058742 | 0.579311 | 0.896692 | Nln |
| ENSMUSG( | 35.59956 | 0.071241 | 0.829381 | 0.969343 | Nlrc3 |
| ENSMUSG( | 15.30372 | -0.20172 | 0.699529 | NA | Nlrc4 |
| ENSMUSG( | 37.09013 | -0.30844 | 0.341352 | 0.778828 | Nlrc5 |
| ENSMUSG( | 34.19006 | -0.17467 | 0.63126 | 0.914073 | Nlrp1b |
| ENSMUSG( | 43.26631 | -0.04719 | 0.880046 | 0.981469 | Nlrp3 |
| ENSMUSG( | 15.3571 | 0.80202 | 0.132785 | NA | Nlrp6 |
| ENSMUSG( | 300.6471 | 0.011894 | 0.930563 | 0.991965 | Nlrx1 |
| ENSMUSG( | 111.3299 | -0.04275 | 0.826607 | 0.968404 | Nmb |
| ENSMUSG( | 57.40223 | -0.6175 | 0.024202 | 0.238178 | Nmbr |
| ENSMUSG( | 513.0357 | -0.08711 | 0.455854 | 0.838601 | Nmd3 |
| ENSMUSG( | 1501.557 | 0.028133 | 0.805702 | 0.962616 | Nme1 |
| ENSMUSG( | 43.25933 | 0.121197 | 0.680685 | 0.931139 | Nme2 |
| ENSMUSG( | 72.39709 | 0.121836 | 0.587215 | 0.900334 | Nme4 |
| ENSMUSG( | 177.1043 | -0.13788 | 0.421989 | 0.823495 | Nme5 |
| ENSMUSG( | 134.5178 | 0.278125 | 0.132158 | 0.544244 | Nme6 |
| ENSMUSG( | 1025.307 | 0.061582 | 0.592324 | 0.902094 | Nme7 |
| ENSMUSG( | 201.8906 | 0.010318 | 0.949521 | 0.993821 | Nmi |
| ENSMUSG( | 168.4385 | 0.105166 | 0.552164 | 0.886141 | Nmnat1 |
| ENSMUSG( | 3655.273 | 0.060659 | 0.543935 | 0.884297 | Nmnat2 |
| ENSMUSG( | 53.66938 | -0.26233 | 0.335645 | 0.777 | Nmnat3 |
| ENSMUSG( | 270.725 | 0.038829 | 0.824159 | 0.966864 | Nmral1 |
| ENSMUSG( | 129.9189 | -0.06514 | 0.744764 | 0.94902 | Nmrk1 |
| ENSMUSG( | 3840.569 | 0.127371 | 0.199746 | 0.640239 | Nmt1 |
| ENSMUSG( | 2652.641 | 0.036931 | 0.66653 | 0.927247 | Nmt2 |
| ENSMUSG( | 9368.527 | -0.06361 | 0.62698 | 0.912825 | Nnat |
| ENSMUSG( | 18.02908 | -0.04871 | 0.909953 | NA | Nnmt |
| ENSMUSG( | 10.65973 | -0.22646 | 0.714867 | NA | Nnt |
| ENSMUSG( | 526.3188 | 0.151022 | 0.215727 | 0.660456 | Noa1 |
| ENSMUSG( | 552.9691 | -0.03997 | 0.71709 | 0.941697 | Nob1 |
| ENSMUSG( | 2406.326 | 0.133278 | 0.165029 | 0.596428 | Noc2l |
| ENSMUSG( | 433.2561 | 0.188407 | 0.130111 | 0.540593 | Noc3l |
| ENSMUSG( | 419.4747 | 0.107596 | 0.557244 | 0.888279 | Noc4l |
| ENSMUSG( | 903.1493 | -0.03484 | 0.793651 | 0.959799 | Noct |
| ENSMUSG( | 243.5321 | -0.1164 | 0.513333 | 0.871762 | Nod1 |
| ENSMUSG( | 15.95999 | -0.73539 | 0.122631 | NA | Nod2 |
| ENSMUSG( | 23.96119 | -0.59515 | 0.202819 | 0.643279 | Nodal |
| ENSMUSG( | 365.94 | -0.47843 | 0.003308 | 0.072126 | Nog |
| ENSMUSG( | 506.5975 | 0.147197 | 0.237781 | 0.685809 | Nol10 |
| ENSMUSG( | 468.7763 | 0.001209 | 0.991942 | 0.999232 | Nol11 |
| ENSMUSG( | 823.6854 | 0.101825 | 0.298592 | 0.747287 | Nol12 |
| ENSMUSG( | 882.0782 | -0.02435 | 0.822964 | 0.96644 | Nol3 |
| ENSMUSG( | 1388.3 | 0.033059 | 0.706529 | 0.939259 | Nol4 |
| ENSMUSG( | 2270.664 | 0.181696 | 0.078079 | 0.432233 | Nol4l |
| ENSMUSG( | 2952.013 | 0.075898 | 0.48074 | 0.855094 | Nol6 |
| ENSMUSG( | 1127.811 | 0.064151 | 0.594132 | 0.902869 | Nol7 |
| ENSMUSG( | 640.4942 | 0.115275 | 0.336338 | 0.777 | Nol8 |
| ENSMUSG( | 607.2052 | 0.113272 | 0.328612 | 0.773148 | Nol9 |
| ENSMUSG( | 1158.434 | 0.116021 | 0.240655 | 0.688247 | Nolc1 |
| ENSMUSG( | 853.8447 | 0.04638 | 0.664209 | 0.926591 | Nom1 |
| ENSMUSG( | 5473.384 | -0.11558 | 0.386754 | 0.805046 | Nomo1 |
| ENSMUSG( | 3468.17 | 0.013406 | 0.878484 | 0.981431 | Nono |
| ENSMUSG( | 813.4089 | -0.07342 | 0.502259 | 0.864924 | Nop10 |
| ENSMUSG( | 644.8658 | 0.169484 | 0.157173 | 0.585516 | Nop14 |
| ENSMUSG( | 333.5234 | 0.192054 | 0.209386 | 0.65137 | Nop16 |
| ENSMUSG( | 570.7333 | 0.150448 | 0.165352 | 0.596691 | Nop2 |
| ENSMUSG( | 3441.134 | 0.029284 | 0.727064 | 0.944223 | Nop53 |
| ENSMUSG( | 1917.407 | 0.085472 | 0.42554 | 0.824898 | Nop56 |
| ENSMUSG( | 797.2025 | 0.088192 | 0.401683 | 0.814923 | Nop58 |
| ENSMUSG( | 338.1695 | 0.190975 | 0.154039 | 0.580901 | Nop9 |
| ENSMUSG( | 1120.546 | 0.105476 | 0.283733 | 0.732203 | Nopchap1 |
| ENSMUSG( | 34785.76 | 0.113516 | 0.15554 | 0.583591 | Norad |
| ENSMUSG( | 888.6835 | 0.174972 | 0.106917 | 0.494634 | Nos1 |
| ENSMUSG( | 1144.182 | 0.380086 | 0.006082 | 0.106072 | Nos1ap |
| ENSMUSG( | 33.78285 | 0.168566 | 0.643613 | 0.917316 | Nos2 |
| ENSMUSG( | 388.1407 | 0.398121 | 0.043456 | 0.319227 | Nos3 |
| ENSMUSG( | 1164.514 | -0.01885 | 0.867232 | 0.976559 | Nosip |
| ENSMUSG( | 232.5353 | 0.276032 | 0.174504 | 0.610304 | Nostrin |
| ENSMUSG( | 1542.44 | -0.12831 | 0.433783 | 0.828541 | Notch1 |
| ENSMUSG( | 1388.972 | -0.30751 | 0.024144 | 0.237772 | Notch2 |
| ENSMUSG( | 760.8927 | -0.15842 | 0.317914 | 0.764145 | Notch3 |
| ENSMUSG( | 398.4602 | 0.641142 | 0.000146 | 0.008459 | Notch4 |
| ENSMUSG( | 159.3534 | 0.107977 | 0.561708 | 0.889052 | Notum |
| ENSMUSG( | 1559.038 | 0.07405 | 0.434785 | 0.829025 | Nova1 |
| ENSMUSG( | 3524.362 | 0.164433 | 0.066084 | 0.397686 | Nova2 |
| ENSMUSG( | 100.2271 | 0.152001 | 0.454746 | 0.837513 | Noxo1 |
| ENSMUSG( | 40.77795 | 0.291613 | 0.358539 | 0.789791 | Noxred1 |
| ENSMUSG( | 296.6626 | 0.465292 | 0.006422 | 0.110055 | Npas1 |
| ENSMUSG( | 1920.613 | 0.200291 | 0.027555 | 0.2548 | Npas2 |
| ENSMUSG( | 756.7756 | -0.00307 | 0.979578 | 0.997428 | Npas3 |
| ENSMUSG( | 558.5984 | -0.59053 | 0.357181 | 0.788702 | Npas4 |
| ENSMUSG( | 546.7853 | -0.03197 | 0.781289 | 0.957558 | Npat |
| ENSMUSG( | 21.29479 | -0.94625 | 0.022587 | 0.229429 | Npb |
| ENSMUSG( | 45.90949 | -0.48066 | 0.151504 | 0.576717 | Npbwr1 |
| ENSMUSG( | 3336.281 | -0.33884 | 0.031166 | 0.268462 | Npc1 |
| ENSMUSG( | 86.5722 | 0.186516 | 0.423104 | 0.823662 | Npc1l1 |
| ENSMUSG( | 5610.883 | -0.14068 | 0.201161 | 0.641362 | Npc2 |
| ENSMUSG( | 9068.627 | -0.1948 | 0.096979 | 0.477375 | Npdc1 |
| ENSMUSG( | 1042.382 | 0.106021 | 0.274163 | 0.724254 | Npepl1 |
| ENSMUSG( | 5337.181 | 0.169891 | 0.035115 | 0.288256 | Npepps |
| ENSMUSG( | 39.96941 | 0.3854 | 0.25983 | 0.709951 | Npff |
| ENSMUSG( | 35.47242 | 0.106568 | 0.742196 | 0.948478 | Npffr1 |
| ENSMUSG( | 644.7389 | 0.032443 | 0.75351 | 0.950753 | Nphp1 |
| ENSMUSG( | 418.9768 | -0.24322 | 0.05248 | 0.35151 | Nphp3 |
| ENSMUSG( | 575.9062 | 0.072902 | 0.553805 | 0.886394 | Nphp4 |
| ENSMUSG( | 301.6568 | 0.173741 | 0.235117 | 0.682364 | Nphs1 |
| ENSMUSG( | 36.65198 | -0.17233 | 0.639475 | 0.916434 | Nphs1os |

| | | | | | |
|---|---|---|---|---|---|
| ENSMUSG( | 3039.873 | 0.011554 | 0.914919 | 0.992407 | Nova2 |
| ENSMUSG( | 96.81048 | 0.31104 | 0.175085 | 0.845305 | Noxo1 |
| ENSMUSG( | 38.33074 | 0.380825 | 0.297161 | 0.898538 | Noxred1 |
| ENSMUSG( | 234.377 | 0.096251 | 0.586727 | 0.944975 | Npas1 |
| ENSMUSG( | 1554.198 | -0.13887 | 0.188169 | 0.85698 | Npas2 |
| ENSMUSG( | 632.4674 | -0.26857 | 0.037941 | 0.6763 | Npas3 |
| ENSMUSG( | 519.1175 | -0.51754 | 0.418264 | 0.917257 | Npas4 |
| ENSMUSG( | 521.6855 | 0.103534 | 0.463056 | 0.923329 | Npat |
| ENSMUSG( | 27.73961 | 0.234807 | 0.539003 | 0.937216 | Npb |
| ENSMUSG( | 48.82852 | 0.038287 | 0.918304 | 0.992529 | Npbwr1 |
| ENSMUSG( | 3734.206 | 0.267832 | 0.12906 | 0.820358 | Npc1 |
| ENSMUSG( | 72.45776 | -0.0499 | 0.853276 | 0.985598 | Npc1l1 |
| ENSMUSG( | 5169.517 | -0.10592 | 0.395473 | 0.917257 | Npc2 |
| ENSMUSG( | 8905.842 | 0.031089 | 0.810606 | 0.978779 | Npdc1 |
| ENSMUSG( | 915.53 | 0.003531 | 0.977784 | 0.996416 | Npepl1 |
| ENSMUSG( | 4497.37 | -0.05227 | 0.614725 | 0.951177 | Npepps |
| ENSMUSG( | 30.93809 | -0.0335 | 0.934714 | 0.99306 | Npff |
| ENSMUSG( | 39.1097 | 0.610223 | 0.063873 | 0.736482 | Npffr1 |
| ENSMUSG( | 576.8633 | -0.02223 | 0.859889 | 0.9861 | Nphp1 |
| ENSMUSG( | 395.6777 | -0.12726 | 0.406277 | 0.917257 | Nphp3 |
| ENSMUSG( | 511.9495 | 0.004552 | 0.972783 | 0.996416 | Nphp4 |
| ENSMUSG( | 250.7223 | -0.09064 | 0.558455 | 0.93957 | Nphs1 |
| ENSMUSG( | 29.96449 | -0.5736 | 0.166308 | 0.840082 | Nphs1os |
| ENSMUSG( | 14.01011 | -0.18977 | 0.733169 | 0.967228 | Nphs2 |
| ENSMUSG( | 134.6392 | 0.009893 | 0.959031 | 0.994798 | Npl |
| ENSMUSG( | 1996.122 | 0.00342 | 0.973521 | 0.996416 | Nploc4 |
| ENSMUSG( | 2689.069 | -0.1396 | 0.311052 | 0.899422 | Npm1 |
| ENSMUSG( | 12.64186 | 0.078254 | 0.889544 | 0.990558 | Npm2 |
| ENSMUSG( | 163.6645 | 0.055454 | 0.770808 | 0.974236 | Npm3 |
| ENSMUSG( | 1375.979 | 0.360811 | 0.047586 | 0.701648 | Npnt |
| ENSMUSG( | 11.45606 | -0.50161 | 0.509983 | 0.929089 | Nppa |
| ENSMUSG( | 217.2619 | 0.268433 | 0.233384 | 0.868507 | Nppc |
| ENSMUSG( | 137.8045 | 0.168984 | 0.399019 | 0.917257 | Npr1 |
| ENSMUSG( | 1979.013 | 0.02924 | 0.804459 | 0.977232 | Npr2 |
| ENSMUSG( | 344.6638 | 0.391803 | 0.018528 | 0.545482 | Npr3 |
| ENSMUSG( | 619.9524 | -0.01009 | 0.948011 | 0.994798 | Nprl2 |
| ENSMUSG( | 852.0015 | 0.187149 | 0.168311 | 0.840222 | Nprl3 |
| ENSMUSG( | 52.50402 | -0.10841 | 0.708652 | 0.964042 | Npsr1 |
| ENSMUSG( | 19559.28 | 0.043745 | 0.768838 | 0.974236 | Nptn |
| ENSMUSG( | 11340.44 | 0.296768 | 0.111104 | 0.807885 | Nptx1 |
| ENSMUSG( | 2314.096 | -0.23483 | 0.117579 | 0.813384 | Nptx2 |
| ENSMUSG( | 39431.59 | 0.109028 | 0.428076 | 0.917257 | Nptxr |
| ENSMUSG( | 4233.994 | -0.10066 | 0.506366 | 0.927825 | Npy |
| ENSMUSG( | 828.4475 | 0.400829 | 0.032195 | 0.634374 | Npy1r |
| ENSMUSG( | 235.6651 | 0.35546 | 0.081394 | 0.77681 | Npy2r |
| ENSMUSG( | 119.9308 | 0.312846 | 0.164234 | 0.838505 | Npy5r |
| ENSMUSG( | 224.2981 | -0.13447 | 0.455334 | 0.923266 | Nqo1 |
| ENSMUSG( | 564.5062 | -0.20693 | 0.209564 | 0.864412 | Nqo2 |
| ENSMUSG( | 2600.782 | 0.148744 | 0.258797 | 0.883971 | Nr1d1 |
| ENSMUSG( | 2551.79 | 0.59036 | 3.92E-07 | 0.000229 | Nr1d2 |
| ENSMUSG( | 1799.863 | 0.050319 | 0.661598 | 0.956205 | Nr1h2 |
| ENSMUSG( | 110.4397 | -0.08258 | 0.707909 | 0.964042 | Nr1h3 |
| ENSMUSG( | 16.93713 | -1.40402 | 0.005189 | 0.296096 | Nr1i3 |
| ENSMUSG( | 256.8575 | 0.003783 | 0.984943 | 0.997067 | Nr2c1 |
| ENSMUSG( | 3766.988 | -0.0659 | 0.538134 | 0.937216 | Nr2c2 |
| ENSMUSG( | 269.1082 | -0.13911 | 0.400351 | 0.917257 | Nr2c2ap |
| ENSMUSG( | 1489.771 | -0.21378 | 0.141312 | 0.830678 | Nr2e1 |
| ENSMUSG( | 3061.59 | 0.09485 | 0.405631 | 0.917257 | Nr2f1 |
| ENSMUSG( | 448.326 | 0.201825 | 0.228468 | 0.868507 | Nr2f2 |
| ENSMUSG( | 1166.084 | 0.122338 | 0.301947 | 0.898538 | Nr2f6 |
| ENSMUSG( | 2589.462 | 0.016155 | 0.891663 | 0.990558 | Nr3c1 |
| ENSMUSG( | 1052.171 | -0.0717 | 0.577939 | 0.943131 | Nr3c2 |
| ENSMUSG( | 2186.604 | -0.50112 | 0.025364 | 0.598637 | Nr4a1 |
| ENSMUSG( | 1175.14 | 0.067916 | 0.554898 | 0.939565 | Nr4a2 |
| ENSMUSG( | 689.0757 | 0.098196 | 0.436661 | 0.919879 | Nr4a3 |
| ENSMUSG( | 179.6878 | -0.09522 | 0.592415 | 0.945026 | Nr6a1 |
| ENSMUSG( | 46.70624 | -0.22868 | 0.48754 | 0.925427 | Nradd |
| ENSMUSG( | 25.50821 | -0.49427 | 0.264545 | 0.886134 | Nrap |
| ENSMUSG( | 1821.579 | 0.074406 | 0.585313 | 0.944975 | Nrarp |
| ENSMUSG( | 2941.082 | -0.17126 | 0.239378 | 0.87431 | Nras |
| ENSMUSG( | 187.092 | 0.021917 | 0.912049 | 0.992196 | Nrbf2 |
| ENSMUSG( | 1594.56 | -0.00395 | 0.975082 | 0.996416 | Nrbp1 |
| ENSMUSG( | 5276.952 | 0.044082 | 0.828906 | 0.980224 | Nrbp2 |
| ENSMUSG( | 12424.49 | 0.0263 | 0.862009 | 0.98632 | Nrcam |
| ENSMUSG( | 4668.303 | -0.09082 | 0.365451 | 0.914516 | Nrdc |
| ENSMUSG( | 349.9127 | 0.191103 | 0.284877 | 0.892412 | Nrde2 |
| ENSMUSG( | 3362.136 | -0.145 | 0.250451 | 0.877932 | Nrep |
| ENSMUSG( | 483.8722 | 0.134464 | 0.350105 | 0.911652 | Nrf1 |
| ENSMUSG( | 310.7741 | 0.059619 | 0.736776 | 0.967344 | Nrg1 |
| ENSMUSG( | 231.3492 | 0.078506 | 0.617797 | 0.952134 | Nrg2 |
| ENSMUSG( | 1147.473 | 0.08101 | 0.650909 | 0.95422 | Nrg3 |
| ENSMUSG( | 11.09099 | -0.2677 | 0.644158 | NA | Nrg4 |
| ENSMUSG( | 49555.42 | -0.20037 | 0.155162 | 0.837724 | Nrgn |
| ENSMUSG( | 689.5887 | 0.17326 | 0.234849 | 0.871042 | Nrip1 |
| ENSMUSG( | 149.7503 | -0.09877 | 0.654801 | 0.955015 | Nrip2 |
| ENSMUSG( | 2613.675 | -0.02315 | 0.85202 | 0.985598 | Nrip3 |
| ENSMUSG( | 144.6835 | 0.029039 | 0.89477 | 0.990558 | Nrm |
| ENSMUSG( | 5672.454 | -0.07306 | 0.575596 | 0.942912 | Nrn1 |
| ENSMUSG( | 844.5595 | 0.339738 | 0.08695 | 0.782278 | Nrp1 |
| ENSMUSG( | 1146.845 | 0.048812 | 0.770554 | 0.974236 | Nrp2 |
| ENSMUSG( | 302.2 | 0.360549 | 0.09566 | 0.796133 | Nrros |
| ENSMUSG( | 28941.98 | -0.01712 | 0.90131 | 0.990605 | Nrsn1 |
| ENSMUSG( | 2482.981 | -0.07937 | 0.59266 | 0.945158 | Nrsn2 |
| ENSMUSG( | 140.5582 | -0.10592 | 0.589844 | 0.944975 | Nrtn |
| ENSMUSG( | 15598.49 | 0.164297 | 0.327128 | 0.905768 | Nrxn1 |
| ENSMUSG( | 20628.95 | 0.130159 | 0.349195 | 0.911652 | Nrxn2 |
| ENSMUSG( | 4354.122 | 0.246182 | 0.098509 | 0.798438 | Nrxn3 |
| ENSMUSG( | 1361.612 | -0.03466 | 0.80479 | 0.977232 | Nsa2 |
| ENSMUSG( | 15.00316 | 0.616915 | 0.34068 | 0.907972 | Nscme3l |
| ENSMUSG( | 4209.554 | 0.089353 | 0.407158 | 0.917257 | Nsd1 |
| ENSMUSG( | 2502.152 | -0.01341 | 0.894725 | 0.990558 | Nsd2 |
| ENSMUSG( | 2321.556 | 0.116441 | 0.315982 | 0.901868 | Nsd3 |
| ENSMUSG( | 1351.519 | -0.08885 | 0.555517 | 0.939565 | Nsdhl |
| ENSMUSG( | 10651.29 | -0.11647 | 0.421228 | 0.917257 | Nsf |
| ENSMUSG( | 1360.256 | -0.02898 | 0.820692 | 0.979198 | Nsfl1c |
| ENSMUSG( | 8673.535 | 0.084863 | 0.517993 | 0.930939 | Nsg1 |
| ENSMUSG( | 18563.74 | 0.092338 | 0.489629 | 0.925858 | Nsg2 |
| ENSMUSG( | 26.79494 | 0.024501 | 0.954621 | 0.994798 | Nsl1 |
| ENSMUSG( | 1077.055 | 0.04452 | 0.740633 | 0.968811 | Nsmaf |
| ENSMUSG( | 317.6973 | -0.2326 | 0.131817 | 0.822912 | Nsmce1 |
| ENSMUSG( | 376.9368 | -0.08218 | 0.65659 | 0.955533 | Nsmce2 |
| ENSMUSG( | 877.175 | 0.147937 | 0.326428 | 0.905371 | Nsmce3 |
| ENSMUSG( | 621.241 | 0.072393 | 0.581976 | 0.944266 | Nsmce4a |
| ENSMUSG( | 40333.75 | 0.099704 | 0.364173 | 0.914223 | Nsmf |
| ENSMUSG( | 396.66 | -0.10886 | 0.493405 | 0.927372 | Nsrp1 |
| ENSMUSG( | 1387.067 | -0.1707 | 0.1371 | 0.828492 | Nsun2 |
| ENSMUSG( | 284.7507 | -0.02377 | 0.887812 | 0.990399 | Nsun3 |
| ENSMUSG( | 294.3611 | -0.0947 | 0.534984 | 0.936475 | Nsun4 |
| ENSMUSG( | 339.0152 | 0.179421 | 0.290001 | 0.896726 | Nsun5 |
| ENSMUSG( | 145.3399 | -0.00733 | 0.969272 | 0.99594 | Nsun6 |
| ENSMUSG( | 263.5576 | -0.23546 | 0.238203 | 0.87259 | Nsun7 |
| ENSMUSG( | 657.2621 | 0.082867 | 0.545084 | 0.938063 | Nt5c |
| ENSMUSG( | 77.09647 | -0.12009 | 0.639999 | 0.953903 | Nt5c1a |
| ENSMUSG( | 2664.87 | 0.067925 | 0.497383 | 0.927825 | Nt5c2 |
| ENSMUSG( | 1065.596 | 0.050444 | 0.698078 | 0.960867 | Nt5c3 |
| ENSMUSG( | 1105.256 | -0.15806 | 0.189684 | 0.857355 | Nt5c3b |
| ENSMUSG( | 61.67725 | 0.051567 | 0.854111 | 0.98563 | Nt5dc1 |
| ENSMUSG( | 348.0142 | 0.171251 | 0.278251 | 0.891468 | Nt5dc2 |
| ENSMUSG( | 2133.703 | -0.11452 | 0.379175 | 0.916995 | Nt5dc3 |
| ENSMUSG( | 258.7371 | 0.075057 | 0.662684 | 0.956205 | Nt5e |
| ENSMUSG( | 1106.569 | -0.19348 | 0.083211 | 0.77681 | Nt5m |
| ENSMUSG( | 1182.998 | -0.10383 | 0.378706 | 0.916995 | Ntan1 |
| ENSMUSG( | 416.6954 | -0.10524 | 0.456784 | 0.923329 | Ntaq1 |
| ENSMUSG( | 54.97556 | -0.05775 | 0.851725 | 0.985591 | Ntf3 |
| ENSMUSG( | 79.87407 | 0.36643 | 0.124901 | 0.81516 | Nthl1 |
| ENSMUSG( | 14504.22 | -0.08623 | 0.546184 | 0.938063 | Ntm |
| ENSMUSG( | 458.4811 | -0.16533 | 0.331974 | 0.907907 | Ntmt1 |
| ENSMUSG( | 143.9168 | 0.080266 | 0.683061 | 0.958678 | Ntmt2 |
| ENSMUSG( | 535.4092 | -0.07147 | 0.675331 | 0.95694 | Ntn1 |
| ENSMUSG( | 446.6419 | 0.071953 | 0.619555 | 0.952896 | Ntn3 |
| ENSMUSG( | 487.9759 | 0.094219 | 0.568395 | 0.941777 | Ntn4 |
| ENSMUSG( | 157.9334 | -0.13756 | 0.503195 | 0.927825 | Ntn5 |
| ENSMUSG( | 726.12 | -0.13875 | 0.345199 | 0.91045 | Ntng1 |
| ENSMUSG( | 1533.667 | 0.146545 | 0.271594 | 0.888062 | Ntng2 |
| ENSMUSG( | 532.7932 | -0.1036 | 0.44619 | 0.921734 | Ntpcr |
| ENSMUSG( | 31.10906 | -0.24266 | 0.508771 | 0.929089 | Ntrk1 |
| ENSMUSG( | 51270.89 | 0.075075 | 0.538316 | 0.937216 | Ntrk2 |
| ENSMUSG( | 3864.266 | 0.221465 | 0.099476 | 0.80042 | Ntrk3 |
| ENSMUSG( | 162.8857 | -0.33525 | 0.149709 | 0.836241 | Nts |
| ENSMUSG( | 342.949 | 0.00991 | 0.958232 | 0.994798 | Ntsr1 |
| ENSMUSG( | 13583.74 | -0.02669 | 0.831612 | 0.980656 | Ntsr2 |
| ENSMUSG( | 1640.242 | -0.01937 | 0.883822 | 0.989644 | Nuak1 |
| ENSMUSG( | 34.90466 | 0.19063 | 0.599665 | 0.947071 | Nuak2 |
| ENSMUSG( | 2130.964 | -0.09004 | 0.400287 | 0.917257 | Nub1 |
| ENSMUSG( | 455.6017 | -0.09452 | 0.541126 | 0.93767 | Nubp1 |
| ENSMUSG( | 762.9991 | -0.20128 | 0.086045 | 0.780251 | Nubp2 |
| ENSMUSG( | 137.32 | -0.23482 | 0.306175 | 0.898538 | Nubpl |
| ENSMUSG( | 3656.271 | 0.191154 | 0.177202 | 0.846973 | Nucb1 |

| | | | | | | | | | | | |
|---|---|---|---|---|---|---|---|---|---|---|---|
| ENSMUSG( | Nudt14 | 164.0442 | -0.08764 | 0.597544 | 0.903694 | ENSMUSG( | Nudt14 | 154.5559 | -0.0004 | 0.9984 | 0.999313 |
| ENSMUSG( | Nudt15 | 125.0412 | -0.02718 | 0.880352 | 0.981469 | ENSMUSG( | Nudt15 | 109.7427 | -0.15885 | 0.474011 | 0.923329 |
| ENSMUSG( | Nudt16 | 321.0437 | -0.15187 | 0.304617 | 0.752574 | ENSMUSG( | Nudt16 | 292.8673 | -0.15772 | 0.363019 | 0.914223 |
| ENSMUSG( | Nudt16l1 | 1259.769 | 0.022662 | 0.852239 | 0.973168 | ENSMUSG( | Nudt16l1 | 1076.589 | -0.17068 | 0.228262 | 0.868507 |
| ENSMUSG( | Nudt17 | 146.2758 | 0.097776 | 0.582525 | 0.897918 | ENSMUSG( | Nudt17 | 123.3647 | -0.13779 | 0.521256 | 0.932489 |
| ENSMUSG( | Nudt18 | 1171.566 | 0.020903 | 0.848511 | 0.972374 | ENSMUSG( | Nudt18 | 1015.271 | -0.12778 | 0.322387 | 0.904023 |
| ENSMUSG( | Nudt19 | 3131.702 | 0.080921 | 0.441294 | 0.832449 | ENSMUSG( | Nudt19 | 2578.811 | -0.2215 | 0.096068 | 0.796133 |
| ENSMUSG( | Nudt2 | 450.4133 | 0.053157 | 0.700995 | 0.93743 | ENSMUSG( | Nudt2 | 354.0787 | -0.41224 | 0.015772 | 0.502615 |
| ENSMUSG( | Nudt21 | 1196.377 | 0.041237 | 0.697463 | 0.935662 | ENSMUSG( | Nudt21 | 1108.041 | 0.086474 | 0.486006 | 0.92541 |
| ENSMUSG( | Nudt22 | 159.033 | 0.201315 | 0.297488 | 0.746042 | ENSMUSG( | Nudt22 | 143.9491 | 0.185527 | 0.348068 | 0.911652 |
| ENSMUSG( | Nudt3 | 13746.91 | 0.054488 | 0.561702 | 0.889052 | ENSMUSG( | Nudt3 | 11810.71 | -0.11908 | 0.372486 | 0.915491 |
| ENSMUSG( | Nudt4 | 7644.8 | 0.059087 | 0.514518 | 0.872055 | ENSMUSG( | Nudt4 | 6762.531 | -0.02824 | 0.837581 | 0.981335 |
| ENSMUSG( | Nudt5 | 388.3784 | 0.062916 | 0.614941 | 0.910234 | ENSMUSG( | Nudt5 | 347.3667 | 0.007474 | 0.958062 | 0.994798 |
| ENSMUSG( | Nudt6 | 101.5964 | -0.13002 | 0.545088 | 0.884712 | ENSMUSG( | Nudt6 | 94.38392 | -0.08598 | 0.718169 | 0.966441 |
| ENSMUSG( | Nudt7 | 200.0514 | 0.497666 | 0.001296 | 0.039151 | ENSMUSG( | Nudt7 | 164.04 | 0.23557 | 0.202324 | 0.864412 |
| ENSMUSG( | Nudt8 | 18.45453 | -0.17573 | 0.692337 | NA | ENSMUSG( | Nudt8 | 17.73471 | 0.00589 | 0.990236 | 0.998104 |
| ENSMUSG( | Nudt9 | 2787.527 | 0.039371 | 0.627084 | 0.912825 | ENSMUSG( | Nudt9 | 2477.825 | -0.03289 | 0.76721 | 0.974236 |
| ENSMUSG( | Nuf2 | 342.7755 | 0.076229 | 0.689005 | 0.933441 | ENSMUSG( | Nuf2 | 297.3039 | -0.07238 | 0.719951 | 0.966887 |
| ENSMUSG( | Nufip1 | 240.6245 | 0.023274 | 0.879729 | 0.981436 | ENSMUSG( | Nufip1 | 211.8018 | -0.09054 | 0.631657 | 0.953251 |
| ENSMUSG( | Nufip2 | 479.1447 | -0.09634 | 0.460339 | 0.840676 | ENSMUSG( | Nufip2 | 487.2928 | 0.212124 | 0.131723 | 0.822912 |
| ENSMUSG( | Numa1 | 2394.15 | 0.070952 | 0.404876 | 0.816396 | ENSMUSG( | Numa1 | 2122.105 | -0.00566 | 0.958646 | 0.994798 |
| ENSMUSG( | Numb | 1403.514 | 0.125066 | 0.170437 | 0.605271 | ENSMUSG( | Numb | 1212.556 | -0.02886 | 0.830069 | 0.980224 |
| ENSMUSG( | Numbl | 2755.721 | 0.131253 | 0.265127 | 0.715962 | ENSMUSG( | Numbl | 2346.478 | -0.06195 | 0.622661 | 0.953234 |
| ENSMUSG( | Nup107 | 374.2117 | 0.05869 | 0.662262 | 0.925946 | ENSMUSG( | Nup107 | 329.9853 | -0.03679 | 0.819815 | 0.979198 |
| ENSMUSG( | Nup133 | 726.0206 | 0.124208 | 0.239343 | 0.687192 | ENSMUSG( | Nup133 | 633.8751 | 0.004567 | 0.970241 | 0.99594 |
| ENSMUSG( | Nup153 | 1128.51 | 0.129413 | 0.234812 | 0.682188 | ENSMUSG( | Nup153 | 1065.222 | 0.227062 | 0.052016 | 0.712974 |
| ENSMUSG( | Nup155 | 493.3186 | -0.05845 | 0.63962 | 0.916434 | ENSMUSG( | Nup155 | 492.9125 | 0.205741 | 0.127114 | 0.817278 |
| ENSMUSG( | Nup160 | 386.8917 | 0.075665 | 0.620312 | 0.911432 | ENSMUSG( | Nup160 | 353.7887 | 0.089185 | 0.583 | 0.944266 |
| ENSMUSG( | Nup188 | 744.6374 | 0.067519 | 0.578818 | 0.896692 | ENSMUSG( | Nup188 | 679.2503 | 0.074018 | 0.567927 | 0.941777 |
| ENSMUSG( | Nup205 | 803.1738 | 0.131044 | 0.232043 | 0.679932 | ENSMUSG( | Nup205 | 722.869 | 0.10028 | 0.412754 | 0.917257 |
| ENSMUSG( | Nup210 | 1040.307 | -0.21229 | 0.070047 | 0.40908 | ENSMUSG( | Nup210 | 1049.055 | 0.091534 | 0.497638 | 0.927825 |
| ENSMUSG( | Nup210l | 30.11755 | 0.106928 | 0.761665 | 0.953201 | ENSMUSG( | Nup210l | 23.23969 | -0.36214 | 0.394124 | 0.917257 |
| ENSMUSG( | Nup214 | 1134.685 | 0.06426 | 0.533119 | 0.879905 | ENSMUSG( | Nup214 | 1057.643 | 0.132395 | 0.302893 | 0.898538 |
| ENSMUSG( | Nup35 | 225.9977 | 0.04194 | 0.782705 | 0.957558 | ENSMUSG( | Nup35 | 204.3071 | 0.00736 | 0.966527 | 0.995475 |
| ENSMUSG( | Nup37 | 75.94856 | 0.131503 | 0.606442 | 0.906481 | ENSMUSG( | Nup37 | 68.64717 | 0.109151 | 0.670948 | 0.95694 |
| ENSMUSG( | Nup42 | 214.8454 | -0.10763 | 0.501877 | 0.86481 | ENSMUSG( | Nup42 | 194.4299 | -0.12736 | 0.460556 | 0.923329 |
| ENSMUSG( | Nup43 | 55.72655 | 0.108589 | 0.68998 | 0.93379 | ENSMUSG( | Nup43 | 51.12272 | 0.13172 | 0.653811 | 0.955015 |
| ENSMUSG( | Nup50 | 1086.495 | 0.034211 | 0.73005 | 0.944956 | ENSMUSG( | Nup50 | 1006.093 | 0.084288 | 0.497736 | 0.927825 |
| ENSMUSG( | Nup54 | 400.932 | 0.15926 | 0.182359 | 0.619591 | ENSMUSG( | Nup54 | 359.5408 | 0.116218 | 0.400891 | 0.917257 |
| ENSMUSG( | Nup58 | 745.4165 | 0.120427 | 0.303496 | 0.751968 | ENSMUSG( | Nup58 | 669.512 | 0.075198 | 0.583934 | 0.944398 |
| ENSMUSG( | Nup62 | 445.9624 | 0.146231 | 0.339136 | 0.777518 | ENSMUSG( | Nup62 | 388.853 | 0.025511 | 0.87216 | 0.988907 |
| ENSMUSG( | Nup62cl | 15.15806 | 0.294171 | 0.546512 | NA | ENSMUSG( | Nup62cl | 11.24224 | -0.28513 | 0.630322 | NA |
| ENSMUSG( | Nup85 | 754.6691 | 0.174387 | 0.126372 | 0.533885 | ENSMUSG( | Nup85 | 683.4039 | 0.159926 | 0.258348 | 0.883971 |
| ENSMUSG( | Nup88 | 1210.266 | 0.071258 | 0.594382 | 0.902872 | ENSMUSG( | Nup88 | 1110.77 | 0.092981 | 0.538093 | 0.937216 |
| ENSMUSG( | Nup93 | 598.6955 | -0.02413 | 0.845535 | 0.97218 | ENSMUSG( | Nup93 | 544.3454 | -0.02822 | 0.839901 | 0.98221 |
| ENSMUSG( | Nup98 | 1005.662 | 0.020373 | 0.840411 | 0.971179 | ENSMUSG( | Nup98 | 938.0885 | 0.088213 | 0.476771 | 0.923329 |
| ENSMUSG( | Nupr1 | 592.6412 | 0.201411 | 0.082252 | 0.442642 | ENSMUSG( | Nupr1 | 488.3207 | -0.08634 | 0.541725 | 0.937841 |
| ENSMUSG( | Nupr2 | 41.63081 | -0.07416 | 0.81185 | 0.964426 | ENSMUSG( | Nupr2 | 44.18155 | 0.354666 | 0.241709 | 0.876528 |
| ENSMUSG( | Nus1 | 2998.912 | -0.02032 | 0.864936 | 0.975565 | ENSMUSG( | Nus1 | 2873.643 | 0.122341 | 0.367143 | 0.915058 |
| ENSMUSG( | Nusap1 | 39.94407 | -0.46055 | 0.160263 | 0.589853 | ENSMUSG( | Nusap1 | 40.33233 | -0.16121 | 0.652833 | 0.954897 |
| ENSMUSG( | Nutf2 | 427.5583 | -0.00866 | 0.942524 | 0.993415 | ENSMUSG( | Nutf2 | 378.2196 | -0.10381 | 0.485285 | 0.92541 |
| ENSMUSG( | Nvl | 1464.53 | 0.022702 | 0.835006 | 0.970426 | ENSMUSG( | Nvl | 1421.171 | 0.200755 | 0.190614 | 0.857801 |
| ENSMUSG( | Nwd1 | 16269.95 | 0.142899 | 0.227793 | 0.674849 | ENSMUSG( | Nwd1 | 13323 | -0.16656 | 0.197555 | 0.862161 |
| ENSMUSG( | Nwd2 | 956.0197 | -0.08997 | 0.383099 | 0.803237 | ENSMUSG( | Nwd2 | 873.2164 | -0.08093 | 0.504647 | 0.927825 |
| ENSMUSG( | Nxf1 | 2108.581 | 0.119159 | 0.300853 | 0.749979 | ENSMUSG( | Nxf1 | 1977.707 | 0.200465 | 0.221394 | 0.866104 |
| ENSMUSG( | Nxf7 | 32.97931 | -0.02364 | 0.952811 | 0.993874 | ENSMUSG( | Nxf7 | 27.49927 | -0.28445 | 0.519319 | 0.931392 |
| ENSMUSG( | Nxn | 962.383 | 0.350334 | 0.000556 | 0.022059 | ENSMUSG( | Nxn | 938.5938 | 0.518509 | 1.83E-05 | 0.005843 |
| ENSMUSG( | Nxpe2 | 11.76067 | -0.28163 | 0.63356 | NA | ENSMUSG( | Nxpe2 | 13.12135 | 0.330965 | 0.542553 | 0.938034 |
| ENSMUSG( | Nxpe3 | 492.134 | -0.31376 | 0.026552 | 0.249087 | ENSMUSG( | Nxpe3 | 542.1063 | 0.248999 | 0.173819 | 0.844317 |
| ENSMUSG( | Nxpe4 | 621.7809 | -0.2419 | 0.062493 | 0.386467 | ENSMUSG( | Nxpe4 | 593.2313 | -0.09321 | 0.566869 | 0.941777 |
| ENSMUSG( | Nxph1 | 787.4561 | -0.57622 | 0.001058 | 0.034188 | ENSMUSG( | Nxph1 | 900.7394 | 0.141581 | 0.490407 | 0.926343 |
| ENSMUSG( | Nxph2 | 38.74297 | -0.45317 | 0.207296 | 0.649561 | ENSMUSG( | Nxph2 | 46.14329 | 0.332865 | 0.395424 | 0.917257 |
| ENSMUSG( | Nxph3 | 1554.497 | -0.10821 | 0.470633 | 0.84674 | ENSMUSG( | Nxph3 | 1544.711 | 0.144091 | 0.361149 | 0.914223 |
| ENSMUSG( | Nxph4 | 253.1777 | -0.75464 | 4.43E-06 | 0.000589 | ENSMUSG( | Nxph4 | 288.1013 | -0.01853 | 0.917255 | 0.992529 |
| ENSMUSG( | Nxt1 | 133.2567 | -0.18 | 0.310739 | 0.758828 | ENSMUSG( | Nxt1 | 129.5693 | 0.005198 | 0.979014 | 0.996468 |
| ENSMUSG( | Nxt2 | 1319.257 | 0.016672 | 0.901964 | 0.985633 | ENSMUSG( | Nxt2 | 1175.985 | -0.0521 | 0.755014 | 0.971098 |
| ENSMUSG( | Nyap1 | 3183.621 | 0.215654 | 0.022447 | 0.228899 | ENSMUSG( | Nyap1 | 2769.545 | 0.089898 | 0.378671 | 0.916995 |
| ENSMUSG( | Nyap2 | 323.0938 | -0.36706 | 0.006111 | 0.106351 | ENSMUSG( | Nyap2 | 317.38 | -0.12531 | 0.389565 | 0.917257 |
| ENSMUSG( | Nynrin | 387.7416 | -0.22672 | 0.085087 | 0.449548 | ENSMUSG( | Nynrin | 400.065 | 0.147359 | 0.375189 | 0.915491 |
| ENSMUSG( | Nyx | 19.15078 | 0.421115 | 0.353163 | NA | ENSMUSG( | Nyx | 18.17997 | 0.537716 | 0.249556 | 0.877904 |
| ENSMUSG( | Oacyl | 60.48842 | -0.64727 | 0.014596 | 0.177737 | ENSMUSG( | Oacyl | 57.48488 | -0.5003 | 0.082079 | 0.77681 |
| ENSMUSG( | Oaf | 876.2935 | -0.14071 | 0.346261 | 0.779153 | ENSMUSG( | Oaf | 949.1431 | 0.347203 | 0.050312 | 0.708648 |
| ENSMUSG( | Oard1 | 388.1092 | -0.0339 | 0.805669 | 0.962616 | ENSMUSG( | Oard1 | 335.4631 | -0.19673 | 0.213713 | 0.864421 |
| ENSMUSG( | Oas1a | 32.48788 | -0.48867 | 0.13694 | 0.55149 | ENSMUSG( | Oas1a | 32.49893 | -0.15876 | 0.656318 | 0.955533 |
| ENSMUSG( | Oas1b | 23.08542 | 0.079701 | 0.834302 | 0.970333 | ENSMUSG( | Oas1b | 21.98164 | 0.242778 | 0.590276 | 0.944975 |
| ENSMUSG( | Oas1c | 38.80898 | 0.032656 | 0.916661 | 0.988856 | ENSMUSG( | Oas1c | 31.8141 | -0.26294 | 0.464234 | 0.923329 |
| ENSMUSG( | Oas2 | 31.6575 | 0.012319 | 0.974616 | 0.997179 | ENSMUSG( | Oas2 | 23.09324 | -0.68456 | 0.163782 | 0.838291 |
| ENSMUSG( | Oas3 | 9.882222 | -0.63101 | 0.347503 | NA | ENSMUSG( | Oas3 | 9.248201 | -0.53522 | 0.447957 | NA |
| ENSMUSG( | Oasl1 | 11.84226 | -0.25211 | 0.657317 | NA | ENSMUSG( | Oasl1 | 10.71842 | -0.19919 | 0.752232 | NA |
| ENSMUSG( | Oasl2 | 205.7395 | -0.30613 | 0.094046 | 0.471556 | ENSMUSG( | Oasl2 | 188.3729 | -0.28435 | 0.159644 | 0.837724 |
| ENSMUSG( | Oat | 4626.955 | 0.139755 | 0.170984 | 0.605536 | ENSMUSG( | Oat | 3788.462 | -0.17201 | 0.161146 | 0.837724 |
| ENSMUSG( | Oaz1 | 7337.269 | 0.057306 | 0.441425 | 0.832452 | ENSMUSG( | Oaz1 | 6285.329 | -0.12326 | 0.245265 | 0.877904 |
| ENSMUSG( | Oaz2 | 4236.903 | 0.068891 | 0.397319 | 0.812127 | ENSMUSG( | Oaz2 | 3640.607 | -0.10303 | 0.33835 | 0.907972 |
| ENSMUSG( | Obi1 | 230.3271 | -0.05833 | 0.723279 | 0.943761 | ENSMUSG( | Obi1 | 213.6699 | -0.01274 | 0.945861 | 0.994798 |
| ENSMUSG( | Obscn | 205.6785 | -0.87275 | 4.86E-07 | 0.000104 | ENSMUSG( | Obscn | 238.2989 | -0.04615 | 0.797473 | 0.97681 |
| ENSMUSG( | Obsl1 | 567.9047 | 0.109128 | 0.332012 | 0.775301 | ENSMUSG( | Obsl1 | 543.7142 | 0.248451 | 0.083597 | 0.77681 |
| ENSMUSG( | Oca2 | 18.92162 | -0.39562 | 0.361835 | NA | ENSMUSG( | Oca2 | 26.09099 | 0.686488 | 0.129919 | 0.820944 |
| ENSMUSG( | Ocel1 | 606.698 | 0.157997 | 0.171392 | 0.605943 | ENSMUSG( | Ocel1 | 563.3357 | 0.21656 | 0.145202 | 0.83339 |
| ENSMUSG( | Ociad1 | 9240.916 | -0.33228 | 0.000988 | 0.032693 | ENSMUSG( | Ociad1 | 9296.302 | -0.02642 | 0.8302 | 0.980224 |
| ENSMUSG( | Ociad2 | 1656.084 | -0.13693 | 0.212627 | 0.6567 | ENSMUSG( | Ociad2 | 1589.219 | 0.016269 | 0.885115 | 0.989688 |
| ENSMUSG( | Ocln | 321.2424 | -0.04122 | 0.876615 | 0.980871 | ENSMUSG( | Ocln | 389.7562 | 0.712232 | 0.149662 | 0.836241 |
| ENSMUSG( | Ocm | 14.39042 | -0.27267 | 0.581705 | NA | ENSMUSG( | Ocm | 15.22848 | 0.166955 | 0.749238 | 0.96998 |
| ENSMUSG( | Ocrl | 2072.77 | 0.210603 | 0.018653 | 0.205542 | ENSMUSG( | Ocrl | 1819.473 | 0.107867 | 0.349684 | 0.911652 |
| ENSMUSG( | Odad1 | 136.753 | -0.0808 | 0.640713 | 0.916434 | ENSMUSG( | Odad1 | 128.2307 | 0.017702 | 0.936291 | 0.993573 |
| ENSMUSG( | Odad2 | 19.23189 | -0.24017 | 0.591971 | NA | ENSMUSG( | Odad2 | 17.88945 | -0.10357 | 0.854268 | 0.98563 |
| ENSMUSG( | Odad3 | 108.6595 | -0.02434 | 0.909095 | 0.98794 | ENSMUSG( | Odad3 | 105.1733 | 0.144906 | 0.510933 | 0.929546 |
| ENSMUSG( | Odad4 | 45.93444 | -0.37501 | 0.223459 | 0.669469 | ENSMUSG( | Odad4 | 49.45499 | 0.14926 | 0.672675 | 0.95694 |
| ENSMUSG( | Odc1 | 2067.932 | -0.06701 | 0.524418 | 0.87583 | ENSMUSG( | Odc1 | 1805.948 | -0.19916 | 0.130522 | 0.822225 |
| ENSMUSG( | Odf2 | 2406.447 | 0.140421 | 0.196053 | 0.635489 | ENSMUSG( | Odf2 | 2174.054 | 0.1163 | 0.390995 | 0.917257 |
| ENSMUSG( | Odf2l | 364.5961 | 0.079922 | 0.523595 | 0.875628 | ENSMUSG( | Odf2l | 319.3429 | -0.03189 | 0.861368 | 0.986281 |
| ENSMUSG( | Odr4 | 2398.968 | -0.02007 | 0.852773 | 0.973168 | ENSMUSG( | Odr4 | 2172.402 | -0.03982 | 0.746958 | 0.969763 |
| ENSMUSG( | Ofd1 | 291.2791 | 3.49E-05 | 0.999813 | 0.99989 | ENSMUSG( | Ofd1 | 252.5936 | -0.15317 | 0.366251 | 0.914703 |
| ENSMUSG( | Oga | 5738.298 | 0.137443 | 0.137104 | 0.551819 | ENSMUSG( | Oga | 4734.707 | -0.14995 | 0.136848 | 0.828492 |
| ENSMUSG( | Ogdh | 9654.122 | 0.140301 | 0.083463 | 0.445586 | ENSMUSG( | Ogdh | 7874.08 | -0.18123 | 0.066 | 0.746371 |
| ENSMUSG( | Ogdhl | 5615.674 | 0.186619 | 0.034773 | 0.286739 | ENSMUSG( | Ogdhl | 4625.888 | -0.10122 | 0.316054 | 0.901868 |
| ENSMUSG( | Ogfod1 | 3032.16 | 0.181527 | 0.04274 | 0.316989 | ENSMUSG( | Ogfod1 | 2609.723 | 0.022056 | 0.841028 | 0.9826 |
| ENSMUSG( | Ogfod2 | 752.2273 | 0.008521 | 0.941634 | 0.993415 | ENSMUSG( | Ogfod2 | 685.0405 | 0.004659 | 0.972795 | 0.996416 |
| ENSMUSG( | Ogfod3 | 283.9544 | -0.39117 | 0.021671 | 0.225182 | ENSMUSG( | Ogfod3 | 299.5371 | 0.064908 | 0.718834 | 0.966668 |
| ENSMUSG( | Ogfr | 1615.409 | 0.088842 | 0.380154 | 0.801757 | ENSMUSG( | Ogfr | 1412.174 | -0.02777 | 0.824362 | 0.979198 |
| ENSMUSG( | Ogfrl1 | 2695.056 | 0.086242 | 0.329471 | 0.773311 | ENSMUSG( | Ogfrl1 | 2577.412 | 0.216743 | 0.117242 | 0.813384 |
| ENSMUSG( | Ogg1 | 200.1112 | 0.080952 | 0.598964 | 0.903801 | ENSMUSG( | Ogg1 | 182.2343 | 0.084861 | 0.633158 | 0.953844 |
| ENSMUSG( | Ogn | 336.9876 | -0.21458 | 0.267382 | 0.717525 | ENSMUSG( | Ogn | 363.6704 | 0.268193 | 0.09866 | 0.798438 |
| ENSMUSG( | Ogt | 6530.254 | 0.038436 | 0.737897 | 0.947329 | ENSMUSG( | Ogt | 6345.55 | 0.215888 | 0.19146 | 0.858412 |
| ENSMUSG( | Oip5 | 13.07207 | 0.101376 | 0.852286 | NA | ENSMUSG( | Oip5 | 17.25875 | 0.981577 | 0.040188 | 0.681706 |
| ENSMUSG( | Oip5os1 | 16087.44 | 0.140792 | 0.132746 | 0.545099 | ENSMUSG( | Oip5os1 | 13743.18 | -0.04314 | 0.711482 | 0.964363 |
| ENSMUSG( | Ola1 | 1610.625 | -0.04246 | 0.716953 | 0.941697 | ENSMUSG( | Ola1 | 1436.803 | -0.10987 | 0.460226 | 0.923329 |
| ENSMUSG( | Olfm1 | 49119.54 | -0.12284 | 0.226052 | 0.672763 | ENSMUSG( | Olfm1 | 45937.09 | -0.04446 | 0.714447 | 0.964986 |
| ENSMUSG( | Olfm2 | 6250.384 | -0.3254 | 0.006856 | 0.11446 | ENSMUSG( | Olfm2 | 6533.597 | 0.090473 | 0.522291 | 0.932681 |
| ENSMUSG( | Olfm3 | 512.0904 | -0.94433 | 2.70E-06 | 0.000405 | ENSMUSG( | Olfm3 | 667.5722 | 0.234306 | 0.298645 | 0.898538 |
| ENSMUSG( | Olfm4 | 42.7271 | 0.132156 | 0.663315 | 0.9265 | ENSMUSG( | Olfm4 | 34.54222 | -0.21719 | 0.531996 | 0.934795 |
| ENSMUSG( | Olfml1 | 709.7437 | -0.12852 | 0.410842 | 0.818585 | ENSMUSG( | Olfml1 | 675.865 | 0.008952 | 0.960624 | 0.995047 |
| ENSMUSG( | Olfml2a | 221.0833 | -0.26048 | 0.215228 | 0.659815 | ENSMUSG( | Olfml2a | 212.11 | -0.08829 | 0.671405 | 0.95694 |
| ENSMUSG( | Olfml2b | 347.245 | 0.16041 | 0.494548 | 0.860776 | ENSMUSG( | Olfml2b | 391.6296 | 0.708874 | 0.007462 | 0.367159 |
| ENSMUSG( | Olfml3 | 1509.104 | -0.27058 | 0.101159 | 0.484996 | ENSMUSG( | Olfml3 | 1541.983 | 0.078029 | 0.700086 | 0.961335 |
| ENSMUSG( | Olig1 | 9658.095 | -0.18742 | 0.089539 | 0.459596 | ENSMUSG( | Olig1 | 8629.857 | -0.24919 | 0.047457 | 0.700936 |
| ENSMUSG( | Olig2 | 1278.812 | -0.59517 | 1.37E-05 | 0.001449 | ENSMUSG( | Olig2 | 1389.313 | -0.02288 | 0.887323 | 0.990266 |
| ENSMUSG( | Oma1 | 263.4406 | 0.068145 | 0.639389 | 0.916434 | ENSMUSG( | Oma1 | 227.546 | -0.09084 | 0.589135 | 0.944975 |
| ENSMUSG( | Omd | 33.54041 | 0.193253 | 0.573566 | 0.893584 | ENSMUSG( | Omd | 37.05948 | 0.671079 | 0.061128 | 0.736111 |
| ENSMUSG( | Omg | 3937.536 | -0.45476 | 0.006737 | 0.113006 | ENSMUSG( | Omg | 4458.351 | 0.206966 | 0.301306 | 0.898538 |
| ENSMUSG( | Omp | 49.65054 | -2.62222 | 0.01154 | 0.154695 | ENSMUSG( | Omp | 108.1038 | 0.819761 | 0.581272 | 0.944266 |
| ENSMUSG( | Onecut2 | 81.09059 | -0.52451 | 0.020523 | 0.216158 | ENSMUSG( | Onecut2 | 79.50093 | -0.26394 | 0.298804 | 0.898538 |
| ENSMUSG( | Opa1 | 4577.166 | 0.090824 | 0.233096 | 0.681351 | ENSMUSG( | Opa1 | 3992.591 | -0.03363 | 0.730439 | 0.967228 |
| ENSMUSG( | Opa3 | 2637.841 | 0.060338 | 0.462635 | 0.842333 | ENSMUSG( | Opa3 | 2252.494 | -0.12924 | 0.220639 | 0.866104 |
| ENSMUSG( | Opalin | 1199.33 | -0.74577 | 1.04E-06 | 0.000197 | ENSMUSG( | Opalin | 1262.729 | -0.23712 | 0.194388 | 0.860096 |
| ENSMUSG( | Opcml | 5380.971 | -0.37148 | 0.016177 | 0.189041 | ENSMUSG( | Opcml | 5918.232 | 0.193501 | 0.286598 | 0.894091 |
| ENSMUSG( | Ophn1 | 1474.951 | 0.071466 | 0.532683 | 0.879905 | ENSMUSG( | Ophn1 | 1350.167 | 0.086513 | 0.5126 | 0.930164 |
| ENSMUSG( | Oplah | 1875.306 | 0.11513 | 0.271711 | 0.722082 | ENSMUSG( | Oplah | 1614.829 | -0.04642 | 0.695835 | 0.960867 |
| ENSMUSG( | Opn3 | 144.0092 | -0.12075 | 0.533152 | 0.879905 | ENSMUSG( | Opn3 | 142.8195 | 0.134151 | 0.526902 | 0.934449 |
| ENSMUSG( | Oprd1 | 230.1327 | -0.20744 | 0.343253 | 0.779153 | ENSMUSG( | Oprd1 | 238.8881 | 0.187764 | 0.47862 | 0.923329 |
| ENSMUSG( | Oprk1 | 212.4665 | -0.79983 | 3.27E-06 | 0.000465 | ENSMUSG( | Oprk1 | 267.0119 | 0.228212 | 0.25375 | 0.880692 |
| ENSMUSG( | Oprl1 | 1052.428 | -0.06374 | 0.640233 | 0.916434 | ENSMUSG( | Oprl1 | 1082.424 | 0.27868 | 0.066903 | 0.751729 |
| ENSMUSG( | Oprm1 | 11.24422 | -0.46656 | 0.4519 | NA | ENSMUSG( | Oprm1 | 12.96103 | 0.245726 | 0.654712 | 0.955015 |
| ENSMUSG( | Optc | 33.48434 | -0.2722 | 0.421421 | 0.823126 | ENSMUSG( | Optc | 33.88062 | 0.049231 | 0.884929 | 0.989688 |
| ENSMUSG( | Optn | 500.6619 | -0.0619 | 0.604914 | 0.905556 | ENSMUSG( | Optn | 453.8398 | -0.07808 | 0.588059 | 0.944975 |
| ENSMUSG( | Or10ad1 | 599.8774 | 0.087601 | 0.466761 | 0.844819 | ENSMUSG( | Or10ad1 | 515.4452 | -0.09114 | 0.593659 | 0.945408 |
| ENSMUSG( | Or2ak4 | 51.08857 | -0.08019 | 0.776394 | 0.955658 | ENSMUSG( | Or2ak4 | 46.82857 | -0.05265 | 0.864733 | 0.986591 |
| ENSMUSG( | Or2y1g | 18.93615 | -1.97517 | 0.00169 | NA | ENSMUSG( | Or2y1g | 26.64285 | -0.13456 | 0.770953 | 0.974236 |
| ENSMUSG( | Or4d1 | 37.09659 | -0.34971 | 0.261005 | 0.710889 | ENSMUSG( | Or4d1 | 28.15855 | -1.0424 | 0.005644 | 0.304216 |
| ENSMUSG( | Or51e2 | 24.79305 | -0.36418 | 0.368323 | 0.795863 | ENSMUSG( | Or51e2 | 25.63155 | 0.036122 | 0.933588 | 0.99306 |
| ENSMUSG( | Or5m3b | 15.85828 | 0.362996 | 0.537246 | NA | ENSMUSG( | Or5m3b | 10.63115 | -0.51438 | 0.390444 | NA |
| ENSMUSG( | Orai1 | 365.5107 | -0.48675 | 0.00242 | 0.057903 | ENSMUSG( | Orai1 | 438.6925 | 0.338025 | 0.073061 | 0.763629 |
| ENSMUSG( | Orai2 | 3447.817 | -0.04205 | 0.68038 | 0.931044 | ENSMUSG( | Orai2 | 3426.219 | 0.204584 | 0.082907 | 0.77681 |
| ENSMUSG( | Orai3 | 735.2552 | 0.021095 | 0.864243 | 0.975358 | ENSMUSG( | Orai3 | 662.065 | -0.00766 | 0.960301 | 0.994831 |
| ENSMUSG( | Orc2 | 933.4692 | 0.184369 | 0.059358 | 0.376382 | ENSMUSG( | Orc2 | 794.0299 | -0.01262 | 0.921239 | 0.992644 |
| ENSMUSG( | Orc3 | 714.2317 | -0.02622 | 0.801961 | 0.962116 | ENSMUSG( | Orc3 | 663.7288 | 0.032032 | 0.798415 | 0.97681 |
| ENSMUSG( | Orc4 | 898.2217 | -0.10818 | 0.340168 | 0.777962 | ENSMUSG( | Orc4 | 886.0747 | 0.116917 | 0.393876 | 0.917257 |
| ENSMUSG( | Orc5 | 447.2922 | -0.31305 | 0.012346 | 0.160458 | ENSMUSG( | Orc5 | 439.2624 | -0.07645 | 0.584715 | 0.944975 |
| ENSMUSG( | Orc6 | 521.621 | 0.039786 | 0.742574 | 0.948478 | ENSMUSG( | Orc6 | 455.1714 | -0.08379 | 0.561047 | 0.940254 |

| | | | | | |
|---|---|---|---|---|---|
| ENSMUSG( | 15.7841 | -0.06931 | 0.897481 | NA | Nphs2 |
| ENSMUSG( | 164.9555 | 0.308194 | 0.086355 | 0.452045 | Npl |
| ENSMUSG( | 2287.747 | 0.125053 | 0.133772 | 0.547197 | Nploc4 |
| ENSMUSG( | 3038.752 | -0.04927 | 0.624472 | 0.91187 | Npm1 |
| ENSMUSG( | 15.44514 | 0.347948 | 0.477253 | NA | Npm2 |
| ENSMUSG( | 184.1042 | 0.135754 | 0.384864 | 0.804031 | Npm3 |
| ENSMUSG( | 1292.646 | -0.0731 | 0.664868 | 0.926824 | Npnt |
| ENSMUSG( | 10.40045 | -1.13139 | 0.147178 | NA | Nppa |
| ENSMUSG( | 178.9428 | -0.61631 | 0.002465 | 0.058361 | Nppc |
| ENSMUSG( | 128.9001 | -0.31248 | 0.101789 | 0.486216 | Npr1 |
| ENSMUSG( | 2204.931 | 0.070424 | 0.414589 | 0.820184 | Npr2 |
| ENSMUSG( | 330.4608 | 0.025491 | 0.871228 | 0.978114 | Npr3 |
| ENSMUSG( | 693.6934 | 0.046891 | 0.73214 | 0.944956 | Nprl2 |
| ENSMUSG( | 877.9907 | 0.008877 | 0.946189 | 0.99369 | Nprl3 |
| ENSMUSG( | 48.28876 | -0.68802 | 0.044126 | 0.322108 | Npsr1 |
| ENSMUSG( | 17488.89 | -0.61425 | 7.85E-07 | 0.000159 | Nptn |
| ENSMUSG( | 10092.01 | -0.31648 | 0.055853 | 0.364301 | Nptx1 |
| ENSMUSG( | 2298.464 | -0.57492 | 5.40E-05 | 0.004212 | Nptx2 |
| ENSMUSG( | 40260.18 | -0.1028 | 0.401805 | 0.814965 | Nptxr |
| ENSMUSG( | 3812.252 | -0.77417 | 1.99E-09 | 8.98E-07 | Npy |
| ENSMUSG( | 701.8671 | -0.34521 | 0.040487 | 0.311082 | Npy1r |
| ENSMUSG( | 205.1565 | -0.31828 | 0.086163 | 0.451605 | Npy2r |
| ENSMUSG( | 105.1711 | -0.34576 | 0.136741 | 0.551176 | Npy5r |
| ENSMUSG( | 259.6419 | 0.034172 | 0.820619 | 0.965888 | Nqo1 |
| ENSMUSG( | 678.3729 | 0.066393 | 0.667116 | 0.92729 | Nqo2 |
| ENSMUSG( | 2903.091 | 0.195688 | 0.105485 | 0.492417 | Nr1d1 |
| ENSMUSG( | 2310.528 | 0.095682 | 0.331383 | 0.77519 | Nr1d2 |
| ENSMUSG( | 2068.322 | 0.176681 | 0.049003 | 0.337938 | Nr1h2 |
| ENSMUSG( | 139.5272 | 0.298335 | 0.093408 | 0.469793 | Nr1h3 |
| ENSMUSG( | 24.33574 | -0.34897 | 0.369175 | 0.796641 | Nr1i3 |
| ENSMUSG( | 256.8529 | -0.28714 | 0.13306 | 0.545688 | Nr2c1 |
| ENSMUSG( | 4317.523 | 0.059733 | 0.519729 | 0.874979 | Nr2c2 |
| ENSMUSG( | 311.6272 | 0.019982 | 0.886709 | 0.983255 | Nr2c2ap |
| ENSMUSG( | 1728.646 | -0.04265 | 0.724847 | 0.944096 | Nr2e1 |
| ENSMUSG( | 3485.297 | 0.195936 | 0.025656 | 0.246969 | Nr2f1 |
| ENSMUSG( | 458.8348 | 0.015147 | 0.928845 | 0.991747 | Nr2f2 |
| ENSMUSG( | 1259.673 | 0.075433 | 0.405514 | 0.81652 | Nr2f6 |
| ENSMUSG( | 2792.646 | -0.03425 | 0.747097 | 0.949526 | Nr3c1 |
| ENSMUSG( | 1148.873 | -0.08446 | 0.412461 | 0.819452 | Nr3c2 |
| ENSMUSG( | 2282.211 | -0.68222 | 7.14E-05 | 0.005192 | Nr4a1 |
| ENSMUSG( | 1312.134 | 0.11763 | 0.210725 | 0.653529 | Nr4a2 |
| ENSMUSG( | 751.8325 | 0.084712 | 0.399132 | 0.813048 | Nr4a3 |
| ENSMUSG( | 189.4328 | -0.2233 | 0.165109 | 0.596585 | Nr6a1 |
| ENSMUSG( | 48.89396 | -0.40268 | 0.179995 | 0.617634 | Nradd |
| ENSMUSG( | 32.64834 | -0.01446 | 0.970467 | 0.996369 | Nrap |
| ENSMUSG( | 2003.444 | 0.084378 | 0.436539 | 0.829373 | Nrarp |
| ENSMUSG( | 3423.33 | 0.006105 | 0.958785 | 0.993874 | Nras |
| ENSMUSG( | 205.7725 | 0.024712 | 0.900805 | 0.985633 | Nrbf2 |
| ENSMUSG( | 1828.127 | 0.120654 | 0.270632 | 0.720561 | Nrbp1 |
| ENSMUSG( | 5895.503 | 0.099089 | 0.590938 | 0.90113 | Nrbp2 |
| ENSMUSG( | 13306.94 | -0.04798 | 0.718618 | 0.941974 | Nrcam |
| ENSMUSG( | 5089.563 | -0.11072 | 0.140487 | 0.557685 | Nrdc |
| ENSMUSG( | 369.218 | 0.075234 | 0.630276 | 0.914073 | Nrde2 |
| ENSMUSG( | 4019.184 | 0.106257 | 0.256176 | 0.706924 | Nrep |
| ENSMUSG( | 542.4099 | 0.197538 | 0.069395 | 0.40782 | Nrf1 |
| ENSMUSG( | 323.7025 | -0.1011 | 0.524414 | 0.87583 | Nrg1 |
| ENSMUSG( | 241.4858 | -0.07078 | 0.61245 | 0.908747 | Nrg2 |
| ENSMUSG( | 1048.247 | -0.49556 | 0.002722 | 0.063172 | Nrg3 |
| ENSMUSG( | 11.61529 | -0.41272 | 0.443418 | NA | Nrg4 |
| ENSMUSG( | 60013.31 | 0.09104 | 0.348595 | 0.78132 | Nrgn |
| ENSMUSG( | 687.2703 | -0.10514 | 0.305581 | 0.753616 | Nrip1 |
| ENSMUSG( | 177.5155 | 0.129982 | 0.507769 | 0.86795 | Nrip2 |
| ENSMUSG( | 3019.309 | 0.124373 | 0.190774 | 0.630215 | Nrip3 |
| ENSMUSG( | 139.9948 | -0.37331 | 0.038611 | 0.302696 | Nrm |
| ENSMUSG( | 5766.661 | -0.31146 | 0.003254 | 0.071142 | Nrn1 |
| ENSMUSG( | 800.3894 | -0.07418 | 0.68547 | 0.932981 | Nrp1 |
| ENSMUSG( | 1083.651 | -0.41883 | 0.020106 | 0.214413 | Nrp2 |
| ENSMUSG( | 291.4668 | -0.00458 | 0.978242 | 0.997428 | Nrros |
| ENSMUSG( | 34606.39 | 0.222345 | 0.03289 | 0.277278 | Nrsn1 |
| ENSMUSG( | 2251.078 | -0.71633 | 1.85E-10 | 1.07E-07 | Nrsn2 |
| ENSMUSG( | 155.2788 | -0.07854 | 0.671496 | 0.928781 | Nrtn |
| ENSMUSG( | 15091.53 | -0.20345 | 0.151458 | 0.576717 | Nrxn1 |
| ENSMUSG( | 20810.85 | -0.11565 | 0.34414 | 0.779153 | Nrxn2 |
| ENSMUSG( | 3825.202 | -0.41876 | 0.001186 | 0.037079 | Nrxn3 |
| ENSMUSG( | 1556.889 | 0.086607 | 0.383621 | 0.803504 | Nsa2 |
| ENSMUSG( | 10.82608 | -0.6796 | 0.246563 | NA | Nscme3l |
| ENSMUSG( | 4389.521 | -0.05975 | 0.521904 | 0.875186 | Nsd1 |
| ENSMUSG( | 2834.504 | 0.075904 | 0.363177 | 0.793211 | Nsd2 |
| ENSMUSG( | 2438.254 | -0.00936 | 0.926851 | 0.991229 | Nsd3 |
| ENSMUSG( | 1460.312 | -0.13766 | 0.332302 | 0.775474 | Nsdhl |
| ENSMUSG( | 12973.4 | 0.181818 | 0.155258 | 0.583155 | Nsf |
| ENSMUSG( | 1535.27 | 0.053388 | 0.589146 | 0.90113 | Nsfl1c |
| ENSMUSG( | 8031.302 | -0.43982 | 4.99E-05 | 0.00395 | Nsg1 |
| ENSMUSG( | 17230.65 | -0.42364 | 0.000252 | 0.012746 | Nsg2 |
| ENSMUSG( | 29.34348 | -0.01625 | 0.965404 | 0.995377 | Nsl1 |
| ENSMUSG( | 1217.346 | 0.126543 | 0.208447 | 0.650319 | Nsmaf |
| ENSMUSG( | 375.8568 | 0.001554 | 0.990543 | 0.999232 | Nsmce1 |
| ENSMUSG( | 448.9543 | 0.160581 | 0.279435 | 0.729114 | Nsmce2 |
| ENSMUSG( | 755.6087 | -0.61655 | 1.91E-06 | 0.000324 | Nsmce3 |
| ENSMUSG( | 680.2453 | 0.069362 | 0.539935 | 0.88315 | Nsmce4a |
| ENSMUSG( | 46491.73 | 0.23259 | 0.006802 | 0.113745 | Nsmf |
| ENSMUSG( | 463.2242 | 0.079927 | 0.524174 | 0.87583 | Nsrp1 |
| ENSMUSG( | 1673.443 | 0.104889 | 0.277619 | 0.72746 | Nsun2 |
| ENSMUSG( | 314.8393 | 0.008356 | 0.955908 | 0.993874 | Nsun3 |
| ENSMUSG( | 349.5936 | 0.13778 | 0.290759 | 0.740751 | Nsun4 |
| ENSMUSG( | 358.9904 | 0.071728 | 0.597564 | 0.903694 | Nsun5 |
| ENSMUSG( | 157.3169 | -0.04331 | 0.790539 | 0.959461 | Nsun6 |
| ENSMUSG( | 317.8613 | 0.056789 | 0.686027 | 0.932981 | Nsun7 |
| ENSMUSG( | 716.925 | 0.066819 | 0.583872 | 0.898666 | Nt5c |
| ENSMUSG( | 92.11023 | 0.106273 | 0.623169 | 0.911599 | Nt5c1a |
| ENSMUSG( | 2925.488 | 0.068944 | 0.381952 | 0.802983 | Nt5c2 |
| ENSMUSG( | 1072.886 | -0.20536 | 0.044529 | 0.324173 | Nt5c3 |
| ENSMUSG( | 1318.843 | 0.088883 | 0.34231 | 0.778828 | Nt5c3b |
| ENSMUSG( | 61.95924 | -0.24164 | 0.3423 | 0.778828 | Nt5dc1 |
| ENSMUSG( | 380.1357 | 0.160095 | 0.238234 | 0.686308 | Nt5dc2 |
| ENSMUSG( | 2626.471 | 0.210833 | 0.076083 | 0.42702 | Nt5dc3 |
| ENSMUSG( | 237.5643 | -0.48362 | 0.01221 | 0.160227 | Nt5e |
| ENSMUSG( | 1334.804 | 0.085368 | 0.337744 | 0.777422 | Nt5m |
| ENSMUSG( | 1346.862 | 0.006026 | 0.947499 | 0.993821 | Ntan1 |
| ENSMUSG( | 463.328 | -0.06217 | 0.62675 | 0.912825 | Ntaq1 |
| ENSMUSG( | 56.19566 | -0.30139 | 0.273385 | 0.72385 | Ntf3 |
| ENSMUSG( | 83.78553 | 0.252095 | 0.283449 | 0.73186 | Nthl1 |
| ENSMUSG( | 15260.7 | -0.21553 | 0.059232 | 0.375733 | Ntm |
| ENSMUSG( | 505.7741 | -0.14947 | 0.348465 | 0.78132 | Ntmt1 |
| ENSMUSG( | 191.1062 | 0.582249 | 0.000381 | 0.017215 | Ntmt2 |
| ENSMUSG( | 522.1964 | -0.45494 | 0.004738 | 0.09051 | Ntn1 |
| ENSMUSG( | 467.8534 | -0.06432 | 0.594952 | 0.903007 | Ntn3 |
| ENSMUSG( | 488.1515 | -0.17373 | 0.226546 | 0.673549 | Ntn4 |
| ENSMUSG( | 173.2394 | -0.15289 | 0.404414 | 0.816132 | Ntn5 |
| ENSMUSG( | 717.1214 | -0.47641 | 8.35E-05 | 0.005713 | Ntng1 |
| ENSMUSG( | 1549.418 | -0.09401 | 0.410539 | 0.818455 | Ntng2 |
| ENSMUSG( | 583.4327 | -0.1073 | 0.355619 | 0.787945 | Ntpcr |
| ENSMUSG( | 23.32465 | -1.89014 | 9.15E-05 | 0.006056 | Ntrk1 |
| ENSMUSG( | 56407.18 | 0.08077 | 0.473566 | 0.848919 | Ntrk2 |
| ENSMUSG( | 3856.502 | -0.04767 | 0.679482 | 0.930983 | Ntrk3 |
| ENSMUSG( | 149.1341 | -0.99162 | 1.58E-06 | 0.000275 | Nts |
| ENSMUSG( | 332.5423 | -0.38225 | 0.027653 | 0.255168 | Ntsr1 |
| ENSMUSG( | 15135.74 | 0.015664 | 0.888674 | 0.983434 | Ntsr2 |
| ENSMUSG( | 1956.273 | 0.21044 | 0.08669 | 0.452707 | Nuak1 |
| ENSMUSG( | 31.61192 | -0.42493 | 0.262699 | 0.713259 | Nuak2 |
| ENSMUSG( | 2458.053 | 0.054239 | 0.536835 | 0.882123 | Nub1 |
| ENSMUSG( | 506.4902 | -0.05994 | 0.62255 | 0.911432 | Nubp1 |
| ENSMUSG( | 933.008 | 0.11409 | 0.241731 | 0.689179 | Nubp2 |
| ENSMUSG( | 149.9321 | -0.23087 | 0.207545 | 0.64962 | Nubpl |
| ENSMUSG( | 3434.475 | -0.26737 | 0.027306 | 0.253597 | Nucb1 |
| ENSMUSG( | 509.8105 | -0.30825 | 0.014893 | 0.179275 | Nucb2 |
| ENSMUSG( | 2806.364 | 0.016897 | 0.851576 | 0.973046 | Nucks1 |
| ENSMUSG( | 3929.832 | -0.0019 | 0.981665 | 0.997512 | Nudc |
| ENSMUSG( | 282.3328 | 0.143518 | 0.295416 | 0.743814 | Nudcd1 |
| ENSMUSG( | 557.6909 | 0.032037 | 0.832521 | 0.970333 | Nudcd2 |
| ENSMUSG( | 3963.35 | 0.005581 | 0.951048 | 0.993821 | Nudcd3 |
| ENSMUSG( | 155.9462 | 0.078607 | 0.659438 | 0.924506 | Nudt1 |
| ENSMUSG( | 397.2792 | 0.039767 | 0.763741 | 0.954013 | Nudt10 |
| ENSMUSG( | 675.9869 | 0.010414 | 0.933161 | 0.99297 | Nudt11 |
| ENSMUSG( | 189.0995 | 0.217502 | 0.275922 | 0.725669 | Nudt12 |
| ENSMUSG( | 520.3273 | 0.024055 | 0.833926 | 0.970333 | Nudt13 |
| ENSMUSG( | 164.0442 | -0.08764 | 0.597544 | 0.903694 | Nudt14 |
| ENSMUSG( | 125.0412 | -0.02718 | 0.880352 | 0.981469 | Nudt15 |
| ENSMUSG( | 321.0437 | -0.15187 | 0.304617 | 0.752574 | Nudt16 |
| ENSMUSG( | 1259.769 | 0.022662 | 0.852239 | 0.973168 | Nudt16l1 |
| ENSMUSG( | 146.2758 | 0.097776 | 0.582525 | 0.897918 | Nudt17 |
| ENSMUSG( | 1171.566 | 0.020903 | 0.848511 | 0.972374 | Nudt18 |
| ENSMUSG( | 3131.702 | 0.080921 | 0.441294 | 0.832449 | Nudt19 |
| ENSMUSG( | 450.4133 | 0.053157 | 0.700995 | 0.93743 | Nudt2 |
| ENSMUSG( | 1196.377 | 0.041237 | 0.697463 | 0.935662 | Nudt21 |
| ENSMUSG( | 159.033 | 0.201315 | 0.297488 | 0.746042 | Nudt22 |
| ENSMUSG( | 13746.91 | 0.054488 | 0.561702 | 0.889052 | Nudt3 |
| ENSMUSG( | 7644.8 | 0.059087 | 0.514518 | 0.872055 | Nudt4 |

| | | | | | |
|---|---|---|---|---|---|
| ENSMUSG( | 507.3759 | -0.03727 | 0.780792 | 0.976106 | Nucb2 |
| ENSMUSG( | 2491.768 | -0.06154 | 0.666707 | 0.956766 | Nucks1 |
| ENSMUSG( | 3240.066 | -0.30808 | 0.005271 | 0.297824 | Nudc |
| ENSMUSG( | 257.2985 | 0.133728 | 0.429895 | 0.917257 | Nudcd1 |
| ENSMUSG( | 496.1043 | -0.04472 | 0.774553 | 0.974931 | Nudcd2 |
| ENSMUSG( | 3640.617 | 0.031153 | 0.776877 | 0.975717 | Nudcd3 |
| ENSMUSG( | 124.9246 | -0.30986 | 0.119061 | 0.813384 | Nudt1 |
| ENSMUSG( | 352.1525 | -0.05207 | 0.758958 | 0.971858 | Nudt10 |
| ENSMUSG( | 619.155 | 0.016789 | 0.926764 | 0.99306 | Nudt11 |
| ENSMUSG( | 166.6036 | 0.129488 | 0.516823 | 0.930939 | Nudt12 |
| ENSMUSG( | 461.4396 | -0.05529 | 0.688157 | 0.959742 | Nudt13 |
| ENSMUSG( | 154.5559 | -0.0004 | 0.9984 | 0.999313 | Nudt14 |
| ENSMUSG( | 109.7427 | -0.15885 | 0.474011 | 0.923329 | Nudt15 |
| ENSMUSG( | 292.8673 | -0.15772 | 0.363019 | 0.914223 | Nudt16 |
| ENSMUSG( | 1076.589 | -0.17068 | 0.228262 | 0.868507 | Nudt16l1 |
| ENSMUSG( | 123.3647 | -0.13779 | 0.521256 | 0.932489 | Nudt17 |
| ENSMUSG( | 1015.271 | -0.12778 | 0.322387 | 0.904023 | Nudt18 |
| ENSMUSG( | 2578.811 | -0.2215 | 0.096068 | 0.796133 | Nudt19 |
| ENSMUSG( | 354.0787 | -0.41224 | 0.015772 | 0.502615 | Nudt2 |
| ENSMUSG( | 1108.041 | 0.086474 | 0.486006 | 0.92541 | Nudt21 |
| ENSMUSG( | 143.9491 | 0.185527 | 0.348068 | 0.911652 | Nudt22 |
| ENSMUSG( | 11810.71 | -0.11908 | 0.372486 | 0.915491 | Nudt3 |
| ENSMUSG( | 6762.531 | -0.02824 | 0.837581 | 0.981335 | Nudt4 |
| ENSMUSG( | 347.3667 | 0.007474 | 0.958062 | 0.994798 | Nudt5 |
| ENSMUSG( | 94.38392 | -0.08598 | 0.718169 | 0.966441 | Nudt6 |
| ENSMUSG( | 164.04 | 0.23557 | 0.202324 | 0.864412 | Nudt7 |
| ENSMUSG( | 17.73471 | 0.00589 | 0.990236 | 0.998104 | Nudt8 |
| ENSMUSG( | 2477.825 | -0.03289 | 0.76721 | 0.974236 | Nudt9 |
| ENSMUSG( | 297.3039 | -0.07238 | 0.719951 | 0.966887 | Nuf2 |
| ENSMUSG( | 211.8018 | -0.09054 | 0.631657 | 0.953251 | Nufip1 |
| ENSMUSG( | 487.2928 | 0.212124 | 0.131723 | 0.822912 | Nufip2 |
| ENSMUSG( | 2122.105 | -0.00566 | 0.958646 | 0.994798 | Numa1 |
| ENSMUSG( | 1212.556 | -0.02886 | 0.830069 | 0.980224 | Numb |
| ENSMUSG( | 2346.478 | -0.06195 | 0.622661 | 0.953234 | Numbl |
| ENSMUSG( | 329.9853 | -0.03679 | 0.819815 | 0.979198 | Nup107 |
| ENSMUSG( | 633.8751 | 0.004567 | 0.970241 | 0.99594 | Nup133 |
| ENSMUSG( | 1065.222 | 0.227062 | 0.052016 | 0.712974 | Nup153 |
| ENSMUSG( | 492.9125 | 0.205741 | 0.127114 | 0.817278 | Nup155 |
| ENSMUSG( | 353.7887 | 0.089185 | 0.583 | 0.944266 | Nup160 |
| ENSMUSG( | 679.2503 | 0.074018 | 0.567927 | 0.941777 | Nup188 |
| ENSMUSG( | 722.869 | 0.10028 | 0.412754 | 0.917257 | Nup205 |
| ENSMUSG( | 1049.055 | 0.091534 | 0.497638 | 0.927825 | Nup210 |
| ENSMUSG( | 23.23969 | -0.36214 | 0.394124 | 0.917257 | Nup210l |
| ENSMUSG( | 1057.643 | 0.132395 | 0.302893 | 0.898538 | Nup214 |
| ENSMUSG( | 204.3071 | 0.00736 | 0.966527 | 0.995475 | Nup35 |
| ENSMUSG( | 68.64717 | 0.109151 | 0.670948 | 0.95694 | Nup37 |
| ENSMUSG( | 194.4299 | -0.12736 | 0.460556 | 0.923329 | Nup42 |
| ENSMUSG( | 51.12272 | 0.13172 | 0.653811 | 0.955015 | Nup43 |
| ENSMUSG( | 1006.093 | 0.084288 | 0.497736 | 0.927825 | Nup50 |
| ENSMUSG( | 359.5408 | 0.116218 | 0.400891 | 0.917257 | Nup54 |
| ENSMUSG( | 669.512 | 0.075198 | 0.583934 | 0.944398 | Nup58 |
| ENSMUSG( | 388.853 | 0.025511 | 0.87216 | 0.988907 | Nup62 |
| ENSMUSG( | 11.24224 | -0.28513 | 0.630322 | NA | Nup62cl |
| ENSMUSG( | 683.4039 | 0.159926 | 0.258348 | 0.883971 | Nup85 |
| ENSMUSG( | 1110.77 | 0.092981 | 0.538093 | 0.937216 | Nup88 |
| ENSMUSG( | 544.3454 | -0.02822 | 0.839901 | 0.98221 | Nup93 |
| ENSMUSG( | 938.0885 | 0.088213 | 0.476771 | 0.923329 | Nup98 |
| ENSMUSG( | 488.3207 | -0.08634 | 0.541725 | 0.937841 | Nupr1 |
| ENSMUSG( | 44.18155 | 0.354666 | 0.241709 | 0.876528 | Nupr2 |
| ENSMUSG( | 2873.643 | 0.122341 | 0.367143 | 0.915058 | Nus1 |
| ENSMUSG( | 40.33233 | -0.16121 | 0.652833 | 0.954897 | Nusap1 |
| ENSMUSG( | 378.2196 | -0.10381 | 0.485285 | 0.92541 | Nutf2 |
| ENSMUSG( | 1421.171 | 0.200755 | 0.190614 | 0.857801 | Nvl |
| ENSMUSG( | 13323 | -0.16656 | 0.197555 | 0.862161 | Nwd1 |
| ENSMUSG( | 873.2164 | -0.08093 | 0.504647 | 0.927825 | Nwd2 |
| ENSMUSG( | 1977.707 | 0.200465 | 0.221394 | 0.866104 | Nxf1 |
| ENSMUSG( | 27.49927 | -0.28445 | 0.519319 | 0.931392 | Nxf7 |
| ENSMUSG( | 938.5938 | 0.518509 | 1.83E-05 | 0.005843 | Nxn |
| ENSMUSG( | 13.12135 | 0.330965 | 0.542553 | 0.938034 | Nxpe2 |
| ENSMUSG( | 542.1063 | 0.248999 | 0.173819 | 0.844317 | Nxpe3 |
| ENSMUSG( | 593.2313 | -0.09321 | 0.566869 | 0.941777 | Nxpe4 |
| ENSMUSG( | 900.7394 | 0.141581 | 0.490407 | 0.926343 | Nxph1 |
| ENSMUSG( | 46.14329 | 0.332865 | 0.395424 | 0.917257 | Nxph2 |
| ENSMUSG( | 1544.711 | 0.144091 | 0.361149 | 0.914223 | Nxph3 |
| ENSMUSG( | 288.1013 | -0.01853 | 0.917255 | 0.992529 | Nxph4 |
| ENSMUSG( | 129.5693 | 0.005198 | 0.979014 | 0.996468 | Nxt1 |
| ENSMUSG( | 1175.985 | -0.0521 | 0.755014 | 0.971098 | Nxt2 |
| ENSMUSG( | 2769.545 | 0.089898 | 0.378671 | 0.916995 | Nyap1 |
| ENSMUSG( | 317.38 | -0.12531 | 0.389565 | 0.917257 | Nyap2 |
| ENSMUSG( | 400.065 | 0.147359 | 0.375189 | 0.915491 | Nynrin |
| ENSMUSG( | 18.17997 | 0.537716 | 0.249556 | 0.877904 | Nyx |
| ENSMUSG( | 57.48488 | -0.5003 | 0.082079 | 0.77681 | Oacyl |
| ENSMUSG( | 949.1431 | 0.347203 | 0.050312 | 0.708648 | Oaf |
| ENSMUSG( | 335.4631 | -0.19673 | 0.213713 | 0.864421 | Oard1 |
| ENSMUSG( | 32.49893 | -0.15876 | 0.656318 | 0.955533 | Oas1a |
| ENSMUSG( | 21.98164 | 0.242778 | 0.590276 | 0.944975 | Oas1b |
| ENSMUSG( | 31.8141 | -0.26294 | 0.464234 | 0.923329 | Oas1c |
| ENSMUSG( | 23.09324 | -0.68456 | 0.163782 | 0.838291 | Oas2 |
| ENSMUSG( | 9.248201 | -0.53522 | 0.447957 | NA | Oas3 |
| ENSMUSG( | 10.71842 | -0.19919 | 0.752232 | NA | Oasl1 |
| ENSMUSG( | 188.3729 | -0.28435 | 0.159644 | 0.837724 | Oasl2 |
| ENSMUSG( | 3788.462 | -0.17201 | 0.161146 | 0.837724 | Oat |
| ENSMUSG( | 6285.329 | -0.12326 | 0.245265 | 0.877904 | Oaz1 |
| ENSMUSG( | 3640.607 | -0.10303 | 0.33835 | 0.907972 | Oaz2 |
| ENSMUSG( | 213.6699 | -0.01274 | 0.945861 | 0.994798 | Obi1 |
| ENSMUSG( | 238.2989 | -0.04615 | 0.797473 | 0.97681 | Obscn |
| ENSMUSG( | 543.7142 | 0.248451 | 0.083597 | 0.77681 | Obsl1 |
| ENSMUSG( | 26.09099 | 0.686488 | 0.129919 | 0.820944 | Oca2 |
| ENSMUSG( | 563.3357 | 0.21656 | 0.145202 | 0.83339 | Ocel1 |
| ENSMUSG( | 9296.302 | -0.02642 | 0.8302 | 0.980224 | Ociad1 |
| ENSMUSG( | 1589.219 | 0.016269 | 0.885115 | 0.989688 | Ociad2 |
| ENSMUSG( | 389.7562 | 0.712232 | 0.149662 | 0.836241 | Ocln |
| ENSMUSG( | 15.22848 | 0.166955 | 0.749238 | 0.96998 | Ocm |
| ENSMUSG( | 1819.473 | 0.107867 | 0.349684 | 0.911652 | Ocrl |
| ENSMUSG( | 128.2307 | 0.017702 | 0.936291 | 0.993573 | Odad1 |
| ENSMUSG( | 17.88945 | -0.10357 | 0.854268 | 0.98563 | Odad2 |
| ENSMUSG( | 105.1733 | 0.144906 | 0.510933 | 0.929546 | Odad3 |
| ENSMUSG( | 49.45499 | 0.14926 | 0.672675 | 0.95694 | Odad4 |
| ENSMUSG( | 1805.948 | -0.19916 | 0.130522 | 0.822225 | Odc1 |
| ENSMUSG( | 2174.054 | 0.1163 | 0.390995 | 0.917257 | Odf2 |
| ENSMUSG( | 319.3429 | -0.03189 | 0.861368 | 0.986281 | Odf2l |
| ENSMUSG( | 2172.402 | -0.03982 | 0.746958 | 0.969763 | Odr4 |
| ENSMUSG( | 252.5936 | -0.15317 | 0.366251 | 0.914703 | Ofd1 |
| ENSMUSG( | 4734.707 | -0.14995 | 0.136848 | 0.828492 | Oga |
| ENSMUSG( | 7874.08 | -0.18123 | 0.066 | 0.746371 | Ogdh |
| ENSMUSG( | 4625.888 | -0.10122 | 0.316054 | 0.901868 | Ogdhl |
| ENSMUSG( | 2609.723 | 0.022056 | 0.841028 | 0.9826 | Ogfod1 |
| ENSMUSG( | 685.0405 | 0.004659 | 0.972795 | 0.996416 | Ogfod2 |
| ENSMUSG( | 299.5371 | 0.064908 | 0.718834 | 0.966668 | Ogfod3 |
| ENSMUSG( | 1412.174 | -0.02777 | 0.824362 | 0.979198 | Ogfr |
| ENSMUSG( | 2577.412 | 0.216743 | 0.117242 | 0.813384 | Ogfrl1 |
| ENSMUSG( | 182.2343 | 0.084861 | 0.633158 | 0.953844 | Ogg1 |
| ENSMUSG( | 363.6704 | 0.268193 | 0.09866 | 0.798438 | Ogn |
| ENSMUSG( | 6345.55 | 0.215888 | 0.19146 | 0.858412 | Ogt |
| ENSMUSG( | 17.25875 | 0.981577 | 0.040188 | 0.681706 | Oip5 |
| ENSMUSG( | 13743.18 | -0.04314 | 0.711482 | 0.964363 | Oip5os1 |
| ENSMUSG( | 1436.803 | -0.10987 | 0.460226 | 0.923329 | Ola1 |
| ENSMUSG( | 45937.09 | -0.04446 | 0.714447 | 0.964986 | Olfm1 |
| ENSMUSG( | 6533.597 | 0.090473 | 0.522291 | 0.932681 | Olfm2 |
| ENSMUSG( | 667.5722 | 0.234306 | 0.298645 | 0.898538 | Olfm3 |
| ENSMUSG( | 34.54222 | -0.21719 | 0.531996 | 0.934795 | Olfm4 |
| ENSMUSG( | 675.865 | 0.008952 | 0.960624 | 0.995047 | Olfml1 |
| ENSMUSG( | 212.11 | -0.08829 | 0.671405 | 0.95694 | Olfml2a |
| ENSMUSG( | 391.6296 | 0.708874 | 0.007462 | 0.367159 | Olfml2b |
| ENSMUSG( | 1541.983 | 0.078029 | 0.700086 | 0.961335 | Olfml3 |
| ENSMUSG( | 8629.857 | -0.24919 | 0.047457 | 0.700936 | Olig1 |
| ENSMUSG( | 1389.313 | -0.02288 | 0.887323 | 0.990266 | Olig2 |
| ENSMUSG( | 227.546 | -0.09084 | 0.589135 | 0.944975 | Oma1 |
| ENSMUSG( | 37.05948 | 0.671079 | 0.061128 | 0.736111 | Omd |
| ENSMUSG( | 4458.351 | 0.206966 | 0.301306 | 0.898538 | Omg |
| ENSMUSG( | 108.1038 | 0.819761 | 0.581272 | 0.944266 | Omp |
| ENSMUSG( | 79.50093 | -0.26394 | 0.298804 | 0.898538 | Onecut2 |
| ENSMUSG( | 3992.591 | -0.03363 | 0.730439 | 0.967228 | Opa1 |
| ENSMUSG( | 2252.494 | -0.12924 | 0.220639 | 0.866104 | Opa3 |
| ENSMUSG( | 1262.729 | -0.23712 | 0.194388 | 0.860096 | Opalin |
| ENSMUSG( | 5918.232 | 0.193501 | 0.286598 | 0.894091 | Opcml |
| ENSMUSG( | 1350.167 | 0.086513 | 0.5126 | 0.930164 | Ophn1 |
| ENSMUSG( | 1614.829 | -0.04642 | 0.695835 | 0.960867 | Oplah |
| ENSMUSG( | 142.8195 | 0.134151 | 0.526902 | 0.934449 | Opn3 |
| ENSMUSG( | 238.8881 | 0.187764 | 0.47862 | 0.923329 | Oprd1 |
| ENSMUSG( | 267.0119 | 0.228212 | 0.25375 | 0.880692 | Oprk1 |
| ENSMUSG( | 1082.424 | 0.27868 | 0.066903 | 0.751729 | Oprl1 |
| ENSMUSG( | 12.96103 | 0.245726 | 0.654712 | 0.955015 | Oprm1 |
| ENSMUSG( | 33.88062 | 0.049231 | 0.884929 | 0.989688 | Optc |
| ENSMUSG( | 453.8398 | -0.07808 | 0.588059 | 0.944975 | Optn |
| ENSMUSG( | 515.4452 | -0.09114 | 0.593659 | 0.945408 | Or10ad1 |
| ENSMUSG( | 46.82857 | -0.05265 | 0.864733 | 0.986591 | Or2ak4 |
| ENSMUSG( | 26.64285 | -0.13456 | 0.770953 | 0.974236 | Or2y1g |

| | | | | | | | | | | | |
|---|---|---|---|---|---|---|---|---|---|---|---|
| ENSMUSG( | Ormdl1 | 178.743 | -0.29643 | 0.115377 | 0.512362 | ENSMUSG( | Ormdl1 | 189.0626 | 0.154847 | 0.494826 | 0.927372 |
| ENSMUSG( | Ormdl2 | 288.5841 | -0.08945 | 0.600504 | 0.90443 | ENSMUSG( | Ormdl2 | 276.0338 | 0.047615 | 0.776669 | 0.975717 |
| ENSMUSG( | Ormdl3 | 2726.992 | -0.14586 | 0.181487 | 0.618685 | ENSMUSG( | Ormdl3 | 2642.344 | 0.037384 | 0.768686 | 0.974236 |
| ENSMUSG( | Os9 | 6640.905 | -0.25841 | 0.023781 | 0.235641 | ENSMUSG( | Os9 | 6896.928 | 0.130872 | 0.329367 | 0.907083 |
| ENSMUSG( | Osbp | 2259.892 | 0.04487 | 0.615054 | 0.910318 | ENSMUSG( | Osbp | 2077.938 | 0.07155 | 0.500194 | 0.927825 |
| ENSMUSG( | Osbp2 | 2814.769 | 0.067074 | 0.545118 | 0.884712 | ENSMUSG( | Osbp2 | 2550.402 | 0.052268 | 0.661143 | 0.956205 |
| ENSMUSG( | Osbpl10 | 697.9335 | 0.060804 | 0.640326 | 0.916434 | ENSMUSG( | Osbpl10 | 617.0427 | -0.01931 | 0.894683 | 0.990558 |
| ENSMUSG( | Osbpl11 | 552.6876 | -0.15639 | 0.162096 | 0.592429 | ENSMUSG( | Osbpl11 | 525.4153 | -0.0251 | 0.854856 | 0.985649 |
| ENSMUSG( | Osbpl1a | 8165.591 | 0.033622 | 0.693424 | 0.934202 | ENSMUSG( | Osbpl1a | 6972.717 | -0.15908 | 0.134528 | 0.823094 |
| ENSMUSG( | Osbpl2 | 1636.717 | 0.052664 | 0.585143 | 0.899171 | ENSMUSG( | Osbpl2 | 1482.623 | 0.034056 | 0.781136 | 0.976106 |
| ENSMUSG( | Osbpl3 | 838.0628 | 0.106725 | 0.394884 | 0.810072 | ENSMUSG( | Osbpl3 | 791.3599 | 0.204499 | 0.090457 | 0.791074 |
| ENSMUSG( | Osbpl5 | 921.2652 | 0.234726 | 0.052779 | 0.352597 | ENSMUSG( | Osbpl5 | 770.3099 | -0.0034 | 0.980092 | 0.996479 |
| ENSMUSG( | Osbpl6 | 1610.204 | 0.227968 | 0.058089 | 0.371677 | ENSMUSG( | Osbpl6 | 1406.024 | 0.109243 | 0.423392 | 0.917257 |
| ENSMUSG( | Osbpl7 | 468.6924 | -0.01718 | 0.896461 | 0.984868 | ENSMUSG( | Osbpl7 | 432.9852 | 0.019616 | 0.891049 | 0.990558 |
| ENSMUSG( | Osbpl8 | 1472.842 | -0.08137 | 0.397586 | 0.812127 | ENSMUSG( | Osbpl8 | 1463.464 | 0.162386 | 0.263531 | 0.886134 |
| ENSMUSG( | Osbpl9 | 1342.379 | 0.060862 | 0.51349 | 0.871762 | ENSMUSG( | Osbpl9 | 1259.766 | 0.14235 | 0.216391 | 0.865167 |
| ENSMUSG( | Oscar | 71.52519 | 0.345514 | 0.243103 | 0.690443 | ENSMUSG( | Oscar | 63.31238 | 0.243936 | 0.452842 | 0.922316 |
| ENSMUSG( | Oscp1 | 1043.654 | -0.05465 | 0.559316 | 0.888638 | ENSMUSG( | Oscp1 | 973.8465 | 0.014905 | 0.893909 | 0.990558 |
| ENSMUSG( | Oser1 | 126.3927 | -0.09644 | 0.613549 | 0.909044 | ENSMUSG( | Oser1 | 122.0184 | 0.077189 | 0.725919 | 0.966887 |
| ENSMUSG( | Osgep | 901.4706 | 0.063623 | 0.539694 | 0.88315 | ENSMUSG( | Osgep | 734.9431 | -0.27303 | 0.035273 | 0.661159 |
| ENSMUSG( | Osgepl1 | 66.50375 | -0.52542 | 0.069375 | 0.40782 | ENSMUSG( | Osgepl1 | 80.91866 | 0.361468 | 0.258958 | 0.883971 |
| ENSMUSG( | Osgin1 | 13.85954 | -0.85039 | 0.094126 | NA | ENSMUSG( | Osgin1 | 19.47046 | 0.506782 | 0.271758 | 0.888132 |
| ENSMUSG( | Osgin2 | 564.3748 | 0.054062 | 0.609591 | 0.9077 | ENSMUSG( | Osgin2 | 501.5826 | -0.02017 | 0.875703 | 0.989644 |
| ENSMUSG( | Osmr | 264.3928 | 0.18326 | 0.321253 | 0.766497 | ENSMUSG( | Osmr | 217.0736 | -0.09893 | 0.665368 | 0.956766 |
| ENSMUSG( | Osr1 | 82.27286 | -0.21162 | 0.369802 | 0.797313 | ENSMUSG( | Osr1 | 74.67211 | -0.2388 | 0.37189 | 0.915491 |
| ENSMUSG( | Ost4 | 1436.581 | -0.0085 | 0.926274 | 0.991229 | ENSMUSG( | Ost4 | 1191.117 | -0.3 | 0.016835 | 0.522883 |
| ENSMUSG( | Ostc | 1214.466 | -0.34832 | 0.006641 | 0.112089 | ENSMUSG( | Ostc | 1289.501 | 0.113745 | 0.421224 | 0.917257 |
| ENSMUSG( | Ostf1 | 575.8882 | -0.17164 | 0.152944 | 0.579346 | ENSMUSG( | Ostf1 | 541.2331 | -0.07567 | 0.568056 | 0.941777 |
| ENSMUSG( | Ostm1 | 2006.76 | -0.35134 | 0.01323 | 0.167129 | ENSMUSG( | Ostm1 | 2168.459 | 0.161799 | 0.289983 | 0.896726 |
| ENSMUSG( | Otoa | 12.77106 | 0.703321 | 0.210623 | NA | ENSMUSG( | Otoa | 10.17899 | 0.336694 | 0.593698 | NA |
| ENSMUSG( | Otof | 675.7485 | -0.09596 | 0.638893 | 0.916418 | ENSMUSG( | Otof | 606.8449 | -0.14621 | 0.560665 | 0.940254 |
| ENSMUSG( | Otogl | 12.40966 | 0.141495 | 0.795696 | NA | ENSMUSG( | Otogl | 11.67635 | 0.236813 | 0.685248 | 0.958678 |
| ENSMUSG( | Otop2 | 34.10886 | -0.73986 | 0.023385 | 0.233882 | ENSMUSG( | Otop2 | 33.55613 | -0.51753 | 0.179243 | 0.849741 |
| ENSMUSG( | Otub1 | 4222.892 | 0.030152 | 0.798999 | 0.960922 | ENSMUSG( | Otub1 | 3600.069 | -0.16885 | 0.243767 | 0.877904 |
| ENSMUSG( | Otub2 | 1002.193 | 0.189809 | 0.083737 | 0.446196 | ENSMUSG( | Otub2 | 868.9703 | 0.055192 | 0.656419 | 0.955533 |
| ENSMUSG( | Otud1 | 412.1051 | -0.17067 | 0.205104 | 0.646229 | ENSMUSG( | Otud1 | 407.5179 | 0.063339 | 0.700895 | 0.96162 |
| ENSMUSG( | Otud3 | 319.2434 | 0.069375 | 0.615422 | 0.910481 | ENSMUSG( | Otud3 | 268.3567 | -0.16682 | 0.303265 | 0.898538 |
| ENSMUSG( | Otud4 | 1307.273 | -0.16632 | 0.07959 | 0.435679 | ENSMUSG( | Otud4 | 1268.39 | 0.020881 | 0.846933 | 0.985027 |
| ENSMUSG( | Otud5 | 2937.928 | 0.142123 | 0.083518 | 0.445612 | ENSMUSG( | Otud5 | 2551.275 | 0.005856 | 0.954898 | 0.994798 |
| ENSMUSG( | Otud6b | 2515.93 | 0.139447 | 0.274499 | 0.724787 | ENSMUSG( | Otud6b | 2118.544 | -0.08964 | 0.532824 | 0.934795 |
| ENSMUSG( | Otud7a | 1489.916 | 0.26882 | 0.02333 | 0.233882 | ENSMUSG( | Otud7a | 1260.247 | 0.069424 | 0.595183 | 0.945701 |
| ENSMUSG( | Otud7b | 2976.92 | -0.00523 | 0.951975 | 0.993821 | ENSMUSG( | Otud7b | 2690.258 | -0.0303 | 0.797115 | 0.97681 |
| ENSMUSG( | Otulin | 684.1343 | 0.027193 | 0.80645 | 0.962804 | ENSMUSG( | Otulin | 589.2056 | -0.14389 | 0.282966 | 0.892412 |
| ENSMUSG( | Otulinl | 672.0144 | -0.3462 | 0.037742 | 0.30023 | ENSMUSG( | Otulinl | 742.3336 | 0.229314 | 0.201073 | 0.86439 |
| ENSMUSG( | Otx1 | 246.243 | -0.01709 | 0.906353 | 0.986998 | ENSMUSG( | Otx1 | 240.4888 | 0.166089 | 0.37538 | 0.915491 |
| ENSMUSG( | Otx2 | 51.47999 | -0.47072 | 0.641347 | 0.916693 | ENSMUSG( | Otx2 | 52.05564 | -0.14627 | 0.868152 | 0.98788 |
| ENSMUSG( | Ovca2 | 293.3432 | 0.058153 | 0.659491 | 0.924506 | ENSMUSG( | Ovca2 | 249.9226 | -0.1279 | 0.446581 | 0.921734 |
| ENSMUSG( | Ovgp1 | 106.9787 | 0.390627 | 0.161477 | 0.59148 | ENSMUSG( | Ovgp1 | 97.62942 | 0.397857 | 0.230956 | 0.868507 |
| ENSMUSG( | Ovol2 | 132.5283 | -0.06429 | 0.797855 | 0.96053 | ENSMUSG( | Ovol2 | 106.0662 | -0.49332 | 0.041138 | 0.687358 |
| ENSMUSG( | Oxa1l | 1872.195 | -0.03572 | 0.693041 | 0.934202 | ENSMUSG( | Oxa1l | 1654.604 | -0.12784 | 0.291819 | 0.897854 |
| ENSMUSG( | Oxct1 | 14931.27 | 0.131116 | 0.168993 | 0.603241 | ENSMUSG( | Oxct1 | 12336.67 | -0.15391 | 0.219906 | 0.866104 |
| ENSMUSG( | Oxld1 | 128.4377 | -0.06371 | 0.732089 | 0.944956 | ENSMUSG( | Oxld1 | 108.7508 | -0.2944 | 0.176588 | 0.846973 |
| ENSMUSG( | Oxnad1 | 600.9802 | 0.066524 | 0.561974 | 0.889052 | ENSMUSG( | Oxnad1 | 516.2541 | -0.10944 | 0.42784 | 0.917257 |
| ENSMUSG( | Oxr1 | 4090.294 | 0.137292 | 0.134805 | 0.54834 | ENSMUSG( | Oxr1 | 3544.441 | -0.00713 | 0.953525 | 0.994798 |
| ENSMUSG( | Oxsm | 328.6633 | 0.046835 | 0.745457 | 0.949158 | ENSMUSG( | Oxsm | 282.5185 | -0.12307 | 0.457844 | 0.923329 |
| ENSMUSG( | Oxsr1 | 4484.675 | -0.05898 | 0.492223 | 0.860297 | ENSMUSG( | Oxsr1 | 3806.194 | -0.27863 | 0.024578 | 0.592201 |
| ENSMUSG( | Oxtr | 270.1516 | -0.06391 | 0.68407 | 0.932484 | ENSMUSG( | Oxtr | 275.3264 | 0.249442 | 0.109633 | 0.806227 |
| ENSMUSG( | P2rx1 | 23.83939 | -1.12174 | 0.005727 | 0.101308 | ENSMUSG( | P2rx1 | 22.00668 | -1.00573 | 0.031406 | 0.631192 |
| ENSMUSG( | P2rx3 | 55.64878 | 0.054978 | 0.828641 | 0.969097 | ENSMUSG( | P2rx3 | 49.50001 | 0.003414 | 0.990711 | 0.998287 |
| ENSMUSG( | P2rx4 | 571.8559 | -0.29964 | 0.01806 | 0.202299 | ENSMUSG( | P2rx4 | 585.1576 | 0.055831 | 0.692222 | 0.960404 |
| ENSMUSG( | P2rx5 | 25.44991 | -0.3527 | 0.363508 | 0.793339 | ENSMUSG( | P2rx5 | 24.69684 | -0.17064 | 0.677588 | 0.957106 |
| ENSMUSG( | P2rx6 | 102.569 | -0.29234 | 0.258014 | 0.709319 | ENSMUSG( | P2rx6 | 100.3989 | -0.04495 | 0.879802 | 0.989644 |
| ENSMUSG( | P2rx7 | 457.7173 | -0.329 | 0.027656 | 0.255168 | ENSMUSG( | P2rx7 | 480.6777 | 0.104168 | 0.542608 | 0.938034 |
| ENSMUSG( | P2ry1 | 65.61439 | -0.62154 | 0.012791 | 0.163486 | ENSMUSG( | P2ry1 | 86.49368 | 0.477162 | 0.043645 | 0.690304 |
| ENSMUSG( | P2ry10b | 10.11119 | -0.21674 | 0.704826 | NA | ENSMUSG( | P2ry10b | 11.32058 | 0.364803 | 0.521886 | NA |
| ENSMUSG( | P2ry12 | 1845.237 | -0.378 | 0.034639 | 0.285926 | ENSMUSG( | P2ry12 | 2111.842 | 0.295253 | 0.159513 | 0.837724 |
| ENSMUSG( | P2ry13 | 667.5492 | -0.59192 | 0.004455 | 0.086961 | ENSMUSG( | P2ry13 | 833.7998 | 0.365315 | 0.13823 | 0.828492 |
| ENSMUSG( | P2ry14 | 111.1355 | -0.13863 | 0.594827 | 0.903007 | ENSMUSG( | P2ry14 | 133.1146 | 0.613547 | 0.042782 | 0.690304 |
| ENSMUSG( | P2ry2 | 27.73503 | 0.009067 | 0.982412 | 0.997512 | ENSMUSG( | P2ry2 | 30.27225 | 0.504383 | 0.263956 | 0.886134 |
| ENSMUSG( | P2ry6 | 156.0238 | -0.13151 | 0.608203 | 0.907154 | ENSMUSG( | P2ry6 | 163.8961 | 0.284097 | 0.279182 | 0.892356 |
| ENSMUSG( | P3h1 | 334.4806 | -0.0397 | 0.778463 | 0.956215 | ENSMUSG( | P3h1 | 339.0512 | 0.263695 | 0.093262 | 0.795363 |
| ENSMUSG( | P3h2 | 173.2876 | 0.186132 | 0.272162 | 0.7223 | ENSMUSG( | P3h2 | 171.5038 | 0.409612 | 0.026782 | 0.603041 |
| ENSMUSG( | P3h3 | 876.6731 | 0.094622 | 0.346527 | 0.779166 | ENSMUSG( | P3h3 | 815.6752 | 0.153615 | 0.356792 | 0.913358 |
| ENSMUSG( | P3h4 | 579.7754 | -0.24093 | 0.058414 | 0.372872 | ENSMUSG( | P3h4 | 603.4273 | 0.155593 | 0.309947 | 0.898538 |
| ENSMUSG( | P4ha1 | 2526.872 | -0.12279 | 0.416393 | 0.820842 | ENSMUSG( | P4ha1 | 2528.933 | 0.150121 | 0.332608 | 0.907907 |
| ENSMUSG( | P4ha2 | 368.1449 | 0.295899 | 0.023119 | 0.233201 | ENSMUSG( | P4ha2 | 380.8407 | 0.61761 | 2.63E-05 | 0.007461 |
| ENSMUSG( | P4ha3 | 350.898 | -0.24616 | 0.091892 | 0.465937 | ENSMUSG( | P4ha3 | 326.5597 | -0.18174 | 0.221925 | 0.866104 |
| ENSMUSG( | P4hb | 5400.44 | -0.36472 | 0.007975 | 0.124256 | ENSMUSG( | P4hb | 5822.229 | 0.144238 | 0.346095 | 0.910966 |
| ENSMUSG( | P4htm | 2406.247 | -0.22593 | 0.080939 | 0.439809 | ENSMUSG( | P4htm | 2561.924 | 0.228202 | 0.150922 | 0.837173 |
| ENSMUSG( | Pa2g4 | 1798.908 | 0.067249 | 0.507873 | 0.86795 | ENSMUSG( | Pa2g4 | 1540.493 | -0.11683 | 0.392665 | 0.917257 |
| ENSMUSG( | Pabir1 | 250.8912 | -0.01098 | 0.947683 | 0.993821 | ENSMUSG( | Pabir1 | 237.8224 | 0.091882 | 0.622338 | 0.953234 |
| ENSMUSG( | Pabir2 | 218.8725 | -0.06259 | 0.679828 | 0.930983 | ENSMUSG( | Pabir2 | 193.4709 | -0.17114 | 0.386094 | 0.917257 |
| ENSMUSG( | Pabpc1 | 11318.29 | 0.079593 | 0.438517 | 0.830637 | ENSMUSG( | Pabpc1 | 9834.715 | -0.05646 | 0.621644 | 0.953199 |
| ENSMUSG( | Pabpc1l | 22.84123 | -0.31236 | 0.451474 | 0.835567 | ENSMUSG( | Pabpc1l | 19.17555 | -0.59068 | 0.199349 | 0.863016 |
| ENSMUSG( | Pabpc4 | 1463.831 | 0.025215 | 0.809424 | 0.963674 | ENSMUSG( | Pabpc4 | 1317.547 | -0.00981 | 0.929622 | 0.99306 |
| ENSMUSG( | Pabpc4l | 10.25778 | -0.09439 | 0.867319 | NA | ENSMUSG( | Pabpc4l | 10.00539 | 0.121661 | 0.841197 | NA |
| ENSMUSG( | Pabpc5 | 11.12523 | -0.36286 | 0.52493 | NA | ENSMUSG( | Pabpc5 | 12.10477 | 0.191142 | 0.746491 | 0.969692 |
| ENSMUSG( | Pabpn1 | 1208.852 | 0.144446 | 0.428396 | 0.826032 | ENSMUSG( | Pabpn1 | 1186.918 | 0.342598 | 0.10654 | 0.804969 |
| ENSMUSG( | Pacc1 | 336.1251 | -0.15369 | 0.42672 | 0.825155 | ENSMUSG( | Pacc1 | 361.4954 | 0.321064 | 0.137746 | 0.828492 |
| ENSMUSG( | Pacrg | 511.1745 | 0.008122 | 0.957859 | 0.993874 | ENSMUSG( | Pacrg | 419.7186 | -0.31237 | 0.027934 | 0.61023 |
| ENSMUSG( | Pacrgl | 215.7422 | 0.093462 | 0.514533 | 0.872055 | ENSMUSG( | Pacrgl | 194.2501 | 0.056902 | 0.746467 | 0.969692 |
| ENSMUSG( | Pacs1 | 2048.459 | 0.21027 | 0.091831 | 0.465904 | ENSMUSG( | Pacs1 | 1721.33 | -0.0166 | 0.904786 | 0.991089 |
| ENSMUSG( | Pacs2 | 13186.09 | -0.02461 | 0.780264 | 0.957491 | ENSMUSG( | Pacs2 | 11392.75 | -0.1855 | 0.084971 | 0.779799 |
| ENSMUSG( | Pacsin1 | 12362.73 | 0.109172 | 0.239233 | 0.687031 | ENSMUSG( | Pacsin1 | 10586.99 | -0.06751 | 0.504405 | 0.927825 |
| ENSMUSG( | Pacsin2 | 1663.155 | 0.036645 | 0.710671 | 0.939759 | ENSMUSG( | Pacsin2 | 1440.022 | -0.11432 | 0.345745 | 0.910877 |
| ENSMUSG( | Pacsin3 | 2147.971 | 0.03653 | 0.663138 | 0.9265 | ENSMUSG( | Pacsin3 | 1882.121 | -0.07975 | 0.497553 | 0.927825 |
| ENSMUSG( | Padi2 | 1984.102 | -0.01848 | 0.84444 | 0.972023 | ENSMUSG( | Padi2 | 1866.049 | 0.071231 | 0.527413 | 0.934466 |
| ENSMUSG( | Padi6 | 40.6829 | 0.227505 | 0.483983 | 0.856501 | ENSMUSG( | Padi6 | 35.0693 | 0.083143 | 0.819566 | 0.979198 |
| ENSMUSG( | Paf1 | 1669.465 | 0.036937 | 0.768021 | 0.954621 | ENSMUSG( | Paf1 | 1493.806 | -0.01873 | 0.900793 | 0.990558 |
| ENSMUSG( | Pafah1b1 | 12593.86 | 0.070795 | 0.364456 | 0.793796 | ENSMUSG( | Pafah1b1 | 10991.61 | -0.05426 | 0.616768 | 0.951636 |
| ENSMUSG( | Pafah1b2 | 3355.894 | 0.016671 | 0.854004 | 0.973491 | ENSMUSG( | Pafah1b2 | 2964.336 | -0.0758 | 0.552769 | 0.939565 |
| ENSMUSG( | Pafah1b3 | 192.2595 | -0.07214 | 0.669265 | 0.928136 | ENSMUSG( | Pafah1b3 | 165.9813 | -0.24603 | 0.209809 | 0.864412 |
| ENSMUSG( | Pafah2 | 441.8403 | 0.134007 | 0.290487 | 0.740483 | ENSMUSG( | Pafah2 | 368.1553 | -0.12803 | 0.379808 | 0.916995 |
| ENSMUSG( | Pag1 | 1035.18 | -0.30931 | 0.025596 | 0.246694 | ENSMUSG( | Pag1 | 1106.258 | 0.167331 | 0.335081 | 0.907907 |
| ENSMUSG( | Pagr1a | 32.34215 | 0.118441 | 0.742883 | 0.948478 | ENSMUSG( | Pagr1a | 28.91376 | 0.08924 | 0.812296 | 0.979148 |
| ENSMUSG( | Paics | 3838.791 | 0.115302 | 0.259618 | 0.709951 | ENSMUSG( | Paics | 3208.727 | -0.13711 | 0.274982 | 0.889874 |
| ENSMUSG( | Paip1 | 2053.597 | 0.031903 | 0.763373 | 0.953781 | ENSMUSG( | Paip1 | 1816.615 | -0.05743 | 0.67616 | 0.95694 |
| ENSMUSG( | Paip2 | 2787.481 | 0.038715 | 0.707305 | 0.93949 | ENSMUSG( | Paip2 | 2406.369 | -0.12168 | 0.331757 | 0.907907 |
| ENSMUSG( | Paip2b | 2327.563 | -0.01153 | 0.901049 | 0.985633 | ENSMUSG( | Paip2b | 2002.326 | -0.18672 | 0.151104 | 0.837173 |
| ENSMUSG( | Pak1 | 7383.858 | 0.154817 | 0.129171 | 0.538324 | ENSMUSG( | Pak1 | 6138.91 | -0.10791 | 0.348623 | 0.911652 |
| ENSMUSG( | Pak1ip1 | 825.3512 | 0.099147 | 0.444025 | 0.833239 | ENSMUSG( | Pak1ip1 | 723.4903 | -0.01238 | 0.933583 | 0.99306 |
| ENSMUSG( | Pak2 | 2360.441 | 0.126966 | 0.260668 | 0.710468 | ENSMUSG( | Pak2 | 2062.746 | 0.007071 | 0.955795 | 0.994798 |
| ENSMUSG( | Pak3 | 1136.855 | 0.097127 | 0.348102 | 0.78132 | ENSMUSG( | Pak3 | 988.3181 | -0.04148 | 0.756697 | 0.971779 |
| ENSMUSG( | Pak4 | 487.5066 | -0.14787 | 0.197385 | 0.637004 | ENSMUSG( | Pak4 | 439.7989 | -0.1794 | 0.168801 | 0.840222 |
| ENSMUSG( | Pak5 | 669.2795 | -0.35295 | 0.000731 | 0.026277 | ENSMUSG( | Pak5 | 606.2658 | -0.37493 | 0.003007 | 0.219289 |
| ENSMUSG( | Pak6 | 5622.984 | 0.183096 | 0.026244 | 0.247758 | ENSMUSG( | Pak6 | 4972.445 | 0.101786 | 0.327156 | 0.905768 |
| ENSMUSG( | Pakap | 103.0999 | 0.2986 | 0.169529 | 0.603608 | ENSMUSG( | Pakap | 90.94988 | 0.213579 | 0.353107 | 0.912263 |
| ENSMUSG( | Palb2 | 29.2244 | -0.04761 | 0.905016 | 0.986449 | ENSMUSG( | Palb2 | 31.21633 | 0.416726 | 0.274927 | 0.889874 |
| ENSMUSG( | Pald1 | 519.433 | -0.10962 | 0.444422 | 0.833239 | ENSMUSG( | Pald1 | 562.5585 | 0.376944 | 0.070049 | 0.757435 |
| ENSMUSG( | Palld | 587.0079 | 0.25103 | 0.128479 | 0.536774 | ENSMUSG( | Palld | 470.4542 | -0.11733 | 0.467439 | 0.923329 |
| ENSMUSG( | Palm | 20210.7 | 0.269982 | 0.002112 | 0.053164 | ENSMUSG( | Palm | 17976.97 | 0.207244 | 0.052677 | 0.71525 |
| ENSMUSG( | Palm3 | 70.16313 | -0.67632 | 0.003081 | 0.068742 | ENSMUSG( | Palm3 | 84.1862 | 0.170046 | 0.482478 | 0.924578 |
| ENSMUSG( | Palmd | 547.4944 | 0.096834 | 0.536158 | 0.881449 | ENSMUSG( | Palmd | 533.8978 | 0.289577 | 0.128265 | 0.818934 |
| ENSMUSG( | Pals1 | 1091.394 | 0.079655 | 0.525287 | 0.876559 | ENSMUSG( | Pals1 | 998.6763 | 0.088064 | 0.538511 | 0.937216 |
| ENSMUSG( | Pals2 | 3180.588 | 0.198868 | 0.079581 | 0.435679 | ENSMUSG( | Pals2 | 2592.012 | -0.1216 | 0.394772 | 0.917257 |
| ENSMUSG( | Pam | 5200.861 | -0.2163 | 0.117156 | 0.516516 | ENSMUSG( | Pam | 5584.635 | 0.258757 | 0.133759 | 0.822912 |
| ENSMUSG( | Pam16 | 632.0082 | -0.01621 | 0.889108 | 0.983434 | ENSMUSG( | Pam16 | 544.1595 | -0.19309 | 0.189309 | 0.85698 |
| ENSMUSG( | Pam16l | 14.54931 | 0.178825 | 0.718946 | NA | ENSMUSG( | Pam16l | 9.872897 | -0.79703 | 0.216634 | NA |
| ENSMUSG( | Pamr1 | 1653.583 | -0.23216 | 0.149609 | 0.573617 | ENSMUSG( | Pamr1 | 1629.63 | 0.010251 | 0.955702 | 0.994798 |
| ENSMUSG( | Pan2 | 750.1519 | 0.135698 | 0.334593 | 0.776386 | ENSMUSG( | Pan2 | 677.3104 | 0.111937 | 0.542009 | 0.937993 |
| ENSMUSG( | Pan3 | 1198.608 | 0.079962 | 0.382429 | 0.803227 | ENSMUSG( | Pan3 | 1134.817 | 0.186611 | 0.109733 | 0.80637 |
| ENSMUSG( | Panct2 | 74.27845 | -0.08374 | 0.712305 | 0.94031 | ENSMUSG( | Panct2 | 70.58836 | 0.040898 | 0.871474 | 0.988779 |
| ENSMUSG( | Pank1 | 1295.911 | 0.174803 | 0.170019 | 0.604759 | ENSMUSG( | Pank1 | 1083.865 | -0.07284 | 0.650877 | 0.95422 |
| ENSMUSG( | Pank2 | 1586.719 | 0.074827 | 0.431621 | 0.827432 | ENSMUSG( | Pank2 | 1439.64 | 0.059881 | 0.627062 | 0.953234 |
| ENSMUSG( | Pank3 | 2324.484 | 0.050487 | 0.685902 | 0.932981 | ENSMUSG( | Pank3 | 2089.979 | 0.010305 | 0.94016 | 0.994334 |
| ENSMUSG( | Pank4 | 901.4028 | 0.100491 | 0.450375 | 0.835344 | ENSMUSG( | Pank4 | 805.6932 | 0.049248 | 0.763633 | 0.974 |
| ENSMUSG( | Panx1 | 634.5144 | -0.34963 | 0.032553 | 0.276297 | ENSMUSG( | Panx1 | 680.8273 | 0.14439 | 0.364057 | 0.914223 |
| ENSMUSG( | Panx2 | 6692.878 | 0.218016 | 0.016291 | 0.18942 | ENSMUSG( | Panx2 | 5920.237 | 0.138308 | 0.219041 | 0.866104 |
| ENSMUSG( | Paox | 241.1106 | -0.02566 | 0.855747 | 0.973813 | ENSMUSG( | Paox | 217.2274 | -0.05304 | 0.749483 | 0.96998 |
| ENSMUSG( | Papln | 91.34283 | -0.08847 | 0.683293 | 0.932153 | ENSMUSG( | Papln | 87.34263 | 0.052411 | 0.845463 | 0.984117 |
| ENSMUSG( | Papola | 3061.006 | 0.121191 | 0.131656 | 0.544205 | ENSMUSG( | Papola | 2741.315 | 0.071034 | 0.545881 | 0.938063 |
| ENSMUSG( | Papolb | 8.28081 | -0.50636 | 0.481772 | NA | ENSMUSG( | Papolb | 7.785588 | -0.4077 | 0.58055 | NA |
| ENSMUSG( | Papolg | 276.1517 | 0.206973 | 0.160035 | 0.589679 | ENSMUSG( | Papolg | 260.2969 | 0.307244 | 0.068183 | 0.75212 |
| ENSMUSG( | Pappa | 14.34333 | -0.1075 | 0.831856 | NA | ENSMUSG( | Pappa | 12.73438 | -0.20713 | 0.719732 | 0.966887 |
| ENSMUSG( | Pappa2 | 81.15466 | -0.71532 | 0.007643 | 0.120942 | ENSMUSG( | Pappa2 | 76.94558 | -0.57823 | 0.03918 | 0.679972 |
| ENSMUSG( | Papss1 | 3486.389 | 0.052877 | 0.526142 | 0.877173 | ENSMUSG( | Papss1 | 3013.726 | -0.10201 | 0.382003 | 0.917257 |
| ENSMUSG( | Papss2 | 1947.953 | 0.096211 | 0.45557 | 0.838412 | ENSMUSG( | Papss2 | 1606.325 | -0.20122 | 0.185742 | 0.855305 |
| ENSMUSG( | Paqr3 | 268.5597 | -0.23931 | 0.222965 | 0.668853 | ENSMUSG( | Paqr3 | 287.7313 | 0.241511 | 0.269487 | 0.887332 |
| ENSMUSG( | Paqr4 | 3370.504 | -0.26169 | 0.01668 | 0.192198 | ENSMUSG( | Paqr4 | 3302.028 | -0.03813 | 0.770903 | 0.974236 |
| ENSMUSG( | Paqr5 | 161.522 | 0.679169 | 0.00311 | 0.069189 | ENSMUSG( | Paqr5 | 199.5219 | 1.350877 | 0.009726 | 0.406458 |
| ENSMUSG( | Paqr6 | 468.3904 | -0.1265 | 0.271702 | 0.722082 | ENSMUSG( | Paqr6 | 428.7528 | -0.11187 | 0.402918 | 0.917257 |
| ENSMUSG( | Paqr7 | 4106.932 | -0.23156 | 0.102781 | 0.488221 | ENSMUSG( | Paqr7 | 4269.639 | 0.157368 | 0.327779 | 0.906255 |
| ENSMUSG( | Paqr8 | 24398.75 | 0.29942 | 0.013064 | 0.165414 | ENSMUSG( | Paqr8 | 21065.04 | 0.154118 | 0.236992 | 0.872173 |
| ENSMUSG( | Paqr9 | 1625.942 | -0.18089 | 0.282704 | 0.73186 | ENSMUSG( | Paqr9 | 1752.549 | 0.301546 | 0.128284 | 0.818934 |
| ENSMUSG( | Pard3 | 874.563 | -0.15465 | 0.162281 | 0.592836 | ENSMUSG( | Pard3 | 804.7196 | -0.12762 | 0.293471 | 0.898538 |
| ENSMUSG( | Pard3b | 450.9795 | 0.229685 | 0.057552 | 0.370143 | ENSMUSG( | Pard3b | 378.67 | -0.00757 | 0.963814 | 0.995433 |
| ENSMUSG( | Pard6a | 640.8458 | -0.00702 | 0.955418 | 0.993874 | ENSMUSG( | Pard6a | 596.0495 | 0.058521 | 0.687528 | 0.959673 |
| ENSMUSG( | Pard6b | 77.03757 | -0.15157 | 0.604733 | 0.905556 | ENSMUSG( | Pard6b | 74.3552 | 0.009101 | 0.976002 | 0.996416 |

| | | | | | |
|---|---|---|---|---|---|
| ENSMUSG( | 388.3784 | 0.062916 | 0.614941 | 0.910234 | Nudt5 |
| ENSMUSG( | 101.5964 | -0.13002 | 0.545088 | 0.884712 | Nudt6 |
| ENSMUSG( | 200.0514 | 0.497666 | 0.001296 | 0.039151 | Nudt7 |
| ENSMUSG( | 18.45453 | -0.17573 | 0.692337 | NA | Nudt8 |
| ENSMUSG( | 2787.527 | 0.039371 | 0.627084 | 0.912825 | Nudt9 |
| ENSMUSG( | 342.7755 | 0.076229 | 0.689005 | 0.933441 | Nuf2 |
| ENSMUSG( | 240.6245 | 0.023274 | 0.879729 | 0.981436 | Nufip1 |
| ENSMUSG( | 479.1447 | -0.09634 | 0.460339 | 0.840676 | Nufip2 |
| ENSMUSG( | 2394.15 | 0.070952 | 0.404876 | 0.816396 | Numa1 |
| ENSMUSG( | 1403.514 | 0.125066 | 0.170437 | 0.605271 | Numb |
| ENSMUSG( | 2755.721 | 0.131253 | 0.265127 | 0.715962 | Numbl |
| ENSMUSG( | 374.2117 | 0.05869 | 0.662262 | 0.925946 | Nup107 |
| ENSMUSG( | 726.0206 | 0.124208 | 0.239343 | 0.687192 | Nup133 |
| ENSMUSG( | 1128.51 | 0.129413 | 0.234812 | 0.682188 | Nup153 |
| ENSMUSG( | 493.3186 | -0.05845 | 0.63962 | 0.916434 | Nup155 |
| ENSMUSG( | 386.8917 | 0.075665 | 0.620312 | 0.911432 | Nup160 |
| ENSMUSG( | 744.6374 | 0.067519 | 0.578818 | 0.896692 | Nup188 |
| ENSMUSG( | 803.1738 | 0.131044 | 0.232043 | 0.679932 | Nup205 |
| ENSMUSG( | 1040.307 | -0.21229 | 0.070047 | 0.40908 | Nup210 |
| ENSMUSG( | 30.11755 | 0.106928 | 0.761665 | 0.953201 | Nup210l |
| ENSMUSG( | 1134.685 | 0.06426 | 0.533119 | 0.879905 | Nup214 |
| ENSMUSG( | 225.9977 | 0.04194 | 0.782705 | 0.957558 | Nup35 |
| ENSMUSG( | 75.94856 | 0.131503 | 0.606442 | 0.906481 | Nup37 |
| ENSMUSG( | 214.8454 | -0.10763 | 0.501877 | 0.86481 | Nup42 |
| ENSMUSG( | 55.72655 | 0.108589 | 0.68998 | 0.93379 | Nup43 |
| ENSMUSG( | 1086.495 | 0.034211 | 0.73005 | 0.944956 | Nup50 |
| ENSMUSG( | 400.932 | 0.15926 | 0.182359 | 0.619591 | Nup54 |
| ENSMUSG( | 745.4165 | 0.120427 | 0.303496 | 0.751968 | Nup58 |
| ENSMUSG( | 445.9624 | 0.146231 | 0.339136 | 0.777518 | Nup62 |
| ENSMUSG( | 15.15806 | 0.294171 | 0.546512 | NA | Nup62cl |
| ENSMUSG( | 754.6691 | 0.174387 | 0.126372 | 0.533885 | Nup85 |
| ENSMUSG( | 1210.266 | 0.071258 | 0.594382 | 0.902872 | Nup88 |
| ENSMUSG( | 598.6955 | -0.02413 | 0.845535 | 0.97218 | Nup93 |
| ENSMUSG( | 1005.662 | 0.020373 | 0.840411 | 0.971179 | Nup98 |
| ENSMUSG( | 592.6412 | 0.201411 | 0.082252 | 0.442642 | Nupr1 |
| ENSMUSG( | 41.63081 | -0.07416 | 0.81185 | 0.964426 | Nupr2 |
| ENSMUSG( | 2998.912 | -0.02032 | 0.864936 | 0.975565 | Nus1 |
| ENSMUSG( | 39.94407 | -0.46055 | 0.160263 | 0.589853 | Nusap1 |
| ENSMUSG( | 427.5583 | -0.00866 | 0.942524 | 0.993415 | Nutf2 |
| ENSMUSG( | 1464.53 | 0.022702 | 0.835006 | 0.970426 | Nvl |
| ENSMUSG( | 16269.95 | 0.142899 | 0.227793 | 0.674849 | Nwd1 |
| ENSMUSG( | 956.0197 | -0.08997 | 0.383099 | 0.803237 | Nwd2 |
| ENSMUSG( | 2108.581 | 0.119159 | 0.300853 | 0.749979 | Nxf1 |
| ENSMUSG( | 32.97931 | -0.02364 | 0.952811 | 0.993874 | Nxf7 |
| ENSMUSG( | 962.383 | 0.350334 | 0.000556 | 0.022059 | Nxn |
| ENSMUSG( | 11.76067 | -0.28163 | 0.63356 | NA | Nxpe2 |
| ENSMUSG( | 492.134 | -0.31376 | 0.026552 | 0.249087 | Nxpe3 |
| ENSMUSG( | 621.7809 | -0.2419 | 0.062493 | 0.386467 | Nxpe4 |
| ENSMUSG( | 787.4561 | -0.57622 | 0.001058 | 0.034188 | Nxph1 |
| ENSMUSG( | 38.74297 | -0.45317 | 0.207296 | 0.649561 | Nxph2 |
| ENSMUSG( | 1554.497 | -0.10821 | 0.470633 | 0.84674 | Nxph3 |
| ENSMUSG( | 253.1777 | -0.75464 | 4.43E-06 | 0.000589 | Nxph4 |
| ENSMUSG( | 133.2567 | -0.18 | 0.310739 | 0.758828 | Nxt1 |
| ENSMUSG( | 1319.257 | 0.016672 | 0.901964 | 0.985633 | Nxt2 |
| ENSMUSG( | 3183.621 | 0.215654 | 0.022447 | 0.228899 | Nyap1 |
| ENSMUSG( | 323.0938 | -0.36706 | 0.006111 | 0.106351 | Nyap2 |
| ENSMUSG( | 387.7416 | -0.22672 | 0.085087 | 0.449548 | Nynrin |
| ENSMUSG( | 19.15078 | 0.421115 | 0.353163 | NA | Nyx |
| ENSMUSG( | 60.48842 | -0.64727 | 0.014596 | 0.177737 | Oacyl |
| ENSMUSG( | 876.2935 | -0.14071 | 0.346261 | 0.779153 | Oaf |
| ENSMUSG( | 388.1092 | -0.0339 | 0.805669 | 0.962616 | Oard1 |
| ENSMUSG( | 32.48788 | -0.48867 | 0.13694 | 0.55149 | Oas1a |
| ENSMUSG( | 23.08542 | 0.079701 | 0.834302 | 0.970333 | Oas1b |
| ENSMUSG( | 38.80898 | 0.032656 | 0.916661 | 0.988856 | Oas1c |
| ENSMUSG( | 31.6575 | 0.012319 | 0.974616 | 0.997179 | Oas2 |
| ENSMUSG( | 9.882222 | -0.63101 | 0.347503 | NA | Oas3 |
| ENSMUSG( | 11.84226 | -0.25211 | 0.657317 | NA | Oasl1 |
| ENSMUSG( | 205.7395 | -0.30613 | 0.094046 | 0.471556 | Oasl2 |
| ENSMUSG( | 4626.955 | 0.139755 | 0.170984 | 0.605536 | Oat |
| ENSMUSG( | 7337.269 | 0.057306 | 0.441425 | 0.832452 | Oaz1 |
| ENSMUSG( | 4236.903 | 0.068891 | 0.397319 | 0.812127 | Oaz2 |
| ENSMUSG( | 230.3271 | -0.05833 | 0.723279 | 0.943761 | Obi1 |
| ENSMUSG( | 205.6785 | -0.87275 | 4.86E-07 | 0.000104 | Obscn |
| ENSMUSG( | 567.9047 | 0.109128 | 0.332012 | 0.775301 | Obsl1 |
| ENSMUSG( | 18.92162 | -0.39562 | 0.361835 | NA | Oca2 |
| ENSMUSG( | 606.698 | 0.157997 | 0.171392 | 0.605943 | Ocel1 |
| ENSMUSG( | 9240.916 | -0.33228 | 0.000988 | 0.032693 | Ociad1 |
| ENSMUSG( | 1656.084 | -0.13693 | 0.212627 | 0.6567 | Ociad2 |
| ENSMUSG( | 321.2424 | -0.04122 | 0.876615 | 0.980871 | Ocln |
| ENSMUSG( | 14.39042 | -0.27267 | 0.581705 | NA | Ocm |
| ENSMUSG( | 2072.77 | 0.210603 | 0.018653 | 0.205542 | Ocrl |
| ENSMUSG( | 136.753 | -0.0808 | 0.640713 | 0.916434 | Odad1 |
| ENSMUSG( | 19.23189 | -0.24017 | 0.591971 | NA | Odad2 |
| ENSMUSG( | 108.6595 | -0.02434 | 0.909095 | 0.98794 | Odad3 |
| ENSMUSG( | 45.93444 | -0.37501 | 0.223459 | 0.669469 | Odad4 |
| ENSMUSG( | 2067.932 | -0.06701 | 0.524418 | 0.87583 | Odc1 |
| ENSMUSG( | 2406.447 | 0.140421 | 0.196053 | 0.635489 | Odf2 |
| ENSMUSG( | 364.5961 | 0.079922 | 0.523595 | 0.875628 | Odf2l |
| ENSMUSG( | 2398.968 | -0.02007 | 0.852773 | 0.973168 | Odr4 |
| ENSMUSG( | 291.2791 | 3.49E-05 | 0.999813 | 0.99989 | Ofd1 |
| ENSMUSG( | 5738.298 | 0.137443 | 0.137104 | 0.551819 | Oga |
| ENSMUSG( | 9654.122 | 0.140301 | 0.083463 | 0.445586 | Ogdh |
| ENSMUSG( | 5615.674 | 0.186619 | 0.034773 | 0.286739 | Ogdhl |
| ENSMUSG( | 3032.16 | 0.181527 | 0.04274 | 0.316989 | Ogfod1 |
| ENSMUSG( | 752.2273 | 0.008521 | 0.941634 | 0.993415 | Ogfod2 |
| ENSMUSG( | 283.9544 | -0.39117 | 0.021671 | 0.225182 | Ogfod3 |
| ENSMUSG( | 1615.409 | 0.088842 | 0.380154 | 0.801757 | Ogfr |
| ENSMUSG( | 2695.056 | 0.086242 | 0.329471 | 0.773311 | Ogfrl1 |
| ENSMUSG( | 200.1112 | 0.080952 | 0.598964 | 0.903801 | Ogg1 |
| ENSMUSG( | 336.9876 | -0.21458 | 0.267382 | 0.717525 | Ogn |
| ENSMUSG( | 6530.254 | 0.038436 | 0.737897 | 0.947329 | Ogt |
| ENSMUSG( | 13.07207 | 0.101376 | 0.852286 | NA | Oip5 |
| ENSMUSG( | 16087.44 | 0.140792 | 0.132746 | 0.545099 | Oip5os1 |
| ENSMUSG( | 1610.625 | -0.04246 | 0.716953 | 0.941697 | Ola1 |
| ENSMUSG( | 49119.54 | -0.12284 | 0.226052 | 0.672763 | Olfm1 |
| ENSMUSG( | 6250.384 | -0.3254 | 0.006856 | 0.11446 | Olfm2 |
| ENSMUSG( | 512.0904 | -0.94433 | 2.70E-06 | 0.000405 | Olfm3 |
| ENSMUSG( | 42.7271 | 0.132156 | 0.663315 | 0.9265 | Olfm4 |
| ENSMUSG( | 709.7437 | -0.12852 | 0.410842 | 0.818585 | Olfml1 |
| ENSMUSG( | 221.0833 | -0.26048 | 0.215228 | 0.659815 | Olfml2a |
| ENSMUSG( | 347.245 | 0.16041 | 0.494548 | 0.860776 | Olfml2b |
| ENSMUSG( | 1509.104 | -0.27058 | 0.101159 | 0.484996 | Olfml3 |
| ENSMUSG( | 9658.095 | -0.18742 | 0.089539 | 0.459596 | Olig1 |
| ENSMUSG( | 1278.812 | -0.59517 | 1.37E-05 | 0.001449 | Olig2 |
| ENSMUSG( | 263.4406 | 0.068145 | 0.639389 | 0.916434 | Oma1 |
| ENSMUSG( | 33.54041 | 0.193253 | 0.573566 | 0.893584 | Omd |
| ENSMUSG( | 3937.536 | -0.45476 | 0.006737 | 0.113006 | Omg |
| ENSMUSG( | 49.65054 | -2.62222 | 0.01154 | 0.154695 | Omp |
| ENSMUSG( | 81.09059 | -0.52451 | 0.020523 | 0.216158 | Onecut2 |
| ENSMUSG( | 4577.166 | 0.090824 | 0.233096 | 0.681351 | Opa1 |
| ENSMUSG( | 2637.841 | 0.060338 | 0.462635 | 0.842333 | Opa3 |
| ENSMUSG( | 1199.33 | -0.74577 | 1.04E-06 | 0.000197 | Opalin |
| ENSMUSG( | 5380.971 | -0.37148 | 0.016177 | 0.189041 | Opcml |
| ENSMUSG( | 1474.951 | 0.071466 | 0.532683 | 0.879905 | Ophn1 |
| ENSMUSG( | 1875.306 | 0.11513 | 0.271711 | 0.722082 | Oplah |
| ENSMUSG( | 144.0092 | -0.12075 | 0.533152 | 0.879905 | Opn3 |
| ENSMUSG( | 230.1327 | -0.20744 | 0.343253 | 0.779153 | Oprd1 |
| ENSMUSG( | 212.4665 | -0.79983 | 3.27E-06 | 0.000465 | Oprk1 |
| ENSMUSG( | 1052.428 | -0.06374 | 0.640233 | 0.916434 | Oprl1 |
| ENSMUSG( | 11.24422 | -0.46656 | 0.4519 | NA | Oprm1 |
| ENSMUSG( | 33.48434 | -0.2722 | 0.421421 | 0.823126 | Optc |
| ENSMUSG( | 500.6619 | -0.0619 | 0.604914 | 0.905556 | Optn |
| ENSMUSG( | 599.8774 | 0.087601 | 0.466761 | 0.844819 | Or10ad1 |
| ENSMUSG( | 51.08857 | -0.08019 | 0.776394 | 0.955658 | Or2ak4 |
| ENSMUSG( | 18.93615 | -1.97517 | 0.00169 | NA | Or2y1g |
| ENSMUSG( | 37.09659 | -0.34971 | 0.261005 | 0.710889 | Or4d1 |
| ENSMUSG( | 24.79305 | -0.36418 | 0.368323 | 0.795863 | Or51e2 |
| ENSMUSG( | 15.85828 | 0.362996 | 0.537246 | NA | Or5m3b |
| ENSMUSG( | 365.5107 | -0.48675 | 0.00242 | 0.057903 | Orai1 |
| ENSMUSG( | 3447.817 | -0.04205 | 0.68038 | 0.931044 | Orai2 |
| ENSMUSG( | 735.2552 | 0.021095 | 0.864243 | 0.975358 | Orai3 |
| ENSMUSG( | 933.4692 | 0.184369 | 0.059358 | 0.376382 | Orc2 |
| ENSMUSG( | 714.2317 | -0.02622 | 0.801961 | 0.962116 | Orc3 |
| ENSMUSG( | 898.2217 | -0.10818 | 0.340168 | 0.777962 | Orc4 |
| ENSMUSG( | 447.2922 | -0.31305 | 0.012346 | 0.160458 | Orc5 |
| ENSMUSG( | 521.621 | 0.039786 | 0.742574 | 0.948478 | Orc6 |
| ENSMUSG( | 178.743 | -0.29643 | 0.115377 | 0.512362 | Ormdl1 |
| ENSMUSG( | 288.5841 | -0.08945 | 0.600504 | 0.90443 | Ormdl2 |
| ENSMUSG( | 2726.992 | -0.14586 | 0.181487 | 0.618685 | Ormdl3 |
| ENSMUSG( | 6640.905 | -0.25841 | 0.023781 | 0.235641 | Os9 |
| ENSMUSG( | 2259.892 | 0.04487 | 0.615054 | 0.910318 | Osbp |
| ENSMUSG( | 2814.769 | 0.067074 | 0.545118 | 0.884712 | Osbp2 |
| ENSMUSG( | 697.9335 | 0.060804 | 0.640326 | 0.916434 | Osbpl10 |
| ENSMUSG( | 552.6876 | -0.15639 | 0.162096 | 0.592429 | Osbpl11 |
| ENSMUSG( | 8165.591 | 0.033622 | 0.693424 | 0.934202 | Osbpl1a |
| ENSMUSG( | 1636.717 | 0.052664 | 0.585143 | 0.899171 | Osbpl2 |
| ENSMUSG( | 838.0628 | 0.106725 | 0.394884 | 0.810072 | Osbpl3 |
| ENSMUSG( | 921.2652 | 0.234726 | 0.052779 | 0.352597 | Osbpl5 |

| | | | | | |
|---|---|---|---|---|---|
| ENSMUSG( | 28.15855 | -1.0424 | 0.005644 | 0.304216 | Or4d1 |
| ENSMUSG( | 25.63155 | 0.036122 | 0.933588 | 0.99306 | Or51e2 |
| ENSMUSG( | 10.63115 | -0.51438 | 0.390444 | NA | Or5m3b |
| ENSMUSG( | 438.6925 | 0.338025 | 0.073061 | 0.763629 | Orai1 |
| ENSMUSG( | 3426.219 | 0.204584 | 0.082907 | 0.77681 | Orai2 |
| ENSMUSG( | 662.065 | -0.00766 | 0.960301 | 0.994831 | Orai3 |
| ENSMUSG( | 794.0299 | -0.01262 | 0.921239 | 0.992644 | Orc2 |
| ENSMUSG( | 663.7288 | 0.032032 | 0.798415 | 0.97681 | Orc3 |
| ENSMUSG( | 886.0747 | 0.116917 | 0.393876 | 0.917257 | Orc4 |
| ENSMUSG( | 439.2624 | -0.07645 | 0.584715 | 0.944975 | Orc5 |
| ENSMUSG( | 455.1714 | -0.08379 | 0.561047 | 0.940254 | Orc6 |
| ENSMUSG( | 189.0626 | 0.154847 | 0.494826 | 0.927372 | Ormdl1 |
| ENSMUSG( | 276.0338 | 0.047615 | 0.776669 | 0.975717 | Ormdl2 |
| ENSMUSG( | 2642.344 | 0.037384 | 0.768686 | 0.974236 | Ormdl3 |
| ENSMUSG( | 6896.928 | 0.130872 | 0.329367 | 0.907083 | Os9 |
| ENSMUSG( | 2077.938 | 0.07155 | 0.500194 | 0.927825 | Osbp |
| ENSMUSG( | 2550.402 | 0.052268 | 0.661143 | 0.956205 | Osbp2 |
| ENSMUSG( | 617.0427 | -0.01931 | 0.894683 | 0.990558 | Osbpl10 |
| ENSMUSG( | 525.4153 | -0.0251 | 0.854856 | 0.985649 | Osbpl11 |
| ENSMUSG( | 6972.717 | -0.15908 | 0.134528 | 0.823094 | Osbpl1a |
| ENSMUSG( | 1482.623 | 0.034056 | 0.781136 | 0.976106 | Osbpl2 |
| ENSMUSG( | 791.3599 | 0.204499 | 0.090457 | 0.791074 | Osbpl3 |
| ENSMUSG( | 770.3099 | -0.0034 | 0.980092 | 0.996479 | Osbpl5 |
| ENSMUSG( | 1406.024 | 0.109243 | 0.423392 | 0.917257 | Osbpl6 |
| ENSMUSG( | 432.9852 | 0.019616 | 0.891049 | 0.990558 | Osbpl7 |
| ENSMUSG( | 1463.464 | 0.162386 | 0.263531 | 0.886134 | Osbpl8 |
| ENSMUSG( | 1259.766 | 0.14235 | 0.216391 | 0.865167 | Osbpl9 |
| ENSMUSG( | 63.31238 | 0.243936 | 0.452842 | 0.922316 | Oscar |
| ENSMUSG( | 973.8465 | 0.014905 | 0.893909 | 0.990558 | Oscp1 |
| ENSMUSG( | 122.0184 | 0.077189 | 0.725919 | 0.966887 | Oser1 |
| ENSMUSG( | 734.9431 | -0.27303 | 0.035273 | 0.661159 | Osgep |
| ENSMUSG( | 80.91866 | 0.361468 | 0.258958 | 0.883971 | Osgepl1 |
| ENSMUSG( | 19.47046 | 0.506782 | 0.271758 | 0.888132 | Osgin1 |
| ENSMUSG( | 501.5826 | -0.02017 | 0.875703 | 0.989644 | Osgin2 |
| ENSMUSG( | 217.0736 | -0.09893 | 0.665368 | 0.956766 | Osmr |
| ENSMUSG( | 74.67211 | -0.2388 | 0.37189 | 0.915491 | Osr1 |
| ENSMUSG( | 1191.117 | -0.3 | 0.016835 | 0.522883 | Ost4 |
| ENSMUSG( | 1289.501 | 0.113745 | 0.421224 | 0.917257 | Ostc |
| ENSMUSG( | 541.2331 | -0.07567 | 0.568056 | 0.941777 | Ostf1 |
| ENSMUSG( | 2168.459 | 0.161799 | 0.289983 | 0.896726 | Ostm1 |
| ENSMUSG( | 10.17899 | 0.336694 | 0.593698 | NA | Otoa |
| ENSMUSG( | 606.8449 | -0.14621 | 0.560665 | 0.940254 | Otof |
| ENSMUSG( | 11.67635 | 0.236813 | 0.685248 | 0.958678 | Otogl |
| ENSMUSG( | 33.55613 | -0.51753 | 0.179243 | 0.849741 | Otop2 |
| ENSMUSG( | 3600.069 | -0.16885 | 0.243767 | 0.877904 | Otub1 |
| ENSMUSG( | 868.9703 | 0.055192 | 0.656419 | 0.955533 | Otub2 |
| ENSMUSG( | 407.5179 | 0.063339 | 0.700895 | 0.96162 | Otud1 |
| ENSMUSG( | 268.3567 | -0.16682 | 0.303265 | 0.898538 | Otud3 |
| ENSMUSG( | 1268.39 | 0.020881 | 0.846933 | 0.985027 | Otud4 |
| ENSMUSG( | 2551.275 | 0.005856 | 0.954898 | 0.994798 | Otud5 |
| ENSMUSG( | 2118.544 | -0.08964 | 0.532824 | 0.934795 | Otud6b |
| ENSMUSG( | 1260.247 | 0.069424 | 0.595183 | 0.945701 | Otud7a |
| ENSMUSG( | 2690.258 | -0.0303 | 0.797115 | 0.97681 | Otud7b |
| ENSMUSG( | 589.2056 | -0.14389 | 0.282966 | 0.892412 | Otulin |
| ENSMUSG( | 742.3336 | 0.229314 | 0.201073 | 0.86439 | Otulinl |
| ENSMUSG( | 240.4888 | 0.166089 | 0.37538 | 0.915491 | Otx1 |
| ENSMUSG( | 52.05564 | -0.14627 | 0.868152 | 0.98788 | Otx2 |
| ENSMUSG( | 249.9226 | -0.1279 | 0.446581 | 0.921734 | Ovca2 |
| ENSMUSG( | 97.62942 | 0.397857 | 0.230956 | 0.868507 | Ovgp1 |
| ENSMUSG( | 106.0662 | -0.49332 | 0.041138 | 0.687358 | Ovol2 |
| ENSMUSG( | 1654.604 | -0.12784 | 0.291819 | 0.897854 | Oxa1l |
| ENSMUSG( | 12336.67 | -0.15391 | 0.219906 | 0.866104 | Oxct1 |
| ENSMUSG( | 108.7508 | -0.2944 | 0.176588 | 0.846973 | Oxld1 |
| ENSMUSG( | 516.2541 | -0.10944 | 0.42784 | 0.917257 | Oxnad1 |
| ENSMUSG( | 3544.441 | -0.00713 | 0.953525 | 0.994798 | Oxr1 |
| ENSMUSG( | 282.5185 | -0.12307 | 0.457844 | 0.923329 | Oxsm |
| ENSMUSG( | 3806.194 | -0.27863 | 0.024578 | 0.592201 | Oxsr1 |
| ENSMUSG( | 275.3264 | 0.249442 | 0.109633 | 0.806227 | Oxtr |
| ENSMUSG( | 22.00668 | -1.00573 | 0.031406 | 0.631192 | P2rx1 |
| ENSMUSG( | 49.50001 | 0.003414 | 0.990711 | 0.998287 | P2rx3 |
| ENSMUSG( | 585.1576 | 0.055831 | 0.692222 | 0.960404 | P2rx4 |
| ENSMUSG( | 24.69684 | -0.17064 | 0.677588 | 0.957106 | P2rx5 |
| ENSMUSG( | 100.3989 | -0.04495 | 0.879802 | 0.989644 | P2rx6 |
| ENSMUSG( | 480.6777 | 0.104168 | 0.542608 | 0.938034 | P2rx7 |
| ENSMUSG( | 86.49368 | 0.477162 | 0.043645 | 0.690304 | P2ry1 |
| ENSMUSG( | 11.32058 | 0.364803 | 0.521886 | NA | P2ry10b |
| ENSMUSG( | 2111.842 | 0.295253 | 0.159513 | 0.837724 | P2ry12 |
| ENSMUSG( | 833.7998 | 0.365315 | 0.13823 | 0.828492 | P2ry13 |
| ENSMUSG( | 133.1146 | 0.613547 | 0.042782 | 0.690304 | P2ry14 |
| ENSMUSG( | 30.27225 | 0.504383 | 0.263956 | 0.886134 | P2ry2 |
| ENSMUSG( | 163.8961 | 0.284097 | 0.279182 | 0.892356 | P2ry6 |
| ENSMUSG( | 339.0512 | 0.263695 | 0.093262 | 0.795363 | P3h1 |
| ENSMUSG( | 171.5038 | 0.409612 | 0.026782 | 0.603041 | P3h2 |
| ENSMUSG( | 815.6752 | 0.153615 | 0.356792 | 0.913358 | P3h3 |
| ENSMUSG( | 603.4273 | 0.155593 | 0.309947 | 0.898538 | P3h4 |
| ENSMUSG( | 2528.933 | 0.150121 | 0.332608 | 0.907907 | P4ha1 |
| ENSMUSG( | 380.8407 | 0.61761 | 2.63E-05 | 0.007461 | P4ha2 |
| ENSMUSG( | 326.5597 | -0.18174 | 0.221925 | 0.866104 | P4ha3 |
| ENSMUSG( | 5822.229 | 0.144238 | 0.346095 | 0.910966 | P4hb |
| ENSMUSG( | 2561.924 | 0.228202 | 0.150922 | 0.837173 | P4htm |
| ENSMUSG( | 1540.493 | -0.11683 | 0.392665 | 0.917257 | Pa2g4 |
| ENSMUSG( | 237.8224 | 0.091882 | 0.622338 | 0.953234 | Pabir1 |
| ENSMUSG( | 193.4709 | -0.17114 | 0.386094 | 0.917257 | Pabir2 |
| ENSMUSG( | 9834.715 | -0.05646 | 0.621644 | 0.953199 | Pabpc1 |
| ENSMUSG( | 19.17555 | -0.59068 | 0.199349 | 0.863016 | Pabpc1l |
| ENSMUSG( | 1317.547 | -0.00981 | 0.929622 | 0.99306 | Pabpc4 |
| ENSMUSG( | 10.00539 | 0.121661 | 0.841197 | NA | Pabpc4l |
| ENSMUSG( | 12.10477 | 0.191142 | 0.746491 | 0.969692 | Pabpc5 |
| ENSMUSG( | 1186.918 | 0.342598 | 0.10654 | 0.804969 | Pabpn1 |
| ENSMUSG( | 361.4954 | 0.321064 | 0.137746 | 0.828492 | Pacc1 |
| ENSMUSG( | 419.7186 | -0.31237 | 0.027934 | 0.61023 | Pacrg |
| ENSMUSG( | 194.2501 | 0.056902 | 0.746467 | 0.969692 | Pacrgl |
| ENSMUSG( | 1721.33 | -0.0166 | 0.904786 | 0.991089 | Pacs1 |
| ENSMUSG( | 11392.75 | -0.1855 | 0.084971 | 0.779799 | Pacs2 |
| ENSMUSG( | 10586.99 | -0.06751 | 0.504405 | 0.927825 | Pacsin1 |
| ENSMUSG( | 1440.022 | -0.11432 | 0.345745 | 0.910877 | Pacsin2 |
| ENSMUSG( | 1882.121 | -0.07975 | 0.497553 | 0.927825 | Pacsin3 |
| ENSMUSG( | 1866.049 | 0.071231 | 0.527413 | 0.934466 | Padi2 |
| ENSMUSG( | 35.0693 | 0.083143 | 0.819566 | 0.979198 | Padi6 |
| ENSMUSG( | 1493.806 | -0.01873 | 0.900793 | 0.990558 | Paf1 |
| ENSMUSG( | 10991.61 | -0.05426 | 0.616768 | 0.951636 | Pafah1b1 |
| ENSMUSG( | 2964.336 | -0.0758 | 0.552769 | 0.939565 | Pafah1b2 |
| ENSMUSG( | 165.9813 | -0.24603 | 0.209809 | 0.864412 | Pafah1b3 |
| ENSMUSG( | 368.1553 | -0.12803 | 0.379808 | 0.916995 | Pafah2 |
| ENSMUSG( | 1106.258 | 0.167331 | 0.335081 | 0.907907 | Pag1 |
| ENSMUSG( | 28.91376 | 0.08924 | 0.812296 | 0.979148 | Pagr1a |
| ENSMUSG( | 3208.727 | -0.13711 | 0.274982 | 0.889874 | Paics |
| ENSMUSG( | 1816.615 | -0.05743 | 0.67616 | 0.95694 | Paip1 |
| ENSMUSG( | 2406.369 | -0.12168 | 0.331757 | 0.907907 | Paip2 |
| ENSMUSG( | 2002.326 | -0.18672 | 0.151104 | 0.837173 | Paip2b |
| ENSMUSG( | 6138.91 | -0.10791 | 0.348623 | 0.911652 | Pak1 |
| ENSMUSG( | 723.4903 | -0.01238 | 0.933583 | 0.99306 | Pak1ip1 |
| ENSMUSG( | 2062.746 | 0.007071 | 0.955795 | 0.994798 | Pak2 |
| ENSMUSG( | 988.3181 | -0.04148 | 0.756697 | 0.971779 | Pak3 |
| ENSMUSG( | 439.7989 | -0.1794 | 0.168801 | 0.840222 | Pak4 |
| ENSMUSG( | 606.2658 | -0.37493 | 0.003007 | 0.219289 | Pak5 |
| ENSMUSG( | 4972.445 | 0.101786 | 0.327156 | 0.905768 | Pak6 |
| ENSMUSG( | 90.94988 | 0.213579 | 0.353107 | 0.912263 | Pakap |
| ENSMUSG( | 31.21633 | 0.416726 | 0.274927 | 0.889874 | Palb2 |
| ENSMUSG( | 562.5585 | 0.376944 | 0.070049 | 0.757435 | Pald1 |
| ENSMUSG( | 470.4542 | -0.11733 | 0.467439 | 0.923329 | Palld |
| ENSMUSG( | 17976.97 | 0.207244 | 0.052677 | 0.71525 | Palm |
| ENSMUSG( | 84.1862 | 0.170046 | 0.482478 | 0.924578 | Palm3 |
| ENSMUSG( | 533.8978 | 0.289577 | 0.128265 | 0.818934 | Palmd |
| ENSMUSG( | 998.6763 | 0.088064 | 0.538511 | 0.937216 | Pals1 |
| ENSMUSG( | 2592.012 | -0.1216 | 0.394772 | 0.917257 | Pals2 |
| ENSMUSG( | 5584.635 | 0.258757 | 0.133759 | 0.822912 | Pam |
| ENSMUSG( | 544.1595 | -0.19309 | 0.189309 | 0.85698 | Pam16 |
| ENSMUSG( | 9.872897 | -0.79703 | 0.216634 | NA | Pam16l |
| ENSMUSG( | 1629.63 | 0.010251 | 0.955702 | 0.994798 | Pamr1 |
| ENSMUSG( | 677.3104 | 0.111937 | 0.542009 | 0.937993 | Pan2 |
| ENSMUSG( | 1134.817 | 0.186611 | 0.109733 | 0.80637 | Pan3 |
| ENSMUSG( | 70.58836 | 0.040898 | 0.871474 | 0.988779 | Panct2 |
| ENSMUSG( | 1083.865 | -0.07284 | 0.650877 | 0.95422 | Pank1 |
| ENSMUSG( | 1439.64 | 0.059881 | 0.627062 | 0.953234 | Pank2 |
| ENSMUSG( | 2089.979 | 0.010305 | 0.94016 | 0.994334 | Pank3 |
| ENSMUSG( | 805.6932 | 0.049248 | 0.763633 | 0.974 | Pank4 |
| ENSMUSG( | 680.8273 | 0.14439 | 0.364057 | 0.914223 | Panx1 |
| ENSMUSG( | 5920.237 | 0.138308 | 0.219041 | 0.866104 | Panx2 |
| ENSMUSG( | 217.2274 | -0.05304 | 0.749483 | 0.96998 | Paox |
| ENSMUSG( | 87.34263 | 0.052411 | 0.845463 | 0.984117 | Papln |
| ENSMUSG( | 2741.315 | 0.071034 | 0.545881 | 0.938063 | Papola |
| ENSMUSG( | 7.785588 | -0.4077 | 0.58055 | NA | Papolb |
| ENSMUSG( | 260.2969 | 0.307244 | 0.068183 | 0.75212 | Papolg |
| ENSMUSG( | 12.73438 | -0.20713 | 0.719732 | 0.966887 | Pappa |
| ENSMUSG( | 76.94558 | -0.57823 | 0.03918 | 0.679972 | Pappa2 |
| ENSMUSG( | 3013.726 | -0.10201 | 0.382003 | 0.917257 | Papss1 |
| ENSMUSG( | 1606.325 | -0.20122 | 0.185742 | 0.855305 | Papss2 |

| | | | | | | | | | | |
|---|---|---|---|---|---|---|---|---|---|---|
| ENSMUSG( | Pard6g | 230.0828 | -0.21226 | 0.147403 | 0.569664 | ENSMUSG( | Pard6g | 218.8697 | -0.08001 | 0.619326 | 0.952814 |
| ENSMUSG( | Parg | 710.7913 | 0.226178 | 0.028792 | 0.258585 | ENSMUSG( | Parg | 646.3934 | 0.219667 | 0.078482 | 0.77321 |
| ENSMUSG( | Park7 | 2769.187 | 0.010189 | 0.91993 | 0.990247 | ENSMUSG( | Park7 | 2317.014 | -0.25001 | 0.06443 | 0.737935 |
| ENSMUSG( | Parl | 936.341 | -0.1313 | 0.188987 | 0.627678 | ENSMUSG( | Parl | 838.2016 | -0.18354 | 0.15453 | 0.837724 |
| ENSMUSG( | Parm1 | 1686.586 | -0.29088 | 0.088493 | 0.456254 | ENSMUSG( | Parm1 | 1800.105 | 0.180465 | 0.340249 | 0.907972 |
| ENSMUSG( | Parn | 889.5421 | -0.02978 | 0.764508 | 0.954055 | ENSMUSG( | Parn | 787.2 | -0.11408 | 0.335295 | 0.907907 |
| ENSMUSG( | Parp1 | 1890.905 | 0.112967 | 0.260408 | 0.710245 | ENSMUSG( | Parp1 | 1635.795 | -0.03516 | 0.767297 | 0.974236 |
| ENSMUSG( | Parp10 | 92.95848 | -0.07741 | 0.701276 | 0.93743 | ENSMUSG( | Parp10 | 96.03824 | 0.28655 | 0.195986 | 0.862161 |
| ENSMUSG( | Parp11 | 558.303 | 0.003641 | 0.974828 | 0.997207 | ENSMUSG( | Parp11 | 491.475 | -0.10016 | 0.457342 | 0.923329 |
| ENSMUSG( | Parp12 | 461.6631 | -0.00675 | 0.958126 | 0.993874 | ENSMUSG( | Parp12 | 423.924 | 0.022761 | 0.88156 | 0.989644 |
| ENSMUSG( | Parp14 | 203.8275 | -0.15448 | 0.351029 | 0.783067 | ENSMUSG( | Parp14 | 192.4911 | -0.0363 | 0.847033 | 0.985027 |
| ENSMUSG( | Parp16 | 89.86136 | 0.064337 | 0.762727 | 0.953519 | ENSMUSG( | Parp16 | 86.48555 | 0.226202 | 0.371046 | 0.915491 |
| ENSMUSG( | Parp2 | 524.9396 | 0.129632 | 0.238836 | 0.6865 | ENSMUSG( | Parp2 | 452.8825 | -0.02085 | 0.888047 | 0.990442 |
| ENSMUSG( | Parp3 | 231.5314 | -0.00859 | 0.958097 | 0.993874 | ENSMUSG( | Parp3 | 197.9519 | -0.19559 | 0.275871 | 0.890441 |
| ENSMUSG( | Parp4 | 331.8213 | 0.108821 | 0.386234 | 0.804623 | ENSMUSG( | Parp4 | 305.9791 | 0.146078 | 0.315325 | 0.901306 |
| ENSMUSG( | Parp6 | 7609.704 | 0.199102 | 0.007734 | 0.122031 | ENSMUSG( | Parp6 | 6537.542 | 0.036264 | 0.704507 | 0.962677 |
| ENSMUSG( | Parp8 | 724.7642 | 0.180103 | 0.088279 | 0.455783 | ENSMUSG( | Parp8 | 621.9549 | 0.012727 | 0.924091 | 0.992904 |
| ENSMUSG( | Parp9 | 221.3365 | -0.02873 | 0.870427 | 0.977906 | ENSMUSG( | Parp9 | 204.9653 | 0.021172 | 0.91208 | 0.992196 |
| ENSMUSG( | Parpbp | 11.15662 | -0.30166 | 0.608633 | NA | ENSMUSG( | Parpbp | 10.42009 | -0.23284 | 0.706895 | NA |
| ENSMUSG( | Pars2 | 179.0451 | 0.165784 | 0.294567 | 0.743754 | ENSMUSG( | Pars2 | 151.5258 | -0.03245 | 0.862245 | 0.986412 |
| ENSMUSG( | Particl | 485.545 | -0.20338 | 0.100653 | 0.48402 | ENSMUSG( | Particl | 445.9144 | -0.18316 | 0.203902 | 0.864412 |
| ENSMUSG( | Parva | 4310.262 | 0.115267 | 0.263744 | 0.714352 | ENSMUSG( | Parva | 3582.726 | -0.15279 | 0.181893 | 0.850579 |
| ENSMUSG( | Parvaos | 15.18422 | 0.357538 | 0.485904 | NA | ENSMUSG( | Parvaos | 14.40969 | 0.408847 | 0.45772 | 0.923329 |
| ENSMUSG( | Parvb | 2076.88 | 0.007635 | 0.938174 | 0.993415 | ENSMUSG( | Parvb | 1763.556 | -0.20261 | 0.082374 | 0.77681 |
| ENSMUSG( | Parvg | 358.6145 | -0.08453 | 0.507416 | 0.86795 | ENSMUSG( | Parvg | 312.7973 | -0.21094 | 0.153477 | 0.837724 |
| ENSMUSG( | Pasd1 | 18.83737 | 0.541537 | 0.214967 | NA | ENSMUSG( | Pasd1 | 18.93963 | 0.727057 | 0.141731 | 0.830709 |
| ENSMUSG( | Pask | 106.3174 | -0.04603 | 0.840546 | 0.971179 | ENSMUSG( | Pask | 115.9476 | 0.445628 | 0.044973 | 0.693959 |
| ENSMUSG( | Pate2 | 22.32177 | 0.343123 | 0.379945 | 0.801757 | ENSMUSG( | Pate2 | 19.52445 | 0.2655 | 0.553593 | 0.939565 |
| ENSMUSG( | Patj | 320.7931 | -0.14995 | 0.349947 | 0.781943 | ENSMUSG( | Patj | 280.7857 | -0.28348 | 0.12116 | 0.813384 |
| ENSMUSG( | Patl1 | 987.8956 | -0.07706 | 0.468287 | 0.845336 | ENSMUSG( | Patl1 | 915.5396 | -0.03204 | 0.810719 | 0.978779 |
| ENSMUSG( | Patz1 | 1051.658 | 0.032491 | 0.743441 | 0.948478 | ENSMUSG( | Patz1 | 951.3341 | 0.010754 | 0.927268 | 0.99306 |
| ENSMUSG( | Pawr | 88.02693 | -0.09095 | 0.706177 | 0.938869 | ENSMUSG( | Pawr | 83.14167 | 0.027155 | 0.915597 | 0.992529 |
| ENSMUSG( | Pax5 | 11.30284 | 0.373624 | 0.517821 | NA | ENSMUSG( | Pax5 | 9.231073 | 0.139313 | 0.838325 | NA |
| ENSMUSG( | Pax6 | 770.059 | -0.00297 | 0.9827 | 0.997522 | ENSMUSG( | Pax6 | 699.0728 | -0.02054 | 0.904958 | 0.991089 |
| ENSMUSG( | Pax6os1 | 24.2327 | 0.572585 | 0.175762 | 0.611006 | ENSMUSG( | Pax6os1 | 18.3458 | 0.037158 | 0.940632 | 0.99445 |
| ENSMUSG( | Paxbp1 | 1640.406 | -0.04004 | 0.717222 | 0.941697 | ENSMUSG( | Paxbp1 | 1606.006 | 0.163425 | 0.162032 | 0.837724 |
| ENSMUSG( | Paxip1 | 2265.373 | 0.213946 | 0.026256 | 0.247758 | ENSMUSG( | Paxip1 | 2021.232 | 0.160528 | 0.184439 | 0.853168 |
| ENSMUSG( | Paxx | 1753.997 | 0.095501 | 0.263741 | 0.714352 | ENSMUSG( | Paxx | 1551.394 | 0.010027 | 0.926859 | 0.99306 |
| ENSMUSG( | Pbdc1 | 434.9551 | 0.179211 | 0.129156 | 0.538324 | ENSMUSG( | Pbdc1 | 349.605 | -0.18969 | 0.199969 | 0.863016 |
| ENSMUSG( | Pbk | 45.82963 | -0.45054 | 0.121413 | 0.525713 | ENSMUSG( | Pbk | 44.28567 | -0.25225 | 0.420072 | 0.917257 |
| ENSMUSG( | Pbld2 | 37.00389 | -0.21095 | 0.512924 | 0.871762 | ENSMUSG( | Pbld2 | 31.38915 | -0.4532 | 0.219921 | 0.866104 |
| ENSMUSG( | Pbrm1 | 1216.762 | -0.01235 | 0.894875 | 0.984512 | ENSMUSG( | Pbrm1 | 1143.975 | 0.077991 | 0.505275 | 0.927825 |
| ENSMUSG( | Pbx1 | 5519.02 | 0.056087 | 0.522936 | 0.875628 | ENSMUSG( | Pbx1 | 4863.154 | -0.04069 | 0.734614 | 0.967318 |
| ENSMUSG( | Pbx2 | 1416.866 | -0.01379 | 0.898202 | 0.985633 | ENSMUSG( | Pbx2 | 1328.933 | 0.066486 | 0.608165 | 0.950113 |
| ENSMUSG( | Pbx3 | 406.032 | -0.24804 | 0.211899 | 0.655541 | ENSMUSG( | Pbx3 | 421.8744 | 0.128342 | 0.584574 | 0.944975 |
| ENSMUSG( | Pbx4 | 360.5205 | 0.099107 | 0.478475 | 0.853567 | ENSMUSG( | Pbx4 | 309.1024 | -0.08009 | 0.621618 | 0.953199 |
| ENSMUSG( | Pbxip1 | 8003.441 | -0.09627 | 0.408213 | 0.818167 | ENSMUSG( | Pbxip1 | 7060.88 | -0.19301 | 0.128726 | 0.819665 |
| ENSMUSG( | Pcbd1 | 118.3927 | 0.17935 | 0.346392 | 0.779153 | ENSMUSG( | Pcbd1 | 102.2878 | 0.012706 | 0.955164 | 0.994798 |
| ENSMUSG( | Pcbd2 | 253.0123 | -0.05622 | 0.705229 | 0.938303 | ENSMUSG( | Pcbd2 | 218.0192 | -0.24238 | 0.204182 | 0.864412 |
| ENSMUSG( | Pcbp1 | 3562.284 | 0.058111 | 0.641262 | 0.916693 | ENSMUSG( | Pcbp1 | 3115.604 | -0.06373 | 0.685879 | 0.958956 |
| ENSMUSG( | Pcbp2 | 8361.06 | 0.054675 | 0.493062 | 0.860552 | ENSMUSG( | Pcbp2 | 7365.816 | -0.04358 | 0.730663 | 0.967228 |
| ENSMUSG( | Pcbp3 | 1024.225 | 0.055315 | 0.723921 | 0.944096 | ENSMUSG( | Pcbp3 | 949.4531 | 0.09752 | 0.646176 | 0.953988 |
| ENSMUSG( | Pcbp4 | 7726.86 | 0.186904 | 0.047977 | 0.334562 | ENSMUSG( | Pcbp4 | 6785.46 | 0.085627 | 0.391728 | 0.917257 |
| ENSMUSG( | Pcca | 1091.371 | 0.071488 | 0.482982 | 0.856216 | ENSMUSG( | Pcca | 908.063 | -0.19772 | 0.093953 | 0.795735 |
| ENSMUSG( | Pccb | 3290.838 | 0.043614 | 0.622251 | 0.911432 | ENSMUSG( | Pccb | 2768.658 | -0.19479 | 0.092968 | 0.79479 |
| ENSMUSG( | Pcdh1 | 6356.206 | -0.19637 | 0.250387 | 0.700322 | ENSMUSG( | Pcdh1 | 6908.38 | 0.309006 | 0.10904 | 0.804969 |
| ENSMUSG( | Pcdh10 | 6965.028 | -0.214 | 0.097864 | 0.478264 | ENSMUSG( | Pcdh10 | 7289.415 | 0.19019 | 0.19662 | 0.862161 |
| ENSMUSG( | Pcdh11x | 310.1821 | -0.49257 | 0.001869 | 0.049292 | ENSMUSG( | Pcdh11x | 375.528 | 0.35212 | 0.074737 | 0.767423 |
| ENSMUSG( | Pcdh12 | 46.70995 | -0.11559 | 0.733412 | 0.944956 | ENSMUSG( | Pcdh12 | 43.34487 | -0.03649 | 0.924015 | 0.992882 |
| ENSMUSG( | Pcdh15 | 678.9386 | -0.19803 | 0.082435 | 0.443042 | ENSMUSG( | Pcdh15 | 690.3736 | 0.127055 | 0.359809 | 0.914223 |
| ENSMUSG( | Pcdh17 | 2624.584 | -0.1957 | 0.151958 | 0.577721 | ENSMUSG( | Pcdh17 | 2804.79 | 0.261631 | 0.098544 | 0.798438 |
| ENSMUSG( | Pcdh18 | 165.0134 | -0.18512 | 0.487328 | 0.858812 | ENSMUSG( | Pcdh18 | 210.4492 | 0.718314 | 0.008613 | 0.391682 |
| ENSMUSG( | Pcdh19 | 737.8118 | -0.10157 | 0.609379 | 0.907634 | ENSMUSG( | Pcdh19 | 823.9659 | 0.461437 | 0.042796 | 0.690304 |
| ENSMUSG( | Pcdh20 | 798.9438 | -0.34246 | 0.049801 | 0.340731 | ENSMUSG( | Pcdh20 | 900.3437 | 0.284117 | 0.150873 | 0.837173 |
| ENSMUSG( | Pcdh7 | 3461.386 | -0.35849 | 0.012563 | 0.161972 | ENSMUSG( | Pcdh7 | 3732.981 | 0.148519 | 0.398989 | 0.917257 |
| ENSMUSG( | Pcdh8 | 1113.824 | -0.31758 | 0.052835 | 0.352814 | ENSMUSG( | Pcdh8 | 1278.356 | 0.350047 | 0.044803 | 0.693959 |
| ENSMUSG( | Pcdh9 | 2836.848 | -0.33905 | 0.033677 | 0.281425 | ENSMUSG( | Pcdh9 | 3191.236 | 0.27971 | 0.139979 | 0.830678 |
| ENSMUSG( | Pcdha1 | 14.56935 | -0.39747 | 0.563022 | NA | ENSMUSG( | Pcdha1 | 17.41268 | 0.427038 | 0.500672 | 0.927825 |
| ENSMUSG( | Pcdha10 | 27.86372 | -0.51488 | 0.279383 | 0.729114 | ENSMUSG( | Pcdha10 | 30.64754 | 0.104345 | 0.814332 | 0.979198 |
| ENSMUSG( | Pcdha11 | 25.40797 | -0.54742 | 0.180674 | 0.617841 | ENSMUSG( | Pcdha11 | 36.93201 | 0.789007 | 0.049379 | 0.708648 |
| ENSMUSG( | Pcdha12 | 91.59458 | -0.3375 | 0.369589 | 0.797172 | ENSMUSG( | Pcdha12 | 89.37836 | -0.09537 | 0.812682 | 0.979148 |
| ENSMUSG( | Pcdha2 | 28.79544 | -0.44728 | 0.332808 | 0.775602 | ENSMUSG( | Pcdha2 | 33.9606 | 0.354934 | 0.44788 | 0.922202 |
| ENSMUSG( | Pcdha3 | 37.20827 | -0.49883 | 0.186569 | 0.623869 | ENSMUSG( | Pcdha3 | 42.74481 | 0.247802 | 0.568512 | 0.941777 |
| ENSMUSG( | Pcdha5 | 50.80137 | -0.65164 | 0.111399 | 0.503454 | ENSMUSG( | Pcdha5 | 63.55202 | 0.346175 | 0.410887 | 0.917257 |
| ENSMUSG( | Pcdha6 | 40.16593 | -0.5791 | 0.105421 | 0.492417 | ENSMUSG( | Pcdha6 | 46.14334 | 0.156869 | 0.654299 | 0.955015 |
| ENSMUSG( | Pcdha7 | 28.43247 | -0.48439 | 0.347703 | 0.780695 | ENSMUSG( | Pcdha7 | 36.07979 | 0.502878 | 0.263264 | 0.885994 |
| ENSMUSG( | Pcdha9 | 56.35446 | -0.61497 | 0.057895 | 0.371023 | ENSMUSG( | Pcdha9 | 65.8707 | 0.197812 | 0.605134 | 0.949082 |
| ENSMUSG( | Pcdhac1 | 39.41171 | -0.20954 | 0.597434 | 0.903694 | ENSMUSG( | Pcdhac1 | 43.27194 | 0.351491 | 0.392422 | 0.917257 |
| ENSMUSG( | Pcdhac2 | 345.3705 | -0.3621 | 0.441165 | 0.832445 | ENSMUSG( | Pcdhac2 | 420.5172 | 0.473606 | 0.428421 | 0.917257 |
| ENSMUSG( | Pcdhb10 | 103.2229 | -0.68416 | 0.001253 | 0.038624 | ENSMUSG( | Pcdhb10 | 115.0023 | -0.01758 | 0.938984 | 0.993898 |
| ENSMUSG( | Pcdhb11 | 71.66782 | -0.65043 | 0.01083 | 0.150145 | ENSMUSG( | Pcdhb11 | 89.36709 | 0.327764 | 0.179955 | 0.850372 |
| ENSMUSG( | Pcdhb12 | 182.0841 | -0.70979 | 0.000184 | 0.009888 | ENSMUSG( | Pcdhb12 | 205.7488 | 0.005614 | 0.977991 | 0.996416 |
| ENSMUSG( | Pcdhb13 | 122.3234 | -0.56803 | 0.016016 | 0.187923 | ENSMUSG( | Pcdhb13 | 139.569 | 0.14123 | 0.545849 | 0.938063 |
| ENSMUSG( | Pcdhb14 | 195.3793 | -0.61636 | 0.002479 | 0.058622 | ENSMUSG( | Pcdhb14 | 220.0633 | 0.06819 | 0.736437 | 0.967344 |
| ENSMUSG( | Pcdhb15 | 94.46829 | -0.44583 | 0.029338 | 0.259647 | ENSMUSG( | Pcdhb15 | 95.57737 | -0.10504 | 0.634759 | 0.953903 |
| ENSMUSG( | Pcdhb16 | 94.0157 | -0.6885 | 0.003412 | 0.073541 | ENSMUSG( | Pcdhb16 | 109.0226 | 0.106348 | 0.687852 | 0.959705 |
| ENSMUSG( | Pcdhb17 | 321.667 | -0.50801 | 0.000845 | 0.029095 | ENSMUSG( | Pcdhb17 | 375.9785 | 0.252226 | 0.131775 | 0.822912 |
| ENSMUSG( | Pcdhb18 | 139.1876 | -0.82731 | 7.94E-05 | 0.005507 | ENSMUSG( | Pcdhb18 | 161.6869 | -0.0154 | 0.950577 | 0.994798 |
| ENSMUSG( | Pcdhb19 | 180.412 | -0.53688 | 0.00128 | 0.039067 | ENSMUSG( | Pcdhb19 | 190.8152 | -0.05205 | 0.786163 | 0.976285 |
| ENSMUSG( | Pcdhb2 | 32.27831 | -0.90905 | 0.028319 | 0.257185 | ENSMUSG( | Pcdhb2 | 29.60837 | -0.83455 | 0.061536 | 0.736111 |
| ENSMUSG( | Pcdhb20 | 311.2079 | -0.54494 | 0.000186 | 0.009972 | ENSMUSG( | Pcdhb20 | 341.1259 | 0.045586 | 0.782207 | 0.976106 |
| ENSMUSG( | Pcdhb21 | 53.37876 | -0.61636 | 0.031856 | 0.272382 | ENSMUSG( | Pcdhb21 | 54.42267 | -0.2121 | 0.492937 | 0.927358 |
| ENSMUSG( | Pcdhb22 | 348.2199 | -0.39581 | 0.025955 | 0.247245 | ENSMUSG( | Pcdhb22 | 375.1176 | 0.120221 | 0.521726 | 0.932571 |
| ENSMUSG( | Pcdhb3 | 136.5871 | -0.38638 | 0.104156 | 0.489971 | ENSMUSG( | Pcdhb3 | 137.4956 | -0.07206 | 0.775758 | 0.975472 |
| ENSMUSG( | Pcdhb4 | 52.36489 | -0.21185 | 0.457968 | 0.839769 | ENSMUSG( | Pcdhb4 | 58.99828 | 0.384122 | 0.194336 | 0.860096 |
| ENSMUSG( | Pcdhb5 | 132.9149 | -0.41419 | 0.089912 | 0.460493 | ENSMUSG( | Pcdhb5 | 143.6165 | 0.122025 | 0.645477 | 0.953988 |
| ENSMUSG( | Pcdhb6 | 65.68541 | -0.36097 | 0.208634 | 0.650496 | ENSMUSG( | Pcdhb6 | 61.41479 | -0.26622 | 0.346997 | 0.911434 |
| ENSMUSG( | Pcdhb7 | 203.2934 | -0.45199 | 0.009884 | 0.141868 | ENSMUSG( | Pcdhb7 | 220.389 | 0.086859 | 0.654416 | 0.955015 |
| ENSMUSG( | Pcdhb8 | 68.88744 | -0.02755 | 0.910312 | 0.9881 | ENSMUSG( | Pcdhb8 | 70.37716 | 0.280604 | 0.288 | 0.894644 |
| ENSMUSG( | Pcdhb9 | 236.895 | -0.64632 | 4.09E-05 | 0.00344 | ENSMUSG( | Pcdhb9 | 263.993 | 0.010001 | 0.954104 | 0.994798 |
| ENSMUSG( | Pcdhga1 | 112.447 | -0.21566 | 0.346531 | 0.779166 | ENSMUSG( | Pcdhga1 | 116.5546 | 0.182945 | 0.498075 | 0.927825 |
| ENSMUSG( | Pcdhga10 | 184.3248 | -0.15579 | 0.430109 | 0.826789 | ENSMUSG( | Pcdhga10 | 178.24 | 0.037712 | 0.867941 | 0.98781 |
| ENSMUSG( | Pcdhga11 | 692.7978 | 0.168372 | 0.333147 | 0.775602 | ENSMUSG( | Pcdhga11 | 676.7007 | 0.35861 | 0.040607 | 0.683847 |
| ENSMUSG( | Pcdhga12 | 568.6135 | 0.111161 | 0.552513 | 0.886141 | ENSMUSG( | Pcdhga12 | 544.2462 | 0.253133 | 0.202179 | 0.864412 |
| ENSMUSG( | Pcdhga2 | 152.5251 | -0.11898 | 0.5482 | 0.885197 | ENSMUSG( | Pcdhga2 | 165.5662 | 0.381919 | 0.093678 | 0.795735 |
| ENSMUSG( | Pcdhga3 | 301.4449 | -0.1571 | 0.448014 | 0.834051 | ENSMUSG( | Pcdhga3 | 312.8059 | 0.227896 | 0.328684 | 0.906727 |
| ENSMUSG( | Pcdhga4 | 204.1931 | -0.13846 | 0.488247 | 0.859092 | ENSMUSG( | Pcdhga4 | 201.8199 | 0.112106 | 0.573079 | 0.942912 |
| ENSMUSG( | Pcdhga5 | 210.1918 | -0.0414 | 0.83688 | 0.971111 | ENSMUSG( | Pcdhga5 | 201.6608 | 0.119341 | 0.576142 | 0.942912 |
| ENSMUSG( | Pcdhga6 | 373.1514 | -0.12028 | 0.386576 | 0.804912 | ENSMUSG( | Pcdhga6 | 347.6491 | -0.04969 | 0.782734 | 0.976106 |
| ENSMUSG( | Pcdhga7 | 339.9122 | -0.17392 | 0.349978 | 0.781943 | ENSMUSG( | Pcdhga7 | 345.658 | 0.155439 | 0.431324 | 0.917257 |
| ENSMUSG( | Pcdhga8 | 257.2998 | 0.038189 | 0.823305 | 0.966603 | ENSMUSG( | Pcdhga8 | 231.2322 | 0.007566 | 0.972348 | 0.996416 |
| ENSMUSG( | Pcdhga9 | 118.8968 | -0.00735 | 0.971216 | 0.996369 | ENSMUSG( | Pcdhga9 | 120.2164 | 0.293281 | 0.211693 | 0.864412 |
| ENSMUSG( | Pcdhgb1 | 200.9363 | -0.13451 | 0.571059 | 0.893319 | ENSMUSG( | Pcdhgb1 | 208.4861 | 0.244181 | 0.286476 | 0.894091 |
| ENSMUSG( | Pcdhgb2 | 168.4484 | -0.419 | 0.028544 | 0.257525 | ENSMUSG( | Pcdhgb2 | 173.1974 | -0.01955 | 0.938146 | 0.993731 |
| ENSMUSG( | Pcdhgb4 | 159.5724 | -0.34108 | 0.087656 | 0.454663 | ENSMUSG( | Pcdhgb4 | 162.3735 | 0.013341 | 0.952533 | 0.994798 |
| ENSMUSG( | Pcdhgb5 | 144.9526 | -0.06718 | 0.746152 | 0.949289 | ENSMUSG( | Pcdhgb5 | 140.1861 | 0.120975 | 0.601316 | 0.947749 |
| ENSMUSG( | Pcdhgb6 | 346.543 | -0.21883 | 0.277453 | 0.727432 | ENSMUSG( | Pcdhgb6 | 341.561 | 0.026842 | 0.894413 | 0.990558 |
| ENSMUSG( | Pcdhgb7 | 242.7267 | -0.07198 | 0.683656 | 0.932154 | ENSMUSG( | Pcdhgb7 | 239.9986 | 0.174475 | 0.385063 | 0.917257 |
| ENSMUSG( | Pcdhgc3 | 7755.958 | 0.0334 | 0.766379 | 0.954173 | ENSMUSG( | Pcdhgc3 | 6793.842 | -0.07902 | 0.503397 | 0.927825 |
| ENSMUSG( | Pcdhgc4 | 232.4103 | -0.38074 | 0.051458 | 0.348059 | ENSMUSG( | Pcdhgc4 | 242.4608 | 0.049348 | 0.832061 | 0.980738 |
| ENSMUSG( | Pcdhgc5 | 2079.941 | -0.37685 | 0.044576 | 0.32426 | ENSMUSG( | Pcdhgc5 | 2181.775 | 0.061978 | 0.76822 | 0.974236 |
| ENSMUSG( | Pced1a | 741.3872 | 0.147088 | 0.167628 | 0.600338 | ENSMUSG( | Pced1a | 672.1877 | 0.136074 | 0.273241 | 0.889657 |
| ENSMUSG( | Pced1b | 197.9943 | -0.09799 | 0.511251 | 0.870859 | ENSMUSG( | Pced1b | 180.3401 | -0.10511 | 0.546343 | 0.938063 |
| ENSMUSG( | Pcf11 | 838.0973 | -0.17154 | 0.095143 | 0.473815 | ENSMUSG( | Pcf11 | 847.0551 | 0.133728 | 0.284318 | 0.892412 |
| ENSMUSG( | Pcgf1 | 240.9821 | 0.080541 | 0.632657 | 0.914073 | ENSMUSG( | Pcgf1 | 211.1443 | -0.02693 | 0.889422 | 0.990558 |
| ENSMUSG( | Pcgf2 | 844.7038 | 0.026827 | 0.888442 | 0.983434 | ENSMUSG( | Pcgf2 | 733.0539 | -0.1214 | 0.567901 | 0.941777 |
| ENSMUSG( | Pcgf3 | 1381.332 | 0.141675 | 0.108585 | 0.497529 | ENSMUSG( | Pcgf3 | 1232.253 | 0.081256 | 0.470799 | 0.923329 |
| ENSMUSG( | Pcgf5 | 554.4493 | 0.018878 | 0.878584 | 0.981431 | ENSMUSG( | Pcgf5 | 543.9534 | 0.221254 | 0.115404 | 0.813375 |
| ENSMUSG( | Pcgf6 | 383.0957 | 0.005827 | 0.965033 | 0.995303 | ENSMUSG( | Pcgf6 | 365.6133 | 0.139692 | 0.40026 | 0.917257 |
| ENSMUSG( | Pcid2 | 1102.471 | 0.195786 | 0.039544 | 0.3066 | ENSMUSG( | Pcid2 | 930.6366 | -0.01802 | 0.873247 | 0.98947 |
| ENSMUSG( | Pcif1 | 2699.419 | 0.108146 | 0.211883 | 0.655541 | ENSMUSG( | Pcif1 | 2418.319 | 0.062233 | 0.554193 | 0.939565 |
| ENSMUSG( | Pck2 | 783.4638 | 0.179595 | 0.100091 | 0.483032 | ENSMUSG( | Pck2 | 689.1969 | 0.085622 | 0.520286 | 0.931964 |
| ENSMUSG( | Pclaf | 40.13826 | -0.07568 | 0.803201 | 0.962169 | ENSMUSG( | Pclaf | 34.31144 | -0.27426 | 0.460001 | 0.923329 |
| ENSMUSG( | Pclo | 7003.077 | -0.57458 | 0.066609 | 0.398819 | ENSMUSG( | Pclo | 8352.767 | 0.255179 | 0.238049 | 0.87259 |
| ENSMUSG( | Pcm1 | 2277.823 | 0.025939 | 0.791289 | 0.959461 | ENSMUSG( | Pcm1 | 2046.939 | -0.01518 | 0.896972 | 0.990558 |
| ENSMUSG( | Pcmt1 | 3754.055 | -0.23578 | 0.00794 | 0.124138 | ENSMUSG( | Pcmt1 | 3682.304 | -0.01244 | 0.911267 | 0.992196 |
| ENSMUSG( | Pcmtd1 | 3028.337 | -0.0299 | 0.77096 | 0.954726 | ENSMUSG( | Pcmtd1 | 2819.408 | 0.029698 | 0.828811 | 0.980224 |
| ENSMUSG( | Pcmtd2 | 1453.908 | -0.02461 | 0.809253 | 0.963595 | ENSMUSG( | Pcmtd2 | 1374.591 | 0.079636 | 0.501251 | 0.927825 |
| ENSMUSG( | Pcna | 958.7802 | 0.062056 | 0.573561 | 0.893584 | ENSMUSG( | Pcna | 835.6733 | -0.06982 | 0.604201 | 0.948139 |
| ENSMUSG( | Pcnp | 2775.029 | -0.0418 | 0.714614 | 0.941144 | ENSMUSG( | Pcnp | 2643.506 | 0.081948 | 0.625086 | 0.953234 |
| ENSMUSG( | Pcnt | 1195.194 | 0.114054 | 0.301388 | 0.749979 | ENSMUSG( | Pcnt | 1132.753 | 0.226952 | 0.105354 | 0.804969 |
| ENSMUSG( | Pcnx1 | 3128.033 | -0.05034 | 0.72514 | 0.944096 | ENSMUSG( | Pcnx1 | 3217.488 | 0.291028 | 0.10184 | 0.801719 |
| ENSMUSG( | Pcnx2 | 3162.741 | -0.11226 | 0.452631 | 0.836145 | ENSMUSG( | Pcnx2 | 3292.533 | 0.268124 | 0.138081 | 0.828492 |
| ENSMUSG( | Pcnx3 | 2127.965 | -0.06688 | 0.601114 | 0.904603 | ENSMUSG( | Pcnx3 | 2173.347 | 0.25708 | 0.123676 | 0.814691 |
| ENSMUSG( | Pcnx4 | 1096.859 | -0.20846 | 0.12693 | 0.534201 | ENSMUSG( | Pcnx4 | 1150.384 | 0.203524 | 0.203039 | 0.864412 |
| ENSMUSG( | Pcolce | 417.9677 | -0.15866 | 0.239204 | 0.687031 | ENSMUSG( | Pcolce | 431.1121 | 0.195606 | 0.16211 | 0.837724 |
| ENSMUSG( | Pcolce2 | 180.0372 | -0.08539 | 0.615521 | 0.910481 | ENSMUSG( | Pcolce2 | 192.3564 | 0.357213 | 0.05636 | 0.723362 |
| ENSMUSG( | Pcp4 | 1646.9 | -0.34363 | 0.024257 | 0.238311 | ENSMUSG( | Pcp4 | 1514.808 | -0.31998 | 0.120764 | 0.813384 |
| ENSMUSG( | Pcp4l1 | 2858.142 | -0.74112 | 9.11E-06 | 0.001083 | ENSMUSG( | Pcp4l1 | 3046.838 | -0.20404 | 0.250007 | 0.877904 |
| ENSMUSG( | Pcsk1 | 883.3353 | -0.69091 | 0.000676 | 0.025081 | ENSMUSG( | Pcsk1 | 1122.714 | 0.334376 | 0.117904 | 0.813384 |
| ENSMUSG( | Pcsk1n | 46432.86 | -0.0769 | 0.363901 | 0.793343 | ENSMUSG( | Pcsk1n | 42247.24 | -0.0815 | 0.483958 | 0.925289 |
| ENSMUSG( | Pcsk2 | 10817.89 | -0.44034 | 0.003594 | 0.075902 | ENSMUSG( | Pcsk2 | 12027.24 | 0.167692 | 0.337653 | 0.907972 |
| ENSMUSG( | Pcsk2os1 | 103.0621 | 0.114554 | 0.59162 | 0.901449 | ENSMUSG( | Pcsk2os1 | 92.25362 | 0.064182 | 0.771408 | 0.974236 |
| ENSMUSG( | Pcsk4 | 166.2986 | 0.052262 | 0.766259 | 0.954173 | ENSMUSG( | Pcsk4 | 167.121 | 0.308807 | 0.146343 | 0.835973 |
| ENSMUSG( | Pcsk5 | 460.0404 | -0.36424 | 0.005278 | 0.096847 | ENSMUSG( | Pcsk5 | 466.9502 | -0.02383 | 0.879081 | 0.989644 |
| ENSMUSG( | Pcsk6 | 1424.319 | -0.10413 | 0.462378 | 0.842104 | ENSMUSG( | Pcsk6 | 1343.308 | 0.000332 | 0.998326 | 0.999313 |
| ENSMUSG( | Pcsk7 | 912.2202 | -0.03985 | 0.752021 | 0.950417 | ENSMUSG( | Pcsk7 | 872.3185 | 0.102625 | 0.443807 | 0.921734 |
| ENSMUSG( | Pcsk9 | 103.6682 | -2.99754 | 7.87E-31 | 1.82E-27 | ENSMUSG( | Pcsk9 | 150.1556 | -0.33112 | 0.084833 | 0.779799 |

| | | | | | |
|---|---|---|---|---|---|
| ENSMUSG( | 1610.204 | 0.227968 | 0.058089 | 0.371677 | Osbpl6 |
| ENSMUSG( | 468.6924 | -0.01718 | 0.896461 | 0.984868 | Osbpl7 |
| ENSMUSG( | 1472.842 | -0.08137 | 0.397586 | 0.812127 | Osbpl8 |
| ENSMUSG( | 1342.379 | 0.060862 | 0.51349 | 0.871762 | Osbpl9 |
| ENSMUSG( | 71.52519 | 0.345514 | 0.243103 | 0.690443 | Oscar |
| ENSMUSG( | 1043.654 | -0.05465 | 0.559316 | 0.888638 | Oscp1 |
| ENSMUSG( | 126.3927 | -0.09644 | 0.613549 | 0.909044 | Oser1 |
| ENSMUSG( | 901.4706 | 0.063623 | 0.539694 | 0.88315 | Osgep |
| ENSMUSG( | 66.50375 | -0.52542 | 0.069375 | 0.40782 | Osgepl1 |
| ENSMUSG( | 13.85954 | -0.85039 | 0.094126 | NA | Osgin1 |
| ENSMUSG( | 564.3748 | 0.054062 | 0.609591 | 0.9077 | Osgin2 |
| ENSMUSG( | 264.3928 | 0.18326 | 0.321253 | 0.766497 | Osmr |
| ENSMUSG( | 82.27286 | -0.21162 | 0.369802 | 0.797313 | Osr1 |
| ENSMUSG( | 1436.581 | -0.0085 | 0.926274 | 0.991229 | Ost4 |
| ENSMUSG( | 1214.466 | -0.34832 | 0.006641 | 0.112089 | Ostc |
| ENSMUSG( | 575.8882 | -0.17164 | 0.152944 | 0.579346 | Ostf1 |
| ENSMUSG( | 2006.76 | -0.35134 | 0.01323 | 0.167129 | Ostm1 |
| ENSMUSG( | 12.77106 | 0.703321 | 0.210623 | NA | Otoa |
| ENSMUSG( | 675.7485 | -0.09596 | 0.638893 | 0.916418 | Otof |
| ENSMUSG( | 12.40966 | 0.141495 | 0.795696 | NA | Otogl |
| ENSMUSG( | 34.10886 | -0.73986 | 0.023385 | 0.233882 | Otop2 |
| ENSMUSG( | 4222.892 | 0.030152 | 0.798999 | 0.960922 | Otub1 |
| ENSMUSG( | 1002.193 | 0.189809 | 0.083737 | 0.446196 | Otub2 |
| ENSMUSG( | 412.1051 | -0.17067 | 0.205104 | 0.646229 | Otud1 |
| ENSMUSG( | 319.2434 | 0.069375 | 0.615422 | 0.910481 | Otud3 |
| ENSMUSG( | 1307.273 | -0.16632 | 0.07959 | 0.435679 | Otud4 |
| ENSMUSG( | 2937.928 | 0.142123 | 0.083518 | 0.445612 | Otud5 |
| ENSMUSG( | 2515.93 | 0.139447 | 0.274499 | 0.724787 | Otud6b |
| ENSMUSG( | 1489.916 | 0.26882 | 0.02333 | 0.233882 | Otud7a |
| ENSMUSG( | 2976.92 | -0.00523 | 0.951975 | 0.993821 | Otud7b |
| ENSMUSG( | 684.1343 | 0.027193 | 0.80645 | 0.962804 | Otulin |
| ENSMUSG( | 672.0144 | -0.3462 | 0.037742 | 0.30023 | Otulinl |
| ENSMUSG( | 246.243 | -0.01709 | 0.906353 | 0.986998 | Otx1 |
| ENSMUSG( | 51.47999 | -0.47072 | 0.641347 | 0.916693 | Otx2 |
| ENSMUSG( | 293.3432 | 0.058153 | 0.659491 | 0.924506 | Ovca2 |
| ENSMUSG( | 106.9787 | 0.390627 | 0.161477 | 0.59148 | Ovgp1 |
| ENSMUSG( | 132.5283 | -0.06429 | 0.797855 | 0.96053 | Ovol2 |
| ENSMUSG( | 1872.195 | -0.03572 | 0.693041 | 0.934202 | Oxa1l |
| ENSMUSG( | 14931.27 | 0.131116 | 0.168993 | 0.603241 | Oxct1 |
| ENSMUSG( | 128.4377 | -0.06371 | 0.732089 | 0.944956 | Oxld1 |
| ENSMUSG( | 600.9802 | 0.066524 | 0.561974 | 0.889052 | Oxnad1 |
| ENSMUSG( | 4090.294 | 0.137292 | 0.134805 | 0.54834 | Oxr1 |
| ENSMUSG( | 328.6633 | 0.046835 | 0.745457 | 0.949158 | Oxsm |
| ENSMUSG( | 4484.675 | -0.05898 | 0.492223 | 0.860297 | Oxsr1 |
| ENSMUSG( | 270.1516 | -0.06391 | 0.68407 | 0.932484 | Oxtr |
| ENSMUSG( | 23.83939 | -1.12174 | 0.005727 | 0.101308 | P2rx1 |
| ENSMUSG( | 55.64878 | 0.054978 | 0.828641 | 0.969097 | P2rx3 |
| ENSMUSG( | 571.8559 | -0.29964 | 0.01806 | 0.202299 | P2rx4 |
| ENSMUSG( | 25.44991 | -0.3527 | 0.363508 | 0.793339 | P2rx5 |
| ENSMUSG( | 102.569 | -0.29234 | 0.258014 | 0.709319 | P2rx6 |
| ENSMUSG( | 457.7173 | -0.329 | 0.027656 | 0.255168 | P2rx7 |
| ENSMUSG( | 65.61439 | -0.62154 | 0.012791 | 0.163486 | P2ry1 |
| ENSMUSG( | 10.11119 | -0.21674 | 0.704826 | NA | P2ry10b |
| ENSMUSG( | 1845.237 | -0.378 | 0.034639 | 0.285926 | P2ry12 |
| ENSMUSG( | 667.5492 | -0.59192 | 0.004455 | 0.086961 | P2ry13 |
| ENSMUSG( | 111.1355 | -0.13863 | 0.594827 | 0.903007 | P2ry14 |
| ENSMUSG( | 27.73503 | 0.009067 | 0.982412 | 0.997512 | P2ry2 |
| ENSMUSG( | 156.0238 | -0.13151 | 0.608203 | 0.907154 | P2ry6 |
| ENSMUSG( | 334.4806 | -0.0397 | 0.778463 | 0.956215 | P3h1 |
| ENSMUSG( | 173.2876 | 0.186132 | 0.272162 | 0.7223 | P3h2 |
| ENSMUSG( | 876.6731 | 0.094622 | 0.346527 | 0.779166 | P3h3 |
| ENSMUSG( | 579.7754 | -0.24093 | 0.058414 | 0.372872 | P3h4 |
| ENSMUSG( | 2526.872 | -0.12279 | 0.416393 | 0.820842 | P4ha1 |
| ENSMUSG( | 368.1449 | 0.295899 | 0.023119 | 0.233201 | P4ha2 |
| ENSMUSG( | 350.898 | -0.24616 | 0.091892 | 0.465937 | P4ha3 |
| ENSMUSG( | 5400.44 | -0.36472 | 0.007975 | 0.124256 | P4hb |
| ENSMUSG( | 2406.247 | -0.22593 | 0.080939 | 0.439809 | P4htm |
| ENSMUSG( | 1798.908 | 0.067249 | 0.507873 | 0.86795 | Pa2g4 |
| ENSMUSG( | 250.8912 | -0.01098 | 0.947683 | 0.993821 | Pabir1 |
| ENSMUSG( | 218.8725 | -0.06259 | 0.679828 | 0.930983 | Pabir2 |
| ENSMUSG( | 11318.29 | 0.079593 | 0.438517 | 0.830637 | Pabpc1 |
| ENSMUSG( | 22.84123 | -0.31236 | 0.451474 | 0.835567 | Pabpc1l |
| ENSMUSG( | 1463.831 | 0.025215 | 0.809424 | 0.963674 | Pabpc4 |
| ENSMUSG( | 10.25778 | -0.09439 | 0.867319 | NA | Pabpc4l |
| ENSMUSG( | 11.12523 | -0.36286 | 0.52493 | NA | Pabpc5 |
| ENSMUSG( | 1208.852 | 0.144446 | 0.428396 | 0.826032 | Pabpn1 |
| ENSMUSG( | 336.1251 | -0.15369 | 0.42672 | 0.825155 | Pacc1 |
| ENSMUSG( | 511.1745 | 0.008122 | 0.957859 | 0.993874 | Pacrg |
| ENSMUSG( | 215.7422 | 0.093462 | 0.514533 | 0.872055 | Pacrgl |
| ENSMUSG( | 2048.459 | 0.21027 | 0.091831 | 0.465904 | Pacs1 |
| ENSMUSG( | 13186.09 | -0.02461 | 0.780264 | 0.957491 | Pacs2 |
| ENSMUSG( | 12362.73 | 0.109172 | 0.239233 | 0.687031 | Pacsin1 |
| ENSMUSG( | 1663.155 | 0.036645 | 0.710671 | 0.939759 | Pacsin2 |
| ENSMUSG( | 2147.971 | 0.03653 | 0.663138 | 0.9265 | Pacsin3 |
| ENSMUSG( | 1984.102 | -0.01848 | 0.84444 | 0.972023 | Padi2 |
| ENSMUSG( | 40.6829 | 0.227505 | 0.483983 | 0.856501 | Padi6 |
| ENSMUSG( | 1669.465 | 0.036937 | 0.768021 | 0.954621 | Paf1 |
| ENSMUSG( | 12593.86 | 0.070795 | 0.364456 | 0.793796 | Pafah1b1 |
| ENSMUSG( | 3355.894 | 0.016671 | 0.854004 | 0.973491 | Pafah1b2 |
| ENSMUSG( | 192.2595 | -0.07214 | 0.669265 | 0.928136 | Pafah1b3 |
| ENSMUSG( | 441.8403 | 0.134007 | 0.290487 | 0.740483 | Pafah2 |
| ENSMUSG( | 1035.18 | -0.30931 | 0.025596 | 0.246694 | Pag1 |
| ENSMUSG( | 32.34215 | 0.118441 | 0.742883 | 0.948478 | Pagr1a |
| ENSMUSG( | 3838.791 | 0.115302 | 0.259618 | 0.709951 | Paics |
| ENSMUSG( | 2053.597 | 0.031903 | 0.763373 | 0.953781 | Paip1 |
| ENSMUSG( | 2787.481 | 0.038715 | 0.707305 | 0.93949 | Paip2 |
| ENSMUSG( | 2327.563 | -0.01153 | 0.901049 | 0.985633 | Paip2b |
| ENSMUSG( | 7383.858 | 0.154817 | 0.129171 | 0.538324 | Pak1 |
| ENSMUSG( | 825.3512 | 0.099147 | 0.444025 | 0.833239 | Pak1ip1 |
| ENSMUSG( | 2360.441 | 0.126966 | 0.260668 | 0.710468 | Pak2 |
| ENSMUSG( | 1136.855 | 0.097127 | 0.348102 | 0.78132 | Pak3 |
| ENSMUSG( | 487.5066 | -0.14787 | 0.197385 | 0.637004 | Pak4 |
| ENSMUSG( | 669.2795 | -0.35295 | 0.000731 | 0.026277 | Pak5 |
| ENSMUSG( | 5622.984 | 0.183096 | 0.026244 | 0.247758 | Pak6 |
| ENSMUSG( | 103.0999 | 0.2986 | 0.169529 | 0.603608 | Pakap |
| ENSMUSG( | 29.2244 | -0.04761 | 0.905016 | 0.986449 | Palb2 |
| ENSMUSG( | 519.433 | -0.10962 | 0.444422 | 0.833239 | Pald1 |
| ENSMUSG( | 587.0079 | 0.25103 | 0.128479 | 0.536774 | Palld |
| ENSMUSG( | 20210.7 | 0.269982 | 0.002112 | 0.053164 | Palm |
| ENSMUSG( | 70.16313 | -0.67632 | 0.003081 | 0.068742 | Palm3 |
| ENSMUSG( | 547.4944 | 0.096834 | 0.536158 | 0.881449 | Palmd |
| ENSMUSG( | 1091.394 | 0.079655 | 0.525287 | 0.876559 | Pals1 |
| ENSMUSG( | 3180.588 | 0.198868 | 0.079581 | 0.435679 | Pals2 |
| ENSMUSG( | 5200.861 | -0.2163 | 0.117156 | 0.516516 | Pam |
| ENSMUSG( | 632.0082 | -0.01621 | 0.889108 | 0.983434 | Pam16 |
| ENSMUSG( | 14.54931 | 0.178825 | 0.718946 | NA | Pam16l |
| ENSMUSG( | 1653.583 | -0.23216 | 0.149609 | 0.573617 | Pamr1 |
| ENSMUSG( | 750.1519 | 0.135698 | 0.334593 | 0.776386 | Pan2 |
| ENSMUSG( | 1198.608 | 0.079962 | 0.382429 | 0.803227 | Pan3 |
| ENSMUSG( | 74.27845 | -0.08374 | 0.712305 | 0.94031 | Panct2 |
| ENSMUSG( | 1295.911 | 0.174803 | 0.170019 | 0.604759 | Pank1 |
| ENSMUSG( | 1586.719 | 0.074827 | 0.431621 | 0.827432 | Pank2 |
| ENSMUSG( | 2324.484 | 0.050487 | 0.685902 | 0.932981 | Pank3 |
| ENSMUSG( | 901.4028 | 0.100491 | 0.450375 | 0.835344 | Pank4 |
| ENSMUSG( | 634.5144 | -0.34963 | 0.032553 | 0.276297 | Panx1 |
| ENSMUSG( | 6692.878 | 0.218016 | 0.016291 | 0.18942 | Panx2 |
| ENSMUSG( | 241.1106 | -0.02566 | 0.855747 | 0.973813 | Paox |
| ENSMUSG( | 91.34283 | -0.08847 | 0.683293 | 0.932153 | Papln |
| ENSMUSG( | 3061.006 | 0.121191 | 0.131656 | 0.544205 | Papola |
| ENSMUSG( | 8.28081 | -0.50636 | 0.481772 | NA | Papolb |
| ENSMUSG( | 276.1517 | 0.206973 | 0.160035 | 0.589679 | Papolg |
| ENSMUSG( | 14.34333 | -0.1075 | 0.831856 | NA | Pappa |
| ENSMUSG( | 81.15466 | -0.71532 | 0.007643 | 0.120942 | Pappa2 |
| ENSMUSG( | 3486.389 | 0.052877 | 0.526142 | 0.877173 | Papss1 |
| ENSMUSG( | 1947.953 | 0.096211 | 0.45557 | 0.838412 | Papss2 |
| ENSMUSG( | 268.5597 | -0.23931 | 0.222965 | 0.668853 | Paqr3 |
| ENSMUSG( | 3370.504 | -0.26169 | 0.01668 | 0.192198 | Paqr4 |
| ENSMUSG( | 161.522 | 0.679169 | 0.00311 | 0.069189 | Paqr5 |
| ENSMUSG( | 468.3904 | -0.1265 | 0.271702 | 0.722082 | Paqr6 |
| ENSMUSG( | 4106.932 | -0.23156 | 0.102781 | 0.488221 | Paqr7 |
| ENSMUSG( | 24398.75 | 0.29942 | 0.013064 | 0.165414 | Paqr8 |
| ENSMUSG( | 1625.942 | -0.18089 | 0.282704 | 0.73186 | Paqr9 |
| ENSMUSG( | 874.563 | -0.15465 | 0.162281 | 0.592836 | Pard3 |
| ENSMUSG( | 450.9795 | 0.229685 | 0.057552 | 0.370143 | Pard3b |
| ENSMUSG( | 640.8458 | -0.00702 | 0.955418 | 0.993874 | Pard6a |
| ENSMUSG( | 77.03757 | -0.15157 | 0.604733 | 0.905556 | Pard6b |
| ENSMUSG( | 230.0828 | -0.21226 | 0.147403 | 0.569664 | Pard6g |
| ENSMUSG( | 710.7913 | 0.226178 | 0.028792 | 0.258585 | Parg |
| ENSMUSG( | 2769.187 | 0.010189 | 0.91993 | 0.990247 | Park7 |
| ENSMUSG( | 936.341 | -0.1313 | 0.188987 | 0.627678 | Parl |
| ENSMUSG( | 1686.586 | -0.29088 | 0.088493 | 0.456254 | Parm1 |
| ENSMUSG( | 889.5421 | -0.02978 | 0.764508 | 0.954055 | Parn |
| ENSMUSG( | 1890.905 | 0.112967 | 0.260408 | 0.710245 | Parp1 |
| ENSMUSG( | 92.95848 | -0.07741 | 0.701276 | 0.93743 | Parp10 |
| ENSMUSG( | 558.303 | 0.003641 | 0.974828 | 0.997207 | Parp11 |
| ENSMUSG( | 461.6631 | -0.00675 | 0.958126 | 0.993874 | Parp12 |
| ENSMUSG( | 203.8275 | -0.15448 | 0.351029 | 0.783067 | Parp14 |
| ENSMUSG( | 89.86136 | 0.064337 | 0.762727 | 0.953519 | Parp16 |
| ENSMUSG( | 287.7313 | 0.241511 | 0.269487 | 0.887332 | Paqr3 |
| ENSMUSG( | 3302.028 | -0.03813 | 0.770903 | 0.974236 | Paqr4 |
| ENSMUSG( | 199.5219 | 1.350877 | 0.009726 | 0.406458 | Paqr5 |
| ENSMUSG( | 428.7528 | -0.11187 | 0.402918 | 0.917257 | Paqr6 |
| ENSMUSG( | 4269.639 | 0.157368 | 0.327779 | 0.906255 | Paqr7 |
| ENSMUSG( | 21065.04 | 0.154118 | 0.236992 | 0.872173 | Paqr8 |
| ENSMUSG( | 1752.549 | 0.301546 | 0.128284 | 0.818934 | Paqr9 |
| ENSMUSG( | 804.7196 | -0.12762 | 0.293471 | 0.898538 | Pard3 |
| ENSMUSG( | 378.67 | -0.00757 | 0.963814 | 0.995433 | Pard3b |
| ENSMUSG( | 596.0495 | 0.058521 | 0.687528 | 0.959673 | Pard6a |
| ENSMUSG( | 74.3552 | 0.009101 | 0.976002 | 0.996416 | Pard6b |
| ENSMUSG( | 218.8697 | -0.08001 | 0.619326 | 0.952814 | Pard6g |
| ENSMUSG( | 646.3934 | 0.219667 | 0.078482 | 0.77321 | Parg |
| ENSMUSG( | 2317.014 | -0.25001 | 0.06443 | 0.737935 | Park7 |
| ENSMUSG( | 838.2016 | -0.18354 | 0.15453 | 0.837724 | Parl |
| ENSMUSG( | 1800.105 | 0.180465 | 0.340249 | 0.907972 | Parm1 |
| ENSMUSG( | 787.2 | -0.11408 | 0.335295 | 0.907907 | Parn |
| ENSMUSG( | 1635.795 | -0.03516 | 0.767297 | 0.974236 | Parp1 |
| ENSMUSG( | 96.03824 | 0.28655 | 0.195986 | 0.862161 | Parp10 |
| ENSMUSG( | 491.475 | -0.10016 | 0.457342 | 0.923329 | Parp11 |
| ENSMUSG( | 423.924 | 0.022761 | 0.88156 | 0.989644 | Parp12 |
| ENSMUSG( | 192.4911 | -0.0363 | 0.847033 | 0.985027 | Parp14 |
| ENSMUSG( | 86.48555 | 0.226202 | 0.371046 | 0.915491 | Parp16 |
| ENSMUSG( | 452.8825 | -0.02085 | 0.888047 | 0.990442 | Parp2 |
| ENSMUSG( | 197.9519 | -0.19559 | 0.275871 | 0.890441 | Parp3 |
| ENSMUSG( | 305.9791 | 0.146078 | 0.315325 | 0.901306 | Parp4 |
| ENSMUSG( | 6537.542 | 0.036264 | 0.704507 | 0.962677 | Parp6 |
| ENSMUSG( | 621.9549 | 0.012727 | 0.924091 | 0.992904 | Parp8 |
| ENSMUSG( | 204.9653 | 0.021172 | 0.91208 | 0.992196 | Parp9 |
| ENSMUSG( | 10.42009 | -0.23284 | 0.706895 | NA | Parpbp |
| ENSMUSG( | 151.5258 | -0.03245 | 0.862245 | 0.986412 | Pars2 |
| ENSMUSG( | 445.9144 | -0.18316 | 0.203902 | 0.864412 | Particl |
| ENSMUSG( | 3582.726 | -0.15279 | 0.181893 | 0.850579 | Parva |
| ENSMUSG( | 14.40969 | 0.408847 | 0.45772 | 0.923329 | Parvaos |
| ENSMUSG( | 1763.556 | -0.20261 | 0.082374 | 0.77681 | Parvb |
| ENSMUSG( | 312.7973 | -0.21094 | 0.153477 | 0.837724 | Parvg |
| ENSMUSG( | 18.93963 | 0.727057 | 0.141731 | 0.830709 | Pasd1 |
| ENSMUSG( | 115.9476 | 0.445628 | 0.044973 | 0.693959 | Pask |
| ENSMUSG( | 19.52445 | 0.2655 | 0.553593 | 0.939565 | Pate2 |
| ENSMUSG( | 280.7857 | -0.28348 | 0.12116 | 0.813384 | Patj |
| ENSMUSG( | 915.5396 | -0.03204 | 0.810719 | 0.978779 | Patl1 |
| ENSMUSG( | 951.3341 | 0.010754 | 0.927268 | 0.99306 | Patz1 |
| ENSMUSG( | 83.14167 | 0.027155 | 0.915597 | 0.992529 | Pawr |
| ENSMUSG( | 9.231073 | 0.139313 | 0.838325 | NA | Pax5 |
| ENSMUSG( | 699.0728 | -0.02054 | 0.904958 | 0.991089 | Pax6 |
| ENSMUSG( | 18.3458 | 0.037158 | 0.940632 | 0.99445 | Pax6os1 |
| ENSMUSG( | 1606.006 | 0.163425 | 0.162032 | 0.837724 | Paxbp1 |
| ENSMUSG( | 2021.232 | 0.160528 | 0.184439 | 0.853168 | Paxip1 |
| ENSMUSG( | 1551.394 | 0.010027 | 0.926859 | 0.99306 | Paxx |
| ENSMUSG( | 349.605 | -0.18969 | 0.199969 | 0.863016 | Pbdc1 |
| ENSMUSG( | 44.28567 | -0.25225 | 0.420072 | 0.917257 | Pbk |
| ENSMUSG( | 31.38915 | -0.4532 | 0.219921 | 0.866104 | Pbld2 |
| ENSMUSG( | 7.898323 | -0.43018 | 0.590094 | NA | Pbp2 |
| ENSMUSG( | 1143.975 | 0.077991 | 0.505275 | 0.927825 | Pbrm1 |
| ENSMUSG( | 4863.154 | -0.04069 | 0.734614 | 0.967318 | Pbx1 |
| ENSMUSG( | 1328.933 | 0.066486 | 0.608165 | 0.950113 | Pbx2 |
| ENSMUSG( | 421.8744 | 0.128342 | 0.584574 | 0.944975 | Pbx3 |
| ENSMUSG( | 309.1024 | -0.08009 | 0.621618 | 0.953199 | Pbx4 |
| ENSMUSG( | 7060.88 | -0.19301 | 0.128726 | 0.819665 | Pbxip1 |
| ENSMUSG( | 102.2878 | 0.012706 | 0.955164 | 0.994798 | Pcbd1 |
| ENSMUSG( | 218.0192 | -0.24238 | 0.204182 | 0.864412 | Pcbd2 |
| ENSMUSG( | 3115.604 | -0.06373 | 0.685879 | 0.958956 | Pcbp1 |
| ENSMUSG( | 7365.816 | -0.04358 | 0.730663 | 0.967228 | Pcbp2 |
| ENSMUSG( | 949.4531 | 0.09752 | 0.646176 | 0.953988 | Pcbp3 |
| ENSMUSG( | 6785.46 | 0.085627 | 0.391728 | 0.917257 | Pcbp4 |
| ENSMUSG( | 908.063 | -0.19772 | 0.093953 | 0.795735 | Pcca |
| ENSMUSG( | 2768.658 | -0.19479 | 0.092968 | 0.79479 | Pccb |
| ENSMUSG( | 6908.38 | 0.309006 | 0.10904 | 0.804969 | Pcdh1 |
| ENSMUSG( | 7289.415 | 0.19019 | 0.19662 | 0.862161 | Pcdh10 |
| ENSMUSG( | 375.528 | 0.35212 | 0.074737 | 0.767423 | Pcdh11x |
| ENSMUSG( | 43.34487 | -0.03649 | 0.924015 | 0.992882 | Pcdh12 |
| ENSMUSG( | 690.3736 | 0.127055 | 0.359809 | 0.914223 | Pcdh15 |
| ENSMUSG( | 2804.79 | 0.261631 | 0.098544 | 0.798438 | Pcdh17 |
| ENSMUSG( | 210.4492 | 0.718314 | 0.008613 | 0.391682 | Pcdh18 |
| ENSMUSG( | 823.9659 | 0.461437 | 0.042796 | 0.690304 | Pcdh19 |
| ENSMUSG( | 900.3437 | 0.284117 | 0.150873 | 0.837173 | Pcdh20 |
| ENSMUSG( | 3732.981 | 0.148519 | 0.398989 | 0.917257 | Pcdh7 |
| ENSMUSG( | 1278.356 | 0.350047 | 0.044803 | 0.693959 | Pcdh8 |
| ENSMUSG( | 3191.236 | 0.27971 | 0.139979 | 0.830678 | Pcdh9 |
| ENSMUSG( | 17.41268 | 0.427038 | 0.500672 | 0.927825 | Pcdha1 |
| ENSMUSG( | 30.64754 | 0.104345 | 0.814332 | 0.979198 | Pcdha10 |
| ENSMUSG( | 36.93201 | 0.789007 | 0.049379 | 0.708648 | Pcdha11 |
| ENSMUSG( | 89.37836 | -0.09537 | 0.812682 | 0.979148 | Pcdha12 |
| ENSMUSG( | 33.9606 | 0.354934 | 0.44788 | 0.922202 | Pcdha2 |
| ENSMUSG( | 42.74481 | 0.247802 | 0.568512 | 0.941777 | Pcdha3 |
| ENSMUSG( | 63.55202 | 0.346175 | 0.410887 | 0.917257 | Pcdha5 |
| ENSMUSG( | 46.14334 | 0.156869 | 0.654299 | 0.955015 | Pcdha6 |
| ENSMUSG( | 36.07979 | 0.502878 | 0.263264 | 0.885994 | Pcdha7 |
| ENSMUSG( | 65.8707 | 0.197812 | 0.605134 | 0.949082 | Pcdha9 |
| ENSMUSG( | 43.27194 | 0.351491 | 0.392422 | 0.917257 | Pcdhac1 |
| ENSMUSG( | 420.5172 | 0.473606 | 0.428421 | 0.917257 | Pcdhac2 |
| ENSMUSG( | 115.0023 | -0.01758 | 0.938984 | 0.993898 | Pcdhb10 |
| ENSMUSG( | 89.36709 | 0.327764 | 0.179955 | 0.850372 | Pcdhb11 |
| ENSMUSG( | 205.7488 | 0.005614 | 0.977991 | 0.996416 | Pcdhb12 |
| ENSMUSG( | 139.569 | 0.14123 | 0.545849 | 0.938063 | Pcdhb13 |
| ENSMUSG( | 220.0633 | 0.06819 | 0.736437 | 0.967344 | Pcdhb14 |
| ENSMUSG( | 95.57737 | -0.10504 | 0.634759 | 0.953903 | Pcdhb15 |
| ENSMUSG( | 109.0226 | 0.106348 | 0.687852 | 0.959705 | Pcdhb16 |
| ENSMUSG( | 375.9785 | 0.252226 | 0.131775 | 0.822912 | Pcdhb17 |
| ENSMUSG( | 161.6869 | -0.0154 | 0.950577 | 0.994798 | Pcdhb18 |
| ENSMUSG( | 190.8152 | -0.05205 | 0.786163 | 0.976285 | Pcdhb19 |
| ENSMUSG( | 29.60837 | -0.83455 | 0.061536 | 0.736111 | Pcdhb2 |
| ENSMUSG( | 341.1259 | 0.045586 | 0.782207 | 0.976106 | Pcdhb20 |
| ENSMUSG( | 54.42267 | -0.2121 | 0.492937 | 0.927358 | Pcdhb21 |
| ENSMUSG( | 375.1176 | 0.120221 | 0.521726 | 0.932571 | Pcdhb22 |
| ENSMUSG( | 137.4956 | -0.07206 | 0.775758 | 0.975472 | Pcdhb3 |
| ENSMUSG( | 58.99828 | 0.384122 | 0.194336 | 0.860096 | Pcdhb4 |
| ENSMUSG( | 143.6165 | 0.122025 | 0.645477 | 0.953988 | Pcdhb5 |
| ENSMUSG( | 61.41479 | -0.26622 | 0.346997 | 0.911434 | Pcdhb6 |
| ENSMUSG( | 220.389 | 0.086859 | 0.654416 | 0.955015 | Pcdhb7 |
| ENSMUSG( | 70.37716 | 0.280604 | 0.288 | 0.894644 | Pcdhb8 |
| ENSMUSG( | 263.993 | 0.010001 | 0.954104 | 0.994798 | Pcdhb9 |
| ENSMUSG( | 116.5546 | 0.182945 | 0.498075 | 0.927825 | Pcdhga1 |
| ENSMUSG( | 178.24 | 0.037712 | 0.867941 | 0.98781 | Pcdhga10 |
| ENSMUSG( | 676.7007 | 0.35861 | 0.040607 | 0.683847 | Pcdhga11 |
| ENSMUSG( | 544.2462 | 0.253133 | 0.202179 | 0.864412 | Pcdhga12 |
| ENSMUSG( | 165.5662 | 0.381919 | 0.093678 | 0.795735 | Pcdhga2 |
| ENSMUSG( | 312.8059 | 0.227896 | 0.328684 | 0.906727 | Pcdhga3 |
| ENSMUSG( | 201.8199 | 0.112106 | 0.573079 | 0.942912 | Pcdhga4 |
| ENSMUSG( | 201.6608 | 0.119341 | 0.576142 | 0.942912 | Pcdhga5 |
| ENSMUSG( | 347.6491 | -0.04969 | 0.782734 | 0.976106 | Pcdhga6 |
| ENSMUSG( | 345.658 | 0.155439 | 0.431324 | 0.917257 | Pcdhga7 |
| ENSMUSG( | 231.2322 | 0.007566 | 0.972348 | 0.996416 | Pcdhga8 |
| ENSMUSG( | 120.2164 | 0.293281 | 0.211693 | 0.864412 | Pcdhga9 |
| ENSMUSG( | 208.4861 | 0.244181 | 0.286476 | 0.894091 | Pcdhgb1 |
| ENSMUSG( | 173.1974 | -0.01955 | 0.938146 | 0.993731 | Pcdhgb2 |
| ENSMUSG( | 162.3735 | 0.013341 | 0.952533 | 0.994798 | Pcdhgb4 |
| ENSMUSG( | 140.1861 | 0.120975 | 0.601316 | 0.947749 | Pcdhgb5 |
| ENSMUSG( | 341.561 | 0.026842 | 0.894413 | 0.990558 | Pcdhgb6 |
| ENSMUSG( | 239.9986 | 0.174475 | 0.385063 | 0.917257 | Pcdhgb7 |
| ENSMUSG( | 6793.842 | -0.07902 | 0.503397 | 0.927825 | Pcdhgc3 |
| ENSMUSG( | 242.4608 | 0.049348 | 0.832061 | 0.980738 | Pcdhgc4 |
| ENSMUSG( | 2181.775 | 0.061978 | 0.76822 | 0.974236 | Pcdhgc5 |
| ENSMUSG( | 672.1877 | 0.136074 | 0.273241 | 0.889657 | Pced1a |
| ENSMUSG( | 180.3401 | -0.10511 | 0.546343 | 0.938063 | Pced1b |
| ENSMUSG( | 847.0551 | 0.133728 | 0.284318 | 0.892412 | Pcf11 |
| ENSMUSG( | 211.1443 | -0.02693 | 0.889422 | 0.990558 | Pcgf1 |
| ENSMUSG( | 733.0539 | -0.1214 | 0.567901 | 0.941777 | Pcgf2 |
| ENSMUSG( | 1232.253 | 0.081256 | 0.470799 | 0.923329 | Pcgf3 |
| ENSMUSG( | 543.9534 | 0.221254 | 0.115404 | 0.813375 | Pcgf5 |
| ENSMUSG( | 365.6133 | 0.139692 | 0.40026 | 0.917257 | Pcgf6 |
| ENSMUSG( | 930.6366 | -0.01802 | 0.873247 | 0.98947 | Pcid2 |
| ENSMUSG( | 2418.319 | 0.062233 | 0.554193 | 0.939565 | Pcif1 |
| ENSMUSG( | 689.1969 | 0.085622 | 0.520286 | 0.931964 | Pck2 |
| ENSMUSG( | 34.31144 | -0.27426 | 0.460001 | 0.923329 | Pclaf |
| ENSMUSG( | 8352.767 | 0.255179 | 0.238049 | 0.87259 | Pclo |
| ENSMUSG( | 2046.939 | -0.01518 | 0.896972 | 0.990558 | Pcm1 |
| ENSMUSG( | 3682.304 | -0.01244 | 0.911267 | 0.992196 | Pcmt1 |
| ENSMUSG( | 2819.408 | 0.029698 | 0.828811 | 0.980224 | Pcmtd1 |
| ENSMUSG( | 1374.591 | 0.079636 | 0.501251 | 0.927825 | Pcmtd2 |
| ENSMUSG( | 835.6733 | -0.06982 | 0.604201 | 0.948139 | Pcna |
| ENSMUSG( | 2643.506 | 0.081948 | 0.625086 | 0.953234 | Pcnp |
| ENSMUSG( | 1132.753 | 0.226952 | 0.105354 | 0.804969 | Pcnt |
| ENSMUSG( | 3217.488 | 0.291028 | 0.10184 | 0.801719 | Pcnx1 |
| ENSMUSG( | 3292.533 | 0.268124 | 0.138081 | 0.828492 | Pcnx2 |
| ENSMUSG( | 2173.347 | 0.25708 | 0.123676 | 0.814691 | Pcnx3 |
| ENSMUSG( | 1150.384 | 0.203524 | 0.203039 | 0.864412 | Pcnx4 |
| ENSMUSG( | 431.1121 | 0.195606 | 0.16211 | 0.837724 | Pcolce |

| | | | | | | | | | | | |
|---|---|---|---|---|---|---|---|---|---|---|---|
| ENSMUSG( | Pctp | 122.8271 | 0.07413 | 0.699361 | 0.936948 | ENSMUSG( | Pctp | 101.8712 | -0.21531 | 0.318958 | 0.903344 |
| ENSMUSG( | Pcx | 5001.462 | 0.143297 | 0.071593 | 0.413189 | ENSMUSG( | Pcx | 4204.088 | -0.08839 | 0.401983 | 0.917257 |
| ENSMUSG( | Pcyox1 | 4365.045 | -0.30527 | 0.031102 | 0.268324 | ENSMUSG( | Pcyox1 | 4812.064 | 0.254128 | 0.123819 | 0.814691 |
| ENSMUSG( | Pcyox1l | 882.734 | -0.3913 | 0.001392 | 0.040902 | ENSMUSG( | Pcyox1l | 969.8247 | 0.171328 | 0.232504 | 0.868507 |
| ENSMUSG( | Pcyt1a | 2575.814 | 0.081095 | 0.388574 | 0.806278 | ENSMUSG( | Pcyt1a | 2197.406 | -0.11096 | 0.309953 | 0.898538 |
| ENSMUSG( | Pcyt1b | 1841.741 | 0.026735 | 0.774615 | 0.955606 | ENSMUSG( | Pcyt1b | 1594.751 | -0.12381 | 0.250097 | 0.877904 |
| ENSMUSG( | Pcyt2 | 3256.163 | 0.092146 | 0.301659 | 0.749979 | ENSMUSG( | Pcyt2 | 2848.345 | -0.02373 | 0.85895 | 0.985884 |
| ENSMUSG( | Pdap1 | 2526.091 | 0.082355 | 0.404456 | 0.816132 | ENSMUSG( | Pdap1 | 2164.059 | -0.0995 | 0.48615 | 0.92541 |
| ENSMUSG( | Pdcd10 | 598.7237 | 0.040659 | 0.727261 | 0.944223 | ENSMUSG( | Pdcd10 | 550.8675 | 0.065895 | 0.613502 | 0.951177 |
| ENSMUSG( | Pdcd11 | 998.1013 | 0.081631 | 0.449174 | 0.834943 | ENSMUSG( | Pdcd11 | 930.9598 | 0.147877 | 0.224203 | 0.867489 |
| ENSMUSG( | Pdcd2 | 223.6109 | -0.15887 | 0.296205 | 0.744386 | ENSMUSG( | Pdcd2 | 206.419 | -0.12357 | 0.501964 | 0.927825 |
| ENSMUSG( | Pdcd2l | 244.5124 | 0.056514 | 0.695274 | 0.934948 | ENSMUSG( | Pdcd2l | 219.3201 | 0.010758 | 0.948216 | 0.994798 |
| ENSMUSG( | Pdcd4 | 1632.806 | 0.036185 | 0.691544 | 0.933881 | ENSMUSG( | Pdcd4 | 1494.38 | 0.04523 | 0.729215 | 0.966913 |
| ENSMUSG( | Pdcd5 | 982.8029 | 0.070105 | 0.498605 | 0.862973 | ENSMUSG( | Pdcd5 | 825.1384 | -0.17339 | 0.192713 | 0.859764 |
| ENSMUSG( | Pdcd6 | 1448.329 | 0.036618 | 0.712247 | 0.94031 | ENSMUSG( | Pdcd6 | 1234.704 | -0.16221 | 0.191461 | 0.858412 |
| ENSMUSG( | Pdcd6ip | 2502.174 | 0.064577 | 0.437955 | 0.830072 | ENSMUSG( | Pdcd6ip | 2252.847 | 0.030923 | 0.793324 | 0.97681 |
| ENSMUSG( | Pdcd7 | 428.8609 | 0.02311 | 0.854356 | 0.973491 | ENSMUSG( | Pdcd7 | 383.6712 | -0.03072 | 0.828317 | 0.980224 |
| ENSMUSG( | Pdcl | 606.3914 | 0.140346 | 0.29012 | 0.739897 | ENSMUSG( | Pdcl | 534.1716 | 0.045729 | 0.731184 | 0.967228 |
| ENSMUSG( | Pdcl3 | 700.529 | -0.06231 | 0.551739 | 0.886141 | ENSMUSG( | Pdcl3 | 633.8383 | -0.08563 | 0.494099 | 0.927372 |
| ENSMUSG( | Pde10a | 3272.502 | -0.33034 | 0.020601 | 0.216558 | ENSMUSG( | Pde10a | 3328.28 | 0.005622 | 0.966665 | 0.995475 |
| ENSMUSG( | Pde11a | 58.99107 | 0.255974 | 0.33619 | 0.777 | ENSMUSG( | Pde11a | 47.1478 | -0.12834 | 0.68323 | 0.958678 |
| ENSMUSG( | Pde12 | 312.5311 | -0.00332 | 0.982344 | 0.997512 | ENSMUSG( | Pde12 | 280.7705 | -0.04952 | 0.7747 | 0.974931 |
| ENSMUSG( | Pde1a | 2668.26 | 0.176233 | 0.168161 | 0.601451 | ENSMUSG( | Pde1a | 2278.967 | -0.00672 | 0.959808 | 0.994798 |
| ENSMUSG( | Pde1b | 5327.85 | -0.41567 | 0.006685 | 0.112486 | ENSMUSG( | Pde1b | 5181.653 | -0.20671 | 0.215004 | 0.865167 |
| ENSMUSG( | Pde1c | 550.7545 | -0.05658 | 0.722972 | 0.943761 | ENSMUSG( | Pde1c | 467.924 | -0.27932 | 0.06714 | 0.75212 |
| ENSMUSG( | Pde2a | 256.5706 | 0.060413 | 0.691494 | 0.933881 | ENSMUSG( | Pde2a | 226.2583 | -0.04428 | 0.810154 | 0.978779 |
| ENSMUSG( | Pde3a | 186.4086 | -0.00171 | 0.991565 | 0.999232 | ENSMUSG( | Pde3a | 179.8192 | 0.163002 | 0.367506 | 0.915217 |
| ENSMUSG( | Pde3b | 576.1687 | -0.00758 | 0.954228 | 0.993874 | ENSMUSG( | Pde3b | 557.8043 | 0.168807 | 0.283577 | 0.892412 |
| ENSMUSG( | Pde4a | 4730.013 | 0.13551 | 0.201219 | 0.641362 | ENSMUSG( | Pde4a | 4269.126 | 0.107563 | 0.394371 | 0.917257 |
| ENSMUSG( | Pde4b | 2270.197 | -0.09312 | 0.282475 | 0.73186 | ENSMUSG( | Pde4b | 2071.673 | -0.09021 | 0.417535 | 0.917257 |
| ENSMUSG( | Pde4c | 100.0719 | 0.543913 | 0.012589 | 0.161972 | ENSMUSG( | Pde4c | 95.7559 | 0.667644 | 0.005252 | 0.297744 |
| ENSMUSG( | Pde4d | 521.791 | 0.196878 | 0.114075 | 0.509818 | ENSMUSG( | Pde4d | 479.9489 | 0.230058 | 0.136324 | 0.827092 |
| ENSMUSG( | Pde4dip | 5415.917 | 0.156399 | 0.04731 | 0.332697 | ENSMUSG( | Pde4dip | 4774.79 | 0.065084 | 0.506126 | 0.927825 |
| ENSMUSG( | Pde5a | 246.9813 | 0.006244 | 0.969219 | 0.996369 | ENSMUSG( | Pde5a | 250.6778 | 0.31206 | 0.082439 | 0.77681 |
| ENSMUSG( | Pde6a | 17.25231 | -0.68963 | 0.126146 | NA | ENSMUSG( | Pde6a | 19.43232 | 0.031676 | 0.943327 | 0.994798 |
| ENSMUSG( | Pde6d | 1501.404 | 0.080951 | 0.47559 | 0.850919 | ENSMUSG( | Pde6d | 1258.382 | -0.16868 | 0.223274 | 0.867404 |
| ENSMUSG( | Pde6h | 10.35306 | -0.14338 | 0.811489 | NA | ENSMUSG( | Pde6h | 11.94258 | 0.447825 | 0.472116 | 0.923329 |
| ENSMUSG( | Pde7a | 859.3986 | 0.0304 | 0.777731 | 0.956168 | ENSMUSG( | Pde7a | 798.6501 | 0.083734 | 0.523858 | 0.932848 |
| ENSMUSG( | Pde7b | 1337.299 | 0.079998 | 0.55276 | 0.886141 | ENSMUSG( | Pde7b | 1133.504 | -0.13481 | 0.296531 | 0.898538 |
| ENSMUSG( | Pde8a | 2034.521 | -0.00049 | 0.995621 | 0.99943 | ENSMUSG( | Pde8a | 1728.044 | -0.21352 | 0.065917 | 0.746371 |
| ENSMUSG( | Pde8b | 3192.966 | 0.050439 | 0.563025 | 0.889738 | ENSMUSG( | Pde8b | 2765.845 | -0.09642 | 0.345917 | 0.910966 |
| ENSMUSG( | Pde9a | 703.4926 | -0.06269 | 0.577556 | 0.895864 | ENSMUSG( | Pde9a | 594.2803 | -0.29469 | 0.025248 | 0.597654 |
| ENSMUSG( | Pdf | 30.71488 | 0.331852 | 0.36208 | 0.79289 | ENSMUSG( | Pdf | 28.61816 | 0.388578 | 0.312884 | 0.900409 |
| ENSMUSG( | Pdgfa | 1766.356 | -0.00426 | 0.967861 | 0.996237 | ENSMUSG( | Pdgfa | 1734.736 | 0.204817 | 0.105987 | 0.804969 |
| ENSMUSG( | Pdgfb | 1104.569 | 0.054714 | 0.709261 | 0.939656 | ENSMUSG( | Pdgfb | 1074.166 | 0.24011 | 0.141973 | 0.830709 |
| ENSMUSG( | Pdgfc | 401.7253 | -0.09278 | 0.483166 | 0.856216 | ENSMUSG( | Pdgfc | 410.9668 | 0.237355 | 0.106364 | 0.804969 |
| ENSMUSG( | Pdgfd | 180.7306 | 0.151331 | 0.39031 | 0.80794 | ENSMUSG( | Pdgfd | 202.6276 | 0.684443 | 0.000301 | 0.047107 |
| ENSMUSG( | Pdgfra | 2675.174 | -0.37299 | 0.027895 | 0.255773 | ENSMUSG( | Pdgfra | 3042.722 | 0.282428 | 0.162287 | 0.837724 |
| ENSMUSG( | Pdgfrb | 1447.848 | -0.29155 | 0.126631 | 0.534187 | ENSMUSG( | Pdgfrb | 1620.225 | 0.309268 | 0.181045 | 0.850372 |
| ENSMUSG( | Pdgfrl | 92.98141 | -0.35783 | 0.139217 | 0.555317 | ENSMUSG( | Pdgfrl | 96.26113 | 0.041804 | 0.864082 | 0.986552 |
| ENSMUSG( | Pdha1 | 9304.314 | 0.049131 | 0.591179 | 0.90113 | ENSMUSG( | Pdha1 | 7922.038 | -0.15247 | 0.253051 | 0.88018 |
| ENSMUSG( | Pdhb | 2990.646 | 0.11148 | 0.265197 | 0.715962 | ENSMUSG( | Pdhb | 2512.494 | -0.12553 | 0.298019 | 0.898538 |
| ENSMUSG( | Pdhx | 1302.369 | 0.129149 | 0.156555 | 0.584667 | ENSMUSG( | Pdhx | 1077.244 | -0.15449 | 0.23248 | 0.868507 |
| ENSMUSG( | Pdia3 | 7929.542 | -0.24987 | 0.089226 | 0.458482 | ENSMUSG( | Pdia3 | 8446.791 | 0.207743 | 0.190712 | 0.857801 |
| ENSMUSG( | Pdia4 | 1859.693 | -0.11882 | 0.442559 | 0.83278 | ENSMUSG( | Pdia4 | 2092.538 | 0.459413 | 0.004834 | 0.287322 |
| ENSMUSG( | Pdia5 | 123.7435 | -0.17982 | 0.32084 | 0.765984 | ENSMUSG( | Pdia5 | 116.8567 | -0.06681 | 0.767722 | 0.974236 |
| ENSMUSG( | Pdia6 | 3921.628 | -0.11332 | 0.473346 | 0.848919 | ENSMUSG( | Pdia6 | 4066.345 | 0.255523 | 0.13306 | 0.822912 |
| ENSMUSG( | Pdik1l | 256.4257 | -0.08961 | 0.520702 | 0.875186 | ENSMUSG( | Pdik1l | 260.9183 | 0.225632 | 0.154682 | 0.837724 |
| ENSMUSG( | Pdk1 | 1867.621 | 0.169386 | 0.06765 | 0.402445 | ENSMUSG( | Pdk1 | 1636.061 | 0.057969 | 0.611343 | 0.950771 |
| ENSMUSG( | Pdk2 | 4305.775 | 0.046265 | 0.594947 | 0.903007 | ENSMUSG( | Pdk2 | 3658.086 | -0.16142 | 0.165986 | 0.839855 |
| ENSMUSG( | Pdk3 | 691.4144 | -0.04424 | 0.693235 | 0.934202 | ENSMUSG( | Pdk3 | 599.8783 | -0.19716 | 0.17264 | 0.843702 |
| ENSMUSG( | Pdk4 | 1079.97 | -0.17481 | 0.190178 | 0.629268 | ENSMUSG( | Pdk4 | 889.2281 | -0.51184 | 0.000368 | 0.054165 |
| ENSMUSG( | Pdlim1 | 113.6233 | -0.1044 | 0.606625 | 0.906529 | ENSMUSG( | Pdlim1 | 105.5188 | -0.05784 | 0.803103 | 0.976937 |
| ENSMUSG( | Pdlim2 | 2301.716 | -0.22877 | 0.027569 | 0.2548 | ENSMUSG( | Pdlim2 | 2082.632 | -0.25528 | 0.078032 | 0.77321 |
| ENSMUSG( | Pdlim3 | 12.80344 | -0.14734 | 0.787591 | NA | ENSMUSG( | Pdlim3 | 12.44068 | 0.04368 | 0.936446 | 0.993573 |
| ENSMUSG( | Pdlim4 | 640.9144 | 0.213835 | 0.070293 | 0.40908 | ENSMUSG( | Pdlim4 | 530.5086 | -0.05858 | 0.67525 | 0.95694 |
| ENSMUSG( | Pdlim5 | 2252.866 | 0.068886 | 0.597216 | 0.903694 | ENSMUSG( | Pdlim5 | 1996.642 | -0.01253 | 0.935252 | 0.993078 |
| ENSMUSG( | Pdlim7 | 1334.73 | 0.219567 | 0.133046 | 0.545688 | ENSMUSG( | Pdlim7 | 1132.64 | 0.01995 | 0.903407 | 0.991089 |
| ENSMUSG( | Pdp1 | 1406.885 | -0.17388 | 0.192029 | 0.630811 | ENSMUSG( | Pdp1 | 1291.056 | -0.15419 | 0.324675 | 0.905257 |
| ENSMUSG( | Pdp2 | 380.2565 | 0.183531 | 0.146086 | 0.567605 | ENSMUSG( | Pdp2 | 338.8032 | 0.125596 | 0.393701 | 0.917257 |
| ENSMUSG( | Pdpk1 | 3032.189 | 0.086667 | 0.337013 | 0.777008 | ENSMUSG( | Pdpk1 | 2680.03 | 0.000937 | 0.992711 | 0.99874 |
| ENSMUSG( | Pdpn | 816.213 | -0.38693 | 0.015462 | 0.183729 | ENSMUSG( | Pdpn | 820.3544 | -0.07513 | 0.636974 | 0.953903 |
| ENSMUSG( | Pdpr | 1071.62 | 0.019053 | 0.871685 | 0.97812 | ENSMUSG( | Pdpr | 943.3268 | -0.08082 | 0.511091 | 0.929571 |
| ENSMUSG( | Pdrg1 | 962.0494 | -0.07118 | 0.478366 | 0.853567 | ENSMUSG( | Pdrg1 | 836.885 | -0.21461 | 0.080009 | 0.776379 |
| ENSMUSG( | Pds5a | 1608.15 | 0.119153 | 0.225671 | 0.672494 | ENSMUSG( | Pds5a | 1458.998 | 0.108506 | 0.33887 | 0.907972 |
| ENSMUSG( | Pds5b | 1916.334 | 0.072483 | 0.406751 | 0.816928 | ENSMUSG( | Pds5b | 1788.462 | 0.138109 | 0.212092 | 0.864412 |
| ENSMUSG( | Pdss1 | 222.2553 | -0.11001 | 0.51997 | 0.874979 | ENSMUSG( | Pdss1 | 220.9102 | 0.149384 | 0.371017 | 0.915491 |
| ENSMUSG( | Pdss2 | 270.971 | 0.072448 | 0.636308 | 0.914873 | ENSMUSG( | Pdss2 | 232.0679 | -0.11729 | 0.523252 | 0.932848 |
| ENSMUSG( | Pdxdc1 | 4755.247 | 0.085284 | 0.317583 | 0.763932 | ENSMUSG( | Pdxdc1 | 3976.956 | -0.16758 | 0.147253 | 0.836241 |
| ENSMUSG( | Pdxk | 5812.211 | 0.072318 | 0.399169 | 0.813048 | ENSMUSG( | Pdxk | 5262.387 | 0.055637 | 0.596643 | 0.946541 |
| ENSMUSG( | Pdxp | 2569.961 | 0.061507 | 0.59592 | 0.903152 | ENSMUSG( | Pdxp | 2231.341 | -0.08028 | 0.570715 | 0.941973 |
| ENSMUSG( | Pdyn | 957.884 | -0.78916 | 9.93E-09 | 3.91E-06 | ENSMUSG( | Pdyn | 1141.345 | 0.092421 | 0.547008 | 0.938063 |
| ENSMUSG( | Pdzd11 | 944.3874 | 0.004835 | 0.961273 | 0.994061 | ENSMUSG( | Pdzd11 | 794.9703 | -0.23635 | 0.05433 | 0.723362 |
| ENSMUSG( | Pdzd2 | 1962.103 | 0.103531 | 0.253187 | 0.70338 | ENSMUSG( | Pdzd2 | 1902.061 | 0.274078 | 0.037315 | 0.673514 |
| ENSMUSG( | Pdzd4 | 7721.36 | 0.182601 | 0.015606 | 0.184634 | ENSMUSG( | Pdzd4 | 6644.648 | 0.023605 | 0.808269 | 0.978622 |
| ENSMUSG( | Pdzd7 | 168.7144 | 0.434575 | 0.029767 | 0.261412 | ENSMUSG( | Pdzd7 | 136.2152 | 0.122981 | 0.616241 | 0.951413 |
| ENSMUSG( | Pdzd8 | 2234.67 | -0.23194 | 0.090986 | 0.463502 | ENSMUSG( | Pdzd8 | 2346.405 | 0.184339 | 0.25771 | 0.883401 |
| ENSMUSG( | Pdzk1 | 19.3414 | 3.37E-06 | 0.999994 | NA | ENSMUSG( | Pdzk1 | 16.3898 | -0.24176 | 0.635311 | 0.953903 |
| ENSMUSG( | Pdzk1ip1 | 19.8319 | -0.02284 | 0.956337 | NA | ENSMUSG( | Pdzk1ip1 | 14.32994 | -0.75811 | 0.168835 | 0.840222 |
| ENSMUSG( | Pdzph1 | 337.8989 | 0.257024 | 0.087663 | 0.454663 | ENSMUSG( | Pdzph1 | 276.1009 | -0.05795 | 0.747132 | 0.969763 |
| ENSMUSG( | Pdzrn3 | 1033.349 | -0.10016 | 0.432221 | 0.82768 | ENSMUSG( | Pdzrn3 | 905.2145 | -0.22243 | 0.122933 | 0.814458 |
| ENSMUSG( | Pdzrn4 | 190.4867 | -0.33793 | 0.037663 | 0.299895 | ENSMUSG( | Pdzrn4 | 186.5109 | -0.11689 | 0.502288 | 0.927825 |
| ENSMUSG( | Pea15a | 49662.94 | -0.01273 | 0.884618 | 0.982647 | ENSMUSG( | Pea15a | 42142.36 | -0.22841 | 0.071301 | 0.761942 |
| ENSMUSG( | Peak1 | 1670.661 | -0.05945 | 0.509964 | 0.869743 | ENSMUSG( | Peak1 | 1618.9 | 0.115158 | 0.295423 | 0.898538 |
| ENSMUSG( | Pear1 | 303.1885 | 0.262326 | 0.120217 | 0.524481 | ENSMUSG( | Pear1 | 300.8117 | 0.493739 | 0.043149 | 0.690304 |
| ENSMUSG( | Pebp1 | 9233.884 | 0.028148 | 0.748809 | 0.949604 | ENSMUSG( | Pebp1 | 7896.133 | -0.16076 | 0.196953 | 0.862161 |
| ENSMUSG( | Pecam1 | 883.768 | 0.130312 | 0.545334 | 0.884774 | ENSMUSG( | Pecam1 | 915.0949 | 0.47393 | 0.270089 | 0.88756 |
| ENSMUSG( | Pecr | 106.8168 | -0.08219 | 0.680431 | 0.931044 | ENSMUSG( | Pecr | 93.58095 | -0.20327 | 0.372366 | 0.915491 |
| ENSMUSG( | Peds1 | 2156.299 | -0.35696 | 0.002789 | 0.064345 | ENSMUSG( | Peds1 | 2267.791 | 0.079722 | 0.562292 | 0.940752 |
| ENSMUSG( | Pef1 | 1195.774 | 0.043159 | 0.667536 | 0.927404 | ENSMUSG( | Pef1 | 1045.333 | -0.0796 | 0.560681 | 0.940254 |
| ENSMUSG( | Peg10 | 649.2691 | 0.09231 | 0.589191 | 0.90113 | ENSMUSG( | Peg10 | 594.8564 | 0.100762 | 0.519773 | 0.931729 |
| ENSMUSG( | Peg12 | 15.93085 | -0.08679 | 0.857551 | NA | ENSMUSG( | Peg12 | 16.62904 | 0.287211 | 0.566224 | 0.941665 |
| ENSMUSG( | Peg13 | 4131.367 | -0.09868 | 0.34019 | 0.777962 | ENSMUSG( | Peg13 | 3778.244 | -0.08768 | 0.466397 | 0.923329 |
| ENSMUSG( | Peg3 | 7853.012 | -0.17012 | 0.084318 | 0.447526 | ENSMUSG( | Peg3 | 7431.243 | -0.05921 | 0.642649 | 0.953903 |
| ENSMUSG( | Peli1 | 914.2104 | 0.074982 | 0.445418 | 0.833239 | ENSMUSG( | Peli1 | 851.4494 | 0.137166 | 0.283957 | 0.892412 |
| ENSMUSG( | Peli2 | 2406.708 | 0.071955 | 0.529817 | 0.878883 | ENSMUSG( | Peli2 | 2128.573 | -0.01608 | 0.912989 | 0.992298 |
| ENSMUSG( | Peli3 | 471.9281 | 0.258855 | 0.070165 | 0.40908 | ENSMUSG( | Peli3 | 383.373 | -0.05305 | 0.747099 | 0.969763 |
| ENSMUSG( | Pelo | 115.0651 | 0.054676 | 0.766817 | 0.954331 | ENSMUSG( | Pelo | 108.5057 | 0.148808 | 0.470794 | 0.923329 |
| ENSMUSG( | Pelp1 | 1275.021 | 0.094821 | 0.342231 | 0.778828 | ENSMUSG( | Pelp1 | 1107.413 | -0.04481 | 0.727292 | 0.966887 |
| ENSMUSG( | Pemt | 82.89085 | 0.044822 | 0.837554 | 0.971111 | ENSMUSG( | Pemt | 68.81474 | -0.20964 | 0.437406 | 0.920144 |
| ENSMUSG( | Penk | 6763.388 | -1.40769 | 0.000113 | 0.007021 | ENSMUSG( | Penk | 8555.92 | -0.13272 | 0.39837 | 0.917257 |
| ENSMUSG( | Pepd | 616.7714 | 0.285642 | 0.014429 | 0.176902 | ENSMUSG( | Pepd | 532.4656 | 0.144339 | 0.298379 | 0.898538 |
| ENSMUSG( | Per1 | 2472.233 | -0.37681 | 0.001599 | 0.044877 | ENSMUSG( | Per1 | 2381.238 | -0.19664 | 0.27474 | 0.889874 |
| ENSMUSG( | Per2 | 930.1725 | 0.055443 | 0.604822 | 0.905556 | ENSMUSG( | Per2 | 1188.755 | 0.895709 | 6.56E-10 | 6.76E-07 |
| ENSMUSG( | Per3 | 1368.009 | 0.180778 | 0.098242 | 0.479534 | ENSMUSG( | Per3 | 1562.023 | 0.744534 | 1.18E-06 | 0.000604 |
| ENSMUSG( | Perm1 | 84.54323 | 0.028982 | 0.904171 | 0.986246 | ENSMUSG( | Perm1 | 75.13714 | -0.07134 | 0.801107 | 0.97681 |
| ENSMUSG( | Perp | 284.9024 | -0.02998 | 0.835272 | 0.970426 | ENSMUSG( | Perp | 309.6235 | 0.448433 | 0.002539 | 0.197651 |
| ENSMUSG( | Pes1 | 1309.697 | 0.167934 | 0.079615 | 0.435679 | ENSMUSG( | Pes1 | 1088.023 | -0.10002 | 0.444167 | 0.921734 |
| ENSMUSG( | Pet100 | 766.213 | 0.046842 | 0.660919 | 0.92497 | ENSMUSG( | Pet100 | 677.345 | -0.04514 | 0.731885 | 0.967228 |
| ENSMUSG( | Pex1 | 542.1516 | 0.183435 | 0.13185 | 0.544244 | ENSMUSG( | Pex1 | 494.7383 | 0.19246 | 0.201622 | 0.864412 |
| ENSMUSG( | Pex10 | 299.404 | -0.01566 | 0.912189 | 0.988483 | ENSMUSG( | Pex10 | 273.1516 | -0.01731 | 0.914383 | 0.992407 |
| ENSMUSG( | Pex11a | 612.8012 | 0.244351 | 0.026097 | 0.247245 | ENSMUSG( | Pex11a | 510.9001 | -0.00472 | 0.971934 | 0.996416 |
| ENSMUSG( | Pex11b | 534.0611 | -0.0031 | 0.981306 | 0.997512 | ENSMUSG( | Pex11b | 482.8045 | -0.03027 | 0.84268 | 0.983099 |
| ENSMUSG( | Pex11g | 50.36415 | 0.171474 | 0.530917 | 0.879289 | ENSMUSG( | Pex11g | 42.39372 | -0.04164 | 0.895113 | 0.990558 |
| ENSMUSG( | Pex12 | 334.1048 | -0.00101 | 0.994559 | 0.999369 | ENSMUSG( | Pex12 | 301.568 | -0.0346 | 0.829388 | 0.980224 |
| ENSMUSG( | Pex13 | 980.5718 | -0.11192 | 0.346916 | 0.779684 | ENSMUSG( | Pex13 | 930.3931 | 0.005758 | 0.966719 | 0.995475 |
| ENSMUSG( | Pex14 | 1694.142 | -0.23929 | 0.039097 | 0.304587 | ENSMUSG( | Pex14 | 1692.874 | 0.040413 | 0.780941 | 0.976106 |
| ENSMUSG( | Pex16 | 1035.354 | 0.046229 | 0.617697 | 0.910872 | ENSMUSG( | Pex16 | 901.3368 | -0.08783 | 0.445585 | 0.921734 |
| ENSMUSG( | Pex19 | 3378.512 | 0.01812 | 0.84822 | 0.972374 | ENSMUSG( | Pex19 | 2862.371 | -0.20105 | 0.082955 | 0.77681 |
| ENSMUSG( | Pex2 | 903.0024 | -0.02594 | 0.821178 | 0.966248 | ENSMUSG( | Pex2 | 842.0441 | 0.044067 | 0.737987 | 0.967709 |
| ENSMUSG( | Pex26 | 357.0299 | 0.255577 | 0.058143 | 0.371733 | ENSMUSG( | Pex26 | 326.6268 | 0.276677 | 0.088913 | 0.788051 |
| ENSMUSG( | Pex3 | 693.9868 | -0.09604 | 0.366077 | 0.794603 | ENSMUSG( | Pex3 | 658.2275 | 0.020879 | 0.862962 | 0.986547 |
| ENSMUSG( | Pex5 | 1448.276 | 0.056613 | 0.591085 | 0.90113 | ENSMUSG( | Pex5 | 1311.468 | 0.03812 | 0.764931 | 0.974236 |
| ENSMUSG( | Pex5l | 3655.328 | -0.00755 | 0.929972 | 0.991848 | ENSMUSG( | Pex5l | 3184.596 | -0.13935 | 0.194745 | 0.860433 |
| ENSMUSG( | Pex6 | 2273.708 | 0.227767 | 0.008262 | 0.126789 | ENSMUSG( | Pex6 | 1891.312 | -0.02433 | 0.826248 | 0.979795 |
| ENSMUSG( | Pex7 | 977.9955 | 0.03212 | 0.785259 | 0.958473 | ENSMUSG( | Pex7 | 825.7148 | -0.20131 | 0.168945 | 0.840222 |
| ENSMUSG( | Pf4 | 64.07624 | 0.06386 | 0.803138 | 0.962169 | ENSMUSG( | Pf4 | 58.71438 | 0.056748 | 0.841131 | 0.9826 |
| ENSMUSG( | Pfas | 1210.45 | 0.080978 | 0.407063 | 0.817163 | ENSMUSG( | Pfas | 1089.002 | 0.046498 | 0.706138 | 0.96335 |
| ENSMUSG( | Pfdn1 | 1438.031 | -0.00702 | 0.955401 | 0.993874 | ENSMUSG( | Pfdn1 | 1235.786 | -0.18792 | 0.247393 | 0.877904 |
| ENSMUSG( | Pfdn2 | 3373.222 | 0.017247 | 0.845024 | 0.972023 | ENSMUSG( | Pfdn2 | 2729.084 | -0.35008 | 0.020221 | 0.563435 |
| ENSMUSG( | Pfdn4 | 252.7105 | 0.051231 | 0.75473 | 0.951007 | ENSMUSG( | Pfdn4 | 215.7508 | -0.14219 | 0.442282 | 0.921554 |
| ENSMUSG( | Pfdn5 | 6003.208 | 0.064516 | 0.454449 | 0.837347 | ENSMUSG( | Pfdn5 | 5070.968 | -0.15996 | 0.215648 | 0.865167 |
| ENSMUSG( | Pfdn6 | 950.2099 | 0.028601 | 0.827746 | 0.968924 | ENSMUSG( | Pfdn6 | 811.8811 | -0.16616 | 0.303628 | 0.898538 |
| ENSMUSG( | Pfkfb1 | 16.93686 | -0.58829 | 0.186367 | NA | ENSMUSG( | Pfkfb1 | 16.62624 | -0.31764 | 0.502707 | 0.927825 |
| ENSMUSG( | Pfkfb2 | 850.5434 | 0.075218 | 0.47431 | 0.849425 | ENSMUSG( | Pfkfb2 | 772.7628 | 0.065876 | 0.651295 | 0.95449 |
| ENSMUSG( | Pfkfb3 | 844.2902 | -0.12048 | 0.22327 | 0.669382 | ENSMUSG( | Pfkfb3 | 772.3692 | -0.10777 | 0.393437 | 0.917257 |
| ENSMUSG( | Pfkfb4 | 727.7577 | -0.0595 | 0.57675 | 0.895432 | ENSMUSG( | Pfkfb4 | 614.0314 | -0.29546 | 0.021307 | 0.575341 |
| ENSMUSG( | Pfkl | 4961.916 | 0.139154 | 0.241286 | 0.689026 | ENSMUSG( | Pfkl | 4071.348 | -0.16752 | 0.237473 | 0.872173 |
| ENSMUSG( | Pfkm | 17880.94 | 0.145088 | 0.096441 | 0.476986 | ENSMUSG( | Pfkm | 14747.85 | -0.14388 | 0.194328 | 0.860096 |
| ENSMUSG( | Pfkp | 7235.397 | 0.183346 | 0.054715 | 0.360135 | ENSMUSG( | Pfkp | 5737.513 | -0.22129 | 0.080784 | 0.77681 |
| ENSMUSG( | Pfn1 | 3485.58 | -0.04068 | 0.674228 | 0.929638 | ENSMUSG( | Pfn1 | 3086.418 | -0.12882 | 0.35188 | 0.912093 |
| ENSMUSG( | Pfn2 | 12544.22 | -0.02922 | 0.758934 | 0.952116 | ENSMUSG( | Pfn2 | 11220.41 | -0.08639 | 0.55514 | 0.939565 |
| ENSMUSG( | Pfn4 | 555.1689 | 0.014344 | 0.908151 | 0.9878 | ENSMUSG( | Pfn4 | 464.887 | -0.24422 | 0.116267 | 0.813384 |
| ENSMUSG( | Pgam1 | 4486.235 | 0.074375 | 0.434122 | 0.828931 | ENSMUSG( | Pgam1 | 3773.456 | -0.16238 | 0.195906 | 0.862161 |
| ENSMUSG( | Pgam2 | 84.1486 | -0.38484 | 0.08423 | 0.447434 | ENSMUSG( | Pgam2 | 76.56359 | -0.41554 | 0.11642 | 0.813384 |
| ENSMUSG( | Pgam5 | 1468.511 | 0.187145 | 0.096508 | 0.476986 | ENSMUSG( | Pgam5 | 1268.321 | 0.036386 | 0.78325 | 0.976106 |
| ENSMUSG( | Pgap1 | 792.0436 | -0.38894 | 0.03806 | 0.300991 | ENSMUSG( | Pgap1 | 930.1895 | 0.353716 | 0.071503 | 0.761942 |
| ENSMUSG( | Pgap2 | 534.5954 | -0.19795 | 0.135591 | 0.549038 | ENSMUSG( | Pgap2 | 498.0959 | -0.12873 | 0.340552 | 0.907972 |

| | | | | | |
|---|---|---|---|---|---|
| ENSMUSG( | 524.9396 | 0.129632 | 0.238836 | 0.6865 | Parp2 |
| ENSMUSG( | 231.5314 | -0.00859 | 0.958097 | 0.993874 | Parp3 |
| ENSMUSG( | 331.8213 | 0.108821 | 0.386234 | 0.804623 | Parp4 |
| ENSMUSG( | 7609.704 | 0.199102 | 0.007734 | 0.122031 | Parp6 |
| ENSMUSG( | 724.7642 | 0.180103 | 0.088279 | 0.455783 | Parp8 |
| ENSMUSG( | 221.3365 | -0.02873 | 0.870427 | 0.977906 | Parp9 |
| ENSMUSG( | 11.15662 | -0.30166 | 0.608633 | NA | Parpbp |
| ENSMUSG( | 179.0451 | 0.165784 | 0.294567 | 0.743754 | Pars2 |
| ENSMUSG( | 485.545 | -0.20338 | 0.100653 | 0.48402 | Particl |
| ENSMUSG( | 4310.262 | 0.115267 | 0.263744 | 0.714352 | Parva |
| ENSMUSG( | 15.18422 | 0.357538 | 0.485904 | NA | Parvaos |
| ENSMUSG( | 2076.88 | 0.007635 | 0.938174 | 0.993415 | Parvb |
| ENSMUSG( | 358.6145 | -0.08453 | 0.507416 | 0.86795 | Parvg |
| ENSMUSG( | 18.83737 | 0.541537 | 0.214967 | NA | Pasd1 |
| ENSMUSG( | 106.3174 | -0.04603 | 0.840546 | 0.971179 | Pask |
| ENSMUSG( | 22.32177 | 0.343123 | 0.379945 | 0.801757 | Pate2 |
| ENSMUSG( | 320.7931 | -0.14995 | 0.349947 | 0.781943 | Patj |
| ENSMUSG( | 987.8956 | -0.07706 | 0.468287 | 0.845336 | Patl1 |
| ENSMUSG( | 1051.658 | 0.032491 | 0.743441 | 0.948478 | Patz1 |
| ENSMUSG( | 88.02693 | -0.09095 | 0.706177 | 0.938869 | Pawr |
| ENSMUSG( | 11.30284 | 0.373624 | 0.517821 | NA | Pax5 |
| ENSMUSG( | 770.059 | -0.00297 | 0.9827 | 0.997522 | Pax6 |
| ENSMUSG( | 24.2327 | 0.572585 | 0.175762 | 0.611006 | Pax6os1 |
| ENSMUSG( | 1640.406 | -0.04004 | 0.717222 | 0.941697 | Paxbp1 |
| ENSMUSG( | 2265.373 | 0.213946 | 0.026256 | 0.247758 | Paxip1 |
| ENSMUSG( | 1753.997 | 0.095501 | 0.263741 | 0.714352 | Paxx |
| ENSMUSG( | 434.9551 | 0.179211 | 0.129156 | 0.538324 | Pbdc1 |
| ENSMUSG( | 45.82963 | -0.45054 | 0.121413 | 0.525713 | Pbk |
| ENSMUSG( | 37.00389 | -0.21095 | 0.512924 | 0.871762 | Pbld2 |
| ENSMUSG( | 1216.762 | -0.01235 | 0.894875 | 0.984512 | Pbrm1 |
| ENSMUSG( | 5519.02 | 0.056087 | 0.522936 | 0.875628 | Pbx1 |
| ENSMUSG( | 1416.866 | -0.01379 | 0.898202 | 0.985633 | Pbx2 |
| ENSMUSG( | 406.032 | -0.24804 | 0.211899 | 0.655541 | Pbx3 |
| ENSMUSG( | 360.5205 | 0.099107 | 0.478475 | 0.853567 | Pbx4 |
| ENSMUSG( | 8003.441 | -0.09627 | 0.408213 | 0.818167 | Pbxip1 |
| ENSMUSG( | 118.3927 | 0.17935 | 0.346392 | 0.779153 | Pcbd1 |
| ENSMUSG( | 253.0123 | -0.05622 | 0.705229 | 0.938303 | Pcbd2 |
| ENSMUSG( | 3562.284 | 0.058111 | 0.641262 | 0.916693 | Pcbp1 |
| ENSMUSG( | 8361.06 | 0.054675 | 0.493062 | 0.860552 | Pcbp2 |
| ENSMUSG( | 1024.225 | 0.055315 | 0.723921 | 0.944096 | Pcbp3 |
| ENSMUSG( | 7726.86 | 0.186904 | 0.047977 | 0.334562 | Pcbp4 |
| ENSMUSG( | 1091.371 | 0.071488 | 0.482982 | 0.856216 | Pcca |
| ENSMUSG( | 3290.838 | 0.043614 | 0.622251 | 0.911432 | Pccb |
| ENSMUSG( | 6356.206 | -0.19637 | 0.250387 | 0.700322 | Pcdh1 |
| ENSMUSG( | 6965.028 | -0.214 | 0.097864 | 0.478264 | Pcdh10 |
| ENSMUSG( | 310.1821 | -0.49257 | 0.001869 | 0.049292 | Pcdh11x |
| ENSMUSG( | 46.70995 | -0.11559 | 0.733412 | 0.944956 | Pcdh12 |
| ENSMUSG( | 678.9386 | -0.19803 | 0.082435 | 0.443042 | Pcdh15 |
| ENSMUSG( | 2624.584 | -0.1957 | 0.151958 | 0.577721 | Pcdh17 |
| ENSMUSG( | 165.0134 | -0.18512 | 0.487328 | 0.858812 | Pcdh18 |
| ENSMUSG( | 737.8118 | -0.10157 | 0.609379 | 0.907634 | Pcdh19 |
| ENSMUSG( | 798.9438 | -0.34246 | 0.049801 | 0.340731 | Pcdh20 |
| ENSMUSG( | 3461.386 | -0.35849 | 0.012563 | 0.161972 | Pcdh7 |
| ENSMUSG( | 1113.824 | -0.31758 | 0.052835 | 0.352814 | Pcdh8 |
| ENSMUSG( | 2836.848 | -0.33905 | 0.033677 | 0.281425 | Pcdh9 |
| ENSMUSG( | 14.56935 | -0.39747 | 0.563022 | NA | Pcdha1 |
| ENSMUSG( | 27.86372 | -0.51488 | 0.279383 | 0.729114 | Pcdha10 |
| ENSMUSG( | 25.40797 | -0.54742 | 0.180674 | 0.617841 | Pcdha11 |
| ENSMUSG( | 91.59458 | -0.3375 | 0.369589 | 0.797172 | Pcdha12 |
| ENSMUSG( | 28.79544 | -0.44728 | 0.332808 | 0.775602 | Pcdha2 |
| ENSMUSG( | 37.20827 | -0.49883 | 0.186569 | 0.623869 | Pcdha3 |
| ENSMUSG( | 50.80137 | -0.65164 | 0.111399 | 0.503454 | Pcdha5 |
| ENSMUSG( | 40.16593 | -0.5791 | 0.105421 | 0.492417 | Pcdha6 |
| ENSMUSG( | 28.43247 | -0.48439 | 0.347703 | 0.780695 | Pcdha7 |
| ENSMUSG( | 56.35446 | -0.61497 | 0.057895 | 0.371023 | Pcdha9 |
| ENSMUSG( | 39.41171 | -0.20954 | 0.597434 | 0.903694 | Pcdhac1 |
| ENSMUSG( | 345.3705 | -0.3621 | 0.441165 | 0.832445 | Pcdhac2 |
| ENSMUSG( | 103.2229 | -0.68416 | 0.001253 | 0.038624 | Pcdhb10 |
| ENSMUSG( | 71.66782 | -0.65043 | 0.01083 | 0.150145 | Pcdhb11 |
| ENSMUSG( | 182.0841 | -0.70979 | 0.000184 | 0.009888 | Pcdhb12 |
| ENSMUSG( | 122.3234 | -0.56803 | 0.016016 | 0.187923 | Pcdhb13 |
| ENSMUSG( | 195.3793 | -0.61636 | 0.002479 | 0.058622 | Pcdhb14 |
| ENSMUSG( | 94.46829 | -0.44583 | 0.029338 | 0.259647 | Pcdhb15 |
| ENSMUSG( | 94.0157 | -0.6885 | 0.003412 | 0.073541 | Pcdhb16 |
| ENSMUSG( | 321.667 | -0.50801 | 0.000845 | 0.029095 | Pcdhb17 |
| ENSMUSG( | 139.1876 | -0.82731 | 7.94E-05 | 0.005507 | Pcdhb18 |
| ENSMUSG( | 180.412 | -0.53688 | 0.00128 | 0.039067 | Pcdhb19 |
| ENSMUSG( | 32.27831 | -0.90905 | 0.028319 | 0.257185 | Pcdhb2 |
| ENSMUSG( | 311.2079 | -0.54494 | 0.000186 | 0.009972 | Pcdhb20 |
| ENSMUSG( | 53.37876 | -0.61636 | 0.031856 | 0.272382 | Pcdhb21 |
| ENSMUSG( | 348.2199 | -0.39581 | 0.025955 | 0.247245 | Pcdhb22 |
| ENSMUSG( | 136.5871 | -0.38638 | 0.104156 | 0.489971 | Pcdhb3 |
| ENSMUSG( | 52.36489 | -0.21185 | 0.457968 | 0.839769 | Pcdhb4 |
| ENSMUSG( | 132.9149 | -0.41419 | 0.089912 | 0.460493 | Pcdhb5 |
| ENSMUSG( | 65.68541 | -0.36097 | 0.208634 | 0.650496 | Pcdhb6 |
| ENSMUSG( | 203.2934 | -0.45199 | 0.009884 | 0.141868 | Pcdhb7 |
| ENSMUSG( | 68.88744 | -0.02755 | 0.910312 | 0.9881 | Pcdhb8 |
| ENSMUSG( | 236.895 | -0.64632 | 4.09E-05 | 0.00344 | Pcdhb9 |
| ENSMUSG( | 112.447 | -0.21566 | 0.346531 | 0.779166 | Pcdhga1 |
| ENSMUSG( | 184.3248 | -0.15579 | 0.430109 | 0.826789 | Pcdhga10 |
| ENSMUSG( | 692.7978 | 0.168372 | 0.333147 | 0.775602 | Pcdhga11 |
| ENSMUSG( | 568.6135 | 0.111161 | 0.552513 | 0.886141 | Pcdhga12 |
| ENSMUSG( | 152.5251 | -0.11898 | 0.5482 | 0.885197 | Pcdhga2 |
| ENSMUSG( | 301.4449 | -0.1571 | 0.448014 | 0.834051 | Pcdhga3 |
| ENSMUSG( | 204.1931 | -0.13846 | 0.488247 | 0.859092 | Pcdhga4 |
| ENSMUSG( | 210.1918 | -0.0414 | 0.83688 | 0.971111 | Pcdhga5 |
| ENSMUSG( | 373.1514 | -0.12028 | 0.386576 | 0.804912 | Pcdhga6 |
| ENSMUSG( | 339.9122 | -0.17392 | 0.349978 | 0.781943 | Pcdhga7 |
| ENSMUSG( | 257.2998 | 0.038189 | 0.823305 | 0.966603 | Pcdhga8 |
| ENSMUSG( | 118.8968 | -0.00735 | 0.971216 | 0.996369 | Pcdhga9 |
| ENSMUSG( | 200.9363 | -0.13451 | 0.571059 | 0.893319 | Pcdhgb1 |
| ENSMUSG( | 168.4484 | -0.419 | 0.028544 | 0.257525 | Pcdhgb2 |
| ENSMUSG( | 159.5724 | -0.34108 | 0.087656 | 0.454663 | Pcdhgb4 |
| ENSMUSG( | 144.9526 | -0.06718 | 0.746152 | 0.949289 | Pcdhgb5 |
| ENSMUSG( | 346.543 | -0.21883 | 0.277453 | 0.727432 | Pcdhgb6 |
| ENSMUSG( | 242.7267 | -0.07198 | 0.683656 | 0.932154 | Pcdhgb7 |
| ENSMUSG( | 7755.958 | 0.0334 | 0.766379 | 0.954173 | Pcdhgc3 |
| ENSMUSG( | 232.4103 | -0.38074 | 0.051458 | 0.348059 | Pcdhgc4 |
| ENSMUSG( | 2079.941 | -0.37685 | 0.044576 | 0.32426 | Pcdhgc5 |
| ENSMUSG( | 741.3872 | 0.147088 | 0.167628 | 0.600338 | Pced1a |
| ENSMUSG( | 197.9943 | -0.09799 | 0.511251 | 0.870859 | Pced1b |
| ENSMUSG( | 838.0973 | -0.17154 | 0.095143 | 0.473815 | Pcf11 |
| ENSMUSG( | 240.9821 | 0.080541 | 0.632657 | 0.914073 | Pcgf1 |
| ENSMUSG( | 844.7038 | 0.026827 | 0.888442 | 0.983434 | Pcgf2 |
| ENSMUSG( | 1381.332 | 0.141675 | 0.108585 | 0.497529 | Pcgf3 |
| ENSMUSG( | 554.4493 | 0.018878 | 0.878584 | 0.981431 | Pcgf5 |
| ENSMUSG( | 383.0957 | 0.005827 | 0.965033 | 0.995303 | Pcgf6 |
| ENSMUSG( | 1102.471 | 0.195786 | 0.039544 | 0.3066 | Pcid2 |
| ENSMUSG( | 2699.419 | 0.108146 | 0.211883 | 0.655541 | Pcif1 |
| ENSMUSG( | 783.4638 | 0.179595 | 0.100091 | 0.483032 | Pck2 |
| ENSMUSG( | 40.13826 | -0.07568 | 0.803201 | 0.962169 | Pclaf |
| ENSMUSG( | 7003.077 | -0.57458 | 0.066609 | 0.398819 | Pclo |
| ENSMUSG( | 2277.823 | 0.025939 | 0.791289 | 0.959461 | Pcm1 |
| ENSMUSG( | 3754.055 | -0.23578 | 0.00794 | 0.124138 | Pcmt1 |
| ENSMUSG( | 3028.337 | -0.0299 | 0.77096 | 0.954726 | Pcmtd1 |
| ENSMUSG( | 1453.908 | -0.02461 | 0.809253 | 0.963595 | Pcmtd2 |
| ENSMUSG( | 958.7802 | 0.062056 | 0.573561 | 0.893584 | Pcna |
| ENSMUSG( | 2775.029 | -0.0418 | 0.714614 | 0.941144 | Pcnp |
| ENSMUSG( | 1195.194 | 0.114054 | 0.301388 | 0.749979 | Pcnt |
| ENSMUSG( | 3128.033 | -0.05034 | 0.72514 | 0.944096 | Pcnx1 |
| ENSMUSG( | 3162.741 | -0.11226 | 0.452631 | 0.836145 | Pcnx2 |
| ENSMUSG( | 2127.965 | -0.06688 | 0.601114 | 0.904603 | Pcnx3 |
| ENSMUSG( | 1096.859 | -0.20846 | 0.12693 | 0.534201 | Pcnx4 |
| ENSMUSG( | 417.9677 | -0.15866 | 0.239204 | 0.687031 | Pcolce |
| ENSMUSG( | 180.0372 | -0.08539 | 0.615521 | 0.910481 | Pcolce2 |
| ENSMUSG( | 1646.9 | -0.34363 | 0.024257 | 0.238311 | Pcp4 |
| ENSMUSG( | 2858.142 | -0.74112 | 9.11E-06 | 0.001083 | Pcp4l1 |
| ENSMUSG( | 883.3353 | -0.69091 | 0.000676 | 0.025081 | Pcsk1 |
| ENSMUSG( | 46432.86 | -0.0769 | 0.363901 | 0.793343 | Pcsk1n |
| ENSMUSG( | 10817.89 | -0.44034 | 0.003594 | 0.075902 | Pcsk2 |
| ENSMUSG( | 103.0621 | 0.114554 | 0.59162 | 0.901449 | Pcsk2os1 |
| ENSMUSG( | 166.2986 | 0.052262 | 0.766259 | 0.954173 | Pcsk4 |
| ENSMUSG( | 460.0404 | -0.36424 | 0.005278 | 0.096847 | Pcsk5 |
| ENSMUSG( | 1424.319 | -0.10413 | 0.462378 | 0.842104 | Pcsk6 |
| ENSMUSG( | 912.2202 | -0.03985 | 0.752021 | 0.950417 | Pcsk7 |
| ENSMUSG( | 103.6682 | -2.99754 | 7.87E-31 | 1.82E-27 | Pcsk9 |
| ENSMUSG( | 122.8271 | 0.07413 | 0.699361 | 0.936948 | Pctp |
| ENSMUSG( | 5001.462 | 0.143297 | 0.071593 | 0.413189 | Pcx |
| ENSMUSG( | 4365.045 | -0.30527 | 0.031102 | 0.268324 | Pcyox1 |
| ENSMUSG( | 882.734 | -0.3913 | 0.001392 | 0.040902 | Pcyox1l |
| ENSMUSG( | 2575.814 | 0.081095 | 0.388574 | 0.806278 | Pcyt1a |
| ENSMUSG( | 1841.741 | 0.026735 | 0.774615 | 0.955606 | Pcyt1b |
| ENSMUSG( | 3256.163 | 0.092146 | 0.301659 | 0.749979 | Pcyt2 |
| ENSMUSG( | 2526.091 | 0.082355 | 0.404456 | 0.816132 | Pdap1 |
| ENSMUSG( | 598.7237 | 0.040659 | 0.727261 | 0.944223 | Pdcd10 |
| ENSMUSG( | 998.1013 | 0.081631 | 0.449174 | 0.834943 | Pdcd11 |
| ENSMUSG( | 223.6109 | -0.15887 | 0.296205 | 0.744386 | Pdcd2 |
| ENSMUSG( | 244.5124 | 0.056514 | 0.695274 | 0.934948 | Pdcd2l |

| | | | | | |
|---|---|---|---|---|---|
| ENSMUSG( | 192.3564 | 0.357213 | 0.05636 | 0.723362 | Pcolce2 |
| ENSMUSG( | 1514.808 | -0.31998 | 0.120764 | 0.813384 | Pcp4 |
| ENSMUSG( | 3046.838 | -0.20404 | 0.250007 | 0.877904 | Pcp4l1 |
| ENSMUSG( | 1122.714 | 0.334376 | 0.117904 | 0.813384 | Pcsk1 |
| ENSMUSG( | 42247.24 | -0.0815 | 0.483958 | 0.925289 | Pcsk1n |
| ENSMUSG( | 12027.24 | 0.167692 | 0.337653 | 0.907972 | Pcsk2 |
| ENSMUSG( | 92.25362 | 0.064182 | 0.771408 | 0.974236 | Pcsk2os1 |
| ENSMUSG( | 167.121 | 0.308807 | 0.146343 | 0.835973 | Pcsk4 |
| ENSMUSG( | 466.9502 | -0.02383 | 0.879081 | 0.989644 | Pcsk5 |
| ENSMUSG( | 1343.308 | 0.000332 | 0.998326 | 0.999313 | Pcsk6 |
| ENSMUSG( | 872.3185 | 0.102625 | 0.443807 | 0.921734 | Pcsk7 |
| ENSMUSG( | 150.1556 | -0.33112 | 0.084833 | 0.779799 | Pcsk9 |
| ENSMUSG( | 101.8712 | -0.21531 | 0.318958 | 0.903344 | Pctp |
| ENSMUSG( | 4204.088 | -0.08839 | 0.401983 | 0.917257 | Pcx |
| ENSMUSG( | 4812.064 | 0.254128 | 0.123819 | 0.814691 | Pcyox1 |
| ENSMUSG( | 969.8247 | 0.171328 | 0.232504 | 0.868507 | Pcyox1l |
| ENSMUSG( | 2197.406 | -0.11096 | 0.309953 | 0.898538 | Pcyt1a |
| ENSMUSG( | 1594.751 | -0.12381 | 0.250097 | 0.877904 | Pcyt1b |
| ENSMUSG( | 2848.345 | -0.02373 | 0.85895 | 0.985884 | Pcyt2 |
| ENSMUSG( | 2164.059 | -0.0995 | 0.48615 | 0.92541 | Pdap1 |
| ENSMUSG( | 550.8675 | 0.065895 | 0.613502 | 0.951177 | Pdcd10 |
| ENSMUSG( | 930.9598 | 0.147877 | 0.224203 | 0.867489 | Pdcd11 |
| ENSMUSG( | 206.419 | -0.12357 | 0.501964 | 0.927825 | Pdcd2 |
| ENSMUSG( | 219.3201 | 0.010758 | 0.948216 | 0.994798 | Pdcd2l |
| ENSMUSG( | 1494.38 | 0.04523 | 0.729215 | 0.966913 | Pdcd4 |
| ENSMUSG( | 825.1384 | -0.17339 | 0.192713 | 0.859764 | Pdcd5 |
| ENSMUSG( | 1234.704 | -0.16221 | 0.191461 | 0.858412 | Pdcd6 |
| ENSMUSG( | 2252.847 | 0.030923 | 0.793324 | 0.97681 | Pdcd6ip |
| ENSMUSG( | 383.6712 | -0.03072 | 0.828317 | 0.980224 | Pdcd7 |
| ENSMUSG( | 534.1716 | 0.045729 | 0.731184 | 0.967228 | Pdcl |
| ENSMUSG( | 633.8383 | -0.08563 | 0.494099 | 0.927372 | Pdcl3 |
| ENSMUSG( | 3328.28 | 0.005622 | 0.966665 | 0.995475 | Pde10a |
| ENSMUSG( | 47.1478 | -0.12834 | 0.68323 | 0.958678 | Pde11a |
| ENSMUSG( | 280.7705 | -0.04952 | 0.7747 | 0.974931 | Pde12 |
| ENSMUSG( | 2278.967 | -0.00672 | 0.959808 | 0.994798 | Pde1a |
| ENSMUSG( | 5181.653 | -0.20671 | 0.215004 | 0.865167 | Pde1b |
| ENSMUSG( | 467.924 | -0.27932 | 0.06714 | 0.75212 | Pde1c |
| ENSMUSG( | 226.2583 | -0.04428 | 0.810154 | 0.978779 | Pde2a |
| ENSMUSG( | 179.8192 | 0.163002 | 0.367506 | 0.915217 | Pde3a |
| ENSMUSG( | 557.8043 | 0.168807 | 0.283577 | 0.892412 | Pde3b |
| ENSMUSG( | 4269.126 | 0.107563 | 0.394371 | 0.917257 | Pde4a |
| ENSMUSG( | 2071.673 | -0.09021 | 0.417535 | 0.917257 | Pde4b |
| ENSMUSG( | 95.7559 | 0.667644 | 0.005252 | 0.297744 | Pde4c |
| ENSMUSG( | 479.9489 | 0.230058 | 0.136324 | 0.827092 | Pde4d |
| ENSMUSG( | 4774.79 | 0.065084 | 0.506126 | 0.927825 | Pde4dip |
| ENSMUSG( | 250.6778 | 0.31206 | 0.082439 | 0.77681 | Pde5a |
| ENSMUSG( | 19.43232 | 0.031676 | 0.943327 | 0.994798 | Pde6a |
| ENSMUSG( | 1258.382 | -0.16868 | 0.223274 | 0.867404 | Pde6d |
| ENSMUSG( | 11.94258 | 0.447825 | 0.472116 | 0.923329 | Pde6h |
| ENSMUSG( | 798.6501 | 0.083734 | 0.523858 | 0.932848 | Pde7a |
| ENSMUSG( | 1133.504 | -0.13481 | 0.296531 | 0.898538 | Pde7b |
| ENSMUSG( | 1728.044 | -0.21352 | 0.065917 | 0.746371 | Pde8a |
| ENSMUSG( | 2765.845 | -0.09642 | 0.345917 | 0.910966 | Pde8b |
| ENSMUSG( | 594.2803 | -0.29469 | 0.025248 | 0.597654 | Pde9a |
| ENSMUSG( | 28.61816 | 0.388578 | 0.312884 | 0.900409 | Pdf |
| ENSMUSG( | 1734.736 | 0.204817 | 0.105987 | 0.804969 | Pdgfa |
| ENSMUSG( | 1074.166 | 0.24011 | 0.141973 | 0.830709 | Pdgfb |
| ENSMUSG( | 410.9668 | 0.237355 | 0.106364 | 0.804969 | Pdgfc |
| ENSMUSG( | 202.6276 | 0.684443 | 0.000301 | 0.047107 | Pdgfd |
| ENSMUSG( | 3042.722 | 0.282428 | 0.162287 | 0.837724 | Pdgfra |
| ENSMUSG( | 1620.225 | 0.309268 | 0.181045 | 0.850372 | Pdgfrb |
| ENSMUSG( | 96.26113 | 0.041804 | 0.864082 | 0.986552 | Pdgfrl |
| ENSMUSG( | 7922.038 | -0.15247 | 0.253051 | 0.88018 | Pdha1 |
| ENSMUSG( | 2512.494 | -0.12553 | 0.298019 | 0.898538 | Pdhb |
| ENSMUSG( | 1077.244 | -0.15449 | 0.23248 | 0.868507 | Pdhx |
| ENSMUSG( | 8446.791 | 0.207743 | 0.190712 | 0.857801 | Pdia3 |
| ENSMUSG( | 2092.538 | 0.459413 | 0.004834 | 0.287322 | Pdia4 |
| ENSMUSG( | 116.8567 | -0.06681 | 0.767722 | 0.974236 | Pdia5 |
| ENSMUSG( | 4066.345 | 0.255523 | 0.13306 | 0.822912 | Pdia6 |
| ENSMUSG( | 260.9183 | 0.225632 | 0.154682 | 0.837724 | Pdik1l |
| ENSMUSG( | 1636.061 | 0.057969 | 0.611343 | 0.950771 | Pdk1 |
| ENSMUSG( | 3658.086 | -0.16142 | 0.165986 | 0.839855 | Pdk2 |
| ENSMUSG( | 599.8783 | -0.19716 | 0.17264 | 0.843702 | Pdk3 |
| ENSMUSG( | 889.2281 | -0.51184 | 0.000368 | 0.054165 | Pdk4 |
| ENSMUSG( | 105.5188 | -0.05784 | 0.803103 | 0.976937 | Pdlim1 |
| ENSMUSG( | 2082.632 | -0.25528 | 0.078032 | 0.77321 | Pdlim2 |
| ENSMUSG( | 12.44068 | 0.04368 | 0.936446 | 0.993573 | Pdlim3 |
| ENSMUSG( | 530.5086 | -0.05858 | 0.67525 | 0.95694 | Pdlim4 |
| ENSMUSG( | 1996.642 | -0.01253 | 0.935252 | 0.993078 | Pdlim5 |
| ENSMUSG( | 1132.64 | 0.01995 | 0.903407 | 0.991089 | Pdlim7 |
| ENSMUSG( | 1291.056 | -0.15419 | 0.324675 | 0.905257 | Pdp1 |
| ENSMUSG( | 338.8032 | 0.125596 | 0.393701 | 0.917257 | Pdp2 |
| ENSMUSG( | 2680.03 | 0.000937 | 0.992711 | 0.99874 | Pdpk1 |
| ENSMUSG( | 820.3544 | -0.07513 | 0.636974 | 0.953903 | Pdpn |
| ENSMUSG( | 943.3268 | -0.08082 | 0.511091 | 0.929571 | Pdpr |
| ENSMUSG( | 836.885 | -0.21461 | 0.080009 | 0.776379 | Pdrg1 |
| ENSMUSG( | 1458.998 | 0.108506 | 0.33887 | 0.907972 | Pds5a |
| ENSMUSG( | 1788.462 | 0.138109 | 0.212092 | 0.864412 | Pds5b |
| ENSMUSG( | 220.9102 | 0.149384 | 0.371017 | 0.915491 | Pdss1 |
| ENSMUSG( | 232.0679 | -0.11729 | 0.523252 | 0.932848 | Pdss2 |
| ENSMUSG( | 3976.956 | -0.16758 | 0.147253 | 0.836241 | Pdxdc1 |
| ENSMUSG( | 5262.387 | 0.055637 | 0.596643 | 0.946541 | Pdxk |
| ENSMUSG( | 2231.341 | -0.08028 | 0.570715 | 0.941973 | Pdxp |
| ENSMUSG( | 1141.345 | 0.092421 | 0.547008 | 0.938063 | Pdyn |
| ENSMUSG( | 794.9703 | -0.23635 | 0.05433 | 0.723362 | Pdzd11 |
| ENSMUSG( | 1902.061 | 0.274078 | 0.037315 | 0.673514 | Pdzd2 |
| ENSMUSG( | 6644.648 | 0.023605 | 0.808269 | 0.978622 | Pdzd4 |
| ENSMUSG( | 136.2152 | 0.122981 | 0.616241 | 0.951413 | Pdzd7 |
| ENSMUSG( | 2346.405 | 0.184339 | 0.25771 | 0.883401 | Pdzd8 |
| ENSMUSG( | 16.3898 | -0.24176 | 0.635311 | 0.953903 | Pdzk1 |
| ENSMUSG( | 14.32994 | -0.75811 | 0.168835 | 0.840222 | Pdzk1ip1 |
| ENSMUSG( | 276.1009 | -0.05795 | 0.747132 | 0.969763 | Pdzph1 |
| ENSMUSG( | 905.2145 | -0.22243 | 0.122933 | 0.814458 | Pdzrn3 |
| ENSMUSG( | 186.5109 | -0.11689 | 0.502288 | 0.927825 | Pdzrn4 |
| ENSMUSG( | 42142.36 | -0.22841 | 0.071301 | 0.761942 | Pea15a |
| ENSMUSG( | 1618.9 | 0.115158 | 0.295423 | 0.898538 | Peak1 |
| ENSMUSG( | 300.8117 | 0.493739 | 0.043149 | 0.690304 | Pear1 |
| ENSMUSG( | 7896.133 | -0.16076 | 0.196953 | 0.862161 | Pebp1 |
| ENSMUSG( | 915.0949 | 0.47393 | 0.270089 | 0.88756 | Pecam1 |
| ENSMUSG( | 93.58095 | -0.20327 | 0.372366 | 0.915491 | Pecr |
| ENSMUSG( | 2267.791 | 0.079722 | 0.562292 | 0.940752 | Peds1 |
| ENSMUSG( | 1045.333 | -0.0796 | 0.560681 | 0.940254 | Pef1 |
| ENSMUSG( | 594.8564 | 0.100762 | 0.519773 | 0.931729 | Peg10 |
| ENSMUSG( | 16.62904 | 0.287211 | 0.566224 | 0.941665 | Peg12 |
| ENSMUSG( | 3778.244 | -0.08768 | 0.466397 | 0.923329 | Peg13 |
| ENSMUSG( | 7431.243 | -0.05921 | 0.642649 | 0.953903 | Peg3 |
| ENSMUSG( | 851.4494 | 0.137166 | 0.283957 | 0.892412 | Peli1 |
| ENSMUSG( | 2128.573 | -0.01608 | 0.912989 | 0.992298 | Peli2 |
| ENSMUSG( | 383.373 | -0.05305 | 0.747099 | 0.969763 | Peli3 |
| ENSMUSG( | 108.5057 | 0.148808 | 0.470794 | 0.923329 | Pelo |
| ENSMUSG( | 1107.413 | -0.04481 | 0.727292 | 0.966887 | Pelp1 |
| ENSMUSG( | 68.81474 | -0.20964 | 0.437406 | 0.920144 | Pemt |
| ENSMUSG( | 8555.92 | -0.13272 | 0.39837 | 0.917257 | Penk |
| ENSMUSG( | 532.4656 | 0.144339 | 0.298379 | 0.898538 | Pepd |
| ENSMUSG( | 2381.238 | -0.19664 | 0.27474 | 0.889874 | Per1 |
| ENSMUSG( | 1188.755 | 0.895709 | 6.56E-10 | 6.76E-07 | Per2 |
| ENSMUSG( | 1562.023 | 0.744534 | 1.18E-06 | 0.000604 | Per3 |
| ENSMUSG( | 75.13714 | -0.07134 | 0.801107 | 0.97681 | Perm1 |
| ENSMUSG( | 309.6235 | 0.448433 | 0.002539 | 0.197651 | Perp |
| ENSMUSG( | 1088.023 | -0.10002 | 0.444167 | 0.921734 | Pes1 |
| ENSMUSG( | 677.345 | -0.04514 | 0.731885 | 0.967228 | Pet100 |
| ENSMUSG( | 494.7383 | 0.19246 | 0.201622 | 0.864412 | Pex1 |
| ENSMUSG( | 273.1516 | -0.01731 | 0.914383 | 0.992407 | Pex10 |
| ENSMUSG( | 510.9001 | -0.00472 | 0.971934 | 0.996416 | Pex11a |
| ENSMUSG( | 482.8045 | -0.03027 | 0.84268 | 0.983099 | Pex11b |
| ENSMUSG( | 42.39372 | -0.04164 | 0.895113 | 0.990558 | Pex11g |
| ENSMUSG( | 301.568 | -0.0346 | 0.829388 | 0.980224 | Pex12 |
| ENSMUSG( | 930.3931 | 0.005758 | 0.966719 | 0.995475 | Pex13 |
| ENSMUSG( | 1692.874 | 0.040413 | 0.780941 | 0.976106 | Pex14 |
| ENSMUSG( | 901.3368 | -0.08783 | 0.445585 | 0.921734 | Pex16 |
| ENSMUSG( | 2862.371 | -0.20105 | 0.082955 | 0.77681 | Pex19 |
| ENSMUSG( | 842.0441 | 0.044067 | 0.737987 | 0.967709 | Pex2 |
| ENSMUSG( | 326.6268 | 0.276677 | 0.088913 | 0.788051 | Pex26 |
| ENSMUSG( | 658.2275 | 0.020879 | 0.862962 | 0.986547 | Pex3 |
| ENSMUSG( | 1311.468 | 0.03812 | 0.764931 | 0.974236 | Pex5 |
| ENSMUSG( | 3184.596 | -0.13935 | 0.194745 | 0.860433 | Pex5l |
| ENSMUSG( | 1891.312 | -0.02433 | 0.826248 | 0.979795 | Pex6 |
| ENSMUSG( | 825.7148 | -0.20131 | 0.168945 | 0.840222 | Pex7 |
| ENSMUSG( | 58.71438 | 0.056748 | 0.841131 | 0.9826 | Pf4 |
| ENSMUSG( | 1089.002 | 0.046498 | 0.706138 | 0.96335 | Pfas |
| ENSMUSG( | 1235.786 | -0.18792 | 0.247393 | 0.877904 | Pfdn1 |
| ENSMUSG( | 2729.084 | -0.35008 | 0.020221 | 0.563435 | Pfdn2 |
| ENSMUSG( | 215.7508 | -0.14219 | 0.442282 | 0.921554 | Pfdn4 |
| ENSMUSG( | 5070.968 | -0.15996 | 0.215648 | 0.865167 | Pfdn5 |
| ENSMUSG( | 811.8811 | -0.16616 | 0.303628 | 0.898538 | Pfdn6 |
| ENSMUSG( | 16.62624 | -0.31764 | 0.502707 | 0.927825 | Pfkfb1 |
| ENSMUSG( | 772.7628 | 0.065876 | 0.651295 | 0.95449 | Pfkfb2 |
| ENSMUSG( | 772.3692 | -0.10777 | 0.393437 | 0.917257 | Pfkfb3 |

| | | | | | | | | | | | |
|---|---|---|---|---|---|---|---|---|---|---|---|
| ENSMUSG( | Pgap3 | 385.399 | -0.37201 | 0.013029 | 0.165126 | ENSMUSG( | Pgap3 | 388.3315 | -0.04811 | 0.786618 | 0.976303 |
| ENSMUSG( | Pgap4 | 2135.274 | -0.226 | 0.099094 | 0.481086 | ENSMUSG( | Pgap4 | 2193.444 | 0.131043 | 0.443071 | 0.921554 |
| ENSMUSG( | Pgap6 | 587.9163 | 0.133119 | 0.213253 | 0.657598 | ENSMUSG( | Pgap6 | 537.5314 | 0.14647 | 0.319106 | 0.903431 |
| ENSMUSG( | Pgbd1 | 60.13261 | 0.155591 | 0.531934 | 0.879687 | ENSMUSG( | Pgbd1 | 52.78059 | 0.04115 | 0.885254 | 0.989688 |
| ENSMUSG( | Pgbd5 | 3797.98 | 0.217716 | 0.048876 | 0.337919 | ENSMUSG( | Pgbd5 | 3123.432 | -0.07149 | 0.561218 | 0.940254 |
| ENSMUSG( | Pgd | 1246.728 | -0.10301 | 0.257839 | 0.709198 | ENSMUSG( | Pgd | 1125.563 | -0.13158 | 0.240173 | 0.875205 |
| ENSMUSG( | Pgf | 113.1449 | -0.35348 | 0.075853 | 0.426666 | ENSMUSG( | Pgf | 115.3821 | -0.01116 | 0.957095 | 0.994798 |
| ENSMUSG( | Pgghg | 534.9775 | 0.05915 | 0.653892 | 0.922914 | ENSMUSG( | Pgghg | 447.702 | -0.19097 | 0.168144 | 0.840222 |
| ENSMUSG( | Pggt1b | 1034.524 | 0.140605 | 0.20799 | 0.650144 | ENSMUSG( | Pggt1b | 939.4327 | 0.127763 | 0.408283 | 0.917257 |
| ENSMUSG( | Pgk1 | 6268.659 | 0.041481 | 0.655084 | 0.923872 | ENSMUSG( | Pgk1 | 5087.005 | -0.31205 | 0.015025 | 0.491964 |
| ENSMUSG( | Pgls | 976.1789 | 0.004184 | 0.965554 | 0.995377 | ENSMUSG( | Pgls | 856.704 | -0.10865 | 0.373208 | 0.915491 |
| ENSMUSG( | Pglyrp1 | 465.7245 | 0.164605 | 0.379935 | 0.801757 | ENSMUSG( | Pglyrp1 | 476.5347 | 0.474795 | 0.036532 | 0.66949 |
| ENSMUSG( | Pgm1 | 2901.51 | 0.106487 | 0.335969 | 0.777 | ENSMUSG( | Pgm1 | 2525.651 | -0.02648 | 0.829882 | 0.980224 |
| ENSMUSG( | Pgm2 | 334.2692 | 0.008942 | 0.945239 | 0.99365 | ENSMUSG( | Pgm2 | 311.573 | 0.078359 | 0.597356 | 0.946609 |
| ENSMUSG( | Pgm2l1 | 6131.367 | 0.157337 | 0.065325 | 0.395951 | ENSMUSG( | Pgm2l1 | 5439.353 | 0.082052 | 0.443905 | 0.921734 |
| ENSMUSG( | Pgm3 | 354.9795 | 0.133451 | 0.310771 | 0.758828 | ENSMUSG( | Pgm3 | 294.3654 | -0.14355 | 0.338239 | 0.907972 |
| ENSMUSG( | Pgm5 | 253.5384 | 0.284438 | 0.053611 | 0.35565 | ENSMUSG( | Pgm5 | 220.5188 | 0.173023 | 0.362882 | 0.914223 |
| ENSMUSG( | Pgp | 3234.641 | 0.097055 | 0.217936 | 0.663176 | ENSMUSG( | Pgp | 2754.568 | -0.09972 | 0.355447 | 0.912555 |
| ENSMUSG( | Pgpep1 | 690.1804 | -0.06057 | 0.599954 | 0.903978 | ENSMUSG( | Pgpep1 | 607.8456 | -0.16553 | 0.190034 | 0.857355 |
| ENSMUSG( | Pgpep1l | 11.7673 | -0.37596 | 0.484768 | NA | ENSMUSG( | Pgpep1l | 13.13164 | 0.232216 | 0.662264 | 0.956205 |
| ENSMUSG( | Pgr | 345.8985 | 0.010926 | 0.94148 | 0.993415 | ENSMUSG( | Pgr | 300.8039 | -0.12829 | 0.422141 | 0.917257 |
| ENSMUSG( | Pgrmc1 | 7357.776 | -0.37368 | 0.00237 | 0.057164 | ENSMUSG( | Pgrmc1 | 8215.94 | 0.229641 | 0.121252 | 0.813384 |
| ENSMUSG( | Pgrmc2 | 3585.612 | -0.24368 | 0.026799 | 0.250822 | ENSMUSG( | Pgrmc2 | 3471.094 | -0.05825 | 0.653464 | 0.955015 |
| ENSMUSG( | Pgs1 | 3702.769 | 0.12941 | 0.10383 | 0.489428 | ENSMUSG( | Pgs1 | 3176.985 | -0.0414 | 0.682049 | 0.958678 |
| ENSMUSG( | Phactr1 | 20391.19 | 0.027843 | 0.772134 | 0.954994 | ENSMUSG( | Phactr1 | 19251.82 | 0.129813 | 0.238243 | 0.87259 |
| ENSMUSG( | Phactr2 | 1331.963 | -0.04475 | 0.685183 | 0.932981 | ENSMUSG( | Phactr2 | 1141.226 | -0.23397 | 0.048741 | 0.706195 |
| ENSMUSG( | Phactr3 | 13159.14 | 0.167142 | 0.046352 | 0.329764 | ENSMUSG( | Phactr3 | 11136.56 | -0.04224 | 0.700427 | 0.961475 |
| ENSMUSG( | Phactr4 | 835.9069 | 0.06703 | 0.516757 | 0.873014 | ENSMUSG( | Phactr4 | 806.9009 | 0.220707 | 0.095967 | 0.796133 |
| ENSMUSG( | Phaf1 | 779.0927 | 0.082363 | 0.559558 | 0.888674 | ENSMUSG( | Phaf1 | 676.9795 | -0.05511 | 0.702537 | 0.962215 |
| ENSMUSG( | Phax | 842.6316 | 0.136565 | 0.289901 | 0.739839 | ENSMUSG( | Phax | 723.8539 | -0.03455 | 0.820253 | 0.979198 |
| ENSMUSG( | Phb1 | 1614.796 | 0.017753 | 0.840604 | 0.971179 | ENSMUSG( | Phb1 | 1388.141 | -0.15642 | 0.201468 | 0.864412 |
| ENSMUSG( | Phb2 | 2113.809 | 0.008665 | 0.927936 | 0.991229 | ENSMUSG( | Phb2 | 1878.376 | -0.06622 | 0.587407 | 0.944975 |
| ENSMUSG( | Phc1 | 2854.476 | 0.196989 | 0.018206 | 0.203238 | ENSMUSG( | Phc1 | 2492.376 | 0.079647 | 0.437258 | 0.920144 |
| ENSMUSG( | Phc2 | 2769.788 | 0.022273 | 0.827381 | 0.968732 | ENSMUSG( | Phc2 | 2477.001 | -0.03131 | 0.782768 | 0.976106 |
| ENSMUSG( | Phc3 | 2080.467 | 0.104197 | 0.322505 | 0.767663 | ENSMUSG( | Phc3 | 1872.828 | 0.068806 | 0.569678 | 0.941777 |
| ENSMUSG( | Pheta1 | 349.8509 | 0.055124 | 0.694178 | 0.9343 | ENSMUSG( | Pheta1 | 326.1749 | 0.126308 | 0.416871 | 0.917257 |
| ENSMUSG( | Pheta2 | 28.80017 | -0.11741 | 0.748002 | 0.949604 | ENSMUSG( | Pheta2 | 24.6715 | -0.29678 | 0.474104 | 0.923329 |
| ENSMUSG( | Phex | 18.34749 | -0.41583 | 0.329147 | NA | ENSMUSG( | Phex | 18.10945 | -0.1591 | 0.723686 | 0.966887 |
| ENSMUSG( | Phf1 | 1075.482 | -0.00853 | 0.941246 | 0.993415 | ENSMUSG( | Phf1 | 1013.681 | 0.088327 | 0.498916 | 0.927825 |
| ENSMUSG( | Phf10 | 2890.142 | 0.032122 | 0.719835 | 0.941986 | ENSMUSG( | Phf10 | 2604.378 | -0.00246 | 0.984598 | 0.996963 |
| ENSMUSG( | Phf11b | 16.36857 | -0.42934 | 0.377626 | NA | ENSMUSG( | Phf11b | 14.52955 | -0.52961 | 0.355182 | 0.912411 |
| ENSMUSG( | Phf11c | 22.62355 | 0.008606 | 0.984406 | 0.99781 | ENSMUSG( | Phf11c | 19.19302 | -0.16743 | 0.734417 | 0.967228 |
| ENSMUSG( | Phf11d | 17.54333 | -0.89663 | 0.063906 | NA | ENSMUSG( | Phf11d | 17.53852 | -0.53015 | 0.284753 | 0.892412 |
| ENSMUSG( | Phf12 | 1487.482 | 0.03228 | 0.745976 | 0.949289 | ENSMUSG( | Phf12 | 1381.074 | 0.085803 | 0.432818 | 0.918434 |
| ENSMUSG( | Phf13 | 337.7536 | -0.75232 | 7.61E-08 | 2.13E-05 | ENSMUSG( | Phf13 | 357.195 | -0.23766 | 0.090648 | 0.791074 |
| ENSMUSG( | Phf14 | 1516.89 | -0.08013 | 0.366647 | 0.794842 | ENSMUSG( | Phf14 | 1466.478 | 0.088333 | 0.435255 | 0.919062 |
| ENSMUSG( | Phf19 | 50.59511 | -0.23978 | 0.367391 | 0.795221 | ENSMUSG( | Phf19 | 54.65553 | 0.243054 | 0.384998 | 0.917257 |
| ENSMUSG( | Phf2 | 2819.064 | 0.113353 | 0.237834 | 0.685809 | ENSMUSG( | Phf2 | 2529.69 | 0.069876 | 0.531739 | 0.934795 |
| ENSMUSG( | Phf20 | 2112.223 | -0.12602 | 0.177376 | 0.614553 | ENSMUSG( | Phf20 | 2021.166 | 0.018795 | 0.876758 | 0.989644 |
| ENSMUSG( | Phf20l1 | 2030.911 | -0.114 | 0.286517 | 0.736031 | ENSMUSG( | Phf20l1 | 1971.562 | 0.069566 | 0.568201 | 0.941777 |
| ENSMUSG( | Phf21a | 2056.935 | 0.129223 | 0.149287 | 0.573248 | ENSMUSG( | Phf21a | 1809.401 | 0.027913 | 0.808955 | 0.97873 |
| ENSMUSG( | Phf21b | 651.3076 | 0.000923 | 0.993524 | 0.999369 | ENSMUSG( | Phf21b | 601.6591 | 0.038632 | 0.77215 | 0.974236 |
| ENSMUSG( | Phf23 | 1127.48 | 0.042952 | 0.678116 | 0.930542 | ENSMUSG( | Phf23 | 1048.943 | 0.097806 | 0.456746 | 0.923329 |
| ENSMUSG( | Phf24 | 12296.43 | 0.105719 | 0.319831 | 0.765305 | ENSMUSG( | Phf24 | 11044.54 | 0.065603 | 0.540411 | 0.937383 |
| ENSMUSG( | Phf3 | 2538.859 | 0.040098 | 0.64528 | 0.917957 | ENSMUSG( | Phf3 | 2315.798 | 0.041889 | 0.710695 | 0.964363 |
| ENSMUSG( | Phf5a | 624.0293 | 0.09087 | 0.403506 | 0.816132 | ENSMUSG( | Phf5a | 529.3645 | -0.1176 | 0.369382 | 0.915491 |
| ENSMUSG( | Phf6 | 451.2021 | 0.247272 | 0.055098 | 0.361231 | ENSMUSG( | Phf6 | 421.3289 | 0.305532 | 0.042364 | 0.690304 |
| ENSMUSG( | Phf7 | 132.7465 | -0.19628 | 0.291244 | 0.741248 | ENSMUSG( | Phf7 | 121.0041 | -0.17566 | 0.45153 | 0.922202 |
| ENSMUSG( | Phf8 | 626.6155 | 0.093455 | 0.42053 | 0.822898 | ENSMUSG( | Phf8 | 575.9045 | 0.115246 | 0.391233 | 0.917257 |
| ENSMUSG( | Phgdh | 5282.412 | 0.149158 | 0.14932 | 0.573248 | ENSMUSG( | Phgdh | 4291.401 | -0.18628 | 0.141498 | 0.830678 |
| ENSMUSG( | Phip | 1108.716 | -0.19579 | 0.157018 | 0.58551 | ENSMUSG( | Phip | 1163.017 | 0.212367 | 0.148884 | 0.836241 |
| ENSMUSG( | Phka1 | 3518.819 | 0.171182 | 0.120783 | 0.524807 | ENSMUSG( | Phka1 | 2906.933 | -0.1106 | 0.37906 | 0.916995 |
| ENSMUSG( | Phka2 | 823.8671 | 0.220759 | 0.158104 | 0.586829 | ENSMUSG( | Phka2 | 722.387 | 0.115807 | 0.541006 | 0.937647 |
| ENSMUSG( | Phkb | 2469.874 | 0.087543 | 0.473982 | 0.849425 | ENSMUSG( | Phkb | 2060.654 | -0.17264 | 0.182987 | 0.850741 |
| ENSMUSG( | Phkg1 | 1862.168 | 0.030008 | 0.790653 | 0.959461 | ENSMUSG( | Phkg1 | 1564.617 | -0.2111 | 0.096318 | 0.796133 |
| ENSMUSG( | Phkg2 | 2418.806 | 0.052227 | 0.559717 | 0.88884 | ENSMUSG( | Phkg2 | 2115.564 | -0.06873 | 0.554235 | 0.939565 |
| ENSMUSG( | Phlda1 | 482.9242 | -0.14798 | 0.351674 | 0.783381 | ENSMUSG( | Phlda1 | 451.1887 | -0.07808 | 0.673685 | 0.95694 |
| ENSMUSG( | Phlda3 | 699.67 | -0.05843 | 0.649066 | 0.920025 | ENSMUSG( | Phlda3 | 592.5187 | -0.28497 | 0.04913 | 0.708275 |
| ENSMUSG( | Phldb1 | 6400.787 | -0.22775 | 0.010127 | 0.143118 | ENSMUSG( | Phldb1 | 6058.489 | -0.10992 | 0.403813 | 0.917257 |
| ENSMUSG( | Phldb2 | 372.4411 | -0.21202 | 0.136785 | 0.551217 | ENSMUSG( | Phldb2 | 394.7733 | 0.220458 | 0.174227 | 0.844317 |
| ENSMUSG( | Phlpp1 | 5612.261 | 0.028814 | 0.740944 | 0.948028 | ENSMUSG( | Phlpp1 | 4761.312 | -0.18251 | 0.102703 | 0.803503 |
| ENSMUSG( | Phlpp2 | 1581.74 | 0.075791 | 0.412173 | 0.819452 | ENSMUSG( | Phlpp2 | 1451.998 | 0.096338 | 0.367581 | 0.915217 |
| ENSMUSG( | Phospho1 | 131.0883 | 0.158672 | 0.418595 | 0.821966 | ENSMUSG( | Phospho1 | 116.5019 | 0.105629 | 0.63506 | 0.953903 |
| ENSMUSG( | Phospho2 | 381.5118 | 0.061921 | 0.65955 | 0.924506 | ENSMUSG( | Phospho2 | 340.4567 | -0.00578 | 0.973192 | 0.996416 |
| ENSMUSG( | Phpt1 | 332.3751 | -0.00846 | 0.949814 | 0.993821 | ENSMUSG( | Phpt1 | 298.8212 | -0.05044 | 0.757535 | 0.971858 |
| ENSMUSG( | Phrf1 | 1650.718 | 0.184515 | 0.043364 | 0.319133 | ENSMUSG( | Phrf1 | 1479.882 | 0.142517 | 0.230493 | 0.868507 |
| ENSMUSG( | Phtf1 | 1319.636 | 0.066336 | 0.548867 | 0.885686 | ENSMUSG( | Phtf1 | 1312.972 | 0.308112 | 0.015734 | 0.502615 |
| ENSMUSG( | Phtf1os | 32.66911 | -0.01671 | 0.960593 | 0.993991 | ENSMUSG( | Phtf1os | 34.4613 | 0.366426 | 0.303546 | 0.898538 |
| ENSMUSG( | Phtf2 | 404.325 | -0.25728 | 0.102686 | 0.488148 | ENSMUSG( | Phtf2 | 440.5589 | 0.263084 | 0.107265 | 0.804969 |
| ENSMUSG( | Phyh | 4270.913 | 0.028371 | 0.811474 | 0.964426 | ENSMUSG( | Phyh | 3562.423 | -0.23907 | 0.069951 | 0.757435 |
| ENSMUSG( | Phyhd1 | 126.2988 | 0.384126 | 0.07424 | 0.422054 | ENSMUSG( | Phyhd1 | 111.7598 | 0.328793 | 0.259301 | 0.883971 |
| ENSMUSG( | Phyhip | 10677.06 | 0.075087 | 0.590791 | 0.90113 | ENSMUSG( | Phyhip | 9344.324 | -0.04158 | 0.781284 | 0.976106 |
| ENSMUSG( | Phyhipl | 4888.165 | -0.05241 | 0.583933 | 0.898666 | ENSMUSG( | Phyhipl | 4160.758 | -0.2623 | 0.042903 | 0.690304 |
| ENSMUSG( | Phykpl | 144.1143 | 0.086413 | 0.625993 | 0.912356 | ENSMUSG( | Phykpl | 127.0939 | 0.000447 | 0.998236 | 0.999313 |
| ENSMUSG( | Pi16 | 20.66511 | -0.14628 | 0.771141 | NA | ENSMUSG( | Pi16 | 18.63298 | -0.19475 | 0.715002 | 0.965063 |
| ENSMUSG( | Pi4k2a | 1929.229 | 0.125632 | 0.234043 | 0.681906 | ENSMUSG( | Pi4k2a | 1656.967 | -0.04208 | 0.726652 | 0.966887 |
| ENSMUSG( | Pi4k2b | 122.0925 | -0.13614 | 0.466318 | 0.844631 | ENSMUSG( | Pi4k2b | 104.1212 | -0.34118 | 0.125735 | 0.81516 |
| ENSMUSG( | Pi4ka | 7979.473 | 0.193896 | 0.0281 | 0.25643 | ENSMUSG( | Pi4ka | 6837.364 | 0.023495 | 0.833853 | 0.981218 |
| ENSMUSG( | Pi4kb | 1339.51 | 0.157437 | 0.133139 | 0.545727 | ENSMUSG( | Pi4kb | 1197.689 | 0.10927 | 0.39704 | 0.917257 |
| ENSMUSG( | Pianp | 6260.826 | -0.41367 | 0.000177 | 0.009611 | ENSMUSG( | Pianp | 6730.866 | 0.09379 | 0.467331 | 0.923329 |
| ENSMUSG( | Pias1 | 1239.668 | 0.150573 | 0.123553 | 0.530552 | ENSMUSG( | Pias1 | 1081.345 | 0.026748 | 0.824401 | 0.979198 |
| ENSMUSG( | Pias2 | 1329.159 | 0.12697 | 0.171464 | 0.605943 | ENSMUSG( | Pias2 | 1201.723 | 0.10246 | 0.422864 | 0.917257 |
| ENSMUSG( | Pias3 | 701.1843 | 0.050411 | 0.691382 | 0.933881 | ENSMUSG( | Pias3 | 662.9296 | 0.155665 | 0.253362 | 0.88018 |
| ENSMUSG( | Pias4 | 631.1969 | 0.134061 | 0.361374 | 0.792797 | ENSMUSG( | Pias4 | 557.1776 | 0.043189 | 0.769443 | 0.974236 |
| ENSMUSG( | Pibf1 | 201.7678 | 0.010078 | 0.945936 | 0.99369 | ENSMUSG( | Pibf1 | 184.1046 | 0.008521 | 0.960974 | 0.99511 |
| ENSMUSG( | Picalm | 4031.344 | 0.106641 | 0.200643 | 0.640661 | ENSMUSG( | Picalm | 3682.649 | 0.11366 | 0.290914 | 0.897019 |
| ENSMUSG( | Pick1 | 527.9048 | 0.074815 | 0.533796 | 0.880095 | ENSMUSG( | Pick1 | 465.9266 | -0.01882 | 0.911112 | 0.992196 |
| ENSMUSG( | Pid1 | 970.6525 | -0.05071 | 0.649385 | 0.920317 | ENSMUSG( | Pid1 | 882.3761 | -0.06267 | 0.655856 | 0.955533 |
| ENSMUSG( | Pidd1 | 78.32187 | 0.196734 | 0.390984 | 0.80828 | ENSMUSG( | Pidd1 | 66.4542 | 0.017756 | 0.94844 | 0.994798 |
| ENSMUSG( | Pierce1 | 67.00374 | -0.12633 | 0.607009 | 0.906685 | ENSMUSG( | Pierce1 | 57.70504 | -0.28702 | 0.296911 | 0.898538 |
| ENSMUSG( | Pierce2 | 140.3554 | -0.18908 | 0.294471 | 0.743674 | ENSMUSG( | Pierce2 | 120.8072 | -0.36579 | 0.069392 | 0.755467 |
| ENSMUSG( | Piezo1 | 398.1419 | 0.180221 | 0.185135 | 0.623005 | ENSMUSG( | Piezo1 | 382.4842 | 0.329588 | 0.093416 | 0.795507 |
| ENSMUSG( | Piezo2 | 76.4377 | -0.30279 | 0.194036 | 0.633126 | ENSMUSG( | Piezo2 | 76.68709 | 0.008059 | 0.973895 | 0.996416 |
| ENSMUSG( | Pif1 | 14.55567 | -0.39621 | 0.484334 | NA | ENSMUSG( | Pif1 | 15.41355 | 0.051702 | 0.930831 | 0.99306 |
| ENSMUSG( | Piga | 250.7622 | -0.11257 | 0.456002 | 0.838683 | ENSMUSG( | Piga | 261.0954 | 0.273264 | 0.121569 | 0.813384 |
| ENSMUSG( | Pigb | 489.457 | -0.14784 | 0.23161 | 0.67949 | ENSMUSG( | Pigb | 510.1978 | 0.241129 | 0.098271 | 0.798438 |
| ENSMUSG( | Pigbos1 | 163.51 | 0.068513 | 0.678374 | 0.930542 | ENSMUSG( | Pigbos1 | 137.6328 | -0.16405 | 0.394203 | 0.917257 |
| ENSMUSG( | Pigc | 495.6872 | -0.02669 | 0.840132 | 0.971179 | ENSMUSG( | Pigc | 495.3007 | 0.23828 | 0.128441 | 0.81925 |
| ENSMUSG( | Pigf | 255.3368 | -0.183 | 0.247905 | 0.696522 | ENSMUSG( | Pigf | 247.0072 | -0.0122 | 0.947364 | 0.994798 |
| ENSMUSG( | Pigg | 595.0037 | -0.18035 | 0.269591 | 0.719729 | ENSMUSG( | Pigg | 600.6912 | 0.127676 | 0.508823 | 0.929089 |
| ENSMUSG( | Pigh | 462.5954 | -0.22196 | 0.071434 | 0.413071 | ENSMUSG( | Pigh | 447.0477 | -0.03956 | 0.774428 | 0.974931 |
| ENSMUSG( | Pigk | 2026.547 | -0.33554 | 0.004419 | 0.086676 | ENSMUSG( | Pigk | 2128.686 | 0.096127 | 0.469392 | 0.923329 |
| ENSMUSG( | Pigl | 163.315 | -0.52581 | 0.00295 | 0.066455 | ENSMUSG( | Pigl | 172.6666 | -0.039 | 0.845145 | 0.983878 |
| ENSMUSG( | Pigm | 397.6124 | -0.23378 | 0.142276 | 0.560124 | ENSMUSG( | Pigm | 428.0484 | 0.256113 | 0.185476 | 0.855305 |
| ENSMUSG( | Pign | 458.5178 | -0.2104 | 0.153032 | 0.579541 | ENSMUSG( | Pign | 456.5023 | 0.058743 | 0.717561 | 0.965964 |
| ENSMUSG( | Pigo | 734.0951 | 0.035377 | 0.820435 | 0.965888 | ENSMUSG( | Pigo | 747.6553 | 0.344818 | 0.048824 | 0.706228 |
| ENSMUSG( | Pigp | 702.4955 | -0.40541 | 0.001026 | 0.033548 | ENSMUSG( | Pigp | 727.0584 | -0.00815 | 0.949222 | 0.994798 |
| ENSMUSG( | Pigq | 6112.635 | -0.00291 | 0.976982 | 0.997428 | ENSMUSG( | Pigq | 5473.734 | -0.05102 | 0.665724 | 0.956766 |
| ENSMUSG( | Pigs | 2550.631 | -0.19014 | 0.165817 | 0.597653 | ENSMUSG( | Pigs | 2603.233 | 0.144903 | 0.368367 | 0.915491 |
| ENSMUSG( | Pigt | 3384.395 | -0.27967 | 0.057891 | 0.371023 | ENSMUSG( | Pigt | 3524.867 | 0.122653 | 0.456379 | 0.923329 |
| ENSMUSG( | Pigu | 814.7961 | -0.36927 | 0.009377 | 0.137017 | ENSMUSG( | Pigu | 879.1926 | 0.143431 | 0.360648 | 0.914223 |
| ENSMUSG( | Pigv | 285.3858 | -0.2725 | 0.086094 | 0.451605 | ENSMUSG( | Pigv | 298.4221 | 0.147469 | 0.468546 | 0.923329 |
| ENSMUSG( | Pigw | 85.27875 | -0.43861 | 0.073057 | 0.417421 | ENSMUSG( | Pigw | 90.92509 | 0.053316 | 0.818123 | 0.979198 |
| ENSMUSG( | Pigx | 351.8054 | -0.27332 | 0.035881 | 0.290992 | ENSMUSG( | Pigx | 366.2283 | 0.120482 | 0.426047 | 0.917257 |
| ENSMUSG( | Pigyl | 657.2767 | -0.2156 | 0.040347 | 0.310226 | ENSMUSG( | Pigyl | 610.8942 | -0.15754 | 0.244119 | 0.877904 |
| ENSMUSG( | Pigz | 266.9531 | -0.51537 | 0.011952 | 0.157992 | ENSMUSG( | Pigz | 275.8739 | -0.09541 | 0.660249 | 0.956205 |
| ENSMUSG( | Pih1d1 | 755.527 | 0.066404 | 0.515297 | 0.872546 | ENSMUSG( | Pih1d1 | 652.6269 | -0.08946 | 0.473855 | 0.923329 |
| ENSMUSG( | Pih1d2 | 54.6152 | 0.094062 | 0.722013 | 0.943535 | ENSMUSG( | Pih1d2 | 49.35969 | 0.073847 | 0.801799 | 0.97681 |
| ENSMUSG( | Pik3ap1 | 270.4479 | -0.21173 | 0.157068 | 0.58551 | ENSMUSG( | Pik3ap1 | 247.5141 | -0.19507 | 0.24061 | 0.875927 |
| ENSMUSG( | Pik3c2a | 991.8835 | 0.119124 | 0.351152 | 0.783067 | ENSMUSG( | Pik3c2a | 982.429 | 0.345942 | 0.031461 | 0.631192 |
| ENSMUSG( | Pik3c2b | 652.1856 | -0.16257 | 0.151084 | 0.576088 | ENSMUSG( | Pik3c2b | 586.0956 | -0.20445 | 0.114669 | 0.811703 |
| ENSMUSG( | Pik3c3 | 1011.547 | 0.232808 | 0.029259 | 0.259574 | ENSMUSG( | Pik3c3 | 848.8135 | 0.007358 | 0.952497 | 0.994798 |
| ENSMUSG( | Pik3ca | 2406.509 | 0.153083 | 0.108184 | 0.497094 | ENSMUSG( | Pik3ca | 2136.027 | 0.078252 | 0.515755 | 0.930939 |
| ENSMUSG( | Pik3cb | 703.7518 | 0.158373 | 0.135375 | 0.548884 | ENSMUSG( | Pik3cb | 614.1547 | 0.034907 | 0.790116 | 0.97681 |
| ENSMUSG( | Pik3cd | 964.4 | 0.172224 | 0.130769 | 0.542059 | ENSMUSG( | Pik3cd | 884.9716 | 0.194538 | 0.138385 | 0.828492 |
| ENSMUSG( | Pik3cg | 192.6332 | 0.228656 | 0.15255 | 0.578935 | ENSMUSG( | Pik3cg | 182.8144 | 0.346022 | 0.065551 | 0.745138 |
| ENSMUSG( | Pik3ip1 | 1377.971 | -0.30661 | 0.039396 | 0.305596 | ENSMUSG( | Pik3ip1 | 1458.44 | 0.144424 | 0.423372 | 0.917257 |
| ENSMUSG( | Pik3r1 | 6554.888 | 0.216665 | 0.008042 | 0.124826 | ENSMUSG( | Pik3r1 | 5498.199 | -0.01486 | 0.897034 | 0.990558 |
| ENSMUSG( | Pik3r2 | 5597.444 | 0.158769 | 0.084568 | 0.448139 | ENSMUSG( | Pik3r2 | 4962.376 | 0.083957 | 0.400799 | 0.917257 |
| ENSMUSG( | Pik3r3 | 1737.551 | -0.21275 | 0.050899 | 0.345939 | ENSMUSG( | Pik3r3 | 1691.265 | -0.01537 | 0.902571 | 0.991089 |
| ENSMUSG( | Pik3r4 | 1001.455 | -0.04153 | 0.716172 | 0.94152 | ENSMUSG( | Pik3r4 | 993.2001 | 0.20226 | 0.183737 | 0.85191 |
| ENSMUSG( | Pik3r5 | 168.8925 | -0.00284 | 0.986324 | 0.998326 | ENSMUSG( | Pik3r5 | 153.3886 | -0.00118 | 0.995533 | 0.999116 |
| ENSMUSG( | Pik3r6 | 142.1159 | 0.139739 | 0.472743 | 0.848388 | ENSMUSG( | Pik3r6 | 130.3873 | 0.177938 | 0.450958 | 0.922202 |
| ENSMUSG( | Pikfyve | 1182.422 | 0.073535 | 0.427964 | 0.825894 | ENSMUSG( | Pikfyve | 1117.29 | 0.173617 | 0.120251 | 0.813384 |
| ENSMUSG( | Pilra | 32.05064 | -0.41224 | 0.39696 | 0.811839 | ENSMUSG( | Pilra | 27.16878 | -0.67276 | 0.173639 | 0.844317 |
| ENSMUSG( | Pim1 | 95.34493 | 0.052812 | 0.816832 | 0.96566 | ENSMUSG( | Pim1 | 87.12679 | 0.063191 | 0.788132 | 0.97637 |
| ENSMUSG( | Pim2 | 921.2436 | 0.085589 | 0.528126 | 0.878224 | ENSMUSG( | Pim2 | 790.4219 | -0.08843 | 0.573478 | 0.942912 |
| ENSMUSG( | Pim3 | 893.3166 | -0.35305 | 0.054183 | 0.357985 | ENSMUSG( | Pim3 | 896.7321 | -0.0482 | 0.820508 | 0.979198 |
| ENSMUSG( | Pimreg | 25.70673 | 0.094848 | 0.816133 | 0.965455 | ENSMUSG( | Pimreg | 20.17729 | -0.35555 | 0.446087 | 0.921734 |
| ENSMUSG( | Pin1 | 2501.079 | 0.001103 | 0.990741 | 0.999232 | ENSMUSG( | Pin1 | 2091.983 | -0.25951 | 0.028538 | 0.614171 |
| ENSMUSG( | Pin4 | 357.2458 | 0.049469 | 0.733261 | 0.944956 | ENSMUSG( | Pin4 | 296.5189 | -0.2362 | 0.199055 | 0.863016 |
| ENSMUSG( | Pink1 | 30765.79 | 0.193724 | 0.017602 | 0.198821 | ENSMUSG( | Pink1 | 25757.58 | -0.04582 | 0.714872 | 0.964986 |
| ENSMUSG( | Pinx1 | 158.0458 | -0.22801 | 0.193788 | 0.633085 | ENSMUSG( | Pinx1 | 142.3963 | -0.27286 | 0.178745 | 0.849041 |
| ENSMUSG( | Pip4k2a | 4285.289 | -0.01458 | 0.865378 | 0.975565 | ENSMUSG( | Pip4k2a | 3747.093 | -0.1381 | 0.268321 | 0.886493 |
| ENSMUSG( | Pip4k2b | 5155.56 | 0.162568 | 0.035734 | 0.290677 | ENSMUSG( | Pip4k2b | 4393.221 | -0.02637 | 0.788828 | 0.976593 |
| ENSMUSG( | Pip4k2c | 4005.075 | -0.08073 | 0.42599 | 0.824898 | ENSMUSG( | Pip4k2c | 3732.321 | -0.01173 | 0.920658 | 0.992644 |
| ENSMUSG( | Pip4p1 | 1639.076 | -0.0109 | 0.907088 | 0.987447 | ENSMUSG( | Pip4p1 | 1516.578 | 0.034198 | 0.749591 | 0.96998 |
| ENSMUSG( | Pip4p2 | 1019.511 | -0.06385 | 0.558088 | 0.888427 | ENSMUSG( | Pip4p2 | 947.1894 | -0.01334 | 0.926932 | 0.99306 |

| | | | | | |
|---|---|---|---|---|---|
| ENSMUSG( | 1632.806 | 0.036185 | 0.691544 | 0.933881 | Pdcd4 |
| ENSMUSG( | 982.8029 | 0.070105 | 0.498605 | 0.862973 | Pdcd5 |
| ENSMUSG( | 1448.329 | 0.036618 | 0.712247 | 0.94031 | Pdcd6 |
| ENSMUSG( | 2502.174 | 0.064577 | 0.437955 | 0.830072 | Pdcd6ip |
| ENSMUSG( | 428.8609 | 0.02311 | 0.854356 | 0.973491 | Pdcd7 |
| ENSMUSG( | 606.3914 | 0.140346 | 0.29012 | 0.739897 | Pdcl |
| ENSMUSG( | 700.529 | -0.06231 | 0.551739 | 0.886141 | Pdcl3 |
| ENSMUSG( | 3272.502 | -0.33034 | 0.020601 | 0.216558 | Pde10a |
| ENSMUSG( | 58.99107 | 0.255974 | 0.33619 | 0.777 | Pde11a |
| ENSMUSG( | 312.5311 | -0.00332 | 0.982344 | 0.997512 | Pde12 |
| ENSMUSG( | 2668.26 | 0.176233 | 0.168161 | 0.601451 | Pde1a |
| ENSMUSG( | 5327.85 | -0.41567 | 0.006685 | 0.112486 | Pde1b |
| ENSMUSG( | 550.7545 | -0.05658 | 0.722972 | 0.943761 | Pde1c |
| ENSMUSG( | 256.5706 | 0.060413 | 0.691494 | 0.933881 | Pde2a |
| ENSMUSG( | 186.4086 | -0.00171 | 0.991565 | 0.999232 | Pde3a |
| ENSMUSG( | 576.1687 | -0.00758 | 0.954228 | 0.993874 | Pde3b |
| ENSMUSG( | 4730.013 | 0.13551 | 0.201219 | 0.641362 | Pde4a |
| ENSMUSG( | 2270.197 | -0.09312 | 0.282475 | 0.73186 | Pde4b |
| ENSMUSG( | 100.0719 | 0.543913 | 0.012589 | 0.161972 | Pde4c |
| ENSMUSG( | 521.791 | 0.196878 | 0.114075 | 0.509818 | Pde4d |
| ENSMUSG( | 5415.917 | 0.156399 | 0.04731 | 0.332697 | Pde4dip |
| ENSMUSG( | 246.9813 | 0.006244 | 0.969219 | 0.996369 | Pde5a |
| ENSMUSG( | 17.25231 | -0.68963 | 0.126146 | NA | Pde6a |
| ENSMUSG( | 1501.404 | 0.080951 | 0.47559 | 0.850919 | Pde6d |
| ENSMUSG( | 10.35306 | -0.14338 | 0.811489 | NA | Pde6h |
| ENSMUSG( | 859.3986 | 0.0304 | 0.777731 | 0.956168 | Pde7a |
| ENSMUSG( | 1337.299 | 0.079998 | 0.55276 | 0.886141 | Pde7b |
| ENSMUSG( | 2034.521 | -0.00049 | 0.995621 | 0.99943 | Pde8a |
| ENSMUSG( | 3192.966 | 0.050439 | 0.563025 | 0.889738 | Pde8b |
| ENSMUSG( | 703.4926 | -0.06269 | 0.577556 | 0.895864 | Pde9a |
| ENSMUSG( | 30.71488 | 0.331852 | 0.36208 | 0.79289 | Pdf |
| ENSMUSG( | 1766.356 | -0.00426 | 0.967861 | 0.996237 | Pdgfa |
| ENSMUSG( | 1104.569 | 0.054714 | 0.709261 | 0.939656 | Pdgfb |
| ENSMUSG( | 401.7253 | -0.09278 | 0.483166 | 0.856216 | Pdgfc |
| ENSMUSG( | 180.7306 | 0.151331 | 0.39031 | 0.80794 | Pdgfd |
| ENSMUSG( | 2675.174 | -0.37299 | 0.027895 | 0.255773 | Pdgfra |
| ENSMUSG( | 1447.848 | -0.29155 | 0.126631 | 0.534187 | Pdgfrb |
| ENSMUSG( | 92.98141 | -0.35783 | 0.139217 | 0.555317 | Pdgfrl |
| ENSMUSG( | 9304.314 | 0.049131 | 0.591179 | 0.90113 | Pdha1 |
| ENSMUSG( | 2990.646 | 0.11148 | 0.265197 | 0.715962 | Pdhb |
| ENSMUSG( | 1302.369 | 0.129149 | 0.156555 | 0.584667 | Pdhx |
| ENSMUSG( | 9.026518 | 1.240792 | 0.05789 | NA | Pdia2 |
| ENSMUSG( | 7929.542 | -0.24987 | 0.089226 | 0.458482 | Pdia3 |
| ENSMUSG( | 1859.693 | -0.11882 | 0.442559 | 0.83278 | Pdia4 |
| ENSMUSG( | 123.7435 | -0.17982 | 0.32084 | 0.765984 | Pdia5 |
| ENSMUSG( | 3921.628 | -0.11332 | 0.473346 | 0.848919 | Pdia6 |
| ENSMUSG( | 256.4257 | -0.08961 | 0.520702 | 0.875186 | Pdik1l |
| ENSMUSG( | 1867.621 | 0.169386 | 0.06765 | 0.402445 | Pdk1 |
| ENSMUSG( | 4305.775 | 0.046265 | 0.594947 | 0.903007 | Pdk2 |
| ENSMUSG( | 691.4144 | -0.04424 | 0.693235 | 0.934202 | Pdk3 |
| ENSMUSG( | 1079.97 | -0.17481 | 0.190178 | 0.629268 | Pdk4 |
| ENSMUSG( | 113.6233 | -0.1044 | 0.606625 | 0.906529 | Pdlim1 |
| ENSMUSG( | 2301.716 | -0.22877 | 0.027569 | 0.2548 | Pdlim2 |
| ENSMUSG( | 12.80344 | -0.14734 | 0.787591 | NA | Pdlim3 |
| ENSMUSG( | 640.9144 | 0.213835 | 0.070293 | 0.40908 | Pdlim4 |
| ENSMUSG( | 2252.866 | 0.068886 | 0.597216 | 0.903694 | Pdlim5 |
| ENSMUSG( | 1334.73 | 0.219567 | 0.133046 | 0.545688 | Pdlim7 |
| ENSMUSG( | 1406.885 | -0.17388 | 0.192029 | 0.630811 | Pdp1 |
| ENSMUSG( | 380.2565 | 0.183531 | 0.146086 | 0.567605 | Pdp2 |
| ENSMUSG( | 3032.189 | 0.086667 | 0.337013 | 0.777008 | Pdpk1 |
| ENSMUSG( | 816.213 | -0.38693 | 0.015462 | 0.183729 | Pdpn |
| ENSMUSG( | 1071.62 | 0.019053 | 0.871685 | 0.97812 | Pdpr |
| ENSMUSG( | 962.0494 | -0.07118 | 0.478366 | 0.853567 | Pdrg1 |
| ENSMUSG( | 1608.15 | 0.119153 | 0.225671 | 0.672494 | Pds5a |
| ENSMUSG( | 1916.334 | 0.072483 | 0.406751 | 0.816928 | Pds5b |
| ENSMUSG( | 222.2553 | -0.11001 | 0.51997 | 0.874979 | Pdss1 |
| ENSMUSG( | 270.971 | 0.072448 | 0.636308 | 0.914873 | Pdss2 |
| ENSMUSG( | 4755.247 | 0.085284 | 0.317583 | 0.763932 | Pdxdc1 |
| ENSMUSG( | 5812.211 | 0.072318 | 0.399169 | 0.813048 | Pdxk |
| ENSMUSG( | 2569.961 | 0.061507 | 0.59592 | 0.903152 | Pdxp |
| ENSMUSG( | 957.884 | -0.78916 | 9.93E-09 | 3.91E-06 | Pdyn |
| ENSMUSG( | 944.3874 | 0.004835 | 0.961273 | 0.994061 | Pdzd11 |
| ENSMUSG( | 1962.103 | 0.103531 | 0.253187 | 0.70338 | Pdzd2 |
| ENSMUSG( | 7721.36 | 0.182601 | 0.015606 | 0.184634 | Pdzd4 |
| ENSMUSG( | 168.7144 | 0.434575 | 0.029767 | 0.261412 | Pdzd7 |
| ENSMUSG( | 2234.67 | -0.23194 | 0.090986 | 0.463502 | Pdzd8 |
| ENSMUSG( | 19.3414 | 3.37E-06 | 0.999994 | NA | Pdzk1 |
| ENSMUSG( | 19.8319 | -0.02284 | 0.956337 | NA | Pdzk1ip1 |
| ENSMUSG( | 337.8989 | 0.257024 | 0.087663 | 0.454663 | Pdzph1 |
| ENSMUSG( | 1033.349 | -0.10016 | 0.432221 | 0.82768 | Pdzrn3 |
| ENSMUSG( | 190.4867 | -0.33793 | 0.037663 | 0.299895 | Pdzrn4 |
| ENSMUSG( | 49662.94 | -0.01273 | 0.884618 | 0.982647 | Pea15a |
| ENSMUSG( | 1670.661 | -0.05945 | 0.509964 | 0.869743 | Peak1 |
| ENSMUSG( | 303.1885 | 0.262326 | 0.120217 | 0.524481 | Pear1 |
| ENSMUSG( | 9233.884 | 0.028148 | 0.748809 | 0.949604 | Pebp1 |
| ENSMUSG( | 883.768 | 0.130312 | 0.545334 | 0.884774 | Pecam1 |
| ENSMUSG( | 106.8168 | -0.08219 | 0.680431 | 0.931044 | Pecr |
| ENSMUSG( | 2156.299 | -0.35696 | 0.002789 | 0.064345 | Peds1 |
| ENSMUSG( | 1195.774 | 0.043159 | 0.667536 | 0.927404 | Pef1 |
| ENSMUSG( | 649.2691 | 0.09231 | 0.589191 | 0.90113 | Peg10 |
| ENSMUSG( | 15.93085 | -0.08679 | 0.857551 | NA | Peg12 |
| ENSMUSG( | 4131.367 | -0.09868 | 0.34019 | 0.777962 | Peg13 |
| ENSMUSG( | 7853.012 | -0.17012 | 0.084318 | 0.447526 | Peg3 |
| ENSMUSG( | 914.2104 | 0.074982 | 0.445418 | 0.833239 | Peli1 |
| ENSMUSG( | 2406.708 | 0.071955 | 0.529817 | 0.878883 | Peli2 |
| ENSMUSG( | 471.9281 | 0.258855 | 0.070165 | 0.40908 | Peli3 |
| ENSMUSG( | 115.0651 | 0.054676 | 0.766817 | 0.954331 | Pelo |
| ENSMUSG( | 1275.021 | 0.094821 | 0.342231 | 0.778828 | Pelp1 |
| ENSMUSG( | 82.89085 | 0.044822 | 0.837554 | 0.971111 | Pemt |
| ENSMUSG( | 6763.388 | -1.40769 | 0.000113 | 0.007021 | Penk |
| ENSMUSG( | 616.7714 | 0.285642 | 0.014429 | 0.176902 | Pepd |
| ENSMUSG( | 2472.233 | -0.37681 | 0.001599 | 0.044877 | Per1 |
| ENSMUSG( | 930.1725 | 0.055443 | 0.604822 | 0.905556 | Per2 |
| ENSMUSG( | 1368.009 | 0.180778 | 0.098242 | 0.479534 | Per3 |
| ENSMUSG( | 84.54323 | 0.028982 | 0.904171 | 0.986246 | Perm1 |
| ENSMUSG( | 284.9024 | -0.02998 | 0.835272 | 0.970426 | Perp |
| ENSMUSG( | 1309.697 | 0.167934 | 0.079615 | 0.435679 | Pes1 |
| ENSMUSG( | 766.213 | 0.046842 | 0.660919 | 0.92497 | Pet100 |
| ENSMUSG( | 542.1516 | 0.183435 | 0.13185 | 0.544244 | Pex1 |
| ENSMUSG( | 299.404 | -0.01566 | 0.912189 | 0.988483 | Pex10 |
| ENSMUSG( | 612.8012 | 0.244351 | 0.026097 | 0.247245 | Pex11a |
| ENSMUSG( | 534.0611 | -0.0031 | 0.981306 | 0.997512 | Pex11b |
| ENSMUSG( | 50.36415 | 0.171474 | 0.530917 | 0.879289 | Pex11g |
| ENSMUSG( | 334.1048 | -0.00101 | 0.994559 | 0.999369 | Pex12 |
| ENSMUSG( | 980.5718 | -0.11192 | 0.346916 | 0.779684 | Pex13 |
| ENSMUSG( | 1694.142 | -0.23929 | 0.039097 | 0.304587 | Pex14 |
| ENSMUSG( | 1035.354 | 0.046229 | 0.617697 | 0.910872 | Pex16 |
| ENSMUSG( | 3378.512 | 0.01812 | 0.84822 | 0.972374 | Pex19 |
| ENSMUSG( | 903.0024 | -0.02594 | 0.821178 | 0.966248 | Pex2 |
| ENSMUSG( | 357.0299 | 0.255577 | 0.058143 | 0.371733 | Pex26 |
| ENSMUSG( | 693.9868 | -0.09604 | 0.366077 | 0.794603 | Pex3 |
| ENSMUSG( | 1448.276 | 0.056613 | 0.591085 | 0.90113 | Pex5 |
| ENSMUSG( | 3655.328 | -0.00755 | 0.929972 | 0.991848 | Pex5l |
| ENSMUSG( | 2273.708 | 0.227767 | 0.008262 | 0.126789 | Pex6 |
| ENSMUSG( | 977.9955 | 0.03212 | 0.785259 | 0.958473 | Pex7 |
| ENSMUSG( | 64.07624 | 0.06386 | 0.803138 | 0.962169 | Pf4 |
| ENSMUSG( | 1210.45 | 0.080978 | 0.407063 | 0.817163 | Pfas |
| ENSMUSG( | 1438.031 | -0.00702 | 0.955401 | 0.993874 | Pfdn1 |
| ENSMUSG( | 3373.222 | 0.017247 | 0.845024 | 0.972023 | Pfdn2 |
| ENSMUSG( | 252.7105 | 0.051231 | 0.75473 | 0.951007 | Pfdn4 |
| ENSMUSG( | 6003.208 | 0.064516 | 0.454449 | 0.837347 | Pfdn5 |
| ENSMUSG( | 950.2099 | 0.028601 | 0.827746 | 0.968924 | Pfdn6 |
| ENSMUSG( | 16.93686 | -0.58829 | 0.186367 | NA | Pfkfb1 |
| ENSMUSG( | 850.5434 | 0.075218 | 0.47431 | 0.849425 | Pfkfb2 |
| ENSMUSG( | 844.2902 | -0.12048 | 0.22327 | 0.669382 | Pfkfb3 |
| ENSMUSG( | 727.7577 | -0.0595 | 0.57675 | 0.895432 | Pfkfb4 |
| ENSMUSG( | 4961.916 | 0.139154 | 0.241286 | 0.689026 | Pfkl |
| ENSMUSG( | 17880.94 | 0.145088 | 0.096441 | 0.476986 | Pfkm |
| ENSMUSG( | 7235.397 | 0.183346 | 0.054715 | 0.360135 | Pfkp |
| ENSMUSG( | 3485.58 | -0.04068 | 0.674228 | 0.929638 | Pfn1 |
| ENSMUSG( | 12544.22 | -0.02922 | 0.758934 | 0.952116 | Pfn2 |
| ENSMUSG( | 555.1689 | 0.014344 | 0.908151 | 0.9878 | Pfn4 |
| ENSMUSG( | 4486.235 | 0.074375 | 0.434122 | 0.828931 | Pgam1 |
| ENSMUSG( | 84.1486 | -0.38484 | 0.08423 | 0.447434 | Pgam2 |
| ENSMUSG( | 1468.511 | 0.187145 | 0.096508 | 0.476986 | Pgam5 |
| ENSMUSG( | 792.0436 | -0.38894 | 0.03806 | 0.300991 | Pgap1 |
| ENSMUSG( | 534.5954 | -0.19795 | 0.135591 | 0.549038 | Pgap2 |
| ENSMUSG( | 385.399 | -0.37201 | 0.013029 | 0.165126 | Pgap3 |
| ENSMUSG( | 2135.274 | -0.226 | 0.099094 | 0.481086 | Pgap4 |
| ENSMUSG( | 587.9163 | 0.133119 | 0.213253 | 0.657598 | Pgap6 |
| ENSMUSG( | 60.13261 | 0.155591 | 0.531934 | 0.879687 | Pgbd1 |
| ENSMUSG( | 3797.98 | 0.217716 | 0.048876 | 0.337919 | Pgbd5 |
| ENSMUSG( | 1246.728 | -0.10301 | 0.257839 | 0.709198 | Pgd |
| ENSMUSG( | 113.1449 | -0.35348 | 0.075853 | 0.426666 | Pgf |
| ENSMUSG( | 534.9775 | 0.05915 | 0.653892 | 0.922914 | Pgghg |
| ENSMUSG( | 1034.524 | 0.140605 | 0.20799 | 0.650144 | Pggt1b |
| ENSMUSG( | 6268.659 | 0.041481 | 0.655084 | 0.923872 | Pgk1 |
| ENSMUSG( | 976.1789 | 0.004184 | 0.965554 | 0.995377 | Pgls |

| | | | | | |
|---|---|---|---|---|---|
| ENSMUSG( | 614.0314 | -0.29546 | 0.021307 | 0.575341 | Pfkfb4 |
| ENSMUSG( | 4071.348 | -0.16752 | 0.237473 | 0.872173 | Pfkl |
| ENSMUSG( | 14747.85 | -0.14388 | 0.194328 | 0.860096 | Pfkm |
| ENSMUSG( | 5737.513 | -0.22129 | 0.080784 | 0.77681 | Pfkp |
| ENSMUSG( | 3086.418 | -0.12882 | 0.35188 | 0.912093 | Pfn1 |
| ENSMUSG( | 11220.41 | -0.08639 | 0.55514 | 0.939565 | Pfn2 |
| ENSMUSG( | 464.887 | -0.24422 | 0.116267 | 0.813384 | Pfn4 |
| ENSMUSG( | 3773.456 | -0.16238 | 0.195906 | 0.862161 | Pgam1 |
| ENSMUSG( | 76.56359 | -0.41554 | 0.11642 | 0.813384 | Pgam2 |
| ENSMUSG( | 1268.321 | 0.036386 | 0.78325 | 0.976106 | Pgam5 |
| ENSMUSG( | 930.1895 | 0.353716 | 0.071503 | 0.761942 | Pgap1 |
| ENSMUSG( | 498.0959 | -0.12873 | 0.340552 | 0.907972 | Pgap2 |
| ENSMUSG( | 388.3315 | -0.04811 | 0.786618 | 0.976303 | Pgap3 |
| ENSMUSG( | 2193.444 | 0.131043 | 0.443071 | 0.921554 | Pgap4 |
| ENSMUSG( | 537.5314 | 0.14647 | 0.319106 | 0.903431 | Pgap6 |
| ENSMUSG( | 52.78059 | 0.04115 | 0.885254 | 0.989688 | Pgbd1 |
| ENSMUSG( | 3123.432 | -0.07149 | 0.561218 | 0.940254 | Pgbd5 |
| ENSMUSG( | 1125.563 | -0.13158 | 0.240173 | 0.875205 | Pgd |
| ENSMUSG( | 115.3821 | -0.01116 | 0.957095 | 0.994798 | Pgf |
| ENSMUSG( | 447.702 | -0.19097 | 0.168144 | 0.840222 | Pgghg |
| ENSMUSG( | 939.4327 | 0.127763 | 0.408283 | 0.917257 | Pggt1b |
| ENSMUSG( | 5087.005 | -0.31205 | 0.015025 | 0.491964 | Pgk1 |
| ENSMUSG( | 856.704 | -0.10865 | 0.373208 | 0.915491 | Pgls |
| ENSMUSG( | 476.5347 | 0.474795 | 0.036532 | 0.66949 | Pglyrp1 |
| ENSMUSG( | 2525.651 | -0.02648 | 0.829882 | 0.980224 | Pgm1 |
| ENSMUSG( | 311.573 | 0.078359 | 0.597356 | 0.946609 | Pgm2 |
| ENSMUSG( | 5439.353 | 0.082052 | 0.443905 | 0.921734 | Pgm2l1 |
| ENSMUSG( | 294.3654 | -0.14355 | 0.338239 | 0.907972 | Pgm3 |
| ENSMUSG( | 220.5188 | 0.173023 | 0.362882 | 0.914223 | Pgm5 |
| ENSMUSG( | 2754.568 | -0.09972 | 0.355447 | 0.912555 | Pgp |
| ENSMUSG( | 607.8456 | -0.16553 | 0.190034 | 0.857355 | Pgpep1 |
| ENSMUSG( | 13.13164 | 0.232216 | 0.662264 | 0.956205 | Pgpep1l |
| ENSMUSG( | 300.8039 | -0.12829 | 0.422141 | 0.917257 | Pgr |
| ENSMUSG( | 8215.94 | 0.229641 | 0.121252 | 0.813384 | Pgrmc1 |
| ENSMUSG( | 3471.094 | -0.05825 | 0.653464 | 0.955015 | Pgrmc2 |
| ENSMUSG( | 3176.985 | -0.0414 | 0.682049 | 0.958678 | Pgs1 |
| ENSMUSG( | 19251.82 | 0.129813 | 0.238243 | 0.87259 | Phactr1 |
| ENSMUSG( | 1141.226 | -0.23397 | 0.048741 | 0.706195 | Phactr2 |
| ENSMUSG( | 11136.56 | -0.04224 | 0.700427 | 0.961475 | Phactr3 |
| ENSMUSG( | 806.9009 | 0.220707 | 0.095967 | 0.796133 | Phactr4 |
| ENSMUSG( | 676.9795 | -0.05511 | 0.702537 | 0.962215 | Phaf1 |
| ENSMUSG( | 723.8539 | -0.03455 | 0.820253 | 0.979198 | Phax |
| ENSMUSG( | 1388.141 | -0.15642 | 0.201468 | 0.864412 | Phb1 |
| ENSMUSG( | 1878.376 | -0.06622 | 0.587407 | 0.944975 | Phb2 |
| ENSMUSG( | 2492.376 | 0.079647 | 0.437258 | 0.920144 | Phc1 |
| ENSMUSG( | 2477.001 | -0.03131 | 0.782768 | 0.976106 | Phc2 |
| ENSMUSG( | 1872.828 | 0.068806 | 0.569678 | 0.941777 | Phc3 |
| ENSMUSG( | 326.1749 | 0.126308 | 0.416871 | 0.917257 | Pheta1 |
| ENSMUSG( | 24.6715 | -0.29678 | 0.474104 | 0.923329 | Pheta2 |
| ENSMUSG( | 18.10945 | -0.1591 | 0.723686 | 0.966887 | Phex |
| ENSMUSG( | 1013.681 | 0.088327 | 0.498916 | 0.927825 | Phf1 |
| ENSMUSG( | 2604.378 | -0.00246 | 0.984598 | 0.996963 | Phf10 |
| ENSMUSG( | 14.52955 | -0.52961 | 0.355182 | 0.912411 | Phf11b |
| ENSMUSG( | 19.19302 | -0.16743 | 0.734417 | 0.967228 | Phf11c |
| ENSMUSG( | 17.53852 | -0.53015 | 0.284753 | 0.892412 | Phf11d |
| ENSMUSG( | 1381.074 | 0.085803 | 0.432818 | 0.918434 | Phf12 |
| ENSMUSG( | 357.195 | -0.23766 | 0.090648 | 0.791074 | Phf13 |
| ENSMUSG( | 1466.478 | 0.088333 | 0.435255 | 0.919062 | Phf14 |
| ENSMUSG( | 54.65553 | 0.243054 | 0.384998 | 0.917257 | Phf19 |
| ENSMUSG( | 2529.69 | 0.069876 | 0.531739 | 0.934795 | Phf2 |
| ENSMUSG( | 2021.166 | 0.018795 | 0.876758 | 0.989644 | Phf20 |
| ENSMUSG( | 1971.562 | 0.069566 | 0.568201 | 0.941777 | Phf20l1 |
| ENSMUSG( | 1809.401 | 0.027913 | 0.808955 | 0.97873 | Phf21a |
| ENSMUSG( | 601.6591 | 0.038632 | 0.77215 | 0.974236 | Phf21b |
| ENSMUSG( | 1048.943 | 0.097806 | 0.456746 | 0.923329 | Phf23 |
| ENSMUSG( | 11044.54 | 0.065603 | 0.540411 | 0.937383 | Phf24 |
| ENSMUSG( | 2315.798 | 0.041889 | 0.710695 | 0.964363 | Phf3 |
| ENSMUSG( | 529.3645 | -0.1176 | 0.369382 | 0.915491 | Phf5a |
| ENSMUSG( | 421.3289 | 0.305532 | 0.042364 | 0.690304 | Phf6 |
| ENSMUSG( | 121.0041 | -0.17566 | 0.45153 | 0.922202 | Phf7 |
| ENSMUSG( | 575.9045 | 0.115246 | 0.391233 | 0.917257 | Phf8 |
| ENSMUSG( | 4291.401 | -0.18628 | 0.141498 | 0.830678 | Phgdh |
| ENSMUSG( | 1163.017 | 0.212367 | 0.148884 | 0.836241 | Phip |
| ENSMUSG( | 2906.933 | -0.1106 | 0.37906 | 0.916995 | Phka1 |
| ENSMUSG( | 722.387 | 0.115807 | 0.541006 | 0.937647 | Phka2 |
| ENSMUSG( | 2060.654 | -0.17264 | 0.182987 | 0.850741 | Phkb |
| ENSMUSG( | 1564.617 | -0.2111 | 0.096318 | 0.796133 | Phkg1 |
| ENSMUSG( | 2115.564 | -0.06873 | 0.554235 | 0.939565 | Phkg2 |
| ENSMUSG( | 451.1887 | -0.07808 | 0.673685 | 0.95694 | Phlda1 |
| ENSMUSG( | 592.5187 | -0.28497 | 0.04913 | 0.708275 | Phlda3 |
| ENSMUSG( | 6058.489 | -0.10992 | 0.403813 | 0.917257 | Phldb1 |
| ENSMUSG( | 394.7733 | 0.220458 | 0.174227 | 0.844317 | Phldb2 |
| ENSMUSG( | 4761.312 | -0.18251 | 0.102703 | 0.803503 | Phlpp1 |
| ENSMUSG( | 1451.998 | 0.096338 | 0.367581 | 0.915217 | Phlpp2 |
| ENSMUSG( | 116.5019 | 0.105629 | 0.63506 | 0.953903 | Phospho1 |
| ENSMUSG( | 340.4567 | -0.00578 | 0.973192 | 0.996416 | Phospho2 |
| ENSMUSG( | 298.8212 | -0.05044 | 0.757535 | 0.971858 | Phpt1 |
| ENSMUSG( | 1479.882 | 0.142517 | 0.230493 | 0.868507 | Phrf1 |
| ENSMUSG( | 1312.972 | 0.308112 | 0.015734 | 0.502615 | Phtf1 |
| ENSMUSG( | 34.4613 | 0.366426 | 0.303546 | 0.898538 | Phtf1os |
| ENSMUSG( | 440.5589 | 0.263084 | 0.107265 | 0.804969 | Phtf2 |
| ENSMUSG( | 3562.423 | -0.23907 | 0.069951 | 0.757435 | Phyh |
| ENSMUSG( | 111.7598 | 0.328793 | 0.259301 | 0.883971 | Phyhd1 |
| ENSMUSG( | 9344.324 | -0.04158 | 0.781284 | 0.976106 | Phyhip |
| ENSMUSG( | 4160.758 | -0.2623 | 0.042903 | 0.690304 | Phyhipl |
| ENSMUSG( | 127.0939 | 0.000447 | 0.998236 | 0.999313 | Phykpl |
| ENSMUSG( | 8.576864 | 0.503791 | 0.508327 | NA | Pi15 |
| ENSMUSG( | 18.63298 | -0.19475 | 0.715002 | 0.965063 | Pi16 |
| ENSMUSG( | 1656.967 | -0.04208 | 0.726652 | 0.966887 | Pi4k2a |
| ENSMUSG( | 104.1212 | -0.34118 | 0.125735 | 0.81516 | Pi4k2b |
| ENSMUSG( | 6837.364 | 0.023495 | 0.833853 | 0.981218 | Pi4ka |
| ENSMUSG( | 1197.689 | 0.10927 | 0.39704 | 0.917257 | Pi4kb |
| ENSMUSG( | 6730.866 | 0.09379 | 0.467331 | 0.923329 | Pianp |
| ENSMUSG( | 1081.345 | 0.026748 | 0.824401 | 0.979198 | Pias1 |
| ENSMUSG( | 1201.723 | 0.10246 | 0.422864 | 0.917257 | Pias2 |
| ENSMUSG( | 662.9296 | 0.155665 | 0.253362 | 0.88018 | Pias3 |
| ENSMUSG( | 557.1776 | 0.043189 | 0.769443 | 0.974236 | Pias4 |
| ENSMUSG( | 184.1046 | 0.008521 | 0.960974 | 0.99511 | Pibf1 |
| ENSMUSG( | 3682.649 | 0.11366 | 0.290914 | 0.897019 | Picalm |
| ENSMUSG( | 465.9266 | -0.01882 | 0.911112 | 0.992196 | Pick1 |
| ENSMUSG( | 882.3761 | -0.06267 | 0.655856 | 0.955533 | Pid1 |
| ENSMUSG( | 66.4542 | 0.017756 | 0.94844 | 0.994798 | Pidd1 |
| ENSMUSG( | 57.70504 | -0.28702 | 0.296911 | 0.898538 | Pierce1 |
| ENSMUSG( | 120.8072 | -0.36579 | 0.069392 | 0.755467 | Pierce2 |
| ENSMUSG( | 382.4842 | 0.329588 | 0.093416 | 0.795507 | Piezo1 |
| ENSMUSG( | 76.68709 | 0.008059 | 0.973895 | 0.996416 | Piezo2 |
| ENSMUSG( | 15.41355 | 0.051702 | 0.930831 | 0.99306 | Pif1 |
| ENSMUSG( | 261.0954 | 0.273264 | 0.121569 | 0.813384 | Piga |
| ENSMUSG( | 510.1978 | 0.241129 | 0.098271 | 0.798438 | Pigb |
| ENSMUSG( | 137.6328 | -0.16405 | 0.394203 | 0.917257 | Pigbos1 |
| ENSMUSG( | 495.3007 | 0.23828 | 0.128441 | 0.81925 | Pigc |
| ENSMUSG( | 247.0072 | -0.0122 | 0.947364 | 0.994798 | Pigf |
| ENSMUSG( | 600.6912 | 0.127676 | 0.508823 | 0.929089 | Pigg |
| ENSMUSG( | 447.0477 | -0.03956 | 0.774428 | 0.974931 | Pigh |
| ENSMUSG( | 2128.686 | 0.096127 | 0.469392 | 0.923329 | Pigk |
| ENSMUSG( | 172.6666 | -0.039 | 0.845145 | 0.983878 | Pigl |
| ENSMUSG( | 428.0484 | 0.256113 | 0.185476 | 0.855305 | Pigm |
| ENSMUSG( | 456.5023 | 0.058743 | 0.717561 | 0.965964 | Pign |
| ENSMUSG( | 747.6553 | 0.344818 | 0.048824 | 0.706228 | Pigo |
| ENSMUSG( | 727.0584 | -0.00815 | 0.949222 | 0.994798 | Pigp |
| ENSMUSG( | 5473.734 | -0.05102 | 0.665724 | 0.956766 | Pigq |
| ENSMUSG( | 2603.233 | 0.144903 | 0.368367 | 0.915491 | Pigs |
| ENSMUSG( | 3524.867 | 0.122653 | 0.456379 | 0.923329 | Pigt |
| ENSMUSG( | 879.1926 | 0.143431 | 0.360648 | 0.914223 | Pigu |
| ENSMUSG( | 298.4221 | 0.147469 | 0.468546 | 0.923329 | Pigv |
| ENSMUSG( | 90.92509 | 0.053316 | 0.818123 | 0.979198 | Pigw |
| ENSMUSG( | 366.2283 | 0.120482 | 0.426047 | 0.917257 | Pigx |
| ENSMUSG( | 610.8942 | -0.15754 | 0.244119 | 0.877904 | Pigyl |
| ENSMUSG( | 275.8739 | -0.09541 | 0.660249 | 0.956205 | Pigz |
| ENSMUSG( | 652.6269 | -0.08946 | 0.473855 | 0.923329 | Pih1d1 |
| ENSMUSG( | 49.35969 | 0.073847 | 0.801799 | 0.97681 | Pih1d2 |
| ENSMUSG( | 247.5141 | -0.19507 | 0.24061 | 0.875927 | Pik3ap1 |
| ENSMUSG( | 982.429 | 0.345942 | 0.031461 | 0.631192 | Pik3c2a |
| ENSMUSG( | 586.0956 | -0.20445 | 0.114669 | 0.811703 | Pik3c2b |
| ENSMUSG( | 848.8135 | 0.007358 | 0.952497 | 0.994798 | Pik3c3 |
| ENSMUSG( | 2136.027 | 0.078252 | 0.515755 | 0.930939 | Pik3ca |
| ENSMUSG( | 614.1547 | 0.034907 | 0.790116 | 0.97681 | Pik3cb |
| ENSMUSG( | 884.9716 | 0.194538 | 0.138385 | 0.828492 | Pik3cd |
| ENSMUSG( | 182.8144 | 0.346022 | 0.065551 | 0.745138 | Pik3cg |
| ENSMUSG( | 1458.44 | 0.144424 | 0.423372 | 0.917257 | Pik3ip1 |
| ENSMUSG( | 5498.199 | -0.01486 | 0.897034 | 0.990558 | Pik3r1 |
| ENSMUSG( | 4962.376 | 0.083957 | 0.400799 | 0.917257 | Pik3r2 |
| ENSMUSG( | 1691.265 | -0.01537 | 0.902571 | 0.991089 | Pik3r3 |
| ENSMUSG( | 993.2001 | 0.20226 | 0.183737 | 0.85191 | Pik3r4 |
| ENSMUSG( | 153.3886 | -0.00118 | 0.995533 | 0.999116 | Pik3r5 |
| ENSMUSG( | 130.3873 | 0.177938 | 0.450958 | 0.922202 | Pik3r6 |
| ENSMUSG( | 1117.29 | 0.173617 | 0.120251 | 0.813384 | Pikfyve |
| ENSMUSG( | 27.16878 | -0.67276 | 0.173639 | 0.844317 | Pilra |

| | | | | | | | | | | | |
|---|---|---|---|---|---|---|---|---|---|---|---|
| ENSMUSG( | Pip5k1a | 1204.917 | 0.001886 | 0.986764 | 0.998619 | ENSMUSG( | Pip5k1a | 1066.258 | -0.08478 | 0.514963 | 0.930939 |
| ENSMUSG( | Pip5k1b | 332.1936 | 0.307895 | 0.091969 | 0.465937 | ENSMUSG( | Pip5k1b | 285.7008 | 0.156942 | 0.435018 | 0.918876 |
| ENSMUSG( | Pip5k1c | 22856.69 | 0.225216 | 0.01092 | 0.150598 | ENSMUSG( | Pip5k1c | 19613.27 | 0.061443 | 0.552684 | 0.939565 |
| ENSMUSG( | Pip5kl1 | 153.351 | 0.198394 | 0.333859 | 0.775602 | ENSMUSG( | Pip5kl1 | 139.1706 | 0.184937 | 0.481283 | 0.924564 |
| ENSMUSG( | Pipox | 57.75046 | -0.4613 | 0.065654 | 0.396911 | ENSMUSG( | Pipox | 62.55317 | 0.05269 | 0.846574 | 0.984821 |
| ENSMUSG( | Pir | 168.1804 | -0.18485 | 0.289039 | 0.739705 | ENSMUSG( | Pir | 154.714 | -0.15191 | 0.422781 | 0.917257 |
| ENSMUSG( | Pirb | 32.59257 | -0.60708 | 0.245513 | 0.693175 | ENSMUSG( | Pirb | 33.29575 | -0.23113 | 0.665415 | 0.956766 |
| ENSMUSG( | Pirt | 12.77829 | -0.97207 | 0.064302 | NA | ENSMUSG( | Pirt | 14.21648 | -0.22755 | 0.656504 | 0.955533 |
| ENSMUSG( | Pisd | 2419.665 | 0.051051 | 0.552683 | 0.886141 | ENSMUSG( | Pisd | 2150.425 | -0.02174 | 0.839746 | 0.982162 |
| ENSMUSG( | Pisd-ps1 | 301.2569 | -0.12475 | 0.417041 | 0.8209 | ENSMUSG( | Pisd-ps1 | 299.2961 | 0.133712 | 0.441125 | 0.921166 |
| ENSMUSG( | Pithd1 | 2236.182 | 0.029138 | 0.749258 | 0.949664 | ENSMUSG( | Pithd1 | 1984.557 | -0.04983 | 0.677164 | 0.957106 |
| ENSMUSG( | Pitpna | 9022.034 | 0.043888 | 0.59026 | 0.90113 | ENSMUSG( | Pitpna | 7742.517 | -0.13292 | 0.237573 | 0.872264 |
| ENSMUSG( | Pitpnb | 3127.509 | 0.072824 | 0.382509 | 0.803227 | ENSMUSG( | Pitpnb | 2692.968 | -0.0932 | 0.473102 | 0.923329 |
| ENSMUSG( | Pitpnc1 | 3245.036 | 0.118688 | 0.214543 | 0.659193 | ENSMUSG( | Pitpnc1 | 2801.112 | -0.03572 | 0.734491 | 0.967228 |
| ENSMUSG( | Pitpnm1 | 10355.28 | 0.198484 | 0.026074 | 0.247245 | ENSMUSG( | Pitpnm1 | 9172.164 | 0.122396 | 0.241218 | 0.876259 |
| ENSMUSG( | Pitpnm2 | 9454.564 | 0.250778 | 0.016452 | 0.190546 | ENSMUSG( | Pitpnm2 | 8849.051 | 0.32293 | 0.005913 | 0.316645 |
| ENSMUSG( | Pitpnm3 | 5957.641 | 0.138361 | 0.104187 | 0.489971 | ENSMUSG( | Pitpnm3 | 5345.645 | 0.096286 | 0.340699 | 0.907972 |
| ENSMUSG( | Pitrm1 | 1482.284 | 0.149904 | 0.093408 | 0.469793 | ENSMUSG( | Pitrm1 | 1261.651 | -0.04099 | 0.724968 | 0.966887 |
| ENSMUSG( | Piwil2 | 33.18198 | 0.256306 | 0.450457 | 0.835401 | ENSMUSG( | Piwil2 | 28.49296 | 0.086303 | 0.815626 | 0.979198 |
| ENSMUSG( | Pja1 | 2605.59 | -0.12362 | 0.231801 | 0.679725 | ENSMUSG( | Pja1 | 2452.79 | -0.02883 | 0.818853 | 0.979198 |
| ENSMUSG( | Pja2 | 6922.792 | -0.05555 | 0.511945 | 0.871327 | ENSMUSG( | Pja2 | 6438.402 | 0.002787 | 0.980538 | 0.996479 |
| ENSMUSG( | Pkd1 | 4873.364 | -0.11527 | 0.221651 | 0.667258 | ENSMUSG( | Pkd1 | 4835.888 | 0.131604 | 0.27049 | 0.888062 |
| ENSMUSG( | Pkd1l3 | 27.67281 | 0.021715 | 0.954038 | 0.993874 | ENSMUSG( | Pkd1l3 | 25.39376 | 0.030127 | 0.945018 | 0.994798 |
| ENSMUSG( | Pkd2 | 2467.59 | -0.0084 | 0.940333 | 0.993415 | ENSMUSG( | Pkd2 | 2343.232 | 0.109664 | 0.387725 | 0.917257 |
| ENSMUSG( | Pkd2l1 | 24.34262 | -0.87156 | 0.03789 | 0.30067 | ENSMUSG( | Pkd2l1 | 30.20111 | 0.167885 | 0.667912 | 0.956766 |
| ENSMUSG( | Pkd2l2 | 46.63829 | -0.57876 | 0.070293 | 0.40908 | ENSMUSG( | Pkd2l2 | 46.76412 | -0.24828 | 0.421295 | 0.917257 |
| ENSMUSG( | Pkdcc | 369.1221 | -0.09748 | 0.517757 | 0.873611 | ENSMUSG( | Pkdcc | 365.1132 | 0.148214 | 0.376224 | 0.915491 |
| ENSMUSG( | Pkdrej | 41.21298 | 0.298361 | 0.415616 | 0.82084 | ENSMUSG( | Pkdrej | 36.56319 | 0.262317 | 0.532271 | 0.934795 |
| ENSMUSG( | Pkhd1l1 | 16.64944 | 0.144179 | 0.765824 | NA | ENSMUSG( | Pkhd1l1 | 17.96453 | 0.593697 | 0.233392 | 0.868507 |
| ENSMUSG( | Pkia | 4203.835 | 0.023709 | 0.820495 | 0.965888 | ENSMUSG( | Pkia | 3724.436 | -0.0606 | 0.685528 | 0.958678 |
| ENSMUSG( | Pkib | 127.9835 | -0.13373 | 0.495663 | 0.861237 | ENSMUSG( | Pkib | 113.6056 | -0.21744 | 0.319357 | 0.903453 |
| ENSMUSG( | Pkig | 3338.024 | 0.075686 | 0.346708 | 0.779432 | ENSMUSG( | Pkig | 2841.615 | -0.1243 | 0.350611 | 0.911652 |
| ENSMUSG( | Pkm | 26282.16 | -0.01809 | 0.842747 | 0.971663 | ENSMUSG( | Pkm | 22531.09 | -0.20218 | 0.081517 | 0.77681 |
| ENSMUSG( | Pkmyt1 | 162.6713 | -0.21272 | 0.253913 | 0.704011 | ENSMUSG( | Pkmyt1 | 157.463 | -0.02394 | 0.895517 | 0.990558 |
| ENSMUSG( | Pkn1 | 3789.106 | 0.059679 | 0.466167 | 0.844453 | ENSMUSG( | Pkn1 | 3293.658 | -0.0759 | 0.443369 | 0.921704 |
| ENSMUSG( | Pkn2 | 1033.842 | 0.057005 | 0.631405 | 0.914073 | ENSMUSG( | Pkn2 | 984.5302 | 0.178622 | 0.188545 | 0.85698 |
| ENSMUSG( | Pkn3 | 37.42968 | 0.125083 | 0.726235 | 0.944223 | ENSMUSG( | Pkn3 | 33.83965 | 0.139014 | 0.736303 | 0.967344 |
| ENSMUSG( | Pknox1 | 685.4738 | 0.122944 | 0.272674 | 0.722557 | ENSMUSG( | Pknox1 | 587.6775 | -0.05272 | 0.713657 | 0.96497 |
| ENSMUSG( | Pknox2 | 1520.112 | 0.161717 | 0.148838 | 0.572605 | ENSMUSG( | Pknox2 | 1269.413 | -0.08784 | 0.505205 | 0.927825 |
| ENSMUSG( | Pkp1 | 16.40002 | 0.242656 | 0.610195 | NA | ENSMUSG( | Pkp1 | 11.26248 | -0.58686 | 0.325524 | NA |
| ENSMUSG( | Pkp2 | 202.2546 | 0.599904 | 0.004895 | 0.092653 | ENSMUSG( | Pkp2 | 167.5606 | 0.366088 | 0.057121 | 0.723362 |
| ENSMUSG( | Pkp4 | 17277.05 | 0.08508 | 0.246547 | 0.694958 | ENSMUSG( | Pkp4 | 14871.1 | -0.07955 | 0.434224 | 0.91869 |
| ENSMUSG( | Pla1a | 52.95319 | 0.151161 | 0.624908 | 0.912167 | ENSMUSG( | Pla1a | 85.18257 | 1.46713 | 1.48E-05 | 0.005074 |
| ENSMUSG( | Pla2g12a | 617.0078 | -0.29477 | 0.017683 | 0.199454 | ENSMUSG( | Pla2g12a | 637.028 | 0.08362 | 0.521381 | 0.932489 |
| ENSMUSG( | Pla2g15 | 935.6799 | -0.22679 | 0.200198 | 0.640302 | ENSMUSG( | Pla2g15 | 1025.479 | 0.307015 | 0.120859 | 0.813384 |
| ENSMUSG( | Pla2g3 | 177.7508 | 0.16596 | 0.324398 | 0.769598 | ENSMUSG( | Pla2g3 | 186.1036 | 0.533179 | 0.003961 | 0.253257 |
| ENSMUSG( | Pla2g4a | 130.7308 | -0.44031 | 0.028732 | 0.258426 | ENSMUSG( | Pla2g4a | 146.434 | 0.201841 | 0.407052 | 0.917257 |
| ENSMUSG( | Pla2g4b | 20.92289 | 0.322651 | 0.472338 | NA | ENSMUSG( | Pla2g4b | 18.54977 | 0.291152 | 0.552078 | 0.939565 |
| ENSMUSG( | Pla2g4e | 417.629 | -0.28303 | 0.04506 | 0.325595 | ENSMUSG( | Pla2g4e | 372.0693 | -0.3503 | 0.021973 | 0.575341 |
| ENSMUSG( | Pla2g5 | 179.8486 | -0.21996 | 0.169398 | 0.603554 | ENSMUSG( | Pla2g5 | 148.2177 | -0.55569 | 0.002578 | 0.198937 |
| ENSMUSG( | Pla2g6 | 718.0013 | 0.013426 | 0.915491 | 0.988634 | ENSMUSG( | Pla2g6 | 633.9232 | -0.07084 | 0.635512 | 0.953903 |
| ENSMUSG( | Pla2g7 | 12888 | -0.08786 | 0.514887 | 0.872306 | ENSMUSG( | Pla2g7 | 11930.77 | -0.04008 | 0.779984 | 0.976106 |
| ENSMUSG( | Pla2r1 | 21.47986 | -0.13035 | 0.752614 | 0.950536 | ENSMUSG( | Pla2r1 | 24.87554 | 0.558916 | 0.207761 | 0.864412 |
| ENSMUSG( | Plaa | 1227.406 | 0.107107 | 0.248152 | 0.696853 | ENSMUSG( | Plaa | 1087.921 | 0.025605 | 0.840592 | 0.982495 |
| ENSMUSG( | Plaat1 | 404.8416 | -0.06469 | 0.674007 | 0.929626 | ENSMUSG( | Plaat1 | 345.2324 | -0.27475 | 0.13557 | 0.825163 |
| ENSMUSG( | Plaat3 | 2719.946 | -0.02536 | 0.828634 | 0.969097 | ENSMUSG( | Plaat3 | 2345.447 | -0.19474 | 0.209383 | 0.864412 |
| ENSMUSG( | Plac8 | 26.54671 | -0.8906 | 0.099388 | 0.48179 | ENSMUSG( | Plac8 | 23.61376 | -1.03044 | 0.058493 | 0.728256 |
| ENSMUSG( | Plac9 | 374.2895 | -0.27911 | 0.065696 | 0.397014 | ENSMUSG( | Plac9 | 358.4139 | -0.11498 | 0.536789 | 0.937108 |
| ENSMUSG( | Plag1 | 83.4431 | -0.04621 | 0.857618 | 0.974667 | ENSMUSG( | Plag1 | 90.21395 | 0.423791 | 0.098547 | 0.798438 |
| ENSMUSG( | Plagl1 | 674.342 | 0.0298 | 0.84479 | 0.972023 | ENSMUSG( | Plagl1 | 633.046 | 0.10452 | 0.614961 | 0.951177 |
| ENSMUSG( | Plagl2 | 203.8513 | -0.44162 | 0.002554 | 0.059956 | ENSMUSG( | Plagl2 | 208.9873 | -0.05661 | 0.745643 | 0.96951 |
| ENSMUSG( | Plat | 1867.393 | -0.38962 | 0.015581 | 0.184473 | ENSMUSG( | Plat | 1899.853 | -0.03953 | 0.819243 | 0.979198 |
| ENSMUSG( | Platr14 | 14.27124 | -0.30942 | 0.543436 | NA | ENSMUSG( | Platr14 | 13.02218 | -0.27621 | 0.625728 | 0.953234 |
| ENSMUSG( | Platr25 | 224.8899 | 0.05833 | 0.695879 | 0.934979 | ENSMUSG( | Platr25 | 211.6547 | 0.142852 | 0.410488 | 0.917257 |
| ENSMUSG( | Plau | 138.9241 | -0.26516 | 0.179197 | 0.616715 | ENSMUSG( | Plau | 154.9808 | 0.325294 | 0.113653 | 0.809623 |
| ENSMUSG( | Plaur | 41.42849 | -0.36833 | 0.246987 | 0.695749 | ENSMUSG( | Plaur | 36.75199 | -0.4722 | 0.157371 | 0.837724 |
| ENSMUSG( | Plb1 | 135.3207 | 0.055889 | 0.752819 | 0.950621 | ENSMUSG( | Plb1 | 117.1767 | -0.10674 | 0.621608 | 0.953199 |
| ENSMUSG( | Plbd1 | 29.57865 | -0.73259 | 0.149184 | 0.573248 | ENSMUSG( | Plbd1 | 29.56168 | -0.45062 | 0.393677 | 0.917257 |
| ENSMUSG( | Plbd2 | 2778.523 | -0.23031 | 0.095627 | 0.475062 | ENSMUSG( | Plbd2 | 3013.179 | 0.272875 | 0.078438 | 0.77321 |
| ENSMUSG( | Plcb1 | 4724.588 | 0.083088 | 0.410857 | 0.818585 | ENSMUSG( | Plcb1 | 4051.806 | -0.0945 | 0.440667 | 0.921166 |
| ENSMUSG( | Plcb2 | 170.829 | -0.20979 | 0.228208 | 0.675678 | ENSMUSG( | Plcb2 | 157.3016 | -0.16882 | 0.382251 | 0.917257 |
| ENSMUSG( | Plcb3 | 2187.298 | 0.07432 | 0.497597 | 0.862196 | ENSMUSG( | Plcb3 | 1893.619 | -0.07642 | 0.530886 | 0.934795 |
| ENSMUSG( | Plcb4 | 1045.574 | 0.039683 | 0.7431 | 0.948478 | ENSMUSG( | Plcb4 | 961.3275 | 0.066896 | 0.639964 | 0.953903 |
| ENSMUSG( | Plcd1 | 635.2078 | -0.02819 | 0.830616 | 0.96974 | ENSMUSG( | Plcd1 | 571.0147 | -0.06204 | 0.672146 | 0.95694 |
| ENSMUSG( | Plcd3 | 764.5742 | 0.151209 | 0.24583 | 0.693547 | ENSMUSG( | Plcd3 | 681.5133 | 0.095652 | 0.475725 | 0.923329 |
| ENSMUSG( | Plcd4 | 1191.983 | 0.207823 | 0.091933 | 0.465937 | ENSMUSG( | Plcd4 | 971.9492 | -0.10389 | 0.42855 | 0.917257 |
| ENSMUSG( | Plce1 | 948.2447 | 0.183771 | 0.20462 | 0.645534 | ENSMUSG( | Plce1 | 906.9296 | 0.307175 | 0.043743 | 0.690304 |
| ENSMUSG( | Plcg1 | 3247.314 | 0.197033 | 0.043231 | 0.318825 | ENSMUSG( | Plcg1 | 2814.955 | 0.05989 | 0.615075 | 0.951177 |
| ENSMUSG( | Plcg2 | 272.4842 | 0.146344 | 0.296031 | 0.744386 | ENSMUSG( | Plcg2 | 245.3396 | 0.114601 | 0.462361 | 0.923329 |
| ENSMUSG( | Plch1 | 265.5551 | -0.01264 | 0.927294 | 0.991229 | ENSMUSG( | Plch1 | 241.5146 | -0.01239 | 0.942353 | 0.994774 |
| ENSMUSG( | Plch2 | 3078.715 | 0.321333 | 0.000787 | 0.027699 | ENSMUSG( | Plch2 | 2608.644 | 0.127684 | 0.306706 | 0.898538 |
| ENSMUSG( | Plcl1 | 760.1925 | 0.043153 | 0.674209 | 0.929638 | ENSMUSG( | Plcl1 | 741.2884 | 0.232691 | 0.049545 | 0.708648 |
| ENSMUSG( | Plcl2 | 1346.658 | 0.310973 | 0.013658 | 0.170671 | ENSMUSG( | Plcl2 | 1245.088 | 0.347037 | 0.023787 | 0.588226 |
| ENSMUSG( | Plcxd1 | 682.5671 | -0.24325 | 0.033457 | 0.280305 | ENSMUSG( | Plcxd1 | 662.7432 | -0.04591 | 0.718936 | 0.96673 |
| ENSMUSG( | Plcxd2 | 2587.782 | 0.24794 | 0.01778 | 0.2 | ENSMUSG( | Plcxd2 | 2098.093 | -0.07854 | 0.487073 | 0.92541 |
| ENSMUSG( | Plcxd3 | 31.47808 | -0.34834 | 0.441123 | 0.832445 | ENSMUSG( | Plcxd3 | 30.294 | -0.15606 | 0.718738 | 0.966668 |
| ENSMUSG( | Pld1 | 503.1961 | 0.276683 | 0.013736 | 0.171117 | ENSMUSG( | Pld1 | 488.6127 | 0.441777 | 0.001853 | 0.15992 |
| ENSMUSG( | Pld2 | 1588.281 | 0.123936 | 0.207033 | 0.649503 | ENSMUSG( | Pld2 | 1397.996 | 0.027236 | 0.803585 | 0.976978 |
| ENSMUSG( | Pld3 | 33142.5 | -0.2085 | 0.096518 | 0.476986 | ENSMUSG( | Pld3 | 32913.98 | 0.050589 | 0.722264 | 0.966887 |
| ENSMUSG( | Pld4 | 751.2501 | -0.22135 | 0.130766 | 0.542059 | ENSMUSG( | Pld4 | 760.0688 | 0.095072 | 0.607916 | 0.950113 |
| ENSMUSG( | Pld5 | 182.7708 | -0.36418 | 0.047227 | 0.332553 | ENSMUSG( | Pld5 | 176.8419 | -0.17466 | 0.364006 | 0.914223 |
| ENSMUSG( | Pld6 | 28.52375 | -0.37732 | 0.291564 | 0.741366 | ENSMUSG( | Pld6 | 26.53745 | -0.28901 | 0.462958 | 0.923329 |
| ENSMUSG( | Plec | 7925.559 | -0.2142 | 0.046607 | 0.330354 | ENSMUSG( | Plec | 8114.406 | 0.130102 | 0.297783 | 0.898538 |
| ENSMUSG( | Plek | 629.0488 | -0.26071 | 0.065889 | 0.397536 | ENSMUSG( | Plek | 579.6914 | -0.22867 | 0.163824 | 0.838291 |
| ENSMUSG( | Plekha1 | 3604.157 | 9.67E-05 | 0.999138 | 0.99989 | ENSMUSG( | Plekha1 | 3311.179 | 0.024872 | 0.810309 | 0.978779 |
| ENSMUSG( | Plekha2 | 577.297 | -0.14332 | 0.22476 | 0.671058 | ENSMUSG( | Plekha2 | 537.9602 | -0.07907 | 0.555522 | 0.939565 |
| ENSMUSG( | Plekha3 | 1242.996 | 0.037521 | 0.703696 | 0.937885 | ENSMUSG( | Plekha3 | 1130.988 | 0.029861 | 0.808952 | 0.97873 |
| ENSMUSG( | Plekha4 | 101.5576 | 0.070994 | 0.749842 | 0.949664 | ENSMUSG( | Plekha4 | 91.3456 | 0.0211 | 0.935095 | 0.993078 |
| ENSMUSG( | Plekha5 | 2471.972 | 0.095835 | 0.377773 | 0.801561 | ENSMUSG( | Plekha5 | 2184.015 | 0.007088 | 0.957093 | 0.994798 |
| ENSMUSG( | Plekha6 | 2809.866 | 0.013126 | 0.915803 | 0.988634 | ENSMUSG( | Plekha6 | 2537.882 | -0.01379 | 0.913266 | 0.992298 |
| ENSMUSG( | Plekha7 | 1022.194 | 0.019771 | 0.845849 | 0.972198 | ENSMUSG( | Plekha7 | 904.1372 | -0.064 | 0.61352 | 0.951177 |
| ENSMUSG( | Plekha8 | 433.0845 | -0.00122 | 0.992489 | 0.999327 | ENSMUSG( | Plekha8 | 411.3098 | 0.124894 | 0.434693 | 0.918742 |
| ENSMUSG( | Plekhb1 | 42141.84 | -0.00227 | 0.979698 | 0.997428 | ENSMUSG( | Plekhb1 | 35558.84 | -0.23557 | 0.054708 | 0.723362 |
| ENSMUSG( | Plekhb2 | 11829.53 | 0.072841 | 0.475813 | 0.850919 | ENSMUSG( | Plekhb2 | 9791.708 | -0.21202 | 0.062485 | 0.736482 |
| ENSMUSG( | Plekhd1 | 151.7425 | -0.39498 | 0.03337 | 0.279991 | ENSMUSG( | Plekhd1 | 127.8547 | -0.69008 | 0.000984 | 0.107109 |
| ENSMUSG( | Plekhf1 | 160.3688 | -0.18856 | 0.313308 | 0.76033 | ENSMUSG( | Plekhf1 | 139.7753 | -0.30911 | 0.150776 | 0.837173 |
| ENSMUSG( | Plekhf2 | 379.4272 | 0.073264 | 0.599555 | 0.903874 | ENSMUSG( | Plekhf2 | 357.5983 | 0.165822 | 0.318216 | 0.902849 |
| ENSMUSG( | Plekhg1 | 1162.533 | 0.230755 | 0.015402 | 0.18329 | ENSMUSG( | Plekhg1 | 1051.985 | 0.215553 | 0.085436 | 0.779799 |
| ENSMUSG( | Plekhg2 | 310.0196 | 0.08855 | 0.53083 | 0.879289 | ENSMUSG( | Plekhg2 | 319.6341 | 0.425154 | 0.026906 | 0.603041 |
| ENSMUSG( | Plekhg3 | 810.7649 | -0.32647 | 0.002319 | 0.05646 | ENSMUSG( | Plekhg3 | 819.1436 | -0.00331 | 0.982113 | 0.996756 |
| ENSMUSG( | Plekhg4 | 101.6073 | 0.963001 | 2.58E-05 | 0.002351 | ENSMUSG( | Plekhg4 | 74.13595 | 0.432796 | 0.233376 | 0.868507 |
| ENSMUSG( | Plekhg5 | 2548.884 | 0.082207 | 0.520533 | 0.875186 | ENSMUSG( | Plekhg5 | 2226.43 | -0.03692 | 0.753143 | 0.970622 |
| ENSMUSG( | Plekhh1 | 945.2187 | -0.43349 | 1.57E-05 | 0.001615 | ENSMUSG( | Plekhh1 | 946.9985 | -0.12455 | 0.434427 | 0.91869 |
| ENSMUSG( | Plekhh2 | 330.7372 | -0.21702 | 0.109956 | 0.500285 | ENSMUSG( | Plekhh2 | 347.0971 | 0.194171 | 0.199848 | 0.863016 |
| ENSMUSG( | Plekhh3 | 405.7217 | -0.12142 | 0.403602 | 0.816132 | ENSMUSG( | Plekhh3 | 408.576 | 0.175854 | 0.324101 | 0.905257 |
| ENSMUSG( | Plekhj1 | 980.4452 | -0.17619 | 0.075808 | 0.426666 | ENSMUSG( | Plekhj1 | 906.2899 | -0.13396 | 0.255565 | 0.883043 |
| ENSMUSG( | Plekhm1 | 1066.026 | 0.257572 | 0.019009 | 0.207571 | ENSMUSG( | Plekhm1 | 913.9964 | 0.094529 | 0.448992 | 0.922202 |
| ENSMUSG( | Plekhm2 | 10372.37 | 0.187652 | 0.024392 | 0.238772 | ENSMUSG( | Plekhm2 | 8359.194 | -0.16675 | 0.107322 | 0.804969 |
| ENSMUSG( | Plekhm3 | 844.2516 | -0.08003 | 0.466438 | 0.844661 | ENSMUSG( | Plekhm3 | 778.859 | -0.04251 | 0.731352 | 0.967228 |
| ENSMUSG( | Plekhn1 | 310.5406 | -0.13782 | 0.348712 | 0.781339 | ENSMUSG( | Plekhn1 | 299.6114 | 0.038579 | 0.851049 | 0.985541 |
| ENSMUSG( | Plekho1 | 1395.057 | -0.04803 | 0.633514 | 0.914073 | ENSMUSG( | Plekho1 | 1255.58 | -0.08513 | 0.46037 | 0.923329 |
| ENSMUSG( | Plekho2 | 1353.054 | -0.21405 | 0.062707 | 0.387171 | ENSMUSG( | Plekho2 | 1389.723 | 0.132769 | 0.320042 | 0.903785 |
| ENSMUSG( | Plekhs1 | 16.03835 | -0.40472 | 0.370292 | NA | ENSMUSG( | Plekhs1 | 15.10709 | -0.27578 | 0.582409 | 0.944266 |
| ENSMUSG( | Plgrkt | 403.1945 | -0.08422 | 0.533237 | 0.879905 | ENSMUSG( | Plgrkt | 399.6798 | 0.16283 | 0.370343 | 0.915491 |
| ENSMUSG( | Plin2 | 356.5948 | 0.000902 | 0.994871 | 0.999369 | ENSMUSG( | Plin2 | 301.7202 | -0.22654 | 0.149778 | 0.836357 |
| ENSMUSG( | Plin3 | 731.0681 | 0.002817 | 0.980087 | 0.997491 | ENSMUSG( | Plin3 | 645.0272 | -0.09001 | 0.464511 | 0.923329 |
| ENSMUSG( | Plin4 | 70.92371 | 1.574707 | 6.02E-06 | 0.000775 | ENSMUSG( | Plin4 | 79.52487 | 1.978387 | 2.17E-10 | 2.37E-07 |
| ENSMUSG( | Plin5 | 17.44454 | 0.119706 | 0.793254 | NA | ENSMUSG( | Plin5 | 17.88315 | 0.446766 | 0.340875 | 0.907972 |
| ENSMUSG( | Plk1 | 91.26287 | 0.09346 | 0.685876 | 0.932981 | ENSMUSG( | Plk1 | 85.82364 | 0.169669 | 0.476889 | 0.923329 |
| ENSMUSG( | Plk2 | 2839.994 | -0.05611 | 0.732006 | 0.944956 | ENSMUSG( | Plk2 | 2544.594 | -0.1069 | 0.521096 | 0.932489 |
| ENSMUSG( | Plk3 | 439.2999 | -0.49166 | 7.64E-05 | 0.005386 | ENSMUSG( | Plk3 | 413.0231 | -0.37721 | 0.030913 | 0.629448 |
| ENSMUSG( | Plk4 | 91.18762 | -0.06929 | 0.765339 | 0.954127 | ENSMUSG( | Plk4 | 92.60171 | 0.252612 | 0.347614 | 0.911652 |
| ENSMUSG( | Plk5 | 326.5313 | -0.07532 | 0.687966 | 0.93324 | ENSMUSG( | Plk5 | 281.6043 | -0.24781 | 0.282707 | 0.892412 |
| ENSMUSG( | Pllp | 3563.962 | -0.72481 | 3.74E-08 | 1.24E-05 | ENSMUSG( | Pllp | 3870.846 | -0.12655 | 0.4452 | 0.921734 |
| ENSMUSG( | Plod1 | 1489.99 | -0.39295 | 0.01026 | 0.144326 | ENSMUSG( | Plod1 | 1659.424 | 0.211701 | 0.24844 | 0.877904 |
| ENSMUSG( | Plod2 | 311.5179 | -0.06308 | 0.670748 | 0.928683 | ENSMUSG( | Plod2 | 310.2069 | 0.195884 | 0.248842 | 0.877904 |
| ENSMUSG( | Plod3 | 1251.904 | -0.11396 | 0.391854 | 0.809274 | ENSMUSG( | Plod3 | 1304.214 | 0.267698 | 0.07433 | 0.766789 |
| ENSMUSG( | Plp1 | 125848 | -0.45502 | 0.000294 | 0.014126 | ENSMUSG( | Plp1 | 136096.4 | 0.07633 | 0.630614 | 0.953251 |
| ENSMUSG( | Plp2 | 201.3654 | -0.49802 | 0.003464 | 0.074411 | ENSMUSG( | Plp2 | 209.024 | -0.07821 | 0.666725 | 0.956766 |
| ENSMUSG( | Plpbp | 1757.903 | -0.00216 | 0.982779 | 0.997522 | ENSMUSG( | Plpbp | 1625.43 | 0.043466 | 0.721444 | 0.966887 |
| ENSMUSG( | Plpp1 | 590.933 | -0.29808 | 0.021308 | 0.222975 | ENSMUSG( | Plpp1 | 606.4987 | 0.062451 | 0.646554 | 0.953988 |
| ENSMUSG( | Plpp2 | 147.9045 | -0.36408 | 0.05753 | 0.370143 | ENSMUSG( | Plpp2 | 156.7959 | 0.106436 | 0.615551 | 0.951177 |
| ENSMUSG( | Plpp3 | 21119.23 | -0.27773 | 0.048819 | 0.337919 | ENSMUSG( | Plpp3 | 20759.29 | -0.0421 | 0.784796 | 0.976231 |
| ENSMUSG( | Plpp4 | 257.9324 | -0.15704 | 0.394686 | 0.810072 | ENSMUSG( | Plpp4 | 262.8961 | 0.170893 | 0.391008 | 0.917257 |
| ENSMUSG( | Plpp5 | 698.5524 | -0.30101 | 0.00498 | 0.093596 | ENSMUSG( | Plpp5 | 698.9982 | -0.01253 | 0.921726 | 0.992644 |
| ENSMUSG( | Plpp6 | 1545.544 | -0.39808 | 0.000716 | 0.025854 | ENSMUSG( | Plpp6 | 1614.981 | 0.027509 | 0.834236 | 0.981237 |
| ENSMUSG( | Plpp7 | 793.5395 | -0.35285 | 0.017267 | 0.196955 | ENSMUSG( | Plpp7 | 853.004 | 0.149206 | 0.385467 | 0.917257 |
| ENSMUSG( | Plppr1 | 337.2408 | -1.04938 | 4.93E-11 | 3.07E-08 | ENSMUSG( | Plppr1 | 417.5301 | 0.026238 | 0.878104 | 0.989644 |
| ENSMUSG( | Plppr2 | 8621.794 | -0.18227 | 0.11486 | 0.510978 | ENSMUSG( | Plppr2 | 8734.73 | 0.129696 | 0.305909 | 0.898538 |
| ENSMUSG( | Plppr3 | 2044.582 | -0.31429 | 0.000179 | 0.009711 | ENSMUSG( | Plppr3 | 2050.298 | -0.02317 | 0.851581 | 0.985541 |
| ENSMUSG( | Plppr4 | 4956.41 | -0.51567 | 0.004665 | 0.089972 | ENSMUSG( | Plppr4 | 6215.79 | 0.427208 | 0.024629 | 0.592568 |
| ENSMUSG( | Plppr5 | 1513.753 | -0.61072 | 2.05E-05 | 0.001993 | ENSMUSG( | Plppr5 | 1763.485 | 0.159424 | 0.336822 | 0.907972 |
| ENSMUSG( | Plrg1 | 663.5968 | 0.206839 | 0.072718 | 0.416435 | ENSMUSG( | Plrg1 | 564.6791 | 0.015649 | 0.908504 | 0.991958 |
| ENSMUSG( | Pls1 | 214.0531 | -0.20211 | 0.183655 | 0.620678 | ENSMUSG( | Pls1 | 205.0488 | -0.04749 | 0.776754 | 0.975717 |
| ENSMUSG( | Pls3 | 3440.344 | -0.01184 | 0.883431 | 0.982465 | ENSMUSG( | Pls3 | 3278.908 | 0.116728 | 0.266227 | 0.886134 |
| ENSMUSG( | Plscr1 | 76.18093 | -0.01483 | 0.95365 | 0.993874 | ENSMUSG( | Plscr1 | 65.8658 | -0.18543 | 0.494417 | 0.927372 |
| ENSMUSG( | Plscr2 | 41.02159 | -0.80997 | 0.006434 | 0.110086 | ENSMUSG( | Plscr2 | 47.58127 | -0.00838 | 0.977409 | 0.996416 |
| ENSMUSG( | Plscr3 | 565.7219 | -0.04465 | 0.749081 | 0.949604 | ENSMUSG( | Plscr3 | 518.5 | -0.02817 | 0.858808 | 0.985884 |

| | | | | | |
|---|---|---|---|---|---|
| ENSMUSG( | 465.7245 | 0.164605 | 0.379935 | 0.801757 | Pglyrp1 |
| ENSMUSG( | 2901.51 | 0.106487 | 0.335969 | 0.777 | Pgm1 |
| ENSMUSG( | 334.2692 | 0.008942 | 0.945239 | 0.99365 | Pgm2 |
| ENSMUSG( | 6131.367 | 0.157337 | 0.065325 | 0.395951 | Pgm2l1 |
| ENSMUSG( | 354.9795 | 0.133451 | 0.310771 | 0.758828 | Pgm3 |
| ENSMUSG( | 253.5384 | 0.284438 | 0.053611 | 0.35565 | Pgm5 |
| ENSMUSG( | 3234.641 | 0.097055 | 0.217936 | 0.663176 | Pgp |
| ENSMUSG( | 690.1804 | -0.06057 | 0.599954 | 0.903978 | Pgpep1 |
| ENSMUSG( | 11.7673 | -0.37596 | 0.484768 | NA | Pgpep1l |
| ENSMUSG( | 345.8985 | 0.010926 | 0.94148 | 0.993415 | Pgr |
| ENSMUSG( | 7357.776 | -0.37368 | 0.00237 | 0.057164 | Pgrmc1 |
| ENSMUSG( | 3585.612 | -0.24368 | 0.026799 | 0.250822 | Pgrmc2 |
| ENSMUSG( | 3702.769 | 0.12941 | 0.10383 | 0.489428 | Pgs1 |
| ENSMUSG( | 20391.19 | 0.027843 | 0.772134 | 0.954994 | Phactr1 |
| ENSMUSG( | 1331.963 | -0.04475 | 0.685183 | 0.932981 | Phactr2 |
| ENSMUSG( | 13159.14 | 0.167142 | 0.046352 | 0.329764 | Phactr3 |
| ENSMUSG( | 835.9069 | 0.06703 | 0.516757 | 0.873014 | Phactr4 |
| ENSMUSG( | 779.0927 | 0.082363 | 0.559558 | 0.888674 | Phaf1 |
| ENSMUSG( | 842.6316 | 0.136565 | 0.289901 | 0.739839 | Phax |
| ENSMUSG( | 1614.796 | 0.017753 | 0.840604 | 0.971179 | Phb1 |
| ENSMUSG( | 2113.809 | 0.008665 | 0.927936 | 0.991229 | Phb2 |
| ENSMUSG( | 2854.476 | 0.196989 | 0.018206 | 0.203238 | Phc1 |
| ENSMUSG( | 2769.788 | 0.022273 | 0.827381 | 0.968732 | Phc2 |
| ENSMUSG( | 2080.467 | 0.104197 | 0.322505 | 0.767663 | Phc3 |
| ENSMUSG( | 349.8509 | 0.055124 | 0.694178 | 0.9343 | Pheta1 |
| ENSMUSG( | 28.80017 | -0.11741 | 0.748002 | 0.949604 | Pheta2 |
| ENSMUSG( | 18.34749 | -0.41583 | 0.329147 | NA | Phex |
| ENSMUSG( | 1075.482 | -0.00853 | 0.941246 | 0.993415 | Phf1 |
| ENSMUSG( | 2890.142 | 0.032122 | 0.719835 | 0.941986 | Phf10 |
| ENSMUSG( | 16.36857 | -0.42934 | 0.377626 | NA | Phf11b |
| ENSMUSG( | 22.62355 | 0.008606 | 0.984406 | 0.99781 | Phf11c |
| ENSMUSG( | 17.54333 | -0.89663 | 0.063906 | NA | Phf11d |
| ENSMUSG( | 1487.482 | 0.03228 | 0.745976 | 0.949289 | Phf12 |
| ENSMUSG( | 337.7536 | -0.75232 | 7.61E-08 | 2.13E-05 | Phf13 |
| ENSMUSG( | 1516.89 | -0.08013 | 0.366647 | 0.794842 | Phf14 |
| ENSMUSG( | 50.59511 | -0.23978 | 0.367391 | 0.795221 | Phf19 |
| ENSMUSG( | 2819.064 | 0.113353 | 0.237834 | 0.685809 | Phf2 |
| ENSMUSG( | 2112.223 | -0.12602 | 0.177376 | 0.614553 | Phf20 |
| ENSMUSG( | 2030.911 | -0.114 | 0.286517 | 0.736031 | Phf20l1 |
| ENSMUSG( | 2056.935 | 0.129223 | 0.149287 | 0.573248 | Phf21a |
| ENSMUSG( | 651.3076 | 0.000923 | 0.993524 | 0.999369 | Phf21b |
| ENSMUSG( | 1127.48 | 0.042952 | 0.678116 | 0.930542 | Phf23 |
| ENSMUSG( | 12296.43 | 0.105719 | 0.319831 | 0.765305 | Phf24 |
| ENSMUSG( | 2538.859 | 0.040098 | 0.64528 | 0.917957 | Phf3 |
| ENSMUSG( | 624.0293 | 0.09087 | 0.403506 | 0.816132 | Phf5a |
| ENSMUSG( | 451.2021 | 0.247272 | 0.055098 | 0.361231 | Phf6 |
| ENSMUSG( | 132.7465 | -0.19628 | 0.291244 | 0.741248 | Phf7 |
| ENSMUSG( | 626.6155 | 0.093455 | 0.42053 | 0.822898 | Phf8 |
| ENSMUSG( | 5282.412 | 0.149158 | 0.14932 | 0.573248 | Phgdh |
| ENSMUSG( | 1108.716 | -0.19579 | 0.157018 | 0.58551 | Phip |
| ENSMUSG( | 3518.819 | 0.171182 | 0.120783 | 0.524807 | Phka1 |
| ENSMUSG( | 823.8671 | 0.220759 | 0.158104 | 0.586829 | Phka2 |
| ENSMUSG( | 2469.874 | 0.087543 | 0.473982 | 0.849425 | Phkb |
| ENSMUSG( | 1862.168 | 0.030008 | 0.790653 | 0.959461 | Phkg1 |
| ENSMUSG( | 2418.806 | 0.052227 | 0.559717 | 0.88884 | Phkg2 |
| ENSMUSG( | 482.9242 | -0.14798 | 0.351674 | 0.783381 | Phlda1 |
| ENSMUSG( | 699.67 | -0.05843 | 0.649066 | 0.920025 | Phlda3 |
| ENSMUSG( | 6400.787 | -0.22775 | 0.010127 | 0.143118 | Phldb1 |
| ENSMUSG( | 372.4411 | -0.21202 | 0.136785 | 0.551217 | Phldb2 |
| ENSMUSG( | 5612.261 | 0.028814 | 0.740944 | 0.948028 | Phlpp1 |
| ENSMUSG( | 1581.74 | 0.075791 | 0.412173 | 0.819452 | Phlpp2 |
| ENSMUSG( | 131.0883 | 0.158672 | 0.418595 | 0.821966 | Phospho1 |
| ENSMUSG( | 381.5118 | 0.061921 | 0.65955 | 0.924506 | Phospho2 |
| ENSMUSG( | 332.3751 | -0.00846 | 0.949814 | 0.993821 | Phpt1 |
| ENSMUSG( | 1650.718 | 0.184515 | 0.043364 | 0.319133 | Phrf1 |
| ENSMUSG( | 1319.636 | 0.066336 | 0.548867 | 0.885686 | Phtf1 |
| ENSMUSG( | 32.66911 | -0.01671 | 0.960593 | 0.993991 | Phtf1os |
| ENSMUSG( | 404.325 | -0.25728 | 0.102686 | 0.488148 | Phtf2 |
| ENSMUSG( | 4270.913 | 0.028371 | 0.811474 | 0.964426 | Phyh |
| ENSMUSG( | 126.2988 | 0.384126 | 0.07424 | 0.422054 | Phyhd1 |
| ENSMUSG( | 10677.06 | 0.075087 | 0.590791 | 0.90113 | Phyhip |
| ENSMUSG( | 4888.165 | -0.05241 | 0.583933 | 0.898666 | Phyhipl |
| ENSMUSG( | 144.1143 | 0.086413 | 0.625993 | 0.912356 | Phykpl |
| ENSMUSG( | 20.66511 | -0.14628 | 0.771141 | NA | Pi16 |
| ENSMUSG( | 1929.229 | 0.125632 | 0.234043 | 0.681906 | Pi4k2a |
| ENSMUSG( | 122.0925 | -0.13614 | 0.466318 | 0.844631 | Pi4k2b |
| ENSMUSG( | 7979.473 | 0.193896 | 0.0281 | 0.25643 | Pi4ka |
| ENSMUSG( | 1339.51 | 0.157437 | 0.133139 | 0.545727 | Pi4kb |
| ENSMUSG( | 6260.826 | -0.41367 | 0.000177 | 0.009611 | Pianp |
| ENSMUSG( | 1239.668 | 0.150573 | 0.123553 | 0.530552 | Pias1 |
| ENSMUSG( | 1329.159 | 0.12697 | 0.171464 | 0.605943 | Pias2 |
| ENSMUSG( | 701.1843 | 0.050411 | 0.691382 | 0.933881 | Pias3 |
| ENSMUSG( | 631.1969 | 0.134061 | 0.361374 | 0.792797 | Pias4 |
| ENSMUSG( | 201.7678 | 0.010078 | 0.945936 | 0.99369 | Pibf1 |
| ENSMUSG( | 4031.344 | 0.106641 | 0.200643 | 0.640661 | Picalm |
| ENSMUSG( | 527.9048 | 0.074815 | 0.533796 | 0.880095 | Pick1 |
| ENSMUSG( | 970.6525 | -0.05071 | 0.649385 | 0.920317 | Pid1 |
| ENSMUSG( | 78.32187 | 0.196734 | 0.390984 | 0.80828 | Pidd1 |
| ENSMUSG( | 67.00374 | -0.12633 | 0.607009 | 0.906685 | Pierce1 |
| ENSMUSG( | 140.3554 | -0.18908 | 0.294471 | 0.743674 | Pierce2 |
| ENSMUSG( | 398.1419 | 0.180221 | 0.185135 | 0.623005 | Piezo1 |
| ENSMUSG( | 76.4377 | -0.30279 | 0.194036 | 0.633126 | Piezo2 |
| ENSMUSG( | 14.55567 | -0.39621 | 0.484334 | NA | Pif1 |
| ENSMUSG( | 250.7622 | -0.11257 | 0.456002 | 0.838683 | Piga |
| ENSMUSG( | 489.457 | -0.14784 | 0.23161 | 0.67949 | Pigb |
| ENSMUSG( | 163.51 | 0.068513 | 0.678374 | 0.930542 | Pigbos1 |
| ENSMUSG( | 495.6872 | -0.02669 | 0.840132 | 0.971179 | Pigc |
| ENSMUSG( | 255.3368 | -0.183 | 0.247905 | 0.696522 | Pigf |
| ENSMUSG( | 595.0037 | -0.18035 | 0.269591 | 0.719729 | Pigg |
| ENSMUSG( | 462.5954 | -0.22196 | 0.071434 | 0.413071 | Pigh |
| ENSMUSG( | 2026.547 | -0.33554 | 0.004419 | 0.086676 | Pigk |
| ENSMUSG( | 163.315 | -0.52581 | 0.00295 | 0.066455 | Pigl |
| ENSMUSG( | 397.6124 | -0.23378 | 0.142276 | 0.560124 | Pigm |
| ENSMUSG( | 458.5178 | -0.2104 | 0.153032 | 0.579541 | Pign |
| ENSMUSG( | 734.0951 | 0.035377 | 0.820435 | 0.965888 | Pigo |
| ENSMUSG( | 702.4955 | -0.40541 | 0.001026 | 0.033548 | Pigp |
| ENSMUSG( | 6112.635 | -0.00291 | 0.976982 | 0.997428 | Pigq |
| ENSMUSG( | 2550.631 | -0.19014 | 0.165817 | 0.597653 | Pigs |
| ENSMUSG( | 3384.395 | -0.27967 | 0.057891 | 0.371023 | Pigt |
| ENSMUSG( | 814.7961 | -0.36927 | 0.009377 | 0.137017 | Pigu |
| ENSMUSG( | 285.3858 | -0.2725 | 0.086094 | 0.451605 | Pigv |
| ENSMUSG( | 85.27875 | -0.43861 | 0.073057 | 0.417421 | Pigw |
| ENSMUSG( | 351.8054 | -0.27332 | 0.035881 | 0.290992 | Pigx |
| ENSMUSG( | 657.2767 | -0.2156 | 0.040347 | 0.310226 | Pigyl |
| ENSMUSG( | 266.9531 | -0.51537 | 0.011952 | 0.157992 | Pigz |
| ENSMUSG( | 755.527 | 0.066404 | 0.515297 | 0.872546 | Pih1d1 |
| ENSMUSG( | 54.6152 | 0.094062 | 0.722013 | 0.943535 | Pih1d2 |
| ENSMUSG( | 270.4479 | -0.21173 | 0.157068 | 0.58551 | Pik3ap1 |
| ENSMUSG( | 991.8835 | 0.119124 | 0.351152 | 0.783067 | Pik3c2a |
| ENSMUSG( | 652.1856 | -0.16257 | 0.151084 | 0.576088 | Pik3c2b |
| ENSMUSG( | 1011.547 | 0.232808 | 0.029259 | 0.259574 | Pik3c3 |
| ENSMUSG( | 2406.509 | 0.153083 | 0.108184 | 0.497094 | Pik3ca |
| ENSMUSG( | 703.7518 | 0.158373 | 0.135375 | 0.548884 | Pik3cb |
| ENSMUSG( | 964.4 | 0.172224 | 0.130769 | 0.542059 | Pik3cd |
| ENSMUSG( | 192.6332 | 0.228656 | 0.15255 | 0.578935 | Pik3cg |
| ENSMUSG( | 1377.971 | -0.30661 | 0.039396 | 0.305596 | Pik3ip1 |
| ENSMUSG( | 6554.888 | 0.216665 | 0.008042 | 0.124826 | Pik3r1 |
| ENSMUSG( | 5597.444 | 0.158769 | 0.084568 | 0.448139 | Pik3r2 |
| ENSMUSG( | 1737.551 | -0.21275 | 0.050899 | 0.345939 | Pik3r3 |
| ENSMUSG( | 1001.455 | -0.04153 | 0.716172 | 0.94152 | Pik3r4 |
| ENSMUSG( | 168.8925 | -0.00284 | 0.986324 | 0.998326 | Pik3r5 |
| ENSMUSG( | 142.1159 | 0.139739 | 0.472743 | 0.848388 | Pik3r6 |
| ENSMUSG( | 1182.422 | 0.073535 | 0.427964 | 0.825894 | Pikfyve |
| ENSMUSG( | 32.05064 | -0.41224 | 0.39696 | 0.811839 | Pilra |
| ENSMUSG( | 95.34493 | 0.052812 | 0.816832 | 0.96566 | Pim1 |
| ENSMUSG( | 921.2436 | 0.085589 | 0.528126 | 0.878224 | Pim2 |
| ENSMUSG( | 893.3166 | -0.35305 | 0.054183 | 0.357985 | Pim3 |
| ENSMUSG( | 25.70673 | 0.094848 | 0.816133 | 0.965455 | Pimreg |
| ENSMUSG( | 2501.079 | 0.001103 | 0.990741 | 0.999232 | Pin1 |
| ENSMUSG( | 357.2458 | 0.049469 | 0.733261 | 0.944956 | Pin4 |
| ENSMUSG( | 30765.79 | 0.193724 | 0.017602 | 0.198821 | Pink1 |
| ENSMUSG( | 158.0458 | -0.22801 | 0.193788 | 0.633085 | Pinx1 |
| ENSMUSG( | 4285.289 | -0.01458 | 0.865378 | 0.975565 | Pip4k2a |
| ENSMUSG( | 5155.56 | 0.162568 | 0.035734 | 0.290677 | Pip4k2b |
| ENSMUSG( | 4005.075 | -0.08073 | 0.42599 | 0.824898 | Pip4k2c |
| ENSMUSG( | 1639.076 | -0.0109 | 0.907088 | 0.987447 | Pip4p1 |
| ENSMUSG( | 1019.511 | -0.06385 | 0.558088 | 0.888427 | Pip4p2 |
| ENSMUSG( | 1204.917 | 0.001886 | 0.986764 | 0.998619 | Pip5k1a |
| ENSMUSG( | 332.1936 | 0.307895 | 0.091969 | 0.465937 | Pip5k1b |
| ENSMUSG( | 22856.69 | 0.225216 | 0.01092 | 0.150598 | Pip5k1c |
| ENSMUSG( | 153.351 | 0.198394 | 0.333859 | 0.775602 | Pip5kl1 |
| ENSMUSG( | 57.75046 | -0.4613 | 0.065654 | 0.396911 | Pipox |
| ENSMUSG( | 168.1804 | -0.18485 | 0.289039 | 0.739705 | Pir |
| ENSMUSG( | 32.59257 | -0.60708 | 0.245513 | 0.693175 | Pirb |
| ENSMUSG( | 12.77829 | -0.97207 | 0.064302 | NA | Pirt |
| ENSMUSG( | 2419.665 | 0.051051 | 0.552683 | 0.886141 | Pisd |
| ENSMUSG( | 301.2569 | -0.12475 | 0.417041 | 0.8209 | Pisd-ps1 |
| ENSMUSG( | 2236.182 | 0.029138 | 0.749258 | 0.949664 | Pithd1 |

| | | | | | |
|---|---|---|---|---|---|
| ENSMUSG( | 87.12679 | 0.063191 | 0.788132 | 0.97637 | Pim1 |
| ENSMUSG( | 790.4219 | -0.08843 | 0.573478 | 0.942912 | Pim2 |
| ENSMUSG( | 896.7321 | -0.0482 | 0.820508 | 0.979198 | Pim3 |
| ENSMUSG( | 20.17729 | -0.35555 | 0.446087 | 0.921734 | Pimreg |
| ENSMUSG( | 2091.983 | -0.25951 | 0.028538 | 0.614171 | Pin1 |
| ENSMUSG( | 296.5189 | -0.2362 | 0.199055 | 0.863016 | Pin4 |
| ENSMUSG( | 25757.58 | -0.04582 | 0.714872 | 0.964986 | Pink1 |
| ENSMUSG( | 142.3963 | -0.27286 | 0.178745 | 0.849041 | Pinx1 |
| ENSMUSG( | 3747.093 | -0.1381 | 0.268321 | 0.886493 | Pip4k2a |
| ENSMUSG( | 4393.221 | -0.02637 | 0.788828 | 0.976593 | Pip4k2b |
| ENSMUSG( | 3732.321 | -0.01173 | 0.920658 | 0.992644 | Pip4k2c |
| ENSMUSG( | 1516.578 | 0.034198 | 0.749591 | 0.96998 | Pip4p1 |
| ENSMUSG( | 947.1894 | -0.01334 | 0.926932 | 0.99306 | Pip4p2 |
| ENSMUSG( | 1066.258 | -0.08478 | 0.514963 | 0.930939 | Pip5k1a |
| ENSMUSG( | 285.7008 | 0.156942 | 0.435018 | 0.918876 | Pip5k1b |
| ENSMUSG( | 19613.27 | 0.061443 | 0.552684 | 0.939565 | Pip5k1c |
| ENSMUSG( | 139.1706 | 0.184937 | 0.481283 | 0.924564 | Pip5kl1 |
| ENSMUSG( | 62.55317 | 0.05269 | 0.846574 | 0.984821 | Pipox |
| ENSMUSG( | 154.714 | -0.15191 | 0.422781 | 0.917257 | Pir |
| ENSMUSG( | 33.29575 | -0.23113 | 0.665415 | 0.956766 | Pirb |
| ENSMUSG( | 14.21648 | -0.22755 | 0.656504 | 0.955533 | Pirt |
| ENSMUSG( | 2150.425 | -0.02174 | 0.839746 | 0.982162 | Pisd |
| ENSMUSG( | 299.2961 | 0.133712 | 0.441125 | 0.921166 | Pisd-ps1 |
| ENSMUSG( | 1984.557 | -0.04983 | 0.677164 | 0.957106 | Pithd1 |
| ENSMUSG( | 7742.517 | -0.13292 | 0.237573 | 0.872264 | Pitpna |
| ENSMUSG( | 2692.968 | -0.0932 | 0.473102 | 0.923329 | Pitpnb |
| ENSMUSG( | 2801.112 | -0.03572 | 0.734491 | 0.967228 | Pitpnc1 |
| ENSMUSG( | 9172.164 | 0.122396 | 0.241218 | 0.876259 | Pitpnm1 |
| ENSMUSG( | 8849.051 | 0.32293 | 0.005913 | 0.316645 | Pitpnm2 |
| ENSMUSG( | 5345.645 | 0.096286 | 0.340699 | 0.907972 | Pitpnm3 |
| ENSMUSG( | 1261.651 | -0.04099 | 0.724968 | 0.966887 | Pitrm1 |
| ENSMUSG( | 28.49296 | 0.086303 | 0.815626 | 0.979198 | Piwil2 |
| ENSMUSG( | 2452.79 | -0.02883 | 0.818853 | 0.979198 | Pja1 |
| ENSMUSG( | 6438.402 | 0.002787 | 0.980538 | 0.996479 | Pja2 |
| ENSMUSG( | 4835.888 | 0.131604 | 0.27049 | 0.888062 | Pkd1 |
| ENSMUSG( | 25.39376 | 0.030127 | 0.945018 | 0.994798 | Pkd1l3 |
| ENSMUSG( | 2343.232 | 0.109664 | 0.387725 | 0.917257 | Pkd2 |
| ENSMUSG( | 30.20111 | 0.167885 | 0.667912 | 0.956766 | Pkd2l1 |
| ENSMUSG( | 46.76412 | -0.24828 | 0.421295 | 0.917257 | Pkd2l2 |
| ENSMUSG( | 365.1132 | 0.148214 | 0.376224 | 0.915491 | Pkdcc |
| ENSMUSG( | 36.56319 | 0.262317 | 0.532271 | 0.934795 | Pkdrej |
| ENSMUSG( | 17.96453 | 0.593697 | 0.233392 | 0.868507 | Pkhd1l1 |
| ENSMUSG( | 3724.436 | -0.0606 | 0.685528 | 0.958678 | Pkia |
| ENSMUSG( | 113.6056 | -0.21744 | 0.319357 | 0.903453 | Pkib |
| ENSMUSG( | 2841.615 | -0.1243 | 0.350611 | 0.911652 | Pkig |
| ENSMUSG( | 22531.09 | -0.20218 | 0.081517 | 0.77681 | Pkm |
| ENSMUSG( | 157.463 | -0.02394 | 0.895517 | 0.990558 | Pkmyt1 |
| ENSMUSG( | 3293.658 | -0.0759 | 0.443369 | 0.921704 | Pkn1 |
| ENSMUSG( | 984.5302 | 0.178622 | 0.188545 | 0.85698 | Pkn2 |
| ENSMUSG( | 33.83965 | 0.139014 | 0.736303 | 0.967344 | Pkn3 |
| ENSMUSG( | 587.6775 | -0.05272 | 0.713657 | 0.96497 | Pknox1 |
| ENSMUSG( | 1269.413 | -0.08784 | 0.505205 | 0.927825 | Pknox2 |
| ENSMUSG( | 11.26248 | -0.58686 | 0.325524 | NA | Pkp1 |
| ENSMUSG( | 167.5606 | 0.366088 | 0.057121 | 0.723362 | Pkp2 |
| ENSMUSG( | 14871.1 | -0.07955 | 0.434224 | 0.91869 | Pkp4 |
| ENSMUSG( | 85.18257 | 1.46713 | 1.48E-05 | 0.005074 | Pla1a |
| ENSMUSG( | 637.028 | 0.08362 | 0.521381 | 0.932489 | Pla2g12a |
| ENSMUSG( | 1025.479 | 0.307015 | 0.120859 | 0.813384 | Pla2g15 |
| ENSMUSG( | 186.1036 | 0.533179 | 0.003961 | 0.253257 | Pla2g3 |
| ENSMUSG( | 146.434 | 0.201841 | 0.407052 | 0.917257 | Pla2g4a |
| ENSMUSG( | 18.54977 | 0.291152 | 0.552078 | 0.939565 | Pla2g4b |
| ENSMUSG( | 372.0693 | -0.3503 | 0.021973 | 0.575341 | Pla2g4e |
| ENSMUSG( | 148.2177 | -0.55569 | 0.002578 | 0.198937 | Pla2g5 |
| ENSMUSG( | 633.9232 | -0.07084 | 0.635512 | 0.953903 | Pla2g6 |
| ENSMUSG( | 11930.77 | -0.04008 | 0.779984 | 0.976106 | Pla2g7 |
| ENSMUSG( | 24.87554 | 0.558916 | 0.207761 | 0.864412 | Pla2r1 |
| ENSMUSG( | 1087.921 | 0.025605 | 0.840592 | 0.982495 | Plaa |
| ENSMUSG( | 345.2324 | -0.27475 | 0.13557 | 0.825163 | Plaat1 |
| ENSMUSG( | 2345.447 | -0.19474 | 0.209383 | 0.864412 | Plaat3 |
| ENSMUSG( | 23.61376 | -1.03044 | 0.058493 | 0.728256 | Plac8 |
| ENSMUSG( | 358.4139 | -0.11498 | 0.536789 | 0.937108 | Plac9 |
| ENSMUSG( | 90.21395 | 0.423791 | 0.098547 | 0.798438 | Plag1 |
| ENSMUSG( | 633.046 | 0.10452 | 0.614961 | 0.951177 | Plagl1 |
| ENSMUSG( | 208.9873 | -0.05661 | 0.745643 | 0.96951 | Plagl2 |
| ENSMUSG( | 1899.853 | -0.03953 | 0.819243 | 0.979198 | Plat |
| ENSMUSG( | 13.02218 | -0.27621 | 0.625728 | 0.953234 | Platr14 |
| ENSMUSG( | 211.6547 | 0.142852 | 0.410488 | 0.917257 | Platr25 |
| ENSMUSG( | 154.9808 | 0.325294 | 0.113653 | 0.809623 | Plau |
| ENSMUSG( | 36.75199 | -0.4722 | 0.157371 | 0.837724 | Plaur |
| ENSMUSG( | 117.1767 | -0.10674 | 0.621608 | 0.953199 | Plb1 |
| ENSMUSG( | 29.56168 | -0.45062 | 0.393677 | 0.917257 | Plbd1 |
| ENSMUSG( | 3013.179 | 0.272875 | 0.078438 | 0.77321 | Plbd2 |
| ENSMUSG( | 4051.806 | -0.0945 | 0.440667 | 0.921166 | Plcb1 |
| ENSMUSG( | 157.3016 | -0.16882 | 0.382251 | 0.917257 | Plcb2 |
| ENSMUSG( | 1893.619 | -0.07642 | 0.530886 | 0.934795 | Plcb3 |
| ENSMUSG( | 961.3275 | 0.066896 | 0.639964 | 0.953903 | Plcb4 |
| ENSMUSG( | 571.0147 | -0.06204 | 0.672146 | 0.95694 | Plcd1 |
| ENSMUSG( | 681.5133 | 0.095652 | 0.475725 | 0.923329 | Plcd3 |
| ENSMUSG( | 971.9492 | -0.10389 | 0.42855 | 0.917257 | Plcd4 |
| ENSMUSG( | 906.9296 | 0.307175 | 0.043743 | 0.690304 | Plce1 |
| ENSMUSG( | 2814.955 | 0.05989 | 0.615075 | 0.951177 | Plcg1 |
| ENSMUSG( | 245.3396 | 0.114601 | 0.462361 | 0.923329 | Plcg2 |
| ENSMUSG( | 241.5146 | -0.01239 | 0.942353 | 0.994774 | Plch1 |
| ENSMUSG( | 2608.644 | 0.127684 | 0.306706 | 0.898538 | Plch2 |
| ENSMUSG( | 741.2884 | 0.232691 | 0.049545 | 0.708648 | Plcl1 |
| ENSMUSG( | 1245.088 | 0.347037 | 0.023787 | 0.588226 | Plcl2 |
| ENSMUSG( | 662.7432 | -0.04591 | 0.718936 | 0.96673 | Plcxd1 |
| ENSMUSG( | 2098.093 | -0.07854 | 0.487073 | 0.92541 | Plcxd2 |
| ENSMUSG( | 30.294 | -0.15606 | 0.718738 | 0.966668 | Plcxd3 |
| ENSMUSG( | 8.617795 | 0.334852 | 0.614834 | NA | Plcz1 |
| ENSMUSG( | 488.6127 | 0.441777 | 0.001853 | 0.15992 | Pld1 |
| ENSMUSG( | 1397.996 | 0.027236 | 0.803585 | 0.976978 | Pld2 |
| ENSMUSG( | 32913.98 | 0.050589 | 0.722264 | 0.966887 | Pld3 |
| ENSMUSG( | 760.0688 | 0.095072 | 0.607916 | 0.950113 | Pld4 |
| ENSMUSG( | 176.8419 | -0.17466 | 0.364006 | 0.914223 | Pld5 |
| ENSMUSG( | 26.53745 | -0.28901 | 0.462958 | 0.923329 | Pld6 |
| ENSMUSG( | 8114.406 | 0.130102 | 0.297783 | 0.898538 | Plec |
| ENSMUSG( | 579.6914 | -0.22867 | 0.163824 | 0.838291 | Plek |
| ENSMUSG( | 3311.179 | 0.024872 | 0.810309 | 0.978779 | Plekha1 |
| ENSMUSG( | 537.9602 | -0.07907 | 0.555522 | 0.939565 | Plekha2 |
| ENSMUSG( | 1130.988 | 0.029861 | 0.808952 | 0.97873 | Plekha3 |
| ENSMUSG( | 91.3456 | 0.0211 | 0.935095 | 0.993078 | Plekha4 |
| ENSMUSG( | 2184.015 | 0.007088 | 0.957093 | 0.994798 | Plekha5 |
| ENSMUSG( | 2537.882 | -0.01379 | 0.913266 | 0.992298 | Plekha6 |
| ENSMUSG( | 904.1372 | -0.064 | 0.61352 | 0.951177 | Plekha7 |
| ENSMUSG( | 411.3098 | 0.124894 | 0.434693 | 0.918742 | Plekha8 |
| ENSMUSG( | 35558.84 | -0.23557 | 0.054708 | 0.723362 | Plekhb1 |
| ENSMUSG( | 9791.708 | -0.21202 | 0.062485 | 0.736482 | Plekhb2 |
| ENSMUSG( | 127.8547 | -0.69008 | 0.000984 | 0.107109 | Plekhd1 |
| ENSMUSG( | 139.7753 | -0.30911 | 0.150776 | 0.837173 | Plekhf1 |
| ENSMUSG( | 357.5983 | 0.165822 | 0.318216 | 0.902849 | Plekhf2 |
| ENSMUSG( | 1051.985 | 0.215553 | 0.085436 | 0.779799 | Plekhg1 |
| ENSMUSG( | 319.6341 | 0.425154 | 0.026906 | 0.603041 | Plekhg2 |
| ENSMUSG( | 819.1436 | -0.00331 | 0.982113 | 0.996756 | Plekhg3 |
| ENSMUSG( | 74.13595 | 0.432796 | 0.233376 | 0.868507 | Plekhg4 |
| ENSMUSG( | 2226.43 | -0.03692 | 0.753143 | 0.970622 | Plekhg5 |
| ENSMUSG( | 946.9985 | -0.12455 | 0.434427 | 0.91869 | Plekhh1 |
| ENSMUSG( | 347.0971 | 0.194171 | 0.199848 | 0.863016 | Plekhh2 |
| ENSMUSG( | 408.576 | 0.175854 | 0.324101 | 0.905257 | Plekhh3 |
| ENSMUSG( | 906.2899 | -0.13396 | 0.255565 | 0.883043 | Plekhj1 |
| ENSMUSG( | 913.9964 | 0.094529 | 0.448992 | 0.922202 | Plekhm1 |
| ENSMUSG( | 8359.194 | -0.16675 | 0.107322 | 0.804969 | Plekhm2 |
| ENSMUSG( | 778.859 | -0.04251 | 0.731352 | 0.967228 | Plekhm3 |
| ENSMUSG( | 299.6114 | 0.038579 | 0.851049 | 0.985541 | Plekhn1 |
| ENSMUSG( | 1255.58 | -0.08513 | 0.46037 | 0.923329 | Plekho1 |
| ENSMUSG( | 1389.723 | 0.132769 | 0.320042 | 0.903785 | Plekho2 |
| ENSMUSG( | 15.10709 | -0.27578 | 0.582409 | 0.944266 | Plekhs1 |
| ENSMUSG( | 399.6798 | 0.16283 | 0.370343 | 0.915491 | Plgrkt |
| ENSMUSG( | 301.7202 | -0.22654 | 0.149778 | 0.836357 | Plin2 |
| ENSMUSG( | 645.0272 | -0.09001 | 0.464511 | 0.923329 | Plin3 |
| ENSMUSG( | 79.52487 | 1.978387 | 2.17E-10 | 2.37E-07 | Plin4 |
| ENSMUSG( | 17.88315 | 0.446766 | 0.340875 | 0.907972 | Plin5 |
| ENSMUSG( | 85.82364 | 0.169669 | 0.476889 | 0.923329 | Plk1 |
| ENSMUSG( | 2544.594 | -0.1069 | 0.521096 | 0.932489 | Plk2 |
| ENSMUSG( | 413.0231 | -0.37721 | 0.030913 | 0.629448 | Plk3 |
| ENSMUSG( | 92.60171 | 0.252612 | 0.347614 | 0.911652 | Plk4 |
| ENSMUSG( | 281.6043 | -0.24781 | 0.282707 | 0.892412 | Plk5 |
| ENSMUSG( | 3870.846 | -0.12655 | 0.4452 | 0.921734 | Pllp |
| ENSMUSG( | 1659.424 | 0.211701 | 0.24844 | 0.877904 | Plod1 |
| ENSMUSG( | 310.2069 | 0.195884 | 0.248842 | 0.877904 | Plod2 |
| ENSMUSG( | 1304.214 | 0.267698 | 0.07433 | 0.766789 | Plod3 |
| ENSMUSG( | 136096.4 | 0.07633 | 0.630614 | 0.953251 | Plp1 |
| ENSMUSG( | 209.024 | -0.07821 | 0.666725 | 0.956766 | Plp2 |
| ENSMUSG( | 1625.43 | 0.043466 | 0.721444 | 0.966887 | Plpbp |
| ENSMUSG( | 606.4987 | 0.062451 | 0.646554 | 0.953988 | Plpp1 |
| ENSMUSG( | 156.7959 | 0.106436 | 0.615551 | 0.951177 | Plpp2 |
| ENSMUSG( | 20759.29 | -0.0421 | 0.784796 | 0.976231 | Plpp3 |
| ENSMUSG( | 262.8961 | 0.170893 | 0.391008 | 0.917257 | Plpp4 |

| | | | | | | | | | | | |
|---|---|---|---|---|---|---|---|---|---|---|---|
| ENSMUSG( | Plscr4 | 135.7151 | 0.087172 | 0.673096 | 0.929007 | ENSMUSG( | Plscr4 | 138.958 | 0.406701 | 0.058018 | 0.728256 |
| ENSMUSG( | Pltp | 3387.373 | -0.06563 | 0.72598 | 0.944223 | ENSMUSG( | Pltp | 3959.469 | 0.60066 | 0.010886 | 0.422321 |
| ENSMUSG( | Plvap | 221.3893 | -0.25489 | 0.113251 | 0.507541 | ENSMUSG( | Plvap | 194.3563 | -0.37532 | 0.031798 | 0.631941 |
| ENSMUSG( | Plxdc1 | 959.9187 | -0.5157 | 0.000446 | 0.01925 | ENSMUSG( | Plxdc1 | 1029.384 | 0.00662 | 0.970083 | 0.99594 |
| ENSMUSG( | Plxdc2 | 1342.258 | -0.21377 | 0.201657 | 0.641362 | ENSMUSG( | Plxdc2 | 1426.762 | 0.235914 | 0.223446 | 0.867404 |
| ENSMUSG( | Plxna1 | 3376.439 | 0.018328 | 0.902821 | 0.986055 | ENSMUSG( | Plxna1 | 3412.641 | 0.30684 | 0.094004 | 0.795735 |
| ENSMUSG( | Plxna2 | 3437.431 | -0.23928 | 0.098713 | 0.480598 | ENSMUSG( | Plxna2 | 3793.029 | 0.312022 | 0.069189 | 0.755467 |
| ENSMUSG( | Plxna3 | 881.7129 | 0.192409 | 0.056188 | 0.36528 | ENSMUSG( | Plxna3 | 839.6141 | 0.309924 | 0.019082 | 0.549166 |
| ENSMUSG( | Plxna4 | 2444.3 | -0.35748 | 0.033595 | 0.28103 | ENSMUSG( | Plxna4 | 2747.978 | 0.265963 | 0.154435 | 0.837724 |
| ENSMUSG( | Plxna4os1 | 19.11579 | -0.08709 | 0.842984 | NA | ENSMUSG( | Plxna4os1 | 15.836 | -0.37265 | 0.445732 | 0.921734 |
| ENSMUSG( | Plxnb1 | 14589.99 | 0.090719 | 0.381346 | 0.802629 | ENSMUSG( | Plxnb1 | 12658.06 | -0.04893 | 0.646459 | 0.953988 |
| ENSMUSG( | Plxnb2 | 3594.584 | -0.01402 | 0.902157 | 0.985633 | ENSMUSG( | Plxnb2 | 3582.253 | 0.238743 | 0.111652 | 0.808384 |
| ENSMUSG( | Plxnb3 | 1107.823 | -0.1066 | 0.356538 | 0.788187 | ENSMUSG( | Plxnb3 | 1122.68 | 0.201207 | 0.267593 | 0.886134 |
| ENSMUSG( | Plxnc1 | 1504.828 | -0.03129 | 0.803037 | 0.962169 | ENSMUSG( | Plxnc1 | 1509.926 | 0.240626 | 0.072113 | 0.763629 |
| ENSMUSG( | Plxnd1 | 4094.574 | 0.077339 | 0.490518 | 0.859387 | ENSMUSG( | Plxnd1 | 3936.048 | 0.227968 | 0.114872 | 0.811703 |
| ENSMUSG( | Pm20d1 | 168.6117 | -0.38028 | 0.052188 | 0.350744 | ENSMUSG( | Pm20d1 | 160.4897 | -0.23302 | 0.267141 | 0.886134 |
| ENSMUSG( | Pm20d2 | 110.8783 | 0.053005 | 0.801917 | 0.962116 | ENSMUSG( | Pm20d2 | 95.146 | -0.14725 | 0.530547 | 0.934795 |
| ENSMUSG( | Pmaip1 | 56.75089 | -0.10616 | 0.7061 | 0.938843 | ENSMUSG( | Pmaip1 | 52.41637 | -0.04498 | 0.880084 | 0.989644 |
| ENSMUSG( | Pmch | 22.78877 | -0.13457 | 0.751461 | 0.950237 | ENSMUSG( | Pmch | 17.68025 | -0.73707 | 0.155214 | 0.837724 |
| ENSMUSG( | Pmel | 36.50002 | 0.03991 | 0.901154 | 0.985633 | ENSMUSG( | Pmel | 31.93924 | -0.05033 | 0.890239 | 0.990558 |
| ENSMUSG( | Pmepa1 | 2394.937 | -0.33415 | 0.006656 | 0.11214 | ENSMUSG( | Pmepa1 | 2471.323 | 0.04718 | 0.744801 | 0.96951 |
| ENSMUSG( | Pmf1 | 150.4441 | -0.11294 | 0.533724 | 0.880095 | ENSMUSG( | Pmf1 | 131.4344 | -0.22855 | 0.248153 | 0.877904 |
| ENSMUSG( | Pml | 389.0303 | 0.040989 | 0.755526 | 0.951395 | ENSMUSG( | Pml | 348.274 | -0.00809 | 0.956935 | 0.994798 |
| ENSMUSG( | Pmm1 | 6632.303 | 0.092401 | 0.279365 | 0.729114 | ENSMUSG( | Pmm1 | 5405.156 | -0.23995 | 0.041596 | 0.690304 |
| ENSMUSG( | Pmm2 | 712.4488 | 0.031776 | 0.781357 | 0.957558 | ENSMUSG( | Pmm2 | 646.3685 | 0.021295 | 0.879629 | 0.989644 |
| ENSMUSG( | Pmp22 | 1271.575 | -0.43463 | 0.002383 | 0.057268 | ENSMUSG( | Pmp22 | 1397.299 | 0.139893 | 0.421336 | 0.917257 |
| ENSMUSG( | Pmpca | 3519.523 | 0.092775 | 0.248338 | 0.697133 | ENSMUSG( | Pmpca | 3052.447 | -0.0477 | 0.628791 | 0.953251 |
| ENSMUSG( | Pmpcb | 1314.544 | 0.090041 | 0.37507 | 0.800786 | ENSMUSG( | Pmpcb | 1114.125 | -0.12111 | 0.272301 | 0.888771 |
| ENSMUSG( | Pms1 | 94.98091 | -0.01044 | 0.960899 | 0.993991 | ENSMUSG( | Pms1 | 83.84736 | -0.10944 | 0.666173 | 0.956766 |
| ENSMUSG( | Pms2 | 243.233 | 0.029819 | 0.89568 | 0.984527 | ENSMUSG( | Pms2 | 371.2944 | -0.05413 | 0.739632 | 0.968611 |
| ENSMUSG( | Pms2 | 414.7957 | -0.01229 | 0.93423 | 0.99323 | ENSMUSG( | Pms2 | 220.2022 | 0.013208 | 0.950423 | 0.994798 |
| ENSMUSG( | Pmvk | 1625.204 | 0.05402 | 0.561442 | 0.889052 | ENSMUSG( | Pmvk | 1347.517 | -0.22996 | 0.058661 | 0.728256 |
| ENSMUSG( | Pnck | 1286.448 | 0.253569 | 0.075827 | 0.426666 | ENSMUSG( | Pnck | 1058.811 | -0.03282 | 0.851304 | 0.985541 |
| ENSMUSG( | Pnisr | 3385.597 | -0.02614 | 0.853752 | 0.973491 | ENSMUSG( | Pnisr | 3642.729 | 0.423496 | 0.040201 | 0.681706 |
| ENSMUSG( | Pnkd | 5792.311 | -0.10383 | 0.287194 | 0.737655 | ENSMUSG( | Pnkd | 5427.692 | -0.01911 | 0.858429 | 0.985884 |
| ENSMUSG( | Pnkp | 762.5934 | 0.007299 | 0.94807 | 0.993821 | ENSMUSG( | Pnkp | 696.794 | 0.016367 | 0.89238 | 0.990558 |
| ENSMUSG( | Pnldc1 | 60.11853 | 0.470081 | 0.065611 | 0.396799 | ENSMUSG( | Pnldc1 | 48.53731 | 0.152646 | 0.63858 | 0.953903 |
| ENSMUSG( | Pnlip | 61.08304 | -0.00125 | 0.99631 | 0.99943 | ENSMUSG( | Pnlip | 46.87665 | -0.5627 | 0.060701 | 0.734317 |
| ENSMUSG( | Pnma1 | 56.16498 | -0.40516 | 0.142998 | 0.561716 | ENSMUSG( | Pnma1 | 51.43636 | -0.38074 | 0.202043 | 0.864412 |
| ENSMUSG( | Pnma2 | 882.1426 | 0.115799 | 0.372442 | 0.798768 | ENSMUSG( | Pnma2 | 768.5459 | -0.00905 | 0.950009 | 0.994798 |
| ENSMUSG( | Pnma3 | 404.1646 | 0.121184 | 0.426693 | 0.825155 | ENSMUSG( | Pnma3 | 343.3315 | -0.07677 | 0.617562 | 0.951952 |
| ENSMUSG( | Pnma8a | 2877.67 | -0.03059 | 0.751435 | 0.950237 | ENSMUSG( | Pnma8a | 2655.956 | 0.00938 | 0.93446 | 0.99306 |
| ENSMUSG( | Pnma8b | 6103.632 | 0.158857 | 0.20234 | 0.642343 | ENSMUSG( | Pnma8b | 5175.286 | -0.04695 | 0.752372 | 0.970493 |
| ENSMUSG( | Pnma8c | 1869.036 | -0.0095 | 0.941514 | 0.993415 | ENSMUSG( | Pnma8c | 1651.286 | -0.10115 | 0.507586 | 0.929089 |
| ENSMUSG( | Pnn | 2018.506 | 0.04501 | 0.68547 | 0.932981 | ENSMUSG( | Pnn | 1947.106 | 0.202603 | 0.22271 | 0.86713 |
| ENSMUSG( | Pno1 | 620.8594 | 0.046209 | 0.702207 | 0.937536 | ENSMUSG( | Pno1 | 524.4287 | -0.18408 | 0.221714 | 0.866104 |
| ENSMUSG( | Pnoc | 326.5301 | -0.71557 | 2.12E-06 | 0.000341 | ENSMUSG( | Pnoc | 336.5415 | -0.28739 | 0.108822 | 0.804969 |
| ENSMUSG( | Pnp | 368.1052 | 0.121851 | 0.370289 | 0.797542 | ENSMUSG( | Pnp | 315.5012 | -0.05209 | 0.719828 | 0.966887 |
| ENSMUSG( | Pnp2 | 22.77906 | -0.46632 | 0.251129 | 0.700723 | ENSMUSG( | Pnp2 | 18.22499 | -0.98402 | 0.048668 | 0.706195 |
| ENSMUSG( | Pnpla1 | 10.30752 | 0.142771 | 0.807096 | NA | ENSMUSG( | Pnpla1 | 8.855757 | -0.09217 | 0.894863 | NA |
| ENSMUSG( | Pnpla2 | 1221.604 | 0.187821 | 0.06607 | 0.397686 | ENSMUSG( | Pnpla2 | 1020.21 | -0.05294 | 0.702125 | 0.962014 |
| ENSMUSG( | Pnpla3 | 100.7158 | -0.01895 | 0.934552 | 0.993231 | ENSMUSG( | Pnpla3 | 97.84796 | 0.180351 | 0.551874 | 0.939565 |
| ENSMUSG( | Pnpla6 | 1782.055 | -0.00774 | 0.957404 | 0.993874 | ENSMUSG( | Pnpla6 | 1777.516 | 0.248134 | 0.117287 | 0.813384 |
| ENSMUSG( | Pnpla7 | 1091.276 | 0.073442 | 0.611849 | 0.908586 | ENSMUSG( | Pnpla7 | 1007.139 | 0.115563 | 0.517314 | 0.930939 |
| ENSMUSG( | Pnpla8 | 5463.382 | 0.150914 | 0.138725 | 0.554336 | ENSMUSG( | Pnpla8 | 4797.707 | 0.047155 | 0.702054 | 0.962014 |
| ENSMUSG( | Pnpo | 755.9631 | -0.09652 | 0.544532 | 0.884649 | ENSMUSG( | Pnpo | 697.9429 | -0.0599 | 0.725208 | 0.966887 |
| ENSMUSG( | Pnpt1 | 788.7652 | 0.103432 | 0.316467 | 0.763626 | ENSMUSG( | Pnpt1 | 731.0112 | 0.15134 | 0.251009 | 0.879208 |
| ENSMUSG( | Pnrc1 | 590.4536 | -0.24862 | 0.073365 | 0.418269 | ENSMUSG( | Pnrc1 | 575.1223 | -0.05015 | 0.760106 | 0.97252 |
| ENSMUSG( | Pnrc2 | 1858.37 | -0.03896 | 0.742764 | 0.948478 | ENSMUSG( | Pnrc2 | 1576.437 | -0.26019 | 0.068984 | 0.754298 |
| ENSMUSG( | Poc1a | 455.6216 | -0.02895 | 0.832877 | 0.970333 | ENSMUSG( | Poc1a | 367.2746 | -0.42156 | 0.012859 | 0.457834 |
| ENSMUSG( | Poc1b | 310.6452 | -0.00612 | 0.966807 | 0.995784 | ENSMUSG( | Poc1b | 293.9848 | 0.089468 | 0.614746 | 0.951177 |
| ENSMUSG( | Poc5 | 792.3596 | 0.089509 | 0.432071 | 0.82768 | ENSMUSG( | Poc5 | 708.3901 | 0.034525 | 0.799671 | 0.97681 |
| ENSMUSG( | Podn | 221.4729 | 0.272065 | 0.126792 | 0.534201 | ENSMUSG( | Podn | 211.97 | 0.406687 | 0.028954 | 0.614171 |
| ENSMUSG( | Podxl | 925.3112 | 0.308309 | 0.373564 | 0.800375 | ENSMUSG( | Podxl | 1111.195 | 0.971733 | 0.036211 | 0.668395 |
| ENSMUSG( | Podxl2 | 6415.244 | -0.08903 | 0.452414 | 0.836145 | ENSMUSG( | Podxl2 | 6199.121 | 0.083511 | 0.497453 | 0.927825 |
| ENSMUSG( | Pofut1 | 1091.257 | -0.14023 | 0.169087 | 0.603241 | ENSMUSG( | Pofut1 | 1041.511 | 0.001195 | 0.992682 | 0.99874 |
| ENSMUSG( | Pofut2 | 1297.777 | -0.30039 | 0.017724 | 0.199784 | ENSMUSG( | Pofut2 | 1357.795 | 0.116059 | 0.422933 | 0.917257 |
| ENSMUSG( | Pogk | 5305.029 | 0.034199 | 0.69149 | 0.933881 | ENSMUSG( | Pogk | 4601.913 | -0.10977 | 0.348033 | 0.911652 |
| ENSMUSG( | Poglut1 | 2136.367 | 0.000709 | 0.994128 | 0.999369 | ENSMUSG( | Poglut1 | 1999.116 | 0.078136 | 0.475912 | 0.923329 |
| ENSMUSG( | Poglut2 | 219.7351 | -0.14077 | 0.451819 | 0.83592 | ENSMUSG( | Poglut2 | 239.604 | 0.373097 | 0.121165 | 0.813384 |
| ENSMUSG( | Poglut3 | 404.72 | -0.06148 | 0.624083 | 0.91187 | ENSMUSG( | Poglut3 | 398.2972 | 0.162427 | 0.282458 | 0.892412 |
| ENSMUSG( | Pogz | 1578.93 | 0.032882 | 0.719563 | 0.941986 | ENSMUSG( | Pogz | 1452.973 | 0.05948 | 0.609997 | 0.950639 |
| ENSMUSG( | Pola1 | 164.6493 | 0.015857 | 0.927027 | 0.991229 | ENSMUSG( | Pola1 | 153.9353 | 0.098894 | 0.625574 | 0.953234 |
| ENSMUSG( | Pola2 | 196.6496 | 0.409351 | 0.019393 | 0.209852 | ENSMUSG( | Pola2 | 167.7725 | 0.236372 | 0.269855 | 0.887508 |
| ENSMUSG( | Polb | 1121.489 | 0.109948 | 0.226939 | 0.674167 | ENSMUSG( | Polb | 1001.657 | 0.0531 | 0.656122 | 0.955533 |
| ENSMUSG( | Pold1 | 218.4983 | 0.401385 | 0.011149 | 0.152217 | ENSMUSG( | Pold1 | 176.7589 | 0.090006 | 0.64974 | 0.95422 |
| ENSMUSG( | Pold2 | 351.3426 | 0.095049 | 0.481769 | 0.855573 | ENSMUSG( | Pold2 | 304.9726 | -0.04459 | 0.772991 | 0.974463 |
| ENSMUSG( | Pold3 | 619.0082 | -0.04419 | 0.726401 | 0.944223 | ENSMUSG( | Pold3 | 549.7107 | -0.11701 | 0.383989 | 0.917257 |
| ENSMUSG( | Pold4 | 304.8972 | 0.058358 | 0.667479 | 0.927404 | ENSMUSG( | Pold4 | 266.0106 | -0.0729 | 0.647928 | 0.95422 |
| ENSMUSG( | Poldip2 | 4423.824 | 0.064371 | 0.404271 | 0.816132 | ENSMUSG( | Poldip2 | 3832.276 | -0.08206 | 0.418384 | 0.917257 |
| ENSMUSG( | Poldip3 | 1605.32 | 0.02746 | 0.769444 | 0.954674 | ENSMUSG( | Poldip3 | 1427.934 | -0.04009 | 0.729076 | 0.966913 |
| ENSMUSG( | Pole | 82.94138 | 0.056166 | 0.811566 | 0.964426 | ENSMUSG( | Pole | 76.19365 | 0.098256 | 0.706489 | 0.963529 |
| ENSMUSG( | Pole2 | 9.073527 | -0.84348 | 0.194892 | NA | ENSMUSG( | Pole2 | 12.47477 | 0.416045 | 0.460972 | 0.923329 |
| ENSMUSG( | Pole3 | 962.614 | 0.135029 | 0.205322 | 0.64654 | ENSMUSG( | Pole3 | 831.9909 | -0.01794 | 0.890345 | 0.990558 |
| ENSMUSG( | Pole4 | 1053.942 | 0.185408 | 0.09474 | 0.472243 | ENSMUSG( | Pole4 | 895.6857 | -0.01477 | 0.917382 | 0.992529 |
| ENSMUSG( | Polg | 2356.599 | 0.082224 | 0.409309 | 0.818258 | ENSMUSG( | Polg | 2191.387 | 0.142846 | 0.296976 | 0.898538 |
| ENSMUSG( | Polg2 | 82.66902 | -0.0719 | 0.748785 | 0.949604 | ENSMUSG( | Polg2 | 74.32983 | -0.11052 | 0.659538 | 0.956205 |
| ENSMUSG( | Polh | 155.5675 | 0.191805 | 0.250757 | 0.700403 | ENSMUSG( | Polh | 151.9409 | 0.383598 | 0.041178 | 0.687358 |
| ENSMUSG( | Poli | 411.9091 | 0.103539 | 0.445072 | 0.833239 | ENSMUSG( | Poli | 363.257 | 0.00756 | 0.960736 | 0.995097 |
| ENSMUSG( | Polk | 240.6796 | -0.16027 | 0.355536 | 0.787945 | ENSMUSG( | Polk | 259.4445 | 0.32509 | 0.120864 | 0.813384 |
| ENSMUSG( | Poll | 348.4316 | 0.046113 | 0.708673 | 0.939519 | ENSMUSG( | Poll | 297.7893 | -0.14759 | 0.32729 | 0.905995 |
| ENSMUSG( | Polm | 236.3724 | -0.02712 | 0.850828 | 0.972638 | ENSMUSG( | Polm | 218.4705 | 0.022144 | 0.903105 | 0.991089 |
| ENSMUSG( | Poln | 12.60981 | -0.13707 | 0.812278 | NA | ENSMUSG( | Poln | 13.27282 | 0.256679 | 0.636656 | 0.953903 |
| ENSMUSG( | Polq | 69.15422 | -0.16948 | 0.462712 | 0.842333 | ENSMUSG( | Polq | 63.44255 | -0.15207 | 0.553865 | 0.939565 |
| ENSMUSG( | Polr1a | 993.5439 | 0.168756 | 0.163267 | 0.594466 | ENSMUSG( | Polr1a | 868.6665 | 0.054139 | 0.69364 | 0.960767 |
| ENSMUSG( | Polr1b | 478.1786 | 0.196898 | 0.134897 | 0.548434 | ENSMUSG( | Polr1b | 429.5823 | 0.16466 | 0.251879 | 0.879902 |
| ENSMUSG( | Polr1c | 459.8444 | -0.04065 | 0.783782 | 0.957645 | ENSMUSG( | Polr1c | 408.7909 | -0.11116 | 0.489751 | 0.925955 |
| ENSMUSG( | Polr1d | 1229.538 | 0.041183 | 0.68709 | 0.933063 | ENSMUSG( | Polr1d | 1071.316 | -0.09097 | 0.44054 | 0.921166 |
| ENSMUSG( | Polr1e | 152.0596 | 0.108546 | 0.546048 | 0.884974 | ENSMUSG( | Polr1e | 126.2454 | -0.15876 | 0.442592 | 0.921554 |
| ENSMUSG( | Polr1f | 293.3986 | -0.0856 | 0.536996 | 0.882123 | ENSMUSG( | Polr1f | 261.0406 | -0.16274 | 0.421678 | 0.917257 |
| ENSMUSG( | Polr1g | 563.4736 | 0.174868 | 0.116218 | 0.514678 | ENSMUSG( | Polr1g | 492.6695 | 0.057576 | 0.669184 | 0.956766 |
| ENSMUSG( | Polr1h | 261.3443 | -0.04044 | 0.795266 | 0.960062 | ENSMUSG( | Polr1h | 240.5177 | -0.01798 | 0.916676 | 0.992529 |
| ENSMUSG( | Polr1has | 83.01392 | -0.03011 | 0.89126 | 0.983819 | ENSMUSG( | Polr1has | 68.21545 | -0.35263 | 0.166773 | 0.840183 |
| ENSMUSG( | Polr2a | 2529.604 | 0.092384 | 0.304766 | 0.752574 | ENSMUSG( | Polr2a | 2358.685 | 0.155475 | 0.178204 | 0.84849 |
| ENSMUSG( | Polr2b | 1735.444 | 0.208765 | 0.017338 | 0.197483 | ENSMUSG( | Polr2b | 1472.735 | 0.012719 | 0.908939 | 0.991958 |
| ENSMUSG( | Polr2c | 951.8225 | -0.0214 | 0.872712 | 0.978753 | ENSMUSG( | Polr2c | 862.1237 | -0.0409 | 0.781925 | 0.976106 |
| ENSMUSG( | Polr2d | 378.1469 | 0.149013 | 0.329083 | 0.773233 | ENSMUSG( | Polr2d | 324.763 | -0.02484 | 0.888102 | 0.990442 |
| ENSMUSG( | Polr2e | 1168.629 | -0.00468 | 0.963301 | 0.994911 | ENSMUSG( | Polr2e | 994.4291 | -0.21122 | 0.083626 | 0.77681 |
| ENSMUSG( | Polr2f | 1098.219 | 0.007058 | 0.948186 | 0.993821 | ENSMUSG( | Polr2f | 906.0328 | -0.30005 | 0.043117 | 0.690304 |
| ENSMUSG( | Polr2g | 1305.09 | 0.043401 | 0.687789 | 0.93324 | ENSMUSG( | Polr2g | 1150.671 | -0.05439 | 0.677702 | 0.957106 |
| ENSMUSG( | Polr2h | 318.3187 | 0.081262 | 0.63779 | 0.91586 | ENSMUSG( | Polr2h | 274.9034 | -0.07712 | 0.688094 | 0.959742 |
| ENSMUSG( | Polr2i | 515.7315 | 0.067489 | 0.561789 | 0.889052 | ENSMUSG( | Polr2i | 436.1749 | -0.15287 | 0.282024 | 0.892412 |
| ENSMUSG( | Polr2j | 591.2538 | 0.035679 | 0.790983 | 0.959461 | ENSMUSG( | Polr2j | 507.5854 | -0.1455 | 0.341975 | 0.909146 |
| ENSMUSG( | Polr2k | 224.7142 | -0.03377 | 0.816134 | 0.965455 | ENSMUSG( | Polr2k | 186.2548 | -0.33192 | 0.059806 | 0.729747 |
| ENSMUSG( | Polr2l | 793.5582 | -0.05442 | 0.634736 | 0.914073 | ENSMUSG( | Polr2l | 670.9668 | -0.29222 | 0.06875 | 0.753052 |
| ENSMUSG( | Polr2m | 9234.436 | 0.131991 | 0.154915 | 0.582458 | ENSMUSG( | Polr2m | 7833.27 | -0.07407 | 0.546282 | 0.938063 |
| ENSMUSG( | Polr3a | 638.2285 | 0.031417 | 0.791863 | 0.959461 | ENSMUSG( | Polr3a | 579.6329 | 0.029248 | 0.854161 | 0.98563 |
| ENSMUSG( | Polr3b | 603.4989 | 0.006905 | 0.951385 | 0.993821 | ENSMUSG( | Polr3b | 552.3989 | 0.021936 | 0.861703 | 0.986307 |
| ENSMUSG( | Polr3c | 652.686 | 0.101961 | 0.320824 | 0.765984 | ENSMUSG( | Polr3c | 591.648 | 0.086546 | 0.480221 | 0.924399 |
| ENSMUSG( | Polr3d | 401.1316 | 0.091838 | 0.467563 | 0.844819 | ENSMUSG( | Polr3d | 355.0569 | 0.01363 | 0.927164 | 0.99306 |
| ENSMUSG( | Polr3e | 809.6964 | -0.00757 | 0.952645 | 0.993874 | ENSMUSG( | Polr3e | 755.4556 | 0.063081 | 0.664872 | 0.956766 |
| ENSMUSG( | Polr3f | 607.2757 | -0.04169 | 0.710362 | 0.939748 | ENSMUSG( | Polr3f | 555.2375 | -0.02975 | 0.819256 | 0.979198 |
| ENSMUSG( | Polr3g | 161.0559 | -0.12103 | 0.502889 | 0.865717 | ENSMUSG( | Polr3g | 154.1042 | 0.003229 | 0.988318 | 0.997862 |
| ENSMUSG( | Polr3gl | 792.0033 | 0.125845 | 0.25597 | 0.706694 | ENSMUSG( | Polr3gl | 712.8202 | 0.084277 | 0.628732 | 0.953251 |
| ENSMUSG( | Polr3h | 2622.53 | 0.044743 | 0.628627 | 0.913658 | ENSMUSG( | Polr3h | 2246.588 | -0.13829 | 0.216757 | 0.865299 |
| ENSMUSG( | Polr3k | 1073.76 | -0.04314 | 0.700262 | 0.937391 | ENSMUSG( | Polr3k | 945.933 | -0.15001 | 0.317545 | 0.902602 |
| ENSMUSG( | Polrmt | 977.7348 | 0.110893 | 0.310293 | 0.758431 | ENSMUSG( | Polrmt | 888.2073 | 0.105588 | 0.399439 | 0.917257 |
| ENSMUSG( | Pom121 | 2363.02 | -0.36699 | 0.039629 | 0.306963 | ENSMUSG( | Pom121 | 2694.064 | 0.295588 | 0.157017 | 0.837724 |
| ENSMUSG( | Pomc | 71.22052 | 0.015532 | 0.946689 | 0.993731 | ENSMUSG( | Pomc | 59.34392 | -0.256 | 0.342907 | 0.909646 |
| ENSMUSG( | Pomgnt1 | 3355.071 | -0.08506 | 0.445433 | 0.833239 | ENSMUSG( | Pomgnt1 | 3234.797 | 0.081408 | 0.538521 | 0.937216 |
| ENSMUSG( | Pomgnt2 | 922.5709 | -0.25449 | 0.063474 | 0.38983 | ENSMUSG( | Pomgnt2 | 1008.858 | 0.276763 | 0.091174 | 0.791889 |
| ENSMUSG( | Pomk | 1094.079 | -0.25092 | 0.085042 | 0.449548 | ENSMUSG( | Pomk | 1134.03 | 0.134844 | 0.417753 | 0.917257 |
| ENSMUSG( | Pomp | 2539.2 | -0.01186 | 0.903464 | 0.986055 | ENSMUSG( | Pomp | 2129.131 | -0.26682 | 0.030695 | 0.628144 |
| ENSMUSG( | Pomt1 | 672.0191 | -0.11399 | 0.457751 | 0.839765 | ENSMUSG( | Pomt1 | 668.6933 | 0.146478 | 0.38234 | 0.917257 |
| ENSMUSG( | Pomt2 | 1186.245 | -0.11785 | 0.414806 | 0.820184 | ENSMUSG( | Pomt2 | 1159.174 | 0.091419 | 0.612614 | 0.951071 |
| ENSMUSG( | Pon2 | 9656.305 | 0.080336 | 0.521973 | 0.875186 | ENSMUSG( | Pon2 | 8518.922 | -0.01235 | 0.929823 | 0.99306 |
| ENSMUSG( | Pon3 | 139.5597 | -0.23901 | 0.24149 | 0.689139 | ENSMUSG( | Pon3 | 128.6282 | -0.18773 | 0.369899 | 0.915491 |
| ENSMUSG( | Pop1 | 228.5752 | 0.271025 | 0.081518 | 0.441102 | ENSMUSG( | Pop1 | 199.6315 | 0.157914 | 0.361685 | 0.914223 |
| ENSMUSG( | Pop4 | 630.6935 | -0.03958 | 0.750837 | 0.950042 | ENSMUSG( | Pop4 | 561.1396 | -0.11463 | 0.424172 | 0.917257 |
| ENSMUSG( | Pop5 | 932.5325 | 0.032515 | 0.777697 | 0.956168 | ENSMUSG( | Pop5 | 848.1128 | 0.023384 | 0.868869 | 0.988097 |
| ENSMUSG( | Pop7 | 254.1215 | -0.07948 | 0.652895 | 0.922632 | ENSMUSG( | Pop7 | 236.694 | -0.02148 | 0.913444 | 0.992298 |
| ENSMUSG( | Popdc2 | 11.0978 | 0.413239 | 0.471139 | NA | ENSMUSG( | Popdc2 | 11.88046 | 0.767898 | 0.185957 | 0.855305 |
| ENSMUSG( | Popdc3 | 27.10604 | 0.032431 | 0.939168 | 0.993415 | ENSMUSG( | Popdc3 | 27.94216 | 0.385248 | 0.360734 | 0.914223 |
| ENSMUSG( | Por | 2953.716 | -0.35197 | 0.003521 | 0.075237 | ENSMUSG( | Por | 3385.048 | 0.317553 | 0.028902 | 0.614171 |
| ENSMUSG( | Porcn | 2318.917 | -0.25752 | 0.031278 | 0.269138 | ENSMUSG( | Porcn | 2370.778 | 0.086805 | 0.485704 | 0.92541 |
| ENSMUSG( | Postn | 82.87729 | -0.25694 | 0.241439 | 0.689139 | ENSMUSG( | Postn | 101.0869 | 0.561227 | 0.017103 | 0.52741 |
| ENSMUSG( | Pot1a | 317.3915 | -0.10197 | 0.478737 | 0.853778 | ENSMUSG( | Pot1a | 312.7037 | 0.135601 | 0.458356 | 0.923329 |
| ENSMUSG( | Pot1b | 182.5412 | 0.170691 | 0.331558 | 0.77519 | ENSMUSG( | Pot1b | 168.7551 | 0.207469 | 0.334347 | 0.907907 |
| ENSMUSG( | Pou2af3 | 83.46781 | 0.075621 | 0.734157 | 0.945382 | ENSMUSG( | Pou2af3 | 72.52444 | -0.07645 | 0.768985 | 0.974236 |
| ENSMUSG( | Pou2f1 | 794.3408 | -0.1694 | 0.147179 | 0.569203 | ENSMUSG( | Pou2f1 | 767.8717 | 0.008953 | 0.945893 | 0.994798 |
| ENSMUSG( | Pou2f2 | 420.0238 | 0.031697 | 0.827282 | 0.968732 | ENSMUSG( | Pou2f2 | 368.9338 | -0.08482 | 0.618405 | 0.952254 |
| ENSMUSG( | Pou3f1 | 1005.143 | 0.135113 | 0.273175 | 0.723531 | ENSMUSG( | Pou3f1 | 878.1947 | 0.01285 | 0.928634 | 0.99306 |
| ENSMUSG( | Pou3f2 | 587.6509 | -0.15275 | 0.148905 | 0.572605 | ENSMUSG( | Pou3f2 | 599.1064 | 0.168569 | 0.22088 | 0.866104 |
| ENSMUSG( | Pou3f3 | 5470.905 | 0.045026 | 0.644934 | 0.91787 | ENSMUSG( | Pou3f3 | 5145.064 | 0.13378 | 0.235925 | 0.87215 |
| ENSMUSG( | Pou3f4 | 195.7541 | -0.31781 | 0.0489 | 0.337919 | ENSMUSG( | Pou3f4 | 199.5386 | 0.032233 | 0.85487 | 0.985649 |
| ENSMUSG( | Pou6f1 | 8807.68 | 0.151537 | 0.072486 | 0.415742 | ENSMUSG( | Pou6f1 | 7813.702 | 0.077626 | 0.476893 | 0.923329 |
| ENSMUSG( | Pou6f2 | 40.57066 | -1.10456 | 0.000452 | 0.019387 | ENSMUSG( | Pou6f2 | 36.14001 | -1.23751 | 0.000562 | 0.072369 |
| ENSMUSG( | Ppa1 | 1232.245 | -0.0156 | 0.894948 | 0.984512 | ENSMUSG( | Ppa1 | 1044.295 | -0.2395 | 0.104159 | 0.804969 |

| | | | | | |
|---|---|---|---|---|---|
| ENSMUSG( | 9022.034 | 0.043888 | 0.59026 | 0.90113 | Pitpna |
| ENSMUSG( | 3127.509 | 0.072824 | 0.382509 | 0.803227 | Pitpnb |
| ENSMUSG( | 3245.036 | 0.118688 | 0.214543 | 0.659193 | Pitpnc1 |
| ENSMUSG( | 10355.28 | 0.198484 | 0.026074 | 0.247245 | Pitpnm1 |
| ENSMUSG( | 9454.564 | 0.250778 | 0.016452 | 0.190546 | Pitpnm2 |
| ENSMUSG( | 5957.641 | 0.138361 | 0.104187 | 0.489971 | Pitpnm3 |
| ENSMUSG( | 1482.284 | 0.149904 | 0.093408 | 0.469793 | Pitrm1 |
| ENSMUSG( | 33.18198 | 0.256306 | 0.450457 | 0.835401 | Piwil2 |
| ENSMUSG( | 2605.59 | -0.12362 | 0.231801 | 0.679725 | Pja1 |
| ENSMUSG( | 6922.792 | -0.05555 | 0.511945 | 0.871327 | Pja2 |
| ENSMUSG( | 4873.364 | -0.11527 | 0.221651 | 0.667258 | Pkd1 |
| ENSMUSG( | 27.67281 | 0.021715 | 0.954038 | 0.993874 | Pkd1l3 |
| ENSMUSG( | 2467.59 | -0.0084 | 0.940333 | 0.993415 | Pkd2 |
| ENSMUSG( | 24.34262 | -0.87156 | 0.03789 | 0.30067 | Pkd2l1 |
| ENSMUSG( | 46.63829 | -0.57876 | 0.070293 | 0.40908 | Pkd2l2 |
| ENSMUSG( | 369.1221 | -0.09748 | 0.517757 | 0.873611 | Pkdcc |
| ENSMUSG( | 41.21298 | 0.298361 | 0.415616 | 0.82084 | Pkdrej |
| ENSMUSG( | 16.64944 | 0.144179 | 0.765824 | NA | Pkhd1l1 |
| ENSMUSG( | 4203.835 | 0.023709 | 0.820495 | 0.965888 | Pkia |
| ENSMUSG( | 127.9835 | -0.13373 | 0.495663 | 0.861237 | Pkib |
| ENSMUSG( | 3338.024 | 0.075686 | 0.346708 | 0.779432 | Pkig |
| ENSMUSG( | 26282.16 | -0.01809 | 0.842747 | 0.971663 | Pkm |
| ENSMUSG( | 162.6713 | -0.21272 | 0.253913 | 0.704011 | Pkmyt1 |
| ENSMUSG( | 3789.106 | 0.059679 | 0.466167 | 0.844453 | Pkn1 |
| ENSMUSG( | 1033.842 | 0.057005 | 0.631405 | 0.914073 | Pkn2 |
| ENSMUSG( | 37.42968 | 0.125083 | 0.726235 | 0.944223 | Pkn3 |
| ENSMUSG( | 685.4738 | 0.122944 | 0.272674 | 0.722557 | Pknox1 |
| ENSMUSG( | 1520.112 | 0.161717 | 0.148838 | 0.572605 | Pknox2 |
| ENSMUSG( | 16.40002 | 0.242656 | 0.610195 | NA | Pkp1 |
| ENSMUSG( | 202.2546 | 0.599904 | 0.004895 | 0.092653 | Pkp2 |
| ENSMUSG( | 17277.05 | 0.08508 | 0.246547 | 0.694958 | Pkp4 |
| ENSMUSG( | 52.95319 | 0.151161 | 0.624908 | 0.912167 | Pla1a |
| ENSMUSG( | 617.0078 | -0.29477 | 0.017683 | 0.199454 | Pla2g12a |
| ENSMUSG( | 935.6799 | -0.22679 | 0.200198 | 0.640302 | Pla2g15 |
| ENSMUSG( | 12.30132 | -0.43888 | 0.444953 | NA | Pla2g2f |
| ENSMUSG( | 177.7508 | 0.16596 | 0.324398 | 0.769598 | Pla2g3 |
| ENSMUSG( | 130.7308 | -0.44031 | 0.028732 | 0.258426 | Pla2g4a |
| ENSMUSG( | 20.92289 | 0.322651 | 0.472338 | NA | Pla2g4b |
| ENSMUSG( | 10.36229 | 0.536967 | 0.37419 | NA | Pla2g4c |
| ENSMUSG( | 417.629 | -0.28303 | 0.04506 | 0.325595 | Pla2g4e |
| ENSMUSG( | 179.8486 | -0.21996 | 0.169398 | 0.603554 | Pla2g5 |
| ENSMUSG( | 718.0013 | 0.013426 | 0.915491 | 0.988634 | Pla2g6 |
| ENSMUSG( | 12888 | -0.08786 | 0.514887 | 0.872306 | Pla2g7 |
| ENSMUSG( | 21.47986 | -0.13035 | 0.752614 | 0.950536 | Pla2r1 |
| ENSMUSG( | 1227.406 | 0.107107 | 0.248152 | 0.696853 | Plaa |
| ENSMUSG( | 404.8416 | -0.06469 | 0.674007 | 0.929626 | Plaat1 |
| ENSMUSG( | 2719.946 | -0.02536 | 0.828634 | 0.969097 | Plaat3 |
| ENSMUSG( | 26.54671 | -0.8906 | 0.099388 | 0.48179 | Plac8 |
| ENSMUSG( | 374.2895 | -0.27911 | 0.065696 | 0.397014 | Plac9 |
| ENSMUSG( | 83.4431 | -0.04621 | 0.857618 | 0.974667 | Plag1 |
| ENSMUSG( | 674.342 | 0.0298 | 0.84479 | 0.972023 | Plagl1 |
| ENSMUSG( | 203.8513 | -0.44162 | 0.002554 | 0.059956 | Plagl2 |
| ENSMUSG( | 1867.393 | -0.38962 | 0.015581 | 0.184473 | Plat |
| ENSMUSG( | 14.27124 | -0.30942 | 0.543436 | NA | Platr14 |
| ENSMUSG( | 224.8899 | 0.05833 | 0.695879 | 0.934979 | Platr25 |
| ENSMUSG( | 138.9241 | -0.26516 | 0.179197 | 0.616715 | Plau |
| ENSMUSG( | 41.42849 | -0.36833 | 0.246987 | 0.695749 | Plaur |
| ENSMUSG( | 135.3207 | 0.055889 | 0.752819 | 0.950621 | Plb1 |
| ENSMUSG( | 29.57865 | -0.73259 | 0.149184 | 0.573248 | Plbd1 |
| ENSMUSG( | 2778.523 | -0.23031 | 0.095627 | 0.475062 | Plbd2 |
| ENSMUSG( | 4724.588 | 0.083088 | 0.410857 | 0.818585 | Plcb1 |
| ENSMUSG( | 170.829 | -0.20979 | 0.228208 | 0.675678 | Plcb2 |
| ENSMUSG( | 2187.298 | 0.07432 | 0.497597 | 0.862196 | Plcb3 |
| ENSMUSG( | 1045.574 | 0.039683 | 0.7431 | 0.948478 | Plcb4 |
| ENSMUSG( | 635.2078 | -0.02819 | 0.830616 | 0.96974 | Plcd1 |
| ENSMUSG( | 764.5742 | 0.151209 | 0.24583 | 0.693547 | Plcd3 |
| ENSMUSG( | 1191.983 | 0.207823 | 0.091933 | 0.465937 | Plcd4 |
| ENSMUSG( | 948.2447 | 0.183771 | 0.20462 | 0.645534 | Plce1 |
| ENSMUSG( | 3247.314 | 0.197033 | 0.043231 | 0.318825 | Plcg1 |
| ENSMUSG( | 272.4842 | 0.146344 | 0.296031 | 0.744386 | Plcg2 |
| ENSMUSG( | 265.5551 | -0.01264 | 0.927294 | 0.991229 | Plch1 |
| ENSMUSG( | 3078.715 | 0.321333 | 0.000787 | 0.027699 | Plch2 |
| ENSMUSG( | 760.1925 | 0.043153 | 0.674209 | 0.929638 | Plcl1 |
| ENSMUSG( | 1346.658 | 0.310973 | 0.013658 | 0.170671 | Plcl2 |
| ENSMUSG( | 682.5671 | -0.24325 | 0.033457 | 0.280305 | Plcxd1 |
| ENSMUSG( | 2587.782 | 0.24794 | 0.01778 | 0.2 | Plcxd2 |
| ENSMUSG( | 31.47808 | -0.34834 | 0.441123 | 0.832445 | Plcxd3 |
| ENSMUSG( | 503.1961 | 0.276683 | 0.013736 | 0.171117 | Pld1 |
| ENSMUSG( | 1588.281 | 0.123936 | 0.207033 | 0.649503 | Pld2 |
| ENSMUSG( | 33142.5 | -0.2085 | 0.096518 | 0.476986 | Pld3 |
| ENSMUSG( | 751.2501 | -0.22135 | 0.130766 | 0.542059 | Pld4 |
| ENSMUSG( | 182.7708 | -0.36418 | 0.047227 | 0.332553 | Pld5 |
| ENSMUSG( | 28.52375 | -0.37732 | 0.291564 | 0.741366 | Pld6 |
| ENSMUSG( | 7925.559 | -0.2142 | 0.046607 | 0.330354 | Plec |
| ENSMUSG( | 629.0488 | -0.26071 | 0.065889 | 0.397536 | Plek |
| ENSMUSG( | 3604.157 | 9.67E-05 | 0.999138 | 0.99989 | Plekha1 |
| ENSMUSG( | 577.297 | -0.14332 | 0.22476 | 0.671058 | Plekha2 |
| ENSMUSG( | 1242.996 | 0.037521 | 0.703696 | 0.937885 | Plekha3 |
| ENSMUSG( | 101.5576 | 0.070994 | 0.749842 | 0.949664 | Plekha4 |
| ENSMUSG( | 2471.972 | 0.095835 | 0.377773 | 0.801561 | Plekha5 |
| ENSMUSG( | 2809.866 | 0.013126 | 0.915803 | 0.988634 | Plekha6 |
| ENSMUSG( | 1022.194 | 0.019771 | 0.845849 | 0.972198 | Plekha7 |
| ENSMUSG( | 433.0845 | -0.00122 | 0.992489 | 0.999327 | Plekha8 |
| ENSMUSG( | 42141.84 | -0.00227 | 0.979698 | 0.997428 | Plekhb1 |
| ENSMUSG( | 11829.53 | 0.072841 | 0.475813 | 0.850919 | Plekhb2 |
| ENSMUSG( | 151.7425 | -0.39498 | 0.03337 | 0.279991 | Plekhd1 |
| ENSMUSG( | 160.3688 | -0.18856 | 0.313308 | 0.76033 | Plekhf1 |
| ENSMUSG( | 379.4272 | 0.073264 | 0.599555 | 0.903874 | Plekhf2 |
| ENSMUSG( | 1162.533 | 0.230755 | 0.015402 | 0.18329 | Plekhg1 |
| ENSMUSG( | 310.0196 | 0.08855 | 0.53083 | 0.879289 | Plekhg2 |
| ENSMUSG( | 810.7649 | -0.32647 | 0.002319 | 0.05646 | Plekhg3 |
| ENSMUSG( | 101.6073 | 0.963001 | 2.58E-05 | 0.002351 | Plekhg4 |
| ENSMUSG( | 2548.884 | 0.082207 | 0.520533 | 0.875186 | Plekhg5 |
| ENSMUSG( | 945.2187 | -0.43349 | 1.57E-05 | 0.001615 | Plekhh1 |
| ENSMUSG( | 330.7372 | -0.21702 | 0.109956 | 0.500285 | Plekhh2 |
| ENSMUSG( | 405.7217 | -0.12142 | 0.403602 | 0.816132 | Plekhh3 |
| ENSMUSG( | 980.4452 | -0.17619 | 0.075808 | 0.426666 | Plekhj1 |
| ENSMUSG( | 1066.026 | 0.257572 | 0.019009 | 0.207571 | Plekhm1 |
| ENSMUSG( | 10372.37 | 0.187652 | 0.024392 | 0.238772 | Plekhm2 |
| ENSMUSG( | 844.2516 | -0.08003 | 0.466438 | 0.844661 | Plekhm3 |
| ENSMUSG( | 310.5406 | -0.13782 | 0.348712 | 0.781339 | Plekhn1 |
| ENSMUSG( | 1395.057 | -0.04803 | 0.633514 | 0.914073 | Plekho1 |
| ENSMUSG( | 1353.054 | -0.21405 | 0.062707 | 0.387171 | Plekho2 |
| ENSMUSG( | 16.03835 | -0.40472 | 0.370292 | NA | Plekhs1 |
| ENSMUSG( | 403.1945 | -0.08422 | 0.533237 | 0.879905 | Plgrkt |
| ENSMUSG( | 356.5948 | 0.000902 | 0.994871 | 0.999369 | Plin2 |
| ENSMUSG( | 731.0681 | 0.002817 | 0.980087 | 0.997491 | Plin3 |
| ENSMUSG( | 70.92371 | 1.574707 | 6.02E-06 | 0.000775 | Plin4 |
| ENSMUSG( | 17.44454 | 0.119706 | 0.793254 | NA | Plin5 |
| ENSMUSG( | 91.26287 | 0.09346 | 0.685876 | 0.932981 | Plk1 |
| ENSMUSG( | 2839.994 | -0.05611 | 0.732006 | 0.944956 | Plk2 |
| ENSMUSG( | 439.2999 | -0.49166 | 7.64E-05 | 0.005386 | Plk3 |
| ENSMUSG( | 91.18762 | -0.06929 | 0.765339 | 0.954127 | Plk4 |
| ENSMUSG( | 326.5313 | -0.07532 | 0.687966 | 0.93324 | Plk5 |
| ENSMUSG( | 3563.962 | -0.72481 | 3.74E-08 | 1.24E-05 | Pllp |
| ENSMUSG( | 1489.99 | -0.39295 | 0.01026 | 0.144326 | Plod1 |
| ENSMUSG( | 311.5179 | -0.06308 | 0.670748 | 0.928683 | Plod2 |
| ENSMUSG( | 1251.904 | -0.11396 | 0.391854 | 0.809274 | Plod3 |
| ENSMUSG( | 125848 | -0.45502 | 0.000294 | 0.014126 | Plp1 |
| ENSMUSG( | 201.3654 | -0.49802 | 0.003464 | 0.074411 | Plp2 |
| ENSMUSG( | 1757.903 | -0.00216 | 0.982779 | 0.997522 | Plpbp |
| ENSMUSG( | 590.933 | -0.29808 | 0.021308 | 0.222975 | Plpp1 |
| ENSMUSG( | 147.9045 | -0.36408 | 0.05753 | 0.370143 | Plpp2 |
| ENSMUSG( | 21119.23 | -0.27773 | 0.048819 | 0.337919 | Plpp3 |
| ENSMUSG( | 257.9324 | -0.15704 | 0.394686 | 0.810072 | Plpp4 |
| ENSMUSG( | 698.5524 | -0.30101 | 0.00498 | 0.093596 | Plpp5 |
| ENSMUSG( | 1545.544 | -0.39808 | 0.000716 | 0.025854 | Plpp6 |
| ENSMUSG( | 793.5395 | -0.35285 | 0.017267 | 0.196955 | Plpp7 |
| ENSMUSG( | 337.2408 | -1.04938 | 4.93E-11 | 3.07E-08 | Plppr1 |
| ENSMUSG( | 8621.794 | -0.18227 | 0.11486 | 0.510978 | Plppr2 |
| ENSMUSG( | 2044.582 | -0.31429 | 0.000179 | 0.009711 | Plppr3 |
| ENSMUSG( | 4956.41 | -0.51567 | 0.004665 | 0.089972 | Plppr4 |
| ENSMUSG( | 1513.753 | -0.61072 | 2.05E-05 | 0.001993 | Plppr5 |
| ENSMUSG( | 663.5968 | 0.206839 | 0.072718 | 0.416435 | Plrg1 |
| ENSMUSG( | 214.0531 | -0.20211 | 0.183655 | 0.620678 | Pls1 |
| ENSMUSG( | 3440.344 | -0.01184 | 0.883431 | 0.982465 | Pls3 |
| ENSMUSG( | 76.18093 | -0.01483 | 0.95365 | 0.993874 | Plscr1 |
| ENSMUSG( | 41.02159 | -0.80997 | 0.006434 | 0.110086 | Plscr2 |
| ENSMUSG( | 565.7219 | -0.04465 | 0.749081 | 0.949604 | Plscr3 |
| ENSMUSG( | 135.7151 | 0.087172 | 0.673096 | 0.929007 | Plscr4 |
| ENSMUSG( | 3387.373 | -0.06563 | 0.72598 | 0.944223 | Pltp |
| ENSMUSG( | 221.3893 | -0.25489 | 0.113251 | 0.507541 | Plvap |
| ENSMUSG( | 959.9187 | -0.5157 | 0.000446 | 0.01925 | Plxdc1 |
| ENSMUSG( | 1342.258 | -0.21377 | 0.201657 | 0.641362 | Plxdc2 |
| ENSMUSG( | 3376.439 | 0.018328 | 0.902821 | 0.986055 | Plxna1 |
| ENSMUSG( | 3437.431 | -0.23928 | 0.098713 | 0.480598 | Plxna2 |
| ENSMUSG( | 881.7129 | 0.192409 | 0.056188 | 0.36528 | Plxna3 |
| ENSMUSG( | 2444.3 | -0.35748 | 0.033595 | 0.28103 | Plxna4 |

| | | | | | |
|---|---|---|---|---|---|
| ENSMUSG( | 698.9982 | -0.01253 | 0.921726 | 0.992644 | Plpp5 |
| ENSMUSG( | 1614.981 | 0.027509 | 0.834236 | 0.981237 | Plpp6 |
| ENSMUSG( | 853.004 | 0.149206 | 0.385467 | 0.917257 | Plpp7 |
| ENSMUSG( | 417.5301 | 0.026238 | 0.878104 | 0.989644 | Plppr1 |
| ENSMUSG( | 8734.73 | 0.129696 | 0.305909 | 0.898538 | Plppr2 |
| ENSMUSG( | 2050.298 | -0.02317 | 0.851581 | 0.985541 | Plppr3 |
| ENSMUSG( | 6215.79 | 0.427208 | 0.024629 | 0.592568 | Plppr4 |
| ENSMUSG( | 1763.485 | 0.159424 | 0.336822 | 0.907972 | Plppr5 |
| ENSMUSG( | 564.6791 | 0.015649 | 0.908504 | 0.991958 | Plrg1 |
| ENSMUSG( | 205.0488 | -0.04749 | 0.776754 | 0.975717 | Pls1 |
| ENSMUSG( | 3278.908 | 0.116728 | 0.266227 | 0.886134 | Pls3 |
| ENSMUSG( | 65.8658 | -0.18543 | 0.494417 | 0.927372 | Plscr1 |
| ENSMUSG( | 47.58127 | -0.00838 | 0.977409 | 0.996416 | Plscr2 |
| ENSMUSG( | 518.5 | -0.02817 | 0.858808 | 0.985884 | Plscr3 |
| ENSMUSG( | 138.958 | 0.406701 | 0.058018 | 0.728256 | Plscr4 |
| ENSMUSG( | 3959.469 | 0.60066 | 0.010886 | 0.422321 | Pltp |
| ENSMUSG( | 194.3563 | -0.37532 | 0.031798 | 0.631941 | Plvap |
| ENSMUSG( | 1029.384 | 0.00662 | 0.970083 | 0.99594 | Plxdc1 |
| ENSMUSG( | 1426.762 | 0.235914 | 0.223446 | 0.867404 | Plxdc2 |
| ENSMUSG( | 3412.641 | 0.30684 | 0.094004 | 0.795735 | Plxna1 |
| ENSMUSG( | 3793.029 | 0.312022 | 0.069189 | 0.755467 | Plxna2 |
| ENSMUSG( | 839.6141 | 0.309924 | 0.019082 | 0.549166 | Plxna3 |
| ENSMUSG( | 2747.978 | 0.265963 | 0.154435 | 0.837724 | Plxna4 |
| ENSMUSG( | 15.836 | -0.37265 | 0.445732 | 0.921734 | Plxna4os1 |
| ENSMUSG( | 12658.06 | -0.04893 | 0.646459 | 0.953988 | Plxnb1 |
| ENSMUSG( | 3582.253 | 0.238743 | 0.111652 | 0.808384 | Plxnb2 |
| ENSMUSG( | 1122.68 | 0.201207 | 0.267593 | 0.886134 | Plxnb3 |
| ENSMUSG( | 1509.926 | 0.240626 | 0.072113 | 0.763629 | Plxnc1 |
| ENSMUSG( | 3936.048 | 0.227968 | 0.114872 | 0.811703 | Plxnd1 |
| ENSMUSG( | 160.4897 | -0.23302 | 0.267141 | 0.886134 | Pm20d1 |
| ENSMUSG( | 95.146 | -0.14725 | 0.530547 | 0.934795 | Pm20d2 |
| ENSMUSG( | 52.41637 | -0.04498 | 0.880084 | 0.989644 | Pmaip1 |
| ENSMUSG( | 17.68025 | -0.73707 | 0.155214 | 0.837724 | Pmch |
| ENSMUSG( | 31.93924 | -0.05033 | 0.890239 | 0.990558 | Pmel |
| ENSMUSG( | 2471.323 | 0.04718 | 0.744801 | 0.96951 | Pmepa1 |
| ENSMUSG( | 131.4344 | -0.22855 | 0.248153 | 0.877904 | Pmf1 |
| ENSMUSG( | 10.42845 | -0.31663 | 0.597674 | NA | Pmfbp1 |
| ENSMUSG( | 348.274 | -0.00809 | 0.956935 | 0.994798 | Pml |
| ENSMUSG( | 5405.156 | -0.23995 | 0.041596 | 0.690304 | Pmm1 |
| ENSMUSG( | 646.3685 | 0.021295 | 0.879629 | 0.989644 | Pmm2 |
| ENSMUSG( | 1397.299 | 0.139893 | 0.421336 | 0.917257 | Pmp22 |
| ENSMUSG( | 3052.447 | -0.0477 | 0.628791 | 0.953251 | Pmpca |
| ENSMUSG( | 1114.125 | -0.12111 | 0.272301 | 0.888771 | Pmpcb |
| ENSMUSG( | 83.84736 | -0.10944 | 0.666173 | 0.956766 | Pms1 |
| ENSMUSG( | 371.2944 | -0.05413 | 0.739632 | 0.968611 | Pms2 |
| ENSMUSG( | 220.2022 | 0.013208 | 0.950423 | 0.994798 | Pms2 |
| ENSMUSG( | 1347.517 | -0.22996 | 0.058661 | 0.728256 | Pmvk |
| ENSMUSG( | 1058.811 | -0.03282 | 0.851304 | 0.985541 | Pnck |
| ENSMUSG( | 3642.729 | 0.423496 | 0.040201 | 0.681706 | Pnisr |
| ENSMUSG( | 5427.692 | -0.01911 | 0.858429 | 0.985884 | Pnkd |
| ENSMUSG( | 696.794 | 0.016367 | 0.89238 | 0.990558 | Pnkp |
| ENSMUSG( | 48.53731 | 0.152646 | 0.63858 | 0.953903 | Pnldc1 |
| ENSMUSG( | 46.87665 | -0.5627 | 0.060701 | 0.734317 | Pnlip |
| ENSMUSG( | 51.43636 | -0.38074 | 0.202043 | 0.864412 | Pnma1 |
| ENSMUSG( | 768.5459 | -0.00905 | 0.950009 | 0.994798 | Pnma2 |
| ENSMUSG( | 343.3315 | -0.07677 | 0.617562 | 0.951952 | Pnma3 |
| ENSMUSG( | 2655.956 | 0.00938 | 0.93446 | 0.99306 | Pnma8a |
| ENSMUSG( | 5175.286 | -0.04695 | 0.752372 | 0.970493 | Pnma8b |
| ENSMUSG( | 1651.286 | -0.10115 | 0.507586 | 0.929089 | Pnma8c |
| ENSMUSG( | 1947.106 | 0.202603 | 0.22271 | 0.86713 | Pnn |
| ENSMUSG( | 524.4287 | -0.18408 | 0.221714 | 0.866104 | Pno1 |
| ENSMUSG( | 336.5415 | -0.28739 | 0.108822 | 0.804969 | Pnoc |
| ENSMUSG( | 315.5012 | -0.05209 | 0.719828 | 0.966887 | Pnp |
| ENSMUSG( | 18.22499 | -0.98402 | 0.048668 | 0.706195 | Pnp2 |
| ENSMUSG( | 8.855757 | -0.09217 | 0.894863 | NA | Pnpla1 |
| ENSMUSG( | 1020.21 | -0.05294 | 0.702125 | 0.962014 | Pnpla2 |
| ENSMUSG( | 97.84796 | 0.180351 | 0.551874 | 0.939565 | Pnpla3 |
| ENSMUSG( | 1777.516 | 0.248134 | 0.117287 | 0.813384 | Pnpla6 |
| ENSMUSG( | 1007.139 | 0.115563 | 0.517314 | 0.930939 | Pnpla7 |
| ENSMUSG( | 4797.707 | 0.047155 | 0.702054 | 0.962014 | Pnpla8 |
| ENSMUSG( | 697.9429 | -0.0599 | 0.725208 | 0.966887 | Pnpo |
| ENSMUSG( | 731.0112 | 0.15134 | 0.251009 | 0.879208 | Pnpt1 |
| ENSMUSG( | 575.1223 | -0.05015 | 0.760106 | 0.97252 | Pnrc1 |
| ENSMUSG( | 1576.437 | -0.26019 | 0.068984 | 0.754298 | Pnrc2 |
| ENSMUSG( | 367.2746 | -0.42156 | 0.012859 | 0.457834 | Poc1a |
| ENSMUSG( | 293.9848 | 0.089468 | 0.614746 | 0.951177 | Poc1b |
| ENSMUSG( | 708.3901 | 0.034525 | 0.799671 | 0.97681 | Poc5 |
| ENSMUSG( | 211.97 | 0.406687 | 0.028954 | 0.614171 | Podn |
| ENSMUSG( | 1111.195 | 0.971733 | 0.036211 | 0.668395 | Podxl |
| ENSMUSG( | 6199.121 | 0.083511 | 0.497453 | 0.927825 | Podxl2 |
| ENSMUSG( | 1041.511 | 0.001195 | 0.992682 | 0.99874 | Pofut1 |
| ENSMUSG( | 1357.795 | 0.116059 | 0.422933 | 0.917257 | Pofut2 |
| ENSMUSG( | 4601.913 | -0.10977 | 0.348033 | 0.911652 | Pogk |
| ENSMUSG( | 1999.116 | 0.078136 | 0.475912 | 0.923329 | Poglut1 |
| ENSMUSG( | 239.604 | 0.373097 | 0.121165 | 0.813384 | Poglut2 |
| ENSMUSG( | 398.2972 | 0.162427 | 0.282458 | 0.892412 | Poglut3 |
| ENSMUSG( | 1452.973 | 0.05948 | 0.609997 | 0.950639 | Pogz |
| ENSMUSG( | 153.9353 | 0.098894 | 0.625574 | 0.953234 | Pola1 |
| ENSMUSG( | 167.7725 | 0.236372 | 0.269855 | 0.887508 | Pola2 |
| ENSMUSG( | 1001.657 | 0.0531 | 0.656122 | 0.955533 | Polb |
| ENSMUSG( | 176.7589 | 0.090006 | 0.64974 | 0.95422 | Pold1 |
| ENSMUSG( | 304.9726 | -0.04459 | 0.772991 | 0.974463 | Pold2 |
| ENSMUSG( | 549.7107 | -0.11701 | 0.383989 | 0.917257 | Pold3 |
| ENSMUSG( | 266.0106 | -0.0729 | 0.647928 | 0.95422 | Pold4 |
| ENSMUSG( | 3832.276 | -0.08206 | 0.418384 | 0.917257 | Poldip2 |
| ENSMUSG( | 1427.934 | -0.04009 | 0.729076 | 0.966913 | Poldip3 |
| ENSMUSG( | 76.19365 | 0.098256 | 0.706489 | 0.963529 | Pole |
| ENSMUSG( | 12.47477 | 0.416045 | 0.460972 | 0.923329 | Pole2 |
| ENSMUSG( | 831.9909 | -0.01794 | 0.890345 | 0.990558 | Pole3 |
| ENSMUSG( | 895.6857 | -0.01477 | 0.917382 | 0.992529 | Pole4 |
| ENSMUSG( | 2191.387 | 0.142846 | 0.296976 | 0.898538 | Polg |
| ENSMUSG( | 74.32983 | -0.11052 | 0.659538 | 0.956205 | Polg2 |
| ENSMUSG( | 151.9409 | 0.383598 | 0.041178 | 0.687358 | Polh |
| ENSMUSG( | 363.257 | 0.00756 | 0.960736 | 0.995097 | Poli |
| ENSMUSG( | 259.4445 | 0.32509 | 0.120864 | 0.813384 | Polk |
| ENSMUSG( | 297.7893 | -0.14759 | 0.32729 | 0.905995 | Poll |
| ENSMUSG( | 218.4705 | 0.022144 | 0.903105 | 0.991089 | Polm |
| ENSMUSG( | 13.27282 | 0.256679 | 0.636656 | 0.953903 | Poln |
| ENSMUSG( | 63.44255 | -0.15207 | 0.553865 | 0.939565 | Polq |
| ENSMUSG( | 868.6665 | 0.054139 | 0.69364 | 0.960767 | Polr1a |
| ENSMUSG( | 429.5823 | 0.16466 | 0.251879 | 0.879902 | Polr1b |
| ENSMUSG( | 408.7909 | -0.11116 | 0.489751 | 0.925955 | Polr1c |
| ENSMUSG( | 1071.316 | -0.09097 | 0.44054 | 0.921166 | Polr1d |
| ENSMUSG( | 126.2454 | -0.15876 | 0.442592 | 0.921554 | Polr1e |
| ENSMUSG( | 261.0406 | -0.16274 | 0.421678 | 0.917257 | Polr1f |
| ENSMUSG( | 492.6695 | 0.057576 | 0.669184 | 0.956766 | Polr1g |
| ENSMUSG( | 240.5177 | -0.01798 | 0.916676 | 0.992529 | Polr1h |
| ENSMUSG( | 68.21545 | -0.35263 | 0.166773 | 0.840183 | Polr1has |
| ENSMUSG( | 2358.685 | 0.155475 | 0.178204 | 0.84849 | Polr2a |
| ENSMUSG( | 1472.735 | 0.012719 | 0.908939 | 0.991958 | Polr2b |
| ENSMUSG( | 862.1237 | -0.0409 | 0.781925 | 0.976106 | Polr2c |
| ENSMUSG( | 324.763 | -0.02484 | 0.888102 | 0.990442 | Polr2d |
| ENSMUSG( | 994.4291 | -0.21122 | 0.083626 | 0.77681 | Polr2e |
| ENSMUSG( | 906.0328 | -0.30005 | 0.043117 | 0.690304 | Polr2f |
| ENSMUSG( | 1150.671 | -0.05439 | 0.677702 | 0.957106 | Polr2g |
| ENSMUSG( | 274.9034 | -0.07712 | 0.688094 | 0.959742 | Polr2h |
| ENSMUSG( | 436.1749 | -0.15287 | 0.282024 | 0.892412 | Polr2i |
| ENSMUSG( | 507.5854 | -0.1455 | 0.341975 | 0.909146 | Polr2j |
| ENSMUSG( | 186.2548 | -0.33192 | 0.059806 | 0.729747 | Polr2k |
| ENSMUSG( | 670.9668 | -0.29222 | 0.06875 | 0.753052 | Polr2l |
| ENSMUSG( | 7833.27 | -0.07407 | 0.546282 | 0.938063 | Polr2m |
| ENSMUSG( | 579.6329 | 0.029248 | 0.854161 | 0.98563 | Polr3a |
| ENSMUSG( | 552.3989 | 0.021936 | 0.861703 | 0.986307 | Polr3b |
| ENSMUSG( | 591.648 | 0.086546 | 0.480221 | 0.924399 | Polr3c |
| ENSMUSG( | 355.0569 | 0.01363 | 0.927164 | 0.99306 | Polr3d |
| ENSMUSG( | 755.4556 | 0.063081 | 0.664872 | 0.956766 | Polr3e |
| ENSMUSG( | 555.2375 | -0.02975 | 0.819256 | 0.979198 | Polr3f |
| ENSMUSG( | 154.1042 | 0.003229 | 0.988318 | 0.997862 | Polr3g |
| ENSMUSG( | 712.8202 | 0.084277 | 0.628732 | 0.953251 | Polr3gl |
| ENSMUSG( | 2246.588 | -0.13829 | 0.216757 | 0.865299 | Polr3h |
| ENSMUSG( | 945.933 | -0.15001 | 0.317545 | 0.902602 | Polr3k |
| ENSMUSG( | 888.2073 | 0.105588 | 0.399439 | 0.917257 | Polrmt |
| ENSMUSG( | 2694.064 | 0.295588 | 0.157017 | 0.837724 | Pom121 |
| ENSMUSG( | 59.34392 | -0.256 | 0.342907 | 0.909646 | Pomc |
| ENSMUSG( | 3234.797 | 0.081408 | 0.538521 | 0.937216 | Pomgnt1 |
| ENSMUSG( | 1008.858 | 0.276763 | 0.091174 | 0.791889 | Pomgnt2 |
| ENSMUSG( | 1134.03 | 0.134844 | 0.417753 | 0.917257 | Pomk |
| ENSMUSG( | 2129.131 | -0.26682 | 0.030695 | 0.628144 | Pomp |
| ENSMUSG( | 668.6933 | 0.146478 | 0.38234 | 0.917257 | Pomt1 |
| ENSMUSG( | 1159.174 | 0.091419 | 0.612614 | 0.951071 | Pomt2 |
| ENSMUSG( | 8518.922 | -0.01235 | 0.929823 | 0.99306 | Pon2 |
| ENSMUSG( | 128.6282 | -0.18773 | 0.369899 | 0.915491 | Pon3 |
| ENSMUSG( | 199.6315 | 0.157914 | 0.361685 | 0.914223 | Pop1 |
| ENSMUSG( | 561.1396 | -0.11463 | 0.424172 | 0.917257 | Pop4 |
| ENSMUSG( | 848.1128 | 0.023384 | 0.868869 | 0.988097 | Pop5 |
| ENSMUSG( | 236.694 | -0.02148 | 0.913444 | 0.992298 | Pop7 |
| ENSMUSG( | 11.88046 | 0.767898 | 0.185957 | 0.855305 | Popdc2 |
| ENSMUSG( | 27.94216 | 0.385248 | 0.360734 | 0.914223 | Popdc3 |

| | | | | | | | | | | |
|---|---|---|---|---|---|---|---|---|---|---|
| ENSMUSG( | Ppa2 | 1225.17 | 0.112356 | 0.254205 | 0.704461 | ENSMUSG( | Ppa2 | 1020.859 | -0.15059 | 0.206962 | 0.864412 |
| ENSMUSG( | Ppan | 489.2214 | -0.12986 | 0.339753 | 0.777939 | ENSMUSG( | Ppan | 431.7109 | -0.22722 | 0.108506 | 0.804969 |
| ENSMUSG( | Ppara | 288.7945 | 0.432922 | 0.00694 | 0.114527 | ENSMUSG( | Ppara | 247.9875 | 0.274471 | 0.124567 | 0.81516 |
| ENSMUSG( | Ppard | 1691.757 | 0.18764 | 0.067271 | 0.40086 | ENSMUSG( | Ppard | 1492.047 | 0.10167 | 0.41216 | 0.917257 |
| ENSMUSG( | Pparg | 110.5834 | -0.10228 | 0.632479 | 0.914073 | ENSMUSG( | Pparg | 102.5038 | -0.06422 | 0.792298 | 0.97681 |
| ENSMUSG( | Ppargc1a | 1317.161 | 0.044658 | 0.651367 | 0.921451 | ENSMUSG( | Ppargc1a | 1158.658 | -0.06077 | 0.625071 | 0.953234 |
| ENSMUSG( | Ppargc1b | 62.19183 | 0.112725 | 0.647544 | 0.919716 | ENSMUSG( | Ppargc1b | 60.343 | 0.289254 | 0.284837 | 0.892412 |
| ENSMUSG( | Ppat | 265.89 | 0.184183 | 0.183754 | 0.620678 | ENSMUSG( | Ppat | 243.5687 | 0.198347 | 0.215417 | 0.865167 |
| ENSMUSG( | Ppcdc | 451.8386 | -0.01197 | 0.915816 | 0.988634 | ENSMUSG( | Ppcdc | 416.7575 | 0.029156 | 0.846923 | 0.985027 |
| ENSMUSG( | Ppcs | 107.2534 | -0.10202 | 0.658889 | 0.924506 | ENSMUSG( | Ppcs | 96.09535 | -0.15275 | 0.539221 | 0.937216 |
| ENSMUSG( | Ppdpf | 2130.708 | 0.0911 | 0.404548 | 0.816132 | ENSMUSG( | Ppdpf | 1783.093 | -0.16019 | 0.180695 | 0.850372 |
| ENSMUSG( | Ppef1 | 15.03243 | -0.36116 | 0.473801 | NA | ENSMUSG( | Ppef1 | 15.63384 | 0.066189 | 0.893938 | 0.990558 |
| ENSMUSG( | Ppef2 | 24.53845 | -0.08564 | 0.821724 | 0.96635 | ENSMUSG( | Ppef2 | 26.49744 | 0.424479 | 0.328435 | 0.906727 |
| ENSMUSG( | Ppfia1 | 2329.201 | -0.06727 | 0.484348 | 0.856665 | ENSMUSG( | Ppfia1 | 2054.762 | -0.16806 | 0.17053 | 0.841739 |
| ENSMUSG( | Ppfia2 | 2102.909 | 0.072504 | 0.46915 | 0.845876 | ENSMUSG( | Ppfia2 | 1781.337 | -0.14509 | 0.277474 | 0.891468 |
| ENSMUSG( | Ppfia3 | 15123.75 | 0.064208 | 0.424737 | 0.824675 | ENSMUSG( | Ppfia3 | 12183.29 | -0.30834 | 0.002634 | 0.202395 |
| ENSMUSG( | Ppfia4 | 3724.345 | 0.105523 | 0.239763 | 0.68758 | ENSMUSG( | Ppfia4 | 3205.952 | -0.0561 | 0.612637 | 0.951071 |
| ENSMUSG( | Ppfibp1 | 1537.428 | 0.012048 | 0.905006 | 0.986449 | ENSMUSG( | Ppfibp1 | 1372.824 | -0.04768 | 0.67892 | 0.957617 |
| ENSMUSG( | Ppfibp2 | 369.3578 | -0.16587 | 0.212385 | 0.65642 | ENSMUSG( | Ppfibp2 | 335.8162 | -0.15992 | 0.36996 | 0.915491 |
| ENSMUSG( | Pphln1 | 842.2058 | -0.06146 | 0.576038 | 0.895114 | ENSMUSG( | Pphln1 | 752.0265 | -0.1262 | 0.341096 | 0.90819 |
| ENSMUSG( | Ppia | 15832.13 | -0.01979 | 0.849212 | 0.972638 | ENSMUSG( | Ppia | 13679.51 | -0.18111 | 0.183118 | 0.850844 |
| ENSMUSG( | Ppib | 4829.897 | -0.20009 | 0.059671 | 0.377567 | ENSMUSG( | Ppib | 4872.506 | 0.101446 | 0.401473 | 0.917257 |
| ENSMUSG( | Ppic | 199.8695 | -0.17273 | 0.372041 | 0.798768 | ENSMUSG( | Ppic | 211.4257 | 0.267525 | 0.231998 | 0.868507 |
| ENSMUSG( | Ppid | 1329.861 | -0.0287 | 0.775422 | 0.955606 | ENSMUSG( | Ppid | 1252.762 | 0.064743 | 0.590349 | 0.944975 |
| ENSMUSG( | Ppie | 313.1766 | 0.04926 | 0.71268 | 0.940652 | ENSMUSG( | Ppie | 280.4473 | -0.00482 | 0.976524 | 0.996416 |
| ENSMUSG( | Ppif | 582.2214 | -0.01484 | 0.910396 | 0.9881 | ENSMUSG( | Ppif | 520.2651 | -0.07038 | 0.642475 | 0.953903 |
| ENSMUSG( | Ppig | 1507.504 | 0.065688 | 0.466923 | 0.844819 | ENSMUSG( | Ppig | 1368.402 | 0.051782 | 0.676475 | 0.95694 |
| ENSMUSG( | Ppih | 165.637 | 0.061887 | 0.752047 | 0.950417 | ENSMUSG( | Ppih | 157.1867 | 0.159923 | 0.47472 | 0.923329 |
| ENSMUSG( | Ppil1 | 519.4482 | -0.04913 | 0.701613 | 0.937441 | ENSMUSG( | Ppil1 | 463.0828 | -0.12349 | 0.488009 | 0.925453 |
| ENSMUSG( | Ppil2 | 1668.345 | 0.202968 | 0.055978 | 0.364712 | ENSMUSG( | Ppil2 | 1467.201 | 0.105778 | 0.375003 | 0.915491 |
| ENSMUSG( | Ppil3 | 313.0348 | -0.02883 | 0.827975 | 0.968966 | ENSMUSG( | Ppil3 | 280.5358 | -0.08214 | 0.595809 | 0.946047 |
| ENSMUSG( | Ppil4 | 944.3626 | 0.072689 | 0.474018 | 0.849425 | ENSMUSG( | Ppil4 | 852.9079 | 0.048118 | 0.714024 | 0.964986 |
| ENSMUSG( | Ppil6 | 121.4489 | 0.333732 | 0.065089 | 0.395697 | ENSMUSG( | Ppil6 | 93.09906 | -0.12933 | 0.588566 | 0.944975 |
| ENSMUSG( | Ppip5k1 | 2920.929 | 0.214035 | 0.04981 | 0.340731 | ENSMUSG( | Ppip5k1 | 2470.695 | 0.006695 | 0.9597 | 0.994798 |
| ENSMUSG( | Ppip5k2 | 577.1014 | -0.09138 | 0.509189 | 0.869008 | ENSMUSG( | Ppip5k2 | 588.7888 | 0.233335 | 0.126985 | 0.817278 |
| ENSMUSG( | Ppl | 24.29392 | 0.08418 | 0.848043 | 0.972374 | ENSMUSG( | Ppl | 21.55089 | 0.023361 | 0.957449 | 0.994798 |
| ENSMUSG( | Ppm1a | 6734.738 | 0.095373 | 0.203694 | 0.643787 | ENSMUSG( | Ppm1a | 5977.262 | 0.020316 | 0.84928 | 0.985321 |
| ENSMUSG( | Ppm1b | 3885.87 | 0.103224 | 0.275969 | 0.725669 | ENSMUSG( | Ppm1b | 3310.725 | -0.09311 | 0.485513 | 0.92541 |
| ENSMUSG( | Ppm1d | 364.706 | 0.002072 | 0.987262 | 0.998777 | ENSMUSG( | Ppm1d | 337.3364 | 0.04043 | 0.789142 | 0.976593 |
| ENSMUSG( | Ppm1e | 4440.623 | 0.139774 | 0.229478 | 0.67735 | ENSMUSG( | Ppm1e | 3919.366 | 0.048115 | 0.726401 | 0.966887 |
| ENSMUSG( | Ppm1f | 2584.838 | 0.226295 | 0.013564 | 0.170154 | ENSMUSG( | Ppm1f | 2220.731 | 0.06805 | 0.571071 | 0.942052 |
| ENSMUSG( | Ppm1g | 2110.578 | -0.10034 | 0.326307 | 0.771301 | ENSMUSG( | Ppm1g | 1988.659 | -0.00196 | 0.98695 | 0.997485 |
| ENSMUSG( | Ppm1h | 2330.798 | 0.146471 | 0.137666 | 0.552903 | ENSMUSG( | Ppm1h | 2099.601 | 0.117421 | 0.314098 | 0.900949 |
| ENSMUSG( | Ppm1j | 21.64896 | -0.71011 | 0.096135 | 0.476417 | ENSMUSG( | Ppm1j | 20.65985 | -0.57101 | 0.191985 | 0.859497 |
| ENSMUSG( | Ppm1k | 1683.806 | 0.088046 | 0.342296 | 0.778828 | ENSMUSG( | Ppm1k | 1482.382 | -0.01324 | 0.915932 | 0.992529 |
| ENSMUSG( | Ppm1l | 3861.726 | -0.43462 | 0.002269 | 0.055678 | ENSMUSG( | Ppm1l | 4227.758 | 0.129677 | 0.445167 | 0.921734 |
| ENSMUSG( | Ppm1m | 1092.926 | 0.106905 | 0.377353 | 0.801213 | ENSMUSG( | Ppm1m | 934.502 | -0.07819 | 0.586662 | 0.944975 |
| ENSMUSG( | Ppme1 | 4227.213 | 0.010842 | 0.924567 | 0.99114 | ENSMUSG( | Ppme1 | 3645.518 | -0.15229 | 0.211568 | 0.864412 |
| ENSMUSG( | Ppox | 829.7584 | 0.14584 | 0.317654 | 0.763989 | ENSMUSG( | Ppox | 763.4113 | 0.174789 | 0.374959 | 0.915491 |
| ENSMUSG( | Ppp1ca | 6751.158 | -0.0647 | 0.447476 | 0.834051 | ENSMUSG( | Ppp1ca | 5821.795 | -0.23254 | 0.030816 | 0.629149 |
| ENSMUSG( | Ppp1cb | 5300.713 | 0.034028 | 0.786405 | 0.958843 | ENSMUSG( | Ppp1cb | 4786.941 | 0.005026 | 0.976824 | 0.996416 |
| ENSMUSG( | Ppp1cc | 1670.96 | -0.11062 | 0.214865 | 0.659433 | ENSMUSG( | Ppp1cc | 1593.07 | 0.022119 | 0.843777 | 0.983462 |
| ENSMUSG( | Ppp1r10 | 1370.207 | 0.175608 | 0.167189 | 0.599823 | ENSMUSG( | Ppp1r10 | 1209.031 | 0.088105 | 0.541247 | 0.937693 |
| ENSMUSG( | Ppp1r11 | 1317.58 | -0.06351 | 0.542625 | 0.883495 | ENSMUSG( | Ppp1r11 | 1124.701 | -0.26788 | 0.090879 | 0.791074 |
| ENSMUSG( | Ppp1r12a | 1990.942 | 0.019854 | 0.813909 | 0.964574 | ENSMUSG( | Ppp1r12a | 1848.749 | 0.074589 | 0.486257 | 0.92541 |
| ENSMUSG( | Ppp1r12b | 6174.255 | 0.176197 | 0.066266 | 0.398381 | ENSMUSG( | Ppp1r12b | 5542.905 | 0.137213 | 0.192567 | 0.859764 |
| ENSMUSG( | Ppp1r12c | 4522.104 | 0.075864 | 0.414632 | 0.820184 | ENSMUSG( | Ppp1r12c | 3973.699 | -0.02886 | 0.787288 | 0.97637 |
| ENSMUSG( | Ppp1r13b | 1898.273 | 0.153154 | 0.166975 | 0.599655 | ENSMUSG( | Ppp1r13b | 1740.01 | 0.173476 | 0.214945 | 0.865167 |
| ENSMUSG( | Ppp1r13l | 141.0344 | 0.002128 | 0.990143 | 0.999144 | ENSMUSG( | Ppp1r13l | 131.7356 | 0.071387 | 0.723869 | 0.966887 |
| ENSMUSG( | Ppp1r14a | 491.2186 | -0.33921 | 0.004553 | 0.088443 | ENSMUSG( | Ppp1r14a | 459.3456 | -0.2516 | 0.070317 | 0.758336 |
| ENSMUSG( | Ppp1r14b | 1150.093 | 0.054611 | 0.58802 | 0.900712 | ENSMUSG( | Ppp1r14b | 1039.053 | 0.023908 | 0.883714 | 0.989644 |
| ENSMUSG( | Ppp1r14c | 466.2906 | 0.069874 | 0.595475 | 0.903058 | ENSMUSG( | Ppp1r14c | 400.0204 | -0.11331 | 0.482032 | 0.924578 |
| ENSMUSG( | Ppp1r15a | 526.5428 | -0.09007 | 0.496144 | 0.861554 | ENSMUSG( | Ppp1r15a | 492.2758 | -0.01081 | 0.942831 | 0.994774 |
| ENSMUSG( | Ppp1r15b | 2104.932 | -0.26187 | 0.060757 | 0.380787 | ENSMUSG( | Ppp1r15b | 2279.736 | 0.242698 | 0.127609 | 0.818199 |
| ENSMUSG( | Ppp1r16a | 4489.704 | 0.137465 | 0.123316 | 0.530552 | ENSMUSG( | Ppp1r16a | 3995.447 | 0.073589 | 0.514952 | 0.930939 |
| ENSMUSG( | Ppp1r16b | 9081.719 | 0.065023 | 0.531025 | 0.879289 | ENSMUSG( | Ppp1r16b | 8054.492 | -0.0095 | 0.941462 | 0.994578 |
| ENSMUSG( | Ppp1r18 | 646.4938 | -0.02065 | 0.841899 | 0.971237 | ENSMUSG( | Ppp1r18 | 586.5756 | -0.03619 | 0.776676 | 0.975717 |
| ENSMUSG( | Ppp1r1a | 4237.8 | 0.021233 | 0.810081 | 0.963864 | ENSMUSG( | Ppp1r1a | 3657.464 | -0.14066 | 0.242993 | 0.877399 |
| ENSMUSG( | Ppp1r1b | 5779.604 | -0.67769 | 0.000686 | 0.025185 | ENSMUSG( | Ppp1r1b | 5886.203 | -0.29581 | 0.151995 | 0.837724 |
| ENSMUSG( | Ppp1r2 | 3296.185 | 0.015572 | 0.876905 | 0.980991 | ENSMUSG( | Ppp1r2 | 2956.96 | -0.03256 | 0.811535 | 0.979107 |
| ENSMUSG( | Ppp1r21 | 2154.062 | 0.121026 | 0.224104 | 0.669929 | ENSMUSG( | Ppp1r21 | 1820.112 | -0.09754 | 0.457897 | 0.923329 |
| ENSMUSG( | Ppp1r26 | 816.8861 | 0.220503 | 0.028134 | 0.256512 | ENSMUSG( | Ppp1r26 | 705.2111 | 0.073663 | 0.5415 | 0.937841 |
| ENSMUSG( | Ppp1r35 | 270.0105 | -0.08713 | 0.566367 | 0.89163 | ENSMUSG( | Ppp1r35 | 245.9915 | -0.08996 | 0.574609 | 0.942912 |
| ENSMUSG( | Ppp1r36 | 54.6368 | -0.37419 | 0.15147 | 0.576717 | ENSMUSG( | Ppp1r36 | 59.09228 | 0.137349 | 0.610938 | 0.950771 |
| ENSMUSG( | Ppp1r36dn | 12.94644 | -0.02934 | 0.955017 | NA | ENSMUSG( | Ppp1r36dn | 11.0863 | -0.23042 | 0.686037 | NA |
| ENSMUSG( | Ppp1r37 | 5855.451 | 0.206197 | 0.011857 | 0.157504 | ENSMUSG( | Ppp1r37 | 5109.27 | 0.088329 | 0.397851 | 0.917257 |
| ENSMUSG( | Ppp1r3b | 66.58213 | 0.081757 | 0.7464 | 0.949289 | ENSMUSG( | Ppp1r3b | 67.9988 | 0.378902 | 0.210822 | 0.864412 |
| ENSMUSG( | Ppp1r3c | 6201.186 | 0.202625 | 0.085523 | 0.450845 | ENSMUSG( | Ppp1r3c | 4889.878 | -0.21709 | 0.129555 | 0.820557 |
| ENSMUSG( | Ppp1r3d | 402.3089 | 0.125552 | 0.333117 | 0.775602 | ENSMUSG( | Ppp1r3d | 338.725 | -0.10972 | 0.477177 | 0.923329 |
| ENSMUSG( | Ppp1r3e | 465.1078 | -0.00961 | 0.937937 | 0.993415 | ENSMUSG( | Ppp1r3e | 417.0857 | -0.06269 | 0.725024 | 0.966887 |
| ENSMUSG( | Ppp1r3f | 209.8516 | 0.114611 | 0.470304 | 0.84674 | ENSMUSG( | Ppp1r3f | 177.1098 | -0.10811 | 0.540371 | 0.937383 |
| ENSMUSG( | Ppp1r3g | 677.1121 | 0.308407 | 0.011612 | 0.1554 | ENSMUSG( | Ppp1r3g | 547.0478 | -0.01898 | 0.885448 | 0.989688 |
| ENSMUSG( | Ppp1r42 | 15.11729 | 0.046918 | 0.921114 | NA | ENSMUSG( | Ppp1r42 | 13.79684 | 0.001609 | 0.997672 | 0.999268 |
| ENSMUSG( | Ppp1r7 | 2887.22 | 0.006018 | 0.942361 | 0.993415 | ENSMUSG( | Ppp1r7 | 2574.391 | -0.05744 | 0.591434 | 0.944975 |
| ENSMUSG( | Ppp1r8 | 977.7155 | 0.04804 | 0.629095 | 0.913838 | ENSMUSG( | Ppp1r8 | 862.7639 | -0.04579 | 0.704343 | 0.962677 |
| ENSMUSG( | Ppp1r9a | 9982.593 | -0.20027 | 0.063066 | 0.388234 | ENSMUSG( | Ppp1r9a | 9982.833 | 0.076582 | 0.516235 | 0.930939 |
| ENSMUSG( | Ppp1r9b | 88729.42 | 0.236378 | 0.005714 | 0.101299 | ENSMUSG( | Ppp1r9b | 78048.11 | 0.141663 | 0.188212 | 0.85698 |
| ENSMUSG( | Ppp2ca | 9199.29 | 0.030755 | 0.70257 | 0.937735 | ENSMUSG( | Ppp2ca | 7879.277 | -0.15325 | 0.212665 | 0.864412 |
| ENSMUSG( | Ppp2cb | 3833.143 | 0.041705 | 0.597597 | 0.903694 | ENSMUSG( | Ppp2cb | 3294.946 | -0.13035 | 0.251803 | 0.879902 |
| ENSMUSG( | Ppp2r1a | 13257.27 | -0.11551 | 0.25096 | 0.700615 | ENSMUSG( | Ppp2r1a | 11747.37 | -0.20126 | 0.100629 | 0.801444 |
| ENSMUSG( | Ppp2r1b | 524.9434 | 0.049056 | 0.663972 | 0.926511 | ENSMUSG( | Ppp2r1b | 458.2748 | -0.08319 | 0.569379 | 0.941777 |
| ENSMUSG( | Ppp2r2a | 1984.782 | -0.04978 | 0.682666 | 0.93198 | ENSMUSG( | Ppp2r2a | 1800.72 | -0.06397 | 0.646139 | 0.953988 |
| ENSMUSG( | Ppp2r2b | 3380.628 | 0.033375 | 0.720044 | 0.941986 | ENSMUSG( | Ppp2r2b | 2962.36 | -0.08223 | 0.503565 | 0.927825 |
| ENSMUSG( | Ppp2r2c | 40760.15 | 0.167458 | 0.034982 | 0.28772 | ENSMUSG( | Ppp2r2c | 35584.55 | 0.049284 | 0.615225 | 0.951177 |
| ENSMUSG( | Ppp2r2d | 1775.843 | 0.042486 | 0.667205 | 0.92729 | ENSMUSG( | Ppp2r2d | 1579.114 | -0.0286 | 0.806521 | 0.978042 |
| ENSMUSG( | Ppp2r3a | 2002.625 | 0.043756 | 0.653635 | 0.922872 | ENSMUSG( | Ppp2r3a | 1749.709 | -0.07869 | 0.504852 | 0.927825 |
| ENSMUSG( | Ppp2r3c | 390.6557 | 0.20904 | 0.106152 | 0.493488 | ENSMUSG( | Ppp2r3c | 329.3428 | -0.00699 | 0.962591 | 0.995284 |
| ENSMUSG( | Ppp2r3d | 1222.194 | 0.046737 | 0.643388 | 0.917267 | ENSMUSG( | Ppp2r3d | 1074.136 | -0.05721 | 0.641463 | 0.953903 |
| ENSMUSG( | Ppp2r5a | 5072.76 | 0.093385 | 0.350947 | 0.782993 | ENSMUSG( | Ppp2r5a | 4291.179 | -0.12356 | 0.328797 | 0.906727 |
| ENSMUSG( | Ppp2r5b | 4888.478 | 0.076161 | 0.368344 | 0.795863 | ENSMUSG( | Ppp2r5b | 4378.719 | 0.027551 | 0.786002 | 0.976285 |
| ENSMUSG( | Ppp2r5c | 9251.915 | 0.0778 | 0.397371 | 0.812127 | ENSMUSG( | Ppp2r5c | 7984.853 | -0.07965 | 0.492943 | 0.927358 |
| ENSMUSG( | Ppp2r5d | 2713.616 | 0.053965 | 0.577394 | 0.895776 | ENSMUSG( | Ppp2r5d | 2350.798 | -0.09451 | 0.432125 | 0.918158 |
| ENSMUSG( | Ppp2r5e | 1839.384 | 0.072798 | 0.429437 | 0.826446 | ENSMUSG( | Ppp2r5e | 1610.245 | -0.04379 | 0.72016 | 0.966887 |
| ENSMUSG( | Ppp3ca | 20068.6 | 0.014034 | 0.895725 | 0.984527 | ENSMUSG( | Ppp3ca | 18085.37 | -0.02041 | 0.90107 | 0.990558 |
| ENSMUSG( | Ppp3cb | 5794.225 | 0.016564 | 0.862103 | 0.975055 | ENSMUSG( | Ppp3cb | 5233.196 | -0.01088 | 0.931409 | 0.99306 |
| ENSMUSG( | Ppp3cc | 638.3212 | 0.087193 | 0.428924 | 0.826446 | ENSMUSG( | Ppp3cc | 576.7286 | 0.061715 | 0.644705 | 0.953903 |
| ENSMUSG( | Ppp3r1 | 10455.95 | -0.00337 | 0.979259 | 0.997428 | ENSMUSG( | Ppp3r1 | 9151.465 | -0.12576 | 0.490498 | 0.926367 |
| ENSMUSG( | Ppp3r2 | 13.72621 | -0.5265 | 0.316973 | NA | ENSMUSG( | Ppp3r2 | 16.16994 | 0.243981 | 0.630715 | 0.953251 |
| ENSMUSG( | Ppp4c | 836.9251 | -0.04279 | 0.671709 | 0.928781 | ENSMUSG( | Ppp4c | 754.083 | -0.07765 | 0.528834 | 0.934795 |
| ENSMUSG( | Ppp4r1 | 1455.748 | 0.177044 | 0.066248 | 0.398381 | ENSMUSG( | Ppp4r1 | 1233.061 | -0.0291 | 0.822984 | 0.979198 |
| ENSMUSG( | Ppp4r1l-ps | 757.9734 | 0.086987 | 0.403008 | 0.815774 | ENSMUSG( | Ppp4r1l-ps | 700.2347 | 0.122292 | 0.364349 | 0.914223 |
| ENSMUSG( | Ppp4r2 | 1495.722 | 0.144608 | 0.150187 | 0.574666 | ENSMUSG( | Ppp4r2 | 1347.158 | 0.107723 | 0.484871 | 0.92541 |
| ENSMUSG( | Ppp4r3a | 1305.371 | -0.0242 | 0.812714 | 0.964574 | ENSMUSG( | Ppp4r3a | 1260.788 | 0.142198 | 0.217759 | 0.865589 |
| ENSMUSG( | Ppp4r3b | 1457.28 | 0.041922 | 0.70064 | 0.93743 | ENSMUSG( | Ppp4r3b | 1391.764 | 0.170736 | 0.194262 | 0.860096 |
| ENSMUSG( | Ppp4r4 | 881.3327 | 0.053071 | 0.634117 | 0.914073 | ENSMUSG( | Ppp4r4 | 782.7061 | -0.02442 | 0.856932 | 0.985884 |
| ENSMUSG( | Ppp5c | 3594.285 | 0.081178 | 0.473119 | 0.848727 | ENSMUSG( | Ppp5c | 3115.645 | -0.06435 | 0.628967 | 0.953251 |
| ENSMUSG( | Ppp6c | 1165.313 | -0.04157 | 0.743666 | 0.948478 | ENSMUSG( | Ppp6c | 1053.002 | -0.0718 | 0.647223 | 0.95422 |
| ENSMUSG( | Ppp6r1 | 8782.448 | 0.244781 | 0.002019 | 0.051507 | ENSMUSG( | Ppp6r1 | 7286.96 | -0.01481 | 0.878657 | 0.989644 |
| ENSMUSG( | Ppp6r2 | 3128.171 | 0.167807 | 0.105693 | 0.492581 | ENSMUSG( | Ppp6r2 | 2745.481 | 0.064478 | 0.55691 | 0.939565 |
| ENSMUSG( | Ppp6r3 | 2521.2 | 0.096148 | 0.265379 | 0.716109 | ENSMUSG( | Ppp6r3 | 2247.916 | 0.034303 | 0.757438 | 0.971858 |
| ENSMUSG( | Pprc1 | 611.7275 | -0.0062 | 0.953872 | 0.993874 | ENSMUSG( | Pprc1 | 592.7569 | 0.17166 | 0.204957 | 0.864412 |
| ENSMUSG( | Ppt1 | 6202.783 | 0.355822 | 0.001506 | 0.043083 | ENSMUSG( | Ppt1 | 4947.743 | -0.0039 | 0.975796 | 0.996416 |
| ENSMUSG( | Ppt2 | 713.6361 | -0.18131 | 0.143114 | 0.561716 | ENSMUSG( | Ppt2 | 722.2431 | 0.125526 | 0.300079 | 0.898538 |
| ENSMUSG( | Pptc7 | 1772.082 | 0.130827 | 0.153625 | 0.580569 | ENSMUSG( | Pptc7 | 1650.711 | 0.188495 | 0.164098 | 0.838291 |
| ENSMUSG( | Ppwd1 | 203.6485 | -0.05224 | 0.737098 | 0.946839 | ENSMUSG( | Ppwd1 | 202.1678 | 0.187614 | 0.285373 | 0.893136 |
| ENSMUSG( | Pqbp1 | 1417.802 | 0.024415 | 0.834843 | 0.970426 | ENSMUSG( | Pqbp1 | 1202.362 | -0.19346 | 0.217498 | 0.865589 |
| ENSMUSG( | Pradc1 | 427.9864 | -0.19621 | 0.096883 | 0.477375 | ENSMUSG( | Pradc1 | 440.7874 | 0.159097 | 0.22695 | 0.868507 |
| ENSMUSG( | Praf2 | 1885.75 | -0.42926 | 0.000122 | 0.007362 | ENSMUSG( | Praf2 | 1917.451 | -0.08322 | 0.541619 | 0.937841 |
| ENSMUSG( | Prag1 | 626.4675 | -0.01852 | 0.889604 | 0.98353 | ENSMUSG( | Prag1 | 615.115 | 0.195058 | 0.180938 | 0.850372 |
| ENSMUSG( | Pramel12 | 294.1254 | 0.185469 | 0.192693 | 0.631789 | ENSMUSG( | Pramel12 | 262.8252 | 0.136175 | 0.408436 | 0.917257 |
| ENSMUSG( | Prc1 | 75.66289 | -0.3494 | 0.150208 | 0.574666 | ENSMUSG( | Prc1 | 67.30508 | -0.42391 | 0.125589 | 0.81516 |
| ENSMUSG( | Prcc | 1026.123 | 0.156844 | 0.173894 | 0.609325 | ENSMUSG( | Prcc | 892.7232 | 0.024732 | 0.858167 | 0.985884 |
| ENSMUSG( | Prcd | 89.23784 | -0.25048 | 0.33715 | 0.777008 | ENSMUSG( | Prcd | 82.22455 | -0.22233 | 0.384981 | 0.917257 |
| ENSMUSG( | Prcp | 393.0558 | -0.36485 | 0.035046 | 0.287963 | ENSMUSG( | Prcp | 396.6486 | -0.03671 | 0.847838 | 0.985242 |
| ENSMUSG( | Prdm1 | 37.5411 | -0.32818 | 0.282363 | 0.73186 | ENSMUSG( | Prdm1 | 40.80373 | 0.195312 | 0.529786 | 0.934795 |
| ENSMUSG( | Prdm10 | 625.5875 | 0.039179 | 0.722654 | 0.943756 | ENSMUSG( | Prdm10 | 596.0516 | 0.16077 | 0.26161 | 0.884926 |
| ENSMUSG( | Prdm11 | 216.8989 | 0.095001 | 0.538403 | 0.88259 | ENSMUSG( | Prdm11 | 189.2154 | -0.01066 | 0.960178 | 0.994798 |
| ENSMUSG( | Prdm15 | 483.4947 | 0.244263 | 0.062259 | 0.386085 | ENSMUSG( | Prdm15 | 444.5234 | 0.271997 | 0.062485 | 0.736482 |
| ENSMUSG( | Prdm16 | 865.4372 | -0.0766 | 0.556056 | 0.887457 | ENSMUSG( | Prdm16 | 764.3286 | -0.17213 | 0.234059 | 0.869201 |
| ENSMUSG( | Prdm16os | 2316.412 | 0.075951 | 0.546048 | 0.884974 | ENSMUSG( | Prdm16os | 1870.993 | -0.2878 | 0.046449 | 0.697814 |
| ENSMUSG( | Prdm2 | 1547.685 | 0.098675 | 0.341969 | 0.778828 | ENSMUSG( | Prdm2 | 1393.941 | 0.069642 | 0.555803 | 0.939565 |
| ENSMUSG( | Prdm4 | 821.3939 | 0.061221 | 0.558593 | 0.888533 | ENSMUSG( | Prdm4 | 702.7533 | -0.12013 | 0.335341 | 0.907907 |
| ENSMUSG( | Prdm5 | 165.0524 | 0.122286 | 0.45188 | 0.835936 | ENSMUSG( | Prdm5 | 160.95 | 0.299149 | 0.104926 | 0.804969 |
| ENSMUSG( | Prdm6 | 39.07229 | -0.09538 | 0.78399 | 0.957689 | ENSMUSG( | Prdm6 | 38.50046 | 0.112639 | 0.752053 | 0.970461 |
| ENSMUSG( | Prdm8 | 612.3544 | 0.265707 | 0.05671 | 0.367404 | ENSMUSG( | Prdm8 | 548.2761 | 0.214358 | 0.130358 | 0.821987 |
| ENSMUSG( | Prdm9 | 74.15447 | 0.065691 | 0.775966 | 0.955606 | ENSMUSG( | Prdm9 | 64.88744 | -0.05364 | 0.843499 | 0.983392 |
| ENSMUSG( | Prdx1 | 7182.158 | 0.061024 | 0.575611 | 0.894836 | ENSMUSG( | Prdx1 | 5948.43 | -0.22425 | 0.075418 | 0.767994 |
| ENSMUSG( | Prdx2 | 6139.3 | -0.002 | 0.982742 | 0.997522 | ENSMUSG( | Prdx2 | 5175.55 | -0.23884 | 0.059178 | 0.729429 |
| ENSMUSG( | Prdx3 | 1744.35 | 0.076334 | 0.419621 | 0.822309 | ENSMUSG( | Prdx3 | 1473.252 | -0.14953 | 0.293911 | 0.898538 |
| ENSMUSG( | Prdx4 | 680.7719 | -0.14371 | 0.356628 | 0.788187 | ENSMUSG( | Prdx4 | 677.2962 | 0.11398 | 0.491466 | 0.926825 |
| ENSMUSG( | Prdx5 | 4961.911 | -0.02894 | 0.768041 | 0.954621 | ENSMUSG( | Prdx5 | 4231.834 | -0.23179 | 0.080754 | 0.77681 |
| ENSMUSG( | Prdx6 | 17638.88 | 0.082631 | 0.442883 | 0.83278 | ENSMUSG( | Prdx6 | 14873.31 | -0.14441 | 0.253186 | 0.88018 |
| ENSMUSG( | Preb | 2002.198 | 0.114901 | 0.231032 | 0.679229 | ENSMUSG( | Preb | 1735.081 | -0.0262 | 0.820878 | 0.979198 |
| ENSMUSG( | Prelid1 | 3267.176 | 0.028298 | 0.743617 | 0.948478 | ENSMUSG( | Prelid1 | 2783.731 | -0.17178 | 0.142814 | 0.831221 |
| ENSMUSG( | Prelid3a | 1341.57 | 0.088518 | 0.325505 | 0.770741 | ENSMUSG( | Prelid3a | 1163.664 | -0.05504 | 0.64891 | 0.95422 |
| ENSMUSG( | Prelid3b | 1761.12 | 0.028739 | 0.791266 | 0.959461 | ENSMUSG( | Prelid3b | 1521.312 | -0.13262 | 0.371471 | 0.915491 |
| ENSMUSG( | Prelp | 1086.806 | -0.16983 | 0.338076 | 0.777422 | ENSMUSG( | Prelp | 1054.923 | 0.026291 | 0.895104 | 0.990558 |
| ENSMUSG( | Prep | 745.1982 | -0.09854 | 0.342638 | 0.778971 | ENSMUSG( | Prep | 709.4481 | 0.024418 | 0.867431 | 0.987764 |

| | | | | | |
|---|---|---|---|---|---|
| ENSMUSG( | 19.11579 | -0.08709 | 0.842984 | NA | Plxna4os1 |
| ENSMUSG( | 14589.99 | 0.090719 | 0.381346 | 0.802629 | Plxnb1 |
| ENSMUSG( | 3594.584 | -0.01402 | 0.902157 | 0.985633 | Plxnb2 |
| ENSMUSG( | 1107.823 | -0.1066 | 0.356538 | 0.788187 | Plxnb3 |
| ENSMUSG( | 1504.828 | -0.03129 | 0.803037 | 0.962169 | Plxnc1 |
| ENSMUSG( | 4094.574 | 0.077339 | 0.490518 | 0.859387 | Plxnd1 |
| ENSMUSG( | 168.6117 | -0.38028 | 0.052188 | 0.350744 | Pm20d1 |
| ENSMUSG( | 110.8783 | 0.053005 | 0.801917 | 0.962116 | Pm20d2 |
| ENSMUSG( | 56.75089 | -0.10616 | 0.7061 | 0.938843 | Pmaip1 |
| ENSMUSG( | 22.78877 | -0.13457 | 0.751461 | 0.950237 | Pmch |
| ENSMUSG( | 36.50002 | 0.03991 | 0.901154 | 0.985633 | Pmel |
| ENSMUSG( | 2394.937 | -0.33415 | 0.006656 | 0.11214 | Pmepa1 |
| ENSMUSG( | 150.4441 | -0.11294 | 0.533724 | 0.880095 | Pmf1 |
| ENSMUSG( | 389.0303 | 0.040989 | 0.755526 | 0.951395 | Pml |
| ENSMUSG( | 6632.303 | 0.092401 | 0.279365 | 0.729114 | Pmm1 |
| ENSMUSG( | 712.4488 | 0.031776 | 0.781357 | 0.957558 | Pmm2 |
| ENSMUSG( | 1271.575 | -0.43463 | 0.002383 | 0.057268 | Pmp22 |
| ENSMUSG( | 3519.523 | 0.092775 | 0.248338 | 0.697133 | Pmpca |
| ENSMUSG( | 1314.544 | 0.090041 | 0.37507 | 0.800786 | Pmpcb |
| ENSMUSG( | 94.98091 | -0.01044 | 0.960899 | 0.993991 | Pms1 |
| ENSMUSG( | 243.233 | 0.029819 | 0.89568 | 0.984527 | Pms2 |
| ENSMUSG( | 414.7957 | -0.01229 | 0.93423 | 0.99323 | Pms2 |
| ENSMUSG( | 1625.204 | 0.05402 | 0.561442 | 0.889052 | Pmvk |
| ENSMUSG( | 1286.448 | 0.253569 | 0.075827 | 0.426666 | Pnck |
| ENSMUSG( | 3385.597 | -0.02614 | 0.853752 | 0.973491 | Pnisr |
| ENSMUSG( | 5792.311 | -0.10383 | 0.287194 | 0.737655 | Pnkd |
| ENSMUSG( | 762.5934 | 0.007299 | 0.94807 | 0.993821 | Pnkp |
| ENSMUSG( | 60.11853 | 0.470081 | 0.065611 | 0.396799 | Pnldc1 |
| ENSMUSG( | 61.08304 | -0.00125 | 0.99631 | 0.99943 | Pnlip |
| ENSMUSG( | 56.16498 | -0.40516 | 0.142998 | 0.561716 | Pnma1 |
| ENSMUSG( | 882.1426 | 0.115799 | 0.372442 | 0.798768 | Pnma2 |
| ENSMUSG( | 404.1646 | 0.121184 | 0.426693 | 0.825155 | Pnma3 |
| ENSMUSG( | 2877.67 | -0.03059 | 0.751435 | 0.950237 | Pnma8a |
| ENSMUSG( | 6103.632 | 0.158857 | 0.20234 | 0.642343 | Pnma8b |
| ENSMUSG( | 1869.036 | -0.0095 | 0.941514 | 0.993415 | Pnma8c |
| ENSMUSG( | 10.46602 | 0.557478 | 0.330671 | NA | Pnmt |
| ENSMUSG( | 2018.506 | 0.04501 | 0.68547 | 0.932981 | Pnn |
| ENSMUSG( | 620.8594 | 0.046209 | 0.702207 | 0.937536 | Pno1 |
| ENSMUSG( | 326.5301 | -0.71557 | 2.12E-06 | 0.000341 | Pnoc |
| ENSMUSG( | 368.1052 | 0.121851 | 0.370289 | 0.797542 | Pnp |
| ENSMUSG( | 22.77906 | -0.46632 | 0.251129 | 0.700723 | Pnp2 |
| ENSMUSG( | 10.30752 | 0.142771 | 0.807096 | NA | Pnpla1 |
| ENSMUSG( | 1221.604 | 0.187821 | 0.06607 | 0.397686 | Pnpla2 |
| ENSMUSG( | 100.7158 | -0.01895 | 0.934552 | 0.993231 | Pnpla3 |
| ENSMUSG( | 1782.055 | -0.00774 | 0.957404 | 0.993874 | Pnpla6 |
| ENSMUSG( | 1091.276 | 0.073442 | 0.611849 | 0.908586 | Pnpla7 |
| ENSMUSG( | 5463.382 | 0.150914 | 0.138725 | 0.554336 | Pnpla8 |
| ENSMUSG( | 755.9631 | -0.09652 | 0.544532 | 0.884649 | Pnpo |
| ENSMUSG( | 788.7652 | 0.103432 | 0.316467 | 0.763626 | Pnpt1 |
| ENSMUSG( | 590.4536 | -0.24862 | 0.073365 | 0.418269 | Pnrc1 |
| ENSMUSG( | 1858.37 | -0.03896 | 0.742764 | 0.948478 | Pnrc2 |
| ENSMUSG( | 455.6216 | -0.02895 | 0.832877 | 0.970333 | Poc1a |
| ENSMUSG( | 310.6452 | -0.00612 | 0.966807 | 0.995784 | Poc1b |
| ENSMUSG( | 792.3596 | 0.089509 | 0.432071 | 0.82768 | Poc5 |
| ENSMUSG( | 221.4729 | 0.272065 | 0.126792 | 0.534201 | Podn |
| ENSMUSG( | 925.3112 | 0.308309 | 0.373564 | 0.800375 | Podxl |
| ENSMUSG( | 6415.244 | -0.08903 | 0.452414 | 0.836145 | Podxl2 |
| ENSMUSG( | 1091.257 | -0.14023 | 0.169087 | 0.603241 | Pofut1 |
| ENSMUSG( | 1297.777 | -0.30039 | 0.017724 | 0.199784 | Pofut2 |
| ENSMUSG( | 5305.029 | 0.034199 | 0.69149 | 0.933881 | Pogk |
| ENSMUSG( | 2136.367 | 0.000709 | 0.994128 | 0.999369 | Poglut1 |
| ENSMUSG( | 219.7351 | -0.14077 | 0.451819 | 0.83592 | Poglut2 |
| ENSMUSG( | 404.72 | -0.06148 | 0.624083 | 0.91187 | Poglut3 |
| ENSMUSG( | 1578.93 | 0.032882 | 0.719563 | 0.941986 | Pogz |
| ENSMUSG( | 164.6493 | 0.015857 | 0.927027 | 0.991229 | Pola1 |
| ENSMUSG( | 196.6496 | 0.409351 | 0.019393 | 0.209852 | Pola2 |
| ENSMUSG( | 1121.489 | 0.109948 | 0.226939 | 0.674167 | Polb |
| ENSMUSG( | 218.4983 | 0.401385 | 0.011149 | 0.152217 | Pold1 |
| ENSMUSG( | 351.3426 | 0.095049 | 0.481769 | 0.855573 | Pold2 |
| ENSMUSG( | 619.0082 | -0.04419 | 0.726401 | 0.944223 | Pold3 |
| ENSMUSG( | 304.8972 | 0.058358 | 0.667479 | 0.927404 | Pold4 |
| ENSMUSG( | 4423.824 | 0.064371 | 0.404271 | 0.816132 | Poldip2 |
| ENSMUSG( | 1605.32 | 0.02746 | 0.769444 | 0.954674 | Poldip3 |
| ENSMUSG( | 82.94138 | 0.056166 | 0.811566 | 0.964426 | Pole |
| ENSMUSG( | 9.073527 | -0.84348 | 0.194892 | NA | Pole2 |
| ENSMUSG( | 962.614 | 0.135029 | 0.205322 | 0.64654 | Pole3 |
| ENSMUSG( | 1053.942 | 0.185408 | 0.09474 | 0.472243 | Pole4 |
| ENSMUSG( | 2356.599 | 0.082224 | 0.409309 | 0.818258 | Polg |
| ENSMUSG( | 82.66902 | -0.0719 | 0.748785 | 0.949604 | Polg2 |
| ENSMUSG( | 155.5675 | 0.191805 | 0.250757 | 0.700403 | Polh |
| ENSMUSG( | 411.9091 | 0.103539 | 0.445072 | 0.833239 | Poli |
| ENSMUSG( | 240.6796 | -0.16027 | 0.355736 | 0.787945 | Polk |
| ENSMUSG( | 348.4316 | 0.046113 | 0.708673 | 0.939519 | Poll |
| ENSMUSG( | 236.3724 | -0.02712 | 0.850828 | 0.972638 | Polm |
| ENSMUSG( | 12.60981 | -0.13707 | 0.812278 | NA | Poln |
| ENSMUSG( | 69.15422 | -0.16948 | 0.462712 | 0.842333 | Polq |
| ENSMUSG( | 993.5439 | 0.168756 | 0.163267 | 0.594466 | Polr1a |
| ENSMUSG( | 478.1786 | 0.196898 | 0.134897 | 0.548434 | Polr1b |
| ENSMUSG( | 459.8444 | -0.04065 | 0.783782 | 0.957645 | Polr1c |
| ENSMUSG( | 1229.538 | 0.041183 | 0.68709 | 0.933063 | Polr1d |
| ENSMUSG( | 152.0596 | 0.108546 | 0.545048 | 0.884974 | Polr1e |
| ENSMUSG( | 293.3986 | -0.0856 | 0.536996 | 0.882123 | Polr1f |
| ENSMUSG( | 563.4736 | 0.174868 | 0.116218 | 0.514678 | Polr1g |
| ENSMUSG( | 261.3443 | -0.04044 | 0.795266 | 0.960062 | Polr1h |
| ENSMUSG( | 83.01392 | -0.03011 | 0.89126 | 0.983819 | Polr1has |
| ENSMUSG( | 2529.604 | 0.092384 | 0.304766 | 0.752574 | Polr2a |
| ENSMUSG( | 1735.444 | 0.208765 | 0.017338 | 0.197483 | Polr2b |
| ENSMUSG( | 951.8225 | -0.0214 | 0.872712 | 0.978753 | Polr2c |
| ENSMUSG( | 378.1469 | 0.149013 | 0.329083 | 0.773233 | Polr2d |
| ENSMUSG( | 1168.629 | -0.00468 | 0.963301 | 0.994911 | Polr2e |
| ENSMUSG( | 1098.219 | 0.007058 | 0.948186 | 0.993821 | Polr2f |
| ENSMUSG( | 1305.09 | 0.043401 | 0.687789 | 0.93324 | Polr2g |
| ENSMUSG( | 318.3187 | 0.081262 | 0.63779 | 0.91586 | Polr2h |
| ENSMUSG( | 515.7315 | 0.067489 | 0.561789 | 0.889052 | Polr2i |
| ENSMUSG( | 591.2538 | 0.035679 | 0.790983 | 0.959461 | Polr2j |
| ENSMUSG( | 224.7142 | -0.03377 | 0.816134 | 0.965455 | Polr2k |
| ENSMUSG( | 793.5582 | -0.05442 | 0.634736 | 0.914073 | Polr2l |
| ENSMUSG( | 9234.436 | 0.131991 | 0.154915 | 0.582458 | Polr2m |
| ENSMUSG( | 638.2285 | 0.031417 | 0.791863 | 0.959461 | Polr3a |
| ENSMUSG( | 603.4989 | 0.006905 | 0.951385 | 0.993821 | Polr3b |
| ENSMUSG( | 652.686 | 0.101961 | 0.320824 | 0.765984 | Polr3c |
| ENSMUSG( | 401.1316 | 0.091838 | 0.467563 | 0.844819 | Polr3d |
| ENSMUSG( | 809.6964 | -0.00757 | 0.952645 | 0.993874 | Polr3e |
| ENSMUSG( | 607.2757 | -0.04169 | 0.710362 | 0.939748 | Polr3f |
| ENSMUSG( | 161.0559 | -0.12103 | 0.502889 | 0.865717 | Polr3g |
| ENSMUSG( | 792.0033 | 0.125845 | 0.25597 | 0.706694 | Polr3gl |
| ENSMUSG( | 2622.53 | 0.044743 | 0.628627 | 0.913658 | Polr3h |
| ENSMUSG( | 1073.76 | -0.04314 | 0.700262 | 0.937391 | Polr3k |
| ENSMUSG( | 977.7348 | 0.110893 | 0.310293 | 0.758431 | Polrmt |
| ENSMUSG( | 2363.02 | -0.36699 | 0.039629 | 0.306963 | Pom121 |
| ENSMUSG( | 71.22052 | 0.015532 | 0.946689 | 0.993731 | Pomc |
| ENSMUSG( | 3355.071 | -0.08506 | 0.445433 | 0.833239 | Pomgnt1 |
| ENSMUSG( | 922.5709 | -0.25449 | 0.063474 | 0.38983 | Pomgnt2 |
| ENSMUSG( | 1094.079 | -0.25092 | 0.085042 | 0.449548 | Pomk |
| ENSMUSG( | 2539.2 | -0.01186 | 0.903464 | 0.986055 | Pomp |
| ENSMUSG( | 672.0191 | -0.11399 | 0.457751 | 0.839765 | Pomt1 |
| ENSMUSG( | 1186.245 | -0.11785 | 0.414806 | 0.820184 | Pomt2 |
| ENSMUSG( | 9656.305 | 0.080336 | 0.521973 | 0.875186 | Pon2 |
| ENSMUSG( | 139.5597 | -0.23901 | 0.24149 | 0.689139 | Pon3 |
| ENSMUSG( | 228.5752 | 0.271025 | 0.081518 | 0.441102 | Pop1 |
| ENSMUSG( | 630.6935 | -0.03958 | 0.750837 | 0.950042 | Pop4 |
| ENSMUSG( | 932.5325 | 0.032515 | 0.777697 | 0.956168 | Pop5 |
| ENSMUSG( | 254.1215 | -0.07948 | 0.652895 | 0.922632 | Pop7 |
| ENSMUSG( | 11.0978 | 0.413239 | 0.471139 | NA | Popdc2 |
| ENSMUSG( | 27.10604 | 0.032431 | 0.939168 | 0.993415 | Popdc3 |
| ENSMUSG( | 2953.716 | -0.35197 | 0.003521 | 0.075237 | Por |
| ENSMUSG( | 2318.917 | -0.25752 | 0.031278 | 0.269138 | Porcn |
| ENSMUSG( | 82.87729 | -0.25694 | 0.241439 | 0.689139 | Postn |
| ENSMUSG( | 317.3915 | -0.10197 | 0.478737 | 0.853778 | Pot1a |
| ENSMUSG( | 182.5412 | 0.170691 | 0.331558 | 0.77519 | Pot1b |
| ENSMUSG( | 8.314335 | -0.87872 | 0.22804 | NA | Pou2af1 |
| ENSMUSG( | 83.46781 | 0.075621 | 0.734157 | 0.945382 | Pou2af3 |
| ENSMUSG( | 794.3408 | -0.1694 | 0.147179 | 0.569203 | Pou2f1 |
| ENSMUSG( | 420.0238 | 0.031697 | 0.827282 | 0.968732 | Pou2f2 |
| ENSMUSG( | 1005.143 | 0.135113 | 0.273175 | 0.723531 | Pou3f1 |
| ENSMUSG( | 587.6509 | -0.15275 | 0.148905 | 0.572605 | Pou3f2 |
| ENSMUSG( | 5470.905 | 0.045026 | 0.644934 | 0.91787 | Pou3f3 |
| ENSMUSG( | 195.7541 | -0.31781 | 0.0489 | 0.337919 | Pou3f4 |
| ENSMUSG( | 8807.68 | 0.151537 | 0.072486 | 0.415742 | Pou6f1 |
| ENSMUSG( | 40.57066 | -1.10456 | 0.000452 | 0.019387 | Pou6f2 |
| ENSMUSG( | 1232.245 | -0.0156 | 0.894948 | 0.984512 | Ppa1 |
| ENSMUSG( | 1225.17 | 0.112356 | 0.254205 | 0.704461 | Ppa2 |
| ENSMUSG( | 489.2214 | -0.12986 | 0.339753 | 0.777939 | Ppan |
| ENSMUSG( | 288.7945 | 0.432922 | 0.00694 | 0.114527 | Ppara |
| ENSMUSG( | 1691.757 | 0.18764 | 0.067271 | 0.40086 | Ppard |
| ENSMUSG( | 110.5834 | -0.10228 | 0.632479 | 0.914073 | Pparg |
| ENSMUSG( | 1317.161 | 0.044658 | 0.651367 | 0.921451 | Ppargc1a |
| ENSMUSG( | 62.19183 | 0.112725 | 0.647544 | 0.919716 | Ppargc1b |

| | | | | | |
|---|---|---|---|---|---|
| ENSMUSG( | 3385.048 | 0.317553 | 0.028902 | 0.614171 | Por |
| ENSMUSG( | 2370.778 | 0.086805 | 0.485704 | 0.92541 | Porcn |
| ENSMUSG( | 101.0869 | 0.561227 | 0.017103 | 0.52741 | Postn |
| ENSMUSG( | 312.7037 | 0.135601 | 0.458356 | 0.923329 | Pot1a |
| ENSMUSG( | 168.7551 | 0.207469 | 0.334347 | 0.907907 | Pot1b |
| ENSMUSG( | 72.52444 | -0.07645 | 0.768985 | 0.974236 | Pou2af3 |
| ENSMUSG( | 767.8717 | 0.008953 | 0.945893 | 0.994798 | Pou2f1 |
| ENSMUSG( | 368.9338 | -0.08482 | 0.618405 | 0.952254 | Pou2f2 |
| ENSMUSG( | 878.1947 | 0.01285 | 0.928634 | 0.99306 | Pou3f1 |
| ENSMUSG( | 599.1064 | 0.168569 | 0.22088 | 0.866104 | Pou3f2 |
| ENSMUSG( | 5145.064 | 0.13378 | 0.235925 | 0.87215 | Pou3f3 |
| ENSMUSG( | 199.5386 | 0.032233 | 0.85487 | 0.985649 | Pou3f4 |
| ENSMUSG( | 7813.702 | 0.077626 | 0.476893 | 0.923329 | Pou6f1 |
| ENSMUSG( | 36.14001 | -1.23751 | 0.000562 | 0.072369 | Pou6f2 |
| ENSMUSG( | 1044.295 | -0.2395 | 0.104159 | 0.804969 | Ppa1 |
| ENSMUSG( | 1020.859 | -0.15059 | 0.206962 | 0.864412 | Ppa2 |
| ENSMUSG( | 431.7109 | -0.22722 | 0.108506 | 0.804969 | Ppan |
| ENSMUSG( | 247.9875 | 0.274471 | 0.124567 | 0.81516 | Ppara |
| ENSMUSG( | 1492.047 | 0.10167 | 0.41216 | 0.917257 | Ppard |
| ENSMUSG( | 102.5038 | -0.06422 | 0.792298 | 0.97681 | Pparg |
| ENSMUSG( | 1158.658 | -0.06077 | 0.625071 | 0.953234 | Ppargc1a |
| ENSMUSG( | 60.343 | 0.289254 | 0.284837 | 0.892412 | Ppargc1b |
| ENSMUSG( | 243.5687 | 0.198347 | 0.215417 | 0.865167 | Ppat |
| ENSMUSG( | 416.7575 | 0.029156 | 0.846923 | 0.985027 | Ppcdc |
| ENSMUSG( | 96.09535 | -0.15275 | 0.539221 | 0.937216 | Ppcs |
| ENSMUSG( | 1783.093 | -0.16019 | 0.180695 | 0.850372 | Ppdpf |
| ENSMUSG( | 15.63384 | 0.066189 | 0.893938 | 0.990558 | Ppef1 |
| ENSMUSG( | 26.49744 | 0.424479 | 0.328435 | 0.906727 | Ppef2 |
| ENSMUSG( | 2054.762 | -0.16806 | 0.17053 | 0.841739 | Ppfia1 |
| ENSMUSG( | 1781.337 | -0.14509 | 0.277474 | 0.891468 | Ppfia2 |
| ENSMUSG( | 12183.29 | -0.30834 | 0.002634 | 0.202395 | Ppfia3 |
| ENSMUSG( | 3205.952 | -0.0561 | 0.612637 | 0.951071 | Ppfia4 |
| ENSMUSG( | 1372.824 | -0.04768 | 0.67892 | 0.957617 | Ppfibp1 |
| ENSMUSG( | 335.8162 | -0.15992 | 0.36996 | 0.915491 | Ppfibp2 |
| ENSMUSG( | 752.0265 | -0.1262 | 0.341096 | 0.90819 | Pphln1 |
| ENSMUSG( | 13679.51 | -0.18111 | 0.183118 | 0.850844 | Ppia |
| ENSMUSG( | 4872.506 | 0.101446 | 0.401473 | 0.917257 | Ppib |
| ENSMUSG( | 211.4257 | 0.267525 | 0.231998 | 0.868507 | Ppic |
| ENSMUSG( | 1252.762 | 0.064743 | 0.590349 | 0.944975 | Ppid |
| ENSMUSG( | 280.4473 | -0.00482 | 0.976524 | 0.996416 | Ppie |
| ENSMUSG( | 520.2651 | -0.07038 | 0.642475 | 0.953903 | Ppif |
| ENSMUSG( | 1368.402 | 0.051782 | 0.676475 | 0.95694 | Ppig |
| ENSMUSG( | 157.1867 | 0.159923 | 0.47472 | 0.923329 | Ppih |
| ENSMUSG( | 463.0828 | -0.12349 | 0.488009 | 0.925453 | Ppil1 |
| ENSMUSG( | 1467.201 | 0.105778 | 0.375003 | 0.915491 | Ppil2 |
| ENSMUSG( | 280.5358 | -0.08214 | 0.595809 | 0.946047 | Ppil3 |
| ENSMUSG( | 852.9079 | 0.048118 | 0.714024 | 0.964986 | Ppil4 |
| ENSMUSG( | 93.09906 | -0.12933 | 0.588566 | 0.944975 | Ppil6 |
| ENSMUSG( | 2470.695 | 0.006695 | 0.9597 | 0.994798 | Ppip5k1 |
| ENSMUSG( | 588.7888 | 0.233335 | 0.126985 | 0.817278 | Ppip5k2 |
| ENSMUSG( | 21.55089 | 0.023361 | 0.957449 | 0.994798 | Ppl |
| ENSMUSG( | 5977.262 | 0.020316 | 0.84928 | 0.985321 | Ppm1a |
| ENSMUSG( | 3310.725 | -0.09311 | 0.485513 | 0.92541 | Ppm1b |
| ENSMUSG( | 337.3364 | 0.04043 | 0.789142 | 0.976593 | Ppm1d |
| ENSMUSG( | 3919.366 | 0.048115 | 0.726401 | 0.966887 | Ppm1e |
| ENSMUSG( | 2220.731 | 0.06805 | 0.571071 | 0.942052 | Ppm1f |
| ENSMUSG( | 1988.659 | -0.00196 | 0.98695 | 0.997485 | Ppm1g |
| ENSMUSG( | 2099.601 | 0.117421 | 0.314098 | 0.900949 | Ppm1h |
| ENSMUSG( | 20.65985 | -0.57101 | 0.191985 | 0.859497 | Ppm1j |
| ENSMUSG( | 1482.382 | -0.01324 | 0.915932 | 0.992529 | Ppm1k |
| ENSMUSG( | 4227.758 | 0.129677 | 0.445167 | 0.921734 | Ppm1l |
| ENSMUSG( | 934.502 | -0.07819 | 0.586662 | 0.944975 | Ppm1m |
| ENSMUSG( | 3645.518 | -0.15229 | 0.211568 | 0.864412 | Ppme1 |
| ENSMUSG( | 763.4113 | 0.174789 | 0.374959 | 0.915491 | Ppox |
| ENSMUSG( | 5821.795 | -0.23254 | 0.030816 | 0.629149 | Ppp1ca |
| ENSMUSG( | 4786.941 | 0.005026 | 0.976824 | 0.996416 | Ppp1cb |
| ENSMUSG( | 1593.07 | 0.022119 | 0.843777 | 0.983462 | Ppp1cc |
| ENSMUSG( | 1209.031 | 0.088105 | 0.541247 | 0.937693 | Ppp1r10 |
| ENSMUSG( | 1124.701 | -0.26788 | 0.090879 | 0.791074 | Ppp1r11 |
| ENSMUSG( | 1848.749 | 0.074589 | 0.486257 | 0.92541 | Ppp1r12a |
| ENSMUSG( | 5542.905 | 0.137213 | 0.192567 | 0.859764 | Ppp1r12b |
| ENSMUSG( | 3973.699 | -0.02886 | 0.787288 | 0.97637 | Ppp1r12c |
| ENSMUSG( | 1740.01 | 0.173476 | 0.214945 | 0.865167 | Ppp1r13b |
| ENSMUSG( | 131.7356 | 0.071387 | 0.723869 | 0.966887 | Ppp1r13l |
| ENSMUSG( | 459.3456 | -0.2516 | 0.070317 | 0.758336 | Ppp1r14a |
| ENSMUSG( | 1039.053 | 0.023908 | 0.883714 | 0.989644 | Ppp1r14b |
| ENSMUSG( | 400.0204 | -0.11331 | 0.482032 | 0.924578 | Ppp1r14c |
| ENSMUSG( | 492.2758 | -0.01081 | 0.942831 | 0.994774 | Ppp1r15a |
| ENSMUSG( | 2279.736 | 0.242698 | 0.127609 | 0.818199 | Ppp1r15b |
| ENSMUSG( | 3995.447 | 0.073589 | 0.514952 | 0.930939 | Ppp1r16a |
| ENSMUSG( | 8054.492 | -0.0095 | 0.941462 | 0.994578 | Ppp1r16b |
| ENSMUSG( | 586.5756 | -0.03619 | 0.776676 | 0.975717 | Ppp1r18 |
| ENSMUSG( | 3657.464 | -0.14066 | 0.242993 | 0.877399 | Ppp1r1a |
| ENSMUSG( | 5886.203 | -0.29581 | 0.151995 | 0.837724 | Ppp1r1b |
| ENSMUSG( | 2956.96 | -0.03256 | 0.811535 | 0.979107 | Ppp1r2 |
| ENSMUSG( | 1820.112 | -0.09754 | 0.457897 | 0.923329 | Ppp1r21 |
| ENSMUSG( | 705.2111 | 0.073663 | 0.5415 | 0.937841 | Ppp1r26 |
| ENSMUSG( | 245.9915 | -0.08996 | 0.574609 | 0.942912 | Ppp1r35 |
| ENSMUSG( | 59.09228 | 0.137349 | 0.610938 | 0.950771 | Ppp1r36 |
| ENSMUSG( | 11.0863 | -0.23042 | 0.686037 | NA | Ppp1r36dn |
| ENSMUSG( | 5109.27 | 0.088329 | 0.397851 | 0.917257 | Ppp1r37 |
| ENSMUSG( | 67.9988 | 0.378902 | 0.210822 | 0.864412 | Ppp1r3b |
| ENSMUSG( | 4889.878 | -0.21709 | 0.129555 | 0.820557 | Ppp1r3c |
| ENSMUSG( | 338.725 | -0.10972 | 0.477177 | 0.923329 | Ppp1r3d |
| ENSMUSG( | 417.0857 | -0.06269 | 0.725024 | 0.966887 | Ppp1r3e |
| ENSMUSG( | 177.1098 | -0.10811 | 0.540371 | 0.937383 | Ppp1r3f |
| ENSMUSG( | 547.0478 | -0.01898 | 0.885448 | 0.989688 | Ppp1r3g |
| ENSMUSG( | 13.79684 | 0.001609 | 0.997672 | 0.999268 | Ppp1r42 |
| ENSMUSG( | 2574.391 | -0.05744 | 0.591434 | 0.944975 | Ppp1r7 |
| ENSMUSG( | 862.7639 | -0.04579 | 0.704343 | 0.962677 | Ppp1r8 |
| ENSMUSG( | 9982.833 | 0.076582 | 0.516235 | 0.930939 | Ppp1r9a |
| ENSMUSG( | 78048.11 | 0.141663 | 0.188212 | 0.85598 | Ppp1r9b |
| ENSMUSG( | 7879.277 | -0.15325 | 0.212665 | 0.864412 | Ppp2ca |
| ENSMUSG( | 3294.946 | -0.13035 | 0.251803 | 0.879902 | Ppp2cb |
| ENSMUSG( | 11747.37 | -0.2016 | 0.100629 | 0.801444 | Ppp2r1a |
| ENSMUSG( | 458.2748 | -0.08319 | 0.569379 | 0.941777 | Ppp2r1b |
| ENSMUSG( | 1800.72 | -0.06397 | 0.646139 | 0.953988 | Ppp2r2a |
| ENSMUSG( | 2962.36 | -0.08223 | 0.503565 | 0.927825 | Ppp2r2b |
| ENSMUSG( | 35584.55 | 0.049284 | 0.615225 | 0.951177 | Ppp2r2c |
| ENSMUSG( | 1579.114 | -0.0286 | 0.806521 | 0.978042 | Ppp2r2d |
| ENSMUSG( | 1749.709 | -0.07869 | 0.504852 | 0.927825 | Ppp2r3a |
| ENSMUSG( | 329.3428 | -0.00699 | 0.962591 | 0.994798 | Ppp2r3c |
| ENSMUSG( | 1074.136 | -0.05721 | 0.641463 | 0.953903 | Ppp2r3d |
| ENSMUSG( | 4291.179 | -0.12356 | 0.328797 | 0.906727 | Ppp2r5a |
| ENSMUSG( | 4378.719 | 0.027551 | 0.786002 | 0.976285 | Ppp2r5b |
| ENSMUSG( | 7984.853 | -0.07965 | 0.492943 | 0.927358 | Ppp2r5c |
| ENSMUSG( | 2350.798 | -0.09451 | 0.432125 | 0.918158 | Ppp2r5d |
| ENSMUSG( | 1610.245 | -0.04379 | 0.72016 | 0.966887 | Ppp2r5e |
| ENSMUSG( | 18085.37 | -0.02041 | 0.90107 | 0.990558 | Ppp3ca |
| ENSMUSG( | 5233.196 | -0.01088 | 0.931409 | 0.99306 | Ppp3cb |
| ENSMUSG( | 576.7286 | 0.061715 | 0.644705 | 0.953903 | Ppp3cc |
| ENSMUSG( | 9151.465 | -0.12576 | 0.490498 | 0.926367 | Ppp3r1 |
| ENSMUSG( | 16.16994 | 0.243981 | 0.630715 | 0.953251 | Ppp3r2 |
| ENSMUSG( | 754.083 | -0.07765 | 0.528834 | 0.934795 | Ppp4c |
| ENSMUSG( | 1233.061 | -0.0291 | 0.822984 | 0.979198 | Ppp4r1 |
| ENSMUSG( | 700.2347 | 0.122292 | 0.364349 | 0.914223 | Ppp4r1l-ps |
| ENSMUSG( | 1347.158 | 0.107723 | 0.484871 | 0.92541 | Ppp4r2 |
| ENSMUSG( | 1260.788 | 0.142198 | 0.217759 | 0.865589 | Ppp4r3a |
| ENSMUSG( | 1391.764 | 0.170736 | 0.194262 | 0.860096 | Ppp4r3b |
| ENSMUSG( | 782.7061 | -0.02442 | 0.856932 | 0.985884 | Ppp4r4 |
| ENSMUSG( | 3115.645 | -0.06435 | 0.628967 | 0.953251 | Ppp5c |
| ENSMUSG( | 1053.002 | -0.0718 | 0.647223 | 0.95422 | Ppp6c |
| ENSMUSG( | 7286.96 | -0.01481 | 0.878657 | 0.989644 | Ppp6r1 |
| ENSMUSG( | 2745.481 | 0.064478 | 0.55691 | 0.939565 | Ppp6r2 |
| ENSMUSG( | 2247.916 | 0.034303 | 0.757438 | 0.971858 | Ppp6r3 |
| ENSMUSG( | 592.7569 | 0.17166 | 0.204957 | 0.864412 | Pprc1 |
| ENSMUSG( | 4947.743 | -0.0039 | 0.975796 | 0.996416 | Ppt1 |
| ENSMUSG( | 722.2431 | 0.125526 | 0.300079 | 0.898538 | Ppt2 |
| ENSMUSG( | 1650.711 | 0.188495 | 0.164098 | 0.838291 | Pptc7 |
| ENSMUSG( | 202.1678 | 0.187614 | 0.285373 | 0.893136 | Ppwd1 |
| ENSMUSG( | 1202.362 | -0.19346 | 0.217498 | 0.865589 | Pqbp1 |
| ENSMUSG( | 440.7874 | 0.159097 | 0.22695 | 0.868507 | Pradc1 |
| ENSMUSG( | 1917.451 | -0.08322 | 0.541619 | 0.937841 | Praf2 |
| ENSMUSG( | 615.115 | 0.195058 | 0.180938 | 0.850372 | Prag1 |
| ENSMUSG( | 262.8252 | 0.136175 | 0.408436 | 0.917257 | Pramel12 |
| ENSMUSG( | 67.30508 | -0.42391 | 0.125589 | 0.81516 | Prc1 |
| ENSMUSG( | 892.7232 | 0.024732 | 0.858167 | 0.985884 | Prcc |
| ENSMUSG( | 82.22455 | -0.22233 | 0.384981 | 0.917257 | Prcd |
| ENSMUSG( | 396.6486 | -0.03671 | 0.847838 | 0.985242 | Prcp |
| ENSMUSG( | 40.80373 | 0.195312 | 0.529786 | 0.934795 | Prdm1 |
| ENSMUSG( | 596.0516 | 0.16077 | 0.26161 | 0.884926 | Prdm10 |
| ENSMUSG( | 189.2154 | -0.01066 | 0.960178 | 0.994798 | Prdm11 |
| ENSMUSG( | 444.5234 | 0.271997 | 0.062485 | 0.736482 | Prdm15 |
| ENSMUSG( | 764.3286 | -0.17213 | 0.234059 | 0.869201 | Prdm16 |
| ENSMUSG( | 1870.993 | -0.2878 | 0.046449 | 0.697814 | Prdm16os |
| ENSMUSG( | 1393.941 | 0.069642 | 0.555803 | 0.939565 | Prdm2 |
| ENSMUSG( | 702.7533 | -0.12013 | 0.335341 | 0.907907 | Prdm4 |
| ENSMUSG( | 160.95 | 0.299149 | 0.104926 | 0.804969 | Prdm5 |

| | | | | | | | | | | | |
|---|---|---|---|---|---|---|---|---|---|---|---|
| ENSMUSG( | Prepl | 4685.756 | 0.068014 | 0.459617 | 0.840338 | ENSMUSG( | Prepl | 4145.401 | -0.01507 | 0.893183 | 0.990558 |
| ENSMUSG( | Prex1 | 15466.15 | 0.025608 | 0.779549 | 0.956885 | ENSMUSG( | Prex1 | 13089.24 | -0.19402 | 0.093757 | 0.795735 |
| ENSMUSG( | Prex2 | 11927.51 | 0.211428 | 0.121179 | 0.525683 | ENSMUSG( | Prex2 | 9901.9 | -0.05086 | 0.714087 | 0.964986 |
| ENSMUSG( | Prg4 | 292.7796 | 0.581713 | 0.000147 | 0.008459 | ENSMUSG( | Prg4 | 256.0207 | 0.486286 | 0.013304 | 0.461493 |
| ENSMUSG( | Prickle1 | 2111.337 | -0.01258 | 0.908613 | 0.987848 | ENSMUSG( | Prickle1 | 2010.76 | 0.115404 | 0.340425 | 0.907972 |
| ENSMUSG( | Prickle2 | 5992.362 | 0.118408 | 0.164135 | 0.5949 | ENSMUSG( | Prickle2 | 5412.192 | 0.095432 | 0.343079 | 0.909646 |
| ENSMUSG( | Prickle3 | 45.00355 | 0.138407 | 0.647795 | 0.919759 | ENSMUSG( | Prickle3 | 45.22548 | 0.421549 | 0.204577 | 0.864412 |
| ENSMUSG( | Prim1 | 127.2928 | 0.136633 | 0.451265 | 0.835567 | ENSMUSG( | Prim1 | 116.2968 | 0.146147 | 0.494137 | 0.927372 |
| ENSMUSG( | Prim2 | 187.1327 | 0.141736 | 0.365494 | 0.794085 | ENSMUSG( | Prim2 | 156.3347 | -0.10162 | 0.581418 | 0.944266 |
| ENSMUSG( | Prima1 | 116.0028 | -0.7694 | 0.000203 | 0.010726 | ENSMUSG( | Prima1 | 126.3402 | -0.14475 | 0.496397 | 0.927825 |
| ENSMUSG( | Primpol | 177.2422 | 0.015877 | 0.925061 | 0.990436 | ENSMUSG( | Primpol | 165.7149 | 0.097033 | 0.595574 | 0.945931 |
| ENSMUSG( | Prkaa1 | 572.2544 | 0.039244 | 0.716137 | 0.94152 | ENSMUSG( | Prkaa1 | 519.3181 | 0.028403 | 0.822977 | 0.979198 |
| ENSMUSG( | Prkaa2 | 578.3166 | 0.083338 | 0.564006 | 0.890139 | ENSMUSG( | Prkaa2 | 535.4439 | 0.127604 | 0.426162 | 0.917257 |
| ENSMUSG( | Prkab1 | 469.4959 | -0.21725 | 0.106073 | 0.493403 | ENSMUSG( | Prkab1 | 444.097 | -0.09363 | 0.555912 | 0.939565 |
| ENSMUSG( | Prkab2 | 1175.554 | 0.033179 | 0.725521 | 0.944223 | ENSMUSG( | Prkab2 | 1059.011 | 0.002408 | 0.983937 | 0.996963 |
| ENSMUSG( | Prkaca | 4501.927 | 0.002178 | 0.985245 | 0.997982 | ENSMUSG( | Prkaca | 3951.991 | -0.0585 | 0.426727 | 0.917257 |
| ENSMUSG( | Prkacb | 9100.655 | 0.02566 | 0.793781 | 0.959799 | ENSMUSG( | Prkacb | 7966.513 | -0.09287 | 0.459551 | 0.923329 |
| ENSMUSG( | Prkag1 | 1485.578 | 0.052241 | 0.560345 | 0.888934 | ENSMUSG( | Prkag1 | 1287.99 | -0.09305 | 0.405147 | 0.917257 |
| ENSMUSG( | Prkag2 | 2634.399 | 0.175984 | 0.039116 | 0.304587 | ENSMUSG( | Prkag2 | 2255.104 | 0.000511 | 0.996194 | 0.999178 |
| ENSMUSG( | Prkar1a | 13770.48 | 0.0234 | 0.760773 | 0.952907 | ENSMUSG( | Prkar1a | 12276.99 | -0.04038 | 0.69607 | 0.960867 |
| ENSMUSG( | Prkar1b | 16443.69 | 0.100284 | 0.378751 | 0.801688 | ENSMUSG( | Prkar1b | 13672.94 | -0.16774 | 0.178952 | 0.849041 |
| ENSMUSG( | Prkar2a | 1589.5 | 0.09452 | 0.305405 | 0.753523 | ENSMUSG( | Prkar2a | 1486.326 | 0.164595 | 0.158884 | 0.837724 |
| ENSMUSG( | Prkar2b | 1013.742 | -0.3035 | 0.010086 | 0.143118 | ENSMUSG( | Prkar2b | 1002.405 | -0.05758 | 0.727994 | 0.966913 |
| ENSMUSG( | Prkca | 4336.61 | 2.76E-05 | 0.999734 | 0.99989 | ENSMUSG( | Prkca | 3898.416 | -0.039 | 0.691329 | 0.960071 |
| ENSMUSG( | Prkcb | 16174.43 | 0.073 | 0.409662 | 0.818258 | ENSMUSG( | Prkcb | 14551.67 | 0.036883 | 0.764973 | 0.974236 |
| ENSMUSG( | Prkcd | 934.6838 | -0.29071 | 0.075523 | 0.425931 | ENSMUSG( | Prkcd | 845.3422 | -0.31995 | 0.088983 | 0.788051 |
| ENSMUSG( | Prkce | 9754.211 | 0.12529 | 0.118329 | 0.520133 | ENSMUSG( | Prkce | 8828.534 | 0.106895 | 0.377891 | 0.916784 |
| ENSMUSG( | Prkcg | 9208.465 | 0.212728 | 0.0437 | 0.320345 | ENSMUSG( | Prkcg | 7956.973 | 0.064708 | 0.680192 | 0.9581 |
| ENSMUSG( | Prkch | 176.0884 | -0.36993 | 0.047529 | 0.333288 | ENSMUSG( | Prkch | 204.0216 | 0.340829 | 0.091452 | 0.791889 |
| ENSMUSG( | Prkci | 1429.979 | 0.060598 | 0.55076 | 0.886038 | ENSMUSG( | Prkci | 1253.856 | -0.05334 | 0.661825 | 0.956205 |
| ENSMUSG( | Prkcq | 944.2275 | -0.01102 | 0.926875 | 0.991229 | ENSMUSG( | Prkcq | 833.4965 | -0.10934 | 0.404104 | 0.917257 |
| ENSMUSG( | Prkcsh | 3360.084 | -0.21514 | 0.090662 | 0.462287 | ENSMUSG( | Prkcsh | 3549.749 | 0.21594 | 0.13944 | 0.830678 |
| ENSMUSG( | Prkcz | 10732.83 | 0.095822 | 0.241598 | 0.689179 | ENSMUSG( | Prkcz | 9634.887 | 0.055283 | 0.592124 | 0.944975 |
| ENSMUSG( | Prkd1 | 1021.229 | 0.193459 | 0.128337 | 0.536641 | ENSMUSG( | Prkd1 | 913.5259 | 0.137297 | 0.380552 | 0.917202 |
| ENSMUSG( | Prkd2 | 212.4043 | 0.068532 | 0.665696 | 0.927054 | ENSMUSG( | Prkd2 | 210.3347 | 0.301978 | 0.101817 | 0.801719 |
| ENSMUSG( | Prkd3 | 1848.734 | 0.089866 | 0.469119 | 0.845876 | ENSMUSG( | Prkd3 | 1609.624 | -0.0414 | 0.759248 | 0.97199 |
| ENSMUSG( | Prkdc | 854.2805 | -0.0866 | 0.439368 | 0.830988 | ENSMUSG( | Prkdc | 806.6718 | 0.022546 | 0.865093 | 0.98675 |
| ENSMUSG( | Prkg1 | 177.6018 | 0.209902 | 0.180171 | 0.617634 | ENSMUSG( | Prkg1 | 171.2052 | 0.352506 | 0.045521 | 0.693959 |
| ENSMUSG( | Prkg2 | 455.2041 | 0.025079 | 0.854887 | 0.973665 | ENSMUSG( | Prkg2 | 406.2438 | -0.03892 | 0.791776 | 0.97681 |
| ENSMUSG( | Prkn | 508.3227 | 0.135512 | 0.417919 | 0.821964 | ENSMUSG( | Prkn | 431.6055 | -0.07726 | 0.719497 | 0.966876 |
| ENSMUSG( | Prkra | 519.0406 | 0.133843 | 0.245811 | 0.693547 | ENSMUSG( | Prkra | 443.1087 | -0.05738 | 0.683736 | 0.958678 |
| ENSMUSG( | Prkrip1 | 447.7661 | -0.00828 | 0.940783 | 0.993415 | ENSMUSG( | Prkrip1 | 390.5372 | 0.121988 | 0.423762 | 0.917257 |
| ENSMUSG( | Prkx | 640.3791 | 0.126427 | 0.296259 | 0.744386 | ENSMUSG( | Prkx | 558.3665 | -1.78E-05 | 0.999889 | 0.999947 |
| ENSMUSG( | Prlr | 83.89142 | 0.464129 | 0.474706 | 0.849485 | ENSMUSG( | Prlr | 92.03954 | 0.888522 | 0.017763 | 0.534632 |
| ENSMUSG( | Prmt1 | 1291.804 | -0.006 | 0.956739 | 0.993874 | ENSMUSG( | Prmt1 | 1139.372 | -0.10403 | 0.445492 | 0.921734 |
| ENSMUSG( | Prmt2 | 1413.942 | 0.094671 | 0.310783 | 0.758828 | ENSMUSG( | Prmt2 | 1229.22 | -0.04058 | 0.717406 | 0.965964 |
| ENSMUSG( | Prmt3 | 598.0676 | 0.171338 | 0.108793 | 0.497919 | ENSMUSG( | Prmt3 | 517.398 | 0.023212 | 0.860525 | 0.986137 |
| ENSMUSG( | Prmt5 | 1968.81 | 0.149431 | 0.12761 | 0.535393 | ENSMUSG( | Prmt5 | 1686.493 | -0.02613 | 0.827861 | 0.980224 |
| ENSMUSG( | Prmt6 | 314.5604 | 0.028658 | 0.827424 | 0.968732 | ENSMUSG( | Prmt6 | 284.5437 | 0.009972 | 0.950033 | 0.994798 |
| ENSMUSG( | Prmt7 | 978.2709 | 0.193856 | 0.100639 | 0.48402 | ENSMUSG( | Prmt7 | 834.7346 | 0.010814 | 0.932793 | 0.99306 |
| ENSMUSG( | Prmt8 | 2672.019 | 0.083596 | 0.403538 | 0.816132 | ENSMUSG( | Prmt8 | 2276.439 | -0.11226 | 0.35276 | 0.912209 |
| ENSMUSG( | Prmt9 | 422.8075 | -0.02925 | 0.827739 | 0.968924 | ENSMUSG( | Prmt9 | 396.7716 | 0.061001 | 0.679482 | 0.957966 |
| ENSMUSG( | Prnp | 33726.91 | -0.3915 | 0.001845 | 0.048742 | ENSMUSG( | Prnp | 35216.36 | 0.030083 | 0.835242 | 0.981237 |
| ENSMUSG( | Prob1 | 710.1887 | -0.03036 | 0.781511 | 0.957558 | ENSMUSG( | Prob1 | 639.6894 | -0.06464 | 0.658842 | 0.956205 |
| ENSMUSG( | Proca1 | 422.8272 | 0.234871 | 0.099811 | 0.482257 | ENSMUSG( | Proca1 | 367.142 | 0.094596 | 0.59415 | 0.945408 |
| ENSMUSG( | Procr | 32.92901 | 0.28518 | 0.416239 | 0.82084 | ENSMUSG( | Procr | 31.83967 | 0.458621 | 0.241651 | 0.876528 |
| ENSMUSG( | Prodh | 3448.32 | 0.212239 | 0.018318 | 0.203658 | ENSMUSG( | Prodh | 2921.126 | 0.011496 | 0.92324 | 0.99272 |
| ENSMUSG( | Prokr2 | 77.65413 | -0.99557 | 0.001014 | 0.033228 | ENSMUSG( | Prokr2 | 100.1715 | 0.161495 | 0.509406 | 0.929089 |
| ENSMUSG( | Prom1 | 1137.145 | 0.075024 | 0.687865 | 0.93324 | ENSMUSG( | Prom1 | 1244.689 | 0.560378 | 0.01361 | 0.466552 |
| ENSMUSG( | Prom2 | 29.05425 | -0.01944 | 0.963445 | 0.994911 | ENSMUSG( | Prom2 | 24.99417 | -0.19245 | 0.664884 | 0.956766 |
| ENSMUSG( | Prorp | 130.4838 | 0.139135 | 0.444346 | 0.833239 | ENSMUSG( | Prorp | 110.0832 | -0.07663 | 0.715925 | 0.965424 |
| ENSMUSG( | Prorsd1 | 94.79561 | 0.286134 | 0.190749 | 0.630215 | ENSMUSG( | Prorsd1 | 85.97378 | 0.246712 | 0.339892 | 0.907972 |
| ENSMUSG( | Pros1 | 830.3952 | -0.0674 | 0.652297 | 0.922188 | ENSMUSG( | Pros1 | 842.1307 | 0.239096 | 0.150525 | 0.837173 |
| ENSMUSG( | Proser1 | 618.4634 | 0.021806 | 0.852205 | 0.973168 | ENSMUSG( | Proser1 | 582.6106 | 0.120324 | 0.391027 | 0.917257 |
| ENSMUSG( | Proser2 | 94.91984 | 0.064262 | 0.768052 | 0.954621 | ENSMUSG( | Proser2 | 95.2815 | 0.350813 | 0.203165 | 0.864412 |
| ENSMUSG( | Proser3 | 80.28365 | 0.113529 | 0.609572 | 0.9077 | ENSMUSG( | Proser3 | 81.43676 | 0.387917 | 0.114419 | 0.811703 |
| ENSMUSG( | Prox1 | 533.9876 | 0.026819 | 0.833724 | 0.970333 | ENSMUSG( | Prox1 | 492.4774 | 0.06896 | 0.658312 | 0.956205 |
| ENSMUSG( | Prox1os | 33.86422 | 0.262606 | 0.446168 | 0.833546 | ENSMUSG( | Prox1os | 25.89245 | -0.21071 | 0.615794 | 0.951306 |
| ENSMUSG( | Prox2 | 82.39059 | -0.00229 | 0.992921 | 0.999327 | ENSMUSG( | Prox2 | 72.76707 | -0.09942 | 0.695797 | 0.960867 |
| ENSMUSG( | Prpf18 | 1082.604 | 0.101756 | 0.307951 | 0.75624 | ENSMUSG( | Prpf18 | 970.009 | 0.054056 | 0.641686 | 0.953903 |
| ENSMUSG( | Prpf19 | 4153.406 | 0.149768 | 0.162449 | 0.593257 | ENSMUSG( | Prpf19 | 3494.652 | -0.07962 | 0.556998 | 0.939565 |
| ENSMUSG( | Prpf3 | 529.611 | 0.125759 | 0.245246 | 0.692812 | ENSMUSG( | Prpf3 | 481.7423 | 0.125157 | 0.391109 | 0.917257 |
| ENSMUSG( | Prpf31 | 977.9445 | 0.064603 | 0.552333 | 0.886141 | ENSMUSG( | Prpf31 | 823.9937 | -0.16853 | 0.198008 | 0.862161 |
| ENSMUSG( | Prpf38a | 541.614 | 0.001457 | 0.991263 | 0.999232 | ENSMUSG( | Prpf38a | 484.1985 | -0.05506 | 0.703409 | 0.962481 |
| ENSMUSG( | Prpf38b | 1662.065 | -0.15491 | 0.10183 | 0.486216 | ENSMUSG( | Prpf38b | 1701.159 | 0.177603 | 0.126275 | 0.81516 |
| ENSMUSG( | Prpf39 | 1505.587 | -0.10165 | 0.469601 | 0.846167 | ENSMUSG( | Prpf39 | 1587.833 | 0.308196 | 0.045888 | 0.695944 |
| ENSMUSG( | Prpf4 | 563.8293 | 0.036794 | 0.743615 | 0.948478 | ENSMUSG( | Prpf4 | 529.4148 | 0.119962 | 0.356183 | 0.913358 |
| ENSMUSG( | Prpf40a | 1367.794 | 0.056758 | 0.567602 | 0.891774 | ENSMUSG( | Prpf40a | 1241.701 | 0.043126 | 0.742105 | 0.969415 |
| ENSMUSG( | Prpf40b | 2125.27 | 0.197967 | 0.152674 | 0.579011 | ENSMUSG( | Prpf40b | 1800.826 | -0.00735 | 0.965155 | 0.995433 |
| ENSMUSG( | Prpf4b | 1872.689 | -0.01969 | 0.857233 | 0.97444 | ENSMUSG( | Prpf4b | 1911.805 | 0.295913 | 0.063202 | 0.736482 |
| ENSMUSG( | Prpf6 | 1435.894 | 0.157797 | 0.150021 | 0.574564 | ENSMUSG( | Prpf6 | 1204.697 | -0.07665 | 0.546053 | 0.938063 |
| ENSMUSG( | Prpf8 | 5486.71 | 0.097911 | 0.265057 | 0.715962 | ENSMUSG( | Prpf8 | 4779.434 | -0.0285 | 0.792873 | 0.97681 |
| ENSMUSG( | Prph | 90.86567 | 0.265131 | 0.261599 | 0.711405 | ENSMUSG( | Prph | 73.80376 | -0.06596 | 0.807412 | 0.978513 |
| ENSMUSG( | Prps1 | 1030.741 | 0.012874 | 0.909557 | 0.988084 | ENSMUSG( | Prps1 | 886.7219 | -0.16033 | 0.280273 | 0.892356 |
| ENSMUSG( | Prps1l3 | 153.952 | -0.00439 | 0.979745 | 0.997428 | ENSMUSG( | Prps1l3 | 136.3989 | -0.08677 | 0.669257 | 0.956766 |
| ENSMUSG( | Prps2 | 769.3521 | -0.06601 | 0.539438 | 0.883094 | ENSMUSG( | Prps2 | 699.5033 | -0.07862 | 0.613918 | 0.951177 |
| ENSMUSG( | Prpsap1 | 1177.047 | -0.05042 | 0.644029 | 0.917536 | ENSMUSG( | Prpsap1 | 1127.642 | 0.096976 | 0.431505 | 0.917315 |
| ENSMUSG( | Prpsap2 | 833.2954 | 0.117454 | 0.25938 | 0.709948 | ENSMUSG( | Prpsap2 | 690.6333 | -0.15976 | 0.204724 | 0.864412 |
| ENSMUSG( | Prr11 | 43.23564 | -0.50758 | 0.102655 | 0.488148 | ENSMUSG( | Prr11 | 42.25001 | -0.27077 | 0.410455 | 0.917257 |
| ENSMUSG( | Prr12 | 2534.61 | 0.058567 | 0.535172 | 0.880733 | ENSMUSG( | Prr12 | 2392.259 | 0.157805 | 0.15838 | 0.837724 |
| ENSMUSG( | Prr13 | 1135.617 | 0.036357 | 0.709686 | 0.939669 | ENSMUSG( | Prr13 | 942.5171 | -0.24813 | 0.06299 | 0.736482 |
| ENSMUSG( | Prr14 | 801.675 | -0.03238 | 0.783531 | 0.957558 | ENSMUSG( | Prr14 | 760.0365 | 0.028223 | 0.854891 | 0.985649 |
| ENSMUSG( | Prr14l | 1262.939 | 0.209691 | 0.054316 | 0.358716 | ENSMUSG( | Prr14l | 1168.397 | 0.253096 | 0.031269 | 0.631113 |
| ENSMUSG( | Prr15 | 14.10023 | -0.07522 | 0.885101 | NA | ENSMUSG( | Prr15 | 12.03821 | -0.31589 | 0.589775 | 0.944975 |
| ENSMUSG( | Prr16 | 174.2096 | -0.25878 | 0.164482 | 0.595363 | ENSMUSG( | Prr16 | 160.8019 | -0.22469 | 0.277231 | 0.891468 |
| ENSMUSG( | Prr18 | 10568.77 | -0.20489 | 0.031337 | 0.269505 | ENSMUSG( | Prr18 | 9091.054 | -0.39298 | 0.001616 | 0.150169 |
| ENSMUSG( | Prr22 | 29.50564 | 0.006016 | 0.987149 | 0.998764 | ENSMUSG( | Prr22 | 27.05451 | 0.003674 | 0.992972 | 0.998813 |
| ENSMUSG( | Prr3 | 758.7714 | 0.028654 | 0.832339 | 0.970333 | ENSMUSG( | Prr3 | 678.6904 | -0.02728 | 0.860653 | 0.986137 |
| ENSMUSG( | Prr32 | 24.94699 | -0.42175 | 0.514026 | 0.872055 | ENSMUSG( | Prr32 | 24.68112 | -0.15196 | 0.795209 | 0.97681 |
| ENSMUSG( | Prr36 | 859.4479 | 0.187258 | 0.109517 | 0.499679 | ENSMUSG( | Prr36 | 716.6753 | -0.06085 | 0.642557 | 0.953903 |
| ENSMUSG( | Prr5 | 630.5375 | 0.535428 | 1.09E-05 | 0.001199 | ENSMUSG( | Prr5 | 502.0012 | 0.195248 | 0.246411 | 0.877904 |
| ENSMUSG( | Prr5l | 557.1306 | -0.28489 | 0.01203 | 0.158657 | ENSMUSG( | Prr5l | 520.2305 | -0.20332 | 0.155556 | 0.837724 |
| ENSMUSG( | Prr7 | 1669.753 | -0.18397 | 0.057306 | 0.369582 | ENSMUSG( | Prr7 | 1681.525 | 0.110056 | 0.359346 | 0.914126 |
| ENSMUSG( | Prrc1 | 592.8008 | 0.011757 | 0.921818 | 0.990436 | ENSMUSG( | Prrc1 | 527.4643 | -0.05675 | 0.669331 | 0.956766 |
| ENSMUSG( | Prrc2a | 17228.63 | 0.204446 | 0.012482 | 0.161449 | ENSMUSG( | Prrc2a | 15189.9 | 0.115776 | 0.302112 | 0.898538 |
| ENSMUSG( | Prrc2b | 14784.71 | 0.058302 | 0.591897 | 0.901716 | ENSMUSG( | Prrc2b | 13812.47 | 0.131356 | 0.348723 | 0.911652 |
| ENSMUSG( | Prrc2c | 4004.133 | -0.07039 | 0.468471 | 0.845415 | ENSMUSG( | Prrc2c | 3968.562 | 0.1703 | 0.121783 | 0.813384 |
| ENSMUSG( | Prrg1 | 117.773 | -0.34238 | 0.096555 | 0.476986 | ENSMUSG( | Prrg1 | 134.8263 | 0.330172 | 0.194392 | 0.860096 |
| ENSMUSG( | Prrg2 | 260.835 | -0.27749 | 0.109357 | 0.499374 | ENSMUSG( | Prrg2 | 261.7752 | 0.021941 | 0.908463 | 0.991958 |
| ENSMUSG( | Prrg3 | 1145.318 | -0.19271 | 0.194534 | 0.633457 | ENSMUSG( | Prrg3 | 1154.885 | 0.109647 | 0.478886 | 0.923329 |
| ENSMUSG( | Prrg4 | 15.10317 | -0.50224 | 0.324077 | NA | ENSMUSG( | Prrg4 | 18.27152 | 0.328742 | 0.494139 | 0.927372 |
| ENSMUSG( | Prrt1 | 4777.747 | -0.0164 | 0.853901 | 0.973491 | ENSMUSG( | Prrt1 | 4347.498 | -0.0203 | 0.848824 | 0.985321 |
| ENSMUSG( | Prrt1b | 11.61736 | -0.36594 | 0.506643 | NA | ENSMUSG( | Prrt1b | 9.566333 | -0.65027 | 0.343519 | NA |
| ENSMUSG( | Prrt2 | 12.62721 | -0.18639 | 0.744981 | NA | ENSMUSG( | Prrt2 | 12.26208 | 0.06448 | 0.91439 | 0.992407 |
| ENSMUSG( | Prrt3 | 1841.835 | -0.27722 | 0.079124 | 0.435105 | ENSMUSG( | Prrt3 | 1934.576 | 0.148867 | 0.425161 | 0.917257 |
| ENSMUSG( | Prrt4 | 114.9997 | -0.54483 | 0.016779 | 0.193019 | ENSMUSG( | Prrt4 | 134.8478 | 0.243982 | 0.320156 | 0.903876 |
| ENSMUSG( | Prrx1 | 460.1502 | -0.04656 | 0.732652 | 0.944956 | ENSMUSG( | Prrx1 | 458.2687 | 0.195642 | 0.216252 | 0.865167 |
| ENSMUSG( | Prrx2 | 63.30033 | -0.28215 | 0.248445 | 0.697313 | ENSMUSG( | Prrx2 | 58.9983 | -0.20626 | 0.435666 | 0.919133 |
| ENSMUSG( | Prss12 | 692.4064 | -0.38498 | 0.001402 | 0.040091 | ENSMUSG( | Prss12 | 738.8571 | 0.096336 | 0.482378 | 0.924578 |
| ENSMUSG( | Prss16 | 20.32367 | 0.069201 | 0.869023 | NA | ENSMUSG( | Prss16 | 18.56747 | 0.051459 | 0.914101 | 0.992407 |
| ENSMUSG( | Prss22 | 34.70903 | -0.02676 | 0.934343 | 0.99323 | ENSMUSG( | Prss22 | 26.47101 | -0.66429 | 0.140066 | 0.831355 |
| ENSMUSG( | Prss23 | 520.7874 | -0.47409 | 0.006279 | 0.108547 | ENSMUSG( | Prss23 | 555.7009 | 0.026945 | 0.889429 | 0.990558 |
| ENSMUSG( | Prss35 | 114.1127 | -0.64354 | 0.001177 | 0.036941 | ENSMUSG( | Prss35 | 133.3857 | 0.146412 | 0.455954 | 0.923329 |
| ENSMUSG( | Prss36 | 239.6814 | 0.055546 | 0.690852 | 0.933881 | ENSMUSG( | Prss36 | 205.9357 | -0.12483 | 0.490326 | 0.926341 |
| ENSMUSG( | Prss3l | 23.36243 | -0.33041 | 0.396681 | 0.811742 | ENSMUSG( | Prss3l | 22.13642 | -0.20162 | 0.634844 | 0.953903 |
| ENSMUSG( | Prss43 | 15.95186 | 0.034756 | 0.941996 | NA | ENSMUSG( | Prss43 | 12.18248 | -0.54125 | 0.349649 | 0.911652 |
| ENSMUSG( | Prss45 | 14.00935 | -0.09136 | 0.860145 | NA | ENSMUSG( | Prss45 | 14.48726 | 0.284806 | 0.602233 | 0.94789 |
| ENSMUSG( | Prss50 | 32.85415 | -0.69344 | 0.04011 | 0.308919 | ENSMUSG( | Prss50 | 37.70643 | 0.029958 | 0.930254 | 0.99306 |
| ENSMUSG( | Prss53 | 26.77481 | -0.0854 | 0.823916 | 0.966648 | ENSMUSG( | Prss53 | 25.70397 | 0.04179 | 0.91803 | 0.992529 |
| ENSMUSG( | Prss56 | 26.45774 | -0.86397 | 0.0419 | 0.315367 | ENSMUSG( | Prss56 | 30.46127 | -0.04127 | 0.912407 | 0.992225 |
| ENSMUSG( | Prss57 | 25.57531 | 0.24642 | 0.542366 | 0.883182 | ENSMUSG( | Prss57 | 21.32855 | 0.010526 | 0.980552 | 0.996479 |
| ENSMUSG( | Prss8 | 11.52107 | -0.10012 | 0.858275 | NA | ENSMUSG( | Prss8 | 10.88962 | 0.010505 | 0.985464 | NA |
| ENSMUSG( | Prtg | 66.21274 | -0.06944 | 0.791313 | 0.959461 | ENSMUSG( | Prtg | 62.87737 | 0.069803 | 0.801938 | 0.97681 |
| ENSMUSG( | Prtn3 | 39.6036 | -0.04146 | 0.893558 | 0.984512 | ENSMUSG( | Prtn3 | 35.34592 | -0.08721 | 0.799928 | 0.97681 |
| ENSMUSG( | Prune1 | 700.3497 | -0.06349 | 0.609797 | 0.907755 | ENSMUSG( | Prune1 | 659.2935 | 0.030715 | 0.818435 | 0.979198 |
| ENSMUSG( | Prune2 | 1788.218 | -0.26795 | 0.007627 | 0.120942 | ENSMUSG( | Prune2 | 1832.898 | 0.084504 | 0.450479 | 0.922202 |
| ENSMUSG( | Prx | 57.21377 | -0.03386 | 0.891223 | 0.983819 | ENSMUSG( | Prx | 48.45643 | -0.25791 | 0.379616 | 0.916995 |
| ENSMUSG( | Prxl2a | 12743.6 | 0.095027 | 0.331733 | 0.775288 | ENSMUSG( | Prxl2a | 11409.31 | 0.044773 | 0.738901 | 0.968305 |
| ENSMUSG( | Prxl2b | 2450.069 | 0.005664 | 0.946087 | 0.99369 | ENSMUSG( | Prxl2b | 1959.334 | -0.40116 | 0.001307 | 0.130907 |
| ENSMUSG( | Prxl2c | 493.6898 | 0.029453 | 0.825608 | 0.967934 | ENSMUSG( | Prxl2c | 451.1934 | 0.041572 | 0.768276 | 0.974236 |
| ENSMUSG( | Psap | 80192.68 | -0.19871 | 0.077425 | 0.429743 | ENSMUSG( | Psap | 80198.79 | 0.078393 | 0.556446 | 0.939565 |
| ENSMUSG( | Psat1 | 10366.95 | 0.139174 | 0.189658 | 0.628577 | ENSMUSG( | Psat1 | 8657.638 | -0.11224 | 0.403572 | 0.917257 |
| ENSMUSG( | Psd | 39238.83 | 0.125815 | 0.152169 | 0.578042 | ENSMUSG( | Psd | 34472.1 | 0.023598 | 0.806705 | 0.97813 |
| ENSMUSG( | Psd2 | 6318.698 | 0.145921 | 0.114452 | 0.510426 | ENSMUSG( | Psd2 | 5001.121 | -0.27045 | 0.028043 | 0.61023 |
| ENSMUSG( | Psd3 | 18983.24 | 0.143602 | 0.167361 | 0.599971 | ENSMUSG( | Psd3 | 16484.46 | 0.010083 | 0.931145 | 0.99306 |
| ENSMUSG( | Psd4 | 62.43151 | -0.24542 | 0.303569 | 0.751968 | ENSMUSG( | Psd4 | 66.72259 | 0.22819 | 0.382698 | 0.917257 |
| ENSMUSG( | Psen1 | 2171.668 | -0.2709 | 0.079181 | 0.435105 | ENSMUSG( | Psen1 | 2315.905 | 0.194729 | 0.2621 | 0.885136 |
| ENSMUSG( | Psen2 | 665.4113 | -0.2883 | 0.020306 | 0.214844 | ENSMUSG( | Psen2 | 709.9074 | 0.180133 | 0.224915 | 0.86838 |
| ENSMUSG( | Psenen | 1112.498 | -0.20279 | 0.109932 | 0.500285 | ENSMUSG( | Psenen | 1084.885 | 0.000783 | 0.995446 | 0.999116 |
| ENSMUSG( | Psg16 | 80.63178 | -0.03214 | 0.900576 | 0.985633 | ENSMUSG( | Psg16 | 72.63532 | -0.05887 | 0.823803 | 0.979198 |
| ENSMUSG( | Psip1 | 3166.483 | 0.025165 | 0.794513 | 0.959799 | ENSMUSG( | Psip1 | 2877.233 | 0.014036 | 0.918535 | 0.992529 |
| ENSMUSG( | Pskh1 | 361.3743 | -0.0863 | 0.598648 | 0.903801 | ENSMUSG( | Pskh1 | 332.7023 | -0.04845 | 0.782755 | 0.976106 |
| ENSMUSG( | Psma1 | 1283.983 | -0.07391 | 0.546149 | 0.884974 | ENSMUSG( | Psma1 | 1145.574 | -0.1395 | 0.303087 | 0.898538 |
| ENSMUSG( | Psma2 | 2069.151 | 0.01571 | 0.87073 | 0.977951 | ENSMUSG( | Psma2 | 1755.622 | -0.20002 | 0.129803 | 0.820802 |
| ENSMUSG( | Psma3 | 2326.959 | 0.051339 | 0.557858 | 0.888427 | ENSMUSG( | Psma3 | 2058.887 | -0.03454 | 0.754005 | 0.970671 |
| ENSMUSG( | Psma4 | 1964.213 | -0.00651 | 0.946388 | 0.99369 | ENSMUSG( | Psma4 | 1736.573 | -0.09893 | 0.468371 | 0.923329 |
| ENSMUSG( | Psma5 | 1214.09 | 0.00945 | 0.936096 | 0.993415 | ENSMUSG( | Psma5 | 1041.241 | -0.17192 | 0.202904 | 0.864412 |
| ENSMUSG( | Psma6 | 1505.507 | 0.02117 | 0.807009 | 0.963044 | ENSMUSG( | Psma6 | 1321.415 | -0.09148 | 0.47249 | 0.923329 |

| | | | | | |
|---|---|---|---|---|---|
| ENSMUSG( | 265.89 | 0.184183 | 0.183754 | 0.620678 | Ppat |
| ENSMUSG( | 451.8386 | -0.01197 | 0.915816 | 0.988634 | Ppcdc |
| ENSMUSG( | 107.2534 | -0.10202 | 0.658889 | 0.924506 | Ppcs |
| ENSMUSG( | 2130.708 | 0.0911 | 0.404548 | 0.816132 | Ppdpf |
| ENSMUSG( | 15.03243 | -0.36116 | 0.473801 | NA | Ppef1 |
| ENSMUSG( | 24.53845 | -0.08564 | 0.821724 | 0.96635 | Ppef2 |
| ENSMUSG( | 2329.201 | -0.06727 | 0.484348 | 0.856665 | Ppfia1 |
| ENSMUSG( | 2102.909 | 0.072504 | 0.46915 | 0.845876 | Ppfia2 |
| ENSMUSG( | 15123.75 | 0.064208 | 0.424737 | 0.824675 | Ppfia3 |
| ENSMUSG( | 3724.345 | 0.105523 | 0.239763 | 0.68758 | Ppfia4 |
| ENSMUSG( | 1537.428 | 0.012048 | 0.905006 | 0.986449 | Ppfibp1 |
| ENSMUSG( | 369.3578 | -0.16587 | 0.212385 | 0.65642 | Ppfibp2 |
| ENSMUSG( | 842.2058 | -0.06146 | 0.576038 | 0.895114 | Pphln1 |
| ENSMUSG( | 15832.13 | -0.01979 | 0.849212 | 0.972638 | Ppia |
| ENSMUSG( | 4829.897 | -0.20009 | 0.059671 | 0.377567 | Ppib |
| ENSMUSG( | 199.8695 | -0.17273 | 0.372041 | 0.798768 | Ppic |
| ENSMUSG( | 1329.861 | -0.0287 | 0.775422 | 0.955606 | Ppid |
| ENSMUSG( | 313.1766 | 0.04926 | 0.71268 | 0.940652 | Ppie |
| ENSMUSG( | 582.2214 | -0.01484 | 0.910396 | 0.9881 | Ppif |
| ENSMUSG( | 1507.504 | 0.065688 | 0.466923 | 0.844819 | Ppig |
| ENSMUSG( | 165.637 | 0.061887 | 0.752047 | 0.950417 | Ppih |
| ENSMUSG( | 519.4482 | -0.04913 | 0.701613 | 0.937441 | Ppil1 |
| ENSMUSG( | 1668.345 | 0.202968 | 0.055978 | 0.364712 | Ppil2 |
| ENSMUSG( | 313.0348 | -0.02883 | 0.827975 | 0.968966 | Ppil3 |
| ENSMUSG( | 944.3626 | 0.072689 | 0.474018 | 0.849425 | Ppil4 |
| ENSMUSG( | 121.4489 | 0.333732 | 0.065089 | 0.395697 | Ppil6 |
| ENSMUSG( | 2920.929 | 0.214035 | 0.04981 | 0.340731 | Ppip5k1 |
| ENSMUSG( | 577.1014 | -0.09138 | 0.509189 | 0.869008 | Ppip5k2 |
| ENSMUSG( | 24.29392 | 0.08418 | 0.848043 | 0.972374 | Ppl |
| ENSMUSG( | 6734.738 | 0.095373 | 0.203694 | 0.643787 | Ppm1a |
| ENSMUSG( | 3885.87 | 0.103224 | 0.275969 | 0.725669 | Ppm1b |
| ENSMUSG( | 364.706 | 0.002072 | 0.987262 | 0.998777 | Ppm1d |
| ENSMUSG( | 4440.623 | 0.139774 | 0.229478 | 0.67735 | Ppm1e |
| ENSMUSG( | 2584.838 | 0.226295 | 0.013564 | 0.170154 | Ppm1f |
| ENSMUSG( | 2110.578 | -0.10034 | 0.326307 | 0.771301 | Ppm1g |
| ENSMUSG( | 2330.798 | 0.146471 | 0.137666 | 0.552903 | Ppm1h |
| ENSMUSG( | 21.64896 | -0.71011 | 0.096135 | 0.476417 | Ppm1j |
| ENSMUSG( | 1683.806 | 0.088046 | 0.342296 | 0.778828 | Ppm1k |
| ENSMUSG( | 3861.726 | -0.43462 | 0.002269 | 0.055678 | Ppm1l |
| ENSMUSG( | 1092.926 | 0.106905 | 0.377353 | 0.801213 | Ppm1m |
| ENSMUSG( | 4227.213 | 0.010842 | 0.924567 | 0.99114 | Ppme1 |
| ENSMUSG( | 829.7584 | 0.14584 | 0.317654 | 0.763989 | Ppox |
| ENSMUSG( | 6751.158 | -0.0647 | 0.447476 | 0.834051 | Ppp1ca |
| ENSMUSG( | 5300.713 | 0.034028 | 0.786405 | 0.958843 | Ppp1cb |
| ENSMUSG( | 1670.96 | -0.11062 | 0.214865 | 0.659433 | Ppp1cc |
| ENSMUSG( | 1370.207 | 0.175608 | 0.167189 | 0.599823 | Ppp1r10 |
| ENSMUSG( | 1317.58 | -0.06351 | 0.542625 | 0.883495 | Ppp1r11 |
| ENSMUSG( | 1990.942 | 0.019854 | 0.813909 | 0.964574 | Ppp1r12a |
| ENSMUSG( | 6174.255 | 0.176197 | 0.066266 | 0.398381 | Ppp1r12b |
| ENSMUSG( | 4522.104 | 0.075864 | 0.414632 | 0.820184 | Ppp1r12c |
| ENSMUSG( | 1898.273 | 0.153154 | 0.166975 | 0.599655 | Ppp1r13b |
| ENSMUSG( | 141.0344 | 0.002128 | 0.990143 | 0.999144 | Ppp1r13l |
| ENSMUSG( | 491.2186 | -0.33921 | 0.004553 | 0.088443 | Ppp1r14a |
| ENSMUSG( | 1150.093 | 0.054611 | 0.58802 | 0.900712 | Ppp1r14b |
| ENSMUSG( | 466.2906 | 0.069874 | 0.595475 | 0.903058 | Ppp1r14c |
| ENSMUSG( | 526.5428 | -0.09007 | 0.496144 | 0.861554 | Ppp1r15a |
| ENSMUSG( | 2104.932 | -0.26187 | 0.060757 | 0.380787 | Ppp1r15b |
| ENSMUSG( | 4489.704 | 0.137465 | 0.123316 | 0.530552 | Ppp1r16a |
| ENSMUSG( | 9081.719 | 0.065023 | 0.531025 | 0.879289 | Ppp1r16b |
| ENSMUSG( | 646.4938 | -0.02065 | 0.841899 | 0.971237 | Ppp1r18 |
| ENSMUSG( | 4237.8 | 0.021233 | 0.810081 | 0.963864 | Ppp1r1a |
| ENSMUSG( | 5779.604 | -0.67769 | 0.000686 | 0.025185 | Ppp1r1b |
| ENSMUSG( | 3296.185 | 0.015572 | 0.876905 | 0.980991 | Ppp1r2 |
| ENSMUSG( | 2154.062 | 0.121026 | 0.224104 | 0.669929 | Ppp1r21 |
| ENSMUSG( | 816.8861 | 0.220503 | 0.028134 | 0.256512 | Ppp1r26 |
| ENSMUSG( | 270.0105 | -0.08713 | 0.566367 | 0.89163 | Ppp1r35 |
| ENSMUSG( | 54.6368 | -0.37419 | 0.15147 | 0.576717 | Ppp1r36 |
| ENSMUSG( | 12.94644 | -0.02934 | 0.955017 | NA | Ppp1r36dn |
| ENSMUSG( | 5855.451 | 0.206197 | 0.011857 | 0.157504 | Ppp1r37 |
| ENSMUSG( | 66.58213 | 0.081757 | 0.7464 | 0.949289 | Ppp1r3b |
| ENSMUSG( | 6201.186 | 0.202625 | 0.085523 | 0.450845 | Ppp1r3c |
| ENSMUSG( | 402.3089 | 0.125552 | 0.333117 | 0.775602 | Ppp1r3d |
| ENSMUSG( | 465.1078 | -0.00961 | 0.937937 | 0.993415 | Ppp1r3e |
| ENSMUSG( | 209.8516 | 0.114611 | 0.470304 | 0.84674 | Ppp1r3f |
| ENSMUSG( | 677.1121 | 0.308407 | 0.011612 | 0.1554 | Ppp1r3g |
| ENSMUSG( | 15.11729 | 0.046918 | 0.921114 | NA | Ppp1r42 |
| ENSMUSG( | 2887.22 | 0.006018 | 0.942361 | 0.993415 | Ppp1r7 |
| ENSMUSG( | 977.7155 | 0.04804 | 0.629095 | 0.913838 | Ppp1r8 |
| ENSMUSG( | 9982.593 | -0.20027 | 0.063066 | 0.388234 | Ppp1r9a |
| ENSMUSG( | 88729.42 | 0.236378 | 0.005714 | 0.101299 | Ppp1r9b |
| ENSMUSG( | 9199.29 | 0.030755 | 0.70257 | 0.937735 | Ppp2ca |
| ENSMUSG( | 3833.143 | 0.041705 | 0.597597 | 0.903694 | Ppp2cb |
| ENSMUSG( | 13257.27 | -0.11551 | 0.25096 | 0.700615 | Ppp2r1a |
| ENSMUSG( | 524.9434 | 0.049056 | 0.663972 | 0.926511 | Ppp2r1b |
| ENSMUSG( | 1984.782 | -0.04978 | 0.682666 | 0.93198 | Ppp2r2a |
| ENSMUSG( | 3380.628 | 0.033375 | 0.720044 | 0.941986 | Ppp2r2b |
| ENSMUSG( | 40760.15 | 0.167458 | 0.034982 | 0.28772 | Ppp2r2c |
| ENSMUSG( | 1775.843 | 0.042486 | 0.667205 | 0.92729 | Ppp2r2d |
| ENSMUSG( | 2002.625 | 0.043756 | 0.653635 | 0.922872 | Ppp2r3a |
| ENSMUSG( | 390.6557 | 0.20904 | 0.106152 | 0.493488 | Ppp2r3c |
| ENSMUSG( | 1222.194 | 0.046737 | 0.643388 | 0.917267 | Ppp2r3d |
| ENSMUSG( | 5072.76 | 0.093385 | 0.350947 | 0.782993 | Ppp2r5a |
| ENSMUSG( | 4888.478 | 0.076161 | 0.368344 | 0.795863 | Ppp2r5b |
| ENSMUSG( | 9251.915 | 0.0778 | 0.397371 | 0.812127 | Ppp2r5c |
| ENSMUSG( | 2713.616 | 0.053965 | 0.577394 | 0.895776 | Ppp2r5d |
| ENSMUSG( | 1839.384 | 0.072798 | 0.429437 | 0.826446 | Ppp2r5e |
| ENSMUSG( | 20068.6 | 0.014034 | 0.895725 | 0.984527 | Ppp3ca |
| ENSMUSG( | 5794.225 | 0.016564 | 0.862103 | 0.975055 | Ppp3cb |
| ENSMUSG( | 638.3212 | 0.087193 | 0.428924 | 0.826446 | Ppp3cc |
| ENSMUSG( | 10455.95 | -0.00337 | 0.979259 | 0.997428 | Ppp3r1 |
| ENSMUSG( | 13.72621 | -0.5265 | 0.316973 | NA | Ppp3r2 |
| ENSMUSG( | 836.9251 | -0.04279 | 0.671709 | 0.928781 | Ppp4c |
| ENSMUSG( | 1455.748 | 0.177044 | 0.066248 | 0.398381 | Ppp4r1 |
| ENSMUSG( | 757.9734 | 0.086987 | 0.403008 | 0.815774 | Ppp4r1l-ps |
| ENSMUSG( | 1495.722 | 0.144608 | 0.150187 | 0.574666 | Ppp4r2 |
| ENSMUSG( | 1305.371 | -0.0242 | 0.812714 | 0.964574 | Ppp4r3a |
| ENSMUSG( | 1457.28 | 0.041922 | 0.70064 | 0.93743 | Ppp4r3b |
| ENSMUSG( | 881.3327 | 0.053071 | 0.634117 | 0.914073 | Ppp4r4 |
| ENSMUSG( | 3594.285 | 0.081178 | 0.473119 | 0.848727 | Ppp5c |
| ENSMUSG( | 1165.313 | -0.04157 | 0.743666 | 0.948478 | Ppp6c |
| ENSMUSG( | 8782.448 | 0.244781 | 0.002019 | 0.051507 | Ppp6r1 |
| ENSMUSG( | 3128.171 | 0.167807 | 0.105693 | 0.492581 | Ppp6r2 |
| ENSMUSG( | 2521.2 | 0.096148 | 0.265379 | 0.716109 | Ppp6r3 |
| ENSMUSG( | 611.7275 | -0.0062 | 0.953872 | 0.993874 | Pprc1 |
| ENSMUSG( | 6202.783 | 0.355822 | 0.001506 | 0.043083 | Ppt1 |
| ENSMUSG( | 713.6361 | -0.18131 | 0.143114 | 0.561716 | Ppt2 |
| ENSMUSG( | 1772.082 | 0.130827 | 0.153625 | 0.580569 | Pptc7 |
| ENSMUSG( | 203.6485 | -0.05224 | 0.737098 | 0.946839 | Ppwd1 |
| ENSMUSG( | 1417.802 | 0.024415 | 0.834843 | 0.970426 | Pqbp1 |
| ENSMUSG( | 427.9864 | -0.19621 | 0.096883 | 0.477375 | Pradc1 |
| ENSMUSG( | 1885.75 | -0.42926 | 0.000122 | 0.007362 | Praf2 |
| ENSMUSG( | 626.4675 | -0.01852 | 0.889604 | 0.98353 | Prag1 |
| ENSMUSG( | 294.1254 | 0.185469 | 0.192693 | 0.631789 | Pramel12 |
| ENSMUSG( | 75.66289 | -0.3494 | 0.150208 | 0.574666 | Prc1 |
| ENSMUSG( | 1026.123 | 0.156844 | 0.173894 | 0.609325 | Prcc |
| ENSMUSG( | 89.23784 | -0.25048 | 0.33715 | 0.777008 | Prcd |
| ENSMUSG( | 393.0558 | -0.36485 | 0.035046 | 0.287963 | Prcp |
| ENSMUSG( | 37.5411 | -0.32818 | 0.282363 | 0.73186 | Prdm1 |
| ENSMUSG( | 625.5875 | 0.039179 | 0.722654 | 0.943756 | Prdm10 |
| ENSMUSG( | 216.8989 | 0.095001 | 0.538403 | 0.88259 | Prdm11 |
| ENSMUSG( | 483.4947 | 0.244263 | 0.062259 | 0.386085 | Prdm15 |
| ENSMUSG( | 865.4372 | -0.0766 | 0.556056 | 0.887457 | Prdm16 |
| ENSMUSG( | 2316.412 | 0.075951 | 0.546048 | 0.884974 | Prdm16os |
| ENSMUSG( | 1547.685 | 0.098675 | 0.341969 | 0.778828 | Prdm2 |
| ENSMUSG( | 821.3939 | 0.061221 | 0.558593 | 0.888533 | Prdm4 |
| ENSMUSG( | 165.0524 | 0.122286 | 0.45188 | 0.835936 | Prdm5 |
| ENSMUSG( | 39.07229 | -0.09538 | 0.78399 | 0.957689 | Prdm6 |
| ENSMUSG( | 612.3544 | 0.265707 | 0.05671 | 0.367404 | Prdm8 |
| ENSMUSG( | 74.15447 | 0.065691 | 0.775966 | 0.955606 | Prdm9 |
| ENSMUSG( | 7182.158 | 0.061024 | 0.575611 | 0.894836 | Prdx1 |
| ENSMUSG( | 6139.3 | -0.002 | 0.982742 | 0.997522 | Prdx2 |
| ENSMUSG( | 1744.35 | 0.076334 | 0.419621 | 0.822309 | Prdx3 |
| ENSMUSG( | 680.7719 | -0.14371 | 0.356628 | 0.788187 | Prdx4 |
| ENSMUSG( | 4961.911 | -0.02894 | 0.768041 | 0.954621 | Prdx5 |
| ENSMUSG( | 17638.88 | 0.082631 | 0.442883 | 0.83278 | Prdx6 |
| ENSMUSG( | 2002.198 | 0.114901 | 0.231032 | 0.679229 | Preb |
| ENSMUSG( | 3267.176 | 0.028298 | 0.743617 | 0.948478 | Prelid1 |
| ENSMUSG( | 1341.57 | 0.088518 | 0.325505 | 0.770741 | Prelid3a |
| ENSMUSG( | 1761.12 | 0.028739 | 0.791266 | 0.959461 | Prelid3b |
| ENSMUSG( | 1086.806 | -0.16983 | 0.338076 | 0.777422 | Prelp |
| ENSMUSG( | 745.1982 | -0.09854 | 0.342638 | 0.778971 | Prep |
| ENSMUSG( | 4685.756 | 0.068014 | 0.459617 | 0.840338 | Prepl |
| ENSMUSG( | 15466.15 | 0.025608 | 0.779549 | 0.956885 | Prex1 |
| ENSMUSG( | 11927.51 | 0.211428 | 0.121179 | 0.525683 | Prex2 |
| ENSMUSG( | 292.7796 | 0.581713 | 0.000147 | 0.008459 | Prg4 |
| ENSMUSG( | 2111.337 | -0.01258 | 0.908613 | 0.987848 | Prickle1 |
| ENSMUSG( | 5992.362 | 0.118408 | 0.164135 | 0.5949 | Prickle2 |
| ENSMUSG( | 45.00355 | 0.138407 | 0.647795 | 0.919759 | Prickle3 |

| | | | | | |
|---|---|---|---|---|---|
| ENSMUSG( | 38.50046 | 0.112639 | 0.752053 | 0.970461 | Prdm6 |
| ENSMUSG( | 548.2761 | 0.214358 | 0.130358 | 0.821987 | Prdm8 |
| ENSMUSG( | 64.88744 | -0.05364 | 0.843499 | 0.983392 | Prdm9 |
| ENSMUSG( | 5948.43 | -0.22425 | 0.075418 | 0.767994 | Prdx1 |
| ENSMUSG( | 5175.55 | -0.23884 | 0.059178 | 0.729429 | Prdx2 |
| ENSMUSG( | 1473.252 | -0.14953 | 0.293911 | 0.898538 | Prdx3 |
| ENSMUSG( | 677.2962 | 0.11398 | 0.491466 | 0.926825 | Prdx4 |
| ENSMUSG( | 4231.834 | -0.23179 | 0.080754 | 0.77681 | Prdx5 |
| ENSMUSG( | 14873.31 | -0.14441 | 0.253186 | 0.88018 | Prdx6 |
| ENSMUSG( | 1735.081 | -0.0262 | 0.820878 | 0.979198 | Preb |
| ENSMUSG( | 2783.731 | -0.17178 | 0.142814 | 0.831221 | Prelid1 |
| ENSMUSG( | 1163.664 | -0.05504 | 0.64891 | 0.95422 | Prelid3a |
| ENSMUSG( | 1521.312 | -0.13262 | 0.371471 | 0.915491 | Prelid3b |
| ENSMUSG( | 1054.923 | 0.026291 | 0.895104 | 0.990558 | Prelp |
| ENSMUSG( | 709.4481 | 0.024418 | 0.867431 | 0.987764 | Prep |
| ENSMUSG( | 4145.401 | -0.01507 | 0.893183 | 0.990558 | Prepl |
| ENSMUSG( | 13089.24 | -0.19402 | 0.093757 | 0.795735 | Prex1 |
| ENSMUSG( | 9901.9 | -0.05086 | 0.714087 | 0.964986 | Prex2 |
| ENSMUSG( | 256.0207 | 0.486286 | 0.013304 | 0.461493 | Prg4 |
| ENSMUSG( | 2010.76 | 0.115404 | 0.340425 | 0.907972 | Prickle1 |
| ENSMUSG( | 5412.192 | 0.095432 | 0.343079 | 0.909646 | Prickle2 |
| ENSMUSG( | 45.22548 | 0.421559 | 0.204577 | 0.864412 | Prickle3 |
| ENSMUSG( | 116.2968 | 0.146147 | 0.494137 | 0.927372 | Prim1 |
| ENSMUSG( | 156.3347 | -0.10162 | 0.581418 | 0.944266 | Prim2 |
| ENSMUSG( | 126.3402 | -0.14475 | 0.496397 | 0.927825 | Prima1 |
| ENSMUSG( | 165.7149 | 0.097033 | 0.595574 | 0.945931 | Primpol |
| ENSMUSG( | 519.3181 | 0.028403 | 0.822977 | 0.979198 | Prkaa1 |
| ENSMUSG( | 535.4439 | 0.127604 | 0.426162 | 0.917257 | Prkaa2 |
| ENSMUSG( | 444.097 | -0.09363 | 0.555912 | 0.939565 | Prkab1 |
| ENSMUSG( | 1059.011 | 0.002408 | 0.983937 | 0.996963 | Prkab2 |
| ENSMUSG( | 3951.991 | -0.10853 | 0.426727 | 0.917257 | Prkaca |
| ENSMUSG( | 7966.513 | -0.09287 | 0.459551 | 0.923329 | Prkacb |
| ENSMUSG( | 1287.99 | -0.09305 | 0.405147 | 0.917257 | Prkag1 |
| ENSMUSG( | 2255.104 | 0.000511 | 0.996194 | 0.999178 | Prkag2 |
| ENSMUSG( | 12276.99 | -0.04038 | 0.69607 | 0.960867 | Prkar1a |
| ENSMUSG( | 13672.94 | -0.16774 | 0.178952 | 0.849041 | Prkar1b |
| ENSMUSG( | 1486.326 | 0.164595 | 0.158884 | 0.837724 | Prkar2a |
| ENSMUSG( | 1002.405 | -0.05758 | 0.727994 | 0.966913 | Prkar2b |
| ENSMUSG( | 3898.416 | -0.039 | 0.691329 | 0.960071 | Prkca |
| ENSMUSG( | 14551.67 | 0.036883 | 0.764973 | 0.974236 | Prkcb |
| ENSMUSG( | 845.3422 | -0.31995 | 0.088983 | 0.788051 | Prkcd |
| ENSMUSG( | 8828.534 | 0.106895 | 0.377891 | 0.916784 | Prkce |
| ENSMUSG( | 7956.973 | 0.064708 | 0.680192 | 0.9581 | Prkcg |
| ENSMUSG( | 204.0216 | 0.340829 | 0.091452 | 0.791889 | Prkch |
| ENSMUSG( | 1253.856 | -0.05334 | 0.661825 | 0.956205 | Prkci |
| ENSMUSG( | 833.4965 | -0.10934 | 0.404104 | 0.917257 | Prkcq |
| ENSMUSG( | 3549.749 | 0.215487 | 0.13944 | 0.830678 | Prkcsh |
| ENSMUSG( | 9634.887 | 0.055283 | 0.592124 | 0.944975 | Prkcz |
| ENSMUSG( | 913.5259 | 0.137297 | 0.380552 | 0.917202 | Prkd1 |
| ENSMUSG( | 210.3347 | 0.301978 | 0.101817 | 0.801719 | Prkd2 |
| ENSMUSG( | 1609.624 | -0.0414 | 0.759248 | 0.97199 | Prkd3 |
| ENSMUSG( | 806.6718 | 0.022546 | 0.865093 | 0.98675 | Prkdc |
| ENSMUSG( | 171.2052 | 0.352506 | 0.045521 | 0.693959 | Prkg1 |
| ENSMUSG( | 406.2438 | -0.03892 | 0.791776 | 0.97681 | Prkg2 |
| ENSMUSG( | 431.6055 | -0.07726 | 0.719497 | 0.966876 | Prkn |
| ENSMUSG( | 443.1087 | -0.05738 | 0.683736 | 0.958678 | Prkra |
| ENSMUSG( | 390.5372 | -0.11988 | 0.423762 | 0.917257 | Prkrip1 |
| ENSMUSG( | 558.3665 | -1.78E-05 | 0.999889 | 0.999947 | Prkx |
| ENSMUSG( | 92.03954 | 0.888522 | 0.017763 | 0.534632 | Prlr |
| ENSMUSG( | 1139.372 | -0.10403 | 0.445492 | 0.921734 | Prmt1 |
| ENSMUSG( | 1229.22 | -0.04058 | 0.717406 | 0.965964 | Prmt2 |
| ENSMUSG( | 517.398 | 0.023212 | 0.860525 | 0.986137 | Prmt3 |
| ENSMUSG( | 1686.493 | -0.02613 | 0.827861 | 0.980224 | Prmt5 |
| ENSMUSG( | 284.5437 | 0.009972 | 0.950033 | 0.994798 | Prmt6 |
| ENSMUSG( | 834.7346 | 0.010814 | 0.932793 | 0.99306 | Prmt7 |
| ENSMUSG( | 2276.439 | -0.11226 | 0.35276 | 0.912209 | Prmt8 |
| ENSMUSG( | 396.7716 | 0.061001 | 0.679482 | 0.957966 | Prmt9 |
| ENSMUSG( | 35216.36 | 0.030083 | 0.835242 | 0.981237 | Prnp |
| ENSMUSG( | 639.6894 | -0.06464 | 0.658842 | 0.956205 | Prob1 |
| ENSMUSG( | 367.142 | 0.094596 | 0.59415 | 0.945408 | Proca1 |
| ENSMUSG( | 31.83967 | 0.458621 | 0.241651 | 0.876528 | Procr |
| ENSMUSG( | 2921.126 | 0.011496 | 0.92324 | 0.99272 | Prodh |
| ENSMUSG( | 100.1715 | 0.161495 | 0.509406 | 0.929089 | Prokr2 |
| ENSMUSG( | 1244.689 | 0.560378 | 0.01361 | 0.466552 | Prom1 |
| ENSMUSG( | 24.99417 | -0.19245 | 0.664884 | 0.956766 | Prom2 |
| ENSMUSG( | 110.0832 | -0.07663 | 0.715925 | 0.965424 | Prorp |
| ENSMUSG( | 85.97378 | 0.246712 | 0.339892 | 0.907972 | Prorsd1 |
| ENSMUSG( | 842.1307 | 0.239096 | 0.150525 | 0.837173 | Pros1 |
| ENSMUSG( | 582.6106 | 0.120324 | 0.391027 | 0.917257 | Proser1 |
| ENSMUSG( | 95.2815 | 0.350813 | 0.203165 | 0.864412 | Proser2 |
| ENSMUSG( | 81.43676 | 0.387917 | 0.114419 | 0.811703 | Proser3 |
| ENSMUSG( | 492.4774 | 0.06896 | 0.658312 | 0.956205 | Prox1 |
| ENSMUSG( | 25.89245 | -0.21071 | 0.615794 | 0.951306 | Prox1os |
| ENSMUSG( | 72.76707 | -0.09942 | 0.695797 | 0.960867 | Prox2 |
| ENSMUSG( | 970.009 | 0.054056 | 0.641686 | 0.953903 | Prpf18 |
| ENSMUSG( | 3494.652 | -0.07962 | 0.556998 | 0.939565 | Prpf19 |
| ENSMUSG( | 481.7423 | 0.125157 | 0.391109 | 0.917257 | Prpf3 |
| ENSMUSG( | 823.9937 | -0.16853 | 0.198008 | 0.862161 | Prpf31 |
| ENSMUSG( | 484.1985 | -0.05506 | 0.703409 | 0.962481 | Prpf38a |
| ENSMUSG( | 1701.159 | 0.177603 | 0.126275 | 0.81516 | Prpf38b |
| ENSMUSG( | 1587.833 | 0.308196 | 0.045888 | 0.695944 | Prpf39 |
| ENSMUSG( | 529.4148 | 0.119962 | 0.356183 | 0.913358 | Prpf4 |
| ENSMUSG( | 1241.701 | 0.043126 | 0.742105 | 0.969415 | Prpf40a |
| ENSMUSG( | 1800.826 | -0.00735 | 0.965155 | 0.995433 | Prpf40b |
| ENSMUSG( | 1911.805 | 0.295913 | 0.063202 | 0.736482 | Prpf4b |
| ENSMUSG( | 1204.697 | -0.07665 | 0.546053 | 0.938063 | Prpf6 |
| ENSMUSG( | 4779.434 | -0.0285 | 0.792473 | 0.97681 | Prpf8 |
| ENSMUSG( | 73.80376 | -0.06596 | 0.807412 | 0.978513 | Prph |
| ENSMUSG( | 886.7219 | -0.16033 | 0.280273 | 0.892356 | Prps1 |
| ENSMUSG( | 136.3989 | -0.08677 | 0.669257 | 0.956766 | Prps1l3 |
| ENSMUSG( | 699.5033 | -0.07862 | 0.613918 | 0.951177 | Prps2 |
| ENSMUSG( | 1127.642 | 0.096976 | 0.431505 | 0.917315 | Prpsap1 |
| ENSMUSG( | 690.6333 | -0.15976 | 0.204724 | 0.864412 | Prpsap2 |
| ENSMUSG( | 42.25001 | -0.27077 | 0.410455 | 0.917257 | Prr11 |
| ENSMUSG( | 2392.259 | 0.157805 | 0.15838 | 0.837724 | Prr12 |
| ENSMUSG( | 942.5171 | -0.24813 | 0.06299 | 0.736482 | Prr13 |
| ENSMUSG( | 760.0365 | 0.082228 | 0.544891 | 0.938063 | Prr14 |
| ENSMUSG( | 1168.397 | 0.253096 | 0.031269 | 0.631113 | Prr14l |
| ENSMUSG( | 12.03821 | -0.31589 | 0.589775 | 0.944975 | Prr15 |
| ENSMUSG( | 160.8019 | -0.22469 | 0.277231 | 0.891468 | Prr16 |
| ENSMUSG( | 9091.054 | -0.39298 | 0.001616 | 0.150169 | Prr18 |
| ENSMUSG( | 27.05451 | 0.003674 | 0.992972 | 0.998813 | Prr22 |
| ENSMUSG( | 678.6904 | -0.02728 | 0.860653 | 0.986137 | Prr3 |
| ENSMUSG( | 24.68112 | -0.15196 | 0.795209 | 0.97681 | Prr32 |
| ENSMUSG( | 14.8462 | -0.22744 | 0.70157 | 0.962014 | Prr35 |
| ENSMUSG( | 716.6753 | -0.06085 | 0.642557 | 0.953903 | Prr36 |
| ENSMUSG( | 502.0012 | 0.195248 | 0.246411 | 0.877904 | Prr5 |
| ENSMUSG( | 520.2305 | -0.20332 | 0.155556 | 0.837724 | Prr5l |
| ENSMUSG( | 1681.525 | 0.110056 | 0.359346 | 0.914126 | Prr7 |
| ENSMUSG( | 527.4643 | -0.05675 | 0.669331 | 0.956766 | Prrc1 |
| ENSMUSG( | 15189.9 | 0.115776 | 0.302112 | 0.898538 | Prrc2a |
| ENSMUSG( | 13812.47 | 0.131356 | 0.348723 | 0.911652 | Prrc2b |
| ENSMUSG( | 3968.562 | 0.1703 | 0.121783 | 0.813384 | Prrc2c |
| ENSMUSG( | 134.8263 | 0.330172 | 0.194392 | 0.860096 | Prrg1 |
| ENSMUSG( | 261.7752 | 0.021941 | 0.908463 | 0.991958 | Prrg2 |
| ENSMUSG( | 1154.885 | 0.109647 | 0.478886 | 0.923329 | Prrg3 |
| ENSMUSG( | 18.27152 | 0.328742 | 0.494139 | 0.927372 | Prrg4 |
| ENSMUSG( | 4347.498 | -0.0203 | 0.848824 | 0.985321 | Prrt1 |
| ENSMUSG( | 9.566333 | -0.65027 | 0.343519 | NA | Prrt1b |
| ENSMUSG( | 12.26208 | 0.06448 | 0.91439 | 0.992407 | Prrt2 |
| ENSMUSG( | 1934.576 | 0.148867 | 0.425161 | 0.917257 | Prrt3 |
| ENSMUSG( | 134.8478 | 0.243982 | 0.320156 | 0.903876 | Prrt4 |
| ENSMUSG( | 458.2687 | 0.195642 | 0.216252 | 0.865167 | Prrx1 |
| ENSMUSG( | 58.9983 | -0.20626 | 0.435666 | 0.919133 | Prrx2 |
| ENSMUSG( | 738.8571 | 0.096336 | 0.482378 | 0.924578 | Prss12 |
| ENSMUSG( | 18.56747 | 0.051459 | 0.914101 | 0.992407 | Prss16 |
| ENSMUSG( | 26.47101 | -0.66429 | 0.144066 | 0.831355 | Prss22 |
| ENSMUSG( | 555.7009 | 0.026945 | 0.889429 | 0.990558 | Prss23 |
| ENSMUSG( | 11.79152 | -0.20515 | 0.738005 | 0.967709 | Prss30 |
| ENSMUSG( | 133.3857 | 0.146412 | 0.455954 | 0.923329 | Prss35 |
| ENSMUSG( | 205.9357 | -0.12483 | 0.490326 | 0.926341 | Prss36 |
| ENSMUSG( | 22.13642 | -0.20162 | 0.634844 | 0.953903 | Prss3l |
| ENSMUSG( | 12.18248 | -0.54125 | 0.349649 | 0.911652 | Prss43 |
| ENSMUSG( | 14.48726 | 0.284806 | 0.602233 | 0.94789 | Prss45 |
| ENSMUSG( | 37.70643 | 0.029958 | 0.930254 | 0.99306 | Prss50 |
| ENSMUSG( | 25.70397 | 0.04179 | 0.91803 | 0.992529 | Prss53 |
| ENSMUSG( | 30.46127 | -0.04127 | 0.912407 | 0.992225 | Prss56 |
| ENSMUSG( | 21.32855 | 0.010526 | 0.980552 | 0.996479 | Prss57 |
| ENSMUSG( | 10.88962 | 0.010505 | 0.985464 | NA | Prss8 |
| ENSMUSG( | 62.87737 | 0.069803 | 0.801938 | 0.97681 | Prtg |
| ENSMUSG( | 35.34592 | -0.08721 | 0.799928 | 0.97681 | Prtn3 |
| ENSMUSG( | 659.2935 | 0.030715 | 0.818435 | 0.979198 | Prune1 |
| ENSMUSG( | 1832.898 | 0.084504 | 0.450479 | 0.922202 | Prune2 |
| ENSMUSG( | 48.45643 | -0.25791 | 0.379616 | 0.916995 | Prx |
| ENSMUSG( | 11409.31 | 0.044773 | 0.738901 | 0.968305 | Prxl2a |
| ENSMUSG( | 1959.334 | -0.40116 | 0.001307 | 0.130907 | Prxl2b |
| ENSMUSG( | 451.1934 | 0.041572 | 0.768276 | 0.974236 | Prxl2c |
| ENSMUSG( | 80198.79 | 0.078393 | 0.556446 | 0.939565 | Psap |

| | | | | | | | | | | | |
|---|---|---|---|---|---|---|---|---|---|---|---|
| ENSMUSG( | Psma7 | 2002.27 | -0.00874 | 0.929826 | 0.991848 | ENSMUSG( | Psma7 | 1785.034 | -0.07263 | 0.529546 | 0.934795 |
| ENSMUSG( | Psma8 | 16.77069 | -0.20192 | 0.682095 | NA | ENSMUSG( | Psma8 | 14.18264 | -0.44363 | 0.38379 | 0.917257 |
| ENSMUSG( | Psmb1 | 3168.439 | 0.012315 | 0.890938 | 0.983764 | ENSMUSG( | Psmb1 | 2716.582 | -0.17089 | 0.167971 | 0.840222 |
| ENSMUSG( | Psmb10 | 1092.976 | -0.08409 | 0.411528 | 0.819317 | ENSMUSG( | Psmb10 | 972.8693 | -0.15847 | 0.21225 | 0.864412 |
| ENSMUSG( | Psmb2 | 1258.613 | -0.07958 | 0.48337 | 0.856304 | ENSMUSG( | Psmb2 | 1093.678 | -0.22898 | 0.107003 | 0.804969 |
| ENSMUSG( | Psmb3 | 1478.187 | -0.01357 | 0.900758 | 0.985633 | ENSMUSG( | Psmb3 | 1260.847 | -0.21584 | 0.14268 | 0.831221 |
| ENSMUSG( | Psmb4 | 2849.735 | 0.012968 | 0.904363 | 0.986246 | ENSMUSG( | Psmb4 | 2447.083 | -0.16663 | 0.249931 | 0.877904 |
| ENSMUSG( | Psmb5 | 1593.619 | 0.006635 | 0.951566 | 0.993821 | ENSMUSG( | Psmb5 | 1348.989 | -0.21697 | 0.12423 | 0.814691 |
| ENSMUSG( | Psmb6 | 2195.013 | -0.03637 | 0.694388 | 0.9343 | ENSMUSG( | Psmb6 | 1878.048 | -0.22979 | 0.060829 | 0.734852 |
| ENSMUSG( | Psmb7 | 2478.936 | 0.043318 | 0.656713 | 0.923929 | ENSMUSG( | Psmb7 | 2098.366 | -0.17664 | 0.142149 | 0.830709 |
| ENSMUSG( | Psmb8 | 154.5277 | -0.23094 | 0.178937 | 0.616298 | ENSMUSG( | Psmb8 | 143.2643 | -0.17049 | 0.391101 | 0.917257 |
| ENSMUSG( | Psmb9 | 63.30072 | -0.50778 | 0.053094 | 0.353671 | ENSMUSG( | Psmb9 | 59.08315 | -0.42129 | 0.155845 | 0.837724 |
| ENSMUSG( | Psmc1 | 3133.23 | 0.039799 | 0.621763 | 0.911432 | ENSMUSG( | Psmc1 | 2702.011 | -0.12235 | 0.268284 | 0.886493 |
| ENSMUSG( | Psmc2 | 1730.743 | 0.049011 | 0.690526 | 0.933881 | ENSMUSG( | Psmc2 | 1488.405 | -0.12256 | 0.380864 | 0.917202 |
| ENSMUSG( | Psmc3 | 3660.379 | 0.103082 | 0.282412 | 0.73186 | ENSMUSG( | Psmc3 | 3099.598 | -0.11053 | 0.389812 | 0.917257 |
| ENSMUSG( | Psmc3ip | 76.48445 | -0.26625 | 0.22651 | 0.673549 | ENSMUSG( | Psmc3ip | 69.56449 | -0.28473 | 0.282919 | 0.892412 |
| ENSMUSG( | Psmc4 | 2068.047 | 0.007482 | 0.943341 | 0.993415 | ENSMUSG( | Psmc4 | 1797.951 | -0.13242 | 0.313669 | 0.900949 |
| ENSMUSG( | Psmc5 | 4554.538 | 0.045562 | 0.60076 | 0.90443 | ENSMUSG( | Psmc5 | 3708.518 | -0.29571 | 0.021543 | 0.575341 |
| ENSMUSG( | Psmc6 | 1394.336 | 0.066438 | 0.529593 | 0.878883 | ENSMUSG( | Psmc6 | 1255.987 | 0.03154 | 0.789247 | 0.976593 |
| ENSMUSG( | Psmd1 | 2308.484 | 0.154198 | 0.082977 | 0.44459 | ENSMUSG( | Psmd1 | 1974.171 | -0.02738 | 0.828678 | 0.980224 |
| ENSMUSG( | Psmd10 | 302.5238 | -0.06167 | 0.632393 | 0.914073 | ENSMUSG( | Psmd10 | 257.6548 | -0.27231 | 0.079371 | 0.77458 |
| ENSMUSG( | Psmd11 | 1429.258 | 0.175171 | 0.057438 | 0.370033 | ENSMUSG( | Psmd11 | 1271.906 | 0.107479 | 0.385359 | 0.917257 |
| ENSMUSG( | Psmd12 | 1228.857 | 0.044474 | 0.71348 | 0.940865 | ENSMUSG( | Psmd12 | 1047.073 | -0.15596 | 0.267598 | 0.886134 |
| ENSMUSG( | Psmd13 | 1555.437 | 0.067114 | 0.553038 | 0.886204 | ENSMUSG( | Psmd13 | 1346.613 | -0.0826 | 0.527832 | 0.934603 |
| ENSMUSG( | Psmd14 | 1397.22 | 0.122104 | 0.23298 | 0.681135 | ENSMUSG( | Psmd14 | 1188.089 | -0.07916 | 0.557541 | 0.939565 |
| ENSMUSG( | Psmd2 | 5602.376 | 0.132672 | 0.157119 | 0.58551 | ENSMUSG( | Psmd2 | 4590.583 | -0.17673 | 0.111885 | 0.808673 |
| ENSMUSG( | Psmd3 | 3134.267 | -0.01314 | 0.870469 | 0.977906 | ENSMUSG( | Psmd3 | 2896.823 | 0.028158 | 0.779135 | 0.976106 |
| ENSMUSG( | Psmd4 | 1920.237 | 0.064193 | 0.53791 | 0.882548 | ENSMUSG( | Psmd4 | 1654.998 | -0.10002 | 0.434273 | 0.91869 |
| ENSMUSG( | Psmd5 | 842.5334 | 0.123992 | 0.305145 | 0.752998 | ENSMUSG( | Psmd5 | 766.6693 | 0.118063 | 0.395795 | 0.917257 |
| ENSMUSG( | Psmd6 | 1150.561 | 0.034714 | 0.721191 | 0.942909 | ENSMUSG( | Psmd6 | 1033.98 | -0.00335 | 0.975963 | 0.996416 |
| ENSMUSG( | Psmd7 | 2335.559 | 0.129418 | 0.136401 | 0.550651 | ENSMUSG( | Psmd7 | 2069.33 | 0.048773 | 0.686509 | 0.95899 |
| ENSMUSG( | Psmd8 | 2763.607 | -0.02156 | 0.802287 | 0.962119 | ENSMUSG( | Psmd8 | 2366.103 | -0.21048 | 0.059823 | 0.729747 |
| ENSMUSG( | Psmd9 | 718.3301 | 0.148841 | 0.223686 | 0.669845 | ENSMUSG( | Psmd9 | 614.6698 | -0.03535 | 0.823488 | 0.979198 |
| ENSMUSG( | Psme1 | 898.3763 | -0.07298 | 0.565231 | 0.890779 | ENSMUSG( | Psme1 | 823.5686 | -0.05977 | 0.673558 | 0.95694 |
| ENSMUSG( | Psme2 | 637.9945 | 0.019034 | 0.859439 | 0.97481 | ENSMUSG( | Psme2 | 570.7483 | -0.0349 | 0.784243 | 0.976166 |
| ENSMUSG( | Psme2b | 21.57686 | -0.34663 | 0.449343 | 0.835057 | ENSMUSG( | Psme2b | 20.71638 | -0.18859 | 0.6866 | 0.95899 |
| ENSMUSG( | Psme3 | 3182.751 | 0.030376 | 0.708651 | 0.939519 | ENSMUSG( | Psme3 | 2805.309 | -0.06642 | 0.550077 | 0.938699 |
| ENSMUSG( | Psme3ip1 | 1199.12 | 0.001094 | 0.991443 | 0.999232 | ENSMUSG( | Psme3ip1 | 1043.519 | -0.13976 | 0.2988 | 0.898538 |
| ENSMUSG( | Psme4 | 1622.425 | 0.039094 | 0.703385 | 0.937735 | ENSMUSG( | Psme4 | 1488.743 | 0.062116 | 0.591863 | 0.944975 |
| ENSMUSG( | Psmf1 | 1045.029 | 0.00113 | 0.991419 | 0.999232 | ENSMUSG( | Psmf1 | 920.4289 | -0.10278 | 0.464266 | 0.923329 |
| ENSMUSG( | Psmg1 | 266.7919 | 0.175608 | 0.257405 | 0.708607 | ENSMUSG( | Psmg1 | 229.4018 | 0.004703 | 0.980073 | 0.996479 |
| ENSMUSG( | Psmg2 | 366.1137 | -0.07336 | 0.593673 | 0.90281 | ENSMUSG( | Psmg2 | 346.8791 | 0.033332 | 0.829634 | 0.980224 |
| ENSMUSG( | Psmg3 | 126.5397 | 0.041418 | 0.833243 | 0.970333 | ENSMUSG( | Psmg3 | 117.6318 | 0.088817 | 0.707891 | 0.964042 |
| ENSMUSG( | Psmg4 | 199.5887 | 0.039821 | 0.792998 | 0.95947 | ENSMUSG( | Psmg4 | 180.278 | -0.00277 | 0.98913 | 0.997893 |
| ENSMUSG( | Pspc1 | 826.2059 | -0.03473 | 0.78075 | 0.957491 | ENSMUSG( | Pspc1 | 718.2042 | -0.18001 | 0.207265 | 0.864412 |
| ENSMUSG( | Psph | 1031.958 | -0.01939 | 0.867087 | 0.976541 | ENSMUSG( | Psph | 889.0025 | -0.1909 | 0.143929 | 0.831311 |
| ENSMUSG( | Psrc1 | 238.9478 | 0.178079 | 0.267199 | 0.717525 | ENSMUSG( | Psrc1 | 190.8574 | -0.18748 | 0.371278 | 0.915491 |
| ENSMUSG( | Pstk | 450.8653 | 0.097464 | 0.467352 | 0.844819 | ENSMUSG( | Pstk | 423.2881 | 0.186642 | 0.208242 | 0.864412 |
| ENSMUSG( | Pstpip1 | 85.97245 | -0.36689 | 0.097304 | 0.477629 | ENSMUSG( | Pstpip1 | 77.87348 | -0.39579 | 0.104461 | 0.804969 |
| ENSMUSG( | Pstpip2 | 148.819 | -0.44852 | 0.023925 | 0.236484 | ENSMUSG( | Pstpip2 | 159.7341 | 0.065856 | 0.738387 | 0.967967 |
| ENSMUSG( | Ptafr | 83.17238 | -0.44998 | 0.203258 | 0.643631 | ENSMUSG( | Ptafr | 104.1778 | 0.489872 | 0.197977 | 0.862161 |
| ENSMUSG( | Ptar1 | 3226.179 | -0.11927 | 0.382734 | 0.803227 | ENSMUSG( | Ptar1 | 2838.033 | -0.22889 | 0.132715 | 0.822912 |
| ENSMUSG( | Ptbp1 | 1252.349 | -0.10213 | 0.306888 | 0.754887 | ENSMUSG( | Ptbp1 | 1237.762 | 0.135124 | 0.259716 | 0.883971 |
| ENSMUSG( | Ptbp2 | 2092.609 | 0.03747 | 0.683122 | 0.93213 | ENSMUSG( | Ptbp2 | 1878.113 | -0.00904 | 0.943524 | 0.994798 |
| ENSMUSG( | Ptbp3 | 537.6181 | -0.04363 | 0.747669 | 0.949604 | ENSMUSG( | Ptbp3 | 542.5892 | 0.238433 | 0.125518 | 0.81516 |
| ENSMUSG( | Ptcd1 | 570.7723 | 0.045294 | 0.75606 | 0.951479 | ENSMUSG( | Ptcd1 | 528.5883 | 0.090558 | 0.581205 | 0.944266 |
| ENSMUSG( | Ptcd2 | 570.106 | 0.137996 | 0.236099 | 0.683188 | ENSMUSG( | Ptcd2 | 492.8843 | -0.01114 | 0.935657 | 0.993387 |
| ENSMUSG( | Ptcd3 | 1068.177 | 0.140451 | 0.174395 | 0.610304 | ENSMUSG( | Ptcd3 | 917.9543 | -0.02132 | 0.860207 | 0.986137 |
| ENSMUSG( | Ptch1 | 3608.294 | 0.063189 | 0.507501 | 0.86795 | ENSMUSG( | Ptch1 | 3402.682 | 0.158188 | 0.171912 | 0.842999 |
| ENSMUSG( | Ptch2 | 51.00197 | -0.79461 | 0.005187 | 0.096057 | ENSMUSG( | Ptch2 | 59.2672 | 0.002111 | 0.993745 | 0.998813 |
| ENSMUSG( | Ptchd1 | 96.2041 | -0.20731 | 0.335126 | 0.776511 | ENSMUSG( | Ptchd1 | 108.9746 | 0.403932 | 0.055406 | 0.723362 |
| ENSMUSG( | Ptchd4 | 105.0897 | 0.039747 | 0.866948 | 0.976541 | ENSMUSG( | Ptchd4 | 97.27171 | 0.092595 | 0.708866 | 0.964042 |
| ENSMUSG( | Ptdss1 | 3963.376 | -0.17513 | 0.169803 | 0.604256 | ENSMUSG( | Ptdss1 | 3855.658 | 0.020575 | 0.882376 | 0.989644 |
| ENSMUSG( | Ptdss2 | 2273.431 | -0.1245 | 0.258086 | 0.70936 | ENSMUSG( | Ptdss2 | 2278.88 | 0.152518 | 0.234002 | 0.86917 |
| ENSMUSG( | Pten | 3560.06 | 0.06226 | 0.560939 | 0.888934 | ENSMUSG( | Pten | 3411.096 | 0.199942 | 0.118688 | 0.813384 |
| ENSMUSG( | Pter | 115.2911 | 0.187929 | 0.334701 | 0.776386 | ENSMUSG( | Pter | 95.0298 | -0.11274 | 0.623104 | 0.953234 |
| ENSMUSG( | Ptgdr | 54.10478 | -0.29967 | 0.251271 | 0.700869 | ENSMUSG( | Ptgdr | 60.54412 | 0.309801 | 0.250494 | 0.877932 |
| ENSMUSG( | Ptgds | 34036.71 | 0.209071 | 0.039882 | 0.308163 | ENSMUSG( | Ptgds | 28994.76 | 0.021494 | 0.859258 | 0.985884 |
| ENSMUSG( | Ptger1 | 123.7763 | -0.02575 | 0.887945 | 0.983434 | ENSMUSG( | Ptger1 | 103.3067 | -0.26797 | 0.259371 | 0.883971 |
| ENSMUSG( | Ptger3 | 103.643 | -0.28006 | 0.22576 | 0.672634 | ENSMUSG( | Ptger3 | 116.0538 | 0.314583 | 0.175697 | 0.845305 |
| ENSMUSG( | Ptger4 | 40.27545 | -0.14829 | 0.651287 | 0.921451 | ENSMUSG( | Ptger4 | 38.52926 | -0.00977 | 0.977156 | 0.996416 |
| ENSMUSG( | Ptges | 410.1336 | -0.0181 | 0.893425 | 0.984512 | ENSMUSG( | Ptges | 395.034 | 0.142587 | 0.338275 | 0.907972 |
| ENSMUSG( | Ptges2 | 1153.054 | 0.0514 | 0.579162 | 0.896692 | ENSMUSG( | Ptges2 | 1016.318 | -0.04268 | 0.703877 | 0.96259 |
| ENSMUSG( | Ptges3 | 1989.158 | 0.056416 | 0.638285 | 0.916154 | ENSMUSG( | Ptges3 | 1727.168 | -0.08814 | 0.584951 | 0.944975 |
| ENSMUSG( | Ptges3l | 38.30628 | -0.08129 | 0.794401 | 0.959799 | ENSMUSG( | Ptges3l | 34.88363 | -0.09387 | 0.783319 | 0.976106 |
| ENSMUSG( | Ptgfr | 51.92709 | 0.491512 | 0.062831 | 0.387645 | ENSMUSG( | Ptgfr | 39.30164 | 0.002305 | 0.994266 | 0.998813 |
| ENSMUSG( | Ptgfrn | 2228.228 | -0.11484 | 0.384448 | 0.804007 | ENSMUSG( | Ptgfrn | 2153.606 | 0.060857 | 0.700161 | 0.961335 |
| ENSMUSG( | Ptgis | 78.39753 | -0.23994 | 0.29502 | 0.743798 | ENSMUSG( | Ptgis | 83.77334 | 0.227386 | 0.329444 | 0.907083 |
| ENSMUSG( | Ptgr1 | 59.4979 | 0.439106 | 0.098495 | 0.480334 | ENSMUSG( | Ptgr1 | 45.96854 | -0.02471 | 0.937349 | 0.993685 |
| ENSMUSG( | Ptgr2 | 4118.417 | 0.002924 | 0.981941 | 0.997512 | ENSMUSG( | Ptgr2 | 3433.466 | -0.2693 | 0.064064 | 0.736482 |
| ENSMUSG( | Ptgr3 | 1398.285 | 0.023884 | 0.797131 | 0.960507 | ENSMUSG( | Ptgr3 | 1243.878 | -0.04982 | 0.695451 | 0.960867 |
| ENSMUSG( | Ptgs1 | 786.5756 | -0.15767 | 0.345722 | 0.779153 | ENSMUSG( | Ptgs1 | 798.6953 | 0.163717 | 0.426603 | 0.917257 |
| ENSMUSG( | Ptgs2 | 228.5216 | -0.44185 | 0.01473 | 0.178407 | ENSMUSG( | Ptgs2 | 264.9398 | 0.284248 | 0.168514 | 0.840222 |
| ENSMUSG( | Pth1r | 427.057 | -0.14325 | 0.274771 | 0.725078 | ENSMUSG( | Pth1r | 406.8921 | -0.00662 | 0.965551 | 0.995433 |
| ENSMUSG( | Pth2r | 51.71923 | -0.66619 | 0.020202 | 0.214698 | ENSMUSG( | Pth2r | 58.43578 | 0.030006 | 0.918752 | 0.992529 |
| ENSMUSG( | Pthlh | 102.86 | -1.08736 | 7.51E-08 | 2.13E-05 | ENSMUSG( | Pthlh | 122.7074 | -0.10341 | 0.672737 | 0.95694 |
| ENSMUSG( | Ptk2 | 3577.802 | 0.119592 | 0.173358 | 0.60863 | ENSMUSG( | Ptk2 | 3179.386 | 0.051547 | 0.631909 | 0.953251 |
| ENSMUSG( | Ptk2b | 8536.608 | 0.20336 | 0.203398 | 0.643631 | ENSMUSG( | Ptk2b | 7070.28 | -0.06727 | 0.700071 | 0.961335 |
| ENSMUSG( | Ptk7 | 495.8339 | -0.40736 | 0.013963 | 0.172788 | ENSMUSG( | Ptk7 | 533.8343 | 0.110547 | 0.569796 | 0.941777 |
| ENSMUSG( | Ptma | 5977.331 | 0.04233 | 0.64352 | 0.917316 | ENSMUSG( | Ptma | 5574.502 | 0.105544 | 0.470276 | 0.923329 |
| ENSMUSG( | Ptms | 22594.1 | -0.00719 | 0.939662 | 0.993415 | ENSMUSG( | Ptms | 19727.97 | -0.13538 | 0.398439 | 0.917257 |
| ENSMUSG( | Ptn | 21841.96 | -0.38297 | 0.004104 | 0.08269 | ENSMUSG( | Ptn | 21745.35 | -0.10221 | 0.464015 | 0.923329 |
| ENSMUSG( | Ptov1 | 4615.962 | 0.054823 | 0.507934 | 0.867961 | ENSMUSG( | Ptov1 | 3979.245 | -0.10788 | 0.319921 | 0.903771 |
| ENSMUSG( | Ptp4a1 | 16.62145 | -0.779 | 0.107269 | NA | ENSMUSG( | Ptp4a1 | 18.49635 | -0.04048 | 0.935485 | 0.993265 |
| ENSMUSG( | Ptp4a1 | 13.06645 | -0.01105 | 0.985393 | NA | ENSMUSG( | Ptp4a1 | 10.79748 | -0.32043 | 0.637216 | NA |
| ENSMUSG( | Ptp4a2 | 6175.966 | -0.10182 | 0.341033 | 0.778828 | ENSMUSG( | Ptp4a2 | 5647.451 | -0.09349 | 0.520702 | 0.932121 |
| ENSMUSG( | Ptp4a3 | 2073.902 | 0.011456 | 0.911087 | 0.988483 | ENSMUSG( | Ptp4a3 | 1871.198 | -0.01646 | 0.885359 | 0.989688 |
| ENSMUSG( | Ptpa | 8010.494 | 0.126108 | 0.08779 | 0.454663 | ENSMUSG( | Ptpa | 6840.074 | -0.05886 | 0.535169 | 0.936508 |
| ENSMUSG( | Ptpdc1 | 1186.821 | -0.08626 | 0.413104 | 0.819541 | ENSMUSG( | Ptpdc1 | 1158.237 | 0.112812 | 0.316049 | 0.901868 |
| ENSMUSG( | Ptpmt1 | 1663.42 | -0.01447 | 0.889096 | 0.983434 | ENSMUSG( | Ptpmt1 | 1477.331 | -0.09064 | 0.470624 | 0.923329 |
| ENSMUSG( | Ptpn1 | 2124.732 | 0.088002 | 0.300809 | 0.749979 | ENSMUSG( | Ptpn1 | 1885.92 | 0.014706 | 0.892284 | 0.990558 |
| ENSMUSG( | Ptpn11 | 13193.96 | 0.085159 | 0.277923 | 0.727904 | ENSMUSG( | Ptpn11 | 11275.32 | -0.10165 | 0.365919 | 0.914623 |
| ENSMUSG( | Ptpn12 | 1317.831 | 0.087295 | 0.346072 | 0.779153 | ENSMUSG( | Ptpn12 | 1176.698 | 0.028812 | 0.800697 | 0.97681 |
| ENSMUSG( | Ptpn13 | 1772.559 | 0.071752 | 0.561892 | 0.889052 | ENSMUSG( | Ptpn13 | 1493.638 | -0.15804 | 0.155827 | 0.837724 |
| ENSMUSG( | Ptpn14 | 441.9212 | 0.127737 | 0.319483 | 0.765296 | ENSMUSG( | Ptpn14 | 396.3623 | 0.089857 | 0.545327 | 0.938063 |
| ENSMUSG( | Ptpn18 | 106.2762 | -0.11724 | 0.554617 | 0.886731 | ENSMUSG( | Ptpn18 | 92.58385 | -0.2486 | 0.273912 | 0.889859 |
| ENSMUSG( | Ptpn2 | 413.236 | 0.039922 | 0.805707 | 0.962616 | ENSMUSG( | Ptpn2 | 371.3216 | -0.00662 | 0.972297 | 0.996416 |
| ENSMUSG( | Ptpn21 | 403.8491 | 0.156501 | 0.183425 | 0.62047 | ENSMUSG( | Ptpn21 | 359.536 | 0.097118 | 0.491491 | 0.926825 |
| ENSMUSG( | Ptpn22 | 43.23365 | 0.070784 | 0.831461 | 0.969889 | ENSMUSG( | Ptpn22 | 36.33939 | -0.14969 | 0.690589 | 0.960071 |
| ENSMUSG( | Ptpn23 | 2079.578 | 0.164367 | 0.2101 | 0.652466 | ENSMUSG( | Ptpn23 | 1741.5 | -0.07848 | 0.60737 | 0.950022 |
| ENSMUSG( | Ptpn3 | 1223.904 | 0.19722 | 0.133613 | 0.546858 | ENSMUSG( | Ptpn3 | 1013.048 | -0.07332 | 0.592266 | 0.944975 |
| ENSMUSG( | Ptpn4 | 2233.277 | -0.07708 | 0.610484 | 0.908189 | ENSMUSG( | Ptpn4 | 2012.875 | -0.11329 | 0.530351 | 0.934795 |
| ENSMUSG( | Ptpn5 | 7043.261 | -0.26532 | 0.006954 | 0.114623 | ENSMUSG( | Ptpn5 | 7109.219 | 0.044212 | 0.698048 | 0.960867 |
| ENSMUSG( | Ptpn6 | 257.6582 | -0.0772 | 0.585513 | 0.899171 | ENSMUSG( | Ptpn6 | 247.0329 | 0.070563 | 0.662864 | 0.956205 |
| ENSMUSG( | Ptpn7 | 32.00888 | -0.85786 | 0.023624 | 0.235222 | ENSMUSG( | Ptpn7 | 32.92422 | -0.45334 | 0.223725 | 0.867404 |
| ENSMUSG( | Ptpn9 | 1434.656 | 0.20341 | 0.019709 | 0.21171 | ENSMUSG( | Ptpn9 | 1212.428 | -0.0075 | 0.944995 | 0.994798 |
| ENSMUSG( | Ptpra | 9573.561 | -0.11513 | 0.348608 | 0.78132 | ENSMUSG( | Ptpra | 9224.653 | 0.051223 | 0.726973 | 0.966887 |
| ENSMUSG( | Ptprb | 1157.032 | 0.207323 | 0.593775 | 0.902869 | ENSMUSG( | Ptprb | 1356.625 | 0.835012 | 0.105103 | 0.804969 |
| ENSMUSG( | Ptprc | 265.2712 | -0.51813 | 0.014487 | 0.177206 | ENSMUSG( | Ptprc | 270.4131 | -0.15113 | 0.522991 | 0.932848 |
| ENSMUSG( | Ptprcap | 15.01423 | -0.7991 | 0.108061 | NA | ENSMUSG( | Ptprcap | 16.29824 | -0.24914 | 0.629937 | 0.953251 |
| ENSMUSG( | Ptprd | 8399.71 | -0.31753 | 0.023185 | 0.233583 | ENSMUSG( | Ptprd | 8816.358 | 0.109661 | 0.474417 | 0.923329 |
| ENSMUSG( | Ptpre | 927.961 | -0.34832 | 0.032154 | 0.273917 | ENSMUSG( | Ptpre | 1005.513 | 0.174806 | 0.344942 | 0.91045 |
| ENSMUSG( | Ptprf | 4317.084 | -0.03865 | 0.763546 | 0.953845 | ENSMUSG( | Ptprf | 4315.606 | 0.224602 | 0.120059 | 0.813384 |
| ENSMUSG( | Ptprg | 3394.971 | -0.02894 | 0.834182 | 0.970333 | ENSMUSG( | Ptprg | 3348.511 | 0.19569 | 0.182516 | 0.850579 |
| ENSMUSG( | Ptprh | 39.39953 | -0.10625 | 0.736014 | 0.946267 | ENSMUSG( | Ptprh | 38.22707 | 0.078146 | 0.811387 | 0.978997 |
| ENSMUSG( | Ptprj | 2734.745 | -0.01102 | 0.939188 | 0.993415 | ENSMUSG( | Ptprj | 2888.008 | 0.393776 | 0.025303 | 0.598154 |
| ENSMUSG( | Ptprk | 1756.344 | -0.1479 | 0.341915 | 0.778828 | ENSMUSG( | Ptprk | 1790.713 | 0.180799 | 0.287521 | 0.894091 |
| ENSMUSG( | Ptprm | 1188.477 | -0.62154 | 4.19E-05 | 0.003503 | ENSMUSG( | Ptprm | 1406.686 | 0.199784 | 0.344287 | 0.910321 |
| ENSMUSG( | Ptprn | 19630.21 | -0.10538 | 0.37882 | 0.801731 | ENSMUSG( | Ptprn | 19421.07 | 0.134429 | 0.325783 | 0.905339 |
| ENSMUSG( | Ptprn2 | 5074.286 | -0.07023 | 0.727201 | 0.944223 | ENSMUSG( | Ptprn2 | 5499.222 | 0.411004 | 0.276772 | 0.890889 |
| ENSMUSG( | Ptpro | 896.6646 | -0.59631 | 0.00041 | 0.018302 | ENSMUSG( | Ptpro | 1040.486 | 0.162307 | 0.294853 | 0.898538 |
| ENSMUSG( | Ptprr | 494.9051 | -0.52131 | 0.00097 | 0.03229 | ENSMUSG( | Ptprr | 562.9252 | 0.167516 | 0.32049 | 0.903887 |
| ENSMUSG( | Ptprs | 18015.22 | 0.008399 | 0.931338 | 0.992327 | ENSMUSG( | Ptprs | 17505.23 | 0.188905 | 0.077105 | 0.770558 |
| ENSMUSG( | Ptprt | 9891.329 | -0.09817 | 0.425204 | 0.82478 | ENSMUSG( | Ptprt | 9279.107 | -0.01144 | 0.927638 | 0.99306 |
| ENSMUSG( | Ptpru | 1861.241 | 0.234163 | 0.065201 | 0.395697 | ENSMUSG( | Ptpru | 1723.304 | 0.280104 | 0.031482 | 0.631192 |
| ENSMUSG( | Ptprv | 83.79861 | -1.18724 | 8.54E-05 | 0.005781 | ENSMUSG( | Ptprv | 93.67729 | -0.39952 | 0.215734 | 0.865167 |
| ENSMUSG( | Ptprz1 | 23315.93 | -0.03481 | 0.788411 | 0.959226 | ENSMUSG( | Ptprz1 | 21401.61 | -0.0124 | 0.928996 | 0.99306 |
| ENSMUSG( | Ptrh1 | 115.6432 | 0.155408 | 0.397455 | 0.812127 | ENSMUSG( | Ptrh1 | 98.89572 | -0.0188 | 0.931461 | 0.99306 |
| ENSMUSG( | Ptrh2 | 295.6317 | 0.017902 | 0.900926 | 0.985633 | ENSMUSG( | Ptrh2 | 262.7744 | -0.06009 | 0.720875 | 0.966887 |
| ENSMUSG( | Ptrhd1 | 425.7713 | -0.01088 | 0.934201 | 0.99323 | ENSMUSG( | Ptrhd1 | 356.949 | -0.27299 | 0.079201 | 0.77458 |
| ENSMUSG( | Pts | 1433.835 | 0.013027 | 0.901805 | 0.985633 | ENSMUSG( | Pts | 1247.905 | -0.12336 | 0.27809 | 0.891468 |
| ENSMUSG( | Pttg1 | 596.0987 | 0.002234 | 0.984461 | 0.99781 | ENSMUSG( | Pttg1 | 509.9832 | -0.18847 | 0.17064 | 0.841762 |
| ENSMUSG( | Pttg1ip | 3829.805 | -0.13564 | 0.296148 | 0.744386 | ENSMUSG( | Pttg1ip | 3805.939 | 0.119349 | 0.395508 | 0.917257 |
| ENSMUSG( | Ptx3 | 12.03456 | 0.250563 | 0.644985 | NA | ENSMUSG( | Ptx3 | 16.52202 | 1.199517 | 0.027273 | 0.60551 |
| ENSMUSG( | Puf60 | 3363.713 | 0.091493 | 0.370532 | 0.797778 | ENSMUSG( | Puf60 | 2815.137 | -0.15806 | 0.20141 | 0.864412 |
| ENSMUSG( | Pum1 | 2592.651 | 0.103359 | 0.206364 | 0.648436 | ENSMUSG( | Pum1 | 2273.148 | -0.00792 | 0.945771 | 0.994798 |
| ENSMUSG( | Pum2 | 6900.384 | 0.043614 | 0.626833 | 0.912825 | ENSMUSG( | Pum2 | 6311.72 | 0.054205 | 0.630929 | 0.953251 |
| ENSMUSG( | Pum3 | 1085.137 | 0.15159 | 0.148013 | 0.571232 | ENSMUSG( | Pum3 | 897.4495 | -0.12957 | 0.306692 | 0.898538 |
| ENSMUSG( | Pura | 1845.154 | -0.01171 | 0.924792 | 0.991152 | ENSMUSG( | Pura | 1617.303 | -0.13029 | 0.409103 | 0.917257 |
| ENSMUSG( | Purb | 34513.99 | 0.073377 | 0.484214 | 0.856577 | ENSMUSG( | Purb | 30564.6 | -0.00865 | 0.951939 | 0.994798 |
| ENSMUSG( | Purg | 733.9863 | 0.143838 | 0.193947 | 0.633085 | ENSMUSG( | Purg | 631.6914 | -0.02523 | 0.868019 | 0.98781 |
| ENSMUSG( | Pus1 | 292.5674 | -0.00385 | 0.980762 | 0.997512 | ENSMUSG( | Pus1 | 268.4948 | 0.02119 | 0.90027 | 0.990558 |
| ENSMUSG( | Pus10 | 299.3864 | 0.025964 | 0.841702 | 0.971179 | ENSMUSG( | Pus10 | 267.8519 | -0.03012 | 0.844039 | 0.983571 |
| ENSMUSG( | Pus3 | 256.6271 | 0.193951 | 0.169845 | 0.604274 | ENSMUSG( | Pus3 | 224.659 | 0.082738 | 0.605624 | 0.949298 |
| ENSMUSG( | Pus7 | 607.329 | 0.126565 | 0.339802 | 0.777939 | ENSMUSG( | Pus7 | 564.0881 | 0.17393 | 0.28343 | 0.892412 |
| ENSMUSG( | Pus7l | 86.80069 | -0.20254 | 0.372396 | 0.798768 | ENSMUSG( | Pus7l | 82.10664 | -0.10008 | 0.694512 | 0.960867 |
| ENSMUSG( | Pusl1 | 765.4575 | 0.113217 | 0.289262 | 0.739725 | ENSMUSG( | Pusl1 | 704.8866 | 0.144795 | 0.253925 | 0.880967 |
| ENSMUSG( | Pvalb | 856.7935 | -0.08165 | 0.632558 | 0.914073 | ENSMUSG( | Pvalb | 682.3404 | -0.51874 | 0.002697 | 0.205145 |

| | | | | | |
|---|---|---|---|---|---|
| ENSMUSG( | 127.2928 | 0.136633 | 0.451265 | 0.835567 | Prim1 |
| ENSMUSG( | 187.1327 | 0.141736 | 0.365494 | 0.794085 | Prim2 |
| ENSMUSG( | 116.0028 | -0.7694 | 0.000203 | 0.010726 | Prima1 |
| ENSMUSG( | 177.2422 | 0.015877 | 0.921556 | 0.990436 | Primpol |
| ENSMUSG( | 572.2544 | 0.039244 | 0.716137 | 0.94152 | Prkaa1 |
| ENSMUSG( | 578.3166 | 0.083338 | 0.564006 | 0.890139 | Prkaa2 |
| ENSMUSG( | 469.4959 | -0.21725 | 0.106073 | 0.493403 | Prkab1 |
| ENSMUSG( | 1175.554 | 0.033179 | 0.725521 | 0.944223 | Prkab2 |
| ENSMUSG( | 4501.927 | 0.002178 | 0.985245 | 0.997982 | Prkaca |
| ENSMUSG( | 9100.655 | 0.02566 | 0.793781 | 0.959799 | Prkacb |
| ENSMUSG( | 1485.578 | 0.052241 | 0.560345 | 0.888934 | Prkag1 |
| ENSMUSG( | 2634.399 | 0.175984 | 0.039116 | 0.304587 | Prkag2 |
| ENSMUSG( | 13770.48 | 0.0234 | 0.760773 | 0.952907 | Prkar1a |
| ENSMUSG( | 16443.69 | 0.100284 | 0.378751 | 0.801688 | Prkar1b |
| ENSMUSG( | 1589.5 | 0.09452 | 0.305405 | 0.753523 | Prkar2a |
| ENSMUSG( | 1013.742 | -0.3035 | 0.010086 | 0.143118 | Prkar2b |
| ENSMUSG( | 4336.61 | 2.76E-05 | 0.999734 | 0.99989 | Prkca |
| ENSMUSG( | 16174.43 | 0.073 | 0.409662 | 0.818258 | Prkcb |
| ENSMUSG( | 934.6838 | -0.29071 | 0.075523 | 0.425931 | Prkcd |
| ENSMUSG( | 9754.211 | 0.12529 | 0.118329 | 0.520133 | Prkce |
| ENSMUSG( | 9208.465 | 0.212728 | 0.0437 | 0.320345 | Prkcg |
| ENSMUSG( | 176.0884 | -0.36993 | 0.047529 | 0.333288 | Prkch |
| ENSMUSG( | 1429.979 | 0.060598 | 0.55076 | 0.886038 | Prkci |
| ENSMUSG( | 944.2275 | -0.01102 | 0.926875 | 0.991229 | Prkcq |
| ENSMUSG( | 3360.084 | -0.21514 | 0.090662 | 0.462287 | Prkcsh |
| ENSMUSG( | 10732.83 | 0.095822 | 0.241598 | 0.689179 | Prkcz |
| ENSMUSG( | 1021.229 | 0.193459 | 0.128337 | 0.536641 | Prkd1 |
| ENSMUSG( | 212.4043 | 0.068532 | 0.665696 | 0.927054 | Prkd2 |
| ENSMUSG( | 1848.734 | 0.089866 | 0.469119 | 0.845876 | Prkd3 |
| ENSMUSG( | 854.2805 | -0.0866 | 0.439368 | 0.830988 | Prkdc |
| ENSMUSG( | 177.6018 | 0.209902 | 0.180171 | 0.617634 | Prkg1 |
| ENSMUSG( | 455.2041 | 0.025079 | 0.854887 | 0.973665 | Prkg2 |
| ENSMUSG( | 508.3227 | 0.135512 | 0.417919 | 0.821964 | Prkn |
| ENSMUSG( | 519.0406 | 0.133843 | 0.245811 | 0.693547 | Prkra |
| ENSMUSG( | 447.7642 | 0.009828 | 0.940783 | 0.993415 | Prkrip1 |
| ENSMUSG( | 640.3791 | 0.126427 | 0.296259 | 0.744386 | Prkx |
| ENSMUSG( | 83.89142 | 0.464129 | 0.474706 | 0.849485 | Prlr |
| ENSMUSG( | 1291.804 | -0.006 | 0.956739 | 0.993874 | Prmt1 |
| ENSMUSG( | 1413.942 | 0.094671 | 0.310783 | 0.758828 | Prmt2 |
| ENSMUSG( | 598.0676 | 0.171338 | 0.108793 | 0.497919 | Prmt3 |
| ENSMUSG( | 1968.81 | 0.149431 | 0.12761 | 0.535393 | Prmt5 |
| ENSMUSG( | 314.5604 | 0.028658 | 0.827424 | 0.968732 | Prmt6 |
| ENSMUSG( | 978.2709 | 0.193856 | 0.100639 | 0.48402 | Prmt7 |
| ENSMUSG( | 2672.019 | 0.083596 | 0.403538 | 0.816132 | Prmt8 |
| ENSMUSG( | 422.8075 | -0.02925 | 0.827739 | 0.968924 | Prmt9 |
| ENSMUSG( | 33726.91 | -0.3915 | 0.001845 | 0.048742 | Prnp |
| ENSMUSG( | 710.1887 | -0.03036 | 0.781511 | 0.957558 | Prob1 |
| ENSMUSG( | 422.8272 | 0.234871 | 0.099811 | 0.482257 | Proca1 |
| ENSMUSG( | 32.92901 | 0.28518 | 0.416239 | 0.82084 | Procr |
| ENSMUSG( | 3448.32 | 0.212239 | 0.018318 | 0.203658 | Prodh |
| ENSMUSG( | 77.65413 | -0.99557 | 0.001014 | 0.033228 | Prokr2 |
| ENSMUSG( | 1137.145 | 0.075024 | 0.687865 | 0.93324 | Prom1 |
| ENSMUSG( | 29.05425 | -0.01944 | 0.963445 | 0.994911 | Prom2 |
| ENSMUSG( | 130.4838 | 0.139135 | 0.444346 | 0.833239 | Prorp |
| ENSMUSG( | 94.79561 | 0.280134 | 0.190749 | 0.630215 | Prorsd1 |
| ENSMUSG( | 830.3952 | -0.0674 | 0.652297 | 0.922188 | Pros1 |
| ENSMUSG( | 618.4634 | 0.021806 | 0.852205 | 0.973168 | Proser1 |
| ENSMUSG( | 94.91984 | 0.064262 | 0.768052 | 0.954621 | Proser2 |
| ENSMUSG( | 80.28365 | 0.113529 | 0.609572 | 0.9077 | Proser3 |
| ENSMUSG( | 533.9876 | 0.026819 | 0.833724 | 0.970333 | Prox1 |
| ENSMUSG( | 33.86422 | 0.262606 | 0.446168 | 0.833546 | Prox1os |
| ENSMUSG( | 82.39059 | -0.00229 | 0.992921 | 0.999327 | Prox2 |
| ENSMUSG( | 1082.604 | 0.101756 | 0.307951 | 0.75624 | Prpf18 |
| ENSMUSG( | 4153.406 | 0.149768 | 0.162449 | 0.593257 | Prpf19 |
| ENSMUSG( | 529.611 | 0.125759 | 0.245246 | 0.692812 | Prpf3 |
| ENSMUSG( | 977.9445 | 0.064603 | 0.552333 | 0.886141 | Prpf31 |
| ENSMUSG( | 541.6164 | 0.001457 | 0.991263 | 0.999232 | Prpf38a |
| ENSMUSG( | 1662.065 | -0.15491 | 0.10183 | 0.486216 | Prpf38b |
| ENSMUSG( | 1505.587 | -0.10165 | 0.469601 | 0.846167 | Prpf39 |
| ENSMUSG( | 563.8293 | 0.036794 | 0.743615 | 0.948478 | Prpf4 |
| ENSMUSG( | 1367.794 | 0.056758 | 0.567602 | 0.891774 | Prpf40a |
| ENSMUSG( | 2125.27 | 0.197967 | 0.152674 | 0.579011 | Prpf40b |
| ENSMUSG( | 1872.689 | -0.01969 | 0.857233 | 0.97444 | Prpf4b |
| ENSMUSG( | 1435.894 | 0.157797 | 0.150021 | 0.574564 | Prpf6 |
| ENSMUSG( | 5486.71 | 0.097911 | 0.265057 | 0.715962 | Prpf8 |
| ENSMUSG( | 90.86567 | 0.265131 | 0.261329 | 0.711405 | Prph |
| ENSMUSG( | 1030.741 | 0.012874 | 0.909557 | 0.988084 | Prps1 |
| ENSMUSG( | 153.952 | -0.00439 | 0.979745 | 0.997428 | Prps1l3 |
| ENSMUSG( | 769.3521 | -0.06601 | 0.539438 | 0.883094 | Prps2 |
| ENSMUSG( | 1177.047 | -0.05042 | 0.644029 | 0.917536 | Prpsap1 |
| ENSMUSG( | 833.2954 | 0.117454 | 0.25938 | 0.709948 | Prpsap2 |
| ENSMUSG( | 43.23564 | -0.50758 | 0.102655 | 0.488148 | Prr11 |
| ENSMUSG( | 2534.61 | 0.058567 | 0.535172 | 0.880733 | Prr12 |
| ENSMUSG( | 1135.617 | 0.036357 | 0.709686 | 0.939669 | Prr13 |
| ENSMUSG( | 801.675 | -0.03238 | 0.783511 | 0.957558 | Prr14 |
| ENSMUSG( | 1262.939 | 0.209691 | 0.054316 | 0.358716 | Prr14l |
| ENSMUSG( | 14.10023 | -0.07522 | 0.885101 | NA | Prr15 |
| ENSMUSG( | 174.2096 | -0.25878 | 0.164482 | 0.595363 | Prr16 |
| ENSMUSG( | 10568.77 | -0.20489 | 0.031337 | 0.269505 | Prr18 |
| ENSMUSG( | 29.50564 | 0.006016 | 0.987149 | 0.998764 | Prr22 |
| ENSMUSG( | 758.7714 | 0.028654 | 0.832339 | 0.970333 | Prr3 |
| ENSMUSG( | 24.94699 | -0.42175 | 0.514026 | 0.872055 | Prr32 |
| ENSMUSG( | 859.4479 | 0.187258 | 0.109517 | 0.499679 | Prr36 |
| ENSMUSG( | 630.5375 | 0.535428 | 1.09E-05 | 0.001199 | Prr5 |
| ENSMUSG( | 557.1306 | -0.28489 | 0.012031 | 0.158657 | Prr5l |
| ENSMUSG( | 1669.753 | -0.18397 | 0.057306 | 0.369582 | Prr7 |
| ENSMUSG( | 592.8008 | 0.011757 | 0.921818 | 0.990436 | Prrc1 |
| ENSMUSG( | 17228.63 | 0.204446 | 0.012482 | 0.161449 | Prrc2a |
| ENSMUSG( | 14784.71 | 0.058302 | 0.591897 | 0.901716 | Prrc2b |
| ENSMUSG( | 4004.133 | -0.07039 | 0.468471 | 0.845415 | Prrc2c |
| ENSMUSG( | 117.773 | -0.34238 | 0.096555 | 0.476986 | Prrg1 |
| ENSMUSG( | 260.835 | -0.27749 | 0.109357 | 0.499374 | Prrg2 |
| ENSMUSG( | 1145.318 | -0.19271 | 0.194534 | 0.633457 | Prrg3 |
| ENSMUSG( | 15.10317 | -0.50224 | 0.324077 | NA | Prrg4 |
| ENSMUSG( | 4777.747 | -0.0164 | 0.853901 | 0.973491 | Prrt1 |
| ENSMUSG( | 11.61736 | -0.36594 | 0.506643 | NA | Prrt1b |
| ENSMUSG( | 12.62721 | -0.18639 | 0.744981 | NA | Prrt2 |
| ENSMUSG( | 1841.835 | -0.27722 | 0.079124 | 0.435105 | Prrt3 |
| ENSMUSG( | 114.9997 | -0.54483 | 0.016779 | 0.193019 | Prrt4 |
| ENSMUSG( | 460.1502 | -0.04656 | 0.732652 | 0.944956 | Prrx1 |
| ENSMUSG( | 63.30033 | -0.28215 | 0.248445 | 0.697313 | Prrx2 |
| ENSMUSG( | 692.4064 | -0.38498 | 0.001402 | 0.040991 | Prss12 |
| ENSMUSG( | 20.32367 | 0.069201 | 0.869023 | NA | Prss16 |
| ENSMUSG( | 34.70903 | -0.02676 | 0.934343 | 0.99323 | Prss22 |
| ENSMUSG( | 520.7874 | -0.47409 | 0.006279 | 0.108547 | Prss23 |
| ENSMUSG( | 114.1127 | -0.64354 | 0.001177 | 0.036941 | Prss35 |
| ENSMUSG( | 239.6814 | 0.055546 | 0.690852 | 0.933881 | Prss36 |
| ENSMUSG( | 23.33243 | -0.33041 | 0.396681 | 0.811742 | Prss3l |
| ENSMUSG( | 15.95186 | 0.034756 | 0.941996 | NA | Prss43 |
| ENSMUSG( | 14.00935 | -0.09136 | 0.860145 | NA | Prss45 |
| ENSMUSG( | 32.85415 | -0.69344 | 0.04011 | 0.308919 | Prss50 |
| ENSMUSG( | 26.77481 | -0.0854 | 0.823916 | 0.966648 | Prss53 |
| ENSMUSG( | 26.45774 | -0.86397 | 0.0419 | 0.315367 | Prss56 |
| ENSMUSG( | 25.57531 | 0.24642 | 0.542366 | 0.883182 | Prss57 |
| ENSMUSG( | 11.52107 | -0.10012 | 0.858275 | NA | Prss8 |
| ENSMUSG( | 66.21274 | -0.06944 | 0.791313 | 0.959461 | Prtg |
| ENSMUSG( | 39.6036 | -0.04146 | 0.893558 | 0.984512 | Prtn3 |
| ENSMUSG( | 700.3497 | -0.06349 | 0.609797 | 0.907755 | Prune1 |
| ENSMUSG( | 1788.218 | -0.26795 | 0.007627 | 0.120942 | Prune2 |
| ENSMUSG( | 57.21377 | -0.03386 | 0.891223 | 0.983819 | Prx |
| ENSMUSG( | 12743.6 | 0.095027 | 0.331733 | 0.775288 | Prxl2a |
| ENSMUSG( | 2450.069 | 0.005664 | 0.946087 | 0.99369 | Prxl2b |
| ENSMUSG( | 493.6898 | 0.029453 | 0.825608 | 0.967934 | Prxl2c |
| ENSMUSG( | 80192.68 | -0.19871 | 0.077425 | 0.429743 | Psap |
| ENSMUSG( | 10366.95 | 0.139174 | 0.189658 | 0.628577 | Psat1 |
| ENSMUSG( | 39238.83 | 0.125815 | 0.152169 | 0.578042 | Psd |
| ENSMUSG( | 6318.698 | 0.145921 | 0.114452 | 0.510426 | Psd2 |
| ENSMUSG( | 18983.24 | 0.143602 | 0.167361 | 0.599971 | Psd3 |
| ENSMUSG( | 62.43151 | -0.24542 | 0.303569 | 0.751968 | Psd4 |
| ENSMUSG( | 2171.668 | -0.2709 | 0.079181 | 0.435105 | Psen1 |
| ENSMUSG( | 665.4113 | -0.2883 | 0.020306 | 0.214844 | Psen2 |
| ENSMUSG( | 1112.498 | -0.20279 | 0.109932 | 0.500285 | Psenen |
| ENSMUSG( | 80.63178 | -0.03214 | 0.900576 | 0.985633 | Psg16 |
| ENSMUSG( | 3166.483 | 0.025165 | 0.794513 | 0.959799 | Psip1 |
| ENSMUSG( | 361.3743 | -0.0863 | 0.598648 | 0.903801 | Pskh1 |
| ENSMUSG( | 1283.983 | -0.07391 | 0.546149 | 0.884974 | Psma1 |
| ENSMUSG( | 2069.151 | 0.01571 | 0.87073 | 0.977951 | Psma2 |
| ENSMUSG( | 2326.959 | 0.051339 | 0.557858 | 0.888427 | Psma3 |
| ENSMUSG( | 1964.213 | -0.00651 | 0.946388 | 0.99369 | Psma4 |
| ENSMUSG( | 1214.09 | 0.00945 | 0.936096 | 0.993415 | Psma5 |
| ENSMUSG( | 1505.507 | 0.02117 | 0.807009 | 0.963044 | Psma6 |
| ENSMUSG( | 2002.27 | -0.00874 | 0.929826 | 0.991848 | Psma7 |
| ENSMUSG( | 16.77069 | -0.20192 | 0.682095 | NA | Psma8 |
| ENSMUSG( | 3168.439 | 0.012315 | 0.890938 | 0.983764 | Psmb1 |
| ENSMUSG( | 1092.976 | -0.08409 | 0.411528 | 0.819317 | Psmb10 |
| ENSMUSG( | 1258.613 | -0.07958 | 0.48337 | 0.856304 | Psmb2 |
| ENSMUSG( | 1478.187 | -0.01357 | 0.900758 | 0.985633 | Psmb3 |
| ENSMUSG( | 2849.735 | 0.012968 | 0.904363 | 0.986246 | Psmb4 |

| | | | | | |
|---|---|---|---|---|---|
| ENSMUSG( | 8657.638 | -0.11224 | 0.403572 | 0.917257 | Psat1 |
| ENSMUSG( | 34472.1 | 0.023696 | 0.806705 | 0.97813 | Psd |
| ENSMUSG( | 5001.121 | -0.27045 | 0.028043 | 0.61023 | Psd2 |
| ENSMUSG( | 16484.46 | 0.010083 | 0.931145 | 0.99306 | Psd3 |
| ENSMUSG( | 66.72259 | 0.22819 | 0.382698 | 0.917257 | Psd4 |
| ENSMUSG( | 2315.905 | 0.194729 | 0.2621 | 0.885136 | Psen1 |
| ENSMUSG( | 709.9074 | 0.180133 | 0.224915 | 0.86838 | Psen2 |
| ENSMUSG( | 1084.885 | 0.000783 | 0.995446 | 0.999116 | Psenen |
| ENSMUSG( | 72.63532 | -0.05887 | 0.823803 | 0.979198 | Psg16 |
| ENSMUSG( | 2877.233 | 0.014036 | 0.918535 | 0.992529 | Psip1 |
| ENSMUSG( | 332.7023 | -0.04845 | 0.782755 | 0.976106 | Pskh1 |
| ENSMUSG( | 1145.574 | -0.1395 | 0.303087 | 0.898538 | Psma1 |
| ENSMUSG( | 1755.622 | -0.20002 | 0.129803 | 0.820802 | Psma2 |
| ENSMUSG( | 2058.887 | -0.03454 | 0.754005 | 0.970671 | Psma3 |
| ENSMUSG( | 1736.573 | -0.09893 | 0.468371 | 0.923329 | Psma4 |
| ENSMUSG( | 1041.241 | -0.17192 | 0.202904 | 0.864412 | Psma5 |
| ENSMUSG( | 1321.415 | -0.09148 | 0.47249 | 0.923329 | Psma6 |
| ENSMUSG( | 1785.034 | -0.07263 | 0.529546 | 0.934795 | Psma7 |
| ENSMUSG( | 14.18264 | -0.44363 | 0.38379 | 0.917257 | Psma8 |
| ENSMUSG( | 2716.582 | -0.17089 | 0.167971 | 0.840222 | Psmb1 |
| ENSMUSG( | 972.8693 | -0.15847 | 0.21225 | 0.864412 | Psmb10 |
| ENSMUSG( | 1093.678 | -0.22898 | 0.107003 | 0.804969 | Psmb2 |
| ENSMUSG( | 1260.847 | -0.21584 | 0.14268 | 0.831221 | Psmb3 |
| ENSMUSG( | 2447.083 | -0.16663 | 0.249931 | 0.877904 | Psmb4 |
| ENSMUSG( | 1348.989 | -0.21697 | 0.12423 | 0.814691 | Psmb5 |
| ENSMUSG( | 1878.048 | -0.22979 | 0.060829 | 0.734852 | Psmb6 |
| ENSMUSG( | 2098.366 | -0.17664 | 0.142149 | 0.830709 | Psmb7 |
| ENSMUSG( | 143.2643 | -0.17049 | 0.391101 | 0.917257 | Psmb8 |
| ENSMUSG( | 59.08315 | -0.42129 | 0.155845 | 0.837724 | Psmb9 |
| ENSMUSG( | 2702.011 | -0.12235 | 0.268284 | 0.886493 | Psmc1 |
| ENSMUSG( | 1488.405 | -0.12256 | 0.380864 | 0.917202 | Psmc2 |
| ENSMUSG( | 3099.598 | -0.11053 | 0.389812 | 0.917257 | Psmc3 |
| ENSMUSG( | 69.56449 | -0.28473 | 0.282919 | 0.892412 | Psmc3ip |
| ENSMUSG( | 1797.951 | -0.13242 | 0.313669 | 0.900949 | Psmc4 |
| ENSMUSG( | 3708.518 | -0.29571 | 0.021543 | 0.575341 | Psmc5 |
| ENSMUSG( | 1255.987 | 0.03154 | 0.789247 | 0.976593 | Psmc6 |
| ENSMUSG( | 1974.171 | -0.02738 | 0.828678 | 0.980224 | Psmd1 |
| ENSMUSG( | 257.6548 | -0.27231 | 0.079371 | 0.77458 | Psmd10 |
| ENSMUSG( | 1271.906 | 0.107479 | 0.385359 | 0.917257 | Psmd11 |
| ENSMUSG( | 1047.073 | -0.15596 | 0.267598 | 0.886134 | Psmd12 |
| ENSMUSG( | 1346.613 | -0.0826 | 0.527832 | 0.934603 | Psmd13 |
| ENSMUSG( | 1188.089 | -0.07916 | 0.557541 | 0.939565 | Psmd14 |
| ENSMUSG( | 4590.583 | -0.17673 | 0.111885 | 0.808673 | Psmd2 |
| ENSMUSG( | 2896.823 | 0.028158 | 0.779135 | 0.976106 | Psmd3 |
| ENSMUSG( | 1654.998 | -0.10002 | 0.434273 | 0.91869 | Psmd4 |
| ENSMUSG( | 766.6693 | 0.118063 | 0.395795 | 0.917257 | Psmd5 |
| ENSMUSG( | 1033.98 | -0.00335 | 0.975963 | 0.996416 | Psmd6 |
| ENSMUSG( | 2069.33 | 0.048773 | 0.686509 | 0.95899 | Psmd7 |
| ENSMUSG( | 2366.103 | -0.21048 | 0.059823 | 0.729747 | Psmd8 |
| ENSMUSG( | 614.6698 | -0.03535 | 0.823488 | 0.979198 | Psmd9 |
| ENSMUSG( | 823.5686 | -0.05977 | 0.673558 | 0.95694 | Psme1 |
| ENSMUSG( | 570.7483 | -0.0349 | 0.784243 | 0.976166 | Psme2 |
| ENSMUSG( | 20.71638 | -0.18859 | 0.6866 | 0.95899 | Psme2b |
| ENSMUSG( | 2805.309 | -0.06642 | 0.550077 | 0.938699 | Psme3 |
| ENSMUSG( | 1043.519 | -0.13976 | 0.2988 | 0.898538 | Psme3ip1 |
| ENSMUSG( | 1488.743 | 0.062116 | 0.591863 | 0.944975 | Psme4 |
| ENSMUSG( | 920.4289 | -0.10278 | 0.464266 | 0.923329 | Psmf1 |
| ENSMUSG( | 229.4018 | 0.004703 | 0.980073 | 0.996479 | Psmg1 |
| ENSMUSG( | 346.8791 | 0.033332 | 0.829634 | 0.980224 | Psmg2 |
| ENSMUSG( | 117.6318 | 0.088817 | 0.707891 | 0.964042 | Psmg3 |
| ENSMUSG( | 180.278 | -0.00277 | 0.98913 | 0.997893 | Psmg4 |
| ENSMUSG( | 718.2042 | -0.18001 | 0.207265 | 0.864412 | Pspc1 |
| ENSMUSG( | 889.0025 | -0.1909 | 0.143929 | 0.831311 | Psph |
| ENSMUSG( | 190.8574 | -0.18748 | 0.371278 | 0.915491 | Psrc1 |
| ENSMUSG( | 423.2881 | 0.186642 | 0.208242 | 0.864412 | Pstk |
| ENSMUSG( | 77.87348 | -0.39579 | 0.104461 | 0.804969 | Pstpip1 |
| ENSMUSG( | 159.7341 | 0.065856 | 0.738387 | 0.967967 | Pstpip2 |
| ENSMUSG( | 104.1778 | 0.489872 | 0.197977 | 0.862161 | Ptafr |
| ENSMUSG( | 2838.033 | -0.22889 | 0.132715 | 0.822912 | Ptar1 |
| ENSMUSG( | 1237.762 | 0.135124 | 0.259716 | 0.883971 | Ptbp1 |
| ENSMUSG( | 1878.113 | -0.00904 | 0.943524 | 0.994798 | Ptbp2 |
| ENSMUSG( | 542.5892 | 0.238433 | 0.125518 | 0.81516 | Ptbp3 |
| ENSMUSG( | 528.5883 | 0.090558 | 0.581205 | 0.944266 | Ptcd1 |
| ENSMUSG( | 492.8843 | -0.01114 | 0.935657 | 0.993387 | Ptcd2 |
| ENSMUSG( | 917.9543 | -0.02132 | 0.860207 | 0.986137 | Ptcd3 |
| ENSMUSG( | 3402.682 | 0.158188 | 0.171912 | 0.842999 | Ptch1 |
| ENSMUSG( | 59.2672 | 0.002111 | 0.993745 | 0.998813 | Ptch2 |
| ENSMUSG( | 108.9746 | 0.403932 | 0.055406 | 0.723362 | Ptchd1 |
| ENSMUSG( | 97.27171 | 0.092595 | 0.708866 | 0.964042 | Ptchd4 |
| ENSMUSG( | 3855.658 | 0.020575 | 0.882376 | 0.989644 | Ptdss1 |
| ENSMUSG( | 2278.88 | 0.152518 | 0.234002 | 0.86917 | Ptdss2 |
| ENSMUSG( | 3411.096 | 0.199942 | 0.118688 | 0.813384 | Pten |
| ENSMUSG( | 95.0298 | -0.11274 | 0.623104 | 0.953234 | Pter |
| ENSMUSG( | 60.54412 | 0.309801 | 0.250494 | 0.877932 | Ptgdr |
| ENSMUSG( | 28994.76 | 0.021494 | 0.859258 | 0.985884 | Ptgds |
| ENSMUSG( | 103.3067 | -0.26797 | 0.259371 | 0.883971 | Ptger1 |
| ENSMUSG( | 9.589884 | -0.15382 | 0.801231 | NA | Ptger2 |
| ENSMUSG( | 116.0538 | 0.314583 | 0.175697 | 0.845305 | Ptger3 |
| ENSMUSG( | 38.52926 | -0.00977 | 0.977156 | 0.996416 | Ptger4 |
| ENSMUSG( | 395.034 | 0.142587 | 0.338275 | 0.907972 | Ptges |
| ENSMUSG( | 1016.318 | -0.04268 | 0.703877 | 0.96259 | Ptges2 |
| ENSMUSG( | 1727.168 | -0.08814 | 0.584951 | 0.944975 | Ptges3 |
| ENSMUSG( | 34.88363 | -0.09387 | 0.783319 | 0.976106 | Ptges3l |
| ENSMUSG( | 39.30164 | 0.002305 | 0.994266 | 0.998813 | Ptgfr |
| ENSMUSG( | 2153.606 | 0.060857 | 0.700161 | 0.961335 | Ptgfrn |
| ENSMUSG( | 6.678136 | -0.45559 | 0.573686 | NA | Ptgir |
| ENSMUSG( | 83.77334 | 0.227386 | 0.329444 | 0.907083 | Ptgis |
| ENSMUSG( | 45.96854 | -0.02471 | 0.937349 | 0.993685 | Ptgr1 |
| ENSMUSG( | 3433.466 | -0.2693 | 0.064064 | 0.736482 | Ptgr2 |
| ENSMUSG( | 1243.878 | -0.04982 | 0.695451 | 0.960867 | Ptgr3 |
| ENSMUSG( | 798.6953 | 0.163717 | 0.426603 | 0.917257 | Ptgs1 |
| ENSMUSG( | 264.9398 | 0.284248 | 0.168514 | 0.840222 | Ptgs2 |
| ENSMUSG( | 406.8921 | -0.00662 | 0.965551 | 0.995433 | Pth1r |
| ENSMUSG( | 58.43578 | 0.030006 | 0.918752 | 0.992529 | Pth2r |
| ENSMUSG( | 122.7074 | -0.10341 | 0.672737 | 0.95694 | Pthlh |
| ENSMUSG( | 3179.386 | 0.051547 | 0.631909 | 0.953251 | Ptk2 |
| ENSMUSG( | 7070.28 | -0.06727 | 0.700071 | 0.961335 | Ptk2b |
| ENSMUSG( | 533.8343 | 0.110547 | 0.569796 | 0.941777 | Ptk7 |
| ENSMUSG( | 5574.502 | 0.105544 | 0.470276 | 0.923329 | Ptma |
| ENSMUSG( | 19727.97 | -0.13538 | 0.398439 | 0.917257 | Ptms |
| ENSMUSG( | 21745.35 | -0.10221 | 0.464015 | 0.923329 | Ptn |
| ENSMUSG( | 3979.245 | -0.10788 | 0.319921 | 0.903771 | Ptov1 |
| ENSMUSG( | 18.49635 | -0.04048 | 0.935485 | 0.993265 | Ptp4a1 |
| ENSMUSG( | 10.79748 | -0.32043 | 0.637216 | NA | Ptp4a1 |
| ENSMUSG( | 5647.451 | -0.09349 | 0.520702 | 0.932121 | Ptp4a2 |
| ENSMUSG( | 1871.198 | -0.01646 | 0.885359 | 0.989688 | Ptp4a3 |
| ENSMUSG( | 6840.074 | -0.05886 | 0.535169 | 0.936508 | Ptpa |
| ENSMUSG( | 1158.237 | 0.112812 | 0.316049 | 0.901868 | Ptpdc1 |
| ENSMUSG( | 1477.331 | -0.09064 | 0.470624 | 0.923329 | Ptpmt1 |
| ENSMUSG( | 1885.92 | 0.014706 | 0.892284 | 0.990558 | Ptpn1 |
| ENSMUSG( | 11275.32 | -0.10165 | 0.365919 | 0.914623 | Ptpn11 |
| ENSMUSG( | 1176.698 | 0.028812 | 0.800697 | 0.97681 | Ptpn12 |
| ENSMUSG( | 1493.638 | -0.15804 | 0.155827 | 0.837724 | Ptpn13 |
| ENSMUSG( | 396.3623 | 0.089857 | 0.545327 | 0.938063 | Ptpn14 |
| ENSMUSG( | 92.58385 | -0.2486 | 0.273912 | 0.889859 | Ptpn18 |
| ENSMUSG( | 371.3216 | -0.00662 | 0.972297 | 0.996416 | Ptpn2 |
| ENSMUSG( | 359.536 | 0.097118 | 0.491491 | 0.926825 | Ptpn21 |
| ENSMUSG( | 36.33939 | -0.14969 | 0.690589 | 0.960071 | Ptpn22 |
| ENSMUSG( | 1741.5 | -0.07848 | 0.60737 | 0.950022 | Ptpn23 |
| ENSMUSG( | 1013.048 | -0.07332 | 0.592266 | 0.944975 | Ptpn3 |
| ENSMUSG( | 2012.875 | -0.11329 | 0.530351 | 0.934795 | Ptpn4 |
| ENSMUSG( | 7109.219 | 0.044212 | 0.698048 | 0.960867 | Ptpn5 |
| ENSMUSG( | 247.0329 | 0.070563 | 0.662864 | 0.956205 | Ptpn6 |
| ENSMUSG( | 32.92422 | -0.45334 | 0.223725 | 0.867404 | Ptpn7 |
| ENSMUSG( | 1212.428 | -0.0075 | 0.944995 | 0.994798 | Ptpn9 |
| ENSMUSG( | 9224.653 | 0.051223 | 0.726973 | 0.966887 | Ptpra |
| ENSMUSG( | 1356.625 | 0.835012 | 0.105103 | 0.804969 | Ptprb |
| ENSMUSG( | 270.4131 | -0.15113 | 0.522991 | 0.932848 | Ptprc |
| ENSMUSG( | 16.29824 | -0.24914 | 0.629937 | 0.953251 | Ptprcap |
| ENSMUSG( | 8816.358 | 0.109661 | 0.474417 | 0.923329 | Ptprd |
| ENSMUSG( | 1005.513 | 0.174806 | 0.344942 | 0.91045 | Ptpre |
| ENSMUSG( | 4315.606 | 0.224602 | 0.120059 | 0.813384 | Ptprf |
| ENSMUSG( | 3348.511 | 0.19569 | 0.182516 | 0.850579 | Ptprg |
| ENSMUSG( | 38.22707 | 0.078146 | 0.811387 | 0.978997 | Ptprh |
| ENSMUSG( | 2888.008 | 0.393776 | 0.025303 | 0.598154 | Ptprj |
| ENSMUSG( | 1790.713 | 0.180799 | 0.287521 | 0.894091 | Ptprk |
| ENSMUSG( | 1406.686 | 0.199784 | 0.344287 | 0.910321 | Ptprm |
| ENSMUSG( | 19421.07 | 0.134429 | 0.325783 | 0.905339 | Ptprn |
| ENSMUSG( | 5499.222 | 0.411004 | 0.276772 | 0.890889 | Ptprn2 |
| ENSMUSG( | 1040.486 | 0.162307 | 0.294853 | 0.898538 | Ptpro |
| ENSMUSG( | 562.9252 | 0.167516 | 0.32049 | 0.903887 | Ptprr |
| ENSMUSG( | 17505.23 | 0.188905 | 0.077105 | 0.770558 | Ptprs |
| ENSMUSG( | 9279.107 | -0.01144 | 0.927638 | 0.99306 | Ptprt |
| ENSMUSG( | 1723.304 | 0.280104 | 0.031482 | 0.631192 | Ptpru |
| ENSMUSG( | 93.67729 | -0.39952 | 0.215734 | 0.865167 | Ptprv |
| ENSMUSG( | 21401.61 | -0.0124 | 0.928996 | 0.99306 | Ptprz1 |
| ENSMUSG( | 98.89572 | -0.0188 | 0.931461 | 0.99306 | Ptrh1 |
| ENSMUSG( | 262.7744 | -0.06009 | 0.720875 | 0.966887 | Ptrh2 |

| | | | | | | | | | | | |
|---|---|---|---|---|---|---|---|---|---|---|---|
| ENSMUSG( | Pvr | 240.7496 | -0.74064 | 2.00E-05 | 0.001952 | ENSMUSG( | Pvr | 255.0205 | -0.21094 | 0.256421 | 0.883082 |
| ENSMUSG( | Pvt1 | 494.9657 | 0.059314 | 0.612147 | 0.908586 | ENSMUSG( | Pvt1 | 440.4456 | -0.00848 | 0.948963 | 0.994798 |
| ENSMUSG( | Pwp1 | 476.4593 | 0.075986 | 0.562366 | 0.889448 | ENSMUSG( | Pwp1 | 414.9164 | -0.05734 | 0.706765 | 0.963529 |
| ENSMUSG( | Pwp2 | 692.9405 | -0.08186 | 0.425762 | 0.824898 | ENSMUSG( | Pwp2 | 635.3258 | -0.06468 | 0.598218 | 0.946676 |
| ENSMUSG( | Pwwp2a | 575.135 | 0.123563 | 0.276932 | 0.727074 | ENSMUSG( | Pwwp2a | 526.0302 | 0.132226 | 0.317614 | 0.902602 |
| ENSMUSG( | Pwwp2b | 314.3647 | -0.07244 | 0.648479 | 0.919997 | ENSMUSG( | Pwwp2b | 289.5551 | -0.04295 | 0.802559 | 0.97681 |
| ENSMUSG( | Pwwp3a | 902.832 | -0.06058 | 0.632367 | 0.914073 | ENSMUSG( | Pwwp3a | 860.7886 | 0.076072 | 0.621766 | 0.953234 |
| ENSMUSG( | Pwwp3b | 58.9325 | 0.057293 | 0.814793 | 0.964668 | ENSMUSG( | Pwwp3b | 56.72454 | 0.215338 | 0.431026 | 0.917257 |
| ENSMUSG( | Pxdc1 | 248.3205 | -0.05596 | 0.699257 | 0.936886 | ENSMUSG( | Pxdc1 | 228.8238 | -0.02381 | 0.883614 | 0.989644 |
| ENSMUSG( | Pxdn | 3211.175 | 0.047574 | 0.664612 | 0.926824 | ENSMUSG( | Pxdn | 2954.579 | 0.076499 | 0.568428 | 0.941777 |
| ENSMUSG( | Pxk | 790.7224 | 0.12236 | 0.303932 | 0.752178 | ENSMUSG( | Pxk | 701.7704 | 0.052063 | 0.729933 | 0.967228 |
| ENSMUSG( | Pxmp2 | 401.0668 | 0.087448 | 0.534971 | 0.880733 | ENSMUSG( | Pxmp2 | 343.2177 | -0.09608 | 0.497118 | 0.927825 |
| ENSMUSG( | Pxmp4 | 944.3619 | 0.010803 | 0.92699 | 0.991229 | ENSMUSG( | Pxmp4 | 783.7721 | -0.2751 | 0.029868 | 0.61992 |
| ENSMUSG( | Pxn | 1427.346 | 0.156532 | 0.20702 | 0.649503 | ENSMUSG( | Pxn | 1211.78 | -0.04311 | 0.75712 | 0.971858 |
| ENSMUSG( | Pxylp1 | 366.6396 | -0.49725 | 0.001146 | 0.036093 | ENSMUSG( | Pxylp1 | 378.3416 | -0.09053 | 0.615139 | 0.951177 |
| ENSMUSG( | Pycard | 156.4359 | -0.04222 | 0.817303 | 0.965736 | ENSMUSG( | Pycard | 133.0262 | -0.24951 | 0.203476 | 0.864412 |
| ENSMUSG( | Pycr1 | 114.6604 | -0.22847 | 0.272265 | 0.7223 | ENSMUSG( | Pycr1 | 98.0756 | -0.44981 | 0.054389 | 0.723362 |
| ENSMUSG( | Pycr2 | 682.1058 | 0.027108 | 0.852471 | 0.973168 | ENSMUSG( | Pycr2 | 589.9886 | -0.12872 | 0.42204 | 0.917257 |
| ENSMUSG( | Pycr3 | 454.8671 | 0.055313 | 0.677464 | 0.930542 | ENSMUSG( | Pycr3 | 400.1503 | -0.04925 | 0.747498 | 0.969776 |
| ENSMUSG( | Pygb | 18375.89 | 0.120622 | 0.225921 | 0.672685 | ENSMUSG( | Pygb | 14981.14 | -0.20609 | 0.087615 | 0.785039 |
| ENSMUSG( | Pygl | 210.9093 | 0.073369 | 0.640057 | 0.916434 | ENSMUSG( | Pygl | 186.8608 | -0.02481 | 0.903982 | 0.991089 |
| ENSMUSG( | Pygm | 4141.233 | 0.07134 | 0.511405 | 0.870898 | ENSMUSG( | Pygm | 3383.486 | -0.25613 | 0.032377 | 0.635121 |
| ENSMUSG( | Pygo1 | 1637.26 | 0.08418 | 0.434192 | 0.828931 | ENSMUSG( | Pygo1 | 1458.288 | 0.016785 | 0.907847 | 0.991958 |
| ENSMUSG( | Pygo2 | 1289.758 | 0.094904 | 0.392858 | 0.809274 | ENSMUSG( | Pygo2 | 1127.133 | -0.02422 | 0.831079 | 0.98051 |
| ENSMUSG( | Pym1 | 463.6441 | -0.11469 | 0.49895 | 0.862973 | ENSMUSG( | Pym1 | 399.1609 | -0.2987 | 0.120225 | 0.813384 |
| ENSMUSG( | Pyroxd1 | 353.5685 | 0.02373 | 0.856515 | 0.974154 | ENSMUSG( | Pyroxd1 | 313.7525 | -0.04412 | 0.768936 | 0.974236 |
| ENSMUSG( | Pyroxd2 | 142.9502 | -0.02363 | 0.893593 | 0.984512 | ENSMUSG( | Pyroxd2 | 129.8098 | -0.02633 | 0.901555 | 0.990784 |
| ENSMUSG( | Pyurf | 835.2062 | -0.03403 | 0.81712 | 0.96566 | ENSMUSG( | Pyurf | 684.6748 | -0.36798 | 0.02892 | 0.614171 |
| ENSMUSG( | Qars1 | 1307.611 | -0.11675 | 0.253912 | 0.704011 | ENSMUSG( | Qars1 | 1262.807 | 0.056567 | 0.638429 | 0.953903 |
| ENSMUSG( | Qdpr | 7792.715 | -0.19066 | 0.027943 | 0.255775 | ENSMUSG( | Qdpr | 6546.782 | -0.45666 | 9.71E-05 | 0.02045 |
| ENSMUSG( | Qki | 25292.53 | 0.025472 | 0.841453 | 0.971179 | ENSMUSG( | Qki | 22932.26 | 0.009899 | 0.954627 | 0.994798 |
| ENSMUSG( | Qng1 | 355.9408 | 0.108526 | 0.475618 | 0.850919 | ENSMUSG( | Qng1 | 282.6632 | -0.30987 | 0.060263 | 0.731553 |
| ENSMUSG( | Qpct | 263.6947 | -0.72134 | 7.56E-05 | 0.005353 | ENSMUSG( | Qpct | 343.3493 | 0.374969 | 0.050535 | 0.708889 |
| ENSMUSG( | Qpctl | 648.614 | -0.42946 | 0.001431 | 0.041661 | ENSMUSG( | Qpctl | 701.1617 | 0.100319 | 0.543312 | 0.938063 |
| ENSMUSG( | Qprt | 28.99353 | -0.02114 | 0.955798 | 0.993874 | ENSMUSG( | Qprt | 22.76373 | -0.51525 | 0.243049 | 0.877399 |
| ENSMUSG( | Qrfpr | 73.01563 | -0.40674 | 0.089655 | 0.45961 | ENSMUSG( | Qrfpr | 86.96327 | 0.37153 | 0.114996 | 0.811921 |
| ENSMUSG( | Qrfprl | 14.37825 | -1.63143 | 0.001798 | NA | ENSMUSG( | Qrfprl | 18.29847 | -0.25175 | 0.587286 | 0.944975 |
| ENSMUSG( | Qrich1 | 1718.357 | 0.053323 | 0.585412 | 0.899171 | ENSMUSG( | Qrich1 | 1503.155 | -0.06657 | 0.572732 | 0.94262 |
| ENSMUSG( | Qrsl1 | 350.6252 | 0.093176 | 0.467306 | 0.844819 | ENSMUSG( | Qrsl1 | 318.845 | 0.082203 | 0.581389 | 0.944266 |
| ENSMUSG( | Qser1 | 872.4875 | 0.022149 | 0.87395 | 0.979336 | ENSMUSG( | Qser1 | 788.302 | -0.00383 | 0.978524 | 0.996459 |
| ENSMUSG( | Qsox1 | 969.5062 | -0.20024 | 0.1808 | 0.617841 | ENSMUSG( | Qsox1 | 1043.817 | 0.281945 | 0.109283 | 0.805353 |
| ENSMUSG( | Qsox2 | 862.4655 | -0.31595 | 0.031117 | 0.268324 | ENSMUSG( | Qsox2 | 922.0275 | 0.164532 | 0.340621 | 0.907972 |
| ENSMUSG( | Qtrt1 | 290.8231 | -0.05139 | 0.762677 | 0.953519 | ENSMUSG( | Qtrt1 | 262.0088 | -0.08332 | 0.675399 | 0.95694 |
| ENSMUSG( | Qtrt2 | 382.8352 | 0.169692 | 0.260158 | 0.710129 | ENSMUSG( | Qtrt2 | 369.4803 | 0.326702 | 0.046795 | 0.700612 |
| ENSMUSG( | R3hcc1 | 1071.092 | -0.01445 | 0.885883 | 0.983008 | ENSMUSG( | R3hcc1 | 931.2313 | -0.15807 | 0.244471 | 0.877904 |
| ENSMUSG( | R3hcc1l | 387.5122 | 0.051186 | 0.67827 | 0.930542 | ENSMUSG( | R3hcc1l | 336.6753 | -0.08408 | 0.578984 | 0.943849 |
| ENSMUSG( | R3hdm1 | 12166.64 | 0.066013 | 0.487192 | 0.858812 | ENSMUSG( | R3hdm1 | 10310.81 | -0.14768 | 0.204221 | 0.864412 |
| ENSMUSG( | R3hdm2 | 13904.49 | 0.132262 | 0.093332 | 0.469793 | ENSMUSG( | R3hdm2 | 12113.64 | 0.005483 | 0.957592 | 0.994798 |
| ENSMUSG( | R3hdm4 | 18738.72 | 0.081832 | 0.313008 | 0.76033 | ENSMUSG( | R3hdm4 | 16253.52 | -0.0591 | 0.565165 | 0.94131 |
| ENSMUSG( | R74862 | 308.9222 | -0.09282 | 0.500346 | 0.863455 | ENSMUSG( | R74862 | 275.2941 | -0.15879 | 0.312251 | 0.900037 |
| ENSMUSG( | Rab10 | 6600.21 | -0.02796 | 0.803758 | 0.962169 | ENSMUSG( | Rab10 | 5857.662 | -0.10761 | 0.443134 | 0.921569 |
| ENSMUSG( | Rab11a | 1746.257 | 0.015951 | 0.873075 | 0.978753 | ENSMUSG( | Rab11a | 1558.774 | -0.04732 | 0.726559 | 0.966887 |
| ENSMUSG( | Rab11b | 5779.091 | 0.021727 | 0.773017 | 0.955272 | ENSMUSG( | Rab11b | 4990.644 | -0.13655 | 0.188422 | 0.85698 |
| ENSMUSG( | Rab11fip1 | 120.074 | 0.22965 | 0.282771 | 0.73186 | ENSMUSG( | Rab11fip1 | 130.2018 | 0.683173 | 0.016201 | 0.512256 |
| ENSMUSG( | Rab11fip2 | 1079.261 | -0.04149 | 0.663156 | 0.9265 | ENSMUSG( | Rab11fip2 | 1010.004 | 0.034519 | 0.769402 | 0.974236 |
| ENSMUSG( | Rab11fip3 | 4502.014 | 0.100532 | 0.267981 | 0.718218 | ENSMUSG( | Rab11fip3 | 4071.194 | 0.080138 | 0.431116 | 0.917257 |
| ENSMUSG( | Rab11fip4 | 13982.86 | 0.139268 | 0.089204 | 0.458482 | ENSMUSG( | Rab11fip4 | 11818.55 | -0.07473 | 0.427419 | 0.917257 |
| ENSMUSG( | Rab11fip5 | 2528.592 | 0.13968 | 0.149887 | 0.574325 | ENSMUSG( | Rab11fip5 | 2223.131 | 0.042551 | 0.732425 | 0.967228 |
| ENSMUSG( | Rab12 | 4674.785 | -0.57942 | 0.000228 | 0.01183 | ENSMUSG( | Rab12 | 3756.78 | -1.08666 | 5.68E-17 | 9.04E-14 |
| ENSMUSG( | Rab13 | 532.046 | 0.082352 | 0.529684 | 0.878883 | ENSMUSG( | Rab13 | 423.3293 | -0.33545 | 0.032378 | 0.635121 |
| ENSMUSG( | Rab14 | 5181.048 | 0.068363 | 0.420715 | 0.822925 | ENSMUSG( | Rab14 | 4589.093 | -0.01432 | 0.905728 | 0.991089 |
| ENSMUSG( | Rab15 | 13304.25 | 0.17832 | 0.01859 | 0.20512 | ENSMUSG( | Rab15 | 11170.28 | -0.05194 | 0.581671 | 0.944266 |
| ENSMUSG( | Rab18 | 3021.173 | -0.00009 | 0.992404 | 0.999327 | ENSMUSG( | Rab18 | 2737.789 | -0.01981 | 0.894428 | 0.990558 |
| ENSMUSG( | Rab1a | 5171.428 | 0.019916 | 0.838964 | 0.971179 | ENSMUSG( | Rab1a | 4501.057 | -0.11751 | 0.416254 | 0.917257 |
| ENSMUSG( | Rab1b | 6186.566 | 0.074195 | 0.341484 | 0.778828 | ENSMUSG( | Rab1b | 5235.859 | -0.14278 | 0.192478 | 0.859726 |
| ENSMUSG( | Rab20 | 19.41741 | -0.56962 | 0.184816 | NA | ENSMUSG( | Rab20 | 18.05924 | -0.48881 | 0.29901 | 0.898538 |
| ENSMUSG( | Rab21 | 4502.941 | 0.046857 | 0.703417 | 0.937735 | ENSMUSG( | Rab21 | 3927.539 | -0.08252 | 0.623094 | 0.953234 |
| ENSMUSG( | Rab22a | 3524.034 | 0.134412 | 0.182606 | 0.620026 | ENSMUSG( | Rab22a | 3082.554 | 0.020526 | 0.856863 | 0.985884 |
| ENSMUSG( | Rab23 | 1171.57 | 0.163353 | 0.143741 | 0.562581 | ENSMUSG( | Rab23 | 1043.362 | 0.095594 | 0.511887 | 0.930005 |
| ENSMUSG( | Rab24 | 1376.759 | 0.017456 | 0.874483 | 0.979498 | ENSMUSG( | Rab24 | 1240.812 | -0.01575 | 0.907535 | 0.991907 |
| ENSMUSG( | Rab26 | 1076.595 | 0.181098 | 0.242551 | 0.690206 | ENSMUSG( | Rab26 | 914.4395 | -0.01692 | 0.925261 | 0.99306 |
| ENSMUSG( | Rab26os | 53.9504 | 0.145001 | 0.643909 | 0.917526 | ENSMUSG( | Rab26os | 47.00802 | 0.037459 | 0.920233 | 0.992644 |
| ENSMUSG( | Rab27a | 118.3851 | 0.162721 | 0.389558 | 0.807428 | ENSMUSG( | Rab27a | 112.7342 | 0.266937 | 0.217653 | 0.865589 |
| ENSMUSG( | Rab27b | 602.0685 | -0.0255 | 0.821631 | 0.966344 | ENSMUSG( | Rab27b | 538.5569 | -0.08914 | 0.57996 | 0.944209 |
| ENSMUSG( | Rab28 | 1693.522 | 0.073495 | 0.429506 | 0.826446 | ENSMUSG( | Rab28 | 1465.705 | -0.07629 | 0.48084 | 0.924548 |
| ENSMUSG( | Rab29 | 215.7953 | 0.240011 | 0.143062 | 0.561716 | ENSMUSG( | Rab29 | 173.4077 | -0.11534 | 0.529092 | 0.934795 |
| ENSMUSG( | Rab2a | 5339.441 | 0.009588 | 0.921552 | 0.990436 | ENSMUSG( | Rab2a | 4746.05 | -0.06557 | 0.65375 | 0.955015 |
| ENSMUSG( | Rab2b | 1051.428 | 0.11683 | 0.229695 | 0.677538 | ENSMUSG( | Rab2b | 898.442 | -0.06921 | 0.574078 | 0.942912 |
| ENSMUSG( | Rab30 | 812.9456 | -0.08399 | 0.465586 | 0.84415 | ENSMUSG( | Rab30 | 774.6724 | 0.041062 | 0.794716 | 0.97681 |
| ENSMUSG( | Rab31 | 7914.087 | 0.076555 | 0.464737 | 0.843568 | ENSMUSG( | Rab31 | 6533.129 | -0.21638 | 0.075307 | 0.767994 |
| ENSMUSG( | Rab32 | 79.34169 | -0.02611 | 0.905214 | 0.98645 | ENSMUSG( | Rab32 | 68.58145 | -0.20406 | 0.437893 | 0.920495 |
| ENSMUSG( | Rab33a | 446.3436 | -0.05962 | 0.754437 | 0.951007 | ENSMUSG( | Rab33a | 381.6012 | -0.25883 | 0.196208 | 0.862161 |
| ENSMUSG( | Rab33b | 1033.053 | -0.06858 | 0.51248 | 0.871762 | ENSMUSG( | Rab33b | 924.0611 | -0.12882 | 0.379442 | 0.916995 |
| ENSMUSG( | Rab34 | 730.3997 | -0.01598 | 0.886513 | 0.983204 | ENSMUSG( | Rab34 | 674.3013 | 0.025493 | 0.861037 | 0.986226 |
| ENSMUSG( | Rab35 | 2485.483 | 0.075065 | 0.350643 | 0.782422 | ENSMUSG( | Rab35 | 2132.635 | -0.0997 | 0.343253 | 0.909646 |
| ENSMUSG( | Rab36 | 1106.423 | 0.019528 | 0.867316 | 0.976559 | ENSMUSG( | Rab36 | 990.0933 | -0.03433 | 0.812731 | 0.979148 |
| ENSMUSG( | Rab37 | 89.17656 | -0.84828 | 0.001773 | 0.04746 | ENSMUSG( | Rab37 | 91.35747 | -0.44203 | 0.117932 | 0.813384 |
| ENSMUSG( | Rab39 | 23.13067 | -0.4096 | 0.314667 | 0.761775 | ENSMUSG( | Rab39 | 24.31024 | 0.038244 | 0.927864 | 0.99306 |
| ENSMUSG( | Rab39b | 583.428 | -0.40736 | 0.000131 | 0.007715 | ENSMUSG( | Rab39b | 599.687 | -0.03614 | 0.827863 | 0.980224 |
| ENSMUSG( | Rab3a | 9995.129 | 0.021837 | 0.874035 | 0.979336 | ENSMUSG( | Rab3a | 8596.364 | -0.15092 | 0.336528 | 0.907972 |
| ENSMUSG( | Rab3b | 871.5621 | -0.80373 | 1.98E-06 | 0.000328 | ENSMUSG( | Rab3b | 954.4823 | -0.17327 | 0.358397 | 0.913759 |
| ENSMUSG( | Rab3c | 7556.126 | 0.221815 | 0.003672 | 0.077033 | ENSMUSG( | Rab3c | 6491.764 | 0.058934 | 0.634965 | 0.953903 |
| ENSMUSG( | Rab3d | 337.5612 | 0.126612 | 0.344942 | 0.779153 | ENSMUSG( | Rab3d | 311.7258 | 0.168223 | 0.281162 | 0.892356 |
| ENSMUSG( | Rab3gap1 | 2292.649 | 0.168472 | 0.042497 | 0.316636 | ENSMUSG( | Rab3gap1 | 2000.701 | 0.049796 | 0.638527 | 0.953903 |
| ENSMUSG( | Rab3gap2 | 2353.918 | -0.00202 | 0.981886 | 0.997512 | ENSMUSG( | Rab3gap2 | 2144.078 | -0.00355 | 0.974141 | 0.996416 |
| ENSMUSG( | Rab3il1 | 589.0316 | 0.132588 | 0.233597 | 0.681508 | ENSMUSG( | Rab3il1 | 506.4268 | -0.03108 | 0.808568 | 0.97873 |
| ENSMUSG( | Rab3ip | 815.5936 | 0.04622 | 0.710392 | 0.939748 | ENSMUSG( | Rab3ip | 727.5878 | -0.01951 | 0.898691 | 0.990558 |
| ENSMUSG( | Rab40b | 563.8609 | -0.24077 | 0.116577 | 0.514987 | ENSMUSG( | Rab40b | 535.9829 | -0.11778 | 0.509308 | 0.929089 |
| ENSMUSG( | Rab40c | 1433.564 | 0.100214 | 0.350056 | 0.781943 | ENSMUSG( | Rab40c | 1240.142 | -0.04612 | 0.712606 | 0.964363 |
| ENSMUSG( | Rab42 | 9.792169 | 0.259178 | 0.664705 | NA | ENSMUSG( | Rab42 | 10.9206 | 0.77414 | 0.183826 | NA |
| ENSMUSG( | Rab43 | 40.86638 | 0.124336 | 0.676799 | 0.930542 | ENSMUSG( | Rab43 | 38.78863 | 0.236026 | 0.465578 | 0.923329 |
| ENSMUSG( | Rab4a | 1423.316 | 0.164172 | 0.187424 | 0.625777 | ENSMUSG( | Rab4a | 1319.924 | 0.209919 | 0.080136 | 0.77681 |
| ENSMUSG( | Rab4b | 131.2653 | -0.21204 | 0.330552 | 0.774568 | ENSMUSG( | Rab4b | 131.4399 | 0.077976 | 0.712846 | 0.96439 |
| ENSMUSG( | Rab5a | 2516.023 | -0.00949 | 0.925674 | 0.991204 | ENSMUSG( | Rab5a | 2215.209 | -0.11395 | 0.43933 | 0.920721 |
| ENSMUSG( | Rab5b | 5947.175 | 0.07317 | 0.375607 | 0.800887 | ENSMUSG( | Rab5b | 5016.538 | -0.15391 | 0.162023 | 0.837724 |
| ENSMUSG( | Rab5c | 7438.58 | 0.020479 | 0.811476 | 0.964426 | ENSMUSG( | Rab5c | 6337.6 | -0.18099 | 0.182721 | 0.850579 |
| ENSMUSG( | Rab5if | 1544.509 | 0.006405 | 0.951484 | 0.993821 | ENSMUSG( | Rab5if | 1412.564 | 0.019698 | 0.866952 | 0.987487 |
| ENSMUSG( | Rab6a | 10390.19 | -0.0243 | 0.806738 | 0.962932 | ENSMUSG( | Rab6a | 9090.577 | -0.14715 | 0.300428 | 0.898538 |
| ENSMUSG( | Rab6b | 39401.76 | 0.155018 | 0.037503 | 0.299216 | ENSMUSG( | Rab6b | 33347.34 | -0.05392 | 0.568211 | 0.941777 |
| ENSMUSG( | Rab7 | 9615.723 | 0.019474 | 0.834423 | 0.970342 | ENSMUSG( | Rab7 | 8269.108 | -0.15367 | 0.26667 | 0.886134 |
| ENSMUSG( | Rab7b | 258.9487 | -0.37909 | 0.016031 | 0.187923 | ENSMUSG( | Rab7b | 219.8222 | -0.64361 | 0.000331 | 0.049413 |
| ENSMUSG( | Rab8a | 1322.734 | 0.144683 | 0.105096 | 0.492392 | ENSMUSG( | Rab8a | 1134.264 | -0.02892 | 0.810124 | 0.978779 |
| ENSMUSG( | Rab8b | 895.718 | 0.074522 | 0.491322 | 0.859873 | ENSMUSG( | Rab8b | 788.2417 | -0.02908 | 0.837198 | 0.981335 |
| ENSMUSG( | Rab9 | 959.3523 | 0.060118 | 0.594324 | 0.902869 | ENSMUSG( | Rab9 | 809.1372 | -0.17268 | 0.219383 | 0.866104 |
| ENSMUSG( | Rab9b | 282.0667 | 0.04324 | 0.775441 | 0.955606 | ENSMUSG( | Rab9b | 251.6227 | -0.02921 | 0.882242 | 0.989644 |
| ENSMUSG( | Rabac1 | 4044.679 | -0.21134 | 0.046236 | 0.329472 | ENSMUSG( | Rabac1 | 4092.991 | 0.099617 | 0.446803 | 0.921867 |
| ENSMUSG( | Rabep1 | 4119.864 | 0.152234 | 0.076426 | 0.4279 | ENSMUSG( | Rabep1 | 3491.695 | -0.05425 | 0.617385 | 0.951952 |
| ENSMUSG( | Rabep2 | 356.9682 | -0.07015 | 0.621891 | 0.911432 | ENSMUSG( | Rabep2 | 350.2148 | 0.145655 | 0.396331 | 0.917257 |
| ENSMUSG( | Rabepk | 549.0588 | -0.01651 | 0.883044 | 0.982422 | ENSMUSG( | Rabepk | 480.4944 | -0.14323 | 0.321175 | 0.904019 |
| ENSMUSG( | Rabgap1 | 2392.574 | 0.125958 | 0.116251 | 0.514678 | ENSMUSG( | Rabgap1 | 2097.433 | 0.015901 | 0.882383 | 0.989644 |
| ENSMUSG( | Rabgap1l | 2060.812 | -0.01759 | 0.850609 | 0.972638 | ENSMUSG( | Rabgap1l | 1906.673 | 0.022744 | 0.885431 | 0.989688 |
| ENSMUSG( | Rabgef1 | 858.006 | -0.01412 | 0.890397 | 0.983687 | ENSMUSG( | Rabgef1 | 761.6151 | -0.09508 | 0.468386 | 0.923329 |
| ENSMUSG( | Rabggta | 981.3219 | 0.039512 | 0.69553 | 0.934979 | ENSMUSG( | Rabggta | 867.1742 | -0.04976 | 0.700697 | 0.961547 |
| ENSMUSG( | Rabggtb | 1152.56 | -0.03058 | 0.753243 | 0.950753 | ENSMUSG( | Rabggtb | 1012.179 | -0.14252 | 0.240147 | 0.875205 |
| ENSMUSG( | Rabif | 805.9563 | 0.000284 | 0.997808 | 0.99989 | ENSMUSG( | Rabif | 717.2828 | -0.07345 | 0.590235 | 0.944975 |
| ENSMUSG( | Rabl2 | 667.8795 | 0.122633 | 0.473547 | 0.848919 | ENSMUSG( | Rabl2 | 583.083 | 0.001269 | 0.994881 | 0.998815 |
| ENSMUSG( | Rabl3 | 156.239 | 0.083038 | 0.632874 | 0.914073 | ENSMUSG( | Rabl3 | 137.9621 | -0.01835 | 0.930996 | 0.99306 |
| ENSMUSG( | Rabl6 | 7470.364 | 0.200653 | 0.012385 | 0.160838 | ENSMUSG( | Rabl6 | 6459.579 | 0.057147 | 0.559005 | 0.939587 |
| ENSMUSG( | Rac1 | 12163.29 | 0.01221 | 0.894651 | 0.984512 | ENSMUSG( | Rac1 | 10565.69 | -0.1306 | 0.336155 | 0.907972 |
| ENSMUSG( | Rac2 | 222.1353 | -0.11548 | 0.509919 | 0.869743 | ENSMUSG( | Rac2 | 187.0178 | -0.37217 | 0.05909 | 0.729429 |
| ENSMUSG( | Rac3 | 622.9792 | 0.108881 | 0.318366 | 0.764234 | ENSMUSG( | Rac3 | 530.2977 | -0.09258 | 0.505543 | 0.927825 |
| ENSMUSG( | Racgap1 | 258.095 | -0.12605 | 0.379796 | 0.801757 | ENSMUSG( | Racgap1 | 255.4273 | 0.107633 | 0.506302 | 0.927825 |
| ENSMUSG( | Rack1 | 9524.308 | 0.087281 | 0.266281 | 0.716441 | ENSMUSG( | Rack1 | 7950.495 | -0.16998 | 0.092846 | 0.79479 |
| ENSMUSG( | Rad1 | 181.6258 | 0.159944 | 0.342561 | 0.778971 | ENSMUSG( | Rad1 | 160.1549 | 0.070369 | 0.703154 | 0.962428 |
| ENSMUSG( | Rad17 | 490.8531 | 0.117623 | 0.309986 | 0.758025 | ENSMUSG( | Rad17 | 446.5678 | 0.109633 | 0.43237 | 0.918158 |
| ENSMUSG( | Rad18 | 122.4259 | -0.15321 | 0.394898 | 0.810072 | ENSMUSG( | Rad18 | 110.4824 | -0.19399 | 0.355164 | 0.912411 |
| ENSMUSG( | Rad21 | 6100.88 | 0.103226 | 0.257552 | 0.708686 | ENSMUSG( | Rad21 | 5199.486 | -0.09037 | 0.416005 | 0.917257 |
| ENSMUSG( | Rad23a | 1941.488 | 0.089219 | 0.361448 | 0.792797 | ENSMUSG( | Rad23a | 1714.265 | -0.00187 | 0.988569 | 0.997862 |
| ENSMUSG( | Rad23b | 3306.688 | 0.10108 | 0.297497 | 0.746042 | ENSMUSG( | Rad23b | 2844.674 | -0.06735 | 0.648996 | 0.95422 |
| ENSMUSG( | Rad50 | 407.8451 | 0.113816 | 0.342707 | 0.778971 | ENSMUSG( | Rad50 | 362.8573 | 0.052085 | 0.720694 | 0.966887 |
| ENSMUSG( | Rad51 | 42.16685 | 0.009577 | 0.973653 | 0.99675 | ENSMUSG( | Rad51 | 37.90488 | -0.0251 | 0.937695 | 0.993685 |
| ENSMUSG( | Rad51ap1 | 35.42372 | -0.45762 | 0.140295 | 0.557263 | ENSMUSG( | Rad51ap1 | 38.02405 | 0.071415 | 0.82702 | 0.980159 |
| ENSMUSG( | Rad51ap2 | 18.89086 | -0.46454 | 0.289578 | NA | ENSMUSG( | Rad51ap2 | 16.65335 | -0.5789 | 0.230551 | 0.868507 |
| ENSMUSG( | Rad51b | 11.33381 | -0.35883 | 0.545938 | NA | ENSMUSG( | Rad51b | 10.62669 | -0.25036 | 0.698346 | NA |
| ENSMUSG( | Rad51c | 67.89532 | 0.135827 | 0.560807 | 0.888934 | ENSMUSG( | Rad51c | 65.95523 | 0.300656 | 0.244136 | 0.877904 |
| ENSMUSG( | Rad51d | 491.5878 | 0.035434 | 0.778271 | 0.956215 | ENSMUSG( | Rad51d | 421.2291 | -0.1443 | 0.299963 | 0.898538 |
| ENSMUSG( | Rad52 | 513.6276 | 0.084904 | 0.494347 | 0.860611 | ENSMUSG( | Rad52 | 470.4162 | 0.101543 | 0.480105 | 0.924276 |
| ENSMUSG( | Rad54b | 31.40736 | -0.43794 | 0.231423 | 0.679437 | ENSMUSG( | Rad54b | 35.15391 | 0.204166 | 0.574342 | 0.942912 |
| ENSMUSG( | Rad54l | 52.47589 | 0.003996 | 0.987693 | 0.998902 | ENSMUSG( | Rad54l | 45.59437 | -0.12378 | 0.686378 | 0.95899 |
| ENSMUSG( | Rad54l2 | 934.3593 | 0.014934 | 0.879399 | 0.981436 | ENSMUSG( | Rad54l2 | 838.1461 | -0.02744 | 0.822775 | 0.979198 |
| ENSMUSG( | Rad9a | 253.2758 | -0.37543 | 0.006336 | 0.108996 | ENSMUSG( | Rad9a | 253.3373 | -0.07174 | 0.669237 | 0.956766 |
| ENSMUSG( | Rad9b | 129.2853 | -0.20848 | 0.265935 | 0.716441 | ENSMUSG( | Rad9b | 130.0894 | 0.083114 | 0.703124 | 0.962428 |
| ENSMUSG( | Radil | 335.5188 | 0.073822 | 0.596918 | 0.903526 | ENSMUSG( | Radil | 304.5742 | 0.068558 | 0.691963 | 0.960393 |
| ENSMUSG( | Rae1 | 765.9093 | -0.09103 | 0.46582 | 0.844222 | ENSMUSG( | Rae1 | 708.8378 | -0.042 | 0.778591 | 0.976106 |
| ENSMUSG( | Raf1 | 2563.066 | 0.0745 | 0.422836 | 0.823633 | ENSMUSG( | Raf1 | 2276.178 | 0.003085 | 0.97702 | 0.996416 |
| ENSMUSG( | Rai1 | 1951.535 | -0.22994 | 0.032869 | 0.277278 | ENSMUSG( | Rai1 | 1950.757 | 0.049976 | 0.687016 | 0.959226 |
| ENSMUSG( | Rai14 | 430.7238 | 0.272612 | 0.028726 | 0.258426 | ENSMUSG( | Rai14 | 397.1689 | 0.310121 | 0.042064 | 0.690304 |
| ENSMUSG( | Rai2 | 142.3415 | 0.010077 | 0.960611 | 0.993991 | ENSMUSG( | Rai2 | 123.6654 | -0.11738 | 0.602801 | 0.94789 |

| | | | | | |
|---|---|---|---|---|---|
| ENSMUSG( | 1593.619 | 0.006635 | 0.951566 | 0.993821 | Psmb5 |
| ENSMUSG( | 2195.013 | -0.03637 | 0.694388 | 0.9343 | Psmb6 |
| ENSMUSG( | 2478.936 | 0.043318 | 0.656713 | 0.923929 | Psmb7 |
| ENSMUSG( | 154.5277 | -0.23094 | 0.178937 | 0.616298 | Psmb8 |
| ENSMUSG( | 63.30072 | -0.50778 | 0.053094 | 0.353671 | Psmb9 |
| ENSMUSG( | 3133.23 | 0.039799 | 0.621763 | 0.911432 | Psmc1 |
| ENSMUSG( | 1730.743 | 0.049011 | 0.690526 | 0.933881 | Psmc2 |
| ENSMUSG( | 3660.379 | 0.103082 | 0.282412 | 0.73186 | Psmc3 |
| ENSMUSG( | 76.48445 | -0.26625 | 0.22651 | 0.673549 | Psmc3ip |
| ENSMUSG( | 2068.047 | 0.007482 | 0.943341 | 0.993415 | Psmc4 |
| ENSMUSG( | 4554.538 | 0.045562 | 0.60076 | 0.90443 | Psmc5 |
| ENSMUSG( | 1394.336 | 0.066438 | 0.529593 | 0.878883 | Psmc6 |
| ENSMUSG( | 2308.484 | 0.154198 | 0.082977 | 0.44459 | Psmd1 |
| ENSMUSG( | 302.5238 | -0.06167 | 0.632393 | 0.914073 | Psmd10 |
| ENSMUSG( | 1429.258 | 0.175171 | 0.057438 | 0.370033 | Psmd11 |
| ENSMUSG( | 1228.857 | 0.044474 | 0.71348 | 0.940865 | Psmd12 |
| ENSMUSG( | 1555.437 | 0.067114 | 0.553038 | 0.886204 | Psmd13 |
| ENSMUSG( | 1397.22 | 0.122104 | 0.23298 | 0.681135 | Psmd14 |
| ENSMUSG( | 5602.376 | 0.132672 | 0.157119 | 0.58551 | Psmd2 |
| ENSMUSG( | 3134.267 | -0.01314 | 0.870469 | 0.977906 | Psmd3 |
| ENSMUSG( | 1920.237 | 0.064193 | 0.53791 | 0.882548 | Psmd4 |
| ENSMUSG( | 842.5334 | 0.123992 | 0.305145 | 0.752998 | Psmd5 |
| ENSMUSG( | 1150.561 | 0.034714 | 0.721191 | 0.942909 | Psmd6 |
| ENSMUSG( | 2335.559 | 0.129418 | 0.136401 | 0.550651 | Psmd7 |
| ENSMUSG( | 2763.607 | -0.02156 | 0.802287 | 0.962119 | Psmd8 |
| ENSMUSG( | 718.3301 | 0.148841 | 0.223686 | 0.669845 | Psmd9 |
| ENSMUSG( | 898.3763 | -0.07298 | 0.565231 | 0.890779 | Psme1 |
| ENSMUSG( | 637.9945 | 0.019034 | 0.859439 | 0.97481 | Psme2 |
| ENSMUSG( | 21.57686 | -0.34663 | 0.449343 | 0.835057 | Psme2b |
| ENSMUSG( | 3182.751 | 0.030376 | 0.708651 | 0.939519 | Psme3 |
| ENSMUSG( | 1199.12 | 0.001094 | 0.991443 | 0.999232 | Psme3ip1 |
| ENSMUSG( | 1622.425 | 0.039094 | 0.703385 | 0.937735 | Psme4 |
| ENSMUSG( | 1045.029 | 0.00113 | 0.991419 | 0.999232 | Psmf1 |
| ENSMUSG( | 266.7919 | 0.175608 | 0.257405 | 0.708607 | Psmg1 |
| ENSMUSG( | 366.1137 | -0.07336 | 0.593673 | 0.90281 | Psmg2 |
| ENSMUSG( | 126.5397 | 0.041418 | 0.833243 | 0.970333 | Psmg3 |
| ENSMUSG( | 199.5887 | 0.039821 | 0.792998 | 0.95947 | Psmg4 |
| ENSMUSG( | 826.2059 | -0.03473 | 0.78075 | 0.957491 | Pspc1 |
| ENSMUSG( | 1031.958 | -0.01939 | 0.867087 | 0.976541 | Psph |
| ENSMUSG( | 238.9478 | 0.178079 | 0.267199 | 0.717525 | Psrc1 |
| ENSMUSG( | 450.8653 | 0.097464 | 0.467352 | 0.844819 | Pstk |
| ENSMUSG( | 85.97245 | -0.36689 | 0.097304 | 0.477629 | Pstpip1 |
| ENSMUSG( | 148.819 | -0.44852 | 0.023925 | 0.236484 | Pstpip2 |
| ENSMUSG( | 83.17238 | -0.44998 | 0.203258 | 0.643631 | Ptafr |
| ENSMUSG( | 3226.179 | -0.11927 | 0.382734 | 0.803227 | Ptar1 |
| ENSMUSG( | 1252.349 | -0.10213 | 0.306888 | 0.754887 | Ptbp1 |
| ENSMUSG( | 2092.609 | 0.03747 | 0.683122 | 0.93213 | Ptbp2 |
| ENSMUSG( | 537.6181 | -0.04363 | 0.747669 | 0.949604 | Ptbp3 |
| ENSMUSG( | 570.7723 | 0.045294 | 0.75606 | 0.951479 | Ptcd1 |
| ENSMUSG( | 570.106 | 0.137996 | 0.236099 | 0.683188 | Ptcd2 |
| ENSMUSG( | 1068.177 | 0.140451 | 0.174395 | 0.610304 | Ptcd3 |
| ENSMUSG( | 3608.294 | 0.063189 | 0.507501 | 0.86795 | Ptch1 |
| ENSMUSG( | 51.00197 | -0.79461 | 0.005187 | 0.096057 | Ptch2 |
| ENSMUSG( | 96.2041 | -0.20731 | 0.335126 | 0.776511 | Ptchd1 |
| ENSMUSG( | 105.0897 | 0.039747 | 0.866948 | 0.976541 | Ptchd4 |
| ENSMUSG( | 3963.376 | -0.17513 | 0.169803 | 0.604256 | Ptdss1 |
| ENSMUSG( | 2273.431 | -0.1245 | 0.258086 | 0.70936 | Ptdss2 |
| ENSMUSG( | 3560.06 | 0.06226 | 0.560939 | 0.888934 | Pten |
| ENSMUSG( | 115.2911 | 0.187929 | 0.334701 | 0.776386 | Pter |
| ENSMUSG( | 54.10478 | -0.29967 | 0.251271 | 0.700869 | Ptgdr |
| ENSMUSG( | 34036.71 | 0.209071 | 0.039882 | 0.308163 | Ptgds |
| ENSMUSG( | 123.7763 | -0.02575 | 0.887945 | 0.983434 | Ptger1 |
| ENSMUSG( | 103.643 | -0.28006 | 0.22576 | 0.672634 | Ptger3 |
| ENSMUSG( | 40.27545 | -0.14829 | 0.651287 | 0.921451 | Ptger4 |
| ENSMUSG( | 410.1336 | -0.0181 | 0.893425 | 0.984512 | Ptges |
| ENSMUSG( | 1153.054 | 0.0514 | 0.579162 | 0.896692 | Ptges2 |
| ENSMUSG( | 1989.158 | 0.056416 | 0.638285 | 0.916154 | Ptges3 |
| ENSMUSG( | 38.30628 | -0.08129 | 0.794401 | 0.959799 | Ptges3l |
| ENSMUSG( | 51.92709 | 0.491512 | 0.062831 | 0.387645 | Ptgfr |
| ENSMUSG( | 2228.228 | -0.11484 | 0.384448 | 0.804007 | Ptgfrn |
| ENSMUSG( | 78.39753 | -0.23994 | 0.29502 | 0.743798 | Ptgis |
| ENSMUSG( | 59.4979 | 0.439106 | 0.098495 | 0.480334 | Ptgr1 |
| ENSMUSG( | 4118.417 | 0.002924 | 0.981941 | 0.997512 | Ptgr2 |
| ENSMUSG( | 1398.285 | 0.023884 | 0.797131 | 0.960507 | Ptgr3 |
| ENSMUSG( | 786.5756 | -0.15767 | 0.345722 | 0.779153 | Ptgs1 |
| ENSMUSG( | 228.5216 | -0.44185 | 0.01473 | 0.178407 | Ptgs2 |
| ENSMUSG( | 427.057 | -0.14325 | 0.274771 | 0.725078 | Pth1r |
| ENSMUSG( | 51.71923 | -0.66619 | 0.020202 | 0.214698 | Pth2r |
| ENSMUSG( | 102.86 | -1.08736 | 7.51E-08 | 2.13E-05 | Pthlh |
| ENSMUSG( | 3577.802 | 0.119592 | 0.173358 | 0.60863 | Ptk2 |
| ENSMUSG( | 8536.608 | 0.20336 | 0.203398 | 0.643631 | Ptk2b |
| ENSMUSG( | 495.8339 | -0.40736 | 0.013963 | 0.172788 | Ptk7 |
| ENSMUSG( | 5977.331 | 0.04233 | 0.64352 | 0.917316 | Ptma |
| ENSMUSG( | 22594.1 | -0.00719 | 0.939662 | 0.993415 | Ptms |
| ENSMUSG( | 21841.96 | -0.38297 | 0.004104 | 0.08269 | Ptn |
| ENSMUSG( | 4615.962 | 0.054823 | 0.507934 | 0.867961 | Ptov1 |
| ENSMUSG( | 16.62145 | -0.779 | 0.107269 | NA | Ptp4a1 |
| ENSMUSG( | 13.06645 | -0.01105 | 0.985393 | NA | Ptp4a1 |
| ENSMUSG( | 6175.966 | -0.10182 | 0.341033 | 0.778828 | Ptp4a2 |
| ENSMUSG( | 2073.902 | 0.011456 | 0.911087 | 0.988483 | Ptp4a3 |
| ENSMUSG( | 8010.494 | 0.126108 | 0.08779 | 0.454663 | Ptpa |
| ENSMUSG( | 1186.821 | -0.08626 | 0.413104 | 0.819541 | Ptpdc1 |
| ENSMUSG( | 1663.42 | -0.01447 | 0.889096 | 0.983434 | Ptpmt1 |
| ENSMUSG( | 2124.732 | 0.088002 | 0.300809 | 0.749979 | Ptpn1 |
| ENSMUSG( | 13193.96 | 0.085159 | 0.277923 | 0.727904 | Ptpn11 |
| ENSMUSG( | 1317.831 | 0.087295 | 0.346072 | 0.779153 | Ptpn12 |
| ENSMUSG( | 1772.559 | 0.071752 | 0.561892 | 0.889052 | Ptpn13 |
| ENSMUSG( | 441.9212 | 0.127737 | 0.319483 | 0.765296 | Ptpn14 |
| ENSMUSG( | 106.2762 | -0.11724 | 0.554617 | 0.886731 | Ptpn18 |
| ENSMUSG( | 413.236 | 0.039922 | 0.805707 | 0.962616 | Ptpn2 |
| ENSMUSG( | 403.8491 | 0.156501 | 0.183425 | 0.62047 | Ptpn21 |
| ENSMUSG( | 43.23365 | 0.070784 | 0.831461 | 0.969889 | Ptpn22 |
| ENSMUSG( | 2079.578 | 0.164367 | 0.2101 | 0.652466 | Ptpn23 |
| ENSMUSG( | 1223.904 | 0.19722 | 0.133613 | 0.546858 | Ptpn3 |
| ENSMUSG( | 2233.277 | -0.07708 | 0.610484 | 0.908189 | Ptpn4 |
| ENSMUSG( | 7043.261 | -0.26532 | 0.006954 | 0.114623 | Ptpn5 |
| ENSMUSG( | 257.6582 | -0.0772 | 0.585513 | 0.899171 | Ptpn6 |
| ENSMUSG( | 32.00888 | -0.85786 | 0.023624 | 0.235222 | Ptpn7 |
| ENSMUSG( | 1434.656 | 0.20341 | 0.019709 | 0.21171 | Ptpn9 |
| ENSMUSG( | 9573.561 | -0.11513 | 0.348608 | 0.78132 | Ptpra |
| ENSMUSG( | 1157.032 | 0.207323 | 0.593775 | 0.902869 | Ptprb |
| ENSMUSG( | 265.2712 | -0.51813 | 0.014487 | 0.177206 | Ptprc |
| ENSMUSG( | 15.01423 | -0.7991 | 0.108061 | NA | Ptprcap |
| ENSMUSG( | 8399.71 | -0.31753 | 0.023185 | 0.233583 | Ptprd |
| ENSMUSG( | 927.961 | -0.34832 | 0.032154 | 0.273917 | Ptpre |
| ENSMUSG( | 4317.084 | -0.03865 | 0.763546 | 0.953845 | Ptprf |
| ENSMUSG( | 3394.971 | -0.02894 | 0.834182 | 0.970333 | Ptprg |
| ENSMUSG( | 39.39953 | -0.10625 | 0.736014 | 0.946267 | Ptprh |
| ENSMUSG( | 2734.745 | -0.01102 | 0.939188 | 0.993415 | Ptprj |
| ENSMUSG( | 1756.344 | -0.1479 | 0.341915 | 0.778828 | Ptprk |
| ENSMUSG( | 1188.477 | -0.62154 | 4.19E-05 | 0.003503 | Ptprm |
| ENSMUSG( | 19630.21 | -0.10538 | 0.37882 | 0.801731 | Ptprn |
| ENSMUSG( | 5074.286 | -0.07023 | 0.727201 | 0.944223 | Ptprn2 |
| ENSMUSG( | 896.6646 | -0.59631 | 0.00041 | 0.018302 | Ptpro |
| ENSMUSG( | 494.9051 | -0.52131 | 0.00097 | 0.03229 | Ptprr |
| ENSMUSG( | 18015.22 | 0.008399 | 0.931338 | 0.992327 | Ptprs |
| ENSMUSG( | 9891.329 | -0.09817 | 0.425204 | 0.82478 | Ptprt |
| ENSMUSG( | 1861.241 | 0.234163 | 0.065201 | 0.395697 | Ptpru |
| ENSMUSG( | 83.79861 | -1.18724 | 8.54E-05 | 0.005781 | Ptprv |
| ENSMUSG( | 23315.93 | -0.03481 | 0.788411 | 0.959226 | Ptprz1 |
| ENSMUSG( | 115.6432 | 0.155408 | 0.397455 | 0.812127 | Ptrh1 |
| ENSMUSG( | 295.6317 | 0.017902 | 0.900926 | 0.985633 | Ptrh2 |
| ENSMUSG( | 425.7713 | -0.01088 | 0.934201 | 0.99323 | Ptrhd1 |
| ENSMUSG( | 1433.835 | 0.013027 | 0.901805 | 0.985633 | Pts |
| ENSMUSG( | 596.0987 | 0.002234 | 0.984461 | 0.99781 | Pttg1 |
| ENSMUSG( | 3829.805 | -0.13564 | 0.296148 | 0.744386 | Pttg1ip |
| ENSMUSG( | 12.03456 | 0.250563 | 0.644985 | NA | Ptx3 |
| ENSMUSG( | 3363.713 | 0.091493 | 0.370532 | 0.797778 | Puf60 |
| ENSMUSG( | 2592.651 | 0.103359 | 0.206364 | 0.648436 | Pum1 |
| ENSMUSG( | 6900.384 | 0.043614 | 0.626833 | 0.912825 | Pum2 |
| ENSMUSG( | 1085.137 | 0.15159 | 0.148013 | 0.571232 | Pum3 |
| ENSMUSG( | 1845.154 | -0.01171 | 0.924792 | 0.991152 | Pura |
| ENSMUSG( | 34513.99 | 0.073377 | 0.484214 | 0.856577 | Purb |
| ENSMUSG( | 733.9863 | 0.143838 | 0.193947 | 0.633085 | Purg |
| ENSMUSG( | 292.5674 | -0.00385 | 0.980762 | 0.997512 | Pus1 |
| ENSMUSG( | 299.3864 | 0.025964 | 0.841702 | 0.971179 | Pus10 |
| ENSMUSG( | 256.6271 | 0.193951 | 0.169845 | 0.604274 | Pus3 |
| ENSMUSG( | 607.329 | 0.126565 | 0.339802 | 0.777939 | Pus7 |
| ENSMUSG( | 86.80069 | -0.20254 | 0.372396 | 0.798768 | Pus7l |
| ENSMUSG( | 765.4575 | 0.113217 | 0.289262 | 0.739725 | Pusl1 |
| ENSMUSG( | 856.7935 | -0.08165 | 0.632558 | 0.914073 | Pvalb |
| ENSMUSG( | 240.7496 | -0.74064 | 2.00E-05 | 0.001952 | Pvr |
| ENSMUSG( | 494.9657 | 0.059314 | 0.612147 | 0.908586 | Pvt1 |
| ENSMUSG( | 476.4593 | 0.075986 | 0.562366 | 0.889448 | Pwp1 |
| ENSMUSG( | 692.9405 | -0.08186 | 0.425762 | 0.824898 | Pwp2 |
| ENSMUSG( | 575.135 | 0.123563 | 0.276932 | 0.727074 | Pwwp2a |
| ENSMUSG( | 314.3647 | -0.07244 | 0.648479 | 0.919997 | Pwwp2b |
| ENSMUSG( | 902.832 | -0.06058 | 0.632367 | 0.914073 | Pwwp3a |

| | | | | | |
|---|---|---|---|---|---|
| ENSMUSG( | 356.949 | -0.27299 | 0.079201 | 0.77458 | Ptrhd1 |
| ENSMUSG( | 1247.905 | -0.12336 | 0.27809 | 0.891468 | Pts |
| ENSMUSG( | 509.9832 | -0.18847 | 0.17064 | 0.841762 | Pttg1 |
| ENSMUSG( | 3805.939 | 0.119349 | 0.395508 | 0.917257 | Pttg1ip |
| ENSMUSG( | 16.52202 | 1.199517 | 0.027273 | 0.60551 | Ptx3 |
| ENSMUSG( | 2815.137 | -0.15806 | 0.20141 | 0.864412 | Puf60 |
| ENSMUSG( | 2273.148 | -0.00792 | 0.945771 | 0.994798 | Pum1 |
| ENSMUSG( | 6311.72 | 0.054205 | 0.630929 | 0.953251 | Pum2 |
| ENSMUSG( | 897.4495 | -0.12957 | 0.306692 | 0.898538 | Pum3 |
| ENSMUSG( | 1617.303 | -0.13029 | 0.409103 | 0.917257 | Pura |
| ENSMUSG( | 30564.6 | -0.00865 | 0.951939 | 0.994798 | Purb |
| ENSMUSG( | 631.6914 | -0.02523 | 0.868019 | 0.98781 | Purg |
| ENSMUSG( | 268.4948 | 0.02119 | 0.90027 | 0.990558 | Pus1 |
| ENSMUSG( | 267.8519 | -0.03012 | 0.844039 | 0.983571 | Pus10 |
| ENSMUSG( | 224.659 | 0.082738 | 0.605624 | 0.949298 | Pus3 |
| ENSMUSG( | 564.0881 | 0.17393 | 0.28343 | 0.892412 | Pus7 |
| ENSMUSG( | 82.10664 | -0.10008 | 0.694512 | 0.960867 | Pus7l |
| ENSMUSG( | 704.8866 | 0.144795 | 0.253925 | 0.880967 | Pusl1 |
| ENSMUSG( | 682.3404 | -0.51874 | 0.002697 | 0.205145 | Pvalb |
| ENSMUSG( | 255.0205 | -0.21094 | 0.256421 | 0.883082 | Pvr |
| ENSMUSG( | 440.4456 | -0.00848 | 0.948963 | 0.994798 | Pvt1 |
| ENSMUSG( | 414.9164 | -0.05734 | 0.706765 | 0.963529 | Pwp1 |
| ENSMUSG( | 635.3258 | -0.06468 | 0.598218 | 0.946676 | Pwp2 |
| ENSMUSG( | 526.0302 | 0.132226 | 0.317614 | 0.902602 | Pwwp2a |
| ENSMUSG( | 289.5551 | -0.04295 | 0.802559 | 0.97681 | Pwwp2b |
| ENSMUSG( | 860.7886 | 0.076072 | 0.621766 | 0.953234 | Pwwp3a |
| ENSMUSG( | 56.72454 | 0.215338 | 0.431026 | 0.917257 | Pwwp3b |
| ENSMUSG( | 228.8238 | -0.02381 | 0.883614 | 0.989644 | Pxdc1 |
| ENSMUSG( | 2954.579 | 0.076499 | 0.568428 | 0.941777 | Pxdn |
| ENSMUSG( | 701.7704 | 0.052063 | 0.729933 | 0.967228 | Pxk |
| ENSMUSG( | 343.2177 | -0.09608 | 0.497118 | 0.927825 | Pxmp2 |
| ENSMUSG( | 783.7721 | -0.2751 | 0.029868 | 0.61992 | Pxmp4 |
| ENSMUSG( | 1211.78 | -0.04311 | 0.75712 | 0.971858 | Pxn |
| ENSMUSG( | 378.3416 | -0.09053 | 0.615139 | 0.951177 | Pxylp1 |
| ENSMUSG( | 133.0262 | -0.24951 | 0.203476 | 0.864412 | Pycard |
| ENSMUSG( | 98.0756 | -0.44981 | 0.054389 | 0.723362 | Pycr1 |
| ENSMUSG( | 589.9886 | -0.12872 | 0.42204 | 0.917257 | Pycr2 |
| ENSMUSG( | 400.1503 | -0.04925 | 0.747498 | 0.969776 | Pycr3 |
| ENSMUSG( | 14981.14 | -0.20609 | 0.087615 | 0.785039 | Pygb |
| ENSMUSG( | 186.8608 | -0.02481 | 0.903982 | 0.991089 | Pygl |
| ENSMUSG( | 3383.486 | -0.25613 | 0.032377 | 0.635121 | Pygm |
| ENSMUSG( | 1458.288 | 0.016785 | 0.907847 | 0.991958 | Pygo1 |
| ENSMUSG( | 1127.133 | -0.02422 | 0.831079 | 0.98051 | Pygo2 |
| ENSMUSG( | 399.1609 | -0.2987 | 0.120225 | 0.813384 | Pym1 |
| ENSMUSG( | 313.7525 | -0.04412 | 0.768936 | 0.974236 | Pyroxd1 |
| ENSMUSG( | 129.8098 | -0.02633 | 0.901555 | 0.990784 | Pyroxd2 |
| ENSMUSG( | 684.6748 | -0.36798 | 0.02892 | 0.614171 | Pyurf |
| ENSMUSG( | 1262.807 | 0.056567 | 0.638429 | 0.953903 | Qars1 |
| ENSMUSG( | 6546.782 | -0.45666 | 9.71E-05 | 0.02045 | Qdpr |
| ENSMUSG( | 22932.26 | 0.009899 | 0.954627 | 0.994798 | Qki |
| ENSMUSG( | 282.6632 | -0.30987 | 0.060263 | 0.731553 | Qng1 |
| ENSMUSG( | 343.3493 | 0.374969 | 0.050535 | 0.708889 | Qpct |
| ENSMUSG( | 701.1617 | 0.100319 | 0.543312 | 0.938063 | Qpctl |
| ENSMUSG( | 22.76373 | -0.51525 | 0.243049 | 0.877399 | Qprt |
| ENSMUSG( | 86.96327 | 0.37153 | 0.114996 | 0.811921 | Qrfpr |
| ENSMUSG( | 18.29847 | -0.25175 | 0.587286 | 0.944975 | Qrfprl |
| ENSMUSG( | 1503.155 | -0.06657 | 0.572732 | 0.94262 | Qrich1 |
| ENSMUSG( | 318.845 | 0.082203 | 0.581389 | 0.944266 | Qrsl1 |
| ENSMUSG( | 788.302 | -0.00383 | 0.978524 | 0.996459 | Qser1 |
| ENSMUSG( | 1043.817 | 0.281945 | 0.109283 | 0.805353 | Qsox1 |
| ENSMUSG( | 922.0275 | 0.164532 | 0.340621 | 0.907972 | Qsox2 |
| ENSMUSG( | 262.0088 | -0.08332 | 0.675399 | 0.95694 | Qtrt1 |
| ENSMUSG( | 369.4803 | 0.326702 | 0.046795 | 0.700612 | Qtrt2 |
| ENSMUSG( | 931.2313 | -0.15807 | 0.244471 | 0.877904 | R3hcc1 |
| ENSMUSG( | 336.6753 | -0.08408 | 0.578984 | 0.943849 | R3hcc1l |
| ENSMUSG( | 10310.81 | -0.14768 | 0.204221 | 0.864412 | R3hdm1 |
| ENSMUSG( | 12113.64 | 0.005483 | 0.957592 | 0.994798 | R3hdm2 |
| ENSMUSG( | 16253.52 | -0.0591 | 0.565165 | 0.94131 | R3hdm4 |
| ENSMUSG( | 275.2941 | -0.15879 | 0.312251 | 0.900037 | R74862 |
| ENSMUSG( | 5857.662 | -0.10761 | 0.443134 | 0.921569 | Rab10 |
| ENSMUSG( | 1558.774 | -0.04732 | 0.726559 | 0.966887 | Rab11a |
| ENSMUSG( | 4990.644 | -0.13655 | 0.188422 | 0.85698 | Rab11b |
| ENSMUSG( | 130.2018 | 0.683173 | 0.016201 | 0.512256 | Rab11fip1 |
| ENSMUSG( | 1010.004 | 0.034519 | 0.769402 | 0.974236 | Rab11fip2 |
| ENSMUSG( | 4071.194 | 0.080138 | 0.431116 | 0.917257 | Rab11fip3 |
| ENSMUSG( | 11818.55 | -0.07473 | 0.427419 | 0.917257 | Rab11fip4 |
| ENSMUSG( | 2223.131 | 0.042551 | 0.732425 | 0.967228 | Rab11fip5 |
| ENSMUSG( | 3756.78 | -1.08666 | 5.68E-17 | 9.04E-14 | Rab12 |
| ENSMUSG( | 423.3293 | -0.33545 | 0.032378 | 0.635121 | Rab13 |
| ENSMUSG( | 4589.093 | -0.01432 | 0.905728 | 0.991089 | Rab14 |
| ENSMUSG( | 11170.28 | -0.05194 | 0.581671 | 0.944266 | Rab15 |
| ENSMUSG( | 2737.789 | -0.01981 | 0.894428 | 0.990558 | Rab18 |
| ENSMUSG( | 4501.057 | -0.11751 | 0.416254 | 0.917257 | Rab1a |
| ENSMUSG( | 5235.859 | -0.14278 | 0.192478 | 0.859726 | Rab1b |
| ENSMUSG( | 18.05924 | -0.48881 | 0.29901 | 0.898538 | Rab20 |
| ENSMUSG( | 3927.539 | -0.08252 | 0.623094 | 0.953234 | Rab21 |
| ENSMUSG( | 3082.554 | 0.020526 | 0.856863 | 0.985884 | Rab22a |
| ENSMUSG( | 1043.362 | 0.095594 | 0.511887 | 0.930005 | Rab23 |
| ENSMUSG( | 1240.812 | -0.01575 | 0.907535 | 0.991907 | Rab24 |
| ENSMUSG( | 914.4395 | -0.01692 | 0.925261 | 0.99306 | Rab26 |
| ENSMUSG( | 47.00802 | 0.037459 | 0.920233 | 0.992644 | Rab26os |
| ENSMUSG( | 112.7342 | 0.266937 | 0.217653 | 0.865589 | Rab27a |
| ENSMUSG( | 538.5569 | -0.08914 | 0.57996 | 0.944209 | Rab27b |
| ENSMUSG( | 1465.705 | -0.07629 | 0.48084 | 0.924548 | Rab28 |
| ENSMUSG( | 173.4077 | -0.11534 | 0.529092 | 0.934795 | Rab29 |
| ENSMUSG( | 4746.05 | -0.06557 | 0.65375 | 0.955015 | Rab2a |
| ENSMUSG( | 898.442 | -0.06921 | 0.574078 | 0.942912 | Rab2b |
| ENSMUSG( | 774.6724 | 0.041062 | 0.794716 | 0.97681 | Rab30 |
| ENSMUSG( | 6533.129 | -0.21638 | 0.075307 | 0.767994 | Rab31 |
| ENSMUSG( | 68.58145 | -0.20406 | 0.437893 | 0.920495 | Rab32 |
| ENSMUSG( | 381.6012 | -0.25883 | 0.196208 | 0.862161 | Rab33a |
| ENSMUSG( | 924.0611 | -0.12882 | 0.379442 | 0.916995 | Rab33b |
| ENSMUSG( | 674.3013 | 0.025493 | 0.861037 | 0.986226 | Rab34 |
| ENSMUSG( | 2132.635 | -0.0997 | 0.343253 | 0.909646 | Rab35 |
| ENSMUSG( | 990.0933 | -0.03433 | 0.812731 | 0.979148 | Rab36 |
| ENSMUSG( | 91.35747 | -0.44203 | 0.117932 | 0.813384 | Rab37 |
| ENSMUSG( | 24.31024 | 0.038244 | 0.927864 | 0.99306 | Rab39 |
| ENSMUSG( | 599.687 | -0.03614 | 0.827863 | 0.980224 | Rab39b |
| ENSMUSG( | 8596.364 | -0.15092 | 0.336528 | 0.907972 | Rab3a |
| ENSMUSG( | 954.4823 | -0.17327 | 0.358397 | 0.913759 | Rab3b |
| ENSMUSG( | 6491.764 | 0.058934 | 0.634965 | 0.953903 | Rab3c |
| ENSMUSG( | 311.7258 | 0.168223 | 0.281162 | 0.892356 | Rab3d |
| ENSMUSG( | 2000.701 | 0.049796 | 0.638527 | 0.953903 | Rab3gap1 |
| ENSMUSG( | 2144.078 | -0.00355 | 0.974141 | 0.996416 | Rab3gap2 |
| ENSMUSG( | 506.4268 | -0.03108 | 0.808568 | 0.97873 | Rab3il1 |
| ENSMUSG( | 727.5878 | -0.01951 | 0.898691 | 0.990558 | Rab3ip |
| ENSMUSG( | 535.9829 | -0.11778 | 0.509308 | 0.929089 | Rab40b |
| ENSMUSG( | 1240.142 | -0.04612 | 0.712606 | 0.964363 | Rab40c |
| ENSMUSG( | 10.9206 | 0.77414 | 0.183826 | NA | Rab42 |
| ENSMUSG( | 38.78863 | 0.236026 | 0.465578 | 0.923329 | Rab43 |
| ENSMUSG( | 1319.924 | 0.209919 | 0.080136 | 0.77681 | Rab4a |
| ENSMUSG( | 131.4399 | 0.077976 | 0.712846 | 0.96439 | Rab4b |
| ENSMUSG( | 2215.209 | -0.11395 | 0.43933 | 0.920721 | Rab5a |
| ENSMUSG( | 5016.538 | -0.15391 | 0.162023 | 0.837724 | Rab5b |
| ENSMUSG( | 6337.6 | -0.18099 | 0.182721 | 0.850579 | Rab5c |
| ENSMUSG( | 1412.564 | 0.019698 | 0.866952 | 0.987487 | Rab5if |
| ENSMUSG( | 9090.577 | -0.14715 | 0.300428 | 0.898538 | Rab6a |
| ENSMUSG( | 33347.34 | -0.05392 | 0.568211 | 0.941777 | Rab6b |
| ENSMUSG( | 8269.108 | -0.15367 | 0.26667 | 0.886134 | Rab7 |
| ENSMUSG( | 219.8222 | -0.64361 | 0.000331 | 0.049413 | Rab7b |
| ENSMUSG( | 1134.264 | -0.02892 | 0.810124 | 0.978779 | Rab8a |
| ENSMUSG( | 788.2417 | -0.02908 | 0.837198 | 0.981335 | Rab8b |
| ENSMUSG( | 809.1372 | -0.17268 | 0.219383 | 0.866104 | Rab9 |
| ENSMUSG( | 251.6227 | -0.02921 | 0.882242 | 0.989644 | Rab9b |
| ENSMUSG( | 4092.991 | 0.099617 | 0.446803 | 0.921867 | Rabac1 |
| ENSMUSG( | 3491.695 | -0.05425 | 0.617385 | 0.951952 | Rabep1 |
| ENSMUSG( | 350.2148 | 0.145655 | 0.396331 | 0.917257 | Rabep2 |
| ENSMUSG( | 480.4944 | -0.14323 | 0.321175 | 0.904019 | Rabepk |
| ENSMUSG( | 2097.433 | 0.015901 | 0.882383 | 0.989644 | Rabgap1 |
| ENSMUSG( | 1906.673 | 0.022744 | 0.885431 | 0.989688 | Rabgap1l |
| ENSMUSG( | 761.6151 | -0.09508 | 0.468386 | 0.923329 | Rabgef1 |
| ENSMUSG( | 867.1742 | -0.04976 | 0.700697 | 0.961547 | Rabggta |
| ENSMUSG( | 1012.179 | -0.14252 | 0.240147 | 0.875205 | Rabggtb |
| ENSMUSG( | 717.2828 | -0.07345 | 0.590235 | 0.944975 | Rabif |
| ENSMUSG( | 583.083 | 0.001269 | 0.994881 | 0.998815 | Rabl2 |
| ENSMUSG( | 137.9621 | -0.01835 | 0.930996 | 0.99306 | Rabl3 |
| ENSMUSG( | 6459.579 | 0.057147 | 0.559005 | 0.939587 | Rabl6 |
| ENSMUSG( | 10565.69 | -0.1306 | 0.336155 | 0.907972 | Rac1 |
| ENSMUSG( | 187.0178 | -0.37217 | 0.05909 | 0.729429 | Rac2 |
| ENSMUSG( | 530.2977 | -0.09258 | 0.505543 | 0.927825 | Rac3 |
| ENSMUSG( | 255.4273 | 0.107633 | 0.506302 | 0.927825 | Racgap1 |
| ENSMUSG( | 7950.495 | -0.16998 | 0.092846 | 0.79479 | Rack1 |
| ENSMUSG( | 160.1549 | 0.070369 | 0.703154 | 0.962428 | Rad1 |
| ENSMUSG( | 446.5678 | 0.109633 | 0.43237 | 0.918158 | Rad17 |
| ENSMUSG( | 110.4824 | -0.19399 | 0.355164 | 0.912411 | Rad18 |
| ENSMUSG( | 5199.486 | -0.09037 | 0.416005 | 0.917257 | Rad21 |
| ENSMUSG( | 1714.265 | -0.00187 | 0.988569 | 0.997862 | Rad23a |
| ENSMUSG( | 2844.674 | -0.06735 | 0.648996 | 0.95422 | Rad23b |

| | | | | | | | | | | | |
|---|---|---|---|---|---|---|---|---|---|---|---|
| ENSMUSG( | Rala | 2664.163 | 0.078279 | 0.449583 | 0.835234 | ENSMUSG( | Rala | 2270.368 | -0.12 | 0.410738 | 0.917257 |
| ENSMUSG( | Ralb | 798.4142 | -0.05006 | 0.63343 | 0.914073 | ENSMUSG( | Ralb | 747.8215 | 0.025009 | 0.847853 | 0.985242 |
| ENSMUSG( | Ralbp1 | 2587.973 | 0.249556 | 0.008025 | 0.124801 | ENSMUSG( | Ralbp1 | 2212.816 | 0.074456 | 0.521856 | 0.932571 |
| ENSMUSG( | Ralgapa1 | 2828.47 | 0.045969 | 0.575065 | 0.894572 | ENSMUSG( | Ralgapa1 | 2620.828 | 0.093511 | 0.356959 | 0.913358 |
| ENSMUSG( | Ralgapa2 | 738.8928 | 0.01232 | 0.905178 | 0.98645 | ENSMUSG( | Ralgapa2 | 698.7667 | 0.116147 | 0.348422 | 0.911652 |
| ENSMUSG( | Ralgapb | 3946.816 | 0.018212 | 0.847732 | 0.972374 | ENSMUSG( | Ralgapb | 3710.917 | 0.109491 | 0.337821 | 0.907972 |
| ENSMUSG( | Ralgds | 4816.448 | 0.19362 | 0.046688 | 0.330693 | ENSMUSG( | Ralgds | 4087.094 | -0.00425 | 0.970871 | 0.996061 |
| ENSMUSG( | Ralgps1 | 2343.527 | 0.125544 | 0.130723 | 0.542059 | ENSMUSG( | Ralgps1 | 2126.883 | 0.115341 | 0.262657 | 0.885357 |
| ENSMUSG( | Ralgps2 | 1331.452 | 0.32121 | 0.000396 | 0.017766 | ENSMUSG( | Ralgps2 | 1179.982 | 0.245247 | 0.08916 | 0.788502 |
| ENSMUSG( | Raly | 2268.213 | 0.011986 | 0.88794 | 0.983434 | ENSMUSG( | Raly | 1967.567 | -0.13438 | 0.220231 | 0.866104 |
| ENSMUSG( | Ralyl | 1194.228 | 0.092829 | 0.444749 | 0.833239 | ENSMUSG( | Ralyl | 1002.098 | -0.14882 | 0.272435 | 0.888825 |
| ENSMUSG( | Ramac | 713.2335 | -0.09143 | 0.45317 | 0.836701 | ENSMUSG( | Ramac | 611.5317 | -0.28424 | 0.046868 | 0.700664 |
| ENSMUSG( | Ramacl | 47.17636 | 0.010565 | 0.971056 | 0.996369 | ENSMUSG( | Ramacl | 39.40933 | -0.25068 | 0.454439 | 0.922444 |
| ENSMUSG( | Ramp1 | 5098.744 | -0.14153 | 0.140381 | 0.557399 | ENSMUSG( | Ramp1 | 4575.466 | -0.18848 | 0.094361 | 0.795735 |
| ENSMUSG( | Ramp2 | 931.3564 | 0.182088 | 0.153294 | 0.57989 | ENSMUSG( | Ramp2 | 1043.253 | 0.705482 | 2.64E-05 | 0.007461 |
| ENSMUSG( | Ramp3 | 543.0218 | -0.54269 | 2.17E-06 | 0.000346 | ENSMUSG( | Ramp3 | 558.0575 | -0.1452 | 0.384054 | 0.917257 |
| ENSMUSG( | Ran | 3177.171 | 0.030031 | 0.79222 | 0.959461 | ENSMUSG( | Ran | 2729.919 | -0.14616 | 0.297215 | 0.898538 |
| ENSMUSG( | Ranbp1 | 687.5898 | -0.01211 | 0.913913 | 0.988483 | ENSMUSG( | Ranbp1 | 607.0443 | -0.11146 | 0.455701 | 0.923329 |
| ENSMUSG( | Ranbp10 | 1238.813 | 0.083063 | 0.456284 | 0.838832 | ENSMUSG( | Ranbp10 | 1136.125 | 0.103837 | 0.396192 | 0.917257 |
| ENSMUSG( | Ranbp17 | 192.9217 | -0.02269 | 0.882639 | 0.982283 | ENSMUSG( | Ranbp17 | 168.674 | -0.14178 | 0.418092 | 0.917257 |
| ENSMUSG( | Ranbp2 | 3075.873 | -0.00426 | 0.970781 | 0.996369 | ENSMUSG( | Ranbp2 | 3012.225 | 0.200698 | 0.138144 | 0.828492 |
| ENSMUSG( | Ranbp3 | 2152.671 | 0.02386 | 0.832115 | 0.970303 | ENSMUSG( | Ranbp3 | 1912.984 | -0.04648 | 0.675065 | 0.95694 |
| ENSMUSG( | Ranbp3l | 567.6107 | 0.1192 | 0.388822 | 0.806585 | ENSMUSG( | Ranbp3l | 499.0592 | 0.014387 | 0.925082 | 0.99306 |
| ENSMUSG( | Ranbp6 | 551.0374 | -0.03538 | 0.749939 | 0.949664 | ENSMUSG( | Ranbp6 | 535.8236 | 0.152937 | 0.268779 | 0.886755 |
| ENSMUSG( | Ranbp9 | 2269.484 | -0.00404 | 0.963838 | 0.995001 | ENSMUSG( | Ranbp9 | 2135.657 | 0.089028 | 0.403765 | 0.917257 |
| ENSMUSG( | Rangap1 | 4842.92 | 0.124494 | 0.257793 | 0.709192 | ENSMUSG( | Rangap1 | 4045.931 | -0.12755 | 0.330467 | 0.907398 |
| ENSMUSG( | Rangrf | 219.5141 | 0.002878 | 0.984832 | 0.997815 | ENSMUSG( | Rangrf | 207.4137 | 0.094062 | 0.601583 | 0.947749 |
| ENSMUSG( | Rap1a | 1427.832 | -0.09139 | 0.450703 | 0.835475 | ENSMUSG( | Rap1a | 1251.041 | -0.21516 | 0.15835 | 0.837724 |
| ENSMUSG( | Rap1b | 1993.957 | 0.027403 | 0.788207 | 0.959226 | ENSMUSG( | Rap1b | 1827.08 | 0.039358 | 0.784552 | 0.976207 |
| ENSMUSG( | Rap1gap | 6071.083 | 0.089554 | 0.506521 | 0.867425 | ENSMUSG( | Rap1gap | 5481.219 | 0.062296 | 0.668323 | 0.956766 |
| ENSMUSG( | Rap1gap2 | 13068.74 | 0.24959 | 0.002598 | 0.060717 | ENSMUSG( | Rap1gap2 | 11154.19 | 0.072213 | 0.521467 | 0.932489 |
| ENSMUSG( | Rap1gds1 | 9922.516 | 0.03043 | 0.719056 | 0.941986 | ENSMUSG( | Rap1gds1 | 8419.653 | -0.18127 | 0.100511 | 0.801444 |
| ENSMUSG( | Rap2a | 4013.421 | 0.134367 | 0.112911 | 0.506708 | ENSMUSG( | Rap2a | 3407.764 | -0.06819 | 0.526667 | 0.934374 |
| ENSMUSG( | Rap2b | 1488.914 | 0.155963 | 0.209189 | 0.651133 | ENSMUSG( | Rap2b | 1337.321 | 0.111194 | 0.503749 | 0.927825 |
| ENSMUSG( | Rap2c | 1619.767 | 0.043768 | 0.767198 | 0.954438 | ENSMUSG( | Rap2c | 1485.829 | 0.057619 | 0.755857 | 0.971371 |
| ENSMUSG( | Rapgef1 | 9755.038 | 0.20834 | 0.007334 | 0.118347 | ENSMUSG( | Rapgef1 | 8538.478 | 0.098838 | 0.32316 | 0.904023 |
| ENSMUSG( | Rapgef2 | 2562.671 | 0.194801 | 0.060544 | 0.379893 | ENSMUSG( | Rapgef2 | 2364.517 | 0.230593 | 0.058101 | 0.728256 |
| ENSMUSG( | Rapgef3 | 2053.936 | 0.169914 | 0.092178 | 0.466065 | ENSMUSG( | Rapgef3 | 1765.339 | 0.01034 | 0.939258 | 0.993898 |
| ENSMUSG( | Rapgef4 | 73724.29 | 0.091952 | 0.32831 | 0.772854 | ENSMUSG( | Rapgef4 | 62072.32 | -0.13925 | 0.220109 | 0.866104 |
| ENSMUSG( | Rapgef5 | 2684.778 | 0.170522 | 0.090631 | 0.462274 | ENSMUSG( | Rapgef5 | 2480.483 | 0.211221 | 0.108277 | 0.804969 |
| ENSMUSG( | Rapgef6 | 790.1025 | -0.04772 | 0.654481 | 0.923227 | ENSMUSG( | Rapgef6 | 789.6634 | 0.214648 | 0.072475 | 0.763629 |
| ENSMUSG( | Rapgefl1 | 5994.341 | 0.226155 | 0.008405 | 0.128255 | ENSMUSG( | Rapgefl1 | 5241.505 | 0.113977 | 0.268262 | 0.886493 |
| ENSMUSG( | Raph1 | 982.7614 | 0.027955 | 0.783109 | 0.957558 | ENSMUSG( | Raph1 | 913.3522 | 0.085565 | 0.471086 | 0.923329 |
| ENSMUSG( | Rapsn | 24.2721 | -0.54167 | 0.174826 | 0.610304 | ENSMUSG( | Rapsn | 22.4751 | -0.52112 | 0.223573 | 0.867404 |
| ENSMUSG( | Rara | 352.6343 | -0.19759 | 0.200297 | 0.640413 | ENSMUSG( | Rara | 307.4257 | -0.34332 | 0.039591 | 0.68111 |
| ENSMUSG( | Rarb | 312.4008 | -0.59057 | 0.003915 | 0.080172 | ENSMUSG( | Rarb | 318.5542 | -0.22376 | 0.297841 | 0.898538 |
| ENSMUSG( | Rarg | 381.7313 | -0.13832 | 0.309721 | 0.757658 | ENSMUSG( | Rarg | 401.611 | 0.268039 | 0.076404 | 0.770504 |
| ENSMUSG( | Rarres1 | 63.68718 | -0.30077 | 0.243012 | 0.690307 | ENSMUSG( | Rarres1 | 74.86075 | 0.417068 | 0.110094 | 0.806572 |
| ENSMUSG( | Rarres2 | 512.5683 | -0.121 | 0.393156 | 0.809274 | ENSMUSG( | Rarres2 | 498.9492 | 0.080604 | 0.649343 | 0.95422 |
| ENSMUSG( | Rars1 | 839.6013 | 0.050836 | 0.620255 | 0.911432 | ENSMUSG( | Rars1 | 715.4305 | -0.14944 | 0.29402 | 0.898538 |
| ENSMUSG( | Rars2 | 373.691 | 0.078651 | 0.576915 | 0.89546 | ENSMUSG( | Rars2 | 334.4121 | 0.028637 | 0.861181 | 0.986226 |
| ENSMUSG( | Rasa1 | 1062.746 | 0.072517 | 0.464474 | 0.843364 | ENSMUSG( | Rasa1 | 1014.248 | 0.201474 | 0.078097 | 0.77321 |
| ENSMUSG( | Rasa2 | 1727.737 | -0.02396 | 0.85287 | 0.973168 | ENSMUSG( | Rasa2 | 1411.228 | -0.36417 | 0.010579 | 0.419247 |
| ENSMUSG( | Rasa3 | 2990.208 | 0.035258 | 0.677405 | 0.930542 | ENSMUSG( | Rasa3 | 2588.673 | -0.11632 | 0.290866 | 0.897019 |
| ENSMUSG( | Rasa4 | 242.0743 | -0.01898 | 0.894443 | 0.984512 | ENSMUSG( | Rasa4 | 222.8747 | 0.016654 | 0.92468 | 0.99306 |
| ENSMUSG( | Rasal1 | 2968.304 | 0.301551 | 0.014309 | 0.17583 | ENSMUSG( | Rasal1 | 2569.154 | 0.165875 | 0.209715 | 0.864412 |
| ENSMUSG( | Rasal2 | 1827.156 | 0.162521 | 0.081881 | 0.442044 | ENSMUSG( | Rasal2 | 1669.816 | 0.172892 | 0.175363 | 0.845305 |
| ENSMUSG( | Rasal3 | 162.683 | -0.08876 | 0.636046 | 0.914873 | ENSMUSG( | Rasal3 | 171.8523 | 0.330302 | 0.126439 | 0.815475 |
| ENSMUSG( | Rasd1 | 145.2491 | -0.13145 | 0.644139 | 0.917612 | ENSMUSG( | Rasd1 | 134.0375 | -0.09347 | 0.731536 | 0.967228 |
| ENSMUSG( | Rasd2 | 3565.632 | -0.50375 | 0.001969 | 0.051087 | ENSMUSG( | Rasd2 | 3710.502 | -0.07765 | 0.564822 | 0.941206 |
| ENSMUSG( | Rasgef1a | 11694.34 | 0.1103 | 0.15713 | 0.58551 | ENSMUSG( | Rasgef1a | 10285.8 | 0.010651 | 0.923043 | 0.992644 |
| ENSMUSG( | Rasgef1c | 1131.193 | 0.063592 | 0.535227 | 0.880733 | ENSMUSG( | Rasgef1c | 961.7382 | -0.13494 | 0.27534 | 0.889984 |
| ENSMUSG( | Rasgrf1 | 7684.617 | -0.02922 | 0.771585 | 0.954769 | ENSMUSG( | Rasgrf1 | 7481.612 | 0.160325 | 0.164479 | 0.838699 |
| ENSMUSG( | Rasgrf2 | 2506.333 | 0.149189 | 0.139428 | 0.555633 | ENSMUSG( | Rasgrf2 | 2195.917 | 0.040031 | 0.759124 | 0.971909 |
| ENSMUSG( | Rasgrp1 | 8572.724 | 0.017971 | 0.839666 | 0.971179 | ENSMUSG( | Rasgrp1 | 7840.66 | 0.0289 | 0.783847 | 0.976154 |
| ENSMUSG( | Rasgrp2 | 1005.209 | -0.36238 | 0.039779 | 0.307538 | ENSMUSG( | Rasgrp2 | 987.0321 | -0.13231 | 0.494559 | 0.927372 |
| ENSMUSG( | Rasgrp3 | 517.4416 | -0.0578 | 0.741048 | 0.948083 | ENSMUSG( | Rasgrp3 | 510.1348 | 0.176542 | 0.424101 | 0.917257 |
| ENSMUSG( | Rasgrp4 | 68.35749 | -0.08574 | 0.725766 | 0.944223 | ENSMUSG( | Rasgrp4 | 71.17763 | 0.285331 | 0.300761 | 0.898538 |
| ENSMUSG( | Rasip1 | 334.4796 | 0.290154 | 0.095238 | 0.474145 | ENSMUSG( | Rasip1 | 334.6666 | 0.538234 | 0.009143 | 0.401114 |
| ENSMUSG( | Rasl10a | 613.2374 | -0.02798 | 0.799541 | 0.961275 | ENSMUSG( | Rasl10a | 551.7278 | -0.06967 | 0.627214 | 0.953234 |
| ENSMUSG( | Rasl10b | 22824.6 | 0.179396 | 0.049256 | 0.33882 | ENSMUSG( | Rasl10b | 20229.53 | 0.103338 | 0.334211 | 0.907907 |
| ENSMUSG( | Rasl11a | 93.60653 | -0.24149 | 0.450232 | 0.835344 | ENSMUSG( | Rasl11a | 74.3339 | -0.69884 | 0.013239 | 0.460134 |
| ENSMUSG( | Rasl11b | 1339.2 | -0.08575 | 0.585472 | 0.899171 | ENSMUSG( | Rasl11b | 1199.659 | -0.14045 | 0.418814 | 0.917257 |
| ENSMUSG( | Rasl12 | 74.31596 | -0.19894 | 0.472785 | 0.848388 | ENSMUSG( | Rasl12 | 84.28986 | 0.425821 | 0.136977 | 0.828492 |
| ENSMUSG( | Rassf1 | 358.3327 | -0.18825 | 0.132097 | 0.544244 | ENSMUSG( | Rassf1 | 337.5502 | -0.07729 | 0.64163 | 0.953903 |
| ENSMUSG( | Rassf10 | 99.80686 | -0.0853 | 0.671357 | 0.928781 | ENSMUSG( | Rassf10 | 96.6867 | 0.078318 | 0.750055 | 0.970113 |
| ENSMUSG( | Rassf2 | 6746.401 | 0.109722 | 0.23889 | 0.686534 | ENSMUSG( | Rassf2 | 5930.006 | 0.00722 | 0.949535 | 0.994798 |
| ENSMUSG( | Rassf3 | 1305.326 | -0.08822 | 0.357166 | 0.788702 | ENSMUSG( | Rassf3 | 1153.584 | -0.17969 | 0.125776 | 0.81516 |
| ENSMUSG( | Rassf4 | 218.8454 | -0.28394 | 0.055366 | 0.361966 | ENSMUSG( | Rassf4 | 215.2517 | -0.03858 | 0.823689 | 0.979198 |
| ENSMUSG( | Rassf5 | 570.8779 | 0.151463 | 0.269696 | 0.719729 | ENSMUSG( | Rassf5 | 476.2843 | -0.1033 | 0.511481 | 0.92971 |
| ENSMUSG( | Rassf6 | 12.5699 | -0.51491 | 0.350109 | NA | ENSMUSG( | Rassf6 | 13.23513 | -0.08218 | 0.884109 | 0.989644 |
| ENSMUSG( | Rassf7 | 18.8349 | 0.454312 | 0.292688 | NA | ENSMUSG( | Rassf7 | 16.75609 | 0.434987 | 0.375952 | 0.915491 |
| ENSMUSG( | Rassf8 | 1317.564 | -0.06227 | 0.580184 | 0.896952 | ENSMUSG( | Rassf8 | 1215.28 | -0.03086 | 0.819119 | 0.979198 |
| ENSMUSG( | Rassf9 | 51.13077 | -0.30026 | 0.393058 | 0.809274 | ENSMUSG( | Rassf9 | 67.43103 | 0.7254 | 0.048215 | 0.706142 |
| ENSMUSG( | Raver2 | 1437.117 | 0.282777 | 0.001324 | 0.039614 | ENSMUSG( | Raver2 | 1234.246 | 0.120982 | 0.315923 | 0.901868 |
| ENSMUSG( | Rb1 | 1903.214 | 0.053304 | 0.698269 | 0.936084 | ENSMUSG( | Rb1 | 1601.281 | -0.18473 | 0.237149 | 0.872173 |
| ENSMUSG( | Rb1cc1 | 2317.882 | 0.110072 | 0.183981 | 0.620925 | ENSMUSG( | Rb1cc1 | 2064.657 | 0.045724 | 0.66153 | 0.956205 |
| ENSMUSG( | Rbak | 401.8835 | -0.27272 | 0.09494 | 0.472949 | ENSMUSG( | Rbak | 400.4053 | 0.003846 | 0.981705 | 0.996756 |
| ENSMUSG( | Rbbp4 | 3060.485 | 0.014134 | 0.878134 | 0.981431 | ENSMUSG( | Rbbp4 | 2762.076 | -0.01559 | 0.8953 | 0.990558 |
| ENSMUSG( | Rbbp5 | 1112.049 | 0.120646 | 0.266482 | 0.716441 | ENSMUSG( | Rbbp5 | 968.2954 | -0.00677 | 0.953658 | 0.994798 |
| ENSMUSG( | Rbbp6 | 2259.568 | -0.09603 | 0.288274 | 0.739023 | ENSMUSG( | Rbbp6 | 2172.11 | 0.057336 | 0.63581 | 0.953903 |
| ENSMUSG( | Rbbp7 | 2636.72 | 0.132698 | 0.156844 | 0.5851 | ENSMUSG( | Rbbp7 | 2255.911 | -0.04844 | 0.691019 | 0.960071 |
| ENSMUSG( | Rbbp8 | 127.9864 | 0.050779 | 0.781037 | 0.957558 | ENSMUSG( | Rbbp8 | 128.1705 | 0.305107 | 0.12935 | 0.820358 |
| ENSMUSG( | Rbbp9 | 1466.417 | 0.005083 | 0.963119 | 0.994911 | ENSMUSG( | Rbbp9 | 1243.248 | -0.2151 | 0.113335 | 0.809623 |
| ENSMUSG( | Rbck1 | 3948.122 | 0.054656 | 0.523706 | 0.875723 | ENSMUSG( | Rbck1 | 3548.778 | 0.016548 | 0.872282 | 0.988944 |
| ENSMUSG( | Rbfa | 734.5637 | -0.05115 | 0.682924 | 0.93205 | ENSMUSG( | Rbfa | 678.6329 | -0.014 | 0.921578 | 0.992644 |
| ENSMUSG( | Rbfaos | 17.8173 | -0.18596 | 0.674566 | NA | ENSMUSG( | Rbfaos | 17.27439 | 0.003396 | 0.99427 | 0.998813 |
| ENSMUSG( | Rbfox1 | 3319.466 | -0.09537 | 0.33641 | 0.777 | ENSMUSG( | Rbfox1 | 3069.107 | -0.05215 | 0.65052 | 0.95422 |
| ENSMUSG( | Rbfox2 | 6857.607 | 0.149791 | 0.086176 | 0.451605 | ENSMUSG( | Rbfox2 | 6052.475 | 0.062553 | 0.552208 | 0.939565 |
| ENSMUSG( | Rbfox3 | 7088.968 | 0.071638 | 0.497786 | 0.862341 | ENSMUSG( | Rbfox3 | 6314.601 | 0.006702 | 0.958057 | 0.994798 |
| ENSMUSG( | Rbis | 394.6232 | -0.06657 | 0.604297 | 0.905469 | ENSMUSG( | Rbis | 375.3456 | 0.049228 | 0.752124 | 0.970461 |
| ENSMUSG( | Rbks | 58.80861 | -0.2691 | 0.303125 | 0.751443 | ENSMUSG( | Rbks | 51.72782 | -0.38885 | 0.181421 | 0.850579 |
| ENSMUSG( | Rbl1 | 89.9802 | -0.20324 | 0.337325 | 0.7773 | ENSMUSG( | Rbl1 | 93.01035 | 0.171416 | 0.434283 | 0.91869 |
| ENSMUSG( | Rbl2 | 2567.277 | 0.081003 | 0.436995 | 0.829501 | ENSMUSG( | Rbl2 | 2189.439 | -0.11365 | 0.357496 | 0.913759 |
| ENSMUSG( | Rbm10 | 1680.565 | 0.230357 | 0.040597 | 0.311337 | ENSMUSG( | Rbm10 | 1407.335 | -0.00571 | 0.965553 | 0.995433 |
| ENSMUSG( | Rbm11 | 110.249 | -0.28679 | 0.140944 | 0.557902 | ENSMUSG( | Rbm11 | 106.8544 | -0.11303 | 0.626837 | 0.953234 |
| ENSMUSG( | Rbm12 | 255.8886 | 0.250356 | 0.074542 | 0.423195 | ENSMUSG( | Rbm12 | 212.558 | -0.01507 | 0.93737 | 0.993685 |
| ENSMUSG( | Rbm12b1 | 143.4049 | -0.1408 | 0.435003 | 0.829093 | ENSMUSG( | Rbm12b1 | 149.1129 | 0.24374 | 0.273073 | 0.889275 |
| ENSMUSG( | Rbm12b2 | 219.4676 | -0.01481 | 0.921497 | 0.990436 | ENSMUSG( | Rbm12b2 | 208.8867 | 0.110988 | 0.5237 | 0.932848 |
| ENSMUSG( | Rbm14 | 429.7648 | 0.103193 | 0.372392 | 0.798768 | ENSMUSG( | Rbm14 | 366.7068 | -0.07539 | 0.63545 | 0.953903 |
| ENSMUSG( | Rbm15 | 153.0125 | -0.24988 | 0.142518 | 0.560815 | ENSMUSG( | Rbm15 | 159.7979 | 0.154278 | 0.405963 | 0.917257 |
| ENSMUSG( | Rbm15b | 1922.799 | -0.02952 | 0.773 | 0.955272 | ENSMUSG( | Rbm15b | 1800.374 | 0.051871 | 0.649787 | 0.95422 |
| ENSMUSG( | Rbm17 | 1533.924 | 0.013161 | 0.892443 | 0.984455 | ENSMUSG( | Rbm17 | 1379.518 | -0.0285 | 0.830642 | 0.98051 |
| ENSMUSG( | Rbm18 | 972.5413 | 0.075419 | 0.496164 | 0.861554 | ENSMUSG( | Rbm18 | 868.0057 | 0.014216 | 0.911839 | 0.992196 |
| ENSMUSG( | Rbm19 | 316.7509 | 0.292111 | 0.041183 | 0.313607 | ENSMUSG( | Rbm19 | 263.2593 | 0.039971 | 0.80483 | 0.977232 |
| ENSMUSG( | Rbm20 | 156.8855 | -0.03727 | 0.8336 | 0.970333 | ENSMUSG( | Rbm20 | 143.8739 | -0.01022 | 0.956847 | 0.994798 |
| ENSMUSG( | Rbm22 | 922.443 | -0.03145 | 0.773111 | 0.955272 | ENSMUSG( | Rbm22 | 871.5122 | 0.076075 | 0.554597 | 0.939565 |
| ENSMUSG( | Rbm24 | 307.4676 | -0.03333 | 0.838077 | 0.971111 | ENSMUSG( | Rbm24 | 276.2797 | -0.07396 | 0.703687 | 0.962481 |
| ENSMUSG( | Rbm25 | 3188.543 | -0.01123 | 0.913756 | 0.988483 | ENSMUSG( | Rbm25 | 3107.44 | 0.175704 | 0.138091 | 0.828492 |
| ENSMUSG( | Rbm26 | 1115.889 | -0.07188 | 0.463057 | 0.842422 | ENSMUSG( | Rbm26 | 1110.32 | 0.180374 | 0.121219 | 0.813384 |
| ENSMUSG( | Rbm27 | 742.9225 | 0.118531 | 0.238165 | 0.686299 | ENSMUSG( | Rbm27 | 687.6461 | 0.156852 | 0.227864 | 0.868507 |
| ENSMUSG( | Rbm28 | 1822.393 | 0.213042 | 0.017376 | 0.197592 | ENSMUSG( | Rbm28 | 1538.179 | -0.00286 | 0.981661 | 0.996756 |
| ENSMUSG( | Rbm3 | 1815.924 | -0.20765 | 0.106913 | 0.494634 | ENSMUSG( | Rbm3 | 1861.601 | 0.135148 | 0.327896 | 0.906255 |
| ENSMUSG( | Rbm33 | 1681.943 | -0.16074 | 0.179718 | 0.617507 | ENSMUSG( | Rbm33 | 1667.73 | 0.091695 | 0.558068 | 0.939565 |
| ENSMUSG( | Rbm34 | 423.2986 | 0.0129 | 0.915645 | 0.988634 | ENSMUSG( | Rbm34 | 385.5593 | 0.002578 | 0.987236 | 0.997487 |
| ENSMUSG( | Rbm38 | 117.3862 | -0.0683 | 0.746494 | 0.949289 | ENSMUSG( | Rbm38 | 115.2875 | 0.155769 | 0.476731 | 0.923329 |
| ENSMUSG( | Rbm39 | 5068.088 | 0.034249 | 0.716877 | 0.941697 | ENSMUSG( | Rbm39 | 4953.64 | 0.228896 | 0.037817 | 0.6763 |
| ENSMUSG( | Rbm3os | 11.21085 | 0.019777 | 0.972693 | NA | ENSMUSG( | Rbm3os | 10.33749 | 0.078491 | 0.897756 | NA |
| ENSMUSG( | Rbm4 | 71.47659 | 0.462333 | 0.052201 | 0.350744 | ENSMUSG( | Rbm4 | 63.23156 | 0.359581 | 0.212236 | 0.864412 |
| ENSMUSG( | Rbm41 | 166.894 | 0.1471 | 0.36257 | 0.792943 | ENSMUSG( | Rbm41 | 151.9539 | 0.160263 | 0.423092 | 0.917257 |
| ENSMUSG( | Rbm43 | 283.5669 | 0.246904 | 0.097184 | 0.47759 | ENSMUSG( | Rbm43 | 237.2107 | 0.007241 | 0.967639 | 0.995833 |
| ENSMUSG( | Rbm45 | 550.4678 | 0.102122 | 0.379031 | 0.801757 | ENSMUSG( | Rbm45 | 463.4912 | -0.12255 | 0.37636 | 0.915491 |
| ENSMUSG( | Rbm46 | 22.92166 | 0.105218 | 0.840306 | 0.971179 | ENSMUSG( | Rbm46 | 20.83712 | 0.079124 | 0.885803 | 0.98977 |
| ENSMUSG( | Rbm47 | 36.90408 | -0.24695 | 0.494523 | 0.860776 | ENSMUSG( | Rbm47 | 35.10248 | -0.09498 | 0.806613 | 0.978086 |
| ENSMUSG( | Rbm48 | 129.4237 | 0.269216 | 0.198784 | 0.63899 | ENSMUSG( | Rbm48 | 119.8696 | 0.310216 | 0.122914 | 0.814458 |
| ENSMUSG( | Rbm4b | 451.3605 | -0.01704 | 0.899455 | 0.985633 | ENSMUSG( | Rbm4b | 394.9434 | -0.14357 | 0.352119 | 0.912093 |
| ENSMUSG( | Rbm5 | 4223.997 | -0.11515 | 0.315602 | 0.762671 | ENSMUSG( | Rbm5 | 4174.915 | 0.122048 | 0.360012 | 0.914223 |
| ENSMUSG( | Rbm6 | 1578.567 | 0.219539 | 0.025763 | 0.247138 | ENSMUSG( | Rbm6 | 1366.31 | 0.077429 | 0.542788 | 0.938044 |
| ENSMUSG( | Rbm7 | 804.1205 | -0.07474 | 0.456424 | 0.838832 | ENSMUSG( | Rbm7 | 767.1583 | 0.055064 | 0.66858 | 0.956766 |
| ENSMUSG( | Rbm8a | 1528.299 | 0.08603 | 0.390259 | 0.80794 | ENSMUSG( | Rbm8a | 1317.589 | -0.07784 | 0.573251 | 0.942912 |
| ENSMUSG( | Rbms1 | 570.3905 | -0.17897 | 0.156061 | 0.58379 | ENSMUSG( | Rbms1 | 558.6919 | 0.03624 | 0.798306 | 0.97681 |
| ENSMUSG( | Rbms2 | 759.4639 | 0.258404 | 0.02647 | 0.24888 | ENSMUSG( | Rbms2 | 631.9937 | 0.005055 | 0.968656 | 0.99594 |
| ENSMUSG( | Rbms3 | 364.1357 | -0.07636 | 0.576752 | 0.895432 | ENSMUSG( | Rbms3 | 338.1973 | -0.01986 | 0.896643 | 0.990558 |
| ENSMUSG( | Rbmx | 1010.286 | -0.24944 | 0.031711 | 0.271573 | ENSMUSG( | Rbmx | 1074.604 | 0.203674 | 0.208152 | 0.864412 |
| ENSMUSG( | Rbmx2 | 145.5921 | 0.177255 | 0.387738 | 0.805835 | ENSMUSG( | Rbmx2 | 124.6101 | -0.01399 | 0.954387 | 0.994798 |
| ENSMUSG( | Rbmxl1 | 432.6553 | 0.001978 | 0.989401 | 0.999111 | ENSMUSG( | Rbmxl1 | 391.2368 | -0.02158 | 0.89428 | 0.990558 |
| ENSMUSG( | Rbp1 | 1463.532 | 0.085313 | 0.526869 | 0.877267 | ENSMUSG( | Rbp1 | 1200.945 | -0.23108 | 0.175474 | 0.845305 |
| ENSMUSG( | Rbp4 | 505.0287 | -0.36823 | 0.019407 | 0.209852 | ENSMUSG( | Rbp4 | 488.7103 | -0.17422 | 0.326501 | 0.905371 |
| ENSMUSG( | Rbpj | 800.1811 | -0.02268 | 0.821756 | 0.96635 | ENSMUSG( | Rbpj | 724.6156 | -0.04242 | 0.731178 | 0.967228 |
| ENSMUSG( | Rbpjl | 41.20613 | 0.394443 | 0.259994 | 0.709951 | ENSMUSG( | Rbpjl | 39.52677 | 0.54447 | 0.167959 | 0.840222 |
| ENSMUSG( | Rbpms | 100.8974 | -0.01885 | 0.938515 | 0.993415 | ENSMUSG( | Rbpms | 119.2541 | 0.677411 | 0.01872 | 0.547826 |
| ENSMUSG( | Rbpms2 | 150.4382 | 0.035712 | 0.830546 | 0.96974 | ENSMUSG( | Rbpms2 | 146.9479 | 0.216478 | 0.264196 | 0.886134 |
| ENSMUSG( | Rbsn | 1574.845 | 0.153484 | 0.099514 | 0.481972 | ENSMUSG( | Rbsn | 1334.348 | -0.05284 | 0.624396 | 0.953234 |
| ENSMUSG( | Rbx1 | 2631.838 | 0.050545 | 0.547148 | 0.885118 | ENSMUSG( | Rbx1 | 2295.429 | -0.07885 | 0.548599 | 0.938063 |
| ENSMUSG( | Rc3h1 | 1015.055 | -0.00588 | 0.960619 | 0.993991 | ENSMUSG( | Rc3h1 | 1013.367 | 0.246115 | 0.062069 | 0.736482 |
| ENSMUSG( | Rc3h2 | 1516.107 | 0.002411 | 0.977858 | 0.997428 | ENSMUSG( | Rc3h2 | 1435.208 | 0.107091 | 0.427353 | 0.917257 |
| ENSMUSG( | Rcan1 | 2211.589 | 0.043445 | 0.607473 | 0.906876 | ENSMUSG( | Rcan1 | 1896.774 | -0.13424 | 0.22009 | 0.866104 |
| ENSMUSG( | Rcan2 | 6293.249 | 0.018089 | 0.820781 | 0.965962 | ENSMUSG( | Rcan2 | 5284.782 | -0.22776 | 0.058749 | 0.728832 |

| | | | | | |
|---|---|---|---|---|---|
| ENSMUSG( | 58.9325 | 0.057293 | 0.814793 | 0.964668 | Pwwp3b |
| ENSMUSG( | 248.3205 | -0.05596 | 0.699257 | 0.936886 | Pxdc1 |
| ENSMUSG( | 3211.175 | 0.047574 | 0.664612 | 0.926824 | Pxdn |
| ENSMUSG( | 790.7224 | 0.12236 | 0.303932 | 0.752178 | Pxk |
| ENSMUSG( | 401.0668 | 0.087448 | 0.534971 | 0.880733 | Pxmp2 |
| ENSMUSG( | 944.3619 | 0.010803 | 0.92699 | 0.991229 | Pxmp4 |
| ENSMUSG( | 1427.346 | 0.156532 | 0.20702 | 0.649503 | Pxn |
| ENSMUSG( | 366.6396 | -0.49725 | 0.001146 | 0.036093 | Pxylp1 |
| ENSMUSG( | 156.4359 | -0.04222 | 0.817303 | 0.965736 | Pycard |
| ENSMUSG( | 114.6604 | -0.22847 | 0.272265 | 0.7223 | Pycr1 |
| ENSMUSG( | 682.1058 | 0.027108 | 0.852471 | 0.973168 | Pycr2 |
| ENSMUSG( | 454.8671 | 0.055313 | 0.677464 | 0.930542 | Pycr3 |
| ENSMUSG( | 18375.89 | 0.120622 | 0.225921 | 0.672685 | Pygb |
| ENSMUSG( | 210.9093 | 0.073369 | 0.640057 | 0.916434 | Pygl |
| ENSMUSG( | 4141.233 | 0.07134 | 0.511405 | 0.870898 | Pygm |
| ENSMUSG( | 1637.26 | 0.08418 | 0.434192 | 0.828931 | Pygo1 |
| ENSMUSG( | 1289.758 | 0.094904 | 0.392858 | 0.809274 | Pygo2 |
| ENSMUSG( | 463.6441 | -0.11469 | 0.49895 | 0.862973 | Pym1 |
| ENSMUSG( | 353.5685 | 0.02373 | 0.856515 | 0.974154 | Pyroxd1 |
| ENSMUSG( | 142.9502 | -0.02363 | 0.893593 | 0.984512 | Pyroxd2 |
| ENSMUSG( | 835.2062 | -0.03403 | 0.81712 | 0.96566 | Pyurf |
| ENSMUSG( | 1307.611 | -0.11675 | 0.253912 | 0.704011 | Qars1 |
| ENSMUSG( | 7792.715 | -0.19066 | 0.027943 | 0.255775 | Qdpr |
| ENSMUSG( | 25292.53 | 0.025472 | 0.841453 | 0.971179 | Qki |
| ENSMUSG( | 355.9408 | 0.108526 | 0.475618 | 0.850919 | Qng1 |
| ENSMUSG( | 263.6947 | -0.72134 | 7.56E-05 | 0.005353 | Qpct |
| ENSMUSG( | 648.614 | -0.42946 | 0.001431 | 0.041661 | Qpctl |
| ENSMUSG( | 28.99353 | -0.02114 | 0.955798 | 0.993874 | Qprt |
| ENSMUSG( | 73.01563 | -0.40674 | 0.089655 | 0.45961 | Qrfpr |
| ENSMUSG( | 14.37825 | -1.63143 | 0.001798 | NA | Qrfprl |
| ENSMUSG( | 1718.357 | 0.053323 | 0.585412 | 0.899171 | Qrich1 |
| ENSMUSG( | 350.6252 | 0.093176 | 0.467306 | 0.844819 | Qrsl1 |
| ENSMUSG( | 872.4875 | 0.022149 | 0.87395 | 0.979336 | Qser1 |
| ENSMUSG( | 969.5062 | -0.20024 | 0.1808 | 0.617841 | Qsox1 |
| ENSMUSG( | 862.4655 | -0.31595 | 0.031117 | 0.268324 | Qsox2 |
| ENSMUSG( | 290.8231 | -0.05139 | 0.762677 | 0.953519 | Qtrt1 |
| ENSMUSG( | 382.8352 | 0.169692 | 0.260158 | 0.710129 | Qtrt2 |
| ENSMUSG( | 1071.092 | -0.01445 | 0.885883 | 0.983008 | R3hcc1 |
| ENSMUSG( | 387.5122 | 0.051186 | 0.67827 | 0.930542 | R3hcc1l |
| ENSMUSG( | 12166.64 | 0.066013 | 0.487192 | 0.858812 | R3hdm1 |
| ENSMUSG( | 13904.49 | 0.132262 | 0.093332 | 0.469793 | R3hdm2 |
| ENSMUSG( | 18738.72 | 0.081832 | 0.313008 | 0.76033 | R3hdm4 |
| ENSMUSG( | 308.9222 | -0.09282 | 0.500346 | 0.863455 | R74862 |
| ENSMUSG( | 6600.21 | -0.02796 | 0.803758 | 0.962169 | Rab10 |
| ENSMUSG( | 1746.257 | 0.015951 | 0.873075 | 0.978753 | Rab11a |
| ENSMUSG( | 5779.091 | 0.021727 | 0.773017 | 0.955272 | Rab11b |
| ENSMUSG( | 120.074 | 0.22965 | 0.282771 | 0.73186 | Rab11fip1 |
| ENSMUSG( | 1079.261 | -0.04149 | 0.663156 | 0.9265 | Rab11fip2 |
| ENSMUSG( | 4502.014 | 0.100532 | 0.267981 | 0.718218 | Rab11fip3 |
| ENSMUSG( | 13982.86 | 0.139268 | 0.089204 | 0.458482 | Rab11fip4 |
| ENSMUSG( | 2528.592 | 0.13968 | 0.149887 | 0.574325 | Rab11fip5 |
| ENSMUSG( | 4674.785 | -0.57942 | 0.000228 | 0.01183 | Rab12 |
| ENSMUSG( | 532.046 | 0.082352 | 0.529684 | 0.878883 | Rab13 |
| ENSMUSG( | 5181.048 | 0.068363 | 0.420715 | 0.822925 | Rab14 |
| ENSMUSG( | 13304.25 | 0.17832 | 0.01859 | 0.20512 | Rab15 |
| ENSMUSG( | 3021.173 | -0.00099 | 0.992404 | 0.999327 | Rab18 |
| ENSMUSG( | 5171.428 | 0.019916 | 0.838964 | 0.971179 | Rab1a |
| ENSMUSG( | 6186.566 | 0.074195 | 0.341484 | 0.778828 | Rab1b |
| ENSMUSG( | 19.41741 | -0.56962 | 0.184816 | NA | Rab20 |
| ENSMUSG( | 4502.941 | 0.046857 | 0.703417 | 0.937735 | Rab21 |
| ENSMUSG( | 3524.034 | 0.134412 | 0.182606 | 0.620026 | Rab22a |
| ENSMUSG( | 1171.57 | 0.163353 | 0.143741 | 0.562581 | Rab23 |
| ENSMUSG( | 1376.759 | 0.017456 | 0.874483 | 0.979498 | Rab24 |
| ENSMUSG( | 1076.595 | 0.181098 | 0.242551 | 0.690206 | Rab26 |
| ENSMUSG( | 53.9504 | 0.145001 | 0.643909 | 0.917526 | Rab26os |
| ENSMUSG( | 118.3851 | 0.162721 | 0.389558 | 0.807428 | Rab27a |
| ENSMUSG( | 602.0685 | -0.0255 | 0.821631 | 0.966344 | Rab27b |
| ENSMUSG( | 1693.522 | 0.073495 | 0.429506 | 0.826446 | Rab28 |
| ENSMUSG( | 215.7953 | 0.240011 | 0.143062 | 0.561716 | Rab29 |
| ENSMUSG( | 5339.441 | 0.009588 | 0.921552 | 0.990436 | Rab2a |
| ENSMUSG( | 1051.428 | 0.11683 | 0.229695 | 0.677538 | Rab2b |
| ENSMUSG( | 812.9456 | -0.08399 | 0.465586 | 0.84415 | Rab30 |
| ENSMUSG( | 7914.087 | 0.076555 | 0.464737 | 0.843568 | Rab31 |
| ENSMUSG( | 79.34169 | -0.02611 | 0.905214 | 0.98645 | Rab32 |
| ENSMUSG( | 446.3436 | -0.05962 | 0.754437 | 0.951007 | Rab33a |
| ENSMUSG( | 1033.053 | -0.06858 | 0.51248 | 0.871762 | Rab33b |
| ENSMUSG( | 730.3997 | -0.01598 | 0.886513 | 0.983204 | Rab34 |
| ENSMUSG( | 2485.483 | 0.075065 | 0.350643 | 0.782422 | Rab35 |
| ENSMUSG( | 1106.423 | 0.019528 | 0.867316 | 0.976559 | Rab36 |
| ENSMUSG( | 89.17656 | -0.84828 | 0.001773 | 0.04746 | Rab37 |
| ENSMUSG( | 23.13067 | -0.4096 | 0.314667 | 0.761775 | Rab39 |
| ENSMUSG( | 583.428 | -0.40736 | 0.000131 | 0.007715 | Rab39b |
| ENSMUSG( | 9995.129 | 0.021837 | 0.874035 | 0.979336 | Rab3a |
| ENSMUSG( | 871.5621 | -0.80373 | 1.98E-06 | 0.000328 | Rab3b |
| ENSMUSG( | 7556.126 | 0.221815 | 0.003672 | 0.077033 | Rab3c |
| ENSMUSG( | 337.5612 | 0.126612 | 0.344942 | 0.779153 | Rab3d |
| ENSMUSG( | 2292.649 | 0.168472 | 0.042497 | 0.316636 | Rab3gap1 |
| ENSMUSG( | 2353.918 | -0.00202 | 0.981886 | 0.997512 | Rab3gap2 |
| ENSMUSG( | 589.0316 | 0.132588 | 0.233597 | 0.681508 | Rab3il1 |
| ENSMUSG( | 815.5936 | 0.04622 | 0.710392 | 0.939748 | Rab3ip |
| ENSMUSG( | 563.8609 | -0.24077 | 0.116577 | 0.514987 | Rab40b |
| ENSMUSG( | 1433.564 | 0.100214 | 0.350056 | 0.781943 | Rab40c |
| ENSMUSG( | 9.792169 | 0.259178 | 0.664705 | NA | Rab42 |
| ENSMUSG( | 40.86638 | 0.124336 | 0.676799 | 0.930542 | Rab43 |
| ENSMUSG( | 9.12132 | -0.52554 | 0.458135 | NA | Rab44 |
| ENSMUSG( | 1423.316 | 0.164172 | 0.187424 | 0.625777 | Rab4a |
| ENSMUSG( | 131.2653 | -0.21204 | 0.330552 | 0.774568 | Rab4b |
| ENSMUSG( | 2516.023 | -0.00949 | 0.925674 | 0.991204 | Rab5a |
| ENSMUSG( | 5947.175 | 0.07317 | 0.375607 | 0.800887 | Rab5b |
| ENSMUSG( | 7438.58 | 0.020479 | 0.811476 | 0.964426 | Rab5c |
| ENSMUSG( | 1544.509 | 0.006405 | 0.951484 | 0.993821 | Rab5if |
| ENSMUSG( | 10390.19 | -0.0243 | 0.806738 | 0.962932 | Rab6a |
| ENSMUSG( | 39401.76 | 0.155018 | 0.037503 | 0.299216 | Rab6b |
| ENSMUSG( | 9615.723 | 0.019474 | 0.834423 | 0.970342 | Rab7 |
| ENSMUSG( | 258.9487 | -0.37909 | 0.016031 | 0.187923 | Rab7b |
| ENSMUSG( | 1322.734 | 0.144683 | 0.105096 | 0.492392 | Rab8a |
| ENSMUSG( | 895.718 | 0.074522 | 0.491322 | 0.859873 | Rab8b |
| ENSMUSG( | 959.3523 | 0.060118 | 0.594324 | 0.902869 | Rab9 |
| ENSMUSG( | 282.0667 | 0.04324 | 0.775441 | 0.955606 | Rab9b |
| ENSMUSG( | 4044.679 | -0.21134 | 0.046236 | 0.329472 | Rabac1 |
| ENSMUSG( | 4119.864 | 0.152234 | 0.076426 | 0.4279 | Rabep1 |
| ENSMUSG( | 356.9682 | -0.07015 | 0.621891 | 0.911432 | Rabep2 |
| ENSMUSG( | 549.0588 | -0.01651 | 0.883044 | 0.982422 | Rabepk |
| ENSMUSG( | 2392.574 | 0.125958 | 0.116251 | 0.514678 | Rabgap1 |
| ENSMUSG( | 2060.812 | -0.01759 | 0.850609 | 0.972638 | Rabgap1l |
| ENSMUSG( | 858.006 | -0.01412 | 0.890397 | 0.983687 | Rabgef1 |
| ENSMUSG( | 981.3219 | 0.039512 | 0.69553 | 0.934979 | Rabggta |
| ENSMUSG( | 1152.56 | -0.03058 | 0.753243 | 0.950753 | Rabggtb |
| ENSMUSG( | 805.9563 | 0.000284 | 0.997808 | 0.99989 | Rabif |
| ENSMUSG( | 667.8795 | 0.122633 | 0.473547 | 0.848919 | Rabl2 |
| ENSMUSG( | 156.239 | 0.083038 | 0.632874 | 0.914073 | Rabl3 |
| ENSMUSG( | 7470.364 | 0.200653 | 0.012385 | 0.160838 | Rabl6 |
| ENSMUSG( | 12163.29 | 0.01221 | 0.894651 | 0.984512 | Rac1 |
| ENSMUSG( | 222.1353 | -0.11548 | 0.509919 | 0.869743 | Rac2 |
| ENSMUSG( | 622.9792 | 0.108881 | 0.318366 | 0.764234 | Rac3 |
| ENSMUSG( | 258.095 | -0.12605 | 0.379796 | 0.801757 | Racgap1 |
| ENSMUSG( | 9524.308 | 0.087281 | 0.266281 | 0.716441 | Rack1 |
| ENSMUSG( | 181.6258 | 0.159944 | 0.342561 | 0.778971 | Rad1 |
| ENSMUSG( | 490.8531 | 0.117623 | 0.309986 | 0.758025 | Rad17 |
| ENSMUSG( | 122.4259 | -0.15321 | 0.394898 | 0.810072 | Rad18 |
| ENSMUSG( | 6100.88 | 0.103226 | 0.257552 | 0.708686 | Rad21 |
| ENSMUSG( | 1941.488 | 0.089219 | 0.361448 | 0.792797 | Rad23a |
| ENSMUSG( | 3306.688 | 0.10108 | 0.297497 | 0.746042 | Rad23b |
| ENSMUSG( | 407.8451 | 0.113816 | 0.342707 | 0.778971 | Rad50 |
| ENSMUSG( | 42.16685 | 0.009577 | 0.973653 | 0.99675 | Rad51 |
| ENSMUSG( | 35.42372 | -0.45762 | 0.140295 | 0.557263 | Rad51ap1 |
| ENSMUSG( | 18.89086 | -0.46454 | 0.289578 | NA | Rad51ap2 |
| ENSMUSG( | 11.33381 | -0.35883 | 0.545938 | NA | Rad51b |
| ENSMUSG( | 67.89532 | 0.135827 | 0.560807 | 0.888934 | Rad51c |
| ENSMUSG( | 491.5878 | 0.035434 | 0.778271 | 0.956215 | Rad51d |
| ENSMUSG( | 513.6276 | 0.084904 | 0.494347 | 0.860611 | Rad52 |
| ENSMUSG( | 31.40736 | -0.43794 | 0.231423 | 0.679437 | Rad54b |
| ENSMUSG( | 52.47589 | 0.003996 | 0.987693 | 0.998902 | Rad54l |
| ENSMUSG( | 934.3593 | 0.014934 | 0.879399 | 0.981436 | Rad54l2 |
| ENSMUSG( | 253.2758 | -0.37543 | 0.006336 | 0.108996 | Rad9a |
| ENSMUSG( | 129.2853 | -0.20848 | 0.265935 | 0.716441 | Rad9b |
| ENSMUSG( | 335.5188 | 0.073822 | 0.596918 | 0.903526 | Radil |
| ENSMUSG( | 765.9093 | -0.09103 | 0.46582 | 0.844222 | Rae1 |
| ENSMUSG( | 2563.066 | 0.0745 | 0.422836 | 0.823633 | Raf1 |
| ENSMUSG( | 1951.535 | -0.22994 | 0.032869 | 0.277278 | Rai1 |
| ENSMUSG( | 430.7238 | 0.272612 | 0.028726 | 0.258426 | Rai14 |
| ENSMUSG( | 142.3415 | 0.010077 | 0.960611 | 0.993991 | Rai2 |
| ENSMUSG( | 2664.163 | 0.078279 | 0.449583 | 0.835234 | Rala |
| ENSMUSG( | 798.4142 | -0.05006 | 0.63343 | 0.914073 | Ralb |
| ENSMUSG( | 2587.973 | 0.249556 | 0.008025 | 0.124801 | Ralbp1 |
| ENSMUSG( | 2828.47 | 0.045969 | 0.575065 | 0.894572 | Ralgapa1 |
| ENSMUSG( | 738.8928 | 0.01232 | 0.905178 | 0.98645 | Ralgapa2 |
| ENSMUSG( | 3946.816 | 0.018212 | 0.847732 | 0.972374 | Ralgapb |

| | | | | | |
|---|---|---|---|---|---|
| ENSMUSG( | 362.8573 | 0.052085 | 0.720694 | 0.966887 | Rad50 |
| ENSMUSG( | 37.90488 | -0.0251 | 0.937695 | 0.993685 | Rad51 |
| ENSMUSG( | 38.02405 | 0.071415 | 0.82702 | 0.980159 | Rad51ap1 |
| ENSMUSG( | 16.65335 | -0.5789 | 0.230551 | 0.868507 | Rad51ap2 |
| ENSMUSG( | 10.62669 | -0.25036 | 0.698346 | NA | Rad51b |
| ENSMUSG( | 65.95523 | 0.300656 | 0.244136 | 0.877904 | Rad51c |
| ENSMUSG( | 421.2291 | -0.1443 | 0.299963 | 0.898538 | Rad51d |
| ENSMUSG( | 470.4162 | 0.101543 | 0.480105 | 0.924276 | Rad52 |
| ENSMUSG( | 35.15391 | 0.204166 | 0.574342 | 0.942912 | Rad54b |
| ENSMUSG( | 45.59437 | -0.12378 | 0.686378 | 0.95899 | Rad54l |
| ENSMUSG( | 838.1461 | -0.02744 | 0.822775 | 0.979198 | Rad54l2 |
| ENSMUSG( | 253.3373 | -0.07174 | 0.669237 | 0.956766 | Rad9a |
| ENSMUSG( | 130.0894 | 0.083114 | 0.703124 | 0.962428 | Rad9b |
| ENSMUSG( | 304.5742 | 0.068558 | 0.691963 | 0.960393 | Radil |
| ENSMUSG( | 708.8378 | -0.042 | 0.778591 | 0.976106 | Rae1 |
| ENSMUSG( | 2276.178 | 0.003085 | 0.97702 | 0.996416 | Raf1 |
| ENSMUSG( | 1950.757 | 0.049976 | 0.687016 | 0.959226 | Rai1 |
| ENSMUSG( | 397.1689 | 0.310121 | 0.042064 | 0.690304 | Rai14 |
| ENSMUSG( | 123.6654 | -0.11738 | 0.602801 | 0.94789 | Rai2 |
| ENSMUSG( | 2270.368 | -0.12 | 0.410738 | 0.917257 | Rala |
| ENSMUSG( | 747.8215 | 0.025009 | 0.847853 | 0.985242 | Ralb |
| ENSMUSG( | 2212.816 | 0.074456 | 0.521856 | 0.932571 | Ralbp1 |
| ENSMUSG( | 2620.828 | 0.093511 | 0.356959 | 0.913358 | Ralgapa1 |
| ENSMUSG( | 698.7667 | 0.116147 | 0.348422 | 0.911652 | Ralgapa2 |
| ENSMUSG( | 3710.917 | 0.109491 | 0.337821 | 0.907972 | Ralgapb |
| ENSMUSG( | 4087.094 | -0.00425 | 0.970871 | 0.996061 | Ralgds |
| ENSMUSG( | 2126.883 | 0.115341 | 0.262657 | 0.885357 | Ralgps1 |
| ENSMUSG( | 1179.982 | 0.245247 | 0.08916 | 0.788502 | Ralgps2 |
| ENSMUSG( | 1967.567 | -0.13438 | 0.220231 | 0.866104 | Raly |
| ENSMUSG( | 1002.098 | -0.14882 | 0.272435 | 0.888825 | Ralyl |
| ENSMUSG( | 611.5317 | -0.28424 | 0.046868 | 0.700664 | Ramac |
| ENSMUSG( | 39.40933 | -0.25068 | 0.454439 | 0.922444 | Ramacl |
| ENSMUSG( | 4575.466 | -0.18848 | 0.094361 | 0.795735 | Ramp1 |
| ENSMUSG( | 1043.253 | 0.705482 | 2.64E-05 | 0.007461 | Ramp2 |
| ENSMUSG( | 558.0575 | -0.1452 | 0.384054 | 0.917257 | Ramp3 |
| ENSMUSG( | 2729.919 | -0.14616 | 0.297215 | 0.898538 | Ran |
| ENSMUSG( | 607.0443 | -0.11146 | 0.455701 | 0.923329 | Ranbp1 |
| ENSMUSG( | 1136.125 | 0.103837 | 0.396192 | 0.917257 | Ranbp10 |
| ENSMUSG( | 168.674 | -0.14178 | 0.418092 | 0.917257 | Ranbp17 |
| ENSMUSG( | 3012.225 | 0.200698 | 0.138144 | 0.828492 | Ranbp2 |
| ENSMUSG( | 1912.984 | -0.04648 | 0.675065 | 0.95694 | Ranbp3 |
| ENSMUSG( | 499.0592 | 0.014387 | 0.925082 | 0.99306 | Ranbp3l |
| ENSMUSG( | 535.8236 | 0.152937 | 0.268779 | 0.886755 | Ranbp6 |
| ENSMUSG( | 2135.657 | 0.089028 | 0.403765 | 0.917257 | Ranbp9 |
| ENSMUSG( | 4045.931 | -0.12755 | 0.330467 | 0.907398 | Rangap1 |
| ENSMUSG( | 207.4137 | 0.094062 | 0.601583 | 0.947749 | Rangrf |
| ENSMUSG( | 1251.041 | -0.21516 | 0.15835 | 0.837724 | Rap1a |
| ENSMUSG( | 1827.08 | 0.039358 | 0.784552 | 0.976207 | Rap1b |
| ENSMUSG( | 5481.219 | 0.062296 | 0.668323 | 0.956766 | Rap1gap |
| ENSMUSG( | 11154.19 | 0.072213 | 0.521467 | 0.932489 | Rap1gap2 |
| ENSMUSG( | 8419.653 | -0.18127 | 0.100511 | 0.801444 | Rap1gds1 |
| ENSMUSG( | 3407.764 | -0.06819 | 0.526667 | 0.934374 | Rap2a |
| ENSMUSG( | 1337.321 | 0.111194 | 0.503749 | 0.927825 | Rap2b |
| ENSMUSG( | 1485.829 | 0.057619 | 0.755857 | 0.971371 | Rap2c |
| ENSMUSG( | 8538.478 | 0.098838 | 0.32316 | 0.904023 | Rapgef1 |
| ENSMUSG( | 2364.517 | 0.230593 | 0.058101 | 0.728256 | Rapgef2 |
| ENSMUSG( | 1765.339 | 0.01034 | 0.939258 | 0.993898 | Rapgef3 |
| ENSMUSG( | 62072.32 | -0.13925 | 0.220109 | 0.866104 | Rapgef4 |
| ENSMUSG( | 2480.483 | 0.211221 | 0.108277 | 0.804969 | Rapgef5 |
| ENSMUSG( | 789.6634 | 0.214648 | 0.072475 | 0.763629 | Rapgef6 |
| ENSMUSG( | 5241.505 | 0.113977 | 0.268262 | 0.886493 | Rapgefl1 |
| ENSMUSG( | 913.3522 | 0.085565 | 0.471086 | 0.923329 | Raph1 |
| ENSMUSG( | 22.4751 | -0.52112 | 0.223573 | 0.867404 | Rapsn |
| ENSMUSG( | 307.4257 | -0.34332 | 0.039591 | 0.68111 | Rara |
| ENSMUSG( | 318.5542 | -0.22376 | 0.297841 | 0.898538 | Rarb |
| ENSMUSG( | 401.611 | 0.268039 | 0.076404 | 0.770504 | Rarg |
| ENSMUSG( | 74.86075 | 0.417068 | 0.110094 | 0.806572 | Rarres1 |
| ENSMUSG( | 498.9492 | 0.080604 | 0.649343 | 0.95422 | Rarres2 |
| ENSMUSG( | 715.4305 | -0.14944 | 0.29402 | 0.898538 | Rars1 |
| ENSMUSG( | 334.4121 | 0.028637 | 0.861181 | 0.986226 | Rars2 |
| ENSMUSG( | 1014.248 | 0.201474 | 0.078097 | 0.77321 | Rasa1 |
| ENSMUSG( | 1411.228 | -0.36417 | 0.010579 | 0.419247 | Rasa2 |
| ENSMUSG( | 2588.673 | -0.11632 | 0.290866 | 0.897019 | Rasa3 |
| ENSMUSG( | 222.8747 | 0.016654 | 0.92468 | 0.99306 | Rasa4 |
| ENSMUSG( | 2569.154 | 0.165875 | 0.209715 | 0.864412 | Rasal1 |
| ENSMUSG( | 1669.816 | 0.172892 | 0.175363 | 0.845305 | Rasal2 |
| ENSMUSG( | 171.8523 | 0.330302 | 0.126439 | 0.815475 | Rasal3 |
| ENSMUSG( | 134.0375 | -0.09347 | 0.731536 | 0.967228 | Rasd1 |
| ENSMUSG( | 3710.502 | -0.07765 | 0.564822 | 0.941206 | Rasd2 |
| ENSMUSG( | 10285.8 | 0.010651 | 0.923043 | 0.992644 | Rasgef1a |
| ENSMUSG( | 961.7382 | -0.13494 | 0.27534 | 0.889984 | Rasgef1c |
| ENSMUSG( | 7481.612 | 0.160325 | 0.164479 | 0.838699 | Rasgrf1 |
| ENSMUSG( | 2195.917 | 0.040031 | 0.759124 | 0.971909 | Rasgrf2 |
| ENSMUSG( | 7840.66 | 0.0289 | 0.783847 | 0.976154 | Rasgrp1 |
| ENSMUSG( | 987.0321 | -0.13231 | 0.494559 | 0.927372 | Rasgrp2 |
| ENSMUSG( | 510.1348 | 0.176542 | 0.424101 | 0.917257 | Rasgrp3 |
| ENSMUSG( | 71.17763 | 0.285331 | 0.300761 | 0.898538 | Rasgrp4 |
| ENSMUSG( | 334.6666 | 0.538234 | 0.009143 | 0.401114 | Rasip1 |
| ENSMUSG( | 551.7278 | -0.06967 | 0.627214 | 0.953234 | Rasl10a |
| ENSMUSG( | 20229.53 | 0.103338 | 0.334211 | 0.907907 | Rasl10b |
| ENSMUSG( | 74.3339 | -0.69884 | 0.013239 | 0.460134 | Rasl11a |
| ENSMUSG( | 1199.659 | -0.14045 | 0.418814 | 0.917257 | Rasl11b |
| ENSMUSG( | 84.28986 | 0.425821 | 0.136977 | 0.828492 | Rasl12 |
| ENSMUSG( | 337.5502 | -0.07729 | 0.64163 | 0.953903 | Rassf1 |
| ENSMUSG( | 96.6867 | 0.078318 | 0.750055 | 0.970113 | Rassf10 |
| ENSMUSG( | 5930.006 | 0.00722 | 0.949535 | 0.994798 | Rassf2 |
| ENSMUSG( | 1153.584 | -0.17969 | 0.125776 | 0.81516 | Rassf3 |
| ENSMUSG( | 215.2517 | -0.03858 | 0.823689 | 0.979198 | Rassf4 |
| ENSMUSG( | 476.2843 | -0.1033 | 0.511481 | 0.92971 | Rassf5 |
| ENSMUSG( | 13.23513 | -0.08218 | 0.884109 | 0.989644 | Rassf6 |
| ENSMUSG( | 16.75609 | 0.434987 | 0.375952 | 0.915491 | Rassf7 |
| ENSMUSG( | 1215.28 | -0.03086 | 0.819119 | 0.979198 | Rassf8 |
| ENSMUSG( | 67.43103 | 0.7254 | 0.048215 | 0.706142 | Rassf9 |
| ENSMUSG( | 1234.246 | 0.120982 | 0.315923 | 0.901868 | Raver2 |
| ENSMUSG( | 1601.281 | -0.18473 | 0.237149 | 0.872173 | Rb1 |
| ENSMUSG( | 2064.657 | 0.045724 | 0.66153 | 0.956205 | Rb1cc1 |
| ENSMUSG( | 400.4053 | 0.003846 | 0.981705 | 0.996756 | Rbak |
| ENSMUSG( | 2762.076 | -0.01559 | 0.8953 | 0.990558 | Rbbp4 |
| ENSMUSG( | 968.2954 | -0.00677 | 0.953658 | 0.994798 | Rbbp5 |
| ENSMUSG( | 2172.11 | 0.057336 | 0.63581 | 0.953903 | Rbbp6 |
| ENSMUSG( | 2255.911 | -0.04844 | 0.691019 | 0.960071 | Rbbp7 |
| ENSMUSG( | 128.1705 | 0.305107 | 0.12935 | 0.820358 | Rbbp8 |
| ENSMUSG( | 1243.248 | -0.2151 | 0.113335 | 0.809623 | Rbbp9 |
| ENSMUSG( | 3548.778 | 0.016548 | 0.872282 | 0.988944 | Rbck1 |
| ENSMUSG( | 678.6329 | -0.014 | 0.921578 | 0.992644 | Rbfa |
| ENSMUSG( | 17.27439 | 0.003396 | 0.99427 | 0.998813 | Rbfaos |
| ENSMUSG( | 3069.107 | -0.05215 | 0.65052 | 0.95422 | Rbfox1 |
| ENSMUSG( | 6052.475 | 0.062553 | 0.552208 | 0.939565 | Rbfox2 |
| ENSMUSG( | 6314.601 | 0.006702 | 0.958057 | 0.994798 | Rbfox3 |
| ENSMUSG( | 375.3456 | 0.049228 | 0.752124 | 0.970461 | Rbis |
| ENSMUSG( | 51.72782 | -0.38885 | 0.181421 | 0.850579 | Rbks |
| ENSMUSG( | 93.01035 | 0.171416 | 0.434283 | 0.91869 | Rbl1 |
| ENSMUSG( | 2189.439 | -0.11365 | 0.357496 | 0.913759 | Rbl2 |
| ENSMUSG( | 1407.335 | -0.00571 | 0.965553 | 0.995433 | Rbm10 |
| ENSMUSG( | 106.8544 | -0.11303 | 0.626837 | 0.953234 | Rbm11 |
| ENSMUSG( | 212.558 | -0.01507 | 0.93737 | 0.993685 | Rbm12 |
| ENSMUSG( | 149.1129 | 0.24374 | 0.273073 | 0.889275 | Rbm12b1 |
| ENSMUSG( | 208.8867 | 0.110988 | 0.5237 | 0.932848 | Rbm12b2 |
| ENSMUSG( | 366.7068 | -0.07539 | 0.63545 | 0.953903 | Rbm14 |
| ENSMUSG( | 159.7979 | 0.154278 | 0.405963 | 0.917257 | Rbm15 |
| ENSMUSG( | 1800.374 | 0.051871 | 0.649787 | 0.95422 | Rbm15b |
| ENSMUSG( | 1379.518 | -0.0285 | 0.830642 | 0.98051 | Rbm17 |
| ENSMUSG( | 868.0057 | 0.014216 | 0.911839 | 0.992196 | Rbm18 |
| ENSMUSG( | 263.2593 | 0.039971 | 0.80483 | 0.977232 | Rbm19 |
| ENSMUSG( | 143.8739 | -0.01022 | 0.956847 | 0.994798 | Rbm20 |
| ENSMUSG( | 871.5122 | 0.076075 | 0.554597 | 0.939565 | Rbm22 |
| ENSMUSG( | 276.2797 | -0.07396 | 0.703687 | 0.962481 | Rbm24 |
| ENSMUSG( | 3107.44 | 0.175704 | 0.138091 | 0.828492 | Rbm25 |
| ENSMUSG( | 1110.32 | 0.180374 | 0.121219 | 0.813384 | Rbm26 |
| ENSMUSG( | 687.6461 | 0.156852 | 0.227864 | 0.868507 | Rbm27 |
| ENSMUSG( | 1538.179 | -0.00286 | 0.981661 | 0.996756 | Rbm28 |
| ENSMUSG( | 1861.601 | 0.135148 | 0.327896 | 0.906255 | Rbm3 |
| ENSMUSG( | 1667.73 | 0.091695 | 0.558068 | 0.939565 | Rbm33 |
| ENSMUSG( | 385.5593 | 0.002578 | 0.987236 | 0.997487 | Rbm34 |
| ENSMUSG( | 115.2875 | 0.155769 | 0.476731 | 0.923329 | Rbm38 |
| ENSMUSG( | 4953.64 | 0.228896 | 0.037817 | 0.6763 | Rbm39 |
| ENSMUSG( | 10.33749 | 0.078491 | 0.897756 | NA | Rbm3os |
| ENSMUSG( | 63.23156 | 0.359581 | 0.212236 | 0.864412 | Rbm4 |
| ENSMUSG( | 151.9539 | 0.160263 | 0.423092 | 0.917257 | Rbm41 |
| ENSMUSG( | 237.2107 | 0.007241 | 0.967639 | 0.995833 | Rbm43 |
| ENSMUSG( | 463.4912 | -0.12255 | 0.37636 | 0.915491 | Rbm45 |
| ENSMUSG( | 20.83712 | 0.079124 | 0.885803 | 0.98977 | Rbm46 |
| ENSMUSG( | 35.10248 | -0.09498 | 0.806613 | 0.978086 | Rbm47 |
| ENSMUSG( | 119.8696 | 0.310216 | 0.122914 | 0.814458 | Rbm48 |
| ENSMUSG( | 394.9434 | -0.14357 | 0.352119 | 0.912093 | Rbm4b |
| ENSMUSG( | 4174.915 | 0.122048 | 0.360012 | 0.914223 | Rbm5 |
| ENSMUSG( | 1366.31 | 0.077429 | 0.542788 | 0.938044 | Rbm6 |
| ENSMUSG( | 767.1583 | 0.055064 | 0.66858 | 0.956766 | Rbm7 |

| | | | | | | | | | | | |
|---|---|---|---|---|---|---|---|---|---|---|---|
| ENSMUSG( | Rcan3 | 2055.902 | -0.03899 | 0.717831 | 0.941697 | ENSMUSG( | Rcan3 | 1788.949 | -0.17896 | 0.151012 | 0.837173 |
| ENSMUSG( | Rcbtb1 | 1568.133 | -0.07741 | 0.39389 | 0.809643 | ENSMUSG( | Rcbtb1 | 1422.845 | -0.08843 | 0.416705 | 0.917257 |
| ENSMUSG( | Rcbtb2 | 965.3181 | -0.00783 | 0.941183 | 0.993415 | ENSMUSG( | Rcbtb2 | 847.3388 | -0.12347 | 0.387993 | 0.917257 |
| ENSMUSG( | Rcc1 | 393.3153 | 0.197743 | 0.139965 | 0.556732 | ENSMUSG( | Rcc1 | 344.8003 | 0.09211 | 0.513112 | 0.930358 |
| ENSMUSG( | Rcc1l | 442.4627 | 0.120914 | 0.381505 | 0.802699 | ENSMUSG( | Rcc1l | 387.7873 | 0.011134 | 0.942575 | 0.994774 |
| ENSMUSG( | Rcc2 | 3545.412 | 0.022834 | 0.795823 | 0.960062 | ENSMUSG( | Rcc2 | 3122.418 | -0.07768 | 0.504805 | 0.927825 |
| ENSMUSG( | Rccd1 | 256.0541 | 0.037806 | 0.793492 | 0.959799 | ENSMUSG( | Rccd1 | 244.3124 | 0.178062 | 0.360289 | 0.914223 |
| ENSMUSG( | Rce1 | 534.0348 | -0.21161 | 0.134414 | 0.547651 | ENSMUSG( | Rce1 | 547.2511 | 0.139224 | 0.396516 | 0.917257 |
| ENSMUSG( | Rchy1 | 869.1336 | 0.05566 | 0.639116 | 0.916434 | ENSMUSG( | Rchy1 | 751.0841 | -0.10325 | 0.462645 | 0.923329 |
| ENSMUSG( | Rcl1 | 231.5864 | 0.141082 | 0.360891 | 0.792797 | ENSMUSG( | Rcl1 | 208.1906 | 0.093241 | 0.624573 | 0.953234 |
| ENSMUSG( | Rcn1 | 1152.01 | -0.2041 | 0.134326 | 0.547472 | ENSMUSG( | Rcn1 | 1187.359 | 0.156241 | 0.282792 | 0.892412 |
| ENSMUSG( | Rcn2 | 9233.355 | -0.07228 | 0.570324 | 0.893192 | ENSMUSG( | Rcn2 | 8765.062 | 0.046588 | 0.736644 | 0.967344 |
| ENSMUSG( | Rcn3 | 341.5626 | -0.11884 | 0.379346 | 0.801757 | ENSMUSG( | Rcn3 | 339.8595 | 0.144707 | 0.407967 | 0.917257 |
| ENSMUSG( | Rcor1 | 574.4313 | 0.089749 | 0.431237 | 0.827055 | ENSMUSG( | Rcor1 | 514.0831 | 0.04511 | 0.743673 | 0.96951 |
| ENSMUSG( | Rcor2 | 386.3448 | -0.10041 | 0.483722 | 0.85636 | ENSMUSG( | Rcor2 | 381.72 | 0.131558 | 0.348675 | 0.911652 |
| ENSMUSG( | Rcor3 | 475.4282 | -0.11383 | 0.375529 | 0.800845 | ENSMUSG( | Rcor3 | 469.992 | 0.12285 | 0.378839 | 0.916995 |
| ENSMUSG( | Rcsd1 | 269.9426 | -0.03915 | 0.801052 | 0.96188 | ENSMUSG( | Rcsd1 | 244.6972 | -0.03925 | 0.83743 | 0.981335 |
| ENSMUSG( | Rcvrn | 10.48894 | 0.302596 | 0.612406 | NA | ENSMUSG( | Rcvrn | 9.116288 | 0.237574 | 0.72153 | NA |
| ENSMUSG( | Rd3 | 16.20069 | -0.85934 | 0.066824 | NA | ENSMUSG( | Rd3 | 23.70237 | 0.575518 | 0.210896 | 0.864412 |
| ENSMUSG( | Rdh10 | 416.6182 | -0.06043 | 0.692105 | 0.934101 | ENSMUSG( | Rdh10 | 425.0249 | 0.260177 | 0.116698 | 0.813384 |
| ENSMUSG( | Rdh11 | 1151.775 | -0.50827 | 0.001191 | 0.037142 | ENSMUSG( | Rdh11 | 1292.415 | 0.13823 | 0.404013 | 0.917257 |
| ENSMUSG( | Rdh12 | 51.54103 | -0.56921 | 0.042543 | 0.316636 | ENSMUSG( | Rdh12 | 58.62344 | 0.116079 | 0.679514 | 0.957966 |
| ENSMUSG( | Rdh13 | 534.9422 | 0.139386 | 0.277196 | 0.727175 | ENSMUSG( | Rdh13 | 496.1068 | 0.186288 | 0.188951 | 0.85698 |
| ENSMUSG( | Rdh14 | 889.0477 | -0.24786 | 0.057619 | 0.370247 | ENSMUSG( | Rdh14 | 911.047 | 0.1028 | 0.468091 | 0.923329 |
| ENSMUSG( | Rdh5 | 178.296 | 0.216566 | 0.212759 | 0.6567 | ENSMUSG( | Rdh5 | 155.5588 | 0.101975 | 0.597092 | 0.946609 |
| ENSMUSG( | Rdm1 | 225.9844 | 0.171688 | 0.258829 | 0.709814 | ENSMUSG( | Rdm1 | 200.1253 | 0.092612 | 0.593386 | 0.945408 |
| ENSMUSG( | Rdx | 5610.252 | -0.04374 | 0.757469 | 0.951513 | ENSMUSG( | Rdx | 5198.222 | 0.004564 | 0.9742 | 0.996416 |
| ENSMUSG( | Rec8 | 97.95878 | 0.574968 | 0.009132 | 0.134662 | ENSMUSG( | Rec8 | 80.26459 | 0.293209 | 0.309071 | 0.898538 |
| ENSMUSG( | Reck | 412.7998 | -0.0156 | 0.901593 | 0.985633 | ENSMUSG( | Reck | 413.5255 | 0.256988 | 0.11339 | 0.809623 |
| ENSMUSG( | Recql | 548.0003 | 0.021294 | 0.852872 | 0.973168 | ENSMUSG( | Recql | 463.09 | -0.20386 | 0.136101 | 0.827092 |
| ENSMUSG( | Recql4 | 87.03752 | 0.382384 | 0.127253 | 0.534725 | ENSMUSG( | Recql4 | 77.1931 | 0.324593 | 0.259823 | 0.883971 |
| ENSMUSG( | Recql5 | 605.5029 | 0.129428 | 0.274094 | 0.724254 | ENSMUSG( | Recql5 | 557.703 | 0.165847 | 0.332872 | 0.907907 |
| ENSMUSG( | Redic1 | 19.76548 | 0.030917 | 0.950809 | NA | ENSMUSG( | Redic1 | 14.45717 | -0.74763 | 0.22183 | 0.866104 |
| ENSMUSG( | Reep1 | 3920.607 | -0.40664 | 0.001767 | 0.047422 | ENSMUSG( | Reep1 | 4167.098 | 0.068893 | 0.650823 | 0.95422 |
| ENSMUSG( | Reep2 | 5649.658 | -0.29706 | 0.00764 | 0.120942 | ENSMUSG( | Reep2 | 5671.299 | 0.000567 | 0.996553 | 0.999178 |
| ENSMUSG( | Reep3 | 2382.317 | -0.18379 | 0.21159 | 0.654835 | ENSMUSG( | Reep3 | 2401.076 | 0.113586 | 0.48323 | 0.92499 |
| ENSMUSG( | Reep4 | 133.3155 | -0.27977 | 0.176973 | 0.61401 | ENSMUSG( | Reep4 | 143.0571 | 0.203011 | 0.334804 | 0.907907 |
| ENSMUSG( | Reep5 | 17669.9 | -0.2659 | 0.036449 | 0.294207 | ENSMUSG( | Reep5 | 17637.03 | 0.012155 | 0.932991 | 0.99306 |
| ENSMUSG( | Reep6 | 510.9878 | -0.15764 | 0.203417 | 0.643631 | ENSMUSG( | Reep6 | 498.218 | 0.047683 | 0.751589 | 0.97035 |
| ENSMUSG( | Rel | 120.1887 | -0.1962 | 0.316717 | 0.763648 | ENSMUSG( | Rel | 112.3105 | -0.10919 | 0.618449 | 0.952254 |
| ENSMUSG( | Rela | 813.4692 | -0.04436 | 0.656452 | 0.923929 | ENSMUSG( | Rela | 744.5553 | -0.0266 | 0.836078 | 0.981237 |
| ENSMUSG( | Relb | 187.9067 | -0.10298 | 0.543076 | 0.883579 | ENSMUSG( | Relb | 174.5357 | -0.04203 | 0.808205 | 0.978622 |
| ENSMUSG( | Relch | 1166.179 | 0.038966 | 0.6961 | 0.935007 | ENSMUSG( | Relch | 1085.646 | 0.099116 | 0.400998 | 0.917257 |
| ENSMUSG( | Rell1 | 635.8793 | -0.40103 | 0.019693 | 0.21168 | ENSMUSG( | Rell1 | 708.098 | 0.206449 | 0.29596 | 0.898538 |
| ENSMUSG( | Rell2 | 7530.962 | 0.076028 | 0.414917 | 0.820184 | ENSMUSG( | Rell2 | 6831.214 | 0.064494 | 0.552405 | 0.939565 |
| ENSMUSG( | Reln | 1713.968 | -0.38249 | 0.006588 | 0.111324 | ENSMUSG( | Reln | 1908.997 | 0.217192 | 0.202764 | 0.864412 |
| ENSMUSG( | Relt | 653.2848 | -0.41788 | 0.000707 | 0.025719 | ENSMUSG( | Relt | 624.3858 | -0.25693 | 0.068585 | 0.753052 |
| ENSMUSG( | Rem1 | 11.64861 | 0.28338 | 0.610838 | NA | ENSMUSG( | Rem1 | 15.26453 | 1.14103 | 0.028389 | 0.613486 |
| ENSMUSG( | Rem2 | 162.5022 | -0.58563 | 0.021806 | 0.225679 | ENSMUSG( | Rem2 | 163.0653 | -0.25985 | 0.319463 | 0.903461 |
| ENSMUSG( | Renbp | 197.7206 | -0.00926 | 0.950998 | 0.993821 | ENSMUSG( | Renbp | 165.5335 | -0.25906 | 0.143964 | 0.831311 |
| ENSMUSG( | Reno1 | 282.5803 | -0.12522 | 0.397602 | 0.812127 | ENSMUSG( | Reno1 | 309.3706 | 0.392195 | 0.05121 | 0.711023 |
| ENSMUSG( | Rep15 | 44.16196 | 0.038352 | 0.899383 | 0.985633 | ENSMUSG( | Rep15 | 32.6071 | -0.62063 | 0.089283 | 0.788502 |
| ENSMUSG( | Repin1 | 1084.698 | 0.030486 | 0.784377 | 0.958003 | ENSMUSG( | Repin1 | 1017.897 | 0.115574 | 0.367102 | 0.915058 |
| ENSMUSG( | Reps1 | 1169.315 | 0.140569 | 0.190216 | 0.629268 | ENSMUSG( | Reps1 | 1011.425 | -0.00681 | 0.958213 | 0.994798 |
| ENSMUSG( | Reps2 | 2570.027 | 0.135807 | 0.097542 | 0.477842 | ENSMUSG( | Reps2 | 2231.913 | -0.00033 | 0.997568 | 0.999268 |
| ENSMUSG( | Rer1 | 2804.112 | -0.18381 | 0.105527 | 0.492417 | ENSMUSG( | Rer1 | 2764.12 | 0.049923 | 0.701231 | 0.961756 |
| ENSMUSG( | Rere | 4401.785 | 0.168082 | 0.033511 | 0.280615 | ENSMUSG( | Rere | 4025.792 | 0.177821 | 0.08406 | 0.777852 |
| ENSMUSG( | Rerg | 108.5945 | 0.174835 | 0.465484 | 0.844064 | ENSMUSG( | Rerg | 102.8104 | 0.265085 | 0.33525 | 0.907907 |
| ENSMUSG( | Resf1 | 317.3522 | 0.078987 | 0.552522 | 0.886141 | ENSMUSG( | Resf1 | 321.864 | 0.37471 | 0.020896 | 0.571913 |
| ENSMUSG( | Resp18 | 1697.968 | -0.38956 | 0.000473 | 0.019871 | ENSMUSG( | Resp18 | 1690.884 | -0.10883 | 0.401717 | 0.917257 |
| ENSMUSG( | Rest | 239.2977 | -0.20566 | 0.141493 | 0.558606 | ENSMUSG( | Rest | 231.1971 | -0.02667 | 0.866388 | 0.987347 |
| ENSMUSG( | Ret | 108.6057 | -0.50374 | 0.033133 | 0.278602 | ENSMUSG( | Ret | 124.177 | 0.193588 | 0.369371 | 0.915491 |
| ENSMUSG( | Retn | 12.13385 | 1.201246 | 0.045284 | NA | ENSMUSG( | Retn | 8.962973 | 0.70563 | 0.3809 | NA |
| ENSMUSG( | Retnlg | 15.10013 | -1.12741 | 0.106763 | NA | ENSMUSG( | Retnlg | 18.71473 | -0.08991 | 0.89134 | 0.990558 |
| ENSMUSG( | Retreg1 | 1250.314 | -0.59114 | 0.000839 | 0.02897 | ENSMUSG( | Retreg1 | 1474.799 | 0.209853 | 0.295734 | 0.898538 |
| ENSMUSG( | Retreg2 | 7643.602 | -0.19388 | 0.12138 | 0.525713 | ENSMUSG( | Retreg2 | 7755.853 | 0.124316 | 0.406982 | 0.917257 |
| ENSMUSG( | Retreg3 | 3163.197 | -0.11138 | 0.394464 | 0.810072 | ENSMUSG( | Retreg3 | 3101.001 | 0.104603 | 0.476743 | 0.923329 |
| ENSMUSG( | Retsat | 813.8298 | 0.176112 | 0.154582 | 0.581712 | ENSMUSG( | Retsat | 786.1883 | 0.336223 | 0.024848 | 0.592568 |
| ENSMUSG( | Rev1 | 837.0076 | -0.10749 | 0.390657 | 0.808144 | ENSMUSG( | Rev1 | 805.8658 | 0.05575 | 0.698975 | 0.961064 |
| ENSMUSG( | Rev3l | 1056.013 | -0.32714 | 0.001932 | 0.050384 | ENSMUSG( | Rev3l | 1064.16 | -0.01601 | 0.891383 | 0.990558 |
| ENSMUSG( | Rex1bd | 1415.482 | 0.036085 | 0.692568 | 0.934101 | ENSMUSG( | Rex1bd | 1175.368 | -0.24589 | 0.05283 | 0.71525 |
| ENSMUSG( | Rexo1 | 1925.782 | 0.118349 | 0.209926 | 0.652297 | ENSMUSG( | Rexo1 | 1759.704 | 0.129327 | 0.278641 | 0.891628 |
| ENSMUSG( | Rexo2 | 1614.488 | -0.00436 | 0.963545 | 0.994922 | ENSMUSG( | Rexo2 | 1407.911 | -0.13883 | 0.305812 | 0.898538 |
| ENSMUSG( | Rexo4 | 1020.784 | 0.110044 | 0.332881 | 0.775602 | ENSMUSG( | Rexo4 | 910.0243 | 0.053998 | 0.706854 | 0.963529 |
| ENSMUSG( | Rexo5 | 42.08372 | 0.186225 | 0.522191 | 0.875186 | ENSMUSG( | Rexo5 | 38.77879 | 0.199728 | 0.57119 | 0.942052 |
| ENSMUSG( | Rfc1 | 1610.319 | 0.164005 | 0.081895 | 0.442044 | ENSMUSG( | Rfc1 | 1325.077 | -0.13033 | 0.248479 | 0.877904 |
| ENSMUSG( | Rfc2 | 1027.898 | 0.125473 | 0.201922 | 0.641612 | ENSMUSG( | Rfc2 | 882.7771 | -0.04115 | 0.722001 | 0.966887 |
| ENSMUSG( | Rfc3 | 109.0265 | -0.24557 | 0.208316 | 0.650319 | ENSMUSG( | Rfc3 | 106.2737 | -0.03916 | 0.856068 | 0.985884 |
| ENSMUSG( | Rfc4 | 87.01623 | 0.244952 | 0.267846 | 0.718054 | ENSMUSG( | Rfc4 | 72.60274 | -0.00908 | 0.970267 | 0.99594 |
| ENSMUSG( | Rfc5 | 228.4094 | 0.103851 | 0.541934 | 0.883182 | ENSMUSG( | Rfc5 | 191.4142 | -0.12602 | 0.498129 | 0.927825 |
| ENSMUSG( | Rfesd | 194.7129 | 0.009498 | 0.95602 | 0.993874 | ENSMUSG( | Rfesd | 176.0202 | -0.02709 | 0.897022 | 0.990558 |
| ENSMUSG( | Rffl | 412.6357 | -0.34501 | 0.00599 | 0.104817 | ENSMUSG( | Rffl | 431.3376 | 0.079168 | 0.60978 | 0.950397 |
| ENSMUSG( | Rfk | 2855.078 | 0.003038 | 0.978415 | 0.997428 | ENSMUSG( | Rfk | 2454.196 | -0.17398 | 0.267284 | 0.886134 |
| ENSMUSG( | Rflna | 23.76651 | 0.495129 | 0.217128 | 0.661995 | ENSMUSG( | Rflna | 25.48625 | 0.869171 | 0.02967 | 0.61992 |
| ENSMUSG( | Rflnb | 653.7944 | 0.089527 | 0.504981 | 0.866883 | ENSMUSG( | Rflnb | 563.6337 | -0.06585 | 0.637694 | 0.953903 |
| ENSMUSG( | Rfng | 3209.64 | 0.025248 | 0.754238 | 0.950981 | ENSMUSG( | Rfng | 2923.37 | 0.024034 | 0.809097 | 0.97873 |
| ENSMUSG( | Rft1 | 418.7687 | -0.3281 | 0.02489 | 0.242767 | ENSMUSG( | Rft1 | 433.6787 | 0.066993 | 0.677739 | 0.957106 |
| ENSMUSG( | Rftn1 | 246.3619 | -0.10944 | 0.449629 | 0.835234 | ENSMUSG( | Rftn1 | 222.7033 | -0.12298 | 0.492473 | 0.927315 |
| ENSMUSG( | Rftn2 | 4322.072 | 0.085679 | 0.487226 | 0.858812 | ENSMUSG( | Rftn2 | 3440.487 | -0.32373 | 0.025691 | 0.599234 |
| ENSMUSG( | Rfwd3 | 616.5993 | 0.183764 | 0.08829 | 0.455783 | ENSMUSG( | Rfwd3 | 543.3939 | 0.089908 | 0.491042 | 0.926604 |
| ENSMUSG( | Rfx1 | 842.2833 | 0.255264 | 0.061163 | 0.381822 | ENSMUSG( | Rfx1 | 702.745 | 0.008525 | 0.956083 | 0.994798 |
| ENSMUSG( | Rfx2 | 167.1448 | -0.00302 | 0.985774 | 0.998326 | ENSMUSG( | Rfx2 | 149.7173 | -0.04652 | 0.813076 | 0.979198 |
| ENSMUSG( | Rfx3 | 1240.154 | 0.061135 | 0.561094 | 0.888934 | ENSMUSG( | Rfx3 | 1134.182 | 0.069273 | 0.591752 | 0.944975 |
| ENSMUSG( | Rfx4 | 1408.766 | 0.104002 | 0.370631 | 0.797829 | ENSMUSG( | Rfx4 | 1192.363 | -0.11123 | 0.377813 | 0.916784 |
| ENSMUSG( | Rfx5 | 683.596 | -0.09188 | 0.406202 | 0.816525 | ENSMUSG( | Rfx5 | 638.8079 | -0.01976 | 0.879023 | 0.989644 |
| ENSMUSG( | Rfx7 | 936.812 | -0.20526 | 0.041335 | 0.314189 | ENSMUSG( | Rfx7 | 901.5244 | -0.04643 | 0.733052 | 0.967228 |
| ENSMUSG( | Rfx8 | 17.43806 | 0.62232 | 0.159117 | NA | ENSMUSG( | Rfx8 | 11.62372 | -0.28405 | 0.626805 | 0.953234 |
| ENSMUSG( | Rfxank | 474.0032 | -0.12512 | 0.275239 | 0.72509 | ENSMUSG( | Rfxank | 453.0933 | 0.021527 | 0.886793 | 0.990118 |
| ENSMUSG( | Rfxap | 578.768 | 0.067381 | 0.569548 | 0.893041 | ENSMUSG( | Rfxap | 510.4819 | -0.03216 | 0.837561 | 0.981335 |
| ENSMUSG( | Rgcc | 1202.868 | 0.008688 | 0.935274 | 0.993331 | ENSMUSG( | Rgcc | 1009.886 | -0.23721 | 0.043274 | 0.690304 |
| ENSMUSG( | Rgl1 | 3279.538 | 0.10499 | 0.242682 | 0.690216 | ENSMUSG( | Rgl1 | 2846.128 | -0.03348 | 0.751416 | 0.970317 |
| ENSMUSG( | Rgl2 | 1959.763 | 0.354935 | 0.042154 | 0.316282 | ENSMUSG( | Rgl2 | 1629.966 | 0.107946 | 0.599965 | 0.947322 |
| ENSMUSG( | Rgl3 | 252.2173 | -0.05005 | 0.731797 | 0.944956 | ENSMUSG( | Rgl3 | 239.9045 | 0.072855 | 0.642768 | 0.953903 |
| ENSMUSG( | Rgma | 3741.574 | -0.23381 | 0.074677 | 0.423665 | ENSMUSG( | Rgma | 4057.669 | 0.270095 | 0.068363 | 0.752306 |
| ENSMUSG( | Rgmb | 2146.085 | -0.4009 | 0.001747 | 0.047136 | ENSMUSG( | Rgmb | 2317.967 | 0.11848 | 0.427037 | 0.917257 |
| ENSMUSG( | Rgp1 | 2229.817 | 0.100169 | 0.253281 | 0.703461 | ENSMUSG( | Rgp1 | 1945.683 | -0.02297 | 0.8248 | 0.979198 |
| ENSMUSG( | Rgs10 | 1052.087 | 0.153863 | 0.174136 | 0.60991 | ENSMUSG( | Rgs10 | 885.4166 | -0.07534 | 0.564577 | 0.941206 |
| ENSMUSG( | Rgs11 | 1197.207 | 0.418063 | 0.244171 | 0.691435 | ENSMUSG( | Rgs11 | 984.9856 | 0.144844 | 0.72518 | 0.966887 |
| ENSMUSG( | Rgs12 | 2411.548 | 0.040725 | 0.685201 | 0.932981 | ENSMUSG( | Rgs12 | 2164.919 | 0.001674 | 0.989843 | 0.997928 |
| ENSMUSG( | Rgs14 | 1055.387 | -0.07952 | 0.464935 | 0.843635 | ENSMUSG( | Rgs14 | 919.1831 | -0.21575 | 0.094741 | 0.796133 |
| ENSMUSG( | Rgs16 | 202.5612 | 0.103534 | 0.535017 | 0.880733 | ENSMUSG( | Rgs16 | 170.6759 | -0.11255 | 0.571455 | 0.942116 |
| ENSMUSG( | Rgs17 | 997.5712 | 0.206256 | 0.088686 | 0.45666 | ENSMUSG( | Rgs17 | 907.7107 | 0.195149 | 0.271156 | 0.888062 |
| ENSMUSG( | Rgs19 | 394.9996 | 0.044242 | 0.725174 | 0.944096 | ENSMUSG( | Rgs19 | 342.8452 | -0.10514 | 0.507149 | 0.928677 |
| ENSMUSG( | Rgs2 | 739.9327 | -0.20854 | 0.245612 | 0.693322 | ENSMUSG( | Rgs2 | 704.6134 | -0.07849 | 0.686523 | 0.95899 |
| ENSMUSG( | Rgs20 | 2172.858 | 0.028567 | 0.779812 | 0.957136 | ENSMUSG( | Rgs20 | 1770.824 | -0.31402 | 0.038752 | 0.679972 |
| ENSMUSG( | Rgs22 | 32.64111 | 0.009499 | 0.977994 | 0.997428 | ENSMUSG( | Rgs22 | 36.92768 | 0.593086 | 0.074365 | 0.766789 |
| ENSMUSG( | Rgs3 | 786.7508 | 0.043539 | 0.693279 | 0.934202 | ENSMUSG( | Rgs3 | 729.3414 | 0.09554 | 0.434134 | 0.91869 |
| ENSMUSG( | Rgs4 | 7170.864 | -0.17433 | 0.165159 | 0.596597 | ENSMUSG( | Rgs4 | 6232.382 | -0.32506 | 0.018558 | 0.545482 |
| ENSMUSG( | Rgs5 | 3573.145 | -0.0126 | 0.971722 | 0.996543 | ENSMUSG( | Rgs5 | 4385.976 | 0.759561 | 0.114542 | 0.811703 |
| ENSMUSG( | Rgs6 | 3041.417 | 0.03762 | 0.656596 | 0.923929 | ENSMUSG( | Rgs6 | 2561.065 | -0.19778 | 0.060146 | 0.731553 |
| ENSMUSG( | Rgs7 | 2112.876 | 0.028862 | 0.789088 | 0.959299 | ENSMUSG( | Rgs7 | 1771.196 | -0.22302 | 0.08479 | 0.779799 |
| ENSMUSG( | Rgs7bp | 5260.462 | -0.00383 | 0.963582 | 0.994922 | ENSMUSG( | Rgs7bp | 4898.186 | 0.056532 | 0.650659 | 0.95422 |
| ENSMUSG( | Rgs8 | 1806.441 | -0.03877 | 0.738299 | 0.947329 | ENSMUSG( | Rgs8 | 1611.071 | -0.10026 | 0.453035 | 0.922316 |
| ENSMUSG( | Rgs9 | 1338.861 | -0.79993 | 0.03567 | 0.290384 | ENSMUSG( | Rgs9 | 1381.078 | -0.36 | 0.098561 | 0.798438 |
| ENSMUSG( | Rgs9bp | 9.082488 | 0.306145 | 0.637247 | NA | ENSMUSG( | Rgs9bp | 9.857764 | 0.747536 | 0.260503 | NA |
| ENSMUSG( | Rgsl1 | 109.1612 | 0.273572 | 0.18097 | 0.618094 | ENSMUSG( | Rgsl1 | 93.06984 | 0.093572 | 0.677292 | 0.957106 |
| ENSMUSG( | Rhbdd1 | 391.896 | -0.49874 | 0.001606 | 0.044935 | ENSMUSG( | Rhbdd1 | 413.6236 | -0.02368 | 0.895427 | 0.990558 |
| ENSMUSG( | Rhbdd2 | 3496.833 | -0.16696 | 0.106611 | 0.494174 | ENSMUSG( | Rhbdd2 | 3388.056 | 0.01776 | 0.882677 | 0.989644 |
| ENSMUSG( | Rhbdd3 | 587.959 | -0.24918 | 0.044666 | 0.324591 | ENSMUSG( | Rhbdd3 | 572.35 | -0.03906 | 0.802636 | 0.97681 |
| ENSMUSG( | Rhbdf1 | 818.6686 | 0.040767 | 0.719581 | 0.941986 | ENSMUSG( | Rhbdf1 | 830.0374 | 0.334451 | 0.019294 | 0.554059 |
| ENSMUSG( | Rhbdf2 | 64.28267 | 0.046546 | 0.865134 | 0.975565 | ENSMUSG( | Rhbdf2 | 73.1562 | 0.630218 | 0.019891 | 0.558472 |
| ENSMUSG( | Rhbdl1 | 174.8843 | 0.099981 | 0.580697 | 0.897294 | ENSMUSG( | Rhbdl1 | 163.3784 | 0.183244 | 0.347936 | 0.911652 |
| ENSMUSG( | Rhbdl2 | 28.54944 | -2.48474 | 2.77E-08 | 9.55E-06 | ENSMUSG( | Rhbdl2 | 43.46046 | -0.0359 | 0.915933 | 0.992529 |
| ENSMUSG( | Rhbdl3 | 967.883 | 0.006249 | 0.95204 | 0.993821 | ENSMUSG( | Rhbdl3 | 966.3944 | 0.257501 | 0.025012 | 0.594476 |
| ENSMUSG( | Rhcg | 45.04572 | 0.071149 | 0.844691 | 0.972023 | ENSMUSG( | Rhcg | 40.41365 | -0.01713 | 0.965127 | 0.995433 |
| ENSMUSG( | Rheb | 2445.083 | 0.053829 | 0.595115 | 0.903007 | ENSMUSG( | Rheb | 2101.584 | -0.12029 | 0.350348 | 0.911652 |
| ENSMUSG( | Rhebl1 | 141.8852 | -0.01271 | 0.94803 | 0.993821 | ENSMUSG( | Rhebl1 | 137.8634 | 0.175312 | 0.404497 | 0.917257 |
| ENSMUSG( | Rhno1 | 369.0489 | 0.135287 | 0.283265 | 0.73186 | ENSMUSG( | Rhno1 | 326.9312 | 0.061655 | 0.669516 | 0.956834 |
| ENSMUSG( | Rho | 29.7152 | 0.731474 | 0.117099 | 0.516406 | ENSMUSG( | Rho | 21.54982 | 0.151243 | 0.739405 | 0.968458 |
| ENSMUSG( | Rhoa | 6411.053 | 0.031956 | 0.754597 | 0.951007 | ENSMUSG( | Rhoa | 5630.671 | -0.07724 | 0.545078 | 0.938063 |
| ENSMUSG( | Rhob | 5997.049 | 0.002556 | 0.982568 | 0.997512 | ENSMUSG( | Rhob | 5267.273 | -0.10555 | 0.432624 | 0.918256 |
| ENSMUSG( | Rhobtb1 | 229.9782 | 0.011842 | 0.937844 | 0.993415 | ENSMUSG( | Rhobtb1 | 228.1464 | 0.259777 | 0.156901 | 0.837724 |
| ENSMUSG( | Rhobtb2 | 3919.969 | 0.05507 | 0.476919 | 0.851851 | ENSMUSG( | Rhobtb2 | 3747.313 | 0.188504 | 0.055016 | 0.723362 |
| ENSMUSG( | Rhobtb3 | 1646.939 | 0.001066 | 0.99444 | 0.999369 | ENSMUSG( | Rhobtb3 | 1454.424 | -0.09474 | 0.556076 | 0.939565 |
| ENSMUSG( | Rhoc | 1143.235 | -0.08207 | 0.459222 | 0.840244 | ENSMUSG( | Rhoc | 1000.743 | -0.2044 | 0.075453 | 0.767994 |
| ENSMUSG( | Rhod | 33.09224 | -0.11796 | 0.715851 | 0.941484 | ENSMUSG( | Rhod | 27.27177 | -0.45403 | 0.23411 | 0.869203 |
| ENSMUSG( | Rhof | 796.2128 | 0.238797 | 0.062632 | 0.38686 | ENSMUSG( | Rhof | 692.8192 | 0.114603 | 0.392122 | 0.917257 |
| ENSMUSG( | Rhog | 1080.685 | -0.18847 | 0.06448 | 0.393628 | ENSMUSG( | Rhog | 996.7728 | -0.15039 | 0.233593 | 0.868507 |
| ENSMUSG( | Rhoh | 33.46448 | 0.046385 | 0.898435 | 0.985633 | ENSMUSG( | Rhoh | 31.40606 | 0.149901 | 0.671147 | 0.95694 |
| ENSMUSG( | Rhoj | 215.0622 | -0.08222 | 0.65932 | 0.924506 | ENSMUSG( | Rhoj | 174.6124 | -0.43681 | 0.035178 | 0.660461 |
| ENSMUSG( | Rhoq | 2429.771 | -0.21619 | 0.086429 | 0.452045 | ENSMUSG( | Rhoq | 2162.025 | -0.29575 | 0.062942 | 0.736482 |
| ENSMUSG( | Rhot1 | 1614.453 | 0.075202 | 0.378048 | 0.801561 | ENSMUSG( | Rhot1 | 1453.383 | 0.041171 | 0.69837 | 0.960908 |
| ENSMUSG( | Rhot2 | 1341.945 | 0.057161 | 0.542001 | 0.883182 | ENSMUSG( | Rhot2 | 1167.09 | -0.07983 | 0.510009 | 0.929089 |
| ENSMUSG( | Rhou | 1997.79 | 0.17288 | 0.129505 | 0.539023 | ENSMUSG( | Rhou | 1634.081 | -0.13581 | 0.275458 | 0.889984 |
| ENSMUSG( | Rhov | 112.1324 | -0.03455 | 0.867056 | 0.976541 | ENSMUSG( | Rhov | 94.71762 | -0.27859 | 0.248848 | 0.877904 |
| ENSMUSG( | Rhpn1 | 398.7293 | 0.091519 | 0.475827 | 0.850919 | ENSMUSG( | Rhpn1 | 354.3943 | 0.019438 | 0.894318 | 0.990558 |
| ENSMUSG( | Rhpn2 | 265.5476 | 0.104008 | 0.441372 | 0.832449 | ENSMUSG( | Rhpn2 | 230.6171 | -0.04431 | 0.800155 | 0.97681 |
| ENSMUSG( | Ribc1 | 11.50801 | 0.028021 | 0.958897 | NA | ENSMUSG( | Ribc1 | 9.559443 | -0.27062 | 0.664467 | NA |

| | | | | | |
|---|---|---|---|---|---|
| ENSMUSG( | 4816.448 | 0.19362 | 0.046688 | 0.330693 | Ralgds |
| ENSMUSG( | 2343.527 | 0.125544 | 0.130723 | 0.542059 | Ralgps1 |
| ENSMUSG( | 1331.452 | 0.32121 | 0.000396 | 0.017766 | Ralgps2 |
| ENSMUSG( | 2268.213 | 0.011986 | 0.88794 | 0.983434 | Raly |
| ENSMUSG( | 1194.228 | 0.092829 | 0.444749 | 0.833239 | Ralyl |
| ENSMUSG( | 713.2335 | -0.09143 | 0.45317 | 0.836701 | Ramac |
| ENSMUSG( | 47.17636 | 0.010565 | 0.971056 | 0.996369 | Ramacl |
| ENSMUSG( | 5098.744 | -0.14153 | 0.140381 | 0.557399 | Ramp1 |
| ENSMUSG( | 931.3564 | 0.182088 | 0.153294 | 0.57989 | Ramp2 |
| ENSMUSG( | 543.0218 | -0.54269 | 2.17E-06 | 0.000346 | Ramp3 |
| ENSMUSG( | 3177.171 | 0.030031 | 0.79222 | 0.959461 | Ran |
| ENSMUSG( | 687.5898 | -0.01211 | 0.913913 | 0.988483 | Ranbp1 |
| ENSMUSG( | 1238.813 | 0.083063 | 0.456284 | 0.838832 | Ranbp10 |
| ENSMUSG( | 192.9217 | -0.02269 | 0.882639 | 0.982283 | Ranbp17 |
| ENSMUSG( | 3075.873 | -0.00426 | 0.970781 | 0.996369 | Ranbp2 |
| ENSMUSG( | 2152.671 | 0.02386 | 0.832115 | 0.970303 | Ranbp3 |
| ENSMUSG( | 567.6107 | 0.1192 | 0.388822 | 0.806585 | Ranbp3l |
| ENSMUSG( | 551.0374 | -0.03538 | 0.749939 | 0.949664 | Ranbp6 |
| ENSMUSG( | 2269.484 | -0.00404 | 0.963838 | 0.995001 | Ranbp9 |
| ENSMUSG( | 4842.92 | 0.124494 | 0.257793 | 0.709192 | Rangap1 |
| ENSMUSG( | 219.5141 | 0.002878 | 0.984832 | 0.997815 | Rangrf |
| ENSMUSG( | 1427.832 | -0.09139 | 0.450703 | 0.835475 | Rap1a |
| ENSMUSG( | 1993.957 | 0.027403 | 0.788207 | 0.959226 | Rap1b |
| ENSMUSG( | 6071.083 | 0.089554 | 0.506521 | 0.867425 | Rap1gap |
| ENSMUSG( | 13068.74 | 0.24959 | 0.002598 | 0.060717 | Rap1gap2 |
| ENSMUSG( | 9922.516 | 0.03043 | 0.719056 | 0.941986 | Rap1gds1 |
| ENSMUSG( | 4013.421 | 0.134367 | 0.112911 | 0.506708 | Rap2a |
| ENSMUSG( | 1488.914 | 0.155963 | 0.209189 | 0.651133 | Rap2b |
| ENSMUSG( | 1619.767 | 0.043768 | 0.767198 | 0.954438 | Rap2c |
| ENSMUSG( | 9755.038 | 0.20834 | 0.007334 | 0.118347 | Rapgef1 |
| ENSMUSG( | 2562.671 | 0.194801 | 0.060544 | 0.379893 | Rapgef2 |
| ENSMUSG( | 2053.936 | 0.169914 | 0.092178 | 0.466065 | Rapgef3 |
| ENSMUSG( | 73724.29 | 0.091952 | 0.32831 | 0.772854 | Rapgef4 |
| ENSMUSG( | 2684.778 | 0.170522 | 0.090631 | 0.462274 | Rapgef5 |
| ENSMUSG( | 790.1025 | -0.04772 | 0.654481 | 0.923227 | Rapgef6 |
| ENSMUSG( | 5994.341 | 0.226155 | 0.008405 | 0.128255 | Rapgefl1 |
| ENSMUSG( | 982.7614 | 0.027955 | 0.783109 | 0.957558 | Raph1 |
| ENSMUSG( | 24.2721 | -0.54167 | 0.174826 | 0.610304 | Rapsn |
| ENSMUSG( | 352.6343 | -0.19759 | 0.200297 | 0.640413 | Rara |
| ENSMUSG( | 312.4008 | -0.59057 | 0.003915 | 0.080172 | Rarb |
| ENSMUSG( | 381.7313 | -0.13832 | 0.309721 | 0.757658 | Rarg |
| ENSMUSG( | 63.68718 | -0.30077 | 0.243012 | 0.690307 | Rarres1 |
| ENSMUSG( | 512.5683 | -0.121 | 0.393156 | 0.809274 | Rarres2 |
| ENSMUSG( | 839.6013 | 0.050836 | 0.620255 | 0.911432 | Rars1 |
| ENSMUSG( | 373.691 | 0.078651 | 0.576915 | 0.89546 | Rars2 |
| ENSMUSG( | 1062.746 | 0.072517 | 0.464474 | 0.843364 | Rasa1 |
| ENSMUSG( | 1727.737 | -0.02396 | 0.85287 | 0.973168 | Rasa2 |
| ENSMUSG( | 2990.208 | 0.035258 | 0.677405 | 0.930542 | Rasa3 |
| ENSMUSG( | 242.0743 | -0.01898 | 0.894443 | 0.984512 | Rasa4 |
| ENSMUSG( | 2968.304 | 0.301551 | 0.014309 | 0.17583 | Rasal1 |
| ENSMUSG( | 1827.156 | 0.162521 | 0.081881 | 0.442044 | Rasal2 |
| ENSMUSG( | 162.683 | -0.08876 | 0.636046 | 0.914873 | Rasal3 |
| ENSMUSG( | 145.2491 | -0.13145 | 0.644139 | 0.917612 | Rasd1 |
| ENSMUSG( | 3565.632 | -0.50375 | 0.001969 | 0.051087 | Rasd2 |
| ENSMUSG( | 11694.34 | 0.1103 | 0.15713 | 0.58551 | Rasgef1a |
| ENSMUSG( | 1131.193 | 0.063592 | 0.535227 | 0.880733 | Rasgef1c |
| ENSMUSG( | 7684.617 | -0.02922 | 0.771585 | 0.954769 | Rasgrf1 |
| ENSMUSG( | 2506.333 | 0.149189 | 0.139428 | 0.555633 | Rasgrf2 |
| ENSMUSG( | 8572.724 | 0.017971 | 0.839666 | 0.971179 | Rasgrp1 |
| ENSMUSG( | 1005.209 | -0.36238 | 0.039779 | 0.307538 | Rasgrp2 |
| ENSMUSG( | 517.4416 | -0.0578 | 0.741048 | 0.948083 | Rasgrp3 |
| ENSMUSG( | 68.35749 | -0.08574 | 0.725766 | 0.944223 | Rasgrp4 |
| ENSMUSG( | 334.4796 | 0.290154 | 0.095238 | 0.474145 | Rasip1 |
| ENSMUSG( | 613.2374 | -0.02798 | 0.799541 | 0.961275 | Rasl10a |
| ENSMUSG( | 22824.6 | 0.179396 | 0.049256 | 0.33882 | Rasl10b |
| ENSMUSG( | 93.60653 | -0.24149 | 0.450232 | 0.835344 | Rasl11a |
| ENSMUSG( | 1339.2 | -0.08575 | 0.585472 | 0.899171 | Rasl11b |
| ENSMUSG( | 74.31596 | -0.19894 | 0.472785 | 0.848388 | Rasl12 |
| ENSMUSG( | 358.3327 | -0.18825 | 0.132097 | 0.544244 | Rassf1 |
| ENSMUSG( | 99.80686 | -0.0853 | 0.671357 | 0.928781 | Rassf10 |
| ENSMUSG( | 6746.401 | 0.109722 | 0.23889 | 0.686534 | Rassf2 |
| ENSMUSG( | 1305.326 | -0.08822 | 0.357166 | 0.788702 | Rassf3 |
| ENSMUSG( | 218.8454 | -0.28394 | 0.055366 | 0.361966 | Rassf4 |
| ENSMUSG( | 570.8779 | 0.151463 | 0.269696 | 0.719729 | Rassf5 |
| ENSMUSG( | 12.5699 | -0.51491 | 0.350109 | NA | Rassf6 |
| ENSMUSG( | 18.8349 | 0.454312 | 0.292688 | NA | Rassf7 |
| ENSMUSG( | 1317.564 | -0.06227 | 0.580184 | 0.896952 | Rassf8 |
| ENSMUSG( | 51.13077 | -0.30026 | 0.393058 | 0.809274 | Rassf9 |
| ENSMUSG( | 1437.117 | 0.282777 | 0.001324 | 0.039614 | Raver2 |
| ENSMUSG( | 1903.214 | 0.053304 | 0.698269 | 0.936084 | Rb1 |
| ENSMUSG( | 2317.882 | 0.110072 | 0.183981 | 0.620925 | Rb1cc1 |
| ENSMUSG( | 401.8835 | -0.27272 | 0.09494 | 0.472949 | Rbak |
| ENSMUSG( | 3060.485 | 0.014134 | 0.878134 | 0.981431 | Rbbp4 |
| ENSMUSG( | 1112.049 | 0.120646 | 0.266482 | 0.716441 | Rbbp5 |
| ENSMUSG( | 2259.568 | -0.09603 | 0.288274 | 0.739023 | Rbbp6 |
| ENSMUSG( | 2636.72 | 0.132698 | 0.156844 | 0.5851 | Rbbp7 |
| ENSMUSG( | 127.9864 | 0.050779 | 0.781037 | 0.957558 | Rbbp8 |
| ENSMUSG( | 1466.417 | 0.005083 | 0.963119 | 0.994911 | Rbbp9 |
| ENSMUSG( | 3948.122 | 0.054656 | 0.523706 | 0.875723 | Rbck1 |
| ENSMUSG( | 734.5637 | -0.05115 | 0.682924 | 0.93205 | Rbfa |
| ENSMUSG( | 17.8173 | -0.18596 | 0.674566 | NA | Rbfaos |
| ENSMUSG( | 3319.466 | -0.09537 | 0.33641 | 0.777 | Rbfox1 |
| ENSMUSG( | 6857.607 | 0.149791 | 0.086176 | 0.451605 | Rbfox2 |
| ENSMUSG( | 7088.968 | 0.071638 | 0.497786 | 0.862341 | Rbfox3 |
| ENSMUSG( | 394.6232 | -0.06657 | 0.604297 | 0.905469 | Rbis |
| ENSMUSG( | 58.80861 | -0.2691 | 0.303125 | 0.751443 | Rbks |
| ENSMUSG( | 89.9802 | -0.20324 | 0.337325 | 0.7773 | Rbl1 |
| ENSMUSG( | 2567.277 | 0.081003 | 0.436995 | 0.829501 | Rbl2 |
| ENSMUSG( | 1680.565 | 0.230357 | 0.040597 | 0.311337 | Rbm10 |
| ENSMUSG( | 110.249 | -0.28679 | 0.140944 | 0.557902 | Rbm11 |
| ENSMUSG( | 255.8886 | 0.250356 | 0.074542 | 0.423195 | Rbm12 |
| ENSMUSG( | 143.4049 | -0.1408 | 0.435003 | 0.829093 | Rbm12b1 |
| ENSMUSG( | 219.4676 | -0.01481 | 0.921497 | 0.990436 | Rbm12b2 |
| ENSMUSG( | 429.7648 | 0.103193 | 0.372392 | 0.798768 | Rbm14 |
| ENSMUSG( | 153.0125 | -0.24988 | 0.142518 | 0.560815 | Rbm15 |
| ENSMUSG( | 1922.799 | -0.02952 | 0.773 | 0.955272 | Rbm15b |
| ENSMUSG( | 1533.924 | 0.013161 | 0.892443 | 0.984455 | Rbm17 |
| ENSMUSG( | 972.5413 | 0.075419 | 0.496164 | 0.861554 | Rbm18 |
| ENSMUSG( | 316.7509 | 0.292111 | 0.041183 | 0.313607 | Rbm19 |
| ENSMUSG( | 156.8855 | -0.03727 | 0.8336 | 0.970333 | Rbm20 |
| ENSMUSG( | 922.443 | -0.03145 | 0.773111 | 0.955272 | Rbm22 |
| ENSMUSG( | 307.4676 | -0.03333 | 0.838077 | 0.971111 | Rbm24 |
| ENSMUSG( | 3188.543 | -0.01123 | 0.913756 | 0.988483 | Rbm25 |
| ENSMUSG( | 1115.889 | -0.07188 | 0.463057 | 0.842422 | Rbm26 |
| ENSMUSG( | 742.9225 | 0.118531 | 0.238165 | 0.686299 | Rbm27 |
| ENSMUSG( | 1822.393 | 0.213042 | 0.017376 | 0.197592 | Rbm28 |
| ENSMUSG( | 1815.924 | -0.20765 | 0.106913 | 0.494634 | Rbm3 |
| ENSMUSG( | 1681.943 | -0.16074 | 0.179718 | 0.617507 | Rbm33 |
| ENSMUSG( | 423.2986 | 0.0129 | 0.915645 | 0.988634 | Rbm34 |
| ENSMUSG( | 117.3862 | -0.0683 | 0.746494 | 0.949289 | Rbm38 |
| ENSMUSG( | 5068.088 | 0.034249 | 0.716877 | 0.941697 | Rbm39 |
| ENSMUSG( | 11.21085 | 0.019777 | 0.972693 | NA | Rbm3os |
| ENSMUSG( | 71.47659 | 0.462333 | 0.052201 | 0.350744 | Rbm4 |
| ENSMUSG( | 166.894 | 0.1471 | 0.36257 | 0.792943 | Rbm41 |
| ENSMUSG( | 283.5669 | 0.246904 | 0.097184 | 0.47759 | Rbm43 |
| ENSMUSG( | 550.4678 | 0.102122 | 0.379031 | 0.801757 | Rbm45 |
| ENSMUSG( | 22.92166 | 0.105218 | 0.840306 | 0.971179 | Rbm46 |
| ENSMUSG( | 36.90408 | -0.24695 | 0.494523 | 0.860776 | Rbm47 |
| ENSMUSG( | 129.4237 | 0.269216 | 0.198784 | 0.63899 | Rbm48 |
| ENSMUSG( | 451.3605 | -0.01704 | 0.899455 | 0.985633 | Rbm4b |
| ENSMUSG( | 4223.997 | -0.11515 | 0.315602 | 0.762671 | Rbm5 |
| ENSMUSG( | 1578.567 | 0.219539 | 0.025763 | 0.247138 | Rbm6 |
| ENSMUSG( | 804.1205 | -0.07474 | 0.456424 | 0.838832 | Rbm7 |
| ENSMUSG( | 1528.299 | 0.08603 | 0.390259 | 0.80794 | Rbm8a |
| ENSMUSG( | 570.3905 | -0.17897 | 0.156061 | 0.58379 | Rbms1 |
| ENSMUSG( | 759.4639 | 0.258404 | 0.02647 | 0.24888 | Rbms2 |
| ENSMUSG( | 364.1357 | -0.07636 | 0.576752 | 0.895432 | Rbms3 |
| ENSMUSG( | 1010.286 | -0.24944 | 0.031711 | 0.271573 | Rbmx |
| ENSMUSG( | 145.5921 | 0.177255 | 0.387738 | 0.805835 | Rbmx2 |
| ENSMUSG( | 432.6553 | 0.001978 | 0.989401 | 0.999111 | Rbmxl1 |
| ENSMUSG( | 1463.532 | 0.085313 | 0.526869 | 0.877267 | Rbp1 |
| ENSMUSG( | 505.0287 | -0.36823 | 0.019407 | 0.209852 | Rbp4 |
| ENSMUSG( | 800.1811 | -0.02268 | 0.821756 | 0.96635 | Rbpj |
| ENSMUSG( | 41.20613 | 0.394443 | 0.259994 | 0.709951 | Rbpjl |
| ENSMUSG( | 100.8974 | -0.01885 | 0.938515 | 0.993415 | Rbpms |
| ENSMUSG( | 150.4382 | 0.035712 | 0.830546 | 0.96974 | Rbpms2 |
| ENSMUSG( | 1574.845 | 0.153484 | 0.099514 | 0.481972 | Rbsn |
| ENSMUSG( | 2631.838 | 0.050545 | 0.547148 | 0.885118 | Rbx1 |
| ENSMUSG( | 1015.055 | -0.00588 | 0.960619 | 0.993991 | Rc3h1 |
| ENSMUSG( | 1516.107 | 0.002411 | 0.977858 | 0.997428 | Rc3h2 |
| ENSMUSG( | 2211.589 | 0.043445 | 0.607473 | 0.906876 | Rcan1 |
| ENSMUSG( | 6293.249 | 0.018089 | 0.820781 | 0.965962 | Rcan2 |
| ENSMUSG( | 2055.902 | -0.03899 | 0.717831 | 0.941697 | Rcan3 |
| ENSMUSG( | 1568.133 | -0.07741 | 0.39389 | 0.809643 | Rcbtb1 |
| ENSMUSG( | 965.3181 | -0.00783 | 0.941183 | 0.993415 | Rcbtb2 |
| ENSMUSG( | 393.3153 | 0.197743 | 0.139965 | 0.556732 | Rcc1 |
| ENSMUSG( | 442.4627 | 0.120914 | 0.381505 | 0.802699 | Rcc1l |
| ENSMUSG( | 3545.412 | 0.022834 | 0.795823 | 0.960062 | Rcc2 |

| | | | | | |
|---|---|---|---|---|---|
| ENSMUSG( | 1317.589 | -0.07784 | 0.573251 | 0.942912 | Rbm8a |
| ENSMUSG( | 558.6919 | 0.03624 | 0.798306 | 0.97681 | Rbms1 |
| ENSMUSG( | 631.9937 | 0.005055 | 0.968656 | 0.99594 | Rbms2 |
| ENSMUSG( | 338.1973 | -0.01986 | 0.896643 | 0.990558 | Rbms3 |
| ENSMUSG( | 1074.604 | 0.203674 | 0.208152 | 0.864412 | Rbmx |
| ENSMUSG( | 124.6101 | -0.01399 | 0.954387 | 0.994798 | Rbmx2 |
| ENSMUSG( | 391.2368 | -0.02158 | 0.89428 | 0.990558 | Rbmxl1 |
| ENSMUSG( | 1200.945 | -0.23108 | 0.175474 | 0.845305 | Rbp1 |
| ENSMUSG( | 488.7103 | -0.17422 | 0.326501 | 0.905371 | Rbp4 |
| ENSMUSG( | 724.6156 | -0.04242 | 0.731178 | 0.967228 | Rbpj |
| ENSMUSG( | 39.52677 | 0.54447 | 0.167959 | 0.840222 | Rbpjl |
| ENSMUSG( | 119.2541 | 0.677411 | 0.01872 | 0.547826 | Rbpms |
| ENSMUSG( | 146.9479 | 0.216478 | 0.264196 | 0.886134 | Rbpms2 |
| ENSMUSG( | 1334.348 | -0.05284 | 0.624396 | 0.953234 | Rbsn |
| ENSMUSG( | 2295.429 | -0.07885 | 0.548599 | 0.938063 | Rbx1 |
| ENSMUSG( | 1013.367 | 0.246115 | 0.062069 | 0.736482 | Rc3h1 |
| ENSMUSG( | 1435.208 | 0.107091 | 0.427353 | 0.917257 | Rc3h2 |
| ENSMUSG( | 1896.774 | -0.13424 | 0.22009 | 0.866104 | Rcan1 |
| ENSMUSG( | 5284.782 | -0.22776 | 0.058749 | 0.728832 | Rcan2 |
| ENSMUSG( | 1788.949 | -0.17896 | 0.151012 | 0.837173 | Rcan3 |
| ENSMUSG( | 1422.845 | -0.08843 | 0.416705 | 0.917257 | Rcbtb1 |
| ENSMUSG( | 847.3388 | -0.12347 | 0.387993 | 0.917257 | Rcbtb2 |
| ENSMUSG( | 344.8003 | 0.09211 | 0.513112 | 0.930358 | Rcc1 |
| ENSMUSG( | 387.7873 | 0.011134 | 0.942575 | 0.994774 | Rcc1l |
| ENSMUSG( | 3122.418 | -0.07768 | 0.504805 | 0.927825 | Rcc2 |
| ENSMUSG( | 244.3124 | 0.178062 | 0.360289 | 0.914223 | Rccd1 |
| ENSMUSG( | 547.2511 | 0.139224 | 0.396516 | 0.917257 | Rce1 |
| ENSMUSG( | 751.0841 | -0.10325 | 0.462645 | 0.923329 | Rchy1 |
| ENSMUSG( | 208.1906 | 0.093241 | 0.624573 | 0.953234 | Rcl1 |
| ENSMUSG( | 1187.359 | 0.156241 | 0.282792 | 0.892412 | Rcn1 |
| ENSMUSG( | 8765.062 | 0.046588 | 0.736644 | 0.967344 | Rcn2 |
| ENSMUSG( | 339.8595 | 0.144707 | 0.407967 | 0.917257 | Rcn3 |
| ENSMUSG( | 514.0831 | 0.04511 | 0.743673 | 0.96951 | Rcor1 |
| ENSMUSG( | 381.72 | 0.131558 | 0.348675 | 0.911652 | Rcor2 |
| ENSMUSG( | 469.992 | 0.12285 | 0.378839 | 0.916995 | Rcor3 |
| ENSMUSG( | 244.6972 | -0.03925 | 0.83743 | 0.981335 | Rcsd1 |
| ENSMUSG( | 9.116288 | 0.237574 | 0.72153 | NA | Rcvrn |
| ENSMUSG( | 23.70237 | 0.575518 | 0.210896 | 0.864412 | Rd3 |
| ENSMUSG( | 425.0249 | 0.260177 | 0.116698 | 0.813384 | Rdh10 |
| ENSMUSG( | 1292.415 | 0.13823 | 0.404013 | 0.917257 | Rdh11 |
| ENSMUSG( | 58.62344 | 0.116079 | 0.679514 | 0.957966 | Rdh12 |
| ENSMUSG( | 496.1068 | 0.186288 | 0.188951 | 0.85698 | Rdh13 |
| ENSMUSG( | 911.047 | 0.1028 | 0.468091 | 0.923329 | Rdh14 |
| ENSMUSG( | 155.5588 | 0.101975 | 0.597092 | 0.946609 | Rdh5 |
| ENSMUSG( | 200.1253 | 0.092612 | 0.593386 | 0.945408 | Rdm1 |
| ENSMUSG( | 5198.222 | 0.004564 | 0.9742 | 0.996416 | Rdx |
| ENSMUSG( | 80.26459 | 0.293209 | 0.309071 | 0.898538 | Rec8 |
| ENSMUSG( | 413.5255 | 0.256988 | 0.11339 | 0.809623 | Reck |
| ENSMUSG( | 463.09 | -0.20386 | 0.136101 | 0.827092 | Recql |
| ENSMUSG( | 77.1931 | 0.324593 | 0.259823 | 0.883971 | Recql4 |
| ENSMUSG( | 557.703 | 0.165847 | 0.332872 | 0.907907 | Recql5 |
| ENSMUSG( | 14.45717 | -0.74763 | 0.22183 | 0.866104 | Redic1 |
| ENSMUSG( | 4167.098 | 0.068893 | 0.650823 | 0.95422 | Reep1 |
| ENSMUSG( | 5671.299 | 0.000567 | 0.996553 | 0.999178 | Reep2 |
| ENSMUSG( | 2401.076 | 0.113586 | 0.48323 | 0.92499 | Reep3 |
| ENSMUSG( | 143.0571 | 0.203011 | 0.334804 | 0.907907 | Reep4 |
| ENSMUSG( | 17637.03 | 0.012155 | 0.932991 | 0.99306 | Reep5 |
| ENSMUSG( | 498.218 | 0.047683 | 0.751589 | 0.97035 | Reep6 |
| ENSMUSG( | 112.3105 | -0.10919 | 0.618449 | 0.952254 | Rel |
| ENSMUSG( | 744.5553 | -0.0266 | 0.836078 | 0.981237 | Rela |
| ENSMUSG( | 174.5357 | -0.04203 | 0.808205 | 0.978622 | Relb |
| ENSMUSG( | 1085.646 | 0.099116 | 0.400998 | 0.917257 | Relch |
| ENSMUSG( | 708.098 | 0.206449 | 0.29596 | 0.898538 | Rell1 |
| ENSMUSG( | 6831.214 | 0.064494 | 0.552405 | 0.939565 | Rell2 |
| ENSMUSG( | 1908.997 | 0.217192 | 0.202764 | 0.864412 | Reln |
| ENSMUSG( | 624.3858 | -0.25693 | 0.068585 | 0.753052 | Relt |
| ENSMUSG( | 15.26453 | 1.14103 | 0.028389 | 0.613486 | Rem1 |
| ENSMUSG( | 163.0653 | -0.25985 | 0.319463 | 0.903461 | Rem2 |
| ENSMUSG( | 165.5335 | -0.25906 | 0.143964 | 0.831311 | Renbp |
| ENSMUSG( | 309.3706 | 0.392195 | 0.05121 | 0.711023 | Reno1 |
| ENSMUSG( | 32.6071 | -0.62063 | 0.089283 | 0.788502 | Rep15 |
| ENSMUSG( | 1017.897 | 0.115574 | 0.367102 | 0.915058 | Repin1 |
| ENSMUSG( | 1011.425 | -0.00681 | 0.958213 | 0.994798 | Reps1 |
| ENSMUSG( | 2231.913 | -0.00033 | 0.997568 | 0.999268 | Reps2 |
| ENSMUSG( | 2764.12 | 0.049923 | 0.701231 | 0.961756 | Rer1 |
| ENSMUSG( | 4025.792 | 0.177821 | 0.08406 | 0.777852 | Rere |
| ENSMUSG( | 102.8104 | 0.265085 | 0.33525 | 0.907907 | Rerg |
| ENSMUSG( | 321.864 | 0.37471 | 0.020896 | 0.571913 | Resf1 |
| ENSMUSG( | 1690.884 | -0.10883 | 0.401717 | 0.917257 | Resp18 |
| ENSMUSG( | 231.1971 | -0.02667 | 0.866388 | 0.987347 | Rest |
| ENSMUSG( | 124.177 | 0.193588 | 0.369371 | 0.915491 | Ret |
| ENSMUSG( | 8.962973 | 0.70563 | 0.3809 | NA | Retn |
| ENSMUSG( | 18.71473 | -0.08991 | 0.89134 | 0.990558 | Retnlg |
| ENSMUSG( | 1474.799 | 0.209853 | 0.295734 | 0.898538 | Retreg1 |
| ENSMUSG( | 7755.853 | 0.124316 | 0.406982 | 0.917257 | Retreg2 |
| ENSMUSG( | 3101.001 | 0.104603 | 0.476743 | 0.923329 | Retreg3 |
| ENSMUSG( | 786.1883 | 0.336223 | 0.024848 | 0.592568 | Retsat |
| ENSMUSG( | 805.8658 | 0.05575 | 0.698975 | 0.961064 | Rev1 |
| ENSMUSG( | 1064.16 | -0.01601 | 0.891383 | 0.990558 | Rev3l |
| ENSMUSG( | 1175.368 | -0.24589 | 0.05283 | 0.71525 | Rex1bd |
| ENSMUSG( | 1759.704 | 0.129327 | 0.278641 | 0.891628 | Rexo1 |
| ENSMUSG( | 1407.911 | -0.13883 | 0.305812 | 0.898538 | Rexo2 |
| ENSMUSG( | 910.0243 | 0.053998 | 0.706854 | 0.963529 | Rexo4 |
| ENSMUSG( | 38.77879 | 0.199728 | 0.57119 | 0.942052 | Rexo5 |
| ENSMUSG( | 1325.077 | -0.13033 | 0.248479 | 0.877904 | Rfc1 |
| ENSMUSG( | 882.7771 | -0.04115 | 0.722001 | 0.966887 | Rfc2 |
| ENSMUSG( | 106.2737 | -0.03916 | 0.856068 | 0.985884 | Rfc3 |
| ENSMUSG( | 72.60274 | -0.00908 | 0.970267 | 0.99594 | Rfc4 |
| ENSMUSG( | 191.4142 | -0.12602 | 0.498129 | 0.927825 | Rfc5 |
| ENSMUSG( | 176.0202 | -0.02709 | 0.897022 | 0.990558 | Rfesd |
| ENSMUSG( | 431.3376 | 0.079168 | 0.60978 | 0.950397 | Rffl |
| ENSMUSG( | 2454.196 | -0.17398 | 0.267284 | 0.886134 | Rfk |
| ENSMUSG( | 25.48625 | 0.869171 | 0.02967 | 0.61992 | Rflna |
| ENSMUSG( | 563.6337 | -0.06585 | 0.637694 | 0.953903 | Rflnb |
| ENSMUSG( | 2923.37 | 0.024034 | 0.809097 | 0.97873 | Rfng |
| ENSMUSG( | 433.6787 | 0.066993 | 0.677739 | 0.957106 | Rft1 |
| ENSMUSG( | 222.7033 | -0.12298 | 0.492473 | 0.927315 | Rftn1 |
| ENSMUSG( | 3440.487 | -0.32373 | 0.025691 | 0.599234 | Rftn2 |
| ENSMUSG( | 543.3939 | 0.089908 | 0.491042 | 0.926604 | Rfwd3 |
| ENSMUSG( | 702.745 | 0.008525 | 0.956083 | 0.994798 | Rfx1 |
| ENSMUSG( | 149.7173 | -0.04652 | 0.813076 | 0.979198 | Rfx2 |
| ENSMUSG( | 1134.182 | 0.069273 | 0.591752 | 0.944975 | Rfx3 |
| ENSMUSG( | 1192.363 | -0.11123 | 0.377813 | 0.916784 | Rfx4 |
| ENSMUSG( | 638.8079 | -0.01976 | 0.879023 | 0.989644 | Rfx5 |
| ENSMUSG( | 901.5244 | -0.04643 | 0.733052 | 0.967228 | Rfx7 |
| ENSMUSG( | 11.62372 | -0.28405 | 0.626805 | 0.953234 | Rfx8 |
| ENSMUSG( | 453.0933 | 0.021527 | 0.886793 | 0.990118 | Rfxank |
| ENSMUSG( | 510.4819 | -0.03216 | 0.837561 | 0.981335 | Rfxap |
| ENSMUSG( | 1009.886 | -0.23721 | 0.043274 | 0.690304 | Rgcc |
| ENSMUSG( | 2846.128 | -0.03348 | 0.751416 | 0.970317 | Rgl1 |
| ENSMUSG( | 1629.966 | 0.107946 | 0.599965 | 0.947322 | Rgl2 |
| ENSMUSG( | 239.9045 | 0.072855 | 0.642768 | 0.953903 | Rgl3 |
| ENSMUSG( | 4057.669 | 0.270095 | 0.068363 | 0.752306 | Rgma |
| ENSMUSG( | 2317.967 | 0.11848 | 0.427037 | 0.917257 | Rgmb |
| ENSMUSG( | 1945.683 | -0.02297 | 0.8248 | 0.979198 | Rgp1 |
| ENSMUSG( | 885.4166 | -0.07534 | 0.564577 | 0.941206 | Rgs10 |
| ENSMUSG( | 984.9856 | 0.144844 | 0.72518 | 0.966887 | Rgs11 |
| ENSMUSG( | 2164.919 | 0.001674 | 0.989843 | 0.997928 | Rgs12 |
| ENSMUSG( | 919.1831 | -0.21575 | 0.094741 | 0.796133 | Rgs14 |
| ENSMUSG( | 170.6759 | -0.11255 | 0.571455 | 0.942116 | Rgs16 |
| ENSMUSG( | 907.7107 | 0.195149 | 0.271156 | 0.888062 | Rgs17 |
| ENSMUSG( | 342.8452 | -0.10514 | 0.507149 | 0.928677 | Rgs19 |
| ENSMUSG( | 704.6134 | -0.07849 | 0.686523 | 0.95899 | Rgs2 |
| ENSMUSG( | 1770.824 | -0.31402 | 0.038752 | 0.679972 | Rgs20 |
| ENSMUSG( | 36.92768 | 0.593086 | 0.074365 | 0.766789 | Rgs22 |
| ENSMUSG( | 729.3414 | 0.09554 | 0.434134 | 0.91869 | Rgs3 |
| ENSMUSG( | 6232.382 | -0.32506 | 0.018558 | 0.545482 | Rgs4 |
| ENSMUSG( | 4385.976 | 0.759561 | 0.114542 | 0.811703 | Rgs5 |
| ENSMUSG( | 2561.065 | -0.19778 | 0.060146 | 0.731553 | Rgs6 |
| ENSMUSG( | 1771.196 | -0.22302 | 0.08479 | 0.779799 | Rgs7 |
| ENSMUSG( | 4898.186 | 0.056532 | 0.650659 | 0.95422 | Rgs7bp |
| ENSMUSG( | 1611.071 | -0.10026 | 0.453035 | 0.922316 | Rgs8 |
| ENSMUSG( | 1381.078 | -0.36 | 0.098561 | 0.798438 | Rgs9 |
| ENSMUSG( | 9.857764 | 0.747536 | 0.260503 | NA | Rgs9bp |
| ENSMUSG( | 93.06984 | 0.093572 | 0.677292 | 0.957106 | Rgsl1 |
| ENSMUSG( | 413.6236 | -0.02368 | 0.895427 | 0.990558 | Rhbdd1 |
| ENSMUSG( | 3388.056 | 0.01776 | 0.882677 | 0.989644 | Rhbdd2 |
| ENSMUSG( | 572.35 | -0.03906 | 0.802636 | 0.97681 | Rhbdd3 |
| ENSMUSG( | 830.0374 | 0.334451 | 0.019294 | 0.554059 | Rhbdf1 |
| ENSMUSG( | 73.1562 | 0.630218 | 0.019891 | 0.558472 | Rhbdf2 |
| ENSMUSG( | 163.3784 | 0.183244 | 0.347936 | 0.911652 | Rhbdl1 |
| ENSMUSG( | 43.46046 | -0.0359 | 0.915933 | 0.992529 | Rhbdl2 |
| ENSMUSG( | 966.3944 | 0.257501 | 0.025012 | 0.594476 | Rhbdl3 |
| ENSMUSG( | 40.41365 | -0.01713 | 0.965127 | 0.995433 | Rhcg |
| ENSMUSG( | 2101.584 | -0.12029 | 0.350348 | 0.911652 | Rheb |
| ENSMUSG( | 137.8634 | 0.175312 | 0.404497 | 0.917257 | Rhebl1 |
| ENSMUSG( | 326.9312 | 0.061655 | 0.669516 | 0.956834 | Rhno1 |
| ENSMUSG( | 21.54982 | 0.151243 | 0.739405 | 0.968458 | Rho |

| | | | | | |
|---|---|---|---|---|---|
| ENSMUSG( | Ric1 | 1406.512 | -0.02821 | 0.787886 | 0.959226 |
| ENSMUSG( | Ric3 | 3659.017 | -0.16255 | 0.181962 | 0.619138 |
| ENSMUSG( | Ric8a | 1879.253 | 0.054123 | 0.543875 | 0.884287 |
| ENSMUSG( | Ric8b | 1083.542 | -0.04324 | 0.747159 | 0.949526 |
| ENSMUSG( | Rictor | 1338.672 | -0.10175 | 0.331251 | 0.77519 |
| ENSMUSG( | Rida | 2581.732 | 0.067874 | 0.603332 | 0.904887 |
| ENSMUSG( | Rif1 | 669.4912 | -0.00437 | 0.971889 | 0.996585 |
| ENSMUSG( | Rigi | 279.3725 | 0.04922 | 0.718879 | 0.941986 |
| ENSMUSG( | Riiad1 | 74.73947 | -0.21436 | 0.345172 | 0.779153 |
| ENSMUSG( | Rilp | 73.26487 | 0.029406 | 0.905342 | 0.986465 |
| ENSMUSG( | Rilpl1 | 1451.06 | 0.186326 | 0.133344 | 0.54617 |
| ENSMUSG( | Rilpl2 | 358.4678 | 0.048478 | 0.732579 | 0.944956 |
| ENSMUSG( | Rimbp2 | 4391.53 | 0.156752 | 0.192987 | 0.632208 |
| ENSMUSG( | Rimbp3 | 37.32234 | -0.05215 | 0.862524 | 0.975055 |
| ENSMUSG( | Rimkla | 332.8722 | -0.36177 | 0.01157 | 0.154961 |
| ENSMUSG( | Rimklb | 1931.429 | 0.084939 | 0.43467 | 0.828978 |
| ENSMUSG( | Rimoc1 | 1702.394 | -0.00145 | 0.990484 | 0.999232 |
| ENSMUSG( | Rims1 | 36657.46 | 0.026924 | 0.770691 | 0.954722 |
| ENSMUSG( | Rims2 | 6674.111 | -0.00463 | 0.958824 | 0.993874 |
| ENSMUSG( | Rims3 | 6037.531 | 0.243366 | 0.006012 | 0.105086 |
| ENSMUSG( | Rims4 | 7216.178 | 0.232318 | 0.022589 | 0.229429 |
| ENSMUSG( | Rin1 | 2248.158 | 0.290003 | 0.010557 | 0.14749 |
| ENSMUSG( | Rin2 | 3184.174 | 0.598016 | 4.29E-12 | 3.49E-09 |
| ENSMUSG( | Rin3 | 145.4296 | 0.414707 | 0.028602 | 0.257732 |
| ENSMUSG( | Ring1 | 829.4811 | 0.081408 | 0.490994 | 0.859739 |
| ENSMUSG( | Rinl | 147.5686 | 0.234935 | 0.257215 | 0.708432 |
| ENSMUSG( | Rint1 | 496.1826 | 0.190016 | 0.091813 | 0.465904 |
| ENSMUSG( | Riok1 | 401.7236 | 0.152889 | 0.219473 | 0.664984 |
| ENSMUSG( | Riok2 | 486.132 | -0.07037 | 0.53172 | 0.879628 |
| ENSMUSG( | Riok3 | 2383.345 | -0.00645 | 0.944127 | 0.993503 |
| ENSMUSG( | Riox1 | 267.6327 | -0.05115 | 0.770456 | 0.954689 |
| ENSMUSG( | Riox2 | 294.6576 | 0.037174 | 0.788253 | 0.959226 |
| ENSMUSG( | Ripk1 | 330.6083 | 0.207239 | 0.124718 | 0.531714 |
| ENSMUSG( | Ripk2 | 184.0438 | 0.197686 | 0.200651 | 0.640661 |
| ENSMUSG( | Ripk3 | 14.34151 | -0.75835 | 0.140899 | NA |
| ENSMUSG( | Ripor1 | 2885.892 | 0.278174 | 0.007551 | 0.120197 |
| ENSMUSG( | Ripor2 | 1464.665 | 0.079889 | 0.407265 | 0.817447 |
| ENSMUSG( | Ripor3 | 17.64042 | -0.22024 | 0.656574 | NA |
| ENSMUSG( | Ripply2 | 40.45021 | -0.18268 | 0.573732 | 0.893584 |
| ENSMUSG( | Rit1 | 983.7058 | 0.134754 | 0.244873 | 0.692492 |
| ENSMUSG( | Rit2 | 1066.441 | -0.01186 | 0.929276 | 0.991848 |
| ENSMUSG( | Rita1 | 508.1288 | 0.120327 | 0.323409 | 0.768488 |
| ENSMUSG( | Rlbp1 | 1964.918 | 0.087527 | 0.49211 | 0.860297 |
| ENSMUSG( | Rlf | 884.7252 | -0.13499 | 0.184574 | 0.621633 |
| ENSMUSG( | Rlig1 | 297.8268 | -0.0418 | 0.791576 | 0.959461 |
| ENSMUSG( | Rlim | 1224.888 | 0.088409 | 0.343151 | 0.779153 |
| ENSMUSG( | Rln1 | 9.27392 | -0.06123 | 0.92786 | NA |
| ENSMUSG( | Rmc1 | 663.0102 | 0.081495 | 0.548893 | 0.885686 |
| ENSMUSG( | Rmdn1 | 174.5017 | -0.02179 | 0.896423 | 0.984868 |
| ENSMUSG( | Rmdn2 | 288.4023 | 0.042127 | 0.767634 | 0.954621 |
| ENSMUSG( | Rmdn3 | 2812.185 | 0.058461 | 0.582516 | 0.897918 |
| ENSMUSG( | Rmi1 | 482.0739 | -0.00801 | 0.954673 | 0.993874 |
| ENSMUSG( | Rmi2 | 11.32506 | -0.42244 | 0.435683 | NA |
| ENSMUSG( | Rmnd1 | 708.5823 | 0.07872 | 0.473605 | 0.848919 |
| ENSMUSG( | Rmnd5a | 3958.512 | 0.091341 | 0.301875 | 0.750166 |
| ENSMUSG( | Rmnd5b | 1183.951 | 7.74E-05 | 0.999398 | 0.99989 |
| ENSMUSG( | Rmst | 110.9629 | 0.680282 | 0.001085 | 0.034804 |
| ENSMUSG( | Rn18s | 176819.4 | -1.17146 | 1.43E-52 | 1.16E-48 |
| ENSMUSG( | Rn7sk | 55.64563 | -1.21053 | 0.001008 | 0.03318 |
| ENSMUSG( | Rnase1 | 45.84547 | -0.28456 | 0.389597 | 0.807428 |
| ENSMUSG( | Rnase4 | 1239.059 | -0.07689 | 0.579092 | 0.896692 |
| ENSMUSG( | Rnase6 | 24.64681 | -1.1296 | 0.013998 | 0.172788 |
| ENSMUSG( | Rnaseh1 | 252.883 | -0.05482 | 0.737786 | 0.947329 |
| ENSMUSG( | Rnaseh2a | 379.1798 | -0.05027 | 0.688434 | 0.933441 |
| ENSMUSG( | Rnaseh2b | 223.2972 | -0.01483 | 0.922397 | 0.99045 |
| ENSMUSG( | Rnaseh2c | 455.6975 | 0.072643 | 0.522697 | 0.875389 |
| ENSMUSG( | Rnasek | 43.89869 | 0.069732 | 0.807444 | 0.963257 |
| ENSMUSG( | Rnasel | 319.2816 | -0.02248 | 0.878598 | 0.981431 |
| ENSMUSG( | Rnaset2a | 9.892792 | -0.67512 | 0.301164 | NA |
| ENSMUSG( | Rnaset2b | 13.13425 | 0.010071 | 0.983775 | NA |
| ENSMUSG( | Rnd1 | 432.7659 | -0.04147 | 0.795778 | 0.960062 |
| ENSMUSG( | Rnd2 | 8047.091 | 0.040311 | 0.693794 | 0.934202 |
| ENSMUSG( | Rnd3 | 430.8951 | -0.25304 | 0.086436 | 0.452045 |
| ENSMUSG( | Rnf10 | 16925.6 | 0.158772 | 0.042109 | 0.316282 |
| ENSMUSG( | Rnf103 | 1572.869 | -0.35118 | 0.021817 | 0.225679 |
| ENSMUSG( | Rnf11 | 7593.461 | 0.050395 | 0.639173 | 0.916434 |
| ENSMUSG( | Rnf111 | 813.0577 | -0.08957 | 0.366194 | 0.794603 |
| ENSMUSG( | Rnf112 | 8939.398 | 0.320285 | 0.056742 | 0.367404 |
| ENSMUSG( | Rnf113a1 | 71.48076 | -0.20895 | 0.424392 | 0.824645 |
| ENSMUSG( | Rnf113a2 | 178.9007 | -0.00997 | 0.950863 | 0.993821 |
| ENSMUSG( | Rnf114 | 3143.314 | 0.097542 | 0.220947 | 0.666476 |
| ENSMUSG( | Rnf115 | 1188.938 | -0.00677 | 0.94612 | 0.99369 |
| ENSMUSG( | Rnf121 | 1048.447 | -0.13059 | 0.267412 | 0.717525 |
| ENSMUSG( | Rnf122 | 450.2316 | -0.60726 | 0.001885 | 0.049559 |
| ENSMUSG( | Rnf123 | 2373.377 | 0.081901 | 0.33556 | 0.776986 |
| ENSMUSG( | Rnf125 | 21.24715 | -0.09528 | 0.842662 | 0.971663 |
| ENSMUSG( | Rnf126 | 2359.546 | -0.00668 | 0.948188 | 0.993821 |
| ENSMUSG( | Rnf128 | 377.5498 | -0.59691 | 0.000323 | 0.015194 |
| ENSMUSG( | Rnf13 | 3924.277 | -0.38179 | 0.011987 | 0.158212 |
| ENSMUSG( | Rnf130 | 5942.738 | -0.25819 | 0.073145 | 0.417454 |
| ENSMUSG( | Rnf135 | 149.573 | 0.175671 | 0.300324 | 0.749479 |
| ENSMUSG( | Rnf138 | 325.441 | -0.01248 | 0.927951 | 0.991229 |
| ENSMUSG( | Rnf138rt1 | 10.33167 | -0.31121 | 0.633239 | NA |
| ENSMUSG( | Rnf139 | 945.9143 | -0.28593 | 0.097858 | 0.478264 |
| ENSMUSG( | Rnf14 | 6824.963 | 0.175569 | 0.044619 | 0.32439 |
| ENSMUSG( | Rnf141 | 3738.408 | -0.01758 | 0.860307 | 0.974842 |
| ENSMUSG( | Rnf144a | 2310.222 | -0.07684 | 0.532963 | 0.879905 |
| ENSMUSG( | Rnf144b | 547.892 | 0.10524 | 0.412938 | 0.819541 |
| ENSMUSG( | Rnf145 | 3476.113 | -0.24034 | 0.070831 | 0.410605 |
| ENSMUSG( | Rnf146 | 1394.785 | 0.121078 | 0.170225 | 0.60523 |
| ENSMUSG( | Rnf149 | 1008.195 | -0.14425 | 0.191172 | 0.630245 |
| ENSMUSG( | Rnf150 | 1448.929 | -0.21821 | 0.124718 | 0.531714 |
| ENSMUSG( | Rnf152 | 296.6349 | 0.014587 | 0.920583 | 0.990307 |
| ENSMUSG( | Rnf157 | 16225.52 | 0.13613 | 0.100776 | 0.484034 |
| ENSMUSG( | Rnf166 | 1035.723 | 0.132542 | 0.173854 | 0.609316 |
| ENSMUSG( | Rnf167 | 2527.152 | -0.21285 | 0.036814 | 0.296095 |
| ENSMUSG( | Rnf168 | 872.8869 | 0.173591 | 0.067052 | 0.400593 |
| ENSMUSG( | Rnf169 | 1160.145 | 0.042784 | 0.709966 | 0.939669 |
| ENSMUSG( | Rnf17 | 18.6777 | 0.687718 | 0.175369 | NA |
| ENSMUSG( | Rnf170 | 822.2009 | -0.12476 | 0.266052 | 0.716441 |
| ENSMUSG( | Rnf180 | 748.728 | 0.163127 | 0.234595 | 0.681922 |
| ENSMUSG( | Rnf181 | 2428.062 | 0.025349 | 0.782687 | 0.957558 |
| ENSMUSG( | Rnf182 | 1043.373 | 0.154642 | 0.238796 | 0.6865 |
| ENSMUSG( | Rnf185 | 596.4889 | -0.00732 | 0.956336 | 0.993874 |
| ENSMUSG( | Rnf187 | 14707.69 | 0.07293 | 0.327933 | 0.772842 |
| ENSMUSG( | Rnf19a | 802.5499 | 0.062101 | 0.558307 | 0.888427 |
| ENSMUSG( | Rnf19b | 1050.885 | 0.016616 | 0.891688 | 0.983979 |
| ENSMUSG( | Rnf2 | 480.3668 | -0.01509 | 0.903559 | 0.986055 |
| ENSMUSG( | Rnf20 | 2978.41 | 0.095171 | 0.256949 | 0.708193 |
| ENSMUSG( | Rnf207 | 187.8333 | 0.034678 | 0.884897 | 0.982815 |
| ENSMUSG( | Rnf208 | 3274.474 | 0.046496 | 0.644535 | 0.917773 |
| ENSMUSG( | Rnf213 | 754.5125 | -0.23962 | 0.077013 | 0.429516 |
| ENSMUSG( | Rnf214 | 1314.487 | 0.080629 | 0.371499 | 0.798768 |
| ENSMUSG( | Rnf215 | 1115.022 | -0.1916 | 0.126817 | 0.534201 |
| ENSMUSG( | Rnf216 | 1505.704 | 0.180611 | 0.045803 | 0.328083 |
| ENSMUSG( | Rnf217 | 559.3215 | 0.003065 | 0.979656 | 0.997428 |
| ENSMUSG( | Rnf220 | 8010.464 | 0.107715 | 0.171117 | 0.605644 |
| ENSMUSG( | Rnf223 | 51.25616 | 0.268389 | 0.314242 | 0.761542 |
| ENSMUSG( | Rnf225 | 25.93104 | -0.22461 | 0.546916 | 0.884974 |
| ENSMUSG( | Rnf227 | 2443.184 | 0.025633 | 0.819811 | 0.965888 |
| ENSMUSG( | Rnf24 | 1327.714 | -0.39187 | 0.009943 | 0.142596 |
| ENSMUSG( | Rnf25 | 994.717 | 0.072469 | 0.605823 | 0.905919 |
| ENSMUSG( | Rnf26 | 369.4785 | -0.26828 | 0.083027 | 0.4446 |
| ENSMUSG( | Rnf31 | 744.675 | 0.048323 | 0.667871 | 0.92763 |
| ENSMUSG( | Rnf32 | 293.431 | 0.093773 | 0.546155 | 0.884974 |
| ENSMUSG( | Rnf34 | 1198.428 | 0.036757 | 0.690946 | 0.933881 |
| ENSMUSG( | Rnf38 | 1687.321 | -0.0377 | 0.664146 | 0.926583 |
| ENSMUSG( | Rnf39 | 114.0808 | -0.04444 | 0.841361 | 0.971179 |
| ENSMUSG( | Rnf4 | 1909.795 | 0.040985 | 0.655206 | 0.923929 |
| ENSMUSG( | Rnf40 | 2193.113 | 0.046753 | 0.605882 | 0.905919 |
| ENSMUSG( | Rnf41 | 2128.035 | 0.119328 | 0.220104 | 0.665863 |
| ENSMUSG( | Rnf43 | 107.7482 | -0.01921 | 0.944499 | 0.993583 |
| ENSMUSG( | Rnf44 | 5941.127 | 0.034416 | 0.665119 | 0.926824 |
| ENSMUSG( | Rnf5 | 2786.764 | -0.01178 | 0.904611 | 0.986275 |
| ENSMUSG( | Rnf6 | 1016.137 | 0.068232 | 0.602143 | 0.904887 |
| ENSMUSG( | Rnf7 | 2014.289 | 0.023169 | 0.822335 | 0.966352 |
| ENSMUSG( | Rnf8 | 206.4838 | -0.00143 | 0.993261 | 0.999369 |
| ENSMUSG( | Rnft1 | 673.3599 | -0.0077 | 0.953807 | 0.993874 |
| ENSMUSG( | Rnft2 | 3694.568 | -0.20294 | 0.100505 | 0.483976 |
| ENSMUSG( | Rngtt | 512.0239 | 0.097917 | 0.416293 | 0.82084 |
| ENSMUSG( | Rnh1 | 1017.013 | -0.00632 | 0.948272 | 0.993821 |
| ENSMUSG( | Rnls | 34.72205 | 0.157569 | 0.653428 | 0.922819 |
| ENSMUSG( | Rnmt | 1158.768 | -0.00365 | 0.975413 | 0.997302 |
| ENSMUSG( | Rnpc3 | 308.0821 | -0.15343 | 0.328848 | 0.773233 |
| ENSMUSG( | Rnpep | 1288.716 | 0.193714 | 0.045525 | 0.327592 |
| ENSMUSG( | Rnpepl1 | 1989.457 | 0.107633 | 0.1972 | 0.636662 |
| ENSMUSG( | Rnps1 | 2461.221 | 0.082441 | 0.498398 | 0.862846 |
| ENSMUSG( | Rny1 | 33.35874 | -1.22743 | 0.022438 | 0.228899 |

| | | | | | |
|---|---|---|---|---|---|
| ENSMUSG( | Ric1 | 1330.529 | 0.080447 | 0.483677 | 0.925155 |
| ENSMUSG( | Ric3 | 3552.63 | 0.027365 | 0.833029 | 0.981118 |
| ENSMUSG( | Ric8a | 1649.616 | -0.05232 | 0.621447 | 0.953199 |
| ENSMUSG( | Ric8b | 982.0038 | -0.06213 | 0.691632 | 0.960161 |
| ENSMUSG( | Rictor | 1330.67 | 0.150013 | 0.207433 | 0.864412 |
| ENSMUSG( | Rida | 2122.165 | -0.24317 | 0.143977 | 0.831311 |
| ENSMUSG( | Rif1 | 657.8351 | 0.211376 | 0.133497 | 0.822912 |
| ENSMUSG( | Rigi | 262.5678 | 0.137151 | 0.394574 | 0.917257 |
| ENSMUSG( | Riiad1 | 66.85445 | -0.26046 | 0.299113 | 0.898538 |
| ENSMUSG( | Rilp | 62.31967 | -0.17261 | 0.536476 | 0.937108 |
| ENSMUSG( | Rilpl1 | 1259.34 | 0.047361 | 0.750411 | 0.970113 |
| ENSMUSG( | Rilpl2 | 309.5368 | -0.10866 | 0.504019 | 0.927825 |
| ENSMUSG( | Rimbp2 | 3943.401 | 0.116903 | 0.371698 | 0.915491 |
| ENSMUSG( | Rimbp3 | 32.33968 | -0.17799 | 0.615286 | 0.951177 |
| ENSMUSG( | Rimkla | 327.0692 | -0.12821 | 0.41892 | 0.917257 |
| ENSMUSG( | Rimklb | 1703.278 | -0.01247 | 0.926384 | 0.99306 |
| ENSMUSG( | Rimoc1 | 1658.011 | 0.184787 | 0.179891 | 0.850372 |
| ENSMUSG( | Rims1 | 30858.74 | -0.21014 | 0.076923 | 0.770558 |
| ENSMUSG( | Rims2 | 5883.86 | -0.10381 | 0.354796 | 0.912411 |
| ENSMUSG( | Rims3 | 4944.087 | -0.05393 | 0.640086 | 0.953903 |
| ENSMUSG( | Rims4 | 6009.326 | -0.01555 | 0.905273 | 0.991089 |
| ENSMUSG( | Rin1 | 1963.585 | 0.179678 | 0.182721 | 0.850579 |
| ENSMUSG( | Rin2 | 2419.322 | 0.132525 | 0.220486 | 0.866104 |
| ENSMUSG( | Rin3 | 138.5996 | 0.539473 | 0.012079 | 0.445877 |
| ENSMUSG( | Ring1 | 757.5356 | 0.089347 | 0.560907 | 0.940254 |
| ENSMUSG( | Rinl | 117.6647 | -0.13566 | 0.567838 | 0.941777 |
| ENSMUSG( | Rint1 | 437.2123 | 0.102528 | 0.476175 | 0.923329 |
| ENSMUSG( | Riok1 | 374.0259 | 0.216763 | 0.153945 | 0.837724 |
| ENSMUSG( | Riok2 | 433.7797 | -0.13892 | 0.321503 | 0.904019 |
| ENSMUSG( | Riok3 | 2171.154 | -0.00799 | 0.942519 | 0.994774 |
| ENSMUSG( | Riox1 | 232.9152 | -0.19157 | 0.30864 | 0.898538 |
| ENSMUSG( | Riox2 | 242.2224 | -0.26829 | 0.107794 | 0.804969 |
| ENSMUSG( | Ripk1 | 294.0754 | 0.147139 | 0.339069 | 0.907972 |
| ENSMUSG( | Ripk2 | 181.0307 | 0.392333 | 0.027004 | 0.603041 |
| ENSMUSG( | Ripk3 | 15.78279 | -0.14557 | 0.778345 | 0.976106 |
| ENSMUSG( | Ripor1 | 2450.859 | 0.08686 | 0.488621 | 0.92554 |
| ENSMUSG( | Ripor2 | 1316.109 | 0.039496 | 0.742958 | 0.969415 |
| ENSMUSG( | Ripor3 | 19.45023 | 0.355246 | 0.475096 | 0.923329 |
| ENSMUSG( | Ripply2 | 32.38548 | -0.63171 | 0.091922 | 0.791889 |
| ENSMUSG( | Rit1 | 846.6067 | -0.03048 | 0.826181 | 0.979795 |
| ENSMUSG( | Rit2 | 905.0449 | -0.23302 | 0.213845 | 0.864421 |
| ENSMUSG( | Rita1 | 454.3396 | 0.07332 | 0.589166 | 0.944975 |
| ENSMUSG( | Rlbp1 | 1639.46 | -0.17253 | 0.218668 | 0.866104 |
| ENSMUSG( | Rlf | 831.9771 | -0.04025 | 0.729072 | 0.966913 |
| ENSMUSG( | Rlig1 | 272.2835 | -0.03297 | 0.843994 | 0.983571 |
| ENSMUSG( | Rlim | 1068.992 | -0.03875 | 0.760728 | 0.972695 |
| ENSMUSG( | Rln1 | 11.267 | 0.70247 | 0.244316 | NA |
| ENSMUSG( | Rmc1 | 578.4697 | -0.04276 | 0.797814 | 0.97681 |
| ENSMUSG( | Rmdn1 | 165.3225 | 0.081901 | 0.650045 | 0.95422 |
| ENSMUSG( | Rmdn2 | 260.7008 | 0.02533 | 0.879662 | 0.989644 |
| ENSMUSG( | Rmdn3 | 2488.373 | -0.0254 | 0.835635 | 0.981237 |
| ENSMUSG( | Rmi1 | 436.4218 | -0.03162 | 0.829739 | 0.980224 |
| ENSMUSG( | Rmi2 | 11.15302 | -0.15517 | 0.789836 | NA |
| ENSMUSG( | Rmnd1 | 666.7584 | 0.16699 | 0.31355 | 0.900883 |
| ENSMUSG( | Rmnd5a | 3527.236 | 0.026672 | 0.807987 | 0.978622 |
| ENSMUSG( | Rmnd5b | 1092.901 | 0.037469 | 0.742686 | 0.969415 |
| ENSMUSG( | Rmst | 89.3287 | 0.366875 | 0.158753 | 0.837724 |
| ENSMUSG( | Rn18s | 144151 | -1.77476 | 2.83E-54 | 4.96E-50 |
| ENSMUSG( | Rn7sk | 42.85587 | -2.20772 | 6.25E-07 | 0.000342 |
| ENSMUSG( | Rnase1 | 38.41039 | -0.52814 | 0.168444 | 0.840222 |
| ENSMUSG( | Rnase4 | 1290.644 | 0.300328 | 0.055683 | 0.723362 |
| ENSMUSG( | Rnase6 | 23.79593 | -0.89478 | 0.050707 | 0.709454 |
| ENSMUSG( | Rnaseh1 | 240.2876 | 0.079836 | 0.648115 | 0.95422 |
| ENSMUSG( | Rnaseh2a | 350.9725 | -0.00979 | 0.946203 | 0.994798 |
| ENSMUSG( | Rnaseh2b | 212.4627 | 0.097094 | 0.590131 | 0.944975 |
| ENSMUSG( | Rnaseh2c | 402.186 | -0.01627 | 0.905626 | 0.991089 |
| ENSMUSG( | Rnasek | 41.01453 | 0.157362 | 0.622882 | 0.953234 |
| ENSMUSG( | Rnasel | 299.3566 | 0.057784 | 0.728606 | 0.966913 |
| ENSMUSG( | Rnaset2a | 10.30302 | -0.2597 | 0.666031 | NA |
| ENSMUSG( | Rnaset2b | 13.48713 | 0.363163 | 0.489319 | 0.925673 |
| ENSMUSG( | Rnd1 | 383.4745 | -0.13194 | 0.466149 | 0.923329 |
| ENSMUSG( | Rnd2 | 6826.876 | -0.17275 | 0.193815 | 0.860096 |
| ENSMUSG( | Rnd3 | 443.3177 | 0.099013 | 0.518907 | 0.930939 |
| ENSMUSG( | Rnf10 | 14793.64 | 0.043165 | 0.650864 | 0.95422 |
| ENSMUSG( | Rnf103 | 1723.837 | 0.199793 | 0.237346 | 0.872173 |
| ENSMUSG( | Rnf11 | 6762.585 | -0.01779 | 0.90819 | 0.991958 |
| ENSMUSG( | Rnf111 | 772.5701 | 0.03063 | 0.797668 | 0.97681 |
| ENSMUSG( | Rnf112 | 7644.412 | 0.1477 | 0.423866 | 0.917257 |
| ENSMUSG( | Rnf113a1 | 64.36444 | -0.23793 | 0.414617 | 0.917257 |
| ENSMUSG( | Rnf113a2 | 160.3664 | -0.06194 | 0.73578 | 0.967344 |
| ENSMUSG( | Rnf114 | 2752.966 | -0.0147 | 0.883355 | 0.989644 |
| ENSMUSG( | Rnf115 | 1110.366 | 0.059826 | 0.65425 | 0.955015 |
| ENSMUSG( | Rnf121 | 1070.656 | 0.198369 | 0.1522 | 0.837724 |
| ENSMUSG( | Rnf122 | 524.8535 | 0.171489 | 0.433314 | 0.918434 |
| ENSMUSG( | Rnf123 | 2164.924 | 0.086631 | 0.415043 | 0.917257 |
| ENSMUSG( | Rnf125 | 23.67422 | 0.486607 | 0.322959 | 0.904023 |
| ENSMUSG( | Rnf126 | 2116.006 | -0.05426 | 0.651697 | 0.954576 |
| ENSMUSG( | Rnf128 | 455.2649 | 0.260298 | 0.209781 | 0.864412 |
| ENSMUSG( | Rnf13 | 4292.033 | 0.168626 | 0.349235 | 0.911652 |
| ENSMUSG( | Rnf130 | 6240.086 | 0.163139 | 0.348277 | 0.911652 |
| ENSMUSG( | Rnf135 | 123.9102 | -0.08344 | 0.684759 | 0.958678 |
| ENSMUSG( | Rnf138 | 313.3417 | 0.133765 | 0.437433 | 0.920144 |
| ENSMUSG( | Rnf138rt1 | 10.2435 | 0.015141 | 0.98353 | NA |
| ENSMUSG( | Rnf139 | 1017.07 | 0.204897 | 0.293182 | 0.898538 |
| ENSMUSG( | Rnf14 | 5752.177 | -0.04479 | 0.679595 | 0.957966 |
| ENSMUSG( | Rnf141 | 3187.94 | -0.21969 | 0.127027 | 0.817278 |
| ENSMUSG( | Rnf144a | 1958.689 | -0.29984 | 0.024389 | 0.590702 |
| ENSMUSG( | Rnf144b | 526.4605 | 0.258688 | 0.142199 | 0.830709 |
| ENSMUSG( | Rnf145 | 3712.593 | 0.223577 | 0.148866 | 0.836241 |
| ENSMUSG( | Rnf146 | 1220.064 | 0.002439 | 0.985295 | 0.997084 |
| ENSMUSG( | Rnf149 | 988.5351 | 0.070192 | 0.567968 | 0.941777 |
| ENSMUSG( | Rnf150 | 1617.515 | 0.359083 | 0.036084 | 0.668395 |
| ENSMUSG( | Rnf152 | 267.3627 | -0.01938 | 0.906763 | 0.991517 |
| ENSMUSG( | Rnf157 | 14313.9 | 0.04722 | 0.654821 | 0.955015 |
| ENSMUSG( | Rnf166 | 897.8973 | -0.01176 | 0.931407 | 0.99306 |
| ENSMUSG( | Rnf167 | 2579.065 | 0.122203 | 0.302712 | 0.898538 |
| ENSMUSG( | Rnf168 | 809.5899 | 0.216845 | 0.126835 | 0.816972 |
| ENSMUSG( | Rnf169 | 1065.594 | 0.069209 | 0.587766 | 0.944975 |
| ENSMUSG( | Rnf17 | 12.5105 | -0.15614 | 0.806272 | 0.977809 |
| ENSMUSG( | Rnf170 | 813.2378 | 0.114016 | 0.368771 | 0.915491 |
| ENSMUSG( | Rnf180 | 661.0725 | 0.068991 | 0.680143 | 0.9581 |
| ENSMUSG( | Rnf181 | 2117.724 | -0.10556 | 0.375202 | 0.915491 |
| ENSMUSG( | Rnf182 | 827.4726 | -0.25875 | 0.094963 | 0.796133 |
| ENSMUSG( | Rnf185 | 524.4714 | -0.10824 | 0.452381 | 0.922316 |
| ENSMUSG( | Rnf187 | 12222.06 | -0.19892 | 0.039933 | 0.681706 |
| ENSMUSG( | Rnf19a | 739.3499 | 0.093579 | 0.422508 | 0.917257 |
| ENSMUSG( | Rnf19b | 882.3416 | -0.23091 | 0.0958 | 0.796133 |
| ENSMUSG( | Rnf2 | 448.3814 | 0.047455 | 0.754896 | 0.971098 |
| ENSMUSG( | Rnf20 | 2627.499 | 0.002397 | 0.981687 | 0.996756 |
| ENSMUSG( | Rnf207 | 163.5082 | -0.11282 | 0.704398 | 0.962677 |
| ENSMUSG( | Rnf208 | 2917.854 | -0.01771 | 0.881395 | 0.989644 |
| ENSMUSG( | Rnf213 | 700.9895 | -0.17173 | 0.275271 | 0.889984 |
| ENSMUSG( | Rnf214 | 1153.144 | -0.02936 | 0.791258 | 0.97681 |
| ENSMUSG( | Rnf215 | 1197 | 0.279686 | 0.074414 | 0.766789 |
| ENSMUSG( | Rnf216 | 1336.538 | 0.108169 | 0.313411 | 0.900883 |
| ENSMUSG( | Rnf217 | 536.3399 | 0.138675 | 0.375897 | 0.915491 |
| ENSMUSG( | Rnf220 | 6935.511 | -0.03886 | 0.709696 | 0.96416 |
| ENSMUSG( | Rnf223 | 56.21831 | 0.718967 | 0.010091 | 0.40957 |
| ENSMUSG( | Rnf225 | 23.30373 | -0.24072 | 0.564181 | 0.941206 |
| ENSMUSG( | Rnf227 | 2118.675 | -0.12441 | 0.395388 | 0.917257 |
| ENSMUSG( | Rnf24 | 1422.904 | 0.106456 | 0.556264 | 0.939565 |
| ENSMUSG( | Rnf25 | 859.5353 | -0.08363 | 0.633683 | 0.953889 |
| ENSMUSG( | Rnf26 | 382.4991 | 0.104173 | 0.542614 | 0.938034 |
| ENSMUSG( | Rnf31 | 671.1844 | 0.020668 | 0.878135 | 0.989644 |
| ENSMUSG( | Rnf32 | 252.6792 | -0.0582 | 0.739017 | 0.968312 |
| ENSMUSG( | Rnf34 | 1075.825 | -0.01026 | 0.938105 | 0.993731 |
| ENSMUSG( | Rnf38 | 1535.989 | -0.04209 | 0.706333 | 0.963467 |
| ENSMUSG( | Rnf39 | 96.80015 | -0.26888 | 0.290334 | 0.897019 |
| ENSMUSG( | Rnf4 | 1679.677 | -0.05964 | 0.582748 | 0.944266 |
| ENSMUSG( | Rnf40 | 1955.898 | -0.01582 | 0.887237 | 0.990263 |
| ENSMUSG( | Rnf41 | 1875.895 | 0.026229 | 0.807256 | 0.978393 |
| ENSMUSG( | Rnf43 | 115.5658 | 0.438144 | 0.106464 | 0.804969 |
| ENSMUSG( | Rnf44 | 5446.319 | 0.052158 | 0.598537 | 0.946859 |
| ENSMUSG( | Rnf5 | 2365.175 | -0.22915 | 0.09007 | 0.790741 |
| ENSMUSG( | Rnf6 | 917.0567 | 0.041099 | 0.776282 | 0.975717 |
| ENSMUSG( | Rnf7 | 1748.628 | -0.12308 | 0.38402 | 0.917257 |
| ENSMUSG( | Rnf8 | 179.4927 | -0.13743 | 0.470888 | 0.923329 |
| ENSMUSG( | Rnft1 | 654.244 | 0.173207 | 0.234718 | 0.871042 |
| ENSMUSG( | Rnft2 | 3700.983 | 0.0806 | 0.579237 | 0.943942 |
| ENSMUSG( | Rngtt | 458.0263 | 0.043533 | 0.758696 | 0.971858 |
| ENSMUSG( | Rnh1 | 889.1932 | -0.1252 | 0.293741 | 0.898538 |
| ENSMUSG( | Rnls | 30.78944 | 0.054287 | 0.888929 | 0.990558 |
| ENSMUSG( | Rnmt | 1087.89 | 0.084277 | 0.538312 | 0.937216 |
| ENSMUSG( | Rnpc3 | 370.7382 | 0.60117 | 0.002889 | 0.217231 |
| ENSMUSG( | Rnpep | 1083.177 | -0.03179 | 0.787793 | 0.97637 |
| ENSMUSG( | Rnpepl1 | 1719.952 | -0.04093 | 0.71662 | 0.965807 |
| ENSMUSG( | Rnps1 | 2158.763 | -0.02771 | 0.829186 | 0.980224 |
| ENSMUSG( | Rny1 | 24.33302 | -2.82741 | 9.48E-06 | 0.003679 |

| | | | | | |
|---|---|---|---|---|---|
| ENSMUSG( | 256.0541 | 0.037806 | 0.793492 | 0.959799 | Rccd1 |
| ENSMUSG( | 534.0348 | -0.21161 | 0.134414 | 0.547651 | Rce1 |
| ENSMUSG( | 869.1336 | 0.05566 | 0.639116 | 0.916434 | Rchy1 |
| ENSMUSG( | 231.5864 | 0.141082 | 0.360891 | 0.792797 | Rcl1 |
| ENSMUSG( | 1152.01 | -0.2041 | 0.134326 | 0.547472 | Rcn1 |
| ENSMUSG( | 9233.355 | -0.07228 | 0.570324 | 0.893192 | Rcn2 |
| ENSMUSG( | 341.5626 | -0.11884 | 0.379346 | 0.801757 | Rcn3 |
| ENSMUSG( | 574.4313 | 0.089749 | 0.431237 | 0.827055 | Rcor1 |
| ENSMUSG( | 386.3448 | -0.10041 | 0.483722 | 0.85636 | Rcor2 |
| ENSMUSG( | 475.4282 | -0.11383 | 0.375529 | 0.800845 | Rcor3 |
| ENSMUSG( | 269.9426 | -0.03915 | 0.801052 | 0.96188 | Rcsd1 |
| ENSMUSG( | 10.48894 | 0.302596 | 0.612406 | NA | Rcvrn |
| ENSMUSG( | 16.20069 | -0.85934 | 0.066824 | NA | Rd3 |
| ENSMUSG( | 416.6182 | -0.06043 | 0.692105 | 0.934101 | Rdh10 |
| ENSMUSG( | 1151.775 | -0.50827 | 0.001191 | 0.037142 | Rdh11 |
| ENSMUSG( | 51.54103 | -0.56921 | 0.042543 | 0.316636 | Rdh12 |
| ENSMUSG( | 534.9422 | 0.139386 | 0.277196 | 0.727175 | Rdh13 |
| ENSMUSG( | 889.0477 | -0.24786 | 0.057619 | 0.370247 | Rdh14 |
| ENSMUSG( | 178.296 | 0.216566 | 0.212759 | 0.6567 | Rdh5 |
| ENSMUSG( | 225.9844 | 0.171688 | 0.258829 | 0.709814 | Rdm1 |
| ENSMUSG( | 5610.252 | -0.04374 | 0.757469 | 0.951513 | Rdx |
| ENSMUSG( | 97.95878 | 0.574968 | 0.009132 | 0.134662 | Rec8 |
| ENSMUSG( | 412.7998 | -0.0156 | 0.901593 | 0.985633 | Reck |
| ENSMUSG( | 548.0003 | 0.021294 | 0.852872 | 0.973168 | Recql |
| ENSMUSG( | 87.03752 | 0.382384 | 0.127253 | 0.534725 | Recql4 |
| ENSMUSG( | 605.5029 | 0.129428 | 0.274094 | 0.724254 | Recql5 |
| ENSMUSG( | 19.76548 | 0.030917 | 0.950809 | NA | Redic1 |
| ENSMUSG( | 3920.607 | -0.40664 | 0.001767 | 0.047422 | Reep1 |
| ENSMUSG( | 5649.658 | -0.29706 | 0.00764 | 0.120942 | Reep2 |
| ENSMUSG( | 2382.317 | -0.18379 | 0.21159 | 0.654835 | Reep3 |
| ENSMUSG( | 133.3155 | -0.27977 | 0.176973 | 0.61401 | Reep4 |
| ENSMUSG( | 17669.9 | -0.2659 | 0.036449 | 0.294207 | Reep5 |
| ENSMUSG( | 510.9878 | -0.15764 | 0.203417 | 0.643631 | Reep6 |
| ENSMUSG( | 120.1887 | -0.1962 | 0.316717 | 0.763648 | Rel |
| ENSMUSG( | 813.4692 | -0.04436 | 0.656452 | 0.923929 | Rela |
| ENSMUSG( | 187.9067 | -0.10298 | 0.543076 | 0.883579 | Relb |
| ENSMUSG( | 1166.179 | 0.038966 | 0.6961 | 0.935007 | Relch |
| ENSMUSG( | 635.8793 | -0.40103 | 0.019693 | 0.21168 | Rell1 |
| ENSMUSG( | 7530.962 | 0.076028 | 0.414917 | 0.820184 | Rell2 |
| ENSMUSG( | 1713.968 | -0.38249 | 0.006588 | 0.111324 | Reln |
| ENSMUSG( | 653.2848 | -0.41788 | 0.000707 | 0.025719 | Relt |
| ENSMUSG( | 11.64861 | 0.28338 | 0.610838 | NA | Rem1 |
| ENSMUSG( | 162.5022 | -0.58563 | 0.021806 | 0.225679 | Rem2 |
| ENSMUSG( | 197.7206 | -0.00926 | 0.950998 | 0.993821 | Renbp |
| ENSMUSG( | 282.5803 | -0.12522 | 0.397602 | 0.812127 | Reno1 |
| ENSMUSG( | 44.16196 | 0.038352 | 0.899383 | 0.985633 | Rep15 |
| ENSMUSG( | 1084.698 | 0.030486 | 0.784377 | 0.958003 | Repin1 |
| ENSMUSG( | 1169.315 | 0.140569 | 0.190216 | 0.629268 | Reps1 |
| ENSMUSG( | 2570.027 | 0.135807 | 0.097542 | 0.477842 | Reps2 |
| ENSMUSG( | 2804.112 | -0.18381 | 0.105527 | 0.492417 | Rer1 |
| ENSMUSG( | 4401.785 | 0.168082 | 0.033511 | 0.280615 | Rere |
| ENSMUSG( | 108.5945 | 0.174835 | 0.465484 | 0.844064 | Rerg |
| ENSMUSG( | 317.3522 | 0.078987 | 0.552522 | 0.886141 | Resf1 |
| ENSMUSG( | 1697.968 | -0.38956 | 0.000473 | 0.019871 | Resp18 |
| ENSMUSG( | 239.2977 | -0.20566 | 0.141493 | 0.558606 | Rest |
| ENSMUSG( | 108.6057 | -0.50374 | 0.033133 | 0.278602 | Ret |
| ENSMUSG( | 12.13385 | 1.201246 | 0.045284 | NA | Retn |
| ENSMUSG( | 15.10013 | -1.12741 | 0.106763 | NA | Retnlg |
| ENSMUSG( | 1250.314 | -0.59114 | 0.000839 | 0.02897 | Retreg1 |
| ENSMUSG( | 7643.602 | -0.19388 | 0.12138 | 0.525713 | Retreg2 |
| ENSMUSG( | 3163.197 | -0.11138 | 0.394464 | 0.810072 | Retreg3 |
| ENSMUSG( | 813.8298 | 0.176112 | 0.154582 | 0.581712 | Retsat |
| ENSMUSG( | 837.0076 | -0.10749 | 0.390657 | 0.808144 | Rev1 |
| ENSMUSG( | 1056.013 | -0.32714 | 0.001932 | 0.050384 | Rev3l |
| ENSMUSG( | 1415.482 | 0.036085 | 0.692568 | 0.934101 | Rex1bd |
| ENSMUSG( | 1925.782 | 0.118349 | 0.209926 | 0.652297 | Rexo1 |
| ENSMUSG( | 1614.488 | -0.00436 | 0.963545 | 0.994922 | Rexo2 |
| ENSMUSG( | 1020.784 | 0.110044 | 0.332881 | 0.775602 | Rexo4 |
| ENSMUSG( | 42.08372 | 0.186225 | 0.522191 | 0.875186 | Rexo5 |
| ENSMUSG( | 1610.319 | 0.164005 | 0.081895 | 0.442044 | Rfc1 |
| ENSMUSG( | 1027.898 | 0.125473 | 0.201922 | 0.641612 | Rfc2 |
| ENSMUSG( | 109.0265 | -0.24557 | 0.208316 | 0.650319 | Rfc3 |
| ENSMUSG( | 87.01623 | 0.244952 | 0.267846 | 0.718054 | Rfc4 |
| ENSMUSG( | 228.4094 | 0.103851 | 0.541934 | 0.883182 | Rfc5 |
| ENSMUSG( | 194.7129 | 0.009498 | 0.95602 | 0.993874 | Rfesd |
| ENSMUSG( | 412.6357 | -0.34501 | 0.00599 | 0.104817 | Rffl |
| ENSMUSG( | 2855.078 | 0.003038 | 0.978415 | 0.997428 | Rfk |
| ENSMUSG( | 23.76651 | 0.495129 | 0.217128 | 0.661995 | Rflna |
| ENSMUSG( | 653.7944 | 0.089527 | 0.504981 | 0.866883 | Rflnb |
| ENSMUSG( | 3209.64 | 0.025248 | 0.754238 | 0.950981 | Rfng |
| ENSMUSG( | 418.7687 | -0.3281 | 0.02489 | 0.242767 | Rft1 |
| ENSMUSG( | 246.3619 | -0.10944 | 0.449629 | 0.835234 | Rftn1 |
| ENSMUSG( | 4322.072 | 0.085679 | 0.487226 | 0.858812 | Rftn2 |
| ENSMUSG( | 616.5993 | 0.183764 | 0.08829 | 0.455783 | Rfwd3 |
| ENSMUSG( | 842.2833 | 0.255264 | 0.061163 | 0.381822 | Rfx1 |
| ENSMUSG( | 167.1448 | -0.00302 | 0.985774 | 0.998326 | Rfx2 |
| ENSMUSG( | 1240.154 | 0.061135 | 0.561094 | 0.888934 | Rfx3 |
| ENSMUSG( | 1408.766 | 0.104002 | 0.370631 | 0.797829 | Rfx4 |
| ENSMUSG( | 683.596 | -0.09188 | 0.406202 | 0.816525 | Rfx5 |
| ENSMUSG( | 936.812 | -0.20526 | 0.041335 | 0.314189 | Rfx7 |
| ENSMUSG( | 17.43806 | 0.62232 | 0.159117 | NA | Rfx8 |
| ENSMUSG( | 474.0032 | -0.12512 | 0.275239 | 0.72509 | Rfxank |
| ENSMUSG( | 578.768 | 0.067381 | 0.569548 | 0.893041 | Rfxap |
| ENSMUSG( | 1202.868 | 0.008688 | 0.935274 | 0.993331 | Rgcc |
| ENSMUSG( | 3279.538 | 0.10499 | 0.242682 | 0.690216 | Rgl1 |
| ENSMUSG( | 1959.763 | 0.354935 | 0.042154 | 0.316282 | Rgl2 |
| ENSMUSG( | 252.2173 | -0.05005 | 0.731797 | 0.944956 | Rgl3 |
| ENSMUSG( | 3741.574 | -0.23381 | 0.074677 | 0.423665 | Rgma |
| ENSMUSG( | 2146.085 | -0.4009 | 0.001747 | 0.047136 | Rgmb |
| ENSMUSG( | 2229.817 | 0.100169 | 0.253281 | 0.703461 | Rgp1 |
| ENSMUSG( | 1052.087 | 0.153863 | 0.174136 | 0.60991 | Rgs10 |
| ENSMUSG( | 1197.207 | 0.418063 | 0.244171 | 0.691435 | Rgs11 |
| ENSMUSG( | 2411.548 | 0.040725 | 0.685201 | 0.932981 | Rgs12 |
| ENSMUSG( | 1055.387 | -0.07952 | 0.464935 | 0.843635 | Rgs14 |
| ENSMUSG( | 202.5612 | 0.103534 | 0.535017 | 0.880733 | Rgs16 |
| ENSMUSG( | 997.5712 | 0.206256 | 0.088686 | 0.45666 | Rgs17 |
| ENSMUSG( | 394.9996 | 0.044242 | 0.725174 | 0.944096 | Rgs19 |
| ENSMUSG( | 739.9327 | -0.20854 | 0.245612 | 0.693322 | Rgs2 |
| ENSMUSG( | 2172.858 | 0.028567 | 0.779812 | 0.957136 | Rgs20 |
| ENSMUSG( | 32.64111 | 0.009499 | 0.977994 | 0.997428 | Rgs22 |
| ENSMUSG( | 786.7508 | 0.043539 | 0.693279 | 0.934202 | Rgs3 |
| ENSMUSG( | 7170.864 | -0.17433 | 0.165159 | 0.596597 | Rgs4 |
| ENSMUSG( | 3573.145 | -0.0126 | 0.971722 | 0.996543 | Rgs5 |
| ENSMUSG( | 3041.417 | 0.03762 | 0.656596 | 0.923929 | Rgs6 |
| ENSMUSG( | 2112.876 | 0.028862 | 0.789088 | 0.959299 | Rgs7 |
| ENSMUSG( | 5260.462 | -0.00383 | 0.963582 | 0.994922 | Rgs7bp |
| ENSMUSG( | 1806.441 | -0.03877 | 0.738299 | 0.947329 | Rgs8 |
| ENSMUSG( | 1338.861 | -0.79993 | 0.03567 | 0.290384 | Rgs9 |
| ENSMUSG( | 9.082488 | 0.306145 | 0.637247 | NA | Rgs9bp |
| ENSMUSG( | 109.1612 | 0.273572 | 0.18097 | 0.618094 | Rgsl1 |
| ENSMUSG( | 391.896 | -0.49874 | 0.001606 | 0.044935 | Rhbdd1 |
| ENSMUSG( | 3496.833 | -0.16696 | 0.106611 | 0.494174 | Rhbdd2 |
| ENSMUSG( | 587.959 | -0.24918 | 0.044666 | 0.324591 | Rhbdd3 |
| ENSMUSG( | 818.6686 | 0.040767 | 0.719581 | 0.941986 | Rhbdf1 |
| ENSMUSG( | 64.28267 | 0.046546 | 0.865134 | 0.975565 | Rhbdf2 |
| ENSMUSG( | 174.8843 | 0.099981 | 0.580697 | 0.897294 | Rhbdl1 |
| ENSMUSG( | 28.54944 | -2.48474 | 2.77E-08 | 9.55E-06 | Rhbdl2 |
| ENSMUSG( | 967.883 | 0.006249 | 0.95204 | 0.993821 | Rhbdl3 |
| ENSMUSG( | 45.04572 | 0.071149 | 0.844691 | 0.972023 | Rhcg |
| ENSMUSG( | 2445.083 | 0.053829 | 0.595115 | 0.903007 | Rheb |
| ENSMUSG( | 141.8852 | -0.01271 | 0.94803 | 0.993821 | Rhebl1 |
| ENSMUSG( | 369.0489 | 0.135287 | 0.283265 | 0.73186 | Rhno1 |
| ENSMUSG( | 29.7152 | 0.731474 | 0.117099 | 0.516406 | Rho |
| ENSMUSG( | 6411.053 | 0.031956 | 0.754597 | 0.951007 | Rhoa |
| ENSMUSG( | 5997.049 | 0.002556 | 0.982568 | 0.997512 | Rhob |
| ENSMUSG( | 229.9782 | 0.011842 | 0.937844 | 0.993415 | Rhobtb1 |
| ENSMUSG( | 3919.969 | 0.05507 | 0.476919 | 0.851851 | Rhobtb2 |
| ENSMUSG( | 1646.939 | 0.001066 | 0.99444 | 0.999369 | Rhobtb3 |
| ENSMUSG( | 1143.235 | -0.08207 | 0.459222 | 0.840244 | Rhoc |
| ENSMUSG( | 33.09224 | -0.11796 | 0.715851 | 0.941484 | Rhod |
| ENSMUSG( | 796.2128 | 0.238797 | 0.062632 | 0.38686 | Rhof |
| ENSMUSG( | 1080.685 | -0.18847 | 0.06448 | 0.393628 | Rhog |
| ENSMUSG( | 33.46448 | 0.046385 | 0.898435 | 0.985633 | Rhoh |
| ENSMUSG( | 215.0622 | -0.08222 | 0.65932 | 0.924506 | Rhoj |
| ENSMUSG( | 2429.771 | -0.21619 | 0.086429 | 0.452045 | Rhoq |
| ENSMUSG( | 1614.453 | 0.075202 | 0.378048 | 0.801561 | Rhot1 |
| ENSMUSG( | 1341.945 | 0.057161 | 0.542001 | 0.883182 | Rhot2 |
| ENSMUSG( | 1997.79 | 0.17288 | 0.129505 | 0.539023 | Rhou |
| ENSMUSG( | 112.1324 | -0.03455 | 0.867056 | 0.976541 | Rhov |
| ENSMUSG( | 398.7293 | 0.091519 | 0.475827 | 0.850919 | Rhpn1 |
| ENSMUSG( | 265.5476 | 0.104008 | 0.441372 | 0.832449 | Rhpn2 |
| ENSMUSG( | 11.50801 | 0.028021 | 0.958897 | NA | Ribc1 |
| ENSMUSG( | 1406.512 | -0.02821 | 0.787886 | 0.959226 | Ric1 |
| ENSMUSG( | 3659.017 | -0.16255 | 0.181962 | 0.619138 | Ric3 |
| ENSMUSG( | 1879.253 | 0.054123 | 0.543875 | 0.884287 | Ric8a |
| ENSMUSG( | 1083.542 | -0.04324 | 0.747159 | 0.949526 | Ric8b |
| ENSMUSG( | 1338.672 | -0.10175 | 0.331251 | 0.77519 | Rictor |
| ENSMUSG( | 2581.732 | 0.067874 | 0.603332 | 0.904887 | Rida |

| | | | | | |
|---|---|---|---|---|---|
| ENSMUSG( | 5630.671 | -0.07724 | 0.545078 | 0.938063 | Rhoa |
| ENSMUSG( | 5267.273 | -0.10555 | 0.432624 | 0.918256 | Rhob |
| ENSMUSG( | 228.1464 | 0.259777 | 0.156901 | 0.837724 | Rhobtb1 |
| ENSMUSG( | 3747.313 | 0.188504 | 0.055016 | 0.723362 | Rhobtb2 |
| ENSMUSG( | 1454.424 | -0.09474 | 0.556076 | 0.939565 | Rhobtb3 |
| ENSMUSG( | 1000.743 | -0.2044 | 0.075453 | 0.767994 | Rhoc |
| ENSMUSG( | 27.27177 | -0.45403 | 0.23411 | 0.869203 | Rhod |
| ENSMUSG( | 692.8192 | 0.114603 | 0.392122 | 0.917257 | Rhof |
| ENSMUSG( | 996.7728 | -0.15039 | 0.233593 | 0.868507 | Rhog |
| ENSMUSG( | 31.40606 | 0.149901 | 0.671147 | 0.95694 | Rhoh |
| ENSMUSG( | 174.6124 | -0.43681 | 0.035178 | 0.660461 | Rhoj |
| ENSMUSG( | 2162.025 | -0.29575 | 0.062942 | 0.736482 | Rhoq |
| ENSMUSG( | 1453.383 | 0.041171 | 0.69837 | 0.960908 | Rhot1 |
| ENSMUSG( | 1167.09 | -0.07983 | 0.510009 | 0.929089 | Rhot2 |
| ENSMUSG( | 1634.081 | -0.13581 | 0.275458 | 0.889984 | Rhou |
| ENSMUSG( | 94.71762 | -0.27859 | 0.248848 | 0.877904 | Rhov |
| ENSMUSG( | 354.3943 | 0.019438 | 0.894318 | 0.990558 | Rhpn1 |
| ENSMUSG( | 230.6171 | -0.04431 | 0.800155 | 0.97681 | Rhpn2 |
| ENSMUSG( | 9.559443 | -0.27062 | 0.664467 | NA | Ribc1 |
| ENSMUSG( | 1330.529 | 0.080447 | 0.483677 | 0.925155 | Ric1 |
| ENSMUSG( | 3552.63 | 0.027365 | 0.833029 | 0.981118 | Ric3 |
| ENSMUSG( | 1649.616 | -0.05232 | 0.621447 | 0.953199 | Ric8a |
| ENSMUSG( | 982.0038 | -0.06213 | 0.691632 | 0.960161 | Ric8b |
| ENSMUSG( | 1330.67 | 0.150013 | 0.207433 | 0.864412 | Rictor |
| ENSMUSG( | 2122.165 | -0.24317 | 0.143977 | 0.831311 | Rida |
| ENSMUSG( | 657.8351 | 0.211376 | 0.133497 | 0.822912 | Rif1 |
| ENSMUSG( | 262.5678 | 0.137151 | 0.394574 | 0.917257 | Rigi |
| ENSMUSG( | 66.85445 | -0.26046 | 0.299113 | 0.898538 | Riiad1 |
| ENSMUSG( | 62.31967 | -0.17261 | 0.536476 | 0.937108 | Rilp |
| ENSMUSG( | 1259.34 | 0.047361 | 0.750411 | 0.970113 | Rilpl1 |
| ENSMUSG( | 309.5368 | -0.10866 | 0.504019 | 0.927825 | Rilpl2 |
| ENSMUSG( | 3943.401 | 0.116903 | 0.371698 | 0.915491 | Rimbp2 |
| ENSMUSG( | 32.33968 | -0.17799 | 0.615286 | 0.951177 | Rimbp3 |
| ENSMUSG( | 327.0692 | -0.12821 | 0.41892 | 0.917257 | Rimkla |
| ENSMUSG( | 1703.278 | -0.01247 | 0.926384 | 0.99306 | Rimklb |
| ENSMUSG( | 1658.011 | 0.184787 | 0.179891 | 0.850372 | Rimoc1 |
| ENSMUSG( | 30858.74 | -0.21014 | 0.076923 | 0.770558 | Rims1 |
| ENSMUSG( | 5883.86 | -0.10381 | 0.354796 | 0.912411 | Rims2 |
| ENSMUSG( | 4944.087 | -0.05393 | 0.640086 | 0.953903 | Rims3 |
| ENSMUSG( | 6009.326 | -0.01555 | 0.905273 | 0.991089 | Rims4 |
| ENSMUSG( | 1963.585 | 0.179678 | 0.182721 | 0.850579 | Rin1 |
| ENSMUSG( | 2419.322 | 0.132525 | 0.220486 | 0.866104 | Rin2 |
| ENSMUSG( | 138.5996 | 0.539473 | 0.012079 | 0.445877 | Rin3 |
| ENSMUSG( | 757.5356 | 0.089347 | 0.560907 | 0.940254 | Ring1 |
| ENSMUSG( | 117.6647 | -0.13566 | 0.567838 | 0.941777 | Rinl |
| ENSMUSG( | 437.2123 | 0.102528 | 0.476175 | 0.923329 | Rint1 |
| ENSMUSG( | 374.0259 | 0.216763 | 0.153945 | 0.837724 | Riok1 |
| ENSMUSG( | 433.7797 | -0.13892 | 0.321503 | 0.904019 | Riok2 |
| ENSMUSG( | 2171.154 | -0.00799 | 0.942519 | 0.994774 | Riok3 |
| ENSMUSG( | 232.9152 | -0.19157 | 0.30864 | 0.898538 | Riox1 |
| ENSMUSG( | 242.2224 | -0.26829 | 0.107794 | 0.804969 | Riox2 |
| ENSMUSG( | 294.0754 | 0.147139 | 0.339069 | 0.907972 | Ripk1 |
| ENSMUSG( | 181.0307 | 0.392333 | 0.027004 | 0.603041 | Ripk2 |
| ENSMUSG( | 15.78279 | -0.14557 | 0.778345 | 0.976106 | Ripk3 |
| ENSMUSG( | 2450.859 | 0.08686 | 0.488621 | 0.92554 | Ripor1 |
| ENSMUSG( | 1316.109 | 0.039496 | 0.742958 | 0.969415 | Ripor2 |
| ENSMUSG( | 19.45023 | 0.355246 | 0.475096 | 0.923329 | Ripor3 |
| ENSMUSG( | 32.38548 | -0.63171 | 0.091922 | 0.791889 | Ripply2 |
| ENSMUSG( | 10.55397 | 0.496262 | 0.425707 | NA | Ripply3 |
| ENSMUSG( | 846.6067 | -0.03048 | 0.826181 | 0.979795 | Rit1 |
| ENSMUSG( | 905.0449 | -0.23302 | 0.213845 | 0.864421 | Rit2 |
| ENSMUSG( | 454.3396 | 0.07332 | 0.589166 | 0.944975 | Rita1 |
| ENSMUSG( | 1639.46 | -0.17253 | 0.218668 | 0.866104 | Rlbp1 |
| ENSMUSG( | 831.9771 | -0.04025 | 0.729072 | 0.966913 | Rlf |
| ENSMUSG( | 272.2835 | -0.03297 | 0.843994 | 0.983571 | Rlig1 |
| ENSMUSG( | 1068.992 | -0.03875 | 0.760728 | 0.972695 | Rlim |
| ENSMUSG( | 11.267 | 0.70247 | 0.244316 | NA | Rln1 |
| ENSMUSG( | 578.4697 | -0.04276 | 0.797814 | 0.97681 | Rmc1 |
| ENSMUSG( | 165.3225 | 0.081901 | 0.650045 | 0.95422 | Rmdn1 |
| ENSMUSG( | 260.7008 | 0.02533 | 0.879662 | 0.989644 | Rmdn2 |
| ENSMUSG( | 2488.373 | -0.0254 | 0.835635 | 0.981237 | Rmdn3 |
| ENSMUSG( | 436.4218 | -0.03162 | 0.829739 | 0.980224 | Rmi1 |
| ENSMUSG( | 11.15302 | -0.15517 | 0.789836 | NA | Rmi2 |
| ENSMUSG( | 666.7584 | 0.16699 | 0.31355 | 0.900883 | Rmnd1 |
| ENSMUSG( | 3527.236 | 0.026672 | 0.807987 | 0.978622 | Rmnd5a |
| ENSMUSG( | 1092.901 | 0.037469 | 0.742686 | 0.969415 | Rmnd5b |
| ENSMUSG( | 89.3287 | 0.366875 | 0.158753 | 0.837724 | Rmst |
| ENSMUSG( | 144151 | -1.77476 | 2.83E-54 | 4.96E-50 | Rn18s |
| ENSMUSG( | 42.85587 | -2.20772 | 6.25E-07 | 0.000342 | Rn7sk |
| ENSMUSG( | 38.41039 | -0.52814 | 0.168444 | 0.840222 | Rnase1 |
| ENSMUSG( | 1290.644 | 0.300328 | 0.055683 | 0.723362 | Rnase4 |
| ENSMUSG( | 23.79593 | -0.89478 | 0.050707 | 0.709454 | Rnase6 |
| ENSMUSG( | 240.2876 | 0.079836 | 0.648115 | 0.95422 | Rnaseh1 |
| ENSMUSG( | 350.9725 | -0.00979 | 0.946203 | 0.994798 | Rnaseh2a |
| ENSMUSG( | 212.4627 | 0.097094 | 0.590131 | 0.944975 | Rnaseh2b |
| ENSMUSG( | 402.186 | -0.01627 | 0.905626 | 0.991089 | Rnaseh2c |
| ENSMUSG( | 41.01453 | 0.157362 | 0.622882 | 0.953234 | Rnasek |
| ENSMUSG( | 299.3566 | 0.057784 | 0.728606 | 0.966913 | Rnasel |
| ENSMUSG( | 10.30302 | -0.2597 | 0.666031 | NA | Rnaset2a |
| ENSMUSG( | 13.48713 | 0.363163 | 0.489319 | 0.925673 | Rnaset2b |
| ENSMUSG( | 383.4745 | -0.13194 | 0.466149 | 0.923329 | Rnd1 |
| ENSMUSG( | 6826.876 | -0.17275 | 0.193815 | 0.860096 | Rnd2 |
| ENSMUSG( | 443.3177 | 0.099013 | 0.518907 | 0.930939 | Rnd3 |
| ENSMUSG( | 14793.64 | 0.043165 | 0.650864 | 0.95422 | Rnf10 |
| ENSMUSG( | 1723.837 | 0.199793 | 0.237346 | 0.872173 | Rnf103 |
| ENSMUSG( | 6762.585 | -0.01779 | 0.90819 | 0.991958 | Rnf11 |
| ENSMUSG( | 772.5701 | 0.03063 | 0.797668 | 0.97681 | Rnf111 |
| ENSMUSG( | 7644.412 | 0.1477 | 0.423866 | 0.917257 | Rnf112 |
| ENSMUSG( | 64.36444 | -0.23793 | 0.414617 | 0.917257 | Rnf113a1 |
| ENSMUSG( | 160.3664 | -0.06194 | 0.73578 | 0.967344 | Rnf113a2 |
| ENSMUSG( | 2752.966 | -0.0147 | 0.883355 | 0.989644 | Rnf114 |
| ENSMUSG( | 1110.366 | 0.059826 | 0.65425 | 0.955015 | Rnf115 |
| ENSMUSG( | 1070.656 | 0.198369 | 0.1522 | 0.837724 | Rnf121 |
| ENSMUSG( | 524.8535 | 0.171489 | 0.433314 | 0.918434 | Rnf122 |
| ENSMUSG( | 2164.924 | 0.086631 | 0.415043 | 0.917257 | Rnf123 |
| ENSMUSG( | 23.67422 | 0.486607 | 0.322959 | 0.904023 | Rnf125 |
| ENSMUSG( | 2116.006 | -0.05426 | 0.651697 | 0.954576 | Rnf126 |
| ENSMUSG( | 455.2649 | 0.260298 | 0.209781 | 0.864412 | Rnf128 |
| ENSMUSG( | 4292.033 | 0.168626 | 0.349235 | 0.911652 | Rnf13 |
| ENSMUSG( | 6240.086 | 0.163139 | 0.348277 | 0.911652 | Rnf130 |
| ENSMUSG( | 123.9102 | -0.08344 | 0.684759 | 0.958678 | Rnf135 |
| ENSMUSG( | 313.3417 | 0.133765 | 0.437433 | 0.920144 | Rnf138 |
| ENSMUSG( | 10.2435 | 0.015141 | 0.98353 | NA | Rnf138rt1 |
| ENSMUSG( | 1017.07 | 0.204897 | 0.293182 | 0.898538 | Rnf139 |
| ENSMUSG( | 5752.177 | -0.04479 | 0.679595 | 0.957966 | Rnf14 |
| ENSMUSG( | 3187.94 | -0.21969 | 0.127027 | 0.817278 | Rnf141 |
| ENSMUSG( | 1958.689 | -0.29984 | 0.024389 | 0.590702 | Rnf144a |
| ENSMUSG( | 526.4605 | 0.258688 | 0.142199 | 0.830709 | Rnf144b |
| ENSMUSG( | 3712.593 | 0.223577 | 0.148866 | 0.836241 | Rnf145 |
| ENSMUSG( | 1220.064 | 0.002439 | 0.985295 | 0.997084 | Rnf146 |
| ENSMUSG( | 988.5351 | 0.070192 | 0.567968 | 0.941777 | Rnf149 |
| ENSMUSG( | 1617.515 | 0.359083 | 0.036084 | 0.668395 | Rnf150 |
| ENSMUSG( | 267.3627 | -0.01938 | 0.906763 | 0.991517 | Rnf152 |
| ENSMUSG( | 14313.9 | 0.04722 | 0.654821 | 0.955015 | Rnf157 |
| ENSMUSG( | 897.8973 | -0.01176 | 0.931407 | 0.99306 | Rnf166 |
| ENSMUSG( | 2579.065 | 0.122203 | 0.302712 | 0.898538 | Rnf167 |
| ENSMUSG( | 809.5899 | 0.216845 | 0.126835 | 0.816972 | Rnf168 |
| ENSMUSG( | 1065.594 | 0.069209 | 0.587766 | 0.944975 | Rnf169 |
| ENSMUSG( | 12.5105 | -0.15614 | 0.806272 | 0.977809 | Rnf17 |
| ENSMUSG( | 813.2378 | 0.114016 | 0.368771 | 0.915491 | Rnf170 |
| ENSMUSG( | 661.0725 | 0.068991 | 0.680143 | 0.9581 | Rnf180 |
| ENSMUSG( | 2117.724 | -0.10556 | 0.375202 | 0.915491 | Rnf181 |
| ENSMUSG( | 827.4726 | -0.25875 | 0.094963 | 0.796133 | Rnf182 |
| ENSMUSG( | 524.4714 | -0.10824 | 0.452381 | 0.922316 | Rnf185 |
| ENSMUSG( | 12222.06 | -0.19892 | 0.039933 | 0.681706 | Rnf187 |
| ENSMUSG( | 739.3499 | 0.093579 | 0.422508 | 0.917257 | Rnf19a |
| ENSMUSG( | 882.3416 | -0.23091 | 0.0958 | 0.796133 | Rnf19b |
| ENSMUSG( | 448.3814 | 0.047455 | 0.754896 | 0.971098 | Rnf2 |
| ENSMUSG( | 2627.499 | 0.002397 | 0.981687 | 0.996756 | Rnf20 |
| ENSMUSG( | 163.5082 | -0.11282 | 0.704398 | 0.962677 | Rnf207 |
| ENSMUSG( | 2917.854 | -0.01771 | 0.881395 | 0.989644 | Rnf208 |
| ENSMUSG( | 700.9895 | -0.17173 | 0.275271 | 0.889984 | Rnf213 |
| ENSMUSG( | 1153.144 | -0.02936 | 0.791258 | 0.97681 | Rnf214 |
| ENSMUSG( | 1197 | 0.279686 | 0.074414 | 0.766789 | Rnf215 |
| ENSMUSG( | 1336.538 | 0.108169 | 0.313411 | 0.900883 | Rnf216 |
| ENSMUSG( | 536.3399 | 0.138675 | 0.375897 | 0.915491 | Rnf217 |
| ENSMUSG( | 6935.511 | -0.03886 | 0.709696 | 0.96416 | Rnf220 |
| ENSMUSG( | 56.21831 | 0.718967 | 0.010091 | 0.40957 | Rnf223 |
| ENSMUSG( | 23.30373 | -0.24072 | 0.564181 | 0.941206 | Rnf225 |
| ENSMUSG( | 2118.675 | -0.12441 | 0.395388 | 0.917257 | Rnf227 |
| ENSMUSG( | 1422.904 | 0.106456 | 0.556264 | 0.939565 | Rnf24 |
| ENSMUSG( | 859.5353 | -0.08363 | 0.633683 | 0.953889 | Rnf25 |
| ENSMUSG( | 382.4991 | 0.104173 | 0.542614 | 0.938034 | Rnf26 |
| ENSMUSG( | 671.1844 | 0.020668 | 0.878135 | 0.989644 | Rnf31 |
| ENSMUSG( | 252.6792 | -0.0582 | 0.739017 | 0.968312 | Rnf32 |
| ENSMUSG( | 1075.825 | -0.01026 | 0.938105 | 0.993731 | Rnf34 |
| ENSMUSG( | 1535.989 | -0.04209 | 0.706333 | 0.963467 | Rnf38 |
| ENSMUSG( | 96.80015 | -0.26888 | 0.290334 | 0.897019 | Rnf39 |

| | | | | | |
|---|---|---|---|---|---|
| ENSMUSG( | Ro60 | 1223.437 | 0.013211 | 0.914348 | 0.988483 |
| ENSMUSG( | Robo1 | 1553.436 | -0.2209 | 0.10403 | 0.489802 |
| ENSMUSG( | Robo2 | 2820.319 | -0.45852 | 0.001692 | 0.046443 |
| ENSMUSG( | Robo3 | 1257.37 | 0.768007 | 0.035793 | 0.290916 |
| ENSMUSG( | Robo4 | 370.1655 | 0.226521 | 0.227535 | 0.674441 |
| ENSMUSG( | Rock1 | 1340.973 | 0.089048 | 0.496646 | 0.86175 |
| ENSMUSG( | Rock2 | 4459.048 | 0.090231 | 0.292802 | 0.742763 |
| ENSMUSG( | Rogdi | 2894.35 | 0.182019 | 0.180156 | 0.617634 |
| ENSMUSG( | Rom1 | 523.9815 | -0.10738 | 0.401601 | 0.814923 |
| ENSMUSG( | Romo1 | 1521.447 | -0.02515 | 0.789634 | 0.959461 |
| ENSMUSG( | Ropn1l | 34.67869 | -0.20708 | 0.533177 | 0.879905 |
| ENSMUSG( | Ror1 | 55.82557 | -0.09773 | 0.718512 | 0.941911 |
| ENSMUSG( | Ror2 | 41.11521 | -0.01995 | 0.948781 | 0.993821 |
| ENSMUSG( | Rora | 2383.532 | 0.090674 | 0.416837 | 0.8209 |
| ENSMUSG( | Rorb | 1655.577 | -0.10691 | 0.342256 | 0.778828 |
| ENSMUSG( | Rorc | 115.4549 | -0.19943 | 0.29407 | 0.743459 |
| ENSMUSG( | Rp2 | 656.2671 | 0.076096 | 0.591181 | 0.90113 |
| ENSMUSG( | Rp9 | 965.1964 | 0.094288 | 0.411814 | 0.81942 |
| ENSMUSG( | Rpa1 | 1823.222 | 0.224627 | 0.033184 | 0.278882 |
| ENSMUSG( | Rpa2 | 340.0576 | 0.039814 | 0.789168 | 0.959299 |
| ENSMUSG( | Rpa3 | 162.1989 | -0.11257 | 0.497084 | 0.861754 |
| ENSMUSG( | Rpain | 291.9784 | -0.00706 | 0.968439 | 0.996369 |
| ENSMUSG( | Rpap1 | 696.3964 | 0.177297 | 0.098915 | 0.480935 |
| ENSMUSG( | Rpap2 | 541.4064 | 0.023565 | 0.852865 | 0.973168 |
| ENSMUSG( | Rpap3 | 559.9078 | 0.159249 | 0.201445 | 0.641362 |
| ENSMUSG( | Rpe | 614.4978 | 0.108449 | 0.36905 | 0.796581 |
| ENSMUSG( | Rpe65 | 85.38052 | 0.211612 | 0.368704 | 0.796356 |
| ENSMUSG( | Rpf1 | 313.4799 | 0.133943 | 0.320487 | 0.765921 |
| ENSMUSG( | Rpf2 | 339.6436 | 0.027815 | 0.837642 | 0.971111 |
| ENSMUSG( | Rpgr | 348.4614 | 0.022486 | 0.879635 | 0.981436 |
| ENSMUSG( | Rpgrip1 | 24.81347 | 0.025564 | 0.947832 | 0.993821 |
| ENSMUSG( | Rpgrip1l | 419.8062 | 0.239539 | 0.05238 | 0.35132 |
| ENSMUSG( | Rph3a | 30550.33 | -0.02913 | 0.689067 | 0.933441 |
| ENSMUSG( | Rph3al | 70.22893 | 0.357479 | 0.144383 | 0.56431 |
| ENSMUSG( | Rpia | 243.5359 | -0.13541 | 0.400813 | 0.81387 |
| ENSMUSG( | Rpl10 | 9448.112 | 0.106306 | 0.267438 | 0.717525 |
| ENSMUSG( | Rpl10a | 4630.183 | 0.044512 | 0.566983 | 0.891644 |
| ENSMUSG( | Rpl10a-ps1 | 104.1193 | -0.03168 | 0.888783 | 0.983434 |
| ENSMUSG( | Rpl11 | 8913.379 | 0.090095 | 0.282387 | 0.73186 |
| ENSMUSG( | Rpl12 | 13276.85 | 0.06961 | 0.409135 | 0.818258 |
| ENSMUSG( | Rpl13 | 16234.77 | 0.079701 | 0.326411 | 0.771327 |
| ENSMUSG( | Rpl13a | 1180.678 | 0.174533 | 0.092866 | 0.468228 |
| ENSMUSG( | Rpl13-ps6 | 17.44141 | -0.23162 | 0.595865 | NA |
| ENSMUSG( | Rpl14 | 8852.132 | 0.072081 | 0.379849 | 0.801757 |
| ENSMUSG( | Rpl14-ps1 | 79.87483 | 0.095171 | 0.661662 | 0.925271 |
| ENSMUSG( | Rpl15 | 4706.748 | 0.034587 | 0.70299 | 0.937735 |
| ENSMUSG( | Rpl17 | 13645.03 | 0.091949 | 0.265785 | 0.716441 |
| ENSMUSG( | Rpl17-ps10 | 93.77396 | 0.122053 | 0.564975 | 0.890735 |
| ENSMUSG( | Rpl17-ps5 | 11.25199 | 0.077398 | 0.887663 | NA |
| ENSMUSG( | Rpl18 | 5629.009 | 0.067575 | 0.430951 | 0.826896 |
| ENSMUSG( | Rpl18a | 10749.58 | 0.067963 | 0.396197 | 0.811615 |
| ENSMUSG( | Rpl18a-ps1 | 13.91912 | -1.20404 | 0.022017 | NA |
| ENSMUSG( | Rpl18-ps1 | 17.2684 | -0.08898 | 0.8414 | NA |
| ENSMUSG( | Rpl19 | 11956.59 | 0.072961 | 0.384918 | 0.80404 |
| ENSMUSG( | Rpl19-ps11 | 97.28339 | 0.075795 | 0.713596 | 0.940866 |
| ENSMUSG( | Rpl21 | 14555.99 | 0.124562 | 0.138474 | 0.553773 |
| ENSMUSG( | Rpl21-ps11 | 27.13102 | 0.322531 | 0.362279 | 0.792943 |
| ENSMUSG( | Rpl21-ps8 | 20.76819 | 0.060458 | 0.885407 | NA |
| ENSMUSG( | Rpl22 | 5360.809 | 0.039158 | 0.65889 | 0.924506 |
| ENSMUSG( | Rpl22l1 | 1301.939 | -0.01419 | 0.906914 | 0.987391 |
| ENSMUSG( | Rpl23 | 9250.528 | 0.061222 | 0.485554 | 0.857637 |
| ENSMUSG( | Rpl23a | 9185.722 | 0.124581 | 0.148484 | 0.571935 |
| ENSMUSG( | Rpl23a-ps3 | 27.29178 | 0.0893 | 0.806267 | 0.962804 |
| ENSMUSG( | Rpl24 | 63.25084 | -0.0607 | 0.804768 | 0.962473 |
| ENSMUSG( | Rpl26 | 11204.03 | 0.132789 | 0.119344 | 0.523036 |
| ENSMUSG( | Rpl27 | 2782.203 | 0.02292 | 0.795692 | 0.960062 |
| ENSMUSG( | Rpl27a | 9031.711 | 0.109851 | 0.235283 | 0.682423 |
| ENSMUSG( | Rpl28 | 9303.741 | 0.039519 | 0.622458 | 0.911432 |
| ENSMUSG( | Rpl28-ps1 | 51.72437 | 0.030068 | 0.910055 | 0.9881 |
| ENSMUSG( | Rpl28-ps3 | 36.85917 | -0.25167 | 0.413841 | 0.819998 |
| ENSMUSG( | Rpl29 | 8351.473 | 0.080927 | 0.328191 | 0.772854 |
| ENSMUSG( | Rpl3 | 18152.41 | 0.06525 | 0.3962 | 0.811615 |
| ENSMUSG( | Rpl30 | 6303.662 | 0.043716 | 0.652744 | 0.922498 |
| ENSMUSG( | Rpl31 | 10957.92 | 0.051224 | 0.506268 | 0.867425 |
| ENSMUSG( | Rpl32 | 7413.378 | 0.089178 | 0.336301 | 0.777 |
| ENSMUSG( | Rpl32l | 28.06241 | 0.04143 | 0.911778 | 0.988483 |
| ENSMUSG( | Rpl34 | 5060.723 | 0.112631 | 0.218502 | 0.664312 |
| ENSMUSG( | Rpl34-ps1 | 50.89771 | 0.240905 | 0.357936 | 0.789465 |
| ENSMUSG( | Rpl35 | 5389.33 | 0.075014 | 0.404499 | 0.816132 |
| ENSMUSG( | Rpl35a | 5971.658 | 0.134699 | 0.17214 | 0.607115 |
| ENSMUSG( | Rpl35rt | 439.1068 | 0.093524 | 0.422241 | 0.823508 |
| ENSMUSG( | Rpl36 | 5695.019 | 0.065087 | 0.523061 | 0.875628 |
| ENSMUSG( | Rpl36a | 3604.278 | 0.069977 | 0.405939 | 0.81652 |
| ENSMUSG( | Rpl36al | 11.5924 | -0.3337 | 0.594309 | NA |
| ENSMUSG( | Rpl36a-ps1 | 30.05313 | -0.32387 | 0.372263 | 0.798768 |
| ENSMUSG( | Rpl36a-ps2 | 50.20572 | 0.141104 | 0.599115 | 0.903801 |
| ENSMUSG( | Rpl36-ps12 | 31.78516 | 0.162213 | 0.648332 | 0.919949 |
| ENSMUSG( | Rpl37 | 9336.886 | 0.132111 | 0.153197 | 0.57989 |
| ENSMUSG( | Rpl37a | 7056.42 | 0.04831 | 0.657894 | 0.92425 |
| ENSMUSG( | Rpl38 | 6567.161 | 0.079353 | 0.364647 | 0.793796 |
| ENSMUSG( | Rpl39 | 3852.453 | 0.116349 | 0.282175 | 0.73186 |
| ENSMUSG( | Rpl3l | 12.53184 | 0.232771 | 0.673503 | NA |
| ENSMUSG( | Rpl3-ps1 | 215.6948 | 0.208005 | 0.176512 | 0.612904 |
| ENSMUSG( | Rpl3-ps2 | 28.08217 | 0.488931 | 0.196843 | 0.636268 |
| ENSMUSG( | Rpl4 | 24701.14 | 0.123511 | 0.11045 | 0.50108 |
| ENSMUSG( | Rpl41 | 18195.86 | 0.074134 | 0.327196 | 0.772056 |
| ENSMUSG( | Rpl5 | 9169.467 | 0.057618 | 0.488504 | 0.859092 |
| ENSMUSG( | Rpl6 | 8608.473 | 0.068739 | 0.411846 | 0.81942 |
| ENSMUSG( | Rpl6l | 28.28501 | -0.05082 | 0.887654 | 0.983411 |
| ENSMUSG( | Rpl7 | 9860.35 | 0.096591 | 0.261441 | 0.711405 |
| ENSMUSG( | Rpl7a | 12120.9 | 0.042766 | 0.555022 | 0.88708 |
| ENSMUSG( | Rpl7a-ps7 | 9.846013 | -0.42251 | 0.471698 | NA |
| ENSMUSG( | Rpl7l1 | 1167.557 | 0.127334 | 0.207376 | 0.649561 |
| ENSMUSG( | Rpl7-ps7 | 21.18984 | -0.10766 | 0.794166 | NA |
| ENSMUSG( | Rpl7-ps9 | 11.83647 | 0.106575 | 0.8432 | NA |
| ENSMUSG( | Rpl8 | 13443.61 | 0.076313 | 0.365302 | 0.794085 |
| ENSMUSG( | Rpl9 | 19180.16 | 0.115639 | 0.249655 | 0.699258 |
| ENSMUSG( | Rpl9-ps6 | 133.172 | -0.1291 | 0.553629 | 0.886216 |
| ENSMUSG( | Rplp0 | 15244.56 | 0.126136 | 0.137309 | 0.552371 |
| ENSMUSG( | Rplp0-ps1 | 13.37509 | -0.16042 | 0.754544 | NA |
| ENSMUSG( | Rplp1 | 28207.83 | 0.089583 | 0.349923 | 0.781943 |
| ENSMUSG( | Rplp2 | 7455.379 | 0.093563 | 0.395532 | 0.810727 |
| ENSMUSG( | Rpn1 | 5680.649 | -0.27735 | 0.038554 | 0.302696 |
| ENSMUSG( | Rpn2 | 5007.381 | -0.18913 | 0.121719 | 0.526333 |
| ENSMUSG( | Rpp14 | 1415.461 | 0.227588 | 0.027214 | 0.253245 |
| ENSMUSG( | Rpp21 | 319.4343 | -0.1456 | 0.320294 | 0.76591 |
| ENSMUSG( | Rpp25 | 300.3855 | -0.0072 | 0.970483 | 0.996369 |
| ENSMUSG( | Rpp25l | 330.2131 | 0.025794 | 0.857291 | 0.97444 |
| ENSMUSG( | Rpp30 | 215.7679 | 0.044752 | 0.789203 | 0.959299 |
| ENSMUSG( | Rpp38 | 95.52977 | -0.12102 | 0.62678 | 0.912825 |
| ENSMUSG( | Rpp40 | 93.88332 | 0.163728 | 0.437067 | 0.829501 |
| ENSMUSG( | Rprd1a | 1718.355 | 0.074154 | 0.428186 | 0.825894 |
| ENSMUSG( | Rprd1b | 558.871 | 0.081165 | 0.492927 | 0.860552 |
| ENSMUSG( | Rprd2 | 1139.335 | 0.095427 | 0.289261 | 0.739725 |
| ENSMUSG( | Rprm | 1209.277 | -0.0384 | 0.799361 | 0.961201 |
| ENSMUSG( | Rprml | 1066.408 | -0.19026 | 0.213407 | 0.65774 |
| ENSMUSG( | Rps10 | 69.69386 | -0.31218 | 0.264637 | 0.71564 |
| ENSMUSG( | Rps10-ps1 | 86.36258 | 0.101784 | 0.660209 | 0.924627 |
| ENSMUSG( | Rps10-ps2 | 40.57311 | 0.362166 | 0.214719 | 0.659193 |
| ENSMUSG( | Rps11 | 8608.344 | 0.08815 | 0.276111 | 0.725738 |
| ENSMUSG( | Rps11-ps1 | 13.36604 | -0.34593 | 0.538146 | NA |
| ENSMUSG( | Rps11-ps2 | 24.13041 | -0.27169 | 0.472387 | 0.848048 |
| ENSMUSG( | Rps12 | 10339.84 | 0.11746 | 0.271532 | 0.722082 |
| ENSMUSG( | Rps13 | 4637.59 | 0.083203 | 0.355 | 0.787423 |
| ENSMUSG( | Rps13-ps1 | 32.01057 | -0.13445 | 0.692634 | 0.934101 |
| ENSMUSG( | Rps13-ps2 | 27.82877 | -0.02597 | 0.94332 | 0.993415 |
| ENSMUSG( | Rps14 | 11618.58 | 0.109926 | 0.144213 | 0.563869 |
| ENSMUSG( | Rps15 | 21208.4 | 0.151801 | 0.042016 | 0.315804 |
| ENSMUSG( | Rps15a | 5104.639 | 0.064187 | 0.462705 | 0.842333 |
| ENSMUSG( | Rps15a-ps3 | 22.46896 | -0.2516 | 0.531043 | 0.879289 |
| ENSMUSG( | Rps15a-ps7 | 10.15164 | -0.52789 | 0.423908 | NA |
| ENSMUSG( | Rps15-ps3 | 18.43288 | 0.450318 | 0.297574 | NA |
| ENSMUSG( | Rps16 | 7988.823 | 0.06768 | 0.434352 | 0.828978 |
| ENSMUSG( | Rps17 | 7049.323 | 0.080369 | 0.400157 | 0.813641 |
| ENSMUSG( | Rps18 | 7710.581 | 0.036948 | 0.640686 | 0.916434 |
| ENSMUSG( | Rps18-ps5 | 47.38534 | 0.162305 | 0.600709 | 0.90443 |
| ENSMUSG( | Rps19 | 8099.286 | 0.086084 | 0.297208 | 0.745805 |
| ENSMUSG( | Rps19bp1 | 393.3012 | 0.061098 | 0.713206 | 0.940734 |
| ENSMUSG( | Rps19-ps3 | 51.4495 | 0.57744 | 0.04647 | 0.330252 |
| ENSMUSG( | Rps19-ps6 | 33.32892 | -0.11144 | 0.72589 | 0.944223 |
| ENSMUSG( | Rps19-ps7 | 16.80112 | 0.117373 | 0.801362 | NA |
| ENSMUSG( | Rps2 | 23946.14 | 0.116562 | 0.157701 | 0.585927 |
| ENSMUSG( | Rps20 | 8486.326 | 0.09918 | 0.293037 | 0.742763 |
| ENSMUSG( | Rps21 | 12031.27 | 0.21712 | 0.039271 | 0.305097 |
| ENSMUSG( | Rps23 | 9400.677 | 0.099867 | 0.221809 | 0.667414 |
| ENSMUSG( | Rps23-ps1 | 177.5093 | 0.221091 | 0.250575 | 0.700403 |
| ENSMUSG( | Rps23-ps2 | 47.89604 | -0.23361 | 0.432465 | 0.82768 |
| ENSMUSG( | Rps24 | 15098.14 | 0.076037 | 0.404458 | 0.816132 |

| | | | | | |
|---|---|---|---|---|---|
| ENSMUSG( | Ro60 | 1148.077 | 0.092021 | 0.588493 | 0.944975 |
| ENSMUSG( | Robo1 | 1581.131 | 0.108792 | 0.458004 | 0.923329 |
| ENSMUSG( | Robo2 | 3294.611 | 0.283614 | 0.088803 | 0.788051 |
| ENSMUSG( | Robo3 | 837.8393 | -0.03862 | 0.925085 | 0.99306 |
| ENSMUSG( | Robo4 | 387.5965 | 0.589257 | 0.010652 | 0.419324 |
| ENSMUSG( | Rock1 | 1285.441 | 0.226344 | 0.111447 | 0.808371 |
| ENSMUSG( | Rock2 | 3822.784 | -0.08718 | 0.500807 | 0.927825 |
| ENSMUSG( | Rogdi | 2461.207 | -0.01417 | 0.930375 | 0.99306 |
| ENSMUSG( | Rom1 | 474.884 | -0.13105 | 0.376644 | 0.915491 |
| ENSMUSG( | Romo1 | 1222.425 | -0.42337 | 0.001626 | 0.150169 |
| ENSMUSG( | Ropn1l | 31.76862 | -0.17756 | 0.635872 | 0.953903 |
| ENSMUSG( | Ror1 | 67.21868 | 0.642592 | 0.012009 | 0.445877 |
| ENSMUSG( | Ror2 | 40.66925 | 0.243169 | 0.478328 | 0.923329 |
| ENSMUSG( | Rora | 2059.462 | -0.06415 | 0.642502 | 0.953903 |
| ENSMUSG( | Rorb | 1539.032 | -0.05035 | 0.716285 | 0.965692 |
| ENSMUSG( | Rorc | 104.0926 | -0.22089 | 0.299196 | 0.898538 |
| ENSMUSG( | Rp2 | 546.7733 | -0.19145 | 0.165864 | 0.839724 |
| ENSMUSG( | Rp9 | 861.765 | 0.034044 | 0.800525 | 0.97681 |
| ENSMUSG( | Rpa1 | 1501.051 | -0.06318 | 0.62713 | 0.953234 |
| ENSMUSG( | Rpa2 | 296.9369 | -0.08993 | 0.594195 | 0.945408 |
| ENSMUSG( | Rpa3 | 154.4536 | 0.016465 | 0.927823 | 0.99306 |
| ENSMUSG( | Rpain | 300.5042 | 0.334988 | 0.073677 | 0.764578 |
| ENSMUSG( | Rpap1 | 611.7476 | 0.080038 | 0.54551 | 0.938063 |
| ENSMUSG( | Rpap2 | 503.1872 | 0.074131 | 0.607815 | 0.950113 |
| ENSMUSG( | Rpap3 | 477.9856 | -0.02441 | 0.864733 | 0.986591 |
| ENSMUSG( | Rpe | 549.1743 | 0.045954 | 0.762234 | 0.973255 |
| ENSMUSG( | Rpe65 | 65.68206 | -0.28214 | 0.299795 | 0.898538 |
| ENSMUSG( | Rpf1 | 273.4468 | 0.010671 | 0.946271 | 0.994798 |
| ENSMUSG( | Rpf2 | 287.3732 | -0.2008 | 0.202866 | 0.864412 |
| ENSMUSG( | Rpgr | 313.8227 | -0.01728 | 0.919819 | 0.992644 |
| ENSMUSG( | Rpgrip1 | 19.09658 | -0.47525 | 0.298182 | 0.898538 |
| ENSMUSG( | Rpgrip1l | 380.0832 | 0.22541 | 0.106122 | 0.804969 |
| ENSMUSG( | Rph3a | 25160.35 | -0.34101 | 0.00086 | 0.09911 |
| ENSMUSG( | Rph3al | 57.16946 | 0.05052 | 0.854303 | 0.98563 |
| ENSMUSG( | Rpia | 219.814 | -0.16882 | 0.321883 | 0.904023 |
| ENSMUSG( | Rpl10 | 8026.839 | -0.09736 | 0.465017 | 0.923329 |
| ENSMUSG( | Rpl10a | 3936.314 | -0.16029 | 0.119724 | 0.813384 |
| ENSMUSG( | Rpl10a-ps1 | 81.41285 | -0.53451 | 0.041116 | 0.687358 |
| ENSMUSG( | Rpl11 | 7545.5 | -0.12515 | 0.314534 | 0.900949 |
| ENSMUSG( | Rpl12 | 11141.75 | -0.17398 | 0.163262 | 0.838291 |
| ENSMUSG( | Rpl13 | 13676.83 | -0.15075 | 0.207999 | 0.864412 |
| ENSMUSG( | Rpl13a | 982.7602 | -0.08754 | 0.513553 | 0.930482 |
| ENSMUSG( | Rpl13-ps6 | 13.86465 | -0.77545 | 0.200989 | 0.86439 |
| ENSMUSG( | Rpl14 | 7498.819 | -0.14213 | 0.258517 | 0.883971 |
| ENSMUSG( | Rpl14-ps1 | 68.68719 | -0.0961 | 0.736842 | 0.967344 |
| ENSMUSG( | Rpl15 | 4011.788 | -0.16441 | 0.201152 | 0.86439 |
| ENSMUSG( | Rpl17 | 11567.2 | -0.11878 | 0.343164 | 0.909646 |
| ENSMUSG( | Rpl17-ps10 | 85.4601 | 0.097891 | 0.719532 | 0.966876 |
| ENSMUSG( | Rpl17-ps5 | 10.85284 | 0.232208 | 0.693766 | NA |
| ENSMUSG( | Rpl18 | 4898.642 | -0.06711 | 0.574924 | 0.942912 |
| ENSMUSG( | Rpl18a | 9106.172 | -0.14644 | 0.246472 | 0.877904 |
| ENSMUSG( | Rpl18a-ps1 | 13.44408 | -0.9988 | 0.095785 | 0.796133 |
| ENSMUSG( | Rpl18-ps1 | 14.34049 | -0.40612 | 0.429538 | 0.917257 |
| ENSMUSG( | Rpl19 | 10131.42 | -0.14067 | 0.267219 | 0.886134 |
| ENSMUSG( | Rpl19-ps11 | 82.87464 | -0.13573 | 0.595987 | 0.946158 |
| ENSMUSG( | Rpl21 | 12192.1 | -0.1199 | 0.335615 | 0.907907 |
| ENSMUSG( | Rpl21-ps11 | 19.25731 | -0.44953 | 0.33436 | 0.907907 |
| ENSMUSG( | Rpl21-ps8 | 15.30812 | -0.6354 | 0.216106 | 0.865167 |
| ENSMUSG( | Rpl22 | 4538.82 | -0.17981 | 0.133674 | 0.822912 |
| ENSMUSG( | Rpl22l1 | 1133.525 | -0.15392 | 0.273873 | 0.889859 |
| ENSMUSG( | Rpl23 | 7785.464 | -0.1745 | 0.174849 | 0.845107 |
| ENSMUSG( | Rpl23a | 7714.98 | -0.11149 | 0.410119 | 0.917257 |
| ENSMUSG( | Rpl23a-ps3 | 22.32642 | -0.24647 | 0.571244 | 0.942052 |
| ENSMUSG( | Rpl24 | 56.37155 | -0.1155 | 0.668175 | 0.956766 |
| ENSMUSG( | Rpl26 | 9374.016 | -0.11426 | 0.362498 | 0.914223 |
| ENSMUSG( | Rpl27 | 2294.604 | -0.28075 | 0.025498 | 0.598732 |
| ENSMUSG( | Rpl27a | 7672.863 | -0.0939 | 0.485298 | 0.92541 |
| ENSMUSG( | Rpl28 | 7776.753 | -0.21905 | 0.072167 | 0.763629 |
| ENSMUSG( | Rpl28-ps1 | 37.88973 | -0.70196 | 0.04113 | 0.687358 |
| ENSMUSG( | Rpl28-ps3 | 30.94784 | -0.55026 | 0.124252 | 0.814691 |
| ENSMUSG( | Rpl29 | 7055.844 | -0.14101 | 0.22533 | 0.868468 |
| ENSMUSG( | Rpl3 | 15707.13 | -0.0849 | 0.475149 | 0.923329 |
| ENSMUSG( | Rpl30 | 5432.701 | -0.12149 | 0.362309 | 0.914223 |
| ENSMUSG( | Rpl31 | 8903.568 | -0.29487 | 0.008659 | 0.391935 |
| ENSMUSG( | Rpl32 | 6277.229 | -0.12572 | 0.326703 | 0.905371 |
| ENSMUSG( | Rpl32l | 23.84906 | -0.18732 | 0.656605 | 0.955533 |
| ENSMUSG( | Rpl34 | 4272.08 | -0.10996 | 0.394661 | 0.917257 |
| ENSMUSG( | Rpl34-ps1 | 38.41152 | -0.31988 | 0.329806 | 0.907083 |
| ENSMUSG( | Rpl35 | 4489.945 | -0.19118 | 0.117747 | 0.813384 |
| ENSMUSG( | Rpl35a | 5045 | -0.08375 | 0.526602 | 0.934374 |
| ENSMUSG( | Rpl35rt | 365.0738 | -0.18703 | 0.273316 | 0.889687 |
| ENSMUSG( | Rpl36 | 4806.028 | -0.16361 | 0.244642 | 0.877904 |
| ENSMUSG( | Rpl36a | 3014.771 | -0.18485 | 0.154313 | 0.837724 |
| ENSMUSG( | Rpl36al | 14.23058 | 0.475465 | 0.4465 | 0.921734 |
| ENSMUSG( | Rpl36a-ps1 | 21.83302 | -1.27255 | 0.006633 | 0.338858 |
| ENSMUSG( | Rpl36a-ps2 | 38.57985 | -0.38667 | 0.242787 | 0.877399 |
| ENSMUSG( | Rpl36-ps12 | 23.11134 | -0.56046 | 0.188738 | 0.85698 |
| ENSMUSG( | Rpl37 | 7727.619 | -0.14844 | 0.25159 | 0.879838 |
| ENSMUSG( | Rpl37a | 5799.108 | -0.26423 | 0.065743 | 0.745828 |
| ENSMUSG( | Rpl38 | 5472.093 | -0.1848 | 0.108139 | 0.804969 |
| ENSMUSG( | Rpl39 | 3252.254 | -0.1063 | 0.414057 | 0.917257 |
| ENSMUSG( | Rpl3l | 11.29231 | 0.181755 | 0.766615 | NA |
| ENSMUSG( | Rpl3-ps1 | 184.9675 | 0.025005 | 0.898749 | 0.990558 |
| ENSMUSG( | Rpl3-ps2 | 18.92255 | -0.35795 | 0.451204 | 0.922202 |
| ENSMUSG( | Rpl4 | 21094.12 | -0.0624 | 0.576961 | 0.942912 |
| ENSMUSG( | Rpl41 | 14874.13 | -0.24988 | 0.029409 | 0.618981 |
| ENSMUSG( | Rpl5 | 7749.991 | -0.1642 | 0.143454 | 0.831311 |
| ENSMUSG( | Rpl6 | 7391.078 | -0.10523 | 0.364549 | 0.914223 |
| ENSMUSG( | Rpl6l | 24.94301 | -0.13243 | 0.744791 | 0.96951 |
| ENSMUSG( | Rpl7 | 8352.565 | -0.11607 | 0.324479 | 0.905257 |
| ENSMUSG( | Rpl7a | 10271.42 | -0.17203 | 0.128241 | 0.818934 |
| ENSMUSG( | Rpl7a-ps7 | 9.359589 | -0.38087 | 0.606796 | NA |
| ENSMUSG( | Rpl7l1 | 1011.096 | -0.01764 | 0.881623 | 0.989644 |
| ENSMUSG( | Rpl7-ps7 | 16.76883 | -0.61874 | 0.21373 | 0.864421 |
| ENSMUSG( | Rpl7-ps9 | 9.447136 | -0.275 | 0.660343 | NA |
| ENSMUSG( | Rpl8 | 11430.48 | -0.12705 | 0.336016 | 0.907913 |
| ENSMUSG( | Rpl9 | 16119.14 | -0.11914 | 0.314083 | 0.900949 |
| ENSMUSG( | Rpl9-ps6 | 95.69907 | -0.98033 | 4.34E-05 | 0.011508 |
| ENSMUSG( | Rplp0 | 12699.45 | -0.13383 | 0.24665 | 0.877904 |
| ENSMUSG( | Rplp0-ps1 | 11.41951 | -0.40402 | 0.501067 | NA |
| ENSMUSG( | Rplp1 | 23374.99 | -0.19154 | 0.163728 | 0.838291 |
| ENSMUSG( | Rplp2 | 6454.631 | -0.05625 | 0.709897 | 0.96416 |
| ENSMUSG( | Rpn1 | 5939.981 | 0.133032 | 0.388519 | 0.917257 |
| ENSMUSG( | Rpn2 | 4958.331 | 0.05975 | 0.672735 | 0.95694 |
| ENSMUSG( | Rpp14 | 1201.422 | 0.027531 | 0.841574 | 0.982981 |
| ENSMUSG( | Rpp21 | 283.253 | -0.23174 | 0.161503 | 0.837724 |
| ENSMUSG( | Rpp25 | 264.6295 | -0.11561 | 0.597913 | 0.946676 |
| ENSMUSG( | Rpp25l | 281.5221 | -0.18117 | 0.302458 | 0.898538 |
| ENSMUSG( | Rpp30 | 198.6732 | 0.063227 | 0.731613 | 0.967228 |
| ENSMUSG( | Rpp38 | 79.82637 | -0.38466 | 0.186819 | 0.856099 |
| ENSMUSG( | Rpp40 | 83.49366 | 0.082479 | 0.745433 | 0.96951 |
| ENSMUSG( | Rprd1a | 1510.253 | -0.02976 | 0.783855 | 0.976154 |
| ENSMUSG( | Rprd1b | 507.2605 | 0.067998 | 0.626425 | 0.953234 |
| ENSMUSG( | Rprd2 | 1054.3 | 0.139478 | 0.206709 | 0.864412 |
| ENSMUSG( | Rprm | 1082.869 | -0.095 | 0.606777 | 0.949772 |
| ENSMUSG( | Rprml | 1004.018 | -0.09531 | 0.581866 | 0.944266 |
| ENSMUSG( | Rps10 | 77.72176 | 0.292034 | 0.312588 | 0.900037 |
| ENSMUSG( | Rps10-ps1 | 68.11706 | -0.34923 | 0.191656 | 0.858764 |
| ENSMUSG( | Rps10-ps2 | 33.0196 | 0.017821 | 0.963064 | 0.995433 |
| ENSMUSG( | Rps11 | 7129.407 | -0.19323 | 0.116084 | 0.813384 |
| ENSMUSG( | Rps11-ps1 | 11.86009 | -0.47469 | 0.423907 | 0.917257 |
| ENSMUSG( | Rps11-ps2 | 17.90396 | -1.00527 | 0.041739 | 0.690304 |
| ENSMUSG( | Rps12 | 8585.264 | -0.15496 | 0.238888 | 0.873304 |
| ENSMUSG( | Rps13 | 3860.518 | -0.18475 | 0.137588 | 0.828492 |
| ENSMUSG( | Rps13-ps1 | 28.99758 | -0.19832 | 0.63646 | 0.953903 |
| ENSMUSG( | Rps13-ps2 | 18.85177 | -1.00493 | 0.039151 | 0.679972 |
| ENSMUSG( | Rps14 | 9892.028 | -0.08581 | 0.512224 | 0.930096 |
| ENSMUSG( | Rps15 | 17480.41 | -0.13782 | 0.245221 | 0.877904 |
| ENSMUSG( | Rps15a | 4354.726 | -0.12929 | 0.254892 | 0.882307 |
| ENSMUSG( | Rps15a-ps3 | 20.69536 | -0.29493 | 0.558302 | 0.93957 |
| ENSMUSG( | Rps15a-ps7 | 9.829559 | -0.30331 | 0.62641 | NA |
| ENSMUSG( | Rps15-ps3 | 12.5536 | -0.37667 | 0.492916 | 0.927358 |
| ENSMUSG( | Rps16 | 6735.102 | -0.16188 | 0.214409 | 0.864421 |
| ENSMUSG( | Rps17 | 6005.625 | -0.1174 | 0.360082 | 0.914223 |
| ENSMUSG( | Rps18 | 6513.687 | -0.1883 | 0.118443 | 0.813384 |
| ENSMUSG( | Rps18-ps5 | 37.67715 | -0.24091 | 0.5091 | 0.929089 |
| ENSMUSG( | Rps19 | 6789.679 | -0.15899 | 0.217292 | 0.865589 |
| ENSMUSG( | Rps19bp1 | 338.1997 | -0.11323 | 0.53867 | 0.937216 |
| ENSMUSG( | Rps19-ps3 | 36.70575 | -0.10062 | 0.771671 | 0.974236 |
| ENSMUSG( | Rps19-ps6 | 29.321 | -0.24989 | 0.50655 | 0.928006 |
| ENSMUSG( | Rps19-ps7 | 11.67415 | -0.84089 | 0.162394 | 0.837724 |
| ENSMUSG( | Rps2 | 20273.87 | -0.09582 | 0.46065 | 0.923329 |
| ENSMUSG( | Rps20 | 7109.101 | -0.14763 | 0.264544 | 0.886134 |
| ENSMUSG( | Rps21 | 10101.56 | -0.01355 | 0.919612 | 0.992644 |
| ENSMUSG( | Rps23 | 7930.679 | -0.12453 | 0.301936 | 0.898538 |
| ENSMUSG( | Rps23-ps1 | 147.6482 | -0.04941 | 0.824344 | 0.979198 |
| ENSMUSG( | Rps23-ps2 | 39.82339 | -0.56773 | 0.133084 | 0.822912 |
| ENSMUSG( | Rps24 | 12951.48 | -0.10037 | 0.439132 | 0.920721 |

| | | | | | |
|---|---|---|---|---|---|
| ENSMUSG( | 669.4912 | -0.00437 | 0.971889 | 0.996585 | Rif1 |
| ENSMUSG( | 279.3725 | 0.04922 | 0.718879 | 0.941986 | Rigi |
| ENSMUSG( | 74.73947 | -0.21436 | 0.345172 | 0.779153 | Riiad1 |
| ENSMUSG( | 73.26487 | 0.029406 | 0.905342 | 0.986465 | Rilp |
| ENSMUSG( | 1451.06 | 0.186326 | 0.133344 | 0.54617 | Rilpl1 |
| ENSMUSG( | 358.4678 | 0.048478 | 0.732579 | 0.944956 | Rilpl2 |
| ENSMUSG( | 4391.53 | 0.156752 | 0.192987 | 0.632208 | Rimbp2 |
| ENSMUSG( | 37.32234 | -0.05215 | 0.862524 | 0.975055 | Rimbp3 |
| ENSMUSG( | 332.8722 | -0.36177 | 0.01157 | 0.154961 | Rimkla |
| ENSMUSG( | 1931.429 | 0.084939 | 0.43467 | 0.828978 | Rimklb |
| ENSMUSG( | 1702.394 | -0.00145 | 0.990484 | 0.999232 | Rimoc1 |
| ENSMUSG( | 36657.46 | 0.026924 | 0.770691 | 0.954722 | Rims1 |
| ENSMUSG( | 6674.111 | -0.00463 | 0.958824 | 0.993874 | Rims2 |
| ENSMUSG( | 6037.531 | 0.243366 | 0.006012 | 0.105086 | Rims3 |
| ENSMUSG( | 7216.178 | 0.232318 | 0.022589 | 0.229429 | Rims4 |
| ENSMUSG( | 2248.158 | 0.290003 | 0.010557 | 0.14749 | Rin1 |
| ENSMUSG( | 3184.174 | 0.598016 | 4.29E-12 | 3.49E-09 | Rin2 |
| ENSMUSG( | 145.4296 | 0.414707 | 0.028602 | 0.257732 | Rin3 |
| ENSMUSG( | 829.4811 | 0.081408 | 0.490994 | 0.859739 | Ring1 |
| ENSMUSG( | 147.5686 | 0.234935 | 0.257215 | 0.708432 | Rinl |
| ENSMUSG( | 496.1826 | 0.190016 | 0.091813 | 0.465904 | Rint1 |
| ENSMUSG( | 401.7236 | 0.152889 | 0.219473 | 0.664984 | Riok1 |
| ENSMUSG( | 486.132 | -0.07037 | 0.53172 | 0.879628 | Riok2 |
| ENSMUSG( | 2383.345 | -0.00645 | 0.944127 | 0.993503 | Riok3 |
| ENSMUSG( | 267.6327 | -0.05115 | 0.770456 | 0.954689 | Riox1 |
| ENSMUSG( | 294.6576 | 0.037174 | 0.788253 | 0.959226 | Riox2 |
| ENSMUSG( | 330.6083 | 0.207239 | 0.124718 | 0.531714 | Ripk1 |
| ENSMUSG( | 184.0438 | 0.197686 | 0.200651 | 0.640661 | Ripk2 |
| ENSMUSG( | 14.34151 | -0.75835 | 0.140899 | NA | Ripk3 |
| ENSMUSG( | 2885.892 | 0.278174 | 0.007551 | 0.120197 | Ripor1 |
| ENSMUSG( | 1464.665 | 0.079889 | 0.407265 | 0.817447 | Ripor2 |
| ENSMUSG( | 17.64042 | -0.22024 | 0.656574 | NA | Ripor3 |
| ENSMUSG( | 40.45021 | -0.18268 | 0.573732 | 0.893584 | Ripply2 |
| ENSMUSG( | 983.7058 | 0.134754 | 0.244873 | 0.692492 | Rit1 |
| ENSMUSG( | 1066.441 | -0.01186 | 0.929276 | 0.991848 | Rit2 |
| ENSMUSG( | 508.1288 | 0.120327 | 0.323409 | 0.768488 | Rita1 |
| ENSMUSG( | 1964.918 | 0.087527 | 0.49211 | 0.860297 | Rlbp1 |
| ENSMUSG( | 884.7252 | -0.13499 | 0.184574 | 0.621633 | Rlf |
| ENSMUSG( | 297.8268 | -0.0418 | 0.791576 | 0.959461 | Rlig1 |
| ENSMUSG( | 1224.888 | 0.088409 | 0.343151 | 0.779153 | Rlim |
| ENSMUSG( | 9.27392 | -0.06123 | 0.92786 | NA | Rln1 |
| ENSMUSG( | 663.0102 | 0.081495 | 0.548893 | 0.885686 | Rmc1 |
| ENSMUSG( | 174.5017 | -0.02179 | 0.896423 | 0.984868 | Rmdn1 |
| ENSMUSG( | 288.4023 | 0.042127 | 0.767634 | 0.954621 | Rmdn2 |
| ENSMUSG( | 2812.185 | 0.058461 | 0.582516 | 0.897918 | Rmdn3 |
| ENSMUSG( | 482.0739 | -0.00801 | 0.954673 | 0.993874 | Rmi1 |
| ENSMUSG( | 11.32506 | -0.42244 | 0.435683 | NA | Rmi2 |
| ENSMUSG( | 708.5823 | 0.07872 | 0.473605 | 0.848919 | Rmnd1 |
| ENSMUSG( | 3958.512 | 0.091341 | 0.301875 | 0.750166 | Rmnd5a |
| ENSMUSG( | 1183.951 | 7.74E-05 | 0.999398 | 0.99989 | Rmnd5b |
| ENSMUSG( | 110.9629 | 0.680282 | 0.001085 | 0.034804 | Rmst |
| ENSMUSG( | 176819.4 | -1.17146 | 1.43E-52 | 1.16E-48 | Rn18s |
| ENSMUSG( | 55.64563 | -1.21053 | 0.001008 | 0.03318 | Rn7sk |
| ENSMUSG( | 45.84547 | -0.28456 | 0.389597 | 0.807428 | Rnase1 |
| ENSMUSG( | 1239.059 | -0.07689 | 0.579092 | 0.896692 | Rnase4 |
| ENSMUSG( | 24.64681 | -1.1296 | 0.013998 | 0.172788 | Rnase6 |
| ENSMUSG( | 252.883 | -0.05482 | 0.737786 | 0.947329 | Rnaseh1 |
| ENSMUSG( | 379.1798 | -0.05027 | 0.688434 | 0.933441 | Rnaseh2a |
| ENSMUSG( | 223.2972 | -0.01483 | 0.922397 | 0.99045 | Rnaseh2b |
| ENSMUSG( | 455.6975 | 0.072643 | 0.522697 | 0.875389 | Rnaseh2c |
| ENSMUSG( | 43.89869 | 0.069732 | 0.807444 | 0.963257 | Rnasek |
| ENSMUSG( | 319.2816 | -0.02248 | 0.878598 | 0.981431 | Rnasel |
| ENSMUSG( | 9.892792 | -0.67512 | 0.301164 | NA | Rnaset2a |
| ENSMUSG( | 13.13425 | 0.010071 | 0.983775 | NA | Rnaset2b |
| ENSMUSG( | 432.7659 | -0.04147 | 0.795778 | 0.960062 | Rnd1 |
| ENSMUSG( | 8047.091 | 0.040311 | 0.693794 | 0.934202 | Rnd2 |
| ENSMUSG( | 430.8951 | -0.25304 | 0.086436 | 0.452045 | Rnd3 |
| ENSMUSG( | 16925.6 | 0.158772 | 0.042109 | 0.316282 | Rnf10 |
| ENSMUSG( | 1572.869 | -0.35118 | 0.021817 | 0.225679 | Rnf103 |
| ENSMUSG( | 7593.461 | 0.050395 | 0.639173 | 0.916434 | Rnf11 |
| ENSMUSG( | 813.0577 | -0.08957 | 0.366194 | 0.794603 | Rnf111 |
| ENSMUSG( | 8939.398 | 0.320285 | 0.056742 | 0.367404 | Rnf112 |
| ENSMUSG( | 71.48076 | -0.20895 | 0.424392 | 0.824645 | Rnf113a1 |
| ENSMUSG( | 178.9007 | -0.00997 | 0.950863 | 0.993821 | Rnf113a2 |
| ENSMUSG( | 3143.314 | 0.097542 | 0.220947 | 0.666476 | Rnf114 |
| ENSMUSG( | 1188.938 | -0.00677 | 0.94612 | 0.99369 | Rnf115 |
| ENSMUSG( | 1048.447 | -0.13059 | 0.267412 | 0.717525 | Rnf121 |
| ENSMUSG( | 450.2316 | -0.60726 | 0.001885 | 0.049559 | Rnf122 |
| ENSMUSG( | 2373.377 | 0.081901 | 0.33556 | 0.776986 | Rnf123 |
| ENSMUSG( | 21.24715 | -0.09528 | 0.842662 | 0.971663 | Rnf125 |
| ENSMUSG( | 2359.546 | -0.00668 | 0.948188 | 0.993821 | Rnf126 |
| ENSMUSG( | 377.5498 | -0.59691 | 0.000323 | 0.015194 | Rnf128 |
| ENSMUSG( | 3924.277 | -0.38179 | 0.011987 | 0.158212 | Rnf13 |
| ENSMUSG( | 5942.738 | -0.25819 | 0.073145 | 0.417454 | Rnf130 |
| ENSMUSG( | 149.573 | 0.175671 | 0.300324 | 0.749479 | Rnf135 |
| ENSMUSG( | 325.441 | -0.01248 | 0.927951 | 0.991229 | Rnf138 |
| ENSMUSG( | 10.33167 | -0.31121 | 0.633239 | NA | Rnf138rt1 |
| ENSMUSG( | 945.9143 | -0.28593 | 0.097858 | 0.478264 | Rnf139 |
| ENSMUSG( | 6824.963 | 0.175569 | 0.044619 | 0.32439 | Rnf14 |
| ENSMUSG( | 3738.408 | -0.01758 | 0.860307 | 0.974842 | Rnf141 |
| ENSMUSG( | 2310.222 | -0.07684 | 0.532963 | 0.879905 | Rnf144a |
| ENSMUSG( | 547.892 | 0.10524 | 0.412938 | 0.819541 | Rnf144b |
| ENSMUSG( | 3476.113 | -0.24034 | 0.070831 | 0.410605 | Rnf145 |
| ENSMUSG( | 1394.785 | 0.121078 | 0.170225 | 0.60523 | Rnf146 |
| ENSMUSG( | 1008.195 | -0.14425 | 0.191172 | 0.630245 | Rnf149 |
| ENSMUSG( | 1448.929 | -0.21821 | 0.124718 | 0.531714 | Rnf150 |
| ENSMUSG( | 296.6349 | 0.014587 | 0.920583 | 0.990307 | Rnf152 |
| ENSMUSG( | 16225.52 | 0.13613 | 0.100776 | 0.484034 | Rnf157 |
| ENSMUSG( | 1035.723 | 0.132542 | 0.173854 | 0.609316 | Rnf166 |
| ENSMUSG( | 2527.152 | -0.21285 | 0.036814 | 0.296095 | Rnf167 |
| ENSMUSG( | 872.8869 | 0.173591 | 0.067052 | 0.400593 | Rnf168 |
| ENSMUSG( | 1160.145 | 0.042784 | 0.709966 | 0.939669 | Rnf169 |
| ENSMUSG( | 18.6777 | 0.687718 | 0.175369 | NA | Rnf17 |
| ENSMUSG( | 822.2009 | -0.12476 | 0.266052 | 0.716441 | Rnf170 |
| ENSMUSG( | 748.728 | 0.163127 | 0.234595 | 0.681922 | Rnf180 |
| ENSMUSG( | 2428.062 | 0.025349 | 0.782687 | 0.957558 | Rnf181 |
| ENSMUSG( | 1043.373 | 0.154642 | 0.238796 | 0.6865 | Rnf182 |
| ENSMUSG( | 596.4889 | -0.00732 | 0.956336 | 0.993874 | Rnf185 |
| ENSMUSG( | 14707.69 | 0.07293 | 0.327933 | 0.772842 | Rnf187 |
| ENSMUSG( | 802.5499 | 0.062101 | 0.558307 | 0.888427 | Rnf19a |
| ENSMUSG( | 1050.885 | 0.016616 | 0.891688 | 0.983979 | Rnf19b |
| ENSMUSG( | 480.3668 | -0.01509 | 0.903559 | 0.986055 | Rnf2 |
| ENSMUSG( | 2978.41 | 0.095171 | 0.256949 | 0.708193 | Rnf20 |
| ENSMUSG( | 187.8333 | 0.034678 | 0.884897 | 0.982815 | Rnf207 |
| ENSMUSG( | 3274.474 | 0.046496 | 0.644535 | 0.917773 | Rnf208 |
| ENSMUSG( | 754.5125 | -0.23962 | 0.077013 | 0.429516 | Rnf213 |
| ENSMUSG( | 1314.487 | 0.080629 | 0.371499 | 0.798768 | Rnf214 |
| ENSMUSG( | 1115.022 | -0.1916 | 0.126817 | 0.534201 | Rnf215 |
| ENSMUSG( | 1505.704 | 0.180611 | 0.045803 | 0.328083 | Rnf216 |
| ENSMUSG( | 559.3215 | 0.003065 | 0.979656 | 0.997428 | Rnf217 |
| ENSMUSG( | 8010.464 | 0.107715 | 0.171117 | 0.605644 | Rnf220 |
| ENSMUSG( | 51.25616 | 0.268389 | 0.314242 | 0.761542 | Rnf223 |
| ENSMUSG( | 25.93104 | -0.22461 | 0.546916 | 0.884974 | Rnf225 |
| ENSMUSG( | 2443.184 | 0.025633 | 0.819811 | 0.965888 | Rnf227 |
| ENSMUSG( | 1327.714 | -0.39187 | 0.009943 | 0.142596 | Rnf24 |
| ENSMUSG( | 994.717 | 0.072469 | 0.605823 | 0.905919 | Rnf25 |
| ENSMUSG( | 369.4785 | -0.26828 | 0.083027 | 0.4446 | Rnf26 |
| ENSMUSG( | 744.675 | 0.048323 | 0.667871 | 0.92763 | Rnf31 |
| ENSMUSG( | 293.431 | 0.093773 | 0.546155 | 0.884974 | Rnf32 |
| ENSMUSG( | 1198.428 | 0.036757 | 0.690946 | 0.933881 | Rnf34 |
| ENSMUSG( | 1687.321 | -0.0377 | 0.664146 | 0.926583 | Rnf38 |
| ENSMUSG( | 114.0808 | -0.04444 | 0.841361 | 0.971179 | Rnf39 |
| ENSMUSG( | 1909.795 | 0.040985 | 0.655206 | 0.923929 | Rnf4 |
| ENSMUSG( | 2193.113 | 0.046753 | 0.605882 | 0.905919 | Rnf40 |
| ENSMUSG( | 2128.035 | 0.119328 | 0.220104 | 0.665863 | Rnf41 |
| ENSMUSG( | 107.7482 | -0.01921 | 0.944499 | 0.993583 | Rnf43 |
| ENSMUSG( | 5941.127 | 0.034416 | 0.665119 | 0.926824 | Rnf44 |
| ENSMUSG( | 2786.764 | -0.01178 | 0.904611 | 0.986275 | Rnf5 |
| ENSMUSG( | 1016.137 | 0.068232 | 0.602143 | 0.904887 | Rnf6 |
| ENSMUSG( | 2014.289 | 0.023169 | 0.822335 | 0.966352 | Rnf7 |
| ENSMUSG( | 206.4838 | -0.00143 | 0.993261 | 0.999369 | Rnf8 |
| ENSMUSG( | 673.3599 | -0.0077 | 0.953807 | 0.993874 | Rnft1 |
| ENSMUSG( | 3694.568 | -0.20294 | 0.100505 | 0.483976 | Rnft2 |
| ENSMUSG( | 512.0239 | 0.097917 | 0.416293 | 0.82084 | Rngtt |
| ENSMUSG( | 1017.013 | -0.00632 | 0.948272 | 0.993821 | Rnh1 |
| ENSMUSG( | 34.72205 | 0.157569 | 0.653428 | 0.922819 | Rnls |
| ENSMUSG( | 1158.768 | -0.00365 | 0.975413 | 0.997302 | Rnmt |
| ENSMUSG( | 308.0821 | -0.15343 | 0.328848 | 0.773233 | Rnpc3 |
| ENSMUSG( | 1288.716 | 0.193714 | 0.045525 | 0.327592 | Rnpep |
| ENSMUSG( | 1989.457 | 0.107633 | 0.1972 | 0.636662 | Rnpepl1 |
| ENSMUSG( | 2461.221 | 0.082441 | 0.498398 | 0.862846 | Rnps1 |
| ENSMUSG( | 33.35874 | -1.22743 | 0.022438 | 0.228899 | Rny1 |
| ENSMUSG( | 21.05549 | -2.01116 | 0.007202 | NA | Rny3 |
| ENSMUSG( | 1223.437 | 0.013211 | 0.914348 | 0.988483 | Ro60 |
| ENSMUSG( | 1553.436 | -0.2209 | 0.10403 | 0.489802 | Robo1 |
| ENSMUSG( | 2820.319 | -0.45852 | 0.001692 | 0.046443 | Robo2 |
| ENSMUSG( | 1257.37 | 0.768007 | 0.035793 | 0.290916 | Robo3 |
| ENSMUSG( | 370.1655 | 0.226521 | 0.227535 | 0.674441 | Robo4 |

| | | | | | |
|---|---|---|---|---|---|
| ENSMUSG( | 1679.677 | -0.05964 | 0.582748 | 0.944266 | Rnf4 |
| ENSMUSG( | 1955.898 | -0.01582 | 0.887237 | 0.990263 | Rnf40 |
| ENSMUSG( | 1875.895 | 0.026229 | 0.807256 | 0.978393 | Rnf41 |
| ENSMUSG( | 115.5658 | 0.438144 | 0.106464 | 0.804969 | Rnf43 |
| ENSMUSG( | 5446.319 | 0.052158 | 0.598537 | 0.946859 | Rnf44 |
| ENSMUSG( | 2365.175 | -0.22915 | 0.09007 | 0.790741 | Rnf5 |
| ENSMUSG( | 917.0567 | 0.041099 | 0.776282 | 0.975717 | Rnf6 |
| ENSMUSG( | 1748.628 | -0.12308 | 0.38402 | 0.917257 | Rnf7 |
| ENSMUSG( | 179.4927 | -0.13743 | 0.470888 | 0.923329 | Rnf8 |
| ENSMUSG( | 654.244 | 0.173207 | 0.234718 | 0.871042 | Rnft1 |
| ENSMUSG( | 3700.983 | 0.0806 | 0.579237 | 0.943942 | Rnft2 |
| ENSMUSG( | 458.0263 | 0.043533 | 0.758696 | 0.971858 | Rngtt |
| ENSMUSG( | 889.1932 | -0.1252 | 0.293741 | 0.898538 | Rnh1 |
| ENSMUSG( | 30.78944 | 0.054287 | 0.888929 | 0.990558 | Rnls |
| ENSMUSG( | 1087.89 | 0.084277 | 0.538312 | 0.937216 | Rnmt |
| ENSMUSG( | 370.7382 | 0.60117 | 0.002889 | 0.217231 | Rnpc3 |
| ENSMUSG( | 1083.177 | -0.03179 | 0.787793 | 0.97637 | Rnpep |
| ENSMUSG( | 1719.952 | -0.04093 | 0.71662 | 0.965807 | Rnpepl1 |
| ENSMUSG( | 2158.763 | -0.02771 | 0.829186 | 0.980224 | Rnps1 |
| ENSMUSG( | 24.33302 | -2.82741 | 9.48E-06 | 0.003679 | Rny1 |
| ENSMUSG( | 1148.077 | 0.092021 | 0.588493 | 0.944975 | Ro60 |
| ENSMUSG( | 1581.131 | 0.108792 | 0.458004 | 0.923329 | Robo1 |
| ENSMUSG( | 3294.611 | 0.283614 | 0.088803 | 0.788051 | Robo2 |
| ENSMUSG( | 837.8393 | -0.03862 | 0.925085 | 0.99306 | Robo3 |
| ENSMUSG( | 387.5965 | 0.589257 | 0.010652 | 0.419324 | Robo4 |
| ENSMUSG( | 1285.441 | 0.226344 | 0.111447 | 0.808371 | Rock1 |
| ENSMUSG( | 3822.784 | -0.08718 | 0.500807 | 0.927825 | Rock2 |
| ENSMUSG( | 2461.207 | -0.01417 | 0.930375 | 0.99306 | Rogdi |
| ENSMUSG( | 474.884 | -0.13105 | 0.376644 | 0.915491 | Rom1 |
| ENSMUSG( | 1222.425 | -0.42337 | 0.001626 | 0.150169 | Romo1 |
| ENSMUSG( | 31.76862 | -0.17756 | 0.635872 | 0.953903 | Ropn1l |
| ENSMUSG( | 67.21868 | 0.642592 | 0.012009 | 0.445877 | Ror1 |
| ENSMUSG( | 40.66925 | 0.243169 | 0.478328 | 0.923329 | Ror2 |
| ENSMUSG( | 2059.462 | -0.06415 | 0.642502 | 0.953903 | Rora |
| ENSMUSG( | 1539.032 | -0.05035 | 0.716285 | 0.965692 | Rorb |
| ENSMUSG( | 104.0926 | -0.22089 | 0.299196 | 0.898538 | Rorc |
| ENSMUSG( | 546.7733 | -0.19145 | 0.165864 | 0.839724 | Rp2 |
| ENSMUSG( | 861.765 | 0.034044 | 0.800525 | 0.97681 | Rp9 |
| ENSMUSG( | 1501.051 | -0.06318 | 0.62713 | 0.953234 | Rpa1 |
| ENSMUSG( | 296.9369 | -0.08993 | 0.594195 | 0.945408 | Rpa2 |
| ENSMUSG( | 154.4536 | 0.016465 | 0.927823 | 0.99306 | Rpa3 |
| ENSMUSG( | 300.5042 | 0.334988 | 0.073677 | 0.764578 | Rpain |
| ENSMUSG( | 611.7476 | 0.080038 | 0.54551 | 0.938063 | Rpap1 |
| ENSMUSG( | 503.1872 | 0.074131 | 0.607815 | 0.950113 | Rpap2 |
| ENSMUSG( | 477.9856 | -0.02441 | 0.864733 | 0.986591 | Rpap3 |
| ENSMUSG( | 549.1743 | 0.045954 | 0.762234 | 0.973255 | Rpe |
| ENSMUSG( | 65.68206 | -0.28214 | 0.299795 | 0.898538 | Rpe65 |
| ENSMUSG( | 273.4468 | 0.010671 | 0.946271 | 0.994798 | Rpf1 |
| ENSMUSG( | 287.3732 | -0.2008 | 0.202866 | 0.864412 | Rpf2 |
| ENSMUSG( | 313.8227 | -0.01728 | 0.919819 | 0.992644 | Rpgr |
| ENSMUSG( | 19.09658 | -0.47525 | 0.298182 | 0.898538 | Rpgrip1 |
| ENSMUSG( | 380.0832 | 0.22541 | 0.106122 | 0.804969 | Rpgrip1l |
| ENSMUSG( | 25160.35 | -0.34101 | 0.00086 | 0.09911 | Rph3a |
| ENSMUSG( | 57.16946 | 0.05052 | 0.854303 | 0.98563 | Rph3al |
| ENSMUSG( | 219.814 | -0.16882 | 0.321883 | 0.904023 | Rpia |
| ENSMUSG( | 8026.839 | -0.09736 | 0.465017 | 0.923329 | Rpl10 |
| ENSMUSG( | 3936.314 | -0.16029 | 0.119724 | 0.813384 | Rpl10a |
| ENSMUSG( | 81.41285 | -0.53451 | 0.041116 | 0.687358 | Rpl10a-ps1 |
| ENSMUSG( | 7545.5 | -0.12515 | 0.314534 | 0.900949 | Rpl11 |
| ENSMUSG( | 11141.75 | -0.17398 | 0.163262 | 0.838291 | Rpl12 |
| ENSMUSG( | 13676.83 | -0.15075 | 0.207999 | 0.864412 | Rpl13 |
| ENSMUSG( | 982.7602 | -0.08754 | 0.513553 | 0.930482 | Rpl13a |
| ENSMUSG( | 13.86465 | -0.77545 | 0.200989 | 0.86439 | Rpl13-ps6 |
| ENSMUSG( | 7498.819 | -0.14213 | 0.258517 | 0.883971 | Rpl14 |
| ENSMUSG( | 68.68719 | -0.0961 | 0.736842 | 0.967344 | Rpl14-ps1 |
| ENSMUSG( | 4011.788 | -0.16441 | 0.201152 | 0.86439 | Rpl15 |
| ENSMUSG( | 11567.2 | -0.11878 | 0.343164 | 0.909646 | Rpl17 |
| ENSMUSG( | 85.4601 | 0.097891 | 0.719532 | 0.966876 | Rpl17-ps10 |
| ENSMUSG( | 8.7837 | 0.474233 | 0.475651 | NA | Rpl17-ps3 |
| ENSMUSG( | 10.85284 | 0.232208 | 0.693766 | NA | Rpl17-ps5 |
| ENSMUSG( | 4898.642 | -0.06711 | 0.574924 | 0.942912 | Rpl18 |
| ENSMUSG( | 9106.172 | -0.14644 | 0.246472 | 0.877904 | Rpl18a |
| ENSMUSG( | 13.44408 | -0.9988 | 0.095785 | 0.796133 | Rpl18a-ps1 |
| ENSMUSG( | 14.34049 | -0.40612 | 0.429538 | 0.917257 | Rpl18-ps1 |
| ENSMUSG( | 10131.42 | -0.14067 | 0.267219 | 0.886134 | Rpl19 |
| ENSMUSG( | 82.87464 | -0.13573 | 0.595987 | 0.946158 | Rpl19-ps11 |
| ENSMUSG( | 12192.1 | -0.1199 | 0.335615 | 0.907907 | Rpl21 |
| ENSMUSG( | 19.25731 | -0.44953 | 0.33436 | 0.907907 | Rpl21-ps11 |
| ENSMUSG( | 15.30812 | -0.6354 | 0.216106 | 0.865167 | Rpl21-ps8 |
| ENSMUSG( | 4538.82 | -0.17981 | 0.133674 | 0.822912 | Rpl22 |
| ENSMUSG( | 1133.525 | -0.15392 | 0.273873 | 0.889859 | Rpl22l1 |
| ENSMUSG( | 7785.464 | -0.1745 | 0.174849 | 0.845107 | Rpl23 |
| ENSMUSG( | 7714.98 | -0.11149 | 0.410119 | 0.917257 | Rpl23a |
| ENSMUSG( | 22.32642 | -0.24647 | 0.571244 | 0.942052 | Rpl23a-ps3 |
| ENSMUSG( | 56.37155 | -0.1155 | 0.668175 | 0.956766 | Rpl24 |
| ENSMUSG( | 9374.016 | -0.11426 | 0.362498 | 0.914223 | Rpl26 |
| ENSMUSG( | 2294.604 | -0.28075 | 0.025498 | 0.598732 | Rpl27 |
| ENSMUSG( | 7672.863 | -0.0939 | 0.485298 | 0.92541 | Rpl27a |
| ENSMUSG( | 7776.753 | -0.21905 | 0.072167 | 0.763629 | Rpl28 |
| ENSMUSG( | 37.88973 | -0.70196 | 0.04113 | 0.687358 | Rpl28-ps1 |
| ENSMUSG( | 30.94784 | -0.55026 | 0.124252 | 0.814691 | Rpl28-ps3 |
| ENSMUSG( | 7055.844 | -0.14101 | 0.22533 | 0.868468 | Rpl29 |
| ENSMUSG( | 15707.13 | -0.0849 | 0.475149 | 0.923329 | Rpl3 |
| ENSMUSG( | 5432.701 | -0.12149 | 0.362309 | 0.914223 | Rpl30 |
| ENSMUSG( | 8903.568 | -0.29487 | 0.008659 | 0.391935 | Rpl31 |
| ENSMUSG( | 6277.229 | -0.12572 | 0.326703 | 0.905371 | Rpl32 |
| ENSMUSG( | 23.84906 | -0.18732 | 0.656605 | 0.955533 | Rpl32l |
| ENSMUSG( | 4272.08 | -0.10996 | 0.394661 | 0.917257 | Rpl34 |
| ENSMUSG( | 38.41152 | -0.31988 | 0.329806 | 0.907083 | Rpl34-ps1 |
| ENSMUSG( | 4489.945 | -0.19118 | 0.117747 | 0.813384 | Rpl35 |
| ENSMUSG( | 5045 | -0.08375 | 0.526602 | 0.934374 | Rpl35a |
| ENSMUSG( | 365.0738 | -0.18703 | 0.273316 | 0.889687 | Rpl35rt |
| ENSMUSG( | 4806.028 | -0.16361 | 0.244642 | 0.877904 | Rpl36 |
| ENSMUSG( | 3014.771 | -0.18485 | 0.154313 | 0.837724 | Rpl36a |
| ENSMUSG( | 14.23058 | 0.475465 | 0.4465 | 0.921734 | Rpl36al |
| ENSMUSG( | 21.83302 | -1.27255 | 0.006633 | 0.338858 | Rpl36a-ps1 |
| ENSMUSG( | 38.57985 | -0.38667 | 0.242787 | 0.877399 | Rpl36a-ps2 |
| ENSMUSG( | 23.11134 | -0.56046 | 0.188738 | 0.85698 | Rpl36-ps12 |
| ENSMUSG( | 7727.619 | -0.14844 | 0.25159 | 0.879838 | Rpl37 |
| ENSMUSG( | 5799.108 | -0.26423 | 0.065743 | 0.745828 | Rpl37a |
| ENSMUSG( | 5472.093 | -0.1848 | 0.108139 | 0.804969 | Rpl38 |
| ENSMUSG( | 3252.254 | -0.1063 | 0.414057 | 0.917257 | Rpl39 |
| ENSMUSG( | 11.29231 | 0.181755 | 0.766615 | NA | Rpl3l |
| ENSMUSG( | 184.9675 | 0.025005 | 0.898749 | 0.990558 | Rpl3-ps1 |
| ENSMUSG( | 18.92255 | -0.35795 | 0.451204 | 0.922202 | Rpl3-ps2 |
| ENSMUSG( | 21094.12 | -0.0624 | 0.576961 | 0.942912 | Rpl4 |
| ENSMUSG( | 14874.13 | -0.24988 | 0.029409 | 0.618981 | Rpl41 |
| ENSMUSG( | 7749.991 | -0.1642 | 0.143454 | 0.831311 | Rpl5 |
| ENSMUSG( | 7391.078 | -0.10523 | 0.364549 | 0.914223 | Rpl6 |
| ENSMUSG( | 24.94301 | -0.13243 | 0.744791 | 0.96951 | Rpl6l |
| ENSMUSG( | 8352.565 | -0.11607 | 0.324479 | 0.905257 | Rpl7 |
| ENSMUSG( | 10271.42 | -0.17203 | 0.128241 | 0.818934 | Rpl7a |
| ENSMUSG( | 9.359589 | -0.38087 | 0.606796 | NA | Rpl7a-ps7 |
| ENSMUSG( | 1011.096 | -0.01764 | 0.881623 | 0.989644 | Rpl7l1 |
| ENSMUSG( | 16.76883 | -0.61874 | 0.21373 | 0.864421 | Rpl7-ps7 |
| ENSMUSG( | 9.447136 | -0.275 | 0.660343 | NA | Rpl7-ps9 |
| ENSMUSG( | 11430.48 | -0.12705 | 0.336016 | 0.907913 | Rpl8 |
| ENSMUSG( | 16119.14 | -0.11914 | 0.314083 | 0.900949 | Rpl9 |
| ENSMUSG( | 95.69907 | -0.98033 | 4.34E-05 | 0.011508 | Rpl9-ps6 |
| ENSMUSG( | 12699.45 | -0.13383 | 0.24665 | 0.877904 | Rplp0 |
| ENSMUSG( | 11.41951 | -0.40402 | 0.501067 | NA | Rplp0-ps1 |
| ENSMUSG( | 23374.99 | -0.19154 | 0.163728 | 0.838291 | Rplp1 |
| ENSMUSG( | 6454.631 | -0.05625 | 0.709897 | 0.96416 | Rplp2 |
| ENSMUSG( | 5939.981 | 0.133032 | 0.388519 | 0.917257 | Rpn1 |
| ENSMUSG( | 4958.331 | 0.05975 | 0.672735 | 0.95694 | Rpn2 |
| ENSMUSG( | 1201.422 | 0.027531 | 0.841574 | 0.982981 | Rpp14 |
| ENSMUSG( | 283.253 | -0.23174 | 0.161503 | 0.837724 | Rpp21 |
| ENSMUSG( | 264.6295 | -0.11561 | 0.597913 | 0.946676 | Rpp25 |
| ENSMUSG( | 281.5221 | -0.18117 | 0.302458 | 0.898538 | Rpp25l |
| ENSMUSG( | 198.6732 | 0.063227 | 0.731613 | 0.967228 | Rpp30 |
| ENSMUSG( | 79.82637 | -0.38466 | 0.186819 | 0.856099 | Rpp38 |
| ENSMUSG( | 83.49366 | 0.082479 | 0.745433 | 0.96951 | Rpp40 |
| ENSMUSG( | 1510.253 | -0.02976 | 0.783855 | 0.976154 | Rprd1a |
| ENSMUSG( | 507.2605 | 0.067998 | 0.626425 | 0.953234 | Rprd1b |
| ENSMUSG( | 1054.3 | 0.139478 | 0.206709 | 0.864412 | Rprd2 |
| ENSMUSG( | 1082.869 | -0.095 | 0.606777 | 0.949772 | Rprm |
| ENSMUSG( | 1004.018 | -0.09531 | 0.581866 | 0.944266 | Rprml |
| ENSMUSG( | 77.72176 | 0.292034 | 0.312588 | 0.900037 | Rps10 |
| ENSMUSG( | 68.11706 | -0.34923 | 0.191656 | 0.858764 | Rps10-ps1 |
| ENSMUSG( | 33.0196 | 0.017821 | 0.963064 | 0.995433 | Rps10-ps2 |
| ENSMUSG( | 7129.407 | -0.19323 | 0.116084 | 0.813384 | Rps11 |
| ENSMUSG( | 11.86009 | -0.47469 | 0.423907 | 0.917257 | Rps11-ps1 |
| ENSMUSG( | 17.90396 | -1.00527 | 0.041739 | 0.690304 | Rps11-ps2 |
| ENSMUSG( | 8585.264 | -0.15496 | 0.238888 | 0.873304 | Rps12 |
| ENSMUSG( | 3860.518 | -0.18475 | 0.137588 | 0.828492 | Rps13 |
| ENSMUSG( | 28.99758 | -0.19832 | 0.63646 | 0.953903 | Rps13-ps1 |
| ENSMUSG( | 18.85177 | -1.00493 | 0.039151 | 0.679972 | Rps13-ps2 |
| ENSMUSG( | 9892.028 | -0.08581 | 0.512224 | 0.930096 | Rps14 |

| | | | | | |
|---|---|---|---|---|---|
| ENSMUSG( | Rps25 | 7550.83 | 0.09394 | 0.277474 | 0.727432 |
| ENSMUSG( | Rps26 | 7108.076 | 0.058553 | 0.50651 | 0.867425 |
| ENSMUSG( | Rps27 | 8939.419 | 0.138326 | 0.101395 | 0.484996 |
| ENSMUSG( | Rps27a | 9610.839 | 0.140958 | 0.138223 | 0.553167 |
| ENSMUSG( | Rps27a-ps2 | 17.08272 | 0.497485 | 0.300245 | NA |
| ENSMUSG( | Rps27a-ps3 | 40.45571 | 0.05555 | 0.86193 | 0.975055 |
| ENSMUSG( | Rps27l | 387.4307 | 0.131544 | 0.310549 | 0.758631 |
| ENSMUSG( | Rps27rt | 59.2002 | 0.082495 | 0.745699 | 0.949242 |
| ENSMUSG( | Rps28 | 5212.486 | 0.080078 | 0.356238 | 0.788108 |
| ENSMUSG( | Rps29 | 14775.33 | 0.08334 | 0.366462 | 0.794761 |
| ENSMUSG( | Rps2-ps10 | 19.56695 | 0.521644 | 0.237196 | NA |
| ENSMUSG( | Rps3 | 15228.93 | 0.085991 | 0.299747 | 0.74879 |
| ENSMUSG( | Rps3a1 | 11177.67 | 0.075612 | 0.365209 | 0.794085 |
| ENSMUSG( | Rps3a2 | 48.29444 | 0.379791 | 0.185545 | 0.623185 |
| ENSMUSG( | Rps3a3 | 83.05172 | 0.055276 | 0.817821 | 0.965785 |
| ENSMUSG( | Rps4x | 12125.74 | 0.096732 | 0.262388 | 0.713005 |
| ENSMUSG( | Rps4x-ps | 29.15287 | -0.02069 | 0.951664 | 0.993821 |
| ENSMUSG( | Rps5 | 9645.875 | 0.07795 | 0.336812 | 0.777 |
| ENSMUSG( | Rps6 | 4998.91 | 0.047151 | 0.577656 | 0.895925 |
| ENSMUSG( | Rps6ka1 | 672.2529 | 0.034436 | 0.742662 | 0.948478 |
| ENSMUSG( | Rps6ka2 | 1826.1 | 0.216588 | 0.055289 | 0.361957 |
| ENSMUSG( | Rps6ka3 | 1398.282 | 0.091373 | 0.376737 | 0.801213 |
| ENSMUSG( | Rps6ka4 | 1305.618 | 0.133363 | 0.181763 | 0.618944 |
| ENSMUSG( | Rps6ka5 | 487.3807 | -0.23264 | 0.098446 | 0.480238 |
| ENSMUSG( | Rps6ka6 | 131.4992 | 0.091963 | 0.63392 | 0.914073 |
| ENSMUSG( | Rps6kb1 | 1230.294 | 0.03085 | 0.742036 | 0.948478 |
| ENSMUSG( | Rps6kb2 | 1203.215 | 0.180817 | 0.271344 | 0.72198 |
| ENSMUSG( | Rps6kc1 | 1278.469 | 0.275813 | 0.008541 | 0.129467 |
| ENSMUSG( | Rps6kl1 | 877.5504 | 0.096999 | 0.497847 | 0.862343 |
| ENSMUSG( | Rps6-ps4 | 115.3357 | 0.06254 | 0.752131 | 0.950417 |
| ENSMUSG( | Rps7 | 9331.072 | 0.085976 | 0.292568 | 0.742521 |
| ENSMUSG( | Rps8 | 13527.92 | 0.150916 | 0.09178 | 0.465904 |
| ENSMUSG( | Rps9 | 11899.87 | 0.057028 | 0.543832 | 0.884287 |
| ENSMUSG( | Rpsa | 30877.5 | 0.094664 | 0.262868 | 0.713397 |
| ENSMUSG( | Rpsa-ps10 | 119.8782 | 0.188827 | 0.312373 | 0.759967 |
| ENSMUSG( | Rpsa-ps12 | 85.76074 | -0.02827 | 0.901515 | 0.985633 |
| ENSMUSG( | Rpsa-ps2 | 36.82623 | -0.08028 | 0.794556 | 0.959799 |
| ENSMUSG( | Rpsa-ps7 | 44.38921 | -0.0732 | 0.80869 | 0.963455 |
| ENSMUSG( | Rpsa-ps9 | 20.26819 | 0.337102 | 0.4496 | NA |
| ENSMUSG( | Rptor | 2423.88 | 0.035782 | 0.742907 | 0.948478 |
| ENSMUSG( | Rpusd1 | 1365.354 | 0.043439 | 0.733438 | 0.944956 |
| ENSMUSG( | Rpusd2 | 80.34787 | -0.084 | 0.708653 | 0.939519 |
| ENSMUSG( | Rpusd3 | 231.5776 | 0.021185 | 0.898696 | 0.985633 |
| ENSMUSG( | Rpusd4 | 209.5995 | 0.10501 | 0.470876 | 0.846742 |
| ENSMUSG( | Rrad | 35.86484 | -0.27325 | 0.403475 | 0.816132 |
| ENSMUSG( | Rraga | 1564.294 | -0.10355 | 0.432934 | 0.828185 |
| ENSMUSG( | Rragb | 952.6701 | 0.122909 | 0.269096 | 0.719305 |
| ENSMUSG( | Rragc | 2057.222 | 0.051462 | 0.561147 | 0.888934 |
| ENSMUSG( | Rragd | 4071.336 | -0.04621 | 0.686674 | 0.933041 |
| ENSMUSG( | Rras | 584.8645 | -0.03231 | 0.782737 | 0.957558 |
| ENSMUSG( | Rras2 | 509.9924 | 0.224351 | 0.070104 | 0.40908 |
| ENSMUSG( | Rrbp1 | 3881.878 | -0.10336 | 0.375538 | 0.800845 |
| ENSMUSG( | Rreb1 | 1623.19 | -0.21704 | 0.232763 | 0.680928 |
| ENSMUSG( | Rrm1 | 630.2704 | 0.065453 | 0.549795 | 0.886008 |
| ENSMUSG( | Rrm2 | 127.9801 | -0.24028 | 0.193985 | 0.633085 |
| ENSMUSG( | Rrm2b | 941.4219 | 0.030224 | 0.8141 | 0.964574 |
| ENSMUSG( | Rrn3 | 1612.062 | 0.100211 | 0.259107 | 0.709814 |
| ENSMUSG( | Rrp1 | 7273.358 | 0.006036 | 0.959993 | 0.993991 |
| ENSMUSG( | Rrp12 | 882.2549 | 0.225721 | 0.0399 | 0.308163 |
| ENSMUSG( | Rrp15 | 224.4091 | 0.058398 | 0.691975 | 0.934101 |
| ENSMUSG( | Rrp1b | 454.4894 | -0.03725 | 0.7485 | 0.949604 |
| ENSMUSG( | Rrp36 | 406.6149 | 0.093997 | 0.447549 | 0.834051 |
| ENSMUSG( | Rrp7a | 2300.825 | 0.031811 | 0.763524 | 0.953845 |
| ENSMUSG( | Rrp8 | 488.6509 | 0.407809 | 0.001567 | 0.044386 |
| ENSMUSG( | Rrp9 | 568.2556 | 0.139656 | 0.232066 | 0.679932 |
| ENSMUSG( | Rrs1 | 480.9433 | 0.07989 | 0.622425 | 0.911432 |
| ENSMUSG( | Rsad1 | 418.5389 | 0.152074 | 0.235966 | 0.683136 |
| ENSMUSG( | Rsad2 | 108.2493 | -0.83916 | 6.86E-05 | 0.005034 |
| ENSMUSG( | Rsbn1 | 437.311 | -0.16025 | 0.242401 | 0.6899 |
| ENSMUSG( | Rsbn1l | 518.1372 | -0.15365 | 0.223884 | 0.669845 |
| ENSMUSG( | Rsf1 | 1908.052 | 0.068265 | 0.492569 | 0.860378 |
| ENSMUSG( | Rskr | 213.748 | 0.561091 | 0.01777 | 0.2 |
| ENSMUSG( | Rsl1 | 101.9491 | -0.0725 | 0.728462 | 0.944393 |
| ENSMUSG( | Rsl1d1 | 1028.406 | -0.00985 | 0.924718 | 0.99114 |
| ENSMUSG( | Rsl24d1 | 457.0078 | 0.169901 | 0.143802 | 0.562581 |
| ENSMUSG( | Rsph1 | 105.5385 | -0.2444 | 0.342227 | 0.778828 |
| ENSMUSG( | Rsph3a | 223.106 | 0.215557 | 0.134248 | 0.547472 |
| ENSMUSG( | Rsph3b | 245.8692 | 0.160319 | 0.246577 | 0.694958 |
| ENSMUSG( | Rsph4a | 64.49937 | -0.48996 | 0.063806 | 0.390932 |
| ENSMUSG( | Rsph9 | 466.8439 | -0.24906 | 0.035852 | 0.290992 |
| ENSMUSG( | Rspo1 | 491.5733 | 0.198936 | 0.098048 | 0.479019 |
| ENSMUSG( | Rspo2 | 372.8102 | -0.37434 | 0.070468 | 0.40908 |
| ENSMUSG( | Rspo3 | 462.6077 | -0.18117 | 0.146777 | 0.56887 |
| ENSMUSG( | Rspry1 | 942.5005 | 0.019796 | 0.856445 | 0.974154 |
| ENSMUSG( | Rsrc1 | 1287.331 | -0.07742 | 0.451112 | 0.835567 |
| ENSMUSG( | Rsrc2 | 1718.325 | -0.00018 | 0.998546 | 0.99989 |
| ENSMUSG( | Rsrp1 | 7244.553 | -0.25563 | 0.111497 | 0.503511 |
| ENSMUSG( | Rsu1 | 1040.253 | 0.003023 | 0.978917 | 0.997428 |
| ENSMUSG( | Rtbdn | 44.32147 | 0.228832 | 0.490977 | 0.859739 |
| ENSMUSG( | Rtca | 1207.996 | -0.03283 | 0.775092 | 0.955606 |
| ENSMUSG( | Rtcb | 3943.704 | -0.10859 | 0.445617 | 0.833239 |
| ENSMUSG( | Rtel1 | 554.9454 | 0.08279 | 0.500319 | 0.863455 |
| ENSMUSG( | Rtf1 | 2017.018 | 0.090878 | 0.370984 | 0.798271 |
| ENSMUSG( | Rtf2 | 1771.525 | 0.082864 | 0.494327 | 0.860611 |
| ENSMUSG( | Rtkn | 1712.773 | 0.040018 | 0.726082 | 0.944223 |
| ENSMUSG( | Rtkn2 | 94.77669 | -0.46553 | 0.029709 | 0.261211 |
| ENSMUSG( | Rtl4 | 49.37716 | 0.44768 | 0.105566 | 0.49243 |
| ENSMUSG( | Rtl5 | 693.8528 | -0.07409 | 0.495252 | 0.860831 |
| ENSMUSG( | Rtl6 | 1554.811 | 0.18015 | 0.04566 | 0.327994 |
| ENSMUSG( | Rtl8a | 1387.807 | -0.04331 | 0.676653 | 0.930542 |
| ENSMUSG( | Rtl8b | 2134.857 | -0.02031 | 0.846433 | 0.972238 |
| ENSMUSG( | Rtl8c | 1982.456 | -0.35997 | 4.05E-05 | 0.003419 |
| ENSMUSG( | Rtl9 | 64.8222 | -0.02238 | 0.925976 | 0.991229 |
| ENSMUSG( | Rtn1 | 36885.99 | -0.54889 | 1.57E-05 | 0.001615 |
| ENSMUSG( | Rtn2 | 5782.651 | 0.183967 | 0.027083 | 0.253045 |
| ENSMUSG( | Rtn3 | 39256.99 | -0.29619 | 0.017573 | 0.198626 |
| ENSMUSG( | Rtn4 | 14326.51 | -0.3074 | 0.031059 | 0.268255 |
| ENSMUSG( | Rtn4ip1 | 372.7052 | 0.109886 | 0.371251 | 0.798633 |
| ENSMUSG( | Rtn4r | 3416.549 | -0.06736 | 0.669453 | 0.928147 |
| ENSMUSG( | Rtn4rl1 | 2128.363 | -0.24985 | 0.09265 | 0.46772 |
| ENSMUSG( | Rtn4rl2 | 1161.824 | -0.01464 | 0.931755 | 0.9924 |
| ENSMUSG( | Rtp1 | 83.94647 | 0.620859 | 0.023673 | 0.23542 |
| ENSMUSG( | Rtp4 | 130.7562 | -0.1249 | 0.58265 | 0.897918 |
| ENSMUSG( | Rtraf | 1388.815 | 0.021218 | 0.830612 | 0.96974 |
| ENSMUSG( | Rttn | 126.4887 | -0.18547 | 0.367089 | 0.795164 |
| ENSMUSG( | Rubcn | 3312.645 | 0.133442 | 0.108785 | 0.497919 |
| ENSMUSG( | Rubcnl | 47.41078 | 0.166385 | 0.605595 | 0.905824 |
| ENSMUSG( | Rufy1 | 1145.637 | 0.080436 | 0.47263 | 0.848298 |
| ENSMUSG( | Rufy2 | 2260.44 | 0.018667 | 0.842189 | 0.971365 |
| ENSMUSG( | Rufy3 | 4709.966 | 0.08388 | 0.397875 | 0.812161 |
| ENSMUSG( | Rundc1 | 1100.519 | -0.19362 | 0.081108 | 0.440221 |
| ENSMUSG( | Rundc3a | 19847.41 | 0.16871 | 0.027099 | 0.253045 |
| ENSMUSG( | Rundc3b | 532.3683 | 0.044073 | 0.792871 | 0.95947 |
| ENSMUSG( | Runx1 | 45.25701 | -0.49279 | 0.079216 | 0.435105 |
| ENSMUSG( | Runx1t1 | 276.6211 | 0.210938 | 0.181152 | 0.618139 |
| ENSMUSG( | Runx2 | 202.8395 | 0.291895 | 0.125531 | 0.53257 |
| ENSMUSG( | Rusc1 | 4147.156 | 0.185183 | 0.169392 | 0.603554 |
| ENSMUSG( | Rusc2 | 3952.421 | 0.247245 | 0.019758 | 0.212093 |
| ENSMUSG( | Rusf1 | 889.1951 | -0.28643 | 0.016703 | 0.192277 |
| ENSMUSG( | Ruvbl1 | 507.1103 | 0.009741 | 0.935194 | 0.99331 |
| ENSMUSG( | Ruvbl2 | 779.0269 | 0.049814 | 0.668781 | 0.927963 |
| ENSMUSG( | Rwdd1 | 783.9091 | 0.091418 | 0.354288 | 0.786661 |
| ENSMUSG( | Rwdd2a | 218.9037 | -0.18714 | 0.339067 | 0.777518 |
| ENSMUSG( | Rwdd2b | 178.6351 | -0.05748 | 0.758823 | 0.952113 |
| ENSMUSG( | Rwdd3 | 106.6672 | -0.2496 | 0.228751 | 0.675959 |
| ENSMUSG( | Rwdd4a | 772.6928 | 0.058942 | 0.59497 | 0.903007 |
| ENSMUSG( | Rxfp1 | 585.5142 | -0.08857 | 0.495018 | 0.860798 |
| ENSMUSG( | Rxfp2 | 46.28717 | -0.23525 | 0.411324 | 0.819213 |
| ENSMUSG( | Rxfp3 | 52.90673 | -0.07779 | 0.806464 | 0.962804 |
| ENSMUSG( | Rxra | 1447.952 | 0.353477 | 0.000359 | 0.016466 |
| ENSMUSG( | Rxrb | 1944.278 | 0.065841 | 0.506858 | 0.867769 |
| ENSMUSG( | Rxrg | 560.8071 | -0.68137 | 0.070443 | 0.40908 |
| ENSMUSG( | Rxylt1 | 846.805 | -0.22896 | 0.038559 | 0.302696 |
| ENSMUSG( | Rybp | 440.8919 | 0.125348 | 0.338327 | 0.777489 |
| ENSMUSG( | Ryk | 1987.312 | -0.07127 | 0.541811 | 0.883182 |
| ENSMUSG( | Ryr1 | 646.1574 | -0.37346 | 0.096313 | 0.476986 |
| ENSMUSG( | Ryr2 | 3746.331 | -0.57193 | 2.20E-07 | 5.18E-05 |
| ENSMUSG( | Ryr3 | 1456.901 | -0.69742 | 2.20E-10 | 1.21E-07 |
| ENSMUSG( | S100a1 | 4949.475 | 3.94E-07 | 0.999997 | 0.999997 |
| ENSMUSG( | S100a10 | 575.4697 | 0.193726 | 0.098764 | 0.480598 |
| ENSMUSG( | S100a11 | 133.0402 | -0.12636 | 0.493117 | 0.860552 |
| ENSMUSG( | S100a13 | 1444.917 | 0.070064 | 0.591528 | 0.901407 |
| ENSMUSG( | S100a16 | 5280.299 | -0.04943 | 0.616451 | 0.910866 |
| ENSMUSG( | S100a3 | 16.92826 | -0.08102 | 0.870912 | NA |
| ENSMUSG( | S100a4 | 18.82059 | -0.37918 | 0.376796 | NA |
| ENSMUSG( | S100a6 | 795.6189 | -0.10711 | 0.343513 | 0.779153 |
| ENSMUSG( | S100a7a | 20.20514 | -0.30393 | 0.47192 | NA |
| ENSMUSG( | S100a8 | 78.40156 | -1.44243 | 0.195438 | 0.635016 |

| | | | | | |
|---|---|---|---|---|---|
| ENSMUSG( | Rps25 | 6321.83 | -0.15436 | 0.211049 | 0.864412 |
| ENSMUSG( | Rps26 | 6002.4 | -0.16709 | 0.179133 | 0.849453 |
| ENSMUSG( | Rps27 | 7365.624 | -0.15457 | 0.216615 | 0.865167 |
| ENSMUSG( | Rps27a | 8028.971 | -0.11002 | 0.387909 | 0.917257 |
| ENSMUSG( | Rps27a-ps2 | 13.17384 | 0.046019 | 0.934237 | 0.99306 |
| ENSMUSG( | Rps27a-ps3 | 28.45485 | -0.77177 | 0.049841 | 0.708648 |
| ENSMUSG( | Rps27l | 311.0137 | -0.24628 | 0.099414 | 0.800296 |
| ENSMUSG( | Rps27rt | 44.62135 | -0.49503 | 0.107218 | 0.804969 |
| ENSMUSG( | Rps28 | 4343.645 | -0.18466 | 0.158545 | 0.837724 |
| ENSMUSG( | Rps29 | 12117.89 | -0.23044 | 0.068143 | 0.75212 |
| ENSMUSG( | Rps2-ps10 | 16.02486 | 0.210503 | 0.671037 | 0.95694 |
| ENSMUSG( | Rps3 | 12961.16 | -0.11348 | 0.378078 | 0.916904 |
| ENSMUSG( | Rps3a1 | 9565.717 | -0.10736 | 0.37299 | 0.915491 |
| ENSMUSG( | Rps3a2 | 35.82344 | -0.19793 | 0.568997 | 0.941777 |
| ENSMUSG( | Rps3a3 | 64.6501 | -0.45047 | 0.111242 | 0.807885 |
| ENSMUSG( | Rps4x | 10103.36 | -0.1655 | 0.176412 | 0.846867 |
| ENSMUSG( | Rps4x-ps | 20.03339 | -1.02233 | 0.021197 | 0.57477 |
| ENSMUSG( | Rps5 | 8099.164 | -0.16295 | 0.179433 | 0.850372 |
| ENSMUSG( | Rps6 | 4278.335 | -0.13798 | 0.220641 | 0.866104 |
| ENSMUSG( | Rps6ka1 | 610.234 | 0.028794 | 0.822572 | 0.979198 |
| ENSMUSG( | Rps6ka2 | 1733.787 | 0.329806 | 0.023426 | 0.585845 |
| ENSMUSG( | Rps6ka3 | 1245.998 | 0.026608 | 0.831063 | 0.98051 |
| ENSMUSG( | Rps6ka4 | 1227.547 | 0.221844 | 0.057314 | 0.723362 |
| ENSMUSG( | Rps6ka5 | 465.807 | -0.08675 | 0.562488 | 0.940761 |
| ENSMUSG( | Rps6ka6 | 111.6675 | -0.12571 | 0.58265 | 0.944266 |
| ENSMUSG( | Rps6kb1 | 1160.847 | 0.12567 | 0.305621 | 0.898538 |
| ENSMUSG( | Rps6kb2 | 1038.609 | 0.026933 | 0.885773 | 0.98977 |
| ENSMUSG( | Rps6kc1 | 1038.669 | -0.04116 | 0.733404 | 0.967228 |
| ENSMUSG( | Rps6kl1 | 756.2537 | -0.06613 | 0.680602 | 0.958395 |
| ENSMUSG( | Rps6-ps4 | 92.34173 | -0.3464 | 0.153373 | 0.837724 |
| ENSMUSG( | Rps7 | 7959.861 | -0.10646 | 0.399685 | 0.917257 |
| ENSMUSG( | Rps8 | 11413.68 | -0.06952 | 0.561197 | 0.940254 |
| ENSMUSG( | Rps9 | 10161.26 | -0.13463 | 0.287268 | 0.894091 |
| ENSMUSG( | Rpsa | 25917.02 | -0.14589 | 0.213607 | 0.864421 |
| ENSMUSG( | Rpsa-ps10 | 97.00768 | -0.16002 | 0.462287 | 0.923329 |
| ENSMUSG( | Rpsa-ps12 | 77.62625 | -0.06414 | 0.794592 | 0.97681 |
| ENSMUSG( | Rpsa-ps2 | 32.13954 | -0.2552 | 0.495078 | 0.927372 |
| ENSMUSG( | Rpsa-ps7 | 41.71516 | -0.01418 | 0.966662 | 0.995475 |
| ENSMUSG( | Rpsa-ps9 | 16.61759 | 0.039186 | 0.938869 | 0.993898 |
| ENSMUSG( | Rptor | 2173.121 | -0.00609 | 0.964998 | 0.995433 |
| ENSMUSG( | Rpusd1 | 1183.754 | -0.10202 | 0.462846 | 0.923329 |
| ENSMUSG( | Rpusd2 | 78.63891 | 0.119433 | 0.610741 | 0.950771 |
| ENSMUSG( | Rpusd3 | 228.5407 | 0.244242 | 0.145028 | 0.83339 |
| ENSMUSG( | Rpusd4 | 182.8193 | -0.01495 | 0.931342 | 0.99306 |
| ENSMUSG( | Rrad | 38.49689 | 0.234683 | 0.498404 | 0.927825 |
| ENSMUSG( | Rraga | 1364.389 | -0.24142 | 0.11755 | 0.813384 |
| ENSMUSG( | Rragb | 849.5525 | 0.05861 | 0.67444 | 0.95694 |
| ENSMUSG( | Rragc | 1841.231 | -0.00213 | 0.986596 | 0.997372 |
| ENSMUSG( | Rragd | 3600.564 | -0.13774 | 0.286726 | 0.894091 |
| ENSMUSG( | Rras | 533.1229 | -0.03367 | 0.80006 | 0.97681 |
| ENSMUSG( | Rras2 | 430.7225 | 0.013623 | 0.92831 | 0.99306 |
| ENSMUSG( | Rrbp1 | 3780.787 | 0.092143 | 0.482403 | 0.924578 |
| ENSMUSG( | Rreb1 | 1662.224 | 0.119122 | 0.582197 | 0.944266 |
| ENSMUSG( | Rrm1 | 556.1001 | -0.02812 | 0.836711 | 0.981335 |
| ENSMUSG( | Rrm2 | 118.6423 | -0.19165 | 0.337964 | 0.907972 |
| ENSMUSG( | Rrm2b | 840.7968 | -0.03322 | 0.819927 | 0.979198 |
| ENSMUSG( | Rrn3 | 1391.49 | -0.05786 | 0.657782 | 0.955974 |
| ENSMUSG( | Rrp1 | 6200.762 | -0.19451 | 0.201758 | 0.864412 |
| ENSMUSG( | Rrp12 | 739.0998 | -0.0065 | 0.959806 | 0.994798 |
| ENSMUSG( | Rrp15 | 195.3486 | -0.07372 | 0.667972 | 0.956766 |
| ENSMUSG( | Rrp1b | 432.0176 | 0.083772 | 0.572028 | 0.942369 |
| ENSMUSG( | Rrp36 | 353.6651 | -0.04617 | 0.774877 | 0.974931 |
| ENSMUSG( | Rrp7a | 2009.731 | -0.09084 | 0.447518 | 0.922202 |
| ENSMUSG( | Rrp8 | 466.4208 | 0.522029 | 0.000115 | 0.022849 |
| ENSMUSG( | Rrp9 | 524.644 | 0.176969 | 0.185735 | 0.855305 |
| ENSMUSG( | Rrs1 | 427.7969 | 0.003064 | 0.986882 | 0.997473 |
| ENSMUSG( | Rsad1 | 381.6531 | 0.160697 | 0.307536 | 0.898538 |
| ENSMUSG( | Rsad2 | 121.1105 | -0.13429 | 0.555146 | 0.939565 |
| ENSMUSG( | Rsbn1 | 440.9311 | 0.131184 | 0.39104 | 0.917257 |
| ENSMUSG( | Rsbn1l | 498.2385 | -0.00044 | 0.997622 | 0.999268 |
| ENSMUSG( | Rsf1 | 1715.263 | 0.027635 | 0.81664 | 0.979198 |
| ENSMUSG( | Rskr | 173.3376 | 0.273723 | 0.313749 | 0.900949 |
| ENSMUSG( | Rsl1 | 100.7579 | 0.172535 | 0.439117 | 0.920721 |
| ENSMUSG( | Rsl1d1 | 924.2111 | -0.04867 | 0.673455 | 0.95694 |
| ENSMUSG( | Rsl24d1 | 393.9193 | 0.009739 | 0.948454 | 0.994798 |
| ENSMUSG( | Rsph1 | 101.4077 | -0.07103 | 0.800568 | 0.97681 |
| ENSMUSG( | Rsph3a | 184.8393 | -0.05642 | 0.754952 | 0.971098 |
| ENSMUSG( | Rsph3b | 211.6495 | -0.00709 | 0.966831 | 0.995532 |
| ENSMUSG( | Rsph4a | 86.50558 | 0.6225 | 0.016519 | 0.519511 |
| ENSMUSG( | Rsph9 | 448.3916 | -0.093 | 0.552538 | 0.939565 |
| ENSMUSG( | Rspo1 | 408.9029 | -0.06412 | 0.65379 | 0.955015 |
| ENSMUSG( | Rspo2 | 416.6938 | 0.236793 | 0.256663 | 0.883082 |
| ENSMUSG( | Rspo3 | 495.084 | 0.273226 | 0.065005 | 0.741314 |
| ENSMUSG( | Rspry1 | 873.3851 | 0.066853 | 0.612803 | 0.951071 |
| ENSMUSG( | Rsrc1 | 1123.318 | -0.21075 | 0.073499 | 0.763629 |
| ENSMUSG( | Rsrc2 | 1667.24 | 0.176315 | 0.12209 | 0.813384 |
| ENSMUSG( | Rsrp1 | 7890.575 | 0.258265 | 0.238179 | 0.87259 |
| ENSMUSG( | Rsu1 | 917.1119 | -0.09499 | 0.439637 | 0.920721 |
| ENSMUSG( | Rtbdn | 34.65142 | -0.22218 | 0.586226 | 0.944975 |
| ENSMUSG( | Rtca | 1029.407 | -0.23915 | 0.066638 | 0.751155 |
| ENSMUSG( | Rtcb | 3684.356 | -0.03093 | 0.825101 | 0.979354 |
| ENSMUSG( | Rtel1 | 499.0028 | 0.051248 | 0.772181 | 0.974236 |
| ENSMUSG( | Rtf1 | 1792.928 | 0.019496 | 0.876772 | 0.989644 |
| ENSMUSG( | Rtf2 | 1503.794 | -0.12616 | 0.373553 | 0.915491 |
| ENSMUSG( | Rtkn | 1516.741 | -0.04152 | 0.740326 | 0.968811 |
| ENSMUSG( | Rtkn2 | 93.14564 | -0.2012 | 0.373948 | 0.915491 |
| ENSMUSG( | Rtl4 | 40.12467 | 0.139026 | 0.665209 | 0.956766 |
| ENSMUSG( | Rtl5 | 631.2659 | -0.08001 | 0.539 | 0.937216 |
| ENSMUSG( | Rtl6 | 1342.619 | 0.033887 | 0.770308 | 0.974236 |
| ENSMUSG( | Rtl8a | 1183.665 | -0.24853 | 0.075103 | 0.767994 |
| ENSMUSG( | Rtl8b | 1814.256 | -0.23258 | 0.0822 | 0.77681 |
| ENSMUSG( | Rtl8c | 1882.058 | -0.22979 | 0.047438 | 0.700936 |
| ENSMUSG( | Rtl9 | 59.41001 | -0.00525 | 0.984165 | 0.996963 |
| ENSMUSG( | Rtn1 | 40669.56 | 0.055032 | 0.708551 | 0.964042 |
| ENSMUSG( | Rtn2 | 5038.939 | 0.061453 | 0.560143 | 0.940119 |
| ENSMUSG( | Rtn3 | 40681.78 | 0.091656 | 0.527747 | 0.934603 |
| ENSMUSG( | Rtn4 | 15350.68 | 0.175245 | 0.27604 | 0.890679 |
| ENSMUSG( | Rtn4ip1 | 309.7629 | -0.15662 | 0.284626 | 0.892412 |
| ENSMUSG( | Rtn4r | 3398.331 | 0.185817 | 0.273351 | 0.889687 |
| ENSMUSG( | Rtn4rl1 | 2185.149 | 0.108648 | 0.504785 | 0.927825 |
| ENSMUSG( | Rtn4rl2 | 1209.208 | 0.355291 | 0.055784 | 0.723362 |
| ENSMUSG( | Rtp1 | 93.18373 | 1.06978 | 1.20E-05 | 0.004289 |
| ENSMUSG( | Rtp4 | 126.4319 | 0.05914 | 0.788476 | 0.97637 |
| ENSMUSG( | Rtraf | 1199.095 | -0.14082 | 0.25653 | 0.883082 |
| ENSMUSG( | Rttn | 120.9417 | -0.01947 | 0.933044 | 0.99306 |
| ENSMUSG( | Rubcn | 2837.839 | -0.04235 | 0.68101 | 0.958395 |
| ENSMUSG( | Rubcnl | 45.93961 | 0.358441 | 0.278732 | 0.891628 |
| ENSMUSG( | Rufy1 | 1021.591 | 0.017017 | 0.903055 | 0.991089 |
| ENSMUSG( | Rufy2 | 2085.843 | 0.053283 | 0.663233 | 0.956359 |
| ENSMUSG( | Rufy3 | 4040.554 | -0.09278 | 0.481786 | 0.924578 |
| ENSMUSG( | Rundc1 | 1050.997 | -0.05117 | 0.676377 | 0.95694 |
| ENSMUSG( | Rundc3a | 16924.05 | -0.0179 | 0.847687 | 0.9852 |
| ENSMUSG( | Rundc3b | 460.6842 | -0.11146 | 0.541784 | 0.937841 |
| ENSMUSG( | Runx1 | 46.19983 | -0.12389 | 0.671069 | 0.95694 |
| ENSMUSG( | Runx1t1 | 252.8492 | 0.205298 | 0.342619 | 0.909646 |
| ENSMUSG( | Runx2 | 183.61 | 0.265223 | 0.209594 | 0.864412 |
| ENSMUSG( | Rusc1 | 3449.717 | -0.07416 | 0.593793 | 0.945408 |
| ENSMUSG( | Rusc2 | 3429.161 | 0.114742 | 0.310959 | 0.899422 |
| ENSMUSG( | Rusf1 | 928.5517 | 0.122734 | 0.38962 | 0.917257 |
| ENSMUSG( | Ruvbl1 | 435.3366 | -0.16786 | 0.238796 | 0.873304 |
| ENSMUSG( | Ruvbl2 | 685.647 | -0.05317 | 0.688962 | 0.959742 |
| ENSMUSG( | Rwdd1 | 695.9658 | 0.012399 | 0.931084 | 0.99306 |
| ENSMUSG( | Rwdd2a | 203.3178 | -0.12978 | 0.538401 | 0.937216 |
| ENSMUSG( | Rwdd2b | 173.314 | 0.126537 | 0.528794 | 0.934795 |
| ENSMUSG( | Rwdd3 | 98.09248 | -0.21958 | 0.343384 | 0.909753 |
| ENSMUSG( | Rwdd4a | 703.922 | 0.051601 | 0.731325 | 0.967228 |
| ENSMUSG( | Rxfp1 | 592.5667 | 0.212801 | 0.109256 | 0.805353 |
| ENSMUSG( | Rxfp2 | 47.14075 | 0.103859 | 0.729228 | 0.966913 |
| ENSMUSG( | Rxfp3 | 53.76095 | 0.252418 | 0.459327 | 0.923329 |
| ENSMUSG( | Rxra | 1281.444 | 0.278313 | 0.017209 | 0.52741 |
| ENSMUSG( | Rxrb | 1721.083 | -0.01451 | 0.894754 | 0.990558 |
| ENSMUSG( | Rxrg | 592.1635 | -0.1888 | 0.293986 | 0.898538 |
| ENSMUSG( | Rxylt1 | 850.3027 | 0.061991 | 0.644664 | 0.953903 |
| ENSMUSG( | Rybp | 430.1585 | 0.312353 | 0.026456 | 0.603041 |
| ENSMUSG( | Ryk | 1923.311 | 0.102325 | 0.427747 | 0.917257 |
| ENSMUSG( | Ryr1 | 660.3353 | -0.02242 | 0.927147 | 0.99306 |
| ENSMUSG( | Ryr2 | 4175.092 | 0.068353 | 0.667955 | 0.956766 |
| ENSMUSG( | Ryr3 | 1647.567 | 0.011421 | 0.931959 | 0.99306 |
| ENSMUSG( | S100a1 | 4119.255 | -0.27722 | 0.043119 | 0.690304 |
| ENSMUSG( | S100a10 | 447.4727 | -0.27911 | 0.077687 | 0.77321 |
| ENSMUSG( | S100a11 | 125.4781 | -0.0214 | 0.917189 | 0.992529 |
| ENSMUSG( | S100a13 | 1190.057 | -0.23485 | 0.096067 | 0.796133 |
| ENSMUSG( | S100a16 | 4283.596 | -0.41575 | 0.001845 | 0.15992 |
| ENSMUSG( | S100a3 | 15.72822 | -0.01353 | 0.979402 | 0.996468 |
| ENSMUSG( | S100a4 | 18.04302 | -0.25549 | 0.592275 | 0.944975 |
| ENSMUSG( | S100a6 | 701.5561 | -0.21373 | 0.155326 | 0.837724 |
| ENSMUSG( | S100a7a | 19.02775 | -0.15059 | 0.753501 | 0.970671 |
| ENSMUSG( | S100a8 | 103.9462 | -0.03335 | 0.969305 | 0.99594 |

| | | | | | |
|---|---|---|---|---|---|
| ENSMUSG( | 1340.973 | 0.089048 | 0.496646 | 0.86175 | Rock1 |
| ENSMUSG( | 4459.048 | 0.090231 | 0.292802 | 0.742763 | Rock2 |
| ENSMUSG( | 2894.35 | 0.182019 | 0.180156 | 0.617634 | Rogdi |
| ENSMUSG( | 523.9815 | -0.10738 | 0.401601 | 0.814923 | Rom1 |
| ENSMUSG( | 1521.447 | -0.02515 | 0.789634 | 0.959461 | Romo1 |
| ENSMUSG( | 34.67869 | -0.20708 | 0.533177 | 0.879905 | Ropn1l |
| ENSMUSG( | 55.82557 | -0.09773 | 0.718512 | 0.941911 | Ror1 |
| ENSMUSG( | 41.11521 | -0.01995 | 0.948781 | 0.993821 | Ror2 |
| ENSMUSG( | 2383.532 | 0.090674 | 0.416837 | 0.8209 | Rora |
| ENSMUSG( | 1655.577 | -0.10691 | 0.342256 | 0.778828 | Rorb |
| ENSMUSG( | 115.4549 | -0.19943 | 0.29407 | 0.743459 | Rorc |
| ENSMUSG( | 656.2671 | 0.076096 | 0.591181 | 0.90113 | Rp2 |
| ENSMUSG( | 965.1964 | 0.094288 | 0.411814 | 0.81942 | Rp9 |
| ENSMUSG( | 1823.222 | 0.224627 | 0.033184 | 0.278882 | Rpa1 |
| ENSMUSG( | 340.0576 | 0.039814 | 0.789168 | 0.959299 | Rpa2 |
| ENSMUSG( | 162.1989 | -0.11257 | 0.497084 | 0.861754 | Rpa3 |
| ENSMUSG( | 291.9784 | -0.00706 | 0.968439 | 0.996369 | Rpain |
| ENSMUSG( | 696.3964 | 0.177297 | 0.098915 | 0.480935 | Rpap1 |
| ENSMUSG( | 541.4064 | 0.023565 | 0.852865 | 0.973168 | Rpap2 |
| ENSMUSG( | 559.9078 | 0.159249 | 0.201445 | 0.641362 | Rpap3 |
| ENSMUSG( | 614.4978 | 0.108449 | 0.36905 | 0.796581 | Rpe |
| ENSMUSG( | 85.38052 | 0.211612 | 0.368704 | 0.796356 | Rpe65 |
| ENSMUSG( | 313.4799 | 0.133943 | 0.320487 | 0.765921 | Rpf1 |
| ENSMUSG( | 339.6436 | 0.027815 | 0.837642 | 0.971111 | Rpf2 |
| ENSMUSG( | 348.4614 | 0.022486 | 0.879635 | 0.981436 | Rpgr |
| ENSMUSG( | 24.81347 | 0.025564 | 0.947832 | 0.993821 | Rpgrip1 |
| ENSMUSG( | 419.8062 | 0.239539 | 0.05238 | 0.35132 | Rpgrip1l |
| ENSMUSG( | 30550.33 | -0.02913 | 0.689067 | 0.933441 | Rph3a |
| ENSMUSG( | 70.22893 | 0.357479 | 0.144383 | 0.56431 | Rph3al |
| ENSMUSG( | 243.5359 | -0.13541 | 0.400813 | 0.81387 | Rpia |
| ENSMUSG( | 9448.112 | 0.106306 | 0.267438 | 0.717525 | Rpl10 |
| ENSMUSG( | 4630.183 | 0.044512 | 0.566983 | 0.891644 | Rpl10a |
| ENSMUSG( | 104.1193 | -0.03168 | 0.888783 | 0.983434 | Rpl10a-ps1 |
| ENSMUSG( | 12.89338 | -0.5378 | 0.311671 | NA | Rpl10-ps3 |
| ENSMUSG( | 8913.379 | 0.090095 | 0.282387 | 0.73186 | Rpl11 |
| ENSMUSG( | 13276.85 | 0.06961 | 0.409135 | 0.818258 | Rpl12 |
| ENSMUSG( | 16234.77 | 0.079701 | 0.326411 | 0.771327 | Rpl13 |
| ENSMUSG( | 1180.678 | 0.174533 | 0.092866 | 0.468228 | Rpl13a |
| ENSMUSG( | 17.44141 | -0.23162 | 0.595865 | NA | Rpl13-ps6 |
| ENSMUSG( | 8852.132 | 0.072081 | 0.379849 | 0.801757 | Rpl14 |
| ENSMUSG( | 79.87483 | 0.095171 | 0.661662 | 0.925271 | Rpl14-ps1 |
| ENSMUSG( | 4706.748 | 0.034587 | 0.70299 | 0.937735 | Rpl15 |
| ENSMUSG( | 13645.03 | 0.091949 | 0.265785 | 0.716441 | Rpl17 |
| ENSMUSG( | 93.77396 | 0.122053 | 0.564975 | 0.890735 | Rpl17-ps10 |
| ENSMUSG( | 11.25199 | 0.077398 | 0.887663 | NA | Rpl17-ps5 |
| ENSMUSG( | 10.2967 | 0.289761 | 0.624409 | NA | Rpl17-ps9 |
| ENSMUSG( | 5629.009 | 0.067575 | 0.430951 | 0.826896 | Rpl18 |
| ENSMUSG( | 10749.58 | 0.067963 | 0.396197 | 0.811615 | Rpl18a |
| ENSMUSG( | 13.91912 | -1.20404 | 0.022017 | NA | Rpl18a-ps1 |
| ENSMUSG( | 17.2684 | -0.08898 | 0.8414 | NA | Rpl18-ps1 |
| ENSMUSG( | 8.714891 | -0.07261 | 0.911722 | NA | Rpl18-ps2 |
| ENSMUSG( | 11956.59 | 0.072961 | 0.384918 | 0.80404 | Rpl19 |
| ENSMUSG( | 97.28339 | 0.075795 | 0.713596 | 0.940866 | Rpl19-ps11 |
| ENSMUSG( | 14555.99 | 0.124562 | 0.138474 | 0.553773 | Rpl21 |
| ENSMUSG( | 27.13102 | 0.322531 | 0.362279 | 0.792943 | Rpl21-ps11 |
| ENSMUSG( | 20.76819 | 0.060458 | 0.885407 | NA | Rpl21-ps8 |
| ENSMUSG( | 5360.809 | 0.039158 | 0.65889 | 0.924506 | Rpl22 |
| ENSMUSG( | 1301.939 | -0.01419 | 0.906914 | 0.987391 | Rpl22l1 |
| ENSMUSG( | 9250.528 | 0.061222 | 0.485554 | 0.857637 | Rpl23 |
| ENSMUSG( | 9185.722 | 0.124581 | 0.148484 | 0.571935 | Rpl23a |
| ENSMUSG( | 27.29178 | 0.0893 | 0.806267 | 0.962804 | Rpl23a-ps3 |
| ENSMUSG( | 63.25084 | -0.0607 | 0.804768 | 0.962473 | Rpl24 |
| ENSMUSG( | 11204.03 | 0.132789 | 0.119344 | 0.523036 | Rpl26 |
| ENSMUSG( | 2782.203 | 0.02292 | 0.795692 | 0.960062 | Rpl27 |
| ENSMUSG( | 9031.711 | 0.109851 | 0.235283 | 0.682423 | Rpl27a |
| ENSMUSG( | 9303.741 | 0.039519 | 0.622458 | 0.911432 | Rpl28 |
| ENSMUSG( | 51.72437 | 0.030068 | 0.910055 | 0.9881 | Rpl28-ps1 |
| ENSMUSG( | 36.85917 | -0.25167 | 0.413841 | 0.819998 | Rpl28-ps3 |
| ENSMUSG( | 8351.473 | 0.080927 | 0.328191 | 0.772854 | Rpl29 |
| ENSMUSG( | 18152.41 | 0.06525 | 0.3962 | 0.811615 | Rpl3 |
| ENSMUSG( | 6303.662 | 0.043716 | 0.652744 | 0.922498 | Rpl30 |
| ENSMUSG( | 10957.92 | 0.051224 | 0.506268 | 0.867425 | Rpl31 |
| ENSMUSG( | 7413.378 | 0.089178 | 0.336301 | 0.777 | Rpl32 |
| ENSMUSG( | 28.06241 | 0.04143 | 0.911778 | 0.988483 | Rpl32l |
| ENSMUSG( | 5060.723 | 0.112631 | 0.218502 | 0.664312 | Rpl34 |
| ENSMUSG( | 50.89771 | 0.240905 | 0.357936 | 0.789465 | Rpl34-ps1 |
| ENSMUSG( | 5389.33 | 0.075014 | 0.404499 | 0.816132 | Rpl35 |
| ENSMUSG( | 5971.658 | 0.134699 | 0.17214 | 0.607115 | Rpl35a |
| ENSMUSG( | 439.1068 | 0.093524 | 0.422241 | 0.823508 | Rpl35rt |
| ENSMUSG( | 5695.019 | 0.065087 | 0.523061 | 0.875628 | Rpl36 |
| ENSMUSG( | 3604.278 | 0.069977 | 0.405939 | 0.81652 | Rpl36a |
| ENSMUSG( | 11.5924 | -0.3337 | 0.594309 | NA | Rpl36al |
| ENSMUSG( | 30.05313 | -0.32387 | 0.372263 | 0.798768 | Rpl36a-ps1 |
| ENSMUSG( | 50.20572 | 0.141104 | 0.599115 | 0.903801 | Rpl36a-ps2 |
| ENSMUSG( | 31.78516 | 0.162213 | 0.648332 | 0.919949 | Rpl36-ps12 |
| ENSMUSG( | 9336.886 | 0.132111 | 0.153197 | 0.57989 | Rpl37 |
| ENSMUSG( | 7056.42 | 0.04831 | 0.657894 | 0.92425 | Rpl37a |
| ENSMUSG( | 6567.161 | 0.079353 | 0.364647 | 0.793796 | Rpl38 |
| ENSMUSG( | 3852.453 | 0.116349 | 0.282175 | 0.73186 | Rpl39 |
| ENSMUSG( | 12.53184 | 0.232771 | 0.673503 | NA | Rpl3l |
| ENSMUSG( | 215.6948 | 0.208005 | 0.176512 | 0.612904 | Rpl3-ps1 |
| ENSMUSG( | 28.08217 | 0.488931 | 0.196843 | 0.636268 | Rpl3-ps2 |
| ENSMUSG( | 24701.14 | 0.123511 | 0.11045 | 0.50108 | Rpl4 |
| ENSMUSG( | 18195.86 | 0.074134 | 0.327196 | 0.772056 | Rpl41 |
| ENSMUSG( | 9169.467 | 0.057618 | 0.488504 | 0.859092 | Rpl5 |
| ENSMUSG( | 9.243582 | 0.207431 | 0.759557 | NA | Rpl5-ps1 |
| ENSMUSG( | 8608.473 | 0.068739 | 0.411846 | 0.81942 | Rpl6 |
| ENSMUSG( | 28.28501 | -0.05082 | 0.887654 | 0.983411 | Rpl6l |
| ENSMUSG( | 9860.35 | 0.096591 | 0.261441 | 0.711405 | Rpl7 |
| ENSMUSG( | 12120.9 | 0.042766 | 0.555022 | 0.88708 | Rpl7a |
| ENSMUSG( | 9.846013 | -0.42251 | 0.471698 | NA | Rpl7a-ps7 |
| ENSMUSG( | 1167.557 | 0.127334 | 0.207376 | 0.649561 | Rpl7l1 |
| ENSMUSG( | 21.18984 | -0.10766 | 0.794166 | NA | Rpl7-ps7 |
| ENSMUSG( | 13.82444 | 0.745177 | 0.133192 | NA | Rpl7-ps8 |
| ENSMUSG( | 11.83647 | 0.106575 | 0.8432 | NA | Rpl7-ps9 |
| ENSMUSG( | 13443.61 | 0.076313 | 0.365302 | 0.794085 | Rpl8 |
| ENSMUSG( | 19180.16 | 0.115639 | 0.249655 | 0.699258 | Rpl9 |
| ENSMUSG( | 133.172 | -0.1291 | 0.553629 | 0.886216 | Rpl9-ps6 |
| ENSMUSG( | 15244.56 | 0.126136 | 0.137309 | 0.552371 | Rplp0 |
| ENSMUSG( | 13.37509 | -0.16042 | 0.754544 | NA | Rplp0-ps1 |
| ENSMUSG( | 28207.83 | 0.089583 | 0.349923 | 0.781943 | Rplp1 |
| ENSMUSG( | 7455.379 | 0.093563 | 0.395532 | 0.810727 | Rplp2 |
| ENSMUSG( | 5680.649 | -0.27735 | 0.038554 | 0.302696 | Rpn1 |
| ENSMUSG( | 5007.381 | -0.18913 | 0.121719 | 0.526333 | Rpn2 |
| ENSMUSG( | 1415.461 | 0.227588 | 0.027214 | 0.253245 | Rpp14 |
| ENSMUSG( | 319.4343 | -0.1456 | 0.320294 | 0.76591 | Rpp21 |
| ENSMUSG( | 300.3855 | -0.0072 | 0.970483 | 0.996369 | Rpp25 |
| ENSMUSG( | 330.2131 | 0.025794 | 0.857291 | 0.97444 | Rpp25l |
| ENSMUSG( | 215.7679 | 0.044752 | 0.789203 | 0.959299 | Rpp30 |
| ENSMUSG( | 95.52977 | -0.12102 | 0.62678 | 0.912825 | Rpp38 |
| ENSMUSG( | 93.88332 | 0.163728 | 0.437067 | 0.829501 | Rpp40 |
| ENSMUSG( | 1718.355 | 0.074154 | 0.428186 | 0.825894 | Rprd1a |
| ENSMUSG( | 558.871 | 0.081165 | 0.492927 | 0.860552 | Rprd1b |
| ENSMUSG( | 1139.335 | 0.095427 | 0.289261 | 0.739725 | Rprd2 |
| ENSMUSG( | 1209.277 | -0.0384 | 0.799361 | 0.961201 | Rprm |
| ENSMUSG( | 1066.408 | -0.19026 | 0.213407 | 0.65774 | Rprml |
| ENSMUSG( | 69.69386 | -0.31218 | 0.264637 | 0.71564 | Rps10 |
| ENSMUSG( | 86.36258 | 0.101784 | 0.660209 | 0.924627 | Rps10-ps1 |
| ENSMUSG( | 40.57311 | 0.362166 | 0.214719 | 0.659193 | Rps10-ps2 |
| ENSMUSG( | 8608.344 | 0.08815 | 0.276111 | 0.725738 | Rps11 |
| ENSMUSG( | 13.36604 | -0.34593 | 0.538146 | NA | Rps11-ps1 |
| ENSMUSG( | 24.13041 | -0.27169 | 0.472387 | 0.848048 | Rps11-ps2 |
| ENSMUSG( | 10339.84 | 0.11746 | 0.271532 | 0.722082 | Rps12 |
| ENSMUSG( | 4637.59 | 0.083203 | 0.355 | 0.787423 | Rps13 |
| ENSMUSG( | 32.01057 | -0.13445 | 0.692634 | 0.934101 | Rps13-ps1 |
| ENSMUSG( | 27.82877 | -0.02597 | 0.94332 | 0.993415 | Rps13-ps2 |
| ENSMUSG( | 11618.58 | 0.109926 | 0.144213 | 0.563869 | Rps14 |
| ENSMUSG( | 21208.4 | 0.151801 | 0.042016 | 0.315804 | Rps15 |
| ENSMUSG( | 5104.639 | 0.064187 | 0.462705 | 0.842333 | Rps15a |
| ENSMUSG( | 22.46896 | -0.2516 | 0.531043 | 0.879289 | Rps15a-ps3 |
| ENSMUSG( | 10.15164 | -0.52789 | 0.423908 | NA | Rps15a-ps7 |
| ENSMUSG( | 18.43288 | 0.450318 | 0.297574 | NA | Rps15-ps3 |
| ENSMUSG( | 7988.823 | 0.06768 | 0.434352 | 0.828978 | Rps16 |
| ENSMUSG( | 7049.323 | 0.080369 | 0.400157 | 0.813641 | Rps17 |
| ENSMUSG( | 7710.581 | 0.036948 | 0.640686 | 0.916434 | Rps18 |
| ENSMUSG( | 47.38534 | 0.162305 | 0.600709 | 0.90443 | Rps18-ps5 |
| ENSMUSG( | 8099.286 | 0.086084 | 0.297208 | 0.745805 | Rps19 |
| ENSMUSG( | 393.3012 | 0.061098 | 0.713206 | 0.940734 | Rps19bp1 |
| ENSMUSG( | 51.4495 | 0.57744 | 0.04647 | 0.330252 | Rps19-ps3 |
| ENSMUSG( | 33.32892 | -0.11144 | 0.72589 | 0.944223 | Rps19-ps6 |
| ENSMUSG( | 16.80112 | 0.117373 | 0.801362 | NA | Rps19-ps7 |
| ENSMUSG( | 23946.14 | 0.116562 | 0.157701 | 0.585927 | Rps2 |
| ENSMUSG( | 8486.326 | 0.09918 | 0.293037 | 0.742763 | Rps20 |
| ENSMUSG( | 12031.27 | 0.21712 | 0.039271 | 0.305097 | Rps21 |
| ENSMUSG( | 9400.677 | 0.099867 | 0.221809 | 0.667414 | Rps23 |
| ENSMUSG( | 177.5093 | 0.221091 | 0.250575 | 0.700403 | Rps23-ps1 |
| ENSMUSG( | 47.89604 | -0.23361 | 0.432465 | 0.82768 | Rps23-ps2 |
| ENSMUSG( | 15098.14 | 0.076037 | 0.404458 | 0.816132 | Rps24 |

| | | | | | |
|---|---|---|---|---|---|
| ENSMUSG( | 17480.41 | -0.13782 | 0.245221 | 0.877904 | Rps15 |
| ENSMUSG( | 4354.726 | -0.12929 | 0.254892 | 0.882307 | Rps15a |
| ENSMUSG( | 20.69536 | -0.29493 | 0.558302 | 0.93957 | Rps15a-ps3 |
| ENSMUSG( | 8.69315 | -0.80423 | 0.226328 | NA | Rps15a-ps5 |
| ENSMUSG( | 9.829559 | -0.30331 | 0.62641 | NA | Rps15a-ps7 |
| ENSMUSG( | 12.5536 | -0.37667 | 0.492916 | 0.927358 | Rps15-ps3 |
| ENSMUSG( | 6735.102 | -0.16188 | 0.214409 | 0.864421 | Rps16 |
| ENSMUSG( | 6005.625 | -0.1174 | 0.360082 | 0.914223 | Rps17 |
| ENSMUSG( | 6513.687 | -0.1883 | 0.118443 | 0.813384 | Rps18 |
| ENSMUSG( | 37.67715 | -0.24091 | 0.5091 | 0.929089 | Rps18-ps5 |
| ENSMUSG( | 6789.679 | -0.15899 | 0.217292 | 0.865589 | Rps19 |
| ENSMUSG( | 338.1997 | -0.11323 | 0.53867 | 0.937216 | Rps19bp1 |
| ENSMUSG( | 36.70575 | -0.10062 | 0.771671 | 0.974236 | Rps19-ps3 |
| ENSMUSG( | 29.321 | -0.24989 | 0.50655 | 0.928006 | Rps19-ps6 |
| ENSMUSG( | 11.67415 | -0.84089 | 0.162394 | 0.837724 | Rps19-ps7 |
| ENSMUSG( | 20273.87 | -0.09582 | 0.46065 | 0.923329 | Rps2 |
| ENSMUSG( | 7109.101 | -0.14763 | 0.264544 | 0.886134 | Rps20 |
| ENSMUSG( | 10101.56 | -0.01355 | 0.919612 | 0.992644 | Rps21 |
| ENSMUSG( | 7930.679 | -0.12453 | 0.301936 | 0.898538 | Rps23 |
| ENSMUSG( | 147.6482 | -0.04941 | 0.824344 | 0.979198 | Rps23-ps1 |
| ENSMUSG( | 39.82339 | -0.56773 | 0.133084 | 0.822912 | Rps23-ps2 |
| ENSMUSG( | 12951.48 | -0.10037 | 0.439132 | 0.920721 | Rps24 |
| ENSMUSG( | 6321.83 | -0.15436 | 0.211049 | 0.864412 | Rps25 |
| ENSMUSG( | 6002.4 | -0.16709 | 0.179133 | 0.849453 | Rps26 |
| ENSMUSG( | 7365.624 | -0.15457 | 0.216615 | 0.865167 | Rps27 |
| ENSMUSG( | 8028.971 | -0.11002 | 0.387909 | 0.917257 | Rps27a |
| ENSMUSG( | 13.17384 | 0.046019 | 0.934237 | 0.99306 | Rps27a-ps2 |
| ENSMUSG( | 28.45485 | -0.77177 | 0.049841 | 0.708648 | Rps27a-ps3 |
| ENSMUSG( | 311.0137 | -0.24628 | 0.099414 | 0.800296 | Rps27l |
| ENSMUSG( | 44.62135 | -0.49503 | 0.107218 | 0.804969 | Rps27rt |
| ENSMUSG( | 4343.645 | -0.18466 | 0.158545 | 0.837724 | Rps28 |
| ENSMUSG( | 12117.89 | -0.23044 | 0.068143 | 0.75212 | Rps29 |
| ENSMUSG( | 16.02486 | 0.210503 | 0.671037 | 0.95694 | Rps2-ps10 |
| ENSMUSG( | 12961.16 | -0.11348 | 0.378078 | 0.916904 | Rps3 |
| ENSMUSG( | 9565.717 | -0.10736 | 0.37299 | 0.915491 | Rps3a1 |
| ENSMUSG( | 35.82344 | -0.19793 | 0.568997 | 0.941777 | Rps3a2 |
| ENSMUSG( | 64.6501 | -0.45047 | 0.111242 | 0.807885 | Rps3a3 |
| ENSMUSG( | 10103.36 | -0.1655 | 0.176412 | 0.846867 | Rps4x |
| ENSMUSG( | 20.03339 | -1.02233 | 0.021197 | 0.57477 | Rps4x-ps |
| ENSMUSG( | 8099.164 | -0.16295 | 0.179433 | 0.850372 | Rps5 |
| ENSMUSG( | 4278.335 | -0.13798 | 0.220641 | 0.866104 | Rps6 |
| ENSMUSG( | 610.234 | 0.028794 | 0.822572 | 0.979198 | Rps6ka1 |
| ENSMUSG( | 1733.787 | 0.329806 | 0.023426 | 0.585845 | Rps6ka2 |
| ENSMUSG( | 1245.998 | 0.026608 | 0.831063 | 0.98051 | Rps6ka3 |
| ENSMUSG( | 1227.547 | 0.221844 | 0.057314 | 0.723362 | Rps6ka4 |
| ENSMUSG( | 465.807 | -0.08675 | 0.562488 | 0.940761 | Rps6ka5 |
| ENSMUSG( | 111.6675 | -0.12571 | 0.58265 | 0.944266 | Rps6ka6 |
| ENSMUSG( | 1160.847 | 0.12567 | 0.305621 | 0.898538 | Rps6kb1 |
| ENSMUSG( | 1038.609 | 0.026933 | 0.885773 | 0.98977 | Rps6kb2 |
| ENSMUSG( | 1038.669 | -0.04116 | 0.733404 | 0.967228 | Rps6kc1 |
| ENSMUSG( | 756.2537 | -0.06613 | 0.680602 | 0.958395 | Rps6kl1 |
| ENSMUSG( | 92.34173 | -0.3464 | 0.153373 | 0.837724 | Rps6-ps4 |
| ENSMUSG( | 7959.861 | -0.10646 | 0.399685 | 0.917257 | Rps7 |
| ENSMUSG( | 11413.68 | -0.06952 | 0.561197 | 0.940254 | Rps8 |
| ENSMUSG( | 10161.26 | -0.13463 | 0.287268 | 0.894091 | Rps9 |
| ENSMUSG( | 25917.02 | -0.14589 | 0.213607 | 0.864421 | Rpsa |
| ENSMUSG( | 97.00768 | -0.16002 | 0.462287 | 0.923329 | Rpsa-ps10 |
| ENSMUSG( | 77.62625 | -0.06414 | 0.794592 | 0.97681 | Rpsa-ps12 |
| ENSMUSG( | 32.13954 | -0.2552 | 0.495078 | 0.927372 | Rpsa-ps2 |
| ENSMUSG( | 41.71516 | -0.01418 | 0.966662 | 0.995475 | Rpsa-ps7 |
| ENSMUSG( | 16.61759 | 0.039186 | 0.938869 | 0.993898 | Rpsa-ps9 |
| ENSMUSG( | 2173.121 | -0.00609 | 0.964998 | 0.995433 | Rptor |
| ENSMUSG( | 1183.754 | -0.10202 | 0.462846 | 0.923329 | Rpusd1 |
| ENSMUSG( | 78.63891 | 0.119433 | 0.610741 | 0.950771 | Rpusd2 |
| ENSMUSG( | 228.5407 | 0.244242 | 0.145028 | 0.83339 | Rpusd3 |
| ENSMUSG( | 182.8193 | -0.01495 | 0.931342 | 0.99306 | Rpusd4 |
| ENSMUSG( | 38.49689 | 0.234683 | 0.498404 | 0.927825 | Rrad |
| ENSMUSG( | 1364.389 | -0.24142 | 0.11755 | 0.813384 | Rraga |
| ENSMUSG( | 849.5525 | 0.05861 | 0.67444 | 0.95694 | Rragb |
| ENSMUSG( | 1841.231 | -0.00213 | 0.986596 | 0.997372 | Rragc |
| ENSMUSG( | 3600.564 | -0.13774 | 0.286726 | 0.894091 | Rragd |
| ENSMUSG( | 533.1229 | -0.03367 | 0.80006 | 0.97681 | Rras |
| ENSMUSG( | 430.7225 | 0.013623 | 0.92831 | 0.99306 | Rras2 |
| ENSMUSG( | 3780.787 | 0.092143 | 0.482403 | 0.924578 | Rrbp1 |
| ENSMUSG( | 1662.224 | 0.119122 | 0.582197 | 0.944266 | Rreb1 |
| ENSMUSG( | 556.1001 | -0.02812 | 0.836711 | 0.981335 | Rrm1 |
| ENSMUSG( | 118.6423 | -0.19165 | 0.337964 | 0.907972 | Rrm2 |
| ENSMUSG( | 840.7968 | -0.03322 | 0.819927 | 0.979198 | Rrm2b |
| ENSMUSG( | 1391.49 | -0.05786 | 0.657782 | 0.955974 | Rrn3 |
| ENSMUSG( | 6200.762 | -0.19451 | 0.201758 | 0.864412 | Rrp1 |
| ENSMUSG( | 739.0998 | -0.0065 | 0.959806 | 0.994798 | Rrp12 |
| ENSMUSG( | 195.3486 | -0.07372 | 0.667972 | 0.956766 | Rrp15 |
| ENSMUSG( | 432.0176 | 0.083772 | 0.572028 | 0.942369 | Rrp1b |
| ENSMUSG( | 353.6651 | -0.04617 | 0.774877 | 0.974931 | Rrp36 |
| ENSMUSG( | 2009.731 | -0.09084 | 0.447518 | 0.922202 | Rrp7a |
| ENSMUSG( | 466.4208 | 0.522029 | 0.000115 | 0.022849 | Rrp8 |
| ENSMUSG( | 524.644 | 0.176969 | 0.185735 | 0.855305 | Rrp9 |
| ENSMUSG( | 427.7969 | 0.003064 | 0.986882 | 0.997473 | Rrs1 |
| ENSMUSG( | 381.6531 | 0.160697 | 0.307536 | 0.898538 | Rsad1 |
| ENSMUSG( | 121.1105 | -0.13429 | 0.555146 | 0.939565 | Rsad2 |
| ENSMUSG( | 440.9311 | 0.131184 | 0.39104 | 0.917257 | Rsbn1 |
| ENSMUSG( | 498.2385 | -0.00044 | 0.997622 | 0.999268 | Rsbn1l |
| ENSMUSG( | 1715.263 | 0.027635 | 0.81664 | 0.979198 | Rsf1 |
| ENSMUSG( | 173.3376 | 0.273723 | 0.313749 | 0.900949 | Rskr |
| ENSMUSG( | 100.7579 | 0.172535 | 0.439117 | 0.920721 | Rsl1 |
| ENSMUSG( | 924.2111 | -0.04867 | 0.673455 | 0.95694 | Rsl1d1 |
| ENSMUSG( | 393.9193 | 0.009739 | 0.948454 | 0.994798 | Rsl24d1 |
| ENSMUSG( | 101.4077 | -0.07103 | 0.800568 | 0.97681 | Rsph1 |
| ENSMUSG( | 184.8393 | -0.05642 | 0.754952 | 0.971098 | Rsph3a |
| ENSMUSG( | 211.6495 | -0.00709 | 0.966831 | 0.995532 | Rsph3b |
| ENSMUSG( | 86.50558 | 0.6225 | 0.016519 | 0.519511 | Rsph4a |
| ENSMUSG( | 448.3916 | -0.093 | 0.552538 | 0.939565 | Rsph9 |
| ENSMUSG( | 408.9029 | -0.06412 | 0.65379 | 0.955015 | Rspo1 |
| ENSMUSG( | 416.6938 | 0.236793 | 0.256663 | 0.883082 | Rspo2 |
| ENSMUSG( | 495.084 | 0.273226 | 0.065005 | 0.741314 | Rspo3 |
| ENSMUSG( | 873.3851 | 0.066853 | 0.612803 | 0.951071 | Rspry1 |
| ENSMUSG( | 1123.318 | -0.21075 | 0.073499 | 0.763629 | Rsrc1 |
| ENSMUSG( | 1667.24 | 0.176315 | 0.12209 | 0.813384 | Rsrc2 |
| ENSMUSG( | 7890.575 | 0.258265 | 0.238179 | 0.87259 | Rsrp1 |
| ENSMUSG( | 917.1119 | -0.09499 | 0.439637 | 0.920721 | Rsu1 |
| ENSMUSG( | 34.65142 | -0.22218 | 0.586226 | 0.944975 | Rtbdn |
| ENSMUSG( | 1029.407 | -0.23915 | 0.066638 | 0.751155 | Rtca |
| ENSMUSG( | 3684.356 | -0.03093 | 0.825101 | 0.979354 | Rtcb |
| ENSMUSG( | 499.0028 | 0.051248 | 0.772181 | 0.974236 | Rtel1 |
| ENSMUSG( | 1792.928 | 0.019496 | 0.876772 | 0.989644 | Rtf1 |
| ENSMUSG( | 1503.794 | -0.12616 | 0.373553 | 0.915491 | Rtf2 |
| ENSMUSG( | 1516.741 | -0.04152 | 0.740326 | 0.968811 | Rtkn |
| ENSMUSG( | 93.14564 | -0.2012 | 0.373948 | 0.915491 | Rtkn2 |
| ENSMUSG( | 40.12467 | 0.139026 | 0.665209 | 0.956766 | Rtl4 |
| ENSMUSG( | 631.2659 | -0.08001 | 0.539 | 0.937216 | Rtl5 |
| ENSMUSG( | 1342.619 | 0.033887 | 0.770308 | 0.974236 | Rtl6 |
| ENSMUSG( | 1183.665 | -0.24853 | 0.075103 | 0.767994 | Rtl8a |
| ENSMUSG( | 1814.256 | -0.23258 | 0.0822 | 0.77681 | Rtl8b |
| ENSMUSG( | 1882.058 | -0.22979 | 0.047438 | 0.700936 | Rtl8c |
| ENSMUSG( | 59.41001 | -0.00525 | 0.984165 | 0.996963 | Rtl9 |
| ENSMUSG( | 40669.56 | 0.055032 | 0.708551 | 0.964042 | Rtn1 |
| ENSMUSG( | 5038.939 | 0.061453 | 0.560143 | 0.940119 | Rtn2 |
| ENSMUSG( | 40681.78 | 0.091656 | 0.527747 | 0.934603 | Rtn3 |
| ENSMUSG( | 15350.68 | 0.175245 | 0.27604 | 0.890679 | Rtn4 |
| ENSMUSG( | 309.7629 | -0.15662 | 0.284626 | 0.892412 | Rtn4ip1 |
| ENSMUSG( | 3398.331 | 0.185817 | 0.273351 | 0.889687 | Rtn4r |
| ENSMUSG( | 2185.149 | 0.108648 | 0.504785 | 0.927825 | Rtn4rl1 |
| ENSMUSG( | 1209.208 | 0.355291 | 0.055784 | 0.723362 | Rtn4rl2 |
| ENSMUSG( | 93.18373 | 1.06978 | 1.20E-05 | 0.004289 | Rtp1 |
| ENSMUSG( | 126.4319 | 0.05914 | 0.788476 | 0.97637 | Rtp4 |
| ENSMUSG( | 1199.095 | -0.14082 | 0.25653 | 0.883082 | Rtraf |
| ENSMUSG( | 120.9417 | -0.01947 | 0.933044 | 0.99306 | Rttn |
| ENSMUSG( | 2837.839 | -0.04235 | 0.68101 | 0.958395 | Rubcn |
| ENSMUSG( | 45.93961 | 0.358441 | 0.278732 | 0.891628 | Rubcnl |
| ENSMUSG( | 1021.591 | 0.017017 | 0.903055 | 0.991089 | Rufy1 |
| ENSMUSG( | 2085.843 | 0.053283 | 0.663233 | 0.956359 | Rufy2 |
| ENSMUSG( | 4040.554 | -0.09278 | 0.481786 | 0.924578 | Rufy3 |
| ENSMUSG( | 1050.997 | -0.05117 | 0.676377 | 0.95694 | Rundc1 |
| ENSMUSG( | 16924.05 | -0.0179 | 0.847687 | 0.9852 | Rundc3a |
| ENSMUSG( | 460.6842 | -0.11146 | 0.541784 | 0.937841 | Rundc3b |
| ENSMUSG( | 46.19983 | -0.12389 | 0.671069 | 0.95694 | Runx1 |
| ENSMUSG( | 252.8492 | 0.205298 | 0.342619 | 0.909646 | Runx1t1 |
| ENSMUSG( | 183.61 | 0.265223 | 0.209594 | 0.864412 | Runx2 |
| ENSMUSG( | 3449.717 | -0.07416 | 0.593793 | 0.945408 | Rusc1 |
| ENSMUSG( | 3429.161 | 0.114742 | 0.310959 | 0.899422 | Rusc2 |
| ENSMUSG( | 928.5517 | 0.122734 | 0.38962 | 0.917257 | Rusf1 |
| ENSMUSG( | 435.3366 | -0.16786 | 0.238796 | 0.873304 | Ruvbl1 |
| ENSMUSG( | 685.647 | -0.05317 | 0.688962 | 0.959742 | Ruvbl2 |
| ENSMUSG( | 695.9658 | 0.012399 | 0.931084 | 0.99306 | Rwdd1 |
| ENSMUSG( | 203.3178 | -0.12978 | 0.538401 | 0.937216 | Rwdd2a |
| ENSMUSG( | 173.314 | 0.126537 | 0.528794 | 0.934795 | Rwdd2b |
| ENSMUSG( | 98.09248 | -0.21958 | 0.343384 | 0.909753 | Rwdd3 |
| ENSMUSG( | 703.922 | 0.051601 | 0.731325 | 0.967228 | Rwdd4a |

| | | | | | | | | | | | |
|---|---|---|---|---|---|---|---|---|---|---|---|
| ENSMUSG( | S100a9 | 143.354 | -1.20172 | 0.27505 | 0.72509 | ENSMUSG( | S100a9 | 174.6839 | -0.14088 | 0.876487 | 0.989644 |
| ENSMUSG( | S100b | 18349.94 | -0.006 | 0.956562 | 0.993874 | ENSMUSG( | S100b | 15329.61 | -0.27146 | 0.056998 | 0.723362 |
| ENSMUSG( | S100pbp | 913.6211 | -0.17727 | 0.150653 | 0.575407 | ENSMUSG( | S100pbp | 898.6258 | 0.050889 | 0.710726 | 0.964363 |
| ENSMUSG( | S1pr1 | 14049.93 | -0.18496 | 0.240883 | 0.68836 | ENSMUSG( | S1pr1 | 13422.59 | -0.0397 | 0.807005 | 0.978239 |
| ENSMUSG( | S1pr2 | 104.9802 | -1.04457 | 0.000164 | 0.009126 | ENSMUSG( | S1pr2 | 124.279 | -0.08296 | 0.780211 | 0.976106 |
| ENSMUSG( | S1pr3 | 367.9207 | -0.15708 | 0.534504 | 0.880719 | ENSMUSG( | S1pr3 | 406.6302 | 0.390463 | 0.371976 | 0.915491 |
| ENSMUSG( | S1pr4 | 16.88229 | -0.19592 | 0.678219 | NA | ENSMUSG( | S1pr4 | 18.90023 | 0.419683 | 0.408944 | 0.917257 |
| ENSMUSG( | S1pr5 | 989.9627 | -0.65897 | 9.55E-07 | 0.000184 | ENSMUSG( | S1pr5 | 987.6164 | -0.33811 | 0.040445 | 0.682882 |
| ENSMUSG( | S2bpcox16 | 17.86966 | 0.311363 | 0.504686 | NA | ENSMUSG( | S2bpcox16 | 14.32371 | -0.04312 | 0.932208 | 0.99306 |
| ENSMUSG( | Saal1 | 286.6273 | 0.012052 | 0.926808 | 0.991229 | ENSMUSG( | Saal1 | 277.6328 | 0.187168 | 0.211146 | 0.864412 |
| ENSMUSG( | Sac3d1 | 414.9438 | -0.02988 | 0.831963 | 0.970194 | ENSMUSG( | Sac3d1 | 368.3311 | -0.10977 | 0.487729 | 0.925427 |
| ENSMUSG( | Sacm1l | 1394.824 | -0.00987 | 0.912898 | 0.988483 | ENSMUSG( | Sacm1l | 1315.245 | 0.086478 | 0.442314 | 0.921554 |
| ENSMUSG( | Sacs | 1937.587 | -0.44497 | 0.000414 | 0.018428 | ENSMUSG( | Sacs | 2188.489 | 0.205305 | 0.155812 | 0.837724 |
| ENSMUSG( | Sae1 | 1990.697 | 0.035121 | 0.686075 | 0.932981 | ENSMUSG( | Sae1 | 1664.018 | -0.22452 | 0.075854 | 0.768524 |
| ENSMUSG( | Safb | 3004.508 | 0.157342 | 0.121478 | 0.525713 | ENSMUSG( | Safb | 2626.481 | 0.039715 | 0.759736 | 0.972189 |
| ENSMUSG( | Safb2 | 2605.44 | 0.099321 | 0.416255 | 0.82084 | ENSMUSG( | Safb2 | 2335.359 | 0.049867 | 0.739831 | 0.968726 |
| ENSMUSG( | Sag | 94.56259 | -0.16291 | 0.522306 | 0.875186 | ENSMUSG( | Sag | 87.55996 | -0.09893 | 0.683101 | 0.958678 |
| ENSMUSG( | Sall1 | 1081.6 | 0.033481 | 0.775863 | 0.955606 | ENSMUSG( | Sall1 | 1081.842 | 0.292386 | 0.019694 | 0.556192 |
| ENSMUSG( | Sall2 | 2431.17 | -0.07612 | 0.393471 | 0.809501 | ENSMUSG( | Sall2 | 2119.547 | -0.2125 | 0.074817 | 0.767423 |
| ENSMUSG( | Sall3 | 436.0629 | 0.116152 | 0.440032 | 0.831851 | ENSMUSG( | Sall3 | 424.2461 | 0.30017 | 0.039297 | 0.679972 |
| ENSMUSG( | Samd1 | 1405.568 | 0.05883 | 0.544966 | 0.884712 | ENSMUSG( | Samd1 | 1246.904 | -0.01996 | 0.864249 | 0.986552 |
| ENSMUSG( | Samd10 | 672.6048 | 0.104345 | 0.34313 | 0.779153 | ENSMUSG( | Samd10 | 549.5003 | -0.21824 | 0.103105 | 0.804311 |
| ENSMUSG( | Samd11 | 51.19845 | -0.23645 | 0.400197 | 0.813641 | ENSMUSG( | Samd11 | 49.97212 | -0.03007 | 0.927304 | 0.99306 |
| ENSMUSG( | Samd12 | 649.3216 | 0.073701 | 0.471331 | 0.847103 | ENSMUSG( | Samd12 | 581.2538 | 0.018704 | 0.886924 | 0.990201 |
| ENSMUSG( | Samd14 | 3112.891 | 0.202648 | 0.023769 | 0.235641 | ENSMUSG( | Samd14 | 2750.934 | 0.117622 | 0.294977 | 0.898538 |
| ENSMUSG( | Samd15 | 52.47748 | -0.14098 | 0.589707 | 0.90113 | ENSMUSG( | Samd15 | 43.7738 | -0.40701 | 0.213853 | 0.864421 |
| ENSMUSG( | Samd3 | 18.59469 | -0.43211 | 0.384444 | NA | ENSMUSG( | Samd3 | 17.54627 | -0.31455 | 0.528148 | 0.934677 |
| ENSMUSG( | Samd4 | 2357.327 | -0.17838 | 0.063289 | 0.388993 | ENSMUSG( | Samd4 | 2015.643 | -0.3848 | 0.003278 | 0.22876 |
| ENSMUSG( | Samd4b | 3740.05 | 0.187642 | 0.018712 | 0.205766 | ENSMUSG( | Samd4b | 3248.428 | 0.055017 | 0.593792 | 0.945408 |
| ENSMUSG( | Samd5 | 674.7524 | 0.096914 | 0.407955 | 0.818167 | ENSMUSG( | Samd5 | 555.2753 | -0.20969 | 0.175939 | 0.845985 |
| ENSMUSG( | Samd8 | 2277.249 | -0.28437 | 0.046767 | 0.330815 | ENSMUSG( | Samd8 | 2403.314 | 0.15377 | 0.35048 | 0.911652 |
| ENSMUSG( | Samd9l | 178.3487 | -0.10405 | 0.529664 | 0.878883 | ENSMUSG( | Samd9l | 190.1359 | 0.34595 | 0.07682 | 0.770558 |
| ENSMUSG( | Samhd1 | 703.2065 | -0.14977 | 0.176914 | 0.613937 | ENSMUSG( | Samhd1 | 656.6446 | -0.07527 | 0.573608 | 0.942912 |
| ENSMUSG( | Samm50 | 2381.677 | 0.166662 | 0.126905 | 0.534201 | ENSMUSG( | Samm50 | 2049.328 | 0.005969 | 0.961256 | 0.99511 |
| ENSMUSG( | Samsn1 | 64.24328 | -0.30388 | 0.227995 | 0.675324 | ENSMUSG( | Samsn1 | 65.55392 | 0.041593 | 0.874099 | 0.989644 |
| ENSMUSG( | Sanbr | 959.7647 | 0.148219 | 0.159905 | 0.589679 | ENSMUSG( | Sanbr | 856.7897 | 0.089229 | 0.464241 | 0.923329 |
| ENSMUSG( | Sap130 | 911.973 | 0.095352 | 0.320874 | 0.765984 | ENSMUSG( | Sap130 | 823.0478 | 0.069743 | 0.574027 | 0.942912 |
| ENSMUSG( | Sap18 | 2343.205 | 0.050062 | 0.590053 | 0.90113 | ENSMUSG( | Sap18 | 1949.664 | -0.22335 | 0.066777 | 0.751729 |
| ENSMUSG( | Sap18b | 13.04516 | 0.607557 | 0.229507 | NA | ENSMUSG( | Sap18b | 12.31714 | 0.665666 | 0.231663 | 0.868507 |
| ENSMUSG( | Sap30 | 189.4004 | 0.142859 | 0.447674 | 0.834051 | ENSMUSG( | Sap30 | 168.5474 | 0.06896 | 0.748659 | 0.96984 |
| ENSMUSG( | Sap30bp | 1344.936 | 0.107388 | 0.294163 | 0.743459 | ENSMUSG( | Sap30bp | 1162.749 | -0.04097 | 0.732855 | 0.967228 |
| ENSMUSG( | Sap30l | 1851.563 | 0.085142 | 0.349077 | 0.781665 | ENSMUSG( | Sap30l | 1574.488 | -0.11911 | 0.34289 | 0.909646 |
| ENSMUSG( | Sapcd1 | 27.13021 | 0.341744 | 0.363478 | 0.793339 | ENSMUSG( | Sapcd1 | 20.9278 | -0.11712 | 0.797446 | 0.97681 |
| ENSMUSG( | Sapcd2 | 82.92602 | -0.30325 | 0.205717 | 0.647154 | ENSMUSG( | Sapcd2 | 80.76471 | -0.11285 | 0.669011 | 0.956766 |
| ENSMUSG( | Sar1a | 4693.773 | 0.017711 | 0.822475 | 0.966352 | ENSMUSG( | Sar1a | 4132.542 | -0.08278 | 0.402278 | 0.917257 |
| ENSMUSG( | Sar1b | 1835.403 | -0.33488 | 0.002199 | 0.054456 | ENSMUSG( | Sar1b | 1817.299 | -0.07702 | 0.587087 | 0.944975 |
| ENSMUSG( | Saraf | 12227.19 | -0.21026 | 0.076163 | 0.42702 | ENSMUSG( | Saraf | 11935.21 | -0.00136 | 0.991903 | 0.99872 |
| ENSMUSG( | Sardh | 1847.585 | 0.008428 | 0.93807 | 0.993415 | ENSMUSG( | Sardh | 1570.909 | -0.19952 | 0.117586 | 0.813384 |
| ENSMUSG( | Sarm1 | 969.1526 | 0.173292 | 0.075628 | 0.426231 | ENSMUSG( | Sarm1 | 862.6621 | 0.112122 | 0.366495 | 0.914703 |
| ENSMUSG( | Sarnp | 789.7536 | 0.139557 | 0.240157 | 0.687899 | ENSMUSG( | Sarnp | 689.2677 | 0.013508 | 0.918845 | 0.992529 |
| ENSMUSG( | Sars1 | 4152.132 | 0.017867 | 0.860991 | 0.975025 | ENSMUSG( | Sars1 | 3673.266 | -0.06921 | 0.557766 | 0.939565 |
| ENSMUSG( | Sars2 | 332.9046 | -0.00315 | 0.984425 | 0.99781 | ENSMUSG( | Sars2 | 304.861 | 0.006526 | 0.969467 | 0.99594 |
| ENSMUSG( | Sart1 | 1666.48 | 0.128546 | 0.222924 | 0.668853 | ENSMUSG( | Sart1 | 1429.815 | -0.04631 | 0.720497 | 0.966887 |
| ENSMUSG( | Sart3 | 1091.204 | 0.212854 | 0.084572 | 0.448139 | ENSMUSG( | Sart3 | 957.3911 | 0.108597 | 0.444711 | 0.921734 |
| ENSMUSG( | Sash1 | 11108.32 | -0.03754 | 0.771197 | 0.954726 | ENSMUSG( | Sash1 | 8471.3 | -0.61711 | 1.05E-05 | 0.003931 |
| ENSMUSG( | Sash3 | 110.2421 | 0.104542 | 0.652438 | 0.922226 | ENSMUSG( | Sash3 | 96.53917 | -0.00498 | 0.98467 | 0.996963 |
| ENSMUSG( | Sass6 | 98.16063 | -0.01273 | 0.948717 | 0.993821 | ENSMUSG( | Sass6 | 98.31176 | 0.262783 | 0.25158 | 0.879838 |
| ENSMUSG( | Sat1 | 2746.183 | 0.053214 | 0.644699 | 0.917782 | ENSMUSG( | Sat1 | 2252.318 | -0.25855 | 0.043056 | 0.690304 |
| ENSMUSG( | Sat2 | 255.0042 | 0.102368 | 0.469377 | 0.846015 | ENSMUSG( | Sat2 | 223.9596 | -0.00183 | 0.991141 | 0.998379 |
| ENSMUSG( | Satb1 | 4213.507 | 0.213737 | 0.021033 | 0.220525 | ENSMUSG( | Satb1 | 3643.07 | 0.06881 | 0.556647 | 0.939565 |
| ENSMUSG( | Satb2 | 824.5242 | 0.148226 | 0.317405 | 0.763823 | ENSMUSG( | Satb2 | 728.7916 | 0.06055 | 0.700524 | 0.961533 |
| ENSMUSG( | Sav1 | 696.6651 | 0.012554 | 0.900963 | 0.985633 | ENSMUSG( | Sav1 | 644.1037 | 0.054021 | 0.654091 | 0.955015 |
| ENSMUSG( | Saxo2 | 73.20205 | 0.039051 | 0.874106 | 0.979336 | ENSMUSG( | Saxo2 | 71.09138 | 0.205399 | 0.429519 | 0.917257 |
| ENSMUSG( | Saxo4 | 37.61986 | -0.18978 | 0.602325 | 0.904887 | ENSMUSG( | Saxo4 | 41.80505 | 0.387755 | 0.223268 | 0.867404 |
| ENSMUSG( | Saysd1 | 229.0593 | -0.26972 | 0.083719 | 0.446196 | ENSMUSG( | Saysd1 | 234.1346 | 0.077513 | 0.63423 | 0.953903 |
| ENSMUSG( | Sbds | 2407.26 | -0.01307 | 0.903167 | 0.986055 | ENSMUSG( | Sbds | 2080.501 | -0.17531 | 0.225578 | 0.868468 |
| ENSMUSG( | Sbf1 | 8948.369 | 0.105445 | 0.198861 | 0.639096 | ENSMUSG( | Sbf1 | 7858.624 | 0.002758 | 0.980363 | 0.996479 |
| ENSMUSG( | Sbf2 | 3464.786 | 0.078758 | 0.387748 | 0.805835 | ENSMUSG( | Sbf2 | 3042.175 | -0.02851 | 0.792386 | 0.97681 |
| ENSMUSG( | Sbk1 | 7748.679 | 0.175641 | 0.041488 | 0.314241 | ENSMUSG( | Sbk1 | 6751.734 | 0.051222 | 0.624366 | 0.953234 |
| ENSMUSG( | Sbk3 | 33.86252 | 0.119795 | 0.731062 | 0.944956 | ENSMUSG( | Sbk3 | 27.94869 | -0.20428 | 0.60363 | 0.948137 |
| ENSMUSG( | Sbno1 | 3983.073 | 0.101966 | 0.214072 | 0.658621 | ENSMUSG( | Sbno1 | 3435.786 | -0.05571 | 0.613379 | 0.951177 |
| ENSMUSG( | Sbno2 | 814.3808 | 0.17913 | 0.119744 | 0.523801 | ENSMUSG( | Sbno2 | 706.5065 | 0.045567 | 0.746771 | 0.969763 |
| ENSMUSG( | Sbsn | 123.9865 | -0.12055 | 0.524934 | 0.87624 | ENSMUSG( | Sbsn | 120.2456 | 0.073135 | 0.722809 | 0.966887 |
| ENSMUSG( | Sbspon | 13.39936 | -0.66712 | 0.199714 | NA | ENSMUSG( | Sbspon | 12.71869 | -0.47734 | 0.389409 | 0.917257 |
| ENSMUSG( | Sc5d | 6184.815 | -0.20888 | 0.124151 | 0.531714 | ENSMUSG( | Sc5d | 5803.054 | -0.11826 | 0.421716 | 0.917257 |
| ENSMUSG( | Scaf1 | 8852.477 | 0.184341 | 0.035346 | 0.289405 | ENSMUSG( | Scaf1 | 7751.625 | 0.075756 | 0.462784 | 0.923329 |
| ENSMUSG( | Scaf11 | 2090.416 | 0.13873 | 0.117952 | 0.518899 | ENSMUSG( | Scaf11 | 1869.736 | 0.087211 | 0.420786 | 0.917257 |
| ENSMUSG( | Scaf4 | 1248.898 | 0.006474 | 0.944804 | 0.993612 | ENSMUSG( | Scaf4 | 1158.589 | 0.055794 | 0.660289 | 0.956205 |
| ENSMUSG( | Scaf8 | 1412.084 | 0.194978 | 0.031153 | 0.268462 | ENSMUSG( | Scaf8 | 1206.79 | 0.018115 | 0.86992 | 0.988097 |
| ENSMUSG( | Scai | 669.7103 | -0.18549 | 0.099051 | 0.481021 | ENSMUSG( | Scai | 706.5908 | 0.236948 | 0.104848 | 0.804969 |
| ENSMUSG( | Scamp1 | 7684.57 | -0.35106 | 0.009405 | 0.13706 | ENSMUSG( | Scamp1 | 8005.67 | 0.058165 | 0.702156 | 0.962014 |
| ENSMUSG( | Scamp2 | 1044.894 | -0.26166 | 0.078769 | 0.434128 | ENSMUSG( | Scamp2 | 1118.98 | 0.215636 | 0.217756 | 0.865589 |
| ENSMUSG( | Scamp3 | 1572.811 | -0.21225 | 0.100555 | 0.483976 | ENSMUSG( | Scamp3 | 1613.974 | 0.140185 | 0.325527 | 0.905339 |
| ENSMUSG( | Scamp5 | 14540.85 | -0.30718 | 0.037092 | 0.297239 | ENSMUSG( | Scamp5 | 15159.62 | 0.101156 | 0.547749 | 0.938063 |
| ENSMUSG( | Scand1 | 1111.632 | -0.06127 | 0.584022 | 0.898666 | ENSMUSG( | Scand1 | 955.5333 | -0.24537 | 0.130536 | 0.822225 |
| ENSMUSG( | Scap | 3424.963 | -0.00999 | 0.944195 | 0.993503 | ENSMUSG( | Scap | 3414.201 | 0.244714 | 0.159274 | 0.837724 |
| ENSMUSG( | Scaper | 856.759 | 0.040156 | 0.724215 | 0.944096 | ENSMUSG( | Scaper | 741.7301 | -0.11259 | 0.369998 | 0.915491 |
| ENSMUSG( | Scara3 | 3158.633 | -0.12079 | 0.378577 | 0.801688 | ENSMUSG( | Scara3 | 3040.66 | 0.045061 | 0.779171 | 0.976106 |
| ENSMUSG( | Scara5 | 25.25554 | 0.500248 | 0.200392 | 0.640592 | ENSMUSG( | Scara5 | 21.95935 | 0.419011 | 0.372043 | 0.915491 |
| ENSMUSG( | Scarb1 | 947.384 | -0.19346 | 0.193318 | 0.632794 | ENSMUSG( | Scarb1 | 994.7108 | 0.221401 | 0.204252 | 0.864412 |
| ENSMUSG( | Scarb2 | 8023.86 | -0.18973 | 0.159564 | 0.589015 | ENSMUSG( | Scarb2 | 7938.843 | 0.056661 | 0.709874 | 0.96416 |
| ENSMUSG( | Scarf1 | 109.7224 | 0.029568 | 0.900381 | 0.985633 | ENSMUSG( | Scarf1 | 114.8942 | 0.424279 | 0.110649 | 0.807582 |
| ENSMUSG( | Scarf2 | 285.9164 | -0.13983 | 0.354087 | 0.786537 | ENSMUSG( | Scarf2 | 307.6024 | 0.334735 | 0.068087 | 0.75212 |
| ENSMUSG( | Sccpdh | 2768.414 | 0.01324 | 0.909246 | 0.98794 | ENSMUSG( | Sccpdh | 2513.07 | 0.002846 | 0.981707 | 0.996756 |
| ENSMUSG( | Scd1 | 43979.41 | 0.148328 | 0.203409 | 0.643631 | ENSMUSG( | Scd1 | 36616.68 | -0.11045 | 0.353945 | 0.912411 |
| ENSMUSG( | Scd2 | 172863.2 | 0.052494 | 0.664909 | 0.926824 | ENSMUSG( | Scd2 | 150056.8 | -0.08768 | 0.481041 | 0.924548 |
| ENSMUSG( | Scd3 | 153.629 | 0.069425 | 0.679518 | 0.930983 | ENSMUSG( | Scd3 | 129.8489 | -0.15764 | 0.422112 | 0.917257 |
| ENSMUSG( | Scd4 | 18.08884 | -0.04756 | 0.923293 | NA | ENSMUSG( | Scd4 | 17.49096 | 0.144324 | 0.789912 | 0.976798 |
| ENSMUSG( | Scel | 92.82818 | -0.03776 | 0.885031 | 0.982829 | ENSMUSG( | Scel | 80.4911 | -0.21387 | 0.4404 | 0.921166 |
| ENSMUSG( | Scfd1 | 769.8936 | 0.119062 | 0.277291 | 0.727292 | ENSMUSG( | Scfd1 | 654.766 | -0.07908 | 0.533868 | 0.935551 |
| ENSMUSG( | Scfd2 | 526.0871 | 0.08641 | 0.45979 | 0.840338 | ENSMUSG( | Scfd2 | 443.3823 | -0.13987 | 0.300302 | 0.898538 |
| ENSMUSG( | Scg2 | 5681.015 | -0.4577 | 0.000912 | 0.03071 | ENSMUSG( | Scg2 | 6141.454 | 0.075469 | 0.630124 | 0.953251 |
| ENSMUSG( | Scg3 | 33489.22 | -0.02824 | 0.813933 | 0.964574 | ENSMUSG( | Scg3 | 30860.36 | 0.006162 | 0.963459 | 0.995433 |
| ENSMUSG( | Scg5 | 6283.85 | -0.37429 | 0.000309 | 0.014712 | ENSMUSG( | Scg5 | 6770.805 | 0.132675 | 0.311422 | 0.900022 |
| ENSMUSG( | Schip1 | 190.954 | 0.058771 | 0.774153 | 0.955606 | ENSMUSG( | Schip1 | 161.2724 | -0.16852 | 0.452267 | 0.922316 |
| ENSMUSG( | Sclt1 | 259.6808 | -0.08684 | 0.552613 | 0.886141 | ENSMUSG( | Sclt1 | 247.9744 | 0.05688 | 0.718587 | 0.966633 |
| ENSMUSG( | Scly | 540.6618 | 0.049048 | 0.652354 | 0.922188 | ENSMUSG( | Scly | 455.3591 | -0.17637 | 0.20377 | 0.864412 |
| ENSMUSG( | Scmh1 | 1932.146 | 0.027803 | 0.760489 | 0.952907 | ENSMUSG( | Scmh1 | 1774.789 | 0.052115 | 0.626094 | 0.953234 |
| ENSMUSG( | Scml2 | 23.6272 | -0.68613 | 0.106446 | 0.494174 | ENSMUSG( | Scml2 | 23.23031 | -0.3873 | 0.413833 | 0.917257 |
| ENSMUSG( | Scml4 | 64.35649 | 0.17615 | 0.490616 | 0.859465 | ENSMUSG( | Scml4 | 57.31961 | 0.092363 | 0.747329 | 0.969763 |
| ENSMUSG( | Scn10a | 10.89894 | 0.447803 | 0.430787 | NA | ENSMUSG( | Scn10a | 12.62623 | 1.000994 | 0.072161 | 0.763629 |
| ENSMUSG( | Scn1a | 2675.637 | -0.34914 | 0.011045 | 0.151501 | ENSMUSG( | Scn1a | 2942.439 | 0.210574 | 0.229587 | 0.868507 |
| ENSMUSG( | Scn1b | 20636.07 | -0.16017 | 0.096574 | 0.476986 | ENSMUSG( | Scn1b | 19487.06 | -0.05168 | 0.638114 | 0.953903 |
| ENSMUSG( | Scn2a | 5943.005 | -0.37119 | 0.017905 | 0.200989 | ENSMUSG( | Scn2a | 6731.05 | 0.272114 | 0.12402 | 0.814691 |
| ENSMUSG( | Scn2b | 7508.646 | -0.18481 | 0.179463 | 0.616974 | ENSMUSG( | Scn2b | 7643.039 | 0.142178 | 0.403506 | 0.917257 |
| ENSMUSG( | Scn3a | 924.329 | -0.42398 | 0.007383 | 0.118765 | ENSMUSG( | Scn3a | 1102.189 | 0.364556 | 0.032663 | 0.637512 |
| ENSMUSG( | Scn3b | 3327.812 | -0.29039 | 0.013031 | 0.165126 | ENSMUSG( | Scn3b | 3620.355 | 0.230281 | 0.075544 | 0.768472 |
| ENSMUSG( | Scn4b | 4762.87 | -1.2151 | 1.44E-09 | 6.68E-07 | ENSMUSG( | Scn4b | 5846.858 | -0.10545 | 0.465006 | 0.923329 |
| ENSMUSG( | Scn5a | 86.52799 | -0.08434 | 0.757071 | 0.951513 | ENSMUSG( | Scn5a | 86.74562 | 0.193123 | 0.4085 | 0.917257 |
| ENSMUSG( | Scn7a | 80.24495 | -0.40775 | 0.135482 | 0.548884 | ENSMUSG( | Scn7a | 94.06721 | 0.343668 | 0.196657 | 0.862161 |
| ENSMUSG( | Scn8a | 9901.71 | -0.22851 | 0.082013 | 0.442387 | ENSMUSG( | Scn8a | 10930.17 | 0.319786 | 0.027041 | 0.603041 |
| ENSMUSG( | Scn9a | 54.62667 | -0.4863 | 0.094426 | 0.471713 | ENSMUSG( | Scn9a | 57.48266 | -0.05264 | 0.869617 | 0.988097 |
| ENSMUSG( | Scnn1a | 225.8279 | -0.15119 | 0.336791 | 0.777 | ENSMUSG( | Scnn1a | 221.5387 | 0.077915 | 0.671181 | 0.95694 |
| ENSMUSG( | Sco1 | 242.5355 | -0.01899 | 0.894354 | 0.984512 | ENSMUSG( | Sco1 | 224.0926 | 0.022286 | 0.905267 | 0.991089 |
| ENSMUSG( | Sco2 | 154.4228 | -0.42795 | 0.024973 | 0.243133 | ENSMUSG( | Sco2 | 161.0146 | -0.00065 | 0.997298 | 0.999268 |
| ENSMUSG( | Scoc | 1904.064 | -0.01608 | 0.885556 | 0.983008 | ENSMUSG( | Scoc | 1736.168 | -0.01901 | 0.896428 | 0.990558 |
| ENSMUSG( | Scp2 | 5842.6 | 0.127156 | 0.235414 | 0.682423 | ENSMUSG( | Scp2 | 4848.632 | -0.14625 | 0.258534 | 0.883971 |
| ENSMUSG( | Scpep1 | 764.3629 | -0.45707 | 0.002215 | 0.054686 | ENSMUSG( | Scpep1 | 818.2422 | 0.048854 | 0.761955 | 0.97318 |
| ENSMUSG( | Scrg1 | 2919.776 | 0.23685 | 0.038664 | 0.302965 | ENSMUSG( | Scrg1 | 2471.854 | 0.033264 | 0.788201 | 0.97637 |
| ENSMUSG( | Scrib | 3655.605 | 0.205835 | 0.015575 | 0.184473 | ENSMUSG( | Scrib | 3024.211 | -0.067 | 0.528542 | 0.934795 |
| ENSMUSG( | Scrn1 | 6468.806 | -0.08105 | 0.367046 | 0.795164 | ENSMUSG( | Scrn1 | 5737.946 | -0.16232 | 0.112318 | 0.809018 |
| ENSMUSG( | Scrn2 | 154.118 | 0.022638 | 0.897501 | 0.985633 | ENSMUSG( | Scrn2 | 138.1755 | -0.02276 | 0.909027 | 0.991958 |
| ENSMUSG( | Scrn3 | 621.9394 | 0.061911 | 0.552061 | 0.886141 | ENSMUSG( | Scrn3 | 540.089 | -0.08419 | 0.558407 | 0.93957 |
| ENSMUSG( | Scrt1 | 8878.176 | 0.17435 | 0.056613 | 0.367011 | ENSMUSG( | Scrt1 | 7865.966 | 0.097855 | 0.40255 | 0.917257 |
| ENSMUSG( | Scrt2 | 624.4872 | -0.02379 | 0.850702 | 0.972638 | ENSMUSG( | Scrt2 | 564.9067 | -0.04968 | 0.742737 | 0.969415 |
| ENSMUSG( | Sctr | 49.61086 | 0.389953 | 0.150858 | 0.575879 | ENSMUSG( | Sctr | 41.80593 | 0.209884 | 0.520425 | 0.93204 |
| ENSMUSG( | Scube1 | 1452.76 | -0.16643 | 0.231869 | 0.679725 | ENSMUSG( | Scube1 | 1360.883 | -0.0764 | 0.627589 | 0.953234 |
| ENSMUSG( | Scube2 | 33.10319 | 0.40092 | 0.253169 | 0.70338 | ENSMUSG( | Scube2 | 40.36367 | 1.094537 | 0.002257 | 0.184738 |
| ENSMUSG( | Scube3 | 144.1454 | -0.99898 | 3.15E-06 | 0.000452 | ENSMUSG( | Scube3 | 162.306 | -0.22863 | 0.209358 | 0.864412 |
| ENSMUSG( | Scx | 43.5097 | 0.167837 | 0.560466 | 0.888934 | ENSMUSG( | Scx | 43.84146 | 0.451272 | 0.153504 | 0.837724 |
| ENSMUSG( | Scyl1 | 2636.189 | 0.158749 | 0.119255 | 0.523036 | ENSMUSG( | Scyl1 | 2306.967 | 0.044924 | 0.696042 | 0.960867 |
| ENSMUSG( | Scyl2 | 714.5817 | -0.03151 | 0.7852 | 0.958473 | ENSMUSG( | Scyl2 | 630.3692 | -0.12546 | 0.344142 | 0.910256 |
| ENSMUSG( | Scyl3 | 642.418 | -0.13364 | 0.276968 | 0.727074 | ENSMUSG( | Scyl3 | 622.6652 | 0.05128 | 0.686095 | 0.958956 |
| ENSMUSG( | Sdad1 | 622.8361 | -0.01206 | 0.919523 | 0.990247 | ENSMUSG( | Sdad1 | 550.193 | -0.1006 | 0.438597 | 0.920721 |
| ENSMUSG( | Sdc1 | 45.79341 | -0.20321 | 0.522055 | 0.875186 | ENSMUSG( | Sdc1 | 53.50579 | 0.497447 | 0.124292 | 0.814691 |
| ENSMUSG( | Sdc2 | 1493.207 | -0.29028 | 0.041838 | 0.315339 | ENSMUSG( | Sdc2 | 1550.486 | 0.103138 | 0.525278 | 0.934374 |
| ENSMUSG( | Sdc3 | 11758.19 | -0.19326 | 0.108879 | 0.49803 | ENSMUSG( | Sdc3 | 11424.5 | 0.001709 | 0.990367 | 0.998129 |
| ENSMUSG( | Sdc4 | 6569.437 | 0.219694 | 0.022649 | 0.22975 | ENSMUSG( | Sdc4 | 6310.361 | 0.358215 | 0.002491 | 0.195662 |
| ENSMUSG( | Sdcbp | 3186.01 | 0.040608 | 0.697043 | 0.935543 | ENSMUSG( | Sdcbp | 2818.58 | -0.04813 | 0.725734 | 0.966887 |
| ENSMUSG( | Sdcbp2 | 22.1147 | -0.19535 | 0.642502 | 0.916865 | ENSMUSG( | Sdcbp2 | 19.87338 | -0.21038 | 0.636346 | 0.953903 |
| ENSMUSG( | Sdccag8 | 481.7705 | 0.151602 | 0.213955 | 0.658384 | ENSMUSG( | Sdccag8 | 426.4209 | 0.074945 | 0.614644 | 0.951177 |
| ENSMUSG( | Sde2 | 397.88 | 0.043041 | 0.734679 | 0.945584 | ENSMUSG( | Sde2 | 358.554 | 0.01234 | 0.93133 | 0.99306 |
| ENSMUSG( | Sdf2 | 1625.839 | -0.29802 | 0.028502 | 0.257525 | ENSMUSG( | Sdf2 | 1641.349 | 0.018162 | 0.906227 | 0.991281 |
| ENSMUSG( | Sdf2l1 | 426.4222 | -0.00267 | 0.986877 | 0.998619 | ENSMUSG( | Sdf2l1 | 463.3734 | 0.466863 | 0.007972 | 0.382597 |
| ENSMUSG( | Sdf4 | 9616.81 | -0.13348 | 0.178045 | 0.615613 | ENSMUSG( | Sdf4 | 9129.662 | -0.01006 | 0.932858 | 0.99306 |
| ENSMUSG( | Sdha | 15335.06 | 0.083414 | 0.333656 | 0.775602 | ENSMUSG( | Sdha | 12677.96 | -0.20376 | 0.05086 | 0.709741 |
| ENSMUSG( | Sdhaf1 | 763.2384 | -0.08937 | 0.427609 | 0.825692 | ENSMUSG( | Sdhaf1 | 657.2031 | -0.26957 | 0.056324 | 0.723362 |
| ENSMUSG( | Sdhaf2 | 1309.985 | 0.054492 | 0.555054 | 0.88708 | ENSMUSG( | Sdhaf2 | 1115.766 | -0.14599 | 0.21032 | 0.864412 |

| | | | | | |
|---|---|---|---|---|---|
| ENSMUSG( | 7550.83 | 0.09394 | 0.277474 | 0.727432 | Rps25 |
| ENSMUSG( | 7108.076 | 0.058553 | 0.50651 | 0.867425 | Rps26 |
| ENSMUSG( | 8939.419 | 0.138326 | 0.101395 | 0.484996 | Rps27 |
| ENSMUSG( | 9610.839 | 0.140958 | 0.138223 | 0.553167 | Rps27a |
| ENSMUSG( | 17.08272 | 0.497485 | 0.300245 | NA | Rps27a-ps2 |
| ENSMUSG( | 40.45571 | 0.05555 | 0.86193 | 0.975055 | Rps27a-ps3 |
| ENSMUSG( | 387.4307 | 0.131544 | 0.310549 | 0.758631 | Rps27l |
| ENSMUSG( | 59.2002 | 0.082495 | 0.745699 | 0.949242 | Rps27rt |
| ENSMUSG( | 5212.486 | 0.080078 | 0.356238 | 0.788108 | Rps28 |
| ENSMUSG( | 14775.33 | 0.08334 | 0.366462 | 0.794761 | Rps29 |
| ENSMUSG( | 19.56695 | 0.521644 | 0.237196 | NA | Rps2-ps10 |
| ENSMUSG( | 15228.93 | 0.085991 | 0.299747 | 0.74879 | Rps3 |
| ENSMUSG( | 11177.67 | 0.075612 | 0.365209 | 0.794085 | Rps3a1 |
| ENSMUSG( | 48.29444 | 0.379791 | 0.185545 | 0.623185 | Rps3a2 |
| ENSMUSG( | 83.05172 | 0.055276 | 0.817821 | 0.965785 | Rps3a3 |
| ENSMUSG( | 12125.74 | 0.096732 | 0.262388 | 0.713005 | Rps4x |
| ENSMUSG( | 29.15287 | -0.02069 | 0.951664 | 0.993821 | Rps4x-ps |
| ENSMUSG( | 9645.875 | 0.07795 | 0.336812 | 0.777 | Rps5 |
| ENSMUSG( | 4998.91 | 0.047151 | 0.577656 | 0.895925 | Rps6 |
| ENSMUSG( | 672.2529 | 0.034436 | 0.742662 | 0.948478 | Rps6ka1 |
| ENSMUSG( | 1826.1 | 0.216588 | 0.055289 | 0.361957 | Rps6ka2 |
| ENSMUSG( | 1398.282 | 0.091373 | 0.376737 | 0.801213 | Rps6ka3 |
| ENSMUSG( | 1305.618 | 0.133363 | 0.181763 | 0.618944 | Rps6ka4 |
| ENSMUSG( | 487.3807 | -0.23264 | 0.098446 | 0.480238 | Rps6ka5 |
| ENSMUSG( | 131.4992 | 0.091963 | 0.63392 | 0.914073 | Rps6ka6 |
| ENSMUSG( | 1230.294 | 0.03085 | 0.742036 | 0.948478 | Rps6kb1 |
| ENSMUSG( | 1203.215 | 0.180817 | 0.271344 | 0.72198 | Rps6kb2 |
| ENSMUSG( | 1278.469 | 0.275813 | 0.008541 | 0.129467 | Rps6kc1 |
| ENSMUSG( | 877.5504 | 0.096999 | 0.497847 | 0.862343 | Rps6kl1 |
| ENSMUSG( | 10.72482 | 0.282369 | 0.62269 | NA | Rps6-ps1 |
| ENSMUSG( | 115.3357 | 0.06254 | 0.752131 | 0.950417 | Rps6-ps4 |
| ENSMUSG( | 9331.072 | 0.085976 | 0.292568 | 0.742521 | Rps7 |
| ENSMUSG( | 13527.92 | 0.150916 | 0.09178 | 0.465904 | Rps8 |
| ENSMUSG( | 11899.87 | 0.057028 | 0.543832 | 0.884287 | Rps9 |
| ENSMUSG( | 30877.5 | 0.094664 | 0.262868 | 0.713397 | Rpsa |
| ENSMUSG( | 119.8782 | 0.188827 | 0.312373 | 0.759967 | Rpsa-ps10 |
| ENSMUSG( | 85.76074 | -0.02827 | 0.901515 | 0.985633 | Rpsa-ps12 |
| ENSMUSG( | 36.82623 | -0.08028 | 0.794556 | 0.959799 | Rpsa-ps2 |
| ENSMUSG( | 44.38921 | -0.0732 | 0.80869 | 0.963455 | Rpsa-ps7 |
| ENSMUSG( | 20.26819 | 0.337102 | 0.4496 | NA | Rpsa-ps9 |
| ENSMUSG( | 2423.88 | 0.035782 | 0.742907 | 0.948478 | Rptor |
| ENSMUSG( | 1365.354 | 0.043439 | 0.733438 | 0.944956 | Rpusd1 |
| ENSMUSG( | 80.34787 | -0.084 | 0.708653 | 0.939519 | Rpusd2 |
| ENSMUSG( | 231.5776 | 0.021185 | 0.898696 | 0.985633 | Rpusd3 |
| ENSMUSG( | 209.5995 | 0.10501 | 0.470876 | 0.846742 | Rpusd4 |
| ENSMUSG( | 35.86484 | -0.27325 | 0.403475 | 0.816132 | Rrad |
| ENSMUSG( | 1564.294 | -0.10355 | 0.432934 | 0.828185 | Rraga |
| ENSMUSG( | 952.6701 | 0.122909 | 0.269096 | 0.719305 | Rragb |
| ENSMUSG( | 2057.222 | 0.051462 | 0.561147 | 0.888934 | Rragc |
| ENSMUSG( | 4071.336 | -0.04621 | 0.686674 | 0.933041 | Rragd |
| ENSMUSG( | 584.8645 | -0.03231 | 0.782737 | 0.957558 | Rras |
| ENSMUSG( | 509.9924 | 0.224351 | 0.070104 | 0.40908 | Rras2 |
| ENSMUSG( | 3881.878 | -0.10336 | 0.375538 | 0.800845 | Rrbp1 |
| ENSMUSG( | 1623.19 | -0.21704 | 0.232763 | 0.680928 | Rreb1 |
| ENSMUSG( | 630.2704 | 0.065453 | 0.549795 | 0.886008 | Rrm1 |
| ENSMUSG( | 127.9801 | -0.24028 | 0.193985 | 0.633085 | Rrm2 |
| ENSMUSG( | 941.4219 | 0.030224 | 0.8141 | 0.964574 | Rrm2b |
| ENSMUSG( | 1612.062 | 0.100211 | 0.259107 | 0.709814 | Rrn3 |
| ENSMUSG( | 7273.358 | 0.006036 | 0.959993 | 0.993991 | Rrp1 |
| ENSMUSG( | 882.2549 | 0.225721 | 0.0399 | 0.308163 | Rrp12 |
| ENSMUSG( | 224.4091 | 0.058398 | 0.691975 | 0.934101 | Rrp15 |
| ENSMUSG( | 454.4894 | -0.03725 | 0.7485 | 0.949604 | Rrp1b |
| ENSMUSG( | 406.6149 | 0.093997 | 0.447549 | 0.834051 | Rrp36 |
| ENSMUSG( | 2300.825 | 0.031811 | 0.763524 | 0.953845 | Rrp7a |
| ENSMUSG( | 488.6509 | 0.407809 | 0.001567 | 0.044386 | Rrp8 |
| ENSMUSG( | 568.2556 | 0.139656 | 0.232066 | 0.679932 | Rrp9 |
| ENSMUSG( | 480.9433 | 0.07989 | 0.622425 | 0.911432 | Rrs1 |
| ENSMUSG( | 418.5389 | 0.152074 | 0.235966 | 0.683136 | Rsad1 |
| ENSMUSG( | 108.2493 | -0.83916 | 6.86E-05 | 0.005034 | Rsad2 |
| ENSMUSG( | 437.311 | -0.16025 | 0.242401 | 0.6899 | Rsbn1 |
| ENSMUSG( | 518.1372 | -0.15365 | 0.223884 | 0.669845 | Rsbn1l |
| ENSMUSG( | 1908.052 | 0.068265 | 0.492569 | 0.860378 | Rsf1 |
| ENSMUSG( | 213.748 | 0.561091 | 0.01777 | 0.2 | Rskr |
| ENSMUSG( | 101.9491 | -0.0725 | 0.728462 | 0.944393 | Rsl1 |
| ENSMUSG( | 1028.406 | -0.00985 | 0.924718 | 0.99114 | Rsl1d1 |
| ENSMUSG( | 457.0078 | 0.169901 | 0.143802 | 0.562581 | Rsl24d1 |
| ENSMUSG( | 105.5385 | -0.2444 | 0.342227 | 0.778828 | Rsph1 |
| ENSMUSG( | 223.106 | 0.215557 | 0.134248 | 0.547472 | Rsph3a |
| ENSMUSG( | 245.8692 | 0.160319 | 0.246577 | 0.694958 | Rsph3b |
| ENSMUSG( | 64.49937 | -0.48996 | 0.063806 | 0.390932 | Rsph4a |
| ENSMUSG( | 466.8439 | -0.24906 | 0.035852 | 0.290992 | Rsph9 |
| ENSMUSG( | 491.5733 | 0.198936 | 0.098048 | 0.479019 | Rspo1 |
| ENSMUSG( | 372.8102 | -0.37434 | 0.070468 | 0.40908 | Rspo2 |
| ENSMUSG( | 462.6077 | -0.18117 | 0.146777 | 0.56887 | Rspo3 |
| ENSMUSG( | 942.5005 | 0.019796 | 0.856445 | 0.974154 | Rspry1 |
| ENSMUSG( | 1287.331 | -0.07742 | 0.451112 | 0.835567 | Rsrc1 |
| ENSMUSG( | 1718.325 | -0.00018 | 0.998546 | 0.99989 | Rsrc2 |
| ENSMUSG( | 7244.553 | -0.25563 | 0.111497 | 0.503511 | Rsrp1 |
| ENSMUSG( | 1040.253 | 0.003023 | 0.978917 | 0.997428 | Rsu1 |
| ENSMUSG( | 44.32147 | 0.228832 | 0.490977 | 0.859739 | Rtbdn |
| ENSMUSG( | 1207.996 | -0.03283 | 0.775092 | 0.955606 | Rtca |
| ENSMUSG( | 3943.704 | -0.10859 | 0.445617 | 0.833239 | Rtcb |
| ENSMUSG( | 554.9454 | 0.08279 | 0.500319 | 0.863455 | Rtel1 |
| ENSMUSG( | 2017.018 | 0.090878 | 0.370984 | 0.798271 | Rtf1 |
| ENSMUSG( | 1771.525 | 0.082864 | 0.494327 | 0.860611 | Rtf2 |
| ENSMUSG( | 1712.773 | 0.040018 | 0.726082 | 0.944223 | Rtkn |
| ENSMUSG( | 94.77669 | -0.46553 | 0.029709 | 0.261211 | Rtkn2 |
| ENSMUSG( | 49.37716 | 0.44768 | 0.105566 | 0.49243 | Rtl4 |
| ENSMUSG( | 693.8528 | -0.07409 | 0.495252 | 0.860831 | Rtl5 |
| ENSMUSG( | 1554.811 | 0.18015 | 0.04566 | 0.327994 | Rtl6 |
| ENSMUSG( | 1387.807 | -0.04331 | 0.676653 | 0.930542 | Rtl8a |
| ENSMUSG( | 2134.857 | -0.02031 | 0.846433 | 0.972238 | Rtl8b |
| ENSMUSG( | 1982.456 | -0.35997 | 4.05E-05 | 0.003419 | Rtl8c |
| ENSMUSG( | 64.8222 | -0.02238 | 0.925976 | 0.991229 | Rtl9 |
| ENSMUSG( | 36885.99 | -0.54889 | 1.57E-05 | 0.001615 | Rtn1 |
| ENSMUSG( | 5782.651 | 0.183967 | 0.027083 | 0.253045 | Rtn2 |
| ENSMUSG( | 39256.99 | -0.29619 | 0.017573 | 0.198626 | Rtn3 |
| ENSMUSG( | 14326.51 | -0.3074 | 0.031059 | 0.268255 | Rtn4 |
| ENSMUSG( | 372.7052 | 0.109886 | 0.371251 | 0.798633 | Rtn4ip1 |
| ENSMUSG( | 3416.549 | -0.06736 | 0.669453 | 0.928147 | Rtn4r |
| ENSMUSG( | 2128.363 | -0.24985 | 0.09265 | 0.46772 | Rtn4rl1 |
| ENSMUSG( | 1161.824 | -0.01464 | 0.931755 | 0.9924 | Rtn4rl2 |
| ENSMUSG( | 83.94647 | 0.620859 | 0.023673 | 0.23542 | Rtp1 |
| ENSMUSG( | 130.7562 | -0.1249 | 0.58265 | 0.897918 | Rtp4 |
| ENSMUSG( | 1388.815 | 0.021218 | 0.830612 | 0.96974 | Rtraf |
| ENSMUSG( | 126.4887 | -0.18547 | 0.367089 | 0.795164 | Rttn |
| ENSMUSG( | 3312.645 | 0.133442 | 0.108785 | 0.497919 | Rubcn |
| ENSMUSG( | 47.41078 | 0.166385 | 0.605595 | 0.905824 | Rubcnl |
| ENSMUSG( | 1145.637 | 0.080436 | 0.47263 | 0.848298 | Rufy1 |
| ENSMUSG( | 2260.44 | 0.018667 | 0.842189 | 0.971365 | Rufy2 |
| ENSMUSG( | 4709.966 | 0.08388 | 0.397875 | 0.812161 | Rufy3 |
| ENSMUSG( | 1100.519 | -0.19362 | 0.081108 | 0.440221 | Rundc1 |
| ENSMUSG( | 19847.41 | 0.16871 | 0.027099 | 0.253045 | Rundc3a |
| ENSMUSG( | 532.3683 | 0.044073 | 0.792871 | 0.95947 | Rundc3b |
| ENSMUSG( | 45.25701 | -0.49279 | 0.079216 | 0.435105 | Runx1 |
| ENSMUSG( | 276.6211 | 0.210938 | 0.181152 | 0.618139 | Runx1t1 |
| ENSMUSG( | 202.8395 | 0.291895 | 0.125531 | 0.53257 | Runx2 |
| ENSMUSG( | 4147.156 | 0.185183 | 0.169392 | 0.603554 | Rusc1 |
| ENSMUSG( | 3952.421 | 0.247245 | 0.019758 | 0.212093 | Rusc2 |
| ENSMUSG( | 889.1951 | -0.28643 | 0.016703 | 0.192277 | Rusf1 |
| ENSMUSG( | 507.1103 | 0.009741 | 0.935194 | 0.99331 | Ruvbl1 |
| ENSMUSG( | 779.0269 | 0.049814 | 0.668781 | 0.927963 | Ruvbl2 |
| ENSMUSG( | 783.9091 | 0.091418 | 0.354288 | 0.786661 | Rwdd1 |
| ENSMUSG( | 218.9037 | -0.18714 | 0.339067 | 0.777518 | Rwdd2a |
| ENSMUSG( | 178.6351 | -0.05748 | 0.758823 | 0.952113 | Rwdd2b |
| ENSMUSG( | 106.6672 | -0.2496 | 0.228751 | 0.675959 | Rwdd3 |
| ENSMUSG( | 772.6928 | 0.058942 | 0.59497 | 0.903007 | Rwdd4a |
| ENSMUSG( | 585.5142 | -0.08857 | 0.495018 | 0.860798 | Rxfp1 |
| ENSMUSG( | 46.28717 | -0.23525 | 0.411324 | 0.819213 | Rxfp2 |
| ENSMUSG( | 52.90673 | -0.07779 | 0.806464 | 0.962804 | Rxfp3 |
| ENSMUSG( | 1447.952 | 0.353477 | 0.000359 | 0.016466 | Rxra |
| ENSMUSG( | 1944.278 | 0.065841 | 0.506858 | 0.867769 | Rxrb |
| ENSMUSG( | 560.8071 | -0.68137 | 0.070443 | 0.40908 | Rxrg |
| ENSMUSG( | 846.805 | -0.22896 | 0.038559 | 0.302696 | Rxylt1 |
| ENSMUSG( | 440.8919 | 0.125348 | 0.338327 | 0.777489 | Rybp |
| ENSMUSG( | 1987.312 | -0.07127 | 0.541811 | 0.883182 | Ryk |
| ENSMUSG( | 646.1574 | -0.37346 | 0.096313 | 0.476986 | Ryr1 |
| ENSMUSG( | 3746.331 | -0.57193 | 2.20E-07 | 5.18E-05 | Ryr2 |
| ENSMUSG( | 1456.901 | -0.69742 | 2.20E-10 | 1.21E-07 | Ryr3 |
| ENSMUSG( | 4949.475 | 3.94E-07 | 0.999997 | 0.999997 | S100a1 |
| ENSMUSG( | 575.4697 | 0.193726 | 0.098764 | 0.480598 | S100a10 |
| ENSMUSG( | 133.0402 | -0.12636 | 0.493117 | 0.860552 | S100a11 |
| ENSMUSG( | 1444.917 | 0.070064 | 0.591528 | 0.901407 | S100a13 |
| ENSMUSG( | 5280.299 | -0.04943 | 0.616451 | 0.910866 | S100a16 |
| ENSMUSG( | 10.82353 | 0.495306 | 0.410886 | NA | S100a2 |
| ENSMUSG( | 16.92826 | -0.08102 | 0.870912 | NA | S100a3 |
| ENSMUSG( | 18.82059 | -0.37918 | 0.376796 | NA | S100a4 |
| ENSMUSG( | 795.6189 | -0.10711 | 0.343513 | 0.779153 | S100a6 |

| | | | | | |
|---|---|---|---|---|---|
| ENSMUSG( | 592.5667 | 0.212801 | 0.109256 | 0.805353 | Rxfp1 |
| ENSMUSG( | 47.14075 | 0.103859 | 0.729228 | 0.966913 | Rxfp2 |
| ENSMUSG( | 53.76095 | 0.252418 | 0.459327 | 0.923329 | Rxfp3 |
| ENSMUSG( | 1281.444 | 0.278313 | 0.017209 | 0.52741 | Rxra |
| ENSMUSG( | 1721.083 | -0.01451 | 0.894754 | 0.990558 | Rxrb |
| ENSMUSG( | 592.1635 | -0.1888 | 0.293986 | 0.898538 | Rxrg |
| ENSMUSG( | 850.3027 | 0.061991 | 0.644664 | 0.953903 | Rxylt1 |
| ENSMUSG( | 430.1585 | 0.312353 | 0.026456 | 0.603041 | Rybp |
| ENSMUSG( | 1923.311 | 0.102325 | 0.427747 | 0.917257 | Ryk |
| ENSMUSG( | 660.3353 | -0.02242 | 0.927147 | 0.99306 | Ryr1 |
| ENSMUSG( | 4175.092 | 0.068353 | 0.667955 | 0.956766 | Ryr2 |
| ENSMUSG( | 1647.567 | 0.011421 | 0.931959 | 0.99306 | Ryr3 |
| ENSMUSG( | 4119.255 | -0.27722 | 0.043119 | 0.690304 | S100a1 |
| ENSMUSG( | 447.4727 | -0.27911 | 0.077687 | 0.77321 | S100a10 |
| ENSMUSG( | 125.4781 | -0.0214 | 0.917189 | 0.992529 | S100a11 |
| ENSMUSG( | 1190.057 | -0.23485 | 0.096067 | 0.796133 | S100a13 |
| ENSMUSG( | 4283.596 | -0.41575 | 0.001845 | 0.15992 | S100a16 |
| ENSMUSG( | 15.72822 | -0.01353 | 0.979402 | 0.996468 | S100a3 |
| ENSMUSG( | 18.04302 | -0.25549 | 0.592275 | 0.944975 | S100a4 |
| ENSMUSG( | 67.71497 | 0.971717 | 0.644829 | 0.953903 | S100a5 |
| ENSMUSG( | 701.5561 | -0.21373 | 0.155326 | 0.837724 | S100a6 |
| ENSMUSG( | 19.02775 | -0.15059 | 0.753501 | 0.970671 | S100a7a |
| ENSMUSG( | 103.9462 | -0.03335 | 0.969305 | 0.99594 | S100a8 |
| ENSMUSG( | 174.6839 | -0.14088 | 0.876487 | 0.989644 | S100a9 |
| ENSMUSG( | 15329.61 | -0.27146 | 0.056998 | 0.723362 | S100b |
| ENSMUSG( | 898.6258 | 0.050889 | 0.710726 | 0.964363 | S100pbp |
| ENSMUSG( | 13422.59 | -0.0397 | 0.807005 | 0.978239 | S1pr1 |
| ENSMUSG( | 124.279 | -0.08296 | 0.780211 | 0.976106 | S1pr2 |
| ENSMUSG( | 406.6302 | 0.390463 | 0.371976 | 0.915491 | S1pr3 |
| ENSMUSG( | 18.90023 | 0.419683 | 0.408944 | 0.917257 | S1pr4 |
| ENSMUSG( | 987.6164 | -0.33811 | 0.040445 | 0.682882 | S1pr5 |
| ENSMUSG( | 14.32371 | -0.04312 | 0.932208 | 0.99306 | S2bpcox16 |
| ENSMUSG( | 277.6328 | 0.187168 | 0.211146 | 0.864412 | Saal1 |
| ENSMUSG( | 368.3311 | -0.10977 | 0.487729 | 0.925427 | Sac3d1 |
| ENSMUSG( | 1315.245 | 0.086478 | 0.442314 | 0.921554 | Sacm1l |
| ENSMUSG( | 2188.489 | 0.205305 | 0.155812 | 0.837724 | Sacs |
| ENSMUSG( | 1664.018 | -0.22452 | 0.075854 | 0.768524 | Sae1 |
| ENSMUSG( | 2626.481 | 0.039715 | 0.759736 | 0.972189 | Safb |
| ENSMUSG( | 2335.359 | 0.049867 | 0.739831 | 0.968726 | Safb2 |
| ENSMUSG( | 87.55996 | -0.09893 | 0.683101 | 0.958678 | Sag |
| ENSMUSG( | 1081.842 | 0.292386 | 0.019694 | 0.556192 | Sall1 |
| ENSMUSG( | 2119.547 | -0.2125 | 0.074817 | 0.767423 | Sall2 |
| ENSMUSG( | 424.2461 | 0.30017 | 0.039297 | 0.679972 | Sall3 |
| ENSMUSG( | 1246.904 | -0.01996 | 0.864249 | 0.986552 | Samd1 |
| ENSMUSG( | 549.5003 | -0.21824 | 0.103105 | 0.804311 | Samd10 |
| ENSMUSG( | 49.97212 | -0.03007 | 0.927304 | 0.99306 | Samd11 |
| ENSMUSG( | 581.2538 | 0.018704 | 0.886924 | 0.990201 | Samd12 |
| ENSMUSG( | 2750.934 | 0.117622 | 0.294977 | 0.898538 | Samd14 |
| ENSMUSG( | 43.7738 | -0.40701 | 0.213853 | 0.864421 | Samd15 |
| ENSMUSG( | 17.54627 | -0.31455 | 0.528148 | 0.934677 | Samd3 |
| ENSMUSG( | 2015.643 | -0.3848 | 0.003278 | 0.22876 | Samd4 |
| ENSMUSG( | 3248.428 | 0.055017 | 0.593792 | 0.945408 | Samd4b |
| ENSMUSG( | 555.2753 | -0.20969 | 0.175939 | 0.845985 | Samd5 |
| ENSMUSG( | 2403.314 | 0.15377 | 0.35048 | 0.911652 | Samd8 |
| ENSMUSG( | 190.1359 | 0.34595 | 0.07682 | 0.770558 | Samd9l |
| ENSMUSG( | 656.6446 | -0.07527 | 0.573608 | 0.942912 | Samhd1 |
| ENSMUSG( | 2049.328 | 0.005969 | 0.961256 | 0.99511 | Samm50 |
| ENSMUSG( | 65.55392 | 0.041593 | 0.874099 | 0.989644 | Samsn1 |
| ENSMUSG( | 856.7897 | 0.089229 | 0.464241 | 0.923329 | Sanbr |
| ENSMUSG( | 823.0478 | 0.069743 | 0.574027 | 0.942912 | Sap130 |
| ENSMUSG( | 1949.664 | -0.22335 | 0.066777 | 0.751729 | Sap18 |
| ENSMUSG( | 12.31714 | 0.665666 | 0.231663 | 0.868507 | Sap18b |
| ENSMUSG( | 168.5474 | 0.06896 | 0.748659 | 0.96984 | Sap30 |
| ENSMUSG( | 1162.749 | -0.04097 | 0.732855 | 0.967228 | Sap30bp |
| ENSMUSG( | 1574.488 | -0.11911 | 0.34289 | 0.909646 | Sap30l |
| ENSMUSG( | 20.9278 | -0.11712 | 0.797446 | 0.97681 | Sapcd1 |
| ENSMUSG( | 80.76471 | -0.11285 | 0.669011 | 0.956766 | Sapcd2 |
| ENSMUSG( | 4132.542 | -0.08278 | 0.402278 | 0.917257 | Sar1a |
| ENSMUSG( | 1817.299 | -0.07702 | 0.587087 | 0.944975 | Sar1b |
| ENSMUSG( | 11935.21 | -0.00136 | 0.991903 | 0.99872 | Saraf |
| ENSMUSG( | 1570.909 | -0.19952 | 0.117586 | 0.813384 | Sardh |
| ENSMUSG( | 862.6621 | 0.112122 | 0.366495 | 0.914703 | Sarm1 |
| ENSMUSG( | 689.2677 | 0.013508 | 0.918845 | 0.992529 | Sarnp |
| ENSMUSG( | 3673.266 | -0.06921 | 0.557766 | 0.939565 | Sars1 |
| ENSMUSG( | 304.861 | 0.006526 | 0.969467 | 0.99594 | Sars2 |
| ENSMUSG( | 1429.815 | -0.04631 | 0.720497 | 0.966887 | Sart1 |
| ENSMUSG( | 957.3911 | 0.108597 | 0.444711 | 0.921734 | Sart3 |
| ENSMUSG( | 8471.3 | -0.61711 | 1.05E-05 | 0.003931 | Sash1 |
| ENSMUSG( | 96.53917 | -0.00498 | 0.98467 | 0.996963 | Sash3 |
| ENSMUSG( | 98.31176 | 0.262783 | 0.25158 | 0.879838 | Sass6 |
| ENSMUSG( | 2252.318 | -0.25855 | 0.043056 | 0.690304 | Sat1 |
| ENSMUSG( | 223.9596 | -0.00183 | 0.991141 | 0.998379 | Sat2 |
| ENSMUSG( | 3643.07 | 0.06881 | 0.556647 | 0.939565 | Satb1 |
| ENSMUSG( | 728.7916 | 0.06055 | 0.700524 | 0.961533 | Satb2 |
| ENSMUSG( | 644.1037 | 0.054021 | 0.654091 | 0.955015 | Sav1 |
| ENSMUSG( | 71.09138 | 0.205399 | 0.429519 | 0.917257 | Saxo2 |
| ENSMUSG( | 11.94229 | 0.761543 | 0.241363 | 0.876259 | Saxo3 |
| ENSMUSG( | 41.80505 | 0.387755 | 0.223268 | 0.867404 | Saxo4 |
| ENSMUSG( | 234.1346 | 0.077513 | 0.63423 | 0.953903 | Saysd1 |
| ENSMUSG( | 2080.501 | -0.17531 | 0.225578 | 0.868468 | Sbds |
| ENSMUSG( | 7858.624 | 0.002758 | 0.980363 | 0.996479 | Sbf1 |
| ENSMUSG( | 3042.175 | -0.02851 | 0.792386 | 0.97681 | Sbf2 |
| ENSMUSG( | 6751.734 | 0.051222 | 0.624366 | 0.953234 | Sbk1 |
| ENSMUSG( | 27.94869 | -0.20428 | 0.60363 | 0.948137 | Sbk3 |
| ENSMUSG( | 3435.786 | -0.05571 | 0.613379 | 0.951177 | Sbno1 |
| ENSMUSG( | 706.5065 | 0.045567 | 0.746771 | 0.969763 | Sbno2 |
| ENSMUSG( | 120.2456 | 0.073135 | 0.722809 | 0.966887 | Sbsn |
| ENSMUSG( | 12.71869 | -0.47734 | 0.389409 | 0.917257 | Sbspon |
| ENSMUSG( | 5803.054 | -0.11826 | 0.421716 | 0.917257 | Sc5d |
| ENSMUSG( | 7751.625 | 0.075756 | 0.462784 | 0.923329 | Scaf1 |
| ENSMUSG( | 1869.736 | 0.087211 | 0.420786 | 0.917257 | Scaf11 |
| ENSMUSG( | 1158.589 | 0.055794 | 0.660289 | 0.956205 | Scaf4 |
| ENSMUSG( | 1206.79 | 0.018115 | 0.86992 | 0.988097 | Scaf8 |
| ENSMUSG( | 706.5908 | 0.236948 | 0.104848 | 0.804969 | Scai |
| ENSMUSG( | 8005.67 | 0.058165 | 0.702156 | 0.962014 | Scamp1 |
| ENSMUSG( | 1118.98 | 0.215636 | 0.217756 | 0.865589 | Scamp2 |
| ENSMUSG( | 1613.974 | 0.140185 | 0.325527 | 0.905339 | Scamp3 |
| ENSMUSG( | 15159.62 | 0.101156 | 0.547749 | 0.938063 | Scamp5 |
| ENSMUSG( | 955.5333 | -0.24537 | 0.130536 | 0.822225 | Scand1 |
| ENSMUSG( | 3414.201 | 0.244714 | 0.159274 | 0.837724 | Scap |
| ENSMUSG( | 741.7301 | -0.11259 | 0.369998 | 0.915491 | Scaper |
| ENSMUSG( | 3040.66 | 0.045061 | 0.779171 | 0.976106 | Scara3 |
| ENSMUSG( | 21.95935 | 0.419011 | 0.372043 | 0.915491 | Scara5 |
| ENSMUSG( | 994.7108 | 0.221401 | 0.204252 | 0.864412 | Scarb1 |
| ENSMUSG( | 7938.843 | 0.056661 | 0.709874 | 0.96416 | Scarb2 |
| ENSMUSG( | 114.8942 | 0.424279 | 0.110649 | 0.807582 | Scarf1 |
| ENSMUSG( | 307.6024 | 0.334735 | 0.068087 | 0.75212 | Scarf2 |
| ENSMUSG( | 2513.07 | 0.002846 | 0.981707 | 0.996756 | Sccpdh |
| ENSMUSG( | 36616.68 | -0.11045 | 0.353945 | 0.912411 | Scd1 |
| ENSMUSG( | 150056.8 | -0.08768 | 0.481041 | 0.924548 | Scd2 |
| ENSMUSG( | 129.8489 | -0.15764 | 0.422112 | 0.917257 | Scd3 |
| ENSMUSG( | 17.49096 | 0.144324 | 0.789912 | 0.976798 | Scd4 |
| ENSMUSG( | 80.4911 | -0.21387 | 0.4404 | 0.921166 | Scel |
| ENSMUSG( | 654.766 | -0.07908 | 0.533868 | 0.935551 | Scfd1 |
| ENSMUSG( | 443.3823 | -0.13987 | 0.300302 | 0.898538 | Scfd2 |
| ENSMUSG( | 6141.454 | 0.075469 | 0.630124 | 0.953251 | Scg2 |
| ENSMUSG( | 30860.36 | 0.006162 | 0.963459 | 0.995433 | Scg3 |
| ENSMUSG( | 6770.805 | 0.132675 | 0.311422 | 0.900022 | Scg5 |
| ENSMUSG( | 161.2724 | -0.16852 | 0.452267 | 0.922316 | Schip1 |
| ENSMUSG( | 247.9744 | 0.05688 | 0.718587 | 0.966633 | Sclt1 |
| ENSMUSG( | 455.3591 | -0.17637 | 0.20377 | 0.864412 | Scly |
| ENSMUSG( | 1774.789 | 0.052115 | 0.626094 | 0.953234 | Scmh1 |
| ENSMUSG( | 23.23031 | -0.3873 | 0.413833 | 0.917257 | Scml2 |
| ENSMUSG( | 57.31961 | 0.092363 | 0.747329 | 0.969763 | Scml4 |
| ENSMUSG( | 12.62623 | 1.000994 | 0.072161 | 0.763629 | Scn10a |
| ENSMUSG( | 2942.439 | 0.210574 | 0.229587 | 0.868507 | Scn1a |
| ENSMUSG( | 19487.06 | -0.05168 | 0.638114 | 0.953903 | Scn1b |
| ENSMUSG( | 6731.05 | 0.272114 | 0.12402 | 0.814691 | Scn2a |
| ENSMUSG( | 7643.039 | 0.142178 | 0.403506 | 0.917257 | Scn2b |
| ENSMUSG( | 1102.189 | 0.364556 | 0.032663 | 0.637512 | Scn3a |
| ENSMUSG( | 3620.355 | 0.230281 | 0.075544 | 0.768472 | Scn3b |
| ENSMUSG( | 5846.858 | -0.10545 | 0.465006 | 0.923329 | Scn4b |
| ENSMUSG( | 86.74562 | 0.193123 | 0.4085 | 0.917257 | Scn5a |
| ENSMUSG( | 94.06721 | 0.343668 | 0.196657 | 0.862161 | Scn7a |
| ENSMUSG( | 10930.17 | 0.319786 | 0.027041 | 0.603041 | Scn8a |
| ENSMUSG( | 57.48266 | -0.05264 | 0.869617 | 0.988097 | Scn9a |
| ENSMUSG( | 221.5387 | 0.077915 | 0.671181 | 0.95694 | Scnn1a |
| ENSMUSG( | 224.0926 | 0.022286 | 0.905267 | 0.991089 | Sco1 |
| ENSMUSG( | 161.0146 | -0.00065 | 0.997298 | 0.999268 | Sco2 |
| ENSMUSG( | 1736.168 | -0.01901 | 0.896428 | 0.990558 | Scoc |
| ENSMUSG( | 4848.632 | -0.14625 | 0.258534 | 0.883971 | Scp2 |
| ENSMUSG( | 818.2422 | 0.048854 | 0.761955 | 0.97318 | Scpep1 |
| ENSMUSG( | 2471.854 | 0.033264 | 0.788201 | 0.97637 | Scrg1 |
| ENSMUSG( | 3024.211 | -0.067 | 0.528542 | 0.934795 | Scrib |
| ENSMUSG( | 5737.946 | -0.16232 | 0.112318 | 0.809018 | Scrn1 |
| ENSMUSG( | 138.1755 | -0.02276 | 0.909027 | 0.991958 | Scrn2 |
| ENSMUSG( | 540.089 | -0.08419 | 0.558407 | 0.93957 | Scrn3 |
| ENSMUSG( | 7865.966 | 0.097855 | 0.40255 | 0.917257 | Scrt1 |

| | | | | | | | | | | | |
|---|---|---|---|---|---|---|---|---|---|---|---|
| ENSMUSG( | Sdhaf3 | 210.0694 | -0.0649 | 0.672573 | 0.929007 | ENSMUSG( | Sdhaf3 | 179.5582 | -0.27418 | 0.149264 | 0.836241 |
| ENSMUSG( | Sdhaf4 | 377.7927 | -0.04776 | 0.746673 | 0.949289 | ENSMUSG( | Sdhaf4 | 325.3038 | -0.2294 | 0.197156 | 0.862161 |
| ENSMUSG( | Sdhb | 3426.044 | 0.048103 | 0.608491 | 0.907154 | ENSMUSG( | Sdhb | 2961.371 | -0.10847 | 0.386213 | 0.917257 |
| ENSMUSG( | Sdhc | 4598.205 | 0.088059 | 0.352243 | 0.783835 | ENSMUSG( | Sdhc | 3831.718 | -0.17576 | 0.174984 | 0.845107 |
| ENSMUSG( | Sdhd | 2297.88 | 0.062967 | 0.508146 | 0.867999 | ENSMUSG( | Sdhd | 2008.693 | -0.05888 | 0.600405 | 0.947732 |
| ENSMUSG( | Sdk1 | 349.4875 | -0.18856 | 0.223823 | 0.669845 | ENSMUSG( | Sdk1 | 370.931 | 0.250854 | 0.10889 | 0.804969 |
| ENSMUSG( | Sdk2 | 1280.155 | 0.049447 | 0.736077 | 0.946267 | ENSMUSG( | Sdk2 | 1334.391 | 0.414555 | 0.017609 | 0.53154 |
| ENSMUSG( | Sdr39u1 | 1397.967 | -0.07575 | 0.510964 | 0.870755 | ENSMUSG( | Sdr39u1 | 1209.072 | -0.23793 | 0.094024 | 0.795735 |
| ENSMUSG( | Sdr42e1 | 270.116 | 0.207875 | 0.225395 | 0.672165 | ENSMUSG( | Sdr42e1 | 237.1035 | 0.100535 | 0.604138 | 0.948139 |
| ENSMUSG( | Sdsl | 217.5833 | 0.019548 | 0.907776 | 0.987798 | ENSMUSG( | Sdsl | 183.8914 | -0.21132 | 0.255243 | 0.882761 |
| ENSMUSG( | Sebox | 75.80356 | 0.080762 | 0.729569 | 0.944764 | ENSMUSG( | Sebox | 68.65487 | 0.075419 | 0.774862 | 0.974931 |
| ENSMUSG( | Sec1 | 52.28947 | -0.11144 | 0.675461 | 0.930049 | ENSMUSG( | Sec1 | 38.05826 | -0.8677 | 0.021472 | 0.575341 |
| ENSMUSG( | Sec11a | 740.9883 | -0.27286 | 0.04254 | 0.316636 | ENSMUSG( | Sec11a | 799.8653 | 0.226236 | 0.16153 | 0.837724 |
| ENSMUSG( | Sec11c | 2341.183 | -0.33249 | 0.00652 | 0.110626 | ENSMUSG( | Sec11c | 2319.832 | -0.07114 | 0.60075 | 0.947749 |
| ENSMUSG( | Sec13 | 1360.22 | 0.048559 | 0.619626 | 0.911432 | ENSMUSG( | Sec13 | 1172.057 | -0.11367 | 0.308263 | 0.898538 |
| ENSMUSG( | Sec14l1 | 7918.992 | 0.034328 | 0.677044 | 0.930542 | ENSMUSG( | Sec14l1 | 7047.198 | -0.03375 | 0.740613 | 0.968811 |
| ENSMUSG( | Sec14l2 | 3254.431 | 0.03954 | 0.666893 | 0.92729 | ENSMUSG( | Sec14l2 | 2750.949 | -0.1856 | 0.159774 | 0.837724 |
| ENSMUSG( | Sec14l3 | 35.97401 | 0.29654 | 0.347002 | 0.7797 | ENSMUSG( | Sec14l3 | 27.81823 | -0.20054 | 0.613089 | 0.951154 |
| ENSMUSG( | Sec14l4 | 10.91318 | -0.3192 | 0.624975 | NA | ENSMUSG( | Sec14l4 | 10.97752 | 0.042045 | 0.945416 | NA |
| ENSMUSG( | Sec14l5 | 1066.517 | -0.47782 | 0.001119 | 0.035587 | ENSMUSG( | Sec14l5 | 1128.954 | 0.001563 | 0.993402 | 0.998813 |
| ENSMUSG( | Sec16a | 1755.517 | 0.040011 | 0.671301 | 0.928781 | ENSMUSG( | Sec16a | 1597.749 | 0.039403 | 0.732834 | 0.967228 |
| ENSMUSG( | Sec16b | 44.4557 | -0.04855 | 0.88813 | 0.983434 | ENSMUSG( | Sec16b | 37.15392 | -0.30231 | 0.437761 | 0.920339 |
| ENSMUSG( | Sec22a | 375.4908 | -0.22436 | 0.105391 | 0.492417 | ENSMUSG( | Sec22a | 381.7071 | 0.100813 | 0.517282 | 0.930939 |
| ENSMUSG( | Sec22b | 1487.867 | 0.095839 | 0.362772 | 0.793121 | ENSMUSG( | Sec22b | 1226.072 | -0.2043 | 0.123973 | 0.814691 |
| ENSMUSG( | Sec22c | 1463.671 | -0.08046 | 0.53473 | 0.880719 | ENSMUSG( | Sec22c | 1469.129 | 0.19737 | 0.174007 | 0.844317 |
| ENSMUSG( | Sec23a | 2398.888 | 0.09883 | 0.264368 | 0.715323 | ENSMUSG( | Sec23a | 2115.232 | 0.004514 | 0.966432 | 0.995475 |
| ENSMUSG( | Sec23b | 1154.941 | 0.097397 | 0.293275 | 0.742763 | ENSMUSG( | Sec23b | 985.9558 | -0.0924 | 0.442606 | 0.921554 |
| ENSMUSG( | Sec23ip | 1030.037 | 0.190341 | 0.100153 | 0.483045 | ENSMUSG( | Sec23ip | 895.0851 | 0.056055 | 0.684846 | 0.958678 |
| ENSMUSG( | Sec24a | 596.8164 | 0.022447 | 0.837698 | 0.971111 | ENSMUSG( | Sec24a | 549.4457 | 0.050224 | 0.687658 | 0.959673 |
| ENSMUSG( | Sec24b | 1377.496 | 0.197394 | 0.038552 | 0.302696 | ENSMUSG( | Sec24b | 1202.79 | 0.07999 | 0.511406 | 0.92971 |
| ENSMUSG( | Sec24c | 2557.8 | 0.209338 | 0.025971 | 0.247245 | ENSMUSG( | Sec24c | 2152.736 | -0.01275 | 0.908539 | 0.991958 |
| ENSMUSG( | Sec24d | 541.5351 | 0.215157 | 0.05639 | 0.366006 | ENSMUSG( | Sec24d | 486.1967 | 0.178186 | 0.17437 | 0.844317 |
| ENSMUSG( | Sec31a | 2514.839 | 0.129593 | 0.179133 | 0.616715 | ENSMUSG( | Sec31a | 2208.527 | 0.025612 | 0.822312 | 0.979198 |
| ENSMUSG( | Sec61a1 | 2951.371 | -0.32604 | 0.010803 | 0.14989 | ENSMUSG( | Sec61a1 | 3101.41 | 0.106304 | 0.481741 | 0.924578 |
| ENSMUSG( | Sec61a2 | 1985.342 | -0.32089 | 0.007791 | 0.12245 | ENSMUSG( | Sec61a2 | 2057.295 | 0.069724 | 0.592094 | 0.944975 |
| ENSMUSG( | Sec61b | 509.0312 | -0.00773 | 0.952601 | 0.993874 | ENSMUSG( | Sec61b | 440.7203 | -0.16825 | 0.268752 | 0.886755 |
| ENSMUSG( | Sec61g | 849.8251 | 0.059805 | 0.611802 | 0.908586 | ENSMUSG( | Sec61g | 705.7668 | -0.22167 | 0.187243 | 0.856372 |
| ENSMUSG( | Sec62 | 7001.663 | -0.27691 | 0.040823 | 0.312481 | ENSMUSG( | Sec62 | 7419.422 | 0.168014 | 0.299724 | 0.898538 |
| ENSMUSG( | Sec63 | 4201.228 | -0.06242 | 0.605586 | 0.905824 | ENSMUSG( | Sec63 | 4076.197 | 0.118767 | 0.385165 | 0.917257 |
| ENSMUSG( | Secisbp2 | 555.5105 | 0.092133 | 0.449646 | 0.835234 | ENSMUSG( | Secisbp2 | 512.644 | 0.123887 | 0.359233 | 0.914061 |
| ENSMUSG( | Secisbp2l | 6057.24 | -0.03285 | 0.715309 | 0.941288 | ENSMUSG( | Secisbp2l | 5511.248 | -0.03683 | 0.74847 | 0.96984 |
| ENSMUSG( | Seh1l | 739.4919 | -0.08208 | 0.499714 | 0.863218 | ENSMUSG( | Seh1l | 676.6287 | -0.07081 | 0.61243 | 0.951071 |
| ENSMUSG( | Sel1l | 6775.232 | -0.21876 | 0.144607 | 0.564508 | ENSMUSG( | Sel1l | 7010.05 | 0.155909 | 0.350008 | 0.911652 |
| ENSMUSG( | Sel1l3 | 2704.361 | -0.10485 | 0.420357 | 0.822856 | ENSMUSG( | Sel1l3 | 2630.314 | 0.088149 | 0.568756 | 0.941777 |
| ENSMUSG( | Selenbp1 | 564.7818 | 0.029058 | 0.837343 | 0.971111 | ENSMUSG( | Selenbp1 | 491.4847 | -0.11242 | 0.484806 | 0.92541 |
| ENSMUSG( | Selenof | 8443.261 | -0.27186 | 0.029477 | 0.260274 | ENSMUSG( | Selenof | 8205.303 | -0.07389 | 0.600758 | 0.947749 |
| ENSMUSG( | Selenoh | 410.5574 | 0.12909 | 0.311619 | 0.759296 | ENSMUSG( | Selenoh | 364.2924 | 0.048595 | 0.774514 | 0.974931 |
| ENSMUSG( | Selenoi | 1651.408 | -0.24481 | 0.114707 | 0.510718 | ENSMUSG( | Selenoi | 1764.861 | 0.222866 | 0.180783 | 0.850372 |
| ENSMUSG( | Selenok | 3747.041 | -0.33059 | 0.005324 | 0.097356 | ENSMUSG( | Selenok | 3760.971 | -0.03308 | 0.823288 | 0.979198 |
| ENSMUSG( | Selenom | 2332.78 | -0.15275 | 0.06512 | 0.395697 | ENSMUSG( | Selenom | 2163.038 | -0.10168 | 0.35802 | 0.913759 |
| ENSMUSG( | Selenon | 1046 | -0.25936 | 0.048466 | 0.336526 | ENSMUSG( | Selenon | 1082.794 | 0.124068 | 0.429434 | 0.917257 |
| ENSMUSG( | Selenoo | 843.2782 | -0.03676 | 0.758064 | 0.951614 | ENSMUSG( | Selenoo | 774.9056 | -0.00588 | 0.964752 | 0.995433 |
| ENSMUSG( | Selenop | 25049.63 | 0.245136 | 0.030418 | 0.265114 | ENSMUSG( | Selenop | 22827.07 | 0.245157 | 0.042999 | 0.690304 |
| ENSMUSG( | Selenos | 1562.918 | -0.27792 | 0.033662 | 0.281425 | ENSMUSG( | Selenos | 1644.978 | 0.150713 | 0.307557 | 0.898538 |
| ENSMUSG( | Selenot | 6491.788 | -0.19496 | 0.149503 | 0.573617 | ENSMUSG( | Selenot | 6368.79 | 0.024801 | 0.875452 | 0.989644 |
| ENSMUSG( | Selenov | 22.42348 | -0.39814 | 0.354459 | 0.786878 | ENSMUSG( | Selenov | 23.17132 | -0.04291 | 0.920678 | 0.992644 |
| ENSMUSG( | Selenow | 50472.46 | 0.053545 | 0.509509 | 0.869462 | ENSMUSG( | Selenow | 42355.66 | -0.19089 | 0.110784 | 0.807582 |
| ENSMUSG( | Sell | 18.63745 | -0.62958 | 0.283771 | NA | ENSMUSG( | Sell | 16.75324 | -0.70722 | 0.231846 | 0.868507 |
| ENSMUSG( | Selplg | 2285.99 | -0.13043 | 0.44796 | 0.834051 | ENSMUSG( | Selplg | 2563.359 | 0.444263 | 0.043211 | 0.690304 |
| ENSMUSG( | Sem1 | 1150.311 | 0.020073 | 0.878042 | 0.981431 | ENSMUSG( | Sem1 | 982.5405 | -0.1763 | 0.220884 | 0.866104 |
| ENSMUSG( | Sema3a | 522.2484 | -0.57261 | 0.003593 | 0.075902 | ENSMUSG( | Sema3a | 604.1308 | 0.173653 | 0.43248 | 0.918256 |
| ENSMUSG( | Sema3b | 146.9287 | -0.00103 | 0.995175 | 0.99943 | ENSMUSG( | Sema3b | 144.9269 | 0.234128 | 0.243001 | 0.877399 |
| ENSMUSG( | Sema3c | 780.1209 | -0.14735 | 0.407525 | 0.817672 | ENSMUSG( | Sema3c | 972.2829 | 0.688213 | 0.001243 | 0.127113 |
| ENSMUSG( | Sema3d | 258.2483 | -0.41868 | 0.024279 | 0.238384 | ENSMUSG( | Sema3d | 327.7597 | 0.526847 | 0.014243 | 0.477949 |
| ENSMUSG( | Sema3e | 737.8312 | -0.68297 | 7.74E-05 | 0.005438 | ENSMUSG( | Sema3e | 904.42 | 0.247669 | 0.216422 | 0.865167 |
| ENSMUSG( | Sema3f | 380.1554 | -0.2689 | 0.080843 | 0.439584 | ENSMUSG( | Sema3f | 398.0428 | 0.147643 | 0.389041 | 0.917257 |
| ENSMUSG( | Sema3g | 124.8698 | 0.197139 | 0.4702 | 0.84674 | ENSMUSG( | Sema3g | 151.276 | 0.90762 | 0.004105 | 0.25867 |
| ENSMUSG( | Sema4a | 2065.235 | -0.29015 | 0.009787 | 0.141238 | ENSMUSG( | Sema4a | 2019.008 | -0.06961 | 0.620207 | 0.952896 |
| ENSMUSG( | Sema4b | 779.6796 | -0.31441 | 0.030823 | 0.266969 | ENSMUSG( | Sema4b | 909.4051 | 0.395068 | 0.016517 | 0.519511 |
| ENSMUSG( | Sema4c | 553.8446 | -0.35246 | 0.043683 | 0.320345 | ENSMUSG( | Sema4c | 597.4871 | 0.162792 | 0.444248 | 0.921734 |
| ENSMUSG( | Sema4d | 3592.141 | -0.33217 | 0.029525 | 0.260413 | ENSMUSG( | Sema4d | 3836.293 | 0.147117 | 0.435474 | 0.919062 |
| ENSMUSG( | Sema4f | 1640.939 | -0.12157 | 0.431272 | 0.827055 | ENSMUSG( | Sema4f | 1663.57 | 0.18842 | 0.228263 | 0.868507 |
| ENSMUSG( | Sema4g | 1526.119 | 0.083728 | 0.37288 | 0.799381 | ENSMUSG( | Sema4g | 1350.349 | 0.001199 | 0.991935 | 0.99872 |
| ENSMUSG( | Sema5a | 1875.656 | -0.44042 | 0.002928 | 0.066146 | ENSMUSG( | Sema5a | 1955.691 | -0.01442 | 0.934142 | 0.99306 |
| ENSMUSG( | Sema5b | 970.1706 | -0.43318 | 0.000758 | 0.027082 | ENSMUSG( | Sema5b | 970.9135 | -0.12824 | 0.383234 | 0.917257 |
| ENSMUSG( | Sema6a | 1760.347 | -0.64238 | 2.13E-05 | 0.002059 | ENSMUSG( | Sema6a | 2037.914 | 0.122236 | 0.47634 | 0.923329 |
| ENSMUSG( | Sema6b | 5314.702 | -0.144 | 0.230759 | 0.679066 | ENSMUSG( | Sema6b | 5437.641 | 0.190785 | 0.173781 | 0.844317 |
| ENSMUSG( | Sema6c | 849.9453 | 0.120735 | 0.427138 | 0.82532 | ENSMUSG( | Sema6c | 805.4999 | 0.233974 | 0.124302 | 0.814691 |
| ENSMUSG( | Sema6d | 3398.607 | -0.16593 | 0.249302 | 0.698744 | ENSMUSG( | Sema6d | 3264.975 | -0.00514 | 0.973042 | 0.996416 |
| ENSMUSG( | Sema7a | 5856.054 | -0.21445 | 0.12481 | 0.531813 | ENSMUSG( | Sema7a | 5753.724 | 0.015875 | 0.92748 | 0.99306 |
| ENSMUSG( | Senp1 | 534.2202 | 0.127704 | 0.37425 | 0.800445 | ENSMUSG( | Senp1 | 493.1422 | 0.163663 | 0.274662 | 0.889874 |
| ENSMUSG( | Senp2 | 3440.458 | 0.075625 | 0.342805 | 0.778971 | ENSMUSG( | Senp2 | 2989.386 | -0.06108 | 0.54438 | 0.938063 |
| ENSMUSG( | Senp3 | 2096.099 | 0.128476 | 0.179183 | 0.616715 | ENSMUSG( | Senp3 | 1865.201 | 0.064181 | 0.555525 | 0.939565 |
| ENSMUSG( | Senp5 | 1353.379 | 0.144133 | 0.11325 | 0.507541 | ENSMUSG( | Senp5 | 1165.321 | -0.01647 | 0.882708 | 0.989644 |
| ENSMUSG( | Senp6 | 2643.679 | 0.154797 | 0.075059 | 0.42422 | ENSMUSG( | Senp6 | 2296.189 | 0.018832 | 0.868515 | 0.988097 |
| ENSMUSG( | Senp7 | 527.7693 | -0.10351 | 0.35786 | 0.789405 | ENSMUSG( | Senp7 | 533.9507 | 0.195961 | 0.136456 | 0.827092 |
| ENSMUSG( | Senp8 | 99.78934 | 0.11073 | 0.584111 | 0.898717 | ENSMUSG( | Senp8 | 87.59884 | -0.00536 | 0.98137 | 0.996756 |
| ENSMUSG( | Sephs1 | 1413.684 | 0.010791 | 0.917319 | 0.989158 | ENSMUSG( | Sephs1 | 1249.228 | -0.08097 | 0.504473 | 0.927825 |
| ENSMUSG( | Sephs2 | 762.4178 | -0.00287 | 0.982304 | 0.997512 | ENSMUSG( | Sephs2 | 666.3116 | -0.13021 | 0.396154 | 0.917257 |
| ENSMUSG( | Sepsecs | 645.5046 | 0.068246 | 0.520869 | 0.875186 | ENSMUSG( | Sepsecs | 578.8518 | 0.019016 | 0.883344 | 0.989644 |
| ENSMUSG( | Septin1 | 72.33332 | 0.005354 | 0.983578 | 0.997738 | ENSMUSG( | Septin1 | 63.77345 | -0.09897 | 0.756655 | 0.971779 |
| ENSMUSG( | Septin10 | 346.9794 | -0.07894 | 0.596021 | 0.903152 | ENSMUSG( | Septin10 | 306.9407 | -0.16708 | 0.272674 | 0.888825 |
| ENSMUSG( | Septin11 | 3729.922 | 0.119485 | 0.124933 | 0.531973 | ENSMUSG( | Septin11 | 3168.936 | -0.08185 | 0.418847 | 0.917257 |
| ENSMUSG( | Septin2 | 37.56581 | -0.36511 | 0.287252 | 0.737687 | ENSMUSG( | Septin2 | 39.65687 | 0.109114 | 0.760987 | 0.972695 |
| ENSMUSG( | Septin3 | 24093.33 | 0.15936 | 0.02776 | 0.255176 | ENSMUSG( | Septin3 | 20394.01 | -0.04923 | 0.613645 | 0.951177 |
| ENSMUSG( | Septin4 | 5794.781 | -0.1201 | 0.170908 | 0.605533 | ENSMUSG( | Septin4 | 5325.167 | -0.0942 | 0.397791 | 0.917257 |
| ENSMUSG( | Septin5 | 10648.92 | 0.15496 | 0.045954 | 0.328356 | ENSMUSG( | Septin5 | 9178.82 | -0.00096 | 0.992016 | 0.99874 |
| ENSMUSG( | Septin6 | 1834.693 | 0.155718 | 0.221218 | 0.666945 | ENSMUSG( | Septin6 | 1491.469 | -0.17544 | 0.208504 | 0.864412 |
| ENSMUSG( | Septin7 | 11505.13 | 0.081785 | 0.41066 | 0.818493 | ENSMUSG( | Septin7 | 9941.632 | -0.07268 | 0.564688 | 0.941206 |
| ENSMUSG( | Septin8 | 14313.72 | 0.10662 | 0.215585 | 0.66027 | ENSMUSG( | Septin8 | 11777.05 | -0.19435 | 0.094998 | 0.796133 |
| ENSMUSG( | Septin9 | 3436.41 | 0.200594 | 0.010737 | 0.149272 | ENSMUSG( | Septin9 | 2717.132 | -0.21019 | 0.042535 | 0.690304 |
| ENSMUSG( | Serac1 | 397.7988 | 0.026766 | 0.826274 | 0.968294 | ENSMUSG( | Serac1 | 378.8384 | 0.159318 | 0.308992 | 0.898538 |
| ENSMUSG( | Serbp1 | 4411.431 | 0.060532 | 0.546904 | 0.884974 | ENSMUSG( | Serbp1 | 3912.768 | -0.01844 | 0.886069 | 0.989877 |
| ENSMUSG( | Serf1 | 425.5475 | 0.141349 | 0.247838 | 0.696522 | ENSMUSG( | Serf1 | 364.5736 | -0.043 | 0.797488 | 0.97681 |
| ENSMUSG( | Serf2 | 4479.643 | 0.048207 | 0.59129 | 0.90113 | ENSMUSG( | Serf2 | 3767.469 | -0.19097 | 0.101613 | 0.801601 |
| ENSMUSG( | Sergef | 593.0487 | 0.063427 | 0.618832 | 0.911432 | ENSMUSG( | Sergef | 510.5353 | -0.10793 | 0.474342 | 0.923329 |
| ENSMUSG( | Serhl | 448.6227 | 0.155696 | 0.238747 | 0.686489 | ENSMUSG( | Serhl | 370.9639 | -0.12165 | 0.37406 | 0.915491 |
| ENSMUSG( | Serinc1 | 21906.19 | -0.5745 | 1.62E-05 | 0.001655 | ENSMUSG( | Serinc1 | 25523.73 | 0.187858 | 0.255349 | 0.882761 |
| ENSMUSG( | Serinc2 | 220.929 | -0.09108 | 0.597573 | 0.903694 | ENSMUSG( | Serinc2 | 224.5199 | 0.224108 | 0.24201 | 0.87661 |
| ENSMUSG( | Serinc3 | 10569.75 | -0.223 | 0.144995 | 0.565206 | ENSMUSG( | Serinc3 | 11778.83 | 0.349941 | 0.047415 | 0.700936 |
| ENSMUSG( | Serinc4 | 21.15114 | -0.14429 | 0.721951 | NA | ENSMUSG( | Serinc4 | 20.08364 | -0.05004 | 0.916468 | 0.992529 |
| ENSMUSG( | Serinc5 | 3485.109 | -0.43562 | 0.003779 | 0.078636 | ENSMUSG( | Serinc5 | 3668.873 | 0.019547 | 0.914713 | 0.992407 |
| ENSMUSG( | Serp1 | 2098.095 | 0.063049 | 0.623054 | 0.911589 | ENSMUSG( | Serp1 | 1841.45 | -0.04863 | 0.76512 | 0.974236 |
| ENSMUSG( | Serp2 | 2803.713 | 0.005261 | 0.953633 | 0.993874 | ENSMUSG( | Serp2 | 2378.806 | -0.21154 | 0.107478 | 0.804969 |
| ENSMUSG( | Serpina3n | 629.6631 | -0.3004 | 0.019782 | 0.212216 | ENSMUSG( | Serpina3n | 665.3259 | 0.14572 | 0.374574 | 0.915491 |
| ENSMUSG( | Serpina9 | 109.0943 | -1.85427 | 0.006405 | 0.109928 | ENSMUSG( | Serpina9 | 144.4245 | -0.23398 | 0.299184 | 0.898538 |
| ENSMUSG( | Serpinb1a | 241.5614 | -0.51406 | 0.001688 | 0.046443 | ENSMUSG( | Serpinb1a | 236.4839 | -0.26889 | 0.097445 | 0.796721 |
| ENSMUSG( | Serpinb1b | 28.48752 | -0.0677 | 0.847145 | 0.972374 | ENSMUSG( | Serpinb1b | 24.64819 | -0.25015 | 0.537121 | 0.937216 |
| ENSMUSG( | Serpinb6a | 1006.784 | -0.00258 | 0.982413 | 0.997512 | ENSMUSG( | Serpinb6a | 851.023 | -0.22746 | 0.059796 | 0.729747 |
| ENSMUSG( | Serpinb6b | 70.37164 | 0.027123 | 0.912536 | 0.988483 | ENSMUSG( | Serpinb6b | 76.75636 | 0.529987 | 0.088013 | 0.786591 |
| ENSMUSG( | Serpinb8 | 160.095 | 0.334941 | 0.152392 | 0.578606 | ENSMUSG( | Serpinb8 | 136.748 | 0.163745 | 0.416782 | 0.917257 |
| ENSMUSG( | Serpinb9 | 287.8047 | -0.2302 | 0.133304 | 0.546146 | ENSMUSG( | Serpinb9 | 282.8059 | 0.004096 | 0.980023 | 0.996479 |
| ENSMUSG( | Serpind1 | 92.72478 | 0.221343 | 0.356406 | 0.788187 | ENSMUSG( | Serpind1 | 82.93713 | 0.177496 | 0.464914 | 0.923329 |
| ENSMUSG( | Serpine1 | 39.47955 | -0.59842 | 0.068432 | 0.405244 | ENSMUSG( | Serpine1 | 44.45583 | 0.093615 | 0.772716 | 0.974463 |
| ENSMUSG( | Serpine2 | 20321.29 | -0.12338 | 0.265956 | 0.716441 | ENSMUSG( | Serpine2 | 18612.59 | -0.10813 | 0.402222 | 0.917257 |
| ENSMUSG( | Serpinf1 | 696.18 | -0.33082 | 0.019528 | 0.210473 | ENSMUSG( | Serpinf1 | 734.291 | 0.109841 | 0.430266 | 0.917257 |
| ENSMUSG( | Serpinf2 | 10.88814 | 0.068003 | 0.904896 | NA | ENSMUSG( | Serpinf2 | 11.40894 | 0.424383 | 0.481219 | NA |
| ENSMUSG( | Serping1 | 385.9028 | 0.01584 | 0.918508 | 0.989954 | ENSMUSG( | Serping1 | 366.5037 | 0.142582 | 0.401418 | 0.917257 |
| ENSMUSG( | Serpinh1 | 1632.187 | -0.20238 | 0.202194 | 0.642173 | ENSMUSG( | Serpinh1 | 1803.786 | 0.345521 | 0.037235 | 0.673514 |
| ENSMUSG( | Serpini1 | 5433.157 | -0.53828 | 0.000336 | 0.015671 | ENSMUSG( | Serpini1 | 6625.142 | 0.335665 | 0.052589 | 0.71525 |
| ENSMUSG( | Sertad1 | 188.6208 | -0.4973 | 0.010648 | 0.14851 | ENSMUSG( | Sertad1 | 195.7735 | -0.08835 | 0.695673 | 0.960867 |
| ENSMUSG( | Sertad2 | 848.6815 | -0.10025 | 0.311745 | 0.759348 | ENSMUSG( | Sertad2 | 803.3746 | 0.00937 | 0.940292 | 0.994334 |
| ENSMUSG( | Sertad3 | 68.66621 | 0.09698 | 0.728569 | 0.944393 | ENSMUSG( | Sertad3 | 66.7476 | 0.272723 | 0.363413 | 0.914223 |
| ENSMUSG( | Sertad4 | 308.2553 | -0.17855 | 0.197657 | 0.637377 | ENSMUSG( | Sertad4 | 298.4128 | -0.00228 | 0.988539 | 0.997862 |
| ENSMUSG( | Sertm1 | 734.6042 | -0.0342 | 0.788088 | 0.959226 | ENSMUSG( | Sertm1 | 660.5229 | -0.0723 | 0.566935 | 0.941777 |
| ENSMUSG( | Sesn1 | 1770.795 | 0.262437 | 0.012317 | 0.160458 | ENSMUSG( | Sesn1 | 1532.383 | 0.123496 | 0.272016 | 0.888477 |
| ENSMUSG( | Sesn2 | 427.8545 | 0.033066 | 0.813027 | 0.964574 | ENSMUSG( | Sesn2 | 390.8553 | 0.037328 | 0.81061 | 0.978779 |
| ENSMUSG( | Sesn3 | 10103.4 | 0.123768 | 0.35738 | 0.788775 | ENSMUSG( | Sesn3 | 8016.134 | -0.28814 | 0.035118 | 0.66005 |
| ENSMUSG( | Sestd1 | 1643.278 | 0.09419 | 0.349598 | 0.78192 | ENSMUSG( | Sestd1 | 1444.215 | -0.00994 | 0.929679 | 0.99306 |
| ENSMUSG( | Set | 4126.544 | 0.081263 | 0.371766 | 0.798768 | ENSMUSG( | Set | 3620.957 | -0.02904 | 0.839191 | 0.981906 |
| ENSMUSG( | Setbp1 | 857.9182 | 0.01662 | 0.868746 | 0.976897 | ENSMUSG( | Setbp1 | 797.806 | 0.07908 | 0.532742 | 0.934795 |
| ENSMUSG( | Setd1a | 607.2741 | 0.007686 | 0.943426 | 0.993415 | ENSMUSG( | Setd1a | 553.0296 | 0.005458 | 0.966352 | 0.995475 |
| ENSMUSG( | Setd1b | 1437.218 | 0.069541 | 0.447573 | 0.834051 | ENSMUSG( | Setd1b | 1386.618 | 0.227605 | 0.039395 | 0.679972 |
| ENSMUSG( | Setd2 | 2700.529 | 0.475671 | 0.000101 | 0.006438 | ENSMUSG( | Setd2 | 2993.96 | 0.935074 | 1.57E-14 | 2.11E-11 |
| ENSMUSG( | Setd3 | 6396.974 | 0.066624 | 0.421761 | 0.823375 | ENSMUSG( | Setd3 | 5517.705 | -0.09374 | 0.379056 | 0.916995 |
| ENSMUSG( | Setd4 | 274.6274 | 0.16486 | 0.339145 | 0.777518 | ENSMUSG( | Setd4 | 247.8871 | 0.13841 | 0.415342 | 0.917257 |
| ENSMUSG( | Setd5 | 3555.786 | 0.002009 | 0.982308 | 0.997512 | ENSMUSG( | Setd5 | 3296.392 | 0.053454 | 0.622822 | 0.953234 |
| ENSMUSG( | Setd6 | 329.0628 | -0.02003 | 0.895854 | 0.984551 | ENSMUSG( | Setd6 | 306.5648 | 0.035585 | 0.829135 | 0.980224 |
| ENSMUSG( | Setd7 | 11199.97 | 0.103344 | 0.37405 | 0.800445 | ENSMUSG( | Setd7 | 9393.132 | -0.13779 | 0.236535 | 0.872173 |
| ENSMUSG( | Setdb1 | 1474.33 | 0.068242 | 0.481273 | 0.855292 | ENSMUSG( | Setdb1 | 1364.597 | 0.114341 | 0.322148 | 0.904023 |
| ENSMUSG( | Setdb2 | 173.5732 | -0.11038 | 0.543112 | 0.883579 | ENSMUSG( | Setdb2 | 148.9169 | -0.29489 | 0.145142 | 0.83339 |
| ENSMUSG( | Setmar | 78.69815 | 0.122789 | 0.633433 | 0.914073 | ENSMUSG( | Setmar | 74.71572 | 0.242179 | 0.38318 | 0.917257 |
| ENSMUSG( | Setx | 2024.763 | 0.146118 | 0.114567 | 0.510543 | ENSMUSG( | Setx | 1855.028 | 0.162482 | 0.12984 | 0.820802 |
| ENSMUSG( | Sez6 | 7101.292 | -0.52825 | 6.73E-08 | 2.02E-05 | ENSMUSG( | Sez6 | 7829.464 | 0.071391 | 0.544198 | 0.938063 |
| ENSMUSG( | Sez6l | 6540.725 | -0.2636 | 0.065416 | 0.396102 | ENSMUSG( | Sez6l | 7129.935 | 0.25895 | 0.121999 | 0.813384 |
| ENSMUSG( | Sez6l2 | 9968.793 | -0.28852 | 0.041679 | 0.315317 | ENSMUSG( | Sez6l2 | 10707.81 | 0.198685 | 0.233986 | 0.86917 |
| ENSMUSG( | Sf1 | 3992.382 | 0.029402 | 0.728731 | 0.944393 | ENSMUSG( | Sf1 | 3632.531 | 0.025703 | 0.802454 | 0.97681 |
| ENSMUSG( | Sf3a1 | 1995.257 | 0.177212 | 0.069353 | 0.40782 | ENSMUSG( | Sf3a1 | 1746.923 | 0.065865 | 0.576311 | 0.942912 |

| | | | | | |
|---|---|---|---|---|---|
| ENSMUSG( | 20.20514 | -0.30393 | 0.47192 | NA | S100a7a |
| ENSMUSG( | 78.40156 | -1.44243 | 0.195438 | 0.635016 | S100a8 |
| ENSMUSG( | 143.354 | -1.20172 | 0.27505 | 0.72509 | S100a9 |
| ENSMUSG( | 18349.94 | -0.006 | 0.956562 | 0.993874 | S100b |
| ENSMUSG( | 913.6211 | -0.17727 | 0.150653 | 0.575407 | S100pbp |
| ENSMUSG( | 14049.93 | -0.18496 | 0.240883 | 0.68836 | S1pr1 |
| ENSMUSG( | 104.9802 | -1.04457 | 0.000164 | 0.009126 | S1pr2 |
| ENSMUSG( | 367.9207 | -0.15708 | 0.534504 | 0.880719 | S1pr3 |
| ENSMUSG( | 16.88229 | -0.19592 | 0.678219 | NA | S1pr4 |
| ENSMUSG( | 989.9627 | -0.65897 | 9.55E-07 | 0.000184 | S1pr5 |
| ENSMUSG( | 17.86966 | 0.311363 | 0.504686 | NA | S2bpcox16 |
| ENSMUSG( | 286.6273 | 0.012052 | 0.926808 | 0.991229 | Saal1 |
| ENSMUSG( | 414.9438 | -0.02988 | 0.831963 | 0.970194 | Sac3d1 |
| ENSMUSG( | 1394.824 | -0.00987 | 0.912898 | 0.988483 | Sacm1l |
| ENSMUSG( | 1937.587 | -0.44497 | 0.000414 | 0.018428 | Sacs |
| ENSMUSG( | 1990.697 | 0.035121 | 0.686075 | 0.932981 | Sae1 |
| ENSMUSG( | 3004.508 | 0.157342 | 0.121478 | 0.525713 | Safb |
| ENSMUSG( | 2605.44 | 0.099321 | 0.416255 | 0.82084 | Safb2 |
| ENSMUSG( | 94.56259 | -0.16291 | 0.522306 | 0.875186 | Sag |
| ENSMUSG( | 1081.6 | 0.033481 | 0.775863 | 0.955606 | Sall1 |
| ENSMUSG( | 2431.17 | -0.07612 | 0.393471 | 0.809501 | Sall2 |
| ENSMUSG( | 436.0629 | 0.116152 | 0.440032 | 0.831851 | Sall3 |
| ENSMUSG( | 1405.568 | 0.05883 | 0.544966 | 0.884712 | Samd1 |
| ENSMUSG( | 672.6048 | 0.104345 | 0.34313 | 0.779153 | Samd10 |
| ENSMUSG( | 51.19845 | -0.23645 | 0.400197 | 0.813641 | Samd11 |
| ENSMUSG( | 649.3216 | 0.073701 | 0.471331 | 0.847103 | Samd12 |
| ENSMUSG( | 3112.891 | 0.202648 | 0.023769 | 0.235641 | Samd14 |
| ENSMUSG( | 52.47748 | -0.14098 | 0.589707 | 0.90113 | Samd15 |
| ENSMUSG( | 18.59469 | -0.43211 | 0.384444 | NA | Samd3 |
| ENSMUSG( | 2357.327 | -0.17838 | 0.063289 | 0.388993 | Samd4 |
| ENSMUSG( | 3740.05 | 0.187642 | 0.018712 | 0.205766 | Samd4b |
| ENSMUSG( | 674.7524 | 0.096914 | 0.407955 | 0.818167 | Samd5 |
| ENSMUSG( | 2277.249 | -0.28437 | 0.046767 | 0.330815 | Samd8 |
| ENSMUSG( | 178.3487 | -0.10405 | 0.529664 | 0.878883 | Samd9l |
| ENSMUSG( | 703.2065 | -0.14977 | 0.176914 | 0.613937 | Samhd1 |
| ENSMUSG( | 2381.677 | 0.166662 | 0.126905 | 0.534201 | Samm50 |
| ENSMUSG( | 64.24328 | -0.30388 | 0.227995 | 0.675324 | Samsn1 |
| ENSMUSG( | 959.7647 | 0.148219 | 0.159905 | 0.589679 | Sanbr |
| ENSMUSG( | 911.973 | 0.095352 | 0.320874 | 0.765984 | Sap130 |
| ENSMUSG( | 2343.205 | 0.050062 | 0.590053 | 0.90113 | Sap18 |
| ENSMUSG( | 13.04516 | 0.607557 | 0.229507 | NA | Sap18b |
| ENSMUSG( | 189.4004 | 0.142859 | 0.447674 | 0.834051 | Sap30 |
| ENSMUSG( | 1344.936 | 0.107388 | 0.294163 | 0.743459 | Sap30bp |
| ENSMUSG( | 1851.563 | 0.085142 | 0.349077 | 0.781665 | Sap30l |
| ENSMUSG( | 27.13021 | 0.341744 | 0.363478 | 0.793339 | Sapcd1 |
| ENSMUSG( | 82.92602 | -0.30325 | 0.205717 | 0.647154 | Sapcd2 |
| ENSMUSG( | 4693.773 | 0.017711 | 0.822475 | 0.966352 | Sar1a |
| ENSMUSG( | 1835.403 | -0.33488 | 0.002199 | 0.054456 | Sar1b |
| ENSMUSG( | 12227.19 | -0.21026 | 0.076163 | 0.42702 | Saraf |
| ENSMUSG( | 1847.585 | 0.008428 | 0.93807 | 0.993415 | Sardh |
| ENSMUSG( | 969.1526 | 0.173292 | 0.075628 | 0.426231 | Sarm1 |
| ENSMUSG( | 789.7536 | 0.139557 | 0.240157 | 0.687899 | Sarnp |
| ENSMUSG( | 4152.132 | 0.017867 | 0.860991 | 0.975025 | Sars1 |
| ENSMUSG( | 332.9046 | -0.00315 | 0.984425 | 0.99781 | Sars2 |
| ENSMUSG( | 1666.48 | 0.128546 | 0.222924 | 0.668853 | Sart1 |
| ENSMUSG( | 1091.204 | 0.212854 | 0.084572 | 0.448139 | Sart3 |
| ENSMUSG( | 11108.32 | -0.03754 | 0.771197 | 0.954726 | Sash1 |
| ENSMUSG( | 110.2421 | 0.104542 | 0.652438 | 0.922226 | Sash3 |
| ENSMUSG( | 98.16063 | -0.01273 | 0.948717 | 0.993821 | Sass6 |
| ENSMUSG( | 2746.183 | 0.053214 | 0.644699 | 0.917782 | Sat1 |
| ENSMUSG( | 255.0042 | 0.102368 | 0.469377 | 0.846015 | Sat2 |
| ENSMUSG( | 4213.507 | 0.213737 | 0.021033 | 0.220525 | Satb1 |
| ENSMUSG( | 824.5242 | 0.148226 | 0.317405 | 0.763823 | Satb2 |
| ENSMUSG( | 696.6651 | 0.012554 | 0.900963 | 0.985633 | Sav1 |
| ENSMUSG( | 73.20205 | 0.039051 | 0.874106 | 0.979336 | Saxo2 |
| ENSMUSG( | 37.61986 | -0.18978 | 0.602325 | 0.904887 | Saxo4 |
| ENSMUSG( | 229.0593 | -0.26972 | 0.083719 | 0.446196 | Saysd1 |
| ENSMUSG( | 2407.26 | -0.01307 | 0.903167 | 0.986055 | Sbds |
| ENSMUSG( | 8948.369 | 0.105445 | 0.198861 | 0.639096 | Sbf1 |
| ENSMUSG( | 3464.786 | 0.078758 | 0.387748 | 0.805835 | Sbf2 |
| ENSMUSG( | 7748.679 | 0.175641 | 0.041488 | 0.314241 | Sbk1 |
| ENSMUSG( | 8.579915 | -0.21591 | 0.738072 | NA | Sbk2 |
| ENSMUSG( | 33.86252 | 0.119795 | 0.731062 | 0.944956 | Sbk3 |
| ENSMUSG( | 3983.073 | 0.101966 | 0.214072 | 0.658621 | Sbno1 |
| ENSMUSG( | 814.3808 | 0.17913 | 0.119744 | 0.523801 | Sbno2 |
| ENSMUSG( | 123.9865 | -0.12055 | 0.524934 | 0.87624 | Sbsn |
| ENSMUSG( | 13.39936 | -0.66712 | 0.199714 | NA | Sbspon |
| ENSMUSG( | 6184.815 | -0.20888 | 0.124151 | 0.531714 | Sc5d |
| ENSMUSG( | 8852.477 | 0.184341 | 0.035346 | 0.289405 | Scaf1 |
| ENSMUSG( | 2090.416 | 0.13873 | 0.117952 | 0.518899 | Scaf11 |
| ENSMUSG( | 1248.898 | 0.006474 | 0.944804 | 0.993612 | Scaf4 |
| ENSMUSG( | 1412.084 | 0.194978 | 0.031153 | 0.268462 | Scaf8 |
| ENSMUSG( | 669.7103 | -0.18549 | 0.099051 | 0.481021 | Scai |
| ENSMUSG( | 7684.57 | -0.35106 | 0.009405 | 0.13706 | Scamp1 |
| ENSMUSG( | 1044.894 | -0.26166 | 0.078769 | 0.434128 | Scamp2 |
| ENSMUSG( | 1572.811 | -0.21225 | 0.100555 | 0.483976 | Scamp3 |
| ENSMUSG( | 14540.85 | -0.30718 | 0.037092 | 0.297239 | Scamp5 |
| ENSMUSG( | 1111.632 | -0.06127 | 0.584022 | 0.898666 | Scand1 |
| ENSMUSG( | 3424.963 | -0.00999 | 0.944195 | 0.993503 | Scap |
| ENSMUSG( | 856.759 | 0.040156 | 0.724215 | 0.944096 | Scaper |
| ENSMUSG( | 3158.633 | -0.12079 | 0.378577 | 0.801688 | Scara3 |
| ENSMUSG( | 25.25554 | 0.500248 | 0.200392 | 0.640592 | Scara5 |
| ENSMUSG( | 947.384 | -0.19346 | 0.193318 | 0.632794 | Scarb1 |
| ENSMUSG( | 8023.86 | -0.18973 | 0.159564 | 0.589015 | Scarb2 |
| ENSMUSG( | 109.7224 | 0.029568 | 0.900381 | 0.985633 | Scarf1 |
| ENSMUSG( | 285.9164 | -0.13983 | 0.354087 | 0.786537 | Scarf2 |
| ENSMUSG( | 8.561478 | 0.240997 | 0.701339 | NA | Scart1 |
| ENSMUSG( | 2768.414 | 0.01324 | 0.909246 | 0.98794 | Sccpdh |
| ENSMUSG( | 43979.41 | 0.148328 | 0.203409 | 0.643631 | Scd1 |
| ENSMUSG( | 172863.2 | 0.052494 | 0.664909 | 0.926824 | Scd2 |
| ENSMUSG( | 153.629 | 0.069425 | 0.679518 | 0.930983 | Scd3 |
| ENSMUSG( | 18.08884 | -0.04756 | 0.923293 | NA | Scd4 |
| ENSMUSG( | 92.82818 | -0.03776 | 0.885031 | 0.982829 | Scel |
| ENSMUSG( | 769.8936 | 0.119062 | 0.277291 | 0.727292 | Scfd1 |
| ENSMUSG( | 526.0871 | 0.08641 | 0.45979 | 0.840338 | Scfd2 |
| ENSMUSG( | 5681.015 | -0.4577 | 0.000912 | 0.03071 | Scg2 |
| ENSMUSG( | 33489.22 | -0.02824 | 0.813933 | 0.964574 | Scg3 |
| ENSMUSG( | 6283.85 | -0.37429 | 0.000309 | 0.014712 | Scg5 |
| ENSMUSG( | 10.51188 | 0.151697 | 0.845398 | NA | Scgn |
| ENSMUSG( | 190.954 | 0.058771 | 0.774153 | 0.955606 | Schip1 |
| ENSMUSG( | 259.6808 | -0.08684 | 0.552613 | 0.886141 | Sclt1 |
| ENSMUSG( | 540.6618 | 0.049048 | 0.652354 | 0.922188 | Scly |
| ENSMUSG( | 1932.146 | 0.027803 | 0.760489 | 0.952907 | Scmh1 |
| ENSMUSG( | 23.6272 | -0.68613 | 0.106446 | 0.494174 | Scml2 |
| ENSMUSG( | 64.35649 | 0.17615 | 0.490616 | 0.859465 | Scml4 |
| ENSMUSG( | 10.89894 | 0.447803 | 0.430787 | NA | Scn10a |
| ENSMUSG( | 2675.637 | -0.34914 | 0.011045 | 0.151501 | Scn1a |
| ENSMUSG( | 20636.07 | -0.16017 | 0.096574 | 0.476986 | Scn1b |
| ENSMUSG( | 5943.005 | -0.37119 | 0.017905 | 0.200989 | Scn2a |
| ENSMUSG( | 7508.646 | -0.18481 | 0.179463 | 0.616974 | Scn2b |
| ENSMUSG( | 924.329 | -0.42398 | 0.007383 | 0.118765 | Scn3a |
| ENSMUSG( | 3327.812 | -0.29039 | 0.013031 | 0.165126 | Scn3b |
| ENSMUSG( | 4762.87 | -1.2151 | 1.44E-09 | 6.68E-07 | Scn4b |
| ENSMUSG( | 86.52799 | -0.08434 | 0.757071 | 0.951513 | Scn5a |
| ENSMUSG( | 80.24495 | -0.40775 | 0.135482 | 0.548884 | Scn7a |
| ENSMUSG( | 9901.71 | -0.22851 | 0.082013 | 0.442387 | Scn8a |
| ENSMUSG( | 54.62667 | -0.4863 | 0.094426 | 0.471713 | Scn9a |
| ENSMUSG( | 225.8279 | -0.15119 | 0.336791 | 0.777 | Scnn1a |
| ENSMUSG( | 242.5355 | -0.01899 | 0.894354 | 0.984512 | Sco1 |
| ENSMUSG( | 154.4228 | -0.42795 | 0.024973 | 0.243133 | Sco2 |
| ENSMUSG( | 1904.064 | -0.01608 | 0.885556 | 0.983008 | Scoc |
| ENSMUSG( | 5842.6 | 0.127156 | 0.235414 | 0.682423 | Scp2 |
| ENSMUSG( | 764.3629 | -0.45707 | 0.002215 | 0.054686 | Scpep1 |
| ENSMUSG( | 2919.776 | 0.23685 | 0.038664 | 0.302965 | Scrg1 |
| ENSMUSG( | 3655.605 | 0.205835 | 0.015575 | 0.184473 | Scrib |
| ENSMUSG( | 6468.806 | -0.08105 | 0.367046 | 0.795164 | Scrn1 |
| ENSMUSG( | 154.118 | 0.022638 | 0.897501 | 0.985633 | Scrn2 |
| ENSMUSG( | 621.9394 | 0.061911 | 0.552061 | 0.886141 | Scrn3 |
| ENSMUSG( | 8878.176 | 0.17435 | 0.056613 | 0.367011 | Scrt1 |
| ENSMUSG( | 624.4872 | -0.02379 | 0.850702 | 0.972638 | Scrt2 |
| ENSMUSG( | 49.61086 | 0.389953 | 0.150858 | 0.575879 | Sctr |
| ENSMUSG( | 1452.76 | -0.16643 | 0.231869 | 0.679725 | Scube1 |
| ENSMUSG( | 33.10319 | 0.40092 | 0.253169 | 0.70338 | Scube2 |
| ENSMUSG( | 144.1454 | -0.99898 | 3.15E-06 | 0.000452 | Scube3 |
| ENSMUSG( | 43.5097 | 0.167837 | 0.560466 | 0.888934 | Scx |
| ENSMUSG( | 2636.189 | 0.158749 | 0.119255 | 0.523036 | Scyl1 |
| ENSMUSG( | 714.5817 | -0.03151 | 0.7852 | 0.958473 | Scyl2 |
| ENSMUSG( | 642.418 | -0.13364 | 0.276968 | 0.727074 | Scyl3 |
| ENSMUSG( | 622.8361 | -0.01206 | 0.919523 | 0.990247 | Sdad1 |
| ENSMUSG( | 45.79341 | -0.20321 | 0.522055 | 0.875186 | Sdc1 |
| ENSMUSG( | 1493.207 | -0.29028 | 0.041838 | 0.315339 | Sdc2 |
| ENSMUSG( | 11758.19 | -0.19326 | 0.108879 | 0.49803 | Sdc3 |
| ENSMUSG( | 6569.437 | 0.219694 | 0.022649 | 0.22975 | Sdc4 |
| ENSMUSG( | 3186.01 | 0.040608 | 0.697043 | 0.935543 | Sdcbp |
| ENSMUSG( | 22.1147 | -0.19535 | 0.642502 | 0.916865 | Sdcbp2 |
| ENSMUSG( | 481.7705 | 0.151602 | 0.213955 | 0.658384 | Sdccag8 |
| ENSMUSG( | 397.88 | 0.043041 | 0.734679 | 0.945584 | Sde2 |
| ENSMUSG( | 1625.839 | -0.29802 | 0.028502 | 0.257525 | Sdf2 |

| | | | | | |
|---|---|---|---|---|---|
| ENSMUSG( | 564.9067 | -0.04968 | 0.742737 | 0.969415 | Scrt2 |
| ENSMUSG( | 41.80593 | 0.209884 | 0.520425 | 0.93204 | Sctr |
| ENSMUSG( | 1360.883 | -0.0764 | 0.627589 | 0.953234 | Scube1 |
| ENSMUSG( | 40.36367 | 1.094537 | 0.002257 | 0.184738 | Scube2 |
| ENSMUSG( | 162.306 | -0.22863 | 0.209358 | 0.864412 | Scube3 |
| ENSMUSG( | 43.84146 | 0.451272 | 0.153504 | 0.837724 | Scx |
| ENSMUSG( | 2306.967 | 0.044924 | 0.696042 | 0.960867 | Scyl1 |
| ENSMUSG( | 630.3692 | -0.12546 | 0.344142 | 0.910256 | Scyl2 |
| ENSMUSG( | 622.6652 | 0.05128 | 0.686095 | 0.958956 | Scyl3 |
| ENSMUSG( | 550.193 | -0.1006 | 0.438597 | 0.920721 | Sdad1 |
| ENSMUSG( | 53.50579 | 0.497447 | 0.124292 | 0.814691 | Sdc1 |
| ENSMUSG( | 1550.486 | 0.103138 | 0.525278 | 0.934374 | Sdc2 |
| ENSMUSG( | 11424.5 | 0.001709 | 0.990367 | 0.998129 | Sdc3 |
| ENSMUSG( | 6310.361 | 0.358215 | 0.002491 | 0.195662 | Sdc4 |
| ENSMUSG( | 2818.58 | -0.04813 | 0.725734 | 0.966887 | Sdcbp |
| ENSMUSG( | 19.87338 | -0.21038 | 0.636346 | 0.953903 | Sdcbp2 |
| ENSMUSG( | 426.4209 | 0.074945 | 0.614644 | 0.951177 | Sdccag8 |
| ENSMUSG( | 358.554 | 0.01234 | 0.93133 | 0.99306 | Sde2 |
| ENSMUSG( | 1641.349 | 0.018162 | 0.906227 | 0.991281 | Sdf2 |
| ENSMUSG( | 463.3734 | 0.466863 | 0.007972 | 0.382597 | Sdf2l1 |
| ENSMUSG( | 9129.662 | -0.01006 | 0.932858 | 0.99306 | Sdf4 |
| ENSMUSG( | 12677.96 | -0.20376 | 0.05086 | 0.709741 | Sdha |
| ENSMUSG( | 657.2031 | -0.26957 | 0.056324 | 0.723362 | Sdhaf1 |
| ENSMUSG( | 1115.766 | -0.14599 | 0.21032 | 0.864412 | Sdhaf2 |
| ENSMUSG( | 179.5582 | -0.27418 | 0.149264 | 0.836241 | Sdhaf3 |
| ENSMUSG( | 325.3038 | -0.2294 | 0.197156 | 0.862161 | Sdhaf4 |
| ENSMUSG( | 2961.371 | -0.10847 | 0.386213 | 0.917257 | Sdhb |
| ENSMUSG( | 3831.718 | -0.17576 | 0.174984 | 0.845107 | Sdhc |
| ENSMUSG( | 2008.693 | -0.05888 | 0.600405 | 0.947732 | Sdhd |
| ENSMUSG( | 370.931 | 0.250854 | 0.10889 | 0.804969 | Sdk1 |
| ENSMUSG( | 1334.391 | 0.414555 | 0.017609 | 0.53154 | Sdk2 |
| ENSMUSG( | 1209.072 | -0.23793 | 0.094024 | 0.795735 | Sdr39u1 |
| ENSMUSG( | 237.1035 | 0.100535 | 0.604138 | 0.948139 | Sdr42e1 |
| ENSMUSG( | 183.8914 | -0.21132 | 0.255243 | 0.882761 | Sdsl |
| ENSMUSG( | 68.65487 | 0.075419 | 0.774862 | 0.974931 | Sebox |
| ENSMUSG( | 38.05826 | -0.8677 | 0.021472 | 0.575341 | Sec1 |
| ENSMUSG( | 799.8653 | 0.226236 | 0.16153 | 0.837724 | Sec11a |
| ENSMUSG( | 2319.832 | -0.07114 | 0.60075 | 0.947749 | Sec11c |
| ENSMUSG( | 1172.057 | -0.11367 | 0.308263 | 0.898538 | Sec13 |
| ENSMUSG( | 7047.198 | -0.03375 | 0.740613 | 0.968811 | Sec14l1 |
| ENSMUSG( | 2750.949 | -0.1856 | 0.159774 | 0.837724 | Sec14l2 |
| ENSMUSG( | 27.81823 | -0.20054 | 0.613089 | 0.951154 | Sec14l3 |
| ENSMUSG( | 10.97752 | 0.042045 | 0.945416 | NA | Sec14l4 |
| ENSMUSG( | 1128.954 | 0.001563 | 0.993402 | 0.998813 | Sec14l5 |
| ENSMUSG( | 1597.749 | 0.039403 | 0.732834 | 0.967228 | Sec16a |
| ENSMUSG( | 37.15392 | -0.30231 | 0.437761 | 0.920339 | Sec16b |
| ENSMUSG( | 381.7071 | 0.100813 | 0.517282 | 0.930939 | Sec22a |
| ENSMUSG( | 1226.072 | -0.2043 | 0.123973 | 0.814691 | Sec22b |
| ENSMUSG( | 1469.129 | 0.19737 | 0.174007 | 0.844317 | Sec22c |
| ENSMUSG( | 2115.232 | 0.004514 | 0.966432 | 0.995475 | Sec23a |
| ENSMUSG( | 985.9558 | -0.0924 | 0.442606 | 0.921554 | Sec23b |
| ENSMUSG( | 895.0851 | 0.056055 | 0.684846 | 0.958678 | Sec23ip |
| ENSMUSG( | 549.4457 | 0.050224 | 0.687658 | 0.959673 | Sec24a |
| ENSMUSG( | 1202.79 | 0.07999 | 0.511406 | 0.92971 | Sec24b |
| ENSMUSG( | 2152.736 | -0.01275 | 0.908539 | 0.991958 | Sec24c |
| ENSMUSG( | 486.1967 | 0.178186 | 0.17437 | 0.844317 | Sec24d |
| ENSMUSG( | 2208.527 | 0.025612 | 0.822312 | 0.979198 | Sec31a |
| ENSMUSG( | 11.13745 | 0.814536 | 0.179191 | NA | Sec31b |
| ENSMUSG( | 3101.41 | 0.106304 | 0.481741 | 0.924578 | Sec61a1 |
| ENSMUSG( | 2057.295 | 0.069724 | 0.592094 | 0.944975 | Sec61a2 |
| ENSMUSG( | 440.7203 | -0.16825 | 0.268752 | 0.886755 | Sec61b |
| ENSMUSG( | 705.7668 | -0.22167 | 0.187243 | 0.856372 | Sec61g |
| ENSMUSG( | 7419.422 | 0.168014 | 0.299724 | 0.898538 | Sec62 |
| ENSMUSG( | 4076.197 | 0.118767 | 0.385165 | 0.917257 | Sec63 |
| ENSMUSG( | 512.644 | 0.123887 | 0.359233 | 0.914061 | Secisbp2 |
| ENSMUSG( | 5511.248 | -0.03683 | 0.74847 | 0.96984 | Secisbp2l |
| ENSMUSG( | 676.6287 | -0.07081 | 0.61243 | 0.951071 | Seh1l |
| ENSMUSG( | 7010.05 | 0.155909 | 0.350008 | 0.911652 | Sel1l |
| ENSMUSG( | 2630.314 | 0.088149 | 0.568756 | 0.941777 | Sel1l3 |
| ENSMUSG( | 491.4847 | -0.11242 | 0.484806 | 0.92541 | Selenbp1 |
| ENSMUSG( | 8205.303 | -0.07389 | 0.600758 | 0.947749 | Selenof |
| ENSMUSG( | 364.2924 | 0.048595 | 0.774514 | 0.974931 | Selenoh |
| ENSMUSG( | 1764.861 | 0.222866 | 0.180783 | 0.850372 | Selenoi |
| ENSMUSG( | 3760.971 | -0.03308 | 0.823288 | 0.979198 | Selenok |
| ENSMUSG( | 2163.038 | -0.10168 | 0.35802 | 0.913759 | Selenom |
| ENSMUSG( | 1082.794 | 0.124068 | 0.429434 | 0.917257 | Selenon |
| ENSMUSG( | 774.9056 | -0.00588 | 0.964752 | 0.995433 | Selenoo |
| ENSMUSG( | 22827.07 | 0.245157 | 0.042999 | 0.690304 | Selenop |
| ENSMUSG( | 1644.978 | 0.150713 | 0.307557 | 0.898538 | Selenos |
| ENSMUSG( | 6368.79 | 0.024801 | 0.875452 | 0.989644 | Selenot |
| ENSMUSG( | 23.17132 | -0.04291 | 0.920678 | 0.992644 | Selenov |
| ENSMUSG( | 42355.66 | -0.19089 | 0.110784 | 0.807582 | Selenow |
| ENSMUSG( | 16.75324 | -0.70722 | 0.231846 | 0.868507 | Sell |
| ENSMUSG( | 2563.359 | 0.444263 | 0.043211 | 0.690304 | Selplg |
| ENSMUSG( | 982.5405 | -0.1763 | 0.220884 | 0.866104 | Sem1 |
| ENSMUSG( | 604.1308 | 0.173653 | 0.43248 | 0.918256 | Sema3a |
| ENSMUSG( | 144.9269 | 0.234128 | 0.243001 | 0.877399 | Sema3b |
| ENSMUSG( | 972.2829 | 0.688213 | 0.001243 | 0.127113 | Sema3c |
| ENSMUSG( | 327.7597 | 0.526847 | 0.014243 | 0.477949 | Sema3d |
| ENSMUSG( | 904.42 | 0.247669 | 0.216422 | 0.865167 | Sema3e |
| ENSMUSG( | 398.0428 | 0.147643 | 0.389041 | 0.917257 | Sema3f |
| ENSMUSG( | 151.276 | 0.90762 | 0.004105 | 0.25867 | Sema3g |
| ENSMUSG( | 2019.008 | -0.06961 | 0.620207 | 0.952896 | Sema4a |
| ENSMUSG( | 909.4051 | 0.395068 | 0.016517 | 0.519511 | Sema4b |
| ENSMUSG( | 597.4871 | 0.162792 | 0.444248 | 0.921734 | Sema4c |
| ENSMUSG( | 3836.293 | 0.147117 | 0.435474 | 0.919062 | Sema4d |
| ENSMUSG( | 1663.57 | 0.18842 | 0.228263 | 0.868507 | Sema4f |
| ENSMUSG( | 1350.349 | 0.001199 | 0.991935 | 0.99872 | Sema4g |
| ENSMUSG( | 1955.691 | -0.01442 | 0.934142 | 0.99306 | Sema5a |
| ENSMUSG( | 970.9135 | -0.12824 | 0.383234 | 0.917257 | Sema5b |
| ENSMUSG( | 2037.914 | 0.122236 | 0.47634 | 0.923329 | Sema6a |
| ENSMUSG( | 5437.641 | 0.190785 | 0.173781 | 0.844317 | Sema6b |
| ENSMUSG( | 805.4999 | 0.233974 | 0.124302 | 0.814691 | Sema6c |
| ENSMUSG( | 3264.975 | -0.00514 | 0.973042 | 0.996416 | Sema6d |
| ENSMUSG( | 5753.724 | 0.015875 | 0.92748 | 0.99306 | Sema7a |
| ENSMUSG( | 493.1422 | 0.163663 | 0.274662 | 0.889874 | Senp1 |
| ENSMUSG( | 2989.386 | -0.06108 | 0.54438 | 0.938063 | Senp2 |
| ENSMUSG( | 1865.201 | 0.064181 | 0.555525 | 0.939565 | Senp3 |
| ENSMUSG( | 1165.321 | -0.01647 | 0.882708 | 0.989644 | Senp5 |
| ENSMUSG( | 2296.189 | 0.018832 | 0.868515 | 0.988097 | Senp6 |
| ENSMUSG( | 533.9507 | 0.195961 | 0.136456 | 0.827092 | Senp7 |
| ENSMUSG( | 87.59884 | -0.00536 | 0.98137 | 0.996756 | Senp8 |
| ENSMUSG( | 1249.228 | -0.08097 | 0.504473 | 0.927825 | Sephs1 |
| ENSMUSG( | 666.3116 | -0.13021 | 0.396154 | 0.917257 | Sephs2 |
| ENSMUSG( | 578.8518 | 0.019016 | 0.883344 | 0.989644 | Sepsecs |
| ENSMUSG( | 63.77345 | -0.09897 | 0.756655 | 0.971779 | Septin1 |
| ENSMUSG( | 306.9407 | -0.16708 | 0.272674 | 0.888825 | Septin10 |
| ENSMUSG( | 3168.936 | -0.08185 | 0.418847 | 0.917257 | Septin11 |
| ENSMUSG( | 39.65687 | 0.109114 | 0.760987 | 0.972695 | Septin2 |
| ENSMUSG( | 20394.01 | -0.04923 | 0.613645 | 0.951177 | Septin3 |
| ENSMUSG( | 5325.167 | -0.0942 | 0.397791 | 0.917257 | Septin4 |
| ENSMUSG( | 9178.82 | -0.00096 | 0.992016 | 0.99874 | Septin5 |
| ENSMUSG( | 1491.469 | -0.17544 | 0.208504 | 0.864412 | Septin6 |
| ENSMUSG( | 9941.632 | -0.07268 | 0.564688 | 0.941206 | Septin7 |
| ENSMUSG( | 11777.05 | -0.19435 | 0.094998 | 0.796133 | Septin8 |
| ENSMUSG( | 2717.132 | -0.21019 | 0.042535 | 0.690304 | Septin9 |
| ENSMUSG( | 378.8384 | 0.159318 | 0.308992 | 0.898538 | Serac1 |
| ENSMUSG( | 3912.768 | -0.01844 | 0.886069 | 0.989877 | Serbp1 |
| ENSMUSG( | 364.5736 | -0.043 | 0.797488 | 0.97681 | Serf1 |
| ENSMUSG( | 3767.469 | -0.19097 | 0.101613 | 0.801601 | Serf2 |
| ENSMUSG( | 510.5353 | -0.10793 | 0.474342 | 0.923329 | Sergef |
| ENSMUSG( | 370.9639 | -0.12165 | 0.37406 | 0.915491 | Serhl |
| ENSMUSG( | 25523.73 | 0.187858 | 0.255349 | 0.882761 | Serinc1 |
| ENSMUSG( | 224.5199 | 0.224108 | 0.24201 | 0.87661 | Serinc2 |
| ENSMUSG( | 11778.83 | 0.349941 | 0.047415 | 0.700936 | Serinc3 |
| ENSMUSG( | 20.08364 | -0.05004 | 0.916468 | 0.992529 | Serinc4 |
| ENSMUSG( | 3668.873 | 0.019547 | 0.914713 | 0.992407 | Serinc5 |
| ENSMUSG( | 1841.45 | -0.04863 | 0.76512 | 0.974236 | Serp1 |
| ENSMUSG( | 2378.806 | -0.21154 | 0.107478 | 0.804969 | Serp2 |
| ENSMUSG( | 665.3259 | 0.14572 | 0.374574 | 0.915491 | Serpina3n |
| ENSMUSG( | 144.4245 | -0.23398 | 0.299184 | 0.898538 | Serpina9 |
| ENSMUSG( | 236.4839 | -0.26889 | 0.097445 | 0.796721 | Serpinb1a |
| ENSMUSG( | 24.64819 | -0.25015 | 0.537121 | 0.937216 | Serpinb1b |
| ENSMUSG( | 851.023 | -0.22746 | 0.059796 | 0.729747 | Serpinb6a |
| ENSMUSG( | 76.75636 | 0.529987 | 0.088013 | 0.786591 | Serpinb6b |
| ENSMUSG( | 136.748 | 0.163745 | 0.416782 | 0.917257 | Serpinb8 |
| ENSMUSG( | 282.8059 | 0.004096 | 0.980023 | 0.996479 | Serpinb9 |
| ENSMUSG( | 82.93713 | 0.177496 | 0.464914 | 0.923329 | Serpind1 |
| ENSMUSG( | 44.45583 | 0.093615 | 0.772716 | 0.974463 | Serpine1 |
| ENSMUSG( | 18612.59 | -0.10813 | 0.402222 | 0.917257 | Serpine2 |
| ENSMUSG( | 734.291 | 0.109841 | 0.430266 | 0.917257 | Serpinf1 |
| ENSMUSG( | 11.40894 | 0.424383 | 0.481219 | NA | Serpinf2 |
| ENSMUSG( | 366.5037 | 0.142582 | 0.401418 | 0.917257 | Serping1 |
| ENSMUSG( | 1803.786 | 0.345521 | 0.037235 | 0.673514 | Serpinh1 |
| ENSMUSG( | 6625.142 | 0.335665 | 0.052589 | 0.71525 | Serpini1 |
| ENSMUSG( | 195.7735 | -0.08835 | 0.695673 | 0.960867 | Sertad1 |
| ENSMUSG( | 803.3746 | 0.00937 | 0.940292 | 0.994334 | Sertad2 |
| ENSMUSG( | 66.7476 | 0.272723 | 0.363413 | 0.914223 | Sertad3 |

| | | | | | |
|---|---|---|---|---|---|
| ENSMUSG( | Sf3a2 | 857.2785 | -0.02886 | 0.817659 | 0.965785 |
| ENSMUSG( | Sf3a3 | 563.4046 | 0.054775 | 0.657212 | 0.924142 |
| ENSMUSG( | Sf3b1 | 7722.841 | -0.03063 | 0.709556 | 0.939669 |
| ENSMUSG( | Sf3b2 | 5993.254 | 0.120493 | 0.143751 | 0.562581 |
| ENSMUSG( | Sf3b3 | 1706.909 | 0.051075 | 0.58708 | 0.900235 |
| ENSMUSG( | Sf3b4 | 662.1542 | 0.018385 | 0.893084 | 0.984512 |
| ENSMUSG( | Sf3b5 | 691.8423 | 0.090159 | 0.450535 | 0.83542 |
| ENSMUSG( | Sf3b6 | 1044.381 | 0.040933 | 0.746399 | 0.949289 |
| ENSMUSG( | Sfi1 | 631.9803 | 0.234813 | 0.146459 | 0.568435 |
| ENSMUSG( | Sfmbt1 | 670.0561 | 0.094405 | 0.376666 | 0.801213 |
| ENSMUSG( | Sfmbt2 | 109.0671 | 0.064488 | 0.742972 | 0.948478 |
| ENSMUSG( | Sfn | 66.07871 | 0.072548 | 0.801325 | 0.961947 |
| ENSMUSG( | Sfpq | 3077.605 | 0.07655 | 0.438197 | 0.830322 |
| ENSMUSG( | Sfr1 | 2180.714 | 0.033587 | 0.728437 | 0.944393 |
| ENSMUSG( | Sfrp1 | 741.7904 | -0.25541 | 0.043206 | 0.318825 |
| ENSMUSG( | Sfrp2 | 76.16831 | -0.57147 | 0.021939 | 0.225738 |
| ENSMUSG( | Sfrp4 | 75.9886 | -0.13281 | 0.568049 | 0.892201 |
| ENSMUSG( | Sfrp5 | 66.59807 | -0.71504 | 0.016303 | 0.189424 |
| ENSMUSG( | Sfswap | 2741.295 | 0.140404 | 0.201617 | 0.641362 |
| ENSMUSG( | Sft2d1 | 754.7177 | -0.1971 | 0.065212 | 0.395697 |
| ENSMUSG( | Sft2d1rt | 23.27855 | -0.17118 | 0.652088 | 0.922053 |
| ENSMUSG( | Sft2d2 | 3733.595 | 0.084187 | 0.474323 | 0.849425 |
| ENSMUSG( | Sft2d3 | 168.8239 | -0.51985 | 0.003384 | 0.073288 |
| ENSMUSG( | Sfta3-ps | 8.575746 | -0.73033 | 0.269411 | NA |
| ENSMUSG( | Sftpc | 29.04238 | -0.3625 | 0.361296 | 0.792797 |
| ENSMUSG( | Sfxn1 | 2883.701 | 0.07869 | 0.375477 | 0.800845 |
| ENSMUSG( | Sfxn2 | 225.0212 | -0.12585 | 0.428222 | 0.825894 |
| ENSMUSG( | Sfxn3 | 2763.052 | 0.120692 | 0.380701 | 0.801984 |
| ENSMUSG( | Sfxn4 | 283.97 | 0.076853 | 0.632848 | 0.914073 |
| ENSMUSG( | Sfxn5 | 26613.72 | 0.053362 | 0.64202 | 0.916865 |
| ENSMUSG( | Sgca | 15.50577 | -0.30884 | 0.518604 | NA |
| ENSMUSG( | Sgcb | 3958.855 | -0.17109 | 0.1278 | 0.535913 |
| ENSMUSG( | Sgcd | 38.03395 | -0.17148 | 0.571563 | 0.893319 |
| ENSMUSG( | Sgce | 1043.463 | -0.34613 | 0.0112 | 0.152581 |
| ENSMUSG( | Sgcg | 11.76858 | 0.164143 | 0.782112 | NA |
| ENSMUSG( | Sgcz | 41.92361 | -0.62935 | 0.046813 | 0.330855 |
| ENSMUSG( | Sgf29 | 250.682 | -0.08335 | 0.640086 | 0.916434 |
| ENSMUSG( | Sgip1 | 3315.914 | -0.09292 | 0.313109 | 0.76033 |
| ENSMUSG( | Sgk1 | 1145.982 | -0.38693 | 0.002288 | 0.055887 |
| ENSMUSG( | Sgk2 | 86.60862 | -0.11413 | 0.595616 | 0.90314 |
| ENSMUSG( | Sgk3 | 500.5776 | -0.05463 | 0.661641 | 0.925271 |
| ENSMUSG( | Sgms1 | 1359.443 | 0.050229 | 0.735721 | 0.946267 |
| ENSMUSG( | Sgms1os1 | 158.1314 | -0.01171 | 0.943244 | 0.993415 |
| ENSMUSG( | Sgms2 | 102.0561 | 0.155902 | 0.4795 | 0.854046 |
| ENSMUSG( | Sgo2a | 37.83177 | -0.42082 | 0.200123 | 0.640302 |
| ENSMUSG( | Sgpl1 | 1853.471 | -0.13407 | 0.313243 | 0.76033 |
| ENSMUSG( | Sgpp1 | 1228.171 | -0.28389 | 0.03739 | 0.298461 |
| ENSMUSG( | Sgpp2 | 442.1645 | -0.32697 | 0.070632 | 0.409748 |
| ENSMUSG( | Sgsh | 387.0399 | -0.26174 | 0.146572 | 0.568485 |
| ENSMUSG( | Sgsm1 | 3251.331 | 0.231719 | 0.047164 | 0.33245 |
| ENSMUSG( | Sgsm2 | 7278.548 | 0.098283 | 0.183535 | 0.620584 |
| ENSMUSG( | Sgsm3 | 1613.896 | 0.223905 | 0.097274 | 0.477629 |
| ENSMUSG( | Sgta | 3601.025 | -0.0152 | 0.87486 | 0.979746 |
| ENSMUSG( | Sgtb | 1999.1 | 0.086292 | 0.42836 | 0.826032 |
| ENSMUSG( | Sh2b1 | 2651.238 | 0.092785 | 0.300457 | 0.749525 |
| ENSMUSG( | Sh2b2 | 170.377 | 0.037148 | 0.816957 | 0.96566 |
| ENSMUSG( | Sh2b3 | 1018.238 | 0.265662 | 0.021995 | 0.226001 |
| ENSMUSG( | Sh2d3c | 574.3668 | -0.14698 | 0.336058 | 0.777 |
| ENSMUSG( | Sh2d4b | 34.28766 | 0.491874 | 0.157565 | 0.585653 |
| ENSMUSG( | Sh2d5 | 2938.025 | 0.009584 | 0.931156 | 0.992271 |
| ENSMUSG( | Sh2d6 | 39.03593 | -0.21408 | 0.484579 | 0.856665 |
| ENSMUSG( | Sh3bgr | 36.89069 | 0.011807 | 0.973248 | 0.99675 |
| ENSMUSG( | Sh3bgrl | 3420.968 | 0.022279 | 0.894281 | 0.984512 |
| ENSMUSG( | Sh3bgrl2 | 569.0737 | -0.05434 | 0.683271 | 0.932153 |
| ENSMUSG( | Sh3bgrl3 | 2134.15 | -0.03785 | 0.719087 | 0.941986 |
| ENSMUSG( | Sh3bp1 | 210.5244 | 0.045415 | 0.813876 | 0.964574 |
| ENSMUSG( | Sh3bp2 | 173.1904 | -0.33131 | 0.068491 | 0.405448 |
| ENSMUSG( | Sh3bp4 | 774.4954 | 0.084384 | 0.40946 | 0.818258 |
| ENSMUSG( | Sh3bp5 | 993.0504 | 0.167263 | 0.12554 | 0.53257 |
| ENSMUSG( | Sh3bp5l | 1597.509 | 0.290947 | 0.005438 | 0.098663 |
| ENSMUSG( | Sh3d19 | 892.3127 | -0.00468 | 0.965943 | 0.995377 |
| ENSMUSG( | Sh3d21 | 65.85281 | 0.146805 | 0.568345 | 0.892252 |
| ENSMUSG( | Sh3gl1 | 1195.306 | 0.135975 | 0.198384 | 0.638387 |
| ENSMUSG( | Sh3gl2 | 5483.866 | 0.180449 | 0.233945 | 0.681906 |
| ENSMUSG( | Sh3gl3 | 3828.663 | 0.070685 | 0.469195 | 0.845876 |
| ENSMUSG( | Sh3glb1 | 6517.826 | -0.02849 | 0.795177 | 0.960062 |
| ENSMUSG( | Sh3glb2 | 5891.219 | 0.03731 | 0.73396 | 0.945306 |
| ENSMUSG( | Sh3kbp1 | 986.8965 | -0.13444 | 0.233586 | 0.681508 |
| ENSMUSG( | Sh3pxd2a | 2720.863 | 0.033921 | 0.72676 | 0.944223 |
| ENSMUSG( | Sh3pxd2b | 1302.73 | -0.01065 | 0.913231 | 0.988483 |
| ENSMUSG( | Sh3rf1 | 1393.059 | 0.144531 | 0.100301 | 0.48338 |
| ENSMUSG( | Sh3rf2 | 82.31049 | -1.27696 | 0.000138 | 0.00806 |
| ENSMUSG( | Sh3rf3 | 1221.754 | 0.290021 | 0.002831 | 0.06505 |
| ENSMUSG( | Sh3tc1 | 111.7832 | 0.015143 | 0.947702 | 0.993821 |
| ENSMUSG( | Sh3tc2 | 172.3508 | 0.158644 | 0.389808 | 0.807597 |
| ENSMUSG( | Sh3yl1 | 171.2579 | 0.03242 | 0.854975 | 0.973665 |
| ENSMUSG( | Shank1 | 28175.6 | 0.377486 | 0.00015 | 0.008579 |
| ENSMUSG( | Shank2 | 11876.65 | 0.170206 | 0.094902 | 0.472904 |
| ENSMUSG( | Shank3 | 15098.78 | 0.226096 | 0.009399 | 0.13706 |
| ENSMUSG( | Sharpin | 892.1834 | -0.02361 | 0.837818 | 0.971111 |
| ENSMUSG( | Shb | 297.4424 | -0.08025 | 0.633913 | 0.914073 |
| ENSMUSG( | Shc1 | 519.4609 | 0.052244 | 0.67839 | 0.930542 |
| ENSMUSG( | Shc2 | 1399.652 | 0.08636 | 0.459087 | 0.840244 |
| ENSMUSG( | Shc3 | 3088.06 | 0.199565 | 0.050316 | 0.342969 |
| ENSMUSG( | Shc4 | 659.6339 | -0.12709 | 0.353935 | 0.786371 |
| ENSMUSG( | Shcbp1 | 22.87614 | 0.234383 | 0.538938 | 0.882849 |
| ENSMUSG( | Shcbp1l | 52.47488 | -0.2686 | 0.316391 | 0.763556 |
| ENSMUSG( | Shd | 1025.81 | 0.037036 | 0.710535 | 0.939748 |
| ENSMUSG( | She | 131.8376 | 0.28654 | 0.241772 | 0.689179 |
| ENSMUSG( | Shf | 1280.648 | 0.12462 | 0.243516 | 0.690987 |
| ENSMUSG( | Shfl | 1888.848 | 0.255167 | 0.143462 | 0.562473 |
| ENSMUSG( | Shh | 83.86347 | -0.40219 | 0.076321 | 0.427609 |
| ENSMUSG( | Shisa2 | 75.16707 | -0.4807 | 0.038206 | 0.301559 |
| ENSMUSG( | Shisa3 | 71.11808 | -1.16666 | 6.16E-06 | 0.000787 |
| ENSMUSG( | Shisa4 | 8104.075 | -0.22822 | 0.042612 | 0.316734 |
| ENSMUSG( | Shisa5 | 2134.033 | -0.36344 | 0.009508 | 0.137946 |
| ENSMUSG( | Shisa6 | 747.8048 | 0.01277 | 0.943438 | 0.993415 |
| ENSMUSG( | Shisa7 | 5701.051 | -0.23811 | 0.023601 | 0.235222 |
| ENSMUSG( | Shisa8 | 226.0881 | -0.7746 | 2.72E-06 | 0.000405 |
| ENSMUSG( | Shisa9 | 2470.807 | -0.3645 | 0.005309 | 0.097191 |
| ENSMUSG( | Shisal1 | 3481.278 | -0.25519 | 0.00816 | 0.125816 |
| ENSMUSG( | Shisal2b | 15.46003 | -0.22363 | 0.646813 | NA |
| ENSMUSG( | Shkbp1 | 394.0266 | 0.187778 | 0.226504 | 0.673549 |
| ENSMUSG( | Shld1 | 167.948 | -0.06797 | 0.692479 | 0.934101 |
| ENSMUSG( | Shld2 | 126.6706 | -0.27441 | 0.143974 | 0.563118 |
| ENSMUSG( | Shld3 | 21.81701 | 0.255192 | 0.547608 | 0.885153 |
| ENSMUSG( | Shmt1 | 57.4764 | -0.35566 | 0.181442 | 0.618684 |
| ENSMUSG( | Shmt2 | 1040.339 | 0.030398 | 0.775239 | 0.955606 |
| ENSMUSG( | Shoc2 | 1004.846 | 0.055073 | 0.561656 | 0.889052 |
| ENSMUSG( | Shpk | 141.6324 | -0.09991 | 0.580308 | 0.896952 |
| ENSMUSG( | Shprh | 1078.638 | 0.037436 | 0.683993 | 0.932457 |
| ENSMUSG( | Shq1 | 101.6036 | 0.281513 | 0.172854 | 0.608361 |
| ENSMUSG( | Shroom1 | 197.7803 | -0.06297 | 0.727014 | 0.944223 |
| ENSMUSG( | Shroom2 | 1809.951 | -0.01995 | 0.843574 | 0.971847 |
| ENSMUSG( | Shroom3 | 158.8227 | -0.30695 | 0.071716 | 0.413564 |
| ENSMUSG( | Shroom4 | 219.2706 | 0.120084 | 0.516161 | 0.872701 |
| ENSMUSG( | Shtn1 | 1405.996 | -0.16927 | 0.064905 | 0.395029 |
| ENSMUSG( | Siae | 1017.818 | -0.26705 | 0.027767 | 0.255176 |
| ENSMUSG( | Siah1a | 431.1111 | -0.06007 | 0.634611 | 0.914073 |
| ENSMUSG( | Siah1b | 86.92589 | -0.29976 | 0.147304 | 0.569415 |
| ENSMUSG( | Siah2 | 333.7083 | -0.27355 | 0.082802 | 0.444372 |
| ENSMUSG( | Siah3 | 295.9917 | 0.217418 | 0.114742 | 0.510733 |
| ENSMUSG( | Sidt1 | 2213.649 | 0.097733 | 0.475867 | 0.850919 |
| ENSMUSG( | Sidt2 | 3438.715 | -0.08585 | 0.496051 | 0.861543 |
| ENSMUSG( | Sigirr | 70.74977 | 0.431238 | 0.104171 | 0.489971 |
| ENSMUSG( | Siglec1 | 27.48616 | -0.28473 | 0.516242 | 0.872701 |
| ENSMUSG( | Siglece | 54.10897 | -0.49742 | 0.123806 | 0.531039 |
| ENSMUSG( | Siglecf | 30.14733 | -0.94137 | 0.013802 | 0.171544 |
| ENSMUSG( | Siglech | 1195.118 | -0.03007 | 0.850015 | 0.972638 |
| ENSMUSG( | Sigmar1 | 1585.367 | -0.21043 | 0.078655 | 0.433892 |
| ENSMUSG( | Sik1 | 386.8891 | -0.70363 | 6.24E-08 | 1.91E-05 |
| ENSMUSG( | Sik2 | 598.3326 | -0.047 | 0.701058 | 0.93743 |
| ENSMUSG( | Sik3 | 4989.571 | 0.106386 | 0.222761 | 0.668614 |
| ENSMUSG( | Sike1 | 1768.504 | 0.052602 | 0.55273 | 0.886141 |
| ENSMUSG( | Sil1 | 874.932 | -0.37194 | 0.006932 | 0.114527 |
| ENSMUSG( | Silc1 | 858.5327 | 0.167267 | 0.123813 | 0.531039 |
| ENSMUSG( | Simc1 | 342.5739 | -0.01139 | 0.929468 | 0.991848 |
| ENSMUSG( | Sin3a | 1120.854 | 0.072225 | 0.428148 | 0.825894 |
| ENSMUSG( | Sin3b | 2561.761 | 0.109155 | 0.312627 | 0.760241 |
| ENSMUSG( | Sinhcaf | 22.32148 | -0.0776 | 0.8645 | 0.975398 |
| ENSMUSG( | Sipa1 | 850.8293 | 0.175814 | 0.105446 | 0.492417 |
| ENSMUSG( | Sipa1l1 | 16066.59 | 0.200011 | 0.048726 | 0.33775 |
| ENSMUSG( | Sipa1l2 | 2120.829 | 0.02868 | 0.75208 | 0.950417 |
| ENSMUSG( | Sipa1l3 | 1867.848 | 0.279915 | 0.013978 | 0.172788 |
| ENSMUSG( | Sirpa | 19012.88 | -0.17954 | 0.186481 | 0.623869 |
| ENSMUSG( | Sirt1 | 320.1077 | -0.03807 | 0.786101 | 0.958774 |
| ENSMUSG( | Sirt2 | 18656.48 | 0.086523 | 0.306404 | 0.754119 |
| ENSMUSG( | Sirt3 | 3281.708 | 0.078929 | 0.33747 | 0.777412 |

| | | | | | |
|---|---|---|---|---|---|
| ENSMUSG( | Sf3a2 | 794.5386 | 0.017037 | 0.905915 | 0.991089 |
| ENSMUSG( | Sf3a3 | 499.6928 | -0.02446 | 0.864282 | 0.986552 |
| ENSMUSG( | Sf3b1 | 7416.017 | 0.119039 | 0.262225 | 0.885219 |
| ENSMUSG( | Sf3b2 | 5094.627 | -0.07924 | 0.456166 | 0.923329 |
| ENSMUSG( | Sf3b3 | 1487.04 | -0.07909 | 0.511139 | 0.929571 |
| ENSMUSG( | Sf3b4 | 564.6193 | -0.18108 | 0.232156 | 0.868507 |
| ENSMUSG( | Sf3b5 | 593.654 | -0.0891 | 0.56095 | 0.940254 |
| ENSMUSG( | Sf3b6 | 906.177 | -0.10752 | 0.451004 | 0.922202 |
| ENSMUSG( | Sfi1 | 561.3122 | 0.165931 | 0.453001 | 0.922316 |
| ENSMUSG( | Sfmbt1 | 569.019 | -0.10908 | 0.393592 | 0.917257 |
| ENSMUSG( | Sfmbt2 | 101.0069 | 0.108116 | 0.611655 | 0.950771 |
| ENSMUSG( | Sfn | 63.99219 | 0.246638 | 0.421761 | 0.917257 |
| ENSMUSG( | Sfpq | 2927.194 | 0.194913 | 0.06485 | 0.740531 |
| ENSMUSG( | Sfr1 | 1901.006 | -0.09907 | 0.452274 | 0.922316 |
| ENSMUSG( | Sfrp1 | 786.4787 | 0.189611 | 0.180548 | 0.850372 |
| ENSMUSG( | Sfrp2 | 80.17526 | -0.10561 | 0.671589 | 0.95694 |
| ENSMUSG( | Sfrp4 | 64.9748 | -0.31146 | 0.238071 | 0.87259 |
| ENSMUSG( | Sfrp5 | 88.73952 | 0.429908 | 0.129442 | 0.820358 |
| ENSMUSG( | Sfswap | 2489.858 | 0.132545 | 0.354168 | 0.912411 |
| ENSMUSG( | Sft2d1 | 722.2958 | -0.04437 | 0.747961 | 0.96984 |
| ENSMUSG( | Sft2d1rt | 23.22672 | 0.074895 | 0.855284 | 0.985649 |
| ENSMUSG( | Sft2d2 | 3363.886 | 0.053789 | 0.662871 | 0.956205 |
| ENSMUSG( | Sft2d3 | 180.0232 | -0.01764 | 0.917997 | 0.992529 |
| ENSMUSG( | Sfta3-ps | 10.25905 | 0.174938 | 0.78107 | NA |
| ENSMUSG( | Sftpc | 28.58889 | -0.1458 | 0.714017 | 0.964986 |
| ENSMUSG( | Sfxn1 | 2540.628 | -0.01928 | 0.87507 | 0.989644 |
| ENSMUSG( | Sfxn2 | 244.5396 | 0.373676 | 0.065676 | 0.745596 |
| ENSMUSG( | Sfxn3 | 2336.977 | -0.09507 | 0.539887 | 0.937378 |
| ENSMUSG( | Sfxn4 | 262.8244 | 0.123074 | 0.549331 | 0.938519 |
| ENSMUSG( | Sfxn5 | 21067.17 | -0.37855 | 0.009911 | 0.407479 |
| ENSMUSG( | Sgca | 18.78828 | 0.507082 | 0.270812 | 0.888062 |
| ENSMUSG( | Sgcb | 3794.254 | -0.01857 | 0.883102 | 0.989644 |
| ENSMUSG( | Sgcd | 39.75941 | 0.220552 | 0.484488 | 0.92541 |
| ENSMUSG( | Sgce | 1105.413 | 0.111533 | 0.474015 | 0.923329 |
| ENSMUSG( | Sgcg | 10.1954 | 0.021642 | 0.973096 | NA |
| ENSMUSG( | Sgcz | 45.50481 | -0.06476 | 0.837901 | 0.981376 |
| ENSMUSG( | Sgf29 | 251.0353 | 0.177792 | 0.299026 | 0.898538 |
| ENSMUSG( | Sgip1 | 3141.202 | 0.021623 | 0.848688 | 0.985321 |
| ENSMUSG( | Sgk1 | 1173.996 | -0.01907 | 0.919932 | 0.992644 |
| ENSMUSG( | Sgk2 | 77.99202 | -0.13512 | 0.580273 | 0.944209 |
| ENSMUSG( | Sgk3 | 488.8632 | 0.139378 | 0.340537 | 0.907972 |
| ENSMUSG( | Sgms1 | 1358.715 | 0.309165 | 0.167557 | 0.840222 |
| ENSMUSG( | Sgms1os1 | 135.9434 | -0.1792 | 0.349741 | 0.911652 |
| ENSMUSG( | Sgms2 | 120.6246 | 0.818057 | 0.001239 | 0.127113 |
| ENSMUSG( | Sgo2a | 35.87567 | -0.28955 | 0.402158 | 0.917257 |
| ENSMUSG( | Sgpl1 | 1841.872 | 0.121054 | 0.391434 | 0.917257 |
| ENSMUSG( | Sgpp1 | 1351.732 | 0.264423 | 0.113103 | 0.809623 |
| ENSMUSG( | Sgpp2 | 517.9735 | 0.398259 | 0.043731 | 0.690304 |
| ENSMUSG( | Sgsh | 409.9238 | 0.189306 | 0.363738 | 0.914223 |
| ENSMUSG( | Sgsm1 | 2683.576 | -0.046 | 0.745286 | 0.96951 |
| ENSMUSG( | Sgsm2 | 6242.584 | -0.07519 | 0.439546 | 0.920721 |
| ENSMUSG( | Sgsm3 | 1323.288 | -0.07428 | 0.610254 | 0.950771 |
| ENSMUSG( | Sgta | 3153.185 | -0.13466 | 0.244996 | 0.877904 |
| ENSMUSG( | Sgtb | 1770.707 | 0.007214 | 0.951136 | 0.994798 |
| ENSMUSG( | Sh2b1 | 2304.093 | -0.0431 | 0.697102 | 0.960867 |
| ENSMUSG( | Sh2b2 | 162.1473 | 0.16322 | 0.372628 | 0.915491 |
| ENSMUSG( | Sh2b3 | 945.8376 | 0.321466 | 0.022255 | 0.577034 |
| ENSMUSG( | Sh2d3c | 505.2573 | -0.24783 | 0.130074 | 0.821082 |
| ENSMUSG( | Sh2d4b | 32.03487 | 0.531576 | 0.177241 | 0.846973 |
| ENSMUSG( | Sh2d5 | 2607.141 | -0.06818 | 0.591287 | 0.944975 |
| ENSMUSG( | Sh2d6 | 31.44313 | -0.58555 | 0.150079 | 0.837086 |
| ENSMUSG( | Sh3bgr | 31.48744 | -0.24153 | 0.584832 | 0.944975 |
| ENSMUSG( | Sh3bgrl | 3055.16 | -0.04087 | 0.847677 | 0.9852 |
| ENSMUSG( | Sh3bgrl2 | 505.4104 | -0.13252 | 0.394454 | 0.917257 |
| ENSMUSG( | Sh3bgrl3 | 1819.344 | -0.24254 | 0.066398 | 0.749411 |
| ENSMUSG( | Sh3bp1 | 197.1545 | 0.120458 | 0.545908 | 0.938063 |
| ENSMUSG( | Sh3bp2 | 170.7252 | -0.06917 | 0.737581 | 0.967686 |
| ENSMUSG( | Sh3bp4 | 698.5528 | 0.059734 | 0.62957 | 0.953251 |
| ENSMUSG( | Sh3bp5 | 846.5868 | -0.02335 | 0.860495 | 0.986137 |
| ENSMUSG( | Sh3bp5l | 1334.593 | 0.054234 | 0.653573 | 0.955015 |
| ENSMUSG( | Sh3d19 | 837.5346 | 0.084305 | 0.502856 | 0.927825 |
| ENSMUSG( | Sh3d21 | 63.76457 | 0.321799 | 0.231231 | 0.868507 |
| ENSMUSG( | Sh3gl1 | 1071.514 | 0.091944 | 0.439596 | 0.920721 |
| ENSMUSG( | Sh3gl2 | 4362.548 | -0.21695 | 0.219738 | 0.866104 |
| ENSMUSG( | Sh3gl3 | 3224.961 | -0.16215 | 0.19359 | 0.860096 |
| ENSMUSG( | Sh3glb1 | 5830.07 | -0.08391 | 0.52133 | 0.932489 |
| ENSMUSG( | Sh3glb2 | 5247.65 | -0.02879 | 0.819818 | 0.979198 |
| ENSMUSG( | Sh3kbp1 | 889.8169 | -0.17089 | 0.185554 | 0.855305 |
| ENSMUSG( | Sh3pxd2a | 2365.026 | -0.10584 | 0.36837 | 0.915491 |
| ENSMUSG( | Sh3pxd2b | 1174.388 | -0.04104 | 0.725527 | 0.966887 |
| ENSMUSG( | Sh3rf1 | 1260.375 | 0.123999 | 0.26678 | 0.886134 |
| ENSMUSG( | Sh3rf2 | 94.62635 | -0.38632 | 0.239803 | 0.874948 |
| ENSMUSG( | Sh3rf3 | 1166.969 | 0.415146 | 0.000856 | 0.09911 |
| ENSMUSG( | Sh3tc1 | 117.1271 | 0.411378 | 0.118226 | 0.813384 |
| ENSMUSG( | Sh3tc2 | 152.1213 | 0.078797 | 0.680639 | 0.958395 |
| ENSMUSG( | Sh3yl1 | 156.1754 | 0.033043 | 0.858801 | 0.985884 |
| ENSMUSG( | Shank1 | 24259.58 | 0.229928 | 0.054446 | 0.723362 |
| ENSMUSG( | Shank2 | 10982 | 0.211429 | 0.090908 | 0.791074 |
| ENSMUSG( | Shank3 | 13648.26 | 0.204192 | 0.086838 | 0.782073 |
| ENSMUSG( | Sharpin | 811.0096 | -0.02959 | 0.819566 | 0.979198 |
| ENSMUSG( | Shb | 271.005 | -0.08694 | 0.652655 | 0.954799 |
| ENSMUSG( | Shc1 | 499.8476 | 0.210375 | 0.178312 | 0.848538 |
| ENSMUSG( | Shc2 | 1238.14 | 0.003297 | 0.979318 | 0.996468 |
| ENSMUSG( | Shc3 | 2514.204 | -0.11999 | 0.302491 | 0.898538 |
| ENSMUSG( | Shc4 | 584.6164 | -0.21988 | 0.181529 | 0.850579 |
| ENSMUSG( | Shcbp1 | 18.34071 | -0.13418 | 0.764969 | 0.974236 |
| ENSMUSG( | Shcbp1l | 60.29077 | 0.409567 | 0.188824 | 0.85698 |
| ENSMUSG( | Shd | 916.3805 | -0.01801 | 0.883134 | 0.989644 |
| ENSMUSG( | She | 161.3246 | 1.002054 | 0.000263 | 0.04337 |
| ENSMUSG( | Shf | 1111.673 | -0.01536 | 0.900578 | 0.990558 |
| ENSMUSG( | Shfl | 1612.587 | 0.074726 | 0.699755 | 0.961335 |
| ENSMUSG( | Shh | 92.37979 | 0.152367 | 0.530431 | 0.934795 |
| ENSMUSG( | Shisa2 | 95.41822 | 0.468462 | 0.05397 | 0.722223 |
| ENSMUSG( | Shisa3 | 88.21764 | -0.07269 | 0.784435 | 0.976196 |
| ENSMUSG( | Shisa4 | 7895.693 | -0.0247 | 0.849223 | 0.985321 |
| ENSMUSG( | Shisa5 | 2238.482 | 0.07005 | 0.660267 | 0.956205 |
| ENSMUSG( | Shisa6 | 817.494 | 0.503463 | 0.016594 | 0.519982 |
| ENSMUSG( | Shisa7 | 5916.879 | 0.146123 | 0.18616 | 0.855305 |
| ENSMUSG( | Shisa8 | 294.0336 | 0.328548 | 0.061572 | 0.736111 |
| ENSMUSG( | Shisa9 | 2656.282 | 0.136212 | 0.341851 | 0.909146 |
| ENSMUSG( | Shisal1 | 3436.542 | -0.01095 | 0.924664 | 0.99306 |
| ENSMUSG( | Shisal2b | 12.39512 | -0.66738 | 0.23324 | 0.868507 |
| ENSMUSG( | Shkbp1 | 342.8972 | 0.051961 | 0.795409 | 0.97681 |
| ENSMUSG( | Shld1 | 152.5085 | -0.07615 | 0.683778 | 0.958678 |
| ENSMUSG( | Shld2 | 115.0769 | -0.27193 | 0.210488 | 0.864412 |
| ENSMUSG( | Shld3 | 16.70483 | -0.26402 | 0.586926 | 0.944975 |
| ENSMUSG( | Shmt1 | 50.87829 | -0.42888 | 0.162111 | 0.837724 |
| ENSMUSG( | Shmt2 | 914.2995 | -0.07682 | 0.525599 | 0.934374 |
| ENSMUSG( | Shoc2 | 924.2032 | 0.07735 | 0.539713 | 0.937378 |
| ENSMUSG( | Shpk | 128.7928 | -0.10684 | 0.582413 | 0.944266 |
| ENSMUSG( | Shprh | 998.0605 | 0.081524 | 0.461591 | 0.923329 |
| ENSMUSG( | Shq1 | 80.8259 | -0.09807 | 0.696989 | 0.960867 |
| ENSMUSG( | Shroom1 | 195.5492 | 0.182011 | 0.363751 | 0.914223 |
| ENSMUSG( | Shroom2 | 1667.235 | 0.008524 | 0.949806 | 0.994798 |
| ENSMUSG( | Shroom3 | 155.4546 | -0.06372 | 0.780575 | 0.976106 |
| ENSMUSG( | Shroom4 | 212.9436 | 0.306721 | 0.161129 | 0.837724 |
| ENSMUSG( | Shtn1 | 1212.278 | -0.34627 | 0.002455 | 0.194608 |
| ENSMUSG( | Siae | 984.336 | -0.07904 | 0.585166 | 0.944975 |
| ENSMUSG( | Siah1a | 398.1376 | -0.02066 | 0.879474 | 0.989644 |
| ENSMUSG( | Siah1b | 78.94867 | -0.34302 | 0.198204 | 0.862161 |
| ENSMUSG( | Siah2 | 322.4708 | -0.09515 | 0.578761 | 0.943611 |
| ENSMUSG( | Siah3 | 263.4386 | 0.158269 | 0.32816 | 0.906255 |
| ENSMUSG( | Sidt1 | 2285.708 | 0.431974 | 0.001438 | 0.139907 |
| ENSMUSG( | Sidt2 | 3420.746 | 0.168419 | 0.27062 | 0.888062 |
| ENSMUSG( | Sigirr | 64.1468 | 0.454519 | 0.256429 | 0.883082 |
| ENSMUSG( | Siglec1 | 25.00227 | -0.27715 | 0.523128 | 0.932848 |
| ENSMUSG( | Siglece | 62.22924 | 0.223693 | 0.465806 | 0.923329 |
| ENSMUSG( | Siglecf | 31.54521 | -0.38606 | 0.329658 | 0.907083 |
| ENSMUSG( | Siglech | 1191.406 | 0.227314 | 0.190261 | 0.857355 |
| ENSMUSG( | Sigmar1 | 1675.463 | 0.219702 | 0.097144 | 0.796721 |
| ENSMUSG( | Sik1 | 398.1204 | -0.27517 | 0.100646 | 0.801444 |
| ENSMUSG( | Sik2 | 551.511 | -0.00977 | 0.94263 | 0.994774 |
| ENSMUSG( | Sik3 | 4375.175 | -0.00274 | 0.979644 | 0.996479 |
| ENSMUSG( | Sike1 | 1565.945 | -0.0313 | 0.782 | 0.976106 |
| ENSMUSG( | Sil1 | 929.5745 | 0.098087 | 0.520553 | 0.932079 |
| ENSMUSG( | Silc1 | 763.8653 | 0.100402 | 0.415552 | 0.917257 |
| ENSMUSG( | Simc1 | 309.1924 | -0.03411 | 0.822741 | 0.979198 |
| ENSMUSG( | Sin3a | 1024.157 | 0.077928 | 0.503183 | 0.927825 |
| ENSMUSG( | Sin3b | 2217.247 | -0.03912 | 0.768344 | 0.974236 |
| ENSMUSG( | Sinhcaf | 27.73306 | 0.75811 | 0.068277 | 0.752208 |
| ENSMUSG( | Sipa1 | 825.0785 | 0.345863 | 0.015371 | 0.495791 |
| ENSMUSG( | Sipa1l1 | 14299.49 | 0.137906 | 0.247642 | 0.877904 |
| ENSMUSG( | Sipa1l2 | 1932.726 | 0.031704 | 0.785978 | 0.976285 |
| ENSMUSG( | Sipa1l3 | 1716.093 | 0.303744 | 0.021421 | 0.575341 |
| ENSMUSG( | Sirpa | 19471.54 | 0.162994 | 0.317182 | 0.902275 |
| ENSMUSG( | Sirt1 | 320.8136 | 0.233665 | 0.115882 | 0.813384 |
| ENSMUSG( | Sirt2 | 15505.16 | -0.18547 | 0.128562 | 0.81925 |
| ENSMUSG( | Sirt3 | 2778.57 | -0.13621 | 0.21151 | 0.864412 |

| | | | | | |
|---|---|---|---|---|---|
| ENSMUSG( | 426.4222 | -0.00267 | 0.986877 | 0.998619 | Sdf2l1 |
| ENSMUSG( | 9616.81 | -0.13348 | 0.178045 | 0.615613 | Sdf4 |
| ENSMUSG( | 15335.06 | 0.083414 | 0.333656 | 0.775602 | Sdha |
| ENSMUSG( | 763.2384 | -0.08937 | 0.427609 | 0.825692 | Sdhaf1 |
| ENSMUSG( | 1309.985 | 0.054492 | 0.555054 | 0.88708 | Sdhaf2 |
| ENSMUSG( | 210.0694 | -0.0649 | 0.672573 | 0.929007 | Sdhaf3 |
| ENSMUSG( | 377.7927 | -0.04776 | 0.746673 | 0.949289 | Sdhaf4 |
| ENSMUSG( | 3426.044 | 0.048103 | 0.608491 | 0.907154 | Sdhb |
| ENSMUSG( | 4598.205 | 0.088059 | 0.352243 | 0.783835 | Sdhc |
| ENSMUSG( | 2297.88 | 0.062967 | 0.508146 | 0.867999 | Sdhd |
| ENSMUSG( | 349.4875 | -0.18856 | 0.223823 | 0.669845 | Sdk1 |
| ENSMUSG( | 1280.155 | 0.049447 | 0.736077 | 0.946267 | Sdk2 |
| ENSMUSG( | 1397.967 | -0.07575 | 0.510964 | 0.870755 | Sdr39u1 |
| ENSMUSG( | 270.116 | 0.207875 | 0.225395 | 0.672165 | Sdr42e1 |
| ENSMUSG( | 217.5833 | 0.019548 | 0.907776 | 0.987798 | Sdsl |
| ENSMUSG( | 75.80356 | 0.080762 | 0.729569 | 0.944764 | Sebox |
| ENSMUSG( | 52.28947 | -0.11144 | 0.675461 | 0.930049 | Sec1 |
| ENSMUSG( | 740.9883 | -0.27286 | 0.04254 | 0.316636 | Sec11a |
| ENSMUSG( | 2341.183 | -0.33249 | 0.00652 | 0.110626 | Sec11c |
| ENSMUSG( | 1360.22 | 0.048559 | 0.619626 | 0.911432 | Sec13 |
| ENSMUSG( | 7918.992 | 0.034328 | 0.677044 | 0.930542 | Sec14l1 |
| ENSMUSG( | 3254.431 | 0.03954 | 0.666893 | 0.92729 | Sec14l2 |
| ENSMUSG( | 35.97401 | 0.29654 | 0.347002 | 0.7797 | Sec14l3 |
| ENSMUSG( | 10.91318 | -0.3192 | 0.624975 | NA | Sec14l4 |
| ENSMUSG( | 1066.517 | -0.47782 | 0.001119 | 0.035587 | Sec14l5 |
| ENSMUSG( | 1755.517 | 0.040011 | 0.671301 | 0.928781 | Sec16a |
| ENSMUSG( | 44.4557 | -0.04855 | 0.88813 | 0.983434 | Sec16b |
| ENSMUSG( | 375.4908 | -0.22436 | 0.105391 | 0.492417 | Sec22a |
| ENSMUSG( | 1487.867 | 0.095839 | 0.362772 | 0.793121 | Sec22b |
| ENSMUSG( | 1463.671 | -0.08046 | 0.53473 | 0.880719 | Sec22c |
| ENSMUSG( | 2398.888 | 0.09883 | 0.264368 | 0.715323 | Sec23a |
| ENSMUSG( | 1154.941 | 0.097397 | 0.293275 | 0.742763 | Sec23b |
| ENSMUSG( | 1030.037 | 0.190341 | 0.100153 | 0.483045 | Sec23ip |
| ENSMUSG( | 596.8164 | 0.022447 | 0.837698 | 0.971111 | Sec24a |
| ENSMUSG( | 1377.496 | 0.197394 | 0.038552 | 0.302696 | Sec24b |
| ENSMUSG( | 2557.8 | 0.209338 | 0.025971 | 0.247245 | Sec24c |
| ENSMUSG( | 541.5351 | 0.215157 | 0.05639 | 0.366006 | Sec24d |
| ENSMUSG( | 2514.839 | 0.129593 | 0.179133 | 0.616715 | Sec31a |
| ENSMUSG( | 2951.371 | -0.32604 | 0.010803 | 0.14989 | Sec61a1 |
| ENSMUSG( | 1985.342 | -0.32089 | 0.007791 | 0.12245 | Sec61a2 |
| ENSMUSG( | 509.0312 | -0.00773 | 0.952601 | 0.993874 | Sec61b |
| ENSMUSG( | 849.8251 | 0.059805 | 0.611802 | 0.908586 | Sec61g |
| ENSMUSG( | 7001.663 | -0.27691 | 0.040823 | 0.312481 | Sec62 |
| ENSMUSG( | 4201.228 | -0.06242 | 0.605586 | 0.905824 | Sec63 |
| ENSMUSG( | 555.5105 | 0.092133 | 0.449646 | 0.835234 | Secisbp2 |
| ENSMUSG( | 6057.24 | -0.03285 | 0.715309 | 0.941288 | Secisbp2l |
| ENSMUSG( | 739.4919 | -0.08208 | 0.499714 | 0.863218 | Seh1l |
| ENSMUSG( | 6775.232 | -0.21876 | 0.144607 | 0.564508 | Sel1l |
| ENSMUSG( | 2704.361 | -0.10485 | 0.420357 | 0.822856 | Sel1l3 |
| ENSMUSG( | 564.7818 | 0.029058 | 0.837343 | 0.971111 | Selenbp1 |
| ENSMUSG( | 8443.261 | -0.27186 | 0.029477 | 0.260274 | Selenof |
| ENSMUSG( | 410.5574 | 0.12909 | 0.311619 | 0.759296 | Selenoh |
| ENSMUSG( | 1651.408 | -0.24481 | 0.114707 | 0.510718 | Selenoi |
| ENSMUSG( | 3747.041 | -0.33059 | 0.005324 | 0.097356 | Selenok |
| ENSMUSG( | 2332.78 | -0.15275 | 0.06512 | 0.395697 | Selenom |
| ENSMUSG( | 1046 | -0.25936 | 0.048466 | 0.336526 | Selenon |
| ENSMUSG( | 843.2782 | -0.03676 | 0.758064 | 0.951614 | Selenoo |
| ENSMUSG( | 25049.63 | 0.245136 | 0.030418 | 0.265114 | Selenop |
| ENSMUSG( | 1562.918 | -0.27792 | 0.033662 | 0.281425 | Selenos |
| ENSMUSG( | 6491.788 | -0.19496 | 0.149503 | 0.573617 | Selenot |
| ENSMUSG( | 22.42348 | -0.39814 | 0.354459 | 0.786878 | Selenov |
| ENSMUSG( | 50472.46 | 0.053545 | 0.509509 | 0.869462 | Selenow |
| ENSMUSG( | 18.63745 | -0.62958 | 0.283771 | NA | Sell |
| ENSMUSG( | 2285.99 | -0.13043 | 0.44796 | 0.834051 | Selplg |
| ENSMUSG( | 1150.311 | 0.020073 | 0.878042 | 0.981431 | Sem1 |
| ENSMUSG( | 522.2484 | -0.57261 | 0.003593 | 0.075902 | Sema3a |
| ENSMUSG( | 146.9287 | -0.00103 | 0.995175 | 0.99943 | Sema3b |
| ENSMUSG( | 780.1209 | -0.14735 | 0.407525 | 0.817672 | Sema3c |
| ENSMUSG( | 258.2483 | -0.41868 | 0.024279 | 0.238384 | Sema3d |
| ENSMUSG( | 737.8312 | -0.68297 | 7.74E-05 | 0.005438 | Sema3e |
| ENSMUSG( | 380.1554 | -0.2689 | 0.080843 | 0.439584 | Sema3f |
| ENSMUSG( | 124.8698 | 0.197139 | 0.4702 | 0.84674 | Sema3g |
| ENSMUSG( | 2065.235 | -0.29015 | 0.009787 | 0.141238 | Sema4a |
| ENSMUSG( | 779.6796 | -0.31441 | 0.030823 | 0.266969 | Sema4b |
| ENSMUSG( | 553.8446 | -0.35246 | 0.043683 | 0.320345 | Sema4c |
| ENSMUSG( | 3592.141 | -0.33217 | 0.029525 | 0.260413 | Sema4d |
| ENSMUSG( | 1640.939 | -0.12157 | 0.431272 | 0.827055 | Sema4f |
| ENSMUSG( | 1526.119 | 0.083728 | 0.37288 | 0.799381 | Sema4g |
| ENSMUSG( | 1875.656 | -0.44042 | 0.002928 | 0.066146 | Sema5a |
| ENSMUSG( | 970.1706 | -0.43318 | 0.000758 | 0.027082 | Sema5b |
| ENSMUSG( | 1760.347 | -0.64238 | 2.13E-05 | 0.002059 | Sema6a |
| ENSMUSG( | 5314.702 | -0.144 | 0.230759 | 0.679066 | Sema6b |
| ENSMUSG( | 849.9453 | 0.120735 | 0.427138 | 0.82532 | Sema6c |
| ENSMUSG( | 3398.607 | -0.16593 | 0.249302 | 0.698744 | Sema6d |
| ENSMUSG( | 5856.054 | -0.21445 | 0.12481 | 0.531813 | Sema7a |
| ENSMUSG( | 534.2202 | 0.127704 | 0.37425 | 0.800445 | Senp1 |
| ENSMUSG( | 3440.458 | 0.075625 | 0.342805 | 0.778971 | Senp2 |
| ENSMUSG( | 2096.099 | 0.128476 | 0.179183 | 0.616715 | Senp3 |
| ENSMUSG( | 1353.379 | 0.144133 | 0.11325 | 0.507541 | Senp5 |
| ENSMUSG( | 2643.679 | 0.154797 | 0.075059 | 0.42422 | Senp6 |
| ENSMUSG( | 527.7693 | -0.10351 | 0.35786 | 0.789405 | Senp7 |
| ENSMUSG( | 99.78934 | 0.11073 | 0.584111 | 0.898717 | Senp8 |
| ENSMUSG( | 1413.684 | 0.010791 | 0.917319 | 0.989158 | Sephs1 |
| ENSMUSG( | 762.4178 | -0.00287 | 0.982304 | 0.997512 | Sephs2 |
| ENSMUSG( | 645.5046 | 0.068246 | 0.520869 | 0.875186 | Sepsecs |
| ENSMUSG( | 72.33332 | 0.005354 | 0.983578 | 0.997738 | Septin1 |
| ENSMUSG( | 346.9794 | -0.07894 | 0.596021 | 0.903152 | Septin10 |
| ENSMUSG( | 3729.922 | 0.119485 | 0.124933 | 0.531973 | Septin11 |
| ENSMUSG( | 37.56581 | -0.36511 | 0.287252 | 0.737687 | Septin2 |
| ENSMUSG( | 24093.33 | 0.15936 | 0.02776 | 0.255176 | Septin3 |
| ENSMUSG( | 5794.781 | -0.1201 | 0.170908 | 0.605533 | Septin4 |
| ENSMUSG( | 10648.92 | 0.15496 | 0.045954 | 0.328356 | Septin5 |
| ENSMUSG( | 1834.693 | 0.155718 | 0.221218 | 0.666945 | Septin6 |
| ENSMUSG( | 11505.13 | 0.081785 | 0.41066 | 0.818493 | Septin7 |
| ENSMUSG( | 14313.72 | 0.10662 | 0.215585 | 0.66027 | Septin8 |
| ENSMUSG( | 3436.41 | 0.200594 | 0.010737 | 0.149272 | Septin9 |
| ENSMUSG( | 397.7988 | 0.026766 | 0.826274 | 0.968294 | Serac1 |
| ENSMUSG( | 4411.431 | 0.060532 | 0.546904 | 0.884974 | Serbp1 |
| ENSMUSG( | 425.5475 | 0.141349 | 0.247838 | 0.696522 | Serf1 |
| ENSMUSG( | 4479.643 | 0.048207 | 0.59129 | 0.90113 | Serf2 |
| ENSMUSG( | 593.0487 | 0.063427 | 0.618832 | 0.911432 | Sergef |
| ENSMUSG( | 448.6227 | 0.155696 | 0.238747 | 0.686489 | Serhl |
| ENSMUSG( | 21906.19 | -0.5745 | 1.62E-05 | 0.001655 | Serinc1 |
| ENSMUSG( | 220.929 | -0.09108 | 0.597573 | 0.903694 | Serinc2 |
| ENSMUSG( | 10569.75 | -0.223 | 0.144995 | 0.565206 | Serinc3 |
| ENSMUSG( | 21.15114 | -0.14429 | 0.721951 | NA | Serinc4 |
| ENSMUSG( | 3485.109 | -0.43562 | 0.003779 | 0.078636 | Serinc5 |
| ENSMUSG( | 2098.095 | 0.063049 | 0.623054 | 0.911589 | Serp1 |
| ENSMUSG( | 2803.713 | 0.005261 | 0.953633 | 0.993874 | Serp2 |
| ENSMUSG( | 629.6631 | -0.3004 | 0.019782 | 0.212216 | Serpina3n |
| ENSMUSG( | 109.0943 | -1.85427 | 0.006405 | 0.109928 | Serpina9 |
| ENSMUSG( | 241.5614 | -0.51406 | 0.001688 | 0.046443 | Serpinb1a |
| ENSMUSG( | 28.48752 | -0.0677 | 0.847145 | 0.972374 | Serpinb1b |
| ENSMUSG( | 1006.784 | -0.00258 | 0.982413 | 0.997512 | Serpinb6a |
| ENSMUSG( | 70.37164 | 0.027123 | 0.912536 | 0.988483 | Serpinb6b |
| ENSMUSG( | 11.03225 | 0.297179 | 0.593614 | NA | Serpinb6c |
| ENSMUSG( | 160.095 | 0.334941 | 0.152392 | 0.578606 | Serpinb8 |
| ENSMUSG( | 287.8047 | -0.2302 | 0.133304 | 0.546146 | Serpinb9 |
| ENSMUSG( | 92.72478 | 0.221343 | 0.356406 | 0.788187 | Serpind1 |
| ENSMUSG( | 39.47955 | -0.59842 | 0.068432 | 0.405244 | Serpine1 |
| ENSMUSG( | 20321.29 | -0.12338 | 0.265956 | 0.716441 | Serpine2 |
| ENSMUSG( | 696.18 | -0.33082 | 0.019528 | 0.210473 | Serpinf1 |
| ENSMUSG( | 10.88814 | 0.068003 | 0.904896 | NA | Serpinf2 |
| ENSMUSG( | 385.9028 | 0.01584 | 0.918508 | 0.989954 | Serping1 |
| ENSMUSG( | 1632.187 | -0.20238 | 0.202194 | 0.642173 | Serpinh1 |
| ENSMUSG( | 5433.157 | -0.53828 | 0.000336 | 0.015671 | Serpini1 |
| ENSMUSG( | 188.6208 | -0.4973 | 0.010648 | 0.14851 | Sertad1 |
| ENSMUSG( | 848.6815 | -0.10025 | 0.311745 | 0.759348 | Sertad2 |
| ENSMUSG( | 68.66621 | 0.09698 | 0.728569 | 0.944393 | Sertad3 |
| ENSMUSG( | 308.2553 | -0.17855 | 0.197657 | 0.637377 | Sertad4 |
| ENSMUSG( | 734.6042 | -0.0342 | 0.788088 | 0.959226 | Sertm1 |
| ENSMUSG( | 1770.795 | 0.262437 | 0.012317 | 0.160458 | Sesn1 |
| ENSMUSG( | 427.8545 | 0.033066 | 0.813027 | 0.964574 | Sesn2 |
| ENSMUSG( | 10103.4 | 0.123768 | 0.35738 | 0.788775 | Sesn3 |
| ENSMUSG( | 1643.278 | 0.09419 | 0.349598 | 0.78192 | Sestd1 |
| ENSMUSG( | 4126.544 | 0.081263 | 0.371766 | 0.798768 | Set |
| ENSMUSG( | 857.9182 | 0.01662 | 0.868746 | 0.976897 | Setbp1 |
| ENSMUSG( | 607.2741 | 0.007686 | 0.943426 | 0.993415 | Setd1a |
| ENSMUSG( | 1437.218 | 0.069541 | 0.447573 | 0.834051 | Setd1b |
| ENSMUSG( | 2700.529 | 0.475671 | 0.000101 | 0.006438 | Setd2 |
| ENSMUSG( | 6396.974 | 0.066624 | 0.421761 | 0.823375 | Setd3 |
| ENSMUSG( | 274.6274 | 0.16486 | 0.339145 | 0.777518 | Setd4 |
| ENSMUSG( | 3555.786 | 0.002009 | 0.982308 | 0.997512 | Setd5 |
| ENSMUSG( | 329.0628 | -0.02003 | 0.895854 | 0.984551 | Setd6 |
| ENSMUSG( | 11199.97 | 0.103344 | 0.37405 | 0.800445 | Setd7 |
| ENSMUSG( | 1474.33 | 0.068242 | 0.481273 | 0.855292 | Setdb1 |
| ENSMUSG( | 173.5732 | -0.11038 | 0.543112 | 0.883579 | Setdb2 |
| ENSMUSG( | 78.69815 | 0.122789 | 0.633433 | 0.914073 | Setmar |

| | | | | | |
|---|---|---|---|---|---|
| ENSMUSG( | 298.4128 | -0.00228 | 0.988539 | 0.997862 | Sertad4 |
| ENSMUSG( | 660.5229 | -0.0723 | 0.566935 | 0.941777 | Sertm1 |
| ENSMUSG( | 1532.383 | 0.123496 | 0.272016 | 0.888477 | Sesn1 |
| ENSMUSG( | 390.8553 | 0.037328 | 0.81061 | 0.978779 | Sesn2 |
| ENSMUSG( | 8016.134 | -0.28814 | 0.035118 | 0.66005 | Sesn3 |
| ENSMUSG( | 1444.215 | -0.00994 | 0.929679 | 0.99306 | Sestd1 |
| ENSMUSG( | 3620.957 | -0.02904 | 0.839191 | 0.981906 | Set |
| ENSMUSG( | 797.806 | 0.07908 | 0.532742 | 0.934795 | Setbp1 |
| ENSMUSG( | 553.0296 | 0.005458 | 0.966352 | 0.995475 | Setd1a |
| ENSMUSG( | 1386.618 | 0.227605 | 0.039395 | 0.679972 | Setd1b |
| ENSMUSG( | 2993.96 | 0.935074 | 1.57E-14 | 2.11E-11 | Setd2 |
| ENSMUSG( | 5517.705 | -0.09374 | 0.379056 | 0.916995 | Setd3 |
| ENSMUSG( | 247.8871 | 0.13841 | 0.415342 | 0.917257 | Setd4 |
| ENSMUSG( | 3296.392 | 0.053454 | 0.622822 | 0.953234 | Setd5 |
| ENSMUSG( | 306.5648 | 0.035585 | 0.829135 | 0.980224 | Setd6 |
| ENSMUSG( | 9393.132 | -0.13779 | 0.236535 | 0.872173 | Setd7 |
| ENSMUSG( | 1364.597 | 0.114341 | 0.322148 | 0.904023 | Setdb1 |
| ENSMUSG( | 148.9169 | -0.29489 | 0.145142 | 0.83339 | Setdb2 |
| ENSMUSG( | 74.71572 | 0.242179 | 0.38318 | 0.917257 | Setmar |
| ENSMUSG( | 1855.028 | 0.162482 | 0.12984 | 0.820802 | Setx |
| ENSMUSG( | 7829.464 | 0.071391 | 0.544198 | 0.938063 | Sez6 |
| ENSMUSG( | 7129.935 | 0.25895 | 0.121999 | 0.813384 | Sez6l |
| ENSMUSG( | 10707.81 | 0.198685 | 0.233986 | 0.86917 | Sez6l2 |
| ENSMUSG( | 3632.531 | 0.025703 | 0.802454 | 0.97681 | Sf1 |
| ENSMUSG( | 1746.923 | 0.065865 | 0.576311 | 0.942912 | Sf3a1 |
| ENSMUSG( | 794.5386 | 0.017037 | 0.905915 | 0.991089 | Sf3a2 |
| ENSMUSG( | 499.6928 | -0.02446 | 0.864282 | 0.986552 | Sf3a3 |
| ENSMUSG( | 7416.017 | 0.119039 | 0.262225 | 0.885219 | Sf3b1 |
| ENSMUSG( | 5094.627 | -0.07924 | 0.456166 | 0.923329 | Sf3b2 |
| ENSMUSG( | 1487.04 | -0.07909 | 0.511139 | 0.929571 | Sf3b3 |
| ENSMUSG( | 564.6193 | -0.18108 | 0.232156 | 0.868507 | Sf3b4 |
| ENSMUSG( | 593.654 | -0.0891 | 0.56095 | 0.940254 | Sf3b5 |
| ENSMUSG( | 906.177 | -0.10752 | 0.451004 | 0.922202 | Sf3b6 |
| ENSMUSG( | 561.3122 | 0.165931 | 0.453001 | 0.922316 | Sfi1 |
| ENSMUSG( | 569.019 | -0.10908 | 0.393592 | 0.917257 | Sfmbt1 |
| ENSMUSG( | 101.0069 | 0.108116 | 0.611655 | 0.950771 | Sfmbt2 |
| ENSMUSG( | 63.99219 | 0.246638 | 0.421761 | 0.917257 | Sfn |
| ENSMUSG( | 2927.194 | 0.194913 | 0.06485 | 0.740531 | Sfpq |
| ENSMUSG( | 1901.006 | -0.09907 | 0.452274 | 0.922316 | Sfr1 |
| ENSMUSG( | 786.4787 | 0.189611 | 0.180548 | 0.850372 | Sfrp1 |
| ENSMUSG( | 80.17526 | -0.10561 | 0.671589 | 0.95694 | Sfrp2 |
| ENSMUSG( | 64.9748 | -0.31146 | 0.238071 | 0.87259 | Sfrp4 |
| ENSMUSG( | 88.73952 | 0.429908 | 0.129442 | 0.820358 | Sfrp5 |
| ENSMUSG( | 2489.858 | 0.132545 | 0.354168 | 0.912411 | Sfswap |
| ENSMUSG( | 722.2958 | -0.04437 | 0.747961 | 0.96984 | Sft2d1 |
| ENSMUSG( | 23.22672 | 0.074895 | 0.855284 | 0.985649 | Sft2d1rt |
| ENSMUSG( | 3363.886 | 0.053789 | 0.662871 | 0.956205 | Sft2d2 |
| ENSMUSG( | 180.0232 | -0.01764 | 0.917997 | 0.992529 | Sft2d3 |
| ENSMUSG( | 10.25905 | 0.174938 | 0.78107 | NA | Sfta3-ps |
| ENSMUSG( | 28.58889 | -0.1458 | 0.714017 | 0.964986 | Sftpc |
| ENSMUSG( | 2540.628 | -0.01928 | 0.87507 | 0.989644 | Sfxn1 |
| ENSMUSG( | 244.5396 | 0.373676 | 0.065676 | 0.745596 | Sfxn2 |
| ENSMUSG( | 2336.977 | -0.09507 | 0.539887 | 0.937378 | Sfxn3 |
| ENSMUSG( | 262.8244 | 0.123074 | 0.549331 | 0.938519 | Sfxn4 |
| ENSMUSG( | 21067.17 | -0.37855 | 0.009911 | 0.407479 | Sfxn5 |
| ENSMUSG( | 18.78828 | 0.507082 | 0.270812 | 0.888062 | Sgca |
| ENSMUSG( | 3794.254 | -0.01857 | 0.883102 | 0.989644 | Sgcb |
| ENSMUSG( | 39.75941 | 0.220552 | 0.484488 | 0.92541 | Sgcd |
| ENSMUSG( | 1105.413 | 0.111533 | 0.474015 | 0.923329 | Sgce |
| ENSMUSG( | 10.1954 | 0.021642 | 0.973096 | NA | Sgcg |
| ENSMUSG( | 45.50481 | -0.06476 | 0.837901 | 0.981376 | Sgcz |
| ENSMUSG( | 251.0353 | 0.177792 | 0.299026 | 0.898538 | Sgf29 |
| ENSMUSG( | 3141.202 | 0.021623 | 0.848688 | 0.985321 | Sgip1 |
| ENSMUSG( | 1173.996 | -0.01907 | 0.919932 | 0.992644 | Sgk1 |
| ENSMUSG( | 77.99202 | -0.13512 | 0.580273 | 0.944209 | Sgk2 |
| ENSMUSG( | 488.8632 | 0.139378 | 0.340537 | 0.907972 | Sgk3 |
| ENSMUSG( | 1358.715 | 0.309165 | 0.167557 | 0.840222 | Sgms1 |
| ENSMUSG( | 135.9434 | -0.1792 | 0.349741 | 0.911652 | Sgms1os1 |
| ENSMUSG( | 120.6246 | 0.818057 | 0.001239 | 0.127113 | Sgms2 |
| ENSMUSG( | 8.574128 | -0.01156 | 0.986569 | NA | Sgo1 |
| ENSMUSG( | 35.87567 | -0.28955 | 0.402158 | 0.917257 | Sgo2a |
| ENSMUSG( | 1841.872 | 0.121054 | 0.391434 | 0.917257 | Sgpl1 |
| ENSMUSG( | 1351.732 | 0.264423 | 0.113103 | 0.809623 | Sgpp1 |
| ENSMUSG( | 517.9735 | 0.398259 | 0.043731 | 0.690304 | Sgpp2 |
| ENSMUSG( | 409.9238 | 0.189306 | 0.363738 | 0.914223 | Sgsh |
| ENSMUSG( | 2683.576 | -0.046 | 0.745286 | 0.96951 | Sgsm1 |
| ENSMUSG( | 6242.584 | -0.07519 | 0.439546 | 0.920721 | Sgsm2 |
| ENSMUSG( | 1323.288 | -0.07428 | 0.610254 | 0.950771 | Sgsm3 |
| ENSMUSG( | 3153.185 | -0.13466 | 0.244996 | 0.877904 | Sgta |
| ENSMUSG( | 1770.707 | 0.007214 | 0.951136 | 0.994798 | Sgtb |
| ENSMUSG( | 2304.093 | -0.0431 | 0.697102 | 0.960867 | Sh2b1 |
| ENSMUSG( | 162.1473 | 0.16322 | 0.372628 | 0.915491 | Sh2b2 |
| ENSMUSG( | 945.8376 | 0.321466 | 0.022255 | 0.577034 | Sh2b3 |
| ENSMUSG( | 505.2573 | -0.24783 | 0.130074 | 0.821082 | Sh2d3c |
| ENSMUSG( | 32.03487 | 0.531576 | 0.177241 | 0.846973 | Sh2d4b |
| ENSMUSG( | 2607.141 | -0.06818 | 0.591287 | 0.944975 | Sh2d5 |
| ENSMUSG( | 31.44313 | -0.58555 | 0.150079 | 0.837086 | Sh2d6 |
| ENSMUSG( | 31.48744 | -0.24153 | 0.584832 | 0.944975 | Sh3bgr |
| ENSMUSG( | 3055.16 | -0.04087 | 0.847677 | 0.9852 | Sh3bgrl |
| ENSMUSG( | 505.4104 | -0.13252 | 0.394454 | 0.917257 | Sh3bgrl2 |
| ENSMUSG( | 1819.344 | -0.24254 | 0.066398 | 0.749411 | Sh3bgrl3 |
| ENSMUSG( | 197.1545 | 0.120458 | 0.545908 | 0.938063 | Sh3bp1 |
| ENSMUSG( | 170.7252 | -0.06917 | 0.737581 | 0.967686 | Sh3bp2 |
| ENSMUSG( | 698.5528 | 0.059734 | 0.62957 | 0.953251 | Sh3bp4 |
| ENSMUSG( | 846.5868 | -0.02335 | 0.860495 | 0.986137 | Sh3bp5 |
| ENSMUSG( | 1334.593 | 0.054234 | 0.653573 | 0.955015 | Sh3bp5l |
| ENSMUSG( | 837.5346 | 0.084305 | 0.502856 | 0.927825 | Sh3d19 |
| ENSMUSG( | 63.76457 | 0.321799 | 0.231231 | 0.868507 | Sh3d21 |
| ENSMUSG( | 1071.514 | 0.091944 | 0.439596 | 0.920721 | Sh3gl1 |
| ENSMUSG( | 4362.548 | -0.21695 | 0.219738 | 0.866104 | Sh3gl2 |
| ENSMUSG( | 3224.961 | -0.16215 | 0.19359 | 0.860096 | Sh3gl3 |
| ENSMUSG( | 5830.07 | -0.08391 | 0.52133 | 0.932489 | Sh3glb1 |
| ENSMUSG( | 5247.65 | -0.02879 | 0.819818 | 0.979198 | Sh3glb2 |
| ENSMUSG( | 889.8169 | -0.17089 | 0.185554 | 0.855305 | Sh3kbp1 |
| ENSMUSG( | 2365.026 | -0.10584 | 0.36837 | 0.915491 | Sh3pxd2a |
| ENSMUSG( | 1174.388 | -0.04104 | 0.725527 | 0.966887 | Sh3pxd2b |
| ENSMUSG( | 1260.375 | 0.123999 | 0.26678 | 0.886134 | Sh3rf1 |
| ENSMUSG( | 94.62635 | -0.38632 | 0.239803 | 0.874948 | Sh3rf2 |
| ENSMUSG( | 1166.969 | 0.415146 | 0.000856 | 0.09911 | Sh3rf3 |
| ENSMUSG( | 117.1271 | 0.411378 | 0.118226 | 0.813384 | Sh3tc1 |
| ENSMUSG( | 152.1213 | 0.078797 | 0.680639 | 0.958395 | Sh3tc2 |
| ENSMUSG( | 156.1754 | 0.033043 | 0.858801 | 0.985884 | Sh3yl1 |
| ENSMUSG( | 24259.58 | 0.229928 | 0.054446 | 0.723362 | Shank1 |
| ENSMUSG( | 10982 | 0.211429 | 0.090908 | 0.791074 | Shank2 |
| ENSMUSG( | 13648.26 | 0.204192 | 0.086838 | 0.782073 | Shank3 |
| ENSMUSG( | 811.0096 | -0.02959 | 0.819566 | 0.979198 | Sharpin |
| ENSMUSG( | 271.005 | -0.08694 | 0.652655 | 0.954799 | Shb |
| ENSMUSG( | 499.8476 | 0.210375 | 0.178312 | 0.848538 | Shc1 |
| ENSMUSG( | 1238.14 | 0.003297 | 0.979318 | 0.996468 | Shc2 |
| ENSMUSG( | 2514.204 | -0.11999 | 0.302491 | 0.898538 | Shc3 |
| ENSMUSG( | 584.6164 | -0.21988 | 0.181529 | 0.850579 | Shc4 |
| ENSMUSG( | 18.34071 | -0.13418 | 0.764969 | 0.974236 | Shcbp1 |
| ENSMUSG( | 60.29077 | 0.409567 | 0.188824 | 0.85698 | Shcbp1l |
| ENSMUSG( | 916.3805 | -0.01801 | 0.883134 | 0.989644 | Shd |
| ENSMUSG( | 161.3246 | 1.002054 | 0.000263 | 0.04337 | She |
| ENSMUSG( | 1111.673 | -0.01536 | 0.900578 | 0.990558 | Shf |
| ENSMUSG( | 1612.587 | 0.074726 | 0.699755 | 0.961335 | Shfl |
| ENSMUSG( | 92.37979 | 0.152367 | 0.530431 | 0.934795 | Shh |
| ENSMUSG( | 95.41822 | 0.468462 | 0.05397 | 0.722223 | Shisa2 |
| ENSMUSG( | 88.21764 | -0.07269 | 0.784435 | 0.976196 | Shisa3 |
| ENSMUSG( | 7895.693 | -0.0247 | 0.849223 | 0.985321 | Shisa4 |
| ENSMUSG( | 2238.482 | 0.07005 | 0.660267 | 0.956205 | Shisa5 |
| ENSMUSG( | 817.494 | 0.503463 | 0.016594 | 0.519982 | Shisa6 |
| ENSMUSG( | 5916.879 | 0.146123 | 0.18616 | 0.855305 | Shisa7 |
| ENSMUSG( | 294.0336 | 0.328548 | 0.061572 | 0.736111 | Shisa8 |
| ENSMUSG( | 2656.282 | 0.136212 | 0.341851 | 0.909146 | Shisa9 |
| ENSMUSG( | 3436.542 | -0.01095 | 0.924664 | 0.99306 | Shisal1 |
| ENSMUSG( | 12.39512 | -0.66738 | 0.23324 | 0.868507 | Shisal2b |
| ENSMUSG( | 342.8972 | 0.051961 | 0.795409 | 0.97681 | Shkbp1 |
| ENSMUSG( | 152.5085 | -0.07615 | 0.683778 | 0.958678 | Shld1 |
| ENSMUSG( | 115.0769 | -0.27193 | 0.210488 | 0.864412 | Shld2 |
| ENSMUSG( | 16.70483 | -0.26402 | 0.586926 | 0.944975 | Shld3 |
| ENSMUSG( | 50.87829 | -0.42888 | 0.162111 | 0.837724 | Shmt1 |
| ENSMUSG( | 914.2995 | -0.07682 | 0.525599 | 0.934374 | Shmt2 |
| ENSMUSG( | 924.2032 | 0.07735 | 0.539713 | 0.937378 | Shoc2 |
| ENSMUSG( | 17.86628 | -2.81639 | 0.096264 | 0.796133 | Shox2 |
| ENSMUSG( | 128.7928 | -0.10684 | 0.582413 | 0.944266 | Shpk |
| ENSMUSG( | 998.0605 | 0.081524 | 0.461591 | 0.923329 | Shprh |
| ENSMUSG( | 80.8259 | -0.09807 | 0.696989 | 0.960867 | Shq1 |
| ENSMUSG( | 195.5492 | 0.182011 | 0.363751 | 0.914223 | Shroom1 |
| ENSMUSG( | 1667.235 | 0.008524 | 0.949806 | 0.994798 | Shroom2 |
| ENSMUSG( | 155.4546 | -0.06372 | 0.780575 | 0.976106 | Shroom3 |
| ENSMUSG( | 212.9436 | 0.306721 | 0.161129 | 0.837724 | Shroom4 |
| ENSMUSG( | 1212.278 | -0.34627 | 0.002455 | 0.194608 | Shtn1 |
| ENSMUSG( | 984.336 | -0.07904 | 0.585166 | 0.944975 | Siae |
| ENSMUSG( | 398.1376 | -0.02066 | 0.879474 | 0.989644 | Siah1a |
| ENSMUSG( | 78.94867 | -0.34302 | 0.198204 | 0.862161 | Siah1b |
| ENSMUSG( | 322.4708 | -0.09515 | 0.578761 | 0.943611 | Siah2 |

| | | | | | | | | | | | |
|---|---|---|---|---|---|---|---|---|---|---|---|
| ENSMUSG( | Sirt4 | 412.1186 | 0.137747 | 0.421294 | 0.823126 | ENSMUSG( | Sirt4 | 404.334 | 0.337499 | 0.16038 | 0.837724 |
| ENSMUSG( | Sirt5 | 302.0701 | -0.1846 | 0.165462 | 0.596928 | ENSMUSG( | Sirt5 | 265.5432 | -0.30049 | 0.062193 | 0.736482 |
| ENSMUSG( | Sirt6 | 364.3144 | -0.01353 | 0.931648 | 0.9924 | ENSMUSG( | Sirt6 | 334.1838 | 0.008659 | 0.958055 | 0.994798 |
| ENSMUSG( | Sirt7 | 1201.929 | 0.105391 | 0.364992 | 0.79407 | ENSMUSG( | Sirt7 | 1157.584 | 0.25763 | 0.09596 | 0.796133 |
| ENSMUSG( | Siva1 | 159.304 | -0.22446 | 0.196645 | 0.635942 | ENSMUSG( | Siva1 | 152.877 | -0.06284 | 0.741231 | 0.968811 |
| ENSMUSG( | Six1 | 21.39449 | -0.62272 | 0.150221 | 0.574666 | ENSMUSG( | Six1 | 27.36948 | 0.418126 | 0.290692 | 0.897019 |
| ENSMUSG( | Six2 | 37.99869 | -0.53257 | 0.085926 | 0.451489 | ENSMUSG( | Six2 | 36.7273 | -0.32462 | 0.336703 | 0.907972 |
| ENSMUSG( | Six3 | 144.6131 | -1.54316 | 0.025033 | 0.243138 | ENSMUSG( | Six3 | 172.5154 | -0.42243 | 0.181138 | 0.850372 |
| ENSMUSG( | Six3os1 | 105.1994 | -0.87172 | 0.172798 | 0.608361 | ENSMUSG( | Six3os1 | 111.1347 | -0.35825 | 0.201064 | 0.86439 |
| ENSMUSG( | Six4 | 17.76594 | 0.38754 | 0.417316 | NA | ENSMUSG( | Six4 | 15.60244 | 0.305588 | 0.558043 | 0.939565 |
| ENSMUSG( | Six5 | 102.4662 | -0.12831 | 0.537561 | 0.882536 | ENSMUSG( | Six5 | 109.1267 | 0.316696 | 0.180472 | 0.850372 |
| ENSMUSG( | Ska2 | 450.0013 | 0.079708 | 0.515682 | 0.872701 | ENSMUSG( | Ska2 | 395.869 | -0.02369 | 0.867764 | 0.98781 |
| ENSMUSG( | Ska3 | 21.48139 | 0.131815 | 0.755151 | 0.951279 | ENSMUSG( | Ska3 | 19.75378 | 0.13014 | 0.781067 | 0.976106 |
| ENSMUSG( | Skap1 | 84.05817 | -0.5222 | 0.023898 | 0.236358 | ENSMUSG( | Skap1 | 93.90965 | 0.114562 | 0.621331 | 0.953199 |
| ENSMUSG( | Skap2 | 484.337 | 0.028406 | 0.856579 | 0.974154 | ENSMUSG( | Skap2 | 412.4274 | -0.17935 | 0.246006 | 0.877904 |
| ENSMUSG( | Ski | 6188.07 | 0.142646 | 0.055168 | 0.361542 | ENSMUSG( | Ski | 5562.784 | 0.105165 | 0.286999 | 0.894091 |
| ENSMUSG( | Skic2 | 2593.307 | 0.230904 | 0.019287 | 0.209368 | ENSMUSG( | Skic2 | 2194.693 | 0.026455 | 0.83823 | 0.981575 |
| ENSMUSG( | Skic3 | 729.3033 | 0.217241 | 0.047265 | 0.332598 | ENSMUSG( | Skic3 | 641.4415 | 0.123754 | 0.308648 | 0.898538 |
| ENSMUSG( | Skic8 | 730.262 | 0.146867 | 0.201372 | 0.641362 | ENSMUSG( | Skic8 | 605.9543 | -0.12215 | 0.361858 | 0.914223 |
| ENSMUSG( | Skida1 | 249.487 | 0.032618 | 0.820182 | 0.965888 | ENSMUSG( | Skida1 | 230.951 | 0.068069 | 0.691334 | 0.960071 |
| ENSMUSG( | Skil | 1154.087 | -0.06914 | 0.474484 | 0.849425 | ENSMUSG( | Skil | 1078.174 | 0.002816 | 0.980199 | 0.996479 |
| ENSMUSG( | Skp1 | 8047.481 | 0.015921 | 0.878628 | 0.981431 | ENSMUSG( | Skp1 | 6870.569 | -0.18039 | 0.226396 | 0.868507 |
| ENSMUSG( | Skp2 | 155.1987 | -0.24224 | 0.145182 | 0.565771 | ENSMUSG( | Skp2 | 153.455 | 0.020464 | 0.917498 | 0.992529 |
| ENSMUSG( | Sla | 288.2735 | 0.005175 | 0.969738 | 0.996369 | ENSMUSG( | Sla | 251.0654 | -0.11305 | 0.521893 | 0.932571 |
| ENSMUSG( | Sla2 | 30.96369 | -0.13752 | 0.689351 | 0.933609 | ENSMUSG( | Sla2 | 23.97104 | -0.66439 | 0.113538 | 0.809623 |
| ENSMUSG( | Slain1 | 1105.929 | -0.06667 | 0.537337 | 0.882504 | ENSMUSG( | Slain1 | 989.3597 | -0.12415 | 0.343916 | 0.910256 |
| ENSMUSG( | Slain2 | 823.7385 | 0.081308 | 0.465977 | 0.844391 | ENSMUSG( | Slain2 | 755.1793 | 0.094322 | 0.485799 | 0.92541 |
| ENSMUSG( | Slamf6 | 14.84478 | -0.26723 | 0.582851 | NA | ENSMUSG( | Slamf6 | 14.54163 | 0.027674 | 0.961569 | 0.99511 |
| ENSMUSG( | Slamf9 | 43.21718 | -0.46411 | 0.124414 | 0.531714 | ENSMUSG( | Slamf9 | 41.17273 | -0.2696 | 0.456348 | 0.923329 |
| ENSMUSG( | Slbp | 291.0976 | 0.047267 | 0.743966 | 0.948497 | ENSMUSG( | Slbp | 265.1169 | 0.041743 | 0.803502 | 0.976978 |
| ENSMUSG( | Slc10a3 | 88.81983 | -0.28171 | 0.29145 | 0.741366 | ENSMUSG( | Slc10a3 | 89.35861 | 0.032585 | 0.90851 | 0.991958 |
| ENSMUSG( | Slc10a4 | 62.5772 | -1.63035 | 1.94E-06 | 0.000324 | ENSMUSG( | Slc10a4 | 73.65823 | -0.51674 | 0.063229 | 0.736482 |
| ENSMUSG( | Slc10a7 | 170.531 | 0.263471 | 0.103745 | 0.489428 | ENSMUSG( | Slc10a7 | 178.1787 | 0.61696 | 0.000962 | 0.10597 |
| ENSMUSG( | Slc11a1 | 172.2215 | -0.18545 | 0.332211 | 0.77543 | ENSMUSG( | Slc11a1 | 187.5973 | 0.331245 | 0.152877 | 0.837724 |
| ENSMUSG( | Slc11a2 | 1605.935 | -0.15711 | 0.179902 | 0.617634 | ENSMUSG( | Slc11a2 | 1580.133 | 0.072007 | 0.600066 | 0.947395 |
| ENSMUSG( | Slc12a2 | 2555.055 | -0.5194 | 0.001086 | 0.034804 | ENSMUSG( | Slc12a2 | 3006.649 | 0.258984 | 0.187017 | 0.856099 |
| ENSMUSG( | Slc12a4 | 1858.321 | -0.07962 | 0.540589 | 0.88315 | ENSMUSG( | Slc12a4 | 1933.574 | 0.293745 | 0.051718 | 0.711848 |
| ENSMUSG( | Slc12a5 | 17153.52 | -0.21914 | 0.069873 | 0.408574 | ENSMUSG( | Slc12a5 | 17494.5 | 0.115482 | 0.406898 | 0.917257 |
| ENSMUSG( | Slc12a6 | 2674.907 | -0.21773 | 0.134254 | 0.547472 | ENSMUSG( | Slc12a6 | 2690.762 | 0.078017 | 0.627524 | 0.953234 |
| ENSMUSG( | Slc12a7 | 311.1661 | -0.45402 | 0.004236 | 0.084545 | ENSMUSG( | Slc12a7 | 351.551 | 0.205201 | 0.253795 | 0.880692 |
| ENSMUSG( | Slc12a8 | 31.21323 | -0.51764 | 0.163643 | 0.594869 | ENSMUSG( | Slc12a8 | 33.62143 | 0.021103 | 0.952082 | 0.994798 |
| ENSMUSG( | Slc12a9 | 1580.25 | -0.2105 | 0.051988 | 0.350064 | ENSMUSG( | Slc12a9 | 1592.964 | 0.090089 | 0.454248 | 0.922444 |
| ENSMUSG( | Slc13a3 | 1526.992 | -0.28465 | 0.047735 | 0.333591 | ENSMUSG( | Slc13a3 | 1478.493 | -0.09568 | 0.480083 | 0.924276 |
| ENSMUSG( | Slc13a4 | 551.6801 | 0.117776 | 0.384189 | 0.803686 | ENSMUSG( | Slc13a4 | 522.6759 | 0.224737 | 0.083455 | 0.77681 |
| ENSMUSG( | Slc13a5 | 1516.84 | -0.16335 | 0.189005 | 0.627678 | ENSMUSG( | Slc13a5 | 1448.97 | -0.01982 | 0.882746 | 0.989644 |
| ENSMUSG( | Slc14a1 | 1065.004 | -0.19779 | 0.161046 | 0.59072 | ENSMUSG( | Slc14a1 | 1058.563 | 0.061885 | 0.692757 | 0.96043 |
| ENSMUSG( | Slc14a2 | 35.55809 | -0.06318 | 0.869563 | 0.977498 | ENSMUSG( | Slc14a2 | 34.86821 | 0.17192 | 0.681168 | 0.958395 |
| ENSMUSG( | Slc15a2 | 2132.172 | 0.133409 | 0.258903 | 0.709814 | ENSMUSG( | Slc15a2 | 2021.797 | 0.242835 | 0.044556 | 0.693697 |
| ENSMUSG( | Slc15a3 | 248.8798 | -0.24193 | 0.194453 | 0.63341 | ENSMUSG( | Slc15a3 | 230.4622 | -0.17758 | 0.402585 | 0.917257 |
| ENSMUSG( | Slc15a4 | 699.7803 | -0.11393 | 0.499649 | 0.863218 | ENSMUSG( | Slc15a4 | 736.1405 | 0.294978 | 0.105915 | 0.804969 |
| ENSMUSG( | Slc16a1 | 2073.007 | -0.57516 | 0.002061 | 0.052152 | ENSMUSG( | Slc16a1 | 2014.819 | -0.34549 | 0.086056 | 0.780251 |
| ENSMUSG( | Slc16a10 | 103.9734 | -0.56505 | 0.012343 | 0.160458 | ENSMUSG( | Slc16a10 | 103.0084 | -0.26323 | 0.311588 | 0.900031 |
| ENSMUSG( | Slc16a11 | 442.49 | 0.255061 | 0.153428 | 0.579959 | ENSMUSG( | Slc16a11 | 405.1911 | 0.271723 | 0.202359 | 0.864412 |
| ENSMUSG( | Slc16a12 | 68.16646 | -0.11298 | 0.636045 | 0.914873 | ENSMUSG( | Slc16a12 | 102.3719 | 1.137687 | 1.51E-05 | 0.005081 |
| ENSMUSG( | Slc16a13 | 205.8076 | -0.13839 | 0.385207 | 0.804048 | ENSMUSG( | Slc16a13 | 207.0429 | 0.159486 | 0.407405 | 0.917257 |
| ENSMUSG( | Slc16a14 | 96.39241 | -0.08693 | 0.77484 | 0.955606 | ENSMUSG( | Slc16a14 | 106.0076 | 0.444612 | 0.142977 | 0.831221 |
| ENSMUSG( | Slc16a2 | 2458.564 | -0.31755 | 0.014276 | 0.175552 | ENSMUSG( | Slc16a2 | 2598.113 | 0.128986 | 0.384197 | 0.917257 |
| ENSMUSG( | Slc16a3 | 200.4832 | -0.2855 | 0.094329 | 0.471647 | ENSMUSG( | Slc16a3 | 206.7248 | 0.091054 | 0.638394 | 0.953903 |
| ENSMUSG( | Slc16a4 | 141.7342 | -0.30641 | 0.279741 | 0.729135 | ENSMUSG( | Slc16a4 | 194.2554 | 0.797516 | 0.009967 | 0.407912 |
| ENSMUSG( | Slc16a6 | 625.6651 | -0.30077 | 0.006319 | 0.108996 | ENSMUSG( | Slc16a6 | 618.9686 | -0.05147 | 0.688872 | 0.959742 |
| ENSMUSG( | Slc16a7 | 833.4757 | -0.64144 | 0.000418 | 0.01851 | ENSMUSG( | Slc16a7 | 953.212 | 0.087404 | 0.651529 | 0.95449 |
| ENSMUSG( | Slc16a8 | 15.89648 | -0.0979 | 0.880423 | NA | ENSMUSG( | Slc16a8 | 18.83129 | 0.566572 | 0.323165 | 0.904023 |
| ENSMUSG( | Slc16a9 | 253.8831 | -0.0977 | 0.522188 | 0.875186 | ENSMUSG( | Slc16a9 | 265.9324 | 0.300166 | 0.064901 | 0.740636 |
| ENSMUSG( | Slc17a5 | 650.0216 | -0.41365 | 0.005539 | 0.099488 | ENSMUSG( | Slc17a5 | 722.9959 | 0.190959 | 0.228195 | 0.868507 |
| ENSMUSG( | Slc17a6 | 667.8916 | -0.62444 | 4.40E-05 | 0.003588 | ENSMUSG( | Slc17a6 | 734.6324 | -0.01273 | 0.943515 | 0.994798 |
| ENSMUSG( | Slc17a7 | 73345.29 | -0.18592 | 0.13153 | 0.543959 | ENSMUSG( | Slc17a7 | 73940.87 | 0.112888 | 0.412316 | 0.917257 |
| ENSMUSG( | Slc17a8 | 103.496 | -0.89001 | 0.000452 | 0.019387 | ENSMUSG( | Slc17a8 | 124.0207 | 0.040561 | 0.862052 | 0.98632 |
| ENSMUSG( | Slc17a9 | 18.26414 | -0.11985 | 0.791033 | NA | ENSMUSG( | Slc17a9 | 23.74967 | 0.83056 | 0.07243 | 0.763629 |
| ENSMUSG( | Slc18a2 | 14.03756 | 0.206922 | 0.674341 | NA | ENSMUSG( | Slc18a2 | 15.65351 | 0.691993 | 0.162349 | 0.837724 |
| ENSMUSG( | Slc18a3 | 38.39513 | -0.87633 | 0.019608 | 0.211048 | ENSMUSG( | Slc18a3 | 47.92127 | 0.153728 | 0.649467 | 0.95422 |
| ENSMUSG( | Slc18b1 | 791.3843 | -0.23988 | 0.078998 | 0.434796 | ENSMUSG( | Slc18b1 | 735.6207 | -0.17854 | 0.220204 | 0.866104 |
| ENSMUSG( | Slc19a1 | 478.9383 | -0.21385 | 0.26175 | 0.711988 | ENSMUSG( | Slc19a1 | 530.9225 | 0.350518 | 0.140732 | 0.830678 |
| ENSMUSG( | Slc19a2 | 368.5087 | -0.38295 | 0.028584 | 0.25772 | ENSMUSG( | Slc19a2 | 429.4222 | 0.340204 | 0.061736 | 0.736228 |
| ENSMUSG( | Slc19a3 | 212.2165 | 0.228764 | 0.366177 | 0.794603 | ENSMUSG( | Slc19a3 | 190.6836 | 0.205561 | 0.692577 | 0.96043 |
| ENSMUSG( | Slc1a1 | 3762.219 | -0.35608 | 0.008183 | 0.125929 | ENSMUSG( | Slc1a1 | 4320.649 | 0.319716 | 0.042566 | 0.690304 |
| ENSMUSG( | Slc1a2 | 501546.3 | 0.124176 | 0.311198 | 0.759247 | ENSMUSG( | Slc1a2 | 420988 | -0.11114 | 0.413338 | 0.917257 |
| ENSMUSG( | Slc1a3 | 71268.34 | -0.01563 | 0.903528 | 0.986055 | ENSMUSG( | Slc1a3 | 65949.67 | 0.030841 | 0.820828 | 0.979198 |
| ENSMUSG( | Slc1a4 | 2962.663 | -0.34215 | 0.034097 | 0.283475 | ENSMUSG( | Slc1a4 | 3153.553 | 0.129867 | 0.472926 | 0.923329 |
| ENSMUSG( | Slc1a5 | 152.4252 | -0.11869 | 0.502014 | 0.864904 | ENSMUSG( | Slc1a5 | 182.9229 | 0.624462 | 0.003785 | 0.245533 |
| ENSMUSG( | Slc1a6 | 242.5852 | -0.06955 | 0.662489 | 0.926183 | ENSMUSG( | Slc1a6 | 235.464 | 0.111718 | 0.515916 | 0.930939 |
| ENSMUSG( | Slc20a1 | 6506.381 | -0.18356 | 0.071541 | 0.413178 | ENSMUSG( | Slc20a1 | 6133.699 | -0.07892 | 0.49992 | 0.927825 |
| ENSMUSG( | Slc20a2 | 2739.25 | -0.17796 | 0.166713 | 0.599311 | ENSMUSG( | Slc20a2 | 2597.897 | -0.05235 | 0.73101 | 0.967228 |
| ENSMUSG( | Slc22a1 | 12.26465 | -1.31872 | 0.018382 | NA | ENSMUSG( | Slc22a1 | 12.35397 | -0.7905 | 0.195456 | 0.861579 |
| ENSMUSG( | Slc22a12 | 15.60621 | 0.143906 | 0.754755 | NA | ENSMUSG( | Slc22a12 | 15.7554 | 0.417239 | 0.392439 | 0.917257 |
| ENSMUSG( | Slc22a13b | 22.31001 | 0.319008 | 0.457454 | 0.839558 | ENSMUSG( | Slc22a13b | 19.78949 | 0.263827 | 0.547066 | 0.938063 |
| ENSMUSG( | Slc22a15 | 275.8905 | -0.26416 | 0.120911 | 0.525221 | ENSMUSG( | Slc22a15 | 277.8505 | 0.040166 | 0.809684 | 0.978779 |
| ENSMUSG( | Slc22a17 | 20967.33 | -0.19397 | 0.127156 | 0.534725 | ENSMUSG( | Slc22a17 | 21764.01 | 0.186542 | 0.200923 | 0.86439 |
| ENSMUSG( | Slc22a18 | 25.32263 | -0.21956 | 0.573767 | 0.893584 | ENSMUSG( | Slc22a18 | 26.09229 | 0.166904 | 0.692429 | 0.960404 |
| ENSMUSG( | Slc22a2 | 40.42754 | 0.027623 | 0.926875 | 0.991229 | ENSMUSG( | Slc22a2 | 38.04742 | 0.12718 | 0.697868 | 0.960867 |
| ENSMUSG( | Slc22a21 | 97.61677 | -0.16845 | 0.430985 | 0.826896 | ENSMUSG( | Slc22a21 | 95.46218 | 0.031003 | 0.889099 | 0.990558 |
| ENSMUSG( | Slc22a23 | 3065.937 | -0.39633 | 0.006337 | 0.108996 | ENSMUSG( | Slc22a23 | 3270.284 | 0.08974 | 0.616969 | 0.951666 |
| ENSMUSG( | Slc22a3 | 83.72509 | -0.63978 | 0.003903 | 0.080172 | ENSMUSG( | Slc22a3 | 91.81973 | -0.03692 | 0.872712 | 0.989213 |
| ENSMUSG( | Slc22a4 | 305.2309 | -0.06939 | 0.643648 | 0.917316 | ENSMUSG( | Slc22a4 | 283.4218 | -0.00338 | 0.985635 | 0.997084 |
| ENSMUSG( | Slc22a5 | 517.5964 | -0.10314 | 0.416792 | 0.8209 | ENSMUSG( | Slc22a5 | 499.9432 | 0.068661 | 0.642332 | 0.953903 |
| ENSMUSG( | Slc22a6 | 316.3686 | -0.11333 | 0.4917 | 0.860249 | ENSMUSG( | Slc22a6 | 328.3897 | 0.264484 | 0.139394 | 0.830678 |
| ENSMUSG( | Slc22a7 | 13.03681 | 0.212773 | 0.709327 | NA | ENSMUSG( | Slc22a7 | 12.24418 | 0.292116 | 0.6365 | 0.953903 |
| ENSMUSG( | Slc22a8 | 1131.078 | -0.01869 | 0.90024 | 0.985633 | ENSMUSG( | Slc22a8 | 1133.191 | 0.25096 | 0.163647 | 0.838291 |
| ENSMUSG( | Slc23a2 | 5407.505 | -0.27471 | 0.028217 | 0.256543 | ENSMUSG( | Slc23a2 | 5774.925 | 0.194117 | 0.211219 | 0.864412 |
| ENSMUSG( | Slc23a3 | 21.51445 | 0.172502 | 0.718784 | 0.941986 | ENSMUSG( | Slc23a3 | 17.47821 | -0.14828 | 0.764242 | 0.974 |
| ENSMUSG( | Slc23a4 | 15.62374 | -0.64291 | 0.189016 | NA | ENSMUSG( | Slc23a4 | 14.15188 | -0.67743 | 0.20667 | 0.864412 |
| ENSMUSG( | Slc24a2 | 9939.44 | -0.62581 | 3.75E-05 | 0.003184 | ENSMUSG( | Slc24a2 | 11901.65 | 0.22537 | 0.203778 | 0.864412 |
| ENSMUSG( | Slc24a3 | 6000.345 | -0.29495 | 0.015572 | 0.184473 | ENSMUSG( | Slc24a3 | 6209.634 | 0.089675 | 0.547436 | 0.938063 |
| ENSMUSG( | Slc24a4 | 434.3032 | -0.22472 | 0.177344 | 0.614553 | ENSMUSG( | Slc24a4 | 516.9867 | 0.518602 | 0.004163 | 0.261354 |
| ENSMUSG( | Slc24a5 | 16.84397 | -0.005 | 0.992998 | NA | ENSMUSG( | Slc24a5 | 16.02929 | 0.105945 | 0.856335 | 0.985884 |
| ENSMUSG( | Slc25a1 | 2039.977 | 0.030177 | 0.760527 | 0.952907 | ENSMUSG( | Slc25a1 | 1743.657 | -0.16214 | 0.183576 | 0.851616 |
| ENSMUSG( | Slc25a10 | 640.6597 | 0.040954 | 0.692433 | 0.934101 | ENSMUSG( | Slc25a10 | 584.739 | 0.045236 | 0.721559 | 0.966887 |
| ENSMUSG( | Slc25a11 | 2468.416 | 0.076322 | 0.482612 | 0.856078 | ENSMUSG( | Slc25a11 | 2133.929 | -0.07653 | 0.542697 | 0.938034 |
| ENSMUSG( | Slc25a12 | 3098.999 | 0.074938 | 0.515195 | 0.872463 | ENSMUSG( | Slc25a12 | 2747.772 | -0.00313 | 0.980474 | 0.996479 |
| ENSMUSG( | Slc25a13 | 56.32447 | -0.29509 | 0.277194 | 0.727175 | ENSMUSG( | Slc25a13 | 57.88539 | 0.064063 | 0.815058 | 0.979198 |
| ENSMUSG( | Slc25a14 | 1164.107 | -0.11035 | 0.418721 | 0.821966 | ENSMUSG( | Slc25a14 | 1195.811 | 0.233507 | 0.113428 | 0.809623 |
| ENSMUSG( | Slc25a15 | 677.3291 | -0.01979 | 0.868115 | 0.97675 | ENSMUSG( | Slc25a15 | 607.7352 | -0.06896 | 0.605375 | 0.94924 |
| ENSMUSG( | Slc25a16 | 1091.312 | 0.007385 | 0.937005 | 0.993415 | ENSMUSG( | Slc25a16 | 1007.862 | 0.043282 | 0.722036 | 0.966887 |
| ENSMUSG( | Slc25a17 | 944.1787 | -0.11422 | 0.268172 | 0.718372 | ENSMUSG( | Slc25a17 | 932.4177 | 0.118488 | 0.34619 | 0.910966 |
| ENSMUSG( | Slc25a18 | 8328.475 | 0.150125 | 0.119597 | 0.523437 | ENSMUSG( | Slc25a18 | 6685.003 | -0.22188 | 0.065045 | 0.741314 |
| ENSMUSG( | Slc25a19 | 1526.857 | 0.076086 | 0.410152 | 0.818258 | ENSMUSG( | Slc25a19 | 1348.422 | -0.01394 | 0.902115 | 0.991012 |
| ENSMUSG( | Slc25a20 | 856.5412 | -0.02414 | 0.8444 | 0.972023 | ENSMUSG( | Slc25a20 | 704.0567 | -0.34186 | 0.015425 | 0.495791 |
| ENSMUSG( | Slc25a21 | 56.15608 | -0.08141 | 0.794335 | 0.959799 | ENSMUSG( | Slc25a21 | 47.08511 | -0.37717 | 0.2464 | 0.877904 |
| ENSMUSG( | Slc25a22 | 6327.008 | 0.156569 | 0.173643 | 0.609084 | ENSMUSG( | Slc25a22 | 5402.907 | -0.0273 | 0.836781 | 0.981335 |
| ENSMUSG( | Slc25a23 | 38360.75 | 0.152656 | 0.056163 | 0.36528 | ENSMUSG( | Slc25a23 | 32765.83 | -0.02992 | 0.758745 | 0.971858 |
| ENSMUSG( | Slc25a24 | 58.24348 | 0.13546 | 0.603401 | 0.904887 | ENSMUSG( | Slc25a24 | 62.88308 | 0.580921 | 0.034815 | 0.658559 |
| ENSMUSG( | Slc25a25 | 1478.979 | -0.11125 | 0.290006 | 0.739839 | ENSMUSG( | Slc25a25 | 1336.796 | -0.13399 | 0.257314 | 0.883401 |
| ENSMUSG( | Slc25a26 | 434.6355 | 0.026852 | 0.817972 | 0.965795 | ENSMUSG( | Slc25a26 | 390.5974 | -0.01473 | 0.914208 | 0.992407 |
| ENSMUSG( | Slc25a27 | 1196.943 | 0.176776 | 0.147684 | 0.57034 | ENSMUSG( | Slc25a27 | 1117.529 | 0.241833 | 0.125432 | 0.81516 |
| ENSMUSG( | Slc25a28 | 1315.937 | -0.14226 | 0.249067 | 0.698455 | ENSMUSG( | Slc25a28 | 1266.56 | 0.023217 | 0.859092 | 0.985884 |
| ENSMUSG( | Slc25a29 | 377.0116 | 0.103495 | 0.450041 | 0.835344 | ENSMUSG( | Slc25a29 | 353.513 | 0.185773 | 0.206917 | 0.864412 |
| ENSMUSG( | Slc25a3 | 13597.08 | -0.01666 | 0.850468 | 0.972638 | ENSMUSG( | Slc25a3 | 11405.41 | -0.26942 | 0.017541 | 0.53154 |
| ENSMUSG( | Slc25a30 | 214.305 | -0.08983 | 0.551989 | 0.886141 | ENSMUSG( | Slc25a30 | 206.8432 | 0.066176 | 0.704013 | 0.962625 |
| ENSMUSG( | Slc25a32 | 460.6421 | 0.007758 | 0.94545 | 0.99365 | ENSMUSG( | Slc25a32 | 451.8063 | 0.211323 | 0.119536 | 0.813384 |
| ENSMUSG( | Slc25a33 | 1405.287 | 0.164477 | 0.108468 | 0.497337 | ENSMUSG( | Slc25a33 | 1243.685 | 0.079272 | 0.544391 | 0.938063 |
| ENSMUSG( | Slc25a34 | 835.9111 | -0.03551 | 0.814407 | 0.964574 | ENSMUSG( | Slc25a34 | 650.57 | -0.54929 | 0.000169 | 0.031167 |
| ENSMUSG( | Slc25a35 | 215.9011 | -0.3564 | 0.03013 | 0.263737 | ENSMUSG( | Slc25a35 | 217.3469 | -0.02991 | 0.882188 | 0.989644 |
| ENSMUSG( | Slc25a36 | 1444.815 | -0.05993 | 0.540146 | 0.88315 | ENSMUSG( | Slc25a36 | 1397.893 | 0.11195 | 0.313001 | 0.900449 |
| ENSMUSG( | Slc25a37 | 1075.657 | -0.20595 | 0.029149 | 0.259574 | ENSMUSG( | Slc25a37 | 1054.357 | 0.015099 | 0.897921 | 0.990558 |
| ENSMUSG( | Slc25a38 | 840.9987 | 0.020361 | 0.845425 | 0.97218 | ENSMUSG( | Slc25a38 | 736.8469 | -0.09763 | 0.450102 | 0.922202 |
| ENSMUSG( | Slc25a39 | 2976.401 | 0.040399 | 0.641373 | 0.916693 | ENSMUSG( | Slc25a39 | 2537.363 | -0.15774 | 0.172851 | 0.843702 |
| ENSMUSG( | Slc25a4 | 18497.62 | 0.016783 | 0.856668 | 0.974154 | ENSMUSG( | Slc25a4 | 15844.34 | -0.16874 | 0.192371 | 0.859726 |
| ENSMUSG( | Slc25a40 | 292.8525 | -0.19116 | 0.217918 | 0.663176 | ENSMUSG( | Slc25a40 | 288.5845 | 0.042416 | 0.799118 | 0.97681 |
| ENSMUSG( | Slc25a42 | 1063.894 | 0.114104 | 0.311409 | 0.759296 | ENSMUSG( | Slc25a42 | 950.7986 | 0.058873 | 0.687795 | 0.959703 |
| ENSMUSG( | Slc25a44 | 4135.622 | -0.1559 | 0.14005 | 0.556902 | ENSMUSG( | Slc25a44 | 4161.558 | 0.134425 | 0.267953 | 0.88628 |
| ENSMUSG( | Slc25a45 | 207.7652 | -0.17927 | 0.315695 | 0.762671 | ENSMUSG( | Slc25a45 | 210.4791 | 0.147084 | 0.519579 | 0.931729 |
| ENSMUSG( | Slc25a46 | 2088.305 | 0.041918 | 0.629871 | 0.913965 | ENSMUSG( | Slc25a46 | 1881.684 | 0.007383 | 0.954719 | 0.994798 |
| ENSMUSG( | Slc25a47 | 181.4928 | 0.060843 | 0.717573 | 0.941697 | ENSMUSG( | Slc25a47 | 161.4045 | -0.00123 | 0.994754 | 0.998813 |
| ENSMUSG( | Slc25a5 | 6278.192 | 0.022773 | 0.794292 | 0.959799 | ENSMUSG( | Slc25a5 | 5418.462 | -0.13926 | 0.246444 | 0.877904 |
| ENSMUSG( | Slc25a51 | 2805.948 | 0.029267 | 0.746265 | 0.949289 | ENSMUSG( | Slc25a51 | 2599.589 | 0.074779 | 0.513043 | 0.930358 |
| ENSMUSG( | Slc25a53 | 604.3846 | -0.08169 | 0.521721 | 0.875186 | ENSMUSG( | Slc25a53 | 567.5263 | 0.001706 | 0.991064 | 0.998379 |
| ENSMUSG( | Slc26a1 | 91.76719 | -0.49639 | 0.030299 | 0.264512 | ENSMUSG( | Slc26a1 | 90.01437 | -0.23289 | 0.354463 | 0.912411 |
| ENSMUSG( | Slc26a10 | 196.1071 | -0.07977 | 0.643727 | 0.917348 | ENSMUSG( | Slc26a10 | 193.0022 | 0.152552 | 0.543776 | 0.938063 |
| ENSMUSG( | Slc26a11 | 220.0801 | -0.22972 | 0.21531 | 0.659922 | ENSMUSG( | Slc26a11 | 227.7079 | 0.152514 | 0.409553 | 0.917257 |
| ENSMUSG( | Slc26a2 | 657.7118 | -0.68914 | 5.19E-09 | 2.10E-06 | ENSMUSG( | Slc26a2 | 780.0504 | 0.156551 | 0.362582 | 0.914223 |
| ENSMUSG( | Slc26a4 | 142.1037 | 0.275625 | 0.158503 | 0.587906 | ENSMUSG( | Slc26a4 | 128.6919 | 0.26109 | 0.215284 | 0.865167 |
| ENSMUSG( | Slc26a5 | 10.86265 | -0.08851 | 0.872049 | NA | ENSMUSG( | Slc26a5 | 9.447181 | -0.20596 | 0.740682 | NA |
| ENSMUSG( | Slc26a6 | 432.1622 | -0.01995 | 0.863298 | 0.975055 | ENSMUSG( | Slc26a6 | 420.1582 | 0.171822 | 0.274959 | 0.889874 |
| ENSMUSG( | Slc26a7 | 180.0366 | -0.25208 | 0.131308 | 0.543457 | ENSMUSG( | Slc26a7 | 185.8256 | 0.115988 | 0.494837 | 0.927372 |
| ENSMUSG( | Slc26a8 | 216.7594 | -0.18335 | 0.273802 | 0.724244 | ENSMUSG( | Slc26a8 | 202.6249 | -0.09946 | 0.550741 | 0.939187 |
| ENSMUSG( | Slc27a1 | 5327.977 | -0.09768 | 0.405437 | 0.81652 | ENSMUSG( | Slc27a1 | 5059.541 | 0.026495 | 0.855661 | 0.985738 |
| ENSMUSG( | Slc27a2 | 129.2622 | -0.22495 | 0.260538 | 0.710468 | ENSMUSG( | Slc27a2 | 134.9881 | 0.18456 | 0.394781 | 0.917257 |
| ENSMUSG( | Slc27a3 | 173.061 | -0.02667 | 0.881766 | 0.98196 | ENSMUSG( | Slc27a3 | 169.6909 | 0.197128 | 0.394875 | 0.917257 |
| ENSMUSG( | Slc27a4 | 3796.692 | -0.27301 | 0.067648 | 0.402445 | ENSMUSG( | Slc27a4 | 3989.029 | 0.152612 | 0.379759 | 0.916995 |

| | | | | | |
|---|---|---|---|---|---|
| ENSMUSG( | 2024.763 | 0.146118 | 0.114567 | 0.510543 | Setx |
| ENSMUSG( | 7101.292 | -0.52825 | 6.73E-08 | 2.02E-05 | Sez6 |
| ENSMUSG( | 6540.725 | -0.2636 | 0.065416 | 0.396102 | Sez6l |
| ENSMUSG( | 9968.793 | -0.28852 | 0.041679 | 0.315317 | Sez6l2 |
| ENSMUSG( | 3992.382 | 0.029402 | 0.728731 | 0.944393 | Sf1 |
| ENSMUSG( | 1995.257 | 0.177212 | 0.069353 | 0.40782 | Sf3a1 |
| ENSMUSG( | 857.2785 | -0.02886 | 0.817659 | 0.965785 | Sf3a2 |
| ENSMUSG( | 563.4046 | 0.054775 | 0.657212 | 0.924142 | Sf3a3 |
| ENSMUSG( | 7722.841 | -0.03063 | 0.709556 | 0.939669 | Sf3b1 |
| ENSMUSG( | 5993.254 | 0.120493 | 0.143751 | 0.562581 | Sf3b2 |
| ENSMUSG( | 1706.909 | 0.051075 | 0.58708 | 0.900235 | Sf3b3 |
| ENSMUSG( | 662.1542 | 0.018385 | 0.893084 | 0.984512 | Sf3b4 |
| ENSMUSG( | 691.8423 | 0.090159 | 0.450535 | 0.83542 | Sf3b5 |
| ENSMUSG( | 1044.381 | 0.040933 | 0.746399 | 0.949289 | Sf3b6 |
| ENSMUSG( | 631.9803 | 0.234813 | 0.146459 | 0.568435 | Sfi1 |
| ENSMUSG( | 670.0561 | 0.094405 | 0.376666 | 0.801213 | Sfmbt1 |
| ENSMUSG( | 109.0671 | 0.064488 | 0.742972 | 0.948478 | Sfmbt2 |
| ENSMUSG( | 66.07871 | 0.072548 | 0.801325 | 0.961947 | Sfn |
| ENSMUSG( | 3077.605 | 0.07655 | 0.438197 | 0.830322 | Sfpq |
| ENSMUSG( | 2180.714 | 0.033587 | 0.728437 | 0.944393 | Sfr1 |
| ENSMUSG( | 741.7904 | -0.25541 | 0.043206 | 0.318825 | Sfrp1 |
| ENSMUSG( | 76.16831 | -0.57147 | 0.021939 | 0.225738 | Sfrp2 |
| ENSMUSG( | 75.9886 | -0.13281 | 0.568049 | 0.892201 | Sfrp4 |
| ENSMUSG( | 66.59807 | -0.71504 | 0.016303 | 0.189424 | Sfrp5 |
| ENSMUSG( | 2741.295 | 0.140404 | 0.201617 | 0.641362 | Sfswap |
| ENSMUSG( | 754.7177 | -0.1971 | 0.065212 | 0.395697 | Sft2d1 |
| ENSMUSG( | 23.27855 | -0.17118 | 0.652088 | 0.922053 | Sft2d1rt |
| ENSMUSG( | 3733.595 | 0.084187 | 0.474323 | 0.849425 | Sft2d2 |
| ENSMUSG( | 168.8239 | -0.51985 | 0.003384 | 0.073288 | Sft2d3 |
| ENSMUSG( | 8.575746 | -0.73033 | 0.269411 | NA | Sfta3-ps |
| ENSMUSG( | 29.04238 | -0.3625 | 0.361296 | 0.792797 | Sftpc |
| ENSMUSG( | 2883.701 | 0.07869 | 0.375477 | 0.800845 | Sfxn1 |
| ENSMUSG( | 225.0212 | -0.12585 | 0.428222 | 0.825894 | Sfxn2 |
| ENSMUSG( | 2763.052 | 0.120692 | 0.380701 | 0.801984 | Sfxn3 |
| ENSMUSG( | 283.97 | 0.076853 | 0.632848 | 0.914073 | Sfxn4 |
| ENSMUSG( | 26613.72 | 0.053362 | 0.64202 | 0.916865 | Sfxn5 |
| ENSMUSG( | 15.50577 | -0.30884 | 0.518604 | NA | Sgca |
| ENSMUSG( | 3958.855 | -0.17109 | 0.1278 | 0.535913 | Sgcb |
| ENSMUSG( | 38.03395 | -0.17148 | 0.571563 | 0.893319 | Sgcd |
| ENSMUSG( | 1043.463 | -0.34613 | 0.0112 | 0.152581 | Sgce |
| ENSMUSG( | 11.76858 | 0.164143 | 0.782112 | NA | Sgcg |
| ENSMUSG( | 41.92361 | -0.62935 | 0.046813 | 0.330855 | Sgcz |
| ENSMUSG( | 250.682 | -0.08335 | 0.640086 | 0.916434 | Sgf29 |
| ENSMUSG( | 3315.914 | -0.09292 | 0.313109 | 0.76033 | Sgip1 |
| ENSMUSG( | 1145.982 | -0.38693 | 0.002288 | 0.055887 | Sgk1 |
| ENSMUSG( | 86.60862 | -0.11413 | 0.595616 | 0.90314 | Sgk2 |
| ENSMUSG( | 500.5776 | -0.05463 | 0.661641 | 0.925271 | Sgk3 |
| ENSMUSG( | 1359.443 | 0.050229 | 0.735721 | 0.946267 | Sgms1 |
| ENSMUSG( | 158.1314 | -0.01171 | 0.943244 | 0.993415 | Sgms1os1 |
| ENSMUSG( | 102.0561 | 0.155902 | 0.4795 | 0.854046 | Sgms2 |
| ENSMUSG( | 37.83177 | -0.42082 | 0.200123 | 0.640302 | Sgo2a |
| ENSMUSG( | 1853.471 | -0.13407 | 0.313243 | 0.76033 | Sgpl1 |
| ENSMUSG( | 1228.171 | -0.28389 | 0.03739 | 0.298461 | Sgpp1 |
| ENSMUSG( | 442.1645 | -0.32697 | 0.070632 | 0.409748 | Sgpp2 |
| ENSMUSG( | 387.0399 | -0.26174 | 0.146572 | 0.568485 | Sgsh |
| ENSMUSG( | 3251.331 | 0.231719 | 0.047164 | 0.33245 | Sgsm1 |
| ENSMUSG( | 7278.548 | 0.098283 | 0.183535 | 0.620584 | Sgsm2 |
| ENSMUSG( | 1613.896 | 0.223905 | 0.097274 | 0.477629 | Sgsm3 |
| ENSMUSG( | 3601.025 | -0.0152 | 0.87486 | 0.979746 | Sgta |
| ENSMUSG( | 1999.1 | 0.086292 | 0.42836 | 0.826032 | Sgtb |
| ENSMUSG( | 2651.238 | 0.092785 | 0.300457 | 0.749525 | Sh2b1 |
| ENSMUSG( | 170.377 | 0.037148 | 0.816957 | 0.96566 | Sh2b2 |
| ENSMUSG( | 1018.238 | 0.265662 | 0.021995 | 0.226001 | Sh2b3 |
| ENSMUSG( | 574.3668 | -0.14698 | 0.336058 | 0.777 | Sh2d3c |
| ENSMUSG( | 34.28766 | 0.491874 | 0.157565 | 0.585653 | Sh2d4b |
| ENSMUSG( | 2938.025 | 0.009584 | 0.931156 | 0.992271 | Sh2d5 |
| ENSMUSG( | 39.03593 | -0.21408 | 0.484579 | 0.856665 | Sh2d6 |
| ENSMUSG( | 36.89069 | 0.011807 | 0.973248 | 0.99675 | Sh3bgr |
| ENSMUSG( | 3420.968 | 0.022279 | 0.894281 | 0.984512 | Sh3bgrl |
| ENSMUSG( | 569.0737 | -0.05434 | 0.683271 | 0.932153 | Sh3bgrl2 |
| ENSMUSG( | 2134.15 | -0.03785 | 0.719087 | 0.941986 | Sh3bgrl3 |
| ENSMUSG( | 210.5244 | 0.045415 | 0.813876 | 0.964574 | Sh3bp1 |
| ENSMUSG( | 173.1904 | -0.33131 | 0.068491 | 0.405448 | Sh3bp2 |
| ENSMUSG( | 774.4954 | 0.084384 | 0.40946 | 0.818258 | Sh3bp4 |
| ENSMUSG( | 993.0504 | 0.167263 | 0.12554 | 0.53257 | Sh3bp5 |
| ENSMUSG( | 1597.509 | 0.290947 | 0.005438 | 0.098663 | Sh3bp5l |
| ENSMUSG( | 892.3127 | -0.00468 | 0.965943 | 0.995377 | Sh3d19 |
| ENSMUSG( | 65.85281 | 0.146805 | 0.568345 | 0.892252 | Sh3d21 |
| ENSMUSG( | 1195.306 | 0.135975 | 0.198384 | 0.638387 | Sh3gl1 |
| ENSMUSG( | 5483.866 | 0.180449 | 0.233945 | 0.681906 | Sh3gl2 |
| ENSMUSG( | 3828.663 | 0.070685 | 0.469195 | 0.845876 | Sh3gl3 |
| ENSMUSG( | 6517.826 | -0.02849 | 0.795177 | 0.960062 | Sh3glb1 |
| ENSMUSG( | 5891.219 | 0.03731 | 0.73396 | 0.945306 | Sh3glb2 |
| ENSMUSG( | 986.8965 | -0.13444 | 0.233586 | 0.681508 | Sh3kbp1 |
| ENSMUSG( | 2720.863 | 0.033921 | 0.72676 | 0.944223 | Sh3pxd2a |
| ENSMUSG( | 1302.73 | -0.01065 | 0.913231 | 0.988483 | Sh3pxd2b |
| ENSMUSG( | 1393.059 | 0.144531 | 0.100301 | 0.48338 | Sh3rf1 |
| ENSMUSG( | 82.31049 | -1.27696 | 0.000138 | 0.00806 | Sh3rf2 |
| ENSMUSG( | 1221.754 | 0.290021 | 0.002831 | 0.06505 | Sh3rf3 |
| ENSMUSG( | 111.7832 | 0.015143 | 0.947702 | 0.993821 | Sh3tc1 |
| ENSMUSG( | 172.3508 | 0.158644 | 0.389808 | 0.807597 | Sh3tc2 |
| ENSMUSG( | 171.2579 | 0.03242 | 0.854975 | 0.973665 | Sh3yl1 |
| ENSMUSG( | 28175.6 | 0.377486 | 0.00015 | 0.008579 | Shank1 |
| ENSMUSG( | 11876.65 | 0.170206 | 0.094902 | 0.472904 | Shank2 |
| ENSMUSG( | 15098.78 | 0.226096 | 0.009399 | 0.13706 | Shank3 |
| ENSMUSG( | 892.1834 | -0.02361 | 0.837818 | 0.971111 | Sharpin |
| ENSMUSG( | 297.4424 | -0.08025 | 0.633913 | 0.914073 | Shb |
| ENSMUSG( | 519.4609 | 0.052244 | 0.67839 | 0.930542 | Shc1 |
| ENSMUSG( | 1399.652 | 0.08636 | 0.459087 | 0.840244 | Shc2 |
| ENSMUSG( | 3088.06 | 0.199565 | 0.050316 | 0.342969 | Shc3 |
| ENSMUSG( | 659.6339 | -0.12709 | 0.353935 | 0.786371 | Shc4 |
| ENSMUSG( | 22.87614 | 0.234383 | 0.538938 | 0.882849 | Shcbp1 |
| ENSMUSG( | 52.47488 | -0.2686 | 0.316391 | 0.763556 | Shcbp1l |
| ENSMUSG( | 1025.81 | 0.037036 | 0.710535 | 0.939748 | Shd |
| ENSMUSG( | 131.8376 | 0.28654 | 0.241772 | 0.689179 | She |
| ENSMUSG( | 1280.648 | 0.12462 | 0.243516 | 0.690987 | Shf |
| ENSMUSG( | 1888.848 | 0.255167 | 0.143462 | 0.562473 | Shfl |
| ENSMUSG( | 83.86347 | -0.40219 | 0.076321 | 0.427609 | Shh |
| ENSMUSG( | 75.16707 | -0.4807 | 0.038206 | 0.301559 | Shisa2 |
| ENSMUSG( | 71.11808 | -1.16666 | 6.16E-06 | 0.000787 | Shisa3 |
| ENSMUSG( | 8104.075 | -0.22822 | 0.042612 | 0.316734 | Shisa4 |
| ENSMUSG( | 2134.033 | -0.36344 | 0.009508 | 0.137946 | Shisa5 |
| ENSMUSG( | 747.8048 | 0.01277 | 0.943438 | 0.993415 | Shisa6 |
| ENSMUSG( | 5701.051 | -0.23811 | 0.023601 | 0.235222 | Shisa7 |
| ENSMUSG( | 226.0881 | -0.7746 | 2.72E-06 | 0.000405 | Shisa8 |
| ENSMUSG( | 2470.807 | -0.3645 | 0.005309 | 0.097191 | Shisa9 |
| ENSMUSG( | 3481.278 | -0.25519 | 0.00816 | 0.125816 | Shisal1 |
| ENSMUSG( | 15.46003 | -0.22363 | 0.646813 | NA | Shisal2b |
| ENSMUSG( | 394.0266 | 0.187778 | 0.226504 | 0.673549 | Shkbp1 |
| ENSMUSG( | 167.948 | -0.06797 | 0.692479 | 0.934101 | Shld1 |
| ENSMUSG( | 126.6706 | -0.27441 | 0.143974 | 0.563118 | Shld2 |
| ENSMUSG( | 21.81701 | 0.255192 | 0.547608 | 0.885153 | Shld3 |
| ENSMUSG( | 57.4764 | -0.35566 | 0.181442 | 0.618684 | Shmt1 |
| ENSMUSG( | 1040.339 | 0.030398 | 0.775239 | 0.955606 | Shmt2 |
| ENSMUSG( | 1004.846 | 0.055073 | 0.561656 | 0.889052 | Shoc2 |
| ENSMUSG( | 141.6324 | -0.09991 | 0.580308 | 0.896952 | Shpk |
| ENSMUSG( | 1078.638 | 0.037436 | 0.683993 | 0.932457 | Shprh |
| ENSMUSG( | 101.6036 | 0.281513 | 0.172854 | 0.608361 | Shq1 |
| ENSMUSG( | 197.7803 | -0.06297 | 0.727014 | 0.944223 | Shroom1 |
| ENSMUSG( | 1809.951 | -0.01995 | 0.843574 | 0.971847 | Shroom2 |
| ENSMUSG( | 158.8227 | -0.30695 | 0.071716 | 0.413564 | Shroom3 |
| ENSMUSG( | 219.2706 | 0.120084 | 0.516161 | 0.872701 | Shroom4 |
| ENSMUSG( | 1405.996 | -0.16927 | 0.064905 | 0.395029 | Shtn1 |
| ENSMUSG( | 1017.818 | -0.26705 | 0.027767 | 0.255176 | Siae |
| ENSMUSG( | 431.1111 | -0.06007 | 0.634611 | 0.914073 | Siah1a |
| ENSMUSG( | 86.92589 | -0.29976 | 0.147304 | 0.569415 | Siah1b |
| ENSMUSG( | 333.7083 | -0.27355 | 0.082802 | 0.444372 | Siah2 |
| ENSMUSG( | 295.9917 | 0.217418 | 0.114742 | 0.510733 | Siah3 |
| ENSMUSG( | 2213.649 | 0.097733 | 0.475867 | 0.850919 | Sidt1 |
| ENSMUSG( | 3438.715 | -0.08585 | 0.496051 | 0.861543 | Sidt2 |
| ENSMUSG( | 70.74977 | 0.431238 | 0.104171 | 0.489971 | Sigirr |
| ENSMUSG( | 27.48616 | -0.28473 | 0.516242 | 0.872701 | Siglec1 |
| ENSMUSG( | 54.10897 | -0.49742 | 0.123806 | 0.531039 | Siglece |
| ENSMUSG( | 30.14733 | -0.94137 | 0.013802 | 0.171544 | Siglecf |
| ENSMUSG( | 1195.118 | -0.03007 | 0.850015 | 0.972638 | Siglech |
| ENSMUSG( | 1585.367 | -0.21043 | 0.078655 | 0.433892 | Sigmar1 |
| ENSMUSG( | 386.8891 | -0.70363 | 6.24E-08 | 1.91E-05 | Sik1 |
| ENSMUSG( | 598.3326 | -0.047 | 0.701058 | 0.93743 | Sik2 |
| ENSMUSG( | 4989.571 | 0.106386 | 0.222761 | 0.668614 | Sik3 |
| ENSMUSG( | 1768.504 | 0.052602 | 0.55273 | 0.886141 | Sike1 |
| ENSMUSG( | 874.932 | -0.37194 | 0.006932 | 0.114527 | Sil1 |
| ENSMUSG( | 858.5327 | 0.167267 | 0.123813 | 0.531039 | Silc1 |
| ENSMUSG( | 342.5739 | -0.01139 | 0.929468 | 0.991848 | Simc1 |
| ENSMUSG( | 1120.854 | 0.072225 | 0.428148 | 0.825894 | Sin3a |
| ENSMUSG( | 2561.761 | 0.109155 | 0.312627 | 0.760241 | Sin3b |
| ENSMUSG( | 22.32148 | -0.0776 | 0.8645 | 0.975398 | Sinhcaf |
| ENSMUSG( | 850.8293 | 0.175814 | 0.105446 | 0.492417 | Sipa1 |
| ENSMUSG( | 16066.59 | 0.200011 | 0.048726 | 0.33775 | Sipa1l1 |

| | | | | | |
|---|---|---|---|---|---|
| ENSMUSG( | 263.4386 | 0.158269 | 0.32816 | 0.906255 | Siah3 |
| ENSMUSG( | 2285.708 | 0.431974 | 0.001438 | 0.139907 | Sidt1 |
| ENSMUSG( | 3420.746 | 0.168419 | 0.27062 | 0.888062 | Sidt2 |
| ENSMUSG( | 64.1468 | 0.454519 | 0.256429 | 0.883082 | Sigirr |
| ENSMUSG( | 25.00227 | -0.27715 | 0.523128 | 0.932848 | Siglec1 |
| ENSMUSG( | 62.22924 | 0.223693 | 0.465806 | 0.923329 | Siglece |
| ENSMUSG( | 31.54521 | -0.38606 | 0.329658 | 0.907083 | Siglecf |
| ENSMUSG( | 1191.406 | 0.227314 | 0.190261 | 0.857355 | Siglech |
| ENSMUSG( | 1675.463 | 0.219702 | 0.097144 | 0.796721 | Sigmar1 |
| ENSMUSG( | 398.1204 | -0.27517 | 0.100646 | 0.801444 | Sik1 |
| ENSMUSG( | 551.511 | -0.00977 | 0.94263 | 0.994774 | Sik2 |
| ENSMUSG( | 4375.175 | -0.00274 | 0.979644 | 0.996479 | Sik3 |
| ENSMUSG( | 1565.945 | -0.0313 | 0.782 | 0.976106 | Sike1 |
| ENSMUSG( | 929.5745 | 0.098087 | 0.520553 | 0.932079 | Sil1 |
| ENSMUSG( | 763.8653 | 0.100402 | 0.415552 | 0.917257 | Silc1 |
| ENSMUSG( | 309.1924 | -0.03411 | 0.822741 | 0.979198 | Simc1 |
| ENSMUSG( | 1024.157 | 0.077928 | 0.503183 | 0.927825 | Sin3a |
| ENSMUSG( | 2217.247 | -0.03912 | 0.768344 | 0.974236 | Sin3b |
| ENSMUSG( | 27.73306 | 0.75811 | 0.068277 | 0.752208 | Sinhcaf |
| ENSMUSG( | 825.0785 | 0.345863 | 0.015371 | 0.495791 | Sipa1 |
| ENSMUSG( | 14299.49 | 0.137906 | 0.247642 | 0.877904 | Sipa1l1 |
| ENSMUSG( | 1932.726 | 0.031704 | 0.785978 | 0.976285 | Sipa1l2 |
| ENSMUSG( | 1716.093 | 0.303744 | 0.021421 | 0.575341 | Sipa1l3 |
| ENSMUSG( | 19471.54 | 0.162994 | 0.317182 | 0.902275 | Sirpa |
| ENSMUSG( | 320.8136 | 0.233665 | 0.115882 | 0.813384 | Sirt1 |
| ENSMUSG( | 15505.16 | -0.18547 | 0.128562 | 0.81925 | Sirt2 |
| ENSMUSG( | 2778.57 | -0.13621 | 0.21151 | 0.864412 | Sirt3 |
| ENSMUSG( | 404.334 | 0.337499 | 0.16038 | 0.837724 | Sirt4 |
| ENSMUSG( | 265.5432 | -0.30049 | 0.062193 | 0.736482 | Sirt5 |
| ENSMUSG( | 334.1838 | 0.008659 | 0.958055 | 0.994798 | Sirt6 |
| ENSMUSG( | 1157.584 | 0.25763 | 0.09596 | 0.796133 | Sirt7 |
| ENSMUSG( | 152.877 | -0.06284 | 0.741231 | 0.968811 | Siva1 |
| ENSMUSG( | 27.36948 | 0.418126 | 0.290692 | 0.897019 | Six1 |
| ENSMUSG( | 36.7273 | -0.32462 | 0.336703 | 0.907972 | Six2 |
| ENSMUSG( | 172.5154 | -0.42243 | 0.181138 | 0.850372 | Six3 |
| ENSMUSG( | 111.1347 | -0.35825 | 0.201064 | 0.86439 | Six3os1 |
| ENSMUSG( | 15.60244 | 0.305588 | 0.558043 | 0.939565 | Six4 |
| ENSMUSG( | 109.1267 | 0.316696 | 0.180472 | 0.850372 | Six5 |
| ENSMUSG( | 395.869 | -0.02369 | 0.867764 | 0.98781 | Ska2 |
| ENSMUSG( | 19.75378 | 0.13014 | 0.781067 | 0.976106 | Ska3 |
| ENSMUSG( | 93.90965 | 0.114562 | 0.621331 | 0.953199 | Skap1 |
| ENSMUSG( | 412.4274 | -0.17935 | 0.246006 | 0.877904 | Skap2 |
| ENSMUSG( | 5562.784 | 0.105165 | 0.286999 | 0.894091 | Ski |
| ENSMUSG( | 2194.693 | 0.026455 | 0.83823 | 0.981575 | Skic2 |
| ENSMUSG( | 641.4415 | 0.123754 | 0.308648 | 0.898538 | Skic3 |
| ENSMUSG( | 605.9543 | -0.12215 | 0.361858 | 0.914223 | Skic8 |
| ENSMUSG( | 230.951 | 0.068069 | 0.691334 | 0.960071 | Skida1 |
| ENSMUSG( | 1078.174 | 0.002816 | 0.980199 | 0.996479 | Skil |
| ENSMUSG( | 6870.569 | -0.18039 | 0.226396 | 0.868507 | Skp1 |
| ENSMUSG( | 153.455 | 0.020464 | 0.917498 | 0.992529 | Skp2 |
| ENSMUSG( | 251.0654 | -0.11305 | 0.521893 | 0.932571 | Sla |
| ENSMUSG( | 23.97104 | -0.66439 | 0.113538 | 0.809623 | Sla2 |
| ENSMUSG( | 989.3597 | -0.12415 | 0.343916 | 0.910256 | Slain1 |
| ENSMUSG( | 755.1793 | 0.094322 | 0.485799 | 0.92541 | Slain2 |
| ENSMUSG( | 14.54163 | 0.027674 | 0.961569 | 0.99511 | Slamf6 |
| ENSMUSG( | 41.17273 | -0.2696 | 0.456348 | 0.923329 | Slamf9 |
| ENSMUSG( | 265.1169 | 0.041743 | 0.803502 | 0.976978 | Slbp |
| ENSMUSG( | 89.35861 | 0.032585 | 0.90851 | 0.991958 | Slc10a3 |
| ENSMUSG( | 73.65823 | -0.51674 | 0.063229 | 0.736482 | Slc10a4 |
| ENSMUSG( | 178.1787 | 0.61696 | 0.000962 | 0.10597 | Slc10a7 |
| ENSMUSG( | 187.5973 | 0.331245 | 0.152877 | 0.837724 | Slc11a1 |
| ENSMUSG( | 1580.133 | 0.072007 | 0.600066 | 0.947395 | Slc11a2 |
| ENSMUSG( | 3006.649 | 0.258984 | 0.187017 | 0.856099 | Slc12a2 |
| ENSMUSG( | 1933.574 | 0.293745 | 0.051718 | 0.711848 | Slc12a4 |
| ENSMUSG( | 17494.5 | 0.115482 | 0.406898 | 0.917257 | Slc12a5 |
| ENSMUSG( | 2690.762 | 0.078017 | 0.627524 | 0.953234 | Slc12a6 |
| ENSMUSG( | 351.551 | 0.205201 | 0.253795 | 0.880692 | Slc12a7 |
| ENSMUSG( | 33.62143 | 0.021103 | 0.952082 | 0.994798 | Slc12a8 |
| ENSMUSG( | 1592.964 | 0.090089 | 0.454248 | 0.922444 | Slc12a9 |
| ENSMUSG( | 1478.493 | -0.09568 | 0.480083 | 0.924276 | Slc13a3 |
| ENSMUSG( | 522.6759 | 0.224737 | 0.083455 | 0.77681 | Slc13a4 |
| ENSMUSG( | 1448.97 | -0.01982 | 0.882746 | 0.989644 | Slc13a5 |
| ENSMUSG( | 1058.563 | 0.061885 | 0.692757 | 0.96043 | Slc14a1 |
| ENSMUSG( | 34.86821 | 0.17192 | 0.681168 | 0.958395 | Slc14a2 |
| ENSMUSG( | 2021.797 | 0.242835 | 0.044556 | 0.693697 | Slc15a2 |
| ENSMUSG( | 230.4622 | -0.17758 | 0.402585 | 0.917257 | Slc15a3 |
| ENSMUSG( | 736.1405 | 0.294978 | 0.105915 | 0.804969 | Slc15a4 |
| ENSMUSG( | 2014.819 | -0.34549 | 0.086056 | 0.780251 | Slc16a1 |
| ENSMUSG( | 103.0084 | -0.26323 | 0.311588 | 0.900031 | Slc16a10 |
| ENSMUSG( | 405.1911 | 0.271723 | 0.202359 | 0.864412 | Slc16a11 |
| ENSMUSG( | 102.3719 | 1.137687 | 1.51E-05 | 0.005081 | Slc16a12 |
| ENSMUSG( | 207.0429 | 0.159486 | 0.407405 | 0.917257 | Slc16a13 |
| ENSMUSG( | 106.0076 | 0.444612 | 0.142977 | 0.831221 | Slc16a14 |
| ENSMUSG( | 2598.113 | 0.128986 | 0.384197 | 0.917257 | Slc16a2 |
| ENSMUSG( | 206.7248 | 0.091054 | 0.638394 | 0.953903 | Slc16a3 |
| ENSMUSG( | 194.2554 | 0.797516 | 0.009967 | 0.407912 | Slc16a4 |
| ENSMUSG( | 618.9686 | -0.05147 | 0.688872 | 0.959742 | Slc16a6 |
| ENSMUSG( | 953.212 | 0.087404 | 0.651529 | 0.95449 | Slc16a7 |
| ENSMUSG( | 18.83129 | 0.566572 | 0.323165 | 0.904023 | Slc16a8 |
| ENSMUSG( | 265.9324 | 0.300166 | 0.064901 | 0.740636 | Slc16a9 |
| ENSMUSG( | 722.9959 | 0.190959 | 0.228195 | 0.868507 | Slc17a5 |
| ENSMUSG( | 734.6324 | -0.01273 | 0.943515 | 0.994798 | Slc17a6 |
| ENSMUSG( | 73940.87 | 0.112888 | 0.412316 | 0.917257 | Slc17a7 |
| ENSMUSG( | 124.0207 | 0.040561 | 0.862052 | 0.98632 | Slc17a8 |
| ENSMUSG( | 23.74967 | 0.83056 | 0.07243 | 0.763629 | Slc17a9 |
| ENSMUSG( | 15.65351 | 0.691993 | 0.162349 | 0.837724 | Slc18a2 |
| ENSMUSG( | 47.92127 | 0.153728 | 0.649467 | 0.95422 | Slc18a3 |
| ENSMUSG( | 735.6207 | -0.17854 | 0.220204 | 0.866104 | Slc18b1 |
| ENSMUSG( | 530.9225 | 0.350518 | 0.140732 | 0.830678 | Slc19a1 |
| ENSMUSG( | 429.4222 | 0.340204 | 0.061736 | 0.736228 | Slc19a2 |
| ENSMUSG( | 190.6836 | 0.205561 | 0.692577 | 0.96043 | Slc19a3 |
| ENSMUSG( | 4320.649 | 0.319716 | 0.042566 | 0.690304 | Slc1a1 |
| ENSMUSG( | 420988 | -0.11114 | 0.413338 | 0.917257 | Slc1a2 |
| ENSMUSG( | 65949.67 | 0.030841 | 0.820828 | 0.979198 | Slc1a3 |
| ENSMUSG( | 3153.553 | 0.129867 | 0.472926 | 0.923329 | Slc1a4 |
| ENSMUSG( | 182.9229 | 0.624462 | 0.003785 | 0.245533 | Slc1a5 |
| ENSMUSG( | 235.464 | 0.111718 | 0.515916 | 0.930939 | Slc1a6 |
| ENSMUSG( | 6133.699 | -0.07892 | 0.49992 | 0.927825 | Slc20a1 |
| ENSMUSG( | 2597.897 | -0.05235 | 0.73101 | 0.967228 | Slc20a2 |
| ENSMUSG( | 12.35397 | -0.7905 | 0.195456 | 0.861579 | Slc22a1 |
| ENSMUSG( | 15.7554 | 0.417239 | 0.392439 | 0.917257 | Slc22a12 |
| ENSMUSG( | 19.78949 | 0.263827 | 0.547066 | 0.938063 | Slc22a13b |
| ENSMUSG( | 277.8505 | 0.040166 | 0.809684 | 0.978779 | Slc22a15 |
| ENSMUSG( | 21764.01 | 0.186542 | 0.200923 | 0.86439 | Slc22a17 |
| ENSMUSG( | 26.09229 | 0.166904 | 0.692429 | 0.960404 | Slc22a18 |
| ENSMUSG( | 38.04742 | 0.12718 | 0.697868 | 0.960867 | Slc22a2 |
| ENSMUSG( | 95.46218 | 0.031003 | 0.889099 | 0.990558 | Slc22a21 |
| ENSMUSG( | 3270.284 | 0.08974 | 0.616969 | 0.951666 | Slc22a23 |
| ENSMUSG( | 91.81973 | -0.03692 | 0.872712 | 0.989213 | Slc22a3 |
| ENSMUSG( | 283.4218 | -0.00338 | 0.985635 | 0.997084 | Slc22a4 |
| ENSMUSG( | 499.9432 | 0.068661 | 0.642332 | 0.953903 | Slc22a5 |
| ENSMUSG( | 328.3897 | 0.264484 | 0.139394 | 0.830678 | Slc22a6 |
| ENSMUSG( | 12.24418 | 0.292116 | 0.6365 | 0.953903 | Slc22a7 |
| ENSMUSG( | 1133.191 | 0.25096 | 0.163647 | 0.838291 | Slc22a8 |
| ENSMUSG( | 5774.925 | 0.194117 | 0.211219 | 0.864412 | Slc23a2 |
| ENSMUSG( | 17.47821 | -0.14828 | 0.764242 | 0.974 | Slc23a3 |
| ENSMUSG( | 14.15188 | -0.67743 | 0.20667 | 0.864412 | Slc23a4 |
| ENSMUSG( | 11901.65 | 0.22537 | 0.203778 | 0.864412 | Slc24a2 |
| ENSMUSG( | 6209.634 | 0.089675 | 0.547436 | 0.938063 | Slc24a3 |
| ENSMUSG( | 516.9867 | 0.518602 | 0.004163 | 0.261354 | Slc24a4 |
| ENSMUSG( | 16.02929 | 0.105945 | 0.856335 | 0.985884 | Slc24a5 |
| ENSMUSG( | 1743.657 | -0.16214 | 0.183576 | 0.851616 | Slc25a1 |
| ENSMUSG( | 584.739 | 0.045236 | 0.721559 | 0.966887 | Slc25a10 |
| ENSMUSG( | 2133.929 | -0.07653 | 0.542697 | 0.938034 | Slc25a11 |
| ENSMUSG( | 2747.772 | -0.00313 | 0.980474 | 0.996479 | Slc25a12 |
| ENSMUSG( | 57.88539 | 0.064063 | 0.815058 | 0.979198 | Slc25a13 |
| ENSMUSG( | 1195.811 | 0.233507 | 0.113428 | 0.809623 | Slc25a14 |
| ENSMUSG( | 607.7352 | -0.06896 | 0.605375 | 0.94924 | Slc25a15 |
| ENSMUSG( | 1007.862 | 0.043282 | 0.722036 | 0.966887 | Slc25a16 |
| ENSMUSG( | 932.4177 | 0.118488 | 0.34619 | 0.910966 | Slc25a17 |
| ENSMUSG( | 6685.003 | -0.22188 | 0.065045 | 0.741314 | Slc25a18 |
| ENSMUSG( | 1348.422 | -0.01394 | 0.902115 | 0.991012 | Slc25a19 |
| ENSMUSG( | 704.0567 | -0.34186 | 0.015425 | 0.495791 | Slc25a20 |
| ENSMUSG( | 47.08511 | -0.37717 | 0.2464 | 0.877904 | Slc25a21 |
| ENSMUSG( | 5402.907 | -0.0273 | 0.836781 | 0.981335 | Slc25a22 |
| ENSMUSG( | 32765.83 | -0.02992 | 0.758745 | 0.971858 | Slc25a23 |
| ENSMUSG( | 62.88308 | 0.580921 | 0.034815 | 0.658559 | Slc25a24 |
| ENSMUSG( | 1336.796 | -0.13399 | 0.257314 | 0.883401 | Slc25a25 |
| ENSMUSG( | 390.5974 | -0.01473 | 0.914208 | 0.992407 | Slc25a26 |
| ENSMUSG( | 1117.529 | 0.241833 | 0.125432 | 0.81516 | Slc25a27 |
| ENSMUSG( | 1266.56 | 0.023217 | 0.859092 | 0.985884 | Slc25a28 |
| ENSMUSG( | 353.513 | 0.185773 | 0.206917 | 0.864412 | Slc25a29 |
| ENSMUSG( | 11405.41 | -0.26942 | 0.017541 | 0.53154 | Slc25a3 |
| ENSMUSG( | 206.8432 | 0.066176 | 0.704013 | 0.962625 | Slc25a30 |
| ENSMUSG( | 451.8063 | 0.211323 | 0.119536 | 0.813384 | Slc25a32 |
| ENSMUSG( | 1243.685 | 0.079272 | 0.544391 | 0.938063 | Slc25a33 |
| ENSMUSG( | 650.57 | -0.54929 | 0.000169 | 0.031167 | Slc25a34 |
| ENSMUSG( | 217.3469 | -0.02991 | 0.882188 | 0.989644 | Slc25a35 |

| | | | | | | | | | | | |
|---|---|---|---|---|---|---|---|---|---|---|---|
| ENSMUSG( | Slc27a5 | 10.22818 | -0.45877 | 0.441059 | NA | ENSMUSG( | Slc27a5 | 8.845932 | -0.56932 | 0.43862 | NA |
| ENSMUSG( | Slc27a6 | 13.52668 | 0.139145 | 0.799234 | NA | ENSMUSG( | Slc27a6 | 13.87906 | 0.500369 | 0.375467 | 0.915491 |
| ENSMUSG( | Slc29a1 | 696.7192 | -0.40507 | 0.001012 | 0.033218 | ENSMUSG( | Slc29a1 | 727.5854 | 0.02354 | 0.874714 | 0.989644 |
| ENSMUSG( | Slc29a2 | 451.4037 | -0.29791 | 0.041752 | 0.315339 | ENSMUSG( | Slc29a2 | 478.1084 | 0.15524 | 0.336758 | 0.907972 |
| ENSMUSG( | Slc29a3 | 837.9821 | -0.25148 | 0.129361 | 0.538561 | ENSMUSG( | Slc29a3 | 947.7123 | 0.367021 | 0.051533 | 0.711358 |
| ENSMUSG( | Slc29a4 | 746.7706 | -0.11118 | 0.289235 | 0.739725 | ENSMUSG( | Slc29a4 | 726.6108 | 0.081452 | 0.505774 | 0.927825 |
| ENSMUSG( | Slc2a1 | 4868.066 | 0.17614 | 0.313265 | 0.76033 | ENSMUSG( | Slc2a1 | 5767.175 | 0.828963 | 0.000333 | 0.049413 |
| ENSMUSG( | Slc2a10 | 81.96992 | -0.68591 | 0.034177 | 0.283507 | ENSMUSG( | Slc2a10 | 88.66366 | -0.09636 | 0.763934 | 0.974 |
| ENSMUSG( | Slc2a12 | 189.1789 | -0.16711 | 0.412882 | 0.819541 | ENSMUSG( | Slc2a12 | 194.4577 | 0.186193 | 0.341394 | 0.908568 |
| ENSMUSG( | Slc2a13 | 3229.66 | -0.63095 | 0.000113 | 0.007021 | ENSMUSG( | Slc2a13 | 3892.365 | 0.238749 | 0.191638 | 0.858764 |
| ENSMUSG( | Slc2a3 | 5200.676 | -0.31549 | 0.013454 | 0.169431 | ENSMUSG( | Slc2a3 | 5503.897 | 0.13411 | 0.351866 | 0.912093 |
| ENSMUSG( | Slc2a4 | 60.9813 | -0.28254 | 0.272045 | 0.7223 | ENSMUSG( | Slc2a4 | 85.70646 | 0.8647 | 0.00032 | 0.049152 |
| ENSMUSG( | Slc2a5 | 283.5716 | -0.1636 | 0.434804 | 0.829025 | ENSMUSG( | Slc2a5 | 281.8726 | 0.105397 | 0.680152 | 0.9581 |
| ENSMUSG( | Slc2a6 | 1048.676 | -0.18981 | 0.158803 | 0.588112 | ENSMUSG( | Slc2a6 | 1086.194 | 0.185366 | 0.197001 | 0.862161 |
| ENSMUSG( | Slc2a8 | 807.6209 | -0.09606 | 0.502092 | 0.864904 | ENSMUSG( | Slc2a8 | 816.8732 | 0.207691 | 0.23349 | 0.868507 |
| ENSMUSG( | Slc2a9 | 61.45888 | -0.26132 | 0.342768 | 0.778971 | ENSMUSG( | Slc2a9 | 60.02907 | -0.02216 | 0.936471 | 0.993573 |
| ENSMUSG( | Slc30a1 | 1213.549 | -0.44406 | 0.00777 | 0.122366 | ENSMUSG( | Slc30a1 | 1343.795 | 0.15373 | 0.425481 | 0.917257 |
| ENSMUSG( | Slc30a10 | 1333.642 | -0.14982 | 0.268776 | 0.719042 | ENSMUSG( | Slc30a10 | 1312.46 | 0.079032 | 0.583552 | 0.944266 |
| ENSMUSG( | Slc30a3 | 4258.699 | -0.06568 | 0.548802 | 0.885686 | ENSMUSG( | Slc30a3 | 4140.233 | 0.12122 | 0.284604 | 0.892412 |
| ENSMUSG( | Slc30a4 | 1894.22 | -0.25032 | 0.109116 | 0.498553 | ENSMUSG( | Slc30a4 | 2106.848 | 0.322453 | 0.064538 | 0.737935 |
| ENSMUSG( | Slc30a5 | 1648.093 | -0.14621 | 0.295907 | 0.744386 | ENSMUSG( | Slc30a5 | 1704.753 | 0.219865 | 0.176911 | 0.846973 |
| ENSMUSG( | Slc30a6 | 399.3667 | -0.37264 | 0.033906 | 0.282701 | ENSMUSG( | Slc30a6 | 437.0563 | 0.183913 | 0.342949 | 0.909646 |
| ENSMUSG( | Slc30a7 | 602.2264 | -0.30245 | 0.102305 | 0.487127 | ENSMUSG( | Slc30a7 | 650.2142 | 0.205686 | 0.312669 | 0.900085 |
| ENSMUSG( | Slc30a9 | 7462.158 | 0.07027 | 0.412755 | 0.819518 | ENSMUSG( | Slc30a9 | 6522.103 | -0.04999 | 0.631216 | 0.953251 |
| ENSMUSG( | Slc31a1 | 1339.243 | -0.14896 | 0.228422 | 0.675849 | ENSMUSG( | Slc31a1 | 1395.438 | 0.236395 | 0.134208 | 0.822912 |
| ENSMUSG( | Slc31a2 | 303.4564 | -0.25409 | 0.097812 | 0.478264 | ENSMUSG( | Slc31a2 | 309.5666 | 0.095045 | 0.615378 | 0.951177 |
| ENSMUSG( | Slc32a1 | 3380.945 | -0.55958 | 2.10E-06 | 0.000341 | ENSMUSG( | Slc32a1 | 3941.207 | 0.201216 | 0.16041 | 0.837724 |
| ENSMUSG( | Slc33a1 | 792.9071 | -0.31298 | 0.032213 | 0.274192 | ENSMUSG( | Slc33a1 | 909.5396 | 0.350374 | 0.026788 | 0.603041 |
| ENSMUSG( | Slc35a1 | 523.9797 | -0.29012 | 0.050888 | 0.345939 | ENSMUSG( | Slc35a1 | 539.1689 | 0.077439 | 0.631286 | 0.953251 |
| ENSMUSG( | Slc35a2 | 1194.304 | -0.24541 | 0.0758 | 0.426666 | ENSMUSG( | Slc35a2 | 1222.047 | 0.102152 | 0.518812 | 0.930939 |
| ENSMUSG( | Slc35a3 | 483.1606 | -0.23124 | 0.110958 | 0.50216 | ENSMUSG( | Slc35a3 | 512.4977 | 0.214432 | 0.153022 | 0.837724 |
| ENSMUSG( | Slc35a4 | 2311.614 | 0.066742 | 0.412504 | 0.819452 | ENSMUSG( | Slc35a4 | 2043.817 | -0.01921 | 0.850711 | 0.985541 |
| ENSMUSG( | Slc35a5 | 1409.43 | -0.30307 | 0.039004 | 0.304448 | ENSMUSG( | Slc35a5 | 1505.32 | 0.170553 | 0.287412 | 0.894091 |
| ENSMUSG( | Slc35b1 | 1512.52 | -0.20803 | 0.112461 | 0.505997 | ENSMUSG( | Slc35b1 | 1540.185 | 0.121759 | 0.413177 | 0.917257 |
| ENSMUSG( | Slc35b2 | 1374.574 | -0.39634 | 0.008635 | 0.130416 | ENSMUSG( | Slc35b2 | 1451.83 | 0.062793 | 0.726333 | 0.966887 |
| ENSMUSG( | Slc35b3 | 390.2104 | -0.26266 | 0.111046 | 0.502387 | ENSMUSG( | Slc35b3 | 408.2447 | 0.155454 | 0.410398 | 0.917257 |
| ENSMUSG( | Slc35b4 | 2012.786 | -0.2713 | 0.040938 | 0.312769 | ENSMUSG( | Slc35b4 | 2084.626 | 0.113855 | 0.444855 | 0.921734 |
| ENSMUSG( | Slc35c1 | 613.5514 | -0.42968 | 0.004665 | 0.089972 | ENSMUSG( | Slc35c1 | 659.439 | 0.085633 | 0.619624 | 0.952896 |
| ENSMUSG( | Slc35c2 | 1829.526 | -0.0808 | 0.466507 | 0.844691 | ENSMUSG( | Slc35c2 | 1809.809 | 0.157138 | 0.278219 | 0.891468 |
| ENSMUSG( | Slc35d1 | 705.6316 | -0.2792 | 0.032817 | 0.277278 | ENSMUSG( | Slc35d1 | 728.4462 | 0.095096 | 0.515439 | 0.930939 |
| ENSMUSG( | Slc35d2 | 24.39934 | 0.194163 | 0.617315 | 0.910866 | ENSMUSG( | Slc35d2 | 24.34592 | 0.443641 | 0.280733 | 0.892356 |
| ENSMUSG( | Slc35d3 | 132.2693 | -1.53665 | 1.08E-06 | 0.000199 | ENSMUSG( | Slc35d3 | 183.4135 | 0.057495 | 0.787988 | 0.97637 |
| ENSMUSG( | Slc35e1 | 2324.86 | -0.19527 | 0.178269 | 0.615871 | ENSMUSG( | Slc35e1 | 2329.785 | 0.089891 | 0.601809 | 0.94789 |
| ENSMUSG( | Slc35e2 | 1457.085 | -0.13561 | 0.239373 | 0.687192 | ENSMUSG( | Slc35e2 | 1526.267 | 0.259608 | 0.038541 | 0.679972 |
| ENSMUSG( | Slc35e3 | 1159.096 | -0.20581 | 0.091614 | 0.465535 | ENSMUSG( | Slc35e3 | 1151.945 | 0.053592 | 0.716471 | 0.965773 |
| ENSMUSG( | Slc35e4 | 1071.784 | -0.20377 | 0.141978 | 0.559166 | ENSMUSG( | Slc35e4 | 1047.919 | 0.013895 | 0.939143 | 0.993898 |
| ENSMUSG( | Slc35f1 | 1899.998 | -0.5303 | 0.000564 | 0.02226 | ENSMUSG( | Slc35f1 | 2189.34 | 0.193467 | 0.264172 | 0.886134 |
| ENSMUSG( | Slc35f2 | 178.8487 | -0.16148 | 0.438802 | 0.830714 | ENSMUSG( | Slc35f2 | 193.7935 | 0.340142 | 0.147626 | 0.836241 |
| ENSMUSG( | Slc35f3 | 873.21 | 0.007121 | 0.955715 | 0.993874 | ENSMUSG( | Slc35f3 | 883.4043 | 0.296084 | 0.03253 | 0.63668 |
| ENSMUSG( | Slc35f4 | 342.5614 | -0.38219 | 0.021452 | 0.22362 | ENSMUSG( | Slc35f4 | 356.4726 | 0.037127 | 0.835751 | 0.981237 |
| ENSMUSG( | Slc35f5 | 825.2311 | -0.20345 | 0.187843 | 0.626399 | ENSMUSG( | Slc35f5 | 834.7183 | 0.108838 | 0.514376 | 0.930939 |
| ENSMUSG( | Slc35f6 | 1471.416 | -0.40691 | 0.010928 | 0.150598 | ENSMUSG( | Slc35f6 | 1659.388 | 0.23365 | 0.197096 | 0.862161 |
| ENSMUSG( | Slc35g1 | 161.7754 | -0.21676 | 0.229495 | 0.67735 | ENSMUSG( | Slc35g1 | 165.1715 | 0.125654 | 0.504225 | 0.927825 |
| ENSMUSG( | Slc35g2 | 393.6253 | -0.46864 | 0.004406 | 0.086676 | ENSMUSG( | Slc35g2 | 418.2827 | 0.013943 | 0.936631 | 0.993573 |
| ENSMUSG( | Slc36a1 | 2492.334 | -0.04674 | 0.76512 | 0.954127 | ENSMUSG( | Slc36a1 | 2618.912 | 0.350452 | 0.061504 | 0.736111 |
| ENSMUSG( | Slc36a2 | 21.02118 | -0.7482 | 0.080144 | NA | ENSMUSG( | Slc36a2 | 30.6086 | 0.619 | 0.105676 | 0.804969 |
| ENSMUSG( | Slc36a4 | 1914.516 | -0.08879 | 0.41701 | 0.8209 | ENSMUSG( | Slc36a4 | 1917.807 | 0.183951 | 0.150424 | 0.837173 |
| ENSMUSG( | Slc37a1 | 190.196 | -0.64343 | 5.88E-05 | 0.004481 | ENSMUSG( | Slc37a1 | 186.3707 | -0.38238 | 0.02708 | 0.603041 |
| ENSMUSG( | Slc37a2 | 254.5773 | -0.38891 | 0.013624 | 0.170376 | ENSMUSG( | Slc37a2 | 260.5085 | -0.0158 | 0.9278 | 0.99306 |
| ENSMUSG( | Slc37a3 | 2054.372 | -0.13236 | 0.282323 | 0.73186 | ENSMUSG( | Slc37a3 | 2006.909 | 0.074179 | 0.612379 | 0.951071 |
| ENSMUSG( | Slc37a4 | 818.1262 | -0.22294 | 0.053721 | 0.355948 | ENSMUSG( | Slc37a4 | 875.3289 | 0.242354 | 0.106186 | 0.804969 |
| ENSMUSG( | Slc38a1 | 4371.796 | -0.26201 | 0.054391 | 0.358917 | ENSMUSG( | Slc38a1 | 4766.934 | 0.260536 | 0.100507 | 0.801444 |
| ENSMUSG( | Slc38a10 | 3052.689 | -0.17054 | 0.1644 | 0.59535 | ENSMUSG( | Slc38a10 | 3058.839 | 0.111229 | 0.469564 | 0.923329 |
| ENSMUSG( | Slc38a11 | 27.90256 | -0.25567 | 0.628882 | 0.913816 | ENSMUSG( | Slc38a11 | 44.34364 | 1.165741 | 0.023313 | 0.585577 |
| ENSMUSG( | Slc38a2 | 3933.262 | -0.34103 | 0.009993 | 0.142653 | ENSMUSG( | Slc38a2 | 4706.873 | 0.437725 | 0.006811 | 0.341866 |
| ENSMUSG( | Slc38a3 | 5447.437 | -0.4295 | 0.002039 | 0.051756 | ENSMUSG( | Slc38a3 | 5509.416 | -0.09364 | 0.543125 | 0.938063 |
| ENSMUSG( | Slc38a4 | 30.47634 | 0.05759 | 0.870992 | 0.977951 | ENSMUSG( | Slc38a4 | 29.15176 | 0.166223 | 0.659624 | 0.956205 |
| ENSMUSG( | Slc38a5 | 135.9027 | -0.33577 | 0.156258 | 0.583988 | ENSMUSG( | Slc38a5 | 175.0346 | 0.632577 | 0.016586 | 0.519982 |
| ENSMUSG( | Slc38a6 | 380.3689 | 0.103122 | 0.407939 | 0.818167 | ENSMUSG( | Slc38a6 | 329.5851 | -0.03814 | 0.801327 | 0.97681 |
| ENSMUSG( | Slc38a7 | 1187.7 | -0.30446 | 0.023622 | 0.235222 | ENSMUSG( | Slc38a7 | 1243.291 | 0.113889 | 0.447192 | 0.922126 |
| ENSMUSG( | Slc38a9 | 742.9435 | -0.13351 | 0.311486 | 0.759296 | ENSMUSG( | Slc38a9 | 778.5875 | 0.26678 | 0.080497 | 0.77681 |
| ENSMUSG( | Slc39a1 | 2303.126 | -0.26268 | 0.029905 | 0.262335 | ENSMUSG( | Slc39a1 | 2263.149 | -0.03037 | 0.827745 | 0.980224 |
| ENSMUSG( | Slc39a10 | 4654.419 | -0.27437 | 0.099008 | 0.480953 | ENSMUSG( | Slc39a10 | 5287.62 | 0.358402 | 0.058106 | 0.728256 |
| ENSMUSG( | Slc39a11 | 693.1575 | -0.23979 | 0.055013 | 0.36102 | ENSMUSG( | Slc39a11 | 693.1457 | 0.042571 | 0.77171 | 0.974236 |
| ENSMUSG( | Slc39a12 | 4690.319 | 0.043181 | 0.72635 | 0.944223 | ENSMUSG( | Slc39a12 | 4320.081 | 0.076522 | 0.577668 | 0.943131 |
| ENSMUSG( | Slc39a13 | 912.8571 | -0.12813 | 0.333486 | 0.775602 | ENSMUSG( | Slc39a13 | 887.7069 | 0.068319 | 0.651421 | 0.95449 |
| ENSMUSG( | Slc39a14 | 920.69 | -0.35493 | 0.018878 | 0.206889 | ENSMUSG( | Slc39a14 | 1003.677 | 0.184594 | 0.267831 | 0.886134 |
| ENSMUSG( | Slc39a2 | 280.5141 | 0.510278 | 0.000896 | 0.030286 | ENSMUSG( | Slc39a2 | 234.5603 | 0.290021 | 0.213641 | 0.864421 |
| ENSMUSG( | Slc39a3 | 3303.588 | -0.18779 | 0.141615 | 0.558606 | ENSMUSG( | Slc39a3 | 3304.647 | 0.090647 | 0.541059 | 0.937647 |
| ENSMUSG( | Slc39a4 | 19.42202 | -0.21525 | 0.69092 | NA | ENSMUSG( | Slc39a4 | 19.17544 | 0.045716 | 0.925853 | 0.99306 |
| ENSMUSG( | Slc39a5 | 10.19489 | -0.47343 | 0.46334 | NA | ENSMUSG( | Slc39a5 | 11.46384 | 0.169054 | 0.782873 | 0.976106 |
| ENSMUSG( | Slc39a6 | 1440.88 | -0.18217 | 0.250649 | 0.700403 | ENSMUSG( | Slc39a6 | 1596.409 | 0.369691 | 0.039289 | 0.679972 |
| ENSMUSG( | Slc39a7 | 159.381 | 0.043057 | 0.833442 | 0.970333 | ENSMUSG( | Slc39a7 | 152.6674 | 0.196322 | 0.396014 | 0.917257 |
| ENSMUSG( | Slc39a8 | 281.6954 | -0.20061 | 0.312133 | 0.759724 | ENSMUSG( | Slc39a8 | 287.5207 | 0.142046 | 0.504707 | 0.927825 |
| ENSMUSG( | Slc39a9 | 1821.324 | -0.24112 | 0.071965 | 0.414057 | ENSMUSG( | Slc39a9 | 1904.175 | 0.165636 | 0.277537 | 0.891468 |
| ENSMUSG( | Slc3a1 | 12.41648 | 0.161396 | 0.77161 | NA | ENSMUSG( | Slc3a1 | 10.37546 | -0.0239 | 0.970459 | NA |
| ENSMUSG( | Slc3a2 | 6078.996 | -0.19641 | 0.135466 | 0.548884 | ENSMUSG( | Slc3a2 | 6329.633 | 0.192866 | 0.213946 | 0.864421 |
| ENSMUSG( | Slc40a1 | 609.2665 | -0.23344 | 0.29412 | 0.743459 | ENSMUSG( | Slc40a1 | 628.3447 | 0.14261 | 0.565563 | 0.941404 |
| ENSMUSG( | Slc41a1 | 4083.488 | -0.04804 | 0.673273 | 0.929007 | ENSMUSG( | Slc41a1 | 3993.822 | 0.155755 | 0.249696 | 0.877904 |
| ENSMUSG( | Slc41a2 | 635.3866 | -0.35747 | 0.026073 | 0.247245 | ENSMUSG( | Slc41a2 | 665.5778 | 0.071063 | 0.696887 | 0.960867 |
| ENSMUSG( | Slc41a3 | 178.2615 | -0.32578 | 0.04306 | 0.31819 | ENSMUSG( | Slc41a3 | 189.4981 | 0.14546 | 0.411845 | 0.917257 |
| ENSMUSG( | Slc43a1 | 14.4067 | -0.13307 | 0.79336 | NA | ENSMUSG( | Slc43a1 | 15.78836 | 0.403023 | 0.450346 | 0.922202 |
| ENSMUSG( | Slc43a2 | 2652.729 | -0.33789 | 0.032439 | 0.275619 | ENSMUSG( | Slc43a2 | 2697.161 | 0.003809 | 0.982975 | 0.996891 |
| ENSMUSG( | Slc43a3 | 58.69387 | -0.25067 | 0.350283 | 0.781943 | ENSMUSG( | Slc43a3 | 66.13488 | 0.35371 | 0.194238 | 0.860096 |
| ENSMUSG( | Slc44a1 | 4615.723 | -0.33175 | 0.012343 | 0.160458 | ENSMUSG( | Slc44a1 | 4864.617 | 0.109165 | 0.521474 | 0.932489 |
| ENSMUSG( | Slc44a2 | 2283.082 | -0.19231 | 0.075197 | 0.424832 | ENSMUSG( | Slc44a2 | 2291.898 | 0.095307 | 0.468497 | 0.923329 |
| ENSMUSG( | Slc44a4 | 15.68069 | -0.52962 | 0.27451 | NA | ENSMUSG( | Slc44a4 | 15.19894 | -0.33681 | 0.55698 | 0.939565 |
| ENSMUSG( | Slc44a5 | 346.8637 | -0.0912 | 0.559155 | 0.888638 | ENSMUSG( | Slc44a5 | 376.676 | 0.399366 | 0.018163 | 0.542002 |
| ENSMUSG( | Slc45a1 | 1639.577 | -0.24202 | 0.0905 | 0.461757 | ENSMUSG( | Slc45a1 | 1779.575 | 0.266685 | 0.092074 | 0.791889 |
| ENSMUSG( | Slc45a3 | 94.09164 | -0.78562 | 0.000673 | 0.025051 | ENSMUSG( | Slc45a3 | 116.2954 | 0.20419 | 0.357246 | 0.913692 |
| ENSMUSG( | Slc45a4 | 1892.213 | 0.043424 | 0.760885 | 0.952907 | ENSMUSG( | Slc45a4 | 1881.529 | 0.287871 | 0.108468 | 0.804969 |
| ENSMUSG( | Slc46a1 | 441.356 | -0.36754 | 0.035826 | 0.290986 | ENSMUSG( | Slc46a1 | 438.0002 | -0.0899 | 0.648199 | 0.95422 |
| ENSMUSG( | Slc46a3 | 258.3382 | -0.2336 | 0.22966 | 0.677538 | ENSMUSG( | Slc46a3 | 287.4154 | 0.343962 | 0.110344 | 0.807051 |
| ENSMUSG( | Slc47a1 | 593.2238 | -0.3299 | 0.02047 | 0.21574 | ENSMUSG( | Slc47a1 | 573.2118 | -0.14757 | 0.389476 | 0.917257 |
| ENSMUSG( | Slc47a2 | 13.71288 | -0.02659 | 0.963124 | NA | ENSMUSG( | Slc47a2 | 11.83785 | -0.17824 | 0.748651 | 0.96984 |
| ENSMUSG( | Slc48a1 | 9695.604 | -0.19622 | 0.06104 | 0.381679 | ENSMUSG( | Slc48a1 | 9299.398 | -0.03986 | 0.740721 | 0.968811 |
| ENSMUSG( | Slc49a3 | 51.98212 | -0.26147 | 0.326352 | 0.771301 | ENSMUSG( | Slc49a3 | 53.20104 | 0.070811 | 0.806105 | 0.977674 |
| ENSMUSG( | Slc49a4 | 4384.194 | -0.18153 | 0.112441 | 0.505997 | ENSMUSG( | Slc49a4 | 4327.808 | 0.056591 | 0.676417 | 0.95694 |
| ENSMUSG( | Slc4a1 | 100.8035 | -1.35499 | 0.081848 | 0.442044 | ENSMUSG( | Slc4a1 | 124.0034 | -0.20016 | 0.78333 | 0.976106 |
| ENSMUSG( | Slc4a10 | 3610.906 | -0.34767 | 0.00623 | 0.107962 | ENSMUSG( | Slc4a10 | 4090.612 | 0.290298 | 0.03708 | 0.673514 |
| ENSMUSG( | Slc4a11 | 30.5734 | 0.045196 | 0.896168 | 0.984747 | ENSMUSG( | Slc4a11 | 27.4373 | 0.034763 | 0.930072 | 0.99306 |
| ENSMUSG( | Slc4a1ap | 514.9285 | 0.220633 | 0.08505 | 0.449548 | ENSMUSG( | Slc4a1ap | 454.9501 | 0.135825 | 0.328646 | 0.906727 |
| ENSMUSG( | Slc4a2 | 2096.097 | -0.15787 | 0.135975 | 0.550005 | ENSMUSG( | Slc4a2 | 1989.115 | -0.03307 | 0.800839 | 0.97681 |
| ENSMUSG( | Slc4a3 | 5857.47 | 0.034618 | 0.736543 | 0.946437 | ENSMUSG( | Slc4a3 | 5496.138 | 0.119483 | 0.30563 | 0.898538 |
| ENSMUSG( | Slc4a4 | 16047.56 | -0.08229 | 0.529967 | 0.878883 | ENSMUSG( | Slc4a4 | 14593.93 | -0.08701 | 0.508882 | 0.929089 |
| ENSMUSG( | Slc4a5 | 76.56393 | 0.015235 | 0.989061 | 0.999111 | ENSMUSG( | Slc4a5 | 83.11518 | 0.46977 | 0.582895 | 0.944266 |
| ENSMUSG( | Slc4a7 | 624.1569 | -0.04436 | 0.750814 | 0.950042 | ENSMUSG( | Slc4a7 | 643.1699 | 0.298944 | 0.056654 | 0.723362 |
| ENSMUSG( | Slc4a8 | 3113.923 | -0.1935 | 0.157468 | 0.585653 | ENSMUSG( | Slc4a8 | 3160.752 | 0.126491 | 0.430515 | 0.917257 |
| ENSMUSG( | Slc50a1 | 971.1051 | -0.15666 | 0.123217 | 0.530407 | ENSMUSG( | Slc50a1 | 964.2827 | 0.097244 | 0.450298 | 0.922202 |
| ENSMUSG( | Slc52a2 | 490.9779 | -0.1713 | 0.267547 | 0.717648 | ENSMUSG( | Slc52a2 | 496.2727 | 0.138077 | 0.425598 | 0.917257 |
| ENSMUSG( | Slc52a3 | 100.9392 | 0.047217 | 0.827767 | 0.968924 | ENSMUSG( | Slc52a3 | 111.5734 | 0.575031 | 0.038989 | 0.679972 |
| ENSMUSG( | Slc5a11 | 28.97694 | -0.49017 | 0.163944 | 0.594869 | ENSMUSG( | Slc5a11 | 31.2143 | 0.072329 | 0.847595 | 0.9852 |
| ENSMUSG( | Slc5a3 | 489.747 | -0.15989 | 0.383147 | 0.803237 | ENSMUSG( | Slc5a3 | 541.1891 | 0.380884 | 0.056476 | 0.723362 |
| ENSMUSG( | Slc5a5 | 585.3984 | -0.56247 | 1.13E-05 | 0.001226 | ENSMUSG( | Slc5a5 | 593.6723 | -0.204 | 0.114961 | 0.811921 |
| ENSMUSG( | Slc5a6 | 822.308 | -0.15186 | 0.206952 | 0.649503 | ENSMUSG( | Slc5a6 | 867.2752 | 0.267306 | 0.051233 | 0.711023 |
| ENSMUSG( | Slc5a7 | 86.05923 | -1.17795 | 3.69E-06 | 0.000504 | ENSMUSG( | Slc5a7 | 116.7659 | 0.201779 | 0.366553 | 0.914703 |
| ENSMUSG( | Slc66a1 | 378.0108 | -0.32335 | 0.045008 | 0.325572 | ENSMUSG( | Slc66a1 | 395.3359 | 0.098886 | 0.553276 | 0.939565 |
| ENSMUSG( | Slc66a2 | 614.0798 | -0.10423 | 0.437252 | 0.829501 | ENSMUSG( | Slc66a2 | 605.102 | 0.12621 | 0.409299 | 0.917257 |
| ENSMUSG( | Slc66a3 | 36.28005 | -0.63326 | 0.051395 | 0.347924 | ENSMUSG( | Slc66a3 | 39.28956 | -0.04841 | 0.884545 | 0.989688 |
| ENSMUSG( | Slc6a1 | 23458.36 | -0.22557 | 0.076387 | 0.427832 | ENSMUSG( | Slc6a1 | 23734.85 | 0.087519 | 0.549131 | 0.938452 |
| ENSMUSG( | Slc6a11 | 16051.14 | -0.33727 | 0.005647 | 0.100551 | ENSMUSG( | Slc6a11 | 15270.83 | -0.20012 | 0.128614 | 0.81925 |
| ENSMUSG( | Slc6a12 | 56.28597 | 0.453209 | 0.084706 | 0.448558 | ENSMUSG( | Slc6a12 | 41.2931 | -0.15433 | 0.625211 | 0.953234 |
| ENSMUSG( | Slc6a13 | 417.418 | 0.060454 | 0.694026 | 0.934202 | ENSMUSG( | Slc6a13 | 335.0154 | -0.31181 | 0.081154 | 0.77681 |
| ENSMUSG( | Slc6a15 | 1256.056 | -0.26156 | 0.067104 | 0.400611 | ENSMUSG( | Slc6a15 | 1437.563 | 0.386221 | 0.013188 | 0.460068 |
| ENSMUSG( | Slc6a16 | 17.08506 | 0.047879 | 0.920043 | NA | ENSMUSG( | Slc6a16 | 16.89701 | 0.279682 | 0.555055 | 0.939565 |
| ENSMUSG( | Slc6a17 | 17312.55 | -0.21511 | 0.140917 | 0.557902 | ENSMUSG( | Slc6a17 | 18285.51 | 0.216072 | 0.193644 | 0.860096 |
| ENSMUSG( | Slc6a19 | 15.19279 | -0.07167 | 0.888825 | NA | ENSMUSG( | Slc6a19 | 13.0738 | -0.17329 | 0.763048 | 0.973795 |
| ENSMUSG( | Slc6a20a | 791.0229 | -0.19342 | 0.132133 | 0.544244 | ENSMUSG( | Slc6a20a | 773.4586 | 0.022757 | 0.891592 | 0.990558 |
| ENSMUSG( | Slc6a6 | 4438.442 | -0.28397 | 0.072071 | 0.414057 | ENSMUSG( | Slc6a6 | 5362.223 | 0.506446 | 0.005429 | 0.303106 |
| ENSMUSG( | Slc6a7 | 2483.011 | 0.017981 | 0.88454 | 0.982647 | ENSMUSG( | Slc6a7 | 2541.854 | 0.336249 | 0.003653 | 0.242109 |
| ENSMUSG( | Slc6a8 | 5112.335 | 0.06993 | 0.410996 | 0.818761 | ENSMUSG( | Slc6a8 | 4652.764 | 0.067552 | 0.509874 | 0.929089 |
| ENSMUSG( | Slc6a9 | 2123.466 | -0.6111 | 5.46E-07 | 0.000113 | ENSMUSG( | Slc6a9 | 2431.304 | 0.111795 | 0.384563 | 0.917257 |
| ENSMUSG( | Slc7a1 | 1395.024 | -0.15426 | 0.374751 | 0.800466 | ENSMUSG( | Slc7a1 | 1582.993 | 0.455298 | 0.018565 | 0.545482 |
| ENSMUSG( | Slc7a10 | 2958.07 | -0.19435 | 0.138603 | 0.554003 | ENSMUSG( | Slc7a10 | 2915.118 | 0.041689 | 0.784913 | 0.976285 |
| ENSMUSG( | Slc7a11 | 3729.834 | -0.01029 | 0.952904 | 0.993874 | ENSMUSG( | Slc7a11 | 3389.001 | -0.016 | 0.923156 | 0.992691 |
| ENSMUSG( | Slc7a14 | 2284.898 | -0.35047 | 0.033961 | 0.282701 | ENSMUSG( | Slc7a14 | 2655.602 | 0.357617 | 0.053227 | 0.716108 |
| ENSMUSG( | Slc7a2 | 2823.02 | 0.019576 | 0.887546 | 0.983411 | ENSMUSG( | Slc7a2 | 2788.28 | 0.244584 | 0.091377 | 0.791889 |
| ENSMUSG( | Slc7a3 | 260.4998 | -0.00194 | 0.991636 | 0.999232 | ENSMUSG( | Slc7a3 | 265.1459 | 0.306223 | 0.099092 | 0.799802 |
| ENSMUSG( | Slc7a4 | 2014.881 | -0.2016 | 0.113274 | 0.507541 | ENSMUSG( | Slc7a4 | 2041.109 | 0.113433 | 0.455257 | 0.923216 |
| ENSMUSG( | Slc7a5 | 4503.642 | 0.088653 | 0.562575 | 0.889465 | ENSMUSG( | Slc7a5 | 4636.487 | 0.418309 | 0.040829 | 0.685722 |
| ENSMUSG( | Slc7a6 | 894.5505 | 0.093282 | 0.418484 | 0.821966 | ENSMUSG( | Slc7a6 | 864.4649 | 0.256915 | 0.038718 | 0.679972 |
| ENSMUSG( | Slc7a6os | 625.4786 | -0.03361 | 0.778952 | 0.956442 | ENSMUSG( | Slc7a6os | 605.2641 | 0.141778 | 0.30652 | 0.898538 |
| ENSMUSG( | Slc7a7 | 106.6451 | 0.053749 | 0.803566 | 0.962169 | ENSMUSG( | Slc7a7 | 98.2321 | 0.103451 | 0.683188 | 0.958678 |
| ENSMUSG( | Slc7a8 | 3583.596 | -0.51833 | 5.15E-05 | 0.004052 | ENSMUSG( | Slc7a8 | 3944.565 | 0.075957 | 0.601856 | 0.94789 |
| ENSMUSG( | Slc8a1 | 3413.869 | -0.30895 | 0.073342 | 0.418269 | ENSMUSG( | Slc8a1 | 3904.634 | 0.348551 | 0.072303 | 0.763629 |
| ENSMUSG( | Slc8a2 | 11966.46 | -0.39584 | 0.004955 | 0.093376 | ENSMUSG( | Slc8a2 | 12900.26 | 0.118755 | 0.459731 | 0.923329 |
| ENSMUSG( | Slc8a3 | 917.7832 | -0.50622 | 3.30E-05 | 0.002906 | ENSMUSG( | Slc8a3 | 969.4783 | -0.03327 | 0.810803 | 0.978779 |
| ENSMUSG( | Slc8b1 | 417.1933 | -0.46569 | 0.002918 | 0.066108 | ENSMUSG( | Slc8b1 | 435.9479 | -0.02516 | 0.881848 | 0.989644 |

| | | | | | |
|---|---|---|---|---|---|
| ENSMUSG( | 2120.829 | 0.02868 | 0.75208 | 0.950417 | Sipa1l2 |
| ENSMUSG( | 1867.848 | 0.279915 | 0.013978 | 0.172788 | Sipa1l3 |
| ENSMUSG( | 19012.88 | -0.17954 | 0.186481 | 0.623869 | Sirpa |
| ENSMUSG( | 320.1077 | -0.03807 | 0.786101 | 0.958774 | Sirt1 |
| ENSMUSG( | 18656.48 | 0.086523 | 0.306404 | 0.754119 | Sirt2 |
| ENSMUSG( | 3281.708 | 0.078929 | 0.33747 | 0.777412 | Sirt3 |
| ENSMUSG( | 412.1186 | 0.137747 | 0.421294 | 0.823126 | Sirt4 |
| ENSMUSG( | 302.0701 | -0.1846 | 0.165462 | 0.596928 | Sirt5 |
| ENSMUSG( | 364.3144 | -0.01353 | 0.931648 | 0.9924 | Sirt6 |
| ENSMUSG( | 1201.929 | 0.105391 | 0.364992 | 0.79407 | Sirt7 |
| ENSMUSG( | 159.304 | -0.22446 | 0.196645 | 0.635942 | Siva1 |
| ENSMUSG( | 21.39449 | -0.62272 | 0.150221 | 0.574666 | Six1 |
| ENSMUSG( | 37.99869 | -0.53257 | 0.085926 | 0.451489 | Six2 |
| ENSMUSG( | 144.6131 | -1.54316 | 0.025033 | 0.243138 | Six3 |
| ENSMUSG( | 105.1994 | -0.87172 | 0.172798 | 0.608361 | Six3os1 |
| ENSMUSG( | 17.76594 | 0.38754 | 0.417316 | NA | Six4 |
| ENSMUSG( | 102.4662 | -0.12831 | 0.537561 | 0.882536 | Six5 |
| ENSMUSG( | 450.0013 | 0.079708 | 0.515682 | 0.872701 | Ska2 |
| ENSMUSG( | 21.48139 | 0.131815 | 0.755151 | 0.951279 | Ska3 |
| ENSMUSG( | 84.05817 | -0.5222 | 0.023898 | 0.236358 | Skap1 |
| ENSMUSG( | 484.337 | 0.028406 | 0.856579 | 0.974154 | Skap2 |
| ENSMUSG( | 6188.07 | 0.142646 | 0.055168 | 0.361542 | Ski |
| ENSMUSG( | 2593.307 | 0.230904 | 0.019287 | 0.209368 | Skic2 |
| ENSMUSG( | 729.3033 | 0.217241 | 0.047265 | 0.332598 | Skic3 |
| ENSMUSG( | 730.262 | 0.146867 | 0.201372 | 0.641362 | Skic8 |
| ENSMUSG( | 249.487 | 0.032618 | 0.820182 | 0.965888 | Skida1 |
| ENSMUSG( | 1154.087 | -0.06914 | 0.474484 | 0.849425 | Skil |
| ENSMUSG( | 8047.481 | 0.015921 | 0.878628 | 0.981431 | Skp1 |
| ENSMUSG( | 155.1987 | -0.24224 | 0.145182 | 0.565771 | Skp2 |
| ENSMUSG( | 288.2735 | 0.005175 | 0.969738 | 0.996369 | Sla |
| ENSMUSG( | 30.96369 | -0.13752 | 0.689351 | 0.933609 | Sla2 |
| ENSMUSG( | 1105.929 | -0.06667 | 0.537337 | 0.882504 | Slain1 |
| ENSMUSG( | 823.7385 | 0.081308 | 0.465977 | 0.844391 | Slain2 |
| ENSMUSG( | 14.84478 | -0.26723 | 0.582851 | NA | Slamf6 |
| ENSMUSG( | 9.997961 | -0.26723 | 0.673749 | NA | Slamf7 |
| ENSMUSG( | 11.68987 | -0.68707 | 0.361178 | NA | Slamf8 |
| ENSMUSG( | 43.21718 | -0.46411 | 0.124414 | 0.531714 | Slamf9 |
| ENSMUSG( | 291.0976 | 0.047267 | 0.743966 | 0.948497 | Slbp |
| ENSMUSG( | 88.81983 | -0.28171 | 0.29145 | 0.741366 | Slc10a3 |
| ENSMUSG( | 62.5772 | -1.63035 | 1.94E-06 | 0.000324 | Slc10a4 |
| ENSMUSG( | 170.531 | 0.263471 | 0.103745 | 0.489428 | Slc10a7 |
| ENSMUSG( | 172.2215 | -0.18545 | 0.332211 | 0.77543 | Slc11a1 |
| ENSMUSG( | 1605.935 | -0.15711 | 0.179902 | 0.617634 | Slc11a2 |
| ENSMUSG( | 2555.055 | -0.5194 | 0.001086 | 0.034804 | Slc12a2 |
| ENSMUSG( | 1858.321 | -0.07962 | 0.540589 | 0.88315 | Slc12a4 |
| ENSMUSG( | 17153.52 | -0.21914 | 0.069873 | 0.408574 | Slc12a5 |
| ENSMUSG( | 2674.907 | -0.21773 | 0.134254 | 0.547472 | Slc12a6 |
| ENSMUSG( | 311.1661 | -0.45402 | 0.004236 | 0.084545 | Slc12a7 |
| ENSMUSG( | 31.21323 | -0.51764 | 0.163643 | 0.594869 | Slc12a8 |
| ENSMUSG( | 1580.25 | -0.2105 | 0.051988 | 0.350064 | Slc12a9 |
| ENSMUSG( | 1526.992 | -0.28465 | 0.047735 | 0.333591 | Slc13a3 |
| ENSMUSG( | 551.6801 | 0.117776 | 0.384189 | 0.803686 | Slc13a4 |
| ENSMUSG( | 1516.84 | -0.16335 | 0.189005 | 0.627678 | Slc13a5 |
| ENSMUSG( | 1065.004 | -0.19779 | 0.161046 | 0.59072 | Slc14a1 |
| ENSMUSG( | 35.55809 | -0.06318 | 0.869563 | 0.977498 | Slc14a2 |
| ENSMUSG( | 2132.172 | 0.133409 | 0.258903 | 0.709814 | Slc15a2 |
| ENSMUSG( | 248.8798 | -0.24193 | 0.194453 | 0.63341 | Slc15a3 |
| ENSMUSG( | 699.7803 | -0.11393 | 0.499649 | 0.863218 | Slc15a4 |
| ENSMUSG( | 2073.007 | -0.57516 | 0.002061 | 0.052152 | Slc16a1 |
| ENSMUSG( | 103.9734 | -0.56505 | 0.012343 | 0.160458 | Slc16a10 |
| ENSMUSG( | 442.49 | 0.255061 | 0.153428 | 0.579959 | Slc16a11 |
| ENSMUSG( | 68.16646 | -0.11298 | 0.636045 | 0.914873 | Slc16a12 |
| ENSMUSG( | 205.8076 | -0.13839 | 0.385207 | 0.804048 | Slc16a13 |
| ENSMUSG( | 96.39241 | -0.08693 | 0.77484 | 0.955606 | Slc16a14 |
| ENSMUSG( | 2458.564 | -0.31755 | 0.014276 | 0.175552 | Slc16a2 |
| ENSMUSG( | 200.4832 | -0.2855 | 0.094329 | 0.471647 | Slc16a3 |
| ENSMUSG( | 141.7342 | -0.30641 | 0.279741 | 0.729135 | Slc16a4 |
| ENSMUSG( | 625.6651 | -0.30077 | 0.006319 | 0.108996 | Slc16a6 |
| ENSMUSG( | 833.4757 | -0.64144 | 0.000418 | 0.01851 | Slc16a7 |
| ENSMUSG( | 15.89648 | -0.0979 | 0.880423 | NA | Slc16a8 |
| ENSMUSG( | 253.8831 | -0.0977 | 0.522188 | 0.875186 | Slc16a9 |
| ENSMUSG( | 650.0216 | -0.41365 | 0.005539 | 0.099488 | Slc17a5 |
| ENSMUSG( | 667.8916 | -0.62444 | 4.40E-05 | 0.003588 | Slc17a6 |
| ENSMUSG( | 73345.29 | -0.18592 | 0.13153 | 0.543959 | Slc17a7 |
| ENSMUSG( | 103.496 | -0.89001 | 0.000452 | 0.019387 | Slc17a8 |
| ENSMUSG( | 18.26414 | -0.11985 | 0.791033 | NA | Slc17a9 |
| ENSMUSG( | 14.03756 | 0.206922 | 0.674341 | NA | Slc18a2 |
| ENSMUSG( | 38.39513 | -0.87633 | 0.019608 | 0.211048 | Slc18a3 |
| ENSMUSG( | 791.3843 | -0.23988 | 0.078998 | 0.434796 | Slc18b1 |
| ENSMUSG( | 478.9383 | -0.21385 | 0.26175 | 0.711988 | Slc19a1 |
| ENSMUSG( | 368.5087 | -0.38295 | 0.028584 | 0.25772 | Slc19a2 |
| ENSMUSG( | 212.2165 | 0.228764 | 0.366177 | 0.794603 | Slc19a3 |
| ENSMUSG( | 3762.219 | -0.35608 | 0.008183 | 0.125929 | Slc1a1 |
| ENSMUSG( | 501546.3 | 0.124176 | 0.311198 | 0.759247 | Slc1a2 |
| ENSMUSG( | 71268.34 | -0.01563 | 0.903528 | 0.986055 | Slc1a3 |
| ENSMUSG( | 2962.663 | -0.34215 | 0.034097 | 0.283475 | Slc1a4 |
| ENSMUSG( | 152.4252 | -0.11869 | 0.502014 | 0.864904 | Slc1a5 |
| ENSMUSG( | 242.5852 | -0.06955 | 0.662489 | 0.926183 | Slc1a6 |
| ENSMUSG( | 6506.381 | -0.18356 | 0.071541 | 0.413178 | Slc20a1 |
| ENSMUSG( | 2739.25 | -0.17796 | 0.166713 | 0.599311 | Slc20a2 |
| ENSMUSG( | 12.26465 | -1.31872 | 0.018382 | NA | Slc22a1 |
| ENSMUSG( | 15.60621 | 0.143906 | 0.754755 | NA | Slc22a12 |
| ENSMUSG( | 22.31001 | 0.319008 | 0.457454 | 0.839558 | Slc22a13b |
| ENSMUSG( | 275.8905 | -0.26416 | 0.120911 | 0.525221 | Slc22a15 |
| ENSMUSG( | 20967.33 | -0.19397 | 0.127156 | 0.534725 | Slc22a17 |
| ENSMUSG( | 25.32263 | -0.21956 | 0.573767 | 0.893584 | Slc22a18 |
| ENSMUSG( | 40.42754 | 0.027623 | 0.926875 | 0.991229 | Slc22a2 |
| ENSMUSG( | 97.61677 | -0.16845 | 0.430985 | 0.826896 | Slc22a21 |
| ENSMUSG( | 3065.937 | -0.39633 | 0.006337 | 0.108996 | Slc22a23 |
| ENSMUSG( | 83.72509 | -0.63978 | 0.003903 | 0.080172 | Slc22a3 |
| ENSMUSG( | 305.2309 | -0.06939 | 0.643648 | 0.917316 | Slc22a4 |
| ENSMUSG( | 517.5964 | -0.10314 | 0.416792 | 0.8209 | Slc22a5 |
| ENSMUSG( | 316.3686 | -0.11333 | 0.4917 | 0.860249 | Slc22a6 |
| ENSMUSG( | 13.03681 | 0.212773 | 0.709327 | NA | Slc22a7 |
| ENSMUSG( | 1131.078 | -0.01869 | 0.90024 | 0.985633 | Slc22a8 |
| ENSMUSG( | 5407.505 | -0.27471 | 0.028217 | 0.256543 | Slc23a2 |
| ENSMUSG( | 21.51445 | 0.172502 | 0.718784 | 0.941986 | Slc23a3 |
| ENSMUSG( | 15.62374 | -0.64291 | 0.189016 | NA | Slc23a4 |
| ENSMUSG( | 9939.44 | -0.62581 | 3.75E-05 | 0.003184 | Slc24a2 |
| ENSMUSG( | 6000.345 | -0.29495 | 0.015572 | 0.184473 | Slc24a3 |
| ENSMUSG( | 434.3032 | -0.22472 | 0.177344 | 0.614553 | Slc24a4 |
| ENSMUSG( | 16.84397 | -0.005 | 0.992998 | NA | Slc24a5 |
| ENSMUSG( | 2039.977 | 0.030177 | 0.760527 | 0.952907 | Slc25a1 |
| ENSMUSG( | 640.6597 | 0.040954 | 0.692433 | 0.934101 | Slc25a10 |
| ENSMUSG( | 2468.416 | 0.076322 | 0.482612 | 0.856078 | Slc25a11 |
| ENSMUSG( | 3098.999 | 0.074938 | 0.515195 | 0.872463 | Slc25a12 |
| ENSMUSG( | 56.32447 | -0.29509 | 0.277194 | 0.727175 | Slc25a13 |
| ENSMUSG( | 1164.107 | -0.11035 | 0.418721 | 0.821966 | Slc25a14 |
| ENSMUSG( | 677.3291 | -0.01979 | 0.868115 | 0.97675 | Slc25a15 |
| ENSMUSG( | 1091.312 | 0.007385 | 0.937005 | 0.993415 | Slc25a16 |
| ENSMUSG( | 944.1787 | -0.11422 | 0.268172 | 0.718372 | Slc25a17 |
| ENSMUSG( | 8328.475 | 0.150125 | 0.119597 | 0.523437 | Slc25a18 |
| ENSMUSG( | 1526.857 | 0.076086 | 0.410152 | 0.818258 | Slc25a19 |
| ENSMUSG( | 856.5412 | -0.02414 | 0.8444 | 0.972023 | Slc25a20 |
| ENSMUSG( | 56.15608 | -0.08141 | 0.794335 | 0.959799 | Slc25a21 |
| ENSMUSG( | 6327.008 | 0.156569 | 0.173643 | 0.609084 | Slc25a22 |
| ENSMUSG( | 38360.75 | 0.152656 | 0.056163 | 0.36528 | Slc25a23 |
| ENSMUSG( | 58.24348 | 0.13546 | 0.603401 | 0.904887 | Slc25a24 |
| ENSMUSG( | 1478.979 | -0.11125 | 0.290006 | 0.739839 | Slc25a25 |
| ENSMUSG( | 434.6355 | 0.026852 | 0.817972 | 0.965795 | Slc25a26 |
| ENSMUSG( | 1196.943 | 0.176776 | 0.147684 | 0.57034 | Slc25a27 |
| ENSMUSG( | 1315.937 | -0.14226 | 0.249067 | 0.698455 | Slc25a28 |
| ENSMUSG( | 377.0116 | 0.103495 | 0.450041 | 0.835344 | Slc25a29 |
| ENSMUSG( | 13597.08 | -0.01666 | 0.850468 | 0.972638 | Slc25a3 |
| ENSMUSG( | 214.305 | -0.08983 | 0.551989 | 0.886141 | Slc25a30 |
| ENSMUSG( | 460.6421 | 0.007758 | 0.94545 | 0.99365 | Slc25a32 |
| ENSMUSG( | 1405.287 | 0.164477 | 0.108468 | 0.497337 | Slc25a33 |
| ENSMUSG( | 835.9111 | -0.03551 | 0.814407 | 0.964574 | Slc25a34 |
| ENSMUSG( | 215.9011 | -0.3564 | 0.03013 | 0.263737 | Slc25a35 |
| ENSMUSG( | 1444.815 | -0.05993 | 0.540146 | 0.88315 | Slc25a36 |
| ENSMUSG( | 1075.657 | -0.20595 | 0.029149 | 0.259574 | Slc25a37 |
| ENSMUSG( | 840.9987 | 0.020361 | 0.845425 | 0.97218 | Slc25a38 |
| ENSMUSG( | 2976.401 | 0.040399 | 0.641373 | 0.916693 | Slc25a39 |
| ENSMUSG( | 18497.62 | 0.016783 | 0.856668 | 0.974154 | Slc25a4 |
| ENSMUSG( | 292.8525 | -0.19116 | 0.217918 | 0.663176 | Slc25a40 |
| ENSMUSG( | 1063.894 | 0.114104 | 0.311409 | 0.759296 | Slc25a42 |
| ENSMUSG( | 4135.622 | -0.1559 | 0.14005 | 0.556902 | Slc25a44 |
| ENSMUSG( | 207.7652 | -0.17927 | 0.315695 | 0.762671 | Slc25a45 |
| ENSMUSG( | 2088.305 | 0.041918 | 0.629871 | 0.913965 | Slc25a46 |
| ENSMUSG( | 181.4928 | 0.060843 | 0.717573 | 0.941697 | Slc25a47 |
| ENSMUSG( | 6278.192 | 0.022773 | 0.794292 | 0.959799 | Slc25a5 |
| ENSMUSG( | 2805.948 | 0.029267 | 0.746265 | 0.949289 | Slc25a51 |
| ENSMUSG( | 604.3846 | -0.08169 | 0.521721 | 0.875186 | Slc25a53 |
| ENSMUSG( | 91.76719 | -0.49639 | 0.030299 | 0.264512 | Slc26a1 |
| ENSMUSG( | 196.1071 | -0.07977 | 0.643727 | 0.917348 | Slc26a10 |
| ENSMUSG( | 220.0801 | -0.22972 | 0.21531 | 0.659922 | Slc26a11 |
| ENSMUSG( | 657.7118 | -0.68914 | 5.19E-09 | 2.10E-06 | Slc26a2 |
| ENSMUSG( | 142.1037 | 0.275625 | 0.158503 | 0.587906 | Slc26a4 |

| | | | | | |
|---|---|---|---|---|---|
| ENSMUSG( | 1397.893 | 0.11195 | 0.313001 | 0.900449 | Slc25a36 |
| ENSMUSG( | 1054.357 | 0.015099 | 0.897921 | 0.990558 | Slc25a37 |
| ENSMUSG( | 736.8469 | -0.09763 | 0.450102 | 0.922202 | Slc25a38 |
| ENSMUSG( | 2537.363 | -0.15774 | 0.172851 | 0.843702 | Slc25a39 |
| ENSMUSG( | 15844.34 | -0.16874 | 0.192371 | 0.859726 | Slc25a4 |
| ENSMUSG( | 288.5845 | 0.042416 | 0.799118 | 0.97681 | Slc25a40 |
| ENSMUSG( | 950.7986 | 0.058873 | 0.687795 | 0.959703 | Slc25a42 |
| ENSMUSG( | 4161.558 | 0.134425 | 0.267953 | 0.88628 | Slc25a44 |
| ENSMUSG( | 210.4791 | 0.147084 | 0.519579 | 0.931729 | Slc25a45 |
| ENSMUSG( | 1881.684 | 0.007383 | 0.954719 | 0.994798 | Slc25a46 |
| ENSMUSG( | 161.4045 | -0.00123 | 0.994754 | 0.998813 | Slc25a47 |
| ENSMUSG( | 5418.462 | -0.13926 | 0.246444 | 0.877904 | Slc25a5 |
| ENSMUSG( | 2599.589 | 0.074779 | 0.513043 | 0.930358 | Slc25a51 |
| ENSMUSG( | 567.5263 | 0.001706 | 0.991064 | 0.998379 | Slc25a53 |
| ENSMUSG( | 90.01437 | -0.23289 | 0.354463 | 0.912411 | Slc26a1 |
| ENSMUSG( | 193.0022 | 0.152552 | 0.543776 | 0.938063 | Slc26a10 |
| ENSMUSG( | 227.7079 | 0.152514 | 0.409553 | 0.917257 | Slc26a11 |
| ENSMUSG( | 780.0504 | 0.156551 | 0.362582 | 0.914223 | Slc26a2 |
| ENSMUSG( | 11.23522 | 1.140904 | 0.080478 | NA | Slc26a3 |
| ENSMUSG( | 128.6919 | 0.26109 | 0.215284 | 0.865167 | Slc26a4 |
| ENSMUSG( | 9.447181 | -0.20596 | 0.740682 | NA | Slc26a5 |
| ENSMUSG( | 420.1582 | 0.171822 | 0.274959 | 0.889874 | Slc26a6 |
| ENSMUSG( | 185.8256 | 0.115988 | 0.494837 | 0.927372 | Slc26a7 |
| ENSMUSG( | 202.6249 | -0.09946 | 0.550741 | 0.939187 | Slc26a8 |
| ENSMUSG( | 5059.541 | 0.026495 | 0.855661 | 0.985738 | Slc27a1 |
| ENSMUSG( | 134.9881 | 0.18456 | 0.394781 | 0.917257 | Slc27a2 |
| ENSMUSG( | 169.6909 | 0.197128 | 0.394875 | 0.917257 | Slc27a3 |
| ENSMUSG( | 3989.029 | 0.152612 | 0.379759 | 0.916995 | Slc27a4 |
| ENSMUSG( | 8.845932 | -0.56932 | 0.43862 | NA | Slc27a5 |
| ENSMUSG( | 13.87906 | 0.500369 | 0.375467 | 0.915491 | Slc27a6 |
| ENSMUSG( | 727.5854 | 0.02354 | 0.874714 | 0.989644 | Slc29a1 |
| ENSMUSG( | 478.1084 | 0.15524 | 0.336758 | 0.907972 | Slc29a2 |
| ENSMUSG( | 947.7123 | 0.367021 | 0.051533 | 0.711358 | Slc29a3 |
| ENSMUSG( | 726.6108 | 0.081452 | 0.505774 | 0.927825 | Slc29a4 |
| ENSMUSG( | 5767.175 | 0.828963 | 0.000333 | 0.049413 | Slc2a1 |
| ENSMUSG( | 88.66366 | -0.09636 | 0.763934 | 0.974 | Slc2a10 |
| ENSMUSG( | 194.4577 | 0.186193 | 0.341394 | 0.908568 | Slc2a12 |
| ENSMUSG( | 3892.365 | 0.238749 | 0.191638 | 0.858764 | Slc2a13 |
| ENSMUSG( | 5503.897 | 0.13411 | 0.351866 | 0.912093 | Slc2a3 |
| ENSMUSG( | 85.70646 | 0.8647 | 0.00032 | 0.049152 | Slc2a4 |
| ENSMUSG( | 281.8726 | 0.105397 | 0.680152 | 0.9581 | Slc2a5 |
| ENSMUSG( | 1086.194 | 0.185366 | 0.197001 | 0.862161 | Slc2a6 |
| ENSMUSG( | 816.8732 | 0.207691 | 0.23349 | 0.868507 | Slc2a8 |
| ENSMUSG( | 60.02907 | -0.02216 | 0.936471 | 0.993573 | Slc2a9 |
| ENSMUSG( | 1343.795 | 0.15373 | 0.425481 | 0.917257 | Slc30a1 |
| ENSMUSG( | 1312.46 | 0.079032 | 0.583552 | 0.944266 | Slc30a10 |
| ENSMUSG( | 4140.233 | 0.12122 | 0.284604 | 0.892412 | Slc30a3 |
| ENSMUSG( | 2106.848 | 0.322453 | 0.064538 | 0.737935 | Slc30a4 |
| ENSMUSG( | 1704.753 | 0.219865 | 0.176911 | 0.846973 | Slc30a5 |
| ENSMUSG( | 437.0563 | 0.183913 | 0.342949 | 0.909646 | Slc30a6 |
| ENSMUSG( | 650.2142 | 0.205686 | 0.312669 | 0.900085 | Slc30a7 |
| ENSMUSG( | 6522.103 | -0.04999 | 0.631216 | 0.953251 | Slc30a9 |
| ENSMUSG( | 1395.438 | 0.236395 | 0.134208 | 0.822912 | Slc31a1 |
| ENSMUSG( | 309.5666 | 0.095045 | 0.615378 | 0.951177 | Slc31a2 |
| ENSMUSG( | 3941.207 | 0.201216 | 0.16041 | 0.837724 | Slc32a1 |
| ENSMUSG( | 909.5396 | 0.350374 | 0.026788 | 0.603041 | Slc33a1 |
| ENSMUSG( | 539.1689 | 0.077439 | 0.631286 | 0.953251 | Slc35a1 |
| ENSMUSG( | 1222.047 | 0.102152 | 0.518812 | 0.930939 | Slc35a2 |
| ENSMUSG( | 512.4977 | 0.214432 | 0.153022 | 0.837724 | Slc35a3 |
| ENSMUSG( | 2043.817 | -0.01921 | 0.850711 | 0.985541 | Slc35a4 |
| ENSMUSG( | 1505.32 | 0.170553 | 0.287412 | 0.894091 | Slc35a5 |
| ENSMUSG( | 1540.185 | 0.121759 | 0.413177 | 0.917257 | Slc35b1 |
| ENSMUSG( | 1451.83 | 0.062793 | 0.726333 | 0.966887 | Slc35b2 |
| ENSMUSG( | 408.2447 | 0.155454 | 0.410398 | 0.917257 | Slc35b3 |
| ENSMUSG( | 2084.626 | 0.113855 | 0.444855 | 0.921734 | Slc35b4 |
| ENSMUSG( | 659.439 | 0.085633 | 0.619624 | 0.952896 | Slc35c1 |
| ENSMUSG( | 1809.809 | 0.157138 | 0.278219 | 0.891468 | Slc35c2 |
| ENSMUSG( | 728.4462 | 0.095096 | 0.515439 | 0.930939 | Slc35d1 |
| ENSMUSG( | 24.34592 | 0.443641 | 0.280733 | 0.892356 | Slc35d2 |
| ENSMUSG( | 183.4135 | 0.057495 | 0.787988 | 0.97637 | Slc35d3 |
| ENSMUSG( | 2329.785 | 0.089891 | 0.601809 | 0.94789 | Slc35e1 |
| ENSMUSG( | 1526.267 | 0.259608 | 0.038541 | 0.679972 | Slc35e2 |
| ENSMUSG( | 1151.945 | 0.053592 | 0.716471 | 0.965773 | Slc35e3 |
| ENSMUSG( | 1047.919 | 0.013895 | 0.939143 | 0.993898 | Slc35e4 |
| ENSMUSG( | 2189.34 | 0.193467 | 0.264172 | 0.886134 | Slc35f1 |
| ENSMUSG( | 193.7935 | 0.340142 | 0.147626 | 0.836241 | Slc35f2 |
| ENSMUSG( | 883.4043 | 0.296084 | 0.03253 | 0.63668 | Slc35f3 |
| ENSMUSG( | 356.4726 | 0.037127 | 0.835751 | 0.981237 | Slc35f4 |
| ENSMUSG( | 834.7183 | 0.108838 | 0.514376 | 0.930939 | Slc35f5 |
| ENSMUSG( | 1659.388 | 0.23365 | 0.197096 | 0.862161 | Slc35f6 |
| ENSMUSG( | 165.1715 | 0.125654 | 0.504225 | 0.927825 | Slc35g1 |
| ENSMUSG( | 418.2827 | 0.013943 | 0.936631 | 0.993573 | Slc35g2 |
| ENSMUSG( | 2618.912 | 0.350452 | 0.061504 | 0.736111 | Slc36a1 |
| ENSMUSG( | 30.6086 | 0.619 | 0.105676 | 0.804969 | Slc36a2 |
| ENSMUSG( | 1917.807 | 0.183951 | 0.150424 | 0.837173 | Slc36a4 |
| ENSMUSG( | 186.3707 | -0.38238 | 0.02708 | 0.603041 | Slc37a1 |
| ENSMUSG( | 260.5085 | -0.0158 | 0.9278 | 0.99306 | Slc37a2 |
| ENSMUSG( | 2006.909 | 0.074179 | 0.612379 | 0.951071 | Slc37a3 |
| ENSMUSG( | 875.3289 | 0.242354 | 0.106186 | 0.804969 | Slc37a4 |
| ENSMUSG( | 4766.934 | 0.260536 | 0.100507 | 0.801444 | Slc38a1 |
| ENSMUSG( | 3058.839 | 0.111229 | 0.469564 | 0.923329 | Slc38a10 |
| ENSMUSG( | 44.34364 | 1.165741 | 0.023313 | 0.585577 | Slc38a11 |
| ENSMUSG( | 4706.873 | 0.437725 | 0.006811 | 0.341866 | Slc38a2 |
| ENSMUSG( | 5509.416 | -0.09364 | 0.543125 | 0.938063 | Slc38a3 |
| ENSMUSG( | 29.15176 | 0.166223 | 0.659624 | 0.956205 | Slc38a4 |
| ENSMUSG( | 175.0346 | 0.632577 | 0.016586 | 0.519982 | Slc38a5 |
| ENSMUSG( | 329.5851 | -0.03814 | 0.801327 | 0.97681 | Slc38a6 |
| ENSMUSG( | 1243.291 | 0.113889 | 0.447192 | 0.922126 | Slc38a7 |
| ENSMUSG( | 9.954861 | 0.44283 | 0.503578 | NA | Slc38a8 |
| ENSMUSG( | 778.5875 | 0.26678 | 0.080497 | 0.77681 | Slc38a9 |
| ENSMUSG( | 2263.149 | -0.03037 | 0.827745 | 0.980224 | Slc39a1 |
| ENSMUSG( | 5287.62 | 0.358402 | 0.058106 | 0.728256 | Slc39a10 |
| ENSMUSG( | 693.1457 | 0.042571 | 0.77171 | 0.974236 | Slc39a11 |
| ENSMUSG( | 4320.081 | 0.076522 | 0.577668 | 0.943131 | Slc39a12 |
| ENSMUSG( | 887.7069 | 0.068319 | 0.651421 | 0.95449 | Slc39a13 |
| ENSMUSG( | 1003.677 | 0.184594 | 0.267831 | 0.886134 | Slc39a14 |
| ENSMUSG( | 234.5603 | 0.290021 | 0.213641 | 0.864421 | Slc39a2 |
| ENSMUSG( | 3304.647 | 0.090647 | 0.541059 | 0.937647 | Slc39a3 |
| ENSMUSG( | 19.17544 | 0.045716 | 0.925853 | 0.99306 | Slc39a4 |
| ENSMUSG( | 11.46384 | 0.169054 | 0.782873 | 0.976106 | Slc39a5 |
| ENSMUSG( | 1596.409 | 0.369691 | 0.039289 | 0.679972 | Slc39a6 |
| ENSMUSG( | 152.6674 | 0.196322 | 0.396014 | 0.917257 | Slc39a7 |
| ENSMUSG( | 287.5207 | 0.142046 | 0.504707 | 0.927825 | Slc39a8 |
| ENSMUSG( | 1904.175 | 0.165636 | 0.277537 | 0.891468 | Slc39a9 |
| ENSMUSG( | 10.37546 | -0.0239 | 0.970459 | NA | Slc3a1 |
| ENSMUSG( | 6329.633 | 0.192866 | 0.213946 | 0.864421 | Slc3a2 |
| ENSMUSG( | 628.3447 | 0.14261 | 0.565563 | 0.941404 | Slc40a1 |
| ENSMUSG( | 3993.822 | 0.155755 | 0.249696 | 0.877904 | Slc41a1 |
| ENSMUSG( | 665.5778 | 0.071063 | 0.696887 | 0.960867 | Slc41a2 |
| ENSMUSG( | 189.4981 | 0.14546 | 0.411845 | 0.917257 | Slc41a3 |
| ENSMUSG( | 15.78836 | 0.403023 | 0.450346 | 0.922202 | Slc43a1 |
| ENSMUSG( | 2697.161 | 0.003809 | 0.982975 | 0.996891 | Slc43a2 |
| ENSMUSG( | 66.13488 | 0.35371 | 0.194238 | 0.860096 | Slc43a3 |
| ENSMUSG( | 4864.617 | 0.109165 | 0.521474 | 0.932489 | Slc44a1 |
| ENSMUSG( | 2291.898 | 0.095307 | 0.468497 | 0.923329 | Slc44a2 |
| ENSMUSG( | 15.19894 | -0.33681 | 0.55698 | 0.939565 | Slc44a4 |
| ENSMUSG( | 376.676 | 0.399366 | 0.018163 | 0.542002 | Slc44a5 |
| ENSMUSG( | 1779.575 | 0.266685 | 0.092074 | 0.791889 | Slc45a1 |
| ENSMUSG( | 116.2954 | 0.20419 | 0.357246 | 0.913692 | Slc45a3 |
| ENSMUSG( | 1881.529 | 0.287871 | 0.108468 | 0.804969 | Slc45a4 |
| ENSMUSG( | 438.0002 | -0.0899 | 0.648199 | 0.95422 | Slc46a1 |
| ENSMUSG( | 287.4154 | 0.343962 | 0.110344 | 0.807051 | Slc46a3 |
| ENSMUSG( | 573.2118 | -0.14757 | 0.389476 | 0.917257 | Slc47a1 |
| ENSMUSG( | 11.83785 | -0.17824 | 0.748651 | 0.96984 | Slc47a2 |
| ENSMUSG( | 9299.398 | -0.03986 | 0.740721 | 0.968811 | Slc48a1 |
| ENSMUSG( | 53.20104 | 0.070811 | 0.806105 | 0.977674 | Slc49a3 |
| ENSMUSG( | 4327.808 | 0.056591 | 0.676417 | 0.95694 | Slc49a4 |
| ENSMUSG( | 124.0034 | -0.20016 | 0.78333 | 0.976106 | Slc4a1 |
| ENSMUSG( | 4090.612 | 0.290298 | 0.03708 | 0.673514 | Slc4a10 |
| ENSMUSG( | 27.4373 | 0.034763 | 0.930072 | 0.99306 | Slc4a11 |
| ENSMUSG( | 454.9501 | 0.135825 | 0.328646 | 0.906727 | Slc4a1ap |
| ENSMUSG( | 1989.115 | -0.03307 | 0.800839 | 0.97681 | Slc4a2 |
| ENSMUSG( | 5496.138 | 0.119483 | 0.30563 | 0.898538 | Slc4a3 |
| ENSMUSG( | 14593.93 | -0.08701 | 0.508882 | 0.929089 | Slc4a4 |
| ENSMUSG( | 83.11518 | 0.46977 | 0.582895 | 0.944266 | Slc4a5 |
| ENSMUSG( | 643.1699 | 0.298944 | 0.056654 | 0.723362 | Slc4a7 |
| ENSMUSG( | 3160.752 | 0.126491 | 0.430515 | 0.917257 | Slc4a8 |
| ENSMUSG( | 964.2827 | 0.097244 | 0.450298 | 0.922202 | Slc50a1 |
| ENSMUSG( | 496.2727 | 0.138077 | 0.425598 | 0.917257 | Slc52a2 |
| ENSMUSG( | 111.5734 | 0.575031 | 0.038989 | 0.679972 | Slc52a3 |
| ENSMUSG( | 31.2143 | 0.072329 | 0.847595 | 0.9852 | Slc5a11 |
| ENSMUSG( | 541.1891 | 0.380884 | 0.056476 | 0.723362 | Slc5a3 |
| ENSMUSG( | 593.6723 | -0.204 | 0.114961 | 0.811921 | Slc5a5 |
| ENSMUSG( | 867.2752 | 0.267306 | 0.051233 | 0.711023 | Slc5a6 |
| ENSMUSG( | 116.7659 | 0.201779 | 0.366553 | 0.914703 | Slc5a7 |
| ENSMUSG( | 395.3359 | 0.098886 | 0.553276 | 0.939565 | Slc66a1 |
| ENSMUSG( | 605.102 | 0.12621 | 0.409299 | 0.917257 | Slc66a2 |
| ENSMUSG( | 39.28956 | -0.04841 | 0.884545 | 0.989688 | Slc66a3 |

| | | | | | | | | | | | |
|---|---|---|---|---|---|---|---|---|---|---|---|
| ENSMUSG( | Slc9a1 | 1830.537 | -0.29641 | 0.020252 | 0.214698 | ENSMUSG( | Slc9a1 | 1935.511 | 0.148931 | 0.310967 | 0.899422 |
| ENSMUSG( | Slc9a2 | 380.8778 | 0.000489 | 0.997127 | 0.999565 | ENSMUSG( | Slc9a2 | 381.5225 | 0.268241 | 0.135993 | 0.826859 |
| ENSMUSG( | Slc9a3 | 25.74454 | 0.590194 | 0.15528 | 0.583155 | ENSMUSG( | Slc9a3 | 21.60044 | 0.420705 | 0.337423 | 0.907972 |
| ENSMUSG( | Slc9a4 | 68.68419 | 0.202853 | 0.414898 | 0.820184 | ENSMUSG( | Slc9a4 | 61.72377 | 0.161851 | 0.574996 | 0.942912 |
| ENSMUSG( | Slc9a5 | 1372.877 | -0.05491 | 0.622125 | 0.911432 | ENSMUSG( | Slc9a5 | 1359.853 | 0.184322 | 0.126506 | 0.815605 |
| ENSMUSG( | Slc9a6 | 3468.971 | -0.3086 | 0.013697 | 0.171028 | ENSMUSG( | Slc9a6 | 3673.313 | 0.139735 | 0.361351 | 0.914223 |
| ENSMUSG( | Slc9a7 | 249.5391 | 0.0039 | 0.984444 | 0.99781 | ENSMUSG( | Slc9a7 | 229.1618 | 0.040607 | 0.857781 | 0.985884 |
| ENSMUSG( | Slc9a8 | 1207.83 | -0.11329 | 0.372061 | 0.798768 | ENSMUSG( | Slc9a8 | 1210.373 | 0.164904 | 0.261678 | 0.884926 |
| ENSMUSG( | Slc9a9 | 793.5239 | -0.21877 | 0.123962 | 0.531451 | ENSMUSG( | Slc9a9 | 817.2281 | 0.1443 | 0.354618 | 0.912411 |
| ENSMUSG( | Slc9b2 | 139.4562 | -0.06584 | 0.764527 | 0.954055 | ENSMUSG( | Slc9b2 | 122.3894 | -0.18676 | 0.473307 | 0.923329 |
| ENSMUSG( | Slco1a4 | 2094.03 | -0.04645 | 0.814788 | 0.964668 | ENSMUSG( | Slco1a4 | 2605.913 | 0.762504 | 0.065189 | 0.74247 |
| ENSMUSG( | Slco1b2 | 16.07486 | -0.44645 | 0.341271 | NA | ENSMUSG( | Slco1b2 | 15.25053 | -0.28112 | 0.586441 | 0.944975 |
| ENSMUSG( | Slco1c1 | 3937.37 | -0.17982 | 0.333526 | 0.775602 | ENSMUSG( | Slco1c1 | 4338.724 | 0.35951 | 0.099364 | 0.800296 |
| ENSMUSG( | Slco2a1 | 92.61515 | -0.3672 | 0.10134 | 0.484996 | ENSMUSG( | Slco2a1 | 91.33031 | -0.10267 | 0.685497 | 0.958678 |
| ENSMUSG( | Slco2b1 | 1334.328 | 0.013181 | 0.938917 | 0.993415 | ENSMUSG( | Slco2b1 | 1406.287 | 0.412501 | 0.067952 | 0.75212 |
| ENSMUSG( | Slco3a1 | 944.8162 | -0.23391 | 0.136271 | 0.550578 | ENSMUSG( | Slco3a1 | 1097.782 | 0.449035 | 0.011634 | 0.440174 |
| ENSMUSG( | Slco4a1 | 162.4304 | -0.52544 | 0.002531 | 0.059664 | ENSMUSG( | Slco4a1 | 157.571 | -0.29346 | 0.162766 | 0.837724 |
| ENSMUSG( | Slco4c1 | 56.68072 | -0.61508 | 0.037856 | 0.30067 | ENSMUSG( | Slco4c1 | 75.04026 | 0.510459 | 0.081539 | 0.77681 |
| ENSMUSG( | Slco5a1 | 63.92293 | -0.83931 | 0.000714 | 0.025843 | ENSMUSG( | Slco5a1 | 78.0326 | 0.111844 | 0.648662 | 0.95422 |
| ENSMUSG( | Slf1 | 338.445 | -0.07747 | 0.575678 | 0.894836 | ENSMUSG( | Slf1 | 335.0616 | 0.166186 | 0.27116 | 0.888062 |
| ENSMUSG( | Slf2 | 1454.01 | 0.00808 | 0.932323 | 0.992515 | ENSMUSG( | Slf2 | 1381.745 | 0.124584 | 0.315144 | 0.901306 |
| ENSMUSG( | Slfn2 | 67.00886 | -0.30381 | 0.243178 | 0.69052 | ENSMUSG( | Slfn2 | 57.41389 | -0.5039 | 0.088062 | 0.78663 |
| ENSMUSG( | Slfn3 | 10.88428 | 0.750173 | 0.229238 | NA | ENSMUSG( | Slfn3 | 11.7741 | 1.17308 | 0.081719 | 0.77681 |
| ENSMUSG( | Slfn4 | 10.08023 | 0.077512 | 0.925698 | NA | ENSMUSG( | Slfn4 | 9.423986 | 0.091442 | 0.914344 | NA |
| ENSMUSG( | Slfn5 | 341.21 | 0.107825 | 0.625565 | 0.912249 | ENSMUSG( | Slfn5 | 349.1766 | 0.428281 | 0.10629 | 0.804969 |
| ENSMUSG( | Slfn8 | 71.71834 | -0.31998 | 0.157502 | 0.585653 | ENSMUSG( | Slfn8 | 64.3907 | -0.38376 | 0.199169 | 0.863016 |
| ENSMUSG( | Slfn9 | 61.00641 | -0.18268 | 0.540577 | 0.88315 | ENSMUSG( | Slfn9 | 54.65649 | -0.25583 | 0.380805 | 0.917202 |
| ENSMUSG( | Slirp | 617.4286 | -0.03367 | 0.757677 | 0.951582 | ENSMUSG( | Slirp | 527.0852 | -0.23799 | 0.127469 | 0.817604 |
| ENSMUSG( | Slit1 | 1772.645 | 0.056575 | 0.769493 | 0.954674 | ENSMUSG( | Slit1 | 1814.201 | 0.377519 | 0.092277 | 0.79275 |
| ENSMUSG( | Slit2 | 1031.993 | -0.38265 | 0.002734 | 0.063356 | ENSMUSG( | Slit2 | 1049.134 | -0.03653 | 0.812034 | 0.979148 |
| ENSMUSG( | Slit3 | 710.5014 | -0.29019 | 0.063717 | 0.390618 | ENSMUSG( | Slit3 | 779.3643 | 0.25709 | 0.191735 | 0.858764 |
| ENSMUSG( | Slitrk1 | 2083.275 | -0.65082 | 0.000159 | 0.008938 | ENSMUSG( | Slitrk1 | 2378.856 | 0.078675 | 0.685324 | 0.958678 |
| ENSMUSG( | Slitrk2 | 830.137 | -0.5148 | 0.004248 | 0.084545 | ENSMUSG( | Slitrk2 | 997.5552 | 0.318492 | 0.114562 | 0.811703 |
| ENSMUSG( | Slitrk3 | 1554.717 | -0.39784 | 0.003409 | 0.073541 | ENSMUSG( | Slitrk3 | 1828.653 | 0.34962 | 0.028706 | 0.614171 |
| ENSMUSG( | Slitrk4 | 537.6 | -0.57024 | 0.000576 | 0.022554 | ENSMUSG( | Slitrk4 | 669.1883 | 0.366602 | 0.065339 | 0.743695 |
| ENSMUSG( | Slitrk5 | 1726.015 | -0.37405 | 0.017523 | 0.198483 | ENSMUSG( | Slitrk5 | 2008.013 | 0.340882 | 0.063845 | 0.736482 |
| ENSMUSG( | Slitrk6 | 19.65093 | -0.94285 | 0.027218 | NA | ENSMUSG( | Slitrk6 | 25.51637 | 0.235962 | 0.559635 | 0.939807 |
| ENSMUSG( | Slk | 2811.602 | -0.07442 | 0.355977 | 0.788009 | ENSMUSG( | Slk | 2645.397 | 0.019623 | 0.84806 | 0.985242 |
| ENSMUSG( | Slmap | 2243.457 | -0.03405 | 0.722541 | 0.943752 | ENSMUSG( | Slmap | 2138.372 | 0.093973 | 0.380896 | 0.917202 |
| ENSMUSG( | Slpi | 22.28206 | -0.3704 | 0.405939 | 0.81652 | ENSMUSG( | Slpi | 22.05155 | -0.09972 | 0.820119 | 0.979198 |
| ENSMUSG( | Sltm | 2390.501 | 0.1312 | 0.166979 | 0.599655 | ENSMUSG( | Sltm | 2091.684 | 0.015032 | 0.898924 | 0.990558 |
| ENSMUSG( | Slu7 | 1574.56 | 0.083873 | 0.385986 | 0.804623 | ENSMUSG( | Slu7 | 1352.127 | -0.08739 | 0.424295 | 0.917257 |
| ENSMUSG( | Slx1b | 523.9121 | -0.15577 | 0.195039 | 0.634458 | ENSMUSG( | Slx1b | 493.5552 | -0.05456 | 0.674747 | 0.95694 |
| ENSMUSG( | Slx4 | 806.2251 | 0.053637 | 0.67784 | 0.930542 | ENSMUSG( | Slx4 | 734.082 | 0.056828 | 0.707501 | 0.964029 |
| ENSMUSG( | Slx4ip | 212.3354 | 0.057386 | 0.700269 | 0.937391 | ENSMUSG( | Slx4ip | 188.9508 | -0.01655 | 0.927856 | 0.99306 |
| ENSMUSG( | Slx9 | 1021.869 | 0.06544 | 0.515157 | 0.872463 | ENSMUSG( | Slx9 | 890.6547 | -0.06535 | 0.59675 | 0.946541 |
| ENSMUSG( | Smad1 | 510.0055 | 0.008486 | 0.939125 | 0.993415 | ENSMUSG( | Smad1 | 468.8307 | 0.025897 | 0.864464 | 0.986591 |
| ENSMUSG( | Smad2 | 851.495 | 0.03897 | 0.716774 | 0.941697 | ENSMUSG( | Smad2 | 771.9011 | 0.022856 | 0.850116 | 0.985538 |
| ENSMUSG( | Smad3 | 1571.889 | 0.104436 | 0.279691 | 0.729119 | ENSMUSG( | Smad3 | 1409.215 | 0.058954 | 0.625814 | 0.953234 |
| ENSMUSG( | Smad4 | 1214.203 | 0.05501 | 0.618215 | 0.911006 | ENSMUSG( | Smad4 | 1122.983 | 0.094791 | 0.436559 | 0.919879 |
| ENSMUSG( | Smad5 | 2343.265 | 0.036669 | 0.778917 | 0.956442 | ENSMUSG( | Smad5 | 2071.393 | -0.05319 | 0.697502 | 0.960867 |
| ENSMUSG( | Smad6 | 60.73465 | 0.077445 | 0.797348 | 0.960507 | ENSMUSG( | Smad6 | 82.94553 | 1.065865 | 0.000101 | 0.020718 |
| ENSMUSG( | Smad7 | 599.0351 | -0.0617 | 0.596442 | 0.903295 | ENSMUSG( | Smad7 | 575.8441 | 0.097364 | 0.464411 | 0.923329 |
| ENSMUSG( | Smad9 | 426.2398 | -0.25097 | 0.04266 | 0.316769 | ENSMUSG( | Smad9 | 402.5652 | -0.14868 | 0.349197 | 0.911652 |
| ENSMUSG( | Smagp | 79.43057 | 0.205279 | 0.384003 | 0.803646 | ENSMUSG( | Smagp | 68.3028 | 0.074554 | 0.800966 | 0.97681 |
| ENSMUSG( | Smap1 | 2198.292 | -0.03945 | 0.64148 | 0.916693 | ENSMUSG( | Smap1 | 1978.602 | -0.07845 | 0.560694 | 0.940254 |
| ENSMUSG( | Smap2 | 3244.89 | 0.086321 | 0.423355 | 0.823662 | ENSMUSG( | Smap2 | 2763.899 | -0.11018 | 0.390778 | 0.917257 |
| ENSMUSG( | Smarca1 | 640.9804 | 0.145439 | 0.175803 | 0.611006 | ENSMUSG( | Smarca1 | 585.9376 | 0.152278 | 0.257148 | 0.883082 |
| ENSMUSG( | Smarca2 | 8357.507 | 0.177968 | 0.030783 | 0.266861 | ENSMUSG( | Smarca2 | 7308.033 | 0.06421 | 0.502489 | 0.927825 |
| ENSMUSG( | Smarca4 | 5302.722 | 0.178408 | 0.039163 | 0.304806 | ENSMUSG( | Smarca4 | 4419.779 | -0.07429 | 0.494183 | 0.927372 |
| ENSMUSG( | Smarca5 | 1578.064 | 0.164937 | 0.15757 | 0.585653 | ENSMUSG( | Smarca5 | 1398.271 | 0.084301 | 0.508995 | 0.929089 |
| ENSMUSG( | Smarcad1 | 703.0274 | 0.034911 | 0.73896 | 0.947768 | ENSMUSG( | Smarcad1 | 653.2215 | 0.086009 | 0.532599 | 0.934795 |
| ENSMUSG( | Smarcal1 | 986.6841 | 0.168252 | 0.087821 | 0.454663 | ENSMUSG( | Smarcal1 | 879.1671 | 0.110475 | 0.382244 | 0.917257 |
| ENSMUSG( | Smarcb1 | 1512.83 | 0.043101 | 0.672059 | 0.9288 | ENSMUSG( | Smarcb1 | 1291.727 | -0.14877 | 0.216841 | 0.865436 |
| ENSMUSG( | Smarcc1 | 2144.678 | 0.043939 | 0.607723 | 0.906878 | ENSMUSG( | Smarcc1 | 1808.23 | -0.18738 | 0.09874 | 0.798438 |
| ENSMUSG( | Smarcc2 | 14106.69 | 0.05154 | 0.503864 | 0.866426 | ENSMUSG( | Smarcc2 | 12196.06 | -0.10099 | 0.299432 | 0.898538 |
| ENSMUSG( | Smarcd1 | 6231.497 | 0.017056 | 0.858766 | 0.974707 | ENSMUSG( | Smarcd1 | 5516.334 | -0.06734 | 0.563011 | 0.941158 |
| ENSMUSG( | Smarcd2 | 525.8302 | 0.081568 | 0.599441 | 0.903801 | ENSMUSG( | Smarcd2 | 546.0174 | 0.435831 | 0.008971 | 0.396825 |
| ENSMUSG( | Smarcd3 | 2990.431 | 0.035981 | 0.672439 | 0.929007 | ENSMUSG( | Smarcd3 | 2728.518 | 0.038906 | 0.716222 | 0.965692 |
| ENSMUSG( | Smarce1 | 1436.151 | -0.01212 | 0.899486 | 0.985633 | ENSMUSG( | Smarce1 | 1250.613 | -0.15006 | 0.225647 | 0.868468 |
| ENSMUSG( | Smc1a | 2055.599 | -0.03949 | 0.691776 | 0.934101 | ENSMUSG( | Smc1a | 1913.771 | 0.025773 | 0.850796 | 0.985541 |
| ENSMUSG( | Smc2 | 225.9491 | -0.05353 | 0.723262 | 0.943761 | ENSMUSG( | Smc2 | 223.6939 | 0.176069 | 0.326754 | 0.905371 |
| ENSMUSG( | Smc2os | 24.15828 | -0.19288 | 0.664948 | 0.926824 | ENSMUSG( | Smc2os | 18.19319 | -0.88776 | 0.092034 | 0.791889 |
| ENSMUSG( | Smc3 | 1756.969 | 0.20069 | 0.029715 | 0.261211 | ENSMUSG( | Smc3 | 1528.613 | 0.073583 | 0.492187 | 0.927315 |
| ENSMUSG( | Smc4 | 344.6831 | 0.086292 | 0.567867 | 0.892002 | ENSMUSG( | Smc4 | 338.1591 | 0.285385 | 0.071726 | 0.762854 |
| ENSMUSG( | Smc5 | 796.3666 | 0.080533 | 0.467139 | 0.844819 | ENSMUSG( | Smc5 | 727.9975 | 0.092524 | 0.473881 | 0.923329 |
| ENSMUSG( | Smc6 | 1272.597 | 0.103617 | 0.271013 | 0.721337 | ENSMUSG( | Smc6 | 1169.855 | 0.125932 | 0.276466 | 0.890889 |
| ENSMUSG( | Smchd1 | 911.3996 | 0.061992 | 0.597355 | 0.903694 | ENSMUSG( | Smchd1 | 822.1357 | 0.036232 | 0.776411 | 0.975717 |
| ENSMUSG( | Smco3 | 106.9802 | -0.02113 | 0.914339 | 0.988483 | ENSMUSG( | Smco3 | 101.1829 | 0.086685 | 0.683375 | 0.958678 |
| ENSMUSG( | Smco4 | 59.73276 | 0.053758 | 0.834544 | 0.970344 | ENSMUSG( | Smco4 | 58.6889 | 0.251709 | 0.35918 | 0.914061 |
| ENSMUSG( | Smcr8 | 804.3925 | 0.164287 | 0.119229 | 0.523036 | ENSMUSG( | Smcr8 | 743.8931 | 0.206723 | 0.100275 | 0.801311 |
| ENSMUSG( | Smdt1 | 3232.963 | -0.08036 | 0.419981 | 0.822442 | ENSMUSG( | Smdt1 | 2712.717 | -0.34134 | 0.020264 | 0.563435 |
| ENSMUSG( | Smg1 | 4079.569 | -0.16946 | 0.062992 | 0.388234 | ENSMUSG( | Smg1 | 4202.346 | 0.185691 | 0.121539 | 0.813384 |
| ENSMUSG( | Smg5 | 3053.834 | 0.005063 | 0.95734 | 0.993874 | ENSMUSG( | Smg5 | 2763.932 | -0.01293 | 0.916343 | 0.992529 |
| ENSMUSG( | Smg6 | 1478.606 | -0.00636 | 0.942622 | 0.993415 | ENSMUSG( | Smg6 | 1326.726 | -0.04978 | 0.659479 | 0.956205 |
| ENSMUSG( | Smg7 | 2661.586 | 0.188628 | 0.021846 | 0.225699 | ENSMUSG( | Smg7 | 2318.182 | 0.062815 | 0.566675 | 0.941777 |
| ENSMUSG( | Smg8 | 310.547 | 0.065992 | 0.693265 | 0.934202 | ENSMUSG( | Smg8 | 299.3386 | 0.233667 | 0.237438 | 0.872173 |
| ENSMUSG( | Smg9 | 1058.63 | 0.087506 | 0.459337 | 0.840244 | ENSMUSG( | Smg9 | 880.2239 | -0.18323 | 0.22079 | 0.866104 |
| ENSMUSG( | Smim1 | 1015.488 | -0.04812 | 0.648711 | 0.920025 | ENSMUSG( | Smim1 | 895.0714 | -0.15178 | 0.233157 | 0.868507 |
| ENSMUSG( | Smim10l1 | 3944.648 | 0.011435 | 0.92664 | 0.991229 | ENSMUSG( | Smim10l1 | 3505.989 | -0.0645 | 0.671997 | 0.95694 |
| ENSMUSG( | Smim10l2a | 927.3868 | 0.095479 | 0.399698 | 0.813337 | ENSMUSG( | Smim10l2a | 776.3873 | -0.15211 | 0.220953 | 0.866104 |
| ENSMUSG( | Smim11 | 282.5267 | -0.03581 | 0.7931 | 0.959505 | ENSMUSG( | Smim11 | 253.9778 | -0.08245 | 0.631442 | 0.953251 |
| ENSMUSG( | Smim12 | 1090.039 | -0.23082 | 0.019161 | 0.208619 | ENSMUSG( | Smim12 | 1016.898 | -0.15747 | 0.161921 | 0.837724 |
| ENSMUSG( | Smim13 | 6416.138 | -0.05646 | 0.523238 | 0.875628 | ENSMUSG( | Smim13 | 5912.296 | -0.02341 | 0.834685 | 0.981237 |
| ENSMUSG( | Smim14 | 3607.516 | -0.04541 | 0.619557 | 0.911432 | ENSMUSG( | Smim14 | 3227.826 | -0.10154 | 0.392514 | 0.917257 |
| ENSMUSG( | Smim15 | 1360.173 | 0.028058 | 0.828281 | 0.968966 | ENSMUSG( | Smim15 | 1146.031 | -0.21068 | 0.185885 | 0.855305 |
| ENSMUSG( | Smim17 | 188.849 | -0.12003 | 0.539703 | 0.88315 | ENSMUSG( | Smim17 | 169.3043 | -0.1809 | 0.411956 | 0.917257 |
| ENSMUSG( | Smim18 | 123.8962 | -0.36933 | 0.060015 | 0.378891 | ENSMUSG( | Smim18 | 127.1588 | -0.01448 | 0.946453 | 0.994798 |
| ENSMUSG( | Smim19 | 761.1088 | -0.22762 | 0.054795 | 0.360194 | ENSMUSG( | Smim19 | 714.5732 | -0.13413 | 0.34381 | 0.910159 |
| ENSMUSG( | Smim20 | 822.1227 | 0.006321 | 0.956959 | 0.993874 | ENSMUSG( | Smim20 | 671.1127 | -0.33353 | 0.00818 | 0.388416 |
| ENSMUSG( | Smim24 | 50.40918 | 0.292976 | 0.291162 | 0.741182 | ENSMUSG( | Smim24 | 42.63725 | 0.079577 | 0.801723 | 0.97681 |
| ENSMUSG( | Smim26 | 305.5624 | -0.31964 | 0.016035 | 0.187923 | ENSMUSG( | Smim26 | 300.2728 | -0.09033 | 0.550982 | 0.939232 |
| ENSMUSG( | Smim27 | 115.9169 | 0.132667 | 0.527547 | 0.877641 | ENSMUSG( | Smim27 | 99.26977 | -0.06431 | 0.795888 | 0.97681 |
| ENSMUSG( | Smim29 | 1744.381 | -0.05319 | 0.527253 | 0.877492 | ENSMUSG( | Smim29 | 1515.255 | -0.19875 | 0.067131 | 0.75212 |
| ENSMUSG( | Smim3 | 858.4561 | 0.75385 | 9.04E-14 | 9.77E-11 | ENSMUSG( | Smim3 | 704.7175 | 0.505114 | 4.40E-05 | 0.01151 |
| ENSMUSG( | Smim30 | 754.6061 | 0.063412 | 0.600731 | 0.90443 | ENSMUSG( | Smim30 | 683.4312 | 0.040373 | 0.793648 | 0.97681 |
| ENSMUSG( | Smim32 | 107.8187 | -0.0497 | 0.800507 | 0.961745 | ENSMUSG( | Smim32 | 102.1007 | 0.084664 | 0.727092 | 0.966887 |
| ENSMUSG( | Smim35 | 18.00039 | -0.32129 | 0.533149 | NA | ENSMUSG( | Smim35 | 17.75893 | -0.10633 | 0.837639 | 0.981335 |
| ENSMUSG( | Smim36 | 204.6323 | 0.011654 | 0.939822 | 0.993415 | ENSMUSG( | Smim36 | 179.3425 | -0.11842 | 0.536338 | 0.93705 |
| ENSMUSG( | Smim38 | 11.35281 | 0.342071 | 0.583125 | NA | ENSMUSG( | Smim38 | 9.825458 | 0.152849 | 0.812664 | NA |
| ENSMUSG( | Smim43 | 694.0499 | 0.141082 | 0.298725 | 0.747452 | ENSMUSG( | Smim43 | 613.5455 | 0.058402 | 0.686223 | 0.95896 |
| ENSMUSG( | Smim45 | 5988.451 | 0.094223 | 0.335707 | 0.777 | ENSMUSG( | Smim45 | 5004.06 | -0.16072 | 0.261164 | 0.884926 |
| ENSMUSG( | Smim5 | 12.53214 | -0.8362 | 0.114317 | NA | ENSMUSG( | Smim5 | 17.86212 | 0.532586 | 0.249593 | 0.877904 |
| ENSMUSG( | Smim7 | 3064.426 | -0.10093 | 0.329474 | 0.773311 | ENSMUSG( | Smim7 | 2837.746 | -0.05178 | 0.657032 | 0.955625 |
| ENSMUSG( | Smim8 | 383.0485 | 0.000662 | 0.996909 | 0.999505 | ENSMUSG( | Smim8 | 322.4966 | -0.24831 | 0.208177 | 0.864412 |
| ENSMUSG( | Smn1 | 304.3244 | 0.093148 | 0.549611 | 0.885944 | ENSMUSG( | Smn1 | 264.514 | -0.04554 | 0.800339 | 0.97681 |
| ENSMUSG( | Smndc1 | 669.8251 | 0.000237 | 0.998575 | 0.99989 | ENSMUSG( | Smndc1 | 599.9739 | -0.05883 | 0.736605 | 0.967344 |
| ENSMUSG( | Smo | 1526.569 | 0.166596 | 0.154429 | 0.581463 | ENSMUSG( | Smo | 1340.58 | 0.0643 | 0.612182 | 0.951071 |
| ENSMUSG( | Smoc1 | 816.5548 | -0.3606 | 0.008303 | 0.127252 | ENSMUSG( | Smoc1 | 929.3532 | 0.293991 | 0.043497 | 0.690304 |
| ENSMUSG( | Smoc2 | 453.6695 | -0.12744 | 0.415047 | 0.820184 | ENSMUSG( | Smoc2 | 468.4066 | 0.237675 | 0.209044 | 0.864412 |
| ENSMUSG( | Smox | 3691.581 | 0.272526 | 0.002176 | 0.054296 | ENSMUSG( | Smox | 3056.603 | 0.008153 | 0.938059 | 0.993731 |
| ENSMUSG( | Smpd1 | 4433.55 | -0.18093 | 0.205666 | 0.64712 | ENSMUSG( | Smpd1 | 4509.998 | 0.144405 | 0.393584 | 0.917257 |
| ENSMUSG( | Smpd2 | 920.4046 | -0.08048 | 0.511762 | 0.871106 | ENSMUSG( | Smpd2 | 853.8553 | -0.02231 | 0.878333 | 0.989644 |
| ENSMUSG( | Smpd3 | 3106.771 | -0.43544 | 0.002915 | 0.066108 | ENSMUSG( | Smpd3 | 3523.466 | 0.225905 | 0.184133 | 0.852618 |
| ENSMUSG( | Smpd4 | 1897.713 | 0.278373 | 0.088494 | 0.456254 | ENSMUSG( | Smpd4 | 1571.267 | 0.012687 | 0.946985 | 0.994798 |
| ENSMUSG( | Smpd5 | 27.8632 | -0.34909 | 0.344178 | 0.779153 | ENSMUSG( | Smpd5 | 32.68699 | 0.403489 | 0.273458 | 0.889703 |
| ENSMUSG( | Smpdl3a | 1949.036 | -0.18719 | 0.204061 | 0.644572 | ENSMUSG( | Smpdl3a | 1898.894 | 0.016352 | 0.918707 | 0.992529 |
| ENSMUSG( | Smpdl3b | 68.25841 | -0.76213 | 0.002378 | 0.057221 | ENSMUSG( | Smpdl3b | 94.76502 | 0.518911 | 0.01889 | 0.548832 |
| ENSMUSG( | Sms | 1728.753 | 0.077258 | 0.490264 | 0.859319 | ENSMUSG( | Sms | 1517.773 | -0.03345 | 0.814892 | 0.979198 |
| ENSMUSG( | Smtn | 279.5371 | 0.149299 | 0.390897 | 0.80826 | ENSMUSG( | Smtn | 294.9043 | 0.542068 | 0.02013 | 0.562379 |
| ENSMUSG( | Smtnl2 | 31.3206 | -0.74941 | 0.029324 | 0.259647 | ENSMUSG( | Smtnl2 | 34.25369 | -0.16448 | 0.64249 | 0.953903 |
| ENSMUSG( | Smu1 | 1238.463 | -0.00096 | 0.993647 | 0.999369 | ENSMUSG( | Smu1 | 1084.035 | -0.12208 | 0.365851 | 0.914623 |
| ENSMUSG( | Smug1 | 322.6168 | 0.030333 | 0.813919 | 0.964574 | ENSMUSG( | Smug1 | 303.4347 | 0.123931 | 0.400572 | 0.917257 |
| ENSMUSG( | Smurf1 | 2114.737 | 0.026809 | 0.78283 | 0.957558 | ENSMUSG( | Smurf1 | 1827.053 | -0.12777 | 0.253156 | 0.88018 |
| ENSMUSG( | Smurf2 | 1444.853 | -0.04617 | 0.637827 | 0.91586 | ENSMUSG( | Smurf2 | 1286.838 | -0.1157 | 0.329866 | 0.907107 |
| ENSMUSG( | Smyd1 | 411.5335 | -0.32433 | 0.090373 | 0.461669 | ENSMUSG( | Smyd1 | 330.3068 | -0.79126 | 0.000106 | 0.021511 |
| ENSMUSG( | Smyd2 | 1590.972 | 0.062149 | 0.561693 | 0.889052 | ENSMUSG( | Smyd2 | 1380.553 | -0.07982 | 0.516264 | 0.930939 |
| ENSMUSG( | Smyd3 | 650.2058 | -0.03813 | 0.76132 | 0.953046 | ENSMUSG( | Smyd3 | 578.2971 | -0.11261 | 0.440898 | 0.921166 |
| ENSMUSG( | Smyd4 | 269.6931 | -0.06103 | 0.653068 | 0.922794 | ENSMUSG( | Smyd4 | 261.3765 | 0.127688 | 0.467033 | 0.923329 |
| ENSMUSG( | Smyd5 | 1095.902 | 0.067877 | 0.504718 | 0.866883 | ENSMUSG( | Smyd5 | 906.6216 | -0.21377 | 0.106292 | 0.804969 |
| ENSMUSG( | Snai1 | 24.74806 | 0.10505 | 0.788686 | 0.959226 | ENSMUSG( | Snai1 | 24.91661 | 0.381409 | 0.352955 | 0.912263 |
| ENSMUSG( | Snai2 | 13.14406 | 0.011461 | 0.982041 | NA | ENSMUSG( | Snai2 | 15.17488 | 0.666873 | 0.195264 | 0.861571 |
| ENSMUSG( | Snap23 | 552.2993 | 0.126145 | 0.312958 | 0.76033 | ENSMUSG( | Snap23 | 485.1647 | 0.024473 | 0.863986 | 0.986552 |
| ENSMUSG( | Snap25 | 28906.59 | 0.008654 | 0.949527 | 0.993821 | ENSMUSG( | Snap25 | 25127.57 | -0.13379 | 0.454939 | 0.922891 |
| ENSMUSG( | Snap29 | 970.8261 | 0.150763 | 0.145624 | 0.566706 | ENSMUSG( | Snap29 | 820.0889 | -0.06999 | 0.614389 | 0.951177 |
| ENSMUSG( | Snap47 | 6671.381 | 0.187855 | 0.259406 | 0.709948 | ENSMUSG( | Snap47 | 5323.53 | -0.19926 | 0.291571 | 0.897854 |
| ENSMUSG( | Snap91 | 8221.906 | 0.200584 | 0.011516 | 0.154494 | ENSMUSG( | Snap91 | 7002.716 | 0.011458 | 0.919985 | 0.992644 |
| ENSMUSG( | Snapc1 | 1000.802 | -0.04125 | 0.693856 | 0.934202 | ENSMUSG( | Snapc1 | 924.99 | -0.00112 | 0.992351 | 0.99874 |
| ENSMUSG( | Snapc2 | 600.4324 | 0.032947 | 0.786976 | 0.959037 | ENSMUSG( | Snapc2 | 526.766 | -0.08182 | 0.577978 | 0.943131 |
| ENSMUSG( | Snapc3 | 475.2525 | 0.166302 | 0.180819 | 0.617841 | ENSMUSG( | Snapc3 | 431.6863 | 0.157407 | 0.326204 | 0.905339 |
| ENSMUSG( | Snapc4 | 1614.515 | 0.291842 | 0.022495 | 0.229049 | ENSMUSG( | Snapc4 | 1360.016 | 0.078291 | 0.602436 | 0.94789 |
| ENSMUSG( | Snapc5 | 686.9606 | 0.105339 | 0.386789 | 0.805046 | ENSMUSG( | Snapc5 | 542.4351 | -0.3293 | 0.01383 | 0.46927 |
| ENSMUSG( | Snapin | 2648.27 | -0.03191 | 0.760966 | 0.952907 | ENSMUSG( | Snapin | 2234.858 | -0.26849 | 0.038 | 0.676422 |
| ENSMUSG( | Snca | 4365.414 | 0.124488 | 0.39318 | 0.809274 | ENSMUSG( | Snca | 3659.854 | -0.11799 | 0.48533 | 0.92541 |
| ENSMUSG( | Sncaip | 238.422 | -0.16311 | 0.345836 | 0.779153 | ENSMUSG( | Sncaip | 207.6944 | -0.3182 | 0.116741 | 0.813384 |
| ENSMUSG( | Sncb | 11116.99 | 0.042196 | 0.791883 | 0.959461 | ENSMUSG( | Sncb | 9188.723 | -0.25286 | 0.186154 | 0.855305 |
| ENSMUSG( | Sncg | 22.40137 | -0.59898 | 0.207551 | 0.64962 | ENSMUSG( | Sncg | 22.45852 | -0.27899 | 0.574337 | 0.942912 |
| ENSMUSG( | Snd1 | 2368.108 | -0.15608 | 0.089236 | 0.458482 | ENSMUSG( | Snd1 | 2227.971 | -0.05748 | 0.608317 | 0.950113 |

| | | | | | |
|---|---|---|---|---|---|
| ENSMUSG( | 10.86265 | -0.08851 | 0.872049 | NA | Slc26a5 |
| ENSMUSG( | 432.1622 | -0.01995 | 0.863298 | 0.975055 | Slc26a6 |
| ENSMUSG( | 180.0366 | -0.25208 | 0.131308 | 0.543457 | Slc26a7 |
| ENSMUSG( | 216.7594 | -0.18335 | 0.273802 | 0.724244 | Slc26a8 |
| ENSMUSG( | 5327.977 | -0.09768 | 0.405437 | 0.81652 | Slc27a1 |
| ENSMUSG( | 129.2622 | -0.22495 | 0.260538 | 0.710468 | Slc27a2 |
| ENSMUSG( | 173.061 | -0.02667 | 0.881766 | 0.98196 | Slc27a3 |
| ENSMUSG( | 3796.692 | -0.27301 | 0.067648 | 0.402445 | Slc27a4 |
| ENSMUSG( | 10.22818 | -0.45877 | 0.441059 | NA | Slc27a5 |
| ENSMUSG( | 13.52668 | 0.139145 | 0.799234 | NA | Slc27a6 |
| ENSMUSG( | 696.7192 | -0.40507 | 0.001012 | 0.033218 | Slc29a1 |
| ENSMUSG( | 451.4037 | -0.29791 | 0.041752 | 0.315339 | Slc29a2 |
| ENSMUSG( | 837.9821 | -0.25148 | 0.129361 | 0.538561 | Slc29a3 |
| ENSMUSG( | 746.7706 | -0.11118 | 0.289235 | 0.739725 | Slc29a4 |
| ENSMUSG( | 4868.066 | 0.17614 | 0.313265 | 0.76033 | Slc2a1 |
| ENSMUSG( | 81.96992 | -0.68591 | 0.034177 | 0.283507 | Slc2a10 |
| ENSMUSG( | 189.1789 | -0.16711 | 0.412882 | 0.819541 | Slc2a12 |
| ENSMUSG( | 3229.66 | -0.63095 | 0.000113 | 0.007021 | Slc2a13 |
| ENSMUSG( | 5200.676 | -0.31549 | 0.013454 | 0.169431 | Slc2a3 |
| ENSMUSG( | 60.9813 | -0.28254 | 0.272045 | 0.7223 | Slc2a4 |
| ENSMUSG( | 283.5716 | -0.1636 | 0.434804 | 0.829025 | Slc2a5 |
| ENSMUSG( | 1048.676 | -0.18981 | 0.158803 | 0.588112 | Slc2a6 |
| ENSMUSG( | 807.6209 | -0.09606 | 0.502092 | 0.864904 | Slc2a8 |
| ENSMUSG( | 61.45888 | -0.26132 | 0.342768 | 0.778971 | Slc2a9 |
| ENSMUSG( | 1213.549 | -0.44406 | 0.00777 | 0.122366 | Slc30a1 |
| ENSMUSG( | 1333.642 | -0.14982 | 0.268776 | 0.719042 | Slc30a10 |
| ENSMUSG( | 4258.699 | -0.06568 | 0.548802 | 0.885686 | Slc30a3 |
| ENSMUSG( | 1894.22 | -0.25032 | 0.109116 | 0.498553 | Slc30a4 |
| ENSMUSG( | 1648.093 | -0.14621 | 0.295907 | 0.744386 | Slc30a5 |
| ENSMUSG( | 399.3667 | -0.37264 | 0.033906 | 0.282701 | Slc30a6 |
| ENSMUSG( | 602.2264 | -0.30245 | 0.102305 | 0.487127 | Slc30a7 |
| ENSMUSG( | 7462.158 | 0.07027 | 0.412755 | 0.819518 | Slc30a9 |
| ENSMUSG( | 1339.243 | -0.14896 | 0.228422 | 0.675849 | Slc31a1 |
| ENSMUSG( | 303.4564 | -0.25409 | 0.097812 | 0.478264 | Slc31a2 |
| ENSMUSG( | 3380.945 | -0.55958 | 2.10E-06 | 0.000341 | Slc32a1 |
| ENSMUSG( | 792.9071 | -0.31298 | 0.032213 | 0.274192 | Slc33a1 |
| ENSMUSG( | 523.9797 | -0.29012 | 0.050888 | 0.345939 | Slc35a1 |
| ENSMUSG( | 1194.304 | -0.24541 | 0.0758 | 0.426666 | Slc35a2 |
| ENSMUSG( | 483.1606 | -0.23124 | 0.110958 | 0.50216 | Slc35a3 |
| ENSMUSG( | 2311.614 | 0.066742 | 0.412504 | 0.819452 | Slc35a4 |
| ENSMUSG( | 1409.43 | -0.30307 | 0.039004 | 0.304448 | Slc35a5 |
| ENSMUSG( | 1512.52 | -0.20803 | 0.112461 | 0.505997 | Slc35b1 |
| ENSMUSG( | 1374.574 | -0.39634 | 0.008635 | 0.130416 | Slc35b2 |
| ENSMUSG( | 390.2104 | -0.26266 | 0.111046 | 0.502387 | Slc35b3 |
| ENSMUSG( | 2012.786 | -0.2713 | 0.040938 | 0.312769 | Slc35b4 |
| ENSMUSG( | 613.5514 | -0.42968 | 0.004665 | 0.089972 | Slc35c1 |
| ENSMUSG( | 1829.526 | -0.0808 | 0.466507 | 0.844691 | Slc35c2 |
| ENSMUSG( | 705.6316 | -0.2792 | 0.032817 | 0.277278 | Slc35d1 |
| ENSMUSG( | 24.39934 | 0.194163 | 0.617315 | 0.910866 | Slc35d2 |
| ENSMUSG( | 132.2693 | -1.53665 | 1.08E-06 | 0.000199 | Slc35d3 |
| ENSMUSG( | 2324.86 | -0.19527 | 0.178269 | 0.615871 | Slc35e1 |
| ENSMUSG( | 1457.085 | -0.13561 | 0.239373 | 0.687192 | Slc35e2 |
| ENSMUSG( | 1159.096 | -0.20581 | 0.091614 | 0.465535 | Slc35e3 |
| ENSMUSG( | 1071.784 | -0.20377 | 0.141978 | 0.559166 | Slc35e4 |
| ENSMUSG( | 1899.998 | -0.5303 | 0.000564 | 0.02226 | Slc35f1 |
| ENSMUSG( | 178.8487 | -0.16148 | 0.438802 | 0.830714 | Slc35f2 |
| ENSMUSG( | 873.21 | 0.007121 | 0.955715 | 0.993874 | Slc35f3 |
| ENSMUSG( | 342.5614 | -0.38219 | 0.021452 | 0.22362 | Slc35f4 |
| ENSMUSG( | 825.2311 | -0.20345 | 0.187843 | 0.626399 | Slc35f5 |
| ENSMUSG( | 1471.416 | -0.40691 | 0.010928 | 0.150598 | Slc35f6 |
| ENSMUSG( | 161.7754 | -0.21676 | 0.229495 | 0.67735 | Slc35g1 |
| ENSMUSG( | 393.6253 | -0.46864 | 0.004406 | 0.086676 | Slc35g2 |
| ENSMUSG( | 2492.334 | -0.04674 | 0.76512 | 0.954127 | Slc36a1 |
| ENSMUSG( | 21.02118 | -0.7482 | 0.080144 | NA | Slc36a2 |
| ENSMUSG( | 1914.516 | -0.08879 | 0.41701 | 0.8209 | Slc36a4 |
| ENSMUSG( | 190.196 | -0.64343 | 5.88E-05 | 0.004481 | Slc37a1 |
| ENSMUSG( | 254.5773 | -0.38891 | 0.013624 | 0.170376 | Slc37a2 |
| ENSMUSG( | 2054.372 | -0.13236 | 0.282323 | 0.73186 | Slc37a3 |
| ENSMUSG( | 818.1262 | -0.22294 | 0.053721 | 0.355948 | Slc37a4 |
| ENSMUSG( | 4371.796 | -0.26201 | 0.054391 | 0.358917 | Slc38a1 |
| ENSMUSG( | 3052.689 | -0.17054 | 0.1644 | 0.59535 | Slc38a10 |
| ENSMUSG( | 27.90256 | -0.25567 | 0.628882 | 0.913816 | Slc38a11 |
| ENSMUSG( | 3933.262 | -0.34103 | 0.009993 | 0.142653 | Slc38a2 |
| ENSMUSG( | 5447.437 | -0.4295 | 0.002039 | 0.051756 | Slc38a3 |
| ENSMUSG( | 30.47634 | 0.05759 | 0.870992 | 0.977951 | Slc38a4 |
| ENSMUSG( | 135.9027 | -0.33577 | 0.156258 | 0.583988 | Slc38a5 |
| ENSMUSG( | 380.3689 | 0.103122 | 0.407939 | 0.818167 | Slc38a6 |
| ENSMUSG( | 1187.7 | -0.30446 | 0.023622 | 0.235222 | Slc38a7 |
| ENSMUSG( | 742.9435 | -0.13351 | 0.311486 | 0.759296 | Slc38a9 |
| ENSMUSG( | 2303.126 | -0.26268 | 0.029905 | 0.262335 | Slc39a1 |
| ENSMUSG( | 4654.419 | -0.27437 | 0.099008 | 0.480953 | Slc39a10 |
| ENSMUSG( | 693.1575 | -0.23979 | 0.055013 | 0.36102 | Slc39a11 |
| ENSMUSG( | 4690.319 | 0.043181 | 0.72635 | 0.944223 | Slc39a12 |
| ENSMUSG( | 912.8571 | -0.12813 | 0.333486 | 0.775602 | Slc39a13 |
| ENSMUSG( | 920.69 | -0.35493 | 0.018878 | 0.206889 | Slc39a14 |
| ENSMUSG( | 280.5141 | 0.510278 | 0.000896 | 0.030286 | Slc39a2 |
| ENSMUSG( | 3303.588 | -0.18779 | 0.141615 | 0.558606 | Slc39a3 |
| ENSMUSG( | 19.42202 | -0.21525 | 0.69092 | NA | Slc39a4 |
| ENSMUSG( | 10.19489 | -0.47343 | 0.46334 | NA | Slc39a5 |
| ENSMUSG( | 1440.88 | -0.18217 | 0.250649 | 0.700403 | Slc39a6 |
| ENSMUSG( | 159.381 | 0.043057 | 0.833442 | 0.970333 | Slc39a7 |
| ENSMUSG( | 281.6954 | -0.20061 | 0.312133 | 0.759724 | Slc39a8 |
| ENSMUSG( | 1821.324 | -0.24112 | 0.071965 | 0.414057 | Slc39a9 |
| ENSMUSG( | 12.41648 | 0.161396 | 0.77161 | NA | Slc3a1 |
| ENSMUSG( | 6078.996 | -0.19641 | 0.135466 | 0.548884 | Slc3a2 |
| ENSMUSG( | 609.2665 | -0.23344 | 0.29412 | 0.743459 | Slc40a1 |
| ENSMUSG( | 4083.488 | -0.04804 | 0.673273 | 0.929007 | Slc41a1 |
| ENSMUSG( | 635.3866 | -0.35747 | 0.026073 | 0.247245 | Slc41a2 |
| ENSMUSG( | 178.2615 | -0.32578 | 0.04306 | 0.31819 | Slc41a3 |
| ENSMUSG( | 14.4067 | -0.13307 | 0.79336 | NA | Slc43a1 |
| ENSMUSG( | 2652.729 | -0.33789 | 0.032439 | 0.275619 | Slc43a2 |
| ENSMUSG( | 58.69387 | -0.25067 | 0.350283 | 0.781943 | Slc43a3 |
| ENSMUSG( | 4615.723 | -0.33175 | 0.012343 | 0.160458 | Slc44a1 |
| ENSMUSG( | 2283.082 | -0.19231 | 0.075197 | 0.424832 | Slc44a2 |
| ENSMUSG( | 15.68069 | -0.52962 | 0.27451 | NA | Slc44a4 |
| ENSMUSG( | 346.8637 | -0.0912 | 0.559155 | 0.888638 | Slc44a5 |
| ENSMUSG( | 1639.577 | -0.24202 | 0.0905 | 0.461757 | Slc45a1 |
| ENSMUSG( | 94.09164 | -0.78562 | 0.000673 | 0.025051 | Slc45a3 |
| ENSMUSG( | 1892.213 | 0.043424 | 0.760885 | 0.952907 | Slc45a4 |
| ENSMUSG( | 441.356 | -0.36754 | 0.035826 | 0.290986 | Slc46a1 |
| ENSMUSG( | 258.3382 | -0.2336 | 0.22966 | 0.677538 | Slc46a3 |
| ENSMUSG( | 593.2238 | -0.3299 | 0.02047 | 0.21574 | Slc47a1 |
| ENSMUSG( | 13.71288 | -0.02659 | 0.963124 | NA | Slc47a2 |
| ENSMUSG( | 9695.604 | -0.19622 | 0.06104 | 0.381679 | Slc48a1 |
| ENSMUSG( | 51.98212 | -0.26147 | 0.326352 | 0.771301 | Slc49a3 |
| ENSMUSG( | 4384.194 | -0.18153 | 0.112441 | 0.505997 | Slc49a4 |
| ENSMUSG( | 100.8035 | -1.35499 | 0.081848 | 0.442044 | Slc4a1 |
| ENSMUSG( | 3610.906 | -0.34767 | 0.00623 | 0.107962 | Slc4a10 |
| ENSMUSG( | 30.5734 | 0.045196 | 0.896168 | 0.984747 | Slc4a11 |
| ENSMUSG( | 514.9285 | 0.220633 | 0.08505 | 0.449548 | Slc4a1ap |
| ENSMUSG( | 2096.097 | -0.15787 | 0.135975 | 0.550005 | Slc4a2 |
| ENSMUSG( | 5857.47 | 0.034618 | 0.736543 | 0.946437 | Slc4a3 |
| ENSMUSG( | 16047.56 | -0.08229 | 0.529967 | 0.878883 | Slc4a4 |
| ENSMUSG( | 76.56393 | 0.015235 | 0.989061 | 0.999111 | Slc4a5 |
| ENSMUSG( | 624.1569 | -0.04436 | 0.750814 | 0.950042 | Slc4a7 |
| ENSMUSG( | 3113.923 | -0.1935 | 0.157468 | 0.585653 | Slc4a8 |
| ENSMUSG( | 971.1051 | -0.15666 | 0.123217 | 0.530407 | Slc50a1 |
| ENSMUSG( | 490.9779 | -0.1713 | 0.267547 | 0.717648 | Slc52a2 |
| ENSMUSG( | 100.9392 | 0.047217 | 0.827767 | 0.968924 | Slc52a3 |
| ENSMUSG( | 28.97694 | -0.49017 | 0.163944 | 0.594869 | Slc5a11 |
| ENSMUSG( | 489.747 | -0.15989 | 0.383147 | 0.803237 | Slc5a3 |
| ENSMUSG( | 585.3984 | -0.56247 | 1.13E-05 | 0.001226 | Slc5a5 |
| ENSMUSG( | 822.308 | -0.15186 | 0.206952 | 0.649503 | Slc5a6 |
| ENSMUSG( | 86.05923 | -1.17795 | 3.69E-06 | 0.000504 | Slc5a7 |
| ENSMUSG( | 378.0108 | -0.32335 | 0.045008 | 0.325572 | Slc66a1 |
| ENSMUSG( | 614.0798 | -0.10423 | 0.437252 | 0.829501 | Slc66a2 |
| ENSMUSG( | 36.28005 | -0.63326 | 0.051395 | 0.347924 | Slc66a3 |
| ENSMUSG( | 23458.36 | -0.22557 | 0.076387 | 0.427832 | Slc6a1 |
| ENSMUSG( | 16051.14 | -0.33727 | 0.005647 | 0.100551 | Slc6a11 |
| ENSMUSG( | 56.28597 | 0.453209 | 0.084706 | 0.448558 | Slc6a12 |
| ENSMUSG( | 417.418 | 0.060454 | 0.694026 | 0.934202 | Slc6a13 |
| ENSMUSG( | 1256.056 | -0.26156 | 0.067104 | 0.400611 | Slc6a15 |
| ENSMUSG( | 17.08506 | 0.047879 | 0.920043 | NA | Slc6a16 |
| ENSMUSG( | 17312.55 | -0.21511 | 0.140917 | 0.557902 | Slc6a17 |
| ENSMUSG( | 15.19279 | -0.07167 | 0.888825 | NA | Slc6a19 |
| ENSMUSG( | 791.0229 | -0.19342 | 0.132133 | 0.544244 | Slc6a20a |
| ENSMUSG( | 4438.442 | -0.28397 | 0.072071 | 0.414057 | Slc6a6 |
| ENSMUSG( | 2483.011 | 0.017981 | 0.88454 | 0.982647 | Slc6a7 |
| ENSMUSG( | 5112.335 | 0.06993 | 0.410996 | 0.818761 | Slc6a8 |
| ENSMUSG( | 2123.466 | -0.6111 | 5.46E-07 | 0.000113 | Slc6a9 |
| ENSMUSG( | 1395.024 | -0.15426 | 0.374751 | 0.800466 | Slc7a1 |
| ENSMUSG( | 2958.07 | -0.19435 | 0.138603 | 0.554003 | Slc7a10 |
| ENSMUSG( | 3729.834 | -0.01029 | 0.952904 | 0.993874 | Slc7a11 |
| ENSMUSG( | 2284.898 | -0.35047 | 0.033961 | 0.282701 | Slc7a14 |
| ENSMUSG( | 2823.02 | 0.019576 | 0.887546 | 0.983411 | Slc7a2 |
| ENSMUSG( | 260.4998 | -0.00194 | 0.991636 | 0.999232 | Slc7a3 |
| ENSMUSG( | 2014.881 | -0.2016 | 0.113274 | 0.507541 | Slc7a4 |
| ENSMUSG( | 4503.642 | 0.088653 | 0.562575 | 0.889465 | Slc7a5 |

| | | | | | |
|---|---|---|---|---|---|
| ENSMUSG( | 23734.85 | 0.087519 | 0.549131 | 0.938452 | Slc6a1 |
| ENSMUSG( | 15270.83 | -0.20012 | 0.128614 | 0.81925 | Slc6a11 |
| ENSMUSG( | 41.2931 | -0.15433 | 0.625211 | 0.953234 | Slc6a12 |
| ENSMUSG( | 335.0154 | -0.31181 | 0.081154 | 0.77681 | Slc6a13 |
| ENSMUSG( | 1437.563 | 0.386221 | 0.013188 | 0.460068 | Slc6a15 |
| ENSMUSG( | 16.89701 | 0.279682 | 0.555055 | 0.939565 | Slc6a16 |
| ENSMUSG( | 18285.51 | 0.216072 | 0.193644 | 0.860096 | Slc6a17 |
| ENSMUSG( | 13.0738 | -0.17329 | 0.763048 | 0.973795 | Slc6a19 |
| ENSMUSG( | 773.4586 | 0.022757 | 0.891592 | 0.990558 | Slc6a20a |
| ENSMUSG( | 5362.223 | 0.506446 | 0.005429 | 0.303106 | Slc6a6 |
| ENSMUSG( | 2541.854 | 0.336249 | 0.003653 | 0.242109 | Slc6a7 |
| ENSMUSG( | 4652.764 | 0.067552 | 0.509874 | 0.929089 | Slc6a8 |
| ENSMUSG( | 2431.304 | 0.111795 | 0.384563 | 0.917257 | Slc6a9 |
| ENSMUSG( | 1582.993 | 0.455298 | 0.018565 | 0.545482 | Slc7a1 |
| ENSMUSG( | 2915.118 | 0.041689 | 0.784913 | 0.976285 | Slc7a10 |
| ENSMUSG( | 3389.001 | -0.016 | 0.923156 | 0.992691 | Slc7a11 |
| ENSMUSG( | 2655.602 | 0.357617 | 0.053227 | 0.716108 | Slc7a14 |
| ENSMUSG( | 2788.28 | 0.244584 | 0.091377 | 0.791889 | Slc7a2 |
| ENSMUSG( | 265.1459 | 0.306223 | 0.099092 | 0.799802 | Slc7a3 |
| ENSMUSG( | 2041.109 | 0.113433 | 0.455257 | 0.923216 | Slc7a4 |
| ENSMUSG( | 4636.487 | 0.418309 | 0.040829 | 0.685722 | Slc7a5 |
| ENSMUSG( | 864.4649 | 0.256915 | 0.038718 | 0.679972 | Slc7a6 |
| ENSMUSG( | 605.2641 | 0.141778 | 0.30652 | 0.898538 | Slc7a6os |
| ENSMUSG( | 98.2321 | 0.103451 | 0.683188 | 0.958678 | Slc7a7 |
| ENSMUSG( | 3944.565 | 0.075957 | 0.601856 | 0.94789 | Slc7a8 |
| ENSMUSG( | 3904.634 | 0.348551 | 0.072303 | 0.763629 | Slc8a1 |
| ENSMUSG( | 12900.26 | 0.118755 | 0.459731 | 0.923329 | Slc8a2 |
| ENSMUSG( | 969.4783 | -0.03327 | 0.810803 | 0.978779 | Slc8a3 |
| ENSMUSG( | 435.9479 | -0.02516 | 0.881848 | 0.989644 | Slc8b1 |
| ENSMUSG( | 1935.511 | 0.148931 | 0.310967 | 0.899422 | Slc9a1 |
| ENSMUSG( | 381.5225 | 0.268241 | 0.135993 | 0.826859 | Slc9a2 |
| ENSMUSG( | 21.60044 | 0.420705 | 0.337423 | 0.907972 | Slc9a3 |
| ENSMUSG( | 61.72377 | 0.161851 | 0.574996 | 0.942912 | Slc9a4 |
| ENSMUSG( | 1359.853 | 0.184322 | 0.126506 | 0.815605 | Slc9a5 |
| ENSMUSG( | 3673.313 | 0.139735 | 0.361351 | 0.914223 | Slc9a6 |
| ENSMUSG( | 229.1618 | 0.040607 | 0.857781 | 0.985884 | Slc9a7 |
| ENSMUSG( | 1210.373 | 0.164904 | 0.261678 | 0.884926 | Slc9a8 |
| ENSMUSG( | 817.2281 | 0.1443 | 0.354618 | 0.912411 | Slc9a9 |
| ENSMUSG( | 122.3894 | -0.18676 | 0.473307 | 0.923329 | Slc9b2 |
| ENSMUSG( | 2605.913 | 0.762504 | 0.065189 | 0.74247 | Slco1a4 |
| ENSMUSG( | 9.631202 | 0.545049 | 0.451868 | NA | Slco1a5 |
| ENSMUSG( | 15.25053 | -0.28112 | 0.586441 | 0.944975 | Slco1b2 |
| ENSMUSG( | 4338.724 | 0.35951 | 0.099364 | 0.800296 | Slco1c1 |
| ENSMUSG( | 91.33031 | -0.10267 | 0.685497 | 0.958678 | Slco2a1 |
| ENSMUSG( | 1406.287 | 0.412501 | 0.067952 | 0.75212 | Slco2b1 |
| ENSMUSG( | 1097.782 | 0.449035 | 0.011634 | 0.440174 | Slco3a1 |
| ENSMUSG( | 157.571 | -0.29346 | 0.162766 | 0.837724 | Slco4a1 |
| ENSMUSG( | 75.04026 | 0.510459 | 0.081539 | 0.77681 | Slco4c1 |
| ENSMUSG( | 78.0326 | 0.111844 | 0.648662 | 0.95422 | Slco5a1 |
| ENSMUSG( | 335.0616 | 0.166186 | 0.27116 | 0.888062 | Slf1 |
| ENSMUSG( | 1381.745 | 0.124584 | 0.315144 | 0.901306 | Slf2 |
| ENSMUSG( | 10.45514 | 0.306211 | 0.737809 | NA | Slfn1 |
| ENSMUSG( | 57.41389 | -0.5039 | 0.088062 | 0.78663 | Slfn2 |
| ENSMUSG( | 11.7741 | 1.17308 | 0.081719 | 0.77681 | Slfn3 |
| ENSMUSG( | 9.423986 | 0.091442 | 0.914344 | NA | Slfn4 |
| ENSMUSG( | 349.1766 | 0.428281 | 0.10629 | 0.804969 | Slfn5 |
| ENSMUSG( | 64.3907 | -0.38376 | 0.199169 | 0.863016 | Slfn8 |
| ENSMUSG( | 54.65649 | -0.25583 | 0.380805 | 0.917202 | Slfn9 |
| ENSMUSG( | 527.0852 | -0.23799 | 0.127469 | 0.817604 | Slirp |
| ENSMUSG( | 1814.201 | 0.377519 | 0.092277 | 0.79275 | Slit1 |
| ENSMUSG( | 1049.134 | -0.03653 | 0.812034 | 0.979148 | Slit2 |
| ENSMUSG( | 779.3643 | 0.25709 | 0.191735 | 0.858764 | Slit3 |
| ENSMUSG( | 2378.856 | 0.078675 | 0.685324 | 0.958678 | Slitrk1 |
| ENSMUSG( | 997.5552 | 0.318492 | 0.114562 | 0.811703 | Slitrk2 |
| ENSMUSG( | 1828.653 | 0.34962 | 0.028706 | 0.614171 | Slitrk3 |
| ENSMUSG( | 669.1883 | 0.366602 | 0.065339 | 0.743695 | Slitrk4 |
| ENSMUSG( | 2008.013 | 0.340882 | 0.063845 | 0.736482 | Slitrk5 |
| ENSMUSG( | 25.51637 | 0.235962 | 0.559635 | 0.939807 | Slitrk6 |
| ENSMUSG( | 2645.397 | 0.019623 | 0.84806 | 0.985242 | Slk |
| ENSMUSG( | 2138.372 | 0.093973 | 0.380896 | 0.917202 | Slmap |
| ENSMUSG( | 22.05155 | -0.09972 | 0.820119 | 0.979198 | Slpi |
| ENSMUSG( | 2091.684 | 0.015032 | 0.898924 | 0.990558 | Sltm |
| ENSMUSG( | 1352.127 | -0.08739 | 0.424295 | 0.917257 | Slu7 |
| ENSMUSG( | 493.5552 | -0.05456 | 0.674747 | 0.95694 | Slx1b |
| ENSMUSG( | 734.082 | 0.056828 | 0.707501 | 0.964029 | Slx4 |
| ENSMUSG( | 188.9508 | -0.01655 | 0.927856 | 0.99306 | Slx4ip |
| ENSMUSG( | 890.6547 | -0.06535 | 0.59675 | 0.946541 | Slx9 |
| ENSMUSG( | 468.8307 | 0.025897 | 0.864464 | 0.986591 | Smad1 |
| ENSMUSG( | 771.9011 | 0.022856 | 0.850116 | 0.985538 | Smad2 |
| ENSMUSG( | 1409.215 | 0.058954 | 0.625814 | 0.953234 | Smad3 |
| ENSMUSG( | 1122.983 | 0.094791 | 0.436559 | 0.919879 | Smad4 |
| ENSMUSG( | 2071.393 | -0.05319 | 0.697502 | 0.960867 | Smad5 |
| ENSMUSG( | 82.94553 | 1.065865 | 0.000101 | 0.020718 | Smad6 |
| ENSMUSG( | 575.8441 | 0.097364 | 0.464411 | 0.923329 | Smad7 |
| ENSMUSG( | 402.5652 | -0.14868 | 0.349197 | 0.911652 | Smad9 |
| ENSMUSG( | 68.3028 | 0.074554 | 0.800966 | 0.97681 | Smagp |
| ENSMUSG( | 1978.602 | -0.07845 | 0.560694 | 0.940254 | Smap1 |
| ENSMUSG( | 2763.899 | -0.11018 | 0.390778 | 0.917257 | Smap2 |
| ENSMUSG( | 585.9376 | 0.152278 | 0.257148 | 0.883082 | Smarca1 |
| ENSMUSG( | 7308.033 | 0.06421 | 0.502489 | 0.927825 | Smarca2 |
| ENSMUSG( | 4419.779 | -0.07429 | 0.494183 | 0.927372 | Smarca4 |
| ENSMUSG( | 1398.271 | 0.084301 | 0.508995 | 0.929089 | Smarca5 |
| ENSMUSG( | 653.2215 | 0.086009 | 0.532599 | 0.934795 | Smarcad1 |
| ENSMUSG( | 879.1671 | 0.110475 | 0.382244 | 0.917257 | Smarcal1 |
| ENSMUSG( | 1291.727 | -0.14877 | 0.216841 | 0.865436 | Smarcb1 |
| ENSMUSG( | 1808.23 | -0.18738 | 0.09874 | 0.798438 | Smarcc1 |
| ENSMUSG( | 12196.06 | -0.10099 | 0.299432 | 0.898538 | Smarcc2 |
| ENSMUSG( | 5516.334 | -0.06734 | 0.563011 | 0.941158 | Smarcd1 |
| ENSMUSG( | 546.0174 | 0.435831 | 0.008971 | 0.396825 | Smarcd2 |
| ENSMUSG( | 2728.518 | 0.038906 | 0.716222 | 0.965692 | Smarcd3 |
| ENSMUSG( | 1250.613 | -0.15006 | 0.225647 | 0.868468 | Smarce1 |
| ENSMUSG( | 1913.771 | 0.025773 | 0.850796 | 0.985541 | Smc1a |
| ENSMUSG( | 223.6939 | 0.176069 | 0.326754 | 0.905371 | Smc2 |
| ENSMUSG( | 18.19319 | -0.88776 | 0.092034 | 0.791889 | Smc2os |
| ENSMUSG( | 1528.613 | 0.073583 | 0.492187 | 0.927315 | Smc3 |
| ENSMUSG( | 338.1591 | 0.285385 | 0.071726 | 0.762854 | Smc4 |
| ENSMUSG( | 727.9975 | 0.092524 | 0.473881 | 0.923329 | Smc5 |
| ENSMUSG( | 1169.855 | 0.125932 | 0.276466 | 0.890889 | Smc6 |
| ENSMUSG( | 822.1357 | 0.036232 | 0.776411 | 0.975717 | Smchd1 |
| ENSMUSG( | 101.1829 | 0.086685 | 0.683375 | 0.958678 | Smco3 |
| ENSMUSG( | 58.6889 | 0.251709 | 0.35918 | 0.914061 | Smco4 |
| ENSMUSG( | 743.8931 | 0.206723 | 0.100275 | 0.801311 | Smcr8 |
| ENSMUSG( | 2712.717 | -0.34134 | 0.020264 | 0.563435 | Smdt1 |
| ENSMUSG( | 4202.346 | 0.185691 | 0.121539 | 0.813384 | Smg1 |
| ENSMUSG( | 2763.932 | -0.01293 | 0.916343 | 0.992529 | Smg5 |
| ENSMUSG( | 1326.726 | -0.04978 | 0.659479 | 0.956205 | Smg6 |
| ENSMUSG( | 2318.182 | 0.062815 | 0.566675 | 0.941777 | Smg7 |
| ENSMUSG( | 299.3386 | 0.233667 | 0.237438 | 0.872173 | Smg8 |
| ENSMUSG( | 880.2239 | -0.18323 | 0.22079 | 0.866104 | Smg9 |
| ENSMUSG( | 895.0714 | -0.15178 | 0.233157 | 0.868507 | Smim1 |
| ENSMUSG( | 3505.989 | -0.0645 | 0.671997 | 0.95694 | Smim10l1 |
| ENSMUSG( | 776.3873 | -0.15211 | 0.220953 | 0.866104 | Smim10l2a |
| ENSMUSG( | 253.9778 | -0.08245 | 0.631442 | 0.953251 | Smim11 |
| ENSMUSG( | 1016.898 | -0.15747 | 0.161921 | 0.837724 | Smim12 |
| ENSMUSG( | 5912.296 | -0.02341 | 0.834685 | 0.981237 | Smim13 |
| ENSMUSG( | 3227.826 | -0.10154 | 0.392514 | 0.917257 | Smim14 |
| ENSMUSG( | 1146.031 | -0.21068 | 0.185885 | 0.855305 | Smim15 |
| ENSMUSG( | 169.3043 | -0.1809 | 0.411956 | 0.917257 | Smim17 |
| ENSMUSG( | 127.1588 | -0.01448 | 0.946453 | 0.994798 | Smim18 |
| ENSMUSG( | 714.5732 | -0.13413 | 0.34381 | 0.910159 | Smim19 |
| ENSMUSG( | 671.1127 | -0.33353 | 0.00818 | 0.388416 | Smim20 |
| ENSMUSG( | 42.63725 | 0.079577 | 0.801723 | 0.97681 | Smim24 |
| ENSMUSG( | 300.2728 | -0.09033 | 0.550982 | 0.939232 | Smim26 |
| ENSMUSG( | 99.26977 | -0.06431 | 0.795888 | 0.97681 | Smim27 |
| ENSMUSG( | 1515.255 | -0.19875 | 0.067131 | 0.75212 | Smim29 |
| ENSMUSG( | 704.7175 | 0.505114 | 4.40E-05 | 0.01151 | Smim3 |
| ENSMUSG( | 683.4312 | 0.040373 | 0.793648 | 0.97681 | Smim30 |
| ENSMUSG( | 102.1007 | 0.084664 | 0.727092 | 0.966887 | Smim32 |
| ENSMUSG( | 17.75893 | -0.10633 | 0.837639 | 0.981335 | Smim35 |
| ENSMUSG( | 179.3425 | -0.11842 | 0.536338 | 0.93705 | Smim36 |
| ENSMUSG( | 9.825458 | 0.152849 | 0.812664 | NA | Smim38 |
| ENSMUSG( | 12.16388 | 0.408707 | 0.504572 | 0.927825 | Smim40 |
| ENSMUSG( | 613.5455 | 0.058402 | 0.686223 | 0.95896 | Smim43 |
| ENSMUSG( | 5004.06 | -0.16072 | 0.261164 | 0.884926 | Smim45 |
| ENSMUSG( | 17.86212 | 0.532586 | 0.249593 | 0.877904 | Smim5 |
| ENSMUSG( | 2837.746 | -0.05178 | 0.657032 | 0.955625 | Smim7 |
| ENSMUSG( | 322.4966 | -0.24831 | 0.208177 | 0.864412 | Smim8 |
| ENSMUSG( | 264.514 | -0.04554 | 0.800339 | 0.97681 | Smn1 |
| ENSMUSG( | 599.9739 | -0.05883 | 0.736605 | 0.967344 | Smndc1 |
| ENSMUSG( | 1340.58 | 0.0643 | 0.612182 | 0.951071 | Smo |
| ENSMUSG( | 929.3532 | 0.293991 | 0.043497 | 0.690304 | Smoc1 |
| ENSMUSG( | 468.4066 | 0.237675 | 0.209044 | 0.864412 | Smoc2 |
| ENSMUSG( | 3056.603 | 0.008153 | 0.938059 | 0.993731 | Smox |
| ENSMUSG( | 4509.998 | 0.144405 | 0.393584 | 0.917257 | Smpd1 |
| ENSMUSG( | 853.8553 | -0.02231 | 0.878333 | 0.989644 | Smpd2 |
| ENSMUSG( | 3523.466 | 0.225905 | 0.184133 | 0.852618 | Smpd3 |
| ENSMUSG( | 1571.267 | 0.012687 | 0.946985 | 0.994798 | Smpd4 |
| ENSMUSG( | 32.68699 | 0.403489 | 0.273458 | 0.889703 | Smpd5 |

| | | | | | | | | | | | |
|---|---|---|---|---|---|---|---|---|---|---|---|
| ENSMUSG( | Sned1 | 1192.579 | -0.11429 | 0.332421 | 0.775474 | ENSMUSG( | Sned1 | 1132.268 | 0.007144 | 0.95952 | 0.994798 |
| ENSMUSG( | Snf8 | 2118.517 | -0.01209 | 0.889834 | 0.983651 | ENSMUSG( | Snf8 | 1828.152 | -0.17576 | 0.134523 | 0.823094 |
| ENSMUSG( | Snhg10 | 68.75871 | -0.36313 | 0.127748 | 0.535835 | ENSMUSG( | Snhg10 | 73.58803 | 0.111888 | 0.658571 | 0.956205 |
| ENSMUSG( | Snhg11 | 40128.48 | 0.332068 | 0.099342 | 0.48179 | ENSMUSG( | Snhg11 | 34374.78 | 0.164175 | 0.445061 | 0.921734 |
| ENSMUSG( | Snhg12 | 727.6099 | -0.35997 | 0.000661 | 0.024836 | ENSMUSG( | Snhg12 | 747.1321 | 0.001135 | 0.993519 | 0.998813 |
| ENSMUSG( | Snhg15 | 90.92269 | -0.3154 | 0.130116 | 0.540593 | ENSMUSG( | Snhg15 | 87.76515 | -0.13262 | 0.552359 | 0.939565 |
| ENSMUSG( | Snhg16 | 88.17861 | -0.48745 | 0.021652 | 0.225182 | ENSMUSG( | Snhg16 | 92.04921 | -0.03016 | 0.903523 | 0.991089 |
| ENSMUSG( | Snhg17 | 239.1649 | -0.24187 | 0.107548 | 0.496444 | ENSMUSG( | Snhg17 | 225.3294 | -0.12451 | 0.508939 | 0.929089 |
| ENSMUSG( | Snhg18 | 116.1184 | -0.11563 | 0.577074 | 0.895621 | ENSMUSG( | Snhg18 | 103.6318 | -0.18446 | 0.412923 | 0.917257 |
| ENSMUSG( | Snhg20 | 481.1187 | -0.17525 | 0.291168 | 0.741182 | ENSMUSG( | Snhg20 | 495.6629 | 0.18298 | 0.396146 | 0.917257 |
| ENSMUSG( | Snhg3 | 137.6462 | 0.013291 | 0.941266 | 0.993415 | ENSMUSG( | Snhg3 | 128.5417 | 0.083868 | 0.70161 | 0.962014 |
| ENSMUSG( | Snhg4 | 158.1807 | -0.03149 | 0.863068 | 0.975055 | ENSMUSG( | Snhg4 | 144.8609 | -0.00959 | 0.959157 | 0.994798 |
| ENSMUSG( | Snhg5 | 238.0322 | -0.11564 | 0.458601 | 0.840224 | ENSMUSG( | Snhg5 | 241.2533 | 0.19656 | 0.248624 | 0.877904 |
| ENSMUSG( | Snhg6 | 792.6509 | 0.152597 | 0.161905 | 0.592398 | ENSMUSG( | Snhg6 | 661.21 | -0.10305 | 0.404781 | 0.917257 |
| ENSMUSG( | Snhg7 | 127.3212 | 0.058862 | 0.758042 | 0.951614 | ENSMUSG( | Snhg7 | 128.7956 | 0.350932 | 0.121686 | 0.813384 |
| ENSMUSG( | Snhg8 | 752.3654 | 0.136741 | 0.190556 | 0.629951 | ENSMUSG( | Snhg8 | 661.4949 | 0.032289 | 0.804785 | 0.977232 |
| ENSMUSG( | Snip1 | 278.8505 | 0.023077 | 0.893509 | 0.984512 | ENSMUSG( | Snip1 | 241.9873 | -0.11374 | 0.548276 | 0.938063 |
| ENSMUSG( | Snn | 10912.6 | -0.02231 | 0.771068 | 0.954726 | ENSMUSG( | Snn | 9971.14 | -0.01454 | 0.888621 | 0.990558 |
| ENSMUSG( | Snora31 | 16.55275 | -0.24886 | 0.667724 | NA | ENSMUSG( | Snora31 | 18.35131 | 0.326577 | 0.548535 | 0.938063 |
| ENSMUSG( | Snora73b | 16.91263 | -0.88734 | 0.095914 | NA | ENSMUSG( | Snora73b | 18.30092 | -0.244 | 0.640254 | 0.953903 |
| ENSMUSG( | Snorc | 17.7495 | 0.03572 | 0.936206 | NA | ENSMUSG( | Snorc | 13.55681 | -0.59575 | 0.300212 | 0.898538 |
| ENSMUSG( | Snord55 | 24.88107 | 0.411051 | 0.287702 | 0.73849 | ENSMUSG( | Snord55 | 21.64958 | 0.28314 | 0.508312 | 0.929089 |
| ENSMUSG( | Snph | 20406.32 | 0.301818 | 0.001406 | 0.041028 | ENSMUSG( | Snph | 18176.48 | 0.243583 | 0.048884 | 0.706521 |
| ENSMUSG( | Snrk | 2062.949 | 0.225619 | 0.011793 | 0.156913 | ENSMUSG( | Snrk | 1735.46 | 0.005599 | 0.957871 | 0.994798 |
| ENSMUSG( | Snrnp200 | 3338.269 | 0.183619 | 0.053082 | 0.353671 | ENSMUSG( | Snrnp200 | 2722.794 | -0.13245 | 0.228344 | 0.868507 |
| ENSMUSG( | Snrnp25 | 410.5475 | -0.01636 | 0.892581 | 0.984474 | ENSMUSG( | Snrnp25 | 345.5562 | -0.26551 | 0.084643 | 0.779799 |
| ENSMUSG( | Snrnp27 | 756.603 | 0.07689 | 0.444303 | 0.833239 | ENSMUSG( | Snrnp27 | 659.6603 | -0.05421 | 0.696143 | 0.960867 |
| ENSMUSG( | Snrnp35 | 278.9328 | 0.023257 | 0.868848 | 0.976897 | ENSMUSG( | Snrnp35 | 233.1556 | -0.23309 | 0.142081 | 0.830709 |
| ENSMUSG( | Snrnp40 | 430.6581 | -0.06735 | 0.653845 | 0.922914 | ENSMUSG( | Snrnp40 | 394.0806 | -0.05982 | 0.725022 | 0.966887 |
| ENSMUSG( | Snrnp48 | 599.4268 | 0.000807 | 0.994629 | 0.999369 | ENSMUSG( | Snrnp48 | 611.4608 | 0.314893 | 0.032233 | 0.634419 |
| ENSMUSG( | Snrnp70 | 8137.617 | 0.109332 | 0.499039 | 0.862973 | ENSMUSG( | Snrnp70 | 7505.544 | 0.1408 | 0.44542 | 0.921734 |
| ENSMUSG( | Snrpa | 1202.808 | -0.00295 | 0.98023 | 0.997512 | ENSMUSG( | Snrpa | 1036.884 | -0.17185 | 0.199084 | 0.863016 |
| ENSMUSG( | Snrpa1 | 539.5899 | -0.01733 | 0.887677 | 0.983411 | ENSMUSG( | Snrpa1 | 486.953 | -0.04259 | 0.7558 | 0.971371 |
| ENSMUSG( | Snrpb | 1306.81 | 0.053888 | 0.610756 | 0.908349 | ENSMUSG( | Snrpb | 1156.22 | -0.03151 | 0.7973 | 0.97681 |
| ENSMUSG( | Snrpb2 | 571.8846 | -0.11348 | 0.359903 | 0.791543 | ENSMUSG( | Snrpb2 | 517.1792 | -0.14231 | 0.32808 | 0.906255 |
| ENSMUSG( | Snrpc | 655.3688 | -0.02733 | 0.808573 | 0.963455 | ENSMUSG( | Snrpc | 568.3927 | -0.17889 | 0.188825 | 0.85698 |
| ENSMUSG( | Snrpd1 | 674.8446 | -0.01324 | 0.917705 | 0.989377 | ENSMUSG( | Snrpd1 | 591.4959 | -0.13655 | 0.400462 | 0.917257 |
| ENSMUSG( | Snrpd2 | 1299.874 | -0.04711 | 0.673312 | 0.929007 | ENSMUSG( | Snrpd2 | 1105.876 | -0.26071 | 0.077457 | 0.772673 |
| ENSMUSG( | Snrpd3 | 1345.725 | -0.02833 | 0.821962 | 0.966352 | ENSMUSG( | Snrpd3 | 1163.407 | -0.18947 | 0.211542 | 0.864412 |
| ENSMUSG( | Snrpe | 561.4303 | 0.072415 | 0.548056 | 0.885153 | ENSMUSG( | Snrpe | 478.219 | -0.13206 | 0.408584 | 0.917257 |
| ENSMUSG( | Snrpf | 554.6091 | -0.04196 | 0.740354 | 0.947823 | ENSMUSG( | Snrpf | 477.9185 | -0.22189 | 0.207339 | 0.864412 |
| ENSMUSG( | Snrpg | 488.9993 | 0.009457 | 0.935855 | 0.993415 | ENSMUSG( | Snrpg | 417.3001 | -0.1939 | 0.203399 | 0.864412 |
| ENSMUSG( | Snrpn | 11973.44 | 0.065848 | 0.459287 | 0.840244 | ENSMUSG( | Snrpn | 10214.18 | -0.12784 | 0.307748 | 0.898538 |
| ENSMUSG( | Snta1 | 1737.273 | 0.108843 | 0.218788 | 0.664559 | ENSMUSG( | Snta1 | 1489.289 | -0.06782 | 0.557764 | 0.939565 |
| ENSMUSG( | Sntb1 | 234.166 | -0.17925 | 0.328391 | 0.772854 | ENSMUSG( | Sntb1 | 221.5754 | -0.08073 | 0.678568 | 0.957506 |
| ENSMUSG( | Sntb2 | 477.9562 | -0.28159 | 0.020096 | 0.214413 | ENSMUSG( | Sntb2 | 444.5003 | -0.21362 | 0.123788 | 0.814691 |
| ENSMUSG( | Sntg1 | 257.55 | -0.22112 | 0.125725 | 0.53257 | ENSMUSG( | Sntg1 | 256.1181 | 0.043497 | 0.785522 | 0.976285 |
| ENSMUSG( | Sntg2 | 51.92733 | -0.16414 | 0.566886 | 0.891644 | ENSMUSG( | Sntg2 | 46.37064 | -0.21724 | 0.481195 | 0.924548 |
| ENSMUSG( | Snu13 | 1934.278 | 0.019511 | 0.832454 | 0.970333 | ENSMUSG( | Snu13 | 1630.349 | -0.21379 | 0.085629 | 0.779799 |
| ENSMUSG( | Snupn | 288.3362 | 0.055043 | 0.705768 | 0.938649 | ENSMUSG( | Snupn | 259.5342 | 0.024662 | 0.881309 | 0.989644 |
| ENSMUSG( | Snw1 | 1635.928 | 0.201878 | 0.029626 | 0.26088 | ENSMUSG( | Snw1 | 1425.949 | 0.077954 | 0.494085 | 0.927372 |
| ENSMUSG( | Snx1 | 1576.479 | 0.008677 | 0.920282 | 0.990247 | ENSMUSG( | Snx1 | 1387.92 | -0.09079 | 0.396611 | 0.917257 |
| ENSMUSG( | Snx10 | 1708.675 | -0.03378 | 0.75685 | 0.951513 | ENSMUSG( | Snx10 | 1609.783 | 0.057918 | 0.688412 | 0.959742 |
| ENSMUSG( | Snx11 | 899.1962 | -0.01027 | 0.922391 | 0.99045 | ENSMUSG( | Snx11 | 824.1043 | 0.008188 | 0.947303 | 0.994798 |
| ENSMUSG( | Snx12 | 1147.596 | -0.0104 | 0.919297 | 0.990247 | ENSMUSG( | Snx12 | 949.9955 | -0.30716 | 0.012091 | 0.445877 |
| ENSMUSG( | Snx13 | 3045.798 | -0.05342 | 0.668825 | 0.927963 | ENSMUSG( | Snx13 | 2840.382 | 0.015918 | 0.907029 | 0.991601 |
| ENSMUSG( | Snx14 | 1398.983 | -0.26181 | 0.091765 | 0.465904 | ENSMUSG( | Snx14 | 1510.3 | 0.236253 | 0.163213 | 0.838291 |
| ENSMUSG( | Snx15 | 863.9605 | -0.14429 | 0.254814 | 0.705183 | ENSMUSG( | Snx15 | 784.2728 | -0.15358 | 0.264289 | 0.886134 |
| ENSMUSG( | Snx16 | 285.7198 | -0.03142 | 0.832685 | 0.970333 | ENSMUSG( | Snx16 | 271.7067 | 0.079765 | 0.662471 | 0.956205 |
| ENSMUSG( | Snx17 | 2283.096 | 0.093513 | 0.270373 | 0.720226 | ENSMUSG( | Snx17 | 1913.409 | -0.15072 | 0.146538 | 0.835973 |
| ENSMUSG( | Snx18 | 1229.086 | -0.09356 | 0.447717 | 0.834051 | ENSMUSG( | Snx18 | 1139.229 | -0.04725 | 0.726994 | 0.966887 |
| ENSMUSG( | Snx19 | 2688.77 | -0.22744 | 0.088434 | 0.456236 | ENSMUSG( | Snx19 | 2768.7 | 0.135544 | 0.373769 | 0.915491 |
| ENSMUSG( | Snx2 | 1116.086 | 0.146494 | 0.147565 | 0.570152 | ENSMUSG( | Snx2 | 978.5033 | 0.036552 | 0.764313 | 0.974 |
| ENSMUSG( | Snx20 | 43.80069 | -0.02782 | 0.928039 | 0.991229 | ENSMUSG( | Snx20 | 36.74471 | -0.24333 | 0.496281 | 0.927825 |
| ENSMUSG( | Snx21 | 1288.994 | 0.090756 | 0.367637 | 0.795314 | ENSMUSG( | Snx21 | 1166.251 | 0.074392 | 0.527816 | 0.934603 |
| ENSMUSG( | Snx22 | 364.572 | -0.21267 | 0.094587 | 0.471772 | ENSMUSG( | Snx22 | 361.2717 | 0.044956 | 0.791926 | 0.97681 |
| ENSMUSG( | Snx24 | 507.5881 | 0.089159 | 0.455077 | 0.837743 | ENSMUSG( | Snx24 | 444.0662 | -0.03353 | 0.82283 | 0.979198 |
| ENSMUSG( | Snx25 | 1553.504 | -0.07746 | 0.571855 | 0.893506 | ENSMUSG( | Snx25 | 1578.744 | 0.233884 | 0.159518 | 0.837724 |
| ENSMUSG( | Snx27 | 5760.704 | 0.100311 | 0.265268 | 0.716034 | ENSMUSG( | Snx27 | 4866.783 | -0.1185 | 0.237087 | 0.872173 |
| ENSMUSG( | Snx29 | 331.7938 | -0.02885 | 0.835624 | 0.970557 | ENSMUSG( | Snx29 | 289.6253 | -0.15168 | 0.319292 | 0.903453 |
| ENSMUSG( | Snx3 | 5353.265 | 0.020246 | 0.816519 | 0.965455 | ENSMUSG( | Snx3 | 4491.061 | -0.22958 | 0.085174 | 0.779799 |
| ENSMUSG( | Snx30 | 1225.318 | -0.05149 | 0.613577 | 0.909044 | ENSMUSG( | Snx30 | 1107.288 | -0.0728 | 0.555794 | 0.939565 |
| ENSMUSG( | Snx32 | 2080.218 | 0.126652 | 0.376937 | 0.801213 | ENSMUSG( | Snx32 | 1860.898 | 0.073453 | 0.69331 | 0.96076 |
| ENSMUSG( | Snx33 | 351.8367 | -0.32245 | 0.012823 | 0.163518 | ENSMUSG( | Snx33 | 366.981 | 0.093964 | 0.559732 | 0.939812 |
| ENSMUSG( | Snx4 | 1212.978 | 0.027155 | 0.783364 | 0.957558 | ENSMUSG( | Snx4 | 1064.527 | -0.08647 | 0.488103 | 0.925453 |
| ENSMUSG( | Snx5 | 5252.799 | 0.050397 | 0.61264 | 0.908818 | ENSMUSG( | Snx5 | 4518.582 | -0.11929 | 0.322309 | 0.904023 |
| ENSMUSG( | Snx6 | 1222.125 | 0.136231 | 0.235575 | 0.682693 | ENSMUSG( | Snx6 | 1050.631 | -0.03222 | 0.797277 | 0.97681 |
| ENSMUSG( | Snx7 | 233.5405 | 0.011581 | 0.939977 | 0.993415 | ENSMUSG( | Snx7 | 205.6207 | -0.09363 | 0.599109 | 0.947045 |
| ENSMUSG( | Snx8 | 726.7861 | 0.075121 | 0.544593 | 0.884649 | ENSMUSG( | Snx8 | 615.6119 | -0.14321 | 0.283313 | 0.892412 |
| ENSMUSG( | Snx9 | 795.5339 | 0.055432 | 0.601441 | 0.904603 | ENSMUSG( | Snx9 | 711.9511 | -0.00087 | 0.994799 | 0.998813 |
| ENSMUSG( | Soat1 | 1516.359 | -0.14929 | 0.309239 | 0.75734 | ENSMUSG( | Soat1 | 1581.024 | 0.237279 | 0.147875 | 0.836241 |
| ENSMUSG( | Sobp | 2768.189 | 0.178626 | 0.045371 | 0.326933 | ENSMUSG( | Sobp | 2572.537 | 0.232484 | 0.023711 | 0.588226 |
| ENSMUSG( | Socs1 | 35.0619 | -0.42401 | 0.23843 | 0.686308 | ENSMUSG( | Socs1 | 29.68354 | -0.65496 | 0.132159 | 0.822912 |
| ENSMUSG( | Socs2 | 406.1417 | -0.05063 | 0.681466 | 0.931595 | ENSMUSG( | Socs2 | 393.4051 | 0.116982 | 0.431272 | 0.917257 |
| ENSMUSG( | Socs3 | 100.4822 | -0.43013 | 0.049626 | 0.340062 | ENSMUSG( | Socs3 | 105.9278 | 0.020208 | 0.92515 | 0.99306 |
| ENSMUSG( | Socs4 | 598.4244 | 0.081015 | 0.541713 | 0.883182 | ENSMUSG( | Socs4 | 553.8267 | 0.116956 | 0.482417 | 0.924578 |
| ENSMUSG( | Socs5 | 2805.199 | 0.067852 | 0.442539 | 0.83278 | ENSMUSG( | Socs5 | 2538.939 | 0.047722 | 0.645866 | 0.953988 |
| ENSMUSG( | Socs6 | 405.7878 | 0.16574 | 0.220121 | 0.665863 | ENSMUSG( | Socs6 | 389.7891 | 0.30048 | 0.050062 | 0.708648 |
| ENSMUSG( | Socs7 | 4912.055 | 0.081883 | 0.343365 | 0.779153 | ENSMUSG( | Socs7 | 4329.842 | -0.01204 | 0.906782 | 0.991517 |
| ENSMUSG( | Sod1 | 6726.248 | -0.03327 | 0.680979 | 0.931244 | ENSMUSG( | Sod1 | 5881.247 | -0.15807 | 0.189103 | 0.85698 |
| ENSMUSG( | Sod2 | 4931.645 | 0.011252 | 0.901562 | 0.985633 | ENSMUSG( | Sod2 | 4253.978 | -0.15336 | 0.245983 | 0.877904 |
| ENSMUSG( | Sod3 | 1316.962 | -0.3448 | 0.025768 | 0.247138 | ENSMUSG( | Sod3 | 1333.947 | -0.01288 | 0.94096 | 0.994558 |
| ENSMUSG( | Son | 7271.246 | 0.032063 | 0.744103 | 0.948497 | ENSMUSG( | Son | 7264.388 | 0.285108 | 0.056567 | 0.723362 |
| ENSMUSG( | Sorbs1 | 4753.807 | 0.240897 | 0.04882 | 0.337919 | ENSMUSG( | Sorbs1 | 3811.458 | -0.12124 | 0.2785 | 0.891628 |
| ENSMUSG( | Sorbs2 | 4462.09 | -0.22583 | 0.006795 | 0.113745 | ENSMUSG( | Sorbs2 | 4328.639 | -0.03674 | 0.737936 | 0.967709 |
| ENSMUSG( | Sorbs3 | 2123.247 | -0.03146 | 0.721397 | 0.943026 | ENSMUSG( | Sorbs3 | 1971.449 | 0.024592 | 0.823518 | 0.979198 |
| ENSMUSG( | Sorcs1 | 1144.699 | -0.38393 | 0.005392 | 0.098038 | ENSMUSG( | Sorcs1 | 1207.155 | 0.068043 | 0.667224 | 0.956766 |
| ENSMUSG( | Sorcs2 | 2879.858 | -0.43881 | 0.001759 | 0.047383 | ENSMUSG( | Sorcs2 | 3026.736 | 0.011446 | 0.941282 | 0.994578 |
| ENSMUSG( | Sorcs3 | 1051.729 | -0.5323 | 0.000506 | 0.020608 | ENSMUSG( | Sorcs3 | 1090.316 | -0.10585 | 0.565715 | 0.941404 |
| ENSMUSG( | Sord | 612.9815 | 0.075984 | 0.480018 | 0.85456 | ENSMUSG( | Sord | 550.2941 | 0.031845 | 0.816032 | 0.979198 |
| ENSMUSG( | Sorl1 | 10673.71 | -0.13385 | 0.323671 | 0.768548 | ENSMUSG( | Sorl1 | 10529.8 | 0.100783 | 0.524773 | 0.933907 |
| ENSMUSG( | Sort1 | 16119.36 | -0.2757 | 0.046809 | 0.330855 | ENSMUSG( | Sort1 | 16710.77 | 0.111841 | 0.496841 | 0.927825 |
| ENSMUSG( | Sos1 | 1218.52 | -0.03034 | 0.733426 | 0.944956 | ENSMUSG( | Sos1 | 1146.916 | 0.060292 | 0.606766 | 0.949772 |
| ENSMUSG( | Sos2 | 1325.502 | -0.00108 | 0.991685 | 0.999232 | ENSMUSG( | Sos2 | 1251.642 | 0.097093 | 0.444741 | 0.921734 |
| ENSMUSG( | Sostdc1 | 105.1762 | -0.38095 | 0.546877 | 0.884974 | ENSMUSG( | Sostdc1 | 112.7499 | 0.097892 | 0.770898 | 0.974236 |
| ENSMUSG( | Sowaha | 3507.855 | 0.048594 | 0.590181 | 0.90113 | ENSMUSG( | Sowaha | 3334.453 | 0.167711 | 0.101245 | 0.801456 |
| ENSMUSG( | Sowahb | 275.8507 | -0.00448 | 0.976514 | 0.997428 | ENSMUSG( | Sowahb | 248.8805 | -0.02529 | 0.886158 | 0.989913 |
| ENSMUSG( | Sowahc | 459.0935 | -0.02459 | 0.848871 | 0.972503 | ENSMUSG( | Sowahc | 423.8995 | 0.010251 | 0.947244 | 0.994798 |
| ENSMUSG( | Sox1 | 565.0058 | -0.00061 | 0.996354 | 0.99943 | ENSMUSG( | Sox1 | 575.3152 | 0.299219 | 0.058134 | 0.728256 |
| ENSMUSG( | Sox10 | 1350.861 | -0.12819 | 0.17072 | 0.605271 | ENSMUSG( | Sox10 | 1292.024 | 0.014719 | 0.894599 | 0.990558 |
| ENSMUSG( | Sox11 | 655.0808 | -0.08311 | 0.526377 | 0.877267 | ENSMUSG( | Sox11 | 669.4272 | 0.233061 | 0.14369 | 0.831311 |
| ENSMUSG( | Sox12 | 984.6323 | -0.10545 | 0.37204 | 0.798768 | ENSMUSG( | Sox12 | 897.0586 | -0.10734 | 0.416396 | 0.917257 |
| ENSMUSG( | Sox13 | 455.5114 | 0.039725 | 0.759167 | 0.952284 | ENSMUSG( | Sox13 | 466.492 | 0.354322 | 0.011328 | 0.433258 |
| ENSMUSG( | Sox17 | 195.4073 | 0.670873 | 0.000397 | 0.017766 | ENSMUSG( | Sox17 | 228.2736 | 1.228708 | 5.23E-08 | 3.67E-05 |
| ENSMUSG( | Sox18 | 319.3858 | 0.317021 | 0.17566 | 0.611006 | ENSMUSG( | Sox18 | 339.7014 | 0.707884 | 0.003815 | 0.246611 |
| ENSMUSG( | Sox2 | 1730.305 | -0.14147 | 0.275171 | 0.72509 | ENSMUSG( | Sox2 | 1677.389 | 0.036307 | 0.837154 | 0.981335 |
| ENSMUSG( | Sox21 | 889.235 | 0.284337 | 0.018488 | 0.204273 | ENSMUSG( | Sox21 | 864.3933 | 0.445162 | 0.012172 | 0.447008 |
| ENSMUSG( | Sox21os1 | 61.61023 | -0.22731 | 0.390322 | 0.80794 | ENSMUSG( | Sox21os1 | 54.40303 | -0.34174 | 0.223671 | 0.867404 |
| ENSMUSG( | Sox2ot | 971.3932 | -0.15321 | 0.162572 | 0.593364 | ENSMUSG( | Sox2ot | 912.3572 | -0.0592 | 0.653489 | 0.955015 |
| ENSMUSG( | Sox3 | 112.2918 | 0.1757 | 0.39745 | 0.812127 | ENSMUSG( | Sox3 | 87.89062 | -0.29053 | 0.231385 | 0.868507 |
| ENSMUSG( | Sox4 | 1279.308 | 0.034572 | 0.770356 | 0.954689 | ENSMUSG( | Sox4 | 1247.669 | 0.218282 | 0.224036 | 0.867404 |
| ENSMUSG( | Sox5 | 1431.256 | -0.02799 | 0.817046 | 0.96566 | ENSMUSG( | Sox5 | 1267.169 | -0.1161 | 0.420336 | 0.917257 |
| ENSMUSG( | Sox6 | 1484.736 | -0.08326 | 0.521799 | 0.875186 | ENSMUSG( | Sox6 | 1257.757 | -0.31033 | 0.01385 | 0.46927 |
| ENSMUSG( | Sox7 | 88.36718 | 0.066416 | 0.816351 | 0.965455 | ENSMUSG( | Sox7 | 102.5687 | 0.698548 | 0.016101 | 0.510946 |
| ENSMUSG( | Sox8 | 1590.031 | -0.23861 | 0.010699 | 0.148964 | ENSMUSG( | Sox8 | 1579.855 | 0.02392 | 0.841625 | 0.982981 |
| ENSMUSG( | Sox9 | 1345.704 | -0.011 | 0.91331 | 0.988483 | ENSMUSG( | Sox9 | 1338.724 | 0.230939 | 0.068179 | 0.75212 |
| ENSMUSG( | Sp1 | 809.4683 | -0.03855 | 0.712957 | 0.94066 | ENSMUSG( | Sp1 | 747.4124 | 0.003617 | 0.976667 | 0.996416 |
| ENSMUSG( | Sp100 | 181.6339 | -0.19051 | 0.370083 | 0.797542 | ENSMUSG( | Sp100 | 167.2545 | -0.14298 | 0.556737 | 0.939565 |
| ENSMUSG( | Sp110 | 73.36795 | -0.4886 | 0.050707 | 0.345138 | ENSMUSG( | Sp110 | 76.78305 | -0.03978 | 0.869492 | 0.988097 |
| ENSMUSG( | Sp140 | 41.19746 | -0.2159 | 0.488586 | 0.859092 | ENSMUSG( | Sp140 | 37.38143 | -0.22645 | 0.509435 | 0.929089 |
| ENSMUSG( | Sp2 | 503.0191 | -0.05852 | 0.610536 | 0.908189 | ENSMUSG( | Sp2 | 432.2983 | -0.23463 | 0.083 | 0.77681 |
| ENSMUSG( | Sp3 | 1291.533 | -0.01393 | 0.880665 | 0.981542 | ENSMUSG( | Sp3 | 1224.05 | 0.095998 | 0.437941 | 0.920495 |
| ENSMUSG( | Sp3os | 436.4977 | 0.024856 | 0.849653 | 0.972638 | ENSMUSG( | Sp3os | 381.5785 | -0.10652 | 0.518334 | 0.930939 |
| ENSMUSG( | Sp4 | 346.3537 | -0.10931 | 0.392945 | 0.809274 | ENSMUSG( | Sp4 | 317.7632 | -0.08952 | 0.532153 | 0.934795 |
| ENSMUSG( | Sp5 | 15.12199 | 0.22387 | 0.632413 | NA | ENSMUSG( | Sp5 | 14.51682 | 0.352155 | 0.484692 | 0.92541 |
| ENSMUSG( | Sp6 | 32.5028 | -0.15919 | 0.625424 | 0.912249 | ENSMUSG( | Sp6 | 29.98195 | -0.08389 | 0.831739 | 0.980656 |
| ENSMUSG( | Sp7 | 52.81902 | -0.59878 | 0.02719 | 0.253245 | ENSMUSG( | Sp7 | 55.27437 | -0.17331 | 0.563233 | 0.941158 |
| ENSMUSG( | Sp8 | 67.94008 | -0.66653 | 0.055208 | 0.361664 | ENSMUSG( | Sp8 | 84.6556 | 0.271924 | 0.445367 | 0.921734 |
| ENSMUSG( | Sp9 | 192.363 | -0.4225 | 0.04314 | 0.318494 | ENSMUSG( | Sp9 | 210.0064 | 0.111956 | 0.621186 | 0.953199 |
| ENSMUSG( | Spa17 | 87.19493 | -0.1598 | 0.501525 | 0.864477 | ENSMUSG( | Spa17 | 80.06849 | -0.15803 | 0.526837 | 0.934449 |
| ENSMUSG( | Spaar | 25.71687 | -0.11731 | 0.756284 | 0.951479 | ENSMUSG( | Spaar | 23.77034 | -0.01683 | 0.977551 | 0.996416 |
| ENSMUSG( | Spaca6 | 720.8754 | 0.111363 | 0.527709 | 0.877801 | ENSMUSG( | Spaca6 | 679.3659 | 0.203781 | 0.297556 | 0.898538 |
| ENSMUSG( | Spag1 | 687.731 | 0.151971 | 0.173057 | 0.608361 | ENSMUSG( | Spag1 | 602.2866 | 0.037139 | 0.80417 | 0.977191 |
| ENSMUSG( | Spag16 | 29.74129 | -0.0246 | 0.953062 | 0.993874 | ENSMUSG( | Spag16 | 30.8896 | 0.359567 | 0.450577 | 0.922202 |
| ENSMUSG( | Spag17 | 14.99672 | -0.28845 | 0.540976 | NA | ENSMUSG( | Spag17 | 20.21921 | 0.79318 | 0.082467 | 0.77681 |
| ENSMUSG( | Spag4 | 136.8533 | 0.129529 | 0.488455 | 0.859092 | ENSMUSG( | Spag4 | 117.305 | -0.03322 | 0.877968 | 0.989644 |
| ENSMUSG( | Spag5 | 795.2436 | 0.526308 | 0.003712 | 0.07748 | ENSMUSG( | Spag5 | 603.3619 | 0.041486 | 0.83854 | 0.981622 |
| ENSMUSG( | Spag6 | 46.07336 | 0.192193 | 0.515766 | 0.872701 | ENSMUSG( | Spag6 | 42.16902 | 0.207737 | 0.50048 | 0.927825 |
| ENSMUSG( | Spag6l | 85.7836 | 0.001173 | 0.995949 | 0.99943 | ENSMUSG( | Spag6l | 89.82328 | 0.386216 | 0.111226 | 0.807885 |
| ENSMUSG( | Spag7 | 995.0441 | 0.034412 | 0.781817 | 0.957558 | ENSMUSG( | Spag7 | 878.8745 | -0.06118 | 0.701985 | 0.962014 |
| ENSMUSG( | Spag8 | 49.29074 | 0.242099 | 0.433529 | 0.828346 | ENSMUSG( | Spag8 | 48.40372 | 0.455619 | 0.162038 | 0.837724 |
| ENSMUSG( | Spag9 | 21686.79 | 0.017046 | 0.877827 | 0.981431 | ENSMUSG( | Spag9 | 19372.82 | -0.03997 | 0.730185 | 0.967228 |
| ENSMUSG( | Sparc | 43174.23 | 0.071239 | 0.541606 | 0.883182 | ENSMUSG( | Sparc | 38891.36 | 0.036527 | 0.770246 | 0.974236 |
| ENSMUSG( | Sparcl1 | 203036.2 | 0.082019 | 0.414464 | 0.82013 | ENSMUSG( | Sparcl1 | 181989 | 0.036045 | 0.761117 | 0.972695 |
| ENSMUSG( | Spart | 1237.495 | 0.027909 | 0.781933 | 0.957558 | ENSMUSG( | Spart | 1073.533 | -0.11886 | 0.353359 | 0.912263 |
| ENSMUSG( | Spast | 1688.402 | -0.04706 | 0.663229 | 0.9265 | ENSMUSG( | Spast | 1634.865 | 0.127124 | 0.298751 | 0.898538 |
| ENSMUSG( | Spata1 | 68.63108 | -0.10806 | 0.64556 | 0.918025 | ENSMUSG( | Spata1 | 64.32267 | -0.00498 | 0.985089 | 0.997084 |
| ENSMUSG( | Spata13 | 467.5335 | -0.30238 | 0.046572 | 0.330304 | ENSMUSG( | Spata13 | 476.0061 | 0.032947 | 0.804904 | 0.977232 |

| | | | | | |
|---|---|---|---|---|---|
| ENSMUSG( | 894.5505 | 0.093282 | 0.418484 | 0.821966 | Slc7a6 |
| ENSMUSG( | 625.4786 | -0.03361 | 0.778952 | 0.956442 | Slc7a6os |
| ENSMUSG( | 106.6451 | 0.053749 | 0.803566 | 0.962169 | Slc7a7 |
| ENSMUSG( | 3583.596 | -0.51833 | 5.15E-05 | 0.004052 | Slc7a8 |
| ENSMUSG( | 3413.869 | -0.30895 | 0.073342 | 0.418269 | Slc8a1 |
| ENSMUSG( | 11966.46 | -0.39584 | 0.004955 | 0.093376 | Slc8a2 |
| ENSMUSG( | 917.7832 | -0.50622 | 3.30E-05 | 0.002906 | Slc8a3 |
| ENSMUSG( | 417.1933 | -0.46569 | 0.002918 | 0.066108 | Slc8b1 |
| ENSMUSG( | 1830.537 | -0.29641 | 0.020252 | 0.214698 | Slc9a1 |
| ENSMUSG( | 380.8778 | 0.000489 | 0.997127 | 0.999565 | Slc9a2 |
| ENSMUSG( | 25.74454 | 0.590194 | 0.15528 | 0.583155 | Slc9a3 |
| ENSMUSG( | 68.68419 | 0.202853 | 0.414898 | 0.820184 | Slc9a4 |
| ENSMUSG( | 1372.877 | -0.05491 | 0.622125 | 0.911432 | Slc9a5 |
| ENSMUSG( | 3468.971 | -0.3086 | 0.013697 | 0.171028 | Slc9a6 |
| ENSMUSG( | 249.5391 | 0.0039 | 0.984444 | 0.99781 | Slc9a7 |
| ENSMUSG( | 1207.83 | -0.11329 | 0.372061 | 0.798768 | Slc9a8 |
| ENSMUSG( | 793.5239 | -0.21877 | 0.123962 | 0.531451 | Slc9a9 |
| ENSMUSG( | 139.4562 | -0.06584 | 0.764527 | 0.954055 | Slc9b2 |
| ENSMUSG( | 2094.03 | -0.04645 | 0.814788 | 0.964668 | Slco1a4 |
| ENSMUSG( | 16.07486 | -0.44645 | 0.341271 | NA | Slco1b2 |
| ENSMUSG( | 3937.37 | -0.17982 | 0.333526 | 0.775602 | Slco1c1 |
| ENSMUSG( | 92.61515 | -0.3672 | 0.10134 | 0.484996 | Slco2a1 |
| ENSMUSG( | 1334.328 | 0.013181 | 0.938917 | 0.993415 | Slco2b1 |
| ENSMUSG( | 944.8162 | -0.23391 | 0.136271 | 0.550578 | Slco3a1 |
| ENSMUSG( | 162.4304 | -0.52544 | 0.002531 | 0.059664 | Slco4a1 |
| ENSMUSG( | 56.68072 | -0.61508 | 0.037856 | 0.30067 | Slco4c1 |
| ENSMUSG( | 63.92293 | -0.83931 | 0.000714 | 0.025843 | Slco5a1 |
| ENSMUSG( | 338.445 | -0.07747 | 0.575678 | 0.894836 | Slf1 |
| ENSMUSG( | 1454.01 | 0.00808 | 0.932323 | 0.992515 | Slf2 |
| ENSMUSG( | 67.00886 | -0.30381 | 0.243178 | 0.69052 | Slfn2 |
| ENSMUSG( | 10.88428 | 0.750173 | 0.229238 | NA | Slfn3 |
| ENSMUSG( | 10.08023 | 0.077512 | 0.925698 | NA | Slfn4 |
| ENSMUSG( | 341.21 | 0.107825 | 0.625565 | 0.912249 | Slfn5 |
| ENSMUSG( | 71.71834 | -0.31998 | 0.157502 | 0.585653 | Slfn8 |
| ENSMUSG( | 61.00641 | -0.18268 | 0.540577 | 0.88315 | Slfn9 |
| ENSMUSG( | 617.4286 | -0.03367 | 0.757677 | 0.951582 | Slirp |
| ENSMUSG( | 1772.645 | 0.056575 | 0.769493 | 0.954674 | Slit1 |
| ENSMUSG( | 1031.993 | -0.38265 | 0.002734 | 0.063356 | Slit2 |
| ENSMUSG( | 710.5014 | -0.29019 | 0.063717 | 0.390618 | Slit3 |
| ENSMUSG( | 2083.275 | -0.65082 | 0.000159 | 0.008938 | Slitrk1 |
| ENSMUSG( | 830.137 | -0.5148 | 0.004248 | 0.084545 | Slitrk2 |
| ENSMUSG( | 1554.717 | -0.39784 | 0.003409 | 0.073541 | Slitrk3 |
| ENSMUSG( | 537.6 | -0.57024 | 0.000576 | 0.022554 | Slitrk4 |
| ENSMUSG( | 1726.015 | -0.37405 | 0.017523 | 0.198483 | Slitrk5 |
| ENSMUSG( | 19.65093 | -0.94285 | 0.027218 | NA | Slitrk6 |
| ENSMUSG( | 2811.602 | -0.07442 | 0.355977 | 0.788009 | Slk |
| ENSMUSG( | 2243.457 | -0.03405 | 0.722541 | 0.943752 | Slmap |
| ENSMUSG( | 8.897533 | 0.841141 | 0.191943 | NA | Sln |
| ENSMUSG( | 22.28206 | -0.3704 | 0.405939 | 0.81652 | Slpi |
| ENSMUSG( | 2390.501 | 0.1312 | 0.166979 | 0.599655 | Sltm |
| ENSMUSG( | 1574.56 | 0.083873 | 0.385986 | 0.804623 | Slu7 |
| ENSMUSG( | 523.9121 | -0.15577 | 0.195039 | 0.634458 | Slx1b |
| ENSMUSG( | 806.2251 | 0.053637 | 0.67784 | 0.930542 | Slx4 |
| ENSMUSG( | 212.3354 | 0.057386 | 0.700269 | 0.937391 | Slx4ip |
| ENSMUSG( | 1021.869 | 0.06544 | 0.515157 | 0.872463 | Slx9 |
| ENSMUSG( | 510.0055 | 0.008486 | 0.939125 | 0.993415 | Smad1 |
| ENSMUSG( | 851.495 | 0.03897 | 0.716774 | 0.941697 | Smad2 |
| ENSMUSG( | 1571.889 | 0.104436 | 0.279691 | 0.729119 | Smad3 |
| ENSMUSG( | 1214.203 | 0.05501 | 0.618215 | 0.911006 | Smad4 |
| ENSMUSG( | 2343.265 | 0.036669 | 0.778917 | 0.956442 | Smad5 |
| ENSMUSG( | 60.73465 | 0.077445 | 0.797348 | 0.960507 | Smad6 |
| ENSMUSG( | 599.0351 | -0.0617 | 0.596442 | 0.903295 | Smad7 |
| ENSMUSG( | 426.2398 | -0.25097 | 0.04266 | 0.316769 | Smad9 |
| ENSMUSG( | 79.43057 | 0.205279 | 0.384003 | 0.803646 | Smagp |
| ENSMUSG( | 2198.292 | -0.03945 | 0.64148 | 0.916693 | Smap1 |
| ENSMUSG( | 3244.89 | 0.086321 | 0.423355 | 0.823662 | Smap2 |
| ENSMUSG( | 640.9804 | 0.145439 | 0.175803 | 0.611006 | Smarca1 |
| ENSMUSG( | 8357.507 | 0.177968 | 0.030783 | 0.266861 | Smarca2 |
| ENSMUSG( | 5302.722 | 0.178408 | 0.039163 | 0.304806 | Smarca4 |
| ENSMUSG( | 1578.064 | 0.164937 | 0.15757 | 0.585653 | Smarca5 |
| ENSMUSG( | 703.0274 | 0.034911 | 0.73896 | 0.947768 | Smarcad1 |
| ENSMUSG( | 986.6841 | 0.168252 | 0.087821 | 0.454663 | Smarcal1 |
| ENSMUSG( | 1512.83 | 0.043101 | 0.672059 | 0.9288 | Smarcb1 |
| ENSMUSG( | 2144.678 | 0.043939 | 0.607723 | 0.906878 | Smarcc1 |
| ENSMUSG( | 14106.69 | 0.05154 | 0.503864 | 0.866426 | Smarcc2 |
| ENSMUSG( | 6231.497 | 0.017056 | 0.858766 | 0.974707 | Smarcd1 |
| ENSMUSG( | 525.8302 | 0.081568 | 0.599441 | 0.903801 | Smarcd2 |
| ENSMUSG( | 2990.431 | 0.035981 | 0.672439 | 0.929007 | Smarcd3 |
| ENSMUSG( | 1436.151 | -0.01212 | 0.899486 | 0.985633 | Smarce1 |
| ENSMUSG( | 2055.599 | -0.03949 | 0.691776 | 0.934101 | Smc1a |
| ENSMUSG( | 225.9491 | -0.05353 | 0.723262 | 0.943761 | Smc2 |
| ENSMUSG( | 24.15828 | -0.19288 | 0.664948 | 0.926824 | Smc2os |
| ENSMUSG( | 1756.969 | 0.20069 | 0.029715 | 0.261211 | Smc3 |
| ENSMUSG( | 344.6831 | 0.086292 | 0.567867 | 0.892002 | Smc4 |
| ENSMUSG( | 796.3666 | 0.080533 | 0.467139 | 0.844819 | Smc5 |
| ENSMUSG( | 1272.597 | 0.103617 | 0.271013 | 0.721337 | Smc6 |
| ENSMUSG( | 911.3996 | 0.061992 | 0.597355 | 0.903694 | Smchd1 |
| ENSMUSG( | 106.9802 | -0.02113 | 0.914339 | 0.988483 | Smco3 |
| ENSMUSG( | 59.73276 | 0.053758 | 0.834544 | 0.970344 | Smco4 |
| ENSMUSG( | 804.3925 | 0.164287 | 0.119229 | 0.523036 | Smcr8 |
| ENSMUSG( | 3232.963 | -0.08036 | 0.419981 | 0.822442 | Smdt1 |
| ENSMUSG( | 4079.569 | -0.16946 | 0.062992 | 0.388234 | Smg1 |
| ENSMUSG( | 3053.834 | 0.005063 | 0.95734 | 0.993874 | Smg5 |
| ENSMUSG( | 1478.606 | -0.00636 | 0.942622 | 0.993415 | Smg6 |
| ENSMUSG( | 2661.586 | 0.188628 | 0.021846 | 0.225699 | Smg7 |
| ENSMUSG( | 310.547 | 0.065992 | 0.693265 | 0.934202 | Smg8 |
| ENSMUSG( | 1058.63 | 0.087506 | 0.459337 | 0.840244 | Smg9 |
| ENSMUSG( | 1015.488 | -0.04812 | 0.648711 | 0.920025 | Smim1 |
| ENSMUSG( | 3944.648 | 0.011435 | 0.92664 | 0.991229 | Smim10l1 |
| ENSMUSG( | 927.3868 | 0.095479 | 0.399698 | 0.813337 | Smim10l2a |
| ENSMUSG( | 282.5267 | -0.03581 | 0.7931 | 0.959505 | Smim11 |
| ENSMUSG( | 1090.039 | -0.23082 | 0.019161 | 0.208619 | Smim12 |
| ENSMUSG( | 6416.138 | -0.05646 | 0.523238 | 0.875628 | Smim13 |
| ENSMUSG( | 3607.516 | -0.04541 | 0.619557 | 0.911432 | Smim14 |
| ENSMUSG( | 1360.173 | 0.028058 | 0.828281 | 0.968966 | Smim15 |
| ENSMUSG( | 188.849 | -0.12003 | 0.539703 | 0.88315 | Smim17 |
| ENSMUSG( | 123.8962 | -0.36933 | 0.060015 | 0.378891 | Smim18 |
| ENSMUSG( | 761.1088 | -0.22762 | 0.054795 | 0.360194 | Smim19 |
| ENSMUSG( | 822.1227 | 0.006321 | 0.956959 | 0.993874 | Smim20 |
| ENSMUSG( | 50.40918 | 0.292976 | 0.291162 | 0.741182 | Smim24 |
| ENSMUSG( | 305.5624 | -0.31964 | 0.016035 | 0.187923 | Smim26 |
| ENSMUSG( | 115.9169 | 0.132667 | 0.527547 | 0.877641 | Smim27 |
| ENSMUSG( | 1744.381 | -0.05319 | 0.527253 | 0.877492 | Smim29 |
| ENSMUSG( | 858.4561 | 0.75385 | 9.04E-14 | 9.77E-11 | Smim3 |
| ENSMUSG( | 754.6061 | 0.063412 | 0.600731 | 0.90443 | Smim30 |
| ENSMUSG( | 107.8187 | -0.0497 | 0.800507 | 0.961745 | Smim32 |
| ENSMUSG( | 18.00039 | -0.32129 | 0.533149 | NA | Smim35 |
| ENSMUSG( | 204.6323 | 0.011654 | 0.939822 | 0.993415 | Smim36 |
| ENSMUSG( | 11.35281 | 0.342071 | 0.583125 | NA | Smim38 |
| ENSMUSG( | 694.0499 | 0.141082 | 0.298725 | 0.747452 | Smim43 |
| ENSMUSG( | 5988.451 | 0.094223 | 0.335707 | 0.777 | Smim45 |
| ENSMUSG( | 12.53214 | -0.8362 | 0.114317 | NA | Smim5 |
| ENSMUSG( | 3064.426 | -0.10093 | 0.329474 | 0.773311 | Smim7 |
| ENSMUSG( | 383.0485 | 0.000662 | 0.996909 | 0.999505 | Smim8 |
| ENSMUSG( | 304.3244 | 0.093148 | 0.549611 | 0.885944 | Smn1 |
| ENSMUSG( | 669.8251 | 0.000237 | 0.998575 | 0.99989 | Smndc1 |
| ENSMUSG( | 1526.569 | 0.166596 | 0.154429 | 0.581463 | Smo |
| ENSMUSG( | 816.5548 | -0.3606 | 0.008303 | 0.127252 | Smoc1 |
| ENSMUSG( | 453.6695 | -0.12744 | 0.415047 | 0.820184 | Smoc2 |
| ENSMUSG( | 3691.581 | 0.272526 | 0.002176 | 0.054296 | Smox |
| ENSMUSG( | 4433.55 | -0.18093 | 0.205666 | 0.64712 | Smpd1 |
| ENSMUSG( | 920.4046 | -0.08048 | 0.511762 | 0.871106 | Smpd2 |
| ENSMUSG( | 3106.771 | -0.43544 | 0.002915 | 0.066108 | Smpd3 |
| ENSMUSG( | 1897.713 | 0.278373 | 0.088494 | 0.456254 | Smpd4 |
| ENSMUSG( | 27.8632 | -0.34909 | 0.344178 | 0.779153 | Smpd5 |
| ENSMUSG( | 1949.036 | -0.18719 | 0.204061 | 0.644572 | Smpdl3a |
| ENSMUSG( | 68.25841 | -0.76213 | 0.002378 | 0.057221 | Smpdl3b |
| ENSMUSG( | 1728.753 | 0.077258 | 0.490264 | 0.859319 | Sms |
| ENSMUSG( | 279.5371 | 0.149299 | 0.390897 | 0.80826 | Smtn |
| ENSMUSG( | 31.3206 | -0.74941 | 0.029324 | 0.259647 | Smtnl2 |
| ENSMUSG( | 1238.463 | -0.00096 | 0.993647 | 0.999369 | Smu1 |
| ENSMUSG( | 322.6168 | 0.030333 | 0.813919 | 0.964574 | Smug1 |
| ENSMUSG( | 2114.737 | 0.026809 | 0.78283 | 0.957558 | Smurf1 |
| ENSMUSG( | 1444.853 | -0.04617 | 0.637827 | 0.91586 | Smurf2 |
| ENSMUSG( | 411.5335 | -0.32433 | 0.090373 | 0.461669 | Smyd1 |
| ENSMUSG( | 1590.972 | 0.062149 | 0.561693 | 0.889052 | Smyd2 |
| ENSMUSG( | 650.2058 | -0.03813 | 0.76132 | 0.953046 | Smyd3 |
| ENSMUSG( | 269.6931 | -0.06103 | 0.653068 | 0.922794 | Smyd4 |
| ENSMUSG( | 1095.902 | 0.067877 | 0.504718 | 0.866883 | Smyd5 |
| ENSMUSG( | 24.74806 | 0.10505 | 0.788686 | 0.959226 | Snai1 |
| ENSMUSG( | 13.14406 | 0.011461 | 0.982041 | NA | Snai2 |
| ENSMUSG( | 11.00518 | -0.32249 | 0.586947 | NA | Snai3 |
| ENSMUSG( | 552.2993 | 0.126145 | 0.312958 | 0.76033 | Snap23 |
| ENSMUSG( | 28906.59 | 0.008654 | 0.949527 | 0.993821 | Snap25 |
| ENSMUSG( | 970.8261 | 0.150763 | 0.145624 | 0.566706 | Snap29 |
| ENSMUSG( | 6671.381 | 0.187855 | 0.259406 | 0.709948 | Snap47 |
| ENSMUSG( | 8221.906 | 0.200584 | 0.011516 | 0.154494 | Snap91 |
| ENSMUSG( | 1000.802 | -0.04125 | 0.693856 | 0.934202 | Snapc1 |

| | | | | | |
|---|---|---|---|---|---|
| ENSMUSG( | 1898.894 | 0.016352 | 0.918707 | 0.992529 | Smpdl3a |
| ENSMUSG( | 94.76502 | 0.518911 | 0.01889 | 0.548832 | Smpdl3b |
| ENSMUSG( | 1517.773 | -0.03345 | 0.814892 | 0.979198 | Sms |
| ENSMUSG( | 294.9043 | 0.542068 | 0.02013 | 0.562379 | Smtn |
| ENSMUSG( | 34.25369 | -0.16448 | 0.64249 | 0.953903 | Smtnl2 |
| ENSMUSG( | 1084.035 | -0.12208 | 0.365851 | 0.914623 | Smu1 |
| ENSMUSG( | 303.4347 | 0.123931 | 0.400572 | 0.917257 | Smug1 |
| ENSMUSG( | 1827.053 | -0.12777 | 0.253156 | 0.88018 | Smurf1 |
| ENSMUSG( | 1286.838 | -0.1157 | 0.329866 | 0.907107 | Smurf2 |
| ENSMUSG( | 330.3068 | -0.79126 | 0.000106 | 0.021511 | Smyd1 |
| ENSMUSG( | 1380.553 | -0.07982 | 0.516264 | 0.930939 | Smyd2 |
| ENSMUSG( | 578.2971 | -0.11261 | 0.440898 | 0.921166 | Smyd3 |
| ENSMUSG( | 261.3765 | 0.127688 | 0.467033 | 0.923329 | Smyd4 |
| ENSMUSG( | 906.6216 | -0.21377 | 0.106292 | 0.804969 | Smyd5 |
| ENSMUSG( | 24.91661 | 0.381409 | 0.352955 | 0.912263 | Snai1 |
| ENSMUSG( | 15.17488 | 0.666873 | 0.195264 | 0.861571 | Snai2 |
| ENSMUSG( | 485.1647 | 0.024473 | 0.863986 | 0.986552 | Snap23 |
| ENSMUSG( | 25127.57 | -0.13379 | 0.454939 | 0.922891 | Snap25 |
| ENSMUSG( | 820.0889 | -0.06999 | 0.614389 | 0.951177 | Snap29 |
| ENSMUSG( | 5323.53 | -0.19926 | 0.291571 | 0.897854 | Snap47 |
| ENSMUSG( | 7002.716 | 0.011458 | 0.919985 | 0.992644 | Snap91 |
| ENSMUSG( | 924.99 | -0.00112 | 0.992351 | 0.99874 | Snapc1 |
| ENSMUSG( | 526.766 | -0.08182 | 0.577978 | 0.943131 | Snapc2 |
| ENSMUSG( | 431.6863 | 0.157407 | 0.326204 | 0.905339 | Snapc3 |
| ENSMUSG( | 1360.016 | 0.078291 | 0.602436 | 0.94789 | Snapc4 |
| ENSMUSG( | 542.4351 | -0.3293 | 0.01383 | 0.46927 | Snapc5 |
| ENSMUSG( | 2234.858 | -0.26849 | 0.038 | 0.676422 | Snapin |
| ENSMUSG( | 3659.854 | -0.11799 | 0.48533 | 0.92541 | Snca |
| ENSMUSG( | 207.6944 | -0.3182 | 0.116741 | 0.813384 | Sncaip |
| ENSMUSG( | 9188.723 | -0.25286 | 0.186154 | 0.855305 | Sncb |
| ENSMUSG( | 22.45852 | -0.27899 | 0.574337 | 0.942912 | Sncg |
| ENSMUSG( | 2227.971 | -0.05748 | 0.608317 | 0.950113 | Snd1 |
| ENSMUSG( | 1132.268 | 0.007144 | 0.95952 | 0.994798 | Sned1 |
| ENSMUSG( | 1828.152 | -0.17576 | 0.134523 | 0.823094 | Snf8 |
| ENSMUSG( | 73.58803 | 0.111888 | 0.658571 | 0.956205 | Snhg10 |
| ENSMUSG( | 34374.78 | 0.164175 | 0.445061 | 0.921734 | Snhg11 |
| ENSMUSG( | 747.1321 | 0.001135 | 0.993519 | 0.998813 | Snhg12 |
| ENSMUSG( | 87.76515 | -0.13262 | 0.552359 | 0.939565 | Snhg15 |
| ENSMUSG( | 92.04921 | -0.03016 | 0.903523 | 0.991089 | Snhg16 |
| ENSMUSG( | 225.3294 | -0.12451 | 0.508939 | 0.929089 | Snhg17 |
| ENSMUSG( | 103.6318 | -0.18446 | 0.412923 | 0.917257 | Snhg18 |
| ENSMUSG( | 495.6629 | 0.18298 | 0.396146 | 0.917257 | Snhg20 |
| ENSMUSG( | 128.5417 | 0.083868 | 0.70161 | 0.962014 | Snhg3 |
| ENSMUSG( | 144.8609 | -0.00959 | 0.959157 | 0.994798 | Snhg4 |
| ENSMUSG( | 241.2533 | 0.19656 | 0.248624 | 0.877904 | Snhg5 |
| ENSMUSG( | 661.21 | -0.10305 | 0.404781 | 0.917257 | Snhg6 |
| ENSMUSG( | 128.7956 | 0.350932 | 0.121686 | 0.813384 | Snhg7 |
| ENSMUSG( | 661.4949 | 0.032289 | 0.804785 | 0.977232 | Snhg8 |
| ENSMUSG( | 241.9873 | -0.11374 | 0.548276 | 0.938063 | Snip1 |
| ENSMUSG( | 9971.14 | -0.01454 | 0.888621 | 0.990558 | Snn |
| ENSMUSG( | 18.35131 | 0.326577 | 0.548535 | 0.938063 | Snora31 |
| ENSMUSG( | 12.10194 | 0.528174 | 0.360995 | 0.914223 | Snora57 |
| ENSMUSG( | 18.30092 | -0.244 | 0.640254 | 0.953903 | Snora73b |
| ENSMUSG( | 13.55681 | -0.59575 | 0.300212 | 0.898538 | Snorc |
| ENSMUSG( | 21.64958 | 0.28314 | 0.508312 | 0.929089 | Snord55 |
| ENSMUSG( | 18176.48 | 0.243583 | 0.048884 | 0.706521 | Snph |
| ENSMUSG( | 1735.46 | 0.005599 | 0.957871 | 0.994798 | Snrk |
| ENSMUSG( | 2722.794 | -0.13245 | 0.228344 | 0.868507 | Snrnp200 |
| ENSMUSG( | 345.5562 | -0.26551 | 0.084643 | 0.779799 | Snrnp25 |
| ENSMUSG( | 659.6603 | -0.05421 | 0.696143 | 0.960867 | Snrnp27 |
| ENSMUSG( | 233.1556 | -0.23309 | 0.142081 | 0.830709 | Snrnp35 |
| ENSMUSG( | 394.0806 | -0.05982 | 0.725022 | 0.966887 | Snrnp40 |
| ENSMUSG( | 611.4608 | 0.314893 | 0.032233 | 0.634419 | Snrnp48 |
| ENSMUSG( | 7505.544 | 0.1408 | 0.44542 | 0.921734 | Snrnp70 |
| ENSMUSG( | 1036.884 | -0.17185 | 0.199084 | 0.863016 | Snrpa |
| ENSMUSG( | 486.953 | -0.04259 | 0.7558 | 0.971371 | Snrpa1 |
| ENSMUSG( | 1156.22 | -0.03151 | 0.7973 | 0.97681 | Snrpb |
| ENSMUSG( | 517.1792 | -0.14231 | 0.32808 | 0.906255 | Snrpb2 |
| ENSMUSG( | 568.3927 | -0.17889 | 0.188825 | 0.85698 | Snrpc |
| ENSMUSG( | 591.4959 | -0.13655 | 0.400462 | 0.917257 | Snrpd1 |
| ENSMUSG( | 1105.876 | -0.26071 | 0.077457 | 0.772673 | Snrpd2 |
| ENSMUSG( | 1163.407 | -0.18947 | 0.211542 | 0.864412 | Snrpd3 |
| ENSMUSG( | 478.219 | -0.13206 | 0.408584 | 0.917257 | Snrpe |
| ENSMUSG( | 477.9185 | -0.22189 | 0.207339 | 0.864412 | Snrpf |
| ENSMUSG( | 417.3001 | -0.1939 | 0.203399 | 0.864412 | Snrpg |
| ENSMUSG( | 10214.18 | -0.12784 | 0.307748 | 0.898538 | Snrpn |
| ENSMUSG( | 1489.289 | -0.06782 | 0.557764 | 0.939565 | Snta1 |
| ENSMUSG( | 221.5754 | -0.08073 | 0.678568 | 0.957506 | Sntb1 |
| ENSMUSG( | 444.5003 | -0.21362 | 0.123788 | 0.814691 | Sntb2 |
| ENSMUSG( | 256.1181 | 0.043497 | 0.785522 | 0.976285 | Sntg1 |
| ENSMUSG( | 46.37064 | -0.21724 | 0.481195 | 0.924548 | Sntg2 |
| ENSMUSG( | 13.00133 | 1.0117 | 0.106689 | 0.804969 | Sntn |
| ENSMUSG( | 1630.349 | -0.21379 | 0.085629 | 0.779799 | Snu13 |
| ENSMUSG( | 259.5342 | 0.024662 | 0.881309 | 0.989644 | Snupn |
| ENSMUSG( | 1425.949 | 0.077954 | 0.494085 | 0.927372 | Snw1 |
| ENSMUSG( | 1387.92 | -0.09079 | 0.396611 | 0.917257 | Snx1 |
| ENSMUSG( | 1609.783 | 0.057918 | 0.688412 | 0.959742 | Snx10 |
| ENSMUSG( | 824.1043 | 0.008188 | 0.947303 | 0.994798 | Snx11 |
| ENSMUSG( | 949.9955 | -0.30716 | 0.012091 | 0.445877 | Snx12 |
| ENSMUSG( | 2840.382 | 0.015918 | 0.907029 | 0.991601 | Snx13 |
| ENSMUSG( | 1510.3 | 0.236253 | 0.163213 | 0.838291 | Snx14 |
| ENSMUSG( | 784.2728 | -0.15358 | 0.264289 | 0.886134 | Snx15 |
| ENSMUSG( | 271.7067 | 0.079765 | 0.662471 | 0.956205 | Snx16 |
| ENSMUSG( | 1913.409 | -0.15072 | 0.146538 | 0.835973 | Snx17 |
| ENSMUSG( | 1139.229 | -0.04725 | 0.726994 | 0.966887 | Snx18 |
| ENSMUSG( | 2768.7 | 0.135544 | 0.373769 | 0.915491 | Snx19 |
| ENSMUSG( | 978.5033 | 0.036552 | 0.764313 | 0.974 | Snx2 |
| ENSMUSG( | 36.74471 | -0.24333 | 0.496281 | 0.927825 | Snx20 |
| ENSMUSG( | 1166.251 | 0.074392 | 0.527816 | 0.934603 | Snx21 |
| ENSMUSG( | 361.2717 | 0.044956 | 0.791926 | 0.97681 | Snx22 |
| ENSMUSG( | 444.0662 | -0.03353 | 0.82283 | 0.979198 | Snx24 |
| ENSMUSG( | 1578.744 | 0.233884 | 0.159518 | 0.837724 | Snx25 |
| ENSMUSG( | 4866.783 | -0.1185 | 0.237087 | 0.872173 | Snx27 |
| ENSMUSG( | 289.6253 | -0.15168 | 0.319292 | 0.903453 | Snx29 |
| ENSMUSG( | 4491.061 | -0.22958 | 0.085174 | 0.779799 | Snx3 |
| ENSMUSG( | 1107.288 | -0.0728 | 0.555794 | 0.939565 | Snx30 |
| ENSMUSG( | 1860.898 | 0.073453 | 0.69331 | 0.96076 | Snx32 |
| ENSMUSG( | 366.981 | 0.093964 | 0.559732 | 0.939812 | Snx33 |
| ENSMUSG( | 1064.527 | -0.08647 | 0.488103 | 0.925453 | Snx4 |
| ENSMUSG( | 4518.582 | -0.11929 | 0.322309 | 0.904023 | Snx5 |
| ENSMUSG( | 1050.631 | -0.03222 | 0.797277 | 0.97681 | Snx6 |
| ENSMUSG( | 205.6207 | -0.09363 | 0.599109 | 0.947045 | Snx7 |
| ENSMUSG( | 615.6119 | -0.14321 | 0.283313 | 0.892412 | Snx8 |
| ENSMUSG( | 711.9511 | -0.00087 | 0.994799 | 0.998813 | Snx9 |
| ENSMUSG( | 1581.024 | 0.237279 | 0.147875 | 0.836241 | Soat1 |
| ENSMUSG( | 2572.537 | 0.232484 | 0.023711 | 0.588226 | Sobp |
| ENSMUSG( | 29.68354 | -0.65496 | 0.132159 | 0.822912 | Socs1 |
| ENSMUSG( | 393.4051 | 0.116982 | 0.431272 | 0.917257 | Socs2 |
| ENSMUSG( | 105.9278 | 0.020208 | 0.92515 | 0.99306 | Socs3 |
| ENSMUSG( | 553.8267 | 0.116956 | 0.482417 | 0.924578 | Socs4 |
| ENSMUSG( | 2538.939 | 0.047722 | 0.645866 | 0.953988 | Socs5 |
| ENSMUSG( | 389.7891 | 0.30048 | 0.050062 | 0.708648 | Socs6 |
| ENSMUSG( | 4329.842 | -0.01204 | 0.906782 | 0.991517 | Socs7 |
| ENSMUSG( | 5881.247 | -0.15807 | 0.189103 | 0.85698 | Sod1 |
| ENSMUSG( | 4253.978 | -0.15336 | 0.245983 | 0.877904 | Sod2 |
| ENSMUSG( | 1333.947 | -0.01288 | 0.94096 | 0.994558 | Sod3 |
| ENSMUSG( | 7264.388 | 0.285108 | 0.056567 | 0.723362 | Son |
| ENSMUSG( | 3811.458 | -0.12124 | 0.2785 | 0.891628 | Sorbs1 |
| ENSMUSG( | 4328.639 | -0.03674 | 0.737936 | 0.967709 | Sorbs2 |
| ENSMUSG( | 1971.449 | 0.024592 | 0.823518 | 0.979198 | Sorbs3 |
| ENSMUSG( | 1207.155 | 0.068043 | 0.667224 | 0.956766 | Sorcs1 |
| ENSMUSG( | 3026.736 | 0.011446 | 0.941282 | 0.994578 | Sorcs2 |
| ENSMUSG( | 1090.316 | -0.10585 | 0.565715 | 0.941404 | Sorcs3 |
| ENSMUSG( | 550.2941 | 0.031845 | 0.816032 | 0.979198 | Sord |
| ENSMUSG( | 10529.8 | 0.100783 | 0.524773 | 0.933907 | Sorl1 |
| ENSMUSG( | 16710.77 | 0.111841 | 0.496841 | 0.927825 | Sort1 |
| ENSMUSG( | 1146.916 | 0.060292 | 0.606766 | 0.949772 | Sos1 |
| ENSMUSG( | 1251.642 | 0.097093 | 0.444741 | 0.921734 | Sos2 |
| ENSMUSG( | 112.7499 | 0.097892 | 0.770898 | 0.974236 | Sostdc1 |
| ENSMUSG( | 3334.453 | 0.167711 | 0.101245 | 0.801456 | Sowaha |
| ENSMUSG( | 248.8805 | -0.02529 | 0.886158 | 0.989913 | Sowahb |
| ENSMUSG( | 423.8995 | 0.010251 | 0.947244 | 0.994798 | Sowahc |
| ENSMUSG( | 575.3152 | 0.299219 | 0.058134 | 0.728256 | Sox1 |
| ENSMUSG( | 1292.024 | 0.014719 | 0.894599 | 0.990558 | Sox10 |
| ENSMUSG( | 669.4272 | 0.233061 | 0.14369 | 0.831311 | Sox11 |
| ENSMUSG( | 897.0586 | -0.10734 | 0.416396 | 0.917257 | Sox12 |
| ENSMUSG( | 466.492 | 0.354322 | 0.011328 | 0.433258 | Sox13 |
| ENSMUSG( | 228.2736 | 1.228708 | 5.23E-08 | 3.67E-05 | Sox17 |
| ENSMUSG( | 339.7014 | 0.707884 | 0.003815 | 0.246611 | Sox18 |
| ENSMUSG( | 1677.389 | 0.036307 | 0.837154 | 0.981335 | Sox2 |
| ENSMUSG( | 864.3933 | 0.445162 | 0.012172 | 0.447008 | Sox21 |
| ENSMUSG( | 54.40303 | -0.34174 | 0.223671 | 0.867404 | Sox21os1 |
| ENSMUSG( | 912.3572 | -0.0592 | 0.653489 | 0.955015 | Sox2ot |
| ENSMUSG( | 87.89062 | -0.29053 | 0.231385 | 0.868507 | Sox3 |
| ENSMUSG( | 1247.669 | 0.218282 | 0.224036 | 0.867404 | Sox4 |
| ENSMUSG( | 1267.169 | -0.1161 | 0.420336 | 0.917257 | Sox5 |
| ENSMUSG( | 1257.757 | -0.31033 | 0.01385 | 0.46927 | Sox6 |
| ENSMUSG( | 102.5687 | 0.698548 | 0.016101 | 0.510946 | Sox7 |

| | | | | | | | | | | | |
|---|---|---|---|---|---|---|---|---|---|---|---|
| ENSMUSG( | Spata17 | 19.51955 | -0.64319 | 0.141171 | NA | ENSMUSG( | Spata17 | 20.40278 | -0.22054 | 0.624226 | 0.953234 |
| ENSMUSG( | Spata18 | 11.96414 | 0.187312 | 0.744952 | NA | ENSMUSG( | Spata18 | 15.31704 | 1.039259 | 0.083963 | 0.77737 |
| ENSMUSG( | Spata19 | 12.10775 | -0.83253 | 0.162861 | NA | ENSMUSG( | Spata19 | 10.5568 | -1.05038 | 0.096462 | NA |
| ENSMUSG( | Spata2 | 1662.077 | 0.127945 | 0.205598 | 0.647118 | ENSMUSG( | Spata2 | 1564.922 | 0.220211 | 0.074232 | 0.766789 |
| ENSMUSG( | Spata21 | 21.72123 | -0.25101 | 0.540648 | 0.88315 | ENSMUSG( | Spata21 | 18.59667 | -0.44994 | 0.332582 | 0.907907 |
| ENSMUSG( | Spata24 | 63.36982 | 0.108306 | 0.68483 | 0.932981 | ENSMUSG( | Spata24 | 55.85449 | 0.036598 | 0.897794 | 0.990558 |
| ENSMUSG( | Spata2l | 1330.279 | 0.048763 | 0.763378 | 0.953781 | ENSMUSG( | Spata2l | 1141.057 | -0.13051 | 0.45294 | 0.922316 |
| ENSMUSG( | Spata3 | 72.68455 | 0.574669 | 0.016984 | 0.194647 | ENSMUSG( | Spata3 | 53.51493 | 0.00821 | 0.977274 | 0.996416 |
| ENSMUSG( | Spata31d1 | 13.64142 | -0.32157 | 0.524261 | NA | ENSMUSG( | Spata31d1 | 12.11084 | -0.3998 | 0.532437 | 0.934795 |
| ENSMUSG( | Spata31f1a | 8.698212 | 0.193221 | 0.753403 | NA | ENSMUSG( | Spata31f1a | 8.433549 | 0.39756 | 0.566219 | NA |
| ENSMUSG( | Spata32 | 11.35299 | -0.54741 | 0.318964 | NA | ENSMUSG( | Spata32 | 11.20488 | -0.29241 | 0.623029 | NA |
| ENSMUSG( | Spata33 | 174.3223 | -0.24287 | 0.131652 | 0.544205 | ENSMUSG( | Spata33 | 159.4427 | -0.23059 | 0.20806 | 0.864412 |
| ENSMUSG( | Spata6 | 119.1921 | -0.64008 | 0.000839 | 0.02897 | ENSMUSG( | Spata6 | 142.6603 | 0.213857 | 0.249457 | 0.877904 |
| ENSMUSG( | Spata6l | 135.6998 | 0.029866 | 0.879186 | 0.981436 | ENSMUSG( | Spata6l | 126.3376 | 0.084955 | 0.691204 | 0.960071 |
| ENSMUSG( | Spata7 | 449.8735 | 0.115876 | 0.384841 | 0.804031 | ENSMUSG( | Spata7 | 393.3133 | -0.0006 | 0.996554 | 0.999178 |
| ENSMUSG( | Spats1 | 94.91823 | -0.18111 | 0.408463 | 0.818167 | ENSMUSG( | Spats1 | 80.32173 | -0.41481 | 0.078986 | 0.77458 |
| ENSMUSG( | Spats2 | 982.7599 | 0.050498 | 0.644286 | 0.917741 | ENSMUSG( | Spats2 | 865.0886 | -0.05247 | 0.684806 | 0.958678 |
| ENSMUSG( | Spats2l | 491.8303 | 0.077918 | 0.587796 | 0.900628 | ENSMUSG( | Spats2l | 422.3951 | -0.09349 | 0.561725 | 0.940388 |
| ENSMUSG( | Spc24 | 38.52409 | -0.18952 | 0.529975 | 0.878883 | ENSMUSG( | Spc24 | 35.88844 | -0.1045 | 0.761242 | 0.972695 |
| ENSMUSG( | Spc25 | 33.7665 | 0.218111 | 0.495103 | 0.860798 | ENSMUSG( | Spc25 | 26.61602 | -0.2107 | 0.585445 | 0.944975 |
| ENSMUSG( | Spcs1 | 2358.504 | -0.37341 | 0.005668 | 0.10081 | ENSMUSG( | Spcs1 | 2445.61 | 0.025128 | 0.872491 | 0.989027 |
| ENSMUSG( | Spcs2 | 3114.474 | -0.27298 | 0.030319 | 0.264512 | ENSMUSG( | Spcs2 | 3131.939 | 0.026806 | 0.856286 | 0.985884 |
| ENSMUSG( | Spcs3 | 2105.839 | -0.19954 | 0.192935 | 0.632208 | ENSMUSG( | Spcs3 | 2223.792 | 0.229326 | 0.172581 | 0.843702 |
| ENSMUSG( | Spdef | 11.84411 | -0.31105 | 0.572305 | NA | ENSMUSG( | Spdef | 10.52052 | -0.4037 | 0.518407 | NA |
| ENSMUSG( | Spdl1 | 9.962538 | 0.470596 | 0.44141 | NA | ENSMUSG( | Spdl1 | 8.761127 | 0.43093 | 0.519224 | NA |
| ENSMUSG( | Spdya | 21.24867 | 0.072432 | 0.869839 | 0.977607 | ENSMUSG( | Spdya | 17.96056 | -0.18928 | 0.711884 | 0.964363 |
| ENSMUSG( | Specc1 | 6520.582 | 0.029277 | 0.78157 | 0.957558 | ENSMUSG( | Specc1 | 5507.877 | -0.19678 | 0.078858 | 0.774307 |
| ENSMUSG( | Specc1l | 2428.58 | 0.082742 | 0.367649 | 0.795314 | ENSMUSG( | Specc1l | 2101.621 | -0.06632 | 0.545998 | 0.938063 |
| ENSMUSG( | Spef1 | 717.978 | 0.153842 | 0.144955 | 0.565187 | ENSMUSG( | Spef1 | 630.9061 | 0.056648 | 0.666064 | 0.956766 |
| ENSMUSG( | Spef1l | 15.78445 | 0.179798 | 0.704553 | NA | ENSMUSG( | Spef1l | 14.52901 | 0.236273 | 0.649999 | 0.95422 |
| ENSMUSG( | Spef2 | 67.22996 | 0.163177 | 0.498123 | 0.862463 | ENSMUSG( | Spef2 | 73.21825 | 0.639666 | 0.024864 | 0.592568 |
| ENSMUSG( | Speg | 6933.652 | 0.064203 | 0.453339 | 0.836798 | ENSMUSG( | Speg | 6224.129 | 0.022497 | 0.820973 | 0.979198 |
| ENSMUSG( | Spen | 2614.076 | 0.014806 | 0.901258 | 0.985633 | ENSMUSG( | Spen | 2477.3 | 0.13002 | 0.347354 | 0.911652 |
| ENSMUSG( | Spg11 | 906.3633 | 0.108305 | 0.38329 | 0.80333 | ENSMUSG( | Spg11 | 797.0079 | 0.013162 | 0.92526 | 0.99306 |
| ENSMUSG( | Spg21 | 769.9646 | 0.009866 | 0.925336 | 0.991204 | ENSMUSG( | Spg21 | 709.375 | 0.03513 | 0.813804 | 0.979198 |
| ENSMUSG( | Spg7 | 3205.406 | 0.055266 | 0.540049 | 0.88315 | ENSMUSG( | Spg7 | 2875.761 | 0.012479 | 0.907022 | 0.991601 |
| ENSMUSG( | Sphk1 | 140.1129 | 0.002704 | 0.990133 | 0.999144 | ENSMUSG( | Sphk1 | 138.9442 | 0.232435 | 0.375624 | 0.915491 |
| ENSMUSG( | Sphk2 | 815.5087 | 0.078085 | 0.487444 | 0.858812 | ENSMUSG( | Sphk2 | 669.8748 | -0.22483 | 0.096289 | 0.796133 |
| ENSMUSG( | Sphkap | 3238.186 | -0.02983 | 0.772273 | 0.955037 | ENSMUSG( | Sphkap | 3101.579 | 0.115531 | 0.396532 | 0.917257 |
| ENSMUSG( | Spi1 | 255.1896 | -0.21747 | 0.106264 | 0.49377 | ENSMUSG( | Spi1 | 236.1419 | -0.16736 | 0.362308 | 0.914223 |
| ENSMUSG( | Spib | 11.51458 | -0.02671 | 0.970238 | NA | ENSMUSG( | Spib | 8.583406 | -0.66069 | 0.39654 | NA |
| ENSMUSG( | Spice1 | 371.4198 | 0.206013 | 0.10359 | 0.489428 | ENSMUSG( | Spice1 | 321.8374 | 0.073149 | 0.668066 | 0.956766 |
| ENSMUSG( | Spidr | 238.2691 | 0.200756 | 0.174733 | 0.610304 | ENSMUSG( | Spidr | 194.054 | -0.12366 | 0.475364 | 0.923329 |
| ENSMUSG( | Spin1 | 4506.788 | 0.137685 | 0.078112 | 0.43227 | ENSMUSG( | Spin1 | 3892.622 | -0.01511 | 0.897943 | 0.990558 |
| ENSMUSG( | Spin2c | 62.30846 | -0.10897 | 0.689005 | 0.933441 | ENSMUSG( | Spin2c | 48.14958 | -0.64761 | 0.039291 | 0.679972 |
| ENSMUSG( | Spin4 | 10.48569 | -0.43291 | 0.461714 | NA | ENSMUSG( | Spin4 | 13.11848 | 0.448297 | 0.423175 | 0.917257 |
| ENSMUSG( | Spindoc | 668.1462 | 0.115638 | 0.332337 | 0.775474 | ENSMUSG( | Spindoc | 574.6959 | -0.0507 | 0.723546 | 0.966887 |
| ENSMUSG( | Spink10 | 100.6465 | 0.066356 | 0.73404 | 0.945306 | ENSMUSG( | Spink10 | 92.43932 | 0.098274 | 0.659296 | 0.956205 |
| ENSMUSG( | Spink8 | 67.38712 | -0.15666 | 0.514764 | 0.872189 | ENSMUSG( | Spink8 | 69.87576 | 0.209408 | 0.414714 | 0.917257 |
| ENSMUSG( | Spint1 | 114.7051 | -0.68851 | 0.006483 | 0.110626 | ENSMUSG( | Spint1 | 139.1868 | 0.223233 | 0.334864 | 0.907907 |
| ENSMUSG( | Spint2 | 661.9823 | -0.27675 | 0.006878 | 0.114527 | ENSMUSG( | Spint2 | 675.0594 | 0.058269 | 0.685461 | 0.958678 |
| ENSMUSG( | Spire1 | 13333.1 | 0.261747 | 0.007222 | 0.117261 | ENSMUSG( | Spire1 | 10725.36 | -0.08849 | 0.495085 | 0.927372 |
| ENSMUSG( | Spire2 | 484.7406 | 0.072779 | 0.59972 | 0.903906 | ENSMUSG( | Spire2 | 443.551 | 0.083559 | 0.574858 | 0.942912 |
| ENSMUSG( | Spmip4 | 19.20393 | -0.08366 | 0.841263 | NA | ENSMUSG( | Spmip4 | 18.05357 | -0.00048 | 0.99917 | 0.999503 |
| ENSMUSG( | Spmip6 | 47.12306 | 0.164041 | 0.590033 | 0.90113 | ENSMUSG( | Spmip6 | 52.08184 | 0.652984 | 0.028946 | 0.614171 |
| ENSMUSG( | Spmip7 | 12.89129 | 0.332643 | 0.538821 | NA | ENSMUSG( | Spmip7 | 12.84963 | 0.586922 | 0.311091 | 0.899422 |
| ENSMUSG( | Spn | 51.83756 | -1.26046 | 0.161976 | 0.592429 | ENSMUSG( | Spn | 44.85033 | -1.59171 | 0.100289 | 0.801311 |
| ENSMUSG( | Spns1 | 2087.597 | -0.12211 | 0.339505 | 0.777792 | ENSMUSG( | Spns1 | 2123.817 | 0.196796 | 0.202011 | 0.864412 |
| ENSMUSG( | Spns2 | 2050.359 | 0.100534 | 0.452307 | 0.836145 | ENSMUSG( | Spns2 | 2056.535 | 0.362209 | 0.018733 | 0.547826 |
| ENSMUSG( | Spns3 | 16.38295 | -0.44053 | 0.45715 | NA | ENSMUSG( | Spns3 | 14.91321 | -0.40701 | 0.5016 | 0.927825 |
| ENSMUSG( | Spo11 | 15.07686 | -0.40561 | 0.438834 | NA | ENSMUSG( | Spo11 | 16.42935 | 0.110083 | 0.857982 | 0.985884 |
| ENSMUSG( | Spocd1 | 18.16394 | 0.85603 | 0.063529 | NA | ENSMUSG( | Spocd1 | 11.8438 | -0.00926 | 0.987487 | 0.99751 |
| ENSMUSG( | Spock1 | 9421.137 | -0.61162 | 9.54E-06 | 0.001107 | ENSMUSG( | Spock1 | 10487.99 | 0.033109 | 0.835678 | 0.981237 |
| ENSMUSG( | Spock2 | 31980.3 | -0.18014 | 0.220911 | 0.666476 | ENSMUSG( | Spock2 | 34327.45 | 0.289025 | 0.086323 | 0.781091 |
| ENSMUSG( | Spock3 | 1501.193 | -0.93538 | 4.12E-10 | 2.09E-07 | ENSMUSG( | Spock3 | 1871.543 | 0.118823 | 0.470713 | 0.923329 |
| ENSMUSG( | Spon1 | 3227.419 | -0.34651 | 0.004259 | 0.084556 | ENSMUSG( | Spon1 | 3502.399 | 0.176053 | 0.19476 | 0.860433 |
| ENSMUSG( | Spon2 | 12.46044 | -0.73216 | 0.210926 | NA | ENSMUSG( | Spon2 | 8.950482 | -1.93108 | 0.006368 | NA |
| ENSMUSG( | Spop | 14570.85 | 0.131112 | 0.155767 | 0.58375 | ENSMUSG( | Spop | 12620.52 | -0.01353 | 0.907843 | 0.991958 |
| ENSMUSG( | Spopl | 297.4219 | -0.11885 | 0.383018 | 0.803237 | ENSMUSG( | Spopl | 280.5761 | -0.01285 | 0.934463 | 0.99306 |
| ENSMUSG( | Spout1 | 1388.601 | 0.08564 | 0.341783 | 0.778828 | ENSMUSG( | Spout1 | 1212.967 | -0.03603 | 0.750678 | 0.970209 |
| ENSMUSG( | Spp1 | 299.6249 | -0.09282 | 0.546485 | 0.884974 | ENSMUSG( | Spp1 | 275.0113 | -0.0819 | 0.620137 | 0.952896 |
| ENSMUSG( | Sppl2a | 1776.096 | -0.22113 | 0.104478 | 0.490485 | ENSMUSG( | Sppl2a | 1801.378 | 0.096829 | 0.510159 | 0.929089 |
| ENSMUSG( | Sppl2b | 1663.461 | -0.13594 | 0.334786 | 0.776386 | ENSMUSG( | Sppl2b | 1690.214 | 0.181824 | 0.259938 | 0.883971 |
| ENSMUSG( | Sppl3 | 2611.859 | -0.32184 | 0.02017 | 0.214666 | ENSMUSG( | Sppl3 | 2693.754 | 0.058052 | 0.72206 | 0.966887 |
| ENSMUSG( | Spr | 1219.632 | 0.102331 | 0.338193 | 0.777422 | ENSMUSG( | Spr | 1009.946 | -0.18067 | 0.134216 | 0.822912 |
| ENSMUSG( | Spred1 | 7735.017 | 0.184367 | 0.124958 | 0.531973 | ENSMUSG( | Spred1 | 6829.403 | 0.095496 | 0.570528 | 0.941872 |
| ENSMUSG( | Spred2 | 3508.628 | 0.115013 | 0.183739 | 0.620678 | ENSMUSG( | Spred2 | 3046.489 | -0.02369 | 0.853856 | 0.98563 |
| ENSMUSG( | Spred3 | 1612.535 | 0.19594 | 0.049188 | 0.338637 | ENSMUSG( | Spred3 | 1416.557 | 0.096133 | 0.407361 | 0.917257 |
| ENSMUSG( | Spring1 | 967.1335 | -0.12405 | 0.269051 | 0.719302 | ENSMUSG( | Spring1 | 957.2207 | 0.118899 | 0.339741 | 0.907972 |
| ENSMUSG( | Sprn | 6810.509 | -0.28901 | 0.028547 | 0.257525 | ENSMUSG( | Sprn | 7257.943 | 0.176285 | 0.2627 | 0.885357 |
| ENSMUSG( | Sprtn | 585.9562 | -0.14541 | 0.180585 | 0.617841 | ENSMUSG( | Sprtn | 539.2614 | -0.11307 | 0.377251 | 0.915912 |
| ENSMUSG( | Spry1 | 223.1707 | -0.21516 | 0.197075 | 0.636662 | ENSMUSG( | Spry1 | 241.0044 | 0.274168 | 0.111355 | 0.808371 |
| ENSMUSG( | Spry2 | 1454.183 | 0.013625 | 0.8995 | 0.985633 | ENSMUSG( | Spry2 | 1447.847 | 0.256023 | 0.0394 | 0.679972 |
| ENSMUSG( | Spry4 | 296.2779 | -0.07404 | 0.573472 | 0.893584 | ENSMUSG( | Spry4 | 289.884 | 0.128206 | 0.399599 | 0.917257 |
| ENSMUSG( | Spryd3 | 3235.14 | 0.043807 | 0.71186 | 0.940028 | ENSMUSG( | Spryd3 | 2767.191 | -0.14338 | 0.286603 | 0.894091 |
| ENSMUSG( | Spryd4 | 137.8506 | 0.148906 | 0.433502 | 0.828346 | ENSMUSG( | Spryd4 | 115.9652 | -0.06731 | 0.766825 | 0.974236 |
| ENSMUSG( | Spryd7 | 895.6481 | -0.10455 | 0.378734 | 0.801688 | ENSMUSG( | Spryd7 | 795.1364 | -0.18928 | 0.226562 | 0.868507 |
| ENSMUSG( | Spsb1 | 660.3151 | 0.13194 | 0.317251 | 0.763823 | ENSMUSG( | Spsb1 | 552.3212 | -0.11245 | 0.424595 | 0.917257 |
| ENSMUSG( | Spsb2 | 99.27235 | 0.105658 | 0.592646 | 0.902094 | ENSMUSG( | Spsb2 | 93.56447 | 0.200711 | 0.357997 | 0.913759 |
| ENSMUSG( | Spsb3 | 1444.216 | 0.047895 | 0.685803 | 0.932981 | ENSMUSG( | Spsb3 | 1239.903 | -0.12493 | 0.333134 | 0.907907 |
| ENSMUSG( | Spsb4 | 213.8145 | -0.19065 | 0.198155 | 0.638387 | ENSMUSG( | Spsb4 | 189.3052 | -0.29869 | 0.155648 | 0.837724 |
| ENSMUSG( | Sptan1 | 34530.87 | 0.041624 | 0.704244 | 0.938221 | ENSMUSG( | Sptan1 | 31479.81 | 0.046104 | 0.722651 | 0.966887 |
| ENSMUSG( | Sptb | 8713.961 | 0.102576 | 0.510321 | 0.870063 | ENSMUSG( | Sptb | 8161.364 | 0.183948 | 0.323275 | 0.904023 |
| ENSMUSG( | Sptbn1 | 56515.73 | -0.00901 | 0.942986 | 0.993415 | ENSMUSG( | Sptbn1 | 52283.17 | 0.036997 | 0.775869 | 0.975517 |
| ENSMUSG( | Sptbn2 | 91830.35 | 0.259828 | 0.074792 | 0.42402 | ENSMUSG( | Sptbn2 | 80880.39 | 0.173014 | 0.309483 | 0.898538 |
| ENSMUSG( | Sptbn4 | 9494.51 | 0.158432 | 0.079189 | 0.435105 | ENSMUSG( | Sptbn4 | 8548.225 | 0.126515 | 0.228851 | 0.868507 |
| ENSMUSG( | Sptbn5 | 161.49 | 0.308899 | 0.054453 | 0.359033 | ENSMUSG( | Sptbn5 | 138.192 | 0.146846 | 0.445852 | 0.921734 |
| ENSMUSG( | Sptlc1 | 1218.667 | -0.19434 | 0.191157 | 0.630245 | ENSMUSG( | Sptlc1 | 1261.935 | 0.181433 | 0.271547 | 0.888062 |
| ENSMUSG( | Sptlc2 | 1227.496 | -0.34891 | 0.022627 | 0.229667 | ENSMUSG( | Sptlc2 | 1325.076 | 0.163281 | 0.361925 | 0.914223 |
| ENSMUSG( | Sptssa | 1055.928 | -0.01012 | 0.938803 | 0.993415 | ENSMUSG( | Sptssa | 976.4832 | 0.026877 | 0.867942 | 0.98781 |
| ENSMUSG( | Sptssb | 48.05952 | -0.15793 | 0.571429 | 0.893319 | ENSMUSG( | Sptssb | 44.64552 | -0.11751 | 0.695888 | 0.960867 |
| ENSMUSG( | Spty2d1 | 257.0652 | -0.14986 | 0.290774 | 0.740751 | ENSMUSG( | Spty2d1 | 256.5783 | 0.118712 | 0.450301 | 0.922202 |
| ENSMUSG( | Spx | 57.00944 | 0.33684 | 0.196568 | 0.635942 | ENSMUSG( | Spx | 48.00135 | 0.100125 | 0.748339 | 0.96984 |
| ENSMUSG( | Sqle | 3159.243 | -0.27527 | 0.089839 | 0.460262 | ENSMUSG( | Sqle | 3182.665 | 0.031987 | 0.857171 | 0.985884 |
| ENSMUSG( | Sqor | 312.3457 | 0.184995 | 0.198788 | 0.63899 | ENSMUSG( | Sqor | 265.0802 | -0.00884 | 0.959048 | 0.994798 |
| ENSMUSG( | Sqstm1 | 9322.693 | -0.0441 | 0.586514 | 0.900023 | ENSMUSG( | Sqstm1 | 8594.908 | -0.0095 | 0.92446 | 0.99306 |
| ENSMUSG( | Sra1 | 810.4626 | 0.047499 | 0.650539 | 0.921301 | ENSMUSG( | Sra1 | 696.7804 | -0.12278 | 0.328135 | 0.906255 |
| ENSMUSG( | Srarp | 99.61566 | 0.347354 | 0.16593 | 0.597821 | ENSMUSG( | Srarp | 127.5865 | 1.148463 | 2.05E-05 | 0.006225 |
| ENSMUSG( | Srbd1 | 356.3932 | 0.069882 | 0.595236 | 0.903007 | ENSMUSG( | Srbd1 | 300.293 | -0.16022 | 0.311725 | 0.900031 |
| ENSMUSG( | Src | 6398.976 | 0.10512 | 0.162871 | 0.593778 | ENSMUSG( | Src | 5452.068 | -0.08821 | 0.36007 | 0.914223 |
| ENSMUSG( | Srcap | 149.1808 | 0.005977 | 0.973951 | 0.99675 | ENSMUSG( | Srcap | 142.0794 | 0.140428 | 0.471837 | 0.923329 |
| ENSMUSG( | Srcin1 | 23075.16 | 0.102563 | 0.183347 | 0.62047 | ENSMUSG( | Srcin1 | 20101.16 | -0.02463 | 0.812608 | 0.979148 |
| ENSMUSG( | Srd5a1 | 880.3824 | -0.02816 | 0.803664 | 0.962169 | ENSMUSG( | Srd5a1 | 800.1278 | -0.03583 | 0.773085 | 0.974463 |
| ENSMUSG( | Srd5a3 | 664.7441 | -0.27914 | 0.031563 | 0.270895 | ENSMUSG( | Srd5a3 | 694.0251 | 0.128083 | 0.394555 | 0.917257 |
| ENSMUSG( | Srebf1 | 2545.469 | -0.19408 | 0.129726 | 0.539528 | ENSMUSG( | Srebf1 | 2844.282 | 0.38089 | 0.014485 | 0.48395 |
| ENSMUSG( | Srebf2 | 10953.85 | -0.17955 | 0.189378 | 0.627932 | ENSMUSG( | Srebf2 | 10600.28 | 0.004069 | 0.979111 | 0.996468 |
| ENSMUSG( | Srek1 | 2120.283 | 0.06089 | 0.590031 | 0.90113 | ENSMUSG( | Srek1 | 1988.085 | 0.137281 | 0.341152 | 0.9082 |
| ENSMUSG( | Srek1ip1 | 378.8756 | -0.22292 | 0.119458 | 0.523291 | ENSMUSG( | Srek1ip1 | 355.8587 | -0.14142 | 0.43502 | 0.918876 |
| ENSMUSG( | Srf | 1685.838 | 0.256422 | 0.026596 | 0.249357 | ENSMUSG( | Srf | 1489.1 | 0.177366 | 0.190786 | 0.857801 |
| ENSMUSG( | Srfbp1 | 271.7191 | -0.00522 | 0.972326 | 0.996585 | ENSMUSG( | Srfbp1 | 248.8702 | 0.007167 | 0.965765 | 0.995433 |
| ENSMUSG( | Srgap1 | 877.3007 | -0.17133 | 0.124538 | 0.531714 | ENSMUSG( | Srgap1 | 823.3667 | -0.08713 | 0.523454 | 0.932848 |
| ENSMUSG( | Srgap2 | 2492.512 | 0.175441 | 0.050191 | 0.342492 | ENSMUSG( | Srgap2 | 2236.41 | 0.136246 | 0.257156 | 0.883082 |
| ENSMUSG( | Srgap3 | 12220.41 | 0.221887 | 0.002452 | 0.058231 | ENSMUSG( | Srgap3 | 10110.29 | -0.04829 | 0.621963 | 0.953234 |
| ENSMUSG( | Srgn | 216.5307 | -0.05749 | 0.74661 | 0.949289 | ENSMUSG( | Srgn | 249.8614 | 0.578691 | 0.00332 | 0.229858 |
| ENSMUSG( | Sri | 2981.537 | 0.018179 | 0.871349 | 0.978114 | ENSMUSG( | Sri | 2543.708 | -0.18067 | 0.205766 | 0.864412 |
| ENSMUSG( | Srl | 297.2951 | 0.134984 | 0.299285 | 0.748131 | ENSMUSG( | Srl | 266.4363 | 0.097587 | 0.546613 | 0.938063 |
| ENSMUSG( | Srm | 1234.933 | -0.10375 | 0.408516 | 0.818167 | ENSMUSG( | Srm | 1178.277 | 0.027524 | 0.855786 | 0.985783 |
| ENSMUSG( | Sros1 | 12.21859 | -0.92322 | 0.094615 | NA | ENSMUSG( | Sros1 | 14.33803 | -0.11254 | 0.854393 | 0.985649 |
| ENSMUSG( | Srp14 | 1774.315 | 0.0078 | 0.94279 | 0.993415 | ENSMUSG( | Srp14 | 1535.831 | -0.14911 | 0.331642 | 0.907907 |
| ENSMUSG( | Srp19 | 924.7947 | 0.045164 | 0.690738 | 0.933881 | ENSMUSG( | Srp19 | 828.9615 | -0.00858 | 0.953611 | 0.994798 |
| ENSMUSG( | Srp54a | 58.70916 | 0.134156 | 0.593051 | 0.902372 | ENSMUSG( | Srp54a | 50.59746 | -0.03228 | 0.910452 | 0.992124 |
| ENSMUSG( | Srp54c | 287.5468 | -0.00159 | 0.991269 | 0.999232 | ENSMUSG( | Srp54c | 239.2424 | -0.27336 | 0.085301 | 0.779799 |
| ENSMUSG( | Srp68 | 1617.879 | 0.149078 | 0.114097 | 0.509818 | ENSMUSG( | Srp68 | 1382.915 | -0.03275 | 0.770425 | 0.974236 |
| ENSMUSG( | Srp72 | 2079.345 | 0.094745 | 0.262999 | 0.713397 | ENSMUSG( | Srp72 | 1903.09 | 0.105198 | 0.363037 | 0.914223 |
| ENSMUSG( | Srp9 | 1823.331 | 0.084638 | 0.384203 | 0.803686 | ENSMUSG( | Srp9 | 1558.624 | -0.10466 | 0.464169 | 0.923329 |
| ENSMUSG( | Srpk1 | 2140.334 | 0.029716 | 0.762245 | 0.953459 | ENSMUSG( | Srpk1 | 1964.454 | 0.051586 | 0.666519 | 0.956766 |
| ENSMUSG( | Srpk2 | 6153.228 | 0.026614 | 0.733988 | 0.945306 | ENSMUSG( | Srpk2 | 5486.816 | -0.03755 | 0.749732 | 0.96998 |
| ENSMUSG( | Srpk3 | 230.0886 | 0.291145 | 0.103708 | 0.489428 | ENSMUSG( | Srpk3 | 197.804 | 0.132167 | 0.594916 | 0.945572 |
| ENSMUSG( | Srpra | 1829.709 | -0.24652 | 0.141957 | 0.559166 | ENSMUSG( | Srpra | 1920.42 | 0.173997 | 0.360378 | 0.914223 |
| ENSMUSG( | Srprb | 1576.833 | -0.33333 | 0.025863 | 0.247245 | ENSMUSG( | Srprb | 1682.677 | 0.14327 | 0.385523 | 0.917257 |
| ENSMUSG( | Srpx | 13.3796 | -0.32358 | 0.511745 | NA | ENSMUSG( | Srpx | 18.84942 | 0.856282 | 0.063613 | 0.736482 |
| ENSMUSG( | Srpx2 | 39.4296 | -0.35489 | 0.269048 | 0.719302 | ENSMUSG( | Srpx2 | 42.44646 | 0.172382 | 0.603892 | 0.948137 |
| ENSMUSG( | Srr | 3809.068 | 0.067029 | 0.437734 | 0.830072 | ENSMUSG( | Srr | 3236.364 | -0.13812 | 0.17614 | 0.846252 |
| ENSMUSG( | Srrd | 547.5234 | -0.12633 | 0.305864 | 0.753712 | ENSMUSG( | Srrd | 509.677 | -0.05753 | 0.679887 | 0.957966 |
| ENSMUSG( | Srrm1 | 4180.862 | 0.059306 | 0.553407 | 0.886216 | ENSMUSG( | Srrm1 | 3825.266 | 0.069428 | 0.576567 | 0.942912 |
| ENSMUSG( | Srrm2 | 11982.08 | 0.111399 | 0.412996 | 0.819541 | ENSMUSG( | Srrm2 | 11567.89 | 0.266964 | 0.122103 | 0.813384 |
| ENSMUSG( | Srrm3 | 2110.758 | 0.223406 | 0.072101 | 0.414057 | ENSMUSG( | Srrm3 | 1798.499 | 0.035453 | 0.779899 | 0.976106 |
| ENSMUSG( | Srrm4 | 1329.016 | 0.037842 | 0.744383 | 0.948758 | ENSMUSG( | Srrm4 | 1227.282 | 0.073386 | 0.60347 | 0.948137 |
| ENSMUSG( | Srrt | 3574.526 | 0.230701 | 0.032828 | 0.277278 | ENSMUSG( | Srrt | 3069.12 | 0.065396 | 0.632907 | 0.953844 |
| ENSMUSG( | Srsf1 | 4187.683 | -0.04235 | 0.700173 | 0.937391 | ENSMUSG( | Srsf1 | 4036.926 | 0.115687 | 0.381577 | 0.917257 |
| ENSMUSG( | Srsf10 | 1277.342 | 0.077674 | 0.450224 | 0.835344 | ENSMUSG( | Srsf10 | 1238.754 | 0.244014 | 0.063774 | 0.736482 |
| ENSMUSG( | Srsf11 | 3554.871 | 0.088297 | 0.435679 | 0.829093 | ENSMUSG( | Srsf11 | 3275.264 | 0.116817 | 0.39274 | 0.917257 |
| ENSMUSG( | Srsf12 | 332.328 | -0.116 | 0.391883 | 0.809274 | ENSMUSG( | Srsf12 | 315.2249 | -0.00731 | 0.967941 | 0.995855 |
| ENSMUSG( | Srsf2 | 2861.269 | -0.22358 | 0.008886 | 0.132538 | ENSMUSG( | Srsf2 | 2983.068 | 0.170387 | 0.190289 | 0.857355 |
| ENSMUSG( | Srsf3 | 2584.598 | -0.00714 | 0.939767 | 0.993415 | ENSMUSG( | Srsf3 | 2419.887 | 0.069377 | 0.531112 | 0.934795 |
| ENSMUSG( | Srsf4 | 2808.714 | 0.058498 | 0.492483 | 0.860378 | ENSMUSG( | Srsf4 | 2485.422 | -0.02736 | 0.811356 | 0.978997 |
| ENSMUSG( | Srsf5 | 5431.989 | 4.58E-05 | 0.999718 | 0.99989 | ENSMUSG( | Srsf5 | 5243.512 | 0.160519 | 0.367582 | 0.915217 |
| ENSMUSG( | Srsf6 | 3013.093 | -0.10126 | 0.271973 | 0.722255 | ENSMUSG( | Srsf6 | 3027.48 | 0.177515 | 0.158924 | 0.837724 |

| | | | | | |
|---|---|---|---|---|---|
| ENSMUSG( | 600.4324 | 0.032947 | 0.786976 | 0.959037 | Snapc2 |
| ENSMUSG( | 475.2525 | 0.166302 | 0.180819 | 0.617841 | Snapc3 |
| ENSMUSG( | 1614.515 | 0.291842 | 0.022495 | 0.229049 | Snapc4 |
| ENSMUSG( | 686.9606 | 0.105339 | 0.386789 | 0.805046 | Snapc5 |
| ENSMUSG( | 2648.27 | -0.03191 | 0.760966 | 0.952907 | Snapin |
| ENSMUSG( | 4365.414 | 0.124488 | 0.39318 | 0.809274 | Snca |
| ENSMUSG( | 238.422 | -0.16311 | 0.345836 | 0.779153 | Sncaip |
| ENSMUSG( | 11116.99 | 0.042196 | 0.791883 | 0.959461 | Sncb |
| ENSMUSG( | 22.40137 | -0.59898 | 0.207551 | 0.64962 | Sncg |
| ENSMUSG( | 2368.108 | -0.15608 | 0.089236 | 0.458482 | Snd1 |
| ENSMUSG( | 1192.579 | -0.11429 | 0.332421 | 0.775474 | Sned1 |
| ENSMUSG( | 2118.517 | -0.01209 | 0.889834 | 0.983651 | Snf8 |
| ENSMUSG( | 68.75871 | -0.36313 | 0.127748 | 0.535835 | Snhg10 |
| ENSMUSG( | 40128.48 | 0.332068 | 0.099342 | 0.48179 | Snhg11 |
| ENSMUSG( | 727.6099 | -0.35997 | 0.000661 | 0.024836 | Snhg12 |
| ENSMUSG( | 90.92269 | -0.3154 | 0.130116 | 0.540593 | Snhg15 |
| ENSMUSG( | 88.17861 | -0.48745 | 0.021652 | 0.225182 | Snhg16 |
| ENSMUSG( | 239.1649 | -0.24187 | 0.107548 | 0.496444 | Snhg17 |
| ENSMUSG( | 116.1184 | -0.11563 | 0.577074 | 0.895621 | Snhg18 |
| ENSMUSG( | 481.1187 | -0.17525 | 0.291168 | 0.741182 | Snhg20 |
| ENSMUSG( | 137.6462 | 0.013291 | 0.941266 | 0.993415 | Snhg3 |
| ENSMUSG( | 158.1807 | -0.03149 | 0.863068 | 0.975055 | Snhg4 |
| ENSMUSG( | 238.0322 | -0.11564 | 0.458601 | 0.840224 | Snhg5 |
| ENSMUSG( | 792.6509 | 0.152597 | 0.161905 | 0.592398 | Snhg6 |
| ENSMUSG( | 127.3212 | 0.058862 | 0.758042 | 0.951614 | Snhg7 |
| ENSMUSG( | 752.3654 | 0.136741 | 0.190556 | 0.629951 | Snhg8 |
| ENSMUSG( | 9.757967 | -0.42883 | 0.470373 | NA | Snhg9 |
| ENSMUSG( | 278.8505 | 0.023077 | 0.893509 | 0.984512 | Snip1 |
| ENSMUSG( | 10912.6 | -0.02231 | 0.771068 | 0.954726 | Snn |
| ENSMUSG( | 16.55275 | -0.24886 | 0.667724 | NA | Snora31 |
| ENSMUSG( | 16.91263 | -0.88734 | 0.095914 | NA | Snora73b |
| ENSMUSG( | 17.7495 | 0.03572 | 0.936206 | NA | Snorc |
| ENSMUSG( | 24.88107 | 0.411051 | 0.287702 | 0.73849 | Snord55 |
| ENSMUSG( | 20406.32 | 0.301818 | 0.001406 | 0.041028 | Snph |
| ENSMUSG( | 2062.949 | 0.225619 | 0.011793 | 0.156913 | Snrk |
| ENSMUSG( | 3338.269 | 0.183619 | 0.053082 | 0.353671 | Snrnp200 |
| ENSMUSG( | 410.5475 | -0.01636 | 0.892581 | 0.984474 | Snrnp25 |
| ENSMUSG( | 756.603 | 0.07689 | 0.444303 | 0.833239 | Snrnp27 |
| ENSMUSG( | 278.9328 | 0.023257 | 0.868848 | 0.976897 | Snrnp35 |
| ENSMUSG( | 430.6581 | -0.06735 | 0.653845 | 0.922914 | Snrnp40 |
| ENSMUSG( | 599.4268 | 0.000807 | 0.994629 | 0.999369 | Snrnp48 |
| ENSMUSG( | 8137.617 | 0.109332 | 0.499039 | 0.862973 | Snrnp70 |
| ENSMUSG( | 1202.808 | -0.00295 | 0.98023 | 0.997512 | Snrpa |
| ENSMUSG( | 539.5899 | -0.01733 | 0.887677 | 0.983411 | Snrpa1 |
| ENSMUSG( | 1306.81 | 0.053888 | 0.610756 | 0.908349 | Snrpb |
| ENSMUSG( | 571.8846 | -0.11348 | 0.359903 | 0.791543 | Snrpb2 |
| ENSMUSG( | 655.3688 | -0.02733 | 0.808573 | 0.963455 | Snrpc |
| ENSMUSG( | 674.8446 | -0.01324 | 0.917705 | 0.989377 | Snrpd1 |
| ENSMUSG( | 1299.874 | -0.04711 | 0.673312 | 0.929007 | Snrpd2 |
| ENSMUSG( | 1345.725 | -0.02833 | 0.821962 | 0.966352 | Snrpd3 |
| ENSMUSG( | 561.4303 | 0.072415 | 0.548056 | 0.885153 | Snrpe |
| ENSMUSG( | 554.6091 | -0.04196 | 0.740354 | 0.947823 | Snrpf |
| ENSMUSG( | 488.9993 | 0.009457 | 0.935855 | 0.993415 | Snrpg |
| ENSMUSG( | 11973.44 | 0.065848 | 0.459287 | 0.840244 | Snrpn |
| ENSMUSG( | 1737.273 | 0.108843 | 0.218788 | 0.664559 | Snta1 |
| ENSMUSG( | 234.166 | -0.17925 | 0.328391 | 0.772854 | Sntb1 |
| ENSMUSG( | 477.9562 | -0.28159 | 0.020096 | 0.214413 | Sntb2 |
| ENSMUSG( | 257.55 | -0.22112 | 0.125725 | 0.53257 | Sntg1 |
| ENSMUSG( | 51.92733 | -0.16414 | 0.566886 | 0.891644 | Sntg2 |
| ENSMUSG( | 1934.278 | 0.019511 | 0.832454 | 0.970333 | Snu13 |
| ENSMUSG( | 288.3362 | 0.055043 | 0.705768 | 0.938649 | Snupn |
| ENSMUSG( | 1635.928 | 0.201878 | 0.029626 | 0.26088 | Snw1 |
| ENSMUSG( | 1576.479 | 0.008677 | 0.920282 | 0.990247 | Snx1 |
| ENSMUSG( | 1708.675 | -0.03378 | 0.75685 | 0.951513 | Snx10 |
| ENSMUSG( | 899.1962 | -0.01027 | 0.922391 | 0.99045 | Snx11 |
| ENSMUSG( | 1147.596 | -0.0104 | 0.919297 | 0.990247 | Snx12 |
| ENSMUSG( | 3045.798 | -0.05342 | 0.668825 | 0.927963 | Snx13 |
| ENSMUSG( | 1398.983 | -0.26181 | 0.091765 | 0.465904 | Snx14 |
| ENSMUSG( | 863.9605 | -0.14429 | 0.254814 | 0.705183 | Snx15 |
| ENSMUSG( | 285.7198 | -0.03142 | 0.832685 | 0.970333 | Snx16 |
| ENSMUSG( | 2283.096 | 0.093513 | 0.270373 | 0.720226 | Snx17 |
| ENSMUSG( | 1229.086 | -0.09356 | 0.447717 | 0.834051 | Snx18 |
| ENSMUSG( | 2688.77 | -0.22744 | 0.088434 | 0.456236 | Snx19 |
| ENSMUSG( | 1116.086 | 0.146494 | 0.147565 | 0.570152 | Snx2 |
| ENSMUSG( | 43.80069 | -0.02782 | 0.928039 | 0.991229 | Snx20 |
| ENSMUSG( | 1288.994 | 0.090756 | 0.367637 | 0.795314 | Snx21 |
| ENSMUSG( | 364.572 | -0.21267 | 0.094587 | 0.471772 | Snx22 |
| ENSMUSG( | 507.5881 | 0.089159 | 0.455077 | 0.837743 | Snx24 |
| ENSMUSG( | 1553.504 | -0.07746 | 0.571855 | 0.893506 | Snx25 |
| ENSMUSG( | 5760.704 | 0.100311 | 0.265268 | 0.716034 | Snx27 |
| ENSMUSG( | 331.7938 | -0.02885 | 0.835624 | 0.970557 | Snx29 |
| ENSMUSG( | 5353.265 | 0.020246 | 0.816519 | 0.965455 | Snx3 |
| ENSMUSG( | 1225.318 | -0.05149 | 0.613577 | 0.909044 | Snx30 |
| ENSMUSG( | 2080.218 | 0.126652 | 0.376937 | 0.801213 | Snx32 |
| ENSMUSG( | 351.8367 | -0.32245 | 0.012823 | 0.163518 | Snx33 |
| ENSMUSG( | 1212.978 | 0.027155 | 0.783364 | 0.957558 | Snx4 |
| ENSMUSG( | 5252.799 | 0.050397 | 0.61264 | 0.908818 | Snx5 |
| ENSMUSG( | 1222.125 | 0.136231 | 0.235575 | 0.682693 | Snx6 |
| ENSMUSG( | 233.5405 | 0.011581 | 0.939977 | 0.993415 | Snx7 |
| ENSMUSG( | 726.7861 | 0.075121 | 0.544593 | 0.884649 | Snx8 |
| ENSMUSG( | 795.5339 | 0.055432 | 0.601441 | 0.904603 | Snx9 |
| ENSMUSG( | 1516.359 | -0.14929 | 0.309239 | 0.75734 | Soat1 |
| ENSMUSG( | 2768.189 | 0.178626 | 0.045371 | 0.326933 | Sobp |
| ENSMUSG( | 35.0619 | -0.42401 | 0.23843 | 0.686308 | Socs1 |
| ENSMUSG( | 406.1417 | -0.05063 | 0.681466 | 0.931595 | Socs2 |
| ENSMUSG( | 100.4822 | -0.43013 | 0.049626 | 0.340062 | Socs3 |
| ENSMUSG( | 598.4244 | 0.081015 | 0.541713 | 0.883182 | Socs4 |
| ENSMUSG( | 2805.199 | 0.067852 | 0.442539 | 0.83278 | Socs5 |
| ENSMUSG( | 405.7878 | 0.16574 | 0.220121 | 0.665863 | Socs6 |
| ENSMUSG( | 4912.055 | 0.081883 | 0.343365 | 0.779153 | Socs7 |
| ENSMUSG( | 6726.248 | -0.03327 | 0.680979 | 0.931244 | Sod1 |
| ENSMUSG( | 4931.645 | 0.011252 | 0.901562 | 0.985633 | Sod2 |
| ENSMUSG( | 1316.962 | -0.3448 | 0.025768 | 0.247138 | Sod3 |
| ENSMUSG( | 7271.246 | 0.032063 | 0.744103 | 0.948497 | Son |
| ENSMUSG( | 4753.807 | 0.240897 | 0.04882 | 0.337919 | Sorbs1 |
| ENSMUSG( | 4462.09 | -0.22583 | 0.006795 | 0.113745 | Sorbs2 |
| ENSMUSG( | 2123.247 | -0.03146 | 0.721397 | 0.943026 | Sorbs3 |
| ENSMUSG( | 1144.699 | -0.38393 | 0.005392 | 0.098038 | Sorcs1 |
| ENSMUSG( | 2879.858 | -0.43881 | 0.001759 | 0.047383 | Sorcs2 |
| ENSMUSG( | 1051.729 | -0.5323 | 0.000506 | 0.020608 | Sorcs3 |
| ENSMUSG( | 612.9815 | 0.075984 | 0.480018 | 0.85456 | Sord |
| ENSMUSG( | 10673.71 | -0.13385 | 0.323671 | 0.768548 | Sorl1 |
| ENSMUSG( | 16119.36 | -0.2757 | 0.046809 | 0.330855 | Sort1 |
| ENSMUSG( | 1218.52 | -0.03034 | 0.733426 | 0.944956 | Sos1 |
| ENSMUSG( | 1325.502 | -0.00108 | 0.991685 | 0.999232 | Sos2 |
| ENSMUSG( | 105.1762 | -0.38095 | 0.546877 | 0.884974 | Sostdc1 |
| ENSMUSG( | 3507.855 | 0.048594 | 0.590181 | 0.90113 | Sowaha |
| ENSMUSG( | 275.8507 | -0.00448 | 0.976514 | 0.997428 | Sowahb |
| ENSMUSG( | 459.0935 | -0.02459 | 0.848871 | 0.972503 | Sowahc |
| ENSMUSG( | 565.0058 | -0.00061 | 0.996354 | 0.99943 | Sox1 |
| ENSMUSG( | 1350.861 | -0.12819 | 0.17072 | 0.605271 | Sox10 |
| ENSMUSG( | 655.0808 | -0.08311 | 0.526377 | 0.877267 | Sox11 |
| ENSMUSG( | 984.6323 | -0.10545 | 0.37204 | 0.798768 | Sox12 |
| ENSMUSG( | 455.5114 | 0.039725 | 0.759167 | 0.952284 | Sox13 |
| ENSMUSG( | 195.4073 | 0.670873 | 0.000397 | 0.017766 | Sox17 |
| ENSMUSG( | 319.3858 | 0.317021 | 0.17566 | 0.611006 | Sox18 |
| ENSMUSG( | 1730.305 | -0.14147 | 0.275171 | 0.72509 | Sox2 |
| ENSMUSG( | 889.235 | 0.284337 | 0.018488 | 0.204273 | Sox21 |
| ENSMUSG( | 61.61023 | -0.22731 | 0.390322 | 0.80794 | Sox21os1 |
| ENSMUSG( | 971.3932 | -0.15321 | 0.162572 | 0.593364 | Sox2ot |
| ENSMUSG( | 112.2918 | 0.1757 | 0.39745 | 0.812127 | Sox3 |
| ENSMUSG( | 1279.308 | 0.034572 | 0.770356 | 0.954689 | Sox4 |
| ENSMUSG( | 1431.256 | -0.02799 | 0.817046 | 0.96566 | Sox5 |
| ENSMUSG( | 1484.736 | -0.08326 | 0.521799 | 0.875186 | Sox6 |
| ENSMUSG( | 88.36718 | 0.066416 | 0.816351 | 0.965455 | Sox7 |
| ENSMUSG( | 1590.031 | -0.23861 | 0.010699 | 0.148964 | Sox8 |
| ENSMUSG( | 1345.704 | -0.011 | 0.91331 | 0.988483 | Sox9 |
| ENSMUSG( | 809.4683 | -0.03855 | 0.712957 | 0.94066 | Sp1 |
| ENSMUSG( | 181.6339 | -0.19051 | 0.370083 | 0.797542 | Sp100 |
| ENSMUSG( | 73.36795 | -0.4886 | 0.050707 | 0.345138 | Sp110 |
| ENSMUSG( | 41.19746 | -0.2159 | 0.488586 | 0.859092 | Sp140 |
| ENSMUSG( | 503.0191 | -0.05852 | 0.610536 | 0.908189 | Sp2 |
| ENSMUSG( | 1291.533 | -0.01393 | 0.880665 | 0.981542 | Sp3 |
| ENSMUSG( | 436.4977 | 0.024856 | 0.849653 | 0.972638 | Sp3os |
| ENSMUSG( | 346.3537 | -0.10931 | 0.392945 | 0.809274 | Sp4 |
| ENSMUSG( | 15.12199 | 0.22387 | 0.632413 | NA | Sp5 |
| ENSMUSG( | 32.5028 | -0.15919 | 0.625424 | 0.912249 | Sp6 |
| ENSMUSG( | 52.81902 | -0.59878 | 0.02719 | 0.253245 | Sp7 |
| ENSMUSG( | 67.94008 | -0.66653 | 0.055208 | 0.361664 | Sp8 |
| ENSMUSG( | 192.363 | -0.4225 | 0.04314 | 0.318494 | Sp9 |
| ENSMUSG( | 87.19493 | -0.1598 | 0.501525 | 0.864477 | Spa17 |
| ENSMUSG( | 25.71687 | -0.11731 | 0.756284 | 0.951479 | Spaar |
| ENSMUSG( | 720.8754 | 0.111363 | 0.527709 | 0.877801 | Spaca6 |
| ENSMUSG( | 687.731 | 0.151971 | 0.173057 | 0.608361 | Spag1 |
| ENSMUSG( | 29.74129 | -0.0246 | 0.953062 | 0.993874 | Spag16 |
| ENSMUSG( | 14.99672 | -0.28845 | 0.540976 | NA | Spag17 |
| ENSMUSG( | 136.8533 | 0.129529 | 0.488455 | 0.859092 | Spag4 |
| ENSMUSG( | 795.2436 | 0.526308 | 0.003712 | 0.07748 | Spag5 |

| | | | | | |
|---|---|---|---|---|---|
| ENSMUSG( | 1579.855 | 0.02392 | 0.841625 | 0.982981 | Sox8 |
| ENSMUSG( | 1338.724 | 0.230939 | 0.068179 | 0.75212 | Sox9 |
| ENSMUSG( | 747.4124 | 0.003617 | 0.976667 | 0.996416 | Sp1 |
| ENSMUSG( | 167.2545 | -0.14298 | 0.556737 | 0.939565 | Sp100 |
| ENSMUSG( | 76.78305 | -0.03978 | 0.869492 | 0.988097 | Sp110 |
| ENSMUSG( | 37.38143 | -0.22645 | 0.509435 | 0.929089 | Sp140 |
| ENSMUSG( | 432.2983 | -0.23463 | 0.083 | 0.77681 | Sp2 |
| ENSMUSG( | 1224.05 | 0.095998 | 0.437941 | 0.920495 | Sp3 |
| ENSMUSG( | 381.5785 | -0.10652 | 0.518334 | 0.930939 | Sp3os |
| ENSMUSG( | 317.7632 | -0.08952 | 0.532153 | 0.934795 | Sp4 |
| ENSMUSG( | 14.51682 | 0.352155 | 0.484692 | 0.92541 | Sp5 |
| ENSMUSG( | 29.98195 | -0.08389 | 0.831739 | 0.980656 | Sp6 |
| ENSMUSG( | 55.27437 | -0.17331 | 0.563233 | 0.941158 | Sp7 |
| ENSMUSG( | 84.6556 | 0.271924 | 0.445367 | 0.921734 | Sp8 |
| ENSMUSG( | 210.0064 | 0.111956 | 0.621186 | 0.953199 | Sp9 |
| ENSMUSG( | 80.06849 | -0.15803 | 0.526837 | 0.934449 | Spa17 |
| ENSMUSG( | 23.77034 | -0.01683 | 0.977551 | 0.996416 | Spaar |
| ENSMUSG( | 679.3659 | 0.203781 | 0.297556 | 0.898538 | Spaca6 |
| ENSMUSG( | 602.2866 | 0.037139 | 0.80417 | 0.977191 | Spag1 |
| ENSMUSG( | 30.8896 | 0.359567 | 0.450577 | 0.922202 | Spag16 |
| ENSMUSG( | 20.21921 | 0.79318 | 0.082467 | 0.77681 | Spag17 |
| ENSMUSG( | 117.305 | -0.03322 | 0.877968 | 0.989644 | Spag4 |
| ENSMUSG( | 603.3619 | 0.041486 | 0.83854 | 0.981622 | Spag5 |
| ENSMUSG( | 42.16902 | 0.207737 | 0.50048 | 0.927825 | Spag6 |
| ENSMUSG( | 89.82328 | 0.386216 | 0.111226 | 0.807885 | Spag6l |
| ENSMUSG( | 878.8745 | -0.06118 | 0.701985 | 0.962014 | Spag7 |
| ENSMUSG( | 48.40372 | 0.455619 | 0.162038 | 0.837724 | Spag8 |
| ENSMUSG( | 19372.82 | -0.03997 | 0.730185 | 0.967228 | Spag9 |
| ENSMUSG( | 38891.36 | 0.036527 | 0.770246 | 0.974236 | Sparc |
| ENSMUSG( | 181989 | 0.036045 | 0.761117 | 0.972695 | Sparcl1 |
| ENSMUSG( | 1073.533 | -0.11886 | 0.353359 | 0.912263 | Spart |
| ENSMUSG( | 1634.865 | 0.127124 | 0.298751 | 0.898538 | Spast |
| ENSMUSG( | 64.32267 | -0.00498 | 0.985089 | 0.997084 | Spata1 |
| ENSMUSG( | 476.0061 | 0.032947 | 0.804904 | 0.977232 | Spata13 |
| ENSMUSG( | 20.40278 | -0.22054 | 0.624226 | 0.953234 | Spata17 |
| ENSMUSG( | 15.31704 | 1.039259 | 0.083963 | 0.77737 | Spata18 |
| ENSMUSG( | 10.5568 | -1.05038 | 0.096462 | NA | Spata19 |
| ENSMUSG( | 1564.922 | 0.220211 | 0.074232 | 0.766789 | Spata2 |
| ENSMUSG( | 18.59667 | -0.44994 | 0.332582 | 0.907907 | Spata21 |
| ENSMUSG( | 55.85449 | 0.036598 | 0.897794 | 0.990558 | Spata24 |
| ENSMUSG( | 1141.057 | -0.13051 | 0.45294 | 0.922316 | Spata2l |
| ENSMUSG( | 53.51493 | 0.00821 | 0.977274 | 0.996416 | Spata3 |
| ENSMUSG( | 12.11084 | -0.3998 | 0.532437 | 0.934795 | Spata31d1e |
| ENSMUSG( | 8.433549 | 0.39756 | 0.566219 | NA | Spata31f1a |
| ENSMUSG( | 11.20488 | -0.29241 | 0.623029 | NA | Spata32 |
| ENSMUSG( | 159.4427 | -0.23059 | 0.20806 | 0.864412 | Spata33 |
| ENSMUSG( | 142.6603 | 0.213857 | 0.249457 | 0.877904 | Spata6 |
| ENSMUSG( | 126.3376 | 0.084955 | 0.691204 | 0.960071 | Spata6l |
| ENSMUSG( | 393.3133 | -0.0006 | 0.996554 | 0.999178 | Spata7 |
| ENSMUSG( | 80.32173 | -0.41481 | 0.078986 | 0.77458 | Spats1 |
| ENSMUSG( | 865.0886 | -0.05247 | 0.684806 | 0.958678 | Spats2 |
| ENSMUSG( | 422.3951 | -0.09349 | 0.561725 | 0.940388 | Spats2l |
| ENSMUSG( | 35.88844 | -0.1045 | 0.761242 | 0.972695 | Spc24 |
| ENSMUSG( | 26.61602 | -0.2107 | 0.585445 | 0.944975 | Spc25 |
| ENSMUSG( | 2445.61 | 0.025128 | 0.872491 | 0.989027 | Spcs1 |
| ENSMUSG( | 3131.939 | 0.026806 | 0.856286 | 0.985884 | Spcs2 |
| ENSMUSG( | 2223.792 | 0.229326 | 0.172581 | 0.843702 | Spcs3 |
| ENSMUSG( | 10.52052 | -0.4037 | 0.518407 | NA | Spdef |
| ENSMUSG( | 8.761127 | 0.43093 | 0.519224 | NA | Spdl1 |
| ENSMUSG( | 17.96056 | -0.18928 | 0.711884 | 0.964363 | Spdya |
| ENSMUSG( | 5507.877 | -0.19678 | 0.078858 | 0.774307 | Specc1 |
| ENSMUSG( | 2101.621 | -0.06632 | 0.545998 | 0.938063 | Specc1l |
| ENSMUSG( | 630.9061 | 0.056648 | 0.666064 | 0.956766 | Spef1 |
| ENSMUSG( | 14.52901 | 0.236273 | 0.649999 | 0.95422 | Spef1l |
| ENSMUSG( | 73.21825 | 0.639666 | 0.024864 | 0.592568 | Spef2 |
| ENSMUSG( | 6224.129 | 0.022497 | 0.820973 | 0.979198 | Speg |
| ENSMUSG( | 2477.3 | 0.13002 | 0.347354 | 0.911652 | Spen |
| ENSMUSG( | 797.0079 | 0.013162 | 0.92526 | 0.99306 | Spg11 |
| ENSMUSG( | 709.375 | 0.03513 | 0.813804 | 0.979198 | Spg21 |
| ENSMUSG( | 2875.761 | 0.012479 | 0.907022 | 0.991601 | Spg7 |
| ENSMUSG( | 138.9442 | 0.232435 | 0.375624 | 0.915491 | Sphk1 |
| ENSMUSG( | 669.8748 | -0.22483 | 0.096289 | 0.796133 | Sphk2 |
| ENSMUSG( | 3101.579 | 0.115531 | 0.396532 | 0.917257 | Sphkap |
| ENSMUSG( | 236.1419 | -0.16736 | 0.362308 | 0.914223 | Spi1 |
| ENSMUSG( | 8.583406 | -0.66069 | 0.39654 | NA | Spib |
| ENSMUSG( | 321.8374 | 0.073149 | 0.668066 | 0.956766 | Spice1 |
| ENSMUSG( | 194.054 | -0.12366 | 0.475364 | 0.923329 | Spidr |
| ENSMUSG( | 3892.622 | -0.01511 | 0.897943 | 0.990558 | Spin1 |
| ENSMUSG( | 48.14958 | -0.64761 | 0.039291 | 0.679972 | Spin2c |
| ENSMUSG( | 13.11848 | 0.448297 | 0.423175 | 0.917257 | Spin4 |
| ENSMUSG( | 574.6959 | -0.0507 | 0.723546 | 0.966887 | Spindoc |
| ENSMUSG( | 92.43932 | 0.098274 | 0.659296 | 0.956205 | Spink10 |
| ENSMUSG( | 69.87576 | 0.209408 | 0.414714 | 0.917257 | Spink8 |
| ENSMUSG( | 139.1868 | 0.223233 | 0.334864 | 0.907907 | Spint1 |
| ENSMUSG( | 675.0594 | 0.058269 | 0.685461 | 0.958678 | Spint2 |
| ENSMUSG( | 10725.36 | -0.08849 | 0.495085 | 0.927372 | Spire1 |
| ENSMUSG( | 443.551 | 0.083559 | 0.574858 | 0.942912 | Spire2 |
| ENSMUSG( | 13.43453 | 0.438013 | 0.470173 | 0.923329 | Spmap2 |
| ENSMUSG( | 18.05357 | -0.00048 | 0.99917 | 0.999503 | Spmip4 |
| ENSMUSG( | 52.08184 | 0.652984 | 0.028946 | 0.614171 | Spmip6 |
| ENSMUSG( | 12.84963 | 0.586922 | 0.311091 | 0.899422 | Spmip7 |
| ENSMUSG( | 44.85033 | -1.59171 | 0.100289 | 0.801311 | Spn |
| ENSMUSG( | 2123.817 | 0.196796 | 0.202011 | 0.864412 | Spns1 |
| ENSMUSG( | 2056.535 | 0.362209 | 0.018733 | 0.547826 | Spns2 |
| ENSMUSG( | 14.91321 | -0.40701 | 0.5016 | 0.927825 | Spns3 |
| ENSMUSG( | 16.42935 | 0.110083 | 0.857982 | 0.985884 | Spo11 |
| ENSMUSG( | 11.8438 | -0.00926 | 0.987487 | 0.99751 | Spocd1 |
| ENSMUSG( | 10487.99 | 0.033109 | 0.835678 | 0.981237 | Spock1 |
| ENSMUSG( | 34327.45 | 0.289025 | 0.086323 | 0.781091 | Spock2 |
| ENSMUSG( | 1871.543 | 0.118823 | 0.470713 | 0.923329 | Spock3 |
| ENSMUSG( | 3502.399 | 0.176053 | 0.19476 | 0.860433 | Spon1 |
| ENSMUSG( | 8.950482 | -1.93108 | 0.006368 | NA | Spon2 |
| ENSMUSG( | 12620.52 | -0.01353 | 0.907843 | 0.991958 | Spop |
| ENSMUSG( | 280.5761 | -0.01285 | 0.934463 | 0.99306 | Spopl |
| ENSMUSG( | 1212.967 | -0.03603 | 0.750678 | 0.970209 | Spout1 |
| ENSMUSG( | 275.0113 | -0.0819 | 0.620137 | 0.952896 | Spp1 |
| ENSMUSG( | 1801.378 | 0.096829 | 0.510159 | 0.929089 | Sppl2a |
| ENSMUSG( | 1690.214 | 0.181824 | 0.259938 | 0.883971 | Sppl2b |
| ENSMUSG( | 2693.754 | 0.058052 | 0.72206 | 0.966887 | Sppl3 |
| ENSMUSG( | 1009.946 | -0.18067 | 0.134216 | 0.822912 | Spr |
| ENSMUSG( | 6829.403 | 0.095496 | 0.570528 | 0.941872 | Spred1 |
| ENSMUSG( | 3046.489 | -0.02369 | 0.853856 | 0.98563 | Spred2 |
| ENSMUSG( | 1416.557 | 0.096133 | 0.407361 | 0.917257 | Spred3 |
| ENSMUSG( | 957.2207 | 0.118899 | 0.339741 | 0.907972 | Spring1 |
| ENSMUSG( | 7257.943 | 0.176285 | 0.2627 | 0.885357 | Sprn |
| ENSMUSG( | 539.2614 | -0.11307 | 0.377251 | 0.915912 | Sprtn |
| ENSMUSG( | 241.0044 | 0.274168 | 0.111355 | 0.808371 | Spry1 |
| ENSMUSG( | 1447.847 | 0.256023 | 0.0394 | 0.679972 | Spry2 |
| ENSMUSG( | 289.884 | 0.128206 | 0.399599 | 0.917257 | Spry4 |
| ENSMUSG( | 2767.191 | -0.14338 | 0.286603 | 0.894091 | Spryd3 |
| ENSMUSG( | 115.9652 | -0.06731 | 0.766825 | 0.974236 | Spryd4 |
| ENSMUSG( | 795.1364 | -0.18928 | 0.226562 | 0.868507 | Spryd7 |
| ENSMUSG( | 552.3212 | -0.11245 | 0.424595 | 0.917257 | Spsb1 |
| ENSMUSG( | 93.56447 | 0.200711 | 0.357997 | 0.913759 | Spsb2 |
| ENSMUSG( | 1239.903 | -0.12493 | 0.333134 | 0.907907 | Spsb3 |
| ENSMUSG( | 189.3052 | -0.29869 | 0.155648 | 0.837724 | Spsb4 |
| ENSMUSG( | 17.01606 | 0.072442 | 0.899403 | 0.990558 | Spta1 |
| ENSMUSG( | 31479.81 | 0.046104 | 0.722651 | 0.966887 | Sptan1 |
| ENSMUSG( | 8161.364 | 0.183948 | 0.323275 | 0.904023 | Sptb |
| ENSMUSG( | 52283.17 | 0.036997 | 0.775869 | 0.975517 | Sptbn1 |
| ENSMUSG( | 80880.39 | 0.173014 | 0.309483 | 0.898538 | Sptbn2 |
| ENSMUSG( | 8548.225 | 0.126515 | 0.228851 | 0.868507 | Sptbn4 |
| ENSMUSG( | 138.192 | 0.146846 | 0.445852 | 0.921734 | Sptbn5 |
| ENSMUSG( | 1261.935 | 0.181433 | 0.271547 | 0.888062 | Sptlc1 |
| ENSMUSG( | 1325.076 | 0.163281 | 0.361925 | 0.914223 | Sptlc2 |
| ENSMUSG( | 976.4832 | 0.026877 | 0.867942 | 0.98781 | Sptssa |
| ENSMUSG( | 44.64552 | -0.11751 | 0.695888 | 0.960867 | Sptssb |
| ENSMUSG( | 256.5783 | 0.118712 | 0.450301 | 0.922202 | Spty2d1 |
| ENSMUSG( | 48.00135 | 0.100125 | 0.748339 | 0.96984 | Spx |
| ENSMUSG( | 3182.665 | 0.031987 | 0.857171 | 0.985884 | Sqle |
| ENSMUSG( | 265.0802 | -0.00884 | 0.959048 | 0.994798 | Sqor |
| ENSMUSG( | 8594.908 | -0.0095 | 0.92446 | 0.99306 | Sqstm1 |
| ENSMUSG( | 696.7804 | -0.12278 | 0.328135 | 0.906255 | Sra1 |
| ENSMUSG( | 127.5865 | 1.148463 | 2.05E-05 | 0.006225 | Srarp |
| ENSMUSG( | 300.293 | -0.16022 | 0.311725 | 0.900031 | Srbd1 |
| ENSMUSG( | 5452.068 | -0.08821 | 0.36007 | 0.914223 | Src |
| ENSMUSG( | 142.0794 | 0.140428 | 0.471837 | 0.923329 | Srcap |
| ENSMUSG( | 20101.16 | -0.02463 | 0.812608 | 0.979148 | Srcin1 |
| ENSMUSG( | 800.1278 | -0.03583 | 0.773085 | 0.974463 | Srd5a1 |
| ENSMUSG( | 694.0251 | 0.128083 | 0.394555 | 0.917257 | Srd5a3 |
| ENSMUSG( | 2844.282 | 0.38089 | 0.014485 | 0.48395 | Srebf1 |
| ENSMUSG( | 10600.28 | 0.004069 | 0.979111 | 0.996468 | Srebf2 |
| ENSMUSG( | 1988.085 | 0.137281 | 0.341152 | 0.9082 | Srek1 |
| ENSMUSG( | 355.8587 | -0.14142 | 0.43502 | 0.918876 | Srek1ip1 |
| ENSMUSG( | 1489.1 | 0.177366 | 0.190786 | 0.857801 | Srf |
| ENSMUSG( | 248.8702 | 0.007167 | 0.965765 | 0.995433 | Srfbp1 |
| ENSMUSG( | 823.3667 | -0.08713 | 0.523454 | 0.932848 | Srgap1 |
| ENSMUSG( | 2236.41 | 0.136246 | 0.257156 | 0.883082 | Srgap2 |

| | | | | | | | | | | | |
|---|---|---|---|---|---|---|---|---|---|---|---|
| ENSMUSG( | Srsf7 | 1985.443 | -0.09891 | 0.387228 | 0.805442 | ENSMUSG( | Srsf7 | 2053.121 | 0.257494 | 0.049911 | 0.708648 |
| ENSMUSG( | Srsf9 | 1326.214 | 0.052154 | 0.575685 | 0.894836 | ENSMUSG( | Srsf9 | 1208.083 | 0.047993 | 0.705657 | 0.96315 |
| ENSMUSG( | Srxn1 | 2399.762 | 0.033167 | 0.711362 | 0.940028 | ENSMUSG( | Srxn1 | 2108.931 | -0.07259 | 0.506165 | 0.927825 |
| ENSMUSG( | Ss18 | 1523.783 | 0.137829 | 0.16631 | 0.598791 | ENSMUSG( | Ss18 | 1386.999 | 0.132968 | 0.267328 | 0.886134 |
| ENSMUSG( | Ss18l1 | 1188.382 | 0.126089 | 0.25092 | 0.700615 | ENSMUSG( | Ss18l1 | 1032.408 | -0.01243 | 0.925519 | 0.99306 |
| ENSMUSG( | Ss18l2 | 876.3597 | -0.05901 | 0.543716 | 0.884287 | ENSMUSG( | Ss18l2 | 762.9973 | -0.1996 | 0.103723 | 0.804969 |
| ENSMUSG( | Ssb | 2111.516 | 0.036524 | 0.723023 | 0.943761 | ENSMUSG( | Ssb | 1899.547 | -0.00341 | 0.978163 | 0.996416 |
| ENSMUSG( | Ssbp1 | 458.7952 | -0.05021 | 0.713809 | 0.940903 | ENSMUSG( | Ssbp1 | 443.4115 | 0.11605 | 0.518574 | 0.930939 |
| ENSMUSG( | Ssbp2 | 1807.136 | 0.036996 | 0.708883 | 0.939519 | ENSMUSG( | Ssbp2 | 1624.007 | -0.0051 | 0.970413 | 0.99594 |
| ENSMUSG( | Ssbp3 | 5143.069 | 0.185345 | 0.041495 | 0.314241 | ENSMUSG( | Ssbp3 | 4395.127 | 0.004388 | 0.970682 | 0.99594 |
| ENSMUSG( | Ssbp4 | 6195.954 | 0.103583 | 0.288592 | 0.739489 | ENSMUSG( | Ssbp4 | 5490.772 | 0.023988 | 0.821769 | 0.979198 |
| ENSMUSG( | Ssc4d | 34.4764 | -0.33372 | 0.311992 | 0.759715 | ENSMUSG( | Ssc4d | 31.7866 | -0.25385 | 0.495194 | 0.927372 |
| ENSMUSG( | Ssc5d | 100.7118 | 0.18025 | 0.403328 | 0.816132 | ENSMUSG( | Ssc5d | 98.16447 | 0.341911 | 0.177548 | 0.847209 |
| ENSMUSG( | Ssh1 | 685.8728 | -0.15141 | 0.218988 | 0.664602 | ENSMUSG( | Ssh1 | 647.3484 | -0.03927 | 0.777056 | 0.975748 |
| ENSMUSG( | Ssh2 | 1252.333 | 0.137128 | 0.183843 | 0.620849 | ENSMUSG( | Ssh2 | 1179.949 | 0.231109 | 0.051732 | 0.711848 |
| ENSMUSG( | Ssh3 | 1439.692 | 0.515884 | 0.003584 | 0.075902 | ENSMUSG( | Ssh3 | 1191.987 | 0.269583 | 0.214607 | 0.864599 |
| ENSMUSG( | Ssna1 | 692.8352 | -0.05156 | 0.642128 | 0.916865 | ENSMUSG( | Ssna1 | 631.8168 | -0.04771 | 0.715341 | 0.965063 |
| ENSMUSG( | Sspn | 1539.899 | -0.10176 | 0.383205 | 0.803254 | ENSMUSG( | Sspn | 1631.838 | 0.317499 | 0.010794 | 0.421367 |
| ENSMUSG( | Sspo | 131.067 | 0.161933 | 0.463917 | 0.842812 | ENSMUSG( | Sspo | 154.1969 | 0.811398 | 0.008906 | 0.394945 |
| ENSMUSG( | Ssr1 | 5488.837 | -0.32596 | 0.018372 | 0.203658 | ENSMUSG( | Ssr1 | 5756.462 | 0.097868 | 0.530541 | 0.934795 |
| ENSMUSG( | Ssr2 | 2480.994 | -0.42387 | 0.001568 | 0.044386 | ENSMUSG( | Ssr2 | 2588.352 | 0.000816 | 0.995656 | 0.999116 |
| ENSMUSG( | Ssr3 | 6190.667 | -0.24428 | 0.069735 | 0.408352 | ENSMUSG( | Ssr3 | 6372.679 | 0.117348 | 0.456239 | 0.923329 |
| ENSMUSG( | Ssr4 | 1624.329 | -0.15255 | 0.122958 | 0.529576 | ENSMUSG( | Ssr4 | 1533.666 | -0.04508 | 0.692464 | 0.960404 |
| ENSMUSG( | Ssrp1 | 2579.672 | 0.143624 | 0.177306 | 0.614553 | ENSMUSG( | Ssrp1 | 2230.064 | -0.00678 | 0.956156 | 0.994798 |
| ENSMUSG( | Sst | 9541.266 | -0.62541 | 5.40E-07 | 0.000113 | ENSMUSG( | Sst | 10130.95 | -0.11794 | 0.446176 | 0.921734 |
| ENSMUSG( | Sstr1 | 308.3147 | -0.29319 | 0.201384 | 0.641362 | ENSMUSG( | Sstr1 | 366.4217 | 0.465021 | 0.074296 | 0.766789 |
| ENSMUSG( | Sstr2 | 901.3595 | -0.36881 | 0.024595 | 0.240469 | ENSMUSG( | Sstr2 | 948.9824 | 0.075133 | 0.660516 | 0.956205 |
| ENSMUSG( | Sstr3 | 995.9529 | -0.06922 | 0.590398 | 0.90113 | ENSMUSG( | Sstr3 | 957.8078 | 0.090572 | 0.485633 | 0.92541 |
| ENSMUSG( | Sstr4 | 146.9692 | 0.246938 | 0.261083 | 0.710889 | ENSMUSG( | Sstr4 | 151.7725 | 0.58155 | 0.022986 | 0.584403 |
| ENSMUSG( | Ssu72 | 1295.719 | -0.07083 | 0.581808 | 0.897862 | ENSMUSG( | Ssu72 | 1143.231 | -0.17136 | 0.254582 | 0.882023 |
| ENSMUSG( | Ssx2ip | 1968.666 | 0.015709 | 0.867801 | 0.976631 | ENSMUSG( | Ssx2ip | 1776.889 | -0.01426 | 0.907262 | 0.991683 |
| ENSMUSG( | St13 | 3428.684 | 0.06179 | 0.551987 | 0.886141 | ENSMUSG( | St13 | 2978.585 | -0.07902 | 0.557007 | 0.939565 |
| ENSMUSG( | St14 | 23.23634 | 0.094543 | 0.828807 | 0.969097 | ENSMUSG( | St14 | 20.74108 | 0.084617 | 0.86328 | 0.986547 |
| ENSMUSG( | St18 | 235.6076 | -0.39564 | 0.011367 | 0.153509 | ENSMUSG( | St18 | 241.6577 | -0.01445 | 0.937685 | 0.993685 |
| ENSMUSG( | St3gal1 | 898.3325 | -0.26415 | 0.075863 | 0.426666 | ENSMUSG( | St3gal1 | 972.5691 | 0.241941 | 0.133785 | 0.822912 |
| ENSMUSG( | St3gal2 | 5739.452 | 0.125174 | 0.188172 | 0.627112 | ENSMUSG( | St3gal2 | 5199.907 | 0.111901 | 0.343667 | 0.910026 |
| ENSMUSG( | St3gal3 | 1902.81 | -0.10099 | 0.318811 | 0.764619 | ENSMUSG( | St3gal3 | 1831.305 | 0.061435 | 0.625131 | 0.953234 |
| ENSMUSG( | St3gal4 | 1558.66 | 0.076412 | 0.509669 | 0.869553 | ENSMUSG( | St3gal4 | 1488.184 | 0.205945 | 0.106665 | 0.804969 |
| ENSMUSG( | St3gal5 | 3970.596 | -0.31613 | 0.036339 | 0.293685 | ENSMUSG( | St3gal5 | 4187.126 | 0.125783 | 0.465291 | 0.923329 |
| ENSMUSG( | St3gal6 | 632.7907 | -0.11271 | 0.506499 | 0.867425 | ENSMUSG( | St3gal6 | 673.7959 | 0.329176 | 0.094826 | 0.796133 |
| ENSMUSG( | St6gal1 | 1089.369 | -0.03559 | 0.801858 | 0.962116 | ENSMUSG( | St6gal1 | 1079.875 | 0.207081 | 0.236357 | 0.872173 |
| ENSMUSG( | St6gal2 | 992.1784 | -0.5115 | 0.002117 | 0.053164 | ENSMUSG( | St6gal2 | 1184.972 | 0.301273 | 0.112939 | 0.809623 |
| ENSMUSG( | St6galnac2 | 143.4971 | 0.327675 | 0.130234 | 0.540948 | ENSMUSG( | St6galnac2 | 144.9107 | 0.604339 | 0.026345 | 0.603041 |
| ENSMUSG( | St6galnac3 | 324.2573 | -0.31984 | 0.071922 | 0.414057 | ENSMUSG( | St6galnac3 | 347.1267 | 0.1648 | 0.389463 | 0.917257 |
| ENSMUSG( | St6galnac4 | 1005.462 | -0.08805 | 0.516902 | 0.873055 | ENSMUSG( | St6galnac4 | 961.9098 | 0.060429 | 0.723357 | 0.966887 |
| ENSMUSG( | St6galnac5 | 3075.251 | -0.25541 | 0.045854 | 0.328083 | ENSMUSG( | St6galnac5 | 3049.296 | 0.00297 | 0.984583 | 0.996963 |
| ENSMUSG( | St6galnac6 | 9275.65 | 0.114911 | 0.155647 | 0.583641 | ENSMUSG( | St6galnac6 | 8167.779 | 0.019036 | 0.847442 | 0.985161 |
| ENSMUSG( | St7 | 562.8721 | 0.020574 | 0.891202 | 0.983819 | ENSMUSG( | St7 | 572.6472 | 0.324972 | 0.044646 | 0.693697 |
| ENSMUSG( | St7l | 706.0134 | -0.14421 | 0.38644 | 0.804732 | ENSMUSG( | St7l | 753.4624 | 0.304941 | 0.074766 | 0.767423 |
| ENSMUSG( | St8sia1 | 972.3591 | -0.19227 | 0.223869 | 0.669845 | ENSMUSG( | St8sia1 | 1011.602 | 0.196338 | 0.287961 | 0.894644 |
| ENSMUSG( | St8sia2 | 203.2396 | -0.61123 | 0.001146 | 0.036093 | ENSMUSG( | St8sia2 | 211.8304 | -0.15929 | 0.381226 | 0.917202 |
| ENSMUSG( | St8sia3 | 4160.835 | -0.61712 | 3.46E-06 | 0.000489 | ENSMUSG( | St8sia3 | 4729.892 | 0.088756 | 0.565871 | 0.941499 |
| ENSMUSG( | St8sia4 | 281.3028 | -0.53058 | 0.017438 | 0.197914 | ENSMUSG( | St8sia4 | 382.2843 | 0.615585 | 0.009305 | 0.403453 |
| ENSMUSG( | St8sia5 | 2100.394 | -0.11171 | 0.394182 | 0.809935 | ENSMUSG( | St8sia5 | 2141.496 | 0.211523 | 0.158809 | 0.837724 |
| ENSMUSG( | St8sia6 | 47.53863 | -0.03646 | 0.90526 | 0.98645 | ENSMUSG( | St8sia6 | 62.03307 | 0.887754 | 0.003658 | 0.242109 |
| ENSMUSG( | Stab1 | 1182.494 | 0.074161 | 0.543503 | 0.884037 | ENSMUSG( | Stab1 | 1195.261 | 0.35699 | 0.039003 | 0.679972 |
| ENSMUSG( | Stab2 | 20.1777 | 0.834651 | 0.050846 | NA | ENSMUSG( | Stab2 | 16.54566 | 0.606268 | 0.203583 | 0.864412 |
| ENSMUSG( | Stac | 27.99314 | 0.566553 | 0.125114 | 0.532139 | ENSMUSG( | Stac | 20.7274 | 0.036625 | 0.932347 | 0.99306 |
| ENSMUSG( | Stac2 | 7014.736 | 0.080555 | 0.363692 | 0.793343 | ENSMUSG( | Stac2 | 5913.916 | -0.14521 | 0.15173 | 0.837724 |
| ENSMUSG( | Stac3 | 92.41391 | 0.095372 | 0.696416 | 0.935166 | ENSMUSG( | Stac3 | 76.42817 | -0.18171 | 0.589285 | 0.944975 |
| ENSMUSG( | Stag1 | 1649.389 | 0.154871 | 0.140989 | 0.557902 | ENSMUSG( | Stag1 | 1453.735 | 0.05774 | 0.679083 | 0.957725 |
| ENSMUSG( | Stag2 | 2742.415 | 0.052084 | 0.658226 | 0.924446 | ENSMUSG( | Stag2 | 2462.645 | 0.007835 | 0.955562 | 0.994798 |
| ENSMUSG( | Stag3 | 68.30358 | 0.258378 | 0.336759 | 0.777 | ENSMUSG( | Stag3 | 59.80424 | 0.14413 | 0.654649 | 0.955015 |
| ENSMUSG( | Stam | 1100.97 | 0.049971 | 0.591037 | 0.90113 | ENSMUSG( | Stam | 993.1888 | 0.01942 | 0.871442 | 0.988779 |
| ENSMUSG( | Stam2 | 711.8037 | -0.04345 | 0.686689 | 0.933041 | ENSMUSG( | Stam2 | 716.6674 | 0.240126 | 0.061741 | 0.736228 |
| ENSMUSG( | Stambp | 649.5498 | -0.02541 | 0.808011 | 0.963389 | ENSMUSG( | Stambp | 574.1114 | -0.11809 | 0.356518 | 0.913358 |
| ENSMUSG( | Stambpl1 | 362.8187 | 0.009677 | 0.944108 | 0.993503 | ENSMUSG( | Stambpl1 | 317.7157 | -0.10058 | 0.513438 | 0.930482 |
| ENSMUSG( | Stap2 | 83.3635 | 0.69684 | 0.006589 | 0.111324 | ENSMUSG( | Stap2 | 69.80522 | 0.511126 | 0.129258 | 0.820358 |
| ENSMUSG( | Star | 58.28263 | 0.072788 | 0.790584 | 0.959461 | ENSMUSG( | Star | 58.1598 | 0.316204 | 0.278222 | 0.891468 |
| ENSMUSG( | Stard10 | 2615.895 | -0.02419 | 0.798652 | 0.960896 | ENSMUSG( | Stard10 | 2273.006 | -0.16936 | 0.222137 | 0.866232 |
| ENSMUSG( | Stard13 | 419.3995 | -0.07468 | 0.619637 | 0.911432 | ENSMUSG( | Stard13 | 405.7609 | 0.107327 | 0.540016 | 0.937383 |
| ENSMUSG( | Stard3 | 1125.333 | -0.2616 | 0.047867 | 0.334081 | ENSMUSG( | Stard3 | 1180.873 | 0.158525 | 0.316391 | 0.901868 |
| ENSMUSG( | Stard3nl | 1239.84 | -0.13189 | 0.235125 | 0.682364 | ENSMUSG( | Stard3nl | 1232.601 | 0.122752 | 0.314105 | 0.900949 |
| ENSMUSG( | Stard4 | 1249.834 | 0.101139 | 0.442242 | 0.832774 | ENSMUSG( | Stard4 | 1009.085 | -0.26067 | 0.055043 | 0.723362 |
| ENSMUSG( | Stard5 | 239.7291 | 0.145664 | 0.325985 | 0.771128 | ENSMUSG( | Stard5 | 205.4227 | -0.03685 | 0.83726 | 0.981335 |
| ENSMUSG( | Stard6 | 93.38516 | 0.239126 | 0.242645 | 0.690216 | ENSMUSG( | Stard6 | 82.79991 | 0.148208 | 0.538517 | 0.937216 |
| ENSMUSG( | Stard7 | 4280.559 | 0.061283 | 0.460549 | 0.840787 | ENSMUSG( | Stard7 | 3683.316 | -0.10697 | 0.339372 | 0.907972 |
| ENSMUSG( | Stard8 | 1089.587 | 0.231535 | 0.027765 | 0.255176 | ENSMUSG( | Stard8 | 939.8696 | 0.086618 | 0.527037 | 0.934449 |
| ENSMUSG( | Stard9 | 878.5893 | -0.44572 | 0.000346 | 0.016028 | ENSMUSG( | Stard9 | 982.5391 | 0.180539 | 0.325013 | 0.905273 |
| ENSMUSG( | Stat1 | 317.0103 | -0.1649 | 0.227383 | 0.67424 | ENSMUSG( | Stat1 | 288.8579 | -0.16122 | 0.290915 | 0.897019 |
| ENSMUSG( | Stat2 | 901.7626 | 0.219522 | 0.039052 | 0.304527 | ENSMUSG( | Stat2 | 785.671 | 0.097248 | 0.465071 | 0.923329 |
| ENSMUSG( | Stat3 | 2026.464 | 0.094269 | 0.263422 | 0.714261 | ENSMUSG( | Stat3 | 1765.901 | -0.03401 | 0.750416 | 0.970113 |
| ENSMUSG( | Stat4 | 11.79411 | -0.26312 | 0.634711 | NA | ENSMUSG( | Stat4 | 9.308739 | -0.74755 | 0.305924 | NA |
| ENSMUSG( | Stat5a | 136.7105 | -0.1988 | 0.359564 | 0.791011 | ENSMUSG( | Stat5a | 121.5527 | -0.28081 | 0.200173 | 0.863016 |
| ENSMUSG( | Stat5b | 1219.619 | 0.157493 | 0.101202 | 0.484996 | ENSMUSG( | Stat5b | 1056.84 | 0.014225 | 0.903934 | 0.991089 |
| ENSMUSG( | Stat6 | 441.0101 | 0.230791 | 0.061252 | 0.381822 | ENSMUSG( | Stat6 | 385.6937 | 0.123109 | 0.428246 | 0.917257 |
| ENSMUSG( | Stau1 | 1988.983 | 0.006222 | 0.940364 | 0.993415 | ENSMUSG( | Stau1 | 1801.246 | -0.0116 | 0.911678 | 0.992196 |
| ENSMUSG( | Stau2 | 5784.531 | 0.085364 | 0.291485 | 0.741366 | ENSMUSG( | Stau2 | 4894.122 | -0.13112 | 0.273438 | 0.889703 |
| ENSMUSG( | Stbd1 | 72.55223 | -0.21653 | 0.334108 | 0.775728 | ENSMUSG( | Stbd1 | 61.62338 | -0.4191 | 0.124795 | 0.81516 |
| ENSMUSG( | Stc1 | 89.08797 | -0.44041 | 0.035653 | 0.290384 | ENSMUSG( | Stc1 | 112.4732 | 0.485559 | 0.019645 | 0.556192 |
| ENSMUSG( | Stc2 | 106.6487 | -0.48259 | 0.026058 | 0.247245 | ENSMUSG( | Stc2 | 116.3015 | 0.0722 | 0.745824 | 0.96951 |
| ENSMUSG( | Steap1 | 32.09738 | 0.150866 | 0.87147 | 0.978114 | ENSMUSG( | Steap1 | 33.98817 | 0.526551 | 0.5769 | 0.942912 |
| ENSMUSG( | Steap2 | 619.27 | -0.27083 | 0.048954 | 0.337919 | ENSMUSG( | Steap2 | 610.0081 | -0.03456 | 0.822892 | 0.979198 |
| ENSMUSG( | Steap3 | 615.2029 | -0.3351 | 0.085527 | 0.450845 | ENSMUSG( | Steap3 | 637.2019 | 0.060315 | 0.757927 | 0.971858 |
| ENSMUSG( | Steap4 | 18.25549 | 0.133967 | 0.788635 | NA | ENSMUSG( | Steap4 | 15.71842 | 0.015409 | 0.977393 | 0.996416 |
| ENSMUSG( | Steep1 | 1994.862 | 0.076236 | 0.559231 | 0.888638 | ENSMUSG( | Steep1 | 1699.649 | -0.12364 | 0.438036 | 0.920503 |
| ENSMUSG( | Stil | 21.58661 | -0.66051 | 0.099679 | 0.482191 | ENSMUSG( | Stil | 23.92634 | -0.01996 | 0.960134 | 0.994798 |
| ENSMUSG( | Stim1 | 3494.602 | -0.35219 | 0.004724 | 0.090358 | ENSMUSG( | Stim1 | 3724.237 | 0.122817 | 0.398339 | 0.917257 |
| ENSMUSG( | Stim2 | 1740.675 | -0.29137 | 0.039663 | 0.307079 | ENSMUSG( | Stim2 | 1938.628 | 0.292912 | 0.085638 | 0.779799 |
| ENSMUSG( | Stimate | 976.115 | -0.11772 | 0.346105 | 0.779153 | ENSMUSG( | Stimate | 961.0896 | 0.111517 | 0.432219 | 0.918158 |
| ENSMUSG( | Sting1 | 241.6493 | -0.33634 | 0.109919 | 0.500285 | ENSMUSG( | Sting1 | 263.1521 | 0.203493 | 0.375701 | 0.915491 |
| ENSMUSG( | Stip1 | 2894.881 | 0.099571 | 0.264727 | 0.715763 | ENSMUSG( | Stip1 | 2482.593 | -0.07637 | 0.530744 | 0.934795 |
| ENSMUSG( | Stk10 | 379.2807 | -0.17162 | 0.268708 | 0.719042 | ENSMUSG( | Stk10 | 328.3648 | -0.34142 | 0.05089 | 0.709741 |
| ENSMUSG( | Stk11 | 3635.018 | 0.097539 | 0.223326 | 0.669382 | ENSMUSG( | Stk11 | 3135.65 | -0.06101 | 0.579044 | 0.943849 |
| ENSMUSG( | Stk11ip | 1324.556 | 0.187823 | 0.059831 | 0.378344 | ENSMUSG( | Stk11ip | 1169.72 | 0.105775 | 0.417841 | 0.917257 |
| ENSMUSG( | Stk16 | 1868.478 | 0.12186 | 0.152768 | 0.579081 | ENSMUSG( | Stk16 | 1603.367 | -0.04766 | 0.659631 | 0.956205 |
| ENSMUSG( | Stk17b | 573.8462 | 0.128904 | 0.325935 | 0.771128 | ENSMUSG( | Stk17b | 538.3143 | 0.199557 | 0.286877 | 0.894091 |
| ENSMUSG( | Stk19 | 871.791 | -0.0966 | 0.375183 | 0.800786 | ENSMUSG( | Stk19 | 807.525 | -0.05017 | 0.712459 | 0.964363 |
| ENSMUSG( | Stk24 | 1226.276 | 0.130029 | 0.188382 | 0.627297 | ENSMUSG( | Stk24 | 1084.712 | 0.04655 | 0.687664 | 0.959673 |
| ENSMUSG( | Stk25 | 6424.568 | 0.177122 | 0.08693 | 0.453311 | ENSMUSG( | Stk25 | 5558.446 | 0.031784 | 0.802828 | 0.97681 |
| ENSMUSG( | Stk26 | 84.25461 | -0.65778 | 0.016836 | 0.193535 | ENSMUSG( | Stk26 | 90.81034 | -0.11531 | 0.683152 | 0.958678 |
| ENSMUSG( | Stk3 | 364.5703 | 0.170632 | 0.178822 | 0.616298 | ENSMUSG( | Stk3 | 330.6191 | 0.147872 | 0.363942 | 0.914223 |
| ENSMUSG( | Stk32a | 173.2101 | -0.24328 | 0.147885 | 0.570981 | ENSMUSG( | Stk32a | 177.5058 | 0.094923 | 0.620998 | 0.953199 |
| ENSMUSG( | Stk32b | 63.76928 | 0.175529 | 0.503279 | 0.86596 | ENSMUSG( | Stk32b | 55.47821 | 0.048336 | 0.865058 | 0.98675 |
| ENSMUSG( | Stk32c | 2532.462 | 0.072704 | 0.551679 | 0.886141 | ENSMUSG( | Stk32c | 2227.023 | -0.02901 | 0.828974 | 0.980224 |
| ENSMUSG( | Stk33 | 31.00508 | -0.25729 | 0.459906 | 0.840338 | ENSMUSG( | Stk33 | 26.04035 | -0.52999 | 0.18203 | 0.850579 |
| ENSMUSG( | Stk35 | 952.1736 | 0.11752 | 0.245261 | 0.692812 | ENSMUSG( | Stk35 | 933.8246 | 0.318592 | 0.007682 | 0.373786 |
| ENSMUSG( | Stk36 | 209.7242 | 0.166375 | 0.314103 | 0.761319 | ENSMUSG( | Stk36 | 190.7985 | 0.162051 | 0.362802 | 0.914223 |
| ENSMUSG( | Stk38 | 1868.827 | 0.062866 | 0.531837 | 0.879628 | ENSMUSG( | Stk38 | 1706.626 | 0.069094 | 0.561664 | 0.940388 |
| ENSMUSG( | Stk38l | 1111.212 | 0.186536 | 0.054731 | 0.360135 | ENSMUSG( | Stk38l | 960.3797 | 0.038264 | 0.741188 | 0.968811 |
| ENSMUSG( | Stk39 | 1862.21 | 0.054165 | 0.630214 | 0.914073 | ENSMUSG( | Stk39 | 1599.094 | -0.12182 | 0.380347 | 0.917202 |
| ENSMUSG( | Stk4 | 1591.408 | 0.214676 | 0.024358 | 0.238636 | ENSMUSG( | Stk4 | 1374.674 | 0.06377 | 0.627369 | 0.953234 |
| ENSMUSG( | Stk40 | 1716.693 | -0.0088 | 0.925578 | 0.991204 | ENSMUSG( | Stk40 | 1528.224 | -0.07693 | 0.507766 | 0.929089 |
| ENSMUSG( | Stkld1 | 12.11701 | -0.13306 | 0.815024 | NA | ENSMUSG( | Stkld1 | 11.41565 | 0.002713 | 0.996594 | NA |
| ENSMUSG( | Stmn1 | 7423.648 | -0.05312 | 0.64023 | 0.916434 | ENSMUSG( | Stmn1 | 6528.274 | -0.16246 | 0.260574 | 0.884586 |
| ENSMUSG( | Stmn2 | 6811.212 | -0.03304 | 0.705561 | 0.938511 | ENSMUSG( | Stmn2 | 5986.601 | -0.141 | 0.209285 | 0.864412 |
| ENSMUSG( | Stmn3 | 9267.23 | -0.27376 | 0.001477 | 0.042635 | ENSMUSG( | Stmn3 | 8663.814 | -0.1952 | 0.149551 | 0.836241 |
| ENSMUSG( | Stmn4 | 5523.158 | -0.5352 | 1.45E-11 | 9.68E-09 | ENSMUSG( | Stmn4 | 5435.993 | -0.27704 | 0.010162 | 0.410151 |
| ENSMUSG( | Stmp1 | 821.4746 | 0.156206 | 0.167815 | 0.60061 | ENSMUSG( | Stmp1 | 694.6854 | -0.06102 | 0.652527 | 0.954799 |
| ENSMUSG( | Stn1 | 218.3767 | -0.07487 | 0.656528 | 0.923929 | ENSMUSG( | Stn1 | 194.9193 | -0.12494 | 0.508804 | 0.929089 |
| ENSMUSG( | Stom | 822.482 | -0.16012 | 0.231179 | 0.679229 | ENSMUSG( | Stom | 836.8862 | 0.164004 | 0.280753 | 0.892356 |
| ENSMUSG( | Stoml1 | 2350.748 | -0.2389 | 0.032844 | 0.277278 | ENSMUSG( | Stoml1 | 2302.524 | -0.01737 | 0.890391 | 0.990558 |
| ENSMUSG( | Stoml2 | 1540.062 | 0.099809 | 0.247715 | 0.696522 | ENSMUSG( | Stoml2 | 1307.303 | -0.1057 | 0.331899 | 0.907907 |
| ENSMUSG( | Stoml3 | 44.65559 | -1.0803 | 0.00202 | 0.051507 | ENSMUSG( | Stoml3 | 65.07711 | 0.412288 | 0.305912 | 0.898538 |
| ENSMUSG( | Ston1 | 280.5966 | 0.175739 | 0.222161 | 0.667925 | ENSMUSG( | Ston1 | 223.9258 | -0.21002 | 0.194979 | 0.861121 |
| ENSMUSG( | Ston2 | 3446.885 | -0.14393 | 0.305533 | 0.753611 | ENSMUSG( | Ston2 | 2932.173 | -0.3628 | 0.037411 | 0.673514 |
| ENSMUSG( | Stox1 | 107.0559 | 0.04604 | 0.819515 | 0.965888 | ENSMUSG( | Stox1 | 94.30974 | -0.03373 | 0.884861 | 0.989688 |
| ENSMUSG( | Stox2 | 4682.302 | 0.22322 | 0.021619 | 0.225182 | ENSMUSG( | Stox2 | 4329.413 | 0.263741 | 0.025931 | 0.601649 |
| ENSMUSG( | Stpg1 | 124.0893 | 0.278302 | 0.203446 | 0.643631 | ENSMUSG( | Stpg1 | 99.18433 | -0.09927 | 0.666573 | 0.956766 |
| ENSMUSG( | Stra6 | 351.9203 | 0.167037 | 0.182645 | 0.620029 | ENSMUSG( | Stra6 | 366.4216 | 0.509723 | 0.000269 | 0.043611 |
| ENSMUSG( | Strada | 779.3253 | 0.120064 | 0.433237 | 0.828207 | ENSMUSG( | Strada | 665.56 | -0.06777 | 0.690281 | 0.960071 |
| ENSMUSG( | Stradb | 1340.824 | 0.039193 | 0.670146 | 0.928485 | ENSMUSG( | Stradb | 1225.438 | 0.044674 | 0.711288 | 0.964363 |
| ENSMUSG( | Strap | 3029.597 | 0.013903 | 0.892639 | 0.984474 | ENSMUSG( | Strap | 2613.457 | -0.1513 | 0.283248 | 0.892412 |
| ENSMUSG( | Strbp | 4047.718 | 0.167523 | 0.122264 | 0.527707 | ENSMUSG( | Strbp | 3467.324 | -0.00712 | 0.955887 | 0.994798 |
| ENSMUSG( | Strc | 19.40802 | 0.630128 | 0.189659 | NA | ENSMUSG( | Strc | 13.9644 | -0.00698 | 0.989257 | 0.997916 |
| ENSMUSG( | Strip1 | 2447.942 | 0.116359 | 0.268754 | 0.719042 | ENSMUSG( | Strip1 | 2151.294 | 0.018093 | 0.881536 | 0.989644 |
| ENSMUSG( | Strip2 | 683.715 | -0.70763 | 2.86E-06 | 0.000417 | ENSMUSG( | Strip2 | 823.9475 | 0.179872 | 0.266276 | 0.886134 |
| ENSMUSG( | Strn | 2172.138 | -0.09638 | 0.361014 | 0.792797 | ENSMUSG( | Strn | 2011.47 | -0.05143 | 0.695977 | 0.960867 |
| ENSMUSG( | Strn3 | 2355.586 | 0.051514 | 0.579814 | 0.89687 | ENSMUSG( | Strn3 | 2084.449 | -0.0347 | 0.773553 | 0.974672 |
| ENSMUSG( | Strn4 | 7881.803 | 0.203579 | 0.038999 | 0.304448 | ENSMUSG( | Strn4 | 6897.454 | 0.093286 | 0.376121 | 0.915491 |
| ENSMUSG( | Stt3a | 1466.927 | -0.39201 | 0.013722 | 0.171117 | ENSMUSG( | Stt3a | 1595.355 | 0.145297 | 0.407 | 0.917257 |
| ENSMUSG( | Stt3b | 3876.276 | -0.2189 | 0.158874 | 0.588112 | ENSMUSG( | Stt3b | 4015.023 | 0.159781 | 0.347713 | 0.911652 |
| ENSMUSG( | Stub1 | 3731.642 | -0.00309 | 0.973366 | 0.99675 | ENSMUSG( | Stub1 | 3270.295 | -0.11906 | 0.298321 | 0.898538 |
| ENSMUSG( | Stum | 10731.41 | -0.01562 | 0.889553 | 0.98353 | ENSMUSG( | Stum | 10393.93 | 0.159756 | 0.228502 | 0.868507 |
| ENSMUSG( | Stx11 | 9.564939 | -0.54883 | 0.39182 | NA | ENSMUSG( | Stx11 | 8.665357 | -0.53365 | 0.460085 | NA |
| ENSMUSG( | Stx12 | 2791.699 | 0.072759 | 0.455818 | 0.838601 | ENSMUSG( | Stx12 | 2390.246 | -0.11139 | 0.391052 | 0.917257 |
| ENSMUSG( | Stx16 | 2093.698 | 0.083452 | 0.367239 | 0.795221 | ENSMUSG( | Stx16 | 1862.709 | 0.012466 | 0.921857 | 0.992644 |
| ENSMUSG( | Stx17 | 861.1405 | 0.034912 | 0.730987 | 0.944956 | ENSMUSG( | Stx17 | 753.5647 | -0.08148 | 0.485761 | 0.92541 |

| | | | | | |
|---|---|---|---|---|---|
| ENSMUSG( | 46.07336 | 0.192193 | 0.515766 | 0.872701 | Spag6 |
| ENSMUSG( | 85.7836 | 0.001173 | 0.995949 | 0.99943 | Spag6l |
| ENSMUSG( | 995.0441 | 0.034412 | 0.781817 | 0.957558 | Spag7 |
| ENSMUSG( | 49.29074 | 0.242099 | 0.433529 | 0.828346 | Spag8 |
| ENSMUSG( | 21686.79 | 0.017046 | 0.877827 | 0.981431 | Spag9 |
| ENSMUSG( | 43174.23 | 0.071239 | 0.541606 | 0.883182 | Sparc |
| ENSMUSG( | 203036.2 | 0.082019 | 0.414464 | 0.82013 | Sparcl1 |
| ENSMUSG( | 1237.495 | 0.027909 | 0.781933 | 0.957558 | Spart |
| ENSMUSG( | 1688.402 | -0.04706 | 0.663229 | 0.9265 | Spast |
| ENSMUSG( | 68.63108 | -0.10806 | 0.64556 | 0.918025 | Spata1 |
| ENSMUSG( | 467.5335 | -0.30238 | 0.046572 | 0.330304 | Spata13 |
| ENSMUSG( | 19.51955 | -0.64319 | 0.141171 | NA | Spata17 |
| ENSMUSG( | 11.96414 | 0.187312 | 0.744952 | NA | Spata18 |
| ENSMUSG( | 12.10775 | -0.83253 | 0.162861 | NA | Spata19 |
| ENSMUSG( | 1662.077 | 0.127945 | 0.205598 | 0.647118 | Spata2 |
| ENSMUSG( | 21.72123 | -0.25101 | 0.540648 | 0.88315 | Spata21 |
| ENSMUSG( | 63.36982 | 0.108306 | 0.68483 | 0.932981 | Spata24 |
| ENSMUSG( | 1330.279 | 0.048763 | 0.763378 | 0.953781 | Spata2l |
| ENSMUSG( | 72.68455 | 0.574669 | 0.016984 | 0.194647 | Spata3 |
| ENSMUSG( | 13.64142 | -0.32157 | 0.524261 | NA | Spata31d1e |
| ENSMUSG( | 8.698212 | 0.193221 | 0.753403 | NA | Spata31f1a |
| ENSMUSG( | 11.35299 | -0.54741 | 0.318964 | NA | Spata32 |
| ENSMUSG( | 174.3223 | -0.24287 | 0.131652 | 0.544205 | Spata33 |
| ENSMUSG( | 119.1921 | -0.64008 | 0.000839 | 0.02897 | Spata6 |
| ENSMUSG( | 135.6998 | 0.029866 | 0.879186 | 0.981436 | Spata6l |
| ENSMUSG( | 449.8735 | 0.115876 | 0.384841 | 0.804031 | Spata7 |
| ENSMUSG( | 94.91823 | -0.18111 | 0.408463 | 0.818167 | Spats1 |
| ENSMUSG( | 982.7599 | 0.050498 | 0.644286 | 0.917741 | Spats2 |
| ENSMUSG( | 491.8303 | 0.077918 | 0.587796 | 0.900628 | Spats2l |
| ENSMUSG( | 38.52409 | -0.18952 | 0.529975 | 0.878883 | Spc24 |
| ENSMUSG( | 33.7665 | 0.218111 | 0.495103 | 0.860798 | Spc25 |
| ENSMUSG( | 2358.504 | -0.37341 | 0.005668 | 0.10081 | Spcs1 |
| ENSMUSG( | 3114.474 | -0.27298 | 0.030319 | 0.264512 | Spcs2 |
| ENSMUSG( | 2105.839 | -0.19954 | 0.192935 | 0.632208 | Spcs3 |
| ENSMUSG( | 11.84411 | -0.31105 | 0.572305 | NA | Spdef |
| ENSMUSG( | 9.962538 | 0.470596 | 0.44141 | NA | Spdl1 |
| ENSMUSG( | 21.24867 | 0.072432 | 0.869839 | 0.977607 | Spdya |
| ENSMUSG( | 6520.582 | 0.029277 | 0.78157 | 0.957558 | Specc1 |
| ENSMUSG( | 2428.58 | 0.082742 | 0.367649 | 0.795314 | Specc1l |
| ENSMUSG( | 717.978 | 0.153842 | 0.144955 | 0.565187 | Spef1 |
| ENSMUSG( | 15.78445 | 0.179798 | 0.704553 | NA | Spef1l |
| ENSMUSG( | 67.22996 | 0.163177 | 0.498123 | 0.862463 | Spef2 |
| ENSMUSG( | 6933.652 | 0.064203 | 0.453339 | 0.836798 | Speg |
| ENSMUSG( | 2614.076 | 0.014806 | 0.901258 | 0.985633 | Spen |
| ENSMUSG( | 906.3633 | 0.108305 | 0.38329 | 0.80333 | Spg11 |
| ENSMUSG( | 769.9646 | 0.009866 | 0.925336 | 0.991204 | Spg21 |
| ENSMUSG( | 3205.406 | 0.055266 | 0.540049 | 0.88315 | Spg7 |
| ENSMUSG( | 140.1129 | 0.002704 | 0.990133 | 0.999144 | Sphk1 |
| ENSMUSG( | 815.5087 | 0.078085 | 0.487444 | 0.858812 | Sphk2 |
| ENSMUSG( | 3238.186 | -0.02983 | 0.772273 | 0.955037 | Sphkap |
| ENSMUSG( | 255.1896 | -0.21747 | 0.106264 | 0.49377 | Spi1 |
| ENSMUSG( | 11.51458 | -0.02671 | 0.970238 | NA | Spib |
| ENSMUSG( | 371.4198 | 0.206013 | 0.10359 | 0.489428 | Spice1 |
| ENSMUSG( | 238.2691 | 0.200756 | 0.174733 | 0.610304 | Spidr |
| ENSMUSG( | 4506.788 | 0.137685 | 0.078112 | 0.43227 | Spin1 |
| ENSMUSG( | 62.30846 | -0.10897 | 0.689005 | 0.933441 | Spin2c |
| ENSMUSG( | 10.48569 | -0.43291 | 0.461714 | NA | Spin4 |
| ENSMUSG( | 668.1462 | 0.115638 | 0.332337 | 0.775474 | Spindoc |
| ENSMUSG( | 100.6465 | 0.066356 | 0.73404 | 0.945306 | Spink10 |
| ENSMUSG( | 67.38712 | -0.15666 | 0.514764 | 0.872189 | Spink8 |
| ENSMUSG( | 114.7051 | -0.68851 | 0.006483 | 0.110626 | Spint1 |
| ENSMUSG( | 661.9823 | -0.27675 | 0.006878 | 0.114527 | Spint2 |
| ENSMUSG( | 13333.1 | 0.261747 | 0.007222 | 0.117261 | Spire1 |
| ENSMUSG( | 484.7406 | 0.072779 | 0.59972 | 0.903906 | Spire2 |
| ENSMUSG( | 9.261084 | -0.1856 | 0.761068 | NA | Spmap1 |
| ENSMUSG( | 19.20393 | -0.08366 | 0.841263 | NA | Spmip4 |
| ENSMUSG( | 47.12306 | 0.164041 | 0.590033 | 0.90113 | Spmip6 |
| ENSMUSG( | 12.89129 | 0.332643 | 0.538821 | NA | Spmip7 |
| ENSMUSG( | 51.83756 | -1.26046 | 0.161976 | 0.592429 | Spn |
| ENSMUSG( | 2087.597 | -0.12211 | 0.339505 | 0.777792 | Spns1 |
| ENSMUSG( | 2050.359 | 0.100534 | 0.452307 | 0.836145 | Spns2 |
| ENSMUSG( | 16.38295 | -0.44053 | 0.45715 | NA | Spns3 |
| ENSMUSG( | 15.07686 | -0.40561 | 0.438834 | NA | Spo11 |
| ENSMUSG( | 18.16394 | 0.85603 | 0.063529 | NA | Spocd1 |
| ENSMUSG( | 9421.137 | -0.61162 | 9.54E-06 | 0.001107 | Spock1 |
| ENSMUSG( | 31980.3 | -0.18014 | 0.220911 | 0.666476 | Spock2 |
| ENSMUSG( | 1501.193 | -0.93538 | 4.12E-10 | 2.09E-07 | Spock3 |
| ENSMUSG( | 3227.419 | -0.34651 | 0.004259 | 0.084556 | Spon1 |
| ENSMUSG( | 12.46044 | -0.73216 | 0.210926 | NA | Spon2 |
| ENSMUSG( | 14570.85 | 0.131112 | 0.155767 | 0.58375 | Spop |
| ENSMUSG( | 297.4219 | -0.11885 | 0.383018 | 0.803237 | Spopl |
| ENSMUSG( | 1388.601 | 0.08564 | 0.341783 | 0.778828 | Spout1 |
| ENSMUSG( | 299.6249 | -0.09282 | 0.546485 | 0.884974 | Spp1 |
| ENSMUSG( | 1776.096 | -0.22113 | 0.104478 | 0.490485 | Sppl2a |
| ENSMUSG( | 1663.461 | -0.13594 | 0.334786 | 0.776386 | Sppl2b |
| ENSMUSG( | 2611.859 | -0.32184 | 0.02017 | 0.214666 | Sppl3 |
| ENSMUSG( | 1219.632 | 0.102331 | 0.338193 | 0.777422 | Spr |
| ENSMUSG( | 7735.017 | 0.184367 | 0.124958 | 0.531973 | Spred1 |
| ENSMUSG( | 3508.628 | 0.115013 | 0.183739 | 0.620678 | Spred2 |
| ENSMUSG( | 1612.535 | 0.19594 | 0.049188 | 0.338637 | Spred3 |
| ENSMUSG( | 967.1335 | -0.12405 | 0.269051 | 0.719302 | Spring1 |
| ENSMUSG( | 6810.509 | -0.28901 | 0.028547 | 0.257525 | Sprn |
| ENSMUSG( | 585.9562 | -0.14541 | 0.180585 | 0.617841 | Sprtn |
| ENSMUSG( | 223.1707 | -0.21516 | 0.197075 | 0.636662 | Spry1 |
| ENSMUSG( | 1454.183 | 0.013625 | 0.8995 | 0.985633 | Spry2 |
| ENSMUSG( | 296.2779 | -0.07404 | 0.573472 | 0.893584 | Spry4 |
| ENSMUSG( | 3235.14 | 0.043807 | 0.71186 | 0.940028 | Spryd3 |
| ENSMUSG( | 137.8506 | 0.148906 | 0.433502 | 0.828346 | Spryd4 |
| ENSMUSG( | 895.6481 | -0.10455 | 0.378734 | 0.801688 | Spryd7 |
| ENSMUSG( | 660.3151 | 0.13194 | 0.317251 | 0.763823 | Spsb1 |
| ENSMUSG( | 99.27235 | 0.105658 | 0.592646 | 0.902094 | Spsb2 |
| ENSMUSG( | 1444.216 | 0.047895 | 0.685803 | 0.932981 | Spsb3 |
| ENSMUSG( | 213.8145 | -0.19065 | 0.198155 | 0.638387 | Spsb4 |
| ENSMUSG( | 34530.87 | 0.041624 | 0.704244 | 0.938221 | Sptan1 |
| ENSMUSG( | 8713.961 | 0.102576 | 0.510321 | 0.870063 | Sptb |
| ENSMUSG( | 56515.73 | -0.00901 | 0.942986 | 0.993415 | Sptbn1 |
| ENSMUSG( | 91830.35 | 0.259828 | 0.074792 | 0.42402 | Sptbn2 |
| ENSMUSG( | 9494.51 | 0.158432 | 0.079189 | 0.435105 | Sptbn4 |
| ENSMUSG( | 161.49 | 0.308899 | 0.054453 | 0.359033 | Sptbn5 |
| ENSMUSG( | 1218.667 | -0.19434 | 0.191157 | 0.630245 | Sptlc1 |
| ENSMUSG( | 1227.496 | -0.34891 | 0.022627 | 0.229667 | Sptlc2 |
| ENSMUSG( | 1055.928 | -0.01012 | 0.938803 | 0.993415 | Sptssa |
| ENSMUSG( | 48.05952 | -0.15793 | 0.571429 | 0.893319 | Sptssb |
| ENSMUSG( | 257.0652 | -0.14986 | 0.290774 | 0.740751 | Spty2d1 |
| ENSMUSG( | 57.00944 | 0.33684 | 0.196568 | 0.635942 | Spx |
| ENSMUSG( | 3159.243 | -0.27527 | 0.089839 | 0.460262 | Sqle |
| ENSMUSG( | 312.3457 | 0.184995 | 0.198788 | 0.63899 | Sqor |
| ENSMUSG( | 9322.693 | -0.0441 | 0.586514 | 0.900023 | Sqstm1 |
| ENSMUSG( | 810.4626 | 0.047499 | 0.650539 | 0.921301 | Sra1 |
| ENSMUSG( | 99.61566 | 0.347354 | 0.16593 | 0.597821 | Srarp |
| ENSMUSG( | 356.3932 | 0.069882 | 0.595236 | 0.903007 | Srbd1 |
| ENSMUSG( | 6398.976 | 0.10512 | 0.162871 | 0.593778 | Src |
| ENSMUSG( | 149.1808 | 0.005977 | 0.973951 | 0.99675 | Srcap |
| ENSMUSG( | 23075.16 | 0.102563 | 0.183347 | 0.62047 | Srcin1 |
| ENSMUSG( | 880.3824 | -0.02816 | 0.803664 | 0.962169 | Srd5a1 |
| ENSMUSG( | 664.7441 | -0.27914 | 0.031563 | 0.270895 | Srd5a3 |
| ENSMUSG( | 2545.469 | -0.19408 | 0.129726 | 0.539528 | Srebf1 |
| ENSMUSG( | 10953.85 | -0.17955 | 0.189378 | 0.627932 | Srebf2 |
| ENSMUSG( | 2120.283 | 0.06089 | 0.590031 | 0.90113 | Srek1 |
| ENSMUSG( | 378.8756 | -0.22292 | 0.119458 | 0.523291 | Srek1ip1 |
| ENSMUSG( | 1685.838 | 0.256422 | 0.026596 | 0.249357 | Srf |
| ENSMUSG( | 271.7191 | -0.00522 | 0.972326 | 0.996585 | Srfbp1 |
| ENSMUSG( | 877.3007 | -0.17133 | 0.124538 | 0.531714 | Srgap1 |
| ENSMUSG( | 2492.512 | 0.175441 | 0.050191 | 0.342492 | Srgap2 |
| ENSMUSG( | 12220.41 | 0.221887 | 0.002452 | 0.058231 | Srgap3 |
| ENSMUSG( | 216.5307 | -0.05749 | 0.74661 | 0.949289 | Srgn |
| ENSMUSG( | 2981.537 | 0.018179 | 0.871349 | 0.978114 | Sri |
| ENSMUSG( | 297.2951 | 0.134984 | 0.299285 | 0.748131 | Srl |
| ENSMUSG( | 1234.933 | -0.10375 | 0.408516 | 0.818167 | Srm |
| ENSMUSG( | 12.21859 | -0.92322 | 0.094615 | NA | Sros1 |
| ENSMUSG( | 1774.315 | 0.0078 | 0.94279 | 0.993415 | Srp14 |
| ENSMUSG( | 924.7947 | 0.045164 | 0.690738 | 0.933881 | Srp19 |
| ENSMUSG( | 58.70916 | 0.134156 | 0.593051 | 0.902372 | Srp54a |
| ENSMUSG( | 287.5468 | -0.00159 | 0.991269 | 0.999232 | Srp54c |
| ENSMUSG( | 1617.879 | 0.149078 | 0.114097 | 0.509818 | Srp68 |
| ENSMUSG( | 2079.345 | 0.094745 | 0.262999 | 0.713397 | Srp72 |
| ENSMUSG( | 1823.331 | 0.084638 | 0.384203 | 0.803686 | Srp9 |
| ENSMUSG( | 2140.334 | 0.029716 | 0.762245 | 0.953459 | Srpk1 |
| ENSMUSG( | 6153.228 | 0.026614 | 0.733988 | 0.945306 | Srpk2 |
| ENSMUSG( | 230.0886 | 0.291145 | 0.103708 | 0.489428 | Srpk3 |
| ENSMUSG( | 1829.709 | -0.24652 | 0.141957 | 0.559166 | Srpra |
| ENSMUSG( | 1576.833 | -0.33333 | 0.025863 | 0.247245 | Srprb |
| ENSMUSG( | 13.3796 | -0.32358 | 0.511745 | NA | Srpx |
| ENSMUSG( | 39.4296 | -0.35489 | 0.269048 | 0.719302 | Srpx2 |
| ENSMUSG( | 3809.068 | 0.067029 | 0.437734 | 0.830072 | Srr |
| ENSMUSG( | 547.5234 | -0.12633 | 0.305864 | 0.753712 | Srrd |
| ENSMUSG( | 4180.862 | 0.059306 | 0.553407 | 0.886216 | Srrm1 |
| ENSMUSG( | 11982.08 | 0.111399 | 0.412996 | 0.819541 | Srrm2 |

| | | | | | |
|---|---|---|---|---|---|
| ENSMUSG( | 10110.29 | -0.04829 | 0.621963 | 0.953234 | Srgap3 |
| ENSMUSG( | 249.8614 | 0.578691 | 0.00332 | 0.229858 | Srgn |
| ENSMUSG( | 2543.708 | -0.18067 | 0.205766 | 0.864412 | Sri |
| ENSMUSG( | 266.4363 | 0.097587 | 0.546613 | 0.938063 | Srl |
| ENSMUSG( | 1178.277 | 0.027524 | 0.855786 | 0.985783 | Srm |
| ENSMUSG( | 9.831852 | 0.771144 | 0.287469 | NA | Srms |
| ENSMUSG( | 14.33803 | -0.11254 | 0.854393 | 0.985649 | Sros1 |
| ENSMUSG( | 1535.831 | -0.14911 | 0.331642 | 0.907907 | Srp14 |
| ENSMUSG( | 828.9615 | -0.00858 | 0.953611 | 0.994798 | Srp19 |
| ENSMUSG( | 50.59746 | -0.03228 | 0.910452 | 0.992124 | Srp54a |
| ENSMUSG( | 239.2424 | -0.27336 | 0.085301 | 0.779799 | Srp54c |
| ENSMUSG( | 1382.915 | -0.03275 | 0.770425 | 0.974236 | Srp68 |
| ENSMUSG( | 1903.09 | 0.105198 | 0.363037 | 0.914223 | Srp72 |
| ENSMUSG( | 1558.624 | -0.10466 | 0.464169 | 0.923329 | Srp9 |
| ENSMUSG( | 1964.454 | 0.051586 | 0.666519 | 0.956766 | Srpk1 |
| ENSMUSG( | 5486.816 | -0.03755 | 0.749732 | 0.96998 | Srpk2 |
| ENSMUSG( | 197.804 | 0.132167 | 0.594916 | 0.945572 | Srpk3 |
| ENSMUSG( | 1920.42 | 0.173997 | 0.360378 | 0.914223 | Srpra |
| ENSMUSG( | 1682.677 | 0.14327 | 0.385523 | 0.917257 | Srprb |
| ENSMUSG( | 18.84942 | 0.856282 | 0.063613 | 0.736482 | Srpx |
| ENSMUSG( | 42.44646 | 0.172382 | 0.603892 | 0.948137 | Srpx2 |
| ENSMUSG( | 3236.364 | -0.13812 | 0.17614 | 0.846252 | Srr |
| ENSMUSG( | 509.677 | -0.05753 | 0.679887 | 0.957966 | Srrd |
| ENSMUSG( | 3825.266 | 0.069428 | 0.576567 | 0.942912 | Srrm1 |
| ENSMUSG( | 11567.89 | 0.266964 | 0.122103 | 0.813384 | Srrm2 |
| ENSMUSG( | 1798.499 | 0.035453 | 0.779899 | 0.976106 | Srrm3 |
| ENSMUSG( | 1227.282 | 0.073386 | 0.60347 | 0.948137 | Srrm4 |
| ENSMUSG( | 3069.12 | 0.065396 | 0.632907 | 0.953844 | Srrt |
| ENSMUSG( | 4036.926 | 0.115687 | 0.381577 | 0.917257 | Srsf1 |
| ENSMUSG( | 1238.754 | 0.244014 | 0.063774 | 0.736482 | Srsf10 |
| ENSMUSG( | 3275.264 | 0.116817 | 0.39274 | 0.917257 | Srsf11 |
| ENSMUSG( | 315.2249 | -0.00731 | 0.967941 | 0.995855 | Srsf12 |
| ENSMUSG( | 2983.068 | 0.170387 | 0.190289 | 0.857355 | Srsf2 |
| ENSMUSG( | 2419.887 | 0.069377 | 0.531112 | 0.934795 | Srsf3 |
| ENSMUSG( | 2485.422 | -0.02736 | 0.811356 | 0.978997 | Srsf4 |
| ENSMUSG( | 5243.512 | 0.160519 | 0.367582 | 0.915217 | Srsf5 |
| ENSMUSG( | 3027.48 | 0.177515 | 0.158924 | 0.837724 | Srsf6 |
| ENSMUSG( | 2053.121 | 0.257494 | 0.049911 | 0.708648 | Srsf7 |
| ENSMUSG( | 1208.083 | 0.047993 | 0.705657 | 0.96315 | Srsf9 |
| ENSMUSG( | 2108.931 | -0.07259 | 0.506165 | 0.927825 | Srxn1 |
| ENSMUSG( | 1386.999 | 0.132968 | 0.267328 | 0.886134 | Ss18 |
| ENSMUSG( | 1032.408 | -0.01243 | 0.925519 | 0.99306 | Ss18l1 |
| ENSMUSG( | 762.9973 | -0.1996 | 0.103723 | 0.804969 | Ss18l2 |
| ENSMUSG( | 1899.547 | -0.00341 | 0.978163 | 0.996416 | Ssb |
| ENSMUSG( | 443.4115 | 0.11605 | 0.518574 | 0.930939 | Ssbp1 |
| ENSMUSG( | 1624.007 | -0.0051 | 0.970413 | 0.99594 | Ssbp2 |
| ENSMUSG( | 4395.127 | 0.004388 | 0.970682 | 0.99594 | Ssbp3 |
| ENSMUSG( | 5490.772 | 0.023988 | 0.821769 | 0.979198 | Ssbp4 |
| ENSMUSG( | 31.7866 | -0.25385 | 0.495194 | 0.927372 | Ssc4d |
| ENSMUSG( | 98.16447 | 0.341911 | 0.177548 | 0.847209 | Ssc5d |
| ENSMUSG( | 647.3484 | -0.03927 | 0.777056 | 0.975748 | Ssh1 |
| ENSMUSG( | 1179.949 | 0.231109 | 0.051732 | 0.711848 | Ssh2 |
| ENSMUSG( | 1191.987 | 0.269583 | 0.214607 | 0.864599 | Ssh3 |
| ENSMUSG( | 631.8168 | -0.04771 | 0.715341 | 0.965063 | Ssna1 |
| ENSMUSG( | 1631.838 | 0.317499 | 0.010794 | 0.421367 | Sspn |
| ENSMUSG( | 154.1969 | 0.811398 | 0.008906 | 0.394945 | Sspo |
| ENSMUSG( | 5756.462 | 0.097868 | 0.530541 | 0.934795 | Ssr1 |
| ENSMUSG( | 2588.352 | 0.000816 | 0.995656 | 0.999116 | Ssr2 |
| ENSMUSG( | 6372.679 | 0.117348 | 0.456239 | 0.923329 | Ssr3 |
| ENSMUSG( | 1533.666 | -0.04508 | 0.692464 | 0.960404 | Ssr4 |
| ENSMUSG( | 2230.064 | -0.00678 | 0.956156 | 0.994798 | Ssrp1 |
| ENSMUSG( | 10130.95 | -0.11794 | 0.446176 | 0.921734 | Sst |
| ENSMUSG( | 366.4217 | 0.465021 | 0.074296 | 0.766789 | Sstr1 |
| ENSMUSG( | 948.9824 | 0.075133 | 0.660516 | 0.956205 | Sstr2 |
| ENSMUSG( | 957.8078 | 0.090572 | 0.485633 | 0.92541 | Sstr3 |
| ENSMUSG( | 151.7725 | 0.58155 | 0.022986 | 0.584403 | Sstr4 |
| ENSMUSG( | 10.97013 | -0.84295 | 0.171414 | NA | Sstr5 |
| ENSMUSG( | 1143.231 | -0.17136 | 0.254582 | 0.882023 | Ssu72 |
| ENSMUSG( | 1776.889 | -0.01426 | 0.907262 | 0.991683 | Ssx2ip |
| ENSMUSG( | 2978.585 | -0.07902 | 0.557007 | 0.939565 | St13 |
| ENSMUSG( | 20.74108 | 0.084617 | 0.86328 | 0.986547 | St14 |
| ENSMUSG( | 241.6577 | -0.01445 | 0.937685 | 0.993685 | St18 |
| ENSMUSG( | 972.5691 | 0.241941 | 0.133785 | 0.822912 | St3gal1 |
| ENSMUSG( | 5199.907 | 0.111901 | 0.343667 | 0.910026 | St3gal2 |
| ENSMUSG( | 1831.305 | 0.061435 | 0.625131 | 0.953234 | St3gal3 |
| ENSMUSG( | 1488.184 | 0.205945 | 0.106665 | 0.804969 | St3gal4 |
| ENSMUSG( | 4187.126 | 0.125783 | 0.465291 | 0.923329 | St3gal5 |
| ENSMUSG( | 673.7959 | 0.329176 | 0.094826 | 0.796133 | St3gal6 |
| ENSMUSG( | 1079.875 | 0.207081 | 0.236357 | 0.872173 | St6gal1 |
| ENSMUSG( | 1184.972 | 0.301273 | 0.112939 | 0.809623 | St6gal2 |
| ENSMUSG( | 144.9107 | 0.604339 | 0.026345 | 0.603041 | St6galnac2 |
| ENSMUSG( | 347.1267 | 0.1648 | 0.389463 | 0.917257 | St6galnac3 |
| ENSMUSG( | 961.9098 | 0.060429 | 0.723357 | 0.966887 | St6galnac4 |
| ENSMUSG( | 3049.296 | 0.00297 | 0.984583 | 0.996963 | St6galnac5 |
| ENSMUSG( | 8167.779 | 0.019036 | 0.847442 | 0.985161 | St6galnac6 |
| ENSMUSG( | 572.6472 | 0.324972 | 0.044646 | 0.693697 | St7 |
| ENSMUSG( | 753.4624 | 0.304941 | 0.074766 | 0.767423 | St7l |
| ENSMUSG( | 1011.602 | 0.196338 | 0.287961 | 0.894644 | St8sia1 |
| ENSMUSG( | 211.8304 | -0.15929 | 0.381226 | 0.917202 | St8sia2 |
| ENSMUSG( | 4729.892 | 0.088756 | 0.565871 | 0.941499 | St8sia3 |
| ENSMUSG( | 382.2843 | 0.615585 | 0.009305 | 0.403453 | St8sia4 |
| ENSMUSG( | 2141.496 | 0.211523 | 0.158809 | 0.837724 | St8sia5 |
| ENSMUSG( | 62.03307 | 0.887754 | 0.003658 | 0.242109 | St8sia6 |
| ENSMUSG( | 1195.261 | 0.35699 | 0.039003 | 0.679972 | Stab1 |
| ENSMUSG( | 16.54566 | 0.606268 | 0.203583 | 0.864412 | Stab2 |
| ENSMUSG( | 20.7274 | 0.036625 | 0.932347 | 0.99306 | Stac |
| ENSMUSG( | 5913.916 | -0.14521 | 0.15173 | 0.837724 | Stac2 |
| ENSMUSG( | 76.42817 | -0.18171 | 0.589285 | 0.944975 | Stac3 |
| ENSMUSG( | 1453.735 | 0.05774 | 0.679083 | 0.957725 | Stag1 |
| ENSMUSG( | 2462.645 | 0.007835 | 0.955562 | 0.994798 | Stag2 |
| ENSMUSG( | 59.80424 | 0.14413 | 0.654649 | 0.955015 | Stag3 |
| ENSMUSG( | 993.1888 | 0.01942 | 0.871442 | 0.988779 | Stam |
| ENSMUSG( | 716.6674 | 0.240126 | 0.061741 | 0.736228 | Stam2 |
| ENSMUSG( | 574.1114 | -0.11809 | 0.356518 | 0.913358 | Stambp |
| ENSMUSG( | 317.7157 | -0.10058 | 0.513438 | 0.930482 | Stambpl1 |
| ENSMUSG( | 69.80522 | 0.511126 | 0.129258 | 0.820358 | Stap2 |
| ENSMUSG( | 58.1598 | 0.316204 | 0.278222 | 0.891468 | Star |
| ENSMUSG( | 2273.006 | -0.16936 | 0.222137 | 0.866232 | Stard10 |
| ENSMUSG( | 405.7609 | 0.107327 | 0.540016 | 0.937383 | Stard13 |
| ENSMUSG( | 1180.873 | 0.158525 | 0.316391 | 0.901868 | Stard3 |
| ENSMUSG( | 1232.601 | 0.122752 | 0.314105 | 0.900949 | Stard3nl |
| ENSMUSG( | 1009.085 | -0.26067 | 0.055043 | 0.723362 | Stard4 |
| ENSMUSG( | 205.4227 | -0.03685 | 0.83726 | 0.981335 | Stard5 |
| ENSMUSG( | 82.79991 | 0.148208 | 0.538517 | 0.937216 | Stard6 |
| ENSMUSG( | 3683.316 | -0.10697 | 0.339372 | 0.907972 | Stard7 |
| ENSMUSG( | 939.8696 | 0.086618 | 0.527037 | 0.934449 | Stard8 |
| ENSMUSG( | 982.5391 | 0.180539 | 0.325013 | 0.905273 | Stard9 |
| ENSMUSG( | 288.8579 | -0.16122 | 0.290915 | 0.897019 | Stat1 |
| ENSMUSG( | 785.671 | 0.097248 | 0.465071 | 0.923329 | Stat2 |
| ENSMUSG( | 1765.901 | -0.03401 | 0.750416 | 0.970113 | Stat3 |
| ENSMUSG( | 9.308739 | -0.74755 | 0.305924 | NA | Stat4 |
| ENSMUSG( | 121.5527 | -0.28081 | 0.200173 | 0.863016 | Stat5a |
| ENSMUSG( | 1056.84 | 0.014225 | 0.903934 | 0.991089 | Stat5b |
| ENSMUSG( | 385.6937 | 0.123109 | 0.428246 | 0.917257 | Stat6 |
| ENSMUSG( | 1801.246 | -0.0116 | 0.911678 | 0.992196 | Stau1 |
| ENSMUSG( | 4894.122 | -0.13112 | 0.273438 | 0.889703 | Stau2 |
| ENSMUSG( | 61.62338 | -0.4191 | 0.124795 | 0.81516 | Stbd1 |
| ENSMUSG( | 112.4732 | 0.485559 | 0.019645 | 0.556192 | Stc1 |
| ENSMUSG( | 116.3015 | 0.0722 | 0.745824 | 0.96951 | Stc2 |
| ENSMUSG( | 33.98817 | 0.526551 | 0.5769 | 0.942912 | Steap1 |
| ENSMUSG( | 610.0081 | -0.03456 | 0.822892 | 0.979198 | Steap2 |
| ENSMUSG( | 637.2019 | 0.060315 | 0.757927 | 0.971858 | Steap3 |
| ENSMUSG( | 15.71842 | 0.015409 | 0.977393 | 0.996416 | Steap4 |
| ENSMUSG( | 1699.649 | -0.12364 | 0.438036 | 0.920503 | Steep1 |
| ENSMUSG( | 23.92634 | -0.01996 | 0.960134 | 0.994798 | Stil |
| ENSMUSG( | 3724.237 | 0.122817 | 0.398339 | 0.917257 | Stim1 |
| ENSMUSG( | 1938.628 | 0.292912 | 0.085638 | 0.779799 | Stim2 |
| ENSMUSG( | 961.0896 | 0.111517 | 0.432219 | 0.918158 | Stimate |
| ENSMUSG( | 263.1521 | 0.203493 | 0.375701 | 0.915491 | Sting1 |
| ENSMUSG( | 2482.593 | -0.07637 | 0.530744 | 0.934795 | Stip1 |
| ENSMUSG( | 328.3648 | -0.34142 | 0.05089 | 0.709741 | Stk10 |
| ENSMUSG( | 3135.65 | -0.06101 | 0.579044 | 0.943849 | Stk11 |
| ENSMUSG( | 1169.72 | 0.105775 | 0.417841 | 0.917257 | Stk11ip |
| ENSMUSG( | 1603.367 | -0.04766 | 0.659631 | 0.956205 | Stk16 |
| ENSMUSG( | 538.3143 | 0.199557 | 0.286877 | 0.894091 | Stk17b |
| ENSMUSG( | 807.525 | -0.05017 | 0.712459 | 0.964363 | Stk19 |
| ENSMUSG( | 1084.712 | 0.04655 | 0.687664 | 0.959673 | Stk24 |
| ENSMUSG( | 5558.446 | 0.031784 | 0.802828 | 0.97681 | Stk25 |
| ENSMUSG( | 90.81034 | -0.11531 | 0.683152 | 0.958678 | Stk26 |
| ENSMUSG( | 330.6191 | 0.147872 | 0.363942 | 0.914223 | Stk3 |
| ENSMUSG( | 177.5058 | 0.094923 | 0.620998 | 0.953199 | Stk32a |
| ENSMUSG( | 55.47821 | 0.048336 | 0.865058 | 0.98675 | Stk32b |
| ENSMUSG( | 2227.023 | -0.02901 | 0.828974 | 0.980224 | Stk32c |
| ENSMUSG( | 26.04035 | -0.52999 | 0.18203 | 0.850579 | Stk33 |
| ENSMUSG( | 933.8246 | 0.318592 | 0.007682 | 0.373786 | Stk35 |
| ENSMUSG( | 190.7985 | 0.162051 | 0.362802 | 0.914223 | Stk36 |
| ENSMUSG( | 1706.626 | 0.069094 | 0.561664 | 0.940388 | Stk38 |
| ENSMUSG( | 960.3797 | 0.038264 | 0.741188 | 0.968811 | Stk38l |

| | | | | | | | | | | | |
|---|---|---|---|---|---|---|---|---|---|---|---|
| ENSMUSG( | Stx18 | 744.9897 | 0.128545 | 0.262517 | 0.713142 | ENSMUSG( | Stx18 | 653.0826 | 0.019559 | 0.883323 | 0.989644 |
| ENSMUSG( | Stx1a | 9737.377 | 0.248631 | 0.01232 | 0.160458 | ENSMUSG( | Stx1a | 8057.694 | -0.0192 | 0.858562 | 0.985884 |
| ENSMUSG( | Stx1b | 22766.15 | 0.120066 | 0.134911 | 0.548434 | ENSMUSG( | Stx1b | 19478.55 | -0.05878 | 0.541596 | 0.937841 |
| ENSMUSG( | Stx2 | 1181.734 | -0.03455 | 0.728673 | 0.944393 | ENSMUSG( | Stx2 | 1063.424 | -0.07133 | 0.535229 | 0.936508 |
| ENSMUSG( | Stx3 | 657.1668 | -0.09203 | 0.503937 | 0.866426 | ENSMUSG( | Stx3 | 627.8141 | 0.04932 | 0.73855 | 0.96799 |
| ENSMUSG( | Stx4a | 2233.277 | 0.042653 | 0.668231 | 0.927864 | ENSMUSG( | Stx4a | 1858.329 | -0.23041 | 0.049269 | 0.708275 |
| ENSMUSG( | Stx5a | 1372.462 | 0.058408 | 0.518193 | 0.873754 | ENSMUSG( | Stx5a | 1239.126 | 0.03334 | 0.784175 | 0.976154 |
| ENSMUSG( | Stx6 | 2640.382 | 0.020983 | 0.810133 | 0.963864 | ENSMUSG( | Stx6 | 2281.417 | -0.13482 | 0.215616 | 0.865167 |
| ENSMUSG( | Stx7 | 2716.109 | 0.038499 | 0.715381 | 0.941306 | ENSMUSG( | Stx7 | 2376.052 | -0.08411 | 0.597309 | 0.946609 |
| ENSMUSG( | Stx8 | 552.5679 | -0.00944 | 0.933781 | 0.993206 | ENSMUSG( | Stx8 | 475.8747 | -0.18316 | 0.191997 | 0.859497 |
| ENSMUSG( | Stxbp1 | 19321.19 | 0.055039 | 0.640665 | 0.916434 | ENSMUSG( | Stxbp1 | 16899.38 | -0.06388 | 0.629469 | 0.953251 |
| ENSMUSG( | Stxbp2 | 1138.514 | 0.090627 | 0.599427 | 0.903801 | ENSMUSG( | Stxbp2 | 1011.151 | 0.014939 | 0.939189 | 0.993898 |
| ENSMUSG( | Stxbp3 | 1090.117 | -0.01941 | 0.865974 | 0.976033 | ENSMUSG( | Stxbp3 | 992.7458 | -0.02266 | 0.866208 | 0.98727 |
| ENSMUSG( | Stxbp4 | 902.0573 | -0.03232 | 0.810467 | 0.964049 | ENSMUSG( | Stxbp4 | 782.3179 | -0.18337 | 0.210679 | 0.864412 |
| ENSMUSG( | Stxbp5 | 1837.022 | 0.122455 | 0.227099 | 0.674223 | ENSMUSG( | Stxbp5 | 1689.861 | 0.150342 | 0.195659 | 0.861796 |
| ENSMUSG( | Stxbp5l | 261.7449 | -0.06733 | 0.754062 | 0.950981 | ENSMUSG( | Stxbp5l | 239.8779 | -0.03539 | 0.879355 | 0.989644 |
| ENSMUSG( | Stxbp6 | 1341.574 | 0.075729 | 0.47111 | 0.846975 | ENSMUSG( | Stxbp6 | 1139.438 | -0.1354 | 0.422442 | 0.917257 |
| ENSMUSG( | Styk1 | 9.856588 | -0.49996 | 0.435306 | NA | ENSMUSG( | Styk1 | 10.78763 | 0.0103 | 0.987591 | NA |
| ENSMUSG( | Styx | 175.7409 | -0.00256 | 0.989102 | 0.999111 | ENSMUSG( | Styx | 177.0884 | 0.286292 | 0.170239 | 0.841364 |
| ENSMUSG( | Styxl2 | 17.71919 | -0.06241 | 0.888368 | NA | ENSMUSG( | Styxl2 | 15.91384 | -0.08937 | 0.852765 | 0.985598 |
| ENSMUSG( | Sub1 | 3915.508 | 0.002549 | 0.978636 | 0.997428 | ENSMUSG( | Sub1 | 3367.682 | -0.17224 | 0.2285 | 0.868507 |
| ENSMUSG( | Sucla2 | 3209.107 | 0.043469 | 0.644015 | 0.917536 | ENSMUSG( | Sucla2 | 2715.34 | -0.17761 | 0.133069 | 0.822912 |
| ENSMUSG( | Suclg1 | 2965.43 | 0.164775 | 0.07719 | 0.429655 | ENSMUSG( | Suclg1 | 2461.073 | -0.10297 | 0.340383 | 0.907972 |
| ENSMUSG( | Suclg2 | 2700.701 | 0.104883 | 0.363155 | 0.793211 | ENSMUSG( | Suclg2 | 2291.397 | -0.10413 | 0.454134 | 0.922444 |
| ENSMUSG( | Suco | 1245.602 | -0.35771 | 0.014739 | 0.178407 | ENSMUSG( | Suco | 1330.107 | 0.122224 | 0.441276 | 0.921205 |
| ENSMUSG( | Suds3 | 5226.226 | 0.146523 | 0.057251 | 0.369376 | ENSMUSG( | Suds3 | 4415.707 | -0.0691 | 0.508142 | 0.929089 |
| ENSMUSG( | Sufu | 506.3471 | 0.058313 | 0.641968 | 0.916865 | ENSMUSG( | Sufu | 462.7715 | 0.069564 | 0.63022 | 0.953251 |
| ENSMUSG( | Sugct | 345.7623 | -0.08253 | 0.552795 | 0.886141 | ENSMUSG( | Sugct | 314.1051 | -0.10434 | 0.55911 | 0.939587 |
| ENSMUSG( | Sugp1 | 1407.23 | 0.005117 | 0.959939 | 0.993991 | ENSMUSG( | Sugp1 | 1256.461 | -0.05272 | 0.637413 | 0.953903 |
| ENSMUSG( | Sugp2 | 2436.774 | 0.2037 | 0.095456 | 0.474828 | ENSMUSG( | Sugp2 | 2205.857 | 0.18913 | 0.241533 | 0.876528 |
| ENSMUSG( | Sugt1 | 1881.144 | 0.045008 | 0.625639 | 0.912249 | ENSMUSG( | Sugt1 | 1674.888 | -0.02341 | 0.838691 | 0.981622 |
| ENSMUSG( | Sulf1 | 564.4021 | -0.3404 | 0.004259 | 0.084556 | ENSMUSG( | Sulf1 | 649.6149 | 0.336694 | 0.011004 | 0.425502 |
| ENSMUSG( | Sulf2 | 4603 | -0.34105 | 0.027975 | 0.255784 | ENSMUSG( | Sulf2 | 5149.848 | 0.264868 | 0.171499 | 0.842999 |
| ENSMUSG( | Sult1a1 | 777.6894 | 0.552178 | 1.13E-05 | 0.001226 | ENSMUSG( | Sult1a1 | 654.8469 | 0.357395 | 0.011326 | 0.433258 |
| ENSMUSG( | Sult2b1 | 213.1968 | -0.01088 | 0.947758 | 0.993821 | ENSMUSG( | Sult2b1 | 194.8613 | -0.00121 | 0.994265 | 0.998813 |
| ENSMUSG( | Sult4a1 | 12698.5 | 0.045251 | 0.650439 | 0.921301 | ENSMUSG( | Sult4a1 | 10955.3 | -0.11501 | 0.303796 | 0.898538 |
| ENSMUSG( | Sult5a1 | 10.87675 | 0.61487 | 0.270618 | NA | ENSMUSG( | Sult5a1 | 8.21758 | 0.14837 | 0.822032 | NA |
| ENSMUSG( | Sumf1 | 997.5517 | -0.19156 | 0.161987 | 0.592429 | ENSMUSG( | Sumf1 | 1004.779 | 0.104152 | 0.484113 | 0.92541 |
| ENSMUSG( | Sumf2 | 517.6879 | -0.14426 | 0.278727 | 0.728705 | ENSMUSG( | Sumf2 | 531.4486 | 0.200706 | 0.166536 | 0.840183 |
| ENSMUSG( | Sumo1 | 1270.629 | 0.134214 | 0.217677 | 0.662824 | ENSMUSG( | Sumo1 | 1134.054 | 0.071798 | 0.621424 | 0.953199 |
| ENSMUSG( | Sumo2 | 1277.388 | 0.109638 | 0.297337 | 0.745872 | ENSMUSG( | Sumo2 | 1118 | -0.00884 | 0.947263 | 0.994798 |
| ENSMUSG( | Sumo3 | 2784.961 | -0.04364 | 0.622879 | 0.911589 | ENSMUSG( | Sumo3 | 2421.766 | -0.18655 | 0.173728 | 0.844317 |
| ENSMUSG( | Sun1 | 3247.904 | -0.15868 | 0.182925 | 0.620462 | ENSMUSG( | Sun1 | 3328.948 | 0.183793 | 0.207575 | 0.864412 |
| ENSMUSG( | Sun2 | 1972.858 | -0.38362 | 0.001252 | 0.038624 | ENSMUSG( | Sun2 | 2109.263 | 0.105579 | 0.478721 | 0.923329 |
| ENSMUSG( | Suox | 592.2316 | 0.101008 | 0.356465 | 0.788187 | ENSMUSG( | Suox | 526.1792 | 0.033338 | 0.795593 | 0.97681 |
| ENSMUSG( | Supt16 | 1976.565 | 0.140695 | 0.112712 | 0.506698 | ENSMUSG( | Supt16 | 1668.012 | -0.08032 | 0.471554 | 0.923329 |
| ENSMUSG( | Supt20 | 1011.093 | 0.099729 | 0.397503 | 0.812127 | ENSMUSG( | Supt20 | 919.1601 | 0.093104 | 0.551373 | 0.939565 |
| ENSMUSG( | Supt3 | 231.5766 | 0.063255 | 0.679896 | 0.930983 | ENSMUSG( | Supt3 | 200.1333 | -0.09948 | 0.592957 | 0.945408 |
| ENSMUSG( | Supt4a | 737.9412 | -0.00076 | 0.994431 | 0.999369 | ENSMUSG( | Supt4a | 624.5225 | -0.22102 | 0.091967 | 0.791889 |
| ENSMUSG( | Supt5 | 4016.438 | 0.177659 | 0.075991 | 0.426896 | ENSMUSG( | Supt5 | 3365.234 | -0.06113 | 0.627589 | 0.953234 |
| ENSMUSG( | Supt6 | 5149.604 | 0.051553 | 0.563466 | 0.889914 | ENSMUSG( | Supt6 | 4646.563 | 0.024385 | 0.819172 | 0.979198 |
| ENSMUSG( | Supt7l | 460.5528 | 0.143254 | 0.247495 | 0.696522 | ENSMUSG( | Supt7l | 400.4407 | 0.0114 | 0.937467 | 0.993685 |
| ENSMUSG( | Supv3l1 | 596.6393 | 0.107728 | 0.336586 | 0.777 | ENSMUSG( | Supv3l1 | 540.7874 | 0.092539 | 0.462199 | 0.923329 |
| ENSMUSG( | Surf1 | 2003.536 | -0.1142 | 0.208899 | 0.650854 | ENSMUSG( | Surf1 | 1836.599 | -0.09779 | 0.385624 | 0.917257 |
| ENSMUSG( | Surf2 | 633.4567 | -0.02016 | 0.852583 | 0.973168 | ENSMUSG( | Surf2 | 574.8249 | -0.0336 | 0.807018 | 0.978239 |
| ENSMUSG( | Surf4 | 3261.118 | -0.27177 | 0.042556 | 0.316636 | ENSMUSG( | Surf4 | 3403.725 | 0.133734 | 0.375241 | 0.915491 |
| ENSMUSG( | Surf6 | 542.8656 | 0.136079 | 0.239978 | 0.687848 | ENSMUSG( | Surf6 | 471.9654 | -0.0008 | 0.995473 | 0.999116 |
| ENSMUSG( | Susd1 | 326.7837 | -0.32814 | 0.023084 | 0.233188 | ENSMUSG( | Susd1 | 338.2248 | 0.059042 | 0.706909 | 0.963529 |
| ENSMUSG( | Susd2 | 276.1329 | -0.41956 | 0.005647 | 0.100551 | ENSMUSG( | Susd2 | 302.5861 | 0.139187 | 0.362624 | 0.914223 |
| ENSMUSG( | Susd3 | 111.1858 | -0.1151 | 0.633669 | 0.914073 | ENSMUSG( | Susd3 | 111.8013 | 0.185833 | 0.464141 | 0.923329 |
| ENSMUSG( | Susd4 | 5192.805 | -0.07828 | 0.509619 | 0.869553 | ENSMUSG( | Susd4 | 5209.417 | 0.197386 | 0.133235 | 0.822912 |
| ENSMUSG( | Susd5 | 327.7146 | -0.29673 | 0.129933 | 0.540113 | ENSMUSG( | Susd5 | 382.5601 | 0.415179 | 0.076508 | 0.770558 |
| ENSMUSG( | Susd6 | 1839.195 | -0.10111 | 0.460966 | 0.841042 | ENSMUSG( | Susd6 | 1858.772 | 0.198958 | 0.22175 | 0.866104 |
| ENSMUSG( | Suv39h1 | 1132.803 | 0.150804 | 0.100765 | 0.484034 | ENSMUSG( | Suv39h1 | 1003.764 | 0.072164 | 0.558746 | 0.939574 |
| ENSMUSG( | Suv39h2 | 91.25038 | -0.17715 | 0.408112 | 0.818167 | ENSMUSG( | Suv39h2 | 85.46782 | -0.10006 | 0.672608 | 0.95694 |
| ENSMUSG( | Suz12 | 1117.679 | 0.261422 | 0.013219 | 0.167118 | ENSMUSG( | Suz12 | 1049.39 | 0.337533 | 0.010897 | 0.422321 |
| ENSMUSG( | Sv2a | 15010.12 | -0.40784 | 0.001739 | 0.047093 | ENSMUSG( | Sv2a | 16282.45 | 0.124872 | 0.404932 | 0.917257 |
| ENSMUSG( | Sv2b | 11013.09 | -0.29395 | 0.064635 | 0.393981 | ENSMUSG( | Sv2b | 12231.03 | 0.283249 | 0.105858 | 0.804969 |
| ENSMUSG( | Sv2c | 1099.058 | -0.73073 | 9.14E-06 | 0.001083 | ENSMUSG( | Sv2c | 1301.201 | 0.117894 | 0.444562 | 0.921734 |
| ENSMUSG( | Svbp | 614.09 | 0.345472 | 0.007129 | 0.116294 | ENSMUSG( | Svbp | 462.6472 | -0.19729 | 0.206716 | 0.864412 |
| ENSMUSG( | Svep1 | 44.85846 | 0.090256 | 0.784443 | 0.958003 | ENSMUSG( | Svep1 | 49.20543 | 0.591872 | 0.059753 | 0.729747 |
| ENSMUSG( | Svil | 644.462 | 0.267202 | 0.017449 | 0.197914 | ENSMUSG( | Svil | 593.8366 | 0.296986 | 0.023377 | 0.585822 |
| ENSMUSG( | Svip | 3764.664 | -0.06517 | 0.560943 | 0.888934 | ENSMUSG( | Svip | 3296.245 | -0.18863 | 0.2 | 0.863016 |
| ENSMUSG( | Svop | 6900.906 | -0.18587 | 0.146494 | 0.568435 | ENSMUSG( | Svop | 7189.859 | 0.203862 | 0.156445 | 0.837724 |
| ENSMUSG( | Swap70 | 1187.076 | 0.15658 | 0.231651 | 0.67949 | ENSMUSG( | Swap70 | 991.494 | -0.08921 | 0.533953 | 0.935607 |
| ENSMUSG( | Swi5 | 3125.763 | 0.026972 | 0.80767 | 0.963257 | ENSMUSG( | Swi5 | 2652.723 | -0.18749 | 0.178766 | 0.849041 |
| ENSMUSG( | Swsap1 | 450.6155 | 0.053447 | 0.662667 | 0.926303 | ENSMUSG( | Swsap1 | 392.87 | -0.08385 | 0.6014 | 0.947749 |
| ENSMUSG( | Swt1 | 213.5422 | 0.089497 | 0.576786 | 0.895432 | ENSMUSG( | Swt1 | 210.344 | 0.305702 | 0.086716 | 0.782073 |
| ENSMUSG( | Syap1 | 838.9283 | -0.01502 | 0.885772 | 0.983008 | ENSMUSG( | Syap1 | 769.6058 | -0.0007 | 0.996285 | 0.999178 |
| ENSMUSG( | Sybu | 1683.072 | 0.056589 | 0.6163 | 0.910866 | ENSMUSG( | Sybu | 1464.433 | -0.081 | 0.580717 | 0.944266 |
| ENSMUSG( | Syce2 | 96.63936 | 0.33718 | 0.101816 | 0.486216 | ENSMUSG( | Syce2 | 81.85056 | 0.125725 | 0.621615 | 0.953199 |
| ENSMUSG( | Sycp2 | 52.81181 | -0.40485 | 0.137909 | 0.552903 | ENSMUSG( | Sycp2 | 55.1141 | 0.040015 | 0.891475 | 0.990558 |
| ENSMUSG( | Syde1 | 513.769 | 0.059638 | 0.636715 | 0.91503 | ENSMUSG( | Syde1 | 465.27 | 0.04034 | 0.761039 | 0.972695 |
| ENSMUSG( | Syde2 | 190.6186 | -0.04386 | 0.772737 | 0.955272 | ENSMUSG( | Syde2 | 171.5516 | -0.07908 | 0.666662 | 0.956766 |
| ENSMUSG( | Syf2 | 1843.279 | 0.078168 | 0.454964 | 0.837725 | ENSMUSG( | Syf2 | 1588.731 | -0.08546 | 0.488081 | 0.925453 |
| ENSMUSG( | Syk | 285.1894 | 0.015442 | 0.913727 | 0.988483 | ENSMUSG( | Syk | 271.4986 | 0.137131 | 0.40431 | 0.917257 |
| ENSMUSG( | Sympk | 6010.714 | 0.253895 | 0.001841 | 0.048742 | ENSMUSG( | Sympk | 5061.831 | 0.037475 | 0.725566 | 0.966887 |
| ENSMUSG( | Syn1 | 18885.94 | 0.129727 | 0.287753 | 0.738505 | ENSMUSG( | Syn1 | 15845.61 | -0.10857 | 0.460538 | 0.923329 |
| ENSMUSG( | Syn2 | 11227.36 | 0.21814 | 0.045711 | 0.328047 | ENSMUSG( | Syn2 | 9109.144 | -0.1119 | 0.387353 | 0.917257 |
| ENSMUSG( | Syn3 | 151.3575 | -0.03173 | 0.861323 | 0.975026 | ENSMUSG( | Syn3 | 139.4634 | 0.021585 | 0.922863 | 0.992644 |
| ENSMUSG( | Syna | 27.62259 | -0.34719 | 0.411907 | 0.81942 | ENSMUSG( | Syna | 26.53388 | -0.16334 | 0.715382 | 0.965063 |
| ENSMUSG( | Sync | 25.28675 | 0.298932 | 0.426282 | 0.824898 | ENSMUSG( | Sync | 20.87841 | 0.027444 | 0.949391 | 0.994798 |
| ENSMUSG( | Syncrip | 1945.139 | 0.007725 | 0.942602 | 0.993415 | ENSMUSG( | Syncrip | 1777.363 | 0.01263 | 0.932402 | 0.99306 |
| ENSMUSG( | Syndig1 | 679.5691 | -0.26105 | 0.027153 | 0.253245 | ENSMUSG( | Syndig1 | 608.3118 | -0.31391 | 0.027663 | 0.61023 |
| ENSMUSG( | Syndig1l | 1936.683 | -0.80124 | 0.047082 | 0.332172 | ENSMUSG( | Syndig1l | 2178.041 | -0.08669 | 0.575481 | 0.942912 |
| ENSMUSG( | Syne1 | 12482.06 | -0.53552 | 7.33E-11 | 4.41E-08 | ENSMUSG( | Syne1 | 13441.16 | -0.00261 | 0.981217 | 0.996756 |
| ENSMUSG( | Syne2 | 849.2467 | -0.31008 | 0.003576 | 0.075902 | ENSMUSG( | Syne2 | 844.557 | -0.03717 | 0.76252 | 0.973539 |
| ENSMUSG( | Syne3 | 74.14336 | -0.08647 | 0.732359 | 0.944956 | ENSMUSG( | Syne3 | 61.32659 | -0.3689 | 0.183038 | 0.850741 |
| ENSMUSG( | Syne4 | 77.45376 | 0.237114 | 0.392201 | 0.809274 | ENSMUSG( | Syne4 | 67.5023 | 0.102232 | 0.767622 | 0.974236 |
| ENSMUSG( | Syngap1 | 12101.86 | 0.244805 | 0.002788 | 0.064345 | ENSMUSG( | Syngap1 | 10636.23 | 0.147113 | 0.127189 | 0.817278 |
| ENSMUSG( | Syngr1 | 19797.21 | -0.1946 | 0.211174 | 0.654296 | ENSMUSG( | Syngr1 | 20529.44 | 0.184753 | 0.298941 | 0.898538 |
| ENSMUSG( | Syngr2 | 558.5693 | -0.24776 | 0.081473 | 0.441102 | ENSMUSG( | Syngr2 | 604.8489 | 0.256433 | 0.179797 | 0.850372 |
| ENSMUSG( | Syngr3 | 5982.466 | -0.45099 | 0.000651 | 0.024622 | ENSMUSG( | Syngr3 | 6332.271 | 0.019755 | 0.89313 | 0.990558 |
| ENSMUSG( | Synj1 | 5993.126 | 0.095214 | 0.317058 | 0.763823 | ENSMUSG( | Synj1 | 5377.015 | 0.052736 | 0.607105 | 0.949917 |
| ENSMUSG( | Synj2 | 1892.756 | 0.122207 | 0.308409 | 0.756439 | ENSMUSG( | Synj2 | 1611.223 | -0.06725 | 0.635155 | 0.953903 |
| ENSMUSG( | Synj2bp | 837.7862 | 0.101899 | 0.374549 | 0.800445 | ENSMUSG( | Synj2bp | 712.7477 | -0.09946 | 0.460108 | 0.923329 |
| ENSMUSG( | Synm | 1444.674 | 0.150401 | 0.185803 | 0.623185 | ENSMUSG( | Synm | 1317.319 | 0.156707 | 0.249469 | 0.877904 |
| ENSMUSG( | Synpo | 4889.488 | 0.186403 | 0.107612 | 0.49648 | ENSMUSG( | Synpo | 4238.119 | 0.050111 | 0.699104 | 0.961139 |
| ENSMUSG( | Synpo2 | 322.8388 | -0.04445 | 0.762639 | 0.953519 | ENSMUSG( | Synpo2 | 330.6712 | 0.280215 | 0.072913 | 0.763629 |
| ENSMUSG( | Synpr | 2312.51 | -0.65149 | 1.09E-05 | 0.001199 | ENSMUSG( | Synpr | 2758.852 | 0.193338 | 0.228766 | 0.868507 |
| ENSMUSG( | Synrg | 2107.337 | -0.00077 | 0.992878 | 0.999327 | ENSMUSG( | Synrg | 1848.866 | -0.11312 | 0.304845 | 0.898538 |
| ENSMUSG( | Syp | 56345.22 | -0.3413 | 0.002372 | 0.057164 | ENSMUSG( | Syp | 58643.87 | 0.064279 | 0.646332 | 0.953988 |
| ENSMUSG( | Sypl1 | 1051.078 | -0.45577 | 0.00733 | 0.118347 | ENSMUSG( | Sypl1 | 1243.03 | 0.3219 | 0.109039 | 0.804969 |
| ENSMUSG( | Sypl2 | 241.5772 | -0.0785 | 0.671695 | 0.928781 | ENSMUSG( | Sypl2 | 242.0828 | 0.191015 | 0.344217 | 0.910274 |
| ENSMUSG( | Sys1 | 1064.636 | -0.1859 | 0.092547 | 0.467345 | ENSMUSG( | Sys1 | 993.287 | -0.11195 | 0.382682 | 0.917257 |
| ENSMUSG( | Syt1 | 41994.51 | -0.55476 | 6.09E-05 | 0.004616 | ENSMUSG( | Syt1 | 47966.52 | 0.150631 | 0.346995 | 0.911434 |
| ENSMUSG( | Syt10 | 39.16819 | -0.18938 | 0.59507 | 0.903007 | ENSMUSG( | Syt10 | 60.86531 | 1.156417 | 0.000143 | 0.027166 |
| ENSMUSG( | Syt11 | 28753.57 | -0.24812 | 0.102166 | 0.486961 | ENSMUSG( | Syt11 | 30614.71 | 0.209114 | 0.220877 | 0.866104 |
| ENSMUSG( | Syt12 | 1581.054 | -0.20375 | 0.152468 | 0.578756 | ENSMUSG( | Syt12 | 1573.649 | 0.063588 | 0.68891 | 0.959742 |
| ENSMUSG( | Syt13 | 10432.59 | -0.20237 | 0.097196 | 0.47759 | ENSMUSG( | Syt13 | 10232.75 | 0.020033 | 0.880933 | 0.989644 |
| ENSMUSG( | Syt14 | 446.2061 | -0.11134 | 0.409852 | 0.818258 | ENSMUSG( | Syt14 | 441.3298 | 0.129424 | 0.365177 | 0.914449 |
| ENSMUSG( | Syt15 | 32.1112 | -0.46073 | 0.202405 | 0.642343 | ENSMUSG( | Syt15 | 33.69396 | 0.005099 | 0.988885 | 0.997886 |
| ENSMUSG( | Syt16 | 4084.272 | 0.060534 | 0.546877 | 0.884974 | ENSMUSG( | Syt16 | 3819.86 | 0.135629 | 0.257063 | 0.883082 |
| ENSMUSG( | Syt17 | 1375.089 | 0.234842 | 0.030619 | 0.266434 | ENSMUSG( | Syt17 | 1225.393 | 0.173275 | 0.205351 | 0.864412 |
| ENSMUSG( | Syt2 | 1251.867 | -0.51705 | 0.000125 | 0.007476 | ENSMUSG( | Syt2 | 1353.635 | 0.027548 | 0.878614 | 0.989644 |
| ENSMUSG( | Syt3 | 2147.621 | -0.34647 | 0.001444 | 0.0419 | ENSMUSG( | Syt3 | 2248.844 | 0.077688 | 0.558312 | 0.93957 |
| ENSMUSG( | Syt4 | 3871.716 | -0.53183 | 0.000496 | 0.020385 | ENSMUSG( | Syt4 | 4576.64 | 0.259014 | 0.135362 | 0.825081 |
| ENSMUSG( | Syt5 | 7515.775 | -0.35316 | 0.000533 | 0.021434 | ENSMUSG( | Syt5 | 7805.125 | 0.047253 | 0.714721 | 0.964986 |
| ENSMUSG( | Syt6 | 973.8257 | -0.63074 | 4.64E-07 | 0.0001 | ENSMUSG( | Syt6 | 1053.798 | -0.07134 | 0.674968 | 0.95694 |
| ENSMUSG( | Syt7 | 16272.81 | 0.007664 | 0.941254 | 0.993415 | ENSMUSG( | Syt7 | 15702.16 | 0.169893 | 0.111561 | 0.808371 |
| ENSMUSG( | Syt9 | 309.5638 | -0.42834 | 0.004676 | 0.089972 | ENSMUSG( | Syt9 | 308.4132 | -0.15406 | 0.389404 | 0.917257 |
| ENSMUSG( | Sytl1 | 33.0629 | 0.485527 | 0.153981 | 0.580901 | ENSMUSG( | Sytl1 | 26.95664 | 0.201812 | 0.650578 | 0.95422 |
| ENSMUSG( | Sytl2 | 2063.971 | -0.08539 | 0.410173 | 0.818258 | ENSMUSG( | Sytl2 | 1823.783 | -0.17458 | 0.161218 | 0.837724 |
| ENSMUSG( | Sytl4 | 87.42623 | 0.374974 | 0.129923 | 0.540113 | ENSMUSG( | Sytl4 | 61.86415 | -0.36637 | 0.184111 | 0.852618 |
| ENSMUSG( | Sytl5 | 271.6189 | -0.05106 | 0.732458 | 0.944956 | ENSMUSG( | Sytl5 | 226.8324 | -0.33266 | 0.059174 | 0.729429 |
| ENSMUSG( | Syvn1 | 2122.086 | -0.31717 | 0.020202 | 0.214698 | ENSMUSG( | Syvn1 | 2308.104 | 0.207844 | 0.155781 | 0.837724 |
| ENSMUSG( | Szrd1 | 2513.198 | 0.034542 | 0.693741 | 0.934202 | ENSMUSG( | Szrd1 | 2094.31 | -0.23446 | 0.033728 | 0.647113 |
| ENSMUSG( | Szt2 | 1641.723 | 0.186721 | 0.076892 | 0.429306 | ENSMUSG( | Szt2 | 1474.631 | 0.148366 | 0.314319 | 0.900949 |
| ENSMUSG( | T2 | 23.06993 | -0.15963 | 0.706993 | 0.93949 | ENSMUSG( | T2 | 24.05094 | 0.206387 | 0.606266 | 0.949497 |
| ENSMUSG( | Tab1 | 808.0672 | -0.07122 | 0.533914 | 0.88018 | ENSMUSG( | Tab1 | 736.6179 | -0.06763 | 0.614692 | 0.951177 |
| ENSMUSG( | Tab2 | 2820.21 | 0.049648 | 0.569771 | 0.893041 | ENSMUSG( | Tab2 | 2584.811 | 0.064881 | 0.570109 | 0.941777 |
| ENSMUSG( | Tab3 | 418.6325 | 0.061779 | 0.597024 | 0.903526 | ENSMUSG( | Tab3 | 381.7072 | 0.058909 | 0.68275 | 0.958678 |
| ENSMUSG( | Tac1 | 752.6145 | -1.17419 | 8.03E-10 | 3.83E-07 | ENSMUSG( | Tac1 | 1032.486 | 0.228911 | 0.151969 | 0.837724 |
| ENSMUSG( | Tac2 | 276.5884 | -0.7397 | 2.02E-06 | 0.000331 | ENSMUSG( | Tac2 | 311.7681 | -0.02847 | 0.86349 | 0.986547 |
| ENSMUSG( | Tacc1 | 7468.359 | -0.08717 | 0.393002 | 0.809274 | ENSMUSG( | Tacc1 | 7195.951 | 0.076936 | 0.511976 | 0.930005 |
| ENSMUSG( | Tacc2 | 4257.454 | 0.102909 | 0.485288 | 0.857408 | ENSMUSG( | Tacc2 | 4023.179 | 0.207155 | 0.187017 | 0.856099 |
| ENSMUSG( | Tacc3 | 269.4067 | 0.06534 | 0.650365 | 0.921301 | ENSMUSG( | Tacc3 | 242.8788 | 0.03686 | 0.816864 | 0.979198 |
| ENSMUSG( | Taco1 | 338.7389 | 0.235974 | 0.077321 | 0.429743 | ENSMUSG( | Taco1 | 281.4915 | -0.02376 | 0.877492 | 0.989644 |
| ENSMUSG( | Tacr1 | 116.5645 | -1.07472 | 0.060297 | 0.379667 | ENSMUSG( | Tacr1 | 148.3748 | 0.081465 | 0.665313 | 0.956766 |
| ENSMUSG( | Tacr3 | 59.53012 | -0.70443 | 0.004438 | 0.086945 | ENSMUSG( | Tacr3 | 75.48154 | 0.297395 | 0.283975 | 0.892412 |
| ENSMUSG( | Tada1 | 562.6349 | -0.02998 | 0.793687 | 0.959799 | ENSMUSG( | Tada1 | 524.8181 | 0.037689 | 0.774834 | 0.974931 |
| ENSMUSG( | Tada2a | 701.8008 | -0.01565 | 0.89812 | 0.985633 | ENSMUSG( | Tada2a | 626.4339 | -0.07901 | 0.596823 | 0.946541 |
| ENSMUSG( | Tada2b | 1063.812 | -0.06708 | 0.489358 | 0.859092 | ENSMUSG( | Tada2b | 993.6245 | 0.00213 | 0.98622 | 0.997188 |
| ENSMUSG( | Tada3 | 930.3544 | -0.02521 | 0.831218 | 0.969889 | ENSMUSG( | Tada3 | 845.7824 | -0.03087 | 0.818847 | 0.979198 |
| ENSMUSG( | Taf1 | 3226.103 | 0.070758 | 0.497524 | 0.862162 | ENSMUSG( | Taf1 | 2776.634 | -0.09557 | 0.381752 | 0.917257 |
| ENSMUSG( | Taf10 | 2201.141 | 0.04523 | 0.613178 | 0.908902 | ENSMUSG( | Taf10 | 1820.61 | -0.24775 | 0.050545 | 0.708889 |
| ENSMUSG( | Taf11 | 793.9232 | -0.00694 | 0.948036 | 0.993821 | ENSMUSG( | Taf11 | 696.5887 | -0.12276 | 0.369911 | 0.915491 |
| ENSMUSG( | Taf12 | 401.362 | 0.006613 | 0.95749 | 0.993874 | ENSMUSG( | Taf12 | 351.048 | -0.11889 | 0.427795 | 0.917257 |

| | | | | | |
|---|---|---|---|---|---|
| ENSMUSG( | 2110.758 | 0.223406 | 0.072101 | 0.414057 | Srrm3 |
| ENSMUSG( | 1329.016 | 0.037842 | 0.744383 | 0.948758 | Srrm4 |
| ENSMUSG( | 3574.526 | 0.230701 | 0.032828 | 0.277278 | Srrt |
| ENSMUSG( | 4187.683 | -0.04235 | 0.700173 | 0.937391 | Srsf1 |
| ENSMUSG( | 1277.342 | 0.077674 | 0.450224 | 0.835344 | Srsf10 |
| ENSMUSG( | 3554.871 | 0.088297 | 0.435679 | 0.829093 | Srsf11 |
| ENSMUSG( | 332.328 | -0.116 | 0.391883 | 0.809274 | Srsf12 |
| ENSMUSG( | 2861.269 | -0.22358 | 0.008886 | 0.132538 | Srsf2 |
| ENSMUSG( | 2584.598 | -0.00714 | 0.939767 | 0.993415 | Srsf3 |
| ENSMUSG( | 2808.714 | 0.058498 | 0.492483 | 0.860378 | Srsf4 |
| ENSMUSG( | 5431.989 | 4.58E-05 | 0.999718 | 0.99989 | Srsf5 |
| ENSMUSG( | 3013.093 | -0.10126 | 0.271973 | 0.722255 | Srsf6 |
| ENSMUSG( | 1985.443 | -0.09891 | 0.387228 | 0.805442 | Srsf7 |
| ENSMUSG( | 1326.214 | 0.052154 | 0.575685 | 0.894836 | Srsf9 |
| ENSMUSG( | 2399.762 | 0.033167 | 0.711362 | 0.940028 | Srxn1 |
| ENSMUSG( | 1523.783 | 0.137829 | 0.16631 | 0.598791 | Ss18 |
| ENSMUSG( | 1188.382 | 0.126089 | 0.25092 | 0.700615 | Ss18l1 |
| ENSMUSG( | 876.3597 | -0.05901 | 0.543716 | 0.884287 | Ss18l2 |
| ENSMUSG( | 2111.516 | 0.036524 | 0.723023 | 0.943761 | Ssb |
| ENSMUSG( | 458.7952 | -0.05021 | 0.713809 | 0.940903 | Ssbp1 |
| ENSMUSG( | 1807.136 | 0.036996 | 0.708883 | 0.939519 | Ssbp2 |
| ENSMUSG( | 5143.069 | 0.185345 | 0.041495 | 0.314241 | Ssbp3 |
| ENSMUSG( | 6195.954 | 0.103583 | 0.288592 | 0.739489 | Ssbp4 |
| ENSMUSG( | 34.4764 | -0.33372 | 0.311992 | 0.759715 | Ssc4d |
| ENSMUSG( | 100.7118 | 0.18025 | 0.403328 | 0.816132 | Ssc5d |
| ENSMUSG( | 685.8728 | -0.15141 | 0.218988 | 0.664602 | Ssh1 |
| ENSMUSG( | 1252.333 | 0.137128 | 0.183843 | 0.620849 | Ssh2 |
| ENSMUSG( | 1439.692 | 0.515884 | 0.003584 | 0.075902 | Ssh3 |
| ENSMUSG( | 692.8352 | -0.05156 | 0.642128 | 0.916865 | Ssna1 |
| ENSMUSG( | 1539.899 | -0.10176 | 0.383205 | 0.803254 | Sspn |
| ENSMUSG( | 131.067 | 0.161933 | 0.463917 | 0.842812 | Sspo |
| ENSMUSG( | 5488.837 | -0.32596 | 0.018372 | 0.203658 | Ssr1 |
| ENSMUSG( | 2480.994 | -0.42387 | 0.001568 | 0.044386 | Ssr2 |
| ENSMUSG( | 6190.667 | -0.24428 | 0.069735 | 0.408352 | Ssr3 |
| ENSMUSG( | 1624.329 | -0.15255 | 0.122958 | 0.529576 | Ssr4 |
| ENSMUSG( | 2579.672 | 0.143624 | 0.177306 | 0.614553 | Ssrp1 |
| ENSMUSG( | 9541.266 | -0.62541 | 5.40E-07 | 0.000113 | Sst |
| ENSMUSG( | 308.3147 | -0.29319 | 0.201384 | 0.641362 | Sstr1 |
| ENSMUSG( | 901.3595 | -0.36881 | 0.024595 | 0.240469 | Sstr2 |
| ENSMUSG( | 995.9529 | -0.06922 | 0.590398 | 0.90113 | Sstr3 |
| ENSMUSG( | 146.9692 | 0.246938 | 0.261083 | 0.710889 | Sstr4 |
| ENSMUSG( | 1295.719 | -0.07083 | 0.581808 | 0.897862 | Ssu72 |
| ENSMUSG( | 1968.666 | 0.015709 | 0.867801 | 0.976631 | Ssx2ip |
| ENSMUSG( | 3428.684 | 0.06179 | 0.551987 | 0.886141 | St13 |
| ENSMUSG( | 23.23634 | 0.094543 | 0.828807 | 0.969097 | St14 |
| ENSMUSG( | 235.6076 | -0.39564 | 0.011367 | 0.153509 | St18 |
| ENSMUSG( | 898.3325 | -0.26415 | 0.075863 | 0.426666 | St3gal1 |
| ENSMUSG( | 5739.452 | 0.125174 | 0.188172 | 0.627112 | St3gal2 |
| ENSMUSG( | 1902.81 | -0.10099 | 0.318811 | 0.764619 | St3gal3 |
| ENSMUSG( | 1558.66 | 0.076412 | 0.509669 | 0.869553 | St3gal4 |
| ENSMUSG( | 3970.596 | -0.31613 | 0.036339 | 0.293685 | St3gal5 |
| ENSMUSG( | 632.7907 | -0.11271 | 0.506499 | 0.867425 | St3gal6 |
| ENSMUSG( | 1089.369 | -0.03559 | 0.801858 | 0.962116 | St6gal1 |
| ENSMUSG( | 992.1784 | -0.5115 | 0.002117 | 0.053164 | St6gal2 |
| ENSMUSG( | 143.4971 | 0.327675 | 0.130234 | 0.540948 | St6galnac2 |
| ENSMUSG( | 324.2573 | -0.31984 | 0.071922 | 0.414057 | St6galnac3 |
| ENSMUSG( | 1005.462 | -0.08805 | 0.516902 | 0.873055 | St6galnac4 |
| ENSMUSG( | 3075.251 | -0.25541 | 0.045854 | 0.328083 | St6galnac5 |
| ENSMUSG( | 9275.65 | 0.114911 | 0.155647 | 0.583641 | St6galnac6 |
| ENSMUSG( | 562.8721 | 0.020574 | 0.891202 | 0.983819 | St7 |
| ENSMUSG( | 706.0134 | -0.14421 | 0.38644 | 0.804732 | St7l |
| ENSMUSG( | 972.3591 | -0.19227 | 0.223869 | 0.669845 | St8sia1 |
| ENSMUSG( | 203.2396 | -0.61123 | 0.001146 | 0.036093 | St8sia2 |
| ENSMUSG( | 4160.835 | -0.61712 | 3.46E-06 | 0.000489 | St8sia3 |
| ENSMUSG( | 281.3028 | -0.53058 | 0.017438 | 0.197914 | St8sia4 |
| ENSMUSG( | 2100.394 | -0.11171 | 0.394182 | 0.809935 | St8sia5 |
| ENSMUSG( | 47.53863 | -0.03646 | 0.90526 | 0.98645 | St8sia6 |
| ENSMUSG( | 1182.494 | 0.074161 | 0.543503 | 0.884037 | Stab1 |
| ENSMUSG( | 20.1777 | 0.834651 | 0.050846 | NA | Stab2 |
| ENSMUSG( | 27.99314 | 0.566553 | 0.125114 | 0.532139 | Stac |
| ENSMUSG( | 7014.736 | 0.080555 | 0.363692 | 0.793343 | Stac2 |
| ENSMUSG( | 92.41391 | 0.095372 | 0.696416 | 0.935166 | Stac3 |
| ENSMUSG( | 1649.389 | 0.154871 | 0.140989 | 0.557902 | Stag1 |
| ENSMUSG( | 2742.415 | 0.052084 | 0.658226 | 0.924446 | Stag2 |
| ENSMUSG( | 68.30358 | 0.258378 | 0.336759 | 0.777 | Stag3 |
| ENSMUSG( | 1100.97 | 0.049971 | 0.591037 | 0.90113 | Stam |
| ENSMUSG( | 711.8037 | -0.04345 | 0.686689 | 0.933041 | Stam2 |
| ENSMUSG( | 649.5498 | -0.02541 | 0.808011 | 0.963389 | Stambp |
| ENSMUSG( | 362.8187 | 0.009677 | 0.944108 | 0.993503 | Stambpl1 |
| ENSMUSG( | 83.3635 | 0.69684 | 0.006589 | 0.111324 | Stap2 |
| ENSMUSG( | 58.28263 | 0.072788 | 0.790584 | 0.959461 | Star |
| ENSMUSG( | 2615.895 | -0.02419 | 0.798652 | 0.960896 | Stard10 |
| ENSMUSG( | 419.3995 | -0.07468 | 0.619637 | 0.911432 | Stard13 |
| ENSMUSG( | 1125.333 | -0.2616 | 0.047867 | 0.334081 | Stard3 |
| ENSMUSG( | 1239.84 | -0.13189 | 0.235125 | 0.682364 | Stard3nl |
| ENSMUSG( | 1249.834 | 0.101139 | 0.442242 | 0.832774 | Stard4 |
| ENSMUSG( | 239.7291 | 0.145664 | 0.325985 | 0.771128 | Stard5 |
| ENSMUSG( | 93.38516 | 0.239126 | 0.242645 | 0.690216 | Stard6 |
| ENSMUSG( | 4280.559 | 0.061283 | 0.460549 | 0.840787 | Stard7 |
| ENSMUSG( | 1089.587 | 0.231535 | 0.027765 | 0.255176 | Stard8 |
| ENSMUSG( | 878.5893 | -0.44572 | 0.000346 | 0.016028 | Stard9 |
| ENSMUSG( | 317.0103 | -0.1649 | 0.227383 | 0.67424 | Stat1 |
| ENSMUSG( | 901.7626 | 0.219522 | 0.039052 | 0.304527 | Stat2 |
| ENSMUSG( | 2026.464 | 0.094269 | 0.263422 | 0.714261 | Stat3 |
| ENSMUSG( | 11.79411 | -0.26312 | 0.634711 | NA | Stat4 |
| ENSMUSG( | 136.7105 | -0.1988 | 0.359564 | 0.791011 | Stat5a |
| ENSMUSG( | 1219.619 | 0.157493 | 0.101202 | 0.484996 | Stat5b |
| ENSMUSG( | 441.0101 | 0.230791 | 0.061252 | 0.381822 | Stat6 |
| ENSMUSG( | 1988.983 | 0.006222 | 0.940364 | 0.993415 | Stau1 |
| ENSMUSG( | 5784.531 | 0.085364 | 0.291485 | 0.741366 | Stau2 |
| ENSMUSG( | 72.55223 | -0.21653 | 0.334108 | 0.775728 | Stbd1 |
| ENSMUSG( | 89.08797 | -0.44041 | 0.035653 | 0.290384 | Stc1 |
| ENSMUSG( | 106.6487 | -0.48259 | 0.026058 | 0.247245 | Stc2 |
| ENSMUSG( | 32.09738 | 0.150866 | 0.87147 | 0.978114 | Steap1 |
| ENSMUSG( | 619.27 | -0.27083 | 0.048954 | 0.337919 | Steap2 |
| ENSMUSG( | 615.2029 | -0.3351 | 0.085527 | 0.450845 | Steap3 |
| ENSMUSG( | 18.25549 | 0.133967 | 0.788635 | NA | Steap4 |
| ENSMUSG( | 1994.862 | 0.076236 | 0.559231 | 0.888638 | Steep1 |
| ENSMUSG( | 21.58661 | -0.66051 | 0.099679 | 0.482191 | Stil |
| ENSMUSG( | 3494.602 | -0.35219 | 0.004724 | 0.090358 | Stim1 |
| ENSMUSG( | 1740.675 | -0.29137 | 0.039663 | 0.307079 | Stim2 |
| ENSMUSG( | 976.115 | -0.11772 | 0.346105 | 0.779153 | Stimate |
| ENSMUSG( | 241.6493 | -0.33634 | 0.109919 | 0.500285 | Sting1 |
| ENSMUSG( | 2894.881 | 0.099571 | 0.264727 | 0.715763 | Stip1 |
| ENSMUSG( | 379.2807 | -0.17162 | 0.268708 | 0.719042 | Stk10 |
| ENSMUSG( | 3635.018 | 0.097539 | 0.223326 | 0.669382 | Stk11 |
| ENSMUSG( | 1324.556 | 0.187823 | 0.059831 | 0.378344 | Stk11ip |
| ENSMUSG( | 1868.478 | 0.12186 | 0.152768 | 0.579081 | Stk16 |
| ENSMUSG( | 573.8462 | 0.128904 | 0.325935 | 0.771128 | Stk17b |
| ENSMUSG( | 871.791 | -0.0966 | 0.375183 | 0.800786 | Stk19 |
| ENSMUSG( | 1226.276 | 0.130029 | 0.188382 | 0.627297 | Stk24 |
| ENSMUSG( | 6424.568 | 0.177122 | 0.08693 | 0.453311 | Stk25 |
| ENSMUSG( | 84.25461 | -0.65778 | 0.016836 | 0.193535 | Stk26 |
| ENSMUSG( | 364.5703 | 0.170632 | 0.178822 | 0.616298 | Stk3 |
| ENSMUSG( | 173.2101 | -0.24328 | 0.147885 | 0.570981 | Stk32a |
| ENSMUSG( | 63.76928 | 0.175529 | 0.503279 | 0.86596 | Stk32b |
| ENSMUSG( | 2532.462 | 0.072704 | 0.551679 | 0.886141 | Stk32c |
| ENSMUSG( | 31.00508 | -0.25729 | 0.459906 | 0.840338 | Stk33 |
| ENSMUSG( | 952.1736 | 0.11752 | 0.245261 | 0.692812 | Stk35 |
| ENSMUSG( | 209.7242 | 0.166375 | 0.314103 | 0.761319 | Stk36 |
| ENSMUSG( | 1868.827 | 0.062866 | 0.531837 | 0.879628 | Stk38 |
| ENSMUSG( | 1111.212 | 0.186536 | 0.054731 | 0.360135 | Stk38l |
| ENSMUSG( | 1862.21 | 0.054165 | 0.630214 | 0.914073 | Stk39 |
| ENSMUSG( | 1591.408 | 0.214676 | 0.024358 | 0.238636 | Stk4 |
| ENSMUSG( | 1716.693 | -0.0088 | 0.925578 | 0.991204 | Stk40 |
| ENSMUSG( | 12.11701 | -0.13306 | 0.815024 | NA | Stkld1 |
| ENSMUSG( | 7423.648 | -0.05312 | 0.64023 | 0.916434 | Stmn1 |
| ENSMUSG( | 6811.212 | -0.03304 | 0.705561 | 0.938511 | Stmn2 |
| ENSMUSG( | 9267.23 | -0.27376 | 0.001477 | 0.042635 | Stmn3 |
| ENSMUSG( | 5523.158 | -0.5352 | 1.45E-11 | 9.68E-09 | Stmn4 |
| ENSMUSG( | 821.4746 | 0.156206 | 0.167815 | 0.60061 | Stmp1 |
| ENSMUSG( | 218.3767 | -0.07487 | 0.656528 | 0.923929 | Stn1 |
| ENSMUSG( | 822.482 | -0.16012 | 0.231179 | 0.679229 | Stom |
| ENSMUSG( | 2350.748 | -0.2389 | 0.032844 | 0.277278 | Stoml1 |
| ENSMUSG( | 1540.062 | 0.099809 | 0.247715 | 0.696522 | Stoml2 |
| ENSMUSG( | 44.65559 | -1.0803 | 0.00202 | 0.051507 | Stoml3 |
| ENSMUSG( | 280.5966 | 0.175739 | 0.222161 | 0.667925 | Ston1 |
| ENSMUSG( | 3446.885 | -0.14393 | 0.305533 | 0.753611 | Ston2 |
| ENSMUSG( | 107.0559 | 0.04604 | 0.819515 | 0.965888 | Stox1 |
| ENSMUSG( | 4682.302 | 0.22322 | 0.021619 | 0.225182 | Stox2 |
| ENSMUSG( | 124.0893 | 0.278302 | 0.203446 | 0.643631 | Stpg1 |
| ENSMUSG( | 351.9203 | 0.167037 | 0.182645 | 0.620029 | Stra6 |
| ENSMUSG( | 779.3253 | 0.120064 | 0.433237 | 0.828207 | Strada |
| ENSMUSG( | 1340.824 | 0.039193 | 0.670146 | 0.928485 | Stradb |
| ENSMUSG( | 3029.597 | 0.013903 | 0.892639 | 0.984474 | Strap |
| ENSMUSG( | 4047.718 | 0.167523 | 0.122264 | 0.527707 | Strbp |
| ENSMUSG( | 19.40802 | 0.630128 | 0.189659 | NA | Strc |
| ENSMUSG( | 2447.942 | 0.116359 | 0.268754 | 0.719042 | Strip1 |

| | | | | | |
|---|---|---|---|---|---|
| ENSMUSG( | 1599.094 | -0.12182 | 0.380347 | 0.917202 | Stk39 |
| ENSMUSG( | 1374.674 | 0.06377 | 0.627369 | 0.953234 | Stk4 |
| ENSMUSG( | 1528.224 | -0.07693 | 0.507766 | 0.929089 | Stk40 |
| ENSMUSG( | 11.41565 | 0.002713 | 0.996594 | NA | Stkld1 |
| ENSMUSG( | 6528.274 | -0.16246 | 0.260574 | 0.884586 | Stmn1 |
| ENSMUSG( | 5986.601 | -0.141 | 0.209285 | 0.864412 | Stmn2 |
| ENSMUSG( | 8663.814 | -0.1952 | 0.149551 | 0.836241 | Stmn3 |
| ENSMUSG( | 5435.993 | -0.27704 | 0.010162 | 0.410151 | Stmn4 |
| ENSMUSG( | 694.6854 | -0.06102 | 0.652527 | 0.954799 | Stmp1 |
| ENSMUSG( | 194.9193 | -0.12494 | 0.508804 | 0.929089 | Stn1 |
| ENSMUSG( | 836.8862 | 0.164004 | 0.280753 | 0.892356 | Stom |
| ENSMUSG( | 2302.524 | -0.01737 | 0.890391 | 0.990558 | Stoml1 |
| ENSMUSG( | 1307.303 | -0.1057 | 0.331899 | 0.907907 | Stoml2 |
| ENSMUSG( | 65.07711 | 0.412288 | 0.305912 | 0.898538 | Stoml3 |
| ENSMUSG( | 223.9258 | -0.21002 | 0.194979 | 0.861121 | Ston1 |
| ENSMUSG( | 2932.173 | -0.3628 | 0.037411 | 0.673514 | Ston2 |
| ENSMUSG( | 94.30974 | -0.03373 | 0.884861 | 0.989688 | Stox1 |
| ENSMUSG( | 4329.413 | 0.263741 | 0.025931 | 0.601649 | Stox2 |
| ENSMUSG( | 99.18433 | -0.09927 | 0.666573 | 0.956766 | Stpg1 |
| ENSMUSG( | 366.4216 | 0.509723 | 0.000269 | 0.043611 | Stra6 |
| ENSMUSG( | 665.56 | -0.06777 | 0.690281 | 0.960071 | Strada |
| ENSMUSG( | 1225.438 | 0.044674 | 0.711288 | 0.964363 | Stradb |
| ENSMUSG( | 2613.457 | -0.1513 | 0.283248 | 0.892412 | Strap |
| ENSMUSG( | 3467.324 | -0.00712 | 0.955887 | 0.994798 | Strbp |
| ENSMUSG( | 13.9644 | -0.00698 | 0.989257 | 0.997916 | Strc |
| ENSMUSG( | 2151.294 | 0.018093 | 0.881536 | 0.989644 | Strip1 |
| ENSMUSG( | 823.9475 | 0.179872 | 0.266276 | 0.886134 | Strip2 |
| ENSMUSG( | 2011.47 | -0.05143 | 0.695977 | 0.960867 | Strn |
| ENSMUSG( | 2084.449 | -0.0347 | 0.773553 | 0.974672 | Strn3 |
| ENSMUSG( | 6897.454 | 0.093286 | 0.376121 | 0.915491 | Strn4 |
| ENSMUSG( | 1595.355 | 0.145297 | 0.407 | 0.917257 | Stt3a |
| ENSMUSG( | 4015.023 | 0.159781 | 0.347713 | 0.911652 | Stt3b |
| ENSMUSG( | 3270.295 | -0.11906 | 0.298321 | 0.898538 | Stub1 |
| ENSMUSG( | 10393.93 | 0.159756 | 0.228502 | 0.868507 | Stum |
| ENSMUSG( | 8.665357 | -0.53365 | 0.460085 | NA | Stx11 |
| ENSMUSG( | 2390.246 | -0.11139 | 0.391052 | 0.917257 | Stx12 |
| ENSMUSG( | 1862.709 | 0.012466 | 0.921857 | 0.992644 | Stx16 |
| ENSMUSG( | 753.5647 | -0.08148 | 0.485761 | 0.92541 | Stx17 |
| ENSMUSG( | 653.0826 | 0.019559 | 0.883323 | 0.989644 | Stx18 |
| ENSMUSG( | 8057.694 | -0.0192 | 0.858562 | 0.985884 | Stx1a |
| ENSMUSG( | 19478.55 | -0.05878 | 0.541596 | 0.937841 | Stx1b |
| ENSMUSG( | 1063.424 | -0.07133 | 0.535229 | 0.936508 | Stx2 |
| ENSMUSG( | 627.8141 | 0.04932 | 0.73855 | 0.96799 | Stx3 |
| ENSMUSG( | 1858.329 | -0.23041 | 0.049269 | 0.708275 | Stx4a |
| ENSMUSG( | 1239.126 | 0.03334 | 0.784175 | 0.976154 | Stx5a |
| ENSMUSG( | 2281.417 | -0.13482 | 0.215616 | 0.865167 | Stx6 |
| ENSMUSG( | 2376.052 | -0.08411 | 0.597309 | 0.946609 | Stx7 |
| ENSMUSG( | 475.8747 | -0.18316 | 0.191997 | 0.859497 | Stx8 |
| ENSMUSG( | 16899.38 | -0.06388 | 0.629469 | 0.953251 | Stxbp1 |
| ENSMUSG( | 1011.151 | 0.014939 | 0.939189 | 0.993898 | Stxbp2 |
| ENSMUSG( | 992.7458 | -0.02266 | 0.866208 | 0.98727 | Stxbp3 |
| ENSMUSG( | 782.3179 | -0.18337 | 0.210679 | 0.864412 | Stxbp4 |
| ENSMUSG( | 1689.861 | 0.150342 | 0.195659 | 0.861796 | Stxbp5 |
| ENSMUSG( | 239.8779 | -0.03539 | 0.879355 | 0.989644 | Stxbp5l |
| ENSMUSG( | 1139.438 | -0.1354 | 0.422442 | 0.917257 | Stxbp6 |
| ENSMUSG( | 10.78763 | 0.0103 | 0.987591 | NA | Styk1 |
| ENSMUSG( | 177.0884 | 0.286292 | 0.170239 | 0.841364 | Styx |
| ENSMUSG( | 15.91384 | -0.08937 | 0.852765 | 0.985598 | Styxl2 |
| ENSMUSG( | 3367.682 | -0.17224 | 0.2285 | 0.868507 | Sub1 |
| ENSMUSG( | 2715.34 | -0.17761 | 0.133069 | 0.822912 | Sucla2 |
| ENSMUSG( | 2461.073 | -0.10297 | 0.340383 | 0.907972 | Suclg1 |
| ENSMUSG( | 2291.397 | -0.10413 | 0.454134 | 0.922444 | Suclg2 |
| ENSMUSG( | 1330.107 | 0.122224 | 0.441276 | 0.921205 | Suco |
| ENSMUSG( | 4415.707 | -0.0691 | 0.508142 | 0.929089 | Suds3 |
| ENSMUSG( | 462.7715 | 0.069564 | 0.63022 | 0.953251 | Sufu |
| ENSMUSG( | 314.1051 | -0.10434 | 0.55911 | 0.939587 | Sugct |
| ENSMUSG( | 1256.461 | -0.05272 | 0.637413 | 0.953903 | Sugp1 |
| ENSMUSG( | 2205.857 | 0.18913 | 0.241533 | 0.876528 | Sugp2 |
| ENSMUSG( | 1674.888 | -0.02341 | 0.838691 | 0.981622 | Sugt1 |
| ENSMUSG( | 649.6149 | 0.336694 | 0.011004 | 0.425502 | Sulf1 |
| ENSMUSG( | 5149.848 | 0.264868 | 0.171499 | 0.842999 | Sulf2 |
| ENSMUSG( | 654.8469 | 0.357395 | 0.011326 | 0.433258 | Sult1a1 |
| ENSMUSG( | 194.8613 | -0.00121 | 0.994265 | 0.998813 | Sult2b1 |
| ENSMUSG( | 10955.3 | -0.11501 | 0.303796 | 0.898538 | Sult4a1 |
| ENSMUSG( | 8.21758 | 0.14837 | 0.822032 | NA | Sult5a1 |
| ENSMUSG( | 1004.779 | 0.104152 | 0.484113 | 0.92541 | Sumf1 |
| ENSMUSG( | 531.4486 | 0.200706 | 0.166536 | 0.840183 | Sumf2 |
| ENSMUSG( | 1134.054 | 0.071798 | 0.621424 | 0.953199 | Sumo1 |
| ENSMUSG( | 1118 | -0.00884 | 0.947263 | 0.994798 | Sumo2 |
| ENSMUSG( | 2421.766 | -0.18655 | 0.173728 | 0.844317 | Sumo3 |
| ENSMUSG( | 3328.948 | 0.183793 | 0.207575 | 0.864412 | Sun1 |
| ENSMUSG( | 2109.263 | 0.105579 | 0.478721 | 0.923329 | Sun2 |
| ENSMUSG( | 526.1792 | 0.033338 | 0.795593 | 0.97681 | Suox |
| ENSMUSG( | 1668.012 | -0.08032 | 0.471554 | 0.923329 | Supt16 |
| ENSMUSG( | 919.1601 | 0.093104 | 0.551373 | 0.939565 | Supt20 |
| ENSMUSG( | 200.1333 | -0.09948 | 0.592957 | 0.945408 | Supt3 |
| ENSMUSG( | 624.5225 | -0.22102 | 0.091967 | 0.791889 | Supt4a |
| ENSMUSG( | 3365.234 | -0.06113 | 0.627589 | 0.953234 | Supt5 |
| ENSMUSG( | 4646.563 | 0.024385 | 0.819172 | 0.979198 | Supt6 |
| ENSMUSG( | 400.4407 | 0.0114 | 0.937467 | 0.993685 | Supt7l |
| ENSMUSG( | 540.7874 | 0.092539 | 0.462199 | 0.923329 | Supv3l1 |
| ENSMUSG( | 1836.599 | -0.09779 | 0.385624 | 0.917257 | Surf1 |
| ENSMUSG( | 574.8249 | -0.0336 | 0.807018 | 0.978239 | Surf2 |
| ENSMUSG( | 3403.725 | 0.133734 | 0.375241 | 0.915491 | Surf4 |
| ENSMUSG( | 471.9654 | -0.0008 | 0.995473 | 0.999116 | Surf6 |
| ENSMUSG( | 338.2248 | 0.059042 | 0.706909 | 0.963529 | Susd1 |
| ENSMUSG( | 302.5861 | 0.139187 | 0.362624 | 0.914223 | Susd2 |
| ENSMUSG( | 111.8013 | 0.185833 | 0.464141 | 0.923329 | Susd3 |
| ENSMUSG( | 5209.417 | 0.197386 | 0.133235 | 0.822912 | Susd4 |
| ENSMUSG( | 382.5601 | 0.415179 | 0.076508 | 0.770558 | Susd5 |
| ENSMUSG( | 1858.772 | 0.198958 | 0.22175 | 0.866104 | Susd6 |
| ENSMUSG( | 1003.764 | 0.072164 | 0.558746 | 0.939574 | Suv39h1 |
| ENSMUSG( | 85.46782 | -0.10006 | 0.672608 | 0.95694 | Suv39h2 |
| ENSMUSG( | 1049.39 | 0.337533 | 0.010897 | 0.422321 | Suz12 |
| ENSMUSG( | 16282.45 | 0.124872 | 0.404932 | 0.917257 | Sv2a |
| ENSMUSG( | 12231.03 | 0.283249 | 0.105858 | 0.804969 | Sv2b |
| ENSMUSG( | 1301.201 | 0.117894 | 0.444562 | 0.921734 | Sv2c |
| ENSMUSG( | 462.6472 | -0.19729 | 0.206716 | 0.864412 | Svbp |
| ENSMUSG( | 49.20543 | 0.591872 | 0.059753 | 0.729747 | Svep1 |
| ENSMUSG( | 593.8366 | 0.296986 | 0.023377 | 0.585822 | Svil |
| ENSMUSG( | 3296.245 | -0.18863 | 0.2 | 0.863016 | Svip |
| ENSMUSG( | 7189.859 | 0.203862 | 0.156445 | 0.837724 | Svop |
| ENSMUSG( | 991.494 | -0.08921 | 0.533953 | 0.935607 | Swap70 |
| ENSMUSG( | 2652.723 | -0.18749 | 0.178766 | 0.849041 | Swi5 |
| ENSMUSG( | 392.87 | -0.08385 | 0.6014 | 0.947749 | Swsap1 |
| ENSMUSG( | 210.344 | 0.305702 | 0.086716 | 0.782073 | Swt1 |
| ENSMUSG( | 769.6058 | -0.0007 | 0.996285 | 0.999178 | Syap1 |
| ENSMUSG( | 1464.433 | -0.081 | 0.580717 | 0.944266 | Sybu |
| ENSMUSG( | 81.85056 | 0.125725 | 0.621615 | 0.953199 | Syce2 |
| ENSMUSG( | 55.1141 | 0.040015 | 0.891475 | 0.990558 | Sycp2 |
| ENSMUSG( | 465.27 | 0.04034 | 0.761039 | 0.972695 | Syde1 |
| ENSMUSG( | 171.5516 | -0.07908 | 0.666662 | 0.956766 | Syde2 |
| ENSMUSG( | 1588.731 | -0.08546 | 0.488081 | 0.925453 | Syf2 |
| ENSMUSG( | 271.4986 | 0.137131 | 0.40431 | 0.917257 | Syk |
| ENSMUSG( | 5061.831 | 0.037475 | 0.725566 | 0.966887 | Sympk |
| ENSMUSG( | 15845.61 | -0.10857 | 0.460538 | 0.923329 | Syn1 |
| ENSMUSG( | 9109.144 | -0.1119 | 0.387353 | 0.917257 | Syn2 |
| ENSMUSG( | 139.4634 | 0.021585 | 0.922863 | 0.992644 | Syn3 |
| ENSMUSG( | 26.53388 | -0.16334 | 0.715382 | 0.965063 | Syna |
| ENSMUSG( | 20.87841 | 0.027444 | 0.949391 | 0.994798 | Sync |
| ENSMUSG( | 1777.363 | 0.01263 | 0.932402 | 0.99306 | Syncrip |
| ENSMUSG( | 608.3118 | -0.31391 | 0.027663 | 0.61023 | Syndig1 |
| ENSMUSG( | 2178.041 | -0.08669 | 0.575481 | 0.942912 | Syndig1l |
| ENSMUSG( | 13441.16 | -0.00261 | 0.981217 | 0.996756 | Syne1 |
| ENSMUSG( | 844.557 | -0.03717 | 0.76252 | 0.973539 | Syne2 |
| ENSMUSG( | 61.32659 | -0.3689 | 0.183038 | 0.850741 | Syne3 |
| ENSMUSG( | 67.5023 | 0.102232 | 0.767622 | 0.974236 | Syne4 |
| ENSMUSG( | 10636.23 | 0.147113 | 0.127189 | 0.817278 | Syngap1 |
| ENSMUSG( | 20529.44 | 0.184753 | 0.298941 | 0.898538 | Syngr1 |
| ENSMUSG( | 604.8489 | 0.256433 | 0.179797 | 0.850372 | Syngr2 |
| ENSMUSG( | 6332.271 | 0.019755 | 0.89313 | 0.990558 | Syngr3 |
| ENSMUSG( | 5377.015 | 0.052736 | 0.607105 | 0.949917 | Synj1 |
| ENSMUSG( | 1611.223 | -0.06725 | 0.635155 | 0.953903 | Synj2 |
| ENSMUSG( | 712.7477 | -0.09946 | 0.460108 | 0.923329 | Synj2bp |
| ENSMUSG( | 1317.319 | 0.156707 | 0.249469 | 0.877904 | Synm |
| ENSMUSG( | 4238.119 | 0.050111 | 0.699104 | 0.961139 | Synpo |
| ENSMUSG( | 330.6712 | 0.280215 | 0.072913 | 0.763629 | Synpo2 |
| ENSMUSG( | 2758.852 | 0.193338 | 0.228766 | 0.868507 | Synpr |
| ENSMUSG( | 1848.866 | -0.11312 | 0.304845 | 0.898538 | Synrg |
| ENSMUSG( | 58643.87 | 0.064279 | 0.646332 | 0.953988 | Syp |
| ENSMUSG( | 1243.03 | 0.3219 | 0.109039 | 0.804969 | Sypl1 |
| ENSMUSG( | 242.0828 | 0.191015 | 0.344217 | 0.910274 | Sypl2 |
| ENSMUSG( | 993.287 | -0.11195 | 0.382682 | 0.917257 | Sys1 |
| ENSMUSG( | 47966.52 | 0.150631 | 0.346995 | 0.911434 | Syt1 |
| ENSMUSG( | 60.86531 | 1.156417 | 0.000143 | 0.027166 | Syt10 |
| ENSMUSG( | 30614.71 | 0.209114 | 0.220877 | 0.866104 | Syt11 |
| ENSMUSG( | 1573.649 | 0.063588 | 0.68891 | 0.959742 | Syt12 |
| ENSMUSG( | 10232.75 | 0.020033 | 0.880933 | 0.989644 | Syt13 |

| | | | | | | | | | | | |
|---|---|---|---|---|---|---|---|---|---|---|---|
| ENSMUSG( | Taf13 | 2011.23 | 0.085048 | 0.522468 | 0.875368 | ENSMUSG( | Taf13 | 1675.092 | -0.18281 | 0.235561 | 0.871807 |
| ENSMUSG( | Taf15 | 1823.062 | 0.061233 | 0.535988 | 0.88142 | ENSMUSG( | Taf15 | 1530.649 | -0.18472 | 0.185662 | 0.855305 |
| ENSMUSG( | Taf1a | 283.4154 | 0.029942 | 0.841263 | 0.971179 | ENSMUSG( | Taf1a | 252.1134 | -0.03326 | 0.850082 | 0.985538 |
| ENSMUSG( | Taf1b | 423.6518 | 0.012841 | 0.927381 | 0.991229 | ENSMUSG( | Taf1b | 369.951 | -0.11165 | 0.428584 | 0.917257 |
| ENSMUSG( | Taf1c | 444.9308 | 0.008131 | 0.950037 | 0.993821 | ENSMUSG( | Taf1c | 399.5748 | -0.02568 | 0.867744 | 0.98781 |
| ENSMUSG( | Taf1d | 322.7163 | -0.1383 | 0.330882 | 0.775123 | ENSMUSG( | Taf1d | 343.1992 | 0.308078 | 0.107872 | 0.804969 |
| ENSMUSG( | Taf2 | 1123.213 | 0.153122 | 0.10788 | 0.49674 | ENSMUSG( | Taf2 | 1026.557 | 0.162885 | 0.148922 | 0.836241 |
| ENSMUSG( | Taf3 | 561.1921 | 0.177484 | 0.164054 | 0.594869 | ENSMUSG( | Taf3 | 528.2463 | 0.260219 | 0.071399 | 0.761942 |
| ENSMUSG( | Taf4 | 844.8979 | 0.13473 | 0.204963 | 0.64616 | ENSMUSG( | Taf4 | 758.687 | 0.089838 | 0.489627 | 0.925858 |
| ENSMUSG( | Taf4b | 79.209 | -0.14165 | 0.511124 | 0.870859 | ENSMUSG( | Taf4b | 77.08003 | 0.062096 | 0.793256 | 0.97681 |
| ENSMUSG( | Taf5 | 138.318 | -0.19553 | 0.258981 | 0.709814 | ENSMUSG( | Taf5 | 139.0977 | 0.096938 | 0.613963 | 0.951177 |
| ENSMUSG( | Taf5l | 484.7078 | -0.19662 | 0.124431 | 0.531714 | ENSMUSG( | Taf5l | 486.0429 | 0.090174 | 0.530291 | 0.934795 |
| ENSMUSG( | Taf6 | 1105.287 | 0.173128 | 0.088074 | 0.455247 | ENSMUSG( | Taf6 | 960.1307 | 0.039589 | 0.734031 | 0.967228 |
| ENSMUSG( | Taf6l | 287.792 | 0.260722 | 0.091755 | 0.465904 | ENSMUSG( | Taf6l | 245.7611 | 0.085019 | 0.603848 | 0.948137 |
| ENSMUSG( | Taf7 | 140.7958 | -0.07449 | 0.675441 | 0.930049 | ENSMUSG( | Taf7 | 133.4573 | 0.045141 | 0.818309 | 0.979198 |
| ENSMUSG( | Taf8 | 560.4475 | 0.064082 | 0.578256 | 0.89641 | ENSMUSG( | Taf8 | 500.5817 | 0.013727 | 0.920421 | 0.992644 |
| ENSMUSG( | Taf9 | 188.1028 | 0.220738 | 0.185769 | 0.623185 | ENSMUSG( | Taf9 | 173.5566 | 0.246778 | 0.172911 | 0.843702 |
| ENSMUSG( | Taf9b | 863.4105 | 0.00888 | 0.939036 | 0.993415 | ENSMUSG( | Taf9b | 760.8906 | -0.09443 | 0.52223 | 0.932681 |
| ENSMUSG( | Tafa1 | 982.3401 | -0.41632 | 0.00133 | 0.039731 | ENSMUSG( | Tafa1 | 998.0109 | -0.07239 | 0.612777 | 0.951071 |
| ENSMUSG( | Tafa2 | 878.837 | -0.10445 | 0.428833 | 0.826446 | ENSMUSG( | Tafa2 | 878.7712 | 0.165258 | 0.304352 | 0.898538 |
| ENSMUSG( | Tafa4 | 22.37713 | -0.69597 | 0.167881 | 0.600712 | ENSMUSG( | Tafa4 | 22.16903 | -0.39176 | 0.404245 | 0.917257 |
| ENSMUSG( | Tafa5 | 4846.339 | -0.07975 | 0.302356 | 0.750682 | ENSMUSG( | Tafa5 | 4487.722 | -0.03301 | 0.752576 | 0.970521 |
| ENSMUSG( | Tafazzin | 1144.442 | -0.10329 | 0.376038 | 0.801127 | ENSMUSG( | Tafazzin | 1154.404 | 0.188607 | 0.229172 | 0.868507 |
| ENSMUSG( | Tagap | 34.92736 | 0.204358 | 0.557068 | 0.888198 | ENSMUSG( | Tagap | 29.69209 | 0.044751 | 0.909248 | 0.991958 |
| ENSMUSG( | Tagap1 | 554.462 | -0.12529 | 0.307846 | 0.756096 | ENSMUSG( | Tagap1 | 550.3891 | 0.12642 | 0.321367 | 0.904019 |
| ENSMUSG( | Tagln | 69.3026 | -0.81724 | 0.001741 | 0.047093 | ENSMUSG( | Tagln | 70.2941 | -0.41616 | 0.133848 | 0.822912 |
| ENSMUSG( | Tagln2 | 415.2036 | 0.188594 | 0.204243 | 0.64477 | ENSMUSG( | Tagln2 | 426.1431 | 0.503703 | 0.003702 | 0.2429 |
| ENSMUSG( | Tagln3 | 6981.378 | 0.101359 | 0.253956 | 0.704011 | ENSMUSG( | Tagln3 | 5560.525 | -0.30321 | 0.013526 | 0.464932 |
| ENSMUSG( | Tal1 | 80.15699 | 0.541968 | 0.028949 | 0.259139 | ENSMUSG( | Tal1 | 75.9027 | 0.659528 | 0.023789 | 0.588226 |
| ENSMUSG( | Taldo1 | 2786.811 | -0.03328 | 0.719412 | 0.941986 | ENSMUSG( | Taldo1 | 2383.39 | -0.22728 | 0.058898 | 0.729034 |
| ENSMUSG( | Tamalin | 1173.787 | 0.005022 | 0.971085 | 0.996369 | ENSMUSG( | Tamalin | 1014.302 | -0.15248 | 0.303458 | 0.898538 |
| ENSMUSG( | Tamm41 | 203.5667 | -0.08536 | 0.611842 | 0.908586 | ENSMUSG( | Tamm41 | 183.9071 | -0.11541 | 0.530126 | 0.934795 |
| ENSMUSG( | Tanc1 | 1132.067 | 0.232525 | 0.011305 | 0.153194 | ENSMUSG( | Tanc1 | 962.7341 | 0.043707 | 0.709143 | 0.964042 |
| ENSMUSG( | Tanc2 | 4269.486 | 0.039829 | 0.672373 | 0.929007 | ENSMUSG( | Tanc2 | 3897.12 | 0.047332 | 0.669269 | 0.956766 |
| ENSMUSG( | Tango2 | 1653.06 | 0.089054 | 0.366911 | 0.795164 | ENSMUSG( | Tango2 | 1449.26 | -0.01967 | 0.862386 | 0.986509 |
| ENSMUSG( | Tango6 | 165.2565 | 0.214711 | 0.186245 | 0.623869 | ENSMUSG( | Tango6 | 143.1862 | 0.078108 | 0.67398 | 0.95694 |
| ENSMUSG( | Tank | 1373.449 | 0.152646 | 0.129577 | 0.539182 | ENSMUSG( | Tank | 1213.048 | 0.062379 | 0.62463 | 0.953234 |
| ENSMUSG( | Taok1 | 4008.648 | 0.007245 | 0.945405 | 0.99365 | ENSMUSG( | Taok1 | 3597.643 | -0.03869 | 0.771477 | 0.974236 |
| ENSMUSG( | Taok2 | 4788.204 | 0.096405 | 0.245196 | 0.692812 | ENSMUSG( | Taok2 | 4267.553 | 0.035845 | 0.735475 | 0.967344 |
| ENSMUSG( | Taok3 | 1937.216 | 0.126598 | 0.22812 | 0.675573 | ENSMUSG( | Taok3 | 1642.067 | -0.08293 | 0.558052 | 0.939565 |
| ENSMUSG( | Tap1 | 163.4242 | -0.53293 | 0.003251 | 0.071142 | ENSMUSG( | Tap1 | 172.4392 | -0.04287 | 0.83669 | 0.981335 |
| ENSMUSG( | Tap2 | 440.3148 | -0.35373 | 0.0145 | 0.17723 | ENSMUSG( | Tap2 | 464.7219 | 0.096055 | 0.554286 | 0.939565 |
| ENSMUSG( | Tapbp | 1261.953 | -0.31873 | 0.016131 | 0.188643 | ENSMUSG( | Tapbp | 1347.282 | 0.157704 | 0.316674 | 0.901868 |
| ENSMUSG( | Tapbpl | 306.3903 | -0.30211 | 0.048463 | 0.336526 | ENSMUSG( | Tapbpl | 307.5568 | 0.003201 | 0.984939 | 0.997067 |
| ENSMUSG( | Tapt1 | 1789.715 | -0.20901 | 0.139815 | 0.55638 | ENSMUSG( | Tapt1 | 1911.468 | 0.250249 | 0.109436 | 0.805353 |
| ENSMUSG( | Tarbp1 | 249.6685 | 0.18686 | 0.251904 | 0.701908 | ENSMUSG( | Tarbp1 | 221.275 | 0.116466 | 0.499477 | 0.927825 |
| ENSMUSG( | Tarbp2 | 448.619 | -0.1212 | 0.334187 | 0.7758 | ENSMUSG( | Tarbp2 | 452.4275 | 0.173649 | 0.24635 | 0.877904 |
| ENSMUSG( | Tardbp | 3920.483 | 0.045776 | 0.548921 | 0.885686 | ENSMUSG( | Tardbp | 3616.318 | 0.081082 | 0.49368 | 0.927372 |
| ENSMUSG( | Tars1 | 767.199 | 0.092247 | 0.363679 | 0.793343 | ENSMUSG( | Tars1 | 692.908 | 0.066394 | 0.584947 | 0.944975 |
| ENSMUSG( | Tars2 | 1032.171 | 0.04059 | 0.718344 | 0.941911 | ENSMUSG( | Tars2 | 915.5492 | -0.03603 | 0.781421 | 0.976106 |
| ENSMUSG( | Tars3 | 707.2526 | 0.090201 | 0.381679 | 0.802879 | ENSMUSG( | Tars3 | 630.0311 | 0.027233 | 0.82479 | 0.979198 |
| ENSMUSG( | Tas1r3 | 11.44139 | -0.22442 | 0.685568 | NA | ENSMUSG( | Tas1r3 | 13.03856 | 0.383584 | 0.482274 | 0.924578 |
| ENSMUSG( | Tasl | 11.17433 | 0.222757 | 0.701634 | NA | ENSMUSG( | Tasl | 11.66277 | 0.607971 | 0.316171 | 0.901868 |
| ENSMUSG( | Tasor | 1006.134 | 0.011225 | 0.929098 | 0.991772 | ENSMUSG( | Tasor | 1011.131 | 0.284042 | 0.041201 | 0.687358 |
| ENSMUSG( | Tasor2 | 681.545 | -0.07905 | 0.602775 | 0.904887 | ENSMUSG( | Tasor2 | 683.3713 | 0.198844 | 0.291885 | 0.897854 |
| ENSMUSG( | Tasp1 | 468.4573 | 0.058383 | 0.641306 | 0.916693 | ENSMUSG( | Tasp1 | 418.265 | -0.00918 | 0.956761 | 0.994798 |
| ENSMUSG( | Tatdn1 | 259.6376 | -0.01394 | 0.929043 | 0.991772 | ENSMUSG( | Tatdn1 | 240.0616 | 0.027038 | 0.870536 | 0.988445 |
| ENSMUSG( | Tatdn2 | 1348.091 | 0.147831 | 0.106544 | 0.494174 | ENSMUSG( | Tatdn2 | 1177.116 | 0.029731 | 0.787339 | 0.97637 |
| ENSMUSG( | Tatdn3 | 399.8851 | -0.07217 | 0.615781 | 0.910647 | ENSMUSG( | Tatdn3 | 372.3165 | 0.002623 | 0.989112 | 0.997893 |
| ENSMUSG( | Tax1bp1 | 4307.833 | 0.003521 | 0.972423 | 0.996585 | ENSMUSG( | Tax1bp1 | 3854.908 | -0.04884 | 0.642357 | 0.953903 |
| ENSMUSG( | Tax1bp3 | 169.3899 | 0.099625 | 0.536928 | 0.882123 | ENSMUSG( | Tax1bp3 | 158.7219 | 0.184799 | 0.310188 | 0.898852 |
| ENSMUSG( | Tbata | 143.8399 | 0.021035 | 0.914546 | 0.988483 | ENSMUSG( | Tbata | 128.488 | -0.03221 | 0.871439 | 0.988779 |
| ENSMUSG( | Tbc1d1 | 707.2336 | 0.246835 | 0.070433 | 0.40908 | ENSMUSG( | Tbc1d1 | 645.761 | 0.25867 | 0.089784 | 0.79071 |
| ENSMUSG( | Tbc1d10a | 801.1787 | 0.18949 | 0.082896 | 0.444486 | ENSMUSG( | Tbc1d10a | 693.7116 | 0.048503 | 0.704922 | 0.962741 |
| ENSMUSG( | Tbc1d10b | 2446.389 | 0.095738 | 0.371684 | 0.798768 | ENSMUSG( | Tbc1d10b | 2115.156 | -0.05693 | 0.671401 | 0.95694 |
| ENSMUSG( | Tbc1d10c | 20.59119 | 0.151963 | 0.745997 | NA | ENSMUSG( | Tbc1d10c | 18.62942 | 0.0883 | 0.86692 | 0.987487 |
| ENSMUSG( | Tbc1d12 | 894.3053 | 0.092321 | 0.486546 | 0.858713 | ENSMUSG( | Tbc1d12 | 795.6655 | 0.019974 | 0.896406 | 0.990558 |
| ENSMUSG( | Tbc1d13 | 1592.843 | 0.103054 | 0.323179 | 0.768415 | ENSMUSG( | Tbc1d13 | 1435.22 | 0.073799 | 0.518872 | 0.930939 |
| ENSMUSG( | Tbc1d14 | 2903.153 | 0.101681 | 0.234461 | 0.681922 | ENSMUSG( | Tbc1d14 | 2601.134 | 0.056235 | 0.587831 | 0.944975 |
| ENSMUSG( | Tbc1d15 | 680.2451 | 0.099255 | 0.336987 | 0.777008 | ENSMUSG( | Tbc1d15 | 618.5763 | 0.093464 | 0.440368 | 0.921166 |
| ENSMUSG( | Tbc1d16 | 3675.947 | 0.032906 | 0.736209 | 0.946267 | ENSMUSG( | Tbc1d16 | 3375.219 | 0.055399 | 0.58517 | 0.944975 |
| ENSMUSG( | Tbc1d17 | 1265.283 | 0.09849 | 0.3433 | 0.779153 | ENSMUSG( | Tbc1d17 | 1126.002 | 0.03322 | 0.777887 | 0.976106 |
| ENSMUSG( | Tbc1d19 | 866.2577 | 0.118244 | 0.255059 | 0.705501 | ENSMUSG( | Tbc1d19 | 774.4202 | 0.06407 | 0.609143 | 0.950335 |
| ENSMUSG( | Tbc1d2 | 183.1029 | 0.372632 | 0.103202 | 0.488596 | ENSMUSG( | Tbc1d2 | 147.5827 | 0.050554 | 0.806768 | 0.978139 |
| ENSMUSG( | Tbc1d20 | 2587.932 | 0.072703 | 0.375272 | 0.800804 | ENSMUSG( | Tbc1d20 | 2319.665 | 0.026119 | 0.801747 | 0.97681 |
| ENSMUSG( | Tbc1d22a | 1256.078 | -0.01159 | 0.908193 | 0.9878 | ENSMUSG( | Tbc1d22a | 1093.17 | -0.14456 | 0.226729 | 0.868507 |
| ENSMUSG( | Tbc1d22b | 1847.089 | 0.124323 | 0.232201 | 0.67996 | ENSMUSG( | Tbc1d22b | 1578.427 | -0.0597 | 0.602574 | 0.94789 |
| ENSMUSG( | Tbc1d23 | 1083.848 | -0.07303 | 0.446415 | 0.83372 | ENSMUSG( | Tbc1d23 | 1066.227 | 0.14528 | 0.197186 | 0.862161 |
| ENSMUSG( | Tbc1d24 | 2104.56 | 0.069211 | 0.53608 | 0.88142 | ENSMUSG( | Tbc1d24 | 1901.855 | 0.048871 | 0.679305 | 0.957928 |
| ENSMUSG( | Tbc1d25 | 261.1708 | -0.00737 | 0.970822 | 0.996369 | ENSMUSG( | Tbc1d25 | 225.0854 | -0.17486 | 0.358985 | 0.914004 |
| ENSMUSG( | Tbc1d2b | 566.9635 | -0.08237 | 0.445973 | 0.833467 | ENSMUSG( | Tbc1d2b | 547.6878 | 0.092044 | 0.502629 | 0.927825 |
| ENSMUSG( | Tbc1d30 | 1794.654 | 0.177463 | 0.178161 | 0.615629 | ENSMUSG( | Tbc1d30 | 1525.757 | -0.018 | 0.898035 | 0.990558 |
| ENSMUSG( | Tbc1d31 | 195.6479 | 0.103595 | 0.546017 | 0.884974 | ENSMUSG( | Tbc1d31 | 190.4242 | 0.292929 | 0.139266 | 0.830678 |
| ENSMUSG( | Tbc1d32 | 392.0395 | -0.16725 | 0.166067 | 0.598098 | ENSMUSG( | Tbc1d32 | 359.62 | -0.15345 | 0.305251 | 0.898538 |
| ENSMUSG( | Tbc1d4 | 377.8854 | 0.782617 | 3.07E-10 | 1.61E-07 | ENSMUSG( | Tbc1d4 | 351.5484 | 0.837034 | 5.33E-06 | 0.002278 |
| ENSMUSG( | Tbc1d5 | 1357.798 | 0.098405 | 0.274078 | 0.724254 | ENSMUSG( | Tbc1d5 | 1162.504 | -0.07867 | 0.489927 | 0.925987 |
| ENSMUSG( | Tbc1d7 | 439.0522 | -0.16476 | 0.180173 | 0.617634 | ENSMUSG( | Tbc1d7 | 384.4735 | -0.29135 | 0.047198 | 0.700856 |
| ENSMUSG( | Tbc1d8 | 720.2574 | -0.27471 | 0.04654 | 0.330304 | ENSMUSG( | Tbc1d8 | 670.8298 | -0.20794 | 0.180002 | 0.850372 |
| ENSMUSG( | Tbc1d8b | 154.789 | 0.139497 | 0.463819 | 0.842812 | ENSMUSG( | Tbc1d8b | 146.6643 | 0.254149 | 0.248639 | 0.877904 |
| ENSMUSG( | Tbc1d9 | 3414.295 | 0.102343 | 0.252946 | 0.703255 | ENSMUSG( | Tbc1d9 | 3046.884 | 0.044401 | 0.66825 | 0.956766 |
| ENSMUSG( | Tbc1d9b | 29583.33 | 0.168146 | 0.036093 | 0.292126 | ENSMUSG( | Tbc1d9b | 24814.93 | -0.06786 | 0.540541 | 0.937383 |
| ENSMUSG( | Tbca | 931.4758 | -0.03486 | 0.748176 | 0.949604 | ENSMUSG( | Tbca | 814.6045 | -0.16421 | 0.311752 | 0.900031 |
| ENSMUSG( | Tbcb | 2609.457 | 0.054668 | 0.523558 | 0.875628 | ENSMUSG( | Tbcb | 2243.545 | -0.11746 | 0.340826 | 0.907972 |
| ENSMUSG( | Tbcc | 294.5883 | -0.11658 | 0.484626 | 0.856665 | ENSMUSG( | Tbcc | 252.9299 | -0.31 | 0.100786 | 0.801456 |
| ENSMUSG( | Tbccd1 | 526.6277 | 0.150308 | 0.195271 | 0.634601 | ENSMUSG( | Tbccd1 | 471.0518 | 0.096434 | 0.491534 | 0.926825 |
| ENSMUSG( | Tbcd | 1436.476 | 0.098165 | 0.29438 | 0.74363 | ENSMUSG( | Tbcd | 1278.841 | 0.034427 | 0.761126 | 0.972695 |
| ENSMUSG( | Tbce | 1346.479 | 0.142374 | 0.326902 | 0.771858 | ENSMUSG( | Tbce | 1191.468 | 0.059097 | 0.76119 | 0.972695 |
| ENSMUSG( | Tbcel | 4881.809 | 0.155141 | 0.161787 | 0.592233 | ENSMUSG( | Tbcel | 4158.227 | -0.03754 | 0.779579 | 0.976106 |
| ENSMUSG( | Tbck | 795.5502 | 0.133262 | 0.271327 | 0.72198 | ENSMUSG( | Tbck | 700.9049 | 0.036266 | 0.799507 | 0.97681 |
| ENSMUSG( | Tbk1 | 906.8704 | 0.047486 | 0.634203 | 0.914073 | ENSMUSG( | Tbk1 | 883.5211 | 0.233863 | 0.061594 | 0.736111 |
| ENSMUSG( | Tbkbp1 | 3146.71 | 0.338851 | 0.000538 | 0.021456 | ENSMUSG( | Tbkbp1 | 2788.727 | 0.266646 | 0.02157 | 0.575341 |
| ENSMUSG( | Tbl1x | 2367.077 | -0.06046 | 0.493885 | 0.860611 | ENSMUSG( | Tbl1x | 2156.68 | -0.06002 | 0.557174 | 0.939565 |
| ENSMUSG( | Tbl1xr1 | 2037.103 | -0.07731 | 0.379732 | 0.801757 | ENSMUSG( | Tbl1xr1 | 1892.678 | -0.02298 | 0.859149 | 0.985884 |
| ENSMUSG( | Tbl2 | 1124.552 | -0.10351 | 0.481837 | 0.855573 | ENSMUSG( | Tbl2 | 1157.126 | 0.244321 | 0.137782 | 0.828492 |
| ENSMUSG( | Tbl3 | 738.028 | 0.066963 | 0.585681 | 0.899171 | ENSMUSG( | Tbl3 | 639.4082 | -0.07759 | 0.541867 | 0.937841 |
| ENSMUSG( | Tbp | 457.3684 | -0.03357 | 0.776422 | 0.955658 | ENSMUSG( | Tbp | 458.1469 | 0.230333 | 0.079481 | 0.77458 |
| ENSMUSG( | Tbpl1 | 1097.625 | 0.097751 | 0.400082 | 0.813641 | ENSMUSG( | Tbpl1 | 968.338 | 0.005132 | 0.970447 | 0.99594 |
| ENSMUSG( | Tbr1 | 2464.431 | 0.079888 | 0.409731 | 0.818258 | ENSMUSG( | Tbr1 | 2252.154 | 0.08705 | 0.427147 | 0.917257 |
| ENSMUSG( | Tbrg1 | 763.5364 | 0.045605 | 0.740572 | 0.947823 | ENSMUSG( | Tbrg1 | 680.4668 | -0.02065 | 0.895649 | 0.990558 |
| ENSMUSG( | Tbrg4 | 995.1325 | 0.113281 | 0.312801 | 0.760322 | ENSMUSG( | Tbrg4 | 856.0599 | -0.05125 | 0.712259 | 0.964363 |
| ENSMUSG( | Tbx1 | 52.71239 | 0.293919 | 0.345164 | 0.779153 | ENSMUSG( | Tbx1 | 54.09724 | 0.608929 | 0.051597 | 0.711677 |
| ENSMUSG( | Tbx15 | 88.16541 | -0.34526 | 0.096884 | 0.477375 | ENSMUSG( | Tbx15 | 98.37883 | 0.23934 | 0.277763 | 0.891468 |
| ENSMUSG( | Tbx18 | 103.2505 | 0.094752 | 0.653314 | 0.922819 | ENSMUSG( | Tbx18 | 97.83981 | 0.203243 | 0.38348 | 0.917257 |
| ENSMUSG( | Tbx2 | 81.51664 | 0.237298 | 0.339312 | 0.777616 | ENSMUSG( | Tbx2 | 94.29523 | 0.842347 | 0.004338 | 0.268942 |
| ENSMUSG( | Tbx3 | 109.4104 | 0.50798 | 0.01697 | 0.194647 | ENSMUSG( | Tbx3 | 131.0528 | 1.139598 | 4.61E-07 | 0.00026 |
| ENSMUSG( | Tbx3os1 | 39.63378 | 0.016294 | 0.968973 | 0.996369 | ENSMUSG( | Tbx3os1 | 44.43106 | 0.584418 | 0.140888 | 0.830678 |
| ENSMUSG( | Tbx4 | 14.41776 | -0.0364 | 0.949325 | NA | ENSMUSG( | Tbx4 | 14.27467 | 0.192476 | 0.729062 | 0.966913 |
| ENSMUSG( | Tbx6 | 31.30228 | 0.365131 | 0.348809 | 0.781448 | ENSMUSG( | Tbx6 | 30.49209 | 0.523946 | 0.226969 | 0.868507 |
| ENSMUSG( | Tbxa2r | 47.83263 | 0.271524 | 0.425656 | 0.824898 | ENSMUSG( | Tbxa2r | 64.46267 | 1.197777 | 0.000396 | 0.057854 |
| ENSMUSG( | Tbxas1 | 196.2275 | -0.19268 | 0.338897 | 0.777518 | ENSMUSG( | Tbxas1 | 179.9779 | -0.15242 | 0.540088 | 0.937383 |
| ENSMUSG( | Tcaf1 | 5926.088 | -0.04489 | 0.57157 | 0.893319 | ENSMUSG( | Tcaf1 | 5392.682 | -0.049 | 0.635162 | 0.953903 |
| ENSMUSG( | Tcaf2 | 28.56838 | 0.206338 | 0.573833 | 0.893584 | ENSMUSG( | Tcaf2 | 25.77211 | 0.194157 | 0.630481 | 0.953251 |
| ENSMUSG( | Tcaim | 598.6119 | 0.109381 | 0.323284 | 0.768415 | ENSMUSG( | Tcaim | 525.3785 | -0.00377 | 0.97878 | 0.996468 |
| ENSMUSG( | Tcap | 108.9179 | 0.112898 | 0.599389 | 0.903801 | ENSMUSG( | Tcap | 98.19475 | 0.078224 | 0.714418 | 0.964986 |
| ENSMUSG( | Tcea1 | 980.0729 | 0.015996 | 0.874302 | 0.979431 | ENSMUSG( | Tcea1 | 874.3511 | -0.04904 | 0.711153 | 0.964363 |
| ENSMUSG( | Tcea2 | 1132.157 | 0.038379 | 0.783149 | 0.957558 | ENSMUSG( | Tcea2 | 1027.01 | 0.026702 | 0.884804 | 0.989688 |
| ENSMUSG( | Tcea3 | 20.21669 | -0.77572 | 0.081324 | NA | ENSMUSG( | Tcea3 | 20.69014 | -0.36673 | 0.411184 | 0.917257 |
| ENSMUSG( | Tceal1 | 365.6721 | -0.07401 | 0.583993 | 0.898666 | ENSMUSG( | Tceal1 | 320.8887 | -0.19921 | 0.307197 | 0.898538 |
| ENSMUSG( | Tceal3 | 2100.694 | 0.000486 | 0.996719 | 0.99943 | ENSMUSG( | Tceal3 | 1764.083 | -0.25103 | 0.120217 | 0.813384 |
| ENSMUSG( | Tceal5 | 874.8867 | -0.03964 | 0.790274 | 0.959461 | ENSMUSG( | Tceal5 | 746.5465 | -0.24669 | 0.213226 | 0.864412 |
| ENSMUSG( | Tceal6 | 784.5488 | -0.09426 | 0.583847 | 0.898666 | ENSMUSG( | Tceal6 | 682.5343 | -0.2433 | 0.205599 | 0.864412 |
| ENSMUSG( | Tceal7 | 25.71728 | 0.368619 | 0.329218 | 0.773233 | ENSMUSG( | Tceal7 | 21.06778 | 0.107783 | 0.824354 | 0.979198 |
| ENSMUSG( | Tceal8 | 843.1963 | 0.06851 | 0.62446 | 0.91187 | ENSMUSG( | Tceal8 | 737.8171 | -0.05476 | 0.768746 | 0.974236 |
| ENSMUSG( | Tceal9 | 805.7515 | -0.03535 | 0.795712 | 0.960062 | ENSMUSG( | Tceal9 | 747.6836 | 0.012222 | 0.946396 | 0.994798 |
| ENSMUSG( | Tceanc | 62.5476 | -0.20088 | 0.448281 | 0.834051 | ENSMUSG( | Tceanc | 55.72363 | -0.26502 | 0.372382 | 0.915491 |
| ENSMUSG( | Tceanc2 | 528.7811 | 0.221652 | 0.042215 | 0.316565 | ENSMUSG( | Tceanc2 | 467.4961 | 0.138513 | 0.291794 | 0.897854 |
| ENSMUSG( | Tcerg1 | 1406.159 | -0.03103 | 0.733809 | 0.945265 | ENSMUSG( | Tcerg1 | 1313.446 | 0.040451 | 0.714013 | 0.964986 |
| ENSMUSG( | Tcerg1l | 285.0913 | 0.178262 | 0.187296 | 0.625476 | ENSMUSG( | Tcerg1l | 247.9674 | 0.040063 | 0.817933 | 0.979198 |
| ENSMUSG( | Tcf12 | 2122.082 | 0.037594 | 0.694616 | 0.9343 | ENSMUSG( | Tcf12 | 1925.224 | 0.025512 | 0.825209 | 0.979354 |
| ENSMUSG( | Tcf15 | 13.31679 | -0.1086 | 0.828707 | NA | ENSMUSG( | Tcf15 | 11.62457 | -0.21952 | 0.698044 | 0.960867 |
| ENSMUSG( | Tcf19 | 161.9924 | -0.06661 | 0.707603 | 0.93949 | ENSMUSG( | Tcf19 | 151.2469 | 0.010979 | 0.953595 | 0.994798 |
| ENSMUSG( | Tcf20 | 2477.008 | 0.06361 | 0.542378 | 0.883182 | ENSMUSG( | Tcf20 | 2250.404 | 0.058178 | 0.62647 | 0.953234 |
| ENSMUSG( | Tcf23 | 11.10511 | -0.12799 | 0.829816 | NA | ENSMUSG( | Tcf23 | 11.04542 | 0.184682 | 0.771395 | NA |
| ENSMUSG( | Tcf24 | 12.50178 | 0.420484 | 0.44178 | NA | ENSMUSG( | Tcf24 | 11.06315 | 0.292826 | 0.645529 | NA |
| ENSMUSG( | Tcf25 | 19991.73 | 0.108112 | 0.187186 | 0.625368 | ENSMUSG( | Tcf25 | 16710.24 | -0.14226 | 0.166914 | 0.840183 |
| ENSMUSG( | Tcf3 | 1349.712 | -0.00849 | 0.925019 | 0.991204 | ENSMUSG( | Tcf3 | 1220.147 | -0.02891 | 0.802769 | 0.97681 |
| ENSMUSG( | Tcf4 | 11946.51 | 0.155978 | 0.12347 | 0.530552 | ENSMUSG( | Tcf4 | 10398.09 | 0.027934 | 0.823591 | 0.979198 |
| ENSMUSG( | Tcf7 | 166.5151 | 0.012996 | 0.949323 | 0.993821 | ENSMUSG( | Tcf7 | 198.084 | 0.70955 | 0.000873 | 0.099327 |
| ENSMUSG( | Tcf7l1 | 338.1443 | -0.18695 | 0.194092 | 0.633181 | ENSMUSG( | Tcf7l1 | 313.8856 | -0.13867 | 0.374469 | 0.915491 |
| ENSMUSG( | Tcf7l2 | 297.1693 | -0.44241 | 0.00221 | 0.054639 | ENSMUSG( | Tcf7l2 | 281.847 | -0.31813 | 0.074964 | 0.76762 |
| ENSMUSG( | Tcfl5 | 53.9241 | -0.32217 | 0.212651 | 0.6567 | ENSMUSG( | Tcfl5 | 48.99604 | -0.34122 | 0.308766 | 0.898538 |
| ENSMUSG( | Tchh | 84.46327 | 0.216377 | 0.360016 | 0.791685 | ENSMUSG( | Tchh | 70.22658 | -0.04418 | 0.864247 | 0.986552 |
| ENSMUSG( | Tchp | 323.6726 | 0.222543 | 0.141462 | 0.558606 | ENSMUSG( | Tchp | 271.8093 | 0.001315 | 0.993958 | 0.998813 |
| ENSMUSG( | Tcim | 89.65769 | 0.280808 | 0.220264 | 0.665898 | ENSMUSG( | Tcim | 77.96004 | 0.181179 | 0.50451 | 0.927825 |
| ENSMUSG( | Tcirg1 | 706.864 | -0.08835 | 0.458446 | 0.840036 | ENSMUSG( | Tcirg1 | 751.0572 | 0.342643 | 0.073659 | 0.764578 |
| ENSMUSG( | Tcn2 | 1166.262 | -0.13386 | 0.39026 | 0.80794 | ENSMUSG( | Tcn2 | 1185.909 | 0.187554 | 0.327067 | 0.905768 |
| ENSMUSG( | Tcof1 | 639.3074 | 0.139798 | 0.212753 | 0.6567 | ENSMUSG( | Tcof1 | 544.5683 | -0.05446 | 0.686037 | 0.958956 |
| ENSMUSG( | Tcp1 | 4658.006 | 0.098646 | 0.317251 | 0.763823 | ENSMUSG( | Tcp1 | 3923.818 | -0.13083 | 0.292208 | 0.897854 |

| | | | | | |
|---|---|---|---|---|---|
| ENSMUSG( | 683.715 | -0.70763 | 2.86E-06 | 0.000417 | Strip2 |
| ENSMUSG( | 2172.138 | -0.09638 | 0.361014 | 0.792797 | Strn |
| ENSMUSG( | 2355.586 | 0.051514 | 0.579814 | 0.89687 | Strn3 |
| ENSMUSG( | 7881.803 | 0.203579 | 0.038999 | 0.304448 | Strn4 |
| ENSMUSG( | 1466.927 | -0.39201 | 0.013722 | 0.171117 | Stt3a |
| ENSMUSG( | 3876.276 | -0.2189 | 0.158874 | 0.588112 | Stt3b |
| ENSMUSG( | 3731.642 | -0.00309 | 0.973366 | 0.99675 | Stub1 |
| ENSMUSG( | 10731.41 | -0.01562 | 0.889553 | 0.98353 | Stum |
| ENSMUSG( | 9.564939 | -0.54883 | 0.39182 | NA | Stx11 |
| ENSMUSG( | 2791.699 | 0.072759 | 0.455818 | 0.838601 | Stx12 |
| ENSMUSG( | 2093.698 | 0.083452 | 0.367239 | 0.795221 | Stx16 |
| ENSMUSG( | 861.1405 | 0.034912 | 0.730987 | 0.944956 | Stx17 |
| ENSMUSG( | 744.9897 | 0.128545 | 0.262517 | 0.713142 | Stx18 |
| ENSMUSG( | 9737.377 | 0.248631 | 0.01232 | 0.160458 | Stx1a |
| ENSMUSG( | 22766.15 | 0.120066 | 0.134911 | 0.548434 | Stx1b |
| ENSMUSG( | 1181.734 | -0.03455 | 0.728673 | 0.944393 | Stx2 |
| ENSMUSG( | 657.1668 | -0.09203 | 0.503937 | 0.866426 | Stx3 |
| ENSMUSG( | 2233.277 | 0.042653 | 0.668231 | 0.927864 | Stx4a |
| ENSMUSG( | 1372.462 | 0.058408 | 0.518193 | 0.873754 | Stx5a |
| ENSMUSG( | 2640.382 | 0.020983 | 0.810133 | 0.963864 | Stx6 |
| ENSMUSG( | 2716.109 | 0.038499 | 0.715381 | 0.941306 | Stx7 |
| ENSMUSG( | 552.5679 | -0.00944 | 0.933781 | 0.993206 | Stx8 |
| ENSMUSG( | 19321.19 | 0.055039 | 0.640665 | 0.916434 | Stxbp1 |
| ENSMUSG( | 1138.514 | 0.090627 | 0.599427 | 0.903801 | Stxbp2 |
| ENSMUSG( | 1090.117 | -0.01941 | 0.865974 | 0.976033 | Stxbp3 |
| ENSMUSG( | 902.0573 | -0.03232 | 0.810467 | 0.964049 | Stxbp4 |
| ENSMUSG( | 1837.022 | 0.122455 | 0.227099 | 0.674223 | Stxbp5 |
| ENSMUSG( | 261.7449 | -0.06733 | 0.754062 | 0.950981 | Stxbp5l |
| ENSMUSG( | 1341.574 | 0.075729 | 0.47111 | 0.846975 | Stxbp6 |
| ENSMUSG( | 9.856588 | -0.49996 | 0.435306 | NA | Styk1 |
| ENSMUSG( | 175.7409 | -0.00256 | 0.989102 | 0.999111 | Styx |
| ENSMUSG( | 17.71919 | -0.06241 | 0.888368 | NA | Styxl2 |
| ENSMUSG( | 3915.508 | 0.002549 | 0.978636 | 0.997428 | Sub1 |
| ENSMUSG( | 3209.107 | 0.043469 | 0.644015 | 0.917536 | Sucla2 |
| ENSMUSG( | 2965.43 | 0.164775 | 0.07719 | 0.429655 | Suclg1 |
| ENSMUSG( | 2700.701 | 0.104883 | 0.363155 | 0.793211 | Suclg2 |
| ENSMUSG( | 1245.602 | -0.35771 | 0.014739 | 0.178407 | Suco |
| ENSMUSG( | 5226.226 | 0.146523 | 0.057251 | 0.369376 | Suds3 |
| ENSMUSG( | 506.3471 | 0.058313 | 0.641968 | 0.916865 | Sufu |
| ENSMUSG( | 345.7623 | -0.08253 | 0.552795 | 0.886141 | Sugct |
| ENSMUSG( | 1407.23 | 0.005117 | 0.959939 | 0.993991 | Sugp1 |
| ENSMUSG( | 2436.774 | 0.2037 | 0.095456 | 0.474828 | Sugp2 |
| ENSMUSG( | 1881.144 | 0.045008 | 0.625639 | 0.912249 | Sugt1 |
| ENSMUSG( | 564.4021 | -0.3404 | 0.004259 | 0.084556 | Sulf1 |
| ENSMUSG( | 4603 | -0.34105 | 0.027975 | 0.255784 | Sulf2 |
| ENSMUSG( | 777.6894 | 0.552178 | 1.13E-05 | 0.001226 | Sult1a1 |
| ENSMUSG( | 213.1968 | -0.01088 | 0.947758 | 0.993821 | Sult2b1 |
| ENSMUSG( | 12698.5 | 0.045251 | 0.650439 | 0.921301 | Sult4a1 |
| ENSMUSG( | 10.87675 | 0.61487 | 0.270618 | NA | Sult5a1 |
| ENSMUSG( | 997.5517 | -0.19156 | 0.161987 | 0.592429 | Sumf1 |
| ENSMUSG( | 517.6879 | -0.14426 | 0.278727 | 0.728705 | Sumf2 |
| ENSMUSG( | 1270.629 | 0.134214 | 0.217677 | 0.662824 | Sumo1 |
| ENSMUSG( | 1277.388 | 0.109638 | 0.297337 | 0.745872 | Sumo2 |
| ENSMUSG( | 2784.961 | -0.04364 | 0.622879 | 0.911589 | Sumo3 |
| ENSMUSG( | 3247.904 | -0.15868 | 0.182925 | 0.620462 | Sun1 |
| ENSMUSG( | 1972.858 | -0.38362 | 0.001252 | 0.038624 | Sun2 |
| ENSMUSG( | 592.2316 | 0.101008 | 0.356465 | 0.788187 | Suox |
| ENSMUSG( | 1976.565 | 0.140695 | 0.112712 | 0.506698 | Supt16 |
| ENSMUSG( | 1011.093 | 0.099729 | 0.397503 | 0.812127 | Supt20 |
| ENSMUSG( | 231.5766 | 0.063255 | 0.679896 | 0.930983 | Supt3 |
| ENSMUSG( | 737.9412 | -0.00076 | 0.994431 | 0.999369 | Supt4a |
| ENSMUSG( | 4016.438 | 0.177659 | 0.075991 | 0.426896 | Supt5 |
| ENSMUSG( | 5149.604 | 0.051553 | 0.563466 | 0.889914 | Supt6 |
| ENSMUSG( | 460.5528 | 0.143254 | 0.247495 | 0.696522 | Supt7l |
| ENSMUSG( | 596.6393 | 0.107728 | 0.336586 | 0.777 | Supv3l1 |
| ENSMUSG( | 2003.536 | -0.1142 | 0.208899 | 0.650854 | Surf1 |
| ENSMUSG( | 633.4567 | -0.02016 | 0.852583 | 0.973168 | Surf2 |
| ENSMUSG( | 3261.118 | -0.27177 | 0.042556 | 0.316636 | Surf4 |
| ENSMUSG( | 542.8656 | 0.136079 | 0.239978 | 0.687848 | Surf6 |
| ENSMUSG( | 326.7837 | -0.32814 | 0.023084 | 0.233188 | Susd1 |
| ENSMUSG( | 276.1329 | -0.41956 | 0.005647 | 0.100551 | Susd2 |
| ENSMUSG( | 111.1858 | -0.1151 | 0.633669 | 0.914073 | Susd3 |
| ENSMUSG( | 5192.805 | -0.07828 | 0.509619 | 0.869553 | Susd4 |
| ENSMUSG( | 327.7146 | -0.29673 | 0.129933 | 0.540113 | Susd5 |
| ENSMUSG( | 1839.195 | -0.10111 | 0.460966 | 0.841042 | Susd6 |
| ENSMUSG( | 1132.803 | 0.150804 | 0.100765 | 0.484034 | Suv39h1 |
| ENSMUSG( | 91.25038 | -0.17715 | 0.408112 | 0.818167 | Suv39h2 |
| ENSMUSG( | 1117.679 | 0.261422 | 0.013219 | 0.167118 | Suz12 |
| ENSMUSG( | 15010.12 | -0.40784 | 0.001739 | 0.047093 | Sv2a |
| ENSMUSG( | 11013.09 | -0.29395 | 0.064635 | 0.393981 | Sv2b |
| ENSMUSG( | 1099.058 | -0.73073 | 9.14E-06 | 0.001083 | Sv2c |
| ENSMUSG( | 614.09 | 0.345472 | 0.007129 | 0.116294 | Svbp |
| ENSMUSG( | 44.85846 | 0.090256 | 0.784443 | 0.958003 | Svep1 |
| ENSMUSG( | 644.462 | 0.267202 | 0.017449 | 0.197914 | Svil |
| ENSMUSG( | 3764.664 | -0.06517 | 0.560943 | 0.888934 | Svip |
| ENSMUSG( | 6900.906 | -0.18587 | 0.146494 | 0.568435 | Svop |
| ENSMUSG( | 1187.076 | 0.15658 | 0.231651 | 0.67949 | Swap70 |
| ENSMUSG( | 3125.763 | 0.026972 | 0.80767 | 0.963257 | Swi5 |
| ENSMUSG( | 450.6155 | 0.053447 | 0.662667 | 0.926303 | Swsap1 |
| ENSMUSG( | 213.5422 | 0.089497 | 0.576786 | 0.895432 | Swt1 |
| ENSMUSG( | 838.9283 | -0.01502 | 0.885772 | 0.983008 | Syap1 |
| ENSMUSG( | 1683.072 | 0.056589 | 0.6163 | 0.910866 | Sybu |
| ENSMUSG( | 96.63936 | 0.33718 | 0.101816 | 0.486216 | Syce2 |
| ENSMUSG( | 52.81181 | -0.40485 | 0.137909 | 0.552903 | Sycp2 |
| ENSMUSG( | 513.769 | 0.059638 | 0.636715 | 0.91503 | Syde1 |
| ENSMUSG( | 190.6186 | -0.04386 | 0.772737 | 0.955272 | Syde2 |
| ENSMUSG( | 1843.279 | 0.078168 | 0.454964 | 0.837725 | Syf2 |
| ENSMUSG( | 285.1894 | 0.015442 | 0.913727 | 0.988483 | Syk |
| ENSMUSG( | 6010.714 | 0.253895 | 0.001841 | 0.048742 | Sympk |
| ENSMUSG( | 18885.94 | 0.129727 | 0.287753 | 0.738505 | Syn1 |
| ENSMUSG( | 11227.36 | 0.21814 | 0.045711 | 0.328047 | Syn2 |
| ENSMUSG( | 151.3575 | -0.03173 | 0.861323 | 0.975026 | Syn3 |
| ENSMUSG( | 27.62259 | -0.34719 | 0.411907 | 0.81942 | Syna |
| ENSMUSG( | 25.28675 | 0.298932 | 0.426282 | 0.824898 | Sync |
| ENSMUSG( | 1945.139 | 0.007725 | 0.942602 | 0.993415 | Syncrip |
| ENSMUSG( | 679.5691 | -0.26105 | 0.027153 | 0.253245 | Syndig1 |
| ENSMUSG( | 1936.683 | -0.80124 | 0.047082 | 0.332172 | Syndig1l |
| ENSMUSG( | 12482.06 | -0.53552 | 7.33E-11 | 4.41E-08 | Syne1 |
| ENSMUSG( | 849.2467 | -0.31008 | 0.003576 | 0.075902 | Syne2 |
| ENSMUSG( | 74.14336 | -0.08647 | 0.732359 | 0.944956 | Syne3 |
| ENSMUSG( | 77.45376 | 0.237114 | 0.392201 | 0.809274 | Syne4 |
| ENSMUSG( | 12101.86 | 0.244805 | 0.002788 | 0.064345 | Syngap1 |
| ENSMUSG( | 19797.21 | -0.1946 | 0.211174 | 0.654296 | Syngr1 |
| ENSMUSG( | 558.5693 | -0.24776 | 0.081473 | 0.441102 | Syngr2 |
| ENSMUSG( | 5982.466 | -0.45099 | 0.000651 | 0.024622 | Syngr3 |
| ENSMUSG( | 5993.126 | 0.095214 | 0.317058 | 0.763823 | Synj1 |
| ENSMUSG( | 1892.756 | 0.122207 | 0.308409 | 0.756439 | Synj2 |
| ENSMUSG( | 837.7862 | 0.101899 | 0.374549 | 0.800445 | Synj2bp |
| ENSMUSG( | 1444.674 | 0.150401 | 0.185803 | 0.623185 | Synm |
| ENSMUSG( | 4889.488 | 0.186403 | 0.107612 | 0.49648 | Synpo |
| ENSMUSG( | 322.8388 | -0.04445 | 0.762639 | 0.953519 | Synpo2 |
| ENSMUSG( | 2312.51 | -0.65149 | 1.09E-05 | 0.001199 | Synpr |
| ENSMUSG( | 2107.337 | -0.00077 | 0.992878 | 0.999327 | Synrg |
| ENSMUSG( | 56345.22 | -0.3413 | 0.002372 | 0.057164 | Syp |
| ENSMUSG( | 1051.078 | -0.45577 | 0.00733 | 0.118347 | Sypl1 |
| ENSMUSG( | 241.5772 | -0.0785 | 0.671695 | 0.928781 | Sypl2 |
| ENSMUSG( | 1064.636 | -0.1859 | 0.092547 | 0.467345 | Sys1 |
| ENSMUSG( | 41994.51 | -0.55476 | 6.09E-05 | 0.004616 | Syt1 |
| ENSMUSG( | 39.16819 | -0.18938 | 0.59507 | 0.903007 | Syt10 |
| ENSMUSG( | 28753.57 | -0.24812 | 0.102166 | 0.486961 | Syt11 |
| ENSMUSG( | 1581.054 | -0.20375 | 0.152468 | 0.578756 | Syt12 |
| ENSMUSG( | 10432.59 | -0.20237 | 0.097196 | 0.47759 | Syt13 |
| ENSMUSG( | 446.2061 | -0.11134 | 0.409852 | 0.818258 | Syt14 |
| ENSMUSG( | 32.1112 | -0.46073 | 0.202405 | 0.642343 | Syt15 |
| ENSMUSG( | 4084.272 | 0.060534 | 0.546877 | 0.884974 | Syt16 |
| ENSMUSG( | 1375.089 | 0.234842 | 0.030619 | 0.266434 | Syt17 |
| ENSMUSG( | 1251.867 | -0.51705 | 0.000125 | 0.007476 | Syt2 |
| ENSMUSG( | 2147.621 | -0.34647 | 0.001444 | 0.0419 | Syt3 |
| ENSMUSG( | 3871.716 | -0.53183 | 0.000496 | 0.020385 | Syt4 |
| ENSMUSG( | 7515.775 | -0.35316 | 0.000533 | 0.021434 | Syt5 |
| ENSMUSG( | 973.8257 | -0.63074 | 4.64E-07 | 0.0001 | Syt6 |
| ENSMUSG( | 16272.81 | 0.007664 | 0.941254 | 0.993415 | Syt7 |
| ENSMUSG( | 309.5638 | -0.42834 | 0.004676 | 0.089972 | Syt9 |
| ENSMUSG( | 33.0629 | 0.485527 | 0.153981 | 0.580901 | Sytl1 |
| ENSMUSG( | 2063.971 | -0.08539 | 0.410173 | 0.818258 | Sytl2 |
| ENSMUSG( | 87.42623 | 0.374974 | 0.129923 | 0.540113 | Sytl4 |
| ENSMUSG( | 271.6189 | -0.05106 | 0.732458 | 0.944956 | Sytl5 |
| ENSMUSG( | 2122.086 | -0.31717 | 0.020202 | 0.214698 | Syvn1 |
| ENSMUSG( | 2513.198 | 0.034542 | 0.693741 | 0.934202 | Szrd1 |
| ENSMUSG( | 1641.723 | 0.186721 | 0.076892 | 0.429306 | Szt2 |
| ENSMUSG( | 23.06993 | -0.15963 | 0.706993 | 0.93949 | T2 |
| ENSMUSG( | 808.0672 | -0.07122 | 0.533914 | 0.88018 | Tab1 |
| ENSMUSG( | 2820.21 | 0.049648 | 0.569771 | 0.893041 | Tab2 |
| ENSMUSG( | 418.6325 | 0.061779 | 0.597024 | 0.903526 | Tab3 |
| ENSMUSG( | 752.6145 | -1.17419 | 8.03E-10 | 3.83E-07 | Tac1 |
| ENSMUSG( | 276.5884 | -0.7397 | 2.02E-06 | 0.000331 | Tac2 |
| ENSMUSG( | 7468.359 | -0.08717 | 0.393002 | 0.809274 | Tacc1 |
| ENSMUSG( | 4257.454 | 0.102909 | 0.485288 | 0.857408 | Tacc2 |

| | | | | | |
|---|---|---|---|---|---|
| ENSMUSG( | 441.3298 | 0.129424 | 0.365177 | 0.914449 | Syt14 |
| ENSMUSG( | 33.69396 | 0.005099 | 0.988885 | 0.997886 | Syt15 |
| ENSMUSG( | 3819.86 | 0.135629 | 0.257063 | 0.883082 | Syt16 |
| ENSMUSG( | 1225.393 | 0.173275 | 0.205351 | 0.864412 | Syt17 |
| ENSMUSG( | 1353.635 | 0.027548 | 0.878614 | 0.989644 | Syt2 |
| ENSMUSG( | 2248.844 | 0.077688 | 0.558312 | 0.93957 | Syt3 |
| ENSMUSG( | 4576.64 | 0.259014 | 0.135362 | 0.825081 | Syt4 |
| ENSMUSG( | 7805.125 | 0.047253 | 0.714721 | 0.964986 | Syt5 |
| ENSMUSG( | 1053.798 | -0.07134 | 0.674968 | 0.95694 | Syt6 |
| ENSMUSG( | 15702.16 | 0.169893 | 0.111561 | 0.808371 | Syt7 |
| ENSMUSG( | 308.4132 | -0.15406 | 0.389404 | 0.917257 | Syt9 |
| ENSMUSG( | 26.95664 | 0.201812 | 0.650578 | 0.95422 | Sytl1 |
| ENSMUSG( | 1823.783 | -0.17458 | 0.161218 | 0.837724 | Sytl2 |
| ENSMUSG( | 61.86415 | -0.36637 | 0.184111 | 0.852618 | Sytl4 |
| ENSMUSG( | 226.8324 | -0.33266 | 0.059174 | 0.729429 | Sytl5 |
| ENSMUSG( | 2308.104 | 0.207844 | 0.155781 | 0.837724 | Syvn1 |
| ENSMUSG( | 2094.31 | -0.23446 | 0.033728 | 0.647113 | Szrd1 |
| ENSMUSG( | 1474.631 | 0.148366 | 0.314319 | 0.900949 | Szt2 |
| ENSMUSG( | 24.05094 | 0.206387 | 0.606266 | 0.949497 | T2 |
| ENSMUSG( | 736.6179 | -0.06763 | 0.614692 | 0.951177 | Tab1 |
| ENSMUSG( | 2584.811 | 0.064881 | 0.570109 | 0.941777 | Tab2 |
| ENSMUSG( | 381.7072 | 0.058909 | 0.68275 | 0.958678 | Tab3 |
| ENSMUSG( | 1032.486 | 0.228911 | 0.151969 | 0.837724 | Tac1 |
| ENSMUSG( | 311.7681 | -0.02847 | 0.86349 | 0.986547 | Tac2 |
| ENSMUSG( | 7195.951 | 0.076936 | 0.511976 | 0.930005 | Tacc1 |
| ENSMUSG( | 4023.179 | 0.207155 | 0.187017 | 0.856099 | Tacc2 |
| ENSMUSG( | 242.8788 | 0.03686 | 0.816864 | 0.979198 | Tacc3 |
| ENSMUSG( | 281.4915 | -0.02376 | 0.877492 | 0.989644 | Taco1 |
| ENSMUSG( | 148.3748 | 0.081465 | 0.665313 | 0.956766 | Tacr1 |
| ENSMUSG( | 75.48154 | 0.297395 | 0.283975 | 0.892412 | Tacr3 |
| ENSMUSG( | 524.8181 | 0.037689 | 0.774834 | 0.974931 | Tada1 |
| ENSMUSG( | 626.4339 | -0.07901 | 0.596823 | 0.946541 | Tada2a |
| ENSMUSG( | 993.6245 | 0.00213 | 0.98622 | 0.997188 | Tada2b |
| ENSMUSG( | 845.7824 | -0.03087 | 0.818847 | 0.979198 | Tada3 |
| ENSMUSG( | 2776.634 | -0.09557 | 0.381752 | 0.917257 | Taf1 |
| ENSMUSG( | 1820.61 | -0.24775 | 0.050545 | 0.708889 | Taf10 |
| ENSMUSG( | 696.5887 | -0.12276 | 0.369911 | 0.915491 | Taf11 |
| ENSMUSG( | 351.048 | -0.11889 | 0.427795 | 0.917257 | Taf12 |
| ENSMUSG( | 1675.092 | -0.18281 | 0.235561 | 0.871807 | Taf13 |
| ENSMUSG( | 1530.649 | -0.18472 | 0.185662 | 0.855305 | Taf15 |
| ENSMUSG( | 252.1134 | -0.03326 | 0.850082 | 0.985538 | Taf1a |
| ENSMUSG( | 369.951 | -0.11165 | 0.428584 | 0.917257 | Taf1b |
| ENSMUSG( | 399.5748 | -0.02568 | 0.867744 | 0.98781 | Taf1c |
| ENSMUSG( | 343.1992 | 0.308078 | 0.107872 | 0.804969 | Taf1d |
| ENSMUSG( | 1026.557 | 0.162885 | 0.148922 | 0.836241 | Taf2 |
| ENSMUSG( | 528.2463 | 0.260219 | 0.071399 | 0.761942 | Taf3 |
| ENSMUSG( | 758.687 | 0.089838 | 0.489627 | 0.925858 | Taf4 |
| ENSMUSG( | 77.08003 | 0.062096 | 0.793256 | 0.97681 | Taf4b |
| ENSMUSG( | 139.0977 | 0.096938 | 0.613963 | 0.951177 | Taf5 |
| ENSMUSG( | 486.0429 | 0.090174 | 0.530291 | 0.934795 | Taf5l |
| ENSMUSG( | 960.1307 | 0.039589 | 0.734031 | 0.967228 | Taf6 |
| ENSMUSG( | 245.7611 | 0.085019 | 0.603848 | 0.948137 | Taf6l |
| ENSMUSG( | 133.4573 | 0.045141 | 0.818309 | 0.979198 | Taf7 |
| ENSMUSG( | 500.5817 | 0.013727 | 0.920421 | 0.992644 | Taf8 |
| ENSMUSG( | 173.5566 | 0.246778 | 0.172911 | 0.843702 | Taf9 |
| ENSMUSG( | 760.8906 | -0.09443 | 0.52223 | 0.932681 | Taf9b |
| ENSMUSG( | 998.0109 | -0.07239 | 0.612777 | 0.951071 | Tafa1 |
| ENSMUSG( | 878.7712 | 0.165258 | 0.304352 | 0.898538 | Tafa2 |
| ENSMUSG( | 22.16903 | -0.39176 | 0.404245 | 0.917257 | Tafa4 |
| ENSMUSG( | 4487.722 | -0.03301 | 0.752576 | 0.970521 | Tafa5 |
| ENSMUSG( | 1154.404 | 0.188607 | 0.229172 | 0.868507 | Tafazzin |
| ENSMUSG( | 29.69209 | 0.044751 | 0.909248 | 0.991958 | Tagap |
| ENSMUSG( | 550.3891 | 0.12642 | 0.321367 | 0.904019 | Tagap1 |
| ENSMUSG( | 70.2941 | -0.41616 | 0.133848 | 0.822912 | Tagln |
| ENSMUSG( | 426.1431 | 0.503703 | 0.003702 | 0.2429 | Tagln2 |
| ENSMUSG( | 5560.525 | -0.30321 | 0.013526 | 0.464932 | Tagln3 |
| ENSMUSG( | 75.9027 | 0.659528 | 0.023789 | 0.588226 | Tal1 |
| ENSMUSG( | 2383.39 | -0.22728 | 0.058898 | 0.729034 | Taldo1 |
| ENSMUSG( | 1014.302 | -0.15248 | 0.303458 | 0.898538 | Tamalin |
| ENSMUSG( | 183.9071 | -0.11541 | 0.530126 | 0.934795 | Tamm41 |
| ENSMUSG( | 962.7341 | 0.043707 | 0.709143 | 0.964042 | Tanc1 |
| ENSMUSG( | 3897.12 | 0.047332 | 0.669269 | 0.956766 | Tanc2 |
| ENSMUSG( | 1449.26 | -0.01967 | 0.862386 | 0.986509 | Tango2 |
| ENSMUSG( | 143.1862 | 0.078108 | 0.67398 | 0.95694 | Tango6 |
| ENSMUSG( | 1213.048 | 0.062379 | 0.62463 | 0.953234 | Tank |
| ENSMUSG( | 3597.643 | -0.03869 | 0.771477 | 0.974236 | Taok1 |
| ENSMUSG( | 4267.553 | 0.035845 | 0.735475 | 0.967344 | Taok2 |
| ENSMUSG( | 1642.067 | -0.08293 | 0.558052 | 0.939565 | Taok3 |
| ENSMUSG( | 172.4392 | -0.04287 | 0.83669 | 0.981335 | Tap1 |
| ENSMUSG( | 464.7219 | 0.096055 | 0.554286 | 0.939565 | Tap2 |
| ENSMUSG( | 1347.282 | 0.157704 | 0.316674 | 0.901868 | Tapbp |
| ENSMUSG( | 307.5568 | 0.003201 | 0.984939 | 0.997067 | Tapbpl |
| ENSMUSG( | 1911.468 | 0.250249 | 0.109436 | 0.805353 | Tapt1 |
| ENSMUSG( | 221.275 | 0.116466 | 0.499477 | 0.927825 | Tarbp1 |
| ENSMUSG( | 452.4275 | 0.173649 | 0.24635 | 0.877904 | Tarbp2 |
| ENSMUSG( | 3616.318 | 0.081082 | 0.49368 | 0.927372 | Tardbp |
| ENSMUSG( | 692.908 | 0.066394 | 0.584947 | 0.944975 | Tars1 |
| ENSMUSG( | 915.5492 | -0.03603 | 0.781421 | 0.976106 | Tars2 |
| ENSMUSG( | 630.0311 | 0.027233 | 0.82479 | 0.979198 | Tars3 |
| ENSMUSG( | 13.03856 | 0.383584 | 0.482274 | 0.924578 | Tas1r3 |
| ENSMUSG( | 11.66277 | 0.607971 | 0.316171 | 0.901868 | Tasl |
| ENSMUSG( | 1011.131 | 0.284042 | 0.041201 | 0.687358 | Tasor |
| ENSMUSG( | 683.3713 | 0.198844 | 0.291885 | 0.897854 | Tasor2 |
| ENSMUSG( | 418.265 | -0.00918 | 0.956761 | 0.994798 | Tasp1 |
| ENSMUSG( | 240.0616 | 0.027038 | 0.870536 | 0.988445 | Tatdn1 |
| ENSMUSG( | 1177.116 | 0.029731 | 0.787339 | 0.97637 | Tatdn2 |
| ENSMUSG( | 372.3165 | 0.002623 | 0.989112 | 0.997893 | Tatdn3 |
| ENSMUSG( | 3854.908 | -0.04884 | 0.642357 | 0.953903 | Tax1bp1 |
| ENSMUSG( | 158.7219 | 0.184799 | 0.310188 | 0.898852 | Tax1bp3 |
| ENSMUSG( | 128.488 | -0.03221 | 0.871439 | 0.988779 | Tbata |
| ENSMUSG( | 645.761 | 0.25867 | 0.089784 | 0.79071 | Tbc1d1 |
| ENSMUSG( | 693.7116 | 0.048503 | 0.704922 | 0.962741 | Tbc1d10a |
| ENSMUSG( | 2115.156 | -0.05693 | 0.671401 | 0.95694 | Tbc1d10b |
| ENSMUSG( | 18.62942 | 0.0883 | 0.86692 | 0.987487 | Tbc1d10c |
| ENSMUSG( | 795.6655 | 0.019974 | 0.896406 | 0.990558 | Tbc1d12 |
| ENSMUSG( | 1435.22 | 0.073799 | 0.518872 | 0.930939 | Tbc1d13 |
| ENSMUSG( | 2601.134 | 0.056235 | 0.587831 | 0.944975 | Tbc1d14 |
| ENSMUSG( | 618.5763 | 0.093464 | 0.440368 | 0.921166 | Tbc1d15 |
| ENSMUSG( | 3375.219 | 0.055399 | 0.58517 | 0.944975 | Tbc1d16 |
| ENSMUSG( | 1126.002 | 0.03322 | 0.777887 | 0.976106 | Tbc1d17 |
| ENSMUSG( | 774.4202 | 0.06407 | 0.609143 | 0.950335 | Tbc1d19 |
| ENSMUSG( | 147.5827 | 0.050554 | 0.806768 | 0.978139 | Tbc1d2 |
| ENSMUSG( | 2319.665 | 0.026119 | 0.801747 | 0.97681 | Tbc1d20 |
| ENSMUSG( | 1093.17 | -0.14456 | 0.226729 | 0.868507 | Tbc1d22a |
| ENSMUSG( | 1578.427 | -0.0597 | 0.602574 | 0.94789 | Tbc1d22b |
| ENSMUSG( | 1066.227 | 0.14528 | 0.197186 | 0.862161 | Tbc1d23 |
| ENSMUSG( | 1901.855 | 0.048871 | 0.679305 | 0.957928 | Tbc1d24 |
| ENSMUSG( | 225.0854 | -0.17486 | 0.358985 | 0.914004 | Tbc1d25 |
| ENSMUSG( | 547.6878 | 0.092044 | 0.502629 | 0.927825 | Tbc1d2b |
| ENSMUSG( | 1525.757 | -0.018 | 0.898035 | 0.990558 | Tbc1d30 |
| ENSMUSG( | 190.4242 | 0.292929 | 0.139266 | 0.830678 | Tbc1d31 |
| ENSMUSG( | 359.62 | -0.15345 | 0.305251 | 0.898538 | Tbc1d32 |
| ENSMUSG( | 351.5484 | 0.837034 | 5.33E-06 | 0.002278 | Tbc1d4 |
| ENSMUSG( | 1162.504 | -0.07867 | 0.489927 | 0.925987 | Tbc1d5 |
| ENSMUSG( | 384.4735 | -0.29135 | 0.047198 | 0.700856 | Tbc1d7 |
| ENSMUSG( | 670.8298 | -0.20794 | 0.180002 | 0.850372 | Tbc1d8 |
| ENSMUSG( | 146.6643 | 0.254149 | 0.248639 | 0.877904 | Tbc1d8b |
| ENSMUSG( | 3046.884 | 0.044401 | 0.66825 | 0.956766 | Tbc1d9 |
| ENSMUSG( | 24814.93 | -0.06786 | 0.540541 | 0.937383 | Tbc1d9b |
| ENSMUSG( | 814.6045 | -0.16421 | 0.311752 | 0.900031 | Tbca |
| ENSMUSG( | 2243.545 | -0.11746 | 0.340826 | 0.907972 | Tbcb |
| ENSMUSG( | 252.9299 | -0.31 | 0.100786 | 0.801456 | Tbcc |
| ENSMUSG( | 471.0518 | 0.096434 | 0.491534 | 0.926825 | Tbccd1 |
| ENSMUSG( | 1278.841 | 0.034427 | 0.761126 | 0.972695 | Tbcd |
| ENSMUSG( | 1191.468 | 0.059097 | 0.76119 | 0.972695 | Tbce |
| ENSMUSG( | 4158.227 | -0.03754 | 0.779579 | 0.976106 | Tbcel |
| ENSMUSG( | 700.9049 | 0.036266 | 0.799507 | 0.97681 | Tbck |
| ENSMUSG( | 883.5211 | 0.233863 | 0.061594 | 0.736111 | Tbk1 |
| ENSMUSG( | 2788.727 | 0.266646 | 0.02157 | 0.575341 | Tbkbp1 |
| ENSMUSG( | 2156.68 | -0.06002 | 0.557174 | 0.939565 | Tbl1x |
| ENSMUSG( | 1892.678 | -0.02298 | 0.859149 | 0.985884 | Tbl1xr1 |
| ENSMUSG( | 1157.126 | 0.244321 | 0.137782 | 0.828492 | Tbl2 |
| ENSMUSG( | 639.4082 | -0.07759 | 0.541867 | 0.937841 | Tbl3 |
| ENSMUSG( | 458.1469 | 0.230333 | 0.079481 | 0.77458 | Tbp |
| ENSMUSG( | 968.338 | 0.005132 | 0.970447 | 0.99594 | Tbpl1 |
| ENSMUSG( | 2252.154 | 0.08705 | 0.427147 | 0.917257 | Tbr1 |
| ENSMUSG( | 680.4668 | -0.02065 | 0.895649 | 0.990558 | Tbrg1 |
| ENSMUSG( | 856.0599 | -0.05125 | 0.712259 | 0.964363 | Tbrg4 |
| ENSMUSG( | 54.09724 | 0.608929 | 0.051597 | 0.711677 | Tbx1 |
| ENSMUSG( | 98.37883 | 0.23934 | 0.277763 | 0.891468 | Tbx15 |
| ENSMUSG( | 97.83981 | 0.203243 | 0.38348 | 0.917257 | Tbx18 |
| ENSMUSG( | 94.29523 | 0.842347 | 0.004338 | 0.268942 | Tbx2 |
| ENSMUSG( | 131.0528 | 1.139598 | 4.61E-07 | 0.00026 | Tbx3 |
| ENSMUSG( | 44.43106 | 0.584418 | 0.140888 | 0.830678 | Tbx3os1 |
| ENSMUSG( | 14.27467 | 0.192476 | 0.729062 | 0.966913 | Tbx4 |
| ENSMUSG( | 30.49209 | 0.523946 | 0.226969 | 0.868507 | Tbx6 |
| ENSMUSG( | 64.46267 | 1.197777 | 0.000396 | 0.057854 | Tbxa2r |
| ENSMUSG( | 179.9779 | -0.15242 | 0.540088 | 0.937383 | Tbxas1 |

| | | | | | | | | | | | |
|---|---|---|---|---|---|---|---|---|---|---|---|
| ENSMUSG( | Tcp11 | 16.11799 | -0.26925 | 0.569215 | NA | ENSMUSG( | Tcp11 | 16.1178 | 0.039371 | 0.936143 | 0.993573 |
| ENSMUSG( | Tcp11l1 | 769.0726 | 0.077679 | 0.521103 | 0.875186 | ENSMUSG( | Tcp11l1 | 650.4929 | -0.14089 | 0.294976 | 0.898538 |
| ENSMUSG( | Tcp11l2 | 717.2032 | 0.073321 | 0.513985 | 0.872055 | ENSMUSG( | Tcp11l2 | 643.0379 | 0.028181 | 0.828329 | 0.980224 |
| ENSMUSG( | Tcta | 1074.415 | -0.06638 | 0.543336 | 0.883854 | ENSMUSG( | Tcta | 962.6835 | -0.11235 | 0.354841 | 0.912411 |
| ENSMUSG( | Tcte1 | 85.49818 | -0.09629 | 0.663241 | 0.9265 | ENSMUSG( | Tcte1 | 83.00488 | 0.104575 | 0.675101 | 0.95694 |
| ENSMUSG( | Tcte2 | 115.4548 | 0.095743 | 0.608571 | 0.907154 | ENSMUSG( | Tcte2 | 95.92595 | -0.18663 | 0.39457 | 0.917257 |
| ENSMUSG( | Tctn1 | 911.7519 | -0.12795 | 0.310885 | 0.758963 | ENSMUSG( | Tctn1 | 889.3269 | 0.077462 | 0.599549 | 0.947071 |
| ENSMUSG( | Tctn2 | 841.7372 | 0.033163 | 0.789417 | 0.959415 | ENSMUSG( | Tctn2 | 822.8206 | 0.232468 | 0.093522 | 0.795507 |
| ENSMUSG( | Tctn3 | 822.5312 | 0.003468 | 0.979541 | 0.997428 | ENSMUSG( | Tctn3 | 808.3919 | 0.21851 | 0.132377 | 0.822912 |
| ENSMUSG( | Tdg | 818.9317 | 0.062104 | 0.528631 | 0.878365 | ENSMUSG( | Tdg | 483.3058 | -1.62333 | 1.03E-32 | 4.49E-29 |
| ENSMUSG( | Tdo2 | 24.17162 | -0.27325 | 0.500149 | 0.863455 | ENSMUSG( | Tdo2 | 23.07877 | -0.07533 | 0.871704 | 0.988779 |
| ENSMUSG( | Tdp1 | 379.5183 | 0.271503 | 0.051124 | 0.346957 | ENSMUSG( | Tdp1 | 335.5537 | 0.188872 | 0.200116 | 0.863016 |
| ENSMUSG( | Tdp2 | 366.5895 | 0.030154 | 0.806723 | 0.962932 | ENSMUSG( | Tdp2 | 346.578 | 0.129363 | 0.382715 | 0.917257 |
| ENSMUSG( | Tdrd1 | 16.38759 | 0.394961 | 0.486763 | NA | ENSMUSG( | Tdrd1 | 13.49668 | 0.128706 | 0.826557 | 0.979952 |
| ENSMUSG( | Tdrd12 | 21.33659 | 0.001757 | 0.996679 | 0.99943 | ENSMUSG( | Tdrd12 | 19.55942 | -0.01446 | 0.97671 | 0.996416 |
| ENSMUSG( | Tdrd3 | 811.7336 | 0.100172 | 0.335417 | 0.776877 | ENSMUSG( | Tdrd3 | 711.3055 | -0.01602 | 0.905652 | 0.991089 |
| ENSMUSG( | Tdrd5 | 34.11885 | 0.13594 | 0.704614 | 0.938303 | ENSMUSG( | Tdrd5 | 32.77228 | 0.303554 | 0.426018 | 0.917257 |
| ENSMUSG( | Tdrd6 | 22.30106 | -0.33878 | 0.405729 | 0.81652 | ENSMUSG( | Tdrd6 | 27.16904 | 0.50125 | 0.199792 | 0.863016 |
| ENSMUSG( | Tdrd7 | 1303.476 | 0.063337 | 0.535115 | 0.880733 | ENSMUSG( | Tdrd7 | 1127.006 | -0.09031 | 0.427976 | 0.917257 |
| ENSMUSG( | Tdrd9 | 17.81763 | -0.19936 | 0.699284 | NA | ENSMUSG( | Tdrd9 | 14.40509 | -0.62975 | 0.272731 | 0.888825 |
| ENSMUSG( | Tdrkh | 1435.89 | 0.131733 | 0.198416 | 0.638387 | ENSMUSG( | Tdrkh | 1210.688 | -0.094 | 0.485881 | 0.92541 |
| ENSMUSG( | Tdrp | 361.8013 | 0.031624 | 0.815353 | 0.965049 | ENSMUSG( | Tdrp | 365.6662 | 0.322553 | 0.042112 | 0.690304 |
| ENSMUSG( | Tead1 | 1505.732 | 0.197775 | 0.122504 | 0.528322 | ENSMUSG( | Tead1 | 1304.425 | 0.057477 | 0.672684 | 0.95694 |
| ENSMUSG( | Tead2 | 265.7043 | 0.226639 | 0.097762 | 0.478264 | ENSMUSG( | Tead2 | 247.8233 | 0.287596 | 0.06938 | 0.755467 |
| ENSMUSG( | Tead3 | 164.5208 | -0.10262 | 0.582229 | 0.897918 | ENSMUSG( | Tead3 | 156.2606 | 0.031248 | 0.870624 | 0.988445 |
| ENSMUSG( | Tead4 | 16.98782 | 0.825328 | 0.090328 | NA | ENSMUSG( | Tead4 | 19.52832 | 1.367332 | 0.023713 | 0.588226 |
| ENSMUSG( | Tec | 82.04782 | -0.29257 | 0.210817 | 0.653564 | ENSMUSG( | Tec | 79.95471 | -0.06153 | 0.809755 | 0.978779 |
| ENSMUSG( | Tecpr1 | 3710.872 | 0.129944 | 0.162797 | 0.593778 | ENSMUSG( | Tecpr1 | 3303.658 | 0.06612 | 0.568959 | 0.941777 |
| ENSMUSG( | Tecpr2 | 2702.697 | -0.04844 | 0.632584 | 0.914073 | ENSMUSG( | Tecpr2 | 2581.276 | 0.089496 | 0.441276 | 0.921205 |
| ENSMUSG( | Tecr | 13766.75 | -0.35143 | 0.000571 | 0.022438 | ENSMUSG( | Tecr | 13791.7 | -0.05521 | 0.648708 | 0.95422 |
| ENSMUSG( | Tecta | 16.18311 | -0.65296 | 0.19315 | NA | ENSMUSG( | Tecta | 19.40725 | 0.246134 | 0.606045 | 0.949497 |
| ENSMUSG( | Tectb | 12.05197 | -0.76224 | 0.20093 | NA | ENSMUSG( | Tectb | 9.707516 | -1.35244 | 0.042169 | NA |
| ENSMUSG( | Tedc1 | 60.13413 | 0.054146 | 0.842865 | 0.97173 | ENSMUSG( | Tedc1 | 55.69321 | 0.124104 | 0.683134 | 0.958678 |
| ENSMUSG( | Tedc2 | 241.8608 | 0.099999 | 0.533259 | 0.879905 | ENSMUSG( | Tedc2 | 211.9336 | -0.01577 | 0.927856 | 0.99306 |
| ENSMUSG( | Teddm2 | 637.1458 | 0.112826 | 0.419124 | 0.821966 | ENSMUSG( | Teddm2 | 483.065 | -0.43854 | 0.009823 | 0.406458 |
| ENSMUSG( | Tef | 12363.67 | 0.10403 | 0.213602 | 0.657797 | ENSMUSG( | Tef | 12181.62 | 0.315728 | 0.001711 | 0.152157 |
| ENSMUSG( | Tefm | 107.6327 | -0.03287 | 0.86702 | 0.976541 | ENSMUSG( | Tefm | 88.8089 | -0.33675 | 0.162929 | 0.837925 |
| ENSMUSG( | Tek | 518.7296 | -0.15 | 0.688974 | 0.933441 | ENSMUSG( | Tek | 628.8999 | 0.623628 | 0.22543 | 0.868468 |
| ENSMUSG( | Tekt1 | 71.7715 | -0.31052 | 0.185354 | 0.623185 | ENSMUSG( | Tekt1 | 71.86855 | -0.01478 | 0.95791 | 0.994798 |
| ENSMUSG( | Tekt2 | 49.93274 | -1.57405 | 5.22E-08 | 1.69E-05 | ENSMUSG( | Tekt2 | 65.79499 | -0.10543 | 0.719973 | 0.966887 |
| ENSMUSG( | Tekt4 | 64.71519 | 0.790128 | 0.004016 | 0.081522 | ENSMUSG( | Tekt4 | 58.78311 | 0.778916 | 0.007575 | 0.370638 |
| ENSMUSG( | Tekt5 | 241.3986 | -0.0925 | 0.570952 | 0.8933 | ENSMUSG( | Tekt5 | 206.9477 | -0.29205 | 0.141504 | 0.830678 |
| ENSMUSG( | Tektip1 | 19.29813 | -0.32075 | 0.50846 | NA | ENSMUSG( | Tektip1 | 19.05262 | -0.04194 | 0.929251 | 0.99306 |
| ENSMUSG( | Telo2 | 597.0896 | 0.261172 | 0.026092 | 0.247245 | ENSMUSG( | Telo2 | 493.1507 | -0.00449 | 0.975165 | 0.996416 |
| ENSMUSG( | Ten1 | 519.844 | -0.08545 | 0.471243 | 0.847103 | ENSMUSG( | Ten1 | 439.4345 | -0.32411 | 0.025932 | 0.601649 |
| ENSMUSG( | Tenm1 | 834.4057 | -0.44386 | 0.004387 | 0.086463 | ENSMUSG( | Tenm1 | 946.0542 | 0.217994 | 0.21036 | 0.864412 |
| ENSMUSG( | Tenm2 | 2258.82 | -0.18066 | 0.186056 | 0.62352 | ENSMUSG( | Tenm2 | 2468.139 | 0.334687 | 0.032755 | 0.637512 |
| ENSMUSG( | Tenm3 | 2086.132 | -0.1775 | 0.24279 | 0.690275 | ENSMUSG( | Tenm3 | 2167.617 | 0.20575 | 0.261495 | 0.884926 |
| ENSMUSG( | Tenm4 | 3607.101 | -0.36749 | 0.001979 | 0.051182 | ENSMUSG( | Tenm4 | 3713.365 | 0.012386 | 0.929117 | 0.99306 |
| ENSMUSG( | Tent2 | 574.4728 | 0.070083 | 0.547854 | 0.885153 | ENSMUSG( | Tent2 | 524.057 | 0.071953 | 0.594682 | 0.945444 |
| ENSMUSG( | Tent4a | 1370.412 | 0.10637 | 0.249765 | 0.699445 | ENSMUSG( | Tent4a | 1246.084 | 0.102541 | 0.350594 | 0.911652 |
| ENSMUSG( | Tent4b | 987.414 | 0.086176 | 0.383906 | 0.803646 | ENSMUSG( | Tent4b | 931.8899 | 0.181596 | 0.121335 | 0.813384 |
| ENSMUSG( | Tent5a | 123.7602 | -0.54765 | 0.002266 | 0.055678 | ENSMUSG( | Tent5a | 125.5847 | -0.19286 | 0.325572 | 0.905339 |
| ENSMUSG( | Tent5c | 179.6612 | -0.25709 | 0.31306 | 0.76033 | ENSMUSG( | Tent5c | 177.1884 | -0.02798 | 0.919645 | 0.992644 |
| ENSMUSG( | Tep1 | 192.6078 | -0.07904 | 0.61089 | 0.908466 | ENSMUSG( | Tep1 | 176.5257 | -0.04817 | 0.813727 | 0.979198 |
| ENSMUSG( | Tepsin | 541.6265 | 0.245737 | 0.044848 | 0.324745 | ENSMUSG( | Tepsin | 477.5734 | 0.158421 | 0.261157 | 0.884926 |
| ENSMUSG( | Terb1 | 32.17371 | 0.336898 | 0.327703 | 0.772692 | ENSMUSG( | Terb1 | 22.53594 | -0.44961 | 0.277186 | 0.891468 |
| ENSMUSG( | Terf1 | 207.2114 | -0.09092 | 0.588968 | 0.90113 | ENSMUSG( | Terf1 | 198.8068 | 0.05889 | 0.752299 | 0.970471 |
| ENSMUSG( | Terf2 | 1018.991 | 0.049816 | 0.649923 | 0.920899 | ENSMUSG( | Terf2 | 887.9988 | -0.08306 | 0.576485 | 0.942912 |
| ENSMUSG( | Terf2ip | 1061.431 | 0.136631 | 0.225618 | 0.67246 | ENSMUSG( | Terf2ip | 899.4158 | -0.07514 | 0.591234 | 0.944975 |
| ENSMUSG( | Tert | 44.32516 | 0.140738 | 0.621775 | 0.911432 | ENSMUSG( | Tert | 36.70847 | -0.13786 | 0.679106 | 0.957725 |
| ENSMUSG( | Tes | 80.62794 | -0.23351 | 0.410223 | 0.818258 | ENSMUSG( | Tes | 85.38502 | 0.208565 | 0.374838 | 0.915491 |
| ENSMUSG( | Tes3-ps | 9.862838 | 0.195671 | 0.735196 | NA | ENSMUSG( | Tes3-ps | 8.916309 | 0.16893 | 0.789936 | NA |
| ENSMUSG( | Tesc | 864.0597 | -0.25504 | 0.061074 | 0.381679 | ENSMUSG( | Tesc | 809.0923 | -0.17346 | 0.268462 | 0.886755 |
| ENSMUSG( | Tesk1 | 3892.084 | 0.162375 | 0.047238 | 0.332553 | ENSMUSG( | Tesk1 | 3455.486 | 0.091715 | 0.364296 | 0.914223 |
| ENSMUSG( | Tesk2 | 287.0617 | 0.218631 | 0.12573 | 0.53257 | ENSMUSG( | Tesk2 | 256.0516 | 0.166932 | 0.316292 | 0.901868 |
| ENSMUSG( | Tet1 | 491.6689 | 0.06356 | 0.646913 | 0.919062 | ENSMUSG( | Tet1 | 460.4913 | 0.135295 | 0.356575 | 0.913358 |
| ENSMUSG( | Tet2 | 1690.567 | -0.39438 | 4.37E-05 | 0.003581 | ENSMUSG( | Tet2 | 1708.355 | -0.06993 | 0.522231 | 0.932681 |
| ENSMUSG( | Tet3 | 1778.761 | -0.0318 | 0.742695 | 0.948478 | ENSMUSG( | Tet3 | 1631.425 | -0.00971 | 0.933322 | 0.99306 |
| ENSMUSG( | Tex10 | 365.7725 | 0.269238 | 0.031798 | 0.272023 | ENSMUSG( | Tex10 | 331.6103 | 0.253431 | 0.095924 | 0.796133 |
| ENSMUSG( | Tex14 | 14.84724 | 0.158548 | 0.738662 | NA | ENSMUSG( | Tex14 | 11.24525 | -0.4025 | 0.48522 | NA |
| ENSMUSG( | Tex15 | 29.74037 | -0.25849 | 0.448158 | 0.834051 | ENSMUSG( | Tex15 | 37.3084 | 0.588798 | 0.084628 | 0.779799 |
| ENSMUSG( | Tex2 | 3751.348 | -0.00733 | 0.946174 | 0.99369 | ENSMUSG( | Tex2 | 3409.693 | -0.01225 | 0.921504 | 0.992644 |
| ENSMUSG( | Tex261 | 2231.036 | -0.12137 | 0.288176 | 0.738889 | ENSMUSG( | Tex261 | 2138.674 | 0.030407 | 0.824335 | 0.979198 |
| ENSMUSG( | Tex264 | 2677.247 | -0.19601 | 0.100718 | 0.484034 | ENSMUSG( | Tex264 | 2694.269 | 0.098132 | 0.489907 | 0.925987 |
| ENSMUSG( | Tex30 | 35.96219 | -0.23429 | 0.538107 | 0.882548 | ENSMUSG( | Tex30 | 40.62562 | 0.368641 | 0.296926 | 0.898538 |
| ENSMUSG( | Tex52 | 64.05448 | -0.69462 | 0.0069 | 0.114527 | ENSMUSG( | Tex52 | 65.0736 | -0.30569 | 0.283788 | 0.892412 |
| ENSMUSG( | Tex9 | 397.2426 | -0.10497 | 0.392237 | 0.809274 | ENSMUSG( | Tex9 | 355.0546 | -0.16472 | 0.25989 | 0.883971 |
| ENSMUSG( | Tfam | 820.9244 | 0.132319 | 0.220295 | 0.665898 | ENSMUSG( | Tfam | 740.1576 | 0.098292 | 0.454861 | 0.922891 |
| ENSMUSG( | Tfap2b | 23.39013 | -0.07359 | 0.854333 | 0.973491 | ENSMUSG( | Tfap2b | 25.99455 | 0.484206 | 0.24779 | 0.877904 |
| ENSMUSG( | Tfap2c | 10.07143 | 0.082829 | 0.885083 | NA | ENSMUSG( | Tfap2c | 10.64793 | 0.478827 | 0.413328 | NA |
| ENSMUSG( | Tfap2d | 14.26992 | -0.62726 | 0.307549 | NA | ENSMUSG( | Tfap2d | 15.47641 | -0.05205 | 0.915018 | 0.992407 |
| ENSMUSG( | Tfap4 | 69.15073 | -0.11187 | 0.629256 | 0.913965 | ENSMUSG( | Tfap4 | 65.35693 | -0.02178 | 0.935228 | 0.993078 |
| ENSMUSG( | Tfb1m | 92.3928 | 0.162094 | 0.478836 | 0.853862 | ENSMUSG( | Tfb1m | 74.30993 | -0.21235 | 0.427161 | 0.917257 |
| ENSMUSG( | Tfb2m | 351.6157 | -0.00943 | 0.945313 | 0.99365 | ENSMUSG( | Tfb2m | 325.2417 | 0.033559 | 0.822125 | 0.979198 |
| ENSMUSG( | Tfcp2 | 611.9003 | 0.107624 | 0.361743 | 0.792797 | ENSMUSG( | Tfcp2 | 563.2864 | 0.138991 | 0.303919 | 0.898538 |
| ENSMUSG( | Tfcp2l1 | 1153.411 | 0.125572 | 0.248751 | 0.697857 | ENSMUSG( | Tfcp2l1 | 988.8776 | -0.05162 | 0.724331 | 0.966887 |
| ENSMUSG( | Tfdp1 | 1426.983 | 0.119791 | 0.191652 | 0.630674 | ENSMUSG( | Tfdp1 | 1314.975 | 0.147935 | 0.225682 | 0.868468 |
| ENSMUSG( | Tfdp2 | 1432.189 | -0.04532 | 0.625965 | 0.912356 | ENSMUSG( | Tfdp2 | 1300.03 | -0.05676 | 0.618057 | 0.952232 |
| ENSMUSG( | Tfe3 | 1327.117 | -0.04519 | 0.621217 | 0.911432 | ENSMUSG( | Tfe3 | 1230.898 | 0.006089 | 0.955968 | 0.994798 |
| ENSMUSG( | Tfeb | 423.0416 | -0.03501 | 0.760491 | 0.952907 | ENSMUSG( | Tfeb | 428.2592 | 0.262184 | 0.059529 | 0.729747 |
| ENSMUSG( | Tfg | 1005.939 | 0.186955 | 0.056513 | 0.366657 | ENSMUSG( | Tfg | 888.0619 | 0.095841 | 0.466257 | 0.923329 |
| ENSMUSG( | Tfip11 | 1343.47 | 0.130016 | 0.339201 | 0.777534 | ENSMUSG( | Tfip11 | 1165.963 | -0.00735 | 0.960213 | 0.994798 |
| ENSMUSG( | Tfpi | 213.0937 | -0.03165 | 0.845369 | 0.97218 | ENSMUSG( | Tfpi | 215.6966 | 0.261795 | 0.139248 | 0.830678 |
| ENSMUSG( | Tfpt | 516.7781 | -0.0031 | 0.980042 | 0.997491 | ENSMUSG( | Tfpt | 448.592 | -0.15466 | 0.336717 | 0.907972 |
| ENSMUSG( | Tfr2 | 231.9736 | 0.325435 | 0.03588 | 0.290992 | ENSMUSG( | Tfr2 | 200.6904 | 0.195734 | 0.272726 | 0.888825 |
| ENSMUSG( | Tfrc | 4133.313 | -0.14963 | 0.286349 | 0.735717 | ENSMUSG( | Tfrc | 4740.128 | 0.481147 | 0.001248 | 0.127113 |
| ENSMUSG( | Tgds | 153.5101 | 0.135698 | 0.413954 | 0.819998 | ENSMUSG( | Tgds | 146.2622 | 0.256215 | 0.160831 | 0.837724 |
| ENSMUSG( | Tgfa | 2155.401 | -0.3269 | 0.007551 | 0.120197 | ENSMUSG( | Tgfa | 2180.469 | -0.00273 | 0.983724 | 0.996963 |
| ENSMUSG( | Tgfb1 | 471.8627 | -0.27204 | 0.084564 | 0.448139 | ENSMUSG( | Tgfb1 | 496.8558 | 0.161954 | 0.346364 | 0.910966 |
| ENSMUSG( | Tgfb1i1 | 831.5445 | 0.04146 | 0.690809 | 0.933881 | ENSMUSG( | Tgfb1i1 | 727.2541 | -0.07534 | 0.551819 | 0.939565 |
| ENSMUSG( | Tgfb2 | 871.308 | -0.04704 | 0.674461 | 0.929638 | ENSMUSG( | Tgfb2 | 866.8825 | 0.201899 | 0.132495 | 0.822912 |
| ENSMUSG( | Tgfb3 | 647.0278 | -0.14241 | 0.205257 | 0.64654 | ENSMUSG( | Tgfb3 | 593.1259 | -0.12691 | 0.330502 | 0.907398 |
| ENSMUSG( | Tgfbi | 318.3878 | -0.215 | 0.127064 | 0.534624 | ENSMUSG( | Tgfbi | 327.3121 | 0.14846 | 0.379159 | 0.916995 |
| ENSMUSG( | Tgfbr1 | 1401.459 | -0.08594 | 0.616429 | 0.910866 | ENSMUSG( | Tgfbr1 | 1471.396 | 0.31396 | 0.103121 | 0.804311 |
| ENSMUSG( | Tgfbr2 | 856.3405 | -0.1898 | 0.261521 | 0.711485 | ENSMUSG( | Tgfbr2 | 891.4774 | 0.203244 | 0.400996 | 0.917257 |
| ENSMUSG( | Tgfbr3 | 399.6452 | -0.23203 | 0.071978 | 0.414057 | ENSMUSG( | Tgfbr3 | 431.8178 | 0.266425 | 0.110043 | 0.806572 |
| ENSMUSG( | Tgfbr3l | 49.45778 | 0.276899 | 0.354855 | 0.78738 | ENSMUSG( | Tgfbr3l | 47.88559 | 0.448755 | 0.169903 | 0.840806 |
| ENSMUSG( | Tgfbrap1 | 2653.8 | 0.093951 | 0.288703 | 0.739578 | ENSMUSG( | Tgfbrap1 | 2341.006 | 0.002502 | 0.980589 | 0.996479 |
| ENSMUSG( | Tgif1 | 53.45826 | -0.47737 | 0.080624 | 0.438981 | ENSMUSG( | Tgif1 | 58.88309 | 0.114255 | 0.679443 | 0.957966 |
| ENSMUSG( | Tgif2 | 82.99375 | -0.19562 | 0.402127 | 0.815103 | ENSMUSG( | Tgif2 | 77.17189 | -0.14795 | 0.564678 | 0.941206 |
| ENSMUSG( | Tgm2 | 418.9951 | 0.195482 | 0.221071 | 0.666624 | ENSMUSG( | Tgm2 | 443.3767 | 0.5861 | 0.003736 | 0.24422 |
| ENSMUSG( | Tgm3 | 96.45144 | 0.055923 | 0.795923 | 0.960062 | ENSMUSG( | Tgm3 | 85.95133 | 0.002243 | 0.992524 | 0.99874 |
| ENSMUSG( | Tgm4 | 21.08925 | 0.03377 | 0.938136 | NA | ENSMUSG( | Tgm4 | 17.96201 | -0.12961 | 0.790732 | 0.97681 |
| ENSMUSG( | Tgm7 | 15.77835 | 0.284889 | 0.564793 | NA | ENSMUSG( | Tgm7 | 14.7159 | 0.320133 | 0.546945 | 0.938063 |
| ENSMUSG( | Tgoln1 | 5353.483 | -0.12198 | 0.389545 | 0.807428 | ENSMUSG( | Tgoln1 | 5319.296 | 0.13147 | 0.411745 | 0.917257 |
| ENSMUSG( | Tgs1 | 641.8414 | 0.163678 | 0.121372 | 0.525713 | ENSMUSG( | Tgs1 | 589.5757 | 0.18462 | 0.146306 | 0.835973 |
| ENSMUSG( | Tgtp1 | 19.92322 | -0.40629 | 0.368893 | NA | ENSMUSG( | Tgtp1 | 21.55854 | 0.14805 | 0.743994 | 0.96951 |
| ENSMUSG( | Tgtp2 | 16.38436 | -0.57098 | 0.216898 | NA | ENSMUSG( | Tgtp2 | 15.72164 | -0.32868 | 0.530227 | 0.934795 |
| ENSMUSG( | Th | 75.2246 | -0.5262 | 0.173037 | 0.608361 | ENSMUSG( | Th | 83.52564 | 0.076664 | 0.928855 | 0.99306 |
| ENSMUSG( | Tha1 | 174.0309 | 0.124202 | 0.46343 | 0.842789 | ENSMUSG( | Tha1 | 159.8435 | 0.153204 | 0.399245 | 0.917257 |
| ENSMUSG( | Thada | 275.88 | 0.212681 | 0.174667 | 0.610304 | ENSMUSG( | Thada | 242.4977 | 0.129145 | 0.500817 | 0.927825 |
| ENSMUSG( | Thap1 | 285.2042 | 0.088434 | 0.496589 | 0.86175 | ENSMUSG( | Thap1 | 243.2328 | -0.10462 | 0.531583 | 0.934795 |
| ENSMUSG( | Thap11 | 638.7743 | 0.107198 | 0.430648 | 0.826896 | ENSMUSG( | Thap11 | 559.1107 | -0.00928 | 0.947628 | 0.994798 |
| ENSMUSG( | Thap12 | 5188.892 | 0.118181 | 0.30141 | 0.749979 | ENSMUSG( | Thap12 | 4416.249 | -0.07911 | 0.547793 | 0.938063 |
| ENSMUSG( | Thap2 | 671.5422 | 0.128963 | 0.251354 | 0.700869 | ENSMUSG( | Thap2 | 622.8681 | 0.172401 | 0.21885 | 0.866104 |
| ENSMUSG( | Thap3 | 343.3462 | -0.10961 | 0.494219 | 0.860611 | ENSMUSG( | Thap3 | 328.4486 | 0.03501 | 0.848253 | 0.985321 |
| ENSMUSG( | Thap4 | 413.9913 | -0.03613 | 0.794694 | 0.959799 | ENSMUSG( | Thap4 | 350.933 | -0.2584 | 0.114845 | 0.811703 |
| ENSMUSG( | Thap6 | 132.9486 | -0.00196 | 0.992057 | 0.999265 | ENSMUSG( | Thap6 | 119.4424 | -0.04871 | 0.816152 | 0.979198 |
| ENSMUSG( | Thap7 | 669.4961 | 0.139265 | 0.193828 | 0.633085 | ENSMUSG( | Thap7 | 570.3061 | -0.05351 | 0.676985 | 0.957106 |
| ENSMUSG( | Thbd | 900.489 | 0.031215 | 0.850865 | 0.972638 | ENSMUSG( | Thbd | 928.4263 | 0.37048 | 0.024077 | 0.588226 |
| ENSMUSG( | Thbs1 | 217.6063 | -1.08742 | 9.48E-07 | 0.000184 | ENSMUSG( | Thbs1 | 259.0496 | -0.10077 | 0.680876 | 0.958395 |
| ENSMUSG( | Thbs2 | 323.8241 | -0.39629 | 0.006941 | 0.114527 | ENSMUSG( | Thbs2 | 361.7188 | 0.217741 | 0.169743 | 0.840806 |
| ENSMUSG( | Thbs3 | 586.0745 | 0.151461 | 0.336647 | 0.777 | ENSMUSG( | Thbs3 | 523.901 | 0.100881 | 0.57669 | 0.942912 |
| ENSMUSG( | Thbs4 | 794.7489 | -0.12426 | 0.530819 | 0.879289 | ENSMUSG( | Thbs4 | 830.4531 | 0.270701 | 0.235779 | 0.871888 |
| ENSMUSG( | Them4 | 700.968 | -0.011 | 0.936864 | 0.993415 | ENSMUSG( | Them4 | 572.8041 | -0.35555 | 0.036428 | 0.66949 |
| ENSMUSG( | Them6 | 1874.953 | -0.46996 | 4.57E-05 | 0.003685 | ENSMUSG( | Them6 | 2010.276 | 0.040964 | 0.767448 | 0.974236 |
| ENSMUSG( | Themis2 | 65.08993 | -0.43186 | 0.10189 | 0.486318 | ENSMUSG( | Themis2 | 68.59467 | 0.031747 | 0.918983 | 0.992529 |
| ENSMUSG( | Thg1l | 208.7484 | -0.03314 | 0.84342 | 0.971847 | ENSMUSG( | Thg1l | 178.8879 | -0.21832 | 0.238831 | 0.873304 |
| ENSMUSG( | Thnsl1 | 402.914 | 0.030937 | 0.791421 | 0.959461 | ENSMUSG( | Thnsl1 | 347.0372 | -0.1351 | 0.389452 | 0.917257 |
| ENSMUSG( | Thnsl2 | 213.3944 | 0.11594 | 0.464196 | 0.843143 | ENSMUSG( | Thnsl2 | 181.8553 | -0.08729 | 0.677456 | 0.957106 |
| ENSMUSG( | Thoc1 | 1037.297 | 0.105752 | 0.341418 | 0.778828 | ENSMUSG( | Thoc1 | 978.0128 | 0.199353 | 0.125543 | 0.81516 |
| ENSMUSG( | Thoc2 | 1151.892 | 0.017853 | 0.878965 | 0.981436 | ENSMUSG( | Thoc2 | 1118.106 | 0.192043 | 0.147269 | 0.836241 |
| ENSMUSG( | Thoc2l | 678.278 | -0.17031 | 0.184633 | 0.621704 | ENSMUSG( | Thoc2l | 712.3339 | 0.239504 | 0.10078 | 0.801456 |
| ENSMUSG( | Thoc3 | 606.3827 | 0.075625 | 0.556032 | 0.887457 | ENSMUSG( | Thoc3 | 507.3462 | -0.1754 | 0.228081 | 0.868507 |
| ENSMUSG( | Thoc5 | 748.5083 | 0.146094 | 0.273787 | 0.724244 | ENSMUSG( | Thoc5 | 632.4956 | -0.07044 | 0.621326 | 0.953199 |
| ENSMUSG( | Thoc6 | 138.7649 | -0.09417 | 0.6263 | 0.912639 | ENSMUSG( | Thoc6 | 123.0435 | -0.16982 | 0.445782 | 0.921734 |
| ENSMUSG( | Thoc7 | 1272.917 | 0.069474 | 0.446352 | 0.833698 | ENSMUSG( | Thoc7 | 1105.125 | -0.07237 | 0.528872 | 0.934795 |
| ENSMUSG( | Thop1 | 1024.84 | 0.08082 | 0.48169 | 0.855573 | ENSMUSG( | Thop1 | 862.2644 | -0.15086 | 0.241025 | 0.876146 |
| ENSMUSG( | Thpo | 243.8679 | 0.369604 | 0.101179 | 0.484996 | ENSMUSG( | Thpo | 188.2991 | -0.09045 | 0.734394 | 0.967228 |
| ENSMUSG( | Thra | 40291.94 | 0.210294 | 0.003482 | 0.074707 | ENSMUSG( | Thra | 34014.68 | -0.00244 | 0.980367 | 0.996479 |
| ENSMUSG( | Thrap3 | 3045.111 | 0.000868 | 0.99171 | 0.999232 | ENSMUSG( | Thrap3 | 2730.937 | -0.04714 | 0.67351 | 0.95694 |
| ENSMUSG( | Thrb | 1100.18 | 0.217606 | 0.018411 | 0.203839 | ENSMUSG( | Thrb | 959.9899 | 0.097609 | 0.403637 | 0.917257 |
| ENSMUSG( | Thrsp | 1319.552 | 0.407935 | 7.47E-05 | 0.005315 | ENSMUSG( | Thrsp | 931.5374 | -0.31517 | 0.006084 | 0.321954 |
| ENSMUSG( | Thsd1 | 193.3952 | -0.14964 | 0.48786 | 0.858824 | ENSMUSG( | Thsd1 | 196.9336 | 0.188877 | 0.487076 | 0.92541 |
| ENSMUSG( | Thsd4 | 532.8169 | -0.21599 | 0.104636 | 0.490944 | ENSMUSG( | Thsd4 | 547.5276 | 0.144126 | 0.353819 | 0.912411 |
| ENSMUSG( | Thsd7a | 862.799 | -0.63535 | 4.94E-05 | 0.00395 | ENSMUSG( | Thsd7a | 987.2684 | 0.094218 | 0.591534 | 0.944975 |
| ENSMUSG( | Thsd7b | 194.4452 | -0.65129 | 0.002837 | 0.065089 | ENSMUSG( | Thsd7b | 228.2235 | 0.160155 | 0.492543 | 0.927315 |

| | | | | | |
|---|---|---|---|---|---|
| ENSMUSG( | 269.4067 | 0.06534 | 0.650365 | 0.921301 | Tacc3 |
| ENSMUSG( | 338.7389 | 0.235974 | 0.077321 | 0.429743 | Taco1 |
| ENSMUSG( | 116.5645 | -1.07472 | 0.060297 | 0.379667 | Tacr1 |
| ENSMUSG( | 59.53012 | -0.70443 | 0.004438 | 0.086945 | Tacr3 |
| ENSMUSG( | 562.6349 | -0.02998 | 0.793687 | 0.959799 | Tada1 |
| ENSMUSG( | 701.8008 | -0.01565 | 0.89812 | 0.985633 | Tada2a |
| ENSMUSG( | 1063.812 | -0.06708 | 0.489358 | 0.859092 | Tada2b |
| ENSMUSG( | 930.3544 | -0.02521 | 0.831218 | 0.969889 | Tada3 |
| ENSMUSG( | 3226.103 | 0.070758 | 0.497524 | 0.862162 | Taf1 |
| ENSMUSG( | 2201.141 | 0.04523 | 0.613178 | 0.908902 | Taf10 |
| ENSMUSG( | 793.9232 | -0.00694 | 0.948036 | 0.993821 | Taf11 |
| ENSMUSG( | 401.362 | 0.006613 | 0.95749 | 0.993874 | Taf12 |
| ENSMUSG( | 2011.23 | 0.085048 | 0.522468 | 0.875368 | Taf13 |
| ENSMUSG( | 1823.062 | 0.061233 | 0.535988 | 0.88142 | Taf15 |
| ENSMUSG( | 283.4154 | 0.029942 | 0.841263 | 0.971179 | Taf1a |
| ENSMUSG( | 423.6518 | 0.012841 | 0.927381 | 0.991229 | Taf1b |
| ENSMUSG( | 444.9308 | 0.008131 | 0.950037 | 0.993821 | Taf1c |
| ENSMUSG( | 322.7163 | -0.1383 | 0.330882 | 0.775123 | Taf1d |
| ENSMUSG( | 1123.213 | 0.153122 | 0.10788 | 0.49674 | Taf2 |
| ENSMUSG( | 561.1921 | 0.177484 | 0.164054 | 0.594869 | Taf3 |
| ENSMUSG( | 844.8979 | 0.13473 | 0.204963 | 0.64616 | Taf4 |
| ENSMUSG( | 79.209 | -0.14165 | 0.511124 | 0.870859 | Taf4b |
| ENSMUSG( | 138.318 | -0.19553 | 0.258981 | 0.709814 | Taf5 |
| ENSMUSG( | 484.7078 | -0.19662 | 0.124431 | 0.531714 | Taf5l |
| ENSMUSG( | 1105.287 | 0.173128 | 0.088074 | 0.455247 | Taf6 |
| ENSMUSG( | 287.792 | 0.260722 | 0.091755 | 0.465904 | Taf6l |
| ENSMUSG( | 140.7958 | -0.07449 | 0.675441 | 0.930049 | Taf7 |
| ENSMUSG( | 560.4475 | 0.064082 | 0.578256 | 0.89641 | Taf8 |
| ENSMUSG( | 188.1028 | 0.220738 | 0.185769 | 0.623185 | Taf9 |
| ENSMUSG( | 863.4105 | 0.00888 | 0.939036 | 0.993415 | Taf9b |
| ENSMUSG( | 982.3401 | -0.41632 | 0.00133 | 0.039731 | Tafa1 |
| ENSMUSG( | 878.837 | -0.10445 | 0.428833 | 0.826446 | Tafa2 |
| ENSMUSG( | 22.37713 | -0.69597 | 0.167881 | 0.600712 | Tafa4 |
| ENSMUSG( | 4846.339 | -0.07975 | 0.302356 | 0.750682 | Tafa5 |
| ENSMUSG( | 1144.442 | -0.10329 | 0.376038 | 0.801127 | Tafazzin |
| ENSMUSG( | 34.92736 | 0.204358 | 0.557068 | 0.888198 | Tagap |
| ENSMUSG( | 554.462 | -0.12529 | 0.307846 | 0.756096 | Tagap1 |
| ENSMUSG( | 69.3026 | -0.81724 | 0.001741 | 0.047093 | Tagln |
| ENSMUSG( | 415.2036 | 0.188594 | 0.204243 | 0.64477 | Tagln2 |
| ENSMUSG( | 6981.378 | 0.101359 | 0.253956 | 0.704011 | Tagln3 |
| ENSMUSG( | 80.15699 | 0.541968 | 0.028949 | 0.259139 | Tal1 |
| ENSMUSG( | 2786.811 | -0.03328 | 0.719412 | 0.941986 | Taldo1 |
| ENSMUSG( | 1173.787 | 0.005022 | 0.971085 | 0.996369 | Tamalin |
| ENSMUSG( | 203.5667 | -0.08536 | 0.611842 | 0.908586 | Tamm41 |
| ENSMUSG( | 1132.067 | 0.232525 | 0.011305 | 0.153194 | Tanc1 |
| ENSMUSG( | 4269.486 | 0.039829 | 0.672373 | 0.929007 | Tanc2 |
| ENSMUSG( | 1653.06 | 0.089054 | 0.366911 | 0.795164 | Tango2 |
| ENSMUSG( | 165.2565 | 0.214711 | 0.186245 | 0.623869 | Tango6 |
| ENSMUSG( | 1373.449 | 0.152646 | 0.129577 | 0.539182 | Tank |
| ENSMUSG( | 4008.648 | 0.007245 | 0.945405 | 0.99365 | Taok1 |
| ENSMUSG( | 4788.204 | 0.096405 | 0.245196 | 0.692812 | Taok2 |
| ENSMUSG( | 1937.216 | 0.126598 | 0.22812 | 0.675573 | Taok3 |
| ENSMUSG( | 163.4242 | -0.53293 | 0.003251 | 0.071142 | Tap1 |
| ENSMUSG( | 440.3148 | -0.35373 | 0.0145 | 0.17723 | Tap2 |
| ENSMUSG( | 1261.953 | -0.31873 | 0.016131 | 0.188643 | Tapbp |
| ENSMUSG( | 306.3903 | -0.30211 | 0.048463 | 0.336526 | Tapbpl |
| ENSMUSG( | 1789.715 | -0.20901 | 0.139815 | 0.55638 | Tapt1 |
| ENSMUSG( | 249.6685 | 0.18686 | 0.251904 | 0.701908 | Tarbp1 |
| ENSMUSG( | 448.619 | -0.1212 | 0.334187 | 0.7758 | Tarbp2 |
| ENSMUSG( | 3920.483 | 0.045776 | 0.548921 | 0.885686 | Tardbp |
| ENSMUSG( | 767.199 | 0.092247 | 0.363679 | 0.793343 | Tars1 |
| ENSMUSG( | 1032.171 | 0.04059 | 0.718344 | 0.941911 | Tars2 |
| ENSMUSG( | 707.2526 | 0.090201 | 0.381679 | 0.802879 | Tars3 |
| ENSMUSG( | 11.44139 | -0.22442 | 0.685568 | NA | Tas1r3 |
| ENSMUSG( | 11.17433 | 0.222757 | 0.701634 | NA | Tasl |
| ENSMUSG( | 1006.134 | 0.011225 | 0.929098 | 0.991772 | Tasor |
| ENSMUSG( | 681.545 | -0.07905 | 0.602775 | 0.904887 | Tasor2 |
| ENSMUSG( | 468.4573 | 0.058383 | 0.641306 | 0.916693 | Tasp1 |
| ENSMUSG( | 259.6376 | -0.01394 | 0.929043 | 0.991772 | Tatdn1 |
| ENSMUSG( | 1348.091 | 0.147831 | 0.106544 | 0.494174 | Tatdn2 |
| ENSMUSG( | 399.8851 | -0.07217 | 0.615781 | 0.910647 | Tatdn3 |
| ENSMUSG( | 4307.833 | 0.003521 | 0.972423 | 0.996585 | Tax1bp1 |
| ENSMUSG( | 169.3899 | 0.099625 | 0.536928 | 0.882123 | Tax1bp3 |
| ENSMUSG( | 143.8399 | 0.021035 | 0.914546 | 0.988483 | Tbata |
| ENSMUSG( | 707.2336 | 0.246835 | 0.070433 | 0.40908 | Tbc1d1 |
| ENSMUSG( | 801.1787 | 0.18949 | 0.082896 | 0.444486 | Tbc1d10a |
| ENSMUSG( | 2446.389 | 0.095738 | 0.371684 | 0.798768 | Tbc1d10b |
| ENSMUSG( | 20.59119 | 0.151963 | 0.745997 | NA | Tbc1d10c |
| ENSMUSG( | 894.3053 | 0.092321 | 0.486546 | 0.858713 | Tbc1d12 |
| ENSMUSG( | 1592.843 | 0.103054 | 0.323179 | 0.768415 | Tbc1d13 |
| ENSMUSG( | 2903.153 | 0.101681 | 0.234461 | 0.681922 | Tbc1d14 |
| ENSMUSG( | 680.2451 | 0.099255 | 0.336987 | 0.777008 | Tbc1d15 |
| ENSMUSG( | 3675.947 | 0.032906 | 0.736209 | 0.946267 | Tbc1d16 |
| ENSMUSG( | 1265.283 | 0.09849 | 0.3433 | 0.779153 | Tbc1d17 |
| ENSMUSG( | 866.2577 | 0.118244 | 0.255059 | 0.705501 | Tbc1d19 |
| ENSMUSG( | 183.1029 | 0.372632 | 0.103202 | 0.488596 | Tbc1d2 |
| ENSMUSG( | 2587.932 | 0.072703 | 0.375272 | 0.800804 | Tbc1d20 |
| ENSMUSG( | 1256.078 | -0.01159 | 0.908193 | 0.9878 | Tbc1d22a |
| ENSMUSG( | 1847.089 | 0.124323 | 0.232201 | 0.67996 | Tbc1d22b |
| ENSMUSG( | 1083.848 | -0.07303 | 0.446415 | 0.83372 | Tbc1d23 |
| ENSMUSG( | 2104.56 | 0.069211 | 0.53608 | 0.88142 | Tbc1d24 |
| ENSMUSG( | 261.1708 | -0.00737 | 0.970822 | 0.996369 | Tbc1d25 |
| ENSMUSG( | 566.9635 | -0.08237 | 0.445973 | 0.833467 | Tbc1d2b |
| ENSMUSG( | 1794.654 | 0.177463 | 0.178161 | 0.615629 | Tbc1d30 |
| ENSMUSG( | 195.6479 | 0.103595 | 0.546017 | 0.884974 | Tbc1d31 |
| ENSMUSG( | 392.0395 | -0.16725 | 0.166067 | 0.598098 | Tbc1d32 |
| ENSMUSG( | 377.8854 | 0.782617 | 3.07E-10 | 1.61E-07 | Tbc1d4 |
| ENSMUSG( | 1357.798 | 0.098405 | 0.274078 | 0.724254 | Tbc1d5 |
| ENSMUSG( | 439.0522 | -0.16476 | 0.180173 | 0.617634 | Tbc1d7 |
| ENSMUSG( | 720.2574 | -0.27471 | 0.04654 | 0.330304 | Tbc1d8 |
| ENSMUSG( | 154.789 | 0.139497 | 0.463819 | 0.842812 | Tbc1d8b |
| ENSMUSG( | 3414.295 | 0.102343 | 0.252946 | 0.703255 | Tbc1d9 |
| ENSMUSG( | 29583.33 | 0.168146 | 0.036093 | 0.292126 | Tbc1d9b |
| ENSMUSG( | 931.4758 | -0.03486 | 0.748176 | 0.949604 | Tbca |
| ENSMUSG( | 2609.457 | 0.054668 | 0.523558 | 0.875628 | Tbcb |
| ENSMUSG( | 294.5883 | -0.11658 | 0.484626 | 0.856665 | Tbcc |
| ENSMUSG( | 526.6277 | 0.150308 | 0.195271 | 0.634601 | Tbccd1 |
| ENSMUSG( | 1436.476 | 0.098165 | 0.29438 | 0.74363 | Tbcd |
| ENSMUSG( | 1346.479 | 0.142374 | 0.326902 | 0.771858 | Tbce |
| ENSMUSG( | 4881.809 | 0.155141 | 0.161787 | 0.592233 | Tbcel |
| ENSMUSG( | 795.5502 | 0.133262 | 0.271327 | 0.72198 | Tbck |
| ENSMUSG( | 906.8704 | 0.047486 | 0.634203 | 0.914073 | Tbk1 |
| ENSMUSG( | 3146.71 | 0.338851 | 0.000538 | 0.021456 | Tbkbp1 |
| ENSMUSG( | 2367.077 | -0.06046 | 0.493885 | 0.860611 | Tbl1x |
| ENSMUSG( | 2037.103 | -0.07731 | 0.379732 | 0.801757 | Tbl1xr1 |
| ENSMUSG( | 1124.552 | -0.10351 | 0.481837 | 0.855573 | Tbl2 |
| ENSMUSG( | 738.028 | 0.066963 | 0.585681 | 0.899171 | Tbl3 |
| ENSMUSG( | 457.3684 | -0.03357 | 0.776422 | 0.955658 | Tbp |
| ENSMUSG( | 1097.625 | 0.097751 | 0.400082 | 0.813641 | Tbpl1 |
| ENSMUSG( | 2464.431 | 0.079888 | 0.409731 | 0.818258 | Tbr1 |
| ENSMUSG( | 763.5364 | 0.045605 | 0.740572 | 0.947823 | Tbrg1 |
| ENSMUSG( | 995.1325 | 0.113281 | 0.312801 | 0.760322 | Tbrg4 |
| ENSMUSG( | 52.71239 | 0.293919 | 0.345164 | 0.779153 | Tbx1 |
| ENSMUSG( | 88.16541 | -0.34526 | 0.096884 | 0.477375 | Tbx15 |
| ENSMUSG( | 103.2505 | 0.094752 | 0.653314 | 0.922819 | Tbx18 |
| ENSMUSG( | 81.51664 | 0.237298 | 0.339312 | 0.777616 | Tbx2 |
| ENSMUSG( | 109.4104 | 0.50798 | 0.01697 | 0.194647 | Tbx3 |
| ENSMUSG( | 39.63378 | 0.016294 | 0.968973 | 0.996369 | Tbx3os1 |
| ENSMUSG( | 14.41776 | -0.0364 | 0.949325 | NA | Tbx4 |
| ENSMUSG( | 31.30228 | 0.365131 | 0.348809 | 0.781448 | Tbx6 |
| ENSMUSG( | 47.83263 | 0.271524 | 0.425656 | 0.824898 | Tbxa2r |
| ENSMUSG( | 196.2275 | -0.19268 | 0.338897 | 0.777518 | Tbxas1 |
| ENSMUSG( | 5926.088 | -0.04489 | 0.57157 | 0.893319 | Tcaf1 |
| ENSMUSG( | 28.56838 | 0.206338 | 0.573833 | 0.893584 | Tcaf2 |
| ENSMUSG( | 598.6119 | 0.109381 | 0.323284 | 0.768415 | Tcaim |
| ENSMUSG( | 108.9179 | 0.112898 | 0.599389 | 0.903801 | Tcap |
| ENSMUSG( | 980.0729 | 0.015996 | 0.874302 | 0.979431 | Tcea1 |
| ENSMUSG( | 1132.157 | 0.038379 | 0.783149 | 0.957558 | Tcea2 |
| ENSMUSG( | 20.21669 | -0.77572 | 0.081324 | NA | Tcea3 |
| ENSMUSG( | 365.6721 | -0.07401 | 0.583993 | 0.898666 | Tceal1 |
| ENSMUSG( | 2100.694 | 0.000486 | 0.996719 | 0.99943 | Tceal3 |
| ENSMUSG( | 874.8867 | -0.03964 | 0.790274 | 0.959461 | Tceal5 |
| ENSMUSG( | 784.5488 | -0.09426 | 0.583847 | 0.898666 | Tceal6 |
| ENSMUSG( | 25.71728 | 0.368619 | 0.329218 | 0.773233 | Tceal7 |
| ENSMUSG( | 843.1963 | 0.06851 | 0.62446 | 0.91187 | Tceal8 |
| ENSMUSG( | 805.7515 | -0.03535 | 0.795712 | 0.960062 | Tceal9 |
| ENSMUSG( | 62.5476 | -0.20088 | 0.448281 | 0.834051 | Tceanc |
| ENSMUSG( | 528.7811 | 0.221652 | 0.042215 | 0.316565 | Tceanc2 |
| ENSMUSG( | 1406.159 | -0.03103 | 0.733809 | 0.945265 | Tcerg1 |
| ENSMUSG( | 285.0913 | 0.178262 | 0.187296 | 0.625476 | Tcerg1l |
| ENSMUSG( | 2122.082 | 0.037594 | 0.694616 | 0.9343 | Tcf12 |
| ENSMUSG( | 13.31679 | -0.1086 | 0.828707 | NA | Tcf15 |
| ENSMUSG( | 161.9924 | -0.06661 | 0.707603 | 0.93949 | Tcf19 |
| ENSMUSG( | 2477.008 | 0.06361 | 0.542378 | 0.883182 | Tcf20 |
| ENSMUSG( | 11.10511 | -0.12799 | 0.829816 | NA | Tcf23 |
| ENSMUSG( | 12.50178 | 0.420484 | 0.44178 | NA | Tcf24 |
| ENSMUSG( | 19991.73 | 0.108112 | 0.187186 | 0.625368 | Tcf25 |
| ENSMUSG( | 1349.712 | -0.00849 | 0.925019 | 0.991204 | Tcf3 |

| | | | | | |
|---|---|---|---|---|---|
| ENSMUSG( | 5392.682 | -0.049 | 0.635162 | 0.953903 | Tcaf1 |
| ENSMUSG( | 25.77211 | 0.194157 | 0.630481 | 0.953251 | Tcaf2 |
| ENSMUSG( | 525.3785 | -0.00377 | 0.97878 | 0.996468 | Tcaim |
| ENSMUSG( | 98.19475 | 0.078224 | 0.714418 | 0.964986 | Tcap |
| ENSMUSG( | 874.3511 | -0.04904 | 0.711153 | 0.964363 | Tcea1 |
| ENSMUSG( | 1027.01 | 0.026702 | 0.884804 | 0.989688 | Tcea2 |
| ENSMUSG( | 20.69014 | -0.36673 | 0.411184 | 0.917257 | Tcea3 |
| ENSMUSG( | 320.8887 | -0.19921 | 0.307197 | 0.898538 | Tceal1 |
| ENSMUSG( | 1764.083 | -0.25103 | 0.120217 | 0.813384 | Tceal3 |
| ENSMUSG( | 746.5465 | -0.24669 | 0.213226 | 0.864412 | Tceal5 |
| ENSMUSG( | 682.5343 | -0.2433 | 0.205599 | 0.864412 | Tceal6 |
| ENSMUSG( | 21.06778 | 0.107783 | 0.824354 | 0.979198 | Tceal7 |
| ENSMUSG( | 737.8171 | -0.05476 | 0.768746 | 0.974236 | Tceal8 |
| ENSMUSG( | 747.6836 | 0.012222 | 0.946396 | 0.994798 | Tceal9 |
| ENSMUSG( | 55.72363 | -0.26502 | 0.372382 | 0.915491 | Tceanc |
| ENSMUSG( | 467.4961 | 0.138513 | 0.291794 | 0.897854 | Tceanc2 |
| ENSMUSG( | 1313.446 | 0.040451 | 0.714013 | 0.964986 | Tcerg1 |
| ENSMUSG( | 247.9674 | 0.040063 | 0.817933 | 0.979198 | Tcerg1l |
| ENSMUSG( | 1925.224 | 0.025512 | 0.825209 | 0.979354 | Tcf12 |
| ENSMUSG( | 11.62457 | -0.21952 | 0.698044 | 0.960867 | Tcf15 |
| ENSMUSG( | 151.2469 | 0.010979 | 0.953595 | 0.994798 | Tcf19 |
| ENSMUSG( | 2250.404 | 0.058178 | 0.62647 | 0.953234 | Tcf20 |
| ENSMUSG( | 11.04542 | 0.184682 | 0.771395 | NA | Tcf23 |
| ENSMUSG( | 11.06315 | 0.292826 | 0.645529 | NA | Tcf24 |
| ENSMUSG( | 16710.24 | -0.14226 | 0.166914 | 0.840183 | Tcf25 |
| ENSMUSG( | 1220.147 | -0.02891 | 0.802769 | 0.97681 | Tcf3 |
| ENSMUSG( | 10398.09 | 0.027934 | 0.823591 | 0.979198 | Tcf4 |
| ENSMUSG( | 198.084 | 0.70955 | 0.000873 | 0.099327 | Tcf7 |
| ENSMUSG( | 313.8856 | -0.13867 | 0.374469 | 0.915491 | Tcf7l1 |
| ENSMUSG( | 281.847 | -0.31813 | 0.074964 | 0.76762 | Tcf7l2 |
| ENSMUSG( | 48.99604 | -0.34122 | 0.308766 | 0.898538 | Tcfl5 |
| ENSMUSG( | 70.22658 | -0.04418 | 0.864247 | 0.986552 | Tchh |
| ENSMUSG( | 271.8093 | 0.001315 | 0.993958 | 0.998813 | Tchp |
| ENSMUSG( | 77.96004 | 0.181179 | 0.50451 | 0.927825 | Tcim |
| ENSMUSG( | 751.0572 | 0.342643 | 0.073659 | 0.764578 | Tcirg1 |
| ENSMUSG( | 1185.909 | 0.187554 | 0.327067 | 0.905768 | Tcn2 |
| ENSMUSG( | 544.5683 | -0.05446 | 0.686037 | 0.958956 | Tcof1 |
| ENSMUSG( | 3923.818 | -0.13083 | 0.292208 | 0.897854 | Tcp1 |
| ENSMUSG( | 9.573714 | 0.165421 | 0.798297 | NA | Tcp10b |
| ENSMUSG( | 16.1178 | 0.039371 | 0.936143 | 0.993573 | Tcp11 |
| ENSMUSG( | 650.4929 | -0.14089 | 0.294976 | 0.898538 | Tcp11l1 |
| ENSMUSG( | 643.0379 | 0.028181 | 0.828329 | 0.980224 | Tcp11l2 |
| ENSMUSG( | 962.6835 | -0.11235 | 0.354841 | 0.912411 | Tcta |
| ENSMUSG( | 83.00488 | 0.104575 | 0.675101 | 0.95694 | Tcte1 |
| ENSMUSG( | 95.92595 | -0.18663 | 0.39457 | 0.917257 | Tcte2 |
| ENSMUSG( | 889.3269 | 0.077462 | 0.599549 | 0.947071 | Tctn1 |
| ENSMUSG( | 822.8206 | 0.232468 | 0.093522 | 0.795507 | Tctn2 |
| ENSMUSG( | 808.3919 | 0.21851 | 0.132377 | 0.822912 | Tctn3 |
| ENSMUSG( | 483.3058 | -1.62333 | 1.03E-32 | 4.49E-29 | Tdg |
| ENSMUSG( | 23.07877 | -0.07533 | 0.871704 | 0.988779 | Tdo2 |
| ENSMUSG( | 335.5537 | 0.188872 | 0.200116 | 0.863016 | Tdp1 |
| ENSMUSG( | 346.578 | 0.129363 | 0.382715 | 0.917257 | Tdp2 |
| ENSMUSG( | 13.49668 | 0.128706 | 0.826557 | 0.979952 | Tdrd1 |
| ENSMUSG( | 19.55942 | -0.01446 | 0.97671 | 0.996416 | Tdrd12 |
| ENSMUSG( | 711.3055 | -0.01602 | 0.905652 | 0.991089 | Tdrd3 |
| ENSMUSG( | 32.77228 | 0.303554 | 0.426018 | 0.917257 | Tdrd5 |
| ENSMUSG( | 27.16904 | 0.50125 | 0.199792 | 0.863016 | Tdrd6 |
| ENSMUSG( | 1127.006 | -0.09031 | 0.427976 | 0.917257 | Tdrd7 |
| ENSMUSG( | 14.40509 | -0.62975 | 0.272731 | 0.888825 | Tdrd9 |
| ENSMUSG( | 1210.688 | -0.094 | 0.485881 | 0.92541 | Tdrkh |
| ENSMUSG( | 365.6662 | 0.322553 | 0.042112 | 0.690304 | Tdrp |
| ENSMUSG( | 1304.425 | 0.057477 | 0.672684 | 0.95694 | Tead1 |
| ENSMUSG( | 247.8233 | 0.287596 | 0.06938 | 0.755467 | Tead2 |
| ENSMUSG( | 156.2606 | 0.031248 | 0.870624 | 0.988445 | Tead3 |
| ENSMUSG( | 19.52832 | 1.367332 | 0.023713 | 0.588226 | Tead4 |
| ENSMUSG( | 79.95471 | -0.06153 | 0.809755 | 0.978779 | Tec |
| ENSMUSG( | 3303.658 | 0.06612 | 0.568959 | 0.941777 | Tecpr1 |
| ENSMUSG( | 2581.276 | 0.089496 | 0.441276 | 0.921205 | Tecpr2 |
| ENSMUSG( | 13791.7 | -0.05521 | 0.648708 | 0.95422 | Tecr |
| ENSMUSG( | 8.862902 | 1.710918 | 0.061282 | NA | Tecrl |
| ENSMUSG( | 19.40725 | 0.246134 | 0.606045 | 0.949497 | Tecta |
| ENSMUSG( | 9.707516 | -1.35244 | 0.042169 | NA | Tectb |
| ENSMUSG( | 55.69321 | 0.124104 | 0.683134 | 0.958678 | Tedc1 |
| ENSMUSG( | 211.9336 | -0.01577 | 0.927856 | 0.99306 | Tedc2 |
| ENSMUSG( | 483.065 | -0.43854 | 0.009823 | 0.406458 | Teddm2 |
| ENSMUSG( | 12181.62 | 0.315728 | 0.001711 | 0.152157 | Tef |
| ENSMUSG( | 88.8089 | -0.33675 | 0.162929 | 0.837925 | Tefm |
| ENSMUSG( | 628.8999 | 0.623628 | 0.22543 | 0.868468 | Tek |
| ENSMUSG( | 71.86855 | -0.01478 | 0.95791 | 0.994798 | Tekt1 |
| ENSMUSG( | 65.79499 | -0.10543 | 0.719973 | 0.966887 | Tekt2 |
| ENSMUSG( | 58.78311 | 0.778916 | 0.007575 | 0.370638 | Tekt4 |
| ENSMUSG( | 206.9477 | -0.29205 | 0.141504 | 0.830678 | Tekt5 |
| ENSMUSG( | 19.05262 | -0.04194 | 0.929251 | 0.99306 | Tektip1 |
| ENSMUSG( | 493.1507 | -0.00449 | 0.975165 | 0.996416 | Telo2 |
| ENSMUSG( | 439.4345 | -0.32411 | 0.025932 | 0.601649 | Ten1 |
| ENSMUSG( | 946.0542 | 0.217994 | 0.21036 | 0.864412 | Tenm1 |
| ENSMUSG( | 2468.139 | 0.334687 | 0.032755 | 0.637512 | Tenm2 |
| ENSMUSG( | 2167.617 | 0.20575 | 0.261495 | 0.884926 | Tenm3 |
| ENSMUSG( | 3713.365 | 0.012386 | 0.929117 | 0.99306 | Tenm4 |
| ENSMUSG( | 524.057 | 0.071953 | 0.594682 | 0.945444 | Tent2 |
| ENSMUSG( | 1246.084 | 0.102541 | 0.350594 | 0.911652 | Tent4a |
| ENSMUSG( | 931.8899 | 0.181596 | 0.121335 | 0.813384 | Tent4b |
| ENSMUSG( | 125.5847 | -0.19286 | 0.325572 | 0.905339 | Tent5a |
| ENSMUSG( | 177.1884 | -0.02798 | 0.919645 | 0.992644 | Tent5c |
| ENSMUSG( | 176.5257 | -0.04817 | 0.813727 | 0.979198 | Tep1 |
| ENSMUSG( | 477.5734 | 0.158421 | 0.261157 | 0.884926 | Tepsin |
| ENSMUSG( | 22.53594 | -0.44961 | 0.277186 | 0.891468 | Terb1 |
| ENSMUSG( | 198.8068 | 0.05889 | 0.752299 | 0.970471 | Terf1 |
| ENSMUSG( | 887.9988 | -0.08306 | 0.576485 | 0.942912 | Terf2 |
| ENSMUSG( | 899.4158 | -0.07514 | 0.591234 | 0.944975 | Terf2ip |
| ENSMUSG( | 36.70847 | -0.13786 | 0.679106 | 0.957725 | Tert |
| ENSMUSG( | 85.38502 | 0.208565 | 0.374838 | 0.915491 | Tes |
| ENSMUSG( | 8.916309 | 0.16893 | 0.789936 | NA | Tes3-ps |
| ENSMUSG( | 809.0923 | -0.17346 | 0.268462 | 0.886755 | Tesc |
| ENSMUSG( | 3455.486 | 0.091715 | 0.364296 | 0.914223 | Tesk1 |
| ENSMUSG( | 256.0516 | 0.166932 | 0.316292 | 0.901868 | Tesk2 |
| ENSMUSG( | 460.4913 | 0.135295 | 0.356575 | 0.913358 | Tet1 |
| ENSMUSG( | 1708.355 | -0.06993 | 0.522231 | 0.932681 | Tet2 |
| ENSMUSG( | 1631.425 | -0.00971 | 0.933322 | 0.99306 | Tet3 |
| ENSMUSG( | 331.6103 | 0.253431 | 0.095924 | 0.796133 | Tex10 |
| ENSMUSG( | 11.24525 | -0.4025 | 0.48522 | NA | Tex14 |
| ENSMUSG( | 37.3084 | 0.588798 | 0.084628 | 0.779799 | Tex15 |
| ENSMUSG( | 3409.693 | -0.01225 | 0.921504 | 0.992644 | Tex2 |
| ENSMUSG( | 2138.674 | 0.030407 | 0.824335 | 0.979198 | Tex261 |
| ENSMUSG( | 2694.269 | 0.098132 | 0.489907 | 0.925987 | Tex264 |
| ENSMUSG( | 40.62562 | 0.368641 | 0.296926 | 0.898538 | Tex30 |
| ENSMUSG( | 65.0736 | -0.30569 | 0.283788 | 0.892412 | Tex52 |
| ENSMUSG( | 355.0546 | -0.16472 | 0.25989 | 0.883971 | Tex9 |
| ENSMUSG( | 740.1576 | 0.098292 | 0.454861 | 0.922891 | Tfam |
| ENSMUSG( | 25.99455 | 0.484206 | 0.24779 | 0.877904 | Tfap2b |
| ENSMUSG( | 10.64793 | 0.478827 | 0.413328 | NA | Tfap2c |
| ENSMUSG( | 15.47641 | -0.05205 | 0.915018 | 0.992407 | Tfap2d |
| ENSMUSG( | 65.35693 | -0.02178 | 0.935228 | 0.993078 | Tfap4 |
| ENSMUSG( | 74.30993 | -0.21235 | 0.427161 | 0.917257 | Tfb1m |
| ENSMUSG( | 325.2417 | 0.033559 | 0.822125 | 0.979198 | Tfb2m |
| ENSMUSG( | 563.2864 | 0.138991 | 0.303919 | 0.898538 | Tfcp2 |
| ENSMUSG( | 988.8776 | -0.05162 | 0.724331 | 0.966887 | Tfcp2l1 |
| ENSMUSG( | 1314.975 | 0.147935 | 0.225682 | 0.868468 | Tfdp1 |
| ENSMUSG( | 1300.03 | -0.05676 | 0.618057 | 0.952232 | Tfdp2 |
| ENSMUSG( | 1230.898 | 0.006089 | 0.955968 | 0.994798 | Tfe3 |
| ENSMUSG( | 428.2592 | 0.262184 | 0.059529 | 0.729747 | Tfeb |
| ENSMUSG( | 888.0619 | 0.095841 | 0.466257 | 0.923329 | Tfg |
| ENSMUSG( | 1165.963 | -0.00735 | 0.960213 | 0.994798 | Tfip11 |
| ENSMUSG( | 215.6966 | 0.261795 | 0.139248 | 0.830678 | Tfpi |
| ENSMUSG( | 448.592 | -0.15466 | 0.336717 | 0.907972 | Tfpt |
| ENSMUSG( | 200.6904 | 0.195734 | 0.272726 | 0.888825 | Tfr2 |
| ENSMUSG( | 4740.128 | 0.481147 | 0.001248 | 0.127113 | Tfrc |
| ENSMUSG( | 146.2622 | 0.256215 | 0.160831 | 0.837724 | Tgds |
| ENSMUSG( | 2180.469 | -0.00273 | 0.983724 | 0.996963 | Tgfa |
| ENSMUSG( | 496.8558 | 0.161954 | 0.346364 | 0.910966 | Tgfb1 |
| ENSMUSG( | 727.2541 | -0.07534 | 0.551819 | 0.939565 | Tgfb1i1 |
| ENSMUSG( | 866.8825 | 0.201899 | 0.132495 | 0.822912 | Tgfb2 |
| ENSMUSG( | 593.1259 | -0.12691 | 0.330502 | 0.907398 | Tgfb3 |
| ENSMUSG( | 327.3121 | 0.14846 | 0.379159 | 0.916995 | Tgfbi |
| ENSMUSG( | 1471.396 | 0.31396 | 0.103121 | 0.804311 | Tgfbr1 |
| ENSMUSG( | 891.4774 | 0.203244 | 0.400996 | 0.917257 | Tgfbr2 |
| ENSMUSG( | 431.8178 | 0.266425 | 0.110043 | 0.806572 | Tgfbr3 |
| ENSMUSG( | 47.88559 | 0.448755 | 0.169903 | 0.840806 | Tgfbr3l |
| ENSMUSG( | 2341.006 | 0.002502 | 0.980589 | 0.996479 | Tgfbrap1 |
| ENSMUSG( | 58.88309 | 0.114255 | 0.679443 | 0.957966 | Tgif1 |
| ENSMUSG( | 77.17189 | -0.14795 | 0.564678 | 0.941206 | Tgif2 |
| ENSMUSG( | 443.3767 | 0.5861 | 0.003736 | 0.24422 | Tgm2 |
| ENSMUSG( | 85.95133 | 0.002243 | 0.992524 | 0.99874 | Tgm3 |
| ENSMUSG( | 17.96201 | -0.12961 | 0.790732 | 0.97681 | Tgm4 |
| ENSMUSG( | 14.7159 | 0.320133 | 0.546945 | 0.938063 | Tgm7 |
| ENSMUSG( | 5319.296 | 0.13147 | 0.411745 | 0.917257 | Tgoln1 |
| ENSMUSG( | 589.5757 | 0.18462 | 0.146306 | 0.835973 | Tgs1 |
| ENSMUSG( | 21.55854 | 0.14805 | 0.743994 | 0.96951 | Tgtp1 |

| | | | | | | | | | | |
|---|---|---|---|---|---|---|---|---|---|---|
| ENSMUSG( | Thtpa | 286.1836 | 0.203587 | 0.135495 | 0.548884 | ENSMUSG( | Thtpa | 240.5885 | -0.01654 | 0.920297 | 0.992644 |
| ENSMUSG( | Thumpd1 | 736.906 | 0.122645 | 0.312599 | 0.760241 | ENSMUSG( | Thumpd1 | 629.7778 | -0.06152 | 0.654289 | 0.955015 |
| ENSMUSG( | Thumpd2 | 136.8343 | 0.032405 | 0.858464 | 0.974683 | ENSMUSG( | Thumpd2 | 120.904 | -0.03269 | 0.886471 | 0.990011 |
| ENSMUSG( | Thumpd3 | 522.9742 | 0.114844 | 0.343684 | 0.779153 | ENSMUSG( | Thumpd3 | 433.9712 | -0.16058 | 0.242348 | 0.877129 |
| ENSMUSG( | Thy1 | 29504.7 | -0.429 | 0.000267 | 0.013312 | ENSMUSG( | Thy1 | 31308.41 | 0.044882 | 0.746114 | 0.969632 |
| ENSMUSG( | Thyn1 | 662.3662 | 0.091337 | 0.446904 | 0.834051 | ENSMUSG( | Thyn1 | 571.5112 | -0.0676 | 0.631192 | 0.953251 |
| ENSMUSG( | Tia1 | 2126.717 | -0.16408 | 0.08572 | 0.451367 | ENSMUSG( | Tia1 | 2291.928 | 0.310611 | 0.026281 | 0.603041 |
| ENSMUSG( | Tial1 | 2365.265 | -0.09627 | 0.289252 | 0.739725 | ENSMUSG( | Tial1 | 2447.948 | 0.262959 | 0.0586 | 0.728256 |
| ENSMUSG( | Tiam1 | 2209.814 | 0.088449 | 0.366077 | 0.794603 | ENSMUSG( | Tiam1 | 2063.532 | 0.15636 | 0.183391 | 0.850984 |
| ENSMUSG( | Tiam2 | 1138.181 | -0.14673 | 0.224775 | 0.671058 | ENSMUSG( | Tiam2 | 1065.276 | -0.06797 | 0.626602 | 0.953234 |
| ENSMUSG( | Ticam1 | 229.4165 | 0.054284 | 0.729284 | 0.944738 | ENSMUSG( | Ticam1 | 196.9371 | -0.11955 | 0.474688 | 0.923329 |
| ENSMUSG( | Ticam2 | 20.70409 | -0.52 | 0.214596 | NA | ENSMUSG( | Ticam2 | 24.55528 | 0.272588 | 0.491175 | 0.926604 |
| ENSMUSG( | Ticrr | 10.17933 | -0.3246 | 0.594872 | NA | ENSMUSG( | Ticrr | 10.20819 | -0.00262 | 0.996881 | NA |
| ENSMUSG( | Tie1 | 563.8738 | 0.090056 | 0.669455 | 0.928147 | ENSMUSG( | Tie1 | 578.6983 | 0.416709 | 0.348956 | 0.911652 |
| ENSMUSG( | Tifa | 372.3953 | 0.160679 | 0.190912 | 0.630245 | ENSMUSG( | Tifa | 326.6787 | 0.054728 | 0.711404 | 0.964363 |
| ENSMUSG( | Tifab | 193.5395 | 0.271981 | 0.136524 | 0.550651 | ENSMUSG( | Tifab | 181.0382 | 0.353873 | 0.053014 | 0.71525 |
| ENSMUSG( | Tigar | 549.5434 | 0.073281 | 0.584382 | 0.898802 | ENSMUSG( | Tigar | 476.6501 | -0.06976 | 0.644759 | 0.953903 |
| ENSMUSG( | Tigd2 | 196.4199 | -0.12452 | 0.40668 | 0.816928 | ENSMUSG( | Tigd2 | 182.1559 | -0.06691 | 0.694278 | 0.960867 |
| ENSMUSG( | Tigd3 | 59.69464 | -0.04132 | 0.89189 | 0.983979 | ENSMUSG( | Tigd3 | 52.5031 | -0.1323 | 0.688149 | 0.959742 |
| ENSMUSG( | Tigd5 | 119.4393 | 0.084546 | 0.708365 | 0.939519 | ENSMUSG( | Tigd5 | 107.6454 | 0.072163 | 0.757198 | 0.971858 |
| ENSMUSG( | Timeless | 146.1563 | 0.319192 | 0.132183 | 0.544244 | ENSMUSG( | Timeless | 130.368 | 0.273572 | 0.231299 | 0.868507 |
| ENSMUSG( | Timm10 | 347.247 | 0.121181 | 0.427082 | 0.82532 | ENSMUSG( | Timm10 | 285.8565 | -0.18601 | 0.353493 | 0.912263 |
| ENSMUSG( | Timm10b | 334.2797 | -0.02929 | 0.860292 | 0.974842 | ENSMUSG( | Timm10b | 308.8277 | 0.019287 | 0.905352 | 0.991089 |
| ENSMUSG( | Timm13 | 1891.355 | -0.02887 | 0.768975 | 0.954674 | ENSMUSG( | Timm13 | 1596.608 | -0.26378 | 0.031864 | 0.632006 |
| ENSMUSG( | Timm17a | 1661.98 | 0.049982 | 0.59704 | 0.903526 | ENSMUSG( | Timm17a | 1489.643 | 0.000657 | 0.995669 | 0.999116 |
| ENSMUSG( | Timm17b | 713.4277 | -0.04202 | 0.722097 | 0.943535 | ENSMUSG( | Timm17b | 613.5564 | -0.22644 | 0.169414 | 0.840806 |
| ENSMUSG( | Timm21 | 529.5998 | -0.01567 | 0.906432 | 0.986998 | ENSMUSG( | Timm21 | 510.0202 | 0.145488 | 0.335467 | 0.907907 |
| ENSMUSG( | Timm22 | 628.2479 | -0.04581 | 0.691109 | 0.933881 | ENSMUSG( | Timm22 | 570.6849 | -0.05443 | 0.673468 | 0.95694 |
| ENSMUSG( | Timm23 | 873.6749 | -0.01942 | 0.847172 | 0.972374 | ENSMUSG( | Timm23 | 758.571 | -0.16812 | 0.183792 | 0.85194 |
| ENSMUSG( | Timm29 | 2154.905 | 0.085417 | 0.31748 | 0.763823 | ENSMUSG( | Timm29 | 1866.506 | -0.06192 | 0.57349 | 0.942912 |
| ENSMUSG( | Timm44 | 1325.137 | 0.11849 | 0.345179 | 0.779153 | ENSMUSG( | Timm44 | 1198.34 | 0.097185 | 0.501917 | 0.927825 |
| ENSMUSG( | Timm50 | 919.3238 | -0.09886 | 0.365067 | 0.79407 | ENSMUSG( | Timm50 | 820.7142 | -0.15634 | 0.215596 | 0.865167 |
| ENSMUSG( | Timm8a1 | 169.4323 | 0.201948 | 0.260599 | 0.710468 | ENSMUSG( | Timm8a1 | 144.3032 | 0.004297 | 0.982416 | 0.996756 |
| ENSMUSG( | Timm8b | 2398.616 | 0.00081 | 0.992978 | 0.999327 | ENSMUSG( | Timm8b | 2011.671 | -0.25315 | 0.049501 | 0.708648 |
| ENSMUSG( | Timm9 | 680.4996 | -0.03183 | 0.817119 | 0.96566 | ENSMUSG( | Timm9 | 620.147 | -0.0343 | 0.834905 | 0.981237 |
| ENSMUSG( | Timmdc1 | 584.566 | -0.02741 | 0.804816 | 0.962473 | ENSMUSG( | Timmdc1 | 540.3079 | 0.014106 | 0.913253 | 0.992298 |
| ENSMUSG( | Timp1 | 36.52868 | -0.23072 | 0.465415 | 0.844064 | ENSMUSG( | Timp1 | 32.01666 | -0.3801 | 0.283365 | 0.892412 |
| ENSMUSG( | Timp2 | 11470.22 | -0.26713 | 0.041776 | 0.315339 | ENSMUSG( | Timp2 | 11840.93 | 0.107148 | 0.474916 | 0.923329 |
| ENSMUSG( | Timp3 | 8397.906 | -0.22462 | 0.056726 | 0.367404 | ENSMUSG( | Timp3 | 8026.364 | -0.07722 | 0.552235 | 0.939565 |
| ENSMUSG( | Timp4 | 4175.523 | 0.07234 | 0.459803 | 0.840338 | ENSMUSG( | Timp4 | 3496.778 | -0.17653 | 0.11811 | 0.813384 |
| ENSMUSG( | Tinagl1 | 249.0223 | 0.910358 | 0.000187 | 0.009972 | ENSMUSG( | Tinagl1 | 269.6928 | 1.2876 | 2.23E-07 | 0.000135 |
| ENSMUSG( | Tincr | 85.23289 | 0.39935 | 0.090352 | 0.461669 | ENSMUSG( | Tincr | 75.4753 | 0.327181 | 0.312406 | 0.900037 |
| ENSMUSG( | Tinf2 | 412.2385 | -0.01769 | 0.898363 | 0.985633 | ENSMUSG( | Tinf2 | 393.6442 | 0.122224 | 0.415835 | 0.917257 |
| ENSMUSG( | Tiparp | 614.2769 | -0.10045 | 0.353326 | 0.785492 | ENSMUSG( | Tiparp | 563.7571 | -0.07603 | 0.557735 | 0.939565 |
| ENSMUSG( | Tipin | 493.6744 | -0.01344 | 0.915296 | 0.988634 | ENSMUSG( | Tipin | 416.3028 | -0.25077 | 0.063408 | 0.736482 |
| ENSMUSG( | Tiprl | 1036.299 | 0.058245 | 0.604053 | 0.905187 | ENSMUSG( | Tiprl | 905.6235 | -0.06686 | 0.640065 | 0.953903 |
| ENSMUSG( | Tirap | 100.8717 | -0.0144 | 0.947214 | 0.993821 | ENSMUSG( | Tirap | 92.35932 | 0.013199 | 0.954241 | 0.994798 |
| ENSMUSG( | Tjap1 | 861.8785 | 0.143318 | 0.220465 | 0.666079 | ENSMUSG( | Tjap1 | 818.5472 | 0.259745 | 0.092138 | 0.791946 |
| ENSMUSG( | Tjp1 | 5639.732 | 0.121307 | 0.304829 | 0.752574 | ENSMUSG( | Tjp1 | 4824.99 | -0.05827 | 0.589501 | 0.944975 |
| ENSMUSG( | Tjp2 | 2380.264 | 0.010223 | 0.923299 | 0.990694 | ENSMUSG( | Tjp2 | 1997.201 | -0.23959 | 0.044337 | 0.693697 |
| ENSMUSG( | Tjp3 | 22.80401 | -0.26411 | 0.532943 | 0.879905 | ENSMUSG( | Tjp3 | 20.84644 | -0.20543 | 0.681013 | 0.958395 |
| ENSMUSG( | Tk1 | 22.94777 | -0.55874 | 0.206685 | 0.648942 | ENSMUSG( | Tk1 | 21.63083 | -0.42568 | 0.323083 | 0.904023 |
| ENSMUSG( | Tk2 | 1381.784 | 0.093892 | 0.379255 | 0.801757 | ENSMUSG( | Tk2 | 1191.843 | -0.06331 | 0.61107 | 0.950771 |
| ENSMUSG( | Tkfc | 528.2319 | 0.082563 | 0.502938 | 0.865717 | ENSMUSG( | Tkfc | 455.2233 | -0.08239 | 0.573957 | 0.942912 |
| ENSMUSG( | Tkt | 6293.67 | -0.076 | 0.363799 | 0.793343 | ENSMUSG( | Tkt | 5739.512 | -0.07315 | 0.489392 | 0.925675 |
| ENSMUSG( | Tktl2 | 9.909634 | 0.454201 | 0.490132 | NA | ENSMUSG( | Tktl2 | 7.493295 | 0.013261 | 0.985931 | NA |
| ENSMUSG( | Tlcd1 | 2284.056 | 0.000618 | 0.995792 | 0.99943 | ENSMUSG( | Tlcd1 | 2211.619 | 0.173878 | 0.193182 | 0.860096 |
| ENSMUSG( | Tlcd2 | 17.48267 | -0.13675 | 0.76445 | NA | ENSMUSG( | Tlcd2 | 16.15216 | -0.06634 | 0.894322 | 0.990558 |
| ENSMUSG( | Tlcd3a | 762.0557 | -0.31029 | 0.024214 | 0.238178 | ENSMUSG( | Tlcd3a | 780.5642 | 0.049723 | 0.74884 | 0.96984 |
| ENSMUSG( | Tlcd3b | 4639.081 | 0.091635 | 0.295159 | 0.743798 | ENSMUSG( | Tlcd3b | 4014.585 | -0.05643 | 0.563579 | 0.941206 |
| ENSMUSG( | Tlcd4 | 2709.936 | -0.17315 | 0.209631 | 0.651841 | ENSMUSG( | Tlcd4 | 2602.181 | -0.0162 | 0.921299 | 0.992644 |
| ENSMUSG( | Tlcd5 | 594.6801 | -0.14765 | 0.183722 | 0.620678 | ENSMUSG( | Tlcd5 | 587.1748 | 0.089786 | 0.478066 | 0.923329 |
| ENSMUSG( | Tle1 | 1448.907 | 0.246888 | 0.004416 | 0.086676 | ENSMUSG( | Tle1 | 1241.702 | 0.078785 | 0.474821 | 0.923329 |
| ENSMUSG( | Tle2 | 746.0895 | 0.386476 | 0.280851 | 0.729868 | ENSMUSG( | Tle2 | 593.4333 | 0.015822 | 0.951438 | 0.994798 |
| ENSMUSG( | Tle3 | 1726.826 | 0.164333 | 0.090201 | 0.461412 | ENSMUSG( | Tle3 | 1559.528 | 0.141852 | 0.224883 | 0.86838 |
| ENSMUSG( | Tle4 | 1433.928 | -0.06807 | 0.44415 | 0.833239 | ENSMUSG( | Tle4 | 1316.603 | -0.04806 | 0.706137 | 0.96335 |
| ENSMUSG( | Tle5 | 29372.25 | 0.037552 | 0.631541 | 0.914073 | ENSMUSG( | Tle5 | 24954.91 | -0.17048 | 0.17802 | 0.848262 |
| ENSMUSG( | Tle6 | 80.53997 | -0.22506 | 0.304565 | 0.752574 | ENSMUSG( | Tle6 | 70.18274 | -0.34437 | 0.192337 | 0.859726 |
| ENSMUSG( | Tlk1 | 2686.373 | 0.067658 | 0.414233 | 0.819998 | ENSMUSG( | Tlk1 | 2441.86 | 0.059432 | 0.583613 | 0.944266 |
| ENSMUSG( | Tlk2 | 1642.546 | 0.063662 | 0.487058 | 0.858812 | ENSMUSG( | Tlk2 | 1465.013 | -0.00053 | 0.996507 | 0.999178 |
| ENSMUSG( | Tll1 | 56.59184 | -0.18212 | 0.515113 | 0.872463 | ENSMUSG( | Tll1 | 60.0792 | 0.248759 | 0.391185 | 0.917257 |
| ENSMUSG( | Tll2 | 29.76221 | -0.44451 | 0.201131 | 0.641362 | ENSMUSG( | Tll2 | 32.3278 | 0.088491 | 0.801703 | 0.97681 |
| ENSMUSG( | Tln1 | 2854.733 | 0.009337 | 0.914837 | 0.988585 | ENSMUSG( | Tln1 | 2551.939 | -0.04582 | 0.653481 | 0.955015 |
| ENSMUSG( | Tln2 | 5005.289 | 0.073471 | 0.409986 | 0.818258 | ENSMUSG( | Tln2 | 4303.489 | -0.09322 | 0.351131 | 0.911901 |
| ENSMUSG( | Tlnrd1 | 946.7176 | -0.02022 | 0.858182 | 0.974672 | ENSMUSG( | Tlnrd1 | 862.4356 | -0.02145 | 0.865807 | 0.987041 |
| ENSMUSG( | Tlr1 | 44.75918 | -0.35673 | 0.255564 | 0.706195 | ENSMUSG( | Tlr1 | 45.96986 | 0.054796 | 0.884037 | 0.989644 |
| ENSMUSG( | Tlr12 | 36.18639 | -0.76258 | 0.043425 | 0.319227 | ENSMUSG( | Tlr12 | 46.31663 | 0.32521 | 0.354991 | 0.912411 |
| ENSMUSG( | Tlr13 | 130.0813 | -0.04747 | 0.822807 | 0.966352 | ENSMUSG( | Tlr13 | 137.2862 | 0.371235 | 0.129272 | 0.820358 |
| ENSMUSG( | Tlr2 | 55.0462 | -0.22492 | 0.446955 | 0.834051 | ENSMUSG( | Tlr2 | 60.11796 | 0.312792 | 0.349106 | 0.911652 |
| ENSMUSG( | Tlr3 | 712.1082 | -0.26349 | 0.15903 | 0.588112 | ENSMUSG( | Tlr3 | 752.365 | 0.177037 | 0.391468 | 0.917257 |
| ENSMUSG( | Tlr4 | 58.36891 | -0.09698 | 0.756608 | 0.951479 | ENSMUSG( | Tlr4 | 63.38124 | 0.395496 | 0.193362 | 0.860096 |
| ENSMUSG( | Tlr5 | 28.33721 | -0.33917 | 0.340057 | 0.777959 | ENSMUSG( | Tlr5 | 32.19617 | 0.331884 | 0.369603 | 0.915491 |
| ENSMUSG( | Tlr6 | 34.622 | -0.56956 | 0.109843 | 0.500285 | ENSMUSG( | Tlr6 | 40.21377 | 0.220036 | 0.5802 | 0.944209 |
| ENSMUSG( | Tlr7 | 198.5941 | 0.077259 | 0.711689 | 0.940028 | ENSMUSG( | Tlr7 | 200.349 | 0.36601 | 0.175674 | 0.845305 |
| ENSMUSG( | Tlr8 | 9.96594 | -0.6858 | 0.359486 | NA | ENSMUSG( | Tlr8 | 12.09992 | 0.193936 | 0.772513 | 0.974454 |
| ENSMUSG( | Tlr9 | 56.11455 | -0.62761 | 0.082607 | 0.443819 | ENSMUSG( | Tlr9 | 63.17229 | 0.074158 | 0.84502 | 0.983863 |
| ENSMUSG( | Tm2d1 | 862.9693 | -0.17538 | 0.288461 | 0.739269 | ENSMUSG( | Tm2d1 | 824.0876 | -0.03236 | 0.857663 | 0.985884 |
| ENSMUSG( | Tm2d2 | 2128.816 | -0.29407 | 0.020226 | 0.214698 | ENSMUSG( | Tm2d2 | 2256.838 | 0.157695 | 0.289876 | 0.896726 |
| ENSMUSG( | Tm2d3 | 997.6178 | -0.31865 | 0.003923 | 0.080248 | ENSMUSG( | Tm2d3 | 1034.322 | 0.06921 | 0.630971 | 0.953251 |
| ENSMUSG( | Tm4sf1 | 352.0994 | -0.20361 | 0.384835 | 0.804031 | ENSMUSG( | Tm4sf1 | 412.2817 | 0.497102 | 0.276765 | 0.890889 |
| ENSMUSG( | Tm4sf5 | 11.00107 | -0.53248 | 0.346784 | NA | ENSMUSG( | Tm4sf5 | 9.669888 | -0.74527 | 0.268555 | NA |
| ENSMUSG( | Tm6sf1 | 459.4952 | -0.45069 | 0.000694 | 0.02534 | ENSMUSG( | Tm6sf1 | 507.7268 | 0.143352 | 0.417208 | 0.917257 |
| ENSMUSG( | Tm6sf2 | 69.91186 | 0.573688 | 0.024354 | 0.238636 | ENSMUSG( | Tm6sf2 | 53.93842 | 0.148834 | 0.647837 | 0.95422 |
| ENSMUSG( | Tm7sf2 | 1023.515 | -0.18478 | 0.114605 | 0.510543 | ENSMUSG( | Tm7sf2 | 942.22 | -0.1499 | 0.270719 | 0.888062 |
| ENSMUSG( | Tm7sf3 | 2032.539 | -0.27797 | 0.047 | 0.331741 | ENSMUSG( | Tm7sf3 | 2065.608 | 0.054268 | 0.735049 | 0.967344 |
| ENSMUSG( | Tm9sf1 | 754.5648 | -0.15759 | 0.33239 | 0.775474 | ENSMUSG( | Tm9sf1 | 804.2061 | 0.292618 | 0.134269 | 0.822912 |
| ENSMUSG( | Tm9sf2 | 3223.976 | -0.42962 | 0.009305 | 0.136095 | ENSMUSG( | Tm9sf2 | 3709.397 | 0.26762 | 0.152652 | 0.837724 |
| ENSMUSG( | Tm9sf3 | 7711.57 | -0.2436 | 0.06952 | 0.408118 | ENSMUSG( | Tm9sf3 | 7819.411 | 0.074594 | 0.631567 | 0.953251 |
| ENSMUSG( | Tm9sf4 | 3393.766 | -0.32309 | 0.013944 | 0.172776 | ENSMUSG( | Tm9sf4 | 3614.344 | 0.145629 | 0.363278 | 0.914223 |
| ENSMUSG( | Tma16 | 289.3086 | -0.2751 | 0.044836 | 0.324745 | ENSMUSG( | Tma16 | 274.0187 | -0.16324 | 0.33494 | 0.907907 |
| ENSMUSG( | Tma7 | 2582.828 | 0.019334 | 0.819795 | 0.965888 | ENSMUSG( | Tma7 | 2239.645 | -0.12923 | 0.293213 | 0.898538 |
| ENSMUSG( | Tmbim1 | 2128.567 | -0.40386 | 0.008821 | 0.132321 | ENSMUSG( | Tmbim1 | 2146.287 | -0.07854 | 0.666061 | 0.956766 |
| ENSMUSG( | Tmbim4 | 1140.839 | -0.226 | 0.057452 | 0.370033 | ENSMUSG( | Tmbim4 | 1171.008 | 0.124715 | 0.368807 | 0.915491 |
| ENSMUSG( | Tmbim6 | 24084.95 | -0.12235 | 0.274717 | 0.725078 | ENSMUSG( | Tmbim6 | 22707.94 | -0.0198 | 0.874135 | 0.989644 |
| ENSMUSG( | Tmc3 | 13.09018 | 0.293535 | 0.564067 | NA | ENSMUSG( | Tmc3 | 12.52334 | 0.457628 | 0.413892 | 0.917257 |
| ENSMUSG( | Tmc4 | 154.2177 | 0.520335 | 0.036459 | 0.294207 | ENSMUSG( | Tmc4 | 132.0013 | 0.350936 | 0.250706 | 0.878324 |
| ENSMUSG( | Tmc6 | 157.2273 | -0.33354 | 0.061973 | 0.38528 | ENSMUSG( | Tmc6 | 161.9996 | 0.055714 | 0.796051 | 0.97681 |
| ENSMUSG( | Tmc7 | 920.0139 | -0.01895 | 0.895441 | 0.984512 | ENSMUSG( | Tmc7 | 908.0158 | 0.209007 | 0.228812 | 0.868507 |
| ENSMUSG( | Tmc8 | 23.83668 | 0.224482 | 0.547278 | 0.885118 | ENSMUSG( | Tmc8 | 23.00856 | 0.386004 | 0.338702 | 0.907972 |
| ENSMUSG( | Tmcc1 | 2295.813 | 0.123319 | 0.228503 | 0.675862 | ENSMUSG( | Tmcc1 | 2246.251 | 0.313862 | 0.003211 | 0.225889 |
| ENSMUSG( | Tmcc2 | 13106.96 | 0.323967 | 0.000273 | 0.013445 | ENSMUSG( | Tmcc2 | 11518.26 | 0.229305 | 0.044662 | 0.693697 |
| ENSMUSG( | Tmcc3 | 1937.075 | -0.05874 | 0.507592 | 0.86795 | ENSMUSG( | Tmcc3 | 1744.618 | -0.09065 | 0.459288 | 0.923329 |
| ENSMUSG( | Tmco1 | 1650.393 | -0.24836 | 0.095882 | 0.475743 | ENSMUSG( | Tmco1 | 1664.646 | 0.058375 | 0.716856 | 0.965861 |
| ENSMUSG( | Tmco3 | 2304.489 | -0.09981 | 0.455663 | 0.838441 | ENSMUSG( | Tmco3 | 2175.409 | 0.007374 | 0.96115 | 0.99511 |
| ENSMUSG( | Tmco4 | 245.7388 | 0.010711 | 0.939792 | 0.993415 | ENSMUSG( | Tmco4 | 222.9113 | 0.002769 | 0.9865 | 0.997372 |
| ENSMUSG( | Tmco5 | 35.14172 | 0.285173 | 0.365321 | 0.794085 | ENSMUSG( | Tmco5 | 27.61011 | -0.12863 | 0.730642 | 0.967228 |
| ENSMUSG( | Tmco6 | 336.3974 | 0.00328 | 0.981072 | 0.997512 | ENSMUSG( | Tmco6 | 320.6091 | 0.140811 | 0.435649 | 0.919133 |
| ENSMUSG( | Tmed1 | 572.8968 | -0.15125 | 0.235238 | 0.682423 | ENSMUSG( | Tmed1 | 552.1312 | 0.018516 | 0.898427 | 0.990558 |
| ENSMUSG( | Tmed10 | 6390.236 | -0.27174 | 0.041821 | 0.315339 | ENSMUSG( | Tmed10 | 6406.037 | 0.018388 | 0.902862 | 0.991089 |
| ENSMUSG( | Tmed2 | 1730.416 | -0.38914 | 0.001791 | 0.047688 | ENSMUSG( | Tmed2 | 1814.462 | 0.044372 | 0.746812 | 0.969763 |
| ENSMUSG( | Tmed3 | 1062.019 | -0.28843 | 0.030556 | 0.266032 | ENSMUSG( | Tmed3 | 1079.213 | 0.044355 | 0.768047 | 0.974236 |
| ENSMUSG( | Tmed4 | 2193.763 | -0.38623 | 0.005236 | 0.096494 | ENSMUSG( | Tmed4 | 2348.956 | 0.107864 | 0.495462 | 0.927641 |
| ENSMUSG( | Tmed5 | 697.0332 | -0.35781 | 0.021052 | 0.22058 | ENSMUSG( | Tmed5 | 841.1898 | 0.443115 | 0.021858 | 0.575341 |
| ENSMUSG( | Tmed7 | 2725.178 | -0.44659 | 0.002259 | 0.055678 | ENSMUSG( | Tmed7 | 3035.068 | 0.164438 | 0.344128 | 0.910256 |
| ENSMUSG( | Tmed8 | 1302.483 | 0.045428 | 0.62035 | 0.911432 | ENSMUSG( | Tmed8 | 1221.919 | 0.128664 | 0.235607 | 0.871807 |
| ENSMUSG( | Tmed9 | 3922.104 | -0.36136 | 0.002448 | 0.058231 | ENSMUSG( | Tmed9 | 4099.034 | 0.058708 | 0.660975 | 0.956205 |
| ENSMUSG( | Tmeff1 | 3167.242 | -0.40787 | 0.000145 | 0.00841 | ENSMUSG( | Tmeff1 | 3395.957 | 0.089538 | 0.530883 | 0.934795 |
| ENSMUSG( | Tmeff2 | 2265.531 | -0.52234 | 0.000331 | 0.015457 | ENSMUSG( | Tmeff2 | 2455.457 | 0.0301 | 0.859737 | 0.98599 |
| ENSMUSG( | Tmem100 | 1146.038 | -0.01013 | 0.949444 | 0.993821 | ENSMUSG( | Tmem100 | 1001.034 | -0.13274 | 0.427763 | 0.917257 |
| ENSMUSG( | Tmem101 | 938.6188 | -0.32608 | 0.012159 | 0.159821 | ENSMUSG( | Tmem101 | 972.9799 | 0.064902 | 0.663976 | 0.956766 |
| ENSMUSG( | Tmem104 | 791.7309 | -0.3595 | 0.049492 | 0.339723 | ENSMUSG( | Tmem104 | 839.3295 | 0.105649 | 0.577305 | 0.942992 |
| ENSMUSG( | Tmem106a | 58.22525 | -0.18523 | 0.46485 | 0.843575 | ENSMUSG( | Tmem106a | 56.96315 | 0.057968 | 0.855691 | 0.985738 |
| ENSMUSG( | Tmem106b | 5025.869 | -0.24303 | 0.057869 | 0.371023 | ENSMUSG( | Tmem106b | 5268.517 | 0.169028 | 0.250235 | 0.877904 |
| ENSMUSG( | Tmem106c | 1102.447 | -0.08079 | 0.478186 | 0.853453 | ENSMUSG( | Tmem106c | 1010.885 | -0.06027 | 0.630624 | 0.953251 |
| ENSMUSG( | Tmem107 | 229.1668 | -0.1433 | 0.322541 | 0.767663 | ENSMUSG( | Tmem107 | 215.974 | -0.04161 | 0.799727 | 0.97681 |
| ENSMUSG( | Tmem108 | 667.1392 | -0.11905 | 0.453878 | 0.836798 | ENSMUSG( | Tmem108 | 764.3043 | 0.507294 | 0.005211 | 0.296373 |
| ENSMUSG( | Tmem109 | 1772.696 | -0.32926 | 0.034803 | 0.286843 | ENSMUSG( | Tmem109 | 1906.749 | 0.168833 | 0.325691 | 0.905339 |
| ENSMUSG( | Tmem11 | 512.562 | -0.01794 | 0.898002 | 0.985633 | ENSMUSG( | Tmem11 | 463.7653 | -0.04535 | 0.793144 | 0.97681 |
| ENSMUSG( | Tmem114 | 62.63847 | -0.36239 | 0.174744 | 0.610304 | ENSMUSG( | Tmem114 | 62.08196 | -0.11681 | 0.676485 | 0.95694 |
| ENSMUSG( | Tmem115 | 1201.762 | -0.3235 | 0.037924 | 0.300779 | ENSMUSG( | Tmem115 | 1329.727 | 0.249966 | 0.147168 | 0.836241 |
| ENSMUSG( | Tmem116 | 66.14251 | -0.32212 | 0.231436 | 0.679437 | ENSMUSG( | Tmem116 | 72.27887 | 0.21984 | 0.399544 | 0.917257 |
| ENSMUSG( | Tmem117 | 196.3338 | -0.90583 | 7.83E-08 | 2.15E-05 | ENSMUSG( | Tmem117 | 249.6266 | 0.186062 | 0.290157 | 0.896947 |
| ENSMUSG( | Tmem119 | 1617.022 | -0.06255 | 0.719026 | 0.941986 | ENSMUSG( | Tmem119 | 1797.281 | 0.480238 | 0.027051 | 0.603041 |
| ENSMUSG( | Tmem120a | 456.1929 | -0.09912 | 0.465231 | 0.843983 | ENSMUSG( | Tmem120a | 444.9107 | 0.104208 | 0.50255 | 0.927825 |
| ENSMUSG( | Tmem120b | 332.724 | -0.27645 | 0.047702 | 0.33359 | ENSMUSG( | Tmem120b | 324.4224 | -0.06417 | 0.667469 | 0.956766 |
| ENSMUSG( | Tmem121 | 492.1134 | -0.25769 | 0.061242 | 0.381822 | ENSMUSG( | Tmem121 | 508.1402 | 0.11795 | 0.470483 | 0.923329 |
| ENSMUSG( | Tmem121b | 2407.174 | -0.2544 | 0.115461 | 0.512397 | ENSMUSG( | Tmem121b | 2484.332 | 0.120415 | 0.492018 | 0.927315 |
| ENSMUSG( | Tmem123 | 692.978 | -0.29598 | 0.099803 | 0.482257 | ENSMUSG( | Tmem123 | 771.9656 | 0.291492 | 0.160819 | 0.837724 |
| ENSMUSG( | Tmem125 | 355.0511 | -0.43499 | 0.005131 | 0.095477 | ENSMUSG( | Tmem125 | 370.9625 | 0.002452 | 0.990861 | 0.998302 |
| ENSMUSG( | Tmem126a | 566.6482 | -0.03783 | 0.754117 | 0.950981 | ENSMUSG( | Tmem126a | 501.8347 | -0.1275 | 0.402921 | 0.917257 |
| ENSMUSG( | Tmem126b | 765.4239 | 0.012096 | 0.906178 | 0.986921 | ENSMUSG( | Tmem126b | 689.6146 | -0.02421 | 0.850275 | 0.985541 |
| ENSMUSG( | Tmem127 | 4974.213 | -0.14954 | 0.216345 | 0.66135 | ENSMUSG( | Tmem127 | 4939.847 | 0.104738 | 0.45807 | 0.923329 |
| ENSMUSG( | Tmem128 | 825.2005 | -0.35627 | 0.011183 | 0.152553 | ENSMUSG( | Tmem128 | 869.6077 | 0.088211 | 0.56971 | 0.941777 |
| ENSMUSG( | Tmem129 | 1103.295 | -0.22087 | 0.105093 | 0.492392 | ENSMUSG( | Tmem129 | 1172.346 | 0.228261 | 0.139625 | 0.830678 |
| ENSMUSG( | Tmem130 | 5743.355 | -0.29118 | 0.018689 | 0.205652 | ENSMUSG( | Tmem130 | 6014.357 | 0.124301 | 0.38274 | 0.917257 |
| ENSMUSG( | Tmem131 | 2614.233 | -0.18944 | 0.131736 | 0.544244 | ENSMUSG( | Tmem131 | 2828.878 | 0.301149 | 0.052772 | 0.71525 |
| ENSMUSG( | Tmem131l | 655.7891 | -0.09123 | 0.480121 | 0.854649 | ENSMUSG( | Tmem131l | 661.151 | 0.201078 | 0.137692 | 0.828492 |
| ENSMUSG( | Tmem132a | 10285.1 | -0.11766 | 0.444598 | 0.833239 | ENSMUSG( | Tmem132a | 10613.66 | 0.23929 | 0.182659 | 0.850579 |

| | | | | | |
|---|---|---|---|---|---|
| ENSMUSG( | 11946.51 | 0.155978 | 0.12347 | 0.530552 | Tcf4 |
| ENSMUSG( | 166.5151 | 0.012996 | 0.949323 | 0.993821 | Tcf7 |
| ENSMUSG( | 338.1443 | -0.18695 | 0.194092 | 0.633181 | Tcf7l1 |
| ENSMUSG( | 297.1693 | -0.44241 | 0.00221 | 0.054639 | Tcf7l2 |
| ENSMUSG( | 53.9241 | -0.32217 | 0.212651 | 0.6567 | Tcfl5 |
| ENSMUSG( | 84.46327 | 0.216377 | 0.360016 | 0.791685 | Tchh |
| ENSMUSG( | 323.6726 | 0.222543 | 0.141462 | 0.558606 | Tchp |
| ENSMUSG( | 89.65769 | 0.280808 | 0.220264 | 0.665898 | Tcim |
| ENSMUSG( | 706.864 | -0.08835 | 0.458446 | 0.840036 | Tcirg1 |
| ENSMUSG( | 1166.262 | -0.13386 | 0.39026 | 0.80794 | Tcn2 |
| ENSMUSG( | 639.3074 | 0.139798 | 0.212753 | 0.6567 | Tcof1 |
| ENSMUSG( | 4658.006 | 0.098646 | 0.317251 | 0.763823 | Tcp1 |
| ENSMUSG( | 16.11799 | -0.26925 | 0.569215 | NA | Tcp11 |
| ENSMUSG( | 769.0726 | 0.077679 | 0.521103 | 0.875186 | Tcp11l1 |
| ENSMUSG( | 717.2032 | 0.073321 | 0.513985 | 0.872055 | Tcp11l2 |
| ENSMUSG( | 1074.415 | -0.06638 | 0.543336 | 0.883854 | Tcta |
| ENSMUSG( | 85.49818 | -0.09629 | 0.663241 | 0.9265 | Tcte1 |
| ENSMUSG( | 115.4548 | 0.095743 | 0.608571 | 0.907154 | Tcte2 |
| ENSMUSG( | 911.7519 | -0.12795 | 0.310885 | 0.758963 | Tctn1 |
| ENSMUSG( | 841.7372 | 0.033163 | 0.789417 | 0.959415 | Tctn2 |
| ENSMUSG( | 822.5312 | 0.003468 | 0.979541 | 0.997428 | Tctn3 |
| ENSMUSG( | 818.9317 | 0.062104 | 0.528631 | 0.878365 | Tdg |
| ENSMUSG( | 24.17162 | -0.27325 | 0.500149 | 0.863455 | Tdo2 |
| ENSMUSG( | 379.5183 | 0.271503 | 0.051124 | 0.346957 | Tdp1 |
| ENSMUSG( | 366.5895 | 0.030154 | 0.806723 | 0.962932 | Tdp2 |
| ENSMUSG( | 16.38759 | 0.394961 | 0.486763 | NA | Tdrd1 |
| ENSMUSG( | 21.33659 | 0.001757 | 0.996679 | 0.99943 | Tdrd12 |
| ENSMUSG( | 811.7336 | 0.100172 | 0.335417 | 0.776877 | Tdrd3 |
| ENSMUSG( | 34.11885 | 0.13594 | 0.704614 | 0.938303 | Tdrd5 |
| ENSMUSG( | 22.30106 | -0.33878 | 0.405729 | 0.81652 | Tdrd6 |
| ENSMUSG( | 1303.476 | 0.063337 | 0.535115 | 0.880733 | Tdrd7 |
| ENSMUSG( | 17.81763 | -0.19936 | 0.699284 | NA | Tdrd9 |
| ENSMUSG( | 1435.89 | 0.131733 | 0.198416 | 0.638387 | Tdrkh |
| ENSMUSG( | 361.8013 | 0.031624 | 0.815353 | 0.965049 | Tdrp |
| ENSMUSG( | 1505.732 | 0.197775 | 0.122504 | 0.528322 | Tead1 |
| ENSMUSG( | 265.7043 | 0.226639 | 0.097762 | 0.478264 | Tead2 |
| ENSMUSG( | 164.5208 | -0.10262 | 0.582229 | 0.897918 | Tead3 |
| ENSMUSG( | 16.98782 | 0.825328 | 0.090328 | NA | Tead4 |
| ENSMUSG( | 82.04782 | -0.29257 | 0.210817 | 0.653564 | Tec |
| ENSMUSG( | 3710.872 | 0.129944 | 0.162797 | 0.593778 | Tecpr1 |
| ENSMUSG( | 2702.697 | -0.04844 | 0.632584 | 0.914073 | Tecpr2 |
| ENSMUSG( | 13766.75 | -0.35143 | 0.000571 | 0.022438 | Tecr |
| ENSMUSG( | 16.18311 | -0.65296 | 0.19315 | NA | Tecta |
| ENSMUSG( | 12.05197 | -0.76224 | 0.20093 | NA | Tectb |
| ENSMUSG( | 60.13413 | 0.054146 | 0.842865 | 0.97173 | Tedc1 |
| ENSMUSG( | 241.8608 | 0.099999 | 0.533259 | 0.879905 | Tedc2 |
| ENSMUSG( | 637.1458 | 0.112826 | 0.419124 | 0.821966 | Teddm2 |
| ENSMUSG( | 11.34315 | -0.17245 | 0.774395 | NA | Teddm3 |
| ENSMUSG( | 12363.67 | 0.10403 | 0.213602 | 0.657797 | Tef |
| ENSMUSG( | 107.6327 | -0.03287 | 0.86702 | 0.976541 | Tefm |
| ENSMUSG( | 518.7296 | -0.15 | 0.688974 | 0.933441 | Tek |
| ENSMUSG( | 71.7715 | -0.31052 | 0.185354 | 0.623185 | Tekt1 |
| ENSMUSG( | 49.93274 | -1.57405 | 5.22E-08 | 1.69E-05 | Tekt2 |
| ENSMUSG( | 64.71519 | 0.790128 | 0.004016 | 0.081522 | Tekt4 |
| ENSMUSG( | 241.3986 | -0.0925 | 0.570952 | 0.8933 | Tekt5 |
| ENSMUSG( | 19.29813 | -0.32075 | 0.50846 | NA | Tektip1 |
| ENSMUSG( | 597.0896 | 0.261172 | 0.026092 | 0.247245 | Telo2 |
| ENSMUSG( | 519.844 | -0.08545 | 0.471243 | 0.847103 | Ten1 |
| ENSMUSG( | 834.4057 | -0.44386 | 0.004387 | 0.086463 | Tenm1 |
| ENSMUSG( | 2258.82 | -0.18066 | 0.186056 | 0.62352 | Tenm2 |
| ENSMUSG( | 2086.132 | -0.1775 | 0.24279 | 0.690275 | Tenm3 |
| ENSMUSG( | 3607.101 | -0.36749 | 0.001979 | 0.051182 | Tenm4 |
| ENSMUSG( | 574.4728 | 0.070083 | 0.547854 | 0.885153 | Tent2 |
| ENSMUSG( | 1370.412 | 0.10637 | 0.249765 | 0.699445 | Tent4a |
| ENSMUSG( | 987.414 | 0.086176 | 0.383906 | 0.803646 | Tent4b |
| ENSMUSG( | 123.7602 | -0.54765 | 0.002266 | 0.055678 | Tent5a |
| ENSMUSG( | 179.6612 | -0.25709 | 0.31306 | 0.76033 | Tent5c |
| ENSMUSG( | 192.6078 | -0.07904 | 0.61089 | 0.908466 | Tep1 |
| ENSMUSG( | 541.6265 | 0.245737 | 0.044848 | 0.324745 | Tepsin |
| ENSMUSG( | 32.17371 | 0.336898 | 0.327703 | 0.772692 | Terb1 |
| ENSMUSG( | 207.2114 | -0.09092 | 0.588968 | 0.90113 | Terf1 |
| ENSMUSG( | 1018.991 | 0.049816 | 0.649923 | 0.920899 | Terf2 |
| ENSMUSG( | 1061.431 | 0.136631 | 0.225618 | 0.67246 | Terf2ip |
| ENSMUSG( | 44.32516 | 0.140738 | 0.621775 | 0.911432 | Tert |
| ENSMUSG( | 80.62794 | -0.23351 | 0.410223 | 0.818258 | Tes |
| ENSMUSG( | 9.862838 | 0.195671 | 0.735196 | NA | Tes3-ps |
| ENSMUSG( | 864.0597 | -0.25504 | 0.061074 | 0.381679 | Tesc |
| ENSMUSG( | 3892.084 | 0.162375 | 0.047238 | 0.332553 | Tesk1 |
| ENSMUSG( | 287.0617 | 0.218631 | 0.12573 | 0.53257 | Tesk2 |
| ENSMUSG( | 491.6689 | 0.06356 | 0.646913 | 0.919062 | Tet1 |
| ENSMUSG( | 1690.567 | -0.39438 | 4.37E-05 | 0.003581 | Tet2 |
| ENSMUSG( | 1778.761 | -0.0318 | 0.742695 | 0.948478 | Tet3 |
| ENSMUSG( | 365.7725 | 0.269238 | 0.031798 | 0.272023 | Tex10 |
| ENSMUSG( | 14.84724 | 0.158548 | 0.738662 | NA | Tex14 |
| ENSMUSG( | 29.74037 | -0.25849 | 0.448158 | 0.834051 | Tex15 |
| ENSMUSG( | 3751.348 | -0.00733 | 0.946174 | 0.99369 | Tex2 |
| ENSMUSG( | 2231.036 | -0.12137 | 0.288176 | 0.738889 | Tex261 |
| ENSMUSG( | 2677.247 | -0.19601 | 0.100718 | 0.484034 | Tex264 |
| ENSMUSG( | 35.96219 | -0.23429 | 0.538107 | 0.882548 | Tex30 |
| ENSMUSG( | 64.05448 | -0.69462 | 0.0069 | 0.114527 | Tex52 |
| ENSMUSG( | 397.2426 | -0.10497 | 0.392237 | 0.809274 | Tex9 |
| ENSMUSG( | 820.9244 | 0.132319 | 0.220295 | 0.665898 | Tfam |
| ENSMUSG( | 23.39013 | -0.07359 | 0.854333 | 0.973491 | Tfap2b |
| ENSMUSG( | 10.07143 | 0.082829 | 0.885083 | NA | Tfap2c |
| ENSMUSG( | 14.26992 | -0.62726 | 0.307549 | NA | Tfap2d |
| ENSMUSG( | 69.15073 | -0.11187 | 0.629256 | 0.913965 | Tfap4 |
| ENSMUSG( | 92.3928 | 0.162094 | 0.478836 | 0.853862 | Tfb1m |
| ENSMUSG( | 351.6157 | -0.00943 | 0.945313 | 0.99365 | Tfb2m |
| ENSMUSG( | 611.9003 | 0.107624 | 0.361743 | 0.792797 | Tfcp2 |
| ENSMUSG( | 1153.411 | 0.125572 | 0.248751 | 0.697857 | Tfcp2l1 |
| ENSMUSG( | 1426.983 | 0.119791 | 0.191652 | 0.630674 | Tfdp1 |
| ENSMUSG( | 1432.189 | -0.04532 | 0.625965 | 0.912356 | Tfdp2 |
| ENSMUSG( | 1327.117 | -0.04519 | 0.621217 | 0.911432 | Tfe3 |
| ENSMUSG( | 423.0416 | -0.03501 | 0.760491 | 0.952907 | Tfeb |
| ENSMUSG( | 1005.939 | 0.186955 | 0.056513 | 0.366657 | Tfg |
| ENSMUSG( | 1343.47 | 0.130016 | 0.339201 | 0.777534 | Tfip11 |
| ENSMUSG( | 213.0937 | -0.03165 | 0.845369 | 0.97218 | Tfpi |
| ENSMUSG( | 516.7781 | -0.0031 | 0.980042 | 0.997491 | Tfpt |
| ENSMUSG( | 231.9736 | 0.325435 | 0.03588 | 0.290992 | Tfr2 |
| ENSMUSG( | 4133.313 | -0.14963 | 0.286349 | 0.735717 | Tfrc |
| ENSMUSG( | 153.5101 | 0.135698 | 0.413954 | 0.819998 | Tgds |
| ENSMUSG( | 2155.401 | -0.3269 | 0.007551 | 0.120197 | Tgfa |
| ENSMUSG( | 471.8627 | -0.27204 | 0.084564 | 0.448139 | Tgfb1 |
| ENSMUSG( | 831.5445 | 0.04146 | 0.690809 | 0.933881 | Tgfb1i1 |
| ENSMUSG( | 871.308 | -0.04704 | 0.674461 | 0.929638 | Tgfb2 |
| ENSMUSG( | 647.0278 | -0.14241 | 0.205257 | 0.64654 | Tgfb3 |
| ENSMUSG( | 318.3878 | -0.215 | 0.127064 | 0.534624 | Tgfbi |
| ENSMUSG( | 1401.459 | -0.08594 | 0.616429 | 0.910866 | Tgfbr1 |
| ENSMUSG( | 856.3405 | -0.1898 | 0.261521 | 0.711485 | Tgfbr2 |
| ENSMUSG( | 399.6452 | -0.23203 | 0.071978 | 0.414057 | Tgfbr3 |
| ENSMUSG( | 49.45778 | 0.276899 | 0.354855 | 0.78738 | Tgfbr3l |
| ENSMUSG( | 2653.8 | 0.093951 | 0.288703 | 0.739578 | Tgfbrap1 |
| ENSMUSG( | 53.45826 | -0.47737 | 0.080624 | 0.438981 | Tgif1 |
| ENSMUSG( | 82.99375 | -0.19562 | 0.402127 | 0.815103 | Tgif2 |
| ENSMUSG( | 418.9951 | 0.195482 | 0.221071 | 0.666624 | Tgm2 |
| ENSMUSG( | 96.45144 | 0.055923 | 0.795923 | 0.960062 | Tgm3 |
| ENSMUSG( | 21.08925 | 0.03377 | 0.938136 | NA | Tgm4 |
| ENSMUSG( | 15.77835 | 0.284889 | 0.564793 | NA | Tgm7 |
| ENSMUSG( | 5353.483 | -0.12198 | 0.389545 | 0.807428 | Tgoln1 |
| ENSMUSG( | 641.8414 | 0.163678 | 0.121372 | 0.525713 | Tgs1 |
| ENSMUSG( | 19.92322 | -0.40629 | 0.368893 | NA | Tgtp1 |
| ENSMUSG( | 16.38436 | -0.57098 | 0.216898 | NA | Tgtp2 |
| ENSMUSG( | 75.2246 | -0.5262 | 0.173037 | 0.608361 | Th |
| ENSMUSG( | 174.0309 | 0.124202 | 0.46343 | 0.842789 | Tha1 |
| ENSMUSG( | 275.88 | 0.212681 | 0.174667 | 0.610304 | Thada |
| ENSMUSG( | 285.2042 | 0.088434 | 0.496589 | 0.86175 | Thap1 |
| ENSMUSG( | 638.7743 | 0.107198 | 0.430648 | 0.826896 | Thap11 |
| ENSMUSG( | 5188.892 | 0.118181 | 0.30141 | 0.749979 | Thap12 |
| ENSMUSG( | 671.5422 | 0.128963 | 0.251354 | 0.700869 | Thap2 |
| ENSMUSG( | 343.3462 | -0.10961 | 0.494219 | 0.860611 | Thap3 |
| ENSMUSG( | 413.9913 | -0.03613 | 0.794694 | 0.959799 | Thap4 |
| ENSMUSG( | 132.9486 | -0.00196 | 0.992057 | 0.999265 | Thap6 |
| ENSMUSG( | 669.4961 | 0.139265 | 0.193828 | 0.633085 | Thap7 |
| ENSMUSG( | 900.489 | 0.031215 | 0.850865 | 0.972638 | Thbd |
| ENSMUSG( | 217.6063 | -1.08742 | 9.48E-07 | 0.000184 | Thbs1 |
| ENSMUSG( | 323.8241 | -0.39629 | 0.006941 | 0.114527 | Thbs2 |
| ENSMUSG( | 586.0745 | 0.151461 | 0.336647 | 0.777 | Thbs3 |
| ENSMUSG( | 794.7489 | -0.12426 | 0.530819 | 0.879289 | Thbs4 |
| ENSMUSG( | 700.968 | -0.011 | 0.936864 | 0.993415 | Them4 |
| ENSMUSG( | 1874.953 | -0.46996 | 4.57E-05 | 0.003685 | Them6 |
| ENSMUSG( | 65.08993 | -0.43186 | 0.10189 | 0.486318 | Themis2 |
| ENSMUSG( | 208.7484 | -0.03314 | 0.84342 | 0.971847 | Thg1l |
| ENSMUSG( | 402.914 | 0.030937 | 0.791421 | 0.959461 | Thnsl1 |
| ENSMUSG( | 213.3944 | 0.11594 | 0.464196 | 0.843143 | Thnsl2 |
| ENSMUSG( | 1037.297 | 0.105752 | 0.341418 | 0.778828 | Thoc1 |
| ENSMUSG( | 1151.892 | 0.017853 | 0.878965 | 0.981436 | Thoc2 |
| ENSMUSG( | 678.278 | -0.17031 | 0.184633 | 0.621704 | Thoc2l |
| ENSMUSG( | 606.3827 | 0.075625 | 0.556032 | 0.887457 | Thoc3 |

| | | | | | |
|---|---|---|---|---|---|
| ENSMUSG( | 15.72164 | -0.32868 | 0.530227 | 0.934795 | Tgtp2 |
| ENSMUSG( | 83.52564 | 0.076664 | 0.928855 | 0.99306 | Th |
| ENSMUSG( | 159.8435 | 0.153204 | 0.399245 | 0.917257 | Tha1 |
| ENSMUSG( | 242.4977 | 0.129145 | 0.500817 | 0.927825 | Thada |
| ENSMUSG( | 243.2328 | -0.10462 | 0.531583 | 0.934795 | Thap1 |
| ENSMUSG( | 559.1107 | -0.00928 | 0.947628 | 0.994798 | Thap11 |
| ENSMUSG( | 4416.249 | -0.07911 | 0.547793 | 0.938063 | Thap12 |
| ENSMUSG( | 622.8681 | 0.172401 | 0.21885 | 0.866104 | Thap2 |
| ENSMUSG( | 328.4486 | 0.03501 | 0.848253 | 0.985321 | Thap3 |
| ENSMUSG( | 350.933 | -0.2584 | 0.114845 | 0.811703 | Thap4 |
| ENSMUSG( | 119.4424 | -0.04871 | 0.816152 | 0.979198 | Thap6 |
| ENSMUSG( | 570.3061 | -0.05351 | 0.676985 | 0.957106 | Thap7 |
| ENSMUSG( | 928.4263 | 0.37048 | 0.024077 | 0.588226 | Thbd |
| ENSMUSG( | 259.0496 | -0.10077 | 0.680876 | 0.958395 | Thbs1 |
| ENSMUSG( | 361.7188 | 0.217741 | 0.169743 | 0.840806 | Thbs2 |
| ENSMUSG( | 523.901 | 0.100881 | 0.57669 | 0.942912 | Thbs3 |
| ENSMUSG( | 830.4531 | 0.270701 | 0.235779 | 0.871888 | Thbs4 |
| ENSMUSG( | 572.8041 | -0.35555 | 0.036428 | 0.66949 | Them4 |
| ENSMUSG( | 2010.276 | 0.040964 | 0.767448 | 0.974236 | Them6 |
| ENSMUSG( | 68.59467 | 0.031747 | 0.918983 | 0.992529 | Themis2 |
| ENSMUSG( | 178.8879 | -0.21832 | 0.238831 | 0.873304 | Thg1l |
| ENSMUSG( | 347.0372 | -0.1351 | 0.389452 | 0.917257 | Thnsl1 |
| ENSMUSG( | 181.8553 | -0.08729 | 0.677456 | 0.957106 | Thnsl2 |
| ENSMUSG( | 978.0128 | 0.199353 | 0.125543 | 0.81516 | Thoc1 |
| ENSMUSG( | 1118.106 | 0.192043 | 0.147269 | 0.836241 | Thoc2 |
| ENSMUSG( | 712.3339 | 0.239504 | 0.10078 | 0.801456 | Thoc2l |
| ENSMUSG( | 507.3462 | -0.1754 | 0.228081 | 0.868507 | Thoc3 |
| ENSMUSG( | 632.4956 | -0.07044 | 0.621326 | 0.953199 | Thoc5 |
| ENSMUSG( | 123.0435 | -0.16982 | 0.445782 | 0.921734 | Thoc6 |
| ENSMUSG( | 1105.125 | -0.07237 | 0.528872 | 0.934795 | Thoc7 |
| ENSMUSG( | 862.2644 | -0.15086 | 0.241025 | 0.876146 | Thop1 |
| ENSMUSG( | 188.2991 | -0.09045 | 0.734394 | 0.967228 | Thpo |
| ENSMUSG( | 34014.68 | -0.00244 | 0.980367 | 0.996479 | Thra |
| ENSMUSG( | 2730.937 | -0.04714 | 0.67351 | 0.95694 | Thrap3 |
| ENSMUSG( | 959.9899 | 0.097609 | 0.403637 | 0.917257 | Thrb |
| ENSMUSG( | 931.5374 | -0.31517 | 0.006084 | 0.321954 | Thrsp |
| ENSMUSG( | 196.9336 | 0.188877 | 0.487076 | 0.92541 | Thsd1 |
| ENSMUSG( | 547.5276 | 0.144126 | 0.353819 | 0.912411 | Thsd4 |
| ENSMUSG( | 987.2684 | 0.094218 | 0.591534 | 0.944975 | Thsd7a |
| ENSMUSG( | 228.2235 | 0.160155 | 0.492543 | 0.927315 | Thsd7b |
| ENSMUSG( | 240.5885 | -0.01654 | 0.920297 | 0.992644 | Thtpa |
| ENSMUSG( | 629.7778 | -0.06152 | 0.654289 | 0.955015 | Thumpd1 |
| ENSMUSG( | 120.904 | -0.03269 | 0.886471 | 0.990011 | Thumpd2 |
| ENSMUSG( | 433.9712 | -0.16058 | 0.242348 | 0.877129 | Thumpd3 |
| ENSMUSG( | 31308.41 | 0.044882 | 0.746114 | 0.969632 | Thy1 |
| ENSMUSG( | 571.5112 | -0.0676 | 0.631192 | 0.953251 | Thyn1 |
| ENSMUSG( | 2291.928 | 0.310611 | 0.026281 | 0.603041 | Tia1 |
| ENSMUSG( | 2447.948 | 0.262959 | 0.0586 | 0.728256 | Tial1 |
| ENSMUSG( | 2063.532 | 0.15636 | 0.183391 | 0.850984 | Tiam1 |
| ENSMUSG( | 1065.276 | -0.06797 | 0.626602 | 0.953234 | Tiam2 |
| ENSMUSG( | 196.9371 | -0.11955 | 0.474688 | 0.923329 | Ticam1 |
| ENSMUSG( | 24.55528 | 0.272588 | 0.491175 | 0.926604 | Ticam2 |
| ENSMUSG( | 10.20819 | -0.00262 | 0.996881 | NA | Ticrr |
| ENSMUSG( | 578.6983 | 0.416709 | 0.348956 | 0.911652 | Tie1 |
| ENSMUSG( | 326.6787 | 0.054728 | 0.711404 | 0.964363 | Tifa |
| ENSMUSG( | 181.0382 | 0.353873 | 0.053014 | 0.71525 | Tifab |
| ENSMUSG( | 476.6501 | -0.06976 | 0.644759 | 0.953903 | Tigar |
| ENSMUSG( | 182.1559 | -0.06691 | 0.694278 | 0.960867 | Tigd2 |
| ENSMUSG( | 52.5031 | -0.1323 | 0.688149 | 0.959742 | Tigd3 |
| ENSMUSG( | 107.6454 | 0.072163 | 0.757198 | 0.971858 | Tigd5 |
| ENSMUSG( | 130.368 | 0.273572 | 0.231299 | 0.868507 | Timeless |
| ENSMUSG( | 285.8565 | -0.18601 | 0.353493 | 0.912263 | Timm10 |
| ENSMUSG( | 308.8277 | 0.019287 | 0.905352 | 0.991089 | Timm10b |
| ENSMUSG( | 1596.608 | -0.26378 | 0.031864 | 0.632006 | Timm13 |
| ENSMUSG( | 1489.643 | 0.000657 | 0.995669 | 0.999116 | Timm17a |
| ENSMUSG( | 613.5564 | -0.22644 | 0.169414 | 0.840806 | Timm17b |
| ENSMUSG( | 510.0202 | 0.145488 | 0.335467 | 0.907907 | Timm21 |
| ENSMUSG( | 570.6849 | -0.05443 | 0.673468 | 0.95694 | Timm22 |
| ENSMUSG( | 758.571 | -0.16812 | 0.183792 | 0.85194 | Timm23 |
| ENSMUSG( | 1866.506 | -0.06192 | 0.57349 | 0.942912 | Timm29 |
| ENSMUSG( | 1198.34 | 0.097185 | 0.501917 | 0.927825 | Timm44 |
| ENSMUSG( | 820.7142 | -0.15634 | 0.215596 | 0.865167 | Timm50 |
| ENSMUSG( | 144.3032 | 0.004297 | 0.982416 | 0.996756 | Timm8a1 |
| ENSMUSG( | 2011.671 | -0.25315 | 0.049501 | 0.708648 | Timm8b |
| ENSMUSG( | 620.147 | -0.0343 | 0.834905 | 0.981237 | Timm9 |
| ENSMUSG( | 540.3079 | 0.014106 | 0.913253 | 0.992298 | Timmdc1 |
| ENSMUSG( | 32.01666 | -0.3801 | 0.283365 | 0.892412 | Timp1 |
| ENSMUSG( | 11840.93 | 0.107148 | 0.474916 | 0.923329 | Timp2 |
| ENSMUSG( | 8026.364 | -0.07722 | 0.552235 | 0.939565 | Timp3 |
| ENSMUSG( | 3496.778 | -0.17653 | 0.11811 | 0.813384 | Timp4 |
| ENSMUSG( | 269.6928 | 1.2876 | 2.23E-07 | 0.000135 | Tinagl1 |
| ENSMUSG( | 75.4753 | 0.327181 | 0.312406 | 0.900037 | Tincr |
| ENSMUSG( | 393.6442 | 0.122224 | 0.415835 | 0.917257 | Tinf2 |
| ENSMUSG( | 563.7571 | -0.07603 | 0.557735 | 0.939565 | Tiparp |
| ENSMUSG( | 416.3028 | -0.25077 | 0.063408 | 0.736482 | Tipin |
| ENSMUSG( | 905.6235 | -0.06686 | 0.640065 | 0.953903 | Tiprl |
| ENSMUSG( | 92.35932 | 0.013199 | 0.954241 | 0.994798 | Tirap |
| ENSMUSG( | 818.5472 | 0.259745 | 0.092138 | 0.791946 | Tjap1 |
| ENSMUSG( | 4824.99 | -0.05827 | 0.589501 | 0.944975 | Tjp1 |
| ENSMUSG( | 1997.201 | -0.23959 | 0.044337 | 0.693697 | Tjp2 |
| ENSMUSG( | 20.84644 | -0.20543 | 0.681013 | 0.958395 | Tjp3 |
| ENSMUSG( | 21.63083 | -0.42568 | 0.323083 | 0.904023 | Tk1 |
| ENSMUSG( | 1191.843 | -0.06331 | 0.61107 | 0.950771 | Tk2 |
| ENSMUSG( | 455.2233 | -0.08239 | 0.573957 | 0.942912 | Tkfc |
| ENSMUSG( | 5739.512 | -0.07315 | 0.489392 | 0.925675 | Tkt |
| ENSMUSG( | 7.493295 | 0.013261 | 0.985931 | NA | Tktl2 |
| ENSMUSG( | 2211.619 | 0.173878 | 0.193182 | 0.860096 | Tlcd1 |
| ENSMUSG( | 16.15216 | -0.06634 | 0.894322 | 0.990558 | Tlcd2 |
| ENSMUSG( | 780.5642 | 0.049723 | 0.74884 | 0.96984 | Tlcd3a |
| ENSMUSG( | 4014.585 | -0.05643 | 0.563579 | 0.941206 | Tlcd3b |
| ENSMUSG( | 2602.181 | -0.0162 | 0.921299 | 0.992644 | Tlcd4 |
| ENSMUSG( | 587.1748 | 0.089786 | 0.478066 | 0.923329 | Tlcd5 |
| ENSMUSG( | 1241.702 | 0.078785 | 0.474821 | 0.923329 | Tle1 |
| ENSMUSG( | 593.4333 | 0.015822 | 0.951438 | 0.994798 | Tle2 |
| ENSMUSG( | 1559.528 | 0.141852 | 0.224883 | 0.86838 | Tle3 |
| ENSMUSG( | 1316.603 | -0.04806 | 0.706137 | 0.96335 | Tle4 |
| ENSMUSG( | 24954.91 | -0.17048 | 0.17802 | 0.848262 | Tle5 |
| ENSMUSG( | 70.18274 | -0.34437 | 0.192337 | 0.859726 | Tle6 |
| ENSMUSG( | 2441.86 | 0.059432 | 0.583613 | 0.944266 | Tlk1 |
| ENSMUSG( | 1465.013 | -0.00053 | 0.996507 | 0.999178 | Tlk2 |
| ENSMUSG( | 60.0792 | 0.248759 | 0.391185 | 0.917257 | Tll1 |
| ENSMUSG( | 32.3278 | 0.088491 | 0.801703 | 0.97681 | Tll2 |
| ENSMUSG( | 2551.939 | -0.04582 | 0.653481 | 0.955015 | Tln1 |
| ENSMUSG( | 4303.489 | -0.09322 | 0.351131 | 0.911901 | Tln2 |
| ENSMUSG( | 862.4356 | -0.02145 | 0.865807 | 0.987041 | Tlnrd1 |
| ENSMUSG( | 45.96986 | 0.054796 | 0.884037 | 0.989644 | Tlr1 |
| ENSMUSG( | 46.31663 | 0.32521 | 0.354991 | 0.912411 | Tlr12 |
| ENSMUSG( | 137.2862 | 0.371235 | 0.129272 | 0.820358 | Tlr13 |
| ENSMUSG( | 60.11796 | 0.312792 | 0.349106 | 0.911652 | Tlr2 |
| ENSMUSG( | 752.365 | 0.177037 | 0.391468 | 0.917257 | Tlr3 |
| ENSMUSG( | 63.38124 | 0.395496 | 0.193362 | 0.860096 | Tlr4 |
| ENSMUSG( | 32.19617 | 0.331884 | 0.369603 | 0.915491 | Tlr5 |
| ENSMUSG( | 40.21377 | 0.220036 | 0.5802 | 0.944209 | Tlr6 |
| ENSMUSG( | 200.349 | 0.36601 | 0.175674 | 0.845305 | Tlr7 |
| ENSMUSG( | 12.09992 | 0.193936 | 0.772513 | 0.974454 | Tlr8 |
| ENSMUSG( | 63.17229 | 0.074158 | 0.84502 | 0.983863 | Tlr9 |
| ENSMUSG( | 824.0876 | -0.03236 | 0.857663 | 0.985884 | Tm2d1 |
| ENSMUSG( | 2256.838 | 0.157695 | 0.289876 | 0.896726 | Tm2d2 |
| ENSMUSG( | 1034.322 | 0.06921 | 0.630971 | 0.953251 | Tm2d3 |
| ENSMUSG( | 412.2817 | 0.497102 | 0.276765 | 0.890889 | Tm4sf1 |
| ENSMUSG( | 9.669888 | -0.74527 | 0.268555 | NA | Tm4sf5 |
| ENSMUSG( | 507.7268 | 0.143352 | 0.417208 | 0.917257 | Tm6sf1 |
| ENSMUSG( | 53.93842 | 0.148834 | 0.647837 | 0.95422 | Tm6sf2 |
| ENSMUSG( | 942.22 | -0.1499 | 0.270719 | 0.888062 | Tm7sf2 |
| ENSMUSG( | 2065.608 | 0.054268 | 0.735049 | 0.967344 | Tm7sf3 |
| ENSMUSG( | 804.2061 | 0.292618 | 0.134269 | 0.822912 | Tm9sf1 |
| ENSMUSG( | 3709.397 | 0.26762 | 0.152652 | 0.837724 | Tm9sf2 |
| ENSMUSG( | 7819.411 | 0.074594 | 0.631567 | 0.953251 | Tm9sf3 |
| ENSMUSG( | 3614.344 | 0.145629 | 0.363278 | 0.914223 | Tm9sf4 |
| ENSMUSG( | 274.0187 | -0.16324 | 0.33494 | 0.907907 | Tma16 |
| ENSMUSG( | 2239.645 | -0.12923 | 0.293213 | 0.898538 | Tma7 |
| ENSMUSG( | 2146.287 | -0.07854 | 0.666061 | 0.956766 | Tmbim1 |
| ENSMUSG( | 1171.008 | 0.124715 | 0.368807 | 0.915491 | Tmbim4 |
| ENSMUSG( | 22707.94 | -0.0198 | 0.874135 | 0.989644 | Tmbim6 |
| ENSMUSG( | 12.52334 | 0.457628 | 0.413892 | 0.917257 | Tmc3 |
| ENSMUSG( | 132.0013 | 0.350936 | 0.250706 | 0.878324 | Tmc4 |
| ENSMUSG( | 161.9996 | 0.055714 | 0.796051 | 0.97681 | Tmc6 |
| ENSMUSG( | 908.0158 | 0.209007 | 0.228812 | 0.868507 | Tmc7 |
| ENSMUSG( | 23.00856 | 0.386004 | 0.338702 | 0.907972 | Tmc8 |
| ENSMUSG( | 2246.251 | 0.313862 | 0.003211 | 0.225889 | Tmcc1 |
| ENSMUSG( | 11518.26 | 0.229305 | 0.044662 | 0.693697 | Tmcc2 |
| ENSMUSG( | 1744.618 | -0.09065 | 0.459288 | 0.923329 | Tmcc3 |
| ENSMUSG( | 1664.646 | 0.058375 | 0.716856 | 0.965861 | Tmco1 |
| ENSMUSG( | 2175.409 | 0.007374 | 0.96115 | 0.99511 | Tmco3 |
| ENSMUSG( | 222.9113 | 0.002769 | 0.9865 | 0.997372 | Tmco4 |
| ENSMUSG( | 27.61011 | -0.12863 | 0.730642 | 0.967228 | Tmco5 |
| ENSMUSG( | 320.6091 | 0.140811 | 0.435649 | 0.919133 | Tmco6 |
| ENSMUSG( | 552.1312 | 0.018516 | 0.898427 | 0.990558 | Tmed1 |

| | | | | | | | | | | | |
|---|---|---|---|---|---|---|---|---|---|---|---|
| ENSMUSG( | Tmem132b | 4492.984 | -0.35666 | 0.01893 | 0.207196 | ENSMUSG( | Tmem132b | 4957.235 | 0.214456 | 0.219161 | 0.866104 |
| ENSMUSG( | Tmem132c | 183.525 | -0.30837 | 0.110394 | 0.50108 | ENSMUSG( | Tmem132c | 198.9485 | 0.217033 | 0.327756 | 0.906255 |
| ENSMUSG( | Tmem132c | 1479.928 | -0.15996 | 0.31749 | 0.763823 | ENSMUSG( | Tmem132c | 1541.687 | 0.227509 | 0.208645 | 0.864412 |
| ENSMUSG( | Tmem132e | 419.6527 | -0.22554 | 0.205988 | 0.647757 | ENSMUSG( | Tmem132e | 438.0812 | 0.176825 | 0.307881 | 0.898538 |
| ENSMUSG( | Tmem134 | 1822.5 | 0.055991 | 0.549245 | 0.885739 | ENSMUSG( | Tmem134 | 1547.01 | -0.15494 | 0.18833 | 0.85698 |
| ENSMUSG( | Tmem135 | 1684.353 | -0.00877 | 0.930049 | 0.991848 | ENSMUSG( | Tmem135 | 1556.667 | 0.034471 | 0.770196 | 0.974236 |
| ENSMUSG( | Tmem138 | 339.7494 | -0.27551 | 0.029728 | 0.261211 | ENSMUSG( | Tmem138 | 353.6402 | 0.12347 | 0.399784 | 0.917257 |
| ENSMUSG( | Tmem140 | 88.1503 | 0.007855 | 0.97275 | 0.996585 | ENSMUSG( | Tmem140 | 93.21862 | 0.430126 | 0.105645 | 0.804969 |
| ENSMUSG( | Tmem141 | 947.7491 | 0.143613 | 0.197793 | 0.637687 | ENSMUSG( | Tmem141 | 788.5162 | -0.11988 | 0.345419 | 0.910581 |
| ENSMUSG( | Tmem143 | 655.0849 | -0.11607 | 0.262636 | 0.713259 | ENSMUSG( | Tmem143 | 571.3099 | -0.25018 | 0.043618 | 0.690304 |
| ENSMUSG( | Tmem144 | 605.5595 | -0.24305 | 0.10559 | 0.49243 | ENSMUSG( | Tmem144 | 626.3785 | 0.132672 | 0.407592 | 0.917257 |
| ENSMUSG( | Tmem145 | 1590.845 | 0.040539 | 0.7922 | 0.959461 | ENSMUSG( | Tmem145 | 1476.841 | 0.093227 | 0.440854 | 0.921166 |
| ENSMUSG( | Tmem147 | 2425.865 | -0.16294 | 0.140949 | 0.557902 | ENSMUSG( | Tmem147 | 2298.359 | -0.04317 | 0.724819 | 0.966887 |
| ENSMUSG( | Tmem14a | 920.0791 | -0.45805 | 6.52E-05 | 0.004839 | ENSMUSG( | Tmem14a | 956.5707 | -0.04338 | 0.759423 | 0.972002 |
| ENSMUSG( | Tmem14c | 1273.969 | -0.2299 | 0.033936 | 0.282701 | ENSMUSG( | Tmem14c | 1232.38 | -0.04738 | 0.697723 | 0.960867 |
| ENSMUSG( | Tmem150a | 415.2757 | 0.255003 | 0.110881 | 0.501952 | ENSMUSG( | Tmem150a | 368.0943 | 0.181565 | 0.31253 | 0.900037 |
| ENSMUSG( | Tmem150b | 226.205 | -0.08326 | 0.56188 | 0.889052 | ENSMUSG( | Tmem150b | 186.0846 | -0.38406 | 0.070201 | 0.757694 |
| ENSMUSG( | Tmem150c | 1781.692 | -0.20517 | 0.074722 | 0.423772 | ENSMUSG( | Tmem150c | 1804.589 | 0.105437 | 0.439203 | 0.920721 |
| ENSMUSG( | Tmem151a | 9244.496 | -0.29858 | 0.025967 | 0.247245 | ENSMUSG( | Tmem151a | 9508.665 | 0.070164 | 0.665294 | 0.956766 |
| ENSMUSG( | Tmem151b | 9230.251 | 0.319131 | 0.00741 | 0.118765 | ENSMUSG( | Tmem151b | 8097.151 | 0.220539 | 0.100158 | 0.801311 |
| ENSMUSG( | Tmem154 | 28.05267 | -0.76781 | 0.036597 | 0.295048 | ENSMUSG( | Tmem154 | 27.33021 | -0.53721 | 0.173518 | 0.844317 |
| ENSMUSG( | Tmem156 | 14.71598 | -0.10893 | 0.822051 | NA | ENSMUSG( | Tmem156 | 11.14825 | -0.67535 | 0.272103 | NA |
| ENSMUSG( | Tmem158 | 1966.229 | -0.7848 | 1.17E-11 | 8.23E-09 | ENSMUSG( | Tmem158 | 2205.841 | -0.08061 | 0.566582 | 0.941777 |
| ENSMUSG( | Tmem160 | 1144.726 | -0.13062 | 0.202615 | 0.642757 | ENSMUSG( | Tmem160 | 1007.608 | -0.2411 | 0.055091 | 0.723362 |
| ENSMUSG( | Tmem161a | 948.631 | -0.1548 | 0.223729 | 0.669845 | ENSMUSG( | Tmem161a | 988.0492 | 0.230347 | 0.103067 | 0.804311 |
| ENSMUSG( | Tmem161b | 536.1984 | -0.27685 | 0.042369 | 0.316636 | ENSMUSG( | Tmem161b | 577.6681 | 0.212566 | 0.163589 | 0.838291 |
| ENSMUSG( | Tmem163 | 1044.964 | -0.50248 | 4.48E-05 | 0.003633 | ENSMUSG( | Tmem163 | 1146.307 | 0.078015 | 0.604737 | 0.948869 |
| ENSMUSG( | Tmem164 | 2177.483 | 0.198245 | 0.027932 | 0.255775 | ENSMUSG( | Tmem164 | 1950.989 | 0.153237 | 0.175218 | 0.845305 |
| ENSMUSG( | Tmem165 | 1017.884 | -0.49584 | 0.000169 | 0.009375 | ENSMUSG( | Tmem165 | 1142.329 | 0.147853 | 0.350069 | 0.911652 |
| ENSMUSG( | Tmem167 | 1490.88 | -0.01838 | 0.867909 | 0.976631 | ENSMUSG( | Tmem167 | 1423.424 | 0.112079 | 0.368113 | 0.915491 |
| ENSMUSG( | Tmem167b | 2464.29 | -0.00061 | 0.995787 | 0.99943 | ENSMUSG( | Tmem167b | 2192.801 | -0.07162 | 0.587729 | 0.944975 |
| ENSMUSG( | Tmem168 | 470.9275 | -0.3469 | 0.022139 | 0.226843 | ENSMUSG( | Tmem168 | 529.087 | 0.271104 | 0.121471 | 0.813384 |
| ENSMUSG( | Tmem169 | 521.5691 | -0.25505 | 0.027785 | 0.255192 | ENSMUSG( | Tmem169 | 535.744 | 0.103194 | 0.450733 | 0.922202 |
| ENSMUSG( | Tmem17 | 201.6398 | -0.51901 | 0.000792 | 0.027736 | ENSMUSG( | Tmem17 | 220.4989 | 0.061999 | 0.715917 | 0.965424 |
| ENSMUSG( | Tmem170 | 166.3601 | 0.147751 | 0.378565 | 0.801688 | ENSMUSG( | Tmem170 | 164.8169 | 0.376078 | 0.042845 | 0.690304 |
| ENSMUSG( | Tmem170b | 2760.598 | -0.02887 | 0.770872 | 0.954726 | ENSMUSG( | Tmem170b | 2515.037 | -0.03103 | 0.804871 | 0.977232 |
| ENSMUSG( | Tmem175 | 1230.265 | -0.197 | 0.12574 | 0.53257 | ENSMUSG( | Tmem175 | 1223.043 | 0.064765 | 0.63592 | 0.953903 |
| ENSMUSG( | Tmem176a | 1099.241 | -0.10729 | 0.394636 | 0.810072 | ENSMUSG( | Tmem176a | 1062.962 | 0.067967 | 0.635041 | 0.953903 |
| ENSMUSG( | Tmem176b | 2405.284 | -0.18195 | 0.102905 | 0.488476 | ENSMUSG( | Tmem176b | 2369.632 | 0.050741 | 0.690158 | 0.960071 |
| ENSMUSG( | Tmem177 | 237.8317 | 0.142737 | 0.37854 | 0.801688 | ENSMUSG( | Tmem177 | 208.2415 | 0.041275 | 0.820454 | 0.979198 |
| ENSMUSG( | Tmem178 | 2063.595 | -0.16825 | 0.266522 | 0.716441 | ENSMUSG( | Tmem178 | 2109.299 | 0.16826 | 0.335891 | 0.907913 |
| ENSMUSG( | Tmem178b | 4235.191 | -0.18487 | 0.179809 | 0.61751 | ENSMUSG( | Tmem178b | 4190.471 | 0.062715 | 0.70001 | 0.961335 |
| ENSMUSG( | Tmem179 | 4328.229 | -0.22139 | 0.064025 | 0.391882 | ENSMUSG( | Tmem179 | 4446.528 | 0.133661 | 0.316979 | 0.902088 |
| ENSMUSG( | Tmem179b | 210.5198 | -0.23723 | 0.126079 | 0.533384 | ENSMUSG( | Tmem179b | 216.1129 | 0.124801 | 0.485998 | 0.92541 |
| ENSMUSG( | Tmem18 | 1358.092 | -0.09114 | 0.433437 | 0.828346 | ENSMUSG( | Tmem18 | 1359.255 | 0.179378 | 0.185881 | 0.855305 |
| ENSMUSG( | Tmem181a | 1996.559 | -0.33678 | 0.017917 | 0.200989 | ENSMUSG( | Tmem181a | 2124.763 | 0.13192 | 0.407477 | 0.917257 |
| ENSMUSG( | Tmem183a | 2291.454 | 0.096369 | 0.273076 | 0.723468 | ENSMUSG( | Tmem183a | 1995.392 | -0.03406 | 0.757381 | 0.971858 |
| ENSMUSG( | Tmem184b | 2708.203 | -0.36999 | 0.00625 | 0.108182 | ENSMUSG( | Tmem184b | 2847.251 | 0.071092 | 0.66236 | 0.956205 |
| ENSMUSG( | Tmem184c | 3291.902 | -0.09784 | 0.419295 | 0.821987 | ENSMUSG( | Tmem184c | 3194.762 | 0.085963 | 0.504095 | 0.927825 |
| ENSMUSG( | Tmem185a | 1453.06 | -0.31938 | 0.014419 | 0.176902 | ENSMUSG( | Tmem185a | 1529.507 | 0.116205 | 0.419426 | 0.917257 |
| ENSMUSG( | Tmem185b | 397.8021 | -0.51773 | 0.002422 | 0.057903 | ENSMUSG( | Tmem185b | 456.0155 | 0.192383 | 0.315719 | 0.901868 |
| ENSMUSG( | Tmem186 | 562.332 | -0.00741 | 0.946025 | 0.99369 | ENSMUSG( | Tmem186 | 515.2351 | 0.011943 | 0.92632 | 0.99306 |
| ENSMUSG( | Tmem19 | 1537.52 | -0.28827 | 0.017549 | 0.198498 | ENSMUSG( | Tmem19 | 1621.503 | 0.147879 | 0.303182 | 0.898538 |
| ENSMUSG( | Tmem191 | 2748.906 | 0.184087 | 0.171729 | 0.606517 | ENSMUSG( | Tmem191 | 2382.467 | 0.041769 | 0.790897 | 0.97681 |
| ENSMUSG( | Tmem192 | 565.2655 | -0.21084 | 0.065526 | 0.396581 | ENSMUSG( | Tmem192 | 555.6488 | 0.020691 | 0.890157 | 0.990558 |
| ENSMUSG( | Tmem196 | 247.443 | -0.57059 | 0.000551 | 0.021914 | ENSMUSG( | Tmem196 | 291.2218 | 0.212719 | 0.251209 | 0.879321 |
| ENSMUSG( | Tmem198 | 2657.502 | -0.12183 | 0.356179 | 0.788108 | ENSMUSG( | Tmem198 | 2554.856 | 0.039259 | 0.768381 | 0.974236 |
| ENSMUSG( | Tmem198b | 591.6915 | -0.4437 | 0.005699 | 0.101138 | ENSMUSG( | Tmem198b | 622.5211 | 0.011182 | 0.950248 | 0.994798 |
| ENSMUSG( | Tmem199 | 508.8638 | -0.13989 | 0.267593 | 0.717651 | ENSMUSG( | Tmem199 | 496.847 | 0.067087 | 0.64343 | 0.953903 |
| ENSMUSG( | Tmem200a | 513.2399 | -0.17429 | 0.355317 | 0.78783 | ENSMUSG( | Tmem200a | 581.7129 | 0.435263 | 0.028605 | 0.614171 |
| ENSMUSG( | Tmem200b | 175.5603 | -0.45615 | 0.00892 | 0.132538 | ENSMUSG( | Tmem200b | 207.1331 | 0.316944 | 0.082643 | 0.77681 |
| ENSMUSG( | Tmem200c | 449.0829 | -0.33703 | 0.064528 | 0.393651 | ENSMUSG( | Tmem200c | 505.7129 | 0.288735 | 0.142624 | 0.831221 |
| ENSMUSG( | Tmem201 | 2883.803 | -0.16668 | 0.184478 | 0.62157 | ENSMUSG( | Tmem201 | 2892.992 | 0.118231 | 0.423351 | 0.917257 |
| ENSMUSG( | Tmem202 | 59.47617 | -0.12128 | 0.625101 | 0.912249 | ENSMUSG( | Tmem202 | 52.26855 | -0.26561 | 0.372882 | 0.915491 |
| ENSMUSG( | Tmem203 | 261.892 | -0.3947 | 0.004659 | 0.089972 | ENSMUSG( | Tmem203 | 265.2252 | -0.05802 | 0.721906 | 0.966887 |
| ENSMUSG( | Tmem204 | 422.5487 | 0.135904 | 0.460089 | 0.840577 | ENSMUSG( | Tmem204 | 415.8183 | 0.352019 | 0.160574 | 0.837724 |
| ENSMUSG( | Tmem205 | 923.4797 | -0.33103 | 0.004086 | 0.082441 | ENSMUSG( | Tmem205 | 916.4093 | -0.0647 | 0.620394 | 0.952896 |
| ENSMUSG( | Tmem208 | 814.8894 | -0.16348 | 0.151608 | 0.576979 | ENSMUSG( | Tmem208 | 774.0014 | -0.03972 | 0.766258 | 0.974236 |
| ENSMUSG( | Tmem209 | 912.4368 | -0.34782 | 0.024127 | 0.237752 | ENSMUSG( | Tmem209 | 990.5167 | 0.176875 | 0.283319 | 0.892412 |
| ENSMUSG( | Tmem212 | 81.42574 | -0.53319 | 0.053817 | 0.356147 | ENSMUSG( | Tmem212 | 90.41718 | 0.104457 | 0.724153 | 0.966887 |
| ENSMUSG( | Tmem214 | 2066.786 | 0.064736 | 0.582164 | 0.897918 | ENSMUSG( | Tmem214 | 1963.393 | 0.183721 | 0.157368 | 0.837724 |
| ENSMUSG( | Tmem215 | 402.6313 | -0.20992 | 0.228611 | 0.675914 | ENSMUSG( | Tmem215 | 406.6127 | 0.098247 | 0.580293 | 0.944209 |
| ENSMUSG( | Tmem216 | 154.4598 | 0.115269 | 0.510074 | 0.869785 | ENSMUSG( | Tmem216 | 137.9257 | 0.068651 | 0.735424 | 0.967344 |
| ENSMUSG( | Tmem217r | 1683.168 | -0.01077 | 0.913848 | 0.988483 | ENSMUSG( | Tmem217r | 1600.837 | 0.113965 | 0.373523 | 0.915491 |
| ENSMUSG( | Tmem218 | 727.2737 | -0.1097 | 0.412315 | 0.819452 | ENSMUSG( | Tmem218 | 642.6669 | -0.20081 | 0.172341 | 0.843502 |
| ENSMUSG( | Tmem219 | 485.0971 | -0.05443 | 0.630524 | 0.914073 | ENSMUSG( | Tmem219 | 433.5509 | -0.11069 | 0.404856 | 0.917257 |
| ENSMUSG( | Tmem220 | 46.36903 | -0.09091 | 0.755456 | 0.951395 | ENSMUSG( | Tmem220 | 41.27411 | -0.12024 | 0.743542 | 0.96951 |
| ENSMUSG( | Tmem221 | 8.954315 | -0.82638 | 0.200934 | NA | ENSMUSG( | Tmem221 | 10.09042 | -0.05921 | 0.926526 | NA |
| ENSMUSG( | Tmem222 | 2705.002 | -0.32201 | 0.004986 | 0.093605 | ENSMUSG( | Tmem222 | 2620.285 | -0.12704 | 0.316589 | 0.901868 |
| ENSMUSG( | Tmem223 | 1217.403 | 0.059974 | 0.542776 | 0.883563 | ENSMUSG( | Tmem223 | 1124.219 | 0.095475 | 0.421618 | 0.917257 |
| ENSMUSG( | Tmem229a | 6780.701 | 0.036271 | 0.797457 | 0.960507 | ENSMUSG( | Tmem229a | 5749.404 | -0.17868 | 0.269937 | 0.887508 |
| ENSMUSG( | Tmem229b | 1225.694 | -0.19008 | 0.065234 | 0.395697 | ENSMUSG( | Tmem229b | 1212.159 | 0.055071 | 0.660913 | 0.956205 |
| ENSMUSG( | Tmem230 | 488.4059 | -0.24203 | 0.105477 | 0.492417 | ENSMUSG( | Tmem230 | 511.9201 | 0.173222 | 0.309107 | 0.898538 |
| ENSMUSG( | Tmem231 | 352.5697 | -0.27197 | 0.088007 | 0.455043 | ENSMUSG( | Tmem231 | 381.0303 | 0.229698 | 0.161492 | 0.837724 |
| ENSMUSG( | Tmem232 | 52.89864 | -0.24683 | 0.44268 | 0.83278 | ENSMUSG( | Tmem232 | 55.25455 | 0.166915 | 0.607289 | 0.949981 |
| ENSMUSG( | Tmem234 | 3147.516 | 0.137667 | 0.148471 | 0.571935 | ENSMUSG( | Tmem234 | 2699.585 | -0.03176 | 0.770307 | 0.974236 |
| ENSMUSG( | Tmem237 | 597.6629 | -0.22262 | 0.136653 | 0.550959 | ENSMUSG( | Tmem237 | 613.0306 | 0.12857 | 0.370859 | 0.915491 |
| ENSMUSG( | Tmem238 | 153.0399 | -0.29761 | 0.100529 | 0.483976 | ENSMUSG( | Tmem238 | 146.7858 | -0.13352 | 0.484512 | 0.92541 |
| ENSMUSG( | Tmem240 | 1576.555 | -0.18692 | 0.069992 | 0.40908 | ENSMUSG( | Tmem240 | 1523.186 | -0.01134 | 0.914621 | 0.992407 |
| ENSMUSG( | Tmem241 | 305.7036 | -0.16922 | 0.241892 | 0.689179 | ENSMUSG( | Tmem241 | 283.9351 | -0.10266 | 0.526847 | 0.934449 |
| ENSMUSG( | Tmem242 | 2395.534 | 0.044625 | 0.636923 | 0.915191 | ENSMUSG( | Tmem242 | 2015.017 | -0.19611 | 0.131533 | 0.822912 |
| ENSMUSG( | Tmem243 | 618.4716 | 0.00721 | 0.950414 | 0.993821 | ENSMUSG( | Tmem243 | 592.8345 | 0.145335 | 0.316399 | 0.901868 |
| ENSMUSG( | Tmem245 | 2275.947 | -0.22812 | 0.117252 | 0.516802 | ENSMUSG( | Tmem245 | 2506.058 | 0.315281 | 0.053336 | 0.717022 |
| ENSMUSG( | Tmem248 | 1499.928 | -0.21826 | 0.099001 | 0.480953 | ENSMUSG( | Tmem248 | 1546.428 | 0.146269 | 0.361686 | 0.914223 |
| ENSMUSG( | Tmem25 | 1710.73 | -0.2238 | 0.077288 | 0.429743 | ENSMUSG( | Tmem25 | 1673.097 | -0.00764 | 0.951129 | 0.994798 |
| ENSMUSG( | Tmem250 | 3312.318 | 0.041586 | 0.604898 | 0.905556 | ENSMUSG( | Tmem250 | 2918.86 | -0.05379 | 0.592355 | 0.945016 |
| ENSMUSG( | Tmem252 | 102.5279 | -0.76407 | 0.007058 | 0.115775 | ENSMUSG( | Tmem252 | 75.64916 | -1.76196 | 2.58E-08 | 2.08E-05 |
| ENSMUSG( | Tmem253 | 15.3743 | -0.40027 | 0.446377 | NA | ENSMUSG( | Tmem253 | 11.75032 | -1.09159 | 0.076715 | 0.770558 |
| ENSMUSG( | Tmem254 | 2667.668 | -0.34445 | 0.001321 | 0.039614 | ENSMUSG( | Tmem254 | 2412.157 | -0.36833 | 0.003311 | 0.229858 |
| ENSMUSG( | Tmem255a | 505.339 | -0.34881 | 0.013612 | 0.170359 | ENSMUSG( | Tmem255a | 541.7848 | 0.135044 | 0.421806 | 0.917257 |
| ENSMUSG( | Tmem255b | 100.9814 | -0.398 | 0.170166 | 0.605149 | ENSMUSG( | Tmem255b | 106.7929 | 0.077071 | 0.799373 | 0.97681 |
| ENSMUSG( | Tmem256 | 1399.368 | 0.067849 | 0.524202 | 0.87583 | ENSMUSG( | Tmem256 | 1168.157 | -0.19513 | 0.145182 | 0.83339 |
| ENSMUSG( | Tmem258 | 785.8924 | -0.16451 | 0.121027 | 0.525487 | ENSMUSG( | Tmem258 | 718.87 | -0.15349 | 0.197594 | 0.862161 |
| ENSMUSG( | Tmem259 | 5500.007 | -0.17193 | 0.078172 | 0.432307 | ENSMUSG( | Tmem259 | 5669.8 | 0.18531 | 0.107348 | 0.804969 |
| ENSMUSG( | Tmem260 | 853.6765 | -0.26835 | 0.025501 | 0.246501 | ENSMUSG( | Tmem260 | 872.0784 | 0.07749 | 0.594994 | 0.94561 |
| ENSMUSG( | Tmem263 | 585.137 | 0.011099 | 0.93152 | 0.992398 | ENSMUSG( | Tmem263 | 521.592 | -0.06148 | 0.701825 | 0.962014 |
| ENSMUSG( | Tmem266 | 342.4012 | -0.06292 | 0.650652 | 0.921301 | ENSMUSG( | Tmem266 | 318.5224 | 0.007498 | 0.962727 | 0.995284 |
| ENSMUSG( | Tmem267 | 349.6496 | -0.73691 | 1.34E-06 | 0.000242 | ENSMUSG( | Tmem267 | 329.6451 | -0.61075 | 0.0171 | 0.52741 |
| ENSMUSG( | Tmem268 | 529.8919 | -0.13895 | 0.251894 | 0.701908 | ENSMUSG( | Tmem268 | 496.7053 | -0.04602 | 0.773714 | 0.974672 |
| ENSMUSG( | Tmem269 | 28.46085 | -0.03971 | 0.916475 | 0.988844 | ENSMUSG( | Tmem269 | 27.69089 | 0.152334 | 0.681879 | 0.958678 |
| ENSMUSG( | Tmem273 | 36.64223 | -0.01718 | 0.960423 | 0.993991 | ENSMUSG( | Tmem273 | 35.45335 | 0.189777 | 0.638883 | 0.953903 |
| ENSMUSG( | Tmem276 | 792.5632 | -0.26719 | 0.012653 | 0.162184 | ENSMUSG( | Tmem276 | 771.1731 | -0.06214 | 0.638146 | 0.953903 |
| ENSMUSG( | Tmem30a | 7753.293 | -0.36823 | 0.020278 | 0.214748 | ENSMUSG( | Tmem30a | 8731.181 | 0.257644 | 0.158159 | 0.837724 |
| ENSMUSG( | Tmem30b | 10.83522 | -0.7022 | 0.218724 | NA | ENSMUSG( | Tmem30b | 11.38543 | -0.15593 | 0.795554 | NA |
| ENSMUSG( | Tmem33 | 4046.054 | -0.2017 | 0.140312 | 0.557263 | ENSMUSG( | Tmem33 | 4010.054 | 0.047809 | 0.767669 | 0.974236 |
| ENSMUSG( | Tmem35a | 2057.866 | -0.4381 | 0.000485 | 0.020239 | ENSMUSG( | Tmem35a | 2195.588 | 0.051859 | 0.727397 | 0.966887 |
| ENSMUSG( | Tmem35b | 95.8924 | 0.027066 | 0.893736 | 0.984512 | ENSMUSG( | Tmem35b | 85.79578 | -0.03806 | 0.874894 | 0.989644 |
| ENSMUSG( | Tmem37 | 53.61466 | 0.102228 | 0.714232 | 0.941016 | ENSMUSG( | Tmem37 | 62.66139 | 0.7501 | 0.012859 | 0.457834 |
| ENSMUSG( | Tmem38a | 4691.985 | -0.22139 | 0.074864 | 0.424133 | ENSMUSG( | Tmem38a | 4783.73 | 0.112775 | 0.432374 | 0.918158 |
| ENSMUSG( | Tmem38b | 590.0769 | 0.001011 | 0.994297 | 0.999369 | ENSMUSG( | Tmem38b | 550.0109 | 0.066069 | 0.662502 | 0.956205 |
| ENSMUSG( | Tmem39a | 251.6376 | -0.19675 | 0.246902 | 0.695632 | ENSMUSG( | Tmem39a | 276.0518 | 0.33962 | 0.09402 | 0.795735 |
| ENSMUSG( | Tmem39b | 261.4599 | -0.50714 | 0.007197 | 0.117084 | ENSMUSG( | Tmem39b | 278.9352 | 0.001352 | 0.994464 | 0.998813 |
| ENSMUSG( | Tmem40 | 85.13151 | -0.04154 | 0.854093 | 0.973491 | ENSMUSG( | Tmem40 | 79.68897 | 0.019309 | 0.936058 | 0.993573 |
| ENSMUSG( | Tmem41a | 307.2106 | -0.26472 | 0.087927 | 0.45492 | ENSMUSG( | Tmem41a | 340.347 | 0.304135 | 0.09075 | 0.791074 |
| ENSMUSG( | Tmem41b | 867.0977 | -0.46017 | 0.001384 | 0.040741 | ENSMUSG( | Tmem41b | 974.6737 | 0.181154 | 0.255223 | 0.882761 |
| ENSMUSG( | Tmem42 | 375.487 | -0.25964 | 0.086507 | 0.452045 | ENSMUSG( | Tmem42 | 371.1208 | -0.01063 | 0.946427 | 0.994798 |
| ENSMUSG( | Tmem43 | 1149.037 | -0.40417 | 0.00734 | 0.118347 | ENSMUSG( | Tmem43 | 1272.596 | 0.187535 | 0.28072 | 0.892356 |
| ENSMUSG( | Tmem44 | 6098.707 | 0.085008 | 0.353964 | 0.786371 | ENSMUSG( | Tmem44 | 5277.514 | -0.06175 | 0.607943 | 0.950113 |
| ENSMUSG( | Tmem45a | 35.44445 | -0.1866 | 0.612008 | 0.908586 | ENSMUSG( | Tmem45a | 28.73202 | -0.57539 | 0.167661 | 0.840222 |
| ENSMUSG( | Tmem47 | 7751.38 | -0.10154 | 0.531759 | 0.879628 | ENSMUSG( | Tmem47 | 7248.364 | -0.02659 | 0.892767 | 0.990558 |
| ENSMUSG( | Tmem50a | 3647.533 | -0.30821 | 0.018789 | 0.206056 | ENSMUSG( | Tmem50a | 3808.138 | 0.102103 | 0.478645 | 0.923329 |
| ENSMUSG( | Tmem50b | 3675.574 | -0.25531 | 0.043522 | 0.31954 | ENSMUSG( | Tmem50b | 3706.865 | 0.05039 | 0.725716 | 0.966887 |
| ENSMUSG( | Tmem51 | 551.4286 | -0.21715 | 0.078588 | 0.433892 | ENSMUSG( | Tmem51 | 473.236 | -0.40833 | 0.005539 | 0.303106 |
| ENSMUSG( | Tmem52 | 47.25145 | 0.770582 | 0.030253 | 0.264384 | ENSMUSG( | Tmem52 | 48.66483 | 1.060357 | 0.001029 | 0.110223 |
| ENSMUSG( | Tmem53 | 75.80147 | -0.51373 | 0.042482 | 0.316636 | ENSMUSG( | Tmem53 | 75.58049 | -0.21743 | 0.40998 | 0.917257 |
| ENSMUSG( | Tmem54 | 74.79574 | 0.03878 | 0.870104 | 0.977767 | ENSMUSG( | Tmem54 | 68.18504 | 0.02847 | 0.914992 | 0.992407 |
| ENSMUSG( | Tmem59 | 7534.545 | -0.27749 | 0.025563 | 0.246694 | ENSMUSG( | Tmem59 | 7689.431 | 0.064735 | 0.641903 | 0.953903 |
| ENSMUSG( | Tmem59l | 10124.67 | -0.37287 | 0.000862 | 0.029481 | ENSMUSG( | Tmem59l | 10516.95 | 0.031817 | 0.809323 | 0.978779 |
| ENSMUSG( | Tmem60 | 486.5879 | -0.43497 | 0.002993 | 0.067057 | ENSMUSG( | Tmem60 | 506.9261 | -0.01417 | 0.928834 | 0.99306 |
| ENSMUSG( | Tmem62 | 1081.18 | -0.20618 | 0.115804 | 0.513769 | ENSMUSG( | Tmem62 | 1073.665 | 0.054069 | 0.716799 | 0.965859 |
| ENSMUSG( | Tmem63a | 1358.471 | -0.79176 | 7.84E-06 | 0.00097 | ENSMUSG( | Tmem63a | 1576.884 | 0.022267 | 0.918881 | 0.992529 |
| ENSMUSG( | Tmem63b | 5588.283 | -0.16389 | 0.165631 | 0.59714 | ENSMUSG( | Tmem63b | 5723.825 | 0.176243 | 0.201183 | 0.86439 |
| ENSMUSG( | Tmem63c | 1145.756 | 0.100633 | 0.480484 | 0.854748 | ENSMUSG( | Tmem63c | 1058.554 | 0.145775 | 0.375339 | 0.915491 |
| ENSMUSG( | Tmem64 | 1253.836 | -0.0888 | 0.497777 | 0.862341 | ENSMUSG( | Tmem64 | 1243.311 | 0.156902 | 0.3045 | 0.898538 |
| ENSMUSG( | Tmem65 | 2707.168 | 0.032178 | 0.750482 | 0.949877 | ENSMUSG( | Tmem65 | 2426.792 | -0.01888 | 0.897927 | 0.990558 |
| ENSMUSG( | Tmem67 | 356.5741 | -0.19207 | 0.150052 | 0.574564 | ENSMUSG( | Tmem67 | 390.9489 | 0.335072 | 0.025635 | 0.599234 |
| ENSMUSG( | Tmem68 | 787.2897 | -0.52788 | 0.001164 | 0.036596 | ENSMUSG( | Tmem68 | 914.7587 | 0.217693 | 0.214251 | 0.864421 |
| ENSMUSG( | Tmem69 | 207.7494 | -0.02204 | 0.888463 | 0.983434 | ENSMUSG( | Tmem69 | 189.9088 | -0.01246 | 0.942219 | 0.994774 |
| ENSMUSG( | Tmem70 | 869.8187 | -0.04924 | 0.639937 | 0.916434 | ENSMUSG( | Tmem70 | 789.2455 | -0.06442 | 0.650231 | 0.95422 |
| ENSMUSG( | Tmem71 | 21.87999 | -0.14375 | 0.725955 | 0.944223 | ENSMUSG( | Tmem71 | 22.6992 | 0.247257 | 0.557809 | 0.939565 |
| ENSMUSG( | Tmem72 | 51.16427 | -0.02953 | 0.975308 | 0.997302 | ENSMUSG( | Tmem72 | 61.43173 | 0.673426 | 0.385964 | 0.917257 |
| ENSMUSG( | Tmem74 | 147.2091 | -0.20194 | 0.341238 | 0.778828 | ENSMUSG( | Tmem74 | 162.9681 | 0.361299 | 0.124345 | 0.814691 |
| ENSMUSG( | Tmem74b | 114.2758 | -0.39382 | 0.077252 | 0.429743 | ENSMUSG( | Tmem74b | 105.6016 | -0.3357 | 0.139619 | 0.830678 |
| ENSMUSG( | Tmem74bc | 17.16618 | 0.670432 | 0.190107 | NA | ENSMUSG( | Tmem74bc | 21.89053 | 1.363797 | 0.004344 | 0.268942 |
| ENSMUSG( | Tmem79 | 27.73437 | -0.55233 | 0.128177 | 0.536641 | ENSMUSG( | Tmem79 | 29.30037 | -0.06279 | 0.868034 | 0.98781 |
| ENSMUSG( | Tmem80 | 1029.534 | 0.064148 | 0.493377 | 0.860611 | ENSMUSG( | Tmem80 | 900.059 | -0.05726 | 0.646415 | 0.953988 |
| ENSMUSG( | Tmem81 | 189.7478 | -0.06234 | 0.689089 | 0.933441 | ENSMUSG( | Tmem81 | 171.39 | -0.09084 | 0.607673 | 0.950113 |
| ENSMUSG( | Tmem82 | 43.98241 | -0.14725 | 0.610595 | 0.908194 | ENSMUSG( | Tmem82 | 37.11854 | -0.38262 | 0.246197 | 0.877904 |

| | | | | | |
|---|---|---|---|---|---|
| ENSMUSG( | 748.5083 | 0.146094 | 0.273787 | 0.724244 | Thoc5 |
| ENSMUSG( | 138.7649 | -0.09417 | 0.6263 | 0.912639 | Thoc6 |
| ENSMUSG( | 1272.917 | 0.069474 | 0.446352 | 0.833698 | Thoc7 |
| ENSMUSG( | 1024.84 | 0.08082 | 0.48169 | 0.855573 | Thop1 |
| ENSMUSG( | 243.8679 | 0.369604 | 0.101179 | 0.484996 | Thpo |
| ENSMUSG( | 40291.94 | 0.210294 | 0.003482 | 0.074707 | Thra |
| ENSMUSG( | 3045.111 | 0.000868 | 0.99171 | 0.999232 | Thrap3 |
| ENSMUSG( | 1100.18 | 0.217606 | 0.018411 | 0.203839 | Thrb |
| ENSMUSG( | 1319.552 | 0.407935 | 7.47E-05 | 0.005315 | Thrsp |
| ENSMUSG( | 193.3952 | -0.14964 | 0.48786 | 0.858824 | Thsd1 |
| ENSMUSG( | 532.8169 | -0.21599 | 0.104636 | 0.490944 | Thsd4 |
| ENSMUSG( | 862.799 | -0.63535 | 4.94E-05 | 0.00395 | Thsd7a |
| ENSMUSG( | 194.4452 | -0.65129 | 0.002837 | 0.065089 | Thsd7b |
| ENSMUSG( | 286.1836 | 0.203587 | 0.135495 | 0.548884 | Thtpa |
| ENSMUSG( | 736.906 | 0.122645 | 0.312599 | 0.760241 | Thumpd1 |
| ENSMUSG( | 136.8343 | 0.032405 | 0.858464 | 0.974683 | Thumpd2 |
| ENSMUSG( | 522.9742 | 0.114844 | 0.343684 | 0.779153 | Thumpd3 |
| ENSMUSG( | 29504.7 | -0.429 | 0.000267 | 0.013312 | Thy1 |
| ENSMUSG( | 662.3662 | 0.091337 | 0.446904 | 0.834051 | Thyn1 |
| ENSMUSG( | 2126.717 | -0.16408 | 0.08572 | 0.451367 | Tia1 |
| ENSMUSG( | 2365.265 | -0.09627 | 0.289252 | 0.739725 | Tial1 |
| ENSMUSG( | 2209.814 | 0.088449 | 0.366077 | 0.794603 | Tiam1 |
| ENSMUSG( | 1138.181 | -0.14673 | 0.224775 | 0.671058 | Tiam2 |
| ENSMUSG( | 229.4165 | 0.054284 | 0.729284 | 0.944738 | Ticam1 |
| ENSMUSG( | 20.70409 | -0.52 | 0.214596 | NA | Ticam2 |
| ENSMUSG( | 10.17933 | -0.3246 | 0.594872 | NA | Ticrr |
| ENSMUSG( | 563.8738 | 0.090056 | 0.669455 | 0.928147 | Tie1 |
| ENSMUSG( | 372.3953 | 0.160679 | 0.190912 | 0.630245 | Tifa |
| ENSMUSG( | 193.5395 | 0.271981 | 0.136524 | 0.550651 | Tifab |
| ENSMUSG( | 549.5434 | 0.073281 | 0.584382 | 0.898802 | Tigar |
| ENSMUSG( | 196.4199 | -0.12452 | 0.40668 | 0.816928 | Tigd2 |
| ENSMUSG( | 59.69464 | -0.04132 | 0.89189 | 0.983979 | Tigd3 |
| ENSMUSG( | 119.4393 | 0.084546 | 0.708365 | 0.939519 | Tigd5 |
| ENSMUSG( | 146.1563 | 0.319192 | 0.132183 | 0.544244 | Timeless |
| ENSMUSG( | 347.247 | 0.121181 | 0.427082 | 0.82532 | Timm10 |
| ENSMUSG( | 334.2797 | -0.02929 | 0.860292 | 0.974842 | Timm10b |
| ENSMUSG( | 1891.355 | -0.02887 | 0.768975 | 0.954674 | Timm13 |
| ENSMUSG( | 1661.98 | 0.049982 | 0.59704 | 0.903526 | Timm17a |
| ENSMUSG( | 713.4277 | -0.04202 | 0.722097 | 0.943535 | Timm17b |
| ENSMUSG( | 529.5998 | -0.01567 | 0.906432 | 0.986998 | Timm21 |
| ENSMUSG( | 628.2479 | -0.04581 | 0.691109 | 0.933881 | Timm22 |
| ENSMUSG( | 873.6749 | -0.01942 | 0.847172 | 0.972374 | Timm23 |
| ENSMUSG( | 2154.905 | 0.085417 | 0.31748 | 0.763823 | Timm29 |
| ENSMUSG( | 1325.137 | 0.11849 | 0.345179 | 0.779153 | Timm44 |
| ENSMUSG( | 919.3238 | -0.09886 | 0.365067 | 0.79407 | Timm50 |
| ENSMUSG( | 169.4323 | 0.201948 | 0.260599 | 0.710468 | Timm8a1 |
| ENSMUSG( | 2398.616 | 0.00081 | 0.992978 | 0.999327 | Timm8b |
| ENSMUSG( | 680.4996 | -0.03183 | 0.817119 | 0.96566 | Timm9 |
| ENSMUSG( | 584.566 | -0.02741 | 0.804816 | 0.962473 | Timmdc1 |
| ENSMUSG( | 36.52868 | -0.23072 | 0.465415 | 0.844064 | Timp1 |
| ENSMUSG( | 11470.22 | -0.26713 | 0.041776 | 0.315339 | Timp2 |
| ENSMUSG( | 8397.906 | -0.22462 | 0.056726 | 0.367404 | Timp3 |
| ENSMUSG( | 4175.523 | 0.07234 | 0.459803 | 0.840338 | Timp4 |
| ENSMUSG( | 249.0223 | 0.910358 | 0.000187 | 0.009972 | Tinagl1 |
| ENSMUSG( | 85.23289 | 0.39935 | 0.090352 | 0.461669 | Tincr |
| ENSMUSG( | 412.2385 | -0.01769 | 0.898363 | 0.985633 | Tinf2 |
| ENSMUSG( | 614.2769 | -0.10045 | 0.353326 | 0.785492 | Tiparp |
| ENSMUSG( | 493.6744 | -0.01344 | 0.915296 | 0.988634 | Tipin |
| ENSMUSG( | 1036.299 | 0.058245 | 0.604053 | 0.905187 | Tiprl |
| ENSMUSG( | 100.8717 | -0.0144 | 0.947214 | 0.993821 | Tirap |
| ENSMUSG( | 861.8785 | 0.143318 | 0.220465 | 0.666079 | Tjap1 |
| ENSMUSG( | 5639.732 | 0.121307 | 0.304829 | 0.752574 | Tjp1 |
| ENSMUSG( | 2380.264 | 0.010223 | 0.923299 | 0.990694 | Tjp2 |
| ENSMUSG( | 22.80401 | -0.26411 | 0.532943 | 0.879905 | Tjp3 |
| ENSMUSG( | 22.94777 | -0.55874 | 0.206685 | 0.648942 | Tk1 |
| ENSMUSG( | 1381.784 | 0.093892 | 0.379255 | 0.801757 | Tk2 |
| ENSMUSG( | 528.2319 | 0.082563 | 0.502938 | 0.865717 | Tkfc |
| ENSMUSG( | 6293.67 | -0.076 | 0.363799 | 0.793343 | Tkt |
| ENSMUSG( | 9.909634 | 0.454201 | 0.490132 | NA | Tktl2 |
| ENSMUSG( | 2284.056 | 0.000618 | 0.995792 | 0.99943 | Tlcd1 |
| ENSMUSG( | 17.48267 | -0.13675 | 0.76445 | NA | Tlcd2 |
| ENSMUSG( | 762.0557 | -0.31029 | 0.024214 | 0.238178 | Tlcd3a |
| ENSMUSG( | 4639.081 | 0.091635 | 0.295159 | 0.743798 | Tlcd3b |
| ENSMUSG( | 2709.936 | -0.17315 | 0.209631 | 0.651841 | Tlcd4 |
| ENSMUSG( | 594.6801 | -0.14765 | 0.183722 | 0.620678 | Tlcd5 |
| ENSMUSG( | 1448.907 | 0.246888 | 0.004416 | 0.086676 | Tle1 |
| ENSMUSG( | 746.0895 | 0.386476 | 0.280851 | 0.729868 | Tle2 |
| ENSMUSG( | 1726.826 | 0.164333 | 0.090201 | 0.461412 | Tle3 |
| ENSMUSG( | 1433.928 | -0.06807 | 0.44415 | 0.833239 | Tle4 |
| ENSMUSG( | 29372.25 | 0.037552 | 0.631541 | 0.914073 | Tle5 |
| ENSMUSG( | 80.53997 | -0.22506 | 0.304565 | 0.752574 | Tle6 |
| ENSMUSG( | 2686.373 | 0.067658 | 0.414233 | 0.819998 | Tlk1 |
| ENSMUSG( | 1642.546 | 0.063662 | 0.487058 | 0.858812 | Tlk2 |
| ENSMUSG( | 56.59184 | -0.18212 | 0.515113 | 0.872463 | Tll1 |
| ENSMUSG( | 29.76221 | -0.44451 | 0.201131 | 0.641362 | Tll2 |
| ENSMUSG( | 2854.733 | 0.009337 | 0.914837 | 0.988585 | Tln1 |
| ENSMUSG( | 5005.289 | 0.073471 | 0.409986 | 0.818258 | Tln2 |
| ENSMUSG( | 946.7176 | -0.02022 | 0.858182 | 0.974672 | Tlnrd1 |
| ENSMUSG( | 44.75918 | -0.35673 | 0.255564 | 0.706195 | Tlr1 |
| ENSMUSG( | 36.18639 | -0.76258 | 0.043425 | 0.319227 | Tlr12 |
| ENSMUSG( | 130.0813 | -0.04747 | 0.822807 | 0.966352 | Tlr13 |
| ENSMUSG( | 55.0462 | -0.22492 | 0.446955 | 0.834051 | Tlr2 |
| ENSMUSG( | 712.1082 | -0.26349 | 0.15903 | 0.588112 | Tlr3 |
| ENSMUSG( | 58.36891 | -0.09698 | 0.756608 | 0.951479 | Tlr4 |
| ENSMUSG( | 28.33721 | -0.33917 | 0.340057 | 0.777959 | Tlr5 |
| ENSMUSG( | 34.622 | -0.56956 | 0.109843 | 0.500285 | Tlr6 |
| ENSMUSG( | 198.5941 | 0.077259 | 0.711689 | 0.940028 | Tlr7 |
| ENSMUSG( | 9.96594 | -0.6858 | 0.359486 | NA | Tlr8 |
| ENSMUSG( | 56.11455 | -0.62761 | 0.082607 | 0.443819 | Tlr9 |
| ENSMUSG( | 862.9693 | -0.17538 | 0.288461 | 0.739269 | Tm2d1 |
| ENSMUSG( | 2128.816 | -0.29407 | 0.020226 | 0.214698 | Tm2d2 |
| ENSMUSG( | 997.6178 | -0.31865 | 0.003923 | 0.080248 | Tm2d3 |
| ENSMUSG( | 352.0994 | -0.20361 | 0.384835 | 0.804031 | Tm4sf1 |
| ENSMUSG( | 11.00107 | -0.53248 | 0.346784 | NA | Tm4sf5 |
| ENSMUSG( | 459.4952 | -0.45069 | 0.000694 | 0.02534 | Tm6sf1 |
| ENSMUSG( | 69.91186 | 0.573688 | 0.024354 | 0.238636 | Tm6sf2 |
| ENSMUSG( | 1023.515 | -0.18478 | 0.114605 | 0.510543 | Tm7sf2 |
| ENSMUSG( | 2032.539 | -0.27797 | 0.047 | 0.331741 | Tm7sf3 |
| ENSMUSG( | 754.5648 | -0.15759 | 0.33239 | 0.775474 | Tm9sf1 |
| ENSMUSG( | 3223.976 | -0.42962 | 0.009305 | 0.136095 | Tm9sf2 |
| ENSMUSG( | 7711.57 | -0.2436 | 0.06952 | 0.408118 | Tm9sf3 |
| ENSMUSG( | 3393.766 | -0.32309 | 0.013944 | 0.172776 | Tm9sf4 |
| ENSMUSG( | 289.3086 | -0.2751 | 0.044836 | 0.324745 | Tma16 |
| ENSMUSG( | 2582.828 | 0.019334 | 0.819795 | 0.965888 | Tma7 |
| ENSMUSG( | 2128.567 | -0.40386 | 0.008821 | 0.132321 | Tmbim1 |
| ENSMUSG( | 1140.839 | -0.226 | 0.057452 | 0.370033 | Tmbim4 |
| ENSMUSG( | 24084.95 | -0.12235 | 0.274717 | 0.725078 | Tmbim6 |
| ENSMUSG( | 13.09018 | 0.293535 | 0.564067 | NA | Tmc3 |
| ENSMUSG( | 154.2177 | 0.520335 | 0.036459 | 0.294207 | Tmc4 |
| ENSMUSG( | 157.2273 | -0.33354 | 0.061973 | 0.38528 | Tmc6 |
| ENSMUSG( | 920.0139 | -0.01895 | 0.895441 | 0.984512 | Tmc7 |
| ENSMUSG( | 23.83668 | 0.224482 | 0.547278 | 0.885118 | Tmc8 |
| ENSMUSG( | 2295.813 | 0.123319 | 0.228503 | 0.675862 | Tmcc1 |
| ENSMUSG( | 13106.96 | 0.323967 | 0.000273 | 0.013445 | Tmcc2 |
| ENSMUSG( | 1937.075 | -0.05874 | 0.507592 | 0.86795 | Tmcc3 |
| ENSMUSG( | 1650.393 | -0.24836 | 0.095882 | 0.475743 | Tmco1 |
| ENSMUSG( | 2304.489 | -0.09981 | 0.455663 | 0.838441 | Tmco3 |
| ENSMUSG( | 245.7388 | 0.010711 | 0.939792 | 0.993415 | Tmco4 |
| ENSMUSG( | 35.14172 | 0.285173 | 0.365321 | 0.794085 | Tmco5 |
| ENSMUSG( | 336.3974 | 0.00328 | 0.981072 | 0.997512 | Tmco6 |
| ENSMUSG( | 572.8968 | -0.15125 | 0.235238 | 0.682423 | Tmed1 |
| ENSMUSG( | 6390.236 | -0.27174 | 0.041821 | 0.315339 | Tmed10 |
| ENSMUSG( | 1730.416 | -0.38914 | 0.001791 | 0.047688 | Tmed2 |
| ENSMUSG( | 1062.019 | -0.28843 | 0.030556 | 0.266032 | Tmed3 |
| ENSMUSG( | 2193.763 | -0.38623 | 0.005236 | 0.096494 | Tmed4 |
| ENSMUSG( | 697.0332 | -0.35781 | 0.021052 | 0.22058 | Tmed5 |
| ENSMUSG( | 2725.178 | -0.44659 | 0.002259 | 0.055678 | Tmed7 |
| ENSMUSG( | 1302.483 | 0.045428 | 0.62035 | 0.911432 | Tmed8 |
| ENSMUSG( | 3922.104 | -0.36136 | 0.002448 | 0.058231 | Tmed9 |
| ENSMUSG( | 3167.242 | -0.40787 | 0.000145 | 0.00841 | Tmeff1 |
| ENSMUSG( | 2265.531 | -0.52234 | 0.000331 | 0.015457 | Tmeff2 |
| ENSMUSG( | 1146.038 | -0.01013 | 0.949444 | 0.993821 | Tmem100 |
| ENSMUSG( | 938.6188 | -0.32608 | 0.012159 | 0.159821 | Tmem101 |
| ENSMUSG( | 791.7309 | -0.3595 | 0.049492 | 0.339723 | Tmem104 |
| ENSMUSG( | 58.22525 | -0.18523 | 0.46485 | 0.843575 | Tmem106a |
| ENSMUSG( | 5025.869 | -0.24303 | 0.057869 | 0.371023 | Tmem106b |
| ENSMUSG( | 1102.447 | -0.08079 | 0.478186 | 0.853453 | Tmem106c |
| ENSMUSG( | 229.1668 | -0.1433 | 0.322541 | 0.767663 | Tmem107 |
| ENSMUSG( | 667.1392 | -0.11905 | 0.453878 | 0.836798 | Tmem108 |
| ENSMUSG( | 1772.696 | -0.32926 | 0.034803 | 0.286843 | Tmem109 |
| ENSMUSG( | 512.562 | -0.01794 | 0.898002 | 0.985633 | Tmem11 |
| ENSMUSG( | 62.63847 | -0.36239 | 0.174744 | 0.610304 | Tmem114 |
| ENSMUSG( | 1201.762 | -0.3235 | 0.037924 | 0.300779 | Tmem115 |
| ENSMUSG( | 66.14251 | -0.32212 | 0.231436 | 0.679437 | Tmem116 |
| ENSMUSG( | 196.3338 | -0.90583 | 7.83E-08 | 2.15E-05 | Tmem117 |
| ENSMUSG( | 1617.022 | -0.06255 | 0.719026 | 0.941986 | Tmem119 |
| ENSMUSG( | 456.1929 | -0.09912 | 0.465231 | 0.843983 | Tmem120a |
| ENSMUSG( | 332.724 | -0.27645 | 0.047702 | 0.33359 | Tmem120b |

| | | | | | |
|---|---|---|---|---|---|
| ENSMUSG( | 6406.037 | 0.018388 | 0.902862 | 0.991089 | Tmed10 |
| ENSMUSG( | 1814.462 | 0.044372 | 0.746812 | 0.969763 | Tmed2 |
| ENSMUSG( | 1079.213 | 0.044355 | 0.768047 | 0.974236 | Tmed3 |
| ENSMUSG( | 2348.956 | 0.107864 | 0.495462 | 0.927641 | Tmed4 |
| ENSMUSG( | 841.1898 | 0.443115 | 0.021858 | 0.575341 | Tmed5 |
| ENSMUSG( | 3035.068 | 0.164438 | 0.344128 | 0.910256 | Tmed7 |
| ENSMUSG( | 1221.919 | 0.128664 | 0.235607 | 0.871807 | Tmed8 |
| ENSMUSG( | 4099.034 | 0.058708 | 0.660975 | 0.956205 | Tmed9 |
| ENSMUSG( | 3395.957 | 0.089538 | 0.530883 | 0.934795 | Tmeff1 |
| ENSMUSG( | 2455.457 | 0.0301 | 0.859737 | 0.98599 | Tmeff2 |
| ENSMUSG( | 1001.034 | -0.13274 | 0.427763 | 0.917257 | Tmem100 |
| ENSMUSG( | 972.9799 | 0.064902 | 0.663976 | 0.956766 | Tmem101 |
| ENSMUSG( | 839.3295 | 0.105649 | 0.577305 | 0.942992 | Tmem104 |
| ENSMUSG( | 56.96315 | 0.057968 | 0.855691 | 0.985738 | Tmem106a |
| ENSMUSG( | 5268.517 | 0.169028 | 0.250235 | 0.877904 | Tmem106b |
| ENSMUSG( | 1010.885 | -0.06027 | 0.630624 | 0.953251 | Tmem106c |
| ENSMUSG( | 215.974 | -0.04161 | 0.799727 | 0.97681 | Tmem107 |
| ENSMUSG( | 764.3043 | 0.507294 | 0.005211 | 0.296373 | Tmem108 |
| ENSMUSG( | 1906.749 | 0.168833 | 0.325691 | 0.905339 | Tmem109 |
| ENSMUSG( | 463.7653 | -0.04535 | 0.793144 | 0.97681 | Tmem11 |
| ENSMUSG( | 62.08196 | -0.11681 | 0.676485 | 0.95694 | Tmem114 |
| ENSMUSG( | 1329.727 | 0.249966 | 0.147168 | 0.836241 | Tmem115 |
| ENSMUSG( | 72.27887 | 0.21984 | 0.399544 | 0.917257 | Tmem116 |
| ENSMUSG( | 249.6266 | 0.186062 | 0.290157 | 0.896947 | Tmem117 |
| ENSMUSG( | 1797.281 | 0.480238 | 0.027051 | 0.603041 | Tmem119 |
| ENSMUSG( | 444.9107 | 0.104208 | 0.50255 | 0.927825 | Tmem120a |
| ENSMUSG( | 324.4224 | -0.06417 | 0.667469 | 0.956766 | Tmem120b |
| ENSMUSG( | 508.1402 | 0.11795 | 0.470483 | 0.923329 | Tmem121 |
| ENSMUSG( | 2484.332 | 0.120415 | 0.492018 | 0.927315 | Tmem121b |
| ENSMUSG( | 771.9656 | 0.291492 | 0.160819 | 0.837724 | Tmem123 |
| ENSMUSG( | 370.9625 | 0.002452 | 0.990861 | 0.998302 | Tmem125 |
| ENSMUSG( | 501.8347 | -0.1275 | 0.402921 | 0.917257 | Tmem126a |
| ENSMUSG( | 689.6146 | -0.02421 | 0.850275 | 0.985541 | Tmem126b |
| ENSMUSG( | 4939.847 | 0.104738 | 0.45807 | 0.923329 | Tmem127 |
| ENSMUSG( | 869.6077 | 0.088211 | 0.56971 | 0.941777 | Tmem128 |
| ENSMUSG( | 1172.346 | 0.228261 | 0.139625 | 0.830678 | Tmem129 |
| ENSMUSG( | 6014.357 | 0.124301 | 0.38274 | 0.917257 | Tmem130 |
| ENSMUSG( | 2828.878 | 0.301149 | 0.052772 | 0.71525 | Tmem131 |
| ENSMUSG( | 661.151 | 0.201078 | 0.137692 | 0.828492 | Tmem131l |
| ENSMUSG( | 10613.66 | 0.23929 | 0.182659 | 0.850579 | Tmem132a |
| ENSMUSG( | 4957.235 | 0.214456 | 0.219161 | 0.866104 | Tmem132b |
| ENSMUSG( | 198.9485 | 0.217033 | 0.327756 | 0.906255 | Tmem132c |
| ENSMUSG( | 1541.687 | 0.227509 | 0.208645 | 0.864412 | Tmem132d |
| ENSMUSG( | 438.0812 | 0.176825 | 0.307881 | 0.898538 | Tmem132e |
| ENSMUSG( | 1547.01 | -0.15494 | 0.18833 | 0.85698 | Tmem134 |
| ENSMUSG( | 1556.667 | 0.034471 | 0.770196 | 0.974236 | Tmem135 |
| ENSMUSG( | 353.6402 | 0.12347 | 0.399784 | 0.917257 | Tmem138 |
| ENSMUSG( | 93.21862 | 0.430126 | 0.105645 | 0.804969 | Tmem140 |
| ENSMUSG( | 788.5162 | -0.11988 | 0.345419 | 0.910581 | Tmem141 |
| ENSMUSG( | 571.3099 | -0.25018 | 0.043618 | 0.690304 | Tmem143 |
| ENSMUSG( | 626.3785 | 0.132672 | 0.407592 | 0.917257 | Tmem144 |
| ENSMUSG( | 1476.841 | 0.093227 | 0.440854 | 0.921166 | Tmem145 |
| ENSMUSG( | 2298.359 | -0.04317 | 0.724819 | 0.966887 | Tmem147 |
| ENSMUSG( | 956.5707 | -0.04338 | 0.759423 | 0.972002 | Tmem14a |
| ENSMUSG( | 1232.38 | -0.04738 | 0.697723 | 0.960867 | Tmem14c |
| ENSMUSG( | 368.0943 | 0.181565 | 0.31253 | 0.900037 | Tmem150a |
| ENSMUSG( | 186.0846 | -0.38406 | 0.070201 | 0.757694 | Tmem150b |
| ENSMUSG( | 1804.589 | 0.105437 | 0.439203 | 0.920721 | Tmem150c |
| ENSMUSG( | 9508.665 | 0.070164 | 0.665294 | 0.956766 | Tmem151a |
| ENSMUSG( | 8097.151 | 0.220539 | 0.100158 | 0.801311 | Tmem151b |
| ENSMUSG( | 27.33021 | -0.53721 | 0.173518 | 0.844317 | Tmem154 |
| ENSMUSG( | 11.14825 | -0.67535 | 0.272103 | NA | Tmem156 |
| ENSMUSG( | 2205.841 | -0.08061 | 0.566582 | 0.941777 | Tmem158 |
| ENSMUSG( | 1007.608 | -0.2411 | 0.055091 | 0.723362 | Tmem160 |
| ENSMUSG( | 988.0492 | 0.230347 | 0.103067 | 0.804311 | Tmem161a |
| ENSMUSG( | 577.6681 | 0.212566 | 0.163589 | 0.838291 | Tmem161b |
| ENSMUSG( | 1146.307 | 0.078015 | 0.604737 | 0.948869 | Tmem163 |
| ENSMUSG( | 1950.989 | 0.153237 | 0.175218 | 0.845305 | Tmem164 |
| ENSMUSG( | 1142.329 | 0.147853 | 0.350069 | 0.911652 | Tmem165 |
| ENSMUSG( | 1423.424 | 0.112079 | 0.368113 | 0.915491 | Tmem167 |
| ENSMUSG( | 2192.801 | -0.07162 | 0.587729 | 0.944975 | Tmem167b |
| ENSMUSG( | 529.087 | 0.271104 | 0.121471 | 0.813384 | Tmem168 |
| ENSMUSG( | 535.744 | 0.103194 | 0.450733 | 0.922202 | Tmem169 |
| ENSMUSG( | 220.4989 | 0.061999 | 0.715917 | 0.965424 | Tmem17 |
| ENSMUSG( | 164.8169 | 0.376078 | 0.042845 | 0.690304 | Tmem170 |
| ENSMUSG( | 2515.037 | -0.03103 | 0.804871 | 0.977232 | Tmem170b |
| ENSMUSG( | 1223.043 | 0.064765 | 0.63592 | 0.953903 | Tmem175 |
| ENSMUSG( | 1062.962 | 0.067967 | 0.635041 | 0.953903 | Tmem176a |
| ENSMUSG( | 2369.632 | 0.050741 | 0.690158 | 0.960071 | Tmem176b |
| ENSMUSG( | 208.2415 | 0.041275 | 0.820454 | 0.979198 | Tmem177 |
| ENSMUSG( | 2109.299 | 0.16826 | 0.335891 | 0.907913 | Tmem178 |
| ENSMUSG( | 4190.471 | 0.062715 | 0.70001 | 0.961335 | Tmem178b |
| ENSMUSG( | 4446.528 | 0.133661 | 0.316979 | 0.902088 | Tmem179 |
| ENSMUSG( | 216.1129 | 0.124801 | 0.485998 | 0.92541 | Tmem179b |
| ENSMUSG( | 1359.255 | 0.179378 | 0.185881 | 0.855305 | Tmem18 |
| ENSMUSG( | 2124.763 | 0.13192 | 0.407477 | 0.917257 | Tmem181a |
| ENSMUSG( | 1995.392 | -0.03406 | 0.757381 | 0.971858 | Tmem183a |
| ENSMUSG( | 2847.251 | 0.071092 | 0.66236 | 0.956205 | Tmem184b |
| ENSMUSG( | 3194.762 | 0.085963 | 0.504095 | 0.927825 | Tmem184c |
| ENSMUSG( | 1529.507 | 0.116205 | 0.419426 | 0.917257 | Tmem185a |
| ENSMUSG( | 456.0155 | 0.192383 | 0.315719 | 0.901868 | Tmem185b |
| ENSMUSG( | 515.2351 | 0.011943 | 0.92632 | 0.99306 | Tmem186 |
| ENSMUSG( | 1621.503 | 0.147879 | 0.303182 | 0.898538 | Tmem19 |
| ENSMUSG( | 2382.467 | 0.041769 | 0.790897 | 0.97681 | Tmem191 |
| ENSMUSG( | 555.6488 | 0.020691 | 0.890157 | 0.990558 | Tmem192 |
| ENSMUSG( | 291.2218 | 0.212719 | 0.251209 | 0.879321 | Tmem196 |
| ENSMUSG( | 2554.856 | 0.039259 | 0.768381 | 0.974236 | Tmem198 |
| ENSMUSG( | 622.5211 | 0.011182 | 0.950248 | 0.994798 | Tmem198b |
| ENSMUSG( | 496.847 | 0.067087 | 0.64343 | 0.953903 | Tmem199 |
| ENSMUSG( | 581.7129 | 0.435263 | 0.028605 | 0.614171 | Tmem200a |
| ENSMUSG( | 207.1331 | 0.316944 | 0.082643 | 0.77681 | Tmem200b |
| ENSMUSG( | 505.7129 | 0.288735 | 0.142624 | 0.831221 | Tmem200c |
| ENSMUSG( | 2892.992 | 0.118231 | 0.423351 | 0.917257 | Tmem201 |
| ENSMUSG( | 52.26855 | -0.26561 | 0.372882 | 0.915491 | Tmem202 |
| ENSMUSG( | 265.2252 | -0.05802 | 0.721906 | 0.966887 | Tmem203 |
| ENSMUSG( | 415.8183 | 0.352019 | 0.160574 | 0.837724 | Tmem204 |
| ENSMUSG( | 916.4093 | -0.0647 | 0.620394 | 0.952896 | Tmem205 |
| ENSMUSG( | 774.0014 | -0.03972 | 0.766258 | 0.974236 | Tmem208 |
| ENSMUSG( | 990.5167 | 0.176875 | 0.283319 | 0.892412 | Tmem209 |
| ENSMUSG( | 90.41718 | 0.104457 | 0.724153 | 0.966887 | Tmem212 |
| ENSMUSG( | 1963.393 | 0.183721 | 0.157368 | 0.837724 | Tmem214 |
| ENSMUSG( | 406.6127 | 0.098247 | 0.580293 | 0.944209 | Tmem215 |
| ENSMUSG( | 137.9257 | 0.068651 | 0.735424 | 0.967344 | Tmem216 |
| ENSMUSG( | 1600.837 | 0.113965 | 0.373523 | 0.915491 | Tmem217rt |
| ENSMUSG( | 642.6669 | -0.20081 | 0.172341 | 0.843502 | Tmem218 |
| ENSMUSG( | 433.5509 | -0.11069 | 0.404856 | 0.917257 | Tmem219 |
| ENSMUSG( | 41.27411 | -0.12024 | 0.743542 | 0.96951 | Tmem220 |
| ENSMUSG( | 10.09042 | -0.05921 | 0.926526 | NA | Tmem221 |
| ENSMUSG( | 2620.285 | -0.12704 | 0.316589 | 0.901868 | Tmem222 |
| ENSMUSG( | 1124.219 | 0.095475 | 0.421618 | 0.917257 | Tmem223 |
| ENSMUSG( | 5749.404 | -0.17868 | 0.269937 | 0.887508 | Tmem229a |
| ENSMUSG( | 1212.159 | 0.055071 | 0.660913 | 0.956205 | Tmem229b |
| ENSMUSG( | 511.9201 | 0.173222 | 0.309107 | 0.898538 | Tmem230 |
| ENSMUSG( | 381.0303 | 0.229698 | 0.161492 | 0.837724 | Tmem231 |
| ENSMUSG( | 55.25455 | 0.166915 | 0.607289 | 0.949981 | Tmem232 |
| ENSMUSG( | 2699.585 | -0.03176 | 0.770307 | 0.974236 | Tmem234 |
| ENSMUSG( | 613.0306 | 0.12857 | 0.370859 | 0.915491 | Tmem237 |
| ENSMUSG( | 146.7858 | -0.13352 | 0.484512 | 0.92541 | Tmem238 |
| ENSMUSG( | 1523.186 | -0.01134 | 0.914621 | 0.992407 | Tmem240 |
| ENSMUSG( | 283.9351 | -0.10266 | 0.526847 | 0.934449 | Tmem241 |
| ENSMUSG( | 2015.017 | -0.19611 | 0.131533 | 0.822912 | Tmem242 |
| ENSMUSG( | 592.8345 | 0.145335 | 0.316399 | 0.901868 | Tmem243 |
| ENSMUSG( | 2506.058 | 0.315281 | 0.053336 | 0.717022 | Tmem245 |
| ENSMUSG( | 1546.428 | 0.146269 | 0.361686 | 0.914223 | Tmem248 |
| ENSMUSG( | 1673.097 | -0.00764 | 0.951129 | 0.994798 | Tmem25 |
| ENSMUSG( | 2918.86 | -0.05379 | 0.592355 | 0.945016 | Tmem250 |
| ENSMUSG( | 75.64916 | -1.76196 | 2.58E-08 | 2.08E-05 | Tmem252 |
| ENSMUSG( | 11.75032 | -1.09159 | 0.076715 | 0.770558 | Tmem253 |
| ENSMUSG( | 2412.157 | -0.36833 | 0.003311 | 0.229858 | Tmem254 |
| ENSMUSG( | 541.7848 | 0.135044 | 0.421806 | 0.917257 | Tmem255a |
| ENSMUSG( | 106.7929 | 0.077071 | 0.799373 | 0.97681 | Tmem255b |
| ENSMUSG( | 1168.157 | -0.19513 | 0.145182 | 0.83339 | Tmem256 |
| ENSMUSG( | 718.87 | -0.15349 | 0.197594 | 0.862161 | Tmem258 |
| ENSMUSG( | 5669.8 | 0.18531 | 0.107348 | 0.804969 | Tmem259 |
| ENSMUSG( | 872.0784 | 0.07749 | 0.594994 | 0.94561 | Tmem260 |
| ENSMUSG( | 521.592 | -0.06148 | 0.701825 | 0.962014 | Tmem263 |
| ENSMUSG( | 318.5224 | 0.007498 | 0.962727 | 0.995284 | Tmem266 |
| ENSMUSG( | 329.6451 | -0.61075 | 0.0171 | 0.52741 | Tmem267 |
| ENSMUSG( | 496.7053 | -0.04602 | 0.773714 | 0.974672 | Tmem268 |
| ENSMUSG( | 27.69089 | 0.152334 | 0.681879 | 0.958678 | Tmem269 |
| ENSMUSG( | 35.45335 | 0.189777 | 0.638883 | 0.953903 | Tmem273 |
| ENSMUSG( | 771.1731 | -0.06214 | 0.638146 | 0.953903 | Tmem276 |
| ENSMUSG( | 8731.181 | 0.257644 | 0.158159 | 0.837724 | Tmem30a |
| ENSMUSG( | 11.38543 | -0.15593 | 0.795554 | NA | Tmem30b |
| ENSMUSG( | 4010.054 | 0.047809 | 0.767669 | 0.974236 | Tmem33 |
| ENSMUSG( | 2195.588 | 0.051859 | 0.727397 | 0.966887 | Tmem35a |
| ENSMUSG( | 85.79578 | -0.03806 | 0.874894 | 0.989644 | Tmem35b |
| ENSMUSG( | 62.66139 | 0.7501 | 0.012859 | 0.457834 | Tmem37 |

| | | | | | | | | | | | |
|---|---|---|---|---|---|---|---|---|---|---|---|
| ENSMUSG( | Tmem86a | 719.1949 | -0.44342 | 0.000935 | 0.031289 | ENSMUSG( | Tmem86a | 818.1291 | 0.229329 | 0.173026 | 0.843788 |
| ENSMUSG( | Tmem86b | 251.3575 | 0.427084 | 0.008192 | 0.12594 | ENSMUSG( | Tmem86b | 209.3995 | 0.192475 | 0.313304 | 0.900878 |
| ENSMUSG( | Tmem87a | 1119.413 | -0.3821 | 0.016201 | 0.189046 | ENSMUSG( | Tmem87a | 1206.826 | 0.131456 | 0.449796 | 0.922202 |
| ENSMUSG( | Tmem87b | 1191.509 | -0.36225 | 0.019919 | 0.213312 | ENSMUSG( | Tmem87b | 1265.441 | 0.106493 | 0.518131 | 0.930939 |
| ENSMUSG( | Tmem88 | 120.4681 | -0.06105 | 0.804259 | 0.962169 | ENSMUSG( | Tmem88 | 141.5598 | 0.615786 | 0.00661 | 0.338858 |
| ENSMUSG( | Tmem88b | 3017.489 | -0.51441 | 1.94E-06 | 0.000324 | ENSMUSG( | Tmem88b | 3057.262 | -0.16724 | 0.219716 | 0.866104 |
| ENSMUSG( | Tmem8b | 4311.502 | -0.06302 | 0.59259 | 0.902094 | ENSMUSG( | Tmem8b | 4219.928 | 0.144182 | 0.30383 | 0.898538 |
| ENSMUSG( | Tmem9 | 2192.361 | -0.36373 | 0.002926 | 0.066146 | ENSMUSG( | Tmem9 | 2297.56 | 0.064187 | 0.644051 | 0.953903 |
| ENSMUSG( | Tmem91 | 417.6507 | -0.19903 | 0.163963 | 0.594869 | ENSMUSG( | Tmem91 | 401.193 | -0.04724 | 0.786163 | 0.976285 |
| ENSMUSG( | Tmem94 | 2875.084 | -0.05826 | 0.616951 | 0.910866 | ENSMUSG( | Tmem94 | 2890.602 | 0.221966 | 0.091905 | 0.791889 |
| ENSMUSG( | Tmem97 | 689.6243 | -0.50931 | 0.001904 | 0.04982 | ENSMUSG( | Tmem97 | 748.0838 | 0.04351 | 0.800544 | 0.97681 |
| ENSMUSG( | Tmem98 | 628.4802 | -0.54649 | 0.001359 | 0.040233 | ENSMUSG( | Tmem98 | 639.4666 | -0.17743 | 0.318573 | 0.903164 |
| ENSMUSG( | Tmem9b | 4510.435 | -0.2603 | 0.045721 | 0.328047 | ENSMUSG( | Tmem9b | 4493.898 | 0.011345 | 0.937546 | 0.993685 |
| ENSMUSG( | Tmf1 | 1282.32 | 0.110468 | 0.260407 | 0.710245 | ENSMUSG( | Tmf1 | 1148.567 | 0.063841 | 0.567749 | 0.941777 |
| ENSMUSG( | Tmie | 360.1258 | -0.21514 | 0.123577 | 0.530552 | ENSMUSG( | Tmie | 329.9965 | -0.2078 | 0.217496 | 0.865589 |
| ENSMUSG( | Tmod1 | 1030.447 | -0.26499 | 0.018931 | 0.207196 | ENSMUSG( | Tmod1 | 966.7987 | -0.17156 | 0.180226 | 0.850372 |
| ENSMUSG( | Tmod2 | 19614.04 | 0.16795 | 0.051513 | 0.34814 | ENSMUSG( | Tmod2 | 16797.41 | -0.00637 | 0.956044 | 0.994798 |
| ENSMUSG( | Tmod3 | 1032.85 | 0.048305 | 0.636365 | 0.914873 | ENSMUSG( | Tmod3 | 973.5961 | 0.143627 | 0.212185 | 0.864412 |
| ENSMUSG( | Tmod4 | 12.07895 | -0.34861 | 0.513639 | NA | ENSMUSG( | Tmod4 | 12.80404 | 0.114567 | 0.829046 | 0.980224 |
| ENSMUSG( | Tmpo | 677.8731 | 0.082896 | 0.432366 | 0.82768 | ENSMUSG( | Tmpo | 675.3683 | 0.324005 | 0.008352 | 0.388416 |
| ENSMUSG( | Tmppe | 84.63354 | 0.072178 | 0.79363 | 0.959799 | ENSMUSG( | Tmppe | 89.87375 | 0.495704 | 0.086589 | 0.782073 |
| ENSMUSG( | Tmprss5 | 35.44228 | -1.2968 | 9.67E-05 | 0.006353 | ENSMUSG( | Tmprss5 | 39.42274 | -0.50428 | 0.126007 | 0.81516 |
| ENSMUSG( | Tmprss6 | 18.20575 | -0.66897 | 0.153499 | NA | ENSMUSG( | Tmprss6 | 17.51833 | -0.46052 | 0.324905 | 0.905257 |
| ENSMUSG( | Tmprss7 | 123.641 | -0.05467 | 0.771176 | 0.954726 | ENSMUSG( | Tmprss7 | 108.6513 | -0.15609 | 0.459136 | 0.923329 |
| ENSMUSG( | Tmprss9 | 17.35452 | -0.28669 | 0.534015 | NA | ENSMUSG( | Tmprss9 | 18.92934 | 0.211219 | 0.639023 | 0.953903 |
| ENSMUSG( | Tmsb10 | 6778.685 | -0.04842 | 0.612462 | 0.908747 | ENSMUSG( | Tmsb10 | 5835.529 | -0.22229 | 0.116213 | 0.813384 |
| ENSMUSG( | Tmsb10b | 61.37838 | 0.003936 | 0.988131 | 0.999109 | ENSMUSG( | Tmsb10b | 48.73443 | -0.44298 | 0.15634 | 0.837724 |
| ENSMUSG( | Tmsb15b1 | 14.75294 | -0.48693 | 0.360247 | NA | ENSMUSG( | Tmsb15b1 | 12.35477 | -0.82334 | 0.163296 | 0.838291 |
| ENSMUSG( | Tmsb4x | 18308.27 | 0.04062 | 0.66474 | 0.926824 | ENSMUSG( | Tmsb4x | 15825.12 | -0.11526 | 0.346455 | 0.910966 |
| ENSMUSG( | Tmt1a | 1068.795 | 0.00736 | 0.953144 | 0.993874 | ENSMUSG( | Tmt1a | 1121.166 | 0.391334 | 0.005603 | 0.303106 |
| ENSMUSG( | Tmtc1 | 2566.031 | -0.38639 | 0.000503 | 0.020541 | ENSMUSG( | Tmtc1 | 2690.962 | 0.048432 | 0.709692 | 0.96416 |
| ENSMUSG( | Tmtc2 | 487.2607 | -0.20499 | 0.242245 | 0.689699 | ENSMUSG( | Tmtc2 | 512.2145 | 0.215475 | 0.291349 | 0.897566 |
| ENSMUSG( | Tmtc3 | 335.3456 | -0.26818 | 0.182151 | 0.619591 | ENSMUSG( | Tmtc3 | 397.6631 | 0.470983 | 0.022532 | 0.581149 |
| ENSMUSG( | Tmtc4 | 336.6084 | -0.06666 | 0.683006 | 0.93205 | ENSMUSG( | Tmtc4 | 349.4996 | 0.301651 | 0.097244 | 0.796721 |
| ENSMUSG( | Tmub1 | 694.6903 | -0.20827 | 0.064062 | 0.391961 | ENSMUSG( | Tmub1 | 702.0572 | 0.101489 | 0.46852 | 0.923329 |
| ENSMUSG( | Tmub2 | 2815.854 | -0.3379 | 0.002511 | 0.059283 | ENSMUSG( | Tmub2 | 2945.174 | 0.081626 | 0.543967 | 0.938063 |
| ENSMUSG( | Tmx1 | 945.1907 | -0.41636 | 0.006326 | 0.108996 | ENSMUSG( | Tmx1 | 1135.733 | 0.389642 | 0.026522 | 0.603041 |
| ENSMUSG( | Tmx2 | 5446.844 | -0.38838 | 0.002 | 0.051341 | ENSMUSG( | Tmx2 | 5588.221 | -0.01726 | 0.905487 | 0.991089 |
| ENSMUSG( | Tmx3 | 1612.806 | -0.41886 | 0.009145 | 0.134728 | ENSMUSG( | Tmx3 | 1855.991 | 0.275703 | 0.132017 | 0.822912 |
| ENSMUSG( | Tmx4 | 5423.419 | -0.41653 | 0.007543 | 0.120197 | ENSMUSG( | Tmx4 | 6111.46 | 0.221978 | 0.209739 | 0.864412 |
| ENSMUSG( | Tnc | 527.6962 | -0.3307 | 0.064244 | 0.392776 | ENSMUSG( | Tnc | 539.6908 | 0.027907 | 0.879219 | 0.989644 |
| ENSMUSG( | Tnfaip1 | 2460.031 | 0.097431 | 0.32474 | 0.770182 | ENSMUSG( | Tnfaip1 | 2148.893 | -0.02091 | 0.851118 | 0.985541 |
| ENSMUSG( | Tnfaip2 | 41.33401 | -0.56472 | 0.057946 | 0.371205 | ENSMUSG( | Tnfaip2 | 36.71438 | -0.63673 | 0.063773 | 0.736482 |
| ENSMUSG( | Tnfaip3 | 104.569 | 0.003832 | 0.984896 | 0.997815 | ENSMUSG( | Tnfaip3 | 118.3591 | 0.587472 | 0.006872 | 0.343883 |
| ENSMUSG( | Tnfaip6 | 220.8315 | -0.83511 | 7.34E-06 | 0.000923 | ENSMUSG( | Tnfaip6 | 277.8405 | 0.216756 | 0.315357 | 0.901306 |
| ENSMUSG( | Tnfaip8 | 584.9865 | -0.09564 | 0.549666 | 0.885944 | ENSMUSG( | Tnfaip8 | 507.8516 | -0.25284 | 0.161616 | 0.837724 |
| ENSMUSG( | Tnfaip8l1 | 151.1227 | -0.05558 | 0.761486 | 0.953106 | ENSMUSG( | Tnfaip8l1 | 137.8226 | -0.04236 | 0.834573 | 0.981237 |
| ENSMUSG( | Tnfaip8l2 | 95.02929 | 0.275638 | 0.18096 | 0.618094 | ENSMUSG( | Tnfaip8l2 | 88.6388 | 0.31778 | 0.228346 | 0.868507 |
| ENSMUSG( | Tnfaip8l3 | 153.063 | 0.244214 | 0.28079 | 0.729868 | ENSMUSG( | Tnfaip8l3 | 132.1507 | 0.07815 | 0.768103 | 0.974236 |
| ENSMUSG( | Tnfrsf10b | 25.97451 | -0.1801 | 0.634779 | 0.914073 | ENSMUSG( | Tnfrsf10b | 30.85393 | 0.5564 | 0.133846 | 0.822912 |
| ENSMUSG( | Tnfrsf11a | 191.0108 | -0.48566 | 0.00365 | 0.076783 | ENSMUSG( | Tnfrsf11a | 206.663 | 0.058042 | 0.770848 | 0.974236 |
| ENSMUSG( | Tnfrsf11b | 102.903 | -0.4501 | 0.128657 | 0.537025 | ENSMUSG( | Tnfrsf11b | 110.7871 | 0.083168 | 0.779356 | 0.976106 |
| ENSMUSG( | Tnfrsf12a | 148.037 | -0.43038 | 0.02878 | 0.258585 | ENSMUSG( | Tnfrsf12a | 138.1658 | -0.33809 | 0.101433 | 0.801456 |
| ENSMUSG( | Tnfrsf13b | 106.5286 | -0.18498 | 0.344191 | 0.779153 | ENSMUSG( | Tnfrsf13b | 95.43402 | -0.23923 | 0.314547 | 0.900949 |
| ENSMUSG( | Tnfrsf13c | 17.33946 | 0.029628 | 0.94731 | NA | ENSMUSG( | Tnfrsf13c | 16.27969 | 0.1358 | 0.796508 | 0.97681 |
| ENSMUSG( | Tnfrsf14 | 20.28251 | 0.086973 | 0.850465 | NA | ENSMUSG( | Tnfrsf14 | 22.57245 | 0.612881 | 0.160751 | 0.837724 |
| ENSMUSG( | Tnfrsf17 | 12.29013 | 0.011335 | 0.9833 | NA | ENSMUSG( | Tnfrsf17 | 12.89798 | 0.401472 | 0.451999 | 0.922316 |
| ENSMUSG( | Tnfrsf19 | 8042.657 | -0.11218 | 0.374503 | 0.800445 | ENSMUSG( | Tnfrsf19 | 7096.04 | -0.20857 | 0.114345 | 0.811703 |
| ENSMUSG( | Tnfrsf1a | 727.7124 | -0.13591 | 0.385275 | 0.804048 | ENSMUSG( | Tnfrsf1a | 763.2966 | 0.267893 | 0.164726 | 0.838743 |
| ENSMUSG( | Tnfrsf1b | 282.1272 | -0.46167 | 0.038036 | 0.300991 | ENSMUSG( | Tnfrsf1b | 306.7505 | 0.090242 | 0.726176 | 0.966887 |
| ENSMUSG( | Tnfrsf21 | 2408.477 | -0.30846 | 0.048928 | 0.337919 | ENSMUSG( | Tnfrsf21 | 2695.234 | 0.290867 | 0.106317 | 0.804969 |
| ENSMUSG( | Tnfrsf22 | 91.72018 | 0.174897 | 0.398135 | 0.812452 | ENSMUSG( | Tnfrsf22 | 74.0516 | -0.1431 | 0.601167 | 0.947749 |
| ENSMUSG( | Tnfrsf23 | 59.48596 | 0.008866 | 0.971306 | 0.996369 | ENSMUSG( | Tnfrsf23 | 49.86155 | -0.23491 | 0.426667 | 0.917257 |
| ENSMUSG( | Tnfrsf25 | 250.4822 | 0.657804 | 0.001742 | 0.047093 | ENSMUSG( | Tnfrsf25 | 185.5714 | 0.125089 | 0.670278 | 0.95694 |
| ENSMUSG( | Tnfrsf4 | 40.46024 | 0.009184 | 0.975355 | 0.997302 | ENSMUSG( | Tnfrsf4 | 32.97697 | -0.35965 | 0.306278 | 0.898538 |
| ENSMUSG( | Tnfrsf8 | 34.39422 | 0.143491 | 0.664038 | 0.926511 | ENSMUSG( | Tnfrsf8 | 30.96188 | 0.108958 | 0.766595 | 0.974236 |
| ENSMUSG( | Tnfsf10 | 72.84979 | -0.14843 | 0.630027 | 0.913965 | ENSMUSG( | Tnfsf10 | 102.8402 | 0.973492 | 0.000514 | 0.068166 |
| ENSMUSG( | Tnfsf12 | 522.4868 | -0.15739 | 0.30116 | 0.749979 | ENSMUSG( | Tnfsf12 | 495.9929 | -0.03717 | 0.814419 | 0.979198 |
| ENSMUSG( | Tnfsf13 | 50.92549 | 0.247029 | 0.424841 | 0.824675 | ENSMUSG( | Tnfsf13 | 42.30647 | -0.00331 | 0.992129 | 0.99874 |
| ENSMUSG( | Tnfsf13b | 35.83161 | -0.69454 | 0.031514 | 0.270738 | ENSMUSG( | Tnfsf13b | 40.86297 | 0.052352 | 0.871618 | 0.988779 |
| ENSMUSG( | Tnfsf9 | 14.10982 | -0.87443 | 0.084803 | NA | ENSMUSG( | Tnfsf9 | 16.53422 | -0.01576 | 0.973746 | 0.996416 |
| ENSMUSG( | Tnfsfm13 | 26.25886 | -0.22168 | 0.551124 | 0.886038 | ENSMUSG( | Tnfsfm13 | 25.39523 | -0.04648 | 0.906592 | 0.991494 |
| ENSMUSG( | Tnik | 4749.996 | -0.00473 | 0.95619 | 0.993874 | ENSMUSG( | Tnik | 4543.407 | 0.133321 | 0.179908 | 0.850372 |
| ENSMUSG( | Tnip1 | 722.0309 | 0.058547 | 0.602202 | 0.904887 | ENSMUSG( | Tnip1 | 618.9676 | -0.11445 | 0.404167 | 0.917257 |
| ENSMUSG( | Tnip2 | 367.8071 | -0.00728 | 0.956815 | 0.993874 | ENSMUSG( | Tnip2 | 349.6929 | 0.124261 | 0.505115 | 0.927825 |
| ENSMUSG( | Tnk2 | 9877.178 | 0.230912 | 0.005492 | 0.0993 | ENSMUSG( | Tnk2 | 8781.976 | 0.165135 | 0.117523 | 0.813384 |
| ENSMUSG( | Tnks | 3405.83 | -0.02845 | 0.789172 | 0.959299 | ENSMUSG( | Tnks | 3170.139 | 0.033975 | 0.782017 | 0.976106 |
| ENSMUSG( | Tnks1bp1 | 2647.744 | 0.018465 | 0.847575 | 0.972374 | ENSMUSG( | Tnks1bp1 | 2490.301 | 0.109329 | 0.308402 | 0.898538 |
| ENSMUSG( | Tnks2 | 4883.683 | 0.006902 | 0.944115 | 0.993503 | ENSMUSG( | Tnks2 | 4504.386 | 0.04161 | 0.720686 | 0.966887 |
| ENSMUSG( | Tnmd | 24.24023 | -0.55405 | 0.159895 | 0.589679 | ENSMUSG( | Tnmd | 25.17003 | -0.11009 | 0.782869 | 0.976106 |
| ENSMUSG( | Tnnc1 | 251.2543 | -0.02181 | 0.891774 | 0.983979 | ENSMUSG( | Tnnc1 | 201.9325 | -0.42082 | 0.026785 | 0.603041 |
| ENSMUSG( | Tnni1 | 259.5544 | -0.10027 | 0.499478 | 0.863218 | ENSMUSG( | Tnni1 | 214.83 | -0.39537 | 0.038758 | 0.679972 |
| ENSMUSG( | Tnni2 | 13.10379 | -0.03805 | 0.942714 | NA | ENSMUSG( | Tnni2 | 12.046 | -0.01171 | 0.984344 | 0.996963 |
| ENSMUSG( | Tnni3 | 26.10511 | -0.28635 | 0.497964 | 0.862372 | ENSMUSG( | Tnni3 | 22.37767 | -0.54491 | 0.240361 | 0.875526 |
| ENSMUSG( | Tnnt1 | 182.9389 | -0.30653 | 0.103162 | 0.48855 | ENSMUSG( | Tnnt1 | 154.287 | -0.57797 | 0.005122 | 0.294733 |
| ENSMUSG( | Tnnt2 | 78.89736 | -0.25905 | 0.392003 | 0.809274 | ENSMUSG( | Tnnt2 | 75.02384 | -0.14025 | 0.630649 | 0.953251 |
| ENSMUSG( | Tnnt3 | 11.67647 | 0.175889 | 0.756775 | NA | ENSMUSG( | Tnnt3 | 9.65547 | -0.19048 | 0.777944 | NA |
| ENSMUSG( | Tnpo1 | 2302.13 | 0.059592 | 0.622514 | 0.911432 | ENSMUSG( | Tnpo1 | 2172.217 | 0.155214 | 0.252157 | 0.879902 |
| ENSMUSG( | Tnpo2 | 5765.069 | 0.139677 | 0.07601 | 0.426896 | ENSMUSG( | Tnpo2 | 4903.754 | -0.05578 | 0.561356 | 0.940254 |
| ENSMUSG( | Tnpo3 | 2163.912 | 0.075576 | 0.389676 | 0.807428 | ENSMUSG( | Tnpo3 | 1928.145 | 0.010602 | 0.927089 | 0.99306 |
| ENSMUSG( | Tnr | 5743.877 | -0.41736 | 0.018375 | 0.203658 | ENSMUSG( | Tnr | 6408.371 | 0.196806 | 0.331903 | 0.907907 |
| ENSMUSG( | Tnrc18 | 4406.372 | -0.00851 | 0.925694 | 0.991204 | ENSMUSG( | Tnrc18 | 4115.293 | 0.064509 | 0.563118 | 0.941158 |
| ENSMUSG( | Tnrc6a | 2670.78 | -0.05646 | 0.525983 | 0.876999 | ENSMUSG( | Tnrc6a | 2504.144 | 0.027089 | 0.80349 | 0.976978 |
| ENSMUSG( | Tnrc6b | 2175.342 | 0.029833 | 0.74517 | 0.949158 | ENSMUSG( | Tnrc6b | 1985.324 | 0.036036 | 0.737561 | 0.967686 |
| ENSMUSG( | Tnrc6c | 2438.062 | -0.21749 | 0.025734 | 0.247138 | ENSMUSG( | Tnrc6c | 2372.149 | -0.0187 | 0.855473 | 0.985738 |
| ENSMUSG( | Tns1 | 2622.007 | 0.084245 | 0.42109 | 0.823126 | ENSMUSG( | Tns1 | 2440.12 | 0.146718 | 0.271563 | 0.888062 |
| ENSMUSG( | Tns2 | 708.8411 | 0.179299 | 0.098725 | 0.480598 | ENSMUSG( | Tns2 | 744.1401 | 0.54946 | 0.001467 | 0.140378 |
| ENSMUSG( | Tns3 | 11815.54 | 0.211388 | 0.047341 | 0.332697 | ENSMUSG( | Tns3 | 9765.288 | -0.06371 | 0.627863 | 0.953234 |
| ENSMUSG( | Tns4 | 13.02429 | 0.010259 | 0.984038 | NA | ENSMUSG( | Tns4 | 11.91243 | 0.024894 | 0.965491 | 0.995433 |
| ENSMUSG( | Tnxb | 235.637 | 0.071755 | 0.658635 | 0.924506 | ENSMUSG( | Tnxb | 282.2118 | 0.774232 | 0.001219 | 0.126346 |
| ENSMUSG( | Tob1 | 701.409 | -0.44465 | 0.005958 | 0.104409 | ENSMUSG( | Tob1 | 843.6551 | 0.374192 | 0.043919 | 0.690976 |
| ENSMUSG( | Tob2 | 855.9641 | -0.42311 | 0.010916 | 0.150598 | ENSMUSG( | Tob2 | 883.9997 | -0.02388 | 0.906664 | 0.991511 |
| ENSMUSG( | Toe1 | 329.3657 | -0.08856 | 0.540772 | 0.88315 | ENSMUSG( | Toe1 | 307.169 | -0.01333 | 0.935879 | 0.993443 |
| ENSMUSG( | Togaram1 | 836.9873 | 0.168527 | 0.084949 | 0.449406 | ENSMUSG( | Togaram1 | 777.7275 | 0.224025 | 0.059024 | 0.729429 |
| ENSMUSG( | Togaram2 | 80.69602 | 0.033067 | 0.89559 | 0.984512 | ENSMUSG( | Togaram2 | 77.82195 | 0.209708 | 0.449707 | 0.922202 |
| ENSMUSG( | Tollip | 5131.69 | 0.112178 | 0.160016 | 0.589679 | ENSMUSG( | Tollip | 4324.212 | -0.11448 | 0.288152 | 0.894794 |
| ENSMUSG( | Tom1 | 810.3748 | 0.023759 | 0.811646 | 0.964426 | ENSMUSG( | Tom1 | 685.8043 | -0.19553 | 0.102392 | 0.802599 |
| ENSMUSG( | Tom1l1 | 1305.211 | 0.11987 | 0.268987 | 0.719302 | ENSMUSG( | Tom1l1 | 1126.147 | -0.04041 | 0.762203 | 0.973255 |
| ENSMUSG( | Tom1l2 | 11342.27 | 0.130567 | 0.095746 | 0.475357 | ENSMUSG( | Tom1l2 | 9503.253 | -0.10962 | 0.256519 | 0.883082 |
| ENSMUSG( | Tomm20 | 8709.712 | -0.17054 | 0.09302 | 0.468859 | ENSMUSG( | Tomm20 | 8429.605 | 0.01107 | 0.925129 | 0.99306 |
| ENSMUSG( | Tomm22 | 1730.96 | -0.00659 | 0.948397 | 0.993821 | ENSMUSG( | Tomm22 | 1501.81 | -0.15558 | 0.209256 | 0.864412 |
| ENSMUSG( | Tomm34 | 1600.949 | 0.068714 | 0.446573 | 0.833726 | ENSMUSG( | Tomm34 | 1368.039 | -0.12086 | 0.318388 | 0.903045 |
| ENSMUSG( | Tomm40 | 1987.93 | -0.03075 | 0.773467 | 0.955447 | ENSMUSG( | Tomm40 | 1790.209 | -0.06545 | 0.605865 | 0.949449 |
| ENSMUSG( | Tomm40l | 3536.155 | 0.107467 | 0.198286 | 0.638387 | ENSMUSG( | Tomm40l | 2882.579 | -0.22232 | 0.051855 | 0.712864 |
| ENSMUSG( | Tomm5 | 653.5988 | 0.018122 | 0.884424 | 0.982647 | ENSMUSG( | Tomm5 | 550.0204 | -0.22249 | 0.119455 | 0.813384 |
| ENSMUSG( | Tomm6os | 157.7836 | 0.074085 | 0.706783 | 0.93949 | ENSMUSG( | Tomm6os | 136.586 | -0.06904 | 0.755113 | 0.971098 |
| ENSMUSG( | Tomm7 | 2137.883 | 0.057933 | 0.587939 | 0.900678 | ENSMUSG( | Tomm7 | 1792.163 | -0.19198 | 0.133076 | 0.822912 |
| ENSMUSG( | Tomm70a | 3080.951 | -0.0455 | 0.658924 | 0.924506 | ENSMUSG( | Tomm70a | 2811.843 | -0.0435 | 0.741207 | 0.968811 |
| ENSMUSG( | Tomt | 12.90196 | -0.51598 | 0.315457 | NA | ENSMUSG( | Tomt | 13.24514 | -0.14079 | 0.794328 | 0.97681 |
| ENSMUSG( | Tonsl | 266.5297 | 0.20329 | 0.175629 | 0.611006 | ENSMUSG( | Tonsl | 232.2153 | 0.079338 | 0.628305 | 0.953251 |
| ENSMUSG( | Top1 | 1834.933 | 0.204611 | 0.05749 | 0.370033 | ENSMUSG( | Top1 | 1692.099 | 0.233142 | 0.067964 | 0.75212 |
| ENSMUSG( | Top1mt | 395.7555 | 0.21606 | 0.078493 | 0.433639 | ENSMUSG( | Top1mt | 345.8406 | 0.097963 | 0.50404 | 0.927825 |
| ENSMUSG( | Top2a | 164.8852 | -0.10984 | 0.550688 | 0.886038 | ENSMUSG( | Top2a | 152.8394 | -0.05002 | 0.801793 | 0.97681 |
| ENSMUSG( | Top2b | 3366.9 | 0.102978 | 0.275567 | 0.725482 | ENSMUSG( | Top2b | 2998.913 | 0.038408 | 0.747815 | 0.96984 |
| ENSMUSG( | Top3a | 179.1504 | -0.05345 | 0.732105 | 0.944956 | ENSMUSG( | Top3a | 162.6891 | -0.04977 | 0.788307 | 0.97637 |
| ENSMUSG( | Top3b | 1169.703 | 0.095037 | 0.403936 | 0.816132 | ENSMUSG( | Top3b | 1037.667 | 0.021843 | 0.87399 | 0.989644 |
| ENSMUSG( | Top6bl | 63.76844 | -0.0406 | 0.888338 | 0.983434 | ENSMUSG( | Top6bl | 64.21178 | 0.236735 | 0.384926 | 0.917257 |
| ENSMUSG( | Topbp1 | 745.7111 | 0.068314 | 0.49973 | 0.863218 | ENSMUSG( | Topbp1 | 697.0285 | 0.137885 | 0.261885 | 0.884926 |
| ENSMUSG( | Topors | 421.4639 | 0.165178 | 0.16206 | 0.592429 | ENSMUSG( | Topors | 379.879 | 0.139795 | 0.325616 | 0.905339 |
| ENSMUSG( | Tor1a | 899.5916 | -0.23688 | 0.038345 | 0.302069 | ENSMUSG( | Tor1a | 914.5454 | 0.092808 | 0.51445 | 0.930939 |
| ENSMUSG( | Tor1aip1 | 3091.648 | 0.030524 | 0.782963 | 0.957558 | ENSMUSG( | Tor1aip1 | 2752.551 | -0.03594 | 0.768384 | 0.974236 |
| ENSMUSG( | Tor1aip2 | 1926.594 | -0.06277 | 0.483701 | 0.85636 | ENSMUSG( | Tor1aip2 | 1837.192 | 0.068982 | 0.530841 | 0.934795 |
| ENSMUSG( | Tor1b | 1104.976 | -0.14464 | 0.3408 | 0.778709 | ENSMUSG( | Tor1b | 1131.747 | 0.198004 | 0.26887 | 0.886755 |
| ENSMUSG( | Tor2a | 810.5673 | -0.21969 | 0.110761 | 0.501952 | ENSMUSG( | Tor2a | 851.4313 | 0.198024 | 0.185325 | 0.855305 |
| ENSMUSG( | Tor3a | 511.3775 | -0.30385 | 0.043132 | 0.318494 | ENSMUSG( | Tor3a | 492.2623 | -0.12733 | 0.429524 | 0.917257 |
| ENSMUSG( | Tor4a | 174.9684 | -0.15194 | 0.38609 | 0.804623 | ENSMUSG( | Tor4a | 169.2527 | 0.044895 | 0.860313 | 0.986137 |
| ENSMUSG( | Tox | 556.7491 | -0.23017 | 0.074938 | 0.42422 | ENSMUSG( | Tox | 492.3807 | -0.33281 | 0.03558 | 0.664751 |
| ENSMUSG( | Tox2 | 767.1108 | 0.10636 | 0.372276 | 0.798768 | ENSMUSG( | Tox2 | 615.8654 | -0.27767 | 0.156277 | 0.837724 |
| ENSMUSG( | Tox3 | 243.0021 | 0.135043 | 0.342525 | 0.778971 | ENSMUSG( | Tox3 | 227.0184 | 0.195595 | 0.245845 | 0.877904 |
| ENSMUSG( | Tox4 | 1477.14 | 0.113178 | 0.201358 | 0.641362 | ENSMUSG( | Tox4 | 1274.146 | -0.04368 | 0.686627 | 0.95899 |
| ENSMUSG( | Tpbg | 647.0324 | -0.66272 | 2.77E-06 | 0.000408 | ENSMUSG( | Tpbg | 781.9286 | 0.221795 | 0.178742 | 0.849041 |
| ENSMUSG( | Tpbgl | 180.0843 | -0.44666 | 0.034124 | 0.283507 | ENSMUSG( | Tpbgl | 204.5283 | 0.223259 | 0.270315 | 0.887887 |
| ENSMUSG( | Tpcn1 | 4341.026 | -0.08253 | 0.45118 | 0.835567 | ENSMUSG( | Tpcn1 | 3973.563 | -0.0664 | 0.583506 | 0.944266 |
| ENSMUSG( | Tpcn2 | 144.3757 | 0.131189 | 0.454187 | 0.837055 | ENSMUSG( | Tpcn2 | 134.7053 | 0.20539 | 0.302254 | 0.898538 |
| ENSMUSG( | Tpd52 | 1914.456 | -0.07443 | 0.462355 | 0.842104 | ENSMUSG( | Tpd52 | 1750.63 | -0.06751 | 0.609598 | 0.950397 |
| ENSMUSG( | Tpd52l1 | 445.5048 | -0.06339 | 0.589636 | 0.90113 | ENSMUSG( | Tpd52l1 | 387.2727 | -0.2156 | 0.159541 | 0.837724 |
| ENSMUSG( | Tpd52l2 | 1596.193 | 0.028916 | 0.745093 | 0.949158 | ENSMUSG( | Tpd52l2 | 1384.733 | -0.11504 | 0.299748 | 0.898538 |
| ENSMUSG( | Tpgs1 | 1156.043 | -0.00083 | 0.993968 | 0.999369 | ENSMUSG( | Tpgs1 | 1013.539 | -0.11644 | 0.376028 | 0.915491 |
| ENSMUSG( | Tpgs2 | 1413.037 | 0.081662 | 0.423243 | 0.823662 | ENSMUSG( | Tpgs2 | 1210.063 | -0.09977 | 0.426129 | 0.917257 |
| ENSMUSG( | Tph2 | 124.6271 | -0.27567 | 0.213303 | 0.657626 | ENSMUSG( | Tph2 | 113.9823 | -0.27374 | 0.275042 | 0.889874 |
| ENSMUSG( | Tpi1 | 12359.55 | 0.0466 | 0.605097 | 0.905594 | ENSMUSG( | Tpi1 | 10267.79 | -0.23049 | 0.060229 | 0.731553 |
| ENSMUSG( | Tpk1 | 77.11349 | -0.21911 | 0.338001 | 0.777422 | ENSMUSG( | Tpk1 | 69.94544 | -0.2305 | 0.355426 | 0.912555 |
| ENSMUSG( | Tpm1 | 3860.947 | -0.02019 | 0.845432 | 0.97218 | ENSMUSG( | Tpm1 | 3448.341 | -0.0799 | 0.516928 | 0.930939 |
| ENSMUSG( | Tpm2 | 167.8259 | -0.2542 | 0.150663 | 0.575407 | ENSMUSG( | Tpm2 | 160.6383 | -0.09526 | 0.631768 | 0.953251 |
| ENSMUSG( | Tpm3 | 3667.807 | 0.040847 | 0.629977 | 0.913965 | ENSMUSG( | Tpm3 | 3172.042 | -0.11332 | 0.311941 | 0.900037 |
| ENSMUSG( | Tpm3-rs7 | 24.27266 | 0.423191 | 0.293097 | 0.742763 | ENSMUSG( | Tpm3-rs7 | 20.28262 | 0.19408 | 0.660539 | 0.956205 |
| ENSMUSG( | Tpm4 | 1881.608 | 0.136729 | 0.266509 | 0.716441 | ENSMUSG( | Tpm4 | 1778.815 | 0.238613 | 0.072463 | 0.763629 |
| ENSMUSG( | Tpmt | 1104.623 | 0.022025 | 0.874946 | 0.979746 | ENSMUSG( | Tpmt | 916.8388 | -0.26388 | 0.090855 | 0.791074 |

| | | | | | |
|---|---|---|---|---|---|
| ENSMUSG( | 492.1134 | -0.25769 | 0.061242 | 0.381822 | Tmem121 |
| ENSMUSG( | 2407.174 | -0.2544 | 0.115461 | 0.512397 | Tmem121b |
| ENSMUSG( | 692.978 | -0.29598 | 0.099803 | 0.482257 | Tmem123 |
| ENSMUSG( | 355.0511 | -0.43499 | 0.005131 | 0.095477 | Tmem125 |
| ENSMUSG( | 566.6482 | -0.03783 | 0.754117 | 0.950981 | Tmem126a |
| ENSMUSG( | 765.4239 | 0.012096 | 0.906178 | 0.986921 | Tmem126b |
| ENSMUSG( | 4974.213 | -0.14954 | 0.216345 | 0.66135 | Tmem127 |
| ENSMUSG( | 825.2005 | -0.35627 | 0.011183 | 0.152553 | Tmem128 |
| ENSMUSG( | 1103.295 | -0.22087 | 0.105093 | 0.492392 | Tmem129 |
| ENSMUSG( | 5743.355 | -0.29118 | 0.018689 | 0.205652 | Tmem130 |
| ENSMUSG( | 2614.233 | -0.18944 | 0.131736 | 0.544244 | Tmem131 |
| ENSMUSG( | 655.7891 | -0.09123 | 0.480121 | 0.854649 | Tmem131l |
| ENSMUSG( | 10285.1 | -0.11766 | 0.444598 | 0.833239 | Tmem132a |
| ENSMUSG( | 4492.984 | -0.35666 | 0.01893 | 0.207196 | Tmem132b |
| ENSMUSG( | 183.525 | -0.30837 | 0.110394 | 0.50108 | Tmem132c |
| ENSMUSG( | 1479.928 | -0.15996 | 0.31749 | 0.763823 | Tmem132d |
| ENSMUSG( | 419.6527 | -0.22554 | 0.205988 | 0.647757 | Tmem132e |
| ENSMUSG( | 1822.5 | 0.055991 | 0.549245 | 0.885739 | Tmem134 |
| ENSMUSG( | 1684.353 | -0.00877 | 0.930049 | 0.991848 | Tmem135 |
| ENSMUSG( | 339.7494 | -0.27551 | 0.029728 | 0.261211 | Tmem138 |
| ENSMUSG( | 88.1503 | 0.007855 | 0.97275 | 0.996585 | Tmem140 |
| ENSMUSG( | 947.7491 | 0.143613 | 0.197793 | 0.637687 | Tmem141 |
| ENSMUSG( | 655.0849 | -0.11607 | 0.262636 | 0.713259 | Tmem143 |
| ENSMUSG( | 605.5595 | -0.24305 | 0.10559 | 0.49243 | Tmem144 |
| ENSMUSG( | 1590.845 | 0.040539 | 0.7922 | 0.959461 | Tmem145 |
| ENSMUSG( | 2425.865 | -0.16294 | 0.140949 | 0.557902 | Tmem147 |
| ENSMUSG( | 920.0791 | -0.45805 | 6.52E-05 | 0.004839 | Tmem14a |
| ENSMUSG( | 1273.969 | -0.2299 | 0.033936 | 0.282701 | Tmem14c |
| ENSMUSG( | 415.2757 | 0.255003 | 0.110881 | 0.501952 | Tmem150a |
| ENSMUSG( | 226.205 | -0.08326 | 0.56188 | 0.889052 | Tmem150b |
| ENSMUSG( | 1781.692 | -0.20517 | 0.074722 | 0.423772 | Tmem150c |
| ENSMUSG( | 9244.496 | -0.29858 | 0.025967 | 0.247245 | Tmem151a |
| ENSMUSG( | 9230.251 | 0.319131 | 0.00741 | 0.118765 | Tmem151b |
| ENSMUSG( | 28.05267 | -0.76781 | 0.036597 | 0.295048 | Tmem154 |
| ENSMUSG( | 14.71598 | -0.10893 | 0.822051 | NA | Tmem156 |
| ENSMUSG( | 1966.229 | -0.7848 | 1.17E-11 | 8.23E-09 | Tmem158 |
| ENSMUSG( | 1144.726 | -0.13062 | 0.202615 | 0.642757 | Tmem160 |
| ENSMUSG( | 948.631 | -0.1548 | 0.223729 | 0.669845 | Tmem161a |
| ENSMUSG( | 536.1984 | -0.27685 | 0.042369 | 0.316636 | Tmem161b |
| ENSMUSG( | 1044.964 | -0.50248 | 4.48E-05 | 0.003633 | Tmem163 |
| ENSMUSG( | 2177.483 | 0.198245 | 0.027932 | 0.255775 | Tmem164 |
| ENSMUSG( | 1017.884 | -0.49584 | 0.000169 | 0.009375 | Tmem165 |
| ENSMUSG( | 1490.88 | -0.01838 | 0.867909 | 0.976631 | Tmem167 |
| ENSMUSG( | 2464.29 | -0.00061 | 0.995787 | 0.99943 | Tmem167b |
| ENSMUSG( | 470.9275 | -0.3469 | 0.022139 | 0.226843 | Tmem168 |
| ENSMUSG( | 521.5691 | -0.25505 | 0.027785 | 0.255192 | Tmem169 |
| ENSMUSG( | 201.6398 | -0.51901 | 0.000792 | 0.027736 | Tmem17 |
| ENSMUSG( | 166.3601 | 0.147751 | 0.378565 | 0.801688 | Tmem170 |
| ENSMUSG( | 2760.598 | -0.02887 | 0.770872 | 0.954726 | Tmem170b |
| ENSMUSG( | 1230.265 | -0.197 | 0.12574 | 0.53257 | Tmem175 |
| ENSMUSG( | 1099.241 | -0.10729 | 0.394636 | 0.810072 | Tmem176a |
| ENSMUSG( | 2405.284 | -0.18195 | 0.102905 | 0.488476 | Tmem176b |
| ENSMUSG( | 237.8317 | 0.142737 | 0.37854 | 0.801688 | Tmem177 |
| ENSMUSG( | 2063.595 | -0.16825 | 0.266522 | 0.716441 | Tmem178 |
| ENSMUSG( | 4235.191 | -0.18487 | 0.179809 | 0.61751 | Tmem178b |
| ENSMUSG( | 4328.229 | -0.22139 | 0.064025 | 0.391882 | Tmem179 |
| ENSMUSG( | 210.5198 | -0.23723 | 0.126079 | 0.533384 | Tmem179b |
| ENSMUSG( | 1358.092 | -0.09114 | 0.433437 | 0.828346 | Tmem18 |
| ENSMUSG( | 1996.559 | -0.33678 | 0.017917 | 0.200989 | Tmem181a |
| ENSMUSG( | 2291.454 | 0.096369 | 0.273076 | 0.723468 | Tmem183a |
| ENSMUSG( | 2708.203 | -0.36999 | 0.00625 | 0.108182 | Tmem184b |
| ENSMUSG( | 3291.902 | -0.09784 | 0.419295 | 0.821987 | Tmem184c |
| ENSMUSG( | 1453.06 | -0.31938 | 0.014419 | 0.176902 | Tmem185a |
| ENSMUSG( | 397.8021 | -0.51773 | 0.002422 | 0.057903 | Tmem185b |
| ENSMUSG( | 562.332 | -0.00741 | 0.946025 | 0.99369 | Tmem186 |
| ENSMUSG( | 1537.52 | -0.28827 | 0.017549 | 0.198498 | Tmem19 |
| ENSMUSG( | 2748.906 | 0.184087 | 0.171729 | 0.606517 | Tmem191 |
| ENSMUSG( | 565.2655 | -0.21084 | 0.065526 | 0.396581 | Tmem192 |
| ENSMUSG( | 247.443 | -0.57059 | 0.000551 | 0.021914 | Tmem196 |
| ENSMUSG( | 2657.502 | -0.12183 | 0.356179 | 0.788108 | Tmem198 |
| ENSMUSG( | 591.6915 | -0.4437 | 0.005699 | 0.101138 | Tmem198b |
| ENSMUSG( | 508.8638 | -0.13989 | 0.267593 | 0.717651 | Tmem199 |
| ENSMUSG( | 513.2399 | -0.17429 | 0.355317 | 0.78783 | Tmem200a |
| ENSMUSG( | 175.5603 | -0.45615 | 0.00892 | 0.132538 | Tmem200b |
| ENSMUSG( | 449.0829 | -0.33703 | 0.064528 | 0.393651 | Tmem200c |
| ENSMUSG( | 2883.803 | -0.16668 | 0.184478 | 0.62157 | Tmem201 |
| ENSMUSG( | 59.47617 | -0.12128 | 0.625101 | 0.912249 | Tmem202 |
| ENSMUSG( | 261.892 | -0.3947 | 0.004659 | 0.089972 | Tmem203 |
| ENSMUSG( | 422.5487 | 0.135904 | 0.460089 | 0.840577 | Tmem204 |
| ENSMUSG( | 923.4797 | -0.33103 | 0.004086 | 0.082441 | Tmem205 |
| ENSMUSG( | 814.8894 | -0.16348 | 0.151608 | 0.576979 | Tmem208 |
| ENSMUSG( | 912.4368 | -0.34782 | 0.024127 | 0.237752 | Tmem209 |
| ENSMUSG( | 81.42574 | -0.53319 | 0.053817 | 0.356147 | Tmem212 |
| ENSMUSG( | 2066.786 | 0.064736 | 0.582164 | 0.897918 | Tmem214 |
| ENSMUSG( | 402.6313 | -0.20992 | 0.228611 | 0.675914 | Tmem215 |
| ENSMUSG( | 154.4598 | 0.115269 | 0.510074 | 0.869785 | Tmem216 |
| ENSMUSG( | 1683.168 | -0.01077 | 0.913848 | 0.988483 | Tmem217rt |
| ENSMUSG( | 727.2737 | -0.1097 | 0.412315 | 0.819452 | Tmem218 |
| ENSMUSG( | 485.0971 | -0.05443 | 0.630524 | 0.914073 | Tmem219 |
| ENSMUSG( | 46.36903 | -0.09091 | 0.755456 | 0.951395 | Tmem220 |
| ENSMUSG( | 8.954315 | -0.82638 | 0.200934 | NA | Tmem221 |
| ENSMUSG( | 2705.002 | -0.32201 | 0.004986 | 0.093605 | Tmem222 |
| ENSMUSG( | 1217.403 | 0.059974 | 0.542776 | 0.883563 | Tmem223 |
| ENSMUSG( | 6780.701 | 0.036271 | 0.797457 | 0.960507 | Tmem229a |
| ENSMUSG( | 1225.694 | -0.19008 | 0.065234 | 0.395697 | Tmem229b |
| ENSMUSG( | 488.4059 | -0.24203 | 0.105477 | 0.492417 | Tmem230 |
| ENSMUSG( | 352.5697 | -0.27197 | 0.088007 | 0.455043 | Tmem231 |
| ENSMUSG( | 52.89864 | -0.24683 | 0.44268 | 0.83278 | Tmem232 |
| ENSMUSG( | 3147.516 | 0.137667 | 0.148471 | 0.571935 | Tmem234 |
| ENSMUSG( | 597.6629 | -0.22262 | 0.136653 | 0.550959 | Tmem237 |
| ENSMUSG( | 153.0399 | -0.29761 | 0.100529 | 0.483976 | Tmem238 |
| ENSMUSG( | 1576.555 | -0.18692 | 0.069992 | 0.40908 | Tmem240 |
| ENSMUSG( | 305.7036 | -0.16922 | 0.241892 | 0.689179 | Tmem241 |
| ENSMUSG( | 2395.534 | 0.044625 | 0.636923 | 0.915191 | Tmem242 |
| ENSMUSG( | 618.4716 | 0.00721 | 0.950414 | 0.993821 | Tmem243 |
| ENSMUSG( | 2275.947 | -0.22812 | 0.117252 | 0.516802 | Tmem245 |
| ENSMUSG( | 1499.928 | -0.21826 | 0.099001 | 0.480953 | Tmem248 |
| ENSMUSG( | 1710.73 | -0.2238 | 0.077288 | 0.429743 | Tmem25 |
| ENSMUSG( | 3312.318 | 0.041586 | 0.604898 | 0.905556 | Tmem250 |
| ENSMUSG( | 102.5279 | -0.76407 | 0.007058 | 0.115775 | Tmem252 |
| ENSMUSG( | 15.3743 | -0.40027 | 0.446377 | NA | Tmem253 |
| ENSMUSG( | 2667.668 | -0.34445 | 0.001321 | 0.039614 | Tmem254 |
| ENSMUSG( | 505.339 | -0.34881 | 0.013612 | 0.170359 | Tmem255a |
| ENSMUSG( | 100.9814 | -0.398 | 0.170166 | 0.605149 | Tmem255b |
| ENSMUSG( | 1399.368 | 0.067849 | 0.524202 | 0.87583 | Tmem256 |
| ENSMUSG( | 785.8924 | -0.16451 | 0.121027 | 0.525487 | Tmem258 |
| ENSMUSG( | 5500.007 | -0.17193 | 0.078172 | 0.432307 | Tmem259 |
| ENSMUSG( | 853.6765 | -0.26835 | 0.025501 | 0.246501 | Tmem260 |
| ENSMUSG( | 585.137 | 0.011099 | 0.93152 | 0.992398 | Tmem263 |
| ENSMUSG( | 342.4012 | -0.06292 | 0.650652 | 0.921301 | Tmem266 |
| ENSMUSG( | 349.6496 | -0.73691 | 1.34E-06 | 0.000242 | Tmem267 |
| ENSMUSG( | 529.8919 | -0.13895 | 0.251894 | 0.701908 | Tmem268 |
| ENSMUSG( | 28.46085 | -0.03971 | 0.916475 | 0.988844 | Tmem269 |
| ENSMUSG( | 36.64223 | -0.01718 | 0.960423 | 0.993991 | Tmem273 |
| ENSMUSG( | 792.5632 | -0.26719 | 0.012653 | 0.162184 | Tmem276 |
| ENSMUSG( | 7753.293 | -0.36823 | 0.020278 | 0.214748 | Tmem30a |
| ENSMUSG( | 10.83522 | -0.7022 | 0.218724 | NA | Tmem30b |
| ENSMUSG( | 11.06606 | 1.122725 | 0.051874 | NA | Tmem30c |
| ENSMUSG( | 4046.054 | -0.2017 | 0.140312 | 0.557263 | Tmem33 |
| ENSMUSG( | 2057.866 | -0.4381 | 0.000485 | 0.020239 | Tmem35a |
| ENSMUSG( | 95.8924 | 0.027066 | 0.893736 | 0.984512 | Tmem35b |
| ENSMUSG( | 53.61466 | 0.102228 | 0.714232 | 0.941016 | Tmem37 |
| ENSMUSG( | 4691.985 | -0.22139 | 0.074864 | 0.424133 | Tmem38a |
| ENSMUSG( | 590.0769 | 0.001011 | 0.994297 | 0.999369 | Tmem38b |
| ENSMUSG( | 251.6376 | -0.19675 | 0.246902 | 0.695632 | Tmem39a |
| ENSMUSG( | 261.4599 | -0.50714 | 0.007197 | 0.117084 | Tmem39b |
| ENSMUSG( | 85.13151 | -0.04154 | 0.854093 | 0.973491 | Tmem40 |
| ENSMUSG( | 307.2106 | -0.26472 | 0.087927 | 0.45492 | Tmem41a |
| ENSMUSG( | 867.0977 | -0.46017 | 0.001384 | 0.040741 | Tmem41b |
| ENSMUSG( | 375.487 | -0.25964 | 0.086507 | 0.452045 | Tmem42 |
| ENSMUSG( | 1149.037 | -0.40417 | 0.00734 | 0.118347 | Tmem43 |
| ENSMUSG( | 6098.707 | 0.085008 | 0.353964 | 0.786371 | Tmem44 |
| ENSMUSG( | 35.44445 | -0.1866 | 0.612008 | 0.908586 | Tmem45a |
| ENSMUSG( | 7751.38 | -0.10154 | 0.531759 | 0.879628 | Tmem47 |
| ENSMUSG( | 3647.533 | -0.30821 | 0.018789 | 0.206056 | Tmem50a |
| ENSMUSG( | 3675.574 | -0.25531 | 0.043522 | 0.31954 | Tmem50b |
| ENSMUSG( | 551.4286 | -0.21715 | 0.078588 | 0.433892 | Tmem51 |
| ENSMUSG( | 47.25145 | 0.770582 | 0.030253 | 0.264384 | Tmem52 |
| ENSMUSG( | 75.80147 | -0.51373 | 0.042482 | 0.316636 | Tmem53 |
| ENSMUSG( | 74.79574 | 0.03878 | 0.870104 | 0.977767 | Tmem54 |
| ENSMUSG( | 7534.545 | -0.27749 | 0.025563 | 0.246694 | Tmem59 |
| ENSMUSG( | 10124.67 | -0.37287 | 0.000862 | 0.029481 | Tmem59l |
| ENSMUSG( | 486.5879 | -0.43497 | 0.002993 | 0.067057 | Tmem60 |
| ENSMUSG( | 1081.18 | -0.20618 | 0.115804 | 0.513769 | Tmem62 |
| ENSMUSG( | 1358.471 | -0.79176 | 7.84E-06 | 0.00097 | Tmem63a |
| ENSMUSG( | 5588.283 | -0.16389 | 0.165631 | 0.59714 | Tmem63b |
| ENSMUSG( | 1145.756 | 0.100633 | 0.480484 | 0.854748 | Tmem63c |
| ENSMUSG( | 1253.836 | -0.0888 | 0.497777 | 0.862341 | Tmem64 |

| | | | | | |
|---|---|---|---|---|---|
| ENSMUSG( | 4783.73 | 0.112775 | 0.432374 | 0.918158 | Tmem38a |
| ENSMUSG( | 550.0109 | 0.066069 | 0.662502 | 0.956205 | Tmem38b |
| ENSMUSG( | 276.0518 | 0.33962 | 0.09402 | 0.795735 | Tmem39a |
| ENSMUSG( | 278.9352 | 0.001352 | 0.994464 | 0.998813 | Tmem39b |
| ENSMUSG( | 79.68897 | 0.019309 | 0.936058 | 0.993573 | Tmem40 |
| ENSMUSG( | 340.347 | 0.304135 | 0.09075 | 0.791074 | Tmem41a |
| ENSMUSG( | 974.6737 | 0.181154 | 0.255223 | 0.882761 | Tmem41b |
| ENSMUSG( | 371.1208 | -0.01063 | 0.946427 | 0.994798 | Tmem42 |
| ENSMUSG( | 1272.596 | 0.187535 | 0.28072 | 0.892356 | Tmem43 |
| ENSMUSG( | 5277.514 | -0.06175 | 0.607943 | 0.950113 | Tmem44 |
| ENSMUSG( | 28.73202 | -0.57539 | 0.167661 | 0.840222 | Tmem45a |
| ENSMUSG( | 7248.364 | -0.02659 | 0.892767 | 0.990558 | Tmem47 |
| ENSMUSG( | 3808.138 | 0.102103 | 0.478645 | 0.923329 | Tmem50a |
| ENSMUSG( | 3706.865 | 0.05039 | 0.725716 | 0.966887 | Tmem50b |
| ENSMUSG( | 473.236 | -0.40833 | 0.005539 | 0.303106 | Tmem51 |
| ENSMUSG( | 48.66483 | 1.060357 | 0.001029 | 0.110223 | Tmem52 |
| ENSMUSG( | 75.58049 | -0.21743 | 0.40998 | 0.917257 | Tmem53 |
| ENSMUSG( | 68.18504 | 0.02847 | 0.914992 | 0.992407 | Tmem54 |
| ENSMUSG( | 7689.431 | 0.064735 | 0.641903 | 0.953903 | Tmem59 |
| ENSMUSG( | 10516.95 | 0.031817 | 0.809323 | 0.978779 | Tmem59l |
| ENSMUSG( | 506.9261 | -0.01417 | 0.928834 | 0.99306 | Tmem60 |
| ENSMUSG( | 1073.665 | 0.054069 | 0.716799 | 0.965859 | Tmem62 |
| ENSMUSG( | 1576.884 | 0.022267 | 0.918881 | 0.992529 | Tmem63a |
| ENSMUSG( | 5723.825 | 0.176243 | 0.201183 | 0.86439 | Tmem63b |
| ENSMUSG( | 1058.554 | 0.145775 | 0.375339 | 0.915491 | Tmem63c |
| ENSMUSG( | 1243.311 | 0.156902 | 0.3045 | 0.898538 | Tmem64 |
| ENSMUSG( | 2426.792 | -0.01888 | 0.897927 | 0.990558 | Tmem65 |
| ENSMUSG( | 390.9489 | 0.335072 | 0.025635 | 0.599234 | Tmem67 |
| ENSMUSG( | 914.7587 | 0.217693 | 0.214251 | 0.864421 | Tmem68 |
| ENSMUSG( | 189.9088 | -0.01246 | 0.942219 | 0.994774 | Tmem69 |
| ENSMUSG( | 789.2455 | -0.06442 | 0.650231 | 0.95422 | Tmem70 |
| ENSMUSG( | 22.6992 | 0.247257 | 0.557809 | 0.939565 | Tmem71 |
| ENSMUSG( | 61.43173 | 0.673426 | 0.385964 | 0.917257 | Tmem72 |
| ENSMUSG( | 162.9681 | 0.361299 | 0.124345 | 0.814691 | Tmem74 |
| ENSMUSG( | 105.6016 | -0.3357 | 0.139619 | 0.830678 | Tmem74b |
| ENSMUSG( | 21.89053 | 1.363797 | 0.004344 | 0.268942 | Tmem74bos |
| ENSMUSG( | 29.30037 | -0.06279 | 0.868034 | 0.98781 | Tmem79 |
| ENSMUSG( | 900.059 | -0.05726 | 0.646415 | 0.953988 | Tmem80 |
| ENSMUSG( | 171.39 | -0.09084 | 0.607673 | 0.950113 | Tmem81 |
| ENSMUSG( | 37.11854 | -0.38262 | 0.246197 | 0.877904 | Tmem82 |
| ENSMUSG( | 818.1291 | 0.229329 | 0.173026 | 0.843788 | Tmem86a |
| ENSMUSG( | 209.3995 | 0.192475 | 0.313304 | 0.900878 | Tmem86b |
| ENSMUSG( | 1206.826 | 0.131456 | 0.449796 | 0.922202 | Tmem87a |
| ENSMUSG( | 1265.441 | 0.106493 | 0.518131 | 0.930939 | Tmem87b |
| ENSMUSG( | 141.5598 | 0.615786 | 0.00661 | 0.338858 | Tmem88 |
| ENSMUSG( | 3057.262 | -0.16724 | 0.219716 | 0.866104 | Tmem88b |
| ENSMUSG( | 4219.928 | 0.144182 | 0.30383 | 0.898538 | Tmem8b |
| ENSMUSG( | 2297.56 | 0.064187 | 0.644051 | 0.953903 | Tmem9 |
| ENSMUSG( | 401.193 | -0.04724 | 0.786163 | 0.976285 | Tmem91 |
| ENSMUSG( | 2890.602 | 0.221966 | 0.091905 | 0.791889 | Tmem94 |
| ENSMUSG( | 748.0838 | 0.04351 | 0.800544 | 0.97681 | Tmem97 |
| ENSMUSG( | 639.4666 | -0.17743 | 0.318573 | 0.903164 | Tmem98 |
| ENSMUSG( | 4493.898 | 0.011345 | 0.937546 | 0.993685 | Tmem9b |
| ENSMUSG( | 1148.567 | 0.063841 | 0.567749 | 0.941777 | Tmf1 |
| ENSMUSG( | 329.9965 | -0.2078 | 0.217496 | 0.865589 | Tmie |
| ENSMUSG( | 966.7987 | -0.17156 | 0.180226 | 0.850372 | Tmod1 |
| ENSMUSG( | 16797.41 | -0.00637 | 0.956044 | 0.994798 | Tmod2 |
| ENSMUSG( | 973.5961 | 0.143627 | 0.212185 | 0.864412 | Tmod3 |
| ENSMUSG( | 12.80404 | 0.114567 | 0.829046 | 0.980224 | Tmod4 |
| ENSMUSG( | 675.3683 | 0.324005 | 0.008352 | 0.388416 | Tmpo |
| ENSMUSG( | 89.87375 | 0.495704 | 0.086589 | 0.782073 | Tmppe |
| ENSMUSG( | 39.42274 | -0.50428 | 0.126007 | 0.81516 | Tmprss5 |
| ENSMUSG( | 17.51833 | -0.46052 | 0.324905 | 0.905257 | Tmprss6 |
| ENSMUSG( | 108.6513 | -0.15609 | 0.459136 | 0.923329 | Tmprss7 |
| ENSMUSG( | 18.92934 | 0.211219 | 0.639023 | 0.953903 | Tmprss9 |
| ENSMUSG( | 5835.529 | -0.22229 | 0.116213 | 0.813384 | Tmsb10 |
| ENSMUSG( | 48.73443 | -0.44298 | 0.15634 | 0.837724 | Tmsb10b |
| ENSMUSG( | 12.35477 | -0.82334 | 0.163296 | 0.838291 | Tmsb15b1 |
| ENSMUSG( | 15825.12 | -0.11526 | 0.346455 | 0.910966 | Tmsb4x |
| ENSMUSG( | 1121.166 | 0.391334 | 0.005603 | 0.303106 | Tmt1a |
| ENSMUSG( | 2690.962 | 0.048432 | 0.709692 | 0.96416 | Tmtc1 |
| ENSMUSG( | 512.2145 | 0.215475 | 0.291349 | 0.897566 | Tmtc2 |
| ENSMUSG( | 397.6631 | 0.470983 | 0.022532 | 0.581149 | Tmtc3 |
| ENSMUSG( | 349.4996 | 0.301651 | 0.097244 | 0.796721 | Tmtc4 |
| ENSMUSG( | 702.0572 | 0.101489 | 0.46852 | 0.923329 | Tmub1 |
| ENSMUSG( | 2945.174 | 0.081626 | 0.543967 | 0.938063 | Tmub2 |
| ENSMUSG( | 1135.733 | 0.389642 | 0.026522 | 0.603041 | Tmx1 |
| ENSMUSG( | 5588.221 | -0.01726 | 0.905487 | 0.991089 | Tmx2 |
| ENSMUSG( | 1855.991 | 0.275703 | 0.132017 | 0.822912 | Tmx3 |
| ENSMUSG( | 6111.46 | 0.221978 | 0.209739 | 0.864412 | Tmx4 |
| ENSMUSG( | 539.6908 | 0.027907 | 0.879219 | 0.989644 | Tnc |
| ENSMUSG( | 2148.893 | -0.02091 | 0.851118 | 0.985541 | Tnfaip1 |
| ENSMUSG( | 36.71438 | -0.63673 | 0.063773 | 0.736482 | Tnfaip2 |
| ENSMUSG( | 118.3591 | 0.587472 | 0.006872 | 0.343883 | Tnfaip3 |
| ENSMUSG( | 277.8405 | 0.216756 | 0.315357 | 0.901306 | Tnfaip6 |
| ENSMUSG( | 507.8516 | -0.25284 | 0.161616 | 0.837724 | Tnfaip8 |
| ENSMUSG( | 137.8226 | -0.04236 | 0.834573 | 0.981237 | Tnfaip8l1 |
| ENSMUSG( | 88.6388 | 0.31778 | 0.228346 | 0.868507 | Tnfaip8l2 |
| ENSMUSG( | 132.1507 | 0.07815 | 0.768103 | 0.974236 | Tnfaip8l3 |
| ENSMUSG( | 30.85393 | 0.5564 | 0.133846 | 0.822912 | Tnfrsf10b |
| ENSMUSG( | 206.663 | 0.058042 | 0.770848 | 0.974236 | Tnfrsf11a |
| ENSMUSG( | 110.7871 | 0.083168 | 0.779356 | 0.976106 | Tnfrsf11b |
| ENSMUSG( | 138.1658 | -0.33809 | 0.101433 | 0.801456 | Tnfrsf12a |
| ENSMUSG( | 95.43402 | -0.23923 | 0.314547 | 0.900949 | Tnfrsf13b |
| ENSMUSG( | 16.27969 | 0.1358 | 0.796508 | 0.97681 | Tnfrsf13c |
| ENSMUSG( | 22.57245 | 0.612881 | 0.160751 | 0.837724 | Tnfrsf14 |
| ENSMUSG( | 12.89798 | 0.401472 | 0.451999 | 0.922316 | Tnfrsf17 |
| ENSMUSG( | 7096.04 | -0.20857 | 0.114345 | 0.811703 | Tnfrsf19 |
| ENSMUSG( | 763.2966 | 0.267893 | 0.164726 | 0.838743 | Tnfrsf1a |
| ENSMUSG( | 306.7505 | 0.090242 | 0.726176 | 0.966887 | Tnfrsf1b |
| ENSMUSG( | 2695.234 | 0.290867 | 0.106317 | 0.804969 | Tnfrsf21 |
| ENSMUSG( | 74.0516 | -0.1431 | 0.601167 | 0.947749 | Tnfrsf22 |
| ENSMUSG( | 49.86155 | -0.23491 | 0.426667 | 0.917257 | Tnfrsf23 |
| ENSMUSG( | 185.5714 | 0.125089 | 0.670278 | 0.95694 | Tnfrsf25 |
| ENSMUSG( | 32.97697 | -0.35965 | 0.306278 | 0.898538 | Tnfrsf4 |
| ENSMUSG( | 30.96188 | 0.108958 | 0.766595 | 0.974236 | Tnfrsf8 |
| ENSMUSG( | 102.8402 | 0.973492 | 0.000514 | 0.068166 | Tnfsf10 |
| ENSMUSG( | 495.9929 | -0.03717 | 0.814419 | 0.979198 | Tnfsf12 |
| ENSMUSG( | 42.30647 | -0.00331 | 0.992129 | 0.99874 | Tnfsf13 |
| ENSMUSG( | 40.86297 | 0.052352 | 0.871618 | 0.988779 | Tnfsf13b |
| ENSMUSG( | 16.53422 | -0.01576 | 0.973746 | 0.996416 | Tnfsf9 |
| ENSMUSG( | 25.39523 | -0.04648 | 0.906592 | 0.991494 | Tnfsfm13 |
| ENSMUSG( | 4543.407 | 0.133321 | 0.179908 | 0.850372 | Tnik |
| ENSMUSG( | 618.9676 | -0.11445 | 0.404167 | 0.917257 | Tnip1 |
| ENSMUSG( | 349.6929 | 0.124261 | 0.505115 | 0.927825 | Tnip2 |
| ENSMUSG( | 8781.976 | 0.165135 | 0.117523 | 0.813384 | Tnk2 |
| ENSMUSG( | 3170.139 | 0.033975 | 0.782017 | 0.976106 | Tnks |
| ENSMUSG( | 2490.301 | 0.109329 | 0.308402 | 0.898538 | Tnks1bp1 |
| ENSMUSG( | 4504.386 | 0.04161 | 0.720686 | 0.966887 | Tnks2 |
| ENSMUSG( | 25.17003 | -0.11009 | 0.782869 | 0.976106 | Tnmd |
| ENSMUSG( | 201.9325 | -0.42082 | 0.026785 | 0.603041 | Tnnc1 |
| ENSMUSG( | 214.83 | -0.39537 | 0.038758 | 0.679972 | Tnni1 |
| ENSMUSG( | 12.046 | -0.01171 | 0.984344 | 0.996963 | Tnni2 |
| ENSMUSG( | 22.37767 | -0.54491 | 0.240361 | 0.875526 | Tnni3 |
| ENSMUSG( | 154.287 | -0.57797 | 0.005122 | 0.294733 | Tnnt1 |
| ENSMUSG( | 75.02384 | -0.14025 | 0.630649 | 0.953251 | Tnnt2 |
| ENSMUSG( | 9.65547 | -0.19048 | 0.777944 | NA | Tnnt3 |
| ENSMUSG( | 2172.217 | 0.155214 | 0.252157 | 0.879902 | Tnpo1 |
| ENSMUSG( | 4903.754 | -0.05578 | 0.561356 | 0.940254 | Tnpo2 |
| ENSMUSG( | 1928.145 | 0.010602 | 0.927089 | 0.99306 | Tnpo3 |
| ENSMUSG( | 6408.371 | 0.196806 | 0.331903 | 0.907907 | Tnr |
| ENSMUSG( | 4115.293 | 0.064509 | 0.563118 | 0.941158 | Tnrc18 |
| ENSMUSG( | 2504.144 | 0.027089 | 0.80349 | 0.976978 | Tnrc6a |
| ENSMUSG( | 1985.324 | 0.036036 | 0.737561 | 0.967686 | Tnrc6b |
| ENSMUSG( | 2372.149 | -0.0187 | 0.855473 | 0.985738 | Tnrc6c |
| ENSMUSG( | 2440.12 | 0.146718 | 0.271563 | 0.888062 | Tns1 |
| ENSMUSG( | 744.1401 | 0.54946 | 0.001467 | 0.140378 | Tns2 |
| ENSMUSG( | 9765.288 | -0.06371 | 0.627863 | 0.953234 | Tns3 |
| ENSMUSG( | 11.91243 | 0.024894 | 0.965491 | 0.995433 | Tns4 |
| ENSMUSG( | 282.2118 | 0.774232 | 0.001219 | 0.126346 | Tnxb |
| ENSMUSG( | 843.6551 | 0.374192 | 0.043919 | 0.690976 | Tob1 |
| ENSMUSG( | 883.9997 | -0.02388 | 0.906664 | 0.991511 | Tob2 |
| ENSMUSG( | 307.169 | -0.01333 | 0.935879 | 0.993443 | Toe1 |
| ENSMUSG( | 777.7275 | 0.224025 | 0.059024 | 0.729429 | Togaram1 |
| ENSMUSG( | 77.82195 | 0.209708 | 0.449707 | 0.922202 | Togaram2 |
| ENSMUSG( | 4324.212 | -0.11448 | 0.288152 | 0.894794 | Tollip |
| ENSMUSG( | 685.8043 | -0.19553 | 0.102392 | 0.802599 | Tom1 |
| ENSMUSG( | 1126.147 | -0.04041 | 0.762203 | 0.973255 | Tom1l1 |
| ENSMUSG( | 9503.253 | -0.10962 | 0.256519 | 0.883082 | Tom1l2 |
| ENSMUSG( | 8429.605 | 0.01107 | 0.925129 | 0.99306 | Tomm20 |
| ENSMUSG( | 1501.81 | -0.15558 | 0.209256 | 0.864412 | Tomm22 |
| ENSMUSG( | 1368.039 | -0.12086 | 0.318388 | 0.903045 | Tomm34 |
| ENSMUSG( | 1790.209 | -0.06545 | 0.605865 | 0.949449 | Tomm40 |
| ENSMUSG( | 2882.579 | -0.22232 | 0.051855 | 0.712864 | Tomm40l |
| ENSMUSG( | 550.0204 | -0.22249 | 0.119455 | 0.813384 | Tomm5 |
| ENSMUSG( | 136.586 | -0.06904 | 0.755113 | 0.971098 | Tomm6os |
| ENSMUSG( | 1792.163 | -0.19198 | 0.133076 | 0.822912 | Tomm7 |
| ENSMUSG( | 2811.843 | -0.0435 | 0.741207 | 0.968811 | Tomm70a |

| | | | | | | | | | | | |
|---|---|---|---|---|---|---|---|---|---|---|---|
| ENSMUSG( | Tpo | 10.57239 | -0.26119 | 0.677965 | NA | ENSMUSG( | Tpo | 12.07303 | 0.313904 | 0.633698 | 0.953889 |
| ENSMUSG( | Tpp1 | 3737.221 | -0.15192 | 0.212027 | 0.655687 | ENSMUSG( | Tpp1 | 3765.095 | 0.140743 | 0.312363 | 0.900037 |
| ENSMUSG( | Tpp2 | 964.01 | 0.014187 | 0.883839 | 0.982647 | ENSMUSG( | Tpp2 | 895.8064 | 0.068129 | 0.566939 | 0.941777 |
| ENSMUSG( | Tppp | 20652.85 | 0.119716 | 0.113442 | 0.508013 | ENSMUSG( | Tppp | 17292.85 | -0.12445 | 0.197522 | 0.862161 |
| ENSMUSG( | Tppp3 | 967.2098 | -0.00257 | 0.980542 | 0.997512 | ENSMUSG( | Tppp3 | 864.0677 | -0.06311 | 0.619939 | 0.952896 |
| ENSMUSG( | Tpr | 3062.372 | 0.198831 | 0.037603 | 0.299568 | ENSMUSG( | Tpr | 2615.867 | 0.01852 | 0.861149 | 0.986226 |
| ENSMUSG( | Tpra1 | 931.5655 | -0.12687 | 0.34461 | 0.779153 | ENSMUSG( | Tpra1 | 981.1563 | 0.285536 | 0.04644 | 0.697814 |
| ENSMUSG( | Tprg1l | 6924.672 | 0.087175 | 0.283381 | 0.73186 | ENSMUSG( | Tprg1l | 5838.129 | -0.14011 | 0.219468 | 0.866104 |
| ENSMUSG( | Tprkb | 5540.846 | 0.082759 | 0.52677 | 0.877267 | ENSMUSG( | Tprkb | 4232.516 | -0.46294 | 0.000443 | 0.061321 |
| ENSMUSG( | Tprn | 631.7621 | -0.4067 | 0.000266 | 0.013312 | ENSMUSG( | Tprn | 635.0463 | -0.08953 | 0.570819 | 0.941973 |
| ENSMUSG( | Tpst1 | 1202.598 | -0.22332 | 0.083872 | 0.446475 | ENSMUSG( | Tpst1 | 1257.945 | 0.183285 | 0.247188 | 0.877904 |
| ENSMUSG( | Tpst2 | 1097.706 | -0.16875 | 0.157464 | 0.585653 | ENSMUSG( | Tpst2 | 1131.886 | 0.191965 | 0.198438 | 0.862367 |
| ENSMUSG( | Tpt1 | 25881.24 | 0.12335 | 0.196186 | 0.635538 | ENSMUSG( | Tpt1 | 22123.73 | -0.06055 | 0.643068 | 0.953903 |
| ENSMUSG( | Tpx2 | 96.86454 | 0.129096 | 0.575497 | 0.894836 | ENSMUSG( | Tpx2 | 88.89313 | 0.154756 | 0.502839 | 0.927825 |
| ENSMUSG( | Tra2a | 1912.281 | 0.155108 | 0.183106 | 0.62047 | ENSMUSG( | Tra2a | 1741.342 | 0.150599 | 0.329087 | 0.907083 |
| ENSMUSG( | Tra2b | 1261.108 | -0.04301 | 0.66118 | 0.92507 | ENSMUSG( | Tra2b | 1185.734 | 0.045428 | 0.703658 | 0.962481 |
| ENSMUSG( | Trabd | 686.0255 | -0.01686 | 0.883902 | 0.982647 | ENSMUSG( | Trabd | 630.1235 | 0.010377 | 0.934073 | 0.99306 |
| ENSMUSG( | Trabd2b | 358.6167 | 0.566908 | 1.77E-05 | 0.001786 | ENSMUSG( | Trabd2b | 305.1603 | 0.403158 | 0.006725 | 0.339232 |
| ENSMUSG( | Tradd | 167.641 | 0.059889 | 0.729393 | 0.944738 | ENSMUSG( | Tradd | 153.5104 | 0.084583 | 0.684574 | 0.958678 |
| ENSMUSG( | Traf1 | 63.09834 | 0.04059 | 0.864135 | 0.975331 | ENSMUSG( | Traf1 | 48.52215 | -0.48952 | 0.092065 | 0.791889 |
| ENSMUSG( | Traf2 | 267.8157 | 0.027427 | 0.836561 | 0.971111 | ENSMUSG( | Traf2 | 244.0524 | 0.034617 | 0.832013 | 0.980738 |
| ENSMUSG( | Traf3 | 7114.629 | 0.364174 | 0.000108 | 0.006836 | ENSMUSG( | Traf3 | 6032.935 | 0.174854 | 0.119071 | 0.813384 |
| ENSMUSG( | Traf3ip1 | 496.0954 | 0.124141 | 0.311029 | 0.759201 | ENSMUSG( | Traf3ip1 | 435.3334 | 0.015253 | 0.917594 | 0.992529 |
| ENSMUSG( | Traf3ip2 | 34.97898 | -0.18383 | 0.554448 | 0.886657 | ENSMUSG( | Traf3ip2 | 30.24744 | -0.31246 | 0.405995 | 0.917257 |
| ENSMUSG( | Traf3ip3 | 49.2679 | -0.04336 | 0.885892 | 0.983008 | ENSMUSG( | Traf3ip3 | 55.32395 | 0.533006 | 0.116714 | 0.813384 |
| ENSMUSG( | Traf4 | 570.9473 | -0.20838 | 0.090501 | 0.461757 | ENSMUSG( | Traf4 | 547.7802 | -0.04971 | 0.717691 | 0.965964 |
| ENSMUSG( | Traf5 | 86.97187 | -0.19067 | 0.387154 | 0.805442 | ENSMUSG( | Traf5 | 88.67286 | 0.162778 | 0.527396 | 0.934466 |
| ENSMUSG( | Traf6 | 643.8057 | -0.10615 | 0.331883 | 0.775288 | ENSMUSG( | Traf6 | 600.5553 | -0.03754 | 0.786042 | 0.976285 |
| ENSMUSG( | Traf7 | 1852.775 | 0.127693 | 0.223915 | 0.669845 | ENSMUSG( | Traf7 | 1624.136 | 0.018142 | 0.900732 | 0.990558 |
| ENSMUSG( | Trafd1 | 1721.501 | 0.057466 | 0.617207 | 0.910866 | ENSMUSG( | Trafd1 | 1587.75 | 0.0918 | 0.471849 | 0.923329 |
| ENSMUSG( | Traip | 633.3302 | -0.01076 | 0.952558 | 0.993874 | ENSMUSG( | Traip | 586.0047 | 0.026335 | 0.903205 | 0.991089 |
| ENSMUSG( | Trak1 | 10926.56 | 0.192742 | 0.016684 | 0.192198 | ENSMUSG( | Trak1 | 9753.324 | 0.137049 | 0.160876 | 0.837724 |
| ENSMUSG( | Trak2 | 36139.13 | 0.007623 | 0.948799 | 0.993821 | ENSMUSG( | Trak2 | 29854.75 | -0.29217 | 0.02939 | 0.618981 |
| ENSMUSG( | Tram1 | 1087.909 | -0.23654 | 0.045865 | 0.328083 | ENSMUSG( | Tram1 | 1152.025 | 0.201368 | 0.174575 | 0.844619 |
| ENSMUSG( | Tram1l1 | 1003.399 | -0.59441 | 0.000293 | 0.014095 | ENSMUSG( | Tram1l1 | 1154.937 | 0.139731 | 0.439566 | 0.920721 |
| ENSMUSG( | Tram2 | 76.40281 | 0.114464 | 0.676236 | 0.930542 | ENSMUSG( | Tram2 | 86.92015 | 0.689576 | 0.011677 | 0.440829 |
| ENSMUSG( | Trank1 | 13543.93 | -0.58955 | 0.001451 | 0.042014 | ENSMUSG( | Trank1 | 15114.17 | 0.050175 | 0.811883 | 0.979148 |
| ENSMUSG( | Trap1 | 2491.805 | 0.051621 | 0.60295 | 0.904887 | ENSMUSG( | Trap1 | 2071.212 | -0.22404 | 0.057054 | 0.723362 |
| ENSMUSG( | Trappc1 | 726.7355 | -0.13019 | 0.288016 | 0.738882 | ENSMUSG( | Trappc1 | 620.6488 | -0.33832 | 0.019626 | 0.556192 |
| ENSMUSG( | Trappc10 | 3634.551 | 0.16756 | 0.05493 | 0.360715 | ENSMUSG( | Trappc10 | 3265.733 | 0.131463 | 0.230367 | 0.868507 |
| ENSMUSG( | Trappc11 | 1493.904 | 0.096038 | 0.263586 | 0.714284 | ENSMUSG( | Trappc11 | 1339.984 | 0.052944 | 0.630997 | 0.953251 |
| ENSMUSG( | Trappc12 | 1247.503 | 0.109022 | 0.376677 | 0.801213 | ENSMUSG( | Trappc12 | 1056.848 | -0.10162 | 0.472827 | 0.923329 |
| ENSMUSG( | Trappc13 | 830.4125 | 0.075361 | 0.469686 | 0.846186 | ENSMUSG( | Trappc13 | 756.1487 | 0.069504 | 0.583163 | 0.944266 |
| ENSMUSG( | Trappc14 | 1565.597 | 0.112282 | 0.399561 | 0.813178 | ENSMUSG( | Trappc14 | 1398.042 | 0.053851 | 0.712746 | 0.96439 |
| ENSMUSG( | Trappc2 | 683.411 | 0.004534 | 0.968633 | 0.996369 | ENSMUSG( | Trappc2 | 591.9624 | -0.15366 | 0.30994 | 0.898538 |
| ENSMUSG( | Trappc2b | 269.7658 | 0.052124 | 0.71782 | 0.941697 | ENSMUSG( | Trappc2b | 241.6658 | -0.01128 | 0.953381 | 0.994798 |
| ENSMUSG( | Trappc2l | 874.1364 | -0.08948 | 0.420048 | 0.82245 | ENSMUSG( | Trappc2l | 770.6641 | -0.19331 | 0.168833 | 0.840222 |
| ENSMUSG( | Trappc3 | 1643.88 | -0.02793 | 0.757154 | 0.951513 | ENSMUSG( | Trappc3 | 1482.262 | -0.06134 | 0.605809 | 0.949445 |
| ENSMUSG( | Trappc4 | 1028.377 | 0.007891 | 0.934886 | 0.993231 | ENSMUSG( | Trappc4 | 891.2054 | -0.14368 | 0.23626 | 0.872173 |
| ENSMUSG( | Trappc5 | 878.0955 | -0.1542 | 0.144472 | 0.564381 | ENSMUSG( | Trappc5 | 820.0419 | -0.08056 | 0.498806 | 0.927825 |
| ENSMUSG( | Trappc6a | 287.6083 | -0.09765 | 0.516306 | 0.872701 | ENSMUSG( | Trappc6a | 253.3853 | -0.21098 | 0.261759 | 0.884926 |
| ENSMUSG( | Trappc6b | 1783.047 | -0.03674 | 0.674587 | 0.929638 | ENSMUSG( | Trappc6b | 1603.73 | -0.07859 | 0.564278 | 0.941206 |
| ENSMUSG( | Trappc8 | 1902.233 | 0.128411 | 0.174672 | 0.610304 | ENSMUSG( | Trappc8 | 1682.142 | 0.042846 | 0.697624 | 0.960867 |
| ENSMUSG( | Trappc9 | 2160.922 | 0.082321 | 0.474677 | 0.849485 | ENSMUSG( | Trappc9 | 1814.472 | -0.1566 | 0.232309 | 0.868507 |
| ENSMUSG( | Trat1 | 15.99858 | -0.4369 | 0.364099 | NA | ENSMUSG( | Trat1 | 16.10575 | -0.0911 | 0.850185 | 0.985541 |
| ENSMUSG( | Trbc2 | 251.5566 | -0.01454 | 0.938061 | 0.993415 | ENSMUSG( | Trbc2 | 209.2731 | -0.2889 | 0.135942 | 0.826837 |
| ENSMUSG( | Trdmt1 | 177.4652 | -0.14584 | 0.371699 | 0.798768 | ENSMUSG( | Trdmt1 | 158.6309 | -0.21015 | 0.24909 | 0.877904 |
| ENSMUSG( | Trem2 | 919.9415 | -0.3573 | 0.023784 | 0.235641 | ENSMUSG( | Trem2 | 941.0491 | 0.005192 | 0.977613 | 0.996416 |
| ENSMUSG( | Treml1 | 9.255316 | -1.01887 | 0.098283 | NA | ENSMUSG( | Treml1 | 8.493739 | -0.90801 | 0.184466 | NA |
| ENSMUSG( | Treml2 | 33.30377 | -0.52706 | 0.11169 | 0.503786 | ENSMUSG( | Treml2 | 33.12185 | -0.20573 | 0.563475 | 0.941206 |
| ENSMUSG( | Treml4 | 24.01 | -2.00112 | 0.005215 | 0.096233 | ENSMUSG( | Treml4 | 23.45106 | -1.64735 | 0.018217 | 0.542424 |
| ENSMUSG( | Trerf1 | 1547.938 | 0.081577 | 0.443535 | 0.833108 | ENSMUSG( | Trerf1 | 1468.735 | 0.191781 | 0.13775 | 0.828492 |
| ENSMUSG( | Trf | 9636.037 | -0.51147 | 0.000532 | 0.021434 | ENSMUSG( | Trf | 10472.27 | 0.046379 | 0.805232 | 0.977427 |
| ENSMUSG( | Trh | 132.9868 | -0.99375 | 0.094221 | 0.471647 | ENSMUSG( | Trh | 150.6367 | -0.22122 | 0.708752 | 0.964042 |
| ENSMUSG( | Trhde | 604.459 | -0.42828 | 0.030665 | 0.266555 | ENSMUSG( | Trhde | 673.0823 | 0.185727 | 0.426879 | 0.917257 |
| ENSMUSG( | Trhr | 71.05024 | -0.11867 | 0.624053 | 0.91187 | ENSMUSG( | Trhr | 81.02267 | 0.464658 | 0.086945 | 0.782278 |
| ENSMUSG( | Trhr2 | 37.44763 | -0.44104 | 0.194606 | 0.633457 | ENSMUSG( | Trhr2 | 37.9574 | -0.0648 | 0.865761 | 0.987041 |
| ENSMUSG( | Triap1 | 362.7045 | 0.184726 | 0.167034 | 0.599655 | ENSMUSG( | Triap1 | 317.5277 | 0.073016 | 0.638076 | 0.953903 |
| ENSMUSG( | Trib1 | 606.7748 | -0.29572 | 0.025597 | 0.246694 | ENSMUSG( | Trib1 | 593.1332 | -0.07344 | 0.588649 | 0.944975 |
| ENSMUSG( | Trib2 | 1789.294 | -0.13088 | 0.238443 | 0.686308 | ENSMUSG( | Trib2 | 1679.307 | -0.04734 | 0.744451 | 0.96951 |
| ENSMUSG( | Trib3 | 12.14063 | 0.023083 | 0.966548 | NA | ENSMUSG( | Trib3 | 14.02915 | 0.611305 | 0.260688 | 0.884675 |
| ENSMUSG( | Tril | 4088.989 | -0.23731 | 0.097192 | 0.47759 | ENSMUSG( | Tril | 3938.443 | -0.06367 | 0.707572 | 0.964029 |
| ENSMUSG( | Trim10 | 22.60445 | -1.48335 | 0.011004 | 0.151333 | ENSMUSG( | Trim10 | 24.02411 | -0.81329 | 0.141449 | 0.830678 |
| ENSMUSG( | Trim11 | 716.8949 | -0.19938 | 0.124733 | 0.531714 | ENSMUSG( | Trim11 | 698.0829 | 0.001991 | 0.988864 | 0.997886 |
| ENSMUSG( | Trim12a | 150.048 | -0.24615 | 0.173713 | 0.609084 | ENSMUSG( | Trim12a | 148.2725 | 0.008985 | 0.966426 | 0.995475 |
| ENSMUSG( | Trim12c | 240.4938 | 0.135767 | 0.425129 | 0.82478 | ENSMUSG( | Trim12c | 192.0783 | -0.25446 | 0.146909 | 0.836241 |
| ENSMUSG( | Trim13 | 181.06 | -0.54013 | 0.007752 | 0.1222 | ENSMUSG( | Trim13 | 211.9249 | 0.236083 | 0.25441 | 0.881624 |
| ENSMUSG( | Trim14 | 20.33043 | -0.06055 | 0.886064 | NA | ENSMUSG( | Trim14 | 17.91409 | -0.1502 | 0.771353 | 0.974236 |
| ENSMUSG( | Trim16 | 74.979 | 0.46713 | 0.105398 | 0.492417 | ENSMUSG( | Trim16 | 70.43173 | 0.56731 | 0.123858 | 0.814691 |
| ENSMUSG( | Trim17 | 258.3395 | 0.204277 | 0.216196 | 0.661017 | ENSMUSG( | Trim17 | 218.3642 | -0.01275 | 0.94356 | 0.994798 |
| ENSMUSG( | Trim2 | 5073.491 | 0.052474 | 0.499109 | 0.862973 | ENSMUSG( | Trim2 | 4424.009 | -0.07589 | 0.495813 | 0.927825 |
| ENSMUSG( | Trim21 | 84.91676 | -0.05691 | 0.798998 | 0.960922 | ENSMUSG( | Trim21 | 78.67608 | 0.0107 | 0.964971 | 0.995433 |
| ENSMUSG( | Trim23 | 850.2359 | -0.00021 | 0.998372 | 0.99989 | ENSMUSG( | Trim23 | 788.6024 | 0.046717 | 0.740963 | 0.968811 |
| ENSMUSG( | Trim24 | 1022.505 | -0.07932 | 0.417936 | 0.821964 | ENSMUSG( | Trim24 | 925.8558 | -0.09639 | 0.415865 | 0.917257 |
| ENSMUSG( | Trim25 | 432.9578 | 0.065119 | 0.675857 | 0.930515 | ENSMUSG( | Trim25 | 373.1327 | -0.08979 | 0.572751 | 0.94262 |
| ENSMUSG( | Trim26 | 768.2918 | -0.1278 | 0.232463 | 0.680606 | ENSMUSG( | Trim26 | 737.6289 | 0.030168 | 0.810297 | 0.978779 |
| ENSMUSG( | Trim27 | 580.612 | 0.123832 | 0.344451 | 0.779153 | ENSMUSG( | Trim27 | 509.5229 | 0.016276 | 0.909396 | 0.991958 |
| ENSMUSG( | Trim28 | 2966.452 | 0.122998 | 0.269142 | 0.719307 | ENSMUSG( | Trim28 | 2568.605 | -0.02274 | 0.852041 | 0.985598 |
| ENSMUSG( | Trim29 | 19.95893 | 0.210761 | 0.657608 | NA | ENSMUSG( | Trim29 | 18.94489 | 0.301034 | 0.526964 | 0.934449 |
| ENSMUSG( | Trim3 | 2962.456 | 0.00972 | 0.944037 | 0.993503 | ENSMUSG( | Trim3 | 2628.218 | -0.06847 | 0.622093 | 0.953234 |
| ENSMUSG( | Trim30a | 109.1775 | 0.020171 | 0.924048 | 0.991077 | ENSMUSG( | Trim30a | 106.7068 | 0.223232 | 0.321368 | 0.904019 |
| ENSMUSG( | Trim30d | 74.33721 | -0.08795 | 0.708617 | 0.939519 | ENSMUSG( | Trim30d | 67.44922 | -0.08489 | 0.753192 | 0.970622 |
| ENSMUSG( | Trim32 | 3094.906 | -0.00042 | 0.997366 | 0.999605 | ENSMUSG( | Trim32 | 2729.209 | -0.09718 | 0.482823 | 0.924615 |
| ENSMUSG( | Trim33 | 1651.857 | 0.042977 | 0.649966 | 0.920899 | ENSMUSG( | Trim33 | 1508.675 | 0.048206 | 0.667719 | 0.956766 |
| ENSMUSG( | Trim34a | 102.5511 | -0.13668 | 0.534697 | 0.880719 | ENSMUSG( | Trim34a | 93.81504 | -0.12975 | 0.568712 | 0.941777 |
| ENSMUSG( | Trim35 | 3379.747 | 0.072005 | 0.438586 | 0.83067 | ENSMUSG( | Trim35 | 2920.764 | -0.08172 | 0.477001 | 0.923329 |
| ENSMUSG( | Trim36 | 167.3625 | -0.38424 | 0.024364 | 0.238636 | ENSMUSG( | Trim36 | 174.6594 | 0.049253 | 0.809142 | 0.97873 |
| ENSMUSG( | Trim37 | 15309.61 | 0.146653 | 0.073626 | 0.41961 | ENSMUSG( | Trim37 | 12823.47 | -0.09467 | 0.353492 | 0.912263 |
| ENSMUSG( | Trim39 | 1016.485 | -0.06612 | 0.611048 | 0.908534 | ENSMUSG( | Trim39 | 946.4585 | -0.00013 | 0.999315 | 0.999503 |
| ENSMUSG( | Trim41 | 2456.098 | -0.00067 | 0.99472 | 0.999369 | ENSMUSG( | Trim41 | 2173.502 | -0.08791 | 0.454088 | 0.922444 |
| ENSMUSG( | Trim43a | 16.2334 | -0.13313 | 0.786032 | NA | ENSMUSG( | Trim43a | 13.39853 | -0.42404 | 0.448015 | 0.922202 |
| ENSMUSG( | Trim44 | 10580.81 | 0.100042 | 0.239512 | 0.687308 | ENSMUSG( | Trim44 | 9423.326 | 0.036014 | 0.745374 | 0.96951 |
| ENSMUSG( | Trim45 | 224.8394 | -0.03191 | 0.822325 | 0.966352 | ENSMUSG( | Trim45 | 188.1783 | -0.28184 | 0.101436 | 0.801456 |
| ENSMUSG( | Trim46 | 2916.592 | 0.152009 | 0.161311 | 0.591331 | ENSMUSG( | Trim46 | 2523.359 | 0.006713 | 0.957652 | 0.994798 |
| ENSMUSG( | Trim47 | 167.0392 | 0.253868 | 0.188272 | 0.627268 | ENSMUSG( | Trim47 | 178.504 | 0.665827 | 0.006124 | 0.322135 |
| ENSMUSG( | Trim5 | 28.72072 | -0.39931 | 0.279935 | 0.729155 | ENSMUSG( | Trim5 | 30.367 | 0.077686 | 0.835991 | 0.981237 |
| ENSMUSG( | Trim54 | 21.35488 | 0.137026 | 0.7928 | 0.95947 | ENSMUSG( | Trim54 | 19.19214 | 0.060806 | 0.896421 | 0.990558 |
| ENSMUSG( | Trim56 | 971.7018 | 0.035324 | 0.797318 | 0.960507 | ENSMUSG( | Trim56 | 778.5881 | -0.3593 | 0.005132 | 0.294733 |
| ENSMUSG( | Trim59 | 304.7082 | -0.47179 | 0.001991 | 0.051331 | ENSMUSG( | Trim59 | 322.0358 | -0.00199 | 0.991119 | 0.998379 |
| ENSMUSG( | Trim6 | 12.54245 | 0.096053 | 0.865094 | NA | ENSMUSG( | Trim6 | 11.46172 | 0.137893 | 0.816165 | 0.979198 |
| ENSMUSG( | Trim62 | 796.499 | -0.04366 | 0.700653 | 0.93743 | ENSMUSG( | Trim62 | 704.9954 | -0.13237 | 0.328974 | 0.906929 |
| ENSMUSG( | Trim65 | 182.9019 | 0.041094 | 0.840081 | 0.971179 | ENSMUSG( | Trim65 | 184.6461 | 0.334428 | 0.121665 | 0.813384 |
| ENSMUSG( | Trim66 | 651.3718 | 0.204877 | 0.118984 | 0.522304 | ENSMUSG( | Trim66 | 551.4396 | 0.006303 | 0.965979 | 0.995436 |
| ENSMUSG( | Trim67 | 244.4612 | 0.000746 | 0.995823 | 0.99943 | ENSMUSG( | Trim67 | 213.1102 | -0.1262 | 0.432562 | 0.918256 |
| ENSMUSG( | Trim68 | 164.2521 | -0.15528 | 0.404395 | 0.816132 | ENSMUSG( | Trim68 | 160.3317 | 0.050197 | 0.827579 | 0.980224 |
| ENSMUSG( | Trim7 | 65.08048 | -0.05518 | 0.815145 | 0.964978 | ENSMUSG( | Trim7 | 61.52348 | 0.0611 | 0.814457 | 0.979198 |
| ENSMUSG( | Trim71 | 24.13394 | -0.50718 | 0.186583 | 0.623869 | ENSMUSG( | Trim71 | 22.73097 | -0.37167 | 0.376802 | 0.915583 |
| ENSMUSG( | Trim72 | 30.96301 | 0.227318 | 0.493522 | 0.860611 | ENSMUSG( | Trim72 | 26.57297 | 0.068234 | 0.85672 | 0.985884 |
| ENSMUSG( | Trim8 | 4405.193 | 0.139909 | 0.104453 | 0.490485 | ENSMUSG( | Trim8 | 4032.592 | 0.151322 | 0.216987 | 0.865589 |
| ENSMUSG( | Trim9 | 11640.63 | 0.103133 | 0.159023 | 0.588112 | ENSMUSG( | Trim9 | 9473.301 | -0.23086 | 0.030257 | 0.623534 |
| ENSMUSG( | Trio | 5182.78 | 0.017547 | 0.844842 | 0.972023 | ENSMUSG( | Trio | 4839.866 | 0.088859 | 0.443958 | 0.921734 |
| ENSMUSG( | Triobp | 830.7504 | -0.00111 | 0.990902 | 0.999232 | ENSMUSG( | Triobp | 757.8503 | 0.003831 | 0.975553 | 0.996416 |
| ENSMUSG( | Trip10 | 259.393 | 0.216185 | 0.128465 | 0.536774 | ENSMUSG( | Trip10 | 245.2957 | 0.32311 | 0.057167 | 0.723362 |
| ENSMUSG( | Trip11 | 1464.764 | 0.092125 | 0.387734 | 0.805835 | ENSMUSG( | Trip11 | 1317.292 | 0.05709 | 0.640591 | 0.953903 |
| ENSMUSG( | Trip12 | 4836.031 | 0.102937 | 0.209698 | 0.651841 | ENSMUSG( | Trip12 | 4293.803 | 0.029407 | 0.786403 | 0.976303 |
| ENSMUSG( | Trip13 | 18.32723 | -0.30891 | 0.469212 | NA | ENSMUSG( | Trip13 | 16.95402 | -0.27388 | 0.566434 | 0.941745 |
| ENSMUSG( | Trip4 | 1211.482 | 0.11909 | 0.231595 | 0.67949 | ENSMUSG( | Trip4 | 1059.775 | 0.000916 | 0.993944 | 0.998813 |
| ENSMUSG( | Trip6 | 260.2593 | -0.00315 | 0.981703 | 0.997512 | ENSMUSG( | Trip6 | 241.3501 | 0.04551 | 0.814843 | 0.979198 |
| ENSMUSG( | Triqk | 303.8246 | 0.007809 | 0.960009 | 0.993991 | ENSMUSG( | Triqk | 260.0869 | -0.19315 | 0.314996 | 0.901306 |
| ENSMUSG( | Trir | 1840.535 | 0.01261 | 0.9217 | 0.990436 | ENSMUSG( | Trir | 1642.344 | -0.05203 | 0.727742 | 0.966887 |
| ENSMUSG( | Trit1 | 365.5762 | 0.067254 | 0.626524 | 0.91272 | ENSMUSG( | Trit1 | 336.9168 | 0.107706 | 0.487997 | 0.925453 |
| ENSMUSG( | Trmo | 72.16338 | -0.13123 | 0.591036 | 0.90113 | ENSMUSG( | Trmo | 65.87742 | -0.14004 | 0.626951 | 0.953234 |
| ENSMUSG( | Trmt1 | 2695.269 | 0.213235 | 0.059088 | 0.375389 | ENSMUSG( | Trmt1 | 2349.24 | 0.091963 | 0.479816 | 0.924025 |
| ENSMUSG( | Trmt10a | 217.6143 | 0.003457 | 0.980794 | 0.997512 | ENSMUSG( | Trmt10a | 194.4016 | -0.06383 | 0.716994 | 0.965864 |
| ENSMUSG( | Trmt10b | 221.8315 | 0.139589 | 0.376203 | 0.801127 | ENSMUSG( | Trmt10b | 199.0685 | 0.097748 | 0.582476 | 0.944266 |
| ENSMUSG( | Trmt10c | 174.5179 | -0.16927 | 0.409632 | 0.818258 | ENSMUSG( | Trmt10c | 168.6658 | 0.008662 | 0.965861 | 0.995433 |
| ENSMUSG( | Trmt11 | 173.302 | -0.08471 | 0.609319 | 0.907627 | ENSMUSG( | Trmt11 | 154.8752 | -0.14581 | 0.464802 | 0.923329 |
| ENSMUSG( | Trmt112 | 419.8923 | 0.136178 | 0.296109 | 0.744386 | ENSMUSG( | Trmt112 | 355.8299 | -0.07744 | 0.610288 | 0.950771 |
| ENSMUSG( | Trmt12 | 172.3865 | 0.102746 | 0.565084 | 0.890735 | ENSMUSG( | Trmt12 | 152.563 | 0.012485 | 0.952442 | 0.994798 |
| ENSMUSG( | Trmt13 | 253.1774 | -0.0553 | 0.704212 | 0.938221 | ENSMUSG( | Trmt13 | 248.9474 | 0.174461 | 0.336437 | 0.907972 |
| ENSMUSG( | Trmt1l | 662.2294 | 0.169817 | 0.135193 | 0.548658 | ENSMUSG( | Trmt1l | 575.6958 | 0.03907 | 0.772 | 0.974236 |
| ENSMUSG( | Trmt2a | 1098.968 | 0.140521 | 0.146882 | 0.56887 | ENSMUSG( | Trmt2a | 965.2859 | 0.039588 | 0.755902 | 0.971371 |
| ENSMUSG( | Trmt2b | 595.0299 | 0.100891 | 0.500416 | 0.863483 | ENSMUSG( | Trmt2b | 481.8282 | -0.25587 | 0.122287 | 0.813384 |
| ENSMUSG( | Trmt44 | 144.7446 | 0.202613 | 0.241681 | 0.689179 | ENSMUSG( | Trmt44 | 133.3244 | 0.244184 | 0.228777 | 0.868507 |
| ENSMUSG( | Trmt5 | 159.12 | -0.13329 | 0.410766 | 0.818585 | ENSMUSG( | Trmt5 | 154.7078 | 0.058274 | 0.758449 | 0.971858 |
| ENSMUSG( | Trmt6 | 711.389 | 0.034195 | 0.771374 | 0.954726 | ENSMUSG( | Trmt6 | 625.6504 | -0.06928 | 0.582913 | 0.944266 |
| ENSMUSG( | Trmt61a | 327.4215 | 0.064675 | 0.705477 | 0.938511 | ENSMUSG( | Trmt61a | 279.7578 | -0.12907 | 0.492279 | 0.927315 |
| ENSMUSG( | Trmt61b | 229.2862 | 0.13134 | 0.390139 | 0.80794 | ENSMUSG( | Trmt61b | 194.0692 | -0.08003 | 0.662078 | 0.956205 |
| ENSMUSG( | Trmt9b | 343.3153 | -0.37302 | 0.003414 | 0.073541 | ENSMUSG( | Trmt9b | 298.7117 | -0.53708 | 0.001422 | 0.139194 |
| ENSMUSG( | Trmu | 424.9134 | -0.02532 | 0.859198 | 0.974707 | ENSMUSG( | Trmu | 410.5788 | 0.14517 | 0.368533 | 0.915491 |
| ENSMUSG( | Trnau1ap | 490.3312 | -0.02411 | 0.824892 | 0.967386 | ENSMUSG( | Trnau1ap | 429.2178 | -0.14108 | 0.299289 | 0.898538 |
| ENSMUSG( | Trnp1 | 8850.231 | 0.026265 | 0.728096 | 0.944355 | ENSMUSG( | Trnp1 | 7900.233 | -0.03398 | 0.740662 | 0.968811 |
| ENSMUSG( | Trnt1 | 594.6637 | 0.083676 | 0.450994 | 0.835567 | ENSMUSG( | Trnt1 | 569.7345 | 0.217217 | 0.09429 | 0.795735 |
| ENSMUSG( | Tro | 3991.982 | 0.071038 | 0.43722 | 0.829501 | ENSMUSG( | Tro | 3724.82 | 0.13998 | 0.241921 | 0.876528 |
| ENSMUSG( | Troap | 11.24407 | -0.68339 | 0.218538 | NA | ENSMUSG( | Troap | 11.53995 | -0.23667 | 0.679831 | 0.957966 |
| ENSMUSG( | Trp53 | 566.3123 | -0.21523 | 0.079775 | 0.436135 | ENSMUSG( | Trp53 | 539.6509 | -0.07469 | 0.595471 | 0.945853 |
| ENSMUSG( | Trp53bp1 | 2120.078 | -0.02108 | 0.831305 | 0.969889 | ENSMUSG( | Trp53bp1 | 1939.768 | -0.00752 | 0.94591 | 0.994798 |
| ENSMUSG( | Trp53bp2 | 2433.354 | 0.242991 | 0.004465 | 0.08705 | ENSMUSG( | Trp53bp2 | 1955.511 | -0.1104 | 0.364525 | 0.914223 |

| | | | | | |
|---|---|---|---|---|---|
| ENSMUSG( | 2707.168 | 0.032178 | 0.750482 | 0.949877 | Tmem65 |
| ENSMUSG( | 356.5741 | -0.19207 | 0.150052 | 0.574564 | Tmem67 |
| ENSMUSG( | 787.2897 | -0.52788 | 0.001164 | 0.036596 | Tmem68 |
| ENSMUSG( | 207.7494 | -0.02204 | 0.888463 | 0.983434 | Tmem69 |
| ENSMUSG( | 869.8187 | -0.04924 | 0.639937 | 0.916434 | Tmem70 |
| ENSMUSG( | 21.87999 | -0.14375 | 0.725955 | 0.944223 | Tmem71 |
| ENSMUSG( | 51.16427 | -0.02953 | 0.975308 | 0.997302 | Tmem72 |
| ENSMUSG( | 147.2091 | -0.20194 | 0.341238 | 0.778828 | Tmem74 |
| ENSMUSG( | 114.2758 | -0.39382 | 0.077252 | 0.429743 | Tmem74b |
| ENSMUSG( | 17.16618 | 0.670432 | 0.190107 | NA | Tmem74bos |
| ENSMUSG( | 27.73437 | -0.55233 | 0.128177 | 0.536641 | Tmem79 |
| ENSMUSG( | 1029.534 | 0.064148 | 0.493377 | 0.860611 | Tmem80 |
| ENSMUSG( | 189.7478 | -0.06234 | 0.689089 | 0.933441 | Tmem81 |
| ENSMUSG( | 43.98241 | -0.14725 | 0.610595 | 0.908194 | Tmem82 |
| ENSMUSG( | 719.1949 | -0.44342 | 0.000935 | 0.031289 | Tmem86a |
| ENSMUSG( | 251.3575 | 0.427084 | 0.008192 | 0.12594 | Tmem86b |
| ENSMUSG( | 1119.413 | -0.3821 | 0.016201 | 0.189046 | Tmem87a |
| ENSMUSG( | 1191.509 | -0.36225 | 0.019919 | 0.213312 | Tmem87b |
| ENSMUSG( | 120.4681 | -0.06105 | 0.804259 | 0.962169 | Tmem88 |
| ENSMUSG( | 3017.489 | -0.51441 | 1.94E-06 | 0.000324 | Tmem88b |
| ENSMUSG( | 4311.502 | -0.06302 | 0.59259 | 0.902094 | Tmem8b |
| ENSMUSG( | 2192.361 | -0.36373 | 0.002926 | 0.066146 | Tmem9 |
| ENSMUSG( | 417.6507 | -0.19903 | 0.163963 | 0.594869 | Tmem91 |
| ENSMUSG( | 2875.084 | -0.05826 | 0.616951 | 0.910866 | Tmem94 |
| ENSMUSG( | 689.6243 | -0.50931 | 0.001904 | 0.04982 | Tmem97 |
| ENSMUSG( | 628.4802 | -0.54649 | 0.001359 | 0.040233 | Tmem98 |
| ENSMUSG( | 4510.435 | -0.2603 | 0.045721 | 0.328047 | Tmem9b |
| ENSMUSG( | 1282.32 | 0.110468 | 0.260407 | 0.710245 | Tmf1 |
| ENSMUSG( | 360.1258 | -0.21514 | 0.123577 | 0.530552 | Tmie |
| ENSMUSG( | 1030.447 | -0.26499 | 0.018931 | 0.207196 | Tmod1 |
| ENSMUSG( | 19614.04 | 0.16795 | 0.051513 | 0.34814 | Tmod2 |
| ENSMUSG( | 1032.85 | 0.048305 | 0.636365 | 0.914873 | Tmod3 |
| ENSMUSG( | 12.07895 | -0.34861 | 0.513639 | NA | Tmod4 |
| ENSMUSG( | 677.8731 | 0.082896 | 0.432366 | 0.82768 | Tmpo |
| ENSMUSG( | 84.63354 | 0.072178 | 0.79363 | 0.959799 | Tmppe |
| ENSMUSG( | 35.44228 | -1.2968 | 9.67E-05 | 0.006353 | Tmprss5 |
| ENSMUSG( | 18.20575 | -0.66897 | 0.153499 | NA | Tmprss6 |
| ENSMUSG( | 123.641 | -0.05467 | 0.771176 | 0.954726 | Tmprss7 |
| ENSMUSG( | 17.35452 | -0.28669 | 0.534015 | NA | Tmprss9 |
| ENSMUSG( | 6778.685 | -0.04842 | 0.612462 | 0.908747 | Tmsb10 |
| ENSMUSG( | 61.37838 | 0.003936 | 0.988131 | 0.999109 | Tmsb10b |
| ENSMUSG( | 14.75294 | -0.48693 | 0.360247 | NA | Tmsb15b1 |
| ENSMUSG( | 18308.27 | 0.04062 | 0.66474 | 0.926824 | Tmsb4x |
| ENSMUSG( | 1068.795 | 0.00736 | 0.953144 | 0.993874 | Tmt1a |
| ENSMUSG( | 2566.031 | -0.38639 | 0.000503 | 0.020541 | Tmtc1 |
| ENSMUSG( | 487.2607 | -0.20499 | 0.242245 | 0.689699 | Tmtc2 |
| ENSMUSG( | 335.3456 | -0.26818 | 0.182151 | 0.619591 | Tmtc3 |
| ENSMUSG( | 336.6084 | -0.06666 | 0.683006 | 0.93205 | Tmtc4 |
| ENSMUSG( | 694.6903 | -0.20827 | 0.064062 | 0.391961 | Tmub1 |
| ENSMUSG( | 2815.854 | -0.3379 | 0.002511 | 0.059283 | Tmub2 |
| ENSMUSG( | 945.1907 | -0.41636 | 0.006326 | 0.108996 | Tmx1 |
| ENSMUSG( | 5446.844 | -0.38838 | 0.002 | 0.051341 | Tmx2 |
| ENSMUSG( | 1612.806 | -0.41886 | 0.009145 | 0.134728 | Tmx3 |
| ENSMUSG( | 5423.419 | -0.41653 | 0.007543 | 0.120197 | Tmx4 |
| ENSMUSG( | 527.6962 | -0.3307 | 0.064244 | 0.392776 | Tnc |
| ENSMUSG( | 2460.031 | 0.097431 | 0.32474 | 0.770182 | Tnfaip1 |
| ENSMUSG( | 41.33401 | -0.56472 | 0.057946 | 0.371205 | Tnfaip2 |
| ENSMUSG( | 104.569 | 0.003832 | 0.984896 | 0.997815 | Tnfaip3 |
| ENSMUSG( | 220.8315 | -0.83511 | 7.34E-06 | 0.000923 | Tnfaip6 |
| ENSMUSG( | 584.9865 | -0.09564 | 0.549666 | 0.885944 | Tnfaip8 |
| ENSMUSG( | 151.1227 | -0.05558 | 0.761486 | 0.953106 | Tnfaip8l1 |
| ENSMUSG( | 95.02929 | 0.275638 | 0.18096 | 0.618094 | Tnfaip8l2 |
| ENSMUSG( | 153.063 | 0.244214 | 0.28079 | 0.729868 | Tnfaip8l3 |
| ENSMUSG( | 25.97451 | -0.1801 | 0.634779 | 0.914073 | Tnfrsf10b |
| ENSMUSG( | 191.0108 | -0.48566 | 0.00365 | 0.076783 | Tnfrsf11a |
| ENSMUSG( | 102.903 | -0.4501 | 0.128657 | 0.537025 | Tnfrsf11b |
| ENSMUSG( | 148.037 | -0.43038 | 0.02878 | 0.258585 | Tnfrsf12a |
| ENSMUSG( | 106.5286 | -0.18498 | 0.344191 | 0.779153 | Tnfrsf13b |
| ENSMUSG( | 17.33946 | 0.029628 | 0.94731 | NA | Tnfrsf13c |
| ENSMUSG( | 20.28251 | 0.086973 | 0.850465 | NA | Tnfrsf14 |
| ENSMUSG( | 12.29013 | 0.011335 | 0.9833 | NA | Tnfrsf17 |
| ENSMUSG( | 8042.657 | -0.11218 | 0.374503 | 0.800445 | Tnfrsf19 |
| ENSMUSG( | 727.7124 | -0.13591 | 0.385275 | 0.804048 | Tnfrsf1a |
| ENSMUSG( | 282.1272 | -0.46167 | 0.038036 | 0.300991 | Tnfrsf1b |
| ENSMUSG( | 2408.477 | -0.30846 | 0.048928 | 0.337919 | Tnfrsf21 |
| ENSMUSG( | 91.72018 | 0.174897 | 0.398135 | 0.812452 | Tnfrsf22 |
| ENSMUSG( | 59.48596 | 0.008866 | 0.971306 | 0.996369 | Tnfrsf23 |
| ENSMUSG( | 250.4822 | 0.657804 | 0.001742 | 0.047093 | Tnfrsf25 |
| ENSMUSG( | 40.46024 | 0.009184 | 0.975355 | 0.997302 | Tnfrsf4 |
| ENSMUSG( | 34.39422 | 0.143491 | 0.664038 | 0.926511 | Tnfrsf8 |
| ENSMUSG( | 72.84979 | -0.14843 | 0.630027 | 0.913965 | Tnfsf10 |
| ENSMUSG( | 522.4868 | -0.15739 | 0.30116 | 0.749979 | Tnfsf12 |
| ENSMUSG( | 50.92549 | 0.247029 | 0.424841 | 0.824675 | Tnfsf13 |
| ENSMUSG( | 35.83161 | -0.69454 | 0.031514 | 0.270738 | Tnfsf13b |
| ENSMUSG( | 14.10982 | -0.87443 | 0.084803 | NA | Tnfsf9 |
| ENSMUSG( | 26.25886 | -0.22168 | 0.551124 | 0.886038 | Tnfsfm13 |
| ENSMUSG( | 4749.996 | -0.00473 | 0.95619 | 0.993874 | Tnik |
| ENSMUSG( | 722.0309 | 0.058547 | 0.602202 | 0.904887 | Tnip1 |
| ENSMUSG( | 367.8071 | -0.00728 | 0.956815 | 0.993874 | Tnip2 |
| ENSMUSG( | 9877.178 | 0.230912 | 0.005492 | 0.0993 | Tnk2 |
| ENSMUSG( | 3405.83 | -0.02845 | 0.789172 | 0.959299 | Tnks |
| ENSMUSG( | 2647.744 | 0.018465 | 0.847575 | 0.972374 | Tnks1bp1 |
| ENSMUSG( | 4883.683 | 0.006902 | 0.944115 | 0.993503 | Tnks2 |
| ENSMUSG( | 24.24023 | -0.55405 | 0.159895 | 0.589679 | Tnmd |
| ENSMUSG( | 251.2543 | -0.02181 | 0.891774 | 0.983979 | Tnnc1 |
| ENSMUSG( | 259.5544 | -0.10027 | 0.499478 | 0.863218 | Tnni1 |
| ENSMUSG( | 13.10379 | -0.03805 | 0.942714 | NA | Tnni2 |
| ENSMUSG( | 26.10511 | -0.28635 | 0.497964 | 0.862372 | Tnni3 |
| ENSMUSG( | 11.09858 | -0.30158 | 0.577598 | NA | Tnni3k |
| ENSMUSG( | 182.9389 | -0.30653 | 0.103162 | 0.48855 | Tnnt1 |
| ENSMUSG( | 78.89736 | -0.25905 | 0.392003 | 0.809274 | Tnnt2 |
| ENSMUSG( | 11.67647 | 0.175889 | 0.756775 | NA | Tnnt3 |
| ENSMUSG( | 2302.13 | 0.059592 | 0.622514 | 0.911432 | Tnpo1 |
| ENSMUSG( | 5765.069 | 0.139677 | 0.07601 | 0.426896 | Tnpo2 |
| ENSMUSG( | 2163.912 | 0.075576 | 0.389676 | 0.807428 | Tnpo3 |
| ENSMUSG( | 5743.877 | -0.41736 | 0.018375 | 0.203658 | Tnr |
| ENSMUSG( | 4406.372 | -0.00851 | 0.925694 | 0.991204 | Tnrc18 |
| ENSMUSG( | 2670.78 | -0.05646 | 0.525983 | 0.876999 | Tnrc6a |
| ENSMUSG( | 2175.342 | 0.029833 | 0.74517 | 0.949158 | Tnrc6b |
| ENSMUSG( | 2438.062 | -0.21749 | 0.025734 | 0.247138 | Tnrc6c |
| ENSMUSG( | 2622.007 | 0.084245 | 0.42109 | 0.823126 | Tns1 |
| ENSMUSG( | 708.8411 | 0.179299 | 0.098725 | 0.480598 | Tns2 |
| ENSMUSG( | 11815.54 | 0.211388 | 0.047341 | 0.332697 | Tns3 |
| ENSMUSG( | 13.02429 | 0.010259 | 0.984038 | NA | Tns4 |
| ENSMUSG( | 235.637 | 0.071755 | 0.658635 | 0.924506 | Tnxb |
| ENSMUSG( | 701.409 | -0.44465 | 0.005958 | 0.104409 | Tob1 |
| ENSMUSG( | 855.9641 | -0.42311 | 0.010916 | 0.150598 | Tob2 |
| ENSMUSG( | 329.3657 | -0.08856 | 0.540772 | 0.88315 | Toe1 |
| ENSMUSG( | 836.9873 | 0.168527 | 0.084949 | 0.449406 | Togaram1 |
| ENSMUSG( | 80.69602 | 0.033067 | 0.89559 | 0.984512 | Togaram2 |
| ENSMUSG( | 5131.69 | 0.112178 | 0.160016 | 0.589679 | Tollip |
| ENSMUSG( | 810.3748 | 0.023759 | 0.811646 | 0.964426 | Tom1 |
| ENSMUSG( | 1305.211 | 0.11987 | 0.268987 | 0.719302 | Tom1l1 |
| ENSMUSG( | 11342.27 | 0.130567 | 0.095746 | 0.475357 | Tom1l2 |
| ENSMUSG( | 8709.712 | -0.17054 | 0.09302 | 0.468859 | Tomm20 |
| ENSMUSG( | 1730.96 | -0.00659 | 0.948397 | 0.993821 | Tomm22 |
| ENSMUSG( | 1600.949 | 0.068714 | 0.446573 | 0.833726 | Tomm34 |
| ENSMUSG( | 1987.93 | -0.03075 | 0.773467 | 0.955447 | Tomm40 |
| ENSMUSG( | 3536.155 | 0.107467 | 0.198286 | 0.638387 | Tomm40l |
| ENSMUSG( | 653.5988 | 0.018122 | 0.884424 | 0.982647 | Tomm5 |
| ENSMUSG( | 157.7836 | 0.074085 | 0.706783 | 0.93949 | Tomm6os |
| ENSMUSG( | 2137.883 | 0.057933 | 0.587939 | 0.900678 | Tomm7 |
| ENSMUSG( | 3080.951 | -0.0455 | 0.658924 | 0.924506 | Tomm70a |
| ENSMUSG( | 12.90196 | -0.51598 | 0.315457 | NA | Tomt |
| ENSMUSG( | 266.5297 | 0.20329 | 0.175629 | 0.611006 | Tonsl |
| ENSMUSG( | 1834.933 | 0.204611 | 0.05749 | 0.370033 | Top1 |
| ENSMUSG( | 395.7555 | 0.21606 | 0.078493 | 0.433639 | Top1mt |
| ENSMUSG( | 164.8852 | -0.10984 | 0.550688 | 0.886038 | Top2a |
| ENSMUSG( | 3366.9 | 0.102978 | 0.275567 | 0.725482 | Top2b |
| ENSMUSG( | 179.1504 | -0.05345 | 0.732105 | 0.944956 | Top3a |
| ENSMUSG( | 1169.703 | 0.095037 | 0.403936 | 0.816132 | Top3b |
| ENSMUSG( | 63.76844 | -0.0406 | 0.888338 | 0.983434 | Top6bl |
| ENSMUSG( | 745.7111 | 0.068314 | 0.49973 | 0.863218 | Topbp1 |
| ENSMUSG( | 421.4639 | 0.165178 | 0.16206 | 0.592429 | Topors |
| ENSMUSG( | 899.5916 | -0.23688 | 0.038345 | 0.302069 | Tor1a |
| ENSMUSG( | 3091.648 | 0.030524 | 0.782963 | 0.957558 | Tor1aip1 |
| ENSMUSG( | 1926.594 | -0.06277 | 0.483701 | 0.85636 | Tor1aip2 |
| ENSMUSG( | 1104.976 | -0.14464 | 0.3408 | 0.778709 | Tor1b |
| ENSMUSG( | 810.5673 | -0.21969 | 0.110761 | 0.501952 | Tor2a |
| ENSMUSG( | 511.3775 | -0.30385 | 0.043132 | 0.318494 | Tor3a |
| ENSMUSG( | 174.9684 | -0.15194 | 0.38609 | 0.804623 | Tor4a |
| ENSMUSG( | 556.7491 | -0.23017 | 0.074938 | 0.42422 | Tox |
| ENSMUSG( | 767.1108 | 0.10636 | 0.372276 | 0.798768 | Tox2 |
| ENSMUSG( | 243.0021 | 0.135043 | 0.342525 | 0.778971 | Tox3 |
| ENSMUSG( | 1477.14 | 0.113178 | 0.201358 | 0.641362 | Tox4 |
| ENSMUSG( | 647.0324 | -0.66272 | 2.77E-06 | 0.000408 | Tpbg |
| ENSMUSG( | 180.0843 | -0.44666 | 0.034124 | 0.283507 | Tpbgl |
| ENSMUSG( | 4341.026 | -0.08253 | 0.45118 | 0.835567 | Tpcn1 |

| | | | | | |
|---|---|---|---|---|---|
| ENSMUSG( | 13.24514 | -0.14079 | 0.794328 | 0.97681 | Tomt |
| ENSMUSG( | 232.2153 | 0.079338 | 0.628305 | 0.953251 | Tonsl |
| ENSMUSG( | 1692.099 | 0.233142 | 0.067964 | 0.75212 | Top1 |
| ENSMUSG( | 345.8406 | 0.097963 | 0.50404 | 0.927825 | Top1mt |
| ENSMUSG( | 152.8394 | -0.05002 | 0.801793 | 0.97681 | Top2a |
| ENSMUSG( | 2998.913 | 0.038408 | 0.747815 | 0.96984 | Top2b |
| ENSMUSG( | 162.6891 | -0.04977 | 0.788307 | 0.97637 | Top3a |
| ENSMUSG( | 1037.667 | 0.021843 | 0.87399 | 0.989644 | Top3b |
| ENSMUSG( | 64.21178 | 0.236735 | 0.384926 | 0.917257 | Top6bl |
| ENSMUSG( | 697.0285 | 0.137885 | 0.261885 | 0.884926 | Topbp1 |
| ENSMUSG( | 379.879 | 0.139795 | 0.325616 | 0.905339 | Topors |
| ENSMUSG( | 914.5454 | 0.092808 | 0.51445 | 0.930939 | Tor1a |
| ENSMUSG( | 2752.551 | -0.03594 | 0.768384 | 0.974236 | Tor1aip1 |
| ENSMUSG( | 1837.192 | 0.068982 | 0.530841 | 0.934795 | Tor1aip2 |
| ENSMUSG( | 1131.747 | 0.198004 | 0.26887 | 0.886755 | Tor1b |
| ENSMUSG( | 851.4313 | 0.198024 | 0.185325 | 0.855305 | Tor2a |
| ENSMUSG( | 492.2623 | -0.12733 | 0.429524 | 0.917257 | Tor3a |
| ENSMUSG( | 169.2527 | 0.044895 | 0.860313 | 0.986137 | Tor4a |
| ENSMUSG( | 492.3807 | -0.33281 | 0.03558 | 0.664751 | Tox |
| ENSMUSG( | 615.8654 | -0.27767 | 0.156277 | 0.837724 | Tox2 |
| ENSMUSG( | 227.0184 | 0.195595 | 0.245845 | 0.877904 | Tox3 |
| ENSMUSG( | 1274.146 | -0.04368 | 0.686627 | 0.95899 | Tox4 |
| ENSMUSG( | 781.9286 | 0.221795 | 0.178742 | 0.849041 | Tpbg |
| ENSMUSG( | 204.5283 | 0.223259 | 0.270315 | 0.887887 | Tpbgl |
| ENSMUSG( | 3973.563 | -0.0664 | 0.583506 | 0.944266 | Tpcn1 |
| ENSMUSG( | 134.7053 | 0.20539 | 0.302254 | 0.898538 | Tpcn2 |
| ENSMUSG( | 1750.63 | -0.06751 | 0.609598 | 0.950397 | Tpd52 |
| ENSMUSG( | 387.2727 | -0.2156 | 0.159541 | 0.837724 | Tpd52l1 |
| ENSMUSG( | 1384.733 | -0.11504 | 0.299748 | 0.898538 | Tpd52l2 |
| ENSMUSG( | 1013.539 | -0.11644 | 0.376028 | 0.915491 | Tpgs1 |
| ENSMUSG( | 1210.063 | -0.09977 | 0.426129 | 0.917257 | Tpgs2 |
| ENSMUSG( | 113.9823 | -0.27374 | 0.275042 | 0.889874 | Tph2 |
| ENSMUSG( | 10267.79 | -0.23049 | 0.060229 | 0.731553 | Tpi1 |
| ENSMUSG( | 69.94544 | -0.2305 | 0.355426 | 0.912555 | Tpk1 |
| ENSMUSG( | 3448.341 | -0.0799 | 0.516928 | 0.930939 | Tpm1 |
| ENSMUSG( | 160.6383 | -0.09526 | 0.631768 | 0.953251 | Tpm2 |
| ENSMUSG( | 3172.042 | -0.11332 | 0.311941 | 0.900037 | Tpm3 |
| ENSMUSG( | 20.28262 | 0.19408 | 0.660539 | 0.956205 | Tpm3-rs7 |
| ENSMUSG( | 1778.815 | 0.238613 | 0.072463 | 0.763629 | Tpm4 |
| ENSMUSG( | 916.8388 | -0.26388 | 0.090855 | 0.791074 | Tpmt |
| ENSMUSG( | 12.07303 | 0.313904 | 0.633698 | 0.953889 | Tpo |
| ENSMUSG( | 3765.095 | 0.140743 | 0.312363 | 0.900037 | Tpp1 |
| ENSMUSG( | 895.8064 | 0.068129 | 0.566939 | 0.941777 | Tpp2 |
| ENSMUSG( | 17292.85 | -0.12445 | 0.197522 | 0.862161 | Tppp |
| ENSMUSG( | 864.0677 | -0.06311 | 0.619939 | 0.952896 | Tppp3 |
| ENSMUSG( | 2615.867 | 0.01852 | 0.861149 | 0.986226 | Tpr |
| ENSMUSG( | 981.1563 | 0.285536 | 0.04644 | 0.697814 | Tpra1 |
| ENSMUSG( | 5838.129 | -0.14011 | 0.219468 | 0.866104 | Tprg1l |
| ENSMUSG( | 4232.516 | -0.46294 | 0.000443 | 0.061321 | Tprkb |
| ENSMUSG( | 635.0463 | -0.08953 | 0.570819 | 0.941973 | Tprn |
| ENSMUSG( | 1257.945 | 0.183285 | 0.247188 | 0.877904 | Tpst1 |
| ENSMUSG( | 1131.886 | 0.191965 | 0.198438 | 0.862367 | Tpst2 |
| ENSMUSG( | 22123.73 | -0.06055 | 0.643068 | 0.953903 | Tpt1 |
| ENSMUSG( | 88.89313 | 0.154756 | 0.502839 | 0.927825 | Tpx2 |
| ENSMUSG( | 1741.342 | 0.150599 | 0.329087 | 0.907083 | Tra2a |
| ENSMUSG( | 1185.734 | 0.045428 | 0.703658 | 0.962481 | Tra2b |
| ENSMUSG( | 630.1235 | 0.010377 | 0.934073 | 0.99306 | Trabd |
| ENSMUSG( | 305.1603 | 0.403158 | 0.006725 | 0.339232 | Trabd2b |
| ENSMUSG( | 153.5104 | 0.084583 | 0.684574 | 0.958678 | Tradd |
| ENSMUSG( | 48.52215 | -0.48952 | 0.092065 | 0.791889 | Traf1 |
| ENSMUSG( | 244.0524 | 0.034617 | 0.832013 | 0.980738 | Traf2 |
| ENSMUSG( | 6032.935 | 0.174854 | 0.119071 | 0.813384 | Traf3 |
| ENSMUSG( | 435.3334 | 0.015253 | 0.917594 | 0.992529 | Traf3ip1 |
| ENSMUSG( | 30.24744 | -0.31246 | 0.405995 | 0.917257 | Traf3ip2 |
| ENSMUSG( | 55.32395 | 0.533006 | 0.116714 | 0.813384 | Traf3ip3 |
| ENSMUSG( | 547.7802 | -0.04971 | 0.717691 | 0.965964 | Traf4 |
| ENSMUSG( | 88.67286 | 0.162778 | 0.527396 | 0.934466 | Traf5 |
| ENSMUSG( | 600.5553 | -0.03754 | 0.786042 | 0.976285 | Traf6 |
| ENSMUSG( | 1624.136 | 0.018142 | 0.900732 | 0.990558 | Traf7 |
| ENSMUSG( | 1587.75 | 0.0918 | 0.471849 | 0.923329 | Trafd1 |
| ENSMUSG( | 586.0047 | 0.026335 | 0.903205 | 0.991089 | Traip |
| ENSMUSG( | 9753.324 | 0.137049 | 0.160876 | 0.837724 | Trak1 |
| ENSMUSG( | 29854.75 | -0.29217 | 0.02939 | 0.618981 | Trak2 |
| ENSMUSG( | 1152.025 | 0.201368 | 0.174575 | 0.844619 | Tram1 |
| ENSMUSG( | 1154.937 | 0.139731 | 0.439566 | 0.920721 | Tram1l1 |
| ENSMUSG( | 86.92015 | 0.689576 | 0.011677 | 0.440829 | Tram2 |
| ENSMUSG( | 15114.17 | 0.050175 | 0.811883 | 0.979148 | Trank1 |
| ENSMUSG( | 2071.212 | -0.22404 | 0.057054 | 0.723362 | Trap1 |
| ENSMUSG( | 620.6488 | -0.33832 | 0.019626 | 0.556192 | Trappc1 |
| ENSMUSG( | 3265.733 | 0.131463 | 0.230367 | 0.868507 | Trappc10 |
| ENSMUSG( | 1339.984 | 0.052944 | 0.630997 | 0.953251 | Trappc11 |
| ENSMUSG( | 1056.848 | -0.10162 | 0.472827 | 0.923329 | Trappc12 |
| ENSMUSG( | 756.1487 | 0.069504 | 0.583163 | 0.944266 | Trappc13 |
| ENSMUSG( | 1398.042 | 0.053851 | 0.712746 | 0.96439 | Trappc14 |
| ENSMUSG( | 591.9624 | -0.15366 | 0.30994 | 0.898538 | Trappc2 |
| ENSMUSG( | 241.6658 | -0.01128 | 0.953381 | 0.994798 | Trappc2b |
| ENSMUSG( | 770.6641 | -0.19331 | 0.168833 | 0.840222 | Trappc2l |
| ENSMUSG( | 1482.262 | -0.06134 | 0.605809 | 0.949445 | Trappc3 |
| ENSMUSG( | 891.2054 | -0.14368 | 0.23626 | 0.872173 | Trappc4 |
| ENSMUSG( | 820.0419 | -0.08056 | 0.498806 | 0.927825 | Trappc5 |
| ENSMUSG( | 253.3853 | -0.21098 | 0.261759 | 0.884926 | Trappc6a |
| ENSMUSG( | 1603.73 | -0.07859 | 0.564278 | 0.941206 | Trappc6b |
| ENSMUSG( | 1682.142 | 0.042846 | 0.697624 | 0.960867 | Trappc8 |
| ENSMUSG( | 1814.472 | -0.1566 | 0.232309 | 0.868507 | Trappc9 |
| ENSMUSG( | 16.10575 | -0.0911 | 0.850185 | 0.985541 | Trat1 |
| ENSMUSG( | 209.2731 | -0.2889 | 0.135942 | 0.826837 | Trbc2 |
| ENSMUSG( | 158.6309 | -0.21015 | 0.24909 | 0.877904 | Trdmt1 |
| ENSMUSG( | 941.0491 | 0.005192 | 0.977613 | 0.996416 | Trem2 |
| ENSMUSG( | 8.493739 | -0.90801 | 0.184466 | NA | Treml1 |
| ENSMUSG( | 33.12185 | -0.20573 | 0.563475 | 0.941206 | Treml2 |
| ENSMUSG( | 23.45106 | -1.64735 | 0.018217 | 0.542424 | Treml4 |
| ENSMUSG( | 1468.735 | 0.191781 | 0.13775 | 0.828492 | Trerf1 |
| ENSMUSG( | 10472.27 | 0.046379 | 0.805232 | 0.977427 | Trf |
| ENSMUSG( | 150.6367 | -0.22122 | 0.708752 | 0.964042 | Trh |
| ENSMUSG( | 673.0823 | 0.185727 | 0.426879 | 0.917257 | Trhde |
| ENSMUSG( | 81.02267 | 0.464658 | 0.086945 | 0.782278 | Trhr |
| ENSMUSG( | 37.9574 | -0.0648 | 0.865761 | 0.987041 | Trhr2 |
| ENSMUSG( | 317.5277 | 0.073016 | 0.638076 | 0.953903 | Triap1 |
| ENSMUSG( | 593.1332 | -0.07344 | 0.588649 | 0.944975 | Trib1 |
| ENSMUSG( | 1679.307 | -0.04734 | 0.744451 | 0.96951 | Trib2 |
| ENSMUSG( | 14.02915 | 0.611305 | 0.260688 | 0.884675 | Trib3 |
| ENSMUSG( | 3938.443 | -0.06367 | 0.707572 | 0.964029 | Tril |
| ENSMUSG( | 24.02411 | -0.81329 | 0.141449 | 0.830678 | Trim10 |
| ENSMUSG( | 698.0829 | 0.001991 | 0.988864 | 0.997886 | Trim11 |
| ENSMUSG( | 148.2725 | 0.008985 | 0.966426 | 0.995475 | Trim12a |
| ENSMUSG( | 192.0783 | -0.25446 | 0.146909 | 0.836241 | Trim12c |
| ENSMUSG( | 211.9249 | 0.236083 | 0.25441 | 0.881624 | Trim13 |
| ENSMUSG( | 17.91409 | -0.1502 | 0.771353 | 0.974236 | Trim14 |
| ENSMUSG( | 70.43173 | 0.56731 | 0.123858 | 0.814691 | Trim16 |
| ENSMUSG( | 218.3642 | -0.01275 | 0.94356 | 0.994798 | Trim17 |
| ENSMUSG( | 4424.009 | -0.07589 | 0.495813 | 0.927825 | Trim2 |
| ENSMUSG( | 78.67608 | 0.0107 | 0.964971 | 0.995433 | Trim21 |
| ENSMUSG( | 788.6024 | 0.046717 | 0.740963 | 0.968811 | Trim23 |
| ENSMUSG( | 925.8558 | -0.09639 | 0.415865 | 0.917257 | Trim24 |
| ENSMUSG( | 373.1327 | -0.08979 | 0.572751 | 0.94262 | Trim25 |
| ENSMUSG( | 737.6289 | 0.030168 | 0.810297 | 0.978779 | Trim26 |
| ENSMUSG( | 509.5229 | 0.016276 | 0.909396 | 0.991958 | Trim27 |
| ENSMUSG( | 2568.605 | -0.02274 | 0.852041 | 0.985598 | Trim28 |
| ENSMUSG( | 18.94489 | 0.301034 | 0.526964 | 0.934449 | Trim29 |
| ENSMUSG( | 2628.218 | -0.06847 | 0.622093 | 0.953234 | Trim3 |
| ENSMUSG( | 106.7068 | 0.223232 | 0.321368 | 0.904019 | Trim30a |
| ENSMUSG( | 9.237617 | -0.06152 | 0.932085 | NA | Trim30b |
| ENSMUSG( | 67.44922 | -0.08489 | 0.753192 | 0.970622 | Trim30d |
| ENSMUSG( | 2729.209 | -0.09718 | 0.482823 | 0.924615 | Trim32 |
| ENSMUSG( | 1508.675 | 0.048206 | 0.667719 | 0.956766 | Trim33 |
| ENSMUSG( | 93.81504 | -0.12975 | 0.568712 | 0.941777 | Trim34a |
| ENSMUSG( | 2920.764 | -0.08172 | 0.477001 | 0.923329 | Trim35 |
| ENSMUSG( | 174.6594 | 0.049253 | 0.809142 | 0.97873 | Trim36 |
| ENSMUSG( | 12823.47 | -0.09467 | 0.353492 | 0.912263 | Trim37 |
| ENSMUSG( | 946.4585 | -0.00013 | 0.999315 | 0.999503 | Trim39 |
| ENSMUSG( | 2173.502 | -0.08791 | 0.454088 | 0.922444 | Trim41 |
| ENSMUSG( | 13.39853 | -0.42404 | 0.448015 | 0.922202 | Trim43a |
| ENSMUSG( | 9423.326 | 0.036014 | 0.745374 | 0.96951 | Trim44 |
| ENSMUSG( | 188.1783 | -0.28184 | 0.101436 | 0.801456 | Trim45 |
| ENSMUSG( | 2523.359 | 0.006713 | 0.957652 | 0.994798 | Trim46 |
| ENSMUSG( | 178.504 | 0.665827 | 0.006124 | 0.322135 | Trim47 |
| ENSMUSG( | 30.367 | 0.077686 | 0.835991 | 0.981237 | Trim5 |
| ENSMUSG( | 19.19214 | 0.060806 | 0.896421 | 0.990558 | Trim54 |
| ENSMUSG( | 778.5881 | -0.3593 | 0.005132 | 0.294733 | Trim56 |
| ENSMUSG( | 322.0358 | -0.00199 | 0.991119 | 0.998379 | Trim59 |
| ENSMUSG( | 11.46172 | 0.137893 | 0.816165 | 0.979198 | Trim6 |
| ENSMUSG( | 704.9954 | -0.13237 | 0.328974 | 0.906929 | Trim62 |
| ENSMUSG( | 184.6461 | 0.334428 | 0.121665 | 0.813384 | Trim65 |
| ENSMUSG( | 551.4396 | 0.006303 | 0.965979 | 0.995436 | Trim66 |
| ENSMUSG( | 213.1102 | -0.1262 | 0.432562 | 0.918256 | Trim67 |
| ENSMUSG( | 160.3317 | 0.050197 | 0.827579 | 0.980224 | Trim68 |
| ENSMUSG( | 61.52348 | 0.0611 | 0.814457 | 0.979198 | Trim7 |
| ENSMUSG( | 22.73097 | -0.37167 | 0.376802 | 0.915583 | Trim71 |

| | | | | | | | | | | | |
|---|---|---|---|---|---|---|---|---|---|---|---|
| ENSMUSG( | Trp53cor1 | 18.98068 | -0.15137 | 0.717645 | NA | ENSMUSG( | Trp53cor1 | 18.56607 | 0.031954 | 0.943839 | 0.994798 |
| ENSMUSG( | Trp53i11 | 2952.332 | -0.27427 | 0.011355 | 0.153509 | ENSMUSG( | Trp53i11 | 2929.817 | -0.01194 | 0.919594 | 0.992644 |
| ENSMUSG( | Trp53i13 | 225.3137 | -0.19608 | 0.206424 | 0.648468 | ENSMUSG( | Trp53i13 | 226.009 | 0.082274 | 0.623665 | 0.953234 |
| ENSMUSG( | Trp53inp1 | 563.1708 | 0.130933 | 0.276068 | 0.725738 | ENSMUSG( | Trp53inp1 | 500.265 | 0.059223 | 0.673231 | 0.95694 |
| ENSMUSG( | Trp53inp2 | 43814.55 | 0.060353 | 0.504141 | 0.866684 | ENSMUSG( | Trp53inp2 | 36201.86 | -0.23215 | 0.051435 | 0.711116 |
| ENSMUSG( | Trp53rka | 510.5424 | 0.042015 | 0.702285 | 0.937536 | ENSMUSG( | Trp53rka | 454.2211 | -0.03524 | 0.816172 | 0.979198 |
| ENSMUSG( | Trp53rkb | 81.28603 | 0.071559 | 0.756463 | 0.951479 | ENSMUSG( | Trp53rkb | 68.84601 | -0.14301 | 0.570487 | 0.941872 |
| ENSMUSG( | Trp63 | 61.90519 | 0.078673 | 0.757515 | 0.951513 | ENSMUSG( | Trp63 | 48.14768 | -0.42289 | 0.161475 | 0.837724 |
| ENSMUSG( | Trp73 | 42.59415 | 0.017098 | 0.955851 | 0.993874 | ENSMUSG( | Trp73 | 41.2589 | 0.224038 | 0.570278 | 0.941823 |
| ENSMUSG( | Trpc1 | 1086.572 | -0.28722 | 0.058936 | 0.37517 | ENSMUSG( | Trpc1 | 1132.299 | 0.118583 | 0.482684 | 0.924615 |
| ENSMUSG( | Trpc3 | 547.2782 | -0.19385 | 0.105798 | 0.492833 | ENSMUSG( | Trpc3 | 557.858 | 0.136676 | 0.334137 | 0.907907 |
| ENSMUSG( | Trpc4 | 470.4879 | 0.07319 | 0.629661 | 0.913965 | ENSMUSG( | Trpc4 | 496.9205 | 0.470093 | 0.003638 | 0.242109 |
| ENSMUSG( | Trpc4ap | 4502.999 | 0.133841 | 0.138477 | 0.553773 | ENSMUSG( | Trpc4ap | 3871.28 | -0.03011 | 0.777432 | 0.975871 |
| ENSMUSG( | Trpc5 | 98.06121 | -0.07042 | 0.780859 | 0.957531 | ENSMUSG( | Trpc5 | 109.4959 | 0.493185 | 0.057014 | 0.723362 |
| ENSMUSG( | Trpc6 | 220.2074 | -0.41331 | 0.04401 | 0.321685 | ENSMUSG( | Trpc6 | 214.5081 | -0.18387 | 0.401661 | 0.917257 |
| ENSMUSG( | Trpc7 | 84.85492 | -0.36834 | 0.115962 | 0.514015 | ENSMUSG( | Trpc7 | 79.76831 | -0.29179 | 0.292285 | 0.897854 |
| ENSMUSG( | Trpm1 | 15.09993 | 0.324307 | 0.507851 | NA | ENSMUSG( | Trpm1 | 13.53829 | 0.291024 | 0.577794 | 0.943131 |
| ENSMUSG( | Trpm2 | 941.5143 | -0.09579 | 0.409577 | 0.818258 | ENSMUSG( | Trpm2 | 936.4804 | 0.159652 | 0.228154 | 0.868507 |
| ENSMUSG( | Trpm3 | 6281.465 | 0.047598 | 0.679157 | 0.930983 | ENSMUSG( | Trpm3 | 5222.031 | -0.22718 | 0.073337 | 0.763629 |
| ENSMUSG( | Trpm4 | 504.1846 | -0.00884 | 0.944295 | 0.993543 | ENSMUSG( | Trpm4 | 467.754 | 0.049587 | 0.755224 | 0.971098 |
| ENSMUSG( | Trpm6 | 37.43886 | -0.43092 | 0.239675 | 0.687561 | ENSMUSG( | Trpm6 | 39.57941 | 0.026695 | 0.94388 | 0.994798 |
| ENSMUSG( | Trpm7 | 1607.243 | -0.15137 | 0.203504 | 0.643689 | ENSMUSG( | Trpm7 | 1636.147 | 0.171391 | 0.22609 | 0.868507 |
| ENSMUSG( | Trps1 | 1129.457 | -0.11631 | 0.351629 | 0.783381 | ENSMUSG( | Trps1 | 1014.004 | -0.1656 | 0.28898 | 0.895783 |
| ENSMUSG( | Trpt1 | 362.8624 | 0.09711 | 0.467432 | 0.844819 | ENSMUSG( | Trpt1 | 310.4733 | -0.08387 | 0.571739 | 0.942247 |
| ENSMUSG( | Trpv2 | 596.039 | -0.14839 | 0.225541 | 0.672427 | ENSMUSG( | Trpv2 | 595.9184 | 0.126525 | 0.387261 | 0.917257 |
| ENSMUSG( | Trpv3 | 29.98779 | -0.64192 | 0.061087 | 0.381679 | ENSMUSG( | Trpv3 | 31.85807 | -0.12105 | 0.733989 | 0.967228 |
| ENSMUSG( | Trpv4 | 87.6897 | 0.137891 | 0.602311 | 0.904887 | ENSMUSG( | Trpv4 | 88.67199 | 0.429069 | 0.125746 | 0.81516 |
| ENSMUSG( | Trpv6 | 83.80126 | 0.225047 | 0.427986 | 0.825894 | ENSMUSG( | Trpv6 | 71.21191 | 0.058771 | 0.8358 | 0.981237 |
| ENSMUSG( | Trrap | 4244.887 | -0.01046 | 0.912164 | 0.988483 | ENSMUSG( | Trrap | 4030.933 | 0.108576 | 0.413632 | 0.917257 |
| ENSMUSG( | Trub1 | 542.4714 | -0.01077 | 0.927335 | 0.991229 | ENSMUSG( | Trub1 | 510.7597 | 0.083558 | 0.505864 | 0.927825 |
| ENSMUSG( | Trub2 | 771.8607 | 0.0414 | 0.687199 | 0.933063 | ENSMUSG( | Trub2 | 673.6173 | -0.08303 | 0.498375 | 0.927825 |
| ENSMUSG( | Tsacc | 31.31727 | -0.10796 | 0.743628 | 0.948478 | ENSMUSG( | Tsacc | 28.45061 | -0.1081 | 0.770539 | 0.974236 |
| ENSMUSG( | Tsbp1 | 99.49761 | 0.46307 | 0.074931 | 0.42422 | ENSMUSG( | Tsbp1 | 79.22985 | 0.109711 | 0.72103 | 0.966887 |
| ENSMUSG( | Tsc1 | 2341.219 | 0.002223 | 0.979583 | 0.997428 | ENSMUSG( | Tsc1 | 2150.305 | 0.026981 | 0.80596 | 0.977674 |
| ENSMUSG( | Tsc2 | 4429.175 | 0.145828 | 0.085812 | 0.451465 | ENSMUSG( | Tsc2 | 3865.387 | 0.025887 | 0.803691 | 0.976978 |
| ENSMUSG( | Tsc22d1 | 8807.104 | 0.15665 | 0.110831 | 0.501952 | ENSMUSG( | Tsc22d1 | 7969.137 | 0.138123 | 0.207138 | 0.864412 |
| ENSMUSG( | Tsc22d2 | 751.0102 | 0.038918 | 0.733108 | 0.944956 | ENSMUSG( | Tsc22d2 | 718.1697 | 0.172195 | 0.165248 | 0.838743 |
| ENSMUSG( | Tsc22d3 | 3399.321 | 0.636438 | 7.44E-08 | 2.13E-05 | ENSMUSG( | Tsc22d3 | 2897.729 | 0.475608 | 0.000289 | 0.04607 |
| ENSMUSG( | Tsc22d4 | 10753.91 | 0.140982 | 0.106391 | 0.494174 | ENSMUSG( | Tsc22d4 | 9071.47 | -0.0806 | 0.518816 | 0.930939 |
| ENSMUSG( | Tsen15 | 411.2511 | -0.11764 | 0.356623 | 0.788187 | ENSMUSG( | Tsen15 | 381.8346 | -0.06175 | 0.667463 | 0.956766 |
| ENSMUSG( | Tsen2 | 386.5867 | 0.029549 | 0.820205 | 0.965888 | ENSMUSG( | Tsen2 | 355.0373 | 0.051921 | 0.726319 | 0.966887 |
| ENSMUSG( | Tsen34 | 1627.783 | 0.055633 | 0.519822 | 0.874979 | ENSMUSG( | Tsen34 | 1387.002 | -0.14291 | 0.209093 | 0.864412 |
| ENSMUSG( | Tsen54 | 217.2787 | -0.03902 | 0.837563 | 0.971111 | ENSMUSG( | Tsen54 | 203.5822 | 0.049545 | 0.776333 | 0.975717 |
| ENSMUSG( | Tsfm | 347.1976 | 0.169988 | 0.218398 | 0.664121 | ENSMUSG( | Tsfm | 290.1746 | -0.08068 | 0.62945 | 0.953251 |
| ENSMUSG( | Tsg101 | 1550.486 | 0.066657 | 0.520631 | 0.875186 | ENSMUSG( | Tsg101 | 1355.541 | -0.05573 | 0.658048 | 0.956123 |
| ENSMUSG( | Tsg101-ps | 11.71665 | 0.04055 | 0.94066 | NA | ENSMUSG( | Tsg101-ps | 10.87538 | 0.082243 | 0.887791 | NA |
| ENSMUSG( | Tsga10 | 483.0961 | 0.033857 | 0.789837 | 0.959461 | ENSMUSG( | Tsga10 | 441.3258 | 0.041074 | 0.778259 | 0.976106 |
| ENSMUSG( | Tshr | 21.5147 | -1.24408 | 0.009598 | 0.138869 | ENSMUSG( | Tshr | 28.19325 | 0.085761 | 0.843057 | 0.983211 |
| ENSMUSG( | Tshz1 | 1223.802 | -0.05204 | 0.595082 | 0.903007 | ENSMUSG( | Tshz1 | 1212.95 | 0.180696 | 0.224032 | 0.867404 |
| ENSMUSG( | Tshz2 | 5080.064 | 0.367539 | 2.67E-05 | 0.002421 | ENSMUSG( | Tshz2 | 3997.799 | -0.03245 | 0.753032 | 0.970622 |
| ENSMUSG( | Tshz3 | 703.8474 | -0.00642 | 0.964691 | 0.995155 | ENSMUSG( | Tshz3 | 631.994 | -0.05359 | 0.694943 | 0.960867 |
| ENSMUSG( | Tsks | 26.6039 | 0.355849 | 0.341354 | 0.778828 | ENSMUSG( | Tsks | 24.86853 | 0.384557 | 0.354708 | 0.912411 |
| ENSMUSG( | Tsku | 151.5043 | -0.39971 | 0.075518 | 0.425931 | ENSMUSG( | Tsku | 179.4497 | 0.372516 | 0.097758 | 0.797222 |
| ENSMUSG( | Tsn | 2852.435 | -0.05505 | 0.572389 | 0.893584 | ENSMUSG( | Tsn | 2613.31 | -0.0392 | 0.724745 | 0.966887 |
| ENSMUSG( | Tsnax | 4074.562 | 0.108144 | 0.245065 | 0.692812 | ENSMUSG( | Tsnax | 3460.477 | -0.09597 | 0.413967 | 0.917257 |
| ENSMUSG( | Tsnaxip1 | 14.49118 | -0.35915 | 0.49792 | NA | ENSMUSG( | Tsnaxip1 | 16.05863 | 0.238836 | 0.654955 | 0.955037 |
| ENSMUSG( | Tspan11 | 391.0281 | -0.24509 | 0.082358 | 0.442878 | ENSMUSG( | Tspan11 | 364.1224 | -0.16336 | 0.395308 | 0.917257 |
| ENSMUSG( | Tspan12 | 1445.828 | -0.00545 | 0.970766 | 0.996369 | ENSMUSG( | Tspan12 | 1309.351 | -0.02571 | 0.879497 | 0.989644 |
| ENSMUSG( | Tspan13 | 5309.805 | -0.50058 | 0.000581 | 0.022718 | ENSMUSG( | Tspan13 | 5969.77 | 0.147628 | 0.376729 | 0.915532 |
| ENSMUSG( | Tspan14 | 2214.5 | -0.13658 | 0.335152 | 0.776511 | ENSMUSG( | Tspan14 | 2126.156 | 0.022951 | 0.893032 | 0.990558 |
| ENSMUSG( | Tspan15 | 1317.528 | -0.3972 | 0.011948 | 0.157992 | ENSMUSG( | Tspan15 | 1372.45 | 0.02203 | 0.904504 | 0.991089 |
| ENSMUSG( | Tspan17 | 1252.263 | 0.048407 | 0.628383 | 0.913637 | ENSMUSG( | Tspan17 | 1115.684 | -0.01148 | 0.938059 | 0.993731 |
| ENSMUSG( | Tspan18 | 305.0088 | -0.35937 | 0.061353 | 0.382309 | ENSMUSG( | Tspan18 | 340.5209 | 0.251125 | 0.241347 | 0.876259 |
| ENSMUSG( | Tspan2 | 8415.604 | -0.57274 | 2.49E-05 | 0.00228 | ENSMUSG( | Tspan2 | 9278.637 | 0.036057 | 0.838676 | 0.981622 |
| ENSMUSG( | Tspan3 | 16331.97 | -0.22367 | 0.064824 | 0.394687 | ENSMUSG( | Tspan3 | 16262.85 | 0.042336 | 0.755631 | 0.971336 |
| ENSMUSG( | Tspan31 | 2842.674 | -0.34109 | 0.005692 | 0.101115 | ENSMUSG( | Tspan31 | 2880.038 | -0.01306 | 0.923008 | 0.992644 |
| ENSMUSG( | Tspan32 | 11.81441 | -0.23937 | 0.666798 | NA | ENSMUSG( | Tspan32 | 9.859046 | -0.47979 | 0.453462 | NA |
| ENSMUSG( | Tspan33 | 975.4536 | -0.23854 | 0.047547 | 0.333288 | ENSMUSG( | Tspan33 | 960.9433 | 0.000398 | 0.997797 | 0.999268 |
| ENSMUSG( | Tspan4 | 444.7257 | -0.36977 | 0.007059 | 0.115775 | ENSMUSG( | Tspan4 | 577.0448 | 0.618515 | 0.000951 | 0.105417 |
| ENSMUSG( | Tspan5 | 6149.485 | -0.2832 | 0.032583 | 0.276411 | ENSMUSG( | Tspan5 | 6431.178 | 0.129431 | 0.369644 | 0.915491 |
| ENSMUSG( | Tspan6 | 711.8286 | -0.30528 | 0.025018 | 0.243138 | ENSMUSG( | Tspan6 | 719.2995 | 0.010848 | 0.942875 | 0.994774 |
| ENSMUSG( | Tspan7 | 109216.5 | -0.17423 | 0.126336 | 0.533885 | ENSMUSG( | Tspan7 | 100675.1 | -0.13747 | 0.305766 | 0.898538 |
| ENSMUSG( | Tspan8 | 40.66659 | -0.15286 | 0.609487 | 0.9077 | ENSMUSG( | Tspan8 | 43.57619 | 0.29296 | 0.348051 | 0.911652 |
| ENSMUSG( | Tspan9 | 2673.555 | -0.28648 | 0.048287 | 0.336144 | ENSMUSG( | Tspan9 | 2427.313 | -0.29168 | 0.117922 | 0.813384 |
| ENSMUSG( | Tspo | 177.8183 | -0.00785 | 0.960846 | 0.993991 | ENSMUSG( | Tspo | 151.5377 | -0.20293 | 0.277994 | 0.891468 |
| ENSMUSG( | Tspoap1 | 18453.8 | 0.217973 | 0.092693 | 0.467791 | ENSMUSG( | Tspoap1 | 15523.92 | -0.00697 | 0.960049 | 0.994798 |
| ENSMUSG( | Tspyl1 | 2558.653 | 0.050017 | 0.677701 | 0.930542 | ENSMUSG( | Tspyl1 | 2214.428 | -0.10204 | 0.464852 | 0.923329 |
| ENSMUSG( | Tspyl2 | 2685.423 | 0.075992 | 0.552697 | 0.886141 | ENSMUSG( | Tspyl2 | 2331.826 | -0.06571 | 0.666438 | 0.956766 |
| ENSMUSG( | Tspyl3 | 489.6837 | 0.202777 | 0.159984 | 0.589679 | ENSMUSG( | Tspyl3 | 412.3729 | -0.01852 | 0.909015 | 0.991958 |
| ENSMUSG( | Tspyl4 | 7163.577 | 0.184072 | 0.038117 | 0.301297 | ENSMUSG( | Tspyl4 | 5889.251 | -0.10996 | 0.306157 | 0.898538 |
| ENSMUSG( | Tspyl5 | 649.8179 | 0.220607 | 0.135236 | 0.548658 | ENSMUSG( | Tspyl5 | 531.4122 | -0.08926 | 0.598934 | 0.947045 |
| ENSMUSG( | Tsr1 | 822.6786 | 0.126944 | 0.218293 | 0.663925 | ENSMUSG( | Tsr1 | 676.1355 | -0.17447 | 0.167878 | 0.840222 |
| ENSMUSG( | Tsr2 | 2202.883 | 0.026628 | 0.775026 | 0.955606 | ENSMUSG( | Tsr2 | 1921.202 | -0.10366 | 0.381924 | 0.917257 |
| ENSMUSG( | Tsr3 | 646.8881 | -0.06095 | 0.633042 | 0.914073 | ENSMUSG( | Tsr3 | 585.0583 | -0.08574 | 0.544884 | 0.938063 |
| ENSMUSG( | Tssc4 | 960.1669 | -0.24212 | 0.013799 | 0.171544 | ENSMUSG( | Tssc4 | 931.457 | -0.05126 | 0.650283 | 0.95422 |
| ENSMUSG( | Tssk4 | 15.00128 | -0.11851 | 0.823919 | NA | ENSMUSG( | Tssk4 | 13.96373 | -0.02714 | 0.963877 | 0.995433 |
| ENSMUSG( | Tst | 3060.526 | 0.064161 | 0.521287 | 0.875186 | ENSMUSG( | Tst | 2437.074 | -0.34698 | 0.003484 | 0.235685 |
| ENSMUSG( | Tstd1 | 18.72951 | 0.333694 | 0.436808 | NA | ENSMUSG( | Tstd1 | 17.19704 | 0.323679 | 0.556733 | 0.939565 |
| ENSMUSG( | Tstd2 | 419.3403 | -0.14026 | 0.23599 | 0.683136 | ENSMUSG( | Tstd2 | 394.9518 | -0.03921 | 0.773751 | 0.974672 |
| ENSMUSG( | Tstd3 | 360.2349 | -0.16904 | 0.209382 | 0.65137 | ENSMUSG( | Tstd3 | 311.0021 | -0.34904 | 0.04524 | 0.693959 |
| ENSMUSG( | Ttbk1 | 8961.314 | 0.31505 | 0.000585 | 0.022791 | ENSMUSG( | Ttbk1 | 7820.823 | 0.202481 | 0.099978 | 0.801311 |
| ENSMUSG( | Ttbk2 | 1722.637 | 0.026196 | 0.800284 | 0.961745 | ENSMUSG( | Ttbk2 | 1548.65 | -0.0138 | 0.914331 | 0.992407 |
| ENSMUSG( | Ttc1 | 895.8716 | 0.03046 | 0.758769 | 0.952113 | ENSMUSG( | Ttc1 | 749.6226 | -0.22544 | 0.062629 | 0.736482 |
| ENSMUSG( | Ttc12 | 140.2214 | -0.20181 | 0.296942 | 0.745676 | ENSMUSG( | Ttc12 | 136.6878 | 0.016308 | 0.941042 | 0.994558 |
| ENSMUSG( | Ttc13 | 1260.946 | -0.21765 | 0.075296 | 0.425242 | ENSMUSG( | Ttc13 | 1315.478 | 0.180047 | 0.174483 | 0.84455 |
| ENSMUSG( | Ttc14 | 2515.257 | -0.07304 | 0.560799 | 0.888934 | ENSMUSG( | Ttc14 | 2628.35 | 0.307886 | 0.061921 | 0.736482 |
| ENSMUSG( | Ttc16 | 46.59223 | 0.226833 | 0.445532 | 0.833239 | ENSMUSG( | Ttc16 | 40.75638 | 0.140418 | 0.683845 | 0.958678 |
| ENSMUSG( | Ttc17 | 1230.134 | -0.15736 | 0.253615 | 0.703909 | ENSMUSG( | Ttc17 | 1247.673 | 0.158579 | 0.343189 | 0.909646 |
| ENSMUSG( | Ttc19 | 1909.689 | -0.29926 | 0.027264 | 0.253564 | ENSMUSG( | Ttc19 | 1990.665 | 0.10625 | 0.468589 | 0.923329 |
| ENSMUSG( | Ttc21a | 56.72654 | -0.20152 | 0.603578 | 0.904887 | ENSMUSG( | Ttc21a | 63.09706 | 0.377022 | 0.260701 | 0.884675 |
| ENSMUSG( | Ttc21b | 305.4589 | 0.171073 | 0.193729 | 0.633085 | ENSMUSG( | Ttc21b | 280.5377 | 0.194222 | 0.19616 | 0.862161 |
| ENSMUSG( | Ttc22 | 28.40091 | -0.85548 | 0.019079 | 0.207968 | ENSMUSG( | Ttc22 | 30.67792 | -0.27605 | 0.466477 | 0.923329 |
| ENSMUSG( | Ttc23 | 263.9384 | 0.096365 | 0.48187 | 0.855573 | ENSMUSG( | Ttc23 | 223.0159 | -0.129 | 0.442915 | 0.921554 |
| ENSMUSG( | Ttc27 | 341.6518 | 0.103365 | 0.479227 | 0.854046 | ENSMUSG( | Ttc27 | 293.6881 | -0.07047 | 0.659375 | 0.956205 |
| ENSMUSG( | Ttc28 | 1055.092 | 0.006113 | 0.951872 | 0.993821 | ENSMUSG( | Ttc28 | 992.7012 | 0.09764 | 0.399292 | 0.917257 |
| ENSMUSG( | Ttc3 | 23001.24 | 0.059739 | 0.530462 | 0.879241 | ENSMUSG( | Ttc3 | 21039.58 | 0.072826 | 0.520154 | 0.931937 |
| ENSMUSG( | Ttc32 | 113.0657 | 0.06786 | 0.725078 | 0.944096 | ENSMUSG( | Ttc32 | 94.07762 | -0.20554 | 0.350892 | 0.911707 |
| ENSMUSG( | Ttc33 | 646.3494 | 0.0978 | 0.412336 | 0.819452 | ENSMUSG( | Ttc33 | 574.9251 | 0.025821 | 0.858369 | 0.985884 |
| ENSMUSG( | Ttc34 | 96.20924 | -0.17763 | 0.394529 | 0.810072 | ENSMUSG( | Ttc34 | 89.32396 | -0.10894 | 0.65743 | 0.955725 |
| ENSMUSG( | Ttc38 | 280.0057 | -0.04665 | 0.732459 | 0.944956 | ENSMUSG( | Ttc38 | 251.376 | -0.0873 | 0.589694 | 0.944975 |
| ENSMUSG( | Ttc39a | 45.24788 | -0.10554 | 0.72131 | 0.942988 | ENSMUSG( | Ttc39a | 40.7752 | -0.14326 | 0.649475 | 0.95422 |
| ENSMUSG( | Ttc39aos1 | 87.9344 | 0.331924 | 0.191187 | 0.630245 | ENSMUSG( | Ttc39aos1 | 71.73281 | 0.012631 | 0.966256 | 0.995475 |
| ENSMUSG( | Ttc39b | 1167.548 | -0.09194 | 0.329586 | 0.773422 | ENSMUSG( | Ttc39b | 1083.1 | -0.03553 | 0.761949 | 0.97318 |
| ENSMUSG( | Ttc39c | 588.072 | 0.035453 | 0.77269 | 0.955272 | ENSMUSG( | Ttc39c | 533.395 | 0.020231 | 0.887836 | 0.990399 |
| ENSMUSG( | Ttc4 | 1064.186 | 0.0119 | 0.913222 | 0.988483 | ENSMUSG( | Ttc4 | 990.2475 | 0.073894 | 0.613221 | 0.951177 |
| ENSMUSG( | Ttc41 | 72.76117 | 0.218055 | 0.364187 | 0.793647 | ENSMUSG( | Ttc41 | 62.57482 | 0.063285 | 0.805334 | 0.977484 |
| ENSMUSG( | Ttc5 | 1187.954 | 0.034912 | 0.70792 | 0.939519 | ENSMUSG( | Ttc5 | 1059.159 | -0.02829 | 0.802604 | 0.97681 |
| ENSMUSG( | Ttc7 | 478.0315 | -0.02517 | 0.833515 | 0.970333 | ENSMUSG( | Ttc7 | 445.2887 | 0.039006 | 0.771833 | 0.974236 |
| ENSMUSG( | Ttc7b | 8486.582 | 0.17542 | 0.027649 | 0.255168 | ENSMUSG( | Ttc7b | 7293.829 | 0.01233 | 0.897772 | 0.990558 |
| ENSMUSG( | Ttc8 | 666.8205 | 0.113306 | 0.299309 | 0.748131 | ENSMUSG( | Ttc8 | 565.4873 | -0.09462 | 0.50058 | 0.927825 |
| ENSMUSG( | Ttc9 | 1621.786 | 0.070611 | 0.482517 | 0.856002 | ENSMUSG( | Ttc9 | 1406.39 | -0.07316 | 0.540913 | 0.93758 |
| ENSMUSG( | Ttc9b | 5763.738 | 0.069986 | 0.397856 | 0.812161 | ENSMUSG( | Ttc9b | 4928.99 | -0.11621 | 0.280475 | 0.892356 |
| ENSMUSG( | Ttc9c | 582.9444 | -0.02901 | 0.794801 | 0.959799 | ENSMUSG( | Ttc9c | 534.0106 | -0.01497 | 0.905857 | 0.991089 |
| ENSMUSG( | Ttf1 | 405.8104 | 0.142571 | 0.336464 | 0.777 | ENSMUSG( | Ttf1 | 363.8434 | 0.106947 | 0.535947 | 0.936952 |
| ENSMUSG( | Ttf2 | 164.8156 | 0.227594 | 0.203216 | 0.643631 | ENSMUSG( | Ttf2 | 135.0012 | -0.06861 | 0.734349 | 0.967228 |
| ENSMUSG( | Tti1 | 502.9245 | 0.070995 | 0.522254 | 0.875186 | ENSMUSG( | Tti1 | 452.0553 | 0.032147 | 0.810455 | 0.978779 |
| ENSMUSG( | Tti2 | 426.1784 | -0.09416 | 0.470119 | 0.846694 | ENSMUSG( | Tti2 | 393.9414 | -0.04567 | 0.758479 | 0.971858 |
| ENSMUSG( | Ttk | 19.00401 | 0.222599 | 0.655518 | NA | ENSMUSG( | Ttk | 16.7071 | 0.116558 | 0.814993 | 0.979198 |
| ENSMUSG( | Ttl | 3198.789 | 0.01415 | 0.870801 | 0.977951 | ENSMUSG( | Ttl | 2711.493 | -0.20259 | 0.062401 | 0.736482 |
| ENSMUSG( | Ttll1 | 740.6954 | 0.016762 | 0.894722 | 0.984512 | ENSMUSG( | Ttll1 | 633.0923 | -0.17821 | 0.266389 | 0.886134 |
| ENSMUSG( | Ttll11 | 1022.347 | 0.093737 | 0.362564 | 0.792943 | ENSMUSG( | Ttll11 | 874.302 | -0.0872 | 0.496986 | 0.927825 |
| ENSMUSG( | Ttll12 | 755.2967 | 0.269523 | 0.016933 | 0.194376 | ENSMUSG( | Ttll12 | 606.4691 | -0.08085 | 0.536382 | 0.93705 |
| ENSMUSG( | Ttll13 | 55.72613 | 0.14951 | 0.560099 | 0.888934 | ENSMUSG( | Ttll13 | 49.70765 | 0.071045 | 0.809155 | 0.97873 |
| ENSMUSG( | Ttll3 | 526.582 | 0.23429 | 0.055027 | 0.36102 | ENSMUSG( | Ttll3 | 468.0397 | 0.1706 | 0.29258 | 0.897982 |
| ENSMUSG( | Ttll4 | 819.4062 | 0.003019 | 0.977725 | 0.997428 | ENSMUSG( | Ttll4 | 797.5115 | 0.189369 | 0.136435 | 0.827092 |
| ENSMUSG( | Ttll5 | 1377.99 | 0.098408 | 0.27652 | 0.726576 | ENSMUSG( | Ttll5 | 1183.416 | -0.07353 | 0.519746 | 0.931729 |
| ENSMUSG( | Ttll6 | 31.31702 | 0.279007 | 0.49224 | 0.860297 | ENSMUSG( | Ttll6 | 22.89333 | -0.36419 | 0.403999 | 0.917257 |
| ENSMUSG( | Ttll7 | 4412.993 | -0.00094 | 0.991877 | 0.999232 | ENSMUSG( | Ttll7 | 3965.992 | -0.043 | 0.709266 | 0.964042 |
| ENSMUSG( | Ttll8 | 70.02563 | 0.011114 | 0.964675 | 0.995155 | ENSMUSG( | Ttll8 | 71.24039 | 0.318206 | 0.250366 | 0.87792 |
| ENSMUSG( | Ttll9 | 45.44423 | 0.237438 | 0.445711 | 0.833239 | ENSMUSG( | Ttll9 | 42.04212 | 0.276764 | 0.410787 | 0.917257 |
| ENSMUSG( | Ttn | 32.89421 | -1.2673 | 0.000173 | 0.00947 | ENSMUSG( | Ttn | 39.21914 | -0.24592 | 0.442512 | 0.921554 |
| ENSMUSG( | Ttpa | 522.8447 | -0.0304 | 0.879279 | 0.981436 | ENSMUSG( | Ttpa | 468.9806 | -0.08483 | 0.680798 | 0.958395 |
| ENSMUSG( | Ttpal | 1667.882 | -0.03289 | 0.728719 | 0.944393 | ENSMUSG( | Ttpal | 1481.159 | -0.10697 | 0.338395 | 0.907972 |
| ENSMUSG( | Ttr | 12964.4 | -0.07646 | 0.943935 | 0.993503 | ENSMUSG( | Ttr | 15441.47 | 0.617921 | 0.534504 | 0.936104 |
| ENSMUSG( | Ttyh1 | 64018.02 | 0.033955 | 0.727246 | 0.944223 | ENSMUSG( | Ttyh1 | 57143.9 | -0.02452 | 0.823605 | 0.979198 |
| ENSMUSG( | Ttyh2 | 1602.417 | -0.6915 | 3.19E-05 | 0.002843 | ENSMUSG( | Ttyh2 | 1954.139 | 0.229095 | 0.253089 | 0.88018 |
| ENSMUSG( | Ttyh3 | 17516.33 | -0.06159 | 0.598286 | 0.903801 | ENSMUSG( | Ttyh3 | 17614.08 | 0.218491 | 0.112041 | 0.808673 |
| ENSMUSG( | Tub | 6515.32 | 0.235108 | 0.015851 | 0.186421 | ENSMUSG( | Tub | 5928.519 | 0.23275 | 0.037638 | 0.674819 |
| ENSMUSG( | Tuba1a | 19507.45 | -0.03368 | 0.654072 | 0.923008 | ENSMUSG( | Tuba1a | 16713.77 | -0.21951 | 0.07145 | 0.761942 |
| ENSMUSG( | Tuba1b | 6321.689 | 0.063779 | 0.460481 | 0.840787 | ENSMUSG( | Tuba1b | 5227.106 | -0.22629 | 0.070039 | 0.757435 |
| ENSMUSG( | Tuba1c | 73.84744 | -0.00351 | 0.989216 | 0.999111 | ENSMUSG( | Tuba1c | 66.0493 | -0.08331 | 0.783422 | 0.976106 |
| ENSMUSG( | Tuba4a | 11043.88 | 0.064224 | 0.550613 | 0.886038 | ENSMUSG( | Tuba4a | 8987.513 | -0.27579 | 0.04837 | 0.706142 |
| ENSMUSG( | Tuba8 | 177.3636 | 0.112219 | 0.621431 | 0.911432 | ENSMUSG( | Tuba8 | 139.3897 | -0.34786 | 0.20777 | 0.864412 |
| ENSMUSG( | Tubb2a | 9978.394 | 0.121865 | 0.272117 | 0.7223 | ENSMUSG( | Tubb2a | 8115.523 | -0.21358 | 0.177092 | 0.846973 |
| ENSMUSG( | Tubb2b | 15006.72 | 0.02221 | 0.859907 | 0.974818 | ENSMUSG( | Tubb2b | 12024.1 | -0.37428 | 0.009125 | 0.401114 |
| ENSMUSG( | Tubb3 | 6740.902 | -0.08718 | 0.555803 | 0.887457 | ENSMUSG( | Tubb3 | 5764.308 | -0.28679 | 0.130071 | 0.821082 |
| ENSMUSG( | Tubb4a | 19548.25 | -0.14622 | 0.275114 | 0.72509 | ENSMUSG( | Tubb4a | 17268.13 | -0.24502 | 0.167437 | 0.840222 |
| ENSMUSG( | Tubb4b | 6387.989 | 0.026914 | 0.785984 | 0.958774 | ENSMUSG( | Tubb4b | 5344.34 | -0.23051 | 0.07537 | 0.767994 |

| | | | | | |
|---|---|---|---|---|---|
| ENSMUSG( | 144.3757 | 0.131189 | 0.454187 | 0.837055 | Tpcn2 |
| ENSMUSG( | 1914.456 | -0.07443 | 0.462355 | 0.842104 | Tpd52 |
| ENSMUSG( | 445.5048 | -0.06339 | 0.589636 | 0.90113 | Tpd52l1 |
| ENSMUSG( | 1596.193 | 0.028916 | 0.745093 | 0.949158 | Tpd52l2 |
| ENSMUSG( | 1156.043 | -0.00083 | 0.993968 | 0.999369 | Tpgs1 |
| ENSMUSG( | 1413.037 | 0.081662 | 0.423243 | 0.823662 | Tpgs2 |
| ENSMUSG( | 124.6271 | -0.27567 | 0.213303 | 0.657626 | Tph2 |
| ENSMUSG( | 12359.55 | 0.0466 | 0.605097 | 0.905594 | Tpi1 |
| ENSMUSG( | 77.11349 | -0.21911 | 0.338001 | 0.777422 | Tpk1 |
| ENSMUSG( | 3860.947 | -0.02019 | 0.845432 | 0.97218 | Tpm1 |
| ENSMUSG( | 167.8259 | -0.2542 | 0.150663 | 0.575407 | Tpm2 |
| ENSMUSG( | 3667.807 | 0.040847 | 0.629977 | 0.913965 | Tpm3 |
| ENSMUSG( | 24.27266 | 0.423191 | 0.293097 | 0.742763 | Tpm3-rs7 |
| ENSMUSG( | 1881.608 | 0.136729 | 0.266509 | 0.716441 | Tpm4 |
| ENSMUSG( | 1104.623 | 0.022025 | 0.874946 | 0.979746 | Tpmt |
| ENSMUSG( | 10.57239 | -0.26119 | 0.677965 | NA | Tpo |
| ENSMUSG( | 3737.221 | -0.15192 | 0.212027 | 0.655687 | Tpp1 |
| ENSMUSG( | 964.01 | 0.014187 | 0.883839 | 0.982647 | Tpp2 |
| ENSMUSG( | 20652.85 | 0.119716 | 0.113442 | 0.508013 | Tppp |
| ENSMUSG( | 967.2098 | -0.00257 | 0.980542 | 0.997512 | Tppp3 |
| ENSMUSG( | 3062.372 | 0.198831 | 0.037603 | 0.299568 | Tpr |
| ENSMUSG( | 931.5655 | -0.12687 | 0.34461 | 0.779153 | Tpra1 |
| ENSMUSG( | 6924.672 | 0.087175 | 0.283381 | 0.73186 | Tprg1l |
| ENSMUSG( | 5540.846 | 0.082759 | 0.52677 | 0.877267 | Tprkb |
| ENSMUSG( | 631.7621 | -0.4067 | 0.000266 | 0.013312 | Tprn |
| ENSMUSG( | 1202.598 | -0.22332 | 0.083872 | 0.446475 | Tpst1 |
| ENSMUSG( | 1097.706 | -0.16875 | 0.157464 | 0.585653 | Tpst2 |
| ENSMUSG( | 25881.24 | 0.12335 | 0.196186 | 0.635538 | Tpt1 |
| ENSMUSG( | 96.86454 | 0.129096 | 0.575497 | 0.894836 | Tpx2 |
| ENSMUSG( | 1912.281 | 0.155108 | 0.183106 | 0.62047 | Tra2a |
| ENSMUSG( | 1261.108 | -0.04301 | 0.66118 | 0.92507 | Tra2b |
| ENSMUSG( | 686.0255 | -0.01686 | 0.883902 | 0.982647 | Trabd |
| ENSMUSG( | 358.6167 | 0.566908 | 1.77E-05 | 0.001786 | Trabd2b |
| ENSMUSG( | 167.641 | 0.059889 | 0.729393 | 0.944738 | Tradd |
| ENSMUSG( | 63.09834 | 0.04059 | 0.864135 | 0.975331 | Traf1 |
| ENSMUSG( | 267.8157 | 0.027427 | 0.836561 | 0.971111 | Traf2 |
| ENSMUSG( | 7114.629 | 0.364174 | 0.000108 | 0.006836 | Traf3 |
| ENSMUSG( | 496.0954 | 0.124141 | 0.311029 | 0.759201 | Traf3ip1 |
| ENSMUSG( | 34.97898 | -0.18383 | 0.554448 | 0.886657 | Traf3ip2 |
| ENSMUSG( | 49.2679 | -0.04336 | 0.885892 | 0.983008 | Traf3ip3 |
| ENSMUSG( | 570.9473 | -0.20838 | 0.090501 | 0.461757 | Traf4 |
| ENSMUSG( | 86.97187 | -0.19067 | 0.387154 | 0.805442 | Traf5 |
| ENSMUSG( | 643.8057 | -0.10615 | 0.331883 | 0.775288 | Traf6 |
| ENSMUSG( | 1852.775 | 0.127693 | 0.223915 | 0.669845 | Traf7 |
| ENSMUSG( | 1721.501 | 0.057466 | 0.617207 | 0.910866 | Trafd1 |
| ENSMUSG( | 633.3302 | -0.01076 | 0.952558 | 0.993874 | Traip |
| ENSMUSG( | 10926.56 | 0.192742 | 0.016684 | 0.192198 | Trak1 |
| ENSMUSG( | 36139.13 | 0.007623 | 0.948799 | 0.993821 | Trak2 |
| ENSMUSG( | 1087.909 | -0.23654 | 0.045865 | 0.328083 | Tram1 |
| ENSMUSG( | 1003.399 | -0.59441 | 0.000293 | 0.014095 | Tram1l1 |
| ENSMUSG( | 76.40281 | 0.114464 | 0.676236 | 0.930542 | Tram2 |
| ENSMUSG( | 13543.93 | -0.58955 | 0.001451 | 0.042014 | Trank1 |
| ENSMUSG( | 2491.805 | 0.051621 | 0.60295 | 0.904887 | Trap1 |
| ENSMUSG( | 726.7355 | -0.13019 | 0.288016 | 0.738882 | Trappc1 |
| ENSMUSG( | 3634.551 | 0.16756 | 0.05493 | 0.360715 | Trappc10 |
| ENSMUSG( | 1493.904 | 0.096038 | 0.263586 | 0.714284 | Trappc11 |
| ENSMUSG( | 1247.503 | 0.109022 | 0.376677 | 0.801213 | Trappc12 |
| ENSMUSG( | 830.4125 | 0.075361 | 0.469686 | 0.846186 | Trappc13 |
| ENSMUSG( | 1565.597 | 0.112282 | 0.399561 | 0.813178 | Trappc14 |
| ENSMUSG( | 683.411 | 0.004534 | 0.968633 | 0.996369 | Trappc2 |
| ENSMUSG( | 269.7658 | 0.052124 | 0.71782 | 0.941697 | Trappc2b |
| ENSMUSG( | 874.1364 | -0.08948 | 0.420048 | 0.82245 | Trappc2l |
| ENSMUSG( | 1643.88 | -0.02793 | 0.757154 | 0.951513 | Trappc3 |
| ENSMUSG( | 1028.377 | 0.007891 | 0.934886 | 0.993231 | Trappc4 |
| ENSMUSG( | 878.0955 | -0.1542 | 0.144472 | 0.564381 | Trappc5 |
| ENSMUSG( | 287.6083 | -0.09765 | 0.516306 | 0.872701 | Trappc6a |
| ENSMUSG( | 1783.047 | -0.03674 | 0.674587 | 0.929638 | Trappc6b |
| ENSMUSG( | 1902.233 | 0.128411 | 0.174672 | 0.610304 | Trappc8 |
| ENSMUSG( | 2160.922 | 0.082321 | 0.474677 | 0.849485 | Trappc9 |
| ENSMUSG( | 11.27029 | 0.579179 | 0.357099 | NA | Trarg1 |
| ENSMUSG( | 15.99858 | -0.4369 | 0.364099 | NA | Trat1 |
| ENSMUSG( | 251.5566 | -0.01454 | 0.938061 | 0.993415 | Trbc2 |
| ENSMUSG( | 177.4652 | -0.14584 | 0.371699 | 0.798768 | Trdmt1 |
| ENSMUSG( | 919.9415 | -0.3573 | 0.023784 | 0.235641 | Trem2 |
| ENSMUSG( | 9.846839 | -1.14383 | 0.09606 | NA | Trem6l |
| ENSMUSG( | 9.255316 | -1.01887 | 0.098283 | NA | Treml1 |
| ENSMUSG( | 33.30377 | -0.52706 | 0.11169 | 0.503786 | Treml2 |
| ENSMUSG( | 24.01 | -2.00112 | 0.005215 | 0.096233 | Treml4 |
| ENSMUSG( | 1547.938 | 0.081577 | 0.443535 | 0.833108 | Trerf1 |
| ENSMUSG( | 9636.037 | -0.51147 | 0.000532 | 0.021434 | Trf |
| ENSMUSG( | 132.9868 | -0.99375 | 0.094221 | 0.471647 | Trh |
| ENSMUSG( | 604.459 | -0.42828 | 0.030665 | 0.266555 | Trhde |
| ENSMUSG( | 71.05024 | -0.11867 | 0.624053 | 0.91187 | Trhr |
| ENSMUSG( | 37.44763 | -0.44104 | 0.194606 | 0.633457 | Trhr2 |
| ENSMUSG( | 362.7045 | 0.184726 | 0.167034 | 0.599655 | Triap1 |
| ENSMUSG( | 606.7748 | -0.29572 | 0.025597 | 0.246694 | Trib1 |
| ENSMUSG( | 1789.294 | -0.13088 | 0.238443 | 0.686308 | Trib2 |
| ENSMUSG( | 12.14063 | 0.023083 | 0.966548 | NA | Trib3 |
| ENSMUSG( | 4088.989 | -0.23731 | 0.097192 | 0.47759 | Tril |
| ENSMUSG( | 22.60445 | -1.48335 | 0.011004 | 0.151333 | Trim10 |
| ENSMUSG( | 716.8949 | -0.19938 | 0.124733 | 0.531714 | Trim11 |
| ENSMUSG( | 150.048 | -0.24615 | 0.173713 | 0.609084 | Trim12a |
| ENSMUSG( | 240.4938 | 0.135767 | 0.425129 | 0.82478 | Trim12c |
| ENSMUSG( | 181.06 | -0.54013 | 0.007752 | 0.1222 | Trim13 |
| ENSMUSG( | 20.33043 | -0.06055 | 0.886064 | NA | Trim14 |
| ENSMUSG( | 74.979 | 0.46713 | 0.105398 | 0.492417 | Trim16 |
| ENSMUSG( | 258.3395 | 0.204277 | 0.216196 | 0.661017 | Trim17 |
| ENSMUSG( | 5073.491 | 0.052474 | 0.499109 | 0.862973 | Trim2 |
| ENSMUSG( | 84.91676 | -0.05691 | 0.798998 | 0.960922 | Trim21 |
| ENSMUSG( | 850.2359 | -0.00021 | 0.998372 | 0.99989 | Trim23 |
| ENSMUSG( | 1022.505 | -0.07932 | 0.417936 | 0.821964 | Trim24 |
| ENSMUSG( | 432.9578 | 0.065119 | 0.675857 | 0.930515 | Trim25 |
| ENSMUSG( | 768.2918 | -0.1278 | 0.232463 | 0.680606 | Trim26 |
| ENSMUSG( | 580.612 | 0.123832 | 0.344451 | 0.779153 | Trim27 |
| ENSMUSG( | 2966.452 | 0.122998 | 0.269142 | 0.719307 | Trim28 |
| ENSMUSG( | 19.95893 | 0.210761 | 0.657608 | NA | Trim29 |
| ENSMUSG( | 2962.456 | 0.00972 | 0.944037 | 0.993503 | Trim3 |
| ENSMUSG( | 109.1775 | 0.020171 | 0.924048 | 0.991077 | Trim30a |
| ENSMUSG( | 74.33721 | -0.08795 | 0.708617 | 0.939519 | Trim30d |
| ENSMUSG( | 3094.906 | -0.00042 | 0.997366 | 0.999605 | Trim32 |
| ENSMUSG( | 1651.857 | 0.042977 | 0.649966 | 0.920899 | Trim33 |
| ENSMUSG( | 102.5511 | -0.13668 | 0.534697 | 0.880719 | Trim34a |
| ENSMUSG( | 3379.747 | 0.072005 | 0.438586 | 0.83067 | Trim35 |
| ENSMUSG( | 167.3625 | -0.38424 | 0.024364 | 0.238636 | Trim36 |
| ENSMUSG( | 15309.61 | 0.146653 | 0.073626 | 0.41961 | Trim37 |
| ENSMUSG( | 1016.485 | -0.06612 | 0.611048 | 0.908534 | Trim39 |
| ENSMUSG( | 2456.098 | -0.00067 | 0.99472 | 0.999369 | Trim41 |
| ENSMUSG( | 16.2334 | -0.13313 | 0.786032 | NA | Trim43a |
| ENSMUSG( | 10580.81 | 0.100042 | 0.239512 | 0.687308 | Trim44 |
| ENSMUSG( | 224.8394 | -0.03191 | 0.822325 | 0.966352 | Trim45 |
| ENSMUSG( | 2916.592 | 0.152009 | 0.161311 | 0.591331 | Trim46 |
| ENSMUSG( | 167.0392 | 0.253868 | 0.188272 | 0.627268 | Trim47 |
| ENSMUSG( | 28.72072 | -0.39931 | 0.279935 | 0.729155 | Trim5 |
| ENSMUSG( | 21.35488 | 0.137026 | 0.7928 | 0.95947 | Trim54 |
| ENSMUSG( | 971.7018 | 0.035324 | 0.797318 | 0.960507 | Trim56 |
| ENSMUSG( | 304.7082 | -0.47179 | 0.001991 | 0.051331 | Trim59 |
| ENSMUSG( | 12.54245 | 0.096053 | 0.865094 | NA | Trim6 |
| ENSMUSG( | 796.499 | -0.04366 | 0.700653 | 0.93743 | Trim62 |
| ENSMUSG( | 182.9019 | 0.041094 | 0.840081 | 0.971179 | Trim65 |
| ENSMUSG( | 651.3718 | 0.204877 | 0.118984 | 0.522304 | Trim66 |
| ENSMUSG( | 244.4612 | 0.000746 | 0.995823 | 0.99943 | Trim67 |
| ENSMUSG( | 164.2521 | -0.15528 | 0.404395 | 0.816132 | Trim68 |
| ENSMUSG( | 65.08048 | -0.05518 | 0.815145 | 0.964978 | Trim7 |
| ENSMUSG( | 24.13394 | -0.50718 | 0.186583 | 0.623869 | Trim71 |
| ENSMUSG( | 30.96301 | 0.227318 | 0.493522 | 0.860611 | Trim72 |
| ENSMUSG( | 4405.193 | 0.139909 | 0.104453 | 0.490485 | Trim8 |
| ENSMUSG( | 11640.63 | 0.103133 | 0.159023 | 0.588112 | Trim9 |
| ENSMUSG( | 5182.78 | 0.017547 | 0.844842 | 0.972023 | Trio |
| ENSMUSG( | 830.7504 | -0.00111 | 0.990902 | 0.999232 | Triobp |
| ENSMUSG( | 259.393 | 0.216185 | 0.128465 | 0.536774 | Trip10 |
| ENSMUSG( | 1464.764 | 0.092125 | 0.387734 | 0.805835 | Trip11 |
| ENSMUSG( | 4836.031 | 0.102937 | 0.209698 | 0.651841 | Trip12 |
| ENSMUSG( | 18.32723 | -0.30891 | 0.469212 | NA | Trip13 |
| ENSMUSG( | 1211.482 | 0.11909 | 0.231595 | 0.67949 | Trip4 |
| ENSMUSG( | 260.2593 | -0.00315 | 0.981703 | 0.997512 | Trip6 |
| ENSMUSG( | 303.8246 | 0.007809 | 0.960009 | 0.993991 | Triqk |
| ENSMUSG( | 1840.535 | 0.01261 | 0.9217 | 0.990436 | Trir |
| ENSMUSG( | 365.5762 | 0.067254 | 0.626524 | 0.91272 | Trit1 |
| ENSMUSG( | 72.16338 | -0.13123 | 0.591036 | 0.90113 | Trmo |
| ENSMUSG( | 2695.269 | 0.213235 | 0.059088 | 0.375389 | Trmt1 |
| ENSMUSG( | 217.6143 | 0.003457 | 0.980794 | 0.997512 | Trmt10a |
| ENSMUSG( | 221.8315 | 0.139589 | 0.376203 | 0.801127 | Trmt10b |
| ENSMUSG( | 174.5179 | -0.16927 | 0.409632 | 0.818258 | Trmt10c |
| ENSMUSG( | 173.302 | -0.08471 | 0.609319 | 0.907627 | Trmt11 |
| ENSMUSG( | 419.8923 | 0.136178 | 0.296109 | 0.744386 | Trmt112 |
| ENSMUSG( | 172.3865 | 0.102746 | 0.565084 | 0.890735 | Trmt12 |
| ENSMUSG( | 253.1774 | -0.0553 | 0.704212 | 0.938221 | Trmt13 |
| ENSMUSG( | 662.2294 | 0.169817 | 0.135193 | 0.548658 | Trmt1l |

| | | | | | |
|---|---|---|---|---|---|
| ENSMUSG( | 26.57297 | 0.068234 | 0.85672 | 0.985884 | Trim72 |
| ENSMUSG( | 4032.592 | 0.151322 | 0.216987 | 0.865589 | Trim8 |
| ENSMUSG( | 9473.301 | -0.23086 | 0.030257 | 0.623534 | Trim9 |
| ENSMUSG( | 4839.866 | 0.088859 | 0.443958 | 0.921734 | Trio |
| ENSMUSG( | 757.8503 | 0.003831 | 0.975553 | 0.996416 | Triobp |
| ENSMUSG( | 245.2957 | 0.32311 | 0.057167 | 0.723362 | Trip10 |
| ENSMUSG( | 1317.292 | 0.05709 | 0.640591 | 0.953903 | Trip11 |
| ENSMUSG( | 4293.803 | 0.029407 | 0.786403 | 0.976303 | Trip12 |
| ENSMUSG( | 16.95402 | -0.27388 | 0.566434 | 0.941745 | Trip13 |
| ENSMUSG( | 1059.775 | 0.000916 | 0.993944 | 0.998813 | Trip4 |
| ENSMUSG( | 241.3501 | 0.04551 | 0.814843 | 0.979198 | Trip6 |
| ENSMUSG( | 260.0869 | -0.19315 | 0.314996 | 0.901306 | Triqk |
| ENSMUSG( | 1642.344 | -0.05203 | 0.727742 | 0.966887 | Trir |
| ENSMUSG( | 336.9168 | 0.107706 | 0.487997 | 0.925453 | Trit1 |
| ENSMUSG( | 65.87742 | -0.14004 | 0.626951 | 0.953234 | Trmo |
| ENSMUSG( | 2349.24 | 0.091963 | 0.479816 | 0.924025 | Trmt1 |
| ENSMUSG( | 194.4016 | -0.06383 | 0.716994 | 0.965864 | Trmt10a |
| ENSMUSG( | 199.0685 | 0.097748 | 0.582476 | 0.944266 | Trmt10b |
| ENSMUSG( | 168.6658 | 0.008662 | 0.965861 | 0.995433 | Trmt10c |
| ENSMUSG( | 154.8752 | -0.14581 | 0.464802 | 0.923329 | Trmt11 |
| ENSMUSG( | 355.8299 | -0.07744 | 0.610288 | 0.950771 | Trmt112 |
| ENSMUSG( | 152.563 | 0.012485 | 0.952442 | 0.994798 | Trmt12 |
| ENSMUSG( | 248.9474 | 0.174461 | 0.336437 | 0.907972 | Trmt13 |
| ENSMUSG( | 575.6958 | 0.03907 | 0.772 | 0.974236 | Trmt1l |
| ENSMUSG( | 965.2859 | 0.039588 | 0.755902 | 0.971371 | Trmt2a |
| ENSMUSG( | 481.8282 | -0.25587 | 0.122287 | 0.813384 | Trmt2b |
| ENSMUSG( | 133.3244 | 0.244184 | 0.228777 | 0.868507 | Trmt44 |
| ENSMUSG( | 154.7078 | 0.058274 | 0.758449 | 0.971858 | Trmt5 |
| ENSMUSG( | 625.6504 | -0.06928 | 0.582913 | 0.944266 | Trmt6 |
| ENSMUSG( | 279.7578 | -0.12907 | 0.492279 | 0.927315 | Trmt61a |
| ENSMUSG( | 194.0692 | -0.08003 | 0.662078 | 0.956205 | Trmt61b |
| ENSMUSG( | 298.7117 | -0.53708 | 0.001422 | 0.139194 | Trmt9b |
| ENSMUSG( | 410.5788 | 0.14517 | 0.368533 | 0.915491 | Trmu |
| ENSMUSG( | 429.2178 | -0.14108 | 0.299289 | 0.898538 | Trnau1ap |
| ENSMUSG( | 7900.233 | -0.03398 | 0.740662 | 0.968811 | Trnp1 |
| ENSMUSG( | 569.7345 | 0.217217 | 0.09429 | 0.795735 | Trnt1 |
| ENSMUSG( | 3724.82 | 0.13998 | 0.241921 | 0.876528 | Tro |
| ENSMUSG( | 11.53995 | -0.23667 | 0.679831 | 0.957966 | Troap |
| ENSMUSG( | 539.6509 | -0.07469 | 0.595471 | 0.945853 | Trp53 |
| ENSMUSG( | 1939.768 | -0.00752 | 0.94591 | 0.994798 | Trp53bp1 |
| ENSMUSG( | 1955.511 | -0.1104 | 0.364525 | 0.914223 | Trp53bp2 |
| ENSMUSG( | 18.56607 | 0.031954 | 0.943839 | 0.994798 | Trp53cor1 |
| ENSMUSG( | 2929.817 | -0.01194 | 0.919594 | 0.992644 | Trp53i11 |
| ENSMUSG( | 226.009 | 0.082274 | 0.623665 | 0.953234 | Trp53i13 |
| ENSMUSG( | 500.265 | 0.059223 | 0.673231 | 0.95694 | Trp53inp1 |
| ENSMUSG( | 36201.86 | -0.23215 | 0.051435 | 0.711116 | Trp53inp2 |
| ENSMUSG( | 454.2211 | -0.03524 | 0.816172 | 0.979198 | Trp53rka |
| ENSMUSG( | 68.84601 | -0.14301 | 0.570487 | 0.941872 | Trp53rkb |
| ENSMUSG( | 48.14768 | -0.42289 | 0.161475 | 0.837724 | Trp63 |
| ENSMUSG( | 41.2589 | 0.224038 | 0.570278 | 0.941823 | Trp73 |
| ENSMUSG( | 1132.299 | 0.118583 | 0.482684 | 0.924615 | Trpc1 |
| ENSMUSG( | 557.858 | 0.136676 | 0.334137 | 0.907907 | Trpc3 |
| ENSMUSG( | 496.9205 | 0.470093 | 0.003638 | 0.242109 | Trpc4 |
| ENSMUSG( | 3871.28 | -0.03011 | 0.777432 | 0.975871 | Trpc4ap |
| ENSMUSG( | 109.4959 | 0.493185 | 0.057014 | 0.723362 | Trpc5 |
| ENSMUSG( | 214.5081 | -0.18387 | 0.401661 | 0.917257 | Trpc6 |
| ENSMUSG( | 79.76831 | -0.29179 | 0.292285 | 0.897854 | Trpc7 |
| ENSMUSG( | 13.53829 | 0.291024 | 0.577794 | 0.943131 | Trpm1 |
| ENSMUSG( | 936.4804 | 0.159652 | 0.228154 | 0.868507 | Trpm2 |
| ENSMUSG( | 5222.031 | -0.22718 | 0.073337 | 0.763629 | Trpm3 |
| ENSMUSG( | 467.754 | 0.049587 | 0.755224 | 0.971098 | Trpm4 |
| ENSMUSG( | 39.57941 | 0.026695 | 0.94388 | 0.994798 | Trpm6 |
| ENSMUSG( | 1636.147 | 0.171391 | 0.22609 | 0.868507 | Trpm7 |
| ENSMUSG( | 1014.004 | -0.1656 | 0.28898 | 0.895783 | Trps1 |
| ENSMUSG( | 310.4733 | -0.08387 | 0.571739 | 0.942247 | Trpt1 |
| ENSMUSG( | 595.9184 | 0.126525 | 0.387261 | 0.917257 | Trpv2 |
| ENSMUSG( | 31.85807 | -0.12105 | 0.733989 | 0.967228 | Trpv3 |
| ENSMUSG( | 88.67199 | 0.429069 | 0.125746 | 0.81516 | Trpv4 |
| ENSMUSG( | 71.21191 | 0.058771 | 0.8358 | 0.981237 | Trpv6 |
| ENSMUSG( | 4030.933 | 0.108576 | 0.413632 | 0.917257 | Trrap |
| ENSMUSG( | 510.7597 | 0.083558 | 0.505864 | 0.927825 | Trub1 |
| ENSMUSG( | 673.6173 | -0.08303 | 0.498375 | 0.927825 | Trub2 |
| ENSMUSG( | 28.45061 | -0.1081 | 0.770539 | 0.974236 | Tsacc |
| ENSMUSG( | 79.22985 | 0.109711 | 0.72103 | 0.966887 | Tsbp1 |
| ENSMUSG( | 2150.305 | 0.026981 | 0.80596 | 0.977674 | Tsc1 |
| ENSMUSG( | 3865.387 | 0.025887 | 0.803691 | 0.976978 | Tsc2 |
| ENSMUSG( | 7969.137 | 0.138123 | 0.207138 | 0.864412 | Tsc22d1 |
| ENSMUSG( | 718.1697 | 0.172195 | 0.165248 | 0.838743 | Tsc22d2 |
| ENSMUSG( | 2897.729 | 0.475608 | 0.000289 | 0.04607 | Tsc22d3 |
| ENSMUSG( | 9071.47 | -0.0806 | 0.518816 | 0.930939 | Tsc22d4 |
| ENSMUSG( | 381.8346 | -0.06175 | 0.667463 | 0.956766 | Tsen15 |
| ENSMUSG( | 355.0373 | 0.051921 | 0.726319 | 0.966887 | Tsen2 |
| ENSMUSG( | 1387.002 | -0.14291 | 0.209093 | 0.864412 | Tsen34 |
| ENSMUSG( | 203.5822 | 0.049545 | 0.776333 | 0.975717 | Tsen54 |
| ENSMUSG( | 290.1746 | -0.08068 | 0.62945 | 0.953251 | Tsfm |
| ENSMUSG( | 1355.541 | -0.05573 | 0.658048 | 0.956123 | Tsg101 |
| ENSMUSG( | 10.87538 | 0.082243 | 0.887791 | NA | Tsg101-ps |
| ENSMUSG( | 441.3258 | 0.041074 | 0.778259 | 0.976106 | Tsga10 |
| ENSMUSG( | 28.19325 | 0.085761 | 0.843057 | 0.983211 | Tshr |
| ENSMUSG( | 1212.95 | 0.180696 | 0.224032 | 0.867404 | Tshz1 |
| ENSMUSG( | 3997.799 | -0.03245 | 0.753032 | 0.970622 | Tshz2 |
| ENSMUSG( | 631.994 | -0.05359 | 0.694943 | 0.960867 | Tshz3 |
| ENSMUSG( | 24.86853 | 0.384557 | 0.354708 | 0.912411 | Tsks |
| ENSMUSG( | 179.4497 | 0.372516 | 0.097758 | 0.797222 | Tsku |
| ENSMUSG( | 2613.31 | -0.0392 | 0.724745 | 0.966887 | Tsn |
| ENSMUSG( | 3460.477 | -0.09597 | 0.413967 | 0.917257 | Tsnax |
| ENSMUSG( | 16.05863 | 0.238836 | 0.654955 | 0.955037 | Tsnaxip1 |
| ENSMUSG( | 364.1224 | -0.16336 | 0.395308 | 0.917257 | Tspan11 |
| ENSMUSG( | 1309.351 | -0.02571 | 0.879497 | 0.989644 | Tspan12 |
| ENSMUSG( | 5969.77 | 0.147628 | 0.376729 | 0.915532 | Tspan13 |
| ENSMUSG( | 2126.156 | 0.022951 | 0.893032 | 0.990558 | Tspan14 |
| ENSMUSG( | 1372.45 | 0.02203 | 0.904504 | 0.991089 | Tspan15 |
| ENSMUSG( | 1115.684 | -0.01148 | 0.938059 | 0.993731 | Tspan17 |
| ENSMUSG( | 340.5209 | 0.251125 | 0.241347 | 0.876259 | Tspan18 |
| ENSMUSG( | 9278.637 | 0.036057 | 0.838676 | 0.981622 | Tspan2 |
| ENSMUSG( | 16262.85 | 0.042336 | 0.755631 | 0.971336 | Tspan3 |
| ENSMUSG( | 2880.038 | -0.01306 | 0.923008 | 0.992644 | Tspan31 |
| ENSMUSG( | 9.859046 | -0.47979 | 0.453462 | NA | Tspan32 |
| ENSMUSG( | 960.9433 | 0.000398 | 0.997797 | 0.999268 | Tspan33 |
| ENSMUSG( | 577.0448 | 0.618515 | 0.000951 | 0.105417 | Tspan4 |
| ENSMUSG( | 6431.178 | 0.129431 | 0.369644 | 0.915491 | Tspan5 |
| ENSMUSG( | 719.2995 | 0.010848 | 0.942875 | 0.994774 | Tspan6 |
| ENSMUSG( | 100675.1 | -0.13747 | 0.305766 | 0.898538 | Tspan7 |
| ENSMUSG( | 43.57619 | 0.29296 | 0.348051 | 0.911652 | Tspan8 |
| ENSMUSG( | 2427.313 | -0.29168 | 0.117922 | 0.813384 | Tspan9 |
| ENSMUSG( | 151.5377 | -0.20293 | 0.277994 | 0.891468 | Tspo |
| ENSMUSG( | 15523.92 | -0.00697 | 0.960049 | 0.994798 | Tspoap1 |
| ENSMUSG( | 2214.428 | -0.10204 | 0.464852 | 0.923329 | Tspyl1 |
| ENSMUSG( | 2331.826 | -0.06571 | 0.666438 | 0.956766 | Tspyl2 |
| ENSMUSG( | 412.3729 | -0.01852 | 0.909015 | 0.991958 | Tspyl3 |
| ENSMUSG( | 5889.251 | -0.10996 | 0.306157 | 0.898538 | Tspyl4 |
| ENSMUSG( | 531.4122 | -0.08926 | 0.598934 | 0.947045 | Tspyl5 |
| ENSMUSG( | 676.1355 | -0.17447 | 0.167878 | 0.840222 | Tsr1 |
| ENSMUSG( | 1921.202 | -0.10366 | 0.381924 | 0.917257 | Tsr2 |
| ENSMUSG( | 585.0583 | -0.08574 | 0.544884 | 0.938063 | Tsr3 |
| ENSMUSG( | 931.457 | -0.05126 | 0.650283 | 0.95422 | Tssc4 |
| ENSMUSG( | 13.96373 | -0.02714 | 0.963877 | 0.995433 | Tssk4 |
| ENSMUSG( | 2437.074 | -0.34698 | 0.003484 | 0.235685 | Tst |
| ENSMUSG( | 17.19704 | 0.323679 | 0.556733 | 0.939565 | Tstd1 |
| ENSMUSG( | 394.9518 | -0.03921 | 0.773751 | 0.974672 | Tstd2 |
| ENSMUSG( | 311.0021 | -0.34904 | 0.04524 | 0.693959 | Tstd3 |
| ENSMUSG( | 7820.823 | 0.202481 | 0.099978 | 0.801311 | Ttbk1 |
| ENSMUSG( | 1548.65 | -0.0138 | 0.914331 | 0.992407 | Ttbk2 |
| ENSMUSG( | 749.6226 | -0.22544 | 0.062629 | 0.736482 | Ttc1 |
| ENSMUSG( | 136.6878 | 0.016308 | 0.941042 | 0.994558 | Ttc12 |
| ENSMUSG( | 1315.478 | 0.180047 | 0.174483 | 0.84455 | Ttc13 |
| ENSMUSG( | 2628.35 | 0.307886 | 0.061921 | 0.736482 | Ttc14 |
| ENSMUSG( | 40.75638 | 0.140418 | 0.683845 | 0.958678 | Ttc16 |
| ENSMUSG( | 1247.673 | 0.158579 | 0.343189 | 0.909646 | Ttc17 |
| ENSMUSG( | 1990.665 | 0.10625 | 0.468589 | 0.923329 | Ttc19 |
| ENSMUSG( | 63.09706 | 0.377022 | 0.260701 | 0.884675 | Ttc21a |
| ENSMUSG( | 280.5377 | 0.194222 | 0.19616 | 0.862161 | Ttc21b |
| ENSMUSG( | 30.67792 | -0.27605 | 0.466477 | 0.923329 | Ttc22 |
| ENSMUSG( | 223.0159 | -0.129 | 0.442915 | 0.921554 | Ttc23 |
| ENSMUSG( | 293.6881 | -0.07047 | 0.659375 | 0.956205 | Ttc27 |
| ENSMUSG( | 992.7012 | 0.09764 | 0.399292 | 0.917257 | Ttc28 |
| ENSMUSG( | 11.11061 | 0.474758 | 0.496562 | NA | Ttc29 |
| ENSMUSG( | 21039.58 | 0.072826 | 0.520154 | 0.931937 | Ttc3 |
| ENSMUSG( | 94.07762 | -0.20554 | 0.350892 | 0.911707 | Ttc32 |
| ENSMUSG( | 574.9251 | 0.025821 | 0.858369 | 0.985884 | Ttc33 |
| ENSMUSG( | 89.32396 | -0.10894 | 0.65743 | 0.955725 | Ttc34 |
| ENSMUSG( | 251.376 | -0.0873 | 0.589694 | 0.944975 | Ttc38 |
| ENSMUSG( | 40.7752 | -0.14326 | 0.649475 | 0.95422 | Ttc39a |
| ENSMUSG( | 71.73281 | 0.012631 | 0.966256 | 0.995475 | Ttc39aos1 |
| ENSMUSG( | 1083.1 | -0.03553 | 0.761949 | 0.97318 | Ttc39b |
| ENSMUSG( | 533.395 | 0.020231 | 0.887836 | 0.990399 | Ttc39c |
| ENSMUSG( | 990.2475 | 0.073894 | 0.613221 | 0.951177 | Ttc4 |
| ENSMUSG( | 62.57482 | 0.063285 | 0.805334 | 0.977484 | Ttc41 |

| | | | | | | | | | | | |
|---|---|---|---|---|---|---|---|---|---|---|---|
| ENSMUSG( | Tubb4b-ps | 31.95892 | 0.107228 | 0.749709 | 0.949664 | ENSMUSG( | Tubb4b-ps | 22.27183 | -0.76244 | 0.074653 | 0.767423 |
| ENSMUSG( | Tubb5 | 14101.56 | 0.029982 | 0.7559 | 0.951479 | ENSMUSG( | Tubb5 | 12118.75 | -0.14361 | 0.274398 | 0.889874 |
| ENSMUSG( | Tubb6 | 79.45986 | -0.22492 | 0.326184 | 0.771128 | ENSMUSG( | Tubb6 | 72.95722 | -0.21411 | 0.405507 | 0.917257 |
| ENSMUSG( | Tubd1 | 109.2068 | 0.19401 | 0.344762 | 0.779153 | ENSMUSG( | Tubd1 | 104.7558 | 0.35097 | 0.145723 | 0.83474 |
| ENSMUSG( | Tube1 | 55.75826 | -0.14029 | 0.582986 | 0.898094 | ENSMUSG( | Tube1 | 55.42655 | 0.122434 | 0.660607 | 0.956205 |
| ENSMUSG( | Tubg1 | 877.9663 | 0.122541 | 0.3462 | 0.779153 | ENSMUSG( | Tubg1 | 739.7641 | -0.10126 | 0.471253 | 0.923329 |
| ENSMUSG( | Tubg2 | 1954.442 | 0.066707 | 0.460235 | 0.840676 | ENSMUSG( | Tubg2 | 1702.295 | -0.06371 | 0.561026 | 0.940254 |
| ENSMUSG( | Tubgcp2 | 1117.293 | 0.053457 | 0.59887 | 0.903801 | ENSMUSG( | Tubgcp2 | 976.6038 | -0.06443 | 0.593555 | 0.945408 |
| ENSMUSG( | Tubgcp3 | 1101.067 | 0.049254 | 0.625768 | 0.912274 | ENSMUSG( | Tubgcp3 | 1022.874 | 0.104937 | 0.388733 | 0.917257 |
| ENSMUSG( | Tubgcp4 | 984.9692 | 0.049761 | 0.632381 | 0.914073 | ENSMUSG( | Tubgcp4 | 884.1967 | 0.009657 | 0.941666 | 0.994578 |
| ENSMUSG( | Tubgcp5 | 570.5782 | 0.051644 | 0.628505 | 0.913637 | ENSMUSG( | Tubgcp5 | 536.912 | 0.147505 | 0.270783 | 0.888062 |
| ENSMUSG( | Tubgcp6 | 1289.862 | 0.205712 | 0.057659 | 0.370247 | ENSMUSG( | Tubgcp6 | 1183.639 | 0.228758 | 0.088853 | 0.788051 |
| ENSMUSG( | Tufm | 1373.521 | 0.064118 | 0.516815 | 0.87302 | ENSMUSG( | Tufm | 1131.517 | -0.2358 | 0.047006 | 0.700664 |
| ENSMUSG( | Tuft1 | 29.85849 | 0.124957 | 0.745428 | 0.949158 | ENSMUSG( | Tuft1 | 26.57214 | 0.070846 | 0.865686 | 0.987041 |
| ENSMUSG( | Tug1 | 2274.685 | -0.24452 | 0.048448 | 0.336526 | ENSMUSG( | Tug1 | 2313.743 | 0.085251 | 0.546052 | 0.938063 |
| ENSMUSG( | Tulp3 | 953.0375 | 0.174594 | 0.083257 | 0.445247 | ENSMUSG( | Tulp3 | 846.7039 | 0.103282 | 0.39396 | 0.917257 |
| ENSMUSG( | Tulp4 | 4983.014 | -0.18742 | 0.045491 | 0.327506 | ENSMUSG( | Tulp4 | 4687.775 | -0.09033 | 0.436807 | 0.919879 |
| ENSMUSG( | Tunar | 315.1437 | 0.022277 | 0.888884 | 0.983434 | ENSMUSG( | Tunar | 273.7736 | -0.12695 | 0.498921 | 0.927825 |
| ENSMUSG( | Tusc1 | 256.1494 | 0.353556 | 0.030028 | 0.26299 | ENSMUSG( | Tusc1 | 217.5733 | 0.157169 | 0.387752 | 0.917257 |
| ENSMUSG( | Tusc2 | 2839.475 | 0.093726 | 0.253093 | 0.70338 | ENSMUSG( | Tusc2 | 2436.927 | -0.07979 | 0.475858 | 0.923329 |
| ENSMUSG( | Tusc3 | 5406.786 | -0.42564 | 0.002644 | 0.061702 | ENSMUSG( | Tusc3 | 5944.471 | 0.147746 | 0.360695 | 0.914223 |
| ENSMUSG( | Tut1 | 569.0288 | 0.109138 | 0.438779 | 0.830714 | ENSMUSG( | Tut1 | 504.2387 | 0.028468 | 0.831321 | 0.980611 |
| ENSMUSG( | Tut4 | 749.6181 | -0.24455 | 0.024323 | 0.238636 | ENSMUSG( | Tut4 | 742.2481 | 0.00856 | 0.947126 | 0.994798 |
| ENSMUSG( | Tut7 | 2497.457 | 0.081707 | 0.338423 | 0.777489 | ENSMUSG( | Tut7 | 2273.827 | 0.079363 | 0.449412 | 0.922202 |
| ENSMUSG( | Tvp23a | 1213.532 | -0.00804 | 0.952873 | 0.993874 | ENSMUSG( | Tvp23a | 1168.133 | 0.152608 | 0.320842 | 0.904019 |
| ENSMUSG( | Tvp23b | 965.0192 | -0.12392 | 0.29875 | 0.747452 | ENSMUSG( | Tvp23b | 925.8748 | 0.028377 | 0.831741 | 0.980656 |
| ENSMUSG( | Twf1 | 2260.672 | 0.0615 | 0.568115 | 0.892217 | ENSMUSG( | Twf1 | 1979.898 | -0.05473 | 0.662135 | 0.956205 |
| ENSMUSG( | Twf2 | 965.4926 | 0.169921 | 0.173348 | 0.60863 | ENSMUSG( | Twf2 | 825.7431 | -0.00553 | 0.965323 | 0.995433 |
| ENSMUSG( | Twist1 | 30.6762 | -0.47533 | 0.164381 | 0.59535 | ENSMUSG( | Twist1 | 29.91954 | -0.27887 | 0.454508 | 0.922444 |
| ENSMUSG( | Twist2 | 21.67519 | -0.00919 | 0.981958 | 0.997512 | ENSMUSG( | Twist2 | 18.69973 | -0.2138 | 0.650095 | 0.95422 |
| ENSMUSG( | Twnk | 409.4289 | 0.176226 | 0.18049 | 0.617841 | ENSMUSG( | Twnk | 361.8469 | 0.098979 | 0.508923 | 0.929089 |
| ENSMUSG( | Twsg1 | 1502.415 | -0.13944 | 0.293514 | 0.742947 | ENSMUSG( | Twsg1 | 1461.991 | 0.054132 | 0.708569 | 0.964042 |
| ENSMUSG( | Txlna | 1614.526 | 0.00835 | 0.926702 | 0.991229 | ENSMUSG( | Txlna | 1513.857 | 0.092014 | 0.443742 | 0.921734 |
| ENSMUSG( | Txlnb | 127.3675 | 0.116168 | 0.600752 | 0.90443 | ENSMUSG( | Txlnb | 98.86785 | -0.37749 | 0.117409 | 0.813384 |
| ENSMUSG( | Txlng | 394.8475 | 0.242209 | 0.046731 | 0.330815 | ENSMUSG( | Txlng | 363.7207 | 0.264706 | 0.085902 | 0.780123 |
| ENSMUSG( | Txn1 | 1566.326 | -0.00476 | 0.964621 | 0.995155 | ENSMUSG( | Txn1 | 1366.868 | -0.13701 | 0.309264 | 0.898538 |
| ENSMUSG( | Txn2 | 5418.1 | 0.051242 | 0.553639 | 0.886216 | ENSMUSG( | Txn2 | 4523.842 | -0.2101 | 0.080311 | 0.77681 |
| ENSMUSG( | Txndc11 | 953.7112 | -0.20118 | 0.204047 | 0.644572 | ENSMUSG( | Txndc11 | 1052.862 | 0.346221 | 0.053102 | 0.715528 |
| ENSMUSG( | Txndc12 | 1451.377 | -0.26289 | 0.0774 | 0.429743 | ENSMUSG( | Txndc12 | 1489.743 | 0.095287 | 0.547711 | 0.938063 |
| ENSMUSG( | Txndc15 | 2374.543 | -0.16143 | 0.205025 | 0.646229 | ENSMUSG( | Txndc15 | 2439.281 | 0.186472 | 0.207138 | 0.864412 |
| ENSMUSG( | Txndc16 | 3239.994 | -0.24601 | 0.072527 | 0.415832 | ENSMUSG( | Txndc16 | 3390.521 | 0.162325 | 0.282866 | 0.892412 |
| ENSMUSG( | Txndc17 | 1011.712 | 0.036148 | 0.748739 | 0.949604 | ENSMUSG( | Txndc17 | 877.7826 | -0.11153 | 0.388039 | 0.917257 |
| ENSMUSG( | Txndc5 | 2306.475 | -0.3198 | 0.028257 | 0.256767 | ENSMUSG( | Txndc5 | 2410.944 | 0.097799 | 0.544789 | 0.938063 |
| ENSMUSG( | Txndc9 | 1023.828 | -0.0421 | 0.66315 | 0.9265 | ENSMUSG( | Txndc9 | 915.4991 | -0.10101 | 0.400988 | 0.917257 |
| ENSMUSG( | Txnip | 869.9134 | -0.19636 | 0.063921 | 0.391395 | ENSMUSG( | Txnip | 843.2703 | -0.00412 | 0.982211 | 0.996756 |
| ENSMUSG( | Txnl1 | 1321.151 | -0.01841 | 0.867404 | 0.976559 | ENSMUSG( | Txnl1 | 1185.473 | -0.06906 | 0.638934 | 0.953903 |
| ENSMUSG( | Txnl4a | 1785.342 | -0.02414 | 0.827325 | 0.968732 | ENSMUSG( | Txnl4a | 1619.843 | -0.03526 | 0.778918 | 0.976106 |
| ENSMUSG( | Txnl4a | 870.285 | 0.230858 | 0.023348 | 0.233882 | ENSMUSG( | Txnl4a | 711.628 | -0.07662 | 0.553002 | 0.939565 |
| ENSMUSG( | Txnl4b | 574.5797 | 0.010246 | 0.925468 | 0.991204 | ENSMUSG( | Txnl4b | 499.3084 | -0.13383 | 0.332126 | 0.907907 |
| ENSMUSG( | Txnrd1 | 1672.595 | 0.067525 | 0.461719 | 0.841564 | ENSMUSG( | Txnrd1 | 1376.867 | -0.23573 | 0.037034 | 0.673514 |
| ENSMUSG( | Txnrd2 | 692.7225 | 0.051637 | 0.619507 | 0.911432 | ENSMUSG( | Txnrd2 | 568.1526 | -0.26094 | 0.038955 | 0.679972 |
| ENSMUSG( | Txnrd3 | 212.7934 | -0.04419 | 0.761083 | 0.952907 | ENSMUSG( | Txnrd3 | 189.4469 | -0.11881 | 0.488968 | 0.925573 |
| ENSMUSG( | Tyk2 | 678.4115 | 0.248305 | 0.027893 | 0.255773 | ENSMUSG( | Tyk2 | 617.6443 | 0.247521 | 0.090688 | 0.791074 |
| ENSMUSG( | Tymp | 26.61389 | -0.27826 | 0.435661 | 0.829093 | ENSMUSG( | Tymp | 26.86272 | 0.005891 | 0.987749 | 0.997659 |
| ENSMUSG( | Tyms | 61.40524 | 0.156811 | 0.541899 | 0.883182 | ENSMUSG( | Tyms | 59.49772 | 0.323099 | 0.247335 | 0.877904 |
| ENSMUSG( | Tyro3 | 7208.514 | -0.2497 | 0.074269 | 0.422054 | ENSMUSG( | Tyro3 | 7785.247 | 0.246124 | 0.146051 | 0.835524 |
| ENSMUSG( | Tyrobp | 1103.575 | -0.03445 | 0.78969 | 0.959461 | ENSMUSG( | Tyrobp | 976.5158 | -0.11955 | 0.366891 | 0.914852 |
| ENSMUSG( | Tysnd1 | 630.7424 | 0.160601 | 0.156684 | 0.584772 | ENSMUSG( | Tysnd1 | 537.982 | -0.02853 | 0.823038 | 0.979198 |
| ENSMUSG( | Tyw1 | 600.5588 | -0.0821 | 0.55853 | 0.888521 | ENSMUSG( | Tyw1 | 612.8491 | 0.24292 | 0.144711 | 0.833302 |
| ENSMUSG( | Tyw3 | 195.5154 | -0.00617 | 0.969081 | 0.996369 | ENSMUSG( | Tyw3 | 186.6753 | 0.13029 | 0.459361 | 0.923329 |
| ENSMUSG( | Tyw5 | 272.4564 | 0.067165 | 0.62559 | 0.912249 | ENSMUSG( | Tyw5 | 262.9484 | 0.229662 | 0.138524 | 0.828492 |
| ENSMUSG( | U2af1 | 1170.093 | -0.01617 | 0.865346 | 0.975565 | ENSMUSG( | U2af1 | 1053.925 | -0.05009 | 0.66565 | 0.956766 |
| ENSMUSG( | U2af2 | 3724.924 | 0.060036 | 0.486852 | 0.858812 | ENSMUSG( | U2af2 | 3172.258 | -0.13986 | 0.208884 | 0.864412 |
| ENSMUSG( | U2surp | 1554.657 | 0.016586 | 0.85237 | 0.973168 | ENSMUSG( | U2surp | 1434.67 | 0.050325 | 0.658879 | 0.956205 |
| ENSMUSG( | Uaca | 547.1519 | -0.04211 | 0.73328 | 0.944956 | ENSMUSG( | Uaca | 568.6758 | 0.323517 | 0.012054 | 0.445877 |
| ENSMUSG( | Uap1 | 1603.347 | 0.082891 | 0.346295 | 0.779153 | ENSMUSG( | Uap1 | 1388.889 | -0.06357 | 0.567014 | 0.941777 |
| ENSMUSG( | Uap1l1 | 423.2286 | 0.138877 | 0.266523 | 0.716441 | ENSMUSG( | Uap1l1 | 366.2856 | -0.00449 | 0.974608 | 0.996416 |
| ENSMUSG( | Uba1 | 8451.89 | 0.095502 | 0.247822 | 0.696522 | ENSMUSG( | Uba1 | 7189.534 | -0.10324 | 0.30307 | 0.898538 |
| ENSMUSG( | Uba1y | 16.07665 | 0.406892 | 0.372138 | NA | ENSMUSG( | Uba1y | 11.38164 | -0.29482 | 0.601799 | NA |
| ENSMUSG( | Uba2 | 1712.299 | 0.135856 | 0.134336 | 0.547472 | ENSMUSG( | Uba2 | 1522.494 | 0.068485 | 0.531661 | 0.934795 |
| ENSMUSG( | Uba3 | 1859.595 | 0.038418 | 0.690961 | 0.933881 | ENSMUSG( | Uba3 | 1597.686 | -0.13762 | 0.270944 | 0.888062 |
| ENSMUSG( | Uba5 | 1635.146 | 0.079462 | 0.368205 | 0.795774 | ENSMUSG( | Uba5 | 1365.337 | -0.17739 | 0.13482 | 0.824049 |
| ENSMUSG( | Uba52 | 466.8862 | -0.01389 | 0.911702 | 0.988483 | ENSMUSG( | Uba52 | 428.2899 | -0.00137 | 0.992671 | 0.99874 |
| ENSMUSG( | Uba52rt | 103.6634 | -0.00467 | 0.981723 | 0.997512 | ENSMUSG( | Uba52rt | 76.57711 | -0.72748 | 0.006739 | 0.339232 |
| ENSMUSG( | Uba6 | 337.309 | 0.090351 | 0.509995 | 0.869743 | ENSMUSG( | Uba6 | 315.0712 | 0.160201 | 0.301047 | 0.898538 |
| ENSMUSG( | Uba7 | 142.1854 | -0.1378 | 0.464765 | 0.843568 | ENSMUSG( | Uba7 | 147.2104 | 0.240895 | 0.302611 | 0.898538 |
| ENSMUSG( | Ubac1 | 1584.305 | 0.076005 | 0.413866 | 0.819998 | ENSMUSG( | Ubac1 | 1381.096 | -0.05265 | 0.644697 | 0.953903 |
| ENSMUSG( | Ubac2 | 1774.622 | -0.22666 | 0.044465 | 0.32391 | ENSMUSG( | Ubac2 | 1771.389 | 0.047744 | 0.711633 | 0.964363 |
| ENSMUSG( | Ubald1 | 2844.124 | 0.051748 | 0.690277 | 0.933881 | ENSMUSG( | Ubald1 | 2442.522 | -0.12276 | 0.380854 | 0.917202 |
| ENSMUSG( | Ubald2 | 397.0479 | 0.039942 | 0.758638 | 0.952113 | ENSMUSG( | Ubald2 | 351.7347 | -0.05046 | 0.75466 | 0.971098 |
| ENSMUSG( | Ubap1 | 1365.485 | 0.080587 | 0.450188 | 0.835344 | ENSMUSG( | Ubap1 | 1211.343 | 0.001053 | 0.993603 | 0.998813 |
| ENSMUSG( | Ubap1l | 92.90351 | 0.306096 | 0.162154 | 0.592508 | ENSMUSG( | Ubap1l | 77.96802 | 0.101877 | 0.694765 | 0.960867 |
| ENSMUSG( | Ubap2 | 2336.01 | 0.166851 | 0.065795 | 0.397169 | ENSMUSG( | Ubap2 | 2021.983 | 0.021471 | 0.847278 | 0.985161 |
| ENSMUSG( | Ubap2l | 2910.542 | 0.015585 | 0.85554 | 0.973813 | ENSMUSG( | Ubap2l | 2621.714 | -0.01783 | 0.873131 | 0.989445 |
| ENSMUSG( | Ubash3b | 783.3579 | -0.10482 | 0.365514 | 0.794085 | ENSMUSG( | Ubash3b | 673.7563 | -0.28262 | 0.024257 | 0.589324 |
| ENSMUSG( | Ubb | 13227.53 | -0.01702 | 0.863665 | 0.975244 | ENSMUSG( | Ubb | 11279.99 | -0.2182 | 0.098374 | 0.798438 |
| ENSMUSG( | Ubc | 1872.765 | -0.03528 | 0.760831 | 0.952907 | ENSMUSG( | Ubc | 1591.227 | -0.2487 | 0.034232 | 0.652059 |
| ENSMUSG( | Ube2a | 904.5902 | 0.041744 | 0.709936 | 0.939669 | ENSMUSG( | Ube2a | 797.6329 | -0.06078 | 0.707911 | 0.964042 |
| ENSMUSG( | Ube2b | 2724.986 | 0.022777 | 0.824784 | 0.967386 | ENSMUSG( | Ube2b | 2460.076 | -0.00631 | 0.961027 | 0.99511 |
| ENSMUSG( | Ube2c | 40.63461 | -0.45753 | 0.1365 | 0.550651 | ENSMUSG( | Ube2c | 47.11221 | 0.238538 | 0.459897 | 0.923329 |
| ENSMUSG( | Ube2d1 | 1005.841 | -0.08924 | 0.383102 | 0.803237 | ENSMUSG( | Ube2d1 | 943.7921 | -0.00732 | 0.956002 | 0.994798 |
| ENSMUSG( | Ube2d2a | 2178.473 | -0.00603 | 0.948697 | 0.993821 | ENSMUSG( | Ube2d2a | 1968.15 | -0.03426 | 0.802956 | 0.976899 |
| ENSMUSG( | Ube2d3 | 3963.737 | -0.0862 | 0.433167 | 0.828207 | ENSMUSG( | Ube2d3 | 3868.676 | 0.10776 | 0.507539 | 0.929089 |
| ENSMUSG( | Ube2d-ps | 797.78 | 0.087551 | 0.415308 | 0.820498 | ENSMUSG( | Ube2d-ps | 678.4669 | -0.1148 | 0.363766 | 0.914223 |
| ENSMUSG( | Ube2e1 | 1536.59 | 0.064578 | 0.613139 | 0.908902 | ENSMUSG( | Ube2e1 | 1345.425 | -0.0562 | 0.732157 | 0.967228 |
| ENSMUSG( | Ube2e2 | 1399.249 | 0.050198 | 0.617 | 0.910866 | ENSMUSG( | Ube2e2 | 1203.795 | -0.12177 | 0.416957 | 0.917257 |
| ENSMUSG( | Ube2e3 | 1731.425 | 0.081145 | 0.440145 | 0.831851 | ENSMUSG( | Ube2e3 | 1566.334 | 0.056279 | 0.701187 | 0.961756 |
| ENSMUSG( | Ube2f | 1551.351 | 0.081836 | 0.344693 | 0.779153 | ENSMUSG( | Ube2f | 1330.139 | -0.09545 | 0.393533 | 0.917257 |
| ENSMUSG( | Ube2g1 | 1197.155 | -0.03948 | 0.712976 | 0.94066 | ENSMUSG( | Ube2g1 | 1130.134 | 0.057177 | 0.694251 | 0.960867 |
| ENSMUSG( | Ube2g2 | 1000.944 | 0.066635 | 0.519869 | 0.874979 | ENSMUSG( | Ube2g2 | 890.6079 | -0.00335 | 0.978711 | 0.996459 |
| ENSMUSG( | Ube2h | 4189.354 | 0.034192 | 0.715847 | 0.941484 | ENSMUSG( | Ube2h | 3678.169 | -0.07578 | 0.529836 | 0.934795 |
| ENSMUSG( | Ube2i | 2138.115 | -0.02637 | 0.79073 | 0.959461 | ENSMUSG( | Ube2i | 2023.835 | 0.085876 | 0.48623 | 0.92541 |
| ENSMUSG( | Ube2j1 | 1848.452 | 0.004457 | 0.965937 | 0.995377 | ENSMUSG( | Ube2j1 | 1640.055 | -0.07679 | 0.591806 | 0.944975 |
| ENSMUSG( | Ube2j2 | 944.2132 | -0.0557 | 0.59365 | 0.90281 | ENSMUSG( | Ube2j2 | 880.0028 | 0.006965 | 0.955652 | 0.994798 |
| ENSMUSG( | Ube2k | 2698.427 | -0.0404 | 0.65847 | 0.924506 | ENSMUSG( | Ube2k | 2417.171 | -0.09401 | 0.47956 | 0.924021 |
| ENSMUSG( | Ube2l3 | 6796.422 | 0.054639 | 0.538296 | 0.88259 | ENSMUSG( | Ube2l3 | 5631.772 | -0.22909 | 0.056646 | 0.723362 |
| ENSMUSG( | Ube2l6 | 320.4565 | -0.43229 | 0.032627 | 0.276639 | ENSMUSG( | Ube2l6 | 312.6121 | -0.21789 | 0.328743 | 0.906727 |
| ENSMUSG( | Ube2m | 6426.709 | 0.004598 | 0.963234 | 0.994911 | ENSMUSG( | Ube2m | 5395.918 | -0.24383 | 0.050314 | 0.708648 |
| ENSMUSG( | Ube2n | 3167.319 | 0.072458 | 0.421461 | 0.823126 | ENSMUSG( | Ube2n | 2719.595 | -0.10217 | 0.406442 | 0.917257 |
| ENSMUSG( | Ube2o | 3996.031 | 0.055837 | 0.62352 | 0.911646 | ENSMUSG( | Ube2o | 3531.271 | -0.03171 | 0.793286 | 0.97681 |
| ENSMUSG( | Ube2q1 | 2983.447 | 0.064339 | 0.480251 | 0.854705 | ENSMUSG( | Ube2q1 | 2576.952 | -0.09196 | 0.420966 | 0.917257 |
| ENSMUSG( | Ube2q2 | 1758.917 | 0.034333 | 0.767104 | 0.954393 | ENSMUSG( | Ube2q2 | 1578.978 | -0.01353 | 0.92459 | 0.99306 |
| ENSMUSG( | Ube2ql1 | 5718.265 | 0.130437 | 0.193727 | 0.633085 | ENSMUSG( | Ube2ql1 | 4852.724 | -0.07311 | 0.518726 | 0.930939 |
| ENSMUSG( | Ube2r2 | 3660.621 | 0.071994 | 0.445082 | 0.833239 | ENSMUSG( | Ube2r2 | 3216.635 | -0.0343 | 0.816977 | 0.979198 |
| ENSMUSG( | Ube2s | 977.1225 | 0.004715 | 0.966738 | 0.995784 | ENSMUSG( | Ube2s | 849.4163 | -0.13818 | 0.304055 | 0.898538 |
| ENSMUSG( | Ube2srt | 24.71777 | -0.18965 | 0.616199 | 0.91085 | ENSMUSG( | Ube2srt | 21.65236 | -0.39467 | 0.430761 | 0.917257 |
| ENSMUSG( | Ube2t | 19.32831 | 0.086012 | 0.844823 | NA | ENSMUSG( | Ube2t | 20.31432 | 0.469039 | 0.32054 | 0.903887 |
| ENSMUSG( | Ube2v1 | 28.90137 | 0.038786 | 0.914507 | 0.988483 | ENSMUSG( | Ube2v1 | 24.93484 | -0.11907 | 0.772993 | 0.974463 |
| ENSMUSG( | Ube2v2 | 657.491 | -0.04852 | 0.700703 | 0.93743 | ENSMUSG( | Ube2v2 | 599.8996 | -0.05317 | 0.757907 | 0.971858 |
| ENSMUSG( | Ube2w | 1083.326 | 0.057311 | 0.59912 | 0.903801 | ENSMUSG( | Ube2w | 965.5292 | -0.01223 | 0.938191 | 0.993731 |
| ENSMUSG( | Ube2z | 2444.681 | 0.089379 | 0.400545 | 0.813717 | ENSMUSG( | Ube2z | 2082.085 | -0.10762 | 0.392001 | 0.917257 |
| ENSMUSG( | Ube3a | 1900.264 | 0.039165 | 0.731472 | 0.944956 | ENSMUSG( | Ube3a | 1796.982 | 0.1393 | 0.417178 | 0.917257 |
| ENSMUSG( | Ube3b | 8020.35 | 0.14234 | 0.091081 | 0.463586 | ENSMUSG( | Ube3b | 7087.429 | 0.05834 | 0.583377 | 0.944266 |
| ENSMUSG( | Ube3c | 2185.918 | 0.16266 | 0.07834 | 0.432941 | ENSMUSG( | Ube3c | 1944.805 | 0.100272 | 0.407612 | 0.917257 |
| ENSMUSG( | Ube3d | 32.57655 | -0.15 | 0.659763 | 0.924524 | ENSMUSG( | Ube3d | 31.68509 | 0.029099 | 0.936444 | 0.993573 |
| ENSMUSG( | Ube4a | 2187.094 | -0.09078 | 0.461887 | 0.841728 | ENSMUSG( | Ube4a | 2169.233 | 0.155639 | 0.26267 | 0.885357 |
| ENSMUSG( | Ube4b | 2800.765 | 0.033089 | 0.737994 | 0.947329 | ENSMUSG( | Ube4b | 2502.454 | -0.02291 | 0.836864 | 0.981335 |
| ENSMUSG( | Ubfd1 | 1361.259 | 0.196178 | 0.039102 | 0.304587 | ENSMUSG( | Ubfd1 | 1192.656 | 0.088576 | 0.418249 | 0.917257 |
| ENSMUSG( | Ubiad1 | 384.3569 | -0.18349 | 0.303675 | 0.752117 | ENSMUSG( | Ubiad1 | 398.9607 | 0.201552 | 0.274137 | 0.889874 |
| ENSMUSG( | Ubl3 | 5303.942 | 0.042016 | 0.626492 | 0.91272 | ENSMUSG( | Ubl3 | 4690.971 | -0.04543 | 0.691357 | 0.960071 |
| ENSMUSG( | Ubl4a | 2266.212 | 0.038632 | 0.640895 | 0.916614 | ENSMUSG( | Ubl4a | 1948.824 | -0.13316 | 0.239735 | 0.874882 |
| ENSMUSG( | Ubl5 | 2745.481 | -0.0348 | 0.70011 | 0.937391 | ENSMUSG( | Ubl5 | 2361.643 | -0.21163 | 0.095325 | 0.796133 |
| ENSMUSG( | Ubl7 | 1561.088 | 0.06311 | 0.525236 | 0.876559 | ENSMUSG( | Ubl7 | 1305.281 | -0.19042 | 0.11617 | 0.813384 |
| ENSMUSG( | Ublcp1 | 686.2223 | 0.120158 | 0.311282 | 0.759247 | ENSMUSG( | Ublcp1 | 613.2388 | 0.061438 | 0.672133 | 0.95694 |
| ENSMUSG( | Ubn1 | 1772.429 | 0.107894 | 0.294221 | 0.743459 | ENSMUSG( | Ubn1 | 1646.894 | 0.164784 | 0.155392 | 0.837724 |
| ENSMUSG( | Ubn2 | 1244.673 | -0.00517 | 0.958251 | 0.993874 | ENSMUSG( | Ubn2 | 1217.465 | 0.193233 | 0.100846 | 0.801456 |
| ENSMUSG( | Ubox5 | 296.2184 | 0.074086 | 0.693118 | 0.934202 | ENSMUSG( | Ubox5 | 264.7862 | 0.015882 | 0.935825 | 0.993443 |
| ENSMUSG( | Ubp1 | 3321.789 | 0.196451 | 0.107622 | 0.49648 | ENSMUSG( | Ubp1 | 2961.122 | 0.133471 | 0.383688 | 0.917257 |
| ENSMUSG( | Ubqln1 | 5620.809 | 0.042389 | 0.672945 | 0.929007 | ENSMUSG( | Ubqln1 | 4862.955 | -0.11068 | 0.391306 | 0.917257 |
| ENSMUSG( | Ubqln2 | 3698.761 | 0.04409 | 0.633886 | 0.914073 | ENSMUSG( | Ubqln2 | 3367.07 | 0.0404 | 0.727505 | 0.966887 |
| ENSMUSG( | Ubqln4 | 7781.489 | 0.141629 | 0.077189 | 0.429655 | ENSMUSG( | Ubqln4 | 6593.631 | -0.0645 | 0.502234 | 0.927825 |
| ENSMUSG( | Ubr1 | 1609.773 | -0.15898 | 0.29993 | 0.749016 | ENSMUSG( | Ubr1 | 1639.003 | 0.164639 | 0.281287 | 0.892412 |
| ENSMUSG( | Ubr2 | 2472.465 | 0.124342 | 0.180785 | 0.617841 | ENSMUSG( | Ubr2 | 2348.888 | 0.240317 | 0.036538 | 0.66949 |
| ENSMUSG( | Ubr3 | 3452.745 | -0.1326 | 0.196455 | 0.635942 | ENSMUSG( | Ubr3 | 3518.548 | 0.189903 | 0.098819 | 0.798438 |
| ENSMUSG( | Ubr4 | 6356.588 | -0.09769 | 0.38012 | 0.801757 | ENSMUSG( | Ubr4 | 6190.076 | 0.097814 | 0.500818 | 0.927825 |
| ENSMUSG( | Ubr5 | 6161.418 | 0.119928 | 0.175457 | 0.610843 | ENSMUSG( | Ubr5 | 5494.057 | 0.060331 | 0.564874 | 0.941206 |
| ENSMUSG( | Ubr7 | 1535.996 | 0.077517 | 0.458172 | 0.839914 | ENSMUSG( | Ubr7 | 1397.726 | 0.071165 | 0.567799 | 0.941777 |
| ENSMUSG( | Ubtd1 | 320.229 | 0.067395 | 0.619897 | 0.911432 | ENSMUSG( | Ubtd1 | 288.9461 | 0.03163 | 0.849574 | 0.985497 |
| ENSMUSG( | Ubtd2 | 477.3357 | 0.105365 | 0.367353 | 0.795221 | ENSMUSG( | Ubtd2 | 430.9161 | 0.078378 | 0.580244 | 0.944209 |
| ENSMUSG( | Ubtf | 3701.928 | 0.095613 | 0.272556 | 0.722478 | ENSMUSG( | Ubtf | 3294.854 | 0.02821 | 0.79779 | 0.97681 |
| ENSMUSG( | Ubxn1 | 3629.23 | 0.014852 | 0.867286 | 0.976559 | ENSMUSG( | Ubxn1 | 3130.646 | -0.14993 | 0.251914 | 0.879902 |
| ENSMUSG( | Ubxn10 | 47.13678 | -0.74727 | 0.010249 | 0.144303 | ENSMUSG( | Ubxn10 | 55.98108 | 0.14233 | 0.661818 | 0.956205 |
| ENSMUSG( | Ubxn11 | 368.9144 | 0.321824 | 0.050212 | 0.342492 | ENSMUSG( | Ubxn11 | 309.53 | 0.10036 | 0.633258 | 0.953844 |
| ENSMUSG( | Ubxn2a | 1744.72 | 0.098501 | 0.315273 | 0.762288 | ENSMUSG( | Ubxn2a | 1497.739 | -0.07593 | 0.545123 | 0.938063 |
| ENSMUSG( | Ubxn2b | 1343.38 | 0.151525 | 0.159026 | 0.588112 | ENSMUSG( | Ubxn2b | 1145.996 | -0.03651 | 0.770696 | 0.974236 |
| ENSMUSG( | Ubxn4 | 2646.019 | 0.005581 | 0.956851 | 0.993874 | ENSMUSG( | Ubxn4 | 2507.143 | 0.117866 | 0.318009 | 0.902849 |
| ENSMUSG( | Ubxn6 | 2830.666 | 0.08444 | 0.324961 | 0.770595 | ENSMUSG( | Ubxn6 | 2440.548 | -0.07414 | 0.487644 | 0.925427 |
| ENSMUSG( | Ubxn7 | 1290.437 | 0.053731 | 0.59086 | 0.90113 | ENSMUSG( | Ubxn7 | 1153.832 | -0.00186 | 0.988181 | 0.997862 |

| | | | | | |
|---|---|---|---|---|---|
| ENSMUSG( | 1098.968 | 0.140521 | 0.146882 | 0.56887 | Trmt2a |
| ENSMUSG( | 595.0299 | 0.100891 | 0.500416 | 0.863483 | Trmt2b |
| ENSMUSG( | 144.7446 | 0.202613 | 0.241681 | 0.689179 | Trmt44 |
| ENSMUSG( | 159.12 | -0.13329 | 0.410766 | 0.818585 | Trmt5 |
| ENSMUSG( | 711.389 | 0.034195 | 0.771374 | 0.954726 | Trmt6 |
| ENSMUSG( | 327.4215 | 0.064675 | 0.705477 | 0.938511 | Trmt61a |
| ENSMUSG( | 229.2862 | 0.13134 | 0.390139 | 0.80794 | Trmt61b |
| ENSMUSG( | 343.3153 | -0.37302 | 0.003414 | 0.073541 | Trmt9b |
| ENSMUSG( | 424.9134 | -0.02532 | 0.859198 | 0.974707 | Trmu |
| ENSMUSG( | 490.3312 | -0.02411 | 0.824892 | 0.967386 | Trnau1ap |
| ENSMUSG( | 8850.231 | 0.026265 | 0.728096 | 0.944355 | Trnp1 |
| ENSMUSG( | 594.6637 | 0.083676 | 0.450994 | 0.835567 | Trnt1 |
| ENSMUSG( | 3991.982 | 0.071038 | 0.43722 | 0.829501 | Tro |
| ENSMUSG( | 11.24407 | -0.68339 | 0.218538 | NA | Troap |
| ENSMUSG( | 566.3123 | -0.21523 | 0.079775 | 0.436135 | Trp53 |
| ENSMUSG( | 2120.078 | -0.02108 | 0.831305 | 0.969889 | Trp53bp1 |
| ENSMUSG( | 2433.354 | 0.242991 | 0.004465 | 0.08705 | Trp53bp2 |
| ENSMUSG( | 18.98068 | -0.15137 | 0.717645 | NA | Trp53cor1 |
| ENSMUSG( | 2952.332 | -0.27427 | 0.011355 | 0.153509 | Trp53i11 |
| ENSMUSG( | 225.3137 | -0.19608 | 0.206424 | 0.648468 | Trp53i13 |
| ENSMUSG( | 563.1708 | 0.130933 | 0.276068 | 0.725738 | Trp53inp1 |
| ENSMUSG( | 43814.55 | 0.060353 | 0.504141 | 0.866684 | Trp53inp2 |
| ENSMUSG( | 510.5424 | 0.042015 | 0.702285 | 0.937536 | Trp53rka |
| ENSMUSG( | 81.28603 | 0.071559 | 0.756463 | 0.951479 | Trp53rkb |
| ENSMUSG( | 61.90519 | 0.078673 | 0.757515 | 0.951513 | Trp63 |
| ENSMUSG( | 42.59415 | 0.017098 | 0.955851 | 0.993874 | Trp73 |
| ENSMUSG( | 1086.572 | -0.28722 | 0.058936 | 0.37517 | Trpc1 |
| ENSMUSG( | 547.2782 | -0.19385 | 0.105798 | 0.492833 | Trpc3 |
| ENSMUSG( | 470.4879 | 0.07319 | 0.629661 | 0.913965 | Trpc4 |
| ENSMUSG( | 4502.999 | 0.133841 | 0.138477 | 0.553773 | Trpc4ap |
| ENSMUSG( | 98.06121 | -0.07042 | 0.780859 | 0.957531 | Trpc5 |
| ENSMUSG( | 220.2074 | -0.41331 | 0.04401 | 0.321685 | Trpc6 |
| ENSMUSG( | 84.85492 | -0.36834 | 0.115962 | 0.514015 | Trpc7 |
| ENSMUSG( | 15.09993 | 0.324307 | 0.507851 | NA | Trpm1 |
| ENSMUSG( | 941.5143 | -0.09579 | 0.409577 | 0.818258 | Trpm2 |
| ENSMUSG( | 6281.465 | 0.047598 | 0.679157 | 0.930983 | Trpm3 |
| ENSMUSG( | 504.1846 | -0.00884 | 0.944295 | 0.993543 | Trpm4 |
| ENSMUSG( | 37.43886 | -0.43092 | 0.239675 | 0.687561 | Trpm6 |
| ENSMUSG( | 1607.243 | -0.15137 | 0.203504 | 0.643689 | Trpm7 |
| ENSMUSG( | 1129.457 | -0.11631 | 0.351629 | 0.783381 | Trps1 |
| ENSMUSG( | 362.8624 | 0.09711 | 0.467432 | 0.844819 | Trpt1 |
| ENSMUSG( | 596.039 | -0.14839 | 0.225541 | 0.672427 | Trpv2 |
| ENSMUSG( | 29.98779 | -0.64192 | 0.061087 | 0.381679 | Trpv3 |
| ENSMUSG( | 87.6897 | 0.137891 | 0.602311 | 0.904887 | Trpv4 |
| ENSMUSG( | 83.80126 | 0.225047 | 0.427986 | 0.825894 | Trpv6 |
| ENSMUSG( | 4244.887 | -0.01046 | 0.912164 | 0.988483 | Trrap |
| ENSMUSG( | 542.4714 | -0.01077 | 0.927335 | 0.991229 | Trub1 |
| ENSMUSG( | 771.8607 | 0.0414 | 0.687199 | 0.933063 | Trub2 |
| ENSMUSG( | 31.31727 | -0.10796 | 0.743628 | 0.948478 | Tsacc |
| ENSMUSG( | 99.49761 | 0.46307 | 0.074931 | 0.42422 | Tsbp1 |
| ENSMUSG( | 2341.219 | 0.002223 | 0.979583 | 0.997428 | Tsc1 |
| ENSMUSG( | 4429.175 | 0.145828 | 0.085812 | 0.451465 | Tsc2 |
| ENSMUSG( | 8807.104 | 0.15665 | 0.110831 | 0.501952 | Tsc22d1 |
| ENSMUSG( | 751.0102 | 0.038918 | 0.733108 | 0.944956 | Tsc22d2 |
| ENSMUSG( | 3399.321 | 0.636438 | 7.44E-08 | 2.13E-05 | Tsc22d3 |
| ENSMUSG( | 10753.91 | 0.140982 | 0.106391 | 0.494174 | Tsc22d4 |
| ENSMUSG( | 411.2511 | -0.11764 | 0.356623 | 0.788187 | Tsen15 |
| ENSMUSG( | 386.5867 | 0.029549 | 0.820205 | 0.965888 | Tsen2 |
| ENSMUSG( | 1627.783 | 0.055633 | 0.519822 | 0.874979 | Tsen34 |
| ENSMUSG( | 217.2787 | -0.03902 | 0.837563 | 0.971111 | Tsen54 |
| ENSMUSG( | 347.1976 | 0.169988 | 0.218398 | 0.664121 | Tsfm |
| ENSMUSG( | 1550.486 | 0.066657 | 0.520631 | 0.875186 | Tsg101 |
| ENSMUSG( | 11.71665 | 0.04055 | 0.94066 | NA | Tsg101-ps |
| ENSMUSG( | 483.0961 | 0.033857 | 0.789837 | 0.959461 | Tsga10 |
| ENSMUSG( | 21.5147 | -1.24408 | 0.009598 | 0.138869 | Tshr |
| ENSMUSG( | 1223.802 | -0.05204 | 0.595082 | 0.903007 | Tshz1 |
| ENSMUSG( | 5080.064 | 0.367539 | 2.67E-05 | 0.002421 | Tshz2 |
| ENSMUSG( | 703.8474 | -0.00642 | 0.964691 | 0.995155 | Tshz3 |
| ENSMUSG( | 26.6039 | 0.355849 | 0.341354 | 0.778828 | Tsks |
| ENSMUSG( | 151.5043 | -0.39971 | 0.075518 | 0.425931 | Tsku |
| ENSMUSG( | 2852.435 | -0.05505 | 0.572389 | 0.893584 | Tsn |
| ENSMUSG( | 4074.562 | 0.108144 | 0.245065 | 0.692812 | Tsnax |
| ENSMUSG( | 14.49118 | -0.35915 | 0.49792 | NA | Tsnaxip1 |
| ENSMUSG( | 391.0281 | -0.24509 | 0.082358 | 0.442878 | Tspan11 |
| ENSMUSG( | 1445.828 | -0.00545 | 0.970766 | 0.996369 | Tspan12 |
| ENSMUSG( | 5309.805 | -0.50058 | 0.000581 | 0.022718 | Tspan13 |
| ENSMUSG( | 2214.5 | -0.13658 | 0.335152 | 0.776511 | Tspan14 |
| ENSMUSG( | 1317.528 | -0.3972 | 0.011948 | 0.157992 | Tspan15 |
| ENSMUSG( | 1252.263 | 0.048407 | 0.628383 | 0.913637 | Tspan17 |
| ENSMUSG( | 305.0088 | -0.35937 | 0.061353 | 0.382309 | Tspan18 |
| ENSMUSG( | 8415.604 | -0.57274 | 2.49E-05 | 0.00228 | Tspan2 |
| ENSMUSG( | 8.402681 | -0.08944 | 0.895221 | NA | Tspan2os |
| ENSMUSG( | 16331.97 | -0.22367 | 0.064824 | 0.394687 | Tspan3 |
| ENSMUSG( | 2842.674 | -0.34109 | 0.005692 | 0.101115 | Tspan31 |
| ENSMUSG( | 11.81441 | -0.23937 | 0.666798 | NA | Tspan32 |
| ENSMUSG( | 975.4536 | -0.23854 | 0.047547 | 0.333288 | Tspan33 |
| ENSMUSG( | 444.7257 | -0.36977 | 0.007059 | 0.115775 | Tspan4 |
| ENSMUSG( | 6149.485 | -0.2832 | 0.032583 | 0.276411 | Tspan5 |
| ENSMUSG( | 711.8286 | -0.30528 | 0.025018 | 0.243138 | Tspan6 |
| ENSMUSG( | 109216.5 | -0.17423 | 0.126336 | 0.533885 | Tspan7 |
| ENSMUSG( | 40.66659 | -0.15286 | 0.649487 | 0.9077 | Tspan8 |
| ENSMUSG( | 2673.555 | -0.28648 | 0.048287 | 0.336144 | Tspan9 |
| ENSMUSG( | 177.8183 | -0.00785 | 0.960846 | 0.993991 | Tspo |
| ENSMUSG( | 18453.8 | 0.217973 | 0.092693 | 0.467791 | Tspoap1 |
| ENSMUSG( | 2558.653 | 0.050017 | 0.677701 | 0.930542 | Tspyl1 |
| ENSMUSG( | 2685.423 | 0.075992 | 0.552697 | 0.886141 | Tspyl2 |
| ENSMUSG( | 489.6837 | 0.202777 | 0.159984 | 0.589679 | Tspyl3 |
| ENSMUSG( | 7163.577 | 0.184072 | 0.038117 | 0.301297 | Tspyl4 |
| ENSMUSG( | 649.8179 | 0.220607 | 0.135236 | 0.548658 | Tspyl5 |
| ENSMUSG( | 822.6786 | 0.126944 | 0.218293 | 0.663925 | Tsr1 |
| ENSMUSG( | 2202.883 | 0.026628 | 0.775026 | 0.955606 | Tsr2 |
| ENSMUSG( | 646.8881 | -0.06095 | 0.633042 | 0.914003 | Tsr3 |
| ENSMUSG( | 960.1669 | -0.24212 | 0.013799 | 0.171544 | Tssc4 |
| ENSMUSG( | 15.00128 | -0.11851 | 0.823919 | NA | Tssk4 |
| ENSMUSG( | 3060.526 | 0.064161 | 0.521287 | 0.875186 | Tst |
| ENSMUSG( | 18.72951 | 0.333694 | 0.436808 | NA | Tstd1 |
| ENSMUSG( | 419.3403 | -0.14026 | 0.23599 | 0.683136 | Tstd2 |
| ENSMUSG( | 360.2349 | -0.16904 | 0.209382 | 0.65137 | Tstd3 |
| ENSMUSG( | 8961.314 | 0.31505 | 0.000585 | 0.022791 | Ttbk1 |
| ENSMUSG( | 1722.637 | 0.026196 | 0.800284 | 0.961745 | Ttbk2 |
| ENSMUSG( | 895.8716 | -0.03046 | 0.758769 | 0.952113 | Ttc1 |
| ENSMUSG( | 140.2214 | -0.20181 | 0.296942 | 0.745676 | Ttc12 |
| ENSMUSG( | 1260.946 | -0.21765 | 0.075296 | 0.425242 | Ttc13 |
| ENSMUSG( | 2515.257 | -0.07304 | 0.560799 | 0.888934 | Ttc14 |
| ENSMUSG( | 46.59223 | 0.226833 | 0.445532 | 0.833239 | Ttc16 |
| ENSMUSG( | 1230.134 | -0.15736 | 0.253615 | 0.703909 | Ttc17 |
| ENSMUSG( | 1909.689 | -0.29926 | 0.027264 | 0.253564 | Ttc19 |
| ENSMUSG( | 56.72654 | -0.20152 | 0.603578 | 0.904887 | Ttc21a |
| ENSMUSG( | 305.4589 | 0.171073 | 0.193729 | 0.633085 | Ttc21b |
| ENSMUSG( | 28.40091 | -0.85548 | 0.019079 | 0.207968 | Ttc22 |
| ENSMUSG( | 263.9384 | 0.096365 | 0.48187 | 0.855573 | Ttc23 |
| ENSMUSG( | 341.6518 | 0.103365 | 0.479227 | 0.854046 | Ttc27 |
| ENSMUSG( | 1055.092 | 0.006113 | 0.951872 | 0.993821 | Ttc28 |
| ENSMUSG( | 23001.24 | 0.059739 | 0.530462 | 0.879241 | Ttc3 |
| ENSMUSG( | 113.0657 | 0.06786 | 0.725078 | 0.944096 | Ttc32 |
| ENSMUSG( | 646.3494 | 0.0978 | 0.412336 | 0.819452 | Ttc33 |
| ENSMUSG( | 96.20924 | -0.17763 | 0.394529 | 0.810072 | Ttc34 |
| ENSMUSG( | 280.0057 | -0.04665 | 0.732459 | 0.944956 | Ttc38 |
| ENSMUSG( | 45.24788 | -0.10554 | 0.72131 | 0.942988 | Ttc39a |
| ENSMUSG( | 87.9344 | 0.331924 | 0.191187 | 0.630245 | Ttc39aos1 |
| ENSMUSG( | 1167.548 | -0.09194 | 0.329586 | 0.773422 | Ttc39b |
| ENSMUSG( | 588.072 | 0.035453 | 0.77269 | 0.955272 | Ttc39c |
| ENSMUSG( | 1064.186 | 0.0119 | 0.913222 | 0.988483 | Ttc4 |
| ENSMUSG( | 72.76117 | 0.218055 | 0.364187 | 0.793647 | Ttc41 |
| ENSMUSG( | 1187.954 | 0.034912 | 0.70792 | 0.939519 | Ttc5 |
| ENSMUSG( | 478.0315 | -0.02517 | 0.833515 | 0.970333 | Ttc7 |
| ENSMUSG( | 8486.582 | 0.17542 | 0.027649 | 0.255168 | Ttc7b |
| ENSMUSG( | 666.8205 | 0.113306 | 0.299309 | 0.748131 | Ttc8 |
| ENSMUSG( | 1621.786 | 0.070611 | 0.482517 | 0.856002 | Ttc9 |
| ENSMUSG( | 5763.738 | 0.069986 | 0.397856 | 0.812161 | Ttc9b |
| ENSMUSG( | 582.9444 | -0.02901 | 0.794801 | 0.959799 | Ttc9c |
| ENSMUSG( | 405.8104 | 0.142571 | 0.336464 | 0.777 | Ttf1 |
| ENSMUSG( | 164.8156 | 0.227594 | 0.203216 | 0.643631 | Ttf2 |
| ENSMUSG( | 502.9245 | 0.070995 | 0.522254 | 0.875186 | Tti1 |
| ENSMUSG( | 426.1784 | -0.09416 | 0.470119 | 0.846694 | Tti2 |
| ENSMUSG( | 19.00401 | 0.222599 | 0.655518 | NA | Ttk |
| ENSMUSG( | 3198.789 | 0.01415 | 0.870801 | 0.977951 | Ttl |
| ENSMUSG( | 740.6954 | 0.016762 | 0.894722 | 0.984512 | Ttll1 |
| ENSMUSG( | 1022.347 | 0.093737 | 0.362564 | 0.792943 | Ttll11 |
| ENSMUSG( | 755.2967 | 0.269523 | 0.016933 | 0.194376 | Ttll12 |
| ENSMUSG( | 55.72613 | 0.14951 | 0.560099 | 0.888934 | Ttll13 |
| ENSMUSG( | 526.582 | 0.23429 | 0.055027 | 0.36102 | Ttll3 |
| ENSMUSG( | 819.4062 | 0.003019 | 0.977725 | 0.997428 | Ttll4 |
| ENSMUSG( | 1377.99 | 0.098408 | 0.27652 | 0.726576 | Ttll5 |
| ENSMUSG( | 31.31702 | 0.279007 | 0.49224 | 0.860297 | Ttll6 |
| ENSMUSG( | 4412.993 | -0.00094 | 0.991877 | 0.999232 | Ttll7 |
| ENSMUSG( | 70.02563 | 0.011114 | 0.964675 | 0.995155 | Ttll8 |
| ENSMUSG( | 45.44423 | 0.237438 | 0.445711 | 0.833239 | Ttll9 |

| | | | | | |
|---|---|---|---|---|---|
| ENSMUSG( | 1059.159 | -0.02829 | 0.802604 | 0.97681 | Ttc5 |
| ENSMUSG( | 445.2887 | 0.039006 | 0.771833 | 0.974236 | Ttc7 |
| ENSMUSG( | 7293.829 | 0.01233 | 0.897772 | 0.990558 | Ttc7b |
| ENSMUSG( | 565.4873 | -0.09462 | 0.50058 | 0.927825 | Ttc8 |
| ENSMUSG( | 1406.39 | -0.07316 | 0.540913 | 0.93758 | Ttc9 |
| ENSMUSG( | 4928.99 | -0.11621 | 0.280475 | 0.892356 | Ttc9b |
| ENSMUSG( | 534.0106 | -0.01497 | 0.905857 | 0.991089 | Ttc9c |
| ENSMUSG( | 363.8434 | 0.106947 | 0.535947 | 0.936952 | Ttf1 |
| ENSMUSG( | 135.0012 | -0.06861 | 0.734349 | 0.967228 | Ttf2 |
| ENSMUSG( | 452.0553 | 0.032147 | 0.810455 | 0.978779 | Tti1 |
| ENSMUSG( | 393.9414 | -0.04567 | 0.758479 | 0.971858 | Tti2 |
| ENSMUSG( | 16.7071 | 0.116558 | 0.814993 | 0.979198 | Ttk |
| ENSMUSG( | 2711.493 | -0.20259 | 0.062401 | 0.736482 | Ttl |
| ENSMUSG( | 633.0923 | -0.17821 | 0.266389 | 0.886134 | Ttll1 |
| ENSMUSG( | 874.302 | -0.0872 | 0.496986 | 0.927825 | Ttll11 |
| ENSMUSG( | 606.4691 | -0.08085 | 0.536382 | 0.93705 | Ttll12 |
| ENSMUSG( | 49.70765 | 0.071045 | 0.809155 | 0.97873 | Ttll13 |
| ENSMUSG( | 468.0397 | 0.1706 | 0.29258 | 0.897982 | Ttll3 |
| ENSMUSG( | 797.5115 | 0.189369 | 0.136435 | 0.827092 | Ttll4 |
| ENSMUSG( | 1183.416 | -0.07353 | 0.519746 | 0.931729 | Ttll5 |
| ENSMUSG( | 22.89333 | -0.36419 | 0.403999 | 0.917257 | Ttll6 |
| ENSMUSG( | 3965.992 | -0.043 | 0.709266 | 0.964042 | Ttll7 |
| ENSMUSG( | 71.24039 | 0.318206 | 0.250366 | 0.87792 | Ttll8 |
| ENSMUSG( | 42.04212 | 0.276764 | 0.410787 | 0.917257 | Ttll9 |
| ENSMUSG( | 39.21914 | -0.24592 | 0.442512 | 0.921554 | Ttn |
| ENSMUSG( | 468.9806 | -0.08483 | 0.680798 | 0.958395 | Ttpa |
| ENSMUSG( | 1481.159 | -0.10697 | 0.338395 | 0.907972 | Ttpal |
| ENSMUSG( | 15441.47 | 0.617921 | 0.534504 | 0.936104 | Ttr |
| ENSMUSG( | 57143.9 | -0.02452 | 0.823605 | 0.979198 | Ttyh1 |
| ENSMUSG( | 1954.139 | 0.229095 | 0.253089 | 0.88018 | Ttyh2 |
| ENSMUSG( | 17614.08 | 0.218491 | 0.112041 | 0.808673 | Ttyh3 |
| ENSMUSG( | 5928.519 | 0.23275 | 0.037638 | 0.674819 | Tub |
| ENSMUSG( | 16713.77 | -0.21951 | 0.07145 | 0.761942 | Tuba1a |
| ENSMUSG( | 5227.106 | -0.22629 | 0.070039 | 0.757435 | Tuba1b |
| ENSMUSG( | 66.0493 | -0.08331 | 0.783422 | 0.976106 | Tuba1c |
| ENSMUSG( | 8987.513 | -0.27579 | 0.04837 | 0.706142 | Tuba4a |
| ENSMUSG( | 139.3897 | -0.34786 | 0.20777 | 0.864412 | Tuba8 |
| ENSMUSG( | 8115.523 | -0.21358 | 0.177092 | 0.846973 | Tubb2a |
| ENSMUSG( | 12024.1 | -0.37428 | 0.009125 | 0.401114 | Tubb2b |
| ENSMUSG( | 5764.308 | -0.28679 | 0.130071 | 0.821082 | Tubb3 |
| ENSMUSG( | 17268.13 | -0.24502 | 0.167437 | 0.840222 | Tubb4a |
| ENSMUSG( | 5344.34 | -0.23051 | 0.07537 | 0.767994 | Tubb4b |
| ENSMUSG( | 22.27183 | -0.76244 | 0.074653 | 0.767423 | Tubb4b-ps1 |
| ENSMUSG( | 12118.75 | -0.14361 | 0.274398 | 0.889874 | Tubb5 |
| ENSMUSG( | 72.95722 | -0.21411 | 0.405507 | 0.917257 | Tubb6 |
| ENSMUSG( | 104.7558 | 0.35097 | 0.145723 | 0.83474 | Tubd1 |
| ENSMUSG( | 55.42655 | 0.122434 | 0.660607 | 0.956205 | Tube1 |
| ENSMUSG( | 739.7641 | -0.10126 | 0.471253 | 0.923329 | Tubg1 |
| ENSMUSG( | 1702.295 | -0.06371 | 0.561026 | 0.940254 | Tubg2 |
| ENSMUSG( | 976.6038 | -0.06443 | 0.593555 | 0.945408 | Tubgcp2 |
| ENSMUSG( | 1022.874 | 0.104937 | 0.388733 | 0.917257 | Tubgcp3 |
| ENSMUSG( | 884.1967 | 0.009657 | 0.941666 | 0.994578 | Tubgcp4 |
| ENSMUSG( | 536.912 | 0.147505 | 0.270783 | 0.888062 | Tubgcp5 |
| ENSMUSG( | 1183.639 | 0.228758 | 0.088853 | 0.788051 | Tubgcp6 |
| ENSMUSG( | 1131.517 | -0.2358 | 0.047006 | 0.700664 | Tufm |
| ENSMUSG( | 26.57214 | 0.070846 | 0.865686 | 0.987041 | Tuft1 |
| ENSMUSG( | 2313.743 | 0.085251 | 0.546052 | 0.938063 | Tug1 |
| ENSMUSG( | 846.7039 | 0.103282 | 0.39396 | 0.917257 | Tulp3 |
| ENSMUSG( | 4687.775 | -0.09033 | 0.436807 | 0.919879 | Tulp4 |
| ENSMUSG( | 273.7736 | -0.12695 | 0.498921 | 0.927825 | Tunar |
| ENSMUSG( | 217.5733 | 0.157169 | 0.387752 | 0.917257 | Tusc1 |
| ENSMUSG( | 2436.927 | -0.07979 | 0.475858 | 0.923329 | Tusc2 |
| ENSMUSG( | 5944.471 | 0.147746 | 0.360695 | 0.914223 | Tusc3 |
| ENSMUSG( | 504.2387 | 0.028468 | 0.831321 | 0.980611 | Tut1 |
| ENSMUSG( | 742.2481 | 0.00856 | 0.947126 | 0.994798 | Tut4 |
| ENSMUSG( | 2273.827 | 0.079363 | 0.449412 | 0.922202 | Tut7 |
| ENSMUSG( | 1168.133 | 0.152608 | 0.320842 | 0.904019 | Tvp23a |
| ENSMUSG( | 925.8748 | 0.028377 | 0.831741 | 0.980656 | Tvp23b |
| ENSMUSG( | 1979.898 | -0.05473 | 0.662135 | 0.956205 | Twf1 |
| ENSMUSG( | 825.7431 | -0.00553 | 0.965323 | 0.995433 | Twf2 |
| ENSMUSG( | 29.91954 | -0.27887 | 0.454508 | 0.922444 | Twist1 |
| ENSMUSG( | 18.69973 | -0.2138 | 0.650095 | 0.95422 | Twist2 |
| ENSMUSG( | 361.8469 | 0.098979 | 0.508923 | 0.929089 | Twnk |
| ENSMUSG( | 1461.991 | 0.054132 | 0.708569 | 0.964042 | Twsg1 |
| ENSMUSG( | 1513.857 | 0.092014 | 0.443742 | 0.921734 | Txlna |
| ENSMUSG( | 98.86785 | -0.37749 | 0.117409 | 0.813384 | Txlnb |
| ENSMUSG( | 363.7207 | 0.264706 | 0.085902 | 0.780123 | Txlng |
| ENSMUSG( | 1366.868 | -0.17408 | 0.309264 | 0.898538 | Txn1 |
| ENSMUSG( | 4523.842 | -0.2101 | 0.080311 | 0.77681 | Txn2 |
| ENSMUSG( | 1052.862 | 0.346221 | 0.053102 | 0.715528 | Txndc11 |
| ENSMUSG( | 1489.743 | 0.095287 | 0.547711 | 0.938063 | Txndc12 |
| ENSMUSG( | 2439.281 | 0.186472 | 0.207138 | 0.864412 | Txndc15 |
| ENSMUSG( | 3390.521 | 0.162325 | 0.282866 | 0.892412 | Txndc16 |
| ENSMUSG( | 877.7826 | -0.11153 | 0.388039 | 0.917257 | Txndc17 |
| ENSMUSG( | 2410.944 | 0.097799 | 0.544789 | 0.938063 | Txndc5 |
| ENSMUSG( | 915.4991 | -0.10101 | 0.400988 | 0.917257 | Txndc9 |
| ENSMUSG( | 843.2703 | -0.00412 | 0.982211 | 0.996756 | Txnip |
| ENSMUSG( | 1185.473 | -0.06906 | 0.638934 | 0.953903 | Txnl1 |
| ENSMUSG( | 1619.843 | -0.03526 | 0.778918 | 0.976106 | Txnl4a |
| ENSMUSG( | 711.628 | -0.07662 | 0.553002 | 0.939565 | Txnl4a |
| ENSMUSG( | 499.3084 | -0.13383 | 0.332126 | 0.907907 | Txnl4b |
| ENSMUSG( | 1376.867 | -0.23573 | 0.037034 | 0.673514 | Txnrd1 |
| ENSMUSG( | 568.1526 | -0.26094 | 0.038955 | 0.679972 | Txnrd2 |
| ENSMUSG( | 189.4469 | -0.11881 | 0.488968 | 0.925573 | Txnrd3 |
| ENSMUSG( | 617.6443 | 0.247521 | 0.090688 | 0.791074 | Tyk2 |
| ENSMUSG( | 26.86272 | 0.005891 | 0.987749 | 0.997659 | Tymp |
| ENSMUSG( | 59.49772 | 0.323099 | 0.247335 | 0.877904 | Tyms |
| ENSMUSG( | 7785.247 | 0.246124 | 0.146051 | 0.835524 | Tyro3 |
| ENSMUSG( | 976.5158 | -0.11955 | 0.366891 | 0.914852 | Tyrobp |
| ENSMUSG( | 537.982 | -0.02853 | 0.823038 | 0.979198 | Tysnd1 |
| ENSMUSG( | 612.8491 | 0.24292 | 0.144711 | 0.833302 | Tyw1 |
| ENSMUSG( | 186.6753 | 0.13029 | 0.459361 | 0.923329 | Tyw3 |
| ENSMUSG( | 262.9484 | 0.229662 | 0.138524 | 0.828492 | Tyw5 |
| ENSMUSG( | 1053.925 | -0.05009 | 0.66565 | 0.956766 | U2af1 |
| ENSMUSG( | 3172.258 | -0.13986 | 0.208884 | 0.864412 | U2af2 |
| ENSMUSG( | 1434.67 | 0.050325 | 0.658879 | 0.956205 | U2surp |
| ENSMUSG( | 568.6758 | 0.323517 | 0.012054 | 0.445877 | Uaca |
| ENSMUSG( | 1388.889 | -0.06357 | 0.567014 | 0.941777 | Uap1 |
| ENSMUSG( | 366.2856 | -0.00449 | 0.974608 | 0.996416 | Uap1l1 |
| ENSMUSG( | 7189.534 | -0.10324 | 0.30307 | 0.898538 | Uba1 |
| ENSMUSG( | 11.38164 | -0.29482 | 0.601799 | NA | Uba1y |
| ENSMUSG( | 1522.494 | 0.068485 | 0.51661 | 0.934795 | Uba2 |
| ENSMUSG( | 1597.686 | -0.13762 | 0.270944 | 0.888062 | Uba3 |
| ENSMUSG( | 1365.337 | -0.17739 | 0.13482 | 0.824049 | Uba5 |
| ENSMUSG( | 428.2899 | -0.00137 | 0.992671 | 0.99874 | Uba52 |
| ENSMUSG( | 76.57711 | -0.72748 | 0.006739 | 0.339232 | Uba52rt |
| ENSMUSG( | 315.0712 | 0.160201 | 0.301047 | 0.898538 | Uba6 |
| ENSMUSG( | 147.2104 | 0.240895 | 0.302611 | 0.898538 | Uba7 |
| ENSMUSG( | 1381.096 | -0.05265 | 0.644697 | 0.953903 | Ubac1 |
| ENSMUSG( | 1771.389 | 0.047744 | 0.711633 | 0.964363 | Ubac2 |
| ENSMUSG( | 2442.522 | -0.12276 | 0.380854 | 0.917202 | Ubald1 |
| ENSMUSG( | 351.7347 | -0.05046 | 0.75466 | 0.971098 | Ubald2 |
| ENSMUSG( | 1211.343 | 0.001053 | 0.993603 | 0.998813 | Ubap1 |
| ENSMUSG( | 77.96802 | 0.101877 | 0.694765 | 0.960867 | Ubap1l |
| ENSMUSG( | 2021.983 | 0.021471 | 0.847278 | 0.985161 | Ubap2 |
| ENSMUSG( | 2621.714 | -0.01783 | 0.873131 | 0.989445 | Ubap2l |
| ENSMUSG( | 673.7563 | -0.26236 | 0.024257 | 0.589324 | Ubash3b |
| ENSMUSG( | 11279.99 | -0.2182 | 0.098374 | 0.798438 | Ubb |
| ENSMUSG( | 1591.227 | -0.2487 | 0.034232 | 0.652059 | Ubc |
| ENSMUSG( | 797.6329 | -0.06078 | 0.707911 | 0.964042 | Ube2a |
| ENSMUSG( | 2460.076 | -0.00631 | 0.961027 | 0.99511 | Ube2b |
| ENSMUSG( | 47.11221 | 0.238538 | 0.459897 | 0.923329 | Ube2c |
| ENSMUSG( | 943.7921 | -0.00732 | 0.956002 | 0.994798 | Ube2d1 |
| ENSMUSG( | 1968.15 | -0.03426 | 0.802956 | 0.976899 | Ube2d2a |
| ENSMUSG( | 3868.676 | 0.10776 | 0.507539 | 0.929089 | Ube2d3 |
| ENSMUSG( | 678.4669 | -0.1148 | 0.363766 | 0.914223 | Ube2d-ps |
| ENSMUSG( | 1345.425 | -0.0562 | 0.732157 | 0.967228 | Ube2e1 |
| ENSMUSG( | 1203.795 | -0.12177 | 0.416957 | 0.917257 | Ube2e2 |
| ENSMUSG( | 1566.334 | 0.056279 | 0.701187 | 0.961756 | Ube2e3 |
| ENSMUSG( | 1330.139 | -0.09545 | 0.393533 | 0.917257 | Ube2f |
| ENSMUSG( | 1130.134 | 0.057177 | 0.694251 | 0.960867 | Ube2g1 |
| ENSMUSG( | 890.6079 | -0.00335 | 0.978711 | 0.996459 | Ube2g2 |
| ENSMUSG( | 3678.169 | -0.07578 | 0.529836 | 0.934795 | Ube2h |
| ENSMUSG( | 2023.835 | 0.085876 | 0.48623 | 0.92541 | Ube2i |
| ENSMUSG( | 1640.055 | -0.07679 | 0.591806 | 0.944975 | Ube2j1 |
| ENSMUSG( | 880.0028 | 0.006965 | 0.955652 | 0.994798 | Ube2j2 |
| ENSMUSG( | 2417.171 | -0.09401 | 0.47956 | 0.924021 | Ube2k |
| ENSMUSG( | 5631.772 | -0.22909 | 0.056646 | 0.723362 | Ube2l3 |
| ENSMUSG( | 312.6121 | -0.21789 | 0.328743 | 0.906727 | Ube2l6 |
| ENSMUSG( | 5395.918 | -0.24383 | 0.050314 | 0.708648 | Ube2m |
| ENSMUSG( | 2719.595 | -0.10217 | 0.406442 | 0.917257 | Ube2n |
| ENSMUSG( | 3531.271 | -0.03171 | 0.793286 | 0.97681 | Ube2o |
| ENSMUSG( | 2576.952 | -0.09196 | 0.420966 | 0.917257 | Ube2q1 |
| ENSMUSG( | 1578.978 | -0.01353 | 0.92459 | 0.99306 | Ube2q2 |
| ENSMUSG( | 4852.724 | -0.07311 | 0.518726 | 0.930939 | Ube2ql1 |
| ENSMUSG( | 3216.635 | -0.0343 | 0.816977 | 0.979198 | Ube2r2 |
| ENSMUSG( | 849.4163 | -0.13818 | 0.304055 | 0.898538 | Ube2s |
| ENSMUSG( | 21.65236 | -0.39467 | 0.430761 | 0.917257 | Ube2srt |

| | | | | | | | | | | | |
|---|---|---|---|---|---|---|---|---|---|---|---|
| ENSMUSG( | Ubxn8 | 909.8195 | -0.29357 | 0.026252 | 0.247758 | ENSMUSG( | Ubxn8 | 937.1648 | 0.075264 | 0.593037 | 0.945408 |
| ENSMUSG( | Uchl1 | 8562.65 | -0.00202 | 0.988911 | 0.999111 | ENSMUSG( | Uchl1 | 7469.803 | -0.13429 | 0.438997 | 0.920721 |
| ENSMUSG( | Uchl1os | 12.50058 | -0.14318 | 0.81501 | NA | ENSMUSG( | Uchl1os | 12.48407 | 0.165003 | 0.791766 | 0.97681 |
| ENSMUSG( | Uchl3 | 507.1148 | 0.110575 | 0.366984 | 0.795164 | ENSMUSG( | Uchl3 | 446.6309 | 0.008079 | 0.960153 | 0.994798 |
| ENSMUSG( | Uchl5 | 869.7079 | 0.120693 | 0.233903 | 0.681906 | ENSMUSG( | Uchl5 | 721.1169 | -0.1578 | 0.236747 | 0.872173 |
| ENSMUSG( | Uck1 | 1163.646 | 0.063121 | 0.48635 | 0.858575 | ENSMUSG( | Uck1 | 1007.511 | -0.08289 | 0.468273 | 0.923329 |
| ENSMUSG( | Uck2 | 1257.739 | 0.10098 | 0.279192 | 0.729109 | ENSMUSG( | Uck2 | 1102.777 | -0.00656 | 0.95453 | 0.994798 |
| ENSMUSG( | Uckl1 | 708.2202 | 0.04242 | 0.708319 | 0.939519 | ENSMUSG( | Uckl1 | 661.6238 | 0.112242 | 0.466665 | 0.923329 |
| ENSMUSG( | Uckl1os | 18.2899 | -0.82673 | 0.074751 | NA | ENSMUSG( | Uckl1os | 22.93098 | 0.207408 | 0.626863 | 0.953234 |
| ENSMUSG( | Ucma | 47.3608 | 0.027059 | 0.926922 | 0.991229 | ENSMUSG( | Ucma | 47.38382 | 0.269697 | 0.403871 | 0.917257 |
| ENSMUSG( | Ucp2 | 1573.918 | 0.111398 | 0.300315 | 0.749479 | ENSMUSG( | Ucp2 | 1564.688 | 0.343921 | 0.008268 | 0.388416 |
| ENSMUSG( | Ucp3 | 276.5557 | -0.35378 | 0.01931 | 0.209368 | ENSMUSG( | Ucp3 | 230.4367 | -0.66945 | 9.60E-05 | 0.02045 |
| ENSMUSG( | Uevld | 311.7434 | -0.04132 | 0.754609 | 0.951007 | ENSMUSG( | Uevld | 282.4699 | -0.06178 | 0.700283 | 0.961427 |
| ENSMUSG( | Ufc1 | 1898.865 | -0.0189 | 0.832846 | 0.970333 | ENSMUSG( | Ufc1 | 1643.665 | -0.17559 | 0.150738 | 0.837173 |
| ENSMUSG( | Ufd1 | 1838.98 | 0.090103 | 0.32922 | 0.773233 | ENSMUSG( | Ufd1 | 1619.497 | -0.00823 | 0.942955 | 0.994774 |
| ENSMUSG( | Ufl1 | 1008.515 | -0.21862 | 0.058327 | 0.372466 | ENSMUSG( | Ufl1 | 1022.97 | 0.102799 | 0.443368 | 0.921704 |
| ENSMUSG( | Ufm1 | 2618.456 | -0.01186 | 0.912573 | 0.988483 | ENSMUSG( | Ufm1 | 2283.567 | -0.14522 | 0.280357 | 0.892356 |
| ENSMUSG( | Ufsp1 | 233.9356 | -0.44406 | 0.002186 | 0.054296 | ENSMUSG( | Ufsp1 | 227.9104 | -0.22396 | 0.183253 | 0.850984 |
| ENSMUSG( | Ufsp2 | 1092.482 | 0.040034 | 0.711532 | 0.940028 | ENSMUSG( | Ufsp2 | 927.2075 | -0.17333 | 0.180512 | 0.850372 |
| ENSMUSG( | Ugcg | 2176.725 | -0.55313 | 0.000488 | 0.02028 | ENSMUSG( | Ugcg | 2513.384 | 0.178727 | 0.323132 | 0.904023 |
| ENSMUSG( | Ugdh | 821.2971 | -0.07583 | 0.479923 | 0.854483 | ENSMUSG( | Ugdh | 751.6179 | -0.06387 | 0.590358 | 0.944975 |
| ENSMUSG( | Uggt1 | 2410.612 | -0.11961 | 0.347565 | 0.780601 | ENSMUSG( | Uggt1 | 2417.149 | 0.158546 | 0.284893 | 0.892412 |
| ENSMUSG( | Uggt2 | 792.9701 | 0.012977 | 0.90699 | 0.987407 | ENSMUSG( | Uggt2 | 741.0736 | 0.090039 | 0.564782 | 0.941206 |
| ENSMUSG( | Ugp2 | 4887.174 | 0.019348 | 0.879844 | 0.981436 | ENSMUSG( | Ugp2 | 4100.766 | -0.22917 | 0.119891 | 0.813384 |
| ENSMUSG( | Ugt1a7c | 31.87806 | -0.00168 | 0.996258 | 0.99943 | ENSMUSG( | Ugt1a7c | 29.71812 | 0.093698 | 0.817774 | 0.979198 |
| ENSMUSG( | Ugt8a | 2586.34 | -0.57782 | 0.001333 | 0.039731 | ENSMUSG( | Ugt8a | 3042.74 | 0.212577 | 0.336956 | 0.907972 |
| ENSMUSG( | Uhmk1 | 9298.296 | 0.062127 | 0.548773 | 0.885686 | ENSMUSG( | Uhmk1 | 8110.268 | -0.06516 | 0.588506 | 0.944975 |
| ENSMUSG( | Uhrf1 | 83.0498 | 0.142534 | 0.558272 | 0.888427 | ENSMUSG( | Uhrf1 | 71.69391 | 0.007412 | 0.976777 | 0.996416 |
| ENSMUSG( | Uhrf2 | 891.7829 | 0.164499 | 0.114793 | 0.510823 | ENSMUSG( | Uhrf2 | 835.7217 | 0.240913 | 0.040313 | 0.682289 |
| ENSMUSG( | Uimc1 | 492.1145 | 0.139164 | 0.268169 | 0.718372 | ENSMUSG( | Uimc1 | 438.1016 | 0.072041 | 0.630734 | 0.953251 |
| ENSMUSG( | Ulk1 | 3128.405 | 0.260867 | 0.006912 | 0.114527 | ENSMUSG( | Ulk1 | 2736.047 | 0.152067 | 0.218683 | 0.866104 |
| ENSMUSG( | Ulk2 | 5093.65 | 0.127174 | 0.203302 | 0.643631 | ENSMUSG( | Ulk2 | 4480.132 | 0.027265 | 0.819869 | 0.979198 |
| ENSMUSG( | Ulk3 | 709.4068 | 0.167994 | 0.261878 | 0.712129 | ENSMUSG( | Ulk3 | 604.2891 | -0.02551 | 0.883736 | 0.989644 |
| ENSMUSG( | Ulk4 | 240.043 | -0.00442 | 0.976577 | 0.997428 | ENSMUSG( | Ulk4 | 223.0076 | 0.05736 | 0.729086 | 0.966913 |
| ENSMUSG( | Umad1 | 484.9143 | -0.04476 | 0.774034 | 0.955606 | ENSMUSG( | Umad1 | 415.1597 | -0.24365 | 0.211229 | 0.864412 |
| ENSMUSG( | Umps | 526.9236 | 0.279545 | 0.024117 | 0.237752 | ENSMUSG( | Umps | 423.9098 | -0.07417 | 0.636144 | 0.953903 |
| ENSMUSG( | Unc119 | 792.9185 | 0.110891 | 0.319362 | 0.765296 | ENSMUSG( | Unc119 | 681.5982 | -0.05945 | 0.667498 | 0.956766 |
| ENSMUSG( | Unc119b | 1564.979 | 0.123095 | 0.154115 | 0.580901 | ENSMUSG( | Unc119b | 1325.806 | -0.08495 | 0.433471 | 0.918564 |
| ENSMUSG( | Unc13a | 12978.02 | 0.092023 | 0.308162 | 0.7563 | ENSMUSG( | Unc13a | 11438.85 | -0.00222 | 0.985271 | 0.997084 |
| ENSMUSG( | Unc13b | 2091.65 | 0.14291 | 0.20743 | 0.649561 | ENSMUSG( | Unc13b | 1779.19 | -0.05339 | 0.689871 | 0.960071 |
| ENSMUSG( | Unc13c | 4941.247 | -0.28349 | 0.106988 | 0.494682 | ENSMUSG( | Unc13c | 4314.397 | -0.42845 | 0.010701 | 0.420304 |
| ENSMUSG( | Unc13d | 54.25233 | 0.010488 | 0.971266 | 0.996369 | ENSMUSG( | Unc13d | 47.76773 | 0.098648 | 0.77859 | 0.976106 |
| ENSMUSG( | Unc45a | 1031.325 | 0.102085 | 0.275986 | 0.725669 | ENSMUSG( | Unc45a | 905.3794 | -0.00253 | 0.982225 | 0.996756 |
| ENSMUSG( | Unc45b | 83.33582 | 0.222087 | 0.325763 | 0.771061 | ENSMUSG( | Unc45b | 75.41512 | 0.216813 | 0.403566 | 0.917257 |
| ENSMUSG( | Unc50 | 1174.237 | -0.2452 | 0.063588 | 0.390386 | ENSMUSG( | Unc50 | 1238.637 | 0.186094 | 0.206735 | 0.864412 |
| ENSMUSG( | Unc5a | 3576.892 | -0.05302 | 0.695907 | 0.934979 | ENSMUSG( | Unc5a | 3764.184 | 0.347235 | 0.021645 | 0.575341 |
| ENSMUSG( | Unc5b | 1002.331 | -0.62879 | 0.000643 | 0.024438 | ENSMUSG( | Unc5b | 1152.84 | 0.117857 | 0.596732 | 0.946541 |
| ENSMUSG( | Unc5c | 948.8395 | -0.64369 | 9.83E-05 | 0.006375 | ENSMUSG( | Unc5c | 1075.309 | 0.062718 | 0.74119 | 0.968811 |
| ENSMUSG( | Unc5d | 1018.459 | -0.2068 | 0.135033 | 0.548656 | ENSMUSG( | Unc5d | 1111.272 | 0.307567 | 0.041948 | 0.690304 |
| ENSMUSG( | Unc79 | 1858.182 | 0.256744 | 0.007728 | 0.122031 | ENSMUSG( | Unc79 | 1747.821 | 0.343098 | 0.00693 | 0.343883 |
| ENSMUSG( | Unc80 | 7743.877 | 0.006434 | 0.942706 | 0.993415 | ENSMUSG( | Unc80 | 7321.158 | 0.111243 | 0.29171 | 0.897854 |
| ENSMUSG( | Unc93b1 | 1200.174 | -0.15128 | 0.256838 | 0.708006 | ENSMUSG( | Unc93b1 | 1213.783 | 0.155061 | 0.408849 | 0.917257 |
| ENSMUSG( | Ung | 52.50206 | -0.03571 | 0.900065 | 0.985633 | ENSMUSG( | Ung | 46.86408 | -0.09634 | 0.756896 | 0.971858 |
| ENSMUSG( | Unk | 755.7728 | -0.03376 | 0.776091 | 0.955606 | ENSMUSG( | Unk | 704.787 | 0.03394 | 0.828025 | 0.980224 |
| ENSMUSG( | Unkl | 621.9606 | 0.074053 | 0.484624 | 0.856665 | ENSMUSG( | Unkl | 557.5582 | 0.033358 | 0.805995 | 0.977674 |
| ENSMUSG( | Uox | 15.79057 | -0.27732 | 0.591149 | NA | ENSMUSG( | Uox | 16.91336 | 0.207138 | 0.668675 | 0.956766 |
| ENSMUSG( | Upb1 | 12.59165 | -0.9829 | 0.110125 | NA | ENSMUSG( | Upb1 | 13.75171 | -0.32403 | 0.538668 | 0.937216 |
| ENSMUSG( | Upf1 | 2140.128 | 0.109434 | 0.2726 | 0.722478 | ENSMUSG( | Upf1 | 1864.198 | -0.01864 | 0.884061 | 0.989644 |
| ENSMUSG( | Upf2 | 1250.791 | 0.157125 | 0.133423 | 0.546357 | ENSMUSG( | Upf2 | 1086.453 | 0.019552 | 0.875008 | 0.989644 |
| ENSMUSG( | Upf3a | 1481.712 | 0.027749 | 0.791007 | 0.959461 | ENSMUSG( | Upf3a | 1292.173 | -0.10477 | 0.430012 | 0.917257 |
| ENSMUSG( | Upf3b | 562.8029 | 0.003031 | 0.977577 | 0.997428 | ENSMUSG( | Upf3b | 541.9227 | 0.159657 | 0.225675 | 0.868468 |
| ENSMUSG( | Upk1a | 28.1291 | 0.397404 | 0.295027 | 0.743798 | ENSMUSG( | Upk1a | 23.66343 | 0.174799 | 0.656003 | 0.955533 |
| ENSMUSG( | Upk1b | 41.54388 | -0.2053 | 0.489688 | 0.859319 | ENSMUSG( | Upk1b | 36.8809 | -0.24961 | 0.465977 | 0.923329 |
| ENSMUSG( | Upp1 | 101.9014 | 0.343621 | 0.107218 | 0.495579 | ENSMUSG( | Upp1 | 88.83322 | 0.211839 | 0.366606 | 0.914703 |
| ENSMUSG( | Upp2 | 1075.056 | 0.138881 | 0.153238 | 0.57989 | ENSMUSG( | Upp2 | 880.6026 | -0.17434 | 0.163635 | 0.838291 |
| ENSMUSG( | Uprt | 131.7125 | 0.093528 | 0.632623 | 0.914073 | ENSMUSG( | Uprt | 111.707 | -0.12194 | 0.570543 | 0.941872 |
| ENSMUSG( | Uqcc1 | 1616.867 | -0.08462 | 0.32327 | 0.768415 | ENSMUSG( | Uqcc1 | 1457.332 | -0.11795 | 0.287362 | 0.894091 |
| ENSMUSG( | Uqcc2 | 2336.624 | -0.02754 | 0.767742 | 0.954621 | ENSMUSG( | Uqcc2 | 1923.954 | -0.34323 | 0.005313 | 0.299238 |
| ENSMUSG( | Uqcc3 | 666.1971 | 0.041448 | 0.685406 | 0.932981 | ENSMUSG( | Uqcc3 | 587.9613 | -0.05367 | 0.690859 | 0.960071 |
| ENSMUSG( | Uqcc4 | 767.9651 | -0.09543 | 0.336548 | 0.777 | ENSMUSG( | Uqcc4 | 703.0109 | -0.08473 | 0.504003 | 0.927825 |
| ENSMUSG( | Uqcc5 | 248.9065 | -0.10353 | 0.486692 | 0.858713 | ENSMUSG( | Uqcc5 | 227.5681 | -0.10817 | 0.564511 | 0.941206 |
| ENSMUSG( | Uqcc6 | 124.0139 | -0.03749 | 0.842713 | 0.971663 | ENSMUSG( | Uqcc6 | 85.53916 | -1.0322 | 1.78E-05 | 0.005763 |
| ENSMUSG( | Uqcr10 | 3631.371 | 0.032511 | 0.695264 | 0.934948 | ENSMUSG( | Uqcr10 | 2995.99 | -0.26846 | 0.021758 | 0.575341 |
| ENSMUSG( | Uqcr11 | 6959.965 | -0.01658 | 0.829221 | 0.969286 | ENSMUSG( | Uqcr11 | 5742.299 | -0.3225 | 0.003759 | 0.244783 |
| ENSMUSG( | Uqcrb | 2102.193 | -0.01406 | 0.884359 | 0.982647 | ENSMUSG( | Uqcrb | 1778.793 | -0.24086 | 0.047108 | 0.700664 |
| ENSMUSG( | Uqcrc1 | 8040.713 | 0.071192 | 0.425086 | 0.82478 | ENSMUSG( | Uqcrc1 | 6736.749 | -0.17743 | 0.145983 | 0.835406 |
| ENSMUSG( | Uqcrc2 | 4932.853 | 0.045114 | 0.64784 | 0.919759 | ENSMUSG( | Uqcrc2 | 4130.17 | -0.20875 | 0.112384 | 0.809134 |
| ENSMUSG( | Uqcrfs1 | 3681.319 | 0.051658 | 0.627347 | 0.912941 | ENSMUSG( | Uqcrfs1 | 2995.076 | -0.2931 | 0.029887 | 0.61992 |
| ENSMUSG( | Uqcrh | 6857.805 | 0.041984 | 0.595449 | 0.903058 | ENSMUSG( | Uqcrh | 5709.645 | -0.22849 | 0.052919 | 0.71525 |
| ENSMUSG( | Uqcrq | 4418.433 | -0.01915 | 0.820535 | 0.965888 | ENSMUSG( | Uqcrq | 3686.785 | -0.29026 | 0.021425 | 0.575341 |
| ENSMUSG( | Urah | 14.53743 | 0.788315 | 0.113429 | NA | ENSMUSG( | Urah | 11.94182 | 0.553082 | 0.335687 | 0.907907 |
| ENSMUSG( | Urb1 | 387.0465 | -0.13667 | 0.295027 | 0.743798 | ENSMUSG( | Urb1 | 384.4697 | 0.124289 | 0.446139 | 0.921734 |
| ENSMUSG( | Urb2 | 310.7218 | -0.16052 | 0.31453 | 0.761558 | ENSMUSG( | Urb2 | 305.7763 | 0.069945 | 0.643386 | 0.953903 |
| ENSMUSG( | Urgcp | 1481.288 | 0.095689 | 0.388567 | 0.806278 | ENSMUSG( | Urgcp | 1295.29 | -0.02071 | 0.864181 | 0.986552 |
| ENSMUSG( | Uri1 | 1987.444 | 0.058684 | 0.538963 | 0.882849 | ENSMUSG( | Uri1 | 1774.063 | -0.00459 | 0.976266 | 0.996416 |
| ENSMUSG( | Urm1 | 1764.173 | 3.54E-05 | 0.999707 | 0.99989 | ENSMUSG( | Urm1 | 1504.954 | -0.19664 | 0.082137 | 0.77681 |
| ENSMUSG( | Urod | 955.9335 | 0.151847 | 0.162797 | 0.593778 | ENSMUSG( | Urod | 814.8968 | -0.04182 | 0.77105 | 0.974236 |
| ENSMUSG( | Uros | 614.6642 | 0.010297 | 0.92821 | 0.991346 | ENSMUSG( | Uros | 555.3216 | -0.01752 | 0.917315 | 0.992529 |
| ENSMUSG( | Usb1 | 258.3057 | -0.02488 | 0.879048 | 0.981436 | ENSMUSG( | Usb1 | 235.0191 | -0.02582 | 0.880015 | 0.989644 |
| ENSMUSG( | Use1 | 2626.748 | 0.065697 | 0.444049 | 0.833239 | ENSMUSG( | Use1 | 2249.994 | -0.11596 | 0.289917 | 0.896726 |
| ENSMUSG( | Usf1 | 1380.874 | 0.110124 | 0.283303 | 0.73186 | ENSMUSG( | Usf1 | 1199.761 | -0.02611 | 0.836484 | 0.981335 |
| ENSMUSG( | Usf2 | 4722.08 | 0.124641 | 0.132646 | 0.544824 | ENSMUSG( | Usf2 | 4108.265 | -0.00815 | 0.944637 | 0.994798 |
| ENSMUSG( | Usf3 | 1803.04 | 0.013973 | 0.898392 | 0.985633 | ENSMUSG( | Usf3 | 1697.723 | 0.109737 | 0.362978 | 0.914223 |
| ENSMUSG( | Ush1g | 12.19892 | -0.36769 | 0.541479 | NA | ENSMUSG( | Ush1g | 13.32856 | 0.148063 | 0.783786 | 0.976154 |
| ENSMUSG( | Ushbp1 | 222.7546 | 0.080559 | 0.604011 | 0.905187 | ENSMUSG( | Ushbp1 | 220.6221 | 0.320168 | 0.228762 | 0.868507 |
| ENSMUSG( | Uso1 | 2770.664 | 0.000223 | 0.997931 | 0.99989 | ENSMUSG( | Uso1 | 2438.421 | -0.10275 | 0.319592 | 0.903535 |
| ENSMUSG( | Usp1 | 895.7783 | 0.075903 | 0.521576 | 0.875186 | ENSMUSG( | Usp1 | 844.7414 | 0.169116 | 0.238577 | 0.872849 |
| ENSMUSG( | Usp10 | 1116.258 | 0.15842 | 0.184568 | 0.621633 | ENSMUSG( | Usp10 | 986.0359 | 0.074103 | 0.572772 | 0.94262 |
| ENSMUSG( | Usp11 | 2549.281 | 0.102017 | 0.398912 | 0.812878 | ENSMUSG( | Usp11 | 2159.911 | -0.10988 | 0.4605 | 0.923329 |
| ENSMUSG( | Usp12 | 1925.552 | 0.040507 | 0.630448 | 0.914073 | ENSMUSG( | Usp12 | 1739.478 | 0.015046 | 0.885992 | 0.989854 |
| ENSMUSG( | Usp13 | 709.6772 | 0.117654 | 0.384036 | 0.803646 | ENSMUSG( | Usp13 | 636.6109 | 0.076105 | 0.563608 | 0.941206 |
| ENSMUSG( | Usp14 | 2008.448 | -0.00404 | 0.970657 | 0.996369 | ENSMUSG( | Usp14 | 1797.083 | -0.0605 | 0.656847 | 0.955567 |
| ENSMUSG( | Usp15 | 1545.004 | 0.087463 | 0.425171 | 0.82478 | ENSMUSG( | Usp15 | 1374.326 | 0.022288 | 0.851432 | 0.985541 |
| ENSMUSG( | Usp16 | 1172.027 | 0.011219 | 0.921494 | 0.990436 | ENSMUSG( | Usp16 | 1066.061 | 0.010572 | 0.934854 | 0.99306 |
| ENSMUSG( | Usp18 | 59.17891 | -0.55919 | 0.045268 | 0.326476 | ENSMUSG( | Usp18 | 60.05878 | -0.18883 | 0.482695 | 0.924615 |
| ENSMUSG( | Usp19 | 3480.881 | 0.251108 | 0.009994 | 0.142653 | ENSMUSG( | Usp19 | 3067.725 | 0.162707 | 0.202653 | 0.864412 |
| ENSMUSG( | Usp2 | 2122.284 | -0.15128 | 0.150723 | 0.575502 | ENSMUSG( | Usp2 | 2357.346 | 0.399334 | 0.00136 | 0.134176 |
| ENSMUSG( | Usp20 | 2202.378 | 0.156177 | 0.12111 | 0.525666 | ENSMUSG( | Usp20 | 1904.122 | 0.011572 | 0.922325 | 0.992644 |
| ENSMUSG( | Usp21 | 1421.322 | 0.130932 | 0.227291 | 0.674223 | ENSMUSG( | Usp21 | 1285.57 | 0.108483 | 0.408075 | 0.917257 |
| ENSMUSG( | Usp22 | 5317.967 | -0.09159 | 0.343242 | 0.779153 | ENSMUSG( | Usp22 | 4866.846 | -0.0761 | 0.497145 | 0.927825 |
| ENSMUSG( | Usp24 | 3615.143 | 0.01823 | 0.845752 | 0.972191 | ENSMUSG( | Usp24 | 3295.807 | 0.021271 | 0.844195 | 0.983576 |
| ENSMUSG( | Usp25 | 981.2639 | 0.135792 | 0.233962 | 0.681906 | ENSMUSG( | Usp25 | 936.7388 | 0.263194 | 0.039817 | 0.681706 |
| ENSMUSG( | Usp27x | 293.5932 | 0.145988 | 0.344412 | 0.779153 | ENSMUSG( | Usp27x | 261.523 | 0.076179 | 0.685127 | 0.958678 |
| ENSMUSG( | Usp28 | 1053.523 | -0.06831 | 0.5242 | 0.87583 | ENSMUSG( | Usp28 | 942.9609 | -0.12188 | 0.339031 | 0.907972 |
| ENSMUSG( | Usp29 | 647.5716 | -0.01851 | 0.895584 | 0.984512 | ENSMUSG( | Usp29 | 593.149 | -0.00551 | 0.972296 | 0.996416 |
| ENSMUSG( | Usp3 | 665.3565 | 0.032605 | 0.754766 | 0.951007 | ENSMUSG( | Usp3 | 600.2282 | 0.003436 | 0.977754 | 0.996416 |
| ENSMUSG( | Usp30 | 1602.817 | -0.22225 | 0.083661 | 0.44608 | ENSMUSG( | Usp30 | 1568.407 | -0.00228 | 0.987923 | 0.997777 |
| ENSMUSG( | Usp31 | 2593.462 | 0.147468 | 0.138528 | 0.553838 | ENSMUSG( | Usp31 | 2320.152 | 0.098849 | 0.383711 | 0.917257 |
| ENSMUSG( | Usp32 | 4894.712 | 0.076313 | 0.38418 | 0.803686 | ENSMUSG( | Usp32 | 4265.862 | -0.05129 | 0.625385 | 0.953234 |
| ENSMUSG( | Usp33 | 2322.052 | -0.0351 | 0.674194 | 0.929638 | ENSMUSG( | Usp33 | 2182.044 | 0.052035 | 0.625126 | 0.953234 |
| ENSMUSG( | Usp34 | 4678.69 | 0.077797 | 0.359483 | 0.791011 | ENSMUSG( | Usp34 | 4235.688 | 0.059361 | 0.555782 | 0.939565 |
| ENSMUSG( | Usp35 | 647.778 | 0.103016 | 0.378367 | 0.801688 | ENSMUSG( | Usp35 | 571.5898 | 0.014162 | 0.917449 | 0.992529 |
| ENSMUSG( | Usp36 | 1243.649 | -0.04599 | 0.674908 | 0.929651 | ENSMUSG( | Usp36 | 1110.475 | -0.10353 | 0.388247 | 0.917257 |
| ENSMUSG( | Usp37 | 682.3558 | 0.141063 | 0.22948 | 0.67735 | ENSMUSG( | Usp37 | 625.055 | 0.154853 | 0.236298 | 0.872173 |
| ENSMUSG( | Usp38 | 658.0862 | 0.065184 | 0.575457 | 0.894836 | ENSMUSG( | Usp38 | 625.0592 | 0.180875 | 0.172995 | 0.843788 |
| ENSMUSG( | Usp39 | 609.3046 | 0.122842 | 0.330402 | 0.774327 | ENSMUSG( | Usp39 | 538.5572 | 0.039669 | 0.782518 | 0.976106 |
| ENSMUSG( | Usp4 | 2754.307 | 0.082209 | 0.317954 | 0.764145 | ENSMUSG( | Usp4 | 2368.547 | -0.08514 | 0.422445 | 0.917257 |
| ENSMUSG( | Usp40 | 834.2363 | 0.274444 | 0.012463 | 0.161344 | ENSMUSG( | Usp40 | 730.6473 | 0.174081 | 0.18578 | 0.855305 |
| ENSMUSG( | Usp42 | 627.9405 | -0.13478 | 0.236209 | 0.683188 | ENSMUSG( | Usp42 | 572.4363 | -0.12642 | 0.362941 | 0.914223 |
| ENSMUSG( | Usp43 | 206.7715 | -0.05265 | 0.762457 | 0.953519 | ENSMUSG( | Usp43 | 175.196 | -0.25855 | 0.20649 | 0.864412 |
| ENSMUSG( | Usp45 | 741.4955 | 0.001069 | 0.991735 | 0.999232 | ENSMUSG( | Usp45 | 699.5754 | 0.099673 | 0.41833 | 0.917257 |
| ENSMUSG( | Usp46 | 4048.619 | 0.047826 | 0.674259 | 0.929638 | ENSMUSG( | Usp46 | 3521.211 | -0.09 | 0.538106 | 0.937216 |
| ENSMUSG( | Usp47 | 2605.184 | 0.192018 | 0.026637 | 0.249596 | ENSMUSG( | Usp47 | 2218.85 | 0.003831 | 0.971112 | 0.996102 |
| ENSMUSG( | Usp48 | 3266.992 | -0.03259 | 0.71921 | 0.941986 | ENSMUSG( | Usp48 | 2977.543 | -0.03108 | 0.780754 | 0.976106 |
| ENSMUSG( | Usp49 | 478.7201 | 0.060753 | 0.628967 | 0.913816 | ENSMUSG( | Usp49 | 442.0345 | 0.094583 | 0.514854 | 0.930939 |
| ENSMUSG( | Usp5 | 4459.453 | 0.102352 | 0.392358 | 0.809274 | ENSMUSG( | Usp5 | 3792.656 | -0.09756 | 0.4665 | 0.923329 |
| ENSMUSG( | Usp51 | 22.85989 | -0.1786 | 0.656054 | 0.923929 | ENSMUSG( | Usp51 | 25.84367 | 0.408692 | 0.342781 | 0.909646 |
| ENSMUSG( | Usp53 | 1140.658 | 0.436846 | 0.000362 | 0.016539 | ENSMUSG( | Usp53 | 1092.512 | 0.561175 | 6.38E-05 | 0.015522 |
| ENSMUSG( | Usp54 | 4231.976 | 0.101427 | 0.258369 | 0.709574 | ENSMUSG( | Usp54 | 3977.401 | 0.18895 | 0.093103 | 0.795363 |
| ENSMUSG( | Usp6nl | 1722.566 | 0.109832 | 0.213502 | 0.65774 | ENSMUSG( | Usp6nl | 1507.58 | -0.00491 | 0.965402 | 0.995433 |
| ENSMUSG( | Usp7 | 4650.58 | 0.088402 | 0.25945 | 0.709948 | ENSMUSG( | Usp7 | 4209.532 | 0.070571 | 0.468257 | 0.923329 |
| ENSMUSG( | Usp8 | 2461.977 | 0.084119 | 0.37158 | 0.798768 | ENSMUSG( | Usp8 | 2096.831 | -0.11416 | 0.325912 | 0.905339 |
| ENSMUSG( | Usp9x | 3366.76 | 0.17039 | 0.171752 | 0.606517 | ENSMUSG( | Usp9x | 2910.676 | 0.025172 | 0.840006 | 0.98221 |
| ENSMUSG( | Uspl1 | 854.2538 | 0.024499 | 0.818161 | 0.965837 | ENSMUSG( | Uspl1 | 794.2979 | 0.083203 | 0.498244 | 0.927825 |
| ENSMUSG( | Ust | 1161.61 | -0.19982 | 0.161181 | 0.591083 | ENSMUSG( | Ust | 1185.595 | 0.137712 | 0.40776 | 0.917257 |
| ENSMUSG( | Utp11 | 478.534 | 0.156309 | 0.16837 | 0.601667 | ENSMUSG( | Utp11 | 439.5697 | 0.174058 | 0.216543 | 0.865167 |
| ENSMUSG( | Utp14a | 496.0494 | 0.238606 | 0.032358 | 0.27508 | ENSMUSG( | Utp14a | 410.8749 | -0.02956 | 0.832648 | 0.980867 |
| ENSMUSG( | Utp14b | 1274.269 | 0.276585 | 0.137788 | 0.552903 | ENSMUSG( | Utp14b | 841.0072 | -0.73893 | 3.70E-08 | 2.70E-05 |
| ENSMUSG( | Utp15 | 560.7019 | -0.02423 | 0.82823 | 0.968966 | ENSMUSG( | Utp15 | 543.7772 | 0.157041 | 0.244178 | 0.877904 |
| ENSMUSG( | Utp18 | 486.1249 | 0.063906 | 0.587427 | 0.900403 | ENSMUSG( | Utp18 | 447.0005 | 0.082198 | 0.591595 | 0.944975 |
| ENSMUSG( | Utp20 | 595.9549 | 0.086571 | 0.430509 | 0.826896 | ENSMUSG( | Utp20 | 562.6705 | 0.18802 | 0.180389 | 0.850372 |
| ENSMUSG( | Utp23 | 188.7645 | 0.295053 | 0.064095 | 0.392013 | ENSMUSG( | Utp23 | 155.4602 | 0.0238 | 0.900838 | 0.990558 |
| ENSMUSG( | Utp25 | 421.7975 | 0.09097 | 0.488668 | 0.859092 | ENSMUSG( | Utp25 | 373.2893 | 0.011979 | 0.932999 | 0.99306 |
| ENSMUSG( | Utp3 | 529.5028 | 0.156495 | 0.242824 | 0.690275 | ENSMUSG( | Utp3 | 467.9618 | 0.067382 | 0.66928 | 0.956766 |
| ENSMUSG( | Utp4 | 550.4278 | 0.176438 | 0.199371 | 0.640054 | ENSMUSG( | Utp4 | 454.0964 | -0.10888 | 0.499422 | 0.927825 |
| ENSMUSG( | Utp6 | 1599.519 | 0.290012 | 0.05701 | 0.368322 | ENSMUSG( | Utp6 | 1336.824 | 0.057264 | 0.761175 | 0.972695 |
| ENSMUSG( | Utrn | 3952.045 | 0.0945 | 0.396373 | 0.811742 | ENSMUSG( | Utrn | 3475.126 | -0.00515 | 0.962098 | 0.995237 |
| ENSMUSG( | Uty | 297.7912 | 0.009084 | 0.95032 | 0.993821 | ENSMUSG( | Uty | 295.602 | 0.251902 | 0.123803 | 0.814691 |

| | | | | | |
|---|---|---|---|---|---|
| ENSMUSG( | 32.89421 | -1.2673 | 0.000173 | 0.00947 | Ttn |
| ENSMUSG( | 522.8447 | -0.0304 | 0.879279 | 0.981436 | Ttpa |
| ENSMUSG( | 1667.882 | -0.03289 | 0.728719 | 0.944393 | Ttpal |
| ENSMUSG( | 12964.4 | -0.07646 | 0.943935 | 0.993503 | Ttr |
| ENSMUSG( | 64018.02 | 0.033955 | 0.727246 | 0.944223 | Ttyh1 |
| ENSMUSG( | 1602.417 | -0.6915 | 3.19E-05 | 0.002843 | Ttyh2 |
| ENSMUSG( | 17516.33 | -0.06159 | 0.598286 | 0.903801 | Ttyh3 |
| ENSMUSG( | 6515.32 | 0.235108 | 0.015851 | 0.186421 | Tub |
| ENSMUSG( | 19507.45 | -0.03368 | 0.654072 | 0.923008 | Tuba1a |
| ENSMUSG( | 6321.689 | 0.063779 | 0.460481 | 0.840787 | Tuba1b |
| ENSMUSG( | 73.84744 | -0.00351 | 0.989216 | 0.999111 | Tuba1c |
| ENSMUSG( | 11043.88 | 0.064224 | 0.550613 | 0.886038 | Tuba4a |
| ENSMUSG( | 177.3636 | 0.112219 | 0.621431 | 0.911432 | Tuba8 |
| ENSMUSG( | 9978.394 | 0.121865 | 0.272117 | 0.7223 | Tubb2a |
| ENSMUSG( | 15006.72 | 0.02221 | 0.859907 | 0.974818 | Tubb2b |
| ENSMUSG( | 6740.902 | -0.08718 | 0.555803 | 0.887457 | Tubb3 |
| ENSMUSG( | 19548.25 | -0.14622 | 0.275114 | 0.72509 | Tubb4a |
| ENSMUSG( | 6387.989 | 0.026914 | 0.785984 | 0.958774 | Tubb4b |
| ENSMUSG( | 31.95892 | 0.107228 | 0.749709 | 0.949664 | Tubb4b-ps1 |
| ENSMUSG( | 14101.56 | 0.029982 | 0.7559 | 0.951479 | Tubb5 |
| ENSMUSG( | 79.45986 | -0.22492 | 0.326184 | 0.771128 | Tubb6 |
| ENSMUSG( | 109.2068 | 0.19401 | 0.344762 | 0.779153 | Tubd1 |
| ENSMUSG( | 55.75826 | -0.14029 | 0.582986 | 0.898094 | Tube1 |
| ENSMUSG( | 877.9663 | 0.122541 | 0.3462 | 0.779153 | Tubg1 |
| ENSMUSG( | 1954.442 | 0.066707 | 0.460235 | 0.840676 | Tubg2 |
| ENSMUSG( | 1117.293 | 0.053457 | 0.59887 | 0.903801 | Tubgcp2 |
| ENSMUSG( | 1101.067 | 0.049254 | 0.625768 | 0.912274 | Tubgcp3 |
| ENSMUSG( | 984.9692 | 0.049761 | 0.632381 | 0.914073 | Tubgcp4 |
| ENSMUSG( | 570.5782 | 0.051644 | 0.628505 | 0.913637 | Tubgcp5 |
| ENSMUSG( | 1289.862 | 0.205712 | 0.057659 | 0.370247 | Tubgcp6 |
| ENSMUSG( | 1373.521 | 0.064118 | 0.516815 | 0.87302 | Tufm |
| ENSMUSG( | 29.85849 | 0.124957 | 0.745428 | 0.949158 | Tuft1 |
| ENSMUSG( | 2274.685 | -0.24452 | 0.048448 | 0.336526 | Tug1 |
| ENSMUSG( | 953.0375 | 0.174594 | 0.083257 | 0.445247 | Tulp3 |
| ENSMUSG( | 4983.014 | -0.18742 | 0.045491 | 0.327506 | Tulp4 |
| ENSMUSG( | 315.1437 | 0.022277 | 0.888884 | 0.983434 | Tunar |
| ENSMUSG( | 256.1494 | 0.353556 | 0.030028 | 0.26299 | Tusc1 |
| ENSMUSG( | 2839.475 | 0.093726 | 0.253093 | 0.70338 | Tusc2 |
| ENSMUSG( | 5406.786 | -0.42564 | 0.002644 | 0.061702 | Tusc3 |
| ENSMUSG( | 569.0288 | 0.109138 | 0.438779 | 0.830714 | Tut1 |
| ENSMUSG( | 749.6181 | -0.24455 | 0.024323 | 0.238636 | Tut4 |
| ENSMUSG( | 2497.457 | 0.081707 | 0.338423 | 0.777489 | Tut7 |
| ENSMUSG( | 1213.532 | -0.00804 | 0.952873 | 0.993874 | Tvp23a |
| ENSMUSG( | 965.0192 | -0.12392 | 0.29875 | 0.747452 | Tvp23b |
| ENSMUSG( | 2260.672 | 0.0615 | 0.568115 | 0.892217 | Twf1 |
| ENSMUSG( | 965.4926 | 0.169921 | 0.173348 | 0.60863 | Twf2 |
| ENSMUSG( | 30.6762 | -0.47533 | 0.164381 | 0.59535 | Twist1 |
| ENSMUSG( | 21.67519 | -0.00919 | 0.981958 | 0.997512 | Twist2 |
| ENSMUSG( | 409.4289 | 0.176226 | 0.18049 | 0.617841 | Twnk |
| ENSMUSG( | 1502.415 | -0.13944 | 0.293514 | 0.742947 | Twsg1 |
| ENSMUSG( | 1614.526 | 0.00835 | 0.926702 | 0.991229 | Txlna |
| ENSMUSG( | 127.3675 | 0.116168 | 0.600752 | 0.90443 | Txlnb |
| ENSMUSG( | 394.8475 | 0.242209 | 0.046731 | 0.330815 | Txlng |
| ENSMUSG( | 1566.326 | -0.00476 | 0.964621 | 0.995155 | Txn1 |
| ENSMUSG( | 5418.1 | 0.051242 | 0.553639 | 0.886216 | Txn2 |
| ENSMUSG( | 953.7112 | -0.20118 | 0.204047 | 0.644572 | Txndc11 |
| ENSMUSG( | 1451.377 | -0.26289 | 0.0774 | 0.429743 | Txndc12 |
| ENSMUSG( | 2374.543 | -0.16143 | 0.205025 | 0.646229 | Txndc15 |
| ENSMUSG( | 3239.994 | -0.24601 | 0.072527 | 0.415832 | Txndc16 |
| ENSMUSG( | 1011.712 | 0.036148 | 0.748739 | 0.949604 | Txndc17 |
| ENSMUSG( | 2306.475 | -0.3198 | 0.028257 | 0.256767 | Txndc5 |
| ENSMUSG( | 1023.828 | -0.0421 | 0.66315 | 0.9265 | Txndc9 |
| ENSMUSG( | 869.9134 | -0.19636 | 0.063921 | 0.391395 | Txnip |
| ENSMUSG( | 1321.151 | -0.01841 | 0.867404 | 0.976559 | Txnl1 |
| ENSMUSG( | 1785.342 | -0.02414 | 0.827325 | 0.968732 | Txnl4a |
| ENSMUSG( | 870.285 | 0.230858 | 0.023348 | 0.233882 | Txnl4a |
| ENSMUSG( | 574.5797 | 0.010246 | 0.925468 | 0.991204 | Txnl4b |
| ENSMUSG( | 1672.595 | 0.067525 | 0.461719 | 0.841564 | Txnrd1 |
| ENSMUSG( | 692.7225 | 0.051637 | 0.619507 | 0.911432 | Txnrd2 |
| ENSMUSG( | 212.7934 | -0.04419 | 0.761083 | 0.952907 | Txnrd3 |
| ENSMUSG( | 678.4115 | 0.248305 | 0.027893 | 0.255773 | Tyk2 |
| ENSMUSG( | 26.61389 | -0.27826 | 0.435661 | 0.829093 | Tymp |
| ENSMUSG( | 61.40524 | 0.156811 | 0.541899 | 0.883182 | Tyms |
| ENSMUSG( | 7208.514 | -0.2497 | 0.074269 | 0.422054 | Tyro3 |
| ENSMUSG( | 1103.575 | -0.03445 | 0.78969 | 0.959461 | Tyrobp |
| ENSMUSG( | 630.7424 | 0.160601 | 0.156684 | 0.584772 | Tysnd1 |
| ENSMUSG( | 600.5588 | -0.0821 | 0.55853 | 0.888521 | Tyw1 |
| ENSMUSG( | 195.5154 | -0.00617 | 0.969081 | 0.996369 | Tyw3 |
| ENSMUSG( | 272.4564 | 0.067165 | 0.62559 | 0.912249 | Tyw5 |
| ENSMUSG( | 1170.093 | -0.01617 | 0.865346 | 0.975565 | U2af1 |
| ENSMUSG( | 3724.924 | 0.060036 | 0.486852 | 0.858812 | U2af2 |
| ENSMUSG( | 1554.657 | 0.016586 | 0.85237 | 0.973168 | U2surp |
| ENSMUSG( | 547.1519 | -0.04211 | 0.73328 | 0.944956 | Uaca |
| ENSMUSG( | 1603.347 | 0.082891 | 0.346295 | 0.779153 | Uap1 |
| ENSMUSG( | 423.2286 | 0.138877 | 0.266523 | 0.716441 | Uap1l1 |
| ENSMUSG( | 8451.89 | 0.095502 | 0.247822 | 0.696522 | Uba1 |
| ENSMUSG( | 16.07665 | 0.406892 | 0.372138 | NA | Uba1y |
| ENSMUSG( | 1712.299 | 0.135856 | 0.134336 | 0.547472 | Uba2 |
| ENSMUSG( | 1859.595 | 0.038418 | 0.690961 | 0.933881 | Uba3 |
| ENSMUSG( | 1635.146 | 0.079462 | 0.368205 | 0.795774 | Uba5 |
| ENSMUSG( | 466.8862 | -0.01389 | 0.911702 | 0.988483 | Uba52 |
| ENSMUSG( | 103.6634 | -0.00467 | 0.981723 | 0.997512 | Uba52rt |
| ENSMUSG( | 337.309 | 0.090351 | 0.509995 | 0.869743 | Uba6 |
| ENSMUSG( | 142.1854 | -0.1378 | 0.464765 | 0.843568 | Uba7 |
| ENSMUSG( | 1584.305 | 0.076005 | 0.413866 | 0.819998 | Ubac1 |
| ENSMUSG( | 1774.622 | -0.22666 | 0.044465 | 0.32391 | Ubac2 |
| ENSMUSG( | 2844.124 | 0.051748 | 0.690277 | 0.933881 | Ubald1 |
| ENSMUSG( | 397.0479 | 0.039942 | 0.758638 | 0.952113 | Ubald2 |
| ENSMUSG( | 1365.485 | 0.080587 | 0.450188 | 0.835344 | Ubap1 |
| ENSMUSG( | 92.90351 | 0.306096 | 0.162154 | 0.592508 | Ubap1l |
| ENSMUSG( | 2336.01 | 0.166851 | 0.065795 | 0.397169 | Ubap2 |
| ENSMUSG( | 2910.542 | 0.015585 | 0.85554 | 0.973813 | Ubap2l |
| ENSMUSG( | 783.3579 | -0.10482 | 0.365514 | 0.794085 | Ubash3b |
| ENSMUSG( | 13227.53 | -0.01702 | 0.863665 | 0.975244 | Ubb |
| ENSMUSG( | 1872.765 | -0.03528 | 0.760831 | 0.952907 | Ubc |
| ENSMUSG( | 904.5902 | 0.041744 | 0.709936 | 0.939669 | Ube2a |
| ENSMUSG( | 2724.986 | 0.022777 | 0.824784 | 0.967386 | Ube2b |
| ENSMUSG( | 40.63461 | -0.45753 | 0.1365 | 0.550651 | Ube2c |
| ENSMUSG( | 1005.841 | -0.08924 | 0.383102 | 0.803237 | Ube2d1 |
| ENSMUSG( | 2178.473 | -0.00603 | 0.948697 | 0.993821 | Ube2d2a |
| ENSMUSG( | 3963.737 | -0.0862 | 0.433167 | 0.828207 | Ube2d3 |
| ENSMUSG( | 797.78 | 0.087551 | 0.415308 | 0.820498 | Ube2d-ps |
| ENSMUSG( | 1536.59 | 0.064578 | 0.613139 | 0.908902 | Ube2e1 |
| ENSMUSG( | 1399.249 | 0.050198 | 0.617 | 0.910866 | Ube2e2 |
| ENSMUSG( | 1731.425 | 0.081145 | 0.440145 | 0.831851 | Ube2e3 |
| ENSMUSG( | 1551.351 | 0.081836 | 0.344693 | 0.779153 | Ube2f |
| ENSMUSG( | 1197.155 | -0.03948 | 0.712976 | 0.94066 | Ube2g1 |
| ENSMUSG( | 1000.944 | 0.066635 | 0.519869 | 0.874979 | Ube2g2 |
| ENSMUSG( | 4189.354 | 0.034192 | 0.715847 | 0.941484 | Ube2h |
| ENSMUSG( | 2138.115 | -0.02637 | 0.79073 | 0.959461 | Ube2i |
| ENSMUSG( | 1848.452 | 0.004457 | 0.965937 | 0.995377 | Ube2j1 |
| ENSMUSG( | 944.2132 | -0.0557 | 0.59365 | 0.90281 | Ube2j2 |
| ENSMUSG( | 2698.427 | -0.0404 | 0.65847 | 0.924506 | Ube2k |
| ENSMUSG( | 6796.422 | 0.054639 | 0.538296 | 0.88259 | Ube2l3 |
| ENSMUSG( | 320.4565 | -0.43229 | 0.032627 | 0.276639 | Ube2l6 |
| ENSMUSG( | 6426.709 | 0.004598 | 0.963234 | 0.994911 | Ube2m |
| ENSMUSG( | 3167.319 | 0.072458 | 0.421461 | 0.823126 | Ube2n |
| ENSMUSG( | 3996.031 | 0.055837 | 0.62352 | 0.911646 | Ube2o |
| ENSMUSG( | 2983.447 | 0.064339 | 0.480251 | 0.854705 | Ube2q1 |
| ENSMUSG( | 1758.917 | 0.034333 | 0.767104 | 0.954393 | Ube2q2 |
| ENSMUSG( | 5718.265 | 0.130437 | 0.193727 | 0.633085 | Ube2ql1 |
| ENSMUSG( | 3660.621 | 0.071994 | 0.445082 | 0.833239 | Ube2r2 |
| ENSMUSG( | 977.1225 | 0.004715 | 0.966738 | 0.995784 | Ube2s |
| ENSMUSG( | 24.71777 | -0.18965 | 0.616199 | 0.91085 | Ube2srt |
| ENSMUSG( | 19.32831 | 0.086012 | 0.844823 | NA | Ube2t |
| ENSMUSG( | 28.90137 | 0.038786 | 0.914507 | 0.988483 | Ube2v1 |
| ENSMUSG( | 657.491 | -0.04852 | 0.700703 | 0.93743 | Ube2v2 |
| ENSMUSG( | 1083.326 | 0.057311 | 0.59912 | 0.903801 | Ube2w |
| ENSMUSG( | 2444.681 | 0.089379 | 0.400545 | 0.813717 | Ube2z |
| ENSMUSG( | 1900.264 | 0.039165 | 0.731472 | 0.944956 | Ube3a |
| ENSMUSG( | 8020.35 | 0.14234 | 0.091081 | 0.463586 | Ube3b |
| ENSMUSG( | 2185.918 | 0.16266 | 0.07834 | 0.432941 | Ube3c |
| ENSMUSG( | 32.57655 | -0.15 | 0.659763 | 0.924524 | Ube3d |
| ENSMUSG( | 2187.094 | -0.09078 | 0.461887 | 0.841728 | Ube4a |
| ENSMUSG( | 2800.765 | 0.033089 | 0.737994 | 0.947329 | Ube4b |
| ENSMUSG( | 1361.259 | 0.196178 | 0.039102 | 0.304587 | Ubfd1 |
| ENSMUSG( | 384.3569 | -0.18349 | 0.303675 | 0.752117 | Ubiad1 |
| ENSMUSG( | 5303.942 | 0.042016 | 0.626492 | 0.91272 | Ubl3 |
| ENSMUSG( | 2266.212 | 0.038632 | 0.640895 | 0.916614 | Ubl4a |
| ENSMUSG( | 2745.481 | -0.0348 | 0.70011 | 0.937391 | Ubl5 |
| ENSMUSG( | 1561.088 | 0.06311 | 0.525236 | 0.876559 | Ubl7 |
| ENSMUSG( | 686.2223 | 0.120158 | 0.311282 | 0.759247 | Ublcp1 |
| ENSMUSG( | 1772.429 | 0.107894 | 0.294221 | 0.743459 | Ubn1 |
| ENSMUSG( | 1244.673 | -0.00517 | 0.958251 | 0.993874 | Ubn2 |
| ENSMUSG( | 296.2184 | 0.074086 | 0.693118 | 0.934202 | Ubox5 |
| ENSMUSG( | 3321.789 | 0.196451 | 0.107622 | 0.49648 | Ubp1 |
| ENSMUSG( | 5620.809 | 0.042389 | 0.672945 | 0.929007 | Ubqln1 |
| ENSMUSG( | 3698.761 | 0.04409 | 0.633886 | 0.914073 | Ubqln2 |

| | | | | | |
|---|---|---|---|---|---|
| ENSMUSG( | 20.31432 | 0.469039 | 0.32054 | 0.903887 | Ube2t |
| ENSMUSG( | 24.93484 | -0.11907 | 0.772993 | 0.974463 | Ube2v1 |
| ENSMUSG( | 599.8996 | -0.05317 | 0.757907 | 0.971858 | Ube2v2 |
| ENSMUSG( | 965.5292 | -0.01223 | 0.938191 | 0.993731 | Ube2w |
| ENSMUSG( | 2082.085 | -0.10762 | 0.392001 | 0.917257 | Ube2z |
| ENSMUSG( | 1796.982 | 0.1393 | 0.417178 | 0.917257 | Ube3a |
| ENSMUSG( | 7087.429 | 0.05834 | 0.583377 | 0.944266 | Ube3b |
| ENSMUSG( | 1944.805 | 0.100272 | 0.407612 | 0.917257 | Ube3c |
| ENSMUSG( | 31.68509 | 0.029099 | 0.936444 | 0.993573 | Ube3d |
| ENSMUSG( | 2169.233 | 0.155639 | 0.26267 | 0.885357 | Ube4a |
| ENSMUSG( | 2502.454 | -0.02291 | 0.836864 | 0.981335 | Ube4b |
| ENSMUSG( | 1192.656 | 0.088576 | 0.418249 | 0.917257 | Ubfd1 |
| ENSMUSG( | 398.9607 | 0.201552 | 0.274137 | 0.889874 | Ubiad1 |
| ENSMUSG( | 4690.971 | -0.04543 | 0.691357 | 0.960071 | Ubl3 |
| ENSMUSG( | 1948.824 | -0.13316 | 0.239735 | 0.874882 | Ubl4a |
| ENSMUSG( | 2361.643 | -0.21163 | 0.095325 | 0.796133 | Ubl5 |
| ENSMUSG( | 1305.281 | -0.19042 | 0.11617 | 0.813384 | Ubl7 |
| ENSMUSG( | 613.2388 | 0.061438 | 0.672133 | 0.95694 | Ublcp1 |
| ENSMUSG( | 1646.894 | 0.164784 | 0.155392 | 0.837724 | Ubn1 |
| ENSMUSG( | 1217.465 | 0.193233 | 0.100846 | 0.801456 | Ubn2 |
| ENSMUSG( | 264.7862 | 0.015882 | 0.935825 | 0.993443 | Ubox5 |
| ENSMUSG( | 2961.122 | 0.133471 | 0.383688 | 0.917257 | Ubp1 |
| ENSMUSG( | 4862.955 | -0.11068 | 0.391306 | 0.917257 | Ubqln1 |
| ENSMUSG( | 3367.07 | 0.0404 | 0.727505 | 0.966887 | Ubqln2 |
| ENSMUSG( | 6593.631 | -0.0645 | 0.502234 | 0.927825 | Ubqln4 |
| ENSMUSG( | 1639.003 | 0.164639 | 0.281287 | 0.892412 | Ubr1 |
| ENSMUSG( | 2348.888 | 0.240317 | 0.036538 | 0.66949 | Ubr2 |
| ENSMUSG( | 3518.548 | 0.189903 | 0.098819 | 0.798438 | Ubr3 |
| ENSMUSG( | 6190.076 | 0.097814 | 0.500818 | 0.927825 | Ubr4 |
| ENSMUSG( | 5494.057 | 0.060331 | 0.564874 | 0.941206 | Ubr5 |
| ENSMUSG( | 1397.726 | 0.071165 | 0.567799 | 0.941777 | Ubr7 |
| ENSMUSG( | 288.9461 | 0.03163 | 0.849574 | 0.985497 | Ubtd1 |
| ENSMUSG( | 430.9161 | 0.078378 | 0.580244 | 0.944209 | Ubtd2 |
| ENSMUSG( | 3294.854 | 0.02821 | 0.79779 | 0.97681 | Ubtf |
| ENSMUSG( | 3130.646 | -0.14993 | 0.251914 | 0.879902 | Ubxn1 |
| ENSMUSG( | 55.98108 | 0.14233 | 0.661818 | 0.956205 | Ubxn10 |
| ENSMUSG( | 309.53 | 0.10036 | 0.633258 | 0.953844 | Ubxn11 |
| ENSMUSG( | 1497.739 | -0.07593 | 0.545123 | 0.938063 | Ubxn2a |
| ENSMUSG( | 1145.996 | -0.03651 | 0.770696 | 0.974236 | Ubxn2b |
| ENSMUSG( | 2507.143 | 0.117866 | 0.318009 | 0.902849 | Ubxn4 |
| ENSMUSG( | 2440.548 | -0.07414 | 0.487644 | 0.925427 | Ubxn6 |
| ENSMUSG( | 1153.832 | -0.00186 | 0.988181 | 0.997862 | Ubxn7 |
| ENSMUSG( | 937.1648 | 0.075264 | 0.593037 | 0.945408 | Ubxn8 |
| ENSMUSG( | 7469.803 | -0.13429 | 0.438997 | 0.920721 | Uchl1 |
| ENSMUSG( | 12.48407 | 0.165003 | 0.791766 | 0.97681 | Uchl1os |
| ENSMUSG( | 446.6309 | 0.008079 | 0.960153 | 0.994798 | Uchl3 |
| ENSMUSG( | 721.1169 | -0.1578 | 0.236747 | 0.872173 | Uchl5 |
| ENSMUSG( | 1007.511 | -0.08289 | 0.468273 | 0.923329 | Uck1 |
| ENSMUSG( | 1102.777 | -0.00656 | 0.95453 | 0.994798 | Uck2 |
| ENSMUSG( | 661.6238 | 0.112242 | 0.466665 | 0.923329 | Uckl1 |
| ENSMUSG( | 22.93098 | 0.207408 | 0.626863 | 0.953234 | Uckl1os |
| ENSMUSG( | 47.38382 | 0.269697 | 0.403871 | 0.917257 | Ucma |
| ENSMUSG( | 8.151249 | -0.15744 | 0.81426 | NA | Ucn |
| ENSMUSG( | 1564.688 | 0.343921 | 0.008268 | 0.388416 | Ucp2 |
| ENSMUSG( | 230.4367 | -0.66945 | 9.60E-05 | 0.02045 | Ucp3 |
| ENSMUSG( | 282.4699 | -0.06178 | 0.700283 | 0.961427 | Uevld |
| ENSMUSG( | 1643.665 | -0.17559 | 0.150738 | 0.837173 | Ufc1 |
| ENSMUSG( | 1619.497 | -0.00823 | 0.942955 | 0.994774 | Ufd1 |
| ENSMUSG( | 1022.97 | 0.102799 | 0.443368 | 0.921704 | Ufl1 |
| ENSMUSG( | 2283.567 | -0.14522 | 0.280357 | 0.892356 | Ufm1 |
| ENSMUSG( | 227.9104 | -0.22396 | 0.183253 | 0.850984 | Ufsp1 |
| ENSMUSG( | 927.2075 | -0.17333 | 0.180512 | 0.850372 | Ufsp2 |
| ENSMUSG( | 2513.384 | 0.178727 | 0.323132 | 0.904023 | Ugcg |
| ENSMUSG( | 751.6179 | -0.06387 | 0.590358 | 0.944975 | Ugdh |
| ENSMUSG( | 2417.149 | 0.158546 | 0.284893 | 0.892412 | Uggt1 |
| ENSMUSG( | 741.0736 | 0.090039 | 0.564782 | 0.941206 | Uggt2 |
| ENSMUSG( | 4100.766 | -0.22917 | 0.119891 | 0.813384 | Ugp2 |
| ENSMUSG( | 29.71812 | 0.093698 | 0.817774 | 0.979198 | Ugt1a7c |
| ENSMUSG( | 3042.74 | 0.212577 | 0.336956 | 0.907972 | Ugt8a |
| ENSMUSG( | 8110.268 | -0.06516 | 0.588506 | 0.944975 | Uhmk1 |
| ENSMUSG( | 71.69391 | 0.007412 | 0.976777 | 0.996416 | Uhrf1 |
| ENSMUSG( | 835.7217 | 0.240913 | 0.040313 | 0.682289 | Uhrf2 |
| ENSMUSG( | 438.1016 | 0.072041 | 0.630734 | 0.953251 | Uimc1 |
| ENSMUSG( | 2736.047 | 0.152067 | 0.218683 | 0.866104 | Ulk1 |
| ENSMUSG( | 4480.132 | 0.027265 | 0.819869 | 0.979198 | Ulk2 |
| ENSMUSG( | 604.2891 | -0.02551 | 0.883736 | 0.989644 | Ulk3 |
| ENSMUSG( | 223.0076 | 0.05736 | 0.729086 | 0.966913 | Ulk4 |
| ENSMUSG( | 415.1597 | -0.24365 | 0.211229 | 0.864412 | Umad1 |
| ENSMUSG( | 423.9098 | -0.07417 | 0.636144 | 0.953903 | Umps |
| ENSMUSG( | 681.5982 | -0.05945 | 0.667498 | 0.956766 | Unc119 |
| ENSMUSG( | 1325.806 | -0.08495 | 0.433471 | 0.918564 | Unc119b |
| ENSMUSG( | 11438.85 | -0.00222 | 0.985271 | 0.997084 | Unc13a |
| ENSMUSG( | 1779.19 | -0.05339 | 0.689871 | 0.960071 | Unc13b |
| ENSMUSG( | 4314.397 | -0.42845 | 0.010701 | 0.420304 | Unc13c |
| ENSMUSG( | 47.76773 | -0.08648 | 0.77859 | 0.976106 | Unc13d |
| ENSMUSG( | 905.3794 | -0.00253 | 0.982225 | 0.996756 | Unc45a |
| ENSMUSG( | 75.41512 | 0.216813 | 0.403566 | 0.917257 | Unc45b |
| ENSMUSG( | 1238.637 | 0.186094 | 0.206735 | 0.864412 | Unc50 |
| ENSMUSG( | 3764.184 | 0.347235 | 0.021645 | 0.575341 | Unc5a |
| ENSMUSG( | 1152.84 | 0.117857 | 0.596732 | 0.946541 | Unc5b |
| ENSMUSG( | 1075.309 | 0.062718 | 0.74119 | 0.968811 | Unc5c |
| ENSMUSG( | 7.994 | 0.717336 | 0.368415 | NA | Unc5cl |
| ENSMUSG( | 1111.272 | 0.307567 | 0.041948 | 0.690304 | Unc5d |
| ENSMUSG( | 1747.821 | 0.343098 | 0.00693 | 0.343883 | Unc79 |
| ENSMUSG( | 7321.158 | 0.111243 | 0.29171 | 0.897854 | Unc80 |
| ENSMUSG( | 1213.783 | 0.155061 | 0.408849 | 0.917257 | Unc93b1 |
| ENSMUSG( | 46.86408 | -0.09634 | 0.756896 | 0.971858 | Ung |
| ENSMUSG( | 704.787 | 0.03394 | 0.828025 | 0.980224 | Unk |
| ENSMUSG( | 557.5582 | 0.033358 | 0.805995 | 0.977674 | Unkl |
| ENSMUSG( | 16.91336 | 0.207138 | 0.668675 | 0.956766 | Uox |
| ENSMUSG( | 13.75171 | -0.32403 | 0.538668 | 0.937216 | Upb1 |
| ENSMUSG( | 1864.198 | -0.01864 | 0.884061 | 0.989644 | Upf1 |
| ENSMUSG( | 1086.453 | 0.019552 | 0.875008 | 0.989644 | Upf2 |
| ENSMUSG( | 1292.173 | -0.10477 | 0.430012 | 0.917257 | Upf3a |
| ENSMUSG( | 541.9227 | 0.159657 | 0.225675 | 0.868468 | Upf3b |
| ENSMUSG( | 23.66343 | 0.177499 | 0.656003 | 0.955533 | Upk1a |
| ENSMUSG( | 36.8809 | -0.24961 | 0.465977 | 0.923329 | Upk1b |
| ENSMUSG( | 88.83322 | 0.211839 | 0.366606 | 0.914703 | Upp1 |
| ENSMUSG( | 880.6026 | -0.17434 | 0.163635 | 0.838291 | Upp2 |
| ENSMUSG( | 111.707 | -0.12194 | 0.570543 | 0.941872 | Uprt |
| ENSMUSG( | 1457.332 | -0.11795 | 0.287362 | 0.894091 | Uqcc1 |
| ENSMUSG( | 1923.954 | -0.34323 | 0.005313 | 0.299238 | Uqcc2 |
| ENSMUSG( | 587.9613 | -0.05367 | 0.690859 | 0.960071 | Uqcc3 |
| ENSMUSG( | 703.0109 | -0.08473 | 0.504003 | 0.927825 | Uqcc4 |
| ENSMUSG( | 227.5681 | -0.10817 | 0.564511 | 0.941206 | Uqcc5 |
| ENSMUSG( | 85.53916 | -1.0322 | 1.78E-05 | 0.005763 | Uqcc6 |
| ENSMUSG( | 2995.99 | -0.26846 | 0.021758 | 0.575341 | Uqcr10 |
| ENSMUSG( | 5742.299 | -0.3225 | 0.003759 | 0.244783 | Uqcr11 |
| ENSMUSG( | 1778.793 | -0.24086 | 0.047108 | 0.700664 | Uqcrb |
| ENSMUSG( | 6736.749 | -0.17743 | 0.145983 | 0.835406 | Uqcrc1 |
| ENSMUSG( | 4130.17 | -0.20875 | 0.112384 | 0.809134 | Uqcrc2 |
| ENSMUSG( | 2995.076 | -0.2931 | 0.029887 | 0.61992 | Uqcrfs1 |
| ENSMUSG( | 5709.645 | -0.22849 | 0.052919 | 0.71525 | Uqcrh |
| ENSMUSG( | 3686.785 | -0.29026 | 0.021425 | 0.575341 | Uqcrq |
| ENSMUSG( | 11.94182 | 0.553082 | 0.335687 | 0.907907 | Urah |
| ENSMUSG( | 384.4697 | 0.124289 | 0.446139 | 0.921734 | Urb1 |
| ENSMUSG( | 305.7763 | 0.069945 | 0.643386 | 0.953903 | Urb2 |
| ENSMUSG( | 1295.29 | -0.02071 | 0.864181 | 0.986552 | Urgcp |
| ENSMUSG( | 1774.063 | -0.00459 | 0.976266 | 0.996416 | Uri1 |
| ENSMUSG( | 1504.954 | -0.19664 | 0.082137 | 0.77681 | Urm1 |
| ENSMUSG( | 814.8968 | -0.04182 | 0.77105 | 0.974236 | Urod |
| ENSMUSG( | 555.3216 | -0.01752 | 0.917315 | 0.992529 | Uros |
| ENSMUSG( | 235.0191 | -0.02582 | 0.880015 | 0.989644 | Usb1 |
| ENSMUSG( | 2249.994 | -0.11596 | 0.289917 | 0.896726 | Use1 |
| ENSMUSG( | 1199.761 | -0.02611 | 0.836484 | 0.981335 | Usf1 |
| ENSMUSG( | 4108.265 | -0.00815 | 0.944637 | 0.994798 | Usf2 |
| ENSMUSG( | 1697.723 | 0.109737 | 0.362978 | 0.914223 | Usf3 |
| ENSMUSG( | 13.32856 | 0.148063 | 0.783786 | 0.976154 | Ush1g |
| ENSMUSG( | 220.6221 | 0.320168 | 0.228762 | 0.868507 | Ushbp1 |
| ENSMUSG( | 2438.421 | -0.10275 | 0.319592 | 0.903535 | Uso1 |
| ENSMUSG( | 844.7414 | 0.169116 | 0.238577 | 0.872849 | Usp1 |
| ENSMUSG( | 986.0359 | 0.074103 | 0.572772 | 0.94262 | Usp10 |
| ENSMUSG( | 2159.911 | -0.10988 | 0.4605 | 0.923329 | Usp11 |
| ENSMUSG( | 1739.478 | 0.015046 | 0.885992 | 0.989854 | Usp12 |
| ENSMUSG( | 636.6109 | 0.076105 | 0.563608 | 0.941206 | Usp13 |
| ENSMUSG( | 1797.083 | -0.0605 | 0.656847 | 0.955567 | Usp14 |
| ENSMUSG( | 1374.326 | 0.022288 | 0.851432 | 0.985541 | Usp15 |
| ENSMUSG( | 1066.061 | 0.010572 | 0.934854 | 0.99306 | Usp16 |
| ENSMUSG( | 60.05878 | -0.18883 | 0.482695 | 0.924615 | Usp18 |
| ENSMUSG( | 3067.725 | 0.162707 | 0.202653 | 0.864412 | Usp19 |
| ENSMUSG( | 2357.346 | 0.399334 | 0.00136 | 0.134176 | Usp2 |
| ENSMUSG( | 1904.122 | 0.011572 | 0.922325 | 0.992644 | Usp20 |
| ENSMUSG( | 1285.57 | 0.108483 | 0.408075 | 0.917257 | Usp21 |
| ENSMUSG( | 4866.846 | -0.0761 | 0.497145 | 0.927825 | Usp22 |
| ENSMUSG( | 3295.807 | 0.021271 | 0.844195 | 0.983576 | Usp24 |
| ENSMUSG( | 936.7388 | 0.263194 | 0.039817 | 0.681706 | Usp25 |
| ENSMUSG( | 261.523 | 0.076179 | 0.685127 | 0.958678 | Usp27x |
| ENSMUSG( | 942.9609 | -0.12188 | 0.339031 | 0.907972 | Usp28 |

| | | | | | | | | | | | |
|---|---|---|---|---|---|---|---|---|---|---|---|
| ENSMUSG( | Uvrag | 930.4071 | 0.117954 | 0.316666 | 0.763648 | ENSMUSG( | Uvrag | 834.3037 | 0.073183 | 0.574226 | 0.942912 |
| ENSMUSG( | Uvssa | 316.1151 | -0.22724 | 0.08153 | 0.441102 | ENSMUSG( | Uvssa | 323.3421 | 0.122211 | 0.486889 | 0.92541 |
| ENSMUSG( | Uxs1 | 1234.576 | -0.40993 | 0.003517 | 0.075237 | ENSMUSG( | Uxs1 | 1336.132 | 0.117202 | 0.474824 | 0.923329 |
| ENSMUSG( | Uxt | 175.414 | -0.12693 | 0.432985 | 0.828185 | ENSMUSG( | Uxt | 148.333 | -0.36469 | 0.048456 | 0.706142 |
| ENSMUSG( | Vac14 | 694.377 | 0.029876 | 0.777846 | 0.956168 | ENSMUSG( | Vac14 | 628.2153 | 0.009001 | 0.941796 | 0.994601 |
| ENSMUSG( | Vamp1 | 2622.86 | -0.02839 | 0.842421 | 0.971564 | ENSMUSG( | Vamp1 | 2284.915 | -0.16304 | 0.282025 | 0.892412 |
| ENSMUSG( | Vamp2 | 39803.92 | 0.103359 | 0.151427 | 0.576717 | ENSMUSG( | Vamp2 | 34107.95 | -0.07302 | 0.522562 | 0.932822 |
| ENSMUSG( | Vamp3 | 1941.376 | 0.104986 | 0.321988 | 0.766908 | ENSMUSG( | Vamp3 | 1742.269 | 0.059882 | 0.63438 | 0.953903 |
| ENSMUSG( | Vamp4 | 2001.699 | 0.068138 | 0.52997 | 0.878883 | ENSMUSG( | Vamp4 | 1815.565 | 0.051599 | 0.702674 | 0.962215 |
| ENSMUSG( | Vamp5 | 114.7248 | -0.01309 | 0.945849 | 0.99369 | ENSMUSG( | Vamp5 | 103.557 | -0.04006 | 0.869341 | 0.988097 |
| ENSMUSG( | Vamp8 | 549.0806 | -0.02791 | 0.800703 | 0.961745 | ENSMUSG( | Vamp8 | 469.9272 | -0.21729 | 0.10865 | 0.804969 |
| ENSMUSG( | Vangl1 | 205.2661 | -0.44356 | 0.017005 | 0.194647 | ENSMUSG( | Vangl1 | 235.771 | 0.26074 | 0.249763 | 0.877904 |
| ENSMUSG( | Vangl2 | 790.4803 | -0.40031 | 0.003126 | 0.069373 | ENSMUSG( | Vangl2 | 848.6244 | 0.10232 | 0.488865 | 0.925573 |
| ENSMUSG( | Vapa | 2733.811 | -0.00939 | 0.932304 | 0.992515 | ENSMUSG( | Vapa | 2383.447 | -0.1436 | 0.380931 | 0.917202 |
| ENSMUSG( | Vapb | 6842.567 | 0.020063 | 0.829271 | 0.969286 | ENSMUSG( | Vapb | 5857.331 | -0.16548 | 0.13386 | 0.822912 |
| ENSMUSG( | Vars1 | 1346.876 | 0.142249 | 0.194874 | 0.634201 | ENSMUSG( | Vars1 | 1169.815 | 0.008266 | 0.949689 | 0.994798 |
| ENSMUSG( | Vars2 | 1015.101 | 0.133288 | 0.153833 | 0.580901 | ENSMUSG( | Vars2 | 913.2913 | 0.099138 | 0.403429 | 0.917257 |
| ENSMUSG( | Vash1 | 2096.347 | -0.04678 | 0.582449 | 0.897918 | ENSMUSG( | Vash1 | 1741.173 | -0.3338 | 0.003114 | 0.221768 |
| ENSMUSG( | Vash2 | 78.33801 | -0.1243 | 0.606506 | 0.906481 | ENSMUSG( | Vash2 | 69.07588 | -0.21437 | 0.428879 | 0.917257 |
| ENSMUSG( | Vasn | 330.3436 | -0.2648 | 0.159822 | 0.589679 | ENSMUSG( | Vasn | 362.9457 | 0.287175 | 0.199534 | 0.863016 |
| ENSMUSG( | Vasp | 564.8695 | -0.07817 | 0.533809 | 0.880095 | ENSMUSG( | Vasp | 568.5691 | 0.208419 | 0.148753 | 0.836241 |
| ENSMUSG( | Vat1 | 1316.547 | -0.06767 | 0.489375 | 0.859092 | ENSMUSG( | Vat1 | 1204.088 | -0.05382 | 0.644148 | 0.953903 |
| ENSMUSG( | Vat1l | 1145.005 | -0.03087 | 0.848487 | 0.972374 | ENSMUSG( | Vat1l | 1011.478 | -0.12839 | 0.465096 | 0.923329 |
| ENSMUSG( | Vav1 | 286.2605 | -0.0383 | 0.796727 | 0.960359 | ENSMUSG( | Vav1 | 266.6919 | 0.034621 | 0.838937 | 0.98174 |
| ENSMUSG( | Vav2 | 683.2186 | 0.035976 | 0.757768 | 0.951582 | ENSMUSG( | Vav2 | 648.1162 | 0.153932 | 0.24288 | 0.877399 |
| ENSMUSG( | Vav3 | 312.2298 | 0.003545 | 0.979885 | 0.997428 | ENSMUSG( | Vav3 | 276.0455 | -0.0973 | 0.577982 | 0.943131 |
| ENSMUSG( | Vax1 | 14.49257 | -0.61305 | 0.211022 | NA | ENSMUSG( | Vax1 | 15.68876 | -0.04087 | 0.934468 | 0.99306 |
| ENSMUSG( | Vbp1 | 908.4149 | -0.08266 | 0.51934 | 0.874979 | ENSMUSG( | Vbp1 | 874.3204 | 0.070034 | 0.657409 | 0.955725 |
| ENSMUSG( | Vcam1 | 2164.857 | -0.24678 | 0.091991 | 0.465937 | ENSMUSG( | Vcam1 | 2180.158 | 0.055186 | 0.728871 | 0.966913 |
| ENSMUSG( | Vcan | 1116.111 | -0.32462 | 0.041787 | 0.315339 | ENSMUSG( | Vcan | 1304.507 | 0.391634 | 0.030639 | 0.628144 |
| ENSMUSG( | Vcf1 | 1235.205 | -0.02381 | 0.808402 | 0.963455 | ENSMUSG( | Vcf1 | 1120.123 | -0.0422 | 0.76324 | 0.973945 |
| ENSMUSG( | Vcf2 | 498.3134 | 0.030045 | 0.812373 | 0.964574 | ENSMUSG( | Vcf2 | 442.3633 | -0.0525 | 0.726964 | 0.966887 |
| ENSMUSG( | Vcl | 1703.896 | 0.25296 | 0.073679 | 0.419767 | ENSMUSG( | Vcl | 1673.949 | 0.452125 | 0.007746 | 0.37555 |
| ENSMUSG( | Vcp | 6301.013 | 0.087561 | 0.321698 | 0.766739 | ENSMUSG( | Vcp | 5446.247 | -0.06461 | 0.557681 | 0.939565 |
| ENSMUSG( | Vcpip1 | 1038.831 | 0.035921 | 0.712843 | 0.94066 | ENSMUSG( | Vcpip1 | 984.7678 | 0.142569 | 0.332858 | 0.907907 |
| ENSMUSG( | Vcpkmt | 518.1018 | -0.11709 | 0.307034 | 0.75508 | ENSMUSG( | Vcpkmt | 453.523 | -0.24086 | 0.066336 | 0.749203 |
| ENSMUSG( | Vdac1 | 10831.62 | 0.020498 | 0.837914 | 0.971111 | ENSMUSG( | Vdac1 | 9340.081 | -0.14481 | 0.309215 | 0.898538 |
| ENSMUSG( | Vdac2 | 4197.165 | -0.01142 | 0.91393 | 0.988483 | ENSMUSG( | Vdac2 | 3664.516 | -0.14129 | 0.301146 | 0.898538 |
| ENSMUSG( | Vdac3 | 2152.719 | -0.00083 | 0.994478 | 0.999369 | ENSMUSG( | Vdac3 | 1851.06 | -0.17791 | 0.262961 | 0.885357 |
| ENSMUSG( | Vegfa | 3395.327 | -0.1003 | 0.378434 | 0.801688 | ENSMUSG( | Vegfa | 3295.233 | 0.084641 | 0.498878 | 0.927825 |
| ENSMUSG( | Vegfb | 5196.293 | 0.018358 | 0.83037 | 0.96974 | ENSMUSG( | Vegfb | 4461.777 | -0.15845 | 0.182711 | 0.850579 |
| ENSMUSG( | Vegfc | 109.9122 | -0.50408 | 0.029042 | 0.259542 | ENSMUSG( | Vegfc | 124.8247 | 0.177739 | 0.443409 | 0.921704 |
| ENSMUSG( | Vegfd | 78.84687 | 0.189016 | 0.453165 | 0.836701 | ENSMUSG( | Vegfd | 83.61095 | 0.574878 | 0.032113 | 0.634374 |
| ENSMUSG( | Veph1 | 16.95004 | 0.572983 | 0.224015 | NA | ENSMUSG( | Veph1 | 12.77843 | 0.044171 | 0.937648 | 0.993685 |
| ENSMUSG( | Vezf1 | 1357.765 | -0.00487 | 0.960836 | 0.993991 | ENSMUSG( | Vezf1 | 1207.35 | -0.07482 | 0.540501 | 0.937383 |
| ENSMUSG( | Vezt | 2219.86 | -0.1197 | 0.290577 | 0.740557 | ENSMUSG( | Vezt | 2213.239 | 0.142832 | 0.288554 | 0.894794 |
| ENSMUSG( | Vgf | 10194.17 | -0.4779 | 0.000119 | 0.007278 | ENSMUSG( | Vgf | 11092.43 | 0.076085 | 0.590189 | 0.944975 |
| ENSMUSG( | Vgll3 | 45.47419 | 0.381507 | 0.248768 | 0.697857 | ENSMUSG( | Vgll3 | 37.65475 | 0.153164 | 0.666919 | 0.956766 |
| ENSMUSG( | Vgll4 | 966.4839 | -0.09569 | 0.322188 | 0.767159 | ENSMUSG( | Vgll4 | 888.6033 | -0.0638 | 0.643586 | 0.953903 |
| ENSMUSG( | Vhl | 3469.838 | 0.057364 | 0.621396 | 0.911432 | ENSMUSG( | Vhl | 2819.888 | -0.28857 | 0.037068 | 0.673514 |
| ENSMUSG( | Vil1 | 27.93865 | 0.361409 | 0.349382 | 0.78176 | ENSMUSG( | Vil1 | 23.45574 | 0.118748 | 0.775457 | 0.975354 |
| ENSMUSG( | Vill | 227.4065 | 0.395349 | 0.096774 | 0.477375 | ENSMUSG( | Vill | 187.9427 | 0.1297 | 0.593266 | 0.945408 |
| ENSMUSG( | Vim | 1921.715 | 0.157211 | 0.139456 | 0.555633 | ENSMUSG( | Vim | 2051.05 | 0.566404 | 3.36E-06 | 0.001549 |
| ENSMUSG( | Vip | 1285.453 | -0.56627 | 0.00126 | 0.038624 | ENSMUSG( | Vip | 1444.762 | 0.099453 | 0.624479 | 0.953234 |
| ENSMUSG( | Vipas39 | 1459.877 | 0.127805 | 0.14615 | 0.567605 | ENSMUSG( | Vipas39 | 1303.216 | 0.071737 | 0.503972 | 0.927825 |
| ENSMUSG( | Vipr1 | 1002.453 | -0.11681 | 0.435331 | 0.829093 | ENSMUSG( | Vipr1 | 992.8651 | 0.129836 | 0.398736 | 0.917257 |
| ENSMUSG( | Vipr2 | 54.9195 | -0.98386 | 0.002198 | 0.054456 | ENSMUSG( | Vipr2 | 65.71883 | -0.0721 | 0.853354 | 0.985598 |
| ENSMUSG( | Virma | 1113.198 | 0.083103 | 0.397509 | 0.812127 | ENSMUSG( | Virma | 1013.797 | 0.083588 | 0.508284 | 0.929089 |
| ENSMUSG( | Vit | 164.968 | -0.04313 | 0.818384 | 0.965837 | ENSMUSG( | Vit | 166.4542 | 0.254402 | 0.202539 | 0.864412 |
| ENSMUSG( | Vkorc1 | 219.1458 | -0.21429 | 0.177788 | 0.615225 | ENSMUSG( | Vkorc1 | 215.6869 | 0.017169 | 0.925816 | 0.99306 |
| ENSMUSG( | Vkorc1l1 | 2358.962 | -0.25657 | 0.027384 | 0.253959 | ENSMUSG( | Vkorc1l1 | 2368.713 | 0.037766 | 0.778884 | 0.976106 |
| ENSMUSG( | Vldlr | 2729.841 | -0.34884 | 0.014474 | 0.177206 | ENSMUSG( | Vldlr | 2858.598 | 0.076964 | 0.629268 | 0.953251 |
| ENSMUSG( | Vma21 | 1351.391 | -0.33804 | 0.020601 | 0.216558 | ENSMUSG( | Vma21 | 1383.647 | 0.017475 | 0.922248 | 0.992644 |
| ENSMUSG( | Vmac | 563.5453 | -0.01731 | 0.879229 | 0.981436 | ENSMUSG( | Vmac | 492.9658 | -0.14045 | 0.283556 | 0.892412 |
| ENSMUSG( | Vmn2r1 | 21.35471 | 0.33989 | 0.454893 | 0.837689 | ENSMUSG( | Vmn2r1 | 24.38504 | 0.882593 | 0.04494 | 0.693959 |
| ENSMUSG( | Vmn2r29 | 90.51452 | -0.16847 | 0.423162 | 0.823662 | ENSMUSG( | Vmn2r29 | 73.64074 | -0.53702 | 0.026995 | 0.603041 |
| ENSMUSG( | Vmn2r57 | 47.29616 | -0.35056 | 0.216902 | 0.661816 | ENSMUSG( | Vmn2r57 | 45.42355 | -0.19951 | 0.531267 | 0.934795 |
| ENSMUSG( | Vmn2r84 | 75.6574 | -0.04604 | 0.866757 | 0.976541 | ENSMUSG( | Vmn2r84 | 69.1076 | -0.01829 | 0.94839 | 0.994798 |
| ENSMUSG( | Vmn2r85 | 10.47293 | 0.430566 | 0.482828 | NA | ENSMUSG( | Vmn2r85 | 8.564871 | 0.172851 | 0.798188 | NA |
| ENSMUSG( | Vmn2r87 | 25.81732 | -0.12483 | 0.760379 | 0.952907 | ENSMUSG( | Vmn2r87 | 23.35766 | -0.10055 | 0.823672 | 0.979198 |
| ENSMUSG( | Vmp1 | 3470.604 | -0.65165 | 1.07E-05 | 0.001199 | ENSMUSG( | Vmp1 | 3943.482 | 0.061463 | 0.722205 | 0.966887 |
| ENSMUSG( | Vnn1 | 27.11492 | -0.59966 | 0.21157 | 0.654835 | ENSMUSG( | Vnn1 | 28.95027 | -0.06053 | 0.893819 | 0.990558 |
| ENSMUSG( | Vopp1 | 3914.122 | -0.24296 | 0.051665 | 0.348878 | ENSMUSG( | Vopp1 | 4030.223 | 0.119449 | 0.396857 | 0.917257 |
| ENSMUSG( | Vps11 | 1752.171 | 0.050295 | 0.632154 | 0.914073 | ENSMUSG( | Vps11 | 1549.47 | -0.03382 | 0.783941 | 0.976154 |
| ENSMUSG( | Vps13a | 1144.891 | -0.14786 | 0.242957 | 0.690307 | ENSMUSG( | Vps13a | 1219.265 | 0.295908 | 0.044733 | 0.693959 |
| ENSMUSG( | Vps13b | 1527.279 | -0.10975 | 0.372496 | 0.798768 | ENSMUSG( | Vps13b | 1465.06 | 0.045531 | 0.752693 | 0.970521 |
| ENSMUSG( | Vps13c | 1590.777 | -0.07119 | 0.587095 | 0.900235 | ENSMUSG( | Vps13c | 1583.255 | 0.184224 | 0.225002 | 0.868422 |
| ENSMUSG( | Vps13d | 4179.849 | -0.11329 | 0.222832 | 0.668701 | ENSMUSG( | Vps13d | 3950.24 | -0.00386 | 0.971426 | 0.996236 |
| ENSMUSG( | Vps16 | 1480.53 | 0.072772 | 0.467604 | 0.844819 | ENSMUSG( | Vps16 | 1322.372 | 0.015652 | 0.902164 | 0.991012 |
| ENSMUSG( | Vps18 | 1154.533 | 0.120662 | 0.385589 | 0.804302 | ENSMUSG( | Vps18 | 996.3691 | -0.03292 | 0.832326 | 0.98075 |
| ENSMUSG( | Vps25 | 34.08543 | 0.373537 | 0.273962 | 0.724254 | ENSMUSG( | Vps25 | 28.8556 | 0.166573 | 0.669959 | 0.95694 |
| ENSMUSG( | Vps26a | 1546.602 | -0.02315 | 0.79864 | 0.960896 | ENSMUSG( | Vps26a | 1376.252 | -0.09626 | 0.442893 | 0.921554 |
| ENSMUSG( | Vps26b | 3462.521 | 0.121865 | 0.135214 | 0.548658 | ENSMUSG( | Vps26b | 2993.564 | -0.02722 | 0.788395 | 0.97637 |
| ENSMUSG( | Vps26c | 1575.603 | 0.104693 | 0.314093 | 0.761319 | ENSMUSG( | Vps26c | 1275.061 | -0.24947 | 0.055062 | 0.723362 |
| ENSMUSG( | Vps28 | 2108.736 | -0.07026 | 0.427402 | 0.82549 | ENSMUSG( | Vps28 | 1820.472 | -0.23752 | 0.043579 | 0.690304 |
| ENSMUSG( | Vps29 | 2381.775 | -0.02148 | 0.801692 | 0.962077 | ENSMUSG( | Vps29 | 2107.678 | -0.11038 | 0.34812 | 0.911652 |
| ENSMUSG( | Vps33a | 2715.772 | 0.128301 | 0.152017 | 0.577721 | ENSMUSG( | Vps33a | 2343.74 | -0.02404 | 0.831382 | 0.980611 |
| ENSMUSG( | Vps33b | 562.9039 | 0.220821 | 0.111536 | 0.503511 | ENSMUSG( | Vps33b | 497.6246 | 0.137789 | 0.417038 | 0.917257 |
| ENSMUSG( | Vps35 | 3363.974 | 0.050479 | 0.622915 | 0.911589 | ENSMUSG( | Vps35 | 3005.033 | -0.00801 | 0.946952 | 0.994798 |
| ENSMUSG( | Vps35l | 2107.021 | 0.103492 | 0.214588 | 0.659193 | ENSMUSG( | Vps35l | 1801.395 | -0.08093 | 0.45555 | 0.923329 |
| ENSMUSG( | Vps36 | 875.2307 | 0.179942 | 0.066763 | 0.399595 | ENSMUSG( | Vps36 | 768.2443 | 0.074721 | 0.532117 | 0.934795 |
| ENSMUSG( | Vps37a | 1847.631 | 0.098355 | 0.343516 | 0.779153 | ENSMUSG( | Vps37a | 1661.665 | 0.060311 | 0.62894 | 0.953251 |
| ENSMUSG( | Vps37b | 1129.875 | -0.15904 | 0.14187 | 0.559166 | ENSMUSG( | Vps37b | 1015.64 | -0.20051 | 0.108333 | 0.804969 |
| ENSMUSG( | Vps37c | 1194.722 | 0.103146 | 0.289435 | 0.739839 | ENSMUSG( | Vps37c | 1017.692 | -0.09201 | 0.434564 | 0.91869 |
| ENSMUSG( | Vps37d | 597.524 | -0.0424 | 0.739433 | 0.947768 | ENSMUSG( | Vps37d | 521.5391 | -0.17749 | 0.219964 | 0.866104 |
| ENSMUSG( | Vps39 | 2425.512 | 0.21545 | 0.017538 | 0.198498 | ENSMUSG( | Vps39 | 2049.539 | 0.005061 | 0.965648 | 0.995433 |
| ENSMUSG( | Vps41 | 4667.629 | 0.054249 | 0.502317 | 0.864924 | ENSMUSG( | Vps41 | 4007.437 | -0.12032 | 0.251242 | 0.879321 |
| ENSMUSG( | Vps45 | 774.1561 | 0.114009 | 0.414347 | 0.819998 | ENSMUSG( | Vps45 | 636.9802 | -0.18638 | 0.217996 | 0.866101 |
| ENSMUSG( | Vps4a | 2917.152 | 0.02198 | 0.790618 | 0.959461 | ENSMUSG( | Vps4a | 2481.919 | -0.18318 | 0.119919 | 0.813384 |
| ENSMUSG( | Vps4b | 917.311 | 0.059985 | 0.580996 | 0.89745 | ENSMUSG( | Vps4b | 867.0928 | 0.162274 | 0.18272 | 0.850579 |
| ENSMUSG( | Vps50 | 1596.547 | 0.096069 | 0.285015 | 0.733568 | ENSMUSG( | Vps50 | 1437.921 | 0.061131 | 0.608272 | 0.950113 |
| ENSMUSG( | Vps51 | 1242.791 | 0.085773 | 0.458294 | 0.839958 | ENSMUSG( | Vps51 | 1080.018 | -0.04964 | 0.700915 | 0.96162 |
| ENSMUSG( | Vps52 | 3353.462 | 0.166942 | 0.043931 | 0.321265 | ENSMUSG( | Vps52 | 2759.931 | -0.12617 | 0.257001 | 0.883082 |
| ENSMUSG( | Vps53 | 1459.775 | 0.134586 | 0.259952 | 0.709951 | ENSMUSG( | Vps53 | 1227.69 | -0.09919 | 0.495129 | 0.927372 |
| ENSMUSG( | Vps54 | 962.5856 | 0.029788 | 0.781671 | 0.957558 | ENSMUSG( | Vps54 | 936.663 | 0.212272 | 0.079427 | 0.77458 |
| ENSMUSG( | Vps72 | 770.5875 | 0.024292 | 0.818735 | 0.965888 | ENSMUSG( | Vps72 | 671.9822 | -0.10686 | 0.407737 | 0.917257 |
| ENSMUSG( | Vps8 | 1524.434 | 0.175883 | 0.066993 | 0.400593 | ENSMUSG( | Vps8 | 1291.015 | -0.02801 | 0.809058 | 0.97873 |
| ENSMUSG( | Vps9d1 | 2825.121 | 0.218386 | 0.012907 | 0.164191 | ENSMUSG( | Vps9d1 | 2405.072 | 0.030798 | 0.789492 | 0.976593 |
| ENSMUSG( | Vrk1 | 383.3551 | -0.03941 | 0.739575 | 0.947768 | ENSMUSG( | Vrk1 | 358.7559 | 0.032532 | 0.823552 | 0.979198 |
| ENSMUSG( | Vrk2 | 69.73264 | -0.15308 | 0.512892 | 0.871762 | ENSMUSG( | Vrk2 | 67.07125 | 0.005469 | 0.982987 | 0.996891 |
| ENSMUSG( | Vrk3 | 1234.633 | 0.036913 | 0.686125 | 0.932981 | ENSMUSG( | Vrk3 | 1080.363 | -0.08135 | 0.475292 | 0.923329 |
| ENSMUSG( | Vsig10 | 124.9721 | -0.40598 | 0.043422 | 0.319227 | ENSMUSG( | Vsig10 | 125.6623 | -0.08211 | 0.703488 | 0.962481 |
| ENSMUSG( | Vsig10l | 140.9465 | 0.219401 | 0.217081 | 0.661995 | ENSMUSG( | Vsig10l | 130.1399 | 0.255786 | 0.195137 | 0.861229 |
| ENSMUSG( | Vsig2 | 76.24601 | 0.394851 | 0.124607 | 0.531714 | ENSMUSG( | Vsig2 | 60.57976 | 0.034567 | 0.904718 | 0.991089 |
| ENSMUSG( | Vsig8 | 16.82013 | -0.99592 | 0.03387 | NA | ENSMUSG( | Vsig8 | 15.41551 | -1.02146 | 0.049965 | 0.708648 |
| ENSMUSG( | Vsir | 2107.435 | -0.07269 | 0.640705 | 0.916434 | ENSMUSG( | Vsir | 2149.116 | 0.248918 | 0.232043 | 0.868507 |
| ENSMUSG( | Vsnl1 | 9044.549 | -0.04728 | 0.730861 | 0.944956 | ENSMUSG( | Vsnl1 | 7779.457 | -0.22614 | 0.201141 | 0.86439 |
| ENSMUSG( | Vstm2a | 2168.757 | -0.63014 | 0.000158 | 0.008882 | ENSMUSG( | Vstm2a | 2658.931 | 0.283139 | 0.141647 | 0.830678 |
| ENSMUSG( | Vstm2b | 1263.824 | -0.26691 | 0.008825 | 0.132321 | ENSMUSG( | Vstm2b | 1317.198 | 0.131622 | 0.272505 | 0.888825 |
| ENSMUSG( | Vstm2l | 4104.932 | -0.46253 | 1.08E-05 | 0.001199 | ENSMUSG( | Vstm2l | 4326.412 | -0.00363 | 0.978151 | 0.996416 |
| ENSMUSG( | Vstm4 | 305.5882 | -0.16127 | 0.364042 | 0.793438 | ENSMUSG( | Vstm4 | 256.4241 | -0.41746 | 0.022905 | 0.584023 |
| ENSMUSG( | Vstm5 | 265.1999 | -0.04575 | 0.763865 | 0.954015 | ENSMUSG( | Vstm5 | 257.7193 | 0.136361 | 0.438822 | 0.920721 |
| ENSMUSG( | Vsx1 | 24.92015 | -0.81721 | 0.044581 | 0.32426 | ENSMUSG( | Vsx1 | 28.57585 | -0.04226 | 0.909562 | 0.991958 |
| ENSMUSG( | Vta1 | 1088.823 | 0.002852 | 0.977605 | 0.997428 | ENSMUSG( | Vta1 | 915.5935 | -0.24188 | 0.050802 | 0.709741 |
| ENSMUSG( | Vti1a | 2088.985 | 0.088572 | 0.341981 | 0.778828 | ENSMUSG( | Vti1a | 1738.857 | -0.17935 | 0.1224 | 0.813384 |
| ENSMUSG( | Vti1b | 5695.538 | 0.005704 | 0.948568 | 0.993821 | ENSMUSG( | Vti1b | 4733.665 | -0.27417 | 0.026734 | 0.603041 |
| ENSMUSG( | Vtn | 3663.583 | -0.03723 | 0.836558 | 0.971111 | ENSMUSG( | Vtn | 3680.577 | 0.242634 | 0.24903 | 0.877904 |
| ENSMUSG( | Vwa1 | 1714.982 | 0.004825 | 0.97615 | 0.997428 | ENSMUSG( | Vwa1 | 1814.396 | 0.413513 | 0.020653 | 0.569724 |
| ENSMUSG( | Vwa3a | 249.1976 | -0.02169 | 0.914028 | 0.988483 | ENSMUSG( | Vwa3a | 220.5603 | -0.0988 | 0.670044 | 0.95694 |
| ENSMUSG( | Vwa3b | 61.66417 | 0.366435 | 0.15199 | 0.577721 | ENSMUSG( | Vwa3b | 49.4922 | 0.05278 | 0.86408 | 0.986552 |
| ENSMUSG( | Vwa5a | 750.1319 | 0.119766 | 0.331059 | 0.77519 | ENSMUSG( | Vwa5a | 629.8062 | -0.11623 | 0.39888 | 0.917257 |
| ENSMUSG( | Vwa5b1 | 363.2073 | 0.267366 | 0.053646 | 0.355738 | ENSMUSG( | Vwa5b1 | 302.146 | 0.029124 | 0.870154 | 0.988232 |
| ENSMUSG( | Vwa5b2 | 3180.797 | 0.195499 | 0.158814 | 0.588112 | ENSMUSG( | Vwa5b2 | 2528.772 | -0.20175 | 0.194578 | 0.860277 |
| ENSMUSG( | Vwa7 | 18.35373 | -0.3119 | 0.480766 | NA | ENSMUSG( | Vwa7 | 17.37516 | -0.18468 | 0.709017 | 0.964042 |
| ENSMUSG( | Vwa8 | 2477.058 | 0.131606 | 0.1446 | 0.564508 | ENSMUSG( | Vwa8 | 2070.469 | -0.11705 | 0.274884 | 0.889874 |
| ENSMUSG( | Vwc2 | 162.3942 | -0.57752 | 0.000881 | 0.029903 | ENSMUSG( | Vwc2 | 169.7424 | -0.11688 | 0.542681 | 0.938034 |
| ENSMUSG( | Vwc2l | 141.19 | -0.29799 | 0.143569 | 0.562581 | ENSMUSG( | Vwc2l | 165.7141 | 0.427201 | 0.03914 | 0.679972 |
| ENSMUSG( | Vwce | 18.51174 | -0.10674 | 0.813982 | NA | ENSMUSG( | Vwce | 17.50558 | -0.01907 | 0.969036 | 0.99594 |
| ENSMUSG( | Vwf | 342.5066 | 0.054669 | 0.79907 | 0.960922 | ENSMUSG( | Vwf | 422.8089 | 0.828837 | 0.002361 | 0.190561 |
| ENSMUSG( | Vxn | 8548.748 | 0.079522 | 0.340817 | 0.778709 | ENSMUSG( | Vxn | 7373.852 | -0.07983 | 0.482302 | 0.924578 |
| ENSMUSG( | Wac | 2392.442 | 0.051856 | 0.578358 | 0.896413 | ENSMUSG( | Wac | 2250.444 | 0.140328 | 0.232854 | 0.868507 |
| ENSMUSG( | Wapl | 1405.823 | 0.097277 | 0.359323 | 0.791011 | ENSMUSG( | Wapl | 1300.736 | 0.137504 | 0.271324 | 0.888062 |
| ENSMUSG( | Wars1 | 2379.772 | 0.164346 | 0.091185 | 0.463787 | ENSMUSG( | Wars1 | 1976.266 | -0.10026 | 0.373898 | 0.915491 |
| ENSMUSG( | Wars2 | 211.3734 | 0.23286 | 0.175011 | 0.610399 | ENSMUSG( | Wars2 | 181.6409 | 0.074946 | 0.68177 | 0.958678 |
| ENSMUSG( | Was | 104.234 | -0.10034 | 0.665798 | 0.927054 | ENSMUSG( | Was | 96.15979 | -0.04688 | 0.853976 | 0.98563 |
| ENSMUSG( | Wasf1 | 4258.219 | -0.01964 | 0.888618 | 0.983434 | ENSMUSG( | Wasf1 | 3817.544 | -0.06986 | 0.678807 | 0.957617 |
| ENSMUSG( | Wasf2 | 1044.703 | 0.042861 | 0.676024 | 0.930542 | ENSMUSG( | Wasf2 | 989.1067 | 0.155372 | 0.309499 | 0.898538 |
| ENSMUSG( | Wasf3 | 11994.29 | 0.066884 | 0.549145 | 0.885723 | ENSMUSG( | Wasf3 | 10096.99 | -0.16723 | 0.222129 | 0.866232 |
| ENSMUSG( | Washc1 | 1167.081 | 0.096486 | 0.394152 | 0.809935 | ENSMUSG( | Washc1 | 986.4632 | -0.12128 | 0.339895 | 0.907972 |
| ENSMUSG( | Washc2 | 5746.333 | 0.048724 | 0.571283 | 0.893319 | ENSMUSG( | Washc2 | 4982.995 | -0.09499 | 0.330967 | 0.907513 |
| ENSMUSG( | Washc3 | 422.6113 | -0.11868 | 0.300346 | 0.749479 | ENSMUSG( | Washc3 | 370.3283 | -0.24065 | 0.086365 | 0.781091 |
| ENSMUSG( | Washc4 | 878.8034 | 0.097611 | 0.342142 | 0.778828 | ENSMUSG( | Washc4 | 808.6608 | 0.127921 | 0.306945 | 0.898538 |
| ENSMUSG( | Washc5 | 1388.308 | 0.117108 | 0.179318 | 0.616869 | ENSMUSG( | Washc5 | 1197.055 | -0.0377 | 0.740269 | 0.968811 |
| ENSMUSG( | Wasl | 1951.069 | 0.07878 | 0.404851 | 0.816396 | ENSMUSG( | Wasl | 1824.485 | 0.148044 | 0.26087 | 0.884853 |
| ENSMUSG( | Wbp1 | 1720.406 | 0.052382 | 0.597901 | 0.90372 | ENSMUSG( | Wbp1 | 1471.636 | -0.13213 | 0.237499 | 0.872173 |
| ENSMUSG( | Wbp11 | 2030.773 | 0.017349 | 0.860349 | 0.974842 | ENSMUSG( | Wbp11 | 1838.236 | -0.00358 | 0.977561 | 0.996416 |
| ENSMUSG( | Wbp1l | 2282.091 | 0.042419 | 0.68203 | 0.931738 | ENSMUSG( | Wbp1l | 2130.314 | 0.113277 | 0.345044 | 0.91045 |
| ENSMUSG( | Wbp2 | 10075.83 | 0.016611 | 0.86126 | 0.975026 | ENSMUSG( | Wbp2 | 8694.714 | -0.14549 | 0.277385 | 0.891468 |

| | | | | | |
|---|---|---|---|---|---|
| ENSMUSG( | 7781.489 | 0.141629 | 0.077189 | 0.429655 | Ubqln4 |
| ENSMUSG( | 1609.773 | -0.15898 | 0.29993 | 0.749016 | Ubr1 |
| ENSMUSG( | 2472.465 | 0.124342 | 0.180785 | 0.617841 | Ubr2 |
| ENSMUSG( | 3452.745 | -0.1326 | 0.196455 | 0.635942 | Ubr3 |
| ENSMUSG( | 6356.588 | -0.09769 | 0.38012 | 0.801757 | Ubr4 |
| ENSMUSG( | 6161.418 | 0.119529 | 0.175457 | 0.610843 | Ubr5 |
| ENSMUSG( | 1535.996 | 0.077517 | 0.458172 | 0.839914 | Ubr7 |
| ENSMUSG( | 320.229 | 0.067395 | 0.619897 | 0.911432 | Ubtd1 |
| ENSMUSG( | 477.3357 | 0.105365 | 0.367353 | 0.795221 | Ubtd2 |
| ENSMUSG( | 3701.928 | 0.095613 | 0.272556 | 0.722478 | Ubtf |
| ENSMUSG( | 3629.23 | 0.014852 | 0.867286 | 0.976559 | Ubxn1 |
| ENSMUSG( | 47.13678 | -0.74727 | 0.010249 | 0.144303 | Ubxn10 |
| ENSMUSG( | 368.9144 | 0.321824 | 0.050212 | 0.342492 | Ubxn11 |
| ENSMUSG( | 1744.72 | 0.098501 | 0.315273 | 0.762288 | Ubxn2a |
| ENSMUSG( | 1343.38 | 0.151525 | 0.159026 | 0.588112 | Ubxn2b |
| ENSMUSG( | 2646.019 | 0.005581 | 0.956851 | 0.993874 | Ubxn4 |
| ENSMUSG( | 2830.666 | 0.08444 | 0.324961 | 0.770595 | Ubxn6 |
| ENSMUSG( | 1290.437 | 0.053731 | 0.59086 | 0.90113 | Ubxn7 |
| ENSMUSG( | 909.8195 | -0.29357 | 0.026252 | 0.247758 | Ubxn8 |
| ENSMUSG( | 8562.65 | -0.00202 | 0.988911 | 0.999111 | Uchl1 |
| ENSMUSG( | 12.50058 | -0.14318 | 0.81501 | NA | Uchl1os |
| ENSMUSG( | 507.1148 | 0.110575 | 0.366984 | 0.795164 | Uchl3 |
| ENSMUSG( | 869.7079 | 0.120693 | 0.233903 | 0.681906 | Uchl5 |
| ENSMUSG( | 1163.646 | 0.063121 | 0.48635 | 0.858575 | Uck1 |
| ENSMUSG( | 1257.739 | 0.10098 | 0.279192 | 0.729109 | Uck2 |
| ENSMUSG( | 708.2202 | 0.04242 | 0.708319 | 0.939519 | Uckl1 |
| ENSMUSG( | 18.2899 | -0.82673 | 0.074751 | NA | Uckl1os |
| ENSMUSG( | 47.3608 | 0.027059 | 0.926922 | 0.991229 | Ucma |
| ENSMUSG( | 1573.918 | 0.111398 | 0.300315 | 0.749479 | Ucp2 |
| ENSMUSG( | 276.5557 | -0.35378 | 0.01931 | 0.209368 | Ucp3 |
| ENSMUSG( | 311.7434 | -0.04132 | 0.754609 | 0.951007 | Uevld |
| ENSMUSG( | 1898.865 | -0.0189 | 0.832846 | 0.970333 | Ufc1 |
| ENSMUSG( | 1838.98 | 0.090103 | 0.32922 | 0.773233 | Ufd1 |
| ENSMUSG( | 1008.515 | -0.21862 | 0.058327 | 0.372466 | Ufl1 |
| ENSMUSG( | 2618.456 | -0.01186 | 0.912573 | 0.988483 | Ufm1 |
| ENSMUSG( | 233.9356 | -0.44406 | 0.002186 | 0.054296 | Ufsp1 |
| ENSMUSG( | 1092.482 | 0.040034 | 0.711532 | 0.940028 | Ufsp2 |
| ENSMUSG( | 2176.725 | -0.55313 | 0.000488 | 0.02028 | Ugcg |
| ENSMUSG( | 821.2971 | -0.07583 | 0.479923 | 0.854483 | Ugdh |
| ENSMUSG( | 2410.612 | -0.11961 | 0.347565 | 0.780601 | Uggt1 |
| ENSMUSG( | 792.9701 | 0.012977 | 0.90699 | 0.987407 | Uggt2 |
| ENSMUSG( | 4887.174 | 0.019348 | 0.879844 | 0.981436 | Ugp2 |
| ENSMUSG( | 31.87806 | -0.00168 | 0.996258 | 0.99943 | Ugt1a7c |
| ENSMUSG( | 2586.34 | -0.57782 | 0.001333 | 0.039731 | Ugt8a |
| ENSMUSG( | 9298.296 | 0.062127 | 0.548773 | 0.885686 | Uhmk1 |
| ENSMUSG( | 83.0498 | 0.142534 | 0.558272 | 0.888427 | Uhrf1 |
| ENSMUSG( | 891.7829 | 0.164499 | 0.114793 | 0.510823 | Uhrf2 |
| ENSMUSG( | 492.1145 | 0.139164 | 0.268169 | 0.718372 | Uimc1 |
| ENSMUSG( | 3128.405 | 0.260867 | 0.006912 | 0.114527 | Ulk1 |
| ENSMUSG( | 5093.65 | 0.127174 | 0.203302 | 0.643631 | Ulk2 |
| ENSMUSG( | 709.4068 | 0.167994 | 0.261878 | 0.712129 | Ulk3 |
| ENSMUSG( | 240.043 | -0.00442 | 0.976577 | 0.997428 | Ulk4 |
| ENSMUSG( | 484.9143 | -0.04476 | 0.774034 | 0.955606 | Umad1 |
| ENSMUSG( | 9.755705 | 0.42837 | 0.482848 | NA | Umodl1 |
| ENSMUSG( | 526.9236 | 0.279545 | 0.024117 | 0.237752 | Umps |
| ENSMUSG( | 792.9185 | 0.110891 | 0.319362 | 0.765296 | Unc119 |
| ENSMUSG( | 1564.979 | 0.123095 | 0.154115 | 0.580901 | Unc119b |
| ENSMUSG( | 12978.02 | 0.092023 | 0.308162 | 0.7563 | Unc13a |
| ENSMUSG( | 2091.65 | 0.14291 | 0.20743 | 0.649561 | Unc13b |
| ENSMUSG( | 4941.247 | -0.28349 | 0.106988 | 0.494682 | Unc13c |
| ENSMUSG( | 54.25233 | 0.010488 | 0.971266 | 0.996369 | Unc13d |
| ENSMUSG( | 1031.325 | 0.102085 | 0.275986 | 0.725669 | Unc45a |
| ENSMUSG( | 83.33582 | 0.222087 | 0.325763 | 0.771061 | Unc45b |
| ENSMUSG( | 1174.237 | -0.2452 | 0.063588 | 0.390386 | Unc50 |
| ENSMUSG( | 3576.892 | -0.05302 | 0.695907 | 0.934979 | Unc5a |
| ENSMUSG( | 1002.331 | -0.62879 | 0.000643 | 0.024438 | Unc5b |
| ENSMUSG( | 948.8395 | -0.64369 | 9.83E-05 | 0.006375 | Unc5c |
| ENSMUSG( | 1018.459 | -0.2068 | 0.135033 | 0.548656 | Unc5d |
| ENSMUSG( | 1858.182 | 0.256744 | 0.007728 | 0.122031 | Unc79 |
| ENSMUSG( | 7743.877 | 0.006434 | 0.942706 | 0.993415 | Unc80 |
| ENSMUSG( | 1200.174 | -0.1528 | 0.256838 | 0.708006 | Unc93b1 |
| ENSMUSG( | 52.50206 | -0.03571 | 0.900065 | 0.985633 | Ung |
| ENSMUSG( | 755.7728 | -0.03376 | 0.776091 | 0.955606 | Unk |
| ENSMUSG( | 621.9606 | 0.074053 | 0.484624 | 0.856665 | Unkl |
| ENSMUSG( | 15.79057 | -0.27732 | 0.591149 | NA | Uox |
| ENSMUSG( | 12.59165 | -0.9829 | 0.110125 | NA | Upb1 |
| ENSMUSG( | 2140.128 | 0.109434 | 0.2726 | 0.722478 | Upf1 |
| ENSMUSG( | 1250.791 | 0.157125 | 0.133423 | 0.546357 | Upf2 |
| ENSMUSG( | 1481.712 | 0.027749 | 0.791007 | 0.959461 | Upf3a |
| ENSMUSG( | 562.8029 | 0.003031 | 0.977577 | 0.997428 | Upf3b |
| ENSMUSG( | 28.1291 | 0.397404 | 0.285739 | 0.734849 | Upk1a |
| ENSMUSG( | 41.54388 | -0.2053 | 0.489688 | 0.859319 | Upk1b |
| ENSMUSG( | 101.9014 | 0.343621 | 0.107218 | 0.495579 | Upp1 |
| ENSMUSG( | 1075.056 | 0.138881 | 0.153238 | 0.57989 | Upp2 |
| ENSMUSG( | 131.7125 | 0.093528 | 0.632623 | 0.914073 | Uprt |
| ENSMUSG( | 1616.867 | -0.08462 | 0.32327 | 0.768415 | Uqcc1 |
| ENSMUSG( | 2336.624 | -0.02754 | 0.767742 | 0.954621 | Uqcc2 |
| ENSMUSG( | 666.1971 | 0.041448 | 0.685406 | 0.932981 | Uqcc3 |
| ENSMUSG( | 767.9651 | -0.09543 | 0.336548 | 0.777 | Uqcc4 |
| ENSMUSG( | 248.9065 | -0.10353 | 0.486692 | 0.858713 | Uqcc5 |
| ENSMUSG( | 124.0139 | -0.03749 | 0.842713 | 0.971663 | Uqcc6 |
| ENSMUSG( | 3631.371 | 0.032511 | 0.695264 | 0.934948 | Uqcr10 |
| ENSMUSG( | 6959.965 | -0.01658 | 0.829221 | 0.969286 | Uqcr11 |
| ENSMUSG( | 2102.193 | -0.01406 | 0.884359 | 0.982647 | Uqcrb |
| ENSMUSG( | 8040.713 | 0.071192 | 0.425086 | 0.82478 | Uqcrc1 |
| ENSMUSG( | 4932.853 | 0.045114 | 0.64784 | 0.919759 | Uqcrc2 |
| ENSMUSG( | 3681.319 | 0.051658 | 0.627347 | 0.912941 | Uqcrfs1 |
| ENSMUSG( | 6857.805 | 0.041984 | 0.595449 | 0.903058 | Uqcrh |
| ENSMUSG( | 4418.433 | -0.01915 | 0.820535 | 0.965888 | Uqcrq |
| ENSMUSG( | 14.53743 | 0.788315 | 0.113429 | NA | Urah |
| ENSMUSG( | 387.0465 | -0.13667 | 0.295027 | 0.743798 | Urb1 |
| ENSMUSG( | 310.7218 | -0.16052 | 0.31453 | 0.761558 | Urb2 |
| ENSMUSG( | 1481.288 | 0.095689 | 0.388567 | 0.806278 | Urgcp |
| ENSMUSG( | 1987.444 | 0.058684 | 0.538963 | 0.882849 | Uri1 |
| ENSMUSG( | 1764.173 | 3.54E-05 | 0.999707 | 0.99989 | Urm1 |
| ENSMUSG( | 955.9335 | 0.151847 | 0.162797 | 0.593778 | Urod |
| ENSMUSG( | 614.6642 | 0.010297 | 0.92821 | 0.991346 | Uros |
| ENSMUSG( | 258.3057 | -0.02488 | 0.879048 | 0.981436 | Usb1 |
| ENSMUSG( | 2626.748 | 0.065697 | 0.444049 | 0.833239 | Use1 |
| ENSMUSG( | 1380.874 | 0.110124 | 0.283303 | 0.73186 | Usf1 |
| ENSMUSG( | 4722.08 | 0.124641 | 0.132646 | 0.544824 | Usf2 |
| ENSMUSG( | 1803.04 | 0.013973 | 0.898392 | 0.985633 | Usf3 |
| ENSMUSG( | 12.19892 | -0.36769 | 0.541479 | NA | Ush1g |
| ENSMUSG( | 222.7546 | 0.080559 | 0.604011 | 0.905187 | Ushbp1 |
| ENSMUSG( | 2770.664 | 0.000223 | 0.997931 | 0.99989 | Uso1 |
| ENSMUSG( | 895.7783 | 0.075903 | 0.521576 | 0.875186 | Usp1 |
| ENSMUSG( | 1116.258 | 0.15842 | 0.184568 | 0.621633 | Usp10 |
| ENSMUSG( | 2549.281 | 0.102017 | 0.398912 | 0.812878 | Usp11 |
| ENSMUSG( | 1925.552 | 0.040507 | 0.630448 | 0.914073 | Usp12 |
| ENSMUSG( | 709.6772 | 0.117654 | 0.384036 | 0.803646 | Usp13 |
| ENSMUSG( | 2008.448 | -0.00404 | 0.970657 | 0.996369 | Usp14 |
| ENSMUSG( | 1545.004 | 0.087463 | 0.425171 | 0.82478 | Usp15 |
| ENSMUSG( | 1172.027 | 0.011219 | 0.921494 | 0.990436 | Usp16 |
| ENSMUSG( | 59.17891 | -0.55919 | 0.045268 | 0.326476 | Usp18 |
| ENSMUSG( | 3480.881 | 0.251108 | 0.009994 | 0.142653 | Usp19 |
| ENSMUSG( | 2122.284 | -0.15128 | 0.150723 | 0.575502 | Usp2 |
| ENSMUSG( | 2202.378 | 0.156177 | 0.12111 | 0.525666 | Usp20 |
| ENSMUSG( | 1421.322 | 0.130932 | 0.227291 | 0.674223 | Usp21 |
| ENSMUSG( | 5317.967 | -0.09159 | 0.343242 | 0.779153 | Usp22 |
| ENSMUSG( | 3615.143 | 0.01823 | 0.845752 | 0.972191 | Usp24 |
| ENSMUSG( | 981.2639 | 0.135792 | 0.233962 | 0.681906 | Usp25 |
| ENSMUSG( | 293.5932 | 0.145988 | 0.344412 | 0.779153 | Usp27x |
| ENSMUSG( | 1053.523 | -0.06831 | 0.5242 | 0.87583 | Usp28 |
| ENSMUSG( | 647.5716 | -0.01851 | 0.895584 | 0.984512 | Usp29 |
| ENSMUSG( | 665.3565 | 0.032605 | 0.754766 | 0.951007 | Usp3 |
| ENSMUSG( | 1602.817 | -0.22225 | 0.083661 | 0.44608 | Usp30 |
| ENSMUSG( | 2593.462 | 0.147468 | 0.138528 | 0.553838 | Usp31 |
| ENSMUSG( | 4894.712 | 0.076313 | 0.38418 | 0.803686 | Usp32 |
| ENSMUSG( | 2322.052 | -0.0351 | 0.674194 | 0.929638 | Usp33 |
| ENSMUSG( | 4678.69 | 0.077797 | 0.359483 | 0.791011 | Usp34 |
| ENSMUSG( | 647.778 | 0.103016 | 0.378367 | 0.801688 | Usp35 |
| ENSMUSG( | 1243.649 | -0.04599 | 0.674908 | 0.929638 | Usp36 |
| ENSMUSG( | 682.3558 | 0.141063 | 0.22948 | 0.67735 | Usp37 |
| ENSMUSG( | 658.0862 | 0.065184 | 0.575457 | 0.894836 | Usp38 |
| ENSMUSG( | 609.3046 | 0.122842 | 0.330402 | 0.774327 | Usp39 |
| ENSMUSG( | 2754.307 | 0.082209 | 0.317954 | 0.764145 | Usp4 |
| ENSMUSG( | 834.2363 | 0.274444 | 0.012463 | 0.161344 | Usp40 |
| ENSMUSG( | 627.9405 | -0.13478 | 0.236209 | 0.683188 | Usp42 |
| ENSMUSG( | 206.7715 | -0.05265 | 0.762457 | 0.953519 | Usp43 |
| ENSMUSG( | 741.4955 | 0.001069 | 0.991735 | 0.999232 | Usp45 |
| ENSMUSG( | 4048.619 | 0.047826 | 0.674259 | 0.929638 | Usp46 |
| ENSMUSG( | 2605.184 | 0.192018 | 0.026637 | 0.249596 | Usp47 |
| ENSMUSG( | 3266.992 | -0.03259 | 0.71921 | 0.941986 | Usp48 |
| ENSMUSG( | 478.7201 | 0.060753 | 0.628967 | 0.913816 | Usp49 |
| ENSMUSG( | 4459.453 | 0.102352 | 0.392358 | 0.809274 | Usp5 |
| ENSMUSG( | 22.85989 | -0.1786 | 0.656054 | 0.923929 | Usp51 |
| ENSMUSG( | 1140.658 | 0.436846 | 0.000362 | 0.016539 | Usp53 |
| ENSMUSG( | 4231.976 | 0.101427 | 0.258369 | 0.709574 | Usp54 |

| | | | | | |
|---|---|---|---|---|---|
| ENSMUSG( | 593.149 | -0.00551 | 0.972296 | 0.996416 | Usp29 |
| ENSMUSG( | 600.2282 | 0.003436 | 0.977754 | 0.996416 | Usp3 |
| ENSMUSG( | 1568.407 | -0.00228 | 0.987923 | 0.997777 | Usp30 |
| ENSMUSG( | 2320.152 | 0.098849 | 0.383711 | 0.917257 | Usp31 |
| ENSMUSG( | 4265.862 | -0.05129 | 0.625385 | 0.953234 | Usp32 |
| ENSMUSG( | 2182.044 | 0.052035 | 0.625126 | 0.953234 | Usp33 |
| ENSMUSG( | 4235.688 | 0.059361 | 0.555782 | 0.939565 | Usp34 |
| ENSMUSG( | 571.5898 | 0.014162 | 0.917449 | 0.992529 | Usp35 |
| ENSMUSG( | 1110.475 | -0.10353 | 0.388247 | 0.917257 | Usp36 |
| ENSMUSG( | 625.055 | 0.154853 | 0.236298 | 0.872173 | Usp37 |
| ENSMUSG( | 625.0592 | 0.180875 | 0.172995 | 0.843788 | Usp38 |
| ENSMUSG( | 538.5572 | 0.039669 | 0.782518 | 0.976106 | Usp39 |
| ENSMUSG( | 2368.547 | -0.08514 | 0.422445 | 0.917257 | Usp4 |
| ENSMUSG( | 730.6473 | 0.174081 | 0.18578 | 0.855305 | Usp40 |
| ENSMUSG( | 572.4363 | -0.12642 | 0.362941 | 0.914223 | Usp42 |
| ENSMUSG( | 175.196 | -0.25855 | 0.20649 | 0.864412 | Usp43 |
| ENSMUSG( | 699.5754 | 0.099673 | 0.41833 | 0.917257 | Usp45 |
| ENSMUSG( | 3521.211 | -0.09 | 0.538106 | 0.937216 | Usp46 |
| ENSMUSG( | 2218.85 | 0.003831 | 0.971112 | 0.996102 | Usp47 |
| ENSMUSG( | 2977.543 | -0.03108 | 0.780754 | 0.976106 | Usp48 |
| ENSMUSG( | 442.0345 | 0.094583 | 0.514854 | 0.930939 | Usp49 |
| ENSMUSG( | 3792.656 | -0.09756 | 0.4665 | 0.923329 | Usp5 |
| ENSMUSG( | 25.84367 | 0.408692 | 0.342781 | 0.909646 | Usp51 |
| ENSMUSG( | 1092.512 | 0.561175 | 6.38E-05 | 0.015522 | Usp53 |
| ENSMUSG( | 3977.401 | 0.18895 | 0.093103 | 0.795363 | Usp54 |
| ENSMUSG( | 1507.58 | -0.00491 | 0.965402 | 0.995433 | Usp6nl |
| ENSMUSG( | 4209.532 | 0.070571 | 0.468257 | 0.923329 | Usp7 |
| ENSMUSG( | 2096.831 | -0.11416 | 0.325912 | 0.905339 | Usp8 |
| ENSMUSG( | 2910.676 | 0.025172 | 0.840006 | 0.98221 | Usp9x |
| ENSMUSG( | 794.2979 | 0.083203 | 0.498244 | 0.927825 | Uspl1 |
| ENSMUSG( | 1185.595 | 0.137712 | 0.40776 | 0.917257 | Ust |
| ENSMUSG( | 439.5697 | 0.174058 | 0.216543 | 0.865167 | Utp11 |
| ENSMUSG( | 410.8749 | -0.02956 | 0.832648 | 0.980867 | Utp14a |
| ENSMUSG( | 841.0072 | -0.73893 | 3.70E-08 | 2.70E-05 | Utp14b |
| ENSMUSG( | 543.7772 | 0.157041 | 0.244178 | 0.877904 | Utp15 |
| ENSMUSG( | 447.0005 | 0.082198 | 0.591595 | 0.944975 | Utp18 |
| ENSMUSG( | 562.6705 | 0.18802 | 0.180389 | 0.850372 | Utp20 |
| ENSMUSG( | 155.4602 | 0.0238 | 0.900838 | 0.990558 | Utp23 |
| ENSMUSG( | 373.2893 | 0.011979 | 0.932999 | 0.99306 | Utp25 |
| ENSMUSG( | 467.9618 | 0.067382 | 0.66928 | 0.956766 | Utp3 |
| ENSMUSG( | 454.0964 | -0.10888 | 0.499422 | 0.927825 | Utp4 |
| ENSMUSG( | 1336.824 | 0.057264 | 0.761175 | 0.972695 | Utp6 |
| ENSMUSG( | 3475.126 | -0.00515 | 0.962098 | 0.995237 | Utrn |
| ENSMUSG( | 295.602 | 0.251902 | 0.123803 | 0.814691 | Uty |
| ENSMUSG( | 834.3037 | 0.073183 | 0.574226 | 0.942912 | Uvrag |
| ENSMUSG( | 323.3421 | 0.122211 | 0.486889 | 0.92541 | Uvssa |
| ENSMUSG( | 1336.132 | 0.117202 | 0.474824 | 0.923329 | Uxs1 |
| ENSMUSG( | 148.333 | -0.36469 | 0.048456 | 0.706142 | Uxt |
| ENSMUSG( | 628.2153 | 0.009001 | 0.941796 | 0.994601 | Vac14 |
| ENSMUSG( | 2284.915 | -0.16304 | 0.282025 | 0.892412 | Vamp1 |
| ENSMUSG( | 34107.95 | -0.07302 | 0.522562 | 0.932822 | Vamp2 |
| ENSMUSG( | 1742.269 | 0.059882 | 0.63438 | 0.953903 | Vamp3 |
| ENSMUSG( | 1815.565 | 0.051599 | 0.702674 | 0.962215 | Vamp4 |
| ENSMUSG( | 103.557 | -0.04006 | 0.869341 | 0.988097 | Vamp5 |
| ENSMUSG( | 469.9272 | -0.21729 | 0.10865 | 0.804969 | Vamp8 |
| ENSMUSG( | 235.771 | 0.26074 | 0.249763 | 0.877904 | Vangl1 |
| ENSMUSG( | 848.6244 | 0.10232 | 0.488865 | 0.925573 | Vangl2 |
| ENSMUSG( | 2383.447 | -0.1436 | 0.380931 | 0.917202 | Vapa |
| ENSMUSG( | 5857.331 | -0.16548 | 0.13386 | 0.822912 | Vapb |
| ENSMUSG( | 1169.815 | 0.008266 | 0.949689 | 0.994798 | Vars1 |
| ENSMUSG( | 913.2913 | 0.099138 | 0.403429 | 0.917257 | Vars2 |
| ENSMUSG( | 1741.173 | -0.3338 | 0.003114 | 0.221768 | Vash1 |
| ENSMUSG( | 69.07588 | -0.21437 | 0.428879 | 0.917257 | Vash2 |
| ENSMUSG( | 362.9457 | 0.287175 | 0.199534 | 0.863016 | Vasn |
| ENSMUSG( | 568.5691 | 0.208419 | 0.148753 | 0.836241 | Vasp |
| ENSMUSG( | 1204.088 | -0.05382 | 0.644148 | 0.953903 | Vat1 |
| ENSMUSG( | 1011.478 | -0.12839 | 0.465096 | 0.923329 | Vat1l |
| ENSMUSG( | 266.6919 | 0.034621 | 0.838937 | 0.98174 | Vav1 |
| ENSMUSG( | 648.1162 | 0.153932 | 0.24288 | 0.877399 | Vav2 |
| ENSMUSG( | 276.0455 | -0.0973 | 0.577982 | 0.943131 | Vav3 |
| ENSMUSG( | 15.68876 | -0.04087 | 0.934468 | 0.99306 | Vax1 |
| ENSMUSG( | 874.3204 | 0.070034 | 0.657409 | 0.955725 | Vbp1 |
| ENSMUSG( | 2180.158 | 0.055186 | 0.728871 | 0.966913 | Vcam1 |
| ENSMUSG( | 1304.507 | 0.391634 | 0.030639 | 0.628144 | Vcan |
| ENSMUSG( | 1120.123 | -0.0422 | 0.76324 | 0.973945 | Vcf1 |
| ENSMUSG( | 442.3633 | -0.0525 | 0.726964 | 0.966887 | Vcf2 |
| ENSMUSG( | 1673.949 | 0.452125 | 0.007746 | 0.37555 | Vcl |
| ENSMUSG( | 5446.247 | -0.06461 | 0.557681 | 0.939565 | Vcp |
| ENSMUSG( | 984.7678 | 0.142569 | 0.332858 | 0.907907 | Vcpip1 |
| ENSMUSG( | 453.523 | -0.24086 | 0.066336 | 0.749203 | Vcpkmt |
| ENSMUSG( | 9340.081 | -0.14481 | 0.309215 | 0.898538 | Vdac1 |
| ENSMUSG( | 3664.516 | -0.14125 | 0.301146 | 0.898538 | Vdac2 |
| ENSMUSG( | 1851.06 | -0.17791 | 0.262961 | 0.885357 | Vdac3 |
| ENSMUSG( | 3295.233 | 0.084641 | 0.498878 | 0.927825 | Vegfa |
| ENSMUSG( | 4461.777 | -0.15845 | 0.182711 | 0.850579 | Vegfb |
| ENSMUSG( | 124.8247 | 0.177739 | 0.443409 | 0.921704 | Vegfc |
| ENSMUSG( | 83.61095 | 0.574878 | 0.032113 | 0.634374 | Vegfd |
| ENSMUSG( | 12.77843 | 0.044171 | 0.937648 | 0.993685 | Veph1 |
| ENSMUSG( | 1207.35 | -0.07482 | 0.540501 | 0.937383 | Vezf1 |
| ENSMUSG( | 2213.239 | 0.142832 | 0.288554 | 0.894794 | Vezt |
| ENSMUSG( | 11092.43 | 0.076085 | 0.590189 | 0.944975 | Vgf |
| ENSMUSG( | 37.65475 | 0.153164 | 0.666919 | 0.956766 | Vgll3 |
| ENSMUSG( | 888.6033 | -0.0638 | 0.643586 | 0.953903 | Vgll4 |
| ENSMUSG( | 2819.888 | -0.28857 | 0.037068 | 0.673514 | Vhl |
| ENSMUSG( | 23.45574 | 0.118748 | 0.775457 | 0.975354 | Vil1 |
| ENSMUSG( | 187.9427 | 0.1297 | 0.593266 | 0.945408 | Vill |
| ENSMUSG( | 2051.05 | 0.566404 | 3.36E-06 | 0.001549 | Vim |
| ENSMUSG( | 1444.762 | 0.099453 | 0.624479 | 0.953234 | Vip |
| ENSMUSG( | 1303.216 | 0.071737 | 0.503972 | 0.927825 | Vipas39 |
| ENSMUSG( | 992.8651 | 0.129836 | 0.398736 | 0.917257 | Vipr1 |
| ENSMUSG( | 65.71883 | -0.0271 | 0.853354 | 0.985598 | Vipr2 |
| ENSMUSG( | 1013.797 | 0.083588 | 0.508284 | 0.929089 | Virma |
| ENSMUSG( | 166.4542 | 0.254402 | 0.202539 | 0.864412 | Vit |
| ENSMUSG( | 215.6869 | 0.017169 | 0.925816 | 0.99306 | Vkorc1 |
| ENSMUSG( | 2368.713 | 0.037766 | 0.778884 | 0.976106 | Vkorc1l1 |
| ENSMUSG( | 2858.598 | 0.076964 | 0.629268 | 0.953251 | Vldlr |
| ENSMUSG( | 1383.647 | 0.017475 | 0.922248 | 0.992644 | Vma21 |
| ENSMUSG( | 492.9658 | -0.14045 | 0.283556 | 0.892412 | Vmac |
| ENSMUSG( | 24.38504 | 0.882593 | 0.04494 | 0.693959 | Vmn2r1 |
| ENSMUSG( | 73.64074 | -0.53702 | 0.026995 | 0.603041 | Vmn2r29 |
| ENSMUSG( | 45.42355 | -0.19951 | 0.531267 | 0.934795 | Vmn2r57 |
| ENSMUSG( | 69.1076 | -0.01829 | 0.94839 | 0.994798 | Vmn2r84 |
| ENSMUSG( | 8.564871 | 0.172851 | 0.798188 | NA | Vmn2r85 |
| ENSMUSG( | 23.35766 | -0.10055 | 0.823672 | 0.979198 | Vmn2r87 |
| ENSMUSG( | 3943.482 | 0.061463 | 0.722205 | 0.966887 | Vmp1 |
| ENSMUSG( | 28.95027 | -0.06053 | 0.893819 | 0.990558 | Vnn1 |
| ENSMUSG( | 4030.223 | 0.119449 | 0.396857 | 0.917257 | Vopp1 |
| ENSMUSG( | 1549.47 | -0.03382 | 0.783941 | 0.976154 | Vps11 |
| ENSMUSG( | 1219.265 | 0.295908 | 0.044733 | 0.693959 | Vps13a |
| ENSMUSG( | 1465.06 | 0.045531 | 0.752693 | 0.970521 | Vps13b |
| ENSMUSG( | 1583.255 | 0.184224 | 0.225002 | 0.868422 | Vps13c |
| ENSMUSG( | 3950.24 | -0.00386 | 0.971426 | 0.996236 | Vps13d |
| ENSMUSG( | 1322.372 | 0.015652 | 0.902164 | 0.991012 | Vps16 |
| ENSMUSG( | 996.3691 | -0.03292 | 0.832326 | 0.98075 | Vps18 |
| ENSMUSG( | 28.8556 | 0.166573 | 0.669959 | 0.95694 | Vps25 |
| ENSMUSG( | 1376.252 | -0.09626 | 0.442893 | 0.921554 | Vps26a |
| ENSMUSG( | 2993.564 | -0.02722 | 0.788395 | 0.97637 | Vps26b |
| ENSMUSG( | 1275.061 | -0.24947 | 0.055062 | 0.723362 | Vps26c |
| ENSMUSG( | 1820.472 | -0.23752 | 0.043579 | 0.690304 | Vps28 |
| ENSMUSG( | 2107.678 | -0.11038 | 0.34812 | 0.911652 | Vps29 |
| ENSMUSG( | 2343.74 | -0.02404 | 0.831382 | 0.980611 | Vps33a |
| ENSMUSG( | 497.6246 | 0.137789 | 0.417038 | 0.917257 | Vps33b |
| ENSMUSG( | 3005.033 | -0.00801 | 0.946952 | 0.994798 | Vps35 |
| ENSMUSG( | 1801.395 | -0.08093 | 0.45555 | 0.923329 | Vps35l |
| ENSMUSG( | 768.2443 | 0.074721 | 0.532117 | 0.934795 | Vps36 |
| ENSMUSG( | 1661.665 | 0.060311 | 0.62894 | 0.953251 | Vps37a |
| ENSMUSG( | 1015.64 | -0.20051 | 0.108333 | 0.804969 | Vps37b |
| ENSMUSG( | 1017.692 | -0.09201 | 0.434564 | 0.91869 | Vps37c |
| ENSMUSG( | 521.5391 | -0.17749 | 0.219964 | 0.866104 | Vps37d |
| ENSMUSG( | 2049.539 | 0.005061 | 0.965648 | 0.995433 | Vps39 |
| ENSMUSG( | 4007.437 | -0.12032 | 0.251242 | 0.879321 | Vps41 |
| ENSMUSG( | 636.9802 | -0.18638 | 0.217996 | 0.866101 | Vps45 |
| ENSMUSG( | 2481.919 | -0.18318 | 0.119919 | 0.813384 | Vps4a |
| ENSMUSG( | 867.0928 | 0.162274 | 0.18272 | 0.850579 | Vps4b |
| ENSMUSG( | 1437.921 | 0.061131 | 0.608272 | 0.950113 | Vps50 |
| ENSMUSG( | 1080.018 | -0.04964 | 0.700915 | 0.96162 | Vps51 |
| ENSMUSG( | 2759.931 | -0.12617 | 0.257001 | 0.883082 | Vps52 |
| ENSMUSG( | 1227.69 | -0.09919 | 0.495129 | 0.927372 | Vps53 |
| ENSMUSG( | 936.663 | 0.212272 | 0.079427 | 0.77458 | Vps54 |
| ENSMUSG( | 671.9822 | -0.10686 | 0.407737 | 0.917257 | Vps72 |
| ENSMUSG( | 1291.015 | -0.02801 | 0.809058 | 0.97873 | Vps8 |
| ENSMUSG( | 2405.072 | 0.030798 | 0.789492 | 0.976593 | Vps9d1 |
| ENSMUSG( | 358.7559 | 0.032532 | 0.823552 | 0.979198 | Vrk1 |
| ENSMUSG( | 67.07125 | 0.005469 | 0.982987 | 0.996891 | Vrk2 |
| ENSMUSG( | 1080.363 | -0.08135 | 0.475292 | 0.923329 | Vrk3 |
| ENSMUSG( | 125.6623 | -0.08211 | 0.703488 | 0.962481 | Vsig10 |
| ENSMUSG( | 130.1399 | 0.255786 | 0.195137 | 0.861229 | Vsig10l |
| ENSMUSG( | 60.57976 | 0.034567 | 0.904718 | 0.991089 | Vsig2 |

| | | | | | | | | | | | |
|---|---|---|---|---|---|---|---|---|---|---|---|
| ENSMUSG( | Wbp4 | 1339.433 | 0.104417 | 0.267298 | 0.717525 | ENSMUSG( | Wbp4 | 1183.384 | 0.015171 | 0.894926 | 0.990558 |
| ENSMUSG( | Wdcp | 221.7327 | 0.274658 | 0.072251 | 0.414687 | ENSMUSG( | Wdcp | 197.5104 | 0.219071 | 0.21747 | 0.865589 |
| ENSMUSG( | Wdfy1 | 548.0839 | -0.2214 | 0.079514 | 0.435569 | ENSMUSG( | Wdfy1 | 732.5622 | 0.786439 | 4.64E-09 | 4.06E-06 |
| ENSMUSG( | Wdfy2 | 1239.918 | -0.01364 | 0.898654 | 0.985633 | ENSMUSG( | Wdfy2 | 1166.319 | 0.079444 | 0.509806 | 0.929089 |
| ENSMUSG( | Wdfy3 | 5148.724 | -0.09293 | 0.374541 | 0.800445 | ENSMUSG( | Wdfy3 | 5082.672 | 0.139501 | 0.261373 | 0.884926 |
| ENSMUSG( | Wdfy4 | 200.1466 | 0.002666 | 0.985755 | 0.998326 | ENSMUSG( | Wdfy4 | 183.9964 | 0.033659 | 0.844449 | 0.983642 |
| ENSMUSG( | Wdhd1 | 170.5862 | 0.113295 | 0.492177 | 0.860297 | ENSMUSG( | Wdhd1 | 156.3821 | 0.131707 | 0.502747 | 0.927825 |
| ENSMUSG( | Wdpcp | 269.6851 | 0.010959 | 0.942308 | 0.993415 | ENSMUSG( | Wdpcp | 233.0321 | -0.14471 | 0.363118 | 0.914223 |
| ENSMUSG( | Wdr1 | 4725.296 | 0.130543 | 0.119958 | 0.523983 | ENSMUSG( | Wdr1 | 3958.835 | -0.11269 | 0.309353 | 0.898538 |
| ENSMUSG( | Wdr11 | 1578.975 | 0.154438 | 0.101911 | 0.486318 | ENSMUSG( | Wdr11 | 1387.328 | 0.054011 | 0.616965 | 0.951666 |
| ENSMUSG( | Wdr12 | 771.5604 | 0.102562 | 0.305977 | 0.753712 | ENSMUSG( | Wdr12 | 680.631 | 0.009025 | 0.942508 | 0.994774 |
| ENSMUSG( | Wdr13 | 15249.63 | 0.178321 | 0.021447 | 0.22362 | ENSMUSG( | Wdr13 | 13288.16 | 0.054051 | 0.575055 | 0.942912 |
| ENSMUSG( | Wdr17 | 681.6332 | -0.02389 | 0.835546 | 0.970557 | ENSMUSG( | Wdr17 | 626.4698 | 0.001366 | 0.991732 | 0.998712 |
| ENSMUSG( | Wdr18 | 1940.504 | 0.105946 | 0.269446 | 0.719729 | ENSMUSG( | Wdr18 | 1640.02 | -0.11147 | 0.345587 | 0.910597 |
| ENSMUSG( | Wdr19 | 609.9653 | -0.04142 | 0.728638 | 0.944393 | ENSMUSG( | Wdr19 | 568.7029 | 0.030896 | 0.824069 | 0.979198 |
| ENSMUSG( | Wdr20 | 489.4655 | 0.020153 | 0.855974 | 0.973813 | ENSMUSG( | Wdr20 | 454.3574 | 0.071431 | 0.58807 | 0.944975 |
| ENSMUSG( | Wdr24 | 597.9034 | 0.108675 | 0.344787 | 0.779153 | ENSMUSG( | Wdr24 | 546.7535 | 0.124668 | 0.353258 | 0.912263 |
| ENSMUSG( | Wdr25 | 257.6559 | -0.03038 | 0.829764 | 0.969582 | ENSMUSG( | Wdr25 | 224.0932 | -0.16798 | 0.295455 | 0.898538 |
| ENSMUSG( | Wdr26 | 4603.176 | 0.079532 | 0.355568 | 0.787945 | ENSMUSG( | Wdr26 | 4147.043 | 0.046658 | 0.67031 | 0.95694 |
| ENSMUSG( | Wdr27 | 52.77515 | 0.344738 | 0.21286 | 0.656885 | ENSMUSG( | Wdr27 | 50.94548 | 0.466638 | 0.169093 | 0.840282 |
| ENSMUSG( | Wdr3 | 430.617 | 0.050851 | 0.693964 | 0.934202 | ENSMUSG( | Wdr3 | 381.7528 | -0.02044 | 0.893271 | 0.990558 |
| ENSMUSG( | Wdr31 | 121.8337 | 0.01658 | 0.928892 | 0.991747 | ENSMUSG( | Wdr31 | 106.7289 | -0.10168 | 0.62673 | 0.953234 |
| ENSMUSG( | Wdr33 | 1696.726 | -0.39209 | 0.005533 | 0.099488 | ENSMUSG( | Wdr33 | 1876.432 | 0.191816 | 0.232267 | 0.868507 |
| ENSMUSG( | Wdr35 | 823.2444 | 0.087907 | 0.396103 | 0.811615 | ENSMUSG( | Wdr35 | 700.1054 | -0.11498 | 0.351975 | 0.912093 |
| ENSMUSG( | Wdr36 | 538.7039 | 0.090857 | 0.448741 | 0.83469 | ENSMUSG( | Wdr36 | 491.5417 | 0.098308 | 0.474886 | 0.923329 |
| ENSMUSG( | Wdr37 | 1377.513 | 0.058933 | 0.540834 | 0.88315 | ENSMUSG( | Wdr37 | 1224.662 | -0.0149 | 0.904298 | 0.991089 |
| ENSMUSG( | Wdr38 | 53.90398 | -0.04056 | 0.880478 | 0.981469 | ENSMUSG( | Wdr38 | 43.71046 | -0.40583 | 0.189117 | 0.85698 |
| ENSMUSG( | Wdr4 | 933.3766 | -0.04196 | 0.703649 | 0.937885 | ENSMUSG( | Wdr4 | 832.424 | -0.10058 | 0.426419 | 0.917257 |
| ENSMUSG( | Wdr41 | 1178.168 | 0.076983 | 0.449835 | 0.835298 | ENSMUSG( | Wdr41 | 993.988 | -0.15065 | 0.222771 | 0.867173 |
| ENSMUSG( | Wdr43 | 779.266 | 0.012533 | 0.900228 | 0.985633 | ENSMUSG( | Wdr43 | 698.6682 | -0.03765 | 0.775517 | 0.975354 |
| ENSMUSG( | Wdr44 | 519.9506 | -0.2309 | 0.130932 | 0.542456 | ENSMUSG( | Wdr44 | 509.4275 | -0.00549 | 0.974845 | 0.996416 |
| ENSMUSG( | Wdr45 | 509.6681 | 0.160962 | 0.219993 | 0.665725 | ENSMUSG( | Wdr45 | 434.8609 | 0.026612 | 0.85857 | 0.985884 |
| ENSMUSG( | Wdr45b | 2531.969 | 0.100577 | 0.236377 | 0.683538 | ENSMUSG( | Wdr45b | 2205.289 | -0.03019 | 0.794478 | 0.97681 |
| ENSMUSG( | Wdr46 | 382.95 | 0.161787 | 0.244271 | 0.691435 | ENSMUSG( | Wdr46 | 321.5978 | -0.07221 | 0.652032 | 0.9547 |
| ENSMUSG( | Wdr47 | 2375.754 | 0.032138 | 0.795028 | 0.959976 | ENSMUSG( | Wdr47 | 2066.971 | -0.10463 | 0.462131 | 0.923329 |
| ENSMUSG( | Wdr48 | 1976.379 | 0.073866 | 0.39243 | 0.809274 | ENSMUSG( | Wdr48 | 1781.218 | 0.044033 | 0.683271 | 0.958678 |
| ENSMUSG( | Wdr5 | 804.4211 | 0.052879 | 0.647971 | 0.919759 | ENSMUSG( | Wdr5 | 691.5756 | -0.11527 | 0.371833 | 0.915491 |
| ENSMUSG( | Wdr53 | 182.6573 | 0.037002 | 0.815174 | 0.964978 | ENSMUSG( | Wdr53 | 163.2986 | -0.02715 | 0.881401 | 0.989644 |
| ENSMUSG( | Wdr54 | 421.8901 | -0.42705 | 0.000272 | 0.013445 | ENSMUSG( | Wdr54 | 452.9026 | 0.074981 | 0.603256 | 0.948137 |
| ENSMUSG( | Wdr55 | 263.2929 | 0.123087 | 0.430745 | 0.826896 | ENSMUSG( | Wdr55 | 220.8264 | -0.12497 | 0.510676 | 0.929501 |
| ENSMUSG( | Wdr59 | 718.6328 | -0.07133 | 0.540397 | 0.88315 | ENSMUSG( | Wdr59 | 663.0399 | -0.0325 | 0.786771 | 0.976303 |
| ENSMUSG( | Wdr5b | 132.9951 | -0.06138 | 0.769107 | 0.954674 | ENSMUSG( | Wdr5b | 106.077 | 0.040869 | 0.855633 | 0.985738 |
| ENSMUSG( | Wdr6 | 3457.435 | 0.029012 | 0.764335 | 0.954055 | ENSMUSG( | Wdr6 | 3185.324 | 0.059176 | 0.638672 | 0.953903 |
| ENSMUSG( | Wdr62 | 65.02871 | 0.070122 | 0.770355 | 0.954689 | ENSMUSG( | Wdr62 | 66.96936 | 0.42486 | 0.161633 | 0.837724 |
| ENSMUSG( | Wdr7 | 3365.864 | 0.07697 | 0.394748 | 0.810072 | ENSMUSG( | Wdr7 | 2972.964 | -0.01051 | 0.921414 | 0.992644 |
| ENSMUSG( | Wdr70 | 541.2576 | -0.01706 | 0.885172 | 0.982851 | ENSMUSG( | Wdr70 | 475.5196 | -0.12404 | 0.371958 | 0.915491 |
| ENSMUSG( | Wdr73 | 815.4853 | 0.044429 | 0.69267 | 0.934101 | ENSMUSG( | Wdr73 | 725.3565 | -0.02195 | 0.863294 | 0.986547 |
| ENSMUSG( | Wdr74 | 581.8326 | 0.052425 | 0.695338 | 0.934957 | ENSMUSG( | Wdr74 | 514.6571 | -0.03592 | 0.813972 | 0.979198 |
| ENSMUSG( | Wdr75 | 595.702 | 0.114305 | 0.323466 | 0.768511 | ENSMUSG( | Wdr75 | 518.6828 | -0.01357 | 0.916874 | 0.992529 |
| ENSMUSG( | Wdr76 | 76.0417 | 0.123295 | 0.584246 | 0.898754 | ENSMUSG( | Wdr76 | 72.16889 | 0.245788 | 0.329501 | 0.907083 |
| ENSMUSG( | Wdr77 | 678.8474 | 0.018737 | 0.856841 | 0.97427 | ENSMUSG( | Wdr77 | 589.3521 | -0.12933 | 0.376629 | 0.915491 |
| ENSMUSG( | Wdr81 | 802.9355 | 0.02205 | 0.850779 | 0.973755 | ENSMUSG( | Wdr81 | 770.5683 | 0.231498 | 0.158669 | 0.837724 |
| ENSMUSG( | Wdr82 | 3506.476 | 0.130535 | 0.126753 | 0.534201 | ENSMUSG( | Wdr82 | 2972.393 | -0.07747 | 0.515569 | 0.930939 |
| ENSMUSG( | Wdr83 | 730.9871 | 0.125779 | 0.295527 | 0.743814 | ENSMUSG( | Wdr83 | 626.4686 | -0.04585 | 0.736215 | 0.967344 |
| ENSMUSG( | Wdr83os | 1080.675 | -0.10709 | 0.265436 | 0.716109 | ENSMUSG( | Wdr83os | 985.2819 | -0.10612 | 0.363876 | 0.914223 |
| ENSMUSG( | Wdr86 | 56.05529 | -0.32255 | 0.43545 | 0.829093 | ENSMUSG( | Wdr86 | 63.83922 | 0.318898 | 0.363262 | 0.914223 |
| ENSMUSG( | Wdr89 | 50.07014 | 0.430221 | 0.108706 | 0.497919 | ENSMUSG( | Wdr89 | 37.58206 | -0.09994 | 0.76081 | 0.972695 |
| ENSMUSG( | Wdr90 | 851.463 | 0.348673 | 0.023687 | 0.23542 | ENSMUSG( | Wdr90 | 731.1944 | 0.19215 | 0.295578 | 0.898538 |
| ENSMUSG( | Wdr91 | 996.2753 | 0.088475 | 0.407474 | 0.817671 | ENSMUSG( | Wdr91 | 922.1569 | 0.135102 | 0.254945 | 0.882307 |
| ENSMUSG( | Wdr93 | 13.89939 | 0.20334 | 0.698095 | NA | ENSMUSG( | Wdr93 | 14.31886 | 0.543482 | 0.326485 | 0.905371 |
| ENSMUSG( | Wdr95 | 27.2281 | 0.307927 | 0.400638 | 0.813717 | ENSMUSG( | Wdr95 | 28.19759 | 0.602028 | 0.142222 | 0.830709 |
| ENSMUSG( | Wdr97 | 207.5352 | -0.06139 | 0.707989 | 0.938734 | ENSMUSG( | Wdr97 | 177.862 | -0.43847 | 0.034763 | 0.624049 |
| ENSMUSG( | Wdsub1 | 326.8595 | 0.087541 | 0.560895 | 0.888934 | ENSMUSG( | Wdsub1 | 287.8332 | -0.00283 | 0.986152 | 0.997188 |
| ENSMUSG( | Wdtc1 | 4453.971 | 0.278625 | 0.000913 | 0.03071 | ENSMUSG( | Wdtc1 | 3836.707 | 0.129122 | 0.235145 | 0.871741 |
| ENSMUSG( | Wee1 | 331.9123 | 0.086416 | 0.528613 | 0.878365 | ENSMUSG( | Wee1 | 332.0046 | 0.33572 | 0.026965 | 0.603041 |
| ENSMUSG( | Wfdc1 | 476.8179 | 0.060883 | 0.701986 | 0.937445 | ENSMUSG( | Wfdc1 | 479.6252 | 0.338558 | 0.093451 | 0.795507 |
| ENSMUSG( | Wfdc18 | 42.45022 | -0.65425 | 0.024084 | 0.237752 | ENSMUSG( | Wfdc18 | 37.95428 | -0.71402 | 0.028224 | 0.611882 |
| ENSMUSG( | Wfdc2 | 23.06007 | 0.583135 | 0.178894 | 0.616298 | ENSMUSG( | Wfdc2 | 24.55169 | 0.95969 | 0.034745 | 0.657974 |
| ENSMUSG( | Wfdc3 | 20.9362 | 0.114931 | 0.775578 | NA | ENSMUSG( | Wfdc3 | 20.29054 | 0.297534 | 0.502678 | 0.927825 |
| ENSMUSG( | Wfikkn1 | 10.1939 | 0.756389 | 0.195304 | NA | ENSMUSG( | Wfikkn1 | 9.930406 | 0.95876 | 0.152973 | NA |
| ENSMUSG( | Wfikkn2 | 136.9045 | 0.475842 | 0.077166 | 0.429655 | ENSMUSG( | Wfikkn2 | 112.3876 | 0.191308 | 0.461798 | 0.923329 |
| ENSMUSG( | Wfs1 | 3688.839 | -0.25405 | 0.050178 | 0.342492 | ENSMUSG( | Wfs1 | 3897.519 | 0.182939 | 0.193554 | 0.860096 |
| ENSMUSG( | Whamm | 641.71 | 0.028689 | 0.793012 | 0.95947 | ENSMUSG( | Whamm | 603.3788 | 0.119949 | 0.332571 | 0.907907 |
| ENSMUSG( | Whrn | 2868.069 | 0.320479 | 0.000453 | 0.019387 | ENSMUSG( | Whrn | 2466.729 | 0.167749 | 0.150169 | 0.837086 |
| ENSMUSG( | Wif1 | 32.9878 | -1.04878 | 0.003109 | 0.069189 | ENSMUSG( | Wif1 | 45.63336 | 0.339809 | 0.288601 | 0.894794 |
| ENSMUSG( | Wipf1 | 1551.808 | -0.09545 | 0.392467 | 0.809274 | ENSMUSG( | Wipf1 | 1389.431 | -0.15063 | 0.291148 | 0.897421 |
| ENSMUSG( | Wipf2 | 2605.43 | 0.171935 | 0.040639 | 0.311461 | ENSMUSG( | Wipf2 | 2309.487 | 0.095079 | 0.378754 | 0.916995 |
| ENSMUSG( | Wipf3 | 4030.503 | 0.159125 | 0.209243 | 0.651175 | ENSMUSG( | Wipf3 | 3706.2 | 0.185301 | 0.178925 | 0.849041 |
| ENSMUSG( | Wipi1 | 1676.832 | 0.002245 | 0.984373 | 0.99781 | ENSMUSG( | Wipi1 | 1396.904 | -0.27139 | 0.022268 | 0.577034 |
| ENSMUSG( | Wipi2 | 4669.091 | 0.096745 | 0.218955 | 0.664602 | ENSMUSG( | Wipi2 | 3960.249 | -0.11008 | 0.266015 | 0.886134 |
| ENSMUSG( | Wiz | 3006.161 | 0.221217 | 0.011931 | 0.157992 | ENSMUSG( | Wiz | 2627.199 | 0.108662 | 0.300496 | 0.898538 |
| ENSMUSG( | Wls | 1485.59 | -0.46019 | 0.004676 | 0.089972 | ENSMUSG( | Wls | 1616.732 | 0.093068 | 0.505299 | 0.927825 |
| ENSMUSG( | Wnk1 | 20068.58 | -0.02171 | 0.839596 | 0.971179 | ENSMUSG( | Wnk1 | 17049.51 | -0.23391 | 0.055121 | 0.723362 |
| ENSMUSG( | Wnk2 | 5707.727 | 0.311714 | 0.000152 | 0.008645 | ENSMUSG( | Wnk2 | 5056.405 | 0.237852 | 0.016175 | 0.512256 |
| ENSMUSG( | Wnk3 | 775.2091 | -0.04555 | 0.696755 | 0.935389 | ENSMUSG( | Wnk3 | 696.4985 | -0.08802 | 0.504709 | 0.927825 |
| ENSMUSG( | Wnk4 | 170.4834 | 0.098511 | 0.605106 | 0.905594 | ENSMUSG( | Wnk4 | 165.2935 | 0.261398 | 0.254416 | 0.881624 |
| ENSMUSG( | Wnt10a | 207.6645 | -0.4036 | 0.032105 | 0.273791 | ENSMUSG( | Wnt10a | 211.4547 | -0.04409 | 0.807009 | 0.978239 |
| ENSMUSG( | Wnt10b | 60.12673 | -0.26073 | 0.33032 | 0.774247 | ENSMUSG( | Wnt10b | 60.88105 | 0.045913 | 0.883656 | 0.989644 |
| ENSMUSG( | Wnt11 | 19.58102 | 0.273251 | 0.558639 | NA | ENSMUSG( | Wnt11 | 17.28466 | 0.225687 | 0.674717 | 0.95694 |
| ENSMUSG( | Wnt16 | 48.55242 | 0.247642 | 0.383014 | 0.803237 | ENSMUSG( | Wnt16 | 35.12372 | -0.45918 | 0.172828 | 0.843702 |
| ENSMUSG( | Wnt2 | 103.8377 | -0.44987 | 0.029125 | 0.259574 | ENSMUSG( | Wnt2 | 122.5068 | 0.31193 | 0.140124 | 0.830678 |
| ENSMUSG( | Wnt2b | 57.12744 | -0.24342 | 0.345158 | 0.779153 | ENSMUSG( | Wnt2b | 56.85329 | 0.028315 | 0.921432 | 0.992644 |
| ENSMUSG( | Wnt3 | 67.39537 | -0.85546 | 0.000732 | 0.026277 | ENSMUSG( | Wnt3 | 69.28729 | -0.39976 | 0.130763 | 0.822884 |
| ENSMUSG( | Wnt4 | 1170.466 | 0.004962 | 0.964315 | 0.995155 | ENSMUSG( | Wnt4 | 1209.305 | 0.349791 | 0.006165 | 0.323304 |
| ENSMUSG( | Wnt5a | 346.5097 | -0.18864 | 0.1603 | 0.589853 | ENSMUSG( | Wnt5a | 357.9228 | 0.174618 | 0.268176 | 0.886493 |
| ENSMUSG( | Wnt5b | 60.46938 | 0.027863 | 0.914118 | 0.988483 | ENSMUSG( | Wnt5b | 64.05395 | 0.453767 | 0.107966 | 0.804969 |
| ENSMUSG( | Wnt6 | 59.9724 | 0.243448 | 0.343409 | 0.779153 | ENSMUSG( | Wnt6 | 62.97359 | 0.587621 | 0.037385 | 0.673514 |
| ENSMUSG( | Wnt7a | 784.5796 | -0.36711 | 0.008899 | 0.132538 | ENSMUSG( | Wnt7a | 813.8311 | 0.033595 | 0.826869 | 0.980159 |
| ENSMUSG( | Wnt7b | 2161.57 | -0.1415 | 0.189054 | 0.627678 | ENSMUSG( | Wnt7b | 2109.366 | 0.061763 | 0.634605 | 0.953903 |
| ENSMUSG( | Wnt9a | 737.4654 | -0.16147 | 0.255681 | 0.706195 | ENSMUSG( | Wnt9a | 685.1612 | -0.09736 | 0.56024 | 0.940131 |
| ENSMUSG( | Wrap53 | 327.6717 | 0.18557 | 0.140691 | 0.557902 | ENSMUSG( | Wrap53 | 287.0394 | 0.080105 | 0.588814 | 0.944975 |
| ENSMUSG( | Wrap73 | 472.7567 | 0.103422 | 0.419875 | 0.822442 | ENSMUSG( | Wrap73 | 428.2032 | 0.094108 | 0.564177 | 0.941206 |
| ENSMUSG( | Wrn | 375.3621 | -0.08187 | 0.491037 | 0.859739 | ENSMUSG( | Wrn | 358.4251 | 0.04662 | 0.750564 | 0.970184 |
| ENSMUSG( | Wrnip1 | 1425.269 | 0.011679 | 0.910876 | 0.988387 | ENSMUSG( | Wrnip1 | 1230.647 | -0.1505 | 0.2501 | 0.877904 |
| ENSMUSG( | Wsb1 | 1590.145 | -0.34055 | 0.007855 | 0.123093 | ENSMUSG( | Wsb1 | 1721.831 | 0.175833 | 0.286133 | 0.893906 |
| ENSMUSG( | Wsb2 | 19599.77 | 0.091523 | 0.237433 | 0.685281 | ENSMUSG( | Wsb2 | 16624.96 | -0.11576 | 0.236434 | 0.872173 |
| ENSMUSG( | Wscd1 | 5353.068 | -0.30043 | 0.014775 | 0.178616 | ENSMUSG( | Wscd1 | 5305.951 | -0.03879 | 0.792286 | 0.97681 |
| ENSMUSG( | Wscd2 | 1828.326 | -0.388 | 0.000673 | 0.025051 | ENSMUSG( | Wscd2 | 1937.002 | 0.076059 | 0.565723 | 0.941404 |
| ENSMUSG( | Wtap | 853.5037 | -0.04125 | 0.683651 | 0.932154 | ENSMUSG( | Wtap | 790.0663 | 0.005231 | 0.964767 | 0.995433 |
| ENSMUSG( | Wtip | 248.0565 | -0.06703 | 0.630964 | 0.914073 | ENSMUSG( | Wtip | 242.7461 | 0.146524 | 0.408333 | 0.917257 |
| ENSMUSG( | Wwc1 | 8443.462 | 0.053223 | 0.536502 | 0.881846 | ENSMUSG( | Wwc1 | 7206.584 | -0.13919 | 0.191369 | 0.858412 |
| ENSMUSG( | Wwc2 | 2428.896 | 0.163253 | 0.093843 | 0.468228 | ENSMUSG( | Wwc2 | 2169.258 | 0.106656 | 0.336803 | 0.907972 |
| ENSMUSG( | Wwox | 557.351 | 0.127137 | 0.259091 | 0.709814 | ENSMUSG( | Wwox | 482.8689 | -0.02235 | 0.880285 | 0.989644 |
| ENSMUSG( | Wwp1 | 2192.668 | 0.025063 | 0.829203 | 0.969286 | ENSMUSG( | Wwp1 | 1900.471 | -0.12455 | 0.38859 | 0.917257 |
| ENSMUSG( | Wwp2 | 1186.847 | 0.09058 | 0.329475 | 0.773311 | ENSMUSG( | Wwp2 | 1027.955 | -0.05547 | 0.625797 | 0.953234 |
| ENSMUSG( | Wwtr1 | 2376.329 | -0.02337 | 0.872722 | 0.978753 | ENSMUSG( | Wwtr1 | 1911.915 | -0.41235 | 0.004618 | 0.279913 |
| ENSMUSG( | Xab2 | 1145.935 | 0.051816 | 0.630812 | 0.914073 | ENSMUSG( | Xab2 | 1019.356 | -0.01762 | 0.890747 | 0.990558 |
| ENSMUSG( | Xaf1 | 107.4586 | -0.26396 | 0.204122 | 0.644638 | ENSMUSG( | Xaf1 | 117.0911 | 0.258549 | 0.251931 | 0.879902 |
| ENSMUSG( | Xbp1 | 2162.307 | -0.32251 | 0.047467 | 0.333152 | ENSMUSG( | Xbp1 | 2575.577 | 0.441288 | 0.01205 | 0.445877 |
| ENSMUSG( | Xdh | 143.8792 | 0.273815 | 0.209673 | 0.651841 | ENSMUSG( | Xdh | 165.8559 | 0.851513 | 0.00106 | 0.111298 |
| ENSMUSG( | Xiap | 3550.986 | -0.0015 | 0.988546 | 0.999109 | ENSMUSG( | Xiap | 3207.861 | -0.02822 | 0.831026 | 0.98051 |
| ENSMUSG( | Xk | 474.9784 | -0.35054 | 0.034171 | 0.283507 | ENSMUSG( | Xk | 531.4999 | 0.257916 | 0.177436 | 0.847137 |
| ENSMUSG( | Xkr4 | 291.5341 | -0.28932 | 0.197168 | 0.636662 | ENSMUSG( | Xkr4 | 331.0019 | 0.351569 | 0.153348 | 0.837724 |
| ENSMUSG( | Xkr6 | 285.9706 | -0.03529 | 0.814112 | 0.964574 | ENSMUSG( | Xkr6 | 250.7826 | -0.16117 | 0.411536 | 0.917257 |
| ENSMUSG( | Xkr7 | 35.25984 | -0.27059 | 0.487236 | 0.858812 | ENSMUSG( | Xkr7 | 30.87644 | -0.35292 | 0.427358 | 0.917257 |
| ENSMUSG( | Xkr8 | 204.2534 | 0.010493 | 0.949154 | 0.993821 | ENSMUSG( | Xkr8 | 200.5488 | 0.21316 | 0.251116 | 0.879321 |
| ENSMUSG( | Xkrx | 64.17304 | -0.50166 | 0.037109 | 0.297239 | ENSMUSG( | Xkrx | 65.86653 | -0.1107 | 0.662927 | 0.956205 |
| ENSMUSG( | Xlr | 29.97477 | 0.539855 | 0.136287 | 0.550578 | ENSMUSG( | Xlr | 26.60617 | 0.481919 | 0.21402 | 0.864421 |
| ENSMUSG( | Xlr3a | 21.2715 | -0.65221 | 0.164084 | 0.594869 | ENSMUSG( | Xlr3a | 23.7002 | -0.03638 | 0.941718 | 0.994578 |
| ENSMUSG( | Xlr3b | 73.56923 | -0.3041 | 0.666566 | 0.927247 | ENSMUSG( | Xlr3b | 52.5537 | -1.24375 | 0.005371 | 0.301579 |
| ENSMUSG( | Xndc1 | 186.8911 | 0.082389 | 0.657328 | 0.924144 | ENSMUSG( | Xndc1 | 182.515 | 0.279571 | 0.146489 | 0.835973 |
| ENSMUSG( | Xntrpc | 11.91503 | -0.02524 | 0.963484 | NA | ENSMUSG( | Xntrpc | 11.49322 | 0.094121 | 0.87553 | 0.989644 |
| ENSMUSG( | Xpa | 211.8907 | 0.107956 | 0.517179 | 0.873295 | ENSMUSG( | Xpa | 189.5146 | 0.055077 | 0.769258 | 0.974236 |
| ENSMUSG( | Xpc | 667.935 | -0.04006 | 0.693983 | 0.934202 | ENSMUSG( | Xpc | 612.0208 | -0.02228 | 0.85531 | 0.985649 |
| ENSMUSG( | Xpnpep1 | 1480.516 | 0.160375 | 0.087468 | 0.454663 | ENSMUSG( | Xpnpep1 | 1287.535 | 0.028713 | 0.796056 | 0.97681 |
| ENSMUSG( | Xpnpep2 | 15.42394 | -0.73354 | 0.136456 | NA | ENSMUSG( | Xpnpep2 | 15.26704 | -0.45942 | 0.372668 | 0.915491 |
| ENSMUSG( | Xpnpep3 | 400.9015 | 0.119148 | 0.328344 | 0.772854 | ENSMUSG( | Xpnpep3 | 367.0997 | 0.138595 | 0.332004 | 0.907907 |
| ENSMUSG( | Xpo1 | 1776.598 | 0.024273 | 0.846483 | 0.972238 | ENSMUSG( | Xpo1 | 1777.875 | 0.278771 | 0.047915 | 0.704961 |
| ENSMUSG( | Xpo4 | 659.7202 | -0.03625 | 0.76628 | 0.954173 | ENSMUSG( | Xpo4 | 588.4321 | -0.10185 | 0.472165 | 0.923329 |
| ENSMUSG( | Xpo5 | 1934.204 | 0.1109 | 0.234563 | 0.681922 | ENSMUSG( | Xpo5 | 1662.958 | -0.05568 | 0.646969 | 0.954033 |
| ENSMUSG( | Xpo6 | 2691.393 | 0.085633 | 0.406731 | 0.816928 | ENSMUSG( | Xpo6 | 2508.52 | -0.15103 | 0.221837 | 0.866104 |
| ENSMUSG( | Xpo7 | 2199.89 | 0.077383 | 0.387159 | 0.806095 | ENSMUSG( | Xpo7 | 1978.77 | -0.01712 | 0.880738 | 0.989644 |
| ENSMUSG( | Xpot | 2851.48 | 0.083824 | 0.319762 | 0.765305 | ENSMUSG( | Xpot | 2569.82 | 0.051764 | 0.626349 | 0.953234 |
| ENSMUSG( | Xpr1 | 5004.807 | -0.12824 | 0.315412 | 0.762441 | ENSMUSG( | Xpr1 | 4972.374 | 0.123711 | 0.39436 | 0.917257 |
| ENSMUSG( | Xrcc1 | 792.2922 | 0.07725 | 0.596203 | 0.903186 | ENSMUSG( | Xrcc1 | 695.9186 | -0.02875 | 0.856183 | 0.985884 |
| ENSMUSG( | Xrcc2 | 261.3117 | 0.063766 | 0.67029 | 0.928525 | ENSMUSG( | Xrcc2 | 226.9874 | -0.07727 | 0.653584 | 0.955015 |
| ENSMUSG( | Xrcc3 | 176.5409 | -0.06303 | 0.734777 | 0.945584 | ENSMUSG( | Xrcc3 | 157.6183 | -0.12681 | 0.535366 | 0.936508 |
| ENSMUSG( | Xrcc4 | 225.1125 | -0.0548 | 0.7296 | 0.944764 | ENSMUSG( | Xrcc4 | 191.6791 | -0.2728 | 0.131837 | 0.822912 |
| ENSMUSG( | Xrcc5 | 651.5512 | -0.10231 | 0.316301 | 0.763453 | ENSMUSG( | Xrcc5 | 567.0088 | -0.24524 | 0.056994 | 0.723362 |
| ENSMUSG( | Xrcc6 | 578.586 | 0.111972 | 0.291104 | 0.741182 | ENSMUSG( | Xrcc6 | 497.9271 | -0.04842 | 0.720489 | 0.966887 |
| ENSMUSG( | Xrn1 | 842.8774 | -0.09113 | 0.368193 | 0.795774 | ENSMUSG( | Xrn1 | 813.3361 | 0.080685 | 0.551875 | 0.939565 |
| ENSMUSG( | Xrn2 | 1181.071 | -0.11544 | 0.239983 | 0.687848 | ENSMUSG( | Xrn2 | 1125.287 | 0.015179 | 0.895471 | 0.990558 |
| ENSMUSG( | Xrra1 | 42.45785 | -0.20267 | 0.513428 | 0.871762 | ENSMUSG( | Xrra1 | 38.75268 | -0.21875 | 0.525651 | 0.934374 |
| ENSMUSG( | Xxylt1 | 878.7225 | -0.41827 | 0.002037 | 0.051756 | ENSMUSG( | Xxylt1 | 925.6483 | 0.035352 | 0.821889 | 0.979198 |
| ENSMUSG( | Xylb | 224.954 | 0.103012 | 0.567612 | 0.891774 | ENSMUSG( | Xylb | 215.4845 | 0.232387 | 0.229391 | 0.868507 |
| ENSMUSG( | Xylt1 | 2581.448 | -0.28314 | 0.041818 | 0.315339 | ENSMUSG( | Xylt1 | 2535.928 | -0.0484 | 0.765046 | 0.974236 |
| ENSMUSG( | Xylt2 | 1030.555 | -0.28378 | 0.065594 | 0.396799 | ENSMUSG( | Xylt2 | 1054.08 | 0.070325 | 0.681264 | 0.958454 |
| ENSMUSG( | Yae1d1 | 880.4933 | 0.013633 | 0.901002 | 0.985633 | ENSMUSG( | Yae1d1 | 777.059 | -0.0814 | 0.506678 | 0.928106 |

| | | | | | |
|---|---|---|---|---|---|
| ENSMUSG( | 1722.566 | 0.109832 | 0.213502 | 0.65774 | Usp6nl |
| ENSMUSG( | 4650.58 | 0.088402 | 0.25945 | 0.709948 | Usp7 |
| ENSMUSG( | 2461.977 | 0.084119 | 0.37158 | 0.798768 | Usp8 |
| ENSMUSG( | 3366.76 | 0.17039 | 0.171752 | 0.606517 | Usp9x |
| ENSMUSG( | 854.2538 | 0.024499 | 0.818161 | 0.965837 | Uspl1 |
| ENSMUSG( | 1161.61 | -0.19982 | 0.161181 | 0.591083 | Ust |
| ENSMUSG( | 478.534 | 0.156309 | 0.16837 | 0.601667 | Utp11 |
| ENSMUSG( | 496.0494 | 0.238606 | 0.032358 | 0.27508 | Utp14a |
| ENSMUSG( | 1274.269 | 0.276585 | 0.137788 | 0.552903 | Utp14b |
| ENSMUSG( | 560.7019 | -0.02423 | 0.82823 | 0.968966 | Utp15 |
| ENSMUSG( | 486.1249 | 0.063906 | 0.587427 | 0.900403 | Utp18 |
| ENSMUSG( | 595.9549 | 0.086571 | 0.430509 | 0.826896 | Utp20 |
| ENSMUSG( | 188.7645 | 0.295053 | 0.064095 | 0.392013 | Utp23 |
| ENSMUSG( | 421.7975 | 0.09097 | 0.488668 | 0.859092 | Utp25 |
| ENSMUSG( | 529.5028 | 0.156495 | 0.242824 | 0.690275 | Utp3 |
| ENSMUSG( | 550.4278 | 0.176438 | 0.199371 | 0.640054 | Utp4 |
| ENSMUSG( | 1599.519 | 0.290012 | 0.05701 | 0.368322 | Utp6 |
| ENSMUSG( | 3952.045 | 0.0945 | 0.396373 | 0.811742 | Utrn |
| ENSMUSG( | 297.7912 | 0.009084 | 0.95032 | 0.993821 | Uty |
| ENSMUSG( | 930.4071 | 0.117954 | 0.316666 | 0.763648 | Uvrag |
| ENSMUSG( | 316.1151 | -0.22724 | 0.08153 | 0.441102 | Uvssa |
| ENSMUSG( | 1234.576 | -0.40993 | 0.003517 | 0.075237 | Uxs1 |
| ENSMUSG( | 175.414 | -0.12693 | 0.432985 | 0.828185 | Uxt |
| ENSMUSG( | 694.377 | 0.029876 | 0.777846 | 0.956168 | Vac14 |
| ENSMUSG( | 2622.86 | -0.02839 | 0.842421 | 0.971564 | Vamp1 |
| ENSMUSG( | 39803.92 | 0.103359 | 0.151427 | 0.576717 | Vamp2 |
| ENSMUSG( | 1941.376 | 0.104986 | 0.321988 | 0.766908 | Vamp3 |
| ENSMUSG( | 2001.699 | 0.068138 | 0.52997 | 0.878883 | Vamp4 |
| ENSMUSG( | 114.7248 | -0.01309 | 0.945849 | 0.99369 | Vamp5 |
| ENSMUSG( | 549.0806 | -0.02791 | 0.800703 | 0.961745 | Vamp8 |
| ENSMUSG( | 205.2661 | -0.44356 | 0.017005 | 0.194647 | Vangl1 |
| ENSMUSG( | 790.4803 | -0.40031 | 0.003126 | 0.069373 | Vangl2 |
| ENSMUSG( | 2733.811 | -0.00939 | 0.932304 | 0.992515 | Vapa |
| ENSMUSG( | 6842.567 | 0.020063 | 0.829271 | 0.969286 | Vapb |
| ENSMUSG( | 1346.876 | 0.142249 | 0.194874 | 0.634201 | Vars1 |
| ENSMUSG( | 1015.101 | 0.133288 | 0.153833 | 0.580901 | Vars2 |
| ENSMUSG( | 2096.347 | -0.04678 | 0.582449 | 0.897918 | Vash1 |
| ENSMUSG( | 78.33801 | -0.1243 | 0.606506 | 0.906481 | Vash2 |
| ENSMUSG( | 330.3436 | -0.2648 | 0.159822 | 0.589679 | Vasn |
| ENSMUSG( | 564.8695 | -0.07817 | 0.533809 | 0.880095 | Vasp |
| ENSMUSG( | 1316.547 | -0.06767 | 0.489375 | 0.859092 | Vat1 |
| ENSMUSG( | 1145.005 | -0.03087 | 0.848487 | 0.972374 | Vat1l |
| ENSMUSG( | 286.2605 | -0.0383 | 0.796727 | 0.960359 | Vav1 |
| ENSMUSG( | 683.2186 | 0.035976 | 0.757768 | 0.951582 | Vav2 |
| ENSMUSG( | 312.2298 | 0.003545 | 0.979885 | 0.997428 | Vav3 |
| ENSMUSG( | 14.49257 | -0.61305 | 0.211022 | NA | Vax1 |
| ENSMUSG( | 908.4149 | -0.08266 | 0.51934 | 0.874979 | Vbp1 |
| ENSMUSG( | 2164.857 | -0.24678 | 0.091991 | 0.465937 | Vcam1 |
| ENSMUSG( | 1116.111 | -0.32462 | 0.041787 | 0.315339 | Vcan |
| ENSMUSG( | 1235.205 | -0.02381 | 0.808402 | 0.963455 | Vcf1 |
| ENSMUSG( | 498.3134 | 0.030045 | 0.812373 | 0.964574 | Vcf2 |
| ENSMUSG( | 1703.896 | 0.25296 | 0.073679 | 0.419767 | Vcl |
| ENSMUSG( | 6301.013 | 0.087561 | 0.321698 | 0.766739 | Vcp |
| ENSMUSG( | 1038.831 | 0.035921 | 0.712843 | 0.94066 | Vcpip1 |
| ENSMUSG( | 518.1018 | -0.11709 | 0.307034 | 0.75508 | Vcpkmt |
| ENSMUSG( | 10831.62 | 0.020498 | 0.837914 | 0.971111 | Vdac1 |
| ENSMUSG( | 4197.165 | -0.01142 | 0.91393 | 0.988483 | Vdac2 |
| ENSMUSG( | 2152.719 | -0.00083 | 0.994478 | 0.999369 | Vdac3 |
| ENSMUSG( | 3395.327 | -0.1003 | 0.378434 | 0.801688 | Vegfa |
| ENSMUSG( | 5196.293 | 0.018358 | 0.83037 | 0.96974 | Vegfb |
| ENSMUSG( | 109.9122 | -0.50408 | 0.029042 | 0.259542 | Vegfc |
| ENSMUSG( | 78.84687 | 0.189016 | 0.453165 | 0.836701 | Vegfd |
| ENSMUSG( | 16.95004 | 0.572983 | 0.224015 | NA | Veph1 |
| ENSMUSG( | 1357.765 | -0.00487 | 0.960836 | 0.993991 | Vezf1 |
| ENSMUSG( | 2219.86 | -0.1197 | 0.290577 | 0.740557 | Vezt |
| ENSMUSG( | 10194.17 | -0.4779 | 0.000119 | 0.007278 | Vgf |
| ENSMUSG( | 45.47419 | 0.381507 | 0.248768 | 0.697857 | Vgll3 |
| ENSMUSG( | 966.4839 | -0.09569 | 0.322188 | 0.767159 | Vgll4 |
| ENSMUSG( | 3469.838 | 0.057364 | 0.621396 | 0.911432 | Vhl |
| ENSMUSG( | 27.93865 | 0.361409 | 0.349382 | 0.78176 | Vil1 |
| ENSMUSG( | 227.4065 | 0.395349 | 0.096774 | 0.477375 | Vill |
| ENSMUSG( | 1921.715 | 0.157211 | 0.139456 | 0.555633 | Vim |
| ENSMUSG( | 9.109177 | -0.0872 | 0.897719 | NA | Vinac1 |
| ENSMUSG( | 1285.453 | -0.56627 | 0.00126 | 0.038624 | Vip |
| ENSMUSG( | 1459.877 | 0.127805 | 0.14615 | 0.567605 | Vipas39 |
| ENSMUSG( | 1002.453 | -0.11681 | 0.435331 | 0.829093 | Vipr1 |
| ENSMUSG( | 54.9195 | -0.98386 | 0.002198 | 0.054456 | Vipr2 |
| ENSMUSG( | 1113.198 | 0.083103 | 0.397509 | 0.812127 | Virma |
| ENSMUSG( | 164.968 | -0.04313 | 0.818384 | 0.965837 | Vit |
| ENSMUSG( | 219.1458 | -0.21429 | 0.177788 | 0.615225 | Vkorc1 |
| ENSMUSG( | 2358.962 | -0.25657 | 0.027384 | 0.253959 | Vkorc1l1 |
| ENSMUSG( | 2729.841 | -0.34884 | 0.014474 | 0.177206 | Vldlr |
| ENSMUSG( | 1351.391 | -0.33804 | 0.020601 | 0.216558 | Vma21 |
| ENSMUSG( | 563.5453 | -0.01731 | 0.879229 | 0.981436 | Vmac |
| ENSMUSG( | 21.35471 | 0.33989 | 0.454893 | 0.837689 | Vmn2r1 |
| ENSMUSG( | 90.51452 | -0.16847 | 0.423162 | 0.823662 | Vmn2r29 |
| ENSMUSG( | 47.29616 | -0.35056 | 0.216902 | 0.661816 | Vmn2r57 |
| ENSMUSG( | 75.6574 | -0.04604 | 0.866757 | 0.976541 | Vmn2r84 |
| ENSMUSG( | 10.47293 | 0.430566 | 0.482828 | NA | Vmn2r85 |
| ENSMUSG( | 25.81732 | -0.12483 | 0.760379 | 0.952907 | Vmn2r87 |
| ENSMUSG( | 3470.604 | -0.65165 | 1.07E-05 | 0.001199 | Vmp1 |
| ENSMUSG( | 27.11492 | -0.59966 | 0.21157 | 0.654835 | Vnn1 |
| ENSMUSG( | 3914.122 | -0.24296 | 0.051665 | 0.348878 | Vopp1 |
| ENSMUSG( | 1752.171 | 0.050295 | 0.632154 | 0.914073 | Vps11 |
| ENSMUSG( | 1144.891 | -0.14786 | 0.242957 | 0.690307 | Vps13a |
| ENSMUSG( | 1527.279 | -0.10975 | 0.372496 | 0.798768 | Vps13b |
| ENSMUSG( | 1590.777 | -0.07119 | 0.587095 | 0.900235 | Vps13c |
| ENSMUSG( | 4179.849 | -0.11329 | 0.222832 | 0.668701 | Vps13d |
| ENSMUSG( | 1480.53 | 0.072772 | 0.467604 | 0.844819 | Vps16 |
| ENSMUSG( | 1154.533 | 0.120662 | 0.385589 | 0.804302 | Vps18 |
| ENSMUSG( | 34.08543 | 0.373537 | 0.273962 | 0.724254 | Vps25 |
| ENSMUSG( | 1546.602 | -0.02315 | 0.79864 | 0.960896 | Vps26a |
| ENSMUSG( | 3462.521 | 0.121865 | 0.135214 | 0.548658 | Vps26b |
| ENSMUSG( | 1575.603 | 0.104693 | 0.314093 | 0.761319 | Vps26c |
| ENSMUSG( | 2108.736 | -0.07026 | 0.427402 | 0.82549 | Vps28 |
| ENSMUSG( | 2381.775 | -0.02148 | 0.801692 | 0.962077 | Vps29 |
| ENSMUSG( | 2715.772 | 0.128301 | 0.152017 | 0.577721 | Vps33a |
| ENSMUSG( | 562.9039 | 0.220821 | 0.111536 | 0.503511 | Vps33b |
| ENSMUSG( | 3363.974 | 0.050479 | 0.622915 | 0.911589 | Vps35 |
| ENSMUSG( | 2107.021 | 0.103492 | 0.214588 | 0.659193 | Vps35l |
| ENSMUSG( | 875.2307 | 0.179942 | 0.066763 | 0.399595 | Vps36 |
| ENSMUSG( | 1847.631 | 0.098355 | 0.343516 | 0.779153 | Vps37a |
| ENSMUSG( | 1129.875 | -0.15904 | 0.14187 | 0.559166 | Vps37b |
| ENSMUSG( | 1194.722 | 0.103146 | 0.289435 | 0.739839 | Vps37c |
| ENSMUSG( | 597.524 | -0.0424 | 0.739433 | 0.947768 | Vps37d |
| ENSMUSG( | 2425.512 | 0.21545 | 0.017538 | 0.198498 | Vps39 |
| ENSMUSG( | 4667.629 | 0.054249 | 0.502317 | 0.864924 | Vps41 |
| ENSMUSG( | 774.1561 | 0.114009 | 0.414347 | 0.819998 | Vps45 |
| ENSMUSG( | 2917.152 | 0.02198 | 0.790618 | 0.959461 | Vps4a |
| ENSMUSG( | 917.311 | 0.059985 | 0.580996 | 0.89745 | Vps4b |
| ENSMUSG( | 1596.547 | 0.096069 | 0.285015 | 0.733568 | Vps50 |
| ENSMUSG( | 1242.791 | 0.085773 | 0.458294 | 0.839958 | Vps51 |
| ENSMUSG( | 3353.462 | 0.166942 | 0.043931 | 0.321265 | Vps52 |
| ENSMUSG( | 1459.775 | 0.134586 | 0.259952 | 0.709951 | Vps53 |
| ENSMUSG( | 962.5856 | 0.029788 | 0.781671 | 0.957558 | Vps54 |
| ENSMUSG( | 770.5875 | 0.024292 | 0.818735 | 0.965888 | Vps72 |
| ENSMUSG( | 1524.434 | 0.175883 | 0.066993 | 0.400593 | Vps8 |
| ENSMUSG( | 2825.121 | 0.218386 | 0.012907 | 0.164191 | Vps9d1 |
| ENSMUSG( | 383.3551 | -0.03941 | 0.739575 | 0.947768 | Vrk1 |
| ENSMUSG( | 69.73264 | -0.15308 | 0.512892 | 0.871762 | Vrk2 |
| ENSMUSG( | 1234.633 | 0.036913 | 0.686125 | 0.932981 | Vrk3 |
| ENSMUSG( | 124.9721 | -0.40598 | 0.043422 | 0.319227 | Vsig10 |
| ENSMUSG( | 140.9465 | 0.219401 | 0.217081 | 0.661995 | Vsig10l |
| ENSMUSG( | 76.24601 | 0.394851 | 0.124607 | 0.531714 | Vsig2 |
| ENSMUSG( | 16.82013 | -0.99592 | 0.03387 | NA | Vsig8 |
| ENSMUSG( | 2107.435 | -0.07269 | 0.640705 | 0.916434 | Vsir |
| ENSMUSG( | 9044.549 | -0.04728 | 0.730861 | 0.944956 | Vsnl1 |
| ENSMUSG( | 2168.757 | -0.63014 | 0.000158 | 0.008882 | Vstm2a |
| ENSMUSG( | 1263.824 | -0.26691 | 0.008825 | 0.132321 | Vstm2b |
| ENSMUSG( | 4104.932 | -0.46253 | 1.08E-05 | 0.001199 | Vstm2l |
| ENSMUSG( | 305.5882 | -0.16127 | 0.364042 | 0.793438 | Vstm4 |
| ENSMUSG( | 265.1999 | -0.04575 | 0.763865 | 0.954015 | Vstm5 |
| ENSMUSG( | 24.92015 | -0.81721 | 0.044581 | 0.32426 | Vsx1 |
| ENSMUSG( | 1088.823 | 0.002852 | 0.977605 | 0.997428 | Vta1 |
| ENSMUSG( | 2088.985 | 0.088572 | 0.341981 | 0.778828 | Vti1a |
| ENSMUSG( | 5695.538 | 0.005704 | 0.948568 | 0.993821 | Vti1b |
| ENSMUSG( | 3663.583 | -0.03723 | 0.836558 | 0.971111 | Vtn |
| ENSMUSG( | 1714.982 | 0.004825 | 0.97615 | 0.997428 | Vwa1 |
| ENSMUSG( | 249.1976 | -0.02169 | 0.914028 | 0.988483 | Vwa3a |
| ENSMUSG( | 61.66417 | 0.366435 | 0.15199 | 0.577721 | Vwa3b |
| ENSMUSG( | 750.1319 | 0.119766 | 0.331059 | 0.77519 | Vwa5a |
| ENSMUSG( | 363.2073 | 0.267366 | 0.053646 | 0.355738 | Vwa5b1 |
| ENSMUSG( | 3180.797 | 0.195499 | 0.158814 | 0.588112 | Vwa5b2 |
| ENSMUSG( | 18.35373 | -0.3119 | 0.480766 | NA | Vwa7 |
| ENSMUSG( | 2477.058 | 0.131606 | 0.1446 | 0.564508 | Vwa8 |
| ENSMUSG( | 162.3942 | -0.57752 | 0.000881 | 0.029903 | Vwc2 |
| ENSMUSG( | 141.19 | -0.29799 | 0.143569 | 0.562581 | Vwc2l |
| ENSMUSG( | 18.51174 | -0.10674 | 0.813982 | NA | Vwce |

| | | | | | |
|---|---|---|---|---|---|
| ENSMUSG( | 15.41551 | -1.02146 | 0.049965 | 0.708648 | Vsig8 |
| ENSMUSG( | 2149.116 | 0.248918 | 0.232043 | 0.868507 | Vsir |
| ENSMUSG( | 7779.457 | -0.22614 | 0.201141 | 0.86439 | Vsnl1 |
| ENSMUSG( | 2658.931 | 0.283139 | 0.141647 | 0.830678 | Vstm2a |
| ENSMUSG( | 1317.198 | 0.131622 | 0.272505 | 0.888825 | Vstm2b |
| ENSMUSG( | 4326.412 | -0.00363 | 0.978151 | 0.996416 | Vstm2l |
| ENSMUSG( | 256.4241 | -0.41746 | 0.022905 | 0.584023 | Vstm4 |
| ENSMUSG( | 257.7193 | 0.136361 | 0.438822 | 0.920721 | Vstm5 |
| ENSMUSG( | 28.57585 | -0.04226 | 0.909562 | 0.991958 | Vsx1 |
| ENSMUSG( | 915.5935 | -0.24188 | 0.050802 | 0.709741 | Vta1 |
| ENSMUSG( | 1738.857 | -0.17935 | 0.1224 | 0.813384 | Vti1a |
| ENSMUSG( | 4733.665 | -0.27417 | 0.026734 | 0.603041 | Vti1b |
| ENSMUSG( | 3680.577 | 0.242634 | 0.24903 | 0.877904 | Vtn |
| ENSMUSG( | 1814.396 | 0.413513 | 0.020653 | 0.569724 | Vwa1 |
| ENSMUSG( | 220.5603 | -0.0988 | 0.670044 | 0.95694 | Vwa3a |
| ENSMUSG( | 49.4922 | 0.05278 | 0.86408 | 0.986552 | Vwa3b |
| ENSMUSG( | 629.8062 | -0.11623 | 0.39888 | 0.917257 | Vwa5a |
| ENSMUSG( | 302.146 | 0.029124 | 0.870154 | 0.988232 | Vwa5b1 |
| ENSMUSG( | 2528.772 | -0.20175 | 0.194578 | 0.860277 | Vwa5b2 |
| ENSMUSG( | 17.37516 | -0.18468 | 0.709017 | 0.964042 | Vwa7 |
| ENSMUSG( | 2070.469 | -0.11705 | 0.274884 | 0.889874 | Vwa8 |
| ENSMUSG( | 169.7424 | -0.11688 | 0.542681 | 0.938034 | Vwc2 |
| ENSMUSG( | 165.7141 | 0.427201 | 0.03914 | 0.679972 | Vwc2l |
| ENSMUSG( | 17.50558 | -0.01907 | 0.969036 | 0.99594 | Vwce |
| ENSMUSG( | 422.8089 | 0.828837 | 0.002361 | 0.190561 | Vwf |
| ENSMUSG( | 7373.852 | -0.07983 | 0.482302 | 0.924578 | Vxn |
| ENSMUSG( | 2250.444 | 0.140328 | 0.232854 | 0.868507 | Wac |
| ENSMUSG( | 1300.736 | 0.137504 | 0.271324 | 0.888062 | Wapl |
| ENSMUSG( | 1976.266 | -0.10026 | 0.373898 | 0.915491 | Wars1 |
| ENSMUSG( | 181.6409 | 0.074946 | 0.68177 | 0.958678 | Wars2 |
| ENSMUSG( | 96.15979 | -0.04688 | 0.853976 | 0.98563 | Was |
| ENSMUSG( | 3817.544 | -0.06986 | 0.678807 | 0.957617 | Wasf1 |
| ENSMUSG( | 989.1067 | 0.155372 | 0.309499 | 0.898538 | Wasf2 |
| ENSMUSG( | 10096.99 | -0.16723 | 0.222129 | 0.866232 | Wasf3 |
| ENSMUSG( | 986.4632 | -0.12128 | 0.339895 | 0.907972 | Washc1 |
| ENSMUSG( | 4982.995 | -0.09499 | 0.330967 | 0.907513 | Washc2 |
| ENSMUSG( | 370.3283 | -0.24065 | 0.086365 | 0.781091 | Washc3 |
| ENSMUSG( | 808.6608 | 0.127921 | 0.306945 | 0.898538 | Washc4 |
| ENSMUSG( | 1197.055 | -0.0377 | 0.740269 | 0.968811 | Washc5 |
| ENSMUSG( | 1824.485 | 0.148044 | 0.26087 | 0.884853 | Wasl |
| ENSMUSG( | 1471.636 | -0.13213 | 0.237499 | 0.872173 | Wbp1 |
| ENSMUSG( | 1838.236 | -0.00358 | 0.977561 | 0.996416 | Wbp11 |
| ENSMUSG( | 2130.314 | 0.113277 | 0.345044 | 0.91045 | Wbp1l |
| ENSMUSG( | 8694.714 | -0.14549 | 0.277385 | 0.891468 | Wbp2 |
| ENSMUSG( | 1183.384 | 0.015171 | 0.894926 | 0.990558 | Wbp4 |
| ENSMUSG( | 197.5104 | 0.219071 | 0.21747 | 0.865589 | Wdcp |
| ENSMUSG( | 732.5622 | 0.786439 | 4.64E-09 | 4.06E-06 | Wdfy1 |
| ENSMUSG( | 1166.319 | 0.079444 | 0.509806 | 0.929089 | Wdfy2 |
| ENSMUSG( | 5082.672 | 0.139501 | 0.261373 | 0.884926 | Wdfy3 |
| ENSMUSG( | 183.9964 | 0.033659 | 0.844449 | 0.983642 | Wdfy4 |
| ENSMUSG( | 156.3821 | 0.131707 | 0.502747 | 0.927825 | Wdhd1 |
| ENSMUSG( | 233.0321 | -0.14471 | 0.363118 | 0.914223 | Wdpcp |
| ENSMUSG( | 3958.835 | -0.11269 | 0.309353 | 0.898538 | Wdr1 |
| ENSMUSG( | 1387.328 | 0.054011 | 0.616965 | 0.951666 | Wdr11 |
| ENSMUSG( | 680.631 | 0.009025 | 0.942508 | 0.994774 | Wdr12 |
| ENSMUSG( | 13288.16 | 0.054051 | 0.575055 | 0.942912 | Wdr13 |
| ENSMUSG( | 626.4698 | 0.001366 | 0.991732 | 0.998712 | Wdr17 |
| ENSMUSG( | 1640.02 | -0.11147 | 0.345587 | 0.910597 | Wdr18 |
| ENSMUSG( | 568.7029 | 0.030896 | 0.824069 | 0.979198 | Wdr19 |
| ENSMUSG( | 454.3574 | 0.071431 | 0.58807 | 0.944975 | Wdr20 |
| ENSMUSG( | 546.7535 | 0.124668 | 0.353258 | 0.912263 | Wdr24 |
| ENSMUSG( | 224.0932 | -0.16798 | 0.295455 | 0.898538 | Wdr25 |
| ENSMUSG( | 4147.043 | 0.046658 | 0.67031 | 0.95694 | Wdr26 |
| ENSMUSG( | 50.94548 | 0.466638 | 0.169093 | 0.840282 | Wdr27 |
| ENSMUSG( | 381.7528 | -0.02044 | 0.893271 | 0.990558 | Wdr3 |
| ENSMUSG( | 106.7289 | -0.10168 | 0.62673 | 0.953234 | Wdr31 |
| ENSMUSG( | 1876.432 | 0.191816 | 0.232267 | 0.868507 | Wdr33 |
| ENSMUSG( | 700.1054 | -0.11498 | 0.351975 | 0.912093 | Wdr35 |
| ENSMUSG( | 491.5417 | 0.098308 | 0.474886 | 0.923329 | Wdr36 |
| ENSMUSG( | 1224.662 | -0.0149 | 0.904298 | 0.991089 | Wdr37 |
| ENSMUSG( | 43.71046 | -0.40583 | 0.189117 | 0.85698 | Wdr38 |
| ENSMUSG( | 832.424 | -0.10058 | 0.426419 | 0.917257 | Wdr4 |
| ENSMUSG( | 993.988 | -0.15065 | 0.222771 | 0.867173 | Wdr41 |
| ENSMUSG( | 698.6682 | -0.03765 | 0.775517 | 0.975354 | Wdr43 |
| ENSMUSG( | 509.4275 | -0.00549 | 0.974845 | 0.996416 | Wdr44 |
| ENSMUSG( | 434.8609 | -0.02612 | 0.85857 | 0.985884 | Wdr45 |
| ENSMUSG( | 2205.289 | -0.03019 | 0.794478 | 0.97681 | Wdr45b |
| ENSMUSG( | 321.5978 | -0.07221 | 0.652032 | 0.9547 | Wdr46 |
| ENSMUSG( | 2066.971 | -0.10463 | 0.462131 | 0.923329 | Wdr47 |
| ENSMUSG( | 1781.218 | 0.044033 | 0.683271 | 0.958678 | Wdr48 |
| ENSMUSG( | 691.5756 | -0.11527 | 0.371833 | 0.915491 | Wdr5 |
| ENSMUSG( | 163.2986 | -0.02715 | 0.881401 | 0.989644 | Wdr53 |
| ENSMUSG( | 452.9026 | 0.074981 | 0.603256 | 0.948137 | Wdr54 |
| ENSMUSG( | 220.8264 | -0.12497 | 0.510676 | 0.929501 | Wdr55 |
| ENSMUSG( | 663.0399 | -0.0325 | 0.786771 | 0.976303 | Wdr59 |
| ENSMUSG( | 106.077 | 0.040869 | 0.855633 | 0.985738 | Wdr5b |
| ENSMUSG( | 3185.324 | 0.059176 | 0.638672 | 0.953903 | Wdr6 |
| ENSMUSG( | 66.96936 | 0.42486 | 0.161633 | 0.837724 | Wdr62 |
| ENSMUSG( | 2972.964 | -0.01051 | 0.921414 | 0.992644 | Wdr7 |
| ENSMUSG( | 475.5196 | -0.12404 | 0.371958 | 0.915491 | Wdr70 |
| ENSMUSG( | 725.3565 | -0.02195 | 0.863294 | 0.986547 | Wdr73 |
| ENSMUSG( | 514.6571 | -0.03592 | 0.813972 | 0.979198 | Wdr74 |
| ENSMUSG( | 518.6828 | -0.01357 | 0.916874 | 0.992529 | Wdr75 |
| ENSMUSG( | 72.16889 | 0.245788 | 0.329501 | 0.907083 | Wdr76 |
| ENSMUSG( | 589.3521 | -0.12933 | 0.376629 | 0.915491 | Wdr77 |
| ENSMUSG( | 770.5683 | 0.231498 | 0.158669 | 0.837724 | Wdr81 |
| ENSMUSG( | 2972.393 | -0.07747 | 0.515569 | 0.930939 | Wdr82 |
| ENSMUSG( | 626.4686 | -0.04585 | 0.736215 | 0.967344 | Wdr83 |
| ENSMUSG( | 985.2819 | -0.10612 | 0.363876 | 0.914223 | Wdr83os |
| ENSMUSG( | 63.83922 | 0.318898 | 0.363262 | 0.914223 | Wdr86 |
| ENSMUSG( | 37.58206 | -0.09994 | 0.76081 | 0.972695 | Wdr89 |
| ENSMUSG( | 731.1944 | 0.19215 | 0.295578 | 0.898538 | Wdr90 |
| ENSMUSG( | 922.1569 | 0.135102 | 0.254945 | 0.882307 | Wdr91 |
| ENSMUSG( | 14.31886 | 0.543482 | 0.326485 | 0.905371 | Wdr93 |
| ENSMUSG( | 28.19759 | 0.602028 | 0.142222 | 0.830709 | Wdr95 |
| ENSMUSG( | 177.862 | 0.43847 | 0.134763 | 0.824049 | Wdr97 |
| ENSMUSG( | 287.8332 | -0.00283 | 0.986152 | 0.997188 | Wdsub1 |
| ENSMUSG( | 3836.707 | 0.129122 | 0.235145 | 0.871741 | Wdtc1 |
| ENSMUSG( | 332.0046 | 0.33572 | 0.026965 | 0.603041 | Wee1 |
| ENSMUSG( | 479.6252 | 0.338558 | 0.093451 | 0.795507 | Wfdc1 |
| ENSMUSG( | 37.95428 | -0.71402 | 0.028224 | 0.611882 | Wfdc18 |
| ENSMUSG( | 24.55169 | 0.95969 | 0.034745 | 0.657974 | Wfdc2 |
| ENSMUSG( | 20.29054 | 0.297534 | 0.502678 | 0.927825 | Wfdc3 |
| ENSMUSG( | 9.930406 | 0.95876 | 0.152973 | NA | Wfikkn1 |
| ENSMUSG( | 112.3876 | 0.191308 | 0.461798 | 0.923329 | Wfikkn2 |
| ENSMUSG( | 3897.519 | 0.182939 | 0.193554 | 0.860096 | Wfs1 |
| ENSMUSG( | 603.3788 | 0.119949 | 0.332571 | 0.907907 | Whamm |
| ENSMUSG( | 2466.729 | 0.167749 | 0.150169 | 0.837086 | Whrn |
| ENSMUSG( | 45.63336 | 0.339809 | 0.288601 | 0.894794 | Wif1 |
| ENSMUSG( | 1389.431 | -0.15063 | 0.291148 | 0.897421 | Wipf1 |
| ENSMUSG( | 2309.487 | 0.095079 | 0.378754 | 0.916995 | Wipf2 |
| ENSMUSG( | 3706.2 | 0.185301 | 0.178925 | 0.849041 | Wipf3 |
| ENSMUSG( | 1396.904 | -0.27139 | 0.022268 | 0.577034 | Wipi1 |
| ENSMUSG( | 3960.249 | -0.11008 | 0.266015 | 0.886134 | Wipi2 |
| ENSMUSG( | 2627.199 | 0.108662 | 0.300496 | 0.898538 | Wiz |
| ENSMUSG( | 1616.732 | 0.093068 | 0.609529 | 0.950397 | Wls |
| ENSMUSG( | 17049.51 | -0.23391 | 0.055121 | 0.723362 | Wnk1 |
| ENSMUSG( | 5056.405 | 0.237852 | 0.016175 | 0.512256 | Wnk2 |
| ENSMUSG( | 696.4985 | -0.08802 | 0.504709 | 0.927825 | Wnk3 |
| ENSMUSG( | 165.2935 | 0.261398 | 0.254416 | 0.881624 | Wnk4 |
| ENSMUSG( | 211.4547 | -0.04409 | 0.807009 | 0.978239 | Wnt10a |
| ENSMUSG( | 60.88105 | 0.045913 | 0.883656 | 0.989644 | Wnt10b |
| ENSMUSG( | 17.28466 | 0.225687 | 0.674717 | 0.95694 | Wnt11 |
| ENSMUSG( | 35.12372 | -0.45918 | 0.172828 | 0.843702 | Wnt16 |
| ENSMUSG( | 122.5068 | 0.31193 | 0.140124 | 0.830678 | Wnt2 |
| ENSMUSG( | 56.85329 | 0.028315 | 0.921432 | 0.992644 | Wnt2b |
| ENSMUSG( | 69.28729 | -0.39976 | 0.130763 | 0.822884 | Wnt3 |
| ENSMUSG( | 1209.305 | 0.349791 | 0.006165 | 0.323304 | Wnt4 |
| ENSMUSG( | 357.9228 | 0.174618 | 0.268176 | 0.886493 | Wnt5a |
| ENSMUSG( | 64.05395 | 0.453767 | 0.107966 | 0.804969 | Wnt5b |
| ENSMUSG( | 62.97359 | 0.587621 | 0.037385 | 0.673514 | Wnt6 |
| ENSMUSG( | 813.8311 | 0.033595 | 0.826869 | 0.980159 | Wnt7a |
| ENSMUSG( | 2109.366 | 0.061763 | 0.634605 | 0.953903 | Wnt7b |
| ENSMUSG( | 685.1612 | -0.09736 | 0.56024 | 0.940131 | Wnt9a |
| ENSMUSG( | 13.64188 | -1.43707 | 0.032712 | 0.637512 | Wnt9b |
| ENSMUSG( | 287.0394 | 0.080105 | 0.588814 | 0.944975 | Wrap53 |
| ENSMUSG( | 428.2032 | 0.094108 | 0.564177 | 0.941206 | Wrap73 |
| ENSMUSG( | 358.4251 | 0.04662 | 0.750564 | 0.970184 | Wrn |
| ENSMUSG( | 1230.647 | -0.1505 | 0.2501 | 0.877904 | Wrnip1 |
| ENSMUSG( | 1721.831 | 0.175833 | 0.286133 | 0.893906 | Wsb1 |
| ENSMUSG( | 16624.96 | -0.11576 | 0.236434 | 0.872173 | Wsb2 |
| ENSMUSG( | 5305.951 | -0.03879 | 0.792286 | 0.97681 | Wscd1 |
| ENSMUSG( | 1937.002 | 0.076059 | 0.565723 | 0.941404 | Wscd2 |
| ENSMUSG( | 790.0663 | 0.005231 | 0.964767 | 0.995433 | Wtap |
| ENSMUSG( | 242.7461 | 0.146524 | 0.408333 | 0.917257 | Wtip |
| ENSMUSG( | 7206.584 | -0.13919 | 0.191369 | 0.858412 | Wwc1 |
| ENSMUSG( | 2169.258 | 0.106656 | 0.336803 | 0.907972 | Wwc2 |
| ENSMUSG( | 482.8689 | -0.02235 | 0.880285 | 0.989644 | Wwox |

| | | | | | | | | | | |
|---|---|---|---|---|---|---|---|---|---|---|---|
| ENSMUSG( | Yaf2 | 3237.423 | 0.069209 | 0.412219 | 0.819452 | ENSMUSG( | Yaf2 | 2826.53 | -0.05655 | 0.663402 | 0.956487 |
| ENSMUSG( | Yap1 | 2414.066 | 0.179589 | 0.098099 | 0.479122 | ENSMUSG( | Yap1 | 2057.918 | -0.01002 | 0.945726 | 0.994798 |
| ENSMUSG( | Yars1 | 2230.034 | 0.11375 | 0.335184 | 0.776511 | ENSMUSG( | Yars1 | 1881.673 | -0.10921 | 0.421034 | 0.917257 |
| ENSMUSG( | Yars2 | 432.8861 | -0.03535 | 0.782241 | 0.957558 | ENSMUSG( | Yars2 | 410.3286 | 0.087205 | 0.589332 | 0.944975 |
| ENSMUSG( | Ybey | 137.6241 | 0.129592 | 0.461913 | 0.841728 | ENSMUSG( | Ybey | 115.2543 | -0.09962 | 0.654547 | 0.955015 |
| ENSMUSG( | Ybx1 | 9103.946 | -0.04524 | 0.611661 | 0.908586 | ENSMUSG( | Ybx1 | 7938.613 | -0.1789 | 0.171815 | 0.842999 |
| ENSMUSG( | Ybx2 | 109.2733 | 0.236201 | 0.392926 | 0.809274 | ENSMUSG( | Ybx2 | 89.97734 | -0.04839 | 0.866351 | 0.987347 |
| ENSMUSG( | Ybx3 | 969.9961 | -0.03484 | 0.736004 | 0.946267 | ENSMUSG( | Ybx3 | 869.5374 | -0.08376 | 0.48887 | 0.925573 |
| ENSMUSG( | Ydjc | 417.6834 | 0.063375 | 0.619007 | 0.911432 | ENSMUSG( | Ydjc | 372.1844 | 0.004684 | 0.974925 | 0.996416 |
| ENSMUSG( | Yeats2 | 1360.485 | 0.022795 | 0.831699 | 0.970026 | ENSMUSG( | Yeats2 | 1250.067 | 0.050908 | 0.698752 | 0.960908 |
| ENSMUSG( | Yeats4 | 989.7163 | 0.010411 | 0.934402 | 0.99323 | ENSMUSG( | Yeats4 | 851.6654 | -0.16445 | 0.288609 | 0.894794 |
| ENSMUSG( | Yes1 | 1089.58 | 0.02167 | 0.878939 | 0.981436 | ENSMUSG( | Yes1 | 962.6183 | -0.0735 | 0.661834 | 0.956205 |
| ENSMUSG( | Yif1a | 578.326 | -0.04155 | 0.724566 | 0.944096 | ENSMUSG( | Yif1a | 533.0381 | -0.00799 | 0.95593 | 0.994798 |
| ENSMUSG( | Yif1b | 1204.34 | -0.2779 | 0.01896 | 0.207297 | ENSMUSG( | Yif1b | 1162.278 | -0.09756 | 0.497627 | 0.927825 |
| ENSMUSG( | Yipf1 | 941.1707 | -0.28671 | 0.03137 | 0.269648 | ENSMUSG( | Yipf1 | 946.4485 | 0.015354 | 0.91313 | 0.992298 |
| ENSMUSG( | Yipf2 | 732.0379 | -0.19891 | 0.094546 | 0.471713 | ENSMUSG( | Yipf2 | 701.4711 | -0.04384 | 0.731057 | 0.967228 |
| ENSMUSG( | Yipf3 | 2232.534 | -0.28006 | 0.021694 | 0.225268 | ENSMUSG( | Yipf3 | 2277.75 | 0.06347 | 0.675418 | 0.95694 |
| ENSMUSG( | Yipf4 | 1061.983 | -0.43288 | 0.009487 | 0.137756 | ENSMUSG( | Yipf4 | 1203.254 | 0.223053 | 0.202796 | 0.864412 |
| ENSMUSG( | Yipf5 | 1382.862 | -0.29164 | 0.063051 | 0.388234 | ENSMUSG( | Yipf5 | 1482.149 | 0.188408 | 0.281615 | 0.892412 |
| ENSMUSG( | Yipf6 | 1541.031 | -0.1523 | 0.262701 | 0.713259 | ENSMUSG( | Yipf6 | 1501.515 | 0.044307 | 0.771369 | 0.974236 |
| ENSMUSG( | Yjefn3 | 269.3969 | 0.035858 | 0.805035 | 0.9626 | ENSMUSG( | Yjefn3 | 245.896 | 0.041389 | 0.798081 | 0.97681 |
| ENSMUSG( | Yju2 | 231.7245 | -0.03803 | 0.802818 | 0.962169 | ENSMUSG( | Yju2 | 209.7599 | -0.05624 | 0.729342 | 0.966991 |
| ENSMUSG( | Yju2b | 282.257 | -0.0165 | 0.915746 | 0.988634 | ENSMUSG( | Yju2b | 253.401 | -0.05807 | 0.731727 | 0.967228 |
| ENSMUSG( | Ykt6 | 4034.448 | 0.040106 | 0.622928 | 0.911589 | ENSMUSG( | Ykt6 | 3428.903 | -0.16671 | 0.112932 | 0.809623 |
| ENSMUSG( | Ylpm1 | 2533.483 | 0.07743 | 0.40963 | 0.818258 | ENSMUSG( | Ylpm1 | 2344.755 | 0.12177 | 0.24065 | 0.875927 |
| ENSMUSG( | Yme1l1 | 1537.591 | 0.017132 | 0.865197 | 0.975565 | ENSMUSG( | Yme1l1 | 1514.001 | 0.232095 | 0.054682 | 0.723362 |
| ENSMUSG( | Yod1 | 299.4168 | -0.03068 | 0.859268 | 0.974707 | ENSMUSG( | Yod1 | 274.0217 | -0.02344 | 0.90046 | 0.990558 |
| ENSMUSG( | Ypel1 | 1335.979 | 0.137044 | 0.146829 | 0.56887 | ENSMUSG( | Ypel1 | 1188.226 | 0.069694 | 0.544684 | 0.938063 |
| ENSMUSG( | Ypel2 | 1064.581 | -0.23971 | 0.015812 | 0.186254 | ENSMUSG( | Ypel2 | 1043.789 | -0.01877 | 0.875014 | 0.989644 |
| ENSMUSG( | Ypel3 | 5314.633 | 0.095859 | 0.383386 | 0.803427 | ENSMUSG( | Ypel3 | 4581.903 | -0.06483 | 0.628091 | 0.953238 |
| ENSMUSG( | Ypel4 | 1120.521 | 0.03516 | 0.839363 | 0.971179 | ENSMUSG( | Ypel4 | 1019.269 | 0.026831 | 0.888775 | 0.990558 |
| ENSMUSG( | Ypel5 | 2016.266 | -0.03234 | 0.795799 | 0.960062 | ENSMUSG( | Ypel5 | 1804.669 | -0.0902 | 0.576539 | 0.942912 |
| ENSMUSG( | Yrdc | 360.4454 | -0.3475 | 0.012464 | 0.161344 | ENSMUSG( | Yrdc | 353.8166 | -0.11535 | 0.461474 | 0.923329 |
| ENSMUSG( | Ythdc1 | 1685.043 | 0.005933 | 0.9521 | 0.993821 | ENSMUSG( | Ythdc1 | 1572.906 | 0.073437 | 0.516356 | 0.930939 |
| ENSMUSG( | Ythdc2 | 519.922 | 0.126695 | 0.331078 | 0.77519 | ENSMUSG( | Ythdc2 | 502.3105 | 0.290135 | 0.049637 | 0.708648 |
| ENSMUSG( | Ythdf1 | 1104.501 | 0.072129 | 0.528768 | 0.878365 | ENSMUSG( | Ythdf1 | 1024.193 | 0.119732 | 0.360862 | 0.914223 |
| ENSMUSG( | Ythdf2 | 548.1396 | -0.08399 | 0.492488 | 0.860378 | ENSMUSG( | Ythdf2 | 545.4991 | 0.169916 | 0.207297 | 0.864412 |
| ENSMUSG( | Ythdf3 | 1239.065 | -0.06691 | 0.511294 | 0.870859 | ENSMUSG( | Ythdf3 | 1215.621 | 0.141853 | 0.25697 | 0.883082 |
| ENSMUSG( | Ywhab | 11158.35 | 0.048437 | 0.530176 | 0.879109 | ENSMUSG( | Ywhab | 9797.26 | -0.05966 | 0.585216 | 0.944975 |
| ENSMUSG( | Ywhae | 23802.87 | 0.02118 | 0.826519 | 0.968372 | ENSMUSG( | Ywhae | 20328.65 | -0.17265 | 0.23104 | 0.868507 |
| ENSMUSG( | Ywhag | 25036.39 | 0.138546 | 0.164466 | 0.595363 | ENSMUSG( | Ywhag | 20510.61 | -0.17124 | 0.169391 | 0.840806 |
| ENSMUSG( | Ywhah | 10261.29 | 0.0046 | 0.972006 | 0.996585 | ENSMUSG( | Ywhah | 8789.819 | -0.18192 | 0.266083 | 0.886134 |
| ENSMUSG( | Ywhaq | 4623.081 | 0.021096 | 0.853759 | 0.973491 | ENSMUSG( | Ywhaq | 4018.365 | -0.12078 | 0.367183 | 0.915058 |
| ENSMUSG( | Ywhaz | 13114.34 | 0.003499 | 0.978052 | 0.997428 | ENSMUSG( | Ywhaz | 11457.57 | -0.12348 | 0.471037 | 0.923329 |
| ENSMUSG( | Yy1 | 2246.418 | 0.068517 | 0.398907 | 0.812878 | ENSMUSG( | Yy1 | 2047.539 | 0.068083 | 0.560559 | 0.940254 |
| ENSMUSG( | Zan | 18.46278 | -0.14571 | 0.769819 | NA | ENSMUSG( | Zan | 18.065 | 0.043829 | 0.933618 | 0.99306 |
| ENSMUSG( | Zap70 | 75.54408 | 0.183658 | 0.427161 | 0.82532 | ENSMUSG( | Zap70 | 63.10465 | -0.06339 | 0.804819 | 0.977232 |
| ENSMUSG( | Zar1l | 32.92196 | 0.186258 | 0.620605 | 0.911432 | ENSMUSG( | Zar1l | 23.36193 | -0.62851 | 0.160542 | 0.837724 |
| ENSMUSG( | Zbed3 | 387.2984 | -0.04379 | 0.73935 | 0.947768 | ENSMUSG( | Zbed3 | 337.9176 | -0.17586 | 0.24646 | 0.877904 |
| ENSMUSG( | Zbed4 | 151.047 | -0.26897 | 0.111928 | 0.50438 | ENSMUSG( | Zbed4 | 150.2923 | -0.0047 | 0.979338 | 0.996468 |
| ENSMUSG( | Zbed5 | 115.8558 | -0.10165 | 0.612713 | 0.908844 | ENSMUSG( | Zbed5 | 100.6377 | -0.254 | 0.257714 | 0.883401 |
| ENSMUSG( | Zbed5-ps | 21.09274 | -0.22444 | 0.578202 | NA | ENSMUSG( | Zbed5-ps | 24.03241 | 0.439904 | 0.308859 | 0.898538 |
| ENSMUSG( | Zbed6 | 415.4686 | -0.17273 | 0.418083 | 0.821966 | ENSMUSG( | Zbed6 | 492.4832 | 0.553027 | 0.036074 | 0.668395 |
| ENSMUSG( | Zbp1 | 15.2785 | -1.07071 | 0.032106 | NA | ENSMUSG( | Zbp1 | 12.61694 | -1.56345 | 0.008276 | 0.388416 |
| ENSMUSG( | Zbtb1 | 425.0851 | -0.10785 | 0.349099 | 0.781665 | ENSMUSG( | Zbtb1 | 416.0651 | 0.102893 | 0.450626 | 0.922202 |
| ENSMUSG( | Zbtb10 | 186.0091 | -0.16037 | 0.294857 | 0.743798 | ENSMUSG( | Zbtb10 | 190.9829 | 0.178791 | 0.289463 | 0.896591 |
| ENSMUSG( | Zbtb11 | 732.1431 | 0.059299 | 0.582476 | 0.897918 | ENSMUSG( | Zbtb11 | 669.6958 | 0.068395 | 0.5943 | 0.945408 |
| ENSMUSG( | Zbtb11os1 | 138.8029 | 0.051372 | 0.76887 | 0.954674 | ENSMUSG( | Zbtb11os1 | 124.5045 | -0.00437 | 0.982803 | 0.996859 |
| ENSMUSG( | Zbtb12 | 232.5905 | -0.11118 | 0.471533 | 0.847331 | ENSMUSG( | Zbtb12 | 202.4877 | -0.26054 | 0.141329 | 0.830678 |
| ENSMUSG( | Zbtb14 | 428.1322 | 0.063244 | 0.611689 | 0.908586 | ENSMUSG( | Zbtb14 | 397.4868 | 0.117504 | 0.40916 | 0.917257 |
| ENSMUSG( | Zbtb16 | 878.6945 | 1.344914 | 1.51E-23 | 2.73E-20 | ENSMUSG( | Zbtb16 | 745.0841 | 1.201249 | 1.61E-15 | 2.35E-12 |
| ENSMUSG( | Zbtb17 | 1167.547 | 0.062218 | 0.562646 | 0.889485 | ENSMUSG( | Zbtb17 | 1014.594 | -0.07187 | 0.543452 | 0.938063 |
| ENSMUSG( | Zbtb18 | 7603.27 | 0.104291 | 0.184186 | 0.621101 | ENSMUSG( | Zbtb18 | 6773.009 | 0.039895 | 0.723726 | 0.966887 |
| ENSMUSG( | Zbtb2 | 293.5644 | 0.137614 | 0.293396 | 0.742763 | ENSMUSG( | Zbtb2 | 272.3555 | 0.185767 | 0.237373 | 0.872173 |
| ENSMUSG( | Zbtb20 | 6029.692 | 0.083886 | 0.550945 | 0.886038 | ENSMUSG( | Zbtb20 | 5280.304 | -0.02848 | 0.837754 | 0.981376 |
| ENSMUSG( | Zbtb21 | 411.8857 | -0.07393 | 0.568214 | 0.892252 | ENSMUSG( | Zbtb21 | 374.5676 | -0.07293 | 0.623685 | 0.953234 |
| ENSMUSG( | Zbtb22 | 891.9516 | 0.032193 | 0.775273 | 0.955606 | ENSMUSG( | Zbtb22 | 815.8076 | 0.044006 | 0.740831 | 0.968811 |
| ENSMUSG( | Zbtb24 | 317.7263 | 0.024482 | 0.8579 | 0.974667 | ENSMUSG( | Zbtb24 | 285.8764 | -0.01679 | 0.922562 | 0.992644 |
| ENSMUSG( | Zbtb25 | 238.4549 | -0.12507 | 0.432199 | 0.82768 | ENSMUSG( | Zbtb25 | 217.9648 | -0.10969 | 0.538499 | 0.937216 |
| ENSMUSG( | Zbtb26 | 165.3895 | 0.071795 | 0.685903 | 0.932981 | ENSMUSG( | Zbtb26 | 160.4885 | 0.249786 | 0.184696 | 0.853871 |
| ENSMUSG( | Zbtb3 | 64.52456 | 0.125147 | 0.671009 | 0.928781 | ENSMUSG( | Zbtb3 | 59.32435 | 0.159254 | 0.569251 | 0.941777 |
| ENSMUSG( | Zbtb33 | 437.4564 | 0.030301 | 0.836303 | 0.971111 | ENSMUSG( | Zbtb33 | 422.0581 | 0.181567 | 0.289098 | 0.895831 |
| ENSMUSG( | Zbtb34 | 400.8558 | 0.02943 | 0.810747 | 0.964272 | ENSMUSG( | Zbtb34 | 374.161 | 0.098977 | 0.477528 | 0.923329 |
| ENSMUSG( | Zbtb37 | 598.9996 | -0.20522 | 0.134554 | 0.547808 | ENSMUSG( | Zbtb37 | 573.0517 | -0.04826 | 0.780203 | 0.976106 |
| ENSMUSG( | Zbtb38 | 753.3175 | -0.034 | 0.768303 | 0.954641 | ENSMUSG( | Zbtb38 | 751.4083 | 0.22463 | 0.09865 | 0.798438 |
| ENSMUSG( | Zbtb39 | 348.9736 | -0.04824 | 0.708113 | 0.939519 | ENSMUSG( | Zbtb39 | 315.7448 | -0.06036 | 0.714667 | 0.964986 |
| ENSMUSG( | Zbtb4 | 6028.978 | 0.139664 | 0.081246 | 0.440604 | ENSMUSG( | Zbtb4 | 5283.499 | 0.030483 | 0.758195 | 0.971858 |
| ENSMUSG( | Zbtb40 | 424.994 | 0.551831 | 3.43E-05 | 0.002956 | ENSMUSG( | Zbtb40 | 395.1123 | 0.603835 | 2.06E-05 | 0.006225 |
| ENSMUSG( | Zbtb41 | 1145.892 | -0.03218 | 0.832804 | 0.970333 | ENSMUSG( | Zbtb41 | 1007.671 | -0.14088 | 0.419536 | 0.917257 |
| ENSMUSG( | Zbtb42 | 29.77021 | 0.275777 | 0.453793 | 0.836798 | ENSMUSG( | Zbtb42 | 31.30322 | 0.665319 | 0.09172 | 0.791889 |
| ENSMUSG( | Zbtb43 | 688.7582 | -0.02148 | 0.855474 | 0.973813 | ENSMUSG( | Zbtb43 | 651.5506 | 0.088546 | 0.478359 | 0.923329 |
| ENSMUSG( | Zbtb44 | 545.5994 | 0.098682 | 0.390141 | 0.80794 | ENSMUSG( | Zbtb44 | 494.4489 | 0.084048 | 0.534467 | 0.936104 |
| ENSMUSG( | Zbtb45 | 748.3788 | 0.131528 | 0.292569 | 0.742521 | ENSMUSG( | Zbtb45 | 666.1719 | 0.068211 | 0.602569 | 0.94789 |
| ENSMUSG( | Zbtb46 | 311.5869 | 0.342681 | 0.013494 | 0.169727 | ENSMUSG( | Zbtb46 | 271.6314 | 0.239347 | 0.170531 | 0.841739 |
| ENSMUSG( | Zbtb47 | 7307.783 | 0.221413 | 0.008911 | 0.132538 | ENSMUSG( | Zbtb47 | 6222.029 | 0.033183 | 0.740814 | 0.968811 |
| ENSMUSG( | Zbtb48 | 287.8644 | -0.12591 | 0.403787 | 0.816132 | ENSMUSG( | Zbtb48 | 277.3931 | 0.045837 | 0.807797 | 0.978622 |
| ENSMUSG( | Zbtb49 | 238.1179 | -0.15188 | 0.37861 | 0.801688 | ENSMUSG( | Zbtb49 | 227.6844 | 0.006725 | 0.973681 | 0.996416 |
| ENSMUSG( | Zbtb5 | 318.886 | -0.0966 | 0.442758 | 0.83278 | ENSMUSG( | Zbtb5 | 308.4735 | 0.079172 | 0.590056 | 0.944975 |
| ENSMUSG( | Zbtb6 | 241.5373 | -0.1136 | 0.428922 | 0.826446 | ENSMUSG( | Zbtb6 | 246.224 | 0.204729 | 0.228339 | 0.868507 |
| ENSMUSG( | Zbtb7a | 3575.93 | 0.127675 | 0.184119 | 0.621101 | ENSMUSG( | Zbtb7a | 3274.58 | 0.140154 | 0.223198 | 0.867404 |
| ENSMUSG( | Zbtb7b | 1301.528 | 0.148686 | 0.124631 | 0.531714 | ENSMUSG( | Zbtb7b | 1158.887 | 0.082574 | 0.497237 | 0.927825 |
| ENSMUSG( | Zbtb7c | 201.3793 | 0.225463 | 0.184949 | 0.62251 | ENSMUSG( | Zbtb7c | 180.513 | 0.191763 | 0.320962 | 0.904019 |
| ENSMUSG( | Zbtb8a | 91.69808 | -0.59941 | 0.010575 | 0.147613 | ENSMUSG( | Zbtb8a | 105.5338 | 0.138759 | 0.59426 | 0.945408 |
| ENSMUSG( | Zbtb8b | 77.36885 | 0.044241 | 0.850788 | 0.972638 | ENSMUSG( | Zbtb8b | 64.89627 | -0.18516 | 0.470556 | 0.923329 |
| ENSMUSG( | Zbtb8os | 456.9884 | 0.282312 | 0.026096 | 0.247245 | ENSMUSG( | Zbtb8os | 355.9802 | -0.1628 | 0.23994 | 0.8752 |
| ENSMUSG( | Zbtb9 | 21.61369 | 0.22638 | 0.595709 | 0.903152 | ENSMUSG( | Zbtb9 | 18.31838 | 0.022526 | 0.96483 | 0.995433 |
| ENSMUSG( | Zc2hc1a | 794.034 | 0.001634 | 0.989501 | 0.999111 | ENSMUSG( | Zc2hc1a | 713.1124 | -0.0485 | 0.764605 | 0.974094 |
| ENSMUSG( | Zc2hc1c | 51.12125 | 0.221289 | 0.420593 | 0.8229 | ENSMUSG( | Zc2hc1c | 48.43672 | 0.327784 | 0.270311 | 0.887887 |
| ENSMUSG( | Zc3h10 | 293.6023 | -0.06045 | 0.655871 | 0.923929 | ENSMUSG( | Zc3h10 | 274.8544 | 0.020759 | 0.892242 | 0.990558 |
| ENSMUSG( | Zc3h11a | 157.8721 | 0.002999 | 0.987527 | 0.998796 | ENSMUSG( | Zc3h11a | 162.3515 | 0.346386 | 0.091919 | 0.791889 |
| ENSMUSG( | Zc3h12a | 47.26298 | 0.071268 | 0.813491 | 0.964574 | ENSMUSG( | Zc3h12a | 45.69879 | 0.266883 | 0.439336 | 0.920721 |
| ENSMUSG( | Zc3h12b | 261.5594 | 0.065595 | 0.664778 | 0.926824 | ENSMUSG( | Zc3h12b | 245.8133 | 0.147432 | 0.401378 | 0.917257 |
| ENSMUSG( | Zc3h12c | 247.646 | -0.07038 | 0.619193 | 0.911432 | ENSMUSG( | Zc3h12c | 244.2193 | 0.160414 | 0.316979 | 0.902088 |
| ENSMUSG( | Zc3h13 | 2237.157 | -0.04668 | 0.7299 | 0.944956 | ENSMUSG( | Zc3h13 | 2161.299 | 0.123681 | 0.397414 | 0.917257 |
| ENSMUSG( | Zc3h14 | 2419.777 | 0.051239 | 0.525544 | 0.876728 | ENSMUSG( | Zc3h14 | 2124.667 | -0.05731 | 0.630464 | 0.953251 |
| ENSMUSG( | Zc3h15 | 1656.247 | 0.127533 | 0.217568 | 0.662824 | ENSMUSG( | Zc3h15 | 1407.006 | -0.07515 | 0.569533 | 0.941777 |
| ENSMUSG( | Zc3h18 | 1259.2 | 0.104442 | 0.289378 | 0.739839 | ENSMUSG( | Zc3h18 | 1124.289 | 0.046724 | 0.690533 | 0.960071 |
| ENSMUSG( | Zc3h3 | 262.6473 | 0.364036 | 0.032884 | 0.277278 | ENSMUSG( | Zc3h3 | 222.7347 | 0.171205 | 0.334921 | 0.907907 |
| ENSMUSG( | Zc3h4 | 2175.925 | 0.045487 | 0.596103 | 0.903152 | ENSMUSG( | Zc3h4 | 1930.456 | -0.03064 | 0.767645 | 0.974236 |
| ENSMUSG( | Zc3h6 | 181.185 | -0.35329 | 0.036692 | 0.295649 | ENSMUSG( | Zc3h6 | 181.878 | -0.05019 | 0.781961 | 0.976106 |
| ENSMUSG( | Zc3h7a | 1235.854 | -0.02019 | 0.848574 | 0.972374 | ENSMUSG( | Zc3h7a | 1164.64 | 0.075474 | 0.530977 | 0.934795 |
| ENSMUSG( | Zc3h7b | 21303.1 | 0.166202 | 0.042911 | 0.317818 | ENSMUSG( | Zc3h7b | 18015.15 | -0.04393 | 0.662716 | 0.956205 |
| ENSMUSG( | Zc3h8 | 134.0041 | -0.0894 | 0.650999 | 0.921451 | ENSMUSG( | Zc3h8 | 128.5392 | 0.06816 | 0.73006 | 0.967228 |
| ENSMUSG( | Zc3hav1 | 319.6076 | -0.0874 | 0.600881 | 0.904492 | ENSMUSG( | Zc3hav1 | 301.4965 | 0.013591 | 0.932601 | 0.99306 |
| ENSMUSG( | Zc3hav1l | 99.56511 | -0.00904 | 0.965503 | 0.995377 | ENSMUSG( | Zc3hav1l | 98.37537 | 0.215057 | 0.357217 | 0.913692 |
| ENSMUSG( | Zc3hc1 | 479.3558 | 0.163491 | 0.286074 | 0.735242 | ENSMUSG( | Zc3hc1 | 396.1828 | -0.11075 | 0.530093 | 0.934795 |
| ENSMUSG( | Zc4h2 | 390.3222 | 0.00903 | 0.945979 | 0.99369 | ENSMUSG( | Zc4h2 | 336.1637 | -0.17141 | 0.345529 | 0.910581 |
| ENSMUSG( | Zcchc10 | 109.3057 | 0.297438 | 0.140647 | 0.557902 | ENSMUSG( | Zcchc10 | 91.88122 | 0.070498 | 0.766557 | 0.974236 |
| ENSMUSG( | Zcchc12 | 895.5168 | -0.27684 | 0.155897 | 0.58375 | ENSMUSG( | Zcchc12 | 828.0105 | -0.23802 | 0.32937 | 0.907083 |
| ENSMUSG( | Zcchc14 | 2832.096 | 0.133713 | 0.135677 | 0.549211 | ENSMUSG( | Zcchc14 | 2607.278 | 0.163915 | 0.132831 | 0.822912 |
| ENSMUSG( | Zcchc17 | 1926.415 | -0.02799 | 0.775928 | 0.955606 | ENSMUSG( | Zcchc17 | 1653.62 | -0.21089 | 0.08857 | 0.786756 |
| ENSMUSG( | Zcchc18 | 5062.938 | -0.009 | 0.932923 | 0.992879 | ENSMUSG( | Zcchc18 | 4611.546 | -0.01137 | 0.92786 | 0.99306 |
| ENSMUSG( | Zcchc2 | 905.1391 | 0.132791 | 0.199448 | 0.640054 | ENSMUSG( | Zcchc2 | 810.3342 | 0.080721 | 0.516158 | 0.930939 |
| ENSMUSG( | Zcchc24 | 11909.21 | 0.06558 | 0.529299 | 0.878733 | ENSMUSG( | Zcchc24 | 10181.84 | -0.12112 | 0.347802 | 0.911652 |
| ENSMUSG( | Zcchc3 | 218.8272 | -0.02092 | 0.895163 | 0.984512 | ENSMUSG( | Zcchc3 | 203.6911 | 0.051435 | 0.779297 | 0.976106 |
| ENSMUSG( | Zcchc4 | 277.8661 | 0.073367 | 0.579089 | 0.896692 | ENSMUSG( | Zcchc4 | 257.7785 | 0.122782 | 0.424822 | 0.917257 |
| ENSMUSG( | Zcchc7 | 1444.636 | 0.095025 | 0.409147 | 0.818258 | ENSMUSG( | Zcchc7 | 1357.956 | 0.180577 | 0.147106 | 0.836241 |
| ENSMUSG( | Zcchc8 | 923.0034 | 0.097819 | 0.378724 | 0.801688 | ENSMUSG( | Zcchc8 | 884.2877 | 0.236588 | 0.076709 | 0.770558 |
| ENSMUSG( | Zcchc9 | 219.2685 | 0.058085 | 0.69877 | 0.936563 | ENSMUSG( | Zcchc9 | 220.9195 | 0.328817 | 0.04828 | 0.706142 |
| ENSMUSG( | Zcrb1 | 1536.941 | -0.01312 | 0.884441 | 0.982647 | ENSMUSG( | Zcrb1 | 1447.345 | 0.078561 | 0.504068 | 0.927825 |
| ENSMUSG( | Zcwpw1 | 240.8784 | -0.01242 | 0.933599 | 0.993194 | ENSMUSG( | Zcwpw1 | 212.932 | -0.10839 | 0.52525 | 0.934374 |
| ENSMUSG( | Zcwpw2 | 20.81335 | 0.120345 | 0.77547 | NA | ENSMUSG( | Zcwpw2 | 19.2153 | 0.154475 | 0.728294 | 0.966913 |
| ENSMUSG( | Zdbf2 | 503.0923 | 0.01291 | 0.927237 | 0.991229 | ENSMUSG( | Zdbf2 | 444.6257 | -0.08056 | 0.55609 | 0.939565 |
| ENSMUSG( | Zdhhc1 | 1685.441 | 0.023906 | 0.811296 | 0.964398 | ENSMUSG( | Zdhhc1 | 1574.151 | 0.095888 | 0.382784 | 0.917257 |
| ENSMUSG( | Zdhhc12 | 267.2717 | -0.43537 | 0.005457 | 0.098899 | ENSMUSG( | Zdhhc12 | 302.958 | 0.22515 | 0.186399 | 0.855939 |
| ENSMUSG( | Zdhhc13 | 579.6708 | -0.22575 | 0.122446 | 0.528223 | ENSMUSG( | Zdhhc13 | 593.6945 | 0.125688 | 0.492979 | 0.927358 |
| ENSMUSG( | Zdhhc14 | 1412.531 | -0.07262 | 0.609891 | 0.907813 | ENSMUSG( | Zdhhc14 | 1513.228 | 0.376839 | 0.02478 | 0.592568 |
| ENSMUSG( | Zdhhc15 | 325.3343 | -0.51084 | 0.010009 | 0.142653 | ENSMUSG( | Zdhhc15 | 347.0179 | -0.00701 | 0.972668 | 0.996416 |
| ENSMUSG( | Zdhhc16 | 1135.068 | -0.09669 | 0.443301 | 0.833108 | ENSMUSG( | Zdhhc16 | 1118.57 | 0.133327 | 0.353119 | 0.912263 |
| ENSMUSG( | Zdhhc17 | 2778.701 | -0.32052 | 0.021772 | 0.225648 | ENSMUSG( | Zdhhc17 | 3041.004 | 0.221649 | 0.15212 | 0.837724 |
| ENSMUSG( | Zdhhc18 | 1917.596 | -0.21192 | 0.051788 | 0.349274 | ENSMUSG( | Zdhhc18 | 1909.475 | 0.054137 | 0.675909 | 0.95694 |
| ENSMUSG( | Zdhhc2 | 1927.731 | -0.17705 | 0.174979 | 0.610399 | ENSMUSG( | Zdhhc2 | 1905.409 | 0.061363 | 0.683818 | 0.958678 |
| ENSMUSG( | Zdhhc20 | 1015.394 | -0.38949 | 0.013989 | 0.172788 | ENSMUSG( | Zdhhc20 | 1208.694 | 0.386902 | 0.022626 | 0.581156 |
| ENSMUSG( | Zdhhc21 | 1422.485 | -0.20207 | 0.105714 | 0.492581 | ENSMUSG( | Zdhhc21 | 1476.833 | 0.179898 | 0.215973 | 0.865167 |
| ENSMUSG( | Zdhhc22 | 1479.627 | -0.23845 | 0.054153 | 0.357985 | ENSMUSG( | Zdhhc22 | 1523.015 | 0.126043 | 0.421319 | 0.917257 |
| ENSMUSG( | Zdhhc23 | 1119.789 | -0.11913 | 0.397898 | 0.812161 | ENSMUSG( | Zdhhc23 | 1150.978 | 0.226964 | 0.105677 | 0.804969 |
| ENSMUSG( | Zdhhc24 | 663.8664 | -0.35709 | 0.001294 | 0.039144 | ENSMUSG( | Zdhhc24 | 701.8191 | 0.096253 | 0.449843 | 0.922202 |
| ENSMUSG( | Zdhhc3 | 4017.056 | -0.03773 | 0.676844 | 0.930542 | ENSMUSG( | Zdhhc3 | 3715.995 | 0.007054 | 0.948033 | 0.994798 |
| ENSMUSG( | Zdhhc4 | 1081.458 | -0.07815 | 0.404221 | 0.816132 | ENSMUSG( | Zdhhc4 | 1016.058 | 0.012171 | 0.913644 | 0.992378 |
| ENSMUSG( | Zdhhc5 | 1423.218 | -0.12659 | 0.433252 | 0.828207 | ENSMUSG( | Zdhhc5 | 1558.187 | 0.386573 | 0.032355 | 0.635121 |
| ENSMUSG( | Zdhhc6 | 1035.847 | -0.12199 | 0.314391 | 0.761558 | ENSMUSG( | Zdhhc6 | 1021.926 | 0.110584 | 0.421929 | 0.917257 |
| ENSMUSG( | Zdhhc7 | 1134.467 | -0.00145 | 0.989887 | 0.999144 | ENSMUSG( | Zdhhc7 | 1085.235 | 0.140795 | 0.302757 | 0.898538 |
| ENSMUSG( | Zdhhc8 | 4730.08 | -0.18332 | 0.155931 | 0.58375 | ENSMUSG( | Zdhhc8 | 4815.04 | 0.143311 | 0.377529 | 0.91645 |
| ENSMUSG( | Zdhhc9 | 2815.717 | -0.4949 | 0.001961 | 0.051054 | ENSMUSG( | Zdhhc9 | 3156.133 | 0.146374 | 0.415707 | 0.917257 |
| ENSMUSG( | Zeb1 | 7260.569 | -0.02187 | 0.84297 | 0.971739 | ENSMUSG( | Zeb1 | 6207.854 | -0.21471 | 0.109285 | 0.805353 |
| ENSMUSG( | Zeb1os1 | 194.0195 | 0.088011 | 0.572227 | 0.893584 | ENSMUSG( | Zeb1os1 | 173.8521 | 0.038331 | 0.827394 | 0.980224 |
| ENSMUSG( | Zeb2 | 5937.546 | 0.015434 | 0.871534 | 0.978114 | ENSMUSG( | Zeb2 | 5287.429 | -0.05305 | 0.684003 | 0.958678 |
| ENSMUSG( | Zeb2os | 237.1339 | -0.12272 | 0.444212 | 0.833239 | ENSMUSG( | Zeb2os | 208.0117 | -0.25175 | 0.180487 | 0.850372 |

| | | | | | |
|---|---|---|---|---|---|
| ENSMUSG( | 342.5066 | 0.054669 | 0.79907 | 0.960922 | Vwf |
| ENSMUSG( | 8548.748 | 0.079522 | 0.340817 | 0.778709 | Vxn |
| ENSMUSG( | 2392.442 | 0.051856 | 0.578358 | 0.896413 | Wac |
| ENSMUSG( | 1405.823 | 0.097277 | 0.359323 | 0.791011 | Wapl |
| ENSMUSG( | 2379.772 | 0.164346 | 0.091185 | 0.463787 | Wars1 |
| ENSMUSG( | 211.3734 | 0.23286 | 0.175011 | 0.610399 | Wars2 |
| ENSMUSG( | 104.234 | -0.10034 | 0.665798 | 0.927054 | Was |
| ENSMUSG( | 4258.219 | -0.01964 | 0.888618 | 0.983434 | Wasf1 |
| ENSMUSG( | 1044.703 | 0.042861 | 0.676024 | 0.930542 | Wasf2 |
| ENSMUSG( | 11994.29 | 0.066884 | 0.549145 | 0.885723 | Wasf3 |
| ENSMUSG( | 1167.081 | 0.096486 | 0.394152 | 0.809935 | Washc1 |
| ENSMUSG( | 5746.333 | 0.048724 | 0.571283 | 0.893319 | Washc2 |
| ENSMUSG( | 422.6113 | -0.11868 | 0.300346 | 0.749479 | Washc3 |
| ENSMUSG( | 878.8034 | 0.097611 | 0.342142 | 0.778828 | Washc4 |
| ENSMUSG( | 1388.308 | 0.117108 | 0.179318 | 0.616869 | Washc5 |
| ENSMUSG( | 1951.069 | 0.07878 | 0.404851 | 0.816396 | Wasl |
| ENSMUSG( | 1720.406 | 0.052382 | 0.597901 | 0.90372 | Wbp1 |
| ENSMUSG( | 2030.773 | 0.017349 | 0.860349 | 0.974842 | Wbp11 |
| ENSMUSG( | 2282.091 | 0.042419 | 0.68203 | 0.931738 | Wbp1l |
| ENSMUSG( | 10075.83 | 0.016611 | 0.86126 | 0.975026 | Wbp2 |
| ENSMUSG( | 1339.433 | 0.104417 | 0.267298 | 0.717525 | Wbp4 |
| ENSMUSG( | 221.7327 | 0.274658 | 0.072251 | 0.414687 | Wdcp |
| ENSMUSG( | 548.0839 | -0.2214 | 0.079514 | 0.435569 | Wdfy1 |
| ENSMUSG( | 1239.918 | -0.01364 | 0.898654 | 0.985633 | Wdfy2 |
| ENSMUSG( | 5148.724 | -0.09293 | 0.374541 | 0.800445 | Wdfy3 |
| ENSMUSG( | 200.1466 | 0.002666 | 0.985755 | 0.998326 | Wdfy4 |
| ENSMUSG( | 170.5862 | 0.113295 | 0.492177 | 0.860297 | Wdhd1 |
| ENSMUSG( | 269.6851 | 0.010959 | 0.942308 | 0.993415 | Wdpcp |
| ENSMUSG( | 4725.296 | 0.130543 | 0.119958 | 0.523983 | Wdr1 |
| ENSMUSG( | 1578.975 | 0.154438 | 0.101911 | 0.486318 | Wdr11 |
| ENSMUSG( | 771.5604 | 0.102562 | 0.305977 | 0.753712 | Wdr12 |
| ENSMUSG( | 15249.63 | 0.178321 | 0.021447 | 0.22362 | Wdr13 |
| ENSMUSG( | 681.6332 | -0.02389 | 0.835546 | 0.970557 | Wdr17 |
| ENSMUSG( | 1940.504 | 0.105946 | 0.269446 | 0.719729 | Wdr18 |
| ENSMUSG( | 609.9653 | -0.04142 | 0.728638 | 0.944393 | Wdr19 |
| ENSMUSG( | 489.4655 | 0.020153 | 0.855974 | 0.973813 | Wdr20 |
| ENSMUSG( | 597.9034 | 0.108675 | 0.344787 | 0.779153 | Wdr24 |
| ENSMUSG( | 257.6559 | -0.03038 | 0.829764 | 0.969582 | Wdr25 |
| ENSMUSG( | 4603.176 | 0.079532 | 0.355568 | 0.787945 | Wdr26 |
| ENSMUSG( | 52.77515 | 0.344738 | 0.21286 | 0.656885 | Wdr27 |
| ENSMUSG( | 430.617 | 0.050851 | 0.693964 | 0.934202 | Wdr3 |
| ENSMUSG( | 121.8337 | 0.01658 | 0.928892 | 0.991747 | Wdr31 |
| ENSMUSG( | 1696.726 | -0.39209 | 0.005533 | 0.099488 | Wdr33 |
| ENSMUSG( | 823.2444 | 0.087907 | 0.396103 | 0.811615 | Wdr35 |
| ENSMUSG( | 538.7039 | 0.090857 | 0.448741 | 0.83469 | Wdr36 |
| ENSMUSG( | 1377.513 | 0.058933 | 0.540834 | 0.88315 | Wdr37 |
| ENSMUSG( | 53.90398 | -0.04056 | 0.880478 | 0.981469 | Wdr38 |
| ENSMUSG( | 933.3766 | -0.04196 | 0.703649 | 0.937885 | Wdr4 |
| ENSMUSG( | 1178.168 | 0.076983 | 0.449835 | 0.835298 | Wdr41 |
| ENSMUSG( | 779.266 | 0.012533 | 0.900228 | 0.985633 | Wdr43 |
| ENSMUSG( | 519.9506 | -0.2309 | 0.130932 | 0.542456 | Wdr44 |
| ENSMUSG( | 509.6681 | 0.160962 | 0.219993 | 0.665725 | Wdr45 |
| ENSMUSG( | 2531.969 | 0.100577 | 0.236377 | 0.683538 | Wdr45b |
| ENSMUSG( | 382.95 | 0.161787 | 0.244271 | 0.691435 | Wdr46 |
| ENSMUSG( | 2375.754 | 0.032138 | 0.795028 | 0.959976 | Wdr47 |
| ENSMUSG( | 1976.379 | 0.073866 | 0.39243 | 0.809274 | Wdr48 |
| ENSMUSG( | 804.4211 | 0.052879 | 0.647971 | 0.919759 | Wdr5 |
| ENSMUSG( | 182.6573 | 0.037002 | 0.815174 | 0.964978 | Wdr53 |
| ENSMUSG( | 421.8901 | -0.42705 | 0.000272 | 0.013445 | Wdr54 |
| ENSMUSG( | 263.2929 | 0.123087 | 0.430745 | 0.826896 | Wdr55 |
| ENSMUSG( | 718.6328 | -0.07133 | 0.540397 | 0.88315 | Wdr59 |
| ENSMUSG( | 112.9951 | -0.06138 | 0.769107 | 0.954674 | Wdr5b |
| ENSMUSG( | 3457.435 | 0.029012 | 0.764335 | 0.954055 | Wdr6 |
| ENSMUSG( | 65.02871 | 0.070122 | 0.770355 | 0.954689 | Wdr62 |
| ENSMUSG( | 3365.864 | 0.07697 | 0.394748 | 0.810072 | Wdr7 |
| ENSMUSG( | 541.2576 | -0.01706 | 0.885172 | 0.982851 | Wdr70 |
| ENSMUSG( | 815.4853 | 0.044429 | 0.69267 | 0.934101 | Wdr73 |
| ENSMUSG( | 581.8326 | 0.052425 | 0.695338 | 0.934957 | Wdr74 |
| ENSMUSG( | 595.702 | 0.114305 | 0.323466 | 0.768511 | Wdr75 |
| ENSMUSG( | 76.0417 | 0.123295 | 0.584246 | 0.898754 | Wdr76 |
| ENSMUSG( | 678.8474 | 0.018737 | 0.856841 | 0.97427 | Wdr77 |
| ENSMUSG( | 802.9355 | 0.08205 | 0.496779 | 0.86175 | Wdr81 |
| ENSMUSG( | 3506.476 | 0.130535 | 0.126753 | 0.534201 | Wdr82 |
| ENSMUSG( | 730.9871 | 0.125779 | 0.295527 | 0.743814 | Wdr83 |
| ENSMUSG( | 1080.675 | -0.10709 | 0.265436 | 0.716109 | Wdr83os |
| ENSMUSG( | 56.05529 | -0.32255 | 0.43545 | 0.829093 | Wdr86 |
| ENSMUSG( | 50.07014 | 0.430221 | 0.108706 | 0.497919 | Wdr89 |
| ENSMUSG( | 851.463 | 0.348673 | 0.023687 | 0.23542 | Wdr90 |
| ENSMUSG( | 996.2753 | 0.088475 | 0.407474 | 0.817671 | Wdr91 |
| ENSMUSG( | 13.89939 | 0.20334 | 0.698095 | NA | Wdr93 |
| ENSMUSG( | 27.2281 | 0.307927 | 0.400638 | 0.813717 | Wdr95 |
| ENSMUSG( | 207.5352 | 0.596195 | 0.007989 | 0.124356 | Wdr97 |
| ENSMUSG( | 326.8595 | 0.087541 | 0.560895 | 0.888934 | Wdsub1 |
| ENSMUSG( | 4453.971 | 0.278625 | 0.000913 | 0.03071 | Wdtc1 |
| ENSMUSG( | 331.9123 | 0.086416 | 0.528613 | 0.878365 | Wee1 |
| ENSMUSG( | 476.8179 | 0.060883 | 0.701986 | 0.937445 | Wfdc1 |
| ENSMUSG( | 42.45022 | -0.65425 | 0.024084 | 0.237752 | Wfdc18 |
| ENSMUSG( | 23.06007 | 0.583135 | 0.178894 | 0.616298 | Wfdc2 |
| ENSMUSG( | 20.9362 | 0.114931 | 0.775578 | NA | Wfdc3 |
| ENSMUSG( | 10.1939 | 0.756389 | 0.195304 | NA | Wfikkn1 |
| ENSMUSG( | 136.9045 | 0.475842 | 0.077166 | 0.429655 | Wfikkn2 |
| ENSMUSG( | 3688.839 | -0.25405 | 0.050178 | 0.342492 | Wfs1 |
| ENSMUSG( | 641.71 | 0.028689 | 0.793012 | 0.95947 | Whamm |
| ENSMUSG( | 2868.069 | 0.320479 | 0.000453 | 0.019387 | Whrn |
| ENSMUSG( | 32.9878 | -1.04878 | 0.003109 | 0.069189 | Wif1 |
| ENSMUSG( | 1551.808 | -0.09545 | 0.392467 | 0.809274 | Wipf1 |
| ENSMUSG( | 2605.43 | 0.171935 | 0.040639 | 0.311461 | Wipf2 |
| ENSMUSG( | 4030.503 | 0.159125 | 0.209243 | 0.651175 | Wipf3 |
| ENSMUSG( | 1676.832 | 0.002245 | 0.984373 | 0.99781 | Wipi1 |
| ENSMUSG( | 4669.091 | 0.096745 | 0.218955 | 0.664602 | Wipi2 |
| ENSMUSG( | 3006.161 | 0.221217 | 0.011931 | 0.157992 | Wiz |
| ENSMUSG( | 1485.59 | -0.46019 | 0.004676 | 0.089972 | Wls |
| ENSMUSG( | 20068.58 | -0.02171 | 0.839596 | 0.971179 | Wnk1 |
| ENSMUSG( | 5707.727 | 0.311714 | 0.000152 | 0.008645 | Wnk2 |
| ENSMUSG( | 775.2091 | -0.04555 | 0.696755 | 0.935389 | Wnk3 |
| ENSMUSG( | 170.4834 | 0.098511 | 0.605106 | 0.905594 | Wnk4 |
| ENSMUSG( | 207.6645 | -0.4036 | 0.032105 | 0.273791 | Wnt10a |
| ENSMUSG( | 60.12673 | -0.26073 | 0.33032 | 0.774247 | Wnt10b |
| ENSMUSG( | 19.58102 | 0.273251 | 0.558639 | NA | Wnt11 |
| ENSMUSG( | 48.55242 | 0.247642 | 0.383014 | 0.803237 | Wnt16 |
| ENSMUSG( | 103.8377 | -0.44987 | 0.029125 | 0.259574 | Wnt2 |
| ENSMUSG( | 57.12744 | -0.24342 | 0.345158 | 0.779153 | Wnt2b |
| ENSMUSG( | 67.39537 | -0.85546 | 0.000732 | 0.026277 | Wnt3 |
| ENSMUSG( | 1170.466 | 0.004962 | 0.964315 | 0.995155 | Wnt4 |
| ENSMUSG( | 346.5097 | -0.18864 | 0.1603 | 0.589853 | Wnt5a |
| ENSMUSG( | 60.46938 | 0.027863 | 0.914118 | 0.988483 | Wnt5b |
| ENSMUSG( | 59.9724 | 0.243448 | 0.343409 | 0.779153 | Wnt6 |
| ENSMUSG( | 784.5756 | -0.36711 | 0.008899 | 0.132538 | Wnt7a |
| ENSMUSG( | 2161.57 | -0.1415 | 0.189054 | 0.627678 | Wnt7b |
| ENSMUSG( | 737.4654 | -0.16147 | 0.255681 | 0.706195 | Wnt9a |
| ENSMUSG( | 327.6717 | 0.18557 | 0.140691 | 0.557902 | Wrap53 |
| ENSMUSG( | 472.7567 | 0.103422 | 0.419875 | 0.822442 | Wrap73 |
| ENSMUSG( | 375.3621 | -0.08187 | 0.491037 | 0.859739 | Wrn |
| ENSMUSG( | 1425.269 | 0.011679 | 0.910876 | 0.988387 | Wrnip1 |
| ENSMUSG( | 1590.145 | -0.34055 | 0.007855 | 0.123093 | Wsb1 |
| ENSMUSG( | 19599.77 | 0.091523 | 0.237433 | 0.685281 | Wsb2 |
| ENSMUSG( | 5353.068 | -0.30043 | 0.014775 | 0.178616 | Wscd1 |
| ENSMUSG( | 1828.326 | -0.388 | 0.000673 | 0.025051 | Wscd2 |
| ENSMUSG( | 853.5037 | -0.04125 | 0.683651 | 0.932154 | Wtap |
| ENSMUSG( | 248.0565 | -0.06703 | 0.630964 | 0.914073 | Wtip |
| ENSMUSG( | 8443.462 | 0.053223 | 0.536502 | 0.881846 | Wwc1 |
| ENSMUSG( | 2428.896 | 0.163253 | 0.092843 | 0.468228 | Wwc2 |
| ENSMUSG( | 557.351 | 0.127137 | 0.259091 | 0.709814 | Wwox |
| ENSMUSG( | 2192.668 | 0.025063 | 0.829203 | 0.969286 | Wwp1 |
| ENSMUSG( | 1186.847 | 0.09058 | 0.329475 | 0.773311 | Wwp2 |
| ENSMUSG( | 2376.329 | -0.02337 | 0.872722 | 0.978753 | Wwtr1 |
| ENSMUSG( | 1145.935 | 0.051816 | 0.630812 | 0.914073 | Xab2 |
| ENSMUSG( | 107.4586 | -0.26396 | 0.204122 | 0.644638 | Xaf1 |
| ENSMUSG( | 2162.307 | -0.32251 | 0.047467 | 0.333152 | Xbp1 |
| ENSMUSG( | 143.8792 | 0.273815 | 0.209673 | 0.651841 | Xdh |
| ENSMUSG( | 3550.986 | -0.0015 | 0.988546 | 0.999109 | Xiap |
| ENSMUSG( | 474.9784 | -0.35054 | 0.034171 | 0.283507 | Xk |
| ENSMUSG( | 291.5341 | -0.28932 | 0.197168 | 0.636662 | Xkr4 |
| ENSMUSG( | 285.9706 | -0.03529 | 0.814112 | 0.964574 | Xkr6 |
| ENSMUSG( | 35.25984 | -0.27059 | 0.487236 | 0.858812 | Xkr7 |
| ENSMUSG( | 204.2534 | 0.010493 | 0.949154 | 0.993821 | Xkr8 |
| ENSMUSG( | 64.17304 | -0.50166 | 0.037109 | 0.297239 | Xkrx |
| ENSMUSG( | 29.97477 | 0.539855 | 0.136287 | 0.550578 | Xlr |
| ENSMUSG( | 21.2715 | -0.65221 | 0.164084 | 0.594869 | Xlr3a |
| ENSMUSG( | 73.56923 | -0.3041 | 0.666566 | 0.927247 | Xlr3b |
| ENSMUSG( | 9.739485 | -0.50724 | 0.481161 | NA | Xlr4a |
| ENSMUSG( | 186.8911 | 0.082389 | 0.657328 | 0.924144 | Xndc1 |
| ENSMUSG( | 11.91503 | -0.02524 | 0.963484 | NA | Xntrpc |
| ENSMUSG( | 211.8907 | 0.107956 | 0.517179 | 0.873295 | Xpa |
| ENSMUSG( | 667.935 | -0.04006 | 0.693983 | 0.934202 | Xpc |
| ENSMUSG( | 1480.516 | 0.160375 | 0.087468 | 0.454663 | Xpnpep1 |
| ENSMUSG( | 15.42394 | -0.73354 | 0.136456 | NA | Xpnpep2 |
| ENSMUSG( | 400.9015 | 0.119148 | 0.328344 | 0.772854 | Xpnpep3 |

| | | | | | |
|---|---|---|---|---|---|
| ENSMUSG( | 1900.471 | -0.12455 | 0.38859 | 0.917257 | Wwp1 |
| ENSMUSG( | 1027.955 | -0.05547 | 0.625797 | 0.953234 | Wwp2 |
| ENSMUSG( | 1911.915 | -0.41235 | 0.004618 | 0.279913 | Wwtr1 |
| ENSMUSG( | 1019.356 | -0.01762 | 0.890747 | 0.990558 | Xab2 |
| ENSMUSG( | 117.0911 | 0.258549 | 0.251931 | 0.879902 | Xaf1 |
| ENSMUSG( | 2575.577 | 0.441288 | 0.01205 | 0.445877 | Xbp1 |
| ENSMUSG( | 165.8559 | 0.851513 | 0.00106 | 0.111298 | Xdh |
| ENSMUSG( | 3207.861 | -0.02822 | 0.831026 | 0.98051 | Xiap |
| ENSMUSG( | 531.4999 | 0.257916 | 0.177436 | 0.847137 | Xk |
| ENSMUSG( | 331.0019 | 0.351569 | 0.153348 | 0.837724 | Xkr4 |
| ENSMUSG( | 250.7826 | -0.16117 | 0.411536 | 0.917257 | Xkr6 |
| ENSMUSG( | 30.87644 | -0.35292 | 0.427358 | 0.917257 | Xkr7 |
| ENSMUSG( | 200.5488 | 0.21316 | 0.251116 | 0.879321 | Xkr8 |
| ENSMUSG( | 65.86653 | -0.1107 | 0.662927 | 0.956205 | Xkrx |
| ENSMUSG( | 26.60617 | 0.481919 | 0.21402 | 0.864421 | Xlr |
| ENSMUSG( | 23.7002 | -0.03638 | 0.941718 | 0.994578 | Xlr3a |
| ENSMUSG( | 52.5537 | -1.24375 | 0.005371 | 0.301579 | Xlr3b |
| ENSMUSG( | 182.515 | 0.279571 | 0.146489 | 0.835973 | Xndc1 |
| ENSMUSG( | 11.49322 | 0.094121 | 0.87553 | 0.989644 | Xntrpc |
| ENSMUSG( | 189.5146 | 0.055077 | 0.769258 | 0.974236 | Xpa |
| ENSMUSG( | 612.0208 | -0.02228 | 0.85531 | 0.985649 | Xpc |
| ENSMUSG( | 1287.535 | 0.028713 | 0.796056 | 0.97681 | Xpnpep1 |
| ENSMUSG( | 15.26704 | -0.45942 | 0.372668 | 0.915491 | Xpnpep2 |
| ENSMUSG( | 367.0997 | 0.138595 | 0.332004 | 0.907907 | Xpnpep3 |
| ENSMUSG( | 1777.875 | 0.278771 | 0.047915 | 0.704961 | Xpo1 |
| ENSMUSG( | 588.4321 | -0.10185 | 0.472165 | 0.923329 | Xpo4 |
| ENSMUSG( | 1662.958 | -0.05568 | 0.646969 | 0.954033 | Xpo5 |
| ENSMUSG( | 2508.52 | 0.15103 | 0.221837 | 0.866104 | Xpo6 |
| ENSMUSG( | 1978.77 | -0.01712 | 0.880738 | 0.989644 | Xpo7 |
| ENSMUSG( | 2569.82 | 0.051764 | 0.626349 | 0.953234 | Xpot |
| ENSMUSG( | 4972.374 | 0.123711 | 0.39436 | 0.917257 | Xpr1 |
| ENSMUSG( | 695.9186 | -0.02875 | 0.856183 | 0.985884 | Xrcc1 |
| ENSMUSG( | 226.9874 | -0.07727 | 0.653584 | 0.955015 | Xrcc2 |
| ENSMUSG( | 157.6183 | -0.12681 | 0.535366 | 0.936508 | Xrcc3 |
| ENSMUSG( | 191.6791 | -0.2728 | 0.131837 | 0.822912 | Xrcc4 |
| ENSMUSG( | 567.0088 | -0.24524 | 0.056994 | 0.723362 | Xrcc5 |
| ENSMUSG( | 497.9271 | -0.04842 | 0.720489 | 0.966887 | Xrcc6 |
| ENSMUSG( | 813.3361 | 0.080685 | 0.551875 | 0.939565 | Xrn1 |
| ENSMUSG( | 1125.287 | 0.015179 | 0.895471 | 0.990558 | Xrn2 |
| ENSMUSG( | 38.75268 | -0.21875 | 0.525651 | 0.934374 | Xrra1 |
| ENSMUSG( | 925.6483 | 0.035352 | 0.821889 | 0.979198 | Xxylt1 |
| ENSMUSG( | 215.4845 | 0.232387 | 0.229391 | 0.868507 | Xylb |
| ENSMUSG( | 2535.928 | -0.0484 | 0.765046 | 0.974236 | Xylt1 |
| ENSMUSG( | 1054.08 | 0.070325 | 0.681264 | 0.958454 | Xylt2 |
| ENSMUSG( | 777.059 | -0.0814 | 0.506678 | 0.928106 | Yae1d1 |
| ENSMUSG( | 2826.53 | -0.05655 | 0.663402 | 0.956487 | Yaf2 |
| ENSMUSG( | 2057.918 | -0.01002 | 0.945726 | 0.994798 | Yap1 |
| ENSMUSG( | 1881.673 | -0.10921 | 0.421034 | 0.917257 | Yars1 |
| ENSMUSG( | 410.3286 | 0.087205 | 0.589332 | 0.944975 | Yars2 |
| ENSMUSG( | 115.2543 | -0.09962 | 0.654547 | 0.955015 | Ybey |
| ENSMUSG( | 7938.613 | -0.1789 | 0.171815 | 0.842999 | Ybx1 |
| ENSMUSG( | 89.97734 | -0.04839 | 0.866351 | 0.987347 | Ybx2 |
| ENSMUSG( | 869.5374 | -0.08376 | 0.48887 | 0.925573 | Ybx3 |
| ENSMUSG( | 372.1844 | 0.004684 | 0.974925 | 0.996416 | Ydjc |
| ENSMUSG( | 1250.067 | 0.050908 | 0.698752 | 0.960908 | Yeats2 |
| ENSMUSG( | 851.6654 | -0.16445 | 0.288609 | 0.894794 | Yeats4 |
| ENSMUSG( | 962.6183 | -0.0735 | 0.661834 | 0.956205 | Yes1 |
| ENSMUSG( | 533.0381 | -0.00799 | 0.95593 | 0.994798 | Yif1a |
| ENSMUSG( | 1162.278 | -0.09756 | 0.497627 | 0.927825 | Yif1b |
| ENSMUSG( | 946.4485 | 0.015354 | 0.91313 | 0.992298 | Yipf1 |
| ENSMUSG( | 701.4711 | -0.04384 | 0.731057 | 0.967228 | Yipf2 |
| ENSMUSG( | 2277.75 | 0.06347 | 0.675418 | 0.95694 | Yipf3 |
| ENSMUSG( | 1203.254 | 0.223053 | 0.202796 | 0.864412 | Yipf4 |
| ENSMUSG( | 1482.149 | 0.188408 | 0.281615 | 0.892412 | Yipf5 |
| ENSMUSG( | 1501.515 | 0.044307 | 0.771369 | 0.974236 | Yipf6 |
| ENSMUSG( | 245.896 | 0.041389 | 0.798081 | 0.97681 | Yjefn3 |
| ENSMUSG( | 209.7599 | -0.05624 | 0.729342 | 0.966991 | Yju2 |
| ENSMUSG( | 253.401 | -0.05807 | 0.731727 | 0.967228 | Yju2b |
| ENSMUSG( | 3428.903 | -0.16671 | 0.112932 | 0.809623 | Ykt6 |
| ENSMUSG( | 2344.755 | 0.12177 | 0.24065 | 0.875927 | Ylpm1 |
| ENSMUSG( | 1514.001 | 0.232095 | 0.054682 | 0.723362 | Yme1l1 |
| ENSMUSG( | 274.0217 | -0.02344 | 0.90046 | 0.990558 | Yod1 |
| ENSMUSG( | 1188.226 | 0.069694 | 0.544684 | 0.938063 | Ypel1 |
| ENSMUSG( | 1043.789 | -0.01877 | 0.875014 | 0.989644 | Ypel2 |
| ENSMUSG( | 4581.903 | -0.06483 | 0.628091 | 0.953238 | Ypel3 |
| ENSMUSG( | 1019.269 | 0.026831 | 0.888775 | 0.990558 | Ypel4 |
| ENSMUSG( | 1804.669 | -0.0902 | 0.576539 | 0.942912 | Ypel5 |
| ENSMUSG( | 353.8166 | -0.11535 | 0.461474 | 0.923329 | Yrdc |
| ENSMUSG( | 1572.906 | 0.073437 | 0.516356 | 0.930939 | Ythdc1 |
| ENSMUSG( | 502.3105 | 0.290135 | 0.049637 | 0.708648 | Ythdc2 |
| ENSMUSG( | 1024.193 | 0.119732 | 0.360862 | 0.914223 | Ythdf1 |
| ENSMUSG( | 545.4991 | 0.169916 | 0.207297 | 0.864412 | Ythdf2 |
| ENSMUSG( | 1215.621 | 0.141853 | 0.25697 | 0.883082 | Ythdf3 |
| ENSMUSG( | 9797.26 | -0.05966 | 0.585216 | 0.944975 | Ywhab |
| ENSMUSG( | 20328.65 | -0.17265 | 0.23104 | 0.868507 | Ywhae |
| ENSMUSG( | 20510.61 | -0.17124 | 0.169391 | 0.840806 | Ywhag |
| ENSMUSG( | 8789.819 | -0.18192 | 0.266083 | 0.886134 | Ywhah |
| ENSMUSG( | 4018.365 | -0.12078 | 0.367183 | 0.915058 | Ywhaq |
| ENSMUSG( | 11457.57 | -0.12348 | 0.471037 | 0.923329 | Ywhaz |
| ENSMUSG( | 2047.539 | 0.068083 | 0.560559 | 0.940254 | Yy1 |
| ENSMUSG( | 18.065 | 0.043829 | 0.933618 | 0.99306 | Zan |
| ENSMUSG( | 63.10465 | -0.06339 | 0.804819 | 0.977232 | Zap70 |
| ENSMUSG( | 23.36193 | -0.62851 | 0.160542 | 0.837724 | Zar1l |
| ENSMUSG( | 8.951335 | 0.658387 | 0.347078 | NA | Zbbx |
| ENSMUSG( | 337.9176 | -0.17586 | 0.24646 | 0.877904 | Zbed3 |
| ENSMUSG( | 150.2923 | -0.0047 | 0.979338 | 0.996468 | Zbed4 |
| ENSMUSG( | 100.6377 | -0.254 | 0.257714 | 0.883401 | Zbed5 |
| ENSMUSG( | 24.03241 | 0.439904 | 0.308859 | 0.898538 | Zbed5-ps |
| ENSMUSG( | 492.4832 | 0.553027 | 0.036074 | 0.668395 | Zbed6 |
| ENSMUSG( | 12.61694 | -1.56345 | 0.008276 | 0.388416 | Zbp1 |
| ENSMUSG( | 416.0651 | 0.102893 | 0.450626 | 0.922202 | Zbtb1 |
| ENSMUSG( | 190.9829 | 0.178791 | 0.289463 | 0.896591 | Zbtb10 |
| ENSMUSG( | 669.6958 | 0.068395 | 0.5943 | 0.945408 | Zbtb11 |
| ENSMUSG( | 124.5045 | -0.00437 | 0.982803 | 0.996859 | Zbtb11os1 |
| ENSMUSG( | 202.4877 | -0.26054 | 0.141329 | 0.830678 | Zbtb12 |
| ENSMUSG( | 397.4868 | 0.117504 | 0.40916 | 0.917257 | Zbtb14 |
| ENSMUSG( | 745.0841 | 1.201249 | 1.61E-15 | 2.35E-12 | Zbtb16 |
| ENSMUSG( | 1014.594 | -0.07187 | 0.543452 | 0.938063 | Zbtb17 |
| ENSMUSG( | 6773.009 | 0.039895 | 0.723726 | 0.966887 | Zbtb18 |
| ENSMUSG( | 272.3555 | 0.185767 | 0.237373 | 0.872173 | Zbtb2 |
| ENSMUSG( | 5280.304 | -0.02848 | 0.837754 | 0.981376 | Zbtb20 |
| ENSMUSG( | 374.5676 | -0.07293 | 0.623685 | 0.953234 | Zbtb21 |
| ENSMUSG( | 815.8076 | 0.044006 | 0.740831 | 0.968811 | Zbtb22 |
| ENSMUSG( | 285.8764 | -0.01679 | 0.922562 | 0.992644 | Zbtb24 |
| ENSMUSG( | 217.9648 | -0.10969 | 0.538499 | 0.937216 | Zbtb25 |
| ENSMUSG( | 160.4885 | 0.249786 | 0.184696 | 0.853871 | Zbtb26 |
| ENSMUSG( | 59.32435 | 0.159254 | 0.569251 | 0.941777 | Zbtb3 |
| ENSMUSG( | 422.0581 | 0.181567 | 0.289098 | 0.895831 | Zbtb33 |
| ENSMUSG( | 374.161 | 0.098977 | 0.477528 | 0.923329 | Zbtb34 |
| ENSMUSG( | 573.0517 | -0.04826 | 0.780203 | 0.976106 | Zbtb37 |
| ENSMUSG( | 751.4083 | 0.22463 | 0.09865 | 0.798438 | Zbtb38 |
| ENSMUSG( | 315.7448 | -0.06036 | 0.714667 | 0.964986 | Zbtb39 |
| ENSMUSG( | 5283.499 | 0.030483 | 0.758195 | 0.971858 | Zbtb4 |
| ENSMUSG( | 395.1123 | 0.603835 | 2.06E-05 | 0.006225 | Zbtb40 |
| ENSMUSG( | 1007.671 | -0.14088 | 0.419536 | 0.917257 | Zbtb41 |
| ENSMUSG( | 31.30322 | 0.665319 | 0.09172 | 0.791889 | Zbtb42 |
| ENSMUSG( | 651.5506 | 0.088546 | 0.478359 | 0.923329 | Zbtb43 |
| ENSMUSG( | 494.4489 | 0.084048 | 0.534467 | 0.936104 | Zbtb44 |
| ENSMUSG( | 666.1719 | 0.068211 | 0.602569 | 0.94789 | Zbtb45 |
| ENSMUSG( | 271.6314 | 0.239347 | 0.170531 | 0.841739 | Zbtb46 |
| ENSMUSG( | 6222.029 | 0.033183 | 0.740814 | 0.968811 | Zbtb47 |
| ENSMUSG( | 277.3931 | 0.045837 | 0.807797 | 0.978622 | Zbtb48 |
| ENSMUSG( | 227.6844 | 0.006725 | 0.973681 | 0.996416 | Zbtb49 |
| ENSMUSG( | 308.4735 | 0.079172 | 0.590056 | 0.944975 | Zbtb5 |
| ENSMUSG( | 246.224 | 0.204729 | 0.228339 | 0.868507 | Zbtb6 |
| ENSMUSG( | 3274.58 | 0.140154 | 0.223198 | 0.867404 | Zbtb7a |
| ENSMUSG( | 1158.887 | 0.082574 | 0.497237 | 0.927825 | Zbtb7b |
| ENSMUSG( | 180.513 | 0.191763 | 0.320962 | 0.904019 | Zbtb7c |
| ENSMUSG( | 105.5338 | 0.138759 | 0.59426 | 0.945408 | Zbtb8a |
| ENSMUSG( | 64.89627 | -0.18516 | 0.470556 | 0.923329 | Zbtb8b |
| ENSMUSG( | 355.9802 | -0.1628 | 0.23994 | 0.8752 | Zbtb8os |
| ENSMUSG( | 18.31838 | 0.022526 | 0.96483 | 0.995433 | Zbtb9 |
| ENSMUSG( | 713.1124 | -0.0485 | 0.764605 | 0.974094 | Zc2hc1a |
| ENSMUSG( | 48.43672 | 0.327784 | 0.270311 | 0.887887 | Zc2hc1c |
| ENSMUSG( | 274.8544 | 0.020759 | 0.892242 | 0.990558 | Zc3h10 |
| ENSMUSG( | 162.3515 | 0.346386 | 0.091919 | 0.791889 | Zc3h11a |
| ENSMUSG( | 45.69879 | 0.266883 | 0.439336 | 0.920721 | Zc3h12a |
| ENSMUSG( | 245.8133 | 0.147432 | 0.401378 | 0.917257 | Zc3h12b |
| ENSMUSG( | 244.2193 | 0.160414 | 0.316979 | 0.902088 | Zc3h12c |
| ENSMUSG( | 2161.299 | 0.123681 | 0.397414 | 0.917257 | Zc3h13 |
| ENSMUSG( | 2124.667 | -0.05731 | 0.630464 | 0.953251 | Zc3h14 |
| ENSMUSG( | 1407.006 | -0.07515 | 0.569533 | 0.941777 | Zc3h15 |
| ENSMUSG( | 1124.289 | 0.046724 | 0.690533 | 0.960071 | Zc3h18 |
| ENSMUSG( | 222.7347 | 0.171205 | 0.334921 | 0.907907 | Zc3h3 |
| ENSMUSG( | 1930.456 | -0.03064 | 0.767645 | 0.974236 | Zc3h4 |
| ENSMUSG( | 181.878 | -0.05019 | 0.781961 | 0.976106 | Zc3h6 |
| ENSMUSG( | 1164.64 | 0.075474 | 0.530977 | 0.934795 | Zc3h7a |
| ENSMUSG( | 18015.15 | -0.04393 | 0.662716 | 0.956205 | Zc3h7b |

| | | | | | | | | | | |
|---|---|---|---|---|---|---|---|---|---|---|
| ENSMUSG( | Zer1 | 5716.165 | 0.226421 | 0.023726 | 0.235641 | ENSMUSG( | Zer1 | 4851.038 | 0.031153 | 0.803211 | 0.976937 |
| ENSMUSG( | Zfand1 | 584.3522 | -0.02332 | 0.833779 | 0.970333 | ENSMUSG( | Zfand1 | 522.4676 | -0.07912 | 0.541189 | 0.937687 |
| ENSMUSG( | Zfand2a | 3498.275 | 0.158129 | 0.053248 | 0.354403 | ENSMUSG( | Zfand2a | 2984.563 | -0.0266 | 0.792062 | 0.97681 |
| ENSMUSG( | Zfand2b | 389.4082 | -0.13253 | 0.274661 | 0.725078 | ENSMUSG( | Zfand2b | 354.1105 | -0.13568 | 0.393461 | 0.917257 |
| ENSMUSG( | Zfand3 | 5229.303 | 0.082171 | 0.382915 | 0.803237 | ENSMUSG( | Zfand3 | 4464.383 | -0.10907 | 0.395991 | 0.917257 |
| ENSMUSG( | Zfand4 | 103.6308 | -0.24802 | 0.269567 | 0.719729 | ENSMUSG( | Zfand4 | 104.4647 | 0.060031 | 0.790894 | 0.97681 |
| ENSMUSG( | Zfand5 | 2516.889 | -0.11225 | 0.333755 | 0.775602 | ENSMUSG( | Zfand5 | 2396.657 | 0.013841 | 0.920475 | 0.992644 |
| ENSMUSG( | Zfand6 | 935.7303 | 0.039221 | 0.677943 | 0.930542 | ENSMUSG( | Zfand6 | 852.4106 | 0.034067 | 0.789463 | 0.976593 |
| ENSMUSG( | Zfas1 | 1330.828 | 0.110356 | 0.236193 | 0.683188 | ENSMUSG( | Zfas1 | 1137.93 | -0.07529 | 0.552304 | 0.939565 |
| ENSMUSG( | Zfat | 164.1491 | 0.079246 | 0.623539 | 0.911646 | ENSMUSG( | Zfat | 143.12 | -0.04746 | 0.80241 | 0.97681 |
| ENSMUSG( | Zfc3h1 | 1222.02 | 0.126388 | 0.294038 | 0.743459 | ENSMUSG( | Zfc3h1 | 1240.636 | 0.413741 | 0.027975 | 0.61023 |
| ENSMUSG( | Zfhx2 | 2297.118 | -0.3748 | 9.77E-05 | 0.006367 | ENSMUSG( | Zfhx2 | 2481.262 | 0.135499 | 0.265763 | 0.886134 |
| ENSMUSG( | Zfhx3 | 168.7045 | -0.59942 | 0.000435 | 0.019019 | ENSMUSG( | Zfhx3 | 208.176 | 0.323952 | 0.048511 | 0.706142 |
| ENSMUSG( | Zfhx4 | 243.228 | -0.50258 | 0.000684 | 0.025185 | ENSMUSG( | Zfhx4 | 286.8299 | 0.269175 | 0.068771 | 0.753052 |
| ENSMUSG( | Zfp1 | 422.9592 | -0.19178 | 0.143667 | 0.562581 | ENSMUSG( | Zfp1 | 406.2037 | -0.03226 | 0.815759 | 0.979198 |
| ENSMUSG( | Zfp1001 | 13.84286 | 0.196383 | 0.692261 | NA | ENSMUSG( | Zfp1001 | 14.08623 | 0.492022 | 0.340209 | 0.907972 |
| ENSMUSG( | Zfp1006 | 307.8814 | 0.022698 | 0.864148 | 0.975331 | ENSMUSG( | Zfp1006 | 299.1705 | 0.204674 | 0.193694 | 0.860096 |
| ENSMUSG( | Zfp1007 | 37.08658 | 0.076483 | 0.814793 | 0.964668 | ENSMUSG( | Zfp1007 | 36.50291 | 0.308416 | 0.369047 | 0.915491 |
| ENSMUSG( | Zfp1008 | 165.4546 | 0.023489 | 0.905456 | 0.986465 | ENSMUSG( | Zfp1008 | 145.2178 | -0.10933 | 0.664995 | 0.956766 |
| ENSMUSG( | Zfp101 | 245.5534 | -0.2545 | 0.078765 | 0.434128 | ENSMUSG( | Zfp101 | 265.0945 | 0.247453 | 0.190631 | 0.857801 |
| ENSMUSG( | Zfp105 | 177.8966 | -0.13478 | 0.427908 | 0.825894 | ENSMUSG( | Zfp105 | 153.192 | -0.31314 | 0.105032 | 0.804969 |
| ENSMUSG( | Zfp106 | 15254.38 | 0.087305 | 0.36365 | 0.793343 | ENSMUSG( | Zfp106 | 12955.77 | -0.11622 | 0.27622 | 0.890749 |
| ENSMUSG( | Zfp108 | 116.6419 | -0.30212 | 0.119868 | 0.523965 | ENSMUSG( | Zfp108 | 122.7005 | 0.139916 | 0.504698 | 0.927825 |
| ENSMUSG( | Zfp109 | 100.5563 | -0.11055 | 0.577806 | 0.895986 | ENSMUSG( | Zfp109 | 92.03925 | -0.09552 | 0.66935 | 0.956766 |
| ENSMUSG( | Zfp11 | 110.773 | -0.33359 | 0.121686 | 0.526331 | ENSMUSG( | Zfp11 | 108.0308 | -0.11803 | 0.613367 | 0.951177 |
| ENSMUSG( | Zfp110 | 322.7807 | 0.01531 | 0.911943 | 0.988483 | ENSMUSG( | Zfp110 | 300.6223 | 0.091255 | 0.639165 | 0.953903 |
| ENSMUSG( | Zfp111 | 179.2192 | 0.035115 | 0.822776 | 0.966352 | ENSMUSG( | Zfp111 | 168.5957 | 0.126261 | 0.484303 | 0.92541 |
| ENSMUSG( | Zfp112 | 229.3429 | -0.16125 | 0.326546 | 0.771368 | ENSMUSG( | Zfp112 | 225.9607 | 0.079583 | 0.660371 | 0.956205 |
| ENSMUSG( | Zfp113 | 181.7314 | 0.040427 | 0.806812 | 0.96295 | ENSMUSG( | Zfp113 | 167.3057 | 0.065709 | 0.726181 | 0.966887 |
| ENSMUSG( | Zfp114 | 78.08989 | -0.00703 | 0.977244 | 0.997428 | ENSMUSG( | Zfp114 | 65.91927 | -0.25333 | 0.352589 | 0.912093 |
| ENSMUSG( | Zfp119a | 65.064 | -0.16735 | 0.5126 | 0.871762 | ENSMUSG( | Zfp119a | 63.58019 | 0.062198 | 0.823465 | 0.979198 |
| ENSMUSG( | Zfp119b | 57.02166 | 0.10991 | 0.667709 | 0.927484 | ENSMUSG( | Zfp119b | 58.0006 | 0.400819 | 0.152001 | 0.837724 |
| ENSMUSG( | Zfp12 | 734.5998 | -0.0241 | 0.823755 | 0.966603 | ENSMUSG( | Zfp12 | 712.9518 | 0.154901 | 0.209154 | 0.864412 |
| ENSMUSG( | Zfp120 | 319.3921 | 0.181152 | 0.156191 | 0.583988 | ENSMUSG( | Zfp120 | 289.4941 | 0.167837 | 0.248555 | 0.877904 |
| ENSMUSG( | Zfp128 | 118.7694 | 0.083763 | 0.665806 | 0.927054 | ENSMUSG( | Zfp128 | 107.7987 | 0.070182 | 0.747377 | 0.969763 |
| ENSMUSG( | Zfp13 | 605.0729 | 0.127257 | 0.335015 | 0.776499 | ENSMUSG( | Zfp13 | 543.8448 | 0.091635 | 0.527748 | 0.934603 |
| ENSMUSG( | Zfp131 | 1514.915 | 0.012997 | 0.882209 | 0.982183 | ENSMUSG( | Zfp131 | 1372.259 | -0.0066 | 0.954725 | 0.994798 |
| ENSMUSG( | Zfp14 | 292.5498 | -0.3169 | 0.029204 | 0.259574 | ENSMUSG( | Zfp14 | 295.1471 | 0.001412 | 0.993637 | 0.998813 |
| ENSMUSG( | Zfp141 | 281.1281 | -0.23106 | 0.075397 | 0.425668 | ENSMUSG( | Zfp141 | 272.751 | -0.04576 | 0.762236 | 0.973255 |
| ENSMUSG( | Zfp142 | 1150.397 | 0.05057 | 0.616044 | 0.91085 | ENSMUSG( | Zfp142 | 1041.478 | 0.033535 | 0.770161 | 0.974236 |
| ENSMUSG( | Zfp143 | 250.6234 | 0.195783 | 0.157276 | 0.585553 | ENSMUSG( | Zfp143 | 221.7358 | 0.121931 | 0.459702 | 0.923329 |
| ENSMUSG( | Zfp146 | 1195.55 | 0.062432 | 0.548163 | 0.885197 | ENSMUSG( | Zfp146 | 1050.66 | -0.04391 | 0.713296 | 0.964766 |
| ENSMUSG( | Zfp148 | 2030.342 | 0.115783 | 0.171964 | 0.606889 | ENSMUSG( | Zfp148 | 1826.308 | 0.078657 | 0.492588 | 0.927315 |
| ENSMUSG( | Zfp157 | 629.5131 | -0.05504 | 0.598888 | 0.903801 | ENSMUSG( | Zfp157 | 577.4439 | -0.0407 | 0.760531 | 0.972688 |
| ENSMUSG( | Zfp160 | 481.5418 | 0.066146 | 0.582936 | 0.898094 | ENSMUSG( | Zfp160 | 468.0773 | 0.24933 | 0.069584 | 0.75601 |
| ENSMUSG( | Zfp169 | 162.0641 | -0.02292 | 0.892703 | 0.984474 | ENSMUSG( | Zfp169 | 158.6683 | 0.176689 | 0.36666 | 0.914703 |
| ENSMUSG( | Zfp174 | 120.5339 | -0.0063 | 0.976041 | 0.997428 | ENSMUSG( | Zfp174 | 106.0666 | -0.08891 | 0.690536 | 0.960071 |
| ENSMUSG( | Zfp180 | 1535.755 | 0.223768 | 0.016419 | 0.190497 | ENSMUSG( | Zfp180 | 1357.13 | 0.142302 | 0.197504 | 0.862161 |
| ENSMUSG( | Zfp182 | 312.8161 | 0.042866 | 0.756216 | 0.951479 | ENSMUSG( | Zfp182 | 296.0985 | 0.14602 | 0.325872 | 0.905339 |
| ENSMUSG( | Zfp184 | 54.96749 | 0.165934 | 0.528558 | 0.878365 | ENSMUSG( | Zfp184 | 52.6818 | 0.327074 | 0.294143 | 0.898538 |
| ENSMUSG( | Zfp185 | 67.02796 | -0.17413 | 0.540145 | 0.88315 | ENSMUSG( | Zfp185 | 66.97687 | 0.080337 | 0.782446 | 0.976106 |
| ENSMUSG( | Zfp189 | 149.6523 | -0.42193 | 0.038249 | 0.301606 | ENSMUSG( | Zfp189 | 135.8954 | -0.42288 | 0.056261 | 0.723362 |
| ENSMUSG( | Zfp2 | 473.222 | 0.133526 | 0.274363 | 0.724547 | ENSMUSG( | Zfp2 | 434.9317 | 0.153651 | 0.269596 | 0.887508 |
| ENSMUSG( | Zfp202 | 134.8965 | -0.0163 | 0.925382 | 0.991204 | ENSMUSG( | Zfp202 | 124.6848 | 0.030947 | 0.878749 | 0.989644 |
| ENSMUSG( | Zfp207 | 2050.861 | 0.071 | 0.431729 | 0.827541 | ENSMUSG( | Zfp207 | 1970.328 | 0.217079 | 0.073448 | 0.763629 |
| ENSMUSG( | Zfp212 | 348.2236 | -0.04572 | 0.73548 | 0.946109 | ENSMUSG( | Zfp212 | 333.3597 | 0.089669 | 0.564468 | 0.941206 |
| ENSMUSG( | Zfp213 | 190.2675 | 0.171932 | 0.28461 | 0.733224 | ENSMUSG( | Zfp213 | 165.9879 | 0.066096 | 0.73353 | 0.967228 |
| ENSMUSG( | Zfp217 | 196.0473 | -0.07343 | 0.646382 | 0.918612 | ENSMUSG( | Zfp217 | 202.0189 | 0.276081 | 0.114598 | 0.811703 |
| ENSMUSG( | Zfp219 | 2192.199 | -0.03304 | 0.707777 | 0.939519 | ENSMUSG( | Zfp219 | 1962.371 | -0.08507 | 0.43013 | 0.917257 |
| ENSMUSG( | Zfp229 | 112.9944 | 0.270492 | 0.24675 | 0.695324 | ENSMUSG( | Zfp229 | 101.4759 | 0.236594 | 0.322771 | 0.904023 |
| ENSMUSG( | Zfp235 | 465.4172 | 0.036943 | 0.78024 | 0.957491 | ENSMUSG( | Zfp235 | 423.6368 | 0.037712 | 0.814024 | 0.979198 |
| ENSMUSG( | Zfp236 | 1751.635 | 0.134362 | 0.177142 | 0.614466 | ENSMUSG( | Zfp236 | 1634.8 | 0.201835 | 0.085088 | 0.779799 |
| ENSMUSG( | Zfp239 | 739.7512 | 0.040398 | 0.702934 | 0.937735 | ENSMUSG( | Zfp239 | 671.1599 | 0.025696 | 0.844234 | 0.983576 |
| ENSMUSG( | Zfp24 | 1706.693 | 0.202271 | 0.053512 | 0.35564 | ENSMUSG( | Zfp24 | 1461.799 | 0.027833 | 0.837425 | 0.981335 |
| ENSMUSG( | Zfp248 | 243.8296 | 0.189005 | 0.190012 | 0.629259 | ENSMUSG( | Zfp248 | 212.5358 | 0.060471 | 0.72045 | 0.966887 |
| ENSMUSG( | Zfp251 | 1528.96 | 0.1645 | 0.089309 | 0.458563 | ENSMUSG( | Zfp251 | 1352.607 | 0.082118 | 0.483378 | 0.92499 |
| ENSMUSG( | Zfp26 | 560.0499 | -0.24796 | 0.038607 | 0.302696 | ENSMUSG( | Zfp26 | 576.5267 | 0.115507 | 0.404187 | 0.917257 |
| ENSMUSG( | Zfp260 | 1211.659 | 0.179903 | 0.05091 | 0.345939 | ENSMUSG( | Zfp260 | 1091.002 | 0.144904 | 0.244524 | 0.877904 |
| ENSMUSG( | Zfp263 | 524.4271 | -0.03452 | 0.809095 | 0.963548 | ENSMUSG( | Zfp263 | 504.3617 | 0.125922 | 0.431414 | 0.917257 |
| ENSMUSG( | Zfp266 | 831.2275 | -0.00731 | 0.950585 | 0.993821 | ENSMUSG( | Zfp266 | 790.3558 | 0.117101 | 0.351661 | 0.912093 |
| ENSMUSG( | Zfp267 | 483.1053 | -0.17637 | 0.127364 | 0.535055 | ENSMUSG( | Zfp267 | 462.3884 | -0.02786 | 0.835585 | 0.981237 |
| ENSMUSG( | Zfp27 | 404.9856 | 0.12147 | 0.324391 | 0.769598 | ENSMUSG( | Zfp27 | 362.9387 | 0.070303 | 0.637125 | 0.953903 |
| ENSMUSG( | Zfp273 | 106.2715 | -0.30532 | 0.148122 | 0.571232 | ENSMUSG( | Zfp273 | 106.3796 | -0.00046 | 0.998385 | 0.999313 |
| ENSMUSG( | Zfp275 | 838.9479 | 0.050504 | 0.661624 | 0.925271 | ENSMUSG( | Zfp275 | 776.0886 | 0.09403 | 0.454037 | 0.922444 |
| ENSMUSG( | Zfp276 | 810.7732 | -0.18307 | 0.213863 | 0.658243 | ENSMUSG( | Zfp276 | 810.2682 | 0.096651 | 0.59356 | 0.945408 |
| ENSMUSG( | Zfp277 | 448.8963 | -0.09521 | 0.498832 | 0.862973 | ENSMUSG( | Zfp277 | 391.0153 | -0.22703 | 0.105371 | 0.804969 |
| ENSMUSG( | Zfp28 | 232.0747 | -0.1814 | 0.232768 | 0.680928 | ENSMUSG( | Zfp28 | 218.83 | -0.06898 | 0.685383 | 0.958678 |
| ENSMUSG( | Zfp280b | 275.9745 | 0.044649 | 0.769693 | 0.954674 | ENSMUSG( | Zfp280b | 257.5452 | 0.12176 | 0.471894 | 0.923329 |
| ENSMUSG( | Zfp280c | 485.015 | -0.26451 | 0.087557 | 0.454663 | ENSMUSG( | Zfp280c | 507.0171 | 0.146923 | 0.334841 | 0.907907 |
| ENSMUSG( | Zfp280d | 800.4414 | -0.0549 | 0.601288 | 0.904603 | ENSMUSG( | Zfp280d | 757.7748 | 0.058236 | 0.641193 | 0.953903 |
| ENSMUSG( | Zfp281 | 552.59 | -0.16772 | 0.132382 | 0.544294 | ENSMUSG( | Zfp281 | 574.7572 | 0.210052 | 0.122787 | 0.814249 |
| ENSMUSG( | Zfp282 | 577.9396 | 0.097341 | 0.458393 | 0.840034 | ENSMUSG( | Zfp282 | 518.734 | 0.056205 | 0.697429 | 0.960867 |
| ENSMUSG( | Zfp286 | 352.0807 | -0.01972 | 0.890368 | 0.983687 | ENSMUSG( | Zfp286 | 311.1936 | -0.10055 | 0.508881 | 0.929089 |
| ENSMUSG( | Zfp287 | 367.1342 | -0.01087 | 0.930215 | 0.991848 | ENSMUSG( | Zfp287 | 324.3551 | -0.09314 | 0.542924 | 0.938063 |
| ENSMUSG( | Zfp292 | 892.1782 | -0.16476 | 0.116482 | 0.514944 | ENSMUSG( | Zfp292 | 850.6014 | -0.02784 | 0.812839 | 0.979148 |
| ENSMUSG( | Zfp296 | 37.23637 | 0.455786 | 0.140906 | 0.557902 | ENSMUSG( | Zfp296 | 34.12942 | 0.496193 | 0.158576 | 0.837724 |
| ENSMUSG( | Zfp3 | 293.5931 | -0.10527 | 0.447505 | 0.834051 | ENSMUSG( | Zfp3 | 266.7145 | -0.11734 | 0.456685 | 0.923329 |
| ENSMUSG( | Zfp30 | 355.2201 | -0.03683 | 0.775837 | 0.955606 | ENSMUSG( | Zfp30 | 346.4061 | 0.155879 | 0.261061 | 0.884926 |
| ENSMUSG( | Zfp300 | 40.80662 | -0.08192 | 0.789715 | 0.959461 | ENSMUSG( | Zfp300 | 41.51251 | 0.24189 | 0.486917 | 0.92541 |
| ENSMUSG( | Zfp316 | 1252.085 | 0.074607 | 0.448232 | 0.834051 | ENSMUSG( | Zfp316 | 1085.316 | -0.06723 | 0.555976 | 0.939565 |
| ENSMUSG( | Zfp317 | 796.7514 | -0.15196 | 0.226003 | 0.672742 | ENSMUSG( | Zfp317 | 759.7717 | -0.01678 | 0.901587 | 0.990784 |
| ENSMUSG( | Zfp318 | 1529.772 | -0.12329 | 0.187985 | 0.626616 | ENSMUSG( | Zfp318 | 1474.375 | 0.04384 | 0.697225 | 0.960867 |
| ENSMUSG( | Zfp319 | 680.8215 | 0.06722 | 0.569768 | 0.893041 | ENSMUSG( | Zfp319 | 627.2959 | 0.105236 | 0.453744 | 0.922316 |
| ENSMUSG( | Zfp322a | 564.2983 | 0.176735 | 0.131878 | 0.544244 | ENSMUSG( | Zfp322a | 520.02 | 0.206156 | 0.172285 | 0.843466 |
| ENSMUSG( | Zfp324 | 252.4222 | 0.245028 | 0.107553 | 0.496444 | ENSMUSG( | Zfp324 | 221.6234 | 0.156724 | 0.379418 | 0.916995 |
| ENSMUSG( | Zfp326 | 657.701 | 0.071012 | 0.533218 | 0.879905 | ENSMUSG( | Zfp326 | 601.2517 | 0.081289 | 0.53274 | 0.934795 |
| ENSMUSG( | Zfp329 | 560.8392 | 0.097049 | 0.360331 | 0.791947 | ENSMUSG( | Zfp329 | 517.6257 | 0.132809 | 0.293449 | 0.898538 |
| ENSMUSG( | Zfp330 | 599.1481 | 0.048981 | 0.677878 | 0.930542 | ENSMUSG( | Zfp330 | 559.8918 | 0.113587 | 0.441151 | 0.921166 |
| ENSMUSG( | Zfp334 | 566.6615 | -0.03593 | 0.778134 | 0.956168 | ENSMUSG( | Zfp334 | 538.0689 | 0.085366 | 0.552417 | 0.939565 |
| ENSMUSG( | Zfp335 | 1458.559 | 0.199121 | 0.060352 | 0.3797 | ENSMUSG( | Zfp335 | 1308.013 | 0.158846 | 0.196195 | 0.862161 |
| ENSMUSG( | Zfp335os | 74.6715 | 0.005237 | 0.983405 | 0.997675 | ENSMUSG( | Zfp335os | 71.82842 | 0.14325 | 0.603541 | 0.948137 |
| ENSMUSG( | Zfp341 | 250.1546 | 0.1696 | 0.285486 | 0.734313 | ENSMUSG( | Zfp341 | 226.8923 | 0.153743 | 0.396117 | 0.917257 |
| ENSMUSG( | Zfp346 | 1006.048 | 0.201992 | 0.047568 | 0.333288 | ENSMUSG( | Zfp346 | 866.3104 | 0.049102 | 0.697011 | 0.960867 |
| ENSMUSG( | Zfp35 | 407.2982 | 0.096759 | 0.431158 | 0.827055 | ENSMUSG( | Zfp35 | 364.7355 | 0.047902 | 0.745697 | 0.96951 |
| ENSMUSG( | Zfp354a | 181.8506 | -0.33014 | 0.086012 | 0.451492 | ENSMUSG( | Zfp354a | 206.4813 | 0.313173 | 0.088314 | 0.786756 |
| ENSMUSG( | Zfp354b | 71.0772 | -0.18928 | 0.495133 | 0.860798 | ENSMUSG( | Zfp354b | 66.92715 | -0.09075 | 0.744624 | 0.96951 |
| ENSMUSG( | Zfp354c | 1236.06 | -0.02532 | 0.797355 | 0.960507 | ENSMUSG( | Zfp354c | 1151.611 | 0.034655 | 0.802608 | 0.97681 |
| ENSMUSG( | Zfp358 | 1172.888 | 0.136261 | 0.139048 | 0.555099 | ENSMUSG( | Zfp358 | 1050.335 | 0.089492 | 0.449677 | 0.922202 |
| ENSMUSG( | Zfp36 | 146.8329 | -0.36015 | 0.060269 | 0.379636 | ENSMUSG( | Zfp36 | 163.8755 | 0.252957 | 0.227901 | 0.868507 |
| ENSMUSG( | Zfp362 | 1204.823 | 0.017739 | 0.854014 | 0.973491 | ENSMUSG( | Zfp362 | 1051.683 | -0.1086 | 0.369814 | 0.915491 |
| ENSMUSG( | Zfp365 | 35872.94 | 0.095298 | 0.31856 | 0.764234 | ENSMUSG( | Zfp365 | 31827.71 | 0.020363 | 0.85099 | 0.985541 |
| ENSMUSG( | Zfp366 | 250.1195 | 0.652813 | 0.006517 | 0.110626 | ENSMUSG( | Zfp366 | 248.5118 | 0.866378 | 0.003178 | 0.224467 |
| ENSMUSG( | Zfp367 | 897.7599 | -0.03567 | 0.781752 | 0.957558 | ENSMUSG( | Zfp367 | 796.8669 | -0.11807 | 0.428667 | 0.917257 |
| ENSMUSG( | Zfp369 | 516.4018 | 0.186727 | 0.1254 | 0.53232 | ENSMUSG( | Zfp369 | 471.7481 | 0.19358 | 0.161298 | 0.837724 |
| ENSMUSG( | Zfp36l1 | 989.9232 | 0.060619 | 0.58978 | 0.90113 | ENSMUSG( | Zfp36l1 | 1038.545 | 0.440831 | 0.001921 | 0.164157 |
| ENSMUSG( | Zfp36l2 | 1034.761 | 0.122523 | 0.344294 | 0.779153 | ENSMUSG( | Zfp36l2 | 1012.775 | 0.314224 | 0.005468 | 0.303106 |
| ENSMUSG( | Zfp37 | 320.3405 | 0.004797 | 0.972042 | 0.996585 | ENSMUSG( | Zfp37 | 308.9929 | 0.159919 | 0.316677 | 0.901868 |
| ENSMUSG( | Zfp382 | 86.89765 | 0.210951 | 0.34237 | 0.778855 | ENSMUSG( | Zfp382 | 78.17076 | 0.203978 | 0.463871 | 0.923329 |
| ENSMUSG( | Zfp383 | 302.9045 | -0.20824 | 0.193538 | 0.633027 | ENSMUSG( | Zfp383 | 316.4407 | 0.195038 | 0.240867 | 0.876146 |
| ENSMUSG( | Zfp384 | 1244.363 | 0.12044 | 0.289641 | 0.739839 | ENSMUSG( | Zfp384 | 1110.553 | 0.063725 | 0.606205 | 0.949497 |
| ENSMUSG( | Zfp385a | 3558.219 | -0.11428 | 0.242873 | 0.690275 | ENSMUSG( | Zfp385a | 3113.154 | -0.24281 | 0.094598 | 0.796133 |
| ENSMUSG( | Zfp385b | 767.3135 | -0.00425 | 0.972807 | 0.996585 | ENSMUSG( | Zfp385b | 706.8793 | 0.021779 | 0.891003 | 0.990558 |
| ENSMUSG( | Zfp385c | 10.13824 | -0.51115 | 0.37775 | NA | ENSMUSG( | Zfp385c | 8.673593 | -0.79985 | 0.237947 | NA |
| ENSMUSG( | Zfp386 | 458.8572 | -0.04456 | 0.70578 | 0.938649 | ENSMUSG( | Zfp386 | 446.5102 | 0.143955 | 0.280091 | 0.892356 |
| ENSMUSG( | Zfp39 | 184.0303 | 0.141468 | 0.475875 | 0.850919 | ENSMUSG( | Zfp39 | 159.0249 | -0.01118 | 0.957399 | 0.994798 |
| ENSMUSG( | Zfp395 | 347.3339 | -0.00464 | 0.972429 | 0.996585 | ENSMUSG( | Zfp395 | 311.9507 | -0.03877 | 0.798892 | 0.97681 |
| ENSMUSG( | Zfp397 | 668.1634 | 0.015122 | 0.897589 | 0.985633 | ENSMUSG( | Zfp397 | 648.0816 | 0.192589 | 0.156919 | 0.837724 |
| ENSMUSG( | Zfp398 | 514.1492 | -0.02083 | 0.85585 | 0.973813 | ENSMUSG( | Zfp398 | 482.0344 | 0.063216 | 0.633232 | 0.953844 |
| ENSMUSG( | Zfp40 | 643.134 | -0.09456 | 0.469193 | 0.845876 | ENSMUSG( | Zfp40 | 624.5981 | 0.08562 | 0.575945 | 0.942912 |
| ENSMUSG( | Zfp407 | 334.2099 | -0.01069 | 0.935378 | 0.993363 | ENSMUSG( | Zfp407 | 288.3991 | -0.17157 | 0.244002 | 0.877904 |
| ENSMUSG( | Zfp408 | 433.6016 | -0.02941 | 0.822706 | 0.966352 | ENSMUSG( | Zfp408 | 385.7132 | -0.09915 | 0.475632 | 0.923329 |
| ENSMUSG( | Zfp41 | 251.9784 | -0.07461 | 0.663573 | 0.9265 | ENSMUSG( | Zfp41 | 248.1059 | 0.156749 | 0.401954 | 0.917257 |
| ENSMUSG( | Zfp410 | 616.1591 | 0.125273 | 0.268088 | 0.718372 | ENSMUSG( | Zfp410 | 547.7336 | 0.062883 | 0.645183 | 0.953988 |
| ENSMUSG( | Zfp414 | 433.6762 | -0.03677 | 0.800599 | 0.961745 | ENSMUSG( | Zfp414 | 379.4076 | -0.15635 | 0.335616 | 0.907907 |
| ENSMUSG( | Zfp418 | 43.99807 | -0.1174 | 0.690423 | 0.933881 | ENSMUSG( | Zfp418 | 44.77688 | 0.213742 | 0.504615 | 0.927825 |
| ENSMUSG( | Zfp420 | 322.0978 | 0.030951 | 0.815764 | 0.965324 | ENSMUSG( | Zfp420 | 306.347 | 0.140395 | 0.396785 | 0.917257 |
| ENSMUSG( | Zfp422 | 590.7254 | 0.020339 | 0.85583 | 0.973813 | ENSMUSG( | Zfp422 | 535.776 | 0.003315 | 0.980683 | 0.996497 |
| ENSMUSG( | Zfp423 | 1009.346 | -0.03433 | 0.792706 | 0.95947 | ENSMUSG( | Zfp423 | 882.5273 | -0.158 | 0.18266 | 0.850579 |
| ENSMUSG( | Zfp426 | 728.3204 | 0.115291 | 0.261977 | 0.712129 | ENSMUSG( | Zfp426 | 619.6006 | -0.08557 | 0.503342 | 0.927825 |
| ENSMUSG( | Zfp428 | 890.7638 | -0.01745 | 0.870899 | 0.977951 | ENSMUSG( | Zfp428 | 737.033 | -0.32015 | 0.040378 | 0.682717 |
| ENSMUSG( | Zfp429 | 52.44877 | -0.14113 | 0.623918 | 0.91187 | ENSMUSG( | Zfp429 | 50.9619 | 0.072713 | 0.81015 | 0.978779 |
| ENSMUSG( | Zfp433 | 18.39064 | 0.256816 | 0.556369 | NA | ENSMUSG( | Zfp433 | 19.17669 | 0.569914 | 0.248485 | 0.877904 |
| ENSMUSG( | Zfp438 | 211.4308 | -0.18853 | 0.231832 | 0.679725 | ENSMUSG( | Zfp438 | 174.8848 | -0.51769 | 0.005929 | 0.316645 |
| ENSMUSG( | Zfp442 | 33.47123 | -0.01816 | 0.955801 | 0.993874 | ENSMUSG( | Zfp442 | 29.61405 | -0.0778 | 0.835009 | 0.981237 |
| ENSMUSG( | Zfp444 | 1047.032 | 0.024396 | 0.81424 | 0.964574 | ENSMUSG( | Zfp444 | 909.9679 | -0.11799 | 0.368041 | 0.915491 |
| ENSMUSG( | Zfp445 | 4811.416 | -0.09714 | 0.349 | 0.781661 | ENSMUSG( | Zfp445 | 4557.212 | 0.014685 | 0.91742 | 0.992529 |
| ENSMUSG( | Zfp446 | 320.7309 | 0.070657 | 0.643553 | 0.917316 | ENSMUSG( | Zfp446 | 295.0415 | 0.105461 | 0.532883 | 0.934795 |
| ENSMUSG( | Zfp449 | 155.8252 | 0.179787 | 0.319767 | 0.765305 | ENSMUSG( | Zfp449 | 137.7436 | 0.088517 | 0.662596 | 0.956205 |
| ENSMUSG( | Zfp451 | 1077.121 | -0.04003 | 0.678573 | 0.930611 | ENSMUSG( | Zfp451 | 1004.165 | 0.025364 | 0.827326 | 0.980224 |
| ENSMUSG( | Zfp454 | 136.9489 | 0.107755 | 0.545107 | 0.884712 | ENSMUSG( | Zfp454 | 124.7191 | 0.095876 | 0.655971 | 0.955533 |
| ENSMUSG( | Zfp455 | 36.79726 | 0.072213 | 0.838225 | 0.971111 | ENSMUSG( | Zfp455 | 35.32375 | 0.19884 | 0.574448 | 0.942912 |
| ENSMUSG( | Zfp456 | 35.56774 | -0.40791 | 0.234192 | 0.68192 | ENSMUSG( | Zfp456 | 34.10173 | -0.23021 | 0.534846 | 0.936328 |
| ENSMUSG( | Zfp457 | 13.92425 | -0.78614 | 0.119923 | NA | ENSMUSG( | Zfp457 | 14.31404 | -0.34872 | 0.493073 | 0.927358 |
| ENSMUSG( | Zfp458 | 60.78613 | -0.24806 | 0.407312 | 0.817447 | ENSMUSG( | Zfp458 | 65.30255 | 0.245333 | 0.394327 | 0.917257 |
| ENSMUSG( | Zfp459 | 56.53197 | 0.063298 | 0.811059 | 0.964318 | ENSMUSG( | Zfp459 | 54.32353 | 0.215529 | 0.497277 | 0.927825 |
| ENSMUSG( | Zfp46 | 2032.089 | -0.04069 | 0.715688 | 0.941484 | ENSMUSG( | Zfp46 | 1929.443 | 0.07751 | 0.502094 | 0.927825 |
| ENSMUSG( | Zfp462 | 1354.034 | -0.16382 | 0.119507 | 0.523326 | ENSMUSG( | Zfp462 | 1298.034 | -0.01499 | 0.900212 | 0.990558 |
| ENSMUSG( | Zfp467 | 979.9189 | 0.145298 | 0.139669 | 0.556069 | ENSMUSG( | Zfp467 | 928.3339 | 0.249758 | 0.030885 | 0.629448 |

| | | | | | |
|---|---|---|---|---|---|
| ENSMUSG( | 1776.598 | 0.024273 | 0.846483 | 0.972238 | Xpo1 |
| ENSMUSG( | 659.7202 | -0.03625 | 0.76628 | 0.954173 | Xpo4 |
| ENSMUSG( | 1934.204 | 0.1109 | 0.234563 | 0.681922 | Xpo5 |
| ENSMUSG( | 2691.393 | 0.085633 | 0.406731 | 0.816928 | Xpo6 |
| ENSMUSG( | 2199.89 | 0.022022 | 0.798169 | 0.96069 | Xpo7 |
| ENSMUSG( | 2851.48 | 0.083824 | 0.319762 | 0.765305 | Xpot |
| ENSMUSG( | 5004.807 | -0.12824 | 0.315412 | 0.762441 | Xpr1 |
| ENSMUSG( | 792.2922 | 0.07725 | 0.596203 | 0.903186 | Xrcc1 |
| ENSMUSG( | 261.3117 | 0.063766 | 0.67029 | 0.928525 | Xrcc2 |
| ENSMUSG( | 176.5409 | -0.06303 | 0.734777 | 0.945584 | Xrcc3 |
| ENSMUSG( | 225.1125 | -0.0548 | 0.7296 | 0.944764 | Xrcc4 |
| ENSMUSG( | 651.1532 | -0.10231 | 0.316301 | 0.763453 | Xrcc5 |
| ENSMUSG( | 578.586 | 0.111972 | 0.291104 | 0.741182 | Xrcc6 |
| ENSMUSG( | 842.8774 | -0.09113 | 0.368193 | 0.795774 | Xrn1 |
| ENSMUSG( | 1181.071 | -0.11544 | 0.239983 | 0.687848 | Xrn2 |
| ENSMUSG( | 42.45785 | -0.20267 | 0.513428 | 0.871762 | Xrra1 |
| ENSMUSG( | 878.7225 | -0.41827 | 0.002037 | 0.051756 | Xxylt1 |
| ENSMUSG( | 224.954 | 0.103012 | 0.567612 | 0.891774 | Xylb |
| ENSMUSG( | 2581.448 | -0.28314 | 0.041818 | 0.315339 | Xylt1 |
| ENSMUSG( | 1030.555 | -0.28378 | 0.065594 | 0.396799 | Xylt2 |
| ENSMUSG( | 880.4933 | 0.013633 | 0.901002 | 0.985633 | Yae1d1 |
| ENSMUSG( | 3237.423 | 0.069209 | 0.412219 | 0.819452 | Yaf2 |
| ENSMUSG( | 2414.066 | 0.179589 | 0.098099 | 0.479122 | Yap1 |
| ENSMUSG( | 2230.034 | 0.11375 | 0.335184 | 0.776511 | Yars1 |
| ENSMUSG( | 432.8861 | -0.03535 | 0.782241 | 0.957558 | Yars2 |
| ENSMUSG( | 137.6241 | 0.129592 | 0.461913 | 0.841728 | Ybey |
| ENSMUSG( | 9103.946 | -0.04524 | 0.611661 | 0.908586 | Ybx1 |
| ENSMUSG( | 109.2733 | 0.236201 | 0.392926 | 0.809274 | Ybx2 |
| ENSMUSG( | 969.9961 | -0.03484 | 0.736004 | 0.946267 | Ybx3 |
| ENSMUSG( | 417.6834 | 0.063375 | 0.619007 | 0.911432 | Ydjc |
| ENSMUSG( | 1360.485 | 0.022795 | 0.831699 | 0.970026 | Yeats2 |
| ENSMUSG( | 989.7163 | 0.010411 | 0.934402 | 0.99323 | Yeats4 |
| ENSMUSG( | 1089.58 | 0.02167 | 0.878939 | 0.981436 | Yes1 |
| ENSMUSG( | 578.326 | -0.04155 | 0.724566 | 0.944096 | Yif1a |
| ENSMUSG( | 1204.34 | -0.2779 | 0.01896 | 0.207297 | Yif1b |
| ENSMUSG( | 941.1707 | -0.28671 | 0.03137 | 0.269648 | Yipf1 |
| ENSMUSG( | 732.0379 | -0.19891 | 0.094546 | 0.471713 | Yipf2 |
| ENSMUSG( | 2232.534 | -0.28006 | 0.021694 | 0.225268 | Yipf3 |
| ENSMUSG( | 1061.983 | -0.43288 | 0.009487 | 0.137756 | Yipf4 |
| ENSMUSG( | 1382.862 | -0.29164 | 0.063051 | 0.388234 | Yipf5 |
| ENSMUSG( | 1541.031 | -0.1523 | 0.262701 | 0.713259 | Yipf6 |
| ENSMUSG( | 269.3969 | 0.035858 | 0.805035 | 0.9626 | Yjefn3 |
| ENSMUSG( | 231.7245 | -0.03803 | 0.802818 | 0.962169 | Yju2 |
| ENSMUSG( | 282.257 | -0.0165 | 0.915746 | 0.988634 | Yju2b |
| ENSMUSG( | 4034.448 | 0.040106 | 0.622928 | 0.911589 | Ykt6 |
| ENSMUSG( | 2533.483 | 0.07743 | 0.40963 | 0.818258 | Ylpm1 |
| ENSMUSG( | 1537.591 | 0.017132 | 0.865197 | 0.975565 | Yme1l1 |
| ENSMUSG( | 299.4168 | -0.03068 | 0.859268 | 0.974707 | Yod1 |
| ENSMUSG( | 1335.979 | 0.137044 | 0.146829 | 0.56887 | Ypel1 |
| ENSMUSG( | 1064.581 | -0.23971 | 0.015812 | 0.186254 | Ypel2 |
| ENSMUSG( | 5314.633 | 0.095859 | 0.383386 | 0.803427 | Ypel3 |
| ENSMUSG( | 1120.521 | 0.03516 | 0.839363 | 0.971179 | Ypel4 |
| ENSMUSG( | 2016.266 | -0.03234 | 0.795799 | 0.960062 | Ypel5 |
| ENSMUSG( | 360.4454 | -0.3475 | 0.012464 | 0.161344 | Yrdc |
| ENSMUSG( | 1685.043 | 0.005933 | 0.9521 | 0.993821 | Ythdc1 |
| ENSMUSG( | 519.922 | 0.126695 | 0.331078 | 0.77519 | Ythdc2 |
| ENSMUSG( | 1104.501 | 0.072129 | 0.528768 | 0.878365 | Ythdf1 |
| ENSMUSG( | 548.1396 | -0.08399 | 0.492488 | 0.860378 | Ythdf2 |
| ENSMUSG( | 1239.065 | -0.06691 | 0.511294 | 0.870859 | Ythdf3 |
| ENSMUSG( | 11158.35 | 0.048437 | 0.530176 | 0.879109 | Ywhab |
| ENSMUSG( | 23802.87 | 0.02118 | 0.826519 | 0.968372 | Ywhae |
| ENSMUSG( | 25036.39 | 0.138546 | 0.164466 | 0.595363 | Ywhag |
| ENSMUSG( | 10261.29 | 0.0046 | 0.972006 | 0.996585 | Ywhah |
| ENSMUSG( | 4623.081 | 0.021096 | 0.853759 | 0.973491 | Ywhaq |
| ENSMUSG( | 13114.34 | 0.003499 | 0.978052 | 0.997428 | Ywhaz |
| ENSMUSG( | 2246.418 | 0.068517 | 0.398907 | 0.812878 | Yy1 |
| ENSMUSG( | 18.46278 | -0.14571 | 0.769819 | NA | Zan |
| ENSMUSG( | 75.54408 | 0.183658 | 0.427161 | 0.82532 | Zap70 |
| ENSMUSG( | 32.92196 | 0.186258 | 0.620605 | 0.911432 | Zar1l |
| ENSMUSG( | 387.2984 | -0.04379 | 0.73935 | 0.947768 | Zbed3 |
| ENSMUSG( | 151.047 | -0.26897 | 0.111928 | 0.50438 | Zbed4 |
| ENSMUSG( | 115.8558 | -0.10165 | 0.612713 | 0.908844 | Zbed5 |
| ENSMUSG( | 21.09274 | -0.22444 | 0.578202 | NA | Zbed5-ps |
| ENSMUSG( | 415.4686 | -0.17273 | 0.418083 | 0.821966 | Zbed6 |
| ENSMUSG( | 15.2785 | -1.07071 | 0.032106 | NA | Zbp1 |
| ENSMUSG( | 425.0851 | -0.10785 | 0.349099 | 0.781665 | Zbtb1 |
| ENSMUSG( | 186.0091 | -0.16037 | 0.294857 | 0.743798 | Zbtb10 |
| ENSMUSG( | 732.1431 | 0.059299 | 0.582476 | 0.897918 | Zbtb11 |
| ENSMUSG( | 138.8029 | 0.051372 | 0.76887 | 0.954674 | Zbtb11os1 |
| ENSMUSG( | 232.5905 | -0.11118 | 0.471533 | 0.847331 | Zbtb12 |
| ENSMUSG( | 428.1322 | 0.063244 | 0.611689 | 0.908586 | Zbtb14 |
| ENSMUSG( | 878.6945 | 1.344914 | 1.51E-23 | 2.73E-20 | Zbtb16 |
| ENSMUSG( | 1167.547 | 0.062218 | 0.562646 | 0.889485 | Zbtb17 |
| ENSMUSG( | 7603.27 | 0.104291 | 0.184186 | 0.621101 | Zbtb18 |
| ENSMUSG( | 293.5644 | 0.137614 | 0.293396 | 0.742763 | Zbtb2 |
| ENSMUSG( | 6029.692 | 0.083886 | 0.550945 | 0.886038 | Zbtb20 |
| ENSMUSG( | 411.8857 | -0.07393 | 0.568214 | 0.892252 | Zbtb21 |
| ENSMUSG( | 891.9516 | 0.032193 | 0.775273 | 0.955606 | Zbtb22 |
| ENSMUSG( | 317.7263 | 0.024482 | 0.8579 | 0.974667 | Zbtb24 |
| ENSMUSG( | 238.4549 | -0.12507 | 0.432199 | 0.82768 | Zbtb25 |
| ENSMUSG( | 165.3895 | 0.071795 | 0.685903 | 0.932981 | Zbtb26 |
| ENSMUSG( | 64.52456 | 0.125147 | 0.671009 | 0.928781 | Zbtb3 |
| ENSMUSG( | 437.4564 | 0.030301 | 0.836303 | 0.971111 | Zbtb33 |
| ENSMUSG( | 400.8558 | 0.02943 | 0.810747 | 0.964272 | Zbtb34 |
| ENSMUSG( | 598.9996 | -0.20522 | 0.134554 | 0.547808 | Zbtb37 |
| ENSMUSG( | 753.3175 | -0.034 | 0.768303 | 0.954641 | Zbtb38 |
| ENSMUSG( | 348.9736 | -0.04824 | 0.708113 | 0.939519 | Zbtb39 |
| ENSMUSG( | 6028.978 | 0.139664 | 0.081246 | 0.440604 | Zbtb4 |
| ENSMUSG( | 424.994 | 0.551831 | 3.43E-05 | 0.002956 | Zbtb40 |
| ENSMUSG( | 1145.892 | -0.03218 | 0.832804 | 0.970333 | Zbtb41 |
| ENSMUSG( | 29.77021 | 0.275777 | 0.453793 | 0.836798 | Zbtb42 |
| ENSMUSG( | 688.7582 | -0.02148 | 0.855474 | 0.973813 | Zbtb43 |
| ENSMUSG( | 545.5994 | 0.098682 | 0.390141 | 0.80794 | Zbtb44 |
| ENSMUSG( | 748.3788 | 0.131528 | 0.292569 | 0.742521 | Zbtb45 |
| ENSMUSG( | 311.5869 | 0.342681 | 0.013494 | 0.169727 | Zbtb46 |
| ENSMUSG( | 7307.783 | 0.221413 | 0.008911 | 0.132538 | Zbtb47 |
| ENSMUSG( | 287.8644 | -0.12591 | 0.403787 | 0.816132 | Zbtb48 |
| ENSMUSG( | 238.1179 | -0.15188 | 0.37861 | 0.801688 | Zbtb49 |
| ENSMUSG( | 318.886 | -0.0966 | 0.442758 | 0.83278 | Zbtb5 |
| ENSMUSG( | 241.5373 | -0.1136 | 0.428922 | 0.826446 | Zbtb6 |
| ENSMUSG( | 3575.93 | 0.127675 | 0.184119 | 0.621101 | Zbtb7a |
| ENSMUSG( | 1301.528 | 0.148686 | 0.124631 | 0.531714 | Zbtb7b |
| ENSMUSG( | 201.3793 | 0.225463 | 0.184949 | 0.62251 | Zbtb7c |
| ENSMUSG( | 91.69808 | -0.59941 | 0.010575 | 0.147613 | Zbtb8a |
| ENSMUSG( | 77.36885 | 0.044241 | 0.850788 | 0.972638 | Zbtb8b |
| ENSMUSG( | 456.9884 | 0.282312 | 0.026096 | 0.247245 | Zbtb8os |
| ENSMUSG( | 21.61369 | 0.22638 | 0.595709 | 0.903152 | Zbtb9 |
| ENSMUSG( | 794.034 | 0.001634 | 0.989501 | 0.999111 | Zc2hc1a |
| ENSMUSG( | 51.12125 | 0.221289 | 0.420593 | 0.8229 | Zc2hc1c |
| ENSMUSG( | 293.6023 | -0.06045 | 0.655871 | 0.923929 | Zc3h10 |
| ENSMUSG( | 157.8721 | 0.002999 | 0.987527 | 0.998796 | Zc3h11a |
| ENSMUSG( | 47.26298 | 0.071268 | 0.813491 | 0.964574 | Zc3h12a |
| ENSMUSG( | 261.5594 | 0.065595 | 0.664778 | 0.926824 | Zc3h12b |
| ENSMUSG( | 247.646 | -0.07038 | 0.619193 | 0.911432 | Zc3h12c |
| ENSMUSG( | 2237.157 | -0.04668 | 0.7299 | 0.944956 | Zc3h13 |
| ENSMUSG( | 2419.777 | 0.051239 | 0.525544 | 0.876728 | Zc3h14 |
| ENSMUSG( | 1656.247 | 0.127533 | 0.217568 | 0.662824 | Zc3h15 |
| ENSMUSG( | 1259.2 | 0.104442 | 0.289378 | 0.739839 | Zc3h18 |
| ENSMUSG( | 262.6473 | 0.364036 | 0.032884 | 0.277278 | Zc3h3 |
| ENSMUSG( | 2175.925 | 0.045487 | 0.596103 | 0.903152 | Zc3h4 |
| ENSMUSG( | 181.185 | -0.35329 | 0.036692 | 0.295649 | Zc3h6 |
| ENSMUSG( | 1235.854 | -0.02019 | 0.848574 | 0.972374 | Zc3h7a |
| ENSMUSG( | 21303.1 | 0.166202 | 0.042911 | 0.317818 | Zc3h7b |
| ENSMUSG( | 134.0041 | -0.0894 | 0.650999 | 0.921451 | Zc3h8 |
| ENSMUSG( | 319.6076 | -0.0874 | 0.600881 | 0.904492 | Zc3hav1 |
| ENSMUSG( | 99.56511 | -0.00904 | 0.965503 | 0.995377 | Zc3hav1l |
| ENSMUSG( | 479.3558 | 0.163491 | 0.286074 | 0.735242 | Zc3hc1 |
| ENSMUSG( | 390.3222 | 0.00903 | 0.945979 | 0.99369 | Zc4h2 |
| ENSMUSG( | 109.3057 | 0.297438 | 0.140647 | 0.557902 | Zcchc10 |
| ENSMUSG( | 895.5168 | -0.27684 | 0.155897 | 0.58375 | Zcchc12 |
| ENSMUSG( | 2832.096 | 0.133713 | 0.135677 | 0.549211 | Zcchc14 |
| ENSMUSG( | 1926.415 | -0.02799 | 0.775928 | 0.955606 | Zcchc17 |
| ENSMUSG( | 5062.938 | -0.009 | 0.932923 | 0.992879 | Zcchc18 |
| ENSMUSG( | 905.1391 | 0.132791 | 0.199448 | 0.640054 | Zcchc2 |
| ENSMUSG( | 11909.21 | 0.06558 | 0.529299 | 0.878733 | Zcchc24 |
| ENSMUSG( | 218.8272 | -0.02092 | 0.895163 | 0.984512 | Zcchc3 |
| ENSMUSG( | 277.8661 | 0.073367 | 0.579089 | 0.896692 | Zcchc4 |
| ENSMUSG( | 1444.636 | 0.095025 | 0.409147 | 0.818258 | Zcchc7 |
| ENSMUSG( | 923.0034 | 0.097819 | 0.378724 | 0.801688 | Zcchc8 |
| ENSMUSG( | 219.2685 | 0.058085 | 0.69877 | 0.936563 | Zcchc9 |
| ENSMUSG( | 1536.941 | -0.01312 | 0.884441 | 0.982647 | Zcrb1 |
| ENSMUSG( | 240.8784 | -0.01242 | 0.933599 | 0.993194 | Zcwpw1 |
| ENSMUSG( | 20.81335 | 0.120345 | 0.77547 | NA | Zcwpw2 |
| ENSMUSG( | 503.0923 | 0.01291 | 0.927237 | 0.991229 | Zdbf2 |
| ENSMUSG( | 1685.441 | 0.023906 | 0.811296 | 0.964398 | Zdhhc1 |
| ENSMUSG( | 267.2717 | -0.43537 | 0.005457 | 0.098899 | Zdhhc12 |
| ENSMUSG( | 579.6708 | -0.22575 | 0.122446 | 0.528223 | Zdhhc13 |
| ENSMUSG( | 1412.531 | -0.07262 | 0.609891 | 0.907813 | Zdhhc14 |

| | | | | | |
|---|---|---|---|---|---|
| ENSMUSG( | 128.5392 | 0.06816 | 0.73006 | 0.967228 | Zc3h8 |
| ENSMUSG( | 301.4965 | 0.013591 | 0.932601 | 0.99306 | Zc3hav1 |
| ENSMUSG( | 98.37537 | 0.215057 | 0.357217 | 0.913692 | Zc3hav1l |
| ENSMUSG( | 396.1828 | -0.11075 | 0.530093 | 0.934795 | Zc3hc1 |
| ENSMUSG( | 336.1637 | -0.17141 | 0.345529 | 0.910581 | Zc4h2 |
| ENSMUSG( | 91.88122 | 0.070498 | 0.766557 | 0.974236 | Zcchc10 |
| ENSMUSG( | 828.0105 | -0.23802 | 0.32937 | 0.907083 | Zcchc12 |
| ENSMUSG( | 2607.278 | 0.163915 | 0.132831 | 0.822912 | Zcchc14 |
| ENSMUSG( | 1653.62 | -0.21089 | 0.08857 | 0.786756 | Zcchc17 |
| ENSMUSG( | 4611.546 | -0.01137 | 0.92786 | 0.99306 | Zcchc18 |
| ENSMUSG( | 810.3342 | 0.080721 | 0.516158 | 0.930939 | Zcchc2 |
| ENSMUSG( | 10181.84 | -0.12112 | 0.347802 | 0.911652 | Zcchc24 |
| ENSMUSG( | 203.6911 | 0.051435 | 0.779297 | 0.976106 | Zcchc3 |
| ENSMUSG( | 257.7785 | 0.122782 | 0.424822 | 0.917257 | Zcchc4 |
| ENSMUSG( | 1357.956 | 0.180577 | 0.147106 | 0.836241 | Zcchc7 |
| ENSMUSG( | 884.2877 | 0.236588 | 0.076709 | 0.770558 | Zcchc8 |
| ENSMUSG( | 220.9195 | 0.328817 | 0.04828 | 0.706142 | Zcchc9 |
| ENSMUSG( | 1447.345 | 0.078561 | 0.504068 | 0.927825 | Zcrb1 |
| ENSMUSG( | 212.932 | -0.10839 | 0.52525 | 0.934374 | Zcwpw1 |
| ENSMUSG( | 19.2153 | 0.154475 | 0.728294 | 0.966913 | Zcwpw2 |
| ENSMUSG( | 444.6257 | -0.08056 | 0.55609 | 0.939565 | Zdbf2 |
| ENSMUSG( | 1574.151 | 0.095888 | 0.382784 | 0.917257 | Zdhhc1 |
| ENSMUSG( | 302.958 | 0.22515 | 0.186399 | 0.855939 | Zdhhc12 |
| ENSMUSG( | 593.6945 | 0.125688 | 0.492979 | 0.927358 | Zdhhc13 |
| ENSMUSG( | 1513.228 | 0.376839 | 0.02478 | 0.592568 | Zdhhc14 |
| ENSMUSG( | 347.0179 | -0.00701 | 0.972668 | 0.996416 | Zdhhc15 |
| ENSMUSG( | 1118.57 | 0.133327 | 0.353119 | 0.912263 | Zdhhc16 |
| ENSMUSG( | 3041.004 | 0.221649 | 0.15212 | 0.837724 | Zdhhc17 |
| ENSMUSG( | 1909.475 | 0.054137 | 0.675909 | 0.95694 | Zdhhc18 |
| ENSMUSG( | 1905.409 | 0.061363 | 0.683818 | 0.958678 | Zdhhc2 |
| ENSMUSG( | 1208.694 | 0.386902 | 0.022626 | 0.581156 | Zdhhc20 |
| ENSMUSG( | 1476.833 | 0.179898 | 0.215973 | 0.865167 | Zdhhc21 |
| ENSMUSG( | 1523.015 | 0.126043 | 0.421319 | 0.917257 | Zdhhc22 |
| ENSMUSG( | 1150.978 | 0.226964 | 0.105677 | 0.804969 | Zdhhc23 |
| ENSMUSG( | 701.8191 | 0.096253 | 0.449843 | 0.922202 | Zdhhc24 |
| ENSMUSG( | 3715.995 | 0.007054 | 0.948033 | 0.994798 | Zdhhc3 |
| ENSMUSG( | 1016.058 | 0.012171 | 0.913644 | 0.992378 | Zdhhc4 |
| ENSMUSG( | 1558.187 | 0.386573 | 0.032355 | 0.635121 | Zdhhc5 |
| ENSMUSG( | 1021.926 | 0.110584 | 0.421929 | 0.917257 | Zdhhc6 |
| ENSMUSG( | 1085.235 | 0.140795 | 0.302757 | 0.898538 | Zdhhc7 |
| ENSMUSG( | 4815.04 | 0.143311 | 0.377529 | 0.91645 | Zdhhc8 |
| ENSMUSG( | 3156.133 | 0.146374 | 0.415707 | 0.917257 | Zdhhc9 |
| ENSMUSG( | 6207.854 | -0.21471 | 0.109285 | 0.805353 | Zeb1 |
| ENSMUSG( | 173.8521 | 0.038331 | 0.827394 | 0.980224 | Zeb1os1 |
| ENSMUSG( | 5287.429 | -0.05305 | 0.684003 | 0.958678 | Zeb2 |
| ENSMUSG( | 208.0117 | -0.25175 | 0.180487 | 0.850372 | Zeb2os |
| ENSMUSG( | 4851.038 | 0.031153 | 0.803211 | 0.976937 | Zer1 |
| ENSMUSG( | 522.4676 | -0.07912 | 0.541189 | 0.937687 | Zfand1 |
| ENSMUSG( | 2984.563 | -0.0266 | 0.792062 | 0.97681 | Zfand2a |
| ENSMUSG( | 354.1105 | -0.13568 | 0.393461 | 0.917257 | Zfand2b |
| ENSMUSG( | 4464.383 | -0.10907 | 0.395991 | 0.917257 | Zfand3 |
| ENSMUSG( | 104.4647 | 0.060031 | 0.790894 | 0.97681 | Zfand4 |
| ENSMUSG( | 2396.657 | 0.013841 | 0.920475 | 0.992644 | Zfand5 |
| ENSMUSG( | 852.4106 | 0.034067 | 0.789463 | 0.976593 | Zfand6 |
| ENSMUSG( | 1137.93 | -0.07529 | 0.552304 | 0.939565 | Zfas1 |
| ENSMUSG( | 143.12 | -0.04746 | 0.80241 | 0.97681 | Zfat |
| ENSMUSG( | 1240.636 | 0.413741 | 0.027975 | 0.61023 | Zfc3h1 |
| ENSMUSG( | 2481.262 | 0.135499 | 0.265763 | 0.886134 | Zfhx2 |
| ENSMUSG( | 208.176 | 0.323952 | 0.048511 | 0.706142 | Zfhx3 |
| ENSMUSG( | 286.8299 | 0.269175 | 0.068771 | 0.753052 | Zfhx4 |
| ENSMUSG( | 406.2037 | -0.03226 | 0.815759 | 0.979198 | Zfp1 |
| ENSMUSG( | 14.08623 | 0.492022 | 0.340209 | 0.907972 | Zfp1001 |
| ENSMUSG( | 299.1705 | 0.204674 | 0.193694 | 0.860096 | Zfp1006 |
| ENSMUSG( | 36.50291 | 0.308416 | 0.369047 | 0.915491 | Zfp1007 |
| ENSMUSG( | 145.2178 | -0.10933 | 0.664995 | 0.956766 | Zfp1008 |
| ENSMUSG( | 265.0945 | 0.247453 | 0.190631 | 0.857801 | Zfp101 |
| ENSMUSG( | 153.192 | -0.31314 | 0.105032 | 0.804969 | Zfp105 |
| ENSMUSG( | 12955.77 | -0.11622 | 0.27622 | 0.890749 | Zfp106 |
| ENSMUSG( | 122.7005 | 0.139916 | 0.504698 | 0.927825 | Zfp108 |
| ENSMUSG( | 92.03925 | -0.09552 | 0.66935 | 0.956766 | Zfp109 |
| ENSMUSG( | 108.0308 | -0.11803 | 0.613367 | 0.951177 | Zfp11 |
| ENSMUSG( | 300.6223 | 0.091255 | 0.639165 | 0.953903 | Zfp110 |
| ENSMUSG( | 168.5957 | 0.126261 | 0.484303 | 0.92541 | Zfp111 |
| ENSMUSG( | 225.9607 | 0.079583 | 0.660371 | 0.956205 | Zfp112 |
| ENSMUSG( | 167.3057 | 0.065709 | 0.726181 | 0.966887 | Zfp113 |
| ENSMUSG( | 65.91927 | -0.25333 | 0.352589 | 0.912093 | Zfp114 |
| ENSMUSG( | 63.58019 | 0.062198 | 0.823465 | 0.979198 | Zfp119a |
| ENSMUSG( | 58.0006 | 0.400819 | 0.152001 | 0.837724 | Zfp119b |
| ENSMUSG( | 712.9518 | 0.154901 | 0.209154 | 0.864412 | Zfp12 |
| ENSMUSG( | 289.4941 | 0.167837 | 0.248555 | 0.877904 | Zfp120 |
| ENSMUSG( | 107.7987 | 0.070182 | 0.747377 | 0.969763 | Zfp128 |
| ENSMUSG( | 543.8448 | 0.091635 | 0.527748 | 0.934603 | Zfp13 |
| ENSMUSG( | 1372.259 | -0.0066 | 0.954725 | 0.994798 | Zfp131 |
| ENSMUSG( | 295.1471 | 0.001412 | 0.993637 | 0.998813 | Zfp14 |
| ENSMUSG( | 272.751 | -0.04576 | 0.762236 | 0.973255 | Zfp141 |
| ENSMUSG( | 1041.478 | 0.033535 | 0.770161 | 0.974236 | Zfp142 |
| ENSMUSG( | 221.7358 | 0.121931 | 0.459702 | 0.923329 | Zfp143 |
| ENSMUSG( | 1050.66 | -0.04391 | 0.713296 | 0.964766 | Zfp146 |
| ENSMUSG( | 1826.308 | 0.078657 | 0.492588 | 0.927315 | Zfp148 |
| ENSMUSG( | 577.4439 | -0.0407 | 0.760531 | 0.972688 | Zfp157 |
| ENSMUSG( | 468.0773 | 0.24933 | 0.069584 | 0.75601 | Zfp160 |
| ENSMUSG( | 158.6683 | 0.176689 | 0.36666 | 0.914703 | Zfp169 |
| ENSMUSG( | 106.0666 | -0.08891 | 0.690536 | 0.960071 | Zfp174 |
| ENSMUSG( | 1357.13 | 0.142302 | 0.197504 | 0.862161 | Zfp180 |
| ENSMUSG( | 296.0985 | 0.14602 | 0.325872 | 0.905339 | Zfp182 |
| ENSMUSG( | 52.6818 | 0.327074 | 0.294143 | 0.898538 | Zfp184 |
| ENSMUSG( | 66.97687 | 0.080337 | 0.782446 | 0.976106 | Zfp185 |
| ENSMUSG( | 135.8954 | -0.42288 | 0.056261 | 0.723362 | Zfp189 |
| ENSMUSG( | 434.9317 | 0.153651 | 0.269596 | 0.887508 | Zfp2 |
| ENSMUSG( | 124.6848 | 0.030947 | 0.878749 | 0.989644 | Zfp202 |
| ENSMUSG( | 1970.328 | 0.217079 | 0.073448 | 0.763629 | Zfp207 |
| ENSMUSG( | 333.3597 | 0.089669 | 0.564468 | 0.941206 | Zfp212 |
| ENSMUSG( | 165.9879 | 0.066096 | 0.73353 | 0.967228 | Zfp213 |
| ENSMUSG( | 202.0189 | 0.276081 | 0.114598 | 0.811703 | Zfp217 |
| ENSMUSG( | 1962.371 | -0.08507 | 0.43013 | 0.917257 | Zfp219 |
| ENSMUSG( | 101.4759 | 0.236594 | 0.322771 | 0.904023 | Zfp229 |
| ENSMUSG( | 423.6368 | 0.037712 | 0.814024 | 0.979198 | Zfp235 |
| ENSMUSG( | 1634.8 | 0.201835 | 0.085088 | 0.779799 | Zfp236 |
| ENSMUSG( | 671.1599 | 0.025696 | 0.844234 | 0.983576 | Zfp239 |
| ENSMUSG( | 1461.799 | 0.027833 | 0.837425 | 0.981335 | Zfp24 |
| ENSMUSG( | 212.5358 | 0.060471 | 0.72045 | 0.966887 | Zfp248 |
| ENSMUSG( | 1352.607 | 0.082118 | 0.483378 | 0.92499 | Zfp251 |
| ENSMUSG( | 576.5267 | 0.115507 | 0.404187 | 0.917257 | Zfp26 |
| ENSMUSG( | 1091.002 | 0.144904 | 0.244524 | 0.877904 | Zfp260 |
| ENSMUSG( | 504.3617 | 0.125922 | 0.431414 | 0.917257 | Zfp263 |
| ENSMUSG( | 790.3558 | 0.117101 | 0.351661 | 0.912093 | Zfp266 |
| ENSMUSG( | 462.3884 | -0.02786 | 0.835585 | 0.981237 | Zfp267 |
| ENSMUSG( | 362.9387 | 0.070303 | 0.637125 | 0.953903 | Zfp27 |
| ENSMUSG( | 106.3796 | -0.00046 | 0.998385 | 0.999313 | Zfp273 |
| ENSMUSG( | 776.0886 | 0.09403 | 0.454037 | 0.922444 | Zfp275 |
| ENSMUSG( | 810.2682 | 0.096651 | 0.59356 | 0.945408 | Zfp276 |
| ENSMUSG( | 391.0153 | -0.22703 | 0.105371 | 0.804969 | Zfp277 |
| ENSMUSG( | 218.83 | -0.06898 | 0.685383 | 0.958678 | Zfp28 |
| ENSMUSG( | 257.5452 | 0.12176 | 0.471894 | 0.923329 | Zfp280b |
| ENSMUSG( | 507.0171 | 0.146923 | 0.334841 | 0.907907 | Zfp280c |
| ENSMUSG( | 757.7748 | 0.058236 | 0.641193 | 0.953903 | Zfp280d |
| ENSMUSG( | 574.7572 | 0.210052 | 0.122787 | 0.814249 | Zfp281 |
| ENSMUSG( | 518.734 | 0.056205 | 0.697429 | 0.960867 | Zfp282 |
| ENSMUSG( | 311.1936 | -0.10055 | 0.508881 | 0.929089 | Zfp286 |
| ENSMUSG( | 324.3551 | -0.09314 | 0.542924 | 0.938063 | Zfp287 |
| ENSMUSG( | 850.6014 | -0.02784 | 0.812839 | 0.979148 | Zfp292 |
| ENSMUSG( | 34.12942 | 0.496193 | 0.158576 | 0.837724 | Zfp296 |
| ENSMUSG( | 266.7145 | -0.11734 | 0.456685 | 0.923329 | Zfp3 |
| ENSMUSG( | 346.4061 | 0.155879 | 0.261061 | 0.884926 | Zfp30 |
| ENSMUSG( | 41.51251 | 0.24189 | 0.486917 | 0.92541 | Zfp300 |
| ENSMUSG( | 1085.316 | -0.06723 | 0.555976 | 0.939565 | Zfp316 |
| ENSMUSG( | 759.7717 | -0.01678 | 0.901587 | 0.990784 | Zfp317 |
| ENSMUSG( | 1474.375 | 0.04384 | 0.697225 | 0.960867 | Zfp318 |
| ENSMUSG( | 627.2959 | 0.105236 | 0.453744 | 0.922316 | Zfp319 |
| ENSMUSG( | 520.02 | 0.206156 | 0.172285 | 0.843466 | Zfp322a |
| ENSMUSG( | 221.6234 | 0.156724 | 0.379418 | 0.916995 | Zfp324 |
| ENSMUSG( | 601.2517 | 0.081289 | 0.53274 | 0.934795 | Zfp326 |
| ENSMUSG( | 517.6257 | 0.132809 | 0.293449 | 0.898538 | Zfp329 |
| ENSMUSG( | 559.8918 | 0.113587 | 0.441151 | 0.921166 | Zfp330 |
| ENSMUSG( | 538.0689 | 0.085366 | 0.552417 | 0.939565 | Zfp334 |
| ENSMUSG( | 1308.013 | 0.158846 | 0.196195 | 0.862161 | Zfp335 |
| ENSMUSG( | 71.82842 | 0.14325 | 0.603541 | 0.948137 | Zfp335os |
| ENSMUSG( | 226.8923 | 0.153743 | 0.396117 | 0.917257 | Zfp341 |
| ENSMUSG( | 866.3104 | 0.049102 | 0.697011 | 0.960867 | Zfp346 |
| ENSMUSG( | 364.7355 | 0.047902 | 0.745697 | 0.96951 | Zfp35 |
| ENSMUSG( | 206.4813 | 0.313173 | 0.088314 | 0.786756 | Zfp354a |
| ENSMUSG( | 66.92715 | -0.09075 | 0.744624 | 0.96951 | Zfp354b |
| ENSMUSG( | 1151.611 | 0.034655 | 0.802608 | 0.97681 | Zfp354c |
| ENSMUSG( | 1050.335 | 0.089492 | 0.449677 | 0.922202 | Zfp358 |
| ENSMUSG( | 163.8755 | 0.252957 | 0.227901 | 0.868507 | Zfp36 |
| ENSMUSG( | 1051.683 | -0.1086 | 0.369814 | 0.915491 | Zfp362 |
| ENSMUSG( | 31827.71 | 0.020363 | 0.85099 | 0.985541 | Zfp365 |
| ENSMUSG( | 248.5118 | 0.866378 | 0.003178 | 0.224467 | Zfp366 |

| | | | | | | | | | | | |
|---|---|---|---|---|---|---|---|---|---|---|---|
| ENSMUSG( | Zfp469 | 204.0971 | -0.71789 | 6.78E-05 | 0.005 | ENSMUSG( | Zfp469 | 227.5456 | -0.0341 | 0.862708 | 0.986547 |
| ENSMUSG( | Zfp472 | 82.49312 | 0.299944 | 0.192238 | 0.631194 | ENSMUSG( | Zfp472 | 71.64307 | 0.184531 | 0.454193 | 0.922444 |
| ENSMUSG( | Zfp473 | 15.753 | -0.8292 | 0.115882 | NA | ENSMUSG( | Zfp473 | 15.38281 | -0.57418 | 0.25985 | 0.883971 |
| ENSMUSG( | Zfp474 | 12.6437 | -0.56829 | 0.283179 | NA | ENSMUSG( | Zfp474 | 13.63389 | 0.020695 | 0.969709 | 0.99594 |
| ENSMUSG( | Zfp488 | 201.1406 | -0.08071 | 0.593873 | 0.902869 | ENSMUSG( | Zfp488 | 192.3186 | 0.069482 | 0.697725 | 0.960867 |
| ENSMUSG( | Zfp493 | 67.15228 | 0.303553 | 0.189371 | 0.627932 | ENSMUSG( | Zfp493 | 57.29872 | 0.129216 | 0.647826 | 0.95422 |
| ENSMUSG( | Zfp503 | 74.31229 | -0.39746 | 0.152208 | 0.578042 | ENSMUSG( | Zfp503 | 89.31862 | 0.393812 | 0.13961 | 0.830678 |
| ENSMUSG( | Zfp507 | 548.1134 | -0.00654 | 0.956252 | 0.993874 | ENSMUSG( | Zfp507 | 492.344 | -0.0399 | 0.779707 | 0.976106 |
| ENSMUSG( | Zfp51 | 91.47198 | -0.10005 | 0.651208 | 0.921451 | ENSMUSG( | Zfp51 | 89.17329 | 0.106701 | 0.646068 | 0.953988 |
| ENSMUSG( | Zfp511 | 459.5985 | 0.083016 | 0.488492 | 0.859092 | ENSMUSG( | Zfp511 | 415.7872 | 0.067454 | 0.6358 | 0.953903 |
| ENSMUSG( | Zfp512 | 1375.889 | 0.141309 | 0.129133 | 0.538324 | ENSMUSG( | Zfp512 | 1161.086 | -0.08095 | 0.495968 | 0.927825 |
| ENSMUSG( | Zfp512b | 2538.798 | 0.090575 | 0.437152 | 0.829501 | ENSMUSG( | Zfp512b | 2225.465 | -0.02156 | 0.869623 | 0.988097 |
| ENSMUSG( | Zfp513 | 813.9625 | 0.056534 | 0.580354 | 0.896952 | ENSMUSG( | Zfp513 | 742.9943 | 0.06268 | 0.611817 | 0.95083 |
| ENSMUSG( | Zfp516 | 754.3463 | -0.1103 | 0.377721 | 0.801561 | ENSMUSG( | Zfp516 | 671.152 | -0.18625 | 0.185791 | 0.855305 |
| ENSMUSG( | Zfp518a | 286.4499 | -0.12363 | 0.385247 | 0.804048 | ENSMUSG( | Zfp518a | 283.9492 | 0.128413 | 0.446457 | 0.921734 |
| ENSMUSG( | Zfp518b | 351.1216 | -0.0897 | 0.517576 | 0.873397 | ENSMUSG( | Zfp518b | 323.0001 | -0.06616 | 0.657157 | 0.955644 |
| ENSMUSG( | Zfp52 | 101.5943 | -0.25575 | 0.186471 | 0.623869 | ENSMUSG( | Zfp52 | 99.36636 | -0.04116 | 0.850546 | 0.985541 |
| ENSMUSG( | Zfp521 | 770.1532 | -0.00748 | 0.955976 | 0.993874 | ENSMUSG( | Zfp521 | 662.641 | -0.17979 | 0.204248 | 0.864412 |
| ENSMUSG( | Zfp523 | 6521.751 | 0.167319 | 0.048964 | 0.337919 | ENSMUSG( | Zfp523 | 5797.466 | 0.099499 | 0.306753 | 0.898538 |
| ENSMUSG( | Zfp524 | 61.57724 | 0.307348 | 0.262526 | 0.713142 | ENSMUSG( | Zfp524 | 51.39929 | 0.046785 | 0.882234 | 0.989644 |
| ENSMUSG( | Zfp526 | 211.9892 | 0.09874 | 0.514467 | 0.872055 | ENSMUSG( | Zfp526 | 196.0357 | 0.154326 | 0.413563 | 0.917257 |
| ENSMUSG( | Zfp53 | 77.82223 | 0.1592 | 0.496602 | 0.86175 | ENSMUSG( | Zfp53 | 73.24239 | 0.254405 | 0.350286 | 0.911652 |
| ENSMUSG( | Zfp532 | 1761.42 | -0.04735 | 0.607754 | 0.906878 | ENSMUSG( | Zfp532 | 1564.541 | -0.12177 | 0.267712 | 0.886134 |
| ENSMUSG( | Zfp536 | 533.6566 | -0.02116 | 0.850113 | 0.972638 | ENSMUSG( | Zfp536 | 503.5746 | 0.075502 | 0.587242 | 0.944975 |
| ENSMUSG( | Zfp54 | 29.9796 | -0.11671 | 0.73832 | 0.947329 | ENSMUSG( | Zfp54 | 26.50805 | -0.21705 | 0.574875 | 0.942912 |
| ENSMUSG( | Zfp541 | 21.93058 | 0.004343 | 0.991592 | 0.999232 | ENSMUSG( | Zfp541 | 20.42952 | 0.039109 | 0.927728 | 0.99306 |
| ENSMUSG( | Zfp551 | 39.14606 | 0.403823 | 0.230858 | 0.679066 | ENSMUSG( | Zfp551 | 36.28829 | 0.481109 | 0.227674 | 0.868507 |
| ENSMUSG( | Zfp553 | 431.4243 | -0.12035 | 0.422705 | 0.823527 | ENSMUSG( | Zfp553 | 394.8766 | -0.10281 | 0.501776 | 0.927825 |
| ENSMUSG( | Zfp558 | 48.64464 | 0.166497 | 0.546238 | 0.884974 | ENSMUSG( | Zfp558 | 41.19736 | -0.03562 | 0.909812 | 0.991958 |
| ENSMUSG( | Zfp560 | 269.0207 | -0.08718 | 0.517341 | 0.873295 | ENSMUSG( | Zfp560 | 230.3768 | -0.28305 | 0.07711 | 0.770558 |
| ENSMUSG( | Zfp563 | 287.3424 | 0.105827 | 0.505294 | 0.866883 | ENSMUSG( | Zfp563 | 266.8727 | 0.148234 | 0.436546 | 0.919879 |
| ENSMUSG( | Zfp566 | 86.03148 | -0.26781 | 0.303102 | 0.751443 | ENSMUSG( | Zfp566 | 82.58485 | -0.09065 | 0.739335 | 0.968458 |
| ENSMUSG( | Zfp568 | 119.667 | -0.05534 | 0.783037 | 0.957558 | ENSMUSG( | Zfp568 | 108.4418 | -0.05143 | 0.829531 | 0.980224 |
| ENSMUSG( | Zfp57 | 366.6327 | 0.069315 | 0.648399 | 0.919965 | ENSMUSG( | Zfp57 | 340.5385 | 0.126755 | 0.439628 | 0.920721 |
| ENSMUSG( | Zfp574 | 965.2709 | 0.000674 | 0.994995 | 0.99943 | ENSMUSG( | Zfp574 | 861.3997 | -0.05744 | 0.638918 | 0.953903 |
| ENSMUSG( | Zfp575 | 263.4251 | -0.10047 | 0.503655 | 0.866308 | ENSMUSG( | Zfp575 | 229.8051 | -0.24074 | 0.192198 | 0.859726 |
| ENSMUSG( | Zfp579 | 589.5476 | 0.130495 | 0.252829 | 0.703255 | ENSMUSG( | Zfp579 | 534.3793 | 0.116646 | 0.449677 | 0.922202 |
| ENSMUSG( | Zfp58 | 72.50337 | -0.1548 | 0.559271 | 0.888638 | ENSMUSG( | Zfp58 | 68.72015 | -0.0378 | 0.883731 | 0.989644 |
| ENSMUSG( | Zfp580 | 751.3001 | -0.06534 | 0.513115 | 0.871762 | ENSMUSG( | Zfp580 | 639.0574 | -0.28168 | 0.038941 | 0.679972 |
| ENSMUSG( | Zfp582 | 103.5598 | 0.349914 | 0.132434 | 0.544369 | ENSMUSG( | Zfp582 | 99.1058 | 0.452436 | 0.096476 | 0.796133 |
| ENSMUSG( | Zfp583 | 123.323 | 0.145323 | 0.42558 | 0.824898 | ENSMUSG( | Zfp583 | 106.0763 | -0.0127 | 0.952972 | 0.994798 |
| ENSMUSG( | Zfp59 | 130.7149 | -0.19506 | 0.293329 | 0.742763 | ENSMUSG( | Zfp59 | 124.2943 | -0.0502 | 0.81953 | 0.979198 |
| ENSMUSG( | Zfp592 | 1158.119 | 0.086812 | 0.348526 | 0.78132 | ENSMUSG( | Zfp592 | 1093.16 | 0.184477 | 0.094279 | 0.795735 |
| ENSMUSG( | Zfp593 | 246.415 | 0.123908 | 0.410177 | 0.818258 | ENSMUSG( | Zfp593 | 206.4627 | -0.123 | 0.492167 | 0.927315 |
| ENSMUSG( | Zfp595 | 96.05183 | 0.071761 | 0.74679 | 0.949289 | ENSMUSG( | Zfp595 | 96.52068 | 0.329103 | 0.168061 | 0.840222 |
| ENSMUSG( | Zfp597 | 260.5256 | -0.09758 | 0.484211 | 0.856577 | ENSMUSG( | Zfp597 | 259.4163 | 0.164083 | 0.300077 | 0.898538 |
| ENSMUSG( | Zfp598 | 1439.891 | 0.200181 | 0.097369 | 0.477629 | ENSMUSG( | Zfp598 | 1313.916 | 0.2041 | 0.161633 | 0.837724 |
| ENSMUSG( | Zfp599 | 54.80962 | -0.07286 | 0.785051 | 0.958473 | ENSMUSG( | Zfp599 | 52.57836 | 0.095978 | 0.744627 | 0.96951 |
| ENSMUSG( | Zfp60 | 774.5585 | -0.02854 | 0.783191 | 0.957558 | ENSMUSG( | Zfp60 | 713.3262 | -0.00263 | 0.984486 | 0.996963 |
| ENSMUSG( | Zfp605 | 335.6208 | 0.105582 | 0.446529 | 0.833726 | ENSMUSG( | Zfp605 | 302.5112 | 0.067203 | 0.678204 | 0.957435 |
| ENSMUSG( | Zfp606 | 448.2568 | -0.05913 | 0.620222 | 0.911432 | ENSMUSG( | Zfp606 | 430.3509 | 0.087928 | 0.531877 | 0.934795 |
| ENSMUSG( | Zfp607a | 29.95055 | -0.19508 | 0.579263 | 0.896692 | ENSMUSG( | Zfp607a | 30.36566 | 0.131072 | 0.72608 | 0.966887 |
| ENSMUSG( | Zfp607b | 411.3892 | 0.024115 | 0.85989 | 0.974818 | ENSMUSG( | Zfp607b | 351.67 | -0.17674 | 0.319195 | 0.903431 |
| ENSMUSG( | Zfp608 | 765.7021 | -0.02679 | 0.823744 | 0.966603 | ENSMUSG( | Zfp608 | 750.6822 | 0.176288 | 0.186414 | 0.855939 |
| ENSMUSG( | Zfp609 | 1738.182 | -0.05442 | 0.59915 | 0.903801 | ENSMUSG( | Zfp609 | 1539.227 | -0.13934 | 0.227748 | 0.868507 |
| ENSMUSG( | Zfp61 | 354.5416 | -0.08996 | 0.590372 | 0.90113 | ENSMUSG( | Zfp61 | 350.6833 | 0.147203 | 0.397008 | 0.917257 |
| ENSMUSG( | Zfp612 | 1164.251 | -0.15723 | 0.141142 | 0.558043 | ENSMUSG( | Zfp612 | 1063.816 | -0.15232 | 0.235471 | 0.871807 |
| ENSMUSG( | Zfp617 | 541.781 | -0.06414 | 0.564212 | 0.890139 | ENSMUSG( | Zfp617 | 501.7442 | -0.02194 | 0.876188 | 0.989644 |
| ENSMUSG( | Zfp618 | 169.4772 | -0.29975 | 0.067768 | 0.402638 | ENSMUSG( | Zfp618 | 149.99 | -0.39804 | 0.041889 | 0.690304 |
| ENSMUSG( | Zfp619 | 110.4338 | -0.12609 | 0.503549 | 0.866217 | ENSMUSG( | Zfp619 | 106.0945 | 0.042466 | 0.84246 | 0.983044 |
| ENSMUSG( | Zfp62 | 975.4054 | -0.09391 | 0.405651 | 0.81652 | ENSMUSG( | Zfp62 | 950.9464 | 0.101728 | 0.42648 | 0.917257 |
| ENSMUSG( | Zfp622 | 925.1379 | 0.125315 | 0.275389 | 0.72513 | ENSMUSG( | Zfp622 | 808.8906 | 0.005519 | 0.968865 | 0.99594 |
| ENSMUSG( | Zfp623 | 184.5651 | -0.25696 | 0.132122 | 0.544244 | ENSMUSG( | Zfp623 | 190.253 | 0.113814 | 0.53868 | 0.937216 |
| ENSMUSG( | Zfp626 | 232.7481 | -0.18782 | 0.19578 | 0.635109 | ENSMUSG( | Zfp626 | 230.3341 | 0.053688 | 0.735257 | 0.967344 |
| ENSMUSG( | Zfp628 | 415.4607 | 0.094283 | 0.424553 | 0.824675 | ENSMUSG( | Zfp628 | 393.0309 | 0.197931 | 0.154637 | 0.837724 |
| ENSMUSG( | Zfp629 | 961.4451 | -0.08773 | 0.387914 | 0.805835 | ENSMUSG( | Zfp629 | 905.4909 | 0.012835 | 0.92015 | 0.992644 |
| ENSMUSG( | Zfp637 | 671.278 | 0.052257 | 0.634626 | 0.914073 | ENSMUSG( | Zfp637 | 598.6034 | -0.00499 | 0.969224 | 0.99594 |
| ENSMUSG( | Zfp638 | 2264.957 | 0.159755 | 0.076664 | 0.429084 | ENSMUSG( | Zfp638 | 2050.342 | 0.143978 | 0.186687 | 0.856099 |
| ENSMUSG( | Zfp639 | 740.2471 | -0.16944 | 0.133882 | 0.547197 | ENSMUSG( | Zfp639 | 715.3419 | 0.00936 | 0.944008 | 0.994798 |
| ENSMUSG( | Zfp64 | 447.1288 | 0.043793 | 0.716958 | 0.941697 | ENSMUSG( | Zfp64 | 410.9436 | 0.072082 | 0.612607 | 0.951071 |
| ENSMUSG( | Zfp641 | 571.6879 | -0.00689 | 0.957044 | 0.993874 | ENSMUSG( | Zfp641 | 504.0325 | -0.09951 | 0.473754 | 0.923329 |
| ENSMUSG( | Zfp644 | 952.333 | 0.011138 | 0.909627 | 0.988084 | ENSMUSG( | Zfp644 | 864.5649 | -0.00147 | 0.989929 | 0.997928 |
| ENSMUSG( | Zfp646 | 724.4437 | -0.09833 | 0.437213 | 0.829501 | ENSMUSG( | Zfp646 | 739.6735 | 0.229666 | 0.110316 | 0.807051 |
| ENSMUSG( | Zfp647 | 72.14534 | 0.320288 | 0.212587 | 0.6567 | ENSMUSG( | Zfp647 | 68.58302 | 0.41859 | 0.140266 | 0.830678 |
| ENSMUSG( | Zfp65 | 216.189 | -0.0189 | 0.902994 | 0.986055 | ENSMUSG( | Zfp65 | 199.8944 | 0.029256 | 0.866795 | 0.987487 |
| ENSMUSG( | Zfp652 | 3663.424 | 0.018691 | 0.854941 | 0.973665 | ENSMUSG( | Zfp652 | 3241.97 | -0.06748 | 0.602933 | 0.947907 |
| ENSMUSG( | Zfp652os | 207.8063 | 0.14931 | 0.331159 | 0.77519 | ENSMUSG( | Zfp652os | 162.6424 | -0.31238 | 0.10126 | 0.801456 |
| ENSMUSG( | Zfp653 | 481.3789 | 0.10493 | 0.470813 | 0.846741 | ENSMUSG( | Zfp653 | 429.6631 | 0.042978 | 0.795725 | 0.97681 |
| ENSMUSG( | Zfp654 | 459.2916 | -0.06838 | 0.631184 | 0.914073 | ENSMUSG( | Zfp654 | 453.7158 | 0.159749 | 0.272816 | 0.888937 |
| ENSMUSG( | Zfp655 | 522.1911 | -0.08301 | 0.51125 | 0.870859 | ENSMUSG( | Zfp655 | 481.6451 | -0.04665 | 0.748315 | 0.96984 |
| ENSMUSG( | Zfp658 | 193.5871 | -0.14882 | 0.331959 | 0.775288 | ENSMUSG( | Zfp658 | 185.457 | -0.00365 | 0.98433 | 0.996963 |
| ENSMUSG( | Zfp660 | 99.5097 | 0.083208 | 0.668376 | 0.927864 | ENSMUSG( | Zfp660 | 93.12671 | 0.147916 | 0.532062 | 0.934795 |
| ENSMUSG( | Zfp661 | 281.5761 | 0.033752 | 0.841326 | 0.971179 | ENSMUSG( | Zfp661 | 250.6258 | -0.02271 | 0.898841 | 0.990558 |
| ENSMUSG( | Zfp664 | 7216.419 | 0.087161 | 0.283153 | 0.73186 | ENSMUSG( | Zfp664 | 6220.052 | -0.07456 | 0.532726 | 0.934795 |
| ENSMUSG( | Zfp667 | 249.0228 | -0.12167 | 0.41681 | 0.8209 | ENSMUSG( | Zfp667 | 233.9431 | -0.03278 | 0.848449 | 0.985321 |
| ENSMUSG( | Zfp668 | 375.6605 | 0.130364 | 0.313424 | 0.760368 | ENSMUSG( | Zfp668 | 344.1721 | 0.151267 | 0.313563 | 0.900883 |
| ENSMUSG( | Zfp672 | 812.9131 | 0.000506 | 0.997107 | 0.999565 | ENSMUSG( | Zfp672 | 748.0306 | 0.032659 | 0.80566 | 0.977674 |
| ENSMUSG( | Zfp677 | 48.85735 | -0.21824 | 0.438692 | 0.830714 | ENSMUSG( | Zfp677 | 41.78866 | -0.41782 | 0.188501 | 0.85698 |
| ENSMUSG( | Zfp68 | 653.6229 | 0.045594 | 0.658983 | 0.924506 | ENSMUSG( | Zfp68 | 615.2179 | 0.139507 | 0.258867 | 0.883971 |
| ENSMUSG( | Zfp687 | 1554.368 | -0.0485 | 0.586953 | 0.900235 | ENSMUSG( | Zfp687 | 1379.74 | -0.12836 | 0.259005 | 0.883971 |
| ENSMUSG( | Zfp688 | 269.9903 | -0.00627 | 0.968097 | 0.996284 | ENSMUSG( | Zfp688 | 248.097 | 0.020992 | 0.902817 | 0.991089 |
| ENSMUSG( | Zfp689 | 160.0231 | -0.00912 | 0.957867 | 0.993874 | ENSMUSG( | Zfp689 | 143.1015 | -0.05128 | 0.797136 | 0.97681 |
| ENSMUSG( | Zfp69 | 48.2738 | -1.75393 | 2.78E-05 | 0.002487 | ENSMUSG( | Zfp69 | 63.21925 | -0.1909 | 0.60144 | 0.947749 |
| ENSMUSG( | Zfp691 | 176.164 | 0.015814 | 0.923507 | 0.990825 | ENSMUSG( | Zfp691 | 170.3682 | 0.2003 | 0.318874 | 0.903344 |
| ENSMUSG( | Zfp692 | 805.8848 | 0.295646 | 0.16097 | 0.590707 | ENSMUSG( | Zfp692 | 674.1085 | 0.060011 | 0.802743 | 0.97681 |
| ENSMUSG( | Zfp697 | 634.0513 | 0.090644 | 0.387463 | 0.805693 | ENSMUSG( | Zfp697 | 557.1738 | -0.01186 | 0.923481 | 0.992815 |
| ENSMUSG( | Zfp7 | 129.0092 | -0.16275 | 0.414015 | 0.819998 | ENSMUSG( | Zfp7 | 139.2139 | 0.312765 | 0.104618 | 0.804969 |
| ENSMUSG( | Zfp703 | 1125.584 | 0.065338 | 0.608507 | 0.907154 | ENSMUSG( | Zfp703 | 1058.619 | 0.151088 | 0.288407 | 0.894794 |
| ENSMUSG( | Zfp704 | 2820.596 | 0.047575 | 0.651379 | 0.921451 | ENSMUSG( | Zfp704 | 2610.392 | 0.09504 | 0.44954 | 0.922202 |
| ENSMUSG( | Zfp706 | 5170.479 | 0.035676 | 0.695719 | 0.934979 | ENSMUSG( | Zfp706 | 4479.537 | -0.11422 | 0.379559 | 0.916995 |
| ENSMUSG( | Zfp707 | 264.5699 | -0.00099 | 0.994711 | 0.999369 | ENSMUSG( | Zfp707 | 244.891 | 0.042181 | 0.798981 | 0.97681 |
| ENSMUSG( | Zfp709 | 156.8672 | -0.01032 | 0.951549 | 0.993821 | ENSMUSG( | Zfp709 | 143.961 | 0.009704 | 0.958689 | 0.994798 |
| ENSMUSG( | Zfp710 | 867.7681 | -0.10657 | 0.290607 | 0.740557 | ENSMUSG( | Zfp710 | 801.166 | -0.06961 | 0.562956 | 0.941158 |
| ENSMUSG( | Zfp711 | 190.4043 | 0.01063 | 0.952133 | 0.993821 | ENSMUSG( | Zfp711 | 181.4412 | 0.13897 | 0.414394 | 0.917257 |
| ENSMUSG( | Zfp712 | 73.57221 | -0.28495 | 0.227283 | 0.674223 | ENSMUSG( | Zfp712 | 74.5234 | 0.049838 | 0.847484 | 0.985161 |
| ENSMUSG( | Zfp715 | 622.2547 | -0.2817 | 0.012662 | 0.162184 | ENSMUSG( | Zfp715 | 647.9858 | 0.114951 | 0.426319 | 0.917257 |
| ENSMUSG( | Zfp719 | 310.6744 | -0.29311 | 0.02569 | 0.247138 | ENSMUSG( | Zfp719 | 309.5247 | -0.02611 | 0.862473 | 0.986545 |
| ENSMUSG( | Zfp72 | 95.01549 | -0.10158 | 0.642326 | 0.916865 | ENSMUSG( | Zfp72 | 92.54952 | 0.071254 | 0.768785 | 0.974236 |
| ENSMUSG( | Zfp729a | 343.777 | -0.06731 | 0.596811 | 0.903526 | ENSMUSG( | Zfp729a | 317.8454 | -0.01655 | 0.913339 | 0.992298 |
| ENSMUSG( | Zfp729b | 176.6054 | -0.04067 | 0.81033 | 0.964028 | ENSMUSG( | Zfp729b | 173.7729 | 0.175632 | 0.350765 | 0.911707 |
| ENSMUSG( | Zfp738 | 663.5617 | -0.21553 | 0.051365 | 0.347869 | ENSMUSG( | Zfp738 | 650.0312 | 0.006589 | 0.966001 | 0.995436 |
| ENSMUSG( | Zfp74 | 397.5129 | 0.072264 | 0.581501 | 0.897767 | ENSMUSG( | Zfp74 | 364.6768 | 0.083526 | 0.608621 | 0.950113 |
| ENSMUSG( | Zfp740 | 3232.785 | 0.011372 | 0.891833 | 0.983979 | ENSMUSG( | Zfp740 | 2914.693 | -0.01843 | 0.854524 | 0.985649 |
| ENSMUSG( | Zfp746 | 767.6551 | 0.137481 | 0.240321 | 0.687899 | ENSMUSG( | Zfp746 | 697.2657 | 0.125267 | 0.387357 | 0.917257 |
| ENSMUSG( | Zfp747 | 103.5219 | -0.09947 | 0.645112 | 0.917934 | ENSMUSG( | Zfp747 | 93.16913 | -0.11746 | 0.635338 | 0.953903 |
| ENSMUSG( | Zfp747l1 | 60.46367 | -0.25544 | 0.31945 | 0.765296 | ENSMUSG( | Zfp747l1 | 61.07285 | 0.077332 | 0.779847 | 0.976106 |
| ENSMUSG( | Zfp748 | 192.3507 | -0.0721 | 0.634344 | 0.914073 | ENSMUSG( | Zfp748 | 191.3618 | 0.178793 | 0.299449 | 0.898538 |
| ENSMUSG( | Zfp758 | 99.66946 | -0.23289 | 0.299846 | 0.748922 | ENSMUSG( | Zfp758 | 111.4476 | 0.341553 | 0.153352 | 0.837724 |
| ENSMUSG( | Zfp759 | 108.1757 | -0.05433 | 0.78204 | 0.957558 | ENSMUSG( | Zfp759 | 103.9288 | 0.109795 | 0.617106 | 0.951739 |
| ENSMUSG( | Zfp760 | 336.8494 | 0.137945 | 0.319138 | 0.765081 | ENSMUSG( | Zfp760 | 315.263 | 0.217832 | 0.269138 | 0.886755 |
| ENSMUSG( | Zfp763 | 281.7909 | 0.005901 | 0.965833 | 0.995377 | ENSMUSG( | Zfp763 | 257.9561 | 0.017523 | 0.91137 | 0.992196 |
| ENSMUSG( | Zfp764 | 84.29132 | -0.16026 | 0.554043 | 0.886397 | ENSMUSG( | Zfp764 | 91.03195 | 0.336051 | 0.228221 | 0.868507 |
| ENSMUSG( | Zfp764l1 | 122.1915 | 0.133958 | 0.484874 | 0.856903 | ENSMUSG( | Zfp764l1 | 114.5822 | 0.200721 | 0.366282 | 0.914703 |
| ENSMUSG( | Zfp768 | 457.0383 | 0.09131 | 0.428148 | 0.825894 | ENSMUSG( | Zfp768 | 401.0197 | -0.01774 | 0.896876 | 0.990558 |
| ENSMUSG( | Zfp770 | 631.5645 | 0.090427 | 0.510874 | 0.870755 | ENSMUSG( | Zfp770 | 569.0417 | 0.051741 | 0.743629 | 0.96951 |
| ENSMUSG( | Zfp771 | 1434.28 | 0.237345 | 0.019354 | 0.209704 | ENSMUSG( | Zfp771 | 1249.169 | 0.11365 | 0.325441 | 0.905339 |
| ENSMUSG( | Zfp772 | 472.4933 | 0.232952 | 0.045104 | 0.325725 | ENSMUSG( | Zfp772 | 389.4586 | -0.05745 | 0.72068 | 0.966887 |
| ENSMUSG( | Zfp773 | 54.31998 | 0.065973 | 0.803205 | 0.962169 | ENSMUSG( | Zfp773 | 54.68742 | 0.342825 | 0.232329 | 0.868507 |
| ENSMUSG( | Zfp775 | 304.4563 | 0.239767 | 0.070147 | 0.40908 | ENSMUSG( | Zfp775 | 257.4352 | 0.028442 | 0.8543 | 0.98563 |
| ENSMUSG( | Zfp777 | 1044.717 | 0.166983 | 0.143621 | 0.562581 | ENSMUSG( | Zfp777 | 901.9268 | 0.017889 | 0.885133 | 0.989688 |
| ENSMUSG( | Zfp78 | 61.11546 | -0.06216 | 0.799458 | 0.961247 | ENSMUSG( | Zfp78 | 54.17324 | -0.12255 | 0.673919 | 0.95694 |
| ENSMUSG( | Zfp780b | 307.7556 | 0.100666 | 0.441031 | 0.832445 | ENSMUSG( | Zfp780b | 282.9843 | 0.122475 | 0.416897 | 0.917257 |
| ENSMUSG( | Zfp781 | 13.15244 | 0.087954 | 0.869129 | NA | ENSMUSG( | Zfp781 | 13.30391 | 0.331054 | 0.554593 | 0.939565 |
| ENSMUSG( | Zfp783 | 236.1604 | 0.01503 | 0.915783 | 0.988634 | ENSMUSG( | Zfp783 | 217.5821 | 0.05094 | 0.796223 | 0.97681 |
| ENSMUSG( | Zfp784 | 166.4579 | 0.113323 | 0.478513 | 0.853567 | ENSMUSG( | Zfp784 | 141.6275 | -0.07566 | 0.68865 | 0.959742 |
| ENSMUSG( | Zfp786 | 72.35443 | -0.0027 | 0.99168 | 0.999232 | ENSMUSG( | Zfp786 | 71.71381 | 0.208051 | 0.485307 | 0.92541 |
| ENSMUSG( | Zfp787 | 587.1858 | 0.043849 | 0.691177 | 0.933881 | ENSMUSG( | Zfp787 | 516.1242 | -0.06027 | 0.643756 | 0.953903 |
| ENSMUSG( | Zfp788 | 724.5691 | 0.056112 | 0.610331 | 0.908189 | ENSMUSG( | Zfp788 | 683.2628 | 0.149548 | 0.233625 | 0.868507 |
| ENSMUSG( | Zfp790 | 289.197 | 0.078951 | 0.561674 | 0.889052 | ENSMUSG( | Zfp790 | 258.6968 | 0.039586 | 0.816779 | 0.979198 |
| ENSMUSG( | Zfp791 | 100.3756 | 0.156342 | 0.447669 | 0.834051 | ENSMUSG( | Zfp791 | 90.65821 | 0.129749 | 0.580012 | 0.944209 |
| ENSMUSG( | Zfp799 | 425.521 | 0.015381 | 0.902069 | 0.985633 | ENSMUSG( | Zfp799 | 384.1661 | -0.0083 | 0.952508 | 0.994798 |
| ENSMUSG( | Zfp800 | 183.6176 | -0.15928 | 0.379585 | 0.801757 | ENSMUSG( | Zfp800 | 184.1504 | 0.132698 | 0.499263 | 0.927825 |
| ENSMUSG( | Zfp804a | 260.3351 | -0.33132 | 0.029496 | 0.260298 | ENSMUSG( | Zfp804a | 249.3769 | -0.16912 | 0.32217 | 0.904023 |
| ENSMUSG( | Zfp808 | 32.59804 | -0.14276 | 0.676136 | 0.930542 | ENSMUSG( | Zfp808 | 30.78662 | -0.04085 | 0.912373 | 0.992225 |
| ENSMUSG( | Zfp809 | 362.9618 | 0.175829 | 0.17048 | 0.605271 | ENSMUSG( | Zfp809 | 325.3788 | 0.135309 | 0.357023 | 0.913389 |
| ENSMUSG( | Zfp81 | 397.9215 | 0.145857 | 0.238135 | 0.686299 | ENSMUSG( | Zfp81 | 372.6247 | 0.215139 | 0.155913 | 0.837724 |
| ENSMUSG( | Zfp810 | 338.8543 | 0.02853 | 0.822276 | 0.966352 | ENSMUSG( | Zfp810 | 334.4981 | 0.249218 | 0.086121 | 0.780433 |
| ENSMUSG( | Zfp811 | 241.4984 | 0.019597 | 0.894927 | 0.984512 | ENSMUSG( | Zfp811 | 243.6416 | 0.298057 | 0.062053 | 0.736482 |
| ENSMUSG( | Zfp820 | 18.36193 | -0.45847 | 0.333836 | NA | ENSMUSG( | Zfp820 | 19.02378 | -0.0035 | 0.994571 | 0.998813 |
| ENSMUSG( | Zfp821 | 1003.284 | 0.133547 | 0.328878 | 0.773233 | ENSMUSG( | Zfp821 | 896.9995 | 0.079988 | 0.562513 | 0.940761 |
| ENSMUSG( | Zfp825 | 116.0219 | 0.126508 | 0.504378 | 0.866883 | ENSMUSG( | Zfp825 | 97.79175 | -0.09852 | 0.64665 | 0.953988 |
| ENSMUSG( | Zfp827 | 1205.85 | 0.050968 | 0.635152 | 0.914296 | ENSMUSG( | Zfp827 | 1095.016 | 0.037692 | 0.761108 | 0.972695 |
| ENSMUSG( | Zfp830 | 204.7322 | -0.16192 | 0.363211 | 0.793211 | ENSMUSG( | Zfp830 | 194.7083 | -0.03323 | 0.863649 | 0.986547 |
| ENSMUSG( | Zfp831 | 605.6453 | 0.039052 | 0.748404 | 0.949604 | ENSMUSG( | Zfp831 | 571.4541 | 0.140747 | 0.396037 | 0.917257 |
| ENSMUSG( | Zfp839 | 297.1059 | 0.052008 | 0.73987 | 0.947823 | ENSMUSG( | Zfp839 | 271.7951 | 0.060567 | 0.735903 | 0.967344 |
| ENSMUSG( | Zfp84 | 365.8888 | 0.120477 | 0.362591 | 0.792943 | ENSMUSG( | Zfp84 | 306.6939 | -0.10936 | 0.487031 | 0.92541 |
| ENSMUSG( | Zfp846 | 519.9683 | 0.035243 | 0.767085 | 0.954393 | ENSMUSG( | Zfp846 | 467.417 | 0.000289 | 0.998288 | 0.999313 |
| ENSMUSG( | Zfp85 | 69.83194 | 0.172991 | 0.495024 | 0.860798 | ENSMUSG( | Zfp85 | 65.50398 | 0.259382 | 0.345163 | 0.91045 |

| | | | | | |
|---|---|---|---|---|---|
| ENSMUSG( | 325.3343 | -0.51084 | 0.010009 | 0.142653 | Zdhhc15 |
| ENSMUSG( | 1135.068 | -0.09669 | 0.443301 | 0.833108 | Zdhhc16 |
| ENSMUSG( | 2778.701 | -0.32052 | 0.021772 | 0.225648 | Zdhhc17 |
| ENSMUSG( | 1917.596 | -0.21192 | 0.051788 | 0.349274 | Zdhhc18 |
| ENSMUSG( | 1927.731 | -0.17705 | 0.174979 | 0.610399 | Zdhhc2 |
| ENSMUSG( | 1015.394 | -0.38949 | 0.013989 | 0.172788 | Zdhhc20 |
| ENSMUSG( | 1422.485 | -0.20207 | 0.105714 | 0.492581 | Zdhhc21 |
| ENSMUSG( | 1479.627 | -0.23845 | 0.054153 | 0.357985 | Zdhhc22 |
| ENSMUSG( | 1119.789 | -0.11913 | 0.397898 | 0.812161 | Zdhhc23 |
| ENSMUSG( | 663.8664 | -0.35709 | 0.001294 | 0.039144 | Zdhhc24 |
| ENSMUSG( | 4017.056 | -0.03773 | 0.676844 | 0.930542 | Zdhhc3 |
| ENSMUSG( | 1081.458 | -0.07815 | 0.404221 | 0.816132 | Zdhhc4 |
| ENSMUSG( | 1423.218 | -0.12659 | 0.433252 | 0.828207 | Zdhhc5 |
| ENSMUSG( | 1035.847 | -0.12199 | 0.314391 | 0.761558 | Zdhhc6 |
| ENSMUSG( | 1134.467 | -0.00145 | 0.989887 | 0.999144 | Zdhhc7 |
| ENSMUSG( | 4730.08 | -0.18332 | 0.155931 | 0.58375 | Zdhhc8 |
| ENSMUSG( | 2815.717 | -0.4949 | 0.001961 | 0.051054 | Zdhhc9 |
| ENSMUSG( | 7260.569 | -0.02187 | 0.84297 | 0.971739 | Zeb1 |
| ENSMUSG( | 194.0195 | 0.088011 | 0.572227 | 0.893584 | Zeb1os1 |
| ENSMUSG( | 5937.546 | 0.015434 | 0.871534 | 0.978114 | Zeb2 |
| ENSMUSG( | 237.1339 | -0.12272 | 0.444212 | 0.833239 | Zeb2os |
| ENSMUSG( | 5716.165 | 0.226421 | 0.023726 | 0.235641 | Zer1 |
| ENSMUSG( | 584.3522 | -0.02332 | 0.833779 | 0.970333 | Zfand1 |
| ENSMUSG( | 3498.275 | 0.158129 | 0.053248 | 0.354403 | Zfand2a |
| ENSMUSG( | 389.4082 | -0.13253 | 0.274661 | 0.725078 | Zfand2b |
| ENSMUSG( | 5229.303 | 0.082171 | 0.382915 | 0.803237 | Zfand3 |
| ENSMUSG( | 103.6308 | -0.24802 | 0.269567 | 0.719729 | Zfand4 |
| ENSMUSG( | 2516.889 | -0.11225 | 0.333755 | 0.775602 | Zfand5 |
| ENSMUSG( | 935.7303 | 0.039221 | 0.677943 | 0.930542 | Zfand6 |
| ENSMUSG( | 1330.828 | 0.110356 | 0.236193 | 0.683188 | Zfas1 |
| ENSMUSG( | 164.1491 | 0.079246 | 0.623539 | 0.911646 | Zfat |
| ENSMUSG( | 1222.02 | 0.126388 | 0.294038 | 0.743459 | Zfc3h1 |
| ENSMUSG( | 2297.118 | -0.3748 | 9.77E-05 | 0.006367 | Zfhx2 |
| ENSMUSG( | 168.7045 | -0.59942 | 0.000435 | 0.019019 | Zfhx3 |
| ENSMUSG( | 243.228 | -0.50258 | 0.000684 | 0.025185 | Zfhx4 |
| ENSMUSG( | 422.9592 | -0.19178 | 0.143667 | 0.562581 | Zfp1 |
| ENSMUSG( | 13.84286 | 0.196383 | 0.692261 | NA | Zfp1001 |
| ENSMUSG( | 307.8814 | 0.022698 | 0.864148 | 0.975331 | Zfp1006 |
| ENSMUSG( | 37.08658 | 0.076483 | 0.814793 | 0.964668 | Zfp1007 |
| ENSMUSG( | 165.4546 | 0.023489 | 0.905456 | 0.986465 | Zfp1008 |
| ENSMUSG( | 245.5534 | -0.2545 | 0.078765 | 0.434128 | Zfp101 |
| ENSMUSG( | 177.8966 | -0.13478 | 0.427908 | 0.825894 | Zfp105 |
| ENSMUSG( | 15254.38 | 0.087305 | 0.36365 | 0.793343 | Zfp106 |
| ENSMUSG( | 116.6419 | -0.30212 | 0.119868 | 0.523965 | Zfp108 |
| ENSMUSG( | 100.5563 | -0.11055 | 0.577806 | 0.895986 | Zfp109 |
| ENSMUSG( | 110.773 | -0.33359 | 0.121686 | 0.526331 | Zfp11 |
| ENSMUSG( | 322.7807 | 0.01531 | 0.911943 | 0.988483 | Zfp110 |
| ENSMUSG( | 179.2192 | 0.035115 | 0.822776 | 0.966352 | Zfp111 |
| ENSMUSG( | 229.3429 | -0.16125 | 0.326546 | 0.771368 | Zfp112 |
| ENSMUSG( | 181.7314 | 0.040427 | 0.806812 | 0.96295 | Zfp113 |
| ENSMUSG( | 78.08989 | -0.00703 | 0.977244 | 0.997428 | Zfp114 |
| ENSMUSG( | 65.064 | -0.16735 | 0.5126 | 0.871762 | Zfp119a |
| ENSMUSG( | 57.02166 | 0.10991 | 0.667709 | 0.927484 | Zfp119b |
| ENSMUSG( | 734.5998 | -0.0241 | 0.823755 | 0.966603 | Zfp12 |
| ENSMUSG( | 319.3921 | 0.181152 | 0.156191 | 0.583988 | Zfp120 |
| ENSMUSG( | 118.7694 | 0.083763 | 0.665806 | 0.927054 | Zfp128 |
| ENSMUSG( | 605.0729 | 0.127257 | 0.335015 | 0.776499 | Zfp13 |
| ENSMUSG( | 1514.915 | 0.012997 | 0.882209 | 0.982183 | Zfp131 |
| ENSMUSG( | 292.5498 | -0.3169 | 0.029204 | 0.259574 | Zfp14 |
| ENSMUSG( | 281.1281 | -0.23106 | 0.075397 | 0.425668 | Zfp141 |
| ENSMUSG( | 1150.397 | 0.05057 | 0.616044 | 0.91085 | Zfp142 |
| ENSMUSG( | 250.6234 | 0.195783 | 0.157276 | 0.585553 | Zfp143 |
| ENSMUSG( | 1195.55 | 0.062432 | 0.548163 | 0.885197 | Zfp146 |
| ENSMUSG( | 2030.342 | 0.115783 | 0.171964 | 0.606889 | Zfp148 |
| ENSMUSG( | 629.5131 | -0.05504 | 0.598888 | 0.903801 | Zfp157 |
| ENSMUSG( | 481.5418 | 0.066146 | 0.582936 | 0.898094 | Zfp160 |
| ENSMUSG( | 162.0641 | -0.02292 | 0.892703 | 0.984474 | Zfp169 |
| ENSMUSG( | 120.5339 | -0.0063 | 0.976041 | 0.997428 | Zfp174 |
| ENSMUSG( | 1535.755 | 0.223768 | 0.016419 | 0.190497 | Zfp180 |
| ENSMUSG( | 312.8161 | 0.042866 | 0.756216 | 0.951479 | Zfp182 |
| ENSMUSG( | 54.96749 | 0.165934 | 0.528558 | 0.878365 | Zfp184 |
| ENSMUSG( | 67.02796 | -0.17413 | 0.540145 | 0.88315 | Zfp185 |
| ENSMUSG( | 149.6523 | -0.42193 | 0.038249 | 0.301606 | Zfp189 |
| ENSMUSG( | 473.222 | 0.133526 | 0.274363 | 0.724547 | Zfp2 |
| ENSMUSG( | 134.8965 | -0.0163 | 0.925382 | 0.991204 | Zfp202 |
| ENSMUSG( | 2050.861 | 0.071 | 0.431729 | 0.827541 | Zfp207 |
| ENSMUSG( | 348.2236 | -0.04572 | 0.73548 | 0.946109 | Zfp212 |
| ENSMUSG( | 190.2675 | 0.171932 | 0.28461 | 0.733224 | Zfp213 |
| ENSMUSG( | 196.0473 | -0.07343 | 0.646382 | 0.918612 | Zfp217 |
| ENSMUSG( | 2192.199 | -0.03304 | 0.707777 | 0.939519 | Zfp219 |
| ENSMUSG( | 112.9944 | 0.270492 | 0.24675 | 0.695324 | Zfp229 |
| ENSMUSG( | 465.4172 | 0.036943 | 0.78024 | 0.957491 | Zfp235 |
| ENSMUSG( | 1751.635 | 0.134362 | 0.177142 | 0.614466 | Zfp236 |
| ENSMUSG( | 739.7512 | 0.040398 | 0.702934 | 0.937735 | Zfp239 |
| ENSMUSG( | 1706.693 | 0.202271 | 0.053512 | 0.35564 | Zfp24 |
| ENSMUSG( | 243.8296 | 0.189005 | 0.190012 | 0.629259 | Zfp248 |
| ENSMUSG( | 1528.96 | 0.1645 | 0.089309 | 0.458563 | Zfp251 |
| ENSMUSG( | 560.0499 | -0.24796 | 0.038607 | 0.302696 | Zfp26 |
| ENSMUSG( | 1211.659 | 0.179903 | 0.05091 | 0.345939 | Zfp260 |
| ENSMUSG( | 524.4271 | -0.03452 | 0.809095 | 0.963548 | Zfp263 |
| ENSMUSG( | 831.2275 | -0.00731 | 0.950585 | 0.993821 | Zfp266 |
| ENSMUSG( | 483.1053 | -0.17637 | 0.127364 | 0.535055 | Zfp267 |
| ENSMUSG( | 404.9856 | 0.12147 | 0.324391 | 0.769598 | Zfp27 |
| ENSMUSG( | 106.2715 | -0.30532 | 0.148122 | 0.571232 | Zfp273 |
| ENSMUSG( | 838.9479 | 0.050504 | 0.661624 | 0.925271 | Zfp275 |
| ENSMUSG( | 810.7732 | -0.18307 | 0.213863 | 0.658243 | Zfp276 |
| ENSMUSG( | 448.8963 | -0.09521 | 0.498832 | 0.862973 | Zfp277 |
| ENSMUSG( | 232.0747 | -0.1814 | 0.232768 | 0.680928 | Zfp28 |
| ENSMUSG( | 275.9745 | 0.044649 | 0.769693 | 0.954674 | Zfp280b |
| ENSMUSG( | 485.015 | -0.26451 | 0.087557 | 0.454663 | Zfp280c |
| ENSMUSG( | 800.4414 | -0.0549 | 0.601288 | 0.904603 | Zfp280d |
| ENSMUSG( | 552.59 | -0.16772 | 0.132382 | 0.544294 | Zfp281 |
| ENSMUSG( | 577.9396 | 0.097341 | 0.458393 | 0.840034 | Zfp282 |
| ENSMUSG( | 352.0807 | -0.01972 | 0.890368 | 0.983687 | Zfp286 |
| ENSMUSG( | 367.1342 | -0.01087 | 0.930215 | 0.991848 | Zfp287 |
| ENSMUSG( | 892.1782 | -0.16476 | 0.116482 | 0.514944 | Zfp292 |
| ENSMUSG( | 37.23637 | 0.455786 | 0.140906 | 0.557902 | Zfp296 |
| ENSMUSG( | 293.5931 | -0.10527 | 0.447505 | 0.834051 | Zfp3 |
| ENSMUSG( | 355.2201 | -0.03683 | 0.775837 | 0.955606 | Zfp30 |
| ENSMUSG( | 40.80662 | -0.08192 | 0.789715 | 0.959461 | Zfp300 |
| ENSMUSG( | 1252.085 | 0.074607 | 0.448232 | 0.834051 | Zfp316 |
| ENSMUSG( | 796.7514 | -0.15196 | 0.226003 | 0.672742 | Zfp317 |
| ENSMUSG( | 1529.772 | -0.12329 | 0.187985 | 0.626616 | Zfp318 |
| ENSMUSG( | 680.8215 | 0.06722 | 0.569768 | 0.893041 | Zfp319 |
| ENSMUSG( | 564.2983 | 0.176735 | 0.131878 | 0.544244 | Zfp322a |
| ENSMUSG( | 252.4222 | 0.245028 | 0.107553 | 0.496444 | Zfp324 |
| ENSMUSG( | 657.701 | 0.071012 | 0.533218 | 0.879905 | Zfp326 |
| ENSMUSG( | 560.8392 | 0.097049 | 0.360331 | 0.791947 | Zfp329 |
| ENSMUSG( | 599.1481 | 0.048981 | 0.677878 | 0.930542 | Zfp330 |
| ENSMUSG( | 566.6615 | -0.03593 | 0.778134 | 0.956168 | Zfp334 |
| ENSMUSG( | 1458.559 | 0.199121 | 0.060352 | 0.3797 | Zfp335 |
| ENSMUSG( | 74.6715 | 0.005237 | 0.983405 | 0.997675 | Zfp335os |
| ENSMUSG( | 250.1546 | 0.1696 | 0.285486 | 0.734313 | Zfp341 |
| ENSMUSG( | 1006.048 | 0.201992 | 0.047568 | 0.333288 | Zfp346 |
| ENSMUSG( | 407.2982 | 0.096759 | 0.431158 | 0.827055 | Zfp35 |
| ENSMUSG( | 181.8506 | -0.33014 | 0.086012 | 0.451492 | Zfp354a |
| ENSMUSG( | 71.0772 | -0.18928 | 0.495133 | 0.860798 | Zfp354b |
| ENSMUSG( | 1236.06 | -0.02532 | 0.797355 | 0.960507 | Zfp354c |
| ENSMUSG( | 1172.888 | 0.136261 | 0.139048 | 0.555099 | Zfp358 |
| ENSMUSG( | 146.8329 | -0.36015 | 0.060269 | 0.379636 | Zfp36 |
| ENSMUSG( | 1204.823 | 0.017739 | 0.854014 | 0.973491 | Zfp362 |
| ENSMUSG( | 35872.94 | 0.095298 | 0.31856 | 0.764234 | Zfp365 |
| ENSMUSG( | 250.1195 | 0.652813 | 0.006517 | 0.110626 | Zfp366 |
| ENSMUSG( | 897.7599 | -0.03567 | 0.781752 | 0.957558 | Zfp367 |
| ENSMUSG( | 516.4018 | 0.186727 | 0.1254 | 0.53232 | Zfp369 |
| ENSMUSG( | 989.9232 | 0.060619 | 0.58978 | 0.90113 | Zfp36l1 |
| ENSMUSG( | 1034.761 | 0.122523 | 0.344294 | 0.779153 | Zfp36l2 |
| ENSMUSG( | 320.3405 | 0.004797 | 0.972042 | 0.996585 | Zfp37 |
| ENSMUSG( | 86.89765 | 0.210951 | 0.34237 | 0.778855 | Zfp382 |
| ENSMUSG( | 302.9045 | -0.20824 | 0.193538 | 0.633027 | Zfp383 |
| ENSMUSG( | 1244.363 | 0.12044 | 0.289641 | 0.739839 | Zfp384 |
| ENSMUSG( | 3558.219 | -0.11428 | 0.242873 | 0.690275 | Zfp385a |
| ENSMUSG( | 767.3135 | -0.00425 | 0.972807 | 0.996585 | Zfp385b |
| ENSMUSG( | 10.13824 | -0.51115 | 0.37775 | NA | Zfp385c |
| ENSMUSG( | 458.8572 | -0.04456 | 0.70578 | 0.938649 | Zfp386 |
| ENSMUSG( | 184.0303 | 0.141468 | 0.475875 | 0.850919 | Zfp39 |
| ENSMUSG( | 347.3339 | -0.00464 | 0.972429 | 0.996585 | Zfp395 |
| ENSMUSG( | 668.1634 | 0.015122 | 0.897589 | 0.985633 | Zfp397 |
| ENSMUSG( | 514.1492 | -0.02083 | 0.85585 | 0.973813 | Zfp398 |
| ENSMUSG( | 643.134 | -0.09456 | 0.469193 | 0.845876 | Zfp40 |
| ENSMUSG( | 334.2099 | -0.01069 | 0.935378 | 0.993363 | Zfp407 |
| ENSMUSG( | 433.6016 | -0.02941 | 0.822706 | 0.966352 | Zfp408 |
| ENSMUSG( | 251.9784 | -0.07461 | 0.663573 | 0.9265 | Zfp41 |
| ENSMUSG( | 616.1591 | 0.125273 | 0.268088 | 0.718372 | Zfp410 |
| ENSMUSG( | 433.6762 | -0.03677 | 0.800599 | 0.961745 | Zfp414 |
| ENSMUSG( | 43.99807 | -0.1174 | 0.690423 | 0.933881 | Zfp418 |
| ENSMUSG( | 322.0978 | 0.030951 | 0.815764 | 0.965324 | Zfp420 |
| ENSMUSG( | 590.7254 | 0.020339 | 0.85583 | 0.973813 | Zfp422 |

| | | | | | |
|---|---|---|---|---|---|
| ENSMUSG( | 796.8669 | -0.11807 | 0.428667 | 0.917257 | Zfp367 |
| ENSMUSG( | 471.7481 | 0.19358 | 0.161298 | 0.837724 | Zfp369 |
| ENSMUSG( | 1038.545 | 0.440831 | 0.001921 | 0.164157 | Zfp36l1 |
| ENSMUSG( | 1012.775 | 0.314224 | 0.005468 | 0.303106 | Zfp36l2 |
| ENSMUSG( | 308.9929 | 0.159919 | 0.316677 | 0.901868 | Zfp37 |
| ENSMUSG( | 78.17076 | 0.203978 | 0.463871 | 0.923329 | Zfp382 |
| ENSMUSG( | 316.4407 | 0.195038 | 0.240867 | 0.876146 | Zfp383 |
| ENSMUSG( | 1110.553 | 0.063725 | 0.606205 | 0.949497 | Zfp384 |
| ENSMUSG( | 3113.154 | -0.24281 | 0.094598 | 0.796133 | Zfp385a |
| ENSMUSG( | 706.8793 | 0.021779 | 0.891003 | 0.990558 | Zfp385b |
| ENSMUSG( | 8.673593 | -0.79985 | 0.237947 | NA | Zfp385c |
| ENSMUSG( | 446.5102 | 0.143955 | 0.280091 | 0.892356 | Zfp386 |
| ENSMUSG( | 159.0249 | -0.01118 | 0.957399 | 0.994798 | Zfp39 |
| ENSMUSG( | 311.9507 | -0.03877 | 0.798892 | 0.97681 | Zfp395 |
| ENSMUSG( | 648.0816 | 0.192589 | 0.156919 | 0.837724 | Zfp397 |
| ENSMUSG( | 482.0344 | 0.063216 | 0.633232 | 0.953844 | Zfp398 |
| ENSMUSG( | 624.5981 | 0.08562 | 0.575945 | 0.942912 | Zfp40 |
| ENSMUSG( | 288.3991 | -0.17157 | 0.244002 | 0.877904 | Zfp407 |
| ENSMUSG( | 385.7132 | -0.09915 | 0.475632 | 0.923329 | Zfp408 |
| ENSMUSG( | 248.1059 | 0.156749 | 0.401954 | 0.917257 | Zfp41 |
| ENSMUSG( | 547.7336 | 0.062883 | 0.645183 | 0.953988 | Zfp410 |
| ENSMUSG( | 379.4076 | -0.15635 | 0.335616 | 0.907907 | Zfp414 |
| ENSMUSG( | 44.77688 | 0.213742 | 0.504615 | 0.927825 | Zfp418 |
| ENSMUSG( | 306.347 | 0.140395 | 0.396785 | 0.917257 | Zfp420 |
| ENSMUSG( | 535.776 | 0.003315 | 0.980683 | 0.996497 | Zfp422 |
| ENSMUSG( | 882.5273 | -0.158 | 0.18266 | 0.850579 | Zfp423 |
| ENSMUSG( | 619.6006 | -0.08557 | 0.503342 | 0.927825 | Zfp426 |
| ENSMUSG( | 737.033 | -0.32015 | 0.040378 | 0.682717 | Zfp428 |
| ENSMUSG( | 50.9619 | 0.072713 | 0.81015 | 0.978779 | Zfp429 |
| ENSMUSG( | 19.17669 | 0.569914 | 0.248485 | 0.877904 | Zfp433 |
| ENSMUSG( | 174.8848 | -0.51769 | 0.005929 | 0.316645 | Zfp438 |
| ENSMUSG( | 29.61405 | -0.0778 | 0.835009 | 0.981237 | Zfp442 |
| ENSMUSG( | 909.9679 | -0.11799 | 0.368041 | 0.915491 | Zfp444 |
| ENSMUSG( | 4557.212 | 0.014685 | 0.91742 | 0.992529 | Zfp445 |
| ENSMUSG( | 295.0415 | 0.105461 | 0.532883 | 0.934795 | Zfp446 |
| ENSMUSG( | 137.7436 | 0.088517 | 0.662596 | 0.956205 | Zfp449 |
| ENSMUSG( | 1004.165 | 0.025364 | 0.827326 | 0.980224 | Zfp451 |
| ENSMUSG( | 124.7191 | 0.095876 | 0.655971 | 0.955533 | Zfp454 |
| ENSMUSG( | 35.32375 | 0.19884 | 0.574448 | 0.942912 | Zfp455 |
| ENSMUSG( | 34.10173 | -0.23021 | 0.534846 | 0.936328 | Zfp456 |
| ENSMUSG( | 14.31404 | -0.34872 | 0.493073 | 0.927358 | Zfp457 |
| ENSMUSG( | 65.30255 | 0.245333 | 0.394327 | 0.917257 | Zfp458 |
| ENSMUSG( | 54.32353 | 0.215529 | 0.497277 | 0.927825 | Zfp459 |
| ENSMUSG( | 1929.443 | 0.07751 | 0.502094 | 0.927825 | Zfp46 |
| ENSMUSG( | 1298.034 | -0.01499 | 0.900212 | 0.990558 | Zfp462 |
| ENSMUSG( | 928.3339 | 0.249758 | 0.030885 | 0.629448 | Zfp467 |
| ENSMUSG( | 227.5456 | -0.0341 | 0.862708 | 0.986547 | Zfp469 |
| ENSMUSG( | 71.64307 | 0.184531 | 0.454193 | 0.922444 | Zfp472 |
| ENSMUSG( | 15.38281 | -0.57418 | 0.25985 | 0.883971 | Zfp473 |
| ENSMUSG( | 13.63389 | 0.020695 | 0.969709 | 0.99594 | Zfp474 |
| ENSMUSG( | 192.3186 | 0.069482 | 0.697725 | 0.960867 | Zfp488 |
| ENSMUSG( | 57.29872 | 0.129216 | 0.647826 | 0.95422 | Zfp493 |
| ENSMUSG( | 89.31862 | 0.393812 | 0.13961 | 0.830678 | Zfp503 |
| ENSMUSG( | 492.344 | -0.0399 | 0.779707 | 0.976106 | Zfp507 |
| ENSMUSG( | 89.17329 | 0.106701 | 0.646068 | 0.953988 | Zfp51 |
| ENSMUSG( | 415.7872 | 0.067454 | 0.6358 | 0.953903 | Zfp511 |
| ENSMUSG( | 1161.086 | -0.08095 | 0.495968 | 0.927825 | Zfp512 |
| ENSMUSG( | 2225.465 | -0.02156 | 0.869623 | 0.988097 | Zfp512b |
| ENSMUSG( | 742.9943 | 0.06268 | 0.611817 | 0.95083 | Zfp513 |
| ENSMUSG( | 671.152 | -0.18625 | 0.185791 | 0.855305 | Zfp516 |
| ENSMUSG( | 283.9492 | 0.128413 | 0.446457 | 0.921734 | Zfp518a |
| ENSMUSG( | 323.0001 | -0.06616 | 0.657157 | 0.955644 | Zfp518b |
| ENSMUSG( | 99.36636 | -0.04116 | 0.850546 | 0.985541 | Zfp52 |
| ENSMUSG( | 662.641 | -0.17979 | 0.204248 | 0.864412 | Zfp521 |
| ENSMUSG( | 5797.466 | 0.099499 | 0.306753 | 0.898538 | Zfp523 |
| ENSMUSG( | 51.39929 | 0.046785 | 0.882234 | 0.989644 | Zfp524 |
| ENSMUSG( | 196.0357 | 0.154326 | 0.413563 | 0.917257 | Zfp526 |
| ENSMUSG( | 73.24239 | 0.254405 | 0.350286 | 0.911652 | Zfp53 |
| ENSMUSG( | 1564.541 | -0.12177 | 0.267712 | 0.886134 | Zfp532 |
| ENSMUSG( | 503.5746 | 0.075502 | 0.587242 | 0.944975 | Zfp536 |
| ENSMUSG( | 26.50805 | -0.21705 | 0.574875 | 0.942912 | Zfp54 |
| ENSMUSG( | 20.42952 | 0.039109 | 0.927728 | 0.99306 | Zfp541 |
| ENSMUSG( | 36.28829 | 0.481109 | 0.227674 | 0.868507 | Zfp551 |
| ENSMUSG( | 394.8766 | -0.10281 | 0.501776 | 0.927825 | Zfp553 |
| ENSMUSG( | 41.19736 | -0.03562 | 0.909812 | 0.991958 | Zfp558 |
| ENSMUSG( | 230.3768 | -0.28305 | 0.07711 | 0.770558 | Zfp560 |
| ENSMUSG( | 266.8727 | 0.148234 | 0.436546 | 0.919879 | Zfp563 |
| ENSMUSG( | 82.58485 | -0.09065 | 0.739335 | 0.968458 | Zfp566 |
| ENSMUSG( | 108.4418 | -0.05143 | 0.829531 | 0.980224 | Zfp568 |
| ENSMUSG( | 340.5385 | 0.126755 | 0.439628 | 0.920721 | Zfp57 |
| ENSMUSG( | 861.3997 | -0.05744 | 0.638918 | 0.953903 | Zfp574 |
| ENSMUSG( | 229.8051 | -0.24074 | 0.192198 | 0.859726 | Zfp575 |
| ENSMUSG( | 534.3793 | 0.116646 | 0.449677 | 0.922202 | Zfp579 |
| ENSMUSG( | 68.72015 | -0.0378 | 0.883731 | 0.989644 | Zfp58 |
| ENSMUSG( | 639.0574 | -0.28168 | 0.038941 | 0.679972 | Zfp580 |
| ENSMUSG( | 99.1058 | 0.452436 | 0.096476 | 0.796133 | Zfp582 |
| ENSMUSG( | 106.0763 | -0.0127 | 0.952972 | 0.994798 | Zfp583 |
| ENSMUSG( | 124.2943 | -0.0502 | 0.81953 | 0.979198 | Zfp59 |
| ENSMUSG( | 1093.16 | 0.184477 | 0.094279 | 0.795735 | Zfp592 |
| ENSMUSG( | 206.4627 | -0.123 | 0.492167 | 0.927315 | Zfp593 |
| ENSMUSG( | 96.52068 | 0.329103 | 0.168061 | 0.840222 | Zfp595 |
| ENSMUSG( | 259.4163 | 0.164083 | 0.300077 | 0.898538 | Zfp597 |
| ENSMUSG( | 1313.916 | 0.2041 | 0.161633 | 0.837724 | Zfp598 |
| ENSMUSG( | 52.57836 | 0.095978 | 0.744627 | 0.96951 | Zfp599 |
| ENSMUSG( | 713.3262 | -0.00263 | 0.984486 | 0.996963 | Zfp60 |
| ENSMUSG( | 302.5112 | 0.067203 | 0.678204 | 0.957435 | Zfp605 |
| ENSMUSG( | 430.3509 | 0.087928 | 0.531877 | 0.934795 | Zfp606 |
| ENSMUSG( | 30.36566 | 0.131072 | 0.72608 | 0.966887 | Zfp607a |
| ENSMUSG( | 351.67 | -0.17674 | 0.319195 | 0.903431 | Zfp607b |
| ENSMUSG( | 750.6822 | 0.176288 | 0.186414 | 0.855939 | Zfp608 |
| ENSMUSG( | 1539.227 | -0.13934 | 0.227748 | 0.868507 | Zfp609 |
| ENSMUSG( | 350.6833 | 0.147203 | 0.397008 | 0.917257 | Zfp61 |
| ENSMUSG( | 1063.816 | -0.15232 | 0.235471 | 0.871807 | Zfp612 |
| ENSMUSG( | 501.7442 | -0.02194 | 0.876188 | 0.989644 | Zfp617 |
| ENSMUSG( | 149.99 | -0.39804 | 0.041889 | 0.690304 | Zfp618 |
| ENSMUSG( | 106.0945 | 0.042466 | 0.84246 | 0.983044 | Zfp619 |
| ENSMUSG( | 950.9464 | 0.101728 | 0.42648 | 0.917257 | Zfp62 |
| ENSMUSG( | 808.8906 | 0.005519 | 0.968865 | 0.99594 | Zfp622 |
| ENSMUSG( | 190.253 | 0.113814 | 0.53868 | 0.937216 | Zfp623 |
| ENSMUSG( | 230.3341 | 0.053688 | 0.735257 | 0.967344 | Zfp626 |
| ENSMUSG( | 393.0309 | 0.197931 | 0.154637 | 0.837724 | Zfp628 |
| ENSMUSG( | 905.4909 | 0.012835 | 0.92015 | 0.992644 | Zfp629 |
| ENSMUSG( | 598.6034 | -0.00499 | 0.969224 | 0.99594 | Zfp637 |
| ENSMUSG( | 2050.342 | 0.143978 | 0.186687 | 0.856099 | Zfp638 |
| ENSMUSG( | 715.3419 | 0.00936 | 0.944008 | 0.994798 | Zfp639 |
| ENSMUSG( | 410.9436 | 0.072082 | 0.612607 | 0.951071 | Zfp64 |
| ENSMUSG( | 504.0325 | -0.09951 | 0.473754 | 0.923329 | Zfp641 |
| ENSMUSG( | 864.5649 | -0.00147 | 0.989929 | 0.997928 | Zfp644 |
| ENSMUSG( | 739.6735 | 0.229666 | 0.110316 | 0.807051 | Zfp646 |
| ENSMUSG( | 68.58302 | 0.41859 | 0.140266 | 0.830678 | Zfp647 |
| ENSMUSG( | 199.8944 | 0.029256 | 0.866795 | 0.987487 | Zfp65 |
| ENSMUSG( | 3241.97 | -0.06748 | 0.602933 | 0.947907 | Zfp652 |
| ENSMUSG( | 162.6424 | -0.31238 | 0.10126 | 0.801456 | Zfp652os |
| ENSMUSG( | 429.6631 | 0.042978 | 0.795725 | 0.97681 | Zfp653 |
| ENSMUSG( | 453.7158 | 0.159749 | 0.272816 | 0.888937 | Zfp654 |
| ENSMUSG( | 481.6451 | -0.04665 | 0.748315 | 0.96984 | Zfp655 |
| ENSMUSG( | 185.457 | -0.00365 | 0.98433 | 0.996963 | Zfp658 |
| ENSMUSG( | 93.12671 | 0.147916 | 0.532062 | 0.934795 | Zfp660 |
| ENSMUSG( | 250.6258 | -0.02271 | 0.898841 | 0.990558 | Zfp661 |
| ENSMUSG( | 6220.052 | -0.07456 | 0.532726 | 0.934795 | Zfp664 |
| ENSMUSG( | 233.9431 | -0.03278 | 0.848449 | 0.985321 | Zfp667 |
| ENSMUSG( | 344.1721 | 0.151267 | 0.313563 | 0.900883 | Zfp668 |
| ENSMUSG( | 748.0306 | 0.032659 | 0.80566 | 0.977674 | Zfp672 |
| ENSMUSG( | 41.78866 | -0.41782 | 0.188501 | 0.85698 | Zfp677 |
| ENSMUSG( | 615.2179 | 0.139507 | 0.258867 | 0.883971 | Zfp68 |
| ENSMUSG( | 1379.74 | -0.12836 | 0.259005 | 0.883971 | Zfp687 |
| ENSMUSG( | 248.097 | 0.020992 | 0.902817 | 0.991089 | Zfp688 |
| ENSMUSG( | 143.1015 | -0.05128 | 0.797136 | 0.97681 | Zfp689 |
| ENSMUSG( | 63.21925 | -0.1909 | 0.60144 | 0.947749 | Zfp69 |
| ENSMUSG( | 170.3682 | 0.2003 | 0.318874 | 0.903344 | Zfp691 |
| ENSMUSG( | 674.1085 | 0.060011 | 0.802743 | 0.97681 | Zfp692 |
| ENSMUSG( | 557.1738 | -0.01186 | 0.923481 | 0.992815 | Zfp697 |
| ENSMUSG( | 139.2139 | 0.312765 | 0.104618 | 0.804969 | Zfp7 |
| ENSMUSG( | 1058.619 | 0.151088 | 0.288407 | 0.894794 | Zfp703 |
| ENSMUSG( | 2610.392 | 0.09504 | 0.44954 | 0.922202 | Zfp704 |
| ENSMUSG( | 4479.537 | -0.11422 | 0.379559 | 0.916995 | Zfp706 |
| ENSMUSG( | 244.891 | 0.042181 | 0.798981 | 0.97681 | Zfp707 |
| ENSMUSG( | 143.961 | 0.009704 | 0.958689 | 0.994798 | Zfp709 |
| ENSMUSG( | 801.166 | -0.06961 | 0.562956 | 0.941158 | Zfp710 |
| ENSMUSG( | 181.4412 | 0.13897 | 0.414394 | 0.917257 | Zfp711 |
| ENSMUSG( | 74.5234 | 0.049838 | 0.847484 | 0.985161 | Zfp712 |
| ENSMUSG( | 647.9858 | 0.114951 | 0.426319 | 0.917257 | Zfp715 |
| ENSMUSG( | 309.5247 | -0.02611 | 0.862473 | 0.986545 | Zfp719 |
| ENSMUSG( | 92.54952 | 0.071254 | 0.768785 | 0.974236 | Zfp72 |
| ENSMUSG( | 317.8454 | -0.01655 | 0.913339 | 0.992298 | Zfp729a |
| ENSMUSG( | 173.7729 | 0.175632 | 0.350765 | 0.911707 | Zfp729b |
| ENSMUSG( | 650.0312 | 0.006589 | 0.966001 | 0.995436 | Zfp738 |
| ENSMUSG( | 364.6768 | 0.083526 | 0.608621 | 0.950113 | Zfp74 |

| | | | | | | | | | | | |
|---|---|---|---|---|---|---|---|---|---|---|---|
| ENSMUSG( | Zfp850 | 114.0402 | 0.073759 | 0.748169 | 0.949604 | ENSMUSG( | Zfp850 | 104.9706 | 0.090779 | 0.702475 | 0.962215 |
| ENSMUSG( | Zfp853 | 485.6879 | 0.217037 | 0.075989 | 0.426896 | ENSMUSG( | Zfp853 | 416.7736 | 0.057434 | 0.682526 | 0.958678 |
| ENSMUSG( | Zfp865 | 1044.171 | 0.02261 | 0.819955 | 0.965888 | ENSMUSG( | Zfp865 | 963.2194 | 0.062674 | 0.638546 | 0.953903 |
| ENSMUSG( | Zfp866 | 589.066 | -0.0044 | 0.970881 | 0.996369 | ENSMUSG( | Zfp866 | 547.2979 | 0.057398 | 0.673344 | 0.95694 |
| ENSMUSG( | Zfp867 | 115.9941 | 0.10802 | 0.564208 | 0.890139 | ENSMUSG( | Zfp867 | 106.1859 | 0.13881 | 0.525907 | 0.934374 |
| ENSMUSG( | Zfp868 | 220.7119 | -0.07684 | 0.598934 | 0.903801 | ENSMUSG( | Zfp868 | 222.6539 | 0.209082 | 0.200852 | 0.86439 |
| ENSMUSG( | Zfp869 | 778.431 | -0.06804 | 0.537989 | 0.882548 | ENSMUSG( | Zfp869 | 695.3436 | -0.12288 | 0.321203 | 0.904019 |
| ENSMUSG( | Zfp87 | 205.5347 | 0.076647 | 0.607257 | 0.906826 | ENSMUSG( | Zfp87 | 195.0541 | 0.185129 | 0.287101 | 0.894091 |
| ENSMUSG( | Zfp870 | 362.0912 | 0.118769 | 0.377853 | 0.801561 | ENSMUSG( | Zfp870 | 318.5612 | 0.02353 | 0.877993 | 0.989644 |
| ENSMUSG( | Zfp871 | 1837.735 | 0.010097 | 0.916755 | 0.988856 | ENSMUSG( | Zfp871 | 1711.439 | 0.073047 | 0.513769 | 0.930586 |
| ENSMUSG( | Zfp873 | 90.87886 | 0.011073 | 0.962473 | 0.994604 | ENSMUSG( | Zfp873 | 85.57332 | 0.122202 | 0.613736 | 0.951177 |
| ENSMUSG( | Zfp874a | 241.7051 | -0.02823 | 0.844029 | 0.971962 | ENSMUSG( | Zfp874a | 239.8002 | 0.215283 | 0.176774 | 0.846973 |
| ENSMUSG( | Zfp874b | 274.5253 | 0.122265 | 0.35844 | 0.789759 | ENSMUSG( | Zfp874b | 265.517 | 0.287018 | 0.05767 | 0.726244 |
| ENSMUSG( | Zfp879 | 67.18048 | 0.086187 | 0.727336 | 0.944244 | ENSMUSG( | Zfp879 | 61.83751 | 0.130781 | 0.62048 | 0.952915 |
| ENSMUSG( | Zfp882 | 135.5352 | 0.159213 | 0.38072 | 0.801984 | ENSMUSG( | Zfp882 | 112.7824 | -0.10391 | 0.626273 | 0.953234 |
| ENSMUSG( | Zfp9 | 369.9101 | 0.096858 | 0.497064 | 0.861754 | ENSMUSG( | Zfp9 | 333.7194 | 0.073762 | 0.651972 | 0.9547 |
| ENSMUSG( | Zfp90 | 403.6699 | -0.21128 | 0.146725 | 0.56887 | ENSMUSG( | Zfp90 | 395.0809 | 0.00635 | 0.967806 | 0.995833 |
| ENSMUSG( | Zfp91 | 5938.344 | 0.005249 | 0.962291 | 0.994543 | ENSMUSG( | Zfp91 | 5312.439 | -0.05043 | 0.740471 | 0.968811 |
| ENSMUSG( | Zfp92 | 41.04488 | 0.395787 | 0.192044 | 0.630811 | ENSMUSG( | Zfp92 | 36.81598 | 0.351324 | 0.286016 | 0.893906 |
| ENSMUSG( | Zfp93 | 223.982 | -0.06488 | 0.702671 | 0.937735 | ENSMUSG( | Zfp93 | 211.9091 | 0.050121 | 0.779644 | 0.976106 |
| ENSMUSG( | Zfp930 | 187.2375 | -0.19186 | 0.248887 | 0.698071 | ENSMUSG( | Zfp930 | 186.3597 | 0.067141 | 0.700103 | 0.961335 |
| ENSMUSG( | Zfp931 | 307.2221 | -0.02902 | 0.889183 | 0.983434 | ENSMUSG( | Zfp931 | 261.1245 | -0.24904 | 0.22868 | 0.868507 |
| ENSMUSG( | Zfp932 | 608.0229 | -0.14334 | 0.28338 | 0.73186 | ENSMUSG( | Zfp932 | 586.8058 | 0.020415 | 0.896242 | 0.990558 |
| ENSMUSG( | Zfp933 | 369.8137 | 0.215547 | 0.121433 | 0.525713 | ENSMUSG( | Zfp933 | 332.1597 | 0.170264 | 0.304104 | 0.898538 |
| ENSMUSG( | Zfp934 | 115.6981 | -0.21294 | 0.316759 | 0.763648 | ENSMUSG( | Zfp934 | 109.9574 | -0.09664 | 0.691256 | 0.960071 |
| ENSMUSG( | Zfp935 | 49.47172 | -0.16392 | 0.545958 | 0.884974 | ENSMUSG( | Zfp935 | 47.79425 | -0.01277 | 0.966057 | 0.995436 |
| ENSMUSG( | Zfp937 | 50.42704 | 0.504996 | 0.073104 | 0.417421 | ENSMUSG( | Zfp937 | 45.647 | 0.449061 | 0.196383 | 0.862161 |
| ENSMUSG( | Zfp938 | 350.5695 | -0.11971 | 0.36705 | 0.795164 | ENSMUSG( | Zfp938 | 340.5444 | 0.058967 | 0.714508 | 0.964986 |
| ENSMUSG( | Zfp939 | 19.06892 | -0.41497 | 0.330208 | NA | ENSMUSG( | Zfp939 | 17.18869 | -0.40109 | 0.406654 | 0.917257 |
| ENSMUSG( | Zfp94 | 236.6167 | -0.1431 | 0.329099 | 0.773233 | ENSMUSG( | Zfp94 | 224.3923 | -0.01314 | 0.940036 | 0.994334 |
| ENSMUSG( | Zfp940 | 425.4895 | -0.12218 | 0.378605 | 0.801688 | ENSMUSG( | Zfp940 | 402.2967 | -0.01083 | 0.941511 | 0.994578 |
| ENSMUSG( | Zfp941 | 322.9533 | -0.16701 | 0.262855 | 0.713397 | ENSMUSG( | Zfp941 | 306.6107 | -0.05042 | 0.771131 | 0.974236 |
| ENSMUSG( | Zfp942 | 306.036 | -0.08463 | 0.559003 | 0.888638 | ENSMUSG( | Zfp942 | 303.7945 | 0.149222 | 0.397312 | 0.917257 |
| ENSMUSG( | Zfp943 | 426.6129 | 0.059516 | 0.686893 | 0.933041 | ENSMUSG( | Zfp943 | 438.6162 | 0.38431 | 0.012958 | 0.458131 |
| ENSMUSG( | Zfp944 | 226.0163 | 0.082195 | 0.622949 | 0.911589 | ENSMUSG( | Zfp944 | 228.5001 | 0.362535 | 0.043782 | 0.690304 |
| ENSMUSG( | Zfp945 | 409.2497 | -0.50895 | 2.23E-05 | 0.002129 | ENSMUSG( | Zfp945 | 448.7485 | 0.073803 | 0.590823 | 0.944975 |
| ENSMUSG( | Zfp946 | 235.8194 | -0.22134 | 0.200223 | 0.640302 | ENSMUSG( | Zfp946 | 239.0397 | 0.091288 | 0.598138 | 0.946676 |
| ENSMUSG( | Zfp948 | 303.4066 | -0.2463 | 0.085708 | 0.451367 | ENSMUSG( | Zfp948 | 285.059 | -0.15946 | 0.333588 | 0.907907 |
| ENSMUSG( | Zfp949 | 402.0725 | 0.175655 | 0.258836 | 0.709814 | ENSMUSG( | Zfp949 | 347.838 | 0.038057 | 0.836337 | 0.981335 |
| ENSMUSG( | Zfp950 | 1066.834 | -0.08689 | 0.44405 | 0.833239 | ENSMUSG( | Zfp950 | 1073.577 | 0.195263 | 0.215647 | 0.865167 |
| ENSMUSG( | Zfp951 | 56.35848 | -0.32278 | 0.202382 | 0.642343 | ENSMUSG( | Zfp951 | 49.78722 | -0.39856 | 0.182437 | 0.850579 |
| ENSMUSG( | Zfp952 | 357.5011 | -0.01334 | 0.914017 | 0.988483 | ENSMUSG( | Zfp952 | 347.6043 | 0.172656 | 0.22747 | 0.868507 |
| ENSMUSG( | Zfp953 | 42.48536 | -0.01278 | 0.965965 | 0.995377 | ENSMUSG( | Zfp953 | 38.47223 | -0.03122 | 0.922868 | 0.992644 |
| ENSMUSG( | Zfp954 | 221.0014 | -0.03546 | 0.848976 | 0.972554 | ENSMUSG( | Zfp954 | 196.2995 | -0.09543 | 0.631374 | 0.953251 |
| ENSMUSG( | Zfp955a | 154.0904 | 0.03669 | 0.841789 | 0.971179 | ENSMUSG( | Zfp955a | 140.6913 | 0.05237 | 0.792667 | 0.97681 |
| ENSMUSG( | Zfp955b | 479.5198 | 0.118989 | 0.30252 | 0.75086 | ENSMUSG( | Zfp955b | 428.8112 | 0.071625 | 0.629254 | 0.953251 |
| ENSMUSG( | Zfp956 | 651.3444 | 0.019706 | 0.870529 | 0.977906 | ENSMUSG( | Zfp956 | 599.6932 | 0.051953 | 0.712416 | 0.964363 |
| ENSMUSG( | Zfp958 | 148.994 | -0.0078 | 0.966134 | 0.995377 | ENSMUSG( | Zfp958 | 147.696 | 0.232356 | 0.233134 | 0.868507 |
| ENSMUSG( | Zfp959 | 119.7587 | 0.001765 | 0.9928 | 0.999327 | ENSMUSG( | Zfp959 | 111.3959 | 0.072711 | 0.726627 | 0.966887 |
| ENSMUSG( | Zfp960 | 22.42254 | -0.16724 | 0.691974 | 0.934101 | ENSMUSG( | Zfp960 | 24.76062 | 0.351802 | 0.410785 | 0.917257 |
| ENSMUSG( | Zfp961 | 337.7569 | 0.09499 | 0.493892 | 0.860611 | ENSMUSG( | Zfp961 | 283.7406 | -0.15059 | 0.39067 | 0.917257 |
| ENSMUSG( | Zfp963 | 88.08923 | -0.28291 | 0.189803 | 0.628929 | ENSMUSG( | Zfp963 | 87.73737 | -0.01349 | 0.953599 | 0.994798 |
| ENSMUSG( | Zfp964 | 87.53645 | -0.22557 | 0.2834 | 0.73186 | ENSMUSG( | Zfp964 | 92.19475 | 0.211423 | 0.358183 | 0.913759 |
| ENSMUSG( | Zfp97 | 25.44673 | -0.13097 | 0.747968 | 0.949604 | ENSMUSG( | Zfp97 | 25.67516 | 0.153887 | 0.708383 | 0.964042 |
| ENSMUSG( | Zfp970 | 249.3874 | 0.042358 | 0.783424 | 0.957558 | ENSMUSG( | Zfp970 | 223.8083 | 0.014097 | 0.940169 | 0.994334 |
| ENSMUSG( | Zfp971 | 157.1719 | 0.139618 | 0.492539 | 0.860378 | ENSMUSG( | Zfp971 | 138.41 | 0.050585 | 0.822622 | 0.979198 |
| ENSMUSG( | Zfp974 | 132.2955 | -0.32027 | 0.082252 | 0.442642 | ENSMUSG( | Zfp974 | 130.2652 | -0.06393 | 0.756686 | 0.971779 |
| ENSMUSG( | Zfp975 | 100.5103 | -0.07197 | 0.720031 | 0.941986 | ENSMUSG( | Zfp975 | 98.8924 | 0.136134 | 0.537916 | 0.937216 |
| ENSMUSG( | Zfp976 | 77.67183 | -0.1299 | 0.608557 | 0.907154 | ENSMUSG( | Zfp976 | 66.75131 | -0.30497 | 0.242824 | 0.877399 |
| ENSMUSG( | Zfp977 | 25.54765 | -0.22056 | 0.565923 | 0.89104 | ENSMUSG( | Zfp977 | 24.78488 | -0.04386 | 0.91559 | 0.992529 |
| ENSMUSG( | Zfp979 | 19.59274 | -0.71028 | 0.100729 | NA | ENSMUSG( | Zfp979 | 22.71536 | 0.051381 | 0.908833 | 0.991958 |
| ENSMUSG( | Zfp982 | 46.7373 | -0.26424 | 0.440298 | 0.831851 | ENSMUSG( | Zfp982 | 42.72833 | -0.28492 | 0.414524 | 0.917257 |
| ENSMUSG( | Zfp983 | 289.4772 | -0.00048 | 0.997245 | 0.999587 | ENSMUSG( | Zfp983 | 268.7589 | 0.051426 | 0.740185 | 0.968811 |
| ENSMUSG( | Zfp984 | 81.12788 | -0.2882 | 0.248527 | 0.697423 | ENSMUSG( | Zfp984 | 84.31172 | 0.087913 | 0.750375 | 0.970113 |
| ENSMUSG( | Zfp991 | 16.73273 | -0.58326 | 0.255126 | NA | ENSMUSG( | Zfp991 | 18.42778 | 0.027548 | 0.954081 | 0.994798 |
| ENSMUSG( | Zfp994 | 111.0117 | -0.24395 | 0.208393 | 0.650319 | ENSMUSG( | Zfp994 | 113.694 | 0.097692 | 0.63686 | 0.953903 |
| ENSMUSG( | Zfp995 | 98.38904 | 0.069092 | 0.746176 | 0.949289 | ENSMUSG( | Zfp995 | 91.20472 | 0.120469 | 0.644803 | 0.953903 |
| ENSMUSG( | Zfpl1 | 377.3263 | -0.0635 | 0.635385 | 0.914296 | ENSMUSG( | Zfpl1 | 352.4038 | 0.007143 | 0.963653 | 0.995433 |
| ENSMUSG( | Zfpm1 | 1866.077 | 0.298117 | 0.007382 | 0.118765 | ENSMUSG( | Zfpm1 | 1650.442 | 0.223134 | 0.098083 | 0.798438 |
| ENSMUSG( | Zfpm2 | 268.6569 | 0.225712 | 0.128066 | 0.536614 | ENSMUSG( | Zfpm2 | 252.7756 | 0.31427 | 0.061579 | 0.736111 |
| ENSMUSG( | Zfr | 13404.11 | 0.175111 | 0.04689 | 0.331252 | ENSMUSG( | Zfr | 11602.14 | 0.032089 | 0.758908 | 0.971858 |
| ENSMUSG( | Zfr2 | 1849.426 | 0.253471 | 0.040161 | 0.309163 | ENSMUSG( | Zfr2 | 1425.722 | -0.22883 | 0.129842 | 0.820802 |
| ENSMUSG( | Zfta | 1955.82 | 0.206521 | 0.0232 | 0.233586 | ENSMUSG( | Zfta | 1753.819 | 0.164042 | 0.118338 | 0.813384 |
| ENSMUSG( | Zftraf1 | 1197.691 | 0.104303 | 0.374107 | 0.800445 | ENSMUSG( | Zftraf1 | 1064.839 | 0.034092 | 0.759376 | 0.972002 |
| ENSMUSG( | Zftraf1 | 3215.067 | 0.166779 | 0.036751 | 0.295976 | ENSMUSG( | Zftraf1 | 2680.568 | -0.08578 | 0.391274 | 0.917257 |
| ENSMUSG( | Zfx | 594.5472 | 0.000841 | 0.993827 | 0.999369 | ENSMUSG( | Zfx | 550.8874 | 0.050152 | 0.693322 | 0.96076 |
| ENSMUSG( | Zfyve1 | 1206.58 | 0.068713 | 0.488172 | 0.859092 | ENSMUSG( | Zfyve1 | 1082.042 | 0.02605 | 0.82532 | 0.979388 |
| ENSMUSG( | Zfyve16 | 590.5183 | -0.05323 | 0.671365 | 0.928781 | ENSMUSG( | Zfyve16 | 542.1366 | -0.02684 | 0.851436 | 0.985541 |
| ENSMUSG( | Zfyve19 | 490.6375 | 0.018338 | 0.870952 | 0.977951 | ENSMUSG( | Zfyve19 | 440.1819 | -0.02239 | 0.869754 | 0.988097 |
| ENSMUSG( | Zfyve21 | 831.547 | 0.070035 | 0.489968 | 0.859319 | ENSMUSG( | Zfyve21 | 736.8009 | -0.01335 | 0.914125 | 0.992407 |
| ENSMUSG( | Zfyve26 | 699.5753 | 0.006584 | 0.957906 | 0.993874 | ENSMUSG( | Zfyve26 | 664.6608 | 0.132282 | 0.402506 | 0.917257 |
| ENSMUSG( | Zfyve27 | 2726.03 | -0.20524 | 0.103773 | 0.489428 | ENSMUSG( | Zfyve27 | 2791.754 | 0.140333 | 0.307434 | 0.898538 |
| ENSMUSG( | Zfyve28 | 1426.708 | -0.20793 | 0.045845 | 0.328083 | ENSMUSG( | Zfyve28 | 1363.464 | -0.06298 | 0.591699 | 0.944975 |
| ENSMUSG( | Zfyve9 | 1839.595 | 0.041669 | 0.720877 | 0.942684 | ENSMUSG( | Zfyve9 | 1648.275 | -0.0079 | 0.953886 | 0.994798 |
| ENSMUSG( | Zgpat | 968.9337 | -5.52E-05 | 0.999704 | 0.99989 | ENSMUSG( | Zgpat | 965.3269 | 0.254839 | 0.257804 | 0.883401 |
| ENSMUSG( | Zgrf1 | 173.5435 | -0.23593 | 0.179998 | 0.617634 | ENSMUSG( | Zgrf1 | 164.5705 | -0.11477 | 0.537757 | 0.937216 |
| ENSMUSG( | Zhx1 | 3182.068 | 0.191475 | 0.113483 | 0.508017 | ENSMUSG( | Zhx1 | 2699.588 | -0.01047 | 0.937134 | 0.993685 |
| ENSMUSG( | Zhx2 | 970.4176 | 0.046299 | 0.657698 | 0.92425 | ENSMUSG( | Zhx2 | 878.033 | 0.023556 | 0.848481 | 0.985321 |
| ENSMUSG( | Zhx3 | 3762.159 | 0.618794 | 1.49E-11 | 9.68E-09 | ENSMUSG( | Zhx3 | 3049.528 | 0.327028 | 0.009996 | 0.408157 |
| ENSMUSG( | Zic1 | 444.4524 | -0.2599 | 0.105665 | 0.492581 | ENSMUSG( | Zic1 | 519.6731 | 0.433921 | 0.013796 | 0.46927 |
| ENSMUSG( | Zic2 | 423.9047 | 0.002824 | 0.982325 | 0.997512 | ENSMUSG( | Zic2 | 448.4564 | 0.412823 | 0.010052 | 0.409511 |
| ENSMUSG( | Zic3 | 102.2077 | 0.144917 | 0.494294 | 0.860611 | ENSMUSG( | Zic3 | 128.4103 | 0.931002 | 6.96E-06 | 0.002837 |
| ENSMUSG( | Zic4 | 151.3492 | -0.24566 | 0.167531 | 0.600122 | ENSMUSG( | Zic4 | 164.5637 | 0.269557 | 0.198167 | 0.862161 |
| ENSMUSG( | Zic5 | 107.4488 | 0.150406 | 0.541503 | 0.883182 | ENSMUSG( | Zic5 | 112.5297 | 0.519769 | 0.029449 | 0.618981 |
| ENSMUSG( | Zik1 | 167.8705 | 0.108475 | 0.516111 | 0.872701 | ENSMUSG( | Zik1 | 155.5944 | 0.153304 | 0.412791 | 0.917257 |
| ENSMUSG( | Zim1 | 112.0786 | -0.08894 | 0.651846 | 0.921871 | ENSMUSG( | Zim1 | 107.6579 | 0.067821 | 0.771106 | 0.974236 |
| ENSMUSG( | Zkscan1 | 945.3398 | -0.03664 | 0.734696 | 0.945584 | ENSMUSG( | Zkscan1 | 875.1176 | 0.011197 | 0.929 | 0.99306 |
| ENSMUSG( | Zkscan14 | 175.9883 | -0.09982 | 0.617311 | 0.910866 | ENSMUSG( | Zkscan14 | 154.6087 | -0.20246 | 0.358633 | 0.913759 |
| ENSMUSG( | Zkscan16 | 261.5279 | -0.19894 | 0.180226 | 0.617634 | ENSMUSG( | Zkscan16 | 268.7601 | 0.162608 | 0.359523 | 0.914223 |
| ENSMUSG( | Zkscan17 | 744.6884 | 0.079508 | 0.442858 | 0.83278 | ENSMUSG( | Zkscan17 | 648.558 | -0.04873 | 0.693521 | 0.960767 |
| ENSMUSG( | Zkscan2 | 1100.044 | -0.58545 | 1.01E-08 | 3.91E-06 | ENSMUSG( | Zkscan2 | 1097.424 | -0.27842 | 0.018255 | 0.542424 |
| ENSMUSG( | Zkscan3 | 1110.401 | 0.060926 | 0.640239 | 0.916434 | ENSMUSG( | Zkscan3 | 1088.214 | 0.265504 | 0.103184 | 0.804311 |
| ENSMUSG( | Zkscan4 | 111.5637 | 0.257225 | 0.201592 | 0.641362 | ENSMUSG( | Zkscan4 | 98.19344 | 0.162727 | 0.475987 | 0.923329 |
| ENSMUSG( | Zkscan5 | 515.8721 | 0.122017 | 0.27932 | 0.729114 | ENSMUSG( | Zkscan5 | 461.009 | 0.068976 | 0.604075 | 0.948139 |
| ENSMUSG( | Zkscan6 | 366.824 | 0.142209 | 0.314502 | 0.761558 | ENSMUSG( | Zkscan6 | 301.3184 | -0.15636 | 0.350554 | 0.911652 |
| ENSMUSG( | Zkscan7 | 153.0997 | -0.10924 | 0.538097 | 0.882548 | ENSMUSG( | Zkscan7 | 148.5212 | 0.087397 | 0.661506 | 0.956205 |
| ENSMUSG( | Zkscan8 | 535.446 | -0.05903 | 0.642354 | 0.916865 | ENSMUSG( | Zkscan8 | 502.4558 | 0.025902 | 0.858732 | 0.985884 |
| ENSMUSG( | Zmat1 | 371.5393 | 0.116206 | 0.340417 | 0.778122 | ENSMUSG( | Zmat1 | 336.8381 | 0.103399 | 0.471254 | 0.923329 |
| ENSMUSG( | Zmat2 | 3929.096 | 0.140673 | 0.154159 | 0.580901 | ENSMUSG( | Zmat2 | 3219.465 | -0.1699 | 0.224068 | 0.867404 |
| ENSMUSG( | Zmat3 | 3987.185 | 0.18973 | 0.023395 | 0.233882 | ENSMUSG( | Zmat3 | 3407.777 | 0.009988 | 0.927316 | 0.99306 |
| ENSMUSG( | Zmat4 | 924.3546 | -0.08429 | 0.570573 | 0.893192 | ENSMUSG( | Zmat4 | 778.9743 | -0.33226 | 0.027873 | 0.61023 |
| ENSMUSG( | Zmat5 | 513.6479 | -0.1951 | 0.116854 | 0.515606 | ENSMUSG( | Zmat5 | 452.6423 | -0.30618 | 0.033773 | 0.647272 |
| ENSMUSG( | Zmiz1 | 4799.292 | 0.069438 | 0.425949 | 0.824898 | ENSMUSG( | Zmiz1 | 4452.875 | 0.12201 | 0.258037 | 0.883684 |
| ENSMUSG( | Zmiz2 | 17493.1 | 0.241003 | 0.004051 | 0.081936 | ENSMUSG( | Zmiz2 | 15234.17 | 0.11936 | 0.242162 | 0.876967 |
| ENSMUSG( | Zmpste24 | 1602.237 | 0.023401 | 0.868555 | 0.976897 | ENSMUSG( | Zmpste24 | 1504.911 | 0.111883 | 0.482462 | 0.924578 |
| ENSMUSG( | Zmym1 | 230.7 | 0.253737 | 0.0848 | 0.448764 | ENSMUSG( | Zmym1 | 203.8446 | 0.172625 | 0.292612 | 0.897982 |
| ENSMUSG( | Zmym2 | 1704.884 | 0.00131 | 0.98776 | 0.998907 | ENSMUSG( | Zmym2 | 1589.292 | 0.065119 | 0.556046 | 0.939565 |
| ENSMUSG( | Zmym3 | 3328.146 | 0.204902 | 0.119543 | 0.523344 | ENSMUSG( | Zmym3 | 2746.074 | -0.0789 | 0.64238 | 0.953903 |
| ENSMUSG( | Zmym4 | 1597.954 | -0.05356 | 0.563326 | 0.889914 | ENSMUSG( | Zmym4 | 1431.081 | -0.10486 | 0.357365 | 0.91373 |
| ENSMUSG( | Zmym5 | 1003.459 | -0.12318 | 0.202524 | 0.642593 | ENSMUSG( | Zmym5 | 981.545 | 0.084174 | 0.48164 | 0.924578 |
| ENSMUSG( | Zmym6 | 819.5051 | -0.47526 | 5.65E-06 | 0.000739 | ENSMUSG( | Zmym6 | 869.9873 | 0.005006 | 0.97495 | 0.996416 |
| ENSMUSG( | Zmynd10 | 99.92058 | 0.059802 | 0.766385 | 0.954173 | ENSMUSG( | Zmynd10 | 97.10567 | 0.234728 | 0.301283 | 0.898538 |
| ENSMUSG( | Zmynd11 | 7828.553 | 0.063148 | 0.431261 | 0.827055 | ENSMUSG( | Zmynd11 | 6938.615 | -0.01754 | 0.87432 | 0.989644 |
| ENSMUSG( | Zmynd15 | 18.23948 | 0.608103 | 0.180853 | NA | ENSMUSG( | Zmynd15 | 17.16934 | 0.692572 | 0.147826 | 0.836241 |
| ENSMUSG( | Zmynd19 | 672.3771 | 0.180603 | 0.160483 | 0.590122 | ENSMUSG( | Zmynd19 | 563.3454 | -0.06171 | 0.711714 | 0.964363 |
| ENSMUSG( | Zmynd8 | 4759.842 | 0.308332 | 0.005195 | 0.096074 | ENSMUSG( | Zmynd8 | 3982.939 | 0.076628 | 0.517762 | 0.930939 |
| ENSMUSG( | Znfx1 | 2176.361 | 0.130015 | 0.155198 | 0.583117 | ENSMUSG( | Znfx1 | 1962.956 | 0.104384 | 0.352878 | 0.912263 |
| ENSMUSG( | Zng1 | 256.6638 | 0.093671 | 0.512674 | 0.871762 | ENSMUSG( | Zng1 | 219.8435 | -0.09673 | 0.581546 | 0.944266 |
| ENSMUSG( | Znhit1 | 1101.213 | 0.009417 | 0.928548 | 0.991641 | ENSMUSG( | Znhit1 | 920.0446 | -0.25514 | 0.046668 | 0.699307 |
| ENSMUSG( | Znhit2 | 913.6878 | 0.114191 | 0.33674 | 0.777 | ENSMUSG( | Znhit2 | 801.4022 | 0.006472 | 0.962338 | 0.995237 |
| ENSMUSG( | Znhit3 | 340.0477 | 0.039261 | 0.824421 | 0.967102 | ENSMUSG( | Znhit3 | 291.8881 | -0.13902 | 0.474565 | 0.923329 |
| ENSMUSG( | Znhit6 | 357.4022 | 0.083118 | 0.500553 | 0.863514 | ENSMUSG( | Znhit6 | 310.7684 | -0.05728 | 0.711468 | 0.964363 |
| ENSMUSG( | Znrf1 | 8400.067 | 0.082116 | 0.306139 | 0.753712 | ENSMUSG( | Znrf1 | 7627.512 | 0.072959 | 0.471944 | 0.923329 |
| ENSMUSG( | Znrf2 | 776.4075 | 0.080559 | 0.414876 | 0.820184 | ENSMUSG( | Znrf2 | 726.7474 | 0.150932 | 0.246776 | 0.877904 |
| ENSMUSG( | Znrf3 | 609.126 | -0.26623 | 0.078979 | 0.434796 | ENSMUSG( | Znrf3 | 635.0989 | 0.138277 | 0.434514 | 0.91869 |
| ENSMUSG( | Zpbp | 35.02303 | -0.31094 | 0.3752 | 0.800786 | ENSMUSG( | Zpbp | 33.26908 | -0.18714 | 0.590399 | 0.944975 |
| ENSMUSG( | Zpr1 | 905.0922 | 0.071797 | 0.5497 | 0.885944 | ENSMUSG( | Zpr1 | 806.6914 | 0.009203 | 0.943666 | 0.994798 |
| ENSMUSG( | Zranb1 | 1714.622 | 0.087521 | 0.380173 | 0.801757 | ENSMUSG( | Zranb1 | 1544.929 | 0.053094 | 0.740165 | 0.968811 |
| ENSMUSG( | Zranb2 | 2325.916 | -0.02404 | 0.813288 | 0.964574 | ENSMUSG( | Zranb2 | 2322.937 | 0.230159 | 0.100541 | 0.801444 |
| ENSMUSG( | Zranb3 | 260.5447 | 0.003519 | 0.981989 | 0.997512 | ENSMUSG( | Zranb3 | 222.8395 | -0.18643 | 0.307414 | 0.898538 |
| ENSMUSG( | Zrsr2 | 903.8007 | -0.0292 | 0.796266 | 0.960088 | ENSMUSG( | Zrsr2 | 832.1852 | -0.00503 | 0.973421 | 0.996416 |
| ENSMUSG( | Zscan12 | 187.6282 | -0.10457 | 0.534478 | 0.880719 | ENSMUSG( | Zscan12 | 176.3039 | -0.01821 | 0.917927 | 0.992529 |
| ENSMUSG( | Zscan18 | 376.7986 | -0.04451 | 0.733164 | 0.944956 | ENSMUSG( | Zscan18 | 328.5279 | -0.1721 | 0.27426 | 0.889874 |
| ENSMUSG( | Zscan2 | 171.6593 | 0.02617 | 0.880068 | 0.981469 | ENSMUSG( | Zscan2 | 154.6079 | -0.01272 | 0.946001 | 0.994798 |
| ENSMUSG( | Zscan20 | 200.6866 | 0.236617 | 0.119327 | 0.523036 | ENSMUSG( | Zscan20 | 168.6934 | 0.021634 | 0.90988 | 0.991958 |
| ENSMUSG( | Zscan21 | 425.6943 | 0.00725 | 0.960556 | 0.993991 | ENSMUSG( | Zscan21 | 395.1055 | 0.058156 | 0.725439 | 0.966887 |
| ENSMUSG( | Zscan22 | 231.6439 | 0.16943 | 0.305145 | 0.752998 | ENSMUSG( | Zscan22 | 206.0064 | 0.09934 | 0.597525 | 0.946609 |
| ENSMUSG( | Zscan25 | 268.8315 | 0.024893 | 0.882424 | 0.982244 | ENSMUSG( | Zscan25 | 246.864 | 0.052015 | 0.781673 | 0.976106 |
| ENSMUSG( | Zscan26 | 2127.517 | -0.04585 | 0.671205 | 0.928781 | ENSMUSG( | Zscan26 | 2055.608 | 0.118854 | 0.385176 | 0.917257 |
| ENSMUSG( | Zscan29 | 272.7885 | 0.112998 | 0.416288 | 0.82084 | ENSMUSG( | Zscan29 | 265.4044 | 0.296474 | 0.059906 | 0.7301 |
| ENSMUSG( | Zscan30 | 27.30813 | 0.504221 | 0.193886 | 0.633085 | ENSMUSG( | Zscan30 | 24.42641 | 0.43987 | 0.330998 | 0.907513 |
| ENSMUSG( | Zswim1 | 468.934 | 0.255813 | 0.080286 | 0.438167 | ENSMUSG( | Zswim1 | 408.0737 | 0.130236 | 0.410019 | 0.917257 |
| ENSMUSG( | Zswim3 | 159.9756 | -0.03983 | 0.813289 | 0.964574 | ENSMUSG( | Zswim3 | 143.0814 | -0.08801 | 0.674908 | 0.95694 |
| ENSMUSG( | Zswim4 | 624.6324 | -0.0023 | 0.985395 | 0.998071 | ENSMUSG( | Zswim4 | 583.0337 | 0.069412 | 0.608217 | 0.950113 |
| ENSMUSG( | Zswim5 | 388.8256 | -0.01194 | 0.926478 | 0.991229 | ENSMUSG( | Zswim5 | 343.4053 | -0.10865 | 0.473713 | 0.923329 |
| ENSMUSG( | Zswim6 | 724.2264 | -0.27097 | 0.025988 | 0.247245 | ENSMUSG( | Zswim6 | 704.7061 | -0.07274 | 0.594676 | 0.945444 |
| ENSMUSG( | Zswim7 | 203.6232 | -0.19969 | 0.195959 | 0.635396 | ENSMUSG( | Zswim7 | 184.7553 | -0.22294 | 0.242353 | 0.877129 |
| ENSMUSG( | Zswim8 | 3793.378 | 0.174073 | 0.145744 | 0.566898 | ENSMUSG( | Zswim8 | 3387.726 | 0.120306 | 0.337504 | 0.907972 |

| | | | | | |
|---|---|---|---|---|---|
| ENSMUSG( | 1009.346 | -0.03433 | 0.792706 | 0.95947 | Zfp423 |
| ENSMUSG( | 728.3204 | 0.115291 | 0.261977 | 0.712129 | Zfp426 |
| ENSMUSG( | 890.7638 | -0.01745 | 0.870899 | 0.977951 | Zfp428 |
| ENSMUSG( | 52.44877 | -0.14113 | 0.623918 | 0.91187 | Zfp429 |
| ENSMUSG( | 18.39064 | 0.256816 | 0.556369 | NA | Zfp433 |
| ENSMUSG( | 211.4308 | -0.18853 | 0.231832 | 0.679725 | Zfp438 |
| ENSMUSG( | 33.47123 | -0.01816 | 0.955801 | 0.993874 | Zfp442 |
| ENSMUSG( | 1047.032 | 0.024396 | 0.81424 | 0.964574 | Zfp444 |
| ENSMUSG( | 4811.416 | -0.09714 | 0.349 | 0.781661 | Zfp445 |
| ENSMUSG( | 320.7309 | 0.070657 | 0.643553 | 0.917316 | Zfp446 |
| ENSMUSG( | 155.8252 | 0.179787 | 0.319767 | 0.765305 | Zfp449 |
| ENSMUSG( | 1077.121 | -0.04003 | 0.678573 | 0.930611 | Zfp451 |
| ENSMUSG( | 136.9489 | 0.107755 | 0.545107 | 0.884712 | Zfp454 |
| ENSMUSG( | 36.79726 | 0.072213 | 0.838225 | 0.971111 | Zfp455 |
| ENSMUSG( | 35.56774 | -0.40791 | 0.234192 | 0.68192 | Zfp456 |
| ENSMUSG( | 13.92425 | -0.78614 | 0.119923 | NA | Zfp457 |
| ENSMUSG( | 60.78613 | -0.24806 | 0.407312 | 0.817447 | Zfp458 |
| ENSMUSG( | 56.53197 | 0.063298 | 0.811059 | 0.964318 | Zfp459 |
| ENSMUSG( | 2032.089 | -0.04069 | 0.715688 | 0.941484 | Zfp46 |
| ENSMUSG( | 1354.034 | -0.16382 | 0.119507 | 0.523326 | Zfp462 |
| ENSMUSG( | 979.9189 | 0.145298 | 0.139669 | 0.556069 | Zfp467 |
| ENSMUSG( | 204.0971 | -0.71789 | 6.78E-05 | 0.005 | Zfp469 |
| ENSMUSG( | 82.49312 | 0.299944 | 0.192238 | 0.631194 | Zfp472 |
| ENSMUSG( | 15.753 | -0.8292 | 0.115882 | NA | Zfp473 |
| ENSMUSG( | 12.6437 | -0.56829 | 0.283179 | NA | Zfp474 |
| ENSMUSG( | 201.1406 | -0.08071 | 0.593873 | 0.902869 | Zfp488 |
| ENSMUSG( | 67.15228 | 0.303553 | 0.189371 | 0.627932 | Zfp493 |
| ENSMUSG( | 74.31229 | -0.39746 | 0.152208 | 0.578042 | Zfp503 |
| ENSMUSG( | 548.1134 | -0.00654 | 0.956252 | 0.993874 | Zfp507 |
| ENSMUSG( | 91.47198 | -0.10005 | 0.651208 | 0.921451 | Zfp51 |
| ENSMUSG( | 459.5985 | 0.083016 | 0.488492 | 0.859092 | Zfp511 |
| ENSMUSG( | 1375.889 | 0.141309 | 0.129133 | 0.538324 | Zfp512 |
| ENSMUSG( | 2538.798 | 0.090575 | 0.437152 | 0.829501 | Zfp512b |
| ENSMUSG( | 813.9625 | 0.056534 | 0.580354 | 0.896952 | Zfp513 |
| ENSMUSG( | 754.3463 | -0.1103 | 0.377721 | 0.801561 | Zfp516 |
| ENSMUSG( | 286.4499 | -0.12363 | 0.385247 | 0.804048 | Zfp518a |
| ENSMUSG( | 351.1216 | -0.0897 | 0.517576 | 0.873397 | Zfp518b |
| ENSMUSG( | 101.5943 | -0.25575 | 0.186471 | 0.623869 | Zfp52 |
| ENSMUSG( | 770.1532 | -0.00748 | 0.955976 | 0.993874 | Zfp521 |
| ENSMUSG( | 6521.751 | 0.167319 | 0.048964 | 0.337919 | Zfp523 |
| ENSMUSG( | 61.57724 | 0.307348 | 0.262526 | 0.713142 | Zfp524 |
| ENSMUSG( | 211.9892 | 0.09874 | 0.514467 | 0.872055 | Zfp526 |
| ENSMUSG( | 77.82223 | 0.1592 | 0.496602 | 0.86175 | Zfp53 |
| ENSMUSG( | 1761.42 | -0.04735 | 0.607754 | 0.906878 | Zfp532 |
| ENSMUSG( | 533.6566 | -0.02116 | 0.850113 | 0.972638 | Zfp536 |
| ENSMUSG( | 29.9796 | -0.11671 | 0.73832 | 0.947329 | Zfp54 |
| ENSMUSG( | 21.93058 | 0.004343 | 0.991592 | 0.999232 | Zfp541 |
| ENSMUSG( | 39.14606 | 0.403823 | 0.230858 | 0.679066 | Zfp551 |
| ENSMUSG( | 431.4243 | -0.12035 | 0.422705 | 0.823527 | Zfp553 |
| ENSMUSG( | 48.64464 | 0.166497 | 0.546238 | 0.884974 | Zfp558 |
| ENSMUSG( | 269.0207 | -0.08718 | 0.517341 | 0.873295 | Zfp560 |
| ENSMUSG( | 287.3424 | 0.105827 | 0.505294 | 0.866883 | Zfp563 |
| ENSMUSG( | 86.03148 | -0.26781 | 0.303102 | 0.751443 | Zfp566 |
| ENSMUSG( | 119.667 | -0.05534 | 0.783037 | 0.957558 | Zfp568 |
| ENSMUSG( | 366.6327 | 0.069315 | 0.648399 | 0.919965 | Zfp57 |
| ENSMUSG( | 965.2709 | 0.000674 | 0.994995 | 0.99943 | Zfp574 |
| ENSMUSG( | 263.4251 | -0.10047 | 0.503655 | 0.866308 | Zfp575 |
| ENSMUSG( | 589.5476 | 0.130495 | 0.252829 | 0.703255 | Zfp579 |
| ENSMUSG( | 72.50337 | -0.1548 | 0.559271 | 0.888638 | Zfp58 |
| ENSMUSG( | 751.3001 | -0.06534 | 0.513115 | 0.871762 | Zfp580 |
| ENSMUSG( | 103.5598 | 0.349914 | 0.132434 | 0.544369 | Zfp582 |
| ENSMUSG( | 123.323 | 0.145323 | 0.42558 | 0.824898 | Zfp583 |
| ENSMUSG( | 130.7149 | -0.19506 | 0.293329 | 0.742763 | Zfp59 |
| ENSMUSG( | 1158.119 | 0.086812 | 0.348526 | 0.78132 | Zfp592 |
| ENSMUSG( | 246.415 | 0.123908 | 0.410177 | 0.818258 | Zfp593 |
| ENSMUSG( | 96.05183 | 0.071761 | 0.74679 | 0.949289 | Zfp595 |
| ENSMUSG( | 260.5256 | -0.09758 | 0.484211 | 0.856577 | Zfp597 |
| ENSMUSG( | 1439.891 | 0.200181 | 0.097369 | 0.477629 | Zfp598 |
| ENSMUSG( | 54.80962 | -0.07286 | 0.785051 | 0.958473 | Zfp599 |
| ENSMUSG( | 774.5585 | -0.02854 | 0.783191 | 0.957558 | Zfp60 |
| ENSMUSG( | 335.6208 | 0.105582 | 0.446529 | 0.833726 | Zfp605 |
| ENSMUSG( | 448.2568 | -0.05913 | 0.620222 | 0.911432 | Zfp606 |
| ENSMUSG( | 29.95055 | -0.19508 | 0.579263 | 0.896692 | Zfp607a |
| ENSMUSG( | 411.3892 | 0.024115 | 0.85989 | 0.974818 | Zfp607b |
| ENSMUSG( | 765.7021 | -0.02679 | 0.823744 | 0.966603 | Zfp608 |
| ENSMUSG( | 1738.182 | -0.05442 | 0.59915 | 0.903801 | Zfp609 |
| ENSMUSG( | 354.5416 | -0.08996 | 0.590372 | 0.90113 | Zfp61 |
| ENSMUSG( | 1164.251 | -0.15723 | 0.141142 | 0.558043 | Zfp612 |
| ENSMUSG( | 541.781 | -0.06414 | 0.564212 | 0.890139 | Zfp617 |
| ENSMUSG( | 169.4772 | -0.29975 | 0.067768 | 0.402638 | Zfp618 |
| ENSMUSG( | 110.4338 | -0.12609 | 0.503549 | 0.866217 | Zfp619 |
| ENSMUSG( | 975.4054 | -0.09391 | 0.405651 | 0.81652 | Zfp62 |
| ENSMUSG( | 925.1379 | 0.125315 | 0.275389 | 0.72513 | Zfp622 |
| ENSMUSG( | 184.5651 | -0.25696 | 0.132122 | 0.544244 | Zfp623 |
| ENSMUSG( | 232.7481 | -0.18782 | 0.19578 | 0.635109 | Zfp626 |
| ENSMUSG( | 415.4607 | 0.094283 | 0.424553 | 0.824675 | Zfp628 |
| ENSMUSG( | 961.4451 | -0.08773 | 0.387914 | 0.805835 | Zfp629 |
| ENSMUSG( | 671.278 | 0.052257 | 0.634626 | 0.914073 | Zfp637 |
| ENSMUSG( | 2264.957 | 0.159755 | 0.076664 | 0.429084 | Zfp638 |
| ENSMUSG( | 740.2471 | -0.16944 | 0.133882 | 0.547197 | Zfp639 |
| ENSMUSG( | 447.1288 | 0.043793 | 0.716958 | 0.941697 | Zfp64 |
| ENSMUSG( | 571.6879 | -0.00689 | 0.957044 | 0.993874 | Zfp641 |
| ENSMUSG( | 952.333 | 0.011138 | 0.909627 | 0.988084 | Zfp644 |
| ENSMUSG( | 724.4437 | -0.09833 | 0.437213 | 0.829501 | Zfp646 |
| ENSMUSG( | 72.14534 | 0.320288 | 0.212587 | 0.6567 | Zfp647 |
| ENSMUSG( | 216.189 | -0.0189 | 0.902994 | 0.986055 | Zfp65 |
| ENSMUSG( | 3663.424 | 0.018691 | 0.854941 | 0.973665 | Zfp652 |
| ENSMUSG( | 207.8063 | 0.14931 | 0.331159 | 0.77519 | Zfp652os |
| ENSMUSG( | 481.3789 | 0.10493 | 0.470813 | 0.846741 | Zfp653 |
| ENSMUSG( | 459.2916 | -0.06838 | 0.631184 | 0.914073 | Zfp654 |
| ENSMUSG( | 522.1911 | -0.08301 | 0.51125 | 0.870859 | Zfp655 |
| ENSMUSG( | 193.5871 | -0.14882 | 0.331959 | 0.775288 | Zfp658 |
| ENSMUSG( | 99.5097 | 0.083208 | 0.668376 | 0.927864 | Zfp660 |
| ENSMUSG( | 281.5761 | 0.033752 | 0.841326 | 0.971179 | Zfp661 |
| ENSMUSG( | 7216.419 | 0.087161 | 0.283153 | 0.73186 | Zfp664 |
| ENSMUSG( | 249.0228 | -0.12167 | 0.41681 | 0.8209 | Zfp667 |
| ENSMUSG( | 375.6605 | 0.130364 | 0.313424 | 0.760368 | Zfp668 |
| ENSMUSG( | 812.9131 | 0.000506 | 0.997107 | 0.999565 | Zfp672 |
| ENSMUSG( | 48.85735 | -0.21824 | 0.438692 | 0.830714 | Zfp677 |
| ENSMUSG( | 653.6229 | 0.045594 | 0.658983 | 0.924506 | Zfp68 |
| ENSMUSG( | 1554.368 | -0.0485 | 0.586953 | 0.900235 | Zfp687 |
| ENSMUSG( | 269.9903 | -0.00627 | 0.968097 | 0.996284 | Zfp688 |
| ENSMUSG( | 160.0231 | -0.00912 | 0.957867 | 0.993874 | Zfp689 |
| ENSMUSG( | 48.2738 | -1.75393 | 2.78E-05 | 0.002487 | Zfp69 |
| ENSMUSG( | 176.164 | 0.015814 | 0.923507 | 0.990825 | Zfp691 |
| ENSMUSG( | 805.8848 | 0.295646 | 0.16097 | 0.590707 | Zfp692 |
| ENSMUSG( | 634.0513 | 0.090644 | 0.387463 | 0.805693 | Zfp697 |
| ENSMUSG( | 129.0092 | -0.16275 | 0.414015 | 0.819998 | Zfp7 |
| ENSMUSG( | 1125.584 | 0.065338 | 0.608507 | 0.907154 | Zfp703 |
| ENSMUSG( | 2820.596 | 0.047575 | 0.651379 | 0.921451 | Zfp704 |
| ENSMUSG( | 5170.479 | 0.035676 | 0.695719 | 0.934979 | Zfp706 |
| ENSMUSG( | 264.5699 | -0.00099 | 0.994711 | 0.999369 | Zfp707 |
| ENSMUSG( | 156.8672 | -0.01032 | 0.951549 | 0.993821 | Zfp709 |
| ENSMUSG( | 867.7681 | -0.10657 | 0.290607 | 0.740557 | Zfp710 |
| ENSMUSG( | 190.4043 | 0.01063 | 0.952133 | 0.993821 | Zfp711 |
| ENSMUSG( | 73.57221 | -0.28495 | 0.227283 | 0.674223 | Zfp712 |
| ENSMUSG( | 622.2547 | -0.2817 | 0.012662 | 0.162184 | Zfp715 |
| ENSMUSG( | 310.6744 | -0.29311 | 0.02569 | 0.247138 | Zfp719 |
| ENSMUSG( | 95.01549 | -0.10158 | 0.642326 | 0.916865 | Zfp72 |
| ENSMUSG( | 343.777 | -0.06731 | 0.596811 | 0.903526 | Zfp729a |
| ENSMUSG( | 176.6054 | -0.04067 | 0.81033 | 0.964028 | Zfp729b |
| ENSMUSG( | 663.5617 | -0.21553 | 0.051365 | 0.347869 | Zfp738 |
| ENSMUSG( | 397.5129 | 0.072264 | 0.581501 | 0.897767 | Zfp74 |
| ENSMUSG( | 3232.785 | 0.011372 | 0.891833 | 0.983979 | Zfp740 |
| ENSMUSG( | 767.6551 | 0.137481 | 0.240321 | 0.687899 | Zfp746 |
| ENSMUSG( | 103.5219 | -0.09947 | 0.645112 | 0.917934 | Zfp747 |
| ENSMUSG( | 60.46367 | -0.25544 | 0.31945 | 0.765296 | Zfp747l1 |
| ENSMUSG( | 192.3507 | -0.0721 | 0.634344 | 0.914073 | Zfp748 |
| ENSMUSG( | 99.66946 | -0.23289 | 0.299846 | 0.748922 | Zfp758 |
| ENSMUSG( | 108.1757 | -0.05433 | 0.78204 | 0.957558 | Zfp759 |
| ENSMUSG( | 336.8494 | 0.137945 | 0.319138 | 0.765081 | Zfp760 |
| ENSMUSG( | 281.7909 | 0.005901 | 0.965833 | 0.995377 | Zfp763 |
| ENSMUSG( | 84.29132 | -0.16026 | 0.554043 | 0.886397 | Zfp764 |
| ENSMUSG( | 122.1915 | 0.133958 | 0.484874 | 0.856903 | Zfp764l1 |
| ENSMUSG( | 457.0383 | 0.09131 | 0.428148 | 0.825894 | Zfp768 |
| ENSMUSG( | 631.5645 | 0.090427 | 0.510874 | 0.870755 | Zfp770 |
| ENSMUSG( | 1434.28 | 0.237345 | 0.019354 | 0.209704 | Zfp771 |
| ENSMUSG( | 472.4933 | 0.232952 | 0.045104 | 0.325725 | Zfp772 |
| ENSMUSG( | 54.31998 | 0.065973 | 0.803205 | 0.962169 | Zfp773 |
| ENSMUSG( | 304.4563 | 0.239767 | 0.070147 | 0.40908 | Zfp775 |
| ENSMUSG( | 1044.717 | 0.166983 | 0.143621 | 0.562581 | Zfp777 |
| ENSMUSG( | 61.11546 | -0.06216 | 0.799458 | 0.961247 | Zfp78 |
| ENSMUSG( | 307.7556 | 0.100666 | 0.441031 | 0.832445 | Zfp780b |
| ENSMUSG( | 13.15244 | 0.087954 | 0.869129 | NA | Zfp781 |
| ENSMUSG( | 236.1604 | 0.01503 | 0.915783 | 0.988634 | Zfp783 |
| ENSMUSG( | 166.4579 | 0.113323 | 0.478513 | 0.853567 | Zfp784 |
| ENSMUSG( | 72.35443 | -0.0027 | 0.99168 | 0.999232 | Zfp786 |
| ENSMUSG( | 587.1858 | 0.043849 | 0.691177 | 0.933881 | Zfp787 |

| | | | | | |
|---|---|---|---|---|---|
| ENSMUSG( | 2914.693 | -0.01843 | 0.854524 | 0.985649 | Zfp740 |
| ENSMUSG( | 697.2657 | 0.125267 | 0.387357 | 0.917257 | Zfp746 |
| ENSMUSG( | 93.16913 | -0.11746 | 0.635338 | 0.953903 | Zfp747 |
| ENSMUSG( | 61.07285 | 0.077332 | 0.779847 | 0.976106 | Zfp747l1 |
| ENSMUSG( | 191.3618 | 0.178793 | 0.299449 | 0.898538 | Zfp748 |
| ENSMUSG( | 111.4476 | 0.341553 | 0.153352 | 0.837724 | Zfp758 |
| ENSMUSG( | 103.9288 | 0.109795 | 0.617106 | 0.951739 | Zfp759 |
| ENSMUSG( | 315.263 | 0.217832 | 0.269138 | 0.886755 | Zfp760 |
| ENSMUSG( | 257.9561 | 0.017523 | 0.91137 | 0.992196 | Zfp763 |
| ENSMUSG( | 91.03195 | 0.336051 | 0.228221 | 0.868507 | Zfp764 |
| ENSMUSG( | 114.5822 | 0.200721 | 0.366282 | 0.914703 | Zfp764l1 |
| ENSMUSG( | 401.0197 | -0.01774 | 0.896876 | 0.990558 | Zfp768 |
| ENSMUSG( | 569.0417 | 0.051741 | 0.743629 | 0.96951 | Zfp770 |
| ENSMUSG( | 1249.169 | 0.11365 | 0.325441 | 0.905339 | Zfp771 |
| ENSMUSG( | 389.4586 | -0.05745 | 0.72068 | 0.966887 | Zfp772 |
| ENSMUSG( | 54.68742 | 0.342825 | 0.232329 | 0.868507 | Zfp773 |
| ENSMUSG( | 257.4352 | 0.028442 | 0.8543 | 0.98563 | Zfp775 |
| ENSMUSG( | 901.9268 | 0.017889 | 0.885133 | 0.989688 | Zfp777 |
| ENSMUSG( | 54.17324 | -0.12255 | 0.673919 | 0.95694 | Zfp78 |
| ENSMUSG( | 282.9843 | 0.122475 | 0.416897 | 0.917257 | Zfp780b |
| ENSMUSG( | 13.30391 | 0.331054 | 0.554593 | 0.939565 | Zfp781 |
| ENSMUSG( | 217.5821 | 0.05094 | 0.796223 | 0.97681 | Zfp783 |
| ENSMUSG( | 141.6275 | -0.07566 | 0.68865 | 0.959742 | Zfp784 |
| ENSMUSG( | 71.71381 | 0.208051 | 0.485307 | 0.92541 | Zfp786 |
| ENSMUSG( | 516.1242 | -0.06027 | 0.643756 | 0.953903 | Zfp787 |
| ENSMUSG( | 683.2628 | 0.149548 | 0.233625 | 0.868507 | Zfp788 |
| ENSMUSG( | 258.6968 | 0.039586 | 0.816779 | 0.979198 | Zfp790 |
| ENSMUSG( | 90.65821 | 0.129749 | 0.580012 | 0.944209 | Zfp791 |
| ENSMUSG( | 384.1661 | -0.0083 | 0.952508 | 0.994798 | Zfp799 |
| ENSMUSG( | 184.1504 | 0.132698 | 0.499263 | 0.927825 | Zfp800 |
| ENSMUSG( | 249.3769 | -0.16912 | 0.32217 | 0.904023 | Zfp804a |
| ENSMUSG( | 30.78662 | -0.04085 | 0.912373 | 0.992225 | Zfp808 |
| ENSMUSG( | 325.3788 | 0.135309 | 0.357023 | 0.913389 | Zfp809 |
| ENSMUSG( | 372.6247 | 0.215139 | 0.155913 | 0.837724 | Zfp81 |
| ENSMUSG( | 334.4981 | 0.249218 | 0.086121 | 0.780433 | Zfp810 |
| ENSMUSG( | 243.6416 | 0.298057 | 0.062053 | 0.736482 | Zfp811 |
| ENSMUSG( | 19.02378 | -0.0035 | 0.994571 | 0.998813 | Zfp820 |
| ENSMUSG( | 896.9995 | 0.079988 | 0.562513 | 0.940761 | Zfp821 |
| ENSMUSG( | 97.79175 | -0.09852 | 0.64665 | 0.953988 | Zfp825 |
| ENSMUSG( | 1095.016 | 0.037692 | 0.761108 | 0.972695 | Zfp827 |
| ENSMUSG( | 194.7083 | -0.03323 | 0.863649 | 0.986547 | Zfp830 |
| ENSMUSG( | 571.4541 | 0.140747 | 0.396037 | 0.917257 | Zfp831 |
| ENSMUSG( | 271.7951 | 0.060567 | 0.735903 | 0.967344 | Zfp839 |
| ENSMUSG( | 306.6939 | -0.10936 | 0.487031 | 0.92541 | Zfp84 |
| ENSMUSG( | 467.417 | 0.000289 | 0.998288 | 0.999313 | Zfp846 |
| ENSMUSG( | 65.50398 | 0.259382 | 0.345163 | 0.91045 | Zfp85 |
| ENSMUSG( | 104.9706 | 0.090779 | 0.702475 | 0.962215 | Zfp850 |
| ENSMUSG( | 416.7736 | 0.057434 | 0.682526 | 0.958678 | Zfp853 |
| ENSMUSG( | 963.2194 | 0.062674 | 0.638546 | 0.953903 | Zfp865 |
| ENSMUSG( | 547.2979 | 0.057398 | 0.673344 | 0.95694 | Zfp866 |
| ENSMUSG( | 106.1859 | 0.13881 | 0.525907 | 0.934374 | Zfp867 |
| ENSMUSG( | 222.6539 | 0.209082 | 0.200852 | 0.86439 | Zfp868 |
| ENSMUSG( | 695.3436 | -0.12288 | 0.321203 | 0.904019 | Zfp869 |
| ENSMUSG( | 195.0541 | 0.185129 | 0.287101 | 0.894091 | Zfp87 |
| ENSMUSG( | 318.5612 | 0.02353 | 0.877993 | 0.989644 | Zfp870 |
| ENSMUSG( | 1711.439 | 0.073047 | 0.513769 | 0.930586 | Zfp871 |
| ENSMUSG( | 85.57332 | 0.122202 | 0.613736 | 0.951177 | Zfp873 |
| ENSMUSG( | 239.8002 | 0.215283 | 0.176774 | 0.846973 | Zfp874a |
| ENSMUSG( | 265.517 | 0.287018 | 0.05767 | 0.726244 | Zfp874b |
| ENSMUSG( | 61.83751 | 0.130781 | 0.62048 | 0.952915 | Zfp879 |
| ENSMUSG( | 112.7824 | -0.10391 | 0.626273 | 0.953234 | Zfp882 |
| ENSMUSG( | 333.7194 | 0.073762 | 0.651972 | 0.9547 | Zfp9 |
| ENSMUSG( | 395.0809 | 0.00635 | 0.967806 | 0.995833 | Zfp90 |
| ENSMUSG( | 5312.439 | -0.05043 | 0.740471 | 0.968811 | Zfp91 |
| ENSMUSG( | 36.81598 | 0.351324 | 0.286016 | 0.893906 | Zfp92 |
| ENSMUSG( | 211.9091 | 0.050121 | 0.779644 | 0.976106 | Zfp93 |
| ENSMUSG( | 186.3597 | 0.067141 | 0.700103 | 0.961335 | Zfp930 |
| ENSMUSG( | 261.1245 | -0.24904 | 0.22868 | 0.868507 | Zfp931 |
| ENSMUSG( | 586.8058 | 0.020415 | 0.896242 | 0.990558 | Zfp932 |
| ENSMUSG( | 332.1597 | 0.170264 | 0.304104 | 0.898538 | Zfp933 |
| ENSMUSG( | 109.9574 | -0.09664 | 0.691256 | 0.960071 | Zfp934 |
| ENSMUSG( | 47.79425 | -0.01277 | 0.966057 | 0.995436 | Zfp935 |
| ENSMUSG( | 45.647 | 0.449061 | 0.196383 | 0.862161 | Zfp937 |
| ENSMUSG( | 340.5444 | 0.058967 | 0.714508 | 0.964986 | Zfp938 |
| ENSMUSG( | 17.18869 | -0.40109 | 0.406654 | 0.917257 | Zfp939 |
| ENSMUSG( | 224.3923 | -0.01314 | 0.940036 | 0.994334 | Zfp94 |
| ENSMUSG( | 402.2967 | -0.01083 | 0.941511 | 0.994578 | Zfp940 |
| ENSMUSG( | 306.6107 | -0.05042 | 0.771131 | 0.974236 | Zfp941 |
| ENSMUSG( | 303.7945 | 0.149222 | 0.397312 | 0.917257 | Zfp942 |
| ENSMUSG( | 438.6162 | 0.38431 | 0.012958 | 0.458131 | Zfp943 |
| ENSMUSG( | 228.5001 | 0.362535 | 0.043782 | 0.690304 | Zfp944 |
| ENSMUSG( | 448.7485 | 0.073803 | 0.590823 | 0.944975 | Zfp945 |
| ENSMUSG( | 239.0397 | 0.091288 | 0.598138 | 0.946676 | Zfp946 |
| ENSMUSG( | 285.059 | -0.15946 | 0.333588 | 0.907907 | Zfp948 |
| ENSMUSG( | 347.838 | 0.038057 | 0.836337 | 0.981335 | Zfp949 |
| ENSMUSG( | 1073.577 | 0.195263 | 0.215647 | 0.865167 | Zfp950 |
| ENSMUSG( | 49.78722 | -0.39856 | 0.182437 | 0.850579 | Zfp951 |
| ENSMUSG( | 347.6043 | 0.172656 | 0.22747 | 0.868507 | Zfp952 |
| ENSMUSG( | 38.47223 | -0.03122 | 0.922868 | 0.992644 | Zfp953 |
| ENSMUSG( | 196.2995 | -0.09543 | 0.631374 | 0.953251 | Zfp954 |
| ENSMUSG( | 140.6913 | 0.05237 | 0.792667 | 0.97681 | Zfp955a |
| ENSMUSG( | 428.8112 | 0.071625 | 0.629254 | 0.953251 | Zfp955b |
| ENSMUSG( | 599.6932 | 0.051953 | 0.712416 | 0.964363 | Zfp956 |
| ENSMUSG( | 147.696 | 0.232356 | 0.233134 | 0.868507 | Zfp958 |
| ENSMUSG( | 111.3959 | 0.072711 | 0.726627 | 0.966887 | Zfp959 |
| ENSMUSG( | 24.76062 | 0.351802 | 0.410785 | 0.917257 | Zfp960 |
| ENSMUSG( | 283.7406 | -0.15059 | 0.39067 | 0.917257 | Zfp961 |
| ENSMUSG( | 87.73737 | -0.01349 | 0.953599 | 0.994798 | Zfp963 |
| ENSMUSG( | 92.19475 | 0.211423 | 0.358183 | 0.913759 | Zfp964 |
| ENSMUSG( | 25.67516 | 0.153887 | 0.708383 | 0.964042 | Zfp97 |
| ENSMUSG( | 223.8083 | 0.014097 | 0.940169 | 0.994334 | Zfp970 |
| ENSMUSG( | 138.41 | 0.050585 | 0.822622 | 0.979198 | Zfp971 |
| ENSMUSG( | 130.2652 | -0.06393 | 0.756686 | 0.971779 | Zfp974 |
| ENSMUSG( | 98.8924 | 0.136134 | 0.537916 | 0.937216 | Zfp975 |
| ENSMUSG( | 66.75131 | -0.30497 | 0.242824 | 0.877399 | Zfp976 |
| ENSMUSG( | 24.78488 | -0.04386 | 0.91559 | 0.992529 | Zfp977 |
| ENSMUSG( | 22.71536 | 0.051381 | 0.908833 | 0.991958 | Zfp979 |
| ENSMUSG( | 42.72833 | -0.28492 | 0.414524 | 0.917257 | Zfp982 |
| ENSMUSG( | 268.7589 | 0.051426 | 0.740185 | 0.968811 | Zfp983 |
| ENSMUSG( | 84.31172 | 0.087913 | 0.750375 | 0.970113 | Zfp984 |
| ENSMUSG( | 18.42778 | 0.027548 | 0.954081 | 0.994798 | Zfp991 |
| ENSMUSG( | 11.1077 | 0.205984 | 0.754786 | NA | Zfp992 |
| ENSMUSG( | 113.694 | 0.097692 | 0.63686 | 0.953903 | Zfp994 |
| ENSMUSG( | 91.20472 | 0.120469 | 0.644803 | 0.953903 | Zfp995 |
| ENSMUSG( | 352.4038 | 0.007143 | 0.963653 | 0.995433 | Zfpl1 |
| ENSMUSG( | 1650.442 | 0.223134 | 0.098083 | 0.798438 | Zfpm1 |
| ENSMUSG( | 252.7756 | 0.31427 | 0.061579 | 0.736111 | Zfpm2 |
| ENSMUSG( | 11602.14 | 0.032089 | 0.758908 | 0.971858 | Zfr |
| ENSMUSG( | 1425.722 | -0.22883 | 0.129842 | 0.820802 | Zfr2 |
| ENSMUSG( | 1753.819 | 0.164042 | 0.118338 | 0.813384 | Zfta |
| ENSMUSG( | 1064.839 | 0.034092 | 0.759376 | 0.972002 | Zftraf1 |
| ENSMUSG( | 2680.568 | -0.08578 | 0.391274 | 0.917257 | Zftraf1 |
| ENSMUSG( | 550.8874 | 0.050152 | 0.693322 | 0.96076 | Zfx |
| ENSMUSG( | 1082.042 | 0.02605 | 0.82532 | 0.979388 | Zfyve1 |
| ENSMUSG( | 542.1366 | -0.02684 | 0.851436 | 0.985541 | Zfyve16 |
| ENSMUSG( | 440.1819 | -0.02239 | 0.869754 | 0.988097 | Zfyve19 |
| ENSMUSG( | 736.8009 | -0.01335 | 0.914125 | 0.992407 | Zfyve21 |
| ENSMUSG( | 664.6608 | 0.132282 | 0.402506 | 0.917257 | Zfyve26 |
| ENSMUSG( | 2791.754 | 0.140333 | 0.307434 | 0.898538 | Zfyve27 |
| ENSMUSG( | 1363.464 | -0.06298 | 0.591699 | 0.944975 | Zfyve28 |
| ENSMUSG( | 1648.275 | -0.0079 | 0.953886 | 0.994798 | Zfyve9 |
| ENSMUSG( | 965.3269 | 0.254839 | 0.257804 | 0.883401 | Zgpat |
| ENSMUSG( | 164.5705 | -0.11477 | 0.537757 | 0.937216 | Zgrf1 |
| ENSMUSG( | 2699.588 | -0.01047 | 0.937134 | 0.993685 | Zhx1 |
| ENSMUSG( | 878.033 | 0.023556 | 0.848481 | 0.985321 | Zhx2 |
| ENSMUSG( | 3049.528 | 0.327028 | 0.009996 | 0.408157 | Zhx3 |
| ENSMUSG( | 519.6731 | 0.433921 | 0.013796 | 0.46927 | Zic1 |
| ENSMUSG( | 448.4564 | 0.412823 | 0.010052 | 0.409511 | Zic2 |
| ENSMUSG( | 128.4103 | 0.931002 | 6.96E-06 | 0.002837 | Zic3 |
| ENSMUSG( | 164.5637 | 0.269557 | 0.198167 | 0.862161 | Zic4 |
| ENSMUSG( | 112.5297 | 0.519769 | 0.029449 | 0.618981 | Zic5 |
| ENSMUSG( | 155.5944 | 0.153304 | 0.412791 | 0.917257 | Zik1 |
| ENSMUSG( | 107.6579 | 0.067821 | 0.771106 | 0.974236 | Zim1 |
| ENSMUSG( | 875.1176 | 0.011197 | 0.929 | 0.99306 | Zkscan1 |
| ENSMUSG( | 154.6087 | -0.20246 | 0.358633 | 0.913759 | Zkscan14 |
| ENSMUSG( | 268.7601 | 0.162608 | 0.359523 | 0.914223 | Zkscan16 |
| ENSMUSG( | 648.558 | -0.04873 | 0.693521 | 0.960767 | Zkscan17 |
| ENSMUSG( | 1097.424 | -0.27842 | 0.018255 | 0.542424 | Zkscan2 |
| ENSMUSG( | 1088.214 | 0.265504 | 0.103184 | 0.804311 | Zkscan3 |
| ENSMUSG( | 98.19344 | 0.162727 | 0.475987 | 0.923329 | Zkscan4 |
| ENSMUSG( | 461.009 | 0.068976 | 0.604075 | 0.948139 | Zkscan5 |
| ENSMUSG( | 301.3184 | -0.15636 | 0.350554 | 0.911652 | Zkscan6 |
| ENSMUSG( | 148.5212 | 0.087397 | 0.661506 | 0.956205 | Zkscan7 |
| ENSMUSG( | 502.4558 | 0.025902 | 0.858732 | 0.985884 | Zkscan8 |
| ENSMUSG( | 336.8381 | 0.103399 | 0.471254 | 0.923329 | Zmat1 |
| ENSMUSG( | 3219.465 | -0.1699 | 0.224068 | 0.867404 | Zmat2 |
| ENSMUSG( | 3407.777 | 0.009988 | 0.927316 | 0.99306 | Zmat3 |
| ENSMUSG( | 778.9743 | -0.33226 | 0.027873 | 0.61023 | Zmat4 |

| | | | | | | | | | | | |
|---|---|---|---|---|---|---|---|---|---|---|---|
| ENSMUSG( | Zswim9 | 616.6812 | -0.03772 | 0.735929 | 0.946267 | ENSMUSG( | Zswim9 | 521.0429 | -0.26967 | 0.042082 | 0.690304 |
| ENSMUSG( | Zup1 | 157.3769 | -0.1823 | 0.311276 | 0.759247 | ENSMUSG( | Zup1 | 166.7686 | 0.261414 | 0.195064 | 0.861121 |
| ENSMUSG( | Zw10 | 499.0257 | 0.171343 | 0.199732 | 0.640239 | ENSMUSG( | Zw10 | 464.0319 | 0.230183 | 0.12234 | 0.813384 |
| ENSMUSG( | Zwilch | 82.15372 | 0.066081 | 0.783441 | 0.957558 | ENSMUSG( | Zwilch | 81.76009 | 0.294592 | 0.245623 | 0.877904 |
| ENSMUSG( | Zwint | 10251.96 | 0.116408 | 0.321343 | 0.766497 | ENSMUSG( | Zwint | 8721.375 | -0.08219 | 0.551587 | 0.939565 |
| ENSMUSG( | Zxdb | 173.4373 | 0.023231 | 0.917125 | 0.989014 | ENSMUSG( | Zxdb | 173.2764 | 0.271163 | 0.280273 | 0.892356 |
| ENSMUSG( | Zxdc | 519.7928 | -0.01236 | 0.927694 | 0.991229 | ENSMUSG( | Zxdc | 466.4322 | -0.05062 | 0.721094 | 0.966887 |
| ENSMUSG( | Zyg11b | 4651.11 | 0.128088 | 0.122834 | 0.529179 | ENSMUSG( | Zyg11b | 3983.705 | -0.04838 | 0.641425 | 0.953903 |
| ENSMUSG( | Zyx | 1708.41 | 0.005346 | 0.953099 | 0.993874 | ENSMUSG( | Zyx | 1505.939 | -0.09064 | 0.425672 | 0.917257 |
| ENSMUSG( | Zzef1 | 1787.866 | -0.11324 | 0.350193 | 0.781943 | ENSMUSG( | Zzef1 | 1785.99 | 0.156069 | 0.314563 | 0.900949 |
| ENSMUSG( | Zzz3 | 2023.366 | 0.012506 | 0.890815 | 0.983764 | ENSMUSG( | Zzz3 | 1839.756 | 0.006758 | 0.95019 | 0.994798 |

| | | | | | |
|---|---|---|---|---|---|
| ENSMUSG( | 724.5691 | 0.056112 | 0.610331 | 0.908189 | Zfp788 |
| ENSMUSG( | 289.197 | 0.078951 | 0.561674 | 0.889052 | Zfp790 |
| ENSMUSG( | 100.3756 | 0.156342 | 0.447669 | 0.834051 | Zfp791 |
| ENSMUSG( | 425.521 | 0.015381 | 0.902069 | 0.985633 | Zfp799 |
| ENSMUSG( | 183.6176 | -0.15928 | 0.379585 | 0.801757 | Zfp800 |
| ENSMUSG( | 260.3351 | -0.33132 | 0.029496 | 0.260298 | Zfp804a |
| ENSMUSG( | 32.59804 | -0.14276 | 0.676136 | 0.930542 | Zfp808 |
| ENSMUSG( | 362.9618 | 0.175829 | 0.17048 | 0.605271 | Zfp809 |
| ENSMUSG( | 397.9215 | 0.145857 | 0.238135 | 0.686299 | Zfp81 |
| ENSMUSG( | 338.8543 | 0.02853 | 0.822276 | 0.966352 | Zfp810 |
| ENSMUSG( | 241.4984 | 0.019597 | 0.894927 | 0.984512 | Zfp811 |
| ENSMUSG( | 18.36193 | -0.45847 | 0.333836 | NA | Zfp820 |
| ENSMUSG( | 1003.284 | 0.133547 | 0.328878 | 0.773233 | Zfp821 |
| ENSMUSG( | 116.0219 | 0.126508 | 0.504378 | 0.866883 | Zfp825 |
| ENSMUSG( | 1205.85 | 0.050968 | 0.635152 | 0.914296 | Zfp827 |
| ENSMUSG( | 204.7322 | -0.16192 | 0.363211 | 0.793211 | Zfp830 |
| ENSMUSG( | 605.6453 | 0.039052 | 0.748404 | 0.949604 | Zfp831 |
| ENSMUSG( | 297.1059 | 0.052008 | 0.73987 | 0.947823 | Zfp839 |
| ENSMUSG( | 365.8888 | 0.120477 | 0.362591 | 0.792943 | Zfp84 |
| ENSMUSG( | 519.9683 | 0.035243 | 0.767085 | 0.954393 | Zfp846 |
| ENSMUSG( | 69.83194 | 0.172991 | 0.495024 | 0.860798 | Zfp85 |
| ENSMUSG( | 114.0402 | 0.073759 | 0.748169 | 0.949604 | Zfp850 |
| ENSMUSG( | 485.6879 | 0.217037 | 0.075989 | 0.426896 | Zfp853 |
| ENSMUSG( | 1044.171 | 0.02261 | 0.819955 | 0.965888 | Zfp865 |
| ENSMUSG( | 589.066 | -0.0044 | 0.970881 | 0.996369 | Zfp866 |
| ENSMUSG( | 115.9941 | 0.10802 | 0.564208 | 0.890139 | Zfp867 |
| ENSMUSG( | 220.7119 | -0.07684 | 0.598934 | 0.903801 | Zfp868 |
| ENSMUSG( | 778.431 | -0.06804 | 0.537989 | 0.882548 | Zfp869 |
| ENSMUSG( | 205.5347 | 0.076647 | 0.607257 | 0.906826 | Zfp87 |
| ENSMUSG( | 362.0912 | 0.118769 | 0.377853 | 0.801561 | Zfp870 |
| ENSMUSG( | 1837.735 | 0.010097 | 0.916755 | 0.988856 | Zfp871 |
| ENSMUSG( | 90.87886 | 0.011073 | 0.962473 | 0.994604 | Zfp873 |
| ENSMUSG( | 241.7051 | -0.02823 | 0.844029 | 0.971962 | Zfp874a |
| ENSMUSG( | 274.5253 | 0.122265 | 0.35844 | 0.789759 | Zfp874b |
| ENSMUSG( | 67.18048 | 0.086187 | 0.727336 | 0.944244 | Zfp879 |
| ENSMUSG( | 135.5352 | 0.159213 | 0.38072 | 0.801984 | Zfp882 |
| ENSMUSG( | 369.9101 | 0.096858 | 0.497064 | 0.861754 | Zfp9 |
| ENSMUSG( | 403.6699 | -0.21128 | 0.146725 | 0.56887 | Zfp90 |
| ENSMUSG( | 5938.344 | 0.005249 | 0.962291 | 0.994543 | Zfp91 |
| ENSMUSG( | 41.04488 | 0.395787 | 0.192044 | 0.630811 | Zfp92 |
| ENSMUSG( | 223.982 | -0.06488 | 0.702671 | 0.937735 | Zfp93 |
| ENSMUSG( | 187.2375 | -0.19186 | 0.248887 | 0.698071 | Zfp930 |
| ENSMUSG( | 307.2221 | -0.02902 | 0.889183 | 0.983434 | Zfp931 |
| ENSMUSG( | 608.0229 | -0.14334 | 0.28338 | 0.73186 | Zfp932 |
| ENSMUSG( | 369.8137 | 0.215547 | 0.121433 | 0.525713 | Zfp933 |
| ENSMUSG( | 115.6981 | -0.21294 | 0.316759 | 0.763648 | Zfp934 |
| ENSMUSG( | 49.47172 | -0.16392 | 0.545958 | 0.884974 | Zfp935 |
| ENSMUSG( | 50.42704 | 0.504996 | 0.073104 | 0.417421 | Zfp937 |
| ENSMUSG( | 350.5695 | -0.11971 | 0.36705 | 0.795164 | Zfp938 |
| ENSMUSG( | 19.06892 | -0.41497 | 0.330208 | NA | Zfp939 |
| ENSMUSG( | 236.6167 | -0.1431 | 0.329099 | 0.773233 | Zfp94 |
| ENSMUSG( | 425.4895 | -0.12218 | 0.378605 | 0.801688 | Zfp940 |
| ENSMUSG( | 322.9533 | -0.16701 | 0.262855 | 0.713397 | Zfp941 |
| ENSMUSG( | 306.036 | -0.08463 | 0.559003 | 0.888638 | Zfp942 |
| ENSMUSG( | 426.6129 | 0.059516 | 0.686893 | 0.933041 | Zfp943 |
| ENSMUSG( | 226.0163 | 0.082195 | 0.622949 | 0.911589 | Zfp944 |
| ENSMUSG( | 409.2497 | -0.50895 | 2.23E-05 | 0.002129 | Zfp945 |
| ENSMUSG( | 235.8194 | -0.22134 | 0.200223 | 0.640302 | Zfp946 |
| ENSMUSG( | 303.4066 | -0.2463 | 0.085708 | 0.451367 | Zfp948 |
| ENSMUSG( | 402.0725 | 0.175655 | 0.258836 | 0.709814 | Zfp949 |
| ENSMUSG( | 1066.834 | -0.08689 | 0.44405 | 0.833239 | Zfp950 |
| ENSMUSG( | 56.35848 | -0.32278 | 0.202382 | 0.642343 | Zfp951 |
| ENSMUSG( | 357.5011 | -0.01334 | 0.914017 | 0.988483 | Zfp952 |
| ENSMUSG( | 42.48536 | -0.01278 | 0.965965 | 0.995377 | Zfp953 |
| ENSMUSG( | 221.0014 | -0.03546 | 0.848976 | 0.972554 | Zfp954 |
| ENSMUSG( | 154.0904 | 0.03669 | 0.841789 | 0.971179 | Zfp955a |
| ENSMUSG( | 479.5198 | 0.118989 | 0.30252 | 0.75086 | Zfp955b |
| ENSMUSG( | 651.3444 | 0.019706 | 0.870529 | 0.977906 | Zfp956 |
| ENSMUSG( | 148.994 | -0.0078 | 0.966134 | 0.995377 | Zfp958 |
| ENSMUSG( | 119.7587 | 0.001765 | 0.9928 | 0.999327 | Zfp959 |
| ENSMUSG( | 22.42254 | -0.16724 | 0.691974 | 0.934101 | Zfp960 |
| ENSMUSG( | 337.7569 | 0.09499 | 0.493892 | 0.860611 | Zfp961 |
| ENSMUSG( | 88.08923 | -0.28291 | 0.189803 | 0.628929 | Zfp963 |
| ENSMUSG( | 87.53645 | -0.22557 | 0.2834 | 0.73186 | Zfp964 |
| ENSMUSG( | 25.44673 | -0.13097 | 0.747968 | 0.949604 | Zfp97 |
| ENSMUSG( | 249.3874 | 0.042358 | 0.783424 | 0.957558 | Zfp970 |
| ENSMUSG( | 157.1719 | 0.139618 | 0.492539 | 0.860378 | Zfp971 |
| ENSMUSG( | 132.2955 | -0.32027 | 0.082252 | 0.442642 | Zfp974 |
| ENSMUSG( | 100.5103 | -0.07197 | 0.720031 | 0.941986 | Zfp975 |
| ENSMUSG( | 77.67183 | -0.1299 | 0.608557 | 0.907154 | Zfp976 |
| ENSMUSG( | 25.54765 | -0.22056 | 0.565923 | 0.89104 | Zfp977 |
| ENSMUSG( | 19.59274 | -0.71028 | 0.100729 | NA | Zfp979 |
| ENSMUSG( | 46.7373 | -0.26424 | 0.440298 | 0.831851 | Zfp982 |
| ENSMUSG( | 289.4772 | -0.00048 | 0.997245 | 0.999587 | Zfp983 |
| ENSMUSG( | 81.12788 | -0.2882 | 0.248527 | 0.697423 | Zfp984 |
| ENSMUSG( | 16.73273 | -0.58326 | 0.255126 | NA | Zfp991 |
| ENSMUSG( | 111.0117 | -0.24395 | 0.208393 | 0.650319 | Zfp994 |
| ENSMUSG( | 98.38904 | 0.069092 | 0.746176 | 0.949289 | Zfp995 |
| ENSMUSG( | 377.3263 | -0.0635 | 0.635385 | 0.914296 | Zfpl1 |
| ENSMUSG( | 1866.077 | 0.298117 | 0.007382 | 0.118765 | Zfpm1 |
| ENSMUSG( | 268.6569 | 0.225712 | 0.128066 | 0.536614 | Zfpm2 |
| ENSMUSG( | 13404.11 | 0.175111 | 0.04689 | 0.331252 | Zfr |
| ENSMUSG( | 1849.426 | 0.253471 | 0.040161 | 0.309163 | Zfr2 |
| ENSMUSG( | 1955.82 | 0.206521 | 0.0232 | 0.233586 | Zfta |
| ENSMUSG( | 1197.691 | 0.104303 | 0.374107 | 0.800445 | Zftraf1 |
| ENSMUSG( | 3215.067 | 0.166779 | 0.036751 | 0.295976 | Zftraf1 |
| ENSMUSG( | 594.5472 | 0.000841 | 0.993827 | 0.999369 | Zfx |
| ENSMUSG( | 1206.58 | 0.068713 | 0.488172 | 0.859092 | Zfyve1 |
| ENSMUSG( | 590.5183 | -0.05323 | 0.671365 | 0.928781 | Zfyve16 |
| ENSMUSG( | 490.6375 | 0.018338 | 0.870952 | 0.977951 | Zfyve19 |
| ENSMUSG( | 831.547 | 0.070035 | 0.489968 | 0.859319 | Zfyve21 |
| ENSMUSG( | 699.5753 | 0.006584 | 0.957906 | 0.993874 | Zfyve26 |
| ENSMUSG( | 2726.03 | -0.20524 | 0.103773 | 0.489428 | Zfyve27 |
| ENSMUSG( | 1426.708 | -0.20793 | 0.045845 | 0.328083 | Zfyve28 |
| ENSMUSG( | 1839.595 | 0.041669 | 0.720877 | 0.942684 | Zfyve9 |
| ENSMUSG( | 968.9337 | -5.52E-05 | 0.999704 | 0.99989 | Zgpat |
| ENSMUSG( | 173.5435 | -0.23593 | 0.179998 | 0.617634 | Zgrf1 |
| ENSMUSG( | 3182.068 | 0.191475 | 0.113483 | 0.508017 | Zhx1 |
| ENSMUSG( | 970.4176 | 0.046299 | 0.657698 | 0.92425 | Zhx2 |
| ENSMUSG( | 3762.159 | 0.618794 | 1.49E-11 | 9.68E-09 | Zhx3 |
| ENSMUSG( | 444.4524 | -0.2599 | 0.105665 | 0.492581 | Zic1 |
| ENSMUSG( | 423.9047 | 0.002824 | 0.982325 | 0.997512 | Zic2 |
| ENSMUSG( | 102.2077 | 0.144917 | 0.494294 | 0.860611 | Zic3 |
| ENSMUSG( | 151.3492 | -0.24566 | 0.167531 | 0.600122 | Zic4 |
| ENSMUSG( | 107.4488 | 0.150406 | 0.541503 | 0.883182 | Zic5 |
| ENSMUSG( | 167.8705 | 0.108475 | 0.516111 | 0.872701 | Zik1 |
| ENSMUSG( | 112.0786 | -0.08894 | 0.651846 | 0.921871 | Zim1 |
| ENSMUSG( | 945.3398 | -0.03664 | 0.734696 | 0.945584 | Zkscan1 |
| ENSMUSG( | 175.9883 | -0.09982 | 0.617311 | 0.910866 | Zkscan14 |
| ENSMUSG( | 261.5279 | -0.19894 | 0.180226 | 0.617634 | Zkscan16 |
| ENSMUSG( | 744.6884 | 0.079508 | 0.442858 | 0.83278 | Zkscan17 |
| ENSMUSG( | 1100.044 | -0.58545 | 1.01E-08 | 3.91E-06 | Zkscan2 |
| ENSMUSG( | 1110.401 | 0.060926 | 0.640239 | 0.916434 | Zkscan3 |
| ENSMUSG( | 111.5637 | 0.257225 | 0.201592 | 0.641362 | Zkscan4 |
| ENSMUSG( | 515.8721 | 0.122017 | 0.27932 | 0.729114 | Zkscan5 |
| ENSMUSG( | 366.824 | 0.142209 | 0.314502 | 0.761558 | Zkscan6 |
| ENSMUSG( | 153.0997 | -0.10924 | 0.538097 | 0.882548 | Zkscan7 |
| ENSMUSG( | 535.446 | -0.05903 | 0.642354 | 0.916865 | Zkscan8 |
| ENSMUSG( | 371.5393 | 0.116206 | 0.340417 | 0.778122 | Zmat1 |
| ENSMUSG( | 3929.096 | 0.140673 | 0.154159 | 0.580901 | Zmat2 |
| ENSMUSG( | 3987.185 | 0.18973 | 0.023395 | 0.233882 | Zmat3 |
| ENSMUSG( | 924.3546 | -0.08429 | 0.570573 | 0.893192 | Zmat4 |
| ENSMUSG( | 513.6479 | -0.1951 | 0.116854 | 0.515606 | Zmat5 |
| ENSMUSG( | 4799.292 | 0.069438 | 0.425949 | 0.824898 | Zmiz1 |
| ENSMUSG( | 17493.1 | 0.241003 | 0.004051 | 0.081936 | Zmiz2 |
| ENSMUSG( | 1602.237 | 0.023401 | 0.868555 | 0.976897 | Zmpste24 |
| ENSMUSG( | 230.7 | 0.253737 | 0.0848 | 0.448764 | Zmym1 |
| ENSMUSG( | 1704.884 | 0.00131 | 0.98776 | 0.998907 | Zmym2 |
| ENSMUSG( | 3328.146 | 0.204902 | 0.119543 | 0.523344 | Zmym3 |
| ENSMUSG( | 1597.954 | -0.05356 | 0.563326 | 0.889914 | Zmym4 |
| ENSMUSG( | 1003.459 | -0.12318 | 0.202524 | 0.642593 | Zmym5 |
| ENSMUSG( | 819.5051 | -0.47526 | 5.65E-06 | 0.000739 | Zmym6 |
| ENSMUSG( | 99.92058 | 0.059802 | 0.766385 | 0.954173 | Zmynd10 |
| ENSMUSG( | 7828.553 | 0.063148 | 0.431261 | 0.827055 | Zmynd11 |
| ENSMUSG( | 18.23948 | 0.608103 | 0.180853 | NA | Zmynd15 |
| ENSMUSG( | 672.3771 | 0.180603 | 0.160483 | 0.590122 | Zmynd19 |
| ENSMUSG( | 4759.842 | 0.308332 | 0.005195 | 0.096074 | Zmynd8 |
| ENSMUSG( | 2176.361 | 0.130015 | 0.155198 | 0.583117 | Znfx1 |
| ENSMUSG( | 256.6638 | 0.093671 | 0.512674 | 0.871762 | Zng1 |
| ENSMUSG( | 1101.213 | 0.009417 | 0.928548 | 0.991641 | Znhit1 |
| ENSMUSG( | 913.6878 | 0.114191 | 0.33674 | 0.777 | Znhit2 |
| ENSMUSG( | 340.0477 | 0.039261 | 0.824421 | 0.967102 | Znhit3 |
| ENSMUSG( | 357.4022 | 0.083118 | 0.500553 | 0.863514 | Znhit6 |
| ENSMUSG( | 8400.067 | 0.082116 | 0.306139 | 0.753712 | Znrf1 |
| ENSMUSG( | 776.4075 | 0.080559 | 0.414876 | 0.820184 | Znrf2 |
| ENSMUSG( | 609.126 | -0.26623 | 0.078979 | 0.434796 | Znrf3 |
| ENSMUSG( | 35.02303 | -0.31094 | 0.3752 | 0.800786 | Zpbp |
| ENSMUSG( | 905.0922 | 0.071797 | 0.5497 | 0.885944 | Zpr1 |

| | | | | | |
|---|---|---|---|---|---|
| ENSMUSG( | 452.6423 | -0.30618 | 0.033773 | 0.647272 | Zmat5 |
| ENSMUSG( | 4452.875 | 0.12201 | 0.258037 | 0.883684 | Zmiz1 |
| ENSMUSG( | 15234.17 | 0.11936 | 0.242162 | 0.876967 | Zmiz2 |
| ENSMUSG( | 1504.911 | 0.111883 | 0.482462 | 0.924578 | Zmpste24 |
| ENSMUSG( | 203.8446 | 0.172625 | 0.292612 | 0.897982 | Zmym1 |
| ENSMUSG( | 1589.292 | 0.065119 | 0.556046 | 0.939565 | Zmym2 |
| ENSMUSG( | 2746.074 | -0.0789 | 0.64238 | 0.953903 | Zmym3 |
| ENSMUSG( | 1431.081 | -0.10486 | 0.357365 | 0.91373 | Zmym4 |
| ENSMUSG( | 981.545 | 0.084174 | 0.48164 | 0.924578 | Zmym5 |
| ENSMUSG( | 869.9873 | 0.005006 | 0.97495 | 0.996416 | Zmym6 |
| ENSMUSG( | 97.10567 | 0.234728 | 0.301283 | 0.898538 | Zmynd10 |
| ENSMUSG( | 6938.615 | -0.01754 | 0.87432 | 0.989644 | Zmynd11 |
| ENSMUSG( | 17.16934 | 0.692572 | 0.147826 | 0.836241 | Zmynd15 |
| ENSMUSG( | 563.3454 | -0.06171 | 0.711714 | 0.964363 | Zmynd19 |
| ENSMUSG( | 3982.939 | 0.076628 | 0.517762 | 0.930939 | Zmynd8 |
| ENSMUSG( | 1962.956 | 0.104384 | 0.352878 | 0.912263 | Znfx1 |
| ENSMUSG( | 219.8435 | -0.09673 | 0.581546 | 0.944266 | Zng1 |
| ENSMUSG( | 920.0446 | -0.25514 | 0.046668 | 0.699307 | Znhit1 |
| ENSMUSG( | 801.4022 | 0.006472 | 0.962338 | 0.995237 | Znhit2 |
| ENSMUSG( | 291.8881 | -0.13902 | 0.474565 | 0.923329 | Znhit3 |
| ENSMUSG( | 310.7684 | -0.05728 | 0.711468 | 0.964363 | Znhit6 |
| ENSMUSG( | 7627.512 | 0.072959 | 0.471944 | 0.923329 | Znrf1 |
| ENSMUSG( | 726.7474 | 0.150932 | 0.246776 | 0.877904 | Znrf2 |
| ENSMUSG( | 635.0989 | 0.138277 | 0.434514 | 0.91869 | Znrf3 |
| ENSMUSG( | 33.26908 | -0.18714 | 0.590399 | 0.944975 | Zpbp |
| ENSMUSG( | 806.6914 | 0.009203 | 0.943666 | 0.994798 | Zpr1 |
| ENSMUSG( | 1544.929 | 0.053094 | 0.740165 | 0.968811 | Zranb1 |
| ENSMUSG( | 2322.937 | 0.230159 | 0.100541 | 0.801444 | Zranb2 |
| ENSMUSG( | 222.8395 | -0.18643 | 0.307414 | 0.898538 | Zranb3 |
| ENSMUSG( | 832.1852 | -0.00503 | 0.973421 | 0.996416 | Zrsr2 |
| ENSMUSG( | 176.3039 | -0.01821 | 0.917927 | 0.992529 | Zscan12 |
| ENSMUSG( | 328.5279 | -0.1721 | 0.27426 | 0.889874 | Zscan18 |
| ENSMUSG( | 154.6079 | -0.01272 | 0.946001 | 0.994798 | Zscan2 |
| ENSMUSG( | 168.6934 | 0.021634 | 0.90988 | 0.991958 | Zscan20 |
| ENSMUSG( | 395.1055 | 0.058156 | 0.725439 | 0.966887 | Zscan21 |
| ENSMUSG( | 206.0064 | 0.09934 | 0.597525 | 0.946609 | Zscan22 |
| ENSMUSG( | 246.864 | 0.052015 | 0.781673 | 0.976106 | Zscan25 |
| ENSMUSG( | 2055.608 | 0.118854 | 0.385176 | 0.917257 | Zscan26 |
| ENSMUSG( | 265.4044 | 0.296474 | 0.059906 | 0.7301 | Zscan29 |
| ENSMUSG( | 24.42641 | 0.43987 | 0.330998 | 0.907513 | Zscan30 |
| ENSMUSG( | 408.0737 | 0.130236 | 0.410019 | 0.917257 | Zswim1 |
| ENSMUSG( | 143.0814 | -0.08801 | 0.674908 | 0.95694 | Zswim3 |
| ENSMUSG( | 583.0337 | 0.069412 | 0.608217 | 0.950113 | Zswim4 |
| ENSMUSG( | 343.4053 | -0.10865 | 0.473713 | 0.923329 | Zswim5 |
| ENSMUSG( | 704.7061 | -0.07274 | 0.594676 | 0.945444 | Zswim6 |
| ENSMUSG( | 184.7553 | -0.22294 | 0.242353 | 0.877129 | Zswim7 |
| ENSMUSG( | 3387.726 | 0.120306 | 0.337504 | 0.907972 | Zswim8 |
| ENSMUSG( | 521.0429 | -0.26967 | 0.042082 | 0.690304 | Zswim9 |
| ENSMUSG( | 166.7686 | 0.261414 | 0.195064 | 0.861121 | Zup1 |
| ENSMUSG( | 464.0319 | 0.230183 | 0.12234 | 0.813384 | Zw10 |
| ENSMUSG( | 81.76009 | 0.294592 | 0.245623 | 0.877904 | Zwilch |
| ENSMUSG( | 8721.375 | -0.08219 | 0.551587 | 0.939565 | Zwint |
| ENSMUSG( | 173.2764 | 0.271163 | 0.280273 | 0.892356 | Zxdb |
| ENSMUSG( | 466.4322 | -0.05062 | 0.721094 | 0.966887 | Zxdc |
| ENSMUSG( | 3983.705 | -0.04838 | 0.641425 | 0.953903 | Zyg11b |
| ENSMUSG( | 1505.939 | -0.09064 | 0.425672 | 0.917257 | Zyx |
| ENSMUSG( | 1785.99 | 0.156069 | 0.314563 | 0.900949 | Zzef1 |
| ENSMUSG( | 1839.756 | 0.006758 | 0.95019 | 0.994798 | Zzz3 |

| | | | | | |
|---|---|---|---|---|---|
| ENSMUSG( | 1714.622 | 0.087521 | 0.380173 | 0.801757 | Zranb1 |
| ENSMUSG( | 2325.916 | -0.02404 | 0.813288 | 0.964574 | Zranb2 |
| ENSMUSG( | 260.5447 | 0.003519 | 0.981989 | 0.997512 | Zranb3 |
| ENSMUSG( | 903.8007 | -0.0292 | 0.796266 | 0.960088 | Zrsr2 |
| ENSMUSG( | 187.6282 | -0.10457 | 0.534478 | 0.880719 | Zscan12 |
| ENSMUSG( | 376.7986 | -0.04451 | 0.733164 | 0.944956 | Zscan18 |
| ENSMUSG( | 171.6593 | 0.02617 | 0.880068 | 0.981469 | Zscan2 |
| ENSMUSG( | 200.6866 | 0.236617 | 0.119327 | 0.523036 | Zscan20 |
| ENSMUSG( | 425.6943 | 0.00725 | 0.960556 | 0.993991 | Zscan21 |
| ENSMUSG( | 231.6439 | 0.16943 | 0.305145 | 0.752998 | Zscan22 |
| ENSMUSG( | 268.8315 | 0.024893 | 0.882424 | 0.982244 | Zscan25 |
| ENSMUSG( | 2127.517 | -0.04585 | 0.671205 | 0.928781 | Zscan26 |
| ENSMUSG( | 272.7885 | 0.112998 | 0.416288 | 0.82084 | Zscan29 |
| ENSMUSG( | 27.30813 | 0.504221 | 0.193886 | 0.633085 | Zscan30 |
| ENSMUSG( | 468.934 | 0.255813 | 0.080286 | 0.438167 | Zswim1 |
| ENSMUSG( | 159.9756 | -0.03983 | 0.813289 | 0.964574 | Zswim3 |
| ENSMUSG( | 624.6324 | -0.0023 | 0.985395 | 0.998071 | Zswim4 |
| ENSMUSG( | 388.8256 | -0.01194 | 0.926478 | 0.991229 | Zswim5 |
| ENSMUSG( | 724.2264 | -0.27097 | 0.025988 | 0.247245 | Zswim6 |
| ENSMUSG( | 203.6232 | -0.19969 | 0.195959 | 0.635396 | Zswim7 |
| ENSMUSG( | 3793.378 | 0.174073 | 0.145744 | 0.566898 | Zswim8 |
| ENSMUSG( | 616.6812 | -0.03772 | 0.735929 | 0.946267 | Zswim9 |
| ENSMUSG( | 157.3769 | -0.1823 | 0.311276 | 0.759247 | Zup1 |
| ENSMUSG( | 499.0257 | 0.171343 | 0.199732 | 0.640239 | Zw10 |
| ENSMUSG( | 82.15372 | 0.066081 | 0.783441 | 0.957558 | Zwilch |
| ENSMUSG( | 10251.96 | 0.116408 | 0.321343 | 0.766497 | Zwint |
| ENSMUSG( | 173.4373 | 0.023231 | 0.917125 | 0.989014 | Zxdb |
| ENSMUSG( | 519.7928 | -0.01236 | 0.927694 | 0.991229 | Zxdc |
| ENSMUSG( | 4651.11 | 0.128088 | 0.122834 | 0.529179 | Zyg11b |
| ENSMUSG( | 1708.41 | 0.005346 | 0.953099 | 0.993874 | Zyx |
| ENSMUSG( | 1787.866 | -0.11324 | 0.350193 | 0.781943 | Zzef1 |
| ENSMUSG( | 2023.366 | 0.012506 | 0.890815 | 0.983764 | Zzz3 |

**K-mean cluster**

Cluster terms

| | Cluster | ID | Descriptior | GeneRatio | BgRatio | pvalue | p.adjust | qvalue | geneID | Count |
|---|---|---|---|---|---|---|---|---|---|---|
| 1 | 2 | GO:009874 | cell-cell adl | 36/531 | 130/8076 | 4.21E-14 | 1.47E-10 | 1.33E-10 | Acvr1/Alca | 36 |
| 2 | 2 | GO:000741 | axon guida | 37/531 | 149/8076 | 7.48E-13 | 8.70E-10 | 7.89E-10 | Alcam/App | 37 |
| 3 | 2 | GO:009748 | neuron prc | 37/531 | 149/8076 | 7.48E-13 | 8.70E-10 | 7.89E-10 | Alcam/App | 37 |
| 4 | 2 | GO:007008 | glycosylatic | 29/531 | 109/8076 | 3.95E-11 | 3.44E-08 | 3.12E-08 | Abca2/Alg1 | 29 |
| 5 | 2 | GO:00091C | glycoprotei | 36/531 | 166/8076 | 1.05E-10 | 7.33E-08 | 6.65E-08 | Abca2/Alg1 | 36 |
| 6 | 2 | GO:000648 | protein gly | 26/531 | 95/8076 | 2.09E-10 | 1.04E-07 | 9.43E-08 | Abca2/Alg1 | 26 |
| 7 | 2 | GO:004341 | macromole | 26/531 | 95/8076 | 2.09E-10 | 1.04E-07 | 9.43E-08 | Abca2/Alg1 | 26 |
| 8 | 2 | GO:004646 | membrane | 19/531 | 57/8076 | 1.53E-09 | 6.68E-07 | 6.06E-07 | Abca2/Abc | 19 |
| 9 | 2 | GO:000664 | membrane | 24/531 | 92/8076 | 3.03E-09 | 1.17E-06 | 1.06E-06 | Abca2/Abc | 24 |
| 10 | 2 | GO:00986C | cell-cell adl | 61/531 | 426/8076 | 3.62E-09 | 1.26E-06 | 1.14E-06 | Acvr1/Alca | 61 |
| 11 | 2 | GO:003497 | response t | 33/531 | 171/8076 | 1.46E-08 | 4.63E-06 | 4.20E-06 | Amfr/App/ | 33 |
| 12 | 2 | GO:000741 | synapse as | 32/531 | 172/8076 | 6.03E-08 | 1.75E-05 | 1.59E-05 | Adgrl2/App | 32 |
| 13 | 2 | GO:000702 | endoplasm | 20/531 | 78/8076 | 8.71E-08 | 2.19E-05 | 1.99E-05 | Atl2/Atl3/E | 20 |
| 14 | 2 | GO:00091C | glycoprotei | 27/531 | 132/8076 | 8.80E-08 | 2.19E-05 | 1.99E-05 | Abca2/Alg1 | 27 |
| 15 | 2 | GO:006156 | axon devel | 53/531 | 381/8076 | 1.11E-07 | 2.45E-05 | 2.22E-05 | Alcam/Aplp | 53 |
| 16 | 2 | GO:003014 | sphingolipi | 14/531 | 40/8076 | 1.12E-07 | 2.45E-05 | 2.22E-05 | Abca2/Abc | 14 |
| 17 | 2 | GO:000715 | homophilic | 18/531 | 66/8076 | 1.40E-07 | 2.75E-05 | 2.49E-05 | Cadm4/Cdl | 18 |
| 18 | 2 | GO:000666 | sphingolipi | 19/531 | 73/8076 | 1.42E-07 | 2.75E-05 | 2.49E-05 | Abca2/Abc | 19 |
| 19 | 2 | GO:000666 | glycolipid r | 15/531 | 47/8076 | 1.61E-07 | 2.80E-05 | 2.54E-05 | Abca2/B4g | 15 |
| 20 | 2 | GO:19035C | liposacchai | 15/531 | 47/8076 | 1.61E-07 | 2.80E-05 | 2.54E-05 | Abca2/B4g | 15 |
| 21 | 2 | GO:00074C | axonogene | 49/531 | 346/8076 | 1.99E-07 | 3.30E-05 | 3.00E-05 | Alcam/Aplp | 49 |
| 22 | 2 | GO:000761 | behavior | 61/531 | 482/8076 | 3.68E-07 | 5.83E-05 | 5.29E-05 | Abca2/Abh | 61 |
| 23 | 2 | GO:000861 | lipid biosyr | 49/531 | 357/8076 | 5.22E-07 | 7.91E-05 | 7.18E-05 | Abca2/Abc | 49 |
| 24 | 2 | GO:000931 | oligosaccha | 10/531 | 23/8076 | 7.28E-07 | 0.000106 | 9.60E-05 | B3galnt1/F | 10 |
| 25 | 2 | GO:00508C | synapse or | 57/531 | 449/8076 | 8.30E-07 | 0.000116 | 0.000105 | Adam10/A | 57 |
| 26 | 2 | GO:000761 | memory | 23/531 | 113/8076 | 8.78E-07 | 0.000118 | 0.000107 | Adcy8/App | 23 |
| 27 | 2 | GO:000649 | protein O-l | 11/531 | 29/8076 | 1.04E-06 | 0.000135 | 0.000122 | B3galnt1/B | 11 |
| 28 | 2 | GO:009956 | synaptic m | 12/531 | 35/8076 | 1.19E-06 | 0.000148 | 0.000134 | Cdh10/Efna | 12 |
| 29 | 2 | GO:000667 | ceramide n | 14/531 | 50/8076 | 2.46E-06 | 0.000296 | 0.000268 | Abca2/B4g | 14 |
| 30 | 2 | GO:000924 | glycolipid b | 10/531 | 26/8076 | 2.82E-06 | 0.000328 | 0.000298 | B4galnt1/F | 10 |
| 31 | 2 | GO:003432 | cell junctio | 44/531 | 325/8076 | 2.97E-06 | 0.000334 | 0.000303 | Ace2/Adgrl | 44 |
| 32 | 2 | GO:004866 | cell morph | 57/531 | 468/8076 | 3.17E-06 | 0.000345 | 0.000313 | Adam10/A | 57 |
| 33 | 2 | GO:005196 | regulation | 21/531 | 106/8076 | 4.14E-06 | 0.000433 | 0.000393 | Adgrl2/App | 21 |
| 34 | 2 | GO:004651 | ceramide b | 10/531 | 27/8076 | 4.22E-06 | 0.000433 | 0.000393 | B4galnt1/C | 10 |
| 35 | 2 | GO:000668 | glycosphing | 10/531 | 28/8076 | 6.18E-06 | 0.000604 | 0.000548 | Abca2/B4g | 10 |
| 36 | 2 | GO:00082C | cholesterol | 17/531 | 76/8076 | 6.23E-06 | 0.000604 | 0.000548 | Abca2/App | 17 |
| 37 | 2 | GO:003096 | endoplasm | 12/531 | 42/8076 | 1.03E-05 | 0.000972 | 0.000881 | Amfr/Dnaj | 12 |
| 38 | 2 | GO:001612 | sterol meta | 17/531 | 79/8076 | 1.08E-05 | 0.000995 | 0.000902 | Abca2/App | 17 |
| 39 | 2 | GO:005196 | regulation | 45/531 | 354/8076 | 1.23E-05 | 0.0011 | 0.000998 | Adgrl2/Bm | 45 |
| 40 | 2 | GO:005089 | cognition | 35/531 | 248/8076 | 1.27E-05 | 0.001106 | 0.001004 | Adcy3/Adc | 35 |
| 41 | 2 | GO:000646 | signal pept | 6/531 | Oct-76 | 1.31E-05 | 0.001118 | 0.001014 | H13/Sec11 | 6 |
| 42 | 2 | GO:000698 | response t | 15/531 | 67/8076 | 2.16E-05 | 0.001795 | 0.001629 | Amfr/Dnaj | 15 |
| 43 | 2 | GO:009905 | presynapse | 12/531 | 45/8076 | 2.23E-05 | 0.001813 | 0.001645 | App/Clstn3 | 12 |
| 44 | 2 | GO:000761 | learning or | 32/531 | 224/8076 | 2.34E-05 | 0.001852 | 0.00168 | Adcy3/Adc | 32 |
| 45 | 2 | GO:005091 | negative ch | 9/531 | 26/8076 | 2.44E-05 | 0.001872 | 0.001698 | Lrtm2/Rob | 9 |
| 46 | 2 | GO:005196 | positive re | 14/531 | 60/8076 | 2.47E-05 | 0.001872 | 0.001698 | Adgrl2/Clst | 14 |
| 47 | 2 | GO:190265 | secondary | 17/531 | 84/8076 | 2.55E-05 | 0.001889 | 0.001714 | Abca2/App | 17 |
| 48 | 2 | GO:000686 | amino acid | 20/531 | 110/8076 | 2.66E-05 | 0.001932 | 0.001752 | Ace2/Cck/C | 20 |
| 49 | 2 | GO:006032 | head devel | 49/531 | 413/8076 | 3.27E-05 | 0.002328 | 0.002112 | Abcb6/Alk/ | 49 |
| 50 | 2 | GO:000742 | brain devel | 46/531 | 384/8076 | 4.53E-05 | 0.003158 | 0.002865 | Abcb6/Alk/ | 46 |
| 51 | 2 | GO:003462 | cellular res | 12/531 | 49/8076 | 5.63E-05 | 0.003849 | 0.003491 | Amfr/Dnaj | 12 |
| 52 | 2 | GO:00365C | ERAD pathy | 16/531 | 81/8076 | 5.99E-05 | 0.004018 | 0.003645 | Amfr/Canx | 16 |
| 53 | 2 | GO:004884 | axon exten | 8/531 | 23/8076 | 6.72E-05 | 0.004263 | 0.003867 | Bmpr2/Dsc | 8 |
| 54 | 2 | GO:190228 | neuron prc | 8/531 | 23/8076 | 6.72E-05 | 0.004263 | 0.003867 | Bmpr2/Dsc | 8 |
| 55 | 2 | GO:190464 | cellular res | 8/531 | 23/8076 | 6.72E-05 | 0.004263 | 0.003867 | Abcc1/App | 8 |
| 56 | 2 | GO:000651 | glycoprotei | 7/531 | 18/8076 | 8.58E-05 | 0.005253 | 0.004766 | Ctsl/Edem3 | 7 |
| 57 | 2 | GO:006041 | ventricular | 7/531 | 18/8076 | 8.58E-05 | 0.005253 | 0.004766 | Acvr1/Bmp | 7 |
| 58 | 2 | GO:005196 | positive re | 33/531 | 251/8076 | 9.46E-05 | 0.005565 | 0.005048 | Adgrl2/Bm | 33 |
| 59 | 2 | GO:190464 | response t | 8/531 | 24/8076 | 9.51E-05 | 0.005565 | 0.005048 | Abcc1/App | 8 |
| 60 | 2 | GO:000803 | neuron rec | 10/531 | 37/8076 | 9.57E-05 | 0.005565 | 0.005048 | App/Cntn5 | 10 |
| 61 | 2 | GO:009917 | presynapse | 12/531 | 52/8076 | 0.000105 | 0.006004 | 0.005446 | App/Clstn3 | 12 |
| 62 | 2 | GO:000693 | chemotaxis | 27/531 | 190/8076 | 0.000111 | 0.006042 | 0.005481 | Abcc1/Ada | 27 |
| 63 | 2 | GO:004233 | taxis | 27/531 | 190/8076 | 0.000111 | 0.006042 | 0.005481 | Abcc1/Ada | 27 |
| 64 | 2 | GO:190161 | organic hy | 34/531 | 264/8076 | 0.000112 | 0.006042 | 0.005481 | Abca2/Alg1 | 34 |
| 65 | 2 | GO:000606 | alcohol me | 26/531 | 180/8076 | 0.000113 | 0.006042 | 0.005481 | Abca2/Alg1 | 26 |
| 66 | 2 | GO:190188 | regulation | 26/531 | 182/8076 | 0.000135 | 0.007159 | 0.006494 | Ace2/Adgrl | 26 |
| 67 | 2 | GO:003596 | response t | 16/531 | 87/8076 | 0.000146 | 0.007545 | 0.006844 | Amfr/Dnajl | 16 |
| 68 | 2 | GO:005092 | regulation | 18/531 | 105/8076 | 0.000147 | 0.007545 | 0.006844 | Adam10/A | 18 |
| 69 | 2 | GO:000157 | ganglioside | 6/531 | 14/8076 | 0.00015 | 0.007562 | 0.00686 | Abca2/B4g | 6 |
| 70 | 2 | GO:00082C | steroid me | 21/531 | 135/8076 | 0.000179 | 0.008896 | 0.00807 | Abca2/App | 21 |
| 71 | 2 | GO:00508C | regulation | 31/531 | 239/8076 | 0.000197 | 0.009678 | 0.00878 | Adam10/A | 31 |
| 72 | 2 | GO:004295 | lipoprotein | 5/531 | Oct-76 | 0.00023 | 0.011006 | 0.009984 | Mia3/Sar1b | 5 |
| 73 | 2 | GO:00715C | cellular res | 5/531 | Oct-76 | 0.00023 | 0.011006 | 0.009984 | Amfr/Eif2a | 5 |
| 74 | 2 | GO:000698 | ER-nucleus | 8/531 | 27/8076 | 0.000241 | 0.011346 | 0.010293 | Amfr/Eif2a | 8 |
| 75 | 2 | GO:004244 | hormone n | 15/531 | 82/8076 | 0.000249 | 0.011581 | 0.010506 | Adam10/C | 15 |
| 76 | 2 | GO:000252 | acute infla | 9/531 | 34/8076 | 0.000255 | 0.011695 | 0.010609 | Acvr1/Aoc3 | 9 |
| 77 | 2 | GO:00508C | regulation | 31/531 | 245/8076 | 0.000308 | 0.013953 | 0.012657 | Adam10/A | 31 |
| 78 | 2 | GO:007097 | protein loc | 12/531 | 58/8076 | 0.000316 | 0.014034 | 0.012731 | Bcap29/Ge | 12 |
| 79 | 2 | GO:00158C | neutral am | 8/531 | 28/8076 | 0.000318 | 0.014034 | 0.012731 | Ace2/Slc1a | 8 |
| 80 | 2 | GO:00083C | associative | 15/531 | 85/8076 | 0.000374 | 0.016316 | 0.014801 | Adcy3/Adra | 15 |
| 81 | 2 | GO:000699 | response t | 5/531 | Nov-76 | 0.0004 | 0.01679 | 0.015231 | Amfr/Eif2a | 5 |
| 82 | 2 | GO:001582 | proline tra | 5/531 | Nov-76 | 0.0004 | 0.01679 | 0.015231 | Ace2/Slc36 | 5 |
| 83 | 2 | GO:004487 | lipoprotein | 5/531 | Nov-76 | 0.0004 | 0.01679 | 0.015231 | Mia3/Sar1b | 5 |
| 84 | 2 | GO:003596 | cellular res | 13/531 | 68/8076 | 0.000408 | 0.016938 | 0.015366 | Amfr/Dnajl | 13 |
| 85 | 2 | GO:190189 | positive re | 16/531 | 95/8076 | 0.000418 | 0.017133 | 0.015542 | Ace2/Adgrl | 16 |
| 86 | 2 | GO:003043 | ubiquitin-d | 12/531 | 61/8076 | 0.000515 | 0.020412 | 0.018518 | Amfr/Canx | 12 |
| 87 | 2 | GO:000319 | endocardia | 6/531 | 17/8076 | 0.00052 | 0.020412 | 0.018518 | Acvr1/Bmp | 6 |
| 88 | 2 | GO:190488 | regulation | 6/531 | 17/8076 | 0.00052 | 0.020412 | 0.018518 | Clstn3/Gric | 6 |
| 89 | 2 | GO:001648 | protein prc | 18/531 | 116/8076 | 0.000525 | 0.020412 | 0.018518 | Adam10/C | 18 |
| 90 | 2 | GO:004801 | ephrin rece | 8/531 | 30/8076 | 0.000533 | 0.020412 | 0.018518 | Efna5/Efnb | 8 |
| 91 | 2 | GO:014096 | multicellula | 8/531 | 30/8076 | 0.000533 | 0.020412 | 0.018518 | Abca3/Aka | 8 |
| 92 | 2 | GO:000194 | vasculature | 40/531 | 356/8076 | 0.000547 | 0.020731 | 0.018806 | Acvr1/Adar | 40 |
| 93 | 2 | GO:00516C | protein ma | 35/531 | 300/8076 | 0.00061 | 0.022821 | 0.020703 | Adam10/Ca | 35 |
| 94 | 2 | GO:000669 | cholesterol | 9/531 | 38/8076 | 0.000625 | 0.022821 | 0.020703 | Dhcr7/Ebp | 9 |
| 95 | 2 | GO:190265 | secondary | 9/531 | 38/8076 | 0.000625 | 0.022821 | 0.020703 | Dhcr7/Ebp | 9 |
| 96 | 2 | GO:000664 | phospholip | 29/531 | 233/8076 | 0.000628 | 0.022821 | 0.020703 | Abca2/Abc | 29 |
| 97 | 2 | GO:000317 | aortic valve | 5/531 | Dec-76 | 0.000648 | 0.023303 | 0.02114 | Bmpr2/Rob | 5 |
| 98 | 2 | GO:19056C | regulation | 8/531 | 31/8076 | 0.000677 | 0.024099 | 0.021862 | App/Clstn3 | 8 |
| 99 | 2 | GO:000156 | blood vess | 38/531 | 337/8076 | 0.0007 | 0.024668 | 0.022378 | Acvr1/Aqp1 | 38 |
| 100 | 2 | GO:004616 | alcohol bio | 14/531 | 82/8076 | 0.00082 | 0.028539 | 0.02589 | Abca2/Dhc | 14 |
| 101 | 2 | GO:000721 | Notch signa | 13/531 | 73/8076 | 0.000829 | 0.028539 | 0.02589 | Adam10/A | 13 |
| 102 | 2 | GO:000328 | ventricular | 8/531 | 32/8076 | 0.000852 | 0.028539 | 0.02589 | Acvr1/Bmp | 8 |
| 103 | 2 | GO:000761 | learning | 20/531 | 141/8076 | 0.000861 | 0.028539 | 0.02589 | Adcy3/Adra | 20 |
| 104 | 2 | GO:000317 | heart valve | 7/531 | 25/8076 | 0.000866 | 0.028539 | 0.02589 | Acvr1/Bmp | 7 |
| 105 | 2 | GO:006041 | cardiac sep | 7/531 | 25/8076 | 0.000866 | 0.028539 | 0.02589 | Acvr1/Bmp | 7 |
| 106 | 2 | GO:004694 | carboxylic a | 24/531 | 183/8076 | 0.000867 | 0.028539 | 0.02589 | Abcc1/Ace2 | 24 |
| 107 | 2 | GO:000004 | transition r | 11/531 | 56/8076 | 0.000886 | 0.028605 | 0.02595 | Abcb6/Atp | 11 |
| 108 | 2 | GO:190589 | regulation | 11/531 | 56/8076 | 0.000886 | 0.028605 | 0.02595 | App/Eif2ak | 11 |
| 109 | 2 | GO:00300C | metal ion t | 50/531 | 486/8076 | 0.000908 | 0.029055 | 0.026358 | Abcb6/Aka | 50 |
| 110 | 2 | GO:001584 | organic aci | 24/531 | 184/8076 | 0.000938 | 0.029461 | 0.026726 | Abcc1/Ace2 | 24 |
| 111 | 2 | GO:005254 | regulation | 24/531 | 184/8076 | 0.000938 | 0.029461 | 0.026726 | App/Aqp1/ | 24 |
| 112 | 2 | GO:190531 | semi-lunar | 5/531 | 13/8076 | 0.000997 | 0.031035 | 0.028154 | Bmpr2/Rob | 5 |
| 113 | 2 | GO:007178 | endoplasm | 6/531 | 19/8076 | 0.001018 | 0.031149 | 0.028258 | Atl2/Atl3/L | 6 |
| 114 | 2 | GO:190486 | inhibitory s | 6/531 | 19/8076 | 0.001018 | 0.031149 | 0.028258 | Clstn3/Gab | 6 |
| 115 | 2 | GO:000715 | heterophili | 8/531 | 33/8076 | 0.001061 | 0.031619 | 0.028683 | Alcam/Fat4 | 8 |
| 116 | 2 | GO:004504 | protein tar | 8/531 | 33/8076 | 0.001061 | 0.031619 | 0.028683 | Get1/Herp | 8 |
| 117 | 2 | GO:190486 | excitatory s | 8/531 | 33/8076 | 0.001061 | 0.031619 | 0.028683 | Clstn3/Gric | 8 |
| 118 | 2 | GO:005077 | regulation | 20/531 | 144/8076 | 0.001125 | 0.033019 | 0.029954 | Bmpr2/Dix | 20 |
| 119 | 2 | GO:001612 | sterol biosy | 9/531 | 41/8076 | 0.001127 | 0.033019 | 0.029954 | Dhcr7/Ebp | 9 |
| 120 | 2 | GO:000716 | enzyme-lin | 48/531 | 468/8076 | 0.001227 | 0.035661 | 0.03235 | Acvr1/Acvr | 48 |
| 121 | 2 | GO:000648 | protein N-l | 9/531 | 42/8076 | 0.001352 | 0.038651 | 0.035063 | Alg10b/Alg | 9 |
| 122 | 2 | GO:000719 | adenylate c | 9/531 | 42/8076 | 0.001352 | 0.038651 | 0.035063 | App/Chrm1 | 9 |
| 123 | 2 | GO:004884 | regulation | 6/531 | 20/8076 | 0.001375 | 0.038979 | 0.035361 | Bmpr2/Dsc | 6 |

Cluster Genes

| | id | symbol | entrez | log2FoldCh | log2FoldCh | logFC.MNK | logFC.MNK | clust |
|---|---|---|---|---|---|---|---|---|
| 2 | ENSMUSG( | 1110059G1 | 66202 | 007348695 | 005661137 | -25775 | 2005 | 1 |
| 40 | ENSMUSG( | Abca7 | 27403 | 005299196 | 01406098C | -494 | -01525 | 1 |
| 507 | ENSMUSG( | Arhgap39 | 223666 | 039403095 | 024364342 | -17125 | 18875 | 1 |
| 521 | ENSMUSG( | Arhgef19 | 213649 | 005785528 | -00783139 | -13125 | 0685 | 1 |
| 549 | ENSMUSG( | Arl6ip4 | 65105 | 011525141 | 008254293 | -112 | 0235 | 1 |
| 588 | ENSMUSG( | Asap2 | 211914 | 030273695 | 019703679 | -1845 | 02275 | 1 |
| 790 | ENSMUSG( | Bicra | 243842 | 015478937 | 02374124C | -10375 | 06775 | 1 |
| 889 | ENSMUSG( | Caap1 | 67770 | 011085232 | 012283571 | -1755 | -01625 | 1 |
| 1303 | ENSMUSG( | Cldn5 | 12741 | -01487777 | 01613669C | -11025 | 09625 | 1 |
| 1658 | ENSMUSG( | Cxxc1 | 74322 | 02423918C | 016229558 | -146 | 1015 | 1 |
| 1686 | ENSMUSG( | Cys1 | 12879 | 033181442 | 063872788 | -14775 | 0035 | 1 |
| 2109 | ENSMUSG( | Efs | 13644 | -01776452 | 001889603 | -14575 | 01 | 1 |
| 2220 | ENSMUSG( | Emsy | 233545 | 018310382 | 030303507 | -116 | 0225 | 1 |
| 2558 | ENSMUSG( | Fndc4 | 64339 | -00089018 | -01012516 | -16225 | 009 | 1 |
| 3054 | ENSMUSG( | Hdgfl2 | 15193 | 017565795 | 010977482 | -15275 | 0295 | 1 |
| 3258 | ENSMUSG( | Ighmbp2 | 20589 | -00305280 | -00182293 | -1265 | 043 | 1 |
| 3514 | ENSMUSG( | Kdm4a | 230674 | 016530823 | 004801227 | -13475 | 01875 | 1 |
| 3563 | ENSMUSG( | Klhl20 | 226541 | -00783044 | 007067091 | -10775 | 09075 | 1 |
| 3644 | ENSMUSG( | Ldlrap1 | 100017 | -04106488 | 014051937 | -13975 | 02925 | 1 |
| 3788 | ENSMUSG( | Lsr | 54135 | 022087308 | 04139643C | -1425 | 124 | 1 |
| 3805 | ENSMUSG( | Ly6g6e | 70274 | -08682348 | -00775057 | -14925 | 01575 | 1 |
| 3915 | ENSMUSG( | Mapk9 | 26420 | 005297017 | -00850602 | -34425 | 107 | 1 |
| 4044 | ENSMUSG( | Mettl22 | 239706 | 004222942 | 003343115 | -21125 | 03625 | 1 |
| 4340 | ENSMUSG( | Myo1f | 17916 | -02704893 | -02974298 | -107 | 061 | 1 |
| 4358 | ENSMUSG( | Naa10 | 56292 | -01215557 | -01296582 | -13275 | 131 | 1 |
| 4687 | ENSMUSG( | Nsmce3 | 66647 | -06165490 | 014793664 | -14225 | 04825 | 1 |
| 4718 | ENSMUSG( | Nucks1 | 98415 | 001689665 | -00615353 | -1605 | 104 | 1 |
| 5133 | ENSMUSG( | Phf8 | 320595 | 009345513 | 011524618 | -12425 | 11675 | 1 |
| 5159 | ENSMUSG( | Pias4 | 59004 | 013406066 | 004318913 | -2255 | 01575 | 1 |
| 5305 | ENSMUSG( | Pno1 | 66249 | 00462093C | -01840808 | -165 | 00625 | 1 |
| 5326 | ENSMUSG( | Poll | 56626 | 004611334 | -01475930 | -3065 | 12175 | 1 |
| 5407 | ENSMUSG( | Ppp1r12c | 232807 | 007586423 | -00288614 | -1075 | 03975 | 1 |
| 5634 | ENSMUSG( | Ptms | 69202 | -00071855 | -01353841 | -1125 | 1145 | 1 |
| 5754 | ENSMUSG( | Rab3b | 69908 | -08037258 | -01732650 | -169 | -00825 | 1 |
| 6008 | ENSMUSG( | Rlim | 19820 | 008840927 | -00387528 | -17575 | 098 | 1 |
| 6017 | ENSMUSG( | Rnaseh2b | 67153 | -00148313 | 009709374 | -13975 | 0625 | 1 |
| 6132 | ENSMUSG( | Rpp25 | 102614 | -00072018 | -01156132 | -1965 | 0645 | 1 |
| 6476 | ENSMUSG( | Sh2b1 | 20399 | 009278462 | -00430972 | -117 | 06975 | 1 |
| 6911 | ENSMUSG( | Spout1 | 227695 | 008564007 | -00360303 | -1785 | -0045 | 1 |
| 7069 | ENSMUSG( | Stxbp2 | 20911 | 009062733 | 001493904 | -10375 | 0815 | 1 |
| 7075 | ENSMUSG( | Sub1 | 20024 | 000254876 | -01722411 | -1465 | 00525 | 1 |
| 7098 | ENSMUSG( | Surf2 | 20931 | -00201622 | -00336037 | -2655 | 143 | 1 |
| 7786 | ENSMUSG( | Ubald1 | 207740 | 005174789 | -01227562 | -148 | 01425 | 1 |
| 7813 | ENSMUSG( | Ube2v1 | 66589 | 00387859 | -01190652 | -12825 | 038 | 1 |
| 7871 | ENSMUSG( | Umad1 | 1E+08 | -00447575 | -02436475 | -101 | 0685 | 1 |
| 7942 | ENSMUSG( | Usp53 | 99526 | 04368463C | 056117473 | -11125 | 02575 | 1 |
| 7948 | ENSMUSG( | Ust | 338362 | -01998171 | 013771219 | -28875 | 107 | 1 |
| 8054 | ENSMUSG( | Vxn | 76982 | 007952154 | -00798263 | -12675 | 01425 | 1 |
| 8084 | ENSMUSG( | Wdr25 | 212198 | -00303776 | -01679766 | -1895 | 0595 | 1 |
| 8158 | ENSMUSG( | Yaf2 | 67057 | 006920916 | -00565472 | -25725 | 03925 | 1 |
| 8194 | ENSMUSG( | Zbed3 | 72114 | -00437876 | -01758561 | -2515 | 04475 | 1 |
| 8233 | ENSMUSG( | Zeb2 | 24136 | 001543388 | -00530480 | -107 | 037 | 1 |
| 8241 | ENSMUSG( | Zfp219 | 69890 | -00330352 | -00850692 | -14575 | 03575 | 1 |
| 8267 | ENSMUSG( | Zgpat | 229007 | -552e-05 | 025483912 | -14325 | 08525 | 1 |
| 8270 | ENSMUSG( | Zkscan1 | 74570 | -00366350 | 001119676 | -155 | 0855 | 1 |
| 8273 | ENSMUSG( | Zmat3 | 22401 | 018973035 | 000998846 | -151 | 07425 | 1 |
| 8286 | ENSMUSG( | Zranb2 | 53861 | -00240385 | 023015928 | -14175 | 048 | 1 |
| 14 | ENSMUSG( | 4933434E2 | 99650 | -01684174 | 014243789 | 01625 | 02075 | 2 |
| 36 | ENSMUSG( | Abca2 | 11305 | -01386696 | 022393465 | 021 | 024 | 2 |
| 37 | ENSMUSG( | Abca3 | 27410 | -01628621 | 01648419C | 037 | 011 | 2 |
| 41 | ENSMUSG( | Abca8b | 27404 | -03174642 | 016751728 | 00725 | 00625 | 2 |
| 45 | ENSMUSG( | Abcb6 | 74104 | -01996104 | -00115545 | 0085 | 005 | 2 |
| 49 | ENSMUSG( | Abcc1 | 17250 | -02911079 | 008215319 | 024 | 00975 | 2 |
| 66 | ENSMUSG( | Abhd12 | 76192 | -02027261 | 003674415 | 0185 | 01075 | 2 |
| 68 | ENSMUSG( | Abhd16a | 193742 | -02976704 | 009864671 | 01775 | -001 | 2 |
| 75 | ENSMUSG( | Abhd6 | 66082 | -02167746 | 019901543 | 016 | 0285 | 2 |
| 109 | ENSMUSG( | Ace2 | 70008 | -02804254 | 043205784 | 00575 | 0285 | 2 |
| 125 | ENSMUSG( | Acp2 | 11432 | -03066847 | 016910729 | 0165 | 006 | 2 |
| 151 | ENSMUSG( | Acvr1 | 11477 | -03279184 | 021171969 | 02525 | 042 | 2 |
| 152 | ENSMUSG( | Acvr1b | 11479 | -02917683 | 018086169 | 046 | 0045 | 2 |
| 154 | ENSMUSG( | Acvr2a | 11480 | -03378873 | 026966002 | 011 | 0125 | 2 |
| 161 | ENSMUSG( | Adam10 | 11487 | -02745081 | 029718305 | 0085 | 01825 | 2 |
| 177 | ENSMUSG( | Adcy3 | 104111 | -01176018 | 034101758 | 0215 | 0045 | 2 |
| 180 | ENSMUSG( | Adcy8 | 11514 | -03270728 | -01869507 | 02725 | -00225 | 2 |
| 197 | ENSMUSG( | Adgrl2 | 99633 | -02374579 | 020017819 | 02175 | 00125 | 2 |
| 208 | ENSMUSG( | Adpgk | 72141 | -03048814 | 02244915C | 024 | 02075 | 2 |
| 209 | ENSMUSG( | Adprh | 11544 | -03072069 | -01036969 | 0145 | 0115 | 2 |
| 213 | ENSMUSG( | Adra1b | 11548 | -02330993 | 011947417 | 036 | 03175 | 2 |
| 240 | ENSMUSG( | Ago3 | 214150 | -01912049 | 012610701 | 032 | 04975 | 2 |
| 243 | ENSMUSG( | Agpat4 | 68262 | -04032792 | 01416628C | 018 | 01025 | 2 |
| 244 | ENSMUSG( | Agpat5 | 52123 | -00585578 | 006777047 | 02 | 023 | 2 |
| 275 | ENSMUSG( | Akap1 | 11640 | -01607199 | 014191475 | 0005 | 02525 | 2 |
| 277 | ENSMUSG( | Akap11 | 219181 | -01425662 | 018812976 | 023 | 00425 | 2 |
| 285 | ENSMUSG( | Akap9 | 100986 | -01928210 | 009234628 | -00025 | 02925 | 2 |
| 295 | ENSMUSG( | Alcam | 11658 | -03073155 | 022001571 | 026 | 04275 | 2 |
| 315 | ENSMUSG( | Alg10b | 380959 | -03920835 | 038718164 | 05975 | 00725 | 2 |
| 318 | ENSMUSG( | Alg2 | 56737 | -05175948 | 009252712 | 0355 | -0035 | 2 |
| 322 | ENSMUSG( | Alk | 11682 | -03934054 | -00555780 | 00525 | 04175 | 2 |
| 337 | ENSMUSG( | Amfr | 23802 | -01284343 | 004776126 | 0305 | 03275 | 2 |
| 408 | ENSMUSG( | Aoc3 | 11754 | -05192728 | 008317176 | 0215 | -0045 | 2 |
| 414 | ENSMUSG( | Ap1s2 | 108012 | -01591518 | 005510966 | 01675 | 013 | 2 |
| 448 | ENSMUSG( | Aplp1 | 11803 | -03541972 | -00281168 | 008 | 03525 | 2 |
| 454 | ENSMUSG( | Apod | 11815 | -03456536 | 019799608 | 0175 | 01675 | 2 |
| 459 | ENSMUSG( | App | 11820 | -04787839 | 017435655 | 0495 | 02075 | 2 |
| 465 | ENSMUSG( | Aqp1 | 11826 | -01463614 | 024669068 | 03075 | 00825 | 2 |
| 579 | ENSMUSG( | Arsb | 11881 | -02830290 | 015657456 | 02625 | 00375 | 2 |
| 597 | ENSMUSG( | Asic1 | 11419 | -01933936 | 014686486 | 01 | 0365 | 2 |
| 609 | ENSMUSG( | Astn1 | 11899 | -02449524 | 013342416 | 00025 | 01425 | 2 |
| 612 | ENSMUSG( | Atad1 | 67979 | -03695240 | 01277218C | 03675 | 0175 | 2 |
| 634 | ENSMUSG( | Atl2 | 56298 | -03325849 | 013589978 | 01075 | -00075 | 2 |
| 635 | ENSMUSG( | Atl3 | 109168 | -01479291 | 015274728 | 0035 | 00575 | 2 |
| 638 | ENSMUSG( | Atp11b | 76295 | -01891557 | 022728308 | 0235 | -00075 | 2 |
| 657 | ENSMUSG( | Atp2c1 | 235574 | -01251876 | 016559249 | 01375 | 027 | 2 |
| 675 | ENSMUSG( | Atp6v0c | 11984 | -05480112 | -00078366 | 022 | 00525 | 2 |
| 695 | ENSMUSG( | Atrn | 11990 | -02846956 | 016091681 | 004 | 025 | 2 |
| 696 | ENSMUSG( | Atrnl1 | 226255 | -04068640 | 017162836 | -00425 | 0295 | 2 |
| 715 | ENSMUSG( | B3galnt1 | 26879 | -04588384 | 01349862C | -0015 | 0455 | 2 |
| 717 | ENSMUSG( | B3galt6 | 117592 | -04312581 | 006303758 | 04475 | -00375 | 2 |
| 720 | ENSMUSG( | B4galnt1 | 14421 | -01400932 | 018283151 | 0375 | 01475 | 2 |
| 756 | ENSMUSG( | Bcap29 | 12033 | -03537886 | 006915373 | 009 | 006 | 2 |
| 797 | ENSMUSG( | Birc6 | 12211 | -02413262 | 01185705C | 00125 | 037 | 2 |
| 809 | ENSMUSG( | Bmpr2 | 12168 | -01574083 | 020886559 | 00175 | 01675 | 2 |
| 845 | ENSMUSG( | Brinp1 | 56710 | -03340845 | 021787396 | 031 | -00275 | 2 |
| 877 | ENSMUSG( | C1qb | 12260 | -00566807 | 012276245 | 017 | 01425 | 2 |
| 881 | ENSMUSG( | C1qtnf4 | 67445 | -03312976 | -00341636 | 024 | 01075 | 2 |
| 925 | ENSMUSG( | Cadm4 | 260299 | -03430885 | -00579685 | 00075 | 01075 | 2 |
| 959 | ENSMUSG( | Canx | 12330 | -01423010 | 010955406 | 016 | 0165 | 2 |
| 982 | ENSMUSG( | Cartpt | 27220 | -05222975 | -01754364 | 01825 | 039 | 2 |
| 1031 | ENSMUSG( | Ccdc134 | 76457 | -01708078 | 037808637 | 041 | 00775 | 2 |
| 1064 | ENSMUSG( | Cck | 12424 | -04973980 | -00710498 | 0145 | 01675 | 2 |
| 1096 | ENSMUSG( | Cd34 | 12490 | -02399328 | 018954314 | -01675 | 0125 | 2 |
| 1101 | ENSMUSG( | Cd55 | 13136 | -04519858 | 014334744 | 02425 | 03875 | 2 |
| 1127 | ENSMUSG( | Cdh10 | 320873 | -04553588 | 019523807 | 00075 | 0185 | 2 |

| | | | | | | | | | | |
|---|---|---|---|---|---|---|---|---|---|---|
| 124 | 2 | GO:003121 | biomineral | 12/531 | 68/8076 | 0.001414 | 0.03977 | 0.036078 | Acvr1/Acvr | 12 |
| 125 | 2 | GO:190589 | positive re | 7/531 | 27/8076 | 0.001426 | 0.03978 | 0.036087 | App/Eif2ak | 7 |
| 126 | 2 | GO:003129 | retinal gan | 5/531 | 14/8076 | 0.001467 | 0.040612 | 0.036842 | Efna5/Epha | 5 |
| 127 | 2 | GO:009917 | regulation | 8/531 | 35/8076 | 0.0016 | 0.043955 | 0.039875 | App/Clstn3 | 8 |
| 128 | 2 | GO:00760 | sensory pe | 24/531 | 192/8076 | 0.0017 | 0.046319 | 0.042019 | Adcy3/Alg1 | 24 |
| 129 | 2 | GO:190113 | carbohydra | 39/531 | 366/8076 | 0.001753 | 0.047392 | 0.042993 | Abca2/Adc | 39 |
| 130 | 2 | GO:000926 | response t | 6/531 | 21/8076 | 0.001819 | 0.048803 | 0.044272 | Arsb/Asic1, | 6 |
| 131 | 3 | GO:000651 | ubiquitin-d | 116/1409 | 421/8076 | 7.84E-08 | 0.000348 | 0.000338 | Agap3/Agt | 116 |
| 132 | 3 | GO:001994 | modificatic | 116/1409 | 428/8076 | 2.05E-07 | 0.000405 | 0.000394 | Agap3/Agt | 116 |
| 133 | 3 | GO:00070 | mitochond | 112/1409 | 412/8076 | 2.85E-07 | 0.000405 | 0.000394 | Acad9/Afg | 112 |
| 134 | 3 | GO:004363 | modificatic | 117/1409 | 437/8076 | 3.65E-07 | 0.000405 | 0.000394 | Agap3/Agt | 117 |
| 135 | 3 | GO:004819 | Golgi vesicl | 69/1409 | 230/8076 | 1.48E-06 | 0.001312 | 0.001274 | Ankfy1/Ap | 69 |
| 136 | 3 | GO:005165 | establishm | 84/1409 | 312/8076 | 1.36E-05 | 0.008925 | 0.008667 | 5730455P1 | 84 |
| 137 | 3 | GO:005164 | organelle l | 108/1409 | 425/8076 | 1.41E-05 | 0.008925 | 0.008667 | 5730455P1 | 108 |
| 138 | 3 | GO:005165 | establishm | 49/1409 | 159/8076 | 2.22E-05 | 0.012317 | 0.011961 | 5730455P1 | 49 |
| 139 | 3 | GO:000683 | mitochond | 38/1409 | 116/8076 | 4.17E-05 | 0.017765 | 0.017251 | Agk/Aip/Bc | 38 |
| 140 | 3 | GO:000701 | microtubul | 119/1409 | 489/8076 | 4.34E-05 | 0.017765 | 0.017251 | Actr10/Agt | 119 |
| 141 | 3 | GO:009902 | vesicle tet | 15/1409 | 30/8076 | 4.40E-05 | 0.017765 | 0.017251 | 5730455P1 | 15 |
| 142 | 3 | GO:000703 | vacuolar tr | 41/1409 | 130/8076 | 5.68E-05 | 0.021002 | 0.020395 | Aktip/Ankf | 41 |
| 143 | 3 | GO:001619 | endosomal | 56/1409 | 196/8076 | 6.59E-05 | 0.022519 | 0.021868 | Ankfy1/Ap | 56 |
| 144 | 3 | GO:001648 | cytosolic tr | 44/1409 | 145/8076 | 8.62E-05 | 0.027329 | 0.026539 | Aktip/Ankf | 44 |
| 145 | 3 | GO:007064 | protein mo | 112/1409 | 464/8076 | 0.000105 | 0.027607 | 0.026809 | Aktip/Anap | 112 |
| 146 | 3 | GO:000616 | purine nuc | 80/1409 | 310/8076 | 0.000107 | 0.027607 | 0.026809 | Acaca/Aca | 80 |
| 147 | 3 | GO:005164 | vesicle loc | 51/1409 | 177/8076 | 0.000109 | 0.027607 | 0.026809 | 5730455P1 | 51 |
| 148 | 3 | GO:004214 | retrograde | 26/1409 | 72/8076 | 0.000112 | 0.027607 | 0.026809 | Ankfy1/Ap | 26 |
| 149 | 3 | GO:000915 | purine ribo | 64/1409 | 237/8076 | 0.00013 | 0.03038 | 0.029501 | Acaca/Aca | 64 |
| 150 | 3 | GO:001969 | ribose pho | 68/1409 | 257/8076 | 0.000158 | 0.034466 | 0.03347 | Acaca/Aca | 68 |
| 151 | 3 | GO:002241 | cellular cor | 83/1409 | 328/8076 | 0.000163 | 0.034466 | 0.03347 | Abce1/Add | 83 |
| 152 | 3 | GO:003254 | mitochond | 36/1409 | 116/8076 | 0.000226 | 0.042667 | 0.041433 | Gatb/Hars1 | 36 |
| 153 | 3 | GO:000925 | ribonucleo | 66/1409 | 251/8076 | 0.000238 | 0.042667 | 0.041433 | Acaca/Aca | 66 |
| 154 | 3 | GO:000911 | nucleotide | 86/1409 | 346/8076 | 0.00024 | 0.042667 | 0.041433 | Acaca/Aca | 86 |
| 155 | 3 | GO:007265 | establishm | 26/1409 | 75/8076 | 0.00024 | 0.042667 | 0.041433 | Agk/Aip/Ap | 26 |
| 156 | 3 | GO:003298 | protein-co | 51/1409 | 183/8076 | 0.00027 | 0.044805 | 0.04351 | Abce1/Add | 51 |
| 157 | 3 | GO:007252 | purine-con | 82/1409 | 328/8076 | 0.000272 | 0.044805 | 0.04351 | Acaca/Aca | 82 |
| 158 | 3 | GO:000675 | nucleoside | 87/1409 | 353/8076 | 0.000299 | 0.046222 | 0.044885 | Acaca/Aca | 87 |
| 159 | 3 | GO:005508 | nucleobase | 95/1409 | 392/8076 | 0.000302 | 0.046222 | 0.044885 | Acaca/Aca | 95 |
| 160 | 5 | GO:003083 | regulation | 32/989 | 103/8076 | 3.11E-07 | 0.001034 | 0.00099 | Actr3/Add1 | 32 |
| 161 | 5 | GO:005169 | actin filam | 15/989 | 30/8076 | 4.89E-07 | 0.001034 | 0.00099 | Add1/Add2 | 15 |
| 162 | 5 | GO:000806 | regulation | 34/989 | 120/8076 | 1.48E-06 | 0.001284 | 0.00123 | Actr3/Add1 | 34 |
| 163 | 5 | GO:003083 | regulation | 34/989 | 120/8076 | 1.48E-06 | 0.001284 | 0.00123 | Actr3/Add1 | 34 |
| 164 | 5 | GO:003083 | negative re | 16/989 | 36/8076 | 1.52E-06 | 0.001284 | 0.00123 | Add1/Add2 | 16 |
| 165 | 5 | GO:003004 | actin filam | 33/989 | 119/8076 | 3.53E-06 | 0.002487 | 0.002382 | Actr3/Add1 | 33 |
| 166 | 5 | GO:003004 | actin filam | 18/989 | 47/8076 | 4.62E-06 | 0.002613 | 0.002502 | Add1/Add2 | 18 |
| 167 | 5 | GO:190274 | positive re | 13/989 | 27/8076 | 4.94E-06 | 0.002613 | 0.002502 | Actr2/Actr | 13 |
| 168 | 5 | GO:003083 | regulation | 17/989 | 44/8076 | 7.38E-06 | 0.003468 | 0.003321 | Add1/Add2 | 17 |
| 169 | 5 | GO:000815 | actin polyn | 36/989 | 140/8076 | 8.60E-06 | 0.003635 | 0.003482 | Actr3/Add1 | 36 |
| 170 | 5 | GO:003083 | negative re | 18/989 | 50/8076 | 1.27E-05 | 0.004215 | 0.004037 | Add1/Add2 | 18 |
| 171 | 5 | GO:190274 | regulation | 15/989 | 37/8076 | 1.28E-05 | 0.004215 | 0.004037 | Actr2/Actr | 15 |
| 172 | 5 | GO:004428 | small mole | 49/989 | 218/8076 | 1.34E-05 | 0.004215 | 0.004037 | Abat/Acaa | 49 |
| 173 | 5 | GO:003278 | monocarbc | 70/989 | 349/8076 | 1.45E-05 | 0.004215 | 0.004037 | Abat/Acaa | 70 |
| 174 | 5 | GO:00904 | organopho | 74/989 | 375/8076 | 1.50E-05 | 0.004215 | 0.004037 | Acss1/Acss | 74 |
| 175 | 5 | GO:001059 | positive re | 11/989 | 22/8076 | 1.72E-05 | 0.004269 | 0.004089 | Actr2/Actr | 11 |
| 176 | 5 | GO:005101 | barbed-en | 11/989 | 22/8076 | 1.72E-05 | 0.004269 | 0.004089 | Add1/Add2 | 11 |
| 177 | 5 | GO:001059 | regulation | 13/989 | 30/8076 | 2.07E-05 | 0.004865 | 0.004659 | Actr2/Actr | 13 |
| 178 | 5 | GO:00070 | mitochond | 24/989 | 83/8076 | 3.53E-05 | 0.007849 | 0.007517 | Acaa2/Apo | 24 |
| 179 | 5 | GO:001605 | organic aci | 37/989 | 156/8076 | 4.50E-05 | 0.009068 | 0.008685 | Abat/Acaa | 37 |
| 180 | 5 | GO:004639 | carboxylic | 37/989 | 156/8076 | 4.50E-05 | 0.009068 | 0.008685 | Abat/Acaa | 37 |
| 181 | 5 | GO:000701 | actin filam | 64/989 | 326/8076 | 6.67E-05 | 0.012828 | 0.012286 | Actr2/Actr | 64 |
| 182 | 5 | GO:001598 | energy der | 49/989 | 233/8076 | 8.33E-05 | 0.015041 | 0.014406 | Acadm/Afg | 49 |
| 183 | 5 | GO:003277 | positive re | 7/989 | Nov-76 | 8.54E-05 | 0.015041 | 0.014406 | Dhfr/Kras/ | 7 |
| 184 | 5 | GO:009758 | lamellipodi | 21/989 | 72/8076 | 9.35E-05 | 0.015045 | 0.014409 | Actr2/Actr | 21 |
| 185 | 5 | GO:004324 | negative re | 20/989 | 67/8076 | 9.56E-05 | 0.015045 | 0.014409 | Add1/Add2 | 20 |
| 186 | 5 | GO:190188 | negative re | 19/989 | 62/8076 | 9.61E-05 | 0.015045 | 0.014409 | Add1/Add2 | 19 |
| 187 | 5 | GO:003227 | regulation | 36/989 | 156/8076 | 0.000104 | 0.015727 | 0.015063 | Actr3/Add1 | 36 |
| 188 | 5 | GO:005125 | protein pol | 43/989 | 201/8076 | 0.00015 | 0.021823 | 0.020901 | Actr3/Add1 | 43 |
| 189 | 5 | GO:000609 | generation | 62/989 | 323/8076 | 0.000169 | 0.023782 | 0.022778 | Acadm/Afg | 62 |
| 190 | 5 | GO:003227 | negative re | 19/989 | 65/8076 | 0.000193 | 0.026329 | 0.025217 | Add1/Add2 | 19 |
| 191 | 5 | GO:003003 | actin cytos | 85/989 | 482/8076 | 0.000245 | 0.032389 | 0.031021 | Actb/Actr2 | 85 |
| 192 | 5 | GO:001969 | ribose pho | 51/989 | 257/8076 | 0.000276 | 0.034753 | 0.033286 | Acaa2/Aco | 51 |
| 193 | 5 | GO:00512 | protein ins | 9/989 | 20/8076 | 0.000284 | 0.034753 | 0.033286 | Hsp90aa1/ | 9 |
| 194 | 5 | GO:000925 | ribonucleo | 50/989 | 251/8076 | 0.000288 | 0.034753 | 0.033286 | Acaa2/Aco | 50 |
| 195 | 5 | GO:000616 | purine nuc | 59/989 | 310/8076 | 0.000305 | 0.035852 | 0.034338 | Acaa2/Aco | 59 |
| 196 | 5 | GO:007064 | protein mo | 22/989 | 84/8076 | 0.000358 | 0.039159 | 0.037506 | Cops2/Cop | 22 |
| 197 | 5 | GO:005508 | nucleobase | 71/989 | 392/8076 | 0.000362 | 0.039159 | 0.037506 | Acaa2/Aco | 71 |
| 198 | 5 | GO:006049 | regulation | 31/989 | 136/8076 | 0.000388 | 0.039159 | 0.037506 | Actr2/Actr | 31 |
| 199 | 5 | GO:000911 | nucleotide | 64/989 | 346/8076 | 0.000391 | 0.039159 | 0.037506 | Acaa2/Aco | 64 |
| 200 | 5 | GO:004277 | ATP synthe | 17/989 | 58/8076 | 0.0004 | 0.039159 | 0.037506 | Afg1l/Coa6 | 17 |
| 201 | 5 | GO:004277 | mitochond | 17/989 | 58/8076 | 0.0004 | 0.039159 | 0.037506 | Afg1l/Coa6 | 17 |
| 202 | 5 | GO:000675 | nucleoside | 65/989 | 353/8076 | 0.000401 | 0.039159 | 0.037506 | Acaa2/Aco | 65 |
| 203 | 5 | GO:004827 | vesicle doc | 16/989 | 53/8076 | 0.000408 | 0.039159 | 0.037506 | Exoc3/Exo | 16 |
| 204 | 5 | GO:190187 | regulation | 20/989 | 74/8076 | 0.00042 | 0.039428 | 0.037763 | Add1/Add2 | 20 |
| 205 | 5 | GO:00324 | melanoson | 6/989 | Oct-76 | 0.00045 | 0.040515 | 0.038805 | Dctn2/Map | 6 |
| 206 | 5 | GO:005187 | pigment gr | 6/989 | Oct-76 | 0.00045 | 0.040515 | 0.038805 | Dctn2/Map | 6 |
| 207 | 5 | GO:000915 | purine ribo | 47/989 | 237/8076 | 0.000481 | 0.042409 | 0.040618 | Acaa2/Aco | 47 |
| 208 | 5 | GO:012003 | positive re | 20/989 | 75/8076 | 0.000508 | 0.043476 | 0.04164 | Actr2/Actr | 20 |
| 209 | 5 | GO:000906 | glutamine | 15/989 | 49/8076 | 0.00052 | 0.043476 | 0.04164 | Aldh5a1/C | 15 |
| 210 | 5 | GO:000038 | spliceosom | 11/989 | 30/8076 | 0.000524 | 0.043476 | 0.04164 | Lsm2/Lsm4 | 11 |
| 211 | 5 | GO:011005 | regulation | 42/989 | 207/8076 | 0.000581 | 0.047248 | 0.045253 | Actr3/Add1 | 42 |
| 212 | 5 | GO:003305 | cellular pig | 12/989 | 35/8076 | 0.000599 | 0.047765 | 0.045749 | Ap1b1/Ap1 | 12 |
| 213 | 7 | GO:004533 | cellular res | 50/916 | 175/8076 | 2.55E-10 | 9.86E-07 | 9.62E-07 | Aco2/Adsl/ | 50 |
| 214 | 7 | GO:000906 | aerobic res | 44/916 | 146/8076 | 4.79E-10 | 9.86E-07 | 9.62E-07 | Aco2/Adsl/ | 44 |
| 215 | 7 | GO:001598 | energy der | 57/916 | 233/8076 | 8.92E-09 | 1.23E-05 | 1.20E-05 | Aco2/Adsl/ | 57 |
| 216 | 7 | GO:000609 | generation | 69/916 | 323/8076 | 9.19E-08 | 9.46E-05 | 9.23E-05 | Aass/Aco2/ | 69 |
| 217 | 7 | GO:000611 | oxidative p | 31/916 | 110/8076 | 9.32E-07 | 0.000768 | 0.00075 | Atp5f1d/At | 31 |
| 218 | 7 | GO:004364 | dicarboxyli | 23/916 | 73/8076 | 2.98E-06 | 0.002044 | 0.001994 | Aasdhppt/ | 23 |
| 219 | 7 | GO:000636 | rRNA proce | 37/916 | 153/8076 | 4.98E-06 | 0.002625 | 0.002561 | Bms1/Ddx | 37 |
| 220 | 7 | GO:000609 | tricarboxyli | 13/916 | 29/8076 | 5.62E-06 | 0.002625 | 0.002561 | Aco2/Dlat/ | 13 |
| 221 | 7 | GO:000911 | nucleotide | 67/916 | 346/8076 | 5.73E-06 | 0.002625 | 0.002561 | Adsl/Ampd | 67 |
| 222 | 7 | GO:000906 | amino acid | 20/916 | 62/8076 | 8.79E-06 | 0.003621 | 0.003533 | Aass/Adhfe | 20 |
| 223 | 7 | GO:000675 | nucleoside | 67/916 | 353/8076 | 1.15E-05 | 0.004316 | 0.004212 | Adsl/Ampd | 67 |
| 224 | 7 | GO:00229 | electron tr | 23/916 | 81/8076 | 2.05E-05 | 0.007034 | 0.006863 | Coq7/Coq | 23 |
| 225 | 7 | GO:001607 | rRNA meta | 38/916 | 169/8076 | 2.26E-05 | 0.007158 | 0.006984 | Bms1/Ddx | 38 |
| 226 | 7 | GO:003447 | ncRNA pro | 54/916 | 274/8076 | 2.82E-05 | 0.00829 | 0.008089 | Ago1/Akt1, | 54 |
| 227 | 7 | GO:000916 | nucleotide | 41/916 | 191/8076 | 3.47E-05 | 0.009537 | 0.009305 | Adsl/Ampd | 41 |
| 228 | 7 | GO:190129 | nucleoside | 41/916 | 192/8076 | 3.95E-05 | 0.010143 | 0.009896 | Adsl/Ampd | 41 |
| 229 | 7 | GO:000926 | ribonucleo | 36/916 | 161/8076 | 4.19E-05 | 0.010143 | 0.009896 | Adsl/Ampd | 36 |
| 230 | 7 | GO:004277 | proton mo | 17/916 | 53/8076 | 4.48E-05 | 0.010244 | 0.009995 | Atp5f1d/At | 17 |
| 231 | 7 | GO:005508 | nucleobase | 70/916 | 392/8076 | 5.94E-05 | 0.012579 | 0.012274 | Adsl/Ampd | 70 |
| 232 | 7 | GO:00990 | vesicle-me | 48/916 | 242/8076 | 6.60E-05 | 0.012579 | 0.012274 | Aak1/Akap | 48 |
| 233 | 7 | GO:000616 | purine nuc | 58/916 | 310/8076 | 6.86E-05 | 0.012579 | 0.012274 | Adsl/Ampd | 58 |
| 234 | 7 | GO:005102 | mRNA tran | 23/916 | 87/8076 | 7.08E-05 | 0.012579 | 0.012274 | Akap8l/Cet | 23 |
| 235 | 7 | GO:000639 | mRNA proc | 65/916 | 359/8076 | 7.13E-05 | 0.012579 | 0.012274 | Acin1/Akap | 65 |
| 236 | 7 | GO:001598 | proton mo | 17/916 | 55/8076 | 7.59E-05 | 0.012579 | 0.012274 | Atp5f1d/At | 17 |
| 237 | 7 | GO:00064 | mRNA exp | 16/916 | 50/8076 | 7.71E-05 | 0.012579 | 0.012274 | Akap8l/Cht | 16 |
| 238 | 7 | GO:00061 | 2-oxogluta | 8/916 | 15/8076 | 8.23E-05 | 0.012579 | 0.012274 | Adhfe1/Dl | 8 |
| 239 | 7 | GO:004639 | ribose pho | 36/916 | 166/8076 | 8.24E-05 | 0.012579 | 0.012274 | Adsl/Ampd | 36 |
| 240 | 7 | GO:001608 | synaptic ve | 11/916 | 27/8076 | 8.79E-05 | 0.012931 | 0.012617 | Cadps2/Er | 11 |
| 241 | 7 | GO:007093 | 3'-UTR-me | 7/916 | Dec-76 | 0.000112 | 0.015456 | 0.01508 | Angel2/Ela | 7 |
| 242 | 7 | GO:001605 | organic aci | 34/916 | 156/8076 | 0.000116 | 0.015456 | 0.01508 | Aasdhppt/ | 34 |
| 243 | 7 | GO:004639 | carboxylic | 34/916 | 156/8076 | 0.000116 | 0.015456 | 0.01508 | Aasdhppt/ | 34 |
| 244 | 7 | GO:000915 | purine ribo | 33/916 | 150/8076 | 0.000121 | 0.015538 | 0.015161 | Adsl/Ampd | 33 |
| 245 | 7 | GO:004225 | ribosome b | 45/916 | 228/8076 | 0.000125 | 0.015629 | 0.01525 | Aatf/Bms1, | 45 |
| 246 | 7 | GO:00229 | respiratory | 20/916 | 75/8076 | 0.000182 | 0.021105 | 0.020593 | Coq7/Coq | 20 |
| 247 | 7 | GO:007252 | purine-con | 59/916 | 328/8076 | 0.000184 | 0.021105 | 0.020593 | Adsl/Ampd | 59 |
| 248 | 7 | GO:000838 | RNA splicin | 57/916 | 314/8076 | 0.000184 | 0.021105 | 0.020593 | Acin1/Aqr/ | 57 |
| 249 | 7 | GO:000616 | purine nuc | 35/916 | 168/8076 | 0.000237 | 0.025188 | 0.024576 | Adsl/Ampd | 35 |

| | | | | | | | | |
|---|---|---|---|---|---|---|---|---|
| 1128 | ENSMUSG | Cdh11 | 12552 | -04670174 | 01504751 | 005 | 004 | 2 |
| 1130 | ENSMUSG | Cdh13 | 12554 | -04328019 | 02905152 | 0685 | 014 | 2 |
| 1134 | ENSMUSG | Cdh20 | 23836 | -00960109 | 00848915 | 024 | 01425 | 2 |
| 1166 | ENSMUSG | Cds1 | 74596 | -02075265 | 00689197 | 03625 | 034 | 2 |
| 1167 | ENSMUSG | Cds2 | 110911 | -01104544 | 01562897 | 02025 | 019 | 2 |
| 1177 | ENSMUSG | Celsr2 | 53883 | -02854693 | 02204603 | 1013 | 01375 | 2 |
| 1178 | ENSMUSG | Cemip2 | 83921 | -02051566 | 02848203 | 0305 | 01275 | 2 |
| 1193 | ENSMUSG | Cers2 | 76893 | -04337496 | 02578012 | 0365 | 048 | 2 |
| 1194 | ENSMUSG | Cers4 | 67260 | -01242778 | 01409280 | 0125 | 03475 | 2 |
| 1236 | ENSMUSG | Chgb | 12653 | -02914347 | 01430165 | 4011 | 02775 | 2 |
| 1260 | ENSMUSG | Chrm1 | 12669 | -02851799 | 02628310 | 1-0175 | 026 | 2 |
| 1296 | ENSMUSG | Clcn6 | 26372 | -02807695 | 00535605 | 8018 | 00825 | 2 |
| 1297 | ENSMUSG | Clcn7 | 26373 | -01498181 | 01721086 | 01925 | 03725 | 2 |
| 1301 | ENSMUSG | Cldn12 | 64945 | -04374259 | 02472717 | 026 | 04 | 2 |
| 1319 | ENSMUSG | Cln6 | 76524 | -03911412 | 00389785 | 102725 | 00125 | 2 |
| 1325 | ENSMUSG | Clptm1 | 56457 | -02375462 | 00666908 | 03025 | 02625 | 2 |
| 1330 | ENSMUSG | Clstn3 | 232370 | -02925228 | 01224891 | 1027 | 021 | 2 |
| 1354 | ENSMUSG | Cnih1 | 12793 | -03508768 | 00087825 | 013 | 00525 | 2 |
| 1363 | ENSMUSG | Cnnm3 | 94218 | -02386614 | 00553026 | 024 | 001 | 2 |
| 1364 | ENSMUSG | Cnnm4 | 94220 | -02372475 | 02539975 | 017 | 02175 | 2 |
| 1375 | ENSMUSG | Cnpy2 | 56530 | -02582889 | -00467625 | -00075 | 01975 | 2 |
| 1377 | ENSMUSG | Cnpy4 | 66455 | -01651065 | 01547312 | 0365 | -009 | 2 |
| 1382 | ENSMUSG | Cntn1 | 12805 | -03111838 | 01000365 | 70155 | -00325 | 2 |
| 1385 | ENSMUSG | Cntn4 | 269784 | -05851128 | 02524038 | 01725 | 0175 | 2 |
| 1386 | ENSMUSG | Cntn5 | 244682 | -01590029 | 01567383 | 006 | 03125 | 2 |
| 1391 | ENSMUSG | Cntnap4 | 170571 | -04710575 | 01457508 | 100575 | 0045 | 2 |
| 1393 | ENSMUSG | Cntnap5b | 241175 | -03557061 | 05285031 | 02375 | 03325 | 2 |
| 1425 | ENSMUSG | Col6a3 | 12835 | -03218421 | -02499400 | 00725 | 026 | 2 |
| 1427 | ENSMUSG | Colgalt1 | 234407 | -02079096 | 02317122 | 20305 | 0105 | 2 |
| 1513 | ENSMUSG | Cpox | 12892 | -03567723 | 00927226 | -0075 | 01625 | 2 |
| 1523 | ENSMUSG | Cpt1c | 78070 | -00698862 | 01430519 | 025 | 034 | 2 |
| 1575 | ENSMUSG | Csmd1 | 94109 | -04115051 | 03481856 | 1012 | 01675 | 2 |
| 1622 | ENSMUSG | Ctsa | 19025 | -02669121 | 00846242 | 702625 | 03975 | 2 |
| 1624 | ENSMUSG | Ctsc | 13032 | -02588978 | 04491121 | 702975 | 04525 | 2 |
| 1625 | ENSMUSG | Ctsd | 13033 | -01395410 | 01254071 | 01875 | 03275 | 2 |
| 1626 | ENSMUSG | Ctsf | 56464 | -02154076 | 02350536 | 102525 | 03825 | 2 |
| 1627 | ENSMUSG | Ctsh | 13036 | -01212901 | 02224895 | 00425 | 03175 | 2 |
| 1628 | ENSMUSG | Ctsl | 13039 | -02484837 | 01974510 | 4042 | 03025 | 2 |
| 1631 | ENSMUSG | Ctsz | 64138 | -02645835 | 00014344 | 70215 | 00725 | 2 |
| 1679 | ENSMUSG | Cyp46a1 | 13116 | -02721626 | 00940439 | 03 | 007 | 2 |
| 1683 | ENSMUSG | Cyp7b1 | 13123 | -02503796 | 01300877 | 10385 | -0015 | 2 |
| 1708 | ENSMUSG | Dag1 | 13138 | -02639336 | 01632195 | 7-00225 | 014 | 2 |
| 1737 | ENSMUSG | Dcc | 13176 | -03227845 | 01762103 | 8-00075 | 013 | 2 |
| 1787 | ENSMUSG | Ddx3x | 13205 | -02323690 | 00589120 | 3-009 | 01175 | 2 |
| 1805 | ENSMUSG | Degs1 | 13244 | -03530334 | 01277063 | -00125 | 02525 | 2 |
| 1842 | ENSMUSG | Dhcr7 | 13360 | -05053828 | -00350150 | 03575 | 0325 | 2 |
| 1850 | ENSMUSG | Dhrs13 | 70451 | -02546465 | 00956326 | 802275 | 0325 | 2 |
| 1882 | ENSMUSG | Dixdc1 | 330938 | -01306457 | 00913232 | 02075 | -001 | 2 |
| 1917 | ENSMUSG | Dnajb12 | 56709 | -03561060 | 01212250 | 700775 | 0145 | 2 |
| 1924 | ENSMUSG | Dnajc10 | 66861 | -02574365 | 02430618 | 031 | 03975 | 2 |
| 1947 | ENSMUSG | Dner | 227325 | -05030833 | 01192978 | 700875 | 03475 | 2 |
| 1963 | ENSMUSG | Dock10 | 210293 | -01868504 | 01587157 | 7-0045 | 00625 | 2 |
| 2013 | ENSMUSG | Dscam | 13508 | -02435814 | 02051782 | -013 | 012 | 2 |
| 2064 | ENSMUSG | Ebp | 13595 | -03832477 | 00137605 | 70515 | 02625 | 2 |
| 2080 | ENSMUSG | Edem3 | 66967 | -02737767 | 02297118 | 02975 | 05225 | 2 |
| 2082 | ENSMUSG | Edil3 | 13612 | -05553130 | 02890960 | 01025 | -001 | 2 |
| 2083 | ENSMUSG | Ednrb | 13618 | -01999833 | 00948563 | 03225 | 00425 | 2 |
| 2103 | ENSMUSG | Efna5 | 13640 | -01418753 | 03578747 | 013 | 00925 | 2 |
| 2105 | ENSMUSG | Efnb2 | 13642 | -03451066 | 03549084 | -01525 | 0165 | 2 |
| 2131 | ENSMUSG | Eif2ak3 | 13666 | -01182422 | 03049009 | 01225 | 007 | 2 |
| 2176 | ENSMUSG | Elapor1 | 229722 | -02176498 | 01168738 | 70085 | 027 | 2 |
| 2202 | ENSMUSG | Emc1 | 230866 | -02655218 | 00952223 | 7047 | -0065 | 2 |
| 2206 | ENSMUSG | Emc4 | 68032 | -05702387 | -01363544 | 03225 | 018 | 2 |
| 2208 | ENSMUSG | Emc7 | 73024 | -02335737 | 00481800 | 043 | 02475 | 2 |
| 2256 | ENSMUSG | Epdr1 | 105298 | -02929217 | 01167116 | 504725 | -00975 | 2 |
| 2259 | ENSMUSG | Epha3 | 13837 | -05570962 | 03824687 | 401225 | 04875 | 2 |
| 2260 | ENSMUSG | Epha4 | 13838 | -04653701 | 02888324 | 102125 | 016 | 2 |
| 2261 | ENSMUSG | Epha5 | 13839 | -02728022 | 02670632 | 4061 | 03275 | 2 |
| 2263 | ENSMUSG | Epha7 | 13841 | -04592621 | 02670282 | 02475 | 0505 | 2 |
| 2266 | ENSMUSG | Ephb3 | 13845 | -03372490 | 01452183 | 016 | 0015 | 2 |
| 2267 | ENSMUSG | Ephb6 | 13848 | -00941131 | 01522028 | 4028 | 01675 | 2 |
| 2283 | ENSMUSG | Erbb3 | 13867 | -06637649 | 02230406 | 503475 | 02925 | 2 |
| 2293 | ENSMUSG | Ergic1 | 67458 | -02885039 | 00686457 | 044 | 00075 | 2 |
| 2294 | ENSMUSG | Ergic2 | 67456 | -01772263 | 00933450 | 302775 | 02925 | 2 |
| 2299 | ENSMUSG | Erlin1 | 226144 | -03403158 | 01318294 | 202875 | 012 | 2 |
| 2300 | ENSMUSG | Erlin2 | 244373 | -01719282 | 00458763 | 803175 | 002 | 2 |
| 2303 | ENSMUSG | Ero1a | 50527 | -02607633 | 02258860 | 202775 | -0085 | 2 |
| 2355 | ENSMUSG | Fa2h | 338521 | -07185125 | 02079641 | 045 | 04225 | 2 |
| 2356 | ENSMUSG | Faah | 14073 | -02168559 | -00184068 | 02725 | 0015 | 2 |
| 2364 | ENSMUSG | Fads3 | 60527 | -02621463 | 01140471 | 03 | -00225 | 2 |
| 2395 | ENSMUSG | Fam171a2 | 217219 | -01861175 | 01938946 | 10215 | 03875 | 2 |
| 2409 | ENSMUSG | Fam234b | 74525 | -01475914 | 00617481 | 024 | 01025 | 2 |
| 2432 | ENSMUSG | Fat4 | 329628 | -06021043 | 01617578 | 0325 | 02975 | 2 |
| 2438 | ENSMUSG | Fbln5 | 23876 | -04066430 | 00058265 | 0085 | 02775 | 2 |
| 2472 | ENSMUSG | Fcgr2b | 14130 | -04478202 | 01452251 | 00575 | 063 | 2 |
| 2538 | ENSMUSG | Flrt2 | 399558 | -04178123 | 04385066 | 2037 | -00225 | 2 |
| 2539 | ENSMUSG | Flrt3 | 71436 | -05526637 | 05709864 | 00675 | 024 | 2 |
| 2576 | ENSMUSG | Frmd4b | 232288 | -01507725 | 01295586 | 102175 | 01025 | 2 |
| 2598 | ENSMUSG | Fuca1 | 71665 | -00815531 | 01044147 | 02 | 026 | 2 |
| 2603 | ENSMUSG | Fut11 | 73068 | -02422486 | 01402246 | 402775 | -0045 | 2 |
| 2604 | ENSMUSG | Fut8 | 53618 | -02794658 | 00164873 | 404625 | 025 | 2 |
| 2605 | ENSMUSG | Fut9 | 14348 | -01887344 | 03411436 | 03625 | 00675 | 2 |
| 2611 | ENSMUSG | Fxyd6 | 59095 | -02323778 | 01504303 | 104875 | 039 | 2 |
| 2617 | ENSMUSG | Fzd3 | 14365 | -02899298 | 01122018 | 7-01025 | 0345 | 2 |
| 2618 | ENSMUSG | G0s2 | 14373 | -05829251 | 04551525 | 202525 | 01275 | 2 |
| 2623 | ENSMUSG | Gaa | 14387 | -01221193 | 02625758 | 0165 | 235514e-1 | 2 |
| 2637 | ENSMUSG | Gabrb2 | 14401 | -07643305 | 04591313 | 704875 | 001 | 2 |
| 2638 | ENSMUSG | Gabrb3 | 14402 | -04510495 | 01911792 | 00725 | 00875 | 2 |
| 2641 | ENSMUSG | Gabrg2 | 14406 | -05757185 | 02424158 | 803825 | -00275 | 2 |
| 2647 | ENSMUSG | Gal3st3 | 545276 | -01775883 | 02881303 | 02775 | 00325 | 2 |
| 2653 | ENSMUSG | Galns | 50917 | -02907258 | 01312700 | 002 | 022 | 2 |
| 2655 | ENSMUSG | Galnt13 | 271786 | -06580519 | 02362486 | 04325 | 0185 | 2 |
| 2656 | ENSMUSG | Galnt14 | 71685 | -02750031 | 00624503 | 00375 | 03275 | 2 |
| 2657 | ENSMUSG | Galnt16 | 108760 | -02824523 | 01303008 | 2036 | 0015 | 2 |
| 2658 | ENSMUSG | Galnt17 | 212996 | -02372913 | 02023709 | 40395 | 00275 | 2 |
| 2659 | ENSMUSG | Galnt2 | 108148 | -02248504 | 01507015 | 702525 | 01825 | 2 |
| 2706 | ENSMUSG | Gde1 | 56209 | -02802556 | -00165237 | 0375 | -01075 | 2 |
| 2711 | ENSMUSG | Gdpd5 | 233552 | -03152461 | 00549866 | 20195 | 01475 | 2 |
| 2713 | ENSMUSG | Gemin4 | 276919 | -04000677 | -00388370 | 06025 | 03625 | 2 |
| 2716 | ENSMUSG | Get1 | 71446 | -01964909 | 00019249 | 802625 | 013 | 2 |
| 2726 | ENSMUSG | Gfra1 | 14585 | -03714765 | -02542688 | 0235 | 03025 | 2 |
| 2734 | ENSMUSG | Ggh | 14590 | -03169205 | 00966657 | 01575 | 00975 | 2 |
| 2749 | ENSMUSG | Gja1 | 14609 | -03372839 | -00680523 | 02 | 0135 | 2 |
| 2757 | ENSMUSG | Gla | 11605 | -05895604 | 01722590 | 20025 | 03075 | 2 |
| 2758 | ENSMUSG | Glb1 | 12091 | -01697828 | 03220389 | 02725 | 01125 | 2 |
| 2761 | ENSMUSG | Glce | 93683 | -04321592 | 02344085 | 80175 | 01025 | 2 |
| 2764 | ENSMUSG | Glg1 | 20340 | -01643768 | 02412255 | 30325 | 02975 | 2 |
| 2838 | ENSMUSG | Golgb1 | 224139 | -02242503 | -00018856 | 0265 | 0175 | 2 |
| 2840 | ENSMUSG | Golm2 | 319996 | -02517051 | 00733459 | 5041 | 0145 | 2 |
| 2843 | ENSMUSG | Golt1b | 66964 | -04188067 | 00982550 | 204925 | -00975 | 2 |
| 2856 | ENSMUSG | Gpat4 | 102247 | -00728616 | 01569480 | 0215 | 0165 | 2 |
| 2861 | ENSMUSG | Gpc1 | 14733 | -01937560 | 01998452 | 1-01025 | 0225 | 2 |
| 2872 | ENSMUSG | Gpm6a | 234267 | -02828105 | 01914147 | 602 | 02275 | 2 |
| 2876 | ENSMUSG | Gpr155 | 68526 | -05561461 | 01940745 | 8-001 | 01925 | 2 |
| 2879 | ENSMUSG | Gpr17 | 574402 | -03279517 | 04894065 | 3-0055 | 06 | 2 |
| 2880 | ENSMUSG | Gpr176 | 381413 | -01407130 | 01508110 | 501875 | 018 | 2 |

| | | | | | | | | | | |
|---|---|---|---|---|---|---|---|---|---|---|
| 250 | 7 | GO:000037 | RNA splicin | 43/916 | 221/8076 | 0.000245 | 0.025188 | 0.024576 | Acin1/Aqr/ | 43 |
| 251 | 7 | GO:000037 | RNA splicin | 43/916 | 221/8076 | 0.000245 | 0.025188 | 0.024576 | Acin1/Aqr/ | 43 |
| 252 | 7 | GO:000039 | mRNA splic | 43/916 | 221/8076 | 0.000245 | 0.025188 | 0.024576 | Acin1/Aqr/ | 43 |
| 253 | 7 | GO:000652 | amino acid | 37/916 | 182/8076 | 0.000267 | 0.026784 | 0.026134 | Aasdhppt/ | 37 |
| 254 | 7 | GO:190160 | alpha-amir | 15/916 | 50/8076 | 0.000287 | 0.02819 | 0.027506 | Aass/Adhfe | 15 |
| 255 | 7 | GO:000640 | RNA expor | 18/916 | 67/8076 | 0.000342 | 0.032748 | 0.031954 | Akap8l/Cht | 18 |
| 256 | 7 | GO:004428 | small mole | 42/916 | 218/8076 | 0.000358 | 0.033529 | 0.032715 | Aasdhppt/ | 42 |
| 257 | 7 | GO:000015 | rRNA modi | 8/916 | 18/8076 | 0.000409 | 0.037471 | 0.036562 | Emg1/Ftsj3 | 8 |
| 258 | 7 | GO:001967 | NAD metal | 10/916 | 27/8076 | 0.000462 | 0.041422 | 0.040416 | Gpd1/Gpd2 | 10 |
| 259 | 7 | GO:007252 | purine-con | 35/916 | 174/8076 | 0.000476 | 0.041758 | 0.040744 | Adsl/Ampd | 35 |
| 260 | 7 | GO:005065 | nucleic aci | 25/916 | 111/8076 | 0.000538 | 0.044323 | 0.043248 | Akap8l/Cet | 25 |
| 261 | 7 | GO:005065 | RNA transp | 25/916 | 111/8076 | 0.000538 | 0.044323 | 0.043248 | Akap8l/Cet | 25 |
| 262 | 7 | GO:005123 | establishm | 25/916 | 111/8076 | 0.000538 | 0.044323 | 0.043248 | Akap8l/Cet | 25 |
| 263 | 7 | GO:001607 | mRNA met | 80/916 | 496/8076 | 0.000577 | 0.046597 | 0.045466 | Acin1/Ago1 | 80 |
| 264 | 7 | GO:000925 | ribonucleo | 46/916 | 251/8076 | 0.000611 | 0.048415 | 0.04724 | Adsl/Ampd | 46 |
| 265 | 8 | GO:004594 | positive reg | Nov-46 | 451/8076 | 3.28E-05 | 0.031501 | 0.029777 | Arid4b/Ccn | 11 |
| 266 | 8 | GO:000012 | negative re | Sep-46 | 329/8076 | 8.04E-05 | 0.038609 | 0.036497 | Arid4b/Epc | 9 |
| 267 | 9 | GO:000317 | heart valve | 8/268 | 22/8076 | 2.83E-07 | 0.000874 | 0.000836 | Emilin1/Mc | 8 |
| 268 | 9 | GO:000317 | heart valve | 8/268 | 25/8076 | 8.77E-07 | 0.001356 | 0.001297 | Emilin1/Mc | 8 |
| 269 | 9 | GO:190267 | negative re | 32/268 | 446/8076 | 2.76E-05 | 0.028418 | 0.027175 | Arid1a/Atf2 | 32 |
| 270 | 9 | GO:004589 | negative re | 31/268 | 444/8076 | 6.24E-05 | 0.048249 | 0.046138 | Arid1a/Atf2 | 31 |
| 271 | 10 | GO:000218 | cytoplasmi | 49/521 | 122/8076 | 2.48E-27 | 9.09E-24 | 9.09E-24 | Dhx36/Eif2 | 49 |
| 272 | 10 | GO:014023 | translation | 29/521 | 46/8076 | 8.91E-24 | 1.63E-20 | 1.63E-20 | Rpl10/Rpl1 | 29 |
| 273 | 10 | GO:014024 | translation | 29/521 | 47/8076 | 2.19E-23 | 2.00E-20 | 2.00E-20 | Rpl10/Rpl1 | 29 |
| 274 | 10 | GO:014024 | translation | 29/521 | 47/8076 | 2.19E-23 | 2.00E-20 | 2.00E-20 | Rpl10/Rpl1 | 29 |
| 275 | 10 | GO:004225 | ribosome b | 35/521 | 228/8076 | 1.19E-06 | 0.000869 | 0.000869 | Ddx28/Ddx | 35 |
| 276 | 10 | GO:003444 | lipid oxidat | 17/521 | 77/8076 | 5.83E-06 | 0.003347 | 0.003347 | Abcd2/Aca | 17 |
| 277 | 10 | GO:004227 | ribosomal s | 17/521 | 78/8076 | 7.02E-06 | 0.003347 | 0.003347 | Gtf2h5/Krr | 17 |
| 278 | 10 | GO:000663 | fatty acid n | 33/521 | 226/8076 | 7.32E-06 | 0.003347 | 0.003347 | Abcd2/Abh | 33 |
| 279 | 10 | GO:001939 | fatty acid c | 16/521 | 75/8076 | 1.74E-05 | 0.00707 | 0.00707 | Abcd2/Aca | 16 |
| 280 | 10 | GO:002261 | ribonucleo | 40/521 | 327/8076 | 5.95E-05 | 0.021747 | 0.021747 | Adar/Ddx2 | 40 |
| 281 | 10 | GO:000636 | rRNA proce | 23/521 | 153/8076 | 0.000114 | 0.037854 | 0.037854 | Ddx47/Dice | 23 |
| 282 | 11 | GO:003422 | monoatom | 96/519 | 486/8076 | 8.70E-25 | 3.15E-21 | 2.66E-21 | Amigo1/An | 96 |
| 283 | 11 | GO:009866 | inorganic ic | 90/519 | 455/8076 | 2.97E-23 | 5.37E-20 | 4.54E-20 | Amigo1/An | 90 |
| 284 | 11 | GO:003000 | metal ion t | 92/519 | 486/8076 | 2.40E-22 | 2.89E-19 | 2.45E-19 | Amigo1/An | 92 |
| 285 | 11 | GO:009865 | monoatom | 81/519 | 427/8076 | 1.00E-19 | 9.09E-17 | 7.68E-17 | Amigo1/An | 81 |
| 286 | 11 | GO:009866 | inorganic c | 80/519 | 423/8076 | 2.12E-19 | 1.54E-16 | 1.30E-16 | Amigo1/An | 80 |
| 287 | 11 | GO:007180 | potassium | 40/519 | 133/8076 | 2.88E-17 | 1.74E-14 | 1.47E-14 | Amigo1/An | 40 |
| 288 | 11 | GO:000681 | potassium | 42/519 | 152/8076 | 1.39E-16 | 7.18E-14 | 6.06E-14 | Amigo1/An | 42 |
| 289 | 11 | GO:000761 | behavior | 80/519 | 482/8076 | 6.65E-16 | 3.01E-13 | 2.54E-13 | Adam22/A | 80 |
| 290 | 11 | GO:004326 | regulation | 65/519 | 350/8076 | 1.92E-15 | 7.73E-13 | 6.54E-13 | Amigo1/An | 65 |
| 291 | 11 | GO:003476 | regulation | 55/519 | 284/8076 | 5.51E-14 | 1.99E-11 | 1.68E-11 | Amigo1/An | 55 |
| 292 | 11 | GO:003476 | regulation | 65/519 | 375/8076 | 6.07E-14 | 2.00E-11 | 1.69E-11 | Acsl1/Acsl5 | 65 |
| 293 | 11 | GO:000741 | synapse as | 41/519 | 172/8076 | 8.33E-14 | 2.51E-11 | 2.12E-11 | Ache/Adgrl | 41 |
| 294 | 11 | GO:005508 | monoatom | 58/519 | 320/8076 | 2.18E-13 | 6.07E-11 | 5.13E-11 | Ank3/Aplp2 | 58 |
| 295 | 11 | GO:005080 | monoatom | 58/519 | 325/8076 | 4.35E-13 | 1.12E-10 | 9.51E-11 | Ank3/Aplp2 | 58 |
| 296 | 11 | GO:000150 | action pote | 30/519 | 102/8076 | 5.82E-13 | 1.40E-10 | 1.19E-10 | Ank3/Atp2 | 30 |
| 297 | 11 | GO:003432 | cell junctio | 57/519 | 325/8076 | 1.50E-12 | 3.37E-10 | 2.85E-10 | Ache/Actn1 | 57 |
| 298 | 11 | GO:009877 | inorganic ic | 52/519 | 280/8076 | 1.58E-12 | 3.37E-10 | 2.85E-10 | Ank3/Aplp2 | 52 |
| 299 | 11 | GO:005080 | synapse or | 69/519 | 449/8076 | 3.90E-12 | 7.70E-10 | 6.50E-10 | Ache/Adgrl | 69 |
| 300 | 11 | GO:004239 | regulation | 55/519 | 314/8076 | 4.04E-12 | 7.70E-10 | 6.50E-10 | Ank3/Atp1 | 55 |
| 301 | 11 | GO:005080 | modulation | 69/519 | 450/8076 | 4.34E-12 | 7.85E-10 | 6.63E-10 | Atp2a2/Atp | 69 |
| 302 | 11 | GO:009917 | regulation | 69/519 | 451/8076 | 4.82E-12 | 8.32E-10 | 7.03E-10 | Atp2a2/Atp | 69 |
| 303 | 11 | GO:003000 | intracellula | 51/519 | 284/8076 | 9.90E-12 | 1.63E-09 | 1.38E-09 | Aplp2/Atp1 | 51 |
| 304 | 11 | GO:009865 | inorganic c | 22/519 | 62/8076 | 1.18E-11 | 1.77E-09 | 1.50E-09 | Atp1a1/Atp | 22 |
| 305 | 11 | GO:009958 | inorganic ic | 22/519 | 62/8076 | 1.18E-11 | 1.77E-09 | 1.50E-09 | Atp1a1/Atp | 22 |
| 306 | 11 | GO:003524 | synaptic tra | 28/519 | 100/8076 | 1.32E-11 | 1.91E-09 | 1.61E-09 | Cacna1a/Ca | 28 |
| 307 | 11 | GO:000687 | intracellula | 51/519 | 288/8076 | 1.70E-11 | 2.37E-09 | 2.01E-09 | Aplp2/Atp1 | 51 |
| 308 | 11 | GO:190189 | positive reg | 27/519 | 95/8076 | 2.10E-11 | 2.81E-09 | 2.38E-09 | Adgrb3/Am | 27 |
| 309 | 11 | GO:009873 | import acro | 30/519 | 117/8076 | 2.80E-11 | 3.62E-09 | 3.06E-09 | Acsl1/Acsl5 | 30 |
| 310 | 11 | GO:001922 | transmissic | 21/519 | 60/8076 | 4.71E-11 | 5.68E-09 | 4.80E-09 | Ank3/Cacn | 21 |
| 311 | 11 | GO:005196 | positive reg | 21/519 | 60/8076 | 4.71E-11 | 5.68E-09 | 4.80E-09 | Adgrb3/Am | 21 |
| 312 | 11 | GO:005196 | regulation | 28/519 | 106/8076 | 6.07E-11 | 7.09E-09 | 5.99E-09 | Adgrb3/Am | 28 |
| 313 | 11 | GO:001972 | cellular hor | 69/519 | 477/8076 | 6.73E-11 | 7.61E-09 | 6.43E-09 | Abcg1/Adc | 69 |
| 314 | 11 | GO:005508 | intracellula | 62/519 | 407/8076 | 7.96E-11 | 8.74E-09 | 7.38E-09 | Abcg1/Adc | 62 |
| 315 | 11 | GO:005196 | regulation | 21/519 | 69/8076 | 9.21E-10 | 9.81E-08 | 8.29E-08 | Cacng2/Ccl | 21 |
| 316 | 11 | GO:006156 | axon devel | 56/519 | 381/8076 | 2.79E-09 | 2.89E-07 | 2.44E-07 | Ache/Amig | 56 |
| 317 | 11 | GO:190188 | regulation | 35/519 | 182/8076 | 3.18E-09 | 3.20E-07 | 2.70E-07 | Adgrb3/Am | 35 |
| 318 | 11 | GO:000761 | learning or | 39/519 | 224/8076 | 7.66E-09 | 7.49E-07 | 6.33E-07 | Adgrb3/Atp | 39 |
| 319 | 11 | GO:005080 | positive reg | 31/519 | 154/8076 | 8.33E-09 | 7.94E-07 | 6.71E-07 | Cacng2/Ccl | 31 |
| 320 | 11 | GO:001922 | neuronal a | 14/519 | 35/8076 | 1.14E-08 | 1.03E-06 | 8.71E-07 | Ank3/Chrn | 14 |
| 321 | 11 | GO:009917 | regulation | 14/519 | 35/8076 | 1.14E-08 | 1.03E-06 | 8.71E-07 | Gpc4/Il1rap | 14 |
| 322 | 11 | GO:190560 | regulation | 13/519 | 31/8076 | 1.94E-08 | 1.71E-06 | 1.45E-06 | Gpc4/Il1rap | 13 |
| 323 | 11 | GO:009860 | cell-cell adl | 58/519 | 426/8076 | 2.55E-08 | 2.19E-06 | 1.85E-06 | Amigo1/An | 58 |
| 324 | 11 | GO:000762 | locomotory | 34/519 | 189/8076 | 3.13E-08 | 2.63E-06 | 2.22E-06 | Adam22/A | 34 |
| 325 | 11 | GO:000740 | axonogene | 50/519 | 346/8076 | 3.71E-08 | 3.05E-06 | 2.58E-06 | Ache/Amig | 50 |
| 326 | 11 | GO:009874 | cell-cell adl | 27/519 | 130/8076 | 3.91E-08 | 3.14E-06 | 2.66E-06 | Amigo1/Ca | 27 |
| 327 | 11 | GO:005089 | cognition | 40/519 | 248/8076 | 4.54E-08 | 3.57E-06 | 3.02E-06 | Adgrb3/Atp | 40 |
| 328 | 11 | GO:009866 | inorganic a | 15/519 | 45/8076 | 6.15E-08 | 4.64E-06 | 3.92E-06 | Ano10/Anc | 15 |
| 329 | 11 | GO:009905 | presynapse | 15/519 | 45/8076 | 6.15E-08 | 4.64E-06 | 3.92E-06 | Cadm1/Gp | 15 |
| 330 | 11 | GO:190207 | response to | 36/519 | 215/8076 | 8.31E-08 | 6.14E-06 | 5.19E-06 | Ache/Amig | 36 |
| 331 | 11 | GO:009890 | regulation | 15/519 | 46/8076 | 8.59E-08 | 6.22E-06 | 5.26E-06 | Ank3/Atp2 | 15 |
| 332 | 11 | GO:003240 | regulation | 34/519 | 198/8076 | 1.02E-07 | 7.27E-06 | 6.15E-06 | Amigo1/An | 34 |
| 333 | 11 | GO:009710 | postsynapt | 14/519 | 41/8076 | 1.21E-07 | 8.41E-06 | 7.11E-06 | C1ql3/Cadr | 14 |
| 334 | 11 | GO:003241 | regulation | 32/519 | 182/8076 | 1.40E-07 | 9.54E-06 | 8.07E-06 | Amigo1/An | 32 |
| 335 | 11 | GO:000681 | calcium ior | 37/519 | 231/8076 | 1.78E-07 | 1.19E-05 | 1.01E-05 | Atp1b1/Atp | 37 |
| 336 | 11 | GO:005196 | regulation | 49/519 | 354/8076 | 2.01E-07 | 1.32E-05 | 1.12E-05 | Acan/Ache | 49 |
| 337 | 11 | GO:199057 | potassium | 11/519 | 26/8076 | 2.22E-07 | 1.43E-05 | 1.21E-05 | Atp1a1/Atp | 11 |
| 338 | 11 | GO:000721 | glutamate | 14/519 | 43/8076 | 2.38E-07 | 1.51E-05 | 1.28E-05 | Atp1a3/Cp | 14 |
| 339 | 11 | GO:005080 | regulation | 38/519 | 245/8076 | 2.83E-07 | 1.77E-05 | 1.49E-05 | Adgrb3/Am | 38 |
| 340 | 11 | GO:004327 | positive reg | 28/519 | 152/8076 | 3.17E-07 | 1.95E-05 | 1.65E-05 | Amigo1/An | 28 |
| 341 | 11 | GO:005148 | regulation | 13/519 | 38/8076 | 3.35E-07 | 2.02E-05 | 1.71E-05 | Atp2b1/Atp | 13 |
| 342 | 11 | GO:002289 | regulation | 32/519 | 189/8076 | 3.42E-07 | 2.03E-05 | 1.72E-05 | Amigo1/An | 32 |
| 343 | 11 | GO:009865 | monoatom | 15/519 | 51/8076 | 3.97E-07 | 2.28E-05 | 1.93E-05 | Ano10/Anc | 15 |
| 344 | 11 | GO:009908 | postsynapt | 15/519 | 51/8076 | 3.97E-07 | 2.28E-05 | 1.93E-05 | C1ql3/Cadr | 15 |
| 345 | 11 | GO:007058 | calcium ior | 31/519 | 181/8076 | 4.03E-07 | 2.28E-05 | 1.93E-05 | Atp1b1/Atp | 31 |
| 346 | 11 | GO:005080 | regulation | 37/519 | 239/8076 | 4.28E-07 | 2.39E-05 | 2.02E-05 | Adgrb3/Am | 37 |
| 347 | 11 | GO:190247 | chloride tra | 12/519 | 33/8076 | 4.46E-07 | 2.45E-05 | 2.07E-05 | Ano10/Anc | 12 |
| 348 | 11 | GO:009917 | presynapse | 15/519 | 52/8076 | 5.25E-07 | 2.80E-05 | 2.37E-05 | Cadm1/Gp | 15 |
| 349 | 11 | GO:005196 | positive reg | 38/519 | 251/8076 | 5.32E-07 | 2.80E-05 | 2.37E-05 | Acan/Ache | 38 |
| 350 | 11 | GO:000681 | sodium ion | 26/519 | 138/8076 | 5.34E-07 | 2.80E-05 | 2.37E-05 | Ank3/Atp1 | 26 |
| 351 | 11 | GO:001569 | inorganic a | 17/519 | 67/8076 | 6.95E-07 | 3.60E-05 | 3.04E-05 | Ano10/Anc | 17 |
| 352 | 11 | GO:000761 | learning | 26/519 | 141/8076 | 8.25E-07 | 4.20E-05 | 3.55E-05 | Adgrb3/Atp | 26 |
| 353 | 11 | GO:005507 | calcium ior | 30/519 | 178/8076 | 8.86E-07 | 4.42E-05 | 3.74E-05 | Atp1b1/Atp | 30 |
| 354 | 11 | GO:003572 | sodium ion | 21/519 | 99/8076 | 8.92E-07 | 4.42E-05 | 3.74E-05 | Ank3/Atp1 | 21 |
| 355 | 11 | GO:000687 | intracellula | 29/519 | 169/8076 | 9.21E-07 | 4.50E-05 | 3.81E-05 | Atp1b1/Atp | 29 |
| 356 | 11 | GO:009709 | presynapti | 7/519 | Nov-76 | 1.14E-06 | 5.53E-05 | 4.67E-05 | Cntn2/Il1ra | 7 |
| 357 | 11 | GO:001583 | amine tran | 19/519 | 85/8076 | 1.28E-06 | 6.09E-05 | 5.14E-05 | Cacna1a/C | 19 |
| 358 | 11 | GO:003476 | positive reg | 23/519 | 118/8076 | 1.31E-06 | 6.16E-05 | 5.21E-05 | Amigo1/An | 23 |
| 359 | 11 | GO:000682 | monoatom | 18/519 | 78/8076 | 1.48E-06 | 6.89E-05 | 5.82E-05 | Ano10/Anc | 18 |
| 360 | 11 | GO:009870 | calcium ior | 9/519 | 20/8076 | 1.55E-06 | 7.09E-05 | 5.99E-05 | Cacna1a/Ca | 9 |
| 361 | 11 | GO:190406 | positive reg | 20/519 | 96/8076 | 2.19E-06 | 9.83E-05 | 8.30E-05 | Amigo1/An | 20 |
| 362 | 11 | GO:005195 | regulation | 18/519 | 80/8076 | 2.20E-06 | 9.83E-05 | 8.30E-05 | Cacna1a/C | 18 |
| 363 | 11 | GO:000688 | intracellula | 7/519 | Dec-76 | 2.60E-06 | 0.000113 | 9.57E-05 | Atp1a1/Atp | 7 |
| 364 | 11 | GO:005196 | positive reg | 11/519 | 32/8076 | 2.60E-06 | 0.000113 | 9.57E-05 | Cacng2/Ccl | 11 |
| 365 | 11 | GO:003053 | adult beha | 24/519 | 132/8076 | 2.84E-06 | 0.000122 | 0.000103 | Adam22/A | 24 |
| 366 | 11 | GO:001095 | regulation | 36/519 | 248/8076 | 2.94E-06 | 0.000125 | 0.000106 | Amigo1/An | 36 |
| 367 | 11 | GO:003476 | positive reg | 27/519 | 160/8076 | 3.06E-06 | 0.000129 | 0.000109 | Acsl1/Acsl5 | 27 |
| 368 | 11 | GO:190207 | cellular res | 24/519 | 133/8076 | 3.26E-06 | 0.000135 | 0.000114 | Ache/Amig | 24 |
| 369 | 11 | GO:004866 | cell morph | 56/519 | 468/8076 | 3.32E-06 | 0.000137 | 0.000115 | Ache/Adgrl | 56 |
| 370 | 11 | GO:000715 | heterophili | 11/519 | 33/8076 | 3.67E-06 | 0.000146 | 0.000123 | Amigo1/Ca | 11 |
| 371 | 11 | GO:007152 | semaphori | 11/519 | 33/8076 | 3.67E-06 | 0.000146 | 0.000123 | Ece1/Ncam | 11 |
| 372 | 11 | GO:190486 | excitatory | 11/519 | 33/8076 | 3.67E-06 | 0.000146 | 0.000123 | C1ql3/Cntr | 11 |
| 373 | 11 | GO:005189 | membrane | 15/519 | 60/8076 | 3.86E-06 | 0.000152 | 0.000128 | Ank3/B2m/ | 15 |
| 374 | 11 | GO:000682 | chloride tra | 14/519 | 53/8076 | 4.02E-06 | 0.000155 | 0.000131 | Ano10/Anc | 14 |
| 375 | 11 | GO:007005 | neuron cell | 14/519 | 53/8076 | 4.02E-06 | 0.000155 | 0.000131 | Atp2a2/Atp | 14 |

| | | | | | | | | |
|---|---|---|---|---|---|---|---|---|
| 2917 | ENSMUSG( | Grid2 | 14804 | -01549669 | 016040221 | 033 | 00725 | 2 |
| 2922 | ENSMUSG( | Grin1 | 14810 | -03065035 | 015884655 | 03875 | -0015 | 2 |
| 2938 | ENSMUSG( | Grm7 | 108073 | -04775824 | 003025568 | -001 | 0055 | 2 |
| 2939 | ENSMUSG( | Grm8 | 14823 | -05664657 | 018071566 | 0005 | 003 | 2 |
| 2941 | ENSMUSG( | Grp | 225642 | -01136808 | 018996915 | 039 | 021 | 2 |
| 3004 | ENSMUSG( | H13 | 14950 | -02521543 | 006890033 | 04025 | -00325 | 2 |
| 3012 | ENSMUSG( | H6pd | 100198 | -03406356 | 026111496 | 01575 | 01725 | 2 |
| 3027 | ENSMUSG( | Hapln4 | 330790 | -03440151 | 001109435 | 0025 | 00825 | 2 |
| 3081 | ENSMUSG( | Herpud1 | 64209 | -02711414 | 045337732 | -0175 | 0225 | 2 |
| 3149 | ENSMUSG( | Hook1 | 77963 | -00591522 | 027583284 | 0375 | 03175 | 2 |
| 3174 | ENSMUSG( | Hsd17b7 | 15490 | -02261943 | 000407600 | 01725 | 01475 | 2 |
| 3202 | ENSMUSG( | Htr1a | 15550 | -01581124 | 028220236 | 02375 | 035 | 2 |
| 3205 | ENSMUSG( | Htr2a | 15558 | -05583869 | 039236803 | -0205 | 03875 | 2 |
| 3206 | ENSMUSG( | Htr5a | 15563 | -03494176 | 027702256 | 02475 | -00225 | 2 |
| 3213 | ENSMUSG( | Hyou1 | 12282 | -02088546 | 024614285 | 03075 | -0015 | 2 |
| 3221 | ENSMUSG( | Icam2 | 15896 | -02873518 | 022025378 | -00925 | 028 | 2 |
| 3234 | ENSMUSG( | Ifngr1 | 15979 | -02940663 | 038923506 | -01075 | 01075 | 2 |
| 3264 | ENSMUSG( | Igsf3 | 78908 | -02808939 | 030979344 | 023 | 03725 | 2 |
| 3265 | ENSMUSG( | Igsf8 | 140559 | -03854321 | 014322396 | -00975 | 01575 | 2 |
| 3274 | ENSMUSG( | Il1rap | 16180 | -04456545 | 038695581 | 0045 | 00925 | 2 |
| 3276 | ENSMUSG( | Il1rapl2 | 60367 | -01018920 | 073346845 | 0405 | 01525 | 2 |
| 3301 | ENSMUSG( | Inpp4b | 234515 | -02286677 | 023336727 | -00575 | 00825 | 2 |
| 3347 | ENSMUSG( | Irf2bp2 | 270110 | -02211219 | -00068424 | 03525 | 0345 | 2 |
| 3357 | ENSMUSG( | Isg15 | 1E+08 | -04639344 | 003545338 | 06375 | 009 | 2 |
| 3359 | ENSMUSG( | Islr2 | 320563 | -01686905 | 028099316 | 01725 | 01425 | 2 |
| 3367 | ENSMUSG( | Itfg1 | 71927 | -03829163 | 015027416 | 04575 | 023 | 2 |
| 3370 | ENSMUSG( | Itga3 | 16400 | -03707203 | 020695054 | 01175 | 047 | 2 |
| 3371 | ENSMUSG( | Itga4 | 16401 | -04483752 | 011540298 | 02975 | 0155 | 2 |
| 3373 | ENSMUSG( | Itga6 | 16403 | -01463507 | 016626151 | -0105 | 02375 | 2 |
| 3448 | ENSMUSG( | Kcna6 | 16494 | -02811702 | 020055471 | 0185 | 00675 | 2 |
| 3458 | ENSMUSG( | Kcnd1 | 16506 | -02012180 | 013514122 | 01575 | 016 | 2 |
| 3479 | ENSMUSG( | Kcnj9 | 16524 | -01233072 | 020151304 | -00175 | 0115 | 2 |
| 3517 | ENSMUSG( | Kdsr | 70750 | -02058212 | 023571337 | 04825 | 0095 | 2 |
| 3547 | ENSMUSG( | Kirrel3 | 67703 | -04036979 | -00713814 | -00375 | 02275 | 2 |
| 3548 | ENSMUSG( | Kit | 16590 | -03708074 | 022073226 | 0145 | -003 | 2 |
| 3595 | ENSMUSG( | L1cam | 16728 | -02929871 | 004299524 | 007 | 026 | 2 |
| 3602 | ENSMUSG( | Lama4 | 16775 | -03457016 | 003287907 | -00625 | 01425 | 2 |
| 3606 | ENSMUSG( | Lamc1 | 226519 | -01820948 | 031736569 | 01475 | 0055 | 2 |
| 3607 | ENSMUSG( | Lamp1 | 16783 | -00904757 | 008970390 | 0125 | 0215 | 2 |
| 3608 | ENSMUSG( | Lamp2 | 16784 | -01967280 | 023956312 | 046 | 001 | 2 |
| 3619 | ENSMUSG( | Laptm4a | 17775 | -02154532 | -00093172 | 02375 | 0375 | 2 |
| 3632 | ENSMUSG( | Lbhd2 | 380787 | -06274024 | 002273833 | 01175 | 009 | 2 |
| 3661 | ENSMUSG( | Lgmn | 19141 | -03488473 | 019166728 | 01125 | 04075 | 2 |
| 3667 | ENSMUSG( | Lifr | 16880 | -00952601 | 033024402 | 011 | 0145 | 2 |
| 3680 | ENSMUSG( | Lingo1 | 235402 | -01229015 | 027318071 | 01475 | 0075 | 2 |
| 3683 | ENSMUSG( | Lipa | 16889 | -03553948 | 020928587 | 049 | 03325 | 2 |
| 3689 | ENSMUSG( | Lman1 | 70361 | -03264822 | 010397748 | 02375 | 032 | 2 |
| 3693 | ENSMUSG( | Lmbrd1 | 68421 | -01747550 | 012496818 | 04225 | 05225 | 2 |
| 3697 | ENSMUSG( | Lmf2 | 105847 | -02460047 | 012009376 | 04675 | -005 | 2 |
| 3698 | ENSMUSG( | Lmln | 239833 | -01662354 | 020802473 | 01575 | 043 | 2 |
| 3705 | ENSMUSG( | Lmtk3 | 381983 | -01161221 | 024418195 | 00225 | 017 | 2 |
| 3706 | ENSMUSG( | Lnpep | 240028 | -02416721 | 017456924 | 0075 | 0175 | 2 |
| 3707 | ENSMUSG( | Lnpk | 69605 | -06083177 | 025537478 | 01475 | 0065 | 2 |
| 3713 | ENSMUSG( | Lpcat2 | 270084 | -03679910 | 006563393 | 01425 | 03275 | 2 |
| 3714 | ENSMUSG( | Lpcat3 | 14792 | -02548623 | 009490941 | 02875 | 01375 | 2 |
| 3716 | ENSMUSG( | Lpgat1 | 226856 | -01689262 | 015635522 | 032 | 005 | 2 |
| 3724 | ENSMUSG( | Lrch2 | 210297 | -01289558 | -00371099 | 014 | 02175 | 2 |
| 3729 | ENSMUSG( | Lrfn4 | 225875 | -01739385 | 012656345 | 011 | 024 | 2 |
| 3730 | ENSMUSG( | Lrfn5 | 238205 | -02618993 | 019184461 | 029 | 00025 | 2 |
| 3734 | ENSMUSG( | Lrp11 | 237253 | -03491062 | 011775856 | 01875 | 049 | 2 |
| 3735 | ENSMUSG( | Lrp1b | 94217 | -04924846 | 028308545 | 008 | 02875 | 2 |
| 3761 | ENSMUSG( | Lrrc8d | 231549 | -03107042 | 018967714 | 02875 | 01625 | 2 |
| 3766 | ENSMUSG( | Lrrn1 | 16979 | -05950989 | 034679905 | 0035 | 025 | 2 |
| 3772 | ENSMUSG( | Lrtm2 | 211187 | -04401519 | 031199912 | -00125 | 0435 | 2 |
| 3806 | ENSMUSG( | Ly6h | 23934 | -02281896 | 004147633 | 043 | 034 | 2 |
| 3831 | ENSMUSG( | Maco1 | 66146 | -02983892 | 024388248 | 04 | -00075 | 2 |
| 3856 | ENSMUSG( | Man1b1 | 227619 | -01629137 | 007817215 | 03475 | 036 | 2 |
| 3860 | ENSMUSG( | Man2b2 | 17160 | -01677040 | 009819542 | 0165 | 0285 | 2 |
| 3862 | ENSMUSG( | Manba | 110173 | -01622314 | 019656113 | 00225 | 03525 | 2 |
| 3864 | ENSMUSG( | Manea | 242362 | -04670867 | 009642028 | -01775 | 04075 | 2 |
| 3865 | ENSMUSG( | Maneal | 215090 | -02871663 | 006186220 | 03075 | 056 | 2 |
| 3923 | ENSMUSG( | Marchf1 | 72925 | -02761193 | -00401537 | 00375 | 03675 | 2 |
| 3946 | ENSMUSG( | Matr3 | 17184 | -01397532 | 012157748 | 02125 | -006 | 2 |
| 4025 | ENSMUSG( | Megf9 | 230316 | -04067493 | 018994408 | 01625 | 009 | 2 |
| 4026 | ENSMUSG( | Meis2 | 17536 | -03287667 | -00649000 | 00925 | 02575 | 2 |
| 4060 | ENSMUSG( | Mfsd3 | 69572 | -02909251 | 004495199 | 0505 | 028 | 2 |
| 4065 | ENSMUSG( | Mgat3 | 17309 | -00750003 | 013942706 | 02925 | 01625 | 2 |
| 4066 | ENSMUSG( | Mgat5 | 107895 | -03470976 | -01090211 | 039 | -001 | 2 |
| 4071 | ENSMUSG( | Mia2 | 338320 | -02569076 | 014606746 | 037 | 01725 | 2 |
| 4072 | ENSMUSG( | Mia3 | 338366 | -02032829 | 024821378 | 0305 | 01175 | 2 |
| 4098 | ENSMUSG( | Minpp1 | 17330 | -01878114 | 029668224 | 007 | 00025 | 2 |
| 4115 | ENSMUSG( | Mme | 17380 | -06971284 | 038412147 | 01825 | 00125 | 2 |
| 4117 | ENSMUSG( | Mmp17 | 23948 | -02446892 | 012877922 | 03075 | 0165 | 2 |
| 4129 | ENSMUSG( | Mogs | 57377 | -02787259 | 021805800 | 05325 | 02225 | 2 |
| 4140 | ENSMUSG( | Moxd1 | 59012 | -03370119 | 000173211 | 015 | 043 | 2 |
| 4268 | ENSMUSG( | Mtdh | 67154 | -03611752 | 023267245 | 049 | 023 | 2 |
| 4314 | ENSMUSG( | Mxra7 | 67622 | -01706586 | 012646555 | 001 | 006 | 2 |
| 4315 | ENSMUSG( | Myadm | 50918 | -04569216 | 02228612 | 02375 | 03025 | 2 |
| 4332 | ENSMUSG( | Myl9 | 98932 | -02225879 | 013243353 | -0075 | 02225 | 2 |
| 4346 | ENSMUSG( | Myoc | 17926 | -07094767 | 028486752 | 02525 | 043 | 2 |
| 4381 | ENSMUSG( | Nalcn | 338370 | -04547965 | 020914365 | 0095 | 03625 | 2 |
| 4414 | ENSMUSG( | Ncam2 | 17968 | -04360606 | 026212896 | 03725 | 01125 | 2 |
| 4443 | ENSMUSG( | Ndfip2 | 76273 | -05200978 | 004427264 | -00975 | 0365 | 2 |
| 4458 | ENSMUSG( | Ndufa4l2 | 407790 | -04234349 | 027116836 | -0005 | 028 | 2 |
| 4503 | ENSMUSG( | Nectin2 | 19294 | -04693316 | 004513399 | 021 | 05175 | 2 |
| 4521 | ENSMUSG( | Nell2 | 54003 | -02606082 | 023188156 | 0265 | 03475 | 2 |
| 4524 | ENSMUSG( | Neo1 | 18007 | -01646704 | 009254089 | 0045 | 02625 | 2 |
| 4531 | ENSMUSG( | Neu4 | 241159 | -02836041 | 018889442 | 0155 | 02675 | 2 |
| 4547 | ENSMUSG( | Nfxl1 | 100978 | -01760322 | 013224090 | 02625 | 0315 | 2 |
| 4554 | ENSMUSG( | Nhlrc3 | 212114 | -03783284 | 004985079 | 0205 | 274766e-1 | 2 |
| 4561 | ENSMUSG( | Nid1 | 18073 | -01958277 | 028250506 | 01225 | 027 | 2 |
| 4562 | ENSMUSG( | Nid2 | 18074 | -03162666 | 010008891 | -00775 | 03525 | 2 |
| 4565 | ENSMUSG( | Nin | 18080 | -02581571 | 005875634 | 00275 | 00975 | 2 |
| 4567 | ENSMUSG( | Nipa1 | 233280 | -02190481 | 015719165 | 018 | 01325 | 2 |
| 4568 | ENSMUSG( | Nipal3 | 74552 | -01939150 | 028259858 | -00725 | 03925 | 2 |
| 4631 | ENSMUSG( | Npc1 | 18145 | -03388377 | 026783223 | 02575 | 0145 | 2 |
| 4649 | ENSMUSG( | Nptxr | 73340 | -01027976 | 010902784 | 01425 | 01275 | 2 |
| 4716 | ENSMUSG( | Nucb1 | 18220 | -02673719 | 019115420 | 04025 | -00575 | 2 |
| 4739 | ENSMUSG( | Nudt8 | 66387 | -01757254 | 000589001 | 0095 | 007 | 2 |
| 4788 | ENSMUSG( | Ogfod3 | 66179 | -03911726 | 006490781 | 0015 | 051 | 2 |
| 4814 | ENSMUSG( | Os9 | 216440 | -02584068 | 013087210 | 04275 | 03525 | 2 |
| 4825 | ENSMUSG( | Osbpl8 | 237542 | -00813727 | 016238634 | 02825 | -00575 | 2 |
| 4831 | ENSMUSG( | Ostm1 | 14628 | -03513366 | 016179928 | -01 | 02775 | 2 |
| 4849 | ENSMUSG( | P2ry12 | 70839 | -03779987 | 029525250 | 00175 | 009 | 2 |
| 4852 | ENSMUSG( | P4htm | 74443 | -02259274 | 022820232 | 055 | 012 | 2 |
| 4880 | ENSMUSG( | Pald1 | 27355 | -01096245 | 037694413 | 01875 | 01725 | 2 |
| 4883 | ENSMUSG( | Palm3 | 74337 | -06763225 | 017004604 | 01375 | 04325 | 2 |
| 4887 | ENSMUSG( | Pam | 18484 | -02162950 | 025875743 | 0085 | 038 | 2 |
| 4894 | ENSMUSG( | Panx1 | 55991 | -03496346 | 014439046 | 0285 | 0195 | 2 |
| 4934 | ENSMUSG( | Pcdh10 | 18526 | -02139950 | 019019014 | 00725 | 033 | 2 |
| 4935 | ENSMUSG( | Pcdh11x | 245578 | -04925699 | 035212001 | -00175 | 04025 | 2 |
| 4937 | ENSMUSG( | Pcdh17 | 219228 | -01957025 | 026163109 | 013 | 01225 | 2 |
| 4940 | ENSMUSG( | Pcdh8 | 18530 | -03175778 | 035004730 | -002 | 0145 | 2 |
| 4941 | ENSMUSG( | Pcdh9 | 211712 | -03390456 | 027971034 | 003 | 00675 | 2 |
| 4943 | ENSMUSG( | Pcdhac2 | 353237 | -03621021 | 047360562 | -00825 | 0105 | 2 |
| 4954 | ENSMUSG( | Pcdhga3 | 93711 | -01570968 | 022789579 | 0205 | 01325 | 2 |

| | | | | | | | | | | |
|---|---|---|---|---|---|---|---|---|---|---|
| 376 | 11 | GO:00076 | sensory pe | 30/519 | 192/8076 | 4.47E-06 | 0.00017 | 0.000144 | Atp2b2/At | 30 |
| 377 | 11 | GO:007124 | cellular res | 21/519 | 109/8076 | 4.60E-06 | 0.000173 | 0.000146 | Ank3/Atp7 | 21 |
| 378 | 11 | GO:014011 | export acr | 12/519 | 40/8076 | 4.64E-06 | 0.000173 | 0.000146 | Abca1/Atp | 12 |
| 379 | 11 | GO:00986 | postsynapt | 11/519 | 34/8076 | 5.12E-06 | 0.000189 | 0.00016 | C1ql3/Lrfn | 11 |
| 380 | 11 | GO:007124 | cellular res | 22/519 | 120/8076 | 6.39E-06 | 0.000234 | 0.000198 | Ank3/Atp7 | 22 |
| 381 | 11 | GO:005507 | potassium | 8/519 | 18/8076 | 6.79E-06 | 0.000246 | 0.000208 | Atp1a1/At | 8 |
| 382 | 11 | GO:000741 | axon guida | 25/519 | 149/8076 | 7.89E-06 | 0.00028 | 0.000237 | Ank3/Cdh4 | 25 |
| 383 | 11 | GO:00974 | neuron pro | 25/519 | 149/8076 | 7.89E-06 | 0.00028 | 0.000237 | Ank3/Cdh4 | 25 |
| 384 | 11 | GO:00440 | regulation | 42/519 | 325/8076 | 9.24E-06 | 0.000325 | 0.000274 | Atp1a1/At | 42 |
| 385 | 11 | GO:00069 | muscle cor | 26/519 | 160/8076 | 9.47E-06 | 0.000329 | 0.000278 | Atp1a1/At | 26 |
| 386 | 11 | GO:00971 | postsynapt | 9/519 | 24/8076 | 9.54E-06 | 0.000329 | 0.000278 | C1ql3/Lrfn | 9 |
| 387 | 11 | GO:00904 | mesenchyr | 10/519 | 30/8076 | 1.03E-05 | 0.000347 | 0.000294 | Cdh2/Erbb | 10 |
| 388 | 11 | GO:19040 | regulation | 31/519 | 210/8076 | 1.03E-05 | 0.000347 | 0.000294 | Amigo1/Ar | 31 |
| 389 | 11 | GO:00216 | nerve deve | 13/519 | 50/8076 | 1.08E-05 | 0.00036 | 0.000305 | Chrnb2/Ec | 13 |
| 390 | 11 | GO:003241 | positive re | 17/519 | 81/8076 | 1.14E-05 | 0.000376 | 0.000318 | Amigo1/Ar | 17 |
| 391 | 11 | GO:00300 | intracellula | 6/519 | Oct-76 | 1.15E-05 | 0.000376 | 0.000318 | Atp1a1/At | 6 |
| 392 | 11 | GO:005192 | positive re | 16/519 | 73/8076 | 1.15E-05 | 0.000376 | 0.000318 | Atp1b1/At | 16 |
| 393 | 11 | GO:00990 | postsynaps | 12/519 | 44/8076 | 1.38E-05 | 0.000443 | 0.000374 | C1ql3/Cdh | 12 |
| 394 | 11 | GO:006007 | regulation | 19/519 | 99/8076 | 1.38E-05 | 0.000443 | 0.000374 | Chrna4/Cn | 19 |
| 395 | 11 | GO:005091 | negative c | 9/519 | 26/8076 | 2.03E-05 | 0.000645 | 0.000545 | Itgav/Nrp2 | 9 |
| 396 | 11 | GO:001972 | calcium-m | 18/519 | 94/8076 | 2.40E-05 | 0.000755 | 0.000638 | Atp1b1/At | 18 |
| 397 | 11 | GO:00100 | response t | 28/519 | 189/8076 | 2.58E-05 | 0.000804 | 0.000679 | Ank3/Atp7 | 28 |
| 398 | 11 | GO:003241 | positive re | 17/519 | 86/8076 | 2.62E-05 | 0.00081 | 0.000684 | Amigo1/Ar | 17 |
| 399 | 11 | GO:003051 | negative re | 7/519 | 16/8076 | 2.99E-05 | 0.000916 | 0.000774 | Cav1/Htra1 | 7 |
| 400 | 11 | GO:00995 | chemical s | 17/519 | 88/8076 | 3.57E-05 | 0.001087 | 0.000919 | Chrna4/Cn | 17 |
| 401 | 11 | GO:00100 | response t | 37/519 | 288/8076 | 3.63E-05 | 0.001095 | 0.000925 | Ank3/Atp2 | 37 |
| 402 | 11 | GO:00515 | response t | 18/519 | 98/8076 | 4.30E-05 | 0.001288 | 0.001088 | Cav1/Crhb | 18 |
| 403 | 11 | GO:005507 | sodium ion | 7/519 | 17/8076 | 4.80E-05 | 0.001424 | 0.001203 | Atp1a1/At | 7 |
| 404 | 11 | GO:001571 | organic ani | 29/519 | 209/8076 | 6.39E-05 | 0.00188 | 0.001588 | Acsl1/Acsl | 29 |
| 405 | 11 | GO:000301 | muscle sys | 30/519 | 220/8076 | 6.65E-05 | 0.001941 | 0.00164 | Atp1a1/At | 30 |
| 406 | 11 | GO:00971 | ammonium | 7/519 | 18/8076 | 7.42E-05 | 0.002149 | 0.001816 | Ache/Atp2 | 7 |
| 407 | 11 | GO:00996 | protein loc | 10/519 | 37/8076 | 7.90E-05 | 0.002233 | 0.001887 | Adam22/C | 10 |
| 408 | 11 | GO:009964 | neurotrans | 10/519 | 37/8076 | 7.90E-05 | 0.002233 | 0.001887 | Adam22/C | 10 |
| 409 | 11 | GO:00355 | calcium-m | 6/519 | 13/8076 | 7.96E-05 | 0.002233 | 0.001887 | Bcap31/Gr | 6 |
| 410 | 11 | GO:19022 | semaphori | 6/519 | 13/8076 | 7.96E-05 | 0.002233 | 0.001887 | Ece1/Nrp1, | 6 |
| 411 | 11 | GO:20012 | regulation | 16/519 | 85/8076 | 8.38E-05 | 0.002334 | 0.001972 | Amigo1/Ar | 16 |
| 412 | 11 | GO:00902 | negative re | 10/519 | 38/8076 | 0.000101 | 0.002794 | 0.002361 | Atp2b4/Ca | 10 |
| 413 | 11 | GO:00481 | regulation | 28/519 | 204/8076 | 0.000103 | 0.002835 | 0.002395 | Atp2b2/Fg | 28 |
| 414 | 11 | GO:00975 | calcium ior | 17/519 | 96/8076 | 0.000112 | 0.003059 | 0.002585 | Cacna1a/C | 17 |
| 415 | 11 | GO:00507 | regulation | 22/519 | 144/8076 | 0.000118 | 0.003201 | 0.002705 | Ache/Amig | 22 |
| 416 | 11 | GO:00140 | regulation | 9/519 | 32/8076 | 0.000129 | 0.003402 | 0.002875 | Chrna4/Ch | 9 |
| 417 | 11 | GO:00860 | membrane | 9/519 | 32/8076 | 0.000129 | 0.003402 | 0.002875 | Atp1a1/At | 9 |
| 418 | 11 | GO:00605 | developme | 27/519 | 196/8076 | 0.000129 | 0.003402 | 0.002875 | Bcan/Cadm | 27 |
| 419 | 11 | GO:00504 | regulation | 11/519 | 47/8076 | 0.000145 | 0.003792 | 0.003204 | Chrna4/Ch | 11 |
| 420 | 11 | GO:004311 | receptor cl | 13/519 | 63/8076 | 0.000147 | 0.003792 | 0.003204 | Cacna1a/C | 13 |
| 421 | 11 | GO:190137 | regulation | 13/519 | 63/8076 | 0.000147 | 0.003792 | 0.003204 | Amigo1/Ar | 13 |
| 422 | 11 | GO:003541 | protein loc | 16/519 | 89/8076 | 0.000148 | 0.0038 | 0.003211 | Adam22/C | 16 |
| 423 | 11 | GO:00519 | synaptic tr | 12/519 | 55/8076 | 0.00015 | 0.003827 | 0.003234 | Cckbr/Clcn | 12 |
| 424 | 11 | GO:000202 | regulation | 7/519 | 20/8076 | 0.000161 | 0.004086 | 0.003453 | Atp1a1/At | 7 |
| 425 | 11 | GO:003051 | regulation | 9/519 | 33/8076 | 0.000167 | 0.0042 | 0.003549 | Cav1/Elapc | 9 |
| 426 | 11 | GO:190352 | regulation | 20/519 | 128/8076 | 0.000175 | 0.004379 | 0.003701 | Atp1a1/At | 20 |
| 427 | 11 | GO:00230 | signal relea | 41/519 | 358/8076 | 0.000193 | 0.004786 | 0.004044 | Abca1/Abc | 41 |
| 428 | 11 | GO:005511 | regulation | 10/519 | 41/8076 | 0.000201 | 0.004892 | 0.004133 | Atp1a1/At | 10 |
| 429 | 11 | GO:190101 | regulation | 10/519 | 41/8076 | 0.000201 | 0.004892 | 0.004133 | Amigo1/Ar | 10 |
| 430 | 11 | GO:20012 | positive re | 10/519 | 41/8076 | 0.000201 | 0.004892 | 0.004133 | Amigo1/Ar | 10 |
| 431 | 11 | GO:000682 | copper ion | 5/519 | Oct-76 | 0.000207 | 0.004914 | 0.004153 | Atp7a/Cp/ | 5 |
| 432 | 11 | GO:00109 | magnesiur | 5/519 | Oct-76 | 0.000207 | 0.004914 | 0.004153 | Ank3/Cnnn | 5 |
| 433 | 11 | GO:003637 | sodium ion | 5/519 | Oct-76 | 0.000207 | 0.004914 | 0.004153 | Atp1a1/At | 5 |
| 434 | 11 | GO:190101 | positive re | 6/519 | 15/8076 | 0.000208 | 0.004914 | 0.004153 | Amigo1/At | 6 |
| 435 | 11 | GO:00069 | smooth mu | 12/519 | 57/8076 | 0.000215 | 0.00503 | 0.00425 | Atp2b1/At | 12 |
| 436 | 11 | GO:009881 | modulatior | 11/519 | 49/8076 | 0.000216 | 0.00503 | 0.00425 | Chrna4/Gri | 11 |
| 437 | 11 | GO:00199 | second-me | 22/519 | 150/8076 | 0.000217 | 0.00503 | 0.00425 | Atp1b1/At | 22 |
| 438 | 11 | GO:00432 | regulation | 14/519 | 74/8076 | 0.000218 | 0.00503 | 0.00425 | Amigo1/Ar | 14 |
| 439 | 11 | GO:000664 | membrane | 16/519 | 92/8076 | 0.000221 | 0.005063 | 0.004278 | Asah1/Asal | 16 |
| 440 | 11 | GO:00509 | neuromusc | 19/519 | 121/8076 | 0.000237 | 0.0054 | 0.004563 | Adcy5/Apl | 19 |
| 441 | 11 | GO:001587 | dopamine | 10/519 | 42/8076 | 0.000249 | 0.005641 | 0.004767 | Chrna4/Ch | 10 |
| 442 | 11 | GO:00516 | detection c | 12/519 | 58/8076 | 0.000256 | 0.005749 | 0.004858 | Atp2b2/At | 12 |
| 443 | 11 | GO:00017 | neural cres | 8/519 | 28/8076 | 0.000272 | 0.005979 | 0.005052 | Erbb4/Folr | 8 |
| 444 | 11 | GO:003164 | regulation | 17/519 | 103/8076 | 0.000273 | 0.005979 | 0.005052 | Chrna4/Ch | 17 |
| 445 | 11 | GO:001404 | dopamine | 9/519 | 35/8076 | 0.000273 | 0.005979 | 0.005052 | Chrna4/Ch | 9 |
| 446 | 11 | GO:00995 | synaptic m | 9/519 | 35/8076 | 0.000273 | 0.005979 | 0.005052 | Cadm1/Gp | 9 |
| 447 | 11 | GO:000961 | response t | 15/519 | 85/8076 | 0.000293 | 0.006383 | 0.005394 | Atp1a1/At | 15 |
| 448 | 11 | GO:00352 | ionotropic | 6/519 | 16/8076 | 0.000314 | 0.006815 | 0.005759 | Atp1a3/Cp | 6 |
| 449 | 11 | GO:00076 | feeding bel | 12/519 | 60/8076 | 0.000357 | 0.007686 | 0.006495 | Aplp2/Atp | 12 |
| 450 | 11 | GO:00451 | neuronal ic | 5/519 | Nov-76 | 0.000359 | 0.00769 | 0.006498 | Ank3/Cntn | 5 |
| 451 | 11 | GO:00485 | developme | 45/519 | 419/8076 | 0.000409 | 0.008705 | 0.007356 | Atp8a2/Bc | 45 |
| 452 | 11 | GO:007127 | cellular res | 12/519 | 61/8076 | 0.000419 | 0.008863 | 0.00749 | Crhbp/Kcnl | 12 |
| 453 | 11 | GO:00163 | calcium-de | 7/519 | 23/8076 | 0.000431 | 0.008967 | 0.007577 | Cdh18/Cdh | 7 |
| 454 | 11 | GO:004884 | axon exten | 7/519 | 23/8076 | 0.000431 | 0.008967 | 0.007577 | Nrp1/Nrp2 | 7 |
| 455 | 11 | GO:19022 | neuron pro | 7/519 | 23/8076 | 0.000431 | 0.008967 | 0.007577 | Nrp1/Nrp2 | 7 |
| 456 | 11 | GO:000742 | brain devel | 42/519 | 384/8076 | 0.000436 | 0.009014 | 0.007617 | Adgrg1/Am | 42 |
| 457 | 11 | GO:00080 | cell recogn | 13/519 | 70/8076 | 0.00044 | 0.00904 | 0.007639 | Amigo1/Ar | 13 |
| 458 | 11 | GO:00069 | regulation | 14/519 | 79/8076 | 0.000442 | 0.009048 | 0.007646 | Atp1a1/At | 14 |
| 459 | 11 | GO:000801 | blood circu | 31/519 | 256/8076 | 0.000449 | 0.009137 | 0.007721 | Atp1a1/At | 31 |
| 460 | 11 | GO:00140 | neural cres | 10/519 | 45/8076 | 0.000454 | 0.009182 | 0.007759 | Cdh2/Erbb | 10 |
| 461 | 11 | GO:000682 | iron ion tra | 8/519 | 30/8076 | 0.000457 | 0.009183 | 0.00776 | Atp7a/B2m | 8 |
| 462 | 11 | GO:00519 | catecholan | 12/519 | 62/8076 | 0.00049 | 0.009791 | 0.008273 | Chrna4/Ch | 12 |
| 463 | 11 | GO:00504 | catecholan | 11/519 | 54/8076 | 0.000529 | 0.010357 | 0.008751 | Chrna4/Ch | 11 |
| 464 | 11 | GO:00996 | regulation | 11/519 | 54/8076 | 0.000529 | 0.010357 | 0.008751 | Cacng2/Ch | 11 |
| 465 | 11 | GO:190442 | positive re | 11/519 | 54/8076 | 0.000529 | 0.010357 | 0.008751 | Atp1b1/Ca | 11 |
| 466 | 11 | GO:00425 | neuron ma | 9/519 | 38/8076 | 0.000529 | 0.010357 | 0.008751 | Adgrb3/An | 9 |
| 467 | 11 | GO:000694 | regulation | 10/519 | 46/8076 | 0.000547 | 0.010655 | 0.009003 | Atp1a1/At | 10 |
| 468 | 11 | GO:00995 | maintenan | 7/519 | 24/8076 | 0.000575 | 0.011051 | 0.009338 | Adgrb3/Ca | 7 |
| 469 | 11 | GO:001074 | macrophag | 5/519 | Dec-76 | 0.000583 | 0.011051 | 0.009338 | Abca5/Abc | 5 |
| 470 | 11 | GO:00108 | regulation | 5/519 | Dec-76 | 0.000583 | 0.011051 | 0.009338 | Atp1b1/At | 5 |
| 471 | 11 | GO:003232 | alanine tra | 5/519 | Dec-76 | 0.000583 | 0.011051 | 0.009338 | Slc38a2/Sl | 5 |
| 472 | 11 | GO:19022 | semaphori | 5/519 | Dec-76 | 0.000583 | 0.011051 | 0.009338 | Ece1/Nrp1, | 5 |
| 473 | 11 | GO:004694 | carboxylic | 24/519 | 183/8076 | 0.000628 | 0.011837 | 0.010002 | Acsl1/Acsl | 24 |
| 474 | 11 | GO:00428 | amide tran | 30/519 | 250/8076 | 0.000644 | 0.012052 | 0.010184 | Abca1/Abc | 30 |
| 475 | 11 | GO:00140 | neural cres | 9/519 | 39/8076 | 0.00065 | 0.012052 | 0.010184 | Cdh2/Erbb | 9 |
| 476 | 11 | GO:000727 | neuromusc | 6/519 | 18/8076 | 0.000653 | 0.012052 | 0.010184 | Cacna1a/C | 6 |
| 477 | 11 | GO:190587 | regulation | 6/519 | 18/8076 | 0.000653 | 0.012052 | 0.010184 | Cdh2/Lrfn1 | 6 |
| 478 | 11 | GO:00439 | cellular cor | 12/519 | 64/8076 | 0.000662 | 0.012122 | 0.010243 | Adgrb3/Ca | 12 |
| 479 | 11 | GO:19035 | regulation | 43/519 | 404/8076 | 0.000663 | 0.012122 | 0.010243 | Abcg1/Acsl | 43 |
| 480 | 11 | GO:006007 | excitatory | 13/519 | 73/8076 | 0.000669 | 0.012174 | 0.010287 | Chrna4/Cn | 13 |
| 481 | 11 | GO:001584 | organic aci | 24/519 | 184/8076 | 0.00068 | 0.012302 | 0.010395 | Acsl1/Acsl | 24 |
| 482 | 11 | GO:001097 | regulation | 44/519 | 417/8076 | 0.00069 | 0.012433 | 0.010506 | Ache/Adgrl | 44 |
| 483 | 11 | GO:000694 | striated mu | 15/519 | 92/8076 | 0.000703 | 0.012602 | 0.010649 | Atp1a1/At | 15 |
| 484 | 11 | GO:00068 | regulation | 11/519 | 56/8076 | 0.000732 | 0.01306 | 0.011036 | Atp6ap1/A | 11 |
| 485 | 11 | GO:19908 | ligand-gate | 7/519 | 25/8076 | 0.000755 | 0.013401 | 0.011324 | Atp1a3/Ch | 7 |
| 486 | 11 | GO:00508 | neuromusc | 10/519 | 48/8076 | 0.000782 | 0.013763 | 0.01163 | Adcy5/Apl | 10 |
| 487 | 11 | GO:000301 | circulatory | 32/519 | 276/8076 | 0.000783 | 0.013763 | 0.01163 | Atp1a1/At | 32 |
| 488 | 11 | GO:00343 | cell junctio | 9/519 | 40/8076 | 0.000791 | 0.013838 | 0.011694 | Adgrb3/Ca | 9 |
| 489 | 11 | GO:00902 | regulation | 19/519 | 133/8076 | 0.000801 | 0.013947 | 0.011785 | Arid4a/Atp | 19 |
| 490 | 11 | GO:19901 | neuron pro | 21/519 | 154/8076 | 0.00081 | 0.014037 | 0.011862 | Bcan/Cdh4 | 21 |
| 491 | 11 | GO:190547 | positive re | 14/519 | 84/8076 | 0.000839 | 0.014467 | 0.012225 | Ank3/Atp2 | 14 |
| 492 | 11 | GO:00217 | developme | 23/519 | 176/8076 | 0.000851 | 0.014596 | 0.012334 | Adgrb3/An | 23 |
| 493 | 11 | GO:00071 | G protein-c | 34/519 | 301/8076 | 0.000865 | 0.014765 | 0.012476 | Abca1/Adc | 34 |
| 494 | 11 | GO:00071 | homophilic | 12/519 | 66/8076 | 0.000882 | 0.014992 | 0.012669 | Amigo1/Ca | 12 |
| 495 | 11 | GO:009007 | foam cell d | 5/519 | 13/8076 | 0.000898 | 0.015124 | 0.01278 | Abca5/Abc | 5 |
| 496 | 11 | GO:00020 | positive re | 6/519 | 19/8076 | 0.000902 | 0.015124 | 0.01278 | Acan/Bcan, | 6 |
| 497 | 11 | GO:19048 | inhibitory s | 6/519 | 19/8076 | 0.000902 | 0.015124 | 0.01278 | Cbln4/Clstr | 6 |
| 498 | 11 | GO:200017 | positive re | 8/519 | 33/8076 | 0.000913 | 0.015232 | 0.012871 | Acan/Adgr | 8 |
| 499 | 11 | GO:00083 | associative | 14/519 | 85/8076 | 0.000947 | 0.015725 | 0.013288 | Atp1a3/Ca | 14 |
| 500 | 11 | GO:005104 | regulation | 44/519 | 424/8076 | 0.000976 | 0.016075 | 0.013584 | Abcg1/Acsl | 44 |
| 501 | 11 | GO:19013 | positive re | 7/519 | 26/8076 | 0.000977 | 0.016075 | 0.013584 | Amigo1/At | 7 |

| | | | | | | | | |
|---|---|---|---|---|---|---|---|---|
| 4960 | ENSMUSG | Pcdhgb2 | 93700 | -04190014 | -00195525 | 01925 | 01175 | 2 |
| 4964 | ENSMUSG | Pcdhgc5 | 93708 | -03768493 | 00619784 | -00575 | 006 | 2 |
| 4982 | ENSMUSG | Pcsk1 | 18548 | -06909070 | 03343760 | 0305 | 04325 | 2 |
| 4984 | ENSMUSG | Pcsk2 | 18549 | -04403430 | 01676918 | 03075 | 001 | 2 |
| 5026 | ENSMUSG | Pdia3 | 14827 | -02498679 | 02077427 | 0365 | 785046e-1 | 2 |
| 5027 | ENSMUSG | Pdia4 | 12304 | -01188216 | 04594132 | 02525 | 00525 | 2 |
| 5029 | ENSMUSG | Pdia6 | 71853 | -01133220 | 02555233 | 032 | 02525 | 2 |
| 5048 | ENSMUSG | Pdyn | 18610 | -07891590 | 00924214 | 0215 | 0395 | 2 |
| 5052 | ENSMUSG | Pdzd8 | 107368 | -02319425 | 01843388 | 00225 | 0125 | 2 |
| 5058 | ENSMUSG | Peds1 | 407243 | -03569576 | 00797223 | 0625 | 043 | 2 |
| 5072 | ENSMUSG | Pex14 | 56273 | -02392949 | 00404133 | 0095 | 01825 | 2 |
| 5097 | ENSMUSG | Pgap1 | 241062 | -03889423 | 03537164 | 0505 | 00975 | 2 |
| 5112 | ENSMUSG | Pgrmc1 | 53328 | -03736779 | 02296413 | 02025 | -00075 | 2 |
| 5128 | ENSMUSG | Phf20l1 | 239510 | -01140038 | 00695657 | 0115 | 02825 | 2 |
| 5165 | ENSMUSG | Pigk | 329777 | -03355381 | 00961272 | 044 | 0165 | 2 |
| 5169 | ENSMUSG | Pigs | 276846 | -01901430 | 01449028 | 038 | -01025 | 2 |
| 5170 | ENSMUSG | Pigt | 78928 | -02796708 | 01226529 | 04675 | -008 | 2 |
| 5220 | ENSMUSG | Plat | 18791 | -03896246 | -00395307 | 03375 | 03475 | 2 |
| 5237 | ENSMUSG | Pld3 | 18807 | -02084983 | 00505892 | 0275 | 012 | 2 |
| 5258 | ENSMUSG | Plod1 | 18822 | -03929467 | 02117009 | 025 | 0075 | 2 |
| 5266 | ENSMUSG | Plpp6 | 74411 | -03980799 | 00275090 | 04425 | 0055 | 2 |
| 5269 | ENSMUSG | Plppr2 | 235044 | -01822733 | 01296964 | 0165 | 03725 | 2 |
| 5271 | ENSMUSG | Plppr4 | 229791 | -05156676 | 04272083 | 00775 | 04975 | 2 |
| 5289 | ENSMUSG | Pm20d1 | 212933 | -03802766 | -02330165 | 02525 | 04375 | 2 |
| 5299 | ENSMUSG | Pnkd | 56695 | -01038330 | -00191146 | 01925 | 02025 | 2 |
| 5348 | ENSMUSG | Pomgnt2 | 215494 | -02544895 | 02767628 | 03875 | -0115 | 2 |
| 5349 | ENSMUSG | Pomk | 74653 | -02509225 | 01348436 | 04075 | 01075 | 2 |
| 5355 | ENSMUSG | Por | 18984 | -03519691 | 03175525 | 00975 | 01375 | 2 |
| 5376 | ENSMUSG | Ppib | 19035 | -02000858 | 01014461 | 04 | -0015 | 2 |
| 5459 | ENSMUSG | Praf2 | 54637 | -04292629 | -00832249 | 027 | 04275 | 2 |
| 5462 | ENSMUSG | Prcp | 72461 | -03648474 | -00367137 | 01525 | 01375 | 2 |
| 5499 | ENSMUSG | Prkcsh | 19089 | -02151417 | 02154871 | 00825 | 007 | 2 |
| 5525 | ENSMUSG | Prpf39 | 328110 | -01016521 | 03081957 | 02675 | -002 | 2 |
| 5542 | ENSMUSG | Prr7 | 432763 | -01839690 | 01100563 | 01375 | 0005 | 2 |
| 5559 | ENSMUSG | Psen1 | 19164 | -02708955 | 01947286 | -01225 | 0335 | 2 |
| 5628 | ENSMUSG | Ptgs1 | 19224 | -01576708 | 01637166 | 01375 | 0305 | 2 |
| 5629 | ENSMUSG | Ptgs2 | 19225 | -04418460 | 02842484 | 06675 | 00675 | 2 |
| 5656 | ENSMUSG | Ptprd | 19266 | -03175302 | 01096613 | -00525 | 007 | 2 |
| 5657 | ENSMUSG | Ptpre | 19267 | -03483160 | 01748059 | 0215 | 00975 | 2 |
| 5660 | ENSMUSG | Ptprk | 19272 | -01478954 | 01807987 | 02325 | 02075 | 2 |
| 5665 | ENSMUSG | Ptprr | 19279 | -05213080 | 01675161 | 014 | 0085 | 2 |
| 5706 | ENSMUSG | Qars1 | 97541 | -01167481 | 00565667 | 0135 | 007 | 2 |
| 5710 | ENSMUSG | Qpctl | 67369 | -04294572 | 01003194 | 01425 | -00075 | 2 |
| 5713 | ENSMUSG | Qsox1 | 104009 | -02002382 | 02819445 | 015 | 03425 | 2 |
| 5714 | ENSMUSG | Qsox2 | 227638 | -03159537 | 01645316 | 03725 | 0135 | 2 |
| 5752 | ENSMUSG | Rab39b | 67790 | -04073642 | -00361391 | 00275 | 00775 | 2 |
| 5871 | ENSMUSG | Rbm15 | 229700 | -02498755 | 01542783 | 0275 | -0045 | 2 |
| 5906 | ENSMUSG | Rcn1 | 19672 | -02041000 | 01562410 | 0395 | -01225 | 2 |
| 5912 | ENSMUSG | Rdh14 | 105014 | -02478608 | 01028002 | -0025 | 0105 | 2 |
| 5915 | ENSMUSG | Reep1 | 52250 | -04066416 | 00688925 | -0035 | 02425 | 2 |
| 5916 | ENSMUSG | Reep2 | 225362 | -02970565 | 00005674 | -005 | 03125 | 2 |
| 5919 | ENSMUSG | Reep6 | 70335 | -01576416 | 00476829 | 018 | 0065 | 2 |
| 5923 | ENSMUSG | Rell1 | 100532 | -04010287 | 02064494 | 0345 | 03275 | 2 |
| 5925 | ENSMUSG | Reln | 19699 | -03824912 | 02171922 | -392523e-1 | 0115 | 2 |
| 5931 | ENSMUSG | Retreg1 | 66270 | -05911356 | 02098533 | 0145 | -002 | 2 |
| 5932 | ENSMUSG | Retreg2 | 227298 | -01938769 | 01243161 | 02825 | 01125 | 2 |
| 5938 | ENSMUSG | Rfc3 | 69263 | -02455736 | -00391637 | 0025 | 03225 | 2 |
| 5952 | ENSMUSG | Rgmb | 68799 | -04009012 | 01184802 | -0075 | 03325 | 2 |
| 5967 | ENSMUSG | Rhbdd2 | 215160 | -01669640 | 00177595 | 011 | 008 | 2 |
| 6033 | ENSMUSG | Rnf130 | 59044 | -02581942 | 01631393 | 051 | 00825 | 2 |
| 6058 | ENSMUSG | Rnft2 | 269695 | -02029433 | 00806003 | 032 | 024 | 2 |
| 6065 | ENSMUSG | Robo1 | 19876 | -02209039 | 01087915 | 01375 | 02825 | 2 |
| 6066 | ENSMUSG | Robo2 | 268902 | -04585168 | 02836139 | 02025 | 0275 | 2 |
| 6208 | ENSMUSG | Rtn4r | 65079 | -00673643 | 01858170 | 0245 | 024 | 2 |
| 6210 | ENSMUSG | Rtn4rl2 | 269295 | -00146364 | 03552906 | 0335 | 022 | 2 |
| 6239 | ENSMUSG | S1pr1 | 13609 | -01849578 | -00396981 | 02125 | 01275 | 2 |
| 6252 | ENSMUSG | Samd8 | 67630 | -02843728 | 01537701 | 051 | 028 | 2 |
| 6262 | ENSMUSG | Sar1b | 66397 | -03348816 | -00770156 | -0035 | 02 | 2 |
| 6263 | ENSMUSG | Saraf | 67887 | -02102624 | -00013563 | 02925 | -00775 | 2 |
| 6279 | ENSMUSG | Sc5d | 235293 | -02088805 | -01182595 | 01025 | 02125 | 2 |
| 6297 | ENSMUSG | Scg2 | 20254 | -04577020 | 00754685 | 0025 | 011 | 2 |
| 6299 | ENSMUSG | Scg5 | 20394 | -03742870 | 01326746 | 031 | 013 | 2 |
| 6307 | ENSMUSG | Scn3b | 235281 | -02903854 | 02302808 | 017 | 02275 | 2 |
| 6309 | ENSMUSG | Scn7a | 20272 | -04077532 | 03436677 | -00325 | 01675 | 2 |
| 6315 | ENSMUSG | Scpep1 | 74617 | -04570745 | 00488538 | 0095 | 01075 | 2 |
| 6334 | ENSMUSG | Sdf4 | 20318 | -01334772 | -00100557 | 01525 | 0245 | 2 |
| 6347 | ENSMUSG | Sec11a | 56529 | -02728595 | 02262356 | 03925 | 02025 | 2 |
| 6348 | ENSMUSG | Sec11c | 66286 | -03324947 | -00711365 | 03125 | 01375 | 2 |
| 6362 | ENSMUSG | Sec61a1 | 53421 | -03260391 | 01063041 | 02975 | 022 | 2 |
| 6366 | ENSMUSG | Sec62 | 69276 | -02769092 | 01680136 | 012 | 01975 | 2 |
| 6370 | ENSMUSG | Sel1l | 20338 | -02187590 | 01559093 | 04475 | -0005 | 2 |
| 6372 | ENSMUSG | Selenof | 93684 | -02718639 | -00738868 | 03225 | -0065 | 2 |
| 6374 | ENSMUSG | Selenoi | 28042 | -02448148 | 02228661 | 0195 | 0035 | 2 |
| 6380 | ENSMUSG | Selenot | 69227 | -01949572 | 00248010 | 031 | -00725 | 2 |
| 6383 | ENSMUSG | Sema4a | 20351 | -02901540 | -00696062 | 011 | 017 | 2 |
| 6390 | ENSMUSG | Sema5b | 20357 | -04331848 | -01282420 | 03725 | 04975 | 2 |
| 6392 | ENSMUSG | Sema6b | 20359 | -01439981 | 01907847 | 01925 | 03225 | 2 |
| 6394 | ENSMUSG | Sema7a | 20361 | -02144458 | 00158747 | 01025 | 01375 | 2 |
| 6404 | ENSMUSG | Septin2 | 18000 | -03651107 | 01091140 | 00525 | 01675 | 2 |
| 6417 | ENSMUSG | Serinc1 | 56442 | -05745015 | 01878578 | 01725 | 0405 | 2 |
| 6420 | ENSMUSG | Serpina3n | 20716 | -03004029 | 01457204 | 01525 | 00375 | 2 |
| 6427 | ENSMUSG | Serpinf1 | 20317 | -03308218 | 01098409 | -00425 | 0255 | 2 |
| 6430 | ENSMUSG | Serpinh1 | 12406 | -02023816 | 03455211 | 032 | 021 | 2 |
| 6431 | ENSMUSG | Serpini1 | 20713 | -05382795 | 03356653 | 06125 | 01325 | 2 |
| 6441 | ENSMUSG | Sez6l2 | 233878 | -02885208 | 01986845 | -0025 | 021 | 2 |
| 6512 | ENSMUSG | Shisa7 | 232813 | -02381117 | 01461229 | 00475 | 037 | 2 |
| 6513 | ENSMUSG | Shisa9 | 72555 | -03644994 | 01362123 | 015 | 0145 | 2 |
| 6531 | ENSMUSG | Sil1 | 81500 | -03719386 | 00980871 | 027 | 0025 | 2 |
| 6548 | ENSMUSG | Slc11a2 | 18174 | -01571120 | 00720065 | 01075 | 0005 | 2 |
| 6562 | ENSMUSG | Slc16a2 | 20502 | -03175542 | 01289859 | 003 | 0105 | 2 |
| 6564 | ENSMUSG | Slc16a7 | 20503 | -06414387 | 00874036 | 0105 | 06025 | 2 |
| 6565 | ENSMUSG | Slc17a5 | 235504 | -04136519 | 01909589 | 06825 | 02775 | 2 |
| 6573 | ENSMUSG | Slc1a1 | 20510 | -03560788 | 03197155 | 0185 | 00575 | 2 |
| 6581 | ENSMUSG | Slc22a17 | 59049 | -01939672 | 01865422 | 01575 | 0275 | 2 |
| 6613 | ENSMUSG | Slc25a35 | 71998 | -03563996 | -00299058 | 0085 | 0305 | 2 |
| 6626 | ENSMUSG | Slc27a2 | 26458 | -02249466 | 01845600 | 009 | 01025 | 2 |
| 6634 | ENSMUSG | Slc2a6 | 227659 | -01898109 | 01853661 | -0055 | 00725 | 2 |
| 6635 | ENSMUSG | Slc30a1 | 22782 | -04440577 | 01537302 | 0385 | 009 | 2 |
| 6636 | ENSMUSG | Slc30a10 | 226781 | -01498208 | 00790323 | 0055 | 0175 | 2 |
| 6641 | ENSMUSG | Slc31a1 | 20529 | -01489622 | 02363951 | 036 | 04625 | 2 |
| 6642 | ENSMUSG | Slc32a1 | 22348 | -05595786 | 02012155 | -0035 | 0105 | 2 |
| 6645 | ENSMUSG | Slc35a3 | 229782 | -02312402 | 02144320 | -0035 | 042 | 2 |
| 6655 | ENSMUSG | Slc36a4 | 234967 | -00887897 | 01839512 | 01425 | 029 | 2 |
| 6657 | ENSMUSG | Slc38a1 | 105727 | -02620121 | 02605362 | -00625 | 01175 | 2 |
| 6662 | ENSMUSG | Slc39a10 | 227059 | -02743741 | 03584024 | -00775 | 01275 | 2 |
| 6667 | ENSMUSG | Slc39a6 | 106957 | -01821696 | 03696905 | -0125 | 02825 | 2 |
| 6694 | ENSMUSG | Slc6a15 | 103098 | -02615596 | 03862209 | 01925 | -00525 | 2 |
| 6695 | ENSMUSG | Slc6a17 | 229706 | -02151062 | 02160719 | 01625 | 031 | 2 |
| 6698 | ENSMUSG | Slc6a7 | 240332 | 00179813 | 03362490 | 03175 | 03625 | 2 |
| 6706 | ENSMUSG | Slc7a3 | 11989 | -00019446 | 03062234 | 0265 | 0185 | 2 |
| 6707 | ENSMUSG | Slc7a4 | 224022 | -02016038 | 01134329 | 01 | 04075 | 2 |
| 6719 | ENSMUSG | Slc9a9 | 331004 | -02187740 | 01442998 | 03175 | 00975 | 2 |
| 6727 | ENSMUSG | Slit2 | 20563 | -03826520 | -00365268 | 00125 | 00975 | 2 |
| 6728 | ENSMUSG | Slit3 | 20564 | -02901885 | 02570900 | 0605 | 044 | 2 |
| 6731 | ENSMUSG | Slitrk3 | 386750 | -03978438 | 03496204 | 0135 | 02875 | 2 |
| 6778 | ENSMUSG | Smpd1 | 20597 | -01809268 | 01444052 | 02875 | 02775 | 2 |

| | | | | | | | | | | |
|---|---|---|---|---|---|---|---|---|---|---|
| 502 | 11 | GO:009917 | postsynaps | 26/519 | 211/8076 | 0.000983 | 0.016106 | 0.01361 | C1ql3/Cadr | 26 |
| 503 | 11 | GO:000834 | adult locon | 13/519 | 76/8076 | 0.000993 | 0.016115 | 0.013617 | Adam22/At | 13 |
| 504 | 11 | GO:006004 | cardiac mu | 13/519 | 76/8076 | 0.000993 | 0.016115 | 0.013617 | Atp1a1/Atp | 13 |
| 505 | 11 | GO:000836 | regulation | 21/519 | 157/8076 | 0.001044 | 0.016872 | 0.014257 | Atp2b2/Atp | 21 |
| 506 | 11 | GO:001046 | regulation | 14/519 | 86/8076 | 0.001066 | 0.017154 | 0.014495 | Aplp2/Cacr | 14 |
| 507 | 11 | GO:001923 | sensory pe | 10/519 | 50/8076 | 0.001094 | 0.017524 | 0.014808 | Cacna1e/C | 10 |
| 508 | 11 | GO:000801 | regulation | 15/519 | 96/8076 | 0.001107 | 0.01758 | 0.014856 | Atp1a1/Atp | 15 |
| 509 | 11 | GO:006029 | long-term s | 15/519 | 96/8076 | 0.001107 | 0.01758 | 0.014856 | Grin2a/Grin | 15 |
| 510 | 11 | GO:190241 | protein loc | 17/519 | 116/8076 | 0.001114 | 0.017618 | 0.014887 | Adam22/C1 | 17 |
| 511 | 11 | GO:200046 | positive reg | 8/519 | 34/8076 | 0.001128 | 0.017754 | 0.015003 | Grin2a/Grin | 8 |
| 512 | 11 | GO:007170 | membrane | 9/519 | 42/8076 | 0.00115 | 0.018027 | 0.015233 | Ank3/Ankle | 9 |
| 513 | 11 | GO:004884 | regulation | 6/519 | 20/8076 | 0.00122 | 0.019037 | 0.016086 | Nrp1/Plxna | 6 |
| 514 | 11 | GO:003134 | negative re | 21/519 | 159/8076 | 0.00123 | 0.019111 | 0.016149 | B2m/Cep97 | 21 |
| 515 | 11 | GO:003064 | regulation | 10/519 | 51/8076 | 0.001284 | 0.019869 | 0.01679 | Atp6ap1/A | 10 |
| 516 | 11 | GO:003223 | positive reg | 5/519 | 14/8076 | 0.001323 | 0.02022 | 0.017086 | Cckbr/Erbb | 5 |
| 517 | 11 | GO:009915 | regulation | 5/519 | 14/8076 | 0.001323 | 0.02022 | 0.017086 | Lrfn1/Lrrc4 | 5 |
| 518 | 11 | GO:000720 | positive reg | 12/519 | 69/8076 | 0.001325 | 0.02022 | 0.017086 | Adcy5/Bcap | 12 |
| 519 | 11 | GO:006223 | protein loc | 11/519 | 60/8076 | 0.001335 | 0.02022 | 0.017086 | Adam22/C1 | 11 |
| 520 | 11 | GO:190353 | protein loc | 11/519 | 60/8076 | 0.001335 | 0.02022 | 0.017086 | Adam22/C1 | 11 |
| 521 | 11 | GO:003237 | positive reg | 9/519 | 43/8076 | 0.001374 | 0.020634 | 0.017436 | Abca1/Abc | 9 |
| 522 | 11 | GO:004886 | stem cell d | 9/519 | 43/8076 | 0.001374 | 0.020634 | 0.017436 | Cdh2/Erbb4 | 9 |
| 523 | 11 | GO:005192 | regulation | 20/519 | 150/8076 | 0.001418 | 0.021212 | 0.017925 | Atp1b1/Atp | 20 |
| 524 | 11 | GO:007265 | protein loc | 48/519 | 484/8076 | 0.001542 | 0.022969 | 0.019409 | Adam22/A | 48 |
| 525 | 11 | GO:000191 | regulation | 6/519 | 21/8076 | 0.001617 | 0.023983 | 0.020266 | Ache/Ece1, | 6 |
| 526 | 11 | GO:003090 | forebrain d | 27/519 | 230/8076 | 0.001645 | 0.024301 | 0.020535 | Adgrg1/Ap | 27 |
| 527 | 11 | GO:000686 | amino acid | 16/519 | 110/8076 | 0.001674 | 0.024631 | 0.020813 | Cacna1a/G | 16 |
| 528 | 11 | GO:001584 | monoamin | 12/519 | 71/8076 | 0.001712 | 0.025086 | 0.021198 | Chrna4/Ch | 12 |
| 529 | 11 | GO:190595 | positive reg | 10/519 | 53/8076 | 0.001746 | 0.025485 | 0.021535 | Abca1/Abc | 10 |
| 530 | 11 | GO:006004 | heart contr | 17/519 | 121/8076 | 0.001788 | 0.025997 | 0.021967 | Atp1a1/Atp | 17 |
| 531 | 11 | GO:006032 | head devel | 42/519 | 413/8076 | 0.001876 | 0.027131 | 0.022926 | Adgrg1/Am | 42 |
| 532 | 11 | GO:004884 | negative re | 5/519 | 15/8076 | 0.001881 | 0.027131 | 0.022926 | Nrp1/Sema | 5 |
| 533 | 11 | GO:006007 | synapse ma | 7/519 | 29/8076 | 0.001959 | 0.028146 | 0.023784 | Adgrb3/Bca | 7 |
| 534 | 11 | GO:190169 | cellular res | 41/519 | 403/8076 | 0.002102 | 0.029983 | 0.025336 | Ache/Adcy | 41 |
| 535 | 11 | GO:008601 | membrane | 6/519 | 22/8076 | 0.002104 | 0.029983 | 0.025336 | Ank3/Cacn | 6 |
| 536 | 11 | GO:005125 | protein cor | 20/519 | 155/8076 | 0.002117 | 0.030052 | 0.025395 | Atl1/B2m/C | 20 |
| 537 | 11 | GO:000689 | receptor-m | 22/519 | 178/8076 | 0.002265 | 0.032029 | 0.027065 | Ache/B2m/ | 22 |
| 538 | 11 | GO:004858 | developme | 24/519 | 201/8076 | 0.002332 | 0.032853 | 0.027761 | Bcan/Cdh4 | 24 |
| 539 | 11 | GO:000188 | receptor re | 8/519 | 38/8076 | 0.002422 | 0.033846 | 0.0286 | Ache/Ece1, | 8 |
| 540 | 11 | GO:004671 | viral entry | 8/519 | 38/8076 | 0.002422 | 0.033846 | 0.0286 | Cav1/Ctsb/ | 8 |
| 541 | 11 | GO:000941 | response to | 19/519 | 146/8076 | 0.002454 | 0.034167 | 0.028872 | Acsl1/Atp1 | 19 |
| 542 | 11 | GO:190382 | organic aci | 14/519 | 94/8076 | 0.002553 | 0.03477 | 0.029381 | Acsl1/Acsl5 | 14 |
| 543 | 11 | GO:190503 | carboxylic a | 14/519 | 94/8076 | 0.002553 | 0.03477 | 0.029381 | Acsl1/Acsl5 | 14 |
| 544 | 11 | GO:000667 | sphingosin | 4/519 | Oct-76 | 0.002592 | 0.03477 | 0.029381 | Asah1/Asal | 4 |
| 545 | 11 | GO:000959 | detection c | 4/519 | Oct-76 | 0.002592 | 0.03477 | 0.029381 | Itgav/Ryr2/ | 4 |
| 546 | 11 | GO:005506 | chloride ion | 4/519 | Oct-76 | 0.002592 | 0.03477 | 0.029381 | Cacna1a/Sl | 4 |
| 547 | 11 | GO:005508 | monoatom | 4/519 | Oct-76 | 0.002592 | 0.03477 | 0.029381 | Cacna1a/Sl | 4 |
| 548 | 11 | GO:009711 | postsynapt | 4/519 | Oct-76 | 0.002592 | 0.03477 | 0.029381 | Cdh2/Nlgn | 4 |
| 549 | 11 | GO:009871 | sodium ion | 4/519 | Oct-76 | 0.002592 | 0.03477 | 0.029381 | Hcn4/Slc12 | 4 |
| 550 | 11 | GO:200080 | regulation | 4/519 | Oct-76 | 0.002592 | 0.03477 | 0.029381 | Cdh2/Nlgn | 4 |
| 551 | 11 | GO:003509 | response to | 5/519 | 16/8076 | 0.002593 | 0.03477 | 0.029381 | B2m/Chrna | 5 |
| 552 | 11 | GO:005145 | regulation | 9/519 | 47/8076 | 0.00264 | 0.035261 | 0.029796 | Atp6v0a1/I | 9 |
| 553 | 11 | GO:000004 | transition r | 10/519 | 56/8076 | 0.002681 | 0.035465 | 0.029969 | Atp7a/B2m | 10 |
| 554 | 11 | GO:008606 | cell commu | 6/519 | 23/8076 | 0.002694 | 0.035465 | 0.029969 | Cacna1c/Ca | 6 |
| 555 | 11 | GO:190464 | cellular res | 6/519 | 23/8076 | 0.002694 | 0.035465 | 0.029969 | Atp1a3/Cac | 6 |
| 556 | 11 | GO:200031 | regulation | 6/519 | 23/8076 | 0.002694 | 0.035465 | 0.029969 | Cacng2/Gsp | 6 |
| 557 | 11 | GO:001585 | organic hyd | 20/519 | 159/8076 | 0.002868 | 0.037353 | 0.031564 | Abca1/Abc | 20 |
| 558 | 11 | GO:000194 | postsynapt | 8/519 | 39/8076 | 0.002879 | 0.037353 | 0.031564 | Cdh2/Glrb/ | 8 |
| 559 | 11 | GO:008600 | cardiac mu | 8/519 | 39/8076 | 0.002879 | 0.037353 | 0.031564 | Ank3/Atp2 | 8 |
| 560 | 11 | GO:009065 | walking be | 8/519 | 39/8076 | 0.002879 | 0.037353 | 0.031564 | Cacna1a/Ca | 8 |
| 561 | 11 | GO:190260 | proton trar | 11/519 | 66/8076 | 0.002952 | 0.03803 | 0.032136 | Atp1a1/Atp | 11 |
| 562 | 11 | GO:007050 | calcium ior | 7/519 | 31/8076 | 0.002952 | 0.03803 | 0.032136 | Cacna1a/Ca | 7 |
| 563 | 11 | GO:004409 | membrane | 9/519 | 48/8076 | 0.003068 | 0.039086 | 0.033028 | Ank3/Ankle | 9 |
| 564 | 11 | GO:004646 | membrane | 10/519 | 57/8076 | 0.00307 | 0.039086 | 0.033028 | Asah1/Asal | 10 |
| 565 | 11 | GO:199077 | protein loc | 32/519 | 300/8076 | 0.00307 | 0.039086 | 0.033028 | Adam22/A | 32 |
| 566 | 11 | GO:000751 | muscle org | 18/519 | 138/8076 | 0.003077 | 0.039086 | 0.033028 | Cav1/Chat/ | 18 |
| 567 | 11 | GO:000689 | endocytosi | 44/519 | 450/8076 | 0.00315 | 0.039871 | 0.033691 | Abca1/Ach | 44 |
| 568 | 11 | GO:001000 | glial cell dif | 21/519 | 172/8076 | 0.003274 | 0.04129 | 0.034891 | Acan/Adam | 21 |
| 569 | 11 | GO:004533 | phospholip | 6/519 | 24/8076 | 0.003401 | 0.042305 | 0.035748 | Abca1/Atp | 6 |
| 570 | 11 | GO:008600 | regulation | 6/519 | 24/8076 | 0.003401 | 0.042305 | 0.035748 | Atp1a1/Atp | 6 |
| 571 | 11 | GO:190464 | response to | 6/519 | 24/8076 | 0.003401 | 0.042305 | 0.035748 | Atp1a3/Cac | 6 |
| 572 | 11 | GO:004440 | entry into h | 8/519 | 40/8076 | 0.003401 | 0.042305 | 0.035748 | Cav1/Ctsb/ | 8 |
| 573 | 11 | GO:003431 | diol metab | 5/519 | 17/8076 | 0.003482 | 0.042732 | 0.036109 | Asah1/Asal | 5 |
| 574 | 11 | GO:006008 | inhibitory p | 5/519 | 17/8076 | 0.003482 | 0.042732 | 0.036109 | Chrna4/Cx3 | 5 |
| 575 | 11 | GO:007086 | regulation | 5/519 | 17/8076 | 0.003482 | 0.042732 | 0.036109 | Bcap31/Cd | 5 |
| 576 | 11 | GO:190488 | regulation | 5/519 | 17/8076 | 0.003482 | 0.042732 | 0.036109 | Lrfn1/Lrrc4 | 5 |
| 577 | 11 | GO:009010 | negative re | 10/519 | 58/8076 | 0.003502 | 0.042824 | 0.036187 | Arid4a/Cav | 10 |
| 578 | 11 | GO:004863 | regulation | 25/519 | 219/8076 | 0.003516 | 0.042853 | 0.036211 | Atp8a2/Cac | 25 |
| 579 | 11 | GO:000301 | heart proce | 17/519 | 129/8076 | 0.003561 | 0.043121 | 0.036438 | Atp1a1/Atp | 17 |
| 580 | 11 | GO:000762 | rhythmic b | 7/519 | 32/8076 | 0.003574 | 0.043121 | 0.036438 | Chat/Chrnb | 7 |
| 581 | 11 | GO:004326 | positive reg | 7/519 | 32/8076 | 0.003574 | 0.043121 | 0.036438 | Amigo1/At | 7 |
| 582 | 11 | GO:009712 | receptor lo | 12/519 | 78/8076 | 0.003858 | 0.04561 | 0.038541 | Adam22/C1 | 12 |
| 583 | 11 | GO:002160 | cranial ner | 4/519 | Nov-76 | 0.003868 | 0.04561 | 0.038541 | Chrnb2/Nr | 4 |
| 584 | 11 | GO:003360 | positive reg | 4/519 | Nov-76 | 0.003868 | 0.04561 | 0.038541 | Chrnb2/Kcn | 4 |
| 585 | 11 | GO:003586 | response to | 4/519 | Nov-76 | 0.003868 | 0.04561 | 0.038541 | Crhbp/Myl | 4 |
| 586 | 11 | GO:003586 | cellular res | 4/519 | Nov-76 | 0.003868 | 0.04561 | 0.038541 | Crhbp/Myl | 4 |
| 587 | 11 | GO:004209 | type 2 imm | 4/519 | Nov-76 | 0.003868 | 0.04561 | 0.038541 | Cd81/Kmt2 | 4 |
| 588 | 11 | GO:004651 | sphingoid r | 4/519 | Nov-76 | 0.003868 | 0.04561 | 0.038541 | Asah1/Asal | 4 |
| 589 | 11 | GO:004298 | amyloid pr | 10/519 | 59/8076 | 0.00398 | 0.046629 | 0.039402 | Abcg1/Efna | 10 |
| 590 | 11 | GO:007190 | negative re | 10/519 | 59/8076 | 0.00398 | 0.046629 | 0.039402 | Cav1/Hmgc | 10 |
| 591 | 11 | GO:001715 | regulation | 8/519 | 41/8076 | 0.003993 | 0.046629 | 0.039402 | Atp2a2/Cac | 8 |
| 592 | 11 | GO:001081 | regulation | 30/519 | 281/8076 | 0.004035 | 0.046961 | 0.039683 | Abcg1/Acsl | 30 |
| 593 | 11 | GO:007141 | cellular res | 38/519 | 380/8076 | 0.004064 | 0.046993 | 0.03971 | Ache/Adcy | 38 |
| 594 | 11 | GO:000664 | triglyceride | 9/519 | 50/8076 | 0.004089 | 0.046993 | 0.03971 | Acsl1/Acsl4 | 9 |
| 595 | 11 | GO:000854 | visual learr | 9/519 | 50/8076 | 0.004089 | 0.046993 | 0.03971 | Atp1a3/Cac | 9 |
| 596 | 11 | GO:003050 | BMP signal | 9/519 | 50/8076 | 0.004089 | 0.046993 | 0.03971 | Cav1/Elapc | 9 |
| 597 | 11 | GO:000663 | acylglycero | 11/519 | 69/8076 | 0.004212 | 0.048228 | 0.040753 | Acsl1/Acsl4 | 11 |
| 598 | 11 | GO:000688 | cell volume | 6/519 | 25/8076 | 0.004236 | 0.048228 | 0.040753 | Clcn3/Kcnn | 6 |
| 599 | 11 | GO:000016 | MAPK casc | 38/519 | 381/8076 | 0.004248 | 0.048228 | 0.040753 | Cav1/Cckbr | 38 |
| 600 | 11 | GO:004206 | gliogenesis | 26/519 | 234/8076 | 0.004253 | 0.048228 | 0.040753 | Acan/Adam | 26 |
| 601 | 11 | GO:003222 | regulation | 7/519 | 33/8076 | 0.00429 | 0.048228 | 0.040753 | Cckbr/Cnr1 | 7 |
| 602 | 11 | GO:003252 | protein exi | 7/519 | 33/8076 | 0.00429 | 0.048228 | 0.040753 | Bcap31/Cd | 7 |
| 603 | 11 | GO:005090 | detection c | 7/519 | 33/8076 | 0.00429 | 0.048228 | 0.040753 | Atp2b2/Atp | 7 |
| 604 | 11 | GO:006106 | muscle stru | 34/519 | 332/8076 | 0.004442 | 0.049785 | 0.042069 | Actn1/Adgr | 34 |
| 605 | 12 | GO:003406 | stress gran | 13/752 | 28/8076 | 3.56E-07 | 0.001459 | 0.001445 | Atxn2l/Cap | 13 |
| 606 | 12 | GO:001607 | mRNA met | 77/752 | 496/8076 | 3.35E-06 | 0.0067 | 0.006638 | Alkbh5/Ape | 77 |
| 607 | 12 | GO:004589 | negative re | 70/752 | 444/8076 | 5.58E-06 | 0.0067 | 0.006638 | Atf7ip/Bptf | 70 |
| 608 | 12 | GO:190267 | negative re | 70/752 | 446/8076 | 6.54E-06 | 0.0067 | 0.006638 | Atf7ip/Bptf | 70 |
| 609 | 12 | GO:003424 | negative re | 31/752 | 146/8076 | 8.81E-06 | 0.007218 | 0.00715 | Apoe/Atca | 31 |
| 610 | 12 | GO:000012 | negative re | 54/752 | 329/8076 | 2.18E-05 | 0.014907 | 0.014768 | Atf7ip/Bptf | 54 |
| 611 | 12 | GO:004477 | mitotic cell | 39/752 | 215/8076 | 3.29E-05 | 0.019241 | 0.019062 | Ambra1/Ar | 39 |
| 612 | 12 | GO:004477 | cell cycle p | 44/752 | 256/8076 | 4.13E-05 | 0.01972 | 0.019536 | Ambra1/Ar | 44 |
| 613 | 12 | GO:001714 | negative re | 27/752 | 129/8076 | 4.33E-05 | 0.01972 | 0.019536 | Caprin1/Cn | 27 |
| 614 | 12 | GO:004341 | macromole | 25/752 | 117/8076 | 5.77E-05 | 0.023622 | 0.023401 | Antkmt/As | 25 |
| 615 | 12 | GO:190280 | positive reg | 11/752 | 32/8076 | 8.78E-05 | 0.032682 | 0.032377 | Ankrd17/A | 11 |
| 616 | 12 | GO:007142 | hematopoi | 8/752 | 19/8076 | 0.000162 | 0.048594 | 0.048142 | Arih2/Ctc1, | 8 |
| 617 | 12 | GO:190198 | regulation | 35/752 | 200/8076 | 0.000173 | 0.048594 | 0.048142 | Ambra1/Ar | 35 |
| 618 | 12 | GO:006021 | hematopoi | 7/752 | 15/8076 | 0.000195 | 0.048594 | 0.048142 | Eif2ak2/Erc | 7 |
| 619 | 12 | GO:190199 | regulation | 30/752 | 163/8076 | 0.000201 | 0.048594 | 0.048142 | Ambra1/Ar | 30 |
| 620 | 12 | GO:003225 | methylatio | 27/752 | 141/8076 | 0.000212 | 0.048594 | 0.048142 | Antkmt/As | 27 |
| 621 | 12 | GO:000626 | DNA replica | 22/752 | 105/8076 | 0.000214 | 0.048594 | 0.048142 | Ankrd17/C | 22 |
| 622 | 12 | GO:190280 | regulation | 22/752 | 105/8076 | 0.000214 | 0.048594 | 0.048142 | Ambra1/Ar | 22 |
| 623 | 14 | GO:190199 | negative re | 10/219 | 70/8076 | 1.67E-05 | 0.048526 | 0.047434 | Inip/Mad1l | 10 |
| 624 | 15 | GO:004308 | regulation | 34/466 | 249/8076 | 2.05E-06 | 0.007585 | 0.007234 | Agap2/Agr | 34 |
| 625 | 15 | GO:000701 | actin filame | 38/466 | 326/8076 | 2.31E-05 | 0.042882 | 0.040899 | Arfgef1/Arl | 38 |
| 626 | 15 | GO:004206 | wound hea | 24/466 | 170/8076 | 3.85E-05 | 0.04547 | 0.043367 | Ahnak/Arfg | 24 |
| 627 | 15 | GO:003523 | tube morpl | 44/466 | 413/8076 | 4.93E-05 | 0.04547 | 0.043367 | Abcc8/Adg | 44 |

| | | | | | | | | |
|---|---|---|---|---|---|---|---|---|
| 6890 | ENSMUSG( | Spcs1 | 69019 | -03734115 | 00251281( | 0405 | -012 | 2 |
| 6891 | ENSMUSG( | Spcs2 | 66624 | -02729829 | 002680605 | 0295 | -0025 | 2 |
| 6892 | ENSMUSG( | Spcs3 | 76687 | -01995445 | 022932563 | 0215 | 01 | 2 |
| 6910 | ENSMUSG( | Spon1 | 233744 | -03465120 | 017605348 | 019 | 01925 | 2 |
| 6912 | ENSMUSG( | Sppl2a | 66552 | -02211252 | 009682858 | -007 | 03175 | 2 |
| 6929 | ENSMUSG( | Sptlc2 | 20773 | -03489094 | 016328093 | -003 | 02225 | 2 |
| 6937 | ENSMUSG( | Srd5a3 | 57357 | -02791411 | 012808327 | 0265 | 01575 | 2 |
| 6952 | ENSMUSG( | Srpra | 67398 | -02465236 | 01739965( | 02575 | 0295 | 2 |
| 6977 | ENSMUSG( | Ssh1 | 231637 | -01514143 | -00392650 | 0185 | 0125 | 2 |
| 6983 | ENSMUSG( | Ssr3 | 67437 | -02442818 | 011734818 | 0445 | -00025 | 2 |
| 6991 | ENSMUSG( | St3gal5 | 20454 | -03161327 | 012578285 | 00575 | 02825 | 2 |
| 6992 | ENSMUSG( | St6galnac5 | 26938 | -02554115 | 000297032 | -0015 | 00575 | 2 |
| 6994 | ENSMUSG( | St8sia3 | 20451 | -06171249 | 008875551 | 019 | 00675 | 2 |
| 7019 | ENSMUSG( | Stim1 | 20866 | -03521888 | 012281657 | 00175 | 00775 | 2 |
| 7021 | ENSMUSG( | Sting1 | 72512 | -03363371 | 020349327 | 0035 | 00675 | 2 |
| 7037 | ENSMUSG( | Stmn3 | 20262 | -02737588 | -01951955 | 017 | 0105 | 2 |
| 7054 | ENSMUSG( | Stt3b | 68292 | -02189015 | 01597809( | 02775 | -0035 | 2 |
| 7099 | ENSMUSG( | Surf4 | 20932 | -02717717 | 013373431 | 035 | -004 | 2 |
| 7109 | ENSMUSG( | Svop | 68666 | -01858713 | 02038621( | -0085 | 01625 | 2 |
| 7125 | ENSMUSG( | Syngr1 | 20972 | -01946042 | 018475341 | 0185 | 02025 | 2 |
| 7126 | ENSMUSG( | Syngr2 | 20973 | -02477635 | 025643338 | -00525 | 02625 | 2 |
| 7127 | ENSMUSG( | Syngr3 | 20974 | -04509859 | 001975508 | -01175 | 0345 | 2 |
| 7137 | ENSMUSG( | Sypl1 | 19027 | -04557739 | 032189956 | 0015 | 029 | 2 |
| 7148 | ENSMUSG( | Syt4 | 20983 | -05318285 | 025901365 | 00275 | 00975 | 2 |
| 7149 | ENSMUSG( | Syt5 | 53420 | -0353164 | 004725285 | 00975 | -0005 | 2 |
| 7196 | ENSMUSG( | Tapt1 | 231225 | -02090142 | 025024883 | 015 | 01275 | 2 |
| 7203 | ENSMUSG( | Tasor2 | 105203 | -00790489 | 019884396 | 0325 | 016 | 2 |
| 7219 | ENSMUSG( | Tbc1d23 | 67581 | -00730347 | 014527965 | 02375 | 0345 | 2 |
| 7242 | ENSMUSG( | Tbl2 | 27368 | -01035133 | 024432081 | 035 | 0145 | 2 |
| 7258 | ENSMUSG( | Tcn2 | 21452 | -01338615 | 018755441 | 01575 | 00075 | 2 |
| 7279 | ENSMUSG( | Tenm2 | 23964 | -01806572 | 033468708 | 02025 | -006 | 2 |
| 7280 | ENSMUSG( | Tenm3 | 23965 | -01775017 | 020574955 | 012 | -00125 | 2 |
| 7286 | ENSMUSG( | Tes | 21753 | -02335090 | 020856459 | 04425 | 03975 | 2 |
| 7291 | ENSMUSG( | Tex264 | 21767 | -01960089 | 009813224 | 01725 | 0005 | 2 |
| 7302 | ENSMUSG( | Tgfbi | 21810 | -02149966 | 014845981 | -00125 | 02925 | 2 |
| 7305 | ENSMUSG( | Tgoln1 | 22134 | -01219775 | 013146989 | 012 | 03275 | 2 |
| 7312 | ENSMUSG( | Thbs4 | 21828 | -01242593 | 027070093 | 01775 | 0285 | 2 |
| 7327 | ENSMUSG( | Thsd7a | 330267 | -06353494 | 009421786 | 00275 | 04375 | 2 |
| 7328 | ENSMUSG( | Thsd7b | 210417 | -06512903 | 01601550( | 0335 | -0005 | 2 |
| 7335 | ENSMUSG( | Tial1 | 21843 | -00962688 | 026295898 | 02 | 005 | 2 |
| 7377 | ENSMUSG( | Tm9sf4 | 99237 | -03230885 | 014562922 | 0515 | 02025 | 2 |
| 7384 | ENSMUSG( | Tmco1 | 68944 | -02483556 | 005837474 | 035 | 02575 | 2 |
| 7387 | ENSMUSG( | Tmed2 | 56334 | -03891420 | 004437241 | 05775 | 00025 | 2 |
| 7389 | ENSMUSG( | Tmed4 | 103694 | -03862276 | 010786445 | 02075 | 0145 | 2 |
| 7394 | ENSMUSG( | Tmeff1 | 230157 | -04078655 | 008953771 | 00525 | 00825 | 2 |
| 7395 | ENSMUSG( | Tmeff2 | 56363 | -05223448 | 003010049 | 01975 | 0205 | 2 |
| 7400 | ENSMUSG( | Tmem115 | 56395 | -03234979 | 024996604 | 01375 | 03025 | 2 |
| 7408 | ENSMUSG( | Tmem131 | 56030 | -01894377 | 030114905 | 0015 | 03875 | 2 |
| 7409 | ENSMUSG( | Tmem132a | 98170 | -01176643 | 023928952 | 02475 | 0425 | 2 |
| 7411 | ENSMUSG( | Tmem132c | 208213 | -03083672 | 021703339 | -0095 | 0185 | 2 |
| 7413 | ENSMUSG( | Tmem132e | 270893 | -02255400 | 01768254( | -0115 | 027 | 2 |
| 7424 | ENSMUSG( | Tmem161b | 72745 | -02768461 | 021256552 | 01125 | 03275 | 2 |
| 7438 | ENSMUSG( | Tmem19 | 67226 | -02882744 | 014787858 | 018 | 03225 | 2 |
| 7443 | ENSMUSG( | Tmem200c | 622645 | -03370287 | 02887346( | 0165 | 03975 | 2 |
| 7445 | ENSMUSG( | Tmem205 | 235043 | -03310287 | -00647031 | 02825 | -0035 | 2 |
| 7454 | ENSMUSG( | Tmem240 | 381582 | -01869214 | -00113404 | 002 | 0265 | 2 |
| 7456 | ENSMUSG( | Tmem245 | 242474 | -02281198 | 031528058 | 0175 | 0185 | 2 |
| 7466 | ENSMUSG( | Tmem33 | 67878 | -02017033 | 004780868 | 02925 | -00925 | 2 |
| 7467 | ENSMUSG( | Tmem35a | 67564 | -04380979 | 005185896 | 02575 | 0115 | 2 |
| 7468 | ENSMUSG( | Tmem38a | 74166 | -02213943 | 011277479 | 02125 | 0015 | 2 |
| 7470 | ENSMUSG( | Tmem41a | 66664 | -02647183 | 030413542 | 05 | 017 | 2 |
| 7476 | ENSMUSG( | Tmem62 | 96957 | -02061838 | 005406922 | 0175 | 03925 | 2 |
| 7477 | ENSMUSG( | Tmem63b | 224807 | -01638906 | 017624346 | 026 | 00225 | 2 |
| 7481 | ENSMUSG( | Tmem68 | 72098 | -05278838 | 021769331 | 001 | 01575 | 2 |
| 7484 | ENSMUSG( | Tmem87a | 211499 | -03821013 | 013145576 | 041 | 00625 | 2 |
| 7486 | ENSMUSG( | Tmem88b | 320587 | -05144104 | -01672435 | 02675 | 0475 | 2 |
| 7487 | ENSMUSG( | Tmem9 | 66241 | -03637271 | 006418708 | 00875 | 044 | 2 |
| 7489 | ENSMUSG( | Tmem97 | 69071 | -05093094 | 004350963 | 05525 | -00475 | 2 |
| 7490 | ENSMUSG( | Tmem9b | 56786 | -02603037 | 001134478 | 028 | 00575 | 2 |
| 7500 | ENSMUSG( | Tmtc3 | 237500 | -02681830 | 047098276 | 03225 | 02925 | 2 |
| 7502 | ENSMUSG( | Tmx1 | 72736 | -04163613 | 03896415( | 00925 | 00925 | 2 |
| 7511 | ENSMUSG( | Tnfrsf21 | 94185 | -03084634 | 029086688 | -0045 | 03375 | 2 |
| 7547 | ENSMUSG( | Tor2a | 30933 | -02196884 | 019802355 | 03625 | 03725 | 2 |
| 7552 | ENSMUSG( | Tpbgl | 1.01E+08 | -04466591 | 022325915 | 006 | 071 | 2 |
| 7573 | ENSMUSG( | Tprn | 97031 | -04066978 | -00895343 | 02 | 007 | 2 |
| 7574 | ENSMUSG( | Tpst1 | 22021 | -02233207 | 01832854( | 0185 | -00375 | 2 |
| 7676 | ENSMUSG( | Tspan31 | 67125 | -03410932 | -00130610 | 01125 | 00875 | 2 |
| 7678 | ENSMUSG( | Tspan4 | 64540 | -03697687 | 061851452 | 03775 | 0375 | 2 |
| 7679 | ENSMUSG( | Tspan5 | 56224 | -02831985 | 012943051 | 02625 | 02725 | 2 |
| 7680 | ENSMUSG( | Tspan6 | 56496 | -03052812 | 001084788 | 0335 | 05225 | 2 |
| 7697 | ENSMUSG( | Ttc19 | 72795 | -02992584 | 010624999 | 004 | 007 | 2 |
| 7759 | ENSMUSG( | Txndc5 | 105245 | -03197994 | 009779856 | 0385 | 196262e-1 | 2 |
| 7769 | ENSMUSG( | Tyro3 | 22174 | -02496970 | 024612432 | 0145 | 00275 | 2 |
| 7770 | ENSMUSG( | Tyw1 | 100929 | -00820983 | 024291991 | 023 | 021 | 2 |
| 7834 | ENSMUSG( | Ubr1 | 22222 | -01589849 | 016463919 | 01175 | 00275 | 2 |
| 7836 | ENSMUSG( | Ubr3 | 68795 | -01325970 | 01899030( | 01325 | 018 | 2 |
| 7858 | ENSMUSG( | Ufl1 | 67490 | -02186243 | 010279939 | 0365 | 00275 | 2 |
| 7880 | ENSMUSG( | Unc5b | 107449 | -06287867 | 011785697 | 00425 | 0445 | 2 |
| 7882 | ENSMUSG( | Unc5d | 210801 | -02067975 | 030756715 | -00125 | 013 | 2 |
| 7927 | ENSMUSG( | Usp30 | 100756 | -02222467 | -00022789 | 0145 | 037 | 2 |
| 7959 | ENSMUSG( | Uxs1 | 67883 | -04099278 | 011720215 | 0475 | -00525 | 2 |
| 7989 | ENSMUSG( | Vezt | 215008 | -01196970 | 014283237 | 0165 | 00725 | 2 |
| 7999 | ENSMUSG( | Vldlr | 22359 | -03488410 | 007696381 | 031 | 0075 | 2 |
| 8001 | ENSMUSG( | Vmp1 | 75909 | -06516539 | 00614628( | 03175 | -005 | 2 |
| 8041 | ENSMUSG( | Vstm2a | 211739 | -06301408 | 028313891 | 035 | -00625 | 2 |
| 8042 | ENSMUSG( | Vstm2b | 58188 | -02669114 | 013162163 | 05075 | -002 | 2 |
| 8087 | ENSMUSG( | Wdr33 | 74320 | -03920949 | 019181588 | 00675 | 0095 | 2 |
| 8100 | ENSMUSG( | Wdr54 | 75659 | -04270549 | 007498061 | -00025 | 01975 | 2 |
| 8113 | ENSMUSG( | Wfs1 | 22393 | -02540461 | 018293927 | 0335 | 023 | 2 |
| 8120 | ENSMUSG( | Wls | 68151 | -04601882 | 009306753 | 047 | 017 | 2 |
| 8148 | ENSMUSG( | Xpr1 | 19775 | -01282358 | 012371061 | 02175 | -00525 | 2 |
| 8169 | ENSMUSG( | Yipf3 | 28064 | -02800569 | 006347012 | 02975 | 048 | 2 |
| 8220 | ENSMUSG( | Zdhhc13 | 243983 | -02257527 | 012568793 | 03625 | 02375 | 2 |
| 8222 | ENSMUSG( | Zdhhc17 | 320150 | -03205245 | 022164947 | 0185 | 00625 | 2 |
| 8226 | ENSMUSG( | Zdhhc22 | 238331 | -02384540 | 012604348 | 04675 | 01775 | 2 |
| 8265 | ENSMUSG( | Zfyve28 | 231125 | -02079343 | -00629757 | 01625 | 023 | 2 |
| 7 | ENSMUSG( | 2310061I0 | 69662 | 003407369 | 00194944( | -017 | 004 | 3 |
| 15 | ENSMUSG( | 5031439G( | 223739 | 010639846 | -00814429 | 00275 | 01875 | 3 |
| 16 | ENSMUSG( | 5730455P1 | 70591 | 007352201 | -00354918 | -02275 | 004 | 3 |
| 25 | ENSMUSG( | Aamp | 227290 | 009390286 | -01204642 | -022 | 009 | 3 |
| 26 | ENSMUSG( | Aar2 | 68295 | 012403455 | -00472997 | -007 | 00575 | 3 |
| 27 | ENSMUSG( | Aars1 | 234734 | 004149507 | -01581661 | -01825 | 003 | 3 |
| 43 | ENSMUSG( | Abcb10 | 56199 | -00590869 | 002356183 | 01475 | 005 | 3 |
| 47 | ENSMUSG( | Abcb8 | 74610 | -00563431 | 007333553 | -01125 | -00425 | 3 |
| 57 | ENSMUSG( | Abce1 | 24015 | 011719052 | -00414415 | -00725 | 021 | 3 |
| 58 | ENSMUSG( | Abcf1 | 224742 | 02191368( | 005868477 | 00525 | 00925 | 3 |
| 59 | ENSMUSG( | Abcf2 | 27407 | 009489989 | 001744608 | -00775 | 00375 | 3 |
| 74 | ENSMUSG( | Abhd5 | 67469 | 012236425 | -01051670 | -00225 | 01275 | 3 |
| 77 | ENSMUSG( | Abi2 | 329165 | -01477399 | 005062748 | -01925 | 00325 | 3 |
| 78 | ENSMUSG( | Abi3 | 66610 | -00619519 | -01134343 | -0005 | -0025 | 3 |
| 87 | ENSMUSG( | Abtb2 | 99382 | 002174711 | -01452054 | -002 | -007 | 3 |
| 90 | ENSMUSG( | Acaca | 107476 | 017054414 | 00037114( | 00825 | 00975 | 3 |
| 94 | ENSMUSG( | Acad9 | 229211 | 00247340( | -01239075 | -0055 | -006 | 3 |
| 97 | ENSMUSG( | Acads | 11409 | 008860309 | -01158546 | 01075 | 0015 | 3 |
| 98 | ENSMUSG( | Acadsb | 66885 | 010829963 | -00523552 | -0085 | -0075 | 3 |
| 103 | ENSMUSG( | Acat1 | 110446 | 000960705 | -03005756 | 003 | 00075 | 3 |

| | | | | | | | | | | |
|---|---|---|---|---|---|---|---|---|---|---|
| 628 | 15 | GO:00300 | actin cytos | 49/466 | 482/8076 | 6.14E-05 | 0.04547 | 0.043367 | Adgrb1/Ag | 49 |
| 629 | 16 | GO:00003 | RNA splicin | 22/249 | 221/8076 | 1.09E-06 | 0.000781 | 0.000781 | Cdk13/Ddx | 22 |
| 630 | 16 | GO:00003 | RNA splicin | 22/249 | 221/8076 | 1.09E-06 | 0.000781 | 0.000781 | Cdk13/Ddx | 22 |
| 631 | 16 | GO:00003 | mRNA splic | 22/249 | 221/8076 | 1.09E-06 | 0.000781 | 0.000781 | Cdk13/Ddx | 22 |
| 632 | 16 | GO:00083 | RNA splicin | 27/249 | 314/8076 | 1.17E-06 | 0.000781 | 0.000781 | Alyref/Cdk | 27 |
| 633 | 16 | GO:00434 | regulation | 15/249 | 127/8076 | 7.58E-06 | 0.004032 | 0.004032 | Hnrnpf/Kh | 15 |
| 634 | 16 | GO:00480 | regulation | 12/249 | 85/8076 | 1.01E-05 | 0.004476 | 0.004476 | Khdrbs2/M | 12 |
| 635 | 16 | GO:00063 | mRNA proc | 26/249 | 359/8076 | 4.08E-05 | 0.015524 | 0.015524 | Alyref/Cdk | 26 |
| 636 | 16 | GO:00506 | regulation | 12/249 | 104/8076 | 7.89E-05 | 0.026247 | 0.026247 | Khdrbs2/M | 12 |
| 637 | 16 | GO:00003 | alternative | 9/249 | 61/8076 | 9.30E-05 | 0.027504 | 0.027504 | Cdk13/Khd | 9 |
| 638 | 16 | GO:00002 | spliceosom | 7/249 | 39/8076 | 0.000161 | 0.042819 | 0.042819 | Ddx39b/Rb | 7 |
| 639 | 17 | GO:000761 | behavior | 29/137 | 482/8076 | 1.41E-09 | 3.33E-06 | 3.00E-06 | Adora2a/A | 29 |
| 640 | 17 | GO:000761 | learning | 14/137 | 141/8076 | 9.67E-08 | 0.000114 | 0.000103 | Crh/Dbh/D | 14 |
| 641 | 17 | GO:00508 | cognition | 18/137 | 248/8076 | 1.71E-07 | 0.000121 | 0.000109 | Arc/Crh/Db | 18 |
| 642 | 17 | GO:000761 | learning or | 17/137 | 224/8076 | 2.04E-07 | 0.000121 | 0.000109 | Arc/Crh/Db | 17 |
| 643 | 17 | GO:000721 | neuropepti | 7/137 | 28/8076 | 3.05E-07 | 0.000144 | 0.00013 | Ecel1/Gpr3 | 7 |
| 644 | 17 | GO:00076 | locomotory | 15/137 | 189/8076 | 6.32E-07 | 0.000249 | 0.000224 | Adora2a/C | 15 |
| 645 | 17 | GO:00485 | positive re | 6/137 | 22/8076 | 1.27E-06 | 0.000377 | 0.00034 | Adora2a/C | 6 |
| 646 | 17 | GO:00083 | associative | 10/137 | 85/8076 | 1.51E-06 | 0.000377 | 0.00034 | Crh/Dbh/D | 10 |
| 647 | 17 | GO:00019 | response t | 5/137 | 13/8076 | 1.51E-06 | 0.000377 | 0.00034 | Adora2a/D | 5 |
| 648 | 17 | GO:00068 | neurotrans | 14/137 | 177/8076 | 1.60E-06 | 0.000377 | 0.00034 | Adora2a/G | 14 |
| 649 | 17 | GO:001407 | response t | 5/137 | 14/8076 | 2.31E-06 | 0.000489 | 0.000441 | Adora2a/D | 5 |
| 650 | 17 | GO:00158 | organic hy | 13/137 | 159/8076 | 2.65E-06 | 0.000489 | 0.000441 | Adora2a/C | 13 |
| 651 | 17 | GO:001584 | monoamin | 9/137 | 71/8076 | 2.73E-06 | 0.000489 | 0.000441 | Adora2a/C | 9 |
| 652 | 17 | GO:00427 | eating beh | 6/137 | 25/8076 | 2.90E-06 | 0.000489 | 0.000441 | Adora2a/D | 6 |
| 653 | 17 | GO:00015 | regulation | 13/137 | 170/8076 | 5.58E-06 | 0.000879 | 0.000792 | Adora2a/G | 13 |
| 654 | 17 | GO:00230 | signal relea | 19/137 | 358/8076 | 8.70E-06 | 0.00119 | 0.001073 | Acvr1c/Ad | 19 |
| 655 | 17 | GO:00519 | catecholan | 8/137 | 62/8076 | 8.74E-06 | 0.00119 | 0.001073 | Adora2a/C | 8 |
| 656 | 17 | GO:004241 | dopamine | 6/137 | 30/8076 | 9.07E-06 | 0.00119 | 0.001073 | Dbh/Drd1/ | 6 |
| 657 | 17 | GO:00071 | G protein- | 17/137 | 301/8076 | 1.21E-05 | 0.001459 | 0.001315 | Actn2/Ado | 17 |
| 658 | 17 | GO:00158 | amine tran | 9/137 | 85/8076 | 1.24E-05 | 0.001459 | 0.001315 | Adora2a/A | 9 |
| 659 | 17 | GO:00071 | homophilic | 8/137 | 66/8076 | 1.40E-05 | 0.001577 | 0.001421 | Cadm2/Cdl | 8 |
| 660 | 17 | GO:00190 | reproducti | 6/137 | 33/8076 | 1.62E-05 | 0.001742 | 0.00157 | Dbh/Drd1/ | 6 |
| 661 | 17 | GO:000761 | memory | 10/137 | 113/8076 | 2.00E-05 | 0.001868 | 0.001683 | Arc/Dbh/D | 10 |
| 662 | 17 | GO:00300 | metal ion t | 22/137 | 486/8076 | 2.02E-05 | 0.001868 | 0.001683 | Actn2/Ado | 22 |
| 663 | 17 | GO:00508 | modulatior | 21/137 | 450/8076 | 2.03E-05 | 0.001868 | 0.001683 | Adora2a/A | 21 |
| 664 | 17 | GO:009917 | regulation | 21/137 | 451/8076 | 2.10E-05 | 0.001868 | 0.001683 | Adora2a/A | 21 |
| 665 | 17 | GO:00019 | synaptic tr | 5/137 | 21/8076 | 2.14E-05 | 0.001868 | 0.001683 | Adora2a/C | 5 |
| 666 | 17 | GO:00065 | catecholan | 6/137 | 35/8076 | 2.31E-05 | 0.001884 | 0.001697 | Dbh/Drd1/ | 6 |
| 667 | 17 | GO:000971 | catechol-c | 6/137 | 35/8076 | 2.31E-05 | 0.001884 | 0.001697 | Dbh/Drd1/ | 6 |
| 668 | 17 | GO:00508 | positive re | 11/137 | 154/8076 | 5.70E-05 | 0.004417 | 0.00398 | Adora2a/A | 11 |
| 669 | 17 | GO:00519 | regulation | 8/137 | 80/8076 | 5.80E-05 | 0.004417 | 0.00398 | Adora2a/A | 8 |
| 670 | 17 | GO:00987 | cell-cell adl | 10/137 | 130/8076 | 6.71E-05 | 0.00483 | 0.004353 | Cadm2/Cdl | 10 |
| 671 | 17 | GO:001587 | dopamine | 6/137 | 42/8076 | 6.78E-05 | 0.00483 | 0.004353 | Drd1/Oprk | 6 |
| 672 | 17 | GO:000727 | synaptic tr | 4/137 | 14/8076 | 6.95E-05 | 0.00483 | 0.004353 | Adora2a/Sl | 4 |
| 673 | 17 | GO:005104 | regulation | 19/137 | 424/8076 | 9.05E-05 | 0.006111 | 0.005506 | Acvr1c/Ad | 19 |
| 674 | 17 | GO:00986 | cell-cell adl | 19/137 | 426/8076 | 9.64E-05 | 0.006324 | 0.005699 | Adora2a/A | 19 |
| 675 | 17 | GO:000681 | sodium ion | 10/137 | 138/8076 | 0.000111 | 0.007085 | 0.006385 | Asic4/Hecw | 10 |
| 676 | 17 | GO:00072 | neurotrans | 10/137 | 140/8076 | 0.000125 | 0.007579 | 0.00683 | Adora2a/G | 10 |
| 677 | 17 | GO:009964 | signal relea | 10/137 | 140/8076 | 0.000125 | 0.007579 | 0.00683 | Adora2a/G | 10 |
| 678 | 17 | GO:00504 | regulation | 6/137 | 47/8076 | 0.000129 | 0.007645 | 0.006889 | Adora2a/C | 6 |
| 679 | 17 | GO:19035 | regulation | 18/137 | 404/8076 | 0.000151 | 0.008727 | 0.007864 | Acvr1c/Ad | 18 |
| 680 | 17 | GO:00420 | regulation | 4/137 | 17/8076 | 0.000159 | 0.008936 | 0.008052 | Drd1/Gpr3 | 4 |
| 681 | 17 | GO:000854 | visual learr | 6/137 | 50/8076 | 0.000184 | 0.010095 | 0.009097 | Dbh/Drd1/ | 6 |
| 682 | 17 | GO:000657 | biogenic ar | 6/137 | 51/8076 | 0.000205 | 0.010846 | 0.009774 | Dbh/Drd1/ | 6 |
| 683 | 17 | GO:00515 | regulation | 8/137 | 96/8076 | 0.000211 | 0.010846 | 0.009774 | Adora2a/G | 8 |
| 684 | 17 | GO:00602 | long-term | 8/137 | 96/8076 | 0.000211 | 0.010846 | 0.009774 | Adora2a/A | 8 |
| 685 | 17 | GO:00076 | visual beha | 6/137 | 52/8076 | 0.000229 | 0.011509 | 0.010371 | Dbh/Drd1/ | 6 |
| 686 | 17 | GO:00420 | regulation | 4/137 | 19/8076 | 0.000252 | 0.012402 | 0.011176 | Drd1/Gpr3 | 4 |
| 687 | 17 | GO:00504 | catecholan | 6/137 | 54/8076 | 0.000283 | 0.013509 | 0.012173 | Adora2a/C | 6 |
| 688 | 17 | GO:00335 | multicellul | 7/137 | 76/8076 | 0.000286 | 0.013509 | 0.012173 | Asic4/Crh/ | 7 |
| 689 | 17 | GO:00507 | regulation | 6/137 | 56/8076 | 0.000346 | 0.016002 | 0.014419 | Adora2a/C | 6 |
| 690 | 17 | GO:00069 | muscle cor | 10/137 | 160/8076 | 0.000372 | 0.016688 | 0.015038 | Actn2/Arg | 10 |
| 691 | 17 | GO:00189 | phenol-cor | 6/137 | 57/8076 | 0.000381 | 0.016688 | 0.015038 | Dbh/Drd1/ | 6 |
| 692 | 17 | GO:000801 | blood circu | 13/137 | 256/8076 | 0.000387 | 0.016688 | 0.015038 | Adora2a/C | 13 |
| 693 | 17 | GO:00305 | adult beha | 9/137 | 132/8076 | 0.000389 | 0.016688 | 0.015038 | Dbh/Drd1/ | 9 |
| 694 | 17 | GO:004692 | regulation | 7/137 | 82/8076 | 0.000457 | 0.01906 | 0.017175 | Adora2a/G | 7 |
| 695 | 17 | GO:00093 | amine met | 6/137 | 59/8076 | 0.00046 | 0.01906 | 0.017175 | Dbh/Drd1/ | 6 |
| 696 | 17 | GO:00076 | feeding bel | 6/137 | 60/8076 | 0.000504 | 0.020526 | 0.018496 | Adora2a/D | 6 |
| 697 | 17 | GO:003422 | monoatom | 19/137 | 486/8076 | 0.000522 | 0.020677 | 0.018632 | Actn2/Ano | 19 |
| 698 | 17 | GO:00421 | neurotrans | 3/137 | Oct-76 | 0.000525 | 0.020677 | 0.018632 | Dbh/Slc5a | 3 |
| 699 | 17 | GO:00420 | gliogenesis | 12/137 | 234/8076 | 0.000598 | 0.023158 | 0.020868 | Adora2a/C | 12 |
| 700 | 17 | GO:00986 | inorganic i | 18/137 | 455/8076 | 0.000644 | 0.024255 | 0.021857 | Actn2/Ano | 18 |
| 701 | 17 | GO:00140 | astrocyte d | 4/137 | 24/8076 | 0.000647 | 0.024255 | 0.021857 | Adora2a/D | 4 |
| 702 | 17 | GO:000761 | mating beh | 4/137 | 25/8076 | 0.00076 | 0.027964 | 0.025199 | Drd1/Ppp1 | 4 |
| 703 | 17 | GO:00986 | inorganic c | 17/137 | 423/8076 | 0.00077 | 0.027964 | 0.025199 | Actn2/Asic | 17 |
| 704 | 17 | GO:000301 | circulatory | 13/137 | 276/8076 | 0.000789 | 0.028253 | 0.025459 | Adora2a/C | 13 |
| 705 | 17 | GO:00990 | vesicle-me | 12/137 | 242/8076 | 0.000805 | 0.028394 | 0.025586 | Arc/Kcnh1/ | 12 |
| 706 | 17 | GO:00986 | monoatom | 17/137 | 427/8076 | 0.000855 | 0.029695 | 0.026759 | Actn2/Asic | 17 |
| 707 | 17 | GO:00332 | regulation | 4/137 | 26/8076 | 0.000887 | 0.03035 | 0.027348 | Drd1/Gpr3 | 4 |
| 708 | 17 | GO:000971 | catechol-c | 3/137 | Dec-76 | 0.000939 | 0.031246 | 0.028156 | Dbh/Gpr37 | 3 |
| 709 | 17 | GO:004242 | catecholan | 3/137 | Dec-76 | 0.000939 | 0.031246 | 0.028156 | Dbh/Gpr37 | 3 |
| 710 | 17 | GO:00071 | adenylate | 7/137 | 95/8076 | 0.001107 | 0.036319 | 0.032727 | Adora2a/C | 7 |
| 711 | 17 | GO:00015 | neurotrans | 4/137 | 28/8076 | 0.001183 | 0.03788 | 0.034134 | Slc17a6/Sl | 4 |
| 712 | 17 | GO:00440 | regulation | 14/137 | 325/8076 | 0.001187 | 0.03788 | 0.034134 | Arg2/Crh/ | 14 |
| 713 | 17 | GO:00425 | fear respor | 5/137 | 48/8076 | 0.001245 | 0.038716 | 0.034888 | Asic4/Crh/ | 5 |
| 714 | 17 | GO:000301 | muscle sys | 11/137 | 220/8076 | 0.001246 | 0.038716 | 0.034888 | Actn2/Arg | 11 |
| 715 | 17 | GO:005104 | positive re | 11/137 | 221/8076 | 0.001292 | 0.039641 | 0.035721 | Adora2a/C | 11 |
| 716 | 17 | GO:190027 | positive re | 4/137 | 29/8076 | 0.001354 | 0.041006 | 0.036951 | Adora2a/D | 4 |
| 717 | 17 | GO:003572 | sodium ion | 7/137 | 99/8076 | 0.00141 | 0.042165 | 0.037995 | Asic4/Hecw | 7 |
| 718 | 17 | GO:00714 | cellular res | 3/137 | 14/8076 | 0.001516 | 0.044755 | 0.040329 | Cd9/Ldlr/S | 3 |
| 719 | 17 | GO:190027 | regulation | 5/137 | 51/8076 | 0.00164 | 0.047815 | 0.043086 | Adora2a/A | 5 |
| 720 | 18 | GO:00091 | glycoprote | 15/139 | 166/8076 | 1.41E-07 | 0.000257 | 0.000255 | Alg11/Alg1 | 15 |
| 721 | 18 | GO:00064 | protein N-l | 8/139 | 42/8076 | 4.53E-07 | 0.000414 | 0.000411 | Alg11/Alg1 | 8 |
| 722 | 18 | GO:00064 | protein gly | 10/139 | 95/8076 | 4.80E-06 | 0.002191 | 0.002175 | Alg11/Alg1 | 10 |
| 723 | 18 | GO:004341 | macromole | 10/139 | 95/8076 | 4.80E-06 | 0.002191 | 0.002175 | Alg11/Alg1 | 10 |
| 724 | 18 | GO:00091 | glycoprote | 11/139 | 132/8076 | 1.55E-05 | 0.005023 | 0.004986 | Alg11/Alg1 | 11 |
| 725 | 18 | GO:00700 | glycosylati | 10/139 | 109/8076 | 1.65E-05 | 0.005023 | 0.004986 | Alg11/Alg1 | 10 |
| 726 | 18 | GO:000924 | glycolipid b | 5/139 | 26/8076 | 6.91E-05 | 0.018034 | 0.017903 | Dpm1/Gpa | 5 |
| 727 | 18 | GO:00065 | GPI anchor | 4/139 | 15/8076 | 9.90E-05 | 0.022609 | 0.022444 | Dpm1/Gpa | 4 |
| 728 | 18 | GO:00065 | GPI anchor | 4/139 | 16/8076 | 0.00013 | 0.026439 | 0.026246 | Dpm1/Gpa | 4 |
| 729 | 18 | GO:003497 | response t | 11/139 | 171/8076 | 0.000167 | 0.028272 | 0.028066 | Calr/Derl1/ | 11 |
| 730 | 18 | GO:19011 | carbohydra | 17/139 | 366/8076 | 0.00017 | 0.028272 | 0.028066 | Alg11/Alg1 | 17 |

| | | | | | | | | |
|---|---|---|---|---|---|---|---|---|
| 104 | ENSMUSG( | Acat2 | 110460 | 004276351 | -02092559 | 01575 | 00575 | 3 |
| 105 | ENSMUSG( | Acbd3 | 170760 | 010839632 | -0111460 | 01625 | 02 | 3 |
| 113 | ENSMUSG( | Aco1 | 11428 | 014227898 | -00741125 | 00825 | 0055 | 3 |
| 116 | ENSMUSG( | Acot11 | 329910 | 023979754 | -01862288 | 017 | 021 | 3 |
| 119 | ENSMUSG( | Acot7 | 70025 | -00019900 | -02847492 | 00675 | 02425 | 3 |
| 138 | ENSMUSG( | Actg1 | 11465 | -00094126 | -01935727 | 0205 | 00925 | 3 |
| 139 | ENSMUSG( | Actl6a | 56456 | 000727791 | 007695368 | 01 | 01375 | 3 |
| 143 | ENSMUSG( | Actn4 | 60595 | -00153492 | 010076025 | 0095 | -007 | 3 |
| 144 | ENSMUSG( | Actr10 | 56444 | 010696186 | -01253654 | 002 | -01025 | 3 |
| 146 | ENSMUSG( | Actr1b | 226977 | 010290425 | -00964591 | 00875 | 0115 | 3 |
| 156 | ENSMUSG( | Acy3 | 71670 | -00635982 | -00981816 | -00275 | 0025 | 3 |
| 158 | ENSMUSG( | Acyp2 | 75572 | 007626606 | -02463807 | -01325 | 016 | 3 |
| 159 | ENSMUSG( | Ada | 11486 | -01384738 | -02301512 | 01025 | 01475 | 3 |
| 171 | ENSMUSG( | Adat2 | 66757 | 005128254 | -02778304 | 004 | 0125 | 3 |
| 183 | ENSMUSG( | Adcyap1r1 | 11517 | -00100842 | 007710361 | 006 | -00175 | 3 |
| 186 | ENSMUSG( | Add3 | 27360 | 010424047 | -01829075 | 002 | -0075 | 3 |
| 198 | ENSMUSG( | Adgrl3 | 319387 | 004369890 | 010468814 | 003 | 01825 | 3 |
| 204 | ENSMUSG( | Adnp | 11538 | 011135110 | -00944223 | -0045 | -0055 | 3 |
| 205 | ENSMUSG( | Ado | 211488 | -00085706 | -00219005 | -0165 | 018 | 3 |
| 211 | ENSMUSG( | Adprs | 100206 | -00634495 | -00197037 | -0185 | -00325 | 3 |
| 227 | ENSMUSG( | Afg3l2 | 69597 | -00447984 | 014668424 | -01425 | -00325 | 3 |
| 228 | ENSMUSG( | Aftph | 216549 | 006164861 | 002253785 | -01 | -003 | 3 |
| 229 | ENSMUSG( | Aga | 11593 | -01405729 | 006096521 | -008 | 01625 | 3 |
| 232 | ENSMUSG( | Agap3 | 213990 | 016799202 | 005168954 | 00375 | 0205 | 3 |
| 236 | ENSMUSG( | Agk | 69923 | 009311694 | -02260093 | -00625 | -00125 | 3 |
| 242 | ENSMUSG( | Agpat3 | 28169 | 004408537 | -00123001 | 015 | 01275 | 3 |
| 248 | ENSMUSG( | Agtpbp1 | 67269 | 017462482 | -00389439 | -01 | 014 | 3 |
| 253 | ENSMUSG( | Ahdc1 | 230793 | 010561841 | 008933926 | 0095 | 00775 | 3 |
| 254 | ENSMUSG( | Ahi1 | 52906 | 015640378 | 002249144 | 00625 | -00125 | 3 |
| 259 | ENSMUSG( | Aida | 108909 | 004775836 | -00511152 | 001 | -00275 | 3 |
| 265 | ENSMUSG( | Aimp1 | 13722 | 001935891 | -01502493 | -00075 | 022 | 3 |
| 267 | ENSMUSG( | Aip | 11632 | 003756640 | -01377073 | -00275 | 003 | 3 |
| 270 | ENSMUSG( | Ak1 | 11636 | 004671233 | -02139233 | -0085 | 00775 | 3 |
| 273 | ENSMUSG( | Ak4 | 11639 | 011311365 | -00358520 | -00175 | -011 | 3 |
| 274 | ENSMUSG( | Ak5 | 229949 | 017747712 | 007480493 | -0005 | 0205 | 3 |
| 281 | ENSMUSG( | Akap6 | 238161 | -01289734 | 007671646 | 00525 | 00625 | 3 |
| 290 | ENSMUSG( | Akt1s1 | 67605 | 013794629 | -00417628 | 003 | -003 | 3 |
| 292 | ENSMUSG( | Akt3 | 23797 | 016320947 | -00070109 | 00375 | 00625 | 3 |
| 293 | ENSMUSG( | Aktip | 14339 | -00716747 | -02474110 | 0215 | 01025 | 3 |
| 297 | ENSMUSG( | Aldh18a1 | 56454 | 006592113 | -01698093 | -024 | 0005 | 3 |
| 301 | ENSMUSG( | Aldh1b1 | 72535 | -00685925 | -01801106 | -01425 | -00275 | 3 |
| 304 | ENSMUSG( | Aldh2 | 11669 | 008193725 | -01030388 | 490654e-1 | -01 | 3 |
| 313 | ENSMUSG( | Aldoc | 11676 | 021073183 | -01125874 | 00225 | -00925 | 3 |
| 325 | ENSMUSG( | Alkbh6 | 233065 | -00304467 | -00959817 | 01825 | 01125 | 3 |
| 332 | ENSMUSG( | Amacr | 17117 | 009347005 | -00338994 | 002 | -0065 | 3 |
| 335 | ENSMUSG( | Amer2 | 72125 | -02406306 | -02340982 | -01125 | 117757e-1 | 3 |
| 346 | ENSMUSG( | Amt | 434437 | 006817978 | -02237788 | 02425 | 00375 | 3 |
| 347 | ENSMUSG( | Amz2 | 13929 | 011820421 | -00817110 | -009 | -0015 | 3 |
| 348 | ENSMUSG( | Anapc1 | 17222 | -00002982 | 010651406 | -006 | -00675 | 3 |
| 351 | ENSMUSG( | Anapc2 | 99152 | 008335586 | -00789793 | -00275 | 0045 | 3 |
| 353 | ENSMUSG( | Anapc5 | 59008 | 012633710 | -00032176 | -01325 | 0085 | 3 |
| 354 | ENSMUSG( | Anapc7 | 56317 | -00506861 | -00915784 | -01075 | 02375 | 3 |
| 357 | ENSMUSG( | Ank2 | 109676 | 003795384 | 011429040 | 00275 | 017 | 3 |
| 360 | ENSMUSG( | Ankfy1 | 11736 | 007088741 | -00801354 | -0005 | -00725 | 3 |
| 368 | ENSMUSG( | Ankrd13d | 68423 | 019013195 | -00750821 | 004 | 392523e-1 | 3 |
| 371 | ENSMUSG( | Ankrd28 | 105522 | -00661703 | -00122164 | -00875 | 00525 | 3 |
| 409 | ENSMUSG( | Aox1 | 11761 | -01609404 | 003976227 | -00625 | 0045 | 3 |
| 413 | ENSMUSG( | Ap1s1 | 11769 | -00952313 | -02967964 | 006 | 01875 | 3 |
| 420 | ENSMUSG( | Ap3b1 | 11774 | 020729734 | 000416378 | -01725 | 00725 | 3 |
| 421 | ENSMUSG( | Ap3b2 | 11775 | 025755807 | -00673506 | -0095 | 01325 | 3 |
| 423 | ENSMUSG( | Ap3m1 | 55946 | 000269176 | -00206698 | 00125 | 00325 | 3 |
| 425 | ENSMUSG( | Ap3s1 | 11777 | 002033197 | -00348205 | 00725 | -00675 | 3 |
| 439 | ENSMUSG( | Apbb2 | 11787 | -00964452 | -00348301 | 00425 | -00275 | 3 |
| 440 | ENSMUSG( | Apc | 11789 | 007658050 | -00782905 | -01225 | 028 | 3 |
| 442 | ENSMUSG( | Apeh | 235606 | 004549780 | -00129825 | 00325 | 024 | 3 |
| 446 | ENSMUSG( | Apip | 56369 | 001813647 | -01344316 | 0165 | 006 | 3 |
| 453 | ENSMUSG( | Apoc3 | 11814 | 015188750 | -08077469 | -025 | 03475 | 3 |
| 458 | ENSMUSG( | Apool | 68117 | -00612960 | -01455780 | 00125 | -00625 | 3 |
| 461 | ENSMUSG( | Appl1 | 72993 | 006477857 | -00080903 | 0015 | 009 | 3 |
| 463 | ENSMUSG( | Aprt | 11821 | -00821487 | -01028192 | -00825 | 0275 | 3 |
| 467 | ENSMUSG( | Aqp4 | 11829 | -01935299 | -01599747 | 0025 | 00325 | 3 |
| 473 | ENSMUSG( | Arcn1 | 213827 | 013013004 | -00371430 | -00575 | 01825 | 3 |
| 477 | ENSMUSG( | Arf4 | 11843 | -00741964 | -00678446 | -00075 | 014 | 3 |
| 484 | ENSMUSG( | Arfgef2 | 99371 | 003320586 | 002981525 | -01275 | -004 | 3 |
| 486 | ENSMUSG( | Arfip1 | 99889 | 012675143 | 001244177 | 0035 | 00775 | 3 |
| 487 | ENSMUSG( | Arfip2 | 76932 | 015319905 | 004491850 | -0155 | 0185 | 3 |
| 496 | ENSMUSG( | Arhgap20 | 244867 | -01480757 | 000168359 | -007 | 02375 | 3 |
| 498 | ENSMUSG( | Arhgap23 | 58996 | -00492813 | -00871040 | -0065 | 007 | 3 |
| 509 | ENSMUSG( | Arhgap44 | 216831 | 016458255 | 004148319 | -00325 | 01225 | 3 |
| 517 | ENSMUSG( | Arhgef11 | 213498 | 014293189 | 003800523 | 00125 | -0025 | 3 |
| 522 | ENSMUSG( | Arhgef2 | 16800 | -00579751 | 002047610 | -006 | 0195 | 3 |
| 524 | ENSMUSG( | Arhgef26 | 622434 | 009660587 | 004669273 | -01 | 007 | 3 |
| 527 | ENSMUSG( | Arhgef40 | 268739 | 004128435 | 006701546 | 0065 | -0015 | 3 |
| 529 | ENSMUSG( | Arhgef7 | 54126 | 015311585 | 000840770 | -00425 | -003 | 3 |
| 538 | ENSMUSG( | Arl1 | 104303 | -00607717 | -00230295 | 00425 | 0215 | 3 |
| 543 | ENSMUSG( | Arl2bp | 107566 | 000301388 | -01790844 | 0075 | -007 | 3 |
| 554 | ENSMUSG( | Armc1 | 74252 | 008875392 | -00044313 | -00325 | 01525 | 3 |
| 555 | ENSMUSG( | Armc10 | 67211 | -01088219 | -00253309 | 00075 | -0025 | 3 |
| 558 | ENSMUSG( | Armc9 | 78795 | 010114453 | 000211495 | 0095 | 0155 | 3 |
| 563 | ENSMUSG( | Armt1 | 73419 | -00124783 | -00629564 | 02075 | 005 | 3 |
| 564 | ENSMUSG( | Arnt2 | 11864 | 008903620 | -01167181 | -0045 | 01275 | 3 |
| 566 | ENSMUSG( | Arpc1b | 11867 | 005890754 | -01106553 | -00075 | -00425 | 3 |
| 577 | ENSMUSG( | Arrdc1 | 215705 | -01499678 | -00088010 | -01 | -009 | 3 |
| 584 | ENSMUSG( | As3mt | 57344 | 004568263 | -01562367 | -00725 | 00975 | 3 |
| 592 | ENSMUSG( | Ascc1 | 69090 | -00645456 | -01052092 | -02175 | 380257e-1 | 3 |
| 601 | ENSMUSG( | Asns | 27053 | 003253986 | -02523317 | 0055 | 02525 | 3 |
| 606 | ENSMUSG( | Aspscr1 | 68938 | 002392347 | 000567366 | -00275 | 00725 | 3 |
| 607 | ENSMUSG( | Asrgl1 | 66514 | 009414942 | -02307863 | 00875 | -0065 | 3 |
| 608 | ENSMUSG( | Ass1 | 11898 | 007052754 | -00509807 | -004 | -008 | 3 |
| 627 | ENSMUSG( | Atg4b | 66615 | 014260661 | 002972269 | 005 | -00525 | 3 |
| 632 | ENSMUSG( | Atic | 108147 | 008718017 | -01082738 | 00075 | 009 | 3 |
| 636 | ENSMUSG( | Atox1 | 11927 | 005825955 | -02481167 | -01025 | 01675 | 3 |
| 643 | ENSMUSG( | Atp13a4 | 224079 | 005566180 | -00892997 | 0135 | 01325 | 3 |
| 645 | ENSMUSG( | Atp1a2 | 98660 | 004328512 | 001094239 | 00125 | -008 | 3 |
| 649 | ENSMUSG( | Atp1b3 | 11933 | -01212729 | 002625234 | -007 | -00825 | 3 |
| 658 | ENSMUSG( | Atp5f1a | 11946 | 015616689 | -01337398 | -00125 | -0095 | 3 |
| 659 | ENSMUSG( | Atp5f1b | 11947 | 009063803 | -01825924 | -0025 | -00025 | 3 |
| 664 | ENSMUSG( | Atp5mg | 27425 | -00326208 | -03567272 | -00625 | 0005 | 3 |
| 668 | ENSMUSG( | Atp5pd | 71679 | 002526645 | -02190317 | -0165 | -00175 | 3 |
| 669 | ENSMUSG( | Atp5pf | 11957 | 002781478 | -02204330 | -025 | 01075 | 3 |
| 674 | ENSMUSG( | Atp6v0a2 | 21871 | 001683169 | -00664042 | 01075 | 01275 | 3 |
| 676 | ENSMUSG( | Atp6v0d1 | 11972 | 001910103 | -01923330 | -0125 | 03 | 3 |
| 677 | ENSMUSG( | Atp6v1a | 11964 | 003920937 | -00777539 | 006 | -0055 | 3 |
| 678 | ENSMUSG( | Atp6v1b2 | 11966 | 014403166 | -0096438 | -006 | 0015 | 3 |
| 689 | ENSMUSG( | Atp9a | 11981 | -01112153 | 016465611 | 0035 | 0025 | 3 |
| 699 | ENSMUSG( | Atxn10 | 54138 | 007045715 | -02081429 | 01875 | 01275 | 3 |
| 728 | ENSMUSG( | Babam1 | 68251 | 005982007 | -01427506 | -01125 | 0225 | 3 |
| 729 | ENSMUSG( | Babam2 | 107976 | 004690833 | -01193903 | 002 | 009 | 3 |
| 736 | ENSMUSG( | Bag5 | 70369 | 001210196 | 000996044 | 01325 | 0195 | 3 |
| 746 | ENSMUSG( | Bbln | 73737 | 011756584 | -02674143 | -01175 | -0075 | 3 |
| 748 | ENSMUSG( | Bbs2 | 67378 | -00347126 | 004963416 | -005 | 003 | 3 |
| 758 | ENSMUSG( | Bcar1 | 12927 | 005123526 | -00027722 | -008 | 003 | 3 |
| 759 | ENSMUSG( | Bcar3 | 29815 | 010945267 | -01827315 | -0235 | -0005 | 3 |
| 763 | ENSMUSG( | Bcat1 | 12035 | 020656919 | 002007602 | -00575 | -01025 | 3 |
| 766 | ENSMUSG( | Bckdha | 12039 | 008553418 | -01897122 | -00875 | 00775 | 3 |
| 767 | ENSMUSG( | Bckdhb | 12040 | 018784148 | -00872217 | -00975 | 01275 | 3 |

| | | | | | | | | |
|---|---|---|---|---|---|---|---|---|
| 769 | ENSMUSG( | Bcl10 | 12042 | -00119158 | -00214011 | 01025 | 02075 | 3 |
| 773 | ENSMUSG( | Bcl2l1 | 12048 | 015171944 | -00059404 | 01475 | 021 | 3 |
| 776 | ENSMUSG( | Bcl7b | 12054 | 00137303C | -01077552 | -00075 | 01325 | 3 |
| 778 | ENSMUSG( | Bcr | 110279 | 003330052 | -00622981 | -00225 | -00375 | 3 |
| 782 | ENSMUSG( | Becn1 | 56208 | 006894009 | -00659904 | 0085 | 0045 | 3 |
| 792 | ENSMUSG( | Bid | 12122 | 004767377 | 006002506 | 018 | 00325 | 3 |
| 799 | ENSMUSG( | Blmh | 104184 | 002873538 | -01502710 | 00475 | 00025 | 3 |
| 801 | ENSMUSG( | Bloc1s2 | 73689 | -01247780 | -02109788 | 00875 | 027 | 3 |
| 804 | ENSMUSG( | Blvrb | 233016 | -00652760 | -01551010 | 00425 | -004 | 3 |
| 805 | ENSMUSG( | Bmerb1 | 67254 | 010323953 | -01829720 | 016 | 012 | 3 |
| 807 | ENSMUSG( | Bmpr1a | 12166 | -00294719 | 010539229 | 00375 | 01675 | 3 |
| 819 | ENSMUSG( | Bok | 51800 | -00035915 | -01168994 | -018 | -0015 | 3 |
| 824 | ENSMUSG( | Borcs5 | 67774 | 005330869 | -00326205 | 00125 | 0065 | 3 |
| 834 | ENSMUSG( | Brap | 72399 | 004351309 | -00428004 | -008 | 01725 | 3 |
| 836 | ENSMUSG( | Brcc3 | 210766 | 006635933 | -00917279 | 02225 | 00625 | 3 |
| 848 | ENSMUSG( | Brix1 | 67832 | 015092059 | -00011746 | 0035 | 00225 | 3 |
| 857 | ENSMUSG( | Bsdc1 | 100383 | -01004727 | -00582776 | -00175 | 02225 | 3 |
| 871 | ENSMUSG( | Bud31 | 231889 | 006549515 | -00990593 | 006 | -0045 | 3 |
| 888 | ENSMUSG( | C9orf72 | 73205 | 007084684 | 004207356 | 00175 | 0075 | 3 |
| 912 | ENSMUSG( | Cacnb4 | 12298 | -00525387 | 000519902 | -0025 | -0055 | 3 |
| 921 | ENSMUSG( | Cad | 69719 | 022037695 | 004368327 | 01925 | 01625 | 3 |
| 938 | ENSMUSG( | Camk1 | 52163 | 002383225 | 000489363 | 01675 | 0105 | 3 |
| 942 | ENSMUSG( | Camk2b | 12323 | 009917048 | -00363437 | -0095 | 008 | 3 |
| 943 | ENSMUSG( | Camk2d | 108058 | 004959782 | -00688226 | 0155 | 02075 | 3 |
| 948 | ENSMUSG( | Camkk2 | 207565 | 012319837 | -00268207 | 00775 | 0095 | 3 |
| 949 | ENSMUSG( | Camkmt | 73582 | 00622185C | -02079851 | -005 | -0115 | 3 |
| 953 | ENSMUSG( | Camsap3 | 69697 | 023503358 | 004447036 | -001 | 02225 | 3 |
| 956 | ENSMUSG( | Cand1 | 71902 | 015022859 | -00251786 | 00325 | -003 | 3 |
| 963 | ENSMUSG( | Capn1 | 12333 | -00565996 | -01173144 | 00875 | 0055 | 3 |
| 965 | ENSMUSG( | Capn2 | 12334 | -00548173 | -01550780 | -00725 | -0045 | 3 |
| 968 | ENSMUSG( | Capns1 | 12336 | 003500929 | -01525243 | -0025 | -003 | 3 |
| 970 | ENSMUSG( | Capza1 | 12340 | 017228607 | 006379791 | 0175 | 01875 | 3 |
| 972 | ENSMUSG( | Capzb | 12345 | 007464481 | -01231691 | -0055 | 01575 | 3 |
| 976 | ENSMUSG( | Carmil1 | 68732 | 014548836 | -00982160 | 0165 | 01475 | 3 |
| 987 | ENSMUSG( | Caskin2 | 140721 | -00945047 | 009264776 | -00675 | -0035 | 3 |
| 1011 | ENSMUSG( | Cbr3 | 109857 | -00496099 | -03225153 | -00225 | -00775 | 3 |
| 1013 | ENSMUSG( | Cbs | 12411 | 024658334 | -01666405 | 0125 | 02575 | 3 |
| 1016 | ENSMUSG( | Cbx5 | 12419 | 006106694 | -00744117 | 009 | -0035 | 3 |
| 1018 | ENSMUSG( | Cby1 | 73739 | 001131833 | -01357901 | -0165 | -0045 | 3 |
| 1029 | ENSMUSG( | Ccdc124 | 234388 | -00565751 | -03272518 | -0125 | 028 | 3 |
| 1032 | ENSMUSG( | Ccdc136 | 232664 | 013536195 | -00968878 | 01225 | -00475 | 3 |
| 1038 | ENSMUSG( | Ccdc184 | 239650 | -01386018 | -03517612 | -0045 | -001 | 3 |
| 1040 | ENSMUSG( | Ccdc22 | 54638 | 038112009 | -00333480 | 0085 | 00575 | 3 |
| 1055 | ENSMUSG( | Ccdc88a | 108686 | 010540904 | 009837648 | -00775 | 02275 | 3 |
| 1059 | ENSMUSG( | Ccdc91 | 67015 | 009539106 | -01518728 | -00625 | -01025 | 3 |
| 1060 | ENSMUSG( | Ccdc92 | 215707 | 009406104 | -01248727 | -00975 | 0145 | 3 |
| 1076 | ENSMUSG( | Ccs | 12460 | -00032621 | -02803059 | 00025 | 01175 | 3 |
| 1080 | ENSMUSG( | Cct2 | 12461 | 001525732 | -02031663 | -009 | -00125 | 3 |
| 1081 | ENSMUSG( | Cct3 | 12462 | 008316818 | -01966444 | -00325 | 0195 | 3 |
| 1083 | ENSMUSG( | Cct5 | 12465 | 011871619 | -00960818 | 0095 | 005 | 3 |
| 1084 | ENSMUSG( | Cct6a | 12466 | 012899533 | -00874675 | 0045 | -00675 | 3 |
| 1086 | ENSMUSG( | Cct8 | 12469 | 011725133 | -02032061 | -01175 | -0045 | 3 |
| 1099 | ENSMUSG( | Cd44 | 12505 | -01295632 | 008216048 | -00475 | -00025 | 3 |
| 1110 | ENSMUSG( | Cdc16 | 69957 | 000635136 | -00678919 | 0145 | 0045 | 3 |
| 1114 | ENSMUSG( | Cdc34 | 216150 | 011923377 | -01071948 | -01275 | 02 | 3 |
| 1119 | ENSMUSG( | Cdc42bpa | 226751 | 017084172 | -00480890 | 008 | -00375 | 3 |
| 1123 | ENSMUSG( | Cdc42ep4 | 56699 | -00100931 | -01567019 | -0115 | 02175 | 3 |
| 1155 | ENSMUSG( | Cdk5r1 | 12569 | -00327329 | 005181693 | -00725 | 0105 | 3 |
| 1157 | ENSMUSG( | Cdk5rap3 | 80280 | 007315118 | -00984187 | 011 | 01725 | 3 |
| 1159 | ENSMUSG( | Cdk9 | 107951 | 02445025C | -00585516 | 01375 | 02025 | 3 |
| 1160 | ENSMUSG( | Cdkal1 | 68916 | -00477261 | -01206868 | -00475 | 01 | 3 |
| 1176 | ENSMUSG( | Celf6 | 76183 | 022563327 | -00041172 | 0115 | 0125 | 3 |
| 1188 | ENSMUSG( | Cep41 | 83922 | -00713493 | 000596733 | 01 | 02275 | 3 |
| 1192 | ENSMUSG( | Cerk | 223753 | 007012351 | -00756805 | -014 | 01275 | 3 |
| 1202 | ENSMUSG( | Cfap300 | 234912 | 019899948 | 006419191 | 002 | 01875 | 3 |
| 1203 | ENSMUSG( | Cfap36 | 216618 | 006354468 | -01400996 | -0145 | 00725 | 3 |
| 1204 | ENSMUSG( | Cfap410 | 67884 | 000538384 | -00011384 | -01625 | 01525 | 3 |
| 1207 | ENSMUSG( | Cfdp1 | 23837 | 003182024 | -02620459 | -013 | 00325 | 3 |
| 1220 | ENSMUSG( | Chchd2 | 14004 | 005342362 | -02666068 | -022 | 01575 | 3 |
| 1223 | ENSMUSG( | Chchd5 | 66170 | -00354548 | -02738599 | 0205 | 01625 | 3 |
| 1227 | ENSMUSG( | Chd3 | 216848 | 022420408 | 003109431 | -008 | 0225 | 3 |
| 1228 | ENSMUSG( | Chd4 | 107932 | -00589892 | -00594796 | -00075 | -002 | 3 |
| 1237 | ENSMUSG( | Chic1 | 12212 | -00309146 | 000532788 | -785046e-1 | -0105 | 3 |
| 1246 | ENSMUSG( | Chmp2a | 68953 | 006317647 | -01626435 | -0205 | 014 | 3 |
| 1249 | ENSMUSG( | Chmp4b | 75608 | 001252484 | -02375237 | -0205 | 00525 | 3 |
| 1253 | ENSMUSG( | Chn1 | 108699 | 003716974 | -01267040 | 00675 | 00675 | 3 |
| 1254 | ENSMUSG( | Chn2 | 69993 | 001670003 | 005767651 | 00525 | -00225 | 3 |
| 1269 | ENSMUSG( | Chuk | 12675 | -00004566 | 009641026 | 005 | 01575 | 3 |
| 1273 | ENSMUSG( | Ciao3 | 67563 | 015119028 | 004586035 | 0185 | 002 | 3 |
| 1274 | ENSMUSG( | Ciapin1 | 109006 | -00207631 | -00234098 | 017 | 016 | 3 |
| 1285 | ENSMUSG( | Ckap5 | 75786 | 016089078 | -01294209 | 00825 | 00425 | 3 |
| 1287 | ENSMUSG( | Ckmt1 | 12716 | -00631939 | -02737418 | -0165 | 01675 | 3 |
| 1299 | ENSMUSG( | Cldn10 | 58187 | -01563305 | -01639098 | -00125 | 005 | 3 |
| 1306 | ENSMUSG( | Clec2l | 665180 | 000907911 | -00462582 | -00075 | 01125 | 3 |
| 1314 | ENSMUSG( | Clip2 | 269713 | -00664275 | -00989526 | -0125 | 018 | 3 |
| 1316 | ENSMUSG( | Clip4 | 78785 | -00278914 | 004802239 | -01225 | 0185 | 3 |
| 1317 | ENSMUSG( | Clmn | 94040 | 00606010C | -00032801 | -011 | 0275 | 3 |
| 1321 | ENSMUSG( | Clns1a | 12729 | 004384211 | -01274320 | 0005 | 01025 | 3 |
| 1323 | ENSMUSG( | Clpb | 20480 | 009751327 | -00364911 | -013 | 01825 | 3 |
| 1327 | ENSMUSG( | Clpx | 270166 | 011766697 | -00317764 | -01925 | 0055 | 3 |
| 1336 | ENSMUSG( | Clvs1 | 74438 | -01256675 | -01491238 | 0015 | -00125 | 3 |
| 1348 | ENSMUSG( | Cmtm4 | 97487 | 002685094 | 005338263 | -01625 | 0165 | 3 |
| 1352 | ENSMUSG( | Cndp2 | 66054 | -00035934 | -01141378 | 017 | 00225 | 3 |
| 1365 | ENSMUSG( | Cnot1 | 234594 | 021735561 | 004914378 | 00025 | 02025 | 3 |
| 1367 | ENSMUSG( | Cnot11 | 52846 | 000974976 | 000950583 | -014 | 021 | 3 |
| 1370 | ENSMUSG( | Cnot4 | 53621 | 003205673 | -00374417 | -00125 | -00425 | 3 |
| 1372 | ENSMUSG( | Cnot7 | 18983 | 000194859 | 005274345 | 01775 | 00575 | 3 |
| 1373 | ENSMUSG( | Cnot9 | 58184 | -00307823 | 000200298 | 0035 | -009 | 3 |
| 1396 | ENSMUSG( | Coa4 | 68185 | 010408735 | -03338707 | 02175 | 01375 | 3 |
| 1398 | ENSMUSG( | Coa7 | 69893 | 013730585 | 002256941 | -01475 | 00475 | 3 |
| 1406 | ENSMUSG( | Cog3 | 338337 | 008179496 | -00128675 | -002 | 0115 | 3 |
| 1407 | ENSMUSG( | Cog4 | 102339 | 016983807 | -00890284 | -00475 | 01825 | 3 |
| 1408 | ENSMUSG( | Cog5 | 238123 | 003345308 | -00325530 | 00025 | -0045 | 3 |
| 1423 | ENSMUSG( | Col6a1 | 12833 | -00485126 | 002201028 | -0125 | 0245 | 3 |
| 1429 | ENSMUSG( | Commd10 | 69456 | 008182012 | -02150886 | -003 | 016 | 3 |
| 1431 | ENSMUSG( | Commd3 | 12238 | -00073211 | 003308912 | 0005 | 0175 | 3 |
| 1433 | ENSMUSG( | Commd6 | 66200 | 000613408 | -01688907 | 002 | 00325 | 3 |
| 1434 | ENSMUSG( | Commd7 | 99311 | 001860724 | -00774232 | -0055 | 006 | 3 |
| 1435 | ENSMUSG( | Commd8 | 27784 | 006891479 | -00665348 | 013 | 00375 | 3 |
| 1443 | ENSMUSG( | Copg1 | 54161 | 013956404 | -01334627 | -004 | 024 | 3 |
| 1447 | ENSMUSG( | Cops3 | 26572 | -00370072 | -00810956 | 0005 | 002 | 3 |
| 1448 | ENSMUSG( | Cops4 | 26891 | 006737589 | -01321668 | -00575 | -0095 | 3 |
| 1453 | ENSMUSG( | Cops8 | 108679 | 002055456 | -01080656 | 00225 | 0105 | 3 |
| 1457 | ENSMUSG( | Coq10a | 210582 | -00524573 | -01152907 | 00825 | 01275 | 3 |
| 1460 | ENSMUSG( | Coq4 | 227683 | 004250792 | -00572313 | 0005 | 0145 | 3 |
| 1468 | ENSMUSG( | Coro1b | 23789 | 003161514 | 002083164 | 0055 | 02275 | 3 |
| 1474 | ENSMUSG( | Cotl1 | 72042 | 012420539 | -01397413 | -01925 | 021 | 3 |
| 1475 | ENSMUSG( | Cox10 | 70383 | -00640457 | 005428475 | 01975 | 00675 | 3 |
| 1486 | ENSMUSG( | Cox5b-ps | 1E+08 | -02375583 | -05699028 | -022 | 01375 | 3 |
| 1488 | ENSMUSG( | Cox6a2 | 12862 | -00506272 | -00970232 | -02175 | -00075 | 3 |
| 1497 | ENSMUSG( | Cpe | 12876 | 009524008 | -00211220 | 00875 | 012 | 3 |
| 1499 | ENSMUSG( | Cpeb3 | 208922 | -00755409 | -00469119 | 0105 | 0015 | 3 |
| 1503 | ENSMUSG( | Cplx3 | 235415 | 019546755 | -01231427 | -01675 | 02575 | 3 |
| 1508 | ENSMUSG( | Cpne5 | 240058 | -02502052 | -02521772 | 001 | 029 | 3 |
| 1511 | ENSMUSG( | Cpne8 | 66871 | -00467890 | -00514831 | 0175 | 018 | 3 |
| 1554 | ENSMUSG( | Crppa | 75847 | -00584638 | -02808011 | -02025 | 00175 | 3 |

| | | | | | | | |
|---|---|---|---|---|---|---|---|
| 1564 | ENSMUSG( | Crym | 12971 | -02780915 | -03052889 | 0105 | 01425 | 3 |
| 1568 | ENSMUSG( | Cs | 12974 | 006461679 | -02289597 | -01075 | -00325 | 3 |
| 1572 | ENSMUSG( | Cse1l | 110750 | 002709181 | 011463526 | -0105 | -002 | 3 |
| 1581 | ENSMUSG( | Csnk1g1 | 214897 | -00208395 | -00367618 | -013 | -0005 | 3 |
| 1596 | ENSMUSG( | Cstf2t | 83410 | -00195402 | -01193491 | -0155 | 02075 | 3 |
| 1597 | ENSMUSG( | Cstf3 | 228410 | -00149764 | 009383492 | 01275 | -0025 | 3 |
| 1598 | ENSMUSG( | Cstpp1 | 228356 | 007441176 | -01785533 | 0165 | 0155 | 3 |
| 1611 | ENSMUSG( | Ctnna1 | 12385 | 015323188 | -00055171 | 014 | 00875 | 3 |
| 1612 | ENSMUSG( | Ctnna2 | 12386 | 019120973 | -01192319 | 0095 | 0015 | 3 |
| 1618 | ENSMUSG( | Ctnnd2 | 18163 | 015327087 | -00849733 | 0055 | -00225 | 3 |
| 1621 | ENSMUSG( | Ctr9 | 22083 | 005497523 | 000365643 | 005 | -00275 | 3 |
| 1632 | ENSMUSG( | Cttn | 13043 | 01237403 | -00229446 | -017 | 009 | 3 |
| 1633 | ENSMUSG( | Cttnbp2 | 30785 | 003063735 | -00315043 | -0125 | 0045 | 3 |
| 1636 | ENSMUSG( | Ctu2 | 66965 | 007144211 | 007316537 | -00425 | -00275 | 3 |
| 1637 | ENSMUSG( | Cuedc2 | 67116 | -00539219 | -02542758 | -0145 | 00575 | 3 |
| 1638 | ENSMUSG( | Cul1 | 26965 | 008513613 | 009215891 | -002 | -002 | 3 |
| 1639 | ENSMUSG( | Cul2 | 71745 | 000538689 | 00051112C | 00275 | 00275 | 3 |
| 1641 | ENSMUSG( | Cul4a | 99375 | 012716413 | -00272876 | -00425 | -007 | 3 |
| 1649 | ENSMUSG( | Cwc15 | 66070 | 005294048 | -01343219 | -00725 | -003 | 3 |
| 1662 | ENSMUSG( | Cyb5b | 66427 | 008099717 | -01415101 | 00375 | -00975 | 3 |
| 1665 | ENSMUSG( | Cyb5r3 | 109754 | -00789120 | -01328120 | 00075 | -0075 | 3 |
| 1666 | ENSMUSG( | Cyb5r4 | 266690 | 017616181 | 002595603 | -004 | 01225 | 3 |
| 1671 | ENSMUSG( | Cyfip2 | 76884 | 013149107 | -00475895 | -0055 | 00925 | 3 |
| 1684 | ENSMUSG( | Cyria | 76820 | 015221022 | -00442511 | 01625 | 00025 | 3 |
| 1688 | ENSMUSG( | Cyth1 | 19157 | 00685438C | 008875491 | -016 | 002 | 3 |
| 1700 | ENSMUSG( | Daam1 | 208846 | -01248124 | -02259926 | -00325 | -0095 | 3 |
| 1702 | ENSMUSG( | Dab1 | 13131 | 007851471 | -00952434 | -0225 | 00575 | 3 |
| 1703 | ENSMUSG( | Dab2 | 13132 | 00234631C | -00178348 | 0095 | 015 | 3 |
| 1725 | ENSMUSG( | Dbnl | 13169 | 011476916 | -01098775 | 00875 | 0125 | 3 |
| 1734 | ENSMUSG( | Dcaf7 | 71833 | 004915607 | -00693799 | 01025 | 0065 | 3 |
| 1735 | ENSMUSG( | Dcaf8 | 98193 | 014456336 | 003020748 | -01325 | 005 | 3 |
| 1740 | ENSMUSG( | Dclk2 | 70762 | 002705544 | -00570214 | -00125 | 00825 | 3 |
| 1744 | ENSMUSG( | Dcp2 | 70640 | 008248905 | -00014435 | 00525 | 0125 | 3 |
| 1748 | ENSMUSG( | Dctn3 | 53598 | 002328285 | -02058543 | -022 | 005 | 3 |
| 1749 | ENSMUSG( | Dctn4 | 67665 | 014286026 | 004929772 | 00925 | 01075 | 3 |
| 1751 | ENSMUSG( | Dctn6 | 22428 | -00360421 | -01518144 | -0165 | 0255 | 3 |
| 1765 | ENSMUSG( | Ddhd1 | 114874 | 002624184 | -00588636 | 00625 | 00525 | 3 |
| 1779 | ENSMUSG( | Ddx20 | 53975 | 007882752 | -01220472 | 00425 | -00275 | 3 |
| 1807 | ENSMUSG( | Dennd10 | 67894 | -01038768 | -01336999 | -02075 | 00525 | 3 |
| 1809 | ENSMUSG( | Dennd1a | 227801 | 01005827C | 007084823 | -00025 | 0055 | 3 |
| 1812 | ENSMUSG( | Dennd4a | 102442 | 000437027 | -00260802 | -02225 | 00075 | 3 |
| 1815 | ENSMUSG( | Dennd5a | 19347 | 007451587 | -01633801 | -00525 | 001 | 3 |
| 1819 | ENSMUSG( | Denr | 68184 | -00196604 | -02066415 | 00175 | 015 | 3 |
| 1820 | ENSMUSG( | Depdc5 | 277854 | 014250443 | 007688749 | 00125 | 00375 | 3 |
| 1827 | ENSMUSG( | Dffa | 13347 | 003391202 | -00362670 | 01775 | 0185 | 3 |
| 1844 | ENSMUSG( | Dhdh | 71755 | 01359104C | 003647719 | 0075 | 003 | 3 |
| 1847 | ENSMUSG( | Dhps | 330817 | 001192068 | 001547113 | -0065 | 00475 | 3 |
| 1853 | ENSMUSG( | Dhrs7b | 216820 | -01925919 | -00104504 | -0025 | 0105 | 3 |
| 1863 | ENSMUSG( | Dhx57 | 106794 | 012350264 | 006169369 | -00475 | 004 | 3 |
| 1864 | ENSMUSG( | Dhx8 | 217207 | 021031569 | 001771967 | -011 | 012 | 3 |
| 1872 | ENSMUSG( | Dip2a | 64451 | 000616943 | -00603392 | 00175 | -00125 | 3 |
| 1873 | ENSMUSG( | Dip2b | 239667 | 011724486 | 007195039 | -01325 | -0035 | 3 |
| 1878 | ENSMUSG( | Diras2 | 68203 | 001566295 | -01938441 | -004 | -006 | 3 |
| 1880 | ENSMUSG( | Dis3l2 | 208718 | 005523106 | -00066617 | 00075 | 00225 | 3 |
| 1888 | ENSMUSG( | Dlg1 | 13383 | -00155874 | -00993466 | -007 | -00125 | 3 |
| 1890 | ENSMUSG( | Dlg3 | 53310 | 013457977 | -00441530 | 01025 | -0055 | 3 |
| 1897 | ENSMUSG( | Dlst | 78920 | 008694647 | -01397656 | -009 | -007 | 3 |
| 1901 | ENSMUSG( | Dmap1 | 66233 | 005555948 | -01193961 | -0035 | -0025 | 3 |
| 1902 | ENSMUSG( | Dmd | 13405 | 002970976 | 006564989 | 00525 | 00925 | 3 |
| 1927 | ENSMUSG( | Dnajc13 | 235567 | 005013282 | 009396901 | 01675 | 012 | 3 |
| 1932 | ENSMUSG( | Dnajc19 | 67713 | -00750995 | -00783382 | -014 | 00125 | 3 |
| 1942 | ENSMUSG( | Dnajc7 | 56354 | 001496878 | -00552378 | -005 | 01475 | 3 |
| 1948 | ENSMUSG( | Dnlz | 52838 | 008577102 | -00556161 | 0005 | -011 | 3 |
| 1949 | ENSMUSG( | Dnm1 | 13429 | 018743544 | -00240846 | -004 | -0035 | 3 |
| 1951 | ENSMUSG( | Dnm2 | 13430 | 003793559 | -00607026 | -00025 | -003 | 3 |
| 1956 | ENSMUSG( | Dnpep | 13437 | 003027039 | -00382933 | 01075 | -00575 | 3 |
| 1965 | ENSMUSG( | Dock2 | 94176 | -00073170 | -01125529 | -01225 | 02025 | 3 |
| 1978 | ENSMUSG( | Dop1a | 320615 | -00028399 | 01216103C | -00175 | 00525 | 3 |
| 1981 | ENSMUSG( | Dpcd | 226162 | -00189808 | -03049220 | -0065 | 00275 | 3 |
| 1985 | ENSMUSG( | Dph2 | 67728 | 000351611 | -01389543 | -004 | 027 | 3 |
| 1987 | ENSMUSG( | Dph6 | 66632 | -00993555 | -01984979 | -0165 | 01525 | 3 |
| 1992 | ENSMUSG( | Dpp3 | 75221 | -00207946 | 001871115 | -0065 | 02175 | 3 |
| 1996 | ENSMUSG( | Dpp8 | 74388 | 013517128 | -00660822 | 00175 | -0045 | 3 |
| 2001 | ENSMUSG( | Dpy30 | 66310 | 00730610C | -01075927 | -00675 | 0215 | 3 |
| 2003 | ENSMUSG( | Dpysl2 | 12934 | -0084202 | -01981495 | 00875 | -001 | 3 |
| 2007 | ENSMUSG( | Dr1 | 13486 | -00569297 | 011174813 | -0105 | -004 | 3 |
| 2010 | ENSMUSG( | Drg1 | 13494 | 000436982 | -00876459 | 004 | 01325 | 3 |
| 2011 | ENSMUSG( | Drg2 | 13495 | 001789283 | -00966574 | 196262e-1 | -00175 | 3 |
| 2012 | ENSMUSG( | Drp2 | 13497 | 012002781 | -01331025 | 0035 | 005 | 3 |
| 2019 | ENSMUSG( | Dtd2 | 328092 | 006391784 | -01221096 | 00225 | 016 | 3 |
| 2020 | ENSMUSG( | Dtna | 13527 | 009185757 | -01672130 | 00375 | 0045 | 3 |
| 2021 | ENSMUSG( | Dtnb | 13528 | 00963960C | 00039353C | -0095 | 02425 | 3 |
| 2031 | ENSMUSG( | Dusp26 | 66959 | 002658292 | -00445697 | 0005 | 02325 | 3 |
| 2032 | ENSMUSG( | Dusp28 | 67446 | 008287808 | -00133021 | -006 | -0095 | 3 |
| 2033 | ENSMUSG( | Dusp3 | 72349 | -00071730 | -00821079 | -004 | 009 | 3 |
| 2034 | ENSMUSG( | Dusp6 | 67603 | -03061999 | -02512964 | 002 | -0005 | 3 |
| 2040 | ENSMUSG( | Dym | 69190 | 004815793 | 004663469 | -0025 | 005 | 3 |
| 2041 | ENSMUSG( | Dync1h1 | 13424 | 00172881C | -00402721 | 012 | 00975 | 3 |
| 2043 | ENSMUSG( | Dync1i2 | 13427 | 018441697 | -00465566 | -006 | 0155 | 3 |
| 2044 | ENSMUSG( | Dync1li1 | 235661 | 007092906 | -00665141 | -00475 | 02275 | 3 |
| 2045 | ENSMUSG( | Dync1li2 | 234663 | -00288285 | -01532083 | 00475 | 01325 | 3 |
| 2052 | ENSMUSG( | Dynlrb1 | 67068 | -00130489 | -04000076 | 002 | 009 | 3 |
| 2055 | ENSMUSG( | Dyrk1a | 13548 | 011640047 | 001997079 | 01025 | -006 | 3 |
| 2057 | ENSMUSG( | Dyrk2 | 69181 | 00092226C | 009414585 | 00825 | 021 | 3 |
| 2058 | ENSMUSG( | Dzank1 | 241688 | 007321066 | -00625144 | 01175 | 0135 | 3 |
| 2076 | ENSMUSG( | Ecpas | 230249 | 007421052 | 006414128 | 00125 | 01025 | 3 |
| 2089 | ENSMUSG( | Eef1akmt1 | 68043 | 015955832 | -03138334 | -00175 | 01275 | 3 |
| 2096 | ENSMUSG( | Eepd1 | 67484 | -02895757 | -03504973 | -00175 | -0015 | 3 |
| 2100 | ENSMUSG( | Efhd2 | 27984 | 013414854 | -01170486 | -01725 | -00425 | 3 |
| 2101 | ENSMUSG( | Efl1 | 101592 | 016449773 | 000770588 | 00925 | 02025 | 3 |
| 2119 | ENSMUSG( | Ehd4 | 98878 | -00061287 | 00796489C | 00275 | 147196e-1 | 3 |
| 2121 | ENSMUSG( | Ehmt1 | 77683 | 012170416 | 005603411 | -0085 | 008 | 3 |
| 2123 | ENSMUSG( | Ei24 | 13663 | -00491534 | -00463315 | 0035 | -00175 | 3 |
| 2129 | ENSMUSG( | Eif2a | 229317 | -00594539 | -01085998 | 003 | 00625 | 3 |
| 2135 | ENSMUSG( | Eif2b3 | 108067 | 003950445 | -00785169 | -00025 | 00175 | 3 |
| 2136 | ENSMUSG( | Eif2b4 | 13667 | 011651797 | -00294938 | -01225 | 02775 | 3 |
| 2137 | ENSMUSG( | Eif2b5 | 224045 | 00825229C | -00988822 | 003 | -0015 | 3 |
| 2138 | ENSMUSG( | Eif2d | 16865 | 011061812 | 006376104 | 00525 | 01625 | 3 |
| 2140 | ENSMUSG( | Eif2s2 | 67204 | 007241014 | -00676034 | 00725 | 02525 | 3 |
| 2142 | ENSMUSG( | Eif2s3y | 26908 | 006036655 | 000064213 | 01025 | -00525 | 3 |
| 2150 | ENSMUSG( | Eif3h | 68135 | 005331002 | -01167708 | 0075 | 02025 | 3 |
| 2151 | ENSMUSG( | Eif3i | 54709 | 00894435C | -01878529 | 0185 | 0155 | 3 |
| 2152 | ENSMUSG( | Eif3j1 | 78655 | 012467006 | -00385351 | -00775 | 02425 | 3 |
| 2160 | ENSMUSG( | Eif4e | 13684 | -00038422 | -00881532 | -003 | -0055 | 3 |
| 2164 | ENSMUSG( | Eif4g1 | 208643 | 013677793 | 004414836 | -0115 | 014 | 3 |
| 2165 | ENSMUSG( | Eif4g2 | 13690 | 009519752 | -00526091 | 0035 | 01775 | 3 |
| 2168 | ENSMUSG( | Eif5 | 217869 | 003788538 | -00879558 | 00225 | -0025 | 3 |
| 2169 | ENSMUSG( | Eif5a | 276770 | 000407775 | -02466151 | 00225 | 011 | 3 |
| 2172 | ENSMUSG( | Eif6 | 16418 | 010449955 | -00475031 | 00825 | 011 | 3 |
| 2190 | ENSMUSG( | Elob | 67673 | -00598158 | -02429811 | -012 | 0155 | 3 |
| 2195 | ENSMUSG( | Elp1 | 230233 | 009921521 | 001929673 | 0135 | 00525 | 3 |
| 2198 | ENSMUSG( | Elp4 | 77766 | 00126545 | -02578735 | -00825 | 0015 | 3 |
| 2199 | ENSMUSG( | Elp5 | 54351 | 006810464 | -01106798 | -00775 | 0155 | 3 |
| 2200 | ENSMUSG( | Elp6 | 72341 | -01235928 | -03795730 | 00375 | -01 | 3 |
| 2224 | ENSMUSG( | Endog | 13804 | -00017195 | -02654358 | -004 | -012 | 3 |
| 2227 | ENSMUSG( | Engase | 217364 | -00368190 | 001206231 | 01325 | 01125 | 3 |

| | | | | | | | | |
|---|---|---|---|---|---|---|---|---|
| 2228 | ENSMUSG( | Enho | 69638 | 006589325 | -02143842 | 157009e-1 | 00125 | 3 |
| 2229 | ENSMUSG( | Eno1 | 13806 | 001752449 | -02080425 | -0035 | 0175 | 3 |
| 2232 | ENSMUSG( | Enoph1 | 67870 | 008243398 | -00408687 | -02 | 00475 | 3 |
| 2234 | ENSMUSG( | Enpp1 | 18605 | -02236288 | -01453837 | 00575 | 00875 | 3 |
| 2243 | ENSMUSG( | Entpd5 | 12499 | -00735132 | 00657526 | 00075 | 0095 | 3 |
| 2248 | ENSMUSG( | Ep400 | 75560 | 009383399 | 008731525 | -01175 | 016 | 3 |
| 2251 | ENSMUSG( | Epb41l2 | 13822 | 009256051 | -01803459 | -00975 | -00175 | 3 |
| 2257 | ENSMUSG( | Epg5 | 1.01E+08 | 008198909 | 006444709 | -00825 | -007 | 3 |
| 2264 | ENSMUSG( | Ephb1 | 270190 | -00021006 | 001202383 | -011 | -00525 | 3 |
| 2269 | ENSMUSG( | Ephx2 | 13850 | -00357672 | -03813684 | 0185 | 01375 | 3 |
| 2274 | ENSMUSG( | Epn2 | 13855 | 001143423 | -00219491 | -02225 | 00425 | 3 |
| 2291 | ENSMUSG( | Erf | 13875 | -01272860 | 002440111 | -016 | -00075 | 3 |
| 2296 | ENSMUSG( | Erh | 13877 | -00464372 | -01899881 | -0125 | 01225 | 3 |
| 2297 | ENSMUSG( | Eri3 | 140546 | 008752116 | -01393079 | 00625 | 0085 | 3 |
| 2324 | ENSMUSG( | Evl | 14026 | 005171796 | -00194845 | -00425 | 0205 | 3 |
| 2328 | ENSMUSG( | Exoc2 | 66482 | -00159644 | -01453128 | 0105 | -00425 | 3 |
| 2332 | ENSMUSG( | Exoc6 | 107371 | 007328655 | -00128505 | 0225 | 00175 | 3 |
| 2333 | ENSMUSG( | Exoc6b | 75914 | -00087144 | -00743246 | -0005 | -007 | 3 |
| 2334 | ENSMUSG( | Exoc7 | 53413 | 013584046 | -00526820 | 00275 | -00775 | 3 |
| 2335 | ENSMUSG( | Exoc8 | 102058 | 029006268 | 003381823 | 00425 | 00675 | 3 |
| 2336 | ENSMUSG( | Exog | 208194 | -00834516 | -00801873 | 01325 | 008 | 3 |
| 2337 | ENSMUSG( | Exosc1 | 66583 | -00155227 | -01419921 | 01925 | 01025 | 3 |
| 2338 | ENSMUSG( | Exosc10 | 50912 | 00456787 | 00528637 | 0095 | 0015 | 3 |
| 2351 | ENSMUSG( | Ezr | 22350 | 011099894 | -0107513 | -009 | -00925 | 3 |
| 2366 | ENSMUSG( | Faf1 | 14084 | -00869997 | -00227188 | 00625 | 01175 | 3 |
| 2374 | ENSMUSG( | Fam107b | 66540 | 005070139 | -01358255 | -01775 | 00425 | 3 |
| 2380 | ENSMUSG( | Fam120a | 218236 | 004858086 | -00713497 | 002 | 012 | 3 |
| 2382 | ENSMUSG( | Fam120c | 207375 | 005537676 | -01718040 | 01125 | 0075 | 3 |
| 2389 | ENSMUSG( | Fam13c | 71721 | -00736449 | 000212032 | 002 | 00525 | 3 |
| 2390 | ENSMUSG( | Fam151b | 73942 | -01636097 | -04037266 | 00975 | -004 | 3 |
| 2391 | ENSMUSG( | Fam162a | 70186 | 001410078 | -01522046 | 001 | -01 | 3 |
| 2397 | ENSMUSG( | Fam174a | 67698 | -02169937 | -00370468 | -00525 | 0055 | 3 |
| 2400 | ENSMUSG( | Fam185a | 330050 | 009856801 | -02245590 | -00425 | -01225 | 3 |
| 2404 | ENSMUSG( | Fam210a | 108654 | 010103574 | 006123589 | -0165 | 00275 | 3 |
| 2407 | ENSMUSG( | Fam219a | 71901 | 002758315 | -01523443 | -00175 | 01425 | 3 |
| 2414 | ENSMUSG( | Fam78b | 226610 | 004807982 | -00423187 | 01725 | 011 | 3 |
| 2417 | ENSMUSG( | Fam91a1 | 210998 | 009423773 | 00586428 | 0065 | -002 | 3 |
| 2424 | ENSMUSG( | Fars2 | 69955 | 005702256 | -01757538 | -0055 | 0015 | 3 |
| 2426 | ENSMUSG( | Farsb | 23874 | 011639557 | -00312496 | -0005 | 02525 | 3 |
| 2427 | ENSMUSG( | Fasn | 14104 | 020569167 | -00039800 | 00075 | 007 | 3 |
| 2430 | ENSMUSG( | Fat1 | 14107 | -01808455 | -00325560 | -0045 | -0025 | 3 |
| 2441 | ENSMUSG( | Fbxl15 | 68431 | 000501933 | -00592314 | 001 | 007 | 3 |
| 2465 | ENSMUSG( | Fbxw11 | 103583 | 009896152 | -00950959 | 0065 | 02225 | 3 |
| 2467 | ENSMUSG( | Fbxw5 | 30839 | 011666829 | -00882387 | -00125 | 00975 | 3 |
| 2474 | ENSMUSG( | Fcho1 | 74015 | 020277015 | -00381808 | 00275 | -00675 | 3 |
| 2486 | ENSMUSG( | Fer | 14158 | 015882689 | -01300845 | -00825 | 014 | 3 |
| 2488 | ENSMUSG( | Fermt3 | 108101 | -00199885 | 012076577 | 0125 | -0045 | 3 |
| 2490 | ENSMUSG( | Fgd1 | 14163 | 007175541 | -00714278 | 00275 | 01275 | 3 |
| 2495 | ENSMUSG( | Fgf12 | 14167 | -00194821 | -00713859 | -00575 | 0125 | 3 |
| 2505 | ENSMUSG( | Fhip2a | 226252 | -00196490 | 009739754 | -004 | -0045 | 3 |
| 2519 | ENSMUSG( | Fkbp15 | 338355 | 006436031 | -00889702 | -00075 | 0165 | 3 |
| 2520 | ENSMUSG( | Fkbp1a | 14225 | -00044402 | -01836530 | 013 | 0225 | 3 |
| 2521 | ENSMUSG( | Fkbp1b | 14226 | 017408681 | 001636596 | 00475 | 01225 | 3 |
| 2531 | ENSMUSG( | Flii | 14248 | 019355758 | 006477371 | -0075 | 00675 | 3 |
| 2552 | ENSMUSG( | Fn3krp | 238024 | 006096088 | -01573691 | 00225 | 02475 | 3 |
| 2561 | ENSMUSG( | Focad | 230393 | 02749948 | 001809074 | -00575 | 0005 | 3 |
| 2567 | ENSMUSG( | Foxk2 | 68837 | 013035281 | 003753533 | 012 | 02 | 3 |
| 2570 | ENSMUSG( | Foxp1 | 108655 | -00947996 | 005291564 | -013 | 00775 | 3 |
| 2573 | ENSMUSG( | Frg1 | 14300 | 003429636 | 004680725 | -004 | 0005 | 3 |
| 2577 | ENSMUSG( | Frmd5 | 228564 | -00497468 | -00399864 | 00375 | 01625 | 3 |
| 2583 | ENSMUSG( | Frs2 | 327826 | 004545539 | -00845235 | -00125 | 0035 | 3 |
| 2586 | ENSMUSG( | Fryl | 72313 | -00087094 | 008935731 | -00375 | 00675 | 3 |
| 2588 | ENSMUSG( | Fsd1 | 240121 | 006107041 | -01808426 | -005 | 0045 | 3 |
| 2589 | ENSMUSG( | Fsd1l | 319636 | -01793306 | 008537786 | -0035 | 01125 | 3 |
| 2600 | ENSMUSG( | Fundc2 | 67391 | -00530549 | -02058130 | -006 | 007 | 3 |
| 2608 | ENSMUSG( | Fxr1 | 14359 | 008240035 | 003920563 | 005 | 01125 | 3 |
| 2609 | ENSMUSG( | Fxr2 | 23879 | 016679289 | -00329192 | -00375 | 025 | 3 |
| 2610 | ENSMUSG( | Fxyd1 | 56188 | -01410217 | -00869550 | -02 | 008 | 3 |
| 2625 | ENSMUSG( | Gab2 | 14389 | 000115921 | -01570173 | -021 | 0015 | 3 |
| 2626 | ENSMUSG( | Gabarapl1 | 57436 | 005597307 | -01581234 | 004 | 021 | 3 |
| 2627 | ENSMUSG( | Gabarapl2 | 93739 | 001637693 | -02143702 | 01225 | 009 | 3 |
| 2646 | ENSMUSG( | Gak | 231580 | 01221671 | 008514129 | 00025 | 0065 | 3 |
| 2650 | ENSMUSG( | Galk1 | 14635 | 003622293 | 011399782 | -0105 | 006 | 3 |
| 2651 | ENSMUSG( | Galk2 | 69976 | -00910028 | -00831189 | -00075 | 02325 | 3 |
| 2654 | ENSMUSG( | Galnt1 | 14423 | -00314766 | -00575461 | 01925 | 00075 | 3 |
| 2667 | ENSMUSG( | Gapvd1 | 66691 | 011542867 | 005669367 | -01075 | 00625 | 3 |
| 2673 | ENSMUSG( | Gart | 14450 | 005523524 | -01254803 | 005 | 005 | 3 |
| 2678 | ENSMUSG( | Gatad1 | 67210 | 00730122 | -01198315 | -0065 | -009 | 3 |
| 2681 | ENSMUSG( | Gatb | 229487 | 009982464 | -01818814 | -00125 | 00275 | 3 |
| 2683 | ENSMUSG( | Gatd1 | 213350 | -01693974 | -01968762 | -01525 | 0025 | 3 |
| 2686 | ENSMUSG( | Gbe1 | 74185 | 013195943 | 004818616 | -0055 | -0025 | 3 |
| 2687 | ENSMUSG( | Gbf1 | 107338 | 016381286 | -00085885 | 01025 | 023 | 3 |
| 2696 | ENSMUSG( | Gcc2 | 70297 | 010082923 | 005141213 | -00825 | 0145 | 3 |
| 2698 | ENSMUSG( | Gclc | 14629 | 010407156 | 005574608 | 00625 | -0005 | 3 |
| 2699 | ENSMUSG( | Gclm | 14630 | -00100506 | -01804866 | 00725 | 0035 | 3 |
| 2700 | ENSMUSG( | Gcn1 | 231659 | 013594064 | 008645348 | -00725 | 01525 | 3 |
| 2703 | ENSMUSG( | Gdap1 | 14545 | 000519124 | 004657448 | 007 | 0005 | 3 |
| 2705 | ENSMUSG( | Gdap2 | 14547 | -00499050 | -00194962 | 005 | -0045 | 3 |
| 2707 | ENSMUSG( | Gdi1 | 14567 | 004323508 | -01407461 | 117757e-1 | 009 | 3 |
| 2712 | ENSMUSG( | Gdpgp1 | 269952 | 011579621 | -00829003 | 01825 | 01075 | 3 |
| 2714 | ENSMUSG( | Gemin5 | 216766 | 003401227 | 006706818 | 0195 | 01175 | 3 |
| 2719 | ENSMUSG( | Gfap | 14580 | 023111434 | -02875228 | -01075 | -00875 | 3 |
| 2720 | ENSMUSG( | Gfer | 11692 | 013117924 | -00535591 | 00025 | 027 | 3 |
| 2724 | ENSMUSG( | Gfod2 | 70575 | 023928439 | -00034647 | 00225 | 00025 | 3 |
| 2725 | ENSMUSG( | Gfpt1 | 14583 | 011298574 | -00057355 | 01275 | 0055 | 3 |
| 2727 | ENSMUSG( | Gfra2 | 14586 | -01400269 | 001472399 | -0115 | 0165 | 3 |
| 2729 | ENSMUSG( | Gga1 | 106039 | 021575913 | 003091256 | 00625 | 0105 | 3 |
| 2730 | ENSMUSG( | Gga2 | 74105 | -00091077 | -00459369 | 002 | 0215 | 3 |
| 2732 | ENSMUSG( | Ggact | 223267 | -00384116 | -02495659 | 01825 | 01575 | 3 |
| 2733 | ENSMUSG( | Ggct | 110175 | -01163305 | -01825466 | -0215 | 0135 | 3 |
| 2739 | ENSMUSG( | Ghdc | 80860 | -00431847 | 007544268 | -00875 | 011 | 3 |
| 2742 | ENSMUSG( | Gid8 | 76425 | 00499972 | -01257500 | 015 | 011 | 3 |
| 2762 | ENSMUSG( | Gldc | 104174 | 006076688 | -01117574 | 0135 | 0015 | 3 |
| 2773 | ENSMUSG( | Glrx3 | 30926 | 005053821 | -02196502 | 00375 | 004 | 3 |
| 2776 | ENSMUSG( | Gls2 | 216456 | -00097570 | -00836635 | -008 | 0145 | 3 |
| 2782 | ENSMUSG( | Gm13889 | 620695 | -03442979 | -03344589 | 015 | 00675 | 3 |
| 2787 | ENSMUSG( | Gm2a | 14667 | -01795268 | -00710989 | 01125 | 00325 | 3 |
| 2788 | ENSMUSG( | Gmds | 218138 | -00168824 | -02967702 | -0005 | 01 | 3 |
| 2790 | ENSMUSG( | Gmeb2 | 229004 | -00377324 | -00556988 | -001 | -0125 | 3 |
| 2793 | ENSMUSG( | Gmppb | 331026 | 00547981 | -00746554 | -006 | 00025 | 3 |
| 2794 | ENSMUSG( | Gmpr | 66355 | -01058095 | -03241150 | -0005 | -00925 | 3 |
| 2795 | ENSMUSG( | Gmpr2 | 105446 | 002213894 | 002556183 | 006 | 0035 | 3 |
| 2796 | ENSMUSG( | Gmps | 229363 | 011602178 | 007264468 | 00475 | -00775 | 3 |
| 2803 | ENSMUSG( | Gnal | 14680 | -02307524 | -00914789 | -01025 | -0015 | 3 |
| 2812 | ENSMUSG( | Gne | 50798 | 013426172 | 002176226 | 007 | -003 | 3 |
| 2833 | ENSMUSG( | Golga3 | 269682 | 016529598 | 004524958 | -00525 | -00875 | 3 |
| 2835 | ENSMUSG( | Golga5 | 27277 | 013352841 | -00289194 | 02 | 01575 | 3 |
| 2836 | ENSMUSG( | Golga7 | 57437 | -00178075 | -01595098 | -013 | -00275 | 3 |
| 2837 | ENSMUSG( | Golga7b | 71146 | 016987589 | -00550772 | -0005 | 0105 | 3 |
| 2846 | ENSMUSG( | Gorasp1 | 74498 | -00216581 | -01166611 | -001 | 00025 | 3 |
| 2849 | ENSMUSG( | Gosr2 | 56494 | 00338537 | -01805759 | -01 | 02725 | 3 |
| 2869 | ENSMUSG( | Gphn | 268566 | 004825342 | -00802705 | 01475 | 006 | 3 |
| 2871 | ENSMUSG( | Gpld1 | 14756 | -01147158 | -00904831 | 001 | 0045 | 3 |
| 2873 | ENSMUSG( | Gpm6b | 14758 | -00328632 | -01080398 | -00125 | -0095 | 3 |
| 2901 | ENSMUSG( | Gpx4 | 625249 | 003775859 | -01705102 | -002 | -01175 | 3 |
| 2904 | ENSMUSG( | Gramd1a | 52857 | -01469024 | 006089323 | 00175 | 00475 | 3 |

| | | | | | | | | |
|---|---|---|---|---|---|---|---|---|
| 2906 | ENSMUSG( | Gramd4 | 223752 | -00480705 | -00678407 | -00075 | -00975 | 3 |
| 2909 | ENSMUSG( | Grb2 | 14784 | 002217347 | -00348456 | 0195 | 018 | 3 |
| 2927 | ENSMUSG( | Grip1 | 74053 | -01797556 | -01152230 | -01 | 002 | 3 |
| 2931 | ENSMUSG( | Grk5 | 14773 | 004416231 | -00529908 | 0005 | 01425 | 3 |
| 2932 | ENSMUSG( | Grk6 | 26385 | 017863021 | 002568744 | 0125 | 01375 | 3 |
| 2942 | ENSMUSG( | Grpel1 | 17713 | 00668906C | -02676036 | -025 | 00775 | 3 |
| 2944 | ENSMUSG( | Grsf1 | 231413 | -00229254 | -00784065 | -0005 | 01675 | 3 |
| 2946 | ENSMUSG( | Grwd1 | 101612 | 01746680€ | 00233669C | 01475 | 00875 | 3 |
| 2951 | ENSMUSG( | Gsk3b | 56637 | 005859465 | 00153129€ | 008 | 0095 | 3 |
| 2952 | ENSMUSG( | Gskip | 66787 | 015110422 | -01603493 | 006 | 00125 | 3 |
| 2954 | ENSMUSG( | Gspt1 | 14852 | 008652364 | -00314833 | 00325 | 0075 | 3 |
| 2955 | ENSMUSG( | Gspt2 | 14853 | -00985683 | 00481986C | -00525 | 00625 | 3 |
| 2957 | ENSMUSG( | Gss | 14854 | -00348419 | -01218582 | -001 | -0095 | 3 |
| 2964 | ENSMUSG( | Gstm7 | 68312 | -01046635 | -01361721 | -009 | -00325 | 3 |
| 2965 | ENSMUSG( | Gsto1 | 14873 | 00949833S | -00551586 | -01025 | 025 | 3 |
| 2981 | ENSMUSG( | Gtf2i | 14886 | 014081147 | 000871975 | -0015 | 009 | 3 |
| 2982 | ENSMUSG( | Gtf3c1 | 233863 | -00402194 | 007279461 | 001 | 00775 | 3 |
| 2983 | ENSMUSG( | Gtf3c2 | 71752 | 011070418 | 00248526C | -0095 | 02175 | 3 |
| 2984 | ENSMUSG( | Gtf3c3 | 98488 | 004305573 | 004868545 | 00125 | -00725 | 3 |
| 2986 | ENSMUSG( | Gtf3c5 | 70239 | 014494168 | 004403073 | -01075 | -0075 | 3 |
| 2996 | ENSMUSG( | Gucy1b1 | 54195 | 001281393 | 002604942 | 196262e-1 | 0045 | 3 |
| 2998 | ENSMUSG( | Guk1 | 14923 | 000564391 | -02635163 | -004 | -0025 | 3 |
| 3007 | ENSMUSG( | H2az2 | 77605 | -00212052 | -01666097 | -00925 | -00775 | 3 |
| 3019 | ENSMUSG( | Hadha | 97212 | 009285955 | -01995880 | -00825 | 001 | 3 |
| 3020 | ENSMUSG( | Hadhb | 231086 | 00950476S | -01928833 | -0165 | -00275 | 3 |
| 3028 | ENSMUSG( | Hars1 | 15115 | 003650002 | -00838094 | 014 | -002 | 3 |
| 3029 | ENSMUSG( | Hars2 | 70791 | 010993498 | 005863414 | 0045 | 001 | 3 |
| 3033 | ENSMUSG( | Hbs1l | 56422 | 010505481 | -00517360 | 0135 | 01625 | 3 |
| 3044 | ENSMUSG( | Hdac2 | 15182 | 008846898 | 000449902 | -00325 | 022 | 3 |
| 3045 | ENSMUSG( | Hdac3 | 15183 | 005614501 | -00521589 | 00925 | -0005 | 3 |
| 3063 | ENSMUSG( | Heatr5b | 320473 | 006421801 | 00377396C | -01675 | 003 | 3 |
| 3068 | ENSMUSG( | Hectd1 | 207304 | 005543563 | 01005140S | -196262e-1 | 01125 | 3 |
| 3070 | ENSMUSG( | Hectd3 | 76608 | 011587628 | 004491672 | -01125 | 0105 | 3 |
| 3071 | ENSMUSG( | Hectd4 | 269700 | 007358608 | 01073910€ | 0155 | 015 | 3 |
| 3075 | ENSMUSG( | Hepacam | 72927 | -00558476 | 001918552 | 00075 | -002 | 3 |
| 3077 | ENSMUSG( | Herc1 | 235439 | -00734926 | 011374301 | -001 | 01075 | 3 |
| 3078 | ENSMUSG( | Herc2 | 15204 | -01468296 | 015592261 | -01 | 01 | 3 |
| 3092 | ENSMUSG( | Hif1an | 319594 | 003114444 | 00401852S | -01725 | 003 | 3 |
| 3112 | ENSMUSG( | Hmgb2 | 97165 | 00063883C | -00744736 | -0145 | 0065 | 3 |
| 3127 | ENSMUSG( | Hnrnpa1 | 15382 | -01202174 | -00732094 | -009 | -005 | 3 |
| 3130 | ENSMUSG( | Hnrnpab | 15384 | 003294432 | 001107708 | -015 | 016 | 3 |
| 3155 | ENSMUSG( | Hpcal1 | 53602 | 012387624 | -01878818 | 00525 | -00875 | 3 |
| 3163 | ENSMUSG( | Hs1bp3 | 58240 | 02262580S | -00102758 | -005 | -00225 | 3 |
| 3169 | ENSMUSG( | Hsd11b1 | 15483 | 000721032 | -01009524 | 014 | 00375 | 3 |
| 3170 | ENSMUSG( | Hsd17b10 | 15108 | 010332905 | -01980723 | -00175 | -00275 | 3 |
| 3185 | ENSMUSG( | Hspa14 | 50497 | 007339752 | -00863283 | -01425 | 0045 | 3 |
| 3188 | ENSMUSG( | Hspa4 | 15525 | 02193366€ | -00734359 | -01925 | 00125 | 3 |
| 3189 | ENSMUSG( | Hspa4l | 18415 | 005842144 | -01238450 | -01325 | 003 | 3 |
| 3191 | ENSMUSG( | Hspa8 | 15481 | 01792331S | -00033779 | 0165 | 006 | 3 |
| 3196 | ENSMUSG( | Hspd1 | 15510 | 01107286C | -00873360 | -01275 | 015 | 3 |
| 3210 | ENSMUSG( | Huwe1 | 59026 | 00400658S | 007689764 | 007 | 00125 | 3 |
| 3218 | ENSMUSG( | Iba57 | 216792 | 001666814 | -02707873 | -00375 | 00875 | 3 |
| 3219 | ENSMUSG( | Ica1 | 15893 | 009432001 | -00605859 | -00025 | 01275 | 3 |
| 3224 | ENSMUSG( | Ide | 15925 | 014547423 | -00973844 | 00625 | 014 | 3 |
| 3225 | ENSMUSG( | Idh1 | 15926 | 00981727€ | -02175647 | -00075 | -00675 | 3 |
| 3236 | ENSMUSG( | Ift122 | 81896 | 004749295 | -01489516 | 012 | 0205 | 3 |
| 3237 | ENSMUSG( | Ift140 | 106633 | 001031211 | 005810848 | 00525 | 02425 | 3 |
| 3245 | ENSMUSG( | Ift80 | 68259 | 00985007C | -01420916 | 01575 | 009 | 3 |
| 3248 | ENSMUSG( | Igbp1 | 18518 | 014491942 | 001153613 | -00025 | 0095 | 3 |
| 3251 | ENSMUSG( | Igf2r | 16004 | -01136230 | 016090772 | 0025 | 002 | 3 |
| 3263 | ENSMUSG( | Igsf21 | 230868 | -00584298 | 010628185 | 0075 | -00575 | 3 |
| 3266 | ENSMUSG( | Igsf9b | 235086 | -00236780 | 00883874€ | -01225 | 02025 | 3 |
| 3269 | ENSMUSG( | Ikbkb | 16150 | 014264153 | 003322251 | -004 | 0105 | 3 |
| 3278 | ENSMUSG( | Ildr2 | 1E+08 | -01471325 | 007594697 | -0055 | 0145 | 3 |
| 3280 | ENSMUSG( | Ilf3 | 16201 | 017653977 | 005719947 | -00875 | -00625 | 3 |
| 3281 | ENSMUSG( | Ilk | 16202 | 00722269S | -00781973 | 011 | 00175 | 3 |
| 3290 | ENSMUSG( | Impa2 | 114663 | -02538723 | -03944003 | -01325 | 02375 | 3 |
| 3291 | ENSMUSG( | Impact | 16210 | 00430000C | 002694107 | -008 | 02 | 3 |
| 3292 | ENSMUSG( | Impdh1 | 23917 | 004862651 | 002912065 | 00175 | 0075 | 3 |
| 3303 | ENSMUSG( | Inpp5b | 16330 | 01014353C | 005764872 | -00925 | 01775 | 3 |
| 3308 | ENSMUSG( | Inpp5k | 19062 | 010443805 | 005382047 | 0085 | 00125 | 3 |
| 3312 | ENSMUSG( | Insyn1 | 319477 | 005340454 | 003903683 | 013 | 0055 | 3 |
| 3313 | ENSMUSG( | Insyn2a | 627214 | -01486694 | -01929047 | 02075 | 01525 | 3 |
| 3316 | ENSMUSG( | Ints11 | 71957 | 017451851 | -00098778 | 00725 | -004 | 3 |
| 3319 | ENSMUSG( | Ints14 | 69882 | 000446123 | 00770734S | 00925 | 00275 | 3 |
| 3325 | ENSMUSG( | Ints6 | 18130 | 000294053 | 000246915 | -00675 | -0085 | 3 |
| 3326 | ENSMUSG( | Ints7 | 77065 | 004060318 | 005677752 | -00425 | 007 | 3 |
| 3334 | ENSMUSG( | Ipo5 | 70572 | 014009905 | -00334653 | 00975 | -0025 | 3 |
| 3338 | ENSMUSG( | Iqcb1 | 320299 | 008436103 | 00867957€ | 005 | 004 | 3 |
| 3356 | ENSMUSG( | Iscu | 66383 | -00585895 | -02352648 | -013 | 00475 | 3 |
| 3362 | ENSMUSG( | Isoc2b | 67441 | -01741162 | -00308641 | -002 | 003 | 3 |
| 3363 | ENSMUSG( | Ist1 | 71955 | -00319630 | 013523138 | -008 | 02075 | 3 |
| 3383 | ENSMUSG( | Itgb8 | 320910 | -00668690 | 005721942 | 01325 | 01425 | 3 |
| 3386 | ENSMUSG( | Itih4 | 16427 | -02030215 | -02709860 | 01125 | 025 | 3 |
| 3391 | ENSMUSG( | Itpa | 16434 | 001603418 | -00633211 | -0085 | -00725 | 3 |
| 3392 | ENSMUSG( | Itpk1 | 217837 | 01672011C | -00035163 | -0125 | 006 | 3 |
| 3396 | ENSMUSG( | Itpr2 | 16439 | -00504637 | -00091082 | 02025 | -00075 | 3 |
| 3401 | ENSMUSG( | Ivd | 56357 | 011056521 | -02025799 | -0145 | 0155 | 3 |
| 3407 | ENSMUSG( | Jakmip1 | 76071 | -00158077 | -02691942 | -01175 | 004 | 3 |
| 3408 | ENSMUSG( | Jakmip2 | 76217 | 00640584S | 00029961€ | 01175 | 00525 | 3 |
| 3409 | ENSMUSG( | Jakmip3 | 74004 | -00030670 | -00248338 | -0025 | 00175 | 3 |
| 3414 | ENSMUSG( | Jmy | 57748 | 00755852 | 010394842 | 0105 | 01325 | 3 |
| 3424 | ENSMUSG( | Jund | 16478 | -00312694 | -01726502 | -014 | 01025 | 3 |
| 3435 | ENSMUSG( | Katnal1 | 231912 | 007574274 | -01248441 | 005 | 025 | 3 |
| 3439 | ENSMUSG( | Kbtbd2 | 210973 | 002665028 | 003163838 | 017 | 01675 | 3 |
| 3496 | ENSMUSG( | Kctd10 | 330171 | 002651584 | -00573481 | 00725 | 022 | 3 |
| 3499 | ENSMUSG( | Kctd13 | 233877 | 010612075 | -01282900 | -0075 | 01575 | 3 |
| 3506 | ENSMUSG( | Kctd5 | 69259 | 007867483 | -01489877 | 002 | -006 | 3 |
| 3507 | ENSMUSG( | Kctd6 | 71393 | 001886037 | 006523591 | -002 | 01775 | 3 |
| 3511 | ENSMUSG( | Kdm1a | 99982 | 020486195 | -00741202 | -011 | 00175 | 3 |
| 3518 | ENSMUSG( | Keap1 | 50868 | 010704382 | -00009382 | 005 | 01225 | 3 |
| 3520 | ENSMUSG( | Khdrbs1 | 20218 | -00320906 | -00949586 | -0115 | 0205 | 3 |
| 3529 | ENSMUSG( | Kif13b | 16554 | -02366449 | -03274091 | -02225 | 0135 | 3 |
| 3534 | ENSMUSG( | Kif21b | 16565 | -00823254 | -01792225 | 00125 | 01875 | 3 |
| 3539 | ENSMUSG( | Kif5a | 16572 | 016992603 | -01256329 | -01075 | 014 | 3 |
| 3540 | ENSMUSG( | Kif5b | 16573 | 00879814C | -01782424 | -01425 | 02125 | 3 |
| 3541 | ENSMUSG( | Kif5c | 16574 | 00564677C | -01166476 | -0085 | 0115 | 3 |
| 3542 | ENSMUSG( | Kifap3 | 16579 | 001408725 | -01155906 | 0055 | -00875 | 3 |
| 3550 | ENSMUSG( | Klc1 | 16593 | 004580084 | -01791939 | 00275 | 0125 | 3 |
| 3557 | ENSMUSG( | Klhdc7a | 242721 | 005478002 | -03134614 | 0195 | 0225 | 3 |
| 3560 | ENSMUSG( | Klhl11 | 217194 | 000181708 | -00339829 | 00075 | 00875 | 3 |
| 3565 | ENSMUSG( | Klhl26 | 234378 | 015023544 | 00439049S | -00025 | 02025 | 3 |
| 3567 | ENSMUSG( | Klhl36 | 234796 | 001522543 | -00441619 | -00775 | 0175 | 3 |
| 3572 | ENSMUSG( | Klhl9 | 242521 | -00646247 | 000676853 | 00425 | 00125 | 3 |
| 3579 | ENSMUSG( | Kpna6 | 16650 | 008740337 | -00149418 | 00075 | 0185 | 3 |
| 3580 | ENSMUSG( | Kpnb1 | 16211 | -00290453 | 004750528 | 588785e-1 | -0085 | 3 |
| 3581 | ENSMUSG( | Kptn | 70394 | -00136619 | -01143316 | -0115 | 011 | 3 |
| 3599 | ENSMUSG( | Lactb2 | 212442 | -00093146 | -00054805 | 00125 | 005 | 3 |
| 3600 | ENSMUSG( | Lama1 | 16772 | -00510507 | 003562932 | 00075 | 02175 | 3 |
| 3610 | ENSMUSG( | Lamtor1 | 66508 | 00511338S | -02135932 | 0015 | 007 | 3 |
| 3612 | ENSMUSG( | Lamtor3 | 56692 | -00913082 | -00105899 | -0025 | -0095 | 3 |
| 3615 | ENSMUSG( | Lancl1 | 14768 | -01052781 | -01625144 | 0155 | -00075 | 3 |
| 3627 | ENSMUSG( | Lars1 | 107045 | 005983464 | -00430735 | 01225 | 014 | 3 |
| 3640 | ENSMUSG( | Ldha | 16828 | 01144743€ | -01946713 | 01125 | 004 | 3 |
| 3656 | ENSMUSG( | Lgalsl | 216551 | 01284501S | 00430591€ | -016 | 004 | 3 |
| 3660 | ENSMUSG( | Lgi4 | 243914 | -00473233 | -01390284 | 00875 | 0165 | 3 |

| | | | | | | | | |
|---|---|---|---|---|---|---|---|---|
| 3670 | ENSMUSG( | Limch1 | 77569 | -00023764 | -02055043 | -027 | 003 | 3 |
| 3673 | ENSMUSG( | Limk1 | 16885 | 012096038 | -01359020 | 00525 | 01975 | 3 |
| 3717 | ENSMUSG( | Lpin1 | 14245 | -00426957 | 003067134 | 00425 | 01025 | 3 |
| 3722 | ENSMUSG( | Lrba | 80877 | 004531792 | -01135723 | -00675 | 01075 | 3 |
| 3733 | ENSMUSG( | Lrp1 | 16971 | -00806681 | 008443478 | 0095 | 00075 | 3 |
| 3736 | ENSMUSG( | Lrp4 | 228357 | 005386422 | -01568455 | -157009e-1 | 01775 | 3 |
| 3740 | ENSMUSG( | Lrpprc | 72416 | 024022077 | -00499468 | 01 | 00475 | 3 |
| 3747 | ENSMUSG( | Lrrc40 | 67144 | 008539243 | -00133534 | 01575 | 02225 | 3 |
| 3752 | ENSMUSG( | Lrrc57 | 66606 | 002503020 | -01315899 | 01075 | -001 | 3 |
| 3765 | ENSMUSG( | Lrrk2 | 66725 | 000408246 | 005357346 | -01275 | 01175 | 3 |
| 3771 | ENSMUSG( | Lrsam1 | 227738 | 013415279 | 005529711 | 0155 | -003 | 3 |
| 3777 | ENSMUSG( | Lsm12 | 268490 | 004890152 | -00150668 | 0045 | 02475 | 3 |
| 3794 | ENSMUSG( | Luc7l2 | 192196 | 000844828 | 002460953 | 01225 | 014 | 3 |
| 3800 | ENSMUSG( | Luzp2 | 233271 | -02284710 | -01429662 | 002 | -001 | 3 |
| 3801 | ENSMUSG( | Lxn | 17035 | 007514614 | -00518198 | 01675 | 0165 | 3 |
| 3811 | ENSMUSG( | Lypd1 | 72585 | -00759591 | 005173479 | 004 | 0235 | 3 |
| 3814 | ENSMUSG( | Lypla2 | 26394 | 003993774 | -01353867 | -00425 | 01975 | 3 |
| 3816 | ENSMUSG( | Lyrm1 | 73919 | 005677534 | 001946132 | -01225 | 01525 | 3 |
| 3820 | ENSMUSG( | Lyrm9 | 66274 | 001370728 | 005362233 | 00725 | 01575 | 3 |
| 3822 | ENSMUSG( | Lysmd2 | 70082 | 008549445 | -00084778 | -00675 | 01325 | 3 |
| 3824 | ENSMUSG( | Lzic | 69151 | 001536482 | -02035352 | -003 | 01575 | 3 |
| 3826 | ENSMUSG( | Lztr1 | 66863 | 014189064 | -00249779 | -01125 | -00525 | 3 |
| 3844 | ENSMUSG( | Maged2 | 80884 | -00467587 | -01266320 | 01775 | 022 | 3 |
| 3845 | ENSMUSG( | Magee1 | 107528 | -01455942 | -00579469 | -001 | 004 | 3 |
| 3847 | ENSMUSG( | Magi1 | 14924 | -00345395 | -00502175 | 009 | 0065 | 3 |
| 3849 | ENSMUSG( | Magi3 | 99470 | 015994028 | 003432204 | -00375 | -00825 | 3 |
| 3852 | ENSMUSG( | Maip1 | 68115 | -00031590 | 007348961 | 00225 | -00525 | 3 |
| 3872 | ENSMUSG( | Map1lc3b | 67443 | 010754324 | -01655394 | 01725 | 02175 | 3 |
| 3877 | ENSMUSG( | Map2k3 | 26397 | 000896984 | 011385549 | -00425 | 018 | 3 |
| 3881 | ENSMUSG( | Map2k7 | 26400 | 001765501 | -00778329 | -0165 | 001 | 3 |
| 3896 | ENSMUSG( | Map4k5 | 399510 | 007909569 | 008818102 | 00275 | -00225 | 3 |
| 3901 | ENSMUSG( | Map7d2 | 78283 | 016816722 | -00413008 | -01 | 02425 | 3 |
| 3903 | ENSMUSG( | Mapk1 | 26413 | 009679213 | -00187741 | 01625 | 014 | 3 |
| 3904 | ENSMUSG( | Mapk10 | 26414 | 014992031 | -00613240 | 005 | 00375 | 3 |
| 3907 | ENSMUSG( | Mapk1ip1l | 218975 | 010492428 | -00190424 | -021 | 011 | 3 |
| 3914 | ENSMUSG( | Mapk8ip3 | 30957 | 019628486 | -00416139 | -00875 | -00275 | 3 |
| 3916 | ENSMUSG( | Mapkap1 | 227743 | 007128777 | -01053551 | -0005 | 0005 | 3 |
| 3924 | ENSMUSG( | Marchf5 | 69104 | 002927599 | 002528128 | 0175 | 0095 | 3 |
| 3927 | ENSMUSG( | Marcks | 17118 | -00758142 | 008957938 | -00325 | 0175 | 3 |
| 3930 | ENSMUSG( | Mark2 | 13728 | 014381716 | 005075757 | -01075 | 90771e-17 | 3 |
| 3942 | ENSMUSG( | Mat2b | 108645 | 002755505 | 004647528 | 02075 | 0095 | 3 |
| 3950 | ENSMUSG( | Mb21d2 | 239796 | 021578473 | 000487887 | -00925 | 0105 | 3 |
| 3952 | ENSMUSG( | Mbd2 | 17191 | -00351188 | -01525232 | -0075 | 027 | 3 |
| 3963 | ENSMUSG( | Mcat | 223722 | 003315899 | -00894455 | -00425 | -002 | 3 |
| 3964 | ENSMUSG( | Mcc | 328949 | 013804814 | -00277330 | 0045 | 0175 | 3 |
| 3978 | ENSMUSG( | Mcrs1 | 51812 | 005391054 | -00431694 | 02125 | 785046e-1 | 3 |
| 3983 | ENSMUSG( | Mcur1 | 76137 | -00200967 | -02488526 | -007 | 0105 | 3 |
| 3986 | ENSMUSG( | Mdga2 | 320772 | -00425010 | 010864182 | -00725 | -00775 | 3 |
| 3988 | ENSMUSG( | Mdh2 | 17448 | 002310720 | -01924059 | 002 | 00325 | 3 |
| 3993 | ENSMUSG( | Mdp1 | 67881 | -00132817 | -00374318 | 01075 | 00775 | 3 |
| 3994 | ENSMUSG( | Me1 | 17436 | 008649259 | -00931914 | -0075 | 01125 | 3 |
| 3999 | ENSMUSG( | Meak7 | 74347 | 023054180 | 001928669 | 01725 | 0115 | 3 |
| 4024 | ENSMUSG( | Megf8 | 269878 | 000208489 | 011424875 | -0045 | 0125 | 3 |
| 4035 | ENSMUSG( | Metap2 | 56307 | 008919629 | -00649389 | 0105 | 00025 | 3 |
| 4037 | ENSMUSG( | Mettl13 | 71449 | 002221087 | -00059483 | -00625 | -0035 | 3 |
| 4038 | ENSMUSG( | Mettl14 | 210529 | 00893236 | -01672721 | -001 | 01775 | 3 |
| 4050 | ENSMUSG( | Mfap3l | 71306 | -00822527 | 006227699 | -01475 | 00525 | 3 |
| 4054 | ENSMUSG( | Mfn1 | 67414 | 022815850 | 001634535 | 018 | -002 | 3 |
| 4055 | ENSMUSG( | Mfn2 | 170731 | 006020463 | -01348447 | 00325 | 00475 | 3 |
| 4068 | ENSMUSG( | Mgrn1 | 17237 | 010633078 | -01224883 | 00975 | -00025 | 3 |
| 4077 | ENSMUSG( | Mical3 | 194401 | -00430599 | 009391233 | -01475 | -00075 | 3 |
| 4086 | ENSMUSG( | Mief1 | 239555 | 00205155 | 012248546 | 00375 | 0085 | 3 |
| 4087 | ENSMUSG( | Mien1 | 103742 | 000598161 | -03248391 | -011 | -0005 | 3 |
| 4091 | ENSMUSG( | Miga2 | 108958 | -00215665 | 011311772 | -0085 | 009 | 3 |
| 4092 | ENSMUSG( | Minar2 | 225583 | 003140242 | -01305204 | 00775 | 0085 | 3 |
| 4113 | ENSMUSG( | Mmaa | 109136 | 021277153 | -00688059 | -002 | -0005 | 3 |
| 4118 | ENSMUSG( | Mms19 | 72199 | 008736388 | -00200173 | 01275 | -0005 | 3 |
| 4119 | ENSMUSG( | Mmut | 17850 | 011151343 | -01946542 | 00425 | -00375 | 3 |
| 4121 | ENSMUSG( | Mob1b | 68473 | -00579503 | -00243304 | 01025 | 00775 | 3 |
| 4126 | ENSMUSG( | Mocs2 | 17434 | 008861609 | -00946597 | 01825 | 00925 | 3 |
| 4127 | ENSMUSG( | Mocs3 | 69372 | 008943807 | 008456284 | -00525 | 01275 | 3 |
| 4131 | ENSMUSG( | Mon1b | 270096 | 009415074 | -00357076 | 0115 | 02425 | 3 |
| 4136 | ENSMUSG( | Morn4 | 226123 | 007113404 | -01592385 | -01175 | 029 | 3 |
| 4154 | ENSMUSG( | Mprip | 26936 | 004957289 | 001507776 | -01525 | 00325 | 3 |
| 4157 | ENSMUSG( | Mpv17l | 93734 | -00111408 | -00439390 | 01325 | 0125 | 3 |
| 4163 | ENSMUSG( | Mri1 | 67873 | -00891806 | 002892710 | 00675 | 00525 | 3 |
| 4168 | ENSMUSG( | Mrpl10 | 107732 | 004447897 | -00955141 | 00425 | 0125 | 3 |
| 4172 | ENSMUSG( | Mrpl14 | 68463 | -00076497 | -03093680 | 0165 | 0005 | 3 |
| 4173 | ENSMUSG( | Mrpl15 | 27395 | 002963236 | -01440626 | -00375 | 018 | 3 |
| 4175 | ENSMUSG( | Mrpl17 | 27397 | 001474868 | -01638078 | -009 | -00675 | 3 |
| 4176 | ENSMUSG( | Mrpl18 | 67681 | 011356004 | 005863994 | 00975 | 004 | 3 |
| 4178 | ENSMUSG( | Mrpl2 | 27398 | -00106528 | -01096462 | 003 | 02075 | 3 |
| 4180 | ENSMUSG( | Mrpl21 | 353242 | 013441547 | -01320550 | 01725 | 005 | 3 |
| 4181 | ENSMUSG( | Mrpl22 | 216767 | -00678388 | -02717964 | 01625 | 014 | 3 |
| 4182 | ENSMUSG( | Mrpl23 | 19935 | 001387778 | -01837839 | 01275 | 0135 | 3 |
| 4183 | ENSMUSG( | Mrpl24 | 67707 | -00119178 | -01025548 | -01075 | 01225 | 3 |
| 4185 | ENSMUSG( | Mrpl28 | 68611 | 001197179 | -01902485 | 0115 | 023 | 3 |
| 4192 | ENSMUSG( | Mrpl37 | 56280 | 015243405 | -00277691 | 0105 | 00925 | 3 |
| 4193 | ENSMUSG( | Mrpl38 | 60441 | 001374501 | -00984400 | -01525 | 01675 | 3 |
| 4202 | ENSMUSG( | Mrpl46 | 67308 | 013010989 | -01461089 | 003 | 01325 | 3 |
| 4204 | ENSMUSG( | Mrpl48 | 52443 | 012897967 | -01067039 | -00125 | -012 | 3 |
| 4207 | ENSMUSG( | Mrpl51 | 66493 | 014943999 | -00873002 | 00725 | -00025 | 3 |
| 4210 | ENSMUSG( | Mrpl55 | 67212 | -00498634 | -02165365 | -0195 | 015 | 3 |
| 4211 | ENSMUSG( | Mrpl57 | 67840 | -01175729 | -02606494 | -0185 | 01775 | 3 |
| 4213 | ENSMUSG( | Mrpl9 | 78523 | 003829262 | -02219836 | -01925 | 01025 | 3 |
| 4217 | ENSMUSG( | Mrps14 | 64659 | 006753348 | -01828850 | -0005 | -0075 | 3 |
| 4229 | ENSMUSG( | Mrps25 | 64658 | 005375749 | -02376905 | -00475 | 019 | 3 |
| 4232 | ENSMUSG( | Mrps28 | 66230 | 004353237 | -00065760 | 0135 | -00125 | 3 |
| 4233 | ENSMUSG( | Mrps30 | 59054 | 018452818 | 004851437 | 001 | 0075 | 3 |
| 4235 | ENSMUSG( | Mrps34 | 79044 | 001255333 | -01444148 | 011 | 01125 | 3 |
| 4240 | ENSMUSG( | Mrps7 | 50529 | 002830382 | -00675416 | -00625 | -002 | 3 |
| 4241 | ENSMUSG( | Mrps9 | 69527 | -00244614 | 003600596 | -00075 | 00475 | 3 |
| 4242 | ENSMUSG( | Mrrf | 67871 | 012104038 | -00296649 | -0145 | 00425 | 3 |
| 4249 | ENSMUSG( | Msi1 | 17690 | -00152533 | -01813938 | -0015 | 00625 | 3 |
| 4253 | ENSMUSG( | Msn | 17698 | -00695451 | -00570676 | -014 | -004 | 3 |
| 4254 | ENSMUSG( | Msra | 110265 | 010736443 | -01390413 | 0055 | 011 | 3 |
| 4255 | ENSMUSG( | Msrb2 | 76467 | 005044742 | -02895619 | -024 | 013 | 3 |
| 4256 | ENSMUSG( | Msto1 | 229524 | -00266824 | -01878626 | -005 | 00975 | 3 |
| 4260 | ENSMUSG( | Mta1 | 116870 | 004428361 | -00634034 | -0035 | 0105 | 3 |
| 4261 | ENSMUSG( | Mta2 | 23942 | 005112526 | 006766741 | -01025 | 02075 | 3 |
| 4272 | ENSMUSG( | Mtfr1l | 76824 | -00191197 | -01594413 | -01325 | -00775 | 3 |
| 4273 | ENSMUSG( | Mtg1 | 212508 | 001170552 | 001043101 | 003 | 021 | 3 |
| 4274 | ENSMUSG( | Mthfd1 | 108156 | 013428312 | -02032604 | 005 | -00325 | 3 |
| 4278 | ENSMUSG( | Mtif2 | 76784 | 015011909 | 006735038 | -00525 | 01175 | 3 |
| 4280 | ENSMUSG( | Mtm1 | 17772 | 012346569 | 007220504 | 0045 | 01975 | 3 |
| 4282 | ENSMUSG( | Mtmr10 | 233315 | 011671033 | 006299449 | 0055 | 0195 | 3 |
| 4283 | ENSMUSG( | Mtmr12 | 268783 | 008995312 | -00463480 | -00475 | -0005 | 3 |
| 4286 | ENSMUSG( | Mtmr3 | 74302 | 010693628 | 005536393 | 00675 | 014 | 3 |
| 4289 | ENSMUSG( | Mtmr9 | 210376 | 010765917 | -00914068 | 015 | 01075 | 3 |
| 4294 | ENSMUSG( | Mtr | 238505 | 012754396 | 001869320 | 01525 | 018 | 3 |
| 4295 | ENSMUSG( | Mtres1 | 67851 | 000535627 | -00638031 | -004 | 00175 | 3 |
| 4296 | ENSMUSG( | Mtrex | 72198 | 023471252 | -00135317 | 00425 | 01625 | 3 |
| 4298 | ENSMUSG( | Mtrfr | 72650 | 022243285 | -00645176 | -02025 | 00925 | 3 |
| 4300 | ENSMUSG( | Mtss1 | 211401 | -00919946 | -01715791 | -0055 | -00125 | 3 |
| 4306 | ENSMUSG( | Mtx3 | 382793 | 009358158 | 002802284 | 003 | -00775 | 3 |
| 4317 | ENSMUSG( | Mycbp | 56309 | 005489500 | 006761263 | -0065 | 0085 | 3 |

| | | | | | | | | |
|---|---|---|---|---|---|---|---|---|
| 4322 | ENSMUSG( | Myh10 | 77579 | -00379861 | -00240848 | -01525 | 0055 | 3 |
| 4325 | ENSMUSG( | Myh9 | 17886 | 010392987 | 009612047 | -01675 | 013 | 3 |
| 4334 | ENSMUSG( | Myo16 | 244281 | 003995942 | 011259284 | 0015 | 0205 | 3 |
| 4335 | ENSMUSG( | Myo18a | 360013 | 013208722 | 000743262 | -01675 | -0035 | 3 |
| 4341 | ENSMUSG( | Myo5a | 17918 | 003954069 | 010246788 | 0055 | -0015 | 3 |
| 4344 | ENSMUSG( | Myo9a | 270163 | -01249968 | 006722700 | -0025 | -00575 | 3 |
| 4347 | ENSMUSG( | Myof | 226101 | -02458632 | 001483549 | -01525 | 014 | 3 |
| 4359 | ENSMUSG( | Naa15 | 74838 | 008510596 | 007942179 | -01325 | 01025 | 3 |
| 4361 | ENSMUSG( | Naa25 | 231713 | -00523373 | 011246211 | -117757e-1 | 00025 | 3 |
| 4363 | ENSMUSG( | Naa35 | 78689 | 012258612 | -00308662 | 008 | 02125 | 3 |
| 4374 | ENSMUSG( | Nadk | 192185 | 014666502 | -00691864 | 01825 | 00475 | 3 |
| 4377 | ENSMUSG( | Naf1 | 234344 | 000801637 | -00987561 | -0005 | 017 | 3 |
| 4379 | ENSMUSG( | Nagk | 56174 | 006973718 | -01194310 | 01525 | 016 | 3 |
| 4383 | ENSMUSG( | Nans | 94181 | 003923450 | -02379245 | 01375 | 012 | 3 |
| 4384 | ENSMUSG( | Nap1l1 | 53605 | 004686011 | -00862940 | -0005 | -00575 | 3 |
| 4398 | ENSMUSG( | Nat2 | 17961 | 018096181 | 005181244 | -0195 | 0055 | 3 |
| 4405 | ENSMUSG( | Naxd | 69225 | -00126149 | -01334981 | -01225 | 01 | 3 |
| 4429 | ENSMUSG( | Ncl | 17975 | -00008982 | -00623157 | -01825 | 0155 | 3 |
| 4440 | ENSMUSG( | Nde1 | 67203 | 028398435 | -00562782 | -023 | 009 | 3 |
| 4441 | ENSMUSG( | Ndel1 | 83431 | 014301259 | -00514657 | -011 | 00025 | 3 |
| 4448 | ENSMUSG( | Ndrg3 | 29812 | 009305680 | -01544926 | -00525 | 014 | 3 |
| 4449 | ENSMUSG( | Ndrg4 | 234593 | -00196186 | -01327122 | 01625 | 007 | 3 |
| 4451 | ENSMUSG( | Ndufa10 | 67273 | 000166723 | -02621159 | 00125 | 00825 | 3 |
| 4457 | ENSMUSG( | Ndufa4 | 17992 | -00780614 | -02465878 | -01375 | -003 | 3 |
| 4459 | ENSMUSG( | Ndufa5 | 68202 | 003952514 | -02437572 | -01125 | 004 | 3 |
| 4462 | ENSMUSG( | Ndufa8 | 68375 | 002412539 | -03000860 | 00375 | 00925 | 3 |
| 4465 | ENSMUSG( | Ndufaf1 | 69702 | -00139887 | -01880950 | 0135 | 01325 | 3 |
| 4467 | ENSMUSG( | Ndufaf3 | 66706 | -00832295 | -00908041 | -0195 | 01225 | 3 |
| 4469 | ENSMUSG( | Ndufaf5 | 69487 | 0085048 | -00617814 | 012 | 00525 | 3 |
| 4470 | ENSMUSG( | Ndufaf6 | 76947 | -01234867 | -01297640 | 0085 | 0165 | 3 |
| 4471 | ENSMUSG( | Ndufaf7 | 73694 | 009028222 | -00453550 | 003 | 01225 | 3 |
| 4478 | ENSMUSG( | Ndufb4 | 68194 | -01142623 | -02353975 | 00425 | -00575 | 3 |
| 4483 | ENSMUSG( | Ndufb9 | 66218 | 005949955 | -01518088 | -01925 | 00175 | 3 |
| 4490 | ENSMUSG( | Ndufs6 | 407785 | -00180029 | -02905994 | -00225 | -01225 | 3 |
| 4494 | ENSMUSG( | Ndufv2 | 72900 | -00159801 | -02048193 | -0035 | -0075 | 3 |
| 4505 | ENSMUSG( | Nedd4 | 17999 | -00929677 | 004964029 | -00175 | 01575 | 3 |
| 4506 | ENSMUSG( | Nedd4l | 83814 | 014407401 | 007869389 | 00075 | -007 | 3 |
| 4510 | ENSMUSG( | Nefm | 18040 | 011530777 | -00725225 | -00725 | -00925 | 3 |
| 4512 | ENSMUSG( | Nek1 | 18004 | 006321810 | -00741064 | -0025 | 00625 | 3 |
| 4513 | ENSMUSG( | Nek4 | 23955 | 011473103 | -00377913 | 01675 | -002 | 3 |
| 4514 | ENSMUSG( | Nek6 | 59126 | 008150130 | -01321371 | -003 | 01475 | 3 |
| 4516 | ENSMUSG( | Nek9 | 217718 | 008777108 | -01562616 | 00675 | -0045 | 3 |
| 4517 | ENSMUSG( | Nelfa | 24116 | 015825727 | -00712803 | -02075 | 0065 | 3 |
| 4518 | ENSMUSG( | Nelfb | 58202 | 012982630 | 001630965 | 00775 | 01625 | 3 |
| 4519 | ENSMUSG( | Nelfcd | 57314 | 014857242 | -00771370 | -009 | 004 | 3 |
| 4522 | ENSMUSG( | Nemf | 66244 | 001885445 | 006259711 | -00275 | 01475 | 3 |
| 4541 | ENSMUSG( | Nfix | 18032 | 005228655 | -01280472 | -00275 | 02775 | 3 |
| 4546 | ENSMUSG( | Nfx1 | 74164 | -00478321 | 005827242 | -014 | 022 | 3 |
| 4550 | ENSMUSG( | Ngef | 53972 | 000456060 | -02845979 | -00325 | -00225 | 3 |
| 4551 | ENSMUSG( | Ngly1 | 59007 | 002415278 | -01135945 | 00475 | 01425 | 3 |
| 4556 | ENSMUSG( | Nhsl1 | 215819 | 031330204 | -01881844 | 00625 | 02825 | 3 |
| 4559 | ENSMUSG( | Niban2 | 227737 | -00584145 | -00639320 | -01 | 01275 | 3 |
| 4563 | ENSMUSG( | Nif3l1 | 65102 | 005808125 | -00796309 | -00725 | -0075 | 3 |
| 4573 | ENSMUSG( | Nisch | 64652 | 015465175 | 007976549 | 0075 | 0145 | 3 |
| 4578 | ENSMUSG( | Nkiras2 | 71966 | 006498170 | -01091848 | -00525 | 022 | 3 |
| 4579 | ENSMUSG( | Nkrf | 77286 | -00265260 | 004361098 | -00125 | -00825 | 3 |
| 4584 | ENSMUSG( | Nln | 75805 | 005874161 | -00756698 | 00675 | 00375 | 3 |
| 4587 | ENSMUSG( | Nme1 | 18102 | 002813322 | -01736361 | -00825 | 01075 | 3 |
| 4588 | ENSMUSG( | Nme7 | 171567 | 006158187 | -01872166 | -00525 | 00075 | 3 |
| 4594 | ENSMUSG( | Nmt2 | 18108 | 003693097 | -01591940 | 00825 | -0045 | 3 |
| 4608 | ENSMUSG( | Nol8 | 70930 | 011527468 | -00294539 | -0085 | 02025 | 3 |
| 4635 | ENSMUSG( | Npepl1 | 228961 | 010602119 | 000353145 | -0075 | 02525 | 3 |
| 4637 | ENSMUSG( | Nphp1 | 53885 | 003244334 | -00222269 | 00325 | -00825 | 3 |
| 4638 | ENSMUSG( | Nphp4 | 260305 | 007290177 | 000455173 | 015 | 02025 | 3 |
| 4640 | ENSMUSG( | Nploc4 | 217365 | 012505293 | 000341977 | -004 | 00425 | 3 |
| 4641 | ENSMUSG( | Npm1 | 18148 | -00492710 | -01396044 | -01475 | 00325 | 3 |
| 4643 | ENSMUSG( | Npr2 | 230103 | 007042399 | 002923957 | -01425 | 0125 | 3 |
| 4655 | ENSMUSG( | Nr2c2 | 22026 | 005973318 | -00659019 | 00475 | -00875 | 3 |
| 4661 | ENSMUSG( | Nrbf2 | 641340 | 002471200 | 002191726 | -00125 | 002 | 3 |
| 4662 | ENSMUSG( | Nrbp1 | 192292 | 012065354 | -00039522 | -0045 | 004 | 3 |
| 4663 | ENSMUSG( | Nrbp2 | 223649 | 009908850 | 004408179 | 00225 | 00975 | 3 |
| 4676 | ENSMUSG( | Nrxn2 | 18190 | -01156529 | 013015935 | 0045 | -00075 | 3 |
| 4679 | ENSMUSG( | Nsd2 | 107823 | 007590439 | -00134120 | 00525 | -0085 | 3 |
| 4692 | ENSMUSG( | Nt5c | 50773 | 006681935 | 008286709 | 0145 | 0155 | 3 |
| 4695 | ENSMUSG( | Nt5c3b | 68106 | 00888826 | -01580599 | -0055 | 0065 | 3 |
| 4703 | ENSMUSG( | Ntmt1 | 66617 | -01494742 | -01653325 | 0005 | 011 | 3 |
| 4705 | ENSMUSG( | Ntng2 | 171171 | -00940087 | 014654519 | 005 | 157009e-1 | 3 |
| 4707 | ENSMUSG( | Ntrk2 | 18212 | 008076974 | 007507508 | 0125 | 0025 | 3 |
| 4712 | ENSMUSG( | Nub1 | 53312 | 005423901 | -00900373 | -00875 | 02275 | 3 |
| 4715 | ENSMUSG( | Nubpl | 76826 | -02308651 | -02348192 | -0065 | -00025 | 3 |
| 4726 | ENSMUSG( | Nudt14 | 66174 | -00876377 | -00004041 | 013 | 0155 | 3 |
| 4740 | ENSMUSG( | Nudt9 | 74167 | 003937119 | -00328934 | -00675 | 012 | 3 |
| 4744 | ENSMUSG( | Numbl | 18223 | 013125323 | -00619479 | 00725 | 001 | 3 |
| 4785 | ENSMUSG( | Ogdh | 18293 | 014030105 | -01812342 | -01325 | -00125 | 3 |
| 4787 | ENSMUSG( | Ogfod1 | 270086 | 018152719 | 002205618 | -00275 | 008 | 3 |
| 4797 | ENSMUSG( | Olig1 | 50914 | -01874198 | -02491920 | -00125 | -00675 | 3 |
| 4815 | ENSMUSG( | Osbp | 76303 | 004486982 | 007155011 | 0005 | 02475 | 3 |
| 4818 | ENSMUSG( | Osbpl11 | 106326 | -01563949 | -00250995 | 013 | 01025 | 3 |
| 4820 | ENSMUSG( | Osbpl2 | 228983 | 005266388 | 003405635 | 012 | 00675 | 3 |
| 4824 | ENSMUSG( | Osbpl7 | 71240 | -00171818 | 001961629 | -0105 | 01225 | 3 |
| 4834 | ENSMUSG( | Otub2 | 68149 | 018980883 | 005519237 | -005 | -004 | 3 |
| 4842 | ENSMUSG( | Oxa1l | 69089 | -00357223 | -01278400 | -00325 | 0005 | 3 |
| 4843 | ENSMUSG( | Oxct1 | 67041 | 013111558 | -01539149 | -0045 | -011 | 3 |
| 4845 | ENSMUSG( | Oxnad1 | 218885 | 006652353 | -01094424 | -00875 | -0075 | 3 |
| 4848 | ENSMUSG( | Oxsr1 | 108737 | -00589780 | -02786296 | 0105 | 0105 | 3 |
| 4853 | ENSMUSG( | Pa2g4 | 18813 | 006724891 | -01168283 | 02025 | 00425 | 3 |
| 4854 | ENSMUSG( | Pabpc1 | 18458 | 007959346 | -00564560 | 00325 | 013 | 3 |
| 4855 | ENSMUSG( | Pabpc4 | 230721 | 002521465 | -00098130 | 01775 | 00625 | 3 |
| 4858 | ENSMUSG( | Pacs2 | 217893 | -00246131 | -01854968 | 0115 | 02425 | 3 |
| 4861 | ENSMUSG( | Pacsin3 | 80708 | 003653017 | -00797480 | 00725 | 0005 | 3 |
| 4864 | ENSMUSG( | Pafah1b1 | 18472 | 007079513 | -00542604 | 00925 | 0015 | 3 |
| 4866 | ENSMUSG( | Pafah1b3 | 18476 | -00721448 | -02460287 | 01625 | 0085 | 3 |
| 4869 | ENSMUSG( | Paics | 67054 | 011530185 | -01371095 | -017 | 0025 | 3 |
| 4870 | ENSMUSG( | Paip1 | 218693 | 003190253 | -00574340 | -015 | 028 | 3 |
| 4875 | ENSMUSG( | Pak3 | 18481 | 009712651 | -00414804 | -00325 | 009 | 3 |
| 4881 | ENSMUSG( | Palld | 72333 | 025103034 | -01173276 | -00325 | -003 | 3 |
| 4892 | ENSMUSG( | Pank2 | 74450 | 007482651 | 005988061 | 00775 | -007 | 3 |
| 4898 | ENSMUSG( | Papss1 | 23971 | 005287745 | -01020079 | 00925 | 02275 | 3 |
| 4903 | ENSMUSG( | Pard3 | 93742 | -01546457 | -01276237 | -0085 | -004 | 3 |
| 4904 | ENSMUSG( | Pard6a | 56513 | -00070210 | 005852108 | -00725 | -001 | 3 |
| 4927 | ENSMUSG( | Pcbp1 | 23983 | 005811108 | -00637337 | 0035 | 024 | 3 |
| 4931 | ENSMUSG( | Pcca | 110821 | 007148846 | -01977156 | -00825 | -009 | 3 |
| 4956 | ENSMUSG( | Pcdhga6 | 93714 | -01202752 | -00496903 | -0165 | 01775 | 3 |
| 4971 | ENSMUSG( | Pcm1 | 18536 | 002593943 | -00151769 | -017 | 0035 | 3 |
| 4975 | ENSMUSG( | Pcna | 18538 | 006205554 | -00698197 | -02425 | 004 | 3 |
| 4983 | ENSMUSG( | Pcsk1n | 30052 | -00769014 | -00815010 | 015 | 012 | 3 |
| 4990 | ENSMUSG( | Pcyt2 | 68671 | 009214588 | -00237303 | 01125 | 0035 | 3 |
| 4991 | ENSMUSG( | Pdap1 | 231887 | 008235544 | -00994974 | -016 | 01725 | 3 |
| 4999 | ENSMUSG( | Pdcl | 67466 | 014034574 | 004572949 | -0125 | 00475 | 3 |
| 5000 | ENSMUSG( | Pdcl3 | 68833 | -00623056 | -00856284 | -013 | 01225 | 3 |
| 5004 | ENSMUSG( | Pde1a | 18573 | 017623262 | -00067162 | 017 | 122663e-1 | 3 |
| 5016 | ENSMUSG( | Pde7b | 29863 | 007999761 | -01348076 | 0105 | -00075 | 3 |
| 5030 | ENSMUSG( | Pdk1 | 228026 | 016938569 | 005796926 | -00125 | -00575 | 3 |
| 5045 | ENSMUSG( | Pdxdc1 | 94184 | 008528387 | -01675760 | 01425 | 026 | 3 |
| 5047 | ENSMUSG( | Pdxp | 57028 | 006150718 | -00802818 | -00875 | 0105 | 3 |
| 5053 | ENSMUSG( | Pdzrn3 | 55983 | -01001573 | -02224331 | 005 | 785046e-1 | 3 |

| | | | | | | | | |
|---|---|---|---|---|---|---|---|---|
| 5055 | ENSMUSG( | Pebp1 | 23980 | 002814781 | -01607614 | 00525 | -00725 | 3 |
| 5073 | ENSMUSG( | Pex16 | 18633 | 004622932 | -00878347 | -022 | 00025 | 3 |
| 5074 | ENSMUSG( | Pex19 | 19298 | 001811985 | -02010494 | 00325 | 02875 | 3 |
| 5077 | ENSMUSG( | Pex3 | 56535 | -00960395 | 002087946 | 0145 | 0145 | 3 |
| 5080 | ENSMUSG( | Pex6 | 224824 | 022776712 | -00243277 | 00825 | 00125 | 3 |
| 5082 | ENSMUSG( | Pfas | 237823 | 008097801 | 004649794 | 01125 | 01425 | 3 |
| 5084 | ENSMUSG( | Pfdn2 | 18637 | 001724728 | -03500750 | 0145 | 01525 | 3 |
| 5087 | ENSMUSG( | Pfdn6 | 14976 | 00286010C | -01661588 | -0035 | 02175 | 3 |
| 5088 | ENSMUSG( | Pfkfb2 | 18640 | 00752182C | 006587614 | 00075 | -00325 | 3 |
| 5092 | ENSMUSG( | Pfkp | 56421 | 018334631 | -02212921 | -004 | 00775 | 3 |
| 5093 | ENSMUSG( | Pfn2 | 18645 | -00292183 | -00863941 | 01425 | -00125 | 3 |
| 5095 | ENSMUSG( | Pgam2 | 56012 | -03848417 | -04155441 | 01175 | 01125 | 3 |
| 5100 | ENSMUSG( | Pgd | 110208 | -01030068 | -01315770 | 00375 | 009 | 3 |
| 5108 | ENSMUSG( | Pgm3 | 109785 | 013345125 | -01435546 | -015 | -0025 | 3 |
| 5113 | ENSMUSG( | Pgrmc2 | 70804 | -02436838 | -00582512 | -013 | 00525 | 3 |
| 5121 | ENSMUSG( | Phb1 | 18673 | 001775314 | -01564163 | -007 | 01175 | 3 |
| 5126 | ENSMUSG( | Phf14 | 75725 | -00801344 | 008833282 | -00725 | 015 | 3 |
| 5161 | ENSMUSG( | Pick1 | 18693 | 007481467 | -00188209 | -00275 | 01875 | 3 |
| 5172 | ENSMUSG( | Pih1d1 | 68845 | 006640439 | -00894553 | -011 | 00275 | 3 |
| 5176 | ENSMUSG( | Pik3ca | 18706 | 01530830C | 007825219 | -00075 | 00675 | 3 |
| 5184 | ENSMUSG( | Pin1 | 23988 | 000110349 | -02595063 | 0145 | -001 | 3 |
| 5191 | ENSMUSG( | Pip5k1a | 18720 | 000188596 | -00847812 | -0125 | 0165 | 3 |
| 5197 | ENSMUSG( | Pitpnb | 56305 | 007282385 | -00932031 | 0025 | 0145 | 3 |
| 5202 | ENSMUSG( | Pitrm1 | 69617 | 014990384 | -00409941 | 0005 | -008 | 3 |
| 5213 | ENSMUSG( | Pkp4 | 227937 | 008507993 | -00795462 | -00875 | -00125 | 3 |
| 5218 | ENSMUSG( | Plaa | 18786 | 010710709 | 002560503 | 00225 | 00925 | 3 |
| 5228 | ENSMUSG( | Plcg1 | 18803 | 019703336 | 005989044 | -012 | -00375 | 3 |
| 5233 | ENSMUSG( | Plcxd2 | 433022 | 024794019 | -00785406 | 01475 | 001 | 3 |
| 5239 | ENSMUSG( | Plek | 56193 | -02607097 | -02286743 | -00375 | 024 | 3 |
| 5242 | ENSMUSG( | Plekha5 | 109135 | 009583547 | 000708761 | -235514e-1 | -00175 | 3 |
| 5244 | ENSMUSG( | Plekha7 | 233765 | 001977103 | -00640034 | 0065 | 001 | 3 |
| 5250 | ENSMUSG( | Plekhg5 | 269608 | 008220715 | -00369232 | -00075 | -0045 | 3 |
| 5253 | ENSMUSG( | Plekho1 | 67220 | -00480323 | -00851261 | 0175 | 01075 | 3 |
| 5262 | ENSMUSG( | Plpbp | 114863 | -00021632 | 004346606 | -0155 | 011 | 3 |
| 5284 | ENSMUSG( | Plxnb1 | 235611 | 00907194 | -00489259 | 00925 | -00575 | 3 |
| 5292 | ENSMUSG( | Pmm1 | 29858 | 009240107 | -02399472 | 00225 | 011 | 3 |
| 5294 | ENSMUSG( | Pmpca | 66865 | 00927753C | -00476970 | -0105 | 00825 | 3 |
| 5295 | ENSMUSG( | Pmpcb | 73078 | 009004095 | -01211065 | -00275 | -0015 | 3 |
| 5297 | ENSMUSG( | Pnck | 93843 | 02535688C | -00328214 | -01425 | 01675 | 3 |
| 5300 | ENSMUSG( | Pnkp | 59047 | 000729937 | 001636738 | -005 | 01525 | 3 |
| 5311 | ENSMUSG( | Pnpla8 | 67452 | 015091374 | 004715547 | 0045 | 0175 | 3 |
| 5312 | ENSMUSG( | Pnpo | 103711 | -00965205 | -00599034 | 00725 | 02525 | 3 |
| 5320 | ENSMUSG( | Pogz | 229584 | 00328821C | 005948033 | -0095 | -00775 | 3 |
| 5323 | ENSMUSG( | Poldip2 | 67811 | 006437067 | -00820574 | -007 | 001 | 3 |
| 5329 | ENSMUSG( | Polr1h | 66136 | -00404419 | -00179831 | -002 | -00875 | 3 |
| 5333 | ENSMUSG( | Polr2d | 69241 | 01490127C | -00248382 | 00325 | 0165 | 3 |
| 5343 | ENSMUSG( | Polr3d | 67065 | 009183848 | 001363043 | 01525 | 00225 | 3 |
| 5360 | ENSMUSG( | Pou6f1 | 19009 | 015153705 | 00776264C | -00075 | 02375 | 3 |
| 5369 | ENSMUSG( | Ppfia2 | 327814 | 007250368 | -01450902 | -0005 | -00475 | 3 |
| 5370 | ENSMUSG( | Ppfia3 | 76787 | 00642080C | -03083373 | -00175 | 003 | 3 |
| 5371 | ENSMUSG( | Ppfia4 | 68507 | 010552302 | -00561010 | -00775 | 00725 | 3 |
| 5372 | ENSMUSG( | Ppfibp1 | 67533 | 001204818 | -00476821 | -01125 | 014 | 3 |
| 5391 | ENSMUSG( | Ppm1e | 320472 | 013977445 | 004811535 | -00125 | -0015 | 3 |
| 5398 | ENSMUSG( | Ppme1 | 72590 | 001084196 | -01522874 | 0045 | 0055 | 3 |
| 5401 | ENSMUSG( | Ppp1cb | 19046 | 003402826 | 000502646 | -016 | -00275 | 3 |
| 5418 | ENSMUSG( | Ppp1r21 | 73825 | 012102644 | -00975417 | -00775 | 01125 | 3 |
| 5423 | ENSMUSG( | Ppp1r7 | 66385 | 000601806 | -00574424 | -002 | 017 | 3 |
| 5428 | ENSMUSG( | Ppp2cb | 19053 | 004170509 | -01303498 | 01025 | 00825 | 3 |
| 5429 | ENSMUSG( | Ppp2r1a | 51792 | -01155099 | -02012612 | 00225 | 01575 | 3 |
| 5431 | ENSMUSG( | Ppp2r2a | 71978 | -00497769 | -00639660 | -01425 | -004 | 3 |
| 5432 | ENSMUSG( | Ppp2r2b | 72930 | 003337505 | -00822318 | 02275 | 002 | 3 |
| 5433 | ENSMUSG( | Ppp2r2c | 269643 | 016745804 | 004928361 | 01525 | 0145 | 3 |
| 5434 | ENSMUSG( | Ppp2r2d | 52432 | 004248587 | -00286005 | -0015 | 00975 | 3 |
| 5435 | ENSMUSG( | Ppp2r5a | 226849 | 009338513 | -01235620 | 00325 | -0075 | 3 |
| 5441 | ENSMUSG( | Ppp3cb | 19056 | 001656419 | -00108761 | 002 | 02575 | 3 |
| 5442 | ENSMUSG( | Ppp3cc | 19057 | 008719268 | 006171453 | -008 | 0005 | 3 |
| 5444 | ENSMUSG( | Ppp4c | 56420 | -00427855 | -00776479 | 00725 | 0105 | 3 |
| 5448 | ENSMUSG( | Ppp4r4 | 74521 | 005307112 | -00244156 | -01175 | 02075 | 3 |
| 5452 | ENSMUSG( | Ppp6r2 | 71474 | 016780733 | 006447762 | 0125 | -002 | 3 |
| 5465 | ENSMUSG( | Prdx2 | 21672 | -00020005 | -02388363 | 0015 | -00925 | 3 |
| 5470 | ENSMUSG( | Preb | 50907 | 011490081 | -00262046 | 01175 | 0055 | 3 |
| 5471 | ENSMUSG( | Prelid3a | 225655 | 00885180C | -00550352 | 01225 | -0025 | 3 |
| 5473 | ENSMUSG( | Prep | 19072 | -00985419 | 002441838 | 00425 | 019 | 3 |
| 5474 | ENSMUSG( | Prepl | 213760 | 006801442 | -00150691 | 011 | -0055 | 3 |
| 5487 | ENSMUSG( | Prkar1a | 19084 | 002339991 | -00403830 | -0015 | 00875 | 3 |
| 5488 | ENSMUSG( | Prkar1b | 19085 | 010028441 | -01677412 | 00925 | 0115 | 3 |
| 5491 | ENSMUSG( | Prkca | 18750 | 276e-05 | -00390025 | -01125 | -00375 | 3 |
| 5495 | ENSMUSG( | Prkcg | 18752 | 021272789 | 006470797 | 01875 | 01475 | 3 |
| 5497 | ENSMUSG( | Prkci | 18759 | 006059803 | -00533401 | 00425 | 00925 | 3 |
| 5500 | ENSMUSG( | Prkcz | 18762 | 009582245 | 005528291 | 00775 | 0095 | 3 |
| 5507 | ENSMUSG( | Prmt1 | 15469 | -00060020 | -01040250 | 01225 | 02375 | 3 |
| 5509 | ENSMUSG( | Prmt3 | 71974 | 017133813 | 00232118C | -011 | 00875 | 3 |
| 5519 | ENSMUSG( | Prpf18 | 67229 | 010175573 | 005405604 | 0065 | 0055 | 3 |
| 5520 | ENSMUSG( | Prpf19 | 28000 | 014976824 | -00796196 | -00225 | 00775 | 3 |
| 5522 | ENSMUSG( | Prpf31 | 68988 | 006460305 | -01685284 | -01375 | 00075 | 3 |
| 5536 | ENSMUSG( | Prpsap2 | 212627 | 011745431 | -01597567 | -002 | -002 | 3 |
| 5547 | ENSMUSG( | Prrt1 | 260297 | -00163954 | -00203008 | -01475 | 0245 | 3 |
| 5550 | ENSMUSG( | Prune1 | 229589 | -00634899 | 003071469 | -0105 | 006 | 3 |
| 5562 | ENSMUSG( | Psma2 | 19166 | 00157095 | -02000181 | -00075 | -00675 | 3 |
| 5564 | ENSMUSG( | Psma4 | 26441 | -00065100 | -00989258 | -00375 | 01875 | 3 |
| 5567 | ENSMUSG( | Psma7 | 26444 | -00087364 | -00726314 | 0055 | 01425 | 3 |
| 5568 | ENSMUSG( | Psmb1 | 19170 | 001231453 | -01708854 | 00075 | 00525 | 3 |
| 5571 | ENSMUSG( | Psmb3 | 26446 | -00135732 | -02158378 | 0095 | 013 | 3 |
| 5573 | ENSMUSG( | Psmb5 | 19173 | 000663497 | -02169721 | 0075 | 00825 | 3 |
| 5577 | ENSMUSG( | Psmc1 | 19179 | 003979926 | -01223472 | 01575 | 0035 | 3 |
| 5578 | ENSMUSG( | Psmc2 | 19181 | 004901131 | -01225612 | 01475 | 0215 | 3 |
| 5580 | ENSMUSG( | Psmc4 | 23996 | 000748168 | -01324239 | 001 | 0045 | 3 |
| 5581 | ENSMUSG( | Psmc5 | 19184 | 00455618C | -02957085 | 011 | 00925 | 3 |
| 5583 | ENSMUSG( | Psmd1 | 70247 | 01541982 | -00273809 | 01175 | 02175 | 3 |
| 5586 | ENSMUSG( | Psmd12 | 66997 | 004447407 | -01559597 | 01775 | 00375 | 3 |
| 5589 | ENSMUSG( | Psmd2 | 21762 | 013267156 | -01767289 | 01025 | 0015 | 3 |
| 5590 | ENSMUSG( | Psmd3 | 22123 | -00131430 | 002815815 | 01125 | 017 | 3 |
| 5593 | ENSMUSG( | Psmd6 | 66413 | 003471395 | -00033544 | -01 | 0065 | 3 |
| 5596 | ENSMUSG( | Psmd9 | 67151 | 014884101 | -00353510 | -01075 | 004 | 3 |
| 5598 | ENSMUSG( | Psme2 | 19188 | 001903384 | -00349006 | 00975 | 00275 | 3 |
| 5600 | ENSMUSG( | Psme3ip1 | 102122 | 000109374 | -01397614 | -01475 | 02275 | 3 |
| 5601 | ENSMUSG( | Psme4 | 103554 | 003909393 | 006211593 | 196262e-1 | 0065 | 3 |
| 5603 | ENSMUSG( | Psmg1 | 56088 | 017560776 | 000470339 | -007 | -0065 | 3 |
| 5604 | ENSMUSG( | Psmg2 | 107047 | -00733552 | 003333215 | 00325 | 372897e-1 | 3 |
| 5606 | ENSMUSG( | Pspc1 | 66645 | -00347274 | -01800071 | -00475 | 01275 | 3 |
| 5607 | ENSMUSG( | Psph | 100678 | -00193924 | -01909044 | -00075 | -00375 | 3 |
| 5610 | ENSMUSG( | Ptbp1 | 19205 | -01021337 | 013512378 | -01625 | 00275 | 3 |
| 5611 | ENSMUSG( | Ptbp2 | 56195 | 003746965 | -00090396 | 0035 | 01775 | 3 |
| 5613 | ENSMUSG( | Ptcd1 | 71799 | 004529433 | 009055775 | -0075 | 0015 | 3 |
| 5614 | ENSMUSG( | Ptcd2 | 68927 | 013799649 | -00111407 | 0155 | 021 | 3 |
| 5623 | ENSMUSG( | Ptgfrn | 19221 | -01148438 | 00608574C | -00925 | 00775 | 3 |
| 5630 | ENSMUSG( | Ptk2 | 14083 | 011959189 | 005154655 | 00525 | -00625 | 3 |
| 5636 | ENSMUSG( | Ptov1 | 84113 | 005482311 | -01078791 | -01125 | 025 | 3 |
| 5639 | ENSMUSG( | Ptp4a3 | 19245 | 001145585 | -00164630 | -016 | 0175 | 3 |
| 5642 | ENSMUSG( | Ptpmt1 | 66461 | -00144748 | -00906351 | 00325 | 0135 | 3 |
| 5643 | ENSMUSG( | Ptpn1 | 19246 | 008800239 | 001470644 | 01675 | 01475 | 3 |
| 5667 | ENSMUSG( | Ptprt | 19281 | -00981715 | -00114430 | 0115 | 0135 | 3 |
| 5669 | ENSMUSG( | Ptprz1 | 19283 | -00348121 | -00124036 | 0045 | 02225 | 3 |
| 5672 | ENSMUSG( | Ptrhd1 | 69709 | -00108838 | -02729856 | -008 | -0055 | 3 |
| 5673 | ENSMUSG( | Pts | 19286 | 001302683 | -01233619 | 01125 | -00025 | 3 |
| 5676 | ENSMUSG( | Pum1 | 80912 | 010335938 | -00079172 | -003 | -00875 | 3 |

| | | | | | | | |
|---|---|---|---|---|---|---|---|
| 5679 | ENSMUSG( | Pura | 19290 | -00117131 | -01302852 | -00525 | -01075 | 3 |
| 5680 | ENSMUSG( | Purb | 19291 | 00733767C | -00086459 | 01475 | 01 | 3 |
| 5686 | ENSMUSG( | Pwp1 | 103136 | 007598552 | -00573364 | -02075 | 0015 | 3 |
| 5689 | ENSMUSG( | Pxdn | 69675 | 004757425 | 007649901 | 00625 | -008 | 3 |
| 5696 | ENSMUSG( | Pycr2 | 69051 | 002710757 | -01287238 | -00075 | 01475 | 3 |
| 5702 | ENSMUSG( | Pym1 | 78428 | -01146879 | -02987033 | 0065 | -002 | 3 |
| 5703 | ENSMUSG( | Pyroxd1 | 232491 | 002372974 | -00441179 | -0115 | 00225 | 3 |
| 5708 | ENSMUSG( | Qki | 19317 | 00254724C | 000989943 | 00725 | 0095 | 3 |
| 5711 | ENSMUSG( | Qrich1 | 69232 | 005332301 | -00665697 | 00575 | 01575 | 3 |
| 5715 | ENSMUSG( | Qtrt1 | 60507 | -00513924 | -00833203 | -00175 | 016 | 3 |
| 5717 | ENSMUSG( | R3hcc1 | 71843 | -00144512 | -01580666 | -01625 | 026 | 3 |
| 5718 | ENSMUSG( | R3hdm1 | 226412 | 006601271 | -01476752 | 01275 | 0015 | 3 |
| 5732 | ENSMUSG( | Rab1b | 76308 | 007419454 | -01427756 | -0045 | 01 | 3 |
| 5734 | ENSMUSG( | Rab22a | 19334 | 013441175 | 002052583 | 0025 | 00975 | 3 |
| 5736 | ENSMUSG( | Rab24 | 19336 | 001745603 | -00157464 | 01075 | 002 | 3 |
| 5745 | ENSMUSG( | Rab31 | 106572 | 007655527 | -02163812 | 00025 | 029 | 3 |
| 5748 | ENSMUSG( | Rab34 | 19376 | -00159779 | 002549274 | -00825 | 0045 | 3 |
| 5749 | ENSMUSG( | Rab35 | 77407 | 007506501 | -00996954 | -00125 | -012 | 3 |
| 5769 | ENSMUSG( | Rab6a | 19346 | -00243032 | -01471517 | 01275 | -00275 | 3 |
| 5770 | ENSMUSG( | Rab6b | 270192 | 015501803 | -00539165 | 01725 | 015 | 3 |
| 5775 | ENSMUSG( | Rabep1 | 54189 | 015223391 | -00542450 | -00875 | 01825 | 3 |
| 5778 | ENSMUSG( | Rabgap1 | 227800 | 01259580C | 001590101 | 01375 | 01675 | 3 |
| 5779 | ENSMUSG( | Rabgap1l | 29809 | -00175911 | 002274384 | 01575 | 0075 | 3 |
| 5780 | ENSMUSG( | Rabgef1 | 56715 | -00141158 | -00950767 | 01975 | -0015 | 3 |
| 5781 | ENSMUSG( | Rabggta | 56187 | 003951199 | -00497618 | -003 | 005 | 3 |
| 5786 | ENSMUSG( | Rabl6 | 227624 | 020065305 | 005714746 | -00925 | -007 | 3 |
| 5787 | ENSMUSG( | Rac1 | 19353 | 001220982 | -01305983 | -01075 | -00075 | 3 |
| 5792 | ENSMUSG( | Rad23a | 19358 | 008921936 | -00018655 | 01125 | -00475 | 3 |
| 5800 | ENSMUSG( | Rala | 56044 | 007827875 | -01200019 | -00275 | 0245 | 3 |
| 5801 | ENSMUSG( | Ralb | 64143 | -00500643 | 002500905 | -0135 | 00375 | 3 |
| 5803 | ENSMUSG( | Ralgapa1 | 56784 | 004596921 | 009351081 | -001 | 014 | 3 |
| 5813 | ENSMUSG( | Ranbp10 | 74334 | 008306332 | 010383715 | 02025 | 0065 | 3 |
| 5827 | ENSMUSG( | Rap2c | 72065 | 004376844 | 005761902 | 00525 | 024 | 3 |
| 5835 | ENSMUSG( | Raph1 | 77300 | 002795474 | 008556545 | -00475 | 0075 | 3 |
| 5844 | ENSMUSG( | Rasgef1a | 70727 | 011030026 | 00106507C | 00725 | -00575 | 3 |
| 5851 | ENSMUSG( | Rassf2 | 215653 | 010972196 | 000722012 | -00475 | 00425 | 3 |
| 5854 | ENSMUSG( | Rb1cc1 | 12421 | 011007247 | 00457236C | -01175 | 00525 | 3 |
| 5858 | ENSMUSG( | Rbbp7 | 245688 | 013269834 | -00484400 | -0055 | 01675 | 3 |
| 5860 | ENSMUSG( | Rbck1 | 24105 | 005465558 | 001654834 | 01075 | 013 | 3 |
| 5872 | ENSMUSG( | Rbm15b | 109095 | -00295151 | 005187096 | 0115 | -00525 | 3 |
| 5873 | ENSMUSG( | Rbm17 | 76938 | 001316085 | -00284980 | 0005 | 00075 | 3 |
| 5893 | ENSMUSG( | Rbsn | 78287 | 015348413 | -00528422 | -0115 | 00625 | 3 |
| 5897 | ENSMUSG( | Rcan1 | 54720 | 004344475 | -01342418 | -012 | 01 | 3 |
| 5908 | ENSMUSG( | Rcor1 | 217864 | 008974929 | 004510985 | 00575 | 002 | 3 |
| 5921 | ENSMUSG( | Rela | 19697 | -00443620 | -00266022 | 00375 | 003 | 3 |
| 5922 | ENSMUSG( | Relch | 227446 | 003896622 | 009911567 | 0115 | 008 | 3 |
| 5949 | ENSMUSG( | Rgl1 | 19731 | 010499007 | -00334831 | 0095 | -005 | 3 |
| 5954 | ENSMUSG( | Rgs10 | 67865 | 015386309 | -00753436 | -0055 | 01775 | 3 |
| 5955 | ENSMUSG( | Rgs12 | 71729 | 004072495 | 000167447 | -00325 | -00375 | 3 |
| 5958 | ENSMUSG( | Rgs19 | 56470 | 004424187 | -01051443 | -02 | 018 | 3 |
| 5970 | ENSMUSG( | Rheb | 19744 | 005382901 | -01202891 | 01625 | 007 | 3 |
| 5972 | ENSMUSG( | Rhob | 11852 | 000255643 | -01055519 | 01825 | 01725 | 3 |
| 5979 | ENSMUSG( | Rhot2 | 214952 | 005716057 | -00798277 | 00775 | 01625 | 3 |
| 5986 | ENSMUSG( | Rictor | 78757 | -01017460 | 015001296 | -0035 | 018 | 3 |
| 5993 | ENSMUSG( | Rims2 | 116838 | -00046261 | -01038064 | -00875 | 00375 | 3 |
| 6000 | ENSMUSG( | Riox1 | 71952 | -00511512 | -01915716 | 0105 | 0005 | 3 |
| 6019 | ENSMUSG( | Rnasel | 24014 | -00224820 | 005778436 | 01425 | 00175 | 3 |
| 6027 | ENSMUSG( | Rnf114 | 81018 | 009754199 | -00147039 | -0135 | 0155 | 3 |
| 6030 | ENSMUSG( | Rnf123 | 84585 | 008190125 | 00866311C | -01 | -0045 | 3 |
| 6039 | ENSMUSG( | Rnf157 | 217340 | 013612997 | 004722006 | 0005 | -0005 | 3 |
| 6047 | ENSMUSG( | Rnf214 | 235315 | 008062892 | -00293604 | -00275 | -006 | 3 |
| 6050 | ENSMUSG( | Rnf25 | 57751 | 007246884 | -00836332 | -0175 | -0005 | 3 |
| 6052 | ENSMUSG( | Rnf31 | 268749 | 004832266 | 002066838 | 0055 | 008 | 3 |
| 6060 | ENSMUSG( | Rnh1 | 107702 | -00063212 | -01251968 | -00825 | 01125 | 3 |
| 6071 | ENSMUSG( | Rp2 | 19889 | 00760964C | -01914526 | -0005 | 00225 | 3 |
| 6074 | ENSMUSG( | Rpa3 | 68240 | -01125723 | 001646535 | -00275 | 01 | 3 |
| 6079 | ENSMUSG( | Rpgr | 19893 | 002248559 | -00172845 | 00025 | 008 | 3 |
| 6082 | ENSMUSG( | Rpia | 19895 | -01354147 | -01688238 | 0005 | 00325 | 3 |
| 6085 | ENSMUSG( | Rpl11 | 67025 | 009009524 | -01251523 | 02025 | 014 | 3 |
| 6100 | ENSMUSG( | Rpl24 | 68193 | -00606955 | -01155048 | 009 | 025 | 3 |
| 6105 | ENSMUSG( | Rpl29 | 19944 | 008092739 | -01410056 | 013 | 0115 | 3 |
| 6120 | ENSMUSG( | Rpl5 | 1.01E+08 | 005761755 | -01641988 | 0195 | 01125 | 3 |
| 6124 | ENSMUSG( | Rpl8 | 26961 | 00763130C | -01270510 | 0195 | 0215 | 3 |
| 6131 | ENSMUSG( | Rpp14 | 67053 | 022758791 | 002753117 | -0225 | 00575 | 3 |
| 6134 | ENSMUSG( | Rprd1a | 225283 | 007415444 | -00297638 | -02025 | 012 | 3 |
| 6150 | ENSMUSG( | Rps21 | 66481 | 021712007 | -00135498 | 002 | 02575 | 3 |
| 6165 | ENSMUSG( | Rps6ka3 | 110651 | 009137322 | 00266076C | 00625 | 0225 | 3 |
| 6169 | ENSMUSG( | Rps6kc1 | 320119 | 027581275 | -00411640 | -00775 | 004 | 3 |
| 6170 | ENSMUSG( | Rps7 | 20115 | 008597645 | -01064591 | 022 | 0025 | 3 |
| 6174 | ENSMUSG( | Rptor | 74370 | 003578212 | -00060900 | -012 | -00675 | 3 |
| 6178 | ENSMUSG( | Rragb | 245670 | 012290901 | 005860993 | -00275 | 00525 | 3 |
| 6179 | ENSMUSG( | Rragc | 54170 | 00514624C | -00021339 | 00175 | 00075 | 3 |
| 6184 | ENSMUSG( | Rrm1 | 20133 | 006545287 | -00281206 | 002 | 01325 | 3 |
| 6188 | ENSMUSG( | Rrp15 | 67223 | 005839801 | -00737152 | -004 | -01125 | 3 |
| 6194 | ENSMUSG( | Rspry1 | 67610 | 001979608 | 006685273 | 00775 | 01175 | 3 |
| 6197 | ENSMUSG( | Rsu1 | 20163 | 00030234C | -00949873 | 022 | 00425 | 3 |
| 6202 | ENSMUSG( | Rtl8b | 553127 | -00203060 | -02325770 | 00425 | 02875 | 3 |
| 6216 | ENSMUSG( | Rundc1 | 217201 | -0193617 | -00511673 | -005 | -00425 | 3 |
| 6224 | ENSMUSG( | Rwdd1 | 66521 | 009141787 | 001239903 | -0025 | 0085 | 3 |
| 6230 | ENSMUSG( | S100a10 | 20194 | 019372602 | -02791128 | -0065 | -01075 | 3 |
| 6233 | ENSMUSG( | S100a16 | 67860 | -00494256 | -04157517 | -00325 | -01025 | 3 |
| 6235 | ENSMUSG( | S100a6 | 20200 | -01071135 | -02137311 | -01275 | 01025 | 3 |
| 6254 | ENSMUSG( | Samhd1 | 56045 | -01497652 | -00752663 | -01675 | 0135 | 3 |
| 6259 | ENSMUSG( | Sap30bp | 57230 | 010738821 | -00409723 | 01475 | 02475 | 3 |
| 6270 | ENSMUSG( | Sash1 | 70097 | -00375421 | -06171139 | -00325 | 00675 | 3 |
| 6275 | ENSMUSG( | Sbds | 66711 | -00130672 | -01753147 | 00175 | -00975 | 3 |
| 6277 | ENSMUSG( | Sbf2 | 319934 | 007875811 | -00285063 | -01275 | 00675 | 3 |
| 6278 | ENSMUSG( | Sbno1 | 243272 | 010196638 | -00557084 | 00425 | 017 | 3 |
| 6290 | ENSMUSG( | Scaper | 244891 | 004015631 | -01125918 | -01025 | 0175 | 3 |
| 6293 | ENSMUSG( | Sccpdh | 109232 | 001323988 | 000284591 | 01675 | 01225 | 3 |
| 6298 | ENSMUSG( | Scg3 | 20255 | -00282361 | 000616237 | -00525 | 02375 | 3 |
| 6311 | ENSMUSG( | Sco1 | 52892 | -00189888 | 002228576 | -01125 | 022 | 3 |
| 6317 | ENSMUSG( | Scrib | 105782 | 02058347C | -00669958 | -00225 | 0185 | 3 |
| 6318 | ENSMUSG( | Scrn1 | 69938 | -00810487 | -01623233 | -01425 | 0055 | 3 |
| 6339 | ENSMUSG( | Sdhaf4 | 68002 | -00477594 | -02294025 | -02 | 0055 | 3 |
| 6349 | ENSMUSG( | Sec13 | 110379 | 004855857 | -01136674 | 006 | -00425 | 3 |
| 6352 | ENSMUSG( | Sec16a | 227648 | 004001076 | 003940274 | -0035 | 02 | 3 |
| 6353 | ENSMUSG( | Sec22b | 20333 | 009583893 | -02042964 | 01025 | -00225 | 3 |
| 6354 | ENSMUSG( | Sec23a | 20334 | 009882988 | 000451379 | 002 | 0195 | 3 |
| 6356 | ENSMUSG( | Sec23ip | 207352 | 019034105 | 005605463 | -0025 | 0155 | 3 |
| 6357 | ENSMUSG( | Sec24a | 77371 | 002244713 | 005022448 | 001 | 0125 | 3 |
| 6393 | ENSMUSG( | Sema6d | 214968 | -01659288 | -00051351 | -019 | 00375 | 3 |
| 6403 | ENSMUSG( | Septin11 | 52398 | 011948459 | -00818472 | -01125 | -00775 | 3 |
| 6415 | ENSMUSG( | Sergef | 27414 | 006342666 | -01079258 | 006 | -0015 | 3 |
| 6433 | ENSMUSG( | Sestd1 | 228071 | 009419013 | -00099440 | 0045 | 02375 | 3 |
| 6458 | ENSMUSG( | Sfxn1 | 14057 | 007869046 | -00192821 | -01425 | 0045 | 3 |
| 6471 | ENSMUSG( | Sgsm1 | 52850 | 023171923 | -00460045 | -0175 | 01925 | 3 |
| 6475 | ENSMUSG( | Sgtb | 218544 | 008629195 | 000721449 | -00125 | 01025 | 3 |
| 6484 | ENSMUSG( | Sh3bp4 | 98402 | 008438373 | 00597341C | 0025 | 0175 | 3 |
| 6491 | ENSMUSG( | Sh3glb1 | 54673 | -00284944 | -00839119 | 0095 | 02225 | 3 |
| 6494 | ENSMUSG( | Sh3pxd2a | 14218 | 003392115 | -01058356 | -02025 | 00975 | 3 |
| 6503 | ENSMUSG( | Sharpin | 106025 | -00236139 | -00295858 | -01625 | 0035 | 3 |
| 6508 | ENSMUSG( | Shf | 435684 | 012461999 | -00153572 | -0065 | 02075 | 3 |
| 6517 | ENSMUSG( | Shpk | 74637 | -00999116 | -01068443 | 01575 | 017 | 3 |
| 6540 | ENSMUSG( | Sirt3 | 64384 | 007892865 | -01362068 | 0115 | 00225 | 3 |
| 6578 | ENSMUSG( | Slc1a6 | 20513 | -00695496 | 011171839 | -011 | 00525 | 3 |

| | | | | | | | | | |
|---|---|---|---|---|---|---|---|---|---|
| 6596 | ENSMUSG( | Slc25a16 | 73132 | 000738474 | 004328168 | 01275 | -00175 | | 3 |
| 6600 | ENSMUSG( | Slc25a20 | 57279 | -00241433 | -03418567 | -0105 | 01225 | | 3 |
| 6622 | ENSMUSG( | Slc25a53 | 67062 | -00816927 | 000170577 | 011 | 0155 | | 3 |
| 6735 | ENSMUSG( | Slmap | 83997 | -00340477 | 009397323 | -0085 | 02375 | | 3 |
| 6739 | ENSMUSG( | Smad3 | 17127 | 010443603 | 005895408 | 007 | 01475 | | 3 |
| 6740 | ENSMUSG( | Smad4 | 17128 | 005501037 | 009479144 | -0155 | 0085 | | 3 |
| 6742 | ENSMUSG( | Smap1 | 98366 | -00394513 | -00784540 | 00925 | 00375 | | 3 |
| 6746 | ENSMUSG( | Smarca4 | 20586 | 017840848 | -00742936 | -01775 | -0035 | | 3 |
| 6749 | ENSMUSG( | Smarcb1 | 20587 | 004310090 | -01487741 | -0035 | -00575 | | 3 |
| 6750 | ENSMUSG( | Smarcc1 | 20588 | 004393860 | -01873787 | 0005 | 0085 | | 3 |
| 6755 | ENSMUSG( | Smc1a | 24061 | -00394885 | 002577275 | -017 | 001 | | 3 |
| 6765 | ENSMUSG( | Smg7 | 226517 | 018862813 | 006281482 | -00225 | -006 | | 3 |
| 6771 | ENSMUSG( | Smim20 | 66278 | 000632093 | -03335267 | 014 | 0035 | | 3 |
| 6774 | ENSMUSG( | Smim8 | 66291 | 000066205 | -02483127 | 00825 | -00025 | | 3 |
| 6784 | ENSMUSG( | Sms | 20603 | 007725802 | -00334480 | -003 | 00325 | | 3 |
| 6787 | ENSMUSG( | Smurf2 | 66313 | -00461734 | -01156985 | -01425 | 004 | | 3 |
| 6790 | ENSMUSG( | Smyd5 | 232187 | 006787726 | -02137705 | -00175 | 0105 | | 3 |
| 6794 | ENSMUSG( | Snap47 | 67826 | 018785469 | -01992611 | 013 | 02425 | | 3 |
| 6796 | ENSMUSG( | Snapin | 20615 | -00319120 | -02684941 | 00075 | 004 | | 3 |
| 6801 | ENSMUSG( | Snf8 | 27681 | -00120863 | -01757586 | 0115 | 001 | | 3 |
| 6805 | ENSMUSG( | Snrk | 20623 | 022561915 | 000559924 | -0075 | 012 | | 3 |
| 6807 | ENSMUSG( | Snrnp27 | 66618 | 007688954 | -00542086 | 00825 | 01225 | | 3 |
| 6814 | ENSMUSG( | Snrpc | 20630 | -00273306 | -01788853 | -00025 | 01875 | | 3 |
| 6821 | ENSMUSG( | Snta1 | 20648 | 010884335 | -00678154 | 00225 | 00375 | | 3 |
| 6825 | ENSMUSG( | Snupn | 66069 | 005504266 | 002466167 | 0115 | 022 | | 3 |
| 6836 | ENSMUSG( | Snx18 | 170625 | -00935582 | -00472489 | -00025 | 02475 | | 3 |
| 6842 | ENSMUSG( | Snx27 | 76742 | 010031132 | -01184958 | -0005 | 00675 | | 3 |
| 6845 | ENSMUSG( | Snx30 | 209131 | -00514944 | -00728025 | -0015 | -01075 | | 3 |
| 6850 | ENSMUSG( | Snx6 | 72183 | 013623111 | -00322247 | -00925 | 014 | | 3 |
| 6859 | ENSMUSG( | Sorbs1 | 20411 | 024089677 | -01212384 | -01175 | 0035 | | 3 |
| 6861 | ENSMUSG( | Sorbs3 | 20410 | -00314553 | 002459218 | 006 | 02125 | | 3 |
| 6868 | ENSMUSG( | Sos1 | 20662 | -00303428 | 006029166 | -00325 | -011 | | 3 |
| 6878 | ENSMUSG( | Spag1 | 26942 | 015197129 | 003713897 | 00475 | 00925 | | 3 |
| 6882 | ENSMUSG( | Sparcl1 | 13602 | 008201864 | 003604535 | 00125 | -0055 | | 3 |
| 6887 | ENSMUSG( | Spata2l | 78779 | 004876337 | -01305124 | -00875 | 02575 | | 3 |
| 6889 | ENSMUSG( | Spats2l | 67198 | 007791787 | -00934933 | -005 | 02725 | | 3 |
| 6893 | ENSMUSG( | Specc1 | 432572 | 002927723 | -01967755 | 0085 | 016 | | 3 |
| 6899 | ENSMUSG( | Spg7 | 234847 | 005526581 | 001247878 | -0025 | -00125 | | 3 |
| 6900 | ENSMUSG( | Sphk2 | 56632 | 007808470 | -02248293 | 01225 | 0015 | | 3 |
| 6904 | ENSMUSG( | Spire2 | 234857 | 007277930 | 008355882 | -00025 | 0175 | | 3 |
| 6914 | ENSMUSG( | Spr | 20751 | 010233123 | -01806677 | 003 | -007 | | 3 |
| 6936 | ENSMUSG( | Srcin1 | 56013 | 010256279 | -00246343 | -019 | 00475 | | 3 |
| 6940 | ENSMUSG( | Srgap1 | 117600 | -01713309 | -00871297 | -0055 | 02 | | 3 |
| 6946 | ENSMUSG( | Srp19 | 66384 | 004516355 | -00085826 | 001 | 02525 | | 3 |
| 6948 | ENSMUSG( | Srp72 | 66661 | 009474471 | 010519752 | -0045 | 01575 | | 3 |
| 6951 | ENSMUSG( | Srpk2 | 20817 | 002661361 | -00375459 | -00025 | 02 | | 3 |
| 6954 | ENSMUSG( | Srr | 27364 | 006702887 | -01381181 | -002 | -0085 | | 3 |
| 6955 | ENSMUSG( | Srrm1 | 51796 | 005930613 | 006942842 | -009 | -00625 | | 3 |
| 6990 | ENSMUSG( | St13 | 70356 | 006179012 | -00790156 | 002 | -00175 | | 3 |
| 6997 | ENSMUSG( | Stac2 | 217154 | 008055456 | -01452101 | -0075 | -006 | | 3 |
| 6999 | ENSMUSG( | Stag2 | 20843 | 005208444 | 000783510 | -00525 | 0055 | | 3 |
| 7000 | ENSMUSG( | Stam | 20844 | 004997121 | 001941980 | 00075 | -00825 | | 3 |
| 7002 | ENSMUSG( | Stambp | 70527 | -00254120 | -01180869 | 0085 | 018 | | 3 |
| 7009 | ENSMUSG( | Stat1 | 20846 | -01649005 | -01612192 | -0075 | -00275 | | 3 |
| 7011 | ENSMUSG( | Stat3 | 20848 | 009426859 | -00340053 | 01125 | -00425 | | 3 |
| 7023 | ENSMUSG( | Stk11 | 20869 | 009753943 | -00610121 | -01575 | 00875 | | 3 |
| 7027 | ENSMUSG( | Stk25 | 59041 | 017712239 | 003178433 | -018 | 014 | | 3 |
| 7036 | ENSMUSG( | Stmn2 | 20257 | -00330371 | -01409992 | -981308e-1 | 008 | | 3 |
| 7045 | ENSMUSG( | Stradb | 227154 | 003919295 | 004467423 | -01525 | 022 | | 3 |
| 7055 | ENSMUSG( | Stub1 | 56424 | -00030856 | -01190575 | 00475 | 021 | | 3 |
| 7060 | ENSMUSG( | Stx18 | 71116 | 012854457 | 001955928 | 0125 | 01475 | | 3 |
| 7063 | ENSMUSG( | Stx2 | 13852 | -00345517 | -00713313 | 00125 | 0255 | | 3 |
| 7065 | ENSMUSG( | Stx6 | 58244 | 002098269 | -01348198 | 0 | -001 | 3 | |
| 7067 | ENSMUSG( | Stx8 | 55943 | -00094431 | -01831577 | 01375 | 0145 | | 3 |
| 7071 | ENSMUSG( | Stxbp4 | 20913 | -00323234 | -01833716 | -01175 | -0015 | | 3 |
| 7084 | ENSMUSG( | Sugt1 | 67955 | 004500750 | -00234068 | 008 | 009 | | 3 |
| 7094 | ENSMUSG( | Supt16 | 114741 | 014069458 | -00803224 | 006 | 02 | | 3 |
| 7106 | ENSMUSG( | Svbp | 69216 | 034547207 | -01972920 | 00175 | 0165 | | 3 |
| 7110 | ENSMUSG( | Swap70 | 20947 | 015658001 | -00892080 | -02125 | 0105 | | 3 |
| 7120 | ENSMUSG( | Syncrip | 56403 | 000772494 | 001263004 | -012 | 01525 | | 3 |
| 7135 | ENSMUSG( | Synrg | 217030 | -00007707 | -01131232 | -024 | 00075 | | 3 |
| 7154 | ENSMUSG( | Sytl5 | 236643 | -00510601 | -03326598 | -005 | -00425 | | 3 |
| 7158 | ENSMUSG( | Tab1 | 66513 | -00712227 | -00676299 | 018 | 004 | | 3 |
| 7160 | ENSMUSG( | Tab3 | 66724 | 006177920 | 005890894 | 02125 | 00675 | | 3 |
| 7163 | ENSMUSG( | Taco1 | 70207 | 023597372 | -00237645 | 004 | -007 | | 3 |
| 7171 | ENSMUSG( | Taf5 | 226182 | -01955295 | 009693768 | -0175 | 0055 | | 3 |
| 7186 | ENSMUSG( | Tanc1 | 66860 | 023252454 | 004370704 | -00125 | 02025 | | 3 |
| 7187 | ENSMUSG( | Tanc2 | 77097 | 003982887 | 004733162 | 00475 | 0235 | | 3 |
| 7188 | ENSMUSG( | Tango2 | 27883 | 008905415 | -00196677 | 01025 | 009 | | 3 |
| 7191 | ENSMUSG( | Taok1 | 216965 | 000724451 | -00386926 | 01775 | 00225 | | 3 |
| 7192 | ENSMUSG( | Taok2 | 381921 | 009640541 | 003584470 | 0005 | 00325 | | 3 |
| 7199 | ENSMUSG( | Tars1 | 110960 | 009224697 | 006639426 | 00775 | -001 | | 3 |
| 7204 | ENSMUSG( | Tatdn1 | 69694 | -00139440 | 002703846 | 01225 | -003 | | 3 |
| 7205 | ENSMUSG( | Tax1bp1 | 52440 | 000352055 | -00488358 | -00075 | -00325 | | 3 |
| 7212 | ENSMUSG( | Tbc1d14 | 100855 | 010168133 | 005623459 | -0005 | -00925 | | 3 |
| 7213 | ENSMUSG( | Tbc1d15 | 66687 | 009925452 | 009346415 | -00225 | 006 | | 3 |
| 7216 | ENSMUSG( | Tbc1d20 | 67231 | 007270313 | 002611876 | 0015 | 01025 | | 3 |
| 7217 | ENSMUSG( | Tbc1d22a | 223754 | -00115890 | -01445563 | -00725 | -00425 | | 3 |
| 7221 | ENSMUSG( | Tbc1d25 | 209815 | -00073695 | -01748569 | 01625 | -00125 | | 3 |
| 7225 | ENSMUSG( | Tbc1d5 | 72238 | 009840511 | -00786680 | -00925 | 02475 | | 3 |
| 7231 | ENSMUSG( | Tbca | 21371 | -00348611 | -01642119 | -0065 | 00525 | | 3 |
| 7234 | ENSMUSG( | Tbcd | 108903 | 009816469 | 003442718 | 011 | 00075 | | 3 |
| 7235 | ENSMUSG( | Tbce | 70430 | 014237359 | 005909694 | -0155 | 0075 | | 3 |
| 7236 | ENSMUSG( | Tbcel | 272589 | 015514123 | -00375399 | 00825 | 004 | | 3 |
| 7243 | ENSMUSG( | Tbl3 | 213773 | 006696305 | -00775908 | 001 | 01425 | | 3 |
| 7246 | ENSMUSG( | Tbrg4 | 21379 | 011328068 | -00512489 | 001 | 00925 | | 3 |
| 7247 | ENSMUSG( | Tcaf1 | 77574 | -00448914 | -00490014 | 196262e-1 | 024 | | 3 |
| 7248 | ENSMUSG( | Tcaim | 382117 | 010938111 | -00037733 | 0005 | -001 | | 3 |
| 7251 | ENSMUSG( | Tceal3 | 594844 | 000048571 | -02510250 | 00225 | 021 | | 3 |
| 7267 | ENSMUSG( | Tdrd3 | 219249 | 010017226 | -00160162 | -00825 | 00675 | | 3 |
| 7269 | ENSMUSG( | Tdrkh | 72634 | 013173321 | -00939967 | 01275 | 00475 | | 3 |
| 7271 | ENSMUSG( | Tecpr1 | 70381 | 012994405 | 006612029 | 0025 | -00075 | | 3 |
| 7272 | ENSMUSG( | Tecpr2 | 104859 | -00484425 | 008949567 | -0025 | 0015 | | 3 |
| 7276 | ENSMUSG( | Telo2 | 71718 | 026117156 | -00044863 | -0025 | 00525 | | 3 |
| 7299 | ENSMUSG( | Tfip11 | 54723 | 013001638 | -00073522 | -01975 | -00075 | | 3 |
| 7303 | ENSMUSG( | Tgfbrap1 | 73122 | 009395070 | 000250161 | -005 | -0015 | | 3 |
| 7313 | ENSMUSG( | Them4 | 75778 | -00110037 | -03555503 | 00025 | -00175 | | 3 |
| 7324 | ENSMUSG( | Thop1 | 50492 | 008082001 | -01508612 | -0035 | -005 | | 3 |
| 7325 | ENSMUSG( | Thra | 21833 | 021029381 | -00024435 | 0095 | -00175 | | 3 |
| 7329 | ENSMUSG( | Thtpa | 105663 | 020358678 | -00165448 | -01875 | 01325 | | 3 |
| 7338 | ENSMUSG( | Tigar | 319801 | 007328129 | -00697645 | 00625 | 0165 | | 3 |
| 7340 | ENSMUSG( | Timm10b | 14356 | -00292854 | 001928716 | 006 | -00025 | | 3 |
| 7342 | ENSMUSG( | Timm17a | 21854 | 004998219 | 000065660 | 01775 | 0195 | | 3 |
| 7345 | ENSMUSG( | Timm22 | 56322 | -00458125 | -00544344 | -0055 | 0245 | | 3 |
| 7346 | ENSMUSG( | Timm23 | 53600 | -00194241 | -01681239 | -00725 | 02125 | | 3 |
| 7349 | ENSMUSG( | Timm50 | 66525 | -00988644 | -01563395 | -00975 | 025 | | 3 |
| 7356 | ENSMUSG( | Tiprl | 226591 | 005824483 | -00668621 | -588785e-1 | 018 | | 3 |
| 7359 | ENSMUSG( | Tk2 | 57813 | 009389187 | -00633145 | 00925 | -0035 | | 3 |
| 7360 | ENSMUSG( | Tkfc | 225913 | 008256347 | -00823942 | 016 | 008 | | 3 |
| 7363 | ENSMUSG( | Tlcd3b | 68952 | 009163461 | -00564313 | -004 | 00425 | | 3 |
| 7367 | ENSMUSG( | Tle4 | 21888 | -00680717 | -00480600 | 00875 | 013 | | 3 |
| 7383 | ENSMUSG( | Tmcc3 | 319880 | -00587398 | -00906487 | -0195 | 0155 | | 3 |
| 7404 | ENSMUSG( | Tmem126a | 66271 | -00378298 | -01275009 | 014 | 024 | | 3 |
| 7415 | ENSMUSG( | Tmem141 | 51875 | 014361304 | -01198845 | -00175 | -00425 | | 3 |
| 7439 | ENSMUSG( | Tmem198 | 319998 | -01218343 | 003925876 | 01 | 0075 | | 3 |
| 7464 | ENSMUSG( | Tmem266 | 244886 | -00629228 | 000749757 | -0115 | -00225 | | 3 |

| | | | | | | | | |
|---|---|---|---|---|---|---|---|---|
| 7491 | ENSMUSG( | Tmf1 | 232286 | 011046844 | 006384057 | 002 | -005 | 3 |
| 7501 | ENSMUSG( | Tmub1 | 64295 | -02082748 | 010148857 | -0135 | 01125 | 3 |
| 7509 | ENSMUSG( | Tnfaip8 | 106869 | -00956360 | -02528409 | -001 | 014 | 3 |
| 7513 | ENSMUSG( | Tnip1 | 57783 | 005854663 | -01144499 | -00675 | 00825 | 3 |
| 7526 | ENSMUSG( | Tns3 | 319939 | 021138772 | -00637056 | -007 | 00725 | 3 |
| 7529 | ENSMUSG( | Tollip | 54473 | 011217813 | -01144751 | 001 | -006 | 3 |
| 7530 | ENSMUSG( | Tom1 | 21968 | 002375922 | -01955308 | -00375 | 0215 | 3 |
| 7532 | ENSMUSG( | Tom1l2 | 216810 | 013056705 | -01096222 | -01375 | -00075 | 3 |
| 7534 | ENSMUSG( | Tomm22 | 223696 | -00065947 | -01555833 | -00775 | 02525 | 3 |
| 7535 | ENSMUSG( | Tomm34 | 67145 | 006871420 | -01208609 | 01275 | 01825 | 3 |
| 7571 | ENSMUSG( | Tprg1l | 67808 | 008717503 | -01401134 | 01125 | 00475 | 3 |
| 7572 | ENSMUSG( | Tprkb | 69786 | 008275903 | -04629446 | -02125 | -196262e-1 | 3 |
| 7583 | ENSMUSG( | Trafd1 | 231712 | 005746623 | 009180031 | 00225 | 01225 | 3 |
| 7589 | ENSMUSG( | Trappc1 | 245828 | -01301894 | -03383223 | 002 | -00225 | 3 |
| 7592 | ENSMUSG( | Trappc12 | 217449 | 010902179 | -01016179 | -005 | 014 | 3 |
| 7594 | ENSMUSG( | Trappc14 | 231807 | 011228166 | 005385051 | -01375 | 01925 | 3 |
| 7596 | ENSMUSG( | Trappc2l | 59005 | -00894810 | -01933079 | 004 | 02075 | 3 |
| 7597 | ENSMUSG( | Trappc3 | 27096 | -00279270 | -00613381 | -01825 | 014 | 3 |
| 7599 | ENSMUSG( | Trappc5 | 66682 | -01541976 | -00805613 | 012 | 0095 | 3 |
| 7601 | ENSMUSG( | Trappc8 | 75964 | 012841099 | 004284565 | -0115 | -004 | 3 |
| 7602 | ENSMUSG( | Trappc9 | 76510 | 008232145 | -01566028 | 00725 | 005 | 3 |
| 7608 | ENSMUSG( | Trim2 | 80890 | 005247444 | -00758946 | 00325 | 00125 | 3 |
| 7613 | ENSMUSG( | Trim28 | 21849 | 012299779 | -00227413 | 0035 | -006 | 3 |
| 7614 | ENSMUSG( | Trim3 | 55992 | 000972039 | -00684669 | 0085 | 015 | 3 |
| 7636 | ENSMUSG( | Trmt10c | 52575 | -01692700 | 000866162 | -004 | 01325 | 3 |
| 7637 | ENSMUSG( | Trmt11 | 73681 | -00847135 | -01458147 | -01575 | 01725 | 3 |
| 7646 | ENSMUSG( | Trp53bp2 | 209456 | 024299050 | -01103999 | -0005 | 0135 | 3 |
| 7659 | ENSMUSG( | Tsc2 | 22084 | 014582825 | 002588705 | -01375 | 00625 | 3 |
| 7666 | ENSMUSG( | Tsg101 | 22088 | 006665678 | -00557329 | -00325 | 0255 | 3 |
| 7681 | ENSMUSG( | Tspan7 | 21912 | -01742294 | -01374689 | 00675 | 007 | 3 |
| 7683 | ENSMUSG( | Tspan9 | 109246 | -02864848 | -02916763 | 011 | -0015 | 3 |
| 7686 | ENSMUSG( | Tspyl4 | 72480 | 018407219 | -01099560 | -0225 | 0015 | 3 |
| 7687 | ENSMUSG( | Tspyl5 | 239364 | 022060656 | -00892632 | -0095 | 02575 | 3 |
| 7688 | ENSMUSG( | Tsr2 | 69499 | 002662828 | -01036636 | -00525 | 00325 | 3 |
| 7691 | ENSMUSG( | Tstd3 | 77032 | -01690371 | -03490417 | -00825 | 00625 | 3 |
| 7694 | ENSMUSG( | Ttc1 | 66827 | 003046038 | -02254393 | -00575 | 02425 | 3 |
| 7700 | ENSMUSG( | Ttc3 | 22129 | 005973886 | 007282623 | -00425 | 02075 | 3 |
| 7712 | ENSMUSG( | Ttl | 69737 | 001414968 | -02025939 | -00325 | -00125 | 3 |
| 7720 | ENSMUSG( | Ttyh1 | 57776 | 003395495 | -00245180 | 0065 | 00025 | 3 |
| 7724 | ENSMUSG( | Tuba1a | 22142 | -00336820 | -02195058 | 01775 | 01625 | 3 |
| 7726 | ENSMUSG( | Tuba4a | 22145 | 006422404 | -02757941 | -0155 | 0205 | 3 |
| 7729 | ENSMUSG( | Tubb2b | 73710 | 002221018 | -03742777 | 01675 | 0085 | 3 |
| 7730 | ENSMUSG( | Tubb3 | 22152 | -00871781 | -02867875 | 01475 | 02275 | 3 |
| 7732 | ENSMUSG( | Tubb4b | 227613 | 002691380 | -02305059 | -00025 | 0175 | 3 |
| 7733 | ENSMUSG( | Tubb5 | 22154 | 002998168 | -01436121 | -00775 | 01425 | 3 |
| 7736 | ENSMUSG( | Tubg2 | 103768 | 006670713 | -00637065 | -01025 | 01625 | 3 |
| 7744 | ENSMUSG( | Tulp4 | 68842 | -01874222 | -00903267 | -00775 | -0045 | 3 |
| 7765 | ENSMUSG( | Txnrd1 | 50493 | 006752546 | -02357323 | -0005 | 022 | 3 |
| 7766 | ENSMUSG( | Txnrd2 | 26462 | 005163698 | -02609390 | -0015 | -0085 | 3 |
| 7773 | ENSMUSG( | U2af1 | 108121 | -00161687 | -00500908 | -014 | 017 | 3 |
| 7778 | ENSMUSG( | Uap1l1 | 227620 | 013887699 | -00044905 | -00875 | -00375 | 3 |
| 7779 | ENSMUSG( | Uba1 | 22201 | 009550242 | -01032363 | -01675 | -00425 | 3 |
| 7782 | ENSMUSG( | Uba5 | 66663 | 007946247 | -01773853 | 00975 | 01275 | 3 |
| 7791 | ENSMUSG( | Ube2a | 22209 | 004174396 | -00607762 | -00375 | -0075 | 3 |
| 7792 | ENSMUSG( | Ube2b | 22210 | 002277663 | -00063091 | 01275 | 02375 | 3 |
| 7798 | ENSMUSG( | Ube2g1 | 67128 | -00394825 | 005717749 | -008 | 015 | 3 |
| 7800 | ENSMUSG( | Ube2h | 22214 | 003419217 | -00757809 | 00825 | 0155 | 3 |
| 7804 | ENSMUSG( | Ube2k | 53323 | -00404013 | -00940061 | -00075 | 01375 | 3 |
| 7805 | ENSMUSG( | Ube2l3 | 22195 | 005463850 | -02290874 | 00475 | -0055 | 3 |
| 7808 | ENSMUSG( | Ube2o | 217342 | 005583722 | -00317063 | 003 | 01925 | 3 |
| 7809 | ENSMUSG( | Ube2q1 | 70093 | 006433940 | -00919648 | -0015 | 0225 | 3 |
| 7816 | ENSMUSG( | Ube2z | 268470 | 008937886 | -01076241 | 00575 | 0095 | 3 |
| 7821 | ENSMUSG( | Ube4b | 63958 | 003308872 | -00229111 | 0105 | 0055 | 3 |
| 7827 | ENSMUSG( | Ubl7 | 69459 | 006310986 | -01904179 | -0195 | 01775 | 3 |
| 7831 | ENSMUSG( | Ubqln1 | 56085 | 004238921 | -01106792 | -00675 | -0025 | 3 |
| 7833 | ENSMUSG( | Ubqln4 | 94232 | 014162903 | -00645017 | 0035 | -0035 | 3 |
| 7837 | ENSMUSG( | Ubr4 | 69116 | -00976913 | 009781373 | 009 | 007 | 3 |
| 7840 | ENSMUSG( | Ubtd2 | 327900 | 010536476 | 007837833 | 012 | 00975 | 3 |
| 7841 | ENSMUSG( | Ubtf | 21429 | 009561264 | 002820956 | -022 | 00125 | 3 |
| 7846 | ENSMUSG( | Ubxn6 | 66530 | 008443969 | -00741437 | 001 | 0185 | 3 |
| 7848 | ENSMUSG( | Uchl1 | 22223 | -00020190 | -01342907 | 00375 | 02025 | 3 |
| 7849 | ENSMUSG( | Uchl3 | 50933 | 011057469 | 000807857 | 01475 | 0085 | 3 |
| 7852 | ENSMUSG( | Uckl1 | 68556 | 004241988 | 011224153 | 02025 | 001 | 3 |
| 7857 | ENSMUSG( | Ufd1 | 22230 | 009010304 | -00082328 | -01325 | 009 | 3 |
| 7859 | ENSMUSG( | Ufm1 | 67890 | -00118633 | -01452217 | -0035 | 022 | 3 |
| 7867 | ENSMUSG( | Uhmk1 | 16589 | 006212704 | -00651612 | -00925 | 02125 | 3 |
| 7870 | ENSMUSG( | Ulk3 | 71742 | 016799381 | -00255101 | -00975 | -0005 | 3 |
| 7874 | ENSMUSG( | Unc119b | 106840 | 012309455 | -00849474 | -011 | -00725 | 3 |
| 7888 | ENSMUSG( | Upf1 | 19704 | 010943362 | -00186421 | 01625 | 00725 | 3 |
| 7893 | ENSMUSG( | Uqcc1 | 56046 | -00846152 | -01179520 | 00375 | 00175 | 3 |
| 7894 | ENSMUSG( | Uqcc2 | 67267 | -00275402 | -03432311 | 0045 | 00275 | 3 |
| 7898 | ENSMUSG( | Uqcrb | 67530 | -00140587 | -02408645 | -006 | -00275 | 3 |
| 7902 | ENSMUSG( | Uqcrh | 66576 | 004198401 | -02284927 | 0085 | 02375 | 3 |
| 7905 | ENSMUSG( | Urb2 | 382038 | -01605179 | 006994530 | -588785e-1 | 00325 | 3 |
| 7908 | ENSMUSG( | Urm1 | 68205 | 354e-05 | -01966424 | 00825 | -005 | 3 |
| 7909 | ENSMUSG( | Urod | 22275 | 015184669 | -00418172 | 011 | 013 | 3 |
| 7914 | ENSMUSG( | Usf2 | 22282 | 012464114 | -00081466 | 0155 | 024 | 3 |
| 7915 | ENSMUSG( | Uso1 | 56041 | 000022252 | -01027534 | 017 | 01625 | 3 |
| 7916 | ENSMUSG( | Usp10 | 22224 | 015841970 | 007410324 | -0165 | 019 | 3 |
| 7917 | ENSMUSG( | Usp11 | 236733 | 010201743 | -01098818 | -0085 | 013 | 3 |
| 7919 | ENSMUSG( | Usp13 | 72607 | 011765357 | 007610498 | 01825 | 0125 | 3 |
| 7920 | ENSMUSG( | Usp14 | 59025 | -00040403 | -00604968 | -00825 | 0255 | 3 |
| 7925 | ENSMUSG( | Usp24 | 329908 | 001823009 | 002127149 | 01775 | 00275 | 3 |
| 7931 | ENSMUSG( | Usp34 | 17847 | 007779720 | 005936124 | 00475 | 00675 | 3 |
| 7935 | ENSMUSG( | Usp4 | 22258 | 008220860 | -00851420 | 00775 | -00025 | 3 |
| 7939 | ENSMUSG( | Usp47 | 74996 | 019201843 | 000383053 | -0025 | 022 | 3 |
| 7941 | ENSMUSG( | Usp5 | 22225 | 010235156 | -00975634 | -01325 | -00025 | 3 |
| 7946 | ENSMUSG( | Usp8 | 84092 | 008411901 | -01141568 | -015 | 029 | 3 |
| 7947 | ENSMUSG( | Usp9x | 22284 | 017039006 | 002517227 | 0185 | 004 | 3 |
| 7968 | ENSMUSG( | Vapb | 56491 | 002006269 | -01654847 | 01 | 01325 | 3 |
| 7969 | ENSMUSG( | Vars1 | 22321 | 014224901 | 000826555 | 01375 | 02025 | 3 |
| 7979 | ENSMUSG( | Vbp1 | 22327 | -00826640 | 007003395 | -0005 | 00175 | 3 |
| 7983 | ENSMUSG( | Vcp | 269523 | 008756053 | -00646079 | 0015 | 0155 | 3 |
| 7995 | ENSMUSG( | Vipas39 | 104799 | 012780452 | 007173731 | 01 | -00225 | 3 |
| 8012 | ENSMUSG( | Vps26b | 69091 | 012186471 | -00272233 | 00475 | 015 | 3 |
| 8013 | ENSMUSG( | Vps26c | 13185 | 010469278 | -02494736 | 01675 | 02725 | 3 |
| 8014 | ENSMUSG( | Vps28 | 66914 | -00702640 | -02375161 | 007 | 012 | 3 |
| 8018 | ENSMUSG( | Vps35 | 65114 | 005047927 | -00080083 | 01825 | -00175 | 3 |
| 8019 | ENSMUSG( | Vps35l | 71517 | 010349202 | -00809344 | 00775 | -002 | 3 |
| 8022 | ENSMUSG( | Vps37b | 330192 | -01590432 | -02005112 | 00825 | -0065 | 3 |
| 8023 | ENSMUSG( | Vps37c | 107305 | 010314615 | -00920080 | 0035 | 01975 | 3 |
| 8027 | ENSMUSG( | Vps45 | 22365 | 011400937 | -01863784 | 0005 | 013 | 3 |
| 8028 | ENSMUSG( | Vps4a | 116733 | 002197966 | -01831816 | -00025 | -392523e-1 | 3 |
| 8031 | ENSMUSG( | Vps51 | 68505 | 008577346 | -00496402 | 01275 | 00025 | 3 |
| 8032 | ENSMUSG( | Vps52 | 224705 | 016694187 | -01261702 | -02075 | 00025 | 3 |
| 8036 | ENSMUSG( | Vps9d1 | 72325 | 021838601 | 003079779 | -0185 | 01675 | 3 |
| 8046 | ENSMUSG( | Vti1b | 53612 | 000570386 | -02741659 | 01 | 0045 | 3 |
| 8057 | ENSMUSG( | Wars1 | 22375 | 016434627 | -01002604 | 009 | -00425 | 3 |
| 8061 | ENSMUSG( | Wasf3 | 245880 | 006688431 | -01672330 | 00175 | 02875 | 3 |
| 8063 | ENSMUSG( | Washc2 | 28006 | 004872393 | -00949942 | -00875 | 02275 | 3 |
| 8066 | ENSMUSG( | Washc5 | 223593 | 011710833 | -00377004 | -011 | 00225 | 3 |
| 8069 | ENSMUSG( | Wbp2 | 22378 | 001661099 | -01454907 | 00025 | 00875 | 3 |
| 8073 | ENSMUSG( | Wdfy2 | 268752 | -00136356 | 007944398 | -0065 | 02475 | 3 |
| 8076 | ENSMUSG( | Wdr11 | 207425 | 015443752 | 005401066 | 011 | 00125 | 3 |
| 8077 | ENSMUSG( | Wdr12 | 57750 | 010256162 | 000902536 | 00925 | -0005 | 3 |
| 8082 | ENSMUSG( | Wdr20 | 69641 | 002015256 | 007143149 | 0095 | 00575 | 3 |

| | | | | | | | |
|---|---|---|---|---|---|---|---|
| 8088 | ENSMUSG( | Wdr35 | 74682 | 00879072€ | -0114977€ | 002 | 0035 | 3 |
| 8091 | ENSMUSG( | Wdr41 | 218460 | 007698274 | -01506532 | -011 | -00275 | 3 |
| 8097 | ENSMUSG( | Wdr47 | 99512 | 003213761 | -01046275 | 00325 | -007 | 3 |
| 8099 | ENSMUSG( | Wdr5 | 140858 | 00528789€ | -01152700 | 00625 | 02625 | 3 |
| 8101 | ENSMUSG( | Wdr59 | 319481 | -00713265 | -00324986 | -0055 | 0015 | 3 |
| 8103 | ENSMUSG( | Wdr7 | 104082 | 00769703C | -00105057 | 0025 | 00025 | 3 |
| 8104 | ENSMUSG( | Wdr73 | 71968 | 00444289C | -00219452 | -0135 | 01325 | 3 |
| 8105 | ENSMUSG( | Wdr75 | 73674 | 01143049€ | -00135684 | -01825 | 00025 | 3 |
| 8106 | ENSMUSG( | Wdr77 | 70465 | 00187374C | -01293301 | 00025 | 00475 | 3 |
| 8109 | ENSMUSG( | Wdr89 | 72338 | 043022123 | -00999373 | -01725 | 00375 | 3 |
| 8118 | ENSMUSG( | Wipi2 | 74781 | 00967453C | -01100845 | -01375 | 0185 | 3 |
| 8124 | ENSMUSG( | Wrnip1 | 78903 | 001167867 | -01504953 | 0075 | 025 | 3 |
| 8139 | ENSMUSG( | Xpc | 22591 | -00400567 | -00222773 | -01025 | -005 | 3 |
| 8140 | ENSMUSG( | Xpnpep1 | 170750 | 016037491 | 002871273 | -00225 | 012 | 3 |
| 8143 | ENSMUSG( | Xpo4 | 57258 | -00362523 | -01018496 | 012 | 00725 | 3 |
| 8144 | ENSMUSG( | Xpo5 | 72322 | 011089983 | -00556803 | 0145 | 01275 | 3 |
| 8147 | ENSMUSG( | Xpot | 73192 | 008382424 | 005176362 | 003 | 0055 | 3 |
| 8149 | ENSMUSG( | Xrcc1 | 22594 | 007724981 | -00287524 | -01575 | 02325 | 3 |
| 8161 | ENSMUSG( | Yars2 | 70120 | -00353464 | 008720477 | -0075 | 004 | 3 |
| 8188 | ENSMUSG( | Ywhae | 22627 | 002118001 | -01726465 | 01075 | 0015 | 3 |
| 8189 | ENSMUSG( | Ywhag | 22628 | 013854567 | -01712404 | -01375 | 0245 | 3 |
| 8190 | ENSMUSG( | Ywhah | 22629 | 000459991 | -01819195 | -01925 | 01375 | 3 |
| 8192 | ENSMUSG( | Ywhaz | 22631 | 000349853 | -01234843 | -01325 | 01725 | 3 |
| 8203 | ENSMUSG( | Zc2hc1a | 67306 | 00016341€ | -00484970 | 0125 | 0245 | 3 |
| 8208 | ENSMUSG( | Zc3h15 | 69082 | 012753327 | -00751510 | -014 | 0285 | 3 |
| 8213 | ENSMUSG( | Zc3h7b | 20286 | 01662023C | -00439280 | 00825 | 01175 | 3 |
| 8214 | ENSMUSG( | Zc3hc1 | 232679 | 016349061 | -01107473 | -004 | 0065 | 3 |
| 8219 | ENSMUSG( | Zcrb1 | 67197 | -00131220 | 00785613 | 01175 | 01225 | 3 |
| 8232 | ENSMUSG( | Zeb1 | 21417 | -00218687 | -02147096 | 01375 | 003 | 3 |
| 8234 | ENSMUSG( | Zer1 | 227693 | 02264211 | 003115264 | 00775 | 003 | 3 |
| 8255 | ENSMUSG( | Zfpl1 | 81909 | -00634976 | 000714283 | 00675 | 0035 | 3 |
| 8266 | ENSMUSG( | Zfyve9 | 230597 | 004166924 | -00078988 | -00575 | -00375 | 3 |
| 8272 | ENSMUSG( | Zmat2 | 66492 | 01406725€ | -01699013 | -00125 | 0025 | 3 |
| 8283 | ENSMUSG( | Znrf1 | 170737 | 008211633 | 007295925 | -0045 | 0025 | 3 |
| 8285 | ENSMUSG( | Zpr1 | 22687 | 00717973C | 000920294 | -00225 | 006 | 3 |
| 8293 | ENSMUSG( | Zyx | 22793 | 000534583 | -00906388 | 011 | 01525 | 3 |
| 8294 | ENSMUSG( | Zzef1 | 195018 | -01132422 | 01560694C | -00025 | 008 | 3 |
| 323 | ENSMUSG( | Alkbh3 | 69113 | -02773351 | -01692520 | -148 | -1725 | 4 |
| 852 | ENSMUSG( | Brpf1 | 78783 | 008083083 | 013987727 | -031 | -17375 | 4 |
| 1021 | ENSMUSG( | Cc2d2a | 231214 | -01250533 | -01313596 | -22275 | -10075 | 4 |
| 1180 | ENSMUSG( | Cenpa | 12615 | 007443373 | 024349858 | -05525 | -20775 | 4 |
| 1232 | ENSMUSG( | Chd9 | 109151 | 008319407 | 01750629€ | -132 | -10125 | 4 |
| 1560 | ENSMUSG( | Cryab | 12955 | -00014402 | -02797423 | -0585 | -14425 | 4 |
| 1796 | ENSMUSG( | Ddx52 | 78394 | 002403183 | 000035532 | -0255 | -29425 | 4 |
| 1870 | ENSMUSG( | Dido1 | 23856 | -01477357 | 008746594 | -25525 | -151 | 4 |
| 2077 | ENSMUSG( | Ecsit | 26940 | 012223074 | -00071565 | -122 | -14825 | 4 |
| 3153 | ENSMUSG( | Hp1bp3 | 15441 | -00212468 | -00547775 | -17675 | -137 | 4 |
| 3395 | ENSMUSG( | Itpr1 | 16438 | -00635925 | 012488281 | -10525 | -129 | 4 |
| 3613 | ENSMUSG( | Lamtor4 | 66096 | 002804363 | -02651511 | -03875 | -15325 | 4 |
| 3762 | ENSMUSG( | Lrrc9 | 78257 | -00079973 | -00751819 | -05575 | -12475 | 4 |
| 3842 | ENSMUSG( | Mag | 17136 | -06278886 | 01039900 | -17575 | -125 | 4 |
| 3960 | ENSMUSG( | Mbp | 17196 | 006492044 | -01351516 | -10325 | -15475 | 4 |
| 3984 | ENSMUSG( | Mdc1 | 240087 | 00481983€ | 020092041 | -097 | -1595 | 4 |
| 4552 | ENSMUSG( | Ngp | 18054 | -14390945 | 010339963 | -246 | -24125 | 4 |
| 4725 | ENSMUSG( | Nudt13 | 67725 | 00240554C | -00552949 | -15725 | -13575 | 4 |
| 4735 | ENSMUSG( | Nudt3 | 56409 | 005448825 | -01190823 | -05 | -17975 | 4 |
| 4860 | ENSMUSG( | Pacsin2 | 23970 | 00366448€ | -01143219 | -0615 | -13775 | 4 |
| 5540 | ENSMUSG( | Prr3 | 75210 | 002865442 | -00272830 | -04775 | -158 | 4 |
| 5661 | ENSMUSG( | Ptprm | 19274 | -06215353 | 019978377 | -16875 | -09675 | 4 |
| 5726 | ENSMUSG( | Rab12 | 19328 | -05794173 | -10866599 | -2015 | -22775 | 4 |
| 5940 | ENSMUSG( | Rfc5 | 72151 | 01038508C | -01260164 | -18525 | -2085 | 4 |
| 6070 | ENSMUSG( | Romo1 | 67067 | -00251489 | -04233690 | -03375 | -14375 | 4 |
| 6237 | ENSMUSG( | S100a9 | 20202 | -12017201 | -01408817 | -18725 | -22175 | 4 |
| 6376 | ENSMUSG( | Selenon | 74777 | -02593586 | 012406844 | -0315 | -15225 | 4 |
| 6419 | ENSMUSG( | Serinc5 | 218442 | -04356236 | 001954745 | -0985 | -1535 | 4 |
| 6421 | ENSMUSG( | Serpinb1a | 66222 | -05140580 | -02688874 | -04275 | -126 | 4 |
| 6568 | ENSMUSG( | Slc17a8 | 216227 | -08900080 | 004056108 | -00975 | -208 | 4 |
| 6608 | ENSMUSG( | Slc25a28 | 246696 | -01422587 | 002321674 | -086 | -118 | 4 |
| 6907 | ENSMUSG( | Spock1 | 20745 | -06116232 | 00331093C | -0525 | -112 | 4 |
| 6975 | ENSMUSG( | Ssbp2 | 66970 | 00369962 | -00051035 | -03025 | -21525 | 4 |
| 7440 | ENSMUSG( | Tmem198b | 73827 | -04437042 | 001118203 | -06 | -12325 | 4 |
| 8171 | ENSMUSG( | Yipf5 | 67180 | -02916362 | 01884081€ | -062 | -1715 | 4 |
| 8179 | ENSMUSG( | Ypel1 | 106369 | 013704363 | 00696937C | -03 | -3085 | 4 |
| 8258 | ENSMUSG( | Zfr2 | 103406 | 025347061 | -02288300 | 0175 | -29125 | 4 |
| 1 | ENSMUSG( | 0610010K1 | 104457 | -00201279 | 019623462 | 021 | -0155 | 5 |
| 5 | ENSMUSG( | 1600014C1 | 72244 | -00291487 | -01526269 | 01575 | -044 | 5 |
| 12 | ENSMUSG( | 4931406CC | 70984 | 01613411€ | -00728305 | 02925 | -018 | 5 |
| 20 | ENSMUSG( | Aaas | 223921 | 003819533 | 003049112 | 04275 | -0225 | 5 |
| 24 | ENSMUSG( | Aamdc | 66273 | 00703162€ | -00878797 | 0165 | -0095 | 5 |
| 34 | ENSMUSG( | Abat | 268860 | 008255567 | -02303170 | 001 | -02025 | 5 |
| 70 | ENSMUSG( | Abhd17b | 226016 | 009667022 | 005865875 | 04025 | -017 | 5 |
| 85 | ENSMUSG( | Abracl | 73112 | 01406431C | 028522715 | 032 | -03825 | 5 |
| 89 | ENSMUSG( | Acaa2 | 52538 | 00192837C | -02373771 | -0035 | -0315 | 5 |
| 92 | ENSMUSG( | Acad12 | 338350 | 001860854 | -01926655 | 01075 | -034 | 5 |
| 93 | ENSMUSG( | Acad8 | 66948 | -00067693 | -01732583 | -00025 | -0145 | 5 |
| 95 | ENSMUSG( | Acadl | 11363 | 012153148 | -01085159 | 017 | -0105 | 5 |
| 96 | ENSMUSG( | Acadm | 11364 | 010652181 | -01096582 | 004 | -00975 | 5 |
| 99 | ENSMUSG( | Acadvl | 11370 | 018211454 | -01296629 | 00125 | -01575 | 5 |
| 115 | ENSMUSG( | Acot1 | 26897 | 015989628 | -03590099 | 023 | -027 | 5 |
| 118 | ENSMUSG( | Acot2 | 171210 | 002401217 | -03490175 | 019 | -02225 | 5 |
| 121 | ENSMUSG( | Acot9 | 56360 | 001915735 | -03610759 | 0135 | -03375 | 5 |
| 135 | ENSMUSG( | Acss1 | 68738 | 017367845 | -01580068 | 01725 | -02275 | 5 |
| 136 | ENSMUSG( | Acss2 | 60525 | 00876202C | -01490731 | 00125 | -015 | 5 |
| 137 | ENSMUSG( | Actb | 11461 | -00791825 | -02379057 | 01675 | -009 | 5 |
| 147 | ENSMUSG( | Actr2 | 66713 | 00168807€ | -00571651 | 02025 | -0155 | 5 |
| 148 | ENSMUSG( | Actr3 | 74117 | 004127998 | -00448644 | 0135 | -04225 | 5 |
| 149 | ENSMUSG( | Actr3b | 242894 | 009088763 | -00282322 | 011 | -0195 | 5 |
| 168 | ENSMUSG( | Adap1 | 231821 | 015436243 | -00066173 | 025 | -02675 | 5 |
| 170 | ENSMUSG( | Adarb1 | 110532 | 000934098 | -00734260 | 01675 | -03825 | 5 |
| 184 | ENSMUSG( | Add1 | 11518 | 010160587 | -01341485 | 00825 | -02275 | 5 |
| 185 | ENSMUSG( | Add2 | 11519 | 014260477 | 008138968 | -588785e-1 | -02475 | 5 |
| 203 | ENSMUSG( | Adk | 11534 | 01846966€ | -02115578 | 01875 | -0035 | 5 |
| 219 | ENSMUSG( | Adss2 | 11566 | -00836751 | -00324350 | 02075 | -02375 | 5 |
| 222 | ENSMUSG( | Afap1l2 | 226250 | -00090600 | -00023172 | 0095 | -0295 | 5 |
| 225 | ENSMUSG( | Afg1l | 215951 | 01596311C | -01232589 | 023 | -03 | 5 |
| 233 | ENSMUSG( | Agbl4 | 78933 | 050546289 | -00532598 | 0495 | -03275 | 5 |
| 237 | ENSMUSG( | Agl | 77559 | 005831292 | -02827396 | 04025 | -022 | 5 |
| 250 | ENSMUSG( | Ahcy | 269378 | 001034244 | -00648183 | 002 | -012 | 5 |
| 251 | ENSMUSG( | Ahcyl1 | 229709 | 007737008 | -02630282 | 0175 | -00575 | 5 |
| 258 | ENSMUSG( | Ahsa2 | 268390 | 010589143 | 007635144 | 01725 | -00575 | 5 |
| 261 | ENSMUSG( | Aifm1 | 26926 | 007903722 | -00516223 | 00375 | -014 | 5 |
| 272 | ENSMUSG( | Ak3 | 56248 | 015766988 | -00373940 | 00475 | -02125 | 5 |
| 276 | ENSMUSG( | Akap10 | 56697 | 00633745€ | 004891795 | 01225 | -0325 | 5 |
| 287 | ENSMUSG( | Akr1b1 | 11677 | -00049592 | -02157120 | 01225 | -0105 | 5 |
| 298 | ENSMUSG( | Aldh1a1 | 11668 | 03389478C | -01935328 | 014 | -01375 | 5 |
| 302 | ENSMUSG( | Aldh1l1 | 107747 | 009374844 | -02438617 | 03 | -01075 | 5 |
| 305 | ENSMUSG( | Aldh3a2 | 11671 | 00736355€ | -01693669 | 0025 | -02175 | 5 |
| 308 | ENSMUSG( | Aldh5a1 | 214579 | 009657983 | -01446554 | 005 | -0485 | 5 |
| 309 | ENSMUSG( | Aldh6a1 | 104776 | 013232628 | -00489646 | 00375 | -0265 | 5 |
| 310 | ENSMUSG( | Aldh7a1 | 110695 | 01267378€ | -01665121 | 01575 | -02875 | 5 |
| 352 | ENSMUSG( | Anapc4 | 52206 | 01495447C | 002819608 | 004 | -0285 | 5 |
| 365 | ENSMUSG( | Ankmy2 | 217473 | -00274596 | -02053022 | 019 | -0065 | 5 |
| 367 | ENSMUSG( | Ankrd13c | 433667 | 002969923 | -00782426 | 025 | -02825 | 5 |
| 380 | ENSMUSG( | Ankrd49 | 56503 | 004313158 | 005816465 | 00125 | -02625 | 5 |
| 405 | ENSMUSG( | Anxa5 | 11747 | 002625258 | -02062425 | 0005 | -03 | 5 |
| 410 | ENSMUSG( | Ap1b1 | 11764 | 016832043 | 00578503€ | 0085 | -0155 | 5 |

| | | | | | | | | |
|---|---|---|---|---|---|---|---|---|
| 411 | ENSMUSG( | Ap1g1 | 11765 | 018070744 | 001973748 | 0155 | -00475 | 5 |
| 412 | ENSMUSG( | Ap1m1 | 11767 | 010665523 | -00726628 | 01925 | -01425 | 5 |
| 415 | ENSMUSG( | Ap2a1 | 11771 | 012137313 | -00795498 | 028 | -027 | 5 |
| 416 | ENSMUSG( | Ap2a2 | 11772 | 018694824 | -01026186 | 022 | -029 | 5 |
| 417 | ENSMUSG( | Ap2b1 | 71770 | 015870312 | -01421864 | 03975 | -01075 | 5 |
| 422 | ENSMUSG( | Ap3d1 | 11776 | 025879533 | -00685408 | 0115 | -04425 | 5 |
| 424 | ENSMUSG( | Ap3m2 | 64933 | 007666678 | -00842012 | 02025 | -00975 | 5 |
| 426 | ENSMUSG( | Ap3s2 | 11778 | 008292520 | 000239895 | 00825 | -017 | 5 |
| 428 | ENSMUSG( | Ap4e1 | 108011 | -00065421 | 006989645 | 012 | -009 | 5 |
| 457 | ENSMUSG( | Apoo | 68316 | -00519468 | -02875849 | -00275 | -031 | 5 |
| 475 | ENSMUSG( | Arf1 | 11840 | -00116044 | -00832777 | 009 | -01775 | 5 |
| 476 | ENSMUSG( | Arf2 | 11841 | -00259384 | -01151982 | -0025 | -02975 | 5 |
| 479 | ENSMUSG( | Arf6 | 11845 | 004681752 | 005671883 | 0005 | -0155 | 5 |
| 480 | ENSMUSG( | Arfgap1 | 228998 | -00893355 | -00964602 | -0005 | -0345 | 5 |
| 488 | ENSMUSG( | Arfrp1 | 76688 | 000797009 | -00854562 | 034 | -011 | 5 |
| 491 | ENSMUSG( | Arhgap1 | 228359 | 012488874 | -01035248 | 00275 | -021 | 5 |
| 493 | ENSMUSG( | Arhgap12 | 75415 | 016294236 | -00903375 | 02725 | -02775 | 5 |
| 510 | ENSMUSG( | Arhgap5 | 11855 | 005297430 | -00838279 | 01525 | -017 | 5 |
| 541 | ENSMUSG( | Arl15 | 218639 | 001813893 | -00337514 | 002 | -0415 | 5 |
| 542 | ENSMUSG( | Arl2 | 56327 | -00353077 | -01781637 | 0255 | -021 | 5 |
| 546 | ENSMUSG( | Arl5a | 75423 | 006680018 | 003600222 | 00725 | -0115 | 5 |
| 547 | ENSMUSG( | Arl6 | 56297 | -00035103 | -00827921 | 03675 | -02175 | 5 |
| 552 | ENSMUSG( | Arl8a | 68724 | -00480061 | -00319692 | 0485 | -035 | 5 |
| 560 | ENSMUSG( | Armcx3 | 71703 | 007798183 | 010149600 | 02575 | -021 | 5 |
| 562 | ENSMUSG( | Armh3 | 71617 | -00624818 | -00566508 | 02175 | -021 | 5 |
| 567 | ENSMUSG( | Arpc2 | 76709 | 007389867 | -00709918 | 02125 | -01825 | 5 |
| 570 | ENSMUSG( | Arpc5 | 67771 | -00375485 | -00874209 | 01825 | -008 | 5 |
| 571 | ENSMUSG( | Arpc5l | 74192 | 000044195 | -01269269 | 00275 | -0365 | 5 |
| 572 | ENSMUSG( | Arpin | 70420 | 006305567 | -00261491 | 0005 | -01225 | 5 |
| 575 | ENSMUSG( | Arrb1 | 109689 | 014514792 | -00678635 | 003 | -019 | 5 |
| 598 | ENSMUSG( | Asic2 | 11418 | 006654116 | 035959582 | 0315 | -02475 | 5 |
| 626 | ENSMUSG( | Atg3 | 67841 | 011237309 | -00153767 | 01975 | -01275 | 5 |
| 629 | ENSMUSG( | Atg5 | 11793 | 017479764 | -00170519 | 0075 | -04575 | 5 |
| 630 | ENSMUSG( | Atg7 | 74244 | 000964850 | -01871434 | -00175 | -01625 | 5 |
| 640 | ENSMUSG( | Atp13a1 | 170759 | -00609830 | 017213044 | 02875 | -01075 | 5 |
| 663 | ENSMUSG( | Atp5me | 11958 | 002553362 | -02515426 | 02175 | -0405 | 5 |
| 666 | ENSMUSG( | Atp5mk | 66477 | -00260023 | -02525628 | 00525 | -0275 | 5 |
| 679 | ENSMUSG( | Atp6v1c1 | 66335 | 006868077 | -01685888 | 01275 | -014 | 5 |
| 682 | ENSMUSG( | Atp6v1f | 66144 | -00274187 | -03593812 | 03375 | -0305 | 5 |
| 709 | ENSMUSG( | Avl9 | 78937 | 000774810 | 011852109 | 00525 | -03475 | 5 |
| 712 | ENSMUSG( | Axl | 26362 | -00138053 | 001428634 | 00425 | -05075 | 5 |
| 718 | ENSMUSG( | B3gat3 | 72727 | 001079399 | -00295037 | 01925 | -015 | 5 |
| 738 | ENSMUSG( | Baiap2 | 108100 | 011770016 | 014120714 | 012 | -0165 | 5 |
| 749 | ENSMUSG( | Bbs4 | 102774 | 000726507 | -02251787 | 037 | -00975 | 5 |
| 750 | ENSMUSG( | Bbs5 | 72569 | 001281445 | 02673243 | 01 | -03575 | 5 |
| 752 | ENSMUSG( | Bbs9 | 319845 | 003376465 | 006993526 | 02375 | -041 | 5 |
| 780 | ENSMUSG( | Bdh1 | 71911 | 007184727 | -00144282 | 0195 | -0045 | 5 |
| 791 | ENSMUSG( | Bicral | 210982 | 009427773 | 001637858 | 00625 | -02425 | 5 |
| 800 | ENSMUSG( | Bloc1s1 | 14533 | 011069411 | -02449587 | 022 | -0015 | 5 |
| 803 | ENSMUSG( | Blvra | 109778 | -00834910 | -03444384 | -00275 | -02175 | 5 |
| 815 | ENSMUSG( | Bnip3l | 12177 | 004091606 | -00765668 | 04625 | -032 | 5 |
| 827 | ENSMUSG( | Borcs8 | 72368 | -00083162 | -01557261 | 01 | -02775 | 5 |
| 828 | ENSMUSG( | Bpgm | 12183 | -00582901 | -03242684 | 0155 | -03525 | 5 |
| 829 | ENSMUSG( | Bphl | 68021 | 002665773 | -03228272 | 012 | -054 | 5 |
| 850 | ENSMUSG( | Brms1l | 52592 | 005105858 | -00558176 | 0235 | -008 | 5 |
| 890 | ENSMUSG( | Cab39 | 12283 | 007135867 | -00250911 | 011 | -023 | 5 |
| 918 | ENSMUSG( | Cactin | 70312 | 007268977 | 005525922 | 01125 | -03125 | 5 |
| 920 | ENSMUSG( | Cacybp | 12301 | 005400824 | 003748410 | 0165 | -02275 | 5 |
| 926 | ENSMUSG( | Cadps | 27062 | 008154339 | -00464308 | 0015 | -01725 | 5 |
| 928 | ENSMUSG( | Calb1 | 12307 | -00066169 | 001135667 | 04125 | -023 | 5 |
| 929 | ENSMUSG( | Calb2 | 12308 | -01641415 | -00587921 | 02225 | -0275 | 5 |
| 950 | ENSMUSG( | Camkv | 235604 | 020466148 | -00007503 | 02375 | -003 | 5 |
| 960 | ENSMUSG( | Cap1 | 12331 | 023877648 | -02017715 | 0125 | -02775 | 5 |
| 961 | ENSMUSG( | Cap2 | 67252 | 002288205 | -01975773 | 01025 | -0325 | 5 |
| 966 | ENSMUSG( | Capn5 | 12337 | 022793498 | -00517767 | 0355 | -01175 | 5 |
| 971 | ENSMUSG( | Capza2 | 12343 | 008405121 | -00815387 | 04025 | -0295 | 5 |
| 975 | ENSMUSG( | Carm1 | 59035 | 002785063 | -00400674 | -001 | -0245 | 5 |
| 977 | ENSMUSG( | Carmil2 | 234695 | 008837117 | -00978707 | 00275 | -03475 | 5 |
| 1009 | ENSMUSG( | Cbr1 | 12408 | -00862988 | -02736020 | 02425 | -01475 | 5 |
| 1014 | ENSMUSG( | Cbx1 | 12412 | 007364042 | -00106059 | 026 | -01825 | 5 |
| 1042 | ENSMUSG( | Ccdc28a | 215814 | -01433239 | -00481599 | 00375 | -0405 | 5 |
| 1085 | ENSMUSG( | Cct7 | 12468 | 018894791 | -02351702 | 012 | -01 | 5 |
| 1087 | ENSMUSG( | Ccz1 | 231874 | 003559343 | -00758127 | 025 | -001 | 5 |
| 1111 | ENSMUSG( | Cdc23 | 52563 | 004105243 | -00009715 | 0225 | -0045 | 5 |
| 1115 | ENSMUSG( | Cdc37 | 12539 | 004050320 | -01076144 | 0145 | -01025 | 5 |
| 1122 | ENSMUSG( | Cdc42ep2 | 104252 | -01510156 | 003588770 | 021 | -024 | 5 |
| 1125 | ENSMUSG( | Cdc5l | 71702 | 019065305 | 000791249 | 00425 | -02575 | 5 |
| 1144 | ENSMUSG( | Cdk11b | 12537 | 008841784 | -00621897 | 0115 | -0075 | 5 |
| 1146 | ENSMUSG( | Cdk14 | 18647 | 011146369 | 001216406 | 0245 | -047 | 5 |
| 1171 | ENSMUSG( | Celf1 | 13046 | -00067320 | 010024169 | 011 | -02 | 5 |
| 1209 | ENSMUSG( | Cfl1 | 12631 | 002854460 | -01833811 | 01625 | -0065 | 5 |
| 1214 | ENSMUSG( | Chac2 | 68044 | -01215382 | -00455001 | -00025 | -01475 | 5 |
| 1234 | ENSMUSG( | Cherp | 27967 | 013859474 | 003085421 | 00025 | -035 | 5 |
| 1239 | ENSMUSG( | Chka | 12660 | 017928300 | 006439573 | 0035 | -01775 | 5 |
| 1251 | ENSMUSG( | Chmp6 | 208092 | 007338647 | -00301868 | 01725 | -01675 | 5 |
| 1256 | ENSMUSG( | Chp1 | 56398 | 005759271 | -01188561 | 00825 | -011 | 5 |
| 1291 | ENSMUSG( | Clcc1 | 229725 | -01673973 | 006121412 | 0295 | -01825 | 5 |
| 1292 | ENSMUSG( | Clcn2 | 12724 | 017622610 | 025576623 | 01625 | -04025 | 5 |
| 1304 | ENSMUSG( | Cldnd1 | 224250 | -02004455 | 000100864 | 04175 | -0195 | 5 |
| 1305 | ENSMUSG( | Clec16a | 74374 | 017600609 | 004574414 | 0115 | -01325 | 5 |
| 1309 | ENSMUSG( | Clic4 | 29876 | -00237914 | -01778970 | 006 | -0215 | 5 |
| 1333 | ENSMUSG( | Cltc | 67300 | 003030582 | -01226196 | 0315 | -0275 | 5 |
| 1340 | ENSMUSG( | Cmbl | 69574 | 001467856 | -03684780 | -0025 | -03325 | 5 |
| 1341 | ENSMUSG( | Cmc1 | 67899 | -00903486 | -02698435 | 02775 | -04725 | 5 |
| 1357 | ENSMUSG( | Cnih4 | 98417 | -00838093 | -00199501 | 0065 | -02125 | 5 |
| 1371 | ENSMUSG( | Cnot6l | 231464 | 004963286 | 018800616 | 02575 | -027 | 5 |
| 1379 | ENSMUSG( | Cnrip1 | 380686 | 004419171 | -00971817 | 006 | -02725 | 5 |
| 1395 | ENSMUSG( | Coa3 | 52469 | -00749186 | -02952930 | 00275 | -01525 | 5 |
| 1397 | ENSMUSG( | Coa6 | 67892 | -01279848 | -01722718 | 003 | -04275 | 5 |
| 1405 | ENSMUSG( | Cog2 | 76332 | -00144257 | -01395648 | -00025 | -02175 | 5 |
| 1430 | ENSMUSG( | Commd2 | 52245 | 000811874 | -00063071 | 02475 | -00825 | 5 |
| 1432 | ENSMUSG( | Commd4 | 66199 | 013349624 | -00662177 | 0225 | -0185 | 5 |
| 1444 | ENSMUSG( | Copg2 | 54160 | 003393195 | -01201639 | 02125 | -0015 | 5 |
| 1446 | ENSMUSG( | Cops2 | 12848 | -00806997 | -00544397 | 0265 | -046 | 5 |
| 1449 | ENSMUSG( | Cops5 | 26754 | 003473303 | -00685030 | 012 | -01725 | 5 |
| 1450 | ENSMUSG( | Cops6 | 26893 | 003394045 | -02221188 | 0095 | -0255 | 5 |
| 1454 | ENSMUSG( | Cops9 | 66915 | 006122284 | -02712469 | 04225 | -04625 | 5 |
| 1464 | ENSMUSG( | Coq8a | 67426 | 009523082 | -00342093 | 00925 | -02975 | 5 |
| 1469 | ENSMUSG( | Coro1c | 23790 | 006825368 | -00670624 | 0135 | -016 | 5 |
| 1473 | ENSMUSG( | Coro7 | 78885 | -00120767 | -00855191 | 00125 | -01875 | 5 |
| 1483 | ENSMUSG( | Cox4i1 | 12857 | 001789391 | -01430897 | 00875 | -01175 | 5 |
| 1485 | ENSMUSG( | Cox5a | 12858 | -00550050 | -01839731 | 00625 | -05 | 5 |
| 1487 | ENSMUSG( | Cox6a1 | 12861 | 000901418 | -02009391 | 02375 | -036 | 5 |
| 1491 | ENSMUSG( | Cox7a1 | 12865 | 012364879 | -03979160 | 04025 | -02975 | 5 |
| 1494 | ENSMUSG( | Cox7c | 12867 | 002672695 | -01554492 | 034 | -036 | 5 |
| 1505 | ENSMUSG( | Cpne2 | 234577 | 006213586 | -02166951 | 017 | -02575 | 5 |
| 1506 | ENSMUSG( | Cpne3 | 70568 | 010277660 | -00341696 | 01625 | -03175 | 5 |
| 1518 | ENSMUSG( | Cpsf3 | 54451 | 002475789 | -00775766 | 014 | -00875 | 5 |
| 1519 | ENSMUSG( | Cpsf4 | 54188 | -00698277 | 010561301 | 014 | -03725 | 5 |
| 1525 | ENSMUSG( | Cptp | 79554 | 002556505 | 008974522 | 04925 | -02325 | 5 |
| 1530 | ENSMUSG( | Cradd | 12905 | -00086093 | -01413819 | 00175 | -0425 | 5 |
| 1531 | ENSMUSG( | Crat | 12908 | 016488267 | -00770786 | 01975 | -0125 | 5 |
| 1532 | ENSMUSG( | Crbn | 58799 | 001136487 | -00405174 | 01775 | -048 | 5 |
| 1546 | ENSMUSG( | Crkl | 12929 | 018680646 | 001419855 | 017 | -0075 | 5 |
| 1563 | ENSMUSG( | Cryl1 | 68631 | -00214812 | -01790257 | 00325 | -02 | 5 |
| 1565 | ENSMUSG( | Cryz | 12972 | 000649342 | -02192227 | 00725 | -03475 | 5 |

| | | | | | | | | |
|---|---|---|---|---|---|---|---|---|
| 1567 | ENSMUSG( | Cryzl2 | 226527 | 00410154 | -03300681 | 0245 | -01975 | 5 |
| 1574 | ENSMUSG( | Csk | 12988 | 006791413 | 001480261 | 0295 | -02425 | 5 |
| 1583 | ENSMUSG( | Csnk1g3 | 70425 | 013131933 | 01351409( | 006 | -014 | 5 |
| 1584 | ENSMUSG( | Csnk2a1 | 12995 | 00076763S | -00954476 | 012 | -01025 | 5 |
| 1585 | ENSMUSG( | Csnk2a2 | 13000 | 013173551 | 001512971 | 00675 | -048 | 5 |
| 1586 | ENSMUSG( | Csnk2b | 13001 | -00120102 | -01162003 | 0195 | -007 | 5 |
| 1589 | ENSMUSG( | Cspp1 | 211660 | 014208828 | 00163968S | 0265 | -01225 | 5 |
| 1594 | ENSMUSG( | Cstf1 | 67337 | -00190623 | -00635444 | 00575 | -028 | 5 |
| 1614 | ENSMUSG( | Ctnnb1 | 12387 | 011669195 | -01119203 | 01975 | -01275 | 5 |
| 1619 | ENSMUSG( | Ctps1 | 51797 | 000021264 | -01058225 | 0165 | -018 | 5 |
| 1620 | ENSMUSG( | Ctps2 | 55936 | 008996417 | 001127697 | 01475 | -034 | 5 |
| 1629 | ENSMUSG( | Ctso | 229445 | 001154495 | 001248042 | 0475 | -03675 | 5 |
| 1640 | ENSMUSG( | Cul3 | 26554 | 007796442 | 000809523 | 016 | -0085 | 5 |
| 1645 | ENSMUSG( | Cul9 | 78309 | 010497376 | 004273459 | 01775 | -0125 | 5 |
| 1646 | ENSMUSG( | Cuta | 67675 | -00896765 | -01293024 | 001 | -04175 | 5 |
| 1648 | ENSMUSG( | Cux1 | 13047 | -00257256 | -01327676 | 02675 | -00975 | 5 |
| 1661 | ENSMUSG( | Cyb5a | 109672 | 00783815C | -01823980 | 04125 | -01475 | 5 |
| 1668 | ENSMUSG( | Cyc1 | 66445 | 003191095 | -01735858 | 0105 | -02975 | 5 |
| 1670 | ENSMUSG( | Cyfip1 | 20430 | 015699673 | -01003585 | 023 | -02275 | 5 |
| 1672 | ENSMUSG( | Cygb | 114886 | -01197439 | -01615129 | 02475 | -0225 | 5 |
| 1673 | ENSMUSG( | Cyld | 74256 | -00578082 | 002723595 | 01425 | -0105 | 5 |
| 1694 | ENSMUSG( | D2hgdh | 98314 | -00573016 | -01246416 | 0445 | -04425 | 5 |
| 1701 | ENSMUSG( | Daam2 | 76441 | 004755291 | -01973573 | 01675 | -01825 | 5 |
| 1721 | ENSMUSG( | Dbi | 13167 | 010949014 | -02416245 | 00475 | -01375 | 5 |
| 1730 | ENSMUSG( | Dcaf11 | 28199 | 015046258 | -00470142 | 01425 | -02075 | 5 |
| 1747 | ENSMUSG( | Dctn2 | 69654 | 006732062 | -01684596 | 015 | -04075 | 5 |
| 1753 | ENSMUSG( | Dcun1d1 | 114893 | 009615713 | -00006220 | 00225 | -02625 | 5 |
| 1754 | ENSMUSG( | Dcun1d2 | 102323 | 017772537 | 003940449 | 0045 | -0425 | 5 |
| 1761 | ENSMUSG( | Ddah1 | 69219 | 000939329 | -03785395 | 0005 | -03025 | 5 |
| 1762 | ENSMUSG( | Ddah2 | 51793 | -00789637 | -00929639 | -00125 | -02625 | 5 |
| 1763 | ENSMUSG( | Ddb1 | 13194 | 014209719 | -00352095 | -00175 | -01375 | 5 |
| 1772 | ENSMUSG( | Ddt | 13202 | -00722746 | -02796211 | 00525 | -046 | 5 |
| 1775 | ENSMUSG( | Ddx17 | 67040 | 001675496 | 013844741 | 01925 | -0035 | 5 |
| 1776 | ENSMUSG( | Ddx18 | 66942 | 009930313 | 004000911 | 0075 | -0215 | 5 |
| 1781 | ENSMUSG( | Ddx23 | 74351 | 011249841 | 006386223 | 01275 | -01975 | 5 |
| 1794 | ENSMUSG( | Ddx50 | 94213 | 00661204C | 017878632 | 003 | -0385 | 5 |
| 1802 | ENSMUSG( | Decr2 | 26378 | 008733407 | -01146829 | 0365 | -019 | 5 |
| 1831 | ENSMUSG( | Dgkb | 217480 | -00651590 | -02182790 | 03575 | -02325 | 5 |
| 1832 | ENSMUSG( | Dgkd | 227333 | 019371467 | 009303012 | 005 | -03875 | 5 |
| 1835 | ENSMUSG( | Dgkh | 380921 | -00214186 | 017549776 | 01525 | -0245 | 5 |
| 1839 | ENSMUSG( | Dglucy | 217830 | -00438749 | -03522317 | 0175 | -0215 | 5 |
| 1840 | ENSMUSG( | Dguok | 27369 | 00002612C | 002629631 | -00175 | -0315 | 5 |
| 1845 | ENSMUSG( | Dhfr | 13361 | 012295497 | -00506388 | 01175 | -0115 | 5 |
| 1855 | ENSMUSG( | Dhx15 | 13204 | 000562738 | 014036171 | -00075 | -01475 | 5 |
| 1858 | ENSMUSG( | Dhx30 | 72831 | 014806732 | 002877657 | 0205 | -0065 | 5 |
| 1866 | ENSMUSG( | Diablo | 66593 | 004382737 | 017485024 | 00325 | -0445 | 5 |
| 1868 | ENSMUSG( | Diaph2 | 54004 | -00168222 | 007512424 | 00525 | -02725 | 5 |
| 1877 | ENSMUSG( | Diras1 | 208666 | 012934126 | -01105999 | 01325 | -008 | 5 |
| 1883 | ENSMUSG( | Dkc1 | 245474 | 012729753 | 00270204C | 0025 | -05075 | 5 |
| 1889 | ENSMUSG( | Dlg2 | 23859 | 011271429 | 007086286 | 00375 | -01875 | 5 |
| 1891 | ENSMUSG( | Dlg4 | 13385 | 01134810C | -00027223 | 02375 | -033 | 5 |
| 1895 | ENSMUSG( | Dlgap3 | 242667 | 023678873 | 017525631 | 00275 | -0275 | 5 |
| 1900 | ENSMUSG( | Dmac2l | 68055 | 00268893 | -01565011 | 00725 | -03275 | 5 |
| 1903 | ENSMUSG( | Dmtn | 13829 | 014989405 | -00114972 | 00425 | -024 | 5 |
| 1912 | ENSMUSG( | Dnaja2 | 56445 | 00853220C | 00040100C | 0045 | -034 | 5 |
| 1925 | ENSMUSG( | Dnajc11 | 230935 | 011519237 | -00588861 | 00875 | -00925 | 5 |
| 1934 | ENSMUSG( | Dnajc24 | 99349 | 014662468 | -03078930 | 005 | -03025 | 5 |
| 1952 | ENSMUSG( | Dnm3 | 103967 | 008058517 | 001479048 | 00275 | -032 | 5 |
| 1966 | ENSMUSG( | Dock3 | 208869 | -00310583 | -00007804 | 0145 | -01125 | 5 |
| 1967 | ENSMUSG( | Dock4 | 238130 | 000795158 | 01355098C | 01975 | -01025 | 5 |
| 1973 | ENSMUSG( | Dok4 | 114255 | 003646208 | 006742429 | 03025 | -00725 | 5 |
| 1986 | ENSMUSG( | Dph5 | 69740 | 004116801 | -00659629 | 017 | -01275 | 5 |
| 1997 | ENSMUSG( | Dpp9 | 224897 | 01627461C | 003402999 | 016 | -0415 | 5 |
| 2015 | ENSMUSG( | Dst | 13518 | -01416738 | 005702883 | 03075 | -0345 | 5 |
| 2027 | ENSMUSG( | Dusp14 | 56405 | -00668532 | -00897175 | 01 | -0205 | 5 |
| 2048 | ENSMUSG( | Dync2i2 | 71820 | 014882059 | -00278893 | 01325 | -0085 | 5 |
| 2050 | ENSMUSG( | Dynll1 | 56455 | 007232523 | -01176077 | 006 | -0195 | 5 |
| 2051 | ENSMUSG( | Dynll2 | 68097 | 005592621 | -01403430 | 01875 | -03975 | 5 |
| 2070 | ENSMUSG( | Ech1 | 51798 | 003695278 | -01755020 | -0025 | -019 | 5 |
| 2071 | ENSMUSG( | Echdc1 | 52665 | 01164405C | -02437819 | 01375 | -02925 | 5 |
| 2074 | ENSMUSG( | Eci1 | 13177 | 01381753C | -01685071 | 0205 | -03175 | 5 |
| 2075 | ENSMUSG( | Eci2 | 23986 | 006677252 | -01344010 | 01925 | -0105 | 5 |
| 2088 | ENSMUSG( | Eef1a2 | 13628 | 01270353C | -00960760 | 02375 | -0305 | 5 |
| 2090 | ENSMUSG( | Eef1akmt2 | 72096 | 001671432 | -00539739 | 014 | -034 | 5 |
| 2094 | ENSMUSG( | Eef2 | 13629 | 009722701 | -01140310 | 009 | -038 | 5 |
| 2097 | ENSMUSG( | Efcab14 | 230648 | -01109032 | -00009590 | 00225 | -0305 | 5 |
| 2108 | ENSMUSG( | Efr3b | 668212 | 018707035 | -00011063 | 00925 | -0335 | 5 |
| 2110 | ENSMUSG( | Eftud2 | 20624 | 013129159 | -00113086 | 00775 | -055 | 5 |
| 2116 | ENSMUSG( | Ehd1 | 13660 | 009454569 | -00321515 | -0005 | -016 | 5 |
| 2118 | ENSMUSG( | Ehd3 | 57440 | 012022438 | -00244282 | 025 | -0355 | 5 |
| 2120 | ENSMUSG( | Ehhadh | 74147 | -00454675 | 004786791 | 00475 | -02925 | 5 |
| 2125 | ENSMUSG( | Eif1 | 20918 | 00482453C | -01386037 | 03825 | -01725 | 5 |
| 2133 | ENSMUSG( | Eif2b1 | 209354 | 010181388 | -00754485 | 002 | -0175 | 5 |
| 2145 | ENSMUSG( | Eif3c | 56347 | 011003185 | -01838919 | 0285 | -007 | 5 |
| 2157 | ENSMUSG( | Eif4a2 | 13682 | 003037655 | -00284874 | 022 | -00375 | 5 |
| 2174 | ENSMUSG( | Elac1 | 114615 | -00634824 | -00133382 | 00275 | -016 | 5 |
| 2179 | ENSMUSG( | Elavl2 | 15569 | -00730255 | -00521535 | 04075 | -031 | 5 |
| 2180 | ENSMUSG( | Elavl3 | 15571 | 00999165C | 010463618 | 0075 | -023 | 5 |
| 2185 | ENSMUSG( | Elmo2 | 140579 | 011592046 | -01570510 | 00975 | -00725 | 5 |
| 2191 | ENSMUSG( | Eloc | 67923 | -00215363 | -01831052 | 0095 | -04175 | 5 |
| 2196 | ENSMUSG( | Elp2 | 58523 | 007033843 | -00528180 | 019 | -004 | 5 |
| 2204 | ENSMUSG( | Emc2 | 66736 | 007988388 | -00806933 | 027 | -00275 | 5 |
| 2209 | ENSMUSG( | Emc8 | 18117 | 002370253 | -01900690 | 023 | -006 | 5 |
| 2222 | ENSMUSG( | Enc1 | 13803 | 016874843 | -01016352 | 02275 | -00475 | 5 |
| 2250 | ENSMUSG( | Epb41l1 | 13821 | 019107774 | 001935781 | 0105 | -013 | 5 |
| 2254 | ENSMUSG( | Epb41l5 | 226352 | 013957052 | 000413596 | 01475 | -00875 | 5 |
| 2277 | ENSMUSG( | Eps15 | 13858 | -00422023 | -01113218 | -00025 | -02325 | 5 |
| 2279 | ENSMUSG( | Eps8 | 13860 | 008686697 | -01024163 | 00225 | -01325 | 5 |
| 2287 | ENSMUSG( | Erc2 | 238988 | -00270606 | 005130202 | 00875 | -03175 | 5 |
| 2308 | ENSMUSG( | Esd | 13885 | 000717263 | -02418511 | -001 | -0395 | 5 |
| 2312 | ENSMUSG( | Etf1 | 225363 | 005885164 | 001368952 | 022 | -002 | 5 |
| 2318 | ENSMUSG( | Etnk1 | 75320 | 000548845 | 005415727 | 00475 | -0355 | 5 |
| 2329 | ENSMUSG( | Exoc3 | 211446 | 019425106 | -01191623 | 00375 | -0145 | 5 |
| 2330 | ENSMUSG( | Exoc4 | 20336 | 005452908 | -02420428 | 019 | -01475 | 5 |
| 2331 | ENSMUSG( | Exoc5 | 105504 | 005664864 | 008620159 | 0215 | -01175 | 5 |
| 2339 | ENSMUSG( | Exosc2 | 227715 | 003165586 | -01385904 | 02525 | -02325 | 5 |
| 2344 | ENSMUSG( | Exosc8 | 69639 | 000079815 | -00160461 | 00825 | -03575 | 5 |
| 2359 | ENSMUSG( | Fabp5 | 16592 | 011924766 | -01240675 | 0395 | -0265 | 5 |
| 2370 | ENSMUSG( | Fahd2a | 68126 | -00691844 | -02346416 | 00775 | -0235 | 5 |
| 2392 | ENSMUSG( | Fam163b | 109349 | -01029557 | 00120597C | 02575 | -0045 | 5 |
| 2402 | ENSMUSG( | Fam20b | 215015 | -00477122 | 008829239 | 01925 | -0085 | 5 |
| 2422 | ENSMUSG( | Farp1 | 223254 | 014375481 | -00227980 | 0325 | -02075 | 5 |
| 2423 | ENSMUSG( | Farp2 | 227377 | 010072099 | 017167183 | 02425 | -042 | 5 |
| 2435 | ENSMUSG( | Fbl | 14113 | 015381451 | -00555452 | 01075 | -0505 | 5 |
| 2436 | ENSMUSG( | Fbll1 | 237730 | -00480957 | -02415296 | 008 | -01925 | 5 |
| 2442 | ENSMUSG( | Fbxl16 | 214931 | 007778116 | 007730268 | 01775 | -02175 | 5 |
| 2450 | ENSMUSG( | Fbxo2 | 230904 | 017287501 | -01454892 | 012 | -03775 | 5 |
| 2451 | ENSMUSG( | Fbxo21 | 231670 | 011118345 | -00386307 | 01025 | -03025 | 5 |
| 2453 | ENSMUSG( | Fbxo3 | 57443 | 014176604 | 002056755 | 0165 | -028 | 5 |
| 2460 | ENSMUSG( | Fbxo44 | 230903 | 00912184C | -00738777 | 043 | -02225 | 5 |
| 2487 | ENSMUSG( | Fermt2 | 218952 | 006346759 | -02217867 | 01 | -0115 | 5 |
| 2492 | ENSMUSG( | Fgd4 | 224014 | 004304729 | -00366259 | 01425 | -00475 | 5 |
| 2493 | ENSMUSG( | Fgd6 | 13998 | 024256248 | -01459269 | 00475 | -01425 | 5 |
| 2494 | ENSMUSG( | Fgf1 | 14164 | 003710486 | -02774168 | -00075 | -03625 | 5 |
| 2496 | ENSMUSG( | Fgf13 | 14168 | 010048324 | -00568961 | 02275 | -009 | 5 |
| 2501 | ENSMUSG( | Fgfr3 | 14184 | 004083874 | 028995169 | 00725 | -03075 | 5 |

| | | | | | | | | |
|---|---|---|---|---|---|---|---|---|
| 2506 | ENSMUSG( | Fhip2b | 239170 | 016394357 | 007452379 | 00075 | -024 | 5 |
| 2507 | ENSMUSG( | Fhit | 14198 | 012004723 | -00992542 | 0025 | -027 | 5 |
| 2513 | ENSMUSG( | Fibp | 58249 | 003039256 | 003153015 | -00075 | -03725 | 5 |
| 2514 | ENSMUSG( | Fig4 | 103199 | 009017100 | 001944898 | 01975 | -0075 | 5 |
| 2522 | ENSMUSG( | Fkbp2 | 14227 | -01969269 | -00987519 | 0325 | -01575 | 5 |
| 2523 | ENSMUSG( | Fkbp3 | 30795 | 004796623 | -00233146 | 0365 | -012 | 5 |
| 2524 | ENSMUSG( | Fkbp4 | 14228 | 020303277 | 004112735 | -00175 | -01975 | 5 |
| 2526 | ENSMUSG( | Fkbp8 | 14232 | 002598184 | -01425011 | 039 | -02325 | 5 |
| 2529 | ENSMUSG( | Flad1 | 319945 | 008846876 | -00097742 | 0035 | -012 | 5 |
| 2542 | ENSMUSG( | Fmn1 | 14260 | 007786611 | 018505563 | 0005 | -02475 | 5 |
| 2545 | ENSMUSG( | Fmnl2 | 71409 | 002710727 | -00780311 | -00275 | -02575 | 5 |
| 2557 | ENSMUSG( | Fndc3b | 72007 | 025937777 | 013330562 | 022 | -054 | 5 |
| 2559 | ENSMUSG( | Fnta | 14272 | 014239587 | -01302154 | 01575 | -0125 | 5 |
| 2560 | ENSMUSG( | Fntb | 110606 | 004141667 | -01056917 | 04025 | -05275 | 5 |
| 2562 | ENSMUSG( | Folh1 | 53320 | 010090658 | 012537653 | 00375 | -0375 | 5 |
| 2574 | ENSMUSG( | Frmd3 | 242506 | -02459966 | -02442989 | 03225 | -01875 | 5 |
| 2581 | ENSMUSG( | Frmpd4 | 333605 | 007421452 | 013549584 | 00875 | -01425 | 5 |
| 2584 | ENSMUSG( | Frs3 | 107971 | 003852344 | -00331503 | 00625 | -0345 | 5 |
| 2591 | ENSMUSG( | Fth1 | 14319 | 012571481 | -01459466 | 024 | -00675 | 5 |
| 2643 | ENSMUSG( | Gad1 | 14415 | -01081803 | -01245953 | 003 | -0165 | 5 |
| 2648 | ENSMUSG( | Galc | 14420 | -00086836 | 009392802 | 01775 | -00525 | 5 |
| 2649 | ENSMUSG( | Gale | 74246 | 00349119 | 012656782 | 00975 | -0245 | 5 |
| 2666 | ENSMUSG( | Gapdh | 14433 | 008304835 | -01985869 | 01325 | -01625 | 5 |
| 2674 | ENSMUSG( | Gas2 | 14453 | -01699634 | -01398937 | 029 | -051 | 5 |
| 2676 | ENSMUSG( | Gas6 | 14456 | -00833762 | 014304506 | 0175 | -0235 | 5 |
| 2697 | ENSMUSG( | Gcdh | 270076 | 016376842 | -00373555 | 025 | -0085 | 5 |
| 2708 | ENSMUSG( | Gdi2 | 14569 | 007370177 | -00731701 | 01825 | -00475 | 5 |
| 2710 | ENSMUSG( | Gdpd2 | 71584 | 028607724 | 004213738 | 00725 | -03075 | 5 |
| 2717 | ENSMUSG( | Get3 | 56495 | -00207291 | -02567680 | 01475 | -0155 | 5 |
| 2721 | ENSMUSG( | Gfm1 | 28030 | 011450500 | -01670149 | 008 | -02925 | 5 |
| 2722 | ENSMUSG( | Gfm2 | 320806 | 012962035 | -00327504 | 0145 | -00525 | 5 |
| 2728 | ENSMUSG( | Gfus | 22122 | 003839492 | -00494348 | -157009e-1 | -05025 | 5 |
| 2735 | ENSMUSG( | Ggps1 | 14593 | 004948722 | 009351395 | -00075 | -01725 | 5 |
| 2755 | ENSMUSG( | Gk | 14933 | 010636603 | -00390803 | 0115 | -00825 | 5 |
| 2763 | ENSMUSG( | Gle1 | 74412 | 008368762 | -00211475 | 02325 | -03125 | 5 |
| 2765 | ENSMUSG( | Glipr2 | 384009 | 025788206 | -03958477 | 02575 | -031 | 5 |
| 2767 | ENSMUSG( | Glo1 | 109801 | 005247838 | -02109024 | 00875 | -013 | 5 |
| 2768 | ENSMUSG( | Glod4 | 67201 | 007807934 | -00935487 | 03125 | -05725 | 5 |
| 2775 | ENSMUSG( | Gls | 14660 | 016427386 | 004981015 | 01325 | -01275 | 5 |
| 2777 | ENSMUSG( | Gltp | 56356 | -00015673 | -02009340 | 002 | -035 | 5 |
| 2778 | ENSMUSG( | Glud1 | 14661 | 011158501 | -01507881 | 01525 | -00675 | 5 |
| 2779 | ENSMUSG( | Glul | 14645 | 024803668 | -00736966 | 02 | -00175 | 5 |
| 2780 | ENSMUSG( | Glyctk | 235582 | -00924358 | -03520756 | 0225 | -048 | 5 |
| 2786 | ENSMUSG( | Gm21992 | 1.03E+08 | 014045918 | 011602005 | 00775 | -0235 | 5 |
| 2799 | ENSMUSG( | Gna13 | 14674 | 017203127 | -00422733 | 01625 | -01975 | 5 |
| 2801 | ENSMUSG( | Gnai2 | 14678 | -00602299 | -01565996 | 02225 | -0385 | 5 |
| 2802 | ENSMUSG( | Gnai3 | 14679 | -00025136 | -00393652 | 0285 | -01375 | 5 |
| 2805 | ENSMUSG( | Gnaq | 14682 | 017276648 | 000515153 | 01575 | -016 | 5 |
| 2806 | ENSMUSG( | Gnas | 14683 | 004487794 | -01329200 | 0275 | -01325 | 5 |
| 2807 | ENSMUSG( | Gnaz | 14687 | 014257230 | 006082517 | 02525 | -02925 | 5 |
| 2816 | ENSMUSG( | Gng13 | 64337 | 003486949 | -02997568 | 0035 | -036 | 5 |
| 2817 | ENSMUSG( | Gng2 | 14702 | 002022471 | -00863664 | 01425 | -03525 | 5 |
| 2820 | ENSMUSG( | Gng5 | 14707 | 011136396 | -01256599 | 01225 | -012 | 5 |
| 2827 | ENSMUSG( | Gnpda2 | 67980 | -00154865 | -00873680 | 0215 | -02025 | 5 |
| 2828 | ENSMUSG( | Gnpnat1 | 54342 | -01175219 | -00340509 | -001 | -03475 | 5 |
| 2829 | ENSMUSG( | Gnptg | 214505 | -01323135 | -00396073 | 0245 | -029 | 5 |
| 2850 | ENSMUSG( | Got1 | 14718 | 004248649 | -01319230 | 196262e-1 | -028 | 5 |
| 2867 | ENSMUSG( | Gpd1l | 333433 | 012062120 | -00457268 | 00925 | -016 | 5 |
| 2887 | ENSMUSG( | Gpr89 | 67549 | -01793243 | -00107251 | 028 | -02025 | 5 |
| 2895 | ENSMUSG( | Gps2 | 56310 | 005104392 | 005149266 | 0435 | -019 | 5 |
| 2899 | ENSMUSG( | Gpx1 | 14775 | -01089274 | -00796690 | -002 | -02775 | 5 |
| 2905 | ENSMUSG( | Gramd1b | 235283 | 004149911 | 015331435 | 02675 | -02375 | 5 |
| 2930 | ENSMUSG( | Grk2 | 110355 | 018090509 | -00817362 | 01325 | -0055 | 5 |
| 2947 | ENSMUSG( | Gsdme | 54722 | -00395254 | -02114406 | 00425 | -013 | 5 |
| 2956 | ENSMUSG( | Gsr | 14782 | -00559258 | -01003869 | 02175 | -0345 | 5 |
| 2958 | ENSMUSG( | Gsta4 | 14860 | -00891407 | -00746023 | 00925 | -01775 | 5 |
| 2959 | ENSMUSG( | Gstk1 | 76263 | 014123503 | -02101527 | 04175 | -05325 | 5 |
| 2960 | ENSMUSG( | Gstm1 | 14862 | 028804837 | -00325136 | 03175 | -035 | 5 |
| 2962 | ENSMUSG( | Gstm4 | 14865 | -01453377 | -02190881 | 0175 | -0295 | 5 |
| 2963 | ENSMUSG( | Gstm5 | 14866 | 023092920 | -01136555 | 0335 | -03075 | 5 |
| 2966 | ENSMUSG( | Gstp1 | 14870 | -00390993 | -02000892 | 007 | -03675 | 5 |
| 2990 | ENSMUSG( | Gtpbp3 | 70359 | 008977754 | 016350085 | 018 | -0175 | 5 |
| 2994 | ENSMUSG( | Gucy1a1 | 60596 | 000254709 | -01895151 | 01125 | -01425 | 5 |
| 2995 | ENSMUSG( | Gucy1a2 | 234889 | -00920536 | -01326035 | 0235 | -026 | 5 |
| 2999 | ENSMUSG( | Gulp1 | 70676 | 017171731 | 010021586 | 02575 | -03525 | 5 |
| 3001 | ENSMUSG( | Gyg1 | 27357 | 004872937 | -01681157 | 02125 | -0175 | 5 |
| 3016 | ENSMUSG( | Hace1 | 209462 | 005187520 | 009363925 | 00175 | -03 | 5 |
| 3021 | ENSMUSG( | Hagh | 14651 | -00489804 | -02132216 | 01325 | -05325 | 5 |
| 3051 | ENSMUSG( | Hddc2 | 69692 | 012049691 | -01171858 | 0025 | -022 | 5 |
| 3089 | ENSMUSG( | Hibadh | 58875 | 010459623 | -01086046 | -00225 | -0215 | 5 |
| 3094 | ENSMUSG( | Hikeshi | 67669 | -00786565 | -01097700 | 00275 | -047 | 5 |
| 3095 | ENSMUSG( | Hint1 | 15254 | 002599321 | -01618508 | -00325 | -01725 | 5 |
| 3097 | ENSMUSG( | Hint3 | 66847 | 002452728 | -01911296 | 02175 | -0175 | 5 |
| 3099 | ENSMUSG( | Hip1r | 29816 | 010769913 | -00640844 | -001 | -01275 | 5 |
| 3105 | ENSMUSG( | Hlcs | 110948 | -02232177 | -03147707 | 04175 | -02425 | 5 |
| 3115 | ENSMUSG( | Hmgcll1 | 208982 | 011261845 | 010111060 | 00775 | -045 | 5 |
| 3117 | ENSMUSG( | Hmgcs1 | 208715 | -00099991 | -04517538 | 04375 | -016 | 5 |
| 3123 | ENSMUSG( | Hmox1 | 15368 | 005049814 | -01249160 | 051 | -018 | 5 |
| 3128 | ENSMUSG( | Hnrnpa2b1 | 53379 | 005264486 | 004912358 | 01075 | -051 | 5 |
| 3129 | ENSMUSG( | Hnrnpa3 | 229279 | 002099888 | 001452757 | 0065 | -029 | 5 |
| 3135 | ENSMUSG( | Hnrnph1 | 59013 | 000333680 | 009103881 | 00525 | -01175 | 5 |
| 3136 | ENSMUSG( | Hnrnph2 | 56258 | -00637412 | -00744026 | 02125 | -0285 | 5 |
| 3139 | ENSMUSG( | Hnrnpl | 15388 | -01852962 | -01466486 | 022 | -034 | 5 |
| 3152 | ENSMUSG( | Hopx | 74318 | 015525636 | -03570037 | 042 | -03475 | 5 |
| 3154 | ENSMUSG( | Hpca | 15444 | -00556379 | -01519048 | 02175 | -02 | 5 |
| 3158 | ENSMUSG( | Hprt1 | 15452 | -00987504 | -01927569 | 01825 | -011 | 5 |
| 3160 | ENSMUSG( | Hras | 15461 | -00423323 | -01903768 | 03825 | -04125 | 5 |
| 3176 | ENSMUSG( | Hsdl1 | 72552 | 002546374 | 000273169 | 01925 | -00575 | 5 |
| 3179 | ENSMUSG( | Hsp90aa1 | 15519 | 011820307 | -01461402 | 02775 | -0125 | 5 |
| 3182 | ENSMUSG( | Hspa12a | 73442 | 010532931 | -00576777 | 02775 | -01825 | 5 |
| 3226 | ENSMUSG( | Idh2 | 269951 | 026178326 | 015304218 | -0005 | -04275 | 5 |
| 3229 | ENSMUSG( | Idh3g | 15929 | 014488412 | -00817729 | 0075 | -0165 | 5 |
| 3230 | ENSMUSG( | Idi1 | 319554 | 002949844 | -05416882 | 01825 | -01375 | 5 |
| 3327 | ENSMUSG( | Ints9 | 210925 | 019855928 | 008566127 | 02225 | -01575 | 5 |
| 3335 | ENSMUSG( | Ipo7 | 233726 | 012994923 | 001279690 | 0195 | -004 | 5 |
| 3354 | ENSMUSG( | Isca1 | 69046 | 001779409 | -00500147 | 0155 | -0105 | 5 |
| 3366 | ENSMUSG( | Itch | 16396 | 013220187 | 014114009 | 04 | -01325 | 5 |
| 3378 | ENSMUSG( | Itgb1bp1 | 16413 | -00310436 | -01734124 | 002 | -0215 | 5 |
| 3399 | ENSMUSG( | Itsn1 | 16443 | 027722848 | -01074538 | 00225 | -01175 | 5 |
| 3400 | ENSMUSG( | Itsn2 | 20403 | 018753026 | 022087439 | 0075 | -03475 | 5 |
| 3415 | ENSMUSG( | Josd1 | 74158 | 014923639 | 003874967 | 01175 | -01675 | 5 |
| 3419 | ENSMUSG( | Jph4 | 319984 | 020549652 | 012582808 | 02475 | -028 | 5 |
| 3436 | ENSMUSG( | Katnb1 | 74187 | 014661310 | -00423696 | 0175 | -0175 | 5 |
| 3468 | ENSMUSG( | Kcnip1 | 70357 | 001546826 | -00313578 | 003 | -018 | 5 |
| 3469 | ENSMUSG( | Kcnip2 | 80906 | 005908755 | -01237580 | 01375 | -04975 | 5 |
| 3470 | ENSMUSG( | Kcnip3 | 56461 | -02201170 | -00823738 | 0265 | -0415 | 5 |
| 3471 | ENSMUSG( | Kcnip4 | 80334 | -00799954 | -00377943 | 00825 | -0295 | 5 |
| 3475 | ENSMUSG( | Kcnj16 | 16517 | 008143785 | -01427969 | 005 | -035 | 5 |
| 3485 | ENSMUSG( | Kcnmb4 | 58802 | 004815508 | 001298591 | 0165 | -03875 | 5 |
| 3502 | ENSMUSG( | Kctd2 | 70382 | 011567121 | -00390444 | 0175 | -04925 | 5 |
| 3505 | ENSMUSG( | Kctd4 | 67516 | -00494913 | 018848206 | 02175 | -01625 | 5 |
| 3522 | ENSMUSG( | Khdrbs3 | 13992 | 014670116 | -01269785 | -0025 | -02475 | 5 |
| 3530 | ENSMUSG( | Kif1a | 16560 | 006678731 | -01351675 | 00675 | -01 | 5 |
| 3531 | ENSMUSG( | Kif1b | 16561 | 005461203 | 004947730 | 007 | -021 | 5 |
| 3554 | ENSMUSG( | Klhdc2 | 69554 | 004074488 | -01116673 | 002 | -0185 | 5 |
| 3566 | ENSMUSG( | Klhl3 | 1.01E+08 | 031954912 | 008746402 | 03025 | -0115 | 5 |

| | | | | | | | |
|---|---|---|---|---|---|---|---|
| 3578 | ENSMUSG( | Kpna4 | 16649 | 005913698 | 005536665 | 02025 | -00425 | 5 |
| 3582 | ENSMUSG( | Kras | 16653 | -00542797 | -00309852 | 02425 | -011 | 5 |
| 3583 | ENSMUSG( | Kri1 | 215194 | 019494134 | 01119651( | 00725 | -04275 | 5 |
| 3611 | ENSMUSG( | Lamtor2 | 83409 | 003381775 | -02066698 | 01225 | -013 | 5 |
| 3614 | ENSMUSG( | Lamtor5 | 68576 | 007388462 | -02065916 | 0125 | -013 | 5 |
| 3616 | ENSMUSG( | Lancl2 | 71835 | -00052560 | -00748778 | 0105 | -01725 | 5 |
| 3618 | ENSMUSG( | Lap3 | 66988 | 01146643( | -03649399 | 02 | -0195 | 5 |
| 3631 | ENSMUSG( | Lbh | 77889 | -01151026 | -02099579 | -00025 | -02125 | 5 |
| 3637 | ENSMUSG( | Lcp1 | 18826 | -02542034 | -03172302 | 01225 | -0315 | 5 |
| 3642 | ENSMUSG( | Ldhd | 52815 | 007023867 | -00136071 | 0095 | -007 | 5 |
| 3651 | ENSMUSG( | Letmd1 | 68614 | -00126593 | 001061034 | 01 | -0155 | 5 |
| 3654 | ENSMUSG( | Lgals8 | 56048 | 003146145 | -00749038 | 0415 | -0205 | 5 |
| 3665 | ENSMUSG( | Lhpp | 76429 | 006643563 | -02210495 | 015 | -032 | 5 |
| 3688 | ENSMUSG( | Llgl1 | 16897 | 003471924 | -01493480 | -0015 | -0215 | 5 |
| 3718 | ENSMUSG( | Lpin2 | 64898 | 024950855 | 007304779 | 01825 | -0105 | 5 |
| 3744 | ENSMUSG( | Lrrc20 | 216011 | 002828237 | 000299558 | 007 | -015 | 5 |
| 3748 | ENSMUSG( | Lrrc47 | 72946 | 007371193 | -00200318 | 014 | -0135 | 5 |
| 3749 | ENSMUSG( | Lrrc49 | 102747 | 004321874 | -00989106 | 001 | -02125 | 5 |
| 3756 | ENSMUSG( | Lrrc73 | 224813 | 010827662 | 004106528 | 04675 | -028 | 5 |
| 3773 | ENSMUSG( | Lsamp | 268890 | -01293884 | 004968828 | 017 | -00575 | 5 |
| 3775 | ENSMUSG( | Lsm1 | 67207 | 000062965 | 003020864 | 02725 | -02 | 5 |
| 3780 | ENSMUSG( | Lsm2 | 27756 | -02572725 | -00324180 | 0375 | -0495 | 5 |
| 3782 | ENSMUSG( | Lsm4 | 50783 | 002452024 | -01943882 | 00075 | -036 | 5 |
| 3786 | ENSMUSG( | Lsm8 | 76522 | 000358198 | 00765898( | 03075 | -033 | 5 |
| 3815 | ENSMUSG( | Lyplal1 | 226791 | 004349523 | -03791706 | 0185 | -005 | 5 |
| 3819 | ENSMUSG( | Lyrm7 | 75530 | 002000479 | -01099533 | 00675 | -016 | 5 |
| 3821 | ENSMUSG( | Lysmd1 | 217779 | 021829476 | 009577669 | 02625 | -033 | 5 |
| 3827 | ENSMUSG( | Lzts1 | 211134 | 019341841 | 00194768( | 0265 | -0105 | 5 |
| 3848 | ENSMUSG( | Magi2 | 50791 | -00058468 | -00413468 | 03475 | -02325 | 5 |
| 3867 | ENSMUSG( | Maoa | 17161 | -00175933 | -00276119 | 03325 | -04625 | 5 |
| 3868 | ENSMUSG( | Maob | 109731 | 025187118 | -02106621 | 01975 | -029 | 5 |
| 3871 | ENSMUSG( | Map1lc3a | 66734 | 001498212 | -02333412 | 007 | -0115 | 5 |
| 3875 | ENSMUSG( | Map2k1 | 26395 | 005882027 | -01145333 | -001 | -0325 | 5 |
| 3876 | ENSMUSG( | Map2k2 | 26396 | 008204951 | -00675040 | 00175 | -02075 | 5 |
| 3878 | ENSMUSG( | Map2k4 | 26398 | 01019680( | -00602324 | 00325 | -0155 | 5 |
| 3920 | ENSMUSG( | Mapre2 | 212307 | 008933234 | -01579631 | 01825 | -0505 | 5 |
| 3941 | ENSMUSG( | Mat2a | 232087 | 01646978( | 024758222 | 00875 | -02975 | 5 |
| 3947 | ENSMUSG( | Mau2 | 74549 | 005672205 | 010741312 | 00875 | -02975 | 5 |
| 3955 | ENSMUSG( | Mblac2 | 72852 | -00480042 | 004437249 | 00575 | -0235 | 5 |
| 3965 | ENSMUSG( | Mccc1 | 72039 | 008052082 | -01470863 | 009 | -019 | 5 |
| 3973 | ENSMUSG( | Mcm4 | 17217 | 004373144 | 007774357 | 0305 | -009 | 5 |
| 3980 | ENSMUSG( | Mcts1 | 68995 | 00059742( | -01190911 | 04075 | -01375 | 5 |
| 4001 | ENSMUSG( | Mecr | 26922 | 008778719 | -01738272 | 00475 | -03275 | 5 |
| 4012 | ENSMUSG( | Med24 | 23989 | 016340088 | 004326197 | 0235 | -027 | 5 |
| 4027 | ENSMUSG( | Memo1 | 76890 | 003343146 | 002704952 | 012 | -02575 | 5 |
| 4031 | ENSMUSG( | Mesd | 67943 | -01814592 | 002575109 | 03575 | -03125 | 5 |
| 4047 | ENSMUSG( | Mettl9 | 59052 | -01107406 | -01290862 | 0025 | -0415 | 5 |
| 4051 | ENSMUSG( | Mff | 75734 | 009979444 | -00763764 | 00125 | -017 | 5 |
| 4073 | ENSMUSG( | Mib1 | 225164 | 005497406 | 003341471 | 014 | -01675 | 5 |
| 4075 | ENSMUSG( | Mical1 | 171580 | -01047976 | -02781443 | 009 | -02775 | 5 |
| 4080 | ENSMUSG( | Micos13 | 224904 | 003947171 | -02586830 | -001 | -03475 | 5 |
| 4082 | ENSMUSG( | Micu2 | 68514 | -00937045 | -02277884 | 00675 | -027 | 5 |
| 4084 | ENSMUSG( | Mid1ip1 | 68041 | 014970523 | -00904970 | -0015 | -0315 | 5 |
| 4088 | ENSMUSG( | Mif | 17319 | -00832819 | -02073369 | 03225 | -037 | 5 |
| 4095 | ENSMUSG( | Mindy3 | 66960 | 003306384 | 000423877 | 00425 | -0135 | 5 |
| 4103 | ENSMUSG( | Mkln1 | 27418 | 01567735( | 01259638( | 0205 | -0175 | 5 |
| 4111 | ENSMUSG( | Mlst8 | 56716 | -00040626 | -00341598 | 0125 | -01575 | 5 |
| 4112 | ENSMUSG( | Mlycd | 56690 | 012404612 | -00904481 | 0075 | -02575 | 5 |
| 4123 | ENSMUSG( | Mob4 | 19070 | 000451873 | -00069976 | 04 | -014 | 5 |
| 4148 | ENSMUSG( | Mpp1 | 17524 | -00196649 | 012509341 | 00925 | -022 | 5 |
| 4149 | ENSMUSG( | Mpp2 | 50997 | 016906245 | 005253183 | 0025 | -0135 | 5 |
| 4151 | ENSMUSG( | Mpp7 | 75739 | -00453679 | -00932351 | 033 | -01425 | 5 |
| 4153 | ENSMUSG( | Mpped2 | 77015 | -02428302 | -02900300 | 02225 | -047 | 5 |
| 4155 | ENSMUSG( | Mpst | 246221 | 003149934 | -02117625 | 00325 | -0375 | 5 |
| 4160 | ENSMUSG( | Mras | 17532 | 009027378 | -01279657 | 006 | -03125 | 5 |
| 4166 | ENSMUSG( | Mroh1 | 223658 | 031197545 | 003436625 | 029 | -0045 | 5 |
| 4167 | ENSMUSG( | Mrpl1 | 94061 | 006420497 | -01124166 | 01625 | -01575 | 5 |
| 4169 | ENSMUSG( | Mrpl11 | 66419 | 002087369 | -01622509 | 02075 | -017 | 5 |
| 4170 | ENSMUSG( | Mrpl12 | 56282 | 001507035 | -01083971 | 00875 | -0195 | 5 |
| 4177 | ENSMUSG( | Mrpl19 | 56284 | 011516065 | 003780686 | 00075 | -0125 | 5 |
| 4179 | ENSMUSG( | Mrpl20 | 66448 | -00277574 | -01624501 | 004 | -03725 | 5 |
| 4194 | ENSMUSG( | Mrpl39 | 27393 | 012397425 | -00757103 | 02775 | -0385 | 5 |
| 4198 | ENSMUSG( | Mrpl42 | 67270 | 007580712 | -01014857 | 00025 | -04175 | 5 |
| 4201 | ENSMUSG( | Mrpl45 | 67036 | 008985004 | -00969875 | -00125 | -0495 | 5 |
| 4203 | ENSMUSG( | Mrpl47 | 74600 | -00310143 | -01309451 | -0025 | -015 | 5 |
| 4206 | ENSMUSG( | Mrpl50 | 28028 | 004772253 | -01474097 | 00625 | -02225 | 5 |
| 4208 | ENSMUSG( | Mrpl52 | 68836 | 003700706 | -01904840 | 0175 | -02775 | 5 |
| 4214 | ENSMUSG( | Mrps10 | 64657 | -00370540 | -00416808 | 02 | -01175 | 5 |
| 4215 | ENSMUSG( | Mrps11 | 67994 | 005471358 | -01921785 | 02975 | -023 | 5 |
| 4225 | ENSMUSG( | Mrps21 | 66292 | -00135498 | -01943291 | -00225 | -02 | 5 |
| 4227 | ENSMUSG( | Mrps23 | 64656 | 002481474 | -01316949 | 00225 | -04775 | 5 |
| 4238 | ENSMUSG( | Mrps5 | 77721 | 001142366 | -00147767 | 023 | -02175 | 5 |
| 4239 | ENSMUSG( | Mrps6 | 121022 | 01929817( | -00032656 | -981308e-1 | -01825 | 5 |
| 4265 | ENSMUSG( | Mtch1 | 56462 | 000076441 | -01854700 | 02575 | -00775 | 5 |
| 4266 | ENSMUSG( | Mtch2 | 56428 | -00573969 | -00592008 | 006 | -03125 | 5 |
| 4271 | ENSMUSG( | Mtfp1 | 67900 | -00294948 | 000240036 | 0155 | -03625 | 5 |
| 4284 | ENSMUSG( | Mtmr14 | 97287 | 013872184 | 006028304 | 01775 | -00775 | 5 |
| 4285 | ENSMUSG( | Mtmr2 | 77116 | 002116085 | 003506579 | 006 | -01475 | 5 |
| 4287 | ENSMUSG( | Mtmr6 | 219135 | 002934978 | 001322838 | 006 | -009 | 5 |
| 4288 | ENSMUSG( | Mtmr7 | 54384 | 000101308 | -00806798 | 031 | -007 | 5 |
| 4293 | ENSMUSG( | Mtpn | 14489 | 000418166 | -01212874 | 0115 | -041 | 5 |
| 4307 | ENSMUSG( | Mul1 | 68350 | -00760760 | 001815442 | 024 | -0105 | 5 |
| 4311 | ENSMUSG( | Mvk | 17855 | -00285010 | -02967597 | 0015 | -0165 | 5 |
| 4312 | ENSMUSG( | Mvp | 78388 | 000394711 | 00510207 | -0005 | -01225 | 5 |
| 4342 | ENSMUSG( | Myo5b | 17919 | 009268215 | 010677975 | -00225 | -01475 | 5 |
| 4343 | ENSMUSG( | Myo6 | 17920 | 007851195 | -02111592 | -00125 | -0125 | 5 |
| 4352 | ENSMUSG( | Mzt1 | 76789 | -00260970 | -00798725 | 03275 | -03725 | 5 |
| 4385 | ENSMUSG( | Nap1l4 | 17955 | 031635671 | -00024089 | 02475 | -01175 | 5 |
| 4390 | ENSMUSG( | Napg | 108123 | 001502037 | 000505647 | 01325 | -04625 | 5 |
| 4392 | ENSMUSG( | Nars1 | 70223 | 008845406 | -00561296 | 0115 | -01575 | 5 |
| 4393 | ENSMUSG( | Nars2 | 244141 | -00908593 | -02417130 | 01475 | -00725 | 5 |
| 4412 | ENSMUSG( | Ncald | 52589 | 006525225 | -01517754 | 02525 | -03275 | 5 |
| 4424 | ENSMUSG( | Nck2 | 17974 | 005012326 | -00599371 | -0005 | -0135 | 5 |
| 4425 | ENSMUSG( | Nckap1 | 50884 | 009901184 | 002412468 | -0015 | -0145 | 5 |
| 4427 | ENSMUSG( | Nckap5l | 380969 | 003578901 | 006204268 | 00925 | -0485 | 5 |
| 4447 | ENSMUSG( | Ndrg2 | 29811 | 020708391 | -02232477 | 0355 | -054 | 5 |
| 4450 | ENSMUSG( | Ndufa1 | 54405 | -00129435 | -01512924 | 01925 | -05075 | 5 |
| 4460 | ENSMUSG( | Ndufa6 | 67130 | -00451000 | -01714795 | -0005 | -019 | 5 |
| 4463 | ENSMUSG( | Ndufa9 | 66108 | 004467305 | -01868732 | 003 | -03275 | 5 |
| 4466 | ENSMUSG( | Ndufaf2 | 75597 | -00337769 | -02789618 | 002 | -03325 | 5 |
| 4473 | ENSMUSG( | Ndufb1 | 1.03E+08 | 001521551 | -02072931 | 01625 | -03375 | 5 |
| 4479 | ENSMUSG( | Ndufb5 | 66046 | -00642162 | -02079472 | -0015 | -03175 | 5 |
| 4481 | ENSMUSG( | Ndufb7 | 66916 | 002132563 | -02091700 | 01575 | -026 | 5 |
| 4484 | ENSMUSG( | Ndufc2 | 68197 | -01999649 | -01788900 | 0065 | -028 | 5 |
| 4485 | ENSMUSG( | Ndufs1 | 227197 | 009330216 | -01773894 | 011 | -0285 | 5 |
| 4486 | ENSMUSG( | Ndufs2 | 226646 | 007064468 | -01854096 | -003 | -0225 | 5 |
| 4487 | ENSMUSG( | Ndufs3 | 68349 | -00524002 | -02021855 | -0005 | -04 | 5 |
| 4501 | ENSMUSG( | Necap2 | 66147 | 007695769 | -01250072 | 016 | -00475 | 5 |
| 4509 | ENSMUSG( | Nefl | 18039 | -00548960 | -02219382 | 01 | -01475 | 5 |
| 4515 | ENSMUSG( | Nek7 | 59125 | -00019870 | -02093654 | -00025 | -01375 | 5 |
| 4534 | ENSMUSG( | Nf1 | 18015 | 003237738 | 001380471 | 008 | -022 | 5 |
| 4537 | ENSMUSG( | Nfat5 | 54446 | -00302863 | -00148866 | 017 | -01575 | 5 |
| 4545 | ENSMUSG( | Nfu1 | 56748 | 000272568 | -01607759 | 0015 | -03675 | 5 |
| 4572 | ENSMUSG( | Nipsnap3b | 66536 | 001291488 | -01249523 | 0175 | -0095 | 5 |
| 4574 | ENSMUSG( | Nit1 | 27045 | 010636642 | -00696194 | 009 | -00975 | 5 |
| 4575 | ENSMUSG( | Nit2 | 52633 | 00316329( | -01454718 | 01925 | -0045 | 5 |

| | | | | | | | | |
|---|---|---|---|---|---|---|---|---|
| 4598 | ENSMUSG( | Noc2l | 57741 | 01332777 | -00030811 | 03525 | -0145 | 5 |
| 4600 | ENSMUSG( | Noc4l | 100608 | 01075962 | -01587055 | 0145 | -0095 | 5 |
| 4617 | ENSMUSG( | Nop53 | 68077 | 00292844 | -00509643 | 0025 | -0265 | 5 |
| 4619 | ENSMUSG( | Nop58 | 55989 | 00881924 | -01931029 | 02675 | -04925 | 5 |
| 4622 | ENSMUSG( | Nos1 | 18125 | 01749718 | -00942540 | 0035 | -0175 | 5 |
| 4623 | ENSMUSG( | Nos1ap | 70729 | 03800860 | 01149653 | 02725 | -014 | 5 |
| 4636 | ENSMUSG( | Npepps | 19155 | 01698914 | -00522671 | 0135 | -00875 | 5 |
| 4652 | ENSMUSG( | Nqo1 | 18104 | 00341718 | -01344659 | 019 | -02125 | 5 |
| 4682 | ENSMUSG( | Nsf | 18195 | 01818177 | -01164677 | 01975 | -01075 | 5 |
| 4697 | ENSMUSG( | Nt5dc3 | 103466 | 02108327 | -01145167 | 001 | -02525 | 5 |
| 4711 | ENSMUSG( | Nuak1 | 77976 | 02104397 | -00193657 | 01375 | -02225 | 5 |
| 4719 | ENSMUSG( | Nudc | 18221 | -00019029 | -03080830 | -00175 | -0235 | 5 |
| 4722 | ENSMUSG( | Nudcd3 | 209586 | 00055811 | 00311527 | 00775 | -00925 | 5 |
| 4728 | ENSMUSG( | Nudt16 | 75686 | -01518707 | -01577177 | 00825 | -034 | 5 |
| 4729 | ENSMUSG( | Nudt16l1 | 66911 | 00226620 | -01706815 | 02875 | -0095 | 5 |
| 4731 | ENSMUSG( | Nudt18 | 213484 | 00209032 | -01277773 | 0045 | -013 | 5 |
| 4732 | ENSMUSG( | Nudt19 | 110959 | 00809214 | -02214995 | 02625 | -0505 | 5 |
| 4733 | ENSMUSG( | Nudt2 | 66401 | 00531567 | -04122362 | 04525 | -037 | 5 |
| 4737 | ENSMUSG( | Nudt5 | 53893 | 00629164 | 00074735 | 003 | -02425 | 5 |
| 4745 | ENSMUSG( | Nup107 | 103468 | 00586902 | -00367882 | -00175 | -04425 | 5 |
| 4746 | ENSMUSG( | Nup133 | 234865 | 01242082 | 00045671 | 0065 | -01875 | 5 |
| 4748 | ENSMUSG( | Nup155 | 170762 | -00584525 | 02057410 | 02375 | -044 | 5 |
| 4758 | ENSMUSG( | Nup54 | 269113 | 01592600 | 01162181 | 02 | -02125 | 5 |
| 4760 | ENSMUSG( | Nup62 | 18226 | 01462313 | 00255113 | 00875 | -043 | 5 |
| 4762 | ENSMUSG( | Nup88 | 19069 | 00712576 | 00929805 | 0195 | -03675 | 5 |
| 4765 | ENSMUSG( | Nutf2 | 68051 | -00086615 | -01038061 | 0105 | -0085 | 5 |
| 4768 | ENSMUSG( | Nwd2 | 319807 | -00899661 | -0080927 | 0215 | -0095 | 5 |
| 4769 | ENSMUSG( | Nxf1 | 53319 | 01191589 | 02004652 | 00175 | -0415 | 5 |
| 4773 | ENSMUSG( | Nxt1 | 56488 | -01800000 | 00051977 | 036 | -0405 | 5 |
| 4777 | ENSMUSG( | Oat | 18242 | 01397554 | -01720109 | 02775 | -02475 | 5 |
| 4782 | ENSMUSG( | Ocrl | 320634 | 02106027 | 01078672 | 02325 | -03825 | 5 |
| 4783 | ENSMUSG( | Odr4 | 226499 | -00200714 | -00398231 | 01975 | -00825 | 5 |
| 4793 | ENSMUSG( | Olfm1 | 56177 | -01228419 | -00444554 | 0065 | -025 | 5 |
| 4801 | ENSMUSG( | Opa1 | 74143 | 00908239 | -00336261 | 01625 | -049 | 5 |
| 4806 | ENSMUSG( | Oplah | 75475 | 01151301 | -00464192 | 00625 | -023 | 5 |
| 4819 | ENSMUSG( | Osbpl1a | 64291 | 00336215 | -01590752 | 01875 | -00675 | 5 |
| 4823 | ENSMUSG( | Osbpl6 | 99031 | 02279678 | 01092430 | -392523e-1 | -02175 | 5 |
| 4833 | ENSMUSG( | Otub1 | 107260 | 00301520 | -01688455 | 03375 | -0155 | 5 |
| 4837 | ENSMUSG( | Otud6b | 72201 | 01394470 | -00896373 | 04225 | -0215 | 5 |
| 4846 | ENSMUSG( | Oxr1 | 170719 | 01372921 | -00071307 | 00875 | -02375 | 5 |
| 4857 | ENSMUSG( | Pacs1 | 107975 | 02102695 | -00165968 | 0125 | -01025 | 5 |
| 4867 | ENSMUSG( | Pafah2 | 100163 | 01340065 | -01280302 | 00275 | -03325 | 5 |
| 4893 | ENSMUSG( | Pank4 | 269614 | 01004913 | 00492482 | 0155 | -04025 | 5 |
| 4899 | ENSMUSG( | Papss2 | 23972 | 00962114 | -02012203 | 027 | -01875 | 5 |
| 4907 | ENSMUSG( | Park7 | 57320 | 00101890 | -02500105 | 0085 | -0155 | 5 |
| 4919 | ENSMUSG( | Paxbp1 | 67367 | -00400412 | 01634252 | 011 | -03175 | 5 |
| 4951 | ENSMUSG( | Pcdhga10 | 93722 | -01557860 | 00377120 | 0325 | -01075 | 5 |
| 4953 | ENSMUSG( | Pcdhga12 | 93724 | 01111605 | 02531330 | 034 | -031 | 5 |
| 4996 | ENSMUSG( | Pdcd5 | 56330 | 00701048 | -01733865 | 02475 | -03 | 5 |
| 5002 | ENSMUSG( | Pde11a | 241489 | 02559739 | -01283389 | 0075 | -01 | 5 |
| 5011 | ENSMUSG( | Pde4d | 238871 | 01968782 | 02300580 | 01875 | -02525 | 5 |
| 5014 | ENSMUSG( | Pde6d | 18582 | 00809508 | -01686760 | 017 | -04775 | 5 |
| 5031 | ENSMUSG( | Pdk2 | 18604 | 00462647 | -01614230 | 01675 | -015 | 5 |
| 5038 | ENSMUSG( | Pdlim7 | 67399 | 02195665 | 00199502 | 00225 | -01775 | 5 |
| 5039 | ENSMUSG( | Pdp1 | 381511 | -01738838 | -01541864 | 00625 | -01575 | 5 |
| 5041 | ENSMUSG( | Pdpk1 | 18607 | 00866674 | 00009365 | 00425 | -03 | 5 |
| 5044 | ENSMUSG( | Pds5b | 100710 | 00724826 | 01381085 | 01825 | -0225 | 5 |
| 5046 | ENSMUSG( | Pdxk | 216134 | 00723182 | 00556371 | 0055 | -0235 | 5 |
| 5057 | ENSMUSG( | Pecr | 111175 | -00821863 | -02032734 | 00625 | -01425 | 5 |
| 5059 | ENSMUSG( | Pef1 | 67898 | 00431593 | -00795957 | 01375 | -03575 | 5 |
| 5066 | ENSMUSG( | Pes1 | 64934 | 01679339 | -01000165 | 0055 | -01825 | 5 |
| 5070 | ENSMUSG( | Pex12 | 103737 | -00010131 | -00346007 | 0175 | -0475 | 5 |
| 5079 | ENSMUSG( | Pex5l | 58869 | -00075472 | -01393499 | -00025 | -013 | 5 |
| 5083 | ENSMUSG( | Pfdn1 | 67199 | -00070192 | -01879212 | 02775 | -0465 | 5 |
| 5085 | ENSMUSG( | Pfdn4 | 109054 | 00512307 | -01421869 | -00325 | -01925 | 5 |
| 5086 | ENSMUSG( | Pfdn5 | 56612 | 00645159 | -01599639 | 02175 | -00275 | 5 |
| 5090 | ENSMUSG( | Pfkl | 18641 | 01391539 | -01675202 | 002 | -02625 | 5 |
| 5091 | ENSMUSG( | Pfkm | 18642 | 01450875 | -01438756 | 007 | -036 | 5 |
| 5094 | ENSMUSG( | Pgam1 | 18648 | 00743746 | -01623796 | 02225 | -04475 | 5 |
| 5096 | ENSMUSG( | Pgam5 | 72542 | 01871448 | 00363857 | 01925 | -041 | 5 |
| 5101 | ENSMUSG( | Pgghg | 212974 | 00591496 | -01909686 | 019 | -004 | 5 |
| 5103 | ENSMUSG( | Pgk1 | 18655 | 00414812 | -03120505 | 0125 | -027 | 5 |
| 5110 | ENSMUSG( | Pgp | 67078 | 00970548 | -00997172 | 003 | -0205 | 5 |
| 5114 | ENSMUSG( | Pgs1 | 74451 | 01294103 | -00413973 | 02375 | -01375 | 5 |
| 5119 | ENSMUSG( | Phaf1 | 234678 | 00823631 | -00551132 | 01825 | -02475 | 5 |
| 5131 | ENSMUSG( | Phf5a | 68479 | 00908701 | -01176036 | 00525 | -0145 | 5 |
| 5134 | ENSMUSG( | Phgdh | 236539 | 01491575 | -01862784 | 02575 | -025 | 5 |
| 5136 | ENSMUSG( | Phka1 | 18679 | 01711820 | -01106007 | 02975 | -0075 | 5 |
| 5137 | ENSMUSG( | Phka2 | 110094 | 02207594 | 01158074 | 015 | -015 | 5 |
| 5138 | ENSMUSG( | Phkb | 102093 | 00875431 | -01726425 | 029 | -025 | 5 |
| 5139 | ENSMUSG( | Phkg1 | 18682 | 00300082 | -02110977 | 0335 | -036 | 5 |
| 5140 | ENSMUSG( | Phkg2 | 68961 | 00522266 | -00687268 | 02475 | -03925 | 5 |
| 5149 | ENSMUSG( | Phpt1 | 75454 | -00084596 | -00504384 | 02075 | -007 | 5 |
| 5154 | ENSMUSG( | Phyhipl | 70911 | -00524053 | -02622980 | 037 | -0435 | 5 |
| 5155 | ENSMUSG( | Pi4k2a | 84095 | 01256316 | -00420830 | 024 | -0035 | 5 |
| 5156 | ENSMUSG( | Pi4ka | 224020 | 01938958 | 00234947 | 0215 | -028 | 5 |
| 5177 | ENSMUSG( | Pik3cb | 74769 | 01583730 | 00349068 | 0135 | -03075 | 5 |
| 5180 | ENSMUSG( | Pik3r1 | 18708 | 02166652 | -00148552 | 0125 | -0285 | 5 |
| 5186 | ENSMUSG( | Pip4k2a | 18718 | -00145769 | -01380992 | -001 | -0235 | 5 |
| 5187 | ENSMUSG( | Pip4k2b | 108083 | 01625677 | -00263747 | 023 | -0365 | 5 |
| 5188 | ENSMUSG( | Pip4k2c | 117150 | -00807324 | -00117266 | 0035 | -03675 | 5 |
| 5189 | ENSMUSG( | Pip4p1 | 219024 | -00109041 | 00341979 | 01125 | -02375 | 5 |
| 5192 | ENSMUSG( | Pip5k1b | 18719 | 03078949 | 01569418 | 0205 | -052 | 5 |
| 5194 | ENSMUSG( | Pir | 69656 | -01848488 | -01519107 | 00375 | -0255 | 5 |
| 5215 | ENSMUSG( | Pla2g4e | 329502 | -02830345 | -03503013 | 02375 | -004 | 5 |
| 5216 | ENSMUSG( | Pla2g6 | 53357 | 00134262 | -00708398 | 00975 | -0265 | 5 |
| 5222 | ENSMUSG( | Plcb1 | 18795 | 00830876 | -00944986 | 004 | -0195 | 5 |
| 5225 | ENSMUSG( | Plcd1 | 18799 | -00281923 | -00620446 | 00225 | -0375 | 5 |
| 5236 | ENSMUSG( | Pld2 | 18806 | 01239358 | 00272356 | 003 | -035 | 5 |
| 5252 | ENSMUSG( | Plekhm2 | 69582 | 01876520 | -01667517 | 0115 | -0395 | 5 |
| 5256 | ENSMUSG( | Plin3 | 66905 | 00028170 | -00900066 | 00575 | -01475 | 5 |
| 5275 | ENSMUSG( | Pls3 | 102866 | -00118385 | 01167277 | 01375 | -01775 | 5 |
| 5290 | ENSMUSG( | Pmch | 110312 | -01345728 | -07370749 | -00175 | -052 | 5 |
| 5318 | ENSMUSG( | Poglut1 | 224143 | 00007090 | 00781361 | 0285 | -02225 | 5 |
| 5319 | ENSMUSG( | Poglut3 | 68304 | -00614819 | 01624272 | 02425 | -00675 | 5 |
| 5362 | ENSMUSG( | Ppa2 | 74776 | 01123563 | -01505893 | 027 | -00525 | 5 |
| 5366 | ENSMUSG( | Ppcs | 106564 | -01020188 | -01527470 | 01175 | -00925 | 5 |
| 5380 | ENSMUSG( | Ppif | 105675 | -00148417 | -00703758 | 01925 | -03425 | 5 |
| 5392 | ENSMUSG( | Ppm1f | 68606 | 02262953 | 00680496 | 021 | -0145 | 5 |
| 5394 | ENSMUSG( | Ppm1h | 319468 | 01464706 | 01174205 | 01825 | -00675 | 5 |
| 5402 | ENSMUSG( | Ppp1cc | 19047 | -01106164 | 00221187 | 022 | -01525 | 5 |
| 5436 | ENSMUSG( | Ppp2r5b | 225849 | 00761608 | 00275507 | 0155 | -0265 | 5 |
| 5437 | ENSMUSG( | Ppp2r5c | 26931 | 00777997 | -00796498 | 00025 | -015 | 5 |
| 5438 | ENSMUSG( | Ppp2r5d | 21770 | 00539652 | -00945109 | 02175 | -0085 | 5 |
| 5439 | ENSMUSG( | Ppp2r5e | 26932 | 00727981 | -00437914 | 002 | -0125 | 5 |
| 5455 | ENSMUSG( | Pptc7 | 320717 | 01308272 | 01884949 | 00525 | -053 | 5 |
| 5466 | ENSMUSG( | Prdx3 | 11757 | 00763342 | -01495291 | 014 | -01575 | 5 |
| 5469 | ENSMUSG( | Prdx6 | 11758 | 00826308 | -01444100 | 005 | -0185 | 5 |
| 5472 | ENSMUSG( | Prelp | 116847 | -01698274 | 00262906 | 0225 | -02675 | 5 |
| 5475 | ENSMUSG( | Prex1 | 277360 | 00256081 | -01940246 | 00375 | -027 | 5 |
| 5480 | ENSMUSG( | Prkaa2 | 108079 | 00833381 | 01276037 | 008 | -01875 | 5 |
| 5481 | ENSMUSG( | Prkab1 | 19079 | -02172476 | -00936265 | 02475 | -0385 | 5 |
| 5485 | ENSMUSG( | Prkag1 | 19082 | 00522407 | -00930495 | 01475 | -0235 | 5 |
| 5492 | ENSMUSG( | Prkcb | 18751 | 00730001 | 00368826 | -0015 | -0475 | 5 |
| 5493 | ENSMUSG( | Prkcd | 18753 | -02907117 | -03199482 | 0115 | -0115 | 5 |
| 5504 | ENSMUSG( | Prkg2 | 19092 | 00250787 | -00389235 | 01525 | -02625 | 5 |

| | | | | | | | | |
|---|---|---|---|---|---|---|---|---|
| 5512 | ENSMUSG( | Prmt8 | 381813 | 008359638 | -01122573 | 036 | -03325 | 5 |
| 5517 | ENSMUSG( | Prorp | 66132 | 013913494 | -00766273 | 01475 | -02925 | 5 |
| 5526 | ENSMUSG( | Prpf4 | 70052 | 003679368 | 011996208 | 00325 | -01725 | 5 |
| 5527 | ENSMUSG( | Prpf40a | 56194 | 005675829 | 004312631 | 01725 | -015 | 5 |
| 5530 | ENSMUSG( | Prpf6 | 68879 | 015779717 | -00766520 | 01825 | -04175 | 5 |
| 5534 | ENSMUSG( | Prps2 | 110639 | -00660098 | -00786238 | 01425 | -025 | 5 |
| 5552 | ENSMUSG( | Prxl2a | 70564 | 009502727 | 004477258 | 03075 | -007 | 5 |
| 5553 | ENSMUSG( | Prxl2b | 66469 | 000566408 | -04011574 | 01325 | -0155 | 5 |
| 5555 | ENSMUSG( | Psat1 | 107272 | 013917441 | -01122381 | 01525 | -02375 | 5 |
| 5561 | ENSMUSG( | Psma1 | 26440 | -00739093 | -01394956 | 0155 | -038 | 5 |
| 5570 | ENSMUSG( | Psmb2 | 26445 | -00795832 | -02289836 | 0515 | -02725 | 5 |
| 5572 | ENSMUSG( | Psmb4 | 19172 | 001296838 | -01666341 | 00875 | -03225 | 5 |
| 5579 | ENSMUSG( | Psmc3 | 19182 | 010308230 | -01105286 | 01 | -0185 | 5 |
| 5582 | ENSMUSG( | Psmc6 | 67089 | 006643755 | 003154018 | 01025 | -007 | 5 |
| 5587 | ENSMUSG( | Psmd13 | 23997 | 006711408 | -00826035 | 01875 | -00925 | 5 |
| 5588 | ENSMUSG( | Psmd14 | 59029 | 012210379 | -00791642 | 02475 | -01425 | 5 |
| 5597 | ENSMUSG( | Psme1 | 19186 | -00729772 | -00597728 | -0025 | -034 | 5 |
| 5605 | ENSMUSG( | Psmg3 | 66506 | 004141801 | 008881702 | 02775 | -016 | 5 |
| 5609 | ENSMUSG( | Ptar1 | 72351 | -01192652 | -02288946 | 00075 | -01825 | 5 |
| 5617 | ENSMUSG( | Ptdss2 | 27388 | -01244969 | 015251846 | 0415 | -01775 | 5 |
| 5622 | ENSMUSG( | Ptges3 | 56351 | 00564161 | -00881422 | 0375 | -01975 | 5 |
| 5626 | ENSMUSG( | Ptgr2 | 77219 | 000292441 | -02693012 | 01725 | -02475 | 5 |
| 5627 | ENSMUSG( | Ptgr3 | 225791 | 002388376 | -00498203 | 00525 | -0155 | 5 |
| 5631 | ENSMUSG( | Ptk2b | 19229 | 020335953 | -00672713 | 0005 | -01375 | 5 |
| 5638 | ENSMUSG( | Ptp4a2 | 19244 | -01018164 | -00934921 | 00375 | -01125 | 5 |
| 5644 | ENSMUSG( | Ptpn11 | 19247 | 008515924 | -01016520 | 013 | -029 | 5 |
| 5650 | ENSMUSG( | Ptpn4 | 19258 | -00770755 | -01132944 | 02375 | -01025 | 5 |
| 5653 | ENSMUSG( | Ptpn9 | 56294 | 020340997 | -00074968 | 0025 | -02475 | 5 |
| 5671 | ENSMUSG( | Ptrh2 | 217057 | 001790219 | -00600859 | 013 | -019 | 5 |
| 5698 | ENSMUSG( | Pygb | 110078 | 012062168 | -02060925 | 01775 | -03225 | 5 |
| 5699 | ENSMUSG( | Pygl | 110095 | 007336874 | -00248056 | 02225 | -009 | 5 |
| 5721 | ENSMUSG( | Rab11b | 19326 | 002172665 | -01365475 | 007 | -01225 | 5 |
| 5728 | ENSMUSG( | Rab14 | 68365 | 006836311 | -00143235 | 0235 | -008 | 5 |
| 5731 | ENSMUSG( | Rab1a | 19324 | 001991628 | -01175113 | -00025 | -023 | 5 |
| 5737 | ENSMUSG( | Rab26 | 328778 | 018109831 | -00169196 | 00875 | -0375 | 5 |
| 5739 | ENSMUSG( | Rab27b | 80718 | -00254999 | -00891412 | 00225 | -04 | 5 |
| 5746 | ENSMUSG( | Rab33a | 19337 | -00596188 | -02588298 | 0175 | -03025 | 5 |
| 5747 | ENSMUSG( | Rab33b | 19338 | -00685784 | -01288169 | 03 | -05175 | 5 |
| 5755 | ENSMUSG( | Rab3c | 67295 | 022181463 | 005893444 | 05425 | -04225 | 5 |
| 5756 | ENSMUSG( | Rab3d | 19340 | 012661229 | 016822280 | 03025 | -0515 | 5 |
| 5763 | ENSMUSG( | Rab4a | 19341 | 016417161 | 020991905 | 0415 | -0265 | 5 |
| 5766 | ENSMUSG( | Rab5b | 19344 | 007316998 | -01539139 | 0055 | -02975 | 5 |
| 5767 | ENSMUSG( | Rab5c | 19345 | 002047905 | -01809867 | 003 | -016 | 5 |
| 5772 | ENSMUSG( | Rab8b | 235442 | 007452215 | -00290773 | 00325 | -0365 | 5 |
| 5773 | ENSMUSG( | Rab9b | 319642 | 004323969 | -00292145 | 02175 | -022 | 5 |
| 5777 | ENSMUSG( | Rabepk | 227746 | -00165135 | -01432256 | 016 | -00975 | 5 |
| 5784 | ENSMUSG( | Rabl2 | 68708 | 012263334 | 000126915 | 0155 | -022 | 5 |
| 5788 | ENSMUSG( | Rac2 | 19354 | -01154801 | -03721671 | -004 | -01875 | 5 |
| 5796 | ENSMUSG( | Rae1 | 66679 | -00910283 | -00419950 | 0185 | -0415 | 5 |
| 5805 | ENSMUSG( | Ralgapb | 228850 | 001821191 | 010949121 | 0045 | -01475 | 5 |
| 5811 | ENSMUSG( | Ran | 19384 | 003003090 | -01461633 | 03125 | -0445 | 5 |
| 5812 | ENSMUSG( | Ranbp1 | 19385 | -00121087 | -01114617 | 024 | -005 | 5 |
| 5817 | ENSMUSG( | Ranbp6 | 240614 | -00353828 | 015293664 | 02325 | -0105 | 5 |
| 5819 | ENSMUSG( | Rangap1 | 19387 | 012449375 | -01275492 | 0205 | -0065 | 5 |
| 5823 | ENSMUSG( | Rap1gap2 | 380711 | 024959005 | 007221259 | 02425 | -00275 | 5 |
| 5824 | ENSMUSG( | Rap1gds1 | 229877 | 003043032 | -01812689 | 02525 | -002 | 5 |
| 5830 | ENSMUSG( | Rapgef3 | 223864 | 016991433 | 001034021 | -0015 | -05325 | 5 |
| 5832 | ENSMUSG( | Rapgef5 | 217944 | 017052175 | 021122134 | 03875 | -02975 | 5 |
| 5834 | ENSMUSG( | Rapgefl1 | 268480 | 022615543 | 011397725 | 00725 | -03075 | 5 |
| 5836 | ENSMUSG( | Rars1 | 104458 | 005083642 | -01494381 | 02125 | -0195 | 5 |
| 5837 | ENSMUSG( | Rars2 | 109093 | 007865068 | 002863718 | 0045 | -03 | 5 |
| 5839 | ENSMUSG( | Rasa2 | 114713 | -00239627 | -03641704 | 00775 | -0415 | 5 |
| 5841 | ENSMUSG( | Rasal1 | 19415 | 030155053 | 016587517 | 03025 | -0125 | 5 |
| 5847 | ENSMUSG( | Rasgrf2 | 19418 | 014918852 | 004003103 | 0105 | -01225 | 5 |
| 5855 | ENSMUSG( | Rbbp4 | 19646 | 001413383 | -00155949 | 01525 | -01025 | 5 |
| 5859 | ENSMUSG( | Rbbp9 | 26450 | 000508291 | -02151029 | 02325 | -018 | 5 |
| 5867 | ENSMUSG( | Rbm10 | 236732 | 023035653 | -00057127 | 00075 | -01725 | 5 |
| 5882 | ENSMUSG( | Rbm39 | 170791 | 003424920 | 022889578 | 0105 | -019 | 5 |
| 5891 | ENSMUSG( | Rbmx2 | 209003 | 017725537 | -00139854 | 01225 | -01925 | 5 |
| 5892 | ENSMUSG( | Rbp1 | 19659 | 008531303 | -02310799 | 03925 | -054 | 5 |
| 5900 | ENSMUSG( | Rcbtb1 | 71330 | -00774076 | -00884332 | 0145 | -03 | 5 |
| 5935 | ENSMUSG( | Rexo2 | 104444 | -00043587 | -01388292 | 02525 | -0135 | 5 |
| 5937 | ENSMUSG( | Rfc2 | 19718 | 012547346 | -00411477 | 0035 | -03975 | 5 |
| 5946 | ENSMUSG( | Rfx3 | 19726 | 006113543 | 006927325 | 001 | -02175 | 5 |
| 5957 | ENSMUSG( | Rgs17 | 56533 | 020625644 | 019514873 | 0335 | -02075 | 5 |
| 5959 | ENSMUSG( | Rgs20 | 58175 | 002856717 | -03140192 | 01275 | -03 | 5 |
| 5962 | ENSMUSG( | Rgs6 | 50779 | 003762016 | -01977805 | -003 | -02725 | 5 |
| 5963 | ENSMUSG( | Rgs7 | 24012 | 002886155 | -02230213 | 01975 | -01975 | 5 |
| 5964 | ENSMUSG( | Rgs7bp | 52882 | -00038320 | 005653225 | -00125 | -032 | 5 |
| 5977 | ENSMUSG( | Rhoq | 104215 | -02161866 | -02957506 | 0175 | -02725 | 5 |
| 5978 | ENSMUSG( | Rhot1 | 59040 | 007520196 | 004117101 | 02275 | -013 | 5 |
| 5987 | ENSMUSG( | Rida | 15473 | 006787409 | -02431686 | 02525 | -0575 | 5 |
| 5990 | ENSMUSG( | Rimbp2 | 231760 | 015675226 | 011690291 | 035 | -021 | 5 |
| 5998 | ENSMUSG( | Rint1 | 72772 | 019001556 | 010252818 | 01725 | -0075 | 5 |
| 6004 | ENSMUSG( | Rit1 | 19769 | 013475448 | -00304797 | 044 | -024 | 5 |
| 6005 | ENSMUSG( | Rit2 | 19762 | -00118562 | -02330234 | 0225 | -0295 | 5 |
| 6009 | ENSMUSG( | Rmc1 | 76482 | 008149498 | -00427561 | 01625 | -0065 | 5 |
| 6010 | ENSMUSG( | Rmdn1 | 66302 | -00217895 | 008190052 | 01025 | -01175 | 5 |
| 6011 | ENSMUSG( | Rmdn2 | 381110 | 004212665 | 002533043 | 0295 | -0565 | 5 |
| 6012 | ENSMUSG( | Rmdn3 | 67809 | 005846069 | -00253972 | 00025 | -02075 | 5 |
| 6025 | ENSMUSG( | Rnf112 | 22671 | 032028496 | 014769998 | 00325 | -02525 | 5 |
| 6035 | ENSMUSG( | Rnf141 | 67150 | -00175794 | -02196853 | 0355 | -02475 | 5 |
| 6036 | ENSMUSG( | Rnf146 | 68031 | 012107832 | 000243893 | 02325 | -002 | 5 |
| 6064 | ENSMUSG( | Ro60 | 20822 | 001321065 | 009202142 | 0105 | -0355 | 5 |
| 6068 | ENSMUSG( | Rock2 | 19878 | 009023113 | -00871764 | -00225 | -02625 | 5 |
| 6069 | ENSMUSG( | Rogdi | 66049 | 018201863 | -00141742 | 002 | -013 | 5 |
| 6072 | ENSMUSG( | Rpa1 | 68275 | 022462677 | -00631776 | -00175 | -02575 | 5 |
| 6077 | ENSMUSG( | Rpe | 66646 | 010844915 | 004595408 | 017 | -00325 | 5 |
| 6101 | ENSMUSG( | Rpl26 | 19941 | 013278917 | -01142649 | 0165 | -0345 | 5 |
| 6102 | ENSMUSG( | Rpl27 | 19942 | 002292041 | -02807481 | 0285 | -0065 | 5 |
| 6106 | ENSMUSG( | Rpl3 | 27367 | 006524979 | -00849036 | 02975 | -0225 | 5 |
| 6121 | ENSMUSG( | Rpl6 | 19988 | 006873948 | -01052288 | 02225 | -0195 | 5 |
| 6125 | ENSMUSG( | Rpl9 | 20005 | 011563894 | -01191395 | 0555 | -03325 | 5 |
| 6143 | ENSMUSG( | Rps15a | 267019 | 006418748 | -01292884 | 03675 | -04325 | 5 |
| 6155 | ENSMUSG( | Rps27 | 57294 | 013832561 | -01545652 | -001 | -021 | 5 |
| 6156 | ENSMUSG( | Rps27a | 78294 | 014095845 | -01100195 | 02625 | -035 | 5 |
| 6168 | ENSMUSG( | Rps6kb1 | 72508 | 003085040 | 012567024 | 014 | -04775 | 5 |
| 6173 | ENSMUSG( | Rpsa | 16785 | 009466377 | -01458902 | 04175 | -02175 | 5 |
| 6182 | ENSMUSG( | Rras2 | 66922 | 022435124 | 001362298 | 00725 | -01125 | 5 |
| 6195 | ENSMUSG( | Rsrc1 | 66880 | -00774218 | -02107510 | 0205 | -015 | 5 |
| 6199 | ENSMUSG( | Rtcb | 28088 | -01085942 | -00309261 | 02575 | -007 | 5 |
| 6222 | ENSMUSG( | Ruvbl1 | 56505 | 000974109 | -01678569 | 00525 | -01425 | 5 |
| 6229 | ENSMUSG( | S100a1 | 20193 | 394e-07 | -02772184 | 0255 | -0275 | 5 |
| 6232 | ENSMUSG( | S100a13 | 20196 | 007006356 | -02348538 | 01325 | -02475 | 5 |
| 6242 | ENSMUSG( | Sacm1l | 83493 | -00098745 | 008647751 | 01625 | -03175 | 5 |
| 6255 | ENSMUSG( | Samm50 | 68653 | 016666179 | 000596858 | -001 | -01475 | 5 |
| 6268 | ENSMUSG( | Sart1 | 20227 | 012854613 | -00463057 | 00725 | -0185 | 5 |
| 6276 | ENSMUSG( | Sbf1 | 77980 | 010544503 | 000275832 | 00775 | -01975 | 5 |
| 6295 | ENSMUSG( | Scfd1 | 76983 | 011906214 | -00790778 | 01675 | -04925 | 5 |
| 6296 | ENSMUSG( | Scfd2 | 212986 | 008640969 | -01398717 | 0375 | -0305 | 5 |
| 6323 | ENSMUSG( | Scyl1 | 78891 | 015874944 | 004492445 | 01425 | -02 | 5 |
| 6346 | ENSMUSG( | Sdsl | 257635 | 001954810 | -02113214 | 0105 | -013 | 5 |
| 6351 | ENSMUSG( | Sec14l2 | 67815 | 00395404 | -01855955 | 0065 | -01925 | 5 |
| 6375 | ENSMUSG( | Selenom | 114679 | -01527498 | -01016790 | 03325 | -03125 | 5 |
| 6395 | ENSMUSG( | Senp2 | 75826 | 007562525 | -00610764 | 0015 | -019 | 5 |
| 6398 | ENSMUSG( | Senp8 | 71599 | 011072954 | -00053620 | 01175 | -01325 | 5 |

| | | | | | | | | |
|---|---|---|---|---|---|---|---|---|
| 6402 | ENSMUSG( | Septin10 | 103080 | -00789361 | -01670802 | 01575 | -04475 | 5 |
| 6405 | ENSMUSG( | Septin3 | 24050 | 015935985 | -00492257 | 014 | -022 | 5 |
| 6407 | ENSMUSG( | Septin5 | 18951 | 015495995 | -00009593 | 00025 | -031 | 5 |
| 6408 | ENSMUSG( | Septin6 | 56526 | 015571760 | -01754388 | 006 | -02225 | 5 |
| 6409 | ENSMUSG( | Septin7 | 235072 | 008178496 | -00726757 | 0035 | -03675 | 5 |
| 6416 | ENSMUSG( | Serhl | 68607 | 01556956 | -01216514 | 02275 | -01 | 5 |
| 6443 | ENSMUSG( | Sf3a1 | 67465 | 017721249 | 006586511 | 00725 | -04675 | 5 |
| 6460 | ENSMUSG( | Sfxn3 | 94280 | 012069213 | -00950681 | 03075 | -0115 | 5 |
| 6474 | ENSMUSG( | Sgta | 52551 | -00151968 | -01346588 | 022 | -014 | 5 |
| 6485 | ENSMUSG( | Sh3bp5 | 24056 | 016726289 | -00233456 | 0265 | -02525 | 5 |
| 6506 | ENSMUSG( | Shc3 | 20418 | 019956474 | -01199868 | -002 | -0205 | 5 |
| 6516 | ENSMUSG( | Shoc2 | 56392 | 005507287 | 007734977 | 03125 | -0355 | 5 |
| 6521 | ENSMUSG( | Shroom2 | 110380 | -00199472 | 000852357 | 008 | -01825 | 5 |
| 6528 | ENSMUSG( | Sik2 | 235344 | -00469989 | -00097738 | 0095 | -047 | 5 |
| 6530 | ENSMUSG( | Sike1 | 66641 | 005260231 | -00312950 | 0115 | -00875 | 5 |
| 6532 | ENSMUSG( | Sin3a | 20466 | 007222466 | 007792759 | 019 | -015 | 5 |
| 6533 | ENSMUSG( | Sin3b | 20467 | 010915465 | -00391183 | 02225 | -011 | 5 |
| 6536 | ENSMUSG( | Sipa1l2 | 244668 | 002868005 | 003170437 | 01975 | -0035 | 5 |
| 6575 | ENSMUSG( | Slc1a3 | 20512 | -00156267 | 003084130 | 025 | -034 | 5 |
| 6590 | ENSMUSG( | Slc25a10 | 27376 | 004095410 | 004523560 | 0225 | -01675 | 5 |
| 6595 | ENSMUSG( | Slc25a15 | 18408 | -00197860 | -00689583 | 01375 | -0335 | 5 |
| 6606 | ENSMUSG( | Slc25a26 | 67582 | 002685221 | -00147349 | 00925 | -02925 | 5 |
| 6615 | ENSMUSG( | Slc25a4 | 11739 | 001678324 | -01687417 | 01775 | -0225 | 5 |
| 6619 | ENSMUSG( | Slc25a46 | 67453 | 004191807 | 000738332 | 03075 | -00825 | 5 |
| 6620 | ENSMUSG( | Slc25a5 | 11740 | 002277258 | -01392595 | 02625 | -0265 | 5 |
| 6637 | ENSMUSG( | Slc30a3 | 22784 | -00656837 | 012121986 | 0195 | -01525 | 5 |
| 6689 | ENSMUSG( | Slc66a2 | 66943 | -01042291 | 012621006 | 03025 | -0325 | 5 |
| 6737 | ENSMUSG( | Slu7 | 193116 | 008387345 | -00873887 | 00175 | -019 | 5 |
| 6759 | ENSMUSG( | Smchd1 | 74355 | 006199240 | 003623240 | -00175 | -03325 | 5 |
| 6775 | ENSMUSG( | Smn1 | 20595 | 009314798 | -00455398 | 046 | -03075 | 5 |
| 6781 | ENSMUSG( | Smpd4 | 77626 | 027837251 | 001268747 | 0125 | -02825 | 5 |
| 6785 | ENSMUSG( | Smu1 | 74255 | -00009550 | -01220765 | 01075 | -05275 | 5 |
| 6791 | ENSMUSG( | Snap23 | 20619 | 012614525 | 002447300 | 00575 | -0145 | 5 |
| 6809 | ENSMUSG( | Snrnp70 | 20637 | 010933212 | 014079972 | 02575 | -03225 | 5 |
| 6810 | ENSMUSG( | Snrpa | 53607 | -00029498 | -01718485 | 02575 | -0525 | 5 |
| 6812 | ENSMUSG( | Snrpb | 20638 | 005388787 | -00315082 | 03275 | -012 | 5 |
| 6815 | ENSMUSG( | Snrpd1 | 20641 | -00132406 | -01365546 | 0315 | -04875 | 5 |
| 6816 | ENSMUSG( | Snrpd2 | 107686 | -00471118 | -02607094 | 026 | -05475 | 5 |
| 6817 | ENSMUSG( | Snrpd3 | 67332 | -00283341 | -01894679 | 02125 | -02175 | 5 |
| 6819 | ENSMUSG( | Snrpf | 69878 | -00419594 | -02218921 | 02175 | -03675 | 5 |
| 6820 | ENSMUSG( | Snrpg | 68011 | 000945696 | -01938977 | 01375 | -0125 | 5 |
| 6827 | ENSMUSG( | Snx1 | 56440 | 000867735 | -00907926 | -0005 | -03125 | 5 |
| 6831 | ENSMUSG( | Snx13 | 217463 | -00534227 | 001591771 | 01475 | -007 | 5 |
| 6840 | ENSMUSG( | Snx24 | 69226 | 008915908 | -00335255 | 005 | -03775 | 5 |
| 6865 | ENSMUSG( | Sord | 20322 | 007598358 | 003184535 | 013 | -04 | 5 |
| 6871 | ENSMUSG( | Sowahb | 78088 | -00044811 | -00252899 | 0015 | -02475 | 5 |
| 6898 | ENSMUSG( | Spg21 | 27965 | 000986635 | 003513044 | 02825 | -023 | 5 |
| 6921 | ENSMUSG( | Spryd4 | 66701 | 014890590 | -00673132 | 00375 | -0295 | 5 |
| 6923 | ENSMUSG( | Sptan1 | 20740 | 004162426 | 004610422 | 007 | -03075 | 5 |
| 6925 | ENSMUSG( | Sptbn1 | 20742 | -00090094 | 003699719 | 019 | -01625 | 5 |
| 6926 | ENSMUSG( | Sptbn2 | 20743 | 025982789 | 017301398 | 0125 | -045 | 5 |
| 6931 | ENSMUSG( | Sqor | 59010 | 018499537 | -00088396 | 00675 | -03225 | 5 |
| 6934 | ENSMUSG( | Srbd1 | 78586 | 006988199 | -01602151 | -00125 | -0505 | 5 |
| 6935 | ENSMUSG( | Src | 20779 | 010512009 | -00882122 | 013 | -03125 | 5 |
| 6943 | ENSMUSG( | Sri | 109552 | 001817883 | -01806709 | 01575 | -025 | 5 |
| 6944 | ENSMUSG( | Srm | 20810 | -01037539 | 002752405 | 01525 | -017 | 5 |
| 6945 | ENSMUSG( | Srp14 | 20813 | 000780006 | -01491135 | 0345 | -01975 | 5 |
| 6949 | ENSMUSG( | Srp9 | 27058 | 008463839 | -01046553 | 05225 | -027 | 5 |
| 6950 | ENSMUSG( | Srpk1 | 20815 | 002971611 | 005158627 | 0275 | -00575 | 5 |
| 6985 | ENSMUSG( | Ssrp1 | 20833 | 014362436 | -00067758 | 0235 | -01625 | 5 |
| 7003 | ENSMUSG( | Stambpl1 | 76630 | 000967672 | -01005817 | 00175 | -0145 | 5 |
| 7008 | ENSMUSG( | Stard7 | 99138 | 006128341 | -01069668 | 015 | -01375 | 5 |
| 7012 | ENSMUSG( | Stat5b | 20851 | 015749347 | 001422476 | 033 | -008 | 5 |
| 7033 | ENSMUSG( | Stk39 | 53416 | 005416536 | -01218185 | 022 | -0125 | 5 |
| 7044 | ENSMUSG( | Strada | 72149 | 012006358 | -00677720 | 00125 | -02425 | 5 |
| 7046 | ENSMUSG( | Strap | 20901 | 001390306 | -01513048 | 012 | -03075 | 5 |
| 7048 | ENSMUSG( | Strip1 | 229707 | 011635934 | 001809282 | 0015 | -02625 | 5 |
| 7050 | ENSMUSG( | Strn | 268980 | -00963847 | -00514255 | 00425 | -01575 | 5 |
| 7051 | ENSMUSG( | Strn3 | 94186 | 005151370 | -00347045 | 00725 | -02225 | 5 |
| 7057 | ENSMUSG( | Stx12 | 100226 | 007275939 | -01113943 | 02175 | -012 | 5 |
| 7058 | ENSMUSG( | Stx16 | 228960 | 008345185 | 001246561 | 0115 | -0215 | 5 |
| 7064 | ENSMUSG( | Stx3 | 20908 | -00920267 | 004932043 | 00975 | -014 | 5 |
| 7066 | ENSMUSG( | Stx7 | 53331 | 003849939 | -00841066 | 013 | -0125 | 5 |
| 7068 | ENSMUSG( | Stxbp1 | 20910 | 005503943 | -00638817 | 0235 | -02275 | 5 |
| 7070 | ENSMUSG( | Stxbp3 | 20912 | -00194068 | -00226564 | 00075 | -01725 | 5 |
| 7073 | ENSMUSG( | Stxbp5l | 207227 | -00673265 | -00353904 | 00525 | -019 | 5 |
| 7074 | ENSMUSG( | Stxbp6 | 217517 | 007572900 | -01353984 | -00225 | -05025 | 5 |
| 7076 | ENSMUSG( | Sucla2 | 20916 | 004346909 | -01776138 | 0015 | -053 | 5 |
| 7087 | ENSMUSG( | Sult4a1 | 29859 | 004525099 | -01150093 | 029 | -02225 | 5 |
| 7093 | ENSMUSG( | Suox | 211389 | 010100752 | 003333841 | 00675 | -025 | 5 |
| 7116 | ENSMUSG( | Sympk | 68188 | 025389465 | 003747518 | 027 | -011 | 5 |
| 7118 | ENSMUSG( | Syn2 | 20965 | 021814006 | -01119030 | 0215 | -0165 | 5 |
| 7124 | ENSMUSG( | Syngap1 | 240057 | 02448047 | 014711303 | 0095 | -0215 | 5 |
| 7153 | ENSMUSG( | Sytl2 | 83671 | -00853876 | -01745772 | 01675 | -02775 | 5 |
| 7182 | ENSMUSG( | Tagln3 | 56370 | 010135863 | -03032077 | 0055 | -01025 | 5 |
| 7200 | ENSMUSG( | Tars2 | 71807 | 004059024 | -00360333 | 02125 | -02975 | 5 |
| 7209 | ENSMUSG( | Tbc1d10b | 68449 | 009573803 | -00569316 | 011 | -036 | 5 |
| 7211 | ENSMUSG( | Tbc1d13 | 70296 | 010305356 | 007379860 | 01175 | -02475 | 5 |
| 7218 | ENSMUSG( | Tbc1d22b | 381085 | 012432300 | -00596999 | 01075 | -008 | 5 |
| 7220 | ENSMUSG( | Tbc1d24 | 224617 | 006921138 | 004887095 | 0145 | -031 | 5 |
| 7223 | ENSMUSG( | Tbc1d32 | 544696 | -01672546 | -01534521 | 03025 | -006 | 5 |
| 7260 | ENSMUSG( | Tcp1 | 21454 | 009864582 | -01308269 | 004 | -02575 | 5 |
| 7261 | ENSMUSG( | Tcp11l1 | 320554 | 007767857 | -01408945 | 0225 | -0015 | 5 |
| 7268 | ENSMUSG( | Tdrd7 | 100121 | 006333668 | -00903079 | 0125 | -01675 | 5 |
| 7284 | ENSMUSG( | Terf2 | 21750 | 004981555 | -00830561 | 03825 | -01625 | 5 |
| 7285 | ENSMUSG( | Terf2ip | 57321 | 013663142 | -00751379 | 0095 | -00825 | 5 |
| 7316 | ENSMUSG( | Thnsl1 | 208967 | 003093747 | -01351014 | 0275 | -0455 | 5 |
| 7320 | ENSMUSG( | Thoc3 | 73666 | 007562512 | -01754009 | 03 | -01 | 5 |
| 7321 | ENSMUSG( | Thoc5 | 107829 | 014609366 | -00704412 | 02225 | -00775 | 5 |
| 7331 | ENSMUSG( | Thumpd3 | 14911 | 011484399 | -01605783 | 0005 | -0155 | 5 |
| 7341 | ENSMUSG( | Timm13 | 30055 | -00288674 | -02637782 | -00225 | -0135 | 5 |
| 7347 | ENSMUSG( | Timm29 | 69773 | 008541688 | -00619241 | 045 | -0195 | 5 |
| 7352 | ENSMUSG( | Timm9 | 30056 | -00318272 | -00343009 | -00175 | -01675 | 5 |
| 7381 | ENSMUSG( | Tmcc1 | 330401 | 012331865 | 031386193 | 018 | -04025 | 5 |
| 7385 | ENSMUSG( | Tmco6 | 71983 | 000328031 | 014081139 | 01425 | -03375 | 5 |
| 7405 | ENSMUSG( | Tmem126b | 68472 | 001209646 | -00242075 | 012 | -044 | 5 |
| 7455 | ENSMUSG( | Tmem243 | 652925 | 000721031 | 014533506 | 01 | -01775 | 5 |
| 7458 | ENSMUSG( | Tmem254 | 66039 | -03444504 | -03683342 | 03325 | -04025 | 5 |
| 7493 | ENSMUSG( | Tmod2 | 50876 | 016795021 | -00063743 | 025 | -0185 | 5 |
| 7517 | ENSMUSG( | Tnpo1 | 238799 | 005959172 | 015521383 | 03075 | -0375 | 5 |
| 7518 | ENSMUSG( | Tnpo2 | 212999 | 013967705 | -00557762 | 001 | -01425 | 5 |
| 7519 | ENSMUSG( | Tnpo3 | 320938 | 007557574 | 001060180 | 022 | -023 | 5 |
| 7527 | ENSMUSG( | Toe1 | 68276 | -00885639 | -00133282 | 01425 | -02975 | 5 |
| 7531 | ENSMUSG( | Tom1l1 | 71943 | 011987011 | -00404080 | 0315 | -0255 | 5 |
| 7536 | ENSMUSG( | Tomm40 | 53333 | -00307530 | -00654527 | 01175 | -01825 | 5 |
| 7538 | ENSMUSG( | Tomm5 | 68512 | 001812191 | -02224868 | 0115 | -018 | 5 |
| 7544 | ENSMUSG( | Tor1aip1 | 208263 | 003052402 | -00359428 | 0125 | -018 | 5 |
| 7557 | ENSMUSG( | Tpgs1 | 110012 | -00008315 | -01164356 | -785046e-1 | -0365 | 5 |
| 7568 | ENSMUSG( | Tppp | 72948 | 011971565 | -01244522 | -00275 | -02475 | 5 |
| 7581 | ENSMUSG( | Traf6 | 22034 | -01061455 | -00375380 | 0175 | -016 | 5 |
| 7588 | ENSMUSG( | Trap1 | 68015 | 005162103 | -02240362 | 00175 | -028 | 5 |
| 7590 | ENSMUSG( | Trappc10 | 216131 | 016755962 | 013146301 | 0275 | -0195 | 5 |
| 7591 | ENSMUSG( | Trappc11 | 320714 | 009603795 | 005294353 | 00475 | -00975 | 5 |
| 7598 | ENSMUSG( | Trappc4 | 60409 | 000789059 | -01436835 | 043 | -026 | 5 |
| 7600 | ENSMUSG( | Trappc6b | 78232 | -00367413 | -00785911 | 00775 | -033 | 5 |
| 7606 | ENSMUSG( | Triap1 | 69076 | 018472628 | 007301645 | 018 | -003 | 5 |

| | | | | | | | | |
|---|---|---|---|---|---|---|---|---|
| 7609 | ENSMUSG( | Trim23 | 81003 | -00002133 | 00467167 | 03475 | -043 | 5 |
| 7625 | ENSMUSG( | Trim66 | 330627 | 020487712 | 00063027 | 01925 | -01475 | 5 |
| 7627 | ENSMUSG( | Trim9 | 94090 | 010313282 | -02308561 | 038 | -01175 | 5 |
| 7628 | ENSMUSG( | Trio | 223435 | 001754723 | 00888585 | 00975 | -00975 | 5 |
| 7653 | ENSMUSG( | Trpm3 | 226025 | 00475983C | -02271780 | -00175 | -03925 | 5 |
| 7670 | ENSMUSG( | Tsnax | 53424 | 010814413 | -00959650 | 00225 | -0125 | 5 |
| 7699 | ENSMUSG( | Ttc27 | 74196 | 01033649 | -00704658 | 0135 | -022 | 5 |
| 7704 | ENSMUSG( | Ttc39c | 72747 | 00354529 | 00202314 | 00525 | -03025 | 5 |
| 7706 | ENSMUSG( | Ttc7b | 104718 | 01754201 | 00123300 | 00875 | -019 | 5 |
| 7718 | ENSMUSG( | Ttpal | 76080 | -00328915 | -01069672 | 02575 | -0185 | 5 |
| 7728 | ENSMUSG( | Tubb2a | 22151 | 012186473 | -02135809 | 00375 | -0135 | 5 |
| 7735 | ENSMUSG( | Tubg1 | 103733 | 012254133 | -01012589 | 0195 | -0045 | 5 |
| 7738 | ENSMUSG( | Tubgcp3 | 259279 | 00492538C | 01049367 | 02275 | -0365 | 5 |
| 7743 | ENSMUSG( | Tulp3 | 22158 | 01745940 | 01032822C | 0305 | -0185 | 5 |
| 7745 | ENSMUSG( | Tusc2 | 80385 | 009372574 | -00797946 | 029 | -00775 | 5 |
| 7750 | ENSMUSG( | Twf1 | 19230 | 00614998 | -00547303 | 0275 | -02325 | 5 |
| 7751 | ENSMUSG( | Twf2 | 23999 | 01699213 | -00055276 | 03125 | -0395 | 5 |
| 7762 | ENSMUSG( | Txnl1 | 53382 | -00184118 | -00690586 | 0335 | -03325 | 5 |
| 7763 | ENSMUSG( | Txnl4a | 27366 | -00241412 | -00352635 | 0375 | -02825 | 5 |
| 7772 | ENSMUSG( | Tyw5 | 68736 | 006716544 | 02296617 | 0435 | -021 | 5 |
| 7775 | ENSMUSG( | U2surp | 67958 | 001658604 | 00503245 | 004 | -04425 | 5 |
| 7787 | ENSMUSG( | Ubap1 | 67123 | 008058671 | 000105341 | -001 | -03525 | 5 |
| 7790 | ENSMUSG( | Ubash3b | 72828 | -01048215 | -02826198 | 02075 | -004 | 5 |
| 7797 | ENSMUSG( | Ube2f | 67921 | 00818359 | -00954479 | -00025 | -0215 | 5 |
| 7810 | ENSMUSG( | Ube2q2 | 109161 | 003433301 | -00135322 | 00675 | -0455 | 5 |
| 7818 | ENSMUSG( | Ube3b | 117146 | 01423398C | 00583396 | 0135 | -025 | 5 |
| 7825 | ENSMUSG( | Ubl4a | 27643 | 00386324 | -01331626 | 01225 | -03025 | 5 |
| 7830 | ENSMUSG( | Ubp1 | 22221 | 01964510 | 01334707 | -00025 | -04425 | 5 |
| 7832 | ENSMUSG( | Ubqln2 | 54609 | 004409007 | 00404003 | 00375 | -015 | 5 |
| 7854 | ENSMUSG( | Ucp3 | 22229 | -03537756 | -06694496 | 01525 | -0155 | 5 |
| 7855 | ENSMUSG( | Uevld | 54122 | -00413190 | -00617755 | 006 | -03125 | 5 |
| 7856 | ENSMUSG( | Ufc1 | 66155 | -00188988 | -01755896 | 0365 | -009 | 5 |
| 7866 | ENSMUSG( | Ugp2 | 216558 | 001934831 | -02291667 | 0175 | -051 | 5 |
| 7868 | ENSMUSG( | Ulk1 | 22241 | 02608669 | 01520665 | 01175 | -0195 | 5 |
| 7872 | ENSMUSG( | Umps | 22247 | 027954484 | -00741655 | 00925 | -02 | 5 |
| 7884 | ENSMUSG( | Unc80 | 329178 | 00064336 | 01112434 | 0045 | -0125 | 5 |
| 7895 | ENSMUSG( | Uqcc3 | 107197 | 00414477 | -00536723 | 04325 | -016 | 5 |
| 7901 | ENSMUSG( | Uqcrfs1 | 66694 | 00516582 | -02931030 | -00325 | -02025 | 5 |
| 7912 | ENSMUSG( | Use1 | 67023 | 00656965 | -01159569 | 023 | -023 | 5 |
| 7921 | ENSMUSG( | Usp15 | 14479 | 00874634 | 00222884 | 01375 | -02525 | 5 |
| 7930 | ENSMUSG( | Usp33 | 170822 | -00350980 | 00520348C | 01775 | -033 | 5 |
| 7932 | ENSMUSG( | Usp35 | 244144 | 010301632 | 00141623 | 00025 | -01725 | 5 |
| 7934 | ENSMUSG( | Usp39 | 28035 | 01228422 | 00396690 | 02775 | -019 | 5 |
| 7938 | ENSMUSG( | Usp46 | 69727 | 00478260 | -00899954 | 011 | -02425 | 5 |
| 7945 | ENSMUSG( | Usp7 | 252870 | 00884016 | 00705713 | 01675 | -01125 | 5 |
| 7978 | ENSMUSG( | Vav3 | 57257 | 000354497 | -00972982 | 0075 | -01625 | 5 |
| 7986 | ENSMUSG( | Vdac1 | 22333 | 00204983 | -01448137 | 012 | -0265 | 5 |
| 7992 | ENSMUSG( | Vill | 22351 | 039534892 | 01297002 | -883177e-1 | -033 | 5 |
| 8003 | ENSMUSG( | Vps11 | 71732 | 00502945 | -00338202 | 02425 | -01025 | 5 |
| 8007 | ENSMUSG( | Vps13d | 230895 | -01132928 | -00038645 | 023 | -007 | 5 |
| 8008 | ENSMUSG( | Vps16 | 80743 | 00727717 | 00156522 | 01125 | -02025 | 5 |
| 8009 | ENSMUSG( | Vps18 | 228545 | 01206616C | -00329155 | -00075 | -024 | 5 |
| 8015 | ENSMUSG( | Vps29 | 56433 | -00214788 | -01103763 | 021 | -01475 | 5 |
| 8016 | ENSMUSG( | Vps33a | 77573 | 012830082 | -00240442 | -00025 | -01775 | 5 |
| 8025 | ENSMUSG( | Vps39 | 269338 | 021545027 | 000506144 | 025 | -01825 | 5 |
| 8026 | ENSMUSG( | Vps41 | 218035 | 00542486 | -01203233 | 02625 | -045 | 5 |
| 8030 | ENSMUSG( | Vps50 | 73288 | 009606947 | 006113102 | -0015 | -018 | 5 |
| 8040 | ENSMUSG( | Vsnl1 | 26950 | -00472816 | -02261386 | 0135 | -013 | 5 |
| 8044 | ENSMUSG( | Vta1 | 66201 | 000285243 | -02418810 | -117757e-1 | -02375 | 5 |
| 8075 | ENSMUSG( | Wdr1 | 22388 | 01305429C | -01126908 | 01875 | -0405 | 5 |
| 8078 | ENSMUSG( | Wdr13 | 73447 | 01783210 | 00540509 | -002 | -0155 | 5 |
| 8094 | ENSMUSG( | Wdr45 | 54636 | 01609623 | -00261184 | 006 | -055 | 5 |
| 8098 | ENSMUSG( | Wdr48 | 67561 | 007386647 | 004403301 | 0085 | -00975 | 5 |
| 8110 | ENSMUSG( | Wdr91 | 101240 | 00884747C | 01351015C | 004 | -0455 | 5 |
| 8117 | ENSMUSG( | Wipi1 | 52639 | 00022449 | -02713861 | 0135 | -046 | 5 |
| 8125 | ENSMUSG( | Wsb2 | 59043 | 009152273 | -01157609 | 02225 | -012 | 5 |
| 8141 | ENSMUSG( | Xpnpep3 | 321003 | 01191476 | 01385954 | 00675 | -045 | 5 |
| 8142 | ENSMUSG( | Xpo1 | 103573 | 002427321 | 027877073 | 0195 | -02975 | 5 |
| 8146 | ENSMUSG( | Xpo7 | 65246 | 00220223 | -00171155 | 002 | -014 | 5 |
| 8205 | ENSMUSG( | Zc3h12b | 547176 | 006559494 | 014743234 | 0065 | -02925 | 5 |
| 8229 | ENSMUSG( | Zdhhc6 | 66980 | -01219864 | 011058403 | 03825 | -02925 | 5 |
| 8249 | ENSMUSG( | Zfp385a | 29813 | -01142762 | -02428142 | 02875 | -018 | 5 |
| 8253 | ENSMUSG( | Zfp532 | 328977 | -00473450 | -01217689 | 009 | -0145 | 5 |
| 8257 | ENSMUSG( | Zfr | 22763 | 017511137 | 003208942 | 003 | -034 | 5 |
| 8259 | ENSMUSG( | Zfyve1 | 217695 | 00687134 | 002605011 | 02775 | -0255 | 5 |
| 8280 | ENSMUSG( | Zmynd8 | 228880 | 03083318C | 007662837 | 01725 | -00775 | 5 |
| 8292 | ENSMUSG( | Zyg11b | 414872 | 01280875C | -00483800 | 00825 | -02 | 5 |
| 3211 | ENSMUSG( | Hyi | 68180 | 051611772 | -00865375 | 522 | -0095 | 6 |
| 5458 | ENSMUSG( | Pradc1 | 73327 | -01962099 | 01590973 | 339 | -08325 | 6 |
| 5914 | ENSMUSG( | Recql5 | 170472 | 01294275C | 016584692 | 50775 | -04625 | 6 |
| 6644 | ENSMUSG( | Slc35a2 | 22232 | -02454060 | 01021519 | 2955 | -02225 | 6 |
| 4 | ENSMUSG( | 1500009L1 | 69784 | 010273324 | -00107497 | -01825 | -0225 | 7 |
| 13 | ENSMUSG( | 4933427D1 | 74477 | 014062691 | -00616856 | -02075 | -02125 | 7 |
| 23 | ENSMUSG( | Aak1 | 269774 | 017650387 | 01144559 | -015 | -02875 | 7 |
| 30 | ENSMUSG( | Aasdhppt | 67618 | 00232696 | 00592644 | -033 | -01625 | 7 |
| 31 | ENSMUSG( | Aass | 30956 | 02409110 | -01099165 | -047 | -02825 | 7 |
| 32 | ENSMUSG( | Aatf | 56321 | 010211612 | -01637736 | -0415 | -04375 | 7 |
| 46 | ENSMUSG( | Abcb7 | 11306 | -00148162 | 00581845C | -006 | -01325 | 7 |
| 60 | ENSMUSG( | Abcf3 | 27406 | 00674170 | -00165499 | -01925 | -01325 | 7 |
| 64 | ENSMUSG( | Abhd10 | 213012 | -01693189 | -01253543 | -01525 | -009 | 7 |
| 65 | ENSMUSG( | Abhd11 | 68758 | -00306899 | -00952897 | -046 | -0385 | 7 |
| 67 | ENSMUSG( | Abhd14b | 76491 | 018046397 | 018588654 | -00225 | -03125 | 7 |
| 82 | ENSMUSG( | Ablim2 | 231148 | -00530262 | -00434394 | -01875 | -0105 | 7 |
| 101 | ENSMUSG( | Acap2 | 78618 | 008076707 | 01446817 | -033 | -0465 | 7 |
| 107 | ENSMUSG( | Acbd6 | 72482 | -00151824 | -01430875 | -0085 | -022 | 7 |
| 111 | ENSMUSG( | Acin1 | 56215 | 00301484 | 00535567 | -0205 | -01725 | 7 |
| 114 | ENSMUSG( | Aco2 | 11429 | 008685167 | -01457630 | -00825 | -016 | 7 |
| 117 | ENSMUSG( | Acot13 | 66834 | 000737957 | -02973141 | -01425 | -009 | 7 |
| 128 | ENSMUSG( | Acsf2 | 264895 | 00348872 | -01048206 | -00825 | -045 | 7 |
| 129 | ENSMUSG( | Acsf3 | 257633 | 003101171 | -02359988 | -0135 | -02425 | 7 |
| 155 | ENSMUSG( | Acy1 | 109652 | -02017787 | -02213756 | -0185 | -016 | 7 |
| 164 | ENSMUSG( | Adam17 | 11491 | -00289995 | 018389893 | -03475 | -00725 | 7 |
| 200 | ENSMUSG( | Adhfe1 | 76187 | 01476731 | -02016225 | -01025 | -015 | 7 |
| 217 | ENSMUSG( | Adsl | 11564 | 00078830 | -01415341 | -0075 | -02175 | 7 |
| 235 | ENSMUSG( | Agfg2 | 231801 | 008717303 | -00360719 | -0245 | -01225 | 7 |
| 238 | ENSMUSG( | Ago1 | 236511 | 008118524 | -00646032 | -02525 | -0055 | 7 |
| 252 | ENSMUSG( | Ahcyl2 | 74340 | 011821407 | -01256616 | -0245 | -0595 | 7 |
| 257 | ENSMUSG( | Ahsa1 | 217737 | 013631992 | -01266724 | -01525 | -02 | 7 |
| 260 | ENSMUSG( | Aif1 | 11629 | 010890417 | -01482360 | -03125 | -013 | 7 |
| 263 | ENSMUSG( | Aifm3 | 72168 | 02997685 | 00077965 | -02225 | -01775 | 7 |
| 280 | ENSMUSG( | Akap5 | 238276 | -00691714 | -0169443 | -04575 | -025 | 7 |
| 284 | ENSMUSG( | Akap8l | 54194 | 02600456 | 01891518 | -02525 | -02675 | 7 |
| 286 | ENSMUSG( | Akr1a1 | 58810 | -00085036 | -01568925 | -01275 | -012 | 7 |
| 289 | ENSMUSG( | Akt1 | 11651 | 00940322 | 000732341 | -00325 | -01925 | 7 |
| 294 | ENSMUSG( | Alad | 17025 | -00249486 | -00975979 | -01725 | -01475 | 7 |
| 306 | ENSMUSG( | Aldh3b1 | 67689 | -00489262 | -03659700 | -0105 | -05575 | 7 |
| 307 | ENSMUSG( | Aldh4a1 | 212647 | 00207818 | -01258685 | -0215 | -01775 | 7 |
| 327 | ENSMUSG( | Alkbh8 | 67667 | 013668004 | 008300402 | -01825 | -039 | 7 |
| 343 | ENSMUSG( | Ampd2 | 109674 | 010142774 | 002440484 | -00625 | -041 | 7 |
| 344 | ENSMUSG( | Ampd3 | 11717 | 000392037 | -01228773 | -00875 | -03825 | 7 |
| 345 | ENSMUSG( | Amph | 218038 | 021936717 | -00293607 | -03825 | -0095 | 7 |
| 349 | ENSMUSG( | Anapc13 | 69010 | -00156268 | -03294356 | -02975 | -0495 | 7 |
| 350 | ENSMUSG( | Anapc16 | 52717 | 006498714 | -01721547 | -04725 | -01475 | 7 |
| 355 | ENSMUSG( | Angel2 | 52477 | 00388218 | 00808492 | -01175 | -02425 | 7 |
| 372 | ENSMUSG( | Ankrd29 | 225187 | 00763864 | -00699949 | -028 | -0465 | 7 |
| 373 | ENSMUSG( | Ankrd33b | 67434 | 008226492 | 013701551 | -0315 | -02925 | 7 |

| | | | | | | | | |
|---|---|---|---|---|---|---|---|---|
| 392 | ENSMUSG( | Ano6 | 105722 | 005372902 | 025346270 | -013 | -023 | 7 |
| 402 | ENSMUSG( | Anxa2 | 12306 | -00094612 | 017941971 | -0105 | -02225 | 7 |
| 404 | ENSMUSG( | Anxa4 | 11746 | 018028683 | 035243582 | -019 | -0545 | 7 |
| 406 | ENSMUSG( | Anxa6 | 11749 | -00438925 | -02076014 | -03025 | -00475 | 7 |
| 407 | ENSMUSG( | Anxa7 | 11750 | 008377939 | 009596112 | -00775 | -03825 | 7 |
| 427 | ENSMUSG( | Ap4b1 | 67489 | 024897467 | -02937817 | -032 | -0185 | 7 |
| 462 | ENSMUSG( | Appl2 | 216190 | 008486394 | 001842166 | -01075 | -04775 | 7 |
| 468 | ENSMUSG( | Aqr | 11834 | 015648811 | 002544867 | -0215 | -018 | 7 |
| 470 | ENSMUSG( | Arap1 | 69710 | -00930404 | -01268215 | -01875 | -007 | 7 |
| 481 | ENSMUSG( | Arfgap2 | 77038 | 005227726 | 000335686 | -01 | -01975 | 7 |
| 482 | ENSMUSG( | Arfgap3 | 66251 | -00059574 | -01919934 | -038 | -00775 | 7 |
| 499 | ENSMUSG( | Arhgap25 | 232201 | 026921865 | -02302327 | -02575 | -019 | 7 |
| 512 | ENSMUSG( | Arhgdia | 192662 | 007845580 | -01567678 | -01775 | -007 | 7 |
| 516 | ENSMUSG( | Arhgef10l | 72754 | 006975037 | 005922181 | -02125 | -0085 | 7 |
| 518 | ENSMUSG( | Arhgef12 | 69632 | 002551132 | -02687201 | -02025 | -01625 | 7 |
| 519 | ENSMUSG( | Arhgef17 | 207212 | 017279362 | 018340175 | -01125 | -03625 | 7 |
| 528 | ENSMUSG( | Arhgef6 | 73341 | 000793223 | -00070547 | -006 | -016 | 7 |
| 536 | ENSMUSG( | Arih1 | 23806 | 008370432 | 006132680 | -012 | -01475 | 7 |
| 557 | ENSMUSG( | Armc8 | 74125 | 008745879 | -00975708 | -0225 | -0145 | 7 |
| 568 | ENSMUSG( | Arpc3 | 56378 | -00451632 | -01937865 | -0115 | -0115 | 7 |
| 600 | ENSMUSG( | Asl | 109900 | 018926168 | -01446936 | -01325 | -01225 | 7 |
| 619 | ENSMUSG( | Atg101 | 68118 | 005369487 | -00989776 | -02825 | -00775 | 7 |
| 660 | ENSMUSG( | Atp5f1d | 66043 | 000997363 | -02165093 | -01425 | -0505 | 7 |
| 667 | ENSMUSG( | Atp5pb | 11950 | 002648077 | -02517480 | -008 | -02425 | 7 |
| 670 | ENSMUSG( | Atp5po | 28080 | 008301721 | -00987087 | -00825 | -01175 | 7 |
| 680 | ENSMUSG( | Atp6v1d | 73834 | -01079604 | -01170638 | -00575 | -01625 | 7 |
| 691 | ENSMUSG( | Atpaf1 | 230649 | -01244684 | -00972439 | -0125 | -015 | 7 |
| 697 | ENSMUSG( | Atrx | 22589 | -00812676 | 009585005 | -038 | -00725 | 7 |
| 705 | ENSMUSG( | Auh | 11992 | 007838002 | -01437561 | -01625 | -033 | 7 |
| 710 | ENSMUSG( | AW209491 | 105351 | 012748634 | -00971145 | -0405 | -026 | 7 |
| 725 | ENSMUSG( | B630019K0 | 102941 | 000440589 | -00813690 | -01425 | -0085 | 7 |
| 732 | ENSMUSG( | Bag1 | 12017 | 005220668 | -01590721 | -01425 | -021 | 7 |
| 733 | ENSMUSG( | Bag2 | 213539 | -00195220 | -01513442 | -027 | -0125 | 7 |
| 737 | ENSMUSG( | Bag6 | 224727 | 017124837 | -00629772 | -01725 | -0255 | 7 |
| 741 | ENSMUSG( | Banf1 | 23825 | -00724674 | -01765423 | -0075 | -0385 | 7 |
| 743 | ENSMUSG( | Bax | 12028 | -00136596 | 000451096 | -01325 | -014 | 7 |
| 744 | ENSMUSG( | Baz1b | 22385 | 020279892 | 016903159 | -0105 | -0325 | 7 |
| 747 | ENSMUSG( | Bbs1 | 52028 | 020091916 | -00050388 | -00725 | -01725 | 7 |
| 760 | ENSMUSG( | Bcas1 | 76960 | -01059750 | -02541770 | -039 | -037 | 7 |
| 761 | ENSMUSG( | Bcas2 | 68183 | -00185703 | -00154896 | -01075 | -01025 | 7 |
| 762 | ENSMUSG( | Bcas3 | 192197 | 003981314 | -00757281 | -007 | -014 | 7 |
| 774 | ENSMUSG( | Bcl2l13 | 94044 | 006487780 | -00688096 | -01 | -03125 | 7 |
| 777 | ENSMUSG( | Bclaf1 | 72567 | 007644994 | 011982070 | -029 | -024 | 7 |
| 779 | ENSMUSG( | Bcs1l | 66821 | 00588339 | -01461105 | -00525 | -0245 | 7 |
| 793 | ENSMUSG( | Bin1 | 30948 | 00372028 | -00129991 | -009 | -03275 | 7 |
| 810 | ENSMUSG( | Bms1 | 213895 | 022846341 | 014857313 | -0455 | -02125 | 7 |
| 833 | ENSMUSG( | Braf | 109880 | -00073847 | 006801681 | -00475 | -0215 | 7 |
| 840 | ENSMUSG( | Brd4 | 57261 | -00951045 | 007460678 | -023 | -032 | 7 |
| 841 | ENSMUSG( | Brd7 | 26992 | 012761924 | -00048679 | -01025 | -021 | 7 |
| 875 | ENSMUSG( | C030006K1 | 223665 | -00530719 | -01124452 | -0395 | -0095 | 7 |
| 878 | ENSMUSG( | C1qbp | 12261 | -00603041 | -00933073 | -01875 | -03125 | 7 |
| 891 | ENSMUSG( | Cab39l | 69008 | 006371299 | -00378421 | -03425 | -03125 | 7 |
| 910 | ENSMUSG( | Cacnb2 | 12296 | 009112267 | 027102537 | -018 | -0255 | 7 |
| 911 | ENSMUSG( | Cacnb3 | 12297 | 018182760 | 004787714 | -012 | -01075 | 7 |
| 927 | ENSMUSG( | Cadps2 | 320405 | 039423565 | 009935294 | -0275 | -047 | 7 |
| 930 | ENSMUSG( | Calcoco1 | 67488 | 004001858 | -00349216 | -048 | -00975 | 7 |
| 939 | ENSMUSG( | Camk1d | 227541 | 002051752 | -00468347 | -003 | -02075 | 7 |
| 945 | ENSMUSG( | Camk2n1 | 66259 | 014001181 | -00560457 | -015 | -0195 | 7 |
| 946 | ENSMUSG( | Camk4 | 12326 | -00285875 | -02104358 | -0055 | -02 | 7 |
| 947 | ENSMUSG( | Camkk1 | 55984 | 003931163 | -00099601 | -0315 | -03025 | 7 |
| 962 | ENSMUSG( | Capg | 12332 | -01944552 | -02350314 | -04175 | -0295 | 7 |
| 974 | ENSMUSG( | Carhsp1 | 52502 | -00029594 | -01843916 | -03475 | -043 | 7 |
| 981 | ENSMUSG( | Cars2 | 71941 | 014508178 | -00624549 | -03325 | -04075 | 7 |
| 992 | ENSMUSG( | Casp8 | 12370 | -00733357 | 018818410 | -0165 | -012 | 7 |
| 1004 | ENSMUSG( | Cbl | 12402 | 016905380 | 009485824 | -028 | -017 | 7 |
| 1012 | ENSMUSG( | Cbr4 | 234309 | 008254614 | -01163873 | -0115 | -0365 | 7 |
| 1026 | ENSMUSG( | Ccdc115 | 69668 | -00381573 | -01434523 | -0145 | -0215 | 7 |
| 1030 | ENSMUSG( | Ccdc127 | 67433 | -00888865 | 002207814 | -01375 | -014 | 7 |
| 1033 | ENSMUSG( | Ccdc137 | 67291 | 012239244 | -00370169 | -01575 | -00725 | 7 |
| 1037 | ENSMUSG( | Ccdc177 | 380768 | 032818694 | 008593181 | -0305 | -01875 | 7 |
| 1049 | ENSMUSG( | Ccdc51 | 66658 | 021366840 | 011201057 | -03875 | -0395 | 7 |
| 1058 | ENSMUSG( | Ccdc90b | 66365 | 008809348 | -01151701 | -01075 | -01125 | 7 |
| 1061 | ENSMUSG( | Ccdc92b | 432582 | 007225236 | 005102673 | -04275 | -02075 | 7 |
| 1066 | ENSMUSG( | Ccm2 | 216527 | 001266160 | -01269126 | -0285 | -0235 | 7 |
| 1073 | ENSMUSG( | Ccny | 67974 | 012940305 | 005977400 | -01475 | -0085 | 7 |
| 1078 | ENSMUSG( | Ccser1 | 232035 | 016682930 | 016578836 | -02525 | -03475 | 7 |
| 1108 | ENSMUSG( | Cda | 72269 | 002395379 | -02371678 | -013 | -05025 | 7 |
| 1109 | ENSMUSG( | Cdc123 | 98828 | 001725250 | -00706618 | -0235 | -0355 | 7 |
| 1112 | ENSMUSG( | Cdc26 | 66440 | -00245983 | -00584606 | -02425 | -00075 | 7 |
| 1117 | ENSMUSG( | Cdc40 | 71713 | 005579206 | 003782248 | -013 | -0135 | 7 |
| 1148 | ENSMUSG( | Cdk17 | 237459 | 007368569 | -00156620 | -022 | -003 | 7 |
| 1149 | ENSMUSG( | Cdk18 | 18557 | -00100175 | -00553328 | -0205 | -00525 | 7 |
| 1161 | ENSMUSG( | Cdkl5 | 382253 | 000400647 | -00692120 | -01275 | -01325 | 7 |
| 1164 | ENSMUSG( | Cdkn2c | 12580 | -00481821 | -02024212 | -0115 | -016 | 7 |
| 1181 | ENSMUSG( | Cenpv | 73139 | 008058807 | -01411309 | -02875 | -0465 | 7 |
| 1182 | ENSMUSG( | Cep112 | 76380 | 005675720 | -01739365 | -027 | -021 | 7 |
| 1185 | ENSMUSG( | Cep135 | 381644 | 007850844 | 019586100 | -016 | -02825 | 7 |
| 1197 | ENSMUSG( | Cetn2 | 26370 | 005919503 | -01461521 | -01475 | -01025 | 7 |
| 1216 | ENSMUSG( | Champ1 | 101994 | 011714817 | -00967284 | -01925 | -00725 | 7 |
| 1242 | ENSMUSG( | Chm | 12662 | -00737486 | 002265218 | -00875 | -015 | 7 |
| 1245 | ENSMUSG( | Chmp1b2 | 67028 | 006835672 | -01804642 | -0235 | -00475 | 7 |
| 1248 | ENSMUSG( | Chmp3 | 66700 | 005154700 | -01348041 | -0235 | -02125 | 7 |
| 1250 | ENSMUSG( | Chmp5 | 76959 | 005786249 | -01910786 | -032 | -00425 | 7 |
| 1257 | ENSMUSG( | Chpf | 74241 | -00290517 | 025198696 | -03625 | -02375 | 7 |
| 1268 | ENSMUSG( | Chtop | 66511 | 000617230 | -01026142 | -015 | -0485 | 7 |
| 1272 | ENSMUSG( | Ciao2b | 68523 | -00889519 | -02235787 | -02 | -0175 | 7 |
| 1275 | ENSMUSG( | Cib1 | 23991 | 014521113 | -01943848 | -03825 | -04175 | 7 |
| 1278 | ENSMUSG( | Cirbp | 12696 | -02748732 | -01292517 | -03525 | -03475 | 7 |
| 1286 | ENSMUSG( | Ckb | 12709 | 003710999 | -02402901 | -00525 | -02025 | 7 |
| 1289 | ENSMUSG( | Clasp2 | 76499 | 007698783 | -00901210 | -04025 | -0155 | 7 |
| 1324 | ENSMUSG( | Clpp | 53895 | 006129706 | -00626444 | -019 | -02175 | 7 |
| 1338 | ENSMUSG( | Clybl | 69634 | 001259593 | -02181664 | -01275 | -017 | 7 |
| 1342 | ENSMUSG( | Cmc2 | 66531 | 005176988 | -01589028 | -022 | -00875 | 7 |
| 1344 | ENSMUSG( | Cmip | 74440 | 026541329 | 009380094 | -00325 | -02825 | 7 |
| 1345 | ENSMUSG( | Cmpk1 | 66588 | 000604777 | -00260256 | -0155 | -017 | 7 |
| 1346 | ENSMUSG( | Cmpk2 | 22169 | -01343790 | -04118136 | -02075 | -01375 | 7 |
| 1374 | ENSMUSG( | Cnp | 12799 | 003155144 | -02127011 | -022 | -05825 | 7 |
| 1380 | ENSMUSG( | Cnst | 226744 | -00790526 | 004264424 | -01525 | -0295 | 7 |
| 1400 | ENSMUSG( | Coasy | 71743 | 017863641 | -00419053 | -006 | -05 | 7 |
| 1428 | ENSMUSG( | Commd1 | 17846 | 005546513 | -02740254 | -034 | -02325 | 7 |
| 1451 | ENSMUSG( | Cops7a | 26894 | 001010942 | -02215015 | -038 | -008 | 7 |
| 1461 | ENSMUSG( | Coq5 | 52064 | 007566602 | -01044610 | -0135 | -0275 | 7 |
| 1463 | ENSMUSG( | Coq7 | 12850 | -00151283 | -01492745 | -0445 | -04875 | 7 |
| 1466 | ENSMUSG( | Coq9 | 67914 | 009137958 | -00719820 | -02275 | -0435 | 7 |
| 1467 | ENSMUSG( | Coro1a | 12721 | -00001375 | -01949406 | -00675 | -02325 | 7 |
| 1471 | ENSMUSG( | Coro2b | 235431 | 004053651 | -01552333 | -014 | -025 | 7 |
| 1472 | ENSMUSG( | Coro6 | 216961 | 040863152 | 000854000 | -01275 | -04275 | 7 |
| 1479 | ENSMUSG( | Cox17 | 12856 | -00379797 | -02001669 | -028 | -00625 | 7 |
| 1489 | ENSMUSG( | Cox6b1 | 110323 | -00168863 | -02862519 | -01025 | -04025 | 7 |
| 1493 | ENSMUSG( | Cox7a2l | 20463 | 004756525 | -01238731 | -0145 | -038 | 7 |
| 1501 | ENSMUSG( | Cplx1 | 12889 | 008043224 | -01513208 | -0215 | -01625 | 7 |
| 1502 | ENSMUSG( | Cplx2 | 12890 | 004105364 | -00455750 | -02875 | -00925 | 7 |
| 1512 | ENSMUSG( | Cpne9 | 211232 | 018887370 | -01031012 | -014 | -02975 | 7 |
| 1524 | ENSMUSG( | Cpt2 | 12896 | 007146671 | -01634852 | -01075 | -03925 | 7 |
| 1545 | ENSMUSG( | Crk | 12928 | 009599071 | -00317469 | -02725 | -027 | 7 |
| 1550 | ENSMUSG( | Crnkl1 | 66877 | 001566060 | 003265520 | -04425 | -04825 | 7 |

| | | | | | | | |
|---|---|---|---|---|---|---|---|
| 1552 | ENSMUSG( | Crocc2 | 381284 | 063943029 | 004720827 -05175 | -05825 | 7 |
| 1556 | ENSMUSG( | Crtc1 | 382056 | 012626672 | 004315551 -01325 | -00725 | 7 |
| 1558 | ENSMUSG( | Crtc3 | 70461 | 004116098 | 000424426 -0275 | -03975 | 7 |
| 1566 | ENSMUSG( | Cryzl1 | 66609 | 002170292 | -01140798 -01925 | -03525 | 7 |
| 1569 | ENSMUSG( | Csad | 246277 | 018157827 | -00042787 -01375 | -01075 | 7 |
| 1590 | ENSMUSG( | Csrp1 | 13007 | 014925095 | -01545769 -01275 | -02825 | 7 |
| 1599 | ENSMUSG( | Ctbp1 | 13016 | 005834727 | -00760444 -01425 | -0065 | 7 |
| 1600 | ENSMUSG( | Ctbp2 | 13017 | 018237148 | 011426083 -03 | -0265 | 7 |
| 1608 | ENSMUSG( | Ctdspl2 | 329506 | 006513314 | 023130266 -032 | -051 | 7 |
| 1615 | ENSMUSG( | Ctnnbip1 | 67087 | 004771652 | -00467378 -01125 | -00825 | 7 |
| 1630 | ENSMUSG( | Ctss | 13040 | -01784959 | 002276701 -0155 | -0325 | 7 |
| 1642 | ENSMUSG( | Cul4b | 72584 | 004649339 | 002169511 -0235 | -0105 | 7 |
| 1652 | ENSMUSG( | Cwc27 | 67285 | 009912344 | -01186517 -03525 | -006 | 7 |
| 1654 | ENSMUSG( | Cx3cr1 | 13051 | -01266929 | 027651925 -03 | -0135 | 7 |
| 1663 | ENSMUSG( | Cyb5d2 | 192986 | -01279843 | 000163748 -0235 | -045 | 7 |
| 1669 | ENSMUSG( | Cycs | 13063 | 002403322 | -02013276 -021 | -042 | 7 |
| 1691 | ENSMUSG( | Czib | 74098 | -01264509 | -00083283 -0245 | -03 | 7 |
| 1695 | ENSMUSG( | D3Ertd751 | 73852 | -00349240 | -01848143 -0215 | -01725 | 7 |
| 1698 | ENSMUSG( | D6Wsu163 | 28040 | 023043899 | 006802884 -00975 | -01675 | 7 |
| 1713 | ENSMUSG( | Dap3 | 65111 | 007312158 | -00958475 -0095 | -017 | 7 |
| 1715 | ENSMUSG( | Dapk3 | 13144 | 017411595 | 000853435 -02375 | -02725 | 7 |
| 1722 | ENSMUSG( | Dbn1 | 56320 | 008519528 | -00085146 -02325 | -0135 | 7 |
| 1726 | ENSMUSG( | Dbr1 | 83703 | 000136233 | 001231462 -0285 | -00225 | 7 |
| 1727 | ENSMUSG( | Dbt | 13171 | 011236189 | -00349790 -01275 | -0335 | 7 |
| 1742 | ENSMUSG( | Dcp1a | 75901 | -00512373 | 013586630 -01025 | -0175 | 7 |
| 1757 | ENSMUSG( | Dcun1d5 | 76863 | -00071924 | 008638686 -03225 | -00975 | 7 |
| 1759 | ENSMUSG( | Dcxr | 67880 | 011201645 | -01344904 -04075 | -00675 | 7 |
| 1766 | ENSMUSG( | Ddhd2 | 72108 | -00199257 | -00130260 -0455 | -02625 | 7 |
| 1767 | ENSMUSG( | Ddi2 | 68817 | 006606666 | 006190932 -00625 | -0155 | 7 |
| 1777 | ENSMUSG( | Ddx19a | 13680 | 006016482 | -00346652 -02625 | -0125 | 7 |
| 1780 | ENSMUSG( | Ddx21 | 56200 | 015074169 | 003847571 -02325 | -021 | 7 |
| 1782 | ENSMUSG( | Ddx24 | 27225 | -01181219 | -00450552 -03325 | -00925 | 7 |
| 1791 | ENSMUSG( | Ddx46 | 212880 | 008115832 | -00332403 -0205 | -005 | 7 |
| 1793 | ENSMUSG( | Ddx5 | 13207 | -00616430 | 016528987 -0135 | -01975 | 7 |
| 1795 | ENSMUSG( | Ddx51 | 69663 | 010254078 | -00442235 -0065 | -0115 | 7 |
| 1797 | ENSMUSG( | Ddx54 | 71990 | 014217033 | -00553381 -0285 | -0205 | 7 |
| 1798 | ENSMUSG( | Ddx55 | 67848 | 008718757 | -00329998 -0515 | -01825 | 7 |
| 1799 | ENSMUSG( | Ddx56 | 52513 | 003867580 | -00971907 -04375 | -0665 | 7 |
| 1801 | ENSMUSG( | Decr1 | 67460 | 005118455 | -02439429 -02075 | -00375 | 7 |
| 1808 | ENSMUSG( | Dennd11 | 243780 | -00204386 | -01086384 -0215 | -0265 | 7 |
| 1811 | ENSMUSG( | Dennd2a | 209773 | 010969299 | -00505459 -03175 | -053 | 7 |
| 1813 | ENSMUSG( | Dennd4b | 229541 | 018520960 | 011957994 -0115 | -02225 | 7 |
| 1818 | ENSMUSG( | Dennd6b | 69440 | 004098543 | 012678147 -00875 | -03075 | 7 |
| 1826 | ENSMUSG( | Desi1 | 28075 | -00752469 | -02561542 -0255 | -02075 | 7 |
| 1828 | ENSMUSG( | Dgat1 | 13350 | -01353060 | -00200395 -01525 | -00625 | 7 |
| 1830 | ENSMUSG( | Dgcr8 | 94223 | 023579080 | 011700668 -021 | -02875 | 7 |
| 1836 | ENSMUSG( | Dgki | 320127 | -01037474 | -02179187 -015 | -01725 | 7 |
| 1846 | ENSMUSG( | Dhodh | 56749 | 009979543 | 018842609 -01425 | -04175 | 7 |
| 1851 | ENSMUSG( | Dhrs4 | 28200 | 012108933 | -02331782 -00875 | -01725 | 7 |
| 1867 | ENSMUSG( | Diaph1 | 13367 | 004313444 | 003464904 -0165 | -02475 | 7 |
| 1874 | ENSMUSG( | Dip2c | 208440 | 009640788 | -00254300 -01125 | -00825 | 7 |
| 1885 | ENSMUSG( | Dlat | 235339 | 009073231 | -01285466 -017 | -01275 | 7 |
| 1886 | ENSMUSG( | Dlc1 | 50768 | 005481506 | -00201216 -0165 | -0305 | 7 |
| 1887 | ENSMUSG( | Dld | 13382 | 009081269 | -00675543 -028 | -0245 | 7 |
| 1894 | ENSMUSG( | Dlgap2 | 244310 | 010279602 | -01827208 -01075 | -031 | 7 |
| 1899 | ENSMUSG( | Dmac2 | 66349 | 008802997 | 007088658 -02725 | -04325 | 7 |
| 1909 | ENSMUSG( | Dnaaf9 | 228602 | 004378292 | -02381820 -0115 | -017 | 7 |
| 1914 | ENSMUSG( | Dnaja4 | 58233 | 013981472 | -00218580 -02925 | -0005 | 7 |
| 1922 | ENSMUSG( | Dnajb6 | 23950 | 004550075 | -01215167 -0235 | -0295 | 7 |
| 1962 | ENSMUSG( | Dock1 | 330662 | 017625002 | -00208255 -01875 | -0365 | 7 |
| 1970 | ENSMUSG( | Dock7 | 67299 | 003886848 | -00183047 -02 | -00725 | 7 |
| 2000 | ENSMUSG( | Dpy19l4 | 381510 | -01903058 | 004108121 -025 | -0155 | 7 |
| 2036 | ENSMUSG( | Dut | 110074 | -00006822 | -02334479 -03725 | -012 | 7 |
| 2039 | ENSMUSG( | Dvl3 | 13544 | 009190535 | -00182504 -014 | -01725 | 7 |
| 2061 | ENSMUSG( | Ears2 | 67417 | 011524634 | -00436821 -01075 | -013 | 7 |
| 2063 | ENSMUSG( | Ebna1bp2 | 69072 | 009636893 | -00550958 -029 | -051 | 7 |
| 2073 | ENSMUSG( | Echs1 | 93747 | 005623395 | -01718351 -0075 | -03475 | 7 |
| 2099 | ENSMUSG( | Efhd1 | 98363 | 008909059 | -02922580 -05 | -011 | 7 |
| 2107 | ENSMUSG( | Efr3a | 76740 | 016459463 | -00095736 -0055 | -03 | 7 |
| 2113 | ENSMUSG( | Egr3 | 13655 | 007071983 | -00816202 -0315 | -0325 | 7 |
| 2158 | ENSMUSG( | Eif4a3 | 192170 | 006334371 | -00908309 -025 | -04375 | 7 |
| 2178 | ENSMUSG( | Elavl1 | 15568 | 006093004 | 000258017 -0085 | -04775 | 7 |
| 2181 | ENSMUSG( | Elavl4 | 15572 | -00391576 | -00169008 -026 | -0085 | 7 |
| 2182 | ENSMUSG( | Elfn1 | 243312 | -00806407 | 014015805 -0175 | -00725 | 7 |
| 2184 | ENSMUSG( | Elmo1 | 140580 | -00870131 | -00931712 -018 | -015 | 7 |
| 2211 | ENSMUSG( | Emg1 | 14791 | 008755762 | 006425306 -0305 | -02625 | 7 |
| 2215 | ENSMUSG( | Eml3 | 225898 | 008763396 | 021203255 -029 | -026 | 7 |
| 2217 | ENSMUSG( | Eml5 | 319670 | -00906887 | 003132885 -02025 | -0235 | 7 |
| 2230 | ENSMUSG( | Eno2 | 13807 | 005148160 | -00429988 -00875 | -0175 | 7 |
| 2249 | ENSMUSG( | Epb41 | 269587 | -00392938 | 004386329 -01725 | -04125 | 7 |
| 2252 | ENSMUSG( | Epb41l3 | 13823 | -02830574 | -00634193 -03075 | -03375 | 7 |
| 2273 | ENSMUSG( | Epn1 | 13854 | 019037121 | 001645821 -0285 | -04425 | 7 |
| 2278 | ENSMUSG( | Eps15l1 | 13859 | 016469229 | 007019803 -014 | -026 | 7 |
| 2285 | ENSMUSG( | Erbin | 59079 | -00866939 | -01193176 -03425 | -00675 | 7 |
| 2286 | ENSMUSG( | Erc1 | 111173 | 016897932 | 003144728 -0045 | -04275 | 7 |
| 2301 | ENSMUSG( | Ermn | 77767 | -01137657 | -01976407 -03925 | -049 | 7 |
| 2313 | ENSMUSG( | Etfa | 110842 | 001092236 | -02711489 -03375 | -01425 | 7 |
| 2314 | ENSMUSG( | Etfb | 110826 | 008078019 | -02236066 -00925 | -02375 | 7 |
| 2315 | ENSMUSG( | Etfdh | 66841 | 009428718 | -02700573 -0135 | -02325 | 7 |
| 2317 | ENSMUSG( | Etl4 | 208618 | 006861201 | 012651555 -01575 | -01025 | 7 |
| 2322 | ENSMUSG( | Evi5 | 14020 | 003035366 | 000455329 -04525 | -01575 | 7 |
| 2358 | ENSMUSG( | Fabp3 | 14077 | -00526063 | -01758572 -00275 | -051 | 7 |
| 2368 | ENSMUSG( | Fah | 14085 | 005800736 | -01685223 -00425 | -026 | 7 |
| 2377 | ENSMUSG( | Fam114a2 | 67726 | 011413350 | 007354788 -0245 | -0155 | 7 |
| 2381 | ENSMUSG( | Fam120b | 67544 | 007341708 | 013997798 -04025 | -0105 | 7 |
| 2384 | ENSMUSG( | Fam131a | 78408 | 005504373 | -02188880 -04 | -04925 | 7 |
| 2385 | ENSMUSG( | Fam131b | 76156 | 004328976 | -00825795 -01475 | -01625 | 7 |
| 2386 | ENSMUSG( | Fam135a | 68187 | -01593760 | 008762503 -0275 | -044 | 7 |
| 2401 | ENSMUSG( | Fam193a | 231128 | 002977501 | -00177485 -0395 | -05225 | 7 |
| 2405 | ENSMUSG( | Fam210b | 67017 | -00480800 | -03382446 -00975 | -03375 | 7 |
| 2429 | ENSMUSG( | Fastkd5 | 380601 | -02779843 | -04510779 -0275 | -032 | 7 |
| 2447 | ENSMUSG( | Fbxl4 | 269514 | -00062058 | 010913160 -023 | -01225 | 7 |
| 2466 | ENSMUSG( | Fbxw2 | 30050 | 004774981 | -00942816 -0365 | -0355 | 7 |
| 2482 | ENSMUSG( | Fdxr | 14149 | 003621828 | -00708036 -03475 | -0195 | 7 |
| 2483 | ENSMUSG( | Fech | 14151 | 001696828 | -01305464 -0065 | -04175 | 7 |
| 2489 | ENSMUSG( | Fez1 | 235180 | 003221491 | -00797809 -02625 | -00175 | 7 |
| 2499 | ENSMUSG( | Fgfr1op2 | 67529 | 002805467 | 004942770 -00875 | -012 | 7 |
| 2517 | ENSMUSG( | Fis1 | 66437 | -00107034 | -02062110 -03575 | -02225 | 7 |
| 2544 | ENSMUSG( | Fmnl1 | 57778 | 018452020 | -00212787 -01925 | -02475 | 7 |
| 2546 | ENSMUSG( | Fmnl3 | 22379 | -00011843 | 031493129 -0255 | -03225 | 7 |
| 2551 | ENSMUSG( | Fn3k | 63828 | -01025602 | -01494984 -021 | -03625 | 7 |
| 2553 | ENSMUSG( | Fnbp1 | 14269 | -00258242 | -02184636 -03925 | -01275 | 7 |
| 2564 | ENSMUSG( | Foxg1 | 15228 | 004416009 | 019248162 -041 | -03325 | 7 |
| 2595 | ENSMUSG( | Ftsj3 | 56095 | 017689126 | 005421799 -008 | -0225 | 7 |
| 2599 | ENSMUSG( | Fundc1 | 72018 | 001733292 | -01230133 -014 | -0135 | 7 |
| 2601 | ENSMUSG( | Fuom | 69064 | -00221861 | -00986331 -033 | -013 | 7 |
| 2607 | ENSMUSG( | Fxn | 14297 | 011399894 | -01475670 -0045 | -05 | 7 |
| 2614 | ENSMUSG( | Fyn | 14360 | -00028835 | -01390921 -013 | -0255 | 7 |
| 2615 | ENSMUSG( | Fyttd1 | 69823 | -00028511 | 00117122 -023 | -05125 | 7 |
| 2624 | ENSMUSG( | Gab1 | 14388 | -01582396 | -00126253 -0265 | -0205 | 7 |
| 2639 | ENSMUSG( | Gabrd | 14403 | -03257159 | -00963295 -03025 | -0465 | 7 |
| 2652 | ENSMUSG( | Galm | 319625 | 008420336 | 014785816 -00575 | -03675 | 7 |
| 2665 | ENSMUSG( | Gap43 | 14432 | -00338438 | -01806196 -028 | -01475 | 7 |
| 2672 | ENSMUSG( | Gars1 | 353172 | 009285750 | -01252379 -00425 | -0165 | 7 |
| 2694 | ENSMUSG( | Gcat | 26912 | 016023371 | -01262925 -01725 | -02 | 7 |
| 2701 | ENSMUSG( | Gcsh | 68133 | 001354139 | -02407913 -01675 | -014 | 7 |

| | | | | | | | | |
|---|---|---|---|---|---|---|---|---|
| 2731 | ENSMUSG( | Gga3 | 260302 | 024836613 | 007405002 | -00725 | -01625 | 7 |
| 2751 | ENSMUSG( | Gjb6 | 14623 | 018418857 | 026326566 | -01725 | -0285 | 7 |
| 2772 | ENSMUSG( | Glrx2 | 69367 | 004693458 | -00010099 | -0285 | -01375 | 7 |
| 2781 | ENSMUSG( | Glyr1 | 74022 | 015858144 | 000834293 | -0255 | -02325 | 7 |
| 2798 | ENSMUSG( | Gna12 | 14673 | 009737112 | -01437523 | -039 | -01 | 7 |
| 2811 | ENSMUSG( | Gnb5 | 14697 | -00628727 | -01416883 | -00925 | -0165 | 7 |
| 2813 | ENSMUSG( | Gng10 | 14700 | 00286263 | -01432419 | -0285 | -04125 | 7 |
| 2815 | ENSMUSG( | Gng12 | 14701 | 012862452 | -01161127 | -0175 | -01575 | 7 |
| 2822 | ENSMUSG( | Gnl1 | 14670 | 011618618 | -00455660 | -00875 | -02525 | 7 |
| 2824 | ENSMUSG( | Gnl3l | 237107 | 002742862 | -01442571 | -01575 | -0065 | 7 |
| 2842 | ENSMUSG( | Golph3l | 229593 | 010720648 | -00030264 | -00425 | -0145 | 7 |
| 2851 | ENSMUSG( | Got2 | 14719 | 002195385 | -01249542 | -00525 | -01775 | 7 |
| 2858 | ENSMUSG( | Gpatch2l | 70373 | 001357335 | -00191677 | -02825 | -005 | 7 |
| 2866 | ENSMUSG( | Gpd1 | 14555 | 020421265 | 003255028 | -02475 | -0145 | 7 |
| 2868 | ENSMUSG( | Gpd2 | 14571 | 013608815 | 000667865 | -014 | -0325 | 7 |
| 2878 | ENSMUSG( | Gpr162 | 14788 | -00705057 | 011232547 | -04325 | -014 | 7 |
| 2911 | ENSMUSG( | Grhpr | 76238 | 008765444 | -02496272 | -0155 | -048 | 7 |
| 2928 | ENSMUSG( | Grip2 | 243547 | 020113852 | 007464758 | -01125 | -01925 | 7 |
| 2929 | ENSMUSG( | Gripap1 | 54645 | 018670320 | -00199884 | -01925 | -01475 | 7 |
| 2935 | ENSMUSG( | Grm3 | 108069 | -01010135 | -02485032 | -016 | -02575 | 7 |
| 2945 | ENSMUSG( | Grtp1 | 66790 | 001695223 | -02070563 | -01625 | -0105 | 7 |
| 2979 | ENSMUSG( | Gtf2h4 | 14885 | 014861186 | 002288273 | -0325 | -00975 | 7 |
| 2991 | ENSMUSG( | Gtpbp4 | 69237 | 018635353 | 002847623 | -02125 | -027 | 7 |
| 2993 | ENSMUSG( | Gtpbp8 | 66067 | 001155434 | 008693135 | -027 | -03275 | 7 |
| 3006 | ENSMUSG( | H2ax | 15270 | -01343886 | 006164799 | -04525 | -0165 | 7 |
| 3008 | ENSMUSG( | H2bc15 | 319187 | -02452626 | -06635984 | -03325 | -01175 | 7 |
| 3018 | ENSMUSG( | Hadh | 15107 | 003600003 | -02198925 | -01075 | -01575 | 7 |
| 3052 | ENSMUSG( | Hddc3 | 68695 | 004572055 | -01700578 | -04375 | -021 | 7 |
| 3056 | ENSMUSG( | Hdhd2 | 76987 | 005079383 | -01978644 | -01825 | -00625 | 7 |
| 3057 | ENSMUSG( | Hdhd3 | 72748 | 008652759 | -01549464 | -0185 | -0475 | 7 |
| 3058 | ENSMUSG( | Hdhd5 | 214932 | 008928729 | 009418389 | -017 | -02 | 7 |
| 3090 | ENSMUSG( | Hibch | 227095 | 010628027 | -01349760 | -01725 | -013 | 7 |
| 3096 | ENSMUSG( | Hint2 | 68917 | -00543779 | -01434734 | -0185 | -02375 | 7 |
| 3102 | ENSMUSG( | Hk1 | 15275 | 005457770 | -00632932 | -02625 | 00025 | 7 |
| 3103 | ENSMUSG( | Hk2 | 15277 | 021536853 | 018370205 | -0385 | -015 | 7 |
| 3104 | ENSMUSG( | Hkdc1 | 216019 | 014971854 | 004506638 | -02425 | -01725 | 7 |
| 3109 | ENSMUSG( | Hmg20a | 66867 | 010721762 | -00137525 | -02025 | -0115 | 7 |
| 3114 | ENSMUSG( | Hmgcl | 15356 | 010162339 | -00031132 | -0145 | -009 | 7 |
| 3124 | ENSMUSG( | Hmox2 | 15369 | -00235993 | -01731334 | -0055 | -0185 | 7 |
| 3126 | ENSMUSG( | Hnrnpa0 | 77134 | -00757356 | -00751088 | -006 | -0315 | 7 |
| 3131 | ENSMUSG( | Hnrnpc | 15381 | 008485360 | -01001578 | -01725 | -04575 | 7 |
| 3137 | ENSMUSG( | Hnrnph3 | 432467 | 010449617 | -01117558 | -00875 | -05425 | 7 |
| 3141 | ENSMUSG( | Hnrnpm | 76936 | 012647698 | 003615563 | -00425 | -0205 | 7 |
| 3143 | ENSMUSG( | Hnrnpu | 51810 | -00509636 | 005042295 | -02025 | -03375 | 7 |
| 3145 | ENSMUSG( | Hnrnpul2 | 68693 | 009358975 | -00739939 | -02725 | -05725 | 7 |
| 3146 | ENSMUSG( | Homer1 | 26556 | 003895564 | -00066429 | -014 | -0195 | 7 |
| 3147 | ENSMUSG( | Homer2 | 26557 | 023606678 | 017798509 | -01325 | -04475 | 7 |
| 3148 | ENSMUSG( | Homer3 | 26558 | 004714542 | 002390728 | -015 | -05475 | 7 |
| 3175 | ENSMUSG( | Hsd17b8 | 14979 | -02510480 | -04943879 | -00875 | -014 | 7 |
| 3192 | ENSMUSG( | Hspa9 | 15526 | 011801091 | -01586965 | -02325 | -00775 | 7 |
| 3194 | ENSMUSG( | Hspb8 | 80888 | 010893057 | -02098369 | -047 | -0435 | 7 |
| 3197 | ENSMUSG( | Hspe1 | 15528 | -00168793 | -02048700 | -02 | -04575 | 7 |
| 3208 | ENSMUSG( | Htra2 | 64704 | 001569671 | -00129368 | -0055 | -01525 | 7 |
| 3209 | ENSMUSG( | Htt | 15194 | 010811470 | 006371787 | -0075 | -02225 | 7 |
| 3215 | ENSMUSG( | Iah1 | 67732 | 001322173 | -00259482 | -016 | -02675 | 7 |
| 3217 | ENSMUSG( | Iars2 | 381314 | 006682464 | 000397116 | -02575 | -03825 | 7 |
| 3227 | ENSMUSG( | Idh3a | 67834 | 000411195 | -02026621 | -00625 | -03575 | 7 |
| 3228 | ENSMUSG( | Idh3b | 170718 | 006091379 | -01835122 | -0075 | -02575 | 7 |
| 3242 | ENSMUSG( | Ift52 | 245866 | -00198427 | -02098242 | -03475 | -02925 | 7 |
| 3243 | ENSMUSG( | Ift57 | 73916 | 006022623 | -01238622 | -02975 | -007 | 7 |
| 3249 | ENSMUSG( | Igdcc4 | 56741 | -00243808 | 002394747 | -0355 | -0355 | 7 |
| 3267 | ENSMUSG( | Igtp | 16145 | -02752486 | -00008824 | -028 | -0265 | 7 |
| 3268 | ENSMUSG( | Ik | 24010 | 010310460 | -00087954 | -02475 | -05575 | 7 |
| 3270 | ENSMUSG( | Ikbkg | 16151 | 010653965 | 002839522 | -01525 | -01475 | 7 |
| 3273 | ENSMUSG( | Il18 | 16173 | 011689434 | -01325331 | -01425 | -021 | 7 |
| 3287 | ENSMUSG( | Immt | 76614 | 016555015 | -01331609 | -041 | -03225 | 7 |
| 3293 | ENSMUSG( | Impdh2 | 23918 | 012023347 | 007034347 | -00525 | -013 | 7 |
| 3294 | ENSMUSG( | Ina | 226180 | 004849243 | -01899128 | -02925 | -05375 | 7 |
| 3296 | ENSMUSG( | Inf2 | 70435 | -00676397 | -02011679 | -02925 | -01275 | 7 |
| 3299 | ENSMUSG( | Inpp1 | 16329 | -00475284 | -00293769 | -02025 | -04525 | 7 |
| 3302 | ENSMUSG( | Inpp5a | 212111 | -00053947 | -01591839 | -0065 | -0195 | 7 |
| 3315 | ENSMUSG( | Ints10 | 70885 | 009511027 | 017970550 | -0185 | -027 | 7 |
| 3317 | ENSMUSG( | Ints12 | 71793 | 006339853 | 016627707 | -046 | -0275 | 7 |
| 3322 | ENSMUSG( | Ints3 | 229543 | 021364697 | 009368793 | -02975 | -02725 | 7 |
| 3330 | ENSMUSG( | Ipcef1 | 320495 | 016703134 | -01308619 | -00525 | -03475 | 7 |
| 3336 | ENSMUSG( | Ipo8 | 320727 | 013330595 | -00537511 | -01175 | -01825 | 7 |
| 3345 | ENSMUSG( | Ireb2 | 64602 | 008184174 | 009543956 | -0145 | -01325 | 7 |
| 3352 | ENSMUSG( | Irs1 | 16367 | 005161199 | 010186728 | -01825 | -0075 | 7 |
| 3355 | ENSMUSG( | Isca2 | 74316 | -00047498 | -00602040 | -02725 | -01825 | 7 |
| 3385 | ENSMUSG( | Itih3 | 16426 | -00826386 | -00024488 | -02425 | -0385 | 7 |
| 3393 | ENSMUSG( | Itpka | 228550 | 011684700 | 005260835 | -02325 | -0385 | 7 |
| 3403 | ENSMUSG( | Iws1 | 73473 | 009830706 | 010777073 | -04025 | -01925 | 7 |
| 3405 | ENSMUSG( | Jak1 | 16451 | 011688246 | 000591255 | -02325 | -00675 | 7 |
| 3406 | ENSMUSG( | Jak2 | 16452 | 013059905 | 005531086 | -01775 | -0115 | 7 |
| 3418 | ENSMUSG( | Jph3 | 57340 | 015422505 | 010963700 | -0025 | -0275 | 7 |
| 3425 | ENSMUSG( | Jup | 16480 | 003346073 | -01009320 | -01125 | -01925 | 7 |
| 3434 | ENSMUSG( | Kat7 | 217127 | 007451870 | 004049405 | -0215 | -034 | 7 |
| 3438 | ENSMUSG( | Kbtbd11 | 74901 | 014503948 | -00746078 | -0245 | -0245 | 7 |
| 3445 | ENSMUSG( | Kcna2 | 16490 | -00356479 | 001576333 | -01225 | -02475 | 7 |
| 3450 | ENSMUSG( | Kcnab2 | 16498 | 015897593 | 009731980 | -0135 | -04125 | 7 |
| 3472 | ENSMUSG( | Kcnj10 | 16513 | 007354660 | -02045252 | -03325 | -03025 | 7 |
| 3492 | ENSMUSG( | Kcnt1 | 227632 | 004631655 | 012285645 | -03 | -0235 | 7 |
| 3501 | ENSMUSG( | Kctd17 | 72844 | -00211863 | -00747167 | -03675 | -00825 | 7 |
| 3523 | ENSMUSG( | Khk | 16548 | -00042034 | -00194062 | -01125 | -0545 | 7 |
| 3537 | ENSMUSG( | Kif3b | 16569 | 023150396 | 001665460 | -0365 | -009 | 7 |
| 3551 | ENSMUSG( | Klc2 | 16594 | 012795254 | -01538318 | -02575 | -0105 | 7 |
| 3569 | ENSMUSG( | Klhl42 | 232539 | 006760079 | 011064772 | -034 | -0385 | 7 |
| 3571 | ENSMUSG( | Klhl8 | 246293 | 016765972 | 006127107 | -0355 | -0315 | 7 |
| 3588 | ENSMUSG( | Ksr1 | 16706 | 037848380 | 026577397 | -03025 | -024 | 7 |
| 3592 | ENSMUSG( | Kxd1 | 75620 | -00303618 | -01817589 | -00425 | -0315 | 7 |
| 3593 | ENSMUSG( | Kyat1 | 70266 | 009855050 | -00083309 | -00475 | -04175 | 7 |
| 3594 | ENSMUSG( | Kyat3 | 229905 | 011461250 | -00747484 | -01925 | -04825 | 7 |
| 3596 | ENSMUSG( | L2hgdh | 217666 | 021425338 | 003448226 | -01275 | -04425 | 7 |
| 3597 | ENSMUSG( | L3hypdh | 67217 | -00604766 | 024695472 | -0325 | -0415 | 7 |
| 3636 | ENSMUSG( | Lcmt1 | 30949 | 010860331 | -01366018 | -00675 | -01475 | 7 |
| 3641 | ENSMUSG( | Ldhb | 16832 | 001857538 | -02540529 | -01325 | -0115 | 7 |
| 3650 | ENSMUSG( | Letm1 | 56384 | 003196916 | -00417928 | -00375 | -01875 | 7 |
| 3676 | ENSMUSG( | Lin7a | 108030 | 022630978 | 002963014 | -00675 | -01075 | 7 |
| 3678 | ENSMUSG( | Lin7c | 22343 | 000145434 | -00468062 | -0305 | -006 | 7 |
| 3684 | ENSMUSG( | Lipe | 16890 | 003786009 | -00618428 | -01025 | -01525 | 7 |
| 3700 | ENSMUSG( | Lmnb1 | 16906 | 007562849 | -01831874 | -01925 | -03625 | 7 |
| 3701 | ENSMUSG( | Lmnb2 | 16907 | 012307452 | 007652330 | -03675 | -049 | 7 |
| 3703 | ENSMUSG( | Lmo7 | 380928 | -02234171 | 009563764 | -03675 | -01475 | 7 |
| 3708 | ENSMUSG( | Lonp1 | 74142 | 014490647 | -01105842 | -024 | -0065 | 7 |
| 3711 | ENSMUSG( | Lpar1 | 14745 | -01730142 | -02734099 | -0225 | -01375 | 7 |
| 3723 | ENSMUSG( | Lrch1 | 380916 | 023563401 | 007189074 | -005 | -0275 | 7 |
| 3757 | ENSMUSG( | Lrrc75a | 192976 | 002117810 | -01358332 | -0245 | -062 | 7 |
| 3781 | ENSMUSG( | Lsm3 | 67678 | -00153591 | -00122980 | -00325 | -035 | 7 |
| 3783 | ENSMUSG( | Lsm5 | 66373 | -00856868 | 003963651 | -03075 | -0195 | 7 |
| 3784 | ENSMUSG( | Lsm6 | 78651 | 003045452 | -01910455 | -01075 | -04875 | 7 |
| 3790 | ENSMUSG( | Lta4h | 16993 | 001972865 | -00902590 | -018 | -0475 | 7 |
| 3828 | ENSMUSG( | Lzts3 | 241638 | 012830759 | 008371518 | -0155 | -01 | 7 |
| 3834 | ENSMUSG( | Macroh2a1 | 26914 | -02159160 | -01131781 | -01375 | -04 | 7 |
| 3869 | ENSMUSG( | Map1a | 17754 | 002821504 | 009477635 | -0425 | -00975 | 7 |
| 3874 | ENSMUSG( | Map2 | 17756 | 009918530 | 000593514 | -047 | -01175 | 7 |
| 3897 | ENSMUSG( | Map6 | 17760 | 010481654 | -00959125 | -0435 | -00625 | 7 |

| | | | | | | | | |
|---|---|---|---|---|---|---|---|---|
| 3905 | ENSMUSG( | Mapk11 | 19094 | 019932146 | -00454200 | -03825 | -026 | 7 |
| 3934 | ENSMUSG( | Mars2 | 212679 | -00501583 | 005209222 | -01325 | -0115 | 7 |
| 3951 | ENSMUSG( | Mbd1 | 17190 | 010098995 | 006379902 | -0335 | -04825 | 7 |
| 3966 | ENSMUSG( | Mccc2 | 78038 | 004561910 | -02765257 | -009 | -0145 | 7 |
| 3967 | ENSMUSG( | Mcee | 73724 | 012106590 | -01933499 | -0125 | -02875 | 7 |
| 3968 | ENSMUSG( | Mcf2l | 17207 | 020618590 | 019325809 | -024 | -016 | 7 |
| 3972 | ENSMUSG( | Mcm3ap | 54387 | 030670142 | 013548442 | -0425 | -0095 | 7 |
| 3987 | ENSMUSG( | Mdh1 | 17449 | 002513098 | -02293389 | -02425 | -0115 | 7 |
| 3995 | ENSMUSG( | Me2 | 107029 | -01942745 | -02893590 | -0385 | -00625 | 7 |
| 3997 | ENSMUSG( | Mea1 | 17256 | 007279816 | -02300504 | -0155 | -032 | 7 |
| 3998 | ENSMUSG( | Meaf6 | 70088 | 010545221 | 005129029 | -02875 | -00225 | 7 |
| 4000 | ENSMUSG( | Mecp2 | 17257 | 009323026 | 001556040 | -0295 | -03675 | 7 |
| 4011 | ENSMUSG( | Med23 | 70208 | -00352107 | 011673507 | -02925 | -00825 | 7 |
| 4036 | ENSMUSG( | Mettl1 | 17299 | 00992382 | -00446439 | -0435 | -05225 | 7 |
| 4039 | ENSMUSG( | Mettl15 | 76894 | 002760647 | -01669789 | -0055 | -02475 | 7 |
| 4040 | ENSMUSG( | Mettl16 | 67493 | 029887668 | -01172936 | -04425 | -0325 | 7 |
| 4067 | ENSMUSG( | Mgll | 23945 | 016741656 | -01045404 | -01075 | -01175 | 7 |
| 4076 | ENSMUSG( | Mical2 | 320878 | 042419481 | 005403595 | -0375 | -046 | 7 |
| 4079 | ENSMUSG( | Micos10 | 433771 | -00140901 | -02069062 | -023 | -00525 | 7 |
| 4081 | ENSMUSG( | Micu1 | 216001 | -00628510 | -02024667 | -03 | -00375 | 7 |
| 4083 | ENSMUSG( | Micu3 | 78506 | 003524523 | 013672347 | -01925 | -04275 | 7 |
| 4085 | ENSMUSG( | Mid2 | 23947 | 020211201 | 007921404 | -0135 | -017 | 7 |
| 4099 | ENSMUSG( | Mios | 252875 | 019431430 | 013655739 | -01825 | -036 | 7 |
| 4100 | ENSMUSG( | Mipep | 70478 | 014738267 | -02575123 | -0065 | -05525 | 7 |
| 4107 | ENSMUSG( | Mlf2 | 30853 | 001600497 | -01861600 | -00425 | -02225 | 7 |
| 4114 | ENSMUSG( | Mmab | 77697 | -00535756 | -02031205 | -005 | -02625 | 7 |
| 4122 | ENSMUSG( | Mob2 | 101513 | 007224904 | 003082192 | -0205 | -03125 | 7 |
| 4125 | ENSMUSG( | Mocs1 | 56738 | -00144332 | 002466158 | -03425 | -0415 | 7 |
| 4135 | ENSMUSG( | Morf4l2 | 56397 | 001885812 | -00950403 | -03625 | -00875 | 7 |
| 4138 | ENSMUSG( | Mospd2 | 76763 | 015307514 | 000126402 | -0335 | -01875 | 7 |
| 4141 | ENSMUSG( | Mpc1 | 55951 | 011642029 | -01176576 | -0265 | -04225 | 7 |
| 4150 | ENSMUSG( | Mpp3 | 13384 | 002114868 | -01343862 | -00725 | -0335 | 7 |
| 4156 | ENSMUSG( | Mpv17 | 17527 | 008161734 | -01278755 | -012 | -017 | 7 |
| 4164 | ENSMUSG( | Mrm1 | 217038 | -00650588 | -02106795 | -019 | -00525 | 7 |
| 4165 | ENSMUSG( | Mrm3 | 67390 | -00371477 | -05105426 | -02825 | -03175 | 7 |
| 4184 | ENSMUSG( | Mrpl27 | 94064 | 008267830 | -01638993 | -027 | -00575 | 7 |
| 4186 | ENSMUSG( | Mrpl3 | 94062 | 008292032 | -00946584 | -005 | -01475 | 7 |
| 4189 | ENSMUSG( | Mrpl33 | 66845 | -00365371 | -02164337 | -021 | -016 | 7 |
| 4197 | ENSMUSG( | Mrpl41 | 107733 | 008257617 | -00501743 | -01375 | -00825 | 7 |
| 4205 | ENSMUSG( | Mrpl49 | 18120 | 005948021 | -01492416 | -0095 | -018 | 7 |
| 4218 | ENSMUSG( | Mrps15 | 66407 | 01278783 | -02189393 | -0295 | -0115 | 7 |
| 4222 | ENSMUSG( | Mrps18b | 66973 | -00172026 | -00887840 | -01025 | -01075 | 7 |
| 4226 | ENSMUSG( | Mrps22 | 64655 | 010358456 | -01198572 | -0045 | -04075 | 7 |
| 4230 | ENSMUSG( | Mrps26 | 99045 | -00283731 | 002325381 | -00375 | -03925 | 7 |
| 4231 | ENSMUSG( | Mrps27 | 218506 | 007689497 | -02032537 | -01075 | -01625 | 7 |
| 4234 | ENSMUSG( | Mrps31 | 57312 | -01310019 | -00354031 | -01625 | -02175 | 7 |
| 4237 | ENSMUSG( | Mrps36 | 66128 | 010206592 | -01535149 | -01575 | -016 | 7 |
| 4243 | ENSMUSG( | Mrs2 | 380836 | 007278190 | 001200575 | -021 | -03325 | 7 |
| 4246 | ENSMUSG( | Mrto4 | 69902 | 005625565 | -01264500 | -024 | -0325 | 7 |
| 4250 | ENSMUSG( | Msi2 | 76626 | 006627852 | -01457191 | -00625 | -034 | 7 |
| 4270 | ENSMUSG( | Mtfmt | 69606 | -01032757 | 000918005 | -026 | -05625 | 7 |
| 4290 | ENSMUSG( | Mto1 | 68291 | 010419783 | -01213664 | -0175 | -009 | 7 |
| 4292 | ENSMUSG( | Mtpap | 67440 | 009847403 | 016528147 | -02475 | -01425 | 7 |
| 4316 | ENSMUSG( | Mybbp1a | 18432 | 010568787 | -00386687 | -0225 | -01675 | 7 |
| 4320 | ENSMUSG( | Myef2 | 17876 | 000855668 | 000331701 | -01025 | -026 | 7 |
| 4324 | ENSMUSG( | Myh14 | 71960 | -00226737 | -02708347 | -00925 | -0155 | 7 |
| 4331 | ENSMUSG( | Myl6b | 216459 | 007437675 | -02544688 | -03075 | -013 | 7 |
| 4336 | ENSMUSG( | Myo1b | 17912 | -01942737 | -00071764 | -02025 | -01775 | 7 |
| 4339 | ENSMUSG( | Myo1e | 71602 | 009454293 | 009960434 | -01275 | -0245 | 7 |
| 4357 | ENSMUSG( | N6amt1 | 67768 | -02272701 | -03284844 | -01125 | -038 | 7 |
| 4375 | ENSMUSG( | Nadk2 | 68646 | 010835589 | -00300792 | -0095 | -0435 | 7 |
| 4376 | ENSMUSG( | Nae1 | 234664 | -00350433 | 000701524 | -0145 | -0215 | 7 |
| 4387 | ENSMUSG( | Napa | 108124 | -00268480 | -01485837 | -00825 | -03175 | 7 |
| 4391 | ENSMUSG( | Naprt | 223646 | -00021468 | -01275344 | -04125 | -035 | 7 |
| 4396 | ENSMUSG( | Nat10 | 98956 | 015878767 | 009557282 | -028 | -02275 | 7 |
| 4406 | ENSMUSG( | Naxe | 246703 | -01252008 | -01587835 | -02425 | -01875 | 7 |
| 4416 | ENSMUSG( | Ncbp1 | 433702 | 009301976 | 004655444 | -006 | -04025 | 7 |
| 4418 | ENSMUSG( | Ncbp3 | 66874 | 010331317 | 005187644 | -02 | -0215 | 7 |
| 4428 | ENSMUSG( | Nckipsd | 80987 | 018051638 | -00571175 | -0095 | -02325 | 7 |
| 4445 | ENSMUSG( | Ndor1 | 78797 | 003546159 | -00405453 | -02275 | -04575 | 7 |
| 4446 | ENSMUSG( | Ndrg1 | 17988 | -00491984 | -00433167 | -02575 | -03825 | 7 |
| 4454 | ENSMUSG( | Ndufa13 | 67184 | -00423889 | -02297437 | -02275 | -0255 | 7 |
| 4455 | ENSMUSG( | Ndufa2 | 17991 | 006239507 | -02500390 | -0185 | -046 | 7 |
| 4456 | ENSMUSG( | Ndufa3 | 66091 | -00395652 | -03663218 | -02875 | -0215 | 7 |
| 4461 | ENSMUSG( | Ndufa7 | 66416 | -00194711 | -03266528 | -04275 | -005 | 7 |
| 4475 | ENSMUSG( | Ndufb11 | 104130 | -00237931 | -01550445 | -0145 | -03125 | 7 |
| 4476 | ENSMUSG( | Ndufb2 | 68198 | 001569637 | -02446089 | -0355 | -01825 | 7 |
| 4477 | ENSMUSG( | Ndufb3 | 66495 | -00826941 | -02645320 | -0175 | -021 | 7 |
| 4480 | ENSMUSG( | Ndufb6 | 230075 | -00516559 | -02127550 | -01325 | -01825 | 7 |
| 4482 | ENSMUSG( | Ndufb8 | 67264 | -00393282 | -02450697 | -01825 | -01475 | 7 |
| 4492 | ENSMUSG( | Ndufs8 | 225887 | -01102802 | -02897140 | -03825 | -01 | 7 |
| 4493 | ENSMUSG( | Ndufv1 | 17995 | 008858731 | -01868307 | -008 | -048 | 7 |
| 4496 | ENSMUSG( | Nebl | 74103 | 013428501 | -00448272 | -0195 | -02225 | 7 |
| 4498 | ENSMUSG( | Necab2 | 117148 | -01124177 | -02215611 | -0075 | -01325 | 7 |
| 4500 | ENSMUSG( | Necap1 | 67602 | 002402360 | -01122760 | -0115 | -02525 | 7 |
| 4508 | ENSMUSG( | Nefh | 380684 | -00373859 | -01169117 | -02925 | -01125 | 7 |
| 4529 | ENSMUSG( | Neu2 | 23956 | -01143397 | -03430656 | -03825 | -004 | 7 |
| 4535 | ENSMUSG( | Nf2 | 18016 | 008034298 | -00839747 | -01 | -012 | 7 |
| 4542 | ENSMUSG( | Nfkb1 | 18033 | -00386620 | 007311038 | -00525 | -03325 | 7 |
| 4543 | ENSMUSG( | Nfrkb | 235134 | -00418027 | 001660227 | -04075 | -0345 | 7 |
| 4544 | ENSMUSG( | Nfs1 | 18041 | 003551874 | -01967948 | -0175 | -006 | 7 |
| 4553 | ENSMUSG( | Nhlrc2 | 66866 | 008393087 | 003675743 | -01 | -01925 | 7 |
| 4560 | ENSMUSG( | Nicn1 | 66257 | 006939932 | -01377887 | -005 | -01425 | 7 |
| 4571 | ENSMUSG( | Nipsnap2 | 14467 | 000196469 | -01929941 | -03375 | -05075 | 7 |
| 4577 | ENSMUSG( | Nkiras1 | 69721 | -00073468 | -01797163 | -0295 | -03025 | 7 |
| 4583 | ENSMUSG( | Nlk | 18099 | 015869359 | 007493342 | -01175 | -0465 | 7 |
| 4585 | ENSMUSG( | Nlrx1 | 270151 | 001189436 | 004486102 | -01525 | -016 | 7 |
| 4592 | ENSMUSG( | Nmral1 | 67824 | 003882894 | 000480275 | -04475 | -0335 | 7 |
| 4601 | ENSMUSG( | Noct | 12457 | -00348370 | -00706242 | -039 | -0345 | 7 |
| 4606 | ENSMUSG( | Nol6 | 230082 | 007589776 | -01244147 | -022 | -008 | 7 |
| 4613 | ENSMUSG( | Nop10 | 66181 | -00734229 | -01787364 | -012 | -0265 | 7 |
| 4614 | ENSMUSG( | Nop14 | 75416 | 016948379 | -00132578 | -019 | -044 | 7 |
| 4618 | ENSMUSG( | Nop56 | 67134 | 008547182 | 002413304 | -01025 | -0335 | 7 |
| 4620 | ENSMUSG( | Nop9 | 67842 | 019097533 | 011735780 | -01875 | -0295 | 7 |
| 4630 | ENSMUSG( | Nova2 | 384569 | 016443299 | 001155442 | -03875 | -005 | 7 |
| 4639 | ENSMUSG( | Npl | 74091 | 030819391 | 000989333 | -0025 | -041 | 7 |
| 4642 | ENSMUSG( | Npm3 | 18150 | 013575384 | 005545411 | -02175 | -0445 | 7 |
| 4644 | ENSMUSG( | Nprl2 | 56032 | 004689147 | -00100888 | -0155 | -01225 | 7 |
| 4653 | ENSMUSG( | Nqo2 | 18105 | 006639344 | -02069285 | -014 | -03375 | 7 |
| 4668 | ENSMUSG( | Nrip2 | 60345 | 012998194 | -00987721 | -04575 | -01475 | 7 |
| 4683 | ENSMUSG( | Nsfl1c | 386649 | 005338792 | -00289816 | -005 | -0275 | 7 |
| 4686 | ENSMUSG( | Nsmaf | 18201 | 012654324 | 004451994 | -0335 | -031 | 7 |
| 4689 | ENSMUSG( | Nsrp1 | 237859 | 007992742 | -01088579 | -02225 | -00275 | 7 |
| 4691 | ENSMUSG( | Nsun4 | 72181 | 013778001 | -00947011 | -018 | -0075 | 7 |
| 4693 | ENSMUSG( | Nt5c1a | 230718 | 010627325 | -01200913 | -0075 | -02625 | 7 |
| 4694 | ENSMUSG( | Nt5c2 | 76952 | 006894448 | 006792486 | -005 | -039 | 7 |
| 4699 | ENSMUSG( | Nt5m | 103850 | 008536789 | -01934784 | -00575 | -0465 | 7 |
| 4700 | ENSMUSG( | Ntan1 | 18203 | 000602586 | -01038253 | -044 | -00825 | 7 |
| 4714 | ENSMUSG( | Nubp2 | 26426 | 011408974 | -02012756 | -02325 | -0105 | 7 |
| 4743 | ENSMUSG( | Numb | 18222 | 012506626 | -00288643 | -024 | -0145 | 7 |
| 4750 | ENSMUSG( | Nup188 | 227699 | 006751927 | 007401806 | -00525 | -02475 | 7 |
| 4753 | ENSMUSG( | Nup214 | 227720 | 006426047 | 013239513 | -02125 | -03975 | 7 |
| 4754 | ENSMUSG( | Nup35 | 69482 | 004194026 | 000735953 | -022 | -0365 | 7 |
| 4755 | ENSMUSG( | Nup37 | 69736 | 013150258 | 010915086 | -02725 | -03425 | 7 |
| 4756 | ENSMUSG( | Nup43 | 69912 | 010858921 | 013171977 | -008 | -0355 | 7 |
| 4757 | ENSMUSG( | Nup50 | 18141 | 003421132 | 008428848 | -02475 | -00825 | 7 |

| | | | | | | | |
|---|---|---|---|---|---|---|---|
| 4759 | ENSMUSG( | Nup58 | 71844 | 012042714 | 007519821 | -01025 | -02675 | 7 |
| 4763 | ENSMUSG( | Nup93 | 71805 | -00241292 | -00282214 | -00625 | -04525 | 7 |
| 4767 | ENSMUSG( | Nwd1 | 319555 | 014289889 | -01665621 | -009 | -0215 | 7 |
| 4786 | ENSMUSG( | Ogdhl | 239017 | 018661901 | -01012171 | -00925 | -02325 | 7 |
| 4802 | ENSMUSG( | Opa3 | 403187 | 006033826 | -01292379 | -02875 | -0125 | 7 |
| 4805 | ENSMUSG( | Ophn1 | 94190 | 007146570 | 008651335 | -025 | -00075 | 7 |
| 4810 | ENSMUSG( | Optn | 71648 | -00618971 | -00780820 | -028 | -01075 | 7 |
| 4816 | ENSMUSG( | Osbp2 | 74309 | 006707387 | 005226810 | -00675 | -02475 | 7 |
| 4827 | ENSMUSG( | Oscp1 | 230751 | -00546491 | 001490466 | -01075 | -04325 | 7 |
| 4836 | ENSMUSG( | Otud5 | 54644 | 014212313 | 000585619 | -0065 | -04725 | 7 |
| 4838 | ENSMUSG( | Otud7a | 170711 | 026881997 | 006942429 | -0175 | -01025 | 7 |
| 4840 | ENSMUSG( | Otulin | 432940 | 002719254 | -01438851 | -04425 | -017 | 7 |
| 4844 | ENSMUSG( | Oxld1 | 66431 | -00637146 | -02944035 | -0375 | -04075 | 7 |
| 4847 | ENSMUSG( | Oxsm | 71147 | 004683515 | -01230698 | -02075 | -03725 | 7 |
| 4859 | ENSMUSG( | Pacsin1 | 23969 | 010917226 | -00675148 | -0155 | -02575 | 7 |
| 4862 | ENSMUSG( | Padi2 | 18600 | -00184760 | 007123139 | -006 | -015 | 7 |
| 4863 | ENSMUSG( | Paf1 | 54624 | 003693742 | -00187255 | -01975 | -0095 | 7 |
| 4865 | ENSMUSG( | Pafah1b2 | 18475 | 001667057 | -00757998 | -0285 | -0235 | 7 |
| 4872 | ENSMUSG( | Pak1 | 18479 | 015481655 | -01079143 | -00725 | -0155 | 7 |
| 4874 | ENSMUSG( | Pak2 | 224105 | 012696577 | 000707107 | -00775 | -012 | 7 |
| 4891 | ENSMUSG( | Pank1 | 75735 | 017480337 | -00728415 | -0455 | -04025 | 7 |
| 4901 | ENSMUSG( | Paqr7 | 71904 | -02315578 | 01573685 | -038 | -036 | 7 |
| 4905 | ENSMUSG( | Pard6b | 58220 | -01515692 | 000910051 | -0145 | -03875 | 7 |
| 4912 | ENSMUSG( | Parp12 | 243771 | -00067527 | 002276137 | -016 | -03275 | 7 |
| 4920 | ENSMUSG( | Paxx | 227622 | 009550096 | 001002672 | -0475 | -016 | 7 |
| 4932 | ENSMUSG( | Pccb | 66904 | 004361371 | -01947852 | -00575 | -0435 | 7 |
| 4958 | ENSMUSG( | Pcdhga8 | 93716 | 003818900 | 000756554 | -011 | -0175 | 7 |
| 4961 | ENSMUSG( | Pcdhgb5 | 93702 | -00671843 | 012097546 | -02375 | -01975 | 7 |
| 4962 | ENSMUSG( | Pcdhgb6 | 93703 | -02188327 | 002684189 | -025 | -03025 | 7 |
| 4966 | ENSMUSG( | Pcgf2 | 22658 | 002682722 | -01214016 | -02325 | -03325 | 7 |
| 4972 | ENSMUSG( | Pcmt1 | 18537 | -02357797 | -00124414 | -03325 | -029 | 7 |
| 4973 | ENSMUSG( | Pcmtd1 | 319263 | -00298962 | 002969831 | -00475 | -03675 | 7 |
| 4985 | ENSMUSG( | Pcx | 18563 | 014329681 | -00883859 | -00825 | -03775 | 7 |
| 4989 | ENSMUSG( | Pcyt1b | 236899 | 002673539 | -01238141 | -037 | -00525 | 7 |
| 4992 | ENSMUSG( | Pdcd10 | 56426 | 004065940 | 006589475 | -00675 | -012 | 7 |
| 4997 | ENSMUSG( | Pdcd6 | 18570 | 003661799 | -01622128 | -0075 | -0535 | 7 |
| 4998 | ENSMUSG( | Pdcd6ip | 18571 | 006457739 | 003092256 | -01875 | -02075 | 7 |
| 5003 | ENSMUSG( | Pde12 | 211948 | -00033157 | -00495223 | -00575 | -01525 | 7 |
| 5010 | ENSMUSG( | Pde4b | 18578 | -00931247 | -00902142 | -02375 | -002 | 7 |
| 5019 | ENSMUSG( | Pde9a | 18585 | -00626890 | -02946886 | -0375 | -03025 | 7 |
| 5023 | ENSMUSG( | Pdha1 | 18597 | 004913133 | -01524694 | -02425 | -01875 | 7 |
| 5024 | ENSMUSG( | Pdhb | 68263 | 011147965 | -01255259 | -01925 | -01675 | 7 |
| 5032 | ENSMUSG( | Pdk3 | 236900 | -00442351 | -01971555 | -007 | -04075 | 7 |
| 5042 | ENSMUSG( | Pdpr | 319518 | 001905269 | -00808162 | -0065 | -02675 | 7 |
| 5051 | ENSMUSG( | Pdzd4 | 245469 | 018260122 | 002360535 | -039 | -02525 | 7 |
| 5078 | ENSMUSG( | Pex5 | 19305 | 005661319 | 003811962 | -03325 | -0235 | 7 |
| 5099 | ENSMUSG( | Pgbd5 | 209966 | 021771605 | -00714898 | -02775 | -01525 | 7 |
| 5102 | ENSMUSG( | Pggt1b | 225467 | 014060521 | 012776309 | -0405 | -0215 | 7 |
| 5115 | ENSMUSG( | Phactr1 | 218194 | 002784310 | 012981295 | -02325 | -0105 | 7 |
| 5116 | ENSMUSG( | Phactr2 | 215789 | -00447534 | -02339652 | -034 | -0125 | 7 |
| 5117 | ENSMUSG( | Phactr3 | 74189 | 016714178 | -00422416 | -023 | -02725 | 7 |
| 5125 | ENSMUSG( | Phf10 | 72057 | 003212181 | -00024553 | -03425 | -04325 | 7 |
| 5129 | ENSMUSG( | Phf24 | 230085 | 010571909 | 006560278 | -0075 | -03125 | 7 |
| 5141 | ENSMUSG( | Phlda1 | 21664 | -01479773 | -00780755 | -025 | -03025 | 7 |
| 5143 | ENSMUSG( | Phldb1 | 102693 | -02277482 | -01099195 | -02875 | -00325 | 7 |
| 5145 | ENSMUSG( | Phlpp1 | 98432 | 002881435 | -01825063 | -0265 | -03125 | 7 |
| 5147 | ENSMUSG( | Phospho1 | 237928 | 015867172 | 010562850 | -03175 | -0315 | 7 |
| 5160 | ENSMUSG( | Picalm | 233489 | 010664104 | 011366044 | -029 | -0585 | 7 |
| 5175 | ENSMUSG( | Pik3c3 | 225326 | 023280833 | 000735820 | -0115 | -016 | 7 |
| 5196 | ENSMUSG( | Pitpna | 18738 | 004388806 | -01329163 | -013 | -0175 | 7 |
| 5205 | ENSMUSG( | Pkd2 | 18764 | -00084004 | 010966427 | -00375 | -0165 | 7 |
| 5207 | ENSMUSG( | Pkib | 18768 | -01337291 | -02174405 | -04775 | -03175 | 7 |
| 5209 | ENSMUSG( | Pkm | 18746 | -00180875 | -02021837 | -00775 | -052 | 7 |
| 5211 | ENSMUSG( | Pkn2 | 109333 | 005700526 | 017862219 | -04325 | -03975 | 7 |
| 5223 | ENSMUSG( | Plcb3 | 18797 | 007431988 | -00764231 | -02125 | -01325 | 7 |
| 5224 | ENSMUSG( | Plcb4 | 18798 | 003968275 | 006689609 | -003 | -02225 | 7 |
| 5230 | ENSMUSG( | Plch2 | 269615 | 032133341 | 012768367 | -02075 | -0395 | 7 |
| 5240 | ENSMUSG( | Plekha1 | 101476 | 967e-05 | 002487198 | -0205 | -00525 | 7 |
| 5243 | ENSMUSG( | Plekha6 | 240753 | 001312644 | -00137922 | -02875 | -01925 | 7 |
| 5245 | ENSMUSG( | Plekhb1 | 27276 | -00022670 | -02355693 | -02 | -008 | 7 |
| 5255 | ENSMUSG( | Plgrkt | 67759 | -00842231 | 016282980 | -03725 | -015 | 7 |
| 5273 | ENSMUSG( | Plrg1 | 53317 | 020683886 | 001564895 | -008 | -0235 | 7 |
| 5274 | ENSMUSG( | Pls1 | 102502 | -02021103 | -00474949 | -03 | -045 | 7 |
| 5293 | ENSMUSG( | Pmm2 | 54128 | 003177568 | 002129452 | -0305 | -0085 | 7 |
| 5302 | ENSMUSG( | Pnma8a | 71691 | -00305862 | 000937991 | -00425 | -0145 | 7 |
| 5330 | ENSMUSG( | Polr2a | 20020 | 009238359 | 015547526 | -01375 | -02175 | 7 |
| 5331 | ENSMUSG( | Polr2b | 231329 | 020876532 | 001271874 | -024 | -0285 | 7 |
| 5332 | ENSMUSG( | Polr2c | 20021 | -00214048 | -00409035 | -03225 | -04425 | 7 |
| 5334 | ENSMUSG( | Polr2e | 66420 | -00046770 | -02112175 | -031 | -021 | 7 |
| 5335 | ENSMUSG( | Polr2g | 67710 | 004340098 | -00543855 | -00325 | -04275 | 7 |
| 5336 | ENSMUSG( | Polr2h | 245841 | 008126224 | -00771157 | -01525 | -03425 | 7 |
| 5363 | ENSMUSG( | Ppan | 235036 | -01298628 | -02272239 | -0425 | -03325 | 7 |
| 5365 | ENSMUSG( | Ppcdc | 66812 | -00119727 | 002915584 | -0165 | -03625 | 7 |
| 5368 | ENSMUSG( | Ppfia1 | 233977 | -00672704 | -01680599 | -031 | -01525 | 7 |
| 5374 | ENSMUSG( | Pphln1 | 223828 | -00614602 | -01262037 | -01025 | -05325 | 7 |
| 5375 | ENSMUSG( | Ppia | 268373 | -00197876 | -01811065 | -01 | -036 | 7 |
| 5382 | ENSMUSG( | Ppih | 66101 | 006188683 | 015992326 | -00525 | -0195 | 7 |
| 5385 | ENSMUSG( | Ppil3 | 70225 | -00288340 | -00821385 | -0355 | -022 | 7 |
| 5389 | ENSMUSG( | Ppm1a | 19042 | 009537336 | 002031614 | -01325 | -007 | 7 |
| 5390 | ENSMUSG( | Ppm1b | 19043 | 010322417 | -00931064 | -01075 | -024 | 7 |
| 5412 | ENSMUSG( | Ppp1r16a | 73062 | 013746464 | 007358933 | -00825 | -03925 | 7 |
| 5413 | ENSMUSG( | Ppp1r16b | 228852 | 006502325 | -00095030 | -0505 | -01625 | 7 |
| 5417 | ENSMUSG( | Ppp1r2 | 66849 | 001557232 | -00325605 | -00525 | -04625 | 7 |
| 5421 | ENSMUSG( | Ppp1r3f | 54646 | 011461095 | -01081082 | -038 | -039 | 7 |
| 5422 | ENSMUSG( | Ppp1r3g | 76487 | 030840738 | -00189789 | -0125 | -0235 | 7 |
| 5425 | ENSMUSG( | Ppp1r9a | 243725 | -02002678 | 007658234 | -031 | -01175 | 7 |
| 5440 | ENSMUSG( | Ppp3ca | 19055 | 001403430 | -00204065 | -014 | -02925 | 7 |
| 5451 | ENSMUSG( | Ppp6r1 | 243819 | 024478062 | -00148123 | -01175 | -01125 | 7 |
| 5453 | ENSMUSG( | Ppp6r3 | 52036 | 009614814 | 003430322 | -009 | -01675 | 7 |
| 5479 | ENSMUSG( | Prkaa1 | 105787 | 003924448 | 002840343 | -016 | -01225 | 7 |
| 5486 | ENSMUSG( | Prkag2 | 108099 | 017598387 | 000051146 | -00625 | -01125 | 7 |
| 5515 | ENSMUSG( | Prodh | 19125 | 021223876 | 001149609 | -0365 | -03225 | 7 |
| 5521 | ENSMUSG( | Prpf3 | 70767 | 012575852 | 012515675 | -01275 | -03575 | 7 |
| 5529 | ENSMUSG( | Prpf4b | 19134 | -00196944 | 029591328 | -01825 | -0545 | 7 |
| 5557 | ENSMUSG( | Psd2 | 74002 | 014592138 | -02704543 | -00925 | -0125 | 7 |
| 5563 | ENSMUSG( | Psma3 | 19167 | 005133922 | -00345378 | -022 | -0075 | 7 |
| 5565 | ENSMUSG( | Psma5 | 26442 | 000944963 | -01719188 | -00975 | -02875 | 7 |
| 5602 | ENSMUSG( | Psmf1 | 228769 | 000113005 | -01027793 | -0325 | -015 | 7 |
| 5615 | ENSMUSG( | Ptcd3 | 69956 | 014045056 | -00213199 | -0045 | -0245 | 7 |
| 5621 | ENSMUSG( | Ptges2 | 96979 | 005140014 | -00426815 | -017 | -00925 | 7 |
| 5640 | ENSMUSG( | Ptpa | 110854 | 012610815 | -00588618 | -006 | -0445 | 7 |
| 5647 | ENSMUSG( | Ptpn2 | 19255 | 003992192 | -00066233 | -03275 | -0345 | 7 |
| 5648 | ENSMUSG( | Ptpn23 | 104831 | 016436683 | -00784761 | -02 | -024 | 7 |
| 5649 | ENSMUSG( | Ptpn3 | 545622 | 019721967 | -00733216 | -00775 | -03425 | 7 |
| 5670 | ENSMUSG( | Ptrh1 | 329384 | 015540782 | -00188025 | -02275 | -03675 | 7 |
| 5678 | ENSMUSG( | Pum3 | 52874 | 015158953 | -01295737 | -0115 | -031 | 7 |
| 5707 | ENSMUSG( | Qdpr | 110391 | -01906619 | -04566645 | -0235 | -03625 | 7 |
| 5724 | ENSMUSG( | Rab11fip4 | 268451 | 013926842 | -00747328 | -01525 | -021 | 7 |
| 5738 | ENSMUSG( | Rab27a | 11891 | 016272118 | 026693674 | -0225 | -031 | 7 |
| 5741 | ENSMUSG( | Rab29 | 226422 | 024001124 | -01153430 | -0305 | -068 | 7 |
| 5771 | ENSMUSG( | Rab8a | 17274 | 014468343 | -00289198 | -014 | -032 | 7 |
| 5782 | ENSMUSG( | Rabggtb | 19352 | -00305797 | -01425214 | -03175 | -019 | 7 |
| 5783 | ENSMUSG( | Rabif | 98710 | 000028392 | -00734464 | -0115 | -02425 | 7 |
| 5791 | ENSMUSG( | Rad21 | 19357 | 010322641 | -00903663 | -0265 | -01825 | 7 |
| 5795 | ENSMUSG( | Radil | 231858 | 007382189 | 006855846 | -02025 | -0085 | 7 |
| 5802 | ENSMUSG( | Ralbp1 | 19765 | 024955618 | 007445646 | -03375 | -031 | 7 |

| | | | | | | | | |
|---|---|---|---|---|---|---|---|---|
| 5804 | ENSMUSG( | Ralgapa2 | 241694 | 001231983 | 011614714 | -01025 | -02225 | 7 |
| 5808 | ENSMUSG( | Raly | 19383 | 001198613 | -01343781 | -0315 | -045 | 7 |
| 5821 | ENSMUSG( | Rap1b | 215449 | 002740302 | 003935842 | -007 | -03625 | 7 |
| 5831 | ENSMUSG( | Rapgef4 | 56508 | 009195234 | -01392509 | -00975 | -01875 | 7 |
| 5833 | ENSMUSG( | Rapgef6 | 192786 | -00477186 | 02146476C | -02725 | -008 | 7 |
| 5852 | ENSMUSG( | Rassf5 | 54354 | 01514626€ | -01033039 | -04175 | -045 | 7 |
| 5857 | ENSMUSG( | Rbbp6 | 19647 | -00960330 | 005733588 | -017 | -0235 | 7 |
| 5862 | ENSMUSG( | Rbfox1 | 268859 | -00953680 | -00521464 | -01975 | -0425 | 7 |
| 5863 | ENSMUSG( | Rbfox2 | 93686 | 014979064 | 00625532S | -04375 | -03 | 7 |
| 5869 | ENSMUSG( | Rbm12b2 | 77604 | -00148106 | 011098788 | -0205 | -044 | 7 |
| 5870 | ENSMUSG( | Rbm14 | 56275 | 010319338 | -00753867 | -035 | -01375 | 7 |
| 5874 | ENSMUSG( | Rbm19 | 74111 | 029211091 | 00399708C | -028 | -011 | 7 |
| 5875 | ENSMUSG( | Rbm22 | 66810 | -00314466 | 007607477 | -04875 | -019 | 7 |
| 5879 | ENSMUSG( | Rbm28 | 68272 | 02130419C | -00028577 | -05125 | -01925 | 7 |
| 5881 | ENSMUSG( | Rbm34 | 52202 | 001290026 | 00025784S | -04775 | -01875 | 7 |
| 5883 | ENSMUSG( | Rbm45 | 241490 | 010212203 | -01225545 | -02175 | -021 | 7 |
| 5902 | ENSMUSG( | Rcc1 | 100088 | 019774316 | 009211014 | -03125 | -01475 | 7 |
| 5905 | ENSMUSG( | Rcl1 | 59028 | 014108205 | 009324105 | -039 | -044 | 7 |
| 5913 | ENSMUSG( | Rdx | 19684 | -00437399 | 00045642 | -01675 | -013 | 7 |
| 5928 | ENSMUSG( | Reps1 | 19707 | 014056881 | -00068116 | -025 | -00325 | 7 |
| 5933 | ENSMUSG( | Retreg3 | 67998 | -01113776 | 010460345 | -01625 | -039 | 7 |
| 5945 | ENSMUSG( | Rftn2 | 74013 | 008567875 | -03237313 | -015 | -03725 | 7 |
| 5956 | ENSMUSG( | Rgs14 | 51791 | -00795213 | -02157530 | -007 | -02 | 7 |
| 5965 | ENSMUSG( | Rgs8 | 67792 | -00387678 | -01002649 | -04175 | -05075 | 7 |
| 5974 | ENSMUSG( | Rhoc | 11853 | -00820667 | -02044024 | -01275 | -01225 | 7 |
| 5994 | ENSMUSG( | Rims3 | 242662 | 024336583 | -00539288 | -0075 | -04 | 7 |
| 5995 | ENSMUSG( | Rims4 | 241770 | 023231833 | -00155453 | -00425 | -01525 | 7 |
| 5999 | ENSMUSG( | Riok3 | 66878 | -00064494 | -00079854 | -00575 | -03875 | 7 |
| 6021 | ENSMUSG( | Rnd2 | 11858 | 00403106C | -01727508 | -0185 | -014 | 7 |
| 6031 | ENSMUSG( | Rnf126 | 70294 | -00066826 | -00542642 | -0355 | -00625 | 7 |
| 6044 | ENSMUSG( | Rnf2 | 19821 | -00150904 | 004745454 | -03675 | -05925 | 7 |
| 6063 | ENSMUSG( | Rnps1 | 19826 | 008244113 | -00277126 | -01575 | -05725 | 7 |
| 6078 | ENSMUSG( | Rpf2 | 67239 | 00278153S | -02007984 | -04425 | -0065 | 7 |
| 6081 | ENSMUSG( | Rph3a | 19894 | -00291252 | -03410072 | -009 | -01575 | 7 |
| 6118 | ENSMUSG( | Rpl39 | 67248 | 011634902 | -01062959 | -03375 | -00525 | 7 |
| 6175 | ENSMUSG( | Rpusd3 | 101122 | 002118537 | 02442420S | -01975 | -01475 | 7 |
| 6185 | ENSMUSG( | Rrm2b | 382985 | 003022358 | -00332150 | -01575 | -0395 | 7 |
| 6187 | ENSMUSG( | Rrp12 | 107094 | 02257213S | -00064958 | -02225 | -01775 | 7 |
| 6189 | ENSMUSG( | Rrp9 | 27966 | 013965562 | 017696913 | -021 | -01225 | 7 |
| 6190 | ENSMUSG( | Rrs1 | 59014 | 00798904S | 000306353 | -042 | -03425 | 7 |
| 6192 | ENSMUSG( | Rsl1d1 | 66409 | -00098547 | -00486683 | -02875 | -02175 | 7 |
| 6207 | ENSMUSG( | Rtn4ip1 | 170728 | 010988615 | -01566200 | -0235 | -02925 | 7 |
| 6218 | ENSMUSG( | Rundc3b | 242819 | 004407291 | -01114634 | -02275 | -00425 | 7 |
| 6223 | ENSMUSG( | Ruvbl2 | 20174 | 004981408 | -00531698 | -0135 | -01875 | 7 |
| 6238 | ENSMUSG( | S100b | 20203 | -00060038 | -02714580 | -01275 | -03125 | 7 |
| 6245 | ENSMUSG( | Safb | 224903 | 015734246 | 003971542 | -01925 | -0345 | 7 |
| 6246 | ENSMUSG( | Safb2 | 224902 | 009932148 | 004986712 | -0065 | -0295 | 7 |
| 6248 | ENSMUSG( | Samd12 | 320679 | 00737012S | 001870415 | -02675 | -00475 | 7 |
| 6264 | ENSMUSG( | Sardh | 192166 | 000842802 | -01995217 | -028 | -02625 | 7 |
| 6266 | ENSMUSG( | Sarnp | 66118 | 013955706 | 001350834 | -02475 | -03675 | 7 |
| 6267 | ENSMUSG( | Sars2 | 71984 | -00031511 | 000652613 | -0145 | -02425 | 7 |
| 6280 | ENSMUSG( | Scaf1 | 233208 | 018434068 | 00757562C | -0305 | -04825 | 7 |
| 6294 | ENSMUSG( | Scd2 | 20250 | 00524942C | -00876777 | -01075 | -0345 | 7 |
| 6320 | ENSMUSG( | Scrn3 | 74616 | 006191061 | -00841929 | -004 | -0225 | 7 |
| 6321 | ENSMUSG( | Scrt1 | 170729 | 017434953 | 009785454 | -0465 | -03375 | 7 |
| 6335 | ENSMUSG( | Sdha | 66945 | 008341362 | -02037571 | -0115 | -0115 | 7 |
| 6340 | ENSMUSG( | Sdhb | 67680 | 00481034C | -01084666 | -013 | -03875 | 7 |
| 6350 | ENSMUSG( | Sec14l1 | 74136 | 003432833 | -00337520 | -0115 | -02125 | 7 |
| 6369 | ENSMUSG( | Seh1l | 72124 | -00820820 | -00708122 | -0205 | -035 | 7 |
| 6377 | ENSMUSG( | Selenoo | 223776 | -00367617 | -00058824 | -023 | -0075 | 7 |
| 6401 | ENSMUSG( | Sepsecs | 211006 | 00682459C | 00190163S | -01775 | -03875 | 7 |
| 6410 | ENSMUSG( | Septin8 | 20362 | 010661951 | -01943522 | -024 | -04375 | 7 |
| 6411 | ENSMUSG( | Septin9 | 53860 | 020059423 | -02101853 | -0065 | -03875 | 7 |
| 6437 | ENSMUSG( | Setd7 | 73251 | 010334413 | -01377930 | -02075 | -01725 | 7 |
| 6447 | ENSMUSG( | Sf3b2 | 319322 | 01204927S | -00792412 | -02025 | -02675 | 7 |
| 6448 | ENSMUSG( | Sf3b3 | 101943 | 00510750S | -00790886 | -0035 | -022 | 7 |
| 6451 | ENSMUSG( | Sf3b6 | 66055 | 004093293 | -01075181 | -044 | -0405 | 7 |
| 6462 | ENSMUSG( | Sfxn5 | 94282 | 005336197 | -03785497 | -01175 | -01825 | 7 |
| 6477 | ENSMUSG( | Sh2b2 | 23921 | 003714838 | 016322005 | -0325 | -064 | 7 |
| 6479 | ENSMUSG( | Sh2d5 | 230863 | 000958396 | -00681776 | -02225 | -022 | 7 |
| 6481 | ENSMUSG( | Sh3bgrl2 | 212531 | -00543409 | -01325247 | -0255 | -0445 | 7 |
| 6482 | ENSMUSG( | Sh3bgrl3 | 73723 | -00378465 | -02425419 | -01725 | -04225 | 7 |
| 6483 | ENSMUSG( | Sh3bp1 | 20401 | 004541457 | 012045828 | -02725 | -027 | 7 |
| 6488 | ENSMUSG( | Sh3gl1 | 20405 | 013597538 | 009194431 | -02025 | -0025 | 7 |
| 6489 | ENSMUSG( | Sh3gl2 | 20404 | 018044925 | -02169542 | -05275 | -01775 | 7 |
| 6490 | ENSMUSG( | Sh3gl3 | 20408 | 007068526 | -01621470 | -01175 | -01875 | 7 |
| 6492 | ENSMUSG( | Sh3glb2 | 227700 | 003730978 | -00287933 | -01875 | -05175 | 7 |
| 6496 | ENSMUSG( | Sh3rf1 | 59009 | 014453072 | 012399886 | -00425 | -019 | 7 |
| 6504 | ENSMUSG( | Shb | 230126 | -00802455 | -00869368 | -0235 | -006 | 7 |
| 6505 | ENSMUSG( | Shc2 | 216148 | 00863598C | 000329714 | -03975 | -01725 | 7 |
| 6529 | ENSMUSG( | Sik3 | 70661 | 010638575 | -00027353 | -00525 | -0435 | 7 |
| 6539 | ENSMUSG( | Sirt2 | 64383 | 008652334 | -01854713 | -01225 | -0505 | 7 |
| 6546 | ENSMUSG( | Slain2 | 75991 | 008130756 | 009432231 | -034 | -021 | 7 |
| 6574 | ENSMUSG( | Slc1a2 | 20511 | 01241762C | -01111368 | -03375 | -032 | 7 |
| 6579 | ENSMUSG( | Slc20a1 | 20515 | -01835603 | -00789182 | -033 | -0085 | 7 |
| 6591 | ENSMUSG( | Slc25a11 | 67863 | 007632156 | -00765261 | -01025 | -0285 | 7 |
| 6592 | ENSMUSG( | Slc25a12 | 78830 | 007493763 | -00031349 | -02875 | -021 | 7 |
| 6603 | ENSMUSG( | Slc25a23 | 66972 | 01526562 | -00299247 | -01175 | -01375 | 7 |
| 6607 | ENSMUSG( | Slc25a27 | 74011 | 017677602 | 024183293 | -01725 | -025 | 7 |
| 6610 | ENSMUSG( | Slc25a3 | 18674 | -00166635 | -02694183 | -01775 | -022 | 7 |
| 6612 | ENSMUSG( | Slc25a33 | 70556 | 016447712 | 007927161 | -021 | -02975 | 7 |
| 6617 | ENSMUSG( | Slc25a42 | 73095 | 0114104 | 00588726S | -01125 | -02425 | 7 |
| 6621 | ENSMUSG( | Slc25a51 | 230125 | 00292665 | 007477896 | -01575 | -016 | 7 |
| 6624 | ENSMUSG( | Slc26a6 | 171429 | -00199493 | 017182191 | -01025 | -02825 | 7 |
| 6640 | ENSMUSG( | Slc30a9 | 109108 | 007026997 | -00499904 | -02425 | -01425 | 7 |
| 6668 | ENSMUSG( | Slc39a7 | 14977 | 004305706 | 019632245 | -0435 | -03025 | 7 |
| 6703 | ENSMUSG( | Slc7a11 | 26570 | -00102932 | -00160009 | -03075 | -0025 | 7 |
| 6734 | ENSMUSG( | Slk | 20874 | -00744247 | 00196229C | -0285 | -02425 | 7 |
| 6743 | ENSMUSG( | Smap2 | 69780 | 00863208C | -01101835 | -03075 | -0135 | 7 |
| 6745 | ENSMUSG( | Smarca2 | 67155 | 01779684S | 006420985 | -02875 | -02 | 7 |
| 6747 | ENSMUSG( | Smarca5 | 93762 | 016493746 | 008430057 | -002 | -03625 | 7 |
| 6752 | ENSMUSG( | Smarcd1 | 83797 | 001705604 | -00673422 | -0175 | -01925 | 7 |
| 6757 | ENSMUSG( | Smc3 | 13006 | 02006899C | 007358251 | -01775 | -01825 | 7 |
| 6760 | ENSMUSG( | Smcr8 | 237782 | 016428652 | 020672331 | -008 | -0305 | 7 |
| 6764 | ENSMUSG( | Smg6 | 103677 | -00063585 | -00497827 | -02675 | -00975 | 7 |
| 6776 | ENSMUSG( | Smndc1 | 76479 | 000023707 | -00588327 | -044 | -057 | 7 |
| 6786 | ENSMUSG( | Smug1 | 71726 | 00303326C | 01239305 | -0245 | -00925 | 7 |
| 6788 | ENSMUSG( | Smyd2 | 226830 | 006214875 | -00798194 | -0065 | -01575 | 7 |
| 6789 | ENSMUSG( | Smyd3 | 69726 | -00381315 | -01126133 | -02725 | -04875 | 7 |
| 6795 | ENSMUSG( | Snap91 | 20616 | 020058426 | 001145751 | -0075 | -015 | 7 |
| 6797 | ENSMUSG( | Snca | 20617 | 012448847 | -01179887 | -02025 | -00325 | 7 |
| 6813 | ENSMUSG( | Snrpb2 | 20639 | -01134800 | -01423116 | -04775 | -01725 | 7 |
| 6822 | ENSMUSG( | Sntb1 | 20649 | -01792518 | -00807342 | -02925 | -011 | 7 |
| 6826 | ENSMUSG( | Snw1 | 66354 | 020187827 | 007795371 | -036 | -0425 | 7 |
| 6830 | ENSMUSG( | Snx12 | 55988 | -00103955 | -03071631 | -008 | -0205 | 7 |
| 6834 | ENSMUSG( | Snx16 | 74718 | -00314176 | 007976458 | -01225 | -012 | 7 |
| 6846 | ENSMUSG( | Snx32 | 225861 | 012665234 | 00734529S | -03475 | -01275 | 7 |
| 6848 | ENSMUSG( | Snx4 | 69150 | 002715477 | -00864745 | -0075 | -01775 | 7 |
| 6849 | ENSMUSG( | Snx5 | 69178 | 005039675 | -01192947 | -008 | -024 | 7 |
| 6851 | ENSMUSG( | Snx7 | 76561 | 001158098 | -00936329 | -0065 | -032 | 7 |
| 6854 | ENSMUSG( | Sobp | 109205 | 01786264 | 023248372 | -025 | -01675 | 7 |
| 6869 | ENSMUSG( | Sos2 | 20663 | -00010834 | 00970932C | -031 | -00575 | 7 |
| 6880 | ENSMUSG( | Spag9 | 70834 | 001704573 | -00399729 | -016 | -019 | 7 |
| 6883 | ENSMUSG( | Spart | 229285 | 002790878 | -01188623 | -0205 | -0085 | 7 |
| 6916 | ENSMUSG( | Spred2 | 114716 | 011501343 | -00236935 | -003 | -01925 | 7 |
| 6920 | ENSMUSG( | Spryd3 | 223918 | 004380706 | -01433785 | -01425 | -03575 | 7 |

| | | | | | | | |
|---|---|---|---|---|---|---|---|
| 6927 | ENSMUSG( | Sptbn4 | 80297 | 015843177 | 012651527 | -01425 | -015 | 7 |
| 6956 | ENSMUSG( | Srrm2 | 75956 | 01113985C | 026696422 | -034 | -032 | 7 |
| 6962 | ENSMUSG( | Srsf12 | 272009 | -01159956 | -00073068 | -01275 | -0535 | 7 |
| 6964 | ENSMUSG( | Srsf3 | 20383 | -00071371 | 00693768€ | -01875 | -02225 | 7 |
| 6965 | ENSMUSG( | Srsf4 | 57317 | 005849751 | -00273581 | -0375 | -0395 | 7 |
| 6970 | ENSMUSG( | Srxn1 | 76650 | 00331671€ | -00725940 | -044 | -01775 | 7 |
| 6974 | ENSMUSG( | Ssbp1 | 381760 | -00502089 | 011604982 | -02525 | -0335 | 7 |
| 6998 | ENSMUSG( | Stag1 | 20842 | 015487084 | 005774047 | -009 | -02975 | 7 |
| 7004 | ENSMUSG( | Stard10 | 56018 | -00241923 | -01693605 | -01 | -02675 | 7 |
| 7010 | ENSMUSG( | Stat2 | 20847 | 021952193 | 00972484 | -03975 | -0345 | 7 |
| 7022 | ENSMUSG( | Stip1 | 20867 | 009957087 | -00763666 | -0255 | -02325 | 7 |
| 7030 | ENSMUSG( | Stk32c | 57740 | 007270364 | -00290068 | -00875 | -04575 | 7 |
| 7032 | ENSMUSG( | Stk38l | 232533 | 018653551 | 003826387 | -02075 | -0315 | 7 |
| 7041 | ENSMUSG( | Stoml2 | 66592 | 009980905 | -01057019 | -04025 | -00925 | 7 |
| 7042 | ENSMUSG( | Ston2 | 108800 | -01439283 | -03627956 | -01575 | -025 | 7 |
| 7047 | ENSMUSG( | Strbp | 20744 | 016752322 | -00071239 | -011 | -053 | 7 |
| 7056 | ENSMUSG( | Stum | 381310 | -00156157 | 015975603 | -0085 | -01725 | 7 |
| 7061 | ENSMUSG( | Stx1a | 20907 | 024863104 | -00192032 | -00725 | -051 | 7 |
| 7062 | ENSMUSG( | Stx1b | 56216 | 012006634 | -00587779 | -01925 | -032 | 7 |
| 7078 | ENSMUSG( | Suclg2 | 20917 | 010488311 | -01041332 | -0215 | -0285 | 7 |
| 7083 | ENSMUSG( | Sugp2 | 234373 | 020369977 | 018913004 | -019 | -04325 | 7 |
| 7096 | ENSMUSG( | Supv3l1 | 338359 | 010772753 | 00925392 | -01575 | -01475 | 7 |
| 7100 | ENSMUSG( | Surf6 | 20935 | 01360788C | -00007970 | -01025 | -0135 | 7 |
| 7108 | ENSMUSG( | Svip | 75744 | -00651658 | -01886271 | -04775 | -0445 | 7 |
| 7114 | ENSMUSG( | Syf2 | 68592 | 00781683C | -00854560 | -0205 | -02075 | 7 |
| 7117 | ENSMUSG( | Syn1 | 20964 | 01297265C | -01085745 | -0255 | -0315 | 7 |
| 7119 | ENSMUSG( | Syn3 | 27204 | -00317345 | 002158508 | -02525 | -0225 | 7 |
| 7121 | ENSMUSG( | Syndig1 | 433485 | -02610517 | -03139109 | -01475 | -01925 | 7 |
| 7123 | ENSMUSG( | Syne2 | 319565 | -03100771 | -00371727 | -0405 | -013 | 7 |
| 7128 | ENSMUSG( | Synj1 | 104015 | 009521367 | 005273553 | -015 | -022 | 7 |
| 7130 | ENSMUSG( | Synj2bp | 24071 | 010189897 | -00994562 | -031 | -0065 | 7 |
| 7132 | ENSMUSG( | Synpo | 104027 | 018640263 | 005011051 | -02475 | -01675 | 7 |
| 7185 | ENSMUSG( | Tamm41 | 68971 | -00853554 | -01154074 | -024 | -023 | 7 |
| 7198 | ENSMUSG( | Tardbp | 230908 | 004577644 | 00810822 | -0155 | -02475 | 7 |
| 7207 | ENSMUSG( | Tbc1d1 | 57915 | 024683493 | 025867017 | -0305 | -044 | 7 |
| 7214 | ENSMUSG( | Tbc1d17 | 233204 | 009849015 | 003322027 | -00725 | -034 | 7 |
| 7226 | ENSMUSG( | Tbc1d7 | 67046 | -01647590 | -02913458 | -008 | -0475 | 7 |
| 7237 | ENSMUSG( | Tbck | 271981 | 013326193 | 003626551 | -00875 | -02125 | 7 |
| 7254 | ENSMUSG( | Tcf20 | 21411 | 006360977 | 005817768 | -02725 | -019 | 7 |
| 7256 | ENSMUSG( | Tcf4 | 21413 | 015597801 | 002793385 | -02225 | -00175 | 7 |
| 7275 | ENSMUSG( | Tefm | 68550 | -00328658 | -03367501 | -006 | -0285 | 7 |
| 7290 | ENSMUSG( | Tex2 | 21763 | -00073321 | -00122458 | -01075 | -015 | 7 |
| 7306 | ENSMUSG( | Tgs1 | 116940 | 016367833 | 018462031 | -0225 | -05425 | 7 |
| 7308 | ENSMUSG( | Tha1 | 71776 | 012420208 | 015320357 | -026 | -01075 | 7 |
| 7315 | ENSMUSG( | Thg1l | 66628 | -00331411 | -02183219 | -03425 | -03475 | 7 |
| 7339 | ENSMUSG( | Timm10 | 30059 | 01211809 | -01860145 | -0155 | -01125 | 7 |
| 7343 | ENSMUSG( | Timm17b | 21855 | -00420199 | -02264387 | -033 | -0185 | 7 |
| 7344 | ENSMUSG( | Timm21 | 67105 | -00156734 | 014548767 | -02 | -017 | 7 |
| 7350 | ENSMUSG( | Timm8a1 | 30058 | 020194828 | 000429671 | -00825 | -03225 | 7 |
| 7351 | ENSMUSG( | Timm8b | 30057 | 000080961 | -02531492 | -03575 | -03975 | 7 |
| 7358 | ENSMUSG( | Tjp2 | 21873 | 001022283 | -02395929 | -04025 | -017 | 7 |
| 7370 | ENSMUSG( | Tln2 | 70549 | 00734705 | -00932234 | -015 | -0075 | 7 |
| 7412 | ENSMUSG( | Tmem132c | 243274 | -01599648 | 02275094 | -036 | -01475 | 7 |
| 7416 | ENSMUSG( | Tmem143 | 70209 | -01160712 | -02501753 | -02575 | -005 | 7 |
| 7418 | ENSMUSG( | Tmem145 | 330485 | 004053932 | 009322661 | -00675 | -01475 | 7 |
| 7437 | ENSMUSG( | Tmem186 | 66690 | -00074126 | 001194271 | -0035 | -03075 | 7 |
| 7479 | ENSMUSG( | Tmem65 | 74868 | 003217758 | -00188808 | -01225 | -01675 | 7 |
| 7492 | ENSMUSG( | Tmod1 | 21916 | -02649905 | -01715589 | -0205 | -01975 | 7 |
| 7494 | ENSMUSG( | Tmod3 | 50875 | 004830508 | 01436265C | -0175 | -01725 | 7 |
| 7514 | ENSMUSG( | Tnk2 | 51789 | 023091217 | 01651348€ | -02675 | -02175 | 7 |
| 7525 | ENSMUSG( | Tns2 | 209039 | 01792989 | 05494600 | -037 | -06325 | 7 |
| 7541 | ENSMUSG( | Top2b | 21974 | 010297841 | 003840754 | -00675 | -029 | 7 |
| 7550 | ENSMUSG( | Tox4 | 268741 | 011317842 | -00436814 | -011 | -0375 | 7 |
| 7555 | ENSMUSG( | Tpd52l1 | 21987 | -00633916 | -02155982 | -01525 | -01275 | 7 |
| 7559 | ENSMUSG( | Tph2 | 216343 | -02756672 | -02737374 | -0345 | -0155 | 7 |
| 7563 | ENSMUSG( | Tpm3 | 59069 | 004084717 | -01133245 | -02275 | -016 | 7 |
| 7570 | ENSMUSG( | Tpr | 108989 | 019883073 | 001851961 | -03675 | -03575 | 7 |
| 7577 | ENSMUSG( | Tra2b | 20462 | -00430144 | 004542848 | -0325 | -05375 | 7 |
| 7582 | ENSMUSG( | Traf7 | 224619 | 01276933 | 001814163 | -02375 | -004 | 7 |
| 7584 | ENSMUSG( | Trak1 | 67095 | 019274192 | 013704893 | -0375 | -029 | 7 |
| 7585 | ENSMUSG( | Trak2 | 70827 | 000762325 | -02921723 | -011 | -0125 | 7 |
| 7593 | ENSMUSG( | Trappc13 | 66975 | 007536125 | 006950413 | -02775 | -0085 | 7 |
| 7612 | ENSMUSG( | Trim26 | 22670 | -01278031 | 003016792 | -03675 | -023 | 7 |
| 7623 | ENSMUSG( | Trim56 | 384309 | 003532418 | -03592978 | -02025 | -0295 | 7 |
| 7631 | ENSMUSG( | Trip12 | 14897 | 01029368 | 002940654 | -026 | -02225 | 7 |
| 7660 | ENSMUSG( | Tsc22d1 | 21807 | 015664954 | 01381226 | -01175 | -02725 | 7 |
| 7661 | ENSMUSG( | Tsc22d2 | 72033 | 003891842 | 017219477 | -039 | -06725 | 7 |
| 7665 | ENSMUSG( | Tsfm | 66399 | 01699884€ | -00806841 | -025 | -00625 | 7 |
| 7669 | ENSMUSG( | Tsn | 22099 | -00550519 | -00392029 | -02425 | -02075 | 7 |
| 7705 | ENSMUSG( | Ttc5 | 219022 | 003491177 | -00282898 | -02225 | -01475 | 7 |
| 7727 | ENSMUSG( | Tuba8 | 53857 | 01122187€ | -03478593 | -03175 | -0535 | 7 |
| 7731 | ENSMUSG( | Tubb4a | 22153 | -01462215 | -02450197 | -0285 | -01675 | 7 |
| 7737 | ENSMUSG( | Tubgcp2 | 74237 | 005345668 | -00644253 | -01025 | -04025 | 7 |
| 7742 | ENSMUSG( | Tufm | 233870 | 006411762 | -02358018 | -018 | -01875 | 7 |
| 7754 | ENSMUSG( | Txn2 | 56551 | 00512418 | -02101035 | -029 | -03825 | 7 |
| 7764 | ENSMUSG( | Txnl4b | 234723 | 001024591 | -01338344 | -04575 | -03 | 7 |
| 7781 | ENSMUSG( | Uba3 | 22200 | 003841832 | -01376212 | -00725 | -021 | 7 |
| 7814 | ENSMUSG( | Ube2v2 | 70620 | -00485175 | -00531675 | -02075 | -0535 | 7 |
| 7815 | ENSMUSG( | Ube2w | 66799 | 005731071 | -00122314 | -02275 | -0105 | 7 |
| 7828 | ENSMUSG( | Ublcp1 | 79560 | 012015814 | 006143835 | -0395 | -03725 | 7 |
| 7838 | ENSMUSG( | Ubr5 | 70790 | 011992757 | 006033081 | -021 | -00825 | 7 |
| 7842 | ENSMUSG( | Ubxn1 | 225896 | 001485237 | -01499347 | -02175 | -00475 | 7 |
| 7843 | ENSMUSG( | Ubxn2a | 217379 | 009850123 | -00759289 | -01375 | -02125 | 7 |
| 7844 | ENSMUSG( | Ubxn2b | 68053 | 01515248C | -00365091 | -011 | -011 | 7 |
| 7863 | ENSMUSG( | Ugdh | 22235 | -00758301 | -00638730 | -04 | -01675 | 7 |
| 7875 | ENSMUSG( | Unc13a | 382018 | 009202343 | -00022202 | -00775 | -022 | 7 |
| 7876 | ENSMUSG( | Unc13b | 22249 | 014291012 | -00533883 | -0185 | -015 | 7 |
| 7877 | ENSMUSG( | Unc13c | 208898 | -02834930 | -04284454 | -0235 | -01675 | 7 |
| 7889 | ENSMUSG( | Upf2 | 326622 | 01571253 | 001955154 | -02475 | -0085 | 7 |
| 7899 | ENSMUSG( | Uqcrc1 | 22273 | 007119152 | -01774331 | -0035 | -02725 | 7 |
| 7900 | ENSMUSG( | Uqcrc2 | 67003 | 004511423 | -02087544 | -00425 | -02275 | 7 |
| 7906 | ENSMUSG( | Urgcp | 72046 | 009568908 | -00207071 | -01325 | -0075 | 7 |
| 7928 | ENSMUSG( | Usp31 | 76179 | 014746838 | 009884877 | -02225 | -007 | 7 |
| 7944 | ENSMUSG( | Usp6nl | 98910 | 010983177 | -00049106 | -0105 | -01075 | 7 |
| 7949 | ENSMUSG( | Utp11 | 67205 | 01563088 | 017405831 | -03275 | -061 | 7 |
| 7951 | ENSMUSG( | Utp15 | 105372 | -00242327 | 015704117 | -032 | -01875 | 7 |
| 7958 | ENSMUSG( | Uvrag | 78610 | 011795423 | 007318308 | -0155 | -03375 | 7 |
| 7960 | ENSMUSG( | Vac14 | 234729 | 00298756€ | 00090007€ | -009 | -02075 | 7 |
| 7970 | ENSMUSG( | Vars2 | 68915 | 013328814 | 009913848 | -0065 | -01275 | 7 |
| 7976 | ENSMUSG( | Vav1 | 22324 | -00382993 | 003462098 | -01175 | -00925 | 7 |
| 8004 | ENSMUSG( | Vps13a | 271564 | -01478606 | 029590752 | -036 | -01325 | 7 |
| 8020 | ENSMUSG( | Vps36 | 70160 | 017994152 | 007472148 | -00525 | -01625 | 7 |
| 8021 | ENSMUSG( | Vps37a | 52348 | 009835508 | 006031091 | -00675 | -01775 | 7 |
| 8033 | ENSMUSG( | Vps53 | 68299 | 013458564 | -00991898 | -01225 | -012 | 7 |
| 8035 | ENSMUSG( | Vps8 | 209018 | 017588257 | -00280114 | -012 | -0185 | 7 |
| 8049 | ENSMUSG( | Vwa5a | 67776 | 011976621 | -01162331 | -00475 | -03275 | 7 |
| 8051 | ENSMUSG( | Vwa8 | 219189 | 013160614 | -01170501 | -015 | -03075 | 7 |
| 8062 | ENSMUSG( | Washc1 | 68767 | 009648558 | -01212828 | -019 | -0065 | 7 |
| 8068 | ENSMUSG( | Wbp11 | 60321 | 00173488C | -00035799 | -021 | -02325 | 7 |
| 8086 | ENSMUSG( | Wdr3 | 269470 | 00508512€ | -00204366 | -0085 | -01075 | 7 |
| 8089 | ENSMUSG( | Wdr36 | 225348 | 009085657 | 009830823 | -0135 | -007 | 7 |
| 8090 | ENSMUSG( | Wdr37 | 207615 | 00589334C | -00149015 | -00975 | -04575 | 7 |
| 8092 | ENSMUSG( | Wdr43 | 72515 | 001253292 | -00376499 | -03325 | -03925 | 7 |
| 8121 | ENSMUSG( | Wnk1 | 232341 | -00217054 | -02339076 | -033 | -011 | 7 |
| 8123 | ENSMUSG( | Wnk3 | 279561 | -00455477 | -00880243 | -0245 | -026 | 7 |
| 8131 | ENSMUSG( | Wwp2 | 66894 | 00905802€ | -00554725 | -01125 | -04775 | 7 |

| | | | | | | | | |
|---|---|---|---|---|---|---|---|---|
| 8176 | ENSMUSG( | Ylpm1 | 56531 | 007743009 | 012176971 | -01975 | -0185 | 7 |
| 8180 | ENSMUSG( | Ypel5 | 383295 | -00323420 | -00902009 | -00625 | -01375 | 7 |
| 8182 | ENSMUSG( | Ythdc1 | 231386 | 000593264 | 00734366( | -0195 | -0505 | 7 |
| 8200 | ENSMUSG( | Zbtb45 | 232879 | 013152752 | 006821077 | -037 | -00975 | 7 |
| 8207 | ENSMUSG( | Zc3h14 | 75553 | 005123868 | -00573136 | -0225 | -046 | 7 |
| 8215 | ENSMUSG( | Zcchc10 | 67966 | 029743805 | 007049831 | -0095 | -02325 | 7 |
| 8244 | ENSMUSG( | Zfp326 | 54367 | 007101242 | 008128931 | -03625 | -05875 | 7 |
| 8250 | ENSMUSG( | Zfp385b | 241494 | -00042493 | 002177913 | -0355 | -03625 | 7 |
| 8251 | ENSMUSG( | Zfp428 | 232969 | -00174453 | -03201548 | -0165 | -01775 | 7 |
| 8262 | ENSMUSG( | Zfyve21 | 68520 | 007003497 | -00133460 | -00575 | -0265 | 7 |
| 8279 | ENSMUSG( | Zmynd11 | 66505 | 006314761 | -00175447 | -04925 | -03275 | 7 |
| 8284 | ENSMUSG( | Znrf2 | 387524 | 00805590( | 015093187 | -04775 | -027 | 7 |
| 189 | ENSMUSG( | Adgra2 | 78560 | 015480941 | 043665202 | -00075 | 2555 | 8 |
| 535 | ENSMUSG( | Arid4b | 94246 | -01608102 | 015291127 | -08575 | 21525 | 8 |
| 1072 | ENSMUSG( | Ccnt2 | 72949 | -00575481 | 016932008 | -04675 | 16975 | 8 |
| 1302 | ENSMUSG( | Cldn3 | 12739 | -06623746 | -05197838 | -13675 | 34725 | 8 |
| 1419 | ENSMUSG( | Col3a1 | 12825 | -02293307 | -01319363 | 01725 | 32075 | 8 |
| 1692 | ENSMUSG( | D030056L2 | 225995 | 011122989 | 001683747 | -05 | 1875 | 8 |
| 2255 | ENSMUSG( | Epc1 | 13831 | 006592521 | 00952400( | 02175 | 3355 | 8 |
| 2440 | ENSMUSG( | Fbn2 | 14119 | -00814054 | 034698929 | -09875 | 17475 | 8 |
| 2534 | ENSMUSG( | Flnc | 68794 | 035894274 | 046809757 | -03325 | 1895 | 8 |
| 2978 | ENSMUSG( | Gtf2h1 | 14884 | 005761305 | -00256192 | 025 | 188 | 8 |
| 3120 | ENSMUSG( | Hmgn2 | 15331 | -00979163 | -00810052 | -0345 | 23375 | 8 |
| 3519 | ENSMUSG( | Khdc4 | 74200 | 000382388 | 032706219 | -08425 | 144 | 8 |
| 3549 | ENSMUSG( | Kl | 16591 | 007962439 | 05385753( | -085 | 14925 | 8 |
| 3647 | ENSMUSG( | Leng1 | 69757 | 023696477 | -01886269 | -09 | 14925 | 8 |
| 3817 | ENSMUSG( | Lyrm2 | 108755 | -00056714 | -01242687 | -04175 | 159 | 8 |
| 3839 | ENSMUSG( | Madd | 228355 | 008489937 | 008680437 | 0405 | 1975 | 8 |
| 4018 | ENSMUSG( | Med8 | 80509 | -04805164 | -01114910 | -08725 | 2115 | 8 |
| 4021 | ENSMUSG( | Mef2d | 17261 | 01218241( | 006253611 | -053 | 1645 | 8 |
| 4419 | ENSMUSG( | Ncdn | 26562 | 006260025 | -00572614 | -07125 | 15675 | 8 |
| 4432 | ENSMUSG( | Ncoa2 | 17978 | 010030476 | 008205046 | -00025 | 17775 | 8 |
| 4472 | ENSMUSG( | Ndufaf8 | 208501 | -00307591 | -04460660 | -04725 | 21 | 8 |
| 4796 | ENSMUSG( | Olfml3 | 99543 | -02705840 | 00780291( | 08575 | 29725 | 8 |
| 4800 | ENSMUSG( | Omp | 18378 | -26222218 | 08197614( | -071 | 4205 | 8 |
| 4980 | ENSMUSG( | Pcolce | 18542 | -01586648 | 019560585 | -00325 | 17975 | 8 |
| 5060 | ENSMUSG( | Peg3 | 18616 | -01701166 | -00592083 | -06125 | 2105 | 8 |
| 5518 | ENSMUSG( | Prorsd1 | 67939 | 028013416 | 024671172 | -0335 | 22575 | 8 |
| 5619 | ENSMUSG( | Pter | 19212 | 018792899 | -01127359 | -0055 | 1625 | 8 |
| 5704 | ENSMUSG( | Pyroxd2 | 74580 | -00236252 | -00263271 | 046 | 1835 | 8 |
| 6015 | ENSMUSG( | Rnase4 | 58809 | -00768947 | 030032769 | 01475 | 2125 | 8 |
| 6396 | ENSMUSG( | Senp3 | 80886 | 012847642 | 006418072 | -01475 | 20425 | 8 |
| 6438 | ENSMUSG( | Setdb1 | 84505 | 006824234 | 011434055 | -0995 | 17775 | 8 |
| 6534 | ENSMUSG( | Sipa1 | 20469 | 017581384 | 03458634( | -026 | 20075 | 8 |
| 6666 | ENSMUSG( | Slc39a3 | 106947 | -01877911 | 009064694 | 00675 | 186 | 8 |
| 6692 | ENSMUSG( | Slc6a12 | 14411 | 045320894 | -01543258 | -03475 | 1755 | 8 |
| 6720 | ENSMUSG( | Slc9b2 | 97086 | -00658360 | -01867617 | -08075 | 263 | 8 |
| 6758 | ENSMUSG( | Smc5 | 226026 | 008053337 | 009252391 | -0445 | 16325 | 8 |
| 6874 | ENSMUSG( | Sox13 | 20668 | 003972458 | 035432193 | -16375 | 2295 | 8 |
| 6938 | ENSMUSG( | Srebf2 | 20788 | -01795469 | 000406948 | -0415 | 16325 | 8 |
| 6976 | ENSMUSG( | Ssbp3 | 72475 | 018534452 | 000438812 | -05925 | 24125 | 8 |
| 7274 | ENSMUSG( | Tef | 21685 | 010403006 | 031572845 | -036 | 17475 | 8 |
| 7334 | ENSMUSG( | Tia1 | 21841 | -01640795 | 031061085 | -0235 | 158 | 8 |
| 7499 | ENSMUSG( | Tmtc1 | 387314 | -03863925 | 004843232 | 0215 | 17675 | 8 |
| 7603 | ENSMUSG( | Trh | 22044 | -09937484 | -02212208 | -13525 | 18 | 8 |
| 7734 | ENSMUSG( | Tubb6 | 67951 | -02249175 | -02141132 | 0005 | 203 | 8 |
| 7853 | ENSMUSG( | Ucp2 | 22228 | 01113976( | 034392069 | 01875 | 15825 | 8 |
| 8193 | ENSMUSG( | Yy1 | 22632 | 006851713 | 0068083 | -16725 | 21175 | 8 |
| 3 | ENSMUSG( | 1110065P2 | 68920 | 009211681 | -02189088 | 00325 | 11925 | 9 |
| 9 | ENSMUSG( | 2410004B1 | 66421 | 014534336 | -01254645 | -07975 | 0785 | 9 |
| 63 | ENSMUSG( | Abcg4 | 192663 | -01607550 | 010567638 | -00675 | 103 | 9 |
| 84 | ENSMUSG( | Abr | 109934 | 016420607 | -00528281 | -0465 | 10425 | 9 |
| 134 | ENSMUSG( | Acsl6 | 216739 | -00120019 | -00030333 | -02325 | 07425 | 9 |
| 214 | ENSMUSG( | Adra2a | 11551 | -01243298 | 032413346 | -03825 | 061 | 9 |
| 220 | ENSMUSG( | Afap1 | 70292 | -00172226 | -01284411 | -031 | 06475 | 9 |
| 226 | ENSMUSG( | Afg3l1 | 114896 | 013692228 | -00761721 | 0175 | 094 | 9 |
| 300 | ENSMUSG( | Aldh1a7 | 26358 | -00878798 | -03676411 | -00975 | 0875 | 9 |
| 312 | ENSMUSG( | Aldoa | 11674 | -00100303 | -02325454 | -0005 | 08775 | 9 |
| 328 | ENSMUSG( | Alox5ap | 11690 | 005038374 | 00241564( | -03125 | 0655 | 9 |
| 336 | ENSMUSG( | Amer3 | 211383 | 007423842 | -00090220 | -0595 | 09125 | 9 |
| 364 | ENSMUSG( | Ankmy1 | 241158 | -00628234 | -00210481 | 0085 | 12625 | 9 |
| 395 | ENSMUSG( | Anp32b | 67628 | 004554494 | 00251164( | -03475 | 09625 | 9 |
| 447 | ENSMUSG( | Aplf | 72103 | 009740398 | 021010216 | -0965 | 1165 | 9 |
| 451 | ENSMUSG( | Apoa2 | 11807 | -00103540 | 002286171 | -093 | 07975 | 9 |
| 531 | ENSMUSG( | Arid1a | 93760 | 007730049 | 008650074 | -02625 | 123 | 9 |
| 545 | ENSMUSG( | Arl4c | 320982 | -00269809 | 010679413 | -048 | 086 | 9 |
| 559 | ENSMUSG( | Armcx2 | 67416 | 011080665 | 005283832 | -019 | 06675 | 9 |
| 581 | ENSMUSG( | Arsj | 271970 | 02017418( | 049781326 | -00375 | 132 | 9 |
| 591 | ENSMUSG( | Asb6 | 72323 | 005213899 | -00264423 | -02075 | 0955 | 9 |
| 617 | ENSMUSG( | Atf2 | 11909 | 005742445 | -00389029 | 01875 | 0945 | 9 |
| 698 | ENSMUSG( | Atxn1 | 20238 | 006408939 | 009210811 | -04625 | 0605 | 9 |
| 701 | ENSMUSG( | Atxn2 | 20239 | 003613375 | 006138874 | -02175 | 061 | 9 |
| 713 | ENSMUSG( | Azi2 | 27215 | 003311505 | 001482867 | -06325 | 074 | 9 |
| 716 | ENSMUSG( | B3galt5 | 93961 | -03111569 | 015937721 | -02075 | 06975 | 9 |
| 723 | ENSMUSG( | B4galt7 | 218271 | -02563059 | 011737328 | -032 | 0965 | 9 |
| 765 | ENSMUSG( | Bccip | 66165 | 000016979 | -00816553 | -03025 | 07875 | 9 |
| 816 | ENSMUSG( | Boc | 117606 | -03035207 | 016318399 | -02925 | 0925 | 9 |
| 818 | ENSMUSG( | Bod1l | 665775 | 006470123 | 005198381 | -02 | 089 | 9 |
| 842 | ENSMUSG( | Brd8 | 78656 | -00442346 | -00365976 | -0335 | 05925 | 9 |
| 865 | ENSMUSG( | Btbd9 | 224671 | 016052201 | 010253504 | -01725 | 0785 | 9 |
| 866 | ENSMUSG( | Btd | 26363 | -04723457 | -00020201 | -02325 | 0765 | 9 |
| 869 | ENSMUSG( | Bub3 | 12237 | -00104962 | -01344402 | -0255 | 074 | 9 |
| 885 | ENSMUSG( | C2cd5 | 74741 | 021869788 | -00683935 | -003 | 1285 | 9 |
| 909 | ENSMUSG( | Cacnb1 | 12295 | 010881757 | -01166368 | 012 | 097 | 9 |
| 919 | ENSMUSG( | Cacul1 | 78832 | 008281375 | 001420729 | -03975 | 08725 | 9 |
| 964 | ENSMUSG( | Capn10 | 23830 | 008705559 | 01031043( | -03575 | 0705 | 9 |
| 979 | ENSMUSG( | Carnmt1 | 67383 | 012262383 | 01149099( | -041 | 082 | 9 |
| 1025 | ENSMUSG( | Ccdc107 | 622404 | -03184415 | -00378017 | -05125 | 1185 | 9 |
| 1046 | ENSMUSG( | Ccdc43 | 52715 | 013144074 | 002256097 | -0405 | 063 | 9 |
| 1107 | ENSMUSG( | Cd99l2 | 171486 | -02432664 | 002357447 | -02725 | 1015 | 9 |
| 1147 | ENSMUSG( | Cdk16 | 18555 | 016203404 | -00119721 | -015 | 08175 | 9 |
| 1175 | ENSMUSG( | Celf5 | 319586 | 025663499 | 004330845 | -0105 | 1135 | 9 |
| 1179 | ENSMUSG( | Cend1 | 57754 | 004182606 | -02194319 | -0835 | 0685 | 9 |
| 1184 | ENSMUSG( | Cep131 | 12009 | 01803150( | -00669053 | -03225 | 06675 | 9 |
| 1206 | ENSMUSG( | Cfap74 | 544678 | 015064165 | 013637525 | -00125 | 07925 | 9 |
| 1235 | ENSMUSG( | Chga | 12652 | -04714968 | -00003817 | 00375 | 0865 | 9 |
| 1258 | ENSMUSG( | Chpf2 | 100910 | -00688735 | 020581619 | -01025 | 095 | 9 |
| 1261 | ENSMUSG( | Chrm2 | 243764 | -05412738 | 041211707 | -04375 | 064 | 9 |
| 1271 | ENSMUSG( | Ciao2a | 68250 | -00017667 | -01760840 | -0045 | 09225 | 9 |
| 1277 | ENSMUSG( | Cic | 71722 | 017103243 | 013162447 | -06075 | 06775 | 9 |
| 1307 | ENSMUSG( | Clgn | 12745 | -03728435 | -00208033 | -03275 | 0605 | 9 |
| 1313 | ENSMUSG( | Clip1 | 56430 | 006698981 | 022782042 | -05425 | 10825 | 9 |
| 1322 | ENSMUSG( | Clock | 12753 | 007875976 | -00443974 | -02075 | 13 | 9 |
| 1332 | ENSMUSG( | Cltb | 74325 | -00756838 | -02840018 | -0615 | 087 | 9 |
| 1339 | ENSMUSG( | Cmas | 12764 | -01015392 | -00839810 | -04125 | 10925 | 9 |
| 1355 | ENSMUSG( | Cnih2 | 12794 | 008625599 | 001462459 | -01625 | 0755 | 9 |
| 1356 | ENSMUSG( | Cnih3 | 72978 | -01387364 | 009607064 | 0165 | 13275 | 9 |
| 1402 | ENSMUSG( | Cobll1 | 319876 | -03386460 | 050440619 | -03075 | 09025 | 9 |
| 1418 | ENSMUSG( | Col25a1 | 77018 | -04710579 | 01141101( | -00825 | 12925 | 9 |
| 1520 | ENSMUSG( | Cpsf6 | 432508 | -00265112 | 037072353 | -013 | 1055 | 9 |
| 1535 | ENSMUSG( | Creg1 | 433375 | -03369178 | 01546665( | -076 | 06425 | 9 |
| 1536 | ENSMUSG( | Creg2 | 263764 | -02902600 | 025807598 | -0675 | 1275 | 9 |
| 1548 | ENSMUSG( | Crls1 | 66586 | 004643629 | 00660957( | -0955 | 129 | 9 |
| 1549 | ENSMUSG( | Crmp1 | 12933 | 008654887 | -01456729 | -02475 | 07125 | 9 |
| 1562 | ENSMUSG( | Crybg3 | 224273 | -00829232 | 012764861 | -00425 | 07275 | 9 |
| 1579 | ENSMUSG( | Csnk1d | 104318 | 017680208 | -00156584 | -03125 | 0665 | 9 |

| | | | | | | | | |
|---|---|---|---|---|---|---|---|---|
| 1644 | ENSMUSG( | Cul7 | 66515 | 008953599 | 004105944 | -046 | 1095 | 9 |
| 1657 | ENSMUSG( | Cxcl14 | 57266 | -01297480 | -00677275 | -033 | 06175 | 9 |
| 1660 | ENSMUSG( | Cyb561d2 | 56368 | -03756388 | -00384775 | -011 | 071 | 9 |
| 1689 | ENSMUSG( | Cyth2 | 19158 | 002306733 | -00454234 | -014 | 0695 | 9 |
| 1705 | ENSMUSG( | Dact2 | 240025 | 007635210 | 000680894 | 01825 | 092 | 9 |
| 1706 | ENSMUSG( | Dact3 | 629378 | 023143904 | 010145603 | -00675 | 08075 | 9 |
| 1729 | ENSMUSG( | Dcaf10 | 242418 | -01212454 | 005888376 | -02325 | 07575 | 9 |
| 1732 | ENSMUSG( | Dcaf5 | 320808 | 009847048 | 000877205 | -055 | 1345 | 9 |
| 1745 | ENSMUSG( | Dcps | 69305 | 020277485 | -00672526 | 0025 | 135 | 9 |
| 1760 | ENSMUSG( | Dda1 | 66498 | 010118803 | 003363730 | 00075 | 0965 | 9 |
| 1843 | ENSMUSG( | Dhdds | 67422 | 010451477 | 003298510 | -066 | 0895 | 9 |
| 1910 | ENSMUSG( | Dnah2 | 327954 | 011119795 | 016905251 | -0815 | 09675 | 9 |
| 2002 | ENSMUSG( | Dpyd | 99586 | -01339212 | -00278188 | 008 | 09725 | 9 |
| 2018 | ENSMUSG( | Dtd1 | 66044 | -01034703 | -01989262 | -034 | 0665 | 9 |
| 2022 | ENSMUSG( | Dtnbp1 | 94245 | 004466431 | -02021864 | -02025 | 0775 | 9 |
| 2066 | ENSMUSG( | Ecd | 70601 | 015674758 | -00413304 | -04425 | 0915 | 9 |
| 2078 | ENSMUSG( | Edc3 | 353190 | 009946378 | 003292877 | 01475 | 0945 | 9 |
| 2091 | ENSMUSG( | Eef1d | 66656 | 014981601 | -00552874 | -00575 | 117 | 9 |
| 2098 | ENSMUSG( | Efemp1 | 216616 | -02824476 | 007617085 | -0255 | 10525 | 9 |
| 2126 | ENSMUSG( | Eif1ad | 69860 | 004646374 | 003874791 | 0025 | 07775 | 9 |
| 2170 | ENSMUSG( | Eif5a2 | 208691 | 004004736 | -00521518 | -01675 | 06875 | 9 |
| 2212 | ENSMUSG( | Emilin1 | 100952 | -04792827 | -00993411 | 01225 | 08025 | 9 |
| 2309 | ENSMUSG( | Ess2 | 27886 | -00075810 | 010378060 | -035 | 072 | 9 |
| 2372 | ENSMUSG( | Faim2 | 72393 | -02396879 | 007059633 | 00475 | 09875 | 9 |
| 2393 | ENSMUSG( | Fam169a | 320557 | -01893593 | 003999164 | -015 | 06725 | 9 |
| 2412 | ENSMUSG( | Fam50a | 108160 | 018103091 | 008727053 | -0375 | 089 | 9 |
| 2416 | ENSMUSG( | Fam8a1 | 97863 | -03280100 | 008580468 | 003 | 0985 | 9 |
| 2456 | ENSMUSG( | Fbxo38 | 107035 | 016280055 | 012982647 | -0185 | 07125 | 9 |
| 2503 | ENSMUSG( | Fhdc1 | 229474 | 002378612 | -01238300 | -0225 | 079 | 9 |
| 2510 | ENSMUSG( | Fhod1 | 234686 | 004094666 | 022235453 | -00525 | 07175 | 9 |
| 2530 | ENSMUSG( | Flcn | 216805 | 001729142 | 017134719 | -018 | 073 | 9 |
| 2566 | ENSMUSG( | Foxk1 | 17425 | 009887434 | 007529993 | -00875 | 089 | 9 |
| 2663 | ENSMUSG( | Ganab | 14376 | -02095994 | 011249335 | -034 | 06375 | 9 |
| 2680 | ENSMUSG( | Gatad2b | 229542 | 001800455 | 008826364 | -01125 | 085 | 9 |
| 2756 | ENSMUSG( | Gkap1 | 56278 | 007399214 | 009406523 | -0275 | 06975 | 9 |
| 2769 | ENSMUSG( | Glra3 | 110304 | -06690234 | 022459971 | -06575 | 122 | 9 |
| 2860 | ENSMUSG( | Gpbp1 | 73274 | 010318823 | 011644196 | -022 | 06625 | 9 |
| 2908 | ENSMUSG( | Grb14 | 50915 | 007740713 | 004091872 | -00925 | 104 | 9 |
| 2910 | ENSMUSG( | Grcc10 | 14790 | -00702326 | -01367030 | 00125 | 09325 | 9 |
| 2940 | ENSMUSG( | Grn | 14824 | -02163312 | 017625278 | 0065 | 15 | 9 |
| 2972 | ENSMUSG( | Gtf2a2 | 235459 | -00413671 | 006396912 | -054 | 07925 | 9 |
| 2989 | ENSMUSG( | Gtpbp2 | 56055 | 024192460 | 007977729 | -00275 | 07825 | 9 |
| 3010 | ENSMUSG( | H2-K1 | 14972 | -03067990 | 016387497 | -03025 | 06425 | 9 |
| 3100 | ENSMUSG( | Hirip3 | 233876 | -00108758 | 016111479 | -045 | 09725 | 9 |
| 3108 | ENSMUSG( | Hmcn1 | 545370 | -00247711 | 022309970 | -03575 | 06275 | 9 |
| 3119 | ENSMUSG( | Hmgn1 | 15312 | 005088943 | -01459625 | -0375 | 0745 | 9 |
| 3122 | ENSMUSG( | Hmgn5 | 50887 | 024054265 | 013593341 | -01725 | 10825 | 9 |
| 3150 | ENSMUSG( | Hook2 | 170833 | 009265860 | 001826542 | -01525 | 09425 | 9 |
| 3159 | ENSMUSG( | Hpx | 15458 | 000891797 | -10354470 | 01875 | 1075 | 9 |
| 3166 | ENSMUSG( | Hs6st1 | 50785 | -01314915 | 026314345 | -018 | 0645 | 9 |
| 3193 | ENSMUSG( | Hspb1 | 15507 | 044595997 | 093851065 | -05425 | 0745 | 9 |
| 3259 | ENSMUSG( | Igkc | 16071 | -00364082 | -04745085 | 00625 | 13325 | 9 |
| 3282 | ENSMUSG( | Ilkap | 67444 | 004436820 | -00661863 | -03275 | 0655 | 9 |
| 3348 | ENSMUSG( | Irf2bpl | 238330 | -01310933 | 000198244 | -02875 | 065 | 9 |
| 3423 | ENSMUSG( | Junb | 16477 | -10254128 | -08074920 | 00925 | 1485 | 9 |
| 3427 | ENSMUSG( | Kank1 | 107351 | 015771184 | -00625510 | -05625 | 112 | 9 |
| 3441 | ENSMUSG( | Kbtbd4 | 67136 | -00181873 | -00387102 | -0095 | 0705 | 9 |
| 3516 | ENSMUSG( | Kdm5b | 75605 | 022307382 | 008819832 | -048 | 12375 | 9 |
| 3535 | ENSMUSG( | Kif2a | 16563 | -00404448 | 006940223 | -0335 | 08375 | 9 |
| 3568 | ENSMUSG( | Klhl4 | 237010 | -00290704 | -00762209 | 01475 | 1135 | 9 |
| 3634 | ENSMUSG( | Lcat | 16816 | -01572168 | 015252307 | -01975 | 07425 | 9 |
| 3639 | ENSMUSG( | Ldb1 | 16825 | -00351392 | 015518718 | -0405 | 07675 | 9 |
| 3648 | ENSMUSG( | Leo1 | 235497 | 015918719 | -03203310 | -002 | 07975 | 9 |
| 3692 | ENSMUSG( | Lmbr1 | 56873 | -02612743 | 020363614 | 0045 | 1335 | 9 |
| 3710 | ENSMUSG( | Lonrf2 | 381338 | 021941137 | -00988935 | -04925 | 13275 | 9 |
| 3721 | ENSMUSG( | Lratd2 | 399603 | -02332026 | 014385410 | 002 | 084 | 9 |
| 3755 | ENSMUSG( | Lrrc7 | 242274 | 009288770 | 016801611 | -08925 | 09775 | 9 |
| 3802 | ENSMUSG( | Ly6a | 110454 | 007268674 | 065108103 | -059 | 081 | 9 |
| 3812 | ENSMUSG( | Lypd6 | 320343 | -01467491 | -02462360 | 0035 | 12825 | 9 |
| 3846 | ENSMUSG( | Magee2 | 272790 | -03212954 | -02913259 | -02825 | 0665 | 9 |
| 3857 | ENSMUSG( | Man2a1 | 17158 | -01573221 | 043491115 | -09675 | 08825 | 9 |
| 3886 | ENSMUSG( | Map3k2 | 26405 | -00101423 | 016784871 | -0585 | 09175 | 9 |
| 3926 | ENSMUSG( | Marchf7 | 57438 | -02304048 | 025676986 | -06625 | 08025 | 9 |
| 3936 | ENSMUSG( | Masp1 | 17174 | -02530833 | 026947874 | -1065 | 1265 | 9 |
| 3959 | ENSMUSG( | Mboat7 | 77582 | -03216301 | 015348748 | 0045 | 0745 | 9 |
| 3991 | ENSMUSG( | Mdm4 | 17248 | -01673382 | 017095653 | -0515 | 0835 | 9 |
| 3992 | ENSMUSG( | Mdn1 | 100019 | -04572215 | 016773874 | -02375 | 0855 | 9 |
| 4006 | ENSMUSG( | Med15 | 94112 | 014922056 | -00864240 | -021 | 0975 | 9 |
| 4013 | ENSMUSG( | Med28 | 66999 | 005346105 | -02154701 | 013 | 093 | 9 |
| 4015 | ENSMUSG( | Med30 | 69790 | -00603020 | -01168991 | 013 | 13325 | 9 |
| 4020 | ENSMUSG( | Mef2c | 17260 | 010498599 | -00328948 | -0215 | 13125 | 9 |
| 4022 | ENSMUSG( | Megf10 | 70417 | -02605030 | 002166078 | -01475 | 11225 | 9 |
| 4023 | ENSMUSG( | Megf6 | 230971 | 052831682 | 083454250 | -01325 | 091 | 9 |
| 4056 | ENSMUSG( | Mfsd1 | 66868 | -00470909 | -00133375 | 01425 | 14975 | 9 |
| 4078 | ENSMUSG( | Micall1 | 27008 | 004423297 | -01094248 | -00925 | 09725 | 9 |
| 4101 | ENSMUSG( | Mipol1 | 73490 | 012806970 | -00442004 | -0505 | 104 | 9 |
| 4109 | ENSMUSG( | Mllt11 | 56772 | -00559710 | -00939105 | -0255 | 10675 | 9 |
| 4133 | ENSMUSG( | Morc2a | 74522 | 009664104 | 002150063 | -01625 | 07725 | 9 |
| 4134 | ENSMUSG( | Morf4l1 | 21761 | 006205293 | -01500824 | -04925 | 084 | 9 |
| 4159 | ENSMUSG( | Mpzl1 | 68481 | -03458646 | 008081211 | -051 | 0815 | 9 |
| 4212 | ENSMUSG( | Mrpl58 | 68572 | -00081698 | -03027776 | -02825 | 0885 | 9 |
| 4247 | ENSMUSG( | Msh2 | 17685 | 005450931 | -00623581 | -0025 | 0925 | 9 |
| 4259 | ENSMUSG( | Mt3 | 17751 | 009119466 | -03320138 | -07425 | 08975 | 9 |
| 4301 | ENSMUSG( | Mtss2 | 244654 | -00413355 | -02948630 | -0035 | 08875 | 9 |
| 4373 | ENSMUSG( | Nacc2 | 67991 | 004705904 | -02686108 | -0155 | 12925 | 9 |
| 4380 | ENSMUSG( | Naglu | 27419 | -04297593 | 014877654 | -02025 | 0905 | 9 |
| 4433 | ENSMUSG( | Ncoa5 | 228869 | 016735863 | 005870431 | 00025 | 0905 | 9 |
| 4603 | ENSMUSG( | Nol11 | 68979 | 000120891 | 010668985 | -0125 | 0975 | 9 |
| 4625 | ENSMUSG( | Nosip | 66394 | -00188499 | -00599538 | -0115 | 08025 | 9 |
| 4627 | ENSMUSG( | Notch1 | 18128 | -01283098 | 059604371 | -04225 | 0585 | 9 |
| 4808 | ENSMUSG( | Oprl1 | 18389 | -00637361 | 027867978 | -079 | 07325 | 9 |
| 4832 | ENSMUSG( | Otof | 83762 | -00959551 | -01462053 | -03075 | 14725 | 9 |
| 4909 | ENSMUSG( | Parm1 | 231440 | -02908763 | 018046478 | -07625 | 10225 | 9 |
| 4916 | ENSMUSG( | Patl1 | 225929 | -00770648 | -00320408 | -0165 | 076 | 9 |
| 4929 | ENSMUSG( | Pcbp3 | 59093 | 005531461 | 009752004 | -04175 | 134 | 9 |
| 4968 | ENSMUSG( | Pcif1 | 228866 | 010814617 | 006223322 | -02575 | 091 | 9 |
| 4988 | ENSMUSG( | Pcyt1a | 13026 | 008109548 | -01109616 | -0065 | 07275 | 9 |
| 5008 | ENSMUSG( | Pde3b | 18576 | -00075787 | 016880652 | -017 | 09 | 9 |
| 5018 | ENSMUSG( | Pde8b | 218461 | 005043895 | -00964190 | -036 | 06 | 9 |
| 5178 | ENSMUSG( | Pik3cd | 18707 | 017222356 | 019453836 | -00175 | 07975 | 9 |
| 5203 | ENSMUSG( | Pja1 | 18744 | -01236194 | -00288268 | -03275 | 05675 | 9 |
| 5204 | ENSMUSG( | Pja2 | 224938 | -00555512 | 000278730 | -05375 | 09825 | 9 |
| 5359 | ENSMUSG( | Pou3f3 | 18993 | 004502627 | 013377996 | -02425 | 08725 | 9 |
| 5386 | ENSMUSG( | Ppil4 | 67418 | 007268946 | 004811824 | -0145 | 077 | 9 |
| 5393 | ENSMUSG( | Ppm1g | 14208 | -01003394 | -00019574 | -0195 | 0955 | 9 |
| 5461 | ENSMUSG( | Prcc | 94315 | 015684372 | 002473224 | 00625 | 101 | 9 |
| 5537 | ENSMUSG( | Prr12 | 233210 | 005856745 | 015780480 | -03825 | 114 | 9 |
| 5560 | ENSMUSG( | Psip1 | 101739 | 002516453 | 001403570 | -07275 | 088 | 9 |
| 5633 | ENSMUSG( | Ptma | 19231 | 004233022 | 010554355 | -0665 | 08225 | 9 |
| 5716 | ENSMUSG( | Qtrt2 | 106248 | 016969242 | 032670207 | -05575 | 0675 | 9 |
| 5789 | ENSMUSG( | Rac3 | 170758 | 010888065 | -00925814 | 02125 | 11325 | 9 |
| 5865 | ENSMUSG( | Rbks | 71336 | -02691042 | -03888514 | -0355 | 067 | 9 |
| 5894 | ENSMUSG( | Rbx1 | 56438 | 005054482 | -00788522 | -0275 | 0865 | 9 |
| 5896 | ENSMUSG( | Rc3h2 | 319817 | 000241137 | 010709075 | 00675 | 09975 | 9 |
| 5904 | ENSMUSG( | Rchy1 | 68098 | 005565986 | -01032453 | 01025 | 08725 | 9 |
| 5939 | ENSMUSG( | Rfc4 | 106344 | 024495198 | -00090773 | -06075 | 085 | 9 |

| | | | | | | | | | |
|---|---|---|---|---|---|---|---|---|---|
| 6026 | ENSMUSG( | Rnf113a2 | 66381 | -00099741 | -00619425 | -029 | 0595 | | 9 |
| 6049 | ENSMUSG( | Rnf220 | 66743 | 010771532 | -00388563 | -02 | 108 | | 9 |
| 6054 | ENSMUSG( | Rnf40 | 233900 | 004675264 | -00158153 | -048 | 0675 | | 9 |
| 6137 | ENSMUSG( | Rps10 | 67097 | -03121807 | 029203363 | 008 | 09325 | | 9 |
| 6177 | ENSMUSG( | Rraga | 68441 | -01035464 | -02414197 | -00825 | 113 | | 9 |
| 6249 | ENSMUSG( | Samd14 | 217125 | 020264816 | 011762163 | -008 | 08875 | | 9 |
| 6251 | ENSMUSG( | Samd5 | 320825 | 009691394 | -02096924 | -06425 | 09275 | | 9 |
| 6257 | ENSMUSG( | Sap130 | 269003 | 009535237 | 006974251 | -05825 | 06625 | | 9 |
| 6306 | ENSMUSG( | Scn3a | 20269 | -04239829 | 036455625 | 0025 | 07375 | | 9 |
| 6343 | ENSMUSG( | Sdk1 | 330222 | -01885615 | 02508540C | -00575 | 076 | | 9 |
| 6373 | ENSMUSG( | Selenoh | 72657 | 012908974 | 004859543 | -054 | 07525 | | 9 |
| 6381 | ENSMUSG( | Selenow | 20364 | 005354527 | -01908869 | -067 | 08125 | | 9 |
| 6414 | ENSMUSG( | Serf2 | 378702 | 00482069C | -01909670 | -0235 | 07 | | 9 |
| 6454 | ENSMUSG( | Sfr1 | 67788 | 00335871C | -00990698 | -02325 | 11325 | | 9 |
| 6463 | ENSMUSG( | Sgcd | 24052 | -01714815 | 022055173 | -031 | 061 | | 9 |
| 6486 | ENSMUSG( | Sh3bp5l | 79566 | 029094746 | 005423368 | -0615 | | 1 9 | |
| 6487 | ENSMUSG( | Sh3d19 | 27059 | -00046764 | 008430454 | -0065 | 08325 | | 9 |
| 6554 | ENSMUSG( | Slc13a3 | 114644 | -02846545 | -00956826 | -0155 | 06325 | | 9 |
| 6648 | ENSMUSG( | Slc35b2 | 73836 | -03963443 | 006279289 | -0095 | 073 | | 9 |
| 6651 | ENSMUSG( | Slc35f3 | 210027 | 000712134 | 029608447 | -0105 | 09975 | | 9 |
| 6652 | ENSMUSG( | Slc35f6 | 74919 | -04069110 | 023364985 | 00775 | 0805 | | 9 |
| 6670 | ENSMUSG( | Slc43a2 | 215113 | -03378868 | 000380882 | -0535 | 06375 | | 9 |
| 6675 | ENSMUSG( | Slc47a1 | 67473 | -03299046 | -01475724 | -024 | 0785 | | 9 |
| 6723 | ENSMUSG( | Slco2b1 | 101488 | 001318148 | 041250071 | -041 | 126 | | 9 |
| 6738 | ENSMUSG( | Smad2 | 17126 | 003896952 | 002285592 | -04625 | 0735 | | 9 |
| 6828 | ENSMUSG( | Snx10 | 71982 | -00337824 | 005791759 | 01375 | 109 | | 9 |
| 6873 | ENSMUSG( | Sox1 | 20664 | -00006117 | 029921896 | 00575 | 11375 | | 9 |
| 6875 | ENSMUSG( | Sox5 | 20678 | -00279917 | -01160968 | -0155 | 09075 | | 9 |
| 6876 | ENSMUSG( | Sox9 | 20682 | -00110024 | 023093881 | -04425 | 09375 | | 9 |
| 6877 | ENSMUSG( | Sp3 | 20687 | -00139322 | 009599805 | -02175 | 07 | | 9 |
| 6897 | ENSMUSG( | Spg11 | 214585 | 010830487 | 001316245 | 00025 | 102 | | 9 |
| 6917 | ENSMUSG( | Spred3 | 101809 | 019594041 | 009613277 | -0425 | 065 | | 9 |
| 6932 | ENSMUSG( | Sqstm1 | 18412 | -00441035 | -00095027 | 02225 | 14025 | | 9 |
| 6981 | ENSMUSG( | Sspn | 16651 | -01017648 | 031749941 | -035 | 05875 | | 9 |
| 6989 | ENSMUSG( | Ssu72 | 68991 | -00708328 | -01713643 | -025 | 07225 | | 9 |
| 6996 | ENSMUSG( | Stab1 | 192187 | 00741610C | 035699016 | -02575 | 06425 | | 9 |
| 7005 | ENSMUSG( | Stard3 | 59045 | -02616001 | 015852487 | -0155 | 083 | | 9 |
| 7043 | ENSMUSG( | Stox2 | 71069 | 022321989 | 026374064 | -00675 | 07225 | | 9 |
| 7082 | ENSMUSG( | Sugp1 | 70616 | 000511665 | -00527156 | -02925 | 08425 | | 9 |
| 7102 | ENSMUSG( | Susd5 | 382111 | -02967290 | 041517862 | 01675 | 0905 | | 9 |
| 7115 | ENSMUSG( | Syk | 20963 | 001544209 | 013713136 | -00275 | 0875 | | 9 |
| 7133 | ENSMUSG( | Synpo2 | 118449 | -00444459 | 028021491 | -03775 | 05775 | | 9 |
| 7245 | ENSMUSG( | Tbr1 | 21375 | 007988815 | 008705003 | -03725 | 06075 | | 9 |
| 7252 | ENSMUSG( | Tceal5 | 331532 | -00396442 | -02466859 | 00975 | 147 | | 9 |
| 7263 | ENSMUSG( | Tdg | 21665 | 006210372 | -16233335 | 005 | 11925 | | 9 |
| 7333 | ENSMUSG( | Thyn1 | 77862 | 009133674 | -00676012 | -0115 | 083 | | 9 |
| 7337 | ENSMUSG( | Tiam2 | 24001 | -01467253 | -00679715 | 01325 | 1435 | | 9 |
| 7366 | ENSMUSG( | Tle3 | 21887 | 016433314 | 014185225 | -026 | 10625 | | 9 |
| 7378 | ENSMUSG( | Tma16 | 66282 | -02750958 | -01632418 | 00175 | 08925 | | 9 |
| 7379 | ENSMUSG( | Tma7 | 66167 | 00193337C | -01292348 | -06075 | 0725 | | 9 |
| 7396 | ENSMUSG( | Tmem106b | 71900 | -02430265 | 016902769 | -00025 | 075 | | 9 |
| 7399 | ENSMUSG( | Tmem114 | 74720 | -03623854 | -01168084 | -03325 | 10025 | | 9 |
| 7406 | ENSMUSG( | Tmem127 | 69470 | -01495353 | 010473778 | 01725 | 0865 | | 9 |
| 7429 | ENSMUSG( | Tmem168 | 101118 | -03468961 | 027110365 | -0115 | 0685 | | 9 |
| 7453 | ENSMUSG( | Tmem237 | 381259 | -02226190 | 012857002 | -02125 | 084 | | 9 |
| 7508 | ENSMUSG( | Tnfaip3 | 21929 | 000383174 | 058747178 | -05775 | 119 | | 9 |
| 7629 | ENSMUSG( | Triobp | 110253 | -00011149 | 000383102 | -00525 | 07975 | | 9 |
| 7633 | ENSMUSG( | Trir | 68544 | 001260957 | -00520325 | -02675 | 129 | | 9 |
| 7641 | ENSMUSG( | Trmt6 | 66926 | 003419469 | -00692832 | -0155 | 106 | | 9 |
| 7657 | ENSMUSG( | Trub1 | 72133 | -00107737 | 008355771 | -0135 | 069 | | 9 |
| 7747 | ENSMUSG( | Tut1 | 70044 | 010913775 | 002846821 | -03325 | 092 | | 9 |
| 7752 | ENSMUSG( | Txlna | 109658 | 000835003 | 00920139C | -0345 | 058 | | 9 |
| 7771 | ENSMUSG( | Tyw3 | 209584 | -00061697 | 013028952 | -025 | 1095 | | 9 |
| 7780 | ENSMUSG( | Uba2 | 50995 | 013585552 | 006848478 | 01325 | 1015 | | 9 |
| 7796 | ENSMUSG( | Ube2e3 | 22193 | 00811450C | 005627919 | -046 | 1085 | | 9 |
| 7847 | ENSMUSG( | Ubxn7 | 224111 | 005373125 | -00018577 | -02325 | 06125 | | 9 |
| 7873 | ENSMUSG( | Unc119 | 22248 | 011089055 | -00594464 | 005 | 09275 | | 9 |
| 7911 | ENSMUSG( | Usb1 | 101985 | -00248751 | -00258238 | -031 | 06825 | | 9 |
| 7936 | ENSMUSG( | Usp43 | 216835 | -00526517 | -02585469 | -011 | 08625 | | 9 |
| 7965 | ENSMUSG( | Vamp8 | 22320 | -00279079 | -02172860 | -02525 | 106 | | 9 |
| 8002 | ENSMUSG( | Vopp1 | 232023 | -02429622 | 011944905 | -002 | 089 | | 9 |
| 8024 | ENSMUSG( | Vps37d | 194309 | -00424013 | -01774876 | -0375 | 0945 | | 9 |
| 8056 | ENSMUSG( | Wapl | 218914 | 009727692 | 013750352 | -0845 | 1355 | | 9 |
| 8070 | ENSMUSG( | Wbp4 | 22380 | 010441724 | 00151708C | 125607e-1 | 0805 | | 9 |
| 8119 | ENSMUSG( | Wiz | 22404 | 02212165C | 010866223 | -0475 | 06175 | | 9 |
| 8133 | ENSMUSG( | Xdh | 22436 | 027381502 | 085151331 | -0065 | 0815 | | 9 |
| 8153 | ENSMUSG( | Xrn1 | 24127 | -00911277 | 008068477 | -02025 | 06575 | | 9 |
| 8156 | ENSMUSG( | Xylb | 102448 | 010301219 | 023238729 | -022 | 07125 | | 9 |
| 8172 | ENSMUSG( | Yipf6 | 77929 | -01523026 | 004430661 | 017 | 08875 | | 9 |
| 8184 | ENSMUSG( | Ythdf1 | 228994 | 00721289C | 011973223 | -0085 | 07625 | | 9 |
| 8197 | ENSMUSG( | Zbtb18 | 30928 | 010429122 | 003989454 | 01175 | 1155 | | 9 |
| 8211 | ENSMUSG( | Zc3h4 | 330474 | 004548695 | -00306442 | -046 | 0905 | | 9 |
| 8225 | ENSMUSG( | Zdhhc20 | 75965 | -03894945 | 038690207 | -02025 | 11425 | | 9 |
| 8248 | ENSMUSG( | Zfp37 | 22696 | 000479703 | 01599189C | -0395 | 0575 | | 9 |
| 8256 | ENSMUSG( | Zfpm1 | 22761 | 02981169C | 022313439 | -01625 | 12975 | | 9 |
| 17 | ENSMUSG( | 6030458C1 | 77877 | -00248171 | -00141871 | 0345 | 03375 | | 10 |
| 54 | ENSMUSG( | Abcd2 | 26874 | -00793676 | -01710385 | 04 | 022 | | 10 |
| 72 | ENSMUSG( | Abhd3 | 106861 | 009520644 | -00377112 | 04875 | 049 | | 10 |
| 73 | ENSMUSG( | Abhd4 | 105501 | 008451827 | -02255822 | 026 | 012 | | 10 |
| 91 | ENSMUSG( | Acad11 | 102632 | 004888939 | -03147915 | 03875 | 0525 | | 10 |
| 120 | ENSMUSG( | Acot8 | 170789 | -01501739 | -01080095 | 02775 | 002 | | 10 |
| 123 | ENSMUSG( | Acox3 | 80911 | -00100051 | -00068609 | 0815 | 035 | | 10 |
| 127 | ENSMUSG( | Acsbg1 | 94180 | 00761819C | -02923948 | 027 | 0035 | | 10 |
| 131 | ENSMUSG( | Acsl3 | 74205 | 003379316 | 004152408 | 0355 | 0245 | | 10 |
| 162 | ENSMUSG( | Adam11 | 11488 | -00148955 | 010015587 | 02225 | 01525 | | 10 |
| 169 | ENSMUSG( | Adar | 56417 | 014397054 | -00213340 | 03875 | -00125 | | 10 |
| 172 | ENSMUSG( | Adck1 | 72113 | 008474031 | -00580632 | 04325 | 00325 | | 10 |
| 176 | ENSMUSG( | Adcy2 | 210044 | -00931993 | -00468142 | 02325 | 01025 | | 10 |
| 191 | ENSMUSG( | Adgrb2 | 230775 | 016666347 | 027199548 | 04125 | 02525 | | 10 |
| 202 | ENSMUSG( | Adipor1 | 72674 | -01229465 | 00098755 | 03275 | 01625 | | 10 |
| 245 | ENSMUSG( | Agps | 228061 | 005102117 | -00178762 | 02575 | 01075 | | 10 |
| 262 | ENSMUSG( | Aifm2 | 71361 | 002715074 | -02114060 | 03075 | 00575 | | 10 |
| 291 | ENSMUSG( | Akt2 | 11652 | 01335691C | -03564295 | 0365 | 0055 | | 10 |
| 311 | ENSMUSG( | Aldh9a1 | 56752 | 009149032 | -01932723 | 04475 | 003 | | 10 |
| 326 | ENSMUSG( | Alkbh7 | 66400 | -00145401 | -01813057 | 04575 | -00325 | | 10 |
| 382 | ENSMUSG( | Ankrd52 | 237615 | 02145385C | 004401202 | 0315 | 01075 | | 10 |
| 418 | ENSMUSG( | Ap2m1 | 11773 | 013338588 | -01356273 | 10925 | 01725 | | 10 |
| 437 | ENSMUSG( | Apbb1 | 11785 | 019157489 | -00032232 | 025 | 00025 | | 10 |
| 471 | ENSMUSG( | Arap2 | 212285 | -00026685 | -00151596 | 02775 | 01675 | | 10 |
| 495 | ENSMUSG( | Arhgap18 | 73910 | -00969054 | -00898143 | 035 | 0375 | | 10 |
| 526 | ENSMUSG( | Arhgef4 | 226970 | 015675488 | 007580531 | 02775 | 02825 | | 10 |
| 544 | ENSMUSG( | Arl3 | 56350 | -00185929 | -01995689 | 0245 | 00425 | | 10 |
| 553 | ENSMUSG( | Arl8b | 67166 | -00784064 | 000774136 | 02925 | 01725 | | 10 |
| 576 | ENSMUSG( | Arrb2 | 216869 | 013036093 | 012111721 | 026 | 01475 | | 10 |
| 611 | ENSMUSG( | Asxl2 | 75302 | -00464381 | 008028781 | 0455 | 008 | | 10 |
| 692 | ENSMUSG( | Atpaf2 | 246782 | 002214452 | -00402729 | 04575 | 008 | | 10 |
| 711 | ENSMUSG( | AW551984 | 244810 | 005358477 | -00555859 | 02225 | 027 | | 10 |
| 719 | ENSMUSG( | B3glct | 381694 | -02824449 | 001096896 | 0495 | 02175 | | 10 |
| 724 | ENSMUSG( | B4gat1 | 108902 | -01897821 | -00297789 | 05375 | 0335 | | 10 |
| 740 | ENSMUSG( | Bak1 | 12018 | 008037206 | -00552357 | 056 | 02075 | | 10 |
| 770 | ENSMUSG( | Bcl11a | 14025 | 001379059 | 002260732 | 0345 | 03 | | 10 |
| 772 | ENSMUSG( | Bcl2 | 12043 | 014120677 | -00176941 | 05625 | -01775 | | 10 |
| 784 | ENSMUSG( | Bet1 | 12068 | 006966813 | -01077291 | 05275 | 037 | | 10 |
| 812 | ENSMUSG( | Bnip1 | 224630 | 003959158 | -01081793 | 03825 | 0225 | | 10 |
| 813 | ENSMUSG( | Bnip2 | 12175 | 000155068 | 008877787 | 03225 | 00775 | | 10 |
| 821 | ENSMUSG( | Bola2 | 66162 | 009787046 | -01093049 | 03775 | 0415 | | 10 |

| | | | | | | | | |
|---|---|---|---|---|---|---|---|---|
| 830 | ENSMUSG( | Bpnt1 | 23827 | 008928732 | -00895676 | 0375 | 04675 | 10 |
| 849 | ENSMUSG( | Brk1 | 101314 | 007028101 | -01669376 | 0405 | 012 | 10 |
| 870 | ENSMUSG( | Bud13 | 215051 | 033233834 | 030888291 | 057 | 03225 | 10 |
| 874 | ENSMUSG( | Bzw2 | 66912 | 007102936 | 002546590 | 023 | 01225 | 10 |
| 894 | ENSMUSG( | Cables2 | 252966 | 003334906 | 000618446 | 037 | 00475 | 10 |
| 896 | ENSMUSG( | Cacfd1 | 381356 | -01489004 | 008482818 | 09275 | 0235 | 10 |
| 957 | ENSMUSG( | Cand2 | 67088 | -00266536 | -02419172 | 04275 | 0155 | 10 |
| 985 | ENSMUSG( | Cask | 12361 | -00798417 | -00183146 | 02625 | 0305 | 10 |
| 996 | ENSMUSG( | Cat | 12359 | 007635117 | -02018869 | 046 | 01375 | 10 |
| 1003 | ENSMUSG( | Cbfb | 12400 | 002030191 | 011390601 | 02 | 01325 | 10 |
| 1005 | ENSMUSG( | Cblb | 208650 | -00054037 | 003661233 | 02575 | 0275 | 10 |
| 1007 | ENSMUSG( | Cbln1 | 12404 | -01773921 | 030006485 | 065 | 016 | 10 |
| 1019 | ENSMUSG( | Cc2d1a | 212139 | 019266267 | 005629386 | 021 | 02175 | 10 |
| 1028 | ENSMUSG( | Ccdc120 | 54648 | 018147657 | 008798387 | 0365 | 0295 | 10 |
| 1047 | ENSMUSG( | Ccdc47 | 67163 | -01650084 | 010368482 | 04675 | -0085 | 10 |
| 1089 | ENSMUSG( | Cd151 | 12476 | -01637008 | 013759270 | 0535 | 024 | 10 |
| 1102 | ENSMUSG( | Cd63 | 12512 | -01240284 | 002214482 | 05375 | -0075 | 10 |
| 1150 | ENSMUSG( | Cdk19 | 78334 | 006058600 | -00133171 | 03925 | 00925 | 10 |
| 1154 | ENSMUSG( | Cdk5 | 12568 | 005589395 | -00195781 | 03025 | -0025 | 10 |
| 1222 | ENSMUSG( | Chchd4 | 72170 | 002239935 | -00798768 | 04025 | 02825 | 10 |
| 1266 | ENSMUSG( | Chst2 | 54371 | 000837116 | 004687575 | 022 | 0145 | 10 |
| 1284 | ENSMUSG( | Ckap4 | 216197 | -01927695 | 016089785 | 08 | 036 | 10 |
| 1315 | ENSMUSG( | Clip3 | 76686 | 011666644 | -00587546 | 03225 | 02075 | 10 |
| 1320 | ENSMUSG( | Cln8 | 26889 | -03132432 | 012205863 | 075 | 0315 | 10 |
| 1328 | ENSMUSG( | Clstn1 | 65945 | -01802925 | 016769363 | 0485 | 0255 | 10 |
| 1334 | ENSMUSG( | Clu | 12759 | 011292539 | 010774832 | 04025 | 01175 | 10 |
| 1349 | ENSMUSG( | Cmtm5 | 67272 | -02224375 | -01628677 | 02925 | 0185 | 10 |
| 1399 | ENSMUSG( | Coa8 | 68020 | 008718833 | -01346710 | 056 | 0465 | 10 |
| 1409 | ENSMUSG( | Cog6 | 67542 | 007928872 | -00387644 | 03575 | 04475 | 10 |
| 1441 | ENSMUSG( | Copb2 | 50797 | 013909144 | -00553153 | 02725 | 01425 | 10 |
| 1442 | ENSMUSG( | Cope | 59042 | 000077918 | -01518332 | 04825 | 02675 | 10 |
| 1458 | ENSMUSG( | Coq2 | 71883 | -00125461 | -00697951 | 03725 | 03875 | 10 |
| 1478 | ENSMUSG( | Cox15 | 226139 | -01189979 | -01444877 | 042 | 006 | 10 |
| 1507 | ENSMUSG( | Cpne4 | 74020 | -00753091 | 003241734 | 04175 | -0055 | 10 |
| 1516 | ENSMUSG( | Cpsf1 | 94230 | 013279887 | -00280317 | 0305 | 0055 | 10 |
| 1517 | ENSMUSG( | Cpsf2 | 51786 | 013480426 | 004169655 | 0345 | -00275 | 10 |
| 1522 | ENSMUSG( | Cpt1a | 12894 | -00414420 | -00431379 | 035 | 01075 | 10 |
| 1526 | ENSMUSG( | Cr1l | 12946 | -00470077 | 008483712 | 03975 | -006 | 10 |
| 1529 | ENSMUSG( | Cracdl | 72097 | 005811454 | 005226203 | 045 | 01625 | 10 |
| 1561 | ENSMUSG( | Crybb1 | 12960 | 032590334 | 005293971 | 03325 | 0315 | 10 |
| 1592 | ENSMUSG( | Cst3 | 13010 | -00406282 | -00698565 | 023 | 02825 | 10 |
| 1593 | ENSMUSG( | Cstb | 13014 | -00329234 | -02250033 | 03225 | 006 | 10 |
| 1606 | ENSMUSG( | Ctdsp1 | 227292 | 011081971 | -00267184 | 0275 | 0265 | 10 |
| 1675 | ENSMUSG( | Cyp2d22 | 56448 | 012367038 | 018898776 | 04375 | 01025 | 10 |
| 1676 | ENSMUSG( | Cyp2j6 | 13110 | -02582148 | 012299300 | 0575 | 001 | 10 |
| 1677 | ENSMUSG( | Cyp2j9 | 74519 | 017770771 | -00486410 | 04675 | 01425 | 10 |
| 1680 | ENSMUSG( | Cyp4f14 | 64385 | 003460761 | -00554885 | 03525 | 01625 | 10 |
| 1690 | ENSMUSG( | Cyth3 | 19159 | 009698494 | 006131565 | 02 | 017 | 10 |
| 1707 | ENSMUSG( | Dad1 | 13135 | -02395175 | -00332414 | 05375 | 0315 | 10 |
| 1716 | ENSMUSG( | Dars1 | 226414 | 004635946 | -02255613 | 03825 | 01575 | 10 |
| 1723 | ENSMUSG( | Dbndd1 | 72185 | 016911812 | 013935541 | 049 | 01775 | 10 |
| 1736 | ENSMUSG( | Dcakd | 68087 | 018552933 | -00694437 | 03225 | 02175 | 10 |
| 1746 | ENSMUSG( | Dctn1 | 13191 | 016956138 | -00771369 | 03225 | 00275 | 10 |
| 1750 | ENSMUSG( | Dctn5 | 59288 | -00174256 | -01142420 | 02075 | 02175 | 10 |
| 1752 | ENSMUSG( | Dctpp1 | 66422 | 002222090 | -01776014 | 0285 | 02125 | 10 |
| 1769 | ENSMUSG( | Ddost | 13200 | -03425995 | 006368070 | 08375 | 0255 | 10 |
| 1785 | ENSMUSG( | Ddx28 | 71986 | 000069583 | -00294131 | 0635 | 01925 | 10 |
| 1792 | ENSMUSG( | Ddx47 | 67755 | 002704474 | 000607632 | 026 | 0085 | 10 |
| 1816 | ENSMUSG( | Dennd5b | 320560 | 006935082 | 007105544 | 02975 | 00575 | 10 |
| 1821 | ENSMUSG( | Deptor | 97998 | 019568329 | 011971890 | 05875 | -0045 | 10 |
| 1861 | ENSMUSG( | Dhx36 | 72162 | 009843751 | 010171137 | 02575 | 01675 | 10 |
| 1869 | ENSMUSG( | Dicer1 | 192119 | 000331747 | 000315244 | 03775 | 047 | 10 |
| 1875 | ENSMUSG( | Dipk1b | 56279 | -02989987 | -00216622 | 0595 | 0015 | 10 |
| 1908 | ENSMUSG( | Dnaaf5 | 433956 | 007757439 | 001074834 | 0245 | 00725 | 10 |
| 1926 | ENSMUSG( | Dnajc12 | 30045 | 021342049 | 007449777 | 0245 | 0115 | 10 |
| 1931 | ENSMUSG( | Dnajc18 | 76594 | -01483977 | 006917254 | 0565 | -0005 | 10 |
| 1945 | ENSMUSG( | Dnal1 | 105000 | 012748896 | -00168559 | 0265 | 029 | 10 |
| 1960 | ENSMUSG( | Doc2a | 13446 | -00062287 | -01308015 | 04575 | -011 | 10 |
| 1961 | ENSMUSG( | Doc2b | 13447 | 014335438 | 004042209 | 06425 | -00625 | 10 |
| 1979 | ENSMUSG( | Dop1b | 70028 | 004824584 | 002128436 | 0255 | 00175 | 10 |
| 1999 | ENSMUSG( | Dpy19l3 | 233115 | -00886143 | 002035347 | 04525 | -00475 | 10 |
| 2016 | ENSMUSG( | Dstn | 56431 | 005912925 | -00624837 | 04575 | 01275 | 10 |
| 2054 | ENSMUSG( | Dynlt3 | 67117 | 008551610 | -00277852 | 06175 | 006 | 10 |
| 2068 | ENSMUSG( | Ece2 | 107522 | 021581849 | 009301496 | 0755 | 03275 | 10 |
| 2087 | ENSMUSG( | Eef1a1 | 13627 | 002992064 | -01648130 | 0375 | 02275 | 10 |
| 2092 | ENSMUSG( | Eef1e1 | 66143 | -00020033 | -01187464 | 05675 | -01025 | 10 |
| 2095 | ENSMUSG( | Eefsec | 65967 | 005054744 | -00832266 | 02 | 02375 | 10 |
| 2128 | ENSMUSG( | Eif1b | 68969 | 012305967 | -00261026 | 06425 | 012 | 10 |
| 2141 | ENSMUSG( | Eif2s3x | 26905 | 008243918 | -01367297 | 0325 | 01075 | 10 |
| 2144 | ENSMUSG( | Eif3b | 27979 | 015184478 | -00165951 | 03525 | 015 | 10 |
| 2146 | ENSMUSG( | Eif3d | 55944 | 002039991 | -01050918 | 04525 | -00225 | 10 |
| 2147 | ENSMUSG( | Eif3e | 16341 | 002069586 | -00489293 | 03475 | 01275 | 10 |
| 2148 | ENSMUSG( | Eif3f | 66085 | 008874275 | -01008246 | 0345 | 01975 | 10 |
| 2154 | ENSMUSG( | Eif3l | 223691 | -00005914 | -01147934 | 05525 | -012 | 10 |
| 2155 | ENSMUSG( | Eif3m | 98221 | 004134825 | -00843073 | 0485 | 0055 | 10 |
| 2207 | ENSMUSG( | Emc6 | 66048 | 004298973 | -00224013 | 04125 | 027 | 10 |
| 2216 | ENSMUSG( | Eml4 | 78798 | 022644801 | 000950159 | 0325 | 03675 | 10 |
| 2244 | ENSMUSG( | Entpd6 | 12497 | -02776120 | 011885216 | 07475 | 013 | 10 |
| 2246 | ENSMUSG( | Eogt | 101351 | -00107719 | 029119178 | 03675 | 032 | 10 |
| 2262 | ENSMUSG( | Epha6 | 13840 | -01578023 | 015804179 | 05475 | 0325 | 10 |
| 2271 | ENSMUSG( | Epm2a | 13853 | 002404945 | -00478353 | 066 | 016 | 10 |
| 2295 | ENSMUSG( | Ergic3 | 66366 | -02445319 | -00321614 | 05725 | 0075 | 10 |
| 2305 | ENSMUSG( | Erp29 | 67397 | -02099412 | -00197642 | 05075 | -00675 | 10 |
| 2320 | ENSMUSG( | Eva1a | 232146 | -00725170 | 011668450 | 02875 | 00575 | 10 |
| 2326 | ENSMUSG( | Exd2 | 97827 | 012204587 | 013685937 | 039 | 0285 | 10 |
| 2342 | ENSMUSG( | Exosc5 | 27998 | 013968102 | -01099454 | 0375 | 01225 | 10 |
| 2350 | ENSMUSG( | Extl3 | 54616 | -00892788 | 010263287 | 03775 | -00925 | 10 |
| 2362 | ENSMUSG( | Fads1 | 76267 | 004181459 | -01320356 | 0495 | -00775 | 10 |
| 2373 | ENSMUSG( | Fam107a | 268709 | 039613802 | 005308533 | 046 | 026 | 10 |
| 2406 | ENSMUSG( | Fam217b | 71532 | 009054932 | -00001929 | 043 | 0245 | 10 |
| 2411 | ENSMUSG( | Fam3c | 27999 | -02958563 | 006889401 | 06425 | 00825 | 10 |
| 2421 | ENSMUSG( | Far1 | 67420 | -00284861 | 005928713 | 07975 | 04175 | 10 |
| 2446 | ENSMUSG( | Fbxl20 | 72194 | -01380365 | 008937792 | 0395 | 03475 | 10 |
| 2449 | ENSMUSG( | Fbxo11 | 225055 | 008667076 | 009928752 | 072 | 046 | 10 |
| 2459 | ENSMUSG( | Fbxo42 | 213499 | 005046358 | -00347763 | 0375 | 026 | 10 |
| 2462 | ENSMUSG( | Fbxo6 | 50762 | -01065339 | -00583636 | 026 | 02225 | 10 |
| 2468 | ENSMUSG( | Fbxw8 | 231672 | 018784587 | -01332357 | 03225 | 036 | 10 |
| 2476 | ENSMUSG( | Fchsd2 | 207278 | 004700356 | -01009012 | 03675 | 005 | 10 |
| 2478 | ENSMUSG( | Fdft1 | 14137 | -01553500 | -02610779 | 028 | 025 | 10 |
| 2516 | ENSMUSG( | Fip1l1 | 66899 | 010172242 | -00006541 | 07025 | 0235 | 10 |
| 2535 | ENSMUSG( | Flot1 | 14251 | 013037059 | -01461456 | 037 | -0015 | 10 |
| 2536 | ENSMUSG( | Flot2 | 14252 | 004792354 | -01465854 | 03575 | 022 | 10 |
| 2547 | ENSMUSG( | Fmo1 | 14261 | 022372444 | 011231129 | 09275 | 02925 | 10 |
| 2587 | ENSMUSG( | Fscn1 | 14086 | 006941619 | -02301286 | 0365 | 01 | 10 |
| 2592 | ENSMUSG( | Ftl1 | 14325 | 009372464 | 001949988 | 0465 | 00975 | 10 |
| 2613 | ENSMUSG( | Fyco1 | 17281 | 011188356 | -01289877 | 05475 | 0075 | 10 |
| 2619 | ENSMUSG( | G3bp1 | 27041 | 010251583 | 001123685 | 038 | 04475 | 10 |
| 2644 | ENSMUSG( | Gad2 | 14417 | -01180288 | 000580548 | 036 | 009 | 10 |
| 2661 | ENSMUSG( | Gamt | 14431 | -00836595 | -03431342 | 053 | 0055 | 10 |
| 2662 | ENSMUSG( | Gan | 209239 | 002814142 | 003969994 | 022 | 0195 | 10 |
| 2670 | ENSMUSG( | Garnl3 | 99326 | 021337829 | 010842598 | 022 | 0225 | 10 |
| 2684 | ENSMUSG( | Gatm | 67092 | -00591193 | -00377411 | 0525 | 03275 | 10 |
| 2693 | ENSMUSG( | Gca | 227960 | 017730674 | -00794360 | 05275 | 005 | 10 |
| 2702 | ENSMUSG( | Gda | 14544 | 003691323 | -00274400 | 07675 | 04125 | 10 |
| 2704 | ENSMUSG( | Gdap1l1 | 228858 | 004118468 | -00805543 | 03925 | 002 | 10 |
| 2741 | ENSMUSG( | Gid4 | 66771 | 017452104 | -00457900 | 07775 | 048 | 10 |

| | | | | | | | |
|---|---|---|---|---|---|---|---|
| 2771 | ENSMUSG( | Glrx | 93692 | -00643239 | -01826978 | 07325 | 00925 | 10 |
| 2791 | ENSMUSG( | Gmfb | 63985 | 000321046 | -00904350 | 05825 | -00525 | 10 |
| 2800 | ENSMUSG( | Gnai1 | 14677 | -00720096 | -00348984 | 02625 | 0075 | 10 |
| 2809 | ENSMUSG( | Gnb2 | 14693 | 001444022 | -01589036 | 02475 | 009 | 10 |
| 2810 | ENSMUSG( | Gnb4 | 14696 | 00067193C | -00562600 | 03925 | -009 | 10 |
| 2818 | ENSMUSG( | Gng3 | 14704 | 006695329 | -01705639 | 02425 | 0235 | 10 |
| 2825 | ENSMUSG( | Gnpat | 14712 | 00937145C | 003113182 | 05125 | 0095 | 10 |
| 2832 | ENSMUSG( | Golga2 | 99412 | 014588747 | -00088040 | 074 | 04075 | 10 |
| 2839 | ENSMUSG( | Golim4 | 73124 | -00308284 | 002347114 | 0245 | 02925 | 10 |
| 2848 | ENSMUSG( | Gosr1 | 53334 | 005569844 | -00765651 | 07025 | 01875 | 10 |
| 2881 | ENSMUSG( | Gpr180 | 58245 | -01113430 | 012356352 | 05975 | -005 | 10 |
| 2902 | ENSMUSG( | Gpx7 | 67305 | -00355869 | 017772478 | 06025 | 015 | 10 |
| 2903 | ENSMUSG( | Gpx8 | 69590 | -00080199 | 017613089 | 0475 | 0275 | 10 |
| 2943 | ENSMUSG( | Grpel2 | 17714 | -01627079 | -01591558 | 0455 | 00725 | 10 |
| 2950 | ENSMUSG( | Gsk3a | 606496 | 01158907C | -00240395 | 03275 | 00825 | 10 |
| 2969 | ENSMUSG( | Gstz1 | 14874 | 017946573 | -00609886 | 02325 | 02525 | 10 |
| 2970 | ENSMUSG( | Gtdc1 | 227835 | 014246355 | 009218665 | 0475 | 01575 | 10 |
| 2980 | ENSMUSG( | Gtf2h5 | 66467 | -00020858 | -01520577 | 039 | -007 | 10 |
| 2985 | ENSMUSG( | Gtf3c4 | 269252 | 023065642 | 004860221 | 0345 | 0005 | 10 |
| 2992 | ENSMUSG( | Gtpbp6 | 107999 | 011606099 | 002086039 | 0585 | 02875 | 10 |
| 3049 | ENSMUSG( | Hdac8 | 70315 | 000635533 | -02554104 | 0235 | 02025 | 10 |
| 3062 | ENSMUSG( | Heatr5a | 320487 | 013731253 | 008915102 | 044 | 05575 | 10 |
| 3083 | ENSMUSG( | Hexa | 15211 | -01026511 | 000956709 | 0335 | -001 | 10 |
| 3084 | ENSMUSG( | Hexb | 15212 | -00535137 | 024648325 | 0485 | 043 | 10 |
| 3086 | ENSMUSG( | Hgh1 | 59053 | -02638151 | -00774217 | 04275 | 01125 | 10 |
| 3107 | ENSMUSG( | Hmbs | 15288 | 000515717 | -00883762 | 03875 | -0035 | 10 |
| 3118 | ENSMUSG( | Hmgcs2 | 15360 | 015523443 | -03702615 | 04525 | 04825 | 10 |
| 3156 | ENSMUSG( | Hpcal4 | 170638 | 027240551 | 008706291 | 0495 | 00975 | 10 |
| 3164 | ENSMUSG( | Hs2st1 | 23908 | -01690330 | 011654122 | 081 | 021 | 10 |
| 3171 | ENSMUSG( | Hsd17b11 | 114664 | -01320530 | 017002929 | 0435 | 00875 | 10 |
| 3173 | ENSMUSG( | Hsd17b4 | 15488 | 015802796 | -01128421 | 08875 | 02875 | 10 |
| 3177 | ENSMUSG( | Hsdl2 | 72479 | 005515051 | -03224362 | 0315 | 025 | 10 |
| 3180 | ENSMUSG( | Hsp90ab1 | 15516 | 010597265 | -01172033 | 02475 | 019 | 10 |
| 3181 | ENSMUSG( | Hsp90b1 | 22027 | -00673692 | 017629117 | 04875 | 02375 | 10 |
| 3200 | ENSMUSG( | Htatip2 | 53415 | -00326938 | -01193548 | 0375 | 0185 | 10 |
| 3216 | ENSMUSG( | Iars1 | 105148 | 005086353 | 00654068 | 028 | 01175 | 10 |
| 3220 | ENSMUSG( | Ica1l | 70375 | 019987759 | -00306013 | 0485 | 03225 | 10 |
| 3235 | ENSMUSG( | Ifngr2 | 15980 | -01664902 | 006435428 | 0535 | 03475 | 10 |
| 3246 | ENSMUSG( | Ift81 | 12589 | 010964207 | -00941904 | 0265 | 02525 | 10 |
| 3271 | ENSMUSG( | Il16 | 16170 | 007793329 | 030184709 | 063 | 03725 | 10 |
| 3337 | ENSMUSG( | Ipo9 | 226432 | 002396329 | -01204786 | 0225 | 01675 | 10 |
| 3387 | ENSMUSG( | Itih5 | 209378 | 011462525 | 018389933 | 0415 | 01775 | 10 |
| 3432 | ENSMUSG( | Kat2a | 14534 | 019414412 | -00187975 | 0655 | -0115 | 10 |
| 3440 | ENSMUSG( | Kbtbd3 | 69149 | -01427731 | -01958409 | 03775 | -0065 | 10 |
| 3442 | ENSMUSG( | Kbtbd7 | 211255 | -00093129 | 023960818 | 037 | -0065 | 10 |
| 3457 | ENSMUSG( | Kcnc4 | 99738 | 005865758 | 021004075 | 0315 | 022 | 10 |
| 3480 | ENSMUSG( | Kcnk1 | 16525 | -02754920 | 020200928 | 07125 | 0195 | 10 |
| 3503 | ENSMUSG( | Kctd21 | 622320 | 023492313 | -02590395 | 04875 | 03325 | 10 |
| 3515 | ENSMUSG( | Kdm5a | 214899 | 001059797 | -00830227 | 042 | 026 | 10 |
| 3526 | ENSMUSG( | Kidins220 | 77480 | -01034085 | 003119122 | 03225 | 0025 | 10 |
| 3528 | ENSMUSG( | Kif13a | 16553 | -00268185 | -01263647 | 066 | -00975 | 10 |
| 3532 | ENSMUSG( | Kif1c | 16562 | -00029532 | -01970808 | 046 | 02625 | 10 |
| 3538 | ENSMUSG( | Kif3c | 16570 | 023362103 | 00805869C | 0305 | 033 | 10 |
| 3545 | ENSMUSG( | Kifc3 | 16582 | 007468483 | 00069809C | 02175 | 025 | 10 |
| 3553 | ENSMUSG( | Klhdc10 | 76788 | 004936424 | -00920428 | 104 | 021 | 10 |
| 3556 | ENSMUSG( | Klhdc4 | 234825 | 005676282 | -00642011 | 02225 | 0155 | 10 |
| 3558 | ENSMUSG( | Klhdc8b | 78267 | 010043207 | 009357071 | 0425 | 01425 | 10 |
| 3576 | ENSMUSG( | Kpna2 | 16647 | 01060244C | 03023438C | 04525 | 00025 | 10 |
| 3585 | ENSMUSG( | Krr1 | 52705 | 002292623 | 00245973C | 029 | 0265 | 10 |
| 3587 | ENSMUSG( | Krtcap2 | 66059 | -01686726 | -01099381 | 04375 | 0075 | 10 |
| 3591 | ENSMUSG( | Ktn1 | 16709 | -00964791 | 009045725 | 0535 | 00825 | 10 |
| 3691 | ENSMUSG( | Lman2l | 214895 | 000171248 | -00485358 | 0745 | 0415 | 10 |
| 3709 | ENSMUSG( | Lonp2 | 66887 | 015075693 | -01366257 | 04025 | 0405 | 10 |
| 3712 | ENSMUSG( | Lpcat1 | 210992 | -01594738 | 007908749 | 04025 | 004 | 10 |
| 3719 | ENSMUSG( | Lpp | 210126 | 009689569 | 003114844 | 0235 | 028 | 10 |
| 3743 | ENSMUSG( | Lrrc14 | 223664 | 00452262C | -01367370 | 03175 | 013 | 10 |
| 3754 | ENSMUSG( | Lrrc59 | 98238 | 002490592 | -00718222 | 04325 | 00525 | 10 |
| 3787 | ENSMUSG( | Lsp1 | 16985 | -03075205 | -03573564 | 055 | 034 | 10 |
| 3789 | ENSMUSG( | Lss | 16987 | -00818968 | -02860445 | 04525 | 006 | 10 |
| 3808 | ENSMUSG( | Lyar | 17089 | 019623956 | 008388405 | 0275 | 018 | 10 |
| 3810 | ENSMUSG( | Lynx1 | 23936 | -00652568 | 009229201 | 03075 | 03125 | 10 |
| 3830 | ENSMUSG( | Macf1 | 11426 | -02550853 | 000379399 | 079 | 01975 | 10 |
| 3833 | ENSMUSG( | Macrod2 | 72899 | 008682753 | -00156295 | 03625 | 01275 | 10 |
| 3858 | ENSMUSG( | Man2a2 | 140481 | -00615272 | 011851228 | 058 | 02975 | 10 |
| 3906 | ENSMUSG( | Mapk14 | 26416 | 015370124 | -00228317 | 034 | 032 | 10 |
| 3908 | ENSMUSG( | Mapk3 | 26417 | 008085072 | -00817211 | 0265 | 023 | 10 |
| 3957 | ENSMUSG( | Mbnl2 | 105559 | -00027464 | 004897349 | 04475 | 0275 | 10 |
| 3958 | ENSMUSG( | Mboat2 | 67216 | -00096626 | 002653346 | 0235 | 018 | 10 |
| 3975 | ENSMUSG( | Mcoln1 | 94178 | -01723524 | 016264464 | 056 | 0285 | 10 |
| 4009 | ENSMUSG( | Med19 | 381379 | 001131597 | -01570677 | 02775 | 018 | 10 |
| 4033 | ENSMUSG( | Metap1 | 75624 | 010164069 | 013854906 | 02875 | 0125 | 10 |
| 4070 | ENSMUSG( | Mgst3 | 66447 | -01485221 | -01301604 | 0585 | 0275 | 10 |
| 4102 | ENSMUSG( | Mix23 | 381045 | 001744136 | -00811476 | 042 | -00025 | 10 |
| 4104 | ENSMUSG( | Mkrn2 | 67027 | 003765992 | -00504726 | 02325 | 02175 | 10 |
| 4106 | ENSMUSG( | Mlec | 109154 | 005308378 | 007547843 | 0485 | 00125 | 10 |
| 4147 | ENSMUSG( | Mpi | 110119 | 008061302 | -00539884 | 02525 | 01375 | 10 |
| 4158 | ENSMUSG( | Mpv17l2 | 234384 | -01535279 | -00952311 | 06825 | 041 | 10 |
| 4171 | ENSMUSG( | Mrpl13 | 68537 | 004899073 | -01765743 | 02575 | 0055 | 10 |
| 4174 | ENSMUSG( | Mrpl16 | 94063 | 005128207 | -02108905 | 02825 | 0185 | 10 |
| 4209 | ENSMUSG( | Mrpl54 | 66047 | -01419821 | -03160745 | 028 | 02475 | 10 |
| 4216 | ENSMUSG( | Mrps12 | 24030 | 012013495 | -00805821 | 0215 | 02575 | 10 |
| 4220 | ENSMUSG( | Mrps17 | 66258 | -00253846 | -00186811 | 05225 | 01125 | 10 |
| 4258 | ENSMUSG( | Mt2 | 17750 | 009189689 | -02155062 | 058 | 02325 | 10 |
| 4348 | ENSMUSG( | Myorg | 329828 | -01352373 | 015463224 | 056 | 024 | 10 |
| 4367 | ENSMUSG( | Naaa | 67111 | -00006408 | 015820014 | 03475 | 00425 | 10 |
| 4388 | ENSMUSG( | Napb | 17957 | 007971688 | -00233142 | 03325 | 01075 | 10 |
| 4400 | ENSMUSG( | Nat8l | 269642 | -00871759 | 002473374 | 0415 | 0305 | 10 |
| 4410 | ENSMUSG( | Nbeal1 | 269198 | -00913837 | 019104154 | 04925 | 0045 | 10 |
| 4430 | ENSMUSG( | Ncln | 103425 | -01767416 | 014094189 | 062 | 00525 | 10 |
| 4452 | ENSMUSG( | Ndufa11 | 69875 | -00776373 | -02774387 | 0295 | 017 | 10 |
| 4497 | ENSMUSG( | Necab1 | 69352 | 000346528 | 002191391 | 028 | 02025 | 10 |
| 4599 | ENSMUSG( | Noc3l | 57753 | 018840738 | 010500884 | 0335 | 0055 | 10 |
| 4611 | ENSMUSG( | Nomo1 | 211548 | -01155846 | 000817489 | 06075 | 02325 | 10 |
| 4657 | ENSMUSG( | Nr2f1 | 13865 | 019593618 | 009484968 | 0375 | 02675 | 10 |
| 4659 | ENSMUSG( | Nr4a3 | 18124 | 008471152 | 00981957C | 0545 | 03875 | 10 |
| 4688 | ENSMUSG( | Nsmf | 56876 | 023258967 | 009970399 | 0475 | -01275 | 10 |
| 4706 | ENSMUSG( | Ntpcr | 66566 | -01073037 | -01035953 | 055 | 03325 | 10 |
| 4717 | ENSMUSG( | Nucb2 | 53322 | -03082467 | -00372666 | 048 | 0035 | 10 |
| 4774 | ENSMUSG( | Nyap1 | 243300 | 021565407 | 008989791 | 054 | 03575 | 10 |
| 4780 | ENSMUSG( | Ociad2 | 433904 | -01369341 | 001626865 | 04075 | 01375 | 10 |
| 4792 | ENSMUSG( | Ola1 | 67059 | -00424636 | -01098654 | 0485 | -014 | 10 |
| 4798 | ENSMUSG( | Oma1 | 67013 | 006814496 | -00908399 | 07975 | 01625 | 10 |
| 4817 | ENSMUSG( | Osbpl10 | 74486 | 006080358 | -00193099 | 02475 | 00925 | 10 |
| 4822 | ENSMUSG( | Osbpl5 | 79196 | 023472629 | -00033952 | 04375 | 02925 | 10 |
| 4828 | ENSMUSG( | Osgep | 66246 | 006362304 | -02730319 | 02625 | 005 | 10 |
| 4882 | ENSMUSG( | Palm | 18483 | 026998196 | 020724383 | 029 | 0285 | 10 |
| 4896 | ENSMUSG( | Paox | 212503 | -00256644 | -00530448 | 03725 | -0035 | 10 |
| 4913 | ENSMUSG( | Pars2 | 230577 | 016578393 | -00324495 | 068 | -01 | 10 |
| 4914 | ENSMUSG( | Parva | 57342 | 011526688 | -01527944 | 03 | 01525 | 10 |
| 4923 | ENSMUSG( | Pbx1 | 18514 | 005608713 | -00406888 | 05125 | 0485 | 10 |
| 4926 | ENSMUSG( | Pcbd2 | 72562 | -00562208 | -02423831 | 04275 | 002 | 10 |
| 4959 | ENSMUSG( | Pcdhgb1 | 93699 | -01345078 | 024418062 | 066 | 021 | 10 |
| 4994 | ENSMUSG( | Pdcd2 | 18567 | -01588669 | -01235731 | 028 | 029 | 10 |
| 5028 | ENSMUSG( | Pdia5 | 72599 | -01798225 | -00668066 | 06175 | -003 | 10 |
| 5034 | ENSMUSG( | Pdlim1 | 54132 | -01044027 | -00578437 | 0345 | 01325 | 10 |
| 5068 | ENSMUSG( | Pex10 | 668173 | -00156586 | -00173146 | 0235 | 00625 | 10 |

| | | | | | | | |
|---|---|---|---|---|---|---|---|
| 5069 | ENSMUSG( | Pex11b | 18632 | -00031014 | -00302730 | 05825 | -0045 | 10 |
| 5071 | ENSMUSG( | Pex13 | 72129 | -01119206 | 00057582( | 04375 | 01625 | 10 |
| 5081 | ENSMUSG( | Pex7 | 18634 | 00321199€ | -02013125 | 045 | 04175 | 10 |
| 5120 | ENSMUSG( | Phax | 56698 | 013656504 | -00345515 | 0215 | 01725 | 10 |
| 5123 | ENSMUSG( | Phc1 | 13619 | 01969893S | 007964714 | 03975 | 016 | 10 |
| 5132 | ENSMUSG( | Phf6 | 70998 | 024727217 | 030553197 | 046 | 005 | 10 |
| 5185 | ENSMUSG( | Pin4 | 69713 | 004946927 | -02362015 | 0405 | 0245 | 10 |
| 5190 | ENSMUSG( | Pip4p2 | 72519 | -00638500 | -00133403 | 0205 | 01375 | 10 |
| 5217 | ENSMUSG( | Pla2g7 | 27226 | -00878566 | -00400753 | 05525 | 00875 | 10 |
| 5227 | ENSMUSG( | Plcd4 | 18802 | 020782337 | -01038871 | 0485 | 0385 | 10 |
| 5287 | ENSMUSG( | Plxnc1 | 54712 | -00312890 | 02406257S | 036 | 01375 | 10 |
| 5307 | ENSMUSG( | Pnp | 18950 | 01218510S | -00520931 | 03725 | -00325 | 10 |
| 5310 | ENSMUSG( | Pnpla7 | 241274 | 00734421S | 011556342 | 05875 | 01575 | 10 |
| 5340 | ENSMUSG( | Polr3a | 218832 | 003141664 | 002924801 | 02675 | 02625 | 10 |
| 5344 | ENSMUSG( | Polr3e | 26939 | -00075661 | 00630808€ | 033 | 03475 | 10 |
| 5352 | ENSMUSG( | Pon2 | 330260 | 008033597 | -00123460 | 0305 | 0005 | 10 |
| 5354 | ENSMUSG( | Pop7 | 74097 | -00794820 | -00214798 | 02575 | 004 | 10 |
| 5400 | ENSMUSG( | Ppp1ca | 19045 | -00647010 | -02325374 | 062 | 022 | 10 |
| 5403 | ENSMUSG( | Ppp1r10 | 52040 | 017560797 | 00881051€ | 032 | 02875 | 10 |
| 5411 | ENSMUSG( | Ppp1r14c | 76142 | 006987424 | -01133087 | 049 | 02425 | 10 |
| 5420 | ENSMUSG( | Ppp1r3d | 228966 | 01255517S | -01097150 | 0325 | -00425 | 10 |
| 5424 | ENSMUSG( | Ppp1r8 | 100336 | 004803951 | -00457851 | 03475 | 003 | 10 |
| 5464 | ENSMUSG( | Prdx1 | 18477 | 006102442 | -02242480 | 02525 | 0015 | 10 |
| 5467 | ENSMUSG( | Prdx4 | 53381 | -01437114 | 01139800€ | 04475 | 00075 | 10 |
| 5476 | ENSMUSG( | Prex2 | 109294 | 02114276€ | -00508645 | 058 | 01825 | 10 |
| 5477 | ENSMUSG( | Prickle1 | 106042 | -00125775 | 01154043€ | 02325 | 024 | 10 |
| 5501 | ENSMUSG( | Prkd1 | 18760 | 019345911 | 013729681 | 02725 | 00775 | 10 |
| 5508 | ENSMUSG( | Prmt2 | 15468 | 009467051 | -00405755 | 047 | -01475 | 10 |
| 5513 | ENSMUSG( | Prmt9 | 102182 | -00292494 | 00610006€ | 0285 | 004 | 10 |
| 5524 | ENSMUSG( | Prpf38b | 66921 | -01549145 | 017760294 | 057 | 019 | 10 |
| 5533 | ENSMUSG( | Prps1l3 | 328099 | -00043924 | -00867704 | 02925 | 016 | 10 |
| 5535 | ENSMUSG( | Prpsap1 | 67763 | -00504162 | 00969759S | 0385 | 04375 | 10 |
| 5584 | ENSMUSG( | Psmd10 | 53380 | -00616726 | -02723093 | 06525 | 03125 | 10 |
| 5595 | ENSMUSG( | Psmd8 | 57296 | -00215611 | -02104779 | 046 | 02925 | 10 |
| 5625 | ENSMUSG( | Ptgr1 | 67103 | 043910584 | -00247144 | 0415 | -00625 | 10 |
| 5675 | ENSMUSG( | Puf60 | 67959 | 00914926S | -01580606 | 02475 | 019 | 10 |
| 5691 | ENSMUSG( | Pxmp2 | 19301 | 008744798 | -00960812 | 0755 | 01375 | 10 |
| 5692 | ENSMUSG( | Pxmp4 | 59038 | 00108033€ | -02750969 | 069 | 0055 | 10 |
| 5701 | ENSMUSG( | Pygo2 | 68911 | 00949035€ | -00242225 | 04125 | 00825 | 10 |
| 5712 | ENSMUSG( | Qrsl1 | 76563 | 009317635 | 00822031€ | 019 | 0195 | 10 |
| 5729 | ENSMUSG( | Rab15 | 104886 | 017831951 | -00519438 | 03325 | 0085 | 10 |
| 5733 | ENSMUSG( | Rab21 | 216344 | 004685674 | -00825190 | 042 | 0155 | 10 |
| 5743 | ENSMUSG( | Rab2b | 76338 | 01168301( | -00692078 | 05025 | -001 | 10 |
| 5757 | ENSMUSG( | Rab3gap1 | 226407 | 01684723S | 004979647 | 02875 | 01625 | 10 |
| 5758 | ENSMUSG( | Rab3gap2 | 98732 | -00020179 | -00035475 | 027 | -0025 | 10 |
| 5759 | ENSMUSG( | Rab3il1 | 74760 | 013258755 | -00310805 | 0545 | 0045 | 10 |
| 5760 | ENSMUSG( | Rab3ip | 216363 | 00462198€ | -00195097 | 06475 | 0075 | 10 |
| 5768 | ENSMUSG( | Rab5if | 67388 | 000640511 | 00196978€ | 041 | 052 | 10 |
| 5790 | ENSMUSG( | Rack1 | 14694 | 008728103 | -01699796 | 04975 | -0115 | 10 |
| 5818 | ENSMUSG( | Ranbp9 | 56705 | -00040354 | 00890278( | 02825 | 00325 | 10 |
| 5825 | ENSMUSG( | Rap2a | 76108 | 013436673 | -00681892 | 029 | 01725 | 10 |
| 5853 | ENSMUSG( | Rassf8 | 71323 | -00622678 | -00308592 | 047 | 00275 | 10 |
| 5884 | ENSMUSG( | Rbm4b | 66704 | -00170363 | -01435672 | 02825 | 01175 | 10 |
| 5898 | ENSMUSG( | Rcan2 | 53901 | 001808885 | -02277646 | 0525 | 015 | 10 |
| 5930 | ENSMUSG( | Rer1 | 67830 | -01838105 | 00499227€ | 06925 | 0165 | 10 |
| 5971 | ENSMUSG( | Rhoa | 11848 | 003195607 | -00772375 | 055 | -006 | 10 |
| 5980 | ENSMUSG( | Rhou | 69581 | 017288031 | -01358056 | 039 | 02525 | 10 |
| 5984 | ENSMUSG( | Ric8a | 101489 | 005412345 | -00523212 | 027 | 033 | 10 |
| 6001 | ENSMUSG( | Ripk1 | 19766 | 02072389( | 014713857 | 031 | 0275 | 10 |
| 6003 | ENSMUSG( | Ripor2 | 193385 | 007988884 | 003949644 | 0295 | 00425 | 10 |
| 6024 | ENSMUSG( | Rnf11 | 29864 | 005039535 | -00177948 | 028 | 015 | 10 |
| 6029 | ENSMUSG( | Rnf121 | 75212 | -01305906 | 01983692€ | 0665 | 021 | 10 |
| 6057 | ENSMUSG( | Rnf7 | 19823 | 002316923 | -01230840 | 036 | 023 | 10 |
| 6083 | ENSMUSG( | Rpl10 | 110954 | 010630641 | -00973586 | 035 | 022 | 10 |
| 6086 | ENSMUSG( | Rpl12 | 269261 | 006960977 | -01739761 | 04725 | 01525 | 10 |
| 6087 | ENSMUSG( | Rpl13 | 270106 | 00797014( | -01507475 | 03975 | 01625 | 10 |
| 6088 | ENSMUSG( | Rpl13a | 22121 | 017453295 | -00875431 | 0425 | 00525 | 10 |
| 6089 | ENSMUSG( | Rpl14 | 67115 | 007208093 | -01421314 | 04775 | 052 | 10 |
| 6090 | ENSMUSG( | Rpl15 | 66480 | 00345872€ | -01644129 | 041 | 01125 | 10 |
| 6091 | ENSMUSG( | Rpl17 | 319195 | 00919489S | -01187835 | 03525 | 0195 | 10 |
| 6092 | ENSMUSG( | Rpl18 | 19899 | 00675751€ | -00671058 | 044 | 015 | 10 |
| 6093 | ENSMUSG( | Rpl18a | 76808 | 00679626S | -01464384 | 02875 | 03425 | 10 |
| 6094 | ENSMUSG( | Rpl19 | 19921 | 007296094 | -01406707 | 06075 | 0125 | 10 |
| 6095 | ENSMUSG( | Rpl21 | 19933 | 012456202 | -01199006 | 0255 | 01225 | 10 |
| 6096 | ENSMUSG( | Rpl22 | 19934 | 003915758 | -01798074 | 029 | 005 | 10 |
| 6097 | ENSMUSG( | Rpl22l1 | 68028 | -00141903 | -01539157 | 05 | 03475 | 10 |
| 6098 | ENSMUSG( | Rpl23 | 65019 | 006122162 | -01744967 | 04275 | 04825 | 10 |
| 6099 | ENSMUSG( | Rpl23a | 268449 | 012458081 | -01114947 | 064 | 01225 | 10 |
| 6103 | ENSMUSG( | Rpl27a | 26451 | 010985102 | -00938970 | 053 | 0125 | 10 |
| 6104 | ENSMUSG( | Rpl28 | 19943 | 003951942 | -02190511 | 043 | 04625 | 10 |
| 6109 | ENSMUSG( | Rpl32 | 19951 | 008917771 | -01257151 | 043 | 028 | 10 |
| 6110 | ENSMUSG( | Rpl34 | 68436 | 01126309€ | -01099618 | 03775 | -00175 | 10 |
| 6111 | ENSMUSG( | Rpl35 | 66489 | 007501367 | -01911807 | 043 | 02775 | 10 |
| 6112 | ENSMUSG( | Rpl35a | 57808 | 01346992€ | -00837483 | 07125 | 03975 | 10 |
| 6113 | ENSMUSG( | Rpl36 | 54217 | 00650866€ | -01636097 | 04475 | 02225 | 10 |
| 6114 | ENSMUSG( | Rpl36a | 19982 | 006997711 | -01848527 | 04075 | 013 | 10 |
| 6116 | ENSMUSG( | Rpl37a | 19981 | 00483102( | -02642291 | 07275 | 04275 | 10 |
| 6117 | ENSMUSG( | Rpl38 | 67671 | 007935257 | -01848029 | 02825 | 005 | 10 |
| 6119 | ENSMUSG( | Rpl4 | 67891 | 012351137 | -00623977 | 037 | 01075 | 10 |
| 6122 | ENSMUSG( | Rpl7 | 19989 | 009659077 | -01160680 | 063 | 02175 | 10 |
| 6123 | ENSMUSG( | Rpl7a | 27176 | 004276612 | -01720320 | 03875 | 01775 | 10 |
| 6128 | ENSMUSG( | Rplp2 | 67186 | 00935631€ | -00562535 | 02375 | 0155 | 10 |
| 6129 | ENSMUSG( | Rpn1 | 103963 | -02773512 | 01330317€ | 061 | 01325 | 10 |
| 6130 | ENSMUSG( | Rpn2 | 20014 | -01891272 | 00597503( | 063 | 01475 | 10 |
| 6138 | ENSMUSG( | Rps11 | 27207 | 008815015 | -01932288 | 02575 | 0295 | 10 |
| 6139 | ENSMUSG( | Rps12 | 20042 | 011745972 | -01549595 | 0265 | 0315 | 10 |
| 6140 | ENSMUSG( | Rps13 | 68052 | 008320274 | -01847532 | 04425 | -0025 | 10 |
| 6141 | ENSMUSG( | Rps14 | 20044 | 01099263S | -00858114 | 0545 | 0205 | 10 |
| 6142 | ENSMUSG( | Rps15 | 20054 | 01518014( | -01378154 | 0305 | 0315 | 10 |
| 6144 | ENSMUSG( | Rps16 | 20055 | 00676798€ | -01618805 | 068 | 0325 | 10 |
| 6145 | ENSMUSG( | Rps17 | 20068 | 00803687€ | -01174021 | 03425 | 00825 | 10 |
| 6146 | ENSMUSG( | Rps18-ps5 | 1E+08 | 016230492 | -02409062 | 02075 | 02825 | 10 |
| 6147 | ENSMUSG( | Rps19 | 20085 | 008608401 | -01589891 | 03075 | 02875 | 10 |
| 6148 | ENSMUSG( | Rps2 | 16898 | 011656155 | -00958211 | 05025 | 01125 | 10 |
| 6149 | ENSMUSG( | Rps20 | 67427 | 00991802S | -01476295 | 10625 | 029 | 10 |
| 6151 | ENSMUSG( | Rps23 | 66475 | 00998666€ | -01245344 | 066 | -0005 | 10 |
| 6152 | ENSMUSG( | Rps24 | 20088 | 007603685 | -01003669 | 0545 | -01725 | 10 |
| 6153 | ENSMUSG( | Rps25 | 75617 | 009394048 | -01543632 | 02425 | 0085 | 10 |
| 6159 | ENSMUSG( | Rps3 | 27050 | 008599105 | -01134804 | 04175 | 0175 | 10 |
| 6160 | ENSMUSG( | Rps4x | 20102 | 009673248 | -01655028 | 056 | -00075 | 10 |
| 6162 | ENSMUSG( | Rps6 | 20104 | 004715083 | -01379847 | 0385 | -002 | 10 |
| 6171 | ENSMUSG( | Rps8 | 20116 | 01509158S | -00695222 | 05325 | 0205 | 10 |
| 6172 | ENSMUSG( | Rps9 | 76846 | 00570283S | -01346307 | 03525 | 01675 | 10 |
| 6211 | ENSMUSG( | Rtraf | 68045 | 002121835 | -01408174 | 0385 | 046 | 10 |
| 6217 | ENSMUSG( | Rundc3a | 51799 | 016871047 | -00179024 | 04625 | 05575 | 10 |
| 6231 | ENSMUSG( | S100a11 | 20195 | -01263637 | -00213976 | 0275 | 03 | 10 |
| 6250 | ENSMUSG( | Samd4b | 233033 | 018764207 | 00550170S | 02775 | 0225 | 10 |
| 6265 | ENSMUSG( | Sarm1 | 237868 | 017329184 | 01121216€ | 03225 | 03525 | 10 |
| 6271 | ENSMUSG( | Sat2 | 69215 | 010236783 | -00018287 | 046 | 01075 | 10 |
| 6300 | ENSMUSG( | Schip1 | 30953 | 005877068 | -01685231 | 03075 | -00225 | 10 |
| 6314 | ENSMUSG( | Scp2 | 20280 | 012715618 | -01462539 | 0475 | 01725 | 10 |
| 6316 | ENSMUSG( | Scrg1 | 20284 | 02368504( | 003326445 | 051 | 02275 | 10 |
| 6319 | ENSMUSG( | Scrn2 | 217140 | 00226376S | -00227610 | 0295 | 027 | 10 |
| 6325 | ENSMUSG( | Scyl3 | 240880 | -01336412 | 005128021 | 04375 | 0095 | 10 |
| 6330 | ENSMUSG( | Sdcbp | 53378 | 004060765 | -00481278 | 031 | 0295 | 10 |
| 6359 | ENSMUSG( | Sec24c | 218811 | 02093380( | -00127456 | 033 | 01975 | 10 |

| | | | | | | | | | |
|---|---|---|---|---|---|---|---|---|---|
| 6364 | ENSMUSG( | Sec61b | 66212 | -00077339 | -01682500 | 051 | 0105 | | 10 |
| 6365 | ENSMUSG( | Sec61g | 20335 | 005980458 | -02216736 | 08775 | 02625 | | 10 |
| 6367 | ENSMUSG( | Sec63 | 140740 | -00624182 | 011876718 | 02925 | -004 | | 10 |
| 6388 | ENSMUSG( | Sema4g | 26456 | 008372820 | 000119926 | 034 | 02475 | | 10 |
| 6400 | ENSMUSG( | Sephs2 | 20768 | -00028732 | -01302088 | 0275 | 0045 | | 10 |
| 6468 | ENSMUSG( | Sgpl1 | 20397 | -01340739 | 012105403 | 038 | 01625 | | 10 |
| 6499 | ENSMUSG( | Sh3yl1 | 24057 | 003242019 | 003304307 | 04375 | 045 | | 10 |
| 6510 | ENSMUSG( | Shisa4 | 77552 | -02282243 | -00246960 | 0475 | 0475 | | 10 |
| 6519 | ENSMUSG( | Shq1 | 72171 | 028151338 | -00980690 | 08175 | 01875 | | 10 |
| 6558 | ENSMUSG( | Slc15a2 | 57738 | 013340924 | 024283456 | 06075 | 00075 | | 10 |
| 6561 | ENSMUSG( | Slc16a10 | 72472 | -05650458 | -02632311 | 06825 | 02525 | | 10 |
| 6589 | ENSMUSG( | Slc25a1 | 13358 | 003017658 | -01621426 | 03575 | 02825 | | 10 |
| 6597 | ENSMUSG( | Slc25a17 | 20524 | -01142209 | 011848842 | 0455 | -017 | | 10 |
| 6601 | ENSMUSG( | Slc25a21 | 217593 | -00814069 | -03771673 | 04875 | -0115 | | 10 |
| 6602 | ENSMUSG( | Slc25a22 | 68267 | 015656919 | -00273013 | 04475 | | 0 10 | |
| 6605 | ENSMUSG( | Slc25a25 | 227731 | -01112540 | -01339943 | 05775 | 007 | | 10 |
| 6625 | ENSMUSG( | Slc27a1 | 26457 | -00976780 | 002649482 | 039 | 00575 | | 10 |
| 6646 | ENSMUSG( | Slc35a4 | 67843 | 006674219 | -00192098 | 05775 | -0005 | | 10 |
| 6709 | ENSMUSG( | Slc7a6 | 330836 | 009328197 | 025691500 | 0375 | 031 | | 10 |
| 6763 | ENSMUSG( | Smg5 | 229512 | 000506340 | -00129276 | 02825 | 02675 | | 10 |
| 6779 | ENSMUSG( | Smpd2 | 20598 | -00804789 | -00223112 | 03825 | 00325 | | 10 |
| 6782 | ENSMUSG( | Smpdl3a | 57319 | -01871879 | 001635243 | 06225 | 01175 | | 10 |
| 6792 | ENSMUSG( | Snap25 | 20614 | 000865375 | -01337934 | 06 | -013 | | 10 |
| 6793 | ENSMUSG( | Snap29 | 67474 | 015076253 | -00699897 | 0415 | 0405 | | 10 |
| 6800 | ENSMUSG( | Snd1 | 56463 | -01560773 | -00574758 | 067 | 01675 | | 10 |
| 6835 | ENSMUSG( | Snx17 | 266781 | 009351280 | -01507168 | 0295 | 02075 | | 10 |
| 6843 | ENSMUSG( | Snx29 | 74478 | -00288451 | -01516820 | 02325 | 01 | | 10 |
| 6844 | ENSMUSG( | Snx3 | 54198 | 002024581 | -02295827 | 0325 | -00375 | | 10 |
| 6855 | ENSMUSG( | Sod1 | 20655 | -00332712 | -01580694 | 04575 | 02275 | | 10 |
| 6922 | ENSMUSG( | Spryd7 | 66674 | -01045527 | -01892837 | 0255 | 00325 | | 10 |
| 6941 | ENSMUSG( | Srgap2 | 14270 | 017544097 | 013624572 | 02775 | 03075 | | 10 |
| 6993 | ENSMUSG( | St7 | 64213 | 002057376 | 032497165 | 05425 | -00125 | | 10 |
| 7015 | ENSMUSG( | Stau2 | 29819 | 008536359 | -01311156 | 026 | 022 | | 10 |
| 7025 | ENSMUSG( | Stk16 | 20872 | 012185998 | -00476601 | 041 | 04175 | | 10 |
| 7034 | ENSMUSG( | Stk4 | 58231 | 021467554 | 006377015 | 024 | 00775 | | 10 |
| 7035 | ENSMUSG( | Stmn1 | 16765 | -00531185 | -01624573 | 0305 | 025 | | 10 |
| 7059 | ENSMUSG( | Stx17 | 67727 | 003491204 | -00814793 | 0385 | 03675 | | 10 |
| 7088 | ENSMUSG( | Sumf2 | 67902 | -01442552 | 020070615 | 0535 | 033 | | 10 |
| 7144 | ENSMUSG( | Syt16 | 238266 | 006053363 | 013562926 | 041 | 02375 | | 10 |
| 7169 | ENSMUSG( | Taf15 | 70439 | 006123254 | -01847161 | 092 | 01975 | | 10 |
| 7174 | ENSMUSG( | Taf6l | 225895 | 026072151 | 008501928 | 0285 | 016 | | 10 |
| 7215 | ENSMUSG( | Tbc1d19 | 67249 | 011824433 | 006407048 | 0355 | 0065 | | 10 |
| 7227 | ENSMUSG( | Tbc1d8 | 54610 | -02747115 | -02079442 | 03075 | -00075 | | 10 |
| 7230 | ENSMUSG( | Tbc1d9b | 76795 | 016814618 | -00678623 | 0345 | 0215 | | 10 |
| 7255 | ENSMUSG( | Tcf25 | 66855 | 010811194 | -01422571 | 02275 | 0165 | | 10 |
| 7264 | ENSMUSG( | Tdp1 | 104884 | 027150269 | 018887230 | 062 | -00775 | | 10 |
| 7293 | ENSMUSG( | Tfb1m | 224481 | 016209397 | -02123504 | 03075 | -00175 | | 10 |
| 7317 | ENSMUSG( | Thnsl2 | 232078 | 011593957 | -00872904 | 05425 | 00975 | | 10 |
| 7318 | ENSMUSG( | Thoc1 | 225160 | 01057524 | 019935313 | 04375 | -006 | | 10 |
| 7322 | ENSMUSG( | Thoc6 | 386612 | -00941716 | -01698153 | 03575 | -0015 | | 10 |
| 7365 | ENSMUSG( | Tle1 | 21885 | 024688818 | 007878543 | 0305 | 00975 | | 10 |
| 7376 | ENSMUSG( | Tm9sf3 | 107358 | -02436023 | 007459396 | 0585 | 0355 | | 10 |
| 7386 | ENSMUSG( | Tmed10 | 68581 | -02717421 | 001838786 | 053 | 00675 | | 10 |
| 7402 | ENSMUSG( | Tmem120a | 215210 | -00991223 | 010420821 | 056 | -00225 | | 10 |
| 7414 | ENSMUSG( | Tmem135 | 72759 | -00087665 | 003447073 | 03725 | -00625 | | 10 |
| 7420 | ENSMUSG( | Tmem14c | 66154 | -02298993 | -00473818 | 0405 | 01175 | | 10 |
| 7441 | ENSMUSG( | Tmem199 | 195040 | -01398939 | 006708704 | 08325 | 03225 | | 10 |
| 7447 | ENSMUSG( | Tmem214 | 68796 | 006473566 | 018372051 | 0525 | -01825 | | 10 |
| 7485 | ENSMUSG( | Tmem87b | 72477 | -03622454 | 010649330 | 088 | 03325 | | 10 |
| 7507 | ENSMUSG( | Tnfaip1 | 21927 | 009743117 | -00209076 | 028 | 0305 | | 10 |
| 7512 | ENSMUSG( | Tnik | 665113 | -00047272 | 013332130 | 034 | 00775 | | 10 |
| 7542 | ENSMUSG( | Top3b | 21976 | 009503656 | 002184303 | 03425 | 01125 | | 10 |
| 7549 | ENSMUSG( | Tox2 | 269389 | 010636034 | -02776694 | 041 | 00825 | | 10 |
| 7553 | ENSMUSG( | Tpcn1 | 252972 | -00825343 | -00663969 | 05575 | 009 | | 10 |
| 7558 | ENSMUSG( | Tpgs2 | 66648 | 008166179 | -00997720 | 04025 | 04075 | | 10 |
| 7561 | ENSMUSG( | Tpk1 | 29807 | -02191146 | -02304996 | 0465 | 01975 | | 10 |
| 7567 | ENSMUSG( | Tpp2 | 22019 | 001418704 | 006812879 | 03475 | 0055 | | 10 |
| 7615 | ENSMUSG( | Trim32 | 69807 | -00004214 | -00971849 | 02625 | 00375 | | 10 |
| 7632 | ENSMUSG( | Trip4 | 56404 | 011909002 | 000091593 | 0225 | 00825 | | 10 |
| 7672 | ENSMUSG( | Tspan14 | 52588 | -01365752 | 002295120 | 0375 | 003 | | 10 |
| 7685 | ENSMUSG( | Tspyl1 | 22110 | 005001692 | -01020378 | 0405 | 04325 | | 10 |
| 7690 | ENSMUSG( | Tst | 22117 | 006416124 | -03469824 | 04 | 007 | | 10 |
| 7702 | ENSMUSG( | Ttc38 | 239570 | -00466483 | -00872982 | 02575 | 006 | | 10 |
| 7713 | ENSMUSG( | Ttll1 | 319953 | 001676233 | -01782124 | 06325 | 00625 | | 10 |
| 7714 | ENSMUSG( | Ttll12 | 223723 | 026952284 | -00808474 | 02325 | 01225 | | 10 |
| 7725 | ENSMUSG( | Tuba1b | 22143 | 006377912 | -02262866 | 034 | -001 | | 10 |
| 7739 | ENSMUSG( | Tubgcp4 | 51885 | 004976144 | 000965724 | 03125 | -00275 | | 10 |
| 7760 | ENSMUSG( | Txndc9 | 98258 | -00420974 | -01010113 | 03175 | 006 | | 10 |
| 7767 | ENSMUSG( | Txnrd3 | 232223 | -00441857 | -01188146 | 02725 | 02725 | | 10 |
| 7776 | ENSMUSG( | Uaca | 72565 | -00421136 | 032351676 | 0545 | 01 | | 10 |
| 7799 | ENSMUSG( | Ube2g2 | 22213 | 006663488 | -00033533 | 0525 | 01875 | | 10 |
| 7801 | ENSMUSG( | Ube2i | 22196 | -00263664 | 008587631 | 03225 | 025 | | 10 |
| 7802 | ENSMUSG( | Ube2j1 | 56228 | 000445748 | -00767881 | 0435 | 05325 | | 10 |
| 7807 | ENSMUSG( | Ube2n | 93765 | 007245782 | -01021705 | 0265 | 0005 | | 10 |
| 7850 | ENSMUSG( | Uchl5 | 56207 | 012069335 | -01578019 | 055 | 037 | | 10 |
| 7851 | ENSMUSG( | Uck1 | 22245 | 006312147 | -00828878 | 04475 | 02375 | | 10 |
| 7861 | ENSMUSG( | Ufsp2 | 192169 | 004003429 | -01733316 | 0625 | 01075 | | 10 |
| 7864 | ENSMUSG( | Uggt1 | 320011 | -01196084 | 015854615 | 0455 | -0135 | | 10 |
| 7878 | ENSMUSG( | Unc45a | 101869 | 010208543 | -00025287 | 04175 | 0285 | | 10 |
| 7891 | ENSMUSG( | Upp1 | 22271 | 034362148 | 021183946 | 057 | 01175 | | 10 |
| 7910 | ENSMUSG( | Uros | 22276 | 001029720 | -00175222 | 029 | 02625 | | 10 |
| 7929 | ENSMUSG( | Usp32 | 237898 | 007631271 | -00512938 | 03225 | 01975 | | 10 |
| 7940 | ENSMUSG( | Usp48 | 170707 | -00325930 | -00310787 | 05675 | 036 | | 10 |
| 7964 | ENSMUSG( | Vamp4 | 53330 | 006813838 | 005159897 | 045 | 0555 | | 10 |
| 7967 | ENSMUSG( | Vapa | 30960 | -00093895 | -01435989 | 04375 | 0025 | | 10 |
| 7974 | ENSMUSG( | Vat1 | 26949 | -00676670 | -00538220 | 0605 | 02 | | 10 |
| 7977 | ENSMUSG( | Vav2 | 22325 | 003597570 | 015393226 | 0255 | 02375 | | 10 |
| 8039 | ENSMUSG( | Vrk3 | 101568 | 003691268 | -00813545 | 032 | 02675 | | 10 |
| 8071 | ENSMUSG( | Wdcp | 238037 | 027465753 | 021907125 | 04575 | 02825 | | 10 |
| 8085 | ENSMUSG( | Wdr26 | 226757 | 007953173 | 004665797 | 0325 | 00175 | | 10 |
| 8102 | ENSMUSG( | Wdr6 | 83669 | 002901241 | 005917600 | 03725 | 02675 | | 10 |
| 8108 | ENSMUSG( | Wdr82 | 77305 | 013053509 | -00774657 | 02425 | 01725 | | 10 |
| 8129 | ENSMUSG( | Wwox | 80707 | 012713657 | -00223497 | 03325 | 00775 | | 10 |
| 8134 | ENSMUSG( | Xiap | 11798 | -00014979 | -00282245 | 026 | 02075 | | 10 |
| 8145 | ENSMUSG( | Xpo6 | 74204 | 008563261 | 015103030 | 049 | 013 | | 10 |
| 8150 | ENSMUSG( | Xrcc4 | 108138 | -00547969 | -02727957 | 03175 | 014 | | 10 |
| 8166 | ENSMUSG( | Yes1 | 22612 | 002167029 | -00735003 | 025 | 027 | | 10 |
| 8167 | ENSMUSG( | Yif1b | 77254 | -02779008 | -00975604 | 03725 | -003 | | 10 |
| 8174 | ENSMUSG( | Yju2 | 72886 | -00380261 | -00562373 | 05075 | 01575 | | 10 |
| 8227 | ENSMUSG( | Zdhhc3 | 69035 | -00377269 | 000705443 | 06425 | 02925 | | 10 |
| 8252 | ENSMUSG( | Zfp512b | 269401 | 009057487 | -00215635 | 06675 | 0325 | | 10 |
| 8275 | ENSMUSG( | Zmpste24 | 230709 | 002340108 | 011188334 | 0395 | -0115 | | 10 |
| 8 | ENSMUSG( | 2410002F2 | 668661 | -02761830 | -01143990 | 02425 | -023 | | 11 |
| 18 | ENSMUSG( | 6820408C1 | 228778 | -01520464 | 012272782 | -0075 | -0285 | | 11 |
| 35 | ENSMUSG( | Abca1 | 11303 | -01955175 | 031516063 | -00525 | -0265 | | 11 |
| 38 | ENSMUSG( | Abca5 | 217265 | -01211934 | 021512130 | -00125 | -032 | | 11 |
| 39 | ENSMUSG( | Abca6 | 76184 | -03237227 | 048962749 | -03625 | -07025 | | 11 |
| 48 | ENSMUSG( | Abcb9 | 56325 | -03368162 | -00664239 | 00875 | -021 | | 11 |
| 61 | ENSMUSG( | Abcg1 | 11307 | -04503070 | 021313270 | -01675 | -03775 | | 11 |
| 100 | ENSMUSG( | Acan | 11595 | -04078297 | 032618879 | -02 | 001 | | 11 |
| 110 | ENSMUSG( | Ache | 11423 | -03322410 | 002857048 | -00775 | -00375 | | 11 |
| 130 | ENSMUSG( | Acsl1 | 14081 | -02316101 | 008960993 | -01425 | -033 | | 11 |
| 132 | ENSMUSG( | Acsl4 | 50790 | -04626666 | 030244055 | 03675 | -04075 | | 11 |
| 133 | ENSMUSG( | Acsl5 | 433256 | -01308072 | 017606607 | -01075 | -0515 | | 11 |
| 141 | ENSMUSG( | Actn1 | 109711 | -01441567 | 013595969 | -00375 | -04725 | | 11 |
| 165 | ENSMUSG( | Adam22 | 11496 | -02984155 | 012208854 | 0075 | -01975 | | 11 |

| | | | | | | | | |
|---|---|---|---|---|---|---|---|---|
| 166 | ENSMUSG( | Adam23 | 23792 | -05052600 | 008386901 | -01825 | -0225 | 11 |
| 178 | ENSMUSG( | Adcy5 | 224129 | -03181793 | 013800625 | -036 | -03375 | 11 |
| 192 | ENSMUSG( | Adgrb3 | 210933 | -01751392 | 022894229 | -01425 | -0145 | 11 |
| 195 | ENSMUSG( | Adgrg1 | 14766 | -03883425 | -00205771 | 002 | -0335 | 11 |
| 269 | ENSMUSG( | Ajm1 | 381353 | -02827834 | 007147735 | -0265 | -031 | 11 |
| 314 | ENSMUSG( | Alg1 | 208211 | -02941925 | 009858082 | -003 | -00125 | 11 |
| 320 | ENSMUSG( | Alg5 | 66248 | -01854079 | 013856011 | -002 | -01025 | 11 |
| 338 | ENSMUSG( | Amigo1 | 229715 | -01946890 | 015277605 | 00375 | -0065 | 11 |
| 358 | ENSMUSG( | Ank3 | 11735 | -02758577 | 009173715 | -015 | -0115 | 11 |
| 363 | ENSMUSG( | Ankle2 | 71782 | -02002344 | 025817855 | -009 | 00375 | 11 |
| 390 | ENSMUSG( | Ano10 | 102566 | -01555502 | 016435554 | 01625 | -0055 | 11 |
| 393 | ENSMUSG( | Ano8 | 382014 | -01066603 | 015066891 | 00775 | -007 | 11 |
| 449 | ENSMUSG( | Aplp2 | 11804 | -02187207 | 007781006 | 0105 | -0135 | 11 |
| 450 | ENSMUSG( | Apmap | 71881 | -03035832 | 018819789 | 022 | -015 | 11 |
| 474 | ENSMUSG( | Arel1 | 68497 | -01100867 | 012331011 | 002 | -021 | 11 |
| 485 | ENSMUSG( | Arfgef3 | 215821 | -00737319 | 010009223 | -0065 | -01075 | 11 |
| 490 | ENSMUSG( | Arglu1 | 234023 | -01347100 | 01972714C | -0095 | -003 | 11 |
| 534 | ENSMUSG( | Arid4a | 238247 | -00780057 | 027874909 | -00725 | -0315 | 11 |
| 539 | ENSMUSG( | Arl10 | 56795 | -03422023 | 018212998 | 0195 | -03725 | 11 |
| 578 | ENSMUSG( | Arsa | 11883 | -01444266 | 015493391 | -0185 | -0105 | 11 |
| 585 | ENSMUSG( | Asah1 | 11886 | -03129248 | 017788497 | -02375 | 003 | 11 |
| 586 | ENSMUSG( | Asah2 | 54447 | -02523599 | -00082748 | 02625 | -01975 | 11 |
| 610 | ENSMUSG( | Astn2 | 56079 | -02531024 | 011044031 | -00075 | -0205 | 11 |
| 633 | ENSMUSG( | Atl1 | 73991 | -04167337 | 011336891 | 0265 | -012 | 11 |
| 637 | ENSMUSG( | Atp11a | 50770 | -01668327 | 022725124 | 01075 | -02 | 11 |
| 639 | ENSMUSG( | Atp11c | 320940 | -03608836 | 035534749 | 023 | -0175 | 11 |
| 644 | ENSMUSG( | Atp1a1 | 11928 | -02424073 | 008902594 | -00125 | -0245 | 11 |
| 646 | ENSMUSG( | Atp1a3 | 232975 | -04245618 | 007518644 | 00625 | -007 | 11 |
| 647 | ENSMUSG( | Atp1b1 | 11931 | -05505797 | 011970009 | -00125 | -0195 | 11 |
| 648 | ENSMUSG( | Atp1b2 | 11932 | -01644274 | 003638388 | 0085 | -00725 | 11 |
| 651 | ENSMUSG( | Atp2a2 | 11938 | -03283185 | 013069563 | 0085 | -0225 | 11 |
| 653 | ENSMUSG( | Atp2b1 | 67972 | -05223947 | 020133446 | -00425 | -023 | 11 |
| 654 | ENSMUSG( | Atp2b2 | 11941 | -03754996 | 005394553 | -011 | -03125 | 11 |
| 655 | ENSMUSG( | Atp2b3 | 320707 | -01684691 | 036131693 | -0065 | -019 | 11 |
| 656 | ENSMUSG( | Atp2b4 | 381290 | -03368110 | 01213177C | 0145 | -012 | 11 |
| 671 | ENSMUSG( | Atp6ap1 | 54411 | -02610466 | 001983165 | 0225 | -0225 | 11 |
| 673 | ENSMUSG( | Atp6v0a1 | 11975 | -0266179 | 010303981 | -0065 | -0075 | 11 |
| 686 | ENSMUSG( | Atp7a | 11977 | -00985993 | 023960205 | -01825 | -037 | 11 |
| 687 | ENSMUSG( | Atp8a1 | 11980 | -02789607 | 019782594 | 00675 | -018 | 11 |
| 688 | ENSMUSG( | Atp8a2 | 50769 | -03316765 | 020158633 | 0235 | -0265 | 11 |
| 694 | ENSMUSG( | Atraid | 381629 | -01902346 | 011745308 | -00125 | -053 | 11 |
| 714 | ENSMUSG( | B2m | 12010 | -03272332 | -00924419 | 0015 | -006 | 11 |
| 722 | ENSMUSG( | B4galt3 | 57370 | -01301007 | 024037182 | 785046e-1 | -042 | 11 |
| 755 | ENSMUSG( | Bcan | 12032 | -00835283 | 010391553 | -00025 | -009 | 11 |
| 757 | ENSMUSG( | Bcap31 | 27061 | -03658458 | -00614203 | -0135 | -01475 | 11 |
| 786 | ENSMUSG( | Bfar | 67118 | -01822197 | 01797190C | 04 | -02475 | 11 |
| 787 | ENSMUSG( | Bgn | 12111 | -02314710 | 033374366 | -00725 | -034 | 11 |
| 798 | ENSMUSG( | Bivm | 246229 | -01405908 | 020949334 | 033 | -0495 | 11 |
| 846 | ENSMUSG( | Brinp2 | 240843 | -00547521 | 038995383 | -00075 | -02175 | 11 |
| 847 | ENSMUSG( | Brinp3 | 215378 | -02331011 | 018881468 | 01225 | -05675 | 11 |
| 863 | ENSMUSG( | Btbd17 | 72014 | -02647005 | -01167485 | 0205 | -01825 | 11 |
| 880 | ENSMUSG( | C1ql3 | 227580 | -03955357 | 042956881 | 02625 | -019 | 11 |
| 883 | ENSMUSG( | C2cd2l | 71764 | -02960283 | 022489745 | 007 | -00175 | 11 |
| 898 | ENSMUSG( | Cacna1a | 12286 | -00233223 | 024372637 | -01 | -0215 | 11 |
| 900 | ENSMUSG( | Cacna1c | 12288 | -03540790 | 015124362 | -0165 | -0255 | 11 |
| 902 | ENSMUSG( | Cacna1e | 12290 | -02742835 | 020965496 | -01325 | -01875 | 11 |
| 906 | ENSMUSG( | Cacna2d1 | 12293 | -04992045 | 020378856 | -00925 | -0265 | 11 |
| 907 | ENSMUSG( | Cacna2d2 | 56808 | -03789513 | -00009347 | -00475 | -038 | 11 |
| 908 | ENSMUSG( | Cacna2d3 | 12294 | -01930576 | 016182209 | 01775 | -02575 | 11 |
| 913 | ENSMUSG( | Cacng2 | 12300 | -00628705 | 02599808C | 00375 | -02425 | 11 |
| 922 | ENSMUSG( | Cadm1 | 54725 | -02046985 | 009694151 | 03475 | -0245 | 11 |
| 973 | ENSMUSG( | Card19 | 68480 | -03348473 | -00861355 | -02525 | -0105 | 11 |
| 984 | ENSMUSG( | Casd1 | 213819 | -04705132 | 01781030C | -045 | -04425 | 11 |
| 997 | ENSMUSG( | Cav1 | 12389 | -02305211 | 030086777 | 0025 | -00325 | 11 |
| 1008 | ENSMUSG( | Cbln4 | 228942 | -01962792 | 02857563C | -03025 | -05425 | 11 |
| 1065 | ENSMUSG( | Cckbr | 12426 | -03675557 | 010029217 | 03275 | -0145 | 11 |
| 1067 | ENSMUSG( | Ccn2 | 14219 | -06086436 | 015040843 | 00025 | -01825 | 11 |
| 1092 | ENSMUSG( | Cd200 | 17470 | -03950362 | 037006468 | 01525 | -016 | 11 |
| 1103 | ENSMUSG( | Cd81 | 12520 | -04207907 | 002589941 | 005 | -02625 | 11 |
| 1106 | ENSMUSG( | Cd93 | 17064 | -02335881 | -00607218 | -0115 | -0425 | 11 |
| 1131 | ENSMUSG( | Cdh18 | 320865 | -04732845 | 021124984 | 01425 | -01375 | 11 |
| 1133 | ENSMUSG( | Cdh2 | 12558 | -01551175 | 02584987C | -235514e-1 | -0025 | 11 |
| 1136 | ENSMUSG( | Cdh4 | 12561 | -04283303 | 003324261 | 0065 | -01825 | 11 |
| 1139 | ENSMUSG( | Cdh7 | 241201 | -02780108 | 020822064 | 009 | -03925 | 11 |
| 1140 | ENSMUSG( | Cdh8 | 12564 | -05070262 | 025588847 | 0155 | -0145 | 11 |
| 1143 | ENSMUSG( | Cdipt | 52858 | -03284208 | 00703270C | 0405 | -0195 | 11 |
| 1190 | ENSMUSG( | Cep97 | 74201 | -01667568 | 004486363 | -012 | -00575 | 11 |
| 1191 | ENSMUSG( | Cept1 | 99712 | -03345345 | 025256988 | 045 | -0295 | 11 |
| 1200 | ENSMUSG( | Cfap20 | 14894 | -01587402 | 00496965C | 00925 | -022 | 11 |
| 1217 | ENSMUSG( | Chat | 12647 | -05177634 | -03421565 | 02125 | -018 | 11 |
| 1230 | ENSMUSG( | Chd6 | 71389 | -02131833 | 003015807 | 0015 | -01 | 11 |
| 1233 | ENSMUSG( | Chdh | 218865 | -01777334 | 032752971 | -0035 | -03975 | 11 |
| 1262 | ENSMUSG( | Chrm3 | 12671 | -02786399 | 02334297C | 00275 | -015 | 11 |
| 1264 | ENSMUSG( | Chrna4 | 11438 | -03713290 | 009923749 | 003 | -00575 | 11 |
| 1265 | ENSMUSG( | Chrnb2 | 11444 | -02666131 | 022679347 | 02725 | -027 | 11 |
| 1280 | ENSMUSG( | Cisd2 | 67006 | -03567150 | 006530451 | 0305 | -0125 | 11 |
| 1293 | ENSMUSG( | Clcn3 | 12725 | -02297240 | 004899023 | -00625 | -02275 | 11 |
| 1294 | ENSMUSG( | Clcn4 | 12727 | -02245611 | 006419517 | 01225 | -008 | 11 |
| 1298 | ENSMUSG( | Cldn1 | 12737 | -03828819 | 028319081 | -00925 | -012 | 11 |
| 1326 | ENSMUSG( | Clptm1l | 218335 | -02392703 | 003438043 | 01 | -02 | 11 |
| 1329 | ENSMUSG( | Clstn2 | 64085 | -03305963 | 002550256 | -0265 | -052 | 11 |
| 1353 | ENSMUSG( | Cnep1r1 | 382030 | -02237529 | 006536757 | 02075 | -01625 | 11 |
| 1361 | ENSMUSG( | Cnnm1 | 83674 | -02951656 | 007621276 | -00975 | 002 | 11 |
| 1362 | ENSMUSG( | Cnnm2 | 94219 | -02193855 | 023576464 | -00325 | -0115 | 11 |
| 1376 | ENSMUSG( | Cnpy3 | 72029 | -03358152 | -00165320 | 0125 | -02125 | 11 |
| 1378 | ENSMUSG( | Cnr1 | 12801 | -05989990 | 027578599 | -00225 | -006 | 11 |
| 1381 | ENSMUSG( | Cntfr | 12804 | -03013179 | 00560350C | 004 | -002 | 11 |
| 1383 | ENSMUSG( | Cntn2 | 21367 | -05773684 | 001467005 | -0115 | -04375 | 11 |
| 1388 | ENSMUSG( | Cntnap1 | 53321 | -02138191 | 003254251 | 0025 | -03 | 11 |
| 1389 | ENSMUSG( | Cntnap2 | 66797 | -04447561 | 01733856C | 00975 | -01425 | 11 |
| 1392 | ENSMUSG( | Cntnap5a | 636808 | -04920342 | 015358412 | 00225 | -002 | 11 |
| 1403 | ENSMUSG( | Coch | 12810 | -04353880 | 015370392 | 0305 | -0125 | 11 |
| 1495 | ENSMUSG( | Cp | 12870 | -02073647 | 028829616 | -00875 | -0065 | 11 |
| 1496 | ENSMUSG( | Cpd | 12874 | -04221297 | 013639252 | -00375 | -01875 | 11 |
| 1500 | ENSMUSG( | Cpeb4 | 67579 | -03049855 | -00555453 | -00125 | -0125 | 11 |
| 1504 | ENSMUSG( | Cpm | 70574 | -02040213 | 076804811 | -00675 | -02075 | 11 |
| 1540 | ENSMUSG( | Crhbp | 12919 | -03988326 | 001013211 | 009 | -03725 | 11 |
| 1555 | ENSMUSG( | Crtac1 | 72832 | -02693273 | 011828556 | 008 | -02475 | 11 |
| 1576 | ENSMUSG( | Csmd2 | 329942 | -04411019 | -00812951 | 02 | -01325 | 11 |
| 1601 | ENSMUSG( | Ctbs | 74245 | -03904337 | 004166492 | 01075 | -02 | 11 |
| 1623 | ENSMUSG( | Ctsb | 13030 | -03242890 | 005813416 | 0145 | -00375 | 11 |
| 1653 | ENSMUSG( | Cx3cl1 | 20312 | -03772966 | 001683095 | 0205 | -00975 | 11 |
| 1664 | ENSMUSG( | Cyb5r1 | 72017 | -01549777 | 015510944 | -00225 | -005 | 11 |
| 1667 | ENSMUSG( | Cybc1 | 217370 | -02106752 | 011277242 | -018 | -02325 | 11 |
| 1682 | ENSMUSG( | Cyp4x1 | 81906 | -03436504 | -01050188 | 03275 | -042 | 11 |
| 1693 | ENSMUSG( | D17H6S53I | 114585 | -03126280 | 012369672 | -0065 | -0175 | 11 |
| 1696 | ENSMUSG( | D430041D( | 241589 | -02590884 | 019616963 | -013 | -0035 | 11 |
| 1697 | ENSMUSG( | D630045J1 | 330286 | -01379477 | 04178100C | 01175 | -00575 | 11 |
| 1710 | ENSMUSG( | Daglb | 231871 | -02891397 | 002435461 | -009 | -0485 | 11 |
| 1741 | ENSMUSG( | Dcn | 13179 | -02086302 | 00402081C | -00875 | -0135 | 11 |
| 1770 | ENSMUSG( | Ddr1 | 12305 | -05174237 | 015033056 | 0095 | -051 | 11 |
| 1833 | ENSMUSG( | Dgke | 56077 | -04217968 | 016030084 | 02075 | -017 | 11 |
| 1852 | ENSMUSG( | Dhrs7 | 66375 | -03416114 | 004091667 | 0075 | -0325 | 11 |
| 1905 | ENSMUSG( | Dmxl1 | 240283 | -00628548 | 026393561 | -008 | -0135 | 11 |
| 1906 | ENSMUSG( | Dmxl2 | 235380 | -01025104 | 01761674C | 001 | -01575 | 11 |
| 1918 | ENSMUSG( | Dnajb14 | 70604 | -01304682 | 014559495 | 024 | -009 | 11 |

| | | | | | | | | | |
|---|---|---|---|---|---|---|---|---|---|
| 1930 | ENSMUSG( | Dnajc16 | 214063 | -02739699 | 02237644 | 02975 | -026 | | 11 |
| 1990 | ENSMUSG( | Dpm3 | 68563 | -02552419 | -00814598 | -00775 | -027 | | 11 |
| 1991 | ENSMUSG( | Dpp10 | 269109 | -02738030 | 03001751 | 0105 | -03175 | | 11 |
| 1994 | ENSMUSG( | Dpp6 | 13483 | -03190461 | 01087676 | 01025 | -0145 | | 11 |
| 2067 | ENSMUSG( | Ece1 | 230857 | -02795697 | 01530822 | 01325 | -0285 | | 11 |
| 2086 | ENSMUSG( | Eed | 13626 | -01375701 | 00856734 | 009 | -0355 | | 11 |
| 2102 | ENSMUSG( | Efna3 | 13638 | -03529964 | 01865383 | -001 | -02475 | | 11 |
| 2111 | ENSMUSG( | Egfr | 13649 | -02074532 | -00603777 | -00325 | -038 | | 11 |
| 2177 | ENSMUSG( | Elapor2 | 231014 | -03208477 | 00885568 | 00625 | -005 | | 11 |
| 2186 | ENSMUSG( | Elmod1 | 270162 | -07026991 | 00647256 | 022 | -013 | | 11 |
| 2187 | ENSMUSG( | Elmod2 | 244548 | -02861929 | 03123629 | -01575 | -048 | | 11 |
| 2192 | ENSMUSG( | Elovl1 | 54325 | -04632490 | 01944695 | 00275 | -01725 | | 11 |
| 2193 | ENSMUSG( | Elovl4 | 83603 | -05373124 | 01428795 | 0325 | -03325 | | 11 |
| 2201 | ENSMUSG( | Emb | 13723 | -02563943 | 01105069 | 01125 | -0355 | | 11 |
| 2203 | ENSMUSG( | Emc10 | 69683 | -02669049 | 00513126 | -01275 | 00575 | | 11 |
| 2223 | ENSMUSG( | Endod1 | 71946 | -04705817 | 02438012 | 01225 | -00325 | | 11 |
| 2233 | ENSMUSG( | Enpep | 13809 | -02425329 | 05545629 | 00025 | -00475 | | 11 |
| 2237 | ENSMUSG( | Enpp5 | 83965 | -04466747 | 00624316 | 01725 | -02325 | | 11 |
| 2238 | ENSMUSG( | Enpp6 | 320981 | -08107221 | 02574138 | 0005 | -07225 | | 11 |
| 2241 | ENSMUSG( | Entpd2 | 12496 | -02313932 | 00864904 | 024 | -03225 | | 11 |
| 2268 | ENSMUSG( | Ephx1 | 13849 | -03472458 | 03034621 | -0125 | -03325 | | 11 |
| 2280 | ENSMUSG( | Eps8l2 | 98845 | -02779643 | -00678017 | 02925 | -039 | | 11 |
| 2282 | ENSMUSG( | Erap1 | 80898 | -02289027 | 00945487 | 01875 | -0385 | | 11 |
| 2284 | ENSMUSG( | Erbb4 | 13869 | -02826758 | 02661923 | 02 | -01275 | | 11 |
| 2302 | ENSMUSG( | Ermp1 | 226090 | -03055147 | 01768084 | 196262e-1 | -02025 | | 11 |
| 2310 | ENSMUSG( | Esyt1 | 23943 | -01740594 | 03863567 | -01775 | -03 | | 11 |
| 2311 | ENSMUSG( | Esyt2 | 52635 | -02440560 | 01963840 | 0245 | -04625 | | 11 |
| 2367 | ENSMUSG( | Faf2 | 76577 | -02631509 | 00833701 | 0 | -01725 | 11 | |
| 2394 | ENSMUSG( | Fam171a1 | 269233 | -01962319 | 02301566 | 003 | -01175 | | 11 |
| 2408 | ENSMUSG( | Fam234a | 106581 | -03449565 | 00175617 | 015 | -0305 | | 11 |
| 2471 | ENSMUSG( | Fcgr1 | 14129 | -03948199 | 02516873 | -0135 | -00025 | | 11 |
| 2497 | ENSMUSG( | Fgf14 | 14169 | -02377134 | -00416023 | 01975 | -03275 | | 11 |
| 2518 | ENSMUSG( | Fitm2 | 228859 | -02435572 | 00994705 | 03225 | -01075 | | 11 |
| 2527 | ENSMUSG( | Fkbp9 | 27055 | -01416210 | -00074258 | -002 | -023 | | 11 |
| 2537 | ENSMUSG( | Flrt1 | 396184 | -01009732 | 02424804 | -01425 | -023 | | 11 |
| 2563 | ENSMUSG( | Folr1 | 14275 | -00489450 | 05461168 | -01025 | -05575 | | 11 |
| 2572 | ENSMUSG( | Fras1 | 231470 | -06475376 | 00735859 | 012 | -00625 | | 11 |
| 2590 | ENSMUSG( | Fstl4 | 320027 | -05434082 | 01373204 | -00375 | -0125 | | 11 |
| 2616 | ENSMUSG( | Fzd2 | 57265 | -00186546 | 07680712 | 02975 | -03875 | | 11 |
| 2629 | ENSMUSG( | Gabbr2 | 242425 | -02520711 | 01314366 | -0095 | -018 | | 11 |
| 2631 | ENSMUSG( | Gabra1 | 14394 | -07127867 | 02746555 | -02275 | -054 | | 11 |
| 2642 | ENSMUSG( | Gabrg3 | 14407 | -03700834 | 02887427 | 026 | -01825 | | 11 |
| 2690 | ENSMUSG( | Gbp5 | 229898 | -00833747 | 05291919 | -02125 | -02525 | | 11 |
| 2738 | ENSMUSG( | Ggt7 | 207182 | -01552657 | 01008538 | 02625 | -01975 | | 11 |
| 2766 | ENSMUSG( | Glmn | 170823 | -01471932 | 00102013 | 00175 | -02775 | | 11 |
| 2770 | ENSMUSG( | Glrb | 14658 | -04873390 | 01989702 | 01975 | -01475 | | 11 |
| 2783 | ENSMUSG( | Gm19410 | 1.01E+08 | -01921995 | 00431770 | 017 | -018 | | 11 |
| 2821 | ENSMUSG( | Gng7 | 14708 | -04492751 | -00894160 | -0105 | -019 | | 11 |
| 2830 | ENSMUSG( | Gns | 75612 | -03027492 | 01867762 | 0185 | -02 | | 11 |
| 2862 | ENSMUSG( | Gpc4 | 14735 | -04635342 | 00941699 | 00375 | -015 | | 11 |
| 2884 | ENSMUSG( | Gpr37l1 | 171469 | -03016171 | -00114430 | 014 | -03625 | | 11 |
| 2912 | ENSMUSG( | Gria1 | 14799 | -01607119 | 01579081 | 0105 | -03275 | | 11 |
| 2913 | ENSMUSG( | Gria2 | 14800 | -03085958 | 03853031 | -00125 | -05875 | | 11 |
| 2914 | ENSMUSG( | Gria3 | 53623 | -01972306 | 01920645 | 02075 | -02725 | | 11 |
| 2915 | ENSMUSG( | Gria4 | 14802 | -03982424 | -00055080 | 02175 | -02125 | | 11 |
| 2916 | ENSMUSG( | Grid1 | 14803 | -02674778 | 01663295 | 00075 | -00975 | | 11 |
| 2919 | ENSMUSG( | Grik2 | 14806 | -02553594 | 01717511 | 01025 | -03625 | | 11 |
| 2923 | ENSMUSG( | Grin2a | 14811 | -03786988 | 01953487 | 00475 | -008 | | 11 |
| 2924 | ENSMUSG( | Grin2b | 14812 | -03788112 | 02088925 | 026 | -014 | | 11 |
| 2933 | ENSMUSG( | Grm1 | 14816 | -06289053 | 02493951 | 017 | -027 | | 11 |
| 2934 | ENSMUSG( | Grm2 | 108068 | -02998499 | 00069647 | -0195 | -03625 | | 11 |
| 2936 | ENSMUSG( | Grm4 | 268934 | -03963022 | 00139888 | -01325 | -01225 | | 11 |
| 2937 | ENSMUSG( | Grm5 | 108071 | -04898618 | 02812662 | -0095 | -003 | | 11 |
| 2949 | ENSMUSG( | Gsg1l | 269994 | -02370966 | 00310368 | -00225 | -0345 | | 11 |
| 2997 | ENSMUSG( | Guf1 | 231279 | -01209372 | 01207642 | -00575 | -0055 | | 11 |
| 3024 | ENSMUSG( | Hapln1 | 12950 | -04097000 | 00461761 | 02275 | -0375 | | 11 |
| 3026 | ENSMUSG( | Hapln3 | 67666 | -02513629 | 02871322 | 03325 | -01575 | | 11 |
| 3037 | ENSMUSG( | Hcn1 | 15165 | -04006795 | 01932440 | -01375 | 006 | | 11 |
| 3040 | ENSMUSG( | Hcn4 | 330953 | -05937792 | 00169409 | 00875 | -00425 | | 11 |
| 3088 | ENSMUSG( | Hhatl | 74770 | -03955541 | 00004330 | 00025 | -03525 | | 11 |
| 3116 | ENSMUSG( | Hmgcr | 15357 | -03535991 | -00109747 | 0195 | -00925 | | 11 |
| 3125 | ENSMUSG( | Hnmt | 140483 | -02737493 | 00684294 | -01375 | -01725 | | 11 |
| 3133 | ENSMUSG( | Hnrnpdl | 50926 | -01659928 | 01517862 | 00575 | -03 | | 11 |
| 3172 | ENSMUSG( | Hsd17b12 | 56348 | -02399152 | -00376260 | 018 | -00625 | | 11 |
| 3207 | ENSMUSG( | Htra1 | 56213 | -02805329 | 00744029 | 01975 | -01675 | | 11 |
| 3222 | ENSMUSG( | Icam5 | 15898 | -00501013 | 02527085 | 00025 | -01775 | | 11 |
| 3260 | ENSMUSG( | Iglon5 | 210094 | -01889580 | 00317912 | -015 | -0095 | | 11 |
| 3262 | ENSMUSG( | Igsf11 | 207683 | -01153588 | 03387130 | -0135 | -02525 | | 11 |
| 3275 | ENSMUSG( | Il1rapl1 | 331461 | -06262915 | 02310165 | 00925 | -0075 | | 11 |
| 3284 | ENSMUSG( | Ilvbl | 216136 | -03277384 | 00830867 | 02575 | -01975 | | 11 |
| 3306 | ENSMUSG( | Inpp5f | 101490 | -02769530 | -00594226 | -01325 | -001 | | 11 |
| 3311 | ENSMUSG( | Insr | 16337 | -01475414 | 02677899 | 0165 | -01375 | | 11 |
| 3375 | ENSMUSG( | Itgam | 16409 | -01577915 | 00694906 | -014 | -02375 | | 11 |
| 3376 | ENSMUSG( | Itgav | 16410 | -01313551 | 01573329 | 0035 | -01425 | | 11 |
| 3379 | ENSMUSG( | Itgb2 | 16414 | -03641230 | -02109453 | -002 | -02875 | | 11 |
| 3389 | ENSMUSG( | Itm2b | 16432 | -02199223 | 01110405 | 00725 | -01325 | | 11 |
| 3390 | ENSMUSG( | Itm2c | 64294 | -03037049 | 00622420 | 02075 | -01875 | | 11 |
| 3410 | ENSMUSG( | Jam2 | 67374 | -01347905 | 00099261 | 0055 | -022 | | 11 |
| 3417 | ENSMUSG( | Jph1 | 57339 | -02295111 | 00413938 | 00125 | -01575 | | 11 |
| 3444 | ENSMUSG( | Kcna1 | 16485 | -03302307 | 05821334 | -02725 | -07025 | | 11 |
| 3446 | ENSMUSG( | Kcna3 | 16491 | -03342720 | 04380037 | -0005 | -01175 | | 11 |
| 3449 | ENSMUSG( | Kcnab1 | 16497 | -02915424 | -01702384 | -002 | -04075 | | 11 |
| 3452 | ENSMUSG( | Kcnb1 | 16500 | -01567479 | 01781858 | -0135 | -03025 | | 11 |
| 3453 | ENSMUSG( | Kcnb2 | 98741 | -04356281 | 01417208 | -03375 | -01225 | | 11 |
| 3454 | ENSMUSG( | Kcnc1 | 16502 | -03347191 | 00663706 | -019 | -04625 | | 11 |
| 3455 | ENSMUSG( | Kcnc2 | 268345 | -02640381 | 01324988 | 03375 | -038 | | 11 |
| 3456 | ENSMUSG( | Kcnc3 | 16504 | -03011633 | 02111624 | -019 | -04525 | | 11 |
| 3459 | ENSMUSG( | Kcnd2 | 16508 | -04837510 | 02241726 | 0075 | -0255 | | 11 |
| 3466 | ENSMUSG( | Kcnh5 | 238271 | -04641369 | 00510360 | -00825 | -056 | | 11 |
| 3467 | ENSMUSG( | Kcnh7 | 170738 | -04642987 | 03218679 | -01075 | -00725 | | 11 |
| 3473 | ENSMUSG( | Kcnj11 | 16514 | -01595635 | 00793190 | -012 | -0265 | | 11 |
| 3476 | ENSMUSG( | Kcnj3 | 16519 | -05012007 | 00439120 | 0015 | -02775 | | 11 |
| 3477 | ENSMUSG( | Kcnj4 | 16520 | -04066707 | 01305498 | -012 | 0005 | | 11 |
| 3478 | ENSMUSG( | Kcnj6 | 16522 | -01072935 | 04799209 | -00725 | -02075 | | 11 |
| 3484 | ENSMUSG( | Kcnma1 | 16531 | -02392936 | 01761909 | 017 | -03525 | | 11 |
| 3489 | ENSMUSG( | Kcnq3 | 110862 | -02700001 | 01058801 | -001 | -0165 | | 11 |
| 3490 | ENSMUSG( | Kcnq5 | 226922 | -03902029 | 02532773 | -0255 | 01125 | | 11 |
| 3493 | ENSMUSG( | Kcnt2 | 240776 | -03637251 | 02857509 | -00175 | -01375 | | 11 |
| 3494 | ENSMUSG( | Kcnv1 | 67498 | -05800820 | 01374293 | -00425 | -05175 | | 11 |
| 3500 | ENSMUSG( | Kctd16 | 383348 | -01281321 | 02722906 | -00625 | -02275 | | 11 |
| 3573 | ENSMUSG( | Kmt2a | 214162 | -05876530 | 01628300 | -011 | -05175 | | 11 |
| 3620 | ENSMUSG( | Large1 | 16795 | -01740265 | 02354727 | 03375 | -03675 | | 11 |
| 3633 | ENSMUSG( | Lbr | 98386 | -04042101 | 02383449 | -0305 | -024 | | 11 |
| 3635 | ENSMUSG( | Lclat1 | 225010 | -03795625 | 01161304 | -0015 | -0095 | | 11 |
| 3645 | ENSMUSG( | Lemd2 | 224640 | -00574611 | 01567657 | 00175 | -0255 | | 11 |
| 3646 | ENSMUSG( | Lemd3 | 380664 | -03560565 | 01750436 | -0065 | -0325 | | 11 |
| 3657 | ENSMUSG( | Lgi1 | 56839 | -03708030 | 01687316 | 02075 | -0555 | | 11 |
| 3659 | ENSMUSG( | Lgi3 | 213469 | -04238574 | 01590179 | -00575 | -0505 | | 11 |
| 3681 | ENSMUSG( | Lingo2 | 242384 | -06381098 | -00836425 | 02325 | -00975 | | 11 |
| 3682 | ENSMUSG( | Lingo3 | 237403 | -05226602 | 00627829 | -03225 | -03775 | | 11 |
| 3694 | ENSMUSG( | Lmbrd2 | 320506 | -04373923 | 01494537 | -00925 | -0045 | | 11 |
| 3704 | ENSMUSG( | Lmtk2 | 231876 | -02835056 | 00644075 | -0125 | -0075 | | 11 |
| 3715 | ENSMUSG( | Lpcat4 | 99010 | -02533665 | 00093271 | 026 | -02 | | 11 |
| 3726 | ENSMUSG( | Lrfn1 | 80749 | -01597234 | 01913910 | 01625 | -02875 | | 11 |
| 3728 | ENSMUSG( | Lrfn3 | 233067 | -01222531 | 02679380 | -0075 | -038 | | 11 |
| 3731 | ENSMUSG( | Lrif1 | 321000 | -01485932 | 02296148 | 017 | -05325 | | 11 |

| | | | | | | | | |
|---|---|---|---|---|---|---|---|---|
| 3739 | ENSMUSG( | Lrpap1 | 16976 | -03063385 | 007820537 | 00825 | -00125 | 11 |
| 3750 | ENSMUSG( | Lrrc4b | 272381 | -02967262 | 025202934 | 00925 | -02675 | 11 |
| 3751 | ENSMUSG( | Lrrc4c | 241568 | -04552469 | 015944021 | 01925 | -03375 | 11 |
| 3758 | ENSMUSG( | Lrrc8a | 241296 | -01279831 | 01573029C | 00225 | -01725 | 11 |
| 3759 | ENSMUSG( | Lrrc8b | 433926 | -04137749 | 01909698S | -002 | -01475 | 11 |
| 3767 | ENSMUSG( | Lrrtm1 | 74342 | -03059053 | 03628382C | 03575 | -031 | 11 |
| 3768 | ENSMUSG( | Lrrtm2 | 107065 | -06299676 | 035033734 | 00975 | -027 | 11 |
| 3769 | ENSMUSG( | Lrrtm3 | 216028 | -04089188 | 03400347E | 014 | -00725 | 11 |
| 3770 | ENSMUSG( | Lrrtm4 | 243499 | -03753734 | 028398544 | 007 | -054 | 11 |
| 3829 | ENSMUSG( | M6pr | 17113 | -03879599 | 006076562 | -0275 | -006 | 11 |
| 3835 | ENSMUSG( | Macroh2a2 | 404634 | -03513820 | -00149639 | -0125 | -0285 | 11 |
| 3853 | ENSMUSG( | Mal2 | 105853 | -04681701 | 013275034 | -0015 | -007 | 11 |
| 3962 | ENSMUSG( | Mcam | 84004 | -06770589 | -00446909 | 008 | -033 | 11 |
| 3979 | ENSMUSG( | Mctp1 | 78771 | -04905108 | 036480947 | 008 | -00575 | 11 |
| 4028 | ENSMUSG( | Men1 | 17283 | -02492502 | 012035113 | -004 | -0355 | 11 |
| 4043 | ENSMUSG( | Mettl21a | 67099 | -03537615 | 019274742 | 015 | -0255 | 11 |
| 4052 | ENSMUSG( | Mfge8 | 17304 | -02037967 | 014323054 | 02825 | -0115 | 11 |
| 4057 | ENSMUSG( | Mfsd10 | 68294 | -02012603 | 01310680S | 0065 | -0035 | 11 |
| 4061 | ENSMUSG( | Mfsd4a | 213006 | -04754553 | 01412332C | -0055 | -00175 | 11 |
| 4090 | ENSMUSG( | Miga1 | 215708 | -01317012 | 01143558C | 00625 | -02325 | 11 |
| 4116 | ENSMUSG( | Mmgt1 | 236792 | -02436066 | 000760357 | 004 | -034 | 11 |
| 4162 | ENSMUSG( | Mrc2 | 17534 | -03646540 | 01703144S | -032 | -01 | 11 |
| 4264 | ENSMUSG( | Mtarc2 | 67247 | -01848044 | 004707744 | -00375 | -00625 | 11 |
| 4333 | ENSMUSG( | Mylk | 107589 | -01839244 | -00693990 | 00575 | -0335 | 11 |
| 4378 | ENSMUSG( | Naga | 17939 | -03398505 | 00633007C | 03375 | -02325 | 11 |
| 4397 | ENSMUSG( | Nat14 | 269854 | -02034292 | -00296188 | 00925 | -04125 | 11 |
| 4407 | ENSMUSG( | Nbas | 71169 | -02115224 | 003159362 | 01975 | -00625 | 11 |
| 4413 | ENSMUSG( | Ncam1 | 17967 | -03083064 | 010589825 | 0365 | -04575 | 11 |
| 4420 | ENSMUSG( | Nceh1 | 320024 | -04487716 | 02232177C | 0365 | -04225 | 11 |
| 4438 | ENSMUSG( | Ncstn | 59287 | -03151278 | 013443894 | 00175 | -02725 | 11 |
| 4439 | ENSMUSG( | Ndc1 | 72787 | -01181535 | 01693830C | 0035 | -0215 | 11 |
| 4442 | ENSMUSG( | Ndfip1 | 65113 | -05452165 | 016533683 | -011 | -02375 | 11 |
| 4502 | ENSMUSG( | Nectin1 | 58235 | -02239915 | 00939303E | -023 | -02875 | 11 |
| 4511 | ENSMUSG( | Negr1 | 320840 | -02810731 | 01476229S | 0155 | -02925 | 11 |
| 4523 | ENSMUSG( | Nenf | 66208 | -02706667 | -01457495 | 01275 | -03575 | 11 |
| 4526 | ENSMUSG( | Neto1 | 246317 | -02954705 | 020571683 | -011 | -01275 | 11 |
| 4527 | ENSMUSG( | Neto2 | 74513 | -05838896 | 02238819E | 02475 | -00925 | 11 |
| 4528 | ENSMUSG( | Neu1 | 18010 | -02432900 | 022250981 | 0345 | -05 | 11 |
| 4536 | ENSMUSG( | Nfasc | 269116 | -02060023 | 018964258 | -02025 | -0195 | 11 |
| 4558 | ENSMUSG( | Niban1 | 63913 | -02873964 | 017687868 | -01775 | -008 | 11 |
| 4580 | ENSMUSG( | Nlgn1 | 192167 | -03532736 | 020372623 | 01475 | -0075 | 11 |
| 4581 | ENSMUSG( | Nlgn2 | 216856 | -01720152 | 013478883 | 0235 | -0095 | 11 |
| 4582 | ENSMUSG( | Nlgn3 | 245537 | -03221298 | 018485581 | -00625 | -014 | 11 |
| 4591 | ENSMUSG( | Nmnat3 | 74080 | -02623345 | 01625585E | -02575 | -01 | 11 |
| 4647 | ENSMUSG( | Nptx1 | 18164 | -03164830 | 029676808 | -02275 | -0095 | 11 |
| 4651 | ENSMUSG( | Npy1r | 18166 | -03452138 | 040082915 | 01625 | -021 | 11 |
| 4670 | ENSMUSG( | Nrm | 106582 | -03733079 | 002903933 | 03625 | -0215 | 11 |
| 4671 | ENSMUSG( | Nrn1 | 68404 | -03114599 | -00730560 | 02875 | -01925 | 11 |
| 4672 | ENSMUSG( | Nrp1 | 18186 | -00741834 | 03397384S | 004 | -03275 | 11 |
| 4673 | ENSMUSG( | Nrp2 | 18187 | -04188276 | 00488117E | 0165 | -0235 | 11 |
| 4675 | ENSMUSG( | Nrxn1 | 18189 | -02034456 | 016429704 | 00625 | -004 | 11 |
| 4677 | ENSMUSG( | Nrxn3 | 18191 | -04187593 | 02461821E | 0205 | -011 | 11 |
| 4685 | ENSMUSG( | Nsg2 | 18197 | -04236386 | 009233794 | -0305 | -01225 | 11 |
| 4698 | ENSMUSG( | Nt5e | 23959 | -04836154 | 00750571S | -01675 | -0345 | 11 |
| 4702 | ENSMUSG( | Ntm | 235106 | -02155317 | -00862338 | 00625 | -0155 | 11 |
| 4752 | ENSMUSG( | Nup210 | 54563 | -02122853 | 009153431 | 0155 | -0335 | 11 |
| 4779 | ENSMUSG( | Ociad1 | 68095 | -03322813 | -00264172 | 0005 | -011 | 11 |
| 4794 | ENSMUSG( | Olfm2 | 244723 | -03253968 | 00904731E | 024 | -01475 | 11 |
| 4799 | ENSMUSG( | Omg | 18377 | -04547581 | 02069664C | 00075 | -04575 | 11 |
| 4804 | ENSMUSG( | Opcml | 330908 | -03714824 | 01935010S | -01725 | -0255 | 11 |
| 4841 | ENSMUSG( | Otulinl | 223433 | -03462046 | 022931438 | 023 | -04375 | 11 |
| 4850 | ENSMUSG( | P4ha1 | 18451 | -01227915 | 015012061 | 00575 | -0025 | 11 |
| 4851 | ENSMUSG( | P4hb | 18453 | -03647181 | 014423751 | 022 | -009 | 11 |
| 4868 | ENSMUSG( | Pag1 | 94212 | -03093052 | 01673312C | -0095 | -00125 | 11 |
| 4877 | ENSMUSG( | Pak5 | 241656 | -03529549 | -03749298 | -00725 | -02275 | 11 |
| 4939 | ENSMUSG( | Pcdh7 | 54216 | -03584869 | 014851862 | -011 | 00425 | 11 |
| 4945 | ENSMUSG( | Pcdhb14 | 93885 | -06163609 | 006818958 | -00075 | -01875 | 11 |
| 4950 | ENSMUSG( | Pcdhb9 | 93880 | -06463183 | 00100010S | 00675 | -0355 | 11 |
| 4957 | ENSMUSG( | Pcdhga7 | 93715 | -01739178 | 015543915 | -00725 | -00475 | 11 |
| 4978 | ENSMUSG( | Pcnx2 | 270109 | -01122647 | 026812438 | -00325 | -00775 | 11 |
| 4986 | ENSMUSG( | Pcyox1 | 66881 | -03052686 | 025412831 | 03775 | -024 | 11 |
| 4987 | ENSMUSG( | Pcyox1l | 240334 | -03912996 | 017132762 | 02075 | -0495 | 11 |
| 5001 | ENSMUSG( | Pde10a | 23984 | -03303448 | 000562245 | -0325 | -03725 | 11 |
| 5022 | ENSMUSG( | Pdgfrb | 18596 | -02915477 | 03092677C | 00525 | -03775 | 11 |
| 5035 | ENSMUSG( | Pdlim3 | 53318 | -01473414 | 00436802S | -00225 | -04425 | 11 |
| 5135 | ENSMUSG( | Phip | 83946 | -01957934 | 021236728 | -013 | -05575 | 11 |
| 5166 | ENSMUSG( | Pign | 27392 | -02103955 | 005874334 | 00475 | -0445 | 11 |
| 5168 | ENSMUSG( | Pigp | 56176 | -04054083 | -00081527 | -004 | -00575 | 11 |
| 5171 | ENSMUSG( | Pigu | 228812 | -03692674 | 014343053 | 0395 | -025 | 11 |
| 5214 | ENSMUSG( | Pla2g15 | 192654 | -02267949 | 030701485 | 00425 | -00175 | 11 |
| 5221 | ENSMUSG( | Plbd2 | 71772 | -02303125 | 027287472 | 021 | -0155 | 11 |
| 5234 | ENSMUSG( | Plcxd3 | 239318 | -03483382 | -01560578 | 02125 | -04025 | 11 |
| 5238 | ENSMUSG( | Plec | 18810 | -02141959 | 013010225 | -024 | -0045 | 11 |
| 5249 | ENSMUSG( | Plekhg3 | 263406 | -03264656 | -00033109 | -033 | -02925 | 11 |
| 5251 | ENSMUSG( | Plekhh2 | 213556 | -02170238 | 019417117 | -00225 | 0025 | 11 |
| 5260 | ENSMUSG( | Plp1 | 18823 | -04550207 | 007632997 | -0055 | -05175 | 11 |
| 5264 | ENSMUSG( | Plpp3 | 67916 | -02777329 | -00420957 | 01525 | -005 | 11 |
| 5265 | ENSMUSG( | Plpp4 | 381925 | -01570449 | 017089311 | 00675 | -027 | 11 |
| 5279 | ENSMUSG( | Plxdc2 | 67448 | -02137725 | 023591375 | -0055 | -005 | 11 |
| 5281 | ENSMUSG( | Plxna2 | 18845 | -02392844 | 031202241 | 01475 | -00975 | 11 |
| 5283 | ENSMUSG( | Plxna4 | 243743 | -03574814 | 026596308 | 00575 | -011 | 11 |
| 5286 | ENSMUSG( | Plxnb3 | 140571 | -01066044 | 020120737 | -00025 | -014 | 11 |
| 5315 | ENSMUSG( | Podxl2 | 319655 | -00890288 | 008351073 | 00525 | -025 | 11 |
| 5347 | ENSMUSG( | Pom121 | 107939 | -03669894 | 029558835 | -0235 | -016 | 11 |
| 5353 | ENSMUSG( | Pon3 | 269823 | -02390080 | -01877289 | 00125 | -01625 | 11 |
| 5356 | ENSMUSG( | Porcn | 53627 | -02575231 | 008680463 | 029 | -01825 | 11 |
| 5357 | ENSMUSG( | Postn | 50706 | -02569387 | 056122688 | 00875 | -0225 | 11 |
| 5377 | ENSMUSG( | Ppic | 19038 | -01727260 | 02675254S | 022 | -0375 | 11 |
| 5388 | ENSMUSG( | Ppip5k2 | 227399 | -00913839 | 023333515 | 006 | -042 | 11 |
| 5396 | ENSMUSG( | Ppm1l | 242083 | -04346225 | 012967733 | -0025 | -02025 | 11 |
| 5490 | ENSMUSG( | Prkar2b | 19088 | -03034999 | -00575829 | 00525 | -00975 | 11 |
| 5496 | ENSMUSG( | Prkch | 18755 | -03699273 | 034082862 | 0285 | -01025 | 11 |
| 5514 | ENSMUSG( | Prnp | 19122 | -03914950 | 003008344 | -01925 | -01375 | 11 |
| 5516 | ENSMUSG( | Prom1 | 19126 | 007502375 | 05603780C | -0055 | -051 | 11 |
| 5651 | ENSMUSG( | Ptpn5 | 19259 | -02653247 | 00442116E | 02375 | -02325 | 11 |
| 5659 | ENSMUSG( | Ptprj | 19271 | -00110239 | 03937755S | 02 | -03725 | 11 |
| 5664 | ENSMUSG( | Ptpro | 19277 | -05963118 | 01623069S | -002 | -02725 | 11 |
| 5709 | ENSMUSG( | Qpct | 70536 | -07213380 | 03749687S | 00325 | 0005 | 11 |
| 5764 | ENSMUSG( | Rab4b | 19342 | -02120378 | 007797648 | 03525 | -02875 | 11 |
| 5798 | ENSMUSG( | Rai1 | 19377 | -02299380 | 004997588 | 0015 | -01825 | 11 |
| 5815 | ENSMUSG( | Ranbp2 | 19386 | -00042634 | 020069763 | 0015 | -019 | 11 |
| 5885 | ENSMUSG( | Rbm5 | 83486 | -01151467 | 012204798 | -0075 | -0185 | 11 |
| 5910 | ENSMUSG( | Rdh11 | 17252 | -05082715 | 013823027 | 00375 | -05 | 11 |
| 6020 | ENSMUSG( | Rnaset2b | 68195 | 001007073 | 036316337 | 0245 | -03525 | 11 |
| 6032 | ENSMUSG( | Rnf13 | 24017 | -03817865 | 016862648 | 0305 | -03025 | 11 |
| 6038 | ENSMUSG( | Rnf150 | 330812 | -02182053 | 035908294 | -008 | -01425 | 11 |
| 6051 | ENSMUSG( | Rnf26 | 213211 | -02682810 | 010417272 | -016 | -03775 | 11 |
| 6205 | ENSMUSG( | Rtn3 | 20168 | -02961895 | 009165643 | -03175 | -0095 | 11 |
| 6209 | ENSMUSG( | Rtn4rl1 | 237847 | -02498528 | 010864832 | 0395 | -0425 | 11 |
| 6227 | ENSMUSG( | Ryr2 | 20191 | -05719317 | 00683525C | -01025 | -03825 | 11 |
| 6243 | ENSMUSG( | Sacs | 50720 | -04449673 | 020530495 | -00725 | -009 | 11 |
| 6284 | ENSMUSG( | Scai | 320271 | -01854862 | 023694781 | 00275 | -03125 | 11 |
| 6285 | ENSMUSG( | Scamp1 | 107767 | -03510600 | 005816504 | -00725 | -0025 | 11 |
| 6292 | ENSMUSG( | Scarb2 | 12492 | -01897253 | 005666131 | 0145 | -0115 | 11 |
| 6304 | ENSMUSG( | Scn2a | 110876 | -03711913 | 027211438 | -0115 | -02425 | 11 |
| 6310 | ENSMUSG( | Scn8a | 20273 | -02285059 | 031978567 | -157009e-1 | -03525 | 11 |

| | | | | | | | |
|---|---|---|---|---|---|---|---|
| 6312 | ENSMUSG( | Sco2 | 1E+08 | -04279521 | -00006499 | 00275 | -0055 | 11 |
| 6379 | ENSMUSG( | Selenos | 109815 | -02779219 | 01507128$ | -02325 | -02625 | 11 |
| 6384 | ENSMUSG( | Sema4b | 20352 | -03144129 | 03950678$ | -01025 | -0015 | 11 |
| 6385 | ENSMUSG( | Sema4c | 20353 | -03524621 | 016279164 | -01725 | -00825 | 11 |
| 6386 | ENSMUSG( | Sema4d | 20354 | -03321667 | 014711657 | 00725 | -006 | 11 |
| 6387 | ENSMUSG( | Sema4f | 20355 | -01215728 | 018842035 | 01375 | -01 | 11 |
| 6418 | ENSMUSG( | Serinc3 | 26943 | -02230019 | 03499410C | -01375 | -0545 | 11 |
| 6425 | ENSMUSG( | Serpinb9 | 20723 | -02301967 | 000409643 | 01975 | -01125 | 11 |
| 6440 | ENSMUSG( | Sez6l | 56747 | -02636021 | 025895022 | 0025 | -03075 | 11 |
| 6455 | ENSMUSG( | Sfrp1 | 20377 | -02554063 | 018961134 | 03 | -02025 | 11 |
| 6526 | ENSMUSG( | Sidt2 | 214597 | -00858475 | 016841938 | -012 | -00925 | 11 |
| 6538 | ENSMUSG( | Sirpa | 19261 | -01795381 | 016299376 | 0135 | -029 | 11 |
| 6549 | ENSMUSG( | Slc12a2 | 20496 | -05193971 | 025898441 | -02525 | -051 | 11 |
| 6551 | ENSMUSG( | Slc12a5 | 57138 | -02191403 | 011548175 | -013 | -0085 | 11 |
| 6553 | ENSMUSG( | Slc12a9 | 83704 | -02104956 | 009008852 | 01725 | -0205 | 11 |
| 6563 | ENSMUSG( | Slc16a3 | 80879 | -02854954 | 00910544C | 01575 | -02075 | 11 |
| 6567 | ENSMUSG( | Slc17a7 | 72961 | -01859155 | 011288811 | 024 | -03575 | 11 |
| 6582 | ENSMUSG( | Slc22a23 | 73102 | -03963254 | 008973952 | -01375 | 00825 | 11 |
| 6585 | ENSMUSG( | Slc23a2 | 54338 | -02747059 | 019411703 | 02175 | -036 | 11 |
| 6587 | ENSMUSG( | Slc24a3 | 94249 | -02949535 | 008967452 | -0375 | -018 | 11 |
| 6588 | ENSMUSG( | Slc24a4 | 238384 | -02247212 | 051860215 | 0125 | -04075 | 11 |
| 6593 | ENSMUSG( | Slc25a13 | 50799 | -02950894 | 006406291 | -015 | -02225 | 11 |
| 6594 | ENSMUSG( | Slc25a14 | 20523 | -01103532 | 023350719 | 019 | -01075 | 11 |
| 6616 | ENSMUSG( | Slc25a40 | 319653 | -01911612 | 004241588 | 011 | -00825 | 11 |
| 6623 | ENSMUSG( | Slc26a2 | 13521 | -06891421 | 015655134 | -01775 | -0505 | 11 |
| 6628 | ENSMUSG( | Slc27a4 | 26569 | -02730134 | 01526122 | 00575 | -017 | 11 |
| 6633 | ENSMUSG( | Slc2a3 | 20527 | -03154892 | 013411015 | -012 | -04025 | 11 |
| 6643 | ENSMUSG( | Slc33a1 | 11416 | -03129760 | 035037397 | 01 | -02075 | 11 |
| 6653 | ENSMUSG( | Slc35g2 | 245020 | -04686416 | 001394283 | 039 | -0495 | 11 |
| 6656 | ENSMUSG( | Slc37a4 | 14385 | -02229411 | 024235433 | 008 | -01275 | 11 |
| 6659 | ENSMUSG( | Slc38a2 | 67760 | -03410269 | 043772514 | -01425 | -03325 | 11 |
| 6660 | ENSMUSG( | Slc38a3 | 76257 | -04294962 | -00936426 | -03275 | -0485 | 11 |
| 6669 | ENSMUSG( | Slc3a2 | 17254 | -01964122 | 019286584 | -00775 | -023 | 11 |
| 6671 | ENSMUSG( | Slc44a1 | 100434 | -03317495 | 01091654$ | -01225 | -0435 | 11 |
| 6678 | ENSMUSG( | Slc4a10 | 94229 | -03476729 | 02902978C | 011 | -01525 | 11 |
| 6684 | ENSMUSG( | Slc4a8 | 59033 | -01934998 | 012649144 | 01625 | -02275 | 11 |
| 6702 | ENSMUSG( | Slc7a10 | 53896 | -01943510 | 00416888$ | -007 | -02275 | 11 |
| 6704 | ENSMUSG( | Slc7a14 | 241919 | -03504688 | 035761653 | 004 | -0205 | 11 |
| 6711 | ENSMUSG( | Slc7a8 | 50934 | -05183273 | 007595686 | -01025 | -024 | 11 |
| 6712 | ENSMUSG( | Slc8a1 | 20541 | -03089525 | 03485509$ | 00075 | -00675 | 11 |
| 6713 | ENSMUSG( | Slc8a2 | 110891 | -03958393 | 011875544 | 014 | -02725 | 11 |
| 6715 | ENSMUSG( | Slc9a1 | 20544 | -02964064 | 01489313C | -01375 | -0145 | 11 |
| 6729 | ENSMUSG( | Slitrk1 | 76965 | -06508155 | 007867507 | 02025 | -03275 | 11 |
| 6730 | ENSMUSG( | Slitrk2 | 245450 | -05148024 | 03184915 | 01 | -002 | 11 |
| 6732 | ENSMUSG( | Slitrk4 | 245446 | -05702372 | 03666020$ | 0175 | -025 | 11 |
| 6733 | ENSMUSG( | Slitrk5 | 75409 | -03740451 | 034088245 | 01625 | -0155 | 11 |
| 6762 | ENSMUSG( | Smg1 | 233789 | -01694550 | 018569054 | -01525 | -002 | 11 |
| 6770 | ENSMUSG( | Smim18 | 72632 | -03693271 | -00144811 | -03775 | -039 | 11 |
| 6783 | ENSMUSG( | Smpdl3b | 100340 | -07621279 | 051891064 | -0125 | -00525 | 11 |
| 6823 | ENSMUSG( | Sntb2 | 20650 | -02815869 | -02136174 | -00825 | -0165 | 11 |
| 6832 | ENSMUSG( | Snx14 | 244962 | -02618061 | 023625308 | 03875 | -0375 | 11 |
| 6837 | ENSMUSG( | Snx19 | 102607 | -02274414 | 01355441$ | -00925 | -00175 | 11 |
| 6841 | ENSMUSG( | Snx25 | 102141 | -00774615 | 02338837C | -004 | -01175 | 11 |
| 6853 | ENSMUSG( | Soat1 | 20652 | -01492868 | 023727907 | 01575 | -035 | 11 |
| 6860 | ENSMUSG( | Sorbs2 | 234214 | -02258252 | -00367440 | -017 | -01025 | 11 |
| 6862 | ENSMUSG( | Sorcs1 | 58178 | -03839298 | 006804281 | -0075 | 00175 | 11 |
| 6863 | ENSMUSG( | Sorcs2 | 81840 | -04388057 | 001144573 | 0015 | -03825 | 11 |
| 6864 | ENSMUSG( | Sorcs3 | 66673 | -05322969 | -01058461 | -005 | -0485 | 11 |
| 6866 | ENSMUSG( | Sorl1 | 20660 | -01338450 | 010078327 | 00675 | -00625 | 11 |
| 6867 | ENSMUSG( | Sort1 | 20661 | -02756990 | 011184123 | 02 | -01275 | 11 |
| 6918 | ENSMUSG( | Sprn | 212518 | -02890143 | 017628486 | -0105 | -01675 | 11 |
| 6928 | ENSMUSG( | Sptlc1 | 268656 | -01943445 | 018143264 | 0035 | -0065 | 11 |
| 6930 | ENSMUSG( | Sqle | 20775 | -02752681 | 003198733 | 0245 | -0295 | 11 |
| 6968 | ENSMUSG( | Srsf7 | 225027 | -00989125 | 02574943$ | -011 | -03775 | 11 |
| 6988 | ENSMUSG( | Sstr2 | 20606 | -03688078 | 007513263 | 00675 | -012 | 11 |
| 7016 | ENSMUSG( | Steap2 | 74051 | -02708306 | -00345610 | 785046e-1 | -018 | 11 |
| 7017 | ENSMUSG( | Steap3 | 68428 | -03350978 | 006031537 | 0095 | -005 | 11 |
| 7020 | ENSMUSG( | Stim2 | 116873 | -02913711 | 029291246 | -00175 | -013 | 11 |
| 7040 | ENSMUSG( | Stom | 13830 | -01601233 | 016400428 | 001 | -01825 | 11 |
| 7085 | ENSMUSG( | Sulf2 | 72043 | -03410463 | 026486817 | 0245 | -0105 | 11 |
| 7092 | ENSMUSG( | Sun2 | 223697 | -03836176 | 010557902 | 0015 | -0115 | 11 |
| 7101 | ENSMUSG( | Susd2 | 71733 | -04195636 | 013918701 | -0015 | -00875 | 11 |
| 7103 | ENSMUSG( | Sv2a | 64051 | -04078378 | 01248723$ | -005 | -006 | 11 |
| 7104 | ENSMUSG( | Sv2b | 64176 | -02939495 | 02832493C | 03075 | -02325 | 11 |
| 7139 | ENSMUSG( | Syt1 | 20979 | -05547635 | 015063093 | 01625 | -01925 | 11 |
| 7141 | ENSMUSG( | Syt11 | 229521 | -02481202 | 020911377 | -00675 | -02725 | 11 |
| 7142 | ENSMUSG( | Syt12 | 171180 | -02037501 | 006358788 | -0195 | -01475 | 11 |
| 7146 | ENSMUSG( | Syt2 | 20980 | -05170516 | 00275477$ | -02275 | -06 | 11 |
| 7147 | ENSMUSG( | Syt3 | 20981 | -03464716 | 007768758 | -00575 | -02225 | 11 |
| 7161 | ENSMUSG( | Tacc1 | 320165 | -00871707 | 007693608 | 00225 | -01125 | 11 |
| 7179 | ENSMUSG( | Tafazzin | 66826 | -01032880 | 018860694 | -01625 | -00575 | 11 |
| 7202 | ENSMUSG( | Tasor | 218850 | 001122478 | 028404185 | 01 | -033 | 11 |
| 7273 | ENSMUSG( | Tecr | 106529 | -03514318 | -00552066 | 03925 | -015 | 11 |
| 7278 | ENSMUSG( | Tenm1 | 23963 | -04438551 | 021799435 | -00325 | -0005 | 11 |
| 7314 | ENSMUSG( | Them6 | 223626 | -04699624 | 00409643C | 011 | -01475 | 11 |
| 7332 | ENSMUSG( | Thy1 | 21838 | -04290017 | 004488152 | -005 | 001 | 11 |
| 7354 | ENSMUSG( | Timp2 | 21858 | -02671324 | 010714754 | 00775 | -00375 | 11 |
| 7380 | ENSMUSG( | Tmbim1 | 69660 | -04038577 | -00785390 | -0345 | -05625 | 11 |
| 7397 | ENSMUSG( | Tmem109 | 68539 | -03292607 | 016883284 | 0035 | -0425 | 11 |
| 7403 | ENSMUSG( | Tmem121k | 94047 | -02544031 | 01204146C | 0125 | -008 | 11 |
| 7407 | ENSMUSG( | Tmem130 | 243339 | -02911783 | 012430107 | -02225 | 00075 | 11 |
| 7410 | ENSMUSG( | Tmem132k | 208151 | -03566649 | 021445635 | -00575 | -0305 | 11 |
| 7417 | ENSMUSG( | Tmem144 | 70652 | -02430501 | 013267195 | 0375 | -02125 | 11 |
| 7421 | ENSMUSG( | Tmem151a | 381199 | -02985811 | 00701643C | 00725 | -02375 | 11 |
| 7427 | ENSMUSG( | Tmem165 | 21982 | -04958415 | 01478526$ | -01 | -02675 | 11 |
| 7431 | ENSMUSG( | Tmem175 | 72392 | -01970003 | 006476452 | -0105 | -0035 | 11 |
| 7434 | ENSMUSG( | Tmem181a | 77106 | -03367841 | 013192011 | 006 | -00725 | 11 |
| 7446 | ENSMUSG( | Tmem209 | 72649 | -03478238 | 01768747C | 003 | -009 | 11 |
| 7448 | ENSMUSG( | Tmem222 | 52174 | -03220065 | -01270368 | 0215 | -05075 | 11 |
| 7461 | ENSMUSG( | Tmem259 | 216157 | -01719325 | 018530993 | 017 | -037 | 11 |
| 7465 | ENSMUSG( | Tmem30a | 69981 | -03682274 | 025764432 | -01025 | -01275 | 11 |
| 7471 | ENSMUSG( | Tmem43 | 74122 | -04041715 | 018753531 | 0075 | -035 | 11 |
| 7488 | ENSMUSG( | Tmem94 | 71947 | -00582579 | 022196584 | -00775 | -0325 | 11 |
| 7503 | ENSMUSG( | Tmx2 | 66958 | -03883801 | -00172634 | -00025 | -0015 | 11 |
| 7504 | ENSMUSG( | Tmx3 | 67988 | -04188612 | 027570284 | 028 | -011 | 11 |
| 7505 | ENSMUSG( | Tmx4 | 52837 | -04165339 | 022197794 | 00725 | -00625 | 11 |
| 7520 | ENSMUSG( | Tnr | 21960 | -04173617 | 01968064$ | 01 | -04125 | 11 |
| 7543 | ENSMUSG( | Tor1a | 30931 | -02368775 | 009280842 | 02525 | -0315 | 11 |
| 7546 | ENSMUSG( | Tor1b | 30934 | -01446359 | 019800365 | 0135 | -0275 | 11 |
| 7548 | ENSMUSG( | Tor3a | 30935 | -03038533 | -01273276 | 01325 | -0465 | 11 |
| 7551 | ENSMUSG( | Tpbg | 21983 | -06627225 | 022179535 | -00975 | -02975 | 11 |
| 7566 | ENSMUSG( | Tpp1 | 12751 | -01519240 | 014074341 | 004 | -0275 | 11 |
| 7586 | ENSMUSG( | Tram1 | 72265 | -02365402 | 020136777 | -02375 | -026 | 11 |
| 7604 | ENSMUSG( | Trhde | 237553 | -04282781 | 018572677 | 01175 | -01175 | 11 |
| 7607 | ENSMUSG( | Tril | 66873 | -02373110 | -00636723 | -00525 | -033 | 11 |
| 7655 | ENSMUSG( | Trpv2 | 22368 | -01483942 | 012652451 | -01125 | -01075 | 11 |
| 7673 | ENSMUSG( | Tspan15 | 70423 | -03971994 | 002203044 | -235514e-1 | -0415 | 11 |
| 7696 | ENSMUSG( | Ttc14 | 67120 | -00730356 | 030788648 | -0125 | -01375 | 11 |
| 7748 | ENSMUSG( | Tut4 | 230594 | -02445529 | 000855975 | -017 | -0525 | 11 |
| 7756 | ENSMUSG( | Txndc15 | 69672 | -01614292 | 01864721$ | 0355 | -0205 | 11 |
| 7761 | ENSMUSG( | Txnip | 56338 | -01963642 | -00041184 | 01325 | -0135 | 11 |
| 7785 | ENSMUSG( | Ubac2 | 68889 | -02266649 | 004774405 | 026 | -0405 | 11 |
| 7820 | ENSMUSG( | Ube4a | 140630 | -00907782 | 015563877 | 0115 | -005 | 11 |
| 7823 | ENSMUSG( | Ubiad1 | 71707 | -01834927 | 020155171 | -004 | 004 | 11 |
| 7860 | ENSMUSG( | Ufsp1 | 70240 | -04440572 | -02239610 | -0075 | -04375 | 11 |
| 7879 | ENSMUSG( | Unc5a | 107448 | -00530225 | 03472348C | 01475 | -024 | 11 |

| | | | | | | | |
|---|---|---|---|---|---|---|---|
| 7881 | ENSMUSG( | Unc5c | 22253 | -06436850 | 006271839 | 00275 | -01575 | 11 |
| 7885 | ENSMUSG( | Unc93b1 | 54445 | -01527961 | 01550609C | -02 | -016 | 11 |
| 7966 | ENSMUSG( | Vangl2 | 93840 | -04003109 | 010231985 | 0085 | -03275 | 11 |
| 7972 | ENSMUSG( | Vasn | 246154 | -02647971 | 028717466 | -03575 | -04875 | 11 |
| 7980 | ENSMUSG( | Vcam1 | 22329 | -02467769 | 005518588 | -01075 | -02725 | 11 |
| 7981 | ENSMUSG( | Vcan | 13003 | -03246226 | 039163352 | -01 | -01125 | 11 |
| 7994 | ENSMUSG( | Vip | 22353 | -05662692 | 009945309 | 02775 | -01025 | 11 |
| 8006 | ENSMUSG( | Vps13c | 320528 | -00711937 | 018422371 | 00575 | -00625 | 11 |
| 8052 | ENSMUSG( | Vwc2l | 320460 | -02979947 | 042720102 | 011 | -014 | 11 |
| 8074 | ENSMUSG( | Wdfy3 | 72145 | -00929297 | 013950088 | 00775 | -017 | 11 |
| 8093 | ENSMUSG( | Wdr44 | 72404 | -02309024 | -00054892 | 015 | -01475 | 11 |
| 8135 | ENSMUSG( | Xk | 22439 | -03505383 | 025791632 | -02975 | -02975 | 11 |
| 8155 | ENSMUSG( | Xxylt1 | 268880 | -04182682 | 003535239 | -01975 | -02275 | 11 |
| 8264 | ENSMUSG( | Zfyve27 | 319740 | -02052407 | 014033288 | -01225 | -05 | 11 |
| 19 | ENSMUSG( | A830018L1 | 320492 | 005631661 | 006841038 | 0055 | 053 | 12 |
| 21 | ENSMUSG( | Aacs | 78894 | 032043727 | 003579243 | 012 | 04575 | 12 |
| 22 | ENSMUSG( | Aagab | 66939 | 005567433 | -00757197 | -006 | 036 | 12 |
| 28 | ENSMUSG( | Aars2 | 224805 | 022611684 | 003738258 | 0025 | 03625 | 12 |
| 53 | ENSMUSG( | Abcd1 | 11666 | -01056995 | 002418521 | 01725 | 04375 | 12 |
| 55 | ENSMUSG( | Abcd3 | 19299 | 008301314 | -01025652 | 022 | 0315 | 12 |
| 69 | ENSMUSG( | Abhd17a | 216169 | 013859745 | -00740458 | -0045 | 05075 | 12 |
| 71 | ENSMUSG( | Abhd17c | 70178 | 006965154 | 001691553 | 0015 | 05175 | 12 |
| 80 | ENSMUSG( | Abl2 | 11352 | -00189989 | 003862387 | -00725 | 02725 | 12 |
| 88 | ENSMUSG( | Acaa1a | 113868 | 002003694 | -00259613 | 009 | 02825 | 12 |
| 106 | ENSMUSG( | Acbd5 | 74159 | 00819902C | -01166490 | 01875 | 03225 | 12 |
| 145 | ENSMUSG( | Actr1a | 54130 | 002624154 | -01379918 | -0045 | 0465 | 12 |
| 163 | ENSMUSG( | Adam15 | 11490 | 000596395 | 01960864C | 02525 | 043 | 12 |
| 167 | ENSMUSG( | Adam9 | 11502 | -00715879 | 013632924 | 01775 | 041 | 12 |
| 187 | ENSMUSG( | Adgb | 215772 | -01208387 | 01020849C | 018 | 05825 | 12 |
| 188 | ENSMUSG( | Adgra1 | 52389 | -00832346 | 022077679 | 023 | 04625 | 12 |
| 199 | ENSMUSG( | Adh5 | 11532 | 001150881 | -01340551 | -008 | 03225 | 12 |
| 210 | ENSMUSG( | Adprm | 66358 | 018375631 | 018601093 | 0035 | 0365 | 12 |
| 216 | ENSMUSG( | Adrm1 | 56436 | -00666697 | 000629444 | 0045 | 0275 | 12 |
| 218 | ENSMUSG( | Adss1 | 11565 | -01587748 | -00199234 | -0095 | 0545 | 12 |
| 224 | ENSMUSG( | Aff4 | 93736 | 010927752 | 008654978 | -0125 | 0365 | 12 |
| 230 | ENSMUSG( | Agap1 | 347722 | 00643973C | 00615022C | 005 | 04675 | 12 |
| 247 | ENSMUSG( | Agt | 11606 | 008456454 | 013200876 | 00475 | 04675 | 12 |
| 256 | ENSMUSG( | Ahnak2 | 1E+08 | -02420238 | -01144469 | -0055 | 046 | 12 |
| 266 | ENSMUSG( | Aimp2 | 231872 | -00261947 | -01134165 | 00725 | 06575 | 12 |
| 278 | ENSMUSG( | Akap12 | 83397 | 005555941 | 004518955 | -0105 | 0455 | 12 |
| 299 | ENSMUSG( | Aldh1a2 | 19378 | 021618792 | -05725324 | 021 | 04525 | 12 |
| 324 | ENSMUSG( | Alkbh5 | 268420 | 011730781 | 010558887 | -00125 | 0595 | 12 |
| 333 | ENSMUSG( | Ambra1 | 228361 | -00291753 | 009642966 | 01 | 06125 | 12 |
| 340 | ENSMUSG( | Amot | 27494 | 022348963 | -01752216 | 01375 | 03875 | 12 |
| 341 | ENSMUSG( | Amotl1 | 75723 | 008285609 | -01654813 | -00575 | 035 | 12 |
| 361 | ENSMUSG( | Ankhd1 | 108857 | -00537063 | 008878509 | 0125 | 03025 | 12 |
| 362 | ENSMUSG( | Ankib1 | 70797 | 003289866 | 010921968 | -0075 | 02825 | 12 |
| 366 | ENSMUSG( | Ankrd13a | 68420 | 002283092 | 01117411C | -00125 | 03125 | 12 |
| 369 | ENSMUSG( | Ankrd17 | 81702 | 012687123 | 007834236 | 0145 | 05225 | 12 |
| 370 | ENSMUSG( | Ankrd27 | 245886 | 019669355 | 003352747 | 0135 | 04075 | 12 |
| 374 | ENSMUSG( | Ankrd34a | 545554 | 009337292 | -00500032 | -01175 | 03925 | 12 |
| 376 | ENSMUSG( | Ankrd40 | 71452 | -00062898 | -02081845 | 0105 | 035 | 12 |
| 377 | ENSMUSG( | Ankrd44 | 329154 | -01100170 | 004351986 | 01225 | 03875 | 12 |
| 379 | ENSMUSG( | Ankrd46 | 68839 | 01279519C | -00729316 | 01625 | 07625 | 12 |
| 388 | ENSMUSG( | Ankzf1 | 52231 | 012609704 | 005729443 | 03075 | 05725 | 12 |
| 394 | ENSMUSG( | Anp32a | 11737 | 010411893 | -01542319 | -0155 | 04425 | 12 |
| 398 | ENSMUSG( | Antkmt | 214917 | 000511736 | -00539102 | 012 | 03 | 12 |
| 429 | ENSMUSG( | Ap4m1 | 11781 | 013130478 | 000576006 | 007 | 02975 | 12 |
| 432 | ENSMUSG( | Ap5z1 | 231855 | 010685312 | 012570157 | 0085 | 05125 | 12 |
| 434 | ENSMUSG( | Apba1 | 319924 | 018948438 | 002003257 | -01025 | 0415 | 12 |
| 435 | ENSMUSG( | Apba2 | 11784 | -02236158 | 006095252 | -0075 | 0385 | 12 |
| 441 | ENSMUSG( | Apc2 | 23805 | 001691926 | -00189595 | -01425 | 03925 | 12 |
| 443 | ENSMUSG( | Apex1 | 11792 | 003430707 | -01084773 | 0135 | 07425 | 12 |
| 455 | ENSMUSG( | Apoe | 11816 | 001809849 | 003244428 | 008 | 02575 | 12 |
| 456 | ENSMUSG( | Apoh | 11818 | 008783468 | -01507379 | 0075 | 04375 | 12 |
| 460 | ENSMUSG( | Appbp2 | 66884 | 000383219 | -00827768 | 03875 | 06275 | 12 |
| 469 | ENSMUSG( | Araf | 11836 | -00849560 | -00363004 | 0265 | 066 | 12 |
| 478 | ENSMUSG( | Arf5 | 11844 | -00478634 | -01136471 | 00725 | 03825 | 12 |
| 504 | ENSMUSG( | Arhgap32 | 330914 | 005199281 | 015775502 | 00325 | 0475 | 12 |
| 505 | ENSMUSG( | Arhgap33 | 233071 | 017295954 | 008125798 | 0135 | 06075 | 12 |
| 523 | ENSMUSG( | Arhgef25 | 52666 | 030091541 | 005141801 | -0035 | 02925 | 12 |
| 537 | ENSMUSG( | Arih2 | 23807 | -00298624 | 004436781 | 0005 | 04275 | 12 |
| 556 | ENSMUSG( | Armc6 | 76813 | -00769168 | -00622760 | 0015 | 0395 | 12 |
| 561 | ENSMUSG( | Armcx4 | 1.01E+08 | -00107594 | 01335834C | 00425 | 0295 | 12 |
| 573 | ENSMUSG( | Arpp19 | 59046 | -00068141 | -01238598 | -01875 | 046 | 12 |
| 593 | ENSMUSG( | Ascc2 | 75452 | 011548806 | -00745972 | -0045 | 06375 | 12 |
| 594 | ENSMUSG( | Ascc3 | 77987 | 008960718 | 009426256 | 0115 | 03525 | 12 |
| 596 | ENSMUSG( | Ash2l | 23808 | -00328355 | -00842663 | -01875 | 04075 | 12 |
| 614 | ENSMUSG( | Atat1 | 73242 | -00228789 | -00963477 | 0015 | 034 | 12 |
| 615 | ENSMUSG( | Atcay | 16467 | -01462256 | 010888438 | -0035 | 0445 | 12 |
| 616 | ENSMUSG( | Ate1 | 11907 | 005972766 | 001714481 | -00775 | 03425 | 12 |
| 618 | ENSMUSG( | Atf7ip | 54343 | 009362206 | 004097328 | 02075 | 0675 | 12 |
| 620 | ENSMUSG( | Atg12 | 67526 | 006851725 | -01929080 | 01425 | 03125 | 12 |
| 652 | ENSMUSG( | Atp2a3 | 53313 | -00209336 | 001211675 | -005 | 03925 | 12 |
| 661 | ENSMUSG( | Atp5f1e | 67126 | 007844314 | -02621713 | 00625 | 03325 | 12 |
| 683 | ENSMUSG( | Atp6v1g1 | 66290 | 001850144 | -00626817 | 0015 | 03475 | 12 |
| 693 | ENSMUSG( | Atpsckmt | 68073 | 007666349 | 00665801C | 0195 | 03075 | 12 |
| 702 | ENSMUSG( | Atxn2l | 233871 | 01205733C | 014306538 | -0095 | 0405 | 12 |
| 703 | ENSMUSG( | Atxn3 | 110616 | 00828491C | 009488763 | -004 | 0455 | 12 |
| 704 | ENSMUSG( | Atxn7l3b | 382423 | 008739964 | -01018173 | 0085 | 043 | 12 |
| 706 | ENSMUSG( | Aup1 | 11993 | -00995896 | 01027722C | -0035 | 036 | 12 |
| 707 | ENSMUSG( | Aurkaip1 | 66077 | -00104117 | -02500552 | 034 | 06675 | 12 |
| 708 | ENSMUSG( | Aven | 74268 | 006127153 | -00499649 | 00275 | 028 | 12 |
| 726 | ENSMUSG( | B9d1 | 27078 | -00637383 | -01327829 | 00625 | 05175 | 12 |
| 739 | ENSMUSG( | Baiap3 | 545192 | 034932223 | 001590941 | 03525 | 07575 | 12 |
| 785 | ENSMUSG( | Bex2 | 12069 | -01373682 | -02629672 | -023 | 053 | 12 |
| 789 | ENSMUSG( | Bicd2 | 76895 | 001579037 | 013925144 | 0225 | 02825 | 12 |
| 795 | ENSMUSG( | Bin3 | 57784 | -01331997 | -00246042 | -01125 | 0445 | 12 |
| 796 | ENSMUSG( | Birc2 | 11797 | -00325654 | -00023127 | 00825 | 06275 | 12 |
| 806 | ENSMUSG( | Bmp2k | 140780 | -01901032 | -00241077 | -00275 | 05875 | 12 |
| 808 | ENSMUSG( | Bmpr1b | 12167 | -00836334 | -02090967 | 0055 | 044 | 12 |
| 825 | ENSMUSG( | Borcs6 | 71923 | 004989137 | -00726282 | 00025 | 031 | 12 |
| 826 | ENSMUSG( | Borcs7 | 66439 | 009523649 | -01110932 | 01575 | 02675 | 12 |
| 832 | ENSMUSG( | Bptf | 207165 | 004375018 | 015923407 | -00175 | 0425 | 12 |
| 858 | ENSMUSG( | Bsg | 12215 | -02400223 | 005223873 | 0005 | 05575 | 12 |
| 867 | ENSMUSG( | Btf3 | 218490 | -00197704 | -02009236 | 00125 | 06325 | 12 |
| 868 | ENSMUSG( | Btf3l4 | 70533 | -02159039 | -00479867 | -0115 | 057 | 12 |
| 879 | ENSMUSG( | C1qc | 12262 | -01600423 | 013079248 | 01975 | 057 | 12 |
| 884 | ENSMUSG( | C2cd4c | 237397 | 025386775 | 000181971 | -01175 | 03125 | 12 |
| 904 | ENSMUSG( | Cacna1h | 58226 | 002370505 | 024468817 | -009 | 0575 | 12 |
| 951 | ENSMUSG( | Camsap1 | 227634 | 013182955 | 002105963 | -0095 | 03 | 12 |
| 955 | ENSMUSG( | Camta2 | 216874 | 026601793 | 009714948 | 01025 | 0315 | 12 |
| 969 | ENSMUSG( | Caprin1 | 53872 | 011810479 | 001834924 | 01825 | 02425 | 12 |
| 989 | ENSMUSG( | Casp3 | 12367 | 00186892C | 000885089 | 01625 | 05425 | 12 |
| 993 | ENSMUSG( | Casq2 | 12373 | -03589443 | -03328919 | 00925 | 055 | 12 |
| 995 | ENSMUSG( | Castor2 | 80909 | 024809606 | 017773038 | -005 | 03525 | 12 |
| 1000 | ENSMUSG( | Cavin3 | 109042 | 018548737 | -02259639 | 017 | 06425 | 12 |
| 1001 | ENSMUSG( | Cbarp | 1.01E+08 | -01380525 | 021207621 | 0065 | 05775 | 12 |
| 1006 | ENSMUSG( | Cbll1 | 104836 | -01590323 | -00012304 | -00775 | 0315 | 12 |
| 1020 | ENSMUSG( | Cc2d1b | 319965 | 006104952 | -00752628 | -01425 | 04 | 12 |
| 1024 | ENSMUSG( | Ccdc106 | 232821 | 007860068 | -00343619 | 0045 | 05275 | 12 |
| 1035 | ENSMUSG( | Ccdc149 | 1.01E+08 | 008236127 | -01446642 | -01175 | 0385 | 12 |
| 1036 | ENSMUSG( | Ccdc167 | 68597 | 009136609 | -00353652 | -00775 | 03075 | 12 |
| 1041 | ENSMUSG( | Ccdc25 | 67179 | 012311537 | -00451486 | 017 | 06075 | 12 |
| 1056 | ENSMUSG( | Ccdc88c | 68339 | -02477187 | -01796933 | -006 | 0475 | 12 |
| 1071 | ENSMUSG( | Ccnt1 | 12455 | -00538961 | 009907549 | 0055 | 0645 | 12 |

| | | | | | | | | |
|---|---|---|---|---|---|---|---|---|
| 1082 | ENSMUSG( | Cct4 | 12464 | 014525029 | -00888137 | 0005 | 03375 | 12 |
| 1095 | ENSMUSG( | Cd2bp2 | 70233 | 004163146 | -01244912 | 02775 | 07525 | 12 |
| 1118 | ENSMUSG( | Cdc42 | 12540 | 001737759 | -01481195 | -01025 | 04075 | 12 |
| 1124 | ENSMUSG( | Cdc42se2 | 72729 | 000547161 | -00866828 | 018 | 0335 | 12 |
| 1137 | ENSMUSG( | Cdh5 | 12562 | -00522886 | 03273492C | -00625 | 04325 | 12 |
| 1152 | ENSMUSG( | Cdk2ap1 | 13445 | 006245184 | -00757998 | 026 | 0525 | 12 |
| 1156 | ENSMUSG( | Cdk5r2 | 12570 | 011922229 | 003954168 | -00575 | 03975 | 12 |
| 1158 | ENSMUSG( | Cdk7 | 12572 | -00708315 | -00031921 | -0005 | 06075 | 12 |
| 1162 | ENSMUSG( | Cdkn1b | 12576 | 010646744 | 015556004 | 00375 | 06025 | 12 |
| 1165 | ENSMUSG( | Cdr2l | 237988 | -00421779 | -00732781 | 0145 | 03075 | 12 |
| 1169 | ENSMUSG( | Cebpz | 12607 | 023438717 | 007342054 | 029 | 076 | 12 |
| 1172 | ENSMUSG( | Celf2 | 14007 | 004878237 | 011886609 | -00875 | 031 | 12 |
| 1174 | ENSMUSG( | Celf4 | 108013 | 016210685 | 016801795 | -0015 | 0345 | 12 |
| 1201 | ENSMUSG( | Cfap298 | 68001 | -00348843 | -01067692 | 024 | 0495 | 12 |
| 1208 | ENSMUSG( | Cfh | 12628 | 007348605 | 027465686 | 01 | 05325 | 12 |
| 1213 | ENSMUSG( | Cgref1 | 68567 | -03454869 | -01464979 | 02975 | 06725 | 12 |
| 1219 | ENSMUSG( | Chchd10 | 103172 | -00265678 | -02789022 | -01075 | 057 | 12 |
| 1229 | ENSMUSG( | Chd5 | 269610 | 029366897 | 003604438 | 033 | 053 | 12 |
| 1252 | ENSMUSG( | Chmp7 | 105513 | 010925259 | -00372787 | 0195 | 0335 | 12 |
| 1270 | ENSMUSG( | Ciao1 | 26371 | -00165003 | -00072514 | -00525 | 05575 | 12 |
| 1288 | ENSMUSG( | Clasp1 | 76707 | 011491755 | -00771629 | 023 | 057 | 12 |
| 1335 | ENSMUSG( | Cluh | 74148 | 011321183 | 005756415 | 02125 | 03325 | 12 |
| 1347 | ENSMUSG( | Cmss1 | 66497 | 004352759 | -02802782 | 03175 | 0495 | 12 |
| 1350 | ENSMUSG( | Cmtr1 | 74157 | 001211477 | 015838553 | -01325 | 0555 | 12 |
| 1359 | ENSMUSG( | Cnn2 | 12798 | 009284796 | 031327257 | 01375 | 077 | 12 |
| 1366 | ENSMUSG( | Cnot10 | 78893 | 003293171 | 01540593C | -0165 | 057 | 12 |
| 1401 | ENSMUSG( | Cobl | 12808 | 023350971 | 003088726 | -0115 | 02925 | 12 |
| 1410 | ENSMUSG( | Cog7 | 233824 | 011971661 | -01980572 | 00675 | 03075 | 12 |
| 1414 | ENSMUSG( | Col18a1 | 12822 | 008434984 | 014206076 | -0125 | 04975 | 12 |
| 1416 | ENSMUSG( | Col1a1 | 12842 | -01662661 | 000016714 | 01125 | 07025 | 12 |
| 1424 | ENSMUSG( | Col6a2 | 12834 | -02824947 | -00025194 | 00725 | 048 | 12 |
| 1438 | ENSMUSG( | Comtd1 | 69156 | -01782153 | 000444047 | -01425 | 04 | 12 |
| 1439 | ENSMUSG( | Copa | 12847 | 005931011 | -01264726 | 0105 | 05575 | 12 |
| 1440 | ENSMUSG( | Copb1 | 70349 | -00525517 | 012541923 | 0045 | 03 | 12 |
| 1445 | ENSMUSG( | Coprs | 66423 | 007755772 | -00446720 | 02375 | 0445 | 12 |
| 1452 | ENSMUSG( | Cops7b | 26895 | 007643682 | -00354119 | -01375 | 03275 | 12 |
| 1465 | ENSMUSG( | Coq8b | 76889 | 00578871C | 004358236 | 024 | 0565 | 12 |
| 1476 | ENSMUSG( | Cox11 | 69802 | -00964044 | 008650061 | 01675 | 05775 | 12 |
| 1477 | ENSMUSG( | Cox14 | 66379 | 001463486 | -02008570 | 01 | 06375 | 12 |
| 1481 | ENSMUSG( | Cox19 | 68033 | 008965224 | -00185889 | 02025 | 051 | 12 |
| 1482 | ENSMUSG( | Cox20 | 66359 | -00586634 | -01604063 | 01425 | 038 | 12 |
| 1509 | ENSMUSG( | Cpne6 | 12891 | 046946436 | 004342994 | 01625 | 03775 | 12 |
| 1514 | ENSMUSG( | Cpped1 | 223978 | -00388867 | -02097822 | 01025 | 0455 | 12 |
| 1515 | ENSMUSG( | Cpq | 54381 | -02074588 | -00421170 | -006 | 02925 | 12 |
| 1521 | ENSMUSG( | Cpsf7 | 269061 | 00112915C | 009333887 | 02625 | 068 | 12 |
| 1528 | ENSMUSG( | Cracd | 320827 | 011582725 | -00028695 | -00225 | 04025 | 12 |
| 1534 | ENSMUSG( | Crebbp | 12914 | 007343943 | 013145934 | 0145 | 07525 | 12 |
| 1537 | ENSMUSG( | Creld1 | 171508 | -02791425 | 011213091 | -01025 | 04625 | 12 |
| 1542 | ENSMUSG( | Crip1 | 12925 | -01306668 | 002411312 | -005 | 06775 | 12 |
| 1543 | ENSMUSG( | Crip2 | 68337 | 001523592 | -01028105 | 02675 | 039 | 12 |
| 1547 | ENSMUSG( | Crlf3 | 54394 | 016098964 | 024159177 | -0205 | 0405 | 12 |
| 1553 | ENSMUSG( | Crot | 74114 | 015153937 | -01286003 | 03025 | 06275 | 12 |
| 1559 | ENSMUSG( | Cry2 | 12953 | 013823236 | 015764962 | 00975 | 0395 | 12 |
| 1571 | ENSMUSG( | Csde1 | 229663 | 000685859 | -00748823 | 314018e-1 | 03625 | 12 |
| 1573 | ENSMUSG( | Csf1r | 12978 | -00890955 | 022361227 | 02175 | 04525 | 12 |
| 1595 | ENSMUSG( | Cstf2 | 108062 | 009737154 | 010128635 | 0125 | 03825 | 12 |
| 1602 | ENSMUSG( | Ctc1 | 68964 | 011241302 | 01885494C | 00775 | 05325 | 12 |
| 1605 | ENSMUSG( | Ctdp1 | 67655 | 004158028 | -00165098 | 00175 | 02675 | 12 |
| 1607 | ENSMUSG( | Ctdspl | 69274 | 00637115C | 00278911C | -00275 | 0645 | 12 |
| 1610 | ENSMUSG( | Ctif | 269037 | 021677085 | 015762625 | 002 | 035 | 12 |
| 1616 | ENSMUSG( | Ctnnbl1 | 66642 | 001980887 | -01936134 | 0135 | 03 | 12 |
| 1617 | ENSMUSG( | Ctnnd1 | 12388 | -00523582 | -01187265 | 00075 | 03 | 12 |
| 1635 | ENSMUSG( | Ctu1 | 233189 | -00494044 | 003781619 | 01075 | 0645 | 12 |
| 1647 | ENSMUSG( | Cutc | 66388 | 012530793 | 001236302 | -00975 | 05975 | 12 |
| 1719 | ENSMUSG( | Dazap1 | 70248 | 00450542C | 00214512C | 0185 | 06625 | 12 |
| 1771 | ENSMUSG( | Ddrgk1 | 77006 | -01199512 | -00436266 | 022 | 0735 | 12 |
| 1773 | ENSMUSG( | Ddx1 | 104721 | 008267321 | -01254678 | 016 | 02875 | 12 |
| 1778 | ENSMUSG( | Ddx19b | 234733 | 005632383 | -00118240 | -00425 | 07175 | 12 |
| 1783 | ENSMUSG( | Ddx25 | 30959 | 018048651 | -00438754 | 0055 | 05575 | 12 |
| 1790 | ENSMUSG( | Ddx42 | 72047 | 004083275 | -00910257 | -01375 | 04725 | 12 |
| 1800 | ENSMUSG( | Ddx6 | 13209 | 008393724 | 000898262 | 0165 | 0275 | 12 |
| 1848 | ENSMUSG( | Dhrs1 | 52585 | 002371036 | -02209721 | 006 | 034 | 12 |
| 1856 | ENSMUSG( | Dhx16 | 69192 | 016676602 | 00437927C | -00125 | 038 | 12 |
| 1857 | ENSMUSG( | Dhx29 | 218629 | 028657427 | 003961538 | 0175 | 0405 | 12 |
| 1865 | ENSMUSG( | Dhx9 | 13211 | 014927902 | -00926025 | 01925 | 04 | 12 |
| 1881 | ENSMUSG( | Disp2 | 214240 | -02693258 | 017387477 | -02125 | 05375 | 12 |
| 1884 | ENSMUSG( | Dkk3 | 50781 | -02512379 | 010049759 | 00425 | 05025 | 12 |
| 1892 | ENSMUSG( | Dlg5 | 71228 | 027270457 | 006417837 | 0215 | 03275 | 12 |
| 1904 | ENSMUSG( | Dmwd | 13401 | 011658773 | -00138331 | 00625 | 02675 | 12 |
| 1911 | ENSMUSG( | Dnaja1 | 15502 | 001238263 | -00187771 | -01425 | 0365 | 12 |
| 1929 | ENSMUSG( | Dnajc15 | 66148 | 003447399 | -00398128 | -0115 | 06675 | 12 |
| 1933 | ENSMUSG( | Dnajc2 | 22791 | 013061987 | -00707600 | -01625 | 037 | 12 |
| 1936 | ENSMUSG( | Dnajc27 | 217378 | 01417620C | -00226210 | -00325 | 0405 | 12 |
| 1939 | ENSMUSG( | Dnajc30 | 66114 | -00265718 | -01848368 | -0005 | 03875 | 12 |
| 1950 | ENSMUSG( | Dnm1l | 74006 | -00258359 | -0079903 | 017 | 043 | 12 |
| 1954 | ENSMUSG( | Dnmt1 | 13433 | 018750441 | 010337254 | 00575 | 0585 | 12 |
| 1955 | ENSMUSG( | Dnmt3a | 13435 | -00048493 | -00647434 | -00775 | 06475 | 12 |
| 1957 | ENSMUSG( | Dnph1 | 381101 | 00860799C | -00298161 | 01575 | 041 | 12 |
| 1969 | ENSMUSG( | Dock6 | 319899 | 018946239 | 022939086 | 01225 | 07225 | 12 |
| 1972 | ENSMUSG( | Dohh | 102115 | -00260636 | -01817854 | 0035 | 0375 | 12 |
| 1974 | ENSMUSG( | Dok6 | 623279 | 010002355 | 01485723C | -002 | 0305 | 12 |
| 1982 | ENSMUSG( | Dpf1 | 29861 | 005420243 | 000927051 | -00175 | 0415 | 12 |
| 1984 | ENSMUSG( | Dph1 | 116905 | 004104998 | 002803428 | 0045 | 056 | 12 |
| 1995 | ENSMUSG( | Dpp7 | 83768 | 008377668 | 01250940C | -01175 | 03025 | 12 |
| 2004 | ENSMUSG( | Dpysl3 | 22240 | 011220484 | 003452363 | 009 | 066 | 12 |
| 2006 | ENSMUSG( | Dpysl5 | 65254 | 014674044 | -00523025 | -0015 | 039 | 12 |
| 2008 | ENSMUSG( | Drap1 | 66556 | 001636791 | -01819971 | -02275 | 0505 | 12 |
| 2025 | ENSMUSG( | Dus2 | 66369 | 000626688 | -02530453 | 01475 | 084 | 12 |
| 2026 | ENSMUSG( | Dus3l | 224907 | 022171705 | 013702156 | -00775 | 058 | 12 |
| 2028 | ENSMUSG( | Dusp15 | 252864 | 001043163 | -00154796 | 00675 | 0265 | 12 |
| 2030 | ENSMUSG( | Dusp23 | 68440 | 016870377 | 010339887 | 00375 | 027 | 12 |
| 2046 | ENSMUSG( | Dync2h1 | 110350 | -01054008 | 011320469 | 01075 | 03525 | 12 |
| 2053 | ENSMUSG( | Dynlt1a | 1E+08 | 00654226C | 006819097 | -022 | 0585 | 12 |
| 2059 | ENSMUSG( | Dzip3 | 224170 | -00472658 | 002862404 | 013 | 0275 | 12 |
| 2084 | ENSMUSG( | Edrf1 | 214764 | 018377558 | 009827071 | 02925 | 06425 | 12 |
| 2093 | ENSMUSG( | Eef1g | 67160 | 009776962 | -01013431 | -004 | 0525 | 12 |
| 2122 | ENSMUSG( | Ehmt2 | 110147 | 016404281 | 004507438 | 016 | 05125 | 12 |
| 2130 | ENSMUSG( | Eif2ak2 | 19106 | 005392304 | -01367624 | 0215 | 04525 | 12 |
| 2134 | ENSMUSG( | Eif2b2 | 217715 | 001372622 | -00965938 | -008 | 02925 | 12 |
| 2139 | ENSMUSG( | Eif2s1 | 13665 | 00847872C | 001823226 | 01375 | 0285 | 12 |
| 2143 | ENSMUSG( | Eif3a | 13669 | 014542849 | -00064812 | 018 | 0335 | 12 |
| 2149 | ENSMUSG( | Eif3g | 53356 | -00160835 | -01158047 | 0145 | 03825 | 12 |
| 2153 | ENSMUSG( | Eif3k | 73830 | 000104772 | -02078717 | -00475 | 03625 | 12 |
| 2156 | ENSMUSG( | Eif4a1 | 13681 | 000681352 | -00392050 | -0065 | 02775 | 12 |
| 2175 | ENSMUSG( | Elac2 | 68626 | 019415077 | -00032410 | 01625 | 0255 | 12 |
| 2194 | ENSMUSG( | Elovl5 | 68801 | -03526431 | -00297751 | 0135 | 066 | 12 |
| 2197 | ENSMUSG( | Elp3 | 74195 | 006675533 | -01918314 | 01025 | 04525 | 12 |
| 2214 | ENSMUSG( | Eml2 | 72205 | 003632339 | 009515223 | 0155 | 03175 | 12 |
| 2219 | ENSMUSG( | Emp2 | 13731 | 007107997 | -03578434 | -0115 | 03625 | 12 |
| 2225 | ENSMUSG( | Endou | 19011 | -02867037 | -01332679 | 02275 | 07825 | 12 |
| 2270 | ENSMUSG( | Ephx4 | 384214 | -02286540 | 00280782C | 01325 | 07125 | 12 |
| 2272 | ENSMUSG( | Epm2aip1 | 77781 | 002668914 | 003166856 | 01675 | 045 | 12 |
| 2276 | ENSMUSG( | Eprs1 | 107508 | 003456026 | -01462960 | 02025 | 034 | 12 |
| 2288 | ENSMUSG( | Ercc2 | 13871 | 01673748C | 001714671 | 0015 | 06 | 12 |
| 2290 | ENSMUSG( | Ercc4 | 50505 | 014358916 | 004777492 | 00075 | 04325 | 12 |
| 2325 | ENSMUSG( | Ewsr1 | 14030 | -00570136 | -00473963 | -0115 | 06475 | 12 |

| | | | | | | | | |
|---|---|---|---|---|---|---|---|---|
| 2343 | ENSMUSG( | Exosc7 | 66446 | 004054423 | -01579311 | -00875 | 03975 | 12 |
| 2345 | ENSMUSG( | Exosc9 | 50911 | 008401467 | -01674830 | 019 | 037 | 12 |
| 2371 | ENSMUSG( | Faim | 23873 | -00285071 | -01228071 | 003 | 0375 | 12 |
| 2376 | ENSMUSG( | Fam114a1 | 68303 | 009627247 | -01042763 | 02425 | 035 | 12 |
| 2378 | ENSMUSG( | Fam117b | 72750 | 003516217 | 001555278 | -0185 | 03675 | 12 |
| 2379 | ENSMUSG( | Fam118a | 73225 | -00195324 | 007269635 | -00025 | 0415 | 12 |
| 2396 | ENSMUSG( | Fam171b | 241520 | -00318675 | 015582597 | 0075 | 03125 | 12 |
| 2403 | ENSMUSG( | Fam20c | 80752 | -01629192 | 028454994 | 0045 | 045 | 12 |
| 2418 | ENSMUSG( | Fam98a | 72722 | 013560803 | 003089142 | 012 | 04525 | 12 |
| 2419 | ENSMUSG( | Fam98b | 68215 | -00090208 | -00687904 | 0085 | 0335 | 12 |
| 2420 | ENSMUSG( | Fam98c | 73833 | -00079566 | -00694169 | -00875 | 0345 | 12 |
| 2425 | ENSMUSG( | Farsa | 66590 | -00061698 | 014028680 | 006 | 02675 | 12 |
| 2434 | ENSMUSG( | Fbh1 | 50755 | 018656477 | 014787486 | -017 | 04025 | 12 |
| 2437 | ENSMUSG( | Fbln1 | 14114 | -01269164 | 021730901 | -00125 | 03725 | 12 |
| 2443 | ENSMUSG( | Fbxl17 | 50758 | 011946837 | -00004182 | 01375 | 06825 | 12 |
| 2444 | ENSMUSG( | Fbxl18 | 231863 | 022123033 | 018464358 | -00225 | 04175 | 12 |
| 2448 | ENSMUSG( | Fbxo10 | 269529 | -00445774 | 013967144 | 005 | 0525 | 12 |
| 2452 | ENSMUSG( | Fbxo22 | 71999 | 009095586 | -00671254 | -009 | 03425 | 12 |
| 2454 | ENSMUSG( | Fbxo30 | 71865 | 008720302 | 000525625 | -0025 | 03275 | 12 |
| 2461 | ENSMUSG( | Fbxo45 | 268882 | 020102524 | 003380561 | 0065 | 02825 | 12 |
| 2484 | ENSMUSG( | Fem1b | 14155 | 025694244 | 011825311 | 0305 | 04825 | 12 |
| 2485 | ENSMUSG( | Fem1c | 240263 | -00608184 | -00510836 | -011 | 05375 | 12 |
| 2509 | ENSMUSG( | Fhl2 | 14200 | 004430691 | -01556328 | 017 | 03325 | 12 |
| 2528 | ENSMUSG( | Fkbpl | 56299 | 009307993 | -01968841 | -0115 | 0345 | 12 |
| 2543 | ENSMUSG( | Fmn2 | 54418 | -00030393 | -00132753 | 0015 | 03825 | 12 |
| 2549 | ENSMUSG( | Fmr1 | 14265 | -00068985 | 012031455 | 011 | 03125 | 12 |
| 2554 | ENSMUSG( | Fnbp1l | 214459 | 013253300 | 019418192 | 01175 | 03975 | 12 |
| 2575 | ENSMUSG( | Frmd4a | 209630 | -00866933 | -01393478 | 004 | 031 | 12 |
| 2579 | ENSMUSG( | Frmpd1 | 666060 | 025495554 | -03386830 | -01425 | 055 | 12 |
| 2593 | ENSMUSG( | Fto | 26383 | 006170768 | -01541213 | 017 | 03525 | 12 |
| 2596 | ENSMUSG( | Fubp1 | 51886 | -00501911 | 009948081 | -0205 | 0555 | 12 |
| 2602 | ENSMUSG( | Fus | 233908 | -00747724 | -00772577 | 0115 | 03375 | 12 |
| 2620 | ENSMUSG( | G3bp2 | 23881 | 003140466 | -00064324 | 007 | 046 | 12 |
| 2630 | ENSMUSG( | Gabpa | 14390 | 010736262 | -00287413 | 02475 | 04175 | 12 |
| 2685 | ENSMUSG( | Gba2 | 230101 | 011006684 | 016680894 | 02075 | 03675 | 12 |
| 2715 | ENSMUSG( | Gemin7 | 69731 | 009967371 | -00220506 | 005 | 0445 | 12 |
| 2744 | ENSMUSG( | Gigyf2 | 227331 | 026575664 | 006169646 | 0195 | 067 | 12 |
| 2745 | ENSMUSG( | Gipc1 | 67903 | 00639224 | -01230027 | 0155 | 0325 | 12 |
| 2784 | ENSMUSG( | Gm20517 | 1.13E+08 | -01901288 | 015496941 | 0105 | 04425 | 12 |
| 2789 | ENSMUSG( | Gmeb1 | 56809 | -00223072 | -00026293 | -00725 | 042 | 12 |
| 2808 | ENSMUSG( | Gnb1 | 14688 | 009791558 | -00997546 | 0165 | 036 | 12 |
| 2819 | ENSMUSG( | Gng4 | 14706 | 002208105 | -00637362 | 01825 | 0425 | 12 |
| 2823 | ENSMUSG( | Gnl3 | 30877 | 007298704 | 010905000 | 00125 | 03225 | 12 |
| 2831 | ENSMUSG( | Golga1 | 76899 | 011233124 | 008307008 | 016 | 0295 | 12 |
| 2834 | ENSMUSG( | Golga4 | 54214 | -00017764 | -00799754 | 00225 | 034 | 12 |
| 2841 | ENSMUSG( | Golph3 | 66629 | 007135073 | -01182599 | 0035 | 0295 | 12 |
| 2844 | ENSMUSG( | Gopc | 94221 | -00506409 | 002958464 | 00675 | 02575 | 12 |
| 2845 | ENSMUSG( | Gorab | 98376 | -01872214 | -00320636 | 0205 | 041 | 12 |
| 2854 | ENSMUSG( | Gpam | 14732 | 018836946 | -00054417 | 00675 | 04 | 12 |
| 2863 | ENSMUSG( | Gpc5 | 103978 | 00266497 | 009361719 | -007 | 03 | 12 |
| 2864 | ENSMUSG( | Gpc6 | 23888 | 019202365 | 017989277 | 01475 | 03375 | 12 |
| 2870 | ENSMUSG( | Gpkow | 209416 | 005393733 | -01101729 | 02525 | 0545 | 12 |
| 2874 | ENSMUSG( | Gpn1 | 74254 | 009405564 | -01487970 | -00225 | 0515 | 12 |
| 2888 | ENSMUSG( | Gprasp1 | 67298 | 009351543 | 007242248 | -0125 | 04525 | 12 |
| 2896 | ENSMUSG( | Gpsm1 | 67839 | 004172633 | -00415421 | -004 | 02925 | 12 |
| 2898 | ENSMUSG( | Gpt2 | 108682 | 034853861 | 011364713 | 03075 | 06325 | 12 |
| 2907 | ENSMUSG( | Grb10 | 14783 | 010475296 | 014152429 | 00475 | 049 | 12 |
| 2918 | ENSMUSG( | Grik1 | 14805 | 001915016 | 023175573 | 00675 | 0765 | 12 |
| 2925 | ENSMUSG( | Grin2d | 14814 | -00426866 | 015096670 | 00125 | 03725 | 12 |
| 2971 | ENSMUSG( | Gtf2a1 | 83602 | -00715222 | 002878599 | -01025 | 03825 | 12 |
| 2973 | ENSMUSG( | Gtf2b | 229906 | 008850066 | -00119791 | 0005 | 061 | 12 |
| 2974 | ENSMUSG( | Gtf2e1 | 74197 | 008888815 | -01867303 | 00175 | 073 | 12 |
| 2987 | ENSMUSG( | Gtpbp1 | 14904 | 014508921 | -00172230 | 00275 | 07625 | 12 |
| 3013 | ENSMUSG( | Habp4 | 56541 | 010702806 | -01128017 | 02175 | 03325 | 12 |
| 3017 | ENSMUSG( | Hacl1 | 56794 | -00146522 | -05208081 | 03775 | 05925 | 12 |
| 3022 | ENSMUSG( | Haghl | 68977 | 001952648 | -00200558 | -01275 | 033 | 12 |
| 3023 | ENSMUSG( | Hap1 | 15114 | 020299028 | 005286421 | -00025 | 029 | 12 |
| 3039 | ENSMUSG( | Hcn3 | 15168 | -01709159 | -00461845 | 007 | 04775 | 12 |
| 3053 | ENSMUSG( | Hdgf | 15191 | 004468619 | -00765242 | -00325 | 057 | 12 |
| 3059 | ENSMUSG( | Hdlbp | 110611 | 0190214 | 001890326 | 01975 | 032 | 12 |
| 3064 | ENSMUSG( | Heatr6 | 217026 | 006268728 | 001176879 | -01875 | 05375 | 12 |
| 3067 | ENSMUSG( | Heca | 380629 | 014159632 | 018983349 | -01375 | 038 | 12 |
| 3074 | ENSMUSG( | Helz | 78455 | 008285597 | 009172482 | -01725 | 0345 | 12 |
| 3091 | ENSMUSG( | Hid1 | 217310 | -00220221 | -00593803 | -00425 | 03775 | 12 |
| 3093 | ENSMUSG( | Higd1a | 56295 | 001094563 | -02016259 | -0155 | 0565 | 12 |
| 3111 | ENSMUSG( | Hmgb1 | 15289 | 006806070 | -00407765 | -00975 | 049 | 12 |
| 3113 | ENSMUSG( | Hmgb3 | 15354 | -01637168 | -01515336 | -00075 | 0495 | 12 |
| 3140 | ENSMUSG( | Hnrnpll | 72692 | -01250468 | -01133127 | -01375 | 036 | 12 |
| 3144 | ENSMUSG( | Hnrnpul1 | 232989 | 005471238 | -00788676 | 00725 | 0405 | 12 |
| 3178 | ENSMUSG( | Hsf1 | 15499 | 019541259 | 009929195 | 02 | 03075 | 12 |
| 3201 | ENSMUSG( | Htatsf1 | 72459 | 014116171 | -00973749 | -0205 | 0385 | 12 |
| 3212 | ENSMUSG( | Hykk | 235386 | 008121911 | -00501488 | 011 | 0415 | 12 |
| 3223 | ENSMUSG( | Icmt | 57295 | -01234339 | 012935752 | -009 | 04075 | 12 |
| 3232 | ENSMUSG( | Iffo1 | 320678 | -01911785 | 002658525 | 00925 | 0445 | 12 |
| 3241 | ENSMUSG( | Ift46 | 76568 | -00354992 | -01869713 | 0165 | 05275 | 12 |
| 3247 | ENSMUSG( | Ift88 | 21821 | 005996196 | 014196234 | -021 | 0535 | 12 |
| 3253 | ENSMUSG( | Igfbp5 | 16011 | 006050862 | 007064162 | 03175 | 0485 | 12 |
| 3254 | ENSMUSG( | Igfbp6 | 16012 | -02593105 | -02127908 | 0055 | 0355 | 12 |
| 3257 | ENSMUSG( | Ighm | 16019 | 039305999 | 005132239 | -01575 | 037 | 12 |
| 3261 | ENSMUSG( | Igsf1 | 209268 | -04090915 | -05413602 | 024 | 07075 | 12 |
| 3283 | ENSMUSG( | Ilrun | 224647 | 016592451 | -00309465 | 03125 | 064 | 12 |
| 3304 | ENSMUSG( | Inpp5d | 16331 | 024990956 | 028126215 | 02825 | 079 | 12 |
| 3331 | ENSMUSG( | Ipo11 | 76582 | 006199654 | -00918154 | -00775 | 0485 | 12 |
| 3332 | ENSMUSG( | Ipo13 | 230673 | -00452068 | -00100887 | -00225 | 0355 | 12 |
| 3333 | ENSMUSG( | Ipo4 | 75751 | 010588115 | -01204603 | 007 | 02975 | 12 |
| 3341 | ENSMUSG( | Iqsec1 | 232227 | 013425161 | 007680356 | -0015 | 04575 | 12 |
| 3351 | ENSMUSG( | Irgq | 210146 | 019523932 | 005158841 | 00425 | 03775 | 12 |
| 3360 | ENSMUSG( | Isoc1 | 66307 | 001191126 | 001048312 | 01125 | 04725 | 12 |
| 3377 | ENSMUSG( | Itgb1 | 16412 | -00980474 | 014814405 | -00725 | 0305 | 12 |
| 3420 | ENSMUSG( | Jpt1 | 15374 | 005038748 | -01247031 | 01075 | 047 | 12 |
| 3421 | ENSMUSG( | Jpt2 | 52009 | -00578889 | -00605374 | 004 | 07575 | 12 |
| 3422 | ENSMUSG( | Jtb | 23922 | -00982635 | 007781297 | 00175 | 0685 | 12 |
| 3431 | ENSMUSG( | Kars1 | 85305 | 004260128 | -02083014 | -00025 | 05275 | 12 |
| 3433 | ENSMUSG( | Kat5 | 81601 | 005805038 | -00478984 | -002 | 0555 | 12 |
| 3481 | ENSMUSG( | Kcnk10 | 72258 | -02064046 | -00415470 | 01925 | 04325 | 12 |
| 3487 | ENSMUSG( | Kcnn3 | 140493 | -02196247 | -01402989 | 0045 | 03225 | 12 |
| 3495 | ENSMUSG( | Kctd1 | 106931 | 008620490 | 012058595 | 02275 | 03 | 12 |
| 3497 | ENSMUSG( | Kctd12 | 239217 | 006225551 | 030370739 | 00125 | 0595 | 12 |
| 3504 | ENSMUSG( | Kctd3 | 226823 | 015228878 | -00208412 | -00325 | 03525 | 12 |
| 3513 | ENSMUSG( | Kdm3b | 277250 | 006568766 | -00266235 | -0125 | 047 | 12 |
| 3524 | ENSMUSG( | Khnyn | 219094 | 004467561 | 012182236 | 00875 | 06925 | 12 |
| 3525 | ENSMUSG( | Khsrp | 16549 | 016842317 | 003318454 | -00575 | 0525 | 12 |
| 3536 | ENSMUSG( | Kif3a | 16568 | 014874742 | 000065312 | -027 | 052 | 12 |
| 3543 | ENSMUSG( | Kifbp | 72320 | 002994580 | -01614080 | -0055 | 041 | 12 |
| 3544 | ENSMUSG( | Kifc2 | 16581 | 014476065 | 001969411 | -01675 | 036 | 12 |
| 3555 | ENSMUSG( | Klhdc3 | 71765 | 018371169 | -00910694 | 0245 | 04875 | 12 |
| 3564 | ENSMUSG( | Klhl22 | 224023 | 016557854 | -00254218 | 0065 | 03 | 12 |
| 3570 | ENSMUSG( | Klhl5 | 71778 | 006025420 | -01650257 | -005 | 062 | 12 |
| 3575 | ENSMUSG( | Kpna1 | 16646 | -00737573 | 014446562 | -02 | 04725 | 12 |
| 3590 | ENSMUSG( | Kti12 | 100087 | 002594051 | 010708976 | -006 | 05225 | 12 |
| 3604 | ENSMUSG( | Lamb1 | 16777 | 004453361 | 030383486 | 01775 | 05525 | 12 |
| 3623 | ENSMUSG( | Larp4 | 207214 | -00583438 | 009770708 | 0085 | 03225 | 12 |
| 3624 | ENSMUSG( | Larp4b | 217980 | 009266947 | -00035889 | 023 | 06875 | 12 |
| 3625 | ENSMUSG( | Larp6 | 67557 | -00539666 | -02734954 | 01075 | 03 | 12 |
| 3626 | ENSMUSG( | Larp7 | 28036 | -01509250 | -01311952 | -02075 | 04875 | 12 |
| 3629 | ENSMUSG( | Las1l | 76130 | 017718561 | -00221867 | -00575 | 052 | 12 |

| | | | | | | | | |
|---|---|---|---|---|---|---|---|---|
| 3638 | ENSMUSG( | Ldah | 68832 | -00162623 | -01941485 | 03475 | 0535 | 12 |
| 3666 | ENSMUSG( | Lias | 79464 | -00458083 | -01433034 | 0035 | 02975 | 12 |
| 3668 | ENSMUSG( | Lig3 | 16882 | 011008358 | -00223993 | 001 | 03625 | 12 |
| 3669 | ENSMUSG( | Lima1 | 65970 | 009278908 | 019348084 | -00025 | 04075 | 12 |
| 3671 | ENSMUSG( | Limd1 | 29806 | -00303085 | -00642079 | 01475 | 06275 | 12 |
| 3674 | ENSMUSG( | Limk2 | 16886 | 015235310 | 003292665 | -013 | 04325 | 12 |
| 3675 | ENSMUSG( | Lims1 | 110829 | 002894976 | 007863840 | -00725 | 03675 | 12 |
| 3677 | ENSMUSG( | Lin7b | 22342 | -00938667 | -00834063 | -00525 | 042 | 12 |
| 3685 | ENSMUSG( | Lipt1 | 623661 | 007616883 | 021586116 | 0115 | 05675 | 12 |
| 3687 | ENSMUSG( | Lix1 | 66643 | 009353319 | -02431945 | -00925 | 038 | 12 |
| 3737 | ENSMUSG( | Lrp6 | 16974 | -00218938 | 018192811 | 01475 | 044 | 12 |
| 3763 | ENSMUSG( | Lrrfip1 | 16978 | 026984544 | -00408215 | -00975 | 06075 | 12 |
| 3764 | ENSMUSG( | Lrrfip2 | 71268 | 006179298 | -00549875 | 0065 | 0285 | 12 |
| 3774 | ENSMUSG( | Lsg1 | 224092 | 009057323 | -00242182 | 00825 | 0385 | 12 |
| 3776 | ENSMUSG( | Lsm11 | 72290 | 032841911 | -00322795 | -01225 | 04325 | 12 |
| 3778 | ENSMUSG( | Lsm14a | 67070 | 003421196 | -00317386 | 00125 | 03075 | 12 |
| 3779 | ENSMUSG( | Lsm14b | 241846 | 012731409 | 003096935 | 00725 | 026 | 12 |
| 3785 | ENSMUSG( | Lsm7 | 66094 | -00241874 | -02492307 | 01 | 02925 | 12 |
| 3791 | ENSMUSG( | Ltbp4 | 108075 | 017113948 | 037186350 | -0075 | 066 | 12 |
| 3793 | ENSMUSG( | Luc7l | 66978 | -00729137 | 011396716 | 039 | 05125 | 12 |
| 3797 | ENSMUSG( | Lurap1 | 68075 | 004706153 | -00878559 | -01675 | 047 | 12 |
| 3832 | ENSMUSG( | Macrod1 | 107227 | 006195236 | -02472137 | 01425 | 03525 | 12 |
| 3840 | ENSMUSG( | Maea | 59003 | 004850207 | -00885962 | 00725 | 0305 | 12 |
| 3841 | ENSMUSG( | Mafg | 17134 | 016598451 | 018973766 | 014 | 07125 | 12 |
| 3859 | ENSMUSG( | Man2b1 | 17159 | -00013389 | 030336954 | 029 | 049 | 12 |
| 3879 | ENSMUSG( | Map2k5 | 23938 | 008090256 | 001572429 | -00525 | 03825 | 12 |
| 3884 | ENSMUSG( | Map3k12 | 26404 | 019760629 | 00369384 | 00825 | 027 | 12 |
| 3890 | ENSMUSG( | Map3k7 | 26409 | 005449364 | -00184527 | 0105 | 036 | 12 |
| 3895 | ENSMUSG( | Map4k4 | 26921 | -00336382 | -02420566 | 00675 | 0655 | 12 |
| 3911 | ENSMUSG( | Mapk8 | 26419 | 000920191 | -01061818 | -0095 | 03475 | 12 |
| 3912 | ENSMUSG( | Mapk8ip1 | 19099 | 013488417 | -00370314 | -011 | 0295 | 12 |
| 3919 | ENSMUSG( | Mapre1 | 13589 | 003428568 | -01959843 | 00375 | 03775 | 12 |
| 3922 | ENSMUSG( | Mapt | 17762 | 008851953 | -01522162 | -01575 | 0435 | 12 |
| 3929 | ENSMUSG( | Mark1 | 226778 | 017772428 | -00034638 | -014 | 039 | 12 |
| 3933 | ENSMUSG( | Mars1 | 216443 | 008191494 | 009471328 | 02675 | 0385 | 12 |
| 3938 | ENSMUSG( | Mast2 | 17776 | 032115207 | 008858829 | 022 | 06075 | 12 |
| 3943 | ENSMUSG( | Matk | 17179 | 021854553 | 008936523 | -0015 | 03625 | 12 |
| 3953 | ENSMUSG( | Mbd3 | 17192 | 005200837 | -01127949 | -022 | 047 | 12 |
| 3954 | ENSMUSG( | Mblac1 | 330216 | -00839325 | -00414366 | 003 | 03125 | 12 |
| 4002 | ENSMUSG( | Med1 | 19014 | 002780703 | 010038416 | -0245 | 0535 | 12 |
| 4016 | ENSMUSG( | Med4 | 67381 | -00258291 | -00197073 | -0055 | 048 | 12 |
| 4029 | ENSMUSG( | Mepce | 231803 | 004440865 | 017137675 | -0145 | 05675 | 12 |
| 4042 | ENSMUSG( | Mettl2 | 52686 | 015805311 | 025906367 | 00925 | 06675 | 12 |
| 4045 | ENSMUSG( | Mettl26 | 68347 | 005776434 | -01516134 | 0035 | 03125 | 12 |
| 4046 | ENSMUSG( | Mettl3 | 56335 | 017649774 | 020711036 | -01625 | 052 | 12 |
| 4105 | ENSMUSG( | Mlc1 | 170790 | 004022741 | -00752270 | 0115 | 05 | 12 |
| 4110 | ENSMUSG( | Mllt6 | 246198 | 019344352 | 007984425 | 0055 | 043 | 12 |
| 4137 | ENSMUSG( | Mospd1 | 70380 | -00133882 | -00067175 | 01925 | 03325 | 12 |
| 4145 | ENSMUSG( | Mphosph6 | 68533 | 023933880 | 020292678 | 00325 | 04225 | 12 |
| 4188 | ENSMUSG( | Mrpl32 | 75398 | -01426461 | -00174004 | 03 | 063 | 12 |
| 4195 | ENSMUSG( | Mrpl4 | 66163 | 001845323 | -00939216 | 01975 | 0585 | 12 |
| 4199 | ENSMUSG( | Mrpl43 | 94067 | -00043986 | -01303487 | -0075 | 05825 | 12 |
| 4200 | ENSMUSG( | Mrpl44 | 69163 | 012938458 | -00301270 | 00925 | 03175 | 12 |
| 4219 | ENSMUSG( | Mrps16 | 66242 | 000167317 | -01266348 | -006 | 04125 | 12 |
| 4221 | ENSMUSG( | Mrps18a | 68565 | 001571540 | -00911397 | -001 | 03825 | 12 |
| 4223 | ENSMUSG( | Mrps18c | 68735 | -00721837 | -02067333 | 0245 | 0505 | 12 |
| 4262 | ENSMUSG( | Mta3 | 116871 | 023136120 | 007420174 | -0035 | 03025 | 12 |
| 4269 | ENSMUSG( | Mterf2 | 74238 | 030747901 | -01352162 | 01525 | 02825 | 12 |
| 4281 | ENSMUSG( | Mtmr1 | 53332 | 013827593 | 000077461 | 0305 | 049 | 12 |
| 4302 | ENSMUSG( | Mturn | 68235 | 016939745 | -00425140 | 00925 | 03275 | 12 |
| 4309 | ENSMUSG( | Mvb12b | 72543 | 014147136 | 010016418 | -003 | 051 | 12 |
| 4310 | ENSMUSG( | Mvd | 192156 | 013801567 | -02736165 | -013 | 04225 | 12 |
| 4321 | ENSMUSG( | Myg1 | 60315 | -00350687 | -00478788 | 016 | 03125 | 12 |
| 4351 | ENSMUSG( | Myt1l | 17933 | -00206335 | 024189130 | 02625 | 057 | 12 |
| 4355 | ENSMUSG( | N4bp2l2 | 381695 | 001691610 | 014238683 | 023 | 05825 | 12 |
| 4362 | ENSMUSG( | Naa30 | 70646 | 000985797 | 012503201 | 02025 | 02975 | 12 |
| 4364 | ENSMUSG( | Naa38 | 78304 | -00808038 | -02111818 | 0015 | 03475 | 12 |
| 4365 | ENSMUSG( | Naa40 | 70999 | -00070840 | 015634569 | -117757e-1 | 03575 | 12 |
| 4366 | ENSMUSG( | Naa50 | 72117 | -00030219 | -00165528 | 005 | 0325 | 12 |
| 4370 | ENSMUSG( | Naca | 17938 | 009616067 | -00965029 | -00425 | 03275 | 12 |
| 4371 | ENSMUSG( | Nacad | 192950 | -01569391 | -00357146 | 00775 | 04825 | 12 |
| 4394 | ENSMUSG( | Nasp | 50927 | 012549797 | 009758372 | 00775 | 06525 | 12 |
| 4402 | ENSMUSG( | Nav1 | 215690 | -01852693 | 011501744 | -00175 | 052 | 12 |
| 4403 | ENSMUSG( | Nav2 | 78286 | 014105378 | 017392546 | 196262e-1 | 0475 | 12 |
| 4415 | ENSMUSG( | Ncan | 13004 | -01968203 | -00854950 | -0035 | 0345 | 12 |
| 4421 | ENSMUSG( | Ncf1 | 17969 | 003343904 | 010495994 | 00025 | 03525 | 12 |
| 4422 | ENSMUSG( | Ncf2 | 17970 | -00078021 | -00658132 | -00075 | 043 | 12 |
| 4426 | ENSMUSG( | Nckap1l | 105855 | 013841197 | 008783009 | -002 | 0345 | 12 |
| 4431 | ENSMUSG( | Ncoa1 | 17977 | 015753784 | 004800304 | 0215 | 06875 | 12 |
| 4444 | ENSMUSG( | Ndn | 17984 | 017527605 | 007133532 | 02525 | 07 | 12 |
| 4504 | ENSMUSG( | Nectin3 | 58998 | -01640175 | 004653953 | -00125 | 0325 | 12 |
| 4538 | ENSMUSG( | Nfia | 18027 | 012085648 | -00083668 | 0115 | 05875 | 12 |
| 4539 | ENSMUSG( | Nfib | 18028 | 015752630 | 017079845 | 00525 | 04275 | 12 |
| 4540 | ENSMUSG( | Nfic | 18029 | 016928077 | 007700277 | -02325 | 06 | 12 |
| 4548 | ENSMUSG( | Nfyc | 18046 | 008483817 | -00131867 | -0075 | 03475 | 12 |
| 4566 | ENSMUSG( | Nip7 | 66164 | 004108959 | -00683375 | 01025 | 0475 | 12 |
| 4576 | ENSMUSG( | Nkain4 | 58237 | 002515900 | -00102886 | 003 | 07025 | 12 |
| 4586 | ENSMUSG( | Nmd3 | 97112 | -00871138 | -01651131 | -00875 | 06125 | 12 |
| 4590 | ENSMUSG( | Nmnat2 | 226518 | 006065870 | -00658260 | 02075 | 065 | 12 |
| 4593 | ENSMUSG( | Nmt1 | 18107 | 012737057 | 006516207 | 0035 | 03175 | 12 |
| 4604 | ENSMUSG( | Nol3 | 78688 | -00243492 | -02253055 | -007 | 03625 | 12 |
| 4612 | ENSMUSG( | Nono | 53610 | 001340621 | -01213503 | 004 | 046 | 12 |
| 4615 | ENSMUSG( | Nop16 | 28126 | 019205438 | 002273311 | -01575 | 0325 | 12 |
| 4633 | ENSMUSG( | Npc2 | 67963 | -01406787 | -01059228 | 004 | 04775 | 12 |
| 4654 | ENSMUSG( | Nr1d2 | 353187 | 009568181 | 059035976 | 02825 | 06475 | 12 |
| 4669 | ENSMUSG( | Nrip3 | 78593 | 012437287 | -00231506 | 026 | 0475 | 12 |
| 4681 | ENSMUSG( | Nsdhl | 18194 | -01376565 | -00888499 | 015 | 04375 | 12 |
| 4690 | ENSMUSG( | Nsun2 | 28114 | 010488932 | -01707042 | 01475 | 0305 | 12 |
| 4723 | ENSMUSG( | Nudt11 | 58242 | 001041438 | 001678945 | 0085 | 03975 | 12 |
| 4724 | ENSMUSG( | Nudt12 | 67993 | 021750161 | 012948823 | -01625 | 03925 | 12 |
| 4734 | ENSMUSG( | Nudt21 | 68219 | 004123721 | 008647355 | 01625 | 08175 | 12 |
| 4741 | ENSMUSG( | Nufip2 | 68564 | -00963352 | 021212393 | 0235 | 045 | 12 |
| 4775 | ENSMUSG( | Nyap2 | 241134 | -03670592 | -01253095 | -0085 | 053 | 12 |
| 4776 | ENSMUSG( | Oard1 | 106821 | -00339048 | -01967267 | -02275 | 0485 | 12 |
| 4784 | ENSMUSG( | Oga | 76055 | 013744332 | -01499546 | 00775 | 02925 | 12 |
| 4812 | ENSMUSG( | Orai2 | 269717 | -00420527 | 020458367 | 01425 | 044 | 12 |
| 4830 | ENSMUSG( | Ostf1 | 20409 | -01716378 | -00756735 | 013 | 03275 | 12 |
| 4835 | ENSMUSG( | Otud4 | 73945 | -01663172 | 002088061 | -01725 | 05125 | 12 |
| 4876 | ENSMUSG( | Pak4 | 70584 | -01478739 | -01794022 | -01175 | 061 | 12 |
| 4878 | ENSMUSG( | Pak6 | 214230 | 018309626 | 010178570 | 0235 | 04025 | 12 |
| 4888 | ENSMUSG( | Pam16 | 66449 | -00162116 | -01930942 | -02625 | 0575 | 12 |
| 4897 | ENSMUSG( | Papola | 18789 | 012119085 | 007103434 | 004 | 035 | 12 |
| 4902 | ENSMUSG( | Paqr9 | 75552 | -01808918 | 030154604 | 00775 | 05575 | 12 |
| 4908 | ENSMUSG( | Parl | 381038 | -01313015 | -01835444 | -014 | 03975 | 12 |
| 4911 | ENSMUSG( | Parp1 | 11545 | 011296699 | -00351629 | -0165 | 056 | 12 |
| 4915 | ENSMUSG( | Parvb | 170736 | 000763539 | -02026075 | 00025 | 0415 | 12 |
| 4918 | ENSMUSG( | Pawr | 114774 | -00909457 | 0027155 | 0135 | 06525 | 12 |
| 4924 | ENSMUSG( | Pbxip1 | 229534 | -00962709 | -01930051 | 02575 | 042 | 12 |
| 4925 | ENSMUSG( | Pcbd1 | 13180 | 017934992 | 001270558 | 00775 | 0345 | 12 |
| 4933 | ENSMUSG( | Pcdh1 | 75599 | -01963700 | 030900619 | -01675 | 04725 | 12 |
| 4942 | ENSMUSG( | Pcdha12 | 192164 | -03374991 | -00953655 | -00425 | 05275 | 12 |
| 4955 | ENSMUSG( | Pcdhga5 | 93713 | -00413978 | 011934063 | 01775 | 061 | 12 |
| 4974 | ENSMUSG( | Pcmtd2 | 245867 | -00246050 | 007963615 | -012 | 039 | 12 |
| 4976 | ENSMUSG( | Pcnp | 76302 | -00418031 | 008194801 | -0205 | 057 | 12 |
| 4995 | ENSMUSG( | Pdcd4 | 18569 | 003618529 | 004523012 | 0185 | 02475 | 12 |
| 5013 | ENSMUSG( | Pde5a | 242202 | 000624394 | 031205966 | -00475 | 04475 | 12 |

| | | | | | | | | |
|---|---|---|---|---|---|---|---|---|
| 5015 | ENSMUSG( | Pde7a | 18583 | 003040025 | 008373363 | 02475 | 0455 | 12 |
| 5036 | ENSMUSG( | Pdlim4 | 30794 | 021383492 | -00585812 | -0125 | 056 | 12 |
| 5037 | ENSMUSG( | Pdlim5 | 56376 | 006888633 | -00125258 | 022 | 0345 | 12 |
| 5049 | ENSMUSG( | Pdzd11 | 72621 | 000483519 | -02363471 | 018 | 0385 | 12 |
| 5063 | ENSMUSG( | Pelp1 | 75273 | 009482106 | -00448074 | 00875 | 03275 | 12 |
| 5104 | ENSMUSG( | Pgls | 66171 | 000418437 | -01086467 | -0005 | 03825 | 12 |
| 5107 | ENSMUSG( | Pgm2l1 | 70974 | 015733743 | 008205171 | 01325 | 04075 | 12 |
| 5118 | ENSMUSG( | Phactr4 | 100169 | 006703007 | 022070650 | 001 | 03825 | 12 |
| 5122 | ENSMUSG( | Phb2 | 12034 | 000866453 | -00662221 | -0115 | 0355 | 12 |
| 5130 | ENSMUSG( | Phf3 | 213109 | 004009809 | 004188861 | -02075 | 0465 | 12 |
| 5146 | ENSMUSG( | Phlpp2 | 244650 | 007579061 | 009633781 | 00775 | 05275 | 12 |
| 5148 | ENSMUSG( | Phospho2 | 73373 | 006192148 | -00057822 | 01675 | 0385 | 12 |
| 5157 | ENSMUSG( | Pi4kb | 107650 | 015743652 | 010927022 | 00225 | 03 | 12 |
| 5162 | ENSMUSG( | Pid1 | 98496 | -00507112 | -00626700 | 00725 | 03475 | 12 |
| 5179 | ENSMUSG( | Pik3ip1 | 216505 | -03066107 | 014442425 | 009 | 05875 | 12 |
| 5206 | ENSMUSG( | Pkia | 18767 | 002370912 | -00606020 | -00825 | 061 | 12 |
| 5208 | ENSMUSG( | Pkig | 18769 | 007568552 | -01242987 | -009 | 045 | 12 |
| 5229 | ENSMUSG( | Plcg2 | 234779 | 014634414 | 011460129 | -00725 | 045 | 12 |
| 5247 | ENSMUSG( | Plekhf2 | 71801 | 007326369 | 016582235 | -0095 | 044 | 12 |
| 5259 | ENSMUSG( | Plod3 | 26433 | -01139580 | 026769771 | 0225 | 07425 | 12 |
| 5282 | ENSMUSG( | Plxna3 | 18846 | 019240874 | 030992448 | 01275 | 04075 | 12 |
| 5301 | ENSMUSG( | Pnma2 | 239157 | 011579928 | -00090549 | 00225 | 05325 | 12 |
| 5321 | ENSMUSG( | Polb | 18970 | 010994787 | 005310034 | 011 | 06775 | 12 |
| 5322 | ENSMUSG( | Pold2 | 18972 | 009504875 | -00445917 | 028 | 08275 | 12 |
| 5337 | ENSMUSG( | Polr2j | 20022 | 003567948 | -01455037 | -0175 | 03975 | 12 |
| 5339 | ENSMUSG( | Polr2m | 28015 | 013199050 | -00740724 | -0035 | 05125 | 12 |
| 5341 | ENSMUSG( | Polr3b | 70428 | 000690469 | 002193555 | 02775 | 04225 | 12 |
| 5358 | ENSMUSG( | Pou3f1 | 18991 | 013511311 | 001285 | -00375 | 0715 | 12 |
| 5361 | ENSMUSG( | Ppa1 | 67895 | -00155962 | -02394969 | 0175 | 03175 | 12 |
| 5395 | ENSMUSG( | Ppm1k | 243382 | 008804598 | -00132425 | 02125 | 0325 | 12 |
| 5410 | ENSMUSG( | Ppp1r14b | 18938 | 005461114 | 002390841 | 00275 | 03175 | 12 |
| 5427 | ENSMUSG( | Ppp2ca | 19052 | 003075507 | -01532472 | -0195 | 04 | 12 |
| 5445 | ENSMUSG( | Ppp4r1 | 70351 | 017704434 | -00290963 | -0055 | 038 | 12 |
| 5446 | ENSMUSG( | Ppp4r2 | 232314 | 014460759 | 010772305 | -00825 | 064 | 12 |
| 5447 | ENSMUSG( | Ppp4r3a | 68734 | -00241984 | 014219775 | 03425 | 0495 | 12 |
| 5463 | ENSMUSG( | Prdm2 | 110593 | 009867520 | 006964245 | 0425 | 0585 | 12 |
| 5478 | ENSMUSG( | Prickle2 | 243548 | 011840822 | 009543215 | 0065 | 034 | 12 |
| 5484 | ENSMUSG( | Prkacb | 18749 | 002566028 | -00928662 | -001 | 029 | 12 |
| 5498 | ENSMUSG( | Prkcq | 18761 | -00110191 | -01093440 | -012 | 03775 | 12 |
| 5506 | ENSMUSG( | Prkra | 23992 | 013384348 | -00573821 | 0145 | 0325 | 12 |
| 5510 | ENSMUSG( | Prmt5 | 27374 | 014943071 | -00261330 | 00175 | 04075 | 12 |
| 5511 | ENSMUSG( | Prmt7 | 214572 | 019385603 | 001081361 | -005 | 044 | 12 |
| 5544 | ENSMUSG( | Prrc2a | 53761 | 020444557 | 011577586 | -0085 | 04825 | 12 |
| 5545 | ENSMUSG( | Prrc2b | 227723 | 005830175 | 013135610 | 021 | 05225 | 12 |
| 5546 | ENSMUSG( | Prrc2c | 226562 | -00703913 | 017030029 | -009 | 04525 | 12 |
| 5551 | ENSMUSG( | Prune2 | 353211 | -02679547 | 008450351 | 002 | 0565 | 12 |
| 5554 | ENSMUSG( | Psap | 19156 | -01987125 | 007839270 | 0255 | 07425 | 12 |
| 5566 | ENSMUSG( | Psma6 | 26443 | 002117005 | -00914796 | 02 | 05325 | 12 |
| 5594 | ENSMUSG( | Psmd7 | 17463 | 012941771 | 004877289 | 00075 | 0435 | 12 |
| 5599 | ENSMUSG( | Psme3 | 19192 | 003037633 | -00664193 | -0095 | 0385 | 12 |
| 5620 | ENSMUSG( | Ptgds | 19215 | 020907125 | 002149351 | -018 | 063 | 12 |
| 5646 | ENSMUSG( | Ptpn13 | 19249 | 007175221 | -01580362 | -00625 | 04175 | 12 |
| 5654 | ENSMUSG( | Ptpra | 19262 | -01151301 | 005122281 | 00625 | 06275 | 12 |
| 5662 | ENSMUSG( | Ptprn | 19275 | -01053842 | 013442887 | -0105 | 03675 | 12 |
| 5677 | ENSMUSG( | Pum2 | 80913 | 004361410 | 005420461 | -00725 | 03225 | 12 |
| 5681 | ENSMUSG( | Purg | 75029 | 014383751 | -00252307 | 01975 | 027 | 12 |
| 5682 | ENSMUSG( | Pus7 | 78697 | 012656464 | 017392985 | -0135 | 04075 | 12 |
| 5690 | ENSMUSG( | Pxk | 218699 | 012236036 | 005206293 | 0355 | 06075 | 12 |
| 5705 | ENSMUSG( | Pyurf | 66459 | -00340274 | -03679752 | 00275 | 077 | 12 |
| 5723 | ENSMUSG( | Rab11fip3 | 215445 | 010053163 | 008013758 | 006 | 06875 | 12 |
| 5730 | ENSMUSG( | Rab18 | 19330 | -00009913 | -00198098 | -00125 | 0495 | 12 |
| 5740 | ENSMUSG( | Rab28 | 100972 | 007349505 | -00762872 | 00025 | 059 | 12 |
| 5750 | ENSMUSG( | Rab36 | 76877 | 001952773 | -00343287 | -001 | 04275 | 12 |
| 5794 | ENSMUSG( | Rad50 | 19360 | 011381618 | 005208480 | -01925 | 039 | 12 |
| 5797 | ENSMUSG( | Raf1 | 110157 | 007450020 | 000308464 | 01125 | 039 | 12 |
| 5814 | ENSMUSG( | Ranbp17 | 66011 | -00226881 | -01417784 | -0135 | 054 | 12 |
| 5816 | ENSMUSG( | Ranbp3 | 71810 | 002386019 | -00464814 | -003 | 052 | 12 |
| 5822 | ENSMUSG( | Rap1gap | 110351 | 008955428 | 006229553 | -0145 | 03725 | 12 |
| 5826 | ENSMUSG( | Rap2b | 74012 | 015596275 | 011119395 | 01175 | 04075 | 12 |
| 5877 | ENSMUSG( | Rbm26 | 74213 | -00718757 | 018037408 | -00925 | 0485 | 12 |
| 5878 | ENSMUSG( | Rbm27 | 225432 | 011853128 | 015685163 | 00075 | 02925 | 12 |
| 5895 | ENSMUSG( | Rc3h1 | 381305 | -00058814 | 024611506 | -01325 | 03525 | 12 |
| 5899 | ENSMUSG( | Rcan3 | 53902 | -00389860 | -01789622 | 02025 | 04875 | 12 |
| 5903 | ENSMUSG( | Rcc2 | 108911 | 002283432 | -00776774 | -01125 | 047 | 12 |
| 5934 | ENSMUSG( | Retsat | 67442 | 017611168 | 033622346 | 02375 | 06325 | 12 |
| 5944 | ENSMUSG( | Rftn1 | 76438 | -01094445 | -01229834 | 02075 | 04525 | 12 |
| 5951 | ENSMUSG( | Rgma | 244058 | -02338083 | 027009531 | 00225 | 0545 | 12 |
| 5981 | ENSMUSG( | Rhpn2 | 52428 | 010400774 | -00443055 | -00825 | 02925 | 12 |
| 5997 | ENSMUSG( | Ring1 | 19763 | 008140822 | 008934734 | -01725 | 05 | 12 |
| 6014 | ENSMUSG( | Rmnd5a | 68477 | 009134064 | 002667190 | 0205 | 02975 | 12 |
| 6034 | ENSMUSG( | Rnf14 | 56736 | 017556895 | -00447856 | -00275 | 0305 | 12 |
| 6045 | ENSMUSG( | Rnf20 | 109331 | 009517073 | 000239719 | 01825 | 04625 | 12 |
| 6053 | ENSMUSG( | Rnf34 | 80751 | 003675654 | -00102559 | 00475 | 04575 | 12 |
| 6061 | ENSMUSG( | Rnmt | 67897 | -00036524 | 008427668 | 024 | 07 | 12 |
| 6062 | ENSMUSG( | Rnpep | 215615 | 019371404 | -00317867 | -00725 | 02925 | 12 |
| 6073 | ENSMUSG( | Rpa2 | 19891 | 003981432 | -00899325 | -01025 | 0645 | 12 |
| 6076 | ENSMUSG( | Rpap3 | 71919 | 015924929 | -00244081 | -00675 | 0315 | 12 |
| 6080 | ENSMUSG( | Rpgrip1l | 244585 | 023953910 | 022541041 | 006 | 04525 | 12 |
| 6107 | ENSMUSG( | Rpl30 | 19946 | 004371619 | -01214923 | 00375 | 04575 | 12 |
| 6108 | ENSMUSG( | Rpl31 | 114641 | 005122417 | -02948708 | 03725 | 06475 | 12 |
| 6126 | ENSMUSG( | Rplp0 | 11837 | 012613563 | -01338302 | 03075 | 043 | 12 |
| 6154 | ENSMUSG( | Rps26 | 27370 | 005855322 | -01670887 | 0165 | 06775 | 12 |
| 6157 | ENSMUSG( | Rps28 | 54127 | 008007783 | -01846647 | -001 | 07175 | 12 |
| 6161 | ENSMUSG( | Rps5 | 20103 | 007795025 | -01629508 | 02275 | 0365 | 12 |
| 6163 | ENSMUSG( | Rps6ka1 | 20111 | 003443619 | 002879425 | -002 | 06825 | 12 |
| 6166 | ENSMUSG( | Rps6ka4 | 56613 | 013336281 | 022184355 | 005 | 0405 | 12 |
| 6167 | ENSMUSG( | Rps6ka5 | 73086 | -02326382 | -00867481 | -0105 | 055 | 12 |
| 6183 | ENSMUSG( | Rrbp1 | 81910 | -01033592 | 009214341 | 03675 | 0565 | 12 |
| 6196 | ENSMUSG( | Rsrc2 | 208606 | -00001830 | 017631530 | -009 | 038 | 12 |
| 6198 | ENSMUSG( | Rtca | 66368 | -00328334 | -02391518 | 0005 | 041 | 12 |
| 6204 | ENSMUSG( | Rtn2 | 20167 | 018396680 | 006145295 | 02 | 05 | 12 |
| 6213 | ENSMUSG( | Rufy1 | 216724 | 00804364 | 001701662 | 02475 | 0375 | 12 |
| 6214 | ENSMUSG( | Rufy2 | 70432 | 001866700 | 005328326 | 02625 | 0345 | 12 |
| 6244 | ENSMUSG( | Sae1 | 56459 | 003512124 | -02245173 | 006 | 07125 | 12 |
| 6260 | ENSMUSG( | Sap30l | 50724 | 008514248 | -01191148 | 005 | 03375 | 12 |
| 6269 | ENSMUSG( | Sart3 | 53890 | 021285380 | 010859702 | 007 | 0705 | 12 |
| 6272 | ENSMUSG( | Satb1 | 20230 | 021373674 | 006880991 | 00375 | 0725 | 12 |
| 6273 | ENSMUSG( | Satb2 | 212712 | 014822635 | 006054988 | 0015 | 0595 | 12 |
| 6282 | ENSMUSG( | Scaf4 | 224432 | 000647419 | 005579416 | -026 | 053 | 12 |
| 6283 | ENSMUSG( | Scaf8 | 106583 | 019497760 | 001811507 | -029 | 05675 | 12 |
| 6289 | ENSMUSG( | Scap | 235623 | -00099898 | 024471416 | 0285 | 05475 | 12 |
| 6291 | ENSMUSG( | Scara3 | 219151 | -01207948 | 004506060 | 00225 | 05925 | 12 |
| 6301 | ENSMUSG( | Scly | 50880 | 004904824 | -01763710 | 0265 | 03775 | 12 |
| 6322 | ENSMUSG( | Scube1 | 64706 | -01664321 | -00763995 | -0105 | 0495 | 12 |
| 6324 | ENSMUSG( | Scyl2 | 213326 | -00315118 | -01254600 | 001 | 031 | 12 |
| 6328 | ENSMUSG( | Sdc3 | 20970 | -01932623 | 000170930 | 01075 | 06625 | 12 |
| 6331 | ENSMUSG( | Sde2 | 208768 | 004304143 | 001233989 | 0045 | 06925 | 12 |
| 6355 | ENSMUSG( | Sec23b | 27054 | 009739680 | -00923976 | -00225 | 03525 | 12 |
| 6358 | ENSMUSG( | Sec24b | 99683 | 019739436 | 007999001 | 00725 | 027 | 12 |
| 6360 | ENSMUSG( | Sec24d | 69608 | 021515724 | 017818597 | -0155 | 04025 | 12 |
| 6361 | ENSMUSG( | Sec31a | 69162 | 012959280 | 002561215 | -00075 | 0275 | 12 |
| 6378 | ENSMUSG( | Selenop | 20363 | 024513603 | 024515746 | -022 | 04525 | 12 |
| 6399 | ENSMUSG( | Sephs1 | 109079 | 001079062 | -00809705 | 01375 | 05325 | 12 |
| 6413 | ENSMUSG( | Serbp1 | 66870 | 006053192 | -00184445 | 01275 | 03175 | 12 |
| 6422 | ENSMUSG( | Serpinb1b | 282663 | -00677008 | -02501453 | -011 | 03925 | 12 |
| 6423 | ENSMUSG( | Serpinb6a | 20719 | -00025762 | -02274634 | 001 | 0725 | 12 |

| | | | | | | | | |
|---|---|---|---|---|---|---|---|---|
| 6426 | ENSMUSG( | Serpine2 | 20720 | -01233807 | -01081315 | 0045 | 03575 | 12 |
| 6432 | ENSMUSG( | Sesn3 | 75747 | 012376845 | -02881430 | 01925 | 0435 | 12 |
| 6434 | ENSMUSG( | Set | 56086 | 008126319 | -00290351 | -00325 | 0425 | 12 |
| 6436 | ENSMUSG( | Setd3 | 52690 | 006662429 | -00937420 | 00625 | 04075 | 12 |
| 6453 | ENSMUSG( | Sfpq | 71514 | 007655039 | 01949127 | -003 | 0365 | 12 |
| 6472 | ENSMUSG( | Sgsm2 | 97761 | 009828283 | -00751940 | -00225 | 04675 | 12 |
| 6478 | ENSMUSG( | Sh2d3c | 27387 | -01469753 | -02478280 | 02025 | 02975 | 12 |
| 6480 | ENSMUSG( | Sh3bgrl | 56726 | 002227862 | -00408722 | 00575 | 0455 | 12 |
| 6493 | ENSMUSG( | Sh3kbp1 | 58194 | -01344404 | -01708882 | -00475 | 0385 | 12 |
| 6507 | ENSMUSG( | Shd | 20420 | 003703578 | -00180118 | -0085 | 04225 | 12 |
| 6518 | ENSMUSG( | Shprh | 268281 | 003743591 | 008152441 | 0155 | 06075 | 12 |
| 6520 | ENSMUSG( | Shroom1 | 71774 | -00629703 | 018201069 | 0185 | 06375 | 12 |
| 6523 | ENSMUSG( | Shtn1 | 71653 | -01692703 | -03462691 | 00575 | 05475 | 12 |
| 6524 | ENSMUSG( | Siae | 22619 | -02670516 | -00790440 | 00825 | 0525 | 12 |
| 6557 | ENSMUSG( | Slc14a1 | 108052 | -01977874 | 006188472 | -00375 | 037 | 12 |
| 6559 | ENSMUSG( | Slc15a4 | 100561 | -01139343 | 02949784C | 0285 | 054 | 12 |
| 6614 | ENSMUSG( | Slc25a36 | 192287 | -00599276 | 011194984 | 022 | 033 | 12 |
| 6658 | ENSMUSG( | Slc38a10 | 72055 | -01705415 | 011122873 | 0045 | 0715 | 12 |
| 6661 | ENSMUSG( | Slc38a7 | 234595 | -03044638 | 011388886 | 003 | 06725 | 12 |
| 6664 | ENSMUSG( | Slc39a12 | 277468 | 00431811C | 007652197 | -01325 | 02875 | 12 |
| 6679 | ENSMUSG( | Slc4a1ap | 20534 | 022063321 | 013582469 | 01275 | 05525 | 12 |
| 6681 | ENSMUSG( | Slc4a3 | 20536 | 003461816 | 011948337 | -0135 | 03075 | 12 |
| 6683 | ENSMUSG( | Slc4a7 | 218756 | -00443562 | 029894381 | 00225 | 04075 | 12 |
| 6741 | ENSMUSG( | Smad5 | 17129 | 003666949 | -00531856 | -01925 | 0515 | 12 |
| 6748 | ENSMUSG( | Smarcal1 | 54380 | 016825216 | 011047492 | -01475 | 04275 | 12 |
| 6802 | ENSMUSG( | Snip1 | 76793 | 002307695 | -01137351 | -0115 | 038 | 12 |
| 6803 | ENSMUSG( | Snn | 20621 | -00223102 | -00145435 | 0205 | 0355 | 12 |
| 6818 | ENSMUSG( | Snrpe | 20643 | 007241455 | -01320559 | 01775 | 04175 | 12 |
| 6829 | ENSMUSG( | Snx11 | 74479 | -00102689 | 000818779 | -0015 | 0275 | 12 |
| 6847 | ENSMUSG( | Snx33 | 235406 | -03224479 | 009396373 | 01025 | 05775 | 12 |
| 6852 | ENSMUSG( | Snx9 | 66616 | 005543218 | -00008679 | -006 | 056 | 12 |
| 6857 | ENSMUSG( | Sod3 | 20657 | -03448005 | -00128754 | 01225 | 06525 | 12 |
| 6888 | ENSMUSG( | Spats2 | 72572 | 005049817 | -00524693 | 00225 | 04725 | 12 |
| 6901 | ENSMUSG( | Sphkap | 77629 | -00298286 | 011553057 | 02875 | 05175 | 12 |
| 6902 | ENSMUSG( | Spin1 | 20729 | 013768538 | -00151092 | -012 | 0355 | 12 |
| 6905 | ENSMUSG( | Spns1 | 73658 | -01221067 | 019679573 | 0065 | 048 | 12 |
| 7006 | ENSMUSG( | Stard3nl | 76205 | -01318943 | 012275171 | 0035 | 03775 | 12 |
| 7024 | ENSMUSG( | Stk11ip | 71728 | 018782337 | 010577541 | -00275 | 05775 | 12 |
| 7031 | ENSMUSG( | Stk38 | 106504 | 006286587 | 006909374 | -009 | 03075 | 12 |
| 7039 | ENSMUSG( | Stn1 | 108689 | -00748659 | -01249442 | 00525 | 047 | 12 |
| 7080 | ENSMUSG( | Suds3 | 71954 | 014652327 | -00691043 | 02425 | 0405 | 12 |
| 7111 | ENSMUSG( | Swi5 | 72931 | 00269717C | -01874878 | 025 | 0435 | 12 |
| 7138 | ENSMUSG( | Sys1 | 66460 | -01859035 | -01119484 | 00625 | 0375 | 12 |
| 7143 | ENSMUSG( | Syt13 | 80976 | -02023663 | 00200327C | 0045 | 044 | 12 |
| 7145 | ENSMUSG( | Syt17 | 110058 | 023484163 | 017327492 | 031 | 04025 | 12 |
| 7151 | ENSMUSG( | Syt7 | 54525 | 000766387 | 016989317 | -00375 | 04025 | 12 |
| 7167 | ENSMUSG( | Taf10 | 24075 | 004523014 | -02477484 | 0225 | 05075 | 12 |
| 7168 | ENSMUSG( | Taf11 | 68776 | -00069433 | -01227603 | -004 | 04175 | 12 |
| 7183 | ENSMUSG( | Taldo1 | 21351 | -00332834 | -02272751 | -0015 | 05375 | 12 |
| 7197 | ENSMUSG( | Tarbp2 | 21357 | -01211982 | 017364911 | 01525 | 07075 | 12 |
| 7201 | ENSMUSG( | Tars3 | 272396 | 009020132 | 002723304 | 00125 | 03175 | 12 |
| 7229 | ENSMUSG( | Tbc1d9 | 71310 | 010234257 | 004440126 | -00375 | 03625 | 12 |
| 7233 | ENSMUSG( | Tbcc | 72726 | -01165754 | -03099988 | 00775 | 03125 | 12 |
| 7240 | ENSMUSG( | Tbl1x | 21372 | -00604573 | -00600183 | 02225 | 03775 | 12 |
| 7241 | ENSMUSG( | Tbl1xr1 | 81004 | -00773128 | -00229754 | -00075 | 035 | 12 |
| 7244 | ENSMUSG( | Tbpl1 | 237336 | 009775131 | 000513214 | -01775 | 05925 | 12 |
| 7250 | ENSMUSG( | Tceal1 | 237052 | -00740102 | -01992070 | 012 | 0585 | 12 |
| 7253 | ENSMUSG( | Tcerg1 | 56070 | -00310338 | 004045105 | -0075 | 03275 | 12 |
| 7265 | ENSMUSG( | Tdp2 | 56196 | 003015421 | 012936327 | 00425 | 06675 | 12 |
| 7283 | ENSMUSG( | Tepsin | 78777 | 024573709 | 015842105 | 02175 | 03725 | 12 |
| 7295 | ENSMUSG( | Tfdp1 | 21781 | 011979095 | 014793465 | 035 | 04725 | 12 |
| 7296 | ENSMUSG( | Tfe3 | 209446 | -00451876 | 000608881 | -01375 | 03175 | 12 |
| 7309 | ENSMUSG( | Thada | 240174 | 021268102 | 012914487 | 0395 | 052 | 12 |
| 7330 | ENSMUSG( | Thumpd1 | 233802 | 012264473 | -00615213 | -004 | 06225 | 12 |
| 7336 | ENSMUSG( | Tiam1 | 21844 | 008844937 | 015635993 | -0115 | 03 | 12 |
| 7353 | ENSMUSG( | Timmdc1 | 76916 | -00274110 | 001410553 | -00925 | 03225 | 12 |
| 7364 | ENSMUSG( | Tlcd4 | 99887 | -01731459 | -00161956 | 01 | 03075 | 12 |
| 7373 | ENSMUSG( | Tm7sf2 | 73166 | -01847815 | -01499043 | 016 | 051 | 12 |
| 7419 | ENSMUSG( | Tmem147 | 69804 | -01629421 | -00431731 | 01975 | 077 | 12 |
| 7423 | ENSMUSG( | Tmem160 | 69094 | -01306223 | -02411049 | 0175 | 041 | 12 |
| 7432 | ENSMUSG( | Tmem177 | 66343 | 014273666 | 004127519 | 00875 | 038 | 12 |
| 7436 | ENSMUSG( | Tmem185a | 236848 | -03193765 | 01162051C | -0035 | 049 | 12 |
| 7450 | ENSMUSG( | Tmem229a | 319832 | 003627098 | -01786777 | -00725 | 04225 | 12 |
| 7460 | ENSMUSG( | Tmem258 | 69038 | -01645118 | -01534914 | 0075 | 06925 | 12 |
| 7473 | ENSMUSG( | Tmem50b | 77975 | -02553108 | 005039029 | 02575 | 0635 | 12 |
| 7482 | ENSMUSG( | Tmem69 | 230657 | -00220414 | -00124628 | 03075 | 044 | 12 |
| 7510 | ENSMUSG( | Tnfaip8l3 | 244882 | 024421442 | 007815036 | -0025 | 032 | 12 |
| 7537 | ENSMUSG( | Tomm40l | 641376 | 010746731 | -02223168 | 0095 | 0295 | 12 |
| 7539 | ENSMUSG( | Tomm7 | 66169 | 005793349 | -01919845 | 01775 | 02975 | 12 |
| 7556 | ENSMUSG( | Tpd52l2 | 66314 | 002891635 | -01150428 | 016 | 067 | 12 |
| 7569 | ENSMUSG( | Tppp3 | 67971 | -00025669 | -00631074 | 00175 | 07475 | 12 |
| 7578 | ENSMUSG( | Trabd | 67976 | -00168579 | 001037709 | -018 | 05225 | 12 |
| 7579 | ENSMUSG( | Traf2 | 22030 | 002742668 | 003461659 | 00425 | 0425 | 12 |
| 7616 | ENSMUSG( | Trim33 | 94093 | 004297735 | 004820593 | -00775 | 0425 | 12 |
| 7618 | ENSMUSG( | Trim36 | 28105 | -03842381 | 004925304 | -00025 | 05825 | 12 |
| 7620 | ENSMUSG( | Trim44 | 80985 | 010004195 | 003601355 | -006 | 027 | 12 |
| 7624 | ENSMUSG( | Trim65 | 338364 | 00410937C | 033442803 | 0275 | 061 | 12 |
| 7638 | ENSMUSG( | Trmt112 | 67674 | 013617818 | -00774443 | 018 | 02625 | 12 |
| 7648 | ENSMUSG( | Trp53rkb | 76367 | 007155857 | -01430056 | 00625 | 031 | 12 |
| 7651 | ENSMUSG( | Trpc4ap | 56407 | 01338413C | -00301058 | 01125 | 02725 | 12 |
| 7663 | ENSMUSG( | Tsc22d4 | 78829 | 014098205 | -00805956 | 026 | 03575 | 12 |
| 7684 | ENSMUSG( | Tspoap1 | 207777 | 021797305 | -00069686 | 02775 | 065 | 12 |
| 7689 | ENSMUSG( | Tssc4 | 56844 | -02421228 | -00512638 | -023 | 05225 | 12 |
| 7693 | ENSMUSG( | Ttbk2 | 140810 | 002619562 | -00137960 | -00375 | 044 | 12 |
| 7701 | ENSMUSG( | Ttc33 | 67515 | 009779996 | 002582143 | 01775 | 07825 | 12 |
| 7708 | ENSMUSG( | Ttc9 | 69480 | 007061078 | -00731578 | 007 | 0335 | 12 |
| 7709 | ENSMUSG( | Ttc9b | 73032 | 00699860C | -01162071 | -01825 | 051 | 12 |
| 7722 | ENSMUSG( | Ttyh3 | 78339 | -00615869 | 021849093 | -01275 | 049 | 12 |
| 7749 | ENSMUSG( | Tut7 | 214290 | 008170729 | 00793627C | 00725 | 0475 | 12 |
| 7753 | ENSMUSG( | Txlng | 353170 | 024220928 | 026470593 | 0035 | 061 | 12 |
| 7774 | ENSMUSG( | U2af2 | 22185 | 006003564 | -01398590 | -0005 | 047 | 12 |
| 7777 | ENSMUSG( | Uap1 | 107652 | 008289099 | -00635679 | 00625 | 045 | 12 |
| 7784 | ENSMUSG( | Ubac1 | 98766 | 007600545 | -00526471 | 003 | 037 | 12 |
| 7788 | ENSMUSG( | Ubap2 | 68926 | 016685093 | 002147067 | -0105 | 0625 | 12 |
| 7789 | ENSMUSG( | Ubap2l | 74383 | 001558463 | -00178299 | -01625 | 0385 | 12 |
| 7794 | ENSMUSG( | Ube2d3 | 66105 | -00862025 | 010775958 | 02625 | 0385 | 12 |
| 7795 | ENSMUSG( | Ube2e2 | 218793 | 005019753 | -01217715 | 02175 | 06 | 12 |
| 7806 | ENSMUSG( | Ube2m | 22192 | 000459775 | -02438349 | 02275 | 06225 | 12 |
| 7822 | ENSMUSG( | Ubfd1 | 28018 | 019617782 | 008857582 | -00025 | 0285 | 12 |
| 7829 | ENSMUSG( | Ubn1 | 170644 | 010789415 | 016478417 | -0145 | 0505 | 12 |
| 7839 | ENSMUSG( | Ubr7 | 66622 | 007751741 | 00711647 | 00925 | 03675 | 12 |
| 7845 | ENSMUSG( | Ubxn4 | 67812 | 000558114 | 011786603 | 024 | 0385 | 12 |
| 7865 | ENSMUSG( | Uggt2 | 66435 | 001297676 | 009003909 | 033 | 06125 | 12 |
| 7886 | ENSMUSG( | Unk | 217331 | -00337637 | 003394047 | 003 | 026 | 12 |
| 7892 | ENSMUSG( | Uprt | 331487 | 009352755 | -01219376 | 02075 | 04375 | 12 |
| 7923 | ENSMUSG( | Usp20 | 74270 | 015617691 | 001157166 | 01 | 0385 | 12 |
| 7924 | ENSMUSG( | Usp22 | 216825 | -00915897 | -00760978 | 02075 | 03975 | 12 |
| 7952 | ENSMUSG( | Utp18 | 217109 | 006390603 | 008219803 | -017 | 04925 | 12 |
| 7962 | ENSMUSG( | Vamp2 | 22318 | 010335881 | -00730241 | 01075 | 054 | 12 |
| 7971 | ENSMUSG( | Vash1 | 238328 | -00467840 | -03337985 | -0175 | 047 | 12 |
| 7975 | ENSMUSG( | Vat1l | 270097 | -00308746 | -01283895 | 01125 | 0305 | 12 |
| 7991 | ENSMUSG( | Vhl | 22346 | 005736355 | -02885686 | 03475 | 06275 | 12 |
| 7996 | ENSMUSG( | Virma | 66185 | 008310309 | 008358833 | 0225 | 04525 | 12 |
| 8037 | ENSMUSG( | Vrk1 | 22367 | -00394128 | 003253178 | 00275 | 0275 | 12 |
| 8045 | ENSMUSG( | Vti1a | 53611 | 008857235 | -01793514 | 03575 | 072 | 12 |
| 8047 | ENSMUSG( | Vtn | 22370 | -00372330 | 024263383 | 0005 | 0565 | 12 |

| | | | | | | | | |
|---|---|---|---|---|---|---|---|---|
| 8050 | ENSMUSG( | Vwa5b2 | 328643 | 019549936 | -02017491 | 02575 | 0655 | 12 |
| 8060 | ENSMUSG( | Wasf2 | 242687 | 004286073 | 015537219 | -00975 | 05775 | 12 |
| 8064 | ENSMUSG( | Washc3 | 67282 | -01186807 | -02406545 | -00425 | 0325 | 12 |
| 8080 | ENSMUSG( | Wdr18 | 216156 | 010594608 | -01114673 | -004 | 03125 | 12 |
| 8096 | ENSMUSG( | Wdr46 | 57315 | 016178733 | -00722095 | -02425 | 054 | 12 |
| 8111 | ENSMUSG( | Wdsub1 | 72137 | 00875407 | -00028250 | -012 | 02825 | 12 |
| 8127 | ENSMUSG( | Wwc1 | 211652 | 005322308 | -01391891 | -00825 | 039 | 12 |
| 8130 | ENSMUSG( | Wwp1 | 107568 | 002506290 | -01245540 | 01875 | 0485 | 12 |
| 8137 | ENSMUSG( | Xkr6 | 219149 | -00352927 | -01611654 | -007 | 0325 | 12 |
| 8152 | ENSMUSG( | Xrcc6 | 14375 | 011197199 | -00484244 | -011 | 05425 | 12 |
| 8154 | ENSMUSG( | Xrn2 | 24128 | -01154378 | 001517914 | -001 | 033 | 12 |
| 8162 | ENSMUSG( | Ybx1 | 22608 | -00452359 | -01789032 | 00425 | 0475 | 12 |
| 8164 | ENSMUSG( | Ydjc | 69101 | 006337477 | 000468431 | -00475 | 0375 | 12 |
| 8168 | ENSMUSG( | Yipf1 | 230584 | -02867082 | 001535425 | 01025 | 04675 | 12 |
| 8173 | ENSMUSG( | Yjefn3 | 234365 | 003585787 | 00413891 | 02175 | 0435 | 12 |
| 8175 | ENSMUSG( | Ykt6 | 56418 | 004010647 | -01667116 | 0235 | 03525 | 12 |
| 8185 | ENSMUSG( | Ythdf2 | 213541 | -00839852 | 016991600 | -00275 | 03175 | 12 |
| 8186 | ENSMUSG( | Ythdf3 | 229096 | -00669131 | 014185295 | 00725 | 06075 | 12 |
| 8187 | ENSMUSG( | Ywhab | 54401 | 004843743 | -00596642 | -0095 | 041 | 12 |
| 8191 | ENSMUSG( | Ywhaq | 22630 | 002109595 | -01207837 | 01525 | 0305 | 12 |
| 8198 | ENSMUSG( | Zbtb20 | 56490 | 008388630 | -00284806 | -00375 | 05175 | 12 |
| 8221 | ENSMUSG( | Zdhhc14 | 224454 | -00726197 | 037683859 | 002 | 06525 | 12 |
| 8224 | ENSMUSG( | Zdhhc2 | 70546 | -01770491 | 006136251 | -017 | 04625 | 12 |
| 8237 | ENSMUSG( | Zfand5 | 22682 | -01122504 | 001384085 | -01275 | 04625 | 12 |
| 8238 | ENSMUSG( | Zfand6 | 65098 | 003922092 | 003406722 | -01525 | 04925 | 12 |
| 8240 | ENSMUSG( | Zfp180 | 210135 | 022376820 | 014230211 | 00075 | 04275 | 12 |
| 8246 | ENSMUSG( | Zfp362 | 230761 | 001773905 | -01086049 | 0075 | 032 | 12 |
| 8274 | ENSMUSG( | Zmiz1 | 328365 | 006943776 | 012200963 | 01275 | 0375 | 12 |
| 8287 | ENSMUSG( | Zscan18 | 232875 | -00445079 | -01720978 | 016 | 043 | 12 |
| 8288 | ENSMUSG( | Zscan26 | 432731 | -00458475 | 011885398 | -01325 | 034 | 12 |
| 8291 | ENSMUSG( | Zwint | 52696 | 011640778 | -00821890 | -007 | 05775 | 12 |
| 10 | ENSMUSG( | 2900026A( | 243219 | 015201412 | 009732492 | 05075 | 0605 | 13 |
| 122 | ENSMUSG( | Acox1 | 11430 | 012957669 | -00565198 | 064 | 053 | 13 |
| 182 | ENSMUSG( | Adcyap1 | 11516 | -05619390 | -01376281 | 05425 | 08925 | 13 |
| 319 | ENSMUSG( | Alg3 | 208624 | -03575634 | -00529538 | 06525 | 08775 | 13 |
| 438 | ENSMUSG( | Apbb1ip | 54519 | -00775499 | -00242511 | 028 | 091 | 13 |
| 464 | ENSMUSG( | Aptx | 66408 | 00109025 | -01824739 | 04075 | 0965 | 13 |
| 530 | ENSMUSG( | Arhgef9 | 236915 | -00145482 | -00570159 | 06775 | 079 | 13 |
| 540 | ENSMUSG( | Arl13b | 68146 | -00123107 | 012100004 | 04725 | 05525 | 13 |
| 622 | ENSMUSG( | Atg14 | 1.01E+08 | 010845740 | 001061220 | 05275 | 06875 | 13 |
| 685 | ENSMUSG( | Atp6v1h | 108664 | -01115597 | -00942763 | 05625 | 11425 | 13 |
| 753 | ENSMUSG( | BC004004 | 80748 | -02916594 | -00030214 | 07175 | 0765 | 13 |
| 802 | ENSMUSG( | Bloc1s4 | 117197 | -01496406 | 000605743 | 029 | 079 | 13 |
| 861 | ENSMUSG( | Btbd1 | 83962 | 007045834 | -02031336 | 03525 | 08975 | 13 |
| 876 | ENSMUSG( | C1qa | 12259 | -00579205 | 017400436 | 047 | 06675 | 13 |
| 886 | ENSMUSG( | C3 | 12266 | -05989094 | -07022089 | 04225 | 06525 | 13 |
| 895 | ENSMUSG( | Cabp1 | 29867 | 009902956 | -00430150 | 0475 | 0815 | 13 |
| 937 | ENSMUSG( | Caly | 68566 | -03736298 | -00083901 | 0205 | 0895 | 13 |
| 958 | ENSMUSG( | Cant1 | 76025 | -01569498 | 022830997 | 046 | 0695 | 13 |
| 988 | ENSMUSG( | Casp1 | 12362 | -02465343 | -02921064 | 09225 | 12225 | 13 |
| 990 | ENSMUSG( | Casp6 | 12368 | 002160908 | -00262290 | 09575 | 11575 | 13 |
| 1068 | ENSMUSG( | Ccn3 | 18133 | -03112766 | 019651381 | 072 | 12825 | 13 |
| 1070 | ENSMUSG( | Ccnl2 | 56036 | 006118626 | 029070337 | 04275 | 07775 | 13 |
| 1215 | ENSMUSG( | Chadl | 214685 | -03216132 | 009603940 | 0375 | 064 | 13 |
| 1218 | ENSMUSG( | Chchd1 | 66121 | 000396047 | -01136288 | 0665 | 0575 | 13 |
| 1241 | ENSMUSG( | Chl1 | 12661 | -03285818 | 023547026 | 0665 | 064 | 13 |
| 1259 | ENSMUSG( | Chpt1 | 212862 | -00405186 | -00030378 | 041 | 081 | 13 |
| 1351 | ENSMUSG( | Cnbp | 12785 | 000402226 | 000379455 | 0445 | 07975 | 13 |
| 1390 | ENSMUSG( | Cntnap3 | 238680 | -04748259 | 035091491 | 066 | 0495 | 13 |
| 1411 | ENSMUSG( | Cog8 | 97484 | 008738804 | -00614244 | 064 | 07725 | 13 |
| 1422 | ENSMUSG( | Col4a5 | 12830 | 020321106 | 026738320 | 11 | 0705 | 13 |
| 1527 | ENSMUSG( | Crabp1 | 12903 | -03018855 | 038922563 | 044 | 0915 | 13 |
| 1588 | ENSMUSG( | Cspg5 | 29873 | -00699028 | 004197743 | 0405 | 0745 | 13 |
| 1655 | ENSMUSG( | Cxadr | 13052 | -04769049 | 014678605 | 0575 | 05575 | 13 |
| 1659 | ENSMUSG( | Cyb561 | 13056 | -02046297 | 016494561 | 0695 | 12775 | 13 |
| 1712 | ENSMUSG( | Dap | 223453 | -00292887 | -01094870 | 08475 | 10175 | 13 |
| 1739 | ENSMUSG( | Dclk1 | 13175 | 009606135 | -00380277 | 04225 | 08375 | 13 |
| 1758 | ENSMUSG( | Dcx | 13193 | -01736497 | 005033154 | 0395 | 1195 | 13 |
| 1824 | ENSMUSG( | Derl2 | 116891 | -01688362 | 032375226 | 11675 | 07375 | 13 |
| 1854 | ENSMUSG( | Dhtkd1 | 209692 | 005612208 | 018501619 | 14025 | 14425 | 13 |
| 1935 | ENSMUSG( | Dnajc25 | 72429 | -03060739 | 017146813 | 084 | 05725 | 13 |
| 1944 | ENSMUSG( | Dnajc9 | 108671 | 000531583 | -01480972 | 05075 | 127 | 13 |
| 2023 | ENSMUSG( | Dtx3 | 80904 | 022380518 | -00250595 | 02825 | 094 | 13 |
| 2127 | ENSMUSG( | Eif1ax | 66235 | 003144899 | -00772166 | 03775 | 119 | 13 |
| 2189 | ENSMUSG( | Eloa | 27224 | 012218123 | 002710353 | 05475 | 108 | 13 |
| 2247 | ENSMUSG( | Ep300 | 328572 | -00403065 | 023913298 | 082 | 13525 | 13 |
| 2327 | ENSMUSG( | Exoc1 | 69940 | 012005837 | 001938970 | 06875 | 09 | 13 |
| 2347 | ENSMUSG( | Ext2 | 14043 | -03099389 | 028112298 | 079 | 0835 | 13 |
| 2348 | ENSMUSG( | Extl1 | 56219 | 008460215 | 026575321 | 07125 | 102 | 13 |
| 2357 | ENSMUSG( | Faap100 | 71885 | 024533145 | 003419046 | 1665 | 124 | 13 |
| 2360 | ENSMUSG( | Fabp7 | 12140 | 020902401 | -04858190 | 04425 | 0935 | 13 |
| 2361 | ENSMUSG( | Fadd | 14082 | 027695506 | 010415280 | 03025 | 092 | 13 |
| 2363 | ENSMUSG( | Fads2 | 56473 | 012072809 | -00680540 | 08275 | 05025 | 13 |
| 2383 | ENSMUSG( | Fam124a | 629059 | 006154788 | -00767259 | 1025 | 08375 | 13 |
| 2470 | ENSMUSG( | Fcf1 | 73736 | -01779152 | -00515879 | 0415 | 084 | 13 |
| 2508 | ENSMUSG( | Fhl1 | 14199 | 005670103 | -00426606 | 081 | 05575 | 13 |
| 2548 | ENSMUSG( | Fmo5 | 14263 | -00487858 | -01638724 | 09275 | 078 | 13 |
| 2565 | ENSMUSG( | Foxj3 | 230700 | -00319117 | -00647029 | 0945 | 16125 | 13 |
| 2582 | ENSMUSG( | Frrs1l | 230235 | -03583832 | 022607901 | 0765 | 03875 | 13 |
| 2612 | ENSMUSG( | Fxyd7 | 57780 | -01642269 | -02548191 | 04675 | 133 | 13 |
| 2636 | ENSMUSG( | Gabrb1 | 14400 | -01072948 | 016483292 | 0725 | 0925 | 13 |
| 2640 | ENSMUSG( | Gabrg1 | 14405 | -02193192 | -00335610 | 11175 | 0835 | 13 |
| 2664 | ENSMUSG( | Ganc | 76051 | -00311705 | -00346260 | 03025 | 0895 | 13 |
| 2746 | ENSMUSG( | Gipc3 | 209047 | -01020817 | 046271743 | 06075 | 08325 | 13 |
| 2875 | ENSMUSG( | Gpr151 | 240239 | 000693858 | 082213832 | 06375 | 0465 | 13 |
| 2900 | ENSMUSG( | Gpx3 | 14778 | -01020041 | 024996858 | 08325 | 06175 | 13 |
| 2926 | ENSMUSG( | Grin3a | 242443 | -02469054 | 014402822 | 0505 | 05375 | 13 |
| 2975 | ENSMUSG( | Gtf2e2 | 68153 | 005553672 | 012212676 | 0485 | 0555 | 13 |
| 3085 | ENSMUSG( | Hexim1 | 192231 | 004112946 | -00025655 | 04425 | 0605 | 13 |
| 3106 | ENSMUSG( | Hltf | 20585 | 007713407 | 013799193 | 06325 | 08625 | 13 |
| 3121 | ENSMUSG( | Hmgn3 | 94353 | 004160608 | 004270712 | 07525 | 15825 | 13 |
| 3203 | ENSMUSG( | Htr1b | 15551 | -06558475 | 038078564 | 085 | 0965 | 13 |
| 3252 | ENSMUSG( | Igfbp4 | 16010 | -01431576 | 010252872 | 06575 | 0825 | 13 |
| 3310 | ENSMUSG( | Insc | 233752 | -04608086 | -02057276 | 02475 | 108 | 13 |
| 3380 | ENSMUSG( | Itgb3 | 16416 | -01460007 | 004440208 | 049 | 06675 | 13 |
| 3398 | ENSMUSG( | Itprid2 | 70599 | 021544768 | 010051391 | 029 | 1005 | 13 |
| 3402 | ENSMUSG( | Ivns1abp | 117198 | -00665371 | 008147312 | 05275 | 103 | 13 |
| 3461 | ENSMUSG( | Kcne5 | 66240 | -02931979 | -03023255 | 09825 | 05625 | 13 |
| 3464 | ENSMUSG( | Kcnh2 | 16511 | -02516046 | 009222656 | 03075 | 072 | 13 |
| 3533 | ENSMUSG( | Kif21a | 16564 | 010135099 | 008476283 | 067 | 0785 | 13 |
| 3586 | ENSMUSG( | Krt12 | 268482 | -01673110 | -02681353 | 05075 | 069 | 13 |
| 3622 | ENSMUSG( | Larp1b | 214048 | 018305551 | 000475561 | 0655 | 129 | 13 |
| 3664 | ENSMUSG( | Lhfpl4 | 269788 | -01481791 | 010502115 | 04025 | 06775 | 13 |
| 3679 | ENSMUSG( | Lin9 | 72568 | 009461055 | 035604968 | 04175 | 08775 | 13 |
| 3745 | ENSMUSG( | Lrrc24 | 378937 | -03674357 | 000790683 | 0355 | 087 | 13 |
| 3803 | ENSMUSG( | Ly6c1 | 17067 | 030694024 | 088145406 | 08725 | 08725 | 13 |
| 3804 | ENSMUSG( | Ly6e | 17069 | -01314702 | 005061928 | 155 | 109 | 13 |
| 3863 | ENSMUSG( | Manbal | 69161 | -02292893 | -01623524 | 10675 | 05925 | 13 |
| 3928 | ENSMUSG( | Marcksl1 | 17357 | 015470787 | -00671324 | 0315 | 10725 | 13 |
| 3944 | ENSMUSG( | Matn2 | 17181 | -00235796 | 048560405 | 06775 | 04675 | 13 |
| 3945 | ENSMUSG( | Matn4 | 17183 | -05499217 | 022633473 | 08325 | 069 | 13 |
| 3961 | ENSMUSG( | Mbtps2 | 270669 | -04102506 | 01075448 | 07125 | 0875 | 13 |
| 3977 | ENSMUSG( | Mcrip2 | 68241 | 003019609 | -01115699 | 07625 | 05275 | 13 |
| 3989 | ENSMUSG( | Mdk | 17242 | -02274474 | -01150239 | 07 | 08925 | 13 |
| 4004 | ENSMUSG( | Med13l | 76199 | 024930242 | 039678705 | 084 | 1595 | 13 |
| 4005 | ENSMUSG( | Med14 | 26896 | 004715404 | 002742747 | 0585 | 0985 | 13 |

| | | | | | | | |
|---|---|---|---|---|---|---|---|
| 4017 | ENSMUSG( | Med6 | 69792 | -01488950 | -01036632 | 04825 | 06275 | 13 |
| 4032 | ENSMUSG( | Met | 17295 | -03102834 | 036656932 | 08425 | 06075 | 13 |
| 4034 | ENSMUSG( | Metap1d | 66559 | -00658840 | 016831274 | 02725 | 10775 | 13 |
| 4059 | ENSMUSG( | Mfsd2a | 76574 | 023214188 | 077886204 | 109 | 1145 | 13 |
| 4063 | ENSMUSG( | Mfsd9 | 211798 | -03060977 | 01919292C | 05125 | 09225 | 13 |
| 4064 | ENSMUSG( | Mgat1 | 17308 | -00482400 | 00902602€ | 091 | 19975 | 13 |
| 4096 | ENSMUSG( | Mindy4 | 330323 | -00261445 | -00144829 | 0745 | 07525 | 13 |
| 4120 | ENSMUSG( | Mnat1 | 17420 | 01569332C | 007820565 | 02875 | 13375 | 13 |
| 4152 | ENSMUSG( | Mpped1 | 223726 | 023614841 | 00387863S | 03975 | 081 | 13 |
| 4196 | ENSMUSG( | Mrpl40 | 18100 | -00937980 | -01431351 | 0745 | 07175 | 13 |
| 4257 | ENSMUSG( | Mt1 | 17748 | 012190202 | -02107704 | 03425 | 099 | 13 |
| 4327 | ENSMUSG( | Myl12a | 67268 | 001665191 | 023054261 | 1075 | 0425 | 13 |
| 4401 | ENSMUSG( | Natd1 | 24083 | 017142058 | 01021138S | 0415 | 07125 | 13 |
| 4411 | ENSMUSG( | Nbr1 | 17966 | -00296085 | -00663729 | 04525 | 07125 | 13 |
| 4549 | ENSMUSG( | Ngb | 64242 | 023958763 | -01915189 | 20125 | 121 | 13 |
| 4605 | ENSMUSG( | Nol4 | 319211 | 003305931 | 006291335 | 10225 | 17825 | 13 |
| 4791 | ENSMUSG( | Ogt | 108155 | 003843635 | 021588754 | 08325 | 0695 | 13 |
| 4936 | ENSMUSG( | Pcdh15 | 11994 | -01980288 | 01270547€ | 094 | 09275 | 13 |
| 5150 | ENSMUSG( | Phrf1 | 101471 | 01845146S | 014251717 | 04975 | 159 | 13 |
| 5167 | ENSMUSG( | Pigo | 56703 | 003537707 | 034481808 | 06475 | 112 | 13 |
| 5296 | ENSMUSG( | Pmvk | 68603 | 005402033 | -02299579 | 0605 | 06125 | 13 |
| 5317 | ENSMUSG( | Pofut2 | 80294 | -03003929 | 01160586C | 023 | 08775 | 13 |
| 5350 | ENSMUSG( | Pomp | 66537 | -00118556 | -02668206 | 10025 | 0855 | 13 |
| 5373 | ENSMUSG( | Ppfibp2 | 19024 | -01658650 | -01599167 | 04975 | 0715 | 13 |
| 5381 | ENSMUSG( | Ppig | 228005 | 006568822 | 005178204 | 03675 | 077 | 13 |
| 5430 | ENSMUSG( | Ppp2r1b | 73699 | 00490562€ | -00831901 | 1005 | 07475 | 13 |
| 5635 | ENSMUSG( | Ptn | 19242 | -03829695 | -01022133 | 1055 | 03525 | 13 |
| 5688 | ENSMUSG( | Pwwp3a | 68114 | -00605771 | 007607201 | 0305 | 07625 | 13 |
| 5774 | ENSMUSG( | Rabac1 | 14470 | -02113395 | 009961721 | 0395 | 0665 | 13 |
| 5845 | ENSMUSG( | Rasgef1c | 74563 | 00635920€ | -01349446 | 071 | 081 | 13 |
| 5889 | ENSMUSG( | Rbms3 | 207181 | -00763580 | -00198558 | 06025 | 06175 | 13 |
| 5989 | ENSMUSG( | Rilpl1 | 75695 | 018632582 | 004736073 | 11425 | 091 | 13 |
| 6007 | ENSMUSG( | Rlf | 109263 | -01349897 | -00402481 | 1175 | 16275 | 13 |
| 6016 | ENSMUSG( | Rnaseh2a | 69724 | -00502721 | -00097866 | 066 | 0575 | 13 |
| 6018 | ENSMUSG( | Rnasek | 52898 | 006973161 | 015736205 | 0445 | 0845 | 13 |
| 6040 | ENSMUSG( | Rnf168 | 70238 | 017359135 | 02168446S | 026 | 13525 | 13 |
| 6056 | ENSMUSG( | Rnf5 | 54197 | -00117846 | -02291502 | 038 | 12925 | 13 |
| 6115 | ENSMUSG( | Rpl37 | 67281 | 01321110S | -01484389 | 04725 | 07075 | 13 |
| 6127 | ENSMUSG( | Rplp1 | 56040 | 008958303 | -01915368 | 1015 | 1005 | 13 |
| 6158 | ENSMUSG( | Rps29 | 20090 | 008333968 | -02304423 | 11325 | 03825 | 13 |
| 6221 | ENSMUSG( | Rusf1 | 233913 | -02864297 | 01227339S | 10325 | 05975 | 13 |
| 6253 | ENSMUSG( | Samd9l | 209086 | -01040488 | 03459498S | 0655 | 15075 | 13 |
| 6274 | ENSMUSG( | Saysd1 | 67509 | -02697167 | 00775133S | 0815 | 141 | 13 |
| 6286 | ENSMUSG( | Scamp2 | 24044 | -02616597 | 021563601 | 02675 | 08725 | 13 |
| 6363 | ENSMUSG( | Sec61a2 | 57743 | -03208875 | 00697235€ | 085 | 05775 | 13 |
| 6429 | ENSMUSG( | Serping1 | 12258 | 00158395€ | 014258192 | 06425 | 1245 | 13 |
| 6470 | ENSMUSG( | Sgsh | 27029 | -02617406 | 01893058C | 12375 | 037 | 13 |
| 6572 | ENSMUSG( | Slc19a2 | 116914 | -03829455 | 034020447 | 062 | 06075 | 13 |
| 6627 | ENSMUSG( | Slc27a3 | 26568 | -00266670 | 01971277€ | 046 | 063 | 13 |
| 6649 | ENSMUSG( | Slc35d3 | 76157 | -15366505 | 005749532 | 04725 | 1725 | 13 |
| 6650 | ENSMUSG( | Slc35f1 | 215085 | -05302959 | 019346722 | 1785 | 139 | 13 |
| 6654 | ENSMUSG( | Slc36a1 | 215335 | -00467412 | 035045242 | 07425 | 0605 | 13 |
| 6696 | ENSMUSG( | Slc6a20a | 102680 | -01934214 | 002275712 | 03125 | 087 | 13 |
| 6756 | ENSMUSG( | Smc2 | 14211 | -00535316 | 017606903 | 0795 | 0665 | 13 |
| 6767 | ENSMUSG( | Smg9 | 71997 | 008750553 | -01832322 | 0255 | 1365 | 13 |
| 6824 | ENSMUSG( | Snu13 | 20826 | 001951083 | -02137859 | 0625 | 14375 | 13 |
| 6881 | ENSMUSG( | Sparc | 20692 | 007123855 | 00365273S | 05475 | 05375 | 13 |
| 6908 | ENSMUSG( | Spock2 | 94214 | -01801408 | 028902482 | 06075 | 06 | 13 |
| 6953 | ENSMUSG( | Srprb | 20818 | -03333288 | 014326998 | 07975 | 04125 | 13 |
| 6980 | ENSMUSG( | Ssna1 | 68475 | -00515634 | -00477062 | 116 | 13425 | 13 |
| 6987 | ENSMUSG( | Sstr1 | 20605 | -02931871 | 046502097 | 11625 | 132 | 13 |
| 6995 | ENSMUSG( | St8sia5 | 225742 | -01117128 | 02115227S | 04825 | 06175 | 13 |
| 7007 | ENSMUSG( | Stard5 | 170460 | 014566371 | -00368450 | 04625 | 07375 | 13 |
| 7086 | ENSMUSG( | Sult1a1 | 20887 | 05521782 | 035739501 | 12375 | 163 | 13 |
| 7136 | ENSMUSG( | Syp | 20977 | -03412984 | 006427937 | 094 | 0535 | 13 |
| 7140 | ENSMUSG( | Syt10 | 54526 | -01893790 | 11564170C | 0865 | 042 | 13 |
| 7165 | ENSMUSG( | Tada1 | 27878 | -00299794 | 003768881 | 09525 | 09425 | 13 |
| 7172 | ENSMUSG( | Taf5l | 102162 | -01966243 | 009017412 | 037 | 10225 | 13 |
| 7177 | ENSMUSG( | Tafa1 | 320265 | -04163156 | -00723898 | 06 | 05525 | 13 |
| 7257 | ENSMUSG( | Tcirg1 | 27060 | -00883500 | 03426426S | 03775 | 076 | 13 |
| 7297 | ENSMUSG( | Tfeb | 21425 | -00350116 | 026218354 | 03975 | 06775 | 13 |
| 7374 | ENSMUSG( | Tm9sf1 | 74140 | -01575892 | 029261778 | 08625 | 056 | 13 |
| 7426 | ENSMUSG( | Tmem164 | 209497 | 01982453C | 01532365C | 056 | 15325 | 13 |
| 7451 | ENSMUSG( | Tmem230 | 70612 | -02420305 | 017322167 | 042 | 08675 | 13 |
| 7452 | ENSMUSG( | Tmem231 | 234740 | -02719733 | 022969842 | 08075 | 06 | 13 |
| 7463 | ENSMUSG( | Tmem263 | 103266 | 001109933 | -00614804 | 0605 | 06525 | 13 |
| 7506 | ENSMUSG( | Tnc | 21923 | -03307045 | 00279067S | 0615 | 05475 | 13 |
| 7545 | ENSMUSG( | Tor1aip2 | 240832 | -00627724 | 006898204 | 0465 | 06425 | 13 |
| 7617 | ENSMUSG( | Trim35 | 66854 | 007200484 | -00817198 | 08675 | 0455 | 13 |
| 7642 | ENSMUSG( | Trmt61a | 328162 | 006467481 | -01290652 | 034 | 10475 | 13 |
| 7645 | ENSMUSG( | Tro | 56191 | 00710375€ | 013997981 | 12775 | 05 | 13 |
| 7652 | ENSMUSG( | Trpc6 | 22068 | -04133117 | -01838710 | 05375 | 06175 | 13 |
| 7671 | ENSMUSG( | Tspan13 | 66109 | -05005769 | 014762847 | 02725 | 079 | 13 |
| 7710 | ENSMUSG( | Ttc9c | 70387 | -00290121 | -00149709 | 05725 | 06975 | 13 |
| 7768 | ENSMUSG( | Tyk2 | 54721 | 024830454 | 024752125 | 04525 | 10875 | 13 |
| 7862 | ENSMUSG( | Ugcg | 22234 | -05531319 | 017872718 | 10325 | 03775 | 13 |
| 7913 | ENSMUSG( | Usf1 | 22278 | 011012431 | -00261069 | 085 | 11125 | 13 |
| 8038 | ENSMUSG( | Vrk2 | 69922 | -01530822 | 000546948 | 06 | 055 | 13 |
| 8043 | ENSMUSG( | Vstm2l | 277432 | -04625298 | -00036297 | 0475 | 065 | 13 |
| 8243 | ENSMUSG( | Zfp324 | 243834 | 024502805 | 015672403 | 1275 | 19775 | 13 |
| 11 | ENSMUSG( | 3110082I1 | 73212 | 005317843 | -00306363 | -06125 | -0345 | 14 |
| 150 | ENSMUSG( | Actr6 | 67019 | 01277017C | 01453637€ | -05425 | -0505 | 14 |
| 174 | ENSMUSG( | Adck5 | 268822 | 005780327 | 00966962C | -063 | -009 | 14 |
| 196 | ENSMUSG( | Adgrl1 | 330814 | 000428528 | 027988428 | -13625 | -046 | 14 |
| 201 | ENSMUSG( | Adi1 | 104923 | 00372934€ | -01799234 | -054 | -04325 | 14 |
| 206 | ENSMUSG( | Adora1 | 11539 | -00123697 | 009393131 | -083 | -0465 | 14 |
| 296 | ENSMUSG( | Aldh16a1 | 69748 | -00168145 | -00176258 | -10175 | -07425 | 14 |
| 329 | ENSMUSG( | Alpl | 11647 | -03275773 | 036416254 | -048 | -0495 | 14 |
| 384 | ENSMUSG( | Ankrd63 | 383787 | -02269625 | -00413988 | -04925 | -03 | 14 |
| 389 | ENSMUSG( | Anln | 68743 | -00772143 | 01259684 | -05025 | -04175 | 14 |
| 400 | ENSMUSG( | Anxa1 | 16952 | -00500307 | -00425639 | -099 | 0115 | 14 |
| 452 | ENSMUSG( | Apoc1 | 11812 | 03686200C | 042715903 | -08375 | -0125 | 14 |
| 492 | ENSMUSG( | Arhgap10 | 78514 | 007335084 | -01243140 | -0665 | -0315 | 14 |
| 508 | ENSMUSG( | Arhgap42 | 71544 | 006843861 | 01165122S | -08325 | -0135 | 14 |
| 587 | ENSMUSG( | Asap1 | 13196 | 022439323 | 002714181 | -0715 | -053 | 14 |
| 589 | ENSMUSG( | Asb13 | 142688 | 003338287 | -00486269 | -057 | -0585 | 14 |
| 604 | ENSMUSG( | Asphd1 | 233879 | -02919782 | -00734022 | -045 | -0555 | 14 |
| 628 | ENSMUSG( | Atg4c | 242557 | 020367617 | 017728508 | -0985 | 015 | 14 |
| 642 | ENSMUSG( | Atp13a3 | 224088 | -02041814 | 030258433 | -0505 | -0565 | 14 |
| 721 | ENSMUSG( | B4galnt4 | 330671 | -00317115 | 025607725 | -176 | -07975 | 14 |
| 745 | ENSMUSG( | Baz2a | 116848 | 01280193€ | 00896025€ | -1195 | -0095 | 14 |
| 781 | ENSMUSG( | Bdnf | 12064 | -05612997 | 009324937 | -1115 | -0615 | 14 |
| 811 | ENSMUSG( | Bmt2 | 101148 | 002310472 | 022879233 | -055 | -0265 | 14 |
| 838 | ENSMUSG( | Brd2 | 14312 | 013910627 | 012621312 | -11325 | -023 | 14 |
| 855 | ENSMUSG( | Brwd1 | 93871 | -01481364 | -00497006 | -08425 | -03275 | 14 |
| 915 | ENSMUSG( | Cacng4 | 54377 | -03286397 | 002003171 | -049 | -022 | 14 |
| 1010 | ENSMUSG( | Cbr2 | 12409 | 026084125 | 028711192 | -0675 | -04175 | 14 |
| 1077 | ENSMUSG( | Ccsap | 73420 | 007467505 | 015761897 | -0995 | -02925 | 14 |
| 1121 | ENSMUSG( | Cdc42ep1 | 104445 | -03121149 | 001958293 | -045 | -063 | 14 |
| 1221 | ENSMUSG( | Chchd3 | 66075 | 00316857S | -01755338 | -062 | -00775 | 14 |
| 1247 | ENSMUSG( | Chmp2b | 68942 | -00826846 | -00261545 | -061 | -00825 | 14 |
| 1290 | ENSMUSG( | Clasrp | 53609 | 007900211 | 01110958S | -07625 | -00675 | 14 |
| 1311 | ENSMUSG( | Clic6 | 209195 | -01664972 | 00425668C | -08025 | -0545 | 14 |
| 1312 | ENSMUSG( | Clint1 | 216705 | 002620847 | 002781585 | -07525 | -091 | 14 |
| 1394 | ENSMUSG( | Cntnap5c | 620292 | 000840708 | 055550242 | -04325 | -07075 | 14 |
| 1412 | ENSMUSG( | Coil | 12812 | 01182998€ | 011886732 | -052 | -03875 | 14 |

| | | | | | | | | |
|---|---|---|---|---|---|---|---|---|
| 1462 | ENSMUSG( | Coq6 | 217707 | -00267517 | -00638001 | -07475 | -0895 | 14 |
| 1541 | ENSMUSG( | Crhr1 | 12921 | -00062032 | 00161243C | -0705 | -04225 | 14 |
| 1557 | ENSMUSG( | Crtc2 | 74343 | -00465573 | -00180120 | -08125 | -01225 | 14 |
| 1578 | ENSMUSG( | Csnk1a1 | 93687 | 003214795 | 005688058 | -09125 | -04275 | 14 |
| 1591 | ENSMUSG( | Csrp2 | 13008 | -00082618 | 026676513 | -05825 | -01175 | 14 |
| 1613 | ENSMUSG( | Ctnna3 | 216033 | -00716287 | -03421252 | -09625 | -08225 | 14 |
| 1650 | ENSMUSG( | Cwc22 | 80744 | 010864748 | 008399765 | -07425 | -03975 | 14 |
| 1651 | ENSMUSG( | Cwc25 | 67480 | 003240377 | 001059746 | -061 | -02425 | 14 |
| 1687 | ENSMUSG( | Cystm1 | 66060 | 003546444 | -02252668 | -1095 | -03225 | 14 |
| 1699 | ENSMUSG( | D8Ertd738 | 101966 | 007178371 | -02903770 | -06525 | -076 | 14 |
| 1724 | ENSMUSG( | Dbndd2 | 52840 | 002057897 | 001774741 | -0885 | -07025 | 14 |
| 1731 | ENSMUSG( | Dcaf13 | 223499 | -00680681 | -00332696 | -05475 | -0175 | 14 |
| 1774 | ENSMUSG( | Ddx10 | 77591 | 021827103 | 00441013C | -12975 | -003 | 14 |
| 1803 | ENSMUSG( | Def6 | 23853 | -01630819 | 007165158 | -10475 | 01225 | 14 |
| 1804 | ENSMUSG( | Def8 | 23854 | 011107582 | -01228427 | -1115 | 00625 | 14 |
| 1814 | ENSMUSG( | Dennd4c | 329877 | 00929784C | -00250554 | -099 | -00925 | 14 |
| 1822 | ENSMUSG( | Dera | 232449 | 003112835 | -01506079 | -103 | -00625 | 14 |
| 1829 | ENSMUSG( | Dgcr2 | 13356 | -02164547 | 017520225 | -059 | -02275 | 14 |
| 1919 | ENSMUSG( | Dnajb2 | 56812 | 004017769 | -01366647 | -05925 | -02575 | 14 |
| 1940 | ENSMUSG( | Dnajc5 | 13002 | 003711611 | -00571093 | -05375 | -01075 | 14 |
| 1953 | ENSMUSG( | Dnmbp | 71972 | -00246341 | -00146955 | -06325 | -02125 | 14 |
| 1968 | ENSMUSG( | Dock5 | 68813 | -01538747 | 012675474 | -04875 | -059 | 14 |
| 1980 | ENSMUSG( | Dpagt1 | 13478 | -01884791 | 029903769 | -057 | -0235 | 14 |
| 2035 | ENSMUSG( | Dusp9 | 75590 | -01258147 | -04392318 | -12775 | -0285 | 14 |
| 2231 | ENSMUSG( | Eno3 | 13808 | 022254736 | -00175004 | -061 | -009 | 14 |
| 2321 | ENSMUSG( | Evi2a | 14017 | -03859523 | -00763076 | -09775 | -091 | 14 |
| 2353 | ENSMUSG( | F13a1 | 74145 | 045248368 | 027212459 | -06275 | -01 | 14 |
| 2369 | ENSMUSG( | Fahd1 | 68636 | -01083145 | -02715953 | -0565 | -034 | 14 |
| 2464 | ENSMUSG( | Fbxo9 | 71538 | 007447779 | -00977306 | -053 | -02325 | 14 |
| 2469 | ENSMUSG( | Fcer1g | 14127 | -01512583 | -00305067 | -11425 | 0085 | 14 |
| 2480 | ENSMUSG( | Fdx1 | 14148 | 00368232C | -00098839 | -08925 | -0635 | 14 |
| 2540 | ENSMUSG( | Flt1 | 14254 | 028235626 | 07667773 | -0865 | 0005 | 14 |
| 2541 | ENSMUSG( | Flywch1 | 224613 | 011704409 | -00196935 | -08075 | -003 | 14 |
| 2568 | ENSMUSG( | Foxo1 | 56458 | 002068922 | -01346055 | -078 | 00275 | 14 |
| 2628 | ENSMUSG( | Gabbr1 | 54393 | -01715756 | -00099925 | -08975 | -00625 | 14 |
| 2633 | ENSMUSG( | Gabra3 | 14396 | -01396363 | 005635347 | -04575 | -04425 | 14 |
| 2634 | ENSMUSG( | Gabra4 | 14397 | -06118525 | 015044303 | -057 | -0675 | 14 |
| 2759 | ENSMUSG( | Glb1l | 74577 | -00068564 | 000093425 | -05525 | -02675 | 14 |
| 2814 | ENSMUSG( | Gng11 | 66066 | -04943354 | -00734042 | -1175 | -03225 | 14 |
| 2859 | ENSMUSG( | Gpatch8 | 237943 | 01255509C | 004618771 | -067 | -0145 | 14 |
| 2882 | ENSMUSG( | Gpr22 | 73010 | -03051428 | 051623511 | -06925 | -01075 | 14 |
| 3009 | ENSMUSG( | H2-D1 | 14964 | -02881521 | 00692507C | -0575 | -0125 | 14 |
| 3011 | ENSMUSG( | H3f3a | 15078 | 000683753 | -01368730 | -154 | -06925 | 14 |
| 3025 | ENSMUSG( | Hapln2 | 73940 | -05209521 | 026966057 | -0765 | -074 | 14 |
| 3036 | ENSMUSG( | Hcls1 | 15163 | 006648406 | -01574655 | -08075 | -049 | 14 |
| 3050 | ENSMUSG( | Hdac9 | 79221 | -00558361 | -02159857 | -08275 | -0855 | 14 |
| 3110 | ENSMUSG( | Hmga1 | 15361 | 002987549 | -00714540 | -0985 | -09775 | 14 |
| 3161 | ENSMUSG( | Hrh1 | 15465 | -03602773 | 01497674C | -06375 | -021 | 14 |
| 3186 | ENSMUSG( | Hspa1a | 193740 | -01062692 | 032553026 | -04825 | -0265 | 14 |
| 3233 | ENSMUSG( | Ifi47 | 15953 | -05724287 | 006907017 | -12325 | -0435 | 14 |
| 3272 | ENSMUSG( | Il17ra | 16172 | -02368302 | -00192606 | -0545 | -044 | 14 |
| 3298 | ENSMUSG( | Inip | 66209 | 002578882 | -00626506 | -077 | -00375 | 14 |
| 3344 | ENSMUSG( | Irak1 | 16179 | 011111206 | -00488733 | -05175 | -06725 | 14 |
| 3346 | ENSMUSG( | Irf2bp1 | 272359 | 005948609 | -00427414 | -10675 | -06425 | 14 |
| 3358 | ENSMUSG( | Isg20l2 | 229504 | -01238273 | 010784467 | -09275 | -026 | 14 |
| 3394 | ENSMUSG( | Itpkc | 233011 | -01930046 | -01094853 | -11225 | -01375 | 14 |
| 3411 | ENSMUSG( | Jam3 | 83964 | -03494336 | 02861971C | -053 | -03625 | 14 |
| 3413 | ENSMUSG( | Jmjd6 | 107817 | -00449967 | 010594547 | -084 | -013 | 14 |
| 3430 | ENSMUSG( | Kank4 | 242553 | -03818009 | 008577469 | -09175 | 0075 | 14 |
| 3462 | ENSMUSG( | Kcnf1 | 382571 | -02236170 | 024411883 | -0455 | -035 | 14 |
| 3474 | ENSMUSG( | Kcnj12 | 16515 | -00435745 | 017273769 | -0705 | -02825 | 14 |
| 3483 | ENSMUSG( | Kcnk4 | 16528 | 014063143 | 021346225 | -0745 | -0545 | 14 |
| 3546 | ENSMUSG( | Kirrel2 | 243911 | 000624991 | 035641262 | -051 | -034 | 14 |
| 3652 | ENSMUSG( | Lgals1 | 16852 | -01554041 | -03121047 | -0555 | -02875 | 14 |
| 3655 | ENSMUSG( | Lgals9 | 16859 | -02315530 | -02309660 | -14075 | -06125 | 14 |
| 3662 | ENSMUSG( | Lhfpl2 | 218454 | -03734652 | -01083133 | -13475 | -05025 | 14 |
| 3672 | ENSMUSG( | Limd2 | 67803 | 003540431 | -00841321 | -051 | -048 | 14 |
| 3742 | ENSMUSG( | Lrrc10b | 278795 | -05846184 | 01783352C | -066 | -034 | 14 |
| 3836 | ENSMUSG( | Mad1l1 | 17120 | -01755447 | -02145776 | -0745 | -0285 | 14 |
| 3837 | ENSMUSG( | Mad2l1 | 56150 | 012363384 | 017304577 | -0555 | -02425 | 14 |
| 3838 | ENSMUSG( | Mad2l1bp | 66591 | 019268882 | 009709619 | -0755 | -028 | 14 |
| 3910 | ENSMUSG( | Mapk7 | 23939 | 000218251 | -00723617 | -09475 | -036 | 14 |
| 3931 | ENSMUSG( | Mark3 | 17169 | 020228282 | 006010324 | -0885 | -056 | 14 |
| 3935 | ENSMUSG( | Mas1 | 17171 | -01319696 | 010192802 | -06475 | -05975 | 14 |
| 3970 | ENSMUSG( | Mchr1 | 207911 | -04192813 | 008718226 | -14825 | -0695 | 14 |
| 3971 | ENSMUSG( | Mcl1 | 17210 | -00149236 | 002399642 | -06175 | -03325 | 14 |
| 3996 | ENSMUSG( | Me3 | 109264 | -00444555 | -00871234 | -0615 | -06225 | 14 |
| 4003 | ENSMUSG( | Med12 | 59024 | 018242975 | 018554294 | -1145 | 0115 | 14 |
| 4048 | ENSMUSG( | Mex3c | 240396 | 000286938 | 003820644 | -1355 | -01725 | 14 |
| 4049 | ENSMUSG( | Mfap1b | 1E+08 | 014757236 | -00065835 | -08325 | -0185 | 14 |
| 4124 | ENSMUSG( | Mobp | 17433 | 018173469 | -00784374 | -09825 | -04925 | 14 |
| 4143 | ENSMUSG( | Mpdu1 | 24070 | -02310967 | -00031054 | -0595 | -0335 | 14 |
| 4190 | ENSMUSG( | Mrpl34 | 94065 | 009490963 | -01155049 | -089 | -05625 | 14 |
| 4263 | ENSMUSG( | Mtap | 66902 | -01531852 | -01554611 | -054 | -012 | 14 |
| 4277 | ENSMUSG( | Mthfsl | 1E+08 | -00326678 | -02278965 | -0745 | -00875 | 14 |
| 4329 | ENSMUSG( | Myl4 | 17896 | -01341396 | -04419542 | -04725 | -062 | 14 |
| 4349 | ENSMUSG( | Mypop | 232934 | 008637922 | -00771820 | -119 | -05625 | 14 |
| 4369 | ENSMUSG( | Nabp2 | 69917 | -00372918 | -01317223 | -06975 | -03375 | 14 |
| 4386 | ENSMUSG( | Nap1l5 | 58243 | 004716508 | -01346288 | -054 | -06575 | 14 |
| 4464 | ENSMUSG( | Ndufab1 | 70316 | 000619551 | -01454225 | -0655 | -0585 | 14 |
| 4488 | ENSMUSG( | Ndufs4 | 17993 | -00584964 | -02194481 | -05325 | -0515 | 14 |
| 4555 | ENSMUSG( | Nhp2 | 52530 | -00904182 | -02530680 | -066 | -02425 | 14 |
| 4597 | ENSMUSG( | Nob1 | 67619 | -00399711 | -00435382 | -05275 | -0275 | 14 |
| 4607 | ENSMUSG( | Nol7 | 70078 | 006415147 | -00731252 | -07325 | -0145 | 14 |
| 4628 | ENSMUSG( | Notch2 | 18129 | -03075139 | -00197055 | -07275 | -0415 | 14 |
| 4656 | ENSMUSG( | Nr2c2ap | 75692 | 001998233 | -01391109 | -06675 | -0225 | 14 |
| 4730 | ENSMUSG( | Nudt17 | 78373 | 009777595 | -01377938 | -07075 | -00675 | 14 |
| 4742 | ENSMUSG( | Numa1 | 101706 | 007095186 | -00056604 | -0545 | -022 | 14 |
| 4809 | ENSMUSG( | Oprm1 | 18390 | -04665634 | 024572637 | -11675 | -00325 | 14 |
| 4826 | ENSMUSG( | Osbpl9 | 100273 | 00608621C | 01423500C | -063 | -0815 | 14 |
| 4871 | ENSMUSG( | Paip2b | 232164 | -00115345 | -01867186 | -102 | -01675 | 14 |
| 5007 | ENSMUSG( | Pde2a | 207728 | 006041335 | -00442836 | -122 | -029 | 14 |
| 5061 | ENSMUSG( | Peli1 | 67245 | 007498187 | 013716587 | -0665 | -06375 | 14 |
| 5105 | ENSMUSG( | Pgm1 | 72157 | 010648727 | -00264843 | -10075 | -00775 | 14 |
| 5291 | ENSMUSG( | Pml | 18854 | 00409894C | -00080916 | -06975 | -0345 | 14 |
| 5298 | ENSMUSG( | Pnisr | 66625 | -00261378 | 042349601 | -11325 | -08575 | 14 |
| 5304 | ENSMUSG( | Pnn | 18949 | 004501033 | 020260274 | -063 | -03075 | 14 |
| 5342 | ENSMUSG( | Polr3c | 74414 | 010196054 | 008654598 | -052 | -04025 | 14 |
| 5346 | ENSMUSG( | Polrmt | 216151 | 011089299 | 010558756 | -0585 | -01 | 14 |
| 5367 | ENSMUSG( | Ppdpf | 66496 | 009109961 | -01601904 | -0775 | -01975 | 14 |
| 5397 | ENSMUSG( | Ppm1m | 67905 | 010690515 | -00781924 | -13925 | -0165 | 14 |
| 5409 | ENSMUSG( | Ppp1r14a | 68458 | -03392097 | -02516001 | -06175 | -04075 | 14 |
| 5502 | ENSMUSG( | Prkdc | 19090 | -00865980 | 00225455C | -0945 | -038 | 14 |
| 5505 | ENSMUSG( | Prkn | 50873 | 013551152 | -00772624 | -08 | -0685 | 14 |
| 5576 | ENSMUSG( | Psmb8 | 16913 | -02309449 | -01704916 | -07375 | -042 | 14 |
| 5608 | ENSMUSG( | Psrc1 | 56742 | 017807946 | -01874795 | -17 | -0885 | 14 |
| 5776 | ENSMUSG( | Rabep2 | 70314 | -00701509 | 014565544 | -06475 | -01 | 14 |
| 5856 | ENSMUSG( | Rbbp5 | 213464 | 012064623 | -00067684 | -06525 | -03525 | 14 |
| 5866 | ENSMUSG( | Rbl2 | 19651 | 008100268 | -01136468 | -099 | -031 | 14 |
| 5920 | ENSMUSG( | Rel | 19696 | -01961961 | -01091888 | -065 | -01475 | 14 |
| 5936 | ENSMUSG( | Rexo4 | 227656 | 011004357 | 005399846 | -05775 | -02625 | 14 |
| 5941 | ENSMUSG( | Rfesd | 218341 | 000949786 | -00270940 | -076 | 00225 | 14 |
| 5947 | ENSMUSG( | Rfxap | 170767 | 00673806 | -00321613 | -06425 | -0375 | 14 |
| 5948 | ENSMUSG( | Rgcc | 66214 | 000868815 | -02372056 | -09575 | -0315 | 14 |
| 5953 | ENSMUSG( | Rgp1 | 242406 | 010016852 | -00229701 | -0765 | -0325 | 14 |
| 5966 | ENSMUSG( | Rgs9 | 19739 | -07999287 | -03600026 | -09675 | -049 | 14 |

| | | | | | | | | |
|---|---|---|---|---|---|---|---|---|
| 5976 | ENSMUSG( | Rhog | 56212 | -01884727 | -01503894 | -06825 | -023 | 14 |
| 6023 | ENSMUSG( | Rnf10 | 50849 | 015877226 | 004316526 | -073 | -025 | 14 |
| 6037 | ENSMUSG( | Rnf149 | 67702 | -01442486 | 007019183 | -0635 | -01225 | 14 |
| 6133 | ENSMUSG( | Rpp30 | 54364 | 004475243 | 006322683 | -06425 | -0225 | 14 |
| 6180 | ENSMUSG( | Rragd | 52187 | -00462080 | -01377362 | -06675 | -0475 | 14 |
| 6215 | ENSMUSG( | Rufy3 | 52822 | 008387974 | -00927840 | -07425 | -01875 | 14 |
| 6234 | ENSMUSG( | S100a4 | 20198 | -03791838 | -02554857 | -071 | -03525 | 14 |
| 6241 | ENSMUSG( | Saal1 | 78935 | 001205237 | 018716791 | -05475 | -04175 | 14 |
| 6258 | ENSMUSG( | Sap18 | 20220 | 005006179 | -02233465 | -06675 | -05025 | 14 |
| 6281 | ENSMUSG( | Scaf11 | 72193 | 013873018 | 008721112 | -0785 | -0785 | 14 |
| 6326 | ENSMUSG( | Sdad1 | 231452 | -00120568 | -01006037 | -11975 | -0185 | 14 |
| 6406 | ENSMUSG( | Septin4 | 18952 | -01200977 | -00942032 | -0445 | -06025 | 14 |
| 6435 | ENSMUSG( | Setd2 | 235626 | 047567097 | 093507410 | -07125 | -04925 | 14 |
| 6444 | ENSMUSG( | Sf3a2 | 20222 | -00288595 | 001703742 | -04775 | -0605 | 14 |
| 6450 | ENSMUSG( | Sf3b5 | 66125 | 009015930 | -00891013 | -093 | -07975 | 14 |
| 6457 | ENSMUSG( | Sft2d2 | 108735 | 008418681 | 005378933 | -09475 | -043 | 14 |
| 6459 | ENSMUSG( | Sfxn2 | 94279 | -01258457 | 037367589 | -10175 | -03275 | 14 |
| 6466 | ENSMUSG( | Sgip1 | 73094 | -00929199 | 002162267 | -05525 | -0405 | 14 |
| 6545 | ENSMUSG( | Slain1 | 105439 | -00666673 | -01241489 | -07125 | -01475 | 14 |
| 6555 | ENSMUSG( | Slc13a4 | 243755 | 011777615 | 022473705 | -1455 | -0615 | 14 |
| 6571 | ENSMUSG( | Slc19a1 | 20509 | -02138455 | 035051834 | -068 | -001 | 14 |
| 6584 | ENSMUSG( | Slc22a5 | 20520 | -01031447 | 006866078 | -054 | -07875 | 14 |
| 6598 | ENSMUSG( | Slc25a18 | 71803 | 015012466 | -02218754 | -059 | -02675 | 14 |
| 6599 | ENSMUSG( | Slc25a19 | 67283 | 007608588 | -00139441 | -05975 | -0275 | 14 |
| 6618 | ENSMUSG( | Slc25a44 | 229517 | -01559047 | 013442541 | -052 | -01175 | 14 |
| 6639 | ENSMUSG( | Slc30a7 | 66500 | -03024529 | 020568602 | -10875 | -0095 | 14 |
| 6665 | ENSMUSG( | Slc39a14 | 213053 | -03549271 | 018459409 | -08075 | 00075 | 14 |
| 6682 | ENSMUSG( | Slc4a4 | 54403 | -00822913 | -00870060 | -05425 | -04175 | 14 |
| 6736 | ENSMUSG( | Sltm | 66660 | 01311996 | 001503185 | -064 | -076 | 14 |
| 6768 | ENSMUSG( | Smim12 | 80284 | -02308200 | -01574677 | -05525 | -04225 | 14 |
| 6769 | ENSMUSG( | Smim13 | 108934 | -00564612 | -00234051 | -06275 | -00525 | 14 |
| 6839 | ENSMUSG( | Snx21 | 101113 | 009075606 | 007439188 | -0545 | -018 | 14 |
| 6858 | ENSMUSG( | Son | 20658 | 003206265 | 028510838 | -05225 | -06075 | 14 |
| 6963 | ENSMUSG( | Srsf2 | 20382 | -02235813 | 017038681 | -04575 | -05975 | 14 |
| 6966 | ENSMUSG( | Srsf5 | 20384 | 458e-05 | 016051948 | -0485 | -0585 | 14 |
| 6969 | ENSMUSG( | Srsf9 | 108014 | 005215404 | 004799258 | -0505 | -0555 | 14 |
| 6971 | ENSMUSG( | Ss18 | 268996 | 013782879 | 013296834 | -078 | -0245 | 14 |
| 7081 | ENSMUSG( | Sugct | 192136 | -00825331 | -01043432 | -104 | 0145 | 14 |
| 7157 | ENSMUSG( | Szt2 | 230676 | 018672066 | 014836638 | -1045 | 0075 | 14 |
| 7166 | ENSMUSG( | Taf1 | 270627 | 007075791 | -00955683 | -11075 | -0895 | 14 |
| 7178 | ENSMUSG( | Tafa5 | 106014 | -00797487 | -00330129 | -09875 | 01875 | 14 |
| 7195 | ENSMUSG( | Tapbpl | 213233 | -03021136 | 000320147 | -0505 | -02125 | 14 |
| 7301 | ENSMUSG( | Tgfb1i1 | 21804 | 004146032 | -00753358 | -061 | -00925 | 14 |
| 7310 | ENSMUSG( | Thbd | 21824 | 003121542 | 037047952 | -054 | -0165 | 14 |
| 7326 | ENSMUSG( | Thrap3 | 230753 | 000086771 | -00471433 | -07575 | -0605 | 14 |
| 7388 | ENSMUSG( | Tmed3 | 66111 | -02884289 | 004435501 | -0925 | -06375 | 14 |
| 7398 | ENSMUSG( | Tmem11 | 216821 | -00179442 | -00453544 | -07425 | -03575 | 14 |
| 7457 | ENSMUSG( | Tmem25 | 71687 | -02238024 | -00076412 | -0745 | -07 | 14 |
| 7469 | ENSMUSG( | Tmem38b | 52076 | 000101123 | 00660692 | -0735 | -0045 | 14 |
| 7474 | ENSMUSG( | Tmem51 | 214359 | -02171504 | -04083308 | -078 | -04525 | 14 |
| 7521 | ENSMUSG( | Tnrc18 | 231861 | -00085131 | 006450938 | -069 | -02825 | 14 |
| 7524 | ENSMUSG( | Tns1 | 21961 | 008424480 | 014671795 | -07775 | -03675 | 14 |
| 7564 | ENSMUSG( | Tpm4 | 326618 | 013672941 | 023861309 | -06925 | -02525 | 14 |
| 7626 | ENSMUSG( | Trim72 | 434246 | 022731837 | 006823440 | -072 | -009 | 14 |
| 7640 | ENSMUSG( | Trmt44 | 78890 | 020261327 | 024418357 | -1005 | -0585 | 14 |
| 7667 | ENSMUSG( | Tsga10 | 211484 | 003385726 | 004107424 | -12775 | -05375 | 14 |
| 7682 | ENSMUSG( | Tspan8 | 216350 | -01528641 | 029296049 | -0625 | -002 | 14 |
| 7707 | ENSMUSG( | Ttc8 | 76260 | 011330614 | -00946241 | -05675 | -03575 | 14 |
| 7758 | ENSMUSG( | Txndc17 | 52700 | 003614830 | -01115330 | -06125 | -0145 | 14 |
| 7803 | ENSMUSG( | Ube2j2 | 140499 | -00557027 | 000696471 | -0625 | -00475 | 14 |
| 7811 | ENSMUSG( | Ube2ql1 | 76980 | 013043651 | -00731129 | -10975 | 012 | 14 |
| 8005 | ENSMUSG( | Vps13b | 666173 | -01097504 | 004553097 | -05125 | -032 | 14 |
| 8055 | ENSMUSG( | Wac | 225131 | 005185608 | 014032838 | -09525 | 0005 | 14 |
| 8159 | ENSMUSG( | Yap1 | 22601 | 017958850 | -00100158 | -07125 | -08525 | 14 |
| 8209 | ENSMUSG( | Zc3h18 | 76014 | 010444246 | 004672416 | -0615 | -01275 | 14 |
| 8278 | ENSMUSG( | Zmym4 | 67785 | -00535600 | -01048635 | -07125 | -00525 | 14 |
| 8282 | ENSMUSG( | Znhit2 | 29805 | 011419107 | 000647231 | -08425 | -00425 | 14 |
| 29 | ENSMUSG( | Aarsd1 | 69684 | 007417213 | 065275582 | 01375 | 0125 | 15 |
| 33 | ENSMUSG( | Aatk | 11302 | 008673757 | 025213384 | 01725 | -01225 | 15 |
| 42 | ENSMUSG( | Abca9 | 217262 | 015766416 | 047873907 | -02 | 0095 | 15 |
| 44 | ENSMUSG( | Abcb1a | 18671 | 011102346 | 078458189 | -022 | 00225 | 15 |
| 50 | ENSMUSG( | Abcc4 | 239273 | 023531565 | 058937613 | 02675 | -009 | 15 |
| 52 | ENSMUSG( | Abcc8 | 20927 | 001766525 | 030948607 | -019 | -01 | 15 |
| 56 | ENSMUSG( | Abcd4 | 19300 | 010635738 | 021854853 | 025 | 0055 | 15 |
| 62 | ENSMUSG( | Abcg2 | 26357 | 005430977 | 046327953 | -00375 | 00075 | 15 |
| 81 | ENSMUSG( | Ablim1 | 226251 | 001956164 | 014950209 | -00875 | 01875 | 15 |
| 102 | ENSMUSG( | Acap3 | 140500 | 020701637 | 008981187 | -011 | 0155 | 15 |
| 126 | ENSMUSG( | Acp6 | 66659 | 006124988 | 023004938 | -00925 | -0085 | 15 |
| 160 | ENSMUSG( | Adal | 75894 | 014902891 | 008805052 | -00525 | 00975 | 15 |
| 175 | ENSMUSG( | Adcy1 | 432530 | 020482496 | 021511059 | 01875 | -00425 | 15 |
| 179 | ENSMUSG( | Adcy6 | 11512 | 022556408 | 033177252 | 0085 | -024 | 15 |
| 181 | ENSMUSG( | Adcy9 | 11515 | -00049713 | 055333860 | -013 | -03475 | 15 |
| 190 | ENSMUSG( | Adgrb1 | 107831 | -00077523 | 020921645 | -00525 | 0055 | 15 |
| 194 | ENSMUSG( | Adgrf5 | 224792 | 017932218 | 075267143 | 01275 | 016 | 15 |
| 223 | ENSMUSG( | Afdn | 17356 | 007592786 | 013087151 | 00225 | 02 | 15 |
| 231 | ENSMUSG( | Agap2 | 216439 | 025007461 | 021084799 | 01025 | 0235 | 15 |
| 234 | ENSMUSG( | Agfg1 | 15463 | 008807230 | 013368308 | -00425 | 005 | 15 |
| 239 | ENSMUSG( | Ago2 | 239528 | 015284323 | 014120187 | 0155 | 0205 | 15 |
| 246 | ENSMUSG( | Agrn | 11603 | 008743613 | 020325777 | 0055 | 01675 | 15 |
| 249 | ENSMUSG( | Ahctf1 | 226747 | -00008993 | 032092428 | -00725 | -014 | 15 |
| 255 | ENSMUSG( | Ahnak | 66395 | 025987246 | 054185595 | -02225 | 0085 | 15 |
| 330 | ENSMUSG( | Als2 | 74018 | 028081928 | 014766105 | -00725 | 01325 | 15 |
| 339 | ENSMUSG( | Amigo2 | 105827 | 009377172 | 051494991 | 00125 | 01275 | 15 |
| 375 | ENSMUSG( | Ankrd34b | 218440 | 012621889 | 013010069 | -00625 | 00825 | 15 |
| 381 | ENSMUSG( | Ankrd50 | 99696 | 015681742 | 012753051 | -0055 | 00175 | 15 |
| 383 | ENSMUSG( | Ankrd54 | 223690 | 004418446 | 017073738 | -01275 | 0245 | 15 |
| 385 | ENSMUSG( | Anks1 | 224650 | 018327051 | 016724266 | -01375 | -012 | 15 |
| 397 | ENSMUSG( | Anpep | 16790 | 010711683 | 058808999 | -01275 | 00625 | 15 |
| 403 | ENSMUSG( | Anxa3 | 11745 | 012059827 | 037663558 | -02725 | -0015 | 15 |
| 483 | ENSMUSG( | Arfgef1 | 211673 | 015771689 | 017360005 | -00325 | 02325 | 15 |
| 497 | ENSMUSG( | Arhgap21 | 71435 | 008787936 | 024646759 | -008 | 001 | 15 |
| 500 | ENSMUSG( | Arhgap26 | 71302 | 022155570 | 008423406 | 00025 | -01125 | 15 |
| 502 | ENSMUSG( | Arhgap29 | 214137 | 014117597 | 033879880 | 0025 | -013 | 15 |
| 503 | ENSMUSG( | Arhgap31 | 12549 | 018926928 | 025231686 | 0285 | -01225 | 15 |
| 506 | ENSMUSG( | Arhgap35 | 232906 | 020354729 | 012873602 | 0115 | 005 | 15 |
| 511 | ENSMUSG( | Arhgap6 | 11856 | -00271753 | 022695848 | -00775 | 00275 | 15 |
| 514 | ENSMUSG( | Arhgef1 | 16801 | 031091914 | 010927168 | -00225 | 001 | 15 |
| 520 | ENSMUSG( | Arhgef18 | 102098 | 015029067 | 015437221 | -00625 | 0065 | 15 |
| 525 | ENSMUSG( | Arhgef28 | 110596 | 008315037 | 012950790 | 00875 | 01575 | 15 |
| 582 | ENSMUSG( | Arvcf | 11877 | 014135734 | 015870303 | -004 | -003 | 15 |
| 590 | ENSMUSG( | Asb3 | 65257 | 027250586 | 012948738 | 00125 | -00075 | 15 |
| 595 | ENSMUSG( | Asf1a | 66403 | 005084541 | 015646098 | 012 | 021 | 15 |
| 624 | ENSMUSG( | Atg2a | 329015 | 025010335 | 008470389 | -01075 | 01675 | 15 |
| 625 | ENSMUSG( | Atg2b | 76559 | 010853251 | 02332311 | -0125 | 01275 | 15 |
| 631 | ENSMUSG( | Atg9a | 245860 | 000019348 | 039854900 | 01625 | 00675 | 15 |
| 641 | ENSMUSG( | Atp13a2 | 74772 | 006126654 | 024628371 | 00975 | 002 | 15 |
| 751 | ENSMUSG( | Bbs7 | 71492 | 014615447 | 009172640 | -00225 | 011 | 15 |
| 754 | ENSMUSG( | Bcam | 57278 | -00844341 | 053079946 | 0155 | 00375 | 15 |
| 764 | ENSMUSG( | Bcat2 | 12036 | 011396114 | 026493568 | 00425 | -018 | 15 |
| 788 | ENSMUSG( | Bicd1 | 12121 | -00552216 | 031138915 | -012 | -01475 | 15 |
| 843 | ENSMUSG( | Brd9 | 105246 | 031053638 | 011000140 | 0195 | 0085 | 15 |
| 853 | ENSMUSG( | Brsk1 | 381979 | 023004186 | 009116963 | 0065 | 005 | 15 |
| 859 | ENSMUSG( | Bsn | 12217 | 013088219 | 026141045 | -02475 | -00775 | 15 |
| 882 | ENSMUSG( | C2cd2 | 207781 | -00455205 | 047371785 | 00325 | -01875 | 15 |
| 887 | ENSMUSG( | C4b | 12268 | 153210721 | 210844048 | 03325 | 0755 | 15 |
| 893 | ENSMUSG( | Cables1 | 63955 | 077354642 | 052338459 | -01675 | -0035 | 15 |

| | | | | | | | | | |
|---|---|---|---|---|---|---|---|---|---|
| 897 | ENSMUSG( | Cachd1 | 320508 | 021872103 | 057828188 | 042 | -0065 | | 15 |
| 899 | ENSMUSG( | Cacna1b | 12287 | -00670667 | 028343681 | -00075 | 00225 | | 15 |
| 903 | ENSMUSG( | Cacna1g | 12291 | 005609389 | 013023482 | 01475 | 00525 | | 15 |
| 917 | ENSMUSG( | Cacng8 | 81905 | -01153337 | 021479291 | -002 | 00375 | | 15 |
| 940 | ENSMUSG( | Camk1g | 215303 | 029161545 | 008055345 | 007 | 0135 | | 15 |
| 978 | ENSMUSG( | Carmil3 | 268747 | 049071953 | 014572911 | 0205 | 01375 | | 15 |
| 980 | ENSMUSG( | Cars1 | 27267 | 012118350 | 014766416 | -00575 | 002 | | 15 |
| 998 | ENSMUSG( | Cavin1 | 19285 | 019451103 | 030983252 | -002 | 0145 | | 15 |
| 999 | ENSMUSG( | Cavin2 | 20324 | 018766672 | 041595630 | -00425 | 0195 | | 15 |
| 1022 | ENSMUSG( | Ccar1 | 67500 | 022543021 | 022770370 | -00125 | 0325 | | 15 |
| 1034 | ENSMUSG( | Ccdc141 | 545428 | 046672313 | 015989065 | -01575 | 022 | | 15 |
| 1051 | ENSMUSG( | Ccdc6 | 76551 | 031093562 | 010985279 | -02125 | 017 | | 15 |
| 1052 | ENSMUSG( | Ccdc85a | 216613 | 029360545 | 023588876 | -011 | 00775 | | 15 |
| 1053 | ENSMUSG( | Ccdc85b | 240514 | 008796307 | 011743083 | 0185 | 196262e-1 | | 15 |
| 1074 | ENSMUSG( | Ccnyl1 | 227210 | -00119485 | 015914233 | -00775 | 02225 | | 15 |
| 1088 | ENSMUSG( | Cd109 | 235505 | 004221544 | 050753141 | 0295 | -00275 | | 15 |
| 1090 | ENSMUSG( | Cd163 | 93671 | 049740348 | 081390734 | 032 | 03575 | | 15 |
| 1116 | ENSMUSG( | Cdc37l1 | 67072 | 004620671 | 022508337 | -01325 | 0165 | | 15 |
| 1120 | ENSMUSG( | Cdc42bpb | 217866 | 016620973 | 016341652 | -0025 | 001 | | 15 |
| 1126 | ENSMUSG( | Cdc73 | 214498 | 012660053 | 022676545 | -022 | -006 | | 15 |
| 1132 | ENSMUSG( | Cdh19 | 227485 | 005867769 | 022275930 | 0015 | 01275 | | 15 |
| 1189 | ENSMUSG( | Cep76 | 225659 | 001793763 | 023898499 | -01775 | 02725 | | 15 |
| 1196 | ENSMUSG( | Cert1 | 68018 | 004870971 | 014437895 | -01575 | -00425 | | 15 |
| 1212 | ENSMUSG( | Cgnl1 | 68178 | 002102069 | 020813116 | -007 | 02625 | | 15 |
| 1226 | ENSMUSG( | Chd2 | 244059 | 019254488 | 008873843 | -01975 | -00225 | | 15 |
| 1231 | ENSMUSG( | Chd8 | 67772 | 006376196 | 014156164 | -0185 | 00475 | | 15 |
| 1240 | ENSMUSG( | Chkb | 12651 | 005127558 | 016712929 | -01425 | -01025 | | 15 |
| 1255 | ENSMUSG( | Chordc1 | 66917 | 011090625 | 024521424 | 01075 | 01225 | | 15 |
| 1282 | ENSMUSG( | Cit | 12704 | 014973630 | 011820580 | -01 | -26986e-16 | | 15 |
| 1337 | ENSMUSG( | Clvs2 | 215890 | 019993676 | 007431510 | -007 | 0015 | | 15 |
| 1343 | ENSMUSG( | Cmc4 | 1.06E+08 | 008128426 | 020131894 | 0085 | 018 | | 15 |
| 1358 | ENSMUSG( | Cnksr2 | 245684 | 016244581 | 028655523 | 00975 | 03075 | | 15 |
| 1369 | ENSMUSG( | Cnot3 | 232791 | 009466973 | 018531186 | -0195 | -00125 | | 15 |
| 1404 | ENSMUSG( | Cog1 | 16834 | 013817936 | 009764018 | -0125 | 01325 | | 15 |
| 1420 | ENSMUSG( | Col4a1 | 12826 | 010587228 | 015954072 | 0045 | 0235 | | 15 |
| 1421 | ENSMUSG( | Col4a2 | 12827 | 033265006 | 023113887 | 02425 | 035 | | 15 |
| 1426 | ENSMUSG( | Colec12 | 140792 | 003887699 | 056299407 | 0015 | -02275 | | 15 |
| 1470 | ENSMUSG( | Coro2a | 107684 | 013468336 | 008618447 | -785046e-1 | -00875 | | 15 |
| 1498 | ENSMUSG( | Cpeb2 | 231207 | -00112157 | 015854272 | 01025 | 018 | | 15 |
| 1510 | ENSMUSG( | Cpne7 | 102278 | 061509665 | 024919347 | 04175 | 00325 | | 15 |
| 1533 | ENSMUSG( | Creb1 | 12912 | 021435768 | 016725100 | -007 | 02475 | | 15 |
| 1551 | ENSMUSG( | Crocc | 230872 | 030024182 | 009021250 | -008 | 02775 | | 15 |
| 1582 | ENSMUSG( | Csnk1g2 | 103236 | 020864528 | 008957576 | -00175 | -00675 | | 15 |
| 1609 | ENSMUSG( | Cth | 107869 | 035137144 | 012240143 | 02275 | 01125 | | 15 |
| 1643 | ENSMUSG( | Cul5 | 75717 | 014389967 | 011943111 | -0085 | -00975 | | 15 |
| 1704 | ENSMUSG( | Dab2ip | 69601 | 020849877 | 009756536 | -00875 | 0035 | | 15 |
| 1711 | ENSMUSG( | Dalrd3 | 67789 | 015291238 | 013331888 | -00525 | -00975 | | 15 |
| 1714 | ENSMUSG( | Dapk1 | 69635 | 022478764 | 017335822 | 00575 | -008 | | 15 |
| 1733 | ENSMUSG( | Dcaf6 | 74106 | 021010959 | 010413266 | -01425 | 02375 | | 15 |
| 1764 | ENSMUSG( | Ddc | 13195 | -00771095 | 032785862 | -01675 | -0065 | | 15 |
| 1768 | ENSMUSG( | Ddn | 13199 | 026844736 | 019305376 | -0035 | -00575 | | 15 |
| 1784 | ENSMUSG( | Ddx27 | 228889 | 028394635 | 009022204 | -02775 | 00175 | | 15 |
| 1810 | ENSMUSG( | Dennd1b | 329260 | 015859572 | 014209308 | -0145 | 021 | | 15 |
| 1817 | ENSMUSG( | Dennd6a | 211922 | 012450666 | 016768923 | -02225 | -00625 | | 15 |
| 1834 | ENSMUSG( | Dgkg | 110197 | 045103956 | 042300486 | 02875 | -02375 | | 15 |
| 1837 | ENSMUSG( | Dgkq | 110524 | 006184451 | 014971642 | 01325 | 00675 | | 15 |
| 1871 | ENSMUSG( | Dimt1 | 66254 | 019702874 | 035966536 | 00175 | -01875 | | 15 |
| 1893 | ENSMUSG( | Dlgap1 | 224997 | 008326882 | 013056961 | -00675 | -0105 | | 15 |
| 1896 | ENSMUSG( | Dlgap4 | 228836 | 019688243 | 010420265 | 011 | 00275 | | 15 |
| 1920 | ENSMUSG( | Dnajb4 | 67035 | 006794340 | 015491224 | 0065 | 01225 | | 15 |
| 1923 | ENSMUSG( | Dnajc1 | 13418 | 008787699 | 020092408 | 01 | -005 | | 15 |
| 1938 | ENSMUSG( | Dnajc3 | 1E+08 | 002620325 | 026789777 | 01425 | -00225 | | 15 |
| 1964 | ENSMUSG( | Dock11 | 75974 | 043925364 | 028550577 | -00025 | -00375 | | 15 |
| 1971 | ENSMUSG( | Dock9 | 105445 | 016810120 | 033437619 | 0055 | -01025 | | 15 |
| 1993 | ENSMUSG( | Dpp4 | 13482 | 015852366 | 018348230 | -00925 | 009 | | 15 |
| 2005 | ENSMUSG( | Dpysl4 | 26757 | 025919877 | 004742076 | -0095 | 001 | | 15 |
| 2014 | ENSMUSG( | Dscaml1 | 114873 | 008386647 | 051867906 | 02475 | 028 | | 15 |
| 2037 | ENSMUSG( | Dvl1 | 13542 | 014160600 | 009677212 | 01325 | -00025 | | 15 |
| 2079 | ENSMUSG( | Edc4 | 234699 | 027157640 | 011183255 | 004 | 0035 | | 15 |
| 2085 | ENSMUSG( | Eea1 | 216238 | 021152163 | 016643141 | 0105 | -00625 | | 15 |
| 2114 | ENSMUSG( | Ehbp1 | 216565 | 019535253 | 010096185 | -016 | 00975 | | 15 |
| 2117 | ENSMUSG( | Ehd2 | 259300 | 031271126 | 063800464 | 0055 | 04575 | | 15 |
| 2132 | ENSMUSG( | Eif2ak4 | 27103 | 008043602 | 013492051 | -013 | 008 | | 15 |
| 2162 | ENSMUSG( | Eif4ebp2 | 13688 | 017782697 | 009525195 | -00625 | 0155 | | 15 |
| 2171 | ENSMUSG( | Eif5b | 226982 | 019731748 | 025216666 | 017 | -0055 | | 15 |
| 2183 | ENSMUSG( | Elfn2 | 207393 | 011115315 | 029060127 | 00125 | 0085 | | 15 |
| 2188 | ENSMUSG( | Eln | 13717 | 006109264 | 025695630 | 00175 | 012 | | 15 |
| 2218 | ENSMUSG( | Eml6 | 237711 | 019800833 | 008733948 | -00175 | -0005 | | 15 |
| 2226 | ENSMUSG( | Eng | 13805 | 006374139 | 073767083 | 00725 | 03475 | | 15 |
| 2240 | ENSMUSG( | Entpd1 | 12495 | 007551665 | 050133275 | -02025 | 02775 | | 15 |
| 2253 | ENSMUSG( | Epb41l4b | 54357 | 020958948 | 017043389 | -0235 | 00425 | | 15 |
| 2265 | ENSMUSG( | Ephb2 | 13844 | -00794421 | 043269377 | -00425 | 0015 | | 15 |
| 2307 | ENSMUSG( | Esam | 69524 | 022408385 | 046560777 | 001 | 00475 | | 15 |
| 2398 | ENSMUSG( | Fam184a | 75906 | 017494424 | 025824267 | -0025 | 01825 | | 15 |
| 2455 | ENSMUSG( | Fbxo31 | 76454 | 019234702 | 012882100 | 007 | 0185 | | 15 |
| 2457 | ENSMUSG( | Fbxo4 | 106052 | 025787490 | 023402111 | 0155 | 0105 | | 15 |
| 2458 | ENSMUSG( | Fbxo41 | 330369 | 015871259 | 020952731 | -003 | 02025 | | 15 |
| 2473 | ENSMUSG( | Fcgrt | 14132 | -00756749 | 028801214 | -00425 | 00125 | | 15 |
| 2475 | ENSMUSG( | Fcho2 | 218503 | 007573369 | 026258950 | -0025 | 004 | | 15 |
| 2477 | ENSMUSG( | Fcsk | 234730 | 027727631 | 018335303 | -002 | -0165 | | 15 |
| 2481 | ENSMUSG( | Fdx2 | 68165 | 056210505 | 021479450 | -01 | -03475 | | 15 |
| 2491 | ENSMUSG( | Fgd2 | 26382 | 002181804 | 024960262 | 022 | -0095 | | 15 |
| 2498 | ENSMUSG( | Fgfr1 | 14182 | 000772730 | 026706289 | -009 | -0015 | | 15 |
| 2504 | ENSMUSG( | Fhip1b | 74349 | 012867771 | 013342559 | -0115 | 00125 | | 15 |
| 2511 | ENSMUSG( | Fhod3 | 225288 | 012481552 | 019780459 | -0125 | -00675 | | 15 |
| 2512 | ENSMUSG( | Fibcd1 | 98970 | 020947466 | 033343816 | -0245 | 02825 | | 15 |
| 2525 | ENSMUSG( | Fkbp5 | 14229 | 083506011 | 083346240 | 02125 | 05575 | | 15 |
| 2532 | ENSMUSG( | Flna | 192176 | 011221315 | 019659734 | 003 | 014 | | 15 |
| 2533 | ENSMUSG( | Flnb | 286940 | 018375949 | 028615974 | -02025 | 005 | | 15 |
| 2550 | ENSMUSG( | Fn1 | 14268 | 020246677 | 063879591 | 02475 | 0245 | | 15 |
| 2556 | ENSMUSG( | Fndc3a | 319448 | 011288441 | 014644004 | 004 | 0185 | | 15 |
| 2580 | ENSMUSG( | Frmpd3 | 245643 | 040811335 | 016595754 | -00375 | -00575 | | 15 |
| 2585 | ENSMUSG( | Fry | 320365 | 002383098 | 016242472 | 01475 | 0195 | | 15 |
| 2606 | ENSMUSG( | Fv1 | 14349 | 028927933 | 035611430 | -01925 | -0015 | | 15 |
| 2669 | ENSMUSG( | Garem1 | 381126 | 007730172 | 015110527 | 021 | 00775 | | 15 |
| 2675 | ENSMUSG( | Gas2l1 | 78926 | 026920473 | 022694958 | 00275 | 03 | | 15 |
| 2723 | ENSMUSG( | Gfod1 | 328232 | 046516901 | 023159979 | 0105 | -0005 | | 15 |
| 2737 | ENSMUSG( | Ggt5 | 23887 | 021002766 | 031102878 | -0105 | -001 | | 15 |
| 2747 | ENSMUSG( | Git1 | 216963 | 029808202 | 013416573 | 001 | -018 | | 15 |
| 2748 | ENSMUSG( | Git2 | 26431 | 003798226 | 018979977 | -0005 | 022 | | 15 |
| 2865 | ENSMUSG( | Gpcpd1 | 74182 | 005553014 | 031242039 | 005 | -00425 | | 15 |
| 2897 | ENSMUSG( | Gpt | 76282 | 016239829 | 022612589 | -00975 | -00125 | | 15 |
| 2920 | ENSMUSG( | Grik3 | 14807 | -00609356 | 040145249 | 0265 | -01025 | | 15 |
| 2921 | ENSMUSG( | Grik5 | 14809 | 011067950 | 039047576 | 01325 | -03375 | | 15 |
| 2948 | ENSMUSG( | Gse1 | 382034 | 012626822 | 022406873 | 02525 | -0105 | | 15 |
| 3042 | ENSMUSG( | Hdac10 | 170787 | 023719935 | 027794885 | 00475 | 0265 | | 15 |
| 3046 | ENSMUSG( | Hdac4 | 208727 | 043345283 | 031167578 | -00225 | -01225 | | 15 |
| 3060 | ENSMUSG( | Heatr1 | 217995 | 024332262 | 013268643 | -0255 | -00275 | | 15 |
| 3072 | ENSMUSG( | Hecw1 | 94253 | -01172527 | 023022729 | -01125 | 0015 | | 15 |
| 3076 | ENSMUSG( | Heph | 15203 | 030025100 | 025595982 | 01025 | -00475 | | 15 |
| 3079 | ENSMUSG( | Herc3 | 73998 | 014988894 | 010236226 | 01425 | 01025 | | 15 |
| 3080 | ENSMUSG( | Herc4 | 67345 | 015966744 | 020515628 | 0155 | 0065 | | 15 |
| 3098 | ENSMUSG( | Hip1 | 215114 | 025653764 | 024882769 | 00625 | 01575 | | 15 |
| 3151 | ENSMUSG( | Hook3 | 320191 | -00009044 | 025938855 | -00725 | -00125 | | 15 |
| 3162 | ENSMUSG( | Hrh3 | 99296 | -01550449 | 052547292 | -01225 | 0305 | | 15 |
| 3183 | ENSMUSG( | Hspa12b | 72630 | 034226853 | 097510657 | -016 | 01425 | | 15 |

| | | | | | | | | | |
|---|---|---|---|---|---|---|---|---|---|
| 3187 | ENSMUSG( | Hspa2 | 15512 | 023083262 | 029302009 | -02225 | 02225 | | 15 |
| 3190 | ENSMUSG( | Hspa5 | 14828 | 005521624 | 046238103 | 0365 | -00725 | | 15 |
| 3198 | ENSMUSG( | Hspg2 | 15530 | -00055100 | 032052685 | 00425 | 02175 | | 15 |
| 3199 | ENSMUSG( | Hsph1 | 15505 | 039293316 | 001501946 | 00575 | 0125 | | 15 |
| 3250 | ENSMUSG( | Igf1r | 16001 | 023249139 | 052494836 | -01025 | 00675 | | 15 |
| 3256 | ENSMUSG( | Ighg2c | 404711 | 060504101 | 059259514 | -01775 | 04675 | | 15 |
| 3300 | ENSMUSG( | Inpp4a | 269180 | 015584351 | 008286030 | 001 | 00225 | | 15 |
| 3305 | ENSMUSG( | Inpp5e | 64436 | 007060028 | 019876313 | 005 | 00875 | | 15 |
| 3321 | ENSMUSG( | Ints2 | 70422 | -00867615 | 022685923 | -01375 | 0225 | | 15 |
| 3329 | ENSMUSG( | Ip6k1 | 27399 | 005542707 | 012404303 | -0005 | 01775 | | 15 |
| 3339 | ENSMUSG( | Iqgap1 | 29875 | -00137184 | 015674387 | -004 | 0085 | | 15 |
| 3340 | ENSMUSG( | Iqgap2 | 544963 | 034652091 | 056542410 | -0135 | -034 | | 15 |
| 3343 | ENSMUSG( | Iqsec3 | 243621 | 028829033 | 009047363 | -00025 | 0045 | | 15 |
| 3349 | ENSMUSG( | Irf3 | 54131 | 017650482 | 032164413 | -02125 | 012 | | 15 |
| 3353 | ENSMUSG( | Irs2 | 384783 | 019250401 | 012458564 | -01725 | 014 | | 15 |
| 3365 | ENSMUSG( | Isyna1 | 71780 | 023142392 | 030602641 | 029 | 03675 | | 15 |
| 3369 | ENSMUSG( | Itga1 | 109700 | -00312418 | 051279089 | -00875 | 0455 | | 15 |
| 3374 | ENSMUSG( | Itga7 | 16404 | 014444510 | 018411933 | 00675 | 196262e-1 | | 15 |
| 3384 | ENSMUSG( | Itih2 | 16425 | 024720086 | 032988428 | 01125 | 025 | | 15 |
| 3388 | ENSMUSG( | Itm2a | 16431 | -01036089 | 069064785 | -00425 | 025 | | 15 |
| 3397 | ENSMUSG( | Itpr3 | 16440 | 037757819 | 081476508 | -0335 | 03525 | | 15 |
| 3412 | ENSMUSG( | Jcad | 240185 | 018213892 | 015525885 | -02925 | -004 | | 15 |
| 3465 | ENSMUSG( | Kcnh3 | 16512 | 010702475 | 025784229 | -0195 | -015 | | 15 |
| 3486 | ENSMUSG( | Kcnn2 | 140492 | 04004885 | 064935537 | 011 | -03625 | | 15 |
| 3488 | ENSMUSG( | Kcnq2 | 16536 | 004575897 | 022262213 | 012 | 017 | | 15 |
| 3498 | ENSMUSG( | Kctd12b | 207474 | 012732484 | 016223867 | 00425 | 0085 | | 15 |
| 3509 | ENSMUSG( | Kctd8 | 243043 | 006869917 | 032355520 | 01325 | 012 | | 15 |
| 3527 | ENSMUSG( | Kif11 | 16551 | 008691796 | 025470519 | 01 | -02325 | | 15 |
| 3584 | ENSMUSG( | Krit1 | 79264 | 010169588 | 035821599 | -017 | 00475 | | 15 |
| 3589 | ENSMUSG( | Ksr2 | 333050 | 016194798 | 009127047 | 0195 | 00625 | | 15 |
| 3603 | ENSMUSG( | Lama5 | 16776 | -00442985 | 033168634 | -0205 | 01075 | | 15 |
| 3605 | ENSMUSG( | Lamb2 | 16779 | -00437866 | 030732648 | -00725 | 02325 | | 15 |
| 3617 | ENSMUSG( | Lancl3 | 236285 | 024722601 | 031228273 | 01875 | 03625 | | 15 |
| 3621 | ENSMUSG( | Larp1 | 73158 | 020280711 | 019418704 | -0075 | -0045 | | 15 |
| 3725 | ENSMUSG( | Lrch3 | 70144 | 009894109 | 012766220 | -02075 | 0075 | | 15 |
| 3727 | ENSMUSG( | Lrfn2 | 70530 | 038655876 | 067071005 | 01425 | -014 | | 15 |
| 3753 | ENSMUSG( | Lrrc58 | 320184 | 019409254 | 014914505 | 024 | 00975 | | 15 |
| 3760 | ENSMUSG( | Lrrc8c | 100604 | 014510211 | 069790354 | 01575 | -01125 | | 15 |
| 3792 | ENSMUSG( | Ltn1 | 78913 | 010809092 | 017065923 | 022 | 01625 | | 15 |
| 3796 | ENSMUSG( | Lum | 17022 | -01429668 | 096605183 | -03675 | -006 | | 15 |
| 3799 | ENSMUSG( | Luzp1 | 269593 | 030702034 | 016610166 | -02725 | -001 | | 15 |
| 3807 | ENSMUSG( | Ly75 | 17076 | 018164640 | 094811678 | 06625 | -0085 | | 15 |
| 3809 | ENSMUSG( | Lyn | 17096 | -00330638 | 023166420 | -0035 | 022 | | 15 |
| 3813 | ENSMUSG( | Lypla1 | 18777 | -00270057 | 020946211 | -01525 | -00775 | | 15 |
| 3823 | ENSMUSG( | Lyst | 17101 | 002947256 | 05687713 | -0065 | -018 | | 15 |
| 3825 | ENSMUSG( | Lztfl1 | 93730 | 012166597 | 017723303 | -0165 | 01025 | | 15 |
| 3861 | ENSMUSG( | Man2c1 | 73744 | 025773351 | 004942345 | -01725 | -003 | | 15 |
| 3880 | ENSMUSG( | Map2k6 | 26399 | 023194683 | 032284765 | -0025 | 0105 | | 15 |
| 3882 | ENSMUSG( | Map3k10 | 269881 | 023501773 | 012522654 | -01825 | -00075 | | 15 |
| 3887 | ENSMUSG( | Map3k20 | 65964 | 004040721 | 044571447 | 033 | -0005 | | 15 |
| 3888 | ENSMUSG( | Map3k4 | 26407 | 019940844 | 011787655 | -02 | -0065 | | 15 |
| 3889 | ENSMUSG( | Map3k5 | 26408 | 038737885 | 003881984 | -018 | 003 | | 15 |
| 3891 | ENSMUSG( | Map3k9 | 338372 | 022553749 | 017822742 | -00525 | 0135 | | 15 |
| 3893 | ENSMUSG( | Map4k2 | 26412 | 033348105 | 027306034 | 035 | 0105 | | 15 |
| 3894 | ENSMUSG( | Map4k3 | 225028 | 026242967 | 022755719 | 006 | -0075 | | 15 |
| 3909 | ENSMUSG( | Mapk4 | 225724 | 038359609 | 020470300 | -002 | 0105 | | 15 |
| 3932 | ENSMUSG( | Mark4 | 232944 | 021059622 | 013882148 | 0075 | 02025 | | 15 |
| 3937 | ENSMUSG( | Mast1 | 56527 | 033506776 | 005978101 | -002 | -00575 | | 15 |
| 3940 | ENSMUSG( | Mast4 | 328329 | 018438109 | 031998781 | -029 | 0205 | | 15 |
| 3948 | ENSMUSG( | Mavs | 228607 | 017704520 | 014479190 | -0165 | 00725 | | 15 |
| 3949 | ENSMUSG( | Max | 17187 | 012475075 | 014048064 | -01075 | 013 | | 15 |
| 3974 | ENSMUSG( | Mcmbp | 210711 | 011588051 | 014529308 | -01 | 017 | | 15 |
| 3982 | ENSMUSG( | Mcub | 66815 | 011101735 | 037918313 | 0355 | -0075 | | 15 |
| 4030 | ENSMUSG( | Mertk | 17289 | 025369764 | 034618028 | -01075 | -02125 | | 15 |
| 4041 | ENSMUSG( | Mettl17 | 52535 | 017324240 | 031504386 | -00025 | -0235 | | 15 |
| 4074 | ENSMUSG( | Mib2 | 76580 | 016804900 | 011061298 | 01075 | -00375 | | 15 |
| 4093 | ENSMUSG( | Mindy1 | 75007 | -00042980 | 024100633 | -015 | -002 | | 15 |
| 4132 | ENSMUSG( | Mon2 | 67074 | 003294658 | 015477083 | 00025 | 02175 | | 15 |
| 4146 | ENSMUSG( | Mphosph8 | 75339 | 035722960 | 025638441 | 01975 | 03125 | | 15 |
| 4161 | ENSMUSG( | Mrc1 | 17533 | 002281785 | 032619255 | 015 | -0005 | | 15 |
| 4251 | ENSMUSG( | Msl1 | 74026 | 024175369 | 017958721 | 0 | 00775 | 15 | |
| 4275 | ENSMUSG( | Mthfd1l | 270685 | 016963701 | 043046894 | -00525 | -03125 | | 15 |
| 4291 | ENSMUSG( | Mtor | 56717 | 011863334 | 012378830 | 0075 | -00575 | | 15 |
| 4297 | ENSMUSG( | Mtrf1l | 108853 | 026233739 | 038430897 | 0335 | 01825 | | 15 |
| 4318 | ENSMUSG( | Mycbp2 | 105689 | -00481146 | 017065970 | -0055 | 0085 | | 15 |
| 4323 | ENSMUSG( | Myh11 | 17880 | 020773148 | 072543601 | -004 | 04075 | | 15 |
| 4328 | ENSMUSG( | Myl12b | 67938 | 016860245 | 019507786 | -01475 | 0105 | | 15 |
| 4337 | ENSMUSG( | Myo1c | 17913 | 000152559 | 032106862 | 00475 | -018 | | 15 |
| 4350 | ENSMUSG( | Myrip | 245049 | 031119623 | 008093755 | 011 | 02475 | | 15 |
| 4382 | ENSMUSG( | Nampt | 59027 | -00593375 | 021709773 | -01725 | 0095 | | 15 |
| 4389 | ENSMUSG( | Napepld | 242864 | 026623285 | 027862056 | -01475 | -022 | | 15 |
| 4409 | ENSMUSG( | Nbea | 26422 | 005286943 | 021752766 | 01675 | 01775 | | 15 |
| 4423 | ENSMUSG( | Nck1 | 17973 | 013776954 | 016035738 | -00275 | -016 | | 15 |
| 4434 | ENSMUSG( | Ncoa7 | 211329 | 005637047 | 017371571 | -004 | 009 | | 15 |
| 4468 | ENSMUSG( | Ndufaf4 | 68493 | 001644715 | 020399791 | -0235 | -00125 | | 15 |
| 4557 | ENSMUSG( | Nhsl2 | 1E+08 | -00140294 | 024564364 | -003 | 009 | | 15 |
| 4569 | ENSMUSG( | Nipbl | 71175 | 007527008 | 019953613 | -02 | -00725 | | 15 |
| 4609 | ENSMUSG( | Nol9 | 74035 | 011327248 | 011647369 | -00575 | 01075 | | 15 |
| 4626 | ENSMUSG( | Nostrin | 329416 | 027603248 | 041074157 | -026 | 01975 | | 15 |
| 4665 | ENSMUSG( | Nrf1 | 18181 | 019753838 | 013446361 | -0055 | 0205 | | 15 |
| 4696 | ENSMUSG( | Nt5dc2 | 70021 | 016009459 | 017125075 | 018 | 01225 | | 15 |
| 4738 | ENSMUSG( | Nudt7 | 67528 | 049766581 | 023556999 | 03225 | 00125 | | 15 |
| 4747 | ENSMUSG( | Nup153 | 218210 | 012941288 | 022706180 | -02975 | -00725 | | 15 |
| 4766 | ENSMUSG( | Nvl | 67459 | 002270228 | 020075484 | -0105 | 0275 | | 15 |
| 4770 | ENSMUSG( | Nxn | 18230 | 035033366 | 051850871 | 018 | 0075 | | 15 |
| 4781 | ENSMUSG( | Ocln | 18260 | -00412183 | 071223228 | -01225 | 0505 | | 15 |
| 4790 | ENSMUSG( | Ogfrl1 | 70155 | 008624204 | 021674286 | -006 | -002 | | 15 |
| 4821 | ENSMUSG( | Osbpl3 | 71720 | 010672456 | 020449857 | -01375 | -00475 | | 15 |
| 4879 | ENSMUSG( | Pakap | 677884 | 029859980 | 021357872 | -01 | 00575 | | 15 |
| 4884 | ENSMUSG( | Palmd | 114301 | 009683362 | 028957693 | -01525 | 0215 | | 15 |
| 4895 | ENSMUSG( | Panx2 | 406218 | 021801605 | 013830819 | -00125 | 015 | | 15 |
| 4906 | ENSMUSG( | Parg | 26430 | 022617809 | 021966693 | -02 | -00275 | | 15 |
| 4952 | ENSMUSG( | Pcdhga11 | 93723 | 016837159 | 035860995 | -008 | -01825 | | 15 |
| 4969 | ENSMUSG( | Pck2 | 74551 | 017959483 | 008562235 | -00175 | 01175 | | 15 |
| 4979 | ENSMUSG( | Pcnx3 | 104401 | -00668752 | 025708023 | 00925 | 0055 | | 15 |
| 4993 | ENSMUSG( | Pdcd11 | 18572 | 008163145 | 014787739 | -02175 | -0035 | | 15 |
| 5009 | ENSMUSG( | Pde4a | 18577 | 013550952 | 010756313 | -00575 | 026 | | 15 |
| 5020 | ENSMUSG( | Pdf | 68023 | 033185211 | 038857837 | 0205 | 02775 | | 15 |
| 5040 | ENSMUSG( | Pdp2 | 382051 | 018353125 | 012559623 | -02225 | 00825 | | 15 |
| 5050 | ENSMUSG( | Pdzd2 | 68070 | 010353058 | 027407831 | 00925 | 032 | | 15 |
| 5056 | ENSMUSG( | Pecam1 | 18613 | 0130312 | 047392957 | -02 | -00125 | | 15 |
| 5062 | ENSMUSG( | Pelo | 105083 | 005467622 | 014880782 | -00625 | -0125 | | 15 |
| 5065 | ENSMUSG( | Pepd | 18624 | 02856416 | 014433910 | -00825 | 01225 | | 15 |
| 5067 | ENSMUSG( | Pex1 | 71382 | 018343498 | 019246039 | 01425 | 02325 | | 15 |
| 5076 | ENSMUSG( | Pex26 | 74043 | 025557729 | 027667696 | 00275 | -008 | | 15 |
| 5109 | ENSMUSG( | Pgm5 | 226041 | 028443823 | 017302281 | -02475 | 006 | | 15 |
| 5152 | ENSMUSG( | Phyhd1 | 227696 | 038412579 | 032879274 | -01475 | 01225 | | 15 |
| 5174 | ENSMUSG( | Pik3c2a | 18704 | 011912449 | 034594172 | 0055 | -00125 | | 15 |
| 5181 | ENSMUSG( | Pik3r2 | 18709 | 015876931 | 008395675 | 001 | 0035 | | 15 |
| 5182 | ENSMUSG( | Pik3r4 | 75669 | -00415272 | 020226043 | -00325 | -01075 | | 15 |
| 5183 | ENSMUSG( | Pikfyve | 18711 | 007353549 | 017361723 | -00225 | 00125 | | 15 |
| 5193 | ENSMUSG( | Pip5k1c | 18717 | 022521605 | 006144269 | -00975 | -0065 | | 15 |
| 5199 | ENSMUSG( | Pitpnm1 | 18739 | 019848407 | 012239636 | 01675 | -00875 | | 15 |
| 5200 | ENSMUSG( | Pitpnm2 | 19679 | 025077765 | 032293009 | 01925 | -021 | | 15 |
| 5201 | ENSMUSG( | Pitpnm3 | 327958 | 013836121 | 009628643 | 01975 | -00125 | | 15 |
| 5212 | ENSMUSG( | Pkp2 | 67451 | 059990435 | 036608848 | 0395 | 0135 | | 15 |

| | | | | | | | | |
|---|---|---|---|---|---|---|---|---|
| 5231 | ENSMUSG( | Plcl1 | 227120 | 004315250 | 023269075 | 00375 | -0145 | 15 |
| 5232 | ENSMUSG( | Plcl2 | 224860 | 031097339 | 034703687 | -0155 | 0065 | 15 |
| 5235 | ENSMUSG( | Pld1 | 18805 | 027668272 | 044177669 | -02675 | -0095 | 15 |
| 5277 | ENSMUSG( | Pltp | 18830 | -00656278 | 060066031 | -02275 | 05225 | 15 |
| 5280 | ENSMUSG( | Plxna1 | 18844 | 001832807 | 030683958 | 01175 | 0225 | 15 |
| 5285 | ENSMUSG( | Plxnb2 | 140570 | -00140157 | 023874258 | -00625 | -0055 | 15 |
| 5288 | ENSMUSG( | Plxnd1 | 67784 | 007733873 | 022796825 | 02125 | 00725 | 15 |
| 5309 | ENSMUSG( | Pnpla6 | 50767 | -00077436 | 024813396 | 01125 | 00075 | 15 |
| 5313 | ENSMUSG( | Pnpt1 | 71701 | 010343199 | 015133999 | 0125 | -003 | 15 |
| 5314 | ENSMUSG( | Podxl | 27205 | 030830859 | 097173299 | 0215 | -03525 | 15 |
| 5364 | ENSMUSG( | Ppat | 231327 | 018418295 | 019834676 | 013 | 03125 | 15 |
| 5384 | ENSMUSG( | Ppil2 | 66053 | 020296752 | 010577780 | 005 | 02475 | 15 |
| 5399 | ENSMUSG( | Ppox | 19044 | 014583981 | 017478865 | -009 | 01075 | 15 |
| 5419 | ENSMUSG( | Ppp1r37 | 232947 | 020619717 | 008832919 | 0035 | -008 | 15 |
| 5426 | ENSMUSG( | Ppp1r9b | 217124 | 023637817 | 014166306 | 012 | 0205 | 15 |
| 5460 | ENSMUSG( | Prag1 | 244418 | -00185201 | 019505820 | -014 | -00575 | 15 |
| 5494 | ENSMUSG( | Prkce | 18754 | 012528966 | 010689547 | 01075 | -0035 | 15 |
| 5503 | ENSMUSG( | Prkg1 | 19091 | 020990220 | 035250607 | 02275 | 001 | 15 |
| 5585 | ENSMUSG( | Psmd11 | 69077 | 017517129 | 010747939 | -00375 | 0155 | 15 |
| 5592 | ENSMUSG( | Psmd5 | 66998 | 012399227 | 011806300 | 0135 | 0195 | 15 |
| 5612 | ENSMUSG( | Ptbp3 | 230257 | -00436286 | 023843285 | 00525 | 02625 | 15 |
| 5618 | ENSMUSG( | Pten | 19211 | 006226047 | 019994189 | -0245 | -005 | 15 |
| 5658 | ENSMUSG( | Ptprf | 19268 | -00386451 | 022460238 | -02225 | -00525 | 15 |
| 5663 | ENSMUSG( | Ptprn2 | 19276 | -00702318 | 041100441 | -0005 | 0245 | 15 |
| 5666 | ENSMUSG( | Ptprs | 19280 | 000839857 | 018890502 | 01 | -00875 | 15 |
| 5668 | ENSMUSG( | Ptpru | 19273 | 023416326 | 028010438 | 01775 | -0065 | 15 |
| 5683 | ENSMUSG( | Pusl1 | 433813 | 011321695 | 014479454 | -0115 | 00525 | 15 |
| 5735 | ENSMUSG( | Rab23 | 19335 | 016335323 | 009559441 | 01225 | -0025 | 15 |
| 5762 | ENSMUSG( | Rab43 | 69834 | 012433616 | 023602561 | 019 | 0105 | 15 |
| 5799 | ENSMUSG( | Rai14 | 75646 | 027261220 | 031012097 | 0005 | -00475 | 15 |
| 5806 | ENSMUSG( | Ralgps1 | 241308 | 012554405 | 011534076 | -0225 | -00225 | 15 |
| 5807 | ENSMUSG( | Ralgps2 | 78255 | 032121003 | 024524688 | 0035 | -002 | 15 |
| 5829 | ENSMUSG( | Rapgef2 | 76089 | 019480099 | 023059331 | -0005 | 004 | 15 |
| 5838 | ENSMUSG( | Rasa1 | 218397 | 007251721 | 020147380 | -01425 | 00125 | 15 |
| 5842 | ENSMUSG( | Rasal2 | 226525 | 016252084 | 017289237 | -0185 | -0065 | 15 |
| 5846 | ENSMUSG( | Rasgrf1 | 19417 | -00292231 | 016032488 | -0025 | 01475 | 15 |
| 5850 | ENSMUSG( | Rasip1 | 69903 | 029015443 | 053823374 | 01275 | 02575 | 15 |
| 5876 | ENSMUSG( | Rbm25 | 67039 | -00112294 | 017570433 | 0165 | 00525 | 15 |
| 5973 | ENSMUSG( | Rhobtb2 | 246710 | 005506998 | 018850441 | 01625 | 01775 | 15 |
| 5975 | ENSMUSG( | Rhof | 23912 | 023879696 | 011460323 | 01075 | 02 | 15 |
| 5991 | ENSMUSG( | Rimoc1 | 106064 | -00014532 | 018478709 | -0095 | 0015 | 15 |
| 5996 | ENSMUSG( | Rin1 | 225870 | 029000252 | 017967837 | -018 | 01775 | 15 |
| 6002 | ENSMUSG( | Ripor1 | 75687 | 027817373 | 008686023 | 0115 | 01075 | 15 |
| 6013 | ENSMUSG( | Rmnd1 | 66084 | 007872022 | 016699009 | 0005 | -012 | 15 |
| 6067 | ENSMUSG( | Rock1 | 19877 | 008904808 | 022634382 | -024 | 0085 | 15 |
| 6164 | ENSMUSG( | Rps6ka2 | 20112 | 021658802 | 032980582 | 032 | -00025 | 15 |
| 6220 | ENSMUSG( | Rusc2 | 100213 | 024724521 | 011474201 | 02 | 00475 | 15 |
| 6256 | ENSMUSG( | Sanbr | 71675 | 014821868 | 008922938 | -00625 | -00925 | 15 |
| 6329 | ENSMUSG( | Sdc4 | 20971 | 021969425 | 035821500 | -02075 | 01525 | 15 |
| 6333 | ENSMUSG( | Sdf2l1 | 64136 | -00026683 | 046686286 | 0285 | -018 | 15 |
| 6344 | ENSMUSG( | Sdk2 | 237979 | 004944655 | 041455536 | 01925 | 0245 | 15 |
| 6382 | ENSMUSG( | Selplg | 20345 | -01304260 | 044426345 | -01575 | -006 | 15 |
| 6412 | ENSMUSG( | Serac1 | 321007 | 002676583 | 015931753 | -02075 | 00225 | 15 |
| 6424 | ENSMUSG( | Serpinb8 | 20725 | 033494121 | 016374476 | -00475 | -0075 | 15 |
| 6428 | ENSMUSG( | Serpinf2 | 18816 | 006800274 | 042438280 | -00425 | 037 | 15 |
| 6452 | ENSMUSG( | Sfn | 55948 | 007254823 | 024663794 | 0025 | 02175 | 15 |
| 6498 | ENSMUSG( | Sh3rf3 | 237353 | 029002078 | 041514596 | 02525 | 02375 | 15 |
| 6500 | ENSMUSG( | Shank1 | 243961 | 037748642 | 022992793 | -00375 | -00425 | 15 |
| 6501 | ENSMUSG( | Shank2 | 210274 | 017020597 | 021142877 | -021 | 00075 | 15 |
| 6502 | ENSMUSG( | Shank3 | 58234 | 022609570 | 020419172 | -00525 | -0085 | 15 |
| 6509 | ENSMUSG( | Shfl | 319278 | 025516749 | 007472628 | -0115 | -0085 | 15 |
| 6511 | ENSMUSG( | Shisa6 | 380702 | 001276978 | 050346290 | 01725 | -0235 | 15 |
| 6525 | ENSMUSG( | Sidt1 | 320007 | 009773311 | 043197449 | 0295 | 0375 | 15 |
| 6535 | ENSMUSG( | Sipa1l1 | 217692 | 020001081 | 013790602 | -0055 | -00475 | 15 |
| 6537 | ENSMUSG( | Sipa1l3 | 74206 | 027991545 | 030374351 | -003 | -01875 | 15 |
| 6541 | ENSMUSG( | Sirt4 | 75387 | 013774668 | 033749895 | 01175 | 036 | 15 |
| 6550 | ENSMUSG( | Slc12a4 | 20498 | -00796153 | 029374491 | -016 | 00925 | 15 |
| 6569 | ENSMUSG( | Slc18a2 | 214084 | 020692191 | 069199291 | -016 | 048 | 15 |
| 6577 | ENSMUSG( | Slc1a5 | 20514 | -01186943 | 062446176 | -01025 | -01675 | 15 |
| 6604 | ENSMUSG( | Slc25a24 | 229731 | 013545950 | 058092142 | 027 | -02075 | 15 |
| 6609 | ENSMUSG( | Slc25a29 | 214663 | 010349455 | 018577253 | -02025 | -005 | 15 |
| 6611 | ENSMUSG( | Slc25a32 | 69906 | 000775780 | 021132272 | -01625 | -011 | 15 |
| 6631 | ENSMUSG( | Slc2a1 | 20525 | 017614039 | 082896284 | -02325 | -02975 | 15 |
| 6686 | ENSMUSG( | Slc5a3 | 53881 | -01598894 | 038088396 | -001 | 00025 | 15 |
| 6701 | ENSMUSG( | Slc7a1 | 11987 | -01542622 | 045529823 | -0015 | 00375 | 15 |
| 6705 | ENSMUSG( | Slc7a2 | 11988 | 001957563 | 024458393 | 00925 | -0145 | 15 |
| 6708 | ENSMUSG( | Slc7a5 | 20539 | 008865300 | 041830944 | -01425 | 01725 | 15 |
| 6721 | ENSMUSG( | Slco1a4 | 28250 | -00464461 | 076250378 | -03525 | 0385 | 15 |
| 6726 | ENSMUSG( | Slit1 | 20562 | 005657457 | 037751866 | 005 | 00325 | 15 |
| 6766 | ENSMUSG( | Smg8 | 74133 | 006599226 | 023366653 | -0065 | 01025 | 15 |
| 6804 | ENSMUSG( | Snph | 241727 | 030181803 | 024358277 | -00525 | -0235 | 15 |
| 6870 | ENSMUSG( | Sowaha | 237761 | 004859394 | 016771069 | -00375 | 003 | 15 |
| 6886 | ENSMUSG( | Spata2 | 263876 | 012794533 | 022021116 | -02375 | 015 | 15 |
| 6906 | ENSMUSG( | Spns2 | 216892 | 010053424 | 036220894 | -003 | 01725 | 15 |
| 6915 | ENSMUSG( | Spred1 | 114715 | 018436683 | 009549571 | -019 | -00075 | 15 |
| 6919 | ENSMUSG( | Spry2 | 24064 | 001362492 | 025602283 | 01525 | -0045 | 15 |
| 6939 | ENSMUSG( | Srek1 | 218543 | 006088994 | 013728067 | -01125 | 0015 | 15 |
| 6961 | ENSMUSG( | Srsf11 | 69207 | 008829737 | 011681675 | 01575 | 01175 | 15 |
| 6978 | ENSMUSG( | Ssh2 | 237860 | 013712807 | 023110906 | -00325 | 0245 | 15 |
| 6979 | ENSMUSG( | Ssh3 | 245857 | 051588357 | 026958301 | 02675 | 0205 | 15 |
| 7013 | ENSMUSG( | Stat6 | 20852 | 023079099 | 012310945 | -007 | -014 | 15 |
| 7029 | ENSMUSG( | Stk3 | 56274 | 017063167 | 014787228 | -00025 | 023 | 15 |
| 7052 | ENSMUSG( | Strn4 | 97387 | 020357921 | 009328623 | 00625 | 0035 | 15 |
| 7072 | ENSMUSG( | Stxbp5 | 78808 | 012245518 | 015034167 | 00975 | 01675 | 15 |
| 7107 | ENSMUSG( | Svil | 225115 | 026720159 | 029698583 | -0025 | 03 | 15 |
| 7170 | ENSMUSG( | Taf2 | 319944 | 015312208 | 016288483 | -002 | -005 | 15 |
| 7176 | ENSMUSG( | Taf9 | 108143 | 022073758 | 024677788 | -0105 | 021 | 15 |
| 7181 | ENSMUSG( | Tagln2 | 21346 | 018859419 | 050370320 | 00775 | 01625 | 15 |
| 7189 | ENSMUSG( | Tango6 | 272538 | 021471139 | 007810842 | -0195 | 0075 | 15 |
| 7224 | ENSMUSG( | Tbc1d4 | 210789 | 078261746 | 083703395 | -025 | -012 | 15 |
| 7228 | ENSMUSG( | Tbc1d8b | 245638 | 013949687 | 025414877 | -02225 | -00025 | 15 |
| 7238 | ENSMUSG( | Tbk1 | 56480 | 004748560 | 023386259 | 00425 | 019 | 15 |
| 7239 | ENSMUSG( | Tbkbp1 | 73174 | 033885123 | 026664577 | -0025 | 02575 | 15 |
| 7266 | ENSMUSG( | Tdrd1 | 83561 | 039496143 | 012870588 | -00675 | -024 | 15 |
| 7270 | ENSMUSG( | Tdrp | 72148 | 003162368 | 032255349 | -00025 | -02375 | 15 |
| 7289 | ENSMUSG( | Tex10 | 269536 | 026923826 | 025343140 | -0045 | 0075 | 15 |
| 7294 | ENSMUSG( | Tfcp2 | 21422 | 010762426 | 013899068 | 01525 | 0035 | 15 |
| 7298 | ENSMUSG( | Tfg | 21787 | 018695490 | 009584102 | -001 | 0015 | 15 |
| 7300 | ENSMUSG( | Tfrc | 22042 | -01496262 | 048114707 | -02275 | 008 | 15 |
| 7304 | ENSMUSG( | Tgm2 | 21817 | 019548197 | 058610032 | -01925 | 0375 | 15 |
| 7319 | ENSMUSG( | Thoc2 | 331401 | 001785250 | 019204284 | -0105 | 0225 | 15 |
| 7355 | ENSMUSG( | Tinagl1 | 94242 | 091035844 | 128760046 | 021 | 01675 | 15 |
| 7382 | ENSMUSG( | Tmcc2 | 68875 | 032396677 | 022930491 | 0065 | 005 | 15 |
| 7392 | ENSMUSG( | Tmed8 | 382620 | 004542789 | 012866418 | 00925 | -00325 | 15 |
| 7422 | ENSMUSG( | Tmem151b | 210573 | 031913129 | 022053900 | -00325 | 0095 | 15 |
| 7478 | ENSMUSG( | Tmem63c | 217733 | 010063340 | 014577541 | -0005 | 01125 | 15 |
| 7495 | ENSMUSG( | Tmpo | 21917 | 008289630 | 032400457 | 00575 | 0375 | 15 |
| 7496 | ENSMUSG( | Tmppe | 1.01E+08 | 00721776 | 049570410 | -023 | 0035 | 15 |
| 7540 | ENSMUSG( | Top1 | 21969 | 020461136 | 023314204 | 01325 | 012 | 15 |
| 7580 | ENSMUSG( | Traf3 | 22031 | 036417374 | 017485367 | -00975 | 00775 | 15 |
| 7622 | ENSMUSG( | Trim47 | 217333 | 025386835 | 066582738 | -0425 | 00325 | 15 |
| 7643 | ENSMUSG( | Trmu | 72026 | -00253238 | 014517021 | -009 | -00875 | 15 |
| 7644 | ENSMUSG( | Trnt1 | 70047 | 008367609 | 021721737 | 0205 | 016 | 15 |
| 7650 | ENSMUSG( | Trpc4 | 22066 | 007319021 | 047009278 | 0345 | 00875 | 15 |
| 7662 | ENSMUSG( | Tsc22d3 | 14605 | 063643812 | 047560815 | 017 | 02825 | 15 |
| 7692 | ENSMUSG( | Ttbk1 | 106763 | 031504965 | 020248051 | -01275 | 008 | 15 |

| | | | | | | | | |
|---|---|---|---|---|---|---|---|---|
| 7698 | ENSMUSG( | Ttc21b | 73668 | 017107317 | 019422213 | 005 | 0005 | 15 |
| 7719 | ENSMUSG( | Ttr | 22139 | -00764594 | 061792102 | 01225 | -009 | 15 |
| 7723 | ENSMUSG( | Tub | 22141 | 023510818 | 023275016 | -01225 | 00625 | 15 |
| 7740 | ENSMUSG( | Tubgcp5 | 233276 | 005164404 | 014750459 | -00775 | 02475 | 15 |
| 7741 | ENSMUSG( | Tubgcp6 | 328580 | 020571178 | 022875826 | -02275 | 005 | 15 |
| 7783 | ENSMUSG( | Uba6 | 231380 | 009035083 | 016020126 | 007 | 006 | 15 |
| 7817 | ENSMUSG( | Ube3a | 22215 | 003916547 | 013929956 | -00725 | -0035 | 15 |
| 7819 | ENSMUSG( | Ube3c | 100763 | 016265978 | 010027154 | 001 | -00425 | 15 |
| 7835 | ENSMUSG( | Ubr2 | 224826 | 012434238 | 024031726 | 02775 | 01325 | 15 |
| 7883 | ENSMUSG( | Unc79 | 217843 | 025674365 | 034309788 | -00125 | -00325 | 15 |
| 7922 | ENSMUSG( | Usp19 | 71472 | 025110791 | 016270682 | 0075 | -003 | 15 |
| 7926 | ENSMUSG( | Usp25 | 30940 | 013579224 | 026319377 | 00675 | 01025 | 15 |
| 7943 | ENSMUSG( | Usp54 | 78787 | 010142741 | 018894961 | 02275 | 0165 | 15 |
| 7953 | ENSMUSG( | Utp20 | 70683 | 008657051 | 018801971 | -02125 | 00125 | 15 |
| 7956 | ENSMUSG( | Utp6 | 216987 | 029001198 | 005726421 | 00575 | 00575 | 15 |
| 7982 | ENSMUSG( | Vcl | 22330 | 025296011 | 045212468 | 00775 | 00425 | 15 |
| 7984 | ENSMUSG( | Vcpip1 | 70675 | 003592120 | 014256939 | -00325 | 002 | 15 |
| 7993 | ENSMUSG( | Vim | 22352 | 015721135 | 056640405 | -03225 | 03975 | 15 |
| 8010 | ENSMUSG( | Vps25 | 28084 | 037353733 | 016657304 | 021 | -009 | 15 |
| 8017 | ENSMUSG( | Vps33b | 233405 | 022082060 | 013778932 | -00525 | 012 | 15 |
| 8029 | ENSMUSG( | Vps4b | 20479 | 005998484 | 016227358 | 007 | -00025 | 15 |
| 8048 | ENSMUSG( | Vwa1 | 246228 | 000482484 | 041351333 | 0205 | 02675 | 15 |
| 8053 | ENSMUSG( | Vwf | 22371 | 005466902 | 082883721 | -00925 | 0555 | 15 |
| 8065 | ENSMUSG( | Washc4 | 319277 | 009761102 | 012792131 | 004 | -003 | 15 |
| 8067 | ENSMUSG( | Wasl | 73178 | 007878038 | 014804358 | -0175 | -0045 | 15 |
| 8107 | ENSMUSG( | Wdr81 | 192652 | 008205028 | 023149751 | -012 | 003 | 15 |
| 8112 | ENSMUSG( | Wdtc1 | 230796 | 027862495 | 012912164 | -02475 | 0175 | 15 |
| 8114 | ENSMUSG( | Whrn | 73750 | 032047949 | 016774913 | 0065 | 00025 | 15 |
| 8115 | ENSMUSG( | Wipf2 | 68524 | 017193519 | 009507920 | -01225 | 0085 | 15 |
| 8116 | ENSMUSG( | Wipf3 | 330319 | 015912541 | 018530137 | 00175 | -01825 | 15 |
| 8122 | ENSMUSG( | Wnk2 | 75607 | 031171379 | 023785224 | -00475 | 019 | 15 |
| 8128 | ENSMUSG( | Wwc2 | 52357 | 016325253 | 010665630 | 012 | 017 | 15 |
| 8183 | ENSMUSG( | Ythdc2 | 240255 | 012669497 | 029013479 | 117757e-1 | 003 | 15 |
| 8196 | ENSMUSG( | Zbtb16 | 235320 | 134491350 | 120124884 | -00575 | 00075 | 15 |
| 8204 | ENSMUSG( | Zc3h11a | 70579 | 000299894 | 034638585 | -02325 | -0215 | 15 |
| 8239 | ENSMUSG( | Zfc3h1 | 216345 | 012638775 | 041374068 | -0185 | 021 | 15 |
| 8254 | ENSMUSG( | Zfp638 | 18139 | 015975497 | 014397836 | -006 | -0105 | 15 |
| 8269 | ENSMUSG( | Zhx3 | 320799 | 061879380 | 032702765 | -038 | 0475 | 15 |
| 8281 | ENSMUSG( | Znfx1 | 98999 | 013001531 | 010438407 | 00225 | 00775 | 15 |
| 8289 | ENSMUSG( | Zswim8 | 268721 | 017407302 | 012030645 | 00375 | -00375 | 15 |
| 8290 | ENSMUSG( | Zw10 | 26951 | 017134306 | 023018288 | 01275 | -0005 | 15 |
| 6 | ENSMUSG( | 1700025G( | 69399 | 006695037 | 005405409 | -01975 | -07125 | 16 |
| 76 | ENSMUSG( | Abi1 | 11308 | 012390816 | 007423639 | -01075 | -11475 | 16 |
| 108 | ENSMUSG( | Ace | 11421 | -02306001 | 031930138 | -02275 | -07 | 16 |
| 157 | ENSMUSG( | Acyp1 | 66204 | 004320876 | -00634569 | -0255 | -06725 | 16 |
| 268 | ENSMUSG( | Ajap1 | 230959 | -02589995 | 031133654 | -02675 | -1135 | 16 |
| 288 | ENSMUSG( | Akr1b10 | 67861 | 016692606 | -00819249 | 03425 | -07675 | 16 |
| 303 | ENSMUSG( | Aldh1l2 | 216188 | 001886187 | -00051041 | -0005 | -0615 | 16 |
| 331 | ENSMUSG( | Alyref | 21681 | 005438382 | -00019176 | -0005 | -11075 | 16 |
| 356 | ENSMUSG( | Ank1 | 11733 | 01000387 | -01426191 | 0035 | -062 | 16 |
| 359 | ENSMUSG( | Ankfn1 | 382543 | -01670312 | 003722174 | 01725 | -058 | 16 |
| 378 | ENSMUSG( | Ankrd45 | 73844 | -00314676 | -00997331 | -04425 | -077 | 16 |
| 399 | ENSMUSG( | Antxr1 | 69538 | -00781112 | -01085535 | -01875 | -081 | 16 |
| 401 | ENSMUSG( | Anxa11 | 11744 | 007026574 | 004827204 | 00475 | -05525 | 16 |
| 419 | ENSMUSG( | Ap2s1 | 232910 | 000640780 | -02496523 | 0345 | -14625 | 16 |
| 431 | ENSMUSG( | Ap5s1 | 69596 | -01295586 | -02656604 | 0545 | -158 | 16 |
| 550 | ENSMUSG( | Arl6ip5 | 65106 | -04415964 | 004039243 | 01725 | -1055 | 16 |
| 551 | ENSMUSG( | Arl6ip6 | 65103 | -03699368 | 020155843 | -0045 | -07875 | 16 |
| 565 | ENSMUSG( | Arpc1a | 56443 | -00086194 | -01516503 | 02 | -05725 | 16 |
| 569 | ENSMUSG( | Arpc4 | 68089 | 003613941 | -00884211 | 0355 | -07175 | 16 |
| 602 | ENSMUSG( | Aspa | 11484 | -01680216 | 025382807 | -02025 | -12225 | 16 |
| 605 | ENSMUSG( | Asphd2 | 72898 | -00325442 | 008398425 | 035 | -06425 | 16 |
| 665 | ENSMUSG( | Atp5mj | 70257 | 006472771 | -01981821 | 01575 | -088 | 16 |
| 672 | ENSMUSG( | Atp6ap2 | 70495 | -05376414 | 017072624 | -011 | -082 | 16 |
| 681 | ENSMUSG( | Atp6v1e1 | 11973 | 002916173 | -01772866 | 03225 | -06925 | 16 |
| 727 | ENSMUSG( | Baalc | 118452 | 004501001 | -02643140 | -00175 | -07075 | 16 |
| 817 | ENSMUSG( | Bod1 | 69556 | 001362846 | -01381376 | 03375 | -06275 | 16 |
| 837 | ENSMUSG( | Brd1 | 223770 | -00144381 | 007672209 | -024 | -13 | 16 |
| 864 | ENSMUSG( | Btbd8 | 1.01E+08 | -01082626 | -00343931 | -042 | -0785 | 16 |
| 872 | ENSMUSG( | Bysl | 53414 | 022481502 | -00355099 | -043 | -101 | 16 |
| 934 | ENSMUSG( | Calm3 | 12315 | 013031328 | -01668976 | 05075 | -1085 | 16 |
| 1045 | ENSMUSG( | Ccdc33 | 382077 | -02535370 | 013248589 | -03525 | -09425 | 16 |
| 1079 | ENSMUSG( | Ccser2 | 72972 | -00023774 | -00402888 | -01025 | -06325 | 16 |
| 1104 | ENSMUSG( | Cd82 | 12521 | -06592022 | -00947510 | -0215 | -09475 | 16 |
| 1135 | ENSMUSG( | Cdh22 | 104010 | 002304050 | 025813123 | -0395 | -1025 | 16 |
| 1145 | ENSMUSG( | Cdk13 | 69562 | 007706406 | -00146965 | -033 | -077 | 16 |
| 1170 | ENSMUSG( | Cebpzos | 68554 | -00882432 | -02398139 | -03125 | -0655 | 16 |
| 1195 | ENSMUSG( | Cers6 | 241447 | -00801455 | 044190585 | 0785 | -21475 | 16 |
| 1198 | ENSMUSG( | Cetn3 | 12626 | 001486774 | 007448720 | 00875 | -0575 | 16 |
| 1210 | ENSMUSG( | Cfl2 | 12632 | -00171215 | -00456719 | 04475 | -1215 | 16 |
| 1295 | ENSMUSG( | Clcn5 | 12728 | -04647872 | 024169757 | 01675 | -0715 | 16 |
| 1300 | ENSMUSG( | Cldn11 | 18417 | -07826651 | 017511214 | -007 | -09575 | 16 |
| 1310 | ENSMUSG( | Clic5 | 224796 | 029587465 | 091965963 | 027 | -0725 | 16 |
| 1331 | ENSMUSG( | Clta | 12757 | 002087398 | -02090783 | 0105 | -139 | 16 |
| 1417 | ENSMUSG( | Col1a2 | 12843 | 000716753 | 029869944 | -03975 | -08175 | 16 |
| 1436 | ENSMUSG( | Commd9 | 76501 | 004348187 | -02311709 | -007 | -0865 | 16 |
| 1455 | ENSMUSG( | Copz1 | 56447 | 002798021 | -01598337 | 02075 | -071 | 16 |
| 1459 | ENSMUSG( | Coq3 | 230027 | 017008579 | -00877130 | -02475 | -081 | 16 |
| 1490 | ENSMUSG( | Cox6c | 12864 | 005450288 | -01741116 | -022 | -06825 | 16 |
| 1492 | ENSMUSG( | Cox7a2 | 12866 | 004305128 | -02289250 | -005 | -077 | 16 |
| 1538 | ENSMUSG( | Creld2 | 76737 | -00012583 | 042623412 | 04425 | -07475 | 16 |
| 1603 | ENSMUSG( | Ctcf | 13018 | 012330895 | 008478325 | 0055 | -0785 | 16 |
| 1604 | ENSMUSG( | Ctdnep1 | 67181 | -00671230 | -00073367 | 00475 | -05425 | 16 |
| 1674 | ENSMUSG( | Cyp20a1 | 77951 | -01463407 | 009297542 | 032 | -08475 | 16 |
| 1681 | ENSMUSG( | Cyp4f16 | 70101 | -03051361 | -00963574 | 03825 | -1135 | 16 |
| 1685 | ENSMUSG( | Cyrib | 223601 | 000217913 | -01613277 | 01575 | -08725 | 16 |
| 1709 | ENSMUSG( | Dagla | 269060 | -03365626 | 009145422 | -03375 | -063 | 16 |
| 1717 | ENSMUSG( | Dars2 | 226539 | 008761771 | -01755265 | -02425 | -0765 | 16 |
| 1786 | ENSMUSG( | Ddx39b | 53817 | 000939630 | -00026862 | 009 | -0915 | 16 |
| 1788 | ENSMUSG( | Ddx3y | 26900 | -00697043 | 019158952 | 00475 | -05625 | 16 |
| 1849 | ENSMUSG( | Dhrs11 | 192970 | 007676615 | -00970602 | 01425 | -05675 | 16 |
| 1913 | ENSMUSG( | Dnaja3 | 83945 | 005399826 | -01041049 | 017 | -08025 | 16 |
| 1937 | ENSMUSG( | Dnajc28 | 246738 | 016728991 | -00125892 | 0115 | -08125 | 16 |
| 1946 | ENSMUSG( | Dnal4 | 54152 | -00288367 | -00620289 | -012 | -11475 | 16 |
| 2024 | ENSMUSG( | Dtymk | 21915 | -00420115 | 000235977 | -00575 | -06975 | 16 |
| 2056 | ENSMUSG( | Dyrk1b | 13549 | -00275881 | -00381228 | 0125 | -13575 | 16 |
| 2210 | ENSMUSG( | Emd | 13726 | -00677858 | -01551536 | -00775 | -06475 | 16 |
| 2304 | ENSMUSG( | Ero1b | 67475 | -01528490 | 024369274 | -00275 | -05775 | 16 |
| 2316 | ENSMUSG( | Ethe1 | 66071 | -00416429 | -00938227 | 0225 | -07125 | 16 |
| 2410 | ENSMUSG( | Fam241b | 69894 | -00440920 | -00954973 | -00425 | -05875 | 16 |
| 2415 | ENSMUSG( | Fam81a | 76886 | 017175252 | -00236000 | 03425 | -08425 | 16 |
| 2445 | ENSMUSG( | Fbxl2 | 72179 | -02268997 | 002011329 | 01975 | -0595 | 16 |
| 2500 | ENSMUSG( | Fgfr2 | 14183 | -01667240 | 010727401 | -02375 | -10125 | 16 |
| 2502 | ENSMUSG( | Fggy | 75578 | -00231100 | -00928480 | -01275 | -057 | 16 |
| 2635 | ENSMUSG( | Gabra5 | 110886 | -02619839 | 028028467 | -01575 | -08325 | 16 |
| 2668 | ENSMUSG( | Gar1 | 68147 | -00608517 | -01823439 | -0005 | -05975 | 16 |
| 2677 | ENSMUSG( | Gas7 | 14457 | 009263548 | -00591546 | 07 | -1235 | 16 |
| 2692 | ENSMUSG( | Gbp9 | 236573 | -04388096 | 006795734 | -02375 | -13375 | 16 |
| 2709 | ENSMUSG( | Gdpd1 | 66569 | -04292384 | 010312195 | 0125 | -0815 | 16 |
| 2736 | ENSMUSG( | Ggt1 | 14598 | -00312431 | 046444387 | -00925 | -08525 | 16 |
| 2753 | ENSMUSG( | Gjc3 | 118446 | -02286362 | -00154448 | -0415 | -09925 | 16 |
| 2826 | ENSMUSG( | Gnpda1 | 26384 | -00204936 | -01659889 | 028 | -0855 | 16 |
| 2967 | ENSMUSG( | Gstt1 | 14871 | -02983579 | -03412858 | 03175 | -06825 | 16 |
| 3003 | ENSMUSG( | Gys1 | 14936 | 009713033 | 001457701 | 01875 | -05675 | 16 |
| 3031 | ENSMUSG( | Hba-a1 | 15122 | -06663103 | -04344143 | 0215 | -16975 | 16 |
| 3032 | ENSMUSG( | Hbb-b1 | 15129 | -07128126 | -05650503 | 02575 | -11175 | 16 |

| | | | | | | | | |
|---|---|---|---|---|---|---|---|---|
| 3043 | ENSMUSG( | Hdac11 | 232232 | 010393311 | -00568696 | 00675 | -06175 | 16 |
| 3132 | ENSMUSG( | Hnrnpd | 11991 | -00328066 | -00788169 | 02575 | -06725 | 16 |
| 3134 | ENSMUSG( | Hnrnpf | 98758 | 009964187 | 007658619 | -00225 | -05825 | 16 |
| 3142 | ENSMUSG( | Hnrnpr | 74326 | 00448568C | 005265772 | -005 | -06625 | 16 |
| 3157 | ENSMUSG( | Hpgds | 54486 | -03243119 | 013405658 | -006 | -08475 | 16 |
| 3168 | ENSMUSG( | Hscb | 100900 | -00353842 | -01762039 | -02075 | -08925 | 16 |
| 3214 | ENSMUSG( | Hypk | 67693 | -02780016 | 009151322 | 04525 | -09475 | 16 |
| 3231 | ENSMUSG( | Ids | 15931 | -00858301 | 013404931 | 00675 | -05375 | 16 |
| 3279 | ENSMUSG( | Ilf2 | 67781 | -00098337 | -00757850 | -00175 | -06575 | 16 |
| 3314 | ENSMUSG( | Ints1 | 68510 | 018299594 | 007150368 | 013 | -0635 | 16 |
| 3350 | ENSMUSG( | Irgm1 | 15944 | -02213552 | -01542488 | -018 | -1155 | 16 |
| 3361 | ENSMUSG( | Isoc2a | 664994 | 002093656 | -03289271 | -039 | -075 | 16 |
| 3382 | ENSMUSG( | Itgb5 | 16419 | 000993159 | 003341915 | -03575 | -076 | 16 |
| 3426 | ENSMUSG( | Kalrn | 545156 | 017468536 | 003767798 | 044 | -12075 | 16 |
| 3451 | ENSMUSG( | Kcnab3 | 16499 | 003953656 | -02369213 | 003 | -06775 | 16 |
| 3460 | ENSMUSG( | Kcnd3 | 56543 | -01259880 | 037892852 | 007 | -0595 | 16 |
| 3508 | ENSMUSG( | Kctd7 | 212919 | 000444922 | -01598114 | 05275 | -149 | 16 |
| 3521 | ENSMUSG( | Khdrbs2 | 170771 | -01406526 | -01748459 | -02875 | -1125 | 16 |
| 3559 | ENSMUSG( | Klhdc9 | 68874 | 00151223C | 009135012 | 00875 | -066 | 16 |
| 3562 | ENSMUSG( | Klhl2 | 77113 | -00303203 | -00292837 | 01725 | -08575 | 16 |
| 3598 | ENSMUSG( | Lactb | 80907 | -00796152 | 000639365 | -00275 | -06125 | 16 |
| 3609 | ENSMUSG( | Lamp5 | 76161 | -02945909 | -00728515 | 0315 | -08675 | 16 |
| 3630 | ENSMUSG( | Lasp1 | 16796 | 013932851 | -00430462 | 027 | -1495 | 16 |
| 3658 | ENSMUSG( | Lgi2 | 246316 | -05130965 | 003595765 | -00075 | -079 | 16 |
| 3699 | ENSMUSG( | Lmna | 16905 | -01970063 | -00334742 | -03275 | -06975 | 16 |
| 3746 | ENSMUSG( | Lrrc4 | 192198 | -01799716 | 015273643 | -01825 | -07225 | 16 |
| 3798 | ENSMUSG( | Lurap1l | 52829 | 026567377 | 016407796 | -016 | -10675 | 16 |
| 3818 | ENSMUSG( | Lyrm4 | 380840 | 001250649 | -02625457 | 05025 | -0925 | 16 |
| 3850 | ENSMUSG( | Magoh | 17149 | -01321419 | -01148582 | -02775 | -0975 | 16 |
| 4007 | ENSMUSG( | Med16 | 216154 | 013764735 | 001323107 | 04175 | -06975 | 16 |
| 4019 | ENSMUSG( | Med9 | 192191 | 005660488 | -01895985 | -01225 | -0765 | 16 |
| 4089 | ENSMUSG( | Mif4gd | 69674 | -01014499 | -01017398 | 0455 | -0955 | 16 |
| 4097 | ENSMUSG( | Mink1 | 50932 | 01305165C | -00212385 | 07525 | -1875 | 16 |
| 4128 | ENSMUSG( | Mog | 17441 | -07525745 | 004516461 | -01475 | -08425 | 16 |
| 4130 | ENSMUSG( | Mon1a | 72825 | 016496662 | -00166762 | -0055 | -06475 | 16 |
| 4142 | ENSMUSG( | Mpc2 | 70456 | -01235591 | -02554083 | 0415 | -07625 | 16 |
| 4224 | ENSMUSG( | Mrps2 | 118451 | 010067754 | -00759131 | -01775 | -07925 | 16 |
| 4228 | ENSMUSG( | Mrps24 | 64660 | -00037587 | -02389196 | -015 | -108 | 16 |
| 4252 | ENSMUSG( | Msl3 | 17692 | 014178537 | 006261038 | 0035 | -079 | 16 |
| 4305 | ENSMUSG( | Mtx2 | 53375 | -00938945 | -00200580 | 04025 | -0725 | 16 |
| 4326 | ENSMUSG( | Myl1 | 17901 | 003187807 | 079578061 | -0105 | -0755 | 16 |
| 4330 | ENSMUSG( | Myl6 | 17904 | -00315192 | -01244999 | 017 | -06425 | 16 |
| 4338 | ENSMUSG( | Myo1d | 338367 | -02271836 | -01809923 | -03775 | -08575 | 16 |
| 4399 | ENSMUSG( | Nat8f1 | 66116 | 004961637 | 001890548 | 06225 | -0935 | 16 |
| 4437 | ENSMUSG( | Ncs1 | 14299 | 019073266 | 009264898 | 03025 | -0745 | 16 |
| 4491 | ENSMUSG( | Ndufs7 | 75406 | 003843548 | -01810794 | -02 | -155 | 16 |
| 4570 | ENSMUSG( | Nipsnap1 | 18082 | -00366120 | -02073398 | -024 | -071 | 16 |
| 4589 | ENSMUSG( | Nmnat1 | 66454 | 010516585 | 019284627 | 013 | -05475 | 16 |
| 4616 | ENSMUSG( | Nop2 | 110109 | 015044777 | -00401603 | -001 | -0715 | 16 |
| 4660 | ENSMUSG( | Nras | 18176 | 000610539 | -01712618 | 04275 | -10075 | 16 |
| 4664 | ENSMUSG( | Nrcam | 319504 | -00479754 | 002630042 | 0325 | -05975 | 16 |
| 4704 | ENSMUSG( | Ntng1 | 80883 | -04764124 | -01387469 | 01875 | -09175 | 16 |
| 4708 | ENSMUSG( | Ntrk3 | 18213 | -00476711 | 022146458 | -00525 | -05725 | 16 |
| 4720 | ENSMUSG( | Nudcd1 | 67429 | 014351785 | 013372782 | -033 | -07625 | 16 |
| 4721 | ENSMUSG( | Nudcd2 | 52653 | 00320373C | -00447228 | -019 | -07475 | 16 |
| 4749 | ENSMUSG( | Nup160 | 59015 | 007566523 | 008918472 | 01425 | -074 | 16 |
| 4751 | ENSMUSG( | Nup205 | 70699 | 013104391 | 010027967 | 018 | -06 | 16 |
| 4761 | ENSMUSG( | Nup85 | 445007 | 017438688 | 015992615 | 012 | -06775 | 16 |
| 4764 | ENSMUSG( | Nup98 | 269966 | 00203733C | 008821269 | 00325 | -06425 | 16 |
| 4811 | ENSMUSG( | Orai1 | 109305 | -04867465 | 033802534 | -01575 | -074 | 16 |
| 4900 | ENSMUSG( | Paqr4 | 76498 | -02616877 | -00381281 | 0085 | -08325 | 16 |
| 4922 | ENSMUSG( | Pbrm1 | 66923 | -00123532 | 007799125 | -00575 | -05325 | 16 |
| 4928 | ENSMUSG( | Pcbp2 | 18521 | 005467496 | -00435836 | -004 | -1185 | 16 |
| 4938 | ENSMUSG( | Pcdh19 | 279653 | -01015671 | 046143671 | 02725 | -07475 | 16 |
| 4946 | ENSMUSG( | Pcdhb17 | 93888 | -05080059 | 025222557 | 0295 | -07075 | 16 |
| 5033 | ENSMUSG( | Pdk4 | 27273 | -01748069 | -05118436 | 0005 | -07725 | 16 |
| 5089 | ENSMUSG( | Pfkfb4 | 270198 | -00595008 | -02954570 | 04025 | -09525 | 16 |
| 5111 | ENSMUSG( | Pgpep1 | 66522 | -00605730 | -01655261 | 04875 | -072 | 16 |
| 5142 | ENSMUSG( | Phlda3 | 27280 | -00584303 | -02849655 | -00925 | -06725 | 16 |
| 5198 | ENSMUSG( | Pitpnc1 | 71795 | 011868781 | -00357192 | 02175 | -07225 | 16 |
| 5226 | ENSMUSG( | Plcd3 | 72469 | 015120892 | 009565234 | -042 | -073 | 16 |
| 5338 | ENSMUSG( | Polr2l | 66491 | -00544194 | -02922216 | -00175 | -1045 | 16 |
| 5345 | ENSMUSG( | Polr3f | 70408 | -00416865 | -00297510 | 032 | -091 | 16 |
| 5378 | ENSMUSG( | Ppid | 67738 | -00286995 | 006474261 | 00625 | -05375 | 16 |
| 5383 | ENSMUSG( | Ppil1 | 68816 | -00491261 | -01234923 | -02975 | -078 | 16 |
| 5443 | ENSMUSG( | Ppp3r1 | 19058 | -00033738 | -01257575 | 0655 | -1265 | 16 |
| 5449 | ENSMUSG( | Ppp5c | 19060 | 008117754 | -00643536 | -0085 | -07525 | 16 |
| 5489 | ENSMUSG( | Prkar2a | 19087 | 00945195C | 016459493 | -0075 | -065 | 16 |
| 5523 | ENSMUSG( | Prpf38a | 230596 | 000145711 | -00550567 | 00625 | -074 | 16 |
| 5531 | ENSMUSG( | Prpf8 | 192159 | 009791114 | -00285002 | -01075 | -06175 | 16 |
| 5556 | ENSMUSG( | Psd | 73728 | 012581488 | 002369569 | -00325 | -0585 | 16 |
| 5558 | ENSMUSG( | Psd3 | 234353 | 014360216 | 001008328 | -022 | -0925 | 16 |
| 5591 | ENSMUSG( | Psmd4 | 19185 | 006419284 | -01000180 | 0175 | -06525 | 16 |
| 5624 | ENSMUSG( | Ptgis | 19223 | -02399441 | 022738603 | 01725 | -07425 | 16 |
| 5684 | ENSMUSG( | Pvalb | 19293 | -00816461 | -05187424 | -025 | -082 | 16 |
| 5700 | ENSMUSG( | Pygm | 19309 | 007133991 | -02561336 | -025 | -087 | 16 |
| 5720 | ENSMUSG( | Rab10 | 19325 | -00279552 | -01076146 | 01225 | -09925 | 16 |
| 5727 | ENSMUSG( | Rab13 | 68328 | 008235188 | -03354455 | 0475 | -11725 | 16 |
| 5753 | ENSMUSG( | Rab3a | 19339 | 002183719 | -01509165 | 02525 | -0615 | 16 |
| 5765 | ENSMUSG( | Rab5a | 271457 | -00094917 | -01139502 | 01175 | -05575 | 16 |
| 5785 | ENSMUSG( | Rabl3 | 67657 | 008303753 | -00183498 | 00975 | -061 | 16 |
| 5809 | ENSMUSG( | Ralyl | 76897 | 009282905 | -01488249 | -00625 | -0635 | 16 |
| 5820 | ENSMUSG( | Rap1a | 109905 | -00913923 | -02151635 | 0085 | -06625 | 16 |
| 5840 | ENSMUSG( | Rasa3 | 19414 | 003525769 | -01163186 | 0045 | -06025 | 16 |
| 5861 | ENSMUSG( | Rbfa | 68731 | -00511462 | -00140016 | -01375 | -09875 | 16 |
| 5864 | ENSMUSG( | Rbfox3 | 52897 | 007163807 | 000670155 | -007 | -06675 | 16 |
| 5888 | ENSMUSG( | Rbm8a | 60365 | 008603044 | -00778356 | -0005 | -0695 | 16 |
| 5890 | ENSMUSG( | Rbmx | 19655 | -02494434 | 020367364 | -0475 | -08025 | 16 |
| 5907 | ENSMUSG( | Rcn2 | 26611 | -00722785 | 00465883C | 04 | -078 | 16 |
| 5942 | ENSMUSG( | Rfk | 54391 | 00030379C | -01739842 | 03275 | -12125 | 16 |
| 5983 | ENSMUSG( | Ric3 | 320360 | -01625480 | 002736456 | -01025 | -073 | 16 |
| 5992 | ENSMUSG( | Rims1 | 116837 | 002692444 | -02101389 | -025 | -1175 | 16 |
| 6022 | ENSMUSG( | Rnd3 | 74194 | -02530423 | 009901301 | 04225 | -06825 | 16 |
| 6181 | ENSMUSG( | Rras | 20130 | -00323113 | -00336693 | 024 | -06125 | 16 |
| 6193 | ENSMUSG( | Rsl24d1 | 225215 | 016990095 | 000973866 | -01775 | -10625 | 16 |
| 6206 | ENSMUSG( | Rtn4 | 68585 | -03074026 | 017524504 | 061 | -17 | 16 |
| 6288 | ENSMUSG( | Scamp5 | 56807 | -03071783 | 010115583 | 024 | -0725 | 16 |
| 6302 | ENSMUSG( | Scn1a | 20265 | -03491355 | 021057384 | -03025 | -065 | 16 |
| 6303 | ENSMUSG( | Scn1b | 20266 | -01601687 | -00516783 | 00475 | -051 | 16 |
| 6345 | ENSMUSG( | Sdr39u1 | 654795 | -00757488 | -02379335 | 007 | -06925 | 16 |
| 6371 | ENSMUSG( | Selenbp1 | 20341 | 002905824 | -01124153 | 005 | -0555 | 16 |
| 6397 | ENSMUSG( | Senp7 | 66315 | -01035132 | 019596144 | 0125 | -0695 | 16 |
| 6445 | ENSMUSG( | Sf3a3 | 75062 | 005477479 | -00244647 | -0115 | -06075 | 16 |
| 6446 | ENSMUSG( | Sf3b1 | 81898 | -00306259 | 011903942 | -019 | -0645 | 16 |
| 6449 | ENSMUSG( | Sf3b4 | 107701 | 001838463 | -01810753 | -0115 | -10875 | 16 |
| 6461 | ENSMUSG( | Sfxn4 | 94281 | 00768534C | 012307424 | 00625 | -16975 | 16 |
| 6522 | ENSMUSG( | Shroom4 | 208431 | 012008422 | 030672096 | 011 | -06225 | 16 |
| 6527 | ENSMUSG( | Sigmar1 | 18391 | -02104303 | 021970196 | 04 | -084 | 16 |
| 6543 | ENSMUSG( | Skp1 | 21402 | 001592111 | -01803923 | -04025 | -095 | 16 |
| 6556 | ENSMUSG( | Slc13a5 | 237831 | -01633512 | -00198198 | 004 | -05325 | 16 |
| 6583 | ENSMUSG( | Slc22a4 | 30805 | -00693904 | -00033757 | 01975 | -1205 | 16 |
| 6680 | ENSMUSG( | Slc4a2 | 20535 | -01578742 | -00330704 | -01225 | -0545 | 16 |
| 6699 | ENSMUSG( | Slc6a8 | 102857 | 006992992 | 006755203 | 0005 | -0525 | 16 |
| 6725 | ENSMUSG( | Slirp | 380773 | -00336668 | -02379920 | 0325 | -08975 | 16 |
| 6772 | ENSMUSG( | Smim26 | 228715 | -03196423 | -00903315 | -026 | -081 | 16 |
| 6798 | ENSMUSG( | Sncb | 104069 | 00421955C | -02528605 | -00175 | -06025 | 16 |

| | | | | | | | |
|---|---|---|---|---|---|---|---|
| 6806 | ENSMUSG( | Snrnp200 | 320632 | 01836187 | -0132448 | -00725 | -067 | 16 |
| 6808 | ENSMUSG( | Snrnp40 | 66585 | -0067345 | -00598188 | 0195 | -0845 | 16 |
| 6811 | ENSMUSG( | Snrpa1 | 68981 | -0017333 | -0042593 | -0225 | -07125 | 16 |
| 6856 | ENSMUSG( | Sod2 | 20656 | 00112519 | -0153360 | -00975 | -07275 | 16 |
| 6885 | ENSMUSG( | Spata1 | 70951 | -0108062 | -00049823 | 04 | -12775 | 16 |
| 6903 | ENSMUSG( | Spire1 | 68166 | 02617474 | -00884900 | -021 | -08325 | 16 |
| 6924 | ENSMUSG( | Sptb | 20741 | 01025758 | 01839475 | -01125 | -079 | 16 |
| 6959 | ENSMUSG( | Srsf1 | 110809 | -00423463 | 01156867 | -01325 | -05925 | 16 |
| 6960 | ENSMUSG( | Srsf10 | 14105 | 00776735 | 02440144 | -0185 | -06675 | 16 |
| 6967 | ENSMUSG( | Srsf6 | 67996 | -0101259 | 01775151 | -05 | -0855 | 16 |
| 6973 | ENSMUSG( | Ssb | 20823 | 00365243 | -0003409 | -0135 | -05725 | 16 |
| 7026 | ENSMUSG( | Stk24 | 223255 | 01300293 | 00465497 | -00475 | -06625 | 16 |
| 7077 | ENSMUSG( | Suclg1 | 56451 | 01647752 | -0102973 | 0055 | -07225 | 16 |
| 7079 | ENSMUSG( | Suco | 226551 | -03577074 | 01222235 | -0135 | -0735 | 16 |
| 7091 | ENSMUSG( | Sun1 | 77053 | -0158679 | 01837929 | 0035 | -05725 | 16 |
| 7095 | ENSMUSG( | Supt20 | 56790 | 00997287 | 00931038 | -042 | -131 | 16 |
| 7208 | ENSMUSG( | Tbc1d10a | 103724 | 01894897 | 00485025 | 01925 | -07125 | 16 |
| 7277 | ENSMUSG( | Ten1 | 69535 | -0085450 | -0324110 | 05175 | -133 | 16 |
| 7287 | ENSMUSG( | Tesc | 57816 | -02550447 | -0173462 | -0115 | -117 | 16 |
| 7372 | ENSMUSG( | Tm2d1 | 94043 | -0175383 | -00323590 | 0195 | -06325 | 16 |
| 7435 | ENSMUSG( | Tmem184 | 223693 | -0369993 | 00710923 | 03675 | -06025 | 16 |
| 7442 | ENSMUSG( | Tmem200 | 77220 | -0174289 | 04352626 | -00725 | -06475 | 16 |
| 7444 | ENSMUSG( | Tmem201 | 230917 | -0166680 | 01182310 | -00225 | -07775 | 16 |
| 7472 | ENSMUSG( | Tmem47 | 192216 | -0101538 | -0026594 | 00825 | -0905 | 16 |
| 7560 | ENSMUSG( | Tpi1 | 21991 | 00465996 | -0230488 | 0245 | -07 | 16 |
| 7565 | ENSMUSG( | Tpmt | 22017 | 00220245 | -0263875 | 045 | -079 | 16 |
| 7576 | ENSMUSG( | Tra2a | 101214 | 01551082 | 01505991 | 008 | -07825 | 16 |
| 7595 | ENSMUSG( | Trappc2 | 66226 | 00045342 | -0153663 | 0325 | -0595 | 16 |
| 7619 | ENSMUSG( | Trim37 | 68729 | 01466530 | -0094666 | 0365 | -13325 | 16 |
| 7647 | ENSMUSG( | Trp53i11 | 277414 | -0274268 | -0011943 | 05325 | -118 | 16 |
| 7649 | ENSMUSG( | Trpc1 | 22063 | -0287219 | 01185832 | -019 | -06 | 16 |
| 7668 | ENSMUSG( | Tshz3 | 243931 | -0006419 | -0053589 | 011 | -1885 | 16 |
| 7674 | ENSMUSG( | Tspan2 | 70747 | -05727435 | 00360569 | 005 | -12225 | 16 |
| 7711 | ENSMUSG( | Tti2 | 234138 | -0094158 | -0045668 | 00275 | -0515 | 16 |
| 7896 | ENSMUSG( | Uqcr10 | 66152 | 00325107 | -0268455 | 06375 | -159 | 16 |
| 7955 | ENSMUSG( | Utp4 | 21771 | 01764383 | -01088847 | 00075 | -09825 | 16 |
| 7987 | ENSMUSG( | Vdac2 | 22334 | -0011420 | -0141286 | 01125 | -0625 | 16 |
| 7988 | ENSMUSG( | Vdac3 | 22335 | -0000830 | -0177911 | 02725 | -078 | 16 |
| 8000 | ENSMUSG( | Vma21 | 67048 | -0338035 | 00174747 | 02625 | -069 | 16 |
| 8058 | ENSMUSG( | Wars2 | 70560 | 02328604 | 00749463 | -005 | -063 | 16 |
| 8095 | ENSMUSG( | Wdr45b | 66840 | 01005773 | -0030194 | 01725 | -06725 | 16 |
| 8126 | ENSMUSG( | Wtap | 60532 | -0041250 | 00052313 | 0275 | -07 | 16 |
| 8157 | ENSMUSG( | Yae1d1 | 67008 | 00136325 | -0081402 | -00725 | -057 | 16 |
| 8206 | ENSMUSG( | Zc3h13 | 67302 | -0046682 | 01236813 | 03275 | -093 | 16 |
| 8235 | ENSMUSG( | Zfand1 | 66361 | -0023319 | -0079116 | -05575 | -09425 | 16 |
| 8271 | ENSMUSG( | Zkscan16 | 1E+08 | -01989448 | 01626083 | 02825 | -0675 | 16 |
| 142 | ENSMUSG( | Actn2 | 11472 | -0723949 | -01078770 | -004 | -0425 | 17 |
| 153 | ENSMUSG( | Acvr1c | 269275 | -0851812 | -01958998 | -015 | -04275 | 17 |
| 207 | ENSMUSG( | Adora2a | 11540 | -1292926 | -01257415 | -09875 | 00775 | 17 |
| 391 | ENSMUSG( | Ano3 | 228432 | -0881266 | 00579459 | -00425 | -04325 | 17 |
| 472 | ENSMUSG( | Arc | 11838 | -1169742 | -09976625 | -01275 | -04225 | 17 |
| 489 | ENSMUSG( | Arg2 | 11847 | -0465434 | -0599602 | 01775 | 0235 | 17 |
| 599 | ENSMUSG( | Asic4 | 241118 | -09360031 | 00366321 | -00125 | 02975 | 17 |
| 923 | ENSMUSG( | Cadm2 | 239857 | -0464773 | 02735166 | -0445 | 00775 | 17 |
| 932 | ENSMUSG( | Calhm2 | 72691 | -0759738 | -01235520 | 00525 | -0095 | 17 |
| 1091 | ENSMUSG( | Cd180 | 17079 | -0888736 | -0240417 | 00475 | 00525 | 17 |
| 1093 | ENSMUSG( | Cd276 | 102657 | -0392522 | -01458628 | -00125 | 0055 | 17 |
| 1105 | ENSMUSG( | Cd9 | 12527 | -0662113 | -0026083 | -00425 | 02275 | 17 |
| 1129 | ENSMUSG( | Cdh12 | 215654 | -07804170 | 01700159 | -00825 | -00525 | 17 |
| 1138 | ENSMUSG( | Cdh6 | 12563 | -0675051 | 01237674 | -04225 | -0115 | 17 |
| 1263 | ENSMUSG( | Chrm4 | 12672 | -07692407 | -00199746 | -02025 | 031 | 17 |
| 1267 | ENSMUSG( | Chsy3 | 78923 | -0655733 | 01095970 | -0065 | 02425 | 17 |
| 1318 | ENSMUSG( | Clmp | 71566 | -0534336 | -0045788 | -05025 | -00175 | 17 |
| 1384 | ENSMUSG( | Cntn3 | 18488 | -0687834 | -0011043 | 00875 | 00575 | 17 |
| 1484 | ENSMUSG( | Cox4i2 | 84682 | -04100081 | -0090923 | -03 | 02025 | 17 |
| 1539 | ENSMUSG( | Crh | 12918 | -07303614 | -0363280 | -00625 | 015 | 17 |
| 1577 | ENSMUSG( | Csmd3 | 239420 | -0869050 | 01070742 | 02 | 015 | 17 |
| 1587 | ENSMUSG( | Cspg4 | 121021 | -07934162 | 01814969 | 00775 | 011 | 17 |
| 1678 | ENSMUSG( | Cyp2s1 | 74134 | -0954304 | -0023155 | -06275 | -056 | 17 |
| 1720 | ENSMUSG( | Dbh | 13166 | -0734067 | -05647508 | -022 | -00325 | 17 |
| 2009 | ENSMUSG( | Drd1 | 13488 | -1292540 | 00297485 | -0815 | 00925 | 17 |
| 2069 | ENSMUSG( | Ecel1 | 13599 | -0760395 | -01046989 | 00525 | 0435 | 17 |
| 2104 | ENSMUSG( | Efnb1 | 13641 | -0513212 | -01621911 | 0015 | -01375 | 17 |
| 2106 | ENSMUSG( | Efnb3 | 13643 | -05308301 | -00361987 | -01475 | 001 | 17 |
| 2236 | ENSMUSG( | Enpp4 | 224794 | -0452032 | 02182591 | -03275 | 02425 | 17 |
| 2242 | ENSMUSG( | Entpd3 | 215446 | -0515272 | -0066492 | 0005 | 01 | 17 |
| 2275 | ENSMUSG( | Epn3 | 71889 | -0791037 | -0744543 | -034 | -0115 | 17 |
| 2431 | ENSMUSG( | Fat3 | 270120 | -0738936 | 01066216 | -0165 | -0065 | 17 |
| 2515 | ENSMUSG( | Filip1 | 70598 | -0600621 | -0153178 | -035 | 03825 | 17 |
| 2688 | ENSMUSG( | Gbp11 | 634650 | -0690180 | -03878875 | -03775 | -01375 | 17 |
| 2750 | ENSMUSG( | Gjb1 | 14618 | -0791393 | -0040904 | -02 | -00825 | 17 |
| 2752 | ENSMUSG( | Gjc2 | 118454 | -0887855 | -00737727 | -018 | -01025 | 17 |
| 2754 | ENSMUSG( | Gjd2 | 14617 | -08044088 | -01088929 | -038 | 053 | 17 |
| 2855 | ENSMUSG( | Gpat3 | 231510 | -0760561 | -0176576 | 00075 | 011 | 17 |
| 2877 | ENSMUSG( | Gpr158 | 241263 | -0570390 | 01234915 | -01625 | -00775 | 17 |
| 2883 | ENSMUSG( | Gpr37 | 14763 | -0587056 | 01365428 | -03025 | -02425 | 17 |
| 2885 | ENSMUSG( | Gpr68 | 238377 | -0461413 | -0217451 | -02825 | 00925 | 17 |
| 2886 | ENSMUSG( | Gpr88 | 64378 | -1269575 | 00192140 | -0155 | -04975 | 17 |
| 2893 | ENSMUSG( | Gprin3 | 243385 | -0743093 | -0180138 | -02025 | 019 | 17 |
| 2953 | ENSMUSG( | Gsn | 227753 | -0478055 | 01626654 | -029 | 031 | 17 |
| 2961 | ENSMUSG( | Gstm2 | 14863 | -0312191 | -0292784 | -02125 | -00125 | 17 |
| 3041 | ENSMUSG( | Hdac1 | 433759 | -0523165 | -0031704 | -01125 | 0015 | 17 |
| 3073 | ENSMUSG( | Hecw2 | 329152 | -0439791 | -0350558 | -008 | -00425 | 17 |
| 3204 | ENSMUSG( | Htr1f | 15557 | -1042420 | 01064665 | -016 | 00775 | 17 |
| 3372 | ENSMUSG( | Itga5 | 16402 | -0548744 | -0166327 | -01125 | 00775 | 17 |
| 3381 | ENSMUSG( | Itgb4 | 192897 | -0518660 | -0258428 | -0505 | 01625 | 17 |
| 3447 | ENSMUSG( | Kcna4 | 16492 | -0651868 | -0007731 | -01675 | 0025 | 17 |
| 3463 | ENSMUSG( | Kcnh1 | 16510 | -04974325 | 00749886 | -0345 | 011 | 17 |
| 3491 | ENSMUSG( | Kcns2 | 16539 | -04889257 | -0052807 | -0155 | 0365 | 17 |
| 3628 | ENSMUSG( | Lars2 | 102436 | -1434285 | -2245513 | 00025 | -0175 | 17 |
| 3643 | ENSMUSG( | Ldlr | 16835 | -1082618 | -0064971 | -0075 | -0015 | 17 |
| 3653 | ENSMUSG( | Lgals3bp | 19039 | -0580500 | -0256886 | -0035 | -03575 | 17 |
| 3695 | ENSMUSG( | Lmcd1 | 30937 | -0406790 | -0722092 | -0425 | 0135 | 17 |
| 4062 | ENSMUSG( | Mfsd6 | 98682 | -04260350 | 01335052 | -0425 | -00175 | 17 |
| 4313 | ENSMUSG( | Mx1 | 17857 | -0688764 | -0797736 | -0615 | -04175 | 17 |
| 4489 | ENSMUSG( | Ndufs5 | 595136 | -1532719 | -0208600 | 01375 | -05025 | 17 |
| 4530 | ENSMUSG( | Neu3 | 50877 | -0530465 | -0377898 | 0135 | -02325 | 17 |
| 4533 | ENSMUSG( | Nexn | 68810 | -0784135 | -0723020 | -08175 | -0315 | 17 |
| 4646 | ENSMUSG( | Nptn | 20320 | -0614246 | 00437448 | -00525 | 00575 | 17 |
| 4648 | ENSMUSG( | Nptx2 | 53324 | -0574922 | -0234825 | -03275 | -03025 | 17 |
| 4650 | ENSMUSG( | Npy | 109648 | -0774166 | -01006637 | 01875 | -01775 | 17 |
| 4666 | ENSMUSG( | Nrg3 | 18183 | -0495559 | 00810095 | -068 | 02075 | 17 |
| 4709 | ENSMUSG( | Nts | 67405 | -0991621 | -0335246 | 0225 | 034 | 17 |
| 4778 | ENSMUSG( | Obscn | 380698 | -0872748 | -00461463 | -02725 | -00125 | 17 |
| 4795 | ENSMUSG( | Olfm3 | 229759 | -0944332 | 02343058 | 01025 | -01675 | 17 |
| 4803 | ENSMUSG( | Opalin | 226115 | -0745772 | -0237118 | -0525 | -002 | 17 |
| 4807 | ENSMUSG( | Oprk1 | 18387 | -0799834 | 02282116 | -09475 | 055 | 17 |
| 4944 | ENSMUSG( | Pcdhb12 | 93883 | -0709790 | 00056141 | -04075 | -003 | 17 |
| 4947 | ENSMUSG( | Pcdhb18 | 93889 | -0827309 | -0015401 | -03975 | 023 | 17 |
| 4948 | ENSMUSG( | Pcdhb20 | 93891 | -0544936 | 00455864 | -0465 | -03725 | 17 |
| 4949 | ENSMUSG( | Pcdhb21 | 93892 | -0616362 | -0212103 | -02075 | -031 | 17 |
| 4970 | ENSMUSG( | Pclo | 26875 | -0574582 | 02551791 | -03475 | -00225 | 17 |
| 4981 | ENSMUSG( | Pcp4l1 | 66425 | -0741119 | -02040437 | -052 | 02175 | 17 |
| 5005 | ENSMUSG( | Pde1b | 18574 | -0415666 | -02067069 | -01325 | 02125 | 17 |
| 5064 | ENSMUSG( | Penk | 18619 | -1407693 | -0132718 | 0075 | -01475 | 17 |
| 5246 | ENSMUSG( | Plekhd1 | 217682 | -03949792 | -0690083 | -01375 | -02 | 17 |

| | | | | | | | | | |
|---|---|---|---|---|---|---|---|---|---|
| 5257 | ENSMUSG( | Pllp | 67801 | -07248056 | -01265464 | -0595 | 0095 | | 17 |
| 5268 | ENSMUSG( | Plppr1 | 272031 | -10493828 | 002623835 | -02825 | 07425 | | 17 |
| 5272 | ENSMUSG( | Plppr5 | 75769 | -06107239 | 015942359 | -056 | 0615 | | 17 |
| 5278 | ENSMUSG( | Plxdc1 | 72324 | -05156953 | 000661961 | -02775 | 015 | | 17 |
| 5306 | ENSMUSG( | Pnoc | 18155 | -07155723 | -02873930 | 00575 | -025 | | 17 |
| 5416 | ENSMUSG( | Ppp1r1b | 19049 | -06776891 | -02958069 | -01425 | 0305 | | 17 |
| 5632 | ENSMUSG( | Ptk7 | 71461 | -04073646 | 011054703 | -02325 | 00825 | | 17 |
| 5637 | ENSMUSG( | Ptp4a1 | 19243 | -07790027 | -00404808 | -0535 | 0075 | | 17 |
| 5655 | ENSMUSG( | Ptprc | 19264 | -05181327 | -01511349 | -0435 | 0095 | | 17 |
| 5685 | ENSMUSG( | Pvr | 52118 | -07406402 | -02109362 | -032 | 00425 | | 17 |
| 5751 | ENSMUSG( | Rab37 | 58222 | -08482772 | -04420271 | -03275 | 017 | | 17 |
| 5843 | ENSMUSG( | Rasd2 | 75141 | -05037539 | -00776485 | -03675 | -0025 | | 17 |
| 5926 | ENSMUSG( | Rem2 | 140743 | -05856347 | -02598483 | -0325 | 023 | | 17 |
| 6203 | ENSMUSG( | Rtn1 | 104001 | -05488943 | 005503177 | -02725 | -0245 | | 17 |
| 6228 | ENSMUSG( | Ryr3 | 20192 | -06974160 | 001142119 | 001 | -02125 | | 17 |
| 6236 | ENSMUSG( | S100a8 | 20201 | -14424343 | -00333478 | -113 | -005 | | 17 |
| 6240 | ENSMUSG( | S1pr5 | 94226 | -06589723 | -03381057 | -0425 | 02725 | | 17 |
| 6308 | ENSMUSG( | Scn4b | 399548 | -12151012 | -01054462 | -071 | 0335 | | 17 |
| 6389 | ENSMUSG( | Sema5a | 20356 | -04404217 | -00144219 | -0265 | -013 | | 17 |
| 6391 | ENSMUSG( | Sema6a | 20358 | -06423790 | 012223559 | 00075 | 004 | | 17 |
| 6439 | ENSMUSG( | Sez6 | 20370 | -05282455 | 007139077 | -019 | -0155 | | 17 |
| 6465 | ENSMUSG( | Sgcz | 244431 | -06293528 | -00647622 | -009 | 041 | | 17 |
| 6497 | ENSMUSG( | Sh3rf2 | 269016 | -12769596 | -03863199 | -02175 | 058 | | 17 |
| 6514 | ENSMUSG( | Shmt1 | 20425 | -03556589 | -04288803 | -036 | 01775 | | 17 |
| 6547 | ENSMUSG( | Slc10a4 | 231290 | -16303508 | -05167442 | -0495 | 04475 | | 17 |
| 6560 | ENSMUSG( | Slc16a1 | 20501 | -05751550 | -03454853 | 001 | 01875 | | 17 |
| 6566 | ENSMUSG( | Slc17a6 | 140919 | -06244357 | -00127263 | -01475 | -0015 | | 17 |
| 6570 | ENSMUSG( | Slc18a3 | 20508 | -08763311 | 015372831 | -00675 | -026 | | 17 |
| 6586 | ENSMUSG( | Slc24a2 | 76376 | -06258080 | 022537047 | -033 | -0075 | | 17 |
| 6632 | ENSMUSG( | Slc2a13 | 239606 | -06309526 | 023874943 | -0195 | 00625 | | 17 |
| 6688 | ENSMUSG( | Slc5a7 | 63993 | -11779474 | 020177857 | -04125 | -01825 | | 17 |
| 6691 | ENSMUSG( | Slc6a11 | 243616 | -03372699 | -02001235 | -03475 | -014 | | 17 |
| 6700 | ENSMUSG( | Slc6a9 | 14664 | -06110973 | 011179451 | -01875 | 00775 | | 17 |
| 6714 | ENSMUSG( | Slc8a3 | 110893 | -05062165 | -00332724 | -0115 | -013 | | 17 |
| 6780 | ENSMUSG( | Smpd3 | 58994 | -04354387 | 022590536 | -05075 | 00575 | | 17 |
| 6799 | ENSMUSG( | Sncg | 20618 | -05989779 | -02789943 | -05225 | 00175 | | 17 |
| 6909 | ENSMUSG( | Spock3 | 72902 | -09353786 | 011882320 | 02425 | -00925 | | 17 |
| 6986 | ENSMUSG( | Sst | 20604 | -06254107 | -01179438 | -0445 | -02 | | 17 |
| 7028 | ENSMUSG( | Stk26 | 70415 | -06577753 | -01153135 | -02625 | 036 | | 17 |
| 7038 | ENSMUSG( | Stmn4 | 56471 | -05352035 | -02770420 | -02325 | -025 | | 17 |
| 7049 | ENSMUSG( | Strip2 | 320609 | -07076330 | 017987191 | -008 | 00875 | | 17 |
| 7105 | ENSMUSG( | Sv2c | 75209 | -07307317 | 011789431 | -022 | 05425 | | 17 |
| 7122 | ENSMUSG( | Syne1 | 64009 | -05355174 | -00026120 | -01675 | -0185 | | 17 |
| 7134 | ENSMUSG( | Synpr | 72003 | -06514886 | 019333761 | -02725 | -01775 | | 17 |
| 7150 | ENSMUSG( | Syt6 | 54524 | -06307372 | -00713387 | -0045 | 0365 | | 17 |
| 7152 | ENSMUSG( | Syt9 | 60510 | -04283364 | -01540604 | -0335 | 0085 | | 17 |
| 7164 | ENSMUSG( | Tacr1 | 21336 | -10747240 | 008146487 | -0775 | -03 | | 17 |
| 7180 | ENSMUSG( | Tagln | 21345 | -08172357 | -04161611 | 002 | 02575 | | 17 |
| 7281 | ENSMUSG( | Tenm4 | 23966 | -03674885 | 001238639 | -0165 | 013 | | 17 |
| 7307 | ENSMUSG( | Th | 21823 | -05261968 | 007666356 | -0195 | 02125 | | 17 |
| 7311 | ENSMUSG( | Thbs1 | 21825 | -10874186 | -01007686 | -02125 | -02525 | | 17 |
| 7425 | ENSMUSG( | Tmem163 | 72160 | -05024838 | 007801488 | -0215 | 03 | | 17 |
| 7483 | ENSMUSG( | Tmem74b | 108832 | -03938189 | -03356981 | -00875 | 035 | | 17 |
| 7587 | ENSMUSG( | Tram1l1 | 229801 | -05944114 | 013973092 | -01725 | -0055 | | 17 |
| 7716 | ENSMUSG( | Ttn | 22138 | -12673018 | -02459212 | -03175 | 01625 | | 17 |
| 7721 | ENSMUSG( | Ttyh2 | 117160 | -06915017 | 022909466 | -0075 | 00275 | | 17 |
| 7990 | ENSMUSG( | Vgf | 381677 | -04778988 | 007608478 | -05225 | 032 | | 17 |
| 8181 | ENSMUSG( | Yrdc | 230734 | -03474954 | -01153472 | -035 | -00225 | | 17 |
| 8231 | ENSMUSG( | Zdhhc9 | 208884 | -04948981 | 014637387 | -01925 | 01325 | | 17 |
| 124 | ENSMUSG( | Acp1 | 11431 | -00367548 | -00890171 | 11725 | -124 | | 18 |
| 241 | ENSMUSG( | Agpat1 | 55979 | -02466861 | -00115283 | 05825 | -01325 | | 18 |
| 316 | ENSMUSG( | Alg11 | 207958 | -04007746 | 029603207 | 0595 | -03175 | | 18 |
| 317 | ENSMUSG( | Alg13 | 67574 | 002738231 | -01708449 | 20275 | -05175 | | 18 |
| 334 | ENSMUSG( | Amdhd2 | 245847 | 000148215 | -00873298 | 082 | -09425 | | 18 |
| 580 | ENSMUSG( | Arsg | 74008 | -06489365 | -00832265 | 0775 | -038 | | 18 |
| 583 | ENSMUSG( | Arxes2 | 76976 | -06180791 | 002948609 | 07075 | -08825 | | 18 |
| 603 | ENSMUSG( | Asph | 65973 | -03916672 | -00827848 | 0585 | -012 | | 18 |
| 650 | ENSMUSG( | Atp23 | 68876 | 002205020 | -00590738 | 072 | -0275 | | 18 |
| 735 | ENSMUSG( | Bag4 | 67384 | 002121610 | 001223456 | 07375 | -06775 | | 18 |
| 831 | ENSMUSG( | Bpnt2 | 242291 | -03832231 | 0136939 | 08225 | -03 | | 18 |
| 844 | ENSMUSG( | Bri3bp | 76809 | -02293000 | 018007205 | 11625 | -01 | | 18 |
| 856 | ENSMUSG( | Bscl2 | 14705 | -02145190 | 016300328 | 0505 | -0185 | | 18 |
| 914 | ENSMUSG( | Cacng3 | 54376 | -03097214 | 011900393 | 08075 | -05025 | | 18 |
| 916 | ENSMUSG( | Cacng7 | 81904 | 001077433 | -00132716 | 066 | -016 | | 18 |
| 933 | ENSMUSG( | Calhm5 | 103511 | -04885562 | 006642116 | 1045 | -018 | | 18 |
| 935 | ENSMUSG( | Calr | 12317 | -01591112 | 011868872 | 11825 | -015 | | 18 |
| 936 | ENSMUSG( | Calu | 12321 | -01622357 | 004106361 | 04425 | -0325 | | 18 |
| 1043 | ENSMUSG( | Ccdc28b | 66264 | -03915645 | 001969841 | 16775 | -06525 | | 18 |
| 1097 | ENSMUSG( | Cd36 | 12491 | -05218303 | -04737844 | 05175 | -017 | | 18 |
| 1098 | ENSMUSG( | Cd38 | 12494 | -03194431 | 003414963 | 06725 | -06625 | | 18 |
| 1100 | ENSMUSG( | Cd47 | 16423 | -05278588 | 013521446 | 07875 | -02 | | 18 |
| 1141 | ENSMUSG( | Cdh9 | 12565 | -04700714 | 021284769 | 05825 | -02725 | | 18 |
| 1238 | ENSMUSG( | Chid1 | 68038 | -02162231 | -00113357 | 062 | -01725 | | 18 |
| 1279 | ENSMUSG( | Cisd1 | 52637 | -01082598 | -00871159 | 14275 | -04975 | | 18 |
| 1437 | ENSMUSG( | Comt | 12846 | -03651780 | -00512122 | 053 | -0625 | | 18 |
| 1656 | ENSMUSG( | Cxcl12 | 20315 | 010666049 | 089779613 | 09675 | -01025 | | 18 |
| 1823 | ENSMUSG( | Derl1 | 67819 | -02974755 | 019806993 | 067 | -00375 | | 18 |
| 1841 | ENSMUSG( | Dhcr24 | 74754 | -03918528 | 008053542 | 09925 | 0045 | | 18 |
| 1876 | ENSMUSG( | Dipk2a | 68861 | -04145444 | 049083938 | 07175 | -01225 | | 18 |
| 1916 | ENSMUSG( | Dnajb11 | 67838 | -01767858 | 015507779 | 059 | -0285 | | 18 |
| 1976 | ENSMUSG( | Dolk | 227697 | -02069625 | 006931913 | 1175 | -08025 | | 18 |
| 1977 | ENSMUSG( | Dolpp1 | 57170 | -03239991 | 024442328 | 1375 | -05525 | | 18 |
| 1989 | ENSMUSG( | Dpm1 | 13480 | -02184520 | 022056981 | 041 | -05525 | | 18 |
| 1998 | ENSMUSG( | Dpy19l1 | 244745 | -02699164 | 013548930 | 06375 | -0415 | | 18 |
| 2062 | ENSMUSG( | Ebag9 | 55960 | -02820313 | 024168896 | 097 | -003 | | 18 |
| 2205 | ENSMUSG( | Emc3 | 66087 | -02260131 | -00020346 | 07175 | -02075 | | 18 |
| 2235 | ENSMUSG( | Enpp2 | 18606 | -02634416 | 046079163 | 08775 | 00475 | | 18 |
| 2292 | ENSMUSG( | Erg28 | 58520 | -05070071 | -00356868 | 0455 | -06825 | | 18 |
| 2298 | ENSMUSG( | Erlec1 | 66753 | -02306786 | 009133436 | 072 | -0025 | | 18 |
| 2306 | ENSMUSG( | Erp44 | 76299 | -02638226 | 024524052 | 0635 | -003 | | 18 |
| 2340 | ENSMUSG( | Exosc3 | 66362 | 020418627 | 001644103 | 07725 | -05225 | | 18 |
| 2341 | ENSMUSG( | Exosc4 | 109075 | 005008404 | -00965702 | 14975 | -04225 | | 18 |
| 2349 | ENSMUSG( | Extl2 | 58193 | -03195108 | 019674345 | 08225 | -01425 | | 18 |
| 2354 | ENSMUSG( | F3 | 14066 | -05461091 | -00981830 | 04475 | -031 | | 18 |
| 2365 | ENSMUSG( | Fads6 | 328035 | -01923602 | 005520613 | 0535 | -068 | | 18 |
| 2413 | ENSMUSG( | Fam76b | 72826 | -02304918 | 028629483 | 1 | 0125 | 18 | |
| 2621 | ENSMUSG( | G6pc3 | 68401 | -01842260 | 011043282 | 07725 | -05825 | | 18 |
| 2671 | ENSMUSG( | Garre1 | 233103 | -00064976 | 003317908 | 05875 | -0665 | | 18 |
| 2785 | ENSMUSG( | Gm20604 | 1.01E+08 | -00623854 | 004983516 | 1395 | -0585 | | 18 |
| 2797 | ENSMUSG( | Gna11 | 14672 | 011307401 | 002815336 | 081 | -09775 | | 18 |
| 2804 | ENSMUSG( | Gnao1 | 14681 | 001382149 | -02134553 | 13975 | -08525 | | 18 |
| 2852 | ENSMUSG( | Gpaa1 | 14731 | -00892962 | 017018426 | 06175 | -00725 | | 18 |
| 3000 | ENSMUSG( | Gusb | 110006 | -01336863 | 013495833 | 0575 | -04075 | | 18 |
| 3002 | ENSMUSG( | Gypa | 14934 | -09082082 | 004591739 | 1005 | -04825 | | 18 |
| 3015 | ENSMUSG( | Hacd3 | 57874 | -03139574 | 004038700 | 0585 | -0105 | | 18 |
| 3034 | ENSMUSG( | Hccs | 15159 | -00264622 | 008744951 | 0605 | -04025 | | 18 |
| 3082 | ENSMUSG( | Herpud2 | 80517 | -00590280 | 007973159 | 1155 | 014 | | 18 |
| 3184 | ENSMUSG( | Hspa13 | 110920 | -03765457 | 042974720 | 0585 | -02725 | | 18 |
| 3239 | ENSMUSG( | Ift22 | 67286 | 010668521 | -01949408 | 0985 | 314018e-1 | | 18 |
| 3286 | ENSMUSG( | Immp2l | 93757 | -03574799 | -02425840 | 0605 | -01925 | | 18 |
| 3288 | ENSMUSG( | Imp3 | 102462 | 000517561 | -00901193 | 08 | -002 | | 18 |
| 3297 | ENSMUSG( | Ing2 | 69260 | -00167058 | -00788722 | 05575 | -02025 | | 18 |
| 3510 | ENSMUSG( | Kdelr1 | 68137 | -04207928 | 010655053 | 17875 | -01775 | | 18 |
| 3561 | ENSMUSG( | Klhl13 | 67455 | -02549369 | -03087178 | 06925 | -00875 | | 18 |
| 3649 | ENSMUSG( | Leprotl1 | 68192 | -04221475 | 013929807 | 08025 | -0455 | | 18 |
| 3663 | ENSMUSG( | Lhfpl3 | 269629 | -04316513 | 019094340 | 071 | -01675 | | 18 |

| | | | | | | | | | |
|---|---|---|---|---|---|---|---|---|---|
| 3690 | ENSMUSG( | Lman2 | 66890 | -00923233 | 016630936 | 0635 | -0295 | | 18 |
| 3720 | ENSMUSG( | Lratd1 | 105005 | 011043255 | 011018396 | 0985 | -068 | | 18 |
| 3851 | ENSMUSG( | Magt1 | 67075 | -00843487 | 013638848 | 07575 | -00675 | | 18 |
| 3855 | ENSMUSG( | Man1a2 | 17156 | -01090010 | 018811325 | 0635 | -0065 | | 18 |
| 3866 | ENSMUSG( | Manf | 74840 | -01393046 | 025903990 | 049 | -0245 | | 18 |
| 3925 | ENSMUSG( | Marchf6 | 223455 | -04644246 | 015069125 | 0445 | -053 | | 18 |
| 3956 | ENSMUSG( | Mbnl1 | 56758 | -01068236 | 010771044 | 20025 | -0365 | | 18 |
| 3969 | ENSMUSG( | Mcfd2 | 193813 | -03262997 | 016962547 | 0555 | -08525 | | 18 |
| 4069 | ENSMUSG( | Mgst1 | 56615 | -00936183 | 003358343 | 09175 | -03575 | | 18 |
| 4108 | ENSMUSG( | Mlip | 69642 | -01784243 | -02785905 | 09775 | -0485 | | 18 |
| 4191 | ENSMUSG( | Mrpl35 | 66223 | 000160733 | -01711918 | 12175 | -056 | | 18 |
| 4279 | ENSMUSG( | Mtif3 | 76366 | 006851022 | -01702600 | 121 | -04825 | | 18 |
| 4319 | ENSMUSG( | Mydgf | 28106 | -03666417 | -00122575 | 054 | -0335 | | 18 |
| 4453 | ENSMUSG( | Ndufa12 | 66414 | 001302280 | -03362337 | 056 | -05725 | | 18 |
| 4624 | ENSMUSG( | Nos3 | 18127 | 039812131 | 081312092 | 05275 | -067 | | 18 |
| 4771 | ENSMUSG( | Nxph1 | 18231 | -05762167 | 014158133 | 08825 | -08725 | | 18 |
| 4772 | ENSMUSG( | Nxph3 | 104079 | -01082079 | 014409124 | 0785 | -0215 | | 18 |
| 4829 | ENSMUSG( | Ostc | 66357 | -03483152 | 011374521 | 106 | 006 | | 18 |
| 4856 | ENSMUSG( | Pabpn1 | 54196 | 014444639 | 034259810 | 07475 | -01875 | | 18 |
| 5017 | ENSMUSG( | Pde8a | 18584 | -00004933 | -02135228 | 05425 | -06325 | | 18 |
| 5098 | ENSMUSG( | Pgap3 | 320655 | -03720136 | -00481095 | 059 | -0125 | | 18 |
| 5153 | ENSMUSG( | Phyhip | 105653 | 007508735 | -00415772 | 11575 | -02775 | | 18 |
| 5164 | ENSMUSG( | Pigg | 433931 | -01803506 | 012767627 | 053 | -01225 | | 18 |
| 5173 | ENSMUSG( | Pik3ap1 | 83490 | -02117258 | -01950650 | 05875 | -02425 | | 18 |
| 5261 | ENSMUSG( | Plp2 | 18824 | -04980161 | -00782098 | 1245 | -089 | | 18 |
| 5316 | ENSMUSG( | Pofut1 | 140484 | -01402261 | 000119501 | 0615 | -02325 | | 18 |
| 5454 | ENSMUSG( | Ppt1 | 19063 | 035582234 | -00039037 | 0975 | -0255 | | 18 |
| 5468 | ENSMUSG( | Prdx5 | 54683 | -00289371 | -02317947 | 06175 | -03875 | | 18 |
| 5482 | ENSMUSG( | Prkab2 | 108097 | 003317872 | 000240791 | 071 | -0545 | | 18 |
| 5616 | ENSMUSG( | Ptdss1 | 19210 | -01751303 | 002057466 | 0535 | -012 | | 18 |
| 5742 | ENSMUSG( | Rab2a | 59021 | 000958761 | -00655740 | 058 | -053 | | 18 |
| 5761 | ENSMUSG( | Rab40c | 224624 | 010021374 | -00461221 | 0875 | -00025 | | 18 |
| 6042 | ENSMUSG( | Rnf181 | 66510 | 002534898 | -01055595 | 11775 | -08 | | 18 |
| 6084 | ENSMUSG( | Rpl10a | 19896 | 004451241 | -01602897 | 057 | -04625 | | 18 |
| 6225 | ENSMUSG( | Rwdd2a | 69519 | -01871375 | -01297752 | 06025 | -0185 | | 18 |
| 6287 | ENSMUSG( | Scamp3 | 24045 | -02122549 | 014018528 | 10625 | -044 | | 18 |
| 6332 | ENSMUSG( | Sdf2 | 20316 | -02980244 | 001816188 | 14275 | -05325 | | 18 |
| 6341 | ENSMUSG( | Sdhc | 66052 | 008805879 | -01757602 | 08975 | -0755 | | 18 |
| 6342 | ENSMUSG( | Sdhd | 66925 | 006296698 | -00588771 | 07975 | -02075 | | 18 |
| 6467 | ENSMUSG( | Sgk2 | 27219 | -01141335 | -01351173 | 11375 | -043 | | 18 |
| 6629 | ENSMUSG( | Slc29a1 | 63959 | -04050744 | 002353974 | 06275 | -04025 | | 18 |
| 6638 | ENSMUSG( | Slc30a6 | 210148 | -03726397 | 018391291 | 07375 | -00075 | | 18 |
| 6677 | ENSMUSG( | Slc4a1 | 20533 | -13549895 | -02001565 | 06 | -0905 | | 18 |
| 6690 | ENSMUSG( | Slc6a1 | 232333 | -02255656 | 008751941 | 08225 | -0275 | | 18 |
| 6761 | ENSMUSG( | Smdt1 | 69029 | -00803617 | -03413418 | 05775 | -05325 | | 18 |
| 6942 | ENSMUSG( | Srgap3 | 259302 | 022188660 | -00482878 | 1655 | -0515 | | 18 |
| 6982 | ENSMUSG( | Ssr1 | 107513 | -03259645 | 009786803 | 05725 | -00725 | | 18 |
| 6984 | ENSMUSG( | Ssr4 | 20832 | -01525484 | -00450757 | 1685 | | -1 18 | |
| 7053 | ENSMUSG( | Stt3a | 16430 | -03920053 | 014529665 | 08975 | -007 | | 18 |
| 7155 | ENSMUSG( | Syvn1 | 74126 | -03171654 | 020784380 | 0545 | -01775 | | 18 |
| 7362 | ENSMUSG( | Tlcd1 | 68385 | 000061783 | 017387809 | 0915 | 00975 | | 18 |
| 7375 | ENSMUSG( | Tm9sf2 | 68059 | -04296157 | 026761996 | 06575 | -02125 | | 18 |
| 7390 | ENSMUSG( | Tmed5 | 73130 | -03578092 | 044311459 | 07925 | -043 | | 18 |
| 7391 | ENSMUSG( | Tmed7 | 66676 | -04465891 | 016443845 | 0725 | 00375 | | 18 |
| 7393 | ENSMUSG( | Tmed9 | 67511 | -03613600 | 005870837 | 0625 | -00675 | | 18 |
| 7401 | ENSMUSG( | Tmem117 | 320709 | -09058322 | 018606183 | 1165 | 01075 | | 18 |
| 7428 | ENSMUSG( | Tmem167 | 66074 | -00183793 | 011207924 | 10125 | -06575 | | 18 |
| 7433 | ENSMUSG( | Tmem178b | 434008 | -01848693 | 006271528 | 10525 | -022 | | 18 |
| 7449 | ENSMUSG( | Tmem223 | 66836 | 00599741 | 009547532 | 05525 | -046 | | 18 |
| 7462 | ENSMUSG( | Tmem260 | 218989 | -02683519 | 007749018 | 06825 | -04775 | | 18 |
| 7475 | ENSMUSG( | Tmem59l | 67937 | -03728707 | 003181748 | 05725 | -00825 | | 18 |
| 7605 | ENSMUSG( | Trhr | 22045 | -01186733 | 046465779 | 05375 | -05125 | | 18 |
| 7639 | ENSMUSG( | Trmt1l | 98685 | 016981671 | 003907010 | 11525 | 01475 | | 18 |
| 7675 | ENSMUSG( | Tspan3 | 56434 | -02236694 | 004233609 | 06975 | -03325 | | 18 |
| 7677 | ENSMUSG( | Tspan33 | 232670 | -02385432 | 000039832 | 04125 | -04775 | | 18 |
| 7695 | ENSMUSG( | Ttc13 | 234875 | -02176470 | 018004734 | 0705 | -0335 | | 18 |
| 7746 | ENSMUSG( | Tusc3 | 80286 | -04256361 | 014774565 | 063 | -0555 | | 18 |
| 7755 | ENSMUSG( | Txndc12 | 66073 | -02628869 | 009528734 | 05 | -01975 | | 18 |
| 7757 | ENSMUSG( | Txndc16 | 70561 | -02460133 | 016232470 | 07025 | 00025 | | 18 |
| 7793 | ENSMUSG( | Ube2d1 | 216080 | -00892423 | -00073153 | 05675 | -0615 | | 18 |
| 7824 | ENSMUSG( | Ubl3 | 24109 | 004201596 | -00454339 | 07275 | -029 | | 18 |
| 7897 | ENSMUSG( | Uqcr11 | 66594 | -00165842 | -03224987 | 0915 | -07475 | | 18 |
| 7918 | ENSMUSG( | Usp12 | 22217 | 004050658 | 001504612 | 056 | -0545 | | 18 |
| 7985 | ENSMUSG( | Vcpkmt | 207965 | -01170885 | -02408590 | 098 | -019 | | 18 |
| 7998 | ENSMUSG( | Vkorc1l1 | 69568 | -02565691 | 003776588 | 05175 | -0255 | | 18 |
| 8011 | ENSMUSG( | Vps26a | 30930 | -00231518 | -00962580 | 0575 | -0385 | | 18 |
| 8170 | ENSMUSG( | Yipf4 | 67864 | -04328786 | 022305302 | 06475 | -06 | | 18 |
| 8202 | ENSMUSG( | Zbtb8os | 67106 | 028231226 | -01627990 | 11 | -0155 | | 18 |
| 8218 | ENSMUSG( | Zcchc8 | 70650 | 009781878 | 023658814 | 07225 | -04475 | | 18 |
| 8263 | ENSMUSG( | Zfyve26 | 211978 | 000658390 | 013228214 | 0555 | -0175 | | 18 |
| 51 | ENSMUSG( | Abcc5 | 27416 | 010355858 | 020650067 | -04025 | 0555 | | 19 |
| 79 | ENSMUSG( | Abl1 | 11350 | -01920943 | 010220750 | -025 | 026 | | 19 |
| 83 | ENSMUSG( | Ablim3 | 319713 | -00201560 | 010595764 | -02225 | 0215 | | 19 |
| 86 | ENSMUSG( | Abraxas2 | 109359 | 006396027 | 002161195 | -034 | 01525 | | 19 |
| 112 | ENSMUSG( | Acly | 104112 | 010622192 | -00822631 | -06825 | 01975 | | 19 |
| 140 | ENSMUSG( | Actl6b | 83766 | 010082496 | -00759567 | -03125 | 01075 | | 19 |
| 173 | ENSMUSG( | Adck2 | 57869 | -01548743 | 029011731 | -05925 | 02775 | | 19 |
| 193 | ENSMUSG( | Adgre1 | 13733 | -02945188 | -00466069 | -07525 | 05525 | | 19 |
| 212 | ENSMUSG( | Adra1a | 11549 | -02086764 | -01879325 | -0635 | 01925 | | 19 |
| 215 | ENSMUSG( | Adra2c | 11553 | -00889638 | 030294728 | -08725 | 049 | | 19 |
| 221 | ENSMUSG( | Afap1l1 | 106877 | -00686517 | 019369628 | -0345 | 0185 | | 19 |
| 271 | ENSMUSG( | Ak2 | 11637 | 016154852 | 000649275 | -02225 | 026 | | 19 |
| 279 | ENSMUSG( | Akap13 | 75547 | -02382188 | 006744461 | -048 | 03175 | | 19 |
| 282 | ENSMUSG( | Akap7 | 432442 | 005967945 | -00861632 | -0175 | 02525 | | 19 |
| 283 | ENSMUSG( | Akap8 | 56399 | 013644969 | 016706656 | -037 | 00625 | | 19 |
| 321 | ENSMUSG( | Alg9 | 102580 | -01996975 | 012011681 | -03025 | 02275 | | 19 |
| 342 | ENSMUSG( | Amotl2 | 56332 | -01253986 | 032792643 | -036 | 00625 | | 19 |
| 386 | ENSMUSG( | Anks1b | 77531 | 004070921 | -00121489 | -01925 | 021 | | 19 |
| 387 | ENSMUSG( | Anks3 | 72615 | 007343989 | 004353173 | -09275 | 0275 | | 19 |
| 396 | ENSMUSG( | Anp32e | 66471 | 001862082 | -00927619 | -038 | 05225 | | 19 |
| 430 | ENSMUSG( | Ap4s1 | 11782 | 005556638 | -01190711 | -0555 | 00075 | | 19 |
| 433 | ENSMUSG( | Apaf1 | 11783 | 033523299 | 013877485 | -03325 | 04875 | | 19 |
| 436 | ENSMUSG( | Apba3 | 57267 | 018765850 | 016049350 | -0345 | 0225 | | 19 |
| 444 | ENSMUSG( | Aph1b | 208117 | -01940977 | 012935232 | -022 | 01425 | | 19 |
| 445 | ENSMUSG( | Api5 | 11800 | 005806081 | -00211777 | -0185 | 02625 | | 19 |
| 466 | ENSMUSG( | Aqp11 | 66333 | 002287273 | 001822947 | -03675 | 0175 | | 19 |
| 494 | ENSMUSG( | Arhgap17 | 70497 | -00833682 | -00063672 | -0345 | 0445 | | 19 |
| 501 | ENSMUSG( | Arhgap27 | 544817 | -00569679 | 017934519 | -02 | 02875 | | 19 |
| 513 | ENSMUSG( | Arhgdib | 11857 | 006773024 | -00986913 | -04925 | 0185 | | 19 |
| 515 | ENSMUSG( | Arhgef10 | 234094 | 003714967 | -00633058 | -03575 | 0005 | | 19 |
| 532 | ENSMUSG( | Arid1b | 239985 | 004033860 | 013873129 | -02225 | 03325 | | 19 |
| 533 | ENSMUSG( | Arid2 | 77044 | 005254025 | 024938182 | -043 | -00225 | | 19 |
| 548 | ENSMUSG( | Arl6ip1 | 54208 | -03394415 | 008709711 | -0335 | 003 | | 19 |
| 574 | ENSMUSG( | Arpp21 | 74100 | -01191098 | -00448402 | -0445 | 02675 | | 19 |
| 613 | ENSMUSG( | Atad2b | 320817 | 002058623 | 016848610 | -057 | 01225 | | 19 |
| 621 | ENSMUSG( | Atg13 | 51897 | 014206087 | 001686428 | -0265 | 01825 | | 19 |
| 623 | ENSMUSG( | Atg16l1 | 77040 | 003328340 | 012478650 | -0475 | 009 | | 19 |
| 662 | ENSMUSG( | Atp5mc1 | 11951 | -00304151 | -02181935 | -03925 | 0005 | | 19 |
| 684 | ENSMUSG( | Atp6v1g2 | 66237 | 002374522 | -01797957 | -04425 | 02875 | | 19 |
| 690 | ENSMUSG( | Atp9b | 50771 | -00075224 | 013699302 | -01825 | 018 | | 19 |
| 700 | ENSMUSG( | Atxn1l | 52335 | 001410099 | -00855600 | -035 | 0215 | | 19 |
| 730 | ENSMUSG( | Bace1 | 23821 | -03147594 | 021871067 | -04575 | 04375 | | 19 |
| 731 | ENSMUSG( | Bad | 12015 | 007930388 | -00298830 | -06 | 012 | | 19 |
| 734 | ENSMUSG( | Bag3 | 29810 | -00837624 | -01279703 | -02725 | 048 | | 19 |
| 742 | ENSMUSG( | Basp1 | 70350 | 001659179 | -00171539 | -05125 | 03775 | | 19 |
| 768 | ENSMUSG( | Bckdk | 12041 | 016558864 | -01065037 | -017 | 0315 | | 19 |

| | | | | | | | | |
|---|---|---|---|---|---|---|---|---|
| 771 | ENSMUSG( | Bcl11b | 58208 | -01526979 | 005746878 | -01825 | 01575 | 19 |
| 775 | ENSMUSG( | Bcl7a | 77045 | 008917499 | -00038441 | -027 | 01075 | 19 |
| 783 | ENSMUSG( | Begain | 380785 | 015539612 | 004482862 | -05375 | -00325 | 19 |
| 794 | ENSMUSG( | Bin2 | 668218 | -01242020 | -01078089 | -022 | 01775 | 19 |
| 814 | ENSMUSG( | Bnip3 | 12176 | 003082326 | -00873256 | -07025 | 0585 | 19 |
| 820 | ENSMUSG( | Bola1 | 69168 | -00612881 | -01559294 | -03375 | 01875 | 19 |
| 822 | ENSMUSG( | Bola3 | 78653 | 007308607 | -02632633 | -04725 | 02875 | 19 |
| 823 | ENSMUSG( | Bop1 | 12181 | 010502889 | 004541255 | -029 | 004 | 19 |
| 835 | ENSMUSG( | Brat1 | 231841 | -00365396 | -00203811 | -032 | 0015 | 19 |
| 839 | ENSMUSG( | Brd3 | 67382 | 005570732 | 001699498 | -05825 | 0135 | 19 |
| 851 | ENSMUSG( | Brox | 71678 | -00275348 | -01375532 | -02225 | 0185 | 19 |
| 854 | ENSMUSG( | Brsk2 | 75770 | 009700771 | 002204803 | -05575 | 0085 | 19 |
| 860 | ENSMUSG( | Btaf1 | 107182 | 005035472 | 005160761 | -0175 | 03275 | 19 |
| 862 | ENSMUSG( | Btbd10 | 68815 | 018775943 | 006066640 | -0245 | 02625 | 19 |
| 873 | ENSMUSG( | Bzw1 | 66882 | 001270934 | -00947468 | -02475 | 01375 | 19 |
| 892 | ENSMUSG( | Cabin1 | 104248 | 013389346 | -00178555 | -05175 | 03 | 19 |
| 901 | ENSMUSG( | Cacna1d | 12289 | 005083886 | 017378750 | -0365 | 005 | 19 |
| 905 | ENSMUSG( | Cacna1i | 239556 | -00858898 | 018195729 | -03575 | 015 | 19 |
| 924 | ENSMUSG( | Cadm3 | 94332 | -02763629 | 019996250 | -0215 | 0115 | 19 |
| 931 | ENSMUSG( | Cald1 | 109624 | -00395001 | 020944542 | -09425 | 04825 | 19 |
| 941 | ENSMUSG( | Camk2a | 12322 | 019567569 | 011277533 | -041 | 03425 | 19 |
| 944 | ENSMUSG( | Camk2g | 12325 | 004085274 | -00614312 | -03675 | 008 | 19 |
| 952 | ENSMUSG( | Camsap2 | 67886 | 010782832 | 011055115 | -034 | 01 | 19 |
| 954 | ENSMUSG( | Camta1 | 100072 | -00140717 | -00765047 | -066 | 047 | 19 |
| 967 | ENSMUSG( | Capn7 | 12339 | 003489255 | 008111507 | -03 | -785046e-1 | 19 |
| 983 | ENSMUSG( | Casc3 | 192160 | 010765314 | 006673041 | -03775 | 0315 | 19 |
| 986 | ENSMUSG( | Caskin1 | 268932 | 012586533 | 006885761 | -03825 | 00075 | 19 |
| 991 | ENSMUSG( | Casp7 | 12369 | -00483603 | 004590912 | -058 | 05675 | 19 |
| 994 | ENSMUSG( | Cast | 12380 | 000940374 | 026172405 | -044 | -00525 | 19 |
| 1002 | ENSMUSG( | Cbfa2t2 | 12396 | 011892265 | 006360952 | -02325 | 00925 | 19 |
| 1015 | ENSMUSG( | Cbx3 | 12417 | 001846053 | -00082591 | -03075 | 01425 | 19 |
| 1017 | ENSMUSG( | Cbx8 | 30951 | -00165203 | -00123617 | -054 | -004 | 19 |
| 1023 | ENSMUSG( | Ccar2 | 219158 | 009461909 | -00214225 | -04275 | 0205 | 19 |
| 1027 | ENSMUSG( | Ccdc12 | 72654 | -00670471 | -02572981 | -0515 | 00475 | 19 |
| 1039 | ENSMUSG( | Ccdc186 | 213993 | 019777950 | 007662524 | -0515 | 01925 | 19 |
| 1044 | ENSMUSG( | Ccdc32 | 269336 | -00050569 | -00850100 | -02925 | 0265 | 19 |
| 1048 | ENSMUSG( | Ccdc50 | 67501 | 005176142 | 015562119 | -046 | 02125 | 19 |
| 1050 | ENSMUSG( | Ccdc59 | 52713 | -01933305 | 009698194 | -03475 | 008 | 19 |
| 1054 | ENSMUSG( | Ccdc85c | 668158 | 018375491 | 008062628 | -0345 | 0255 | 19 |
| 1057 | ENSMUSG( | Ccdc9 | 243846 | 010187239 | -00644532 | -0435 | 01 | 19 |
| 1062 | ENSMUSG( | Ccdc93 | 70829 | 019916592 | 006870558 | -04225 | 00775 | 19 |
| 1063 | ENSMUSG( | Ccdc97 | 52132 | 005644989 | -00877630 | -0325 | 00675 | 19 |
| 1069 | ENSMUSG( | Ccnk | 12454 | 012813650 | 009099486 | -04225 | -0045 | 19 |
| 1075 | ENSMUSG( | Ccp110 | 101565 | -02546929 | -00272964 | -0295 | 0375 | 19 |
| 1094 | ENSMUSG( | Cd2ap | 12488 | -02067611 | 021791429 | -03775 | 01975 | 19 |
| 1113 | ENSMUSG( | Cdc27 | 217232 | 002736143 | 012311473 | -021 | 0345 | 19 |
| 1142 | ENSMUSG( | Cdip1 | 66626 | 011586466 | -00806144 | -033 | 014 | 19 |
| 1151 | ENSMUSG( | Cdk20 | 105278 | 018824418 | 007751180 | -0605 | 0205 | 19 |
| 1153 | ENSMUSG( | Cdk4 | 12567 | -00800254 | -00871721 | -05375 | -003 | 19 |
| 1163 | ENSMUSG( | Cdkn2aip | 70925 | -01211363 | 009070243 | -03475 | 01225 | 19 |
| 1168 | ENSMUSG( | Cdv3 | 321022 | -00149280 | -00999011 | -03 | 003 | 19 |
| 1173 | ENSMUSG( | Celf3 | 78784 | 021154123 | 001548100 | -0245 | 02175 | 19 |
| 1183 | ENSMUSG( | Cep128 | 75216 | -00007968 | -01259590 | -05125 | -007 | 19 |
| 1186 | ENSMUSG( | Cep170 | 545389 | -00483175 | 006390904 | -032 | 00275 | 19 |
| 1187 | ENSMUSG( | Cep170b | 217882 | 028909283 | 010799856 | -02675 | 01875 | 19 |
| 1199 | ENSMUSG( | Cetn4 | 207175 | -00270226 | -01679543 | -0455 | -005 | 19 |
| 1205 | ENSMUSG( | Cfap418 | 67157 | 010580507 | 003606045 | -0365 | 0125 | 19 |
| 1211 | ENSMUSG( | Cgn | 70737 | 031037005 | 012937072 | -0555 | 05875 | 19 |
| 1224 | ENSMUSG( | Chchd6 | 66098 | -01564125 | -02450048 | -0265 | 0195 | 19 |
| 1225 | ENSMUSG( | Chchd7 | 66433 | -00448924 | 016604407 | -02675 | 04475 | 19 |
| 1243 | ENSMUSG( | Chml | 12663 | -03713981 | -00755912 | -065 | 006 | 19 |
| 1244 | ENSMUSG( | Chmp1a | 234852 | 004332706 | -00825687 | -01975 | 02325 | 19 |
| 1276 | ENSMUSG( | Cibar1 | 68099 | -00490830 | 010069793 | -02625 | 035 | 19 |
| 1281 | ENSMUSG( | Cisd3 | 217149 | -00014421 | -01174483 | -01975 | 03775 | 19 |
| 1283 | ENSMUSG( | Ciz1 | 68379 | 027008135 | 012478186 | -072 | 045 | 19 |
| 1308 | ENSMUSG( | Clic1 | 114584 | -00501897 | 009724701 | -027 | 0205 | 19 |
| 1360 | ENSMUSG( | Cnn3 | 71994 | 015204613 | -00704204 | -04225 | 0395 | 19 |
| 1368 | ENSMUSG( | Cnot2 | 72068 | 013753911 | 015109735 | -05425 | -00025 | 19 |
| 1387 | ENSMUSG( | Cntn6 | 53870 | -01183720 | 015822194 | -012 | 025 | 19 |
| 1413 | ENSMUSG( | Col12a1 | 12816 | -00076871 | 01301681 | -0455 | 028 | 19 |
| 1415 | ENSMUSG( | Col19a1 | 12823 | 005936237 | 008161687 | -03025 | 004 | 19 |
| 1456 | ENSMUSG( | Copz2 | 56358 | 010264518 | -02114849 | -05225 | 00325 | 19 |
| 1480 | ENSMUSG( | Cox18 | 231430 | -00351265 | 014956863 | -0715 | 05 | 19 |
| 1570 | ENSMUSG( | Csdc2 | 105859 | 016724737 | -01125835 | -02525 | 015 | 19 |
| 1580 | ENSMUSG( | Csnk1e | 27373 | 006549481 | 007149579 | -01825 | 0195 | 19 |
| 1634 | ENSMUSG( | Cttnbp2nl | 80281 | -00795645 | -01476885 | -03625 | 04225 | 19 |
| 1718 | ENSMUSG( | Daxx | 13163 | 008051541 | 001579353 | -0435 | -00325 | 19 |
| 1728 | ENSMUSG( | Dcaf1 | 321006 | 010058172 | 004210775 | -0485 | 01875 | 19 |
| 1738 | ENSMUSG( | Dchs1 | 233651 | -01389931 | 021751751 | -02675 | 023 | 19 |
| 1743 | ENSMUSG( | Dcp1b | 319618 | 003558197 | 007872126 | -0385 | -001 | 19 |
| 1755 | ENSMUSG( | Dcun1d3 | 233805 | 015881731 | 026497365 | -023 | 03375 | 19 |
| 1756 | ENSMUSG( | Dcun1d4 | 100737 | 009072330 | 005078928 | -027 | 0335 | 19 |
| 1789 | ENSMUSG( | Ddx41 | 72935 | 013820146 | -00726173 | -033 | 02025 | 19 |
| 1806 | ENSMUSG( | Dek | 110052 | 002950494 | 008893848 | -05825 | 02775 | 19 |
| 1825 | ENSMUSG( | Des | 13346 | -02548464 | 018320706 | -041 | -00925 | 19 |
| 1838 | ENSMUSG( | Dgkz | 104418 | 027073320 | 006083504 | -078 | 01175 | 19 |
| 1859 | ENSMUSG( | Dhx32 | 101437 | -00313678 | -00558630 | -02675 | 0095 | 19 |
| 1860 | ENSMUSG( | Dhx35 | 71715 | -00175555 | -00436901 | -046 | -006 | 19 |
| 1862 | ENSMUSG( | Dhx38 | 64340 | 015249759 | 005875100 | -02225 | 0265 | 19 |
| 1879 | ENSMUSG( | Dis3 | 72662 | -00673814 | -01030922 | -03325 | 04725 | 19 |
| 1898 | ENSMUSG( | Dmac1 | 66928 | 005863038 | -01085288 | -02725 | 036 | 19 |
| 1907 | ENSMUSG( | Dnaaf10 | 103784 | 000467841 | 012489899 | -02375 | 00525 | 19 |
| 1915 | ENSMUSG( | Dnajb1 | 81489 | 015068175 | 017663557 | -02925 | 00375 | 19 |
| 1921 | ENSMUSG( | Dnajb5 | 56323 | -00038121 | -02365318 | -026 | 01675 | 19 |
| 1928 | ENSMUSG( | Dnajc14 | 74330 | -01440003 | 007280555 | -07825 | 05975 | 19 |
| 1943 | ENSMUSG( | Dnajc8 | 68598 | 004585948 | -01166904 | -043 | 01225 | 19 |
| 1958 | ENSMUSG( | Dnttip1 | 76233 | 00452112 | 000513590 | -04625 | 016 | 19 |
| 1959 | ENSMUSG( | Dnttip2 | 99480 | -02909013 | 010338003 | -022 | 008 | 19 |
| 1975 | ENSMUSG( | Dok7 | 231134 | -01336236 | -00174055 | -03975 | -0045 | 19 |
| 1983 | ENSMUSG( | Dpf2 | 19708 | 018280537 | 008168864 | -03975 | 03425 | 19 |
| 1988 | ENSMUSG( | Dph7 | 67228 | -00647101 | 001233796 | -03125 | 03425 | 19 |
| 2017 | ENSMUSG( | Dstyk | 213452 | 012218808 | 003907604 | -033 | 04325 | 19 |
| 2029 | ENSMUSG( | Dusp19 | 68082 | 002792433 | -01429694 | -045 | 016 | 19 |
| 2038 | ENSMUSG( | Dvl2 | 13543 | -00028511 | 019818623 | -05375 | 04075 | 19 |
| 2042 | ENSMUSG( | Dync1i1 | 13426 | 008763212 | -01644973 | -025 | 03675 | 19 |
| 2047 | ENSMUSG( | Dync2i1 | 217935 | 002820037 | 005045311 | -054 | 044 | 19 |
| 2049 | ENSMUSG( | Dync2li1 | 213575 | 001241999 | -01337267 | -06575 | 0295 | 19 |
| 2060 | ENSMUSG( | E2f3 | 13557 | -00208234 | 004563818 | -04075 | 02775 | 19 |
| 2065 | ENSMUSG( | Ebpl | 68177 | -01038112 | -01405316 | -0425 | 02175 | 19 |
| 2072 | ENSMUSG( | Echdc2 | 52430 | -00883542 | 019299171 | -0645 | 032 | 19 |
| 2081 | ENSMUSG( | Edf1 | 59022 | -00449930 | -02133056 | -02925 | 0465 | 19 |
| 2112 | ENSMUSG( | Egln1 | 112405 | 009822140 | -00131851 | -0335 | 02975 | 19 |
| 2115 | ENSMUSG( | Ehbp1l1 | 114601 | -00882548 | -00060959 | -02525 | 01025 | 19 |
| 2124 | ENSMUSG( | Eid2 | 386655 | -00938693 | -03026212 | -09225 | 055 | 19 |
| 2159 | ENSMUSG( | Eif4b | 75705 | 194e-05 | -00861861 | -04225 | 03775 | 19 |
| 2161 | ENSMUSG( | Eif4e2 | 26987 | 009177946 | 002915936 | -0365 | 04125 | 19 |
| 2163 | ENSMUSG( | Eif4enif1 | 74203 | 009905909 | 008288396 | -018 | 03175 | 19 |
| 2166 | ENSMUSG( | Eif4g3 | 230861 | 008380824 | -00441357 | -05525 | 03675 | 19 |
| 2167 | ENSMUSG( | Eif4h | 22384 | 015088340 | -00374384 | -06575 | 037 | 19 |
| 2173 | ENSMUSG( | Eipr1 | 380752 | 007640484 | -00138848 | -023 | 02375 | 19 |
| 2213 | ENSMUSG( | Eml1 | 68519 | 000737199 | -01738014 | -054 | 01075 | 19 |
| 2221 | ENSMUSG( | Enah | 13800 | 018030705 | 001543530 | -0195 | 0165 | 19 |
| 2239 | ENSMUSG( | Ensa | 56205 | 009567829 | -01114371 | -0425 | 00975 | 19 |
| 2245 | ENSMUSG( | Eny2 | 223527 | -00400662 | 003993791 | -04725 | -00875 | 19 |
| 2258 | ENSMUSG( | Epha10 | 230735 | -01481031 | 016385834 | -03925 | 03025 | 19 |
| 2281 | ENSMUSG( | Eral1 | 57837 | 010412286 | 001492648 | -0535 | 03025 | 19 |

| | | | | | | | | |
|---|---|---|---|---|---|---|---|---|
| 2289 | ENSMUSG( | Ercc3 | 13872 | 023113368 | -00003338 | -0265 | 0075 | 19 |
| 2319 | ENSMUSG( | Etnppl | 71760 | 05942135C | 017609641 | -0535 | 0205 | 19 |
| 2323 | ENSMUSG( | Evi5l | 213027 | 00806689C | -00294667 | -016 | 02775 | 19 |
| 2346 | ENSMUSG( | Exph5 | 320051 | -00254158 | -01080558 | -06125 | 037 | 19 |
| 2352 | ENSMUSG( | F11r | 16456 | -02056336 | 041394581 | -0355 | 05275 | 19 |
| 2375 | ENSMUSG( | Fam110b | 242297 | 024298244 | 01268382C | -03275 | 0135 | 19 |
| 2387 | ENSMUSG( | Fam135b | 70363 | -00152427 | 025199478 | -04025 | 00775 | 19 |
| 2388 | ENSMUSG( | Fam136a | 66488 | -00175802 | -01697680 | -02175 | 03475 | 19 |
| 2399 | ENSMUSG( | Fam184b | 58227 | -02979720 | -02048662 | -02425 | 03775 | 19 |
| 2428 | ENSMUSG( | Fastkd2 | 75619 | 00757052C | 002881486 | -05225 | 007 | 19 |
| 2433 | ENSMUSG( | Faxc | 76132 | 012715928 | 01146103C | -039 | -0055 | 19 |
| 2439 | ENSMUSG( | Fbn1 | 14118 | -03288104 | -02067270 | -03575 | 05 | 19 |
| 2463 | ENSMUSG( | Fbxo7 | 69754 | -02964530 | 007471929 | -02975 | 0315 | 19 |
| 2479 | ENSMUSG( | Fdps | 110196 | -00828427 | -03279059 | -03575 | 012 | 19 |
| 2555 | ENSMUSG( | Fndc10 | 230991 | -02838551 | 006920318 | -0285 | 01825 | 19 |
| 2569 | ENSMUSG( | Foxo3 | 56484 | 037156136 | 009762763 | -02175 | 0315 | 19 |
| 2571 | ENSMUSG( | Foxred1 | 235169 | 012176811 | -00819469 | -03875 | 005 | 19 |
| 2578 | ENSMUSG( | Frmd8 | 67457 | -00933247 | -01321417 | -02875 | 02875 | 19 |
| 2594 | ENSMUSG( | Ftsj1 | 54632 | 00461506C | -01209692 | -0605 | 0005 | 19 |
| 2597 | ENSMUSG( | Fubp3 | 320267 | 002347325 | -00236746 | -03825 | 021 | 19 |
| 2622 | ENSMUSG( | G6pdx | 14381 | 009406997 | -01187989 | -0265 | 00325 | 19 |
| 2645 | ENSMUSG( | Gadd45gip | 102060 | 003287601 | -01928412 | -06025 | 0285 | 19 |
| 2660 | ENSMUSG( | Galt | 14430 | 017090997 | 004885131 | -02475 | 027 | 19 |
| 2679 | ENSMUSG( | Gatad2a | 234366 | -00115954 | -00309247 | -04225 | 003 | 19 |
| 2682 | ENSMUSG( | Gatc | 384281 | 011321238 | 004502112 | -03225 | -001 | 19 |
| 2689 | ENSMUSG( | Gbp4 | 17472 | -01291792 | 066188289 | -06425 | 0135 | 19 |
| 2691 | ENSMUSG( | Gbp7 | 229900 | -01210473 | -00588425 | -0305 | 02675 | 19 |
| 2695 | ENSMUSG( | Gcc1 | 74375 | 02410980C | 008391299 | -0375 | 05675 | 19 |
| 2718 | ENSMUSG( | Get4 | 67604 | -00686761 | -00220073 | -04475 | -00675 | 19 |
| 2740 | ENSMUSG( | Ghitm | 66092 | 006202513 | -01609033 | -0505 | 02525 | 19 |
| 2743 | ENSMUSG( | Gigyf1 | 57330 | 011206122 | 002464205 | -0325 | 01275 | 19 |
| 2760 | ENSMUSG( | Glcci1 | 170772 | -00317082 | 027426932 | -0235 | 02075 | 19 |
| 2774 | ENSMUSG( | Glrx5 | 73046 | 001303795 | -02197792 | -0335 | 028 | 19 |
| 2792 | ENSMUSG( | Gmppa | 69080 | 024561111 | 005503734 | -0305 | 0555 | 19 |
| 2847 | ENSMUSG( | Gorasp2 | 70231 | -01997199 | -00573302 | -0255 | 00475 | 19 |
| 2853 | ENSMUSG( | Gpalpp1 | 67467 | 007938827 | 003020705 | -08 | 04975 | 19 |
| 2857 | ENSMUSG( | Gpatch2 | 67769 | 008792964 | -00235361 | -085 | 04575 | 19 |
| 2889 | ENSMUSG( | Gprasp2 | 245607 | 008178883 | 007763717 | -0265 | 046 | 19 |
| 2890 | ENSMUSG( | Gprc5b | 64297 | -00633202 | -01923695 | -022 | 02625 | 19 |
| 2891 | ENSMUSG( | Gprin1 | 26913 | 022095987 | -00147728 | -03325 | 01625 | 19 |
| 2892 | ENSMUSG( | Gprin2 | 432839 | -02586354 | -05736892 | -022 | 03725 | 19 |
| 2894 | ENSMUSG( | Gps1 | 209318 | 004754932 | -00895573 | -064 | 05875 | 19 |
| 2968 | ENSMUSG( | Gstt2 | 14872 | -00248239 | 055859986 | -04725 | 01775 | 19 |
| 2976 | ENSMUSG( | Gtf2f1 | 98053 | 00716731C | -01131915 | -0295 | 0265 | 19 |
| 2977 | ENSMUSG( | Gtf2f2 | 68705 | 005351535 | -00769627 | -03225 | 046 | 19 |
| 2988 | ENSMUSG( | Gtpbp10 | 207704 | 009509588 | -00173211 | -0515 | 02275 | 19 |
| 3005 | ENSMUSG( | H1f10 | 243529 | 007881521 | -01100590 | -05425 | 00575 | 19 |
| 3014 | ENSMUSG( | Hacd2 | 70757 | -01643210 | 016128424 | -0205 | 0355 | 19 |
| 3030 | ENSMUSG( | Hax1 | 23897 | 005863652 | 000909879 | -02375 | 028 | 19 |
| 3035 | ENSMUSG( | Hcfc1 | 15161 | 006090415 | 004953864 | -03225 | 0095 | 19 |
| 3038 | ENSMUSG( | Hcn2 | 15166 | 00081444 | 015538481 | -0315 | 00125 | 19 |
| 3047 | ENSMUSG( | Hdac5 | 15184 | 020530432 | 019213969 | -04925 | -00875 | 19 |
| 3048 | ENSMUSG( | Hdac6 | 15185 | 02094587C | -00283435 | -01975 | 01975 | 19 |
| 3055 | ENSMUSG( | Hdgfl3 | 29877 | 005418989 | 002659958 | -03875 | 009 | 19 |
| 3061 | ENSMUSG( | Heatr3 | 234549 | -00808861 | 009452683 | -045 | -00875 | 19 |
| 3066 | ENSMUSG( | Hebp2 | 56016 | -00891055 | -05714933 | -0635 | -0015 | 19 |
| 3069 | ENSMUSG( | Hectd2 | 226098 | 006846511 | 006763772 | -04075 | 02525 | 19 |
| 3087 | ENSMUSG( | Hgs | 15239 | 019298646 | -00745769 | -02825 | 02225 | 19 |
| 3101 | ENSMUSG( | Hivep2 | 15273 | 009543152 | 016420871 | -0635 | 0465 | 19 |
| 3138 | ENSMUSG( | Hnrnpk | 15387 | -00144577 | -01068938 | -02925 | 00675 | 19 |
| 3165 | ENSMUSG( | Hs3st4 | 628779 | 008055311 | 007750802 | -0295 | -00175 | 19 |
| 3167 | ENSMUSG( | Hsbp1 | 68196 | 006181054 | -01597412 | -06275 | 009 | 19 |
| 3195 | ENSMUSG( | Hspbp1 | 66245 | 004170402 | -01823796 | -02525 | 012 | 19 |
| 3238 | ENSMUSG( | Ift172 | 67661 | 021623708 | -00236425 | -03575 | 00225 | 19 |
| 3240 | ENSMUSG( | Ift27 | 67042 | 009295983 | -01511854 | -0365 | 05075 | 19 |
| 3244 | ENSMUSG( | Ift74 | 67694 | 001591088 | -01189956 | -043 | 055 | 19 |
| 3255 | ENSMUSG( | Igfbp7 | 29817 | -01958779 | 040304697 | -0245 | 035 | 19 |
| 3277 | ENSMUSG( | Il6st | 16195 | -03794826 | 00868886C | -04175 | 046 | 19 |
| 3285 | ENSMUSG( | Immp1l | 66541 | 012169258 | -01361426 | -0465 | 04325 | 19 |
| 3289 | ENSMUSG( | Impa1 | 55980 | -00346335 | -00979711 | -02425 | 005 | 19 |
| 3295 | ENSMUSG( | Inafm2 | 1E+08 | -01419633 | 007816445 | -03925 | 0045 | 19 |
| 3307 | ENSMUSG( | Inpp5j | 170835 | 019380999 | -00276067 | -0285 | 01875 | 19 |
| 3309 | ENSMUSG( | Inppl1 | 16332 | 001346998 | 006064612 | -02425 | 03625 | 19 |
| 3318 | ENSMUSG( | Ints13 | 71177 | 011606273 | 009152152 | -02775 | 02325 | 19 |
| 3320 | ENSMUSG( | Ints15 | 231868 | 001241475 | 001148981 | -0455 | 03125 | 19 |
| 3323 | ENSMUSG( | Ints4 | 101861 | 01516163C | -00334506 | -041 | 03225 | 19 |
| 3324 | ENSMUSG( | Ints5 | 109077 | 015521273 | 007865074 | -02425 | 01725 | 19 |
| 3328 | ENSMUSG( | Intu | 380614 | -00654642 | 017182928 | -05725 | 04575 | 19 |
| 3342 | ENSMUSG( | Iqsec2 | 245666 | 014610244 | 015811723 | -04 | 004 | 19 |
| 3364 | ENSMUSG( | Isy1 | 57905 | 004551587 | -00564958 | -02775 | 02375 | 19 |
| 3368 | ENSMUSG( | Itfg2 | 101142 | 008035512 | -01067943 | -052 | -00575 | 19 |
| 3404 | ENSMUSG( | Jagn1 | 67767 | -03520155 | 012419989 | -0565 | 0115 | 19 |
| 3416 | ENSMUSG( | Josd2 | 66124 | -00194358 | 001203499 | -02 | 0295 | 19 |
| 3428 | ENSMUSG( | Kank2 | 235041 | 005623799 | -01532882 | -02575 | 0235 | 19 |
| 3429 | ENSMUSG( | Kank3 | 80880 | 027522842 | 043250237 | -0375 | 03825 | 19 |
| 3437 | ENSMUSG( | Kazn | 71529 | -00714056 | -00035548 | -05225 | 008 | 19 |
| 3443 | ENSMUSG( | Kcmf1 | 74287 | -00058102 | -00955234 | -032 | 055 | 19 |
| 3482 | ENSMUSG( | Kcnk2 | 16526 | -00676725 | 01844565C | -023 | 01125 | 19 |
| 3512 | ENSMUSG( | Kdm2a | 225876 | 004155195 | 002269087 | -02625 | -157009e-1 | 19 |
| 3552 | ENSMUSG( | Klc4 | 74764 | 00903990C | -00830682 | -04075 | 003 | 19 |
| 3574 | ENSMUSG( | Kndc1 | 76484 | 016732981 | 011311573 | -028 | 0055 | 19 |
| 3577 | ENSMUSG( | Kpna3 | 16648 | 010713323 | -00117366 | -029 | 02925 | 19 |
| 3601 | ENSMUSG( | Lama2 | 16773 | -01407352 | -00137481 | -04525 | -00625 | 19 |
| 3686 | ENSMUSG( | Lipt2 | 67164 | 00136728C | -01622213 | -0485 | 02025 | 19 |
| 3696 | ENSMUSG( | Lmf1 | 76483 | -01815413 | 012723927 | -01375 | 0195 | 19 |
| 3702 | ENSMUSG( | Lmo4 | 16911 | 000047227 | -00638327 | -04025 | 0235 | 19 |
| 3732 | ENSMUSG( | Lrig1 | 16206 | -01381616 | 017238433 | -02875 | 00475 | 19 |
| 3738 | ENSMUSG( | Lrp8 | 16975 | -00271703 | -00690374 | -03875 | -003 | 19 |
| 3741 | ENSMUSG( | Lrrc1 | 214345 | 000955747 | -00426405 | -033 | -0015 | 19 |
| 3795 | ENSMUSG( | Luc7l3 | 67684 | -01203004 | 02728448C | -0265 | 0215 | 19 |
| 3843 | ENSMUSG( | Maged1 | 94275 | -01001113 | -00042931 | -01475 | 02725 | 19 |
| 3854 | ENSMUSG( | Malsu1 | 75593 | 002878869 | -01193004 | -0235 | 02725 | 19 |
| 3870 | ENSMUSG( | Map1b | 17755 | 004823164 | 00613715C | -03225 | 00675 | 19 |
| 3873 | ENSMUSG( | Map1s | 270058 | 020726265 | 011965276 | -03525 | 02675 | 19 |
| 3883 | ENSMUSG( | Map3k11 | 26403 | -01003976 | 005953234 | -02775 | 014 | 19 |
| 3885 | ENSMUSG( | Map3k13 | 71751 | -00061115 | 004085733 | -05375 | 02975 | 19 |
| 3892 | ENSMUSG( | Map4 | 17758 | -00030030 | -02439199 | -021 | 03225 | 19 |
| 3898 | ENSMUSG( | Map6d1 | 208158 | -01104566 | -02703787 | -04775 | -00325 | 19 |
| 3899 | ENSMUSG( | Map7 | 17761 | -00211819 | 011017032 | -0135 | 025 | 19 |
| 3900 | ENSMUSG( | Map7d1 | 245877 | 009072125 | 004681767 | -0235 | 02775 | 19 |
| 3902 | ENSMUSG( | Map9 | 213582 | 015496811 | 003781914 | -034 | 012 | 19 |
| 3913 | ENSMUSG( | Mapk8ip2 | 60597 | -00242217 | 012964811 | -03775 | 03825 | 19 |
| 3917 | ENSMUSG( | Mapkapk2 | 17164 | -00419327 | -01178209 | -028 | 047 | 19 |
| 3918 | ENSMUSG( | Mapkbp1 | 26390 | 019849126 | 008046636 | -036 | 03025 | 19 |
| 3921 | ENSMUSG( | Mapre3 | 100732 | -00237819 | -01354709 | -046 | 05025 | 19 |
| 3939 | ENSMUSG( | Mast3 | 546071 | 018598399 | -00445776 | -036 | 0105 | 19 |
| 3976 | ENSMUSG( | Mcrip1 | 192173 | 003306529 | -01723847 | -02575 | 01425 | 19 |
| 3981 | ENSMUSG( | Mcu | 215999 | 005921678 | -00223470 | -0215 | 0145 | 19 |
| 3985 | ENSMUSG( | Mdga1 | 74762 | 008654932 | 024951648 | -02375 | 03075 | 19 |
| 3990 | ENSMUSG( | Mdm1 | 17245 | 018827519 | 019316988 | -02625 | 04975 | 19 |
| 4008 | ENSMUSG( | Med17 | 234959 | 017486878 | 002889871 | -044 | 046 | 19 |
| 4010 | ENSMUSG( | Med22 | 20933 | 005541613 | -01563437 | -04475 | 058 | 19 |
| 4014 | ENSMUSG( | Med29 | 67224 | 033574485 | 007437812 | -06225 | 0025 | 19 |
| 4053 | ENSMUSG( | Mfhas1 | 52065 | 019896375 | 015025501 | -0395 | 037 | 19 |
| 4058 | ENSMUSG( | Mfsd11 | 69900 | -01791421 | 028064625 | -09275 | 02575 | 19 |
| 4094 | ENSMUSG( | Mindy2 | 235461 | -00348922 | -01296071 | -03125 | 00625 | 19 |

| | | | | | | | | |
|---|---|---|---|---|---|---|---|---|
| 4139 | ENSMUSG( | Mov10 | 17454 | 02606106( | 001740262 | -0445 | 00475 | 19 |
| 4144 | ENSMUSG( | Mpdz | 17475 | 00908249( | 002070818 | -04675 | 00075 | 19 |
| 4187 | ENSMUSG( | Mrpl30 | 107734 | 013252345 | -0115951 | -01975 | 036 | 19 |
| 4236 | ENSMUSG( | Mrps35 | 232536 | 011409995 | -0053444 | -042 | 01175 | 19 |
| 4244 | ENSMUSG( | Mrtfa | 223701 | 039267114 | 016771692 | -03425 | 01525 | 19 |
| 4245 | ENSMUSG( | Mrtfb | 239719 | 011558616 | 007535643 | -03625 | 027 | 19 |
| 4248 | ENSMUSG( | Msh6 | 17688 | 023344067 | 011353386 | -0275 | 02725 | 19 |
| 4267 | ENSMUSG( | Mtcl1 | 68617 | -0193708 | -01862779 | -05125 | 018 | 19 |
| 4276 | ENSMUSG( | Mthfd2l | 665563 | -0007360 | -0023671 | -04975 | -004 | 19 |
| 4299 | ENSMUSG( | Mtrr | 210009 | 023923667 | 013042308 | -06675 | 05875 | 19 |
| 4303 | ENSMUSG( | Mtus2 | 77521 | 022196675 | 014646321 | -0405 | 02175 | 19 |
| 4304 | ENSMUSG( | Mtx1 | 17827 | -0001258 | -00441998 | -022 | 01225 | 19 |
| 4308 | ENSMUSG( | Mvb12a | 73711 | -0093427 | 003639768 | -04025 | 0265 | 19 |
| 4345 | ENSMUSG( | Myo9b | 17925 | 017870994 | 004510125 | -03375 | 019 | 19 |
| 4353 | ENSMUSG( | Mzt2 | 72083 | -0007461 | -0289809 | -0425 | 02825 | 19 |
| 4354 | ENSMUSG( | N4bp1 | 80750 | 00824330( | 008379779 | -0315 | 007 | 19 |
| 4356 | ENSMUSG( | N4bp3 | 212706 | 005893958 | 009454994 | -03925 | -006 | 19 |
| 4360 | ENSMUSG( | Naa20 | 67877 | -0025525 | 004228341 | -0355 | 0185 | 19 |
| 4368 | ENSMUSG( | Nab2 | 17937 | -02617499 | -0351307 | -04275 | 009 | 19 |
| 4372 | ENSMUSG( | Nacc1 | 66830 | 013424384 | -0013183 | -0885 | 051 | 19 |
| 4395 | ENSMUSG( | Nat1 | 17960 | -0141696 | -0281558 | -036 | 009 | 19 |
| 4404 | ENSMUSG( | Nav3 | 260315 | 010103762 | 001367413 | -02175 | 034 | 19 |
| 4408 | ENSMUSG( | Nbdy | 70123 | 007549449 | -0116453 | -0425 | 049 | 19 |
| 4417 | ENSMUSG( | Ncbp2 | 68092 | 005080814 | -00727283 | -0295 | 028 | 19 |
| 4435 | ENSMUSG( | Ncor1 | 20185 | 012046255 | 002742795 | -04925 | 04325 | 19 |
| 4436 | ENSMUSG( | Ncor2 | 20602 | -0031813 | 015268141 | -01925 | 03625 | 19 |
| 4474 | ENSMUSG( | Ndufb10 | 68342 | -0005537 | -0190560 | -03975 | 00525 | 19 |
| 4495 | ENSMUSG( | Ndufv3 | 78330 | 000680169 | -02375582 | -074 | 00625 | 19 |
| 4499 | ENSMUSG( | Necab3 | 56846 | 005365619 | -0049236 | -0285 | 02075 | 19 |
| 4507 | ENSMUSG( | Nedd8 | 18002 | -0014971 | -0336735 | -0325 | 029 | 19 |
| 4520 | ENSMUSG( | Nelfe | 27632 | 00818462( | -01514793 | -063 | 025 | 19 |
| 4525 | ENSMUSG( | Nes | 18008 | 007434613 | 050505822 | -0575 | 04075 | 19 |
| 4532 | ENSMUSG( | Neurl4 | 216860 | 013389596 | 004333576 | -03325 | 01825 | 19 |
| 4564 | ENSMUSG( | Nifk | 67949 | 008152017 | -0189979 | -03575 | 01775 | 19 |
| 4596 | ENSMUSG( | Noa1 | 56412 | 015102176 | 001793661 | -03175 | 03475 | 19 |
| 4602 | ENSMUSG( | Nol10 | 217431 | 014719731 | -0039771 | -03025 | 0225 | 19 |
| 4610 | ENSMUSG( | Nolc1 | 70769 | 011602133 | -0041608 | -0605 | 00225 | 19 |
| 4621 | ENSMUSG( | Nopchap1 | 28109 | 010547643 | -0076283 | -0645 | 038 | 19 |
| 4629 | ENSMUSG( | Nova1 | 664883 | 007404973 | -0020352 | -02275 | 025 | 19 |
| 4632 | ENSMUSG( | Npc1l1 | 237636 | 018651585 | -0049898 | -03425 | 00975 | 19 |
| 4634 | ENSMUSG( | Npdc1 | 18146 | -0194796 | 003108882 | -01875 | 0145 | 19 |
| 4645 | ENSMUSG( | Nprl3 | 17168 | 000887734 | 018714925 | -036 | -00375 | 19 |
| 4658 | ENSMUSG( | Nr3c1 | 14815 | -00342471 | 001615484 | -029 | 02 | 19 |
| 4667 | ENSMUSG( | Nrgn | 64011 | 009103987 | -02003686 | -0545 | 01975 | 19 |
| 4674 | ENSMUSG( | Nrsn1 | 22360 | 022234508 | -0017118 | -053 | 06 | 19 |
| 4678 | ENSMUSG( | Nsd1 | 18193 | -00597543 | 008935291 | -01425 | 0275 | 19 |
| 4680 | ENSMUSG( | Nsd3 | 234135 | -0009358 | 011644057 | -04075 | 01 | 19 |
| 4684 | ENSMUSG( | Nsg1 | 18196 | -04398210 | 008486317 | -0685 | 0395 | 19 |
| 4701 | ENSMUSG( | Ntaq1 | 76773 | -00621666 | -01052432 | -04475 | 0215 | 19 |
| 4710 | ENSMUSG( | Ntsr2 | 18217 | 001566428 | -0026689 | -071 | 022 | 19 |
| 4713 | ENSMUSG( | Nubp1 | 26425 | -0059943 | -0094519 | -03325 | 0115 | 19 |
| 4727 | ENSMUSG( | Nudt15 | 214254 | -0027180 | -01588500 | -046 | 0105 | 19 |
| 4736 | ENSMUSG( | Nudt4 | 71207 | 00590873( | -00282407 | -0225 | 011 | 19 |
| 4789 | ENSMUSG( | Ogfr | 72075 | 008884195 | -00277714 | -02525 | 01025 | 19 |
| 4813 | ENSMUSG( | Ormdl3 | 66612 | -01458629 | 003738386 | -03475 | 02875 | 19 |
| 4839 | ENSMUSG( | Otud7b | 229603 | -0005234 | -0030300 | -032 | -0005 | 19 |
| 4873 | ENSMUSG( | Pak1ip1 | 68083 | 009914692 | -0012377 | -0315 | 008 | 19 |
| 4885 | ENSMUSG( | Pals1 | 56217 | 007965478 | 008806362 | -0345 | -001 | 19 |
| 4886 | ENSMUSG( | Pals2 | 56524 | 01988677( | -0121595 | -03525 | 05025 | 19 |
| 4889 | ENSMUSG( | Pan2 | 103135 | 013569785 | 011193652 | -026 | 01475 | 19 |
| 4890 | ENSMUSG( | Pan3 | 72587 | 007996238 | 018661087 | -03175 | 02375 | 19 |
| 4910 | ENSMUSG( | Parn | 74108 | -0029783 | -0114077 | -025 | 026 | 19 |
| 4917 | ENSMUSG( | Patz1 | 56218 | 003249102 | 001075412 | -0625 | 00825 | 19 |
| 4921 | ENSMUSG( | Pbdc1 | 67683 | 017921062 | -0189687 | -02675 | 02425 | 19 |
| 4930 | ENSMUSG( | Pcbp4 | 59092 | 018690418 | 00856272( | -0475 | 027 | 19 |
| 4963 | ENSMUSG( | Pcdhgc3 | 93706 | 003339958 | -00790233 | -0275 | 01025 | 19 |
| 4965 | ENSMUSG( | Pcf11 | 74737 | -0171538 | 013372768 | -04425 | 039 | 19 |
| 4967 | ENSMUSG( | Pcid2 | 234069 | 019578574 | -0018019 | -0235 | 01225 | 19 |
| 4977 | ENSMUSG( | Pcnt | 18541 | 011405383 | 02269515( | -033 | 0295 | 19 |
| 5006 | ENSMUSG( | Pde1c | 18575 | -0056580 | -0279315 | -058 | 05525 | 19 |
| 5012 | ENSMUSG( | Pde4dip | 83679 | 015639925 | 006508412 | -071 | 02575 | 19 |
| 5021 | ENSMUSG( | Pdgfra | 18595 | -0372991 | 028242787 | -02725 | 0365 | 19 |
| 5025 | ENSMUSG( | Pdhx | 27402 | 012914888 | -0154493 | -02775 | 00775 | 19 |
| 5043 | ENSMUSG( | Pds5a | 71521 | 011915273 | 010850623 | -0565 | 03425 | 19 |
| 5054 | ENSMUSG( | Peak1 | 244895 | -00594532 | 011515788 | -02275 | 00375 | 19 |
| 5075 | ENSMUSG( | Pex2 | 19302 | -0025935 | 004406706 | -044 | 009 | 19 |
| 5106 | ENSMUSG( | Pgm2 | 66681 | 000894227 | 007835872 | -02525 | 00425 | 19 |
| 5124 | ENSMUSG( | Phc2 | 54383 | 002227315 | -0031314 | -02025 | 03775 | 19 |
| 5127 | ENSMUSG( | Phf2 | 18676 | 011335313 | 006987588 | -0355 | 04125 | 19 |
| 5144 | ENSMUSG( | Phldb2 | 208177 | -02120193 | 022045816 | -03675 | 024 | 19 |
| 5151 | ENSMUSG( | Phyh | 16922 | 002837129 | -0239069 | -02575 | 04325 | 19 |
| 5158 | ENSMUSG( | Pianp | 319352 | -0413665 | 009378991 | -052 | 0455 | 19 |
| 5163 | ENSMUSG( | Pigb | 55981 | -0147837 | 024112866 | -06525 | 03925 | 19 |
| 5195 | ENSMUSG( | Pithd1 | 66193 | 002913838 | -0049832 | -02675 | 0105 | 19 |
| 5210 | ENSMUSG( | Pkn1 | 320795 | 005967946 | -0075898 | -02275 | 019 | 19 |
| 5219 | ENSMUSG( | Plaat3 | 225845 | -0025357 | -0194741 | -02025 | 03475 | 19 |
| 5241 | ENSMUSG( | Plekha2 | 83436 | -0143316 | -0079066 | -037 | 01175 | 19 |
| 5248 | ENSMUSG( | Plekhg1 | 213783 | 023075512 | 021555311 | -05625 | 0155 | 19 |
| 5254 | ENSMUSG( | Plekho2 | 102595 | -0214053 | 013276883 | -02625 | 0155 | 19 |
| 5263 | ENSMUSG( | Plpp1 | 19012 | -0298079 | 006245125 | -018 | 0285 | 19 |
| 5267 | ENSMUSG( | Plpp7 | 227721 | -03528544 | 014920649 | -02825 | 00975 | 19 |
| 5270 | ENSMUSG( | Plppr3 | 216152 | -0314287 | -0023169 | -01925 | 04075 | 19 |
| 5276 | ENSMUSG( | Plscr3 | 70310 | -0044652 | -0028170 | -03025 | 02075 | 19 |
| 5303 | ENSMUSG( | Pnma8b | 434128 | 015885679 | -0046946 | -04625 | 01525 | 19 |
| 5308 | ENSMUSG( | Pnpla2 | 66853 | 018782112 | -0052941 | -04725 | 008 | 19 |
| 5324 | ENSMUSG( | Poldip3 | 73826 | 00274596( | -0040089 | -04125 | 01275 | 19 |
| 5325 | ENSMUSG( | Polg | 18975 | 00822235( | 014284562 | -03 | 0425 | 19 |
| 5327 | ENSMUSG( | Polr1c | 20016 | -0040646 | -0111159 | -04275 | 013 | 19 |
| 5328 | ENSMUSG( | Polr1d | 20018 | 004118263 | -0090970 | -07375 | 025 | 19 |
| 5351 | ENSMUSG( | Pomt2 | 217734 | -0117845 | 009141916 | -02675 | 02725 | 19 |
| 5379 | ENSMUSG( | Ppie | 56031 | 004926029 | -0004820 | -06875 | 00475 | 19 |
| 5387 | ENSMUSG( | Ppip5k1 | 327655 | 021403488 | 000669504 | -026 | 0025 | 19 |
| 5404 | ENSMUSG( | Ppp1r11 | 76497 | -0063510 | -02678818 | -03 | 0185 | 19 |
| 5405 | ENSMUSG( | Ppp1r12a | 17931 | 001985402 | 007458925 | -01925 | 0155 | 19 |
| 5406 | ENSMUSG( | Ppp1r12b | 329251 | 017619659 | 013721317 | -02125 | 0265 | 19 |
| 5408 | ENSMUSG( | Ppp1r13b | 21981 | 015315361 | 017347562 | -03275 | 023 | 19 |
| 5414 | ENSMUSG( | Ppp1r18 | 76448 | -0020651 | -00361871 | -05625 | 0175 | 19 |
| 5415 | ENSMUSG( | Ppp1r1a | 58200 | 002123274 | -0140659 | -026 | 0255 | 19 |
| 5450 | ENSMUSG( | Ppp6c | 67857 | -0041572 | -0071797 | -017 | 0265 | 19 |
| 5456 | ENSMUSG( | Ppwd1 | 238831 | -0052239 | 01876136( | -0385 | 02425 | 19 |
| 5457 | ENSMUSG( | Pqbp1 | 54633 | 002441524 | -01934597 | -0355 | 0065 | 19 |
| 5483 | ENSMUSG( | Prkaca | 18747 | 000217788 | -0108529 | -022 | 02475 | 19 |
| 5528 | ENSMUSG( | Prpf40b | 54614 | 019796746 | -0007349 | -068 | 03925 | 19 |
| 5532 | ENSMUSG( | Prph | 19132 | 026513069 | -0065956 | -02725 | 008 | 19 |
| 5538 | ENSMUSG( | Prr14 | 233895 | -0032383 | 008222794 | -0315 | 0315 | 19 |
| 5539 | ENSMUSG( | Prr18 | 320111 | -02048898 | -0392979 | -08225 | 05675 | 19 |
| 5541 | ENSMUSG( | Prr36 | 73072 | 01872576( | -0060854 | -042 | 0385 | 19 |
| 5543 | ENSMUSG( | Prrc1 | 73137 | 001175732 | -0056748 | -01975 | 0175 | 19 |
| 5548 | ENSMUSG( | Prrt2 | 69017 | -0186390 | 006448022 | -04475 | -00725 | 19 |
| 5549 | ENSMUSG( | Prrt3 | 210673 | -0277215 | 014886656 | -06725 | 004 | 19 |
| 5569 | ENSMUSG( | Psmb10 | 19171 | -0084094 | -0158470 | -05275 | -0005 | 19 |
| 5574 | ENSMUSG( | Psmb6 | 19175 | -0036372 | -0229785 | -034 | 0115 | 19 |
| 5575 | ENSMUSG( | Psmb7 | 19177 | 004331773 | -01766375 | -021 | 029 | 19 |
| 5641 | ENSMUSG( | Ptpdc1 | 218232 | -0086264 | 011281232 | -04225 | 0455 | 19 |
| 5645 | ENSMUSG( | Ptpn12 | 19248 | 008729453 | 002881227 | -02075 | 03575 | 19 |
| 5652 | ENSMUSG( | Ptpn6 | 15170 | -0077196 | 007056347 | -03075 | 016 | 19 |

| | | | | | | | | |
|---|---|---|---|---|---|---|---|---|
| 5674 | ENSMUSG( | Pttg1ip | 108705 | -01356419 | 011934913 | -04225 | 04 | 19 |
| 5687 | ENSMUSG( | Pwp2 | 110816 | -00818567 | -00646821 | -0955 | 05525 | 19 |
| 5693 | ENSMUSG( | Pxn | 19303 | 015653169 | -00431125 | -02125 | 02275 | 19 |
| 5694 | ENSMUSG( | Pycard | 66824 | -00422165 | -02495072 | -02575 | 0125 | 19 |
| 5695 | ENSMUSG( | Pycr1 | 209027 | -02284706 | -04498123 | -061 | 04575 | 19 |
| 5697 | ENSMUSG( | Pycr3 | 66194 | 005531279 | -00492486 | -0345 | 01975 | 19 |
| 5719 | ENSMUSG( | R3hdm2 | 71750 | 013226178 | 000548334 | -0385 | 03825 | 19 |
| 5722 | ENSMUSG( | Rab11fip2 | 74998 | -00414934 | 003451935 | -0265 | 0095 | 19 |
| 5725 | ENSMUSG( | Rab11fip5 | 52055 | 01396802 | 004255065 | -02425 | 0315 | 19 |
| 5744 | ENSMUSG( | Rab30 | 75985 | -00839946 | 004106159 | -031 | 0075 | 19 |
| 5793 | ENSMUSG( | Rad23b | 19359 | 010108006 | -00673462 | -038 | 00525 | 19 |
| 5810 | ENSMUSG( | Ramac | 67148 | -00914259 | -02842415 | -039 | 0495 | 19 |
| 5828 | ENSMUSG( | Rapgef1 | 107746 | 020833993 | 009883810 | -027 | 0405 | 19 |
| 5848 | ENSMUSG( | Rasgrp1 | 19419 | 001797114 | 002890029 | -04575 | 0275 | 19 |
| 5849 | ENSMUSG( | Rasgrp2 | 19395 | -03623822 | -01323126 | -0205 | 04075 | 19 |
| 5868 | ENSMUSG( | Rbm12 | 75710 | 025035626 | -00150740 | -0295 | 0305 | 19 |
| 5880 | ENSMUSG( | Rbm3 | 19652 | -02076514 | 013514807 | -02375 | 01575 | 19 |
| 5886 | ENSMUSG( | Rbm6 | 19654 | 021953880 | 007742902 | -01975 | 0215 | 19 |
| 5887 | ENSMUSG( | Rbm7 | 67010 | -00747390 | 005506427 | -06025 | 01175 | 19 |
| 5901 | ENSMUSG( | Rcbtb2 | 105670 | -00078319 | -01234734 | -02275 | 02725 | 19 |
| 5909 | ENSMUSG( | Rcsd1 | 226594 | -00391506 | -00392458 | -07975 | 05425 | 19 |
| 5911 | ENSMUSG( | Rdh13 | 108841 | 013938551 | 018628806 | -03275 | -001 | 19 |
| 5917 | ENSMUSG( | Reep3 | 28193 | -01837914 | 011358568 | -03 | 0065 | 19 |
| 5918 | ENSMUSG( | Reep5 | 13476 | -02659035 | 001215506 | -0145 | 02975 | 19 |
| 5924 | ENSMUSG( | Rell2 | 225392 | 007602757 | 006449371 | -049 | 02525 | 19 |
| 5927 | ENSMUSG( | Renbp | 19703 | -00092591 | -02590580 | -05675 | 02825 | 19 |
| 5929 | ENSMUSG( | Reps2 | 194590 | 013580655 | -00003261 | -0265 | 00575 | 19 |
| 5943 | ENSMUSG( | Rft1 | 328370 | -03281019 | 006699315 | -06175 | 04725 | 19 |
| 5950 | ENSMUSG( | Rgl2 | 19732 | 035493544 | 010794598 | -033 | 00575 | 19 |
| 5960 | ENSMUSG( | Rgs3 | 50780 | 004353854 | 009554023 | -03175 | 02125 | 19 |
| 5961 | ENSMUSG( | Rgs4 | 19736 | -01743283 | -03250610 | -046 | 0315 | 19 |
| 5968 | ENSMUSG( | Rhbdf1 | 13650 | 004076719 | 03344511 | -03425 | 021 | 19 |
| 5969 | ENSMUSG( | Rhbdl3 | 246104 | 000624853 | 025750114 | -065 | 0165 | 19 |
| 5982 | ENSMUSG( | Ric1 | 226089 | -00282074 | 008044734 | -0285 | 02175 | 19 |
| 5985 | ENSMUSG( | Ric8b | 237422 | -00432446 | -00621274 | -01625 | 02875 | 19 |
| 5988 | ENSMUSG( | Rif1 | 51869 | -00043651 | 021137602 | -06025 | 0315 | 19 |
| 6006 | ENSMUSG( | Rlbp1 | 19771 | 008752683 | -01725260 | -04975 | -005 | 19 |
| 6028 | ENSMUSG( | Rnf115 | 67845 | -00067670 | 005982614 | -09425 | 0325 | 19 |
| 6041 | ENSMUSG( | Rnf170 | 77733 | -01247624 | 011401601 | -02075 | 00225 | 19 |
| 6043 | ENSMUSG( | Rnf185 | 193670 | -00073244 | -01082398 | -0335 | 01025 | 19 |
| 6046 | ENSMUSG( | Rnf213 | 672511 | -02396247 | -01717306 | -02925 | 02875 | 19 |
| 6048 | ENSMUSG( | Rnf216 | 108086 | 018061083 | 010816855 | -058 | 02625 | 19 |
| 6055 | ENSMUSG( | Rnf41 | 67588 | 011932791 | 002622940 | -07025 | 019 | 19 |
| 6059 | ENSMUSG( | Rngtt | 24018 | 009791650 | 004353346 | -059 | 01825 | 19 |
| 6075 | ENSMUSG( | Rpap1 | 68925 | 017729741 | 008003836 | -02975 | 01025 | 19 |
| 6135 | ENSMUSG( | Rprd1b | 70470 | 008116487 | 006799810 | -03525 | 0335 | 19 |
| 6136 | ENSMUSG( | Rprd2 | 75137 | 009542667 | 013947784 | -04 | 0045 | 19 |
| 6176 | ENSMUSG( | Rpusd4 | 71989 | 010500991 | -00149485 | -025 | 00475 | 19 |
| 6186 | ENSMUSG( | Rrp1 | 18114 | 000603601 | -01945130 | -06525 | 02 | 19 |
| 6191 | ENSMUSG( | Rsf1 | 233532 | 006826476 | 002763540 | -028 | 0115 | 19 |
| 6200 | ENSMUSG( | Rtf1 | 76246 | 009087770 | 00194955 | -09 | 04525 | 19 |
| 6201 | ENSMUSG( | Rtkn | 20166 | 004001825 | -00415175 | -0355 | 0205 | 19 |
| 6212 | ENSMUSG( | Rubcn | 1.01E+08 | 013344241 | -00423500 | -028 | 03025 | 19 |
| 6219 | ENSMUSG( | Rusc1 | 72296 | 018518285 | -00741590 | -0335 | 049 | 19 |
| 6226 | ENSMUSG( | Rxrb | 20182 | 006584092 | -00145075 | -036 | 0095 | 19 |
| 6247 | ENSMUSG( | Sall2 | 50524 | -00761175 | -02124991 | -061 | 02975 | 19 |
| 6261 | ENSMUSG( | Sar1a | 20224 | 001771054 | -00827775 | -04675 | 04425 | 19 |
| 6305 | ENSMUSG( | Scn2b | 72821 | -01848133 | 014217764 | -05525 | 05625 | 19 |
| 6313 | ENSMUSG( | Scoc | 56367 | -00160790 | -00190114 | -04975 | 02225 | 19 |
| 6327 | ENSMUSG( | Sdc2 | 15529 | -02902767 | 010313813 | -036 | -00225 | 19 |
| 6336 | ENSMUSG( | Sdhaf1 | 68332 | -00893744 | -02695659 | -032 | 0195 | 19 |
| 6337 | ENSMUSG( | Sdhaf2 | 66072 | 005449177 | -01459878 | -057 | 001 | 19 |
| 6338 | ENSMUSG( | Sdhaf3 | 71238 | -00649014 | -02741779 | -0805 | 0235 | 19 |
| 6368 | ENSMUSG( | Secisbp2l | 70354 | -00328474 | -00368251 | -0305 | 029 | 19 |
| 6442 | ENSMUSG( | Sf1 | 22668 | 002940151 | 002570343 | -04875 | 0515 | 19 |
| 6456 | ENSMUSG( | Sfswap | 231769 | 014040367 | 013254524 | -01975 | 0345 | 19 |
| 6464 | ENSMUSG( | Sgce | 20392 | -03461282 | 011153316 | -0365 | 013 | 19 |
| 6469 | ENSMUSG( | Sgpp1 | 81535 | -02838900 | 026442254 | -02975 | 0245 | 19 |
| 6473 | ENSMUSG( | Sgsm3 | 105835 | 022390505 | -00742792 | -02575 | 028 | 19 |
| 6495 | ENSMUSG( | Sh3pxd2b | 268396 | -00106534 | -00410390 | -0235 | 02575 | 19 |
| 6515 | ENSMUSG( | Shmt2 | 108037 | 003039794 | -00768183 | -0325 | 00525 | 19 |
| 6542 | ENSMUSG( | Sirt5 | 68346 | -01845979 | -03004902 | -0275 | 00675 | 19 |
| 6544 | ENSMUSG( | Skp2 | 27401 | -02422384 | 002046403 | -0255 | 044 | 19 |
| 6552 | ENSMUSG( | Slc12a6 | 107723 | -02177269 | 007801717 | -02675 | -003 | 19 |
| 6576 | ENSMUSG( | Slc1a4 | 55963 | -03421536 | 012986664 | -04325 | 02325 | 19 |
| 6580 | ENSMUSG( | Slc20a2 | 20516 | -01779605 | -00523455 | -0635 | 007 | 19 |
| 6630 | ENSMUSG( | Slc29a2 | 13340 | -02979137 | 015524032 | -024 | 02775 | 19 |
| 6647 | ENSMUSG( | Slc35a5 | 74102 | -03030747 | 017055306 | -0675 | 03225 | 19 |
| 6663 | ENSMUSG( | Slc39a11 | 69806 | -02397927 | 004257127 | -07125 | 0455 | 19 |
| 6672 | ENSMUSG( | Slc44a2 | 68682 | -01923074 | 009530663 | -036 | 05125 | 19 |
| 6673 | ENSMUSG( | Slc45a1 | 242773 | -02420237 | 026668485 | -0425 | 0325 | 19 |
| 6674 | ENSMUSG( | Slc45a4 | 106068 | 004342379 | 028787052 | -04325 | -00625 | 19 |
| 6676 | ENSMUSG( | Slc49a4 | 224132 | -01815273 | 005659091 | -04925 | 04325 | 19 |
| 6685 | ENSMUSG( | Slc52a2 | 52710 | -01712966 | 013807702 | -016 | 021 | 19 |
| 6687 | ENSMUSG( | Slc5a6 | 330064 | -01518629 | 026730586 | -0225 | 01575 | 19 |
| 6693 | ENSMUSG( | Slc6a13 | 14412 | 006045377 | -03118096 | -0745 | 05825 | 19 |
| 6697 | ENSMUSG( | Slc6a6 | 21366 | -02839675 | 050644553 | -038 | 0345 | 19 |
| 6710 | ENSMUSG( | Slc7a6os | 66432 | -00336099 | 014177781 | -0525 | 04325 | 19 |
| 6716 | ENSMUSG( | Slc9a5 | 277973 | -00549098 | 018432239 | -05725 | 012 | 19 |
| 6717 | ENSMUSG( | Slc9a6 | 236794 | -03086030 | 013973470 | -02525 | 0185 | 19 |
| 6718 | ENSMUSG( | Slc9a7 | 236727 | 000389964 | 004060702 | -027 | 002 | 19 |
| 6722 | ENSMUSG( | Slco1c1 | 58807 | -01798153 | 035950980 | -031 | 0065 | 19 |
| 6724 | ENSMUSG( | Slco3a1 | 108116 | -02339051 | 044903492 | -03025 | 03125 | 19 |
| 6744 | ENSMUSG( | Smarca1 | 93761 | 014543887 | 015227757 | -03525 | 023 | 19 |
| 6751 | ENSMUSG( | Smarcc2 | 68094 | 005153952 | -01009872 | -0305 | 002 | 19 |
| 6753 | ENSMUSG( | Smarcd3 | 66993 | 003598125 | 003890612 | -03625 | 03075 | 19 |
| 6754 | ENSMUSG( | Smarce1 | 57376 | -00121228 | -01500647 | -04575 | 01225 | 19 |
| 6773 | ENSMUSG( | Smim7 | 66818 | -01009327 | -00517786 | -0225 | 039 | 19 |
| 6777 | ENSMUSG( | Smoc1 | 64075 | -03606049 | 029399128 | -066 | 00425 | 19 |
| 6833 | ENSMUSG( | Snx15 | 69024 | -01442944 | -01535826 | -034 | 0195 | 19 |
| 6838 | ENSMUSG( | Snx2 | 67804 | 014649367 | 003655248 | -021 | 014 | 19 |
| 6872 | ENSMUSG( | Sowahc | 268301 | -00245875 | 001025149 | -0465 | 001 | 19 |
| 6879 | ENSMUSG( | Spag7 | 216873 | 003441173 | -00611809 | -03125 | 02675 | 19 |
| 6884 | ENSMUSG( | Spast | 50850 | -00470629 | 012712393 | -01775 | 02075 | 19 |
| 6894 | ENSMUSG( | Specc1l | 74392 | 008274199 | -00663231 | -028 | 0085 | 19 |
| 6895 | ENSMUSG( | Speg | 11790 | 006420333 | 002249690 | -02525 | 02925 | 19 |
| 6896 | ENSMUSG( | Spen | 56381 | 001480624 | 013002037 | -04175 | 00125 | 19 |
| 6913 | ENSMUSG( | Sppl2b | 73218 | -01359403 | 018182353 | -02 | 02975 | 19 |
| 6933 | ENSMUSG( | Sra1 | 24068 | 004749859 | -01227819 | -057 | 02975 | 19 |
| 6947 | ENSMUSG( | Srp68 | 217337 | 014907784 | -00327454 | -02275 | 01675 | 19 |
| 6957 | ENSMUSG( | Srrm3 | 58212 | 022340569 | 003545349 | -03675 | 0085 | 19 |
| 6958 | ENSMUSG( | Srrt | 83701 | 023070118 | 006539567 | -02625 | 013 | 19 |
| 6972 | ENSMUSG( | Ss18l1 | 269397 | 012608868 | -00124263 | -03275 | 01425 | 19 |
| 7001 | ENSMUSG( | Stam2 | 56324 | -00434496 | 024012629 | -039 | -00675 | 19 |
| 7014 | ENSMUSG( | Stau1 | 20853 | 000622186 | -00115994 | -0225 | 0355 | 19 |
| 7018 | ENSMUSG( | Steep1 | 77644 | 007623552 | -01236400 | -0785 | 0495 | 19 |
| 7089 | ENSMUSG( | Sumo1 | 22218 | 013421438 | 007179831 | -032 | 00975 | 19 |
| 7090 | ENSMUSG( | Sumo2 | 170930 | 010963784 | -00088442 | -01925 | 02225 | 19 |
| 7097 | ENSMUSG( | Surf1 | 20930 | -01141995 | -00977910 | -02375 | 00975 | 19 |
| 7112 | ENSMUSG( | Syap1 | 67043 | -00150191 | -00006972 | -02825 | -001 | 19 |
| 7113 | ENSMUSG( | Sybu | 319613 | 005658909 | -00810035 | -0165 | 03075 | 19 |
| 7129 | ENSMUSG( | Synj2 | 20975 | 012220734 | -00672544 | -052 | -00575 | 19 |
| 7131 | ENSMUSG( | Synm | 233335 | 015040102 | 015670741 | -03475 | 03325 | 19 |
| 7156 | ENSMUSG( | Szrd1 | 213491 | 003454220 | -02344645 | -0955 | 02075 | 19 |
| 7159 | ENSMUSG( | Tab2 | 68652 | 004964837 | 006488054 | -033 | 02725 | 19 |
| 7162 | ENSMUSG( | Tacc2 | 57752 | 010290943 | 020715459 | -042 | 008 | 19 |

| | | | | | | | | |
|---|---|---|---|---|---|---|---|---|
| 7173 | ENSMUSG( | Taf6 | 21343 | 017312830 | 003958879 | -0305 | 0035 | 19 |
| 7175 | ENSMUSG( | Taf8 | 63856 | 006408235 | 001372747 | -022 | 03675 | 19 |
| 7184 | ENSMUSG( | Tamalin | 56149 | 000502176 | -01524763 | -0295 | 04125 | 19 |
| 7190 | ENSMUSG( | Tank | 21353 | 015264564 | 006237869 | -04325 | 0185 | 19 |
| 7193 | ENSMUSG( | Taok3 | 330177 | 012659833 | -00829328 | -02425 | 0275 | 19 |
| 7194 | ENSMUSG( | Tapbp | 21356 | -03187348 | 015770435 | -032 | 01425 | 19 |
| 7206 | ENSMUSG( | Tax1bp3 | 76281 | 009962489 | 018479871 | -0485 | -002 | 19 |
| 7210 | ENSMUSG( | Tbc1d12 | 209478 | 009232068 | 001997388 | -03125 | 0215 | 19 |
| 7222 | ENSMUSG( | Tbc1d30 | 74694 | 017746265 | -00180020 | -06325 | 016 | 19 |
| 7232 | ENSMUSG( | Tbcb | 66411 | 005466844 | -01174612 | -023 | 03525 | 19 |
| 7249 | ENSMUSG( | Tcea1 | 21399 | 001599621 | -00490378 | -03025 | 03275 | 19 |
| 7259 | ENSMUSG( | Tcof1 | 21453 | 013979756 | -00544596 | -0865 | 01575 | 19 |
| 7262 | ENSMUSG( | Tctn3 | 67590 | 000346753 | 021851013 | -063 | 0445 | 19 |
| 7282 | ENSMUSG( | Tent4b | 214627 | 008617624 | 018159633 | -058 | 02775 | 19 |
| 7288 | ENSMUSG( | Tesk1 | 21754 | 016237468 | 009171490 | -04375 | 007 | 19 |
| 7292 | ENSMUSG( | Tfam | 21780 | 013231942 | 009829245 | -0435 | -00125 | 19 |
| 7323 | ENSMUSG( | Thoc7 | 66231 | 006947378 | -00723704 | -0655 | 0235 | 19 |
| 7348 | ENSMUSG( | Timm44 | 21856 | 011849043 | 009718547 | -01875 | 01725 | 19 |
| 7357 | ENSMUSG( | Tjp1 | 21872 | 012130729 | -00582654 | -02125 | 018 | 19 |
| 7361 | ENSMUSG( | Tkt | 21881 | -00760001 | -00731535 | -0285 | 00525 | 19 |
| 7368 | ENSMUSG( | Tle5 | 14797 | 003755216 | -01704806 | -0665 | 0305 | 19 |
| 7369 | ENSMUSG( | Tln1 | 21894 | 000933662 | -00458178 | -02425 | 02 | 19 |
| 7371 | ENSMUSG( | Tlnrd1 | 80889 | -00202200 | -00214483 | -0375 | 014 | 19 |
| 7430 | ENSMUSG( | Tmem169 | 271711 | -02550451 | 010319393 | -03975 | 022 | 19 |
| 7459 | ENSMUSG( | Tmem256 | 69186 | 006784885 | -01951274 | -04175 | 00825 | 19 |
| 7480 | ENSMUSG( | Tmem67 | 329795 | -01920733 | 033507222 | -0305 | 01125 | 19 |
| 7497 | ENSMUSG( | Tmsb10 | 19240 | -00484204 | -02222894 | -02475 | 03025 | 19 |
| 7498 | ENSMUSG( | Tmsb4x | 19241 | 004061971 | -01152625 | -0415 | 00825 | 19 |
| 7515 | ENSMUSG( | Tnks | 21951 | -00284522 | 003397478 | -0235 | 01025 | 19 |
| 7516 | ENSMUSG( | Tnks1bp1 | 228140 | 001846468 | 010932932 | -022 | 01975 | 19 |
| 7522 | ENSMUSG( | Tnrc6b | 213988 | 002983308 | 003603594 | -02625 | 04775 | 19 |
| 7523 | ENSMUSG( | Tnrc6c | 217351 | -02174931 | -00187048 | -044 | 01325 | 19 |
| 7528 | ENSMUSG( | Togaram1 | 328108 | 016852667 | 022402542 | -0515 | -00725 | 19 |
| 7533 | ENSMUSG( | Tomm20 | 67952 | -01705431 | 001106970 | -03975 | 00375 | 19 |
| 7554 | ENSMUSG( | Tpd52 | 21985 | -00744274 | -00675054 | -02225 | 029 | 19 |
| 7562 | ENSMUSG( | Tpm1 | 22003 | -00201909 | -00798969 | -05225 | 0445 | 19 |
| 7575 | ENSMUSG( | Tpt1 | 22070 | 012334958 | -00605469 | -0215 | 0315 | 19 |
| 7610 | ENSMUSG( | Trim24 | 21848 | -00793211 | -00963932 | -06 | 03975 | 19 |
| 7611 | ENSMUSG( | Trim25 | 217069 | 006511908 | -00897868 | -03875 | 02225 | 19 |
| 7621 | ENSMUSG( | Trim46 | 360213 | 015200884 | 000671329 | -035 | 0175 | 19 |
| 7630 | ENSMUSG( | Trip11 | 109181 | 009212513 | 005708964 | -0225 | 03725 | 19 |
| 7634 | ENSMUSG( | Trit1 | 66966 | 006725445 | 010770605 | -02175 | 024 | 19 |
| 7635 | ENSMUSG( | Trmt1 | 212528 | 021323496 | 009196329 | -0315 | 0375 | 19 |
| 7654 | ENSMUSG( | Trpm7 | 58800 | -01513667 | 017139081 | -02075 | 001 | 19 |
| 7658 | ENSMUSG( | Tsc1 | 64930 | 000222278 | 002698098 | -03925 | 008 | 19 |
| 7664 | ENSMUSG( | Tsen34 | 66078 | 005563283 | -01429115 | -03225 | 02175 | 19 |
| 7703 | ENSMUSG( | Ttc39b | 69863 | -00919410 | -00355297 | -023 | 02 | 19 |
| 7715 | ENSMUSG( | Ttll7 | 70892 | -00009356 | -00430025 | -0275 | 035 | 19 |
| 7717 | ENSMUSG( | Ttpa | 50500 | -00303986 | -00848306 | -04725 | -00475 | 19 |
| 7812 | ENSMUSG( | Ube2r2 | 67615 | 007199433 | -00342959 | -02625 | 0485 | 19 |
| 7826 | ENSMUSG( | Ubl5 | 66177 | -00348013 | -02116295 | -03175 | 009 | 19 |
| 7869 | ENSMUSG( | Ulk2 | 29869 | 012717375 | 002726529 | -02325 | 004 | 19 |
| 7887 | ENSMUSG( | Unkl | 74154 | 007405308 | 003335792 | -04075 | 045 | 19 |
| 7890 | ENSMUSG( | Upf3b | 68134 | 000303091 | 015965749 | -02825 | 005 | 19 |
| 7903 | ENSMUSG( | Uqcrq | 22272 | -00191469 | -02902636 | -05825 | 009 | 19 |
| 7904 | ENSMUSG( | Urb1 | 207932 | -01366722 | 012428850 | -02825 | 0185 | 19 |
| 7907 | ENSMUSG( | Uri1 | 19777 | 005868405 | -00045902 | -06375 | 01125 | 19 |
| 7933 | ENSMUSG( | Usp38 | 74841 | 006518351 | 018087457 | -019 | 03325 | 19 |
| 7937 | ENSMUSG( | Usp45 | 77593 | 000106946 | 009967262 | -09325 | 04075 | 19 |
| 7950 | ENSMUSG( | Utp14a | 72554 | 023860604 | -00295638 | -0385 | 00075 | 19 |
| 7954 | ENSMUSG( | Utp3 | 65961 | 015649494 | 006738239 | -0485 | 007 | 19 |
| 7957 | ENSMUSG( | Utrn | 22288 | 009449982 | -00051541 | -02075 | 013 | 19 |
| 7961 | ENSMUSG( | Vamp1 | 22317 | -00283907 | -01630447 | -028 | 02925 | 19 |
| 7973 | ENSMUSG( | Vasp | 22323 | -00781697 | 020841875 | -03725 | 036 | 19 |
| 7997 | ENSMUSG( | Vkorc1 | 27973 | -02142892 | 001716925 | -04975 | 0575 | 19 |
| 8034 | ENSMUSG( | Vps54 | 245944 | 002978831 | 021227179 | -054 | 00325 | 19 |
| 8059 | ENSMUSG( | Wasf1 | 83767 | -00196443 | -00698584 | -02225 | 01275 | 19 |
| 8079 | ENSMUSG( | Wdr17 | 244484 | -00238913 | 000136589 | -03275 | 01275 | 19 |
| 8081 | ENSMUSG( | Wdr19 | 213081 | -00414239 | 003089594 | -0535 | -00775 | 19 |
| 8083 | ENSMUSG( | Wdr24 | 268933 | 010867532 | 012466760 | -04875 | 02375 | 19 |
| 8132 | ENSMUSG( | Xab2 | 67439 | 005181604 | -00176185 | -02875 | 04 | 19 |
| 8136 | ENSMUSG( | Xkr4 | 497097 | -02893175 | 035156932 | -0235 | 04625 | 19 |
| 8138 | ENSMUSG( | Xkr7 | 228787 | -02705874 | -03529185 | -043 | 03675 | 19 |
| 8160 | ENSMUSG( | Yars1 | 107271 | 011374954 | -01092104 | -035 | 055 | 19 |
| 8163 | ENSMUSG( | Ybx3 | 56449 | -00348426 | -00837583 | -021 | 0345 | 19 |
| 8165 | ENSMUSG( | Yeats4 | 64050 | 001041086 | -01644495 | -019 | 025 | 19 |
| 8177 | ENSMUSG( | Yme1l1 | 27377 | 001713235 | 023209463 | -023 | 01875 | 19 |
| 8178 | ENSMUSG( | Yod1 | 226418 | -00306766 | -00234386 | -0295 | 02025 | 19 |
| 8195 | ENSMUSG( | Zbtb11 | 271377 | 005929891 | 006839501 | -04 | 00775 | 19 |
| 8199 | ENSMUSG( | Zbtb38 | 245007 | -00339959 | 022463030 | -0265 | 00075 | 19 |
| 8201 | ENSMUSG( | Zbtb7a | 16969 | 012767474 | 014015391 | -038 | 0135 | 19 |
| 8210 | ENSMUSG( | Zc3h3 | 223642 | 036403628 | 017120518 | -0455 | 0105 | 19 |
| 8212 | ENSMUSG( | Zc3h6 | 78751 | -03532921 | -00501867 | -021 | 03375 | 19 |
| 8216 | ENSMUSG( | Zcchc17 | 619605 | -00279881 | -02108851 | -03225 | 02475 | 19 |
| 8217 | ENSMUSG( | Zcchc18 | 66995 | -00089954 | -00113728 | -03475 | 01375 | 19 |
| 8223 | ENSMUSG( | Zdhhc18 | 503610 | -02119169 | 005413714 | -0395 | 031 | 19 |
| 8228 | ENSMUSG( | Zdhhc5 | 228136 | -01265855 | 038657271 | -04875 | 00625 | 19 |
| 8230 | ENSMUSG( | Zdhhc8 | 27801 | -01833164 | 014331112 | -036 | 01375 | 19 |
| 8236 | ENSMUSG( | Zfand2b | 68818 | -01325271 | -01356816 | -07575 | 0215 | 19 |
| 8242 | ENSMUSG( | Zfp280d | 235469 | -00548994 | 005823622 | -086 | 06525 | 19 |
| 8245 | ENSMUSG( | Zfp346 | 26919 | 020199175 | 004910245 | -029 | 00275 | 19 |
| 8247 | ENSMUSG( | Zfp36l2 | 12193 | 012252321 | 031422374 | -03125 | 0335 | 19 |
| 8260 | ENSMUSG( | Zfyve16 | 218441 | -00532276 | -00268350 | -051 | 00075 | 19 |
| 8261 | ENSMUSG( | Zfyve19 | 72008 | 001833772 | -00223948 | -02025 | 03025 | 19 |
| 8268 | ENSMUSG( | Zhx2 | 387609 | 004629852 | 002355619 | -04125 | 03625 | 19 |
| 8276 | ENSMUSG( | Zmym2 | 76007 | 000130955 | 006511947 | -0445 | 02725 | 19 |
| 8277 | ENSMUSG( | Zmym3 | 56364 | 020490218 | -00789010 | -02025 | 02775 | 19 |
| 264 | ENSMUSG( | Aig1 | 66253 | -03558616 | 012703878 | 17725 | 515 | 20 |
| 1544 | ENSMUSG( | Cript | 56724 | 003303120 | -01129170 | 04875 | 388 | 20 |
| 1941 | ENSMUSG( | Dnajc6 | 72685 | 014591898 | 017163438 | 20125 | 2645 | 20 |
| 2632 | ENSMUSG( | Gabra2 | 14395 | 154941682 | 170375342 | 3405 | 271 | 20 |
| 3065 | ENSMUSG( | Hebp1 | 15199 | -00744953 | -01800990 | 35225 | 332 | 20 |
| 4595 | ENSMUSG( | Nnt | 18115 | -02264577 | -04051466 | 44975 | 59625 | 20 |
| 7656 | ENSMUSG( | Trrap | 100683 | -00104645 | 010857585 | 38025 | 39875 | 20 |
| 7963 | ENSMUSG( | Vamp3 | 22319 | 010498560 | 005988207 | 00875 | 4725 | 20 |
| 8072 | ENSMUSG( | Wdfy1 | 69368 | -02213996 | 078643852 | 0135 | 85 | 20 |
| 8151 | ENSMUSG( | Xrcc5 | 22596 | -01023073 | -02452432 | 199 | 27525 | 20 |

**GSEA MNK1KO**

** GSEA mouse MSigDB canonical pathways: m2.cp.v2023.1.Mm.symbols*

| Description | setSize | enrichmen | NES | pvalue | p.adjust | qvalue | rank | leading_ed | core_enrichment | |
|---|---|---|---|---|---|---|---|---|---|---|
| REACTOME_GPCR_LIGAND_BINDING | 239 | -0.76494 | -2.0347 | 3.10E-20 | 5.13E-17 | 4.67E-17 | 1564 | tags=31% | list=10% | signal=28% |
| REACTOME_CLASS_A_1_RHODOPSIN_LIKE_RECEPTORS | 187 | -0.75923 | -1.99368 | 4.29E-15 | 3.56E-12 | 3.23E-12 | 1564 | tags=29% | list=10% | signal=26% |
| REACTOME_TRANSPORT_OF_SMALL_MOLECULES | 548 | -0.61232 | -1.67738 | 3.50E-12 | 1.93E-09 | 1.76E-09 | 3032 | tags=40% | list=19% | signal=33% |
| WP_GPCRS_NONODORANT | 172 | -0.72862 | -1.9067 | 5.38E-11 | 2.03E-08 | 1.85E-08 | 1575 | tags=34% | list=10% | signal=31% |
| REACTOME_SIGNALING_BY_GPCR | 438 | -0.62388 | -1.6973 | 6.13E-11 | 2.03E-08 | 1.85E-08 | 1959 | tags=24% | list=12% | signal=21% |
| REACTOME_SLC_MEDIATED_TRANSMEMBRANE_TRANSPORT | 188 | -0.70571 | -1.85293 | 7.79E-10 | 2.15E-07 | 1.96E-07 | 2537 | tags=47% | list=16% | signal=40% |
| REACTOME_PEPTIDE_LIGAND_BINDING_RECEPTORS | 104 | -0.74402 | -1.88995 | 5.22E-08 | 1.24E-05 | 1.12E-05 | 1564 | tags=26% | list=10% | signal=24% |
| WP_GPCRS_CLASS_A_RHODOPSINLIKE | 104 | -0.74213 | -1.88515 | 6.94E-08 | 1.44E-05 | 1.31E-05 | 1553 | tags=30% | list=10% | signal=27% |
| WP_GPCRS_OTHER | 48 | -0.82258 | -1.93327 | 3.09E-07 | 5.70E-05 | 5.18E-05 | 825 | tags=33% | list=5% | signal=32% |
| REACTOME_G_ALPHA_I_SIGNALLING_EVENTS | 181 | -0.66758 | -1.75088 | 5.50E-07 | 9.12E-05 | 8.29E-05 | 1837 | tags=25% | list=11% | signal=22% |
| REACTOME_TRANSPORT_OF_INORGANIC_CATIONS_ANIONS_AND_AMINO_ACIDS_ | 90 | -0.73091 | -1.83497 | 1.35E-06 | 0.000204 | 0.000185 | 2299 | tags=46% | list=14% | signal=39% |
| WP_SPINAL_CORD_INJURY | 84 | -0.72846 | -1.81183 | 3.90E-06 | 0.000538 | 0.000489 | 2076 | tags=38% | list=13% | signal=33% |
| REACTOME_MAJOR_PATHWAY_OF_RRNA_PROCESSING_IN_THE_NUCLEOLUS_AND | 169 | 0.611335 | 1.974087 | 8.52E-06 | 0.00104 | 0.000945 | 4038 | tags=64% | list=25% | signal=48% |
| WP_GPCRS_ODORANT | 69 | -0.74026 | -1.80699 | 8.78E-06 | 0.00104 | 0.000945 | 2107 | tags=35% | list=13% | signal=30% |
| REACTOME_TRANSLATION | 218 | 0.565882 | 1.931388 | 9.85E-06 | 0.001089 | 0.00099 | 4038 | tags=54% | list=25% | signal=41% |
| REACTOME_O_LINKED_GLYCOSYLATION | 90 | -0.69723 | -1.75043 | 2.21E-05 | 0.002289 | 0.002081 | 2473 | tags=43% | list=15% | signal=37% |
| REACTOME_NEURONAL_SYSTEM | 308 | -0.56796 | -1.52715 | 3.04E-05 | 0.002963 | 0.002693 | 2231 | tags=32% | list=14% | signal=28% |
| REACTOME_GLYCOSAMINOGLYCAN_METABOLISM | 117 | -0.66003 | -1.69134 | 4.16E-05 | 0.003828 | 0.00348 | 3348 | tags=51% | list=21% | signal=41% |
| REACTOME_G_ALPHA_S_SIGNALLING_EVENTS | 97 | -0.68916 | -1.7366 | 6.80E-05 | 0.005934 | 0.005395 | 1026 | tags=19% | list=6% | signal=17% |
| REACTOME_ACTIVATION_OF_C3_AND_C5 | 5 | 0.967203 | 1.765886 | 7.65E-05 | 0.006342 | 0.005766 | 65 | tags=40% | list=0% | signal=40% |
| REACTOME_CARDIAC_CONDUCTION | 88 | -0.69141 | -1.73269 | 9.93E-05 | 0.00784 | 0.007128 | 2448 | tags=43% | list=15% | signal=37% |
| REACTOME_EUKARYOTIC_TRANSLATION_INITIATION | 111 | 0.656225 | 2.099887 | 0.000116 | 0.00867 | 0.007882 | 4226 | tags=70% | list=26% | signal=52% |
| REACTOME_AMINE_LIGAND_BINDING_RECEPTORS | 31 | -0.79755 | -1.78021 | 0.00012 | 0.00867 | 0.007882 | 1553 | tags=55% | list=10% | signal=50% |
| REACTOME_MUSCLE_CONTRACTION | 142 | -0.62406 | -1.61887 | 0.00016 | 0.01108 | 0.010073 | 2659 | tags=34% | list=16% | signal=28% |
| REACTOME_INITIAL_TRIGGERING_OF_COMPLEMENT | 15 | 0.856439 | 2.040583 | 0.000177 | 0.011709 | 0.010645 | 65 | tags=20% | list=0% | signal=20% |
| REACTOME_POST_TRANSLATIONAL_MODIFICATION_SYNTHESIS_OF_GPI_ANCHORE | 63 | -0.70377 | -1.70639 | 0.000202 | 0.012907 | 0.011735 | 3674 | tags=62% | list=23% | signal=48% |
| REACTOME_ADHERENS_JUNCTIONS_INTERACTIONS | 31 | -0.78458 | -1.75125 | 0.000239 | 0.014142 | 0.012857 | 870 | tags=52% | list=5% | signal=49% |
| REACTOME_TRANSPORT_OF_VITAMINS_NUCLEOSIDES_AND_RELATED_MOLECULES | 34 | -0.77686 | -1.73976 | 0.000237 | 0.014142 | 0.012857 | 2537 | tags=59% | list=16% | signal=50% |
| WP_MONOAMINE_GPCRS | 30 | -0.799 | -1.77352 | 0.000252 | 0.014426 | 0.013116 | 1553 | tags=43% | list=10% | signal=39% |
| REACTOME_SPHINGOLIPID_METABOLISM | 71 | -0.69298 | -1.69423 | 0.000267 | 0.014754 | 0.013413 | 2961 | tags=54% | list=18% | signal=44% |
| REACTOME_VOLTAGE_GATED_POTASSIUM_CHANNELS | 37 | -0.75893 | -1.7219 | 0.000376 | 0.0201 | 0.018274 | 1743 | tags=46% | list=11% | signal=41% |
| REACTOME_FORMATION_OF_A_POOL_OF_FREE_40S_SUBUNITS | 93 | 0.653601 | 2.023701 | 0.000397 | 0.020593 | 0.018722 | 4226 | tags=72% | list=26% | signal=53% |
| REACTOME_CLASS_B_2_SECRETIN_FAMILY_RECEPTORS | 41 | -0.75864 | -1.74385 | 0.00042 | 0.021079 | 0.019164 | 1466 | tags=29% | list=9% | signal=27% |
| WP_SEROTONIN_AND_ANXIETY | 16 | -0.87126 | -1.75497 | 0.000438 | 0.021363 | 0.019423 | 453 | tags=44% | list=3% | signal=43% |
| REACTOME_POTASSIUM_CHANNELS | 86 | -0.66335 | -1.65258 | 0.000491 | 0.023272 | 0.021158 | 2974 | tags=44% | list=18% | signal=36% |
| REACTOME_NONSENSE_MEDIATED_DECAY_NMD_INDEPENDENT_OF_THE_EXON_J | 86 | 0.652216 | 2.029911 | 0.000517 | 0.0238 | 0.021638 | 4038 | tags=74% | list=25% | signal=56% |
| WP_CYTOPLASMIC_RIBOSOMAL_PROTEINS | 77 | 0.686664 | 2.090461 | 0.000574 | 0.025725 | 0.023388 | 3722 | tags=81% | list=23% | signal=62% |
| REACTOME_RECEPTOR_TYPE_TYROSINE_PROTEIN_PHOSPHATASES | 20 | -0.82835 | -1.71741 | 0.00072 | 0.031434 | 0.028578 | 1097 | tags=45% | list=7% | signal=42% |
| BIOCARTA_CLASSIC_PATHWAY | 9 | 0.893306 | 1.902312 | 0.000794 | 0.03156 | 0.028693 | 7 | tags=11% | list=0% | signal=11% |
| REACTOME_O_GLYCOSYLATION_OF_TSR_DOMAIN_CONTAINING_PROTEINS | 39 | -0.74443 | -1.69427 | 0.000799 | 0.03156 | 0.028693 | 1071 | tags=36% | list=7% | signal=34% |
| REACTOME_EPH_EPHRIN_MEDIATED_REPULSION_OF_CELLS | 34 | -0.75211 | -1.68434 | 0.000756 | 0.03156 | 0.028693 | 2496 | tags=47% | list=15% | signal=40% |
| REACTOME_IMMUNOREGULATORY_INTERACTIONS_BETWEEN_A_LYMPHOID_AND_ | 65 | -0.67995 | -1.65293 | 0.000775 | 0.03156 | 0.028693 | 2578 | tags=38% | list=16% | signal=32% |
| REACTOME_LYSOSPHINGOLIPID_AND_LPA_RECEPTORS | 14 | -0.86384 | -1.71173 | 0.00096 | 0.037031 | 0.033667 | 698 | tags=57% | list=4% | signal=55% |
| REACTOME_REGULATION_OF_INSULIN_LIKE_GROWTH_FACTOR_IGF_TRANSPORT_/ | 90 | -0.64734 | -1.62518 | 0.000998 | 0.037601 | 0.034185 | 2553 | tags=47% | list=16% | signal=40% |
| REACTOME_CELL_CELL_JUNCTION_ORGANIZATION | 39 | -0.73706 | -1.67749 | 0.001122 | 0.040425 | 0.036752 | 870 | tags=59% | list=5% | signal=56% |
| REACTOME_CELL_CELL_COMMUNICATION | 77 | -0.65259 | -1.60409 | 0.001116 | 0.040425 | 0.036752 | 870 | tags=40% | list=5% | signal=38% |
| REACTOME_NONSENSE_MEDIATED_DECAY_NMD | 106 | 0.595153 | 1.915194 | 0.001415 | 0.047889 | 0.043538 | 4226 | tags=68% | list=26% | signal=50% |
| REACTOME_HEPARAN_SULFATE_HEPARIN_HS_GAG_METABOLISM | 51 | -0.71029 | -1.67923 | 0.0014 | 0.047889 | 0.043538 | 2480 | tags=53% | list=15% | signal=45% |
| REACTOME_TACHYKININ_RECEPTORS_BIND_TACHYKININS | 4 | -0.96919 | -1.56686 | 0.001407 | 0.047889 | 0.043538 | 288 | tags=75% | list=2% | signal=74% |
| REACTOME_ION_CHANNEL_TRANSPORT | 139 | -0.59815 | -1.54776 | 0.001503 | 0.049833 | 0.045306 | 3032 | tags=43% | list=19% | signal=35% |
| REACTOME_GLUCAGON_TYPE_LIGAND_RECEPTORS | 25 | -0.78801 | -1.70913 | 0.001661 | 0.053993 | 0.049088 | 594 | tags=20% | list=4% | signal=19% |
| REACTOME_CHONDROITIN_SULFATE_DERMATAN_SULFATE_METABOLISM | 47 | -0.70489 | -1.64735 | 0.001713 | 0.054612 | 0.04965 | 2328 | tags=51% | list=14% | signal=44% |
| BIOCARTA_LECTIN_PATHWAY | 7 | 0.89652 | 1.849818 | 0.001894 | 0.057677 | 0.052437 | 7 | tags=14% | list=0% | signal=14% |
| BIOCARTA_COMP_PATHWAY | 12 | 0.81206 | 1.830206 | 0.001885 | 0.057677 | 0.052437 | 7 | tags=8% | list=0% | signal=8% |
| WP_CALCIUM_REGULATION_IN_CARDIAC_CELLS | 132 | -0.58825 | -1.52137 | 0.001913 | 0.057677 | 0.052437 | 1870 | tags=22% | list=12% | signal=20% |
| REACTOME_CELL_JUNCTION_ORGANIZATION | 62 | -0.66391 | -1.60852 | 0.002709 | 0.080217 | 0.072929 | 2966 | tags=45% | list=18% | signal=37% |
| REACTOME_ION_HOMEOSTASIS | 48 | -0.68751 | -1.61583 | 0.002792 | 0.081218 | 0.073839 | 2550 | tags=46% | list=16% | signal=39% |
| REACTOME_SUMOYLATION_OF_INTRACELLULAR_RECEPTORS | 22 | 0.706008 | 1.810097 | 0.002922 | 0.082109 | 0.07465 | 1887 | tags=45% | list=12% | signal=40% |
| WP_INSULIN_SIGNALING | 153 | 0.503352 | 1.565741 | 0.002902 | 0.082109 | 0.07465 | 3059 | tags=42% | list=19% | signal=35% |
| REACTOME_LDL_CLEARANCE | 15 | -0.82199 | -1.63938 | 0.002973 | 0.082144 | 0.074681 | 1358 | tags=40% | list=8% | signal=37% |
| BIOCARTA_EGFR_SMRTE_PATHWAY | 11 | 0.850222 | 1.882286 | 0.003259 | 0.088202 | 0.080189 | 987 | tags=45% | list=6% | signal=43% |
| REACTOME_REGULATION_OF_LOCALIZATION_OF_FOXO_TRANSCRIPTION_FACTORS | 12 | 0.791175 | 1.783135 | 0.003298 | 0.088202 | 0.080189 | 2316 | tags=58% | list=14% | signal=50% |
| REACTOME_PLASMA_LIPOPROTEIN_ASSEMBLY_REMODELING_AND_CLEARANCE | 52 | -0.67321 | -1.59532 | 0.003553 | 0.093517 | 0.085021 | 1453 | tags=35% | list=9% | signal=32% |
| REACTOME_ACTIVATION_OF_THE_MRNA_UPON_BINDING_OF_THE_CAP_BINDING_ | 58 | 0.665777 | 1.985881 | 0.003751 | 0.097163 | 0.088336 | 4226 | tags=71% | list=26% | signal=52% |
| REACTOME_GLYCOSPHINGOLIPID_METABOLISM | 38 | -0.7189 | -1.63228 | 0.004112 | 0.104899 | 0.095369 | 2961 | tags=58% | list=18% | signal=47% |
| REACTOME_SEROTONIN_RECEPTORS | 10 | -0.86864 | -1.62339 | 0.00423 | 0.106264 | 0.09661 | 851 | tags=70% | list=5% | signal=66% |
| REACTOME_SYNTHESIS_SECRETION_AND_DEACYLATION_OF_GHRELIN | 9 | -0.8721 | -1.60531 | 0.004494 | 0.111202 | 0.101099 | 1350 | tags=78% | list=8% | signal=71% |
| WP_SEROTONIN_AND_ANXIETYRELATED_EVENTS | 12 | -0.8387 | -1.6323 | 0.004825 | 0.116101 | 0.105553 | 1607 | tags=50% | list=10% | signal=45% |
| REACTOME_PLASMA_LIPOPROTEIN_CLEARANCE | 30 | -0.72904 | -1.61822 | 0.004832 | 0.116101 | 0.105553 | 1358 | tags=23% | list=8% | signal=21% |
| REACTOME_SYNTHESIS_OF_GLYCOSYLPHOSPHATIDYLINOSITOL_GPI | 17 | -0.78727 | -1.58899 | 0.00515 | 0.121974 | 0.110893 | 3083 | tags=82% | list=19% | signal=67% |
| REACTOME_REGULATION_OF_SIGNALING_BY_CBL | 21 | 0.699501 | 1.780813 | 0.00526 | 0.122828 | 0.111669 | 1338 | tags=43% | list=8% | signal=39% |
| BIOCARTA_AMAN_PATHWAY | 9 | -0.86802 | -1.59779 | 0.005338 | 0.122925 | 0.111757 | 1091 | tags=56% | list=7% | signal=52% |
| REACTOME_CHONDROITIN_SULFATE_BIOSYNTHESIS | 18 | -0.79542 | -1.63138 | 0.005682 | 0.129047 | 0.117323 | 2182 | tags=61% | list=14% | signal=53% |
| BIOCARTA_TSP1_PATHWAY | 7 | -0.90599 | -1.57104 | 0.006008 | 0.13462 | 0.122389 | 49 | tags=43% | list=0% | signal=43% |
| BIOCARTA_CK1_PATHWAY | 15 | -0.80138 | -1.59825 | 0.006413 | 0.141774 | 0.128893 | 936 | tags=27% | list=6% | signal=25% |
| REACTOME_REGULATION_OF_GENE_EXPRESSION_BY_HYPOXIA_INDUCIBLE_FACTO | 6 | 0.89323 | 1.747473 | 0.00667 | 0.145504 | 0.132284 | 996 | tags=33% | list=6% | signal=31% |
| REACTOME_TRANSPORT_OF_NUCLEOTIDE_SUGARS | 9 | -0.8611 | -1.58507 | 0.00685 | 0.147489 | 0.13409 | 2105 | tags=89% | list=13% | signal=77% |
| WP_SPHINGOLIPID_METABOLISM_OVERVIEW | 24 | -0.73487 | -1.58418 | 0.007384 | 0.156961 | 0.1427 | 1844 | tags=62% | list=11% | signal=55% |
| REACTOME_INTERLEUKIN_3_INTERLEUKIN_5_AND_GM_CSF_SIGNALING | 35 | 0.608729 | 1.706878 | 0.007515 | 0.15772 | 0.143391 | 2556 | tags=40% | list=16% | signal=34% |
| BIOCARTA_STAT3_PATHWAY | 8 | 0.812469 | 1.70006 | 0.007742 | 0.158473 | 0.144076 | 3034 | tags=100% | list=19% | signal=81% |
| REACTOME_CLASS_C_3_METABOTROPIC_GLUTAMATE_PHEROMONE_RECEPTORS | 11 | -0.84645 | -1.61821 | 0.007698 | 0.158473 | 0.144076 | 1558 | tags=73% | list=10% | signal=66% |
| BIOCARTA_PAR1_PATHWAY | 17 | 0.716215 | 1.785835 | 0.008237 | 0.159906 | 0.145379 | 4111 | tags=82% | list=25% | signal=61% |
| BIOCARTA_ACH_PATHWAY | 12 | 0.752215 | 1.695327 | 0.008054 | 0.159906 | 0.145379 | 2147 | tags=50% | list=13% | signal=43% |
| REACTOME_SYNTHESIS_OF_VERY_LONG_CHAIN_FATTY_ACYL_COAS | 18 | -0.78232 | -1.6045 | 0.008288 | 0.159906 | 0.145379 | 1720 | tags=44% | list=11% | signal=40% |
| REACTOME_TRANSPORT_OF_BILE_SALTS_AND_ORGANIC_ACIDS_METAL_IONS_ANI | 58 | -0.6476 | -1.55347 | 0.008189 | 0.159906 | 0.145379 | 2447 | tags=50% | list=15% | signal=43% |
| REACTOME_PROTEIN_PROTEIN_INTERACTIONS_AT_SYNAPSES | 66 | -0.63155 | -1.53678 | 0.008294 | 0.159906 | 0.145379 | 1206 | tags=35% | list=7% | signal=32% |
| REACTOME_CDC42_GTPASE_CYCLE | 94 | 0.56412 | 1.743605 | 0.00862 | 0.164274 | 0.14935 | 1324 | tags=32% | list=8% | signal=29% |
| WP_OXIDATIVE_DAMAGE_RESPONSE | 37 | 0.60762 | 1.714861 | 0.008946 | 0.168555 | 0.153242 | 579 | tags=14% | list=4% | signal=13% |
| BIOCARTA_RAS_PATHWAY | 21 | 0.678549 | 1.727474 | 0.009052 | 0.168623 | 0.153303 | 3662 | tags=62% | list=23% | signal=48% |
| REACTOME_NUCLEOTIDE_LIKE_PURINERGIC_RECEPTORS | 13 | -0.83761 | -1.64489 | 0.009599 | 0.169425 | 0.154033 | 1742 | tags=46% | list=11% | signal=41% |
| WP_GPCRS_CLASS_C_METABOTROPIC_GLUTAMATE_PHEROMONE | 13 | -0.83747 | -1.64461 | 0.009599 | 0.169425 | 0.154033 | 1558 | tags=62% | list=10% | signal=56% |
| REACTOME_PHOSPHORYLATION_OF_CD3_AND_TCR_ZETA_CHAINS | 11 | -0.8392 | -1.60435 | 0.009319 | 0.169425 | 0.154033 | 1132 | tags=45% | list=7% | signal=42% |
| REACTOME_BIOSYNTHESIS_OF_THE_N_GLYCAN_PRECURSOR_DOLICHOL_LIPID_LIN | 70 | -0.62305 | -1.52348 | 0.009606 | 0.169425 | 0.154033 | 2816 | tags=46% | list=17% | signal=38% |
| REACTOME_G_ALPHA_Q_SIGNALLING_EVENTS | 149 | -0.5529 | -1.4429 | 0.009222 | 0.169425 | 0.154033 | 2290 | tags=26% | list=14% | signal=22% |
| REACTOME_AMINO_ACID_TRANSPORT_ACROSS_THE_PLASMA_MEMBRANE | 29 | -0.70761 | -1.55822 | 0.009812 | 0.169462 | 0.154066 | 2161 | tags=41% | list=13% | signal=36% |

| | | | | | | | | | | |
|---|---|---|---|---|---|---|---|---|---|---|
| REACTOME_HS_GAG_BIOSYNTHESIS | 29 | -0.70705 | -1.55698 | 0.009812 | 0.169462 | 0.154066 | 2378 | tags=45% | list=15% | signal=38% |
| REACTOME_INTERLEUKIN_20_FAMILY_SIGNALING | 10 | 0.767281 | 1.682248 | 0.010195 | 0.172491 | 0.15682 | 2303 | tags=60% | list=14% | signal=51% |
| REACTOME_GAP_JUNCTION_ASSEMBLY | 20 | -0.7569 | -1.56927 | 0.010094 | 0.172491 | 0.15682 | 101 | tags=25% | list=1% | signal=25% |
| REACTOME_RECEPTOR_MEDIATED_MITOPHAGY | 11 | 0.803386 | 1.778595 | 0.010678 | 0.178834 | 0.162587 | 1831 | tags=55% | list=11% | signal=48% |
| WP_MISMATCH_REPAIR | 9 | 0.809347 | 1.723521 | 0.010839 | 0.179716 | 0.163389 | 918 | tags=44% | list=6% | signal=42% |
| REACTOME_O_LINKED_GLYCOSYLATION_OF_MUCINS | 41 | -0.6737 | -1.54859 | 0.011359 | 0.18647 | 0.169529 | 2223 | tags=41% | list=14% | signal=36% |
| REACTOME_NR1H2_NR1H3_REGULATE_GENE_EXPRESSION_TO_CONTROL_BILE_AC | 6 | 0.856995 | 1.676584 | 0.012073 | 0.18725 | 0.170238 | 1196 | tags=67% | list=7% | signal=62% |
| WP_SPHINGOLIPID_METABOLISM_INTEGRATED_PATHWAY | 25 | -0.73014 | -1.58362 | 0.012106 | 0.18725 | 0.170238 | 3216 | tags=64% | list=20% | signal=51% |
| REACTOME_OTHER_SEMAPHORIN_INTERACTIONS | 14 | -0.79557 | -1.57645 | 0.011893 | 0.18725 | 0.170238 | 2001 | tags=57% | list=12% | signal=50% |
| REACTOME_ELASTIC_FIBRE_FORMATION | 34 | -0.67772 | -1.51775 | 0.011766 | 0.18725 | 0.170238 | 1852 | tags=44% | list=11% | signal=39% |
| WP_DRAVET_SYNDROME_SCN1AA1783V_POINT_MUTATION_MODEL | 71 | -0.60955 | -1.49026 | 0.011986 | 0.18725 | 0.170238 | 1986 | tags=37% | list=12% | signal=32% |
| REACTOME_EXTRINSIC_PATHWAY_OF_FIBRIN_CLOT_FORMATION | 3 | -0.9597 | -1.46178 | 0.011787 | 0.18725 | 0.170238 | 292 | tags=67% | list=2% | signal=65% |
| REACTOME_STIMULI_SENSING_CHANNELS | 76 | -0.59125 | -1.44454 | 0.012197 | 0.18725 | 0.170238 | 2734 | tags=34% | list=17% | signal=29% |
| REACTOME_PEPTIDE_HORMONE_METABOLISM | 37 | -0.67656 | -1.53503 | 0.012703 | 0.191462 | 0.174067 | 1435 | tags=32% | list=9% | signal=30% |
| REACTOME_CHYLOMICRON_CLEARANCE | 4 | -0.93333 | -1.5089 | 0.012686 | 0.191462 | 0.174067 | 941 | tags=50% | list=6% | signal=47% |
| REACTOME_CHOLESTEROL_BIOSYNTHESIS | 26 | -0.72174 | -1.56884 | 0.013431 | 0.200623 | 0.182397 | 2258 | tags=50% | list=14% | signal=43% |
| WP_NUCLEOTIDE_GPCRS | 10 | -0.83151 | -1.55399 | 0.013742 | 0.20163 | 0.183312 | 1742 | tags=40% | list=11% | signal=36% |
| BIOCARTA_VIP_PATHWAY | 24 | -0.71101 | -1.53274 | 0.013646 | 0.20163 | 0.183312 | 1481 | tags=21% | list=9% | signal=19% |
| REACTOME_CARNITINE_METABOLISM | 13 | 0.787627 | 1.827166 | 0.014382 | 0.202638 | 0.184228 | 1429 | tags=46% | list=9% | signal=42% |
| REACTOME_SIGNALING_BY_ERYTHROPOIETIN | 16 | 0.710269 | 1.724318 | 0.014073 | 0.202638 | 0.184228 | 1621 | tags=56% | list=10% | signal=51% |
| BIOCARTA_HCMV_PATHWAY | 17 | 0.690911 | 1.722741 | 0.01438 | 0.202638 | 0.184228 | 3034 | tags=59% | list=19% | signal=48% |
| REACTOME_MITOCHONDRIAL_FATTY_ACID_BETA_OXIDATION | 34 | 0.577869 | 1.614505 | 0.014057 | 0.202638 | 0.184228 | 3842 | tags=44% | list=24% | signal=34% |
| REACTOME_PEPTIDE_HORMONE_BIOSYNTHESIS | 6 | -0.89452 | -1.52155 | 0.014422 | 0.202638 | 0.184228 | 354 | tags=50% | list=2% | signal=49% |
| BIOCARTA_RANBP2_PATHWAY | 12 | 0.726863 | 1.63819 | 0.014645 | 0.204046 | 0.185508 | 2135 | tags=58% | list=13% | signal=51% |
| REACTOME_SYNTHESIS_OF_KETONE_BODIES | 8 | 0.767376 | 1.605705 | 0.014833 | 0.204937 | 0.186319 | 3499 | tags=62% | list=22% | signal=49% |
| REACTOME_ACTIVATION_OF_MATRIX_METALLOPROTEINASES | 15 | -0.76975 | -1.53518 | 0.015313 | 0.20983 | 0.190767 | 2295 | tags=60% | list=14% | signal=52% |
| REACTOME_REDUCTION_OF_CYTOSOLIC_CA_LEVELS | 14 | -0.78615 | -1.55778 | 0.01545 | 0.209966 | 0.19089 | 2125 | tags=57% | list=13% | signal=50% |
| REACTOME_BRANCHED_CHAIN_AMINO_ACID_CATABOLISM | 20 | 0.697298 | 1.789395 | 0.01565 | 0.210248 | 0.191147 | 3459 | tags=75% | list=21% | signal=59% |
| WP_CHOLESTEROL_METABOLISM_WITH_BLOCH_AND_KANDUTSCHRUSSELL_PATHW | 47 | -0.64504 | -1.50749 | 0.015724 | 0.210248 | 0.191147 | 2394 | tags=43% | list=15% | signal=36% |
| REACTOME_RESOLUTION_OF_AP_SITES_VIA_THE_MULTIPLE_NUCLEOTIDE_PATCH_ | 24 | 0.615974 | 1.602306 | 0.015866 | 0.210336 | 0.191226 | 2178 | tags=42% | list=13% | signal=36% |
| REACTOME_CD28_CO_STIMULATION | 32 | 0.579826 | 1.589897 | 0.015985 | 0.210336 | 0.191226 | 2852 | tags=41% | list=18% | signal=34% |
| REACTOME_PLATELET_CALCIUM_HOMEOSTASIS | 26 | -0.71667 | -1.55781 | 0.016176 | 0.211185 | 0.191999 | 2203 | tags=50% | list=14% | signal=43% |
| REACTOME_SYNTHESIS_OF_IP3_AND_IP4_IN_THE_CYTOSOL | 27 | 0.60057 | 1.586475 | 0.016903 | 0.218951 | 0.199059 | 3346 | tags=56% | list=21% | signal=44% |
| REACTOME_ACTIVATION_OF_CASPASES_THROUGH_APOPTOSOME_MEDIATED_CLE | 6 | 0.819224 | 1.602691 | 0.017571 | 0.223976 | 0.203627 | 218 | tags=17% | list=1% | signal=16% |
| BIOCARTA_ACTINY_PATHWAY | 10 | -0.82177 | -1.53579 | 0.017566 | 0.223976 | 0.203627 | 8 | tags=20% | list=0% | signal=20% |
| REACTOME_SPHINGOLIPID_DE_NOVO_BIOSYNTHESIS | 33 | -0.66904 | -1.49517 | 0.017697 | 0.223976 | 0.203627 | 3216 | tags=55% | list=20% | signal=44% |
| REACTOME_FORMATION_OF_THE_EARLY_ELONGATION_COMPLEX | 32 | 0.569013 | 1.560247 | 0.018434 | 0.231538 | 0.210502 | 4829 | tags=56% | list=30% | signal=40% |
| WP_OXIDATIVE_PHOSPHORYLATION | 32 | -0.68647 | -1.53772 | 0.018843 | 0.234899 | 0.213558 | 1486 | tags=12% | list=9% | signal=11% |
| REACTOME_NUCLEOTIDE_BIOSYNTHESIS | 12 | 0.708685 | 1.59722 | 0.01927 | 0.23492 | 0.213577 | 3615 | tags=75% | list=22% | signal=58% |
| REACTOME_ORGANIC_ANION_TRANSPORTERS | 8 | -0.84095 | -1.5117 | 0.019048 | 0.23492 | 0.213577 | 664 | tags=62% | list=4% | signal=60% |
| REACTOME_TRANSMISSION_ACROSS_CHEMICAL_SYNAPSES | 194 | -0.51541 | -1.35397 | 0.01917 | 0.23492 | 0.213577 | 2142 | tags=26% | list=13% | signal=23% |
| REACTOME_FLT3_SIGNALING | 29 | 0.672053 | 1.79733 | 0.019732 | 0.235365 | 0.213982 | 2330 | tags=38% | list=14% | signal=33% |
| REACTOME_ARMS_MEDIATED_ACTIVATION | 5 | -0.91534 | -1.51066 | 0.0197 | 0.235365 | 0.213982 | 390 | tags=40% | list=2% | signal=39% |
| WP_GPCRS_PEPTIDE | 35 | -0.65547 | -1.47256 | 0.019727 | 0.235365 | 0.213982 | 3192 | tags=46% | list=20% | signal=37% |
| REACTOME_DOWNSTREAM_SIGNAL_TRANSDUCTION | 27 | 0.594921 | 1.571552 | 0.020005 | 0.23573 | 0.214313 | 2559 | tags=52% | list=16% | signal=44% |
| WP_GLYCOGEN_METABOLISM | 32 | 0.562403 | 1.542123 | 0.020189 | 0.23573 | 0.214313 | 5321 | tags=78% | list=33% | signal=52% |
| REACTOME_ALTERNATIVE_COMPLEMENT_ACTIVATION | 3 | 0.921909 | 1.492655 | 0.020062 | 0.23573 | 0.214313 | 65 | tags=33% | list=0% | signal=33% |
| BIOCARTA_PLC_PATHWAY | 9 | 0.773599 | 1.647396 | 0.020629 | 0.239181 | 0.217451 | 3358 | tags=78% | list=21% | signal=62% |
| REACTOME_TRANSCRIPTIONAL_REGULATION_OF_GRANULOPOIESIS | 6 | 0.809519 | 1.583703 | 0.020918 | 0.240852 | 0.21897 | 104 | tags=33% | list=1% | signal=33% |
| REACTOME_KETONE_BODY_METABOLISM | 9 | 0.769591 | 1.638859 | 0.0214 | 0.2447 | 0.222469 | 3499 | tags=67% | list=22% | signal=52% |
| REACTOME_VEGFR2_MEDIATED_VASCULAR_PERMEABILITY | 29 | 0.66182 | 1.769962 | 0.021657 | 0.245942 | 0.223598 | 2852 | tags=48% | list=18% | signal=40% |
| REACTOME_COMPLEMENT_CASCADE | 29 | 0.654376 | 1.750053 | 0.02294 | 0.258744 | 0.235236 | 65 | tags=10% | list=0% | signal=10% |
| WP_TCA_CYCLE | 30 | 0.591846 | 1.60264 | 0.023427 | 0.262451 | 0.238607 | 4015 | tags=67% | list=25% | signal=50% |
| REACTOME_HSF1_DEPENDENT_TRANSACTIVATION | 23 | 0.654117 | 1.690333 | 0.02525 | 0.267983 | 0.243636 | 2800 | tags=48% | list=17% | signal=40% |
| BIOCARTA_PYK2_PATHWAY | 25 | 0.644178 | 1.668444 | 0.025376 | 0.267983 | 0.243636 | 3518 | tags=56% | list=22% | signal=44% |
| REACTOME_RNA_POLYMERASE_II_PRE_TRANSCRIPTION_EVENTS | 75 | 0.538881 | 1.65469 | 0.025238 | 0.267983 | 0.243636 | 4829 | tags=56% | list=30% | signal=39% |
| REACTOME_ESTROGEN_DEPENDENT_NUCLEAR_EVENTS_DOWNSTREAM_OF_ESR_I | 9 | 0.763439 | 1.62576 | 0.024099 | 0.267983 | 0.243636 | 3034 | tags=78% | list=19% | signal=63% |
| REACTOME_RHOJ_GTPASE_CYCLE | 16 | 0.663507 | 1.610793 | 0.025013 | 0.267983 | 0.243636 | 2403 | tags=44% | list=15% | signal=37% |
| BIOCARTA_VDR_PATHWAY | 21 | 0.629417 | 1.60239 | 0.024303 | 0.267983 | 0.243636 | 2327 | tags=38% | list=14% | signal=33% |
| REACTOME_FOXO_MEDIATED_TRANSCRIPTION | 21 | 0.628401 | 1.599805 | 0.024858 | 0.267983 | 0.243636 | 4957 | tags=67% | list=31% | signal=46% |
| REACTOME_SODIUM_CALCIUM_EXCHANGERS | 12 | -0.79208 | -1.54156 | 0.025133 | 0.267983 | 0.243636 | 1019 | tags=58% | list=6% | signal=55% |
| REACTOME_NEDDYLATION | 213 | 0.402943 | 1.354303 | 0.025288 | 0.267983 | 0.243636 | 4459 | tags=41% | list=28% | signal=30% |
| REACTOME_MRNA_SPLICING | 199 | 0.397208 | 1.317084 | 0.026045 | 0.273311 | 0.24848 | 4601 | tags=41% | list=28% | signal=29% |
| BIOCARTA_CACAM_PATHWAY | 6 | 0.796982 | 1.559178 | 0.026608 | 0.27746 | 0.252252 | 1254 | tags=50% | list=8% | signal=46% |
| REACTOME_ASSOCIATION_OF_TRIC_CCT_WITH_TARGET_PROTEINS_DURING_BIOS' | 9 | 0.756947 | 1.611933 | 0.027184 | 0.281693 | 0.256101 | 3875 | tags=78% | list=24% | signal=59% |
| REACTOME_VLDLR_INTERNALISATION_AND_DEGRADATION | 16 | -0.75554 | -1.52187 | 0.027997 | 0.288318 | 0.262124 | 2 | tags=12% | list=0% | signal=13% |
| BIOCARTA_IL7_PATHWAY | 15 | 0.672427 | 1.602148 | 0.028486 | 0.291539 | 0.265053 | 2422 | tags=53% | list=15% | signal=45% |
| REACTOME_FORMATION_OF_RNA_POL_II_ELONGATION_COMPLEX | 54 | 0.578696 | 1.750125 | 0.029054 | 0.293731 | 0.267045 | 3197 | tags=44% | list=20% | signal=36% |
| REACTOME_METABOLISM_OF_CARBOHYDRATES | 247 | -0.4921 | -1.31016 | 0.028926 | 0.293731 | 0.267045 | 2507 | tags=28% | list=16% | signal=24% |
| REACTOME_PROCESSIVE_SYNTHESIS_ON_THE_LAGGING_STRAND | 14 | 0.677261 | 1.585291 | 0.029785 | 0.294063 | 0.267347 | 452 | tags=21% | list=3% | signal=21% |
| BIOCARTA_PTEN_PATHWAY | 16 | 0.648655 | 1.574738 | 0.029688 | 0.294063 | 0.267347 | 3202 | tags=56% | list=20% | signal=45% |
| REACTOME_RUNX1_INTERACTS_WITH_CO_FACTORS_WHOSE_PRECISE_EFFECT_ON | 35 | 0.532101 | 1.492012 | 0.029672 | 0.294063 | 0.267347 | 3944 | tags=37% | list=24% | signal=28% |
| REACTOME_GLOBAL_GENOME_NUCLEOTIDE_EXCISION_REPAIR_GG_NER | 80 | 0.482483 | 1.470897 | 0.029796 | 0.294063 | 0.267347 | 4670 | tags=48% | list=29% | signal=34% |
| WP_EICOSANOID_METABOLISM_VIA_CYTOCHROME_P450_MONOOXYGENASES | 4 | 0.879589 | 1.521561 | 0.030652 | 0.300716 | 0.273396 | 108 | tags=25% | list=1% | signal=25% |
| WP_G13_SIGNALING_PATHWAY | 38 | 0.610862 | 1.748656 | 0.031045 | 0.302783 | 0.275275 | 1257 | tags=32% | list=8% | signal=29% |
| WP_TRANSLATION_FACTORS | 48 | 0.60348 | 1.7881 | 0.032055 | 0.304068 | 0.276443 | 3794 | tags=54% | list=24% | signal=42% |
| REACTOME_ACTIVATION_OF_IRF3_IRF7_MEDIATED_BY_TBK1_IKK_EPSILON | 17 | 0.638418 | 1.591854 | 0.031816 | 0.304068 | 0.276443 | 2303 | tags=35% | list=14% | signal=30% |
| BIOCARTA_RAC1_PATHWAY | 21 | 0.616992 | 1.57076 | 0.032099 | 0.304068 | 0.276443 | 1999 | tags=43% | list=12% | signal=38% |
| REACTOME_MISMATCH_REPAIR | 14 | 0.669369 | 1.566818 | 0.032514 | 0.304068 | 0.276443 | 584 | tags=21% | list=4% | signal=21% |
| REACTOME_P2Y_RECEPTORS | 9 | -0.815 | -1.50021 | 0.032077 | 0.304068 | 0.276443 | 1316 | tags=56% | list=8% | signal=51% |
| REACTOME_INTERLEUKIN_2_FAMILY_SIGNALING | 26 | 0.559828 | 1.461128 | 0.032238 | 0.304068 | 0.276443 | 2293 | tags=38% | list=14% | signal=33% |
| REACTOME_RESPONSE_TO_ELEVATED_PLATELET_CYTOSOLIC_CA2 | 108 | -0.54619 | -1.39132 | 0.032644 | 0.304068 | 0.276443 | 1884 | tags=20% | list=12% | signal=18% |
| REACTOME_RAC1_GTPASE_CYCLE | 168 | 0.420037 | 1.354256 | 0.032615 | 0.304068 | 0.276443 | 2092 | tags=32% | list=13% | signal=28% |
| REACTOME_AKT_PHOSPHORYLATES_TARGETS_IN_THE_CYTOSOL | 11 | 0.714765 | 1.5824 | 0.03313 | 0.304339 | 0.276689 | 2330 | tags=55% | list=14% | signal=47% |
| REACTOME_FORMATION_OF_INCISION_COMPLEX_IN_GG_NER | 42 | 0.540172 | 1.557166 | 0.033016 | 0.304339 | 0.276689 | 2917 | tags=40% | list=18% | signal=33% |
| REACTOME_ESTABLISHMENT_OF_SISTER_CHROMATID_COHESION | 10 | 0.696315 | 1.526655 | 0.033224 | 0.304339 | 0.276689 | 3071 | tags=60% | list=19% | signal=49% |
| BIOCARTA_PTDINS_PATHWAY | 14 | 0.662147 | 1.549914 | 0.034332 | 0.312019 | 0.283672 | 3960 | tags=57% | list=25% | signal=43% |
| REACTOME_SIALIC_ACID_METABOLISM | 31 | -0.67009 | -1.4957 | 0.034439 | 0.312019 | 0.283672 | 2816 | tags=55% | list=17% | signal=45% |
| REACTOME_TRANSCRIPTIONAL_REGULATION_BY_E2F6 | 23 | 0.615783 | 1.591272 | 0.035495 | 0.319843 | 0.290785 | 2881 | tags=52% | list=18% | signal=43% |
| REACTOME_GAP_JUNCTION_TRAFFICKING_AND_REGULATION | 32 | -0.65568 | -1.46875 | 0.036709 | 0.328991 | 0.299101 | 101 | tags=16% | list=1% | signal=16% |
| REACTOME_LGI_ADAM_INTERACTIONS | 14 | -0.76931 | -1.52442 | 0.037147 | 0.331128 | 0.301045 | 971 | tags=50% | list=6% | signal=47% |
| REACTOME_WNT_LIGAND_BIOGENESIS_AND_TRAFFICKING | 20 | -0.73142 | -1.51646 | 0.037552 | 0.332949 | 0.3027 | 2400 | tags=45% | list=15% | signal=38% |
| WP_METAPATHWAY_BIOTRANSFORMATION | 97 | -0.54681 | -1.37789 | 0.038674 | 0.341072 | 0.310085 | 1904 | tags=34% | list=12% | signal=30% |
| REACTOME_RETROGRADE_NEUROTROPHIN_SIGNALLING | 11 | -0.79105 | -1.51231 | 0.040435 | 0.352853 | 0.320795 | 390 | tags=18% | list=2% | signal=18% |
| REACTOME_MAP2K_AND_MAPK_ACTIVATION | 35 | 0.509013 | 1.427274 | 0.040332 | 0.352853 | 0.320795 | 3575 | tags=43% | list=22% | signal=33% |
| REACTOME_HSP90_CHAPERONE_CYCLE_FOR_STEROID_HORMONE_RECEPTORS_SH | 47 | 0.565606 | 1.688386 | 0.041376 | 0.355612 | 0.323304 | 3184 | tags=36% | list=20% | signal=29% |
| REACTOME_SYNTHESIS_OF_PIPS_AT_THE_PLASMA_MEMBRANE | 48 | 0.560123 | 1.659634 | 0.046266 | 0.355612 | 0.323304 | 3774 | tags=56% | list=23% | signal=43% |
| WP_COMPLEMENT_ACTIVATION_CLASSICAL_PATHWAY | 13 | 0.712064 | 1.651872 | 0.041705 | 0.355612 | 0.323304 | 7 | tags=8% | list=0% | signal=8% |

| | | | | | | | | | | |
|---|---|---|---|---|---|---|---|---|---|---|
| BIOCARTA_PGC1A_PATHWAY | 13 | 0.706616 | 1.639234 | 0.044353 | 0.355612 | 0.323304 | 1254 | tags=31% | list=8% | signal=28% |
| REACTOME_ATTENUATION_PHASE | 13 | 0.698354 | 1.620066 | 0.045681 | 0.355612 | 0.323304 | 2686 | tags=54% | list=17% | signal=45% |
| BIOCARTA_EIF_PATHWAY | 9 | 0.723847 | 1.541447 | 0.042237 | 0.355612 | 0.323304 | 3637 | tags=78% | list=23% | signal=60% |
| REACTOME_ERYTHROPOIETIN_ACTIVATES_PHOSPHOINOSITIDE_3_KINASE_PI3K | 11 | 0.685007 | 1.516521 | 0.045649 | 0.355612 | 0.323304 | 1621 | tags=64% | list=10% | signal=57% |
| REACTOME_SYNTHESIS_OF_PIPS_AT_THE_GOLGI_MEMBRANE | 15 | 0.636135 | 1.515677 | 0.046086 | 0.355612 | 0.323304 | 4029 | tags=60% | list=25% | signal=45% |
| REACTOME_REGULATION_OF_LIPID_METABOLISM_BY_PPARALPHA | 16 | 0.623657 | 1.51405 | 0.043246 | 0.355612 | 0.323304 | 3471 | tags=50% | list=22% | signal=39% |
| BIOCARTA_DREAM_PATHWAY | 12 | -0.77667 | -1.51156 | 0.042169 | 0.355612 | 0.323304 | 1204 | tags=33% | list=7% | signal=31% |
| REACTOME_NA_CL_DEPENDENT_NEUROTRANSMITTER_TRANSPORTERS | 13 | -0.76673 | -1.5057 | 0.046757 | 0.355612 | 0.323304 | 2078 | tags=46% | list=13% | signal=40% |
| REACTOME_ERBB2_REGULATES_CELL_MOTILITY | 11 | -0.78669 | -1.50398 | 0.045101 | 0.355612 | 0.323304 | 2241 | tags=45% | list=14% | signal=39% |
| REACTOME_GAP_FILLING_DNA_REPAIR_SYNTHESIS_AND_LIGATION_IN_GG_NER | 21 | 0.588178 | 1.497404 | 0.04438 | 0.355612 | 0.323304 | 1801 | tags=33% | list=11% | signal=30% |
| REACTOME_PLASMA_LIPOPROTEIN_REMODELING | 18 | -0.72899 | -1.49513 | 0.042017 | 0.355612 | 0.323304 | 3529 | tags=61% | list=22% | signal=48% |
| REACTOME_NEGATIVE_REGULATION_OF_FLT3 | 14 | 0.63834 | 1.494189 | 0.046155 | 0.355612 | 0.323304 | 2069 | tags=29% | list=13% | signal=25% |
| REACTOME_ERBB2_ACTIVATES_PTK6_SIGNALING | 8 | -0.82266 | -1.47882 | 0.045531 | 0.355612 | 0.323304 | 2241 | tags=62% | list=14% | signal=54% |
| WP_NOVEL_JUNDMP1_PATHWAY | 24 | -0.68454 | -1.47567 | 0.046605 | 0.355612 | 0.323304 | 142 | tags=12% | list=1% | signal=12% |
| REACTOME_A_TETRASACCHARIDE_LINKER_SEQUENCE_IS_REQUIRED_FOR_GAG_SY | 24 | -0.68439 | -1.47536 | 0.046605 | 0.355612 | 0.323304 | 2182 | tags=50% | list=14% | signal=43% |
| REACTOME_DERMATAN_SULFATE_BIOSYNTHESIS | 10 | -0.78818 | -1.47303 | 0.046326 | 0.355612 | 0.323304 | 2328 | tags=70% | list=14% | signal=60% |
| REACTOME_CITRIC_ACID_CYCLE_TCA_CYCLE | 22 | 0.564643 | 1.447658 | 0.045967 | 0.355612 | 0.323304 | 4015 | tags=64% | list=25% | signal=48% |
| REACTOME_GLYOXYLATE_METABOLISM_AND_GLYCINE_DEGRADATION | 24 | 0.552169 | 1.436332 | 0.043213 | 0.355612 | 0.323304 | 3846 | tags=46% | list=24% | signal=35% |
| WP_DYSREGULATED_MIRNA_TARGETING_IN_INSULINPI3KAKT_SIGNALING | 26 | 0.549803 | 1.434963 | 0.042253 | 0.355612 | 0.323304 | 3752 | tags=69% | list=23% | signal=53% |
| REACTOME_INTRA_GOLGI_TRAFFIC | 34 | 0.507553 | 1.418051 | 0.043862 | 0.355612 | 0.323304 | 4282 | tags=53% | list=27% | signal=39% |
| REACTOME_SIGNALING_BY_SCF_KIT | 35 | 0.49122 | 1.377383 | 0.046537 | 0.355612 | 0.323304 | 2303 | tags=34% | list=14% | signal=29% |
| REACTOME_NEUROTRANSMITTER_RECEPTORS_AND_POSTSYNAPTIC_SIGNAL_TRAN | 130 | -0.52222 | -1.34547 | 0.044968 | 0.355612 | 0.323304 | 2219 | tags=30% | list=14% | signal=26% |
| REACTOME_S_PHASE | 143 | 0.420012 | 1.325071 | 0.045985 | 0.355612 | 0.323304 | 3371 | tags=27% | list=21% | signal=22% |
| REACTOME_ORGANELLE_BIOGENESIS_AND_MAINTENANCE | 186 | 0.388607 | 1.254201 | 0.041948 | 0.355612 | 0.323304 | 4219 | tags=41% | list=26% | signal=31% |
| REACTOME_NEUTROPHIL_DEGRANULATION | 422 | -0.44831 | -1.21836 | 0.045226 | 0.355612 | 0.323304 | 3525 | tags=31% | list=22% | signal=25% |
| BIOCARTA_AKT_PATHWAY | 19 | 0.614544 | 1.537374 | 0.047314 | 0.356575 | 0.32418 | 2781 | tags=47% | list=17% | signal=39% |
| REACTOME_CD28_DEPENDENT_PI3K_AKT_SIGNALING | 21 | 0.583494 | 1.485479 | 0.047171 | 0.356575 | 0.32418 | 2147 | tags=38% | list=13% | signal=33% |
| WP_GPCRS_SMALL_LIGAND | 14 | -0.75858 | -1.50316 | 0.047548 | 0.35672 | 0.324311 | 219 | tags=21% | list=1% | signal=21% |
| REACTOME_SULFUR_AMINO_ACID_METABOLISM | 22 | 0.561265 | 1.438998 | 0.048251 | 0.360363 | 0.327623 | 3348 | tags=41% | list=21% | signal=32% |
| BIOCARTA_CCR3_PATHWAY | 15 | 0.629522 | 1.499921 | 0.048875 | 0.360367 | 0.327627 | 3508 | tags=60% | list=22% | signal=47% |
| REACTOME_TRANSLOCATION_OF_ZAP_70_TO_IMMUNOLOGICAL_SYNAPSE | 8 | -0.81955 | -1.47324 | 0.048904 | 0.360367 | 0.327627 | 929 | tags=50% | list=6% | signal=47% |
| REACTOME_TRANSFERRIN_ENDOCYTOSIS_AND_RECYCLING | 26 | -0.67347 | -1.46391 | 0.048493 | 0.360367 | 0.327627 | 2993 | tags=50% | list=19% | signal=41% |
| REACTOME_SYNTHESIS_OF_PIPS_AT_THE_LATE_ENDOSOME_MEMBRANE | 10 | 0.677222 | 1.484793 | 0.049183 | 0.360818 | 0.328037 | 3623 | tags=60% | list=22% | signal=47% |
| REACTOME_PHASE_3_RAPID_REPOLARISATION | 6 | -0.86678 | -1.47436 | 0.05045 | 0.364186 | 0.331099 | 2063 | tags=67% | list=13% | signal=58% |
| REACTOME_P75NTR_REGULATES_AXONOGENESIS | 9 | -0.79863 | -1.47008 | 0.050489 | 0.364186 | 0.331099 | 2274 | tags=67% | list=14% | signal=57% |
| REACTOME_PYRUVATE_METABOLISM | 21 | 0.572005 | 1.456229 | 0.05052 | 0.364186 | 0.331099 | 2536 | tags=43% | list=16% | signal=36% |
| REACTOME_RNA_POLYMERASE_II_PROMOTER_ESCAPE | 44 | 0.494777 | 1.454055 | 0.050009 | 0.364186 | 0.331099 | 4814 | tags=50% | list=30% | signal=35% |
| REACTOME_PYRUVATE_METABOLISM_AND_CITRIC_ACID_TCA_CYCLE | 45 | 0.551727 | 1.632641 | 0.051722 | 0.364199 | 0.331111 | 3271 | tags=49% | list=20% | signal=39% |
| REACTOME_REGULATION_OF_PYRUVATE_DEHYDROGENASE_PDH_COMPLEX | 15 | 0.625501 | 1.490341 | 0.051667 | 0.364199 | 0.331111 | 3109 | tags=60% | list=19% | signal=48% |
| REACTOME_RECOGNITION_OF_DNA_DAMAGE_BY_PCNA_CONTAINING_REPLICATI( | 29 | 0.550339 | 1.471818 | 0.051266 | 0.364199 | 0.331111 | 1801 | tags=31% | list=11% | signal=28% |
| REACTOME_G_PROTEIN_ACTIVATION | 21 | -0.70107 | -1.47065 | 0.051771 | 0.364199 | 0.331111 | 594 | tags=14% | list=4% | signal=14% |
| REACTOME_ROS_AND_RNS_PRODUCTION_IN_PHAGOCYTES | 30 | -0.65537 | -1.45469 | 0.051414 | 0.364199 | 0.331111 | 1231 | tags=33% | list=8% | signal=31% |
| REACTOME_DNA_DOUBLE_STRAND_BREAK_RESPONSE | 56 | 0.467124 | 1.385873 | 0.05206 | 0.364199 | 0.331111 | 4097 | tags=38% | list=25% | signal=28% |
| REACTOME_TBC_RABGAPS | 34 | 0.495754 | 1.385084 | 0.052023 | 0.364199 | 0.331111 | 3523 | tags=50% | list=22% | signal=39% |
| REACTOME_MAP3K8_TPL2_DEPENDENT_MAPK1_3_ACTIVATION | 13 | 0.672102 | 1.559167 | 0.054551 | 0.367662 | 0.33426 | 2798 | tags=38% | list=17% | signal=32% |
| REACTOME_MITOPHAGY | 28 | 0.584136 | 1.546251 | 0.054201 | 0.367662 | 0.33426 | 1831 | tags=32% | list=11% | signal=29% |
| BIOCARTA_MTOR_PATHWAY | 19 | 0.608195 | 1.521493 | 0.053129 | 0.367662 | 0.33426 | 2781 | tags=47% | list=17% | signal=39% |
| REACTOME_EPHA_MEDIATED_GROWTH_CONE_COLLAPSE | 18 | -0.71375 | -1.46386 | 0.053221 | 0.367662 | 0.33426 | 711 | tags=33% | list=4% | signal=32% |
| WP_IL3_SIGNALING_PATHWAY | 94 | 0.471385 | 1.456976 | 0.053805 | 0.367662 | 0.33426 | 3059 | tags=41% | list=19% | signal=34% |
| REACTOME_ACTIVATION_OF_G_PROTEIN_GATED_POTASSIUM_CHANNELS | 26 | -0.66578 | -1.44721 | 0.053735 | 0.367662 | 0.33426 | 2116 | tags=27% | list=13% | signal=23% |
| BIOCARTA_EIF4_PATHWAY | 21 | 0.566438 | 1.442056 | 0.05387 | 0.367662 | 0.33426 | 3524 | tags=52% | list=22% | signal=41% |
| WP_TGFBETA_SIGNALING_PATHWAY | 46 | -0.60339 | -1.409 | 0.054414 | 0.367662 | 0.33426 | 1834 | tags=20% | list=11% | signal=17% |
| REACTOME_PROCESSING_OF_CAPPED_INTRON_CONTAINING_PRE_MRNA | 264 | 0.366659 | 1.26473 | 0.054495 | 0.367662 | 0.33426 | 4610 | tags=40% | list=29% | signal=29% |
| REACTOME_DUAL_INCISION_IN_GG_NER | 40 | 0.583543 | 1.685534 | 0.056658 | 0.374261 | 0.340259 | 2178 | tags=38% | list=13% | signal=33% |
| REACTOME_RHOU_GTPASE_CYCLE | 39 | 0.582561 | 1.681816 | 0.056559 | 0.374261 | 0.340259 | 2852 | tags=46% | list=18% | signal=38% |
| REACTOME_VXPX_CARGO_TARGETING_TO_CILIUM | 20 | 0.611536 | 1.569313 | 0.055783 | 0.374261 | 0.340259 | 2424 | tags=40% | list=15% | signal=34% |
| REACTOME_GLYCOGEN_SYNTHESIS | 5 | 0.803761 | 1.467479 | 0.056645 | 0.374261 | 0.340259 | 2778 | tags=80% | list=17% | signal=66% |
| REACTOME_KEAP1_NFE2L2_PATHWAY | 74 | 0.433372 | 1.323078 | 0.056288 | 0.374261 | 0.340259 | 5771 | tags=55% | list=36% | signal=36% |
| REACTOME_RNA_POLYMERASE_II_TRANSCRIBES_SNRNA_GENES | 71 | 0.456277 | 1.386373 | 0.057047 | 0.375335 | 0.341235 | 4580 | tags=46% | list=28% | signal=33% |
| REACTOME_GPER1_SIGNALING | 26 | -0.66079 | -1.43635 | 0.057667 | 0.377913 | 0.343579 | 936 | tags=15% | list=6% | signal=15% |
| REACTOME_SIGNALING_BY_ACTIVIN | 13 | -0.7498 | -1.47245 | 0.058824 | 0.379433 | 0.344961 | 1236 | tags=38% | list=8% | signal=36% |
| REACTOME_TRANSCRIPTION_COUPLED_NUCLEOTIDE_EXCISION_REPAIR_TC_NER | 75 | 0.476852 | 1.464224 | 0.059129 | 0.379433 | 0.344961 | 5703 | tags=64% | list=35% | signal=42% |
| REACTOME_FGFR3B_LIGAND_BINDING_AND_ACTIVATION | 6 | -0.85888 | -1.46093 | 0.059459 | 0.379433 | 0.344961 | 540 | tags=33% | list=3% | signal=32% |
| WP_CYTOKINES_AND_INFLAMMATORY_RESPONSE | 9 | -0.78739 | -1.44938 | 0.058632 | 0.379433 | 0.344961 | 1834 | tags=44% | list=11% | signal=39% |
| WP_GENE_REGULATORY_NETWORK_MODELLING_SOMITOGENESIS | 8 | -0.8055 | -1.44798 | 0.059022 | 0.379433 | 0.344961 | 1454 | tags=38% | list=9% | signal=34% |
| REACTOME_CYP2E1_REACTIONS | 3 | -0.93048 | -1.41727 | 0.059501 | 0.379433 | 0.344961 | 53 | tags=67% | list=0% | signal=66% |
| WP_ALZHEIMERS_DISEASE | 69 | -0.56355 | -1.37565 | 0.059158 | 0.379433 | 0.344961 | 2125 | tags=29% | list=13% | signal=25% |
| REACTOME_N_GLYCAN_ANTENNAE_ELONGATION_IN_THE_MEDIAL_TRANS_GOLGI | 23 | -0.67523 | -1.43388 | 0.060241 | 0.38268 | 0.347913 | 3206 | tags=52% | list=20% | signal=42% |
| REACTOME_TP53_REGULATES_TRANSCRIPTION_OF_DNA_REPAIR_GENES | 43 | 0.567929 | 1.648693 | 0.06124 | 0.386071 | 0.350996 | 3558 | tags=44% | list=22% | signal=35% |
| REACTOME_INWARDLY_RECTIFYING_K_CHANNELS | 31 | -0.63985 | -1.42821 | 0.061224 | 0.386071 | 0.350996 | 2116 | tags=23% | list=13% | signal=20% |
| BIOCARTA_TRKA_PATHWAY | 12 | -0.75666 | -1.47262 | 0.061747 | 0.38779 | 0.352559 | 390 | tags=17% | list=2% | signal=16% |
| REACTOME_RAF_ACTIVATION | 34 | 0.482187 | 1.347181 | 0.062223 | 0.389305 | 0.353936 | 5228 | tags=71% | list=32% | signal=48% |
| BIOCARTA_NDKDYNAMIN_PATHWAY | 13 | 0.651106 | 1.51046 | 0.062533 | 0.389776 | 0.354365 | 3432 | tags=54% | list=21% | signal=42% |
| REACTOME_SUMOYLATION_OF_TRANSCRIPTION_COFACTORS | 38 | 0.542657 | 1.553413 | 0.063127 | 0.390539 | 0.355058 | 3944 | tags=45% | list=24% | signal=34% |
| BIOCARTA_S1P_PATHWAY | 6 | -0.85428 | -1.45309 | 0.063063 | 0.390539 | 0.355058 | 2258 | tags=67% | list=14% | signal=57% |
| REACTOME_ACETYLCHOLINE_BINDING_AND_DOWNSTREAM_EVENTS | 10 | -0.7702 | -1.43943 | 0.063898 | 0.39238 | 0.356731 | 2307 | tags=60% | list=14% | signal=51% |
| REACTOME_HIGHLY_CALCIUM_PERMEABLE_POSTSYNAPTIC_NICOTINIC_ACETYLCH( | 10 | -0.7702 | -1.43943 | 0.063898 | 0.39238 | 0.356731 | 2307 | tags=60% | list=14% | signal=51% |
| REACTOME_RNA_POLYMERASE_I_TRANSCRIPTION_TERMINATION | 29 | 0.521791 | 1.39547 | 0.064163 | 0.392557 | 0.356892 | 3005 | tags=34% | list=19% | signal=28% |
| WP_IL9_SIGNALING_PATHWAY | 22 | 0.54414 | 1.395093 | 0.064844 | 0.395265 | 0.359354 | 3034 | tags=59% | list=19% | signal=48% |
| REACTOME_FOXO_MEDIATED_TRANSCRIPTION_OF_CELL_CYCLE_GENES | 8 | 0.735222 | 1.538423 | 0.066015 | 0.39541 | 0.359486 | 2350 | tags=50% | list=15% | signal=43% |
| WP_MAPK_CASCADE | 28 | 0.560476 | 1.483623 | 0.068166 | 0.39541 | 0.359486 | 3970 | tags=54% | list=25% | signal=40% |
| REACTOME_JOSEPHIN_DOMAIN_DUBS | 11 | 0.669315 | 1.481779 | 0.069638 | 0.39541 | 0.359486 | 3801 | tags=64% | list=24% | signal=49% |
| REACTOME_MRNA_CAPPING | 28 | 0.55615 | 1.472172 | 0.06944 | 0.39541 | 0.359486 | 4829 | tags=54% | list=30% | signal=38% |
| REACTOME_RAB_REGULATION_OF_TRAFFICKING | 112 | 0.454553 | 1.455466 | 0.066219 | 0.39541 | 0.359486 | 3355 | tags=46% | list=21% | signal=37% |
| BIOCARTA_MAPK_PATHWAY | 76 | 0.471245 | 1.453207 | 0.067623 | 0.39541 | 0.359486 | 1253 | tags=24% | list=8% | signal=22% |
| REACTOME_PROLONGED_ERK_ACTIVATION_EVENTS | 14 | -0.73067 | -1.44785 | 0.066865 | 0.39541 | 0.359486 | 390 | tags=14% | list=2% | signal=14% |
| BIOCARTA_ALTERNATIVE_PATHWAY | 5 | 0.789226 | 1.440941 | 0.067538 | 0.39541 | 0.359486 | 1433 | tags=40% | list=9% | signal=36% |
| REACTOME_CELLULAR_RESPONSE_TO_HEAT_STRESS | 81 | 0.473646 | 1.440661 | 0.067765 | 0.39541 | 0.359486 | 3092 | tags=36% | list=19% | signal=29% |
| BIOCARTA_ION_PATHWAY | 5 | 0.788995 | 1.44052 | 0.067538 | 0.39541 | 0.359486 | 3358 | tags=60% | list=21% | signal=48% |
| REACTOME_SURFACTANT_METABOLISM | 15 | -0.722 | -1.43995 | 0.067747 | 0.39541 | 0.359486 | 3478 | tags=67% | list=22% | signal=52% |
| REACTOME_FRUCTOSE_CATABOLISM | 5 | 0.788246 | 1.439152 | 0.067538 | 0.39541 | 0.359486 | 132 | tags=20% | list=1% | signal=20% |
| BIOCARTA_BTG2_PATHWAY | 9 | -0.78175 | -1.439 | 0.066775 | 0.39541 | 0.359486 | 2026 | tags=44% | list=13% | signal=39% |
| REACTOME_REVERSIBLE_HYDRATION_OF_CARBON_DIOXIDE | 10 | -0.76735 | -1.4341 | 0.06869 | 0.39541 | 0.359486 | 378 | tags=30% | list=2% | signal=29% |
| WP_OXIDATION_BY_CYTOCHROME_P450 | 23 | -0.67054 | -1.42391 | 0.069612 | 0.39541 | 0.359486 | 2595 | tags=39% | list=16% | signal=33% |
| REACTOME_TRANSPORT_OF_ORGANIC_ANIONS | 10 | -0.76177 | -1.42366 | 0.06869 | 0.39541 | 0.359486 | 2466 | tags=60% | list=15% | signal=51% |
| REACTOME_ACYL_CHAIN_REMODELLING_OF_PI | 10 | -0.76121 | -1.42262 | 0.06869 | 0.39541 | 0.359486 | 2489 | tags=50% | list=15% | signal=42% |
| REACTOME_CYCLIN_A_CDK2_ASSOCIATED_EVENTS_AT_S_PHASE_ENTRY | 71 | 0.446502 | 1.356673 | 0.065607 | 0.39541 | 0.359486 | 3000 | tags=25% | list=19% | signal=21% |
| REACTOME_SUMOYLATION | 151 | 0.409708 | 1.299306 | 0.065299 | 0.39541 | 0.359486 | 3471 | tags=36% | list=22% | signal=29% |

| | | | | | | | | | | |
|---|---|---|---|---|---|---|---|---|---|---|
| REACTOME_EXTRACELLULAR_MATRIX_ORGANIZATION | 207 | -0.47787 | -1.25987 | 0.069255 | 0.39541 | 0.359486 | 2321 | tags=33% | list=14% | signal=28% |
| WP_EICOSANOID_METABOLISM_VIA_LIPOXYGENASES_LOX | 21 | 0.546203 | 1.390541 | 0.070124 | 0.396813 | 0.360762 | 2174 | tags=24% | list=13% | signal=21% |
| REACTOME_POSITIVE_EPIGENETIC_REGULATION_OF_RRNA_EXPRESSION | 29 | 0.510308 | 1.364759 | 0.070637 | 0.398356 | 0.362165 | 2459 | tags=28% | list=15% | signal=23% |
| WP_MATRIX_METALLOPROTEINASES | 18 | -0.69496 | -1.42534 | 0.072829 | 0.407942 | 0.37088 | 2295 | tags=50% | list=14% | signal=43% |
| REACTOME_GABA_B_RECEPTOR_ACTIVATION | 39 | -0.60652 | -1.3804 | 0.072773 | 0.407942 | 0.37088 | 2116 | tags=28% | list=13% | signal=25% |
| REACTOME_INOSITOL_PHOSPHATE_METABOLISM | 46 | 0.496559 | 1.464803 | 0.073094 | 0.408046 | 0.370974 | 2630 | tags=37% | list=16% | signal=31% |
| REACTOME_HSF1_ACTIVATION | 8 | 0.720945 | 1.50855 | 0.07335 | 0.4081 | 0.371023 | 2709 | tags=62% | list=17% | signal=52% |
| WP_IL2_SIGNALING_PATHWAY | 73 | 0.469623 | 1.432695 | 0.073607 | 0.408163 | 0.371081 | 2835 | tags=40% | list=18% | signal=33% |
| REACTOME_MACROAUTOPHAGY | 112 | 0.452275 | 1.448172 | 0.073953 | 0.408713 | 0.371581 | 4710 | tags=50% | list=29% | signal=36% |
| REACTOME_NECTIN_NECL_TRANS_HETERODIMERIZATION | 7 | -0.81669 | -1.4162 | 0.074468 | 0.410193 | 0.372926 | 2966 | tags=100% | list=18% | signal=82% |
| REACTOME_ESR_MEDIATED_SIGNALING | 121 | 0.383257 | 1.236663 | 0.074932 | 0.411384 | 0.374009 | 2437 | tags=26% | list=15% | signal=22% |
| REACTOME_PCNA_DEPENDENT_LONG_PATCH_BASE_EXCISION_REPAIR | 20 | 0.586268 | 1.504472 | 0.07597 | 0.413455 | 0.375892 | 1974 | tags=35% | list=12% | signal=31% |
| REACTOME_RECYCLING_OF_EIF2_GDP | 8 | 0.716698 | 1.499663 | 0.075795 | 0.413455 | 0.375892 | 3652 | tags=75% | list=23% | signal=58% |
| REACTOME_DNA_STRAND_ELONGATION | 23 | 0.546494 | 1.41222 | 0.076058 | 0.413455 | 0.375892 | 1784 | tags=22% | list=11% | signal=19% |
| REACTOME_MITOCHONDRIAL_FATTY_ACID_BETA_OXIDATION_OF_SATURATED_FA | 9 | 0.68674 | 1.462427 | 0.07732 | 0.414556 | 0.376893 | 4472 | tags=89% | list=28% | signal=64% |
| REACTOME_TRAFFICKING_AND_PROCESSING_OF_ENDOSOMAL_TLR | 11 | -0.75285 | -1.43927 | 0.07776 | 0.414556 | 0.376893 | 3337 | tags=73% | list=21% | signal=58% |
| REACTOME_CS_DS_DEGRADATION | 13 | -0.72796 | -1.42957 | 0.076923 | 0.414556 | 0.376893 | 3535 | tags=69% | list=22% | signal=54% |
| WP_STATIN_PATHWAY | 16 | -0.70649 | -1.42306 | 0.077713 | 0.414556 | 0.376893 | 3529 | tags=62% | list=22% | signal=49% |
| WP_IL7_SIGNALING_PATHWAY | 42 | 0.488499 | 1.408207 | 0.077619 | 0.414556 | 0.376893 | 3960 | tags=43% | list=25% | signal=32% |
| REACTOME_ION_TRANSPORT_BY_P_TYPE_ATPASES | 48 | -0.58214 | -1.36816 | 0.077201 | 0.414556 | 0.376893 | 2559 | tags=46% | list=16% | signal=39% |
| REACTOME_CELLULAR_RESPONSE_TO_HYPOXIA | 68 | 0.525585 | 1.576529 | 0.078395 | 0.415266 | 0.377538 | 2552 | tags=21% | list=16% | signal=17% |
| REACTOME_WNT_MEDIATED_ACTIVATION_OF_DVL | 8 | 0.713397 | 1.492755 | 0.07824 | 0.415266 | 0.377538 | 2743 | tags=50% | list=17% | signal=42% |
| REACTOME_DOPAMINE_RECEPTORS | 4 | -0.87902 | -1.42109 | 0.079096 | 0.417362 | 0.379444 | 22 | tags=75% | list=0% | signal=75% |
| REACTOME_PLATELET_HOMEOSTASIS | 71 | -0.54368 | -1.32922 | 0.079545 | 0.417362 | 0.379444 | 2203 | tags=25% | list=14% | signal=22% |
| REACTOME_METABOLISM_OF_POLYAMINES | 56 | 0.436718 | 1.295666 | 0.079353 | 0.417362 | 0.379444 | 3371 | tags=27% | list=21% | signal=21% |
| REACTOME_TIE2_SIGNALING | 17 | 0.597937 | 1.490918 | 0.081169 | 0.417453 | 0.379527 | 2388 | tags=35% | list=15% | signal=30% |
| REACTOME_INTERLEUKIN_RECEPTOR_SHC_SIGNALING | 16 | 0.608445 | 1.477121 | 0.08125 | 0.417453 | 0.379527 | 1613 | tags=44% | list=10% | signal=39% |
| REACTOME_REGULATION_OF_RAS_BY_GAPS | 65 | 0.48564 | 1.464694 | 0.081037 | 0.417453 | 0.379527 | 2552 | tags=25% | list=16% | signal=21% |
| REACTOME_MISMATCH_REPAIR_MMR_DIRECTED_BY_MSH2_MSH3_MUTSBETA | 9 | 0.683616 | 1.455774 | 0.079897 | 0.417453 | 0.379527 | 452 | tags=22% | list=3% | signal=22% |
| BIOCARTA_CPSF_PATHWAY | 5 | 0.779798 | 1.423729 | 0.08061 | 0.417453 | 0.379527 | 1521 | tags=40% | list=9% | signal=36% |
| REACTOME_SHC1_EVENTS_IN_ERBB2_SIGNALING | 12 | -0.72975 | -1.42025 | 0.081325 | 0.417453 | 0.379527 | 2241 | tags=50% | list=14% | signal=43% |
| REACTOME_FORMATION_OF_TC_NER_PRE_INCISION_COMPLEX | 52 | 0.452006 | 1.381828 | 0.081228 | 0.417453 | 0.379527 | 5703 | tags=60% | list=35% | signal=39% |
| REACTOME_MIRO_GTPASE_CYCLE | 7 | 0.726762 | 1.499553 | 0.082192 | 0.417689 | 0.379742 | 4315 | tags=86% | list=27% | signal=63% |
| REACTOME_NUCLEOTIDE_EXCISION_REPAIR | 107 | 0.461622 | 1.473225 | 0.081968 | 0.417689 | 0.379742 | 5703 | tags=60% | list=35% | signal=39% |
| REACTOME_SYNTHESIS_OF_PE | 12 | 0.643223 | 1.449683 | 0.08284 | 0.417689 | 0.379742 | 669 | tags=25% | list=4% | signal=24% |
| REACTOME_REGULATION_OF_TP53_ACTIVITY_THROUGH_ASSOCIATION_WITH_CO | 12 | 0.642624 | 1.448334 | 0.08284 | 0.417689 | 0.379742 | 2147 | tags=50% | list=13% | signal=43% |
| REACTOME_CHYLOMICRON_ASSEMBLY | 6 | -0.84072 | -1.43003 | 0.082883 | 0.417689 | 0.379742 | 781 | tags=50% | list=5% | signal=48% |
| REACTOME_PHYSIOLOGICAL_FACTORS | 5 | -0.86286 | -1.42405 | 0.082873 | 0.417689 | 0.379742 | 308 | tags=60% | list=2% | signal=59% |
| REACTOME_LAGGING_STRAND_SYNTHESIS | 19 | 0.563186 | 1.408894 | 0.08447 | 0.424398 | 0.385841 | 4670 | tags=58% | list=29% | signal=41% |
| REACTOME_TETRAHYDROBIOPTERIN_BH4_SYNTHESIS_RECYCLING_SALVAGE_AND_ | 9 | 0.679132 | 1.446225 | 0.085052 | 0.425019 | 0.386406 | 2797 | tags=56% | list=17% | signal=46% |
| REACTOME_POLYMERASE_SWITCHING | 14 | 0.61029 | 1.428529 | 0.085106 | 0.425019 | 0.386406 | 4670 | tags=71% | list=29% | signal=51% |
| REACTOME_BASE_EXCISION_REPAIR | 38 | 0.507303 | 1.452206 | 0.087275 | 0.432258 | 0.392987 | 2289 | tags=32% | list=14% | signal=27% |
| REACTOME_PROCESSING_AND_ACTIVATION_OF_SUMO | 10 | 0.649068 | 1.423067 | 0.087766 | 0.432258 | 0.392987 | 3240 | tags=60% | list=20% | signal=48% |
| REACTOME_COHESIN_LOADING_ONTO_CHROMATIN | 10 | 0.64399 | 1.411934 | 0.087766 | 0.432258 | 0.392987 | 4533 | tags=80% | list=28% | signal=58% |
| REACTOME_PHASE_2_PLATEAU_PHASE | 12 | -0.72316 | -1.40741 | 0.087349 | 0.432258 | 0.392987 | 2063 | tags=58% | list=13% | signal=51% |
| REACTOME_REGULATED_PROTEOLYSIS_OF_P75NTR | 10 | -0.74686 | -1.39581 | 0.087859 | 0.432258 | 0.392987 | 3156 | tags=70% | list=20% | signal=56% |
| REACTOME_PEROXISOMAL_PROTEIN_IMPORT | 59 | 0.495569 | 1.477803 | 0.088597 | 0.432823 | 0.3935 | 3582 | tags=41% | list=22% | signal=32% |
| REACTOME_NR1H2_AND_NR1H3_MEDIATED_SIGNALING | 12 | 0.63224 | 1.424932 | 0.088757 | 0.432823 | 0.3935 | 1196 | tags=33% | list=7% | signal=31% |
| REACTOME_ACROSOME_REACTION_AND_SPERM_OOCYTE_MEMBRANE_BINDING | 3 | -0.9115 | -1.38837 | 0.088292 | 0.432823 | 0.3935 | 151 | tags=67% | list=1% | signal=66% |
| REACTOME_SYNTHESIS_OF_BILE_ACIDS_AND_BILE_SALTS_VIA_24_HYDROXYCHOLE | 10 | -0.74511 | -1.39252 | 0.089457 | 0.434955 | 0.395438 | 921 | tags=50% | list=6% | signal=47% |
| WP_OSTEOCLAST_SIGNALING | 11 | -0.7396 | -1.41394 | 0.090202 | 0.437296 | 0.397567 | 2150 | tags=45% | list=13% | signal=39% |
| REACTOME_METABOLISM_OF_COFACTORS | 14 | 0.60066 | 1.405989 | 0.091185 | 0.440774 | 0.400729 | 3129 | tags=50% | list=19% | signal=40% |
| REACTOME_FCGAMMA_RECEPTOR_FCGR_DEPENDENT_PHAGOCYTOSIS | 76 | 0.445418 | 1.373563 | 0.09156 | 0.441297 | 0.401205 | 3034 | tags=42% | list=19% | signal=34% |
| WP_MACROPHAGE_MARKERS | 10 | -0.74133 | -1.38546 | 0.092652 | 0.445266 | 0.404813 | 1356 | tags=70% | list=8% | signal=64% |
| REACTOME_FERTILIZATION | 11 | -0.73538 | -1.40589 | 0.093313 | 0.447145 | 0.406522 | 481 | tags=27% | list=3% | signal=26% |
| REACTOME_DNA_REPAIR | 278 | 0.33478 | 1.181394 | 0.093646 | 0.44745 | 0.406798 | 4542 | tags=37% | list=28% | signal=27% |
| REACTOME_IRAK1_RECRUITS_IKK_COMPLEX | 14 | 0.598381 | 1.400655 | 0.094225 | 0.447636 | 0.406968 | 4227 | tags=57% | list=26% | signal=42% |
| REACTOME_SIGNALING_BY_MST1 | 4 | -0.86615 | -1.40028 | 0.094162 | 0.447636 | 0.406968 | 950 | tags=50% | list=6% | signal=47% |
| REACTOME_PURINE_RIBONUCLEOSIDE_MONOPHOSPHATE_BIOSYNTHESIS | 9 | 0.667263 | 1.42095 | 0.095361 | 0.450699 | 0.409752 | 3295 | tags=67% | list=20% | signal=53% |
| REACTOME_MITOCHONDRIAL_TRANSLATION | 88 | 0.428062 | 1.326664 | 0.095413 | 0.450699 | 0.409752 | 3899 | tags=36% | list=24% | signal=28% |
| BIOCARTA_CTL_PATHWAY | 7 | -0.80641 | -1.39836 | 0.095745 | 0.450979 | 0.410007 | 2076 | tags=57% | list=13% | signal=50% |
| REACTOME_REGULATION_OF_FOXO_TRANSCRIPTIONAL_ACTIVITY_BY_ACETYLATIO | 6 | 0.748742 | 1.464803 | 0.096197 | 0.451826 | 0.410777 | 85 | tags=17% | list=1% | signal=17% |
| REACTOME_LIGAND_RECEPTOR_INTERACTIONS | 6 | -0.82886 | -1.40986 | 0.097297 | 0.454707 | 0.413396 | 2186 | tags=67% | list=14% | signal=58% |
| REACTOME_REGULATION_OF_CHOLESTEROL_BIOSYNTHESIS_BY_SREBP_SREBF | 7 | -0.80604 | -1.39772 | 0.097518 | 0.454707 | 0.413396 | 2258 | tags=57% | list=14% | signal=49% |
| REACTOME_REGULATION_OF_INNATE_IMMUNE_RESPONSES_TO_CYTOSOLIC_DNA | 12 | 0.61975 | 1.39678 | 0.097633 | 0.454707 | 0.413396 | 1379 | tags=33% | list=9% | signal=31% |
| REACTOME_SIGNALING_BY_HIPPO | 20 | 0.560836 | 1.439208 | 0.100408 | 0.458845 | 0.417159 | 4490 | tags=50% | list=28% | signal=36% |
| BIOCARTA_EXTRINSIC_PATHWAY | 5 | -0.85529 | -1.41156 | 0.101289 | 0.458845 | 0.417159 | 1542 | tags=40% | list=10% | signal=36% |
| REACTOME_N_GLYCAN_TRIMMING_AND_ELONGATION_IN_THE_CIS_GOLGI | 5 | -0.85312 | -1.40797 | 0.101289 | 0.458845 | 0.417159 | 535 | tags=60% | list=3% | signal=58% |
| REACTOME_MUSCARINIC_ACETYLCHOLINE_RECEPTORS | 5 | -0.85209 | -1.40628 | 0.101289 | 0.458845 | 0.417159 | 1439 | tags=80% | list=9% | signal=73% |
| REACTOME_PHASE_0_RAPID_DEPOLARISATION | 8 | -0.775 | -1.39315 | 0.10118 | 0.458845 | 0.417159 | 809 | tags=62% | list=5% | signal=59% |
| REACTOME_TRYPTOPHAN_CATABOLISM | 8 | -0.77499 | -1.39312 | 0.10118 | 0.458845 | 0.417159 | 64 | tags=25% | list=0% | signal=25% |
| REACTOME_CELLULAR_HEXOSE_TRANSPORT | 11 | -0.72628 | -1.38848 | 0.101089 | 0.458845 | 0.417159 | 2517 | tags=55% | list=16% | signal=46% |
| BIOCARTA_UCALPAIN_PATHWAY | 12 | -0.71287 | -1.38739 | 0.099398 | 0.458845 | 0.417159 | 121 | tags=25% | list=1% | signal=25% |
| REACTOME_NUCLEOTIDE_SALVAGE | 21 | 0.539701 | 1.373989 | 0.100746 | 0.458845 | 0.417159 | 3145 | tags=43% | list=19% | signal=35% |
| REACTOME_INSULIN_RECEPTOR_RECYCLING | 24 | -0.63539 | -1.36972 | 0.099867 | 0.458845 | 0.417159 | 2917 | tags=42% | list=18% | signal=34% |
| BIOCARTA_HSP27_PATHWAY | 12 | 0.616426 | 1.389289 | 0.10355 | 0.466539 | 0.424153 | 3569 | tags=42% | list=22% | signal=32% |
| REACTOME_SIGNALLING_TO_RAS | 12 | 0.614186 | 1.384242 | 0.10355 | 0.466539 | 0.424153 | 1253 | tags=33% | list=8% | signal=31% |
| REACTOME_HIGHLY_CALCIUM_PERMEABLE_NICOTINIC_ACETYLCHOLINE_RECEPTO | 9 | -0.75728 | -1.39396 | 0.104235 | 0.467083 | 0.424648 | 2307 | tags=56% | list=14% | signal=48% |
| BIOCARTA_NKT_PATHWAY | 14 | -0.69503 | -1.37723 | 0.104012 | 0.467083 | 0.424648 | 3453 | tags=64% | list=21% | signal=51% |
| WP_CHOLESTEROL_BIOSYNTHESIS | 15 | -0.68761 | -1.37137 | 0.104566 | 0.467304 | 0.424848 | 2056 | tags=53% | list=13% | signal=47% |
| BIOCARTA_FOSB_PATHWAY | 5 | -0.8517 | -1.40563 | 0.104972 | 0.467861 | 0.425355 | 968 | tags=60% | list=6% | signal=56% |
| REACTOME_KERATAN_SULFATE_DEGRADATION | 12 | -0.70349 | -1.36914 | 0.105422 | 0.468604 | 0.42603 | 3575 | tags=58% | list=22% | signal=45% |
| REACTOME_DUAL_INCISION_IN_TC_NER | 63 | 0.49936 | 1.501891 | 0.107425 | 0.469366 | 0.426724 | 2459 | tags=32% | list=15% | signal=27% |
| REACTOME_COBALAMIN_CBL_METABOLISM | 7 | 0.701103 | 1.446608 | 0.109589 | 0.469366 | 0.426724 | 3318 | tags=71% | list=21% | signal=57% |
| REACTOME_SIGNALING_BY_NODAL | 6 | 0.732429 | 1.43289 | 0.10962 | 0.469366 | 0.426724 | 3034 | tags=67% | list=19% | signal=54% |
| REACTOME_DNA_REPLICATION_INITIATION | 8 | 0.677471 | 1.417583 | 0.114914 | 0.469366 | 0.426724 | 4166 | tags=62% | list=26% | signal=46% |
| REACTOME_POLB_DEPENDENT_LONG_PATCH_BASE_EXCISION_REPAIR | 8 | 0.67437 | 1.411093 | 0.114914 | 0.469366 | 0.426724 | 2178 | tags=50% | list=13% | signal=43% |
| BIOCARTA_TCYTOTOXIC_PATHWAY | 5 | -0.84939 | -1.40182 | 0.108656 | 0.469366 | 0.426724 | 2076 | tags=80% | list=13% | signal=70% |
| WP_SYNTHESIS_AND_DEGRADATION_OF_KETONE_BODIES | 5 | 0.763655 | 1.394255 | 0.108932 | 0.469366 | 0.426724 | 3499 | tags=80% | list=22% | signal=63% |
| REACTOME_RUNX3_REGULATES_YAP1_MEDIATED_TRANSCRIPTION | 5 | 0.761948 | 1.391139 | 0.115468 | 0.469366 | 0.426724 | 1198 | tags=60% | list=7% | signal=56% |
| REACTOME_ERYTHROPOIETIN_ACTIVATES_RAS | 9 | 0.649169 | 1.382418 | 0.113402 | 0.469366 | 0.426724 | 1613 | tags=44% | list=10% | signal=40% |
| REACTOME_METABOLISM_OF_ANGIOTENSINOGEN_TO_ANGIOTENSINS | 9 | -0.74966 | -1.37993 | 0.112378 | 0.469366 | 0.426724 | 1435 | tags=56% | list=9% | signal=51% |
| REACTOME_ASSEMBLY_OF_ACTIVE_LPL_AND_LIPC_LIPASE_COMPLEXES | 9 | -0.74956 | -1.37975 | 0.112378 | 0.469366 | 0.426724 | 3022 | tags=78% | list=19% | signal=63% |
| REACTOME_GLYCOGEN_METABOLISM | 17 | 0.552662 | 1.378027 | 0.113636 | 0.469366 | 0.426724 | 5321 | tags=76% | list=33% | signal=51% |
| REACTOME_ACYL_CHAIN_REMODELLING_OF_PG | 11 | -0.72031 | -1.37706 | 0.107309 | 0.469366 | 0.426724 | 2589 | tags=55% | list=16% | signal=46% |
| BIOCARTA_ERBB4_PATHWAY | 7 | -0.79269 | -1.37457 | 0.109929 | 0.469366 | 0.426724 | 1769 | tags=43% | list=11% | signal=38% |
| REACTOME_MISCELLANEOUS_TRANSPORT_AND_BINDING_EVENTS | 19 | -0.6657 | -1.37279 | 0.107093 | 0.469366 | 0.426724 | 2577 | tags=63% | list=16% | signal=53% |
| REACTOME_THE_ACTIVATION_OF_ARYLSULFATASES | 8 | -0.76257 | -1.3708 | 0.111298 | 0.469366 | 0.426724 | 3391 | tags=75% | list=21% | signal=59% |

| | | | | | | | | | | |
|---|---|---|---|---|---|---|---|---|---|---|
| REACTOME_PI3K_AKT_ACTIVATION | 8 | -0.76246 | -1.37061 | 0.111298 | 0.469366 | 0.426724 | 390 | tags=25% | list=2% | signal=24% |
| REACTOME_DEGRADATION_OF_CYSTEINE_AND_HOMOCYSTEINE | 13 | 0.588925 | 1.366209 | 0.109145 | 0.469366 | 0.426724 | 4373 | tags=54% | list=27% | signal=39% |
| REACTOME_GROWTH_HORMONE_RECEPTOR_SIGNALING | 10 | 0.622869 | 1.365626 | 0.109043 | 0.469366 | 0.426724 | 3034 | tags=60% | list=19% | signal=49% |
| REACTOME_NEGATIVE_REGULATORS_OF_DDX58_IFIH1_SIGNALING | 14 | 0.583286 | 1.365321 | 0.115502 | 0.469366 | 0.426724 | 1359 | tags=21% | list=8% | signal=20% |
| BIOCARTA_PITX2_PATHWAY | 13 | 0.587604 | 1.363145 | 0.115044 | 0.469366 | 0.426724 | 3960 | tags=54% | list=25% | signal=41% |
| BIOCARTA_ETS_PATHWAY | 17 | -0.67493 | -1.36224 | 0.113833 | 0.469366 | 0.426724 | 391 | tags=24% | list=2% | signal=23% |
| REACTOME_WNT5A_DEPENDENT_INTERNALIZATION_OF_FZD4 | 14 | 0.581463 | 1.361054 | 0.115502 | 0.469366 | 0.426724 | 2395 | tags=43% | list=15% | signal=37% |
| REACTOME_PD_1_SIGNALING | 10 | -0.72795 | -1.36046 | 0.110224 | 0.469366 | 0.426724 | 929 | tags=40% | list=6% | signal=38% |
| WP_SIGNAL_TRANSDUCTION_OF_S1P_RECEPTOR | 21 | -0.64815 | -1.35963 | 0.107629 | 0.469366 | 0.426724 | 156 | tags=14% | list=1% | signal=14% |
| WP_HYPERTROPHY_MODEL | 16 | -0.67454 | -1.35872 | 0.11437 | 0.469366 | 0.426724 | 20 | tags=12% | list=0% | signal=12% |
| REACTOME_PHASE_4_RESTING_MEMBRANE_POTENTIAL | 14 | -0.68554 | -1.35843 | 0.114413 | 0.469366 | 0.426724 | 3276 | tags=64% | list=20% | signal=51% |
| REACTOME_DEADENYLATION_DEPENDENT_MRNA_DECAY | 51 | 0.44558 | 1.34872 | 0.10959 | 0.469366 | 0.426724 | 3529 | tags=35% | list=22% | signal=28% |
| BIOCARTA_GH_PATHWAY | 25 | 0.518092 | 1.341876 | 0.107438 | 0.469366 | 0.426724 | 3508 | tags=44% | list=22% | signal=34% |
| REACTOME_NOTCH_HLH_TRANSCRIPTION_PATHWAY | 22 | 0.520915 | 1.335546 | 0.114504 | 0.469366 | 0.426724 | 1236 | tags=32% | list=8% | signal=29% |
| REACTOME_HATS_ACETYLATE_HISTONES | 44 | 0.448435 | 1.317865 | 0.114027 | 0.469366 | 0.426724 | 4498 | tags=43% | list=28% | signal=31% |
| WP_WNT_SIGNALING_PATHWAY_WP403 | 57 | -0.54843 | -1.31155 | 0.113716 | 0.469366 | 0.426724 | 2400 | tags=26% | list=15% | signal=22% |
| WP_ENDOCHONDRAL_OSSIFICATION | 55 | -0.54462 | -1.3045 | 0.11462 | 0.469366 | 0.426724 | 1994 | tags=25% | list=12% | signal=22% |
| REACTOME_SELECTIVE_AUTOPHAGY | 57 | 0.416417 | 1.240545 | 0.107968 | 0.469366 | 0.426724 | 4710 | tags=44% | list=29% | signal=31% |
| REACTOME_DNA_DOUBLE_STRAND_BREAK_REPAIR | 139 | 0.37762 | 1.198037 | 0.112236 | 0.469366 | 0.426724 | 3327 | tags=29% | list=21% | signal=24% |
| REACTOME_INTERACTION_BETWEEN_L1_AND_ANKYRINS | 9 | 0.647695 | 1.379278 | 0.115979 | 0.469967 | 0.42727 | 2083 | tags=44% | list=13% | signal=39% |
| WP_INFLAMMATORY_RESPONSE_PATHWAY | 22 | -0.63733 | -1.34617 | 0.116216 | 0.469967 | 0.42727 | 2742 | tags=45% | list=17% | signal=38% |
| BIOCARTA_IFNA_PATHWAY | 7 | 0.689818 | 1.423323 | 0.121005 | 0.470056 | 0.427351 | 1448 | tags=43% | list=9% | signal=39% |
| REACTOME_MTOR_SIGNALLING | 40 | 0.492323 | 1.422049 | 0.121319 | 0.470056 | 0.427351 | 4677 | tags=57% | list=29% | signal=41% |
| REACTOME_SUMO_IS_PROTEOLYTICALLY_PROCESSED | 6 | 0.726795 | 1.421867 | 0.120805 | 0.470056 | 0.427351 | 3240 | tags=83% | list=20% | signal=67% |
| WP_EUKARYOTIC_TRANSCRIPTION_INITIATION | 40 | 0.489598 | 1.41418 | 0.122807 | 0.470056 | 0.427351 | 5442 | tags=55% | list=34% | signal=37% |
| REACTOME_STING_MEDIATED_INDUCTION_OF_HOST_IMMUNE_RESPONSES | 6 | 0.722215 | 1.412907 | 0.123043 | 0.470056 | 0.427351 | 1379 | tags=67% | list=9% | signal=61% |
| REACTOME_TRANSPORT_AND_SYNTHESIS_OF_PAPS | 6 | -0.81303 | -1.38294 | 0.120721 | 0.470056 | 0.427351 | 2105 | tags=67% | list=13% | signal=58% |
| REACTOME_CATION_COUPLED_CHLORIDE_COTRANSPORTERS | 5 | -0.83652 | -1.38059 | 0.119705 | 0.470056 | 0.427351 | 2153 | tags=80% | list=13% | signal=69% |
| REACTOME_RECYCLING_OF_BILE_ACIDS_AND_SALTS | 8 | 0.659557 | 1.380097 | 0.122249 | 0.470056 | 0.427351 | 2918 | tags=50% | list=18% | signal=41% |
| REACTOME_ADRENOCEPTORS | 9 | -0.74485 | -1.37108 | 0.12215 | 0.470056 | 0.427351 | 3794 | tags=89% | list=24% | signal=68% |
| REACTOME_TRANSPORT_OF_NUCLEOSIDES_AND_FREE_PURINE_AND_PYRIMIDINE_ | 8 | -0.75762 | -1.36191 | 0.118044 | 0.470056 | 0.427351 | 2934 | tags=50% | list=18% | signal=41% |
| REACTOME_ATTACHMENT_OF_GPI_ANCHOR_TO_UPAR | 7 | -0.78422 | -1.35988 | 0.120567 | 0.470056 | 0.427351 | 2298 | tags=100% | list=14% | signal=86% |
| WP_GPCRS_ORPHAN | 16 | -0.67324 | -1.35609 | 0.118768 | 0.470056 | 0.427351 | 563 | tags=56% | list=3% | signal=54% |
| WP_GLUTATHIONE_AND_ONECARBON_METABOLISM | 28 | 0.511531 | 1.35406 | 0.122881 | 0.470056 | 0.427351 | 4139 | tags=50% | list=26% | signal=37% |
| REACTOME_NRIF_SIGNALS_CELL_DEATH_FROM_THE_NUCLEUS | 14 | -0.68264 | -1.35267 | 0.118871 | 0.470056 | 0.427351 | 2496 | tags=57% | list=15% | signal=48% |
| REACTOME_INTRAFLAGELLAR_TRANSPORT | 45 | 0.457001 | 1.352332 | 0.119418 | 0.470056 | 0.427351 | 4311 | tags=44% | list=27% | signal=33% |
| REACTOME_BLOOD_GROUP_SYSTEMS_BIOSYNTHESIS | 14 | -0.67885 | -1.34516 | 0.118871 | 0.470056 | 0.427351 | 3082 | tags=50% | list=19% | signal=40% |
| BIOCARTA_CXCR4_PATHWAY | 18 | 0.543843 | 1.34444 | 0.121528 | 0.470056 | 0.427351 | 3508 | tags=56% | list=22% | signal=44% |
| REACTOME_RETROGRADE_TRANSPORT_AT_THE_TRANS_GOLGI_NETWORK | 45 | 0.453997 | 1.343444 | 0.122855 | 0.470056 | 0.427351 | 3706 | tags=49% | list=23% | signal=38% |
| REACTOME_DETOXIFICATION_OF_REACTIVE_OXYGEN_SPECIES | 30 | -0.59731 | -1.32583 | 0.122108 | 0.470056 | 0.427351 | 2075 | tags=20% | list=13% | signal=17% |
| BIOCARTA_HDAC_PATHWAY | 22 | 0.512951 | 1.315129 | 0.122137 | 0.470056 | 0.427351 | 2147 | tags=50% | list=13% | signal=43% |
| REACTOME_COMPLEX_I_BIOGENESIS | 50 | -0.54938 | -1.29661 | 0.117788 | 0.470056 | 0.427351 | 12 | tags=32% | list=0% | signal=32% |
| REACTOME_NEUROTRANSMITTER_RELEASE_CYCLE | 50 | -0.54792 | -1.29316 | 0.121394 | 0.470056 | 0.427351 | 855 | tags=14% | list=5% | signal=13% |
| REACTOME_CARGO_TRAFFICKING_TO_THE_PERICILIARY_MEMBRANE | 39 | 0.447392 | 1.291592 | 0.121951 | 0.470056 | 0.427351 | 2485 | tags=36% | list=15% | signal=30% |
| REACTOME_MAPK6_MAPK4_SIGNALING | 72 | 0.418984 | 1.290016 | 0.122903 | 0.470056 | 0.427351 | 2852 | tags=24% | list=18% | signal=20% |
| REACTOME_ETHANOL_OXIDATION | 6 | 0.718312 | 1.405271 | 0.12528 | 0.471151 | 0.428346 | 3792 | tags=67% | list=23% | signal=51% |
| BIOCARTA_BLYMPHOCYTE_PATHWAY | 7 | -0.78001 | -1.35259 | 0.124113 | 0.471151 | 0.428346 | 2076 | tags=57% | list=13% | signal=50% |
| REACTOME_TYPE_I_HEMIDESMOSOME_ASSEMBLY | 7 | -0.77734 | -1.34796 | 0.125887 | 0.471151 | 0.428346 | 3601 | tags=100% | list=22% | signal=78% |
| WP_GPCRS_CLASS_B_SECRETINLIKE | 19 | -0.65123 | -1.34294 | 0.125174 | 0.471151 | 0.428346 | 1882 | tags=42% | list=12% | signal=37% |
| REACTOME_CYTOCHROME_C_MEDIATED_APOPTOTIC_RESPONSE | 11 | 0.60158 | 1.331822 | 0.125348 | 0.471151 | 0.428346 | 3034 | tags=27% | list=19% | signal=22% |
| WP_MECP2_AND_ASSOCIATED_RETT_SYNDROME | 43 | -0.56835 | -1.31479 | 0.12561 | 0.471151 | 0.428346 | 1218 | tags=28% | list=8% | signal=26% |
| REACTOME_EPH_EPHRIN_SIGNALING | 64 | -0.52969 | -1.28813 | 0.125 | 0.471151 | 0.428346 | 2023 | tags=25% | list=13% | signal=22% |
| WP_MICRORNAS_IN_CARDIOMYOCYTE_HYPERTROPHY | 78 | 0.417236 | 1.270031 | 0.124239 | 0.471151 | 0.428346 | 2835 | tags=36% | list=18% | signal=30% |
| REACTOME_SUMOYLATION_OF_CHROMATIN_ORGANIZATION_PROTEINS | 57 | 0.403599 | 1.202359 | 0.124112 | 0.471151 | 0.428346 | 2035 | tags=25% | list=13% | signal=22% |
| REACTOME_REGULATION_OF_PTEN_STABILITY_AND_ACTIVITY | 67 | 0.424888 | 1.267833 | 0.126292 | 0.471604 | 0.428758 | 3371 | tags=28% | list=21% | signal=23% |
| REACTOME_YAP1_AND_WWTR1_TAZ_STIMULATED_GENE_EXPRESSION | 7 | 0.687507 | 1.418557 | 0.127854 | 0.472393 | 0.429476 | 1198 | tags=43% | list=7% | signal=40% |
| REACTOME_APEX1_INDEPENDENT_RESOLUTION_OF_AP_SITES_VIA_THE_SINGLE_N | 7 | 0.686509 | 1.416497 | 0.127854 | 0.472393 | 0.429476 | 4891 | tags=86% | list=30% | signal=60% |
| BIOCARTA_THELPER_PATHWAY | 6 | -0.80499 | -1.36925 | 0.127928 | 0.472393 | 0.429476 | 2076 | tags=83% | list=13% | signal=73% |
| REACTOME_CROSS_PRESENTATION_OF_PARTICULATE_EXOGENOUS_ANTIGENS_PH | 7 | -0.77665 | -1.34675 | 0.12766 | 0.472393 | 0.429476 | 78 | tags=57% | list=0% | signal=57% |
| REACTOME_RNA_POLYMERASE_III_TRANSCRIPTION | 34 | 0.464057 | 1.296527 | 0.127273 | 0.472393 | 0.429476 | 3195 | tags=29% | list=20% | signal=24% |
| REACTOME_DNA_DAMAGE_RECOGNITION_IN_GG_NER | 38 | 0.462416 | 1.323713 | 0.129811 | 0.478281 | 0.434829 | 4385 | tags=42% | list=27% | signal=31% |
| REACTOME_SIGNALING_BY_VEGF | 99 | 0.404283 | 1.260068 | 0.130491 | 0.479721 | 0.436137 | 2852 | tags=39% | list=18% | signal=33% |
| WP_OMEGA9_FATTY_ACID_SYNTHESIS | 13 | -0.67399 | -1.32358 | 0.131222 | 0.48134 | 0.437609 | 1233 | tags=31% | list=8% | signal=28% |
| REACTOME_FRUCTOSE_METABOLISM | 6 | 0.710006 | 1.389021 | 0.134228 | 0.481478 | 0.437735 | 132 | tags=17% | list=1% | signal=17% |
| REACTOME_RAB_GEFS_EXCHANGE_GTP_FOR_GDP_ON_RABS | 86 | 0.440466 | 1.370876 | 0.133376 | 0.481478 | 0.437735 | 3355 | tags=47% | list=21% | signal=37% |
| REACTOME_ANDROGEN_BIOSYNTHESIS | 4 | -0.84255 | -1.36214 | 0.131827 | 0.481478 | 0.437735 | 1720 | tags=75% | list=11% | signal=67% |
| REACTOME_MISCELLANEOUS_SUBSTRATES | 4 | -0.8411 | -1.35979 | 0.13371 | 0.481478 | 0.437735 | 1449 | tags=50% | list=9% | signal=46% |
| WP_MIR222_IN_EXERCISEINDUCED_CARDIAC_GROWTH | 4 | 0.78353 | 1.355393 | 0.133758 | 0.481478 | 0.437735 | 2439 | tags=75% | list=15% | signal=64% |
| REACTOME_TP53_REGULATES_TRANSCRIPTION_OF_ADDITIONAL_CELL_CYCLE_GEN | 18 | -0.65352 | -1.34035 | 0.134454 | 0.481478 | 0.437735 | 343 | tags=17% | list=2% | signal=16% |
| REACTOME_ACTIVATION_OF_BAD_AND_TRANSLOCATION_TO_MITOCHONDRIA | 12 | 0.586579 | 1.32202 | 0.133136 | 0.481478 | 0.437735 | 4268 | tags=42% | list=26% | signal=31% |
| REACTOME_CYTOSOLIC_SENSORS_OF_PATHOGEN_ASSOCIATED_DNA | 31 | 0.482527 | 1.316826 | 0.133028 | 0.481478 | 0.437735 | 2445 | tags=35% | list=15% | signal=30% |
| REACTOME_MRNA_DECAY_BY_3_TO_5_EXORIBONUCLEASE | 12 | 0.58402 | 1.316252 | 0.133136 | 0.481478 | 0.437735 | 3962 | tags=50% | list=25% | signal=38% |
| REACTOME_METABOLISM_OF_FAT_SOLUBLE_VITAMINS | 35 | -0.58588 | -1.31622 | 0.132484 | 0.481478 | 0.437735 | 2130 | tags=29% | list=13% | signal=25% |
| REACTOME_RMTS_METHYLATE_HISTONE_ARGININES | 39 | 0.455348 | 1.314562 | 0.134211 | 0.481478 | 0.437735 | 5818 | tags=54% | list=36% | signal=35% |
| REACTOME_FORMATION_OF_APOPTOSOME | 9 | 0.633774 | 1.349634 | 0.136598 | 0.483535 | 0.439606 | 3034 | tags=33% | list=19% | signal=27% |
| BIOCARTA_REELIN_PATHWAY | 7 | -0.7727 | -1.3399 | 0.136525 | 0.483535 | 0.439606 | 920 | tags=43% | list=6% | signal=40% |
| REACTOME_POLYMERASE_SWITCHING_ON_THE_C_STRAND_OF_THE_TELOMERE | 24 | 0.512858 | 1.334076 | 0.135458 | 0.483535 | 0.439606 | 4670 | tags=54% | list=29% | signal=39% |
| REACTOME_KSRP_KHSRP_BINDS_AND_DESTABILIZES_MRNA | 14 | 0.565724 | 1.324213 | 0.136778 | 0.483535 | 0.439606 | 3962 | tags=43% | list=25% | signal=32% |
| WP_APOPTOSIS | 73 | 0.401499 | 1.224867 | 0.135914 | 0.483535 | 0.439606 | 2422 | tags=23% | list=15% | signal=20% |
| REACTOME_ASPARAGINE_N_LINKED_GLYCOSYLATION | 247 | -0.44055 | -1.17291 | 0.136364 | 0.483535 | 0.439606 | 3111 | tags=31% | list=19% | signal=26% |
| REACTOME_TRIGLYCERIDE_CATABOLISM | 4 | 0.780027 | 1.349332 | 0.138004 | 0.486042 | 0.441884 | 2383 | tags=75% | list=15% | signal=64% |
| REACTOME_DEGRADATION_OF_DVL | 55 | 0.404963 | 1.201439 | 0.138073 | 0.486042 | 0.441884 | 5771 | tags=49% | list=36% | signal=32% |
| REACTOME_G1_S_TRANSITION | 103 | 0.416409 | 1.325658 | 0.138557 | 0.48671 | 0.442492 | 2552 | tags=22% | list=16% | signal=19% |
| REACTOME_ARYL_HYDROCARBON_RECEPTOR_SIGNALLING | 7 | 0.672246 | 1.387068 | 0.139269 | 0.488179 | 0.443827 | 2058 | tags=57% | list=13% | signal=50% |
| REACTOME_MET_RECEPTOR_ACTIVATION | 5 | -0.82036 | -1.35392 | 0.139963 | 0.488545 | 0.44416 | 2563 | tags=80% | list=16% | signal=67% |
| BIOCARTA_ERK5_PATHWAY | 14 | -0.66578 | -1.31927 | 0.139673 | 0.488545 | 0.44416 | 8 | tags=14% | list=0% | signal=14% |
| REACTOME_PTK6_REGULATES_RHO_GTPASES_RAS_GTPASE_AND_MAP_KINASES | 12 | 0.576524 | 1.29936 | 0.142012 | 0.493618 | 0.448772 | 2556 | tags=42% | list=16% | signal=35% |
| REACTOME_MITOCHONDRIAL_BIOGENESIS | 12 | 0.575564 | 1.297196 | 0.142012 | 0.493618 | 0.448772 | 3856 | tags=58% | list=24% | signal=44% |
| REACTOME_NADE_MODULATES_DEATH_SIGNALLING | 6 | -0.79369 | -1.35003 | 0.142342 | 0.493731 | 0.448875 | 2558 | tags=50% | list=16% | signal=42% |
| REACTOME_N_GLYCAN_ANTENNAE_ELONGATION | 15 | -0.65885 | -1.314 | 0.142857 | 0.4941 | 0.44921 | 2198 | tags=47% | list=14% | signal=40% |
| WP_TYPE_II_INTERFERON_SIGNALING_IFNG | 27 | -0.59848 | -1.3025 | 0.143045 | 0.4941 | 0.44921 | 2714 | tags=44% | list=17% | signal=37% |
| REACTOME_ACTIVATED_NTRK2_SIGNALS_THROUGH_FRS2_AND_FRS3 | 5 | -0.81782 | -1.34972 | 0.14733 | 0.4975 | 0.452302 | 307 | tags=40% | list=2% | signal=39% |
| REACTOME_ASSEMBLY_AND_CELL_SURFACE_PRESENTATION_OF_NMDA_RECEPTO | 5 | -0.8174 | -1.34902 | 0.14733 | 0.4975 | 0.452302 | 1860 | tags=60% | list=12% | signal=53% |
| REACTOME_TRANSCRIPTIONAL_ACTIVATION_OF_MITOCHONDRIAL_BIOGENESIS | 10 | 0.602505 | 1.32098 | 0.146277 | 0.4975 | 0.452302 | 3856 | tags=70% | list=24% | signal=53% |
| REACTOME_PYRIMIDINE_SALVAGE | 10 | 0.600421 | 1.316411 | 0.146277 | 0.4975 | 0.452302 | 3936 | tags=60% | list=24% | signal=45% |
| REACTOME_ACTIVATED_TAK1_MEDIATES_P38_MAPK_ACTIVATION | 23 | 0.505932 | 1.307401 | 0.145098 | 0.4975 | 0.452302 | 4744 | tags=57% | list=29% | signal=40% |
| WP_OXIDATIVE_STRESS_RESPONSE | 26 | -0.5985 | -1.30094 | 0.146789 | 0.4975 | 0.452302 | 1122 | tags=15% | list=7% | signal=14% |
| REACTOME_REGULATION_OF_HSF1_MEDIATED_HEAT_SHOCK_RESPONSE | 62 | 0.431262 | 1.290092 | 0.146981 | 0.4975 | 0.452302 | 3092 | tags=32% | list=19% | signal=26% |

| | | | | | | | | | | |
|---|---|---|---|---|---|---|---|---|---|---|
| WP_P38_MAPK_SIGNALING_PATHWAY | 33 | 0.46247 | 1.282851 | 0.145455 | 0.4975 | 0.452302 | 1649 | tags=27% | list=10% | signal=25% |
| REACTOME_RECYCLING_PATHWAY_OF_L1 | 32 | 0.467202 | 1.281079 | 0.146226 | 0.4975 | 0.452302 | 3034 | tags=38% | list=19% | signal=31% |
| BIOCARTA_P38MAPK_PATHWAY | 32 | 0.466569 | 1.279343 | 0.146226 | 0.4975 | 0.452302 | 1628 | tags=25% | list=10% | signal=23% |
| REACTOME_SIGNALING_BY_NTRKS | 74 | -0.51125 | -1.25276 | 0.145289 | 0.4975 | 0.452302 | 961 | tags=9% | list=6% | signal=9% |
| REACTOME_TP53_REGULATES_TRANSCRIPTION_OF_CELL_CYCLE_GENES | 35 | -0.57504 | -1.29186 | 0.147771 | 0.497975 | 0.452734 | 486 | tags=9% | list=3% | signal=8% |
| REACTOME_NEGATIVE_REGULATION_OF_ACTIVITY_OF_TFAP2_AP_2_FAMILY_TRAN | 7 | 0.666746 | 1.375719 | 0.150685 | 0.504718 | 0.458864 | 2758 | tags=43% | list=17% | signal=36% |
| REACTOME_ER_PHAGOSOME_PATHWAY | 20 | -0.63409 | -1.31467 | 0.150209 | 0.504718 | 0.458864 | 3112 | tags=50% | list=19% | signal=40% |
| REACTOME_MITOTIC_TELOPHASE_CYTOKINESIS | 13 | 0.563453 | 1.307118 | 0.150442 | 0.504718 | 0.458864 | 4533 | tags=62% | list=28% | signal=44% |
| REACTOME_GRB2_EVENTS_IN_ERBB2_SIGNALING | 9 | -0.72234 | -1.32964 | 0.151466 | 0.506311 | 0.460312 | 1767 | tags=67% | list=11% | signal=59% |
| BIOCARTA_CALCINEURIN_PATHWAY | 17 | 0.536318 | 1.337274 | 0.152597 | 0.508512 | 0.462313 | 1254 | tags=24% | list=8% | signal=22% |
| REACTOME_ACETYLCHOLINE_NEUROTRANSMITTER_RELEASE_CYCLE | 17 | -0.64703 | -1.30593 | 0.152738 | 0.508512 | 0.462313 | 272 | tags=18% | list=2% | signal=17% |
| REACTOME_RUNX1_REGULATES_GENES_INVOLVED_IN_MEGAKARYOCYTE_DIFFERE | 41 | -0.55687 | -1.28005 | 0.153086 | 0.508652 | 0.46244 | 436 | tags=12% | list=3% | signal=12% |
| REACTOME_ROLE_OF_LAT2_NTAL_LAB_ON_CALCIUM_MOBILIZATION | 12 | 0.572192 | 1.289596 | 0.153846 | 0.510154 | 0.463806 | 1257 | tags=33% | list=8% | signal=31% |
| REACTOME_FORMYL_PEPTIDE_RECEPTORS_BIND_FORMYL_PEPTIDES_AND_MANY_ | 5 | -0.80883 | -1.33489 | 0.154696 | 0.510929 | 0.46451 | 399 | tags=40% | list=2% | signal=39% |
| BIOCARTA_IL3_PATHWAY | 14 | -0.65658 | -1.30104 | 0.154532 | 0.510929 | 0.46451 | 49 | tags=14% | list=0% | signal=14% |
| REACTOME_TRANSCRIPTIONAL_REGULATION_BY_MECP2 | 5 | -0.80468 | -1.32804 | 0.156538 | 0.51496 | 0.468175 | 1051 | tags=40% | list=7% | signal=37% |
| REACTOME_PLATELET_ACTIVATION_SIGNALING_AND_AGGREGATION | 224 | -0.44047 | -1.16702 | 0.156413 | 0.51496 | 0.468175 | 1907 | tags=18% | list=12% | signal=16% |
| REACTOME_P38MAPK_EVENTS | 7 | 0.664028 | 1.370111 | 0.157534 | 0.515029 | 0.468238 | 1253 | tags=43% | list=8% | signal=40% |
| REACTOME_NONCANONICAL_ACTIVATION_OF_NOTCH3 | 7 | -0.75928 | -1.31664 | 0.157801 | 0.515029 | 0.468238 | 3301 | tags=86% | list=20% | signal=68% |
| REACTOME_RNA_POLYMERASE_I_TRANSCRIPTION | 44 | 0.420385 | 1.235431 | 0.157303 | 0.515029 | 0.468238 | 3005 | tags=39% | list=19% | signal=32% |
| WP_KIT_RECEPTOR_SIGNALING_PATHWAY | 66 | 0.412262 | 1.231014 | 0.15745 | 0.515029 | 0.468238 | 3821 | tags=42% | list=24% | signal=33% |
| BIOCARTA_IL12_PATHWAY | 11 | 0.589862 | 1.305881 | 0.158774 | 0.517186 | 0.470199 | 3492 | tags=64% | list=22% | signal=50% |
| BIOCARTA_FMLP_PATHWAY | 29 | 0.485857 | 1.299368 | 0.159483 | 0.51792 | 0.470867 | 3553 | tags=45% | list=22% | signal=35% |
| REACTOME_RET_SIGNALING | 36 | 0.436639 | 1.231305 | 0.159624 | 0.51792 | 0.470867 | 2388 | tags=36% | list=15% | signal=31% |
| WP_WNT_SIGNALING_PATHWAY_AND_PLURIPOTENCY | 88 | -0.48986 | -1.2276 | 0.160398 | 0.519415 | 0.472225 | 1777 | tags=12% | list=11% | signal=11% |
| REACTOME_NGF_STIMULATED_TRANSCRIPTION | 6 | -0.77851 | -1.32421 | 0.162162 | 0.524103 | 0.476488 | 638 | tags=50% | list=4% | signal=48% |
| WP_OSTEOBLAST_SIGNALING | 5 | -0.7998 | -1.31998 | 0.163904 | 0.526511 | 0.478677 | 3237 | tags=100% | list=20% | signal=80% |
| BIOCARTA_INFLAM_PATHWAY | 9 | -0.71475 | -1.31566 | 0.164495 | 0.526511 | 0.478677 | 3388 | tags=44% | list=21% | signal=35% |
| REACTOME_PROSTACYCLIN_SIGNALLING_THROUGH_PROSTACYCLIN_RECEPTOR | 16 | -0.64162 | -1.2924 | 0.164223 | 0.526511 | 0.478677 | 594 | tags=19% | list=4% | signal=18% |
| REACTOME_HDMS_DEMETHYLATE_HISTONES | 29 | 0.479891 | 1.283412 | 0.163793 | 0.526511 | 0.478677 | 5136 | tags=55% | list=32% | signal=38% |
| REACTOME_RHO_GTPASES_ACTIVATE_PAKS | 22 | 0.495439 | 1.270231 | 0.164122 | 0.526511 | 0.478677 | 2852 | tags=45% | list=18% | signal=37% |
| REACTOME_GLYCOPROTEIN_HORMONES | 4 | -0.82017 | -1.32595 | 0.165725 | 0.526674 | 0.478825 | 2429 | tags=50% | list=15% | signal=42% |
| WP_BLOOD_CLOTTING_CASCADE | 10 | -0.70246 | -1.31282 | 0.166134 | 0.526674 | 0.478825 | 2970 | tags=40% | list=18% | signal=33% |
| REACTOME_RAB_GERANYLGERANYLATION | 58 | -0.5217 | -1.25145 | 0.165308 | 0.526674 | 0.478825 | 914 | tags=10% | list=6% | signal=10% |
| REACTOME_AUF1_HNRNP_D0_BINDS_AND_DESTABILIZES_MRNA | 54 | 0.407212 | 1.231514 | 0.165866 | 0.526674 | 0.478825 | 2552 | tags=22% | list=16% | signal=19% |
| REACTOME_UCH_PROTEINASES | 81 | 0.395499 | 1.202964 | 0.165949 | 0.526674 | 0.478825 | 5786 | tags=47% | list=36% | signal=30% |
| REACTOME_SIGNALING_BY_EGFR | 42 | 0.426953 | 1.230786 | 0.166667 | 0.527354 | 0.479443 | 2303 | tags=38% | list=14% | signal=33% |
| REACTOME_SYNTHESIS_OF_PYROPHOSPHATES_IN_THE_CYTOSOL | 10 | 0.584131 | 1.280694 | 0.167553 | 0.528143 | 0.48016 | 4376 | tags=60% | list=27% | signal=44% |
| REACTOME_SYNTHESIS_OF_PA | 29 | -0.57864 | -1.27421 | 0.167532 | 0.528143 | 0.48016 | 1329 | tags=34% | list=8% | signal=32% |
| REACTOME_UNBLOCKING_OF_NMDA_RECEPTORS_GLUTAMATE_BINDING_AND_AC | 22 | -0.60916 | -1.28667 | 0.168919 | 0.530904 | 0.48267 | 1986 | tags=32% | list=12% | signal=28% |
| REACTOME_SIGNALLING_TO_ERKS | 26 | -0.58732 | -1.27666 | 0.169069 | 0.530904 | 0.48267 | 390 | tags=8% | list=2% | signal=8% |
| BIOCARTA_DEATH_PATHWAY | 30 | 0.470357 | 1.273664 | 0.169643 | 0.531697 | 0.483392 | 1531 | tags=13% | list=9% | signal=12% |
| REACTOME_REACTIONS_SPECIFIC_TO_THE_COMPLEX_N_GLYCAN_SYNTHESIS_PATI | 7 | -0.75297 | -1.30569 | 0.170213 | 0.532477 | 0.484101 | 3098 | tags=71% | list=19% | signal=58% |
| WP_ONECARBON_METABOLISM | 26 | 0.49124 | 1.282117 | 0.171548 | 0.532788 | 0.484384 | 4139 | tags=54% | list=26% | signal=40% |
| REACTOME_PECAM1_INTERACTIONS | 12 | 0.564472 | 1.272196 | 0.171598 | 0.532788 | 0.484384 | 2303 | tags=33% | list=14% | signal=29% |
| REACTOME_ECM_PROTEOGLYCANS | 42 | -0.54599 | -1.25934 | 0.171569 | 0.532788 | 0.484384 | 3843 | tags=52% | list=24% | signal=40% |
| WP_ONECARBON_METABOLISM_AND_RELATED_PATHWAYS | 46 | 0.403148 | 1.189248 | 0.171429 | 0.532788 | 0.484384 | 2531 | tags=26% | list=16% | signal=22% |
| WP_P53_SIGNALING | 62 | -0.51466 | -1.24692 | 0.172811 | 0.535553 | 0.486897 | 820 | tags=23% | list=5% | signal=22% |
| BIOCARTA_EPHA4_PATHWAY | 8 | -0.72122 | -1.29647 | 0.173693 | 0.536281 | 0.48756 | 528 | tags=25% | list=3% | signal=24% |
| BIOCARTA_AT1R_PATHWAY | 25 | 0.486493 | 1.260035 | 0.173554 | 0.536281 | 0.48756 | 5404 | tags=68% | list=33% | signal=45% |
| WP_INTEGRINMEDIATED_CELL_ADHESION | 96 | 0.420304 | 1.309672 | 0.175669 | 0.538372 | 0.48946 | 1735 | tags=22% | list=11% | signal=20% |
| BIOCARTA_CDMAC_PATHWAY | 14 | -0.6472 | -1.28245 | 0.175334 | 0.538372 | 0.48946 | 1481 | tags=14% | list=9% | signal=13% |
| REACTOME_RHOBTB3_ATPASE_CYCLE | 10 | 0.580718 | 1.27321 | 0.175532 | 0.538372 | 0.48946 | 4954 | tags=60% | list=31% | signal=42% |
| REACTOME_METABOLISM_OF_STEROIDS | 82 | -0.48618 | -1.20741 | 0.175028 | 0.538372 | 0.48946 | 3316 | tags=34% | list=21% | signal=27% |
| WP_BIOGENIC_AMINE_SYNTHESIS | 12 | -0.65996 | -1.28442 | 0.177711 | 0.543625 | 0.494236 | 669 | tags=50% | list=4% | signal=48% |
| REACTOME_SYNTHESIS_OF_UDP_N_ACETYL_GLUCOSAMINE | 8 | 0.629101 | 1.316371 | 0.178484 | 0.544151 | 0.494714 | 3442 | tags=50% | list=21% | signal=39% |
| REACTOME_PHOSPHATE_BOND_HYDROLYSIS_BY_NTPDASE_PROTEINS | 6 | -0.76814 | -1.30658 | 0.18018 | 0.544151 | 0.494714 | 2718 | tags=67% | list=17% | signal=55% |
| REACTOME_GAMMA_CARBOXYLATION_HYPUSINE_FORMATION_AND_ARYLSULFAT | 31 | -0.57213 | -1.27705 | 0.178571 | 0.544151 | 0.494714 | 3638 | tags=39% | list=23% | signal=30% |
| BIOCARTA_CARDIACEGF_PATHWAY | 18 | -0.62058 | -1.27278 | 0.179272 | 0.544151 | 0.494714 | 2500 | tags=28% | list=15% | signal=24% |
| REACTOME_CROSSLINKING_OF_COLLAGEN_FIBRILS | 15 | -0.63688 | -1.27019 | 0.179676 | 0.544151 | 0.494714 | 1852 | tags=27% | list=11% | signal=24% |
| REACTOME_GABA_RECEPTOR_ACTIVATION | 52 | -0.52209 | -1.23721 | 0.179976 | 0.544151 | 0.494714 | 1697 | tags=33% | list=11% | signal=29% |
| WP_TOLLLIKE_RECEPTOR_SIGNALING | 32 | 0.450216 | 1.234501 | 0.179245 | 0.544151 | 0.494714 | 1788 | tags=31% | list=11% | signal=28% |
| REACTOME_SYNTHESIS_OF_IP2_IP_AND_INS_IN_THE_CYTOSOL | 11 | 0.573091 | 1.268751 | 0.181058 | 0.545809 | 0.496222 | 2630 | tags=55% | list=16% | signal=46% |
| REACTOME_PTEN_REGULATION | 110 | 0.376472 | 1.194498 | 0.181666 | 0.546646 | 0.496982 | 2694 | tags=25% | list=17% | signal=21% |
| BIOCARTA_DSP_PATHWAY | 9 | -0.70245 | -1.29303 | 0.18241 | 0.547352 | 0.497624 | 961 | tags=22% | list=6% | signal=21% |
| BIOCARTA_TALL1_PATHWAY | 13 | 0.55229 | 1.281221 | 0.182891 | 0.547352 | 0.497624 | 987 | tags=15% | list=6% | signal=14% |
| REACTOME_REGULATED_NECROSIS | 46 | 0.396427 | 1.169422 | 0.182857 | 0.547352 | 0.497624 | 5490 | tags=46% | list=34% | signal=30% |
| REACTOME_COENZYME_A_BIOSYNTHESIS | 8 | 0.625368 | 1.308559 | 0.183374 | 0.547809 | 0.49804 | 3764 | tags=50% | list=23% | signal=38% |
| BIOCARTA_IL10_PATHWAY | 9 | -0.70149 | -1.29127 | 0.184039 | 0.548807 | 0.498947 | 116 | tags=22% | list=1% | signal=22% |
| REACTOME_LINOLEIC_ACID_LA_METABOLISM | 7 | -0.74235 | -1.28727 | 0.18617 | 0.554166 | 0.503819 | 1233 | tags=57% | list=8% | signal=53% |
| REACTOME_EUKARYOTIC_TRANSLATION_ELONGATION | 5 | 0.711322 | 1.298707 | 0.187364 | 0.556719 | 0.50614 | 3794 | tags=80% | list=24% | signal=61% |
| REACTOME_ADVANCED_GLYCOSYLATION_ENDPRODUCT_RECEPTOR_SIGNALING | 5 | -0.78577 | -1.29683 | 0.187845 | 0.557151 | 0.506533 | 399 | tags=40% | list=2% | signal=39% |
| REACTOME_RUNX3_REGULATES_CDKN1A_TRANSCRIPTION | 5 | -0.78247 | -1.29138 | 0.191529 | 0.559752 | 0.508897 | 1834 | tags=60% | list=11% | signal=53% |
| REACTOME_TP53_REGULATES_TRANSCRIPTION_OF_GENES_INVOLVED_IN_G2_CEL | 7 | -0.73938 | -1.28214 | 0.191489 | 0.559752 | 0.508897 | 486 | tags=43% | list=3% | signal=42% |
| REACTOME_TRANSLESION_SYNTHESIS_BY_POLH | 18 | 0.514243 | 1.271265 | 0.190972 | 0.559752 | 0.508897 | 3239 | tags=50% | list=20% | signal=40% |
| REACTOME_TNFS_BIND_THEIR_PHYSIOLOGICAL_RECEPTORS | 15 | 0.533275 | 1.2706 | 0.19195 | 0.559752 | 0.508897 | 39 | tags=7% | list=0% | signal=7% |
| REACTOME_RAF_INDEPENDENT_MAPK1_3_ACTIVATION | 21 | -0.60101 | -1.26074 | 0.192098 | 0.559752 | 0.508897 | 1148 | tags=19% | list=7% | signal=18% |
| REACTOME_ADP_SIGNALLING_THROUGH_P2Y_PURINOCEPTOR_12 | 19 | -0.61083 | -1.25964 | 0.190542 | 0.559752 | 0.508897 | 1316 | tags=16% | list=8% | signal=15% |
| REACTOME_ACTIVATION_OF_THE_PRE_REPLICATIVE_COMPLEX | 30 | 0.464787 | 1.258582 | 0.191964 | 0.559752 | 0.508897 | 1830 | tags=27% | list=11% | signal=24% |
| WP_GLYCOLYSIS_AND_GLUCONEOGENESIS | 39 | 0.417602 | 1.20559 | 0.190244 | 0.559752 | 0.508897 | 3746 | tags=44% | list=23% | signal=34% |
| REACTOME_NUCLEAR_EVENTS_MEDIATED_BY_NFE2L2 | 53 | 0.384491 | 1.164994 | 0.191083 | 0.559752 | 0.508897 | 5771 | tags=49% | list=36% | signal=32% |
| REACTOME_DEGRADATION_OF_AXIN | 53 | 0.382627 | 1.159345 | 0.191083 | 0.559752 | 0.508897 | 2552 | tags=21% | list=16% | signal=18% |
| REACTOME_SYNTHESIS_OF_PIPS_AT_THE_ER_MEMBRANE | 5 | 0.705772 | 1.288575 | 0.1939 | 0.56008 | 0.509196 | 1851 | tags=40% | list=11% | signal=35% |
| BIOCARTA_STEM_PATHWAY | 3 | -0.84515 | -1.28731 | 0.193858 | 0.56008 | 0.509196 | 391 | tags=67% | list=2% | signal=65% |
| BIOCARTA_TOB1_PATHWAY | 10 | -0.68861 | -1.28693 | 0.193291 | 0.56008 | 0.509196 | 3078 | tags=60% | list=19% | signal=49% |
| REACTOME_HS_GAG_DEGRADATION | 20 | -0.60742 | -1.25937 | 0.193324 | 0.56008 | 0.509196 | 3179 | tags=55% | list=20% | signal=44% |
| WP_SPLICING_FACTOR_NOVA_REGULATED_SYNAPTIC_PROTEINS | 41 | -0.53712 | -1.23466 | 0.192593 | 0.56008 | 0.509196 | 1710 | tags=32% | list=11% | signal=28% |
| REACTOME_JNK_C_JUN_KINASES_PHOSPHORYLATION_AND_ACTIVATION_MEDIATI | 22 | 0.482763 | 1.237731 | 0.194656 | 0.560313 | 0.509408 | 4744 | tags=55% | list=29% | signal=39% |
| REACTOME_TRANSCRIPTIONAL_REGULATION_BY_TP53 | 289 | 0.30225 | 1.08021 | 0.194597 | 0.560313 | 0.509408 | 2486 | tags=23% | list=15% | signal=20% |
| REACTOME_PURINE_SALVAGE | 12 | 0.553785 | 1.24811 | 0.195266 | 0.561094 | 0.510118 | 3145 | tags=42% | list=19% | signal=34% |
| BIOCARTA_STRESS_PATHWAY | 23 | 0.483223 | 1.24872 | 0.196078 | 0.561973 | 0.510917 | 1235 | tags=26% | list=8% | signal=24% |
| WP_WNT_SIGNALING_IN_KIDNEY_DISEASE | 40 | -0.53751 | -1.22812 | 0.19625 | 0.561973 | 0.510917 | 2933 | tags=28% | list=18% | signal=23% |
| REACTOME_RHOV_GTPASE_CYCLE | 36 | 0.427397 | 1.205242 | 0.197183 | 0.562702 | 0.511579 | 2559 | tags=33% | list=16% | signal=28% |
| WP_IL5_SIGNALING_PATHWAY | 66 | 0.394311 | 1.177412 | 0.19685 | 0.562702 | 0.511579 | 3960 | tags=44% | list=25% | signal=33% |
| REACTOME_CREB1_PHOSPHORYLATION_THROUGH_NMDA_RECEPTOR_MEDIATED_ | 5 | 0.701258 | 1.280333 | 0.200436 | 0.564215 | 0.512955 | 3050 | tags=60% | list=19% | signal=49% |
| REACTOME_SERINE_BIOSYNTHESIS | 9 | -0.69364 | -1.27682 | 0.198697 | 0.564215 | 0.512955 | 585 | tags=33% | list=4% | signal=32% |
| REACTOME_PI3K_EVENTS_IN_ERBB2_SIGNALING | 12 | -0.65207 | -1.26908 | 0.198795 | 0.564215 | 0.512955 | 2398 | tags=50% | list=15% | signal=43% |
| REACTOME_FGFR2_LIGAND_BINDING_AND_ACTIVATION | 14 | -0.63793 | -1.26408 | 0.199108 | 0.564215 | 0.512955 | 540 | tags=21% | list=3% | signal=21% |

| | | | | | | | | | | |
|---|---|---|---|---|---|---|---|---|---|---|
| WP_OMEGA3_OMEGA6_FATTY_ACID_SYNTHESIS | 15 | -0.6237 | -1.2439 | 0.200295 | 0.564215 | 0.512955 | 1327 | tags=33% | list=8% | signal=31% |
| REACTOME_ACYL_CHAIN_REMODELLING_OF_PE | 19 | -0.60241 | -1.24226 | 0.200278 | 0.564215 | 0.512955 | 2489 | tags=37% | list=15% | signal=31% |
| REACTOME_VASOPRESSIN_REGULATES_RENAL_WATER_HOMEOSTASIS_VIA_AQUA | 28 | -0.56586 | -1.23772 | 0.199739 | 0.564215 | 0.512955 | 936 | tags=14% | list=6% | signal=13% |
| REACTOME_PI_METABOLISM | 73 | 0.385622 | 1.17643 | 0.2 | 0.564215 | 0.512955 | 3465 | tags=49% | list=21% | signal=39% |
| REACTOME_BETA_OXIDATION_OF_PRISTANOYL_COA | 7 | 0.629827 | 1.299543 | 0.200913 | 0.5646 | 0.513306 | 2633 | tags=57% | list=16% | signal=48% |
| REACTOME_ACYL_CHAIN_REMODELLING_OF_PS | 15 | -0.62311 | -1.24273 | 0.201767 | 0.565655 | 0.514265 | 2489 | tags=47% | list=15% | signal=40% |
| REACTOME_NONHOMOLOGOUS_END_JOINING_NHEJ | 48 | 0.396716 | 1.17546 | 0.202312 | 0.565655 | 0.514265 | 4097 | tags=31% | list=25% | signal=23% |
| REACTOME_TELOMERE_MAINTENANCE | 48 | 0.395694 | 1.172434 | 0.202312 | 0.565655 | 0.514265 | 4670 | tags=44% | list=29% | signal=31% |
| BIOCARTA_IL22BP_PATHWAY | 9 | 0.597235 | 1.271825 | 0.203608 | 0.568321 | 0.516688 | 2293 | tags=67% | list=14% | signal=57% |
| REACTOME_MITOTIC_G1_PHASE_AND_G1_S_TRANSITION | 127 | 0.343658 | 1.099383 | 0.204897 | 0.570956 | 0.519084 | 2552 | tags=21% | list=16% | signal=18% |
| REACTOME_NEGATIVE_FEEDBACK_REGULATION_OF_MAPK_PATHWAY | 6 | 0.659066 | 1.289364 | 0.205817 | 0.572557 | 0.520539 | 3508 | tags=67% | list=22% | signal=52% |
| REACTOME_MTORC1_MEDIATED_SIGNALLING | 24 | 0.467793 | 1.216849 | 0.207171 | 0.57536 | 0.523088 | 4677 | tags=50% | list=29% | signal=36% |
| REACTOME_PTK6_REGULATES_RTKS_AND_THEIR_EFFECTORS_AKT1_AND_DOK1 | 8 | 0.609132 | 1.274586 | 0.207824 | 0.575874 | 0.523555 | 2147 | tags=38% | list=13% | signal=33% |
| REACTOME_ENDOSOMAL_VACUOLAR_PATHWAY | 11 | -0.66023 | -1.26221 | 0.208398 | 0.575874 | 0.523555 | 4503 | tags=82% | list=28% | signal=59% |
| BIOCARTA_BIOPEPTIDES_PATHWAY | 27 | 0.47649 | 1.258703 | 0.208333 | 0.575874 | 0.523555 | 3508 | tags=44% | list=22% | signal=35% |
| WP_FAS_PATHWAY_AND_STRESS_INDUCTION_OF_HSP_REGULATION | 34 | 0.431851 | 1.206547 | 0.209091 | 0.576827 | 0.524421 | 4810 | tags=50% | list=30% | signal=35% |
| REACTOME_PROPIONYL_COA_CATABOLISM | 5 | 0.697441 | 1.273364 | 0.211329 | 0.580105 | 0.527402 | 4888 | tags=100% | list=30% | signal=70% |
| REACTOME_RUNX3_REGULATES_P14_ARF | 5 | 0.697385 | 1.273261 | 0.211329 | 0.580105 | 0.527402 | 57 | tags=20% | list=0% | signal=20% |
| REACTOME_TELOMERE_C_STRAND_LAGGING_STRAND_SYNTHESIS | 31 | 0.446631 | 1.218865 | 0.211009 | 0.580105 | 0.527402 | 4670 | tags=45% | list=29% | signal=32% |
| WP_MITOCHONDRIAL_GENE_EXPRESSION | 19 | 0.503394 | 1.259316 | 0.212014 | 0.580873 | 0.5281 | 5107 | tags=53% | list=32% | signal=36% |
| BIOCARTA_41BB_PATHWAY | 14 | 0.537145 | 1.257318 | 0.212766 | 0.580873 | 0.5281 | 1095 | tags=21% | list=7% | signal=20% |
| REACTOME_NUCLEAR_EVENTS_KINASE_AND_TRANSCRIPTION_FACTOR_ACTIVATIO | 27 | -0.56984 | -1.24018 | 0.212598 | 0.580873 | 0.5281 | 1894 | tags=19% | list=12% | signal=16% |
| REACTOME_AQUAPORIN_MEDIATED_TRANSPORT | 31 | -0.55404 | -1.23666 | 0.21301 | 0.580873 | 0.5281 | 936 | tags=13% | list=6% | signal=12% |
| REACTOME_SIGNALING_BY_NTRK1_TRKA | 63 | -0.49935 | -1.21074 | 0.214039 | 0.582721 | 0.52978 | 961 | tags=8% | list=6% | signal=7% |
| REACTOME_BETA_OXIDATION_OF_DECANOYL_COA_TO_OCTANOYL_COA_COA | 6 | 0.65405 | 1.279551 | 0.214765 | 0.582787 | 0.52984 | 4472 | tags=83% | list=28% | signal=60% |
| REACTOME_APOPTOTIC_CLEAVAGE_OF_CELL_ADHESION_PROTEINS | 8 | 0.603626 | 1.263064 | 0.215159 | 0.582787 | 0.52984 | 2537 | tags=38% | list=16% | signal=32% |
| REACTOME_THE_NLRP3_INFLAMMASOME | 9 | -0.68541 | -1.26166 | 0.214984 | 0.582787 | 0.52984 | 2417 | tags=44% | list=15% | signal=38% |
| REACTOME_REELIN_SIGNALLING_PATHWAY | 5 | -0.76226 | -1.25802 | 0.21547 | 0.582787 | 0.52984 | 2653 | tags=60% | list=16% | signal=50% |
| REACTOME_MULTIFUNCTIONAL_ANION_EXCHANGERS | 6 | -0.74368 | -1.26497 | 0.216216 | 0.583449 | 0.530442 | 2955 | tags=67% | list=18% | signal=54% |
| REACTOME_NETRIN_1_SIGNALING | 21 | 0.479319 | 1.220267 | 0.216418 | 0.583449 | 0.530442 | 2559 | tags=38% | list=16% | signal=32% |
| BIOCARTA_MALATEX_PATHWAY | 8 | 0.602147 | 1.259969 | 0.217604 | 0.584531 | 0.531426 | 5962 | tags=88% | list=37% | signal=55% |
| REACTOME_SLBP_DEPENDENT_PROCESSING_OF_REPLICATION_DEPENDENT_HISTC | 11 | 0.551212 | 1.220315 | 0.21727 | 0.584531 | 0.531426 | 117 | tags=9% | list=1% | signal=9% |
| REACTOME_REGULATION_OF_TNFR1_SIGNALING | 47 | 0.381144 | 1.137751 | 0.217877 | 0.584531 | 0.531426 | 2692 | tags=26% | list=17% | signal=21% |
| REACTOME_ZBP1_DAI_MEDIATED_INDUCTION_OF_TYPE_I_IFNS | 13 | 0.520343 | 1.207109 | 0.218289 | 0.58469 | 0.53157 | 2308 | tags=38% | list=14% | signal=33% |
| WP_EBV_LMP1_SIGNALING | 18 | 0.506305 | 1.251641 | 0.21875 | 0.58498 | 0.531834 | 3034 | tags=39% | list=19% | signal=32% |
| REACTOME_ORGANIC_CATION_ANION_ZWITTERION_TRANSPORT | 11 | -0.65197 | -1.24642 | 0.222395 | 0.593185 | 0.539293 | 4349 | tags=45% | list=27% | signal=33% |
| REACTOME_SENESCENCE_ASSOCIATED_SECRETORY_PHENOTYPE_SASP | 40 | 0.401768 | 1.160487 | 0.222772 | 0.593185 | 0.539293 | 3050 | tags=30% | list=19% | signal=24% |
| REACTOME_NRAGE_SIGNALS_DEATH_THROUGH_JNK | 51 | 0.375643 | 1.137029 | 0.222892 | 0.593185 | 0.539293 | 1298 | tags=25% | list=8% | signal=24% |
| REACTOME_TRANSPORT_OF_FATTY_ACIDS | 4 | -0.78618 | -1.271 | 0.224105 | 0.594362 | 0.540364 | 1634 | tags=75% | list=10% | signal=67% |
| BIOCARTA_ARENRF2_PATHWAY | 18 | -0.60277 | -1.23626 | 0.22409 | 0.594362 | 0.540364 | 721 | tags=11% | list=4% | signal=11% |
| REACTOME_ANTIGEN_PRESENTATION_FOLDING_ASSEMBLY_AND_PEPTIDE_LOADII | 27 | -0.56094 | -1.2208 | 0.224409 | 0.594362 | 0.540364 | 3112 | tags=52% | list=19% | signal=42% |
| WP_APOPTOSIS_MODULATION_BY_HSP70 | 16 | 0.505878 | 1.228118 | 0.225 | 0.594976 | 0.540922 | 1211 | tags=19% | list=8% | signal=17% |
| REACTOME_HIGHLY_SODIUM_PERMEABLE_POSTSYNAPTIC_ACETYLCHOLINE_NICOT | 4 | -0.78537 | -1.2697 | 0.225989 | 0.595587 | 0.541477 | 1747 | tags=75% | list=11% | signal=67% |
| REACTOME_HDL_REMODELING | 6 | -0.74041 | -1.25942 | 0.227027 | 0.595587 | 0.541477 | 3117 | tags=50% | list=19% | signal=40% |
| BIOCARTA_RECK_PATHWAY | 9 | -0.68047 | -1.25257 | 0.226384 | 0.595587 | 0.541477 | 1939 | tags=78% | list=12% | signal=68% |
| REACTOME_PHASE_1_INACTIVATION_OF_FAST_NA_CHANNELS | 7 | -0.71833 | -1.24562 | 0.22695 | 0.595587 | 0.541477 | 2361 | tags=57% | list=15% | signal=49% |
| REACTOME_SIGNALING_BY_NOTCH3 | 19 | -0.59356 | -1.22402 | 0.226704 | 0.595587 | 0.541477 | 2496 | tags=53% | list=15% | signal=45% |
| REACTOME_RIPK1_MEDIATED_REGULATED_NECROSIS | 30 | 0.442208 | 1.19744 | 0.227679 | 0.596352 | 0.542173 | 6025 | tags=57% | list=37% | signal=36% |
| BIOCARTA_EICOSANOID_PATHWAY | 18 | -0.60031 | -1.23122 | 0.229692 | 0.600677 | 0.546105 | 1327 | tags=44% | list=8% | signal=41% |
| BIOCARTA_PELP1_PATHWAY | 5 | 0.686304 | 1.25303 | 0.230937 | 0.601768 | 0.547096 | 3628 | tags=80% | list=22% | signal=62% |
| BIOCARTA_RAB_PATHWAY | 11 | 0.547101 | 1.211214 | 0.231198 | 0.601768 | 0.547096 | 4701 | tags=45% | list=29% | signal=32% |
| WP_ESC_PLURIPOTENCY_PATHWAYS | 104 | -0.45428 | -1.15396 | 0.230601 | 0.601768 | 0.547096 | 2894 | tags=25% | list=18% | signal=21% |
| REACTOME_G_BETA_GAMMA_SIGNALLING_THROUGH_CDC42 | 18 | -0.59662 | -1.22364 | 0.232493 | 0.60299 | 0.548208 | 594 | tags=17% | list=4% | signal=16% |
| WP_TRIACYLGLYCERIDE_SYNTHESIS | 19 | -0.5903 | -1.2173 | 0.232267 | 0.60299 | 0.548208 | 1696 | tags=37% | list=11% | signal=33% |
| REACTOME_ENDOSOMAL_SORTING_COMPLEX_REQUIRED_FOR_TRANSPORT_ESCR | 29 | 0.450134 | 1.203832 | 0.232759 | 0.60299 | 0.548208 | 4767 | tags=55% | list=30% | signal=39% |
| REACTOME_ORGANIC_CATION_TRANSPORT | 7 | -0.7149 | -1.23967 | 0.234043 | 0.605371 | 0.550372 | 3686 | tags=57% | list=23% | signal=44% |
| REACTOME_NEUREXINS_AND_NEUROLIGINS | 31 | -0.54677 | -1.22045 | 0.234694 | 0.60611 | 0.551044 | 1206 | tags=32% | list=7% | signal=30% |
| BIOCARTA_ERYTH_PATHWAY | 9 | -0.6772 | -1.24656 | 0.236156 | 0.607632 | 0.552427 | 2933 | tags=44% | list=18% | signal=36% |
| BIOCARTA_GSK3_PATHWAY | 20 | 0.476146 | 1.221878 | 0.236749 | 0.607632 | 0.552427 | 2781 | tags=40% | list=17% | signal=33% |
| REACTOME_GLYCOGEN_BREAKDOWN_GLYCOGENOLYSIS | 13 | 0.517016 | 1.199393 | 0.235988 | 0.607632 | 0.552427 | 5321 | tags=77% | list=33% | signal=52% |
| REACTOME_GLI3_IS_PROCESSED_TO_GLI3R_BY_THE_PROTEASOME | 56 | 0.376981 | 1.118437 | 0.236486 | 0.607632 | 0.552427 | 5771 | tags=46% | list=36% | signal=30% |
| WP_GLUCURONIDATION | 7 | 0.596738 | 1.231268 | 0.237443 | 0.60847 | 0.55319 | 4127 | tags=57% | list=26% | signal=43% |
| REACTOME_INTERFERON_SIGNALING | 54 | 0.368781 | 1.115287 | 0.23871 | 0.610773 | 0.555283 | 2928 | tags=35% | list=18% | signal=29% |
| BIOCARTA_CREB_PATHWAY | 20 | 0.467852 | 1.200593 | 0.240283 | 0.61385 | 0.558081 | 3358 | tags=55% | list=21% | signal=44% |
| BIOCARTA_SLRP_PATHWAY | 4 | -0.77177 | -1.2477 | 0.244821 | 0.620663 | 0.564275 | 2182 | tags=75% | list=14% | signal=65% |
| REACTOME_CHYLOMICRON_REMODELING | 4 | -0.77052 | -1.24568 | 0.244821 | 0.620663 | 0.564275 | 1453 | tags=50% | list=9% | signal=46% |
| REACTOME_REGULATION_OF_IFNA_IFNB_SIGNALING | 8 | 0.580123 | 1.213884 | 0.244499 | 0.620663 | 0.564275 | 2551 | tags=50% | list=16% | signal=42% |
| REACTOME_REGULATION_OF_CYTOSKELETAL_REMODELING_AND_CELL_SPREADIN | 8 | 0.576242 | 1.205765 | 0.244499 | 0.620663 | 0.564275 | 2208 | tags=38% | list=14% | signal=32% |
| BIOCARTA_CERAMIDE_PATHWAY | 22 | 0.452782 | 1.160865 | 0.244275 | 0.620663 | 0.564275 | 3848 | tags=45% | list=24% | signal=35% |
| REACTOME_SYNTHESIS_OF_ACTIVE_UBIQUITIN_ROLES_OF_E1_AND_E2_ENZYMES | 30 | 0.433714 | 1.174441 | 0.25 | 0.632824 | 0.575331 | 4958 | tags=50% | list=31% | signal=35% |
| REACTOME_THYROXINE_BIOSYNTHESIS | 5 | -0.74958 | -1.2371 | 0.252302 | 0.636707 | 0.578862 | 237 | tags=40% | list=1% | signal=39% |
| REACTOME_TERMINATION_OF_TRANSLESION_DNA_SYNTHESIS | 31 | 0.427709 | 1.167227 | 0.252294 | 0.636707 | 0.578862 | 1801 | tags=26% | list=11% | signal=23% |
| BIOCARTA_SUMO_PATHWAY | 6 | 0.631833 | 1.236089 | 0.252796 | 0.636986 | 0.579115 | 2474 | tags=50% | list=15% | signal=42% |
| REACTOME_RNA_POLYMERASE_III_TRANSCRIPTION_INITIATION_FROM_TYPE_1_PF | 27 | 0.447998 | 1.183437 | 0.254167 | 0.639466 | 0.58137 | 3823 | tags=30% | list=24% | signal=23% |
| REACTOME_CA2_ACTIVATED_K_CHANNELS | 6 | 0.630259 | 1.233008 | 0.255034 | 0.640675 | 0.582469 | 98 | tags=17% | list=1% | signal=17% |
| REACTOME_ACTIVATION_OF_NMDA_RECEPTORS_AND_POSTSYNAPTIC_EVENTS | 36 | -0.52518 | -1.18443 | 0.25602 | 0.641211 | 0.582956 | 1986 | tags=25% | list=12% | signal=22% |
| BIOCARTA_IGF1R_PATHWAY | 22 | 0.449152 | 1.151557 | 0.255725 | 0.641211 | 0.582956 | 3584 | tags=50% | list=22% | signal=39% |
| REACTOME_XENOBIOTICS | 9 | -0.66135 | -1.21738 | 0.257329 | 0.642547 | 0.584171 | 1489 | tags=22% | list=9% | signal=20% |
| REACTOME_MOLECULES_ASSOCIATED_WITH_ELASTIC_FIBRES | 28 | -0.54278 | -1.18723 | 0.25718 | 0.642547 | 0.584171 | 3313 | tags=39% | list=21% | signal=31% |
| REACTOME_SEPARATION_OF_SISTER_CHROMATIDS | 177 | 0.325119 | 1.072736 | 0.258284 | 0.643963 | 0.585458 | 5777 | tags=45% | list=36% | signal=29% |
| REACTOME_UBIQUITIN_MEDIATED_DEGRADATION_OF_PHOSPHORYLATED_CDC25/ | 50 | 0.364714 | 1.100645 | 0.258824 | 0.644338 | 0.585799 | 5771 | tags=44% | list=36% | signal=28% |
| REACTOME_UTILIZATION_OF_KETONE_BODIES | 3 | 0.742808 | 1.202675 | 0.259875 | 0.645987 | 0.587298 | 3499 | tags=67% | list=22% | signal=52% |
| REACTOME_SHC1_EVENTS_IN_ERBB4_SIGNALING | 9 | -0.66031 | -1.21546 | 0.260586 | 0.646785 | 0.588023 | 1767 | tags=78% | list=11% | signal=69% |
| BIOCARTA_MITOCHONDRIA_PATHWAY | 19 | 0.472268 | 1.18145 | 0.261484 | 0.648043 | 0.589167 | 3848 | tags=21% | list=24% | signal=16% |
| REACTOME_SODIUM_PROTON_EXCHANGERS | 9 | -0.65879 | -1.21266 | 0.262215 | 0.648315 | 0.589414 | 2054 | tags=44% | list=13% | signal=39% |
| REACTOME_DCC_MEDIATED_ATTRACTIVE_SIGNALING | 9 | 0.563136 | 1.19921 | 0.262887 | 0.648315 | 0.589414 | 2559 | tags=44% | list=16% | signal=37% |
| REACTOME_RAP1_SIGNALLING | 15 | 0.494695 | 1.178677 | 0.263158 | 0.648315 | 0.589414 | 2118 | tags=33% | list=13% | signal=29% |
| REACTOME_ESTROGEN_DEPENDENT_GENE_EXPRESSION | 60 | 0.375164 | 1.107076 | 0.263158 | 0.648315 | 0.589414 | 2359 | tags=20% | list=15% | signal=17% |
| REACTOME_IRS_MEDIATED_SIGNALLING | 33 | 0.410527 | 1.138765 | 0.263636 | 0.64853 | 0.58961 | 1735 | tags=24% | list=11% | signal=22% |
| REACTOME_CTLA4_INHIBITORY_SIGNALING | 20 | 0.455651 | 1.169285 | 0.265018 | 0.650962 | 0.591821 | 4737 | tags=60% | list=29% | signal=42% |
| REACTOME_HEMOSTASIS | 436 | -0.39863 | -1.08418 | 0.269347 | 0.660617 | 0.600599 | 1907 | tags=16% | list=12% | signal=14% |
| WP_METHYLATION | 8 | -0.67717 | -1.21728 | 0.271501 | 0.663936 | 0.603616 | 1572 | tags=25% | list=10% | signal=23% |
| WP_PRIMARY_FOCAL_SEGMENTAL_GLOMERULOSCLEROSIS_FSGS | 68 | -0.47254 | -1.15383 | 0.27138 | 0.663936 | 0.603616 | 2072 | tags=18% | list=13% | signal=15% |
| BIOCARTA_IL17_PATHWAY | 3 | -0.79035 | -1.20383 | 0.272553 | 0.664548 | 0.604173 | 3388 | tags=100% | list=21% | signal=79% |
| REACTOME_INTEGRIN_CELL_SURFACE_INTERACTIONS | 65 | -0.47593 | -1.15696 | 0.272414 | 0.664548 | 0.604173 | 2578 | tags=31% | list=16% | signal=26% |
| REACTOME_RHO_GTPASES_ACTIVATE_WASPS_AND_WAVES | 32 | 0.411413 | 1.128104 | 0.273585 | 0.666085 | 0.60557 | 3352 | tags=41% | list=21% | signal=32% |
| REACTOME_ELECTRIC_TRANSMISSION_ACROSS_GAP_JUNCTIONS | 4 | -0.75609 | -1.22236 | 0.276836 | 0.667898 | 0.607219 | 1246 | tags=50% | list=8% | signal=46% |
| REACTOME_ACTIVATION_OF_THE_AP_1_FAMILY_OF_TRANSCRIPTION_FACTORS | 10 | -0.65188 | -1.2183 | 0.277955 | 0.667898 | 0.607219 | 49 | tags=20% | list=0% | signal=20% |

| | | | | | | | | | | |
|---|---|---|---|---|---|---|---|---|---|---|
| REACTOME_MITOCHONDRIAL_PROTEIN_IMPORT | 7 | 0.578268 | 1.193159 | 0.276256 | 0.667898 | 0.607219 | 4624 | tags=57% | list=29% | signal=41% |
| REACTOME_PRESYNAPTIC_FUNCTION_OF_KAINATE_RECEPTORS | 19 | -0.57214 | -1.17984 | 0.275382 | 0.667898 | 0.607219 | 594 | tags=16% | list=4% | signal=15% |
| WP_TYROBP_CAUSAL_NETWORK_IN_MICROGLIA | 52 | -0.48937 | -1.15968 | 0.27652 | 0.667898 | 0.607219 | 4653 | tags=58% | list=29% | signal=41% |
| REACTOME_PINK1_PRKN_MEDIATED_MITOPHAGY | 21 | 0.455453 | 1.159506 | 0.276119 | 0.667898 | 0.607219 | 1831 | tags=24% | list=11% | signal=21% |
| REACTOME_CELL_EXTRACELLULAR_MATRIX_INTERACTIONS | 18 | 0.468558 | 1.158327 | 0.277778 | 0.667898 | 0.607219 | 2208 | tags=28% | list=14% | signal=24% |
| REACTOME_IRON_UPTAKE_AND_TRANSPORT | 49 | -0.48971 | -1.15515 | 0.277844 | 0.667898 | 0.607219 | 2993 | tags=37% | list=19% | signal=30% |
| REACTOME_REGULATION_OF_PLK1_ACTIVITY_AT_G2_M_TRANSITION | 84 | 0.358646 | 1.106113 | 0.27619 | 0.667898 | 0.607219 | 3004 | tags=30% | list=19% | signal=24% |
| REACTOME_REGULATION_OF_TP53_ACTIVITY | 148 | 0.343402 | 1.06963 | 0.278435 | 0.668084 | 0.607387 | 2363 | tags=27% | list=15% | signal=23% |
| REACTOME_SCF_BETA_TRCP_MEDIATED_DEGRADATION_OF_EMI1 | 9 | 0.5534 | 1.178476 | 0.280928 | 0.669311 | 0.608503 | 3000 | tags=33% | list=19% | signal=27% |
| REACTOME_DSCAM_INTERACTIONS | 9 | 0.553217 | 1.178086 | 0.280928 | 0.669311 | 0.608503 | 1253 | tags=33% | list=8% | signal=31% |
| REACTOME_EPHB_MEDIATED_FORWARD_SIGNALING | 36 | -0.51545 | -1.16248 | 0.281369 | 0.669311 | 0.608503 | 1347 | tags=17% | list=8% | signal=15% |
| REACTOME_DEADENYLATION_OF_MRNA | 25 | 0.438863 | 1.13667 | 0.280992 | 0.669311 | 0.608503 | 3524 | tags=36% | list=22% | signal=28% |
| BIOCARTA_RACCYCD_PATHWAY | 25 | 0.438711 | 1.136276 | 0.280992 | 0.669311 | 0.608503 | 3508 | tags=44% | list=22% | signal=34% |
| REACTOME_CHROMATIN_MODIFYING_ENZYMES | 155 | 0.342942 | 1.070796 | 0.280513 | 0.669311 | 0.608503 | 4182 | tags=42% | list=26% | signal=31% |
| REACTOME_ROLE_OF_PHOSPHOLIPIDS_IN_PHAGOCYTOSIS | 20 | 0.444033 | 1.139471 | 0.282686 | 0.671479 | 0.610475 | 1282 | tags=35% | list=8% | signal=32% |
| BIOCARTA_TFF_PATHWAY | 21 | 0.450959 | 1.148065 | 0.283582 | 0.672162 | 0.611095 | 3034 | tags=38% | list=19% | signal=31% |
| REACTOME_DECTIN_1_MEDIATED_NONCANONICAL_NF_KB_SIGNALING | 56 | 0.36214 | 1.074406 | 0.283784 | 0.672162 | 0.611095 | 5771 | tags=45% | list=36% | signal=29% |
| REACTOME_SIGNALING_BY_NOTCH1 | 18 | 0.462507 | 1.143367 | 0.284722 | 0.673423 | 0.612241 | 2214 | tags=39% | list=14% | signal=34% |
| BIOCARTA_CTBP1_PATHWAY | 8 | 0.554989 | 1.161293 | 0.286064 | 0.674619 | 0.613329 | 1887 | tags=38% | list=12% | signal=33% |
| REACTOME_DNA_REPLICATION_PRE_INITIATION | 96 | 0.359531 | 1.120302 | 0.286449 | 0.674619 | 0.613329 | 2680 | tags=22% | list=17% | signal=18% |
| REACTOME_COLLAGEN_CHAIN_TRIMERIZATION | 36 | 0.389092 | 1.097225 | 0.286385 | 0.674619 | 0.613329 | 660 | tags=14% | list=4% | signal=13% |
| REACTOME_LIPID_PARTICLE_ORGANIZATION | 3 | -0.77597 | -1.18193 | 0.287908 | 0.676351 | 0.614904 | 3620 | tags=100% | list=22% | signal=78% |
| REACTOME_RHOB_GTPASE_CYCLE | 68 | 0.352997 | 1.05884 | 0.288 | 0.676351 | 0.614904 | 1231 | tags=22% | list=8% | signal=20% |
| REACTOME_OPIOID_SIGNALLING | 68 | -0.46847 | -1.14388 | 0.289624 | 0.679202 | 0.617496 | 1777 | tags=18% | list=11% | signal=16% |
| REACTOME_EPIGENETIC_REGULATION_OF_GENE_EXPRESSION | 66 | 0.350072 | 1.045315 | 0.291339 | 0.681297 | 0.6194 | 2459 | tags=18% | list=15% | signal=15% |
| REACTOME_SIGNALING_BY_MET | 66 | 0.346892 | 1.03582 | 0.291339 | 0.681297 | 0.6194 | 2556 | tags=35% | list=16% | signal=29% |
| REACTOME_FATTY_ACYL_COA_BIOSYNTHESIS | 28 | -0.53162 | -1.16282 | 0.292428 | 0.681921 | 0.619968 | 1720 | tags=32% | list=11% | signal=29% |
| REACTOME_PROCESSIVE_SYNTHESIS_ON_THE_C_STRAND_OF_THE_TELOMERE | 17 | 0.450283 | 1.122752 | 0.292208 | 0.681921 | 0.619968 | 452 | tags=12% | list=3% | signal=11% |
| REACTOME_INTERLEUKIN_23_SIGNALING | 5 | 0.659665 | 1.204394 | 0.296296 | 0.682144 | 0.62017 | 2293 | tags=80% | list=14% | signal=69% |
| REACTOME_TRANSPORT_OF_RCBL_WITHIN_THE_BODY | 6 | -0.7074 | -1.20326 | 0.297297 | 0.682144 | 0.62017 | 2320 | tags=67% | list=14% | signal=57% |
| REACTOME_GLUCURONIDATION | 5 | -0.72877 | -1.20274 | 0.296501 | 0.682144 | 0.62017 | 3998 | tags=60% | list=25% | signal=45% |
| REACTOME_CALCITONIN_LIKE_LIGAND_RECEPTORS | 6 | -0.70592 | -1.20074 | 0.297297 | 0.682144 | 0.62017 | 2689 | tags=67% | list=17% | signal=56% |
| REACTOME_RUNX2_REGULATES_BONE_DEVELOPMENT | 6 | 0.611735 | 1.196769 | 0.297539 | 0.682144 | 0.62017 | 5296 | tags=67% | list=33% | signal=45% |
| BIOCARTA_LYMPHOCYTE_PATHWAY | 8 | -0.66372 | -1.1931 | 0.293423 | 0.682144 | 0.62017 | 2076 | tags=88% | list=13% | signal=76% |
| BIOCARTA_NGF_PATHWAY | 19 | -0.56643 | -1.16807 | 0.293463 | 0.682144 | 0.62017 | 390 | tags=16% | list=2% | signal=15% |
| WP_EDA_SIGNALING_IN_HAIR_FOLLICLE_DEVELOPMENT | 15 | -0.58399 | -1.16469 | 0.297496 | 0.682144 | 0.62017 | 1151 | tags=20% | list=7% | signal=19% |
| WP_BMP_SIGNALING_PATHWAY_IN_EYELID_DEVELOPMENT | 19 | -0.56216 | -1.15927 | 0.297636 | 0.682144 | 0.62017 | 2467 | tags=47% | list=15% | signal=40% |
| WP_TRANSCRIPTIONAL_ACTIVATION_BY_NFE2L2_IN_RESPONSE_TO_PHYTOCHEMI | 13 | 0.497678 | 1.154532 | 0.294985 | 0.682144 | 0.62017 | 4135 | tags=38% | list=26% | signal=29% |
| REACTOME_SIGNALING_BY_TGFB_FAMILY_MEMBERS | 95 | -0.44313 | -1.11902 | 0.294053 | 0.682144 | 0.62017 | 3078 | tags=26% | list=19% | signal=21% |
| REACTOME_ANCHORING_OF_THE_BASAL_BODY_TO_THE_PLASMA_MEMBRANE | 95 | 0.350837 | 1.085794 | 0.297872 | 0.682144 | 0.62017 | 3004 | tags=28% | list=19% | signal=23% |
| REACTOME_HDR_THROUGH_HOMOLOGOUS_RECOMBINATION_HRR | 54 | 0.357833 | 1.082179 | 0.296774 | 0.682144 | 0.62017 | 1801 | tags=19% | list=11% | signal=17% |
| REACTOME_SENSING_OF_DNA_DOUBLE_STRAND_BREAKS | 6 | 0.611072 | 1.195473 | 0.299776 | 0.685557 | 0.623274 | 4752 | tags=83% | list=29% | signal=59% |
| REACTOME_SUMO_IS_TRANSFERRED_FROM_E1_TO_E2_UBE2I_UBC9 | 5 | 0.654391 | 1.194764 | 0.300654 | 0.685672 | 0.623378 | 1887 | tags=40% | list=12% | signal=35% |
| REACTOME_SUMOYLATION_OF_TRANSCRIPTION_FACTORS | 15 | 0.478533 | 1.14017 | 0.30031 | 0.685672 | 0.623378 | 3167 | tags=33% | list=20% | signal=27% |
| BIOCARTA_TH1TH2_PATHWAY | 9 | -0.64306 | -1.18371 | 0.301303 | 0.686184 | 0.623843 | 3418 | tags=67% | list=21% | signal=53% |
| WP_MYOMETRIAL_RELAXATION_AND_CONTRACTION_PATHWAYS | 135 | -0.42552 | -1.10081 | 0.301706 | 0.686184 | 0.623843 | 1870 | tags=16% | list=12% | signal=15% |
| REACTOME_ABACAVIR_ADME | 5 | -0.72378 | -1.19451 | 0.305709 | 0.690477 | 0.627746 | 330 | tags=40% | list=2% | signal=39% |
| WP_ESTROGEN_METABOLISM | 5 | 0.653296 | 1.192766 | 0.305011 | 0.690477 | 0.627746 | 166 | tags=40% | list=1% | signal=40% |
| BIOCARTA_RARRXR_PATHWAY | 6 | 0.609368 | 1.192139 | 0.304251 | 0.690477 | 0.627746 | 104 | tags=17% | list=1% | signal=17% |
| WP_MIR302367_PROMOTING_CARDIOMYOCYTE_PROLIFERATION | 3 | 0.723228 | 1.170972 | 0.305613 | 0.690477 | 0.627746 | 986 | tags=33% | list=6% | signal=31% |
| REACTOME_ABC_TRANSPORTERS_IN_LIPID_HOMEOSTASIS | 14 | -0.58748 | -1.16412 | 0.306092 | 0.690477 | 0.627746 | 4262 | tags=64% | list=26% | signal=47% |
| REACTOME_REGULATION_OF_MRNA_STABILITY_BY_PROTEINS_THAT_BIND_AU_RI | 82 | 0.349321 | 1.065261 | 0.304762 | 0.690477 | 0.627746 | 5830 | tags=49% | list=36% | signal=31% |
| REACTOME_VITAMIN_B5_PANTOTHENATE_METABOLISM | 13 | 0.493948 | 1.145878 | 0.306785 | 0.691099 | 0.628312 | 3764 | tags=38% | list=23% | signal=30% |
| REACTOME_SIGNALING_BY_BMP | 25 | -0.53658 | -1.16379 | 0.307895 | 0.692659 | 0.62973 | 5196 | tags=64% | list=32% | signal=43% |
| REACTOME_SYNAPTIC_ADHESION_LIKE_MOLECULES | 20 | -0.55691 | -1.15464 | 0.308762 | 0.693326 | 0.630337 | 2671 | tags=45% | list=17% | signal=38% |
| BIOCARTA_CHEMICAL_PATHWAY | 18 | 0.44965 | 1.111584 | 0.309028 | 0.693326 | 0.630337 | 3599 | tags=39% | list=22% | signal=30% |
| REACTOME_EPHRIN_SIGNALING | 18 | -0.56101 | -1.1506 | 0.310924 | 0.695699 | 0.632493 | 1052 | tags=22% | list=7% | signal=21% |
| REACTOME_DNA_REPLICATION | 123 | 0.33875 | 1.100974 | 0.310716 | 0.695699 | 0.632493 | 3371 | tags=24% | list=21% | signal=19% |
| BIOCARTA_D4GDI_PATHWAY | 10 | 0.519135 | 1.138191 | 0.31383 | 0.700711 | 0.63705 | 3492 | tags=30% | list=22% | signal=24% |
| REACTOME_INTRINSIC_PATHWAY_FOR_APOPTOSIS | 37 | 0.385737 | 1.088651 | 0.31401 | 0.700711 | 0.63705 | 4268 | tags=38% | list=26% | signal=28% |
| BIOCARTA_ERBB3_PATHWAY | 6 | -0.69948 | -1.18978 | 0.317117 | 0.700817 | 0.637147 | 197 | tags=50% | list=1% | signal=49% |
| BIOCARTA_NOTCH_PATHWAY | 6 | -0.69686 | -1.18533 | 0.317117 | 0.700817 | 0.637147 | 4015 | tags=67% | list=25% | signal=50% |
| WP_MIR127_IN_MESENDODERM_DIFFERENTIATION | 6 | -0.69655 | -1.1848 | 0.317117 | 0.700817 | 0.637147 | 1116 | tags=50% | list=7% | signal=47% |
| BIOCARTA_MONOCYTE_PATHWAY | 9 | -0.63432 | -1.16762 | 0.315961 | 0.700817 | 0.637147 | 2858 | tags=89% | list=18% | signal=73% |
| WP_HEDGEHOG_SIGNALING_PATHWAY | 21 | -0.55139 | -1.15665 | 0.317439 | 0.700817 | 0.637147 | 1151 | tags=24% | list=7% | signal=22% |
| REACTOME_KERATAN_SULFATE_KERATIN_METABOLISM | 32 | -0.50879 | -1.1397 | 0.316456 | 0.700817 | 0.637147 | 3575 | tags=47% | list=22% | signal=37% |
| BIOCARTA_PCAF_PATHWAY | 9 | 0.534371 | 1.137953 | 0.31701 | 0.700817 | 0.637147 | 4723 | tags=67% | list=29% | signal=47% |
| REACTOME_ANTIGEN_PROCESSING_CROSS_PRESENTATION | 74 | -0.45994 | -1.12703 | 0.316686 | 0.700817 | 0.637147 | 2728 | tags=20% | list=17% | signal=17% |
| REACTOME_BETA_OXIDATION_OF_LAUROYL_COA_TO_DECANOYL_COA_COA | 5 | 0.642163 | 1.172439 | 0.318083 | 0.701024 | 0.637335 | 4472 | tags=80% | list=28% | signal=58% |
| REACTOME_ADRENALINE_NORADRENALINE_INHIBITS_INSULIN_SECRETION | 21 | -0.54959 | -1.15289 | 0.318801 | 0.701024 | 0.637335 | 884 | tags=29% | list=5% | signal=27% |
| REACTOME_GABA_SYNTHESIS_RELEASE_REUPTAKE_AND_DEGRADATION | 19 | -0.55171 | -1.13771 | 0.318498 | 0.701024 | 0.637335 | 765 | tags=21% | list=5% | signal=20% |
| BIOCARTA_ERK_PATHWAY | 25 | -0.53284 | -1.15568 | 0.319737 | 0.702151 | 0.638359 | 2337 | tags=28% | list=14% | signal=24% |
| REACTOME_SIGNALING_BY_NUCLEAR_RECEPTORS | 160 | 0.335416 | 1.063116 | 0.321743 | 0.70469 | 0.640668 | 2437 | tags=26% | list=15% | signal=23% |
| WP_AMINO_ACID_METABOLISM | 78 | 0.339991 | 1.034904 | 0.321739 | 0.70469 | 0.640668 | 3129 | tags=42% | list=19% | signal=34% |
| REACTOME_ELEVATION_OF_CYTOSOLIC_CA2_LEVELS | 12 | -0.59804 | -1.16391 | 0.322289 | 0.704954 | 0.640908 | 2684 | tags=83% | list=17% | signal=70% |
| REACTOME_NEPHRIN_FAMILY_INTERACTIONS | 8 | -0.65246 | -1.17286 | 0.325464 | 0.705384 | 0.641299 | 590 | tags=25% | list=4% | signal=24% |
| REACTOME_BETA_OXIDATION_OF_OCTANOYL_COA_TO_HEXANOYL_COA | 5 | 0.639939 | 1.168378 | 0.324619 | 0.705384 | 0.641299 | 4472 | tags=80% | list=28% | signal=58% |
| BIOCARTA_IL4_PATHWAY | 10 | 0.515593 | 1.130426 | 0.324468 | 0.705384 | 0.641299 | 2147 | tags=40% | list=13% | signal=35% |
| REACTOME_CLEAVAGE_OF_THE_DAMAGED_PYRIMIDINE | 8 | 0.539657 | 1.129212 | 0.325183 | 0.705384 | 0.641299 | 4891 | tags=62% | list=30% | signal=44% |
| BIOCARTA_GCR_PATHWAY | 15 | 0.468674 | 1.116679 | 0.325077 | 0.705384 | 0.641299 | 2147 | tags=47% | list=13% | signal=40% |
| REACTOME_TRANSCRIPTIONAL_REGULATION_BY_THE_AP_2_TFAP2_FAMILY_OF_T | 18 | 0.441964 | 1.092583 | 0.322917 | 0.705384 | 0.641299 | 3426 | tags=28% | list=21% | signal=22% |
| REACTOME_BETA_CATENIN_PHOSPHORYLATION_CASCADE | 17 | 0.435605 | 1.086152 | 0.324675 | 0.705384 | 0.641299 | 5694 | tags=76% | list=35% | signal=50% |
| REACTOME_FACTORS_INVOLVED_IN_MEGAKARYOCYTE_DEVELOPMENT_AND_PLAT | 106 | 0.340495 | 1.095706 | 0.326031 | 0.705691 | 0.641578 | 3184 | tags=32% | list=20% | signal=26% |
| BIOCARTA_NOS1_PATHWAY | 19 | -0.54827 | -1.13063 | 0.326843 | 0.706526 | 0.642337 | 1054 | tags=21% | list=7% | signal=20% |
| REACTOME_HISTIDINE_CATABOLISM | 4 | -0.72177 | -1.16687 | 0.327684 | 0.707421 | 0.643151 | 988 | tags=50% | list=6% | signal=47% |
| REACTOME_BETA_OXIDATION_OF_HEXANOYL_COA_TO_BUTANOYL_COA | 5 | 0.637181 | 1.163343 | 0.328976 | 0.708367 | 0.644011 | 4472 | tags=80% | list=28% | signal=58% |
| BIOCARTA_PPARG_PATHWAY | 7 | 0.549296 | 1.13338 | 0.328767 | 0.708367 | 0.644011 | 104 | tags=14% | list=1% | signal=14% |
| REACTOME_PLASMA_LIPOPROTEIN_ASSEMBLY | 13 | -0.58508 | -1.14898 | 0.330317 | 0.710331 | 0.645796 | 2394 | tags=38% | list=15% | signal=33% |
| WP_MAPK_SIGNALING_PATHWAY | 151 | -0.41668 | -1.08468 | 0.331217 | 0.711344 | 0.646718 | 531 | tags=8% | list=3% | signal=8% |
| REACTOME_RECRUITMENT_OF_NUMA_TO_MITOTIC_CENTROSOMES | 84 | 0.326814 | 1.00794 | 0.333333 | 0.714963 | 0.650008 | 5001 | tags=46% | list=31% | signal=32% |
| REACTOME_INHIBITION_OF_REPLICATION_INITIATION_OF_DAMAGED_DNA_BY_RB | 10 | 0.51163 | 1.121736 | 0.335106 | 0.716911 | 0.651779 | 5627 | tags=80% | list=35% | signal=52% |
| REACTOME_CARGO_RECOGNITION_FOR_CLATHRIN_MEDIATED_ENDOCYTOSIS | 92 | -0.43719 | -1.09712 | 0.334807 | 0.716911 | 0.651779 | 941 | tags=11% | list=6% | signal=10% |
| REACTOME_SIGNALING_BY_NTRK2_TRKB | 8 | -0.64734 | -1.16367 | 0.335582 | 0.717003 | 0.651863 | 307 | tags=25% | list=2% | signal=25% |
| BIOCARTA_MRP_PATHWAY | 4 | -0.71438 | -1.15492 | 0.3371 | 0.719248 | 0.653903 | 2805 | tags=50% | list=17% | signal=41% |
| REACTOME_SPRY_REGULATION_OF_FGF_SIGNALING | 16 | 0.451137 | 1.095223 | 0.3375 | 0.719248 | 0.653903 | 3034 | tags=38% | list=19% | signal=30% |
| REACTOME_PTK6_PROMOTES_HIF1A_STABILIZATION | 5 | -0.71472 | -1.17956 | 0.338858 | 0.720291 | 0.654851 | 2241 | tags=60% | list=14% | signal=52% |
| REACTOME_INTEGRIN_SIGNALING | 24 | 0.414119 | 1.077229 | 0.338645 | 0.720291 | 0.654851 | 2781 | tags=38% | list=17% | signal=31% |
| REACTOME_CREB3_FACTORS_ACTIVATE_GENES | 5 | -0.71122 | -1.17378 | 0.3407 | 0.723278 | 0.657567 | 2115 | tags=60% | list=13% | signal=52% |

| | | | | | | | | | | |
|---|---|---|---|---|---|---|---|---|---|---|
| WP_ADIPOGENESIS_GENES | 110 | -0.42347 | -1.08176 | 0.341622 | 0.723383 | 0.657662 | 1869 | tags=16% | list=12% | signal=15% |
| REACTOME_SIGNALING_BY_RECEPTOR_TYROSINE_KINASES | 372 | -0.39158 | -1.05984 | 0.341439 | 0.723383 | 0.657662 | 2068 | tags=15% | list=13% | signal=13% |
| BIOCARTA_CTLA4_PATHWAY | 9 | 0.52566 | 1.119404 | 0.342784 | 0.724127 | 0.658339 | 2303 | tags=33% | list=14% | signal=29% |
| REACTOME_ACTIVATED_NOTCH1_TRANSMITS_SIGNAL_TO_THE_NUCLEUS | 10 | 0.506797 | 1.11114 | 0.343085 | 0.724127 | 0.658339 | 3441 | tags=50% | list=21% | signal=39% |
| REACTOME_CARBOXYTERMINAL_POST_TRANSLATIONAL_MODIFICATIONS_OF_TUB | 21 | 0.426757 | 1.086453 | 0.343284 | 0.724127 | 0.658339 | 3561 | tags=29% | list=22% | signal=22% |
| REACTOME_ACTIVATION_OF_RAC1_DOWNSTREAM_OF_NMDARS | 6 | 0.592301 | 1.158748 | 0.344519 | 0.724323 | 0.658517 | 1408 | tags=33% | list=9% | signal=30% |
| BIOCARTA_VOBESITY_PATHWAY | 6 | 0.5921 | 1.158356 | 0.344519 | 0.724323 | 0.658517 | 104 | tags=17% | list=1% | signal=17% |
| BIOCARTA_IL2_PATHWAY | 21 | -0.53447 | -1.12118 | 0.344687 | 0.724323 | 0.658517 | 49 | tags=10% | list=0% | signal=10% |
| REACTOME_RHO_GTPASE_EFFECTORS | 234 | 0.296953 | 1.014055 | 0.346184 | 0.726548 | 0.66054 | 3034 | tags=27% | list=19% | signal=22% |
| REACTOME_PEROXISOMAL_LIPID_METABOLISM | 25 | 0.425798 | 1.102833 | 0.347107 | 0.727565 | 0.661465 | 4434 | tags=48% | list=27% | signal=35% |
| REACTOME_CAMK_IV_MEDIATED_PHOSPHORYLATION_OF_CREB | 5 | 0.628633 | 1.147736 | 0.348584 | 0.728809 | 0.662596 | 1408 | tags=40% | list=9% | signal=37% |
| REACTOME_IKK_COMPLEX_RECRUITMENT_MEDIATED_BY_RIP1 | 23 | 0.412734 | 1.066565 | 0.34902 | 0.728809 | 0.662596 | 2308 | tags=22% | list=14% | signal=19% |
| REACTOME_NOD1_2_SIGNALING_PATHWAY | 31 | 0.389875 | 1.063976 | 0.348624 | 0.728809 | 0.662596 | 2308 | tags=26% | list=14% | signal=22% |
| REACTOME_INSULIN_RECEPTOR_SIGNALLING_CASCADE | 38 | 0.375403 | 1.074628 | 0.35122 | 0.73248 | 0.665934 | 3034 | tags=32% | list=19% | signal=26% |
| REACTOME_ANTIMICROBIAL_PEPTIDES | 19 | -0.53652 | -1.10639 | 0.351878 | 0.732931 | 0.666343 | 4160 | tags=37% | list=26% | signal=27% |
| REACTOME_TANDEM_PORE_DOMAIN_POTASSIUM_CHANNELS | 8 | -0.63582 | -1.14296 | 0.354132 | 0.734856 | 0.668093 | 3276 | tags=88% | list=20% | signal=70% |
| REACTOME_GP1B_IX_V_ACTIVATION_SIGNALLING | 11 | 0.495457 | 1.09688 | 0.35376 | 0.734856 | 0.668093 | 2116 | tags=27% | list=13% | signal=24% |
| REACTOME_LYSINE_CATABOLISM | 11 | 0.493527 | 1.092608 | 0.35376 | 0.734856 | 0.668093 | 3109 | tags=55% | list=19% | signal=44% |
| REACTOME_HYDROLYSIS_OF_LPC | 6 | -0.68849 | -1.1711 | 0.354955 | 0.735644 | 0.66881 | 2800 | tags=67% | list=17% | signal=55% |
| REACTOME_SPERM_MOTILITY_AND_TAXES | 5 | -0.70348 | -1.16101 | 0.355433 | 0.735715 | 0.668874 | 481 | tags=40% | list=3% | signal=39% |
| REACTOME_ENERGY_DEPENDENT_REGULATION_OF_MTOR_BY_LKB1_AMPK | 28 | 0.400363 | 1.059793 | 0.355932 | 0.73583 | 0.668979 | 6132 | tags=75% | list=38% | signal=47% |
| REACTOME_DEGRADATION_OF_BETA_CATENIN_BY_THE_DESTRUCTION_COMPLEX | 78 | 0.327667 | 0.997389 | 0.356522 | 0.736131 | 0.669252 | 5771 | tags=55% | list=36% | signal=36% |
| REACTOME_NEUROTRANSMITTER_CLEARANCE | 6 | -0.68619 | -1.16718 | 0.358559 | 0.737709 | 0.670687 | 626 | tags=33% | list=4% | signal=32% |
| WP_ELECTRON_TRANSPORT_CHAIN | 65 | -0.45034 | -1.09475 | 0.358621 | 0.737709 | 0.670687 | 1486 | tags=5% | list=9% | signal=4% |
| REACTOME_MET_PROMOTES_CELL_MOTILITY | 31 | 0.387403 | 1.05723 | 0.357798 | 0.737709 | 0.670687 | 1783 | tags=32% | list=11% | signal=29% |
| BIOCARTA_FBW7_PATHWAY | 8 | 0.522397 | 1.093095 | 0.359413 | 0.738423 | 0.671336 | 2798 | tags=50% | list=17% | signal=41% |
| BIOCARTA_IGF1_PATHWAY | 22 | -0.52752 | -1.11423 | 0.360811 | 0.740377 | 0.673112 | 420 | tags=9% | list=3% | signal=9% |
| REACTOME_GLUTAMATE_NEUROTRANSMITTER_RELEASE_CYCLE | 23 | -0.52375 | -1.1122 | 0.362784 | 0.740759 | 0.67346 | 855 | tags=22% | list=5% | signal=21% |
| REACTOME_EGFR_DOWNREGULATION | 27 | 0.401491 | 1.060584 | 0.3625 | 0.740759 | 0.67346 | 2276 | tags=33% | list=14% | signal=29% |
| REACTOME_ANTIVIRAL_MECHANISM_BY_IFN_STIMULATED_GENES | 31 | 0.384708 | 1.049875 | 0.362385 | 0.740759 | 0.67346 | 3754 | tags=42% | list=23% | signal=32% |
| REACTOME_RHO_GTPASES_ACTIVATE_FORMINS | 127 | 0.313099 | 1.001623 | 0.362319 | 0.740759 | 0.67346 | 3024 | tags=20% | list=19% | signal=16% |
| REACTOME_INSERTION_OF_TAIL_ANCHORED_PROTEINS_INTO_THE_ENDOPLASMIC | 14 | -0.55729 | -1.1043 | 0.364042 | 0.741333 | 0.673982 | 573 | tags=29% | list=4% | signal=28% |
| REACTOME_TELOMERE_C_STRAND_SYNTHESIS_INITIATION | 12 | 0.48322 | 1.089071 | 0.363905 | 0.741333 | 0.673982 | 4422 | tags=50% | list=27% | signal=36% |
| REACTOME_PROCESSING_OF_CAPPED_INTRONLESS_PRE_MRNA | 28 | 0.39788 | 1.053218 | 0.364407 | 0.741333 | 0.673982 | 2857 | tags=29% | list=18% | signal=24% |
| REACTOME_ADP_SIGNALLING_THROUGH_P2Y_PURINOCEPTOR_1 | 22 | -0.52334 | -1.1054 | 0.364865 | 0.741355 | 0.674002 | 1327 | tags=18% | list=8% | signal=17% |
| REACTOME_ARACHIDONATE_PRODUCTION_FROM_DAG | 5 | -0.69791 | -1.15183 | 0.366483 | 0.741781 | 0.674389 | 2196 | tags=80% | list=14% | signal=69% |
| REACTOME_WNT5A_DEPENDENT_INTERNALIZATION_OF_FZD2_FZD5_AND_ROR2 | 12 | 0.481833 | 1.085946 | 0.366864 | 0.741781 | 0.674389 | 2105 | tags=33% | list=13% | signal=29% |
| BIOCARTA_SET_PATHWAY | 9 | 0.508785 | 1.083468 | 0.365979 | 0.741781 | 0.674389 | 6381 | tags=67% | list=40% | signal=40% |
| REACTOME_RND2_GTPASE_CYCLE | 39 | 0.362104 | 1.045371 | 0.365854 | 0.741781 | 0.674389 | 1233 | tags=23% | list=8% | signal=21% |
| WP_ESTROGEN_SIGNALING | 73 | 0.326276 | 0.995382 | 0.368 | 0.743172 | 0.675654 | 3704 | tags=44% | list=23% | signal=34% |
| REACTOME_AURKA_ACTIVATION_BY_TPX2 | 70 | 0.327033 | 0.983447 | 0.368852 | 0.743987 | 0.676395 | 3004 | tags=29% | list=19% | signal=23% |
| REACTOME_DEX_H_BOX_HELICASES_ACTIVATE_TYPE_I_IFN_AND_INFLAMMATORY | 7 | 0.537614 | 1.109277 | 0.369863 | 0.744215 | 0.676602 | 2445 | tags=43% | list=15% | signal=36% |
| REACTOME_CYTOCHROME_P450_ARRANGED_BY_SUBSTRATE_TYPE | 33 | -0.49246 | -1.10054 | 0.369565 | 0.744215 | 0.676602 | 1904 | tags=18% | list=12% | signal=16% |
| REACTOME_AGGREPHAGY | 30 | 0.385756 | 1.044577 | 0.370536 | 0.744654 | 0.677001 | 4615 | tags=40% | list=29% | signal=29% |
| REACTOME_RESOLUTION_OF_SISTER_CHROMATID_COHESION | 114 | 0.315221 | 0.992341 | 0.371429 | 0.744654 | 0.677001 | 5777 | tags=45% | list=36% | signal=29% |
| REACTOME_PCP_CE_PATHWAY | 84 | 0.321387 | 0.991201 | 0.371429 | 0.744654 | 0.677001 | 2880 | tags=21% | list=18% | signal=18% |
| REACTOME_PROLACTIN_RECEPTOR_SIGNALING | 7 | 0.53657 | 1.107122 | 0.372146 | 0.745191 | 0.67749 | 4359 | tags=57% | list=27% | signal=42% |
| REACTOME_FATTY_ACIDS_BOUND_TO_GPR40_FFAR1_REGULATE_INSULIN_SECRET | 6 | 0.578049 | 1.130868 | 0.373602 | 0.747204 | 0.679319 | 3970 | tags=67% | list=25% | signal=50% |
| BIOCARTA_EPO_PATHWAY | 19 | -0.52714 | -1.08705 | 0.374131 | 0.74736 | 0.679461 | 49 | tags=11% | list=0% | signal=11% |
| BIOCARTA_MEF2D_PATHWAY | 15 | 0.445865 | 1.062333 | 0.374613 | 0.747423 | 0.679518 | 3432 | tags=40% | list=21% | signal=32% |
| REACTOME_LECTIN_PATHWAY_OF_COMPLEMENT_ACTIVATION | 4 | -0.69712 | -1.12702 | 0.376648 | 0.750579 | 0.682388 | 2911 | tags=75% | list=18% | signal=61% |
| REACTOME_INTERLEUKIN_7_SIGNALING | 13 | 0.472056 | 1.095092 | 0.377581 | 0.750635 | 0.682439 | 1276 | tags=38% | list=8% | signal=35% |
| BIOCARTA_ERAD_PATHWAY | 14 | -0.55167 | -1.09315 | 0.377415 | 0.750635 | 0.682439 | 2081 | tags=43% | list=13% | signal=37% |
| REACTOME_FCERI_MEDIATED_MAPK_ACTIVATION | 28 | -0.49972 | -1.09304 | 0.37859 | 0.751739 | 0.683443 | 49 | tags=7% | list=0% | signal=7% |
| REACTOME_SIGNAL_REGULATORY_PROTEIN_FAMILY_INTERACTIONS | 7 | -0.65203 | -1.13066 | 0.382979 | 0.753146 | 0.684721 | 313 | tags=29% | list=2% | signal=28% |
| REACTOME_ACTIVATION_OF_THE_PHOTOTRANSDUCTION_CASCADE | 3 | 0.695227 | 1.125637 | 0.380457 | 0.753146 | 0.684721 | 4177 | tags=67% | list=26% | signal=49% |
| REACTOME_METABOLISM_OF_AMINE_DERIVED_HORMONES | 11 | -0.58053 | -1.10984 | 0.382582 | 0.753146 | 0.684721 | 237 | tags=55% | list=1% | signal=54% |
| BIOCARTA_VITCB_PATHWAY | 10 | -0.59168 | -1.10578 | 0.383387 | 0.753146 | 0.684721 | 1208 | tags=30% | list=7% | signal=28% |
| BIOCARTA_PRION_PATHWAY | 10 | -0.59058 | -1.10373 | 0.383387 | 0.753146 | 0.684721 | 625 | tags=30% | list=4% | signal=29% |
| REACTOME_CYTOSOLIC_SULFONATION_OF_SMALL_MOLECULES | 15 | -0.54463 | -1.0862 | 0.381443 | 0.753146 | 0.684721 | 2229 | tags=47% | list=14% | signal=40% |
| BIOCARTA_GABA_PATHWAY | 10 | 0.489362 | 1.072916 | 0.382979 | 0.753146 | 0.684721 | 6 | tags=10% | list=0% | signal=10% |
| WP_SELENIUM_METABOLISM_SELENOPROTEINS | 42 | -0.46337 | -1.06879 | 0.379902 | 0.753146 | 0.684721 | 2230 | tags=19% | list=14% | signal=16% |
| REACTOME_MITOTIC_SPINDLE_CHECKPOINT | 107 | 0.306127 | 0.976977 | 0.380952 | 0.753146 | 0.684721 | 6054 | tags=48% | list=38% | signal=30% |
| REACTOME_GLUCAGON_SIGNALING_IN_METABOLIC_REGULATION | 5 | -0.68535 | -1.13109 | 0.384899 | 0.754265 | 0.685739 | 936 | tags=40% | list=6% | signal=38% |
| REACTOME_PKA_ACTIVATION_IN_GLUCAGON_SIGNALLING | 5 | -0.68535 | -1.13109 | 0.384899 | 0.754265 | 0.685739 | 936 | tags=40% | list=6% | signal=38% |
| WP_NUCLEAR_RECEPTORS | 31 | 0.379306 | 1.035133 | 0.385321 | 0.754265 | 0.685739 | 1196 | tags=26% | list=7% | signal=24% |
| REACTOME_SEMA3A_PAK_DEPENDENT_AXON_REPULSION | 14 | 0.452771 | 1.05982 | 0.386018 | 0.754738 | 0.686169 | 2852 | tags=50% | list=18% | signal=41% |
| REACTOME_REGULATION_BY_C_FLIP | 9 | 0.502992 | 1.07113 | 0.386598 | 0.754982 | 0.68639 | 1531 | tags=22% | list=9% | signal=20% |
| BIOCARTA_NEUTROPHIL_PATHWAY | 7 | -0.64908 | -1.12555 | 0.388298 | 0.756164 | 0.687465 | 2858 | tags=86% | list=18% | signal=71% |
| REACTOME_SYNTHESIS_OF_15_EICOSATETRAENOIC_ACID_DERIVATIVES | 4 | -0.69276 | -1.11996 | 0.387947 | 0.756164 | 0.687465 | 2754 | tags=50% | list=17% | signal=41% |
| REACTOME_CYTOPROTECTION_BY_HMOX1 | 46 | 0.346246 | 1.021393 | 0.388571 | 0.756164 | 0.687465 | 1149 | tags=13% | list=7% | signal=12% |
| BIOCARTA_ARF_PATHWAY | 15 | 0.439404 | 1.04694 | 0.390093 | 0.758234 | 0.689348 | 1621 | tags=33% | list=10% | signal=30% |
| REACTOME_ACYL_CHAIN_REMODELLING_OF_PC | 20 | -0.51985 | -1.07781 | 0.392211 | 0.76146 | 0.69228 | 2589 | tags=40% | list=16% | signal=34% |
| REACTOME_STRIATED_MUSCLE_CONTRACTION | 31 | -0.48835 | -1.09004 | 0.392857 | 0.761821 | 0.692609 | 2618 | tags=16% | list=16% | signal=14% |
| REACTOME_SIGNALING_BY_ERBB4 | 31 | -0.48612 | -1.08507 | 0.395408 | 0.765872 | 0.696292 | 2496 | tags=29% | list=15% | signal=25% |
| REACTOME_TIGHT_JUNCTION_INTERACTIONS | 6 | -0.66804 | -1.13631 | 0.396396 | 0.766891 | 0.697218 | 2542 | tags=67% | list=16% | signal=56% |
| BIOCARTA_ACETAMINOPHEN_PATHWAY | 3 | -0.7336 | -1.11739 | 0.397313 | 0.767768 | 0.698015 | 847 | tags=100% | list=5% | signal=95% |
| BIOCARTA_NFAT_PATHWAY | 42 | 0.355008 | 1.023388 | 0.403226 | 0.778287 | 0.707578 | 3628 | tags=43% | list=22% | signal=33% |
| REACTOME_ASPIRIN_ADME | 7 | -0.64303 | -1.11506 | 0.404255 | 0.778461 | 0.707737 | 2805 | tags=29% | list=17% | signal=24% |
| REACTOME_BICARBONATE_TRANSPORTERS | 9 | -0.59308 | -1.0917 | 0.403909 | 0.778461 | 0.707737 | 2259 | tags=44% | list=14% | signal=38% |
| BIOCARTA_FXR_PATHWAY | 4 | -0.68577 | -1.10868 | 0.40678 | 0.780778 | 0.709844 | 36 | tags=50% | list=0% | signal=50% |
| REACTOME_PRESYNAPTIC_DEPOLARIZATION_AND_CALCIUM_CHANNEL_OPENING | 11 | -0.5663 | -1.08264 | 0.407465 | 0.780778 | 0.709844 | 2408 | tags=36% | list=15% | signal=31% |
| BIOCARTA_CDC42RAC_PATHWAY | 7 | 0.521252 | 1.075516 | 0.406393 | 0.780778 | 0.709844 | 1242 | tags=43% | list=8% | signal=40% |
| WP_HOMOLOGOUS_RECOMBINATION | 12 | 0.46815 | 1.055108 | 0.408284 | 0.780778 | 0.709844 | 502 | tags=25% | list=3% | signal=24% |
| REACTOME_DNA_DAMAGE_BYPASS | 47 | 0.338627 | 1.010834 | 0.407821 | 0.780778 | 0.709844 | 3239 | tags=34% | list=20% | signal=27% |
| REACTOME_DOPAMINE_NEUROTRANSMITTER_RELEASE_CYCLE | 23 | 0.388898 | 1.004969 | 0.407843 | 0.780778 | 0.709844 | 2385 | tags=39% | list=15% | signal=33% |
| WP_SREBF_AND_MIR33_IN_CHOLESTEROL_AND_LIPID_HOMEOSTASIS | 11 | -0.56495 | -1.08006 | 0.40902 | 0.781285 | 0.710304 | 2258 | tags=36% | list=14% | signal=31% |
| REACTOME_NUCLEAR_RECEPTOR_TRANSCRIPTION_PATHWAY | 38 | 0.355649 | 1.018083 | 0.409756 | 0.78179 | 0.710763 | 1196 | tags=24% | list=7% | signal=22% |
| REACTOME_FGFR1B_LIGAND_BINDING_AND_ACTIVATION | 6 | 0.55974 | 1.095048 | 0.41387 | 0.782859 | 0.711735 | 295 | tags=17% | list=2% | signal=16% |
| REACTOME_MOLYBDENUM_COFACTOR_BIOSYNTHESIS | 5 | 0.599228 | 1.09405 | 0.411765 | 0.782859 | 0.711735 | 5889 | tags=80% | list=36% | signal=51% |
| REACTOME_ER_QUALITY_CONTROL_COMPARTMENT_ERQC | 4 | 0.631628 | 1.092623 | 0.414013 | 0.782859 | 0.711735 | 1338 | tags=25% | list=8% | signal=23% |
| BIOCARTA_CFTR_PATHWAY | 9 | -0.58951 | -1.08514 | 0.412052 | 0.782859 | 0.711735 | 936 | tags=33% | list=6% | signal=31% |
| REACTOME_REGULATION_OF_FZD_BY_UBIQUITINATION | 18 | -0.52185 | -1.0703 | 0.414566 | 0.782859 | 0.711735 | 2658 | tags=39% | list=16% | signal=33% |
| REACTOME_INFLAMMASOMES | 12 | -0.54982 | -1.07007 | 0.414157 | 0.782859 | 0.711735 | 2561 | tags=42% | list=16% | signal=35% |
| WP_FATTY_ACID_OXIDATION | 10 | 0.482232 | 1.057282 | 0.412234 | 0.782859 | 0.711735 | 4653 | tags=60% | list=29% | signal=43% |
| REACTOME_TRANSPORT_TO_THE_GOLGI_AND_SUBSEQUENT_MODIFICATION | 166 | -0.39485 | -1.03492 | 0.413866 | 0.782859 | 0.711735 | 1435 | tags=14% | list=9% | signal=13% |
| REACTOME_REGULATION_OF_PTEN_GENE_TRANSCRIPTION | 41 | 0.348713 | 1.008845 | 0.411458 | 0.782859 | 0.711735 | 2258 | tags=24% | list=14% | signal=21% |
| REACTOME_SENSORY_PERCEPTION_OF_TASTE | 5 | 0.594362 | 1.085165 | 0.418301 | 0.783672 | 0.712474 | 1853 | tags=40% | list=11% | signal=35% |

| | | | | | | | | | | |
|---|---|---|---|---|---|---|---|---|---|---|
| REACTOME_FORMATION_OF_XYLULOSE_5_PHOSPHATE | 5 | 0.594298 | 1.085049 | 0.418301 | 0.783672 | 0.712474 | 4950 | tags=60% | list=31% | signal=42% |
| REACTOME_PLATELET_ADHESION_TO_EXPOSED_COLLAGEN | 9 | -0.58674 | -1.08003 | 0.418567 | 0.783672 | 0.712474 | 2624 | tags=44% | list=16% | signal=37% |
| REACTOME_CGMP_EFFECTS | 10 | -0.5776 | -1.07947 | 0.420128 | 0.783672 | 0.712474 | 1026 | tags=40% | list=6% | signal=37% |
| REACTOME_SHC1_EVENTS_IN_EGFR_SIGNALING | 9 | -0.58488 | -1.07661 | 0.420195 | 0.783672 | 0.712474 | 2241 | tags=67% | list=14% | signal=57% |
| REACTOME_UNFOLDED_PROTEIN_RESPONSE_UPR | 11 | -0.56308 | -1.07648 | 0.416796 | 0.783672 | 0.712474 | 2427 | tags=45% | list=15% | signal=39% |
| BIOCARTA_HBX_PATHWAY | 7 | 0.515687 | 1.064034 | 0.420091 | 0.783672 | 0.712474 | 3712 | tags=43% | list=23% | signal=33% |
| WP_LEPTININSULIN_SIGNALING_OVERLAP | 15 | 0.423712 | 1.009552 | 0.417957 | 0.783672 | 0.712474 | 2781 | tags=47% | list=17% | signal=39% |
| REACTOME_INTERFERON_ALPHA_BETA_SIGNALING | 15 | 0.423489 | 1.009019 | 0.417957 | 0.783672 | 0.712474 | 2551 | tags=33% | list=16% | signal=28% |
| REACTOME_DAP12_SIGNALING | 24 | 0.383294 | 0.997046 | 0.418327 | 0.783672 | 0.712474 | 1257 | tags=21% | list=8% | signal=19% |
| REACTOME_CLEC7A_DECTIN_1_SIGNALING | 91 | 0.310218 | 0.97407 | 0.42 | 0.783672 | 0.712474 | 2798 | tags=22% | list=17% | signal=18% |
| REACTOME_RNA_POLYMERASE_III_TRANSCRIPTION_INITIATION_FROM_TYPE_3_PI | 26 | 0.387505 | 1.011373 | 0.422594 | 0.786624 | 0.715158 | 3823 | tags=27% | list=24% | signal=21% |
| REACTOME_SIGNALING_BY_FGFR4 | 33 | 0.360916 | 1.001149 | 0.422727 | 0.786624 | 0.715158 | 1616 | tags=21% | list=10% | signal=19% |
| REACTOME_SHC_RELATED_EVENTS_TRIGGERED_BY_IGF1R | 6 | -0.65713 | -1.11775 | 0.425225 | 0.786856 | 0.71537 | 420 | tags=33% | list=3% | signal=32% |
| BIOCARTA_BCR_PATHWAY | 31 | -0.47814 | -1.06726 | 0.423469 | 0.786856 | 0.71537 | 49 | tags=13% | list=0% | signal=13% |
| BIOCARTA_CCR5_PATHWAY | 14 | -0.52833 | -1.0469 | 0.424963 | 0.786856 | 0.71537 | 49 | tags=14% | list=0% | signal=14% |
| REACTOME_REGULATION_OF_INSULIN_SECRETION | 55 | -0.43595 | -1.04421 | 0.424561 | 0.786856 | 0.71537 | 1620 | tags=15% | list=10% | signal=13% |
| REACTOME_TNFR2_NON_CANONICAL_NF_KB_PATHWAY | 83 | 0.31167 | 0.960995 | 0.424528 | 0.786856 | 0.71537 | 2552 | tags=16% | list=16% | signal=13% |
| REACTOME_CDK_MEDIATED_PHOSPHORYLATION_AND_REMOVAL_OF_CDC6 | 71 | 0.315047 | 0.957254 | 0.42623 | 0.787836 | 0.71626 | 5771 | tags=44% | list=36% | signal=28% |
| REACTOME_FGFRL1_MODULATION_OF_FGFR1_SIGNALING | 10 | 0.477866 | 1.047711 | 0.428191 | 0.789701 | 0.717956 | 1659 | tags=30% | list=10% | signal=27% |
| REACTOME_METABOLISM_OF_NITRIC_OXIDE_NOS3_ACTIVATION_AND_REGULATIO | 16 | 0.417338 | 1.013169 | 0.428125 | 0.789701 | 0.717956 | 2797 | tags=38% | list=17% | signal=31% |
| REACTOME_ARACHIDONIC_ACID_METABOLISM | 36 | -0.45723 | -1.03119 | 0.429658 | 0.791525 | 0.719614 | 3814 | tags=39% | list=24% | signal=30% |
| WP_REGULATION_OF_ACTIN_CYTOSKELETON | 136 | 0.31211 | 0.982939 | 0.430769 | 0.792043 | 0.720085 | 2947 | tags=34% | list=18% | signal=28% |
| REACTOME_TP53_REGULATES_METABOLIC_GENES | 69 | 0.314619 | 0.948326 | 0.430894 | 0.792043 | 0.720085 | 4732 | tags=38% | list=29% | signal=27% |
| REACTOME_ACETYLCHOLINE_REGULATES_INSULIN_SECRETION | 7 | 0.511648 | 1.055699 | 0.431507 | 0.792232 | 0.720256 | 3970 | tags=57% | list=25% | signal=43% |
| REACTOME_P130CAS_LINKAGE_TO_MAPK_SIGNALING_FOR_INTEGRINS | 12 | 0.460941 | 1.038859 | 0.431953 | 0.792232 | 0.720256 | 3575 | tags=42% | list=22% | signal=32% |
| WP_EGFR1_SIGNALING_PATHWAY | 169 | 0.315592 | 1.01909 | 0.43263 | 0.792597 | 0.720588 | 2619 | tags=33% | list=16% | signal=28% |
| REACTOME_SIGNAL_AMPLIFICATION | 29 | -0.47665 | -1.04962 | 0.433766 | 0.793802 | 0.721684 | 1327 | tags=17% | list=8% | signal=16% |
| REACTOME_REGULATION_OF_EXPRESSION_OF_SLITS_AND_ROBOS | 7 | -0.62474 | -1.08334 | 0.434397 | 0.79408 | 0.721937 | 1871 | tags=57% | list=12% | signal=51% |
| REACTOME_THE_PHOTOTRANSDUCTION_CASCADE | 23 | 0.382199 | 0.987657 | 0.435294 | 0.794843 | 0.722631 | 5624 | tags=57% | list=35% | signal=37% |
| REACTOME_VITAMIN_D_CALCIFEROL_METABOLISM | 4 | 0.618125 | 1.069265 | 0.437367 | 0.79775 | 0.725274 | 3167 | tags=50% | list=20% | signal=40% |
| REACTOME_SIGNALING_BY_ROBO_RECEPTORS | 20 | -0.50275 | -1.04236 | 0.438108 | 0.798224 | 0.725704 | 2130 | tags=30% | list=13% | signal=26% |
| REACTOME_INTERLEUKIN_4_AND_INTERLEUKIN_13_SIGNALING | 10 | 0.473468 | 1.038067 | 0.43883 | 0.798651 | 0.726093 | 2293 | tags=40% | list=14% | signal=34% |
| REACTOME_MRNA_SPLICING_MINOR_PATHWAY | 48 | 0.333786 | 0.989001 | 0.439306 | 0.798651 | 0.726093 | 2986 | tags=23% | list=18% | signal=19% |
| BIOCARTA_ACE2_PATHWAY | 11 | 0.46319 | 1.025445 | 0.440111 | 0.798703 | 0.72614 | 1669 | tags=18% | list=10% | signal=16% |
| REACTOME_RNA_POLYMERASE_II_TRANSCRIPTION_TERMINATION | 61 | 0.321749 | 0.950838 | 0.440299 | 0.798703 | 0.72614 | 5281 | tags=48% | list=33% | signal=32% |
| REACTOME_ALPHA_LINOLENIC_OMEGA3_AND_LINOLEIC_OMEGA6_ACID_METABO | 12 | -0.53497 | -1.04116 | 0.441265 | 0.799582 | 0.726939 | 1233 | tags=42% | list=8% | signal=39% |
| REACTOME_RELEASE_OF_HH_NP_FROM_THE_SECRETING_CELL | 6 | -0.64306 | -1.09382 | 0.443243 | 0.801515 | 0.728697 | 2186 | tags=33% | list=14% | signal=29% |
| REACTOME_AMPLIFICATION_OF_SIGNAL_FROM_THE_KINETOCHORES | 90 | 0.308779 | 0.957684 | 0.443299 | 0.801515 | 0.728697 | 5859 | tags=46% | list=36% | signal=29% |
| REACTOME_ACYL_CHAIN_REMODELING_OF_CL | 5 | -0.65693 | -1.08418 | 0.443831 | 0.801602 | 0.728776 | 1327 | tags=60% | list=8% | signal=55% |
| REACTOME_CREB_PHOSPHORYLATION | 7 | 0.498939 | 1.029476 | 0.445205 | 0.803211 | 0.730238 | 759 | tags=29% | list=5% | signal=27% |
| REACTOME_TRIF_MEDIATED_PROGRAMMED_CELL_DEATH | 9 | 0.485531 | 1.033947 | 0.445876 | 0.803547 | 0.730543 | 1531 | tags=22% | list=9% | signal=20% |
| REACTOME_ESTROGEN_STIMULATED_SIGNALING_THROUGH_PRKCZ | 5 | 0.572681 | 1.045582 | 0.446623 | 0.804019 | 0.730972 | 3034 | tags=60% | list=19% | signal=49% |
| BIOCARTA_BCELLSURVIVAL_PATHWAY | 16 | -0.51802 | -1.04344 | 0.447214 | 0.804209 | 0.731146 | 49 | tags=12% | list=0% | signal=12% |
| REACTOME_SUMOYLATION_OF_DNA_REPLICATION_PROTEINS | 41 | 0.33834 | 0.978834 | 0.447917 | 0.8046 | 0.731501 | 3167 | tags=32% | list=20% | signal=26% |
| REACTOME_PRE_NOTCH_EXPRESSION_AND_PROCESSING | 6 | 0.53538 | 1.047392 | 0.449664 | 0.805724 | 0.732522 | 30 | tags=17% | list=0% | signal=17% |
| REACTOME_NERVOUS_SYSTEM_DEVELOPMENT | 253 | -0.38256 | -1.01886 | 0.449741 | 0.805724 | 0.732522 | 2241 | tags=20% | list=14% | signal=17% |
| REACTOME_APOPTOTIC_FACTOR_MEDIATED_RESPONSE | 16 | 0.407217 | 0.988598 | 0.45 | 0.805724 | 0.732522 | 3711 | tags=25% | list=23% | signal=19% |
| REACTOME_REGULATION_OF_TP53_ACTIVITY_THROUGH_PHOSPHORYLATION | 86 | 0.302876 | 0.942651 | 0.45283 | 0.809916 | 0.736334 | 2575 | tags=27% | list=16% | signal=23% |
| REACTOME_METABOLISM_OF_PORPHYRINS | 20 | -0.4957 | -1.02774 | 0.453408 | 0.810075 | 0.736479 | 4786 | tags=45% | list=30% | signal=32% |
| REACTOME_REGULATION_OF_KIT_SIGNALING | 13 | 0.434495 | 1.007958 | 0.454277 | 0.810755 | 0.737097 | 1616 | tags=23% | list=10% | signal=21% |
| WP_HISTONE_MODIFICATIONS | 5 | -0.64471 | -1.06403 | 0.458564 | 0.811122 | 0.737431 | 262 | tags=20% | list=2% | signal=20% |
| REACTOME_NUCLEAR_SIGNALING_BY_ERBB4 | 11 | -0.54567 | -1.04319 | 0.455677 | 0.811122 | 0.737431 | 2496 | tags=55% | list=15% | signal=46% |
| BIOCARTA_CELL2CELL_PATHWAY | 9 | -0.56172 | -1.03398 | 0.457655 | 0.811122 | 0.737431 | 121 | tags=22% | list=1% | signal=22% |
| WP_HEME_BIOSYNTHESIS | 9 | -0.56127 | -1.03315 | 0.459283 | 0.811122 | 0.737431 | 627 | tags=22% | list=4% | signal=21% |
| REACTOME_THE_CANONICAL_RETINOID_CYCLE_IN_RODS_TWILIGHT_VISION | 15 | -0.5142 | -1.02551 | 0.458027 | 0.811122 | 0.737431 | 2269 | tags=27% | list=14% | signal=23% |
| WP_LUNG_FIBROSIS | 42 | -0.44192 | -1.01931 | 0.458333 | 0.811122 | 0.737431 | 2820 | tags=24% | list=17% | signal=20% |
| REACTOME_MAPK1_ERK2_ACTIVATION | 7 | 0.489595 | 1.010197 | 0.456621 | 0.811122 | 0.737431 | 3034 | tags=71% | list=19% | signal=58% |
| REACTOME_DOWNSTREAM_SIGNALING_OF_ACTIVATED_FGFR4 | 19 | 0.398964 | 0.998069 | 0.45583 | 0.811122 | 0.737431 | 1087 | tags=21% | list=7% | signal=20% |
| REACTOME_TRAF6_MEDIATED_INDUCTION_OF_TAK1_COMPLEX_WITHIN_TLR4_CC | 16 | 0.401294 | 0.974219 | 0.459375 | 0.811122 | 0.737431 | 1494 | tags=19% | list=9% | signal=17% |
| REACTOME_ACTIVATION_OF_NF_KAPPAB_IN_B_CELLS | 63 | 0.310203 | 0.932975 | 0.458647 | 0.811122 | 0.737431 | 3371 | tags=24% | list=21% | signal=19% |
| REACTOME_ADENYLATE_CYCLASE_ACTIVATING_PATHWAY | 10 | -0.55944 | -1.04553 | 0.460064 | 0.811427 | 0.737708 | 1697 | tags=40% | list=11% | signal=36% |
| REACTOME_SEMAPHORIN_INTERACTIONS | 51 | -0.42852 | -1.01308 | 0.460526 | 0.811427 | 0.737708 | 2001 | tags=22% | list=12% | signal=19% |
| REACTOME_RHOBTB1_GTPASE_CYCLE | 22 | 0.382948 | 0.981819 | 0.461832 | 0.8119 | 0.738138 | 4496 | tags=55% | list=28% | signal=39% |
| REACTOME_TNFR1_INDUCED_NFKAPPAB_SIGNALING_PATHWAY | 32 | 0.353859 | 0.970288 | 0.462264 | 0.8119 | 0.738138 | 2692 | tags=25% | list=17% | signal=21% |
| WP_CHEMOKINE_SIGNALING_PATHWAY | 138 | 0.295055 | 0.940469 | 0.461538 | 0.8119 | 0.738138 | 2556 | tags=28% | list=16% | signal=23% |
| BIOCARTA_TERC_PATHWAY | 6 | 0.519975 | 1.017254 | 0.463087 | 0.812485 | 0.73867 | 5627 | tags=67% | list=35% | signal=43% |
| REACTOME_PI5P_REGULATES_TP53_ACETYLATION | 9 | 0.47166 | 1.004409 | 0.463918 | 0.813082 | 0.739212 | 1035 | tags=22% | list=6% | signal=21% |
| REACTOME_VLDL_CLEARANCE | 3 | -0.69565 | -1.05959 | 0.466411 | 0.816435 | 0.74226 | 920 | tags=67% | list=6% | signal=63% |
| REACTOME_HEME_DEGRADATION | 8 | -0.58309 | -1.04817 | 0.467116 | 0.816435 | 0.74226 | 5405 | tags=62% | list=33% | signal=42% |
| WP_REGULATION_OF_CARDIAC_HYPERTROPHY_BY_MIR208 | 3 | 0.642154 | 1.039705 | 0.467775 | 0.816435 | 0.74226 | 5779 | tags=100% | list=36% | signal=64% |
| WP_WHITE_FAT_CELL_DIFFERENTIATION | 30 | -0.45891 | -1.01862 | 0.467866 | 0.816435 | 0.74226 | 11 | tags=20% | list=0% | signal=20% |
| REACTOME_TRANSLESION_SYNTHESIS_BY_Y_FAMILY_DNA_POLYMERASES_BYPASS | 38 | 0.339113 | 0.970747 | 0.468293 | 0.816435 | 0.74226 | 2775 | tags=32% | list=17% | signal=26% |
| REACTOME_VISUAL_PHOTOTRANSDUCTION | 64 | -0.41974 | -1.02074 | 0.469037 | 0.816793 | 0.742586 | 2269 | tags=17% | list=14% | signal=15% |
| REACTOME_FANCONI_ANEMIA_PATHWAY | 36 | 0.34683 | 0.978045 | 0.469484 | 0.816793 | 0.742586 | 1604 | tags=17% | list=10% | signal=15% |
| BIOCARTA_CB1R_PATHWAY | 6 | -0.63265 | -1.07612 | 0.472072 | 0.818692 | 0.744312 | 1759 | tags=50% | list=11% | signal=45% |
| REACTOME_IRF3_MEDIATED_INDUCTION_OF_TYPE_I_IFN | 4 | 0.598137 | 1.034689 | 0.471338 | 0.818692 | 0.744312 | 1132 | tags=50% | list=7% | signal=47% |
| WP_DISTAL_CONVOLUTED_TUBULE_1_DCT1_CELL | 5 | 0.55635 | 1.015765 | 0.472767 | 0.818692 | 0.744312 | 5287 | tags=80% | list=33% | signal=54% |
| REACTOME_MITOTIC_G2_G2_M_PHASES | 177 | 0.30597 | 1.009556 | 0.471698 | 0.818692 | 0.744312 | 5040 | tags=39% | list=31% | signal=27% |
| BIOCARTA_ARAP_PATHWAY | 11 | 0.453231 | 1.003398 | 0.473538 | 0.818692 | 0.744312 | 942 | tags=18% | list=6% | signal=17% |
| REACTOME_TNFR1_INDUCED_PROAPOPTOTIC_SIGNALING | 22 | 0.379887 | 0.973974 | 0.473282 | 0.818692 | 0.744312 | 2692 | tags=27% | list=17% | signal=23% |
| REACTOME_BIOSYNTHESIS_OF_EPA_DERIVED_SPMS | 5 | -0.63813 | -1.05316 | 0.480663 | 0.818765 | 0.744379 | 847 | tags=40% | list=5% | signal=38% |
| REACTOME_CELL_SURFACE_INTERACTIONS_AT_THE_VASCULAR_WALL | 74 | -0.41364 | -1.01357 | 0.476731 | 0.818765 | 0.744379 | 3677 | tags=32% | list=23% | signal=25% |
| REACTOME_FORMATION_OF_THE_BETA_CATENIN_TCF_TRANSACTIVATING_COMPI | 23 | -0.47582 | -1.01041 | 0.476573 | 0.818765 | 0.744379 | 1245 | tags=17% | list=8% | signal=16% |
| REACTOME_ADENYLATE_CYCLASE_INHIBITORY_PATHWAY | 13 | -0.51404 | -1.00948 | 0.47813 | 0.818765 | 0.744379 | 1697 | tags=62% | list=11% | signal=55% |
| REACTOME_SULFIDE_OXIDATION_TO_SULFATE | 5 | 0.551853 | 1.007553 | 0.479303 | 0.818765 | 0.744379 | 5803 | tags=80% | list=36% | signal=51% |
| BIOCARTA_PRC2_PATHWAY | 10 | 0.45937 | 1.007159 | 0.478723 | 0.818765 | 0.744379 | 3944 | tags=60% | list=24% | signal=45% |
| REACTOME_PRE_NOTCH_PROCESSING_IN_GOLGI | 5 | 0.551496 | 1.006903 | 0.481481 | 0.818765 | 0.744379 | 30 | tags=20% | list=0% | signal=20% |
| REACTOME_CALCINEURIN_ACTIVATES_NFAT | 10 | 0.457575 | 1.003222 | 0.481383 | 0.818765 | 0.744379 | 1254 | tags=20% | list=8% | signal=18% |
| WP_HYPOXIADEPENDENT_DIFFERENTIATION_OF_MYOBLASTS | 10 | 0.457324 | 1.002673 | 0.481383 | 0.818765 | 0.744379 | 2330 | tags=30% | list=14% | signal=26% |
| REACTOME_DOWNSTREAM_SIGNALING_OF_ACTIVATED_FGFR3 | 21 | 0.382328 | 0.973345 | 0.477612 | 0.818765 | 0.744379 | 1087 | tags=19% | list=7% | signal=18% |
| REACTOME_REGULATION_OF_RUNX3_EXPRESSION_AND_ACTIVITY | 52 | 0.318231 | 0.972863 | 0.478528 | 0.818765 | 0.744379 | 2552 | tags=19% | list=16% | signal=16% |
| REACTOME_PROTEIN_LOCALIZATION | 93 | 0.31002 | 0.959893 | 0.478723 | 0.818765 | 0.744379 | 3582 | tags=33% | list=22% | signal=26% |
| REACTOME_SMOOTH_MUSCLE_CONTRACTION | 31 | 0.348633 | 0.951425 | 0.477064 | 0.818765 | 0.744379 | 2715 | tags=26% | list=17% | signal=22% |
| REACTOME_CELLULAR_SENESCENCE | 89 | 0.302852 | 0.941176 | 0.474227 | 0.818765 | 0.744379 | 3675 | tags=36% | list=23% | signal=28% |
| REACTOME_APC_C_CDH1_MEDIATED_DEGRADATION_OF_CDC20_AND_OTHER_AP | 72 | 0.303334 | 0.933938 | 0.47619 | 0.818765 | 0.744379 | 5771 | tags=42% | list=36% | signal=27% |
| REACTOME_HOMOLOGY_DIRECTED_REPAIR | 111 | 0.291179 | 0.931759 | 0.48 | 0.818765 | 0.744379 | 4097 | tags=32% | list=25% | signal=24% |
| WP_RETINOL_METABOLISM | 29 | 0.354687 | 0.94857 | 0.482759 | 0.820096 | 0.745589 | 132 | tags=10% | list=1% | signal=10% |
| REACTOME_TRAF6_MEDIATED_NF_KB_ACTIVATION | 14 | -0.50135 | -0.99345 | 0.484398 | 0.821199 | 0.746592 | 1481 | tags=14% | list=9% | signal=13% |

| | | | | | | | | | |
|---|---|---|---|---|---|---|---|---|---|
| REACTOME_INTERLEUKIN_6_SIGNALING | 8 | 0.472635 | 0.988971 | 0.484108 | 0.821199 | 0.746592 | 2303 tags=62% | list=14% | signal=54% |
| REACTOME_SYNTHESIS_OF_DOLICHYL_PHOSPHATE | 6 | -0.61545 | -1.04686 | 0.486486 | 0.82264 | 0.747902 | 2616 tags=50% | list=16% | signal=42% |
| REACTOME_SCAVENGING_BY_CLASS_A_RECEPTORS | 7 | -0.59383 | -1.02973 | 0.487589 | 0.82264 | 0.747902 | 2911 tags=57% | list=18% | signal=47% |
| REACTOME_SENSORY_PERCEPTION | 70 | -0.41019 | -1.003 | 0.4875 | 0.82264 | 0.747902 | 2269 tags=17% | list=14% | signal=15% |
| REACTOME_DEUBIQUITINATION | 228 | 0.285154 | 0.983365 | 0.488626 | 0.82264 | 0.747902 | 3469 tags=29% | list=21% | signal=23% |
| REACTOME_SYNTHESIS_OF_PIPS_AT_THE_EARLY_ENDOSOME_MEMBRANE | 15 | 0.399438 | 0.951716 | 0.486068 | 0.82264 | 0.747902 | 3623 tags=60% | list=22% | signal=47% |
| REACTOME_EXTRA_NUCLEAR_ESTROGEN_SIGNALING | 60 | 0.315024 | 0.929608 | 0.488722 | 0.82264 | 0.747902 | 2437 tags=32% | list=15% | signal=27% |
| REACTOME_FCERI_MEDIATED_NF_KB_ACTIVATION | 73 | 0.299109 | 0.912501 | 0.488 | 0.82264 | 0.747902 | 4744 tags=33% | list=29% | signal=23% |
| REACTOME_VLDL_ASSEMBLY | 3 | -0.68741 | -1.04703 | 0.489443 | 0.822829 | 0.748074 | 781 tags=67% | list=5% | signal=63% |
| BIOCARTA_IRES_PATHWAY | 6 | 0.500105 | 0.978381 | 0.489933 | 0.822829 | 0.748074 | 2398 tags=33% | list=15% | signal=28% |
| REACTOME_GOLGI_ASSOCIATED_VESICLE_BIOGENESIS | 54 | 0.319058 | 0.964912 | 0.490323 | 0.822829 | 0.748074 | 2342 tags=30% | list=15% | signal=25% |
| REACTOME_LEUKOTRIENE_RECEPTORS | 4 | -0.64411 | -1.04133 | 0.497175 | 0.824056 | 0.749189 | 5749 tags=100% | list=36% | signal=64% |
| BIOCARTA_EEA1_PATHWAY | 5 | -0.62925 | -1.03851 | 0.491713 | 0.824056 | 0.749189 | 2993 tags=60% | list=19% | signal=49% |
| BIOCARTA_TCRA_PATHWAY | 5 | -0.62688 | -1.03459 | 0.493554 | 0.824056 | 0.749189 | 875 tags=40% | list=5% | signal=38% |
| BIOCARTA_ATRBRCA_PATHWAY | 18 | -0.49139 | -1.00782 | 0.494398 | 0.824056 | 0.749189 | 1703 tags=17% | list=11% | signal=15% |
| BIOCARTA_EPONFKB_PATHWAY | 10 | -0.53823 | -1.00589 | 0.496805 | 0.824056 | 0.749189 | 1832 tags=30% | list=11% | signal=27% |
| REACTOME_GENERATION_OF_SECOND_MESSENGER_MOLECULES | 19 | -0.4877 | -1.00572 | 0.496523 | 0.824056 | 0.749189 | 929 tags=16% | list=6% | signal=15% |
| REACTOME_COLLAGEN_FORMATION | 73 | -0.40964 | -1.00222 | 0.493729 | 0.824056 | 0.749189 | 2245 tags=29% | list=14% | signal=25% |
| REACTOME_G_ALPHA_Z_SIGNALLING_EVENTS | 31 | -0.44603 | -0.99557 | 0.497449 | 0.824056 | 0.749189 | 1697 tags=16% | list=11% | signal=14% |
| WP_ID_SIGNALING_PATHWAY | 43 | -0.43021 | -0.99522 | 0.497561 | 0.824056 | 0.749189 | 3452 tags=26% | list=21% | signal=20% |
| REACTOME_DOWNREGULATION_OF_ERBB2_SIGNALING | 24 | -0.46154 | -0.99496 | 0.499334 | 0.824056 | 0.749189 | 2241 tags=21% | list=14% | signal=18% |
| REACTOME_NUCLEOTIDE_CATABOLISM | 31 | -0.44523 | -0.99378 | 0.5 | 0.824056 | 0.749189 | 2718 tags=26% | list=17% | signal=22% |
| BIOCARTA_LONGEVITY_PATHWAY | 14 | 0.408401 | 0.95596 | 0.49848 | 0.824056 | 0.749189 | 2147 tags=43% | list=13% | signal=37% |
| REACTOME_POST_NMDA_RECEPTOR_ACTIVATION_EVENTS | 16 | 0.393344 | 0.954921 | 0.496875 | 0.824056 | 0.749189 | 3069 tags=38% | list=19% | signal=30% |
| REACTOME_PARACETAMOL_ADME | 16 | 0.392756 | 0.953493 | 0.496875 | 0.824056 | 0.749189 | 829 tags=12% | list=5% | signal=12% |
| REACTOME_RHOBTB_GTPASE_CYCLE | 34 | 0.340769 | 0.952074 | 0.5 | 0.824056 | 0.749189 | 4497 tags=53% | list=28% | signal=38% |
| REACTOME_SUMOYLATION_OF_SUMOYLATION_PROTEINS | 33 | 0.339085 | 0.940593 | 0.495455 | 0.824056 | 0.749189 | 3167 tags=33% | list=20% | signal=27% |
| BIOCARTA_FAS_PATHWAY | 27 | 0.355059 | 0.93793 | 0.5 | 0.824056 | 0.749189 | 4810 tags=48% | list=30% | signal=34% |
| REACTOME_SUMOYLATION_OF_DNA_DAMAGE_RESPONSE_AND_REPAIR_PROTEIN | 72 | 0.298598 | 0.919357 | 0.492063 | 0.824056 | 0.749189 | 2674 tags=28% | list=17% | signal=23% |
| REACTOME_SODIUM_COUPLED_SULPHATE_DI_AND_TRI_CARBOXYLATE_TRANSPOI | 3 | -0.6787 | -1.03377 | 0.50096 | 0.824662 | 0.74974 | 2447 tags=67% | list=15% | signal=57% |
| BIOCARTA_IL6_PATHWAY | 21 | -0.47082 | -0.98766 | 0.501362 | 0.824662 | 0.74974 | 1148 tags=10% | list=7% | signal=9% |
| REACTOME_ERYTHROCYTES_TAKE_UP_CARBON_DIOXIDE_AND_RELEASE_OXYGEN | 9 | 0.458418 | 0.976211 | 0.502577 | 0.825023 | 0.750068 | 833 tags=22% | list=5% | signal=21% |
| REACTOME_SIGNALING_BY_FGFR3 | 36 | 0.341614 | 0.963337 | 0.502347 | 0.825023 | 0.750068 | 1616 tags=19% | list=10% | signal=18% |
| REACTOME_SIGNALING_BY_ALK | 15 | -0.4961 | -0.98941 | 0.503682 | 0.826018 | 0.750973 | 1407 tags=27% | list=9% | signal=24% |
| REACTOME_CASPASE_MEDIATED_CLEAVAGE_OF_CYTOSKELETAL_PROTEINS | 12 | -0.50787 | -0.98841 | 0.504518 | 0.826572 | 0.751477 | 2634 tags=25% | list=16% | signal=21% |
| BIOCARTA_HER2_PATHWAY | 22 | -0.47561 | -1.00458 | 0.508108 | 0.831632 | 0.756077 | 2241 tags=27% | list=14% | signal=24% |
| REACTOME_TERMINAL_PATHWAY_OF_COMPLEMENT | 4 | 0.570493 | 0.986869 | 0.509554 | 0.833176 | 0.757481 | 4543 tags=50% | list=28% | signal=36% |
| REACTOME_WAX_AND_PLASMALOGEN_BIOSYNTHESIS | 4 | -0.62865 | -1.01632 | 0.516008 | 0.835108 | 0.759237 | 2018 tags=75% | list=12% | signal=66% |
| REACTOME_DEFENSINS | 4 | -0.6277 | -1.01479 | 0.519774 | 0.835108 | 0.759237 | 5015 tags=75% | list=31% | signal=52% |
| REACTOME_FGFR2B_LIGAND_BINDING_AND_ACTIVATION | 8 | -0.56125 | -1.00891 | 0.512648 | 0.835108 | 0.759237 | 332 tags=25% | list=2% | signal=24% |
| REACTOME_RUNX3_REGULATES_WNT_SIGNALING | 5 | -0.60917 | -1.00536 | 0.517495 | 0.835108 | 0.759237 | 2564 tags=40% | list=16% | signal=34% |
| REACTOME_TRANSCRIPTIONAL_REGULATION_BY_VENTX | 5 | -0.60917 | -1.00536 | 0.517495 | 0.835108 | 0.759237 | 2564 tags=40% | list=16% | signal=34% |
| REACTOME_BIOSYNTHESIS_OF_E_SERIES_18_S_RESOLVINS | 4 | 0.567576 | 0.981823 | 0.515924 | 0.835108 | 0.759237 | 4812 tags=50% | list=30% | signal=35% |
| REACTOME_THROMBOXANE_SIGNALLING_THROUGH_TP_RECEPTOR | 21 | -0.4675 | -0.98067 | 0.512262 | 0.835108 | 0.759237 | 594 tags=10% | list=4% | signal=9% |
| REACTOME_FORMATION_OF_FIBRIN_CLOT_CLOTTING_CASCADE | 24 | -0.45461 | -0.98001 | 0.516644 | 0.835108 | 0.759237 | 2690 tags=21% | list=17% | signal=17% |
| REACTOME_GLUCONEOGENESIS | 28 | -0.44719 | -0.97814 | 0.519582 | 0.835108 | 0.759237 | 2297 tags=18% | list=14% | signal=15% |
| REACTOME_NF_KB_IS_ACTIVATED_AND_SIGNALS_SURVIVAL | 13 | -0.4975 | -0.97698 | 0.511312 | 0.835108 | 0.759237 | 1900 tags=31% | list=12% | signal=27% |
| REACTOME_CARGO_CONCENTRATION_IN_THE_ER | 30 | -0.43852 | -0.97335 | 0.51671 | 0.835108 | 0.759237 | 2444 tags=47% | list=15% | signal=40% |
| WP_HYPOXIADEPENDENT_PROLIFERATION_OF_MYOBLASTS | 14 | -0.48633 | -0.96368 | 0.518574 | 0.835108 | 0.759237 | 2689 tags=43% | list=17% | signal=36% |
| REACTOME_VEGFR2_MEDIATED_CELL_PROLIFERATION | 12 | 0.426368 | 0.96094 | 0.514793 | 0.835108 | 0.759237 | 3724 tags=50% | list=23% | signal=38% |
| REACTOME_RND3_GTPASE_CYCLE | 38 | 0.332021 | 0.950443 | 0.517073 | 0.835108 | 0.759237 | 2664 tags=39% | list=17% | signal=33% |
| REACTOME_SUMOYLATION_OF_RNA_BINDING_PROTEINS | 45 | 0.31594 | 0.934911 | 0.514286 | 0.835108 | 0.759237 | 3396 tags=29% | list=21% | signal=23% |
| REACTOME_TRANSCRIPTIONAL_REGULATION_BY_SMALL_RNAS | 32 | 0.338092 | 0.927055 | 0.518868 | 0.835108 | 0.759237 | 2035 tags=25% | list=13% | signal=22% |
| REACTOME_RHOF_GTPASE_CYCLE | 40 | 0.317136 | 0.91603 | 0.519802 | 0.835108 | 0.759237 | 3240 tags=40% | list=20% | signal=32% |
| REACTOME_TRANSPORT_OF_MATURE_MRNAS_DERIVED_FROM_INTRONLESS_TRA | 41 | 0.31461 | 0.910183 | 0.515625 | 0.835108 | 0.759237 | 3035 tags=32% | list=19% | signal=26% |
| REACTOME_GRB2_SOS_PROVIDES_LINKAGE_TO_MAPK_SIGNALING_FOR_INTEGRIN | 12 | 0.425611 | 0.959233 | 0.52071 | 0.835757 | 0.759828 | 1783 tags=25% | list=11% | signal=22% |
| REACTOME_AFLATOXIN_ACTIVATION_AND_DETOXIFICATION | 8 | -0.55426 | -0.99633 | 0.527825 | 0.835806 | 0.759872 | 2351 tags=50% | list=15% | signal=43% |
| REACTOME_G_BETA_GAMMA_SIGNALLING_THROUGH_PI3KGAMMA | 23 | -0.46268 | -0.98252 | 0.522088 | 0.835806 | 0.759872 | 594 tags=9% | list=4% | signal=8% |
| REACTOME_SYNTHESIS_OF_LEUKOTRIENES_LT_AND_EOXINS_EX | 12 | -0.50177 | -0.97655 | 0.524096 | 0.835806 | 0.759872 | 2359 tags=25% | list=15% | signal=21% |
| REACTOME_ASSEMBLY_OF_COLLAGEN_FIBRILS_AND_OTHER_MULTIMERIC_STRUC | 43 | -0.41757 | -0.96598 | 0.526829 | 0.835806 | 0.759872 | 2578 tags=30% | list=16% | signal=25% |
| REACTOME_AKT_PHOSPHORYLATES_TARGETS_IN_THE_NUCLEUS | 10 | 0.436362 | 0.956713 | 0.526596 | 0.835806 | 0.759872 | 2629 tags=80% | list=16% | signal=67% |
| REACTOME_MET_RECEPTOR_RECYCLING | 10 | 0.436031 | 0.955989 | 0.526596 | 0.835806 | 0.759872 | 2556 tags=40% | list=16% | signal=34% |
| REACTOME_SIGNAL_ATTENUATION | 8 | 0.456309 | 0.954809 | 0.523227 | 0.835806 | 0.759872 | 3445 tags=50% | list=21% | signal=39% |
| REACTOME_ENOS_ACTIVATION | 13 | 0.408728 | 0.948182 | 0.525074 | 0.835806 | 0.759872 | 2797 tags=38% | list=17% | signal=32% |
| REACTOME_POSTMITOTIC_NUCLEAR_PORE_COMPLEX_NPC_REFORMATION | 22 | 0.369561 | 0.947498 | 0.526718 | 0.835806 | 0.759872 | 2135 tags=27% | list=13% | signal=24% |
| REACTOME_SUMOYLATION_OF_UBIQUITINYLATION_PROTEINS | 37 | 0.334525 | 0.944117 | 0.52657 | 0.835806 | 0.759872 | 3167 tags=32% | list=20% | signal=26% |
| REACTOME_RHOQ_GTPASE_CYCLE | 58 | 0.315674 | 0.941592 | 0.524476 | 0.835806 | 0.759872 | 1279 tags=22% | list=8% | signal=21% |
| REACTOME_PHOSPHOLIPID_METABOLISM | 175 | 0.283937 | 0.940496 | 0.528302 | 0.835806 | 0.759872 | 2007 tags=24% | list=12% | signal=21% |
| REACTOME_SIGNALING_BY_TYPE_1_INSULIN_LIKE_GROWTH_FACTOR_1_RECEPTOI | 38 | 0.326515 | 0.934684 | 0.526829 | 0.835806 | 0.759872 | 1735 tags=24% | list=11% | signal=21% |
| REACTOME_DEACTIVATION_OF_THE_BETA_CATENIN_TRANSACTIVATING_COMPLEX | 31 | 0.333572 | 0.910323 | 0.527523 | 0.835806 | 0.759872 | 2147 tags=23% | list=13% | signal=20% |
| REACTOME_DOWNSTREAM_TCR_SIGNALING | 83 | 0.290626 | 0.896109 | 0.528302 | 0.835806 | 0.759872 | 2798 tags=25% | list=17% | signal=21% |
| BIOCARTA_CIRCADIAN_PATHWAY | 5 | -0.59886 | -0.98835 | 0.530387 | 0.835914 | 0.75997 | 636 tags=40% | list=4% | signal=38% |
| REACTOME_CHOLINE_CATABOLISM | 5 | -0.59854 | -0.98782 | 0.530387 | 0.835914 | 0.75997 | 907 tags=40% | list=6% | signal=38% |
| REACTOME_INTEGRATION_OF_ENERGY_METABOLISM | 69 | -0.39887 | -0.97365 | 0.530148 | 0.835914 | 0.75997 | 1620 tags=14% | list=10% | signal=13% |
| REACTOME_RA_BIOSYNTHESIS_PATHWAY | 12 | -0.50024 | -0.97357 | 0.53012 | 0.835914 | 0.75997 | 1524 tags=25% | list=9% | signal=23% |
| REACTOME_MET_ACTIVATES_PTPN11 | 5 | -0.59394 | -0.98023 | 0.532228 | 0.83802 | 0.761884 | 2563 tags=60% | list=16% | signal=50% |
| REACTOME_METABOLISM_OF_LIPIDS | 487 | -0.36432 | -0.99283 | 0.538616 | 0.842357 | 0.765828 | 2854 tags=23% | list=18% | signal=20% |
| REACTOME_DEVELOPMENTAL_BIOLOGY | 325 | -0.36573 | -0.98526 | 0.542477 | 0.842357 | 0.765828 | 2023 tags=17% | list=13% | signal=15% |
| REACTOME_BIOSYNTHESIS_OF_MARESINS | 3 | 0.605683 | 0.980656 | 0.546778 | 0.842357 | 0.765828 | 2372 tags=33% | list=15% | signal=28% |
| REACTOME_TNFR1_MEDIATED_CERAMIDE_PRODUCTION | 5 | -0.59126 | -0.97581 | 0.535912 | 0.842357 | 0.765828 | 550 tags=60% | list=3% | signal=58% |
| REACTOME_FGFR1C_LIGAND_BINDING_AND_ACTIVATION | 7 | -0.56191 | -0.97438 | 0.544326 | 0.842357 | 0.765828 | 540 tags=29% | list=3% | signal=28% |
| WP_COMPREHENSIVE_IL17A_SIGNALING | 88 | -0.38861 | -0.97386 | 0.544248 | 0.842357 | 0.765828 | 2820 tags=19% | list=17% | signal=16% |
| REACTOME_NEGATIVE_REGULATION_OF_TCF_DEPENDENT_SIGNALING_BY_WNT_L | 5 | -0.58988 | -0.97352 | 0.539595 | 0.842357 | 0.765828 | 6625 tags=100% | list=41% | signal=59% |
| BIOCARTA_INSULIN_PATHWAY | 22 | -0.45928 | -0.97008 | 0.540541 | 0.842357 | 0.765828 | 49 tags=9% | list=0% | signal=9% |
| BIOCARTA_CDK5_PATHWAY | 11 | -0.5073 | -0.96984 | 0.547434 | 0.842357 | 0.765828 | 1900 tags=18% | list=12% | signal=16% |
| WP_ROBO4_AND_VEGF_SIGNALING_PATHWAYS_CROSSTALK | 6 | -0.57014 | -0.96978 | 0.545946 | 0.842357 | 0.765828 | 662 tags=33% | list=4% | signal=32% |
| REACTOME_EGFR_TRANSACTIVATION_BY_GASTRIN | 6 | -0.5689 | -0.96768 | 0.547748 | 0.842357 | 0.765828 | 5824 tags=83% | list=36% | signal=53% |
| REACTOME_MAPK_TARGETS_NUCLEAR_EVENTS_MEDIATED_BY_MAP_KINASES | 28 | -0.43835 | -0.95881 | 0.544386 | 0.842357 | 0.765828 | 961 tags=11% | list=6% | signal=10% |
| WP_EXERCISEINDUCED_CIRCADIAN_REGULATION | 45 | -0.41152 | -0.95823 | 0.540508 | 0.842357 | 0.765828 | 1499 tags=18% | list=9% | signal=16% |
| REACTOME_DOWNREGULATION_OF_ERBB2_ERBB3_SIGNALING | 12 | -0.49181 | -0.95716 | 0.548193 | 0.842357 | 0.765828 | 1495 tags=17% | list=9% | signal=15% |
| REACTOME_REGULATION_OF_BACH1_ACTIVITY | 11 | 0.428652 | 0.948981 | 0.543175 | 0.842357 | 0.765828 | 4824 tags=55% | list=30% | signal=38% |
| REACTOME_COMMON_PATHWAY_OF_FIBRIN_CLOT_FORMATION | 12 | 0.418469 | 0.943136 | 0.54142 | 0.842357 | 0.765828 | 113 tags=8% | list=1% | signal=8% |
| REACTOME_INACTIVATION_OF_CSF3_G_CSF_SIGNALING | 17 | 0.374472 | 0.93372 | 0.542208 | 0.842357 | 0.765828 | 1613 tags=18% | list=10% | signal=16% |
| REACTOME_SYNTHESIS_OF_DIPHTHAMIDE_EEF2 | 7 | 0.449955 | 0.928406 | 0.547945 | 0.842357 | 0.765828 | 6271 tags=71% | list=39% | signal=44% |
| REACTOME_NEGATIVE_REGULATION_OF_MET_ACTIVITY | 21 | 0.363463 | 0.925316 | 0.544776 | 0.842357 | 0.765828 | 3031 tags=33% | list=19% | signal=27% |
| REACTOME_MET_ACTIVATES_PI3K_AKT_SIGNALING | 6 | 0.472976 | 0.925307 | 0.541387 | 0.842357 | 0.765828 | 1087 tags=33% | list=7% | signal=31% |
| WP_EPO_RECEPTOR_SIGNALING | 25 | 0.353767 | 0.916269 | 0.541322 | 0.842357 | 0.765828 | 3724 tags=56% | list=23% | signal=43% |
| REACTOME_TRANSCRIPTIONAL_REGULATION_BY_RUNX2 | 62 | 0.304919 | 0.912143 | 0.537313 | 0.842357 | 0.765828 | 5771 tags=42% | list=36% | signal=27% |

| | | | | | | | | | | |
|---|---|---|---|---|---|---|---|---|---|---|
| WP_BURN_WOUND_HEALING | 16 | 0.375641 | 0.911941 | 0.546875 | 0.842357 | 0.765828 | 718 | tags=19% | list=4% | signal=18% |
| REACTOME_BUTYRATE_RESPONSE_FACTOR_1_BRF1_BINDS_AND_DESTABILIZES_M | 16 | 0.375175 | 0.91081 | 0.546875 | 0.842357 | 0.765828 | 4660 | tags=44% | list=29% | signal=31% |
| REACTOME_DAP12_INTERACTIONS | 27 | 0.343432 | 0.907213 | 0.545833 | 0.842357 | 0.765828 | 1257 | tags=19% | list=8% | signal=17% |
| REACTOME_DOWNSTREAM_SIGNALING_EVENTS_OF_B_CELL_RECEPTOR_BCR | 77 | 0.296852 | 0.903727 | 0.537815 | 0.842357 | 0.765828 | 2798 | tags=19% | list=17% | signal=16% |
| BIOCARTA_CD40_PATHWAY | 13 | -0.48068 | -0.94395 | 0.54902 | 0.842847 | 0.766273 | 81 | tags=31% | list=1% | signal=31% |
| REACTOME_FGFR2C_LIGAND_BINDING_AND_ACTIVATION | 8 | -0.53655 | -0.9645 | 0.549747 | 0.843183 | 0.766578 | 540 | tags=38% | list=3% | signal=36% |
| REACTOME_RHO_GTPASES_ACTIVATE_NADPH_OXIDASES | 20 | -0.45875 | -0.95114 | 0.550765 | 0.843963 | 0.767288 | 78 | tags=15% | list=0% | signal=15% |
| REACTOME_RESPIRATORY_ELECTRON_TRANSPORT | 86 | -0.38822 | -0.96717 | 0.552455 | 0.844992 | 0.768223 | 12 | tags=27% | list=0% | signal=27% |
| REACTOME_TRANSCRIPTIONAL_REGULATION_BY_RUNX3 | 76 | 0.29221 | 0.901107 | 0.552 | 0.844992 | 0.768223 | 2552 | tags=25% | list=16% | signal=21% |
| REACTOME_TRP_CHANNELS | 21 | -0.4504 | -0.94481 | 0.553134 | 0.845249 | 0.768457 | 4680 | tags=57% | list=29% | signal=41% |
| REACTOME_TRANSPORT_OF_GAMMA_CARBOXYLATED_PROTEIN_PRECURSORS_FR | 3 | -0.64167 | -0.97736 | 0.556622 | 0.84534 | 0.76854 | 5217 | tags=100% | list=32% | signal=68% |
| BIOCARTA_CREM_PATHWAY | 3 | 0.598273 | 0.968659 | 0.559252 | 0.84534 | 0.76854 | 1632 | tags=33% | list=10% | signal=30% |
| REACTOME_COPI_MEDIATED_ANTEROGRADE_TRANSPORT | 88 | -0.38355 | -0.9612 | 0.560841 | 0.84534 | 0.76854 | 1353 | tags=11% | list=8% | signal=10% |
| REACTOME_VITAMIN_C_ASCORBATE_METABOLISM | 7 | -0.5524 | -0.9579 | 0.558511 | 0.84534 | 0.76854 | 1208 | tags=57% | list=7% | signal=53% |
| REACTOME_METAL_ION_SLC_TRANSPORTERS | 19 | -0.45952 | -0.9476 | 0.55911 | 0.84534 | 0.76854 | 3697 | tags=74% | list=23% | signal=57% |
| REACTOME_G0_AND_EARLY_G1 | 17 | -0.4688 | -0.94619 | 0.554755 | 0.84534 | 0.76854 | 262 | tags=41% | list=2% | signal=41% |
| BIOCARTA_G2_PATHWAY | 22 | -0.44498 | -0.93988 | 0.559459 | 0.84534 | 0.76854 | 1703 | tags=32% | list=11% | signal=29% |
| REACTOME_SEMA3A_PLEXIN_REPULSION_SIGNALING_BY_INHIBITING_INTEGRIN_A | 9 | 0.435329 | 0.927042 | 0.559278 | 0.84534 | 0.76854 | 676 | tags=22% | list=4% | signal=21% |
| REACTOME_C_TYPE_LECTIN_RECEPTORS_CLRS | 107 | 0.287247 | 0.916722 | 0.559524 | 0.84534 | 0.76854 | 2852 | tags=21% | list=18% | signal=18% |
| WP_MIR193A_AND_MVP_IN_COLON_CANCER_METASTASIS | 6 | 0.466904 | 0.913429 | 0.55481 | 0.84534 | 0.76854 | 3752 | tags=50% | list=23% | signal=38% |
| REACTOME_TCR_SIGNALING | 101 | 0.288745 | 0.913101 | 0.555556 | 0.84534 | 0.76854 | 2852 | tags=28% | list=18% | signal=23% |
| REACTOME_MRNA_DECAY_BY_5_TO_3_EXORIBONUCLEASE | 14 | 0.389838 | 0.91251 | 0.556231 | 0.84534 | 0.76854 | 3529 | tags=29% | list=22% | signal=22% |
| BIOCARTA_IGF1MTOR_PATHWAY | 18 | 0.368009 | 0.909759 | 0.555556 | 0.84534 | 0.76854 | 3960 | tags=61% | list=25% | signal=46% |
| REACTOME_COPI_INDEPENDENT_GOLGI_TO_ER_RETROGRADE_TRAFFIC | 43 | 0.312514 | 0.907227 | 0.56044 | 0.84534 | 0.76854 | 4311 | tags=35% | list=27% | signal=26% |
| REACTOME_REGULATION_OF_TP53_ACTIVITY_THROUGH_ACETYLATION | 29 | 0.336684 | 0.900423 | 0.560345 | 0.84534 | 0.76854 | 2147 | tags=31% | list=13% | signal=27% |
| REACTOME_MET_ACTIVATES_RAP1_AND_RAC1 | 11 | 0.422687 | 0.935776 | 0.562674 | 0.846564 | 0.769653 | 555 | tags=18% | list=3% | signal=18% |
| BIOCARTA_CARM1_PATHWAY | 11 | 0.422436 | 0.935221 | 0.562674 | 0.846564 | 0.769653 | 759 | tags=18% | list=5% | signal=17% |
| BIOCARTA_NTHI_PATHWAY | 20 | -0.45525 | -0.94388 | 0.563282 | 0.846711 | 0.769786 | 81 | tags=20% | list=1% | signal=20% |
| REACTOME_TRANSCRIPTIONAL_REGULATION_BY_NPAS4 | 4 | 0.546269 | 0.944966 | 0.566879 | 0.847272 | 0.770297 | 2058 | tags=50% | list=13% | signal=44% |
| REACTOME_CA_DEPENDENT_EVENTS | 27 | -0.43408 | -0.94471 | 0.565617 | 0.847272 | 0.770297 | 1777 | tags=26% | list=11% | signal=23% |
| REACTOME_PKA_MEDIATED_PHOSPHORYLATION_OF_CREB | 17 | -0.46464 | -0.9378 | 0.564841 | 0.847272 | 0.770297 | 1697 | tags=24% | list=11% | signal=21% |
| REACTOME_INITIATION_OF_NUCLEAR_ENVELOPE_NE_REFORMATION | 17 | -0.46426 | -0.93703 | 0.566282 | 0.847272 | 0.770297 | 6117 | tags=76% | list=38% | signal=48% |
| REACTOME_HEME_BIOSYNTHESIS | 12 | -0.48135 | -0.93681 | 0.569277 | 0.847272 | 0.770297 | 3763 | tags=42% | list=23% | signal=32% |
| BIOCARTA_CSK_PATHWAY | 12 | -0.48126 | -0.93664 | 0.569277 | 0.847272 | 0.770297 | 936 | tags=25% | list=6% | signal=24% |
| BIOCARTA_GLEEVEC_PATHWAY | 23 | -0.44011 | -0.93459 | 0.567604 | 0.847272 | 0.770297 | 49 | tags=9% | list=0% | signal=9% |
| REACTOME_SYNDECAN_INTERACTIONS | 13 | -0.47489 | -0.93259 | 0.568627 | 0.847272 | 0.770297 | 1835 | tags=62% | list=11% | signal=55% |
| REACTOME_PROCESSING_OF_INTRONLESS_PRE_MRNAS | 19 | 0.365512 | 0.914383 | 0.568905 | 0.847272 | 0.770297 | 2857 | tags=37% | list=18% | signal=30% |
| BIOCARTA_NO2IL12_PATHWAY | 7 | 0.442012 | 0.912018 | 0.568493 | 0.847272 | 0.770297 | 1613 | tags=29% | list=10% | signal=26% |
| REACTOME_DOWNSTREAM_SIGNALING_OF_ACTIVATED_FGFR2 | 24 | 0.348704 | 0.907068 | 0.565737 | 0.847272 | 0.770297 | 1087 | tags=17% | list=7% | signal=16% |
| REACTOME_SMAC_XIAP_REGULATED_APOPTOTIC_RESPONSE | 5 | -0.57385 | -0.94707 | 0.570902 | 0.847855 | 0.770827 | 2374 | tags=60% | list=15% | signal=51% |
| REACTOME_GAMMA_CARBOXYLATION_TRANSPORT_AND_AMINO_TERMINAL_CLE | 5 | -0.57176 | -0.94363 | 0.572744 | 0.847855 | 0.770827 | 5217 | tags=80% | list=32% | signal=54% |
| REACTOME_COLLAGEN_DEGRADATION | 40 | -0.4109 | -0.93883 | 0.5725 | 0.847855 | 0.770827 | 2578 | tags=40% | list=16% | signal=34% |
| BIOCARTA_ATM_PATHWAY | 19 | -0.45486 | -0.938 | 0.571627 | 0.847855 | 0.770827 | 1898 | tags=21% | list=12% | signal=19% |
| REACTOME_PROTEIN_METHYLATION | 19 | 0.36493 | 0.912929 | 0.572438 | 0.847855 | 0.770827 | 1837 | tags=32% | list=11% | signal=28% |
| REACTOME_COSTIMULATION_BY_THE_CD28_FAMILY | 53 | 0.299213 | 0.906605 | 0.573248 | 0.847855 | 0.770827 | 3418 | tags=34% | list=21% | signal=27% |
| REACTOME_ACTIVATION_OF_APC_C_AND_APC_C_CDC20_MEDIATED_DEGRADATIO | 75 | 0.28816 | 0.884823 | 0.572581 | 0.847855 | 0.770827 | 5771 | tags=40% | list=36% | signal=26% |
| BIOCARTA_CBL_PATHWAY | 11 | -0.48905 | -0.93496 | 0.575428 | 0.848411 | 0.771331 | 2653 | tags=36% | list=16% | signal=30% |
| WP_GDNFRET_SIGNALING_AXIS | 18 | -0.45383 | -0.93078 | 0.577031 | 0.848411 | 0.771331 | 1728 | tags=33% | list=11% | signal=30% |
| BIOCARTA_PLCE_PATHWAY | 11 | -0.48662 | -0.9303 | 0.576983 | 0.848411 | 0.771331 | 936 | tags=18% | list=6% | signal=17% |
| WP_ERBB_SIGNALING_PATHWAY | 42 | -0.40303 | -0.92961 | 0.577206 | 0.848411 | 0.771331 | 2468 | tags=21% | list=15% | signal=18% |
| REACTOME_SHC_MEDIATED_CASCADE_FGFR4 | 12 | -0.47712 | -0.92858 | 0.575301 | 0.848411 | 0.771331 | 540 | tags=33% | list=3% | signal=32% |
| REACTOME_MAPK3_ERK1_ACTIVATION | 8 | 0.432148 | 0.904252 | 0.574572 | 0.848411 | 0.771331 | 2717 | tags=50% | list=17% | signal=42% |
| REACTOME_REGULATION_OF_TP53_ACTIVITY_THROUGH_METHYLATION | 18 | 0.362511 | 0.896168 | 0.576389 | 0.848411 | 0.771331 | 1338 | tags=22% | list=8% | signal=20% |
| REACTOME_GLYCOLYSIS | 56 | 0.306035 | 0.907953 | 0.581081 | 0.85335 | 0.775822 | 2035 | tags=21% | list=13% | signal=19% |
| REACTOME_P75_NTR_RECEPTOR_MEDIATED_SIGNALLING | 84 | -0.3823 | -0.95085 | 0.58194 | 0.85371 | 0.776149 | 2631 | tags=20% | list=16% | signal=17% |
| REACTOME_PHASE_II_CONJUGATION_OF_COMPOUNDS | 64 | -0.3888 | -0.94549 | 0.583716 | 0.85371 | 0.776149 | 2351 | tags=23% | list=15% | signal=20% |
| REACTOME_FGFR3_LIGAND_BINDING_AND_ACTIVATION | 10 | -0.49306 | -0.92148 | 0.583067 | 0.85371 | 0.776149 | 540 | tags=40% | list=3% | signal=39% |
| REACTOME_SYNTHESIS_OF_PROSTAGLANDINS_PG_AND_THROMBOXANES_TX | 10 | -0.49142 | -0.91841 | 0.583067 | 0.85371 | 0.776149 | 4093 | tags=60% | list=25% | signal=45% |
| REACTOME_FGFR2_ALTERNATIVE_SPLICING | 17 | 0.35647 | 0.888834 | 0.584416 | 0.85371 | 0.776149 | 5442 | tags=53% | list=34% | signal=35% |
| REACTOME_REGULATION_OF_NF_KAPPA_B_SIGNALING | 16 | 0.361225 | 0.876945 | 0.584375 | 0.85371 | 0.776149 | 2939 | tags=25% | list=18% | signal=20% |
| REACTOME_BETA_OXIDATION_OF_VERY_LONG_CHAIN_FATTY_ACIDS | 11 | 0.411812 | 0.911701 | 0.584958 | 0.853751 | 0.776186 | 4434 | tags=45% | list=27% | signal=33% |
| REACTOME_ANTIGEN_PROCESSING_UBIQUITINATION_PROTEASOME_DEGRADATIC | 283 | 0.268412 | 0.955143 | 0.588499 | 0.854423 | 0.776798 | 4484 | tags=37% | list=28% | signal=27% |
| WP_G_PROTEIN_SIGNALING_PATHWAYS | 84 | -0.38081 | -0.94715 | 0.589744 | 0.854423 | 0.776798 | 1777 | tags=17% | list=11% | signal=15% |
| REACTOME_PTK6_REGULATES_PROTEINS_INVOLVED_IN_RNA_PROCESSING | 4 | 0.538715 | 0.931898 | 0.590234 | 0.854423 | 0.776798 | 3513 | tags=50% | list=22% | signal=39% |
| BIOCARTA_GPCR_PATHWAY | 28 | -0.42312 | -0.92551 | 0.587467 | 0.854423 | 0.776798 | 936 | tags=7% | list=6% | signal=7% |
| REACTOME_KERATAN_SULFATE_BIOSYNTHESIS | 26 | -0.42503 | -0.92388 | 0.591088 | 0.854423 | 0.776798 | 3575 | tags=54% | list=22% | signal=42% |
| REACTOME_GDP_FUCOSE_BIOSYNTHESIS | 6 | -0.5407 | -0.91971 | 0.590991 | 0.854423 | 0.776798 | 610 | tags=33% | list=4% | signal=32% |
| REACTOME_DISSOLUTION_OF_FIBRIN_CLOT | 10 | -0.48923 | -0.91432 | 0.586262 | 0.854423 | 0.776798 | 2970 | tags=50% | list=18% | signal=41% |
| REACTOME_RIBAVIRIN_ADME | 10 | -0.48745 | -0.911 | 0.591054 | 0.854423 | 0.776798 | 2249 | tags=20% | list=14% | signal=17% |
| REACTOME_L1CAM_INTERACTIONS | 62 | 0.29901 | 0.894468 | 0.589552 | 0.854423 | 0.776798 | 3575 | tags=37% | list=22% | signal=29% |
| BIOCARTA_RHO_PATHWAY | 20 | 0.346387 | 0.888892 | 0.586572 | 0.854423 | 0.776798 | 2643 | tags=35% | list=16% | signal=29% |
| REACTOME_PKMTS_METHYLATE_HISTONE_LYSINES | 50 | 0.292538 | 0.882831 | 0.588235 | 0.854423 | 0.776798 | 4182 | tags=42% | list=26% | signal=31% |
| REACTOME_TRANSPORT_OF_MATURE_TRANSCRIPT_TO_CYTOPLASM | 76 | 0.28452 | 0.877393 | 0.592 | 0.854997 | 0.777319 | 5309 | tags=50% | list=33% | signal=34% |
| WP_ALANINE_AND_ASPARTATE_METABOLISM | 13 | 0.382254 | 0.886766 | 0.59292 | 0.855538 | 0.777811 | 1421 | tags=31% | list=9% | signal=28% |
| BIOCARTA_KERATINOCYTE_PATHWAY | 43 | 0.301711 | 0.875864 | 0.593407 | 0.855538 | 0.777811 | 1282 | tags=19% | list=8% | signal=17% |
| REACTOME_NEGATIVE_REGULATION_OF_THE_PI3K_AKT_NETWORK | 92 | -0.37484 | -0.94067 | 0.59558 | 0.857925 | 0.779981 | 2933 | tags=25% | list=18% | signal=21% |
| REACTOME_NCAM1_INTERACTIONS | 16 | -0.45374 | -0.91396 | 0.596774 | 0.858154 | 0.78019 | 1091 | tags=31% | list=7% | signal=29% |
| WP_ACETYLCHOLINE_SYNTHESIS | 6 | -0.53669 | -0.9129 | 0.596396 | 0.858154 | 0.78019 | 669 | tags=33% | list=4% | signal=32% |
| REACTOME_COPI_DEPENDENT_GOLGI_TO_ER_RETROGRADE_TRAFFIC | 90 | -0.37501 | -0.94147 | 0.598895 | 0.858735 | 0.780718 | 1360 | tags=11% | list=8% | signal=10% |
| REACTOME_RESPIRATORY_ELECTRON_TRANSPORT_ATP_SYNTHESIS_BY_CHEMIOSM | 93 | -0.374 | -0.94064 | 0.601322 | 0.858735 | 0.780718 | 1718 | tags=28% | list=11% | signal=25% |
| REACTOME_SIGNAL_TRANSDUCTION_BY_L1 | 19 | -0.44345 | -0.91447 | 0.598053 | 0.858735 | 0.780718 | 1053 | tags=32% | list=7% | signal=30% |
| WP_GLYCEROLIPIDS_AND_GLYCEROPHOSPHOLIPIDS | 22 | -0.43167 | -0.91176 | 0.6 | 0.858735 | 0.780718 | 2334 | tags=36% | list=14% | signal=31% |
| REACTOME_OTHER_INTERLEUKIN_SIGNALING | 15 | -0.45604 | -0.90951 | 0.600884 | 0.858735 | 0.780718 | 1294 | tags=40% | list=8% | signal=37% |
| REACTOME_ACYL_CHAIN_REMODELING_OF_DAG_AND_TAG | 6 | -0.53459 | -0.90931 | 0.6 | 0.858735 | 0.780718 | 444 | tags=33% | list=3% | signal=32% |
| BIOCARTA_SAM68_PATHWAY | 6 | 0.453211 | 0.88664 | 0.599553 | 0.858735 | 0.780718 | 3724 | tags=50% | list=23% | signal=38% |
| BIOCARTA_MTA3_PATHWAY | 19 | 0.348381 | 0.871528 | 0.600707 | 0.858735 | 0.780718 | 4311 | tags=37% | list=27% | signal=27% |
| REACTOME_FCERI_MEDIATED_CA_2_MOBILIZATION | 25 | 0.336768 | 0.872241 | 0.603306 | 0.860827 | 0.78262 | 1254 | tags=12% | list=8% | signal=11% |
| REACTOME_PI_3K_CASCADE_FGFR4 | 13 | 0.376041 | 0.872353 | 0.60472 | 0.862103 | 0.783779 | 1087 | tags=23% | list=7% | signal=22% |
| REACTOME_COPII_MEDIATED_VESICLE_TRANSPORT | 65 | 0.292611 | 0.882516 | 0.606061 | 0.863272 | 0.784842 | 1075 | tags=15% | list=7% | signal=14% |
| BIOCARTA_TCAPOPTOSIS_PATHWAY | 3 | -0.61049 | -0.92987 | 0.608445 | 0.865925 | 0.787254 | 3453 | tags=67% | list=21% | signal=52% |
| REACTOME_PROCESSING_OF_DNA_DOUBLE_STRAND_BREAK_ENDS | 69 | 0.287392 | 0.866256 | 0.609756 | 0.867046 | 0.788274 | 4710 | tags=38% | list=29% | signal=27% |
| REACTOME_ZINC_INFLUX_INTO_CELLS_BY_THE_SLC39_GENE_FAMILY | 8 | 0.421034 | 0.880997 | 0.611247 | 0.868421 | 0.789524 | 55 | tags=12% | list=0% | signal=12% |
| REACTOME_RHO_GTPASE_CYCLE | 411 | 0.267473 | 1.017387 | 0.618839 | 0.870259 | 0.791195 | 1324 | tags=18% | list=8% | signal=17% |
| REACTOME_ACTIVATION_OF_KAINATE_RECEPTORS_UPON_GLUTAMATE_BINDING | 30 | -0.40769 | -0.90494 | 0.616967 | 0.870259 | 0.791195 | 1201 | tags=17% | list=7% | signal=15% |
| REACTOME_CLASS_I_PEROXISOMAL_MEMBRANE_PROTEIN_IMPORT | 19 | -0.4377 | -0.90261 | 0.613352 | 0.870259 | 0.791195 | 3328 | tags=32% | list=21% | signal=25% |
| REACTOME_GLUCAGON_LIKE_PEPTIDE_1_GLP1_REGULATES_INSULIN_SECRETION | 29 | -0.40774 | -0.89788 | 0.616883 | 0.870259 | 0.791195 | 1620 | tags=10% | list=10% | signal=9% |
| REACTOME_TRAFFICKING_OF_AMPA_RECEPTORS | 29 | -0.40683 | -0.89587 | 0.618182 | 0.870259 | 0.791195 | 2044 | tags=28% | list=13% | signal=24% |
| BIOCARTA_TOLL_PATHWAY | 24 | -0.4153 | -0.89528 | 0.61518 | 0.870259 | 0.791195 | 49 | tags=8% | list=0% | signal=8% |

| | | | | | | | | | | |
|---|---|---|---|---|---|---|---|---|---|---|
| WP_BDNF_PATHWAY | 14 | -0.45144 | -0.89454 | 0.616642 | 0.870259 | 0.791195 | 1900 | tags=29% | list=12% | signal=25% |
| REACTOME_SYNTHESIS_OF_PI | 5 | 0.489587 | 0.893871 | 0.618736 | 0.870259 | 0.791195 | 1127 | tags=60% | list=7% | signal=56% |
| BIOCARTA_RNA_PATHWAY | 9 | -0.48219 | -0.88759 | 0.617264 | 0.870259 | 0.791195 | 3202 | tags=56% | list=20% | signal=45% |
| REACTOME_CASPASE_ACTIVATION_VIA_DEATH_RECEPTORS_IN_THE_PRESENCE_O | 14 | 0.375624 | 0.879238 | 0.613982 | 0.870259 | 0.791195 | 1531 | tags=14% | list=9% | signal=13% |
| REACTOME_RHO_GTPASES_ACTIVATE_RHOTEKIN_AND_RHOPHILINS | 6 | 0.442171 | 0.865042 | 0.61745 | 0.870259 | 0.791195 | 5940 | tags=83% | list=37% | signal=53% |
| BIOCARTA_TNFR1_PATHWAY | 27 | 0.321904 | 0.850346 | 0.616667 | 0.870259 | 0.791195 | 4622 | tags=41% | list=29% | signal=29% |
| WP_PEROXIREDOXIN_2_INDUCED_OVARIAN_FAILURE | 6 | 0.440759 | 0.862279 | 0.619687 | 0.870712 | 0.791607 | 7292 | tags=67% | list=45% | signal=37% |
| REACTOME_PYROPTOSIS | 16 | 0.355276 | 0.862503 | 0.621875 | 0.873047 | 0.79373 | 4392 | tags=38% | list=27% | signal=27% |
| REACTOME_SUMOYLATION_OF_IMMUNE_RESPONSE_PROTEINS | 11 | 0.397605 | 0.880247 | 0.623955 | 0.875227 | 0.795711 | 3167 | tags=45% | list=20% | signal=37% |
| REACTOME_CA2_PATHWAY | 42 | -0.39319 | -0.9069 | 0.627451 | 0.875601 | 0.796052 | 2936 | tags=19% | list=18% | signal=16% |
| REACTOME_P75NTR_SIGNALS_VIA_NF_KB | 16 | -0.44342 | -0.89318 | 0.6261 | 0.875601 | 0.796052 | 1900 | tags=25% | list=12% | signal=22% |
| BIOCARTA_TPO_PATHWAY | 25 | -0.40916 | -0.88742 | 0.627632 | 0.875601 | 0.796052 | 49 | tags=8% | list=0% | signal=8% |
| BIOCARTA_CYTOKINE_PATHWAY | 5 | 0.484502 | 0.884588 | 0.625272 | 0.875601 | 0.796052 | 5152 | tags=40% | list=32% | signal=27% |
| WP_EICOSANOID_LIPID_SYNTHESIS_MAP | 9 | -0.47963 | -0.88288 | 0.627036 | 0.875601 | 0.796052 | 2489 | tags=33% | list=15% | signal=28% |
| BIOCARTA_NFKB_PATHWAY | 20 | 0.335151 | 0.860058 | 0.628975 | 0.875601 | 0.796052 | 2308 | tags=30% | list=14% | signal=26% |
| REACTOME_GLUTAMATE_AND_GLUTAMINE_METABOLISM | 12 | 0.381253 | 0.859261 | 0.627219 | 0.875601 | 0.796052 | 4083 | tags=58% | list=25% | signal=44% |
| BIOCARTA_EDG1_PATHWAY | 20 | 0.334148 | 0.857486 | 0.628975 | 0.875601 | 0.796052 | 3358 | tags=45% | list=21% | signal=36% |
| REACTOME_MET_ACTIVATES_PTK2_SIGNALING | 20 | 0.333021 | 0.854594 | 0.628975 | 0.875601 | 0.796052 | 1783 | tags=35% | list=11% | signal=31% |
| REACTOME_MITOTIC_PROMETAPHASE | 185 | 0.312251 | 1.010976 | 0.630173 | 0.876533 | 0.796899 | 5001 | tags=39% | list=31% | signal=27% |
| REACTOME_DARPP_32_EVENTS | 17 | -0.43883 | -0.88571 | 0.634006 | 0.878834 | 0.798991 | 936 | tags=24% | list=6% | signal=22% |
| BIOCARTA_SRCRPTP_PATHWAY | 10 | 0.401968 | 0.881306 | 0.635638 | 0.878834 | 0.798991 | 4401 | tags=40% | list=27% | signal=29% |
| REACTOME_SHC_MEDIATED_CASCADE_FGFR3 | 14 | -0.44406 | -0.87992 | 0.634473 | 0.878834 | 0.798991 | 540 | tags=36% | list=3% | signal=35% |
| REACTOME_RUNX3_REGULATES_NOTCH_SIGNALING | 9 | 0.412939 | 0.879361 | 0.636598 | 0.878834 | 0.798991 | 1773 | tags=33% | list=11% | signal=30% |
| REACTOME_BIOSYNTHESIS_OF_SPECIALIZED_PRORESOLVING_MEDIATORS_SPMS | 11 | -0.45763 | -0.87489 | 0.636081 | 0.878834 | 0.798991 | 2927 | tags=27% | list=18% | signal=22% |
| REACTOME_HEDGEHOG_OFF_STATE | 102 | 0.275438 | 0.870846 | 0.636364 | 0.878834 | 0.798991 | 2552 | tags=22% | list=16% | signal=18% |
| REACTOME_SEROTONIN_NEUROTRANSMITTER_RELEASE_CYCLE | 18 | 0.346547 | 0.856702 | 0.635417 | 0.878834 | 0.798991 | 2869 | tags=44% | list=18% | signal=37% |
| REACTOME_SIGNALING_BY_CSF3_G_CSF | 21 | 0.33616 | 0.855807 | 0.634328 | 0.878834 | 0.798991 | 2303 | tags=19% | list=14% | signal=16% |
| REACTOME_HEDGEHOG_ON_STATE | 79 | 0.275618 | 0.846744 | 0.635593 | 0.878834 | 0.798991 | 2552 | tags=27% | list=16% | signal=22% |
| WP_FOCAL_ADHESION_PI3KAKTMTOR_SIGNALING_PATHWAY | 274 | -0.35134 | -0.93934 | 0.63843 | 0.879167 | 0.799293 | 2270 | tags=16% | list=14% | signal=14% |
| BIOCARTA_BARR_MAPK_PATHWAY | 14 | -0.44277 | -0.87738 | 0.637444 | 0.879167 | 0.799293 | 183 | tags=14% | list=1% | signal=14% |
| REACTOME_POST_TRANSCRIPTIONAL_SILENCING_BY_SMALL_RNAS | 7 | -0.49684 | -0.86155 | 0.638298 | 0.879167 | 0.799293 | 2442 | tags=57% | list=15% | signal=49% |
| REACTOME_BIOLOGICAL_OXIDATIONS | 127 | -0.35794 | -0.92213 | 0.642015 | 0.879209 | 0.799332 | 2229 | tags=19% | list=14% | signal=16% |
| BIOCARTA_RANMS_PATHWAY | 9 | 0.412163 | 0.87771 | 0.639175 | 0.879209 | 0.799332 | 2135 | tags=22% | list=13% | signal=19% |
| BIOCARTA_EIF2_PATHWAY | 9 | 0.411467 | 0.876227 | 0.641753 | 0.879209 | 0.799332 | 5741 | tags=78% | list=36% | signal=50% |
| WP_IRON_HOMEOSTASIS | 10 | -0.46323 | -0.86572 | 0.642173 | 0.879209 | 0.799332 | 351 | tags=30% | list=2% | signal=29% |
| REACTOME_SNRNP_ASSEMBLY | 48 | 0.292046 | 0.865326 | 0.641618 | 0.879209 | 0.799332 | 5768 | tags=52% | list=36% | signal=34% |
| REACTOME_PI_3K_CASCADE_FGFR3 | 15 | 0.3567 | 0.849885 | 0.640867 | 0.879209 | 0.799332 | 1087 | tags=20% | list=7% | signal=19% |
| REACTOME_NCAM_SIGNALING_FOR_NEURITE_OUT_GROWTH | 33 | 0.304332 | 0.84419 | 0.640909 | 0.879209 | 0.799332 | 905 | tags=21% | list=6% | signal=20% |
| REACTOME_INTRINSIC_PATHWAY_OF_FIBRIN_CLOT_FORMATION | 14 | -0.43938 | -0.87064 | 0.643388 | 0.880146 | 0.800184 | 2690 | tags=50% | list=17% | signal=42% |
| REACTOME_DRUG_ADME | 55 | -0.37071 | -0.88793 | 0.645614 | 0.881736 | 0.801629 | 2351 | tags=16% | list=15% | signal=14% |
| BIOCARTA_TNFR2_PATHWAY | 17 | -0.43302 | -0.87399 | 0.645533 | 0.881736 | 0.801629 | 1715 | tags=18% | list=11% | signal=16% |
| REACTOME_PROTEIN_REPAIR | 6 | -0.50456 | -0.85824 | 0.646847 | 0.882256 | 0.802102 | 1081 | tags=33% | list=7% | signal=31% |
| REACTOME_METALLOPROTEASE_DUBS | 23 | 0.331328 | 0.856199 | 0.647059 | 0.882256 | 0.802102 | 4723 | tags=39% | list=29% | signal=28% |
| REACTOME_HORMONE_LIGAND_BINDING_RECEPTORS | 3 | -0.58625 | -0.89296 | 0.648752 | 0.882388 | 0.802222 | 5420 | tags=100% | list=34% | signal=66% |
| REACTOME_EGFR_INTERACTS_WITH_PHOSPHOLIPASE_C_GAMMA | 5 | -0.53566 | -0.88405 | 0.64825 | 0.882388 | 0.802222 | 2241 | tags=40% | list=14% | signal=34% |
| REACTOME_BIOTIN_TRANSPORT_AND_METABOLISM | 11 | -0.45268 | -0.86542 | 0.648523 | 0.882388 | 0.802222 | 2537 | tags=27% | list=16% | signal=23% |
| REACTOME_MITOCHONDRIAL_UNCOUPLING | 5 | -0.53046 | -0.87546 | 0.651934 | 0.882671 | 0.802479 | 1718 | tags=60% | list=11% | signal=54% |
| REACTOME_DOWNSTREAM_SIGNALING_OF_ACTIVATED_FGFR1 | 26 | -0.40026 | -0.87004 | 0.652687 | 0.882671 | 0.802479 | 1248 | tags=19% | list=8% | signal=18% |
| REACTOME_CHROMOSOME_MAINTENANCE | 94 | 0.280762 | 0.867792 | 0.652174 | 0.882671 | 0.802479 | 4523 | tags=28% | list=28% | signal=20% |
| REACTOME_TRAFFICKING_OF_GLUR2_CONTAINING_AMPA_RECEPTORS | 16 | -0.42969 | -0.86552 | 0.652493 | 0.882671 | 0.802479 | 2044 | tags=38% | list=13% | signal=33% |
| REACTOME_IRAK2_MEDIATED_ACTIVATION_OF_TAK1_COMPLEX | 10 | 0.391347 | 0.858019 | 0.651596 | 0.882671 | 0.802479 | 4744 | tags=50% | list=29% | signal=35% |
| REACTOME_PYRIMIDINE_CATABOLISM | 10 | -0.45865 | -0.85718 | 0.65016 | 0.882671 | 0.802479 | 1474 | tags=50% | list=9% | signal=45% |
| REACTOME_INTERLEUKIN_1_PROCESSING | 6 | 0.425543 | 0.832512 | 0.651007 | 0.882671 | 0.802479 | 1320 | tags=17% | list=8% | signal=15% |
| WP_NEURAL_CREST_DIFFERENTIATION | 76 | -0.36682 | -0.89621 | 0.653364 | 0.882866 | 0.802657 | 2028 | tags=20% | list=13% | signal=17% |
| REACTOME_BETA_DEFENSINS | 3 | -0.58235 | -0.88701 | 0.654511 | 0.883696 | 0.803411 | 5015 | tags=100% | list=31% | signal=69% |
| REACTOME_PROTEIN_UBIQUITINATION | 62 | 0.288928 | 0.86431 | 0.656716 | 0.884266 | 0.803929 | 4958 | tags=42% | list=31% | signal=29% |
| REACTOME_G_PROTEIN_BETA_GAMMA_SIGNALLING | 30 | -0.38925 | -0.864 | 0.658098 | 0.884266 | 0.803929 | 594 | tags=10% | list=4% | signal=10% |
| REACTOME_OAS_ANTIVIRAL_RESPONSE | 5 | 0.472433 | 0.862551 | 0.657952 | 0.884266 | 0.803929 | 2528 | tags=20% | list=16% | signal=17% |
| REACTOME_TNF_RECEPTOR_SUPERFAMILY_TNFSF_MEMBERS_MEDIATING_NON_C | 14 | -0.43394 | -0.85988 | 0.656761 | 0.884266 | 0.803929 | 3202 | tags=36% | list=20% | signal=29% |
| REACTOME_HYALURONAN_UPTAKE_AND_DEGRADATION | 12 | -0.43656 | -0.84964 | 0.658133 | 0.884266 | 0.803929 | 3535 | tags=33% | list=22% | signal=26% |
| REACTOME_HDACS_DEACETYLATE_HISTONES | 21 | 0.331925 | 0.845025 | 0.656716 | 0.884266 | 0.803929 | 1477 | tags=29% | list=9% | signal=26% |
| WP_DELTANOTCH_SIGNALING_PATHWAY | 80 | -0.35982 | -0.89053 | 0.659193 | 0.884973 | 0.804572 | 2496 | tags=28% | list=15% | signal=23% |
| BIOCARTA_EGF_PATHWAY | 28 | -0.39839 | -0.87142 | 0.660574 | 0.885009 | 0.804605 | 49 | tags=11% | list=0% | signal=11% |
| BIOCARTA_PARKIN_PATHWAY | 8 | -0.4747 | -0.85332 | 0.661046 | 0.885009 | 0.804605 | 374 | tags=25% | list=2% | signal=24% |
| REACTOME_ZINC_TRANSPORTERS | 10 | -0.45536 | -0.85102 | 0.661342 | 0.885009 | 0.804605 | 3084 | tags=70% | list=19% | signal=57% |
| REACTOME_DNA_DAMAGE_TELOMERE_STRESS_INDUCED_SENESCENCE | 24 | 0.320591 | 0.833938 | 0.661355 | 0.885009 | 0.804605 | 4710 | tags=46% | list=29% | signal=33% |
| REACTOME_PI3K_EVENTS_IN_ERBB4_SIGNALING | 6 | -0.49642 | -0.84439 | 0.664865 | 0.888272 | 0.807572 | 1767 | tags=33% | list=11% | signal=30% |
| REACTOME_CAM_PDE_1_ACTIVATION | 6 | -0.49635 | -0.84428 | 0.664865 | 0.888272 | 0.807572 | 720 | tags=33% | list=4% | signal=32% |
| REACTOME_CREATINE_METABOLISM | 8 | -0.47133 | -0.84726 | 0.666105 | 0.889212 | 0.808426 | 765 | tags=50% | list=5% | signal=48% |
| REACTOME_RHOD_GTPASE_CYCLE | 51 | -0.37136 | -0.87795 | 0.671053 | 0.890012 | 0.809154 | 1250 | tags=14% | list=8% | signal=13% |
| REACTOME_SYNTHESIS_OF_EPOXY_EET_AND_DIHYDROXYEICOSATRIENOIC_ACIDS_ | 3 | -0.56665 | -0.86311 | 0.671785 | 0.890012 | 0.809154 | 1489 | tags=67% | list=9% | signal=61% |
| WP_PPAR_SIGNALING_PATHWAY | 54 | 0.285266 | 0.862718 | 0.670968 | 0.890012 | 0.809154 | 2943 | tags=31% | list=18% | signal=26% |
| BIOCARTA_NUCLEARRS_PATHWAY | 17 | -0.42396 | -0.85569 | 0.670029 | 0.890012 | 0.809154 | 505 | tags=35% | list=3% | signal=34% |
| WP_NUCLEAR_RECEPTORS_IN_LIPID_METABOLISM_AND_TOXICITY | 17 | -0.42395 | -0.85568 | 0.670029 | 0.890012 | 0.809154 | 505 | tags=35% | list=3% | signal=34% |
| REACTOME_DOWNREGULATION_OF_ERBB4_SIGNALING | 9 | 0.39953 | 0.850807 | 0.667526 | 0.890012 | 0.809154 | 1977 | tags=33% | list=12% | signal=29% |
| BIOCARTA_DNAFRAGMENT_PATHWAY | 9 | 0.399133 | 0.849962 | 0.670103 | 0.890012 | 0.809154 | 3744 | tags=22% | list=23% | signal=17% |
| REACTOME_INCRETIN_SYNTHESIS_SECRETION_AND_INACTIVATION | 4 | 0.481799 | 0.833442 | 0.66879 | 0.890012 | 0.809154 | 5322 | tags=75% | list=33% | signal=50% |
| REACTOME_ERYTHROCYTES_TAKE_UP_OXYGEN_AND_RELEASE_CARBON_DIOXIDE | 6 | -0.48962 | -0.83282 | 0.672072 | 0.890012 | 0.809154 | 5258 | tags=67% | list=33% | signal=45% |
| REACTOME_DISASSEMBLY_OF_THE_DESTRUCTION_COMPLEX_AND_RECRUITMENT | 26 | 0.315125 | 0.822465 | 0.669456 | 0.890012 | 0.809154 | 5694 | tags=58% | list=35% | signal=37% |
| BIOCARTA_WNT_PATHWAY | 21 | -0.40436 | -0.84824 | 0.673025 | 0.890562 | 0.809654 | 91 | tags=10% | list=1% | signal=9% |
| WP_MICROGLIA_PATHOGEN_PHAGOCYTOSIS_PATHWAY | 39 | -0.38491 | -0.87602 | 0.675031 | 0.891794 | 0.810774 | 2858 | tags=23% | list=18% | signal=19% |
| WP_HEART_DEVELOPMENT | 30 | -0.38319 | -0.85054 | 0.674807 | 0.891794 | 0.810774 | 4015 | tags=40% | list=25% | signal=30% |
| REACTOME_IL_6_TYPE_CYTOKINE_RECEPTOR_LIGAND_INTERACTIONS | 13 | -0.4297 | -0.84385 | 0.675716 | 0.891989 | 0.81095 | 1148 | tags=23% | list=7% | signal=21% |
| BIOCARTA_P53_PATHWAY | 15 | -0.42742 | -0.85244 | 0.677467 | 0.893588 | 0.812404 | 4305 | tags=60% | list=27% | signal=44% |
| REACTOME_APOPTOTIC_CLEAVAGE_OF_CELLULAR_PROTEINS | 34 | -0.38355 | -0.85896 | 0.679028 | 0.893801 | 0.812598 | 1777 | tags=18% | list=11% | signal=16% |
| REACTOME_ASSEMBLY_OF_THE_PRE_REPLICATIVE_COMPLEX | 81 | 0.273177 | 0.830906 | 0.679245 | 0.893801 | 0.812598 | 2680 | tags=19% | list=17% | signal=16% |
| REACTOME_PI_3K_CASCADE_FGFR1 | 17 | 0.329978 | 0.822778 | 0.678571 | 0.893801 | 0.812598 | 1087 | tags=18% | list=7% | signal=16% |
| REACTOME_FRS_MEDIATED_FGFR4_SIGNALING | 14 | -0.42167 | -0.83555 | 0.680535 | 0.894078 | 0.81285 | 540 | tags=29% | list=3% | signal=28% |
| BIOCARTA_HES_PATHWAY | 10 | -0.44408 | -0.82994 | 0.680511 | 0.894078 | 0.81285 | 1454 | tags=40% | list=9% | signal=36% |
| REACTOME_STABILIZATION_OF_P53 | 54 | 0.280628 | 0.84869 | 0.683871 | 0.89633 | 0.814898 | 2552 | tags=19% | list=16% | signal=16% |
| WP_NODLIKE_RECEPTOR_NLR_SIGNALING_PATHWAY | 9 | 0.392105 | 0.834995 | 0.68299 | 0.89633 | 0.814898 | 4061 | tags=33% | list=25% | signal=25% |
| REACTOME_OVARIAN_TUMOR_DOMAIN_PROTEASES | 31 | 0.29973 | 0.81797 | 0.683486 | 0.89633 | 0.814898 | 2709 | tags=26% | list=17% | signal=22% |
| REACTOME_INTERLEUKIN_35_SIGNALLING | 6 | 0.409552 | 0.801227 | 0.684564 | 0.89653 | 0.815079 | 2293 | tags=50% | list=14% | signal=43% |
| REACTOME_NITRIC_OXIDE_STIMULATES_GUANYLATE_CYCLASE | 13 | -0.42488 | -0.83438 | 0.686275 | 0.898061 | 0.816471 | 1026 | tags=15% | list=6% | signal=14% |
| BIOCARTA_TCR_PATHWAY | 37 | -0.37993 | -0.86201 | 0.68805 | 0.899675 | 0.817938 | 49 | tags=5% | list=0% | signal=5% |
| REACTOME_OLFACTORY_SIGNALING_PATHWAY | 4 | -0.51726 | -0.83625 | 0.689266 | 0.899844 | 0.818092 | 1235 | tags=50% | list=8% | signal=46% |
| REACTOME_INSULIN_PROCESSING | 10 | 0.371025 | 0.813464 | 0.68883 | 0.899844 | 0.818092 | 2424 | tags=40% | list=15% | signal=34% |
| REACTOME_LAMININ_INTERACTIONS | 11 | -0.43053 | -0.82307 | 0.690513 | 0.900764 | 0.818928 | 3843 | tags=55% | list=24% | signal=42% |

| | | | | | | | | | | |
|---|---|---|---|---|---|---|---|---|---|---|
| WP_AFLATOXIN_B1_METABOLISM | 4 | -0.51586 | -0.83397 | 0.691149 | 0.900884 | 0.819038 | 1371 | tags=50% | list=8% | signal=46% |
| BIOCARTA_TUBBY_PATHWAY | 5 | -0.50705 | -0.83683 | 0.692449 | 0.901163 | 0.819291 | 1439 | tags=40% | list=9% | signal=36% |
| REACTOME_MITOCHONDRIAL_CALCIUM_ION_TRANSPORT | 16 | -0.41392 | -0.83375 | 0.692082 | 0.901163 | 0.819291 | 3915 | tags=31% | list=24% | signal=24% |
| REACTOME_VITAMIN_B2_RIBOFLAVIN_METABOLISM | 6 | -0.47808 | -0.8132 | 0.693694 | 0.902074 | 0.820119 | 3253 | tags=50% | list=20% | signal=40% |
| REACTOME_PHOSPHOLIPASE_C_MEDIATED_CASCADE_FGFR4 | 8 | -0.45347 | -0.81516 | 0.694772 | 0.902768 | 0.820751 | 540 | tags=25% | list=3% | signal=24% |
| REACTOME_BUTYROPHILIN_BTN_FAMILY_INTERACTIONS | 3 | 0.521967 | 0.845112 | 0.696466 | 0.90329 | 0.821225 | 2738 | tags=33% | list=17% | signal=28% |
| REACTOME_CONVERSION_FROM_APC_C_CDC20_TO_APC_C_CDH1_IN_LATE_ANAF | 20 | 0.316258 | 0.811576 | 0.696113 | 0.90329 | 0.821225 | 5175 | tags=40% | list=32% | signal=27% |
| BIOCARTA_HSWI_SNF_PATHWAY | 10 | 0.366265 | 0.803027 | 0.696809 | 0.90329 | 0.821225 | 5692 | tags=50% | list=35% | signal=32% |
| REACTOME_INNATE_IMMUNE_SYSTEM | 760 | -0.34371 | -0.94448 | 0.703704 | 0.905152 | 0.822918 | 2987 | tags=19% | list=19% | signal=17% |
| REACTOME_DEGRADATION_OF_THE_EXTRACELLULAR_MATRIX | 82 | -0.35096 | -0.87159 | 0.703456 | 0.905152 | 0.822918 | 2295 | tags=30% | list=14% | signal=26% |
| REACTOME_G_PROTEIN_MEDIATED_EVENTS | 38 | -0.37942 | -0.86148 | 0.70138 | 0.905152 | 0.822918 | 1777 | tags=21% | list=11% | signal=19% |
| BIOCARTA_BARRESTIN_SRC_PATHWAY | 17 | -0.41202 | -0.83159 | 0.698847 | 0.905152 | 0.822918 | 183 | tags=12% | list=1% | signal=12% |
| REACTOME_AZATHIOPRINE_ADME | 19 | -0.40089 | -0.82671 | 0.699583 | 0.905152 | 0.822918 | 1346 | tags=21% | list=8% | signal=19% |
| BIOCARTA_LIS1_PATHWAY | 14 | -0.41241 | -0.81721 | 0.702823 | 0.905152 | 0.822918 | 2199 | tags=21% | list=14% | signal=19% |
| REACTOME_SMAD2_SMAD3_SMAD4_HETEROTRIMER_REGULATES_TRANSCRIPTIOI | 25 | 0.312418 | 0.809174 | 0.702479 | 0.905152 | 0.822918 | 3313 | tags=36% | list=21% | signal=29% |
| REACTOME_TGF_BETA_RECEPTOR_SIGNALING_IN_EMT_EPITHELIAL_TO_MESENCH | 15 | 0.337997 | 0.805324 | 0.702786 | 0.905152 | 0.822918 | 2133 | tags=27% | list=13% | signal=23% |
| REACTOME_TP53_REGULATES_TRANSCRIPTION_OF_CELL_DEATH_GENES | 15 | 0.335743 | 0.799954 | 0.702786 | 0.905152 | 0.822918 | 2684 | tags=20% | list=17% | signal=17% |
| REACTOME_TRAIL_SIGNALING | 5 | 0.437512 | 0.798795 | 0.703704 | 0.905152 | 0.822918 | 1531 | tags=20% | list=9% | signal=18% |
| REACTOME_PHASE_I_FUNCTIONALIZATION_OF_COMPOUNDS | 60 | -0.34667 | -0.83891 | 0.704258 | 0.905162 | 0.822927 | 1945 | tags=13% | list=12% | signal=12% |
| REACTOME_VEGF_LIGAND_RECEPTOR_INTERACTIONS | 8 | -0.44701 | -0.80354 | 0.70489 | 0.905274 | 0.823028 | 3376 | tags=38% | list=21% | signal=30% |
| REACTOME_AMPK_INHIBITS_CHREBP_TRANSCRIPTIONAL_ACTIVATION_ACTIVITY | 5 | 0.436415 | 0.796791 | 0.705882 | 0.905846 | 0.823549 | 626 | tags=20% | list=4% | signal=19% |
| REACTOME_FC_EPSILON_RECEPTOR_FCERI_SIGNALING | 115 | 0.270447 | 0.851724 | 0.708333 | 0.907586 | 0.825131 | 2798 | tags=23% | list=17% | signal=20% |
| REACTOME_SHC_MEDIATED_CASCADE_FGFR1 | 16 | -0.4007 | -0.80713 | 0.708211 | 0.907586 | 0.825131 | 540 | tags=50% | list=3% | signal=48% |
| REACTOME_TNF_SIGNALING | 56 | 0.280646 | 0.832627 | 0.709459 | 0.908327 | 0.825804 | 2692 | tags=23% | list=17% | signal=19% |
| REACTOME_SIGNALING_BY_INSULIN_RECEPTOR | 61 | -0.34438 | -0.83393 | 0.714286 | 0.910067 | 0.827386 | 2917 | tags=21% | list=18% | signal=18% |
| REACTOME_THE_ROLE_OF_GTSE1_IN_G2_M_PROGRESSION_AFTER_G2_CHECKPOI | 69 | 0.273917 | 0.82564 | 0.715447 | 0.910067 | 0.827386 | 2552 | tags=17% | list=16% | signal=15% |
| BIOCARTA_PDGF_PATHWAY | 29 | -0.37257 | -0.82043 | 0.714286 | 0.910067 | 0.827386 | 49 | tags=7% | list=0% | signal=7% |
| REACTOME_NON_INTEGRIN_MEMBRANE_ECM_INTERACTIONS | 29 | -0.37152 | -0.81813 | 0.715584 | 0.910067 | 0.827386 | 1845 | tags=21% | list=11% | signal=18% |
| REACTOME_N_GLYCAN_TRIMMING_IN_THE_ER_AND_CALNEXIN_CALRETICULIN_C\ | 15 | -0.40899 | -0.81569 | 0.715758 | 0.910067 | 0.827386 | 3111 | tags=33% | list=19% | signal=27% |
| REACTOME_REGULATION_OF_PTEN_LOCALIZATION | 9 | 0.380361 | 0.809987 | 0.713918 | 0.910067 | 0.827386 | 2250 | tags=22% | list=14% | signal=19% |
| WP_MITOCHONDRIAL_LONG_CHAIN_FATTY_ACID_BETAOXIDATION | 16 | -0.39906 | -0.80383 | 0.71261 | 0.910067 | 0.827386 | 1233 | tags=12% | list=8% | signal=12% |
| REACTOME_HYALURONAN_BIOSYNTHESIS_AND_EXPORT | 5 | 0.43022 | 0.785481 | 0.714597 | 0.910067 | 0.827386 | 2914 | tags=40% | list=18% | signal=33% |
| WP_ACE_INHIBITOR_PATHWAY | 6 | 0.39596 | 0.774638 | 0.711409 | 0.910067 | 0.827386 | 718 | tags=17% | list=4% | signal=16% |
| BIOCARTA_AKAP13_PATHWAY | 9 | -0.43187 | -0.79496 | 0.716612 | 0.910455 | 0.827739 | 1837 | tags=22% | list=11% | signal=20% |
| BIOCARTA_FCER1_PATHWAY | 36 | -0.37094 | -0.83658 | 0.718631 | 0.910507 | 0.827787 | 49 | tags=6% | list=0% | signal=6% |
| REACTOME_LYSOSOME_VESICLE_BIOGENESIS | 34 | -0.36878 | -0.82587 | 0.719949 | 0.910507 | 0.827787 | 1435 | tags=15% | list=9% | signal=13% |
| REACTOME_SYNTHESIS_OF_16_20_HYDROXYEICOSATETRAENOIC_ACIDS_HETE | 3 | -0.53049 | -0.80803 | 0.71977 | 0.910507 | 0.827787 | 1449 | tags=33% | list=9% | signal=30% |
| REACTOME_REGULATION_OF_TLR_BY_ENDOGENOUS_LIGAND | 14 | -0.40636 | -0.80521 | 0.719168 | 0.910507 | 0.827787 | 6759 | tags=71% | list=42% | signal=42% |
| REACTOME_REPRESSION_OF_WNT_TARGET_GENES | 10 | 0.361929 | 0.793521 | 0.718085 | 0.910507 | 0.827787 | 935 | tags=30% | list=6% | signal=28% |
| REACTOME_SYNTHESIS_OF_BILE_ACIDS_AND_BILE_SALTS_VIA_27_HYDROXYCHOLE | 9 | -0.42876 | -0.78924 | 0.718241 | 0.910507 | 0.827787 | 277 | tags=22% | list=2% | signal=22% |
| REACTOME_METABOLISM_OF_VITAMINS_AND_COFACTORS | 152 | -0.33949 | -0.88568 | 0.726027 | 0.911877 | 0.829032 | 2538 | tags=16% | list=16% | signal=14% |
| WP_OXIDATIVE_STRESS_AND_REDOX_PATHWAY | 75 | -0.34589 | -0.84458 | 0.728929 | 0.911877 | 0.829032 | 3566 | tags=31% | list=22% | signal=24% |
| REACTOME_RND1_GTPASE_CYCLE | 40 | 0.280431 | 0.810009 | 0.727723 | 0.911877 | 0.829032 | 1488 | tags=22% | list=9% | signal=20% |
| REACTOME_PLATELET_AGGREGATION_PLUG_FORMATION | 32 | 0.294472 | 0.807448 | 0.721698 | 0.911877 | 0.829032 | 2781 | tags=28% | list=17% | signal=23% |
| BIOCARTA_PEPI_PATHWAY | 3 | -0.52893 | -0.80565 | 0.723608 | 0.911877 | 0.829032 | 2002 | tags=67% | list=12% | signal=58% |
| REACTOME_ACTIVATION_OF_BH3_ONLY_PROTEINS | 18 | 0.32392 | 0.800766 | 0.729167 | 0.911877 | 0.829032 | 4268 | tags=44% | list=26% | signal=33% |
| REACTOME_RHO_GTPASES_ACTIVATE_IQGAPS | 25 | 0.308455 | 0.798908 | 0.727273 | 0.911877 | 0.829032 | 2280 | tags=20% | list=14% | signal=17% |
| REACTOME_P75NTR_RECRUITS_SIGNALLING_COMPLEXES | 13 | -0.40662 | -0.79852 | 0.723982 | 0.911877 | 0.829032 | 1900 | tags=23% | list=12% | signal=20% |
| REACTOME_FRS_MEDIATED_FGFR3_SIGNALING | 16 | -0.39521 | -0.79606 | 0.722874 | 0.911877 | 0.829032 | 540 | tags=31% | list=3% | signal=30% |
| REACTOME_GOLGI_CISTERNAE_PERICENTRIOLAR_STACK_REORGANIZATION | 10 | 0.360605 | 0.790618 | 0.726064 | 0.911877 | 0.829032 | 3034 | tags=40% | list=19% | signal=33% |
| REACTOME_POLO_LIKE_KINASE_MEDIATED_EVENTS | 5 | -0.47833 | -0.78942 | 0.729282 | 0.911877 | 0.829032 | 3152 | tags=60% | list=20% | signal=48% |
| BIOCARTA_ASBCELL_PATHWAY | 4 | -0.48495 | -0.78401 | 0.723164 | 0.911877 | 0.829032 | 5638 | tags=75% | list=35% | signal=49% |
| REACTOME_HEME_SIGNALING | 9 | -0.42473 | -0.78182 | 0.728013 | 0.911877 | 0.829032 | 3573 | tags=33% | list=22% | signal=26% |
| REACTOME_TRIGLYCERIDE_BIOSYNTHESIS | 8 | -0.43062 | -0.77409 | 0.725126 | 0.911877 | 0.829032 | 444 | tags=25% | list=3% | signal=24% |
| REACTOME_INTERLEUKIN_21_SIGNALING | 5 | 0.423419 | 0.773065 | 0.72549 | 0.911877 | 0.829032 | 2293 | tags=40% | list=14% | signal=34% |
| BIOCARTA_SHH_PATHWAY | 15 | -0.40204 | -0.80181 | 0.730486 | 0.912695 | 0.829775 | 2186 | tags=20% | list=14% | signal=17% |
| REACTOME_CLATHRIN_MEDIATED_ENDOCYTOSIS | 131 | -0.33657 | -0.86942 | 0.732409 | 0.91304 | 0.830089 | 941 | tags=8% | list=6% | signal=7% |
| REACTOME_SYNTHESIS_OF_5_EICOSATETRAENOIC_ACIDS | 5 | -0.47372 | -0.78181 | 0.732965 | 0.91304 | 0.830089 | 3226 | tags=40% | list=20% | signal=32% |
| BIOCARTA_IFNG_PATHWAY | 5 | -0.47081 | -0.77701 | 0.732965 | 0.91304 | 0.830089 | 2714 | tags=60% | list=17% | signal=50% |
| REACTOME_FATTY_ACIDS | 6 | -0.44426 | -0.75566 | 0.731532 | 0.91304 | 0.830089 | 1489 | tags=50% | list=9% | signal=45% |
| WP_FATTY_ACID_BETAOXIDATION | 31 | 0.288427 | 0.787124 | 0.733945 | 0.913574 | 0.830575 | 3436 | tags=42% | list=21% | signal=33% |
| BIOCARTA_INTRINSIC_PATHWAY | 13 | 0.336864 | 0.781469 | 0.734513 | 0.913596 | 0.830594 | 1669 | tags=15% | list=10% | signal=14% |
| REACTOME_TRANSPORT_OF_THE_SLBP_DEPENDANT_MATURE_MRNA | 34 | 0.286185 | 0.799571 | 0.736364 | 0.915119 | 0.831979 | 5309 | tags=50% | list=33% | signal=34% |
| REACTOME_EFFECTS_OF_PIP2_HYDROLYSIS | 25 | -0.36533 | -0.79236 | 0.736842 | 0.915119 | 0.831979 | 2196 | tags=40% | list=14% | signal=35% |
| REACTOME_FREE_FATTY_ACIDS_REGULATE_INSULIN_SECRETION | 9 | -0.42132 | -0.77555 | 0.737785 | 0.915604 | 0.832421 | 576 | tags=33% | list=4% | signal=32% |
| REACTOME_CALNEXIN_CALRETICULIN_CYCLE | 7 | -0.45115 | -0.78232 | 0.739362 | 0.916875 | 0.833576 | 3111 | tags=57% | list=19% | signal=46% |
| REACTOME_ACTIVATION_OF_SMO | 12 | -0.40038 | -0.77922 | 0.740964 | 0.917489 | 0.834134 | 2186 | tags=25% | list=14% | signal=22% |
| REACTOME_DOWNREGULATION_OF_TGF_BETA_RECEPTOR_SIGNALING | 16 | -0.38587 | -0.77725 | 0.740469 | 0.917489 | 0.834134 | 3078 | tags=50% | list=19% | signal=41% |
| REACTOME_PREDNISONE_ADME | 4 | 0.444548 | 0.769002 | 0.7431 | 0.919447 | 0.835914 | 8969 | tags=100% | list=56% | signal=44% |
| REACTOME_HDR_THROUGH_SINGLE_STRAND_ANNEALING_SSA | 34 | 0.284216 | 0.79407 | 0.745455 | 0.921119 | 0.837434 | 4710 | tags=47% | list=29% | signal=33% |
| REACTOME_CHK1_CHK2_CDS1_MEDIATED_INACTIVATION_OF_CYCLIN_B_CDK1_CC | 12 | 0.348004 | 0.784324 | 0.745562 | 0.921119 | 0.837434 | 4330 | tags=25% | list=27% | signal=18% |
| REACTOME_THE_FATTY_ACID_CYCLING_MODEL | 4 | -0.47029 | -0.7603 | 0.749529 | 0.925331 | 0.841263 | 994 | tags=50% | list=6% | signal=47% |
| REACTOME_TGF_BETA_RECEPTOR_SIGNALING_ACTIVATES_SMADS | 35 | -0.35997 | -0.80869 | 0.751592 | 0.926511 | 0.842336 | 5206 | tags=51% | list=32% | signal=35% |
| REACTOME_CD28_DEPENDENT_VAV1_PATHWAY | 11 | 0.355481 | 0.786992 | 0.752089 | 0.926511 | 0.842336 | 2852 | tags=27% | list=18% | signal=22% |
| REACTOME_SHC_MEDIATED_CASCADE_FGFR2 | 17 | -0.38296 | -0.77294 | 0.752161 | 0.926511 | 0.842336 | 540 | tags=53% | list=3% | signal=51% |
| REACTOME_NEGATIVE_REGULATION_OF_MAPK_PATHWAY | 42 | -0.36151 | -0.83383 | 0.754902 | 0.92813 | 0.843808 | 961 | tags=19% | list=6% | signal=18% |
| WP_NOTCH_SIGNALING_PATHWAY | 44 | -0.3555 | -0.82436 | 0.758495 | 0.92813 | 0.843808 | 2625 | tags=30% | list=16% | signal=25% |
| WP_ALPHA_6_BETA_4_INTEGRIN_SIGNALING_PATHWAY | 63 | -0.32972 | -0.79945 | 0.758343 | 0.92813 | 0.843808 | 2929 | tags=27% | list=18% | signal=22% |
| BIOCARTA_IL2RB_PATHWAY | 33 | -0.35413 | -0.79141 | 0.755754 | 0.92813 | 0.843808 | 49 | tags=9% | list=0% | signal=9% |
| REACTOME_PI_3K_CASCADE_FGFR2 | 18 | 0.318562 | 0.78752 | 0.756944 | 0.92813 | 0.843808 | 1087 | tags=17% | list=7% | signal=16% |
| REACTOME_CRMPS_IN_SEMA3A_SIGNALING | 15 | 0.321654 | 0.766385 | 0.758514 | 0.92813 | 0.843808 | 2144 | tags=27% | list=13% | signal=23% |
| REACTOME_SEMA4D_INDUCED_CELL_MIGRATION_AND_GROWTH_CONE_COLLAPS | 11 | -0.39913 | -0.76305 | 0.754277 | 0.92813 | 0.843808 | 1495 | tags=27% | list=9% | signal=25% |
| BIOCARTA_BARRESTIN_PATHWAY | 11 | -0.39585 | -0.75677 | 0.755832 | 0.92813 | 0.843808 | 183 | tags=18% | list=1% | signal=18% |
| REACTOME_INTERLEUKIN_9_SIGNALING | 4 | 0.433167 | 0.749315 | 0.757962 | 0.92813 | 0.843808 | 2293 | tags=50% | list=14% | signal=43% |
| WP_EICOSANOID_METABOLISM_VIA_CYCLOOXYGENASES_COX | 23 | 0.301782 | 0.77985 | 0.760784 | 0.930222 | 0.84571 | 2174 | tags=26% | list=13% | signal=23% |
| REACTOME_SIGNALING_BY_ERBB2 | 41 | -0.35299 | -0.8114 | 0.762963 | 0.930513 | 0.845975 | 2398 | tags=17% | list=15% | signal=15% |
| REACTOME_G2_M_DNA_DAMAGE_CHECKPOINT | 71 | 0.258978 | 0.78689 | 0.762295 | 0.930513 | 0.845975 | 4710 | tags=35% | list=29% | signal=25% |
| REACTOME_METABOLISM_OF_STEROID_HORMONES | 19 | -0.37787 | -0.77923 | 0.764951 | 0.930513 | 0.845975 | 3086 | tags=26% | list=19% | signal=21% |
| REACTOME_CHEMOKINE_RECEPTORS_BIND_CHEMOKINES | 21 | -0.36835 | -0.77269 | 0.764305 | 0.930513 | 0.845975 | 1390 | tags=38% | list=9% | signal=35% |
| REACTOME_TRANSPORT_OF_CONNEXONS_TO_THE_PLASMA_MEMBRANE | 13 | -0.39297 | -0.77171 | 0.764706 | 0.930513 | 0.845975 | 4216 | tags=46% | list=26% | signal=34% |
| REACTOME_NOREPINEPHRINE_NEUROTRANSMITTER_RELEASE_CYCLE | 18 | -0.37004 | -0.75895 | 0.764706 | 0.930513 | 0.845975 | 272 | tags=17% | list=2% | signal=16% |
| BIOCARTA_TEL_PATHWAY | 17 | 0.303153 | 0.755892 | 0.762987 | 0.930513 | 0.845975 | 2422 | tags=29% | list=15% | signal=25% |
| REACTOME_SIGNALING_BY_THE_B_CELL_RECEPTOR_BCR | 100 | 0.263354 | 0.825263 | 0.766667 | 0.931916 | 0.84725 | 2798 | tags=20% | list=17% | signal=17% |
| WP_SIDS_SUSCEPTIBILITY_PATHWAYS | 46 | -0.34616 | -0.80834 | 0.771463 | 0.93338 | 0.848581 | 1747 | tags=17% | list=11% | signal=16% |
| WP_TRYPTOPHAN_METABOLISM | 33 | -0.34763 | -0.77689 | 0.768542 | 0.93338 | 0.848581 | 1489 | tags=15% | list=9% | signal=14% |
| REACTOME_TELOMERE_EXTENSION_BY_TELOMERASE | 20 | 0.301883 | 0.774687 | 0.770318 | 0.93338 | 0.848581 | 2287 | tags=25% | list=14% | signal=21% |
| REACTOME_FORMATION_OF_SENESCENCE_ASSOCIATED_HETEROCHROMATIN_FOC | 11 | 0.346949 | 0.768102 | 0.771588 | 0.93338 | 0.848581 | 5918 | tags=73% | list=37% | signal=46% |
| REACTOME_RHO_GTPASES_ACTIVATE_KTN1 | 11 | 0.346786 | 0.76774 | 0.771588 | 0.93338 | 0.848581 | 2963 | tags=36% | list=18% | signal=30% |

| | | | | | | | | | | |
|---|---|---|---|---|---|---|---|---|---|---|
| REACTOME_UREA_CYCLE | 6 | 0.373235 | 0.730178 | 0.771812 | 0.93338 | 0.848581 | 3836 | tags=33% | list=24% | signal=25% |
| BIOCARTA_P35ALZHEIMERS_PATHWAY | 6 | 0.372953 | 0.729628 | 0.771812 | 0.93338 | 0.848581 | 5548 | tags=67% | list=34% | signal=44% |
| REACTOME_G2_M_DNA_REPLICATION_CHECKPOINT | 5 | -0.43998 | -0.72614 | 0.773481 | 0.934716 | 0.849796 | 5695 | tags=80% | list=35% | signal=52% |
| REACTOME_PHOSPHOLIPASE_C_MEDIATED_CASCADE_FGFR1 | 12 | -0.38349 | -0.74634 | 0.774096 | 0.934779 | 0.849853 | 540 | tags=17% | list=3% | signal=16% |
| BIOCARTA_AKAPCENTROSOME_PATHWAY | 12 | -0.38256 | -0.74454 | 0.775602 | 0.935916 | 0.850887 | 1541 | tags=17% | list=10% | signal=15% |
| REACTOME_APOPTOTIC_EXECUTION_PHASE | 43 | -0.34933 | -0.80811 | 0.776829 | 0.936034 | 0.850994 | 1777 | tags=14% | list=11% | signal=12% |
| REACTOME_BBSOME_MEDIATED_CARGO_TARGETING_TO_CILIUM | 13 | -0.38364 | -0.7534 | 0.776772 | 0.936034 | 0.850994 | 517 | tags=15% | list=3% | signal=15% |
| REACTOME_RHOC_GTPASE_CYCLE | 71 | -0.32523 | -0.79515 | 0.782955 | 0.936113 | 0.851065 | 1242 | tags=10% | list=8% | signal=9% |
| REACTOME_THROMBIN_SIGNALLING_THROUGH_PROTEINASE_ACTIVATED_RECEPT | 27 | -0.35487 | -0.77233 | 0.78084 | 0.936113 | 0.851065 | 1542 | tags=11% | list=10% | signal=10% |
| BIOCARTA_MET_PATHWAY | 33 | -0.34411 | -0.76901 | 0.780051 | 0.936113 | 0.851065 | 49 | tags=12% | list=0% | signal=12% |
| BIOCARTA_AGR_PATHWAY | 31 | -0.34173 | -0.76278 | 0.781888 | 0.936113 | 0.851065 | 2903 | tags=23% | list=18% | signal=19% |
| WP_ETHANOL_METABOLISM_RESULTING_IN_PRODUCTION_OF_ROS_BY_CYP2E1 | 9 | 0.354166 | 0.754204 | 0.778351 | 0.936113 | 0.851065 | 5179 | tags=67% | list=32% | signal=45% |
| REACTOME_FRS_MEDIATED_FGFR1_SIGNALING | 18 | -0.35994 | -0.73823 | 0.782913 | 0.936113 | 0.851065 | 540 | tags=33% | list=3% | signal=32% |
| BIOCARTA_FIBRINOLYSIS_PATHWAY | 5 | -0.43706 | -0.72131 | 0.780847 | 0.936113 | 0.851065 | 2970 | tags=80% | list=18% | signal=65% |
| REACTOME_PHENYLALANINE_METABOLISM | 5 | -0.43582 | -0.71928 | 0.782689 | 0.936113 | 0.851065 | 1396 | tags=40% | list=9% | signal=37% |
| BIOCARTA_PKC_PATHWAY | 7 | 0.347998 | 0.718035 | 0.783105 | 0.936113 | 0.851065 | 3358 | tags=43% | list=21% | signal=34% |
| REACTOME_PTK6_REGULATES_CELL_CYCLE | 5 | -0.43437 | -0.71687 | 0.782689 | 0.936113 | 0.851065 | 4688 | tags=80% | list=29% | signal=57% |
| REACTOME_G2_PHASE | 5 | -0.4322 | -0.7133 | 0.782689 | 0.936113 | 0.851065 | 4688 | tags=80% | list=29% | signal=57% |
| BIOCARTA_MCALPAIN_PATHWAY | 18 | -0.35908 | -0.73645 | 0.784314 | 0.936882 | 0.851765 | 2291 | tags=39% | list=14% | signal=33% |
| REACTOME_MYOGENESIS | 20 | -0.36576 | -0.75834 | 0.785814 | 0.937382 | 0.852219 | 586 | tags=25% | list=4% | signal=24% |
| REACTOME_ESTROGEN_BIOSYNTHESIS | 3 | 0.461432 | 0.7471 | 0.785863 | 0.937382 | 0.852219 | 1584 | tags=33% | list=10% | signal=30% |
| REACTOME_MITOTIC_METAPHASE_AND_ANAPHASE | 214 | 0.259379 | 0.873556 | 0.787234 | 0.938342 | 0.853093 | 5782 | tags=43% | list=36% | signal=28% |
| REACTOME_DRUG_MEDIATED_INHIBITION_OF_CDK4_CDK6_ACTIVITY | 5 | -0.42923 | -0.7084 | 0.788214 | 0.938835 | 0.85354 | 3394 | tags=80% | list=21% | signal=63% |
| REACTOME_CELL_DEATH_SIGNALLING_VIA_NRAGE_NRIF_AND_NADE | 69 | -0.32118 | -0.784 | 0.789534 | 0.939637 | 0.85427 | 2631 | tags=19% | list=16% | signal=16% |
| REACTOME_DIGESTION_OF_DIETARY_CARBOHYDRATE | 3 | 0.458541 | 0.74242 | 0.790021 | 0.939637 | 0.85427 | 4052 | tags=67% | list=25% | signal=50% |
| REACTOME_TRANSLESION_SYNTHESIS_BY_POLK | 16 | 0.30076 | 0.730153 | 0.790625 | 0.939682 | 0.854311 | 1784 | tags=25% | list=11% | signal=22% |
| BIOCARTA_INTEGRIN_PATHWAY | 33 | -0.33973 | -0.75923 | 0.795396 | 0.944676 | 0.858851 | 121 | tags=6% | list=1% | signal=6% |
| BIOCARTA_CTCF_PATHWAY | 23 | 0.288172 | 0.74468 | 0.796078 | 0.944809 | 0.858972 | 2330 | tags=26% | list=14% | signal=22% |
| REACTOME_SIGNALING_BY_PDGF | 51 | -0.33205 | -0.78502 | 0.797847 | 0.944879 | 0.859035 | 1193 | tags=10% | list=7% | signal=9% |
| REACTOME_RHOBTB2_GTPASE_CYCLE | 22 | 0.292683 | 0.750395 | 0.79771 | 0.944879 | 0.859035 | 4497 | tags=55% | list=28% | signal=39% |
| REACTOME_MET_INTERACTS_WITH_TNS_PROTEINS | 5 | 0.390021 | 0.712087 | 0.797386 | 0.944879 | 0.859035 | 560 | tags=20% | list=3% | signal=19% |
| REACTOME_RAC2_GTPASE_CYCLE | 85 | -0.31918 | -0.79602 | 0.798889 | 0.945437 | 0.859543 | 2468 | tags=18% | list=15% | signal=15% |
| REACTOME_PROGRAMMED_CELL_DEATH | 122 | 0.250149 | 0.81018 | 0.8 | 0.946077 | 0.860125 | 4398 | tags=37% | list=27% | signal=27% |
| REACTOME_PHENYLALANINE_AND_TYROSINE_METABOLISM | 7 | 0.337293 | 0.695947 | 0.80137 | 0.946347 | 0.86037 | 3777 | tags=57% | list=23% | signal=44% |
| REACTOME_RELAXIN_RECEPTORS | 5 | -0.41627 | -0.687 | 0.801105 | 0.946347 | 0.86037 | 4759 | tags=80% | list=29% | signal=56% |
| WP_FOCAL_ADHESION | 174 | -0.3204 | -0.83914 | 0.803778 | 0.946556 | 0.86056 | 2578 | tags=17% | list=16% | signal=15% |
| REACTOME_FORMATION_OF_THE_CORNIFIED_ENVELOPE | 27 | 0.283655 | 0.749308 | 0.804167 | 0.946556 | 0.86056 | 58 | tags=4% | list=0% | signal=4% |
| REACTOME_KERATINIZATION | 27 | 0.283655 | 0.749308 | 0.804167 | 0.946556 | 0.86056 | 58 | tags=4% | list=0% | signal=4% |
| REACTOME_MET_ACTIVATES_RAS_SIGNALING | 9 | -0.39047 | -0.71875 | 0.802932 | 0.946556 | 0.86056 | 5646 | tags=56% | list=35% | signal=36% |
| REACTOME_INTERLEUKIN_27_SIGNALING | 8 | 0.340777 | 0.713063 | 0.804401 | 0.946556 | 0.86056 | 2293 | tags=38% | list=14% | signal=32% |
| REACTOME_INTRACELLULAR_SIGNALING_BY_SECOND_MESSENGERS | 242 | 0.253341 | 0.861168 | 0.810811 | 0.946707 | 0.860698 | 2330 | tags=22% | list=14% | signal=19% |
| REACTOME_G2_M_CHECKPOINTS | 141 | 0.251887 | 0.797319 | 0.806452 | 0.946707 | 0.860698 | 4736 | tags=31% | list=29% | signal=22% |
| REACTOME_SCF_SKP2_MEDIATED_DEGRADATION_OF_P27_P21 | 59 | 0.264623 | 0.789117 | 0.808511 | 0.946707 | 0.860698 | 2798 | tags=19% | list=17% | signal=15% |
| WP_IL1_SIGNALING_PATHWAY | 34 | 0.273736 | 0.764789 | 0.809091 | 0.946707 | 0.860698 | 3034 | tags=32% | list=19% | signal=26% |
| WP_FIBRIN_COMPLEMENT_RECEPTOR_3_SIGNALING_PATHWAY | 34 | 0.271138 | 0.757533 | 0.809091 | 0.946707 | 0.860698 | 2308 | tags=26% | list=14% | signal=23% |
| REACTOME_NEGATIVE_REGULATION_OF_FGFR4_SIGNALING | 24 | 0.290648 | 0.756049 | 0.808765 | 0.946707 | 0.860698 | 3034 | tags=29% | list=19% | signal=24% |
| BIOCARTA_CASPASE_PATHWAY | 20 | 0.289261 | 0.742297 | 0.809187 | 0.946707 | 0.860698 | 2649 | tags=15% | list=16% | signal=13% |
| BIOCARTA_MAL_PATHWAY | 17 | -0.35913 | -0.72485 | 0.808357 | 0.946707 | 0.860698 | 266 | tags=12% | list=2% | signal=12% |
| BIOCARTA_RB_PATHWAY | 12 | -0.37169 | -0.72339 | 0.810241 | 0.946707 | 0.860698 | 1703 | tags=42% | list=11% | signal=37% |
| REACTOME_METHYLATION | 12 | -0.37167 | -0.72334 | 0.810241 | 0.946707 | 0.860698 | 2099 | tags=17% | list=13% | signal=15% |
| BIOCARTA_AKAP95_PATHWAY | 10 | -0.38218 | -0.71425 | 0.809904 | 0.946707 | 0.860698 | 936 | tags=40% | list=6% | signal=38% |
| BIOCARTA_SODD_PATHWAY | 9 | 0.33533 | 0.714091 | 0.811856 | 0.94726 | 0.8612 | 1531 | tags=22% | list=9% | signal=20% |
| REACTOME_APC_C_CDC20_MEDIATED_DEGRADATION_OF_CYCLIN_B | 24 | 0.288384 | 0.750162 | 0.812749 | 0.947636 | 0.861542 | 5175 | tags=33% | list=32% | signal=23% |
| REACTOME_CHAPERONIN_MEDIATED_PROTEIN_FOLDING | 35 | -0.33103 | -0.74367 | 0.816561 | 0.951411 | 0.864974 | 594 | tags=9% | list=4% | signal=8% |
| REACTOME_APC_C_MEDIATED_DEGRADATION_OF_CELL_CYCLE_PROTEINS | 85 | 0.248196 | 0.763039 | 0.823529 | 0.953432 | 0.866811 | 5771 | tags=39% | list=36% | signal=25% |
| REACTOME_DDX58_IFIH1_MEDIATED_INDUCTION_OF_INTERFERON_ALPHA_BETA | 33 | 0.271223 | 0.752348 | 0.822727 | 0.953432 | 0.866811 | 2308 | tags=24% | list=14% | signal=21% |
| REACTOME_NICOTINATE_METABOLISM | 27 | -0.33738 | -0.73426 | 0.824147 | 0.953432 | 0.866811 | 2279 | tags=30% | list=14% | signal=25% |
| REACTOME_PHOSPHOLIPASE_C_MEDIATED_CASCADE_FGFR2 | 13 | -0.36709 | -0.7209 | 0.820513 | 0.953432 | 0.866811 | 540 | tags=15% | list=3% | signal=15% |
| REACTOME_INTERLEUKIN_12_FAMILY_SIGNALING | 10 | 0.328607 | 0.720464 | 0.821809 | 0.953432 | 0.866811 | 2293 | tags=40% | list=14% | signal=34% |
| REACTOME_HYALURONAN_METABOLISM | 17 | -0.35341 | -0.7133 | 0.821326 | 0.953432 | 0.866811 | 3535 | tags=29% | list=22% | signal=23% |
| REACTOME_HYPUSINE_SYNTHESIS_FROM_EIF5A_LYSINE | 5 | 0.379268 | 0.692454 | 0.823529 | 0.953432 | 0.866811 | 10023 | tags=100% | list=62% | signal=38% |
| REACTOME_REGULATION_OF_IFNG_SIGNALING | 8 | -0.38272 | -0.68797 | 0.824621 | 0.953432 | 0.866811 | 2421 | tags=50% | list=15% | signal=43% |
| REACTOME_ALPHA_OXIDATION_OF_PHYTANATE | 6 | -0.39601 | -0.6736 | 0.823423 | 0.953432 | 0.866811 | 3316 | tags=50% | list=21% | signal=40% |
| REACTOME_PHOSPHATE_BOND_HYDROLYSIS_BY_NUDT_PROTEINS | 7 | 0.324653 | 0.669868 | 0.821918 | 0.953432 | 0.866811 | 5501 | tags=57% | list=34% | signal=38% |
| REACTOME_ATORVASTATIN_ADME | 4 | -0.40284 | -0.65126 | 0.819209 | 0.953432 | 0.866811 | 5405 | tags=75% | list=33% | signal=50% |
| REACTOME_SIGNALING_BY_NOTCH | 38 | -0.32821 | -0.7452 | 0.825596 | 0.953894 | 0.867232 | 2625 | tags=34% | list=16% | signal=29% |
| REACTOME_ABC_FAMILY_PROTEINS_MEDIATED_TRANSPORT | 93 | -0.31118 | -0.78263 | 0.829295 | 0.957268 | 0.870298 | 3588 | tags=31% | list=22% | signal=24% |
| REACTOME_NUCLEOTIDE_BINDING_DOMAIN_LEUCINE_RICH_REPEAT_CONTAININ( | 43 | 0.262391 | 0.761719 | 0.82967 | 0.957268 | 0.870298 | 2422 | tags=26% | list=15% | signal=22% |
| BIOCARTA_PLATELETAPP_PATHWAY | 10 | -0.37357 | -0.69817 | 0.830671 | 0.957755 | 0.870742 | 2970 | tags=30% | list=18% | signal=24% |
| WP_FACTORS_AND_PATHWAYS_AFFECTING_INSULINLIKE_GROWTH_FACTOR_IGF1 | 27 | -0.3311 | -0.7206 | 0.835958 | 0.960512 | 0.873248 | 1931 | tags=22% | list=12% | signal=20% |
| BIOCARTA_BAD_PATHWAY | 23 | -0.33765 | -0.71701 | 0.835341 | 0.960512 | 0.873248 | 1376 | tags=13% | list=9% | signal=12% |
| REACTOME_FRS_MEDIATED_FGFR2_SIGNALING | 19 | -0.34532 | -0.7121 | 0.835883 | 0.960512 | 0.873248 | 540 | tags=47% | list=3% | signal=46% |
| REACTOME_BASE_EXCISION_REPAIR_AP_SITE_FORMATION | 11 | 0.319738 | 0.707861 | 0.835655 | 0.960512 | 0.873248 | 4891 | tags=45% | list=30% | signal=32% |
| REACTOME_CDC6_ASSOCIATION_WITH_THE_ORC_ORIGIN_COMPLEX | 7 | -0.4029 | -0.69866 | 0.835106 | 0.960512 | 0.873248 | 944 | tags=43% | list=6% | signal=40% |
| REACTOME_GOLGI_TO_ER_RETROGRADE_TRANSPORT | 122 | -0.31259 | -0.80229 | 0.838188 | 0.961727 | 0.874352 | 1360 | tags=9% | list=8% | signal=8% |
| REACTOME_APC_CDC20_MEDIATED_DEGRADATION_OF_NEK2A | 26 | 0.277039 | 0.723061 | 0.841004 | 0.961727 | 0.874352 | 5175 | tags=35% | list=32% | signal=24% |
| REACTOME_COBALAMIN_CBL_VITAMIN_B12_TRANSPORT_AND_METABOLISM | 13 | -0.36129 | -0.7095 | 0.840121 | 0.961727 | 0.874352 | 975 | tags=23% | list=6% | signal=22% |
| WP_FOLIC_ACID_NETWORK | 20 | 0.275835 | 0.707845 | 0.840989 | 0.961727 | 0.874352 | 3910 | tags=40% | list=24% | signal=30% |
| REACTOME_INTERLEUKIN_2_SIGNALING | 8 | 0.32522 | 0.68051 | 0.838631 | 0.961727 | 0.874352 | 2084 | tags=25% | list=13% | signal=22% |
| REACTOME_SMALL_INTERFERING_RNA_SIRNA_BIOGENESIS | 8 | 0.321038 | 0.671759 | 0.841076 | 0.961727 | 0.874352 | 2816 | tags=50% | list=17% | signal=41% |
| REACTOME_VITAMIN_B1_THIAMIN_METABOLISM | 5 | -0.38708 | -0.63883 | 0.839779 | 0.961727 | 0.874352 | 1197 | tags=40% | list=7% | signal=37% |
| REACTOME_CYTOKINE_SIGNALING_IN_IMMUNE_SYSTEM | 331 | 0.249631 | 0.889379 | 0.8457 | 0.963691 | 0.876138 | 2566 | tags=21% | list=16% | signal=18% |
| REACTOME_MHC_CLASS_II_ANTIGEN_PRESENTATION | 122 | -0.31063 | -0.79727 | 0.84466 | 0.963691 | 0.876138 | 1064 | tags=20% | list=7% | signal=19% |
| REACTOME_PRC2_METHYLATES_HISTONES_AND_DNA | 36 | 0.259944 | 0.733031 | 0.84507 | 0.963691 | 0.876138 | 1943 | tags=11% | list=12% | signal=10% |
| WP_APOE_AND_MIR146_IN_INFLAMMATION_AND_ATHEROSCLEROSIS | 8 | 0.314829 | 0.658767 | 0.843521 | 0.963691 | 0.876138 | 1788 | tags=25% | list=11% | signal=22% |
| WP_HFE_EFFECT_ON_HEPCIDIN_PRODUCTION | 5 | -0.37941 | -0.62618 | 0.845304 | 0.963691 | 0.876138 | 4696 | tags=80% | list=29% | signal=57% |
| REACTOME_METABOLISM_OF_AMINO_ACIDS_AND_DERIVATIVES | 214 | 0.254205 | 0.85613 | 0.851064 | 0.963694 | 0.876141 | 4140 | tags=38% | list=26% | signal=29% |
| REACTOME_TOLL_LIKE_RECEPTOR_CASCADES | 138 | -0.3095 | -0.80077 | 0.846318 | 0.963694 | 0.876141 | 961 | tags=7% | list=6% | signal=7% |
| REACTOME_DEATH_RECEPTOR_SIGNALING | 137 | -0.309 | -0.79999 | 0.847548 | 0.963694 | 0.876141 | 2631 | tags=15% | list=16% | signal=13% |
| REACTOME_GLYCEROPHOSPHOLIPID_BIOSYNTHESIS | 103 | -0.30149 | -0.76592 | 0.854486 | 0.963694 | 0.876141 | 2801 | tags=25% | list=17% | signal=21% |
| REACTOME_GPVI_MEDIATED_ACTIVATION_CASCADE | 31 | 0.269754 | 0.736163 | 0.848624 | 0.963694 | 0.876141 | 2781 | tags=26% | list=17% | signal=21% |
| REACTOME_SWITCHING_OF_ORIGINS_TO_A_POST_REPLICATIVE_STATE | 89 | 0.236147 | 0.733875 | 0.85567 | 0.963694 | 0.876141 | 5921 | tags=43% | list=37% | signal=27% |
| WP_PARKINSONS_DISEASE | 38 | 0.253171 | 0.724728 | 0.84878 | 0.963694 | 0.876141 | 1253 | tags=13% | list=8% | signal=12% |
| REACTOME_COLLAGEN_BIOSYNTHESIS_AND_MODIFYING_ENZYMES | 64 | -0.29301 | -0.71254 | 0.854358 | 0.963694 | 0.876141 | 2811 | tags=25% | list=17% | signal=21% |
| REACTOME_ACTIVATION_OF_ATR_IN_RESPONSE_TO_REPLICATION_STRESS | 34 | 0.25479 | 0.711857 | 0.854545 | 0.963694 | 0.876141 | 1830 | tags=21% | list=11% | signal=18% |
| REACTOME_SIGNALING_BY_RETINOIC_ACID | 31 | -0.31534 | -0.70387 | 0.855867 | 0.963694 | 0.876141 | 753 | tags=29% | list=5% | signal=28% |
| REACTOME_GAB1_SIGNALOSOME | 13 | -0.357 | -0.70107 | 0.852187 | 0.963694 | 0.876141 | 2398 | tags=31% | list=15% | signal=26% |
| REACTOME_IONOTROPIC_ACTIVITY_OF_KAINATE_RECEPTORS | 12 | 0.305998 | 0.689653 | 0.85503 | 0.963694 | 0.876141 | 2179 | tags=33% | list=13% | signal=29% |

| | | | | | | | | | | |
|---|---|---|---|---|---|---|---|---|---|---|
| REACTOME_METABOLISM_OF_FOLATE_AND_PTERINES | 16 | -0.34227 | -0.68943 | 0.85044 | 0.963694 | 0.876141 | 4527 | tags=44% | list=28% | signal=32% |
| REACTOME_DOWNREGULATION_OF_SMAD2_3_SMAD4_TRANSCRIPTIONAL_ACTIVI | 22 | -0.32422 | -0.68481 | 0.854054 | 0.963694 | 0.876141 | 262 | tags=32% | list=2% | signal=31% |
| REACTOME_ERK_MAPK_TARGETS | 19 | 0.272965 | 0.682863 | 0.855124 | 0.963694 | 0.876141 | 3050 | tags=32% | list=19% | signal=26% |
| BIOCARTA_CDC25_PATHWAY | 8 | -0.36499 | -0.6561 | 0.853288 | 0.963694 | 0.876141 | 1183 | tags=25% | list=7% | signal=23% |
| BIOCARTA_WNT_LRP6_PATHWAY | 3 | -0.42804 | -0.65197 | 0.852207 | 0.963694 | 0.876141 | 9236 | tags=100% | list=57% | signal=43% |
| REACTOME_HUR_ELAVL1_BINDS_AND_STABILIZES_MRNA | 7 | 0.308489 | 0.636515 | 0.856164 | 0.963694 | 0.876141 | 5070 | tags=71% | list=31% | signal=49% |
| REACTOME_RAC3_GTPASE_CYCLE | 88 | -0.30341 | -0.76035 | 0.860619 | 0.964126 | 0.876534 | 1250 | tags=15% | list=8% | signal=14% |
| REACTOME_GENE_SILENCING_BY_RNA | 41 | 0.255666 | 0.739655 | 0.859375 | 0.964126 | 0.876534 | 2095 | tags=24% | list=13% | signal=21% |
| REACTOME_NEGATIVE_REGULATION_OF_FGFR3_SIGNALING | 26 | 0.273719 | 0.714397 | 0.857741 | 0.964126 | 0.876534 | 3034 | tags=27% | list=19% | signal=22% |
| REACTOME_GLUTATHIONE_CONJUGATION | 30 | 0.260867 | 0.706394 | 0.857143 | 0.964126 | 0.876534 | 2333 | tags=20% | list=14% | signal=17% |
| REACTOME_GASTRIN_CREB_SIGNALLING_PATHWAY_VIA_PKC_AND_MAPK | 14 | 0.28732 | 0.672541 | 0.860182 | 0.964126 | 0.876534 | 3050 | tags=36% | list=19% | signal=29% |
| REACTOME_TRIGLYCERIDE_METABOLISM | 12 | -0.34132 | -0.66429 | 0.85994 | 0.964126 | 0.876534 | 444 | tags=17% | list=3% | signal=16% |
| REACTOME_PASSIVE_TRANSPORT_BY_AQUAPORINS | 6 | -0.36993 | -0.62924 | 0.859459 | 0.964126 | 0.876534 | 10175 | tags=100% | list=63% | signal=37% |
| REACTOME_ER_TO_GOLGI_ANTEROGRADE_TRANSPORT | 138 | -0.30294 | -0.7838 | 0.86873 | 0.964594 | 0.87696 | 1435 | tags=12% | list=9% | signal=11% |
| REACTOME_TOLL_LIKE_RECEPTOR_4_TLR4_CASCADE | 114 | -0.29836 | -0.76503 | 0.871245 | 0.964594 | 0.87696 | 961 | tags=4% | list=6% | signal=4% |
| REACTOME_MYD88_INDEPENDENT_TLR4_CASCADE | 99 | 0.238365 | 0.742937 | 0.866667 | 0.964594 | 0.87696 | 2308 | tags=22% | list=14% | signal=19% |
| REACTOME_NUCLEAR_PORE_COMPLEX_NPC_DISASSEMBLY | 31 | 0.260835 | 0.711823 | 0.866972 | 0.964594 | 0.87696 | 2035 | tags=19% | list=13% | signal=17% |
| REACTOME_INHIBITION_OF_THE_PROTEOLYTIC_ACTIVITY_OF_APC_C_REQUIRED_F | 21 | 0.277512 | 0.706498 | 0.873134 | 0.964594 | 0.87696 | 5175 | tags=38% | list=32% | signal=26% |
| REACTOME_OXIDATIVE_STRESS_INDUCED_SENESCENCE | 37 | -0.30542 | -0.69295 | 0.871698 | 0.964594 | 0.87696 | 49 | tags=11% | list=0% | signal=11% |
| BIOCARTA_EFP_PATHWAY | 13 | -0.35225 | -0.69174 | 0.865762 | 0.964594 | 0.87696 | 5695 | tags=54% | list=35% | signal=35% |
| BIOCARTA_G1_PATHWAY | 25 | -0.31419 | -0.68145 | 0.869737 | 0.964594 | 0.87696 | 5206 | tags=52% | list=32% | signal=35% |
| REACTOME_PENTOSE_PHOSPHATE_PATHWAY | 13 | -0.34552 | -0.67853 | 0.873303 | 0.964594 | 0.87696 | 5431 | tags=54% | list=34% | signal=36% |
| WP_PROSTAGLANDIN_SYNTHESIS_AND_REGULATION | 25 | -0.31273 | -0.67828 | 0.875 | 0.964594 | 0.87696 | 3814 | tags=36% | list=24% | signal=28% |
| REACTOME_ERKS_ARE_INACTIVATED | 13 | -0.34484 | -0.67719 | 0.874811 | 0.964594 | 0.87696 | 961 | tags=15% | list=6% | signal=14% |
| REACTOME_ENDOGENOUS_STEROLS | 20 | -0.32462 | -0.67304 | 0.867872 | 0.964594 | 0.87696 | 921 | tags=20% | list=6% | signal=19% |
| REACTOME_SYNTHESIS_OF_BILE_ACIDS_AND_BILE_SALTS | 24 | -0.31144 | -0.67138 | 0.868176 | 0.964594 | 0.87696 | 921 | tags=21% | list=6% | signal=20% |
| REACTOME_INTERLEUKIN_6_FAMILY_SIGNALING | 18 | -0.32562 | -0.66783 | 0.866947 | 0.964594 | 0.87696 | 1869 | tags=17% | list=12% | signal=15% |
| REACTOME_PLATELET_SENSITIZATION_BY_LDL | 16 | 0.27299 | 0.662737 | 0.871875 | 0.964594 | 0.87696 | 5548 | tags=69% | list=34% | signal=45% |
| REACTOME_PURINE_CATABOLISM | 17 | -0.32622 | -0.65842 | 0.871758 | 0.964594 | 0.87696 | 745 | tags=12% | list=5% | signal=11% |
| REACTOME_TERMINATION_OF_O_GLYCAN_BIOSYNTHESIS | 10 | -0.35147 | -0.65686 | 0.864217 | 0.964594 | 0.87696 | 3082 | tags=40% | list=19% | signal=32% |
| REACTOME_BH3_ONLY_PROTEINS_ASSOCIATE_WITH_AND_INACTIVATE_ANTI_APC | 7 | -0.37614 | -0.65226 | 0.874113 | 0.964594 | 0.87696 | 1057 | tags=29% | list=7% | signal=27% |
| BIOCARTA_CHREBP_PATHWAY | 9 | -0.35247 | -0.6488 | 0.874593 | 0.964594 | 0.87696 | 936 | tags=100% | list=6% | signal=94% |
| REACTOME_TP53_REGULATES_TRANSCRIPTION_OF_GENES_INVOLVED_IN_G1_CEL | 10 | -0.3461 | -0.64682 | 0.870607 | 0.964594 | 0.87696 | 5880 | tags=80% | list=36% | signal=51% |
| REACTOME_HDR_THROUGH_MMEJ_ALT_NHEJ | 12 | -0.33105 | -0.64429 | 0.870482 | 0.964594 | 0.87696 | 641 | tags=25% | list=4% | signal=24% |
| BIOCARTA_SKP2E2F_PATHWAY | 9 | 0.297319 | 0.633147 | 0.871134 | 0.964594 | 0.87696 | 2798 | tags=33% | list=17% | signal=28% |
| BIOCARTA_BBCELL_PATHWAY | 3 | -0.41549 | -0.63286 | 0.867562 | 0.964594 | 0.87696 | 5638 | tags=67% | list=35% | signal=43% |
| BIOCARTA_TERT_PATHWAY | 6 | -0.35541 | -0.60453 | 0.872072 | 0.964594 | 0.87696 | 1703 | tags=83% | list=11% | signal=75% |
| REACTOME_TRANS_GOLGI_NETWORK_VESICLE_BUDDING | 70 | -0.29005 | -0.70924 | 0.880682 | 0.967 | 0.879147 | 1435 | tags=11% | list=9% | signal=10% |
| REACTOME_INTERCONVERSION_OF_NUCLEOTIDE_DI_AND_TRIPHOSPHATES | 25 | 0.273329 | 0.707932 | 0.880165 | 0.967 | 0.879147 | 5396 | tags=36% | list=33% | signal=24% |
| REACTOME_REGULATION_OF_APC_C_ACTIVATORS_BETWEEN_G1_S_AND_EARLY_/ | 35 | 0.244636 | 0.68596 | 0.880184 | 0.967 | 0.879147 | 5175 | tags=34% | list=32% | signal=23% |
| REACTOME_HOMOLOGOUS_DNA_PAIRING_AND_STRAND_EXCHANGE | 25 | -0.31012 | -0.67261 | 0.880263 | 0.967 | 0.879147 | 641 | tags=20% | list=4% | signal=19% |
| REACTOME_SYNTHESIS_OF_BILE_ACIDS_AND_BILE_SALTS_VIA_7ALPHA_HYDROXY( | 15 | -0.33455 | -0.66722 | 0.879234 | 0.967 | 0.879147 | 277 | tags=33% | list=2% | signal=33% |
| REACTOME_GAMMA_CARBOXYLATION_OF_PROTEIN_PRECURSORS | 3 | -0.40047 | -0.60998 | 0.879079 | 0.967 | 0.879147 | 9681 | tags=67% | list=60% | signal=27% |
| BIOCARTA_NKCELLS_PATHWAY | 15 | -0.3315 | -0.66113 | 0.883652 | 0.967624 | 0.879714 | 1001 | tags=20% | list=6% | signal=19% |
| REACTOME_ONCOGENE_INDUCED_SENESCENCE | 16 | -0.3216 | -0.6478 | 0.882698 | 0.967624 | 0.879714 | 3551 | tags=31% | list=22% | signal=24% |
| REACTOME_HDL_ASSEMBLY | 7 | -0.36848 | -0.63896 | 0.884752 | 0.967624 | 0.879714 | 2394 | tags=29% | list=15% | signal=24% |
| REACTOME_ASSEMBLY_OF_THE_ORC_COMPLEX_AT_THE_ORIGIN_OF_REPLICATIOI | 8 | -0.35007 | -0.62929 | 0.883642 | 0.967624 | 0.879714 | 944 | tags=38% | list=6% | signal=35% |
| REACTOME_RHO_GTPASES_ACTIVATE_ROCKS | 5 | 0.340352 | 0.621403 | 0.884532 | 0.967624 | 0.879714 | 4111 | tags=40% | list=25% | signal=30% |
| REACTOME_TP53_REGULATES_TRANSCRIPTION_OF_GENES_INVOLVED_IN_CYTOCH | 7 | 0.294984 | 0.60865 | 0.883562 | 0.967624 | 0.879714 | 5846 | tags=71% | list=36% | signal=46% |
| WP_DOPAMINERGIC_NEUROGENESIS | 21 | -0.30919 | -0.64858 | 0.886921 | 0.968719 | 0.880709 | 2922 | tags=43% | list=18% | signal=35% |
| REACTOME_MICRORNA_MIRNA_BIOGENESIS | 6 | -0.34144 | -0.58077 | 0.886486 | 0.968719 | 0.880709 | 3271 | tags=50% | list=20% | signal=40% |
| BIOCARTA_RELA_PATHWAY | 14 | 0.275535 | 0.644955 | 0.887538 | 0.968754 | 0.880742 | 2308 | tags=29% | list=14% | signal=25% |
| WP_SELENIUM_MICRONUTRIENT_NETWORK | 18 | 0.26225 | 0.648311 | 0.888889 | 0.968953 | 0.880922 | 2738 | tags=28% | list=17% | signal=23% |
| REACTOME_SEMA4D_MEDIATED_INHIBITION_OF_CELL_ATTACHMENT_AND_MIGR/ | 8 | -0.34393 | -0.61825 | 0.888702 | 0.968953 | 0.880922 | 2419 | tags=25% | list=15% | signal=21% |
| REACTOME_CELL_CYCLE_CHECKPOINTS | 260 | 0.247795 | 0.858538 | 0.894737 | 0.971983 | 0.883677 | 4737 | tags=31% | list=29% | signal=22% |
| REACTOME_SIGNALING_BY_FGFR | 67 | -0.28562 | -0.69595 | 0.892816 | 0.971983 | 0.883677 | 3036 | tags=16% | list=19% | signal=13% |
| REACTOME_PHOSPHORYLATION_OF_THE_APC_C | 20 | 0.253077 | 0.649442 | 0.893993 | 0.971983 | 0.883677 | 5175 | tags=35% | list=32% | signal=24% |
| BIOCARTA_AGPCR_PATHWAY | 10 | -0.33693 | -0.62969 | 0.896166 | 0.971983 | 0.883677 | 936 | tags=10% | list=6% | signal=9% |
| BIOCARTA_MPR_PATHWAY | 18 | -0.30583 | -0.62725 | 0.896359 | 0.971983 | 0.883677 | 1183 | tags=11% | list=7% | signal=10% |
| REACTOME_PEXOPHAGY | 10 | 0.282149 | 0.618605 | 0.896277 | 0.971983 | 0.883677 | 1831 | tags=20% | list=11% | signal=18% |
| BIOCARTA_FREE_PATHWAY | 7 | -0.35522 | -0.61597 | 0.89539 | 0.971983 | 0.883677 | 6353 | tags=86% | list=39% | signal=52% |
| REACTOME_MRNA_EDITING | 4 | -0.35929 | -0.58086 | 0.894539 | 0.971983 | 0.883677 | 3170 | tags=50% | list=20% | signal=40% |
| REACTOME_REGULATION_OF_RUNX1_EXPRESSION_AND_ACTIVITY | 7 | 0.288846 | 0.595985 | 0.89726 | 0.972325 | 0.883988 | 3752 | tags=29% | list=23% | signal=22% |
| REACTOME_SIGNALING_BY_FGFR2 | 57 | 0.240624 | 0.716842 | 0.899329 | 0.972502 | 0.884149 | 3034 | tags=25% | list=19% | signal=20% |
| BIOCARTA_HIVNEF_PATHWAY | 52 | -0.28994 | -0.68707 | 0.898689 | 0.972502 | 0.884149 | 1777 | tags=13% | list=11% | signal=12% |
| REACTOME_HEDGEHOG_LIGAND_BIOGENESIS | 61 | -0.28007 | -0.67821 | 0.89977 | 0.972502 | 0.884149 | 2638 | tags=15% | list=16% | signal=12% |
| WP_FATTY_ACID_BIOSYNTHESIS | 22 | -0.29693 | -0.62718 | 0.898649 | 0.972502 | 0.884149 | 1233 | tags=9% | list=8% | signal=8% |
| REACTOME_MAPK_FAMILY_SIGNALING_CASCADES | 267 | 0.253017 | 0.882681 | 0.902439 | 0.972805 | 0.884424 | 1659 | tags=18% | list=10% | signal=17% |
| WP_PURINE_METABOLISM | 151 | 0.235579 | 0.747092 | 0.912281 | 0.972805 | 0.884424 | 3394 | tags=27% | list=21% | signal=22% |
| REACTOME_ORC1_REMOVAL_FROM_CHROMATIN | 68 | 0.239007 | 0.716918 | 0.912 | 0.972805 | 0.884424 | 2798 | tags=18% | list=17% | signal=15% |
| REACTOME_METABOLISM_OF_NUCLEOTIDES | 84 | 0.222467 | 0.686118 | 0.914286 | 0.972805 | 0.884424 | 3713 | tags=32% | list=23% | signal=25% |
| REACTOME_SIGNALING_BY_FGFR1 | 44 | -0.28853 | -0.66905 | 0.90534 | 0.972805 | 0.884424 | 2851 | tags=18% | list=18% | signal=15% |
| REACTOME_SIGNALING_BY_PTK6 | 44 | -0.28656 | -0.6645 | 0.908981 | 0.972805 | 0.884424 | 2241 | tags=16% | list=14% | signal=14% |
| REACTOME_INTERLEUKIN_17_SIGNALING | 58 | -0.2768 | -0.66398 | 0.909197 | 0.972805 | 0.884424 | 961 | tags=5% | list=6% | signal=5% |
| REACTOME_REGULATION_OF_TP53_DEGRADATION | 35 | 0.235306 | 0.659799 | 0.912442 | 0.972805 | 0.884424 | 2250 | tags=20% | list=14% | signal=17% |
| REACTOME_NEGATIVE_REGULATION_OF_FGFR2_SIGNALING | 29 | 0.239474 | 0.640447 | 0.913793 | 0.972805 | 0.884424 | 3034 | tags=24% | list=19% | signal=20% |
| REACTOME_NICOTINAMIDE_SALVAGING | 16 | 0.261582 | 0.635042 | 0.903125 | 0.972805 | 0.884424 | 902 | tags=12% | list=6% | signal=12% |
| BIOCARTA_SPRY_PATHWAY | 16 | 0.259154 | 0.629146 | 0.909375 | 0.972805 | 0.884424 | 3724 | tags=31% | list=23% | signal=24% |
| BIOCARTA_ECM_PATHWAY | 21 | -0.29717 | -0.62337 | 0.914169 | 0.972805 | 0.884424 | 445 | tags=10% | list=3% | signal=9% |
| BIOCARTA_STATHMIN_PATHWAY | 14 | -0.31331 | -0.62085 | 0.915305 | 0.972805 | 0.884424 | 936 | tags=29% | list=6% | signal=27% |
| REACTOME_DEPOLYMERISATION_OF_THE_NUCLEAR_LAMINA | 12 | 0.274444 | 0.618537 | 0.914201 | 0.972805 | 0.884424 | 247 | tags=8% | list=2% | signal=8% |
| REACTOME_ALPHA_PROTEIN_KINASE_1_SIGNALING_PATHWAY | 11 | 0.278964 | 0.617592 | 0.913649 | 0.972805 | 0.884424 | 4744 | tags=45% | list=29% | signal=32% |
| REACTOME_INTERLEUKIN_15_SIGNALING | 11 | 0.278685 | 0.616973 | 0.913649 | 0.972805 | 0.884424 | 2293 | tags=18% | list=14% | signal=16% |
| BIOCARTA_NPP1_PATHWAY | 9 | -0.33269 | -0.6124 | 0.90228 | 0.972805 | 0.884424 | 887 | tags=22% | list=5% | signal=21% |
| BIOCARTA_GATA3_PATHWAY | 10 | -0.32555 | -0.60842 | 0.907348 | 0.972805 | 0.884424 | 936 | tags=30% | list=6% | signal=28% |
| BIOCARTA_MCM_PATHWAY | 13 | -0.30789 | -0.60464 | 0.912519 | 0.972805 | 0.884424 | 3203 | tags=38% | list=20% | signal=31% |
| REACTOME_CONDENSATION_OF_PROMETAPHASE_CHROMOSOMES | 8 | 0.287753 | 0.602113 | 0.914425 | 0.972805 | 0.884424 | 1567 | tags=12% | list=10% | signal=11% |
| WP_PTF1A_RELATED_REGULATORY_PATHWAY | 8 | 0.287531 | 0.601648 | 0.914425 | 0.972805 | 0.884424 | 4723 | tags=38% | list=29% | signal=27% |
| REACTOME_RHO_GTPASES_ACTIVATE_CIT | 7 | -0.34603 | -0.60004 | 0.900709 | 0.972805 | 0.884424 | 3903 | tags=43% | list=24% | signal=33% |
| WP_LEPTIN_AND_ADIPONECTIN | 8 | -0.33088 | -0.59479 | 0.91231 | 0.972805 | 0.884424 | 3044 | tags=38% | list=19% | signal=30% |
| REACTOME_APOPTOSIS_INDUCED_DNA_FRAGMENTATION | 9 | 0.276043 | 0.58784 | 0.912371 | 0.972805 | 0.884424 | 3744 | tags=22% | list=23% | signal=17% |
| REACTOME_PHOSPHORYLATION_OF_EMI1 | 6 | 0.284298 | 0.556187 | 0.914989 | 0.972805 | 0.884424 | 11556 | tags=100% | list=72% | signal=28% |
| REACTOME_EICOSANOIDS | 6 | -0.32223 | -0.54811 | 0.908108 | 0.972805 | 0.884424 | 3814 | tags=50% | list=24% | signal=38% |
| REACTOME_G1_S_DNA_DAMAGE_CHECKPOINTS | 65 | 0.235646 | 0.710709 | 0.916667 | 0.973493 | 0.88505 | 2552 | tags=17% | list=16% | signal=14% |
| REACTOME_OREXIN_AND_NEUROPEPTIDES_FF_AND_QRFP_BIND_TO_THEIR_RESPI | 5 | -0.32839 | -0.54197 | 0.917127 | 0.973493 | 0.88505 | 2505 | tags=40% | list=16% | signal=34% |
| REACTOME_METABOLISM_OF_WATER_SOLUBLE_VITAMINS_AND_COFACTORS | 104 | -0.2807 | -0.71304 | 0.918033 | 0.973831 | 0.885357 | 3812 | tags=25% | list=24% | signal=19% |
| REACTOME_RHOA_GTPASE_CYCLE | 138 | -0.28322 | -0.73278 | 0.919957 | 0.975249 | 0.886646 | 1277 | tags=10% | list=8% | signal=9% |
| REACTOME_RHOG_GTPASE_CYCLE | 72 | -0.2737 | -0.66831 | 0.928082 | 0.976769 | 0.888028 | 1250 | tags=31% | list=8% | signal=28% |

| | | | | | | | | | | |
|---|---|---|---|---|---|---|---|---|---|---|
| REACTOME_ASYMMETRIC_LOCALIZATION_OF_PCP_PROTEINS | 60 | 0.223879 | 0.660647 | 0.924812 | 0.976769 | 0.888028 | 2552 | tags=15% | list=16% | signal=13% |
| REACTOME_TRANSCRIPTIONAL_ACTIVITY_OF_SMAD2_SMAD3_SMAD4_HETEROTRI | 38 | 0.224814 | 0.643553 | 0.926829 | 0.976769 | 0.888028 | 3313 | tags=37% | list=21% | signal=29% |
| REACTOME_SEALING_OF_THE_NUCLEAR_ENVELOPE_NE_BY_ESCRT_III | 21 | 0.250136 | 0.636804 | 0.925373 | 0.976769 | 0.888028 | 4557 | tags=38% | list=28% | signal=27% |
| REACTOME_BILE_ACID_AND_BILE_SALT_METABOLISM | 28 | -0.28675 | -0.62721 | 0.924282 | 0.976769 | 0.888028 | 921 | tags=21% | list=6% | signal=20% |
| BIOCARTA_IL1R_PATHWAY | 25 | -0.28191 | -0.61143 | 0.922368 | 0.976769 | 0.888028 | 3202 | tags=24% | list=20% | signal=19% |
| REACTOME_CASPASE_ACTIVATION_VIA_EXTRINSIC_APOPTOTIC_SIGNALLING_PATH | 18 | 0.246295 | 0.608869 | 0.923611 | 0.976769 | 0.888028 | 1531 | tags=11% | list=9% | signal=10% |
| REACTOME_BINDING_AND_UPTAKE_OF_LIGANDS_BY_SCAVENGER_RECEPTORS | 23 | -0.2818 | -0.59841 | 0.92905 | 0.976769 | 0.888028 | 3913 | tags=39% | list=24% | signal=30% |
| BIOCARTA_PTC1_PATHWAY | 11 | 0.26244 | 0.58101 | 0.927577 | 0.976769 | 0.888028 | 4401 | tags=36% | list=27% | signal=26% |
| BIOCARTA_P27_PATHWAY | 11 | 0.262381 | 0.580879 | 0.927577 | 0.976769 | 0.888028 | 2798 | tags=27% | list=17% | signal=23% |
| REACTOME_ASPARTATE_AND_ASPARAGINE_METABOLISM | 11 | -0.29676 | -0.56733 | 0.926905 | 0.976769 | 0.888028 | 4007 | tags=45% | list=25% | signal=34% |
| REACTOME_SEROTONIN_AND_MELATONIN_BIOSYNTHESIS | 3 | -0.35543 | -0.54137 | 0.928983 | 0.976769 | 0.888028 | 4930 | tags=67% | list=31% | signal=46% |
| BIOCARTA_PLCD_PATHWAY | 5 | -0.325 | -0.53637 | 0.922652 | 0.976769 | 0.888028 | 1998 | tags=40% | list=12% | signal=35% |
| WP_EICOSANOID_SYNTHESIS | 15 | -0.30104 | -0.6004 | 0.930781 | 0.977968 | 0.889119 | 3814 | tags=27% | list=24% | signal=20% |
| REACTOME_G_ALPHA_12_13_SIGNALLING_EVENTS | 74 | -0.27097 | -0.66399 | 0.931896 | 0.97852 | 0.88962 | 594 | tags=9% | list=4% | signal=9% |
| REACTOME_BETA_CATENIN_INDEPENDENT_WNT_SIGNALING | 119 | 0.220953 | 0.716767 | 0.935897 | 0.979511 | 0.890521 | 2880 | tags=21% | list=18% | signal=17% |
| WP_IL6_SIGNALING_PATHWAY | 96 | -0.26874 | -0.67809 | 0.933848 | 0.979511 | 0.890521 | 638 | tags=8% | list=4% | signal=8% |
| REACTOME_MYD88_MAL_TIRAP_CASCADE_INITIATED_ON_PLASMA_MEMBRANE | 83 | -0.26804 | -0.66601 | 0.936384 | 0.979511 | 0.890521 | 961 | tags=6% | list=6% | signal=6% |
| REACTOME_RHOH_GTPASE_CYCLE | 35 | -0.27619 | -0.62047 | 0.935032 | 0.979511 | 0.890521 | 3075 | tags=23% | list=19% | signal=19% |
| REACTOME_INTERFERON_GAMMA_SIGNALING | 14 | -0.29191 | -0.57842 | 0.934621 | 0.979511 | 0.890521 | 2421 | tags=29% | list=15% | signal=24% |
| WP_PENTOSE_PHOSPHATE_PATHWAY | 7 | -0.31306 | -0.54287 | 0.93617 | 0.979511 | 0.890521 | 3755 | tags=57% | list=23% | signal=44% |
| BIOCARTA_PS1_PATHWAY | 12 | -0.29644 | -0.57694 | 0.938253 | 0.980847 | 0.891736 | 1769 | tags=25% | list=11% | signal=22% |
| REACTOME_UB_SPECIFIC_PROCESSING_PROTEASES | 159 | 0.235331 | 0.743022 | 0.94 | 0.981289 | 0.892138 | 3371 | tags=28% | list=21% | signal=22% |
| REACTOME_SIGNALING_BY_TGF_BETA_RECEPTOR_COMPLEX | 71 | -0.26518 | -0.64833 | 0.944318 | 0.981289 | 0.892138 | 4696 | tags=38% | list=29% | signal=27% |
| REACTOME_RAS_PROCESSING | 23 | -0.27605 | -0.5862 | 0.941098 | 0.981289 | 0.892138 | 432 | tags=13% | list=3% | signal=13% |
| REACTOME_RHO_GTPASES_ACTIVATE_PKNS | 22 | -0.27105 | -0.5725 | 0.944595 | 0.981289 | 0.892138 | 764 | tags=9% | list=5% | signal=9% |
| WP_KENNEDY_PATHWAY | 14 | -0.28302 | -0.56082 | 0.943536 | 0.981289 | 0.892138 | 3136 | tags=29% | list=19% | signal=23% |
| REACTOME_MITOCHONDRIAL_IRON_SULFUR_CLUSTER_BIOGENESIS | 12 | -0.28657 | -0.55773 | 0.942771 | 0.981289 | 0.892138 | 11522 | tags=100% | list=71% | signal=29% |
| BIOCARTA_MELANOCYTE_PATHWAY | 7 | -0.30583 | -0.53032 | 0.939716 | 0.981289 | 0.892138 | 2933 | tags=29% | list=18% | signal=23% |
| WP_NONHOMOLOGOUS_END_JOINING | 7 | 0.254381 | 0.524872 | 0.940639 | 0.981289 | 0.892138 | 2864 | tags=29% | list=18% | signal=24% |
| REACTOME_PROSTANOID_LIGAND_RECEPTORS | 6 | 0.267025 | 0.522395 | 0.944072 | 0.981289 | 0.892138 | 1675 | tags=17% | list=10% | signal=15% |
| REACTOME_ACTIVATION_OF_NIMA_KINASES_NEK9_NEK6_NEK7 | 5 | 0.28428 | 0.519029 | 0.943355 | 0.981289 | 0.892138 | 11556 | tags=100% | list=72% | signal=28% |
| REACTOME_APOPTOSIS | 94 | 0.221239 | 0.683814 | 0.945652 | 0.981773 | 0.892577 | 4111 | tags=33% | list=25% | signal=25% |
| WP_MECHANISMS_ASSOCIATED_WITH_PLURIPOTENCY | 257 | 0.224608 | 0.776988 | 0.948718 | 0.982451 | 0.893194 | 2507 | tags=24% | list=16% | signal=20% |
| REACTOME_FATTY_ACID_METABOLISM | 136 | -0.26436 | -0.68389 | 0.948773 | 0.982451 | 0.893194 | 1720 | tags=11% | list=11% | signal=10% |
| REACTOME_SEMA4D_IN_SEMAPHORIN_SIGNALING | 15 | -0.2845 | -0.56739 | 0.946981 | 0.982451 | 0.893194 | 2419 | tags=20% | list=15% | signal=17% |
| REACTOME_PREGNENOLONE_BIOSYNTHESIS | 11 | 0.246449 | 0.545606 | 0.949861 | 0.982451 | 0.893194 | 3651 | tags=36% | list=23% | signal=28% |
| REACTOME_INTERACTION_WITH_CUMULUS_CELLS_AND_THE_ZONA_PELLUCIDA | 3 | -0.34142 | -0.52004 | 0.948177 | 0.982451 | 0.893194 | 2826 | tags=67% | list=18% | signal=55% |
| REACTOME_GLUTATHIONE_SYNTHESIS_AND_RECYCLING | 11 | 0.234274 | 0.518654 | 0.949861 | 0.982451 | 0.893194 | 3734 | tags=27% | list=23% | signal=21% |
| REACTOME_GLUCOSE_METABOLISM | 72 | -0.25755 | -0.62887 | 0.955479 | 0.987646 | 0.897917 | 1762 | tags=10% | list=11% | signal=9% |
| WP_HYPOXIADEPENDENT_SELFRENEWAL_OF_MYOBLASTS | 9 | 0.248714 | 0.529641 | 0.956186 | 0.987761 | 0.898021 | 12131 | tags=100% | list=75% | signal=25% |
| REACTOME_KINESINS | 51 | 0.21447 | 0.649178 | 0.957831 | 0.9879 | 0.898148 | 2963 | tags=22% | list=18% | signal=18% |
| WP_OVARIAN_INFERTILITY | 22 | -0.26183 | -0.55305 | 0.958108 | 0.9879 | 0.898148 | 4917 | tags=36% | list=30% | signal=25% |
| REACTOME_RELEASE_OF_APOPTOTIC_FACTORS_FROM_THE_MITOCHONDRIA | 7 | -0.2823 | -0.48953 | 0.957447 | 0.9879 | 0.898148 | 2374 | tags=57% | list=15% | signal=49% |
| REACTOME_ANTIGEN_ACTIVATES_B_CELL_RECEPTOR_BCR_LEADING_TO_GENERAT | 22 | -0.25591 | -0.54053 | 0.959459 | 0.988679 | 0.898856 | 3190 | tags=27% | list=20% | signal=22% |
| REACTOME_AMINO_ACIDS_REGULATE_MTORC1 | 51 | -0.25339 | -0.59906 | 0.960526 | 0.988908 | 0.899064 | 1339 | tags=18% | list=8% | signal=16% |
| REACTOME_RUNX1_REGULATES_TRANSCRIPTION_OF_GENES_INVOLVED_IN_DIFFE | 63 | -0.2427 | -0.58846 | 0.960875 | 0.988908 | 0.899064 | 748 | tags=8% | list=5% | signal=8% |
| WP_WNT_SIGNALING_PATHWAY_WP539 | 101 | -0.25306 | -0.64177 | 0.963816 | 0.991319 | 0.901257 | 1571 | tags=7% | list=10% | signal=6% |
| REACTOME_SIGNALING_BY_HEDGEHOG | 133 | -0.25945 | -0.67036 | 0.964744 | 0.991658 | 0.901565 | 1282 | tags=11% | list=8% | signal=10% |
| REACTOME_CROSS_PRESENTATION_OF_SOLUBLE_EXOGENOUS_ANTIGENS_ENDOS | 46 | 0.215457 | 0.635578 | 0.965714 | 0.992041 | 0.901913 | 2552 | tags=17% | list=16% | signal=15% |
| REACTOME_DECTIN_2_FAMILY | 6 | -0.27958 | -0.47556 | 0.967568 | 0.993329 | 0.903084 | 2624 | tags=50% | list=16% | signal=42% |
| BIOCARTA_PML_PATHWAY | 15 | -0.26426 | -0.52703 | 0.969072 | 0.993644 | 0.90337 | 4053 | tags=40% | list=25% | signal=30% |
| REACTOME_GAP_JUNCTION_DEGRADATION | 11 | -0.25194 | -0.48164 | 0.968896 | 0.993644 | 0.90337 | 3667 | tags=27% | list=23% | signal=21% |
| REACTOME_SYNTHESIS_OF_PC | 25 | -0.23404 | -0.5076 | 0.969737 | 0.993711 | 0.903431 | 1692 | tags=16% | list=10% | signal=14% |
| REACTOME_NUCLEAR_ENVELOPE_BREAKDOWN | 46 | 0.210451 | 0.620812 | 0.971429 | 0.994829 | 0.904448 | 2035 | tags=15% | list=13% | signal=13% |
| REACTOME_TAK1_DEPENDENT_IKK_AND_NF_KAPPA_B_ACTIVATION | 37 | -0.23249 | -0.52749 | 0.973585 | 0.996422 | 0.905896 | 2598 | tags=14% | list=16% | signal=11% |
| REACTOME_CD209_DC_SIGN_SIGNALING | 16 | 0.225057 | 0.546369 | 0.975 | 0.997255 | 0.906653 | 3508 | tags=25% | list=22% | signal=20% |
| REACTOME_TOLL_LIKE_RECEPTOR_9_TLR9_CASCADE | 93 | -0.23204 | -0.5836 | 0.977974 | 0.999064 | 0.908297 | 961 | tags=4% | list=6% | signal=4% |
| REACTOME_EICOSANOID_LIGAND_BINDING_RECEPTORS | 10 | -0.25964 | -0.48525 | 0.977636 | 0.999064 | 0.908297 | 6251 | tags=70% | list=39% | signal=43% |
| REACTOME_M_PHASE | 343 | 0.263746 | 0.950991 | 0.984771 | 1 | 0.909149 | 4965 | tags=35% | list=31% | signal=25% |
| REACTOME_MEMBRANE_TRAFFICKING | 525 | -0.27249 | -0.74531 | 0.981982 | 1 | 0.909149 | 1471 | tags=8% | list=9% | signal=8% |
| REACTOME_ADAPTIVE_IMMUNE_SYSTEM | 623 | -0.26502 | -0.72715 | 0.99 | 1 | 0.909149 | 2899 | tags=16% | list=18% | signal=13% |
| REACTOME_VESICLE_MEDIATED_TRANSPORT | 548 | -0.26367 | -0.72228 | 0.987 | 1 | 0.909149 | 1471 | tags=8% | list=9% | signal=8% |
| REACTOME_CELLULAR_RESPONSE_TO_CHEMICAL_STRESS | 148 | 0.22097 | 0.68828 | 0.981481 | 1 | 0.909149 | 2798 | tags=20% | list=17% | signal=16% |
| REACTOME_SIGNALING_BY_WNT | 219 | -0.24687 | -0.65284 | 0.988482 | 1 | 0.909149 | 1593 | tags=11% | list=10% | signal=10% |
| REACTOME_SIGNALING_BY_INTERLEUKINS | 230 | -0.23967 | -0.63532 | 0.991658 | 1 | 0.909149 | 1500 | tags=5% | list=9% | signal=5% |
| REACTOME_INTRA_GOLGI_AND_RETROGRADE_GOLGI_TO_ER_TRAFFIC | 177 | -0.24137 | -0.6311 | 0.982086 | 1 | 0.909149 | 1360 | tags=7% | list=8% | signal=6% |
| REACTOME_MITOTIC_PROPHASE | 85 | 0.186013 | 0.571866 | 1 | 1 | 0.909149 | 3546 | tags=21% | list=22% | signal=17% |
| REACTOME_INTERLEUKIN_1_FAMILY_SIGNALING | 108 | -0.21747 | -0.55396 | 0.991295 | 1 | 0.909149 | 719 | tags=4% | list=4% | signal=4% |
| REACTOME_THE_CITRIC_ACID_TCA_CYCLE_AND_RESPIRATORY_ELECTRON_TRANSF | 138 | -0.21391 | -0.55345 | 0.998933 | 1 | 0.909149 | 12 | tags=4% | list=0% | signal=4% |
| REACTOME_CLASS_I_MHC_MEDIATED_ANTIGEN_PROCESSING_PRESENTATION | 325 | -0.20311 | -0.54718 | 1 | 1 | 0.909149 | 2728 | tags=13% | list=17% | signal=11% |
| REACTOME_TCF_DEPENDENT_SIGNALING_IN_RESPONSE_TO_WNT | 144 | -0.21059 | -0.54665 | 0.996819 | 1 | 0.909149 | 2658 | tags=10% | list=16% | signal=9% |
| REACTOME_TRANSCRIPTIONAL_REGULATION_BY_RUNX1 | 147 | -0.20892 | -0.54518 | 0.998946 | 1 | 0.909149 | 748 | tags=3% | list=5% | signal=3% |
| REACTOME_CYCLIN_D_ASSOCIATED_EVENTS_IN_G1 | 42 | 0.184734 | 0.532538 | 0.989247 | 1 | 0.909149 | 3752 | tags=24% | list=23% | signal=18% |
| WP_G1_TO_S_CELL_CYCLE_CONTROL | 58 | 0.17729 | 0.52882 | 1 | 1 | 0.909149 | 2435 | tags=16% | list=15% | signal=13% |
| REACTOME_DEPOSITION_OF_NEW_CENPA_CONTAINING_NUCLEOSOMES_AT_THE_ | 46 | 0.178469 | 0.526468 | 0.988571 | 1 | 0.909149 | 13271 | tags=100% | list=82% | signal=18% |
| REACTOME_E3_UBIQUITIN_LIGASES_UBIQUITINATE_TARGET_PROTEINS | 42 | 0.170382 | 0.491164 | 0.994624 | 1 | 0.909149 | 6163 | tags=48% | list=38% | signal=30% |
| REACTOME_RESOLUTION_OF_D_LOOP_STRUCTURES | 31 | -0.21901 | -0.48885 | 0.983418 | 1 | 0.909149 | 641 | tags=19% | list=4% | signal=19% |
| WP_PROTEASOME_DEGRADATION | 47 | 0.162752 | 0.485829 | 1 | 1 | 0.909149 | 2552 | tags=15% | list=16% | signal=13% |
| REACTOME_CONDENSATION_OF_PROPHASE_CHROMOSOMES | 30 | 0.178292 | 0.482792 | 0.991071 | 1 | 0.909149 | 13271 | tags=100% | list=82% | signal=18% |
| BIOCARTA_TGFB_PATHWAY | 17 | -0.23463 | -0.47356 | 0.991354 | 1 | 0.909149 | 5570 | tags=53% | list=35% | signal=35% |
| WP_GLUTATHIONE_METABOLISM | 17 | -0.22817 | -0.46052 | 0.995677 | 1 | 0.909149 | 3389 | tags=24% | list=21% | signal=19% |
| REACTOME_CYCLIN_A_B1_B2_ASSOCIATED_EVENTS_DURING_G2_M_TRANSITION | 25 | 0.176748 | 0.457783 | 1 | 1 | 0.909149 | 13295 | tags=100% | list=82% | signal=18% |
| REACTOME_SYNTHESIS_OF_LIPOXINS_LX | 5 | -0.273 | -0.45055 | 0.981584 | 1 | 0.909149 | 2359 | tags=40% | list=15% | signal=34% |
| REACTOME_ACTIVATION_OF_THE_TFAP2_AP_2_FAMILY_OF_TRANSCRIPTION_FACT | 9 | 0.210784 | 0.448869 | 0.981959 | 1 | 0.909149 | 12743 | tags=100% | list=79% | signal=21% |
| REACTOME_TRISTETRAPROLIN_TTP_ZFP36_BINDS_AND_DESTABILIZES_MRNA | 16 | 0.183475 | 0.445422 | 0.99375 | 1 | 0.909149 | 4959 | tags=38% | list=31% | signal=26% |
| REACTOME_EUKARYOTIC_TRANSLATION_TERMINATION | 6 | 0.226273 | 0.442669 | 0.986577 | 1 | 0.909149 | 5352 | tags=67% | list=33% | signal=45% |
| BIOCARTA_CELLCYCLE_PATHWAY | 22 | -0.20816 | -0.43968 | 0.993243 | 1 | 0.909149 | 5880 | tags=59% | list=36% | signal=38% |
| REACTOME_SYNTHESIS_OF_12_EICOSATETRAENOIC_ACID_DERIVATIVES | 5 | 0.227062 | 0.414561 | 0.986928 | 1 | 0.909149 | 5489 | tags=60% | list=34% | signal=40% |
| REACTOME_DIGESTION_AND_ABSORPTION | 7 | -0.23871 | -0.41394 | 0.987589 | 1 | 0.909149 | 4365 | tags=43% | list=27% | signal=31% |
| REACTOME_DISPLACEMENT_OF_DNA_GLYCOSYLASE_BY_APEX1 | 9 | -0.21779 | -0.40089 | 0.993485 | 1 | 0.909149 | 12631 | tags=100% | list=78% | signal=22% |
| REACTOME_DIGESTION | 5 | -0.24271 | -0.40056 | 0.990792 | 1 | 0.909149 | 4292 | tags=40% | list=27% | signal=29% |
| REACTOME_NUCLEAR_ENVELOPE_NE_REASSEMBLY | 60 | -0.14692 | -0.35554 | 1 | 1 | 0.909149 | 5103 | tags=32% | list=32% | signal=22% |
| WP_MIRNAS_AND_TFS_IN_IPS_CELL_GENERATION | 6 | -0.20258 | -0.34459 | 0.998198 | 1 | 0.909149 | 2922 | tags=33% | list=18% | signal=27% |

**GSEA MNK2KO**

** GSEA mouse MSigDB canonical pathways: m2.cp.v2023.1.Mm.symbols*

| Description | setSize | enrichmen | NES | pvalue | p.adjust | qvalue | rank | leading_ed | core_enrichment | |
|---|---|---|---|---|---|---|---|---|---|---|
| BIOCARTA_COMP_PATHWAY | 12 | 0.957093 | 1.928782 | 0.000128 | 0.106115 | 0.104235 | 2 | tags=8% | list=0% | signal=8% |
| BIOCARTA_CLASSIC_PATHWAY | 9 | 0.97169 | 1.892386 | 0.000105 | 0.106115 | 0.104235 | 2 | tags=11% | list=0% | signal=11% |
| REACTOME_INTEGRIN_CELL_SURFACE_INTERACTIONS | 66 | 0.812892 | 1.898586 | 0.000266 | 0.117568 | 0.115486 | 385 | tags=12% | list=2% | signal=12% |
| BIOCARTA_LECTIN_PATHWAY | 7 | 0.977095 | 1.852214 | 0.000283 | 0.117568 | 0.115486 | 2 | tags=14% | list=0% | signal=14% |
| REACTOME_COMPLEMENT_CASCADE | 28 | 0.877839 | 1.90018 | 0.000393 | 0.130464 | 0.128153 | 2 | tags=4% | list=0% | signal=4% |
| WP_ELECTRON_TRANSPORT_CHAIN | 65 | -0.80998 | -2.12544 | 0.001237 | 0.186755 | 0.183447 | 1024 | tags=72% | list=7% | signal=68% |
| REACTOME_BINDING_AND_UPTAKE_OF_LIGANDS_BY_SCAVENGER_ | 22 | 0.900864 | 1.912186 | 0.000742 | 0.186755 | 0.183447 | 57 | tags=9% | list=0% | signal=9% |
| WP_COMPLEMENT_ACTIVATION_CLASSICAL_PATHWAY | 13 | 0.943318 | 1.908718 | 0.000791 | 0.186755 | 0.183447 | 2 | tags=8% | list=0% | signal=8% |
| REACTOME_INITIAL_TRIGGERING_OF_COMPLEMENT | 14 | 0.92181 | 1.877779 | 0.001193 | 0.186755 | 0.183447 | 2 | tags=7% | list=0% | signal=7% |
| WP_GPCRS_NONODORANT | 169 | 0.678417 | 1.799947 | 0.000991 | 0.186755 | 0.183447 | 1219 | tags=17% | list=8% | signal=16% |
| WP_FOCAL_ADHESION_PI3KAKTMTOR_SIGNALING_PATHWAY | 274 | 0.615001 | 1.70623 | 0.001019 | 0.186755 | 0.183447 | 523 | tags=7% | list=3% | signal=7% |
| REACTOME_RESPIRATORY_ELECTRON_TRANSPORT | 86 | -0.8302 | -2.22419 | 0.001435 | 0.198613 | 0.195095 | 1630 | tags=73% | list=10% | signal=66% |
| REACTOME_RESPIRATORY_ELECTRON_TRANSPORT_ATP_SYNTHESIS_ | 93 | -0.7881 | -2.10852 | 0.001724 | 0.220248 | 0.216347 | 1630 | tags=70% | list=10% | signal=63% |
| WP_OXIDATIVE_PHOSPHORYLATION | 32 | -0.87831 | -2.20414 | 0.002266 | 0.250876 | 0.246432 | 999 | tags=81% | list=6% | signal=76% |
| REACTOME_ACTIVATION_OF_C3_AND_C5 | 5 | 0.976286 | 1.749213 | 0.002208 | 0.250876 | 0.246432 | 2 | tags=20% | list=0% | signal=20% |
| REACTOME_COMPLEX_I_BIOGENESIS | 50 | -0.85187 | -2.19495 | 0.003182 | 0.290983 | 0.285828 | 1630 | tags=78% | list=10% | signal=70% |
| REACTOME_THE_CITRIC_ACID_TCA_CYCLE_AND_RESPIRATORY_ELE( | 137 | -0.76191 | -2.05354 | 0.003329 | 0.290983 | 0.285828 | 1656 | tags=66% | list=11% | signal=60% |
| REACTOME_REGULATION_OF_INSULIN_LIKE_GROWTH_FACTOR_IGF | 87 | 0.724697 | 1.767174 | 0.003144 | 0.290983 | 0.285828 | 1499 | tags=24% | list=10% | signal=22% |
| REACTOME_BUTYROPHILIN_BTN_FAMILY_INTERACTIONS | 3 | 0.98702 | 1.591995 | 0.003023 | 0.290983 | 0.285828 | 62 | tags=33% | list=0% | signal=33% |
| WP_CHOLESTEROL_BIOSYNTHESIS | 15 | -0.92548 | -2.1251 | 0.004094 | 0.304912 | 0.299512 | 353 | tags=53% | list=2% | signal=52% |
| REACTOME_CLEAVAGE_OF_THE_DAMAGED_PYRIMIDINE | 8 | -0.9583 | -1.92845 | 0.003733 | 0.304912 | 0.299512 | 11 | tags=25% | list=0% | signal=25% |
| REACTOME_TRANSLATION | 216 | -0.69354 | -1.91363 | 0.004435 | 0.304912 | 0.299512 | 3298 | tags=69% | list=21% | signal=55% |
| WP_WNT_SIGNALING_IN_KIDNEY_DISEASE | 41 | 0.811981 | 1.812807 | 0.004589 | 0.304912 | 0.299512 | 275 | tags=15% | list=2% | signal=14% |
| WP_MECHANISMS_ASSOCIATED_WITH_PLURIPOTENCY | 256 | 0.593244 | 1.638278 | 0.00446 | 0.304912 | 0.299512 | 891 | tags=12% | list=6% | signal=12% |
| REACTOME_NEURONAL_SYSTEM | 306 | 0.574214 | 1.617885 | 0.004088 | 0.304912 | 0.299512 | 2248 | tags=28% | list=14% | signal=25% |
| REACTOME_NONSENSE_MEDIATED_DECAY_NMD_INDEPENDENT_O | 84 | -0.74437 | -1.9764 | 0.005796 | 0.331994 | 0.326113 | 3443 | tags=83% | list=22% | signal=65% |
| WP_SIDS_SUSCEPTIBILITY_PATHWAYS | 44 | 0.784697 | 1.764039 | 0.005628 | 0.331994 | 0.326113 | 500 | tags=9% | list=3% | signal=9% |
| REACTOME_RELEASE_OF_HH_NP_FROM_THE_SECRETING_CELL | 5 | 0.968308 | 1.734919 | 0.00539 | 0.331994 | 0.326113 | 40 | tags=40% | list=0% | signal=40% |
| BIOCARTA_SLRP_PATHWAY | 4 | 0.961502 | 1.649234 | 0.005718 | 0.331994 | 0.326113 | 274 | tags=50% | list=2% | signal=49% |
| REACTOME_AMINO_ACID_TRANSPORT_ACROSS_THE_PLASMA_MEI | 28 | 0.824371 | 1.784441 | 0.006409 | 0.346671 | 0.340531 | 1588 | tags=46% | list=10% | signal=42% |
| WP_GPCRS_ODORANT | 67 | 0.723903 | 1.701834 | 0.00647 | 0.346671 | 0.340531 | 631 | tags=13% | list=4% | signal=13% |
| REACTOME_BASE_EXCISION_REPAIR_AP_SITE_FORMATION | 10 | -0.93566 | -1.95219 | 0.007881 | 0.409054 | 0.401809 | 11 | tags=20% | list=0% | signal=20% |
| REACTOME_DISPLACEMENT_OF_DNA_GLYCOSYLASE_BY_APEX1 | 9 | -0.93985 | -1.93279 | 0.009191 | 0.462595 | 0.454401 | 11 | tags=22% | list=0% | signal=22% |
| REACTOME_APOPTOSIS_INDUCED_DNA_FRAGMENTATION | 9 | -0.93357 | -1.91989 | 0.010862 | 0.474767 | 0.466358 | 63 | tags=22% | list=0% | signal=22% |
| REACTOME_GRB2_SOS_PROVIDES_LINKAGE_TO_MAPK_SIGNALING_ | 12 | 0.904026 | 1.821839 | 0.010438 | 0.474767 | 0.466358 | 101 | tags=17% | list=1% | signal=17% |
| REACTOME_PLATELET_AGGREGATION_PLUG_FORMATION | 32 | 0.811356 | 1.788314 | 0.010421 | 0.474767 | 0.466358 | 542 | tags=16% | list=3% | signal=15% |
| REACTOME_GABA_RECEPTOR_ACTIVATION | 52 | 0.754712 | 1.742396 | 0.01084 | 0.474767 | 0.466358 | 685 | tags=13% | list=4% | signal=13% |
| REACTOME_SLC_MEDIATED_TRANSMEMBRANE_TRANSPORT | 186 | 0.608101 | 1.621763 | 0.010388 | 0.474767 | 0.466358 | 1997 | tags=27% | list=13% | signal=24% |
| REACTOME_BASE_EXCISION_REPAIR | 37 | -0.81875 | -2.0775 | 0.012031 | 0.50512 | 0.496172 | 11 | tags=8% | list=0% | signal=8% |
| REACTOME_GLYCOSAMINOGLYCAN_METABOLISM | 116 | 0.637568 | 1.623063 | 0.012164 | 0.50512 | 0.496172 | 3973 | tags=54% | list=25% | signal=41% |
| REACTOME_P130CAS_LINKAGE_TO_MAPK_SIGNALING_FOR_INTEGF | 12 | 0.89986 | 1.813442 | 0.012742 | 0.516205 | 0.507061 | 101 | tags=17% | list=1% | signal=17% |
| REACTOME_FORMATION_OF_A_POOL_OF_FREE_40S_SUBUNITS | 91 | -0.75485 | -2.02933 | 0.015113 | 0.528176 | 0.518821 | 3471 | tags=86% | list=22% | signal=67% |
| REACTOME_EUKARYOTIC_TRANSLATION_INITIATION | 109 | -0.70884 | -1.88595 | 0.014568 | 0.528176 | 0.518821 | 3734 | tags=79% | list=24% | signal=61% |
| REACTOME_NOTCH_HLH_TRANSCRIPTION_PATHWAY | 22 | 0.846147 | 1.796042 | 0.015263 | 0.528176 | 0.518821 | 489 | tags=27% | list=3% | signal=26% |
| REACTOME_EICOSANOID_LIGAND_BINDING_RECEPTORS | 10 | 0.915046 | 1.794109 | 0.014391 | 0.528176 | 0.518821 | 20 | tags=10% | list=0% | signal=10% |
| REACTOME_PROSTANOID_LIGAND_RECEPTORS | 6 | 0.951432 | 1.756421 | 0.014147 | 0.528176 | 0.518821 | 20 | tags=17% | list=0% | signal=17% |
| WP_NEURAL_CREST_DIFFERENTIATION | 76 | 0.700408 | 1.668062 | 0.014912 | 0.528176 | 0.518821 | 624 | tags=20% | list=4% | signal=19% |
| REACTOME_CELL_SURFACE_INTERACTIONS_AT_THE_VASCULAR_W/ | 73 | 0.686876 | 1.632208 | 0.01479 | 0.528176 | 0.518821 | 391 | tags=10% | list=2% | signal=9% |
| REACTOME_MITOCHONDRIAL_TRANSLATION | 88 | -0.69185 | -1.8589 | 0.017029 | 0.533668 | 0.524215 | 3229 | tags=69% | list=21% | signal=55% |
| BIOCARTA_GABA_PATHWAY | 10 | 0.911082 | 1.786338 | 0.016956 | 0.533668 | 0.524215 | 4 | tags=10% | list=0% | signal=10% |
| BIOCARTA_NPP1_PATHWAY | 10 | 0.910822 | 1.785828 | 0.016956 | 0.533668 | 0.524215 | 194 | tags=20% | list=1% | signal=20% |
| REACTOME_RUNX3_REGULATES_NOTCH_SIGNALING | 9 | 0.913256 | 1.778586 | 0.016802 | 0.533668 | 0.524215 | 392 | tags=33% | list=2% | signal=33% |
| WP_GPCRS_CLASS_A_RHODOPSINLIKE | 101 | 0.661371 | 1.644155 | 0.016482 | 0.533668 | 0.524215 | 2541 | tags=35% | list=16% | signal=29% |
| REACTOME_TP53_REGULATES_METABOLIC_GENES | 69 | -0.72722 | -1.89958 | 0.019031 | 0.556197 | 0.546345 | 2418 | tags=54% | list=15% | signal=46% |
| REACTOME_NONSENSE_MEDIATED_DECAY_NMD | 104 | -0.67674 | -1.82415 | 0.018503 | 0.556197 | 0.546345 | 3443 | tags=73% | list=22% | signal=57% |
| REACTOME_CELLULAR_HEXOSE_TRANSPORT | 11 | 0.913428 | 1.808328 | 0.019495 | 0.556197 | 0.546345 | 59 | tags=18% | list=0% | signal=18% |
| REACTOME_REGULATION_BY_C_FLIP | 9 | 0.911208 | 1.774597 | 0.019139 | 0.556197 | 0.546345 | 37 | tags=11% | list=0% | signal=11% |
| WP_SYNTHESIS_AND_DEGRADATION_OF_KETONE_BODIES | 5 | -0.9252 | -1.69569 | 0.020018 | 0.556197 | 0.546345 | 1071 | tags=60% | list=7% | signal=56% |
| REACTOME_TRAIL_SIGNALING | 5 | 0.944455 | 1.692181 | 0.018812 | 0.556197 | 0.546345 | 37 | tags=20% | list=0% | signal=20% |
| WP_ESC_PLURIPOTENCY_PATHWAYS | 105 | 0.648158 | 1.629273 | 0.020091 | 0.556197 | 0.546345 | 937 | tags=16% | list=6% | signal=15% |
| REACTOME_DNA_DAMAGE_TELOMERE_STRESS_INDUCED_SENESCE | 24 | -0.82608 | -1.97894 | 0.022592 | 0.562561 | 0.552596 | 63 | tags=17% | list=0% | signal=17% |
| WP_CYTOPLASMIC_RIBOSOMAL_PROTEINS | 77 | -0.73574 | -1.9484 | 0.021345 | 0.562561 | 0.552596 | 3441 | tags=90% | list=22% | signal=70% |
| REACTOME_ACTIVATION_OF_THE_MRNA_UPON_BINDING_OF_THE_ | 58 | -0.69557 | -1.79868 | 0.023031 | 0.562561 | 0.552596 | 2672 | tags=62% | list=17% | signal=52% |
| REACTOME_SYNTHESIS_OF_KETONE_BODIES | 8 | -0.87592 | -1.76267 | 0.022844 | 0.562561 | 0.552596 | 114 | tags=38% | list=1% | signal=37% |
| REACTOME_KERATAN_SULFATE_KERATIN_METABOLISM | 32 | 0.782512 | 1.724739 | 0.022246 | 0.562561 | 0.552596 | 2132 | tags=50% | list=14% | signal=43% |
| REACTOME_PHENYLALANINE_METABOLISM | 5 | -0.92133 | -1.68861 | 0.021523 | 0.562561 | 0.552596 | 954 | tags=40% | list=6% | signal=38% |
| REACTOME_PRE_NOTCH_PROCESSING_IN_GOLGI | 5 | 0.941803 | 1.687429 | 0.021478 | 0.562561 | 0.552596 | 310 | tags=60% | list=2% | signal=59% |
| REACTOME_EXTRACELLULAR_MATRIX_ORGANIZATION | 208 | 0.576642 | 1.564587 | 0.02274 | 0.562561 | 0.552596 | 1345 | tags=20% | list=9% | signal=18% |
| REACTOME_MITOCHONDRIAL_FATTY_ACID_BETA_OXIDATION | 34 | -0.77862 | -1.95192 | 0.023833 | 0.566158 | 0.55613 | 2282 | tags=85% | list=15% | signal=73% |
| WP_TGFBETA_SIGNALING_PATHWAY | 46 | 0.731008 | 1.645989 | 0.02386 | 0.566158 | 0.55613 | 392 | tags=11% | list=2% | signal=11% |
| REACTOME_CHEMOKINE_RECEPTORS_BIND_CHEMOKINES | 19 | -0.84764 | -1.96585 | 0.046435 | 0.572489 | 0.562349 | 14 | tags=11% | list=0% | signal=11% |
| REACTOME_CHOLESTEROL_BIOSYNTHESIS | 26 | -0.80706 | -1.95343 | 0.035682 | 0.572489 | 0.562349 | 560 | tags=35% | list=4% | signal=33% |
| REACTOME_RECYCLING_PATHWAY_OF_L1 | 32 | -0.76484 | -1.91939 | 0.035562 | 0.572489 | 0.562349 | 3130 | tags=75% | list=20% | signal=60% |
| WP_TCA_CYCLE | 30 | -0.75856 | -1.90438 | 0.03108 | 0.572489 | 0.562349 | 2331 | tags=77% | list=15% | signal=65% |
| REACTOME_GLYOXYLATE_METABOLISM_AND_GLYCINE_DEGRADATI | 24 | -0.77854 | -1.86504 | 0.033302 | 0.572489 | 0.562349 | 1986 | tags=67% | list=13% | signal=58% |
| REACTOME_AKT_PHOSPHORYLATES_TARGETS_IN_THE_CYTOSOL | 11 | -0.87032 | -1.86204 | 0.055603 | 0.572489 | 0.562349 | 77 | tags=36% | list=0% | signal=36% |
| REACTOME_FORMATION_OF_SENESCENCE_ASSOCIATED_HETEROCH | 11 | -0.86518 | -1.85103 | 0.062437 | 0.572489 | 0.562349 | 63 | tags=27% | list=0% | signal=27% |
| REACTOME_THE_ROLE_OF_GTSE1_IN_G2_M_PROGRESSION_AFTER_ | 69 | -0.70624 | -1.84478 | 0.032597 | 0.572489 | 0.562349 | 3105 | tags=61% | list=20% | signal=49% |
| WP_CHOLESTEROL_METABOLISM_WITH_BLOCH_AND_KANDUTSCH | 47 | -0.71317 | -1.83491 | 0.061611 | 0.572489 | 0.562349 | 560 | tags=26% | list=4% | signal=25% |
| WP_AMINO_ACID_METABOLISM | 76 | -0.68836 | -1.82817 | 0.038276 | 0.572489 | 0.562349 | 1565 | tags=41% | list=10% | signal=37% |
| REACTOME_TRANSPORT_OF_CONNEXONS_TO_THE_PLASMA_MEM | 13 | -0.82148 | -1.81921 | 0.083938 | 0.572489 | 0.562349 | 1568 | tags=62% | list=10% | signal=55% |
| REACTOME_SEALING_OF_THE_NUCLEAR_ENVELOPE_NE_BY_ESCRT_ | 21 | -0.76068 | -1.79647 | 0.057203 | 0.572489 | 0.562349 | 1568 | tags=57% | list=10% | signal=52% |
| BIOCARTA_EGFR_SMRTE_PATHWAY | 11 | 0.90569 | 1.793008 | 0.027871 | 0.572489 | 0.562349 | 229 | tags=27% | list=1% | signal=27% |
| REACTOME_INTRAFLAGELLAR_TRANSPORT | 44 | -0.70381 | -1.78781 | 0.067797 | 0.572489 | 0.562349 | 1994 | tags=45% | list=13% | signal=40% |
| REACTOME_GAP_JUNCTION_TRAFFICKING_AND_REGULATION | 32 | -0.71208 | -1.78697 | 0.049646 | 0.572489 | 0.562349 | 1662 | tags=47% | list=11% | signal=42% |
| BIOCARTA_RECK_PATHWAY | 9 | -0.86671 | -1.78237 | 0.032217 | 0.572489 | 0.562349 | 589 | tags=33% | list=4% | signal=32% |
| REACTOME_RAB_GERANYLGERANYLATION | 57 | -0.69306 | -1.78119 | 0.061576 | 0.572489 | 0.562349 | 2548 | tags=33% | list=16% | signal=28% |

| | | | | | | | | | | |
|---|---|---|---|---|---|---|---|---|---|---|
| REACTOME_E3_UBIQUITIN_LIGASES_UBIQUITINATE_TARGET_PROTI | 41 | -0.70123 | -1.76968 | 0.070588 | 0.572489 | 0.562349 | 1068 | tags=22% | list=7% | signal=21% |
| REACTOME_ELEVATION_OF_CYTOSOLIC_CA2_LEVELS | 12 | 0.870835 | 1.75495 | 0.034024 | 0.572489 | 0.562349 | 55 | tags=8% | list=0% | signal=8% |
| REACTOME_INTEGRIN_SIGNALING | 24 | 0.818442 | 1.750521 | 0.028444 | 0.572489 | 0.562349 | 101 | tags=8% | list=1% | signal=8% |
| REACTOME_EPIGENETIC_REGULATION_OF_GENE_EXPRESSION | 63 | -0.66501 | -1.74865 | 0.06326 | 0.572489 | 0.562349 | 1084 | tags=19% | list=7% | signal=18% |
| REACTOME_GAP_JUNCTION_DEGRADATION | 11 | -0.81664 | -1.74719 | 0.091777 | 0.572489 | 0.562349 | 931 | tags=73% | list=6% | signal=68% |
| REACTOME_ACTIVATION_OF_MATRIX_METALLOPROTEINASES | 16 | -0.75497 | -1.74661 | 0.059917 | 0.572489 | 0.562349 | 29 | tags=12% | list=0% | signal=12% |
| BIOCARTA_HES_PATHWAY | 10 | 0.886998 | 1.739117 | 0.037515 | 0.572489 | 0.562349 | 465 | tags=30% | list=3% | signal=29% |
| BIOCARTA_AMAN_PATHWAY | 7 | 0.916151 | 1.736686 | 0.028219 | 0.572489 | 0.562349 | 1221 | tags=86% | list=8% | signal=79% |
| REACTOME_KINESINS | 50 | -0.67318 | -1.73454 | 0.070048 | 0.572489 | 0.562349 | 1966 | tags=38% | list=13% | signal=33% |
| WP_MATRIX_METALLOPROTEINASES | 19 | -0.74444 | -1.72651 | 0.062635 | 0.572489 | 0.562349 | 1089 | tags=16% | list=7% | signal=15% |
| REACTOME_KETONE_BODY_METABOLISM | 9 | -0.83944 | -1.72631 | 0.05102 | 0.572489 | 0.562349 | 1692 | tags=44% | list=11% | signal=40% |
| REACTOME_GP1B_IX_V_ACTIVATION_SIGNALLING | 11 | 0.871466 | 1.725254 | 0.052124 | 0.572489 | 0.562349 | 364 | tags=18% | list=2% | signal=18% |
| BIOCARTA_P53_PATHWAY | 15 | -0.75066 | -1.72368 | 0.066116 | 0.572489 | 0.562349 | 523 | tags=40% | list=3% | signal=39% |
| REACTOME_PRE_NOTCH_EXPRESSION_AND_PROCESSING | 6 | 0.933506 | 1.723328 | 0.03429 | 0.572489 | 0.562349 | 310 | tags=50% | list=2% | signal=49% |
| REACTOME_PYRUVATE_METABOLISM_AND_CITRIC_ACID_TCA_CYCL | 44 | -0.67775 | -1.7216 | 0.077482 | 0.572489 | 0.562349 | 1656 | tags=59% | list=11% | signal=53% |
| REACTOME_G_PROTEIN_BETA_GAMMA_SIGNALLING | 30 | -0.68564 | -1.72131 | 0.068075 | 0.572489 | 0.562349 | 3418 | tags=63% | list=22% | signal=50% |
| REACTOME_CHAPERONIN_MEDIATED_PROTEIN_FOLDING | 35 | -0.68687 | -1.72088 | 0.071759 | 0.572489 | 0.562349 | 3418 | tags=71% | list=22% | signal=56% |
| REACTOME_CITRIC_ACID_CYCLE_TCA_CYCLE | 21 | -0.72727 | -1.71758 | 0.067797 | 0.572489 | 0.562349 | 1092 | tags=95% | list=7% | signal=89% |
| REACTOME_PEROXISOMAL_PROTEIN_IMPORT | 59 | -0.65936 | -1.71552 | 0.068966 | 0.572489 | 0.562349 | 2634 | tags=41% | list=17% | signal=34% |
| REACTOME_PINK1_PRKN_MEDIATED_MITOPHAGY | 21 | -0.72598 | -1.71453 | 0.067797 | 0.572489 | 0.562349 | 1234 | tags=52% | list=8% | signal=48% |
| WP_GPCRS_SMALL_LIGAND | 13 | 0.846463 | 1.712741 | 0.062016 | 0.572489 | 0.562349 | 20 | tags=8% | list=0% | signal=8% |
| BIOCARTA_PS1_PATHWAY | 12 | 0.847552 | 1.70803 | 0.067698 | 0.572489 | 0.562349 | 131 | tags=17% | list=1% | signal=17% |
| REACTOME_SCF_SKP2_MEDIATED_DEGRADATION_OF_P27_P21 | 59 | -0.65644 | -1.70791 | 0.068966 | 0.572489 | 0.562349 | 3335 | tags=59% | list=21% | signal=47% |
| REACTOME_ADENYLATE_CYCLASE_ACTIVATING_PATHWAY | 10 | 0.870755 | 1.707269 | 0.060952 | 0.572489 | 0.562349 | 645 | tags=30% | list=4% | signal=29% |
| REACTOME_CASPASE_ACTIVATION_VIA_DEATH_RECEPTORS_IN_THI | 14 | 0.834567 | 1.70006 | 0.060665 | 0.572489 | 0.562349 | 37 | tags=7% | list=0% | signal=7% |
| REACTOME_PROTEIN_LOCALIZATION | 93 | -0.63509 | -1.69915 | 0.028546 | 0.572489 | 0.562349 | 2873 | tags=44% | list=18% | signal=36% |
| WP_OMEGA3_OMEGA6_FATTY_ACID_SYNTHESIS | 15 | -0.73939 | -1.69779 | 0.076446 | 0.572489 | 0.562349 | 1368 | tags=27% | list=9% | signal=24% |
| REACTOME_KERATAN_SULFATE_BIOSYNTHESIS | 26 | 0.786486 | 1.696954 | 0.039439 | 0.572489 | 0.562349 | 2132 | tags=50% | list=14% | signal=43% |
| REACTOME_CONDENSATION_OF_PROPHASE_CHROMOSOMES | 27 | -0.68935 | -1.69427 | 0.071429 | 0.572489 | 0.562349 | 2028 | tags=37% | list=13% | signal=32% |
| WP_SPHINGOLIPID_METABOLISM_INTEGRATED_PATHWAY | 25 | 0.787419 | 1.693123 | 0.050204 | 0.572489 | 0.562349 | 548 | tags=20% | list=3% | signal=19% |
| BIOCARTA_HSP27_PATHWAY | 12 | 0.840076 | 1.692963 | 0.073501 | 0.572489 | 0.562349 | 43 | tags=8% | list=0% | signal=8% |
| WP_GDNFRET_SIGNALING_AXIS | 18 | 0.825016 | 1.69172 | 0.049149 | 0.572489 | 0.562349 | 1421 | tags=50% | list=9% | signal=46% |
| REACTOME_UBIQUITIN_MEDIATED_DEGRADATION_OF_PHOSPHORY | 50 | -0.65619 | -1.69076 | 0.07971 | 0.572489 | 0.562349 | 3335 | tags=62% | list=21% | signal=49% |
| REACTOME_PLATELET_ADHESION_TO_EXPOSED_COLLAGEN | 9 | 0.867966 | 1.690382 | 0.070313 | 0.572489 | 0.562349 | 364 | tags=22% | list=2% | signal=22% |
| REACTOME_ADENYLATE_CYCLASE_INHIBITORY_PATHWAY | 13 | 0.834883 | 1.68931 | 0.077519 | 0.572489 | 0.562349 | 645 | tags=23% | list=4% | signal=22% |
| WP_ACE_INHIBITOR_PATHWAY | 6 | 0.914494 | 1.68823 | 0.058321 | 0.572489 | 0.562349 | 541 | tags=33% | list=3% | signal=32% |
| REACTOME_PROSTACYCLIN_SIGNALLING_THROUGH_PROSTACYCLIN | 16 | -0.72972 | -1.6882 | 0.078512 | 0.572489 | 0.562349 | 3418 | tags=81% | list=22% | signal=64% |
| REACTOME_DOWNREGULATION_OF_ERBB2_ERBB3_SIGNALING | 12 | -0.7817 | -1.6866 | 0.065979 | 0.572489 | 0.562349 | 676 | tags=42% | list=4% | signal=40% |
| REACTOME_PHENYLALANINE_AND_TYROSINE_METABOLISM | 7 | -0.85036 | -1.68637 | 0.038712 | 0.572489 | 0.562349 | 954 | tags=43% | list=6% | signal=40% |
| REACTOME_SPHINGOLIPID_DE_NOVO_BIOSYNTHESIS | 33 | 0.764651 | 1.686266 | 0.033207 | 0.572489 | 0.562349 | 548 | tags=18% | list=3% | signal=18% |
| REACTOME_G_BETA_GAMMA_SIGNALLING_THROUGH_CDC42 | 18 | -0.73011 | -1.68615 | 0.067653 | 0.572489 | 0.562349 | 3418 | tags=78% | list=22% | signal=61% |
| BIOCARTA_NEUTROPHIL_PATHWAY | 7 | 0.887225 | 1.681854 | 0.063872 | 0.572489 | 0.562349 | 102 | tags=14% | list=1% | signal=14% |
| REACTOME_APC_C_CDH1_MEDIATED_DEGRADATION_OF_CDC20_A | 72 | -0.64148 | -1.67937 | 0.066838 | 0.572489 | 0.562349 | 3554 | tags=60% | list=23% | signal=46% |
| REACTOME_TP53_REGULATES_TRANSCRIPTION_OF_GENES_INVOLV | 10 | -0.8048 | -1.67915 | 0.0587 | 0.572489 | 0.562349 | 41 | tags=20% | list=0% | signal=20% |
| REACTOME_DEGRADATION_OF_CYSTEINE_AND_HOMOCYSTEINE | 13 | -0.75799 | -1.67861 | 0.063786 | 0.572489 | 0.562349 | 1199 | tags=31% | list=8% | signal=28% |
| REACTOME_APOPTOTIC_CLEAVAGE_OF_CELL_ADHESION_PROTEINS | 7 | 0.88456 | 1.676801 | 0.065868 | 0.572489 | 0.562349 | 18 | tags=14% | list=0% | signal=14% |
| WP_SPHINGOLIPID_METABOLISM_OVERVIEW | 24 | 0.783317 | 1.675393 | 0.045045 | 0.572489 | 0.562349 | 548 | tags=17% | list=3% | signal=16% |
| WP_SIGNAL_TRANSDUCTION_OF_S1P_RECEPTOR | 21 | -0.70935 | -1.67526 | 0.072034 | 0.572489 | 0.562349 | 941 | tags=19% | list=6% | signal=18% |
| REACTOME_CA2_ACTIVATED_K_CHANNELS | 6 | 0.907336 | 1.675015 | 0.052632 | 0.572489 | 0.562349 | 72 | tags=17% | list=0% | signal=17% |
| REACTOME_ACTIVATION_OF_APC_C_AND_APC_C_CDC20_MEDIATE | 75 | -0.63361 | -1.67433 | 0.069948 | 0.572489 | 0.562349 | 3348 | tags=55% | list=21% | signal=43% |
| REACTOME_DECTIN_1_MEDIATED_NONCANONICAL_NF_KB_SIGNAL | 56 | -0.65317 | -1.67344 | 0.079012 | 0.572489 | 0.562349 | 3335 | tags=62% | list=21% | signal=49% |
| REACTOME_FORMATION_OF_THE_CORNIFIED_ENVELOPE | 25 | 0.778068 | 1.673019 | 0.045936 | 0.572489 | 0.562349 | 160 | tags=8% | list=1% | signal=8% |
| REACTOME_KERATINIZATION | 25 | 0.778068 | 1.673019 | 0.045936 | 0.572489 | 0.562349 | 160 | tags=8% | list=1% | signal=8% |
| REACTOME_BRANCHED_CHAIN_AMINO_ACID_CATABOLISM | 20 | -0.7117 | -1.67161 | 0.076433 | 0.572489 | 0.562349 | 2280 | tags=55% | list=15% | signal=47% |
| REACTOME_REGULATION_OF_GENE_EXPRESSION_BY_HYPOXIA_IND | 6 | 0.905398 | 1.671437 | 0.052632 | 0.572489 | 0.562349 | 392 | tags=33% | list=2% | signal=33% |
| REACTOME_MITOCHONDRIAL_FATTY_ACID_BETA_OXIDATION_OF_S | 9 | -0.81103 | -1.66789 | 0.061224 | 0.572489 | 0.562349 | 2976 | tags=100% | list=19% | signal=81% |
| BIOCARTA_BLYMPHOCYTE_PATHWAY | 7 | 0.879666 | 1.667524 | 0.067864 | 0.572489 | 0.562349 | 102 | tags=14% | list=1% | signal=14% |
| REACTOME_ASSEMBLY_OF_ACTIVE_LPL_AND_LIPC_LIPASE_COMPLE | 7 | -0.83963 | -1.66509 | 0.039912 | 0.572489 | 0.562349 | 46 | tags=29% | list=0% | signal=29% |
| REACTOME_ACYL_CHAIN_REMODELLING_OF_PI | 8 | -0.82661 | -1.66344 | 0.051896 | 0.572489 | 0.562349 | 264 | tags=38% | list=2% | signal=37% |
| BIOCARTA_FREE_PATHWAY | 7 | 0.877391 | 1.663212 | 0.067864 | 0.572489 | 0.562349 | 62 | tags=14% | list=0% | signal=14% |
| WP_EICOSANOID_LIPID_SYNTHESIS_MAP | 9 | -0.80827 | -1.66221 | 0.061224 | 0.572489 | 0.562349 | 48 | tags=22% | list=0% | signal=22% |
| REACTOME_GLUCONEOGENESIS | 27 | -0.67598 | -1.66141 | 0.082949 | 0.572489 | 0.562349 | 2821 | tags=59% | list=18% | signal=49% |
| REACTOME_KERATAN_SULFATE_DEGRADATION | 12 | 0.822761 | 1.658068 | 0.087041 | 0.572489 | 0.562349 | 2033 | tags=50% | list=13% | signal=44% |
| REACTOME_BIOTIN_TRANSPORT_AND_METABOLISM | 11 | -0.77426 | -1.65651 | 0.07438 | 0.572489 | 0.562349 | 1160 | tags=64% | list=7% | signal=59% |
| BIOCARTA_LYMPHOCYTE_PATHWAY | 8 | 0.849515 | 1.655853 | 0.083832 | 0.572489 | 0.562349 | 102 | tags=12% | list=1% | signal=12% |
| REACTOME_SULFUR_AMINO_ACID_METABOLISM | 21 | -0.70055 | -1.65448 | 0.074153 | 0.572489 | 0.562349 | 1199 | tags=29% | list=8% | signal=26% |
| WP_PTF1A_RELATED_REGULATORY_PATHWAY | 8 | 0.84732 | 1.651574 | 0.087824 | 0.572489 | 0.562349 | 124 | tags=12% | list=1% | signal=12% |
| REACTOME_RAB_GEFS_EXCHANGE_GTP_FOR_GDP_ON_RABS | 86 | -0.61638 | -1.65135 | 0.059556 | 0.572489 | 0.562349 | 1439 | tags=17% | list=9% | signal=16% |
| REACTOME_REGULATION_OF_PTEN_STABILITY_AND_ACTIVITY | 67 | -0.63147 | -1.65115 | 0.072864 | 0.572489 | 0.562349 | 3335 | tags=55% | list=21% | signal=44% |
| REACTOME_DETOXIFICATION_OF_REACTIVE_OXYGEN_SPECIES | 30 | -0.65759 | -1.65089 | 0.08216 | 0.572489 | 0.562349 | 1920 | tags=53% | list=12% | signal=47% |
| WP_HFE_EFFECT_ON_HEPCIDIN_PRODUCTION | 5 | 0.92126 | 1.650622 | 0.031856 | 0.572489 | 0.562349 | 488 | tags=40% | list=3% | signal=39% |
| REACTOME_CASPASE_ACTIVATION_VIA_EXTRINSIC_APOPTOTIC_SIG | 18 | 0.804381 | 1.649407 | 0.064272 | 0.572489 | 0.562349 | 37 | tags=6% | list=0% | signal=6% |
| REACTOME_ECM_PROTEOGLYCANS | 42 | 0.732736 | 1.6487 | 0.025403 | 0.572489 | 0.562349 | 607 | tags=14% | list=4% | signal=14% |
| WP_FATTY_ACID_BETAOXIDATION | 30 | -0.65611 | -1.64718 | 0.08216 | 0.572489 | 0.562349 | 2976 | tags=53% | list=19% | signal=43% |
| REACTOME_GLI3_IS_PROCESSED_TO_GLI3R_BY_THE_PROTEASOME | 56 | -0.64267 | -1.64653 | 0.08642 | 0.572489 | 0.562349 | 3355 | tags=62% | list=21% | signal=49% |
| BIOCARTA_CTL_PATHWAY | 7 | 0.86817 | 1.645732 | 0.075848 | 0.572489 | 0.562349 | 102 | tags=14% | list=1% | signal=14% |
| WP_BLOOD_CLOTTING_CASCADE | 9 | 0.843383 | 1.642506 | 0.089844 | 0.572489 | 0.562349 | 73 | tags=11% | list=0% | signal=11% |
| REACTOME_NA_CL_DEPENDENT_NEUROTRANSMITTER_TRANSPORT | 13 | 0.809423 | 1.637793 | 0.100775 | 0.572489 | 0.562349 | 262 | tags=23% | list=2% | signal=23% |
| REACTOME_AQUAPORIN_MEDIATED_TRANSPORT | 29 | -0.66028 | -1.6366 | 0.084507 | 0.572489 | 0.562349 | 3927 | tags=69% | list=25% | signal=52% |
| BIOCARTA_PLCD_PATHWAY | 5 | 0.913161 | 1.636112 | 0.053465 | 0.572489 | 0.562349 | 136 | tags=20% | list=1% | signal=20% |
| BIOCARTA_MONOCYTE_PATHWAY | 9 | 0.839313 | 1.634581 | 0.09375 | 0.572489 | 0.562349 | 102 | tags=11% | list=1% | signal=11% |
| REACTOME_G_BETA_GAMMA_SIGNALLING_THROUGH_PI3KGAMM/ | 23 | -0.68396 | -1.63198 | 0.0837 | 0.572489 | 0.562349 | 3418 | tags=61% | list=22% | signal=48% |
| BIOCARTA_DEATH_PATHWAY | 30 | 0.74464 | 1.631006 | 0.057292 | 0.572489 | 0.562349 | 37 | tags=3% | list=0% | signal=3% |
| REACTOME_IMMUNOREGULATORY_INTERACTIONS_BETWEEN_A_LY | 62 | 0.701645 | 1.628284 | 0.026577 | 0.572489 | 0.562349 | 604 | tags=8% | list=4% | signal=8% |
| WP_WNT_SIGNALING_PATHWAY_WP403 | 57 | 0.704318 | 1.62781 | 0.026487 | 0.572489 | 0.562349 | 275 | tags=9% | list=2% | signal=9% |
| REACTOME_TERMINATION_OF_O_GLYCAN_BIOSYNTHESIS | 10 | 0.830065 | 1.627489 | 0.104762 | 0.572489 | 0.562349 | 1438 | tags=40% | list=9% | signal=36% |
| BIOCARTA_TCYTOTOXIC_PATHWAY | 5 | 0.905765 | 1.62286 | 0.057426 | 0.572489 | 0.562349 | 102 | tags=20% | list=1% | signal=20% |
| REACTOME_MAJOR_PATHWAY_OF_RRNA_PROCESSING_IN_THE_NU | 167 | -0.59809 | -1.62272 | 0.038645 | 0.572489 | 0.562349 | 3339 | tags=55% | list=21% | signal=44% |
| REACTOME_CDK_MEDIATED_PHOSPHORYLATION_AND_REMOVAL_( | 71 | -0.62118 | -1.61923 | 0.079897 | 0.572489 | 0.562349 | 3554 | tags=58% | list=23% | signal=45% |
| REACTOME_G1_S_DNA_DAMAGE_CHECKPOINTS | 65 | -0.61687 | -1.6187 | 0.089109 | 0.572489 | 0.562349 | 3335 | tags=52% | list=21% | signal=41% |

| | | | | | | | | | |
|---|---|---|---|---|---|---|---|---|---|
| BIOCARTA_MELANOCYTE_PATHWAY | 7 | 0.853207 | 1.617367 | 0.083832 | 0.572489 | 0.562349 | 1291 tags=43% | list=8% | signal=39% |
| REACTOME_CTLA4_INHIBITORY_SIGNALING | 20 | -0.68835 | -1.61678 | 0.091295 | 0.572489 | 0.562349 | 1889 tags=50% | list=12% | signal=44% |
| REACTOME_STABILIZATION_OF_P53 | 54 | -0.62675 | -1.61625 | 0.089372 | 0.572489 | 0.562349 | 3335 tags=59% | list=21% | signal=47% |
| REACTOME_GAP_JUNCTION_ASSEMBLY | 20 | -0.68788 | -1.61565 | 0.091295 | 0.572489 | 0.562349 | 1568 tags=40% | list=10% | signal=36% |
| REACTOME_APOPTOTIC_CLEAVAGE_OF_CELLULAR_PROTEINS | 34 | 0.73258 | 1.614404 | 0.059441 | 0.572489 | 0.562349 | 97 tags=6% | list=1% | signal=6% |
| BIOCARTA_THELPER_PATHWAY | 6 | 0.874171 | 1.613791 | 0.072125 | 0.572489 | 0.562349 | 102 tags=17% | list=1% | signal=17% |
| REACTOME_REGULATION_OF_RUNX3_EXPRESSION_AND_ACTIVITY | 52 | -0.63331 | -1.61344 | 0.088235 | 0.572489 | 0.562349 | 3335 tags=65% | list=21% | signal=52% |
| REACTOME_N_GLYCAN_ANTENNAE_ELONGATION_IN_THE_MEDIAL | 23 | 0.757781 | 1.612663 | 0.069343 | 0.572489 | 0.562349 | 2916 tags=70% | list=19% | signal=57% |
| REACTOME_MAP2K_AND_MAPK_ACTIVATION | 35 | 0.731445 | 1.612245 | 0.054386 | 0.572489 | 0.562349 | 325 tags=14% | list=2% | signal=14% |
| REACTOME_KEAP1_NFE2L2_PATHWAY | 74 | -0.61012 | -1.61007 | 0.083117 | 0.572489 | 0.562349 | 3355 tags=51% | list=21% | signal=41% |
| BIOCARTA_EEA1_PATHWAY | 5 | 0.898353 | 1.60958 | 0.063366 | 0.572489 | 0.562349 | 140 tags=20% | list=1% | signal=20% |
| REACTOME_EICOSANOIDS | 6 | 0.871644 | 1.609125 | 0.072125 | 0.572489 | 0.562349 | 180 tags=17% | list=1% | signal=16% |
| REACTOME_N_GLYCAN_ANTENNAE_ELONGATION | 15 | 0.787468 | 1.607994 | 0.108108 | 0.572489 | 0.562349 | 2019 tags=60% | list=13% | signal=52% |
| REACTOME_SUMOYLATION_OF_DNA_DAMAGE_RESPONSE_AND_RE | 71 | -0.61674 | -1.60766 | 0.079897 | 0.572489 | 0.562349 | 11 tags=4% | list=0% | signal=4% |
| WP_BURN_WOUND_HEALING | 16 | 0.788455 | 1.60587 | 0.096525 | 0.572489 | 0.562349 | 65 tags=6% | list=0% | signal=6% |
| WP_GLYCOGEN_METABOLISM | 32 | -0.63938 | -1.60453 | 0.084309 | 0.572489 | 0.562349 | 2935 tags=59% | list=19% | signal=48% |
| REACTOME_VEGF_LIGAND_RECEPTOR_INTERACTIONS | 8 | 0.821721 | 1.601676 | 0.105788 | 0.572489 | 0.562349 | 79 tags=12% | list=1% | signal=12% |
| REACTOME_APC_C_MEDIATED_DEGRADATION_OF_CELL_CYCLE_PR | 85 | -0.60163 | -1.5983 | 0.075419 | 0.572489 | 0.562349 | 3554 tags=53% | list=23% | signal=41% |
| WP_DELTANOTCH_SIGNALING_PATHWAY | 80 | 0.6659 | 1.597474 | 0.036324 | 0.572489 | 0.562349 | 2978 tags=40% | list=19% | signal=33% |
| REACTOME_VASOPRESSIN_REGULATES_RENAL_WATER_HOMEOSTA | 26 | -0.6597 | -1.59676 | 0.089862 | 0.572489 | 0.562349 | 3927 tags=73% | list=25% | signal=55% |
| WP_GENE_REGULATORY_NETWORK_MODELLING_SOMITOGENESIS | 8 | 0.818972 | 1.596318 | 0.10978 | 0.572489 | 0.562349 | 1720 tags=50% | list=11% | signal=45% |
| REACTOME_SYNDECAN_INTERACTIONS | 13 | 0.788598 | 1.595657 | 0.125969 | 0.572489 | 0.562349 | 179 tags=15% | list=1% | signal=15% |
| REACTOME_INTRINSIC_PATHWAY_OF_FIBRIN_CLOT_FORMATION | 14 | 0.783245 | 1.595515 | 0.123288 | 0.572489 | 0.562349 | 781 tags=14% | list=5% | signal=14% |
| REACTOME_ACTIVATION_OF_NF_KAPPAB_IN_B_CELLS | 62 | -0.6088 | -1.59504 | 0.093137 | 0.572489 | 0.562349 | 3335 tags=56% | list=21% | signal=45% |
| WP_PENTOSE_PHOSPHATE_PATHWAY | 7 | -0.8043 | -1.59502 | 0.057884 | 0.572489 | 0.562349 | 2564 tags=86% | list=16% | signal=72% |
| REACTOME_FATTY_ACIDS | 6 | 0.863843 | 1.594724 | 0.081871 | 0.572489 | 0.562349 | 1177 tags=50% | list=7% | signal=46% |
| REACTOME_O_LINKED_GLYCOSYLATION_OF_MUCINS | 41 | 0.71388 | 1.59379 | 0.057192 | 0.572489 | 0.562349 | 3936 tags=71% | list=25% | signal=53% |
| REACTOME_AMINE_LIGAND_BINDING_RECEPTORS | 30 | 0.727138 | 1.592671 | 0.065972 | 0.572489 | 0.562349 | 2310 tags=47% | list=15% | signal=40% |
| BIOCARTA_VITCB_PATHWAY | 10 | 0.811357 | 1.59081 | 0.118095 | 0.572489 | 0.562349 | 59 tags=10% | list=0% | signal=10% |
| REACTOME_PROPIONYL_COA_CATABOLISM | 5 | -0.86783 | -1.59054 | 0.050302 | 0.572489 | 0.562349 | 1389 tags=80% | list=9% | signal=73% |
| REACTOME_EPH_EPHRIN_MEDIATED_REPULSION_OF_CELLS | 34 | 0.72149 | 1.589965 | 0.068182 | 0.572489 | 0.562349 | 1720 tags=38% | list=11% | signal=34% |
| REACTOME_PURINE_CATABOLISM | 17 | 0.77981 | 1.586104 | 0.101727 | 0.572489 | 0.562349 | 62 tags=6% | list=0% | signal=6% |
| REACTOME_NUCLEAR_EVENTS_MEDIATED_BY_NFE2L2 | 53 | -0.61759 | -1.58537 | 0.093976 | 0.572489 | 0.562349 | 3355 tags=66% | list=21% | signal=52% |
| WP_PROTEASOME_DEGRADATION | 47 | -0.61606 | -1.58507 | 0.099526 | 0.572489 | 0.562349 | 3292 tags=57% | list=21% | signal=46% |
| WP_FAS_PATHWAY_AND_STRESS_INDUCTION_OF_HSP_REGULATIO | 34 | 0.718904 | 1.584265 | 0.06993 | 0.572489 | 0.562349 | 490 tags=9% | list=3% | signal=9% |
| BIOCARTA_PGC1A_PATHWAY | 13 | 0.782509 | 1.583335 | 0.127907 | 0.572489 | 0.562349 | 661 tags=23% | list=4% | signal=22% |
| WP_GPCRS_ORPHAN | 16 | 0.777247 | 1.583041 | 0.117761 | 0.572489 | 0.562349 | 414 tags=19% | list=3% | signal=18% |
| REACTOME_CALCITONIN_LIKE_LIGAND_RECEPTORS | 6 | 0.857408 | 1.582844 | 0.08577 | 0.572489 | 0.562349 | 66 tags=17% | list=0% | signal=17% |
| WP_NUCLEOTIDE_GPCRS | 10 | 0.807269 | 1.582794 | 0.125714 | 0.572489 | 0.562349 | 93 tags=10% | list=1% | signal=10% |
| REACTOME_RAS_PROCESSING | 23 | -0.66328 | -1.58264 | 0.096916 | 0.572489 | 0.562349 | 2489 tags=57% | list=16% | signal=48% |
| REACTOME_NEUROTRANSMITTER_RECEPTORS_AND_POSTSYNAPTIC | 128 | 0.620057 | 1.582293 | 0.031835 | 0.572489 | 0.562349 | 1069 tags=13% | list=7% | signal=12% |
| REACTOME_ETHANOL_OXIDATION | 6 | -0.83866 | -1.58163 | 0.051125 | 0.572489 | 0.562349 | 1623 tags=83% | list=10% | signal=75% |
| REACTOME_METABOLISM_OF_POLYAMINES | 55 | -0.61229 | -1.57973 | 0.095122 | 0.572489 | 0.562349 | 3554 tags=62% | list=23% | signal=48% |
| REACTOME_SYNTHESIS_OF_PIPS_AT_THE_LATE_ENDOSOME_MEME | 10 | 0.805225 | 1.578786 | 0.127619 | 0.572489 | 0.562349 | 1603 tags=40% | list=10% | signal=36% |
| REACTOME_VLDLR_INTERNALISATION_AND_DEGRADATION | 16 | -0.68233 | -1.57855 | 0.10124 | 0.572489 | 0.562349 | 2489 tags=69% | list=16% | signal=58% |
| BIOCARTA_CB1R_PATHWAY | 5 | 0.880789 | 1.578111 | 0.075248 | 0.572489 | 0.562349 | 1204 tags=40% | list=8% | signal=37% |
| BIOCARTA_GSK3_PATHWAY | 20 | 0.759258 | 1.575782 | 0.096045 | 0.572489 | 0.562349 | 131 tags=10% | list=1% | signal=10% |
| WP_BMP_SIGNALING_PATHWAY_IN_EYELID_DEVELOPMENT | 19 | 0.756224 | 1.575696 | 0.103896 | 0.572489 | 0.562349 | 1421 tags=32% | list=9% | signal=29% |
| REACTOME_PROTEIN_UBIQUITINATION | 61 | -0.60277 | -1.57545 | 0.093137 | 0.572489 | 0.562349 | 2489 tags=28% | list=16% | signal=24% |
| REACTOME_TRANSCRIPTIONAL_REGULATION_BY_RUNX2 | 62 | -0.60109 | -1.57485 | 0.098039 | 0.572489 | 0.562349 | 3355 tags=58% | list=21% | signal=46% |
| WP_NOTCH_SIGNALING_PATHWAY | 44 | 0.699973 | 1.573577 | 0.056027 | 0.572489 | 0.562349 | 2481 tags=39% | list=16% | signal=33% |
| BIOCARTA_MALATEX_PATHWAY | 8 | -0.78181 | -1.57328 | 0.05988 | 0.572489 | 0.562349 | 3434 tags=100% | list=22% | signal=78% |
| REACTOME_METABOLISM_OF_AMINO_ACIDS_AND_DERIVATIVES | 208 | -0.57224 | -1.57116 | 0.087227 | 0.572489 | 0.562349 | 2079 tags=37% | list=13% | signal=33% |
| REACTOME_COPI_DEPENDENT_GOLGI_TO_ER_RETROGRADE_TRAFF | 89 | -0.58351 | -1.57076 | 0.088398 | 0.572489 | 0.562349 | 2950 tags=43% | list=19% | signal=35% |
| REACTOME_KSRP_KHSRP_BINDS_AND_DESTABILIZES_MRNA | 14 | -0.69506 | -1.57059 | 0.09165 | 0.572489 | 0.562349 | 3659 tags=64% | list=23% | signal=49% |
| REACTOME_RHO_GTPASES_ACTIVATE_KTN1 | 11 | -0.7339 | -1.57017 | 0.092975 | 0.572489 | 0.562349 | 3395 tags=91% | list=22% | signal=71% |
| REACTOME_DEGRADATION_OF_DVL | 55 | -0.60813 | -1.56901 | 0.095122 | 0.572489 | 0.562349 | 3355 tags=58% | list=21% | signal=46% |
| REACTOME_BETA_OXIDATION_OF_HEXANOYL_COA_TO_BUTANOYL | 5 | -0.85503 | -1.56709 | 0.060362 | 0.572489 | 0.562349 | 2282 tags=100% | list=15% | signal=85% |
| REACTOME_VITAMIN_C_ASCORBATE_METABOLISM | 7 | 0.826073 | 1.565932 | 0.0998 | 0.572489 | 0.562349 | 59 tags=14% | list=0% | signal=14% |
| REACTOME_MITOPHAGY | 28 | -0.6357 | -1.56421 | 0.100703 | 0.572489 | 0.562349 | 3161 tags=54% | list=20% | signal=43% |
| REACTOME_PEXOPHAGY | 10 | -0.74861 | -1.56192 | 0.08805 | 0.572489 | 0.562349 | 2709 tags=50% | list=17% | signal=41% |
| BIOCARTA_ATM_PATHWAY | 19 | -0.67293 | -1.56067 | 0.112311 | 0.572489 | 0.562349 | 523 tags=21% | list=3% | signal=20% |
| BIOCARTA_P38MAPK_PATHWAY | 32 | 0.707643 | 1.55972 | 0.086957 | 0.572489 | 0.562349 | 1518 tags=25% | list=10% | signal=23% |
| REACTOME_PKA_MEDIATED_PHOSPHORYLATION_OF_CREB | 17 | 0.766436 | 1.558901 | 0.115163 | 0.572489 | 0.562349 | 645 tags=18% | list=4% | signal=17% |
| BIOCARTA_FXR_PATHWAY | 4 | 0.907988 | 1.557444 | 0.061753 | 0.572489 | 0.562349 | 652 tags=50% | list=4% | signal=48% |
| WP_OXIDATIVE_DAMAGE_RESPONSE | 37 | 0.705324 | 1.557207 | 0.073427 | 0.572489 | 0.562349 | 600 tags=11% | list=4% | signal=10% |
| WP_GPCRS_OTHER | 48 | 0.684859 | 1.556975 | 0.056799 | 0.572489 | 0.562349 | 1998 tags=29% | list=13% | signal=26% |
| REACTOME_DEPOSITION_OF_NEW_CENPA_CONTAINING_NUCLEOS | 43 | -0.61593 | -1.55536 | 0.101695 | 0.572489 | 0.562349 | 1961 tags=23% | list=12% | signal=20% |
| REACTOME_FORMATION_OF_FIBRIN_CLOT_CLOTTING_CASCADE | 24 | 0.727072 | 1.555095 | 0.099099 | 0.572489 | 0.562349 | 2242 tags=33% | list=14% | signal=29% |
| REACTOME_TBC_RABGAPS | 34 | -0.61995 | -1.55413 | 0.106977 | 0.572489 | 0.562349 | 2507 tags=35% | list=16% | signal=30% |
| REACTOME_BETA_OXIDATION_OF_DECANOYL_COA_TO_OCTANOYL | 6 | -0.82356 | -1.55315 | 0.055215 | 0.572489 | 0.562349 | 2777 tags=100% | list=18% | signal=82% |
| REACTOME_ACETYLCHOLINE_NEUROTRANSMITTER_RELEASE_CYCLE | 17 | -0.67225 | -1.55149 | 0.108108 | 0.572489 | 0.562349 | 1851 tags=35% | list=12% | signal=31% |
| BIOCARTA_WNT_LRP6_PATHWAY | 3 | 0.961209 | 1.550365 | 0.026306 | 0.572489 | 0.562349 | 131 tags=33% | list=1% | signal=33% |
| REACTOME_RAB_REGULATION_OF_TRAFFICKING | 112 | -0.5811 | -1.54916 | 0.080925 | 0.572489 | 0.562349 | 1321 tags=17% | list=8% | signal=16% |
| REACTOME_ASSEMBLY_OF_THE_PRE_REPLICATIVE_COMPLEX | 81 | -0.57755 | -1.5486 | 0.090186 | 0.572489 | 0.562349 | 3554 tags=53% | list=23% | signal=41% |
| WP_INFLAMMATORY_RESPONSE_PATHWAY | 22 | 0.729096 | 1.547588 | 0.106618 | 0.572489 | 0.562349 | 2287 tags=32% | list=15% | signal=27% |
| REACTOME_P2Y_RECEPTORS | 9 | 0.79434 | 1.546995 | 0.128906 | 0.572489 | 0.562349 | 93 tags=11% | list=1% | signal=11% |
| REACTOME_RETROGRADE_NEUROTROPHIN_SIGNALLING | 11 | -0.723 | -1.54685 | 0.103306 | 0.572489 | 0.562349 | 3130 tags=73% | list=20% | signal=58% |
| WP_P38_MAPK_SIGNALING_PATHWAY | 33 | 0.701392 | 1.546762 | 0.095652 | 0.572489 | 0.562349 | 1518 tags=27% | list=10% | signal=25% |
| REACTOME_CELLULAR_RESPONSE_TO_CHEMICAL_STRESS | 148 | -0.57363 | -1.54658 | 0.074928 | 0.572489 | 0.562349 | 2063 tags=37% | list=13% | signal=33% |
| REACTOME_DISASSEMBLY_OF_THE_DESTRUCTION_COMPLEX_AND_ | 26 | 0.716387 | 1.545706 | 0.102113 | 0.572489 | 0.562349 | 131 tags=8% | list=1% | signal=8% |
| REACTOME_NOREPINEPHRINE_NEUROTRANSMITTER_RELEASE_CYC | 18 | -0.66892 | -1.54484 | 0.114165 | 0.572489 | 0.562349 | 1851 tags=33% | list=12% | signal=29% |
| REACTOME_ADP_SIGNALLING_THROUGH_P2Y_PURINOCEPTOR_12 | 19 | -0.66589 | -1.54433 | 0.114471 | 0.572489 | 0.562349 | 3418 tags=68% | list=22% | signal=54% |
| REACTOME_CROSS_PRESENTATION_OF_SOLUBLE_EXOGENOUS_AN | 46 | -0.59873 | -1.54366 | 0.106635 | 0.572489 | 0.562349 | 3335 tags=61% | list=21% | signal=48% |
| REACTOME_GLYCOGEN_METABOLISM | 17 | -0.66883 | -1.5436 | 0.108108 | 0.572489 | 0.562349 | 2200 tags=53% | list=14% | signal=46% |
| REACTOME_XENOBIOTICS | 9 | 0.791691 | 1.541836 | 0.128906 | 0.572489 | 0.562349 | 1451 tags=56% | list=9% | signal=50% |
| REACTOME_YAP1_AND_WWTR1_TAZ_STIMULATED_GENE_EXPRESS | 7 | -0.77742 | -1.54171 | 0.071856 | 0.572489 | 0.562349 | 76 tags=29% | list=0% | signal=28% |
| REACTOME_BETA_OXIDATION_OF_LAUROYL_COA_TO_DECANOYL_( | 5 | -0.84087 | -1.54113 | 0.062374 | 0.572489 | 0.562349 | 1169 tags=100% | list=7% | signal=93% |
| BIOCARTA_CALCINEURIN_PATHWAY | 17 | -0.66759 | -1.54073 | 0.110187 | 0.572489 | 0.562349 | 591 tags=12% | list=4% | signal=11% |
| BIOCARTA_BAD_PATHWAY | 23 | 0.723968 | 1.540704 | 0.109489 | 0.572489 | 0.562349 | 1021 tags=17% | list=7% | signal=16% |

| | | | | | | | | | | |
|---|---|---|---|---|---|---|---|---|---|---|
| REACTOME_CROSSLINKING_OF_COLLAGEN_FIBRILS | 15 | -0.67054 | -1.53971 | 0.11157 | 0.572489 | 0.562349 | 31 | tags=13% | list=0% | signal=13% |
| REACTOME_PHASE_4_RESTING_MEMBRANE_POTENTIAL | 14 | 0.754305 | 1.536562 | 0.146771 | 0.572489 | 0.562349 | 2017 | tags=36% | list=13% | signal=31% |
| REACTOME_COPI_MEDIATED_ANTEROGRADE_TRANSPORT | 89 | -0.57007 | -1.53457 | 0.09116 | 0.572489 | 0.562349 | 3169 | tags=49% | list=20% | signal=40% |
| REACTOME_MITOCHONDRIAL_IRON_SULFUR_CLUSTER_BIOGENESIS | 12 | -0.71118 | -1.53443 | 0.103093 | 0.572489 | 0.562349 | 4000 | tags=75% | list=25% | signal=56% |
| REACTOME_SIGNALING_BY_BMP | 25 | 0.713307 | 1.533768 | 0.106007 | 0.572489 | 0.562349 | 1537 | tags=28% | list=10% | signal=25% |
| BIOCARTA_GCR_PATHWAY | 15 | 0.750469 | 1.532444 | 0.146718 | 0.572489 | 0.562349 | 65 | tags=7% | list=0% | signal=7% |
| REACTOME_BETA_OXIDATION_OF_OCTANOYL_COA_TO_HEXANOYL_ | 5 | -0.83593 | -1.53208 | 0.064386 | 0.572489 | 0.562349 | 1169 | tags=100% | list=7% | signal=93% |
| BIOCARTA_TFF_PATHWAY | 21 | 0.735056 | 1.531907 | 0.109434 | 0.572489 | 0.562349 | 27 | tags=5% | list=0% | signal=5% |
| REACTOME_PRESYNAPTIC_FUNCTION_OF_KAINATE_RECEPTORS | 19 | -0.66018 | -1.53108 | 0.116631 | 0.572489 | 0.562349 | 3418 | tags=79% | list=22% | signal=62% |
| WP_ADIPOGENESIS_GENES | 109 | 0.607229 | 1.529584 | 0.039877 | 0.572489 | 0.562349 | 1291 | tags=18% | list=8% | signal=17% |
| BIOCARTA_EPONFKB_PATHWAY | 10 | -0.73271 | -1.52875 | 0.09434 | 0.572489 | 0.562349 | 41 | tags=20% | list=0% | signal=20% |
| REACTOME_HEPARAN_SULFATE_HEPARIN_HS_GAG_METABOLISM | 50 | 0.668138 | 1.528086 | 0.069728 | 0.572489 | 0.562349 | 3973 | tags=60% | list=25% | signal=45% |
| REACTOME_TRANSPORT_OF_INORGANIC_CATIONS_ANIONS_AND_A | 89 | 0.627255 | 1.527695 | 0.045313 | 0.572489 | 0.562349 | 2351 | tags=36% | list=15% | signal=31% |
| REACTOME_MET_ACTIVATES_PTK2_SIGNALING | 20 | 0.736046 | 1.527608 | 0.122411 | 0.572489 | 0.562349 | 2195 | tags=35% | list=14% | signal=30% |
| REACTOME_SEROTONIN_NEUROTRANSMITTER_RELEASE_CYCLE | 18 | -0.66074 | -1.52596 | 0.116279 | 0.572489 | 0.562349 | 4040 | tags=67% | list=26% | signal=50% |
| REACTOME_SIGNALING_BY_NOTCH | 38 | 0.686574 | 1.525347 | 0.09282 | 0.572489 | 0.562349 | 1487 | tags=29% | list=9% | signal=26% |
| WP_IL3_SIGNALING_PATHWAY | 95 | 0.618154 | 1.524172 | 0.049422 | 0.572489 | 0.562349 | 365 | tags=6% | list=2% | signal=6% |
| REACTOME_ASSOCIATION_OF_TRIC_CCT_WITH_TARGET_PROTEINS_ | 9 | -0.74072 | -1.52327 | 0.091837 | 0.572489 | 0.562349 | 3264 | tags=89% | list=21% | signal=70% |
| REACTOME_AZATHIOPRINE_ADME | 19 | 0.731059 | 1.523262 | 0.139147 | 0.572489 | 0.562349 | 122 | tags=11% | list=1% | signal=10% |
| WP_TRYPTOPHAN_METABOLISM | 33 | -0.61461 | -1.52317 | 0.112412 | 0.572489 | 0.562349 | 2300 | tags=42% | list=15% | signal=36% |
| REACTOME_MRNA_DECAY_BY_3_TO_5_EXORIBONUCLEASE | 12 | -0.7056 | -1.52241 | 0.109278 | 0.572489 | 0.562349 | 3683 | tags=75% | list=23% | signal=57% |
| REACTOME_CARBOXYTERMINAL_POST_TRANSLATIONAL_MODIFICA | 21 | -0.64456 | -1.52226 | 0.114407 | 0.572489 | 0.562349 | 1568 | tags=38% | list=10% | signal=34% |
| REACTOME_DOWNSTREAM_SIGNALING_EVENTS_OF_B_CELL_RECEP | 76 | -0.5723 | -1.51994 | 0.119481 | 0.572489 | 0.562349 | 3554 | tags=57% | list=23% | signal=44% |
| REACTOME_PYRUVATE_METABOLISM | 21 | -0.64352 | -1.5198 | 0.116525 | 0.572489 | 0.562349 | 1638 | tags=52% | list=10% | signal=47% |
| REACTOME_RND1_GTPASE_CYCLE | 40 | 0.682501 | 1.519253 | 0.086957 | 0.572489 | 0.562349 | 279 | tags=8% | list=2% | signal=7% |
| REACTOME_SWITCHING_OF_ORIGINS_TO_A_POST_REPLICATIVE_ST | 89 | -0.56416 | -1.51867 | 0.093923 | 0.572489 | 0.562349 | 3554 | tags=51% | list=23% | signal=39% |
| REACTOME_COLLAGEN_BIOSYNTHESIS_AND_MODIFYING_ENZYMES | 63 | 0.65566 | 1.517852 | 0.055838 | 0.572489 | 0.562349 | 3382 | tags=54% | list=22% | signal=43% |
| REACTOME_DOPAMINE_NEUROTRANSMITTER_RELEASE_CYCLE | 23 | -0.63611 | -1.51781 | 0.121145 | 0.572489 | 0.562349 | 4040 | tags=57% | list=26% | signal=42% |
| REACTOME_DOWNREGULATION_OF_ERBB2_SIGNALING | 24 | -0.63303 | -1.51647 | 0.118568 | 0.572489 | 0.562349 | 2057 | tags=38% | list=13% | signal=33% |
| REACTOME_APC_C_CDC20_MEDIATED_DEGRADATION_OF_CYCLIN_ | 24 | -0.63252 | -1.51525 | 0.118568 | 0.572489 | 0.562349 | 3348 | tags=46% | list=21% | signal=36% |
| BIOCARTA_NOTCH_PATHWAY | 6 | 0.820466 | 1.514646 | 0.11501 | 0.572489 | 0.562349 | 1487 | tags=50% | list=9% | signal=45% |
| WP_INSULIN_SIGNALING | 153 | 0.57949 | 1.514543 | 0.038836 | 0.572489 | 0.562349 | 1021 | tags=14% | list=7% | signal=13% |
| REACTOME_EPHA_MEDIATED_GROWTH_CONE_COLLAPSE | 18 | 0.737725 | 1.512727 | 0.141777 | 0.572489 | 0.562349 | 1720 | tags=44% | list=11% | signal=40% |
| BIOCARTA_GH_PATHWAY | 25 | 0.703427 | 1.512522 | 0.121908 | 0.572489 | 0.562349 | 1518 | tags=20% | list=10% | signal=18% |
| WP_APOPTOSIS | 71 | 0.635897 | 1.508472 | 0.050489 | 0.572489 | 0.562349 | 1576 | tags=14% | list=10% | signal=13% |
| REACTOME_APC_CDC20_MEDIATED_DEGRADATION_OF_NEK2A | 26 | -0.62297 | -1.50787 | 0.131336 | 0.572489 | 0.562349 | 3348 | tags=46% | list=21% | signal=36% |
| REACTOME_PLASMA_LIPOPROTEIN_REMODELING | 15 | -0.65662 | -1.50774 | 0.121901 | 0.572489 | 0.562349 | 46 | tags=13% | list=0% | signal=13% |
| REACTOME_REGULATION_OF_LOCALIZATION_OF_FOXO_TRANSCRIP | 12 | -0.69841 | -1.50689 | 0.11134 | 0.572489 | 0.562349 | 2365 | tags=42% | list=15% | signal=35% |
| REACTOME_RUNX3_REGULATES_WNT_SIGNALING | 5 | 0.840953 | 1.506737 | 0.10297 | 0.572489 | 0.562349 | 82 | tags=20% | list=1% | signal=20% |
| REACTOME_TRANSCRIPTIONAL_REGULATION_BY_VENTX | 5 | 0.840953 | 1.506737 | 0.10297 | 0.572489 | 0.562349 | 82 | tags=20% | list=1% | signal=20% |
| REACTOME_METABOLISM_OF_ANGIOTENSINOGEN_TO_ANGIOTENS | 9 | 0.772955 | 1.505346 | 0.140625 | 0.572489 | 0.562349 | 3306 | tags=78% | list=21% | signal=61% |
| REACTOME_SPHINGOLIPID_METABOLISM | 71 | 0.634426 | 1.504983 | 0.063518 | 0.572489 | 0.562349 | 1951 | tags=27% | list=12% | signal=24% |
| REACTOME_OLFACTORY_SIGNALING_PATHWAY | 4 | -0.86095 | -1.50236 | 0.061876 | 0.572489 | 0.562349 | 2142 | tags=75% | list=14% | signal=65% |
| REACTOME_SURFACTANT_METABOLISM | 15 | -0.65398 | -1.50168 | 0.121901 | 0.572489 | 0.562349 | 28 | tags=13% | list=0% | signal=13% |
| REACTOME_PROTEIN_METHYLATION | 19 | -0.64732 | -1.50127 | 0.12527 | 0.572489 | 0.562349 | 3520 | tags=63% | list=22% | signal=49% |
| REACTOME_TETRAHYDROBIOPTERIN_BH4_SYNTHESIS_RECYCLING_S | 9 | 0.770653 | 1.500864 | 0.140625 | 0.572489 | 0.562349 | 65 | tags=11% | list=0% | signal=11% |
| REACTOME_DSCAM_INTERACTIONS | 9 | 0.769344 | 1.498315 | 0.146484 | 0.572489 | 0.562349 | 141 | tags=11% | list=1% | signal=11% |
| WP_WNT_SIGNALING_PATHWAY_AND_PLURIPOTENCY | 89 | 0.615093 | 1.498075 | 0.054688 | 0.572489 | 0.562349 | 392 | tags=8% | list=2% | signal=8% |
| REACTOME_SYNAPTIC_ADHESION_LIKE_MOLECULES | 20 | 0.721368 | 1.497145 | 0.146893 | 0.572489 | 0.562349 | 2031 | tags=50% | list=13% | signal=44% |
| REACTOME_GAMMA_CARBOXYLATION_TRANSPORT_AND_AMINO_T | 5 | 0.835139 | 1.496319 | 0.106931 | 0.572489 | 0.562349 | 2094 | tags=80% | list=13% | signal=69% |
| REACTOME_HS_GAG_BIOSYNTHESIS | 28 | 0.689651 | 1.492825 | 0.123478 | 0.572489 | 0.562349 | 3973 | tags=68% | list=25% | signal=51% |
| REACTOME_UCH_PROTEINASES | 81 | -0.55666 | -1.49258 | 0.132626 | 0.572489 | 0.562349 | 3335 | tags=49% | list=21% | signal=39% |
| REACTOME_RHO_GTPASES_ACTIVATE_CIT | 7 | -0.75258 | -1.49246 | 0.093812 | 0.572489 | 0.562349 | 3395 | tags=71% | list=22% | signal=56% |
| REACTOME_ORC1_REMOVAL_FROM_CHROMATIN | 68 | -0.57062 | -1.49156 | 0.120301 | 0.572489 | 0.562349 | 3355 | tags=51% | list=21% | signal=41% |
| REACTOME_PHASE_I_FUNCTIONALIZATION_OF_COMPOUNDS | 59 | 0.644871 | 1.48985 | 0.075503 | 0.572489 | 0.562349 | 359 | tags=8% | list=2% | signal=8% |
| REACTOME_MHC_CLASS_II_ANTIGEN_PRESENTATION | 121 | -0.55421 | -1.48901 | 0.08596 | 0.572489 | 0.562349 | 2389 | tags=36% | list=15% | signal=31% |
| REACTOME_SYNTHESIS_OF_PI | 5 | 0.830826 | 1.488592 | 0.108911 | 0.572489 | 0.562349 | 1767 | tags=60% | list=11% | signal=53% |
| REACTOME_ENDOSOMAL_SORTING_COMPLEX_REQUIRED_FOR_TRA | 29 | -0.60046 | -1.48832 | 0.13615 | 0.572489 | 0.562349 | 1670 | tags=48% | list=11% | signal=43% |
| BIOCARTA_CIRCADIAN_PATHWAY | 5 | 0.830639 | 1.488257 | 0.108911 | 0.572489 | 0.562349 | 1042 | tags=40% | list=7% | signal=37% |
| REACTOME_REACTIONS_SPECIFIC_TO_THE_COMPLEX_N_GLYCAN_S | 7 | 0.78459 | 1.487295 | 0.125749 | 0.572489 | 0.562349 | 2916 | tags=71% | list=19% | signal=58% |
| WP_WNT_SIGNALING_PATHWAY_WP539 | 101 | 0.59796 | 1.486516 | 0.047766 | 0.572489 | 0.562349 | 661 | tags=12% | list=4% | signal=11% |
| REACTOME_UTILIZATION_OF_KETONE_BODIES | 3 | -0.92171 | -1.48548 | 0.058248 | 0.572489 | 0.562349 | 1071 | tags=67% | list=7% | signal=62% |
| BIOCARTA_INSULIN_PATHWAY | 22 | 0.698839 | 1.483365 | 0.154412 | 0.572489 | 0.562349 | 1518 | tags=27% | list=10% | signal=25% |
| REACTOME_FCERI_MEDIATED_NF_KB_ACTIVATION | 72 | -0.56643 | -1.48288 | 0.125964 | 0.572489 | 0.562349 | 3335 | tags=50% | list=21% | signal=40% |
| WP_LUNG_FIBROSIS | 42 | 0.658576 | 1.481837 | 0.098976 | 0.572489 | 0.562349 | 1712 | tags=24% | list=11% | signal=21% |
| REACTOME_MOLECULES_ASSOCIATED_WITH_ELASTIC_FIBRES | 28 | 0.684284 | 1.481208 | 0.133913 | 0.572489 | 0.562349 | 2101 | tags=36% | list=13% | signal=31% |
| REACTOME_SYNTHESIS_OF_PA | 28 | -0.60191 | -1.48106 | 0.142857 | 0.572489 | 0.562349 | 48 | tags=7% | list=0% | signal=7% |
| REACTOME_O_LINKED_GLYCOSYLATION | 89 | 0.607492 | 1.479562 | 0.060938 | 0.572489 | 0.562349 | 3646 | tags=47% | list=23% | signal=36% |
| REACTOME_TNFR1_INDUCED_PROAPOPTOTIC_SIGNALING | 22 | 0.696988 | 1.479436 | 0.15625 | 0.572489 | 0.562349 | 426 | tags=18% | list=3% | signal=18% |
| REACTOME_GABA_B_RECEPTOR_ACTIVATION | 39 | 0.661986 | 1.478893 | 0.115318 | 0.572489 | 0.562349 | 200 | tags=8% | list=1% | signal=8% |
| REACTOME_REGULATION_OF_INSULIN_SECRETION | 55 | 0.642136 | 1.478741 | 0.087838 | 0.572489 | 0.562349 | 579 | tags=15% | list=4% | signal=14% |
| BIOCARTA_TUBBY_PATHWAY | 5 | 0.824872 | 1.477923 | 0.114851 | 0.572489 | 0.562349 | 1141 | tags=60% | list=7% | signal=56% |
| REACTOME_PROTEIN_PROTEIN_INTERACTIONS_AT_SYNAPSES | 66 | 0.632389 | 1.477004 | 0.078203 | 0.572489 | 0.562349 | 2031 | tags=50% | list=13% | signal=44% |
| REACTOME_COPI_INDEPENDENT_GOLGI_TO_ER_RETROGRADE_TRA | 43 | -0.58466 | -1.4764 | 0.130751 | 0.572489 | 0.562349 | 3455 | tags=60% | list=22% | signal=47% |
| BIOCARTA_RACCYCD_PATHWAY | 25 | -0.61595 | -1.47604 | 0.142202 | 0.572489 | 0.562349 | 377 | tags=20% | list=2% | signal=20% |
| WP_MITOCHONDRIAL_LONG_CHAIN_FATTY_ACID_BETAOXIDATION | 16 | -0.63779 | -1.47551 | 0.123967 | 0.572489 | 0.562349 | 2976 | tags=62% | list=19% | signal=51% |
| WP_FACTORS_AND_PATHWAYS_AFFECTING_INSULINLIKE_GROWTH | 27 | 0.685921 | 1.475148 | 0.149648 | 0.572489 | 0.562349 | 849 | tags=19% | list=5% | signal=18% |
| REACTOME_PROTEIN_REPAIR | 6 | -0.78057 | -1.47207 | 0.09407 | 0.572489 | 0.562349 | 2533 | tags=67% | list=16% | signal=56% |
| REACTOME_CLEC7A_DECTIN_1_SIGNALING | 90 | -0.54572 | -1.4718 | 0.125 | 0.572489 | 0.562349 | 3554 | tags=51% | list=23% | signal=40% |
| WP_EICOSANOID_METABOLISM_VIA_CYTOCHROME_P450_MONOO | 4 | -0.84343 | -1.47179 | 0.06986 | 0.572489 | 0.562349 | 95 | tags=50% | list=1% | signal=50% |
| REACTOME_FORMATION_OF_THE_BETA_CATENIN_TCF_TRANSACTI | 23 | 0.691206 | 1.470982 | 0.153285 | 0.572489 | 0.562349 | 1384 | tags=30% | list=9% | signal=28% |
| REACTOME_G_PROTEIN_ACTIVATION | 21 | -0.62237 | -1.46983 | 0.135593 | 0.572489 | 0.562349 | 3927 | tags=67% | list=25% | signal=50% |
| REACTOME_CAM_PDE_1_ACTIVATION | 6 | -0.7789 | -1.46892 | 0.09407 | 0.572489 | 0.562349 | 2019 | tags=83% | list=13% | signal=73% |
| REACTOME_GLYCOGEN_BREAKDOWN_GLYCOGENOLYSIS | 13 | -0.6632 | -1.46868 | 0.12963 | 0.572489 | 0.562349 | 2200 | tags=69% | list=14% | signal=60% |
| BIOCARTA_CELLCYCLE_PATHWAY | 22 | -0.62344 | -1.4678 | 0.133188 | 0.572489 | 0.562349 | 234 | tags=18% | list=1% | signal=18% |
| REACTOME_CA_DEPENDENT_EVENTS | 27 | 0.682453 | 1.467689 | 0.149648 | 0.572489 | 0.562349 | 645 | tags=11% | list=4% | signal=11% |
| REACTOME_RUNX1_REGULATES_TRANSCRIPTION_OF_GENES_INVO | 63 | -0.55809 | -1.4675 | 0.126521 | 0.572489 | 0.562349 | 3335 | tags=52% | list=21% | signal=41% |
| WP_MYOMETRIAL_RELAXATION_AND_CONTRACTION_PATHWAYS | 134 | 0.570054 | 1.466815 | 0.050305 | 0.572489 | 0.562349 | 200 | tags=5% | list=1% | signal=5% |
| REACTOME_ER_QUALITY_CONTROL_COMPARTMENT_ERQC | 4 | -0.84052 | -1.46672 | 0.06986 | 0.572489 | 0.562349 | 2489 | tags=75% | list=16% | signal=63% |
| REACTOME_SYNTHESIS_OF_PC | 25 | 0.68173 | 1.465869 | 0.15371 | 0.572489 | 0.562349 | 361 | tags=8% | list=2% | signal=8% |

| | | | | | | | | | | |
|---|---|---|---|---|---|---|---|---|---|---|
| REACTOME_SYNTHESIS_OF_ACTIVE_UBIQUITIN_ROLES_OF_E1_AND | 30 | -0.58352 | -1.46494 | 0.143192 | 0.572489 | 0.562349 | 3141 | tags=40% | list=20% | signal=32% |
| REACTOME_PEROXISOMAL_LIPID_METABOLISM | 25 | -0.6112 | -1.46465 | 0.144495 | 0.572489 | 0.562349 | 3881 | tags=60% | list=25% | signal=45% |
| REACTOME_TRANSMISSION_ACROSS_CHEMICAL_SYNAPSES | 192 | 0.543751 | 1.463955 | 0.050074 | 0.572489 | 0.562349 | 1291 | tags=14% | list=8% | signal=13% |
| REACTOME_AKT_PHOSPHORYLATES_TARGETS_IN_THE_NUCLEUS | 10 | -0.70056 | -1.46166 | 0.125786 | 0.572489 | 0.562349 | 885 | tags=30% | list=6% | signal=28% |
| REACTOME_SYNTHESIS_OF_EPOXY_EET_AND_DIHYDROXYEICOSATR | 3 | -0.90675 | -1.46138 | 0.054767 | 0.572489 | 0.562349 | 95 | tags=67% | list=1% | signal=66% |
| REACTOME_TNFR2_NON_CANONICAL_NF_KB_PATHWAY | 83 | -0.54739 | -1.46114 | 0.136612 | 0.572489 | 0.562349 | 3335 | tags=45% | list=21% | signal=35% |
| REACTOME_GOLGI_TO_ER_RETROGRADE_TRANSPORT | 121 | -0.5434 | -1.45998 | 0.103152 | 0.572489 | 0.562349 | 2950 | tags=43% | list=19% | signal=35% |
| WP_FOCAL_ADHESION | 174 | 0.5475 | 1.459725 | 0.048889 | 0.572489 | 0.562349 | 2306 | tags=26% | list=15% | signal=23% |
| REACTOME_SIALIC_ACID_METABOLISM | 31 | 0.662665 | 1.45657 | 0.141115 | 0.572489 | 0.562349 | 2252 | tags=42% | list=14% | signal=36% |
| REACTOME_G_ALPHA_S_SIGNALLING_EVENTS | 95 | 0.590397 | 1.455733 | 0.070093 | 0.572489 | 0.562349 | 137 | tags=4% | list=1% | signal=4% |
| REACTOME_SHC_RELATED_EVENTS_TRIGGERED_BY_IGF1R | 6 | 0.788093 | 1.454883 | 0.136452 | 0.572489 | 0.562349 | 146 | tags=17% | list=1% | signal=17% |
| REACTOME_SYNTHESIS_OF_16_20_HYDROXYEICOSATETRAENOIC_A | 3 | 0.901736 | 1.454439 | 0.084479 | 0.572489 | 0.562349 | 180 | tags=33% | list=1% | signal=33% |
| BIOCARTA_IL17_PATHWAY | 3 | 0.901479 | 1.454023 | 0.084479 | 0.572489 | 0.562349 | 142 | tags=33% | list=1% | signal=33% |
| BIOCARTA_ERYTH_PATHWAY | 8 | 0.745859 | 1.453809 | 0.155689 | 0.572489 | 0.562349 | 874 | tags=25% | list=6% | signal=24% |
| WP_ESTROGEN_METABOLISM | 5 | 0.811294 | 1.453596 | 0.120792 | 0.572489 | 0.562349 | 221 | tags=20% | list=1% | signal=20% |
| REACTOME_SYNTHESIS_OF_DIPHTHAMIDE_EEF2 | 7 | -0.73229 | -1.45222 | 0.113772 | 0.572489 | 0.562349 | 3178 | tags=71% | list=20% | signal=57% |
| REACTOME_GLUTAMATE_AND_GLUTAMINE_METABOLISM | 12 | -0.67249 | -1.45097 | 0.134021 | 0.572489 | 0.562349 | 3909 | tags=75% | list=25% | signal=56% |
| REACTOME_BETA_OXIDATION_OF_PRISTANOYL_COA | 7 | -0.73102 | -1.4497 | 0.115768 | 0.572489 | 0.562349 | 3543 | tags=71% | list=23% | signal=55% |
| REACTOME_PENTOSE_PHOSPHATE_PATHWAY | 13 | -0.65432 | -1.44902 | 0.131687 | 0.572489 | 0.562349 | 4449 | tags=85% | list=28% | signal=61% |
| WP_ID_SIGNALING_PATHWAY | 42 | 0.643214 | 1.447272 | 0.131399 | 0.572489 | 0.562349 | 245 | tags=10% | list=2% | signal=9% |
| REACTOME_G2_M_DNA_REPLICATION_CHECKPOINT | 5 | 0.807617 | 1.447008 | 0.128713 | 0.572489 | 0.562349 | 349 | tags=20% | list=2% | signal=20% |
| REACTOME_LYSINE_CATABOLISM | 11 | -0.67616 | -1.44664 | 0.13843 | 0.572489 | 0.562349 | 4142 | tags=64% | list=26% | signal=47% |
| REACTOME_INTEGRATION_OF_ENERGY_METABOLISM | 69 | 0.611528 | 1.443941 | 0.095868 | 0.572489 | 0.562349 | 579 | tags=13% | list=4% | signal=13% |
| REACTOME_THROMBIN_SIGNALLING_THROUGH_PROTEINASE_ACTI | 27 | -0.58715 | -1.44309 | 0.147465 | 0.572489 | 0.562349 | 3418 | tags=59% | list=22% | signal=46% |
| REACTOME_RAC1_GTPASE_CYCLE | 168 | 0.542836 | 1.442275 | 0.050595 | 0.572489 | 0.562349 | 1624 | tags=20% | list=10% | signal=18% |
| REACTOME_CYP2E1_REACTIONS | 3 | 0.894183 | 1.442256 | 0.088409 | 0.572489 | 0.562349 | 1177 | tags=67% | list=7% | signal=62% |
| REACTOME_DEACTIVATION_OF_THE_BETA_CATENIN_TRANSACTIVA | 31 | 0.656034 | 1.441995 | 0.15331 | 0.572489 | 0.562349 | 483 | tags=16% | list=3% | signal=16% |
| REACTOME_REGULATION_OF_HSF1_MEDIATED_HEAT_SHOCK_RESP | 60 | 0.621948 | 1.439594 | 0.097479 | 0.572489 | 0.562349 | 972 | tags=13% | list=6% | signal=13% |
| REACTOME_SIGNALING_BY_HIPPO | 20 | -0.61265 | -1.43896 | 0.138004 | 0.572489 | 0.562349 | 1223 | tags=35% | list=8% | signal=32% |
| REACTOME_SIGNALING_BY_ERBB2 | 41 | -0.57011 | -1.43879 | 0.152941 | 0.572489 | 0.562349 | 2057 | tags=29% | list=13% | signal=26% |
| REACTOME_METABOLISM_OF_STEROIDS | 80 | -0.53916 | -1.43813 | 0.143236 | 0.572489 | 0.562349 | 623 | tags=14% | list=4% | signal=13% |
| BIOCARTA_EXTRINSIC_PATHWAY | 5 | 0.802392 | 1.437647 | 0.136634 | 0.572489 | 0.562349 | 2009 | tags=80% | list=13% | signal=70% |
| REACTOME_SYNTHESIS_OF_UDP_N_ACETYL_GLUCOSAMINE | 8 | -0.71434 | -1.43752 | 0.117764 | 0.572489 | 0.562349 | 3924 | tags=75% | list=25% | signal=56% |
| REACTOME_CYTOCHROME_P450_ARRANGED_BY_SUBSTRATE_TYPE | 32 | 0.651781 | 1.436594 | 0.156522 | 0.572489 | 0.562349 | 2642 | tags=34% | list=17% | signal=29% |
| BIOCARTA_PPARG_PATHWAY | 7 | 0.756721 | 1.434466 | 0.147705 | 0.572489 | 0.562349 | 1263 | tags=43% | list=8% | signal=39% |
| REACTOME_REGULATION_OF_TP53_ACTIVITY_THROUGH_ASSOCIAT | 12 | -0.66469 | -1.43413 | 0.148454 | 0.572489 | 0.562349 | 764 | tags=25% | list=5% | signal=24% |
| REACTOME_REGULATION_OF_RAS_BY_GAPS | 65 | -0.54622 | -1.43332 | 0.138614 | 0.572489 | 0.562349 | 3355 | tags=54% | list=21% | signal=43% |
| REACTOME_PLATELET_ACTIVATION_SIGNALING_AND_AGGREGATIO | 220 | 0.527721 | 1.432594 | 0.039706 | 0.572489 | 0.562349 | 364 | tags=6% | list=2% | signal=6% |
| BIOCARTA_VOBESITY_PATHWAY | 5 | 0.799492 | 1.43245 | 0.136634 | 0.572489 | 0.562349 | 229 | tags=20% | list=1% | signal=20% |
| REACTOME_IRAK2_MEDIATED_ACTIVATION_OF_TAK1_COMPLEX | 10 | -0.68639 | -1.43209 | 0.140461 | 0.572489 | 0.562349 | 814 | tags=40% | list=5% | signal=38% |
| BIOCARTA_BARRESTIN_SRC_PATHWAY | 17 | -0.62043 | -1.43188 | 0.141372 | 0.572489 | 0.562349 | 3558 | tags=53% | list=23% | signal=41% |
| REACTOME_RND3_GTPASE_CYCLE | 38 | 0.643917 | 1.430576 | 0.155867 | 0.572489 | 0.562349 | 2565 | tags=42% | list=16% | signal=35% |
| REACTOME_REGULATION_OF_INNATE_IMMUNE_RESPONSES_TO_C | 12 | -0.66252 | -1.42946 | 0.148454 | 0.572489 | 0.562349 | 878 | tags=33% | list=6% | signal=31% |
| REACTOME_GLUCAGON_TYPE_LIGAND_RECEPTORS | 25 | -0.59638 | -1.42913 | 0.151376 | 0.572489 | 0.562349 | 3949 | tags=60% | list=25% | signal=45% |
| WP_PEROXIREDOXIN_2_INDUCED_OVARIAN_FAILURE | 6 | -0.75679 | -1.42723 | 0.110429 | 0.572489 | 0.562349 | 1093 | tags=33% | list=7% | signal=31% |
| REACTOME_CYTOPROTECTION_BY_HMOX1 | 46 | -0.55326 | -1.42644 | 0.14455 | 0.572489 | 0.562349 | 1426 | tags=30% | list=9% | signal=28% |
| REACTOME_NR1H2_NR1H3_REGULATE_GENE_EXPRESSION_TO_COI | 6 | 0.77151 | 1.42427 | 0.138402 | 0.572489 | 0.562349 | 1183 | tags=33% | list=8% | signal=31% |
| REACTOME_PKMTS_METHYLATE_HISTONE_LYSINES | 48 | 0.625467 | 1.421952 | 0.127367 | 0.572489 | 0.562349 | 281 | tags=6% | list=2% | signal=6% |
| REACTOME_GLYCEROPHOSPHOLIPID_BIOSYNTHESIS | 101 | 0.571566 | 1.4209 | 0.077042 | 0.572489 | 0.562349 | 361 | tags=5% | list=2% | signal=5% |
| REACTOME_TCF_DEPENDENT_SIGNALING_IN_RESPONSE_TO_WNT | 144 | 0.548808 | 1.417212 | 0.067692 | 0.572489 | 0.562349 | 582 | tags=8% | list=4% | signal=8% |
| REACTOME_TRIGLYCERIDE_CATABOLISM | 4 | -0.81151 | -1.41609 | 0.101796 | 0.572489 | 0.562349 | 2722 | tags=75% | list=17% | signal=62% |
| REACTOME_REGULATION_OF_PYRUVATE_DEHYDROGENASE_PDH_C | 15 | -0.6156 | -1.41356 | 0.146694 | 0.572489 | 0.562349 | 1638 | tags=47% | list=10% | signal=42% |
| WP_TYROBP_CAUSAL_NETWORK_IN_MICROGLIA | 52 | 0.611202 | 1.411077 | 0.141414 | 0.572489 | 0.562349 | 284 | tags=8% | list=2% | signal=8% |
| REACTOME_G_ALPHA_Q_SIGNALLING_EVENTS | 147 | 0.543093 | 1.408211 | 0.070444 | 0.572489 | 0.562349 | 462 | tags=5% | list=3% | signal=5% |
| REACTOME_L1CAM_INTERACTIONS | 63 | -0.53521 | -1.40736 | 0.143552 | 0.572489 | 0.562349 | 3161 | tags=52% | list=20% | signal=42% |
| REACTOME_HYPUSINE_SYNTHESIS_FROM_EIF5A_LYSINE | 5 | -0.76743 | -1.40654 | 0.108652 | 0.572489 | 0.562349 | 1423 | tags=60% | list=9% | signal=55% |
| REACTOME_MITOCHONDRIAL_PROTEIN_IMPORT | 7 | -0.70879 | -1.40561 | 0.127745 | 0.572489 | 0.562349 | 4361 | tags=86% | list=28% | signal=62% |
| REACTOME_INTERCONVERSION_OF_NUCLEOTIDE_DI_AND_TRIPHOS | 25 | -0.58655 | -1.40558 | 0.15367 | 0.572489 | 0.562349 | 2012 | tags=36% | list=13% | signal=31% |
| REACTOME_ASSEMBLY_AND_CELL_SURFACE_PRESENTATION_OF_N | 5 | 0.784433 | 1.40547 | 0.142574 | 0.572489 | 0.562349 | 3359 | tags=80% | list=21% | signal=63% |
| REACTOME_SIGNALING_BY_RETINOIC_ACID | 31 | -0.55876 | -1.40541 | 0.156542 | 0.572489 | 0.562349 | 1638 | tags=39% | list=10% | signal=35% |
| REACTOME_TRANSLOCATION_OF_ZAP_70_TO_IMMUNOLOGICAL_S | 8 | -0.69823 | -1.40509 | 0.129741 | 0.572489 | 0.562349 | 3604 | tags=62% | list=23% | signal=48% |
| REACTOME_HEMOSTASIS | 429 | 0.483353 | 1.404219 | 0.035679 | 0.572489 | 0.562349 | 391 | tags=5% | list=2% | signal=5% |
| REACTOME_INSERTION_OF_TAIL_ANCHORED_PROTEINS_INTO_THE | 14 | -0.62086 | -1.40294 | 0.156823 | 0.572489 | 0.562349 | 2788 | tags=50% | list=18% | signal=41% |
| WP_MICRORNAS_IN_CARDIOMYOCYTE_HYPERTROPHY | 78 | 0.583978 | 1.398608 | 0.104167 | 0.572489 | 0.562349 | 1021 | tags=15% | list=7% | signal=14% |
| REACTOME_RESPONSE_TO_ELEVATED_PLATELET_CYTOSOLIC_CA2 | 104 | 0.55911 | 1.398592 | 0.090769 | 0.572489 | 0.562349 | 317 | tags=8% | list=2% | signal=8% |
| REACTOME_SIGNALING_BY_PDGF | 51 | 0.605016 | 1.391326 | 0.146712 | 0.572489 | 0.562349 | 2694 | tags=33% | list=17% | signal=28% |
| REACTOME_RAC3_GTPASE_CYCLE | 88 | 0.571631 | 1.390761 | 0.105181 | 0.572489 | 0.562349 | 1665 | tags=20% | list=11% | signal=18% |
| WP_MIR193A_AND_MVP_IN_COLON_CANCER_METASTASIS | 6 | -0.73736 | -1.39059 | 0.139059 | 0.572489 | 0.562349 | 234 | tags=33% | list=1% | signal=33% |
| REACTOME_MISCELLANEOUS_SUBSTRATES | 4 | 0.807725 | 1.385465 | 0.133466 | 0.572489 | 0.562349 | 2642 | tags=75% | list=17% | signal=62% |
| REACTOME_CREB1_PHOSPHORYLATION_THROUGH_NMDA_RECEPT | 5 | 0.770705 | 1.380872 | 0.154455 | 0.572489 | 0.562349 | 1291 | tags=40% | list=8% | signal=37% |
| BIOCARTA_FIBRINOLYSIS_PATHWAY | 5 | 0.769497 | 1.378708 | 0.154455 | 0.572489 | 0.562349 | 2242 | tags=60% | list=14% | signal=51% |
| REACTOME_DNA_REPLICATION_PRE_INITIATION | 96 | -0.51375 | -1.37108 | 0.156425 | 0.572489 | 0.562349 | 3554 | tags=45% | list=23% | signal=35% |
| REACTOME_PASSIVE_TRANSPORT_BY_AQUAPORINS | 5 | -0.74776 | -1.37048 | 0.146881 | 0.572489 | 0.562349 | 2536 | tags=40% | list=16% | signal=34% |
| REACTOME_FRUCTOSE_CATABOLISM | 5 | -0.74675 | -1.36863 | 0.152918 | 0.572489 | 0.562349 | 1177 | tags=60% | list=7% | signal=56% |
| REACTOME_EUKARYOTIC_TRANSLATION_ELONGATION | 5 | -0.74624 | -1.3677 | 0.152918 | 0.572489 | 0.562349 | 3990 | tags=100% | list=25% | signal=75% |
| REACTOME_SEPARATION_OF_SISTER_CHROMATIDS | 175 | -0.5073 | -1.3648 | 0.116564 | 0.572489 | 0.562349 | 3348 | tags=43% | list=21% | signal=34% |
| REACTOME_C_TYPE_LECTIN_RECEPTORS_CLRS | 105 | -0.50869 | -1.36328 | 0.148688 | 0.572489 | 0.562349 | 3554 | tags=49% | list=23% | signal=38% |
| REACTOME_TRANSPORT_OF_SMALL_MOLECULES | 541 | 0.460767 | 1.363041 | 0.032763 | 0.572489 | 0.562349 | 3031 | tags=32% | list=19% | signal=26% |
| REACTOME_GAMMA_CARBOXYLATION_OF_PROTEIN_PRECURSORS | 3 | 0.84344 | 1.360411 | 0.125737 | 0.572489 | 0.562349 | 1477 | tags=67% | list=9% | signal=60% |
| REACTOME_STIMULI_SENSING_CHANNELS | 74 | 0.569609 | 1.357336 | 0.136143 | 0.572489 | 0.562349 | 3082 | tags=39% | list=20% | signal=32% |
| REACTOME_CARDIAC_CONDUCTION | 88 | 0.548947 | 1.335572 | 0.147567 | 0.572489 | 0.562349 | 1771 | tags=16% | list=11% | signal=14% |
| REACTOME_INCRETIN_SYNTHESIS_SECRETION_AND_INACTIVATION | 4 | -0.76536 | -1.33556 | 0.149701 | 0.572489 | 0.562349 | 112 | tags=50% | list=1% | signal=50% |
| REACTOME_ION_CHANNEL_TRANSPORT | 138 | 0.516714 | 1.335201 | 0.110092 | 0.572489 | 0.562349 | 3011 | tags=36% | list=19% | signal=29% |
| WP_METAPATHWAY_BIOTRANSFORMATION | 94 | 0.541446 | 1.331586 | 0.140187 | 0.572489 | 0.562349 | 1229 | tags=13% | list=8% | signal=12% |
| REACTOME_FATTY_ACID_METABOLISM | 136 | -0.49237 | -1.32703 | 0.153623 | 0.572489 | 0.562349 | 2976 | tags=42% | list=19% | signal=34% |
| REACTOME_INTRA_GOLGI_AND_RETROGRADE_GOLGI_TO_ER_TRAF | 176 | -0.4917 | -1.32597 | 0.141538 | 0.572489 | 0.562349 | 3169 | tags=40% | list=20% | signal=33% |
| REACTOME_BIOSYNTHESIS_OF_MARESINS | 3 | -0.81961 | -1.32094 | 0.148073 | 0.572489 | 0.562349 | 95 | tags=67% | list=1% | signal=66% |
| WP_MIR302367_PROMOTING_CARDIOMYOCYTE_PROLIFERATION | 3 | 0.818867 | 1.320777 | 0.151277 | 0.572489 | 0.562349 | 1296 | tags=33% | list=8% | signal=31% |
| REACTOME_PHOSPHOLIPID_METABOLISM | 174 | 0.494856 | 1.319368 | 0.108148 | 0.572489 | 0.562349 | 459 | tags=5% | list=3% | signal=5% |
| REACTOME_VLDL_ASSEMBLY | 3 | 0.812954 | 1.311239 | 0.155206 | 0.572489 | 0.562349 | 2940 | tags=100% | list=19% | signal=81% |

| | | | | | | | | | | |
|---|---|---|---|---|---|---|---|---|---|---|
| REACTOME_MEMBRANE_TRAFFICKING | 524 | -0.43119 | -1.30263 | 0.123552 | 0.572489 | 0.562349 | 3130 | tags=34% | list=20% | signal=28% |
| REACTOME_MITOTIC_G2_G2_M_PHASES | 177 | -0.47955 | -1.3009 | 0.151515 | 0.572489 | 0.562349 | 3355 | tags=41% | list=21% | signal=33% |
| REACTOME_CLASS_A_1_RHODOPSIN_LIKE_RECEPTORS | 180 | 0.478781 | 1.274554 | 0.141369 | 0.572489 | 0.562349 | 1410 | tags=15% | list=9% | signal=14% |
| REACTOME_GPCR_LIGAND_BINDING | 232 | 0.462558 | 1.268067 | 0.138081 | 0.572489 | 0.562349 | 776 | tags=8% | list=5% | signal=7% |
| REACTOME_SIGNALING_BY_RECEPTOR_TYROSINE_KINASES | 373 | 0.439322 | 1.261539 | 0.11236 | 0.572489 | 0.562349 | 1603 | tags=14% | list=10% | signal=13% |
| REACTOME_CHROMATIN_MODIFYING_ENZYMES | 153 | 0.480535 | 1.255916 | 0.152568 | 0.572489 | 0.562349 | 1038 | tags=9% | list=7% | signal=9% |
| REACTOME_SIGNALING_BY_WNT | 220 | 0.462052 | 1.254324 | 0.144118 | 0.572489 | 0.562349 | 483 | tags=6% | list=3% | signal=6% |
| REACTOME_SIGNALING_BY_GPCR | 431 | 0.431352 | 1.252619 | 0.114325 | 0.572489 | 0.562349 | 784 | tags=7% | list=5% | signal=7% |
| REACTOME_NEUREXINS_AND_NEUROLIGINS | 31 | 0.652412 | 1.434034 | 0.158537 | 0.576213 | 0.566006 | 2767 | tags=58% | list=18% | signal=48% |
| REACTOME_MET_PROMOTES_CELL_MOTILITY | 31 | 0.652264 | 1.433707 | 0.158537 | 0.576213 | 0.566006 | 2195 | tags=26% | list=14% | signal=22% |
| BIOCARTA_PITX2_PATHWAY | 13 | 0.743654 | 1.504716 | 0.162791 | 0.57927 | 0.56901 | 1384 | tags=38% | list=9% | signal=35% |
| BIOCARTA_CD40_PATHWAY | 13 | 0.743049 | 1.503491 | 0.166667 | 0.57927 | 0.56901 | 600 | tags=23% | list=4% | signal=22% |
| REACTOME_SIGNALING_BY_ACTIVIN | 13 | 0.734559 | 1.486312 | 0.176357 | 0.57927 | 0.56901 | 2918 | tags=54% | list=19% | signal=44% |
| REACTOME_NITRIC_OXIDE_STIMULATES_GUANYLATE_CYCLASE | 13 | 0.732844 | 1.482844 | 0.176357 | 0.57927 | 0.56901 | 1076 | tags=23% | list=7% | signal=22% |
| BIOCARTA_ARF_PATHWAY | 15 | 0.725782 | 1.482033 | 0.175676 | 0.57927 | 0.56901 | 96 | tags=7% | list=1% | signal=7% |
| REACTOME_REPRESSION_OF_WNT_TARGET_GENES | 10 | 0.754826 | 1.479971 | 0.161905 | 0.57927 | 0.56901 | 82 | tags=10% | list=1% | signal=10% |
| WP_SELENIUM_MICRONUTRIENT_NETWORK | 18 | 0.721673 | 1.479812 | 0.162571 | 0.57927 | 0.56901 | 285 | tags=11% | list=2% | signal=11% |
| REACTOME_CASPASE_MEDIATED_CLEAVAGE_OF_CYTOSKELETAL_PF | 12 | 0.733766 | 1.478723 | 0.172147 | 0.57927 | 0.56901 | 97 | tags=8% | list=1% | signal=8% |
| REACTOME_NUCLEOTIDE_LIKE_PURINERGIC_RECEPTORS | 13 | 0.729289 | 1.475651 | 0.180233 | 0.57927 | 0.56901 | 93 | tags=8% | list=1% | signal=8% |
| REACTOME_REGULATION_OF_TLR_BY_ENDOGENOUS_LIGAND | 14 | 0.719572 | 1.46581 | 0.176125 | 0.57927 | 0.56901 | 105 | tags=7% | list=1% | signal=7% |
| REACTOME_NR1H2_AND_NR1H3_MEDIATED_SIGNALING | 12 | 0.726605 | 1.46429 | 0.179884 | 0.57927 | 0.56901 | 1183 | tags=33% | list=8% | signal=31% |
| REACTOME_GPER1_SIGNALING | 26 | 0.667918 | 1.441127 | 0.163732 | 0.57927 | 0.56901 | 101 | tags=8% | list=1% | signal=8% |
| REACTOME_HS_GAG_DEGRADATION | 20 | 0.691787 | 1.435753 | 0.184557 | 0.57927 | 0.56901 | 4566 | tags=80% | list=29% | signal=57% |
| BIOCARTA_HDAC_PATHWAY | 21 | 0.687586 | 1.432977 | 0.184906 | 0.57927 | 0.56901 | 1021 | tags=29% | list=7% | signal=27% |
| WP_SREBF_AND_MIR33_IN_CHOLESTEROL_AND_LIPID_HOMEOSTA | 11 | 0.723181 | 1.431692 | 0.183398 | 0.57927 | 0.56901 | 1736 | tags=36% | list=11% | signal=32% |
| REACTOME_NON_INTEGRIN_MEMBRANE_ECM_INTERACTIONS | 28 | 0.661159 | 1.431151 | 0.170435 | 0.57927 | 0.56901 | 364 | tags=11% | list=2% | signal=10% |
| BIOCARTA_CELL2CELL_PATHWAY | 9 | 0.73389 | 1.429266 | 0.177734 | 0.57927 | 0.56901 | 188 | tags=11% | list=1% | signal=11% |
| REACTOME_PLATELET_CALCIUM_HOMEOSTASIS | 26 | 0.656055 | 1.41553 | 0.176056 | 0.57927 | 0.56901 | 55 | tags=4% | list=0% | signal=4% |
| REACTOME_DNA_DAMAGE_RECOGNITION_IN_GG_NER | 38 | -0.55653 | -1.41207 | 0.160093 | 0.57927 | 0.56901 | 3355 | tags=45% | list=21% | signal=35% |
| REACTOME_TANDEM_PORE_DOMAIN_POTASSIUM_CHANNELS | 8 | 0.718942 | 1.401343 | 0.183633 | 0.57927 | 0.56901 | 3160 | tags=62% | list=20% | signal=50% |
| REACTOME_SODIUM_PROTON_EXCHANGERS | 9 | 0.719164 | 1.400586 | 0.181641 | 0.57927 | 0.56901 | 2896 | tags=67% | list=18% | signal=54% |
| REACTOME_SENESCENCE_ASSOCIATED_SECRETORY_PHENOTYPE_SA | 40 | -0.55388 | -1.39997 | 0.168618 | 0.57927 | 0.56901 | 676 | tags=38% | list=4% | signal=36% |
| REACTOME_G_ALPHA_Z_SIGNALLING_EVENTS | 31 | 0.636874 | 1.39988 | 0.181185 | 0.57927 | 0.56901 | 645 | tags=16% | list=4% | signal=15% |
| REACTOME_NUCLEOTIDE_CATABOLISM | 31 | 0.63592 | 1.397783 | 0.181185 | 0.57927 | 0.56901 | 375 | tags=6% | list=2% | signal=6% |
| REACTOME_LYSOSOME_VESICLE_BIOGENESIS | 34 | -0.55044 | -1.37987 | 0.176744 | 0.57927 | 0.56901 | 3431 | tags=44% | list=22% | signal=35% |
| REACTOME_TP53_REGULATES_TRANSCRIPTION_OF_CELL_CYCLE_GE | 35 | -0.55055 | -1.37935 | 0.178241 | 0.57927 | 0.56901 | 523 | tags=9% | list=3% | signal=8% |
| WP_FATTY_ACID_BIOSYNTHESIS | 22 | -0.58585 | -1.37931 | 0.170306 | 0.57927 | 0.56901 | 2227 | tags=50% | list=14% | signal=43% |
| REACTOME_CYCLIN_D_ASSOCIATED_EVENTS_IN_G1 | 42 | -0.54642 | -1.37755 | 0.175481 | 0.57927 | 0.56901 | 2489 | tags=31% | list=16% | signal=26% |
| REACTOME_ADP_SIGNALLING_THROUGH_P2Y_PURINOCEPTOR_1 | 22 | -0.58355 | -1.37389 | 0.170306 | 0.57927 | 0.56901 | 3418 | tags=59% | list=22% | signal=46% |
| REACTOME_RECEPTOR_MEDIATED_MITOPHAGY | 11 | -0.64125 | -1.37194 | 0.163223 | 0.57927 | 0.56901 | 3161 | tags=64% | list=20% | signal=51% |
| BIOCARTA_G1_PATHWAY | 25 | -0.57058 | -1.3673 | 0.167431 | 0.57927 | 0.56901 | 234 | tags=20% | list=1% | signal=20% |
| REACTOME_ION_TRANSPORT_BY_P_TYPE_ATPASES | 48 | 0.601297 | 1.367002 | 0.165232 | 0.57927 | 0.56901 | 3011 | tags=40% | list=19% | signal=32% |
| REACTOME_INHIBITION_OF_THE_PROTEOLYTIC_ACTIVITY_OF_APC_ | 21 | -0.57862 | -1.36652 | 0.182203 | 0.57927 | 0.56901 | 3348 | tags=43% | list=21% | signal=34% |
| BIOCARTA_MPR_PATHWAY | 18 | -0.59108 | -1.36507 | 0.175476 | 0.57927 | 0.56901 | 2970 | tags=44% | list=19% | signal=36% |
| REACTOME_COSTIMULATION_BY_THE_CD28_FAMILY | 52 | -0.53502 | -1.36304 | 0.164216 | 0.57927 | 0.56901 | 2866 | tags=37% | list=18% | signal=30% |
| REACTOME_SENSORY_PERCEPTION_OF_TASTE | 5 | 0.760717 | 1.362978 | 0.162376 | 0.57927 | 0.56901 | 55 | tags=20% | list=0% | signal=20% |
| REACTOME_CD28_DEPENDENT_PI3K_AKT_SIGNALING | 21 | -0.57679 | -1.36219 | 0.184322 | 0.57927 | 0.56901 | 1769 | tags=19% | list=11% | signal=17% |
| REACTOME_CHONDROITIN_SULFATE_DERMATAN_SULFATE_METAB | 47 | 0.601612 | 1.36186 | 0.172414 | 0.57927 | 0.56901 | 4566 | tags=57% | list=29% | signal=41% |
| REACTOME_REGULATION_OF_APC_C_ACTIVATORS_BETWEEN_G1_S | 35 | -0.54331 | -1.36122 | 0.185185 | 0.57927 | 0.56901 | 3513 | tags=40% | list=22% | signal=31% |
| BIOCARTA_G2_PATHWAY | 22 | -0.57741 | -1.35943 | 0.179039 | 0.57927 | 0.56901 | 2249 | tags=27% | list=14% | signal=23% |
| REACTOME_CD28_CO_STIMULATION | 32 | -0.54132 | -1.35845 | 0.168618 | 0.57927 | 0.56901 | 1769 | tags=22% | list=11% | signal=19% |
| REACTOME_HYALURONAN_BIOSYNTHESIS_AND_EXPORT | 5 | 0.75769 | 1.357553 | 0.162376 | 0.57927 | 0.56901 | 1056 | tags=40% | list=7% | signal=37% |
| REACTOME_PLATELET_SENSITIZATION_BY_LDL | 16 | -0.58604 | -1.35579 | 0.177686 | 0.57927 | 0.56901 | 3800 | tags=62% | list=24% | signal=47% |
| REACTOME_CD28_DEPENDENT_VAV1_PATHWAY | 11 | -0.63292 | -1.35412 | 0.171488 | 0.57927 | 0.56901 | 2708 | tags=36% | list=17% | signal=30% |
| WP_ENDOCHONDRAL_OSSIFICATION | 53 | 0.587507 | 1.353256 | 0.183986 | 0.57927 | 0.56901 | 994 | tags=21% | list=6% | signal=20% |
| REACTOME_N_GLYCAN_TRIMMING_AND_ELONGATION_IN_THE_CIS | 5 | 0.754983 | 1.352704 | 0.162376 | 0.57927 | 0.56901 | 1176 | tags=60% | list=7% | signal=56% |
| REACTOME_DOWNSTREAM_TCR_SIGNALING | 82 | -0.50817 | -1.35199 | 0.167568 | 0.57927 | 0.56901 | 3335 | tags=49% | list=21% | signal=39% |
| REACTOME_AMPK_INHIBITS_CHREBP_TRANSCRIPTIONAL_ACTIVATIO | 5 | 0.753159 | 1.349436 | 0.166337 | 0.57927 | 0.56901 | 2856 | tags=40% | list=18% | signal=33% |
| REACTOME_REGULATION_OF_BACH1_ACTIVITY | 11 | -0.62799 | -1.34357 | 0.181818 | 0.57927 | 0.56901 | 3355 | tags=45% | list=21% | signal=36% |
| REACTOME_PD_1_SIGNALING | 10 | -0.64288 | -1.34132 | 0.176101 | 0.57927 | 0.56901 | 2272 | tags=70% | list=14% | signal=60% |
| REACTOME_REGULATION_OF_EXPRESSION_OF_SLITS_AND_ROBOS | 7 | -0.67568 | -1.33995 | 0.169661 | 0.57927 | 0.56901 | 676 | tags=43% | list=4% | signal=41% |
| REACTOME_SCF_BETA_TRCP_MEDIATED_DEGRADATION_OF_EMI1 | 9 | -0.6504 | -1.33754 | 0.177551 | 0.57927 | 0.56901 | 2489 | tags=56% | list=16% | signal=47% |
| REACTOME_BETA_OXIDATION_OF_VERY_LONG_CHAIN_FATTY_ACID | 11 | -0.62489 | -1.33693 | 0.183884 | 0.57927 | 0.56901 | 3881 | tags=55% | list=25% | signal=41% |
| REACTOME_TRANSPORT_OF_BILE_SALTS_AND_ORGANIC_ACIDS_MI | 58 | 0.577185 | 1.336339 | 0.172819 | 0.57927 | 0.56901 | 2603 | tags=29% | list=17% | signal=25% |
| REACTOME_GLYCOGEN_SYNTHESIS | 5 | -0.72678 | -1.33202 | 0.181087 | 0.57927 | 0.56901 | 987 | tags=60% | list=6% | signal=56% |
| REACTOME_ACTIVATION_OF_RAC1_DOWNSTREAM_OF_NMDARS | 6 | -0.70486 | -1.3293 | 0.165644 | 0.57927 | 0.56901 | 3520 | tags=50% | list=22% | signal=39% |
| REACTOME_ANTIGEN_PROCESSING_CROSS_PRESENTATION | 74 | -0.50309 | -1.32763 | 0.163636 | 0.57927 | 0.56901 | 3335 | tags=47% | list=21% | signal=37% |
| BIOCARTA_PELP1_PATHWAY | 5 | 0.740408 | 1.32659 | 0.178218 | 0.57927 | 0.56901 | 1263 | tags=40% | list=8% | signal=37% |
| REACTOME_HEDGEHOG_ON_STATE | 78 | -0.50178 | -1.32555 | 0.174603 | 0.57927 | 0.56901 | 3355 | tags=47% | list=21% | signal=37% |
| REACTOME_DEGRADATION_OF_AXIN | 53 | -0.51563 | -1.32363 | 0.180723 | 0.57927 | 0.56901 | 3335 | tags=60% | list=21% | signal=48% |
| REACTOME_PTK6_REGULATES_RTKS_AND_THEIR_EFFECTORS_AKT1 | 8 | -0.65764 | -1.32342 | 0.177645 | 0.57927 | 0.56901 | 2489 | tags=50% | list=16% | signal=42% |
| REACTOME_FRUCTOSE_METABOLISM | 6 | -0.69932 | -1.31885 | 0.167689 | 0.57927 | 0.56901 | 1177 | tags=50% | list=7% | signal=46% |
| REACTOME_SENSORY_PERCEPTION | 67 | 0.558149 | 1.312162 | 0.175497 | 0.57927 | 0.56901 | 179 | tags=4% | list=1% | signal=4% |
| WP_REGULATION_OF_CARDIAC_HYPERTROPHY_BY_MIR208 | 3 | -0.81115 | -1.3073 | 0.160243 | 0.57927 | 0.56901 | 1268 | tags=67% | list=8% | signal=61% |
| REACTOME_CYCLIN_A_CDK2_ASSOCIATED_EVENTS_AT_S_PHASE_EI | 71 | -0.49821 | -1.29868 | 0.18299 | 0.57927 | 0.56901 | 1879 | tags=35% | list=12% | signal=31% |
| REACTOME_DNA_REPLICATION | 123 | -0.47935 | -1.28626 | 0.172414 | 0.57927 | 0.56901 | 3554 | tags=44% | list=23% | signal=34% |
| REACTOME_CELLULAR_RESPONSE_TO_HEAT_STRESS | 79 | 0.536826 | 1.284324 | 0.176282 | 0.57927 | 0.56901 | 972 | tags=13% | list=6% | signal=12% |
| REACTOME_CELL_CYCLE_CHECKPOINTS | 256 | -0.44729 | -1.27218 | 0.167192 | 0.57927 | 0.56901 | 3348 | tags=33% | list=21% | signal=26% |
| REACTOME_G2_M_CHECKPOINTS | 138 | -0.46904 | -1.26486 | 0.178161 | 0.57927 | 0.56901 | 2489 | tags=36% | list=16% | signal=31% |
| REACTOME_MITOTIC_METAPHASE_AND_ANAPHASE | 212 | -0.45966 | -1.26208 | 0.18125 | 0.57927 | 0.56901 | 3348 | tags=40% | list=21% | signal=32% |
| REACTOME_M_PHASE | 337 | -0.43022 | -1.26178 | 0.171233 | 0.57927 | 0.56901 | 3210 | tags=35% | list=20% | signal=29% |
| REACTOME_DNA_REPAIR | 272 | -0.43417 | -1.24489 | 0.169329 | 0.57927 | 0.56901 | 2489 | tags=17% | list=16% | signal=14% |
| REACTOME_MUSCLE_CONTRACTION | 139 | 0.479837 | 1.240899 | 0.169207 | 0.57927 | 0.56901 | 1818 | tags=17% | list=12% | signal=15% |
| REACTOME_RHO_GTPASE_CYCLE | 410 | 0.413957 | 1.198843 | 0.160714 | 0.57927 | 0.56901 | 1665 | tags=16% | list=11% | signal=15% |
| WP_MONOAMINE_GPCRS | 29 | 0.645227 | 1.402784 | 0.185764 | 0.579988 | 0.569715 | 2310 | tags=45% | list=15% | signal=38% |
| WP_PPAR_SIGNALING_PATHWAY | 52 | -0.51466 | -1.31115 | 0.186275 | 0.580491 | 0.570209 | 2976 | tags=37% | list=19% | signal=30% |
| REACTOME_ERKS_ARE_INACTIVATED | 13 | -0.60528 | -1.34041 | 0.187243 | 0.582416 | 0.5721 | 3800 | tags=77% | list=24% | signal=58% |
| BIOCARTA_RAC1_PATHWAY | 21 | 0.680622 | 1.418463 | 0.188679 | 0.583387 | 0.573053 | 1021 | tags=19% | list=7% | signal=18% |
| REACTOME_PHOSPHORYLATION_OF_THE_APC_C | 20 | -0.56972 | -1.33812 | 0.18896 | 0.583387 | 0.573053 | 3348 | tags=40% | list=21% | signal=32% |
| REACTOME_CONVERSION_FROM_APC_C_CDC20_TO_APC_C_CDH1_ | 20 | -0.56595 | -1.32928 | 0.18896 | 0.583387 | 0.573053 | 3513 | tags=55% | list=22% | signal=43% |

| | | | | | | | | | |
|---|---|---|---|---|---|---|---|---|---|
| REACTOME_TRANSLESION_SYNTHESIS_BY_POLH | 18 | -0.5745 | -1.32679 | 0.188161 | 0.583387 | 0.573053 | 3984 | tags=56% | list=25% | signal=42% |
| REACTOME_WNT5A_DEPENDENT_INTERNALIZATION_OF_FZD4 | 14 | -0.59515 | -1.34484 | 0.189409 | 0.58369 | 0.573351 | 2283 | tags=50% | list=15% | signal=43% |
| REACTOME_EXTRA_NUCLEAR_ESTROGEN_SIGNALING | 60 | 0.562773 | 1.302624 | 0.189916 | 0.584167 | 0.57382 | 146 | tags=5% | list=1% | signal=5% |
| REACTOME_THE_CANONICAL_RETINOID_CYCLE_IN_RODS_TWILIGHT | 15 | 0.705679 | 1.440983 | 0.19305 | 0.585676 | 0.575302 | 125 | tags=7% | list=1% | signal=7% |
| REACTOME_TRANSLESION_SYNTHESIS_BY_POLK | 16 | -0.56994 | -1.31856 | 0.192149 | 0.585676 | 0.575302 | 3984 | tags=50% | list=25% | signal=37% |
| REACTOME_CAMK_IV_MEDIATED_PHOSPHORYLATION_OF_CREB | 5 | -0.71342 | -1.30755 | 0.191147 | 0.585676 | 0.575302 | 3520 | tags=60% | list=22% | signal=47% |
| REACTOME_OREXIN_AND_NEUROPEPTIDES_FF_AND_QRFP_BIND_T | 5 | 0.725086 | 1.299137 | 0.192079 | 0.585676 | 0.575302 | 2823 | tags=60% | list=18% | signal=49% |
| REACTOME_UREA_CYCLE | 5 | -0.70809 | -1.29778 | 0.193159 | 0.585676 | 0.575302 | 4476 | tags=80% | list=29% | signal=57% |
| REACTOME_ESTROGEN_STIMULATED_SIGNALING_THROUGH_PRKCZ | 5 | -0.7045 | -1.29119 | 0.193159 | 0.585676 | 0.575302 | 377 | tags=60% | list=2% | signal=59% |
| REACTOME_TRANSPORT_OF_GAMMA_CARBOXYLATED_PROTEIN_PI | 3 | 0.797019 | 1.285537 | 0.192534 | 0.585676 | 0.575302 | 2094 | tags=67% | list=13% | signal=58% |
| REACTOME_MUSCARINIC_ACETYLCHOLINE_RECEPTORS | 4 | 0.746085 | 1.279735 | 0.193227 | 0.585676 | 0.575302 | 1141 | tags=50% | list=7% | signal=46% |
| REACTOME_GABA_SYNTHESIS_RELEASE_REUPTAKE_AND_DEGRADA | 19 | -0.57755 | -1.33946 | 0.194384 | 0.587041 | 0.576643 | 3473 | tags=58% | list=22% | signal=45% |
| BIOCARTA_S1P_PATHWAY | 6 | -0.68302 | -1.28811 | 0.194274 | 0.587041 | 0.576643 | 57 | tags=33% | list=0% | signal=33% |
| REACTOME_GLUTATHIONE_CONJUGATION | 30 | -0.52391 | -1.3153 | 0.194836 | 0.587336 | 0.576932 | 3137 | tags=50% | list=20% | signal=40% |
| BIOCARTA_EICOSANOID_PATHWAY | 18 | 0.700063 | 1.435499 | 0.198488 | 0.588729 | 0.578301 | 359 | tags=11% | list=2% | signal=11% |
| REACTOME_REGULATION_OF_KIT_SIGNALING | 13 | 0.705351 | 1.427213 | 0.197674 | 0.588729 | 0.578301 | 284 | tags=15% | list=2% | signal=15% |
| REACTOME_ACYL_CHAIN_REMODELLING_OF_PS | 14 | 0.69998 | 1.4259 | 0.195695 | 0.588729 | 0.578301 | 7 | tags=7% | list=0% | signal=7% |
| REACTOME_THROMBOXANE_SIGNALLING_THROUGH_TP_RECEPTOF | 21 | 0.673017 | 1.402614 | 0.196226 | 0.588729 | 0.578301 | 20 | tags=5% | list=0% | signal=5% |
| REACTOME_G_PROTEIN_MEDIATED_EVENTS | 38 | 0.611217 | 1.357929 | 0.197898 | 0.588729 | 0.578301 | 200 | tags=5% | list=1% | signal=5% |
| REACTOME_MTOR_SIGNALLING | 40 | -0.52757 | -1.33347 | 0.196721 | 0.588729 | 0.578301 | 2451 | tags=32% | list=16% | signal=27% |
| REACTOME_INTERLEUKIN_1_FAMILY_SIGNALING | 108 | -0.47 | -1.26712 | 0.19774 | 0.588729 | 0.578301 | 3355 | tags=41% | list=21% | signal=32% |
| REACTOME_SIGNALING_BY_VEGF | 99 | 0.50938 | 1.258988 | 0.198438 | 0.588729 | 0.578301 | 79 | tags=3% | list=1% | signal=3% |
| REACTOME_BIOLOGICAL_OXIDATIONS | 126 | 0.476762 | 1.218667 | 0.196621 | 0.588729 | 0.578301 | 359 | tags=5% | list=2% | signal=5% |
| REACTOME_NEPHRIN_FAMILY_INTERACTIONS | 8 | 0.685069 | 1.335319 | 0.199601 | 0.590975 | 0.580507 | 2206 | tags=38% | list=14% | signal=32% |
| BIOCARTA_FOSB_PATHWAY | 5 | 0.716209 | 1.283232 | 0.2 | 0.591103 | 0.580633 | 202 | tags=20% | list=1% | signal=20% |
| REACTOME_METABOLISM_OF_COFACTORS | 14 | 0.693122 | 1.411928 | 0.203523 | 0.593726 | 0.58321 | 65 | tags=7% | list=0% | signal=7% |
| BIOCARTA_BARR_MAPK_PATHWAY | 14 | -0.59121 | -1.33593 | 0.201629 | 0.593726 | 0.58321 | 3558 | tags=50% | list=23% | signal=39% |
| REACTOME_PHASE_0_RAPID_DEPOLARISATION | 8 | 0.682004 | 1.329343 | 0.201597 | 0.593726 | 0.58321 | 1631 | tags=38% | list=10% | signal=34% |
| REACTOME_ASPIRIN_ADME | 7 | 0.696171 | 1.319685 | 0.203593 | 0.593726 | 0.58321 | 2597 | tags=43% | list=17% | signal=36% |
| REACTOME_ARYL_HYDROCARBON_RECEPTOR_SIGNALLING | 7 | 0.695874 | 1.319122 | 0.203593 | 0.593726 | 0.58321 | 169 | tags=14% | list=1% | signal=14% |
| REACTOME_COLLAGEN_DEGRADATION | 40 | -0.51267 | -1.29581 | 0.203747 | 0.593726 | 0.58321 | 29 | tags=8% | list=0% | signal=8% |
| REACTOME_CATION_COUPLED_CHLORIDE_COTRANSPORTERS | 5 | 0.715134 | 1.281306 | 0.20198 | 0.593726 | 0.58321 | 3018 | tags=80% | list=19% | signal=65% |
| REACTOME_FC_EPSILON_RECEPTOR_FCERI_SIGNALING | 114 | -0.46882 | -1.25052 | 0.202899 | 0.593726 | 0.58321 | 3554 | tags=45% | list=23% | signal=35% |
| REACTOME_JOSEPHIN_DOMAIN_DUBS | 11 | -0.61539 | -1.31661 | 0.204545 | 0.594769 | 0.584234 | 676 | tags=36% | list=4% | signal=35% |
| REACTOME_RNA_POLYMERASE_III_TRANSCRIPTION_INITIATION_FR | 26 | -0.53103 | -1.28533 | 0.205069 | 0.594769 | 0.584234 | 2109 | tags=31% | list=13% | signal=27% |
| REACTOME_DOPAMINE_RECEPTORS | 4 | 0.737483 | 1.264981 | 0.205179 | 0.594769 | 0.584234 | 1559 | tags=25% | list=10% | signal=23% |
| REACTOME_ESTROGEN_DEPENDENT_NUCLEAR_EVENTS_DOWNSTR | 9 | -0.6324 | -1.30052 | 0.206122 | 0.595425 | 0.584878 | 77 | tags=22% | list=0% | signal=22% |
| WP_OSTEOBLAST_SIGNALING | 5 | 0.70506 | 1.263257 | 0.205941 | 0.595425 | 0.584878 | 2578 | tags=40% | list=16% | signal=33% |
| REACTOME_SYNTHESIS_OF_LEUKOTRIENES_LT_AND_EOXINS_EX | 12 | 0.687393 | 1.385269 | 0.206963 | 0.596816 | 0.586245 | 180 | tags=8% | list=1% | signal=8% |
| WP_OXIDATIVE_STRESS_RESPONSE | 26 | -0.53 | -1.28283 | 0.207373 | 0.596962 | 0.586388 | 1529 | tags=58% | list=10% | signal=52% |
| BIOCARTA_RARRXR_PATHWAY | 6 | 0.706143 | 1.303597 | 0.208577 | 0.598387 | 0.587788 | 1594 | tags=67% | list=10% | signal=60% |
| REACTOME_TRANSCRIPTIONAL_REGULATION_OF_GRANULOPOIESIS | 6 | -0.66732 | -1.25851 | 0.208589 | 0.598387 | 0.587788 | 552 | tags=33% | list=4% | signal=32% |
| REACTOME_NONHOMOLOGOUS_END_JOINING_NHEJ | 45 | -0.5117 | -1.30505 | 0.209135 | 0.598918 | 0.58831 | 2609 | tags=29% | list=17% | signal=24% |
| REACTOME_COBALAMIN_CBL_METABOLISM | 7 | -0.64506 | -1.27923 | 0.209581 | 0.599163 | 0.58855 | 572 | tags=71% | list=4% | signal=69% |
| BIOCARTA_MCM_PATHWAY | 13 | 0.693267 | 1.402763 | 0.21124 | 0.60287 | 0.592191 | 142 | tags=8% | list=1% | signal=8% |
| REACTOME_PRESYNAPTIC_DEPOLARIZATION_AND_CALCIUM_CHAN | 11 | 0.691855 | 1.369675 | 0.212355 | 0.604535 | 0.593827 | 2058 | tags=45% | list=13% | signal=40% |
| REACTOME_GLUCAGON_SIGNALING_IN_METABOLIC_REGULATION | 5 | -0.69227 | -1.26878 | 0.21328 | 0.604535 | 0.593827 | 3701 | tags=100% | list=24% | signal=76% |
| REACTOME_PKA_ACTIVATION_IN_GLUCAGON_SIGNALLING | 5 | -0.69227 | -1.26878 | 0.21328 | 0.604535 | 0.593827 | 3701 | tags=100% | list=24% | signal=76% |
| REACTOME_ESR_MEDIATED_SIGNALING | 118 | 0.480879 | 1.221061 | 0.21319 | 0.604535 | 0.593827 | 538 | tags=7% | list=3% | signal=7% |
| REACTOME_CS_DS_DEGRADATION | 13 | 0.689362 | 1.394861 | 0.215116 | 0.605575 | 0.594849 | 2910 | tags=46% | list=19% | signal=38% |
| REACTOME_NEGATIVE_REGULATION_OF_FGFR4_SIGNALING | 24 | -0.53267 | -1.27605 | 0.214765 | 0.605575 | 0.594849 | 2834 | tags=50% | list=18% | signal=41% |
| REACTOME_CHROMOSOME_MAINTENANCE | 91 | -0.46762 | -1.25714 | 0.21547 | 0.605575 | 0.594849 | 2134 | tags=18% | list=14% | signal=15% |
| REACTOME_POTASSIUM_CHANNELS | 86 | 0.507706 | 1.237995 | 0.215289 | 0.605575 | 0.594849 | 2838 | tags=30% | list=18% | signal=25% |
| REACTOME_TRANSCRIPTIONAL_REGULATION_BY_TP53 | 289 | -0.41042 | -1.19285 | 0.214058 | 0.605575 | 0.594849 | 2489 | tags=26% | list=16% | signal=22% |
| BIOCARTA_CCR3_PATHWAY | 15 | -0.5633 | -1.29347 | 0.216942 | 0.608684 | 0.597902 | 4077 | tags=60% | list=26% | signal=44% |
| REACTOME_CLASS_I_PEROXISOMAL_MEMBRANE_PROTEIN_IMPORT | 19 | -0.55471 | -1.28648 | 0.218143 | 0.610459 | 0.599646 | 2873 | tags=42% | list=18% | signal=34% |
| REACTOME_NEGATIVE_REGULATION_OF_FGFR2_SIGNALING | 29 | -0.5076 | -1.25816 | 0.21831 | 0.610459 | 0.599646 | 2834 | tags=45% | list=18% | signal=37% |
| REACTOME_THE_ACTIVATION_OF_ARYLSULFATASES | 8 | 0.672652 | 1.311116 | 0.219561 | 0.610966 | 0.600144 | 3901 | tags=75% | list=25% | signal=56% |
| REACTOME_DRUG_ADME | 55 | 0.560461 | 1.290656 | 0.219595 | 0.610966 | 0.600144 | 221 | tags=5% | list=1% | signal=5% |
| REACTOME_RECYCLING_OF_BILE_ACIDS_AND_SALTS | 7 | 0.676079 | 1.281598 | 0.219561 | 0.610966 | 0.600144 | 4545 | tags=71% | list=29% | signal=51% |
| REACTOME_PRC2_METHYLATES_HISTONES_AND_DNA | 33 | -0.51831 | -1.28453 | 0.220141 | 0.611461 | 0.60063 | 1888 | tags=27% | list=12% | signal=24% |
| WP_DISTAL_CONVOLUTED_TUBULE_1_DCT1_CELL | 5 | -0.68315 | -1.25207 | 0.221328 | 0.611817 | 0.60098 | 2412 | tags=60% | list=15% | signal=51% |
| REACTOME_CELLULAR_SENESCENCE | 89 | -0.46488 | -1.2514 | 0.220994 | 0.611817 | 0.60098 | 300 | tags=7% | list=2% | signal=7% |
| REACTOME_SIGNALING_BY_HEDGEHOG | 132 | 0.464784 | 1.196595 | 0.221374 | 0.611817 | 0.60098 | 408 | tags=5% | list=3% | signal=4% |
| REACTOME_TP53_REGULATES_TRANSCRIPTION_OF_CELL_DEATH_G | 15 | 0.675065 | 1.37847 | 0.222008 | 0.61255 | 0.6017 | 135 | tags=7% | list=1% | signal=7% |
| REACTOME_DERMATAN_SULFATE_BIOSYNTHESIS | 10 | 0.689353 | 1.351599 | 0.222857 | 0.612857 | 0.602002 | 3561 | tags=60% | list=23% | signal=46% |
| REACTOME_ER_TO_GOLGI_ANTEROGRADE_TRANSPORT | 139 | -0.44611 | -1.20285 | 0.222543 | 0.612857 | 0.602002 | 3672 | tags=45% | list=23% | signal=34% |
| REACTOME_ASYMMETRIC_LOCALIZATION_OF_PCP_PROTEINS | 60 | 0.550154 | 1.273415 | 0.223529 | 0.61369 | 0.60282 | 275 | tags=7% | list=2% | signal=7% |
| REACTOME_FRS_MEDIATED_FGFR4_SIGNALING | 14 | -0.57306 | -1.29491 | 0.224033 | 0.613774 | 0.602902 | 2767 | tags=36% | list=18% | signal=29% |
| REACTOME_SIGNALING_BY_TGFB_FAMILY_MEMBERS | 95 | 0.496561 | 1.224362 | 0.224299 | 0.613774 | 0.602902 | 2649 | tags=32% | list=17% | signal=26% |
| REACTOME_ONCOGENE_INDUCED_SENESCENCE | 16 | 0.678988 | 1.382915 | 0.225869 | 0.617053 | 0.606123 | 1020 | tags=25% | list=6% | signal=23% |
| BIOCARTA_TNFR2_PATHWAY | 17 | 0.677309 | 1.377621 | 0.226488 | 0.617727 | 0.606785 | 600 | tags=18% | list=4% | signal=17% |
| REACTOME_SMAD2_SMAD3_SMAD4_HETEROTRIMER_REGULATES_ | 25 | -0.52129 | -1.2492 | 0.227064 | 0.618285 | 0.607333 | 676 | tags=12% | list=4% | signal=12% |
| REACTOME_G1_S_TRANSITION | 103 | -0.45923 | -1.23404 | 0.22792 | 0.618587 | 0.60763 | 1901 | tags=28% | list=12% | signal=25% |
| REACTOME_PTEN_REGULATION | 110 | -0.46197 | -1.23167 | 0.227666 | 0.618587 | 0.60763 | 3335 | tags=42% | list=21% | signal=33% |
| REACTOME_GOLGI_CISTERNAE_PERICENTRIOLAR_STACK_REORGAN | 10 | -0.6007 | -1.25332 | 0.228512 | 0.619181 | 0.608213 | 4398 | tags=70% | list=28% | signal=50% |
| BIOCARTA_MTA3_PATHWAY | 19 | 0.666126 | 1.387963 | 0.230056 | 0.622349 | 0.611326 | 43 | tags=5% | list=0% | signal=5% |
| REACTOME_ADRENOCEPTORS | 9 | 0.676586 | 1.317665 | 0.230469 | 0.622453 | 0.611428 | 2310 | tags=44% | list=15% | signal=38% |
| REACTOME_EFFECTS_OF_PIP2_HYDROLYSIS | 25 | 0.625361 | 1.344663 | 0.231449 | 0.622556 | 0.611529 | 1558 | tags=20% | list=10% | signal=18% |
| WP_TYPE_II_INTERFERON_SIGNALING_IFNG | 25 | 0.625255 | 1.344435 | 0.231449 | 0.622556 | 0.611529 | 1174 | tags=20% | list=7% | signal=19% |
| REACTOME_SEROTONIN_RECEPTORS | 10 | 0.677275 | 1.327917 | 0.232381 | 0.622556 | 0.611529 | 2095 | tags=60% | list=13% | signal=52% |
| REACTOME_RUNX1_REGULATES_GENES_INVOLVED_IN_MEGAKARY | 38 | -0.49586 | -1.25815 | 0.232019 | 0.622556 | 0.611529 | 2900 | tags=37% | list=18% | signal=30% |
| REACTOME_POLO_LIKE_KINASE_MEDIATED_EVENTS | 5 | 0.696801 | 1.248459 | 0.231683 | 0.622556 | 0.611529 | 349 | tags=20% | list=2% | signal=20% |
| REACTOME_FORMATION_OF_TC_NER_PRE_INCISION_COMPLEX | 52 | -0.48528 | -1.2363 | 0.232843 | 0.62279 | 0.611758 | 3355 | tags=37% | list=21% | signal=29% |
| REACTOME_TRIGLYCERIDE_BIOSYNTHESIS | 8 | 0.656148 | 1.278947 | 0.233533 | 0.623631 | 0.612584 | 3440 | tags=50% | list=22% | signal=39% |
| REACTOME_SEMA3A_PLEXIN_REPULSION_SIGNALING_BY_INHIBITIN | 9 | 0.673498 | 1.311652 | 0.234375 | 0.624875 | 0.613806 | 2254 | tags=67% | list=14% | signal=57% |
| REACTOME_SYNTHESIS_OF_PIPS_AT_THE_EARLY_ENDOSOME_MEM | 15 | 0.665265 | 1.358458 | 0.235521 | 0.626487 | 0.61539 | 2011 | tags=33% | list=13% | signal=29% |
| REACTOME_SIGNAL_AMPLIFICATION | 29 | 0.614517 | 1.336018 | 0.236111 | 0.626487 | 0.61539 | 20 | tags=3% | list=0% | signal=3% |
| WP_MIR127_IN_MESENDODERM_DIFFERENTIATION | 6 | 0.67311 | 1.242615 | 0.235867 | 0.626487 | 0.61539 | 3885 | tags=67% | list=25% | signal=50% |
| REACTOME_ACTIVATION_OF_BAD_AND_TRANSLOCATION_TO_MITO | 12 | -0.58835 | -1.26943 | 0.237113 | 0.627142 | 0.616034 | 3753 | tags=58% | list=24% | signal=44% |

| | | | | | | | | | |
|---|---|---|---|---|---|---|---|---|---|
| REACTOME_GLUCOSE_METABOLISM | 70 | -0.46428 | -1.21315 | 0.236776 | 0.627142 | 0.616034 | 2491 | tags=34% | list=16% | signal=29% |
| REACTOME_PYROPTOSIS | 16 | -0.53277 | -1.23256 | 0.237603 | 0.627439 | 0.616325 | 1445 | tags=62% | list=9% | signal=57% |
| REACTOME_MITOCHONDRIAL_BIOGENESIS | 12 | -0.58811 | -1.2689 | 0.239175 | 0.629279 | 0.618133 | 1529 | tags=50% | list=10% | signal=45% |
| REACTOME_THYROXINE_BIOSYNTHESIS | 5 | -0.67156 | -1.23082 | 0.239437 | 0.629279 | 0.618133 | 4428 | tags=60% | list=28% | signal=43% |
| REACTOME_GLYCOLYSIS | 55 | -0.47222 | -1.21835 | 0.239024 | 0.629279 | 0.618133 | 1640 | tags=27% | list=10% | signal=25% |
| BIOCARTA_HCMV_PATHWAY | 17 | 0.667593 | 1.357858 | 0.241843 | 0.631605 | 0.620417 | 1291 | tags=24% | list=8% | signal=22% |
| REACTOME_REGULATION_OF_TP53_ACTIVITY_THROUGH_ACETYLAT | 29 | -0.49523 | -1.2275 | 0.241784 | 0.631605 | 0.620417 | 190 | tags=14% | list=1% | signal=14% |
| REACTOME_SPRY_REGULATION_OF_FGF_SIGNALING | 16 | -0.52923 | -1.22435 | 0.241736 | 0.631605 | 0.620417 | 2834 | tags=56% | list=18% | signal=46% |
| REACTOME_BETA_CATENIN_PHOSPHORYLATION_CASCADE | 17 | -0.52947 | -1.22196 | 0.241164 | 0.631605 | 0.620417 | 4073 | tags=59% | list=26% | signal=44% |
| REACTOME_A_TETRASACCHARIDE_LINKER_SEQUENCE_IS_REQUIREI | 24 | 0.63234 | 1.352477 | 0.243243 | 0.631754 | 0.620564 | 4566 | tags=62% | list=29% | signal=44% |
| REACTOME_CA2_PATHWAY | 42 | 0.577855 | 1.300209 | 0.242321 | 0.631754 | 0.620564 | 82 | tags=5% | list=1% | signal=5% |
| REACTOME_BIOSYNTHESIS_OF_SPECIALIZED_PRORESOLVING_MEDI | 11 | -0.5837 | -1.24881 | 0.243802 | 0.631754 | 0.620564 | 1552 | tags=27% | list=10% | signal=25% |
| REACTOME_ZINC_INFLUX_INTO_CELLS_BY_THE_SLC39_GENE_FAMI | 8 | 0.640488 | 1.248423 | 0.243513 | 0.631754 | 0.620564 | 4982 | tags=75% | list=32% | signal=51% |
| REACTOME_RUNX3_REGULATES_P14_ARF | 5 | 0.690542 | 1.237244 | 0.243564 | 0.631754 | 0.620564 | 392 | tags=40% | list=2% | signal=39% |
| REACTOME_BETA_CATENIN_INDEPENDENT_WNT_SIGNALING | 119 | 0.463845 | 1.180079 | 0.24424 | 0.631903 | 0.620711 | 275 | tags=5% | list=2% | signal=5% |
| REACTOME_SIGNALING_BY_NOTCH1 | 18 | 0.65725 | 1.34771 | 0.247637 | 0.633715 | 0.62249 | 392 | tags=17% | list=2% | signal=16% |
| REACTOME_CHONDROITIN_SULFATE_BIOSYNTHESIS | 18 | 0.655005 | 1.343106 | 0.251418 | 0.633715 | 0.62249 | 3195 | tags=50% | list=20% | signal=40% |
| REACTOME_PARACETAMOL_ADME | 16 | 0.657869 | 1.339902 | 0.252896 | 0.633715 | 0.62249 | 221 | tags=12% | list=1% | signal=12% |
| REACTOME_ENOS_ACTIVATION | 13 | 0.650544 | 1.316317 | 0.246124 | 0.633715 | 0.62249 | 65 | tags=8% | list=0% | signal=8% |
| REACTOME_RIPK1_MEDIATED_REGULATED_NECROSIS | 30 | 0.600233 | 1.314707 | 0.251736 | 0.633715 | 0.62249 | 37 | tags=3% | list=0% | signal=3% |
| REACTOME_BLOOD_GROUP_SYSTEMS_BIOSYNTHESIS | 12 | 0.642007 | 1.293806 | 0.251451 | 0.633715 | 0.62249 | 3013 | tags=58% | list=19% | signal=47% |
| REACTOME_SYNTHESIS_SECRETION_AND_DEACYLATION_OF_GHREL | 9 | 0.661989 | 1.289238 | 0.248047 | 0.633715 | 0.62249 | 2170 | tags=44% | list=14% | signal=38% |
| REACTOME_ERYTHROPOIETIN_ACTIVATES_RAS | 9 | 0.657358 | 1.280219 | 0.25 | 0.633715 | 0.62249 | 2572 | tags=44% | list=16% | signal=37% |
| REACTOME_ZINC_TRANSPORTERS | 10 | 0.65117 | 1.276735 | 0.253333 | 0.633715 | 0.62249 | 4982 | tags=80% | list=32% | signal=55% |
| BIOCARTA_PCAF_PATHWAY | 9 | 0.645919 | 1.257942 | 0.251953 | 0.633715 | 0.62249 | 1955 | tags=44% | list=12% | signal=39% |
| REACTOME_CARNITINE_METABOLISM | 13 | -0.56197 | -1.24452 | 0.253086 | 0.633715 | 0.62249 | 174 | tags=23% | list=1% | signal=23% |
| REACTOME_RECYCLING_OF_EIF2_GDP | 8 | -0.61506 | -1.23773 | 0.251497 | 0.633715 | 0.62249 | 4879 | tags=88% | list=31% | signal=60% |
| WP_NONHOMOLOGOUS_END_JOINING | 7 | -0.6221 | -1.2337 | 0.251497 | 0.633715 | 0.62249 | 1414 | tags=71% | list=9% | signal=65% |
| BIOCARTA_TERT_PATHWAY | 6 | 0.667994 | 1.23317 | 0.245614 | 0.633715 | 0.62249 | 2962 | tags=50% | list=19% | signal=41% |
| BIOCARTA_ARAP_PATHWAY | 11 | -0.57565 | -1.23159 | 0.25 | 0.633715 | 0.62249 | 3218 | tags=55% | list=20% | signal=43% |
| REACTOME_TRANSPORT_OF_RCBL_WITHIN_THE_BODY | 6 | 0.666539 | 1.230485 | 0.247563 | 0.633715 | 0.62249 | 5241 | tags=100% | list=33% | signal=67% |
| REACTOME_FORMATION_OF_THE_EARLY_ELONGATION_COMPLEX | 32 | -0.48819 | -1.22513 | 0.248244 | 0.633715 | 0.62249 | 3980 | tags=41% | list=25% | signal=30% |
| REACTOME_EPH_EPHRIN_SIGNALING | 64 | 0.52387 | 1.220137 | 0.252508 | 0.633715 | 0.62249 | 1829 | tags=28% | list=12% | signal=25% |
| REACTOME_N_GLYCAN_TRIMMING_IN_THE_ER_AND_CALNEXIN_C/ | 15 | -0.53065 | -1.21849 | 0.252066 | 0.633715 | 0.62249 | 2489 | tags=47% | list=16% | signal=39% |
| WP_FATTY_ACID_OXIDATION | 10 | -0.58057 | -1.21131 | 0.247379 | 0.633715 | 0.62249 | 2777 | tags=70% | list=18% | signal=58% |
| REACTOME_APOPTOSIS | 94 | 0.488789 | 1.202085 | 0.246106 | 0.633715 | 0.62249 | 97 | tags=3% | list=1% | signal=3% |
| REACTOME_S_PHASE | 143 | -0.43045 | -1.156 | 0.252874 | 0.633715 | 0.62249 | 3404 | tags=39% | list=22% | signal=31% |
| BIOCARTA_41BB_PATHWAY | 14 | 0.641347 | 1.306462 | 0.254403 | 0.633883 | 0.622655 | 1955 | tags=21% | list=12% | signal=19% |
| REACTOME_TCR_SIGNALING | 99 | -0.44801 | -1.20873 | 0.254144 | 0.633883 | 0.622655 | 3335 | tags=41% | list=21% | signal=33% |
| WP_PURINE_METABOLISM | 150 | 0.441799 | 1.155 | 0.254545 | 0.633883 | 0.622655 | 375 | tags=3% | list=2% | signal=3% |
| REACTOME_WNT5A_DEPENDENT_INTERNALIZATION_OF_FZD2_FZD | 12 | 0.634745 | 1.279171 | 0.255319 | 0.634858 | 0.623613 | 61 | tags=8% | list=0% | signal=8% |
| REACTOME_LAMININ_INTERACTIONS | 11 | 0.647726 | 1.282312 | 0.256757 | 0.636432 | 0.625159 | 4903 | tags=82% | list=31% | signal=56% |
| REACTOME_HEDGEHOG_LIGAND_BIOGENESIS | 60 | 0.534967 | 1.238263 | 0.257143 | 0.636432 | 0.625159 | 348 | tags=5% | list=2% | signal=5% |
| REACTOME_HSF1_ACTIVATION | 8 | -0.61083 | -1.22922 | 0.257485 | 0.636432 | 0.625159 | 4120 | tags=75% | list=26% | signal=55% |
| REACTOME_MET_INTERACTS_WITH_TNS_PROTEINS | 5 | 0.676819 | 1.212657 | 0.257426 | 0.636432 | 0.625159 | 1913 | tags=40% | list=12% | signal=35% |
| REACTOME_MITOCHONDRIAL_CALCIUM_ION_TRANSPORT | 16 | -0.51341 | -1.18776 | 0.258264 | 0.636465 | 0.625191 | 1661 | tags=38% | list=11% | signal=34% |
| REACTOME_ORGANELLE_BIOGENESIS_AND_MAINTENANCE | 185 | -0.41742 | -1.13918 | 0.258258 | 0.636465 | 0.625191 | 3096 | tags=34% | list=20% | signal=28% |
| REACTOME_RHO_GTPASES_ACTIVATE_WASPS_AND_WAVES | 32 | -0.4816 | -1.20859 | 0.259953 | 0.638101 | 0.626799 | 2844 | tags=56% | list=18% | signal=46% |
| REACTOME_INTRA_GOLGI_TRAFFIC | 34 | -0.48005 | -1.20342 | 0.260465 | 0.638101 | 0.626799 | 3119 | tags=38% | list=20% | signal=31% |
| REACTOME_RECRUITMENT_OF_NUMA_TO_MITOTIC_CENTROSOME | 84 | -0.44473 | -1.18081 | 0.260388 | 0.638101 | 0.626799 | 3676 | tags=44% | list=23% | signal=34% |
| REACTOME_MACROAUTOPHAGY | 112 | -0.43954 | -1.17177 | 0.260116 | 0.638101 | 0.626799 | 2907 | tags=41% | list=19% | signal=34% |
| REACTOME_MISCELLANEOUS_TRANSPORT_AND_BINDING_EVENTS | 19 | 0.636095 | 1.32539 | 0.269017 | 0.638155 | 0.626851 | 3406 | tags=58% | list=22% | signal=45% |
| REACTOME_NCAM1_INTERACTIONS | 16 | 0.643619 | 1.310877 | 0.26834 | 0.638155 | 0.626851 | 2694 | tags=38% | list=17% | signal=31% |
| REACTOME_REGULATION_OF_LIPID_METABOLISM_BY_PPARALPHA | 16 | 0.643437 | 1.310506 | 0.27027 | 0.638155 | 0.626851 | 3196 | tags=56% | list=20% | signal=45% |
| WP_NUCLEAR_RECEPTORS | 31 | 0.589149 | 1.294979 | 0.261324 | 0.638155 | 0.626851 | 1920 | tags=23% | list=12% | signal=20% |
| BIOCARTA_STATHMIN_PATHWAY | 14 | 0.618895 | 1.260723 | 0.268102 | 0.638155 | 0.626851 | 859 | tags=14% | list=5% | signal=14% |
| REACTOME_SODIUM_CALCIUM_EXCHANGERS | 12 | 0.624681 | 1.258889 | 0.26499 | 0.638155 | 0.626851 | 152 | tags=8% | list=1% | signal=8% |
| REACTOME_ERYTHROPOIETIN_ACTIVATES_PHOSPHOINOSITIDE_3_K | 11 | 0.635095 | 1.257308 | 0.266409 | 0.638155 | 0.626851 | 4464 | tags=55% | list=28% | signal=39% |
| WP_EXERCISEINDUCED_CIRCADIAN_REGULATION | 45 | 0.556655 | 1.249835 | 0.264505 | 0.638155 | 0.626851 | 787 | tags=18% | list=5% | signal=17% |
| REACTOME_ACTIVATED_NOTCH1_TRANSMITS_SIGNAL_TO_THE_NU | 10 | 0.631678 | 1.238516 | 0.266667 | 0.638155 | 0.626851 | 124 | tags=10% | list=1% | signal=10% |
| REACTOME_SYNTHESIS_OF_BILE_ACIDS_AND_BILE_SALTS_VIA_27_H | 9 | 0.63492 | 1.23652 | 0.261719 | 0.638155 | 0.626851 | 4731 | tags=67% | list=30% | signal=47% |
| BIOCARTA_PRION_PATHWAY | 10 | 0.629285 | 1.233824 | 0.270476 | 0.638155 | 0.626851 | 851 | tags=20% | list=5% | signal=19% |
| BIOCARTA_IL4_PATHWAY | 10 | 0.627378 | 1.230086 | 0.270476 | 0.638155 | 0.626851 | 4119 | tags=50% | list=26% | signal=37% |
| REACTOME_DOWNREGULATION_OF_ERBB4_SIGNALING | 9 | -0.59084 | -1.21505 | 0.267347 | 0.638155 | 0.626851 | 2526 | tags=56% | list=16% | signal=47% |
| REACTOME_MITOCHONDRIAL_UNCOUPLING | 5 | -0.66124 | -1.21191 | 0.265594 | 0.638155 | 0.626851 | 32 | tags=40% | list=0% | signal=40% |
| REACTOME_SYNTHESIS_OF_LIPOXINS_LX | 5 | 0.671645 | 1.203386 | 0.269307 | 0.638155 | 0.626851 | 4600 | tags=60% | list=29% | signal=42% |
| REACTOME_FLT3_SIGNALING | 29 | -0.48536 | -1.20305 | 0.265258 | 0.638155 | 0.626851 | 676 | tags=14% | list=4% | signal=13% |
| BIOCARTA_IFNA_PATHWAY | 7 | 0.634412 | 1.202612 | 0.265469 | 0.638155 | 0.626851 | 3398 | tags=71% | list=22% | signal=56% |
| REACTOME_NUCLEAR_ENVELOPE_NE_REASSEMBLY | 60 | -0.46075 | -1.19978 | 0.262899 | 0.638155 | 0.626851 | 2083 | tags=35% | list=13% | signal=30% |
| REACTOME_ESTROGEN_DEPENDENT_GENE_EXPRESSION | 57 | -0.46663 | -1.19926 | 0.261084 | 0.638155 | 0.626851 | 1888 | tags=35% | list=12% | signal=31% |
| REACTOME_ACTIVATION_OF_BH3_ONLY_PROTEINS | 18 | -0.51914 | -1.19893 | 0.266385 | 0.638155 | 0.626851 | 4136 | tags=61% | list=26% | signal=45% |
| WP_ERBB_SIGNALING_PATHWAY | 41 | -0.4731 | -1.19397 | 0.263529 | 0.638155 | 0.626851 | 377 | tags=17% | list=2% | signal=17% |
| REACTOME_RECOGNITION_OF_DNA_DAMAGE_BY_PCNA_CONTAINI | 29 | -0.4809 | -1.19198 | 0.269953 | 0.638155 | 0.626851 | 4509 | tags=45% | list=29% | signal=32% |
| REACTOME_PCP_CE_PATHWAY | 84 | 0.48768 | 1.187585 | 0.266771 | 0.638155 | 0.626851 | 275 | tags=6% | list=2% | signal=6% |
| REACTOME_ENERGY_DEPENDENT_REGULATION_OF_MTOR_BY_LKB | 28 | -0.48112 | -1.18384 | 0.262295 | 0.638155 | 0.626851 | 2418 | tags=29% | list=15% | signal=24% |
| REACTOME_RNA_POLYMERASE_III_TRANSCRIPTION | 34 | -0.47139 | -1.18171 | 0.267442 | 0.638155 | 0.626851 | 2109 | tags=26% | list=13% | signal=23% |
| REACTOME_ANTIGEN_PROCESSING_UBIQUITINATION_PROTEASOM | 281 | -0.37786 | -1.09426 | 0.266026 | 0.638155 | 0.626851 | 3367 | tags=33% | list=21% | signal=26% |
| REACTOME_IONOTROPIC_ACTIVITY_OF_KAINATE_RECEPTORS | 12 | 0.622194 | 1.253877 | 0.272727 | 0.642553 | 0.631172 | 1533 | tags=33% | list=10% | signal=30% |
| REACTOME_THE_PHOTOTRANSDUCTION_CASCADE | 21 | -0.51926 | -1.22633 | 0.273305 | 0.643002 | 0.631613 | 2273 | tags=52% | list=14% | signal=45% |
| WP_PRIMARY_FOCAL_SEGMENTAL_GLOMERULOSCLEROSIS_FSGS | 67 | 0.514842 | 1.210349 | 0.274834 | 0.645686 | 0.634249 | 1728 | tags=25% | list=11% | signal=23% |
| REACTOME_SIGNAL_ATTENUATION | 8 | 0.620548 | 1.209556 | 0.275449 | 0.646216 | 0.63477 | 2572 | tags=50% | list=16% | signal=42% |
| WP_GPCRS_PEPTIDE | 35 | 0.571005 | 1.258604 | 0.277193 | 0.648475 | 0.636989 | 5416 | tags=66% | list=34% | signal=43% |
| REACTOME_DNA_DOUBLE_STRAND_BREAK_RESPONSE | 53 | -0.45233 | -1.16114 | 0.277108 | 0.648475 | 0.636989 | 2609 | tags=28% | list=17% | signal=24% |
| REACTOME_TRAFFICKING_AND_PROCESSING_OF_ENDOSOMAL_TLR | 11 | 0.626572 | 1.240433 | 0.277992 | 0.649431 | 0.637927 | 3953 | tags=55% | list=25% | signal=41% |
| REACTOME_METAL_ION_SLC_TRANSPORTERS | 19 | 0.626437 | 1.305266 | 0.280148 | 0.649731 | 0.638222 | 4982 | tags=79% | list=32% | signal=54% |
| REACTOME_ENDOGENOUS_STEROLS | 20 | 0.626585 | 1.300431 | 0.280603 | 0.649731 | 0.638222 | 229 | tags=10% | list=1% | signal=10% |
| REACTOME_COHESIN_LOADING_ONTO_CHROMATIN | 10 | 0.622124 | 1.219784 | 0.28 | 0.649731 | 0.638222 | 3172 | tags=60% | list=20% | signal=48% |
| REACTOME_GAP_FILLING_DNA_REPAIR_SYNTHESIS_AND_LIGATION_ | 21 | -0.50945 | -1.20315 | 0.279661 | 0.649731 | 0.638222 | 3984 | tags=43% | list=25% | signal=32% |
| REACTOME_GLYCOPROTEIN_HORMONES | 3 | 0.735013 | 1.185526 | 0.280943 | 0.649731 | 0.638222 | 2387 | tags=67% | list=15% | signal=57% |
| REACTOME_PLATELET_HOMEOSTASIS | 71 | 0.499365 | 1.184591 | 0.28013 | 0.649731 | 0.638222 | 65 | tags=3% | list=0% | signal=3% |

| | | | | | | | | | | |
|---|---|---|---|---|---|---|---|---|---|---|
| REACTOME_SIGNALING_BY_FGFR4 | 33 | -0.4749 | -1.17695 | 0.278689 | 0.649731 | 0.638222 | 2834 | tags=39% | list=18% | signal=32% |
| REACTOME_NEGATIVE_REGULATION_OF_MAPK_PATHWAY | 42 | -0.46471 | -1.17157 | 0.28125 | 0.649731 | 0.638222 | 2948 | tags=55% | list=19% | signal=45% |
| REACTOME_TRP_CHANNELS | 21 | 0.622136 | 1.296574 | 0.284906 | 0.651475 | 0.639936 | 2976 | tags=43% | list=19% | signal=35% |
| REACTOME_ANTIGEN_ACTIVATES_B_CELL_RECEPTOR_BCR_LEADING | 22 | 0.610503 | 1.295861 | 0.288603 | 0.651475 | 0.639936 | 55 | tags=5% | list=0% | signal=5% |
| BIOCARTA_TOLL_PATHWAY | 24 | 0.603191 | 1.290131 | 0.284685 | 0.651475 | 0.639936 | 1955 | tags=25% | list=12% | signal=22% |
| BIOCARTA_TEL_PATHWAY | 17 | 0.632619 | 1.286723 | 0.287908 | 0.651475 | 0.639936 | 146 | tags=6% | list=1% | signal=6% |
| WP_GLYCEROLIPIDS_AND_GLYCEROPHOSPHOLIPIDS | 22 | 0.604974 | 1.284126 | 0.290441 | 0.651475 | 0.639936 | 2347 | tags=23% | list=15% | signal=19% |
| REACTOME_SIGNALING_BY_ERYTHROPOIETIN | 16 | 0.621258 | 1.265334 | 0.291506 | 0.651475 | 0.639936 | 3601 | tags=44% | list=23% | signal=34% |
| REACTOME_METABOLISM_OF_NITRIC_OXIDE_NOS3_ACTIVATION_A | 16 | 0.618022 | 1.258743 | 0.295367 | 0.651475 | 0.639936 | 65 | tags=6% | list=0% | signal=6% |
| REACTOME_BUTYRATE_RESPONSE_FACTOR_1_BRF1_BINDS_AND_D | 16 | 0.615343 | 1.253288 | 0.295367 | 0.651475 | 0.639936 | 159 | tags=6% | list=1% | signal=6% |
| WP_GLYCOLYSIS_AND_GLUCONEOGENESIS | 38 | 0.563236 | 1.251329 | 0.290718 | 0.651475 | 0.639936 | 59 | tags=5% | list=0% | signal=5% |
| WP_APOPTOSIS_MODULATION_BY_HSP70 | 16 | 0.614358 | 1.251282 | 0.297297 | 0.651475 | 0.639936 | 3832 | tags=44% | list=24% | signal=33% |
| BIOCARTA_CARM1_PATHWAY | 11 | 0.612836 | 1.213241 | 0.291506 | 0.651475 | 0.639936 | 1291 | tags=45% | list=8% | signal=42% |
| REACTOME_RHOQ_GTPASE_CYCLE | 58 | 0.523882 | 1.212929 | 0.283557 | 0.651475 | 0.639936 | 1313 | tags=21% | list=8% | signal=19% |
| REACTOME_TRANSPORT_OF_ORGANIC_ANIONS | 9 | 0.618137 | 1.203835 | 0.289063 | 0.651475 | 0.639936 | 3697 | tags=67% | list=24% | signal=51% |
| REACTOME_ABC_TRANSPORTERS_IN_LIPID_HOMEOSTASIS | 13 | 0.592578 | 1.199028 | 0.296512 | 0.651475 | 0.639936 | 3418 | tags=62% | list=22% | signal=48% |
| REACTOME_TRANSPORT_OF_NUCLEOTIDE_SUGARS | 9 | 0.612148 | 1.192171 | 0.292969 | 0.651475 | 0.639936 | 5311 | tags=89% | list=34% | signal=59% |
| REACTOME_BICARBONATE_TRANSPORTERS | 9 | 0.611417 | 1.190748 | 0.296875 | 0.651475 | 0.639936 | 1833 | tags=33% | list=12% | signal=29% |
| WP_ALPHA_6_BETA_4_INTEGRIN_SIGNALING_PATHWAY | 63 | 0.513456 | 1.188651 | 0.294416 | 0.651475 | 0.639936 | 340 | tags=5% | list=2% | signal=5% |
| BIOCARTA_PLATELETAPP_PATHWAY | 10 | 0.605959 | 1.18809 | 0.291429 | 0.651475 | 0.639936 | 4733 | tags=60% | list=30% | signal=42% |
| REACTOME_CALNEXIN_CALRETICULIN_CYCLE | 7 | -0.59781 | -1.18553 | 0.297405 | 0.651475 | 0.639936 | 676 | tags=43% | list=4% | signal=41% |
| REACTOME_FRS_MEDIATED_FGFR2_SIGNALING | 19 | -0.51074 | -1.1845 | 0.289417 | 0.651475 | 0.639936 | 2772 | tags=32% | list=18% | signal=26% |
| REACTOME_TRYPTOPHAN_CATABOLISM | 8 | 0.605587 | 1.180394 | 0.289421 | 0.651475 | 0.639936 | 1918 | tags=38% | list=12% | signal=33% |
| REACTOME_NF_KB_IS_ACTIVATED_AND_SIGNALS_SURVIVAL | 13 | -0.53161 | -1.17728 | 0.296296 | 0.651475 | 0.639936 | 676 | tags=23% | list=4% | signal=22% |
| REACTOME_PYRIMIDINE_CATABOLISM | 10 | -0.56346 | -1.17561 | 0.291405 | 0.651475 | 0.639936 | 959 | tags=20% | list=6% | signal=19% |
| REACTOME_PURINE_SALVAGE | 12 | -0.54451 | -1.17483 | 0.284536 | 0.651475 | 0.639936 | 3401 | tags=42% | list=22% | signal=33% |
| REACTOME_TRANSCRIPTIONAL_ACTIVATION_OF_MITOCHONDRIAL_ | 10 | -0.56288 | -1.1744 | 0.291405 | 0.651475 | 0.639936 | 1529 | tags=50% | list=10% | signal=45% |
| REACTOME_SCAVENGING_BY_CLASS_A_RECEPTORS | 6 | 0.634659 | 1.171632 | 0.28655 | 0.651475 | 0.639936 | 3721 | tags=50% | list=24% | signal=38% |
| REACTOME_ORGANIC_ANION_TRANSPORTERS | 8 | -0.58169 | -1.17058 | 0.291417 | 0.651475 | 0.639936 | 716 | tags=25% | list=5% | signal=24% |
| REACTOME_REGULATION_OF_CHOLESTEROL_BIOSYNTHESIS_BY_SRI | 7 | 0.616895 | 1.169407 | 0.287425 | 0.651475 | 0.639936 | 1615 | tags=29% | list=10% | signal=26% |
| BIOCARTA_MCALPAIN_PATHWAY | 18 | -0.50581 | -1.16815 | 0.289641 | 0.651475 | 0.639936 | 3701 | tags=67% | list=24% | signal=51% |
| REACTOME_ATTACHMENT_OF_GPI_ANCHOR_TO_UPAR | 7 | 0.613469 | 1.162913 | 0.291417 | 0.651475 | 0.639936 | 3537 | tags=71% | list=23% | signal=55% |
| WP_TRANSLATION_FACTORS | 47 | -0.45187 | -1.16262 | 0.28436 | 0.651475 | 0.639936 | 4670 | tags=64% | list=30% | signal=45% |
| REACTOME_LECTIN_PATHWAY_OF_COMPLEMENT_ACTIVATION | 4 | 0.6773 | 1.161751 | 0.292829 | 0.651475 | 0.639936 | 1662 | tags=25% | list=11% | signal=22% |
| REACTOME_G2_M_DNA_DAMAGE_CHECKPOINT | 68 | -0.44433 | -1.16144 | 0.283208 | 0.651475 | 0.639936 | 2609 | tags=24% | list=17% | signal=20% |
| REACTOME_ACTIVATION_OF_THE_TFAP2_AP_2_FAMILY_OF_TRANS | 8 | 0.593089 | 1.156034 | 0.295409 | 0.651475 | 0.639936 | 1701 | tags=25% | list=11% | signal=22% |
| REACTOME_FGFR2_ALTERNATIVE_SPLICING | 17 | -0.50051 | -1.15512 | 0.295218 | 0.651475 | 0.639936 | 2615 | tags=41% | list=17% | signal=34% |
| REACTOME_INHIBITION_OF_REPLICATION_INITIATION_OF_DAMAGI | 10 | -0.55353 | -1.1549 | 0.297694 | 0.651475 | 0.639936 | 1776 | tags=70% | list=11% | signal=62% |
| REACTOME_SIGNALING_BY_FGFR2 | 57 | -0.44622 | -1.14679 | 0.283251 | 0.651475 | 0.639936 | 2834 | tags=37% | list=18% | signal=30% |
| REACTOME_IRS_MEDIATED_SIGNALLING | 33 | -0.46269 | -1.14668 | 0.297424 | 0.651475 | 0.639936 | 534 | tags=15% | list=3% | signal=15% |
| REACTOME_RHO_GTPASES_ACTIVATE_FORMINS | 126 | -0.42039 | -1.13848 | 0.293447 | 0.651475 | 0.639936 | 2866 | tags=32% | list=18% | signal=26% |
| REACTOME_CELL_CYCLE | 543 | -0.35476 | -1.07902 | 0.295276 | 0.651475 | 0.639936 | 2878 | tags=31% | list=18% | signal=26% |
| REACTOME_COLLAGEN_CHAIN_TRIMERIZATION | 35 | 0.559045 | 1.232243 | 0.3 | 0.654307 | 0.642718 | 3579 | tags=43% | list=23% | signal=33% |
| BIOCARTA_LONGEVITY_PATHWAY | 14 | 0.581659 | 1.184873 | 0.30137 | 0.654307 | 0.642718 | 2223 | tags=36% | list=14% | signal=31% |
| REACTOME_REGULATED_PROTEOLYSIS_OF_P75NTR | 10 | 0.600022 | 1.17645 | 0.300952 | 0.654307 | 0.642718 | 4576 | tags=60% | list=29% | signal=43% |
| REACTOME_PEPTIDE_HORMONE_BIOSYNTHESIS | 5 | 0.652881 | 1.169767 | 0.30099 | 0.654307 | 0.642718 | 2387 | tags=60% | list=15% | signal=51% |
| REACTOME_PROCESSING_OF_DNA_DOUBLE_STRAND_BREAK_ENDS | 66 | -0.44209 | -1.1532 | 0.301746 | 0.654307 | 0.642718 | 1270 | tags=14% | list=8% | signal=13% |
| REACTOME_CREB_PHOSPHORYLATION | 7 | 0.605958 | 1.148674 | 0.301397 | 0.654307 | 0.642718 | 1291 | tags=29% | list=8% | signal=26% |
| REACTOME_CYTOSOLIC_SENSORS_OF_PATHOGEN_ASSOCIATED_DN | 31 | -0.45342 | -1.14046 | 0.301402 | 0.654307 | 0.642718 | 3135 | tags=39% | list=20% | signal=31% |
| REACTOME_PEPTIDE_LIGAND_BINDING_RECEPTORS | 100 | -0.4212 | -1.13726 | 0.302198 | 0.654434 | 0.642842 | 70 | tags=3% | list=0% | signal=3% |
| BIOCARTA_TOB1_PATHWAY | 10 | 0.597832 | 1.172155 | 0.302857 | 0.655007 | 0.643405 | 4633 | tags=80% | list=30% | signal=56% |
| REACTOME_SLBP_DEPENDENT_PROCESSING_OF_REPLICATION_DEP | 11 | -0.54088 | -1.15719 | 0.303719 | 0.655722 | 0.644108 | 5319 | tags=73% | list=34% | signal=48% |
| REACTOME_MITOTIC_G1_PHASE_AND_G1_S_TRANSITION | 127 | -0.41174 | -1.11942 | 0.303977 | 0.655722 | 0.644108 | 2063 | tags=28% | list=13% | signal=24% |
| REACTOME_RHO_GTPASES_ACTIVATE_PAKS | 22 | -0.49238 | -1.15923 | 0.305677 | 0.658533 | 0.646869 | 2708 | tags=41% | list=17% | signal=34% |
| REACTOME_NUCLEAR_RECEPTOR_TRANSCRIPTION_PATHWAY | 38 | 0.554667 | 1.232293 | 0.30648 | 0.659408 | 0.647728 | 1770 | tags=18% | list=11% | signal=16% |
| REACTOME_METABOLISM_OF_CARBOHYDRATES | 242 | 0.402659 | 1.105112 | 0.306902 | 0.659461 | 0.64778 | 2183 | tags=21% | list=14% | signal=19% |
| REACTOME_GLOBAL_GENOME_NUCLEOTIDE_EXCISION_REPAIR_GG | 80 | -0.42228 | -1.12635 | 0.307692 | 0.660306 | 0.64861 | 3153 | tags=29% | list=20% | signal=23% |
| BIOCARTA_P35ALZHEIMERS_PATHWAY | 6 | -0.61582 | -1.16137 | 0.308793 | 0.661814 | 0.650091 | 3139 | tags=50% | list=20% | signal=40% |
| BIOCARTA_STRESS_PATHWAY | 23 | 0.594729 | 1.265665 | 0.310219 | 0.664013 | 0.652251 | 1999 | tags=26% | list=13% | signal=23% |
| REACTOME_MTORC1_MEDIATED_SIGNALLING | 24 | -0.47507 | -1.13807 | 0.310962 | 0.664746 | 0.652972 | 2451 | tags=46% | list=16% | signal=39% |
| REACTOME_REGULATION_OF_FZD_BY_UBIQUITINATION | 18 | 0.601379 | 1.233146 | 0.311909 | 0.665059 | 0.653279 | 937 | tags=17% | list=6% | signal=16% |
| REACTOME_IRAK1_RECRUITS_IKK_COMPLEX | 14 | -0.51836 | -1.17131 | 0.311609 | 0.665059 | 0.653279 | 2489 | tags=29% | list=16% | signal=24% |
| BIOCARTA_NFKB_PATHWAY | 20 | 0.603375 | 1.252261 | 0.312618 | 0.665618 | 0.653828 | 425 | tags=10% | list=3% | signal=10% |
| REACTOME_THE_FATTY_ACID_CYCLING_MODEL | 4 | -0.67437 | -1.17679 | 0.313373 | 0.665618 | 0.653828 | 32 | tags=25% | list=0% | signal=25% |
| REACTOME_TRANSCRIPTIONAL_REGULATION_BY_RUNX1 | 143 | -0.4101 | -1.10136 | 0.313218 | 0.665618 | 0.653828 | 3340 | tags=41% | list=21% | signal=33% |
| BIOCARTA_IGF1_PATHWAY | 22 | 0.584077 | 1.239769 | 0.316176 | 0.666458 | 0.654653 | 2170 | tags=32% | list=14% | signal=27% |
| BIOCARTA_IGF1MTOR_PATHWAY | 18 | 0.599179 | 1.228633 | 0.31569 | 0.666458 | 0.654653 | 2170 | tags=28% | list=14% | signal=24% |
| WP_FIBRIN_COMPLEMENT_RECEPTOR_3_SIGNALING_PATHWAY | 34 | 0.552336 | 1.217196 | 0.314685 | 0.666458 | 0.654653 | 971 | tags=9% | list=6% | signal=8% |
| REACTOME_LDL_CLEARANCE | 15 | -0.50148 | -1.15151 | 0.316116 | 0.666458 | 0.654653 | 2285 | tags=53% | list=15% | signal=46% |
| REACTOME_NEGATIVE_REGULATION_OF_ACTIVITY_OF_TFAP2_AP_2 | 6 | 0.616048 | 1.137274 | 0.315789 | 0.666458 | 0.654653 | 5184 | tags=67% | list=33% | signal=45% |
| REACTOME_INSULIN_RECEPTOR_RECYCLING | 25 | -0.47038 | -1.12719 | 0.31422 | 0.666458 | 0.654653 | 3303 | tags=48% | list=21% | signal=38% |
| REACTOME_INTERLEUKIN_RECEPTOR_SHC_SIGNALING | 16 | 0.590878 | 1.203458 | 0.316602 | 0.66651 | 0.654704 | 4553 | tags=56% | list=29% | signal=40% |
| REACTOME_PLASMA_LIPOPROTEIN_ASSEMBLY | 12 | 0.583665 | 1.17623 | 0.317215 | 0.666954 | 0.65514 | 3122 | tags=58% | list=20% | signal=47% |
| REACTOME_SEMA4D_INDUCED_CELL_MIGRATION_AND_GROWTH_ | 11 | -0.5239 | -1.12087 | 0.318182 | 0.667298 | 0.655478 | 411 | tags=64% | list=3% | signal=62% |
| BIOCARTA_BIOPEPTIDES_PATHWAY | 27 | -0.45527 | -1.11896 | 0.317972 | 0.667298 | 0.655478 | 3624 | tags=48% | list=23% | signal=37% |
| REACTOME_IL_6_TYPE_CYTOKINE_RECEPTOR_LIGAND_INTERACTIOI | 12 | 0.579767 | 1.168375 | 0.319149 | 0.66764 | 0.655814 | 1067 | tags=25% | list=7% | signal=23% |
| REACTOME_PHYSIOLOGICAL_FACTORS | 5 | 0.641304 | 1.149024 | 0.318812 | 0.66764 | 0.655814 | 4091 | tags=60% | list=26% | signal=44% |
| REACTOME_TP53_REGULATES_TRANSCRIPTION_OF_DNA_REPAIR_G | 43 | -0.43719 | -1.104 | 0.319613 | 0.667769 | 0.655941 | 1863 | tags=23% | list=12% | signal=21% |
| REACTOME_ROLE_OF_LAT2_NTAL_LAB_ON_CALCIUM_MOBILIZATIC | 12 | 0.575762 | 1.160304 | 0.321083 | 0.669158 | 0.657306 | 4553 | tags=50% | list=29% | signal=36% |
| REACTOME_PLASMA_LIPOPROTEIN_CLEARANCE | 28 | -0.4485 | -1.10358 | 0.320843 | 0.669158 | 0.657306 | 2489 | tags=39% | list=16% | signal=33% |
| REACTOME_RHOBTB1_GTPASE_CYCLE | 22 | 0.575064 | 1.220638 | 0.323529 | 0.669569 | 0.657709 | 1831 | tags=23% | list=12% | signal=20% |
| BIOCARTA_INTRINSIC_PATHWAY | 13 | 0.576933 | 1.167372 | 0.323643 | 0.669569 | 0.657709 | 4733 | tags=69% | list=30% | signal=48% |
| REACTOME_COMMON_PATHWAY_OF_FIBRIN_CLOT_FORMATION | 12 | 0.570007 | 1.148706 | 0.323017 | 0.669569 | 0.657709 | 2242 | tags=42% | list=14% | signal=36% |
| REACTOME_HDL_REMODELING | 5 | 0.638345 | 1.143723 | 0.322772 | 0.669569 | 0.657709 | 3799 | tags=60% | list=24% | signal=46% |
| WP_STATIN_PATHWAY | 14 | 0.560266 | 1.141295 | 0.324853 | 0.669569 | 0.657709 | 3799 | tags=50% | list=24% | signal=38% |
| BIOCARTA_PRC2_PATHWAY | 10 | 0.579781 | 1.136763 | 0.325714 | 0.669569 | 0.657709 | 4856 | tags=70% | list=31% | signal=48% |
| REACTOME_RHO_GTPASES_ACTIVATE_PKNS | 22 | -0.48174 | -1.1342 | 0.325328 | 0.669569 | 0.657709 | 3405 | tags=55% | list=22% | signal=43% |
| BIOCARTA_CTBP1_PATHWAY | 8 | -0.55991 | -1.12675 | 0.325349 | 0.669569 | 0.657709 | 3097 | tags=62% | list=20% | signal=50% |
| REACTOME_NONCANONICAL_ACTIVATION_OF_NOTCH3 | 7 | 0.587098 | 1.112922 | 0.325349 | 0.669569 | 0.657709 | 3746 | tags=71% | list=24% | signal=54% |
| REACTOME_ACYL_CHAIN_REMODELLING_OF_PE | 17 | -0.48084 | -1.10973 | 0.324324 | 0.669569 | 0.657709 | 525 | tags=29% | list=3% | signal=28% |

| | | | | | | | | | | |
|---|---|---|---|---|---|---|---|---|---|---|
| REACTOME_RESOLUTION_OF_SISTER_CHROMATID_COHESION | 112 | -0.40874 | -1.08967 | 0.323699 | 0.669569 | 0.657709 | 2866 | tags=29% | list=18% | signal=24% |
| REACTOME_GLUCAGON_LIKE_PEPTIDE_1_GLP1_REGULATES_INSULI | 29 | 0.556012 | 1.208822 | 0.326389 | 0.670126 | 0.658256 | 233 | tags=7% | list=1% | signal=7% |
| REACTOME_RECEPTOR_TYPE_TYROSINE_PROTEIN_PHOSPHATASES | 20 | 0.584837 | 1.213786 | 0.327684 | 0.671125 | 0.659238 | 1235 | tags=45% | list=8% | signal=42% |
| REACTOME_SELECTIVE_AUTOPHAGY | 57 | -0.43073 | -1.10698 | 0.327586 | 0.671125 | 0.659238 | 1662 | tags=49% | list=11% | signal=44% |
| REACTOME_TRANSCRIPTIONAL_REGULATION_BY_E2F6 | 23 | 0.577524 | 1.22905 | 0.328467 | 0.671901 | 0.66 | 4856 | tags=61% | list=31% | signal=42% |
| REACTOME_NETRIN_1_SIGNALING | 21 | 0.57228 | 1.192671 | 0.330189 | 0.674592 | 0.662643 | 313 | tags=10% | list=2% | signal=9% |
| WP_FOLIC_ACID_NETWORK | 20 | 0.570891 | 1.184841 | 0.333333 | 0.676028 | 0.664053 | 62 | tags=5% | list=0% | signal=5% |
| BIOCARTA_TCRA_PATHWAY | 5 | -0.63233 | -1.15892 | 0.331992 | 0.676028 | 0.664053 | 2272 | tags=100% | list=14% | signal=86% |
| BIOCARTA_HSWI_SNF_PATHWAY | 10 | -0.53222 | -1.11044 | 0.333333 | 0.676028 | 0.664053 | 1295 | tags=60% | list=8% | signal=55% |
| REACTOME_INACTIVATION_OF_CSF3_G_CSF_SIGNALING | 17 | -0.47398 | -1.09391 | 0.33264 | 0.676028 | 0.664053 | 972 | tags=35% | list=6% | signal=33% |
| BIOCARTA_CDK5_PATHWAY | 11 | -0.51 | -1.09113 | 0.332645 | 0.676028 | 0.664053 | 3558 | tags=73% | list=23% | signal=56% |
| REACTOME_TRANSCRIPTION_COUPLED_NUCLEOTIDE_EXCISION_REI | 75 | -0.41257 | -1.09021 | 0.331606 | 0.676028 | 0.664053 | 3153 | tags=28% | list=20% | signal=22% |
| REACTOME_MITOTIC_PROPHASE | 81 | -0.40952 | -1.09806 | 0.334218 | 0.676994 | 0.665003 | 2834 | tags=27% | list=18% | signal=22% |
| BIOCARTA_RELA_PATHWAY | 14 | 0.55079 | 1.12199 | 0.334638 | 0.67702 | 0.665028 | 3519 | tags=43% | list=22% | signal=33% |
| REACTOME_MULTIFUNCTIONAL_ANION_EXCHANGERS | 6 | 0.601222 | 1.109904 | 0.335283 | 0.677499 | 0.665499 | 4347 | tags=83% | list=28% | signal=60% |
| REACTOME_PROCESSING_OF_INTRONLESS_PRE_MRNAS | 20 | 0.5677 | 1.178219 | 0.3371 | 0.677753 | 0.665748 | 4098 | tags=50% | list=26% | signal=37% |
| REACTOME_MITOTIC_TELOPHASE_CYTOKINESIS | 13 | 0.563014 | 1.139208 | 0.337209 | 0.677753 | 0.665748 | 3460 | tags=54% | list=22% | signal=42% |
| WP_ESTROGEN_SIGNALING | 73 | 0.475789 | 1.130606 | 0.336601 | 0.677753 | 0.665748 | 532 | tags=7% | list=3% | signal=7% |
| REACTOME_MAP3K8_TPL2_DEPENDENT_MAPK1_3_ACTIVATION | 13 | -0.50357 | -1.11518 | 0.337449 | 0.677753 | 0.665748 | 3063 | tags=38% | list=20% | signal=31% |
| REACTOME_NEGATIVE_REGULATION_OF_FGFR3_SIGNALING | 26 | -0.45522 | -1.10184 | 0.336406 | 0.677753 | 0.665748 | 2834 | tags=46% | list=18% | signal=38% |
| REACTOME_OPIOID_SIGNALLING | 68 | 0.489612 | 1.14915 | 0.338308 | 0.678054 | 0.666044 | 200 | tags=4% | list=1% | signal=4% |
| REACTOME_PROGRAMMED_CELL_DEATH | 122 | 0.432331 | 1.108232 | 0.338415 | 0.678054 | 0.666044 | 97 | tags=2% | list=1% | signal=2% |
| REACTOME_SYNTHESIS_OF_PIPS_AT_THE_GOLGI_MEMBRANE | 15 | 0.553152 | 1.129525 | 0.339768 | 0.679946 | 0.667902 | 2783 | tags=47% | list=18% | signal=38% |
| REACTOME_PHOSPHATE_BOND_HYDROLYSIS_BY_NTPDASE_PROTEI | 6 | 0.600273 | 1.108152 | 0.341131 | 0.681851 | 0.669773 | 3007 | tags=50% | list=19% | signal=40% |
| REACTOME_TRANSCRIPTIONAL_REGULATION_BY_MECP2 | 5 | -0.62327 | -1.14232 | 0.346076 | 0.681925 | 0.669846 | 221 | tags=40% | list=1% | signal=39% |
| WP_ALZHEIMERS_DISEASE | 69 | 0.481149 | 1.136089 | 0.343802 | 0.681925 | 0.669846 | 2077 | tags=20% | list=13% | signal=18% |
| REACTOME_P38MAPK_EVENTS | 7 | -0.57058 | -1.13152 | 0.347305 | 0.681925 | 0.669846 | 1828 | tags=71% | list=12% | signal=63% |
| REACTOME_NEGATIVE_REGULATION_OF_TCF_DEPENDENT_SIGNALI | 5 | 0.622863 | 1.115983 | 0.344554 | 0.681925 | 0.669846 | 5645 | tags=80% | list=36% | signal=51% |
| REACTOME_CRMPS_IN_SEMA3A_SIGNALING | 15 | 0.546334 | 1.115603 | 0.341699 | 0.681925 | 0.669846 | 2386 | tags=40% | list=15% | signal=34% |
| REACTOME_SYNTHESIS_OF_BILE_ACIDS_AND_BILE_SALTS_VIA_24_I | 9 | 0.571207 | 1.112438 | 0.34375 | 0.681925 | 0.669846 | 4187 | tags=56% | list=27% | signal=41% |
| REACTOME_METALLOPROTEASE_DUBS | 23 | -0.46243 | -1.1034 | 0.345815 | 0.681925 | 0.669846 | 3554 | tags=48% | list=23% | signal=37% |
| REACTOME_PI_3K_CASCADE_FGFR4 | 13 | -0.49374 | -1.09342 | 0.347737 | 0.681925 | 0.669846 | 2767 | tags=31% | list=18% | signal=25% |
| REACTOME_SIGNALING_BY_INSULIN_RECEPTOR | 62 | -0.4163 | -1.0907 | 0.345588 | 0.681925 | 0.669846 | 3073 | tags=29% | list=20% | signal=23% |
| REACTOME_FOXO_MEDIATED_TRANSCRIPTION | 21 | -0.46177 | -1.09056 | 0.347458 | 0.681925 | 0.669846 | 2365 | tags=29% | list=15% | signal=24% |
| REACTOME_NUCLEOTIDE_SALVAGE | 21 | -0.46177 | -1.09056 | 0.347458 | 0.681925 | 0.669846 | 3794 | tags=48% | list=24% | signal=36% |
| REACTOME_INTERLEUKIN_27_SIGNALING | 8 | 0.557294 | 1.086262 | 0.347305 | 0.681925 | 0.669846 | 5115 | tags=75% | list=33% | signal=51% |
| REACTOME_PLASMA_LIPOPROTEIN_ASSEMBLY_REMODELING_AND_ | 49 | -0.4235 | -1.08624 | 0.347722 | 0.681925 | 0.669846 | 1227 | tags=29% | list=8% | signal=26% |
| REACTOME_ALPHA_PROTEIN_KINASE_1_SIGNALING_PATHWAY | 11 | -0.50184 | -1.07367 | 0.345041 | 0.681925 | 0.669846 | 676 | tags=27% | list=4% | signal=26% |
| REACTOME_TRANSCRIPTIONAL_ACTIVITY_OF_SMAD2_SMAD3_SMA | 38 | -0.42104 | -1.06831 | 0.343387 | 0.681925 | 0.669846 | 676 | tags=8% | list=4% | signal=8% |
| REACTOME_TRANSLESION_SYNTHESIS_BY_Y_FAMILY_DNA_POLYME | 38 | -0.4206 | -1.06719 | 0.343387 | 0.681925 | 0.669846 | 3985 | tags=39% | list=25% | signal=30% |
| WP_GPCRS_CLASS_B_SECRETINLIKE | 19 | 0.560476 | 1.167828 | 0.348794 | 0.682131 | 0.670048 | 4297 | tags=47% | list=27% | signal=34% |
| REACTOME_SIGNALING_BY_SCF_KIT | 35 | 0.528235 | 1.164331 | 0.349123 | 0.682131 | 0.670048 | 316 | tags=9% | list=2% | signal=8% |
| REACTOME_PHASE_2_PLATEAU_PHASE | 12 | 0.550812 | 1.110024 | 0.350097 | 0.682131 | 0.670048 | 3083 | tags=33% | list=20% | signal=27% |
| REACTOME_INSULIN_PROCESSING | 10 | -0.52723 | -1.10002 | 0.350105 | 0.682131 | 0.670048 | 1480 | tags=30% | list=9% | signal=27% |
| REACTOME_MRNA_DECAY_BY_5_TO_3_EXORIBONUCLEASE | 14 | -0.48566 | -1.09742 | 0.350305 | 0.682131 | 0.670048 | 1430 | tags=21% | list=9% | signal=19% |
| REACTOME_RAF_ACTIVATION | 34 | -0.4255 | -1.06668 | 0.348837 | 0.682131 | 0.670048 | 3800 | tags=53% | list=24% | signal=40% |
| REACTOME_INTRINSIC_PATHWAY_FOR_APOPTOSIS | 37 | -0.41773 | -1.05995 | 0.351163 | 0.682999 | 0.670901 | 2839 | tags=49% | list=18% | signal=40% |
| REACTOME_ARMS_MEDIATED_ACTIVATION | 5 | -0.61688 | -1.13062 | 0.352113 | 0.684046 | 0.671929 | 2483 | tags=80% | list=16% | signal=67% |
| REACTOME_CREATINE_METABOLISM | 8 | -0.54724 | -1.10125 | 0.353293 | 0.684079 | 0.671962 | 722 | tags=50% | list=5% | signal=48% |
| REACTOME_SHC_MEDIATED_CASCADE_FGFR4 | 12 | -0.50481 | -1.08917 | 0.352577 | 0.684079 | 0.671962 | 1909 | tags=25% | list=12% | signal=22% |
| REACTOME_ASSEMBLY_OF_COLLAGEN_FIBRILS_AND_OTHER_MULT | 42 | -0.42182 | -1.06343 | 0.353365 | 0.684079 | 0.671962 | 31 | tags=5% | list=0% | signal=5% |
| REACTOME_ACTIVATION_OF_NIMA_KINASES_NEK9_NEK6_NEK7 | 5 | -0.61573 | -1.1285 | 0.354125 | 0.684751 | 0.672622 | 813 | tags=80% | list=5% | signal=76% |
| REACTOME_RHOBTB_GTPASE_CYCLE | 34 | 0.52287 | 1.152261 | 0.354895 | 0.684985 | 0.672852 | 1831 | tags=26% | list=12% | signal=23% |
| REACTOME_REGULATED_NECROSIS | 46 | 0.507358 | 1.142404 | 0.355172 | 0.684985 | 0.672852 | 37 | tags=2% | list=0% | signal=2% |
| REACTOME_JNK_C_JUN_KINASES_PHOSPHORYLATION_AND_ACTIVA | 22 | -0.46413 | -1.09273 | 0.355895 | 0.684985 | 0.672852 | 4174 | tags=59% | list=27% | signal=43% |
| REACTOME_VESICLE_MEDIATED_TRANSPORT | 546 | -0.33525 | -1.02021 | 0.355731 | 0.684985 | 0.672852 | 2292 | tags=25% | list=15% | signal=22% |
| REACTOME_REGULATION_OF_PTEN_LOCALIZATION | 9 | -0.52954 | -1.08899 | 0.357143 | 0.686591 | 0.674429 | 676 | tags=33% | list=4% | signal=32% |
| BIOCARTA_IL1R_PATHWAY | 25 | 0.543018 | 1.167609 | 0.358657 | 0.686733 | 0.674569 | 1955 | tags=24% | list=12% | signal=21% |
| BIOCARTA_NTHI_PATHWAY | 20 | 0.548504 | 1.13838 | 0.359699 | 0.686733 | 0.674569 | 4633 | tags=55% | list=30% | signal=39% |
| WP_GLUTATHIONE_METABOLISM | 17 | 0.555252 | 1.129362 | 0.358925 | 0.686733 | 0.674569 | 1027 | tags=18% | list=7% | signal=17% |
| REACTOME_ION_HOMEOSTASIS | 48 | 0.496569 | 1.128912 | 0.359725 | 0.686733 | 0.674569 | 55 | tags=2% | list=0% | signal=2% |
| REACTOME_ACYL_CHAIN_REMODELLING_OF_PC | 18 | -0.46739 | -1.07941 | 0.359408 | 0.686733 | 0.674569 | 264 | tags=11% | list=2% | signal=11% |
| REACTOME_MRNA_CAPPING | 28 | -0.43557 | -1.07177 | 0.358314 | 0.686733 | 0.674569 | 3443 | tags=32% | list=22% | signal=25% |
| REACTOME_REGULATION_OF_PLK1_ACTIVITY_AT_G2_M_TRANSITIC | 84 | -0.39832 | -1.05758 | 0.360111 | 0.686733 | 0.674569 | 3063 | tags=35% | list=20% | signal=28% |
| REACTOME_SUMOYLATION | 147 | -0.39059 | -1.05417 | 0.361032 | 0.686911 | 0.674744 | 552 | tags=4% | list=4% | signal=4% |
| REACTOME_SIGNALING_BY_FGFR3 | 36 | -0.41426 | -1.04406 | 0.360656 | 0.686911 | 0.674744 | 2834 | tags=36% | list=18% | signal=30% |
| REACTOME_ACYL_CHAIN_REMODELING_OF_CL | 5 | -0.61326 | -1.12398 | 0.362173 | 0.688295 | 0.676103 | 820 | tags=40% | list=5% | signal=38% |
| REACTOME_PHASE_1_INACTIVATION_OF_FAST_NA_CHANNELS | 7 | 0.57482 | 1.089648 | 0.363273 | 0.689597 | 0.677382 | 1408 | tags=29% | list=9% | signal=26% |
| REACTOME_SUMOYLATION_OF_SUMOYLATION_PROTEINS | 32 | 0.522348 | 1.15131 | 0.365217 | 0.690141 | 0.677917 | 4965 | tags=56% | list=32% | signal=39% |
| BIOCARTA_CERAMIDE_PATHWAY | 22 | 0.539348 | 1.144828 | 0.365809 | 0.690141 | 0.677917 | 178 | tags=5% | list=1% | signal=5% |
| REACTOME_SUMOYLATION_OF_CHROMATIN_ORGANIZATION_PRO1 | 54 | 0.490372 | 1.126098 | 0.365646 | 0.690141 | 0.677917 | 4965 | tags=52% | list=32% | signal=36% |
| REACTOME_EUKARYOTIC_TRANSLATION_TERMINATION | 6 | -0.58461 | -1.10252 | 0.366053 | 0.690141 | 0.677917 | 733 | tags=67% | list=5% | signal=64% |
| BIOCARTA_NDKDYNAMIN_PATHWAY | 13 | -0.48339 | -1.07048 | 0.364198 | 0.690141 | 0.677917 | 4187 | tags=54% | list=27% | signal=40% |
| REACTOME_DUAL_INCISION_IN_GG_NER | 40 | -0.41764 | -1.0556 | 0.36534 | 0.690141 | 0.677917 | 5220 | tags=45% | list=33% | signal=30% |
| REACTOME_NRAGE_SIGNALS_DEATH_THROUGH_JNK | 51 | -0.4189 | -1.07102 | 0.366748 | 0.690667 | 0.678434 | 467 | tags=8% | list=3% | signal=8% |
| BIOCARTA_IGF1R_PATHWAY | 22 | 0.539173 | 1.144456 | 0.367647 | 0.69127 | 0.679026 | 1518 | tags=14% | list=10% | signal=12% |
| BIOCARTA_WNT_PATHWAY | 21 | 0.547394 | 1.140807 | 0.367925 | 0.69127 | 0.679026 | 131 | tags=5% | list=1% | signal=5% |
| REACTOME_RUNX3_REGULATES_YAP1_MEDIATED_TRANSCRIPTION | 5 | 0.60597 | 1.085717 | 0.368317 | 0.69127 | 0.679026 | 5238 | tags=60% | list=33% | signal=40% |
| REACTOME_EGFR_TRANSACTIVATION_BY_GASTRIN | 6 | -0.58376 | -1.10091 | 0.370143 | 0.692776 | 0.680505 | 377 | tags=67% | list=2% | signal=65% |
| REACTOME_HDL_ASSEMBLY | 6 | 0.586261 | 1.082285 | 0.37037 | 0.692776 | 0.680505 | 3122 | tags=67% | list=20% | signal=53% |
| REACTOME_TRAF6_MEDIATED_INDUCTION_OF_TAK1_COMPLEX_W | 16 | -0.45783 | -1.05918 | 0.369835 | 0.692776 | 0.680505 | 814 | tags=44% | list=5% | signal=42% |
| WP_G13_SIGNALING_PATHWAY | 38 | 0.513958 | 1.141849 | 0.371278 | 0.692877 | 0.680604 | 2399 | tags=29% | list=15% | signal=25% |
| REACTOME_TRANSPORT_OF_FATTY_ACIDS | 4 | 0.643489 | 1.103756 | 0.37251 | 0.692877 | 0.680604 | 3419 | tags=50% | list=22% | signal=39% |
| REACTOME_REGULATION_OF_NF_KAPPA_B_SIGNALING | 16 | -0.45678 | -1.05675 | 0.371901 | 0.692877 | 0.680604 | 2489 | tags=44% | list=16% | signal=37% |
| REACTOME_SIGNALING_BY_NTRK1_TRKA | 63 | -0.39904 | -1.04929 | 0.372263 | 0.692877 | 0.680604 | 3800 | tags=48% | list=24% | signal=36% |
| REACTOME_TAK1_DEPENDENT_IKK_AND_NF_KAPPA_B_ACTIVATIOI | 37 | -0.40656 | -1.0316 | 0.372093 | 0.692877 | 0.680604 | 2489 | tags=27% | list=16% | signal=23% |
| REACTOME_NUCLEOTIDE_EXCISION_REPAIR | 107 | -0.38667 | -1.03709 | 0.373563 | 0.694059 | 0.681765 | 3153 | tags=27% | list=20% | signal=22% |
| REACTOME_SUMOYLATION_OF_INTRACELLULAR_RECEPTORS | 22 | 0.530141 | 1.125285 | 0.375 | 0.694497 | 0.682196 | 229 | tags=9% | list=1% | signal=9% |
| BIOCARTA_RAS_PATHWAY | 21 | 0.530936 | 1.106507 | 0.375472 | 0.694497 | 0.682196 | 1021 | tags=10% | list=7% | signal=9% |
| BIOCARTA_UCALPAIN_PATHWAY | 12 | -0.49662 | -1.0715 | 0.375258 | 0.694497 | 0.682196 | 3634 | tags=58% | list=23% | signal=45% |

| | | | | | | | | | | |
|---|---|---|---|---|---|---|---|---|---|---|
| REACTOME_RHO_GTPASE_EFFECTORS | 233 | -0.36311 | -1.00652 | 0.375394 | 0.694497 | 0.682196 | 2866 | tags=33% | list=18% | signal=27% |
| BIOCARTA_PDGF_PATHWAY | 29 | 0.515789 | 1.121374 | 0.376736 | 0.695287 | 0.682972 | 2306 | tags=28% | list=15% | signal=24% |
| REACTOME_SIGNALING_BY_NUCLEAR_RECEPTORS | 157 | 0.402311 | 1.056853 | 0.376682 | 0.695287 | 0.682972 | 693 | tags=7% | list=4% | signal=7% |
| REACTOME_VOLTAGE_GATED_POTASSIUM_CHANNELS | 37 | 0.515043 | 1.137105 | 0.377622 | 0.695378 | 0.683061 | 4511 | tags=59% | list=29% | signal=42% |
| REACTOME_ALTERNATIVE_COMPLEMENT_ACTIVATION | 3 | 0.685003 | 1.104863 | 0.37721 | 0.695378 | 0.683061 | 1121 | tags=33% | list=7% | signal=31% |
| REACTOME_P75NTR_RECRUITS_SIGNALLING_COMPLEXES | 13 | -0.47811 | -1.05879 | 0.378601 | 0.696407 | 0.684072 | 676 | tags=38% | list=4% | signal=37% |
| REACTOME_FORMATION_OF_XYLULOSE_5_PHOSPHATE | 5 | -0.60564 | -1.11 | 0.380282 | 0.697953 | 0.685591 | 2307 | tags=60% | list=15% | signal=51% |
| REACTOME_RUNX3_REGULATES_CDKN1A_TRANSCRIPTION | 5 | 0.598335 | 1.072037 | 0.380198 | 0.697953 | 0.685591 | 4628 | tags=80% | list=29% | signal=56% |
| REACTOME_RHO_GTPASES_ACTIVATE_NADPH_OXIDASES | 20 | -0.44495 | -1.04508 | 0.382166 | 0.700637 | 0.688227 | 2990 | tags=30% | list=19% | signal=24% |
| REACTOME_FRS_MEDIATED_FGFR1_SIGNALING | 18 | -0.44685 | -1.03197 | 0.382664 | 0.700777 | 0.688364 | 2772 | tags=33% | list=18% | signal=27% |
| REACTOME_POST_TRANSLATIONAL_MODIFICATION_SYNTHESIS_OF_ | 60 | 0.47364 | 1.096313 | 0.384874 | 0.702501 | 0.690057 | 2147 | tags=25% | list=14% | signal=22% |
| BIOCARTA_SUMO_PATHWAY | 6 | -0.57264 | -1.07994 | 0.384458 | 0.702501 | 0.690057 | 1155 | tags=33% | list=7% | signal=31% |
| REACTOME_VITAMIN_B1_THIAMIN_METABOLISM | 5 | 0.597878 | 1.071219 | 0.384158 | 0.702501 | 0.690057 | 926 | tags=20% | list=6% | signal=19% |
| REACTOME_NUCLEAR_PORE_COMPLEX_NPC_DISASSEMBLY | 30 | 0.507655 | 1.111932 | 0.387153 | 0.70299 | 0.690538 | 4466 | tags=47% | list=28% | signal=33% |
| REACTOME_SODIUM_COUPLED_SULPHATE_DI_AND_TRI_CARBOXYL | 3 | 0.679164 | 1.095445 | 0.387033 | 0.70299 | 0.690538 | 545 | tags=33% | list=3% | signal=32% |
| WP_DRAVET_SYNDROME_SCN1AA1783V_POINT_MUTATION_MODE | 71 | 0.459848 | 1.090849 | 0.387622 | 0.70299 | 0.690538 | 2672 | tags=32% | list=17% | signal=27% |
| REACTOME_RHOG_GTPASE_CYCLE | 72 | 0.456311 | 1.085669 | 0.386623 | 0.70299 | 0.690538 | 471 | tags=10% | list=3% | signal=9% |
| REACTOME_MICRORNA_MIRNA_BIOGENESIS | 6 | 0.578524 | 1.068002 | 0.385965 | 0.70299 | 0.690538 | 1940 | tags=50% | list=12% | signal=44% |
| REACTOME_CELLULAR_RESPONSES_TO_STIMULI | 422 | -0.33653 | -1.0052 | 0.387681 | 0.70299 | 0.690538 | 1890 | tags=25% | list=12% | signal=22% |
| REACTOME_VXPX_CARGO_TARGETING_TO_CILIUM | 20 | -0.43467 | -1.02093 | 0.388535 | 0.70377 | 0.691304 | 1480 | tags=15% | list=9% | signal=14% |
| BIOCARTA_ETS_PATHWAY | 17 | 0.531009 | 1.080051 | 0.389635 | 0.703873 | 0.691406 | 3196 | tags=59% | list=20% | signal=47% |
| BIOCARTA_ERBB4_PATHWAY | 6 | 0.575277 | 1.062008 | 0.389864 | 0.703873 | 0.691406 | 2899 | tags=50% | list=18% | signal=41% |
| REACTOME_RNA_POLYMERASE_III_TRANSCRIPTION_INITIATION_FR | 27 | -0.42531 | -1.04533 | 0.389401 | 0.703873 | 0.691406 | 2109 | tags=41% | list=13% | signal=35% |
| REACTOME_CELL_DEATH_SIGNALLING_VIA_NRAGE_NRIF_AND_NAD | 69 | -0.39326 | -1.02723 | 0.390428 | 0.704127 | 0.691655 | 676 | tags=9% | list=4% | signal=8% |
| REACTOME_SIGNALING_BY_CSF3_G_CSF | 21 | -0.43865 | -1.03595 | 0.391949 | 0.704575 | 0.692095 | 2562 | tags=33% | list=16% | signal=28% |
| WP_G1_TO_S_CELL_CYCLE_CONTROL | 58 | -0.40056 | -1.0358 | 0.391626 | 0.704575 | 0.692095 | 523 | tags=5% | list=3% | signal=5% |
| REACTOME_ALPHA_LINOLENIC_OMEGA3_AND_LINOLEIC_OMEGA6_ | 12 | -0.47998 | -1.03561 | 0.391753 | 0.704575 | 0.692095 | 3881 | tags=50% | list=25% | signal=38% |
| WP_MACROPHAGE_MARKERS | 10 | 0.538927 | 1.056661 | 0.392381 | 0.704589 | 0.692109 | 57 | tags=10% | list=0% | signal=10% |
| REACTOME_INTERLEUKIN_6_FAMILY_SIGNALING | 17 | 0.527767 | 1.073458 | 0.393474 | 0.705789 | 0.693287 | 2152 | tags=29% | list=14% | signal=25% |
| REACTOME_GLYCOSPHINGOLIPID_METABOLISM | 38 | 0.496698 | 1.103502 | 0.394046 | 0.706051 | 0.693545 | 4386 | tags=63% | list=28% | signal=46% |
| BIOCARTA_CTCF_PATHWAY | 23 | 0.509906 | 1.085151 | 0.39781 | 0.709263 | 0.6967 | 3601 | tags=52% | list=23% | signal=40% |
| WP_EICOSANOID_METABOLISM_VIA_CYCLOOXYGENASES_COX | 23 | 0.509517 | 1.084323 | 0.39781 | 0.709263 | 0.6967 | 20 | tags=4% | list=0% | signal=4% |
| REACTOME_ACROSOME_REACTION_AND_SPERM_OOCYTE_MEMBR | 3 | 0.668951 | 1.078972 | 0.398821 | 0.709263 | 0.6967 | 3757 | tags=67% | list=24% | signal=51% |
| WP_CALCIUM_REGULATION_IN_CARDIAC_CELLS | 131 | 0.415153 | 1.067368 | 0.398462 | 0.709263 | 0.6967 | 1191 | tags=10% | list=8% | signal=9% |
| BIOCARTA_SODD_PATHWAY | 9 | 0.531767 | 1.035627 | 0.398438 | 0.709263 | 0.6967 | 7273 | tags=89% | list=46% | signal=48% |
| REACTOME_HOMOLOGY_DIRECTED_REPAIR | 106 | -0.37819 | -1.01136 | 0.398827 | 0.709263 | 0.6967 | 2609 | tags=17% | list=17% | signal=14% |
| REACTOME_AURKA_ACTIVATION_BY_TPX2 | 70 | -0.38698 | -1.01117 | 0.397985 | 0.709263 | 0.6967 | 3020 | tags=34% | list=19% | signal=28% |
| REACTOME_ANTIGEN_PRESENTATION_FOLDING_ASSEMBLY_AND_P | 27 | 0.508891 | 1.094426 | 0.399648 | 0.709963 | 0.697387 | 3907 | tags=33% | list=25% | signal=25% |
| WP_G_PROTEIN_SIGNALING_PATHWAYS | 84 | 0.442676 | 1.077994 | 0.400936 | 0.71149 | 0.698888 | 200 | tags=4% | list=1% | signal=4% |
| WP_LEPTININSULIN_SIGNALING_OVERLAP | 15 | 0.505859 | 1.032954 | 0.401544 | 0.711809 | 0.699201 | 4464 | tags=47% | list=28% | signal=33% |
| REACTOME_MET_ACTIVATES_PI3K_AKT_SIGNALING | 6 | 0.568286 | 1.049101 | 0.403509 | 0.7143 | 0.701648 | 3601 | tags=33% | list=23% | signal=26% |
| REACTOME_INTERLEUKIN_4_AND_INTERLEUKIN_13_SIGNALING | 10 | 0.532712 | 1.044476 | 0.40381 | 0.7143 | 0.701648 | 4601 | tags=60% | list=29% | signal=42% |
| REACTOME_DOWNSTREAM_SIGNALING_OF_ACTIVATED_FGFR1 | 26 | 0.505852 | 1.091447 | 0.40493 | 0.715519 | 0.702845 | 1518 | tags=15% | list=10% | signal=14% |
| REACTOME_ACTIVATION_OF_G_PROTEIN_GATED_POTASSIUM_CHA | 26 | 0.504919 | 1.089434 | 0.40669 | 0.715708 | 0.70303 | 71 | tags=4% | list=0% | signal=4% |
| WP_WHITE_FAT_CELL_DIFFERENTIATION | 29 | 0.489066 | 1.063275 | 0.409722 | 0.715708 | 0.70303 | 2004 | tags=34% | list=13% | signal=30% |
| BIOCARTA_RHO_PATHWAY | 20 | 0.511699 | 1.061994 | 0.40678 | 0.715708 | 0.70303 | 2517 | tags=30% | list=16% | signal=25% |
| REACTOME_WNT_LIGAND_BIOGENESIS_AND_TRAFFICKING | 21 | 0.507971 | 1.058647 | 0.411321 | 0.715708 | 0.70303 | 2701 | tags=29% | list=17% | signal=24% |
| WP_HEART_DEVELOPMENT | 30 | 0.483102 | 1.058153 | 0.413194 | 0.715708 | 0.70303 | 1537 | tags=33% | list=10% | signal=30% |
| REACTOME_NADE_MODULATES_DEATH_SIGNALLING | 6 | -0.5609 | -1.05781 | 0.413088 | 0.715708 | 0.70303 | 1952 | tags=50% | list=12% | signal=44% |
| BIOCARTA_MAL_PATHWAY | 17 | 0.519355 | 1.056349 | 0.40691 | 0.715708 | 0.70303 | 1295 | tags=24% | list=8% | signal=22% |
| REACTOME_TP53_REGULATES_TRANSCRIPTION_OF_GENES_INVOLV | 7 | -0.53141 | -1.05385 | 0.411178 | 0.715708 | 0.70303 | 4311 | tags=71% | list=27% | signal=52% |
| REACTOME_REGULATION_OF_RUNX1_EXPRESSION_AND_ACTIVITY | 7 | -0.53102 | -1.05307 | 0.411178 | 0.715708 | 0.70303 | 1876 | tags=43% | list=12% | signal=38% |
| REACTOME_METABOLISM_OF_VITAMINS_AND_COFACTORS | 150 | 0.399484 | 1.044376 | 0.410606 | 0.715708 | 0.70303 | 423 | tags=4% | list=3% | signal=4% |
| REACTOME_FGFR2C_LIGAND_BINDING_AND_ACTIVATION | 8 | -0.51569 | -1.03776 | 0.409182 | 0.715708 | 0.70303 | 1909 | tags=25% | list=12% | signal=22% |
| REACTOME_TRAFFICKING_OF_GLUR2_CONTAINING_AMPA_RECEPT | 16 | 0.507989 | 1.034635 | 0.407336 | 0.715708 | 0.70303 | 1152 | tags=19% | list=7% | signal=17% |
| REACTOME_INTERLEUKIN_6_SIGNALING | 8 | 0.529186 | 1.031475 | 0.407186 | 0.715708 | 0.70303 | 2152 | tags=38% | list=14% | signal=32% |
| REACTOME_AGGREPHAGY | 30 | -0.40616 | -1.01968 | 0.408451 | 0.715708 | 0.70303 | 1568 | tags=43% | list=10% | signal=39% |
| REACTOME_APOPTOTIC_FACTOR_MEDIATED_RESPONSE | 16 | -0.43557 | -1.00769 | 0.413223 | 0.715708 | 0.70303 | 2834 | tags=38% | list=18% | signal=31% |
| REACTOME_INSULIN_RECEPTOR_SIGNALLING_CASCADE | 38 | -0.39659 | -1.00625 | 0.408353 | 0.715708 | 0.70303 | 534 | tags=8% | list=3% | signal=8% |
| REACTOME_CELL_EXTRACELLULAR_MATRIX_INTERACTIONS | 18 | -0.43362 | -1.00143 | 0.412262 | 0.715708 | 0.70303 | 933 | tags=39% | list=6% | signal=37% |
| BIOCARTA_PML_PATHWAY | 15 | -0.4361 | -1.00139 | 0.411157 | 0.715708 | 0.70303 | 552 | tags=27% | list=4% | signal=26% |
| REACTOME_PI_3K_CASCADE_FGFR2 | 18 | -0.43164 | -0.99684 | 0.412262 | 0.715708 | 0.70303 | 2772 | tags=28% | list=18% | signal=23% |
| REACTOME_INWARDLY_RECTIFYING_K_CHANNELS | 31 | 0.479509 | 1.053984 | 0.414634 | 0.717403 | 0.704696 | 71 | tags=3% | list=0% | signal=3% |
| REACTOME_DEGRADATION_OF_BETA_CATENIN_BY_THE_DESTRUCT | 78 | -0.3799 | -1.00358 | 0.415344 | 0.717645 | 0.704933 | 3355 | tags=56% | list=21% | signal=45% |
| WP_ALANINE_AND_ASPARTATE_METABOLISM | 13 | -0.44594 | -0.98756 | 0.415638 | 0.717645 | 0.704933 | 3417 | tags=62% | list=22% | signal=48% |
| REACTOME_TRANSCRIPTIONAL_REGULATION_BY_RUNX3 | 76 | 0.445774 | 1.061636 | 0.418152 | 0.72053 | 0.707768 | 722 | tags=9% | list=5% | signal=9% |
| REACTOME_IRF3_MEDIATED_INDUCTION_OF_TYPE_I_IFN | 4 | 0.617724 | 1.059562 | 0.418327 | 0.72053 | 0.707768 | 724 | tags=50% | list=5% | signal=48% |
| REACTOME_TRANSPORT_OF_THE_SLBP_DEPENDANT_MATURE_MR | 33 | 0.480224 | 1.059026 | 0.41913 | 0.72053 | 0.707768 | 4931 | tags=52% | list=31% | signal=35% |
| REACTOME_MAPK_FAMILY_SIGNALING_CASCADES | 268 | 0.370634 | 1.028734 | 0.42029 | 0.72053 | 0.707768 | 661 | tags=6% | list=4% | signal=6% |
| REACTOME_HYALURONAN_METABOLISM | 17 | 0.500229 | 1.017447 | 0.420345 | 0.72053 | 0.707768 | 5438 | tags=65% | list=35% | signal=42% |
| REACTOME_NRIF_SIGNALS_CELL_DEATH_FROM_THE_NUCLEUS | 14 | -0.44337 | -1.00187 | 0.419552 | 0.72053 | 0.707768 | 676 | tags=29% | list=4% | signal=27% |
| REACTOME_TRISTETRAPROLIN_TTP_ZFP36_BINDS_AND_DESTABILIZ | 16 | -0.43009 | -0.99499 | 0.419421 | 0.72053 | 0.707768 | 3659 | tags=56% | list=23% | signal=43% |
| REACTOME_GLUCURONIDATION | 5 | -0.58704 | -1.07592 | 0.422535 | 0.722598 | 0.709799 | 3559 | tags=60% | list=23% | signal=46% |
| REACTOME_OTHER_SEMAPHORIN_INTERACTIONS | 14 | 0.499914 | 1.018354 | 0.422701 | 0.722598 | 0.709799 | 2925 | tags=36% | list=19% | signal=29% |
| BIOCARTA_AKAP95_PATHWAY | 10 | 0.518899 | 1.017393 | 0.422857 | 0.722598 | 0.709799 | 1739 | tags=40% | list=11% | signal=36% |
| REACTOME_TRANSCRIPTIONAL_REGULATION_BY_SMALL_RNAS | 31 | 0.475372 | 1.044889 | 0.423345 | 0.722689 | 0.709888 | 4466 | tags=48% | list=28% | signal=35% |
| WP_MAPK_CASCADE | 28 | 0.488305 | 1.056991 | 0.424348 | 0.722986 | 0.71018 | 679 | tags=11% | list=4% | signal=10% |
| REACTOME_BIOSYNTHESIS_OF_THE_N_GLYCAN_PRECURSOR_DOLIC | 70 | 0.446325 | 1.054395 | 0.424793 | 0.722986 | 0.71018 | 2815 | tags=31% | list=18% | signal=26% |
| REACTOME_METABOLISM_OF_FAT_SOLUBLE_VITAMINS | 34 | 0.474716 | 1.046142 | 0.424825 | 0.722986 | 0.71018 | 4566 | tags=44% | list=29% | signal=31% |
| REACTOME_PROLONGED_ERK_ACTIVATION_EVENTS | 14 | -0.43849 | -0.99084 | 0.425662 | 0.723669 | 0.710851 | 3558 | tags=50% | list=23% | signal=39% |
| WP_KIT_RECEPTOR_SIGNALING_PATHWAY | 66 | 0.449505 | 1.049862 | 0.427621 | 0.725514 | 0.712663 | 2426 | tags=23% | list=15% | signal=19% |
| BIOCARTA_ARENRF2_PATHWAY | 18 | 0.487571 | 0.999779 | 0.427221 | 0.725514 | 0.712663 | 2134 | tags=33% | list=14% | signal=29% |
| REACTOME_UNBLOCKING_OF_NMDA_RECEPTORS_GLUTAMATE_BI | 22 | 0.484783 | 1.029007 | 0.428309 | 0.725644 | 0.712791 | 2031 | tags=36% | list=13% | signal=32% |
| REACTOME_MET_RECEPTOR_RECYCLING | 10 | 0.511616 | 1.003114 | 0.428571 | 0.725644 | 0.712791 | 1299 | tags=20% | list=8% | signal=18% |
| REACTOME_HATS_ACETYLATE_HISTONES | 41 | 0.477724 | 1.066554 | 0.433276 | 0.727113 | 0.714234 | 550 | tags=7% | list=4% | signal=7% |
| REACTOME_HYDROLYSIS_OF_LPC | 5 | -0.57508 | -1.054 | 0.438632 | 0.727113 | 0.714234 | 264 | tags=40% | list=2% | signal=39% |
| BIOCARTA_FAS_PATHWAY | 27 | 0.486063 | 1.045332 | 0.431338 | 0.727113 | 0.714234 | 178 | tags=4% | list=1% | signal=4% |
| REACTOME_SIGNALING_BY_MET | 66 | 0.44523 | 1.039876 | 0.434276 | 0.727113 | 0.714234 | 2599 | tags=26% | list=17% | signal=22% |
| BIOCARTA_FMLP_PATHWAY | 29 | 0.477772 | 1.038722 | 0.430556 | 0.727113 | 0.714234 | 679 | tags=14% | list=4% | signal=13% |
| REACTOME_ROS_AND_RNS_PRODUCTION_IN_PHAGOCYTES | 31 | 0.470485 | 1.034148 | 0.437282 | 0.727113 | 0.714234 | 65 | tags=3% | list=0% | signal=3% |

| | | | | | | | | | | |
|---|---|---|---|---|---|---|---|---|---|---|
| REACTOME_TNFR1_INDUCED_NFKAPPAB_SIGNALING_PATHWAY | 32 | 0.468965 | 1.033649 | 0.436522 | 0.727113 | 0.714234 | 426 | tags=9% | list=3% | signal=9% |
| REACTOME_ALPHA_OXIDATION_OF_PHYTANATE | 6 | -0.54515 | -1.02809 | 0.437628 | 0.727113 | 0.714234 | 3555 | tags=67% | list=23% | signal=52% |
| REACTOME_PHOSPHOLIPASE_C_MEDIATED_CASCADE_FGFR4 | 8 | -0.5009 | -1.008 | 0.437126 | 0.727113 | 0.714234 | 1909 | tags=25% | list=12% | signal=22% |
| BIOCARTA_FBW7_PATHWAY | 8 | 0.515892 | 1.005562 | 0.431138 | 0.727113 | 0.714234 | 2960 | tags=62% | list=19% | signal=51% |
| REACTOME_ROLE_OF_PHOSPHOLIPIDS_IN_PHAGOCYTOSIS | 20 | 0.480518 | 0.99728 | 0.433145 | 0.727113 | 0.714234 | 4808 | tags=55% | list=31% | signal=38% |
| WP_IRON_HOMEOSTASIS | 10 | 0.507651 | 0.99534 | 0.434286 | 0.727113 | 0.714234 | 3571 | tags=50% | list=23% | signal=39% |
| REACTOME_LGI_ADAM_INTERACTIONS | 14 | 0.487747 | 0.993568 | 0.436399 | 0.727113 | 0.714234 | 2882 | tags=43% | list=18% | signal=35% |
| REACTOME_SIGNALLING_TO_RAS | 12 | -0.45815 | -0.98849 | 0.435052 | 0.727113 | 0.714234 | 1828 | tags=25% | list=12% | signal=22% |
| REACTOME_CYTOSOLIC_SULFONATION_OF_SMALL_MOLECULES | 15 | 0.481212 | 0.982625 | 0.438224 | 0.727113 | 0.714234 | 3971 | tags=53% | list=25% | signal=40% |
| REACTOME_RA_BIOSYNTHESIS_PATHWAY | 12 | -0.45535 | -0.98246 | 0.437113 | 0.727113 | 0.714234 | 1177 | tags=33% | list=7% | signal=31% |
| REACTOME_MITOTIC_SPINDLE_CHECKPOINT | 106 | -0.36564 | -0.97782 | 0.43695 | 0.727113 | 0.714234 | 3455 | tags=42% | list=22% | signal=33% |
| WP_CHEMOKINE_SIGNALING_PATHWAY | 138 | -0.36114 | -0.9739 | 0.433908 | 0.727113 | 0.714234 | 467 | tags=4% | list=3% | signal=4% |
| REACTOME_DOWNSTREAM_SIGNALING_OF_ACTIVATED_FGFR4 | 19 | -0.41958 | -0.97309 | 0.434125 | 0.727113 | 0.714234 | 2767 | tags=26% | list=18% | signal=22% |
| REACTOME_DNA_DAMAGE_BYPASS | 47 | -0.37686 | -0.96963 | 0.436019 | 0.727113 | 0.714234 | 3985 | tags=43% | list=25% | signal=32% |
| WP_OMEGA9_FATTY_ACID_SYNTHESIS | 13 | -0.43483 | -0.96295 | 0.438272 | 0.727113 | 0.714234 | 2819 | tags=38% | list=18% | signal=32% |
| REACTOME_APOPTOTIC_EXECUTION_PHASE | 43 | 0.463901 | 1.040004 | 0.449915 | 0.730507 | 0.717568 | 97 | tags=5% | list=1% | signal=5% |
| REACTOME_COLLAGEN_FORMATION | 72 | 0.435248 | 1.035556 | 0.445351 | 0.730507 | 0.717568 | 3382 | tags=50% | list=22% | signal=39% |
| REACTOME_CLASS_B_2_SECRETIN_FAMILY_RECEPTORS | 41 | 0.463296 | 1.034341 | 0.448873 | 0.730507 | 0.717568 | 66 | tags=2% | list=0% | signal=2% |
| REACTOME_RHOA_GTPASE_CYCLE | 137 | 0.400698 | 1.033813 | 0.448171 | 0.730507 | 0.717568 | 1649 | tags=17% | list=10% | signal=15% |
| BIOCARTA_MRP_PATHWAY | 4 | 0.601222 | 1.031257 | 0.448207 | 0.730507 | 0.717568 | 4291 | tags=75% | list=27% | signal=55% |
| REACTOME_VISUAL_PHOTOTRANSDUCTION | 61 | 0.443525 | 1.029081 | 0.447811 | 0.730507 | 0.717568 | 179 | tags=3% | list=1% | signal=3% |
| REACTOME_VEGFR2_MEDIATED_VASCULAR_PERMEABILITY | 29 | 0.470346 | 1.022576 | 0.444444 | 0.730507 | 0.717568 | 65 | tags=3% | list=0% | signal=3% |
| WP_RETINOL_METABOLISM | 29 | 0.468738 | 1.019081 | 0.449653 | 0.730507 | 0.717568 | 1217 | tags=17% | list=8% | signal=16% |
| REACTOME_NEUROTRANSMITTER_CLEARANCE | 6 | -0.53971 | -1.01783 | 0.449898 | 0.730507 | 0.717568 | 5617 | tags=83% | list=36% | signal=54% |
| WP_IL5_SIGNALING_PATHWAY | 66 | 0.435371 | 1.01685 | 0.449251 | 0.730507 | 0.717568 | 2285 | tags=18% | list=15% | signal=16% |
| REACTOME_INTERACTION_BETWEEN_L1_AND_ANKYRINS | 10 | -0.48148 | -1.00458 | 0.444444 | 0.730507 | 0.717568 | 1464 | tags=30% | list=9% | signal=27% |
| BIOCARTA_GLEEVEC_PATHWAY | 23 | 0.46959 | 0.999354 | 0.445255 | 0.730507 | 0.717568 | 1021 | tags=9% | list=7% | signal=8% |
| REACTOME_REGULATION_OF_IFNA_IFNB_SIGNALING | 8 | 0.511695 | 0.997382 | 0.443114 | 0.730507 | 0.717568 | 3398 | tags=50% | list=22% | signal=39% |
| BIOCARTA_IL10_PATHWAY | 9 | -0.47922 | -0.98551 | 0.44898 | 0.730507 | 0.717568 | 2346 | tags=56% | list=15% | signal=47% |
| BIOCARTA_BTG2_PATHWAY | 9 | -0.47844 | -0.9839 | 0.44898 | 0.730507 | 0.717568 | 3898 | tags=67% | list=25% | signal=50% |
| WP_ONECARBON_METABOLISM | 26 | -0.40113 | -0.97091 | 0.447005 | 0.730507 | 0.717568 | 3396 | tags=42% | list=22% | signal=33% |
| REACTOME_SIGNALLING_TO_ERKS | 26 | -0.40055 | -0.9695 | 0.447005 | 0.730507 | 0.717568 | 2948 | tags=38% | list=19% | signal=31% |
| REACTOME_RHOJ_GTPASE_CYCLE | 16 | 0.475267 | 0.967991 | 0.447876 | 0.730507 | 0.717568 | 2825 | tags=44% | list=18% | signal=36% |
| REACTOME_TERMINATION_OF_TRANSLESION_DNA_SYNTHESIS | 31 | -0.38375 | -0.96522 | 0.443925 | 0.730507 | 0.717568 | 3985 | tags=35% | list=25% | signal=27% |
| REACTOME_MAPK_TARGETS_NUCLEAR_EVENTS_MEDIATED_BY_MA | 28 | -0.39187 | -0.96423 | 0.442623 | 0.730507 | 0.717568 | 3800 | tags=57% | list=24% | signal=43% |
| REACTOME_CLASS_I_MHC_MEDIATED_ANTIGEN_PROCESSING_PRES | 323 | -0.32546 | -0.94988 | 0.444444 | 0.730507 | 0.717568 | 3367 | tags=31% | list=21% | signal=25% |
| REACTOME_DRUG_MEDIATED_INHIBITION_OF_CDK4_CDK6_ACTIVIT | 5 | -0.56494 | -1.03542 | 0.450704 | 0.731074 | 0.718125 | 2337 | tags=60% | list=15% | signal=51% |
| BIOCARTA_VIP_PATHWAY | 24 | 0.466286 | 0.997314 | 0.452252 | 0.731442 | 0.718486 | 1576 | tags=12% | list=10% | signal=11% |
| REACTOME_NEGATIVE_REGULATORS_OF_DDX58_IFIH1_SIGNALING | 14 | -0.42341 | -0.95677 | 0.452138 | 0.731442 | 0.718486 | 676 | tags=36% | list=4% | signal=34% |
| REACTOME_ERK_MAPK_TARGETS | 19 | -0.40708 | -0.9441 | 0.451404 | 0.731442 | 0.718486 | 3800 | tags=58% | list=24% | signal=44% |
| WP_GLUCURONIDATION | 7 | -0.50512 | -1.00171 | 0.453094 | 0.73209 | 0.719123 | 3559 | tags=57% | list=23% | signal=44% |
| REACTOME_ACTIVATION_OF_THE_PHOTOTRANSDUCTION_CASCADI | 3 | -0.659 | -1.06209 | 0.454361 | 0.732652 | 0.719675 | 2142 | tags=100% | list=14% | signal=86% |
| REACTOME_ACYL_CHAIN_REMODELLING_OF_PG | 10 | -0.47337 | -0.98765 | 0.454927 | 0.732652 | 0.719675 | 48 | tags=10% | list=0% | signal=10% |
| REACTOME_RNA_POLYMERASE_II_PROMOTER_ESCAPE | 44 | -0.37428 | -0.95074 | 0.455206 | 0.732652 | 0.719675 | 2539 | tags=34% | list=16% | signal=29% |
| REACTOME_DUAL_INCISION_IN_TC_NER | 63 | -0.35761 | -0.94033 | 0.454988 | 0.732652 | 0.719675 | 2489 | tags=43% | list=16% | signal=36% |
| BIOCARTA_HBX_PATHWAY | 7 | -0.49923 | -0.99003 | 0.457086 | 0.734347 | 0.721339 | 609 | tags=43% | list=4% | signal=41% |
| BIOCARTA_DNAFRAGMENT_PATHWAY | 9 | -0.47346 | -0.97366 | 0.457143 | 0.734347 | 0.721339 | 5980 | tags=67% | list=38% | signal=41% |
| REACTOME_MAPK1_ERK2_ACTIVATION | 7 | 0.528508 | 1.001858 | 0.459082 | 0.736038 | 0.723 | 751 | tags=29% | list=5% | signal=27% |
| REACTOME_DARPP_32_EVENTS | 17 | 0.473642 | 0.963369 | 0.458733 | 0.736038 | 0.723 | 137 | tags=6% | list=1% | signal=6% |
| REACTOME_SYNTHESIS_OF_PE | 12 | 0.478957 | 0.965218 | 0.460348 | 0.736091 | 0.723053 | 3914 | tags=58% | list=25% | signal=44% |
| BIOCARTA_BARRESTIN_PATHWAY | 11 | -0.45047 | -0.96378 | 0.460744 | 0.736091 | 0.723053 | 3130 | tags=45% | list=20% | signal=36% |
| REACTOME_ANTIMICROBIAL_PEPTIDES | 18 | -0.41247 | -0.95258 | 0.460888 | 0.736091 | 0.723053 | 3183 | tags=28% | list=20% | signal=22% |
| REACTOME_TRAF6_MEDIATED_NF_KB_ACTIVATION | 14 | -0.41676 | -0.94175 | 0.460285 | 0.736091 | 0.723053 | 2314 | tags=29% | list=15% | signal=24% |
| BIOCARTA_PTC1_PATHWAY | 11 | 0.494395 | 0.978761 | 0.46139 | 0.736185 | 0.723145 | 994 | tags=18% | list=6% | signal=17% |
| BIOCARTA_EGF_PATHWAY | 28 | 0.468434 | 1.013976 | 0.462609 | 0.737421 | 0.72436 | 1955 | tags=18% | list=12% | signal=16% |
| BIOCARTA_TNFR1_PATHWAY | 27 | 0.468919 | 1.008462 | 0.466549 | 0.738037 | 0.724964 | 178 | tags=4% | list=1% | signal=4% |
| REACTOME_ATTENUATION_PHASE | 12 | 0.474079 | 0.955388 | 0.466151 | 0.738037 | 0.724964 | 3832 | tags=50% | list=24% | signal=38% |
| REACTOME_RETROGRADE_TRANSPORT_AT_THE_TRANS_GOLGI_NET | 45 | -0.37427 | -0.95454 | 0.463942 | 0.738037 | 0.724964 | 3287 | tags=38% | list=21% | signal=30% |
| REACTOME_INTERFERON_ALPHA_BETA_SIGNALING | 15 | 0.466448 | 0.952478 | 0.465251 | 0.738037 | 0.724964 | 3398 | tags=40% | list=22% | signal=31% |
| REACTOME_SIGNALING_BY_NTRKS | 74 | -0.35984 | -0.9496 | 0.464935 | 0.738037 | 0.724964 | 3800 | tags=45% | list=24% | signal=34% |
| REACTOME_GENERATION_OF_SECOND_MESSENGER_MOLECULES | 18 | -0.40955 | -0.94583 | 0.465116 | 0.738037 | 0.724964 | 2272 | tags=33% | list=14% | signal=29% |
| REACTOME_SHC_MEDIATED_CASCADE_FGFR2 | 17 | -0.4089 | -0.9437 | 0.465696 | 0.738037 | 0.724964 | 377 | tags=24% | list=2% | signal=23% |
| REACTOME_FORMATION_OF_INCISION_COMPLEX_IN_GG_NER | 42 | -0.37426 | -0.94353 | 0.466346 | 0.738037 | 0.724964 | 3355 | tags=24% | list=21% | signal=19% |
| REACTOME_TRANSCRIPTIONAL_REGULATION_BY_THE_AP_2_TFAP2 | 17 | 0.46857 | 0.953053 | 0.46833 | 0.738743 | 0.725658 | 5184 | tags=53% | list=33% | signal=36% |
| REACTOME_MITOTIC_PROMETAPHASE | 183 | -0.34739 | -0.94974 | 0.467456 | 0.738743 | 0.725658 | 3259 | tags=31% | list=21% | signal=25% |
| REACTOME_RHO_GTPASES_ACTIVATE_IQGAPS | 25 | -0.39563 | -0.94807 | 0.46789 | 0.738743 | 0.725658 | 1568 | tags=48% | list=10% | signal=43% |
| REACTOME_SYNTHESIS_OF_12_EICOSATETRAENOIC_ACID_DERIVAT | 5 | -0.55716 | -1.02116 | 0.468813 | 0.738803 | 0.725717 | 3501 | tags=60% | list=22% | signal=47% |
| REACTOME_RELAXIN_RECEPTORS | 4 | 0.586876 | 1.00665 | 0.472112 | 0.739464 | 0.726366 | 3871 | tags=50% | list=25% | signal=38% |
| WP_IL6_SIGNALING_PATHWAY | 97 | 0.40606 | 1.004319 | 0.47352 | 0.739464 | 0.726366 | 751 | tags=7% | list=5% | signal=7% |
| REACTOME_TRANSPORT_OF_MATURE_MRNAS_DERIVED_FROM_IN | 40 | 0.45007 | 1.001859 | 0.473043 | 0.739464 | 0.726366 | 5136 | tags=50% | list=33% | signal=34% |
| REACTOME_SYNTHESIS_OF_DOLICHYL_PHOSPHATE | 6 | -0.5292 | -0.99802 | 0.470348 | 0.739464 | 0.726366 | 157 | tags=17% | list=1% | signal=17% |
| REACTOME_RHOH_GTPASE_CYCLE | 35 | 0.44923 | 0.990189 | 0.473684 | 0.739464 | 0.726366 | 841 | tags=11% | list=5% | signal=11% |
| REACTOME_SEMA3A_PAK_DEPENDENT_AXON_REPULSION | 14 | 0.466256 | 0.94979 | 0.473581 | 0.739464 | 0.726366 | 2386 | tags=43% | list=15% | signal=36% |
| REACTOME_P75_NTR_RECEPTOR_MEDIATED_SIGNALLING | 84 | -0.35657 | -0.94674 | 0.473684 | 0.739464 | 0.726366 | 676 | tags=8% | list=4% | signal=8% |
| BIOCARTA_ERAD_PATHWAY | 14 | 0.464463 | 0.946137 | 0.473581 | 0.739464 | 0.726366 | 3935 | tags=71% | list=25% | signal=54% |
| REACTOME_FRS_MEDIATED_FGFR3_SIGNALING | 16 | -0.40017 | -0.92579 | 0.47314 | 0.739464 | 0.726366 | 1909 | tags=31% | list=12% | signal=27% |
| REACTOME_P75NTR_SIGNALS_VIA_NF_KB | 16 | -0.39977 | -0.92485 | 0.47314 | 0.739464 | 0.726366 | 676 | tags=19% | list=4% | signal=18% |
| REACTOME_CREB3_FACTORS_ACTIVATE_GENES | 5 | 0.561147 | 1.005407 | 0.475248 | 0.741208 | 0.728079 | 5155 | tags=60% | list=33% | signal=40% |
| BIOCARTA_KERATINOCYTE_PATHWAY | 43 | 0.448416 | 1.005288 | 0.47708 | 0.742175 | 0.729029 | 1999 | tags=21% | list=13% | signal=18% |
| BIOCARTA_TCR_PATHWAY | 37 | 0.447939 | 0.988954 | 0.477273 | 0.742175 | 0.729029 | 1955 | tags=19% | list=12% | signal=17% |
| REACTOME_SYNTHESIS_OF_BILE_ACIDS_AND_BILE_SALTS | 23 | 0.444155 | 0.945224 | 0.478102 | 0.742175 | 0.729029 | 3137 | tags=39% | list=20% | signal=31% |
| WP_MECP2_AND_ASSOCIATED_RETT_SYNDROME | 43 | -0.37113 | -0.93718 | 0.476998 | 0.742175 | 0.729029 | 1569 | tags=19% | list=10% | signal=17% |
| BIOCARTA_CHEMICAL_PATHWAY | 18 | -0.40294 | -0.93057 | 0.477801 | 0.742175 | 0.729029 | 4077 | tags=39% | list=26% | signal=29% |
| REACTOME_SEMA4D_IN_SEMAPHORIN_SIGNALING | 15 | -0.40248 | -0.92419 | 0.479339 | 0.7434 | 0.730233 | 411 | tags=47% | list=3% | signal=45% |
| REACTOME_MOLYBDENUM_COFACTOR_BIOSYNTHESIS | 5 | -0.55009 | -1.0082 | 0.480885 | 0.744562 | 0.731374 | 846 | tags=40% | list=5% | signal=38% |
| REACTOME_DOWNSTREAM_SIGNALING_OF_ACTIVATED_FGFR2 | 24 | -0.3886 | -0.93091 | 0.480984 | 0.744562 | 0.731374 | 2772 | tags=25% | list=18% | signal=21% |
| REACTOME_SIGNALING_BY_RHO_GTPASES_MIRO_GTPASES_AND_R | 600 | 0.332579 | 0.991091 | 0.482119 | 0.74493 | 0.731735 | 1665 | tags=13% | list=11% | signal=12% |
| REACTOME_ACTIVATION_OF_NMDA_RECEPTORS_AND_POSTSYNAP | 36 | 0.443278 | 0.978751 | 0.481739 | 0.74493 | 0.731735 | 2031 | tags=28% | list=13% | signal=24% |
| WP_COMPREHENSIVE_IL17A_SIGNALING | 88 | 0.415825 | 1.011689 | 0.483516 | 0.74501 | 0.731814 | 101 | tags=2% | list=1% | signal=2% |
| REACTOME_DEVELOPMENTAL_BIOLOGY | 324 | 0.350155 | 0.993058 | 0.482955 | 0.74501 | 0.731814 | 1649 | tags=15% | list=10% | signal=13% |

| | | | | | | | | | | |
|---|---|---|---|---|---|---|---|---|---|---|
| REACTOME_POSTMITOTIC_NUCLEAR_PORE_COMPLEX_NPC_REFORI | 22 | 0.436257 | 0.926004 | 0.483456 | 0.74501 | 0.731814 | 4965 | tags=55% | list=32% | signal=37% |
| BIOCARTA_INFLAM_PATHWAY | 8 | 0.493419 | 0.961759 | 0.48503 | 0.746649 | 0.733424 | 3571 | tags=62% | list=23% | signal=48% |
| REACTOME_REGULATION_OF_TP53_DEGRADATION | 35 | -0.36728 | -0.92018 | 0.486111 | 0.747621 | 0.734378 | 1337 | tags=17% | list=9% | signal=16% |
| REACTOME_INTERLEUKIN_2_FAMILY_SIGNALING | 26 | 0.453984 | 0.979535 | 0.487676 | 0.748071 | 0.734821 | 4955 | tags=50% | list=32% | signal=34% |
| REACTOME_HEME_DEGRADATION | 8 | -0.47869 | -0.96331 | 0.487026 | 0.748071 | 0.734821 | 2152 | tags=50% | list=14% | signal=43% |
| REACTOME_PURINE_RIBONUCLEOSIDE_MONOPHOSPHATE_BIOSYN | 9 | -0.45907 | -0.94407 | 0.487755 | 0.748071 | 0.734821 | 2302 | tags=44% | list=15% | signal=38% |
| WP_DOPAMINERGIC_NEUROGENESIS | 21 | 0.447151 | 0.931893 | 0.490566 | 0.751688 | 0.738374 | 5991 | tags=76% | list=38% | signal=47% |
| REACTOME_ELECTRIC_TRANSMISSION_ACROSS_GAP_JUNCTIONS | 4 | 0.578262 | 0.991874 | 0.492032 | 0.752546 | 0.739216 | 4993 | tags=75% | list=32% | signal=51% |
| WP_IL2_SIGNALING_PATHWAY | 73 | 0.416917 | 0.990712 | 0.49183 | 0.752546 | 0.739216 | 2278 | tags=19% | list=15% | signal=16% |
| WP_SPINAL_CORD_INJURY | 84 | 0.406168 | 0.989091 | 0.49766 | 0.75476 | 0.741391 | 867 | tags=8% | list=6% | signal=8% |
| REACTOME_ADVANCED_GLYCOSYLATION_ENDPRODUCT_RECEPTOR | 5 | -0.5388 | -0.98751 | 0.49497 | 0.75476 | 0.741391 | 435 | tags=60% | list=3% | signal=58% |
| WP_ACETYLCHOLINE_SYNTHESIS | 6 | -0.51881 | -0.97843 | 0.494888 | 0.75476 | 0.741391 | 1565 | tags=67% | list=10% | signal=60% |
| REACTOME_SUMOYLATION_OF_DNA_REPLICATION_PROTEINS | 40 | 0.435932 | 0.970389 | 0.497391 | 0.75476 | 0.741391 | 3743 | tags=38% | list=24% | signal=29% |
| REACTOME_METABOLISM_OF_RNA | 575 | -0.31104 | -0.94544 | 0.498024 | 0.75476 | 0.741391 | 3443 | tags=37% | list=22% | signal=30% |
| BIOCARTA_AKAP13_PATHWAY | 9 | -0.45603 | -0.93783 | 0.495918 | 0.75476 | 0.741391 | 3395 | tags=56% | list=22% | signal=44% |
| BIOCARTA_IL12_PATHWAY | 10 | 0.478279 | 0.937751 | 0.495238 | 0.75476 | 0.741391 | 1955 | tags=40% | list=12% | signal=35% |
| REACTOME_CALCINEURIN_ACTIVATES_NFAT | 10 | -0.44484 | -0.92813 | 0.496855 | 0.75476 | 0.741391 | 3753 | tags=70% | list=24% | signal=53% |
| WP_PARKINSONS_DISEASE | 38 | -0.35947 | -0.91208 | 0.49652 | 0.75476 | 0.741391 | 1582 | tags=32% | list=10% | signal=28% |
| REACTOME_NEGATIVE_REGULATION_OF_MET_ACTIVITY | 21 | -0.38243 | -0.90317 | 0.497881 | 0.75476 | 0.741391 | 2489 | tags=38% | list=16% | signal=32% |
| REACTOME_HSP90_CHAPERONE_CYCLE_FOR_STEROID_HORMONE_ | 46 | 0.435294 | 0.980139 | 0.501724 | 0.756371 | 0.742973 | 254 | tags=4% | list=2% | signal=4% |
| REACTOME_BILE_ACID_AND_BILE_SALT_METABOLISM | 27 | 0.446258 | 0.959727 | 0.5 | 0.756371 | 0.742973 | 4731 | tags=56% | list=30% | signal=39% |
| REACTOME_CARGO_CONCENTRATION_IN_THE_ER | 30 | 0.436472 | 0.956017 | 0.501736 | 0.756371 | 0.742973 | 2593 | tags=33% | list=17% | signal=28% |
| REACTOME_RELEASE_OF_APOPTOTIC_FACTORS_FROM_THE_MITOC | 7 | -0.47836 | -0.94864 | 0.500998 | 0.756371 | 0.742973 | 2643 | tags=57% | list=17% | signal=48% |
| REACTOME_PHOSPHATE_BOND_HYDROLYSIS_BY_NUDT_PROTEINS | 7 | -0.47829 | -0.9485 | 0.500998 | 0.756371 | 0.742973 | 4949 | tags=71% | list=32% | signal=49% |
| REACTOME_TRANSPORT_OF_NUCLEOSIDES_AND_FREE_PURINE_AN | 8 | -0.4683 | -0.94238 | 0.502994 | 0.756371 | 0.742973 | 1988 | tags=50% | list=13% | signal=44% |
| REACTOME_ORGANIC_CATION_ANION_ZWITTERION_TRANSPORT | 11 | 0.468626 | 0.927745 | 0.501931 | 0.756371 | 0.742973 | 1750 | tags=18% | list=11% | signal=16% |
| REACTOME_PCNA_DEPENDENT_LONG_PATCH_BASE_EXCISION_REP | 20 | -0.38115 | -0.89522 | 0.503185 | 0.756371 | 0.742973 | 5424 | tags=50% | list=35% | signal=33% |
| REACTOME_LYSOSPHINGOLIPID_AND_LPA_RECEPTORS | 14 | -0.39154 | -0.88475 | 0.503055 | 0.756371 | 0.742973 | 515 | tags=14% | list=3% | signal=14% |
| BIOCARTA_VDR_PATHWAY | 21 | 0.440341 | 0.917702 | 0.503774 | 0.756571 | 0.74317 | 1594 | tags=24% | list=10% | signal=21% |
| BIOCARTA_INTEGRIN_PATHWAY | 33 | 0.433175 | 0.955271 | 0.504348 | 0.75675 | 0.743345 | 1955 | tags=27% | list=12% | signal=24% |
| REACTOME_SUMOYLATION_OF_RNA_BINDING_PROTEINS | 44 | 0.43294 | 0.973271 | 0.511036 | 0.757208 | 0.743796 | 4965 | tags=55% | list=32% | signal=37% |
| BIOCARTA_FCER1_PATHWAY | 36 | 0.430065 | 0.949578 | 0.507826 | 0.757208 | 0.743796 | 1955 | tags=17% | list=12% | signal=15% |
| REACTOME_TRAFFICKING_OF_AMPA_RECEPTORS | 29 | 0.435763 | 0.947389 | 0.510417 | 0.757208 | 0.743796 | 1458 | tags=21% | list=9% | signal=19% |
| REACTOME_MAPK3_ERK1_ACTIVATION | 8 | 0.483394 | 0.942218 | 0.510978 | 0.757208 | 0.743796 | 751 | tags=25% | list=5% | signal=24% |
| WP_GLUTATHIONE_AND_ONECARBON_METABOLISM | 28 | 0.435094 | 0.941809 | 0.506087 | 0.757208 | 0.743796 | 1187 | tags=11% | list=8% | signal=10% |
| REACTOME_CONDENSATION_OF_PROMETAPHASE_CHROMOSOMES | 8 | 0.482695 | 0.940856 | 0.510978 | 0.757208 | 0.743796 | 3103 | tags=38% | list=20% | signal=30% |
| REACTOME_HDMS_DEMETHYLATE_HISTONES | 28 | 0.432663 | 0.936547 | 0.506087 | 0.757208 | 0.743796 | 1173 | tags=14% | list=7% | signal=13% |
| REACTOME_ESTABLISHMENT_OF_SISTER_CHROMATID_COHESION | 10 | 0.472737 | 0.926885 | 0.506667 | 0.757208 | 0.743796 | 3140 | tags=40% | list=20% | signal=32% |
| REACTOME_AFLATOXIN_ACTIVATION_AND_DETOXIFICATION | 8 | -0.46042 | -0.92654 | 0.510978 | 0.757208 | 0.743796 | 1694 | tags=38% | list=11% | signal=33% |
| REACTOME_DISSOLUTION_OF_FIBRIN_CLOT | 9 | -0.44977 | -0.92495 | 0.506122 | 0.757208 | 0.743796 | 3245 | tags=44% | list=21% | signal=35% |
| REACTOME_FORMATION_OF_APOPTOSOME | 9 | -0.44844 | -0.92221 | 0.508163 | 0.757208 | 0.743796 | 2834 | tags=44% | list=18% | signal=36% |
| REACTOME_NEUROTRANSMITTER_RELEASE_CYCLE | 50 | -0.34973 | -0.90111 | 0.509662 | 0.757208 | 0.743796 | 1851 | tags=22% | list=12% | signal=19% |
| BIOCARTA_NKT_PATHWAY | 14 | 0.43881 | 0.893882 | 0.510763 | 0.757208 | 0.743796 | 3149 | tags=36% | list=20% | signal=29% |
| REACTOME_ER_PHAGOSOME_PATHWAY | 20 | -0.37983 | -0.89214 | 0.507431 | 0.757208 | 0.743796 | 835 | tags=20% | list=5% | signal=19% |
| REACTOME_MRNA_EDITING | 4 | -0.56508 | -0.98606 | 0.526946 | 0.7589 | 0.745458 | 5865 | tags=75% | list=37% | signal=47% |
| REACTOME_TACHYKININ_RECEPTORS_BIND_TACHYKININS | 4 | 0.569509 | 0.976861 | 0.51992 | 0.7589 | 0.745458 | 5771 | tags=75% | list=37% | signal=47% |
| REACTOME_SIGNALING_BY_MST1 | 4 | 0.56576 | 0.97043 | 0.527888 | 0.7589 | 0.745458 | 6822 | tags=100% | list=43% | signal=57% |
| REACTOME_ERYTHROCYTES_TAKE_UP_OXYGEN_AND_RELEASE_CAF | 6 | -0.51113 | -0.96395 | 0.515337 | 0.7589 | 0.745458 | 675 | tags=33% | list=4% | signal=32% |
| REACTOME_CHYLOMICRON_ASSEMBLY | 5 | 0.537566 | 0.963157 | 0.514851 | 0.7589 | 0.745458 | 2776 | tags=40% | list=18% | signal=33% |
| REACTOME_REGULATION_OF_MRNA_STABILITY_BY_PROTEINS_THA | 82 | 0.392387 | 0.947837 | 0.526899 | 0.7589 | 0.745458 | 159 | tags=2% | list=1% | signal=2% |
| BIOCARTA_BCR_PATHWAY | 31 | 0.42762 | 0.939929 | 0.519164 | 0.7589 | 0.745458 | 1955 | tags=16% | list=12% | signal=14% |
| REACTOME_NCAM_SIGNALING_FOR_NEURITE_OUT_GROWTH | 34 | 0.424335 | 0.935118 | 0.526224 | 0.7589 | 0.745458 | 2787 | tags=26% | list=18% | signal=22% |
| BIOCARTA_STAT3_PATHWAY | 8 | 0.478161 | 0.932019 | 0.51497 | 0.7589 | 0.745458 | 4464 | tags=50% | list=28% | signal=36% |
| BIOCARTA_CDC25_PATHWAY | 8 | 0.475828 | 0.927471 | 0.516966 | 0.7589 | 0.745458 | 349 | tags=12% | list=2% | signal=12% |
| BIOCARTA_PYK2_PATHWAY | 25 | 0.430094 | 0.924797 | 0.515901 | 0.7589 | 0.745458 | 178 | tags=4% | list=1% | signal=4% |
| REACTOME_CYTOCHROME_C_MEDIATED_APOPTOTIC_RESPONSE | 11 | -0.42686 | -0.91326 | 0.520661 | 0.7589 | 0.745458 | 2834 | tags=36% | list=18% | signal=30% |
| REACTOME_SIGNALING_BY_THE_B_CELL_RECEPTOR_BCR | 99 | -0.33698 | -0.90917 | 0.516575 | 0.7589 | 0.745458 | 2063 | tags=31% | list=13% | signal=27% |
| REACTOME_INTERLEUKIN_12_FAMILY_SIGNALING | 10 | 0.462621 | 0.907051 | 0.527619 | 0.7589 | 0.745458 | 5115 | tags=60% | list=33% | signal=40% |
| BIOCARTA_RAB_PATHWAY | 11 | -0.42384 | -0.9068 | 0.524793 | 0.7589 | 0.745458 | 2750 | tags=55% | list=18% | signal=45% |
| REACTOME_THE_NLRP3_INFLAMMASOME | 9 | -0.43881 | -0.90241 | 0.526531 | 0.7589 | 0.745458 | 3003 | tags=44% | list=19% | signal=36% |
| REACTOME_ACTIVATION_OF_THE_AP_1_FAMILY_OF_TRANSCRIPTIC | 10 | -0.43237 | -0.9021 | 0.522013 | 0.7589 | 0.745458 | 3539 | tags=90% | list=23% | signal=70% |
| REACTOME_SERINE_BIOSYNTHESIS | 9 | -0.43848 | -0.90173 | 0.526531 | 0.7589 | 0.745458 | 2477 | tags=44% | list=16% | signal=37% |
| WP_OSTEOCLAST_SIGNALING | 11 | 0.453801 | 0.898397 | 0.519305 | 0.7589 | 0.745458 | 2967 | tags=36% | list=19% | signal=30% |
| REACTOME_NUCLEAR_SIGNALING_BY_ERBB4 | 11 | 0.452918 | 0.896649 | 0.523166 | 0.7589 | 0.745458 | 3746 | tags=45% | list=24% | signal=35% |
| BIOCARTA_SPRY_PATHWAY | 16 | 0.439522 | 0.895187 | 0.523166 | 0.7589 | 0.745458 | 2152 | tags=31% | list=14% | signal=27% |
| REACTOME_EPHRIN_SIGNALING | 18 | 0.436308 | 0.894663 | 0.52741 | 0.7589 | 0.745458 | 1313 | tags=22% | list=8% | signal=20% |
| REACTOME_METHYLATION | 12 | -0.4139 | -0.89302 | 0.521649 | 0.7589 | 0.745458 | 4286 | tags=58% | list=27% | signal=42% |
| REACTOME_ADRENALINE_NORADRENALINE_INHIBITS_INSULIN_SEC | 21 | 0.427837 | 0.891642 | 0.526415 | 0.7589 | 0.745458 | 542 | tags=14% | list=3% | signal=14% |
| REACTOME_HEDGEHOG_OFF_STATE | 102 | -0.33069 | -0.88848 | 0.522857 | 0.7589 | 0.745458 | 2063 | tags=36% | list=13% | signal=32% |
| BIOCARTA_IL7_PATHWAY | 15 | 0.434444 | 0.887127 | 0.527027 | 0.7589 | 0.745458 | 1263 | tags=20% | list=8% | signal=18% |
| REACTOME_PHASE_II_CONJUGATION_OF_COMPOUNDS | 64 | -0.33837 | -0.88365 | 0.519802 | 0.7589 | 0.745458 | 3137 | tags=39% | list=20% | signal=31% |
| REACTOME_GLUTAMATE_NEUROTRANSMITTER_RELEASE_CYCLE | 23 | -0.36995 | -0.88272 | 0.517621 | 0.7589 | 0.745458 | 1851 | tags=30% | list=12% | signal=27% |
| BIOCARTA_MEF2D_PATHWAY | 15 | -0.38389 | -0.8815 | 0.520661 | 0.7589 | 0.745458 | 4077 | tags=47% | list=26% | signal=35% |
| REACTOME_INITIATION_OF_NUCLEAR_ENVELOPE_NE_REFORMATIC | 17 | -0.38084 | -0.87894 | 0.513514 | 0.7589 | 0.745458 | 2299 | tags=35% | list=15% | signal=30% |
| REACTOME_SYNTHESIS_OF_BILE_ACIDS_AND_BILE_SALTS_VIA_7ALF | 14 | 0.431432 | 0.878851 | 0.518591 | 0.7589 | 0.745458 | 4731 | tags=50% | list=30% | signal=35% |
| REACTOME_CARGO_TRAFFICKING_TO_THE_PERICILIARY_MEMBRAN | 39 | -0.34945 | -0.87725 | 0.527316 | 0.7589 | 0.745458 | 1480 | tags=18% | list=9% | signal=16% |
| REACTOME_DDX58_IFIH1_MEDIATED_INDUCTION_OF_INTERFERON | 33 | -0.3516 | -0.87135 | 0.512881 | 0.7589 | 0.745458 | 2489 | tags=30% | list=16% | signal=26% |
| WP_HYPOXIADEPENDENT_PROLIFERATION_OF_MYOBLASTS | 14 | -0.3847 | -0.86928 | 0.519348 | 0.7589 | 0.745458 | 2100 | tags=21% | list=13% | signal=19% |
| REACTOME_ACTIVATION_OF_KAINATE_RECEPTORS_UPON_GLUTAM | 30 | -0.34157 | -0.85751 | 0.528169 | 0.7589 | 0.745458 | 3520 | tags=67% | list=22% | signal=52% |
| BIOCARTA_PTEN_PATHWAY | 16 | 0.435344 | 0.886679 | 0.528958 | 0.759376 | 0.745926 | 2223 | tags=31% | list=14% | signal=27% |
| BIOCARTA_TGFB_PATHWAY | 17 | 0.432875 | 0.880452 | 0.52975 | 0.759858 | 0.746399 | 3375 | tags=41% | list=21% | signal=32% |
| REACTOME_G2_PHASE | 5 | 0.529348 | 0.948432 | 0.534653 | 0.765582 | 0.752022 | 5962 | tags=80% | list=38% | signal=50% |
| REACTOME_ASPARTATE_AND_ASPARAGINE_METABOLISM | 11 | -0.4116 | -0.88061 | 0.535124 | 0.765582 | 0.752022 | 2821 | tags=36% | list=18% | signal=30% |
| REACTOME_TGF_BETA_RECEPTOR_SIGNALING_IN_EMT_EPITHELIAL | 15 | -0.37639 | -0.86428 | 0.535124 | 0.765582 | 0.752022 | 1596 | tags=33% | list=10% | signal=30% |
| REACTOME_FORMATION_OF_RNA_POL_II_ELONGATION_COMPLEX | 54 | -0.34273 | -0.88384 | 0.536232 | 0.766507 | 0.75293 | 1863 | tags=20% | list=12% | signal=18% |
| REACTOME_MIRO_GTPASE_CYCLE | 7 | -0.46176 | -0.91573 | 0.536926 | 0.766839 | 0.753257 | 1234 | tags=43% | list=8% | signal=40% |
| WP_MIR222_IN_EXERCISEINDUCED_CARDIAC_GROWTH | 4 | -0.55777 | -0.97331 | 0.538922 | 0.768369 | 0.754759 | 221 | tags=50% | list=1% | signal=49% |
| WP_DYSREGULATED_MIRNA_TARGETING_IN_INSULINPI3KAKT_SIGN | 26 | 0.421039 | 0.908451 | 0.538732 | 0.768369 | 0.754759 | 2010 | tags=23% | list=13% | signal=20% |
| WP_NOVEL_JUNDMP1_PATHWAY | 24 | -0.36656 | -0.87812 | 0.541387 | 0.771221 | 0.757561 | 3898 | tags=46% | list=25% | signal=35% |
| REACTOME_REGULATION_OF_IFNG_SIGNALING | 8 | 0.461935 | 0.900392 | 0.542914 | 0.772072 | 0.758397 | 5144 | tags=50% | list=33% | signal=34% |

| | | | | | | | | | | |
|---|---|---|---|---|---|---|---|---|---|---|
| WP_LEPTIN_AND_ADIPONECTIN | 8 | 0.461924 | 0.900371 | 0.542914 | 0.772072 | 0.758397 | 427 | tags=12% | list=3% | signal=12% |
| BIOCARTA_CK1_PATHWAY | 15 | -0.37302 | -0.85653 | 0.543388 | 0.772086 | 0.75841 | 2610 | tags=40% | list=17% | signal=33% |
| REACTOME_PHASE_3_RAPID_REPOLARISATION | 6 | 0.509399 | 0.940392 | 0.545809 | 0.77288 | 0.75919 | 4276 | tags=50% | list=27% | signal=36% |
| REACTOME_PROCESSING_OF_CAPPED_INTRONLESS_PRE_MRNA | 29 | 0.41481 | 0.901836 | 0.545139 | 0.77288 | 0.75919 | 4098 | tags=34% | list=26% | signal=26% |
| REACTOME_COPII_MEDIATED_VESICLE_TRANSPORT | 65 | -0.32794 | -0.86053 | 0.544554 | 0.77288 | 0.75919 | 2880 | tags=29% | list=18% | signal=24% |
| REACTOME_TELOMERE_EXTENSION_BY_TELOMERASE | 20 | -0.36372 | -0.85429 | 0.545648 | 0.77288 | 0.75919 | 2134 | tags=35% | list=14% | signal=30% |
| WP_HYPOXIADEPENDENT_DIFFERENTIATION_OF_MYOBLASTS | 10 | 0.447684 | 0.877763 | 0.546667 | 0.773436 | 0.759736 | 124 | tags=10% | list=1% | signal=10% |
| REACTOME_RHO_GTPASES_ACTIVATE_ROCKS | 5 | -0.50616 | -0.92767 | 0.547284 | 0.77365 | 0.759946 | 3395 | tags=80% | list=22% | signal=63% |
| BIOCARTA_NO2IL12_PATHWAY | 6 | 0.508704 | 0.939108 | 0.547758 | 0.773662 | 0.759958 | 4464 | tags=67% | list=28% | signal=48% |
| WP_HYPOXIADEPENDENT_SELFRENEWAL_OF_MYOBLASTS | 9 | 0.454115 | 0.884399 | 0.548828 | 0.774514 | 0.760796 | 465 | tags=22% | list=3% | signal=22% |
| REACTOME_PI_METABOLISM | 74 | 0.393785 | 0.938361 | 0.552674 | 0.775782 | 0.76204 | 4441 | tags=38% | list=28% | signal=27% |
| BIOCARTA_ALTERNATIVE_PATHWAY | 5 | 0.521668 | 0.934673 | 0.552475 | 0.775782 | 0.76204 | 1121 | tags=20% | list=7% | signal=19% |
| REACTOME_STING_MEDIATED_INDUCTION_OF_HOST_IMMUNE_RE | 6 | 0.50347 | 0.929446 | 0.551657 | 0.775782 | 0.76204 | 724 | tags=33% | list=5% | signal=32% |
| REACTOME_ACETYLCHOLINE_BINDING_AND_DOWNSTREAM_EVENT | 8 | 0.454041 | 0.885006 | 0.552894 | 0.775782 | 0.76204 | 6221 | tags=50% | list=40% | signal=30% |
| REACTOME_HIGHLY_CALCIUM_PERMEABLE_POSTSYNAPTIC_NICOTI | 8 | 0.454041 | 0.885006 | 0.552894 | 0.775782 | 0.76204 | 6221 | tags=50% | list=40% | signal=30% |
| REACTOME_INFLAMMASOMES | 12 | -0.40054 | -0.86421 | 0.552577 | 0.775782 | 0.76204 | 3003 | tags=33% | list=19% | signal=27% |
| REACTOME_NUCLEAR_EVENTS_KINASE_AND_TRANSCRIPTION_FACT | 27 | -0.34843 | -0.85636 | 0.552995 | 0.775782 | 0.76204 | 4097 | tags=52% | list=26% | signal=38% |
| REACTOME_MET_RECEPTOR_ACTIVATION | 5 | 0.520478 | 0.93254 | 0.554455 | 0.777173 | 0.763407 | 6308 | tags=80% | list=40% | signal=48% |
| REACTOME_RMTS_METHYLATE_HISTONE_ARGININES | 37 | -0.33678 | -0.85455 | 0.555814 | 0.778421 | 0.764633 | 3340 | tags=46% | list=21% | signal=36% |
| REACTOME_SUMOYLATION_OF_UBIQUITINYLATION_PROTEINS | 36 | 0.412444 | 0.91067 | 0.556522 | 0.778755 | 0.764961 | 4965 | tags=50% | list=32% | signal=34% |
| REACTOME_PI_3K_CASCADE_FGFR1 | 17 | -0.36233 | -0.83622 | 0.559252 | 0.781917 | 0.768067 | 2772 | tags=29% | list=18% | signal=24% |
| REACTOME_EXTRINSIC_PATHWAY_OF_FIBRIN_CLOT_FORMATION | 3 | 0.596812 | 0.962617 | 0.559921 | 0.782195 | 0.76834 | 1587 | tags=33% | list=10% | signal=30% |
| REACTOME_SYNTHESIS_OF_15_EICOSATETRAENOIC_ACID_DERIVAT | 4 | -0.53038 | -0.92553 | 0.560878 | 0.782216 | 0.76836 | 2697 | tags=50% | list=17% | signal=41% |
| REACTOME_TELOMERE_C_STRAND_LAGGING_STRAND_SYNTHESIS | 31 | -0.33317 | -0.838 | 0.560748 | 0.782216 | 0.76836 | 4509 | tags=45% | list=29% | signal=32% |
| REACTOME_FACTORS_INVOLVED_IN_MEGAKARYOCYTE_DEVELOPM | 104 | -0.32218 | -0.86843 | 0.5625 | 0.783162 | 0.76929 | 1786 | tags=23% | list=11% | signal=21% |
| WP_MITOCHONDRIAL_GENE_EXPRESSION | 19 | 0.404407 | 0.842637 | 0.562152 | 0.783162 | 0.76929 | 4204 | tags=47% | list=27% | signal=35% |
| REACTOME_POLYMERASE_SWITCHING_ON_THE_C_STRAND_OF_TH | 24 | -0.36013 | -0.86273 | 0.563758 | 0.784152 | 0.770262 | 4509 | tags=46% | list=29% | signal=33% |
| REACTOME_SHC_MEDIATED_CASCADE_FGFR3 | 14 | -0.37024 | -0.83661 | 0.564155 | 0.784152 | 0.770262 | 377 | tags=21% | list=2% | signal=21% |
| REACTOME_LIPID_PARTICLE_ORGANIZATION | 3 | 0.595911 | 0.961164 | 0.565815 | 0.785012 | 0.771107 | 6348 | tags=100% | list=40% | signal=60% |
| REACTOME_SHC_MEDIATED_CASCADE_FGFR1 | 16 | -0.36633 | -0.84749 | 0.566116 | 0.785012 | 0.771107 | 377 | tags=25% | list=2% | signal=24% |
| BIOCARTA_CDMAC_PATHWAY | 14 | -0.36982 | -0.83568 | 0.566191 | 0.785012 | 0.771107 | 4077 | tags=43% | list=26% | signal=32% |
| REACTOME_TRANS_GOLGI_NETWORK_VESICLE_BUDDING | 70 | -0.32564 | -0.8509 | 0.566751 | 0.785132 | 0.771225 | 3049 | tags=34% | list=19% | signal=28% |
| REACTOME_CDC42_GTPASE_CYCLE | 93 | 0.375801 | 0.923606 | 0.569207 | 0.785876 | 0.771955 | 1722 | tags=22% | list=11% | signal=19% |
| REACTOME_GENE_SILENCING_BY_RNA | 40 | 0.403764 | 0.898783 | 0.568696 | 0.785876 | 0.771955 | 3743 | tags=38% | list=24% | signal=29% |
| REACTOME_TRIF_MEDIATED_PROGRAMMED_CELL_DEATH | 9 | 0.441085 | 0.859023 | 0.570313 | 0.785876 | 0.771955 | 3865 | tags=44% | list=25% | signal=34% |
| REACTOME_METABOLISM_OF_FOLATE_AND_PTERINES | 16 | -0.36399 | -0.84208 | 0.568182 | 0.785876 | 0.771955 | 4217 | tags=56% | list=27% | signal=41% |
| REACTOME_HYALURONAN_UPTAKE_AND_DEGRADATION | 12 | 0.415731 | 0.837803 | 0.5706 | 0.785876 | 0.771955 | 5254 | tags=58% | list=33% | signal=39% |
| REACTOME_DEPOLYMERISATION_OF_THE_NUCLEAR_LAMINA | 12 | 0.415646 | 0.83763 | 0.5706 | 0.785876 | 0.771955 | 279 | tags=8% | list=2% | signal=8% |
| REACTOME_SIGNALING_BY_ERBB4 | 31 | -0.32718 | -0.82294 | 0.570093 | 0.785876 | 0.771955 | 2526 | tags=23% | list=16% | signal=19% |
| REACTOME_INTERLEUKIN_35_SIGNALLING | 6 | 0.49336 | 0.910783 | 0.57115 | 0.785982 | 0.77206 | 5115 | tags=67% | list=33% | signal=45% |
| REACTOME_REGULATION_OF_FOXO_TRANSCRIPTIONAL_ACTIVITY_I | 6 | 0.491346 | 0.907064 | 0.573099 | 0.786709 | 0.772774 | 2223 | tags=50% | list=14% | signal=43% |
| REACTOME_REGULATION_OF_CYTOSKELETAL_REMODELING_AND_( | 8 | -0.43242 | -0.87019 | 0.572854 | 0.786709 | 0.772774 | 933 | tags=38% | list=6% | signal=35% |
| REACTOME_SMALL_INTERFERING_RNA_SIRNA_BIOGENESIS | 8 | 0.444776 | 0.866946 | 0.572854 | 0.786709 | 0.772774 | 1940 | tags=38% | list=12% | signal=33% |
| REACTOME_HIGHLY_SODIUM_PERMEABLE_POSTSYNAPTIC_ACETYL( | 3 | 0.587554 | 0.947684 | 0.575639 | 0.787754 | 0.773801 | 5030 | tags=67% | list=32% | signal=45% |
| WP_P53_SIGNALING | 62 | 0.390847 | 0.907026 | 0.575758 | 0.787754 | 0.773801 | 876 | tags=10% | list=6% | signal=9% |
| WP_SELENIUM_METABOLISM_SELENOPROTEINS | 42 | -0.33768 | -0.8513 | 0.574519 | 0.787754 | 0.773801 | 3599 | tags=40% | list=23% | signal=31% |
| REACTOME_ACTIVATED_TAK1_MEDIATES_P38_MAPK_ACTIVATION | 23 | -0.34796 | -0.83026 | 0.57489 | 0.787754 | 0.773801 | 2489 | tags=26% | list=16% | signal=22% |
| REACTOME_PTK6_REGULATES_RHO_GTPASES_RAS_GTPASE_AND_N | 12 | -0.38633 | -0.83354 | 0.57732 | 0.788592 | 0.774624 | 3655 | tags=50% | list=23% | signal=38% |
| REACTOME_NEGATIVE_REGULATION_OF_FLT3 | 14 | 0.40314 | 0.821219 | 0.577299 | 0.788592 | 0.774624 | 521 | tags=21% | list=3% | signal=21% |
| REACTOME_HIGHLY_CALCIUM_PERMEABLE_NICOTINIC_ACETYLCHO | 7 | 0.472999 | 0.896633 | 0.578842 | 0.788739 | 0.774769 | 6221 | tags=57% | list=40% | signal=35% |
| REACTOME_STRIATED_MUSCLE_CONTRACTION | 29 | 0.395443 | 0.85973 | 0.578125 | 0.788739 | 0.774769 | 1153 | tags=14% | list=7% | signal=13% |
| REACTOME_SIGNALING_BY_EGFR | 42 | -0.33509 | -0.84478 | 0.579327 | 0.788739 | 0.774769 | 2004 | tags=26% | list=13% | signal=23% |
| REACTOME_DNA_STRAND_ELONGATION | 23 | -0.34549 | -0.82437 | 0.579295 | 0.788739 | 0.774769 | 4509 | tags=52% | list=29% | signal=37% |
| REACTOME_TRANSPORT_OF_VITAMINS_NUCLEOSIDES_AND_RELAT | 33 | 0.393631 | 0.868065 | 0.58087 | 0.790192 | 0.776195 | 4141 | tags=52% | list=26% | signal=38% |
| REACTOME_PHOSPHOLIPASE_C_MEDIATED_CASCADE_FGFR2 | 13 | -0.37876 | -0.83877 | 0.582305 | 0.791496 | 0.777476 | 2772 | tags=23% | list=18% | signal=19% |
| BIOCARTA_CBL_PATHWAY | 11 | 0.412769 | 0.817165 | 0.583012 | 0.791809 | 0.777784 | 2306 | tags=36% | list=15% | signal=31% |
| REACTOME_CHYLOMICRON_REMODELING | 3 | 0.582264 | 0.939152 | 0.583497 | 0.791821 | 0.777795 | 5969 | tags=67% | list=38% | signal=41% |
| BIOCARTA_ERK_PATHWAY | 25 | -0.35067 | -0.84032 | 0.584862 | 0.793026 | 0.778979 | 377 | tags=16% | list=2% | signal=16% |
| WP_ETHANOL_METABOLISM_RESULTING_IN_PRODUCTION_OF_RO: | 9 | 0.431645 | 0.840638 | 0.585938 | 0.793835 | 0.779774 | 2134 | tags=33% | list=14% | signal=29% |
| BIOCARTA_AT1R_PATHWAY | 25 | 0.384156 | 0.826021 | 0.586572 | 0.794048 | 0.779983 | 178 | tags=4% | list=1% | signal=4% |
| REACTOME_RHOF_GTPASE_CYCLE | 40 | 0.391935 | 0.872451 | 0.587826 | 0.79445 | 0.780378 | 2278 | tags=25% | list=15% | signal=21% |
| REACTOME_RUNX1_INTERACTS_WITH_CO_FACTORS_WHOSE_PREC | 35 | 0.394422 | 0.869383 | 0.587719 | 0.79445 | 0.780378 | 1586 | tags=20% | list=10% | signal=18% |
| WP_CYTOKINES_AND_INFLAMMATORY_RESPONSE | 8 | 0.43323 | 0.844441 | 0.588822 | 0.79515 | 0.781065 | 3571 | tags=50% | list=23% | signal=39% |
| REACTOME_RHOB_GTPASE_CYCLE | 68 | 0.386724 | 0.907666 | 0.590381 | 0.795719 | 0.781624 | 2334 | tags=26% | list=15% | signal=23% |
| WP_MIRNAS_AND_TFS_IN_IPS_CELL_GENERATION | 6 | -0.47183 | -0.88982 | 0.591002 | 0.795719 | 0.781624 | 2277 | tags=67% | list=14% | signal=57% |
| REACTOME_AMPLIFICATION_OF_SIGNAL_FROM_THE_KINETOCHORI | 89 | -0.31378 | -0.84466 | 0.59116 | 0.795719 | 0.781624 | 2866 | tags=39% | list=18% | signal=32% |
| REACTOME_DEADENYLATION_OF_MRNA | 25 | 0.382158 | 0.821724 | 0.590106 | 0.795719 | 0.781624 | 2629 | tags=28% | list=17% | signal=23% |
| REACTOME_DEATH_RECEPTOR_SIGNALING | 137 | 0.357556 | 0.922507 | 0.596037 | 0.797293 | 0.78317 | 1704 | tags=15% | list=11% | signal=13% |
| REACTOME_PTK6_REGULATES_PROTEINS_INVOLVED_IN_RNA_PROC | 4 | 0.52583 | 0.901939 | 0.59761 | 0.797293 | 0.78317 | 337 | tags=25% | list=2% | signal=24% |
| REACTOME_ACTIVATION_OF_CASPASES_THROUGH_APOPTOSOME_ | 6 | -0.47011 | -0.88658 | 0.595092 | 0.797293 | 0.78317 | 2684 | tags=50% | list=17% | signal=41% |
| REACTOME_AUF1_HNRNP_D0_BINDS_AND_DESTABILIZES_MRNA | 54 | 0.385846 | 0.886063 | 0.596939 | 0.797293 | 0.78317 | 43 | tags=2% | list=0% | signal=2% |
| REACTOME_UB_SPECIFIC_PROCESSING_PROTEASES | 159 | -0.31694 | -0.86207 | 0.59403 | 0.797293 | 0.78317 | 3105 | tags=32% | list=20% | signal=26% |
| REACTOME_HEME_SIGNALING | 8 | 0.430585 | 0.839285 | 0.59481 | 0.797293 | 0.78317 | 483 | tags=12% | list=3% | signal=12% |
| WP_EUKARYOTIC_TRANSCRIPTION_INITIATION | 40 | -0.32599 | -0.82396 | 0.59719 | 0.797293 | 0.78317 | 3458 | tags=32% | list=22% | signal=25% |
| REACTOME_RHOBTB2_GTPASE_CYCLE | 22 | -0.3483 | -0.82002 | 0.59607 | 0.797293 | 0.78317 | 1607 | tags=32% | list=10% | signal=29% |
| REACTOME_CHK1_CHK2_CDS1_MEDIATED_INACTIVATION_OF_CYCL | 12 | -0.37881 | -0.81733 | 0.593814 | 0.797293 | 0.78317 | 2365 | tags=50% | list=15% | signal=43% |
| REACTOME_RAP1_SIGNALLING | 15 | 0.393962 | 0.804462 | 0.596525 | 0.797293 | 0.78317 | 690 | tags=13% | list=4% | signal=13% |
| BIOCARTA_HER2_PATHWAY | 22 | 0.374153 | 0.794181 | 0.597426 | 0.797293 | 0.78317 | 4553 | tags=45% | list=29% | signal=32% |
| REACTOME_G_ALPHA_12_13_SIGNALLING_EVENTS | 74 | 0.37956 | 0.904464 | 0.599676 | 0.798437 | 0.784294 | 841 | tags=9% | list=5% | signal=9% |
| REACTOME_CELLULAR_RESPONSE_TO_HYPOXIA | 68 | 0.381875 | 0.896283 | 0.600332 | 0.798437 | 0.784294 | 23 | tags=1% | list=0% | signal=1% |
| REACTOME_NGF_STIMULATED_TRANSCRIPTION | 6 | 0.474099 | 0.875225 | 0.60039 | 0.798437 | 0.784294 | 1295 | tags=33% | list=8% | signal=31% |
| WP_TRIACYLGLYCERIDE_SYNTHESIS | 18 | 0.394938 | 0.809833 | 0.599244 | 0.798437 | 0.784294 | 5807 | tags=50% | list=37% | signal=32% |
| REACTOME_ELASTIC_FIBRE_FORMATION | 34 | 0.380867 | 0.839326 | 0.601399 | 0.799138 | 0.784983 | 2101 | tags=35% | list=13% | signal=31% |
| REACTOME_TRANSFERRIN_ENDOCYTOSIS_AND_RECYCLING | 27 | 0.381586 | 0.820642 | 0.602113 | 0.799448 | 0.785287 | 1147 | tags=15% | list=7% | signal=14% |
| BIOCARTA_EPO_PATHWAY | 19 | -0.35049 | -0.81286 | 0.604752 | 0.80231 | 0.788099 | 3558 | tags=37% | list=23% | signal=29% |
| REACTOME_CHOLINE_CATABOLISM | 4 | -0.50707 | -0.88485 | 0.606786 | 0.803751 | 0.789514 | 1522 | tags=50% | list=10% | signal=45% |
| REACTOME_TP53_REGULATES_TRANSCRIPTION_OF_ADDITIONAL_CI | 18 | 0.391384 | 0.802545 | 0.606805 | 0.803751 | 0.789514 | 1842 | tags=28% | list=12% | signal=25% |
| REACTOME_RAF_INDEPENDENT_MAPK1_3_ACTIVATION | 21 | -0.34227 | -0.80833 | 0.608051 | 0.804759 | 0.790504 | 1876 | tags=33% | list=12% | signal=29% |
| REACTOME_SYNTHESIS_OF_VERY_LONG_CHAIN_FATTY_ACYL_COAS | 18 | -0.35304 | -0.81532 | 0.608879 | 0.805214 | 0.790951 | 654 | tags=11% | list=4% | signal=11% |
| REACTOME_METABOLISM_OF_LIPIDS | 484 | 0.320143 | 0.935902 | 0.614227 | 0.805385 | 0.791119 | 556 | tags=4% | list=4% | signal=4% |

| | | | | | | | | | | |
|---|---|---|---|---|---|---|---|---|---|---|
| BIOCARTA_CREM_PATHWAY | 3 | 0.555543 | 0.896053 | 0.612967 | 0.805385 | 0.791119 | 483 | tags=33% | list=3% | signal=32% |
| REACTOME_RAC2_GTPASE_CYCLE | 85 | 0.365002 | 0.891055 | 0.613354 | 0.805385 | 0.791119 | 3108 | tags=29% | list=20% | signal=24% |
| REACTOME_NUCLEAR_ENVELOPE_BREAKDOWN | 45 | 0.384101 | 0.862407 | 0.610922 | 0.805385 | 0.791119 | 3743 | tags=33% | list=24% | signal=25% |
| REACTOME_PTK6_REGULATES_CELL_CYCLE | 5 | -0.46728 | -0.85642 | 0.61167 | 0.805385 | 0.791119 | 234 | tags=40% | list=1% | signal=39% |
| REACTOME_REELIN_SIGNALLING_PATHWAY | 5 | -0.46633 | -0.85469 | 0.613682 | 0.805385 | 0.791119 | 1397 | tags=60% | list=9% | signal=55% |
| REACTOME_CELL_CELL_JUNCTION_ORGANIZATION | 39 | 0.37647 | 0.841042 | 0.611015 | 0.805385 | 0.791119 | 2053 | tags=31% | list=13% | signal=27% |
| REACTOME_DCC_MEDIATED_ATTRACTIVE_SIGNALING | 9 | -0.40571 | -0.83435 | 0.612245 | 0.805385 | 0.791119 | 2708 | tags=44% | list=17% | signal=37% |
| BIOCARTA_EPHA4_PATHWAY | 8 | 0.418435 | 0.815603 | 0.61477 | 0.805385 | 0.791119 | 1913 | tags=38% | list=12% | signal=33% |
| REACTOME_POLB_DEPENDENT_LONG_PATCH_BASE_EXCISION_REP | 8 | 0.417946 | 0.81465 | 0.616766 | 0.805385 | 0.791119 | 476 | tags=12% | list=3% | signal=12% |
| WP_GPCRS_CLASS_C_METABOTROPIC_GLUTAMATE_PHEROMONE | 13 | -0.36729 | -0.81339 | 0.611111 | 0.805385 | 0.791119 | 795 | tags=15% | list=5% | signal=15% |
| REACTOME_DOWNREGULATION_OF_SMAD2_3_SMAD4_TRANSCRIP | 22 | -0.34269 | -0.80681 | 0.611354 | 0.805385 | 0.791119 | 2834 | tags=23% | list=18% | signal=19% |
| REACTOME_PROCESSIVE_SYNTHESIS_ON_THE_C_STRAND_OF_THE_ | 17 | -0.34805 | -0.80326 | 0.615385 | 0.805385 | 0.791119 | 4509 | tags=53% | list=29% | signal=38% |
| REACTOME_PEPTIDE_HORMONE_METABOLISM | 36 | -0.31603 | -0.79648 | 0.615925 | 0.805385 | 0.791119 | 154 | tags=11% | list=1% | signal=11% |
| WP_SEROTONIN_AND_ANXIETYRELATED_EVENTS | 12 | 0.394004 | 0.794016 | 0.615087 | 0.805385 | 0.791119 | 2767 | tags=33% | list=18% | signal=27% |
| BIOCARTA_IL2_PATHWAY | 21 | -0.33501 | -0.79118 | 0.616525 | 0.805385 | 0.791119 | 3558 | tags=33% | list=23% | signal=26% |
| REACTOME_FERTILIZATION | 10 | 0.404957 | 0.79399 | 0.619048 | 0.807728 | 0.793421 | 5984 | tags=70% | list=38% | signal=43% |
| REACTOME_DIGESTION | 5 | -0.46243 | -0.84754 | 0.62173 | 0.808055 | 0.793742 | 4367 | tags=60% | list=28% | signal=43% |
| REACTOME_RHOU_GTPASE_CYCLE | 39 | 0.373772 | 0.835016 | 0.621343 | 0.808055 | 0.793742 | 2466 | tags=28% | list=16% | signal=24% |
| REACTOME_RHOBTB3_ATPASE_CYCLE | 10 | -0.39853 | -0.83151 | 0.620545 | 0.808055 | 0.793742 | 3893 | tags=50% | list=25% | signal=38% |
| REACTOME_SYNTHESIS_OF_PYROPHOSPHATES_IN_THE_CYTOSOL | 10 | 0.404902 | 0.793882 | 0.620952 | 0.808055 | 0.793742 | 1328 | tags=20% | list=8% | signal=18% |
| REACTOME_SYNTHESIS_OF_GLYCOSYLPHOSPHATIDYLINOSITOL_GPI | 17 | 0.389027 | 0.791266 | 0.619962 | 0.808055 | 0.793742 | 2147 | tags=24% | list=14% | signal=20% |
| WP_BDNF_PATHWAY | 14 | 0.384828 | 0.783916 | 0.622309 | 0.808175 | 0.79386 | 1518 | tags=21% | list=10% | signal=19% |
| REACTOME_RUNX2_REGULATES_BONE_DEVELOPMENT | 6 | 0.457772 | 0.845084 | 0.623782 | 0.809447 | 0.795109 | 4115 | tags=50% | list=26% | signal=37% |
| REACTOME_BH3_ONLY_PROTEINS_ASSOCIATE_WITH_AND_INACTIV | 7 | 0.437345 | 0.829045 | 0.62475 | 0.809447 | 0.795109 | 1557 | tags=14% | list=10% | signal=13% |
| REACTOME_POST_TRANSCRIPTIONAL_SILENCING_BY_SMALL_RNAS | 7 | 0.437041 | 0.828469 | 0.62475 | 0.809447 | 0.795109 | 1733 | tags=29% | list=11% | signal=25% |
| REACTOME_POLYMERASE_SWITCHING | 14 | -0.34766 | -0.7856 | 0.625255 | 0.809468 | 0.79513 | 5424 | tags=57% | list=35% | signal=37% |
| REACTOME_INTERLEUKIN_23_SIGNALING | 5 | 0.479036 | 0.85829 | 0.625743 | 0.809469 | 0.795131 | 751 | tags=20% | list=5% | signal=19% |
| REACTOME_TRANSPORT_OF_MATURE_TRANSCRIPT_TO_CYTOPLASI | 75 | 0.36766 | 0.875943 | 0.626623 | 0.809483 | 0.795145 | 4485 | tags=39% | list=29% | signal=28% |
| REACTOME_EGFR_DOWNREGULATION | 27 | -0.32011 | -0.78676 | 0.626728 | 0.809483 | 0.795145 | 2489 | tags=30% | list=16% | signal=25% |
| REACTOME_INNATE_IMMUNE_SYSTEM | 748 | 0.309058 | 0.935453 | 0.628719 | 0.810354 | 0.796 | 1163 | tags=6% | list=7% | signal=6% |
| REACTOME_O_GLYCOSYLATION_OF_TSR_DOMAIN_CONTAINING_PI | 38 | 0.361995 | 0.804237 | 0.628722 | 0.810354 | 0.796 | 1761 | tags=21% | list=11% | signal=19% |
| BIOCARTA_RANBP2_PATHWAY | 12 | -0.36723 | -0.79233 | 0.628866 | 0.810354 | 0.796 | 1971 | tags=33% | list=13% | signal=29% |
| REACTOME_SUMOYLATION_OF_TRANSCRIPTION_COFACTORS | 38 | 0.360134 | 0.800101 | 0.630473 | 0.811795 | 0.797416 | 4965 | tags=58% | list=32% | signal=40% |
| BIOCARTA_LIS1_PATHWAY | 14 | -0.34604 | -0.78194 | 0.631365 | 0.812313 | 0.797925 | 4025 | tags=50% | list=26% | signal=37% |
| REACTOME_DNA_DOUBLE_STRAND_BREAK_REPAIR | 134 | -0.30739 | -0.82585 | 0.632948 | 0.812536 | 0.798143 | 2489 | tags=16% | list=16% | signal=14% |
| REACTOME_HUR_ELAVL1_BINDS_AND_STABILIZES_MRNA | 7 | 0.435403 | 0.825365 | 0.632735 | 0.812536 | 0.798143 | 1967 | tags=29% | list=13% | signal=25% |
| REACTOME_INTERLEUKIN_17_SIGNALING | 58 | -0.31183 | -0.80636 | 0.633005 | 0.812536 | 0.798143 | 4174 | tags=47% | list=27% | signal=34% |
| REACTOME_SMOOTH_MUSCLE_CONTRACTION | 30 | 0.360851 | 0.790382 | 0.635417 | 0.815002 | 0.800566 | 1818 | tags=17% | list=12% | signal=15% |
| REACTOME_NERVOUS_SYSTEM_DEVELOPMENT | 254 | 0.327453 | 0.901803 | 0.638767 | 0.815479 | 0.801034 | 2710 | tags=24% | list=17% | signal=20% |
| BIOCARTA_MAPK_PATHWAY | 76 | 0.364748 | 0.868668 | 0.638574 | 0.815479 | 0.801034 | 2123 | tags=25% | list=14% | signal=22% |
| REACTOME_APEX1_INDEPENDENT_RESOLUTION_OF_AP_SITES_VIA_ | 7 | 0.432486 | 0.819835 | 0.638723 | 0.815479 | 0.801034 | 5631 | tags=57% | list=36% | signal=37% |
| REACTOME_GRB2_EVENTS_IN_ERBB2_SIGNALING | 9 | -0.39369 | -0.80962 | 0.638776 | 0.815479 | 0.801034 | 377 | tags=22% | list=2% | signal=22% |
| REACTOME_PI3K_AKT_ACTIVATION | 8 | 0.411274 | 0.801644 | 0.638723 | 0.815479 | 0.801034 | 3601 | tags=38% | list=23% | signal=29% |
| REACTOME_SIGNALING_BY_PTK6 | 44 | -0.31429 | -0.79836 | 0.639225 | 0.815479 | 0.801034 | 2825 | tags=36% | list=18% | signal=30% |
| REACTOME_SIGNALING_BY_NOTCH3 | 19 | 0.367086 | 0.764874 | 0.636364 | 0.815479 | 0.801034 | 1487 | tags=32% | list=9% | signal=29% |
| REACTOME_CELL_CELL_COMMUNICATION | 76 | 0.363759 | 0.866312 | 0.640194 | 0.816088 | 0.801633 | 2279 | tags=29% | list=15% | signal=25% |
| REACTOME_VITAMIN_D_CALCIFEROL_METABOLISM | 3 | 0.542217 | 0.87456 | 0.644401 | 0.816569 | 0.802105 | 6202 | tags=67% | list=39% | signal=40% |
| REACTOME_RHOC_GTPASE_CYCLE | 71 | 0.358299 | 0.849955 | 0.648208 | 0.816569 | 0.802105 | 2334 | tags=27% | list=15% | signal=23% |
| REACTOME_REGULATION_OF_TNFR1_SIGNALING | 47 | 0.370286 | 0.838209 | 0.646552 | 0.816569 | 0.802105 | 426 | tags=9% | list=3% | signal=8% |
| REACTOME_TNFR1_MEDIATED_CERAMIDE_PRODUCTION | 5 | -0.44988 | -0.82454 | 0.645875 | 0.816569 | 0.802105 | 406 | tags=40% | list=3% | signal=39% |
| REACTOME_SHC1_EVENTS_IN_EGFR_SIGNALING | 9 | -0.39136 | -0.80482 | 0.644898 | 0.816569 | 0.802105 | 377 | tags=22% | list=2% | signal=22% |
| REACTOME_RNA_POLYMERASE_II_PRE_TRANSCRIPTION_EVENTS | 75 | -0.30175 | -0.79738 | 0.645078 | 0.816569 | 0.802105 | 1863 | tags=24% | list=12% | signal=21% |
| REACTOME_RND2_GTPASE_CYCLE | 39 | 0.35516 | 0.793437 | 0.64716 | 0.816569 | 0.802105 | 2565 | tags=31% | list=16% | signal=26% |
| BIOCARTA_EIF_PATHWAY | 9 | 0.40432 | 0.787422 | 0.642578 | 0.816569 | 0.802105 | 291 | tags=11% | list=2% | signal=11% |
| REACTOME_MET_ACTIVATES_RAS_SIGNALING | 9 | 0.399995 | 0.778999 | 0.648438 | 0.816569 | 0.802105 | 2569 | tags=44% | list=16% | signal=37% |
| REACTOME_CD209_DC_SIGN_SIGNALING | 16 | -0.33634 | -0.77811 | 0.642562 | 0.816569 | 0.802105 | 2610 | tags=31% | list=17% | signal=26% |
| REACTOME_IKK_COMPLEX_RECRUITMENT_MEDIATED_BY_RIP1 | 23 | -0.3232 | -0.77119 | 0.647577 | 0.816569 | 0.802105 | 676 | tags=26% | list=4% | signal=25% |
| REACTOME_RNA_POLYMERASE_I_TRANSCRIPTION_TERMINATION | 29 | -0.30986 | -0.76804 | 0.647887 | 0.816569 | 0.802105 | 805 | tags=34% | list=5% | signal=33% |
| REACTOME_REGULATION_OF_SIGNALING_BY_CBL | 21 | -0.32503 | -0.76761 | 0.648305 | 0.816569 | 0.802105 | 2489 | tags=24% | list=16% | signal=20% |
| REACTOME_PI3K_EVENTS_IN_ERBB2_SIGNALING | 12 | 0.380854 | 0.767516 | 0.644101 | 0.816569 | 0.802105 | 6209 | tags=58% | list=40% | signal=35% |
| BIOCARTA_EDG1_PATHWAY | 20 | 0.367825 | 0.763393 | 0.642185 | 0.816569 | 0.802105 | 4863 | tags=40% | list=31% | signal=28% |
| WP_EDA_SIGNALING_IN_HAIR_FOLLICLE_DEVELOPMENT | 15 | 0.372564 | 0.760769 | 0.644788 | 0.816569 | 0.802105 | 2799 | tags=27% | list=18% | signal=22% |
| REACTOME_TERMINAL_PATHWAY_OF_COMPLEMENT | 4 | 0.495018 | 0.849089 | 0.651394 | 0.819671 | 0.805153 | 2952 | tags=25% | list=19% | signal=20% |
| REACTOME_NEGATIVE_REGULATION_OF_THE_PI3K_AKT_NETWORK | 91 | 0.351124 | 0.85911 | 0.653125 | 0.819713 | 0.805194 | 891 | tags=9% | list=6% | signal=8% |
| REACTOME_RNA_POLYMERASE_II_TRANSCRIPTION_TERMINATION | 62 | 0.366123 | 0.849649 | 0.654882 | 0.819713 | 0.805194 | 4098 | tags=34% | list=26% | signal=25% |
| REACTOME_NEGATIVE_FEEDBACK_REGULATION_OF_MAPK_PATHW | 6 | -0.44479 | -0.83882 | 0.654397 | 0.819713 | 0.805194 | 3558 | tags=67% | list=23% | signal=52% |
| BIOCARTA_ERBB3_PATHWAY | 6 | 0.443099 | 0.817997 | 0.653021 | 0.819713 | 0.805194 | 6209 | tags=67% | list=40% | signal=40% |
| REACTOME_FGFR1C_LIGAND_BINDING_AND_ACTIVATION | 7 | -0.40916 | -0.81142 | 0.654691 | 0.819713 | 0.805194 | 1909 | tags=29% | list=12% | signal=25% |
| REACTOME_VEGFR2_MEDIATED_CELL_PROLIFERATION | 12 | -0.36009 | -0.77694 | 0.653608 | 0.819713 | 0.805194 | 1662 | tags=25% | list=11% | signal=22% |
| BIOCARTA_BCELLSURVIVAL_PATHWAY | 16 | 0.375541 | 0.764875 | 0.65444 | 0.819713 | 0.805194 | 3601 | tags=38% | list=23% | signal=29% |
| BIOCARTA_SKP2E2F_PATHWAY | 9 | 0.395692 | 0.770618 | 0.65625 | 0.820807 | 0.806268 | 2960 | tags=44% | list=19% | signal=36% |
| WP_EPO_RECEPTOR_SIGNALING | 25 | 0.344634 | 0.74104 | 0.657244 | 0.821431 | 0.806881 | 1518 | tags=16% | list=10% | signal=14% |
| REACTOME_EPHB_MEDIATED_FORWARD_SIGNALING | 36 | -0.30072 | -0.75789 | 0.65808 | 0.821857 | 0.8073 | 2909 | tags=47% | list=19% | signal=39% |
| WP_REGULATION_OF_ACTIN_CYTOSKELETON | 135 | 0.336509 | 0.867969 | 0.664134 | 0.822615 | 0.808044 | 2578 | tags=21% | list=16% | signal=18% |
| REACTOME_WAX_AND_PLASMALOGEN_BIOSYNTHESIS | 4 | 0.48742 | 0.836056 | 0.663347 | 0.822615 | 0.808044 | 5807 | tags=50% | list=37% | signal=32% |
| REACTOME_ABACAVIR_ADME | 5 | 0.45797 | 0.820544 | 0.663366 | 0.822615 | 0.808044 | 3905 | tags=40% | list=25% | signal=30% |
| REACTOME_AMINO_ACIDS_REGULATE_MTORC1 | 51 | -0.30697 | -0.78484 | 0.660147 | 0.822615 | 0.808044 | 2674 | tags=33% | list=17% | signal=28% |
| REACTOME_ERBB2_ACTIVATES_PTK6_SIGNALING | 8 | 0.399811 | 0.779302 | 0.660679 | 0.822615 | 0.808044 | 6209 | tags=75% | list=40% | signal=45% |
| REACTOME_SEMA4D_MEDIATED_INHIBITION_OF_CELL_ATTACHMEI | 8 | 0.396728 | 0.773292 | 0.662675 | 0.822615 | 0.808044 | 1299 | tags=25% | list=8% | signal=23% |
| REACTOME_SIGNALING_BY_TYPE_1_INSULIN_LIKE_GROWTH_FACTC | 38 | 0.345852 | 0.768372 | 0.661996 | 0.822615 | 0.808044 | 624 | tags=8% | list=4% | signal=8% |
| REACTOME_TELOMERE_MAINTENANCE | 48 | -0.29896 | -0.7673 | 0.660333 | 0.822615 | 0.808044 | 4509 | tags=38% | list=29% | signal=27% |
| WP_TOLLLIKE_RECEPTOR_SIGNALING | 32 | 0.348083 | 0.767213 | 0.662609 | 0.822615 | 0.808044 | 4735 | tags=44% | list=30% | signal=31% |
| BIOCARTA_IL3_PATHWAY | 14 | -0.33868 | -0.7653 | 0.663951 | 0.822615 | 0.808044 | 3558 | tags=36% | list=23% | signal=28% |
| REACTOME_UNFOLDED_PROTEIN_RESPONSE_UPR | 11 | 0.381757 | 0.755771 | 0.664093 | 0.822615 | 0.808044 | 3223 | tags=36% | list=21% | signal=29% |
| WP_ONECARBON_METABOLISM_AND_RELATED_PATHWAYS | 44 | 0.352087 | 0.79151 | 0.665535 | 0.823113 | 0.808534 | 1187 | tags=11% | list=8% | signal=11% |
| REACTOME_CARGO_RECOGNITION_FOR_CLATHRIN_MEDIATED_ENI | 91 | -0.2929 | -0.78743 | 0.665746 | 0.823113 | 0.808534 | 3130 | tags=38% | list=20% | signal=31% |
| BIOCARTA_ACE2_PATHWAY | 11 | 0.377715 | 0.747768 | 0.666023 | 0.823113 | 0.808534 | 4733 | tags=64% | list=30% | signal=44% |
| REACTOME_COBALAMIN_CBL_VITAMIN_B12_TRANSPORT_AND_ME | 13 | -0.35059 | -0.77639 | 0.666667 | 0.823296 | 0.808713 | 572 | tags=15% | list=4% | signal=15% |
| WP_OVARIAN_INFERTILITY | 22 | 0.347406 | 0.737409 | 0.667279 | 0.823441 | 0.808855 | 5032 | tags=45% | list=32% | signal=31% |
| REACTOME_SUMO_IS_TRANSFERRED_FROM_E1_TO_E2_UBE2I_UBC | 5 | -0.44282 | -0.81159 | 0.668008 | 0.823728 | 0.809137 | 536 | tags=100% | list=3% | signal=97% |

| | | | | | | | | | |
|---|---|---|---|---|---|---|---|---|---|
| BIOCARTA_AGPCR_PATHWAY | 10 | -0.38102 | -0.79497 | 0.67086 | 0.826018 | 0.811386 | 4077 | tags=60% | list=26% | signal=44% |
| REACTOME_PROCESSING_AND_ACTIVATION_OF_SUMO | 10 | -0.38004 | -0.79293 | 0.67086 | 0.826018 | 0.811386 | 1155 | tags=30% | list=7% | signal=28% |
| REACTOME_PHOSPHORYLATION_OF_CD3_AND_TCR_ZETA_CHAINS | 11 | -0.36536 | -0.78168 | 0.671488 | 0.826178 | 0.811544 | 2272 | tags=55% | list=14% | signal=47% |
| BIOCARTA_CACAM_PATHWAY | 6 | -0.43153 | -0.81382 | 0.672802 | 0.82676 | 0.812116 | 844 | tags=50% | list=5% | signal=47% |
| REACTOME_METABOLISM_OF_AMINE_DERIVED_HORMONES | 10 | -0.37894 | -0.79063 | 0.672956 | 0.82676 | 0.812116 | 4810 | tags=50% | list=31% | signal=35% |
| BIOCARTA_CHREBP_PATHWAY | 9 | -0.37302 | -0.76712 | 0.67551 | 0.829285 | 0.814596 | 2610 | tags=56% | list=17% | signal=46% |
| WP_EICOSANOID_SYNTHESIS | 15 | 0.357561 | 0.730133 | 0.677606 | 0.831244 | 0.81652 | 5332 | tags=47% | list=34% | signal=31% |
| REACTOME_MRNA_SPLICING_MINOR_PATHWAY | 48 | 0.344652 | 0.783541 | 0.681583 | 0.835506 | 0.820706 | 378 | tags=6% | list=2% | signal=6% |
| REACTOME_FOXO_MEDIATED_TRANSCRIPTION_OF_CELL_CYCLE_GE | 8 | 0.388864 | 0.757963 | 0.682635 | 0.835574 | 0.820774 | 2918 | tags=38% | list=19% | signal=31% |
| REACTOME_METABOLISM_OF_PORPHYRINS | 20 | -0.32131 | -0.75468 | 0.683652 | 0.835574 | 0.820774 | 2152 | tags=40% | list=14% | signal=35% |
| REACTOME_DOWNREGULATION_OF_TGF_BETA_RECEPTOR_SIGNAL | 16 | 0.359625 | 0.732458 | 0.683398 | 0.835574 | 0.820774 | 208 | tags=6% | list=1% | signal=6% |
| BIOCARTA_PLCE_PATHWAY | 11 | 0.368784 | 0.730087 | 0.683398 | 0.835574 | 0.820774 | 2277 | tags=27% | list=14% | signal=23% |
| WP_MAPK_SIGNALING_PATHWAY | 151 | 0.329352 | 0.860132 | 0.691489 | 0.835927 | 0.82112 | 1496 | tags=11% | list=10% | signal=10% |
| REACTOME_CLATHRIN_MEDIATED_ENDOCYTOSIS | 130 | -0.29539 | -0.79974 | 0.6875 | 0.835927 | 0.82112 | 3130 | tags=39% | list=20% | signal=32% |
| BIOCARTA_SAM68_PATHWAY | 6 | -0.41774 | -0.78781 | 0.689162 | 0.835927 | 0.82112 | 2834 | tags=50% | list=18% | signal=41% |
| REACTOME_LIGAND_RECEPTOR_INTERACTIONS | 5 | 0.43632 | 0.781755 | 0.691089 | 0.835927 | 0.82112 | 4070 | tags=60% | list=26% | signal=44% |
| BIOCARTA_IRES_PATHWAY | 6 | 0.420871 | 0.776961 | 0.690058 | 0.835927 | 0.82112 | 1497 | tags=17% | list=10% | signal=15% |
| REACTOME_IRON_UPTAKE_AND_TRANSPORT | 50 | -0.29096 | -0.74969 | 0.68599 | 0.835927 | 0.82112 | 2024 | tags=24% | list=13% | signal=21% |
| REACTOME_REGULATION_OF_TP53_ACTIVITY_THROUGH_METHYLA | 18 | -0.32293 | -0.7458 | 0.689218 | 0.835927 | 0.82112 | 676 | tags=33% | list=4% | signal=32% |
| BIOCARTA_CXCR4_PATHWAY | 18 | -0.32195 | -0.74353 | 0.689218 | 0.835927 | 0.82112 | 4077 | tags=33% | list=26% | signal=25% |
| REACTOME_DOWNSTREAM_SIGNALING_OF_ACTIVATED_FGFR3 | 21 | -0.31399 | -0.74154 | 0.690678 | 0.835927 | 0.82112 | 1909 | tags=24% | list=12% | signal=21% |
| BIOCARTA_PTDINS_PATHWAY | 14 | 0.363308 | 0.740079 | 0.690802 | 0.835927 | 0.82112 | 4496 | tags=43% | list=29% | signal=31% |
| BIOCARTA_AGR_PATHWAY | 30 | -0.293 | -0.7356 | 0.687793 | 0.835927 | 0.82112 | 2143 | tags=23% | list=14% | signal=20% |
| BIOCARTA_PAR1_PATHWAY | 17 | 0.360823 | 0.733899 | 0.68906 | 0.835927 | 0.82112 | 1021 | tags=18% | list=7% | signal=17% |
| REACTOME_DOWNSTREAM_SIGNAL_TRANSDUCTION | 27 | 0.337831 | 0.726542 | 0.690141 | 0.835927 | 0.82112 | 3601 | tags=37% | list=23% | signal=29% |
| REACTOME_SIGNALING_BY_ALK | 15 | 0.35153 | 0.717818 | 0.685328 | 0.835927 | 0.82112 | 2278 | tags=20% | list=15% | signal=17% |
| WP_NUCLEAR_RECEPTORS_IN_LIPID_METABOLISM_AND_TOXICITY | 16 | 0.351995 | 0.716918 | 0.689189 | 0.835927 | 0.82112 | 2741 | tags=38% | list=17% | signal=31% |
| BIOCARTA_HIVNEF_PATHWAY | 52 | 0.343934 | 0.794038 | 0.693603 | 0.837872 | 0.823031 | 2763 | tags=17% | list=18% | signal=14% |
| REACTOME_SEROTONIN_AND_MELATONIN_BIOSYNTHESIS | 3 | 0.515083 | 0.830794 | 0.695481 | 0.839291 | 0.824425 | 1312 | tags=33% | list=8% | signal=31% |
| REACTOME_METABOLISM_OF_NUCLEOTIDES | 84 | 0.334127 | 0.813659 | 0.695788 | 0.839291 | 0.824425 | 375 | tags=2% | list=2% | signal=2% |
| REACTOME_SIGNAL_REGULATORY_PROTEIN_FAMILY_INTERACTION | 6 | -0.41545 | -0.7835 | 0.699387 | 0.839365 | 0.824497 | 3176 | tags=50% | list=20% | signal=40% |
| REACTOME_LINOLEIC_ACID_LA_METABOLISM | 7 | -0.39089 | -0.77518 | 0.698603 | 0.839365 | 0.824497 | 2154 | tags=29% | list=14% | signal=25% |
| REACTOME_SIGNALING_BY_FGFR1 | 44 | 0.33812 | 0.760111 | 0.697793 | 0.839365 | 0.824497 | 624 | tags=7% | list=4% | signal=7% |
| BIOCARTA_TRKA_PATHWAY | 12 | -0.34701 | -0.74871 | 0.698969 | 0.839365 | 0.824497 | 377 | tags=33% | list=2% | signal=33% |
| REACTOME_WNT_MEDIATED_ACTIVATION_OF_DVL | 8 | 0.379819 | 0.740332 | 0.696607 | 0.839365 | 0.824497 | 4357 | tags=50% | list=28% | signal=36% |
| REACTOME_SUMOYLATION_OF_TRANSCRIPTION_FACTORS | 14 | 0.359065 | 0.731436 | 0.69863 | 0.839365 | 0.824497 | 3975 | tags=43% | list=25% | signal=32% |
| BIOCARTA_NUCLEARRS_PATHWAY | 16 | 0.348274 | 0.70934 | 0.696911 | 0.839365 | 0.824497 | 2741 | tags=38% | list=17% | signal=31% |
| REACTOME_DEFENSINS | 3 | -0.50729 | -0.81758 | 0.701826 | 0.841097 | 0.826199 | 2272 | tags=67% | list=14% | signal=57% |
| REACTOME_TNF_SIGNALING | 56 | 0.342017 | 0.792695 | 0.701843 | 0.841097 | 0.826199 | 426 | tags=7% | list=3% | signal=7% |
| REACTOME_ADHERENS_JUNCTIONS_INTERACTIONS | 31 | 0.329266 | 0.723742 | 0.703833 | 0.842874 | 0.827944 | 4842 | tags=55% | list=31% | signal=38% |
| REACTOME_INTERLEUKIN_9_SIGNALING | 4 | -0.45379 | -0.79187 | 0.704591 | 0.843174 | 0.828239 | 4739 | tags=50% | list=30% | signal=35% |
| BIOCARTA_TCAPOPTOSIS_PATHWAY | 3 | 0.507974 | 0.819327 | 0.705305 | 0.84342 | 0.828481 | 1314 | tags=33% | list=8% | signal=31% |
| REACTOME_MAPK6_MAPK4_SIGNALING | 72 | 0.336686 | 0.801055 | 0.707993 | 0.84598 | 0.830996 | 648 | tags=4% | list=4% | signal=4% |
| REACTOME_NEDDYLATION | 211 | -0.28868 | -0.79148 | 0.708464 | 0.84598 | 0.830996 | 3117 | tags=35% | list=20% | signal=28% |
| REACTOME_FATTY_ACYL_COA_BIOSYNTHESIS | 28 | -0.29981 | -0.73772 | 0.709602 | 0.846347 | 0.831356 | 883 | tags=21% | list=6% | signal=20% |
| REACTOME_OXIDATIVE_STRESS_INDUCED_SENESCENCE | 37 | 0.329183 | 0.726766 | 0.70979 | 0.846347 | 0.831356 | 876 | tags=14% | list=6% | signal=13% |
| REACTOME_SIGNALING_BY_NTRK2_TRKB | 8 | -0.37929 | -0.76327 | 0.712575 | 0.848134 | 0.833111 | 2767 | tags=38% | list=18% | signal=31% |
| REACTOME_SYNTHESIS_OF_PIPS_AT_THE_PLASMA_MEMBRANE | 49 | 0.329892 | 0.750565 | 0.712821 | 0.848134 | 0.833111 | 4357 | tags=37% | list=28% | signal=27% |
| REACTOME_SIGNAL_TRANSDUCTION_BY_L1 | 19 | 0.332934 | 0.693714 | 0.71243 | 0.848134 | 0.833111 | 3048 | tags=32% | list=19% | signal=25% |
| REACTOME_GOLGI_ASSOCIATED_VESICLE_BIOGENESIS | 54 | -0.29031 | -0.74864 | 0.714976 | 0.850089 | 0.835032 | 3049 | tags=37% | list=19% | signal=30% |
| WP_MICROGLIA_PATHOGEN_PHAGOCYTOSIS_PATHWAY | 39 | 0.324963 | 0.725975 | 0.716007 | 0.850706 | 0.835638 | 4100 | tags=38% | list=26% | signal=28% |
| REACTOME_POST_NMDA_RECEPTOR_ACTIVATION_EVENTS | 16 | -0.31311 | -0.72437 | 0.716942 | 0.851209 | 0.836131 | 3701 | tags=44% | list=24% | signal=33% |
| REACTOME_G_ALPHA_I_SIGNALLING_EVENTS | 178 | 0.315587 | 0.84109 | 0.718101 | 0.851975 | 0.836884 | 645 | tags=6% | list=4% | signal=5% |
| REACTOME_INTERLEUKIN_1_PROCESSING | 6 | 0.405182 | 0.747999 | 0.721248 | 0.854488 | 0.839353 | 4667 | tags=50% | list=30% | signal=35% |
| BIOCARTA_CPSF_PATHWAY | 6 | 0.404399 | 0.746552 | 0.721248 | 0.854488 | 0.839353 | 3938 | tags=50% | list=25% | signal=37% |
| WP_EGFR1_SIGNALING_PATHWAY | 169 | 0.310684 | 0.824292 | 0.725782 | 0.855126 | 0.839979 | 2426 | tags=16% | list=15% | signal=14% |
| REACTOME_HORMONE_LIGAND_BINDING_RECEPTORS | 3 | 0.493282 | 0.795631 | 0.722986 | 0.855126 | 0.839979 | 7682 | tags=67% | list=49% | signal=34% |
| WP_OXIDATIVE_STRESS_AND_REDOX_PATHWAY | 75 | 0.327 | 0.779073 | 0.728896 | 0.855126 | 0.839979 | 1056 | tags=9% | list=7% | signal=9% |
| BIOCARTA_TSP1_PATHWAY | 7 | -0.38041 | -0.75439 | 0.722555 | 0.855126 | 0.839979 | 3532 | tags=71% | list=22% | signal=55% |
| REACTOME_DIGESTION_AND_ABSORPTION | 7 | -0.37927 | -0.75214 | 0.726547 | 0.855126 | 0.839979 | 4367 | tags=43% | list=28% | signal=31% |
| REACTOME_FGFR2B_LIGAND_BINDING_AND_ACTIVATION | 8 | -0.3729 | -0.75042 | 0.728543 | 0.855126 | 0.839979 | 373 | tags=38% | list=2% | signal=37% |
| BIOCARTA_IFNG_PATHWAY | 5 | 0.415402 | 0.744276 | 0.726733 | 0.855126 | 0.839979 | 809 | tags=20% | list=5% | signal=19% |
| REACTOME_MYD88_MAL_TIRAP_CASCADE_INITIATED_ON_PLASMA | 83 | -0.27871 | -0.74396 | 0.729508 | 0.855126 | 0.839979 | 4045 | tags=39% | list=26% | signal=29% |
| REACTOME_SENSING_OF_DNA_DOUBLE_STRAND_BREAKS | 6 | 0.401176 | 0.740604 | 0.729045 | 0.855126 | 0.839979 | 5755 | tags=67% | list=37% | signal=42% |
| BIOCARTA_SET_PATHWAY | 9 | -0.35891 | -0.7381 | 0.728571 | 0.855126 | 0.839979 | 5918 | tags=89% | list=38% | signal=55% |
| REACTOME_FCGAMMA_RECEPTOR_FCGR_DEPENDENT_PHAGOCYT( | 76 | -0.27662 | -0.73465 | 0.724675 | 0.855126 | 0.839979 | 3003 | tags=33% | list=19% | signal=27% |
| REACTOME_ASSEMBLY_OF_THE_ORC_COMPLEX_AT_THE_ORIGIN_( | 8 | 0.369229 | 0.719691 | 0.728543 | 0.855126 | 0.839979 | 2850 | tags=25% | list=18% | signal=20% |
| WP_HEDGEHOG_SIGNALING_PATHWAY | 20 | 0.335452 | 0.696206 | 0.728814 | 0.855126 | 0.839979 | 5016 | tags=55% | list=32% | signal=37% |
| REACTOME_ACTIVATION_OF_IRF3_IRF7_MEDIATED_BY_TBK1_IKK_E | 17 | 0.340428 | 0.692417 | 0.729367 | 0.855126 | 0.839979 | 971 | tags=24% | list=6% | signal=22% |
| REACTOME_NICOTINAMIDE_SALVAGING | 16 | 0.33744 | 0.687273 | 0.727799 | 0.855126 | 0.839979 | 2706 | tags=25% | list=17% | signal=21% |
| REACTOME_TYPE_I_HEMIDESMOSOME_ASSEMBLY | 7 | -0.37722 | -0.74807 | 0.730539 | 0.855213 | 0.840065 | 445 | tags=29% | list=3% | signal=28% |
| REACTOME_PI5P_REGULATES_TP53_ACETYLATION | 9 | -0.35795 | -0.73613 | 0.730612 | 0.855213 | 0.840065 | 1558 | tags=22% | list=10% | signal=20% |
| REACTOME_METABOLISM_OF_WATER_SOLUBLE_VITAMINS_AND_C | 103 | 0.317832 | 0.795173 | 0.731183 | 0.855229 | 0.840081 | 926 | tags=7% | list=6% | signal=6% |
| REACTOME_PYRIMIDINE_SALVAGE | 10 | -0.35338 | -0.73731 | 0.731656 | 0.855229 | 0.840081 | 4769 | tags=50% | list=30% | signal=35% |
| BIOCARTA_AKT_PATHWAY | 19 | -0.30652 | -0.71088 | 0.732181 | 0.855241 | 0.840093 | 2684 | tags=26% | list=17% | signal=22% |
| REACTOME_REGULATION_OF_TP53_ACTIVITY_THROUGH_PHOSPHC | 86 | -0.27425 | -0.73475 | 0.734072 | 0.856246 | 0.841079 | 4183 | tags=30% | list=27% | signal=22% |
| REACTOME_CGMP_EFFECTS | 10 | -0.35199 | -0.7344 | 0.733753 | 0.856246 | 0.841079 | 1796 | tags=20% | list=11% | signal=18% |
| REACTOME_SIGNALING_BY_INTERLEUKINS | 229 | -0.29125 | -0.80565 | 0.736508 | 0.856588 | 0.841415 | 3554 | tags=31% | list=23% | signal=24% |
| REACTOME_FORMYL_PEPTIDE_RECEPTORS_BIND_FORMYL_PEPTIDE | 5 | -0.40542 | -0.74305 | 0.736419 | 0.856588 | 0.841415 | 4523 | tags=80% | list=29% | signal=57% |
| WP_HYPERTROPHY_MODEL | 16 | -0.30755 | -0.71152 | 0.735537 | 0.856588 | 0.841415 | 1594 | tags=25% | list=10% | signal=22% |
| REACTOME_INTERFERON_GAMMA_SIGNALING | 14 | 0.33578 | 0.684003 | 0.735812 | 0.856588 | 0.841415 | 809 | tags=14% | list=5% | signal=14% |
| REACTOME_PECAM1_INTERACTIONS | 12 | 0.33318 | 0.671442 | 0.736944 | 0.856588 | 0.841415 | 2493 | tags=25% | list=16% | signal=21% |
| BIOCARTA_ASBCELL_PATHWAY | 4 | -0.43928 | -0.76655 | 0.738523 | 0.857636 | 0.842444 | 2272 | tags=75% | list=14% | signal=64% |
| REACTOME_SHC1_EVENTS_IN_ERBB2_SIGNALING | 12 | 0.330908 | 0.666863 | 0.738878 | 0.857636 | 0.842444 | 4445 | tags=50% | list=28% | signal=36% |
| REACTOME_GASTRIN_CREB_SIGNALLING_PATHWAY_VIA_PKC_AND_ | 14 | 0.332551 | 0.677425 | 0.741683 | 0.86029 | 0.845052 | 1291 | tags=14% | list=8% | signal=13% |
| REACTOME_HDACS_DEACETYLATE_HISTONES | 21 | 0.324216 | 0.675689 | 0.743396 | 0.861676 | 0.846413 | 1417 | tags=10% | list=9% | signal=9% |
| REACTOME_MET_ACTIVATES_PTPN11 | 5 | 0.4073 | 0.72976 | 0.744554 | 0.862416 | 0.84714 | 1299 | tags=20% | list=8% | signal=18% |
| REACTOME_TRANSCRIPTIONAL_REGULATION_BY_NPAS4 | 4 | -0.43083 | -0.7518 | 0.746507 | 0.864075 | 0.84877 | 895 | tags=75% | list=6% | signal=71% |
| WP_KENNEDY_PATHWAY | 14 | -0.31466 | -0.71102 | 0.747454 | 0.864569 | 0.849255 | 1548 | tags=21% | list=10% | signal=19% |
| REACTOME_ESTROGEN_BIOSYNTHESIS | 3 | 0.472479 | 0.762077 | 0.750491 | 0.866875 | 0.85152 | 1775 | tags=33% | list=11% | signal=30% |

| | | | | | | | | | | |
|---|---|---|---|---|---|---|---|---|---|---|
| WP_TRANSCRIPTIONAL_ACTIVATION_BY_NFE2L2_IN_RESPONSE_TO | 13 | 0.328764 | 0.665224 | 0.75 | 0.866875 | 0.85152 | 1579 | tags=23% | list=10% | signal=21% |
| REACTOME_POSITIVE_EPIGENETIC_REGULATION_OF_RRNA_EXPRES | 29 | 0.302886 | 0.658502 | 0.751736 | 0.867709 | 0.85234 | 1857 | tags=24% | list=12% | signal=21% |
| REACTOME_DEGRADATION_OF_THE_EXTRACELLULAR_MATRIX | 82 | 0.316705 | 0.765022 | 0.754747 | 0.869975 | 0.854566 | 2195 | tags=24% | list=14% | signal=21% |
| REACTOME_DECTIN_2_FAMILY | 5 | 0.399443 | 0.715681 | 0.754455 | 0.869975 | 0.854566 | 3839 | tags=60% | list=24% | signal=45% |
| WP_AFLATOXIN_B1_METABOLISM | 4 | 0.437262 | 0.750022 | 0.758964 | 0.873877 | 0.858398 | 359 | tags=25% | list=2% | signal=24% |
| REACTOME_SHC1_EVENTS_IN_ERBB4_SIGNALING | 9 | -0.34881 | -0.71732 | 0.759184 | 0.873877 | 0.858398 | 377 | tags=22% | list=2% | signal=22% |
| REACTOME_SIGNALING_BY_NODAL | 6 | 0.390675 | 0.721217 | 0.760234 | 0.87448 | 0.85899 | 4628 | tags=50% | list=29% | signal=35% |
| REACTOME_FATTY_ACIDS_BOUND_TO_GPR40_FFAR1_REGULATE_IN | 6 | -0.38076 | -0.71807 | 0.762781 | 0.876802 | 0.861272 | 3170 | tags=50% | list=20% | signal=40% |
| REACTOME_PI_3K_CASCADE_FGFR3 | 15 | -0.30294 | -0.69561 | 0.764463 | 0.877654 | 0.862108 | 2767 | tags=27% | list=18% | signal=22% |
| BIOCARTA_MITOCHONDRIA_PATHWAY | 19 | -0.29456 | -0.68314 | 0.764579 | 0.877654 | 0.862108 | 1093 | tags=47% | list=7% | signal=44% |
| REACTOME_TP53_REGULATES_TRANSCRIPTION_OF_GENES_INVOLV | 7 | -0.36465 | -0.72315 | 0.766467 | 0.87907 | 0.8635 | 4037 | tags=43% | list=26% | signal=32% |
| REACTOME_FGFR3B_LIGAND_BINDING_AND_ACTIVATION | 6 | -0.37905 | -0.71484 | 0.766871 | 0.87907 | 0.8635 | 1909 | tags=33% | list=12% | signal=29% |
| REACTOME_CELL_JUNCTION_ORGANIZATION | 62 | 0.316087 | 0.733533 | 0.767677 | 0.879387 | 0.863811 | 2279 | tags=31% | list=15% | signal=26% |
| REACTOME_NEUTROPHIL_DEGRANULATION | 413 | 0.291059 | 0.841631 | 0.768276 | 0.879467 | 0.863889 | 2984 | tags=20% | list=19% | signal=17% |
| REACTOME_DEUBIQUITINATION | 228 | -0.27809 | -0.77271 | 0.77044 | 0.880125 | 0.864535 | 3505 | tags=32% | list=22% | signal=25% |
| REACTOME_CHYLOMICRON_CLEARANCE | 3 | 0.455997 | 0.735492 | 0.770138 | 0.880125 | 0.864535 | 5969 | tags=67% | list=38% | signal=41% |
| BIOCARTA_CYTOKINE_PATHWAY | 5 | 0.391666 | 0.701749 | 0.770297 | 0.880125 | 0.864535 | 3571 | tags=60% | list=23% | signal=46% |
| REACTOME_ADAPTIVE_IMMUNE_SYSTEM | 614 | -0.26798 | -0.81821 | 0.772358 | 0.880245 | 0.864654 | 2562 | tags=24% | list=16% | signal=21% |
| REACTOME_TOLL_LIKE_RECEPTOR_4_TLR4_CASCADE | 114 | 0.301229 | 0.765365 | 0.780822 | 0.880245 | 0.864654 | 971 | tags=8% | list=6% | signal=7% |
| REACTOME_ATORVASTATIN_ADME | 4 | 0.429092 | 0.736008 | 0.7749 | 0.880245 | 0.864654 | 4291 | tags=50% | list=27% | signal=36% |
| REACTOME_TIGHT_JUNCTION_INTERACTIONS | 6 | 0.384782 | 0.710339 | 0.77193 | 0.880245 | 0.864654 | 748 | tags=17% | list=5% | signal=16% |
| REACTOME_ERYTHROCYTES_TAKE_UP_CARBON_DIOXIDE_AND_REL | 9 | -0.34393 | -0.70729 | 0.77551 | 0.880245 | 0.864654 | 3918 | tags=33% | list=25% | signal=25% |
| REACTOME_RNA_POLYMERASE_II_TRANSCRIBES_SNRNA_GENES | 71 | -0.26836 | -0.69954 | 0.780928 | 0.880245 | 0.864654 | 1838 | tags=15% | list=12% | signal=14% |
| REACTOME_FGFR2_LIGAND_BINDING_AND_ACTIVATION | 14 | -0.30574 | -0.69088 | 0.771894 | 0.880245 | 0.864654 | 2772 | tags=29% | list=18% | signal=24% |
| REACTOME_PI3K_EVENTS_IN_ERBB4_SIGNALING | 6 | 0.373369 | 0.689269 | 0.773879 | 0.880245 | 0.864654 | 6209 | tags=67% | list=40% | signal=40% |
| REACTOME_VITAMIN_B5_PANTOTHENATE_METABOLISM | 13 | -0.31058 | -0.6878 | 0.77572 | 0.880245 | 0.864654 | 335 | tags=31% | list=2% | signal=30% |
| BIOCARTA_CREB_PATHWAY | 20 | -0.2917 | -0.68514 | 0.77707 | 0.880245 | 0.864654 | 4077 | tags=35% | list=26% | signal=26% |
| REACTOME_INTERFERON_SIGNALING | 52 | 0.296703 | 0.684996 | 0.781145 | 0.880245 | 0.864654 | 1029 | tags=13% | list=7% | signal=13% |
| REACTOME_SIGNALING_BY_ROBO_RECEPTORS | 20 | -0.29089 | -0.68323 | 0.779193 | 0.880245 | 0.864654 | 676 | tags=30% | list=4% | signal=29% |
| BIOCARTA_GPCR_PATHWAY | 28 | -0.27746 | -0.68273 | 0.777518 | 0.880245 | 0.864654 | 3701 | tags=39% | list=24% | signal=30% |
| REACTOME_ACYL_CHAIN_REMODELING_OF_DAG_AND_TAG | 6 | 0.369062 | 0.681318 | 0.779727 | 0.880245 | 0.864654 | 1006 | tags=17% | list=6% | signal=16% |
| WP_ROBO4_AND_VEGF_SIGNALING_PATHWAYS_CROSSTALK | 6 | 0.368399 | 0.680095 | 0.779727 | 0.880245 | 0.864654 | 2348 | tags=33% | list=15% | signal=28% |
| WP_SPLICING_FACTOR_NOVA_REGULATED_SYNAPTIC_PROTEINS | 41 | 0.299536 | 0.668737 | 0.779896 | 0.880245 | 0.864654 | 4516 | tags=51% | list=29% | signal=37% |
| REACTOME_TGF_BETA_RECEPTOR_SIGNALING_ACTIVATES_SMADS | 35 | 0.302801 | 0.667432 | 0.773684 | 0.880245 | 0.864654 | 3375 | tags=37% | list=21% | signal=29% |
| BIOCARTA_ERK5_PATHWAY | 14 | 0.319118 | 0.650061 | 0.774951 | 0.880245 | 0.864654 | 1518 | tags=21% | list=10% | signal=19% |
| WP_PROSTAGLANDIN_SYNTHESIS_AND_REGULATION | 25 | 0.291446 | 0.626672 | 0.777385 | 0.880245 | 0.864654 | 2705 | tags=28% | list=17% | signal=23% |
| REACTOME_FCERI_MEDIATED_CA_2_MOBILIZATION | 25 | 0.290805 | 0.625295 | 0.779152 | 0.880245 | 0.864654 | 1596 | tags=16% | list=10% | signal=14% |
| REACTOME_SIGNALING_BY_TGF_BETA_RECEPTOR_COMPLEX | 71 | -0.26602 | -0.69344 | 0.78866 | 0.886909 | 0.871199 | 2009 | tags=17% | list=13% | signal=15% |
| REACTOME_NUCLEOTIDE_BIOSYNTHESIS | 12 | -0.31815 | -0.68644 | 0.787629 | 0.886909 | 0.871199 | 2302 | tags=33% | list=15% | signal=28% |
| REACTOME_LAGGING_STRAND_SYNTHESIS | 19 | -0.2864 | -0.66421 | 0.788337 | 0.886909 | 0.871199 | 5424 | tags=53% | list=35% | signal=34% |
| REACTOME_TRANSPORT_AND_SYNTHESIS_OF_PAPS | 6 | -0.37176 | -0.7011 | 0.791411 | 0.889346 | 0.873593 | 3731 | tags=50% | list=24% | signal=38% |
| REACTOME_REVERSIBLE_HYDRATION_OF_CARBON_DIOXIDE | 10 | -0.33039 | -0.68933 | 0.792453 | 0.889346 | 0.873593 | 2260 | tags=30% | list=14% | signal=26% |
| BIOCARTA_CASPASE_PATHWAY | 20 | -0.28227 | -0.66299 | 0.791932 | 0.889346 | 0.873593 | 5431 | tags=50% | list=35% | signal=33% |
| BIOCARTA_TH1TH2_PATHWAY | 9 | 0.335822 | 0.65402 | 0.792969 | 0.889346 | 0.873593 | 5260 | tags=56% | list=33% | signal=37% |
| REACTOME_HSF1_DEPENDENT_TRANSACTIVATION | 22 | 0.297218 | 0.630879 | 0.794118 | 0.890033 | 0.874268 | 2417 | tags=27% | list=15% | signal=23% |
| BIOCARTA_NKCELLS_PATHWAY | 15 | -0.28943 | -0.6646 | 0.795455 | 0.890931 | 0.87515 | 3558 | tags=40% | list=23% | signal=31% |
| REACTOME_TNFS_BIND_THEIR_PHYSIOLOGICAL_RECEPTORS | 15 | 0.312421 | 0.637957 | 0.797297 | 0.892393 | 0.876586 | 7292 | tags=67% | list=46% | signal=36% |
| REACTOME_SUMO_IS_PROTEOLYTICALLY_PROCESSED | 6 | -0.36719 | -0.69248 | 0.799591 | 0.894357 | 0.878516 | 2951 | tags=33% | list=19% | signal=27% |
| REACTOME_TNF_RECEPTOR_SUPERFAMILY_TNFSF_MEMBERS_MED | 14 | 0.304348 | 0.619975 | 0.800391 | 0.89465 | 0.878803 | 600 | tags=7% | list=4% | signal=7% |
| REACTOME_RHOV_GTPASE_CYCLE | 36 | 0.287078 | 0.633863 | 0.801739 | 0.895554 | 0.879691 | 3679 | tags=33% | list=23% | signal=26% |
| REACTOME_BBSOME_MEDIATED_CARGO_TARGETING_TO_CILIUM | 13 | 0.30429 | 0.615703 | 0.802326 | 0.895607 | 0.879743 | 5843 | tags=62% | list=37% | signal=39% |
| BIOCARTA_BBCELL_PATHWAY | 3 | -0.4434 | -0.71461 | 0.803245 | 0.896031 | 0.88016 | 2272 | tags=67% | list=14% | signal=57% |
| REACTOME_OTHER_INTERLEUKIN_SIGNALING | 15 | -0.28716 | -0.65939 | 0.805785 | 0.898261 | 0.88235 | 2272 | tags=27% | list=14% | signal=23% |
| REACTOME_SIGNALING_BY_FGFR | 67 | 0.297439 | 0.699253 | 0.807947 | 0.898744 | 0.882825 | 1600 | tags=13% | list=10% | signal=12% |
| REACTOME_FGFR1B_LIGAND_BINDING_AND_ACTIVATION | 6 | -0.36218 | -0.68304 | 0.809816 | 0.898744 | 0.882825 | 373 | tags=33% | list=2% | signal=33% |
| REACTOME_ANCHORING_OF_THE_BASAL_BODY_TO_THE_PLASMA_ | 95 | -0.25423 | -0.67889 | 0.808333 | 0.898744 | 0.882825 | 3096 | tags=27% | list=20% | signal=22% |
| REACTOME_TELOMERE_C_STRAND_SYNTHESIS_INITIATION | 12 | -0.31067 | -0.67031 | 0.810309 | 0.898744 | 0.882825 | 4333 | tags=50% | list=28% | signal=36% |
| REACTOME_CYCLIN_A_B1_B2_ASSOCIATED_EVENTS_DURING_G2_M | 25 | -0.27248 | -0.65296 | 0.809633 | 0.898744 | 0.882825 | 1771 | tags=20% | list=11% | signal=18% |
| BIOCARTA_PLC_PATHWAY | 9 | 0.328908 | 0.640556 | 0.810547 | 0.898744 | 0.882825 | 4496 | tags=33% | list=29% | signal=24% |
| WP_EICOSANOID_METABOLISM_VIA_LIPOXYGENASES_LOX | 21 | 0.297368 | 0.619737 | 0.807547 | 0.898744 | 0.882825 | 4600 | tags=38% | list=29% | signal=27% |
| BIOCARTA_MET_PATHWAY | 33 | 0.276818 | 0.61046 | 0.808696 | 0.898744 | 0.882825 | 1955 | tags=21% | list=12% | signal=19% |
| REACTOME_SEMAPHORIN_INTERACTIONS | 51 | 0.278932 | 0.641448 | 0.81113 | 0.89879 | 0.88287 | 2528 | tags=25% | list=16% | signal=21% |
| WP_MISMATCH_REPAIR | 9 | -0.33175 | -0.68224 | 0.812245 | 0.898827 | 0.882906 | 4291 | tags=56% | list=27% | signal=40% |
| BIOCARTA_DSP_PATHWAY | 9 | -0.33127 | -0.68125 | 0.812245 | 0.898827 | 0.882906 | 2834 | tags=67% | list=18% | signal=55% |
| REACTOME_RHOD_GTPASE_CYCLE | 51 | 0.278218 | 0.639804 | 0.814503 | 0.900126 | 0.884182 | 2014 | tags=24% | list=13% | signal=21% |
| WP_OXIDATION_BY_CYTOCHROME_P450 | 22 | 0.28768 | 0.610632 | 0.814338 | 0.900126 | 0.884182 | 2892 | tags=27% | list=18% | signal=22% |
| BIOCARTA_DREAM_PATHWAY | 12 | -0.30833 | -0.66524 | 0.818557 | 0.904004 | 0.887992 | 3701 | tags=58% | list=24% | signal=45% |
| REACTOME_MISMATCH_REPAIR_MMR_DIRECTED_BY_MSH2_MSH3 | 9 | -0.32787 | -0.67427 | 0.822449 | 0.904694 | 0.888669 | 5424 | tags=56% | list=35% | signal=36% |
| REACTOME_TOLL_LIKE_RECEPTOR_9_TLR9_CASCADE | 93 | -0.25006 | -0.66903 | 0.821727 | 0.904694 | 0.888669 | 4045 | tags=38% | list=26% | signal=28% |
| REACTOME_GLUTATHIONE_SYNTHESIS_AND_RECYCLING | 11 | -0.3083 | -0.65961 | 0.820248 | 0.904694 | 0.888669 | 3137 | tags=45% | list=20% | signal=36% |
| REACTOME_DNA_REPLICATION_INITIATION | 8 | 0.324487 | 0.632481 | 0.822355 | 0.904694 | 0.888669 | 6708 | tags=62% | list=43% | signal=36% |
| REACTOME_SUMOYLATION_OF_IMMUNE_RESPONSE_PROTEINS | 11 | 0.316639 | 0.626856 | 0.822394 | 0.904694 | 0.888669 | 3365 | tags=36% | list=21% | signal=29% |
| BIOCARTA_SHH_PATHWAY | 15 | 0.297799 | 0.608099 | 0.820463 | 0.904694 | 0.888669 | 2075 | tags=20% | list=13% | signal=17% |
| REACTOME_ARACHIDONIC_ACID_METABOLISM | 36 | 0.276613 | 0.610757 | 0.824348 | 0.906182 | 0.890131 | 180 | tags=3% | list=1% | signal=3% |
| REACTOME_RNA_POLYMERASE_I_TRANSCRIPTION | 44 | -0.25099 | -0.63756 | 0.825666 | 0.907031 | 0.890965 | 3708 | tags=30% | list=24% | signal=23% |
| REACTOME_PREGNENOLONE_BIOSYNTHESIS | 11 | -0.30546 | -0.65352 | 0.828512 | 0.909344 | 0.893237 | 451 | tags=27% | list=3% | signal=27% |
| REACTOME_PHOSPHOLIPASE_C_MEDIATED_CASCADE_FGFR1 | 12 | -0.2999 | -0.64706 | 0.828866 | 0.909344 | 0.893237 | 2772 | tags=25% | list=18% | signal=21% |
| REACTOME_INTRACELLULAR_SIGNALING_BY_SECOND_MESSENGERS | 241 | 0.2847 | 0.781995 | 0.831625 | 0.911769 | 0.895619 | 798 | tags=5% | list=5% | signal=5% |
| BIOCARTA_TERC_PATHWAY | 6 | 0.341226 | 0.62993 | 0.832359 | 0.911971 | 0.895817 | 3439 | tags=33% | list=22% | signal=26% |
| BIOCARTA_P27_PATHWAY | 11 | 0.309521 | 0.612763 | 0.835907 | 0.915255 | 0.899043 | 2960 | tags=45% | list=19% | signal=37% |
| REACTOME_PREDNISONE_ADME | 4 | 0.395219 | 0.677907 | 0.838645 | 0.917648 | 0.901394 | 4291 | tags=50% | list=27% | signal=36% |
| REACTOME_ERBB2_REGULATES_CELL_MOTILITY | 11 | 0.306123 | 0.606037 | 0.839768 | 0.917668 | 0.901413 | 6209 | tags=64% | list=40% | signal=39% |
| BIOCARTA_CSK_PATHWAY | 12 | 0.298255 | 0.601059 | 0.839458 | 0.917668 | 0.901413 | 1263 | tags=17% | list=8% | signal=15% |
| BIOCARTA_AKAPCENTROSOME_PATHWAY | 12 | 0.297906 | 0.600355 | 0.841393 | 0.91819 | 0.901927 | 3083 | tags=33% | list=20% | signal=27% |
| REACTOME_FGFR3_LIGAND_BINDING_AND_ACTIVATION | 10 | 0.304543 | 0.59711 | 0.841905 | 0.91819 | 0.901927 | 357 | tags=10% | list=2% | signal=10% |
| REACTOME_OVARIAN_TUMOR_DOMAIN_PROTEASES | 31 | 0.26813 | 0.589363 | 0.841463 | 0.91819 | 0.901927 | 798 | tags=13% | list=5% | signal=12% |
| REACTOME_GAMMA_CARBOXYLATION_HYPUSINE_FORMATION_AN | 31 | -0.25141 | -0.63236 | 0.843458 | 0.918678 | 0.902405 | 1423 | tags=32% | list=9% | signal=29% |
| BIOCARTA_EIF4_PATHWAY | 21 | 0.283518 | 0.590871 | 0.843396 | 0.918678 | 0.902405 | 3881 | tags=29% | list=25% | signal=22% |
| REACTOME_TOLL_LIKE_RECEPTOR_CASCADES | 138 | 0.280456 | 0.724705 | 0.848624 | 0.923699 | 0.907337 | 2359 | tags=15% | list=15% | signal=13% |
| REACTOME_FANCONI_ANEMIA_PATHWAY | 35 | -0.24701 | -0.61887 | 0.849537 | 0.924087 | 0.907719 | 676 | tags=34% | list=4% | signal=33% |

| | | | | | | | | | | |
|---|---|---|---|---|---|---|---|---|---|---|
| REACTOME_ABC_FAMILY_PROTEINS_MEDIATED_TRANSPORT | 92 | 0.278718 | 0.684103 | 0.852025 | 0.924351 | 0.907978 | 348 | tags=3% | list=2% | signal=3% |
| REACTOME_SULFIDE_OXIDATION_TO_SULFATE | 5 | -0.35369 | -0.64825 | 0.853119 | 0.924351 | 0.907978 | 10153 | tags=100% | list=65% | signal=35% |
| REACTOME_DEADENYLATION_DEPENDENT_MRNA_DECAY | 51 | -0.24647 | -0.63015 | 0.850856 | 0.924351 | 0.907978 | 3734 | tags=33% | list=24% | signal=25% |
| REACTOME_HDR_THROUGH_HOMOLOGOUS_RECOMBINATION_HRI | 52 | -0.24512 | -0.62448 | 0.852941 | 0.924351 | 0.907978 | 4683 | tags=35% | list=30% | signal=24% |
| REACTOME_BIOSYNTHESIS_OF_EPA_DERIVED_SPMS | 5 | 0.345568 | 0.619154 | 0.851485 | 0.924351 | 0.907978 | 2233 | tags=40% | list=14% | signal=34% |
| BIOCARTA_RB_PATHWAY | 12 | 0.291166 | 0.586773 | 0.852998 | 0.924351 | 0.907978 | 349 | tags=8% | list=2% | signal=8% |
| BIOCARTA_ECM_PATHWAY | 21 | -0.25953 | -0.61293 | 0.853814 | 0.924501 | 0.908125 | 3634 | tags=43% | list=23% | signal=33% |
| REACTOME_SYNTHESIS_OF_IP2_IP_AND_INS_IN_THE_CYTOSOL | 11 | -0.2922 | -0.62515 | 0.855372 | 0.925585 | 0.90919 | 919 | tags=27% | list=6% | signal=26% |
| BIOCARTA_TALL1_PATHWAY | 13 | 0.286246 | 0.579193 | 0.856589 | 0.926299 | 0.909891 | 600 | tags=8% | list=4% | signal=7% |
| REACTOME_CYTOKINE_SIGNALING_IN_IMMUNE_SYSTEM | 328 | -0.25657 | -0.75069 | 0.857627 | 0.926818 | 0.910401 | 3585 | tags=27% | list=23% | signal=21% |
| REACTOME_ORGANIC_CATION_TRANSPORT | 7 | -0.32858 | -0.65161 | 0.860279 | 0.929079 | 0.912623 | 10548 | tags=100% | list=67% | signal=33% |
| REACTOME_ACETYLCHOLINE_REGULATES_INSULIN_SECRETION | 7 | -0.32674 | -0.64797 | 0.862275 | 0.930241 | 0.913764 | 3170 | tags=43% | list=20% | signal=34% |
| REACTOME_VLDL_CLEARANCE | 3 | 0.400342 | 0.645724 | 0.862475 | 0.930241 | 0.913764 | 5203 | tags=67% | list=33% | signal=45% |
| REACTOME_NUCLEOTIDE_BINDING_DOMAIN_LEUCINE_RICH_REPEA | 43 | -0.24269 | -0.61285 | 0.866828 | 0.934329 | 0.91778 | 3003 | tags=23% | list=19% | signal=19% |
| REACTOME_MET_ACTIVATES_RAP1_AND_RAC1 | 11 | 0.287456 | 0.56908 | 0.868726 | 0.935161 | 0.918597 | 1665 | tags=18% | list=11% | signal=16% |
| REACTOME_ACTIVATION_OF_SMO | 11 | 0.284597 | 0.563421 | 0.868726 | 0.935161 | 0.918597 | 4602 | tags=45% | list=29% | signal=32% |
| REACTOME_TIE2_SIGNALING | 16 | -0.26499 | -0.61305 | 0.869835 | 0.935456 | 0.918887 | 2744 | tags=19% | list=17% | signal=15% |
| REACTOME_INOSITOL_PHOSPHATE_METABOLISM | 46 | 0.266829 | 0.600811 | 0.87069 | 0.935456 | 0.918887 | 2057 | tags=22% | list=13% | signal=19% |
| REACTOME_GROWTH_HORMONE_RECEPTOR_SIGNALING | 10 | 0.293786 | 0.576019 | 0.870476 | 0.935456 | 0.918887 | 2365 | tags=20% | list=15% | signal=17% |
| REACTOME_TRANSPORT_TO_THE_GOLGI_AND_SUBSEQUENT_MOD | 167 | -0.24596 | -0.66734 | 0.873112 | 0.93723 | 0.920629 | 1852 | tags=22% | list=12% | signal=20% |
| BIOCARTA_EIF2_PATHWAY | 9 | -0.31 | -0.63751 | 0.873469 | 0.93723 | 0.920629 | 4041 | tags=56% | list=26% | signal=41% |
| REACTOME_NECTIN_NECL_TRANS_HETERODIMERIZATION | 7 | 0.314637 | 0.596437 | 0.874251 | 0.937464 | 0.920859 | 6928 | tags=86% | list=44% | signal=48% |
| REACTOME_MRNA_SPLICING | 199 | -0.24366 | -0.67109 | 0.876161 | 0.938905 | 0.922275 | 2967 | tags=29% | list=19% | signal=24% |
| REACTOME_ACTIVATED_NTRK2_SIGNALS_THROUGH_FRS2_AND_FR | 5 | 0.339096 | 0.607559 | 0.877228 | 0.939442 | 0.922802 | 5164 | tags=60% | list=33% | signal=40% |
| BIOCARTA_GATA3_PATHWAY | 10 | -0.28874 | -0.60243 | 0.878407 | 0.939616 | 0.922973 | 3701 | tags=50% | list=24% | signal=38% |
| REACTOME_NICOTINATE_METABOLISM | 26 | 0.250309 | 0.540077 | 0.878521 | 0.939616 | 0.922973 | 2941 | tags=19% | list=19% | signal=16% |
| REACTOME_ACTIVATION_OF_ATR_IN_RESPONSE_TO_REPLICATION_ | 34 | 0.25371 | 0.559107 | 0.879371 | 0.939919 | 0.923271 | 3483 | tags=24% | list=22% | signal=18% |
| WP_INTEGRINMEDIATED_CELL_ADHESION | 96 | 0.26504 | 0.655697 | 0.880435 | 0.939983 | 0.923333 | 2494 | tags=23% | list=16% | signal=19% |
| REACTOME_FCERI_MEDIATED_MAPK_ACTIVATION | 28 | -0.24369 | -0.59962 | 0.880562 | 0.939983 | 0.923333 | 4208 | tags=39% | list=27% | signal=29% |
| REACTOME_INTERLEUKIN_3_INTERLEUKIN_5_AND_GM_CSF_SIGNAI | 35 | -0.23357 | -0.5852 | 0.881944 | 0.940854 | 0.924189 | 2555 | tags=23% | list=16% | signal=19% |
| WP_METHYLATION | 7 | -0.30826 | -0.61131 | 0.884232 | 0.942579 | 0.925883 | 848 | tags=29% | list=5% | signal=27% |
| BIOCARTA_D4GDI_PATHWAY | 10 | -0.2846 | -0.5938 | 0.884696 | 0.942579 | 0.925883 | 4822 | tags=60% | list=31% | signal=42% |
| REACTOME_SNRNP_ASSEMBLY | 47 | 0.257952 | 0.583921 | 0.886207 | 0.943583 | 0.92687 | 4495 | tags=36% | list=29% | signal=26% |
| BIOCARTA_PARKIN_PATHWAY | 8 | -0.30389 | -0.61153 | 0.888224 | 0.945124 | 0.928384 | 3538 | tags=38% | list=23% | signal=29% |
| REACTOME_REGULATION_OF_TP53_ACTIVITY | 148 | -0.24642 | -0.66437 | 0.89049 | 0.945258 | 0.928514 | 1769 | tags=12% | list=11% | signal=11% |
| REACTOME_FREE_FATTY_ACIDS_REGULATE_INSULIN_SECRETION | 9 | 0.29867 | 0.581665 | 0.890625 | 0.945258 | 0.928514 | 1855 | tags=22% | list=12% | signal=20% |
| REACTOME_MYOGENESIS | 20 | -0.24721 | -0.58064 | 0.889597 | 0.945258 | 0.928514 | 2168 | tags=20% | list=14% | signal=17% |
| REACTOME_ACTIVATION_OF_THE_PRE_REPLICATIVE_COMPLEX | 30 | 0.250497 | 0.548671 | 0.890625 | 0.945258 | 0.928514 | 5649 | tags=40% | list=36% | signal=26% |
| BIOCARTA_NGF_PATHWAY | 19 | -0.24834 | -0.57595 | 0.892009 | 0.945989 | 0.929232 | 3558 | tags=32% | list=23% | signal=24% |
| BIOCARTA_IL6_PATHWAY | 21 | 0.259921 | 0.541694 | 0.892453 | 0.945989 | 0.929232 | 1955 | tags=19% | list=12% | signal=17% |
| BIOCARTA_PEPI_PATHWAY | 3 | 0.381214 | 0.614872 | 0.89391 | 0.946929 | 0.930156 | 2415 | tags=33% | list=15% | signal=28% |
| BIOCARTA_IL22BP_PATHWAY | 9 | 0.290842 | 0.566421 | 0.896484 | 0.949051 | 0.93224 | 751 | tags=11% | list=5% | signal=11% |
| REACTOME_ASPARAGINE_N_LINKED_GLYCOSYLATION | 248 | -0.23888 | -0.673 | 0.902516 | 0.950149 | 0.93332 | 2535 | tags=25% | list=16% | signal=21% |
| REACTOME_PHOSPHORYLATION_OF_EMI1 | 6 | -0.32454 | -0.61206 | 0.903885 | 0.950149 | 0.93332 | 10611 | tags=100% | list=68% | signal=32% |
| REACTOME_COENZYME_A_BIOSYNTHESIS | 8 | -0.29846 | -0.60062 | 0.904192 | 0.950149 | 0.93332 | 11021 | tags=100% | list=70% | signal=30% |
| BIOCARTA_CFTR_PATHWAY | 9 | -0.28953 | -0.59541 | 0.9 | 0.950149 | 0.93332 | 3701 | tags=44% | list=24% | signal=34% |
| REACTOME_P75NTR_REGULATES_AXONOGENESIS | 9 | -0.28799 | -0.59225 | 0.9 | 0.950149 | 0.93332 | 494 | tags=22% | list=3% | signal=22% |
| WP_HEME_BIOSYNTHESIS | 9 | -0.28572 | -0.58759 | 0.904082 | 0.950149 | 0.93332 | 3762 | tags=56% | list=24% | signal=42% |
| REACTOME_ZBP1_DAI_MEDIATED_INDUCTION_OF_TYPE_I_IFNS | 13 | -0.26504 | -0.58695 | 0.899177 | 0.950149 | 0.93332 | 2677 | tags=31% | list=17% | signal=26% |
| WP_SEROTONIN_AND_ANXIETY | 16 | -0.25304 | -0.5854 | 0.904959 | 0.950149 | 0.93332 | 3829 | tags=50% | list=24% | signal=38% |
| REACTOME_MISMATCH_REPAIR | 14 | -0.25837 | -0.58382 | 0.90224 | 0.950149 | 0.93332 | 5424 | tags=57% | list=35% | signal=37% |
| REACTOME_RIBAVIRIN_ADME | 10 | -0.27653 | -0.57695 | 0.901468 | 0.950149 | 0.93332 | 5042 | tags=60% | list=32% | signal=41% |
| REACTOME_GDP_FUCOSE_BIOSYNTHESIS | 6 | 0.31064 | 0.573466 | 0.904483 | 0.950149 | 0.93332 | 1926 | tags=33% | list=12% | signal=29% |
| BIOCARTA_NOS1_PATHWAY | 19 | 0.249479 | 0.519822 | 0.899814 | 0.950149 | 0.93332 | 2903 | tags=26% | list=18% | signal=21% |
| BIOCARTA_MTOR_PATHWAY | 19 | 0.244225 | 0.508876 | 0.903525 | 0.950149 | 0.93332 | 3609 | tags=26% | list=23% | signal=20% |
| REACTOME_REGULATION_OF_PTEN_GENE_TRANSCRIPTION | 41 | 0.248346 | 0.554452 | 0.906412 | 0.951075 | 0.934228 | 746 | tags=7% | list=5% | signal=7% |
| REACTOME_FGFRL1_MODULATION_OF_FGFR1_SIGNALING | 10 | -0.27297 | -0.56954 | 0.907757 | 0.951537 | 0.934682 | 2772 | tags=20% | list=18% | signal=16% |
| REACTOME_INTERLEUKIN_20_FAMILY_SIGNALING | 10 | 0.275854 | 0.540861 | 0.908571 | 0.951537 | 0.934682 | 4809 | tags=40% | list=31% | signal=28% |
| BIOCARTA_TPO_PATHWAY | 25 | 0.242864 | 0.522212 | 0.908127 | 0.951537 | 0.934682 | 1955 | tags=16% | list=12% | signal=14% |
| BIOCARTA_EFP_PATHWAY | 13 | -0.25874 | -0.57298 | 0.909465 | 0.951872 | 0.935012 | 2337 | tags=62% | list=15% | signal=52% |
| REACTOME_TRIGLYCERIDE_METABOLISM | 12 | -0.27071 | -0.58407 | 0.91134 | 0.953234 | 0.93635 | 2722 | tags=25% | list=17% | signal=21% |
| WP_HISTONE_MODIFICATIONS | 5 | 0.322739 | 0.578251 | 0.912871 | 0.954134 | 0.937233 | 10638 | tags=100% | list=68% | signal=32% |
| REACTOME_RET_SIGNALING | 36 | -0.22503 | -0.56713 | 0.913349 | 0.954134 | 0.937233 | 2767 | tags=25% | list=18% | signal=21% |
| REACTOME_REDUCTION_OF_CYTOSOLIC_CA_LEVELS | 14 | -0.25104 | -0.56727 | 0.91446 | 0.954694 | 0.937784 | 3520 | tags=36% | list=22% | signal=28% |
| BIOCARTA_CDC42RAC_PATHWAY | 7 | -0.29108 | -0.57725 | 0.916168 | 0.955276 | 0.938355 | 1727 | tags=43% | list=11% | signal=38% |
| BIOCARTA_REELIN_PATHWAY | 7 | -0.29064 | -0.57638 | 0.916168 | 0.955276 | 0.938355 | 3628 | tags=43% | list=23% | signal=33% |
| BIOCARTA_ACETAMINOPHEN_PATHWAY | 3 | 0.364741 | 0.588303 | 0.921415 | 0.960144 | 0.943137 | 4275 | tags=67% | list=27% | signal=49% |
| BIOCARTA_CTLA4_PATHWAY | 8 | -0.28996 | -0.5835 | 0.922156 | 0.960206 | 0.943198 | 1876 | tags=62% | list=12% | signal=55% |
| REACTOME_HDR_THROUGH_MMEJ_ALT_NHEJ | 12 | 0.261245 | 0.526475 | 0.922631 | 0.960206 | 0.943198 | 3762 | tags=25% | list=24% | signal=19% |
| REACTOME_MYD88_INDEPENDENT_TLR4_CASCADE | 99 | -0.21973 | -0.59284 | 0.925414 | 0.962423 | 0.945376 | 4045 | tags=36% | list=26% | signal=27% |
| REACTOME_GAB1_SIGNALOSOME | 13 | -0.24918 | -0.55182 | 0.925926 | 0.962423 | 0.945376 | 3388 | tags=38% | list=22% | signal=30% |
| BIOCARTA_ACH_PATHWAY | 12 | 0.260611 | 0.525196 | 0.926499 | 0.962423 | 0.945376 | 3601 | tags=25% | list=23% | signal=19% |
| BIOCARTA_CCR5_PATHWAY | 14 | -0.2438 | -0.5509 | 0.928717 | 0.964124 | 0.947047 | 4077 | tags=43% | list=26% | signal=32% |
| BIOCARTA_IL2RB_PATHWAY | 33 | 0.229424 | 0.505942 | 0.930435 | 0.965304 | 0.948206 | 4553 | tags=30% | list=29% | signal=22% |
| REACTOME_SPERM_MOTILITY_AND_TAXES | 5 | -0.3051 | -0.55917 | 0.93159 | 0.965899 | 0.94879 | 10916 | tags=100% | list=70% | signal=31% |
| REACTOME_OAS_ANTIVIRAL_RESPONSE | 4 | 0.334692 | 0.574087 | 0.932271 | 0.966002 | 0.948892 | 10450 | tags=100% | list=67% | signal=33% |
| REACTOME_SMAC_XIAP_REGULATED_APOPTOTIC_RESPONSE | 5 | -0.29808 | -0.54631 | 0.933602 | 0.966176 | 0.949062 | 2218 | tags=40% | list=14% | signal=34% |
| BIOCARTA_NFAT_PATHWAY | 42 | 0.238772 | 0.537252 | 0.933447 | 0.966176 | 0.949062 | 2752 | tags=24% | list=18% | signal=20% |
| BIOCARTA_PKC_PATHWAY | 7 | 0.271725 | 0.515092 | 0.938124 | 0.969751 | 0.952574 | 1576 | tags=14% | list=10% | signal=13% |
| REACTOME_INTERLEUKIN_15_SIGNALING | 11 | 0.254811 | 0.504453 | 0.938224 | 0.969751 | 0.952574 | 4955 | tags=36% | list=32% | signal=25% |
| REACTOME_RESOLUTION_OF_AP_SITES_VIA_THE_MULTIPLE_NUCLE | 24 | -0.21958 | -0.52601 | 0.939597 | 0.969963 | 0.952783 | 5424 | tags=46% | list=35% | signal=30% |
| REACTOME_NOD1_2_SIGNALING_PATHWAY | 31 | 0.23107 | 0.507903 | 0.939024 | 0.969963 | 0.952783 | 679 | tags=10% | list=4% | signal=9% |
| WP_IL7_SIGNALING_PATHWAY | 42 | 0.233216 | 0.52475 | 0.940273 | 0.970058 | 0.952876 | 2572 | tags=21% | list=16% | signal=18% |
| REACTOME_BIOSYNTHESIS_OF_E_SERIES_18_S_RESOLVINS | 4 | 0.313975 | 0.538552 | 0.944223 | 0.971672 | 0.95446 | 921 | tags=25% | list=6% | signal=24% |
| REACTOME_SYNTHESIS_OF_5_EICOSATETRAENOIC_ACIDS | 5 | -0.28854 | -0.52882 | 0.943662 | 0.971672 | 0.95446 | 11176 | tags=60% | list=71% | signal=17% |
| BIOCARTA_ION_PATHWAY | 5 | -0.28631 | -0.52474 | 0.943662 | 0.971672 | 0.95446 | 11211 | tags=40% | list=71% | signal=11% |
| BIOCARTA_SRCRPTP_PATHWAY | 10 | 0.259637 | 0.509064 | 0.944762 | 0.971672 | 0.95446 | 11630 | tags=100% | list=74% | signal=26% |
| REACTOME_GPVI_MEDIATED_ACTIVATION_CASCADE | 31 | 0.22743 | 0.499903 | 0.944251 | 0.971672 | 0.95446 | 3911 | tags=29% | list=25% | signal=22% |
| REACTOME_METABOLISM_OF_STEROID_HORMONES | 19 | -0.22434 | -0.52029 | 0.946004 | 0.972347 | 0.955124 | 3829 | tags=32% | list=24% | signal=24% |
| WP_EBV_LMP1_SIGNALING | 18 | -0.2242 | -0.51777 | 0.94926 | 0.97509 | 0.957819 | 4045 | tags=56% | list=26% | signal=41% |

| | | | | | | | | | | |
|---|---|---|---|---|---|---|---|---|---|---|
| BIOCARTA_RANMS_PATHWAY | 9 | -0.25927 | -0.53318 | 0.95102 | 0.975692 | 0.958409 | 3067 | tags=44% | list=20% | signal=36% |
| REACTOME_ARACHIDONATE_PRODUCTION_FROM_DAG | 5 | 0.294222 | 0.527157 | 0.950495 | 0.975692 | 0.958409 | 1558 | tags=20% | list=10% | signal=18% |
| REACTOME_PROCESSING_OF_CAPPED_INTRON_CONTAINING_PRE_ | 264 | 0.235822 | 0.654126 | 0.952035 | 0.975882 | 0.958596 | 2070 | tags=13% | list=13% | signal=11% |
| REACTOME_SYNTHESIS_OF_PROSTAGLANDINS_PG_AND_THROMBO | 10 | 0.2518 | 0.493698 | 0.952381 | 0.975882 | 0.958596 | 4443 | tags=50% | list=28% | signal=36% |
| REACTOME_INTERLEUKIN_7_SIGNALING | 13 | -0.23269 | -0.5153 | 0.954733 | 0.977518 | 0.960204 | 3761 | tags=38% | list=24% | signal=29% |
| BIOCARTA_ACTINY_PATHWAY | 10 | -0.24369 | -0.50845 | 0.955975 | 0.977518 | 0.960204 | 1542 | tags=30% | list=10% | signal=27% |
| WP_IL9_SIGNALING_PATHWAY | 22 | -0.21215 | -0.49947 | 0.956332 | 0.977518 | 0.960204 | 3558 | tags=36% | list=23% | signal=28% |
| REACTOME_CLASS_C_3_METABOTROPIC_GLUTAMATE_PHEROMON | 11 | 0.248369 | 0.491699 | 0.955598 | 0.977518 | 0.960204 | 1922 | tags=18% | list=12% | signal=16% |
| REACTOME_RHO_GTPASES_ACTIVATE_RHOTEKIN_AND_RHOPHILINS | 6 | -0.27517 | -0.51894 | 0.961145 | 0.980275 | 0.962912 | 5812 | tags=67% | list=37% | signal=42% |
| REACTOME_HEME_BIOSYNTHESIS | 12 | -0.23447 | -0.50588 | 0.960825 | 0.980275 | 0.962912 | 3762 | tags=33% | list=24% | signal=25% |
| REACTOME_INTERLEUKIN_2_SIGNALING | 8 | 0.247367 | 0.482161 | 0.96008 | 0.980275 | 0.962912 | 3591 | tags=25% | list=23% | signal=19% |
| WP_BIOGENIC_AMINE_SYNTHESIS | 11 | 0.242128 | 0.479344 | 0.96139 | 0.980275 | 0.962912 | 1312 | tags=9% | list=8% | signal=8% |
| REACTOME_VITAMIN_B2_RIBOFLAVIN_METABOLISM | 6 | 0.257567 | 0.47549 | 0.966862 | 0.982838 | 0.965429 | 3855 | tags=33% | list=25% | signal=25% |
| REACTOME_SYNTHESIS_OF_IP3_AND_IP4_IN_THE_CYTOSOL | 27 | 0.220237 | 0.473644 | 0.966549 | 0.982838 | 0.965429 | 2057 | tags=19% | list=13% | signal=16% |
| WP_NODLIKE_RECEPTOR_NLR_SIGNALING_PATHWAY | 9 | 0.234839 | 0.457354 | 0.966797 | 0.982838 | 0.965429 | 4314 | tags=33% | list=27% | signal=24% |
| REACTOME_DAP12_INTERACTIONS | 26 | 0.205598 | 0.443607 | 0.964789 | 0.982838 | 0.965429 | 5521 | tags=50% | list=35% | signal=32% |
| BIOCARTA_CARDIACEGF_PATHWAY | 18 | 0.20808 | 0.426673 | 0.965974 | 0.982838 | 0.965429 | 5003 | tags=44% | list=32% | signal=30% |
| WP_IL1_SIGNALING_PATHWAY | 34 | -0.20644 | -0.51753 | 0.969767 | 0.985189 | 0.967738 | 4045 | tags=32% | list=26% | signal=24% |
| REACTOME_SYNTHESIS_OF_PIPS_AT_THE_ER_MEMBRANE | 5 | -0.26063 | -0.47768 | 0.971831 | 0.986279 | 0.968809 | 1847 | tags=40% | list=12% | signal=35% |
| REACTOME_HDR_THROUGH_SINGLE_STRAND_ANNEALING_SSA | 34 | 0.209192 | 0.461001 | 0.972028 | 0.986279 | 0.968809 | 6388 | tags=53% | list=41% | signal=31% |
| REACTOME_PROCESSIVE_SYNTHESIS_ON_THE_LAGGING_STRAND | 14 | -0.20572 | -0.46486 | 0.973523 | 0.987193 | 0.969707 | 12478 | tags=43% | list=79% | signal=9% |
| REACTOME_HISTIDINE_CATABOLISM | 3 | 0.307636 | 0.496195 | 0.97446 | 0.98754 | 0.970048 | 5257 | tags=67% | list=33% | signal=44% |
| REACTOME_DAP12_SIGNALING | 24 | -0.19462 | -0.46622 | 0.975391 | 0.987881 | 0.970383 | 3257 | tags=21% | list=21% | signal=17% |
| REACTOME_EGFR_INTERACTS_WITH_PHOSPHOLIPASE_C_GAMMA | 5 | 0.275628 | 0.493843 | 0.976238 | 0.988136 | 0.970633 | 4496 | tags=40% | list=29% | signal=29% |
| REACTOME_GENERIC_TRANSCRIPTION_PATHWAY | 729 | 0.245956 | 0.74193 | 0.977893 | 0.988759 | 0.971245 | 1882 | tags=12% | list=12% | signal=11% |
| REACTOME_CDC6_ASSOCIATION_WITH_THE_ORC_ORIGIN_COMPLE | 7 | 0.234807 | 0.445108 | 0.978044 | 0.988759 | 0.971245 | 12019 | tags=100% | list=77% | signal=23% |
| REACTOME_PROLACTIN_RECEPTOR_SIGNALING | 7 | 0.233289 | 0.442231 | 0.98004 | 0.990174 | 0.972635 | 4464 | tags=43% | list=28% | signal=31% |
| REACTOME_ANTIVIRAL_MECHANISM_BY_IFN_STIMULATED_GENES | 29 | 0.19546 | 0.424949 | 0.980903 | 0.990443 | 0.9729 | 724 | tags=7% | list=5% | signal=7% |
| BIOCARTA_ATRBRCA_PATHWAY | 18 | -0.19226 | -0.44402 | 0.983087 | 0.992046 | 0.974474 | 12690 | tags=100% | list=81% | signal=19% |
| BIOCARTA_RNA_PATHWAY | 9 | 0.223289 | 0.434861 | 0.984375 | 0.992742 | 0.975158 | 4576 | tags=33% | list=29% | signal=24% |
| REACTOME_DIGESTION_OF_DIETARY_CARBOHYDRATE | 3 | -0.29168 | -0.47009 | 0.985801 | 0.993578 | 0.975978 | 11126 | tags=100% | list=71% | signal=29% |
| WP_APOE_AND_MIR146_IN_INFLAMMATION_AND_ATHEROSCLERC | 8 | -0.24616 | -0.49537 | 0.99002 | 0.995799 | 0.97816 | 11842 | tags=50% | list=75% | signal=12% |
| REACTOME_INTERLEUKIN_21_SIGNALING | 5 | 0.250392 | 0.448627 | 0.990099 | 0.995799 | 0.97816 | 2158 | tags=20% | list=14% | signal=17% |
| REACTOME_LEUKOTRIENE_RECEPTORS | 4 | -0.25495 | -0.44489 | 0.99002 | 0.995799 | 0.97816 | 11703 | tags=50% | list=75% | signal=13% |
| REACTOME_G0_AND_EARLY_G1 | 17 | 0.177093 | 0.3602 | 0.990403 | 0.995799 | 0.97816 | 4911 | tags=35% | list=31% | signal=24% |
| WP_HOMOLOGOUS_RECOMBINATION | 12 | -0.19219 | -0.41466 | 0.991753 | 0.99595 | 0.978309 | 12690 | tags=50% | list=81% | signal=10% |
| REACTOME_RESOLUTION_OF_D_LOOP_STRUCTURES | 29 | 0.174863 | 0.380168 | 0.991319 | 0.99595 | 0.978309 | 5915 | tags=45% | list=38% | signal=28% |
| REACTOME_CROSS_PRESENTATION_OF_PARTICULATE_EXOGENOUS | 7 | -0.20399 | -0.40454 | 0.994012 | 0.99721 | 0.979547 | 4937 | tags=57% | list=31% | signal=39% |
| REACTOME_ENDOSOMAL_VACUOLAR_PATHWAY | 11 | 0.188596 | 0.373365 | 0.994208 | 0.99721 | 0.979547 | 6522 | tags=55% | list=42% | signal=32% |
| WP_MRNA_PROCESSING | 418 | -0.20486 | -0.61052 | 0.99639 | 0.997591 | 0.979921 | 3029 | tags=26% | list=19% | signal=21% |
| REACTOME_ANDROGEN_BIOSYNTHESIS | 4 | 0.243742 | 0.418084 | 0.996016 | 0.997591 | 0.979921 | 11878 | tags=100% | list=76% | signal=24% |
| REACTOME_DEX_H_BOX_HELICASES_ACTIVATE_TYPE_I_IFN_AND_IN | 7 | 0.198277 | 0.375861 | 0.996008 | 0.997591 | 0.979921 | 4576 | tags=29% | list=29% | signal=20% |
| REACTOME_PTK6_PROMOTES_HIF1A_STABILIZATION | 5 | 0.215669 | 0.386414 | 0.99802 | 0.998198 | 0.980517 | 12319 | tags=100% | list=78% | signal=22% |
| REACTOME_HOMOLOGOUS_DNA_PAIRING_AND_STRAND_EXCHAN( | 24 | 0.143855 | 0.307683 | 0.998198 | 0.998198 | 0.980517 | 6235 | tags=46% | list=40% | signal=28% |
